## Supplementary material for "Maleic acid adjuvates BCG to induce CD4^+^CD8^+^ T cells and sterilize tuberculosis": Table S1

PanPep_cluster2

| Peptide | CDR3 | Score |
| --- | --- | --- |
| TQSPALI | ASGELGGSSYEQY | 0.6759249 |
| EFAAAFPQ | ASGELGGSSYEQY | 0.7171448 |
| LLSGNLDVL | ASGELGGSSYEQY | 0.5961815 |
| GPANLGNYNI | ASGELGGSSYEQY | 0.5356473 |
| TLGPTLP | ASGELGGSSYEQY | 0.65381336 |
| VYAYSSCCK | ASGELGGSSYEQY | 0.68268 |
| APAPAPAPNI | ASGELGGSSYEQY | 0.7155036 |
| PGGPVPIP | ASGELGGSSYEQY | 0.6864308 |
| FLTVALEP | ASGELGGSSYEQY | 0.5076953 |
| SQYFGANL | ASGELGGSSYEQY | 0.6648581 |
| GGVKPAA | ASGELGGSSYEQY | 0.6038815 |
| GGTGGAGGAGGW | ASGELGGSSYEQY | 0.6369971 |
| GVGGVAGLGGAGG | ASGELGGSSYEQY | 0.65658027 |
| VLLVLATTI | ASGELGGSSYEQY | 0.631425 |
| IGIFAAML | ASGELGGSSYEQY | 0.62358475 |
| AVGDLID | ASGELGGSSYEQY | 0.57269853 |
| NISPLPL | ASGELGGSSYEQY | 0.6624925 |
| VTAVFVAI | ASGELGGSSYEQY | 0.68347955 |
| SGDDNDAAL | ASGELGGSSYEQY | 0.682304 |
| AVAALAHS | ASGELGGSSYEQY | 0.70183986 |
| GGVTVMATP | ASGELGGSSYEQY | 0.6973531 |
| AGLAANVRD | ASGELGGSSYEQY | 0.6539254 |
| TNRPDLIDPALLRPG | ASGELGGSSYEQY | 0.2780354 |
| LANLGVTAL | ASGELGGSSYEQY | 0.34008244 |
| TGERIEL | ASGELGGSSYEQY | 0.56993234 |
| NGEELAA | ASGELGGSSYEQY | 0.5783057 |
| VLSLVWG | ASGELGGSSYEQY | 0.33822635 |
| IGYALWL | ASGELGGSSYEQY | 0.64419794 |
| LSLLLAPL | ASGELGGSSYEQY | 0.61624056 |
| TLDEALK | ASGELGGSSYEQY | 0.5723225 |
| TDGAGTAEGG | ASGELGGSSYEQY | 0.70076734 |
| NAPGGAGGNGG | ASGELGGSSYEQY | 0.6168668 |
| VSVFMGMGGNF | ASGELGGSSYEQY | 0.49777126 |
| LASYLAAL | ASGELGGSSYEQY | 0.523142 |
| LVGGTPAL | ASGELGGSSYEQY | 0.664073 |
| VRNADVL | ASGELGGSSYEQY | 0.4388168 |
| TCSSPFLVSAM | ASGELGGSSYEQY | 0.67916906 |
| VHITDAE | ASGELGGSSYEQY | 0.46849486 |
| EYAAPSF | ASGELGGSSYEQY | 0.68018174 |
| VVDAFHGGL | ASGELGGSSYEQY | 0.4374906 |
| ASVVDPQ | ASGELGGSSYEQY | 0.6787029 |
| SYNEFGGLS | ASGELGGSSYEQY | 0.63421226 |
| DPDGSVY | ASGELGGSSYEQY | 0.728996 |
| MVSTIHATE | ASGELGGSSYEQY | 0.5880126 |
| LAAGVADPV | ASGELGGSSYEQY | 0.6452134 |
| TKEQIAA | ASGELGGSSYEQY | 0.50820434 |
| SLASPAAL | ASGELGGSSYEQY | 0.7113445 |
| IGLVFYGV | ASGELGGSSYEQY | 0.4676926 |
| FLLDASNL | ASGELGGSSYEQY | 0.6193912 |

| | | |
|---|---|---|
| YQVGNFP | ASGELGGSSYEQY | 0.36777276 |
| ESLLSQST | ASGELGGSSYEQY | 0.65627 |
| LFTVGFAAVG | ASGELGGSSYEQY | 0.6777487 |
| ATPPTPT | ASGELGGSSYEQY | 0.42211998 |
| TDLATLG | ASGELGGSSYEQY | 0.7069544 |
| SAGGTGR | ASGELGGSSYEQY | 0.6196387 |
| GFSNSGS | ASGELGGSSYEQY | 0.543461 |
| GGNGGSGGTSV | ASGELGGSSYEQY | 0.70509124 |
| GTSVATGGAGN | ASGELGGSSYEQY | 0.60881305 |
| NGGNGGSGGTS | ASGELGGSSYEQY | 0.7143199 |
| AVVKFPH | ASGELGGSSYEQY | 0.5976421 |
| TPPPPPP | ASGELGGSSYEQY | 0.60166985 |
| AAPAAAAAPA | ASGELGGSSYEQY | 0.6507252 |
| IALNEDL | ASGELGGSSYEQY | 0.59889513 |
| IVNVSSM | ASGELGGSSYEQY | 0.52656275 |
| TGIPPAPR | ASGELGGSSYEQY | 0.70209295 |
| FDLGGGTF | ASGELGGSSYEQY | 0.8573632 |
| FDLGGGTFDVS | ASGELGGSSYEQY | 0.85335463 |
| NEPTAAA | ASGELGGSSYEQY | 0.69441015 |
| NEPTAAALA | ASGELGGSSYEQY | 0.6887909 |
| GSIINISSI | ASGELGGSSYEQY | 0.7770362 |
| VAPGNAGTA | ASGELGGSSYEQY | 0.8396752 |
| TGGGASLE | ASGELGGSSYEQY | 0.88836294 |
| VAVYNHYKRI | ASGELGGSSYEQY | 0.8355307 |
| TDPAPATT | ASGELGGSSYEQY | 0.67971355 |
| TNRPDILDPALLRPG | ASGELGGSSYEQY | 0.8583324 |
| LTTTTTS | ASGELGGSSYEQY | 0.81862915 |
| PTGSGKTL | ASGELGGSSYEQY | 0.7960743 |
| TGSGKTL | ASGELGGSSYEQY | 0.852952 |
| TIGHVDHGK | ASGELGGSSYEQY | 0.82306206 |
| TIGHVDHGKTT | ASGELGGSSYEQY | 0.7517328 |
| GHVDHGKTT | ASGELGGSSYEQY | 0.7224124 |
| AQFEHTLL | ASGELGGSSYEQY | 0.8513397 |
| IVNQDEL | ASGELGGSSYEQY | 0.87120855 |
| TNGSQFF | ASGELGGSSYEQY | 0.8636012 |
| TGKGIVE | ASGELGGSSYEQY | 0.8796552 |
| LISNASDA | ASGELGGSSYEQY | 0.8673164 |
| AELRTLGELTKIAKAHGAQ | ASGELGGSSYEQY | 0.8462416 |
| APNAVAAAEPSW | ASGELGGSSYEQY | 0.34653744 |
| APQVAGAATAVL | ASGELGGSSYEQY | 0.5630412 |
| ARRDPAAAAAILPTDA | ASGELGGSSYEQY | 0.7167666 |
| DAAARDVLAEAGLAEYF | ASGELGGSSYEQY | 0.7619904 |
| DDQIVLSAKANLAKVEQ | ASGELGGSSYEQY | 0.34099746 |
| DELLPDTGEDSDEDGH | ASGELGGSSYEQY | 0.8182892 |
| ELPEAPANRATVPQD | ASGELGGSSYEQY | 0.80178595 |
| FGQSKIERVTGIG | ASGELGGSSYEQY | 0.7137245 |
| GAGGTGSTAGGAGGAGGAGGLYA | ASGELGGSSYEQY | 0.8698364 |
| GHGGTGAAGGNGGTGGAGGSGIDG | ASGELGGSSYEQY | 0.76415807 |
| GNGGNGGAGGEGGPGGAGGAGA | ASGELGGSSYEQY | 0.6371534 |
| GSTAGNGGNGGNATNGTIAGQPAG | ASGELGGSSYEQY | 0.7616954 |
| GTGGDGGNGGNGGTGG | ASGELGGSSYEQY | 0.8768631 |

| | | |
|---|---|---|
| GVRRPAGIPADAPV | ASGELGGSSYEQY | 0.7066145 |
| GYRQVTDAHLLTLARR | ASGELGGSSYEQY | 0.8795733 |
| IAKMGQDGHDRG | ASGELGGSSYEQY | 0.7336253 |
| KAEPSIEIQYT | ASGELGGSSYEQY | 0.88143754 |
| KPVRIRMAADEDTVAVT | ASGELGGSSYEQY | 0.60490364 |
| MPPRIAGMRLLVIKPEP | ASGELGGSSYEQY | 0.8640988 |
| MKFYAVALFRSRGVPPTR | ASGELGGSSYEQY | 0.8071181 |
| NAGGFGILGAVAHSP | ASGELGGSSYEQY | 0.8992108 |
| NEGREITLAMAEM | ASGELGGSSYEQY | 0.80376434 |
| NELVEWSHPLGHRWRWE | ASGELGGSSYEQY | 0.8694681 |
| NLGAGGTGGNGGNPG | ASGELGGSSYEQY | 0.8382913 |
| NTEWPKWETVMAADPVD | ASGELGGSSYEQY | 0.8768378 |
| PASAAPAETQP | ASGELGGSSYEQY | 0.64356434 |
| QQSSDGKNGVRAAEITGE | ASGELGGSSYEQY | 0.80207616 |
| RAPDAVTTSWWKE | ASGELGGSSYEQY | 0.6438755 |
| SGFGNVGAHTSGWFNQ | ASGELGGSSYEQY | 0.8092553 |
| SGVAGAVGQAASVGGLKV | ASGELGGSSYEQY | 0.84903497 |
| SNGGAGTGSAGGNGGT | ASGELGGSSYEQY | 0.24347113 |
| TGGAGGEGGAAG | ASGELGGSSYEQY | 0.88427365 |
| TGGDGGAAGAGGAPGHG | ASGELGGSSYEQY | 0.55939794 |
| TMPPLLAAIANAGRHGV | ASGELGGSSYEQY | 0.6764305 |
| TSSTAGNSGGA | ASGELGGSSYEQY | 0.8665968 |
| VPATVPVGILC | ASGELGGSSYEQY | 0.8061519 |
| VQRPDSGGVVLTGRIS | ASGELGGSSYEQY | 0.86638737 |
| WTPNPVPATKGVPAGLVQAT | ASGELGGSSYEQY | 0.7577207 |
| YALKRALAKFSDRFDVVII | ASGELGGSSYEQY | 0.8282973 |
| YLIGSVIV | ASGELGGSSYEQY | 0.82881963 |
| YGQLVAVID | ASGELGGSSYEQY | 0.5144826 |
| VTVGGMTLPA | ASGELGGSSYEQY | 0.62889177 |
| VLALPAV | ASGELGGSSYEQY | 0.8806102 |
| VIPPTLNYET | ASGELGGSSYEQY | 0.64844 |
| VGMGGMHPSSG | ASGELGGSSYEQY | 0.6910795 |
| VGLAGGAATA | ASGELGGSSYEQY | 0.59306425 |
| VGGAGGAGGSGP | ASGELGGSSYEQY | 0.5262803 |
| VFAAFMV | ASGELGGSSYEQY | 0.5963479 |
| TLYDKLR | ASGELGGSSYEQY | 0.67848015 |
| TLAVFTVL | ASGELGGSSYEQY | 0.69381255 |
| TGSVQPARV | ASGELGGSSYEQY | 0.66803384 |
| TGGGAGSGGT | ASGELGGSSYEQY | 0.48813576 |
| SHLETSL | ASGELGGSSYEQY | 0.67105436 |
| SFTTGMT | ASGELGGSSYEQY | 0.6161414 |
| QVEVAKRMGVRQPTVS | ASGELGGSSYEQY | 0.6121515 |
| QGSGGPALAM | ASGELGGSSYEQY | 0.6364397 |
| MLAPSNP | ASGELGGSSYEQY | 0.6408221 |
| LVNNAAI | ASGELGGSSYEQY | 0.5758089 |
| LTGEINTLA | ASGELGGSSYEQY | 0.54536414 |
| LSIFGPV | ASGELGGSSYEQY | 0.6562295 |
| LLALLANL | ASGELGGSSYEQY | 0.67185354 |
| LISNASDALDKLR | ASGELGGSSYEQY | 0.67547274 |
| LINEAVP | ASGELGGSSYEQY | 0.7023078 |
| LGYAFSGL | ASGELGGSSYEQY | 0.68538296 |

| | | |
|---|---|---|
| LDFVAVAL | ASGELGGSSYEQY | 0.67660487 |
| LAQPGAE | ASGELGGSSYEQY | 0.34518862 |
| ISNLGAVTGF | ASGELGGSSYEQY | 0.42152628 |
| HQEDPGRI | ASGELGGSSYEQY | 0.5612578 |
| GTPGEPTP | ASGELGGSSYEQY | 0.5987636 |
| GFFNSGS | ASGELGGSSYEQY | 0.36343184 |
| GAGGGGLTAGAN | ASGELGGSSYEQY | 0.6470556 |
| FVDALVNM | ASGELGGSSYEQY | 0.5832022 |
| FLTVAPEL | ASGELGGSSYEQY | 0.55627215 |
| FETLTAL | ASGELGGSSYEQY | 0.682171 |
| DVINPIEL | ASGELGGSSYEQY | 0.5888402 |
| APVSGAPV | ASGELGGSSYEQY | 0.5254615 |
| ALVGGLAW | ASGELGGSSYEQY | 0.5330229 |
| ALAAAAATAI | ASGELGGSSYEQY | 0.66712505 |
| AFGLGLLL | ASGELGGSSYEQY | 0.4660921 |
| AFFAAASL | ASGELGGSSYEQY | 0.6714042 |
| AAAALPAFSPP | ASGELGGSSYEQY | 0.4890643 |
| VVLAPAL | ASGELGGSSYEQY | 0.68955326 |
| TIGHVDHGKTTL | ASGELGGSSYEQY | 0.45459086 |
| LISNASDALDK | ASGELGGSSYEQY | 0.67709094 |
| LISNASDAL | ASGELGGSSYEQY | 0.6247588 |
| ISNASDALDK | ASGELGGSSYEQY | 0.7159968 |
| NQAETLVYQTE | ASGELGGSSYEQY | 0.6002543 |
| TGGTGGTAGAG | ASGELGGSSYEQY | 0.6451512 |
| TGSTAGGAGGA | ASGELGGSSYEQY | 0.5585809 |
| DPLAVPVAAGPV | ASGELGGSSYEQY | 0.6859846 |
| GGTGGTPGNGGN | ASGELGGSSYEQY | 0.42348403 |
| PAGLAGVDPATL | ASGELGGSSYEQY | 0.6062934 |
| GNNLVVEAPHAGASV | ASGELGGSSYEQY | 0.44670463 |
| KVTRIMSASTLEQPA | ASGELGGSSYEQY | 0.64628786 |
| LGFALPIALPRRTVV | ASGELGGSSYEQY | 0.66370046 |
| TRLGNLTARKLVVLV | ASGELGGSSYEQY | 0.3899907 |
| WSQQRRLIYFPSAGP | ASGELGGSSYEQY | 0.6767258 |
| GPNGNPDPMAAAVDIR | ASGELGGSSYEQY | 0.57752836 |
| TQRCVSVADLRAAASR | ASGELGGSSYEQY | 0.54105425 |
| QFVLDVAEHPVEVLTETV | ASGELGGSSYEQY | 0.692176 |
| GGAGGTGGAGGNGGTQAGDG | ASGELGGSSYEQY | 0.649912 |
| PLLATLRAIAVDVDAKPAQV | ASGELGGSSYEQY | 0.68428695 |
| APKFRTGGYMIDFWGVGYQVAKRM | ASGELGGSSYEQY | 0.5796566 |
| GAGGAGGAGGAAGGPGGTGGTGGN | ASGELGGSSYEQY | 0.58665407 |
| IGALGLLATDVAPAGSVAFGVALA | ASGELGGSSYEQY | 0.64761215 |
| AGGAGGIGGNGGAGGAGGVLMGNG | ASGELGGSSYEQY | 0.59597445 |
| ADELIGLL | ASGELGGSSYEQY | 0.54887456 |
| ADRALKA | ASGELGGSSYEQY | 0.6899879 |
| AGAVGQVGGAGGA | ASGELGGSSYEQY | 0.78730106 |
| AGGGGGLGGAP | ASGELGGSSYEQY | 0.79730946 |
| AGGLLYGNGGAG | ASGELGGSSYEQY | 0.72569454 |
| AGLESIGL | ASGELGGSSYEQY | 0.7396352 |
| AGLKTGII | ASGELGGSSYEQY | 0.7601129 |
| ASLAALLL | ASGELGGSSYEQY | 0.80076325 |
| ASVINISSI | ASGELGGSSYEQY | 0.7959901 |

| | | |
|---|---|---|
| ATPPGPA | ASGELGGSSYEQY | 0.7933221 |
| ATVASALV | ASGELGGSSYEQY | 0.74678916 |
| ATVASLIL | ASGELGGSSYEQY | 0.78384745 |
| AVNGPVLIRH | ASGELGGSSYEQY | 0.7636118 |
| DGAAGAVGI | ASGELGGSSYEQY | 0.77023304 |
| DPLLHGI | ASGELGGSSYEQY | 0.7888522 |
| ELGAALGAL | ASGELGGSSYEQY | 0.78237504 |
| ELGRTQAV | ASGELGGSSYEQY | 0.7606105 |
| FVAGGYLS | ASGELGGSSYEQY | 0.7613349 |
| GASGGAGGAGGW | ASGELGGSSYEQY | 0.7823928 |
| GGAGAGGDGGAGGI | ASGELGGSSYEQY | 0.79277 |
| GTGHNGSQPGMGGNG | ASGELGGSSYEQY | 0.7862972 |
| GYAAASASAS | ASGELGGSSYEQY | 0.8040296 |
| HQVDVAAVQQG | ASGELGGSSYEQY | 0.7916132 |
| IGAAIAHT | ASGELGGSSYEQY | 0.7844068 |
| IGMIEVAQL | ASGELGGSSYEQY | 0.4618693 |
| IRLAPEHTSAMD | ASGELGGSSYEQY | 0.6938093 |
| IYNVSVL | ASGELGGSSYEQY | 0.7346697 |
| KQGFLGM | ASGELGGSSYEQY | 0.7698945 |
| LESVITQL | ASGELGGSSYEQY | 0.4302985 |
| LGLGALVVV | ASGELGGSSYEQY | 0.79353964 |
| LIGNGGNGGSGGI | ASGELGGSSYEQY | 0.77154315 |
| LNAINAPTL | ASGELGGSSYEQY | 0.73371613 |
| LQLLNTGV | ASGELGGSSYEQY | 0.79066706 |
| LQTSIEF | ASGELGGSSYEQY | 0.75669503 |
| MSFAVSEL | ASGELGGSSYEQY | 0.7423243 |
| NTGALGL | ASGELGGSSYEQY | 0.7701225 |
| PPGAGGAGGAGTT | ASGELGGSSYEQY | 0.7964246 |
| QLVGLGI | ASGELGGSSYEQY | 0.6986567 |
| RFLEQVHQL | ASGELGGSSYEQY | 0.7953477 |
| RGVGYVLR | ASGELGGSSYEQY | 0.77940536 |
| SATGVFAGVI | ASGELGGSSYEQY | 0.7998049 |
| SLPAITLP | ASGELGGSSYEQY | 0.71414983 |
| STALAFTV | ASGELGGSSYEQY | 0.7925686 |
| SVLTVGLGK | ASGELGGSSYEQY | 0.7750815 |
| TALAGAGSGI | ASGELGGSSYEQY | 0.8053682 |
| TGLPFTGI | ASGELGGSSYEQY | 0.77038455 |
| VAEHVTV | ASGELGGSSYEQY | 0.7938493 |
| VDIVLNSL | ASGELGGSSYEQY | 0.78760505 |
| VLTGGGAVL | ASGELGGSSYEQY | 0.79073274 |
| VNVGLAYGI | ASGELGGSSYEQY | 0.67696846 |
| VSSAAVGAV | ASGELGGSSYEQY | 0.7670547 |
| VTSVYEAGV | ASGELGGSSYEQY | 0.7239281 |
| WVALDRGA | ASGELGGSSYEQY | 0.7808776 |
| YLVGSVVI | ASGELGGSSYEQY | 0.7931581 |
| GPPGTGKTLL | ASGELGGSSYEQY | 0.51593816 |
| GVGMTKF | ASGELGGSSYEQY | 0.7891644 |
| IGIGGSDL | ASGELGGSSYEQY | 0.74704146 |
| IGTLLAEL | ASGELGGSSYEQY | 0.76066375 |
| IYTSGTT | ASGELGGSSYEQY | 0.79363066 |
| LSFPAGV | ASGELGGSSYEQY | 0.76344335 |

| | | |
|---|---|---|
| LYTSGTT | ASGELGGSSYEQY | 0.78939575 |
| VGAFPFGF | ASGELGGSSYEQY | 0.75712514 |
| VGAGTVEF | ASGELGGSSYEQY | 0.77096534 |
| VGKSTLLL | ASGELGGSSYEQY | 0.78649104 |
| AALAHAGGGIV | ASGELGGSSYEQY | 0.57206494 |
| AALVAVSLL | ASGELGGSSYEQY | 0.73241067 |
| AAVSRAVAAL | ASGELGGSSYEQY | 0.8009317 |
| AGGSSGIGGF | ASGELGGSSYEQY | 0.69741756 |
| AGLTVANANG | ASGELGGSSYEQY | 0.7306535 |
| AGLVGAVL | ASGELGGSSYEQY | 0.57444966 |
| ALSQAFTV | ASGELGGSSYEQY | 0.5330366 |
| ALTFSVSL | ASGELGGSSYEQY | 0.6028844 |
| ALTGAALVAL | ASGELGGSSYEQY | 0.72855234 |
| ASFSIGAGVL | ASGELGGSSYEQY | 0.7467778 |
| DGSVYKI | ASGELGGSSYEQY | 0.70136905 |
| EQLAEIV | ASGELGGSSYEQY | 0.58439577 |
| FAVSELTVM | ASGELGGSSYEQY | 0.66704655 |
| GAAPLAAGL | ASGELGGSSYEQY | 0.6163825 |
| GADGGDGGSGGAGG | ASGELGGSSYEQY | 0.63319427 |
| GFGLDTGGAGG | ASGELGGSSYEQY | 0.69708014 |
| GLASGGLL | ASGELGGSSYEQY | 0.66806555 |
| GRIAGAAVLL | ASGELGGSSYEQY | 0.59192467 |
| GSAGSLNTSF | ASGELGGSSYEQY | 0.6040712 |
| GTGQIGI | ASGELGGSSYEQY | 0.6534003 |
| GTLIALIL | ASGELGGSSYEQY | 0.7099682 |
| IAATMGFVVG | ASGELGGSSYEQY | 0.6947212 |
| IFAQALL | ASGELGGSSYEQY | 0.7534167 |
| IGAAVALGF | ASGELGGSSYEQY | 0.71818215 |
| IGLTPLAL | ASGELGGSSYEQY | 0.72866964 |
| ILGSNTSTL | ASGELGGSSYEQY | 0.27640945 |
| IVFSLTVL | ASGELGGSSYEQY | 0.40580684 |
| LASKATGF | ASGELGGSSYEQY | 0.64430594 |
| LASVALIL | ASGELGGSSYEQY | 0.63170916 |
| LFAVGLTL | ASGELGGSSYEQY | 0.34606305 |
| LFNGGIGTYI | ASGELGGSSYEQY | 0.7096013 |
| LTLGAGII | ASGELGGSSYEQY | 0.619964 |
| LVAAALGDI | ASGELGGSSYEQY | 0.54484904 |
| NTAFVQLGI | ASGELGGSSYEQY | 0.6983731 |
| QLFSGAEL | ASGELGGSSYEQY | 0.60425735 |
| SAATFITL | ASGELGGSSYEQY | 0.5625294 |
| SNYGLANLGV | ASGELGGSSYEQY | 0.60073394 |
| SYGGAYAVM | ASGELGGSSYEQY | 0.7255254 |
| TTGVGGLTLG | ASGELGGSSYEQY | 0.44828218 |
| TTQTIYI | ASGELGGSSYEQY | 0.70610714 |
| VAAGAGLGAVI | ASGELGGSSYEQY | 0.5569589 |
| VAAVVTLL | ASGELGGSSYEQY | 0.72847307 |
| VGFLGGLI | ASGELGGSSYEQY | 0.49418652 |
| VGVIPILL | ASGELGGSSYEQY | 0.7038097 |
| VIFASSLI | ASGELGGSSYEQY | 0.6314566 |
| VKGGDGGAAAG | ASGELGGSSYEQY | 0.7569674 |
| VTKTFTNI | ASGELGGSSYEQY | 0.61378837 |

| | | |
|---|---|---|
| VVAAAILL | ASGELGGSSYEQY | 0.6901263 |
| YAALDVEL | ASGELGGSSYEQY | 0.64854383 |
| LASVALLL | ASGELGGSSYEQY | 0.7286267 |
| LDAVHRV | ASGELGGSSYEQY | 0.42363173 |
| QGLSSSIF | ASGELGGSSYEQY | 0.62972695 |
| RGTFANIRL | ASGELGGSSYEQY | 0.42494535 |
| RQLWWGHRI | ASGELGGSSYEQY | 0.68282735 |
| YGPPGTGKTLL | ASGELGGSSYEQY | 0.69483995 |
| AALVGAPANVA | ASGELGGSSYEQY | 0.3913956 |
| AATFRRTE | ASGELGGSSYEQY | 0.70840204 |
| AGALGGGI | ASGELGGSSYEQY | 0.6268487 |
| AGGAGGAGGIGAG | ASGELGGSSYEQY | 0.6006105 |
| AGSGALGL | ASGELGGSSYEQY | 0.7283913 |
| AQIADQI | ASGELGGSSYEQY | 0.6304085 |
| ASAVAVTYL | ASGELGGSSYEQY | 0.7282052 |
| ASTVLGGTGG | ASGELGGSSYEQY | 0.61889875 |
| AVGGEPAK | ASGELGGSSYEQY | 0.5942885 |
| CEQGTFGGTC | ASGELGGSSYEQY | 0.7055676 |
| DEGKAPAG | ASGELGGSSYEQY | 0.39419436 |
| DTVVFPAL | ASGELGGSSYEQY | 0.5320619 |
| FEGITMAG | ASGELGGSSYEQY | 0.73062843 |
| FPLVVQD | ASGELGGSSYEQY | 0.6566684 |
| GAFTTPQ | ASGELGGSSYEQY | 0.79712415 |
| GFGGFGTAGGVGGA | ASGELGGSSYEQY | 0.40581167 |
| GFLAQFL | ASGELGGSSYEQY | 0.45877108 |
| GGAGGAGGNAGVA | ASGELGGSSYEQY | 0.63055515 |
| GGVGTAGAGGAGGAGG | ASGELGGSSYEQY | 0.72194266 |
| GNGGGGDAGNAGSGGN | ASGELGGSSYEQY | 0.793216 |
| GTGGAGGAGNDAGS | ASGELGGSSYEQY | 0.69402176 |
| GTTNTGWFNS | ASGELGGSSYEQY | 0.4850543 |
| IFMGPLFTL | ASGELGGSSYEQY | 0.69476867 |
| ILAHAIEAI | ASGELGGSSYEQY | 0.51423174 |
| IRSAIREAL | ASGELGGSSYEQY | 0.61418074 |
| ITFAGGG | ASGELGGSSYEQY | 0.7306266 |
| ITGMSGHQEV | ASGELGGSSYEQY | 0.61156213 |
| IVAGLACTL | ASGELGGSSYEQY | 0.46957412 |
| IVGGGLAGA | ASGELGGSSYEQY | 0.5261656 |
| IVTGVLNI | ASGELGGSSYEQY | 0.74153364 |
| LAQQGCI | ASGELGGSSYEQY | 0.7575454 |
| LCLHVEN | ASGELGGSSYEQY | 0.741547 |
| LDAISDGS | ASGELGGSSYEQY | 0.8014767 |
| LGGFAAML | ASGELGGSSYEQY | 0.7331995 |
| LGGKNPMI | ASGELGGSSYEQY | 0.68892217 |
| LGPKGVAAIL | ASGELGGSSYEQY | 0.2490146 |
| LMVGVSI | ASGELGGSSYEQY | 0.3890002 |
| LVGAAFVL | ASGELGGSSYEQY | 0.49505574 |
| LVLAVAPL | ASGELGGSSYEQY | 0.6486236 |
| MAAIAVIDAA | ASGELGGSSYEQY | 0.24088608 |
| NNGINAELQ | ASGELGGSSYEQY | 0.7242155 |
| RGEPLVLVHG | ASGELGGSSYEQY | 0.54881144 |
| RGGAGGAAGLI | ASGELGGSSYEQY | 0.45536083 |

| | | |
|---|---|---|
| RSYRMHPAV | ASGELGGSSYEQY | 0.7772026 |
| SAAHLLAGQLA | ASGELGGSSYEQY | 0.49028075 |
| SAAVGSY | ASGELGGSSYEQY | 0.46751362 |
| SPHAVLHRI | ASGELGGSSYEQY | 0.52466273 |
| SQIAKVL | ASGELGGSSYEQY | 0.79532874 |
| TALGHLVP | ASGELGGSSYEQY | 0.37952465 |
| TAPTTSGAA | ASGELGGSSYEQY | 0.73794127 |
| TDGDDGNGGSPGT | ASGELGGSSYEQY | 0.6715369 |
| TDGLGEN | ASGELGGSSYEQY | 0.7562025 |
| TGGPPLPGQA | ASGELGGSSYEQY | 0.36092317 |
| TIPAITI | ASGELGGSSYEQY | 0.7226426 |
| TIPGFTI | ASGELGGSSYEQY | 0.5701927 |
| TSIASGALA | ASGELGGSSYEQY | 0.8187829 |
| TVTLPGAAI | ASGELGGSSYEQY | 0.5888723 |
| VASGLSAALP | ASGELGGSSYEQY | 0.73568845 |
| VGAGVIGI | ASGELGGSSYEQY | 0.67671394 |
| VGPKRFGV | ASGELGGSSYEQY | 0.7792523 |
| VNVAFPD | ASGELGGSSYEQY | 0.35396966 |
| VSMTNSGVS | ASGELGGSSYEQY | 0.5769682 |
| VTDVLGDLSV | ASGELGGSSYEQY | 0.49045765 |
| VVVVGAGFAGL | ASGELGGSSYEQY | 0.6300752 |
| VYTVGTVV | ASGELGGSSYEQY | 0.688336 |
| WTTFSGP | ASGELGGSSYEQY | 0.36050335 |
| YDELVEAF | ASGELGGSSYEQY | 0.76563525 |
| YGTGGAGGNGG | ASGELGGSSYEQY | 0.65873206 |
| YTNVMAV | ASGELGGSSYEQY | 0.446091 |
| FLEQVHQL | ASGELGGSSYEQY | 0.7384335 |
| GLSLLAVAV | ASGELGGSSYEQY | 0.5900092 |
| LEQVHQL | ASGELGGSSYEQY | 0.70194566 |
| LNAAGAI | ASGELGGSSYEQY | 0.48642173 |
| PAVHASL | ASGELGGSSYEQY | 0.6578998 |
| TELEDDA | ASGELGGSSYEQY | 0.6841085 |
| TPPGPPN | ASGELGGSSYEQY | 0.42970714 |
| TPPPPPPP | ASGELGGSSYEQY | 0.44581735 |
| TQSPALI | AWSLGGAETLY | 0.7543733 |
| EFAAAFPQ | AWSLGGAETLY | 0.7170559 |
| LLSGNLDVL | AWSLGGAETLY | 0.7348478 |
| GPANLGNYNI | AWSLGGAETLY | 0.60151136 |
| TLGPTLP | AWSLGGAETLY | 0.59081185 |
| VYAYSSCCK | AWSLGGAETLY | 0.64553237 |
| APAPAPAPNI | AWSLGGAETLY | 0.7145263 |
| PGGPVPIP | AWSLGGAETLY | 0.73492265 |
| FLTVALEP | AWSLGGAETLY | 0.7056658 |
| SQYFGANL | AWSLGGAETLY | 0.5328977 |
| GGVKPAA | AWSLGGAETLY | 0.6860498 |
| GGTGGAGGAGGW | AWSLGGAETLY | 0.66669786 |
| GVGGVAGLGGAGG | AWSLGGAETLY | 0.69608605 |
| VLLVLATTI | AWSLGGAETLY | 0.64418626 |
| IGIFAAML | AWSLGGAETLY | 0.6704186 |
| AVGDLID | AWSLGGAETLY | 0.5427755 |
| NISPLPL | AWSLGGAETLY | 0.5038372 |

| | | |
|---|---|---|
| VTAVFVAI | AWSLGGAETLY | 0.6982434 |
| SGDDNDAAL | AWSLGGAETLY | 0.72823566 |
| AVAALAHS | AWSLGGAETLY | 0.7217651 |
| GGVTVMATP | AWSLGGAETLY | 0.7214037 |
| AGLAANVRD | AWSLGGAETLY | 0.7315917 |
| TNRPDLIDPALLRPG | AWSLGGAETLY | 0.6543316 |
| LANLGVTAL | AWSLGGAETLY | 0.32853293 |
| TGERIEL | AWSLGGAETLY | 0.418128 |
| NGEELAA | AWSLGGAETLY | 0.47361428 |
| VLSLVWG | AWSLGGAETLY | 0.65322447 |
| IGYALWL | AWSLGGAETLY | 0.25505278 |
| LSLLLAPL | AWSLGGAETLY | 0.6650482 |
| TLDEALK | AWSLGGAETLY | 0.5655662 |
| TDGAGTAEGG | AWSLGGAETLY | 0.5869409 |
| NAPGGAGGNGG | AWSLGGAETLY | 0.7227955 |
| VSVFMGMGGNF | AWSLGGAETLY | 0.65618885 |
| LASYLAAL | AWSLGGAETLY | 0.4775341 |
| LVGGTPAL | AWSLGGAETLY | 0.46357882 |
| VRNADVL | AWSLGGAETLY | 0.6803169 |
| TCSSPFLVSAM | AWSLGGAETLY | 0.42355335 |
| VHITDAE | AWSLGGAETLY | 0.7334179 |
| EYAAPSF | AWSLGGAETLY | 0.39062643 |
| VVDAFHGGL | AWSLGGAETLY | 0.7354697 |
| ASVVDPQ | AWSLGGAETLY | 0.4953081 |
| SYNEFGGLS | AWSLGGAETLY | 0.72371995 |
| DPDGSVY | AWSLGGAETLY | 0.709913 |
| MVSTIHATE | AWSLGGAETLY | 0.7483419 |
| LAAGVADPV | AWSLGGAETLY | 0.6745479 |
| TKEQIAA | AWSLGGAETLY | 0.72056174 |
| SLASPAAL | AWSLGGAETLY | 0.6719129 |
| IGLVFYGV | AWSLGGAETLY | 0.74138725 |
| FLLDASNL | AWSLGGAETLY | 0.52310187 |
| YQVGNFP | AWSLGGAETLY | 0.6549805 |
| ESLLSQST | AWSLGGAETLY | 0.347277 |
| LFTVGFAAVG | AWSLGGAETLY | 0.65861905 |
| ATPPTPT | AWSLGGAETLY | 0.71817905 |
| TDLATLG | AWSLGGAETLY | 0.28167486 |
| SAGGTGR | AWSLGGAETLY | 0.64138275 |
| GFSNSGS | AWSLGGAETLY | 0.57890165 |
| GGNGGSGGTSV | AWSLGGAETLY | 0.5970551 |
| GTSVATGGAGN | AWSLGGAETLY | 0.7044814 |
| NGGNGGSGGTS | AWSLGGAETLY | 0.6736917 |
| AVVKFPH | AWSLGGAETLY | 0.676674 |
| TPPPPPP | AWSLGGAETLY | 0.44986808 |
| AAPAAAAAPA | AWSLGGAETLY | 0.53818583 |
| IALNEDL | AWSLGGAETLY | 0.63656455 |
| IVNVSSM | AWSLGGAETLY | 0.4687038 |
| TGIPPAPR | AWSLGGAETLY | 0.3908596 |
| FDLGGGTF | AWSLGGAETLY | 0.7395636 |
| FDLGGGTFDVS | AWSLGGAETLY | 0.6507982 |
| NEPTAAA | AWSLGGAETLY | 0.78856295 |

| | | |
|---|---|---|
| NEPTAAALA | AWSLGGAETLY | 0.44363618 |
| GSIINISSI | AWSLGGAETLY | 0.42393923 |
| VAPGNAGTA | AWSLGGAETLY | 0.5563357 |
| TGGGASLE | AWSLGGAETLY | 0.7728876 |
| VAVYNHYKRI | AWSLGGAETLY | 0.77189314 |
| TDPAPATT | AWSLGGAETLY | 0.679011 |
| TNRPDILDPALLRPG | AWSLGGAETLY | 0.4532454 |
| LTTTTTS | AWSLGGAETLY | 0.5176933 |
| PTGSGKTL | AWSLGGAETLY | 0.49801487 |
| TGSGKTL | AWSLGGAETLY | 0.5545837 |
| TIGHVDHGK | AWSLGGAETLY | 0.65226007 |
| TIGHVDHGKTT | AWSLGGAETLY | 0.6255274 |
| GHVDHGKTT | AWSLGGAETLY | 0.48294616 |
| AQFEHTLL | AWSLGGAETLY | 0.52312976 |
| IVNQDEL | AWSLGGAETLY | 0.6182555 |
| TNGSQFF | AWSLGGAETLY | 0.65794885 |
| TGKGIVE | AWSLGGAETLY | 0.5543967 |
| LISNASDA | AWSLGGAETLY | 0.800318 |
| AELRTLGELTKIAKAHGAQ | AWSLGGAETLY | 0.6982763 |
| APNAVAAAEPSW | AWSLGGAETLY | 0.68824375 |
| APQVAGAATAVL | AWSLGGAETLY | 0.17878775 |
| ARRDPAAAAAILPTDA | AWSLGGAETLY | 0.24496175 |
| DAAARDVLAEAGLAEYF | AWSLGGAETLY | 0.54511344 |
| DDQIVLSAKANLAKVEQ | AWSLGGAETLY | 0.55169797 |
| DELLPDTGEDSDEDGH | AWSLGGAETLY | 0.26123196 |
| ELPEAPANRATVPQD | AWSLGGAETLY | 0.7168899 |
| FGQSKIERVTGIG | AWSLGGAETLY | 0.51436406 |
| GAGGTGSTAGGAGGAGGAGGLYA | AWSLGGAETLY | 0.463759 |
| GHGGTGAAGGNGGTGGAGGSGIDG | AWSLGGAETLY | 0.6821548 |
| GNGGNGGAGGEGGPGGAGGAGA | AWSLGGAETLY | 0.53186244 |
| GSTAGNGGNGGNATNGTIAGQPAG | AWSLGGAETLY | 0.50926185 |
| GTGGDGGNGGNGGTGG | AWSLGGAETLY | 0.48924035 |
| GVRRPAGIPADAPV | AWSLGGAETLY | 0.6677704 |
| GYRQVTDAHLLTLARR | AWSLGGAETLY | 0.2916657 |
| IAKMGQDGHDRG | AWSLGGAETLY | 0.6355498 |
| KAEPSIEIQYT | AWSLGGAETLY | 0.43160653 |
| KPVRIRMAADEDTVAVT | AWSLGGAETLY | 0.73056096 |
| MPPRIAGMRLLVIKPEP | AWSLGGAETLY | 0.3651142 |
| MKFYAVALFRSRGVPPTR | AWSLGGAETLY | 0.6639485 |
| NAGGFGILGAVAHSP | AWSLGGAETLY | 0.5709059 |
| NEGREITLAMAEM | AWSLGGAETLY | 0.8074824 |
| NELVEWSHPLGHRWRWE | AWSLGGAETLY | 0.5021624 |
| NLGAGGTGGNGGNPG | AWSLGGAETLY | 0.63982654 |
| NTEWPKWETVMAADPVD | AWSLGGAETLY | 0.5738432 |
| PASAAPAETQP | AWSLGGAETLY | 0.7458211 |
| QQSSDGKNGVRAAEITGE | AWSLGGAETLY | 0.40877095 |
| RAPDAVTTSWWKE | AWSLGGAETLY | 0.49654394 |
| SGFGNVGAHTSGWFNQ | AWSLGGAETLY | 0.3027524 |
| SGVAGAVGQAASVGGLKV | AWSLGGAETLY | 0.6443817 |
| SNGGAGTGSAGGNGGT | AWSLGGAETLY | 0.6574705 |
| TGGAGGEGGAAG | AWSLGGAETLY | 0.3693965 |

| | | |
|---|---|---|
| TGGDGGAAGAGGAPGHG | AWSLGGAETLY | 0.60613585 |
| TMPPLLAAIANAGRHGV | AWSLGGAETLY | 0.6699121 |
| TSSTAGNSGGA | AWSLGGAETLY | 0.5102726 |
| VPATVPVGILC | AWSLGGAETLY | 0.7248836 |
| VQRPDSGGVVLTGRIS | AWSLGGAETLY | 0.44939768 |
| WTPNPVPATKGVPAGLVQAT | AWSLGGAETLY | 0.73707867 |
| YALKRALAKFSDRFDVVII | AWSLGGAETLY | 0.5878622 |
| YLIGSVIV | AWSLGGAETLY | 0.39070666 |
| YGQLVAVID | AWSLGGAETLY | 0.6655391 |
| VTVGGMTLPA | AWSLGGAETLY | 0.36577404 |
| VLALPAV | AWSLGGAETLY | 0.42294082 |
| VIPPTLNYET | AWSLGGAETLY | 0.7348631 |
| VGMGGMHPSSG | AWSLGGAETLY | 0.77890265 |
| VGLAGGAATA | AWSLGGAETLY | 0.7958977 |
| VGGAGGAGGSGP | AWSLGGAETLY | 0.6723653 |
| VFAAFMV | AWSLGGAETLY | 0.5674906 |
| TLYDKLR | AWSLGGAETLY | 0.69163525 |
| TLAVFTVL | AWSLGGAETLY | 0.7917924 |
| TGSVQPARV | AWSLGGAETLY | 0.8110999 |
| TGGGAGSGGT | AWSLGGAETLY | 0.7547299 |
| SHLETSL | AWSLGGAETLY | 0.49084663 |
| SFTTGMT | AWSLGGAETLY | 0.771542 |
| QVEVAKRMGVRQPTVS | AWSLGGAETLY | 0.6967282 |
| QGSGGPALAM | AWSLGGAETLY | 0.7286808 |
| MLAPSNP | AWSLGGAETLY | 0.75027597 |
| LVNNAAI | AWSLGGAETLY | 0.77196014 |
| LTGEINTLA | AWSLGGAETLY | 0.6911912 |
| LSIFGPV | AWSLGGAETLY | 0.6411891 |
| LLALLANL | AWSLGGAETLY | 0.74469554 |
| LISNASDALDKLR | AWSLGGAETLY | 0.7990627 |
| LINEAVP | AWSLGGAETLY | 0.7859197 |
| LGYAFSGL | AWSLGGAETLY | 0.81489635 |
| LDFVAVAL | AWSLGGAETLY | 0.81129265 |
| LAQPGAE | AWSLGGAETLY | 0.7959122 |
| ISNLGAVTGF | AWSLGGAETLY | 0.22928753 |
| HQEDPGRI | AWSLGGAETLY | 0.40551358 |
| GTPGEPTP | AWSLGGAETLY | 0.5929482 |
| GFFNSGS | AWSLGGAETLY | 0.70294374 |
| GAGGGGLTAGAN | AWSLGGAETLY | 0.27677238 |
| FVDALVNM | AWSLGGAETLY | 0.7424488 |
| FLTVAPEL | AWSLGGAETLY | 0.6407382 |
| FETLTAL | AWSLGGAETLY | 0.5826436 |
| DVINPIEL | AWSLGGAETLY | 0.80680335 |
| APVSGAPV | AWSLGGAETLY | 0.68828857 |
| ALVGGLAW | AWSLGGAETLY | 0.57104635 |
| ALAAAAATAI | AWSLGGAETLY | 0.5388803 |
| AFGLGLLL | AWSLGGAETLY | 0.80212766 |
| AFFAAASL | AWSLGGAETLY | 0.45385942 |
| AAAALPAFSPP | AWSLGGAETLY | 0.7985841 |
| VVLAPAL | AWSLGGAETLY | 0.5627283 |
| TIGHVDHGKTTL | AWSLGGAETLY | 0.8146547 |

| | | |
|---|---|---|
| LISNASDALDK | AWSLGGAETLY | 0.41303784 |
| LISNASDAL | AWSLGGAETLY | 0.7909936 |
| ISNASDALDK | AWSLGGAETLY | 0.72990036 |
| NQAETLVYQTE | AWSLGGAETLY | 0.84105384 |
| TGGTGGTAGAG | AWSLGGAETLY | 0.7019644 |
| TGSTAGGAGGA | AWSLGGAETLY | 0.742298 |
| DPLAVPVAAGPV | AWSLGGAETLY | 0.6859374 |
| GGTGGTPGNGGN | AWSLGGAETLY | 0.8076118 |
| PAGLAGVDPATL | AWSLGGAETLY | 0.41568792 |
| GNNLVVEAPHAGASV | AWSLGGAETLY | 0.7105004 |
| KVTRIMSASTLEQPA | AWSLGGAETLY | 0.38245454 |
| LGFALPIALPRRTVV | AWSLGGAETLY | 0.74536896 |
| TRLGNLTARKLVVLV | AWSLGGAETLY | 0.7742115 |
| WSQQRRLIYFPSAGP | AWSLGGAETLY | 0.28385875 |
| GPNGNPDPMAAAVDIR | AWSLGGAETLY | 0.79653347 |
| TQRCVSVADLRAAASR | AWSLGGAETLY | 0.65448743 |
| QFVLDVAEHPVEVLTETV | AWSLGGAETLY | 0.546937 |
| GGAGGTGGAGGNGGTQAGDG | AWSLGGAETLY | 0.79743165 |
| PLLATLRAIAVDVDAKPAQV | AWSLGGAETLY | 0.71164894 |
| APKFRTGGYMIDFWGVGYQVAKRM | AWSLGGAETLY | 0.81052566 |
| GAGGAGGAGGAAGGPGGTGGTGGN | AWSLGGAETLY | 0.63752353 |
| IGALGLLATDVAPAGSVAFGVALA | AWSLGGAETLY | 0.6898767 |
| AGGAGGIGGNGGAGGAGGVLMGNG | AWSLGGAETLY | 0.76754403 |
| ADELIGLL | AWSLGGAETLY | 0.34708127 |
| ADRALKA | AWSLGGAETLY | 0.4625926 |
| AGAVGQVGGAGGA | AWSLGGAETLY | 0.8177812 |
| AGGGGGLGGAP | AWSLGGAETLY | 0.5714943 |
| AGGLLYGNGGAG | AWSLGGAETLY | 0.6315521 |
| AGLESIGL | AWSLGGAETLY | 0.5027931 |
| AGLKTGII | AWSLGGAETLY | 0.4630885 |
| ASLAALLL | AWSLGGAETLY | 0.54045475 |
| ASVINISSI | AWSLGGAETLY | 0.6190168 |
| ATPPGPA | AWSLGGAETLY | 0.6173167 |
| ATVASALV | AWSLGGAETLY | 0.5948849 |
| ATVASLIL | AWSLGGAETLY | 0.47229177 |
| AVNGPVLIRH | AWSLGGAETLY | 0.5594473 |
| DGAAGAVGI | AWSLGGAETLY | 0.50697696 |
| DPLLHGI | AWSLGGAETLY | 0.5405114 |
| ELGAALGAL | AWSLGGAETLY | 0.5821449 |
| ELGRTQAV | AWSLGGAETLY | 0.5767794 |
| FVAGGYLS | AWSLGGAETLY | 0.5442624 |
| GASGGAGGAGGW | AWSLGGAETLY | 0.5236009 |
| GGAGAGGDGGAGGI | AWSLGGAETLY | 0.58667994 |
| GTGHNGSQPGMGGNG | AWSLGGAETLY | 0.58767605 |
| GYAAASASAS | AWSLGGAETLY | 0.5532195 |
| HQVDVAAVQQG | AWSLGGAETLY | 0.64915246 |
| IGAAIAHT | AWSLGGAETLY | 0.60514194 |
| IGMIEVAQL | AWSLGGAETLY | 0.608959 |
| IRLAPEHTSAMD | AWSLGGAETLY | 0.33723825 |
| IYNVSVL | AWSLGGAETLY | 0.40430868 |
| KQGFLGM | AWSLGGAETLY | 0.5369042 |

| | | |
|---|---|---|
| LESVITQL | AWSLGGAETLY | 0.5209091 |
| LGLGALVVV | AWSLGGAETLY | 0.4019623 |
| LIGNGGNGGSGGI | AWSLGGAETLY | 0.59851384 |
| LNAINAPTL | AWSLGGAETLY | 0.5147222 |
| LQLLNTGV | AWSLGGAETLY | 0.4970958 |
| LQTSIEF | AWSLGGAETLY | 0.5984818 |
| MSFAVSEL | AWSLGGAETLY | 0.52927285 |
| NTGALGL | AWSLGGAETLY | 0.4975196 |
| PPGAGGAGGAGTT | AWSLGGAETLY | 0.5109689 |
| QLVGLGI | AWSLGGAETLY | 0.62380636 |
| RFLEQVHQL | AWSLGGAETLY | 0.41044706 |
| RGVGYVLR | AWSLGGAETLY | 0.5837768 |
| SATGVFAGVI | AWSLGGAETLY | 0.5189223 |
| SLPAITLP | AWSLGGAETLY | 0.61666673 |
| STALAFTV | AWSLGGAETLY | 0.4157549 |
| SVLTVGLGK | AWSLGGAETLY | 0.59248036 |
| TALAGAGSGI | AWSLGGAETLY | 0.5278168 |
| TGLPFTGI | AWSLGGAETLY | 0.65694517 |
| VAEHVTV | AWSLGGAETLY | 0.5121318 |
| VDIVLNSL | AWSLGGAETLY | 0.5577484 |
| VLTGGGAVL | AWSLGGAETLY | 0.5294342 |
| VNVGLAYGI | AWSLGGAETLY | 0.6165957 |
| VSSAAVGAV | AWSLGGAETLY | 0.41488835 |
| VTSVYEAGV | AWSLGGAETLY | 0.52201587 |
| WVALDRGA | AWSLGGAETLY | 0.4421568 |
| YLVGSVVI | AWSLGGAETLY | 0.5711353 |
| GPPGTGKTLL | AWSLGGAETLY | 0.58565116 |
| GVGMTKF | AWSLGGAETLY | 0.43365684 |
| IGIGGSDL | AWSLGGAETLY | 0.62931424 |
| IGTLLAEL | AWSLGGAETLY | 0.56152236 |
| IYTSGTT | AWSLGGAETLY | 0.47858012 |
| LSFPAGV | AWSLGGAETLY | 0.62309176 |
| LYTSGTT | AWSLGGAETLY | 0.5532625 |
| VGAFPFGF | AWSLGGAETLY | 0.62569904 |
| VGAGTVEF | AWSLGGAETLY | 0.5597651 |
| VGKSTLLL | AWSLGGAETLY | 0.537275 |
| AALAHAGGGIV | AWSLGGAETLY | 0.60298884 |
| AALVAVSLL | AWSLGGAETLY | 0.5140071 |
| AAVSRAVAAL | AWSLGGAETLY | 0.48793596 |
| AGGSSGIGGF | AWSLGGAETLY | 0.6195993 |
| AGLTVANANG | AWSLGGAETLY | 0.85891676 |
| AGLVGAVL | AWSLGGAETLY | 0.8854948 |
| ALSQAFTV | AWSLGGAETLY | 0.7943381 |
| ALTFSVSL | AWSLGGAETLY | 0.75302446 |
| ALTGAALVAL | AWSLGGAETLY | 0.81575286 |
| ASFSIGAGVL | AWSLGGAETLY | 0.88174754 |
| DGSVYKI | AWSLGGAETLY | 0.876758 |
| EQLAEIV | AWSLGGAETLY | 0.86375296 |
| FAVSELTVM | AWSLGGAETLY | 0.7299158 |
| GAAPLAAGL | AWSLGGAETLY | 0.8464284 |
| GADGGDGGSGGAGG | AWSLGGAETLY | 0.80831254 |

| | | |
|---|---|---|
| GFGLDTGGAGG | AWSLGGAETLY | 0.81474423 |
| GLASGGLL | AWSLGGAETLY | 0.8654416 |
| GRIAGAAVLL | AWSLGGAETLY | 0.8368879 |
| GSAGSLNTSF | AWSLGGAETLY | 0.80536765 |
| GTGQIGI | AWSLGGAETLY | 0.8203517 |
| GTLIALIL | AWSLGGAETLY | 0.83739865 |
| IAATMGFVVG | AWSLGGAETLY | 0.8564521 |
| IFAQALL | AWSLGGAETLY | 0.8435092 |
| IGAAVALGF | AWSLGGAETLY | 0.8960171 |
| IGLTPLAL | AWSLGGAETLY | 0.87743664 |
| ILGSNTSTL | AWSLGGAETLY | 0.8568765 |
| IVFSLTVL | AWSLGGAETLY | 0.37552357 |
| LASKATGF | AWSLGGAETLY | 0.7285737 |
| LASVALIL | AWSLGGAETLY | 0.8267275 |
| LFAVGLTL | AWSLGGAETLY | 0.828925 |
| LFNGGIGTYI | AWSLGGAETLY | 0.4731509 |
| LTLGAGII | AWSLGGAETLY | 0.86655486 |
| LVAAALGDI | AWSLGGAETLY | 0.81860757 |
| NTAFVQLGI | AWSLGGAETLY | 0.7519189 |
| QLFSGAEL | AWSLGGAETLY | 0.8682306 |
| SAATFITL | AWSLGGAETLY | 0.78371364 |
| SNYGLANLGV | AWSLGGAETLY | 0.7889589 |
| SYGGAYAVM | AWSLGGAETLY | 0.8229777 |
| TTGVGGLTLG | AWSLGGAETLY | 0.8774344 |
| TTQTIYI | AWSLGGAETLY | 0.74259716 |
| VAAGAGLGAVI | AWSLGGAETLY | 0.8701103 |
| VAAVVTLL | AWSLGGAETLY | 0.8289732 |
| VGFLGGLI | AWSLGGAETLY | 0.88461125 |
| VGVIPILL | AWSLGGAETLY | 0.6620824 |
| VIFASSLI | AWSLGGAETLY | 0.87509835 |
| VKGGDGGAAAG | AWSLGGAETLY | 0.83310854 |
| VTKTFTNI | AWSLGGAETLY | 0.8960333 |
| VVAAAILL | AWSLGGAETLY | 0.82083094 |
| YAALDVEL | AWSLGGAETLY | 0.84697497 |
| LASVALLL | AWSLGGAETLY | 0.8222196 |
| LDAVHRV | AWSLGGAETLY | 0.8740994 |
| QGLSSSIF | AWSLGGAETLY | 0.67920053 |
| RGTFANIRL | AWSLGGAETLY | 0.82733625 |
| RQLWWGHRI | AWSLGGAETLY | 0.73183465 |
| YGPPGTGKTLL | AWSLGGAETLY | 0.8528558 |
| AALVGAPANVA | AWSLGGAETLY | 0.8662149 |
| AATFRRTE | AWSLGGAETLY | 0.5815617 |
| AGALGGGI | AWSLGGAETLY | 0.8607426 |
| AGGAGGAGGIGAG | AWSLGGAETLY | 0.81702065 |
| AGSGALGL | AWSLGGAETLY | 0.7525817 |
| AQIADQI | AWSLGGAETLY | 0.8809989 |
| ASAVAVTYL | AWSLGGAETLY | 0.81402266 |
| ASTVLGGTGG | AWSLGGAETLY | 0.8784495 |
| AVGGEPAK | AWSLGGAETLY | 0.82425135 |
| CEQGTFGGTC | AWSLGGAETLY | 0.8211484 |
| DEGKAPAG | AWSLGGAETLY | 0.8687191 |

| | | |
|---|---|---|
| DTVVFPAL | AWSLGGAETLY | 0.70508814 |
| FEGITMAG | AWSLGGAETLY | 0.7795354 |
| FPLVVQD | AWSLGGAETLY | 0.88725996 |
| GAFTTPQ | AWSLGGAETLY | 0.6123185 |
| GFGGFGTAGGVGGA | AWSLGGAETLY | 0.81334186 |
| GFLAQFL | AWSLGGAETLY | 0.25722194 |
| GGAGGAGGNAGVA | AWSLGGAETLY | 0.25141016 |
| GGVGTAGAGGAGGAGG | AWSLGGAETLY | 0.4260937 |
| GNGGGGDAGNAGSGGN | AWSLGGAETLY | 0.7062457 |
| GTGGAGGAGNDAGS | AWSLGGAETLY | 0.8615347 |
| GTTNTGWFNS | AWSLGGAETLY | 0.6798138 |
| IFMGPLFTL | AWSLGGAETLY | 0.2713912 |
| ILAHAIEAI | AWSLGGAETLY | 0.5846808 |
| IRSAIREAL | AWSLGGAETLY | 0.3673743 |
| ITFAGGG | AWSLGGAETLY | 0.4394211 |
| ITGMSGHQEV | AWSLGGAETLY | 0.6258718 |
| IVAGLACTL | AWSLGGAETLY | 0.5433545 |
| IVGGGLAGA | AWSLGGAETLY | 0.37190986 |
| IVTGVLNI | AWSLGGAETLY | 0.29416725 |
| LAQQGCI | AWSLGGAETLY | 0.70037603 |
| LCLHVEN | AWSLGGAETLY | 0.7709515 |
| LDAISDGS | AWSLGGAETLY | 0.72464514 |
| LGGFAAML | AWSLGGAETLY | 0.817299 |
| LGGKNPMI | AWSLGGAETLY | 0.7179448 |
| LGPKGVAAIL | AWSLGGAETLY | 0.7165998 |
| LMVGVSI | AWSLGGAETLY | 0.11557934 |
| LVGAAFVL | AWSLGGAETLY | 0.18569055 |
| LVLAVAPL | AWSLGGAETLY | 0.36078763 |
| MAAIAVIDAA | AWSLGGAETLY | 0.45305854 |
| NNGINAELQ | AWSLGGAETLY | 0.18112113 |
| RGEPLVLVHG | AWSLGGAETLY | 0.68622994 |
| RGGAGGAAGLI | AWSLGGAETLY | 0.3414428 |
| RSYRMHPAV | AWSLGGAETLY | 0.28173727 |
| SAAHLLAGQLA | AWSLGGAETLY | 0.7309544 |
| SAAVGSY | AWSLGGAETLY | 0.32262194 |
| SPHAVLHRI | AWSLGGAETLY | 0.2746715 |
| SQIAKVL | AWSLGGAETLY | 0.37843293 |
| TALGHLVP | AWSLGGAETLY | 0.7148702 |
| TAPTTSGAA | AWSLGGAETLY | 0.22930185 |
| TDGDDGNGGSPGT | AWSLGGAETLY | 0.70597434 |
| TDGLGEN | AWSLGGAETLY | 0.52235544 |
| TGGPPLPGQA | AWSLGGAETLY | 0.70595646 |
| TIPAITI | AWSLGGAETLY | 0.20573449 |
| TIPGFTI | AWSLGGAETLY | 0.7046362 |
| TSIASGALA | AWSLGGAETLY | 0.46290255 |
| TVTLPGAAI | AWSLGGAETLY | 0.8407756 |
| VASGLSAALP | AWSLGGAETLY | 0.379744 |
| VGAGVIGI | AWSLGGAETLY | 0.560941 |
| VGPKRFGV | AWSLGGAETLY | 0.58333373 |
| VNVAFPD | AWSLGGAETLY | 0.81349957 |
| VSMTNSGVS | AWSLGGAETLY | 0.16346262 |

| | | |
|---|---|---|
| VTDVLGDLSV | AWSLGGAETLY | 0.3289979 |
| VVVVGAGFAGL | AWSLGGAETLY | 0.22762905 |
| VYTVGTVV | AWSLGGAETLY | 0.42755002 |
| WTTFSGP | AWSLGGAETLY | 0.64430165 |
| YDELVEAF | AWSLGGAETLY | 0.12858649 |
| YGTGGAGGNGG | AWSLGGAETLY | 0.57506675 |
| YTNVMAV | AWSLGGAETLY | 0.3980727 |
| FLEQVHQL | AWSLGGAETLY | 0.26553965 |
| GLSLLAVAV | AWSLGGAETLY | 0.704328 |
| LEQVHQL | AWSLGGAETLY | 0.56553614 |
| LNAAGAI | AWSLGGAETLY | 0.74481046 |
| PAVHASL | AWSLGGAETLY | 0.49759647 |
| TELEDDA | AWSLGGAETLY | 0.45139915 |
| TPPGPPN | AWSLGGAETLY | 0.46734333 |
| TPPPPPPP | AWSLGGAETLY | 0.23939638 |
| TQSPALI | ASSQVPGQGGEGQNTLY | 0.35362315 |
| EFAAAFPQ | ASSQVPGQGGEGQNTLY | 0.7695546 |
| LLSGNLDVL | ASSQVPGQGGEGQNTLY | 0.53582025 |
| GPANLGNYNI | ASSQVPGQGGEGQNTLY | 0.7175914 |
| TLGPTLP | ASSQVPGQGGEGQNTLY | 0.22816649 |
| VYAYSSCCK | ASSQVPGQGGEGQNTLY | 0.22971755 |
| APAPAPAPNI | ASSQVPGQGGEGQNTLY | 0.37319887 |
| PGGPVPIP | ASSQVPGQGGEGQNTLY | 0.6168556 |
| FLTVALEP | ASSQVPGQGGEGQNTLY | 0.6912917 |
| SQYFGANL | ASSQVPGQGGEGQNTLY | 0.6566492 |
| GGVKPAA | ASSQVPGQGGEGQNTLY | 0.22884218 |
| GGTGGAGGAGGW | ASSQVPGQGGEGQNTLY | 0.37414038 |
| GVGGVAGLGGAGG | ASSQVPGQGGEGQNTLY | 0.248708 |
| VLLVLATTI | ASSQVPGQGGEGQNTLY | 0.30187467 |
| IGIFAAML | ASSQVPGQGGEGQNTLY | 0.40757325 |
| AVGDLID | ASSQVPGQGGEGQNTLY | 0.2598353 |
| NISPLPL | ASSQVPGQGGEGQNTLY | 0.2617308 |
| VTAVFVAI | ASSQVPGQGGEGQNTLY | 0.2367298 |
| SGDDNDAAL | ASSQVPGQGGEGQNTLY | 0.5746635 |
| AVAALAHS | ASSQVPGQGGEGQNTLY | 0.6649606 |
| GGVTVMATP | ASSQVPGQGGEGQNTLY | 0.5285393 |
| AGLAANVRD | ASSQVPGQGGEGQNTLY | 0.7376867 |
| TNRPDLIDPALLRPG | ASSQVPGQGGEGQNTLY | 0.5985345 |
| LANLGVTAL | ASSQVPGQGGEGQNTLY | 0.53265905 |
| TGERIEL | ASSQVPGQGGEGQNTLY | 0.1423712 |
| NGEELAA | ASSQVPGQGGEGQNTLY | 0.1664084 |
| VLSLVWG | ASSQVPGQGGEGQNTLY | 0.20515719 |
| IGYALWL | ASSQVPGQGGEGQNTLY | 0.37328368 |
| LSLLLAPL | ASSQVPGQGGEGQNTLY | 0.14869498 |
| TLDEALK | ASSQVPGQGGEGQNTLY | 0.5065112 |
| TDGAGTAEGG | ASSQVPGQGGEGQNTLY | 0.2900619 |
| NAPGGAGGNGG | ASSQVPGQGGEGQNTLY | 0.20895113 |
| VSVFMGMGGNF | ASSQVPGQGGEGQNTLY | 0.66257274 |
| LASYLAAL | ASSQVPGQGGEGQNTLY | 0.2621417 |
| LVGGTPAL | ASSQVPGQGGEGQNTLY | 0.21492153 |
| VRNADVL | ASSQVPGQGGEGQNTLY | 0.24197073 |

| | | |
|---|---|---|
| TCSSPFLVSAM | ASSQVPGQGGEGQNTLY | 0.68891543 |
| VHITDAE | ASSQVPGQGGEGQNTLY | 0.17328438 |
| EYAAPSF | ASSQVPGQGGEGQNTLY | 0.51816857 |
| VVDAFHGGL | ASSQVPGQGGEGQNTLY | 0.38081372 |
| ASVVDPQ | ASSQVPGQGGEGQNTLY | 0.48198164 |
| SYNEFGGLS | ASSQVPGQGGEGQNTLY | 0.17690653 |
| DPDGSVY | ASSQVPGQGGEGQNTLY | 0.598816 |
| MVSTIHATE | ASSQVPGQGGEGQNTLY | 0.33358192 |
| LAAGVADPV | ASSQVPGQGGEGQNTLY | 0.7827283 |
| TKEQIAA | ASSQVPGQGGEGQNTLY | 0.23911954 |
| SLASPAAL | ASSQVPGQGGEGQNTLY | 0.33432564 |
| IGLVFYGV | ASSQVPGQGGEGQNTLY | 0.25018322 |
| FLLDASNL | ASSQVPGQGGEGQNTLY | 0.5501617 |
| YQVGNFP | ASSQVPGQGGEGQNTLY | 0.16288677 |
| ESLLSQST | ASSQVPGQGGEGQNTLY | 0.25815362 |
| LFTVGFAAVG | ASSQVPGQGGEGQNTLY | 0.18654083 |
| ATPPTPT | ASSQVPGQGGEGQNTLY | 0.35280663 |
| TDLATLG | ASSQVPGQGGEGQNTLY | 0.59473044 |
| SAGGTGR | ASSQVPGQGGEGQNTLY | 0.15281416 |
| GFSNSGS | ASSQVPGQGGEGQNTLY | 0.5837333 |
| GGNGGSGGTSV | ASSQVPGQGGEGQNTLY | 0.26710543 |
| GTSVATGGAGN | ASSQVPGQGGEGQNTLY | 0.21877958 |
| NGGNGGSGGTS | ASSQVPGQGGEGQNTLY | 0.5269263 |
| AVVKFPH | ASSQVPGQGGEGQNTLY | 0.20445934 |
| TPPPPPP | ASSQVPGQGGEGQNTLY | 0.73269844 |
| AAPAAAAAPA | ASSQVPGQGGEGQNTLY | 0.21052277 |
| IALNEDL | ASSQVPGQGGEGQNTLY | 0.42590642 |
| IVNVSSM | ASSQVPGQGGEGQNTLY | 0.45752162 |
| TGIPPAPR | ASSQVPGQGGEGQNTLY | 0.1433429 |
| FDLGGGTF | ASSQVPGQGGEGQNTLY | 0.20473793 |
| FDLGGGTFDVS | ASSQVPGQGGEGQNTLY | 0.66609216 |
| NEPTAAA | ASSQVPGQGGEGQNTLY | 0.74188817 |
| NEPTAAALA | ASSQVPGQGGEGQNTLY | 0.7211041 |
| GSIINISSI | ASSQVPGQGGEGQNTLY | 0.6866056 |
| VAPGNAGTA | ASSQVPGQGGEGQNTLY | 0.6342169 |
| TGGGASLE | ASSQVPGQGGEGQNTLY | 0.6939305 |
| VAVYNHYKRI | ASSQVPGQGGEGQNTLY | 0.7407352 |
| TDPAPATT | ASSQVPGQGGEGQNTLY | 0.75224876 |
| TNRPDILDPALLRPG | ASSQVPGQGGEGQNTLY | 0.7406559 |
| LTTTTTS | ASSQVPGQGGEGQNTLY | 0.6211107 |
| PTGSGKTL | ASSQVPGQGGEGQNTLY | 0.74746203 |
| TGSGKTL | ASSQVPGQGGEGQNTLY | 0.7109902 |
| TIGHVDHGK | ASSQVPGQGGEGQNTLY | 0.71658623 |
| TIGHVDHGKTT | ASSQVPGQGGEGQNTLY | 0.7329822 |
| GHVDHGKTT | ASSQVPGQGGEGQNTLY | 0.7374888 |
| AQFEHTLL | ASSQVPGQGGEGQNTLY | 0.71227396 |
| IVNQDEL | ASSQVPGQGGEGQNTLY | 0.6720841 |
| TNGSQFF | ASSQVPGQGGEGQNTLY | 0.74656963 |
| TGKGIVE | ASSQVPGQGGEGQNTLY | 0.75507855 |
| LISNASDA | ASSQVPGQGGEGQNTLY | 0.74960494 |
| AELRTLGELTKIAKAHGAQ | ASSQVPGQGGEGQNTLY | 0.7650386 |

| | | |
|---|---|---|
| APNAVAAAEPSW | ASSQVPGQGGEGQNTLY | 0.7504807 |
| APQVAGAATAVL | ASSQVPGQGGEGQNTLY | 0.751577 |
| ARRDPAAAAAILPTDA | ASSQVPGQGGEGQNTLY | 0.36414486 |
| DAAARDVLAEAGLAEYF | ASSQVPGQGGEGQNTLY | 0.5905552 |
| DDQIVLSAKANLAKVEQ | ASSQVPGQGGEGQNTLY | 0.66109645 |
| DELLPDTGEDSDEDGH | ASSQVPGQGGEGQNTLY | 0.71525013 |
| ELPEAPANRATVPQD | ASSQVPGQGGEGQNTLY | 0.39148492 |
| FGQSKIERVTGIG | ASSQVPGQGGEGQNTLY | 0.7007332 |
| GAGGTGSTAGGAGGAGGAGGLYA | ASSQVPGQGGEGQNTLY | 0.6992713 |
| GHGGTGAAGGNGGTGGAGGSGIDG | ASSQVPGQGGEGQNTLY | 0.65773773 |
| GNGGNGGAGGEGGPGGAGGAGA | ASSQVPGQGGEGQNTLY | 0.75992954 |
| GSTAGNGGNGGNATNGTIAGQPAG | ASSQVPGQGGEGQNTLY | 0.6988455 |
| GTGGDGGNGGNGGTGG | ASSQVPGQGGEGQNTLY | 0.6444792 |
| GVRRPAGIPADAPV | ASSQVPGQGGEGQNTLY | 0.6817932 |
| GYRQVTDAHLLTLARR | ASSQVPGQGGEGQNTLY | 0.76172817 |
| IAKMGQDGHDRG | ASSQVPGQGGEGQNTLY | 0.59462625 |
| KAEPSIEIQYT | ASSQVPGQGGEGQNTLY | 0.75061667 |
| KPVRIRMAADEDTVAVT | ASSQVPGQGGEGQNTLY | 0.6408479 |
| MPPRIAGMRLLVIKPEP | ASSQVPGQGGEGQNTLY | 0.75555253 |
| MKFYAVALFRSRGVPPTR | ASSQVPGQGGEGQNTLY | 0.5610323 |
| NAGGFGILGAVAHSP | ASSQVPGQGGEGQNTLY | 0.74827087 |
| NEGREITLAMAEM | ASSQVPGQGGEGQNTLY | 0.7090315 |
| NELVEWSHPLGHRWRWE | ASSQVPGQGGEGQNTLY | 0.76765156 |
| NLGAGGTGGNGGNPG | ASSQVPGQGGEGQNTLY | 0.6988675 |
| NTEWPKWETVMAADPVD | ASSQVPGQGGEGQNTLY | 0.746978 |
| PASAAPAETQP | ASSQVPGQGGEGQNTLY | 0.7168462 |
| QQSSDGKNGVRAAEITGE | ASSQVPGQGGEGQNTLY | 0.7347107 |
| RAPDAVTTSWWKE | ASSQVPGQGGEGQNTLY | 0.56694543 |
| SGFGNVGAHTSGWFNQ | ASSQVPGQGGEGQNTLY | 0.7085861 |
| SGVAGAVGQAASVGGLKV | ASSQVPGQGGEGQNTLY | 0.59691024 |
| SNGGAGTGSAGGNGGT | ASSQVPGQGGEGQNTLY | 0.7278558 |
| TGGAGGEGGAAG | ASSQVPGQGGEGQNTLY | 0.7395233 |
| TGGDGGAAGAGGAPGHG | ASSQVPGQGGEGQNTLY | 0.3364006 |
| TMPPLLAAIANAGRHGV | ASSQVPGQGGEGQNTLY | 0.75288016 |
| TSSTAGNSGGA | ASSQVPGQGGEGQNTLY | 0.6437884 |
| VPATVPVGILC | ASSQVPGQGGEGQNTLY | 0.6217249 |
| VQRPDSGGVVLTGRIS | ASSQVPGQGGEGQNTLY | 0.7517011 |
| WTPNPVPATKGVPAGLVQAT | ASSQVPGQGGEGQNTLY | 0.72730803 |
| YALKRALAKFSDRFDVVII | ASSQVPGQGGEGQNTLY | 0.75048625 |
| YLIGSVIV | ASSQVPGQGGEGQNTLY | 0.6981725 |
| YGQLVAVID | ASSQVPGQGGEGQNTLY | 0.70892394 |
| VTVGGMTLPA | ASSQVPGQGGEGQNTLY | 0.7312552 |
| VLALPAV | ASSQVPGQGGEGQNTLY | 0.3826019 |
| VIPPTLNYET | ASSQVPGQGGEGQNTLY | 0.6376947 |
| VGMGGMHPSSG | ASSQVPGQGGEGQNTLY | 0.7574365 |
| VGLAGGAATA | ASSQVPGQGGEGQNTLY | 0.78949845 |
| VGGAGGAGGSGP | ASSQVPGQGGEGQNTLY | 0.80319697 |
| VFAAFMV | ASSQVPGQGGEGQNTLY | 0.678115 |
| TLYDKLR | ASSQVPGQGGEGQNTLY | 0.58304465 |
| TLAVFTVL | ASSQVPGQGGEGQNTLY | 0.72265756 |
| TGSVQPARV | ASSQVPGQGGEGQNTLY | 0.8334147 |

| | | |
|---|---|---|
| TGGGAGSGGT | ASSQVPGQGGEGQNTLY | 0.8277485 |
| SHLETSL | ASSQVPGQGGEGQNTLY | 0.7867265 |
| SFTTGMT | ASSQVPGQGGEGQNTLY | 0.5871645 |
| QVEVAKRMGVRQPTVS | ASSQVPGQGGEGQNTLY | 0.8056035 |
| QGSGGPALAM | ASSQVPGQGGEGQNTLY | 0.7294314 |
| MLAPSNP | ASSQVPGQGGEGQNTLY | 0.7355791 |
| LVNNAAI | ASSQVPGQGGEGQNTLY | 0.8043339 |
| LTGEINTLA | ASSQVPGQGGEGQNTLY | 0.8058868 |
| LSIFGPV | ASSQVPGQGGEGQNTLY | 0.7516209 |
| LLALLANL | ASSQVPGQGGEGQNTLY | 0.68651533 |
| LISNASDALDKLR | ASSQVPGQGGEGQNTLY | 0.7930046 |
| LINEAVP | ASSQVPGQGGEGQNTLY | 0.81374776 |
| LGYAFSGL | ASSQVPGQGGEGQNTLY | 0.808723 |
| LDFVAVAL | ASSQVPGQGGEGQNTLY | 0.85825026 |
| LAQPGAE | ASSQVPGQGGEGQNTLY | 0.8319134 |
| ISNLGAVTGF | ASSQVPGQGGEGQNTLY | 0.8034242 |
| HQEDPGRI | ASSQVPGQGGEGQNTLY | 0.2373516 |
| GTPGEPTP | ASSQVPGQGGEGQNTLY | 0.44487253 |
| GFFNSGS | ASSQVPGQGGEGQNTLY | 0.68051696 |
| GAGGGGLTAGAN | ASSQVPGQGGEGQNTLY | 0.72904944 |
| FVDALVNM | ASSQVPGQGGEGQNTLY | 0.32764482 |
| FLTVAPEL | ASSQVPGQGGEGQNTLY | 0.7907932 |
| FETLTAL | ASSQVPGQGGEGQNTLY | 0.71746624 |
| DVINPIEL | ASSQVPGQGGEGQNTLY | 0.6588409 |
| APVSGAPV | ASSQVPGQGGEGQNTLY | 0.836715 |
| ALVGGLAW | ASSQVPGQGGEGQNTLY | 0.7242187 |
| ALAAAAATAI | ASSQVPGQGGEGQNTLY | 0.6936817 |
| AFGLGLLL | ASSQVPGQGGEGQNTLY | 0.68240833 |
| AFFAAASL | ASSQVPGQGGEGQNTLY | 0.86169755 |
| AAAALPAFSPP | ASSQVPGQGGEGQNTLY | 0.5426024 |
| VVLAPAL | ASSQVPGQGGEGQNTLY | 0.82321316 |
| TIGHVDHGKTTL | ASSQVPGQGGEGQNTLY | 0.6969588 |
| LISNASDALDK | ASSQVPGQGGEGQNTLY | 0.850247 |
| LISNASDAL | ASSQVPGQGGEGQNTLY | 0.500505 |
| ISNASDALDK | ASSQVPGQGGEGQNTLY | 0.817519 |
| NQAETLVYQTE | ASSQVPGQGGEGQNTLY | 0.72862744 |
| TGGTGGTAGAG | ASSQVPGQGGEGQNTLY | 0.87461734 |
| TGSTAGGAGGA | ASSQVPGQGGEGQNTLY | 0.72852993 |
| DPLAVPVAAGPV | ASSQVPGQGGEGQNTLY | 0.79056275 |
| GGTGGTPGNGGN | ASSQVPGQGGEGQNTLY | 0.72756314 |
| PAGLAGVDPATL | ASSQVPGQGGEGQNTLY | 0.80526954 |
| GNNLVVEAPHAGASV | ASSQVPGQGGEGQNTLY | 0.47283024 |
| KVTRIMSASTLEQPA | ASSQVPGQGGEGQNTLY | 0.7492224 |
| LGFALPIALPRRTVV | ASSQVPGQGGEGQNTLY | 0.61189383 |
| TRLGNLTARKLVVLV | ASSQVPGQGGEGQNTLY | 0.78454375 |
| WSQQRRLIYFPSAGP | ASSQVPGQGGEGQNTLY | 0.7900461 |
| GPNGNPDPMAAAVDIR | ASSQVPGQGGEGQNTLY | 0.3586459 |
| TQRCVSVADLRAAASR | ASSQVPGQGGEGQNTLY | 0.8414644 |
| QFVLDVAEHPVEVLTETV | ASSQVPGQGGEGQNTLY | 0.6995429 |
| GGAGGTGGAGGNGGTQAGDG | ASSQVPGQGGEGQNTLY | 0.610828 |
| PLLATLRAIAVDVDAKPAQV | ASSQVPGQGGEGQNTLY | 0.83671385 |

| | | |
|---|---|---|
| APKFRTGGYMIDFWGVGYQVAKRM | ASSQVPGQGGEGQNTLY | 0.7548015 |
| GAGGAGGAGGAAGGPGGTGGTGGN | ASSQVPGQGGEGQNTLY | 0.8381127 |
| IGALGLLATDVAPAGSVAFGVALA | ASSQVPGQGGEGQNTLY | 0.7427553 |
| AGGAGGIGGNGGAGGAGGVLMGNG | ASSQVPGQGGEGQNTLY | 0.7574494 |
| ADELIGLL | ASSQVPGQGGEGQNTLY | 0.8152034 |
| ADRALKA | ASSQVPGQGGEGQNTLY | 0.49810007 |
| AGAVGQVGGAGGA | ASSQVPGQGGEGQNTLY | 0.61037993 |
| AGGGGGLGGAP | ASSQVPGQGGEGQNTLY | 0.8487328 |
| AGGLLYGNGGAG | ASSQVPGQGGEGQNTLY | 0.764146 |
| AGLESIGL | ASSQVPGQGGEGQNTLY | 0.79668045 |
| AGLKTGII | ASSQVPGQGGEGQNTLY | 0.6942625 |
| ASLAALLL | ASSQVPGQGGEGQNTLY | 0.619671 |
| ASVINISSI | ASSQVPGQGGEGQNTLY | 0.7450213 |
| ATPPGPA | ASSQVPGQGGEGQNTLY | 0.7703741 |
| ATVASALV | ASSQVPGQGGEGQNTLY | 0.79635936 |
| ATVASLIL | ASSQVPGQGGEGQNTLY | 0.77389574 |
| AVNGPVLIRH | ASSQVPGQGGEGQNTLY | 0.5606788 |
| DGAAGAVGI | ASSQVPGQGGEGQNTLY | 0.7695232 |
| DPLLHGI | ASSQVPGQGGEGQNTLY | 0.70946395 |
| ELGAALGAL | ASSQVPGQGGEGQNTLY | 0.7360357 |
| ELGRTQAV | ASSQVPGQGGEGQNTLY | 0.7518517 |
| FVAGGYLS | ASSQVPGQGGEGQNTLY | 0.69624376 |
| GASGGAGGAGGW | ASSQVPGQGGEGQNTLY | 0.69941604 |
| GGAGAGGDGGAGGI | ASSQVPGQGGEGQNTLY | 0.686314 |
| GTGHNGSQPGMGGNG | ASSQVPGQGGEGQNTLY | 0.7459364 |
| GYAAASASAS | ASSQVPGQGGEGQNTLY | 0.78088665 |
| HQVDVAAVQQG | ASSQVPGQGGEGQNTLY | 0.77888596 |
| IGAAIAHT | ASSQVPGQGGEGQNTLY | 0.78533316 |
| IGMIEVAQL | ASSQVPGQGGEGQNTLY | 0.77863705 |
| IRLAPEHTSAMD | ASSQVPGQGGEGQNTLY | 0.75842917 |
| IYNVSVL | ASSQVPGQGGEGQNTLY | 0.34476098 |
| KQGFLGM | ASSQVPGQGGEGQNTLY | 0.41173053 |
| LESVITQL | ASSQVPGQGGEGQNTLY | 0.65252644 |
| LGLGALVVV | ASSQVPGQGGEGQNTLY | 0.67475575 |
| LIGNGGNGGSGGI | ASSQVPGQGGEGQNTLY | 0.3303984 |
| LNAINAPTL | ASSQVPGQGGEGQNTLY | 0.7504773 |
| LQLLNTGV | ASSQVPGQGGEGQNTLY | 0.69724125 |
| LQTSIEF | ASSQVPGQGGEGQNTLY | 0.61493206 |
| MSFAVSEL | ASSQVPGQGGEGQNTLY | 0.7871968 |
| NTGALGL | ASSQVPGQGGEGQNTLY | 0.7118434 |
| PPGAGGAGGAGTT | ASSQVPGQGGEGQNTLY | 0.5404367 |
| QLVGLGI | ASSQVPGQGGEGQNTLY | 0.53285587 |
| RFLEQVHQL | ASSQVPGQGGEGQNTLY | 0.72721076 |
| RGVGYVLR | ASSQVPGQGGEGQNTLY | 0.45681095 |
| SATGVFAGVI | ASSQVPGQGGEGQNTLY | 0.7733209 |
| SLPAITLP | ASSQVPGQGGEGQNTLY | 0.53920877 |
| STALAFTV | ASSQVPGQGGEGQNTLY | 0.7687136 |
| SVLTVGLGK | ASSQVPGQGGEGQNTLY | 0.47979057 |
| TALAGAGSGI | ASSQVPGQGGEGQNTLY | 0.7645606 |
| TGLPFTGI | ASSQVPGQGGEGQNTLY | 0.7409129 |
| VAEHVTV | ASSQVPGQGGEGQNTLY | 0.8057263 |

| | | |
|---|---|---|
| VDIVLNSL | ASSQVPGQGGEGQNTLY | 0.7049931 |
| VLTGGGAVL | ASSQVPGQGGEGQNTLY | 0.71781206 |
| VNVGLAYGI | ASSQVPGQGGEGQNTLY | 0.5342998 |
| VSSAAVGAV | ASSQVPGQGGEGQNTLY | 0.79436463 |
| VTSVYEAGV | ASSQVPGQGGEGQNTLY | 0.5626885 |
| WVALDRGA | ASSQVPGQGGEGQNTLY | 0.683608 |
| YLVGSVVI | ASSQVPGQGGEGQNTLY | 0.46253318 |
| GPPGTGKTLL | ASSQVPGQGGEGQNTLY | 0.7398577 |
| GVGMTKF | ASSQVPGQGGEGQNTLY | 0.76628375 |
| IGIGGSDL | ASSQVPGQGGEGQNTLY | 0.47508112 |
| IGTLLAEL | ASSQVPGQGGEGQNTLY | 0.77300775 |
| IYTSGTT | ASSQVPGQGGEGQNTLY | 0.71266115 |
| LSFPAGV | ASSQVPGQGGEGQNTLY | 0.6548095 |
| LYTSGTT | ASSQVPGQGGEGQNTLY | 0.7867379 |
| VGAFPFGF | ASSQVPGQGGEGQNTLY | 0.71862406 |
| VGAGTVEF | ASSQVPGQGGEGQNTLY | 0.79147583 |
| VGKSTLLL | ASSQVPGQGGEGQNTLY | 0.59764934 |
| AALAHAGGGIV | ASSQVPGQGGEGQNTLY | 0.64918727 |
| AALVAVSLL | ASSQVPGQGGEGQNTLY | 0.72514194 |
| AAVSRAVAAL | ASSQVPGQGGEGQNTLY | 0.5326789 |
| AGGSSGIGGF | ASSQVPGQGGEGQNTLY | 0.48882687 |
| AGLTVANANG | ASSQVPGQGGEGQNTLY | 0.7881141 |
| AGLVGAVL | ASSQVPGQGGEGQNTLY | 0.7795154 |
| ALSQAFTV | ASSQVPGQGGEGQNTLY | 0.82000923 |
| ALTFSVSL | ASSQVPGQGGEGQNTLY | 0.72694844 |
| ALTGAALVAL | ASSQVPGQGGEGQNTLY | 0.6604677 |
| ASFSIGAGVL | ASSQVPGQGGEGQNTLY | 0.74579644 |
| DGSVYKI | ASSQVPGQGGEGQNTLY | 0.8090003 |
| EQLAEIV | ASSQVPGQGGEGQNTLY | 0.81459844 |
| FAVSELTVM | ASSQVPGQGGEGQNTLY | 0.791405 |
| GAAPLAAGL | ASSQVPGQGGEGQNTLY | 0.6105856 |
| GADGGDGGSGGAGG | ASSQVPGQGGEGQNTLY | 0.74218297 |
| GFGLDTGGAGG | ASSQVPGQGGEGQNTLY | 0.73793113 |
| GLASGGLL | ASSQVPGQGGEGQNTLY | 0.72314 |
| GRIAGAAVLL | ASSQVPGQGGEGQNTLY | 0.7769494 |
| GSAGSLNTSF | ASSQVPGQGGEGQNTLY | 0.76668084 |
| GTGQIGI | ASSQVPGQGGEGQNTLY | 0.6780504 |
| GTLIALIL | ASSQVPGQGGEGQNTLY | 0.6974021 |
| IAATMGFVVG | ASSQVPGQGGEGQNTLY | 0.7666203 |
| IFAQALL | ASSQVPGQGGEGQNTLY | 0.79593945 |
| IGAAVALGF | ASSQVPGQGGEGQNTLY | 0.7903404 |
| IGLTPLAL | ASSQVPGQGGEGQNTLY | 0.8136127 |
| ILGSNTSTL | ASSQVPGQGGEGQNTLY | 0.8043698 |
| IVFSLTVL | ASSQVPGQGGEGQNTLY | 0.7865821 |
| LASKATGF | ASSQVPGQGGEGQNTLY | 0.24426238 |
| LASVALIL | ASSQVPGQGGEGQNTLY | 0.40077382 |
| LFAVGLTL | ASSQVPGQGGEGQNTLY | 0.71537113 |
| LFNGGIGTYI | ASSQVPGQGGEGQNTLY | 0.7244014 |
| LTLGAGII | ASSQVPGQGGEGQNTLY | 0.47615978 |
| LVAAALGDI | ASSQVPGQGGEGQNTLY | 0.7691942 |
| NTAFVQLGI | ASSQVPGQGGEGQNTLY | 0.7047368 |

| | | |
|---|---|---|
| QLFSGAEL | ASSQVPGQGGEGQNTLY | 0.66774887 |
| SAATFITL | ASSQVPGQGGEGQNTLY | 0.80007267 |
| SNYGLANLGV | ASSQVPGQGGEGQNTLY | 0.72136295 |
| SYGGAYAVM | ASSQVPGQGGEGQNTLY | 0.6628072 |
| TTGVGGLTLG | ASSQVPGQGGEGQNTLY | 0.6410297 |
| TTQTIYI | ASSQVPGQGGEGQNTLY | 0.7950358 |
| VAAGAGLGAVI | ASSQVPGQGGEGQNTLY | 0.47949937 |
| VAAVVTLL | ASSQVPGQGGEGQNTLY | 0.78539443 |
| VGFLGGLI | ASSQVPGQGGEGQNTLY | 0.58970016 |
| VGVIPILL | ASSQVPGQGGEGQNTLY | 0.8074329 |
| VIFASSLI | ASSQVPGQGGEGQNTLY | 0.5294595 |
| VKGGDGGAAAG | ASSQVPGQGGEGQNTLY | 0.792934 |
| VTKTFTNI | ASSQVPGQGGEGQNTLY | 0.7546661 |
| VVAAAILL | ASSQVPGQGGEGQNTLY | 0.826125 |
| YAALDVEL | ASSQVPGQGGEGQNTLY | 0.74040616 |
| LASVALLL | ASSQVPGQGGEGQNTLY | 0.7683166 |
| LDAVHRV | ASSQVPGQGGEGQNTLY | 0.6488496 |
| QGLSSSIF | ASSQVPGQGGEGQNTLY | 0.80967534 |
| RGTFANIRL | ASSQVPGQGGEGQNTLY | 0.6237945 |
| RQLWWGHRI | ASSQVPGQGGEGQNTLY | 0.7458155 |
| YGPPGTGKTLL | ASSQVPGQGGEGQNTLY | 0.4510346 |
| AALVGAPANVA | ASSQVPGQGGEGQNTLY | 0.7799448 |
| AATFRRTE | ASSQVPGQGGEGQNTLY | 0.7874923 |
| AGALGGGI | ASSQVPGQGGEGQNTLY | 0.62242043 |
| AGGAGGAGGIGAG | ASSQVPGQGGEGQNTLY | 0.7983676 |
| AGSGALGL | ASSQVPGQGGEGQNTLY | 0.7782781 |
| AQIADQI | ASSQVPGQGGEGQNTLY | 0.6964415 |
| ASAVAVTYL | ASSQVPGQGGEGQNTLY | 0.8109937 |
| ASTVLGGTGG | ASSQVPGQGGEGQNTLY | 0.7144002 |
| AVGGEPAK | ASSQVPGQGGEGQNTLY | 0.8065703 |
| CEQGTFGGTC | ASSQVPGQGGEGQNTLY | 0.7084347 |
| DEGKAPAG | ASSQVPGQGGEGQNTLY | 0.6982112 |
| DTVVFPAL | ASSQVPGQGGEGQNTLY | 0.7802149 |
| FEGITMAG | ASSQVPGQGGEGQNTLY | 0.7414434 |
| FPLVVQD | ASSQVPGQGGEGQNTLY | 0.64164424 |
| GAFTTPQ | ASSQVPGQGGEGQNTLY | 0.80668044 |
| GFGGFGTAGGVGGA | ASSQVPGQGGEGQNTLY | 0.8385132 |
| GFLAQFL | ASSQVPGQGGEGQNTLY | 0.83471215 |
| GGAGGAGGNAGVA | ASSQVPGQGGEGQNTLY | 0.7879031 |
| GGVGTAGAGGAGGAGG | ASSQVPGQGGEGQNTLY | 0.7472036 |
| GNGGGGDAGNAGSGGN | ASSQVPGQGGEGQNTLY | 0.78371054 |
| GTGGAGGAGNDAGS | ASSQVPGQGGEGQNTLY | 0.82599145 |
| GTTNTGWFNS | ASSQVPGQGGEGQNTLY | 0.849761 |
| IFMGPLFTL | ASSQVPGQGGEGQNTLY | 0.8356924 |
| ILAHAIEAI | ASSQVPGQGGEGQNTLY | 0.67141265 |
| IRSAIREAL | ASSQVPGQGGEGQNTLY | 0.8453171 |
| ITFAGGG | ASSQVPGQGGEGQNTLY | 0.803988 |
| ITGMSGHQEV | ASSQVPGQGGEGQNTLY | 0.80165255 |
| IVAGLACTL | ASSQVPGQGGEGQNTLY | 0.82730305 |
| IVGGGLAGA | ASSQVPGQGGEGQNTLY | 0.8222266 |
| IVTGVLNI | ASSQVPGQGGEGQNTLY | 0.7789384 |

| | | |
|---|---|---|
| LAQQGCI | ASSQVPGQGGEGQNTLY | 0.6859153 |
| LCLHVEN | ASSQVPGQGGEGQNTLY | 0.8171383 |
| LDAISDGS | ASSQVPGQGGEGQNTLY | 0.84635246 |
| LGGFAAML | ASSQVPGQGGEGQNTLY | 0.84538835 |
| LGGKNPMI | ASSQVPGQGGEGQNTLY | 0.84514177 |
| LGPKGVAAIL | ASSQVPGQGGEGQNTLY | 0.8489719 |
| LMVGVSI | ASSQVPGQGGEGQNTLY | 0.83633727 |
| LVGAAFVL | ASSQVPGQGGEGQNTLY | 0.34724498 |
| LVLAVAPL | ASSQVPGQGGEGQNTLY | 0.5582156 |
| MAAIAVIDAA | ASSQVPGQGGEGQNTLY | 0.6757338 |
| NNGINAELQ | ASSQVPGQGGEGQNTLY | 0.7849109 |
| RGEPLVLVHG | ASSQVPGQGGEGQNTLY | 0.30037564 |
| RGGAGGAAGLI | ASSQVPGQGGEGQNTLY | 0.7351448 |
| RSYRMHPAV | ASSQVPGQGGEGQNTLY | 0.7973113 |
| SAAHLLAGQLA | ASSQVPGQGGEGQNTLY | 0.75090295 |
| SAAVGSY | ASSQVPGQGGEGQNTLY | 0.84498006 |
| SPHAVLHRI | ASSQVPGQGGEGQNTLY | 0.7974819 |
| SQIAKVL | ASSQVPGQGGEGQNTLY | 0.6735511 |
| TALGHLVP | ASSQVPGQGGEGQNTLY | 0.72441226 |
| TAPTTSGAA | ASSQVPGQGGEGQNTLY | 0.8378267 |
| TDGDDGNGGSPGT | ASSQVPGQGGEGQNTLY | 0.6394099 |
| TDGLGEN | ASSQVPGQGGEGQNTLY | 0.8491002 |
| TGGPPLPGQA | ASSQVPGQGGEGQNTLY | 0.47171527 |
| TIPAITI | ASSQVPGQGGEGQNTLY | 0.84452975 |
| TIPGFTI | ASSQVPGQGGEGQNTLY | 0.5983546 |
| TSIASGALA | ASSQVPGQGGEGQNTLY | 0.8457448 |
| TVTLPGAAI | ASSQVPGQGGEGQNTLY | 0.82601523 |
| VASGLSAALP | ASSQVPGQGGEGQNTLY | 0.86055225 |
| VGAGVIGI | ASSQVPGQGGEGQNTLY | 0.808453 |
| VGPKRFGV | ASSQVPGQGGEGQNTLY | 0.84044266 |
| VNVAFPD | ASSQVPGQGGEGQNTLY | 0.78015274 |
| VSMTNSGVS | ASSQVPGQGGEGQNTLY | 0.8382292 |
| VTDVLGDLSV | ASSQVPGQGGEGQNTLY | 0.6471081 |
| VVVVGAGFAGL | ASSQVPGQGGEGQNTLY | 0.81185853 |
| VYTVGTVV | ASSQVPGQGGEGQNTLY | 0.5682292 |
| WTTFSGP | ASSQVPGQGGEGQNTLY | 0.8222121 |
| YDELVEAF | ASSQVPGQGGEGQNTLY | 0.8385784 |
| YGTGGAGGNGG | ASSQVPGQGGEGQNTLY | 0.25317022 |
| YTNVMAV | ASSQVPGQGGEGQNTLY | 0.8366461 |
| FLEQVHQL | ASSQVPGQGGEGQNTLY | 0.7164231 |
| GLSLLAVAV | ASSQVPGQGGEGQNTLY | 0.6805556 |
| LEQVHQL | ASSQVPGQGGEGQNTLY | 0.84918827 |
| LNAAGAI | ASSQVPGQGGEGQNTLY | 0.81549925 |
| PAVHASL | ASSQVPGQGGEGQNTLY | 0.8455405 |
| TELEDDA | ASSQVPGQGGEGQNTLY | 0.758129 |
| TPPGPPN | ASSQVPGQGGEGQNTLY | 0.76443124 |
| TPPPPPPP | ASSQVPGQGGEGQNTLY | 0.813234 |
| TQSPALI | ASSQTGDSGNTLY | 0.36786497 |
| EFAAAFPQ | ASSQTGDSGNTLY | 0.6272416 |
| LLSGNLDVL | ASSQTGDSGNTLY | 0.8542174 |
| GPANLGNYNI | ASSQTGDSGNTLY | 0.51655024 |

| | | |
|---|---|---|
| TLGPTLP | ASSQTGDSGNTLY | 0.6447729 |
| VYAYSSCCK | ASSQTGDSGNTLY | 0.43980816 |
| APAPAPAPNI | ASSQTGDSGNTLY | 0.42251077 |
| PGGPVPIP | ASSQTGDSGNTLY | 0.4891985 |
| FLTVALEP | ASSQTGDSGNTLY | 0.57023287 |
| SQYFGANL | ASSQTGDSGNTLY | 0.6460241 |
| GGVKPAA | ASSQTGDSGNTLY | 0.5865379 |
| GGTGGAGGAGGW | ASSQTGDSGNTLY | 0.42901212 |
| GVGGVAGLGGAGG | ASSQTGDSGNTLY | 0.59265596 |
| VLLVLATTI | ASSQTGDSGNTLY | 0.50143206 |
| IGIFAAML | ASSQTGDSGNTLY | 0.4900331 |
| AVGDLID | ASSQTGDSGNTLY | 0.5169224 |
| NISPLPL | ASSQTGDSGNTLY | 0.4759203 |
| VTAVFVAI | ASSQTGDSGNTLY | 0.44914874 |
| SGDDNDAAL | ASSQTGDSGNTLY | 0.42331082 |
| AVAALAHS | ASSQTGDSGNTLY | 0.5528904 |
| GGVTVMATP | ASSQTGDSGNTLY | 0.5732404 |
| AGLAANVRD | ASSQTGDSGNTLY | 0.5601308 |
| TNRPDLIDPALLRPG | ASSQTGDSGNTLY | 0.6206504 |
| LANLGVTAL | ASSQTGDSGNTLY | 0.57430047 |
| TGERIEL | ASSQTGDSGNTLY | 0.46806163 |
| NGEELAA | ASSQTGDSGNTLY | 0.34740087 |
| VLSLVWG | ASSQTGDSGNTLY | 0.387724 |
| IGYALWL | ASSQTGDSGNTLY | 0.46283102 |
| LSLLLAPL | ASSQTGDSGNTLY | 0.5016596 |
| TLDEALK | ASSQTGDSGNTLY | 0.33857802 |
| TDGAGTAEGG | ASSQTGDSGNTLY | 0.58247197 |
| NAPGGAGGNGG | ASSQTGDSGNTLY | 0.49495733 |
| VSVFMGMGGNF | ASSQTGDSGNTLY | 0.47075903 |
| LASYLAAL | ASSQTGDSGNTLY | 0.6071409 |
| LVGGTPAL | ASSQTGDSGNTLY | 0.47373125 |
| VRNADVL | ASSQTGDSGNTLY | 0.41075322 |
| TCSSPFLVSAM | ASSQTGDSGNTLY | 0.43875447 |
| VHITDAE | ASSQTGDSGNTLY | 0.5871494 |
| EYAAPSF | ASSQTGDSGNTLY | 0.3815355 |
| VVDAFHGGL | ASSQTGDSGNTLY | 0.5914732 |
| ASVVDPQ | ASSQTGDSGNTLY | 0.42647296 |
| SYNEFGGLS | ASSQTGDSGNTLY | 0.5931573 |
| DPDGSVY | ASSQTGDSGNTLY | 0.39427072 |
| MVSTIHATE | ASSQTGDSGNTLY | 0.5642294 |
| LAAGVADPV | ASSQTGDSGNTLY | 0.5352534 |
| TKEQIAA | ASSQTGDSGNTLY | 0.6436329 |
| SLASPAAL | ASSQTGDSGNTLY | 0.4567489 |
| IGLVFYGV | ASSQTGDSGNTLY | 0.5404002 |
| FLLDASNL | ASSQTGDSGNTLY | 0.4684838 |
| YQVGNFP | ASSQTGDSGNTLY | 0.6410343 |
| ESLLSQST | ASSQTGDSGNTLY | 0.42115614 |
| LFTVGFAAVG | ASSQTGDSGNTLY | 0.43569288 |
| ATPPTPT | ASSQTGDSGNTLY | 0.4108104 |
| TDLATLG | ASSQTGDSGNTLY | 0.4832144 |
| SAGGTGR | ASSQTGDSGNTLY | 0.5693964 |

| | | |
|---|---|---|
| GFSNSGS | ASSQTGDSGNTLY | 0.4958111 |
| GGNGGSGGTSV | ASSQTGDSGNTLY | 0.52139705 |
| GTSVATGGAGN | ASSQTGDSGNTLY | 0.5864327 |
| NGGNGGSGGTS | ASSQTGDSGNTLY | 0.4265217 |
| AVVKFPH | ASSQTGDSGNTLY | 0.54608774 |
| TPPPPPP | ASSQTGDSGNTLY | 0.56456506 |
| AAPAAAAAPA | ASSQTGDSGNTLY | 0.5187462 |
| IALNEDL | ASSQTGDSGNTLY | 0.39702982 |
| IVNVSSM | ASSQTGDSGNTLY | 0.4735625 |
| TGIPPAPR | ASSQTGDSGNTLY | 0.5319963 |
| FDLGGGTF | ASSQTGDSGNTLY | 0.51862156 |
| FDLGGGTFDVS | ASSQTGDSGNTLY | 0.4381552 |
| NEPTAAA | ASSQTGDSGNTLY | 0.59914905 |
| NEPTAAALA | ASSQTGDSGNTLY | 0.8168879 |
| GSIINISSI | ASSQTGDSGNTLY | 0.83609384 |
| VAPGNAGTA | ASSQTGDSGNTLY | 0.71075463 |
| TGGGASLE | ASSQTGDSGNTLY | 0.6969827 |
| VAVYNHYKRI | ASSQTGDSGNTLY | 0.7458174 |
| TDPAPATT | ASSQTGDSGNTLY | 0.8206274 |
| TNRPDILDPALLRPG | ASSQTGDSGNTLY | 0.8483798 |
| LTTTTTS | ASSQTGDSGNTLY | 0.82579005 |
| PTGSGKTL | ASSQTGDSGNTLY | 0.73987126 |
| TGSGKTL | ASSQTGDSGNTLY | 0.8343625 |
| TIGHVDHGK | ASSQTGDSGNTLY | 0.80369353 |
| TIGHVDHGKTT | ASSQTGDSGNTLY | 0.7807332 |
| GHVDHGKTT | ASSQTGDSGNTLY | 0.8102079 |
| AQFEHTLL | ASSQTGDSGNTLY | 0.82009065 |
| IVNQDEL | ASSQTGDSGNTLY | 0.74851465 |
| TNGSQFF | ASSQTGDSGNTLY | 0.7705149 |
| TGKGIVE | ASSQTGDSGNTLY | 0.803931 |
| LISNASDA | ASSQTGDSGNTLY | 0.8240366 |
| AELRTLGELTKIAKAHGAQ | ASSQTGDSGNTLY | 0.8155518 |
| APNAVAAAEPSW | ASSQTGDSGNTLY | 0.8472978 |
| APQVAGAATAVL | ASSQTGDSGNTLY | 0.82792115 |
| ARRDPAAAAAILPTDA | ASSQTGDSGNTLY | 0.8230369 |
| DAAARDVLAEAGLAEYF | ASSQTGDSGNTLY | 0.5527532 |
| DDQIVLSAKANLAKVEQ | ASSQTGDSGNTLY | 0.6578875 |
| DELLPDTGEDSDEDGH | ASSQTGDSGNTLY | 0.81277686 |
| ELPEAPANRATVPQD | ASSQTGDSGNTLY | 0.7598816 |
| FGQSKIERVTGIG | ASSQTGDSGNTLY | 0.5132099 |
| GAGGTGSTAGGAGGAGGAGGLYA | ASSQTGDSGNTLY | 0.82817125 |
| GHGGTGAAGGNGGTGGAGGSGIDG | ASSQTGDSGNTLY | 0.78963035 |
| GNGGNGGAGGEGGPGGAGGAGA | ASSQTGDSGNTLY | 0.75782704 |
| GSTAGNGGNGGNATNGTIAGQPAG | ASSQTGDSGNTLY | 0.8204978 |
| GTGGDGGNGGNGGTGG | ASSQTGDSGNTLY | 0.7774973 |
| GVRRPAGIPADAPV | ASSQTGDSGNTLY | 0.7568194 |
| GYRQVTDAHLLTLARR | ASSQTGDSGNTLY | 0.7995068 |
| IAKMGQDGHDRG | ASSQTGDSGNTLY | 0.83390576 |
| KAEPSIEIQYT | ASSQTGDSGNTLY | 0.74503744 |
| KPVRIRMAADEDTVAVT | ASSQTGDSGNTLY | 0.8343892 |
| MPPRIAGMRLLVIKPEP | ASSQTGDSGNTLY | 0.7559031 |

| | | |
|---|---|---|
| MKFYAVALFRSRGVPPTR | ASSQTGDSGNTLY | 0.8343424 |
| NAGGFGILGAVAHSP | ASSQTGDSGNTLY | 0.7017582 |
| NEGREITLAMAEM | ASSQTGDSGNTLY | 0.8352325 |
| NELVEWSHPLGHRWRWE | ASSQTGDSGNTLY | 0.78668225 |
| NLGAGGTGGNGGNPG | ASSQTGDSGNTLY | 0.84951675 |
| NTEWPKWETVMAADPVD | ASSQTGDSGNTLY | 0.77263314 |
| PASAAPAETQP | ASSQTGDSGNTLY | 0.8348598 |
| QQSSDGKNGVRAAEITGE | ASSQTGDSGNTLY | 0.82037187 |
| RAPDAVTTSWWKE | ASSQTGDSGNTLY | 0.8487447 |
| SGFGNVGAHTSGWFNQ | ASSQTGDSGNTLY | 0.7128058 |
| SGVAGAVGQAASVGGLKV | ASSQTGDSGNTLY | 0.77534294 |
| SNGGAGTGSAGGNGGT | ASSQTGDSGNTLY | 0.6951683 |
| TGGAGGEGGAAG | ASSQTGDSGNTLY | 0.8047054 |
| TGGDGGAAGAGGAPGHG | ASSQTGDSGNTLY | 0.8323984 |
| TMPPLLAAIANAGRHGV | ASSQTGDSGNTLY | 0.43423843 |
| TSSTAGNSGGA | ASSQTGDSGNTLY | 0.8363197 |
| VPATVPVGILC | ASSQTGDSGNTLY | 0.7012248 |
| VQRPDSGGVVLTGRIS | ASSQTGDSGNTLY | 0.7351258 |
| WTPNPVPATKGVPAGLVQAT | ASSQTGDSGNTLY | 0.83024967 |
| YALKRALAKFSDRFDVVII | ASSQTGDSGNTLY | 0.8175431 |
| YLIGSVIV | ASSQTGDSGNTLY | 0.81864595 |
| YGQLVAVID | ASSQTGDSGNTLY | 0.7841192 |
| VTVGGMTLPA | ASSQTGDSGNTLY | 0.7953353 |
| VLALPAV | ASSQTGDSGNTLY | 0.8017273 |
| VIPPTLNYET | ASSQTGDSGNTLY | 0.61751735 |
| VGMGGMHPSSG | ASSQTGDSGNTLY | 0.7609387 |
| VGLAGGAATA | ASSQTGDSGNTLY | 0.8456664 |
| VGGAGGAGGSGP | ASSQTGDSGNTLY | 0.764233 |
| VFAAFMV | ASSQTGDSGNTLY | 0.84218514 |
| TLYDKLR | ASSQTGDSGNTLY | 0.6499955 |
| TLAVFTVL | ASSQTGDSGNTLY | 0.55507904 |
| TGSVQPARV | ASSQTGDSGNTLY | 0.68864 |
| TGGGAGSGGT | ASSQTGDSGNTLY | 0.8323642 |
| SHLETSL | ASSQTGDSGNTLY | 0.82252264 |
| SFTTGMT | ASSQTGDSGNTLY | 0.78138494 |
| QVEVAKRMGVRQPTVS | ASSQTGDSGNTLY | 0.4913125 |
| QGSGGPALAM | ASSQTGDSGNTLY | 0.69978607 |
| MLAPSNP | ASSQTGDSGNTLY | 0.63297975 |
| LVNNAAI | ASSQTGDSGNTLY | 0.6598344 |
| LTGEINTLA | ASSQTGDSGNTLY | 0.760719 |
| LSIFGPV | ASSQTGDSGNTLY | 0.781 |
| LLALLANL | ASSQTGDSGNTLY | 0.6495424 |
| LISNASDALDKLR | ASSQTGDSGNTLY | 0.6321164 |
| LINEAVP | ASSQTGDSGNTLY | 0.7440797 |
| LGYAFSGL | ASSQTGDSGNTLY | 0.7844111 |
| LDFVAVAL | ASSQTGDSGNTLY | 0.7300619 |
| LAQPGAE | ASSQTGDSGNTLY | 0.85004544 |
| ISNLGAVTGF | ASSQTGDSGNTLY | 0.81839275 |
| HQEDPGRI | ASSQTGDSGNTLY | 0.811982 |
| GTPGEPTP | ASSQTGDSGNTLY | 0.18877512 |
| GFFNSGS | ASSQTGDSGNTLY | 0.35808787 |

| | | |
|---|---|---|
| GAGGGGLTAGAN | ASSQTGDSGNTLY | 0.6295289 |
| FVDALVNM | ASSQTGDSGNTLY | 0.67688155 |
| FLTVAPEL | ASSQTGDSGNTLY | 0.28587648 |
| FETLTAL | ASSQTGDSGNTLY | 0.7797698 |
| DVINPIEL | ASSQTGDSGNTLY | 0.5919846 |
| APVSGAPV | ASSQTGDSGNTLY | 0.55866385 |
| ALVGGLAW | ASSQTGDSGNTLY | 0.8057376 |
| ALAAAAATAI | ASSQTGDSGNTLY | 0.640617 |
| AFGLGLLL | ASSQTGDSGNTLY | 0.61559814 |
| AFFAAASL | ASSQTGDSGNTLY | 0.5667282 |
| AAAALPAFSPP | ASSQTGDSGNTLY | 0.81648004 |
| VVLAPAL | ASSQTGDSGNTLY | 0.3513991 |
| TIGHVDHGKTTL | ASSQTGDSGNTLY | 0.7590374 |
| LISNASDALDK | ASSQTGDSGNTLY | 0.60122484 |
| LISNASDAL | ASSQTGDSGNTLY | 0.8213239 |
| ISNASDALDK | ASSQTGDSGNTLY | 0.43564892 |
| NQAETLVYQTE | ASSQTGDSGNTLY | 0.80234706 |
| TGGTGGTAGAG | ASSQTGDSGNTLY | 0.7215219 |
| TGSTAGGAGGA | ASSQTGDSGNTLY | 0.86722803 |
| DPLAVPVAAGPV | ASSQTGDSGNTLY | 0.6756697 |
| GGTGGTPGNGGN | ASSQTGDSGNTLY | 0.7207606 |
| PAGLAGVDPATL | ASSQTGDSGNTLY | 0.6659054 |
| GNNLVVEAPHAGASV | ASSQTGDSGNTLY | 0.81945044 |
| KVTRIMSASTLEQPA | ASSQTGDSGNTLY | 0.4283749 |
| LGFALPIALPRRTVV | ASSQTGDSGNTLY | 0.6801449 |
| TRLGNLTARKLVVLV | ASSQTGDSGNTLY | 0.4410355 |
| WSQQRRLIYFPSAGP | ASSQTGDSGNTLY | 0.7362726 |
| GPNGNPDPMAAAVDIR | ASSQTGDSGNTLY | 0.7841354 |
| TQRCVSVADLRAAASR | ASSQTGDSGNTLY | 0.31654018 |
| QFVLDVAEHPVEVLTETV | ASSQTGDSGNTLY | 0.7508943 |
| GGAGGTGGAGGNGGTQAGDG | ASSQTGDSGNTLY | 0.7352337 |
| PLLATLRAIAVDVDAKPAQV | ASSQTGDSGNTLY | 0.5804024 |
| APKFRTGGYMIDFWGVGYQVAKRM | ASSQTGDSGNTLY | 0.82812655 |
| GAGGAGGAGGAAGGPGGTGGTGGN | ASSQTGDSGNTLY | 0.67333066 |
| IGALGLLATDVAPAGSVAFGVALA | ASSQTGDSGNTLY | 0.833622 |
| AGGAGGIGGNGGAGGAGGVLMGNG | ASSQTGDSGNTLY | 0.67037606 |
| ADELIGLL | ASSQTGDSGNTLY | 0.6354747 |
| ADRALKA | ASSQTGDSGNTLY | 0.7769452 |
| AGAVGQVGGAGGA | ASSQTGDSGNTLY | 0.52191067 |
| AGGGGGLGGAP | ASSQTGDSGNTLY | 0.52036047 |
| AGGLLYGNGGAG | ASSQTGDSGNTLY | 0.837731 |
| AGLESIGL | ASSQTGDSGNTLY | 0.82012093 |
| AGLKTGII | ASSQTGDSGNTLY | 0.85257375 |
| ASLAALLL | ASSQTGDSGNTLY | 0.464282 |
| ASVINISSI | ASSQTGDSGNTLY | 0.5690968 |
| ATPPGPA | ASSQTGDSGNTLY | 0.63882506 |
| ATVASALV | ASSQTGDSGNTLY | 0.77271056 |
| ATVASLIL | ASSQTGDSGNTLY | 0.86324394 |
| AVNGPVLIRH | ASSQTGDSGNTLY | 0.8335147 |
| DGAAGAVGI | ASSQTGDSGNTLY | 0.5215517 |
| DPLLHGI | ASSQTGDSGNTLY | 0.745453 |

| | | |
|---|---|---|
| ELGAALGAL | ASSQTGDSGNTLY | 0.6368358 |
| ELGRTQAV | ASSQTGDSGNTLY | 0.7158308 |
| FVAGGYLS | ASSQTGDSGNTLY | 0.65298164 |
| GASGGAGGAGGW | ASSQTGDSGNTLY | 0.6665447 |
| GGAGAGGDGGAGGI | ASSQTGDSGNTLY | 0.5370493 |
| GTGHNGSQPGMGGNG | ASSQTGDSGNTLY | 0.45693472 |
| GYAAASASAS | ASSQTGDSGNTLY | 0.6616552 |
| HQVDVAAVQQG | ASSQTGDSGNTLY | 0.8063033 |
| IGAAIAHT | ASSQTGDSGNTLY | 0.79866207 |
| IGMIEVAQL | ASSQTGDSGNTLY | 0.81742704 |
| IRLAPEHTSAMD | ASSQTGDSGNTLY | 0.8072903 |
| IYNVSVL | ASSQTGDSGNTLY | 0.6966388 |
| KQGFLGM | ASSQTGDSGNTLY | 0.18176526 |
| LESVITQL | ASSQTGDSGNTLY | 0.3321168 |
| LGLGALVVV | ASSQTGDSGNTLY | 0.5304728 |
| LIGNGGNGGSGGI | ASSQTGDSGNTLY | 0.5784656 |
| LNAINAPTL | ASSQTGDSGNTLY | 0.19230719 |
| LQLLNTGV | ASSQTGDSGNTLY | 0.7134765 |
| LQTSIEF | ASSQTGDSGNTLY | 0.68701386 |
| MSFAVSEL | ASSQTGDSGNTLY | 0.59875154 |
| NTGALGL | ASSQTGDSGNTLY | 0.8082182 |
| PPGAGGAGGAGTT | ASSQTGDSGNTLY | 0.6292918 |
| QLVGLGI | ASSQTGDSGNTLY | 0.40366095 |
| RFLEQVHQL | ASSQTGDSGNTLY | 0.44768244 |
| RGVGYVLR | ASSQTGDSGNTLY | 0.80922425 |
| SATGVFAGVI | ASSQTGDSGNTLY | 0.39720148 |
| SLPAITLP | ASSQTGDSGNTLY | 0.81198895 |
| STALAFTV | ASSQTGDSGNTLY | 0.42398962 |
| SVLTVGLGK | ASSQTGDSGNTLY | 0.82705295 |
| TALAGAGSGI | ASSQTGDSGNTLY | 0.43385792 |
| TGLPFTGI | ASSQTGDSGNTLY | 0.7971679 |
| VAEHVTV | ASSQTGDSGNTLY | 0.7269908 |
| VDIVLNSL | ASSQTGDSGNTLY | 0.87228465 |
| VLTGGGAVL | ASSQTGDSGNTLY | 0.605652 |
| VNVGLAYGI | ASSQTGDSGNTLY | 0.73688394 |
| VSSAAVGAV | ASSQTGDSGNTLY | 0.48351204 |
| VTSVYEAGV | ASSQTGDSGNTLY | 0.83492076 |
| WVALDRGA | ASSQTGDSGNTLY | 0.4266941 |
| YLVGSVVI | ASSQTGDSGNTLY | 0.5369728 |
| GPPGTGKTLL | ASSQTGDSGNTLY | 0.34585604 |
| GVGMTKF | ASSQTGDSGNTLY | 0.5686121 |
| IGIGGSDL | ASSQTGDSGNTLY | 0.8431481 |
| IGTLLAEL | ASSQTGDSGNTLY | 0.2761051 |
| IYTSGTT | ASSQTGDSGNTLY | 0.6946434 |
| LSFPAGV | ASSQTGDSGNTLY | 0.60954756 |
| LYTSGTT | ASSQTGDSGNTLY | 0.49082404 |
| VGAFPFGF | ASSQTGDSGNTLY | 0.6975176 |
| VGAGTVEF | ASSQTGDSGNTLY | 0.6372023 |
| VGKSTLLL | ASSQTGDSGNTLY | 0.7836243 |
| AALAHAGGGIV | ASSQTGDSGNTLY | 0.45831794 |
| AALVAVSLL | ASSQTGDSGNTLY | 0.5789232 |

| | | |
|---|---|---|
| AAVSRAVAAL | ASSQTGDSGNTLY | 0.7129903 |
| AGGSSGIGGF | ASSQTGDSGNTLY | 0.42213678 |
| AGLTVANANG | ASSQTGDSGNTLY | 0.402218 |
| AGLVGAVL | ASSQTGDSGNTLY | 0.844651 |
| ALSQAFTV | ASSQTGDSGNTLY | 0.12102461 |
| ALTFSVSL | ASSQTGDSGNTLY | 0.120107874 |
| ALTGAALVAL | ASSQTGDSGNTLY | 0.11689159 |
| ASFSIGAGVL | ASSQTGDSGNTLY | 0.12022079 |
| DGSVYKI | ASSQTGDSGNTLY | 0.11785372 |
| EQLAEIV | ASSQTGDSGNTLY | 0.12722808 |
| FAVSELTVM | ASSQTGDSGNTLY | 0.12063443 |
| GAAPLAAGL | ASSQTGDSGNTLY | 0.120147124 |
| GADGGDGGSGGAGG | ASSQTGDSGNTLY | 0.12161384 |
| GFGLDTGGAGG | ASSQTGDSGNTLY | 0.1201645 |
| GLASGGLL | ASSQTGDSGNTLY | 0.11799664 |
| GRIAGAAVLL | ASSQTGDSGNTLY | 0.12074881 |
| GSAGSLNTSF | ASSQTGDSGNTLY | 0.119513474 |
| GTGQIGI | ASSQTGDSGNTLY | 0.12150877 |
| GTLIALIL | ASSQTGDSGNTLY | 0.11663465 |
| IAATMGFVVG | ASSQTGDSGNTLY | 0.12004847 |
| IFAQALL | ASSQTGDSGNTLY | 0.11958432 |
| IGAAVALGF | ASSQTGDSGNTLY | 0.12098101 |
| IGLTPLAL | ASSQTGDSGNTLY | 0.12006547 |
| ILGSNTSTL | ASSQTGDSGNTLY | 0.12836663 |
| IVFSLTVL | ASSQTGDSGNTLY | 0.123353295 |
| LASKATGF | ASSQTGDSGNTLY | 0.11797252 |
| LASVALIL | ASSQTGDSGNTLY | 0.11587732 |
| LFAVGLTL | ASSQTGDSGNTLY | 0.117791854 |
| LFNGGIGTYI | ASSQTGDSGNTLY | 0.11880723 |
| LTLGAGII | ASSQTGDSGNTLY | 0.12080458 |
| LVAAALGDI | ASSQTGDSGNTLY | 0.115052216 |
| NTAFVQLGI | ASSQTGDSGNTLY | 0.12406293 |
| QLFSGAEL | ASSQTGDSGNTLY | 0.12071767 |
| SAATFITL | ASSQTGDSGNTLY | 0.11907896 |
| SNYGLANLGV | ASSQTGDSGNTLY | 0.124352746 |
| SYGGAYAVM | ASSQTGDSGNTLY | 0.11734278 |
| TTGVGGLTLG | ASSQTGDSGNTLY | 0.11923845 |
| TTQTIYI | ASSQTGDSGNTLY | 0.11860249 |
| VAAGAGLGAVI | ASSQTGDSGNTLY | 0.124570265 |
| VAAVVTLL | ASSQTGDSGNTLY | 0.11803724 |
| VGFLGGLI | ASSQTGDSGNTLY | 0.12921199 |
| VGVIPILL | ASSQTGDSGNTLY | 0.11732788 |
| VIFASSLI | ASSQTGDSGNTLY | 0.13513978 |
| VKGGDGGAAAG | ASSQTGDSGNTLY | 0.1206302 |
| VTKTFTNI | ASSQTGDSGNTLY | 0.121175 |
| VVAAAILL | ASSQTGDSGNTLY | 0.12009845 |
| YAALDVEL | ASSQTGDSGNTLY | 0.13390407 |
| LASVALLL | ASSQTGDSGNTLY | 0.12338643 |
| LDAVHRV | ASSQTGDSGNTLY | 0.12665784 |
| QGLSSSIF | ASSQTGDSGNTLY | 0.11874008 |
| RGTFANIRL | ASSQTGDSGNTLY | 0.120641835 |

| | | |
|---|---|---|
| RQLWWGHRI | ASSQTGDSGNTLY | 0.11749982 |
| YGPPGTGKTLL | ASSQTGDSGNTLY | 0.12018728 |
| AALVGAPANVA | ASSQTGDSGNTLY | 0.11673095 |
| AATFRRTE | ASSQTGDSGNTLY | 0.12124827 |
| AGALGGGI | ASSQTGDSGNTLY | 0.1208088 |
| AGGAGGAGGIGAG | ASSQTGDSGNTLY | 0.11581913 |
| AGSGALGL | ASSQTGDSGNTLY | 0.11890824 |
| AQIADQI | ASSQTGDSGNTLY | 0.11609442 |
| ASAVAVTYL | ASSQTGDSGNTLY | 0.118432075 |
| ASTVLGGTGG | ASSQTGDSGNTLY | 0.119994886 |
| AVGGEPAK | ASSQTGDSGNTLY | 0.118850894 |
| CEQGTFGGTC | ASSQTGDSGNTLY | 0.11839665 |
| DEGKAPAG | ASSQTGDSGNTLY | 0.11610047 |
| DTVVFPAL | ASSQTGDSGNTLY | 0.11754788 |
| FEGITMAG | ASSQTGDSGNTLY | 0.12519868 |
| FPLVVQD | ASSQTGDSGNTLY | 0.11671409 |
| GAFTTPQ | ASSQTGDSGNTLY | 0.1176158 |
| GFGGFGTAGGVGGA | ASSQTGDSGNTLY | 0.12399458 |
| GFLAQFL | ASSQTGDSGNTLY | 0.8304093 |
| GGAGGAGGNAGVA | ASSQTGDSGNTLY | 0.79189515 |
| GGVGTAGAGGAGGAGG | ASSQTGDSGNTLY | 0.68736714 |
| GNGGGGDAGNAGSGGN | ASSQTGDSGNTLY | 0.6910156 |
| GTGGAGGAGNDAGS | ASSQTGDSGNTLY | 0.7391081 |
| GTTNTGWFNS | ASSQTGDSGNTLY | 0.84297454 |
| IFMGPLFTL | ASSQTGDSGNTLY | 0.83832973 |
| ILAHAIEAI | ASSQTGDSGNTLY | 0.8316825 |
| IRSAIREAL | ASSQTGDSGNTLY | 0.7048637 |
| ITFAGGG | ASSQTGDSGNTLY | 0.8585181 |
| ITGMSGHQEV | ASSQTGDSGNTLY | 0.79427767 |
| IVAGLACTL | ASSQTGDSGNTLY | 0.7770047 |
| IVGGGLAGA | ASSQTGDSGNTLY | 0.8320558 |
| IVTGVLNI | ASSQTGDSGNTLY | 0.812673 |
| LAQQGCI | ASSQTGDSGNTLY | 0.7581053 |
| LCLHVEN | ASSQTGDSGNTLY | 0.71607745 |
| LDAISDGS | ASSQTGDSGNTLY | 0.8258628 |
| LGGFAAML | ASSQTGDSGNTLY | 0.8425155 |
| LGGKNPMI | ASSQTGDSGNTLY | 0.8282237 |
| LGPKGVAAIL | ASSQTGDSGNTLY | 0.88794035 |
| LMVGVSI | ASSQTGDSGNTLY | 0.8503094 |
| LVGAAFVL | ASSQTGDSGNTLY | 0.8313726 |
| LVLAVAPL | ASSQTGDSGNTLY | 0.37372965 |
| MAAIAVIDAA | ASSQTGDSGNTLY | 0.63613987 |
| NNGINAELQ | ASSQTGDSGNTLY | 0.73893666 |
| RGEPLVLVHG | ASSQTGDSGNTLY | 0.77463233 |
| RGGAGGAAGLI | ASSQTGDSGNTLY | 0.30829772 |
| RSYRMHPAV | ASSQTGDSGNTLY | 0.8275384 |
| SAAHLLAGQLA | ASSQTGDSGNTLY | 0.81295145 |
| SAAVGSY | ASSQTGDSGNTLY | 0.7185477 |
| SPHAVLHRI | ASSQTGDSGNTLY | 0.84850836 |
| SQIAKVL | ASSQTGDSGNTLY | 0.7394276 |
| TALGHLVP | ASSQTGDSGNTLY | 0.7008573 |

| | | |
|---|---|---|
| TAPTTSGAA | ASSQTGDSGNTLY | 0.7748263 |
| TDGDDGNGGSPGT | ASSQTGDSGNTLY | 0.88122654 |
| TDGLGEN | ASSQTGDSGNTLY | 0.69520694 |
| TGGPPLPGQA | ASSQTGDSGNTLY | 0.856044 |
| TIPAITI | ASSQTGDSGNTLY | 0.76787835 |
| TIPGFTI | ASSQTGDSGNTLY | 0.87744987 |
| TSIASGALA | ASSQTGDSGNTLY | 0.6292223 |
| TVTLPGAAI | ASSQTGDSGNTLY | 0.84600985 |
| VASGLSAALP | ASSQTGDSGNTLY | 0.7574314 |
| VGAGVIGI | ASSQTGDSGNTLY | 0.88549936 |
| VGPKRFGV | ASSQTGDSGNTLY | 0.77346796 |
| VNVAFPD | ASSQTGDSGNTLY | 0.857882 |
| VSMTNSGVS | ASSQTGDSGNTLY | 0.78426933 |
| VTDVLGDLSV | ASSQTGDSGNTLY | 0.81961316 |
| VVVVGAGFAGL | ASSQTGDSGNTLY | 0.5823909 |
| VYTVGTVV | ASSQTGDSGNTLY | 0.7818407 |
| WTTFSGP | ASSQTGDSGNTLY | 0.72834706 |
| YDELVEAF | ASSQTGDSGNTLY | 0.8251536 |
| YGTGGAGGNGG | ASSQTGDSGNTLY | 0.8373503 |
| YTNVMAV | ASSQTGDSGNTLY | 0.23869726 |
| FLEQVHQL | ASSQTGDSGNTLY | 0.86340666 |
| GLSLLAVAV | ASSQTGDSGNTLY | 0.5646848 |
| LEQVHQL | ASSQTGDSGNTLY | 0.6824037 |
| LNAAGAI | ASSQTGDSGNTLY | 0.8456801 |
| PAVHASL | ASSQTGDSGNTLY | 0.8165354 |
| TELEDDA | ASSQTGDSGNTLY | 0.8471639 |
| TPPGPPN | ASSQTGDSGNTLY | 0.7396542 |
| TPPPPPPP | ASSQTGDSGNTLY | 0.81807387 |
| TQSPALI | ASSRADRGGNTEVF | 0.84844524 |
| EFAAAFPQ | ASSRADRGGNTEVF | 0.40826872 |
| LLSGNLDVL | ASSRADRGGNTEVF | 0.6968514 |
| GPANLGNYNI | ASSRADRGGNTEVF | 0.86566126 |
| TLGPTLP | ASSRADRGGNTEVF | 0.8753685 |
| VYAYSSCCK | ASSRADRGGNTEVF | 0.8823898 |
| APAPAPAPNI | ASSRADRGGNTEVF | 0.82560253 |
| PGGPVPIP | ASSRADRGGNTEVF | 0.7372794 |
| FLTVALEP | ASSRADRGGNTEVF | 0.8313168 |
| SQYFGANL | ASSRADRGGNTEVF | 0.88612115 |
| GGVKPAA | ASSRADRGGNTEVF | 0.8737154 |
| GGTGGAGGAGGW | ASSRADRGGNTEVF | 0.8660788 |
| GVGGVAGLGGAGG | ASSRADRGGNTEVF | 0.52862096 |
| VLLVLATTI | ASSRADRGGNTEVF | 0.8619423 |
| IGIFAAML | ASSRADRGGNTEVF | 0.8204664 |
| AVGDLID | ASSRADRGGNTEVF | 0.8248087 |
| NISPLPL | ASSRADRGGNTEVF | 0.871385 |
| VTAVFVAI | ASSRADRGGNTEVF | 0.82784647 |
| SGDDNDAAL | ASSRADRGGNTEVF | 0.8482288 |
| AVAALAHS | ASSRADRGGNTEVF | 0.8201817 |
| GGVTVMATP | ASSRADRGGNTEVF | 0.8636434 |
| AGLAANVRD | ASSRADRGGNTEVF | 0.8520571 |
| TNRPDLIDPALLRPG | ASSRADRGGNTEVF | 0.8112001 |

| | | |
|---|---|---|
| LANLGVTAL | ASSRADRGGNTEVF | 0.90159523 |
| TGERIEL | ASSRADRGGNTEVF | 0.8911965 |
| NGEELAA | ASSRADRGGNTEVF | 0.8832816 |
| VLSLVWG | ASSRADRGGNTEVF | 0.21093555 |
| IGYALWL | ASSRADRGGNTEVF | 0.6443517 |
| LSLLLAPL | ASSRADRGGNTEVF | 0.8129028 |
| TLDEALK | ASSRADRGGNTEVF | 0.84111404 |
| TDGAGTAEGG | ASSRADRGGNTEVF | 0.45761526 |
| NAPGGAGGNGG | ASSRADRGGNTEVF | 0.875567 |
| VSVFMGMGGNF | ASSRADRGGNTEVF | 0.8082098 |
| LASYLAAL | ASSRADRGGNTEVF | 0.7815179 |
| LVGGTPAL | ASSRADRGGNTEVF | 0.8910842 |
| VRNADVL | ASSRADRGGNTEVF | 0.8407624 |
| TCSSPFLVSAM | ASSRADRGGNTEVF | 0.7354474 |
| VHITDAE | ASSRADRGGNTEVF | 0.78669184 |
| EYAAPSF | ASSRADRGGNTEVF | 0.88575643 |
| VVDAFHGGL | ASSRADRGGNTEVF | 0.6738087 |
| ASVVDPQ | ASSRADRGGNTEVF | 0.8797157 |
| SYNEFGGLS | ASSRADRGGNTEVF | 0.81346655 |
| DPDGSVY | ASSRADRGGNTEVF | 0.8943334 |
| MVSTIHATE | ASSRADRGGNTEVF | 0.48645693 |
| LAAGVADPV | ASSRADRGGNTEVF | 0.8824016 |
| TKEQIAA | ASSRADRGGNTEVF | 0.8526734 |
| SLASPAAL | ASSRADRGGNTEVF | 0.9053217 |
| IGLVFYGV | ASSRADRGGNTEVF | 0.8246663 |
| FLLDASNL | ASSRADRGGNTEVF | 0.8151468 |
| YQVGNFP | ASSRADRGGNTEVF | 0.72437286 |
| ESLLSQST | ASSRADRGGNTEVF | 0.8857219 |
| LFTVGFAAVG | ASSRADRGGNTEVF | 0.66432184 |
| ATPPTPT | ASSRADRGGNTEVF | 0.79545605 |
| TDLATLG | ASSRADRGGNTEVF | 0.7292789 |
| SAGGTGR | ASSRADRGGNTEVF | 0.85654914 |
| GFSNSGS | ASSRADRGGNTEVF | 0.87009466 |
| GGNGGSGGTSV | ASSRADRGGNTEVF | 0.47152328 |
| GTSVATGGAGN | ASSRADRGGNTEVF | 0.8722595 |
| NGGNGGSGGTS | ASSRADRGGNTEVF | 0.808707 |
| AVVKFPH | ASSRADRGGNTEVF | 0.6393676 |
| TPPPPPP | ASSRADRGGNTEVF | 0.89351976 |
| AAPAAAAAPA | ASSRADRGGNTEVF | 0.78479916 |
| IALNEDL | ASSRADRGGNTEVF | 0.893686 |
| IVNVSSM | ASSRADRGGNTEVF | 0.82700527 |
| TGIPPAPR | ASSRADRGGNTEVF | 0.8175088 |
| FDLGGGTF | ASSRADRGGNTEVF | 0.8649866 |
| FDLGGGTFDVS | ASSRADRGGNTEVF | 0.7865262 |
| NEPTAAA | ASSRADRGGNTEVF | 0.75280225 |
| NEPTAAALA | ASSRADRGGNTEVF | 0.8946314 |
| GSIINISSI | ASSRADRGGNTEVF | 0.8277946 |
| VAPGNAGTA | ASSRADRGGNTEVF | 0.8494824 |
| TGGGASLE | ASSRADRGGNTEVF | 0.7729864 |
| VAVYNHYKRI | ASSRADRGGNTEVF | 0.6605923 |
| TDPAPATT | ASSRADRGGNTEVF | 0.77356017 |

| | | |
|---|---|---|
| TNRPDILDPALLRPG | ASSRADRGGNTEVF | 0.84907734 |
| LTTTTTS | ASSRADRGGNTEVF | 0.8447386 |
| PTGSGKTL | ASSRADRGGNTEVF | 0.84023166 |
| TGSGKTL | ASSRADRGGNTEVF | 0.46951827 |
| TIGHVDHGK | ASSRADRGGNTEVF | 0.8280173 |
| TIGHVDHGKTT | ASSRADRGGNTEVF | 0.77614707 |
| GHVDHGKTT | ASSRADRGGNTEVF | 0.8069501 |
| AQFEHTLL | ASSRADRGGNTEVF | 0.82048273 |
| IVNQDEL | ASSRADRGGNTEVF | 0.77274317 |
| TNGSQFF | ASSRADRGGNTEVF | 0.8044671 |
| TGKGIVE | ASSRADRGGNTEVF | 0.7125716 |
| LISNASDA | ASSRADRGGNTEVF | 0.8188478 |
| AELRTLGELTKIAKAHGAQ | ASSRADRGGNTEVF | 0.8077575 |
| APNAVAAAEPSW | ASSRADRGGNTEVF | 0.76148295 |
| APQVAGAATAVL | ASSRADRGGNTEVF | 0.866189 |
| ARRDPAAAAAILPTDA | ASSRADRGGNTEVF | 0.8510263 |
| DAAARDVLAEAGLAEYF | ASSRADRGGNTEVF | 0.83390784 |
| DDQIVLSAKANLAKVEQ | ASSRADRGGNTEVF | 0.18625787 |
| DELLPDTGEDSDEDGH | ASSRADRGGNTEVF | 0.62340426 |
| ELPEAPANRATVPQD | ASSRADRGGNTEVF | 0.7211869 |
| FGQSKIERVTGIG | ASSRADRGGNTEVF | 0.8090255 |
| GAGGTGSTAGGAGGAGGAGGLYA | ASSRADRGGNTEVF | 0.29507494 |
| GHGGTGAAGGNGGTGGAGGSGIDG | ASSRADRGGNTEVF | 0.83988154 |
| GNGGNGGAGGEGGPGGAGGAGA | ASSRADRGGNTEVF | 0.7506418 |
| GSTAGNGGNGGNATNGTIAGQPAG | ASSRADRGGNTEVF | 0.76974416 |
| GTGGDGGNGGNGGTGG | ASSRADRGGNTEVF | 0.8547485 |
| GVRRPAGIPADAPV | ASSRADRGGNTEVF | 0.80080414 |
| GYRQVTDAHLLTLARR | ASSRADRGGNTEVF | 0.65371835 |
| IAKMGQDGHDRG | ASSRADRGGNTEVF | 0.6755356 |
| KAEPSIEIQYT | ASSRADRGGNTEVF | 0.8615314 |
| KPVRIRMAADEDTVAVT | ASSRADRGGNTEVF | 0.65247416 |
| MPPRIAGMRLLVIKPEP | ASSRADRGGNTEVF | 0.85714805 |
| MKFYAVALFRSRGVPPTR | ASSRADRGGNTEVF | 0.76593804 |
| NAGGFGILGAVAHSP | ASSRADRGGNTEVF | 0.8593504 |
| NEGREITLAMAEM | ASSRADRGGNTEVF | 0.36657485 |
| NELVEWSHPLGHRWRWE | ASSRADRGGNTEVF | 0.85167354 |
| NLGAGGTGGNGGNPG | ASSRADRGGNTEVF | 0.8139007 |
| NTEWPKWETVMAADPVD | ASSRADRGGNTEVF | 0.8662295 |
| PASAAPAETQP | ASSRADRGGNTEVF | 0.73984885 |
| QQSSDGKNGVRAAEITGE | ASSRADRGGNTEVF | 0.7931504 |
| RAPDAVTTSWWKE | ASSRADRGGNTEVF | 0.67543626 |
| SGFGNVGAHTSGWFNQ | ASSRADRGGNTEVF | 0.8574035 |
| SGVAGAVGQAASVGGLKV | ASSRADRGGNTEVF | 0.621907 |
| SNGGAGTGSAGGNGGT | ASSRADRGGNTEVF | 0.6330142 |
| TGGAGGEGGAAG | ASSRADRGGNTEVF | 0.6912583 |
| TGGDGGAAGAGGAPGHG | ASSRADRGGNTEVF | 0.7745153 |
| TMPPLLAAIANAGRHGV | ASSRADRGGNTEVF | 0.8427882 |
| TSSTAGNSGGA | ASSRADRGGNTEVF | 0.3539902 |
| VPATVPVGILC | ASSRADRGGNTEVF | 0.82809186 |
| VQRPDSGGVVLTGRIS | ASSRADRGGNTEVF | 0.75119734 |
| WTPNPVPATKGVPAGLVQAT | ASSRADRGGNTEVF | 0.48110276 |

| | | |
|---|---|---|
| YALKRALAKFSDRFDVVII | ASSRADRGGNTEVF | 0.8413135 |
| YLIGSVIV | ASSRADRGGNTEVF | 0.71170914 |
| YGQLVAVID | ASSRADRGGNTEVF | 0.84106326 |
| VTVGGMTLPA | ASSRADRGGNTEVF | 0.75704944 |
| VLALPAV | ASSRADRGGNTEVF | 0.7701987 |
| VIPPTLNYET | ASSRADRGGNTEVF | 0.808097 |
| VGMGGMHPSSG | ASSRADRGGNTEVF | 0.61972344 |
| VGLAGGAATA | ASSRADRGGNTEVF | 0.6695138 |
| VGGAGGAGGSGP | ASSRADRGGNTEVF | 0.8609259 |
| VFAAFMV | ASSRADRGGNTEVF | 0.72882104 |
| TLYDKLR | ASSRADRGGNTEVF | 0.7754083 |
| TLAVFTVL | ASSRADRGGNTEVF | 0.6385982 |
| TGSVQPARV | ASSRADRGGNTEVF | 0.57742137 |
| TGGGAGSGGT | ASSRADRGGNTEVF | 0.66154087 |
| SHLETSL | ASSRADRGGNTEVF | 0.7523367 |
| SFTTGMT | ASSRADRGGNTEVF | 0.7819594 |
| QVEVAKRMGVRQPTVS | ASSRADRGGNTEVF | 0.7390462 |
| QGSGGPALAM | ASSRADRGGNTEVF | 0.5739961 |
| MLAPSNP | ASSRADRGGNTEVF | 0.7079335 |
| LVNNAAI | ASSRADRGGNTEVF | 0.66973925 |
| LTGEINTLA | ASSRADRGGNTEVF | 0.65409815 |
| LSIFGPV | ASSRADRGGNTEVF | 0.7140287 |
| LLALLANL | ASSRADRGGNTEVF | 0.7203953 |
| LISNASDALDKLR | ASSRADRGGNTEVF | 0.6256006 |
| LINEAVP | ASSRADRGGNTEVF | 0.56068754 |
| LGYAFSGL | ASSRADRGGNTEVF | 0.7111033 |
| LDFVAVAL | ASSRADRGGNTEVF | 0.75526893 |
| LAQPGAE | ASSRADRGGNTEVF | 0.7326671 |
| ISNLGAVTGF | ASSRADRGGNTEVF | 0.7726604 |
| HQEDPGRI | ASSRADRGGNTEVF | 0.75039214 |
| GTPGEPTP | ASSRADRGGNTEVF | 0.73103166 |
| GFFNSGS | ASSRADRGGNTEVF | 0.25596523 |
| GAGGGGLTAGAN | ASSRADRGGNTEVF | 0.37063682 |
| FVDALVNM | ASSRADRGGNTEVF | 0.5694592 |
| FLTVAPEL | ASSRADRGGNTEVF | 0.6713735 |
| FETLTAL | ASSRADRGGNTEVF | 0.301056 |
| DVINPIEL | ASSRADRGGNTEVF | 0.6993832 |
| APVSGAPV | ASSRADRGGNTEVF | 0.62302655 |
| ALVGGLAW | ASSRADRGGNTEVF | 0.59022975 |
| ALAAAAATAI | ASSRADRGGNTEVF | 0.7534596 |
| AFGLGLLL | ASSRADRGGNTEVF | 0.6416824 |
| AFFAAASL | ASSRADRGGNTEVF | 0.5721837 |
| AAAALPAFSPP | ASSRADRGGNTEVF | 0.57583463 |
| VVLAPAL | ASSRADRGGNTEVF | 0.7726686 |
| TIGHVDHGKTTL | ASSRADRGGNTEVF | 0.37830418 |
| LISNASDALDK | ASSRADRGGNTEVF | 0.73442185 |
| LISNASDAL | ASSRADRGGNTEVF | 0.6352043 |
| ISNASDALDK | ASSRADRGGNTEVF | 0.7495463 |
| NQAETLVYQTE | ASSRADRGGNTEVF | 0.4695652 |
| TGGTGGTAGAG | ASSRADRGGNTEVF | 0.7453731 |
| TGSTAGGAGGA | ASSRADRGGNTEVF | 0.6744597 |

| | | |
|---|---|---|
| DPLAVPVAAGPV | ASSRADRGGNTEVF | 0.7843491 |
| GGTGGTPGNGGN | ASSRADRGGNTEVF | 0.6407797 |
| PAGLAGVDPATL | ASSRADRGGNTEVF | 0.71905744 |
| GNNLVVEAPHAGASV | ASSRADRGGNTEVF | 0.6040609 |
| KVTRIMSASTLEQPA | ASSRADRGGNTEVF | 0.76442397 |
| LGFALPIALPRRTVV | ASSRADRGGNTEVF | 0.44606504 |
| TRLGNLTARKLVVLV | ASSRADRGGNTEVF | 0.6535542 |
| WSQQRRLIYFPSAGP | ASSRADRGGNTEVF | 0.49624586 |
| GPNGNPDPMAAAVDIR | ASSRADRGGNTEVF | 0.7099868 |
| TQRCVSVADLRAAASR | ASSRADRGGNTEVF | 0.7366705 |
| QFVLDVAEHPVEVLTETV | ASSRADRGGNTEVF | 0.4041791 |
| GGAGGTGGAGGNGGTQAGDG | ASSRADRGGNTEVF | 0.76522684 |
| PLLATLRAIAVDVDAKPAQV | ASSRADRGGNTEVF | 0.6712643 |
| APKFRTGGYMIDFWGVGYQVAKRM | ASSRADRGGNTEVF | 0.5950847 |
| GAGGAGGAGGAAGGPGGTGGTGGN | ASSRADRGGNTEVF | 0.7487376 |
| IGALGLLATDVAPAGSVAFGVALA | ASSRADRGGNTEVF | 0.6588055 |
| AGGAGGIGGNGGAGGAGGVLMGNG | ASSRADRGGNTEVF | 0.75616515 |
| ADELIGLL | ASSRADRGGNTEVF | 0.6169555 |
| ADRALKA | ASSRADRGGNTEVF | 0.7122124 |
| AGAVGQVGGAGGA | ASSRADRGGNTEVF | 0.73365545 |
| AGGGGGLGGAP | ASSRADRGGNTEVF | 0.5557227 |
| AGGLLYGNGGAG | ASSRADRGGNTEVF | 0.537812 |
| AGLESIGL | ASSRADRGGNTEVF | 0.7589937 |
| AGLKTGII | ASSRADRGGNTEVF | 0.88333285 |
| ASLAALLL | ASSRADRGGNTEVF | 0.90508187 |
| ASVINISSI | ASSRADRGGNTEVF | 0.7698089 |
| ATPPGPA | ASSRADRGGNTEVF | 0.70601606 |
| ATVASALV | ASSRADRGGNTEVF | 0.82531226 |
| ATVASLIL | ASSRADRGGNTEVF | 0.8930635 |
| AVNGPVLIRH | ASSRADRGGNTEVF | 0.91125715 |
| DGAAGAVGI | ASSRADRGGNTEVF | 0.87722194 |
| DPLLHGI | ASSRADRGGNTEVF | 0.71094245 |
| ELGAALGAL | ASSRADRGGNTEVF | 0.8617065 |
| ELGRTQAV | ASSRADRGGNTEVF | 0.83925545 |
| FVAGGYLS | ASSRADRGGNTEVF | 0.81292284 |
| GASGGAGGAGGW | ASSRADRGGNTEVF | 0.88158834 |
| GGAGAGGDGGAGGI | ASSRADRGGNTEVF | 0.8668472 |
| GTGHNGSQPGMGGNG | ASSRADRGGNTEVF | 0.8063998 |
| GYAAASASAS | ASSRADRGGNTEVF | 0.7994102 |
| HQVDVAAVQQG | ASSRADRGGNTEVF | 0.8737291 |
| IGAAIAHT | ASSRADRGGNTEVF | 0.8894237 |
| IGMIEVAQL | ASSRADRGGNTEVF | 0.8861476 |
| IRLAPEHTSAMD | ASSRADRGGNTEVF | 0.90429115 |
| IYNVSVL | ASSRADRGGNTEVF | 0.8929224 |
| KQGFLGM | ASSRADRGGNTEVF | 0.8832154 |
| LESVITQL | ASSRADRGGNTEVF | 0.25147182 |
| LGLGALVVV | ASSRADRGGNTEVF | 0.55233485 |
| LIGNGGNGGSGGI | ASSRADRGGNTEVF | 0.7641728 |
| LNAINAPTL | ASSRADRGGNTEVF | 0.82232577 |
| LQLLNTGV | ASSRADRGGNTEVF | 0.48276454 |
| LQTSIEF | ASSRADRGGNTEVF | 0.85818267 |

| | | |
|---|---|---|
| MSFAVSEL | ASSRADRGGNTEVF | 0.7800734 |
| NTGALGL | ASSRADRGGNTEVF | 0.72609353 |
| PPGAGGAGGAGTT | ASSRADRGGNTEVF | 0.8878304 |
| QLVGLGI | ASSRADRGGNTEVF | 0.8099816 |
| RFLEQVHQL | ASSRADRGGNTEVF | 0.7300416 |
| RGVGYVLR | ASSRADRGGNTEVF | 0.8134483 |
| SATGVFAGVI | ASSRADRGGNTEVF | 0.8990403 |
| SLPAITLP | ASSRADRGGNTEVF | 0.74336666 |
| STALAFTV | ASSRADRGGNTEVF | 0.883993 |
| SVLTVGLGK | ASSRADRGGNTEVF | 0.8244128 |
| TALAGAGSGI | ASSRADRGGNTEVF | 0.8993408 |
| TGLPFTGI | ASSRADRGGNTEVF | 0.591758 |
| VAEHVTV | ASSRADRGGNTEVF | 0.88810855 |
| VDIVLNSL | ASSRADRGGNTEVF | 0.8464309 |
| VLTGGGAVL | ASSRADRGGNTEVF | 0.9143264 |
| VNVGLAYGI | ASSRADRGGNTEVF | 0.8225534 |
| VSSAAVGAV | ASSRADRGGNTEVF | 0.875783 |
| VTSVYEAGV | ASSRADRGGNTEVF | 0.833221 |
| WVALDRGA | ASSRADRGGNTEVF | 0.9096416 |
| YLVGSVVI | ASSRADRGGNTEVF | 0.6400049 |
| GPPGTGKTLL | ASSRADRGGNTEVF | 0.83857477 |
| GVGMTKF | ASSRADRGGNTEVF | 0.5679311 |
| IGIGGSDL | ASSRADRGGNTEVF | 0.8554113 |
| IGTLLAEL | ASSRADRGGNTEVF | 0.8767334 |
| IYTSGTT | ASSRADRGGNTEVF | 0.36392957 |
| LSFPAGV | ASSRADRGGNTEVF | 0.8947476 |
| LYTSGTT | ASSRADRGGNTEVF | 0.79809815 |
| VGAFPFGF | ASSRADRGGNTEVF | 0.75746894 |
| VGAGTVEF | ASSRADRGGNTEVF | 0.89508903 |
| VGKSTLLL | ASSRADRGGNTEVF | 0.8221381 |
| AALAHAGGGIV | ASSRADRGGNTEVF | 0.8963816 |
| AALVAVSLL | ASSRADRGGNTEVF | 0.83235407 |
| AAVSRAVAAL | ASSRADRGGNTEVF | 0.8500222 |
| AGGSSGIGGF | ASSRADRGGNTEVF | 0.8628749 |
| AGLTVANANG | ASSRADRGGNTEVF | 0.6153341 |
| AGLVGAVL | ASSRADRGGNTEVF | 0.75200343 |
| ALSQAFTV | ASSRADRGGNTEVF | 0.9021814 |
| ALTFSVSL | ASSRADRGGNTEVF | 0.64069873 |
| ALTGAALVAL | ASSRADRGGNTEVF | 0.6600195 |
| ASFSIGAGVL | ASSRADRGGNTEVF | 0.5920738 |
| DGSVYKI | ASSRADRGGNTEVF | 0.5741616 |
| EQLAEIV | ASSRADRGGNTEVF | 0.59804726 |
| FAVSELTVM | ASSRADRGGNTEVF | 0.64969707 |
| GAAPLAAGL | ASSRADRGGNTEVF | 0.6653733 |
| GADGGDGGSGGAGG | ASSRADRGGNTEVF | 0.64820135 |
| GFGLDTGGAGG | ASSRADRGGNTEVF | 0.5943377 |
| GLASGGLL | ASSRADRGGNTEVF | 0.65059537 |
| GRIAGAAVLL | ASSRADRGGNTEVF | 0.63292044 |
| GSAGSLNTSF | ASSRADRGGNTEVF | 0.626769 |
| GTGQIGI | ASSRADRGGNTEVF | 0.6387842 |
| GTLIALIL | ASSRADRGGNTEVF | 0.6495865 |

| | | |
|---|---|---|
| IAATMGFVVG | ASSRADRGGNTEVF | 0.61154217 |
| IFAQALL | ASSRADRGGNTEVF | 0.6091511 |
| IGAAVALGF | ASSRADRGGNTEVF | 0.64005685 |
| IGLTPLAL | ASSRADRGGNTEVF | 0.64938444 |
| ILGSNTSTL | ASSRADRGGNTEVF | 0.6443868 |
| IVFSLTVL | ASSRADRGGNTEVF | 0.6661875 |
| LASKATGF | ASSRADRGGNTEVF | 0.6507175 |
| LASVALIL | ASSRADRGGNTEVF | 0.6489815 |
| LFAVGLTL | ASSRADRGGNTEVF | 0.4517712 |
| LFNGGIGTYI | ASSRADRGGNTEVF | 0.55281997 |
| LTLGAGII | ASSRADRGGNTEVF | 0.6318147 |
| LVAAALGDI | ASSRADRGGNTEVF | 0.6203679 |
| NTAFVQLGI | ASSRADRGGNTEVF | 0.43381026 |
| QLFSGAEL | ASSRADRGGNTEVF | 0.6512493 |
| SAATFITL | ASSRADRGGNTEVF | 0.6279936 |
| SNYGLANLGV | ASSRADRGGNTEVF | 0.6046524 |
| SYGGAYAVM | ASSRADRGGNTEVF | 0.6503657 |
| TTGVGGLTLG | ASSRADRGGNTEVF | 0.6199312 |
| TTQTIYI | ASSRADRGGNTEVF | 0.60180444 |
| VAAGAGLGAVI | ASSRADRGGNTEVF | 0.61737466 |
| VAAVVTLL | ASSRADRGGNTEVF | 0.6574185 |
| VGFLGGLI | ASSRADRGGNTEVF | 0.57014143 |
| VGVIPILL | ASSRADRGGNTEVF | 0.65690565 |
| VIFASSLI | ASSRADRGGNTEVF | 0.60800934 |
| VKGGDGGAAAG | ASSRADRGGNTEVF | 0.6595453 |
| VTKTFTNI | ASSRADRGGNTEVF | 0.56193554 |
| VVAAAILL | ASSRADRGGNTEVF | 0.6526848 |
| YAALDVEL | ASSRADRGGNTEVF | 0.6207646 |
| LASVALLL | ASSRADRGGNTEVF | 0.6682704 |
| LDAVHRV | ASSRADRGGNTEVF | 0.6098888 |
| QGLSSSIF | ASSRADRGGNTEVF | 0.6542145 |
| RGTFANIRL | ASSRADRGGNTEVF | 0.6483969 |
| RQLWWGHRI | ASSRADRGGNTEVF | 0.6625166 |
| YGPPGTGKTLL | ASSRADRGGNTEVF | 0.5675874 |
| AALVGAPANVA | ASSRADRGGNTEVF | 0.62227803 |
| AATFRRTE | ASSRADRGGNTEVF | 0.583016 |
| AGALGGGI | ASSRADRGGNTEVF | 0.6413397 |
| AGGAGGAGGIGAG | ASSRADRGGNTEVF | 0.6485724 |
| AGSGALGL | ASSRADRGGNTEVF | 0.41498518 |
| AQIADQI | ASSRADRGGNTEVF | 0.65937907 |
| ASAVAVTYL | ASSRADRGGNTEVF | 0.57400656 |
| ASTVLGGTGG | ASSRADRGGNTEVF | 0.59306186 |
| AVGGEPAK | ASSRADRGGNTEVF | 0.65373856 |
| CEQGTFGGTC | ASSRADRGGNTEVF | 0.63732123 |
| DEGKAPAG | ASSRADRGGNTEVF | 0.64888346 |
| DTVVFPAL | ASSRADRGGNTEVF | 0.62363786 |
| FEGITMAG | ASSRADRGGNTEVF | 0.6307346 |
| FPLVVQD | ASSRADRGGNTEVF | 0.64373523 |
| GAFTTPQ | ASSRADRGGNTEVF | 0.5429762 |
| GFGGFGTAGGVGGA | ASSRADRGGNTEVF | 0.5877981 |
| GFLAQFL | ASSRADRGGNTEVF | 0.6610985 |

| | | |
|---|---|---|
| GGAGGAGGNAGVA | ASSRADRGGNTEVF | 0.7668655 |
| GGVGTAGAGGAGGAGG | ASSRADRGGNTEVF | 0.78428006 |
| GNGGGGDAGNAGSGGN | ASSRADRGGNTEVF | 0.6793456 |
| GTGGAGGAGNDAGS | ASSRADRGGNTEVF | 0.61168295 |
| GTTNTGWFNS | ASSRADRGGNTEVF | 0.65944344 |
| IFMGPLFTL | ASSRADRGGNTEVF | 0.7718053 |
| ILAHAIEAI | ASSRADRGGNTEVF | 0.78449064 |
| IRSAIREAL | ASSRADRGGNTEVF | 0.78237224 |
| ITFAGGG | ASSRADRGGNTEVF | 0.5130834 |
| ITGMSGHQEV | ASSRADRGGNTEVF | 0.75778437 |
| IVAGLACTL | ASSRADRGGNTEVF | 0.7045882 |
| IVGGGLAGA | ASSRADRGGNTEVF | 0.71453154 |
| IVTGVLNI | ASSRADRGGNTEVF | 0.70806587 |
| LAQQGCI | ASSRADRGGNTEVF | 0.6157431 |
| LCLHVEN | ASSRADRGGNTEVF | 0.69724816 |
| LDAISDGS | ASSRADRGGNTEVF | 0.52954173 |
| LGGFAAML | ASSRADRGGNTEVF | 0.5209857 |
| LGGKNPMI | ASSRADRGGNTEVF | 0.73828804 |
| LGPKGVAAIL | ASSRADRGGNTEVF | 0.72041905 |
| LMVGVSI | ASSRADRGGNTEVF | 0.7765556 |
| LVGAAFVL | ASSRADRGGNTEVF | 0.77572227 |
| LVLAVAPL | ASSRADRGGNTEVF | 0.696236 |
| MAAIAVIDAA | ASSRADRGGNTEVF | 0.21698219 |
| NNGINAELQ | ASSRADRGGNTEVF | 0.52070284 |
| RGEPLVLVHG | ASSRADRGGNTEVF | 0.6272613 |
| RGGAGGAAGLI | ASSRADRGGNTEVF | 0.73180133 |
| RSYRMHPAV | ASSRADRGGNTEVF | 0.23103742 |
| SAAHLLAGQLA | ASSRADRGGNTEVF | 0.7378607 |
| SAAVGSY | ASSRADRGGNTEVF | 0.72857815 |
| SPHAVLHRI | ASSRADRGGNTEVF | 0.6933037 |
| SQIAKVL | ASSRADRGGNTEVF | 0.77047235 |
| TALGHLVP | ASSRADRGGNTEVF | 0.7125943 |
| TAPTTSGAA | ASSRADRGGNTEVF | 0.52717966 |
| TDGDDGNGGSPGT | ASSRADRGGNTEVF | 0.48485023 |
| TDGLGEN | ASSRADRGGNTEVF | 0.78045416 |
| TGGPPLPGQA | ASSRADRGGNTEVF | 0.42323864 |
| TIPAITI | ASSRADRGGNTEVF | 0.7714158 |
| TIPGFTI | ASSRADRGGNTEVF | 0.4947968 |
| TSIASGALA | ASSRADRGGNTEVF | 0.7716304 |
| TVTLPGAAI | ASSRADRGGNTEVF | 0.42465854 |
| VASGLSAALP | ASSRADRGGNTEVF | 0.76826733 |
| VGAGVIGI | ASSRADRGGNTEVF | 0.7400819 |
| VGPKRFGV | ASSRADRGGNTEVF | 0.78574955 |
| VNVAFPD | ASSRADRGGNTEVF | 0.6169749 |
| VSMTNSGVS | ASSRADRGGNTEVF | 0.72726214 |
| VTDVLGDLSV | ASSRADRGGNTEVF | 0.4119509 |
| VVVVGAGFAGL | ASSRADRGGNTEVF | 0.7836373 |
| VYTVGTVV | ASSRADRGGNTEVF | 0.5883913 |
| WTTFSGP | ASSRADRGGNTEVF | 0.6131142 |
| YDELVEAF | ASSRADRGGNTEVF | 0.4397849 |
| YGTGGAGGNGG | ASSRADRGGNTEVF | 0.7022298 |

| | | |
|---|---|---|
| YTNVMAV | ASSRADRGGNTEVF | 0.7824899 |
| FLEQVHQL | ASSRADRGGNTEVF | 0.5954226 |
| GLSLLAVAV | ASSRADRGGNTEVF | 0.7762227 |
| LEQVHQL | ASSRADRGGNTEVF | 0.7468877 |
| LNAAGAI | ASSRADRGGNTEVF | 0.4810232 |
| PAVHASL | ASSRADRGGNTEVF | 0.7372122 |
| TELEDDA | ASSRADRGGNTEVF | 0.51091987 |
| TPPGPPN | ASSRADRGGNTEVF | 0.76345026 |
| TPPPPPPP | ASSRADRGGNTEVF | 0.48755836 |
| TQSPALI | ASSSTGGALEQY | 0.7254584 |
| EFAAAFPQ | ASSSTGGALEQY | 0.741212 |
| LLSGNLDVL | ASSSTGGALEQY | 0.5822791 |
| GPANLGNYNI | ASSSTGGALEQY | 0.4413796 |
| TLGPTLP | ASSSTGGALEQY | 0.7822666 |
| VYAYSSCCK | ASSSTGGALEQY | 0.34343952 |
| APAPAPAPNI | ASSSTGGALEQY | 0.6921863 |
| PGGPVPIP | ASSSTGGALEQY | 0.14647067 |
| FLTVALEP | ASSSTGGALEQY | 0.15608715 |
| SQYFGANL | ASSSTGGALEQY | 0.21770915 |
| GGVKPAA | ASSSTGGALEQY | 0.33182198 |
| GGTGGAGGAGGW | ASSSTGGALEQY | 0.7769319 |
| GVGGVAGLGGAGG | ASSSTGGALEQY | 0.309201 |
| VLLVLATTI | ASSSTGGALEQY | 0.16014904 |
| IGIFAAML | ASSSTGGALEQY | 0.33466482 |
| AVGDLID | ASSSTGGALEQY | 0.17327303 |
| NISPLPL | ASSSTGGALEQY | 0.22706121 |
| VTAVFVAI | ASSSTGGALEQY | 0.21917179 |
| SGDDNDAAL | ASSSTGGALEQY | 0.20829815 |
| AVAALAHS | ASSSTGGALEQY | 0.21250364 |
| GGVTVMATP | ASSSTGGALEQY | 0.16264527 |
| AGLAANVRD | ASSSTGGALEQY | 0.32656455 |
| TNRPDLIDPALLRPG | ASSSTGGALEQY | 0.5162612 |
| LANLGVTAL | ASSSTGGALEQY | 0.43458277 |
| TGERIEL | ASSSTGGALEQY | 0.33212447 |
| NGEELAA | ASSSTGGALEQY | 0.37212053 |
| VLSLVWG | ASSSTGGALEQY | 0.2618695 |
| IGYALWL | ASSSTGGALEQY | 0.10510325 |
| LSLLLAPL | ASSSTGGALEQY | 0.121967055 |
| TLDEALK | ASSSTGGALEQY | 0.21402286 |
| TDGAGTAEGG | ASSSTGGALEQY | 0.17926699 |
| NAPGGAGGNGG | ASSSTGGALEQY | 0.11357724 |
| VSVFMGMGGNF | ASSSTGGALEQY | 0.26962572 |
| LASYLAAL | ASSSTGGALEQY | 0.19051248 |
| LVGGTPAL | ASSSTGGALEQY | 0.18698144 |
| VRNADVL | ASSSTGGALEQY | 0.47196627 |
| TCSSPFLVSAM | ASSSTGGALEQY | 0.18336001 |
| VHITDAE | ASSSTGGALEQY | 0.15692113 |
| EYAAPSF | ASSSTGGALEQY | 0.17217551 |
| VVDAFHGGL | ASSSTGGALEQY | 0.37002137 |
| ASVVDPQ | ASSSTGGALEQY | 0.14758211 |
| SYNEFGGLS | ASSSTGGALEQY | 0.32672477 |

| | | |
|---|---|---|
| DPDGSVY | ASSSTGGALEQY | 0.16876456 |
| MVSTIHATE | ASSSTGGALEQY | 0.42984456 |
| LAAGVADPV | ASSSTGGALEQY | 0.14353293 |
| TKEQIAA | ASSSTGGALEQY | 0.27926248 |
| SLASPAAL | ASSSTGGALEQY | 0.2081672 |
| IGLVFYGV | ASSSTGGALEQY | 0.727678 |
| FLLDASNL | ASSSTGGALEQY | 0.17393295 |
| YQVGNFP | ASSSTGGALEQY | 0.20240508 |
| ESLLSQST | ASSSTGGALEQY | 0.18467633 |
| LFTVGFAAVG | ASSSTGGALEQY | 0.80183107 |
| ATPPTPT | ASSSTGGALEQY | 0.124070436 |
| TDLATLG | ASSSTGGALEQY | 0.19327687 |
| SAGGTGR | ASSSTGGALEQY | 0.13565727 |
| GFSNSGS | ASSSTGGALEQY | 0.20727871 |
| GGNGGSGGTSV | ASSSTGGALEQY | 0.29518297 |
| GTSVATGGAGN | ASSSTGGALEQY | 0.10664797 |
| NGGNGGSGGTS | ASSSTGGALEQY | 0.25515142 |
| AVVKFPH | ASSSTGGALEQY | 0.13373776 |
| TPPPPPP | ASSSTGGALEQY | 0.14916033 |
| AAPAAAAAPA | ASSSTGGALEQY | 0.27470368 |
| IALNEDL | ASSSTGGALEQY | 0.21252847 |
| IVNVSSM | ASSSTGGALEQY | 0.37760943 |
| TGIPPAPR | ASSSTGGALEQY | 0.16804041 |
| FDLGGGTF | ASSSTGGALEQY | 0.18878391 |
| FDLGGGTFDVS | ASSSTGGALEQY | 0.24830844 |
| NEPTAAA | ASSSTGGALEQY | 0.14194727 |
| NEPTAAALA | ASSSTGGALEQY | 0.17225796 |
| GSIINISSI | ASSSTGGALEQY | 0.37020773 |
| VAPGNAGTA | ASSSTGGALEQY | 0.62795806 |
| TGGGASLE | ASSSTGGALEQY | 0.8350463 |
| VAVYNHYKRI | ASSSTGGALEQY | 0.42586476 |
| TDPAPATT | ASSSTGGALEQY | 0.3531052 |
| TNRPDILDPALLRPG | ASSSTGGALEQY | 0.63162553 |
| LTTTTTS | ASSSTGGALEQY | 0.76902515 |
| PTGSGKTL | ASSSTGGALEQY | 0.8595004 |
| TGSGKTL | ASSSTGGALEQY | 0.7590261 |
| TIGHVDHGK | ASSSTGGALEQY | 0.3541668 |
| TIGHVDHGKTT | ASSSTGGALEQY | 0.6300664 |
| GHVDHGKTT | ASSSTGGALEQY | 0.4477436 |
| AQFEHTLL | ASSSTGGALEQY | 0.5555355 |
| IVNQDEL | ASSSTGGALEQY | 0.7061622 |
| TNGSQFF | ASSSTGGALEQY | 0.6518786 |
| TGKGIVE | ASSSTGGALEQY | 0.40326375 |
| LISNASDA | ASSSTGGALEQY | 0.42461836 |
| AELRTLGELTKIAKAHGAQ | ASSSTGGALEQY | 0.76529235 |
| APNAVAAAEPSW | ASSSTGGALEQY | 0.81773263 |
| APQVAGAATAVL | ASSSTGGALEQY | 0.7962356 |
| ARRDPAAAAAILPTDA | ASSSTGGALEQY | 0.84671223 |
| DAAARDVLAEAGLAEYF | ASSSTGGALEQY | 0.7450706 |
| DDQIVLSAKANLAKVEQ | ASSSTGGALEQY | 0.784047 |
| DELLPDTGEDSDEDGH | ASSSTGGALEQY | 0.1241147 |

| | | |
|---|---|---|
| ELPEAPANRATVPQD | ASSSTGGALEQY | 0.21934704 |
| FGQSKIERVTGIG | ASSSTGGALEQY | 0.5307257 |
| GAGGTGSTAGGAGGAGGAGGLYA | ASSSTGGALEQY | 0.5732312 |
| GHGGTGAAGGNGGTGGAGGSGIDG | ASSSTGGALEQY | 0.21193832 |
| GNGGNGGAGGEGGPGGAGGAGA | ASSSTGGALEQY | 0.7447183 |
| GSTAGNGGNGGNATNGTIAGQPAG | ASSSTGGALEQY | 0.43637428 |
| GTGGDGGNGGNGGTGG | ASSSTGGALEQY | 0.32720298 |
| GVRRPAGIPADAPV | ASSSTGGALEQY | 0.8035643 |
| GYRQVTDAHLLTLARR | ASSSTGGALEQY | 0.42588204 |
| IAKMGQDGHDRG | ASSSTGGALEQY | 0.3400371 |
| KAEPSIEIQYT | ASSSTGGALEQY | 0.47590187 |
| KPVRIRMAADEDTVAVT | ASSSTGGALEQY | 0.7242235 |
| MPPRIAGMRLLVIKPEP | ASSSTGGALEQY | 0.23291567 |
| MKFYAVALFRSRGVPPTR | ASSSTGGALEQY | 0.7513397 |
| NAGGFGILGAVAHSP | ASSSTGGALEQY | 0.5802332 |
| NEGREITLAMAEM | ASSSTGGALEQY | 0.7450379 |
| NELVEWSHPLGHRWRWE | ASSSTGGALEQY | 0.2751415 |
| NLGAGGTGGNGGNPG | ASSSTGGALEQY | 0.7453245 |
| NTEWPKWETVMAADPVD | ASSSTGGALEQY | 0.61936224 |
| PASAAPAETQP | ASSSTGGALEQY | 0.86873 |
| QQSSDGKNGVRAAEITGE | ASSSTGGALEQY | 0.4878406 |
| RAPDAVTTSWWKE | ASSSTGGALEQY | 0.6218999 |
| SGFGNVGAHTSGWFNQ | ASSSTGGALEQY | 0.6912099 |
| SGVAGAVGQAASVGGLKV | ASSSTGGALEQY | 0.83446264 |
| SNGGAGTGSAGGNGGT | ASSSTGGALEQY | 0.18698819 |
| TGGAGGEGGAAG | ASSSTGGALEQY | 0.5045378 |
| TGGDGGAAGAGGAPGHG | ASSSTGGALEQY | 0.21366665 |
| TMPPLLAAIANAGRHGV | ASSSTGGALEQY | 0.6056942 |
| TSSTAGNSGGA | ASSSTGGALEQY | 0.7163689 |
| VPATVPVGILC | ASSSTGGALEQY | 0.15865189 |
| VQRPDSGGVVLTGRIS | ASSSTGGALEQY | 0.6509111 |
| WTPNPVPATKGVPAGLVQAT | ASSSTGGALEQY | 0.55154467 |
| YALKRALAKFSDRFDVVII | ASSSTGGALEQY | 0.42601916 |
| YLIGSVIV | ASSSTGGALEQY | 0.79881614 |
| YGQLVAVID | ASSSTGGALEQY | 0.6479045 |
| VTVGGMTLPA | ASSSTGGALEQY | 0.793251 |
| VLALPAV | ASSSTGGALEQY | 0.6508714 |
| VIPPTLNYET | ASSSTGGALEQY | 0.5133883 |
| VGMGGMHPSSG | ASSSTGGALEQY | 0.6666239 |
| VGLAGGAATA | ASSSTGGALEQY | 0.32100523 |
| VGGAGGAGGSGP | ASSSTGGALEQY | 0.4691809 |
| VFAAFMV | ASSSTGGALEQY | 0.7961705 |
| TLYDKLR | ASSSTGGALEQY | 0.45117128 |
| TLAVFTVL | ASSSTGGALEQY | 0.71485233 |
| TGSVQPARV | ASSSTGGALEQY | 0.5164665 |
| TGGGAGSGGT | ASSSTGGALEQY | 0.4408146 |
| SHLETSL | ASSSTGGALEQY | 0.5788232 |
| SFTTGMT | ASSSTGGALEQY | 0.6462749 |
| QVEVAKRMGVRQPTVS | ASSSTGGALEQY | 0.7219794 |
| QGSGGPALAM | ASSSTGGALEQY | 0.55039597 |
| MLAPSNP | ASSSTGGALEQY | 0.42942357 |

| | | |
|---|---|---|
| LVNNAAI | ASSSTGGALEQY | 0.6057557 |
| LTGEINTLA | ASSSTGGALEQY | 0.41144148 |
| LSIFGPV | ASSSTGGALEQY | 0.5398391 |
| LLALLANL | ASSSTGGALEQY | 0.5755373 |
| LISNASDALDKLR | ASSSTGGALEQY | 0.6064321 |
| LINEAVP | ASSSTGGALEQY | 0.5516402 |
| LGYAFSGL | ASSSTGGALEQY | 0.39024135 |
| LDFVAVAL | ASSSTGGALEQY | 0.6479051 |
| LAQPGAE | ASSSTGGALEQY | 0.62363005 |
| ISNLGAVTGF | ASSSTGGALEQY | 0.48619992 |
| HQEDPGRI | ASSSTGGALEQY | 0.68057346 |
| GTPGEPTP | ASSSTGGALEQY | 0.6157713 |
| GFFNSGS | ASSSTGGALEQY | 0.51521975 |
| GAGGGGLTAGAN | ASSSTGGALEQY | 0.23379205 |
| FVDALVNM | ASSSTGGALEQY | 0.28657418 |
| FLTVAPEL | ASSSTGGALEQY | 0.44973594 |
| FETLTAL | ASSSTGGALEQY | 0.50840354 |
| DVINPIEL | ASSSTGGALEQY | 0.23445445 |
| APVSGAPV | ASSSTGGALEQY | 0.6703044 |
| ALVGGLAW | ASSSTGGALEQY | 0.450997 |
| ALAAAAATAI | ASSSTGGALEQY | 0.49345803 |
| AFGLGLLL | ASSSTGGALEQY | 0.6927952 |
| AFFAAASL | ASSSTGGALEQY | 0.4892508 |
| AAAALPAFSPP | ASSSTGGALEQY | 0.48664623 |
| VVLAPAL | ASSSTGGALEQY | 0.42522234 |
| TIGHVDHGKTTL | ASSSTGGALEQY | 0.68881667 |
| LISNASDALDK | ASSSTGGALEQY | 0.28579426 |
| LISNASDAL | ASSSTGGALEQY | 0.63578427 |
| ISNASDALDK | ASSSTGGALEQY | 0.42687774 |
| NQAETLVYQTE | ASSSTGGALEQY | 0.63024294 |
| TGGTGGTAGAG | ASSSTGGALEQY | 0.34628814 |
| TGSTAGGAGGA | ASSSTGGALEQY | 0.6028768 |
| DPLAVPVAAGPV | ASSSTGGALEQY | 0.5444692 |
| GGTGGTPGNGGN | ASSSTGGALEQY | 0.74255365 |
| PAGLAGVDPATL | ASSSTGGALEQY | 0.50262165 |
| GNNLVVEAPHAGASV | ASSSTGGALEQY | 0.56108963 |
| KVTRIMSASTLEQPA | ASSSTGGALEQY | 0.46890056 |
| LGFALPIALPRRTVV | ASSSTGGALEQY | 0.7165931 |
| TRLGNLTARKLVVLV | ASSSTGGALEQY | 0.32639614 |
| WSQQRRLIYFPSAGP | ASSSTGGALEQY | 0.4999152 |
| GPNGNPDPMAAAVDIR | ASSSTGGALEQY | 0.43132555 |
| TQRCVSVADLRAAASR | ASSSTGGALEQY | 0.6377592 |
| QFVLDVAEHPVEVLTETV | ASSSTGGALEQY | 0.49186972 |
| GGAGGTGGAGGNGGTQAGDG | ASSSTGGALEQY | 0.2825237 |
| PLLATLRAIAVDVDAKPAQV | ASSSTGGALEQY | 0.63774943 |
| APKFRTGGYMIDFWGVGYQVAKRM | ASSSTGGALEQY | 0.46841544 |
| GAGGAGGAGGAAGGPGGTGGTGGN | ASSSTGGALEQY | 0.38293964 |
| IGALGLLATDVAPAGSVAFGVALA | ASSSTGGALEQY | 0.64320135 |
| AGGAGGIGGNGGAGGAGGVLMGNG | ASSSTGGALEQY | 0.5612562 |
| ADELIGLL | ASSSTGGALEQY | 0.6282276 |
| ADRALKA | ASSSTGGALEQY | 0.55847764 |

| | | |
|---|---|---|
| AGAVGQVGGAGGA | ASSSTGGALEQY | 0.5178752 |
| AGGGGGLGGAP | ASSSTGGALEQY | 0.66860324 |
| AGGLLYGNGGAG | ASSSTGGALEQY | 0.48831415 |
| AGLESIGL | ASSSTGGALEQY | 0.4008425 |
| AGLKTGII | ASSSTGGALEQY | 0.6391216 |
| ASLAALLL | ASSSTGGALEQY | 0.8201394 |
| ASVINISSI | ASSSTGGALEQY | 0.836749 |
| ATPPGPA | ASSSTGGALEQY | 0.7807907 |
| ATVASALV | ASSSTGGALEQY | 0.6996838 |
| ATVASLIL | ASSSTGGALEQY | 0.78084683 |
| AVNGPVLIRH | ASSSTGGALEQY | 0.8279007 |
| DGAAGAVGI | ASSSTGGALEQY | 0.84364086 |
| DPLLHGI | ASSSTGGALEQY | 0.8323867 |
| ELGAALGAL | ASSSTGGALEQY | 0.68348366 |
| ELGRTQAV | ASSSTGGALEQY | 0.83935255 |
| FVAGGYLS | ASSSTGGALEQY | 0.79473287 |
| GASGGAGGAGGW | ASSSTGGALEQY | 0.79237944 |
| GGAGAGGDGGAGGI | ASSSTGGALEQY | 0.79951894 |
| GTGHNGSQPGMGGNG | ASSSTGGALEQY | 0.815593 |
| GYAAASASAS | ASSSTGGALEQY | 0.7948537 |
| HQVDVAAVQQG | ASSSTGGALEQY | 0.7046471 |
| IGAAIAHT | ASSSTGGALEQY | 0.82609165 |
| IGMIEVAQL | ASSSTGGALEQY | 0.82878304 |
| IRLAPEHTSAMD | ASSSTGGALEQY | 0.826136 |
| IYNVSVL | ASSSTGGALEQY | 0.85226953 |
| KQGFLGM | ASSSTGGALEQY | 0.8398306 |
| LESVITQL | ASSSTGGALEQY | 0.8232555 |
| LGLGALVVV | ASSSTGGALEQY | 0.3366166 |
| LIGNGGNGGSGGI | ASSSTGGALEQY | 0.5720236 |
| LNAINAPTL | ASSSTGGALEQY | 0.734568 |
| LQLLNTGV | ASSSTGGALEQY | 0.7776204 |
| LQTSIEF | ASSSTGGALEQY | 0.3437971 |
| MSFAVSEL | ASSSTGGALEQY | 0.7807435 |
| NTGALGL | ASSSTGGALEQY | 0.8059686 |
| PPGAGGAGGAGTT | ASSSTGGALEQY | 0.7372861 |
| QLVGLGI | ASSSTGGALEQY | 0.839296 |
| RFLEQVHQL | ASSSTGGALEQY | 0.7881694 |
| RGVGYVLR | ASSSTGGALEQY | 0.6636455 |
| SATGVFAGVI | ASSSTGGALEQY | 0.6896748 |
| SLPAITLP | ASSSTGGALEQY | 0.84423935 |
| STALAFTV | ASSSTGGALEQY | 0.55066955 |
| SVLTVGLGK | ASSSTGGALEQY | 0.8372493 |
| TALAGAGSGI | ASSSTGGALEQY | 0.6921188 |
| TGLPFTGI | ASSSTGGALEQY | 0.837729 |
| VAEHVTV | ASSSTGGALEQY | 0.5633397 |
| VDIVLNSL | ASSSTGGALEQY | 0.83510584 |
| VLTGGGAVL | ASSSTGGALEQY | 0.797984 |
| VNVGLAYGI | ASSSTGGALEQY | 0.8564793 |
| VSSAAVGAV | ASSSTGGALEQY | 0.7587311 |
| VTSVYEAGV | ASSSTGGALEQY | 0.8273622 |
| WVALDRGA | ASSSTGGALEQY | 0.7502704 |

| | | |
|---|---|---|
| YLVGSVVI | ASSSTGGALEQY | 0.8339498 |
| GPPGTGKTLL | ASSSTGGALEQY | 0.6107923 |
| GVGMTKF | ASSSTGGALEQY | 0.7854743 |
| IGIGGSDL | ASSSTGGALEQY | 0.63470346 |
| IGTLLAEL | ASSSTGGALEQY | 0.8118118 |
| IYTSGTT | ASSSTGGALEQY | 0.83245873 |
| LSFPAGV | ASSSTGGALEQY | 0.4143532 |
| LYTSGTT | ASSSTGGALEQY | 0.8492254 |
| VGAFPFGF | ASSSTGGALEQY | 0.75078654 |
| VGAGTVEF | ASSSTGGALEQY | 0.70485824 |
| VGKSTLLL | ASSSTGGALEQY | 0.83976 |
| AALAHAGGGIV | ASSSTGGALEQY | 0.8093369 |
| AALVAVSLL | ASSSTGGALEQY | 0.8399562 |
| AAVSRAVAAL | ASSSTGGALEQY | 0.72843057 |
| AGGSSGIGGF | ASSSTGGALEQY | 0.792323 |
| AGLTVANANG | ASSSTGGALEQY | 0.8169285 |
| AGLVGAVL | ASSSTGGALEQY | 0.7248856 |
| ALSQAFTV | ASSSTGGALEQY | 0.64317423 |
| ALTFSVSL | ASSSTGGALEQY | 0.8454606 |
| ALTGAALVAL | ASSSTGGALEQY | 0.80398816 |
| ASFSIGAGVL | ASSSTGGALEQY | 0.865355 |
| DGSVYKI | ASSSTGGALEQY | 0.71448255 |
| EQLAEIV | ASSSTGGALEQY | 0.60203993 |
| FAVSELTVM | ASSSTGGALEQY | 0.7331431 |
| GAAPLAAGL | ASSSTGGALEQY | 0.8517974 |
| GADGGDGGSGGAGG | ASSSTGGALEQY | 0.8644489 |
| GFGLDTGGAGG | ASSSTGGALEQY | 0.8160369 |
| GLASGGLL | ASSSTGGALEQY | 0.5333375 |
| GRIAGAAVLL | ASSSTGGALEQY | 0.8052778 |
| GSAGSLNTSF | ASSSTGGALEQY | 0.7180468 |
| GTGQIGI | ASSSTGGALEQY | 0.7494934 |
| GTLIALIL | ASSSTGGALEQY | 0.8257201 |
| IAATMGFVVG | ASSSTGGALEQY | 0.76738966 |
| IFAQALL | ASSSTGGALEQY | 0.6899966 |
| IGAAVALGF | ASSSTGGALEQY | 0.65646803 |
| IGLTPLAL | ASSSTGGALEQY | 0.81268466 |
| ILGSNTSTL | ASSSTGGALEQY | 0.81508607 |
| IVFSLTVL | ASSSTGGALEQY | 0.81677437 |
| LASKATGF | ASSSTGGALEQY | 0.87613547 |
| LASVALIL | ASSSTGGALEQY | 0.8505055 |
| LFAVGLTL | ASSSTGGALEQY | 0.86260986 |
| LFNGGIGTYI | ASSSTGGALEQY | 0.21700442 |
| LTLGAGII | ASSSTGGALEQY | 0.41137582 |
| LVAAALGDI | ASSSTGGALEQY | 0.6871971 |
| NTAFVQLGI | ASSSTGGALEQY | 0.71326864 |
| QLFSGAEL | ASSSTGGALEQY | 0.44292897 |
| SAATFITL | ASSSTGGALEQY | 0.8053755 |
| SNYGLANLGV | ASSSTGGALEQY | 0.67799896 |
| SYGGAYAVM | ASSSTGGALEQY | 0.63691396 |
| TTGVGGLTLG | ASSSTGGALEQY | 0.8416542 |
| TTQTIYI | ASSSTGGALEQY | 0.7052062 |

| | | |
|---|---|---|
| VAAGAGLGAVI | ASSSTGGALEQY | 0.62673604 |
| VAAVVTLL | ASSSTGGALEQY | 0.61499393 |
| VGFLGGLI | ASSSTGGALEQY | 0.83218277 |
| VGVIPILL | ASSSTGGALEQY | 0.4514993 |
| VIFASSLI | ASSSTGGALEQY | 0.8256626 |
| VKGGDGGAAAG | ASSSTGGALEQY | 0.6233343 |
| VTKTFTNI | ASSSTGGALEQY | 0.8486093 |
| VVAAAILL | ASSSTGGALEQY | 0.43543833 |
| YAALDVEL | ASSSTGGALEQY | 0.8443857 |
| LASVALLL | ASSSTGGALEQY | 0.7712921 |
| LDAVHRV | ASSSTGGALEQY | 0.8860105 |
| QGLSSSIF | ASSSTGGALEQY | 0.7149317 |
| RGTFANIRL | ASSSTGGALEQY | 0.7548746 |
| RQLWWGHRI | ASSSTGGALEQY | 0.6154326 |
| YGPPGTGKTLL | ASSSTGGALEQY | 0.86130357 |
| AALVGAPANVA | ASSSTGGALEQY | 0.3889792 |
| AATFRRTE | ASSSTGGALEQY | 0.7543954 |
| AGALGGGI | ASSSTGGALEQY | 0.4259022 |
| AGGAGGAGGIGAG | ASSSTGGALEQY | 0.8139035 |
| AGSGALGL | ASSSTGGALEQY | 0.8194947 |
| AQIADQI | ASSSTGGALEQY | 0.3305482 |
| ASAVAVTYL | ASSSTGGALEQY | 0.82720995 |
| ASTVLGGTGG | ASSSTGGALEQY | 0.7819346 |
| AVGGEPAK | ASSSTGGALEQY | 0.6364188 |
| CEQGTFGGTC | ASSSTGGALEQY | 0.868148 |
| DEGKAPAG | ASSSTGGALEQY | 0.76874185 |
| DTVVFPAL | ASSSTGGALEQY | 0.8713173 |
| FEGITMAG | ASSSTGGALEQY | 0.7725078 |
| FPLVVQD | ASSSTGGALEQY | 0.67738163 |
| GAFTTPQ | ASSSTGGALEQY | 0.7945057 |
| GFGGFGTAGGVGGA | ASSSTGGALEQY | 0.5745384 |
| GFLAQFL | ASSSTGGALEQY | 0.5949546 |
| GGAGGAGGNAGVA | ASSSTGGALEQY | 0.8653481 |
| GGVGTAGAGGAGGAGG | ASSSTGGALEQY | 0.89551973 |
| GNGGGGDAGNAGSGGN | ASSSTGGALEQY | 0.8803779 |
| GTGGAGGAGNDAGS | ASSSTGGALEQY | 0.8027861 |
| GTTNTGWFNS | ASSSTGGALEQY | 0.8061657 |
| IFMGPLFTL | ASSSTGGALEQY | 0.83769166 |
| ILAHAIEAI | ASSSTGGALEQY | 0.89994824 |
| IRSAIREAL | ASSSTGGALEQY | 0.8882947 |
| ITFAGGG | ASSSTGGALEQY | 0.8908638 |
| ITGMSGHQEV | ASSSTGGALEQY | 0.8270882 |
| IVAGLACTL | ASSSTGGALEQY | 0.8643157 |
| IVGGGLAGA | ASSSTGGALEQY | 0.8584359 |
| IVTGVLNI | ASSSTGGALEQY | 0.82561886 |
| LAQQGCI | ASSSTGGALEQY | 0.88384444 |
| LCLHVEN | ASSSTGGALEQY | 0.8761042 |
| LDAISDGS | ASSSTGGALEQY | 0.82964844 |
| LGGFAAML | ASSSTGGALEQY | 0.8793577 |
| LGGKNPMI | ASSSTGGALEQY | 0.8575128 |
| LGPKGVAAIL | ASSSTGGALEQY | 0.88106537 |

| | | |
|---|---|---|
| LMVGVSI | ASSSTGGALEQY | 0.87438667 |
| LVGAAFVL | ASSSTGGALEQY | 0.91594714 |
| LVLAVAPL | ASSSTGGALEQY | 0.90051675 |
| MAAIAVIDAA | ASSSTGGALEQY | 0.89689267 |
| NNGINAELQ | ASSSTGGALEQY | 0.55482244 |
| RGEPLVLVHG | ASSSTGGALEQY | 0.751933 |
| RGGAGGAAGLI | ASSSTGGALEQY | 0.84806526 |
| RSYRMHPAV | ASSSTGGALEQY | 0.8508838 |
| SAAHLLAGQLA | ASSSTGGALEQY | 0.6719713 |
| SAAVGSY | ASSSTGGALEQY | 0.8724958 |
| SPHAVLHRI | ASSSTGGALEQY | 0.87014496 |
| SQIAKVL | ASSSTGGALEQY | 0.77651167 |
| TALGHLVP | ASSSTGGALEQY | 0.8918259 |
| TAPTTSGAA | ASSSTGGALEQY | 0.81963986 |
| TDGDDGNGGSPGT | ASSSTGGALEQY | 0.8355768 |
| TDGLGEN | ASSSTGGALEQY | 0.849262 |
| TGGPPLPGQA | ASSSTGGALEQY | 0.90300715 |
| TIPAITI | ASSSTGGALEQY | 0.80860883 |
| TIPGFTI | ASSSTGGALEQY | 0.8835772 |
| TSIASGALA | ASSSTGGALEQY | 0.8177042 |
| TVTLPGAAI | ASSSTGGALEQY | 0.90787446 |
| VASGLSAALP | ASSSTGGALEQY | 0.785499 |
| VGAGVIGI | ASSSTGGALEQY | 0.8975283 |
| VGPKRFGV | ASSSTGGALEQY | 0.8532553 |
| VNVAFPD | ASSSTGGALEQY | 0.91212046 |
| VSMTNSGVS | ASSSTGGALEQY | 0.8645927 |
| VTDVLGDLSV | ASSSTGGALEQY | 0.8898834 |
| VVVVGAGFAGL | ASSSTGGALEQY | 0.85180056 |
| VYTVGTVV | ASSSTGGALEQY | 0.8877002 |
| WTTFSGP | ASSSTGGALEQY | 0.7225396 |
| YDELVEAF | ASSSTGGALEQY | 0.8684397 |
| YGTGGAGGNGG | ASSSTGGALEQY | 0.6831974 |
| YTNVMAV | ASSSTGGALEQY | 0.87674034 |
| FLEQVHQL | ASSSTGGALEQY | 0.89094687 |
| GLSLLAVAV | ASSSTGGALEQY | 0.6595475 |
| LEQVHQL | ASSSTGGALEQY | 0.904239 |
| LNAAGAI | ASSSTGGALEQY | 0.8025901 |
| PAVHASL | ASSSTGGALEQY | 0.8518063 |
| TELEDDA | ASSSTGGALEQY | 0.8988043 |
| TPPGPPN | ASSSTGGALEQY | 0.8381145 |
| TPPPPPPP | ASSSTGGALEQY | 0.9078961 |
| TQSPALI | AWTGLANSDYT | 0.8325243 |
| EFAAAFPQ | AWTGLANSDYT | 0.82871103 |
| LLSGNLDVL | AWTGLANSDYT | 0.88230467 |
| GPANLGNYNI | AWTGLANSDYT | 0.6556657 |
| TLGPTLP | AWTGLANSDYT | 0.7932651 |
| VYAYSSCCK | AWTGLANSDYT | 0.9101958 |
| APAPAPAPNI | AWTGLANSDYT | 0.5603507 |
| PGGPVPIP | AWTGLANSDYT | 0.63633966 |
| FLTVALEP | AWTGLANSDYT | 0.5265975 |
| SQYFGANL | AWTGLANSDYT | 0.4361561 |

| | | |
|---|---|---|
| GGVKPAA | AWTGLANSDYT | 0.5811773 |
| GGTGGAGGAGGW | AWTGLANSDYT | 0.6038519 |
| GVGGVAGLGGAGG | AWTGLANSDYT | 0.64647734 |
| VLLVLATTI | AWTGLANSDYT | 0.5579753 |
| IGIFAAML | AWTGLANSDYT | 0.4178742 |
| AVGDLID | AWTGLANSDYT | 0.62577003 |
| NISPLPL | AWTGLANSDYT | 0.5917401 |
| VTAVFVAI | AWTGLANSDYT | 0.54871094 |
| SGDDNDAAL | AWTGLANSDYT | 0.598811 |
| AVAALAHS | AWTGLANSDYT | 0.59079456 |
| GGVTVMATP | AWTGLANSDYT | 0.5641097 |
| AGLAANVRD | AWTGLANSDYT | 0.43661824 |
| TNRPDLIDPALLRPG | AWTGLANSDYT | 0.5993708 |
| LANLGVTAL | AWTGLANSDYT | 0.61363685 |
| TGERIEL | AWTGLANSDYT | 0.61407065 |
| NGEELAA | AWTGLANSDYT | 0.64749146 |
| VLSLVWG | AWTGLANSDYT | 0.62223315 |
| IGYALWL | AWTGLANSDYT | 0.574176 |
| LSLLLAPL | AWTGLANSDYT | 0.320498 |
| TLDEALK | AWTGLANSDYT | 0.34976825 |
| TDGAGTAEGG | AWTGLANSDYT | 0.5615061 |
| NAPGGAGGNGG | AWTGLANSDYT | 0.51814735 |
| VSVFMGMGGNF | AWTGLANSDYT | 0.3521946 |
| LASYLAAL | AWTGLANSDYT | 0.6212133 |
| LVGGTPAL | AWTGLANSDYT | 0.5875485 |
| VRNADVL | AWTGLANSDYT | 0.5533294 |
| TCSSPFLVSAM | AWTGLANSDYT | 0.63484836 |
| VHITDAE | AWTGLANSDYT | 0.5441288 |
| EYAAPSF | AWTGLANSDYT | 0.47506088 |
| VVDAFHGGL | AWTGLANSDYT | 0.4916454 |
| ASVVDPQ | AWTGLANSDYT | 0.64153993 |
| SYNEFGGLS | AWTGLANSDYT | 0.3860854 |
| DPDGSVY | AWTGLANSDYT | 0.62679935 |
| MVSTIHATE | AWTGLANSDYT | 0.5355974 |
| LAAGVADPV | AWTGLANSDYT | 0.63706845 |
| TKEQIAA | AWTGLANSDYT | 0.38057676 |
| SLASPAAL | AWTGLANSDYT | 0.5871956 |
| IGLVFYGV | AWTGLANSDYT | 0.53858703 |
| FLLDASNL | AWTGLANSDYT | 0.6561881 |
| YQVGNFP | AWTGLANSDYT | 0.5111476 |
| ESLLSQST | AWTGLANSDYT | 0.5999977 |
| LFTVGFAAVG | AWTGLANSDYT | 0.5193824 |
| ATPPTPT | AWTGLANSDYT | 0.64238405 |
| TDLATLG | AWTGLANSDYT | 0.46143964 |
| SAGGTGR | AWTGLANSDYT | 0.55829906 |
| GFSNSGS | AWTGLANSDYT | 0.46475703 |
| GGNGGSGGTSV | AWTGLANSDYT | 0.59848213 |
| GTSVATGGAGN | AWTGLANSDYT | 0.5641214 |
| NGGNGGSGGTS | AWTGLANSDYT | 0.37668943 |
| AVVKFPH | AWTGLANSDYT | 0.64426935 |
| TPPPPPP | AWTGLANSDYT | 0.54253536 |

| | | |
|---|---|---|
| AAPAAAAAPA | AWTGLANSDYT | 0.44289848 |
| IALNEDL | AWTGLANSDYT | 0.6252803 |
| IVNVSSM | AWTGLANSDYT | 0.5969523 |
| TGIPPAPR | AWTGLANSDYT | 0.6086171 |
| FDLGGGTF | AWTGLANSDYT | 0.5467584 |
| FDLGGGTFDVS | AWTGLANSDYT | 0.60512245 |
| NEPTAAA | AWTGLANSDYT | 0.6147259 |
| NEPTAAALA | AWTGLANSDYT | 0.556671 |
| GSIINISSI | AWTGLANSDYT | 0.5067842 |
| VAPGNAGTA | AWTGLANSDYT | 0.611901 |
| TGGGASLE | AWTGLANSDYT | 0.74493974 |
| VAVYNHYKRI | AWTGLANSDYT | 0.7578368 |
| TDPAPATT | AWTGLANSDYT | 0.6641263 |
| TNRPDILDPALLRPG | AWTGLANSDYT | 0.63087666 |
| LTTTTTS | AWTGLANSDYT | 0.6396639 |
| PTGSGKTL | AWTGLANSDYT | 0.76646686 |
| TGSGKTL | AWTGLANSDYT | 0.7430135 |
| TIGHVDHGK | AWTGLANSDYT | 0.73235834 |
| TIGHVDHGKTT | AWTGLANSDYT | 0.65497464 |
| GHVDHGKTT | AWTGLANSDYT | 0.74212927 |
| AQFEHTLL | AWTGLANSDYT | 0.6980841 |
| IVNQDEL | AWTGLANSDYT | 0.66253674 |
| TNGSQFF | AWTGLANSDYT | 0.7396964 |
| TGKGIVE | AWTGLANSDYT | 0.74415517 |
| LISNASDA | AWTGLANSDYT | 0.6717638 |
| AELRTLGELTKIAKAHGAQ | AWTGLANSDYT | 0.72762954 |
| APNAVAAAEPSW | AWTGLANSDYT | 0.6647336 |
| APQVAGAATAVL | AWTGLANSDYT | 0.7246669 |
| ARRDPAAAAAILPTDA | AWTGLANSDYT | 0.7069872 |
| DAAARDVLAEAGLAEYF | AWTGLANSDYT | 0.7744003 |
| DDQIVLSAKANLAKVEQ | AWTGLANSDYT | 0.75636697 |
| DELLPDTGEDSDEDGH | AWTGLANSDYT | 0.75459963 |
| ELPEAPANRATVPQD | AWTGLANSDYT | 0.4473709 |
| FGQSKIERVTGIG | AWTGLANSDYT | 0.6127414 |
| GAGGTGSTAGGAGGAGGAGGLYA | AWTGLANSDYT | 0.71434176 |
| GHGGTGAAGGNGGTGGAGGSGIDG | AWTGLANSDYT | 0.7001382 |
| GNGGNGGAGGEGGPGGAGGAGA | AWTGLANSDYT | 0.5397418 |
| GSTAGNGGNGGNATNGTIAGQPAG | AWTGLANSDYT | 0.742488 |
| GTGGDGGNGGNGGTGG | AWTGLANSDYT | 0.70049936 |
| GVRRPAGIPADAPV | AWTGLANSDYT | 0.6501996 |
| GYRQVTDAHLLTLARR | AWTGLANSDYT | 0.72872865 |
| IAKMGQDGHDRG | AWTGLANSDYT | 0.67484987 |
| KAEPSIEIQYT | AWTGLANSDYT | 0.71368814 |
| KPVRIRMAADEDTVAVT | AWTGLANSDYT | 0.68599164 |
| MPPRIAGMRLLVIKPEP | AWTGLANSDYT | 0.76195854 |
| MKFYAVALFRSRGVPPTR | AWTGLANSDYT | 0.63270235 |
| NAGGFGILGAVAHSP | AWTGLANSDYT | 0.7395959 |
| NEGREITLAMAEM | AWTGLANSDYT | 0.6513723 |
| NELVEWSHPLGHRWRWE | AWTGLANSDYT | 0.76394814 |
| NLGAGGTGGNGGNPG | AWTGLANSDYT | 0.61662966 |
| NTEWPKWETVMAADPVD | AWTGLANSDYT | 0.7436482 |

| | | |
|---|---|---|
| PASAAPAETQP | AWTGLANSDYT | 0.69661784 |
| QQSSDGKNGVRAAEITGE | AWTGLANSDYT | 0.77350146 |
| RAPDAVTTSWWKE | AWTGLANSDYT | 0.70767456 |
| SGFGNVGAHTSGWFNQ | AWTGLANSDYT | 0.7432094 |
| SGVAGAVGQAASVGGLKV | AWTGLANSDYT | 0.7159168 |
| SNGGAGTGSAGGNGGT | AWTGLANSDYT | 0.7523539 |
| TGGAGGEGGAAG | AWTGLANSDYT | 0.576213 |
| TGGDGGAAGAGGAPGHG | AWTGLANSDYT | 0.7151053 |
| TMPPLLAAIANAGRHGV | AWTGLANSDYT | 0.6055385 |
| TSSTAGNSGGA | AWTGLANSDYT | 0.74009657 |
| VPATVPVGILC | AWTGLANSDYT | 0.7431064 |
| VQRPDSGGVVLTGRIS | AWTGLANSDYT | 0.5830494 |
| WTPNPVPATKGVPAGLVQAT | AWTGLANSDYT | 0.7580287 |
| YALKRALAKFSDRFDVVII | AWTGLANSDYT | 0.69437647 |
| YLIGSVIV | AWTGLANSDYT | 0.69756085 |
| YGQLVAVID | AWTGLANSDYT | 0.7633471 |
| VTVGGMTLPA | AWTGLANSDYT | 0.7060522 |
| VLALPAV | AWTGLANSDYT | 0.7650477 |
| VIPPTLNYET | AWTGLANSDYT | 0.70289415 |
| VGMGGMHPSSG | AWTGLANSDYT | 0.69743425 |
| VGLAGGAATA | AWTGLANSDYT | 0.7428064 |
| VGGAGGAGGSGP | AWTGLANSDYT | 0.57909733 |
| VFAAFMV | AWTGLANSDYT | 0.6288827 |
| TLYDKLR | AWTGLANSDYT | 0.76290035 |
| TLAVFTVL | AWTGLANSDYT | 0.8270166 |
| TGSVQPARV | AWTGLANSDYT | 0.83214784 |
| TGGGAGSGGT | AWTGLANSDYT | 0.67621124 |
| SHLETSL | AWTGLANSDYT | 0.70689857 |
| SFTTGMT | AWTGLANSDYT | 0.7801948 |
| QVEVAKRMGVRQPTVS | AWTGLANSDYT | 0.84853506 |
| QGSGGPALAM | AWTGLANSDYT | 0.8607359 |
| MLAPSNP | AWTGLANSDYT | 0.83732545 |
| LVNNAAI | AWTGLANSDYT | 0.77371275 |
| LTGEINTLA | AWTGLANSDYT | 0.84085155 |
| LSIFGPV | AWTGLANSDYT | 0.8130913 |
| LLALLANL | AWTGLANSDYT | 0.79696476 |
| LISNASDALDKLR | AWTGLANSDYT | 0.85511154 |
| LINEAVP | AWTGLANSDYT | 0.8247261 |
| LGYAFSGL | AWTGLANSDYT | 0.7695812 |
| LDFVAVAL | AWTGLANSDYT | 0.7757759 |
| LAQPGAE | AWTGLANSDYT | 0.8315784 |
| ISNLGAVTGF | AWTGLANSDYT | 0.8499764 |
| HQEDPGRI | AWTGLANSDYT | 0.8437769 |
| GTPGEPTP | AWTGLANSDYT | 0.8775527 |
| GFFNSGS | AWTGLANSDYT | 0.84687525 |
| GAGGGGLTAGAN | AWTGLANSDYT | 0.83351487 |
| FVDALVNM | AWTGLANSDYT | 0.48295093 |
| FLTVAPEL | AWTGLANSDYT | 0.6114853 |
| FETLTAL | AWTGLANSDYT | 0.807143 |
| DVINPIEL | AWTGLANSDYT | 0.77108514 |
| APVSGAPV | AWTGLANSDYT | 0.46826893 |

| | | |
|---|---|---|
| ALVGGLAW | AWTGLANSDYT | 0.842957 |
| ALAAAAATAI | AWTGLANSDYT | 0.80601615 |
| AFGLGLLL | AWTGLANSDYT | 0.73894286 |
| AFFAAASL | AWTGLANSDYT | 0.85412055 |
| AAAALPAFSPP | AWTGLANSDYT | 0.7726957 |
| VVLAPAL | AWTGLANSDYT | 0.77351975 |
| TIGHVDHGKTTL | AWTGLANSDYT | 0.8209357 |
| LISNASDALDK | AWTGLANSDYT | 0.8737951 |
| LISNASDAL | AWTGLANSDYT | 0.7820576 |
| ISNASDALDK | AWTGLANSDYT | 0.86158895 |
| NQAETLVYQTE | AWTGLANSDYT | 0.81697285 |
| TGGTGGTAGAG | AWTGLANSDYT | 0.86148876 |
| TGSTAGGAGGA | AWTGLANSDYT | 0.7068014 |
| DPLAVPVAAGPV | AWTGLANSDYT | 0.8499966 |
| GGTGGTPGNGGN | AWTGLANSDYT | 0.79520476 |
| PAGLAGVDPATL | AWTGLANSDYT | 0.87971413 |
| GNNLVVEAPHAGASV | AWTGLANSDYT | 0.80031335 |
| KVTRIMSASTLEQPA | AWTGLANSDYT | 0.86425626 |
| LGFALPIALPRRTVV | AWTGLANSDYT | 0.83233404 |
| TRLGNLTARKLVVLV | AWTGLANSDYT | 0.8474118 |
| WSQQRRLIYFPSAGP | AWTGLANSDYT | 0.6795411 |
| GPNGNPDPMAAAVDIR | AWTGLANSDYT | 0.77831835 |
| TQRCVSVADLRAAASR | AWTGLANSDYT | 0.7329649 |
| QFVLDVAEHPVEVLTETV | AWTGLANSDYT | 0.8186287 |
| GGAGGTGGAGGNGGTQAGDG | AWTGLANSDYT | 0.84241664 |
| PLLATLRAIAVDVDAKPAQV | AWTGLANSDYT | 0.5101387 |
| APKFRTGGYMIDFWGVGYQVAKRM | AWTGLANSDYT | 0.8657744 |
| GAGGAGGAGGAAGGPGGTGGTGGN | AWTGLANSDYT | 0.7156817 |
| IGALGLLATDVAPAGSVAFGVALA | AWTGLANSDYT | 0.77040553 |
| AGGAGGIGGNGGAGGAGGVLMGNG | AWTGLANSDYT | 0.8563846 |
| ADELIGLL | AWTGLANSDYT | 0.79591286 |
| ADRALKA | AWTGLANSDYT | 0.8309809 |
| AGAVGQVGGAGGA | AWTGLANSDYT | 0.7951727 |
| AGGGGGLGGAP | AWTGLANSDYT | 0.80372566 |
| AGGLLYGNGGAG | AWTGLANSDYT | 0.83354545 |
| AGLESIGL | AWTGLANSDYT | 0.54837865 |
| AGLKTGII | AWTGLANSDYT | 0.7630976 |
| ASLAALLL | AWTGLANSDYT | 0.8646828 |
| ASVINISSI | AWTGLANSDYT | 0.6207267 |
| ATPPGPA | AWTGLANSDYT | 0.74647677 |
| ATVASALV | AWTGLANSDYT | 0.39384872 |
| ATVASLIL | AWTGLANSDYT | 0.40707242 |
| AVNGPVLIRH | AWTGLANSDYT | 0.46265605 |
| DGAAGAVGI | AWTGLANSDYT | 0.5804882 |
| DPLLHGI | AWTGLANSDYT | 0.7533012 |
| ELGAALGAL | AWTGLANSDYT | 0.67550397 |
| ELGRTQAV | AWTGLANSDYT | 0.40512726 |
| FVAGGYLS | AWTGLANSDYT | 0.6070361 |
| GASGGAGGAGGW | AWTGLANSDYT | 0.50242984 |
| GGAGAGGDGGAGGI | AWTGLANSDYT | 0.51309687 |
| GTGHNGSQPGMGGNG | AWTGLANSDYT | 0.47577265 |

| | | |
|---|---|---|
| GYAAASASAS | AWTGLANSDYT | 0.46624184 |
| HQVDVAAVQQG | AWTGLANSDYT | 0.40541583 |
| IGAAIAHT | AWTGLANSDYT | 0.32594606 |
| IGMIEVAQL | AWTGLANSDYT | 0.5417818 |
| IRLAPEHTSAMD | AWTGLANSDYT | 0.608022 |
| IYNVSVL | AWTGLANSDYT | 0.5939566 |
| KQGFLGM | AWTGLANSDYT | 0.6284679 |
| LESVITQL | AWTGLANSDYT | 0.6746632 |
| LGLGALVVV | AWTGLANSDYT | 0.44661903 |
| LIGNGGNGGSGGI | AWTGLANSDYT | 0.2422729 |
| LNAINAPTL | AWTGLANSDYT | 0.28464004 |
| LQLLNTGV | AWTGLANSDYT | 0.41367736 |
| LQTSIEF | AWTGLANSDYT | 0.4794409 |
| MSFAVSEL | AWTGLANSDYT | 0.24278803 |
| NTGALGL | AWTGLANSDYT | 0.55997103 |
| PPGAGGAGGAGTT | AWTGLANSDYT | 0.4313384 |
| QLVGLGI | AWTGLANSDYT | 0.4771607 |
| RFLEQVHQL | AWTGLANSDYT | 0.6846508 |
| RGVGYVLR | AWTGLANSDYT | 0.4963867 |
| SATGVFAGVI | AWTGLANSDYT | 0.31986707 |
| SLPAITLP | AWTGLANSDYT | 0.36103576 |
| STALAFTV | AWTGLANSDYT | 0.64058137 |
| SVLTVGLGK | AWTGLANSDYT | 0.31188786 |
| TALAGAGSGI | AWTGLANSDYT | 0.6417503 |
| TGLPFTGI | AWTGLANSDYT | 0.34484744 |
| VAEHVTV | AWTGLANSDYT | 0.6389591 |
| VDIVLNSL | AWTGLANSDYT | 0.33428648 |
| VLTGGGAVL | AWTGLANSDYT | 0.61798716 |
| VNVGLAYGI | AWTGLANSDYT | 0.5978977 |
| VSSAAVGAV | AWTGLANSDYT | 0.73954064 |
| VTSVYEAGV | AWTGLANSDYT | 0.44336677 |
| WVALDRGA | AWTGLANSDYT | 0.61815923 |
| YLVGSVVI | AWTGLANSDYT | 0.35124922 |
| GPPGTGKTLL | AWTGLANSDYT | 0.74413717 |
| GVGMTKF | AWTGLANSDYT | 0.37987193 |
| IGIGGSDL | AWTGLANSDYT | 0.363203 |
| IGTLLAEL | AWTGLANSDYT | 0.30840623 |
| IYTSGTT | AWTGLANSDYT | 0.39853588 |
| LSFPAGV | AWTGLANSDYT | 0.6488044 |
| LYTSGTT | AWTGLANSDYT | 0.36203 |
| VGAFPFGF | AWTGLANSDYT | 0.5556075 |
| VGAGTVEF | AWTGLANSDYT | 0.5593362 |
| VGKSTLLL | AWTGLANSDYT | 0.36074317 |
| AALAHAGGGIV | AWTGLANSDYT | 0.51399386 |
| AALVAVSLL | AWTGLANSDYT | 0.4453048 |
| AAVSRAVAAL | AWTGLANSDYT | 0.5755459 |
| AGGSSGIGGF | AWTGLANSDYT | 0.35405746 |
| AGLTVANANG | AWTGLANSDYT | 0.48812497 |
| AGLVGAVL | AWTGLANSDYT | 0.47739866 |
| ALSQAFTV | AWTGLANSDYT | 0.38610664 |
| ALTFSVSL | AWTGLANSDYT | 0.3408718 |

| | | |
|---|---|---|
| ALTGAALVAL | AWTGLANSDYT | 0.6773366 |
| ASFSIGAGVL | AWTGLANSDYT | 0.62958324 |
| DGSVYKI | AWTGLANSDYT | 0.6651236 |
| EQLAEIV | AWTGLANSDYT | 0.59425247 |
| FAVSELTVM | AWTGLANSDYT | 0.56645864 |
| GAAPLAAGL | AWTGLANSDYT | 0.59487563 |
| GADGGDGGSGGAGG | AWTGLANSDYT | 0.65147585 |
| GFGLDTGGAGG | AWTGLANSDYT | 0.6572609 |
| GLASGGLL | AWTGLANSDYT | 0.63427603 |
| GRIAGAAVLL | AWTGLANSDYT | 0.5613963 |
| GSAGSLNTSF | AWTGLANSDYT | 0.6321483 |
| GTGQIGI | AWTGLANSDYT | 0.61337084 |
| GTLIALIL | AWTGLANSDYT | 0.60627776 |
| IAATMGFVVG | AWTGLANSDYT | 0.6273581 |
| IFAQALL | AWTGLANSDYT | 0.64079 |
| IGAAVALGF | AWTGLANSDYT | 0.59352046 |
| IGLTPLAL | AWTGLANSDYT | 0.60610783 |
| ILGSNTSTL | AWTGLANSDYT | 0.6077251 |
| IVFSLTVL | AWTGLANSDYT | 0.62886935 |
| LASKATGF | AWTGLANSDYT | 0.61602634 |
| LASVALIL | AWTGLANSDYT | 0.6626066 |
| LFAVGLTL | AWTGLANSDYT | 0.6481772 |
| LFNGGIGTYI | AWTGLANSDYT | 0.6373884 |
| LTLGAGII | AWTGLANSDYT | 0.3826941 |
| LVAAALGDI | AWTGLANSDYT | 0.5146867 |
| NTAFVQLGI | AWTGLANSDYT | 0.60229015 |
| QLFSGAEL | AWTGLANSDYT | 0.61918736 |
| SAATFITL | AWTGLANSDYT | 0.45411175 |
| SNYGLANLGV | AWTGLANSDYT | 0.6493201 |
| SYGGAYAVM | AWTGLANSDYT | 0.59211844 |
| TTGVGGLTLG | AWTGLANSDYT | 0.58456075 |
| TTQTIYI | AWTGLANSDYT | 0.64256966 |
| VAAGAGLGAVI | AWTGLANSDYT | 0.60002446 |
| VAAVVTLL | AWTGLANSDYT | 0.5995957 |
| VGFLGGLI | AWTGLANSDYT | 0.5757333 |
| VGVIPILL | AWTGLANSDYT | 0.64373326 |
| VIFASSLI | AWTGLANSDYT | 0.50841135 |
| VKGGDGGAAAG | AWTGLANSDYT | 0.6424089 |
| VTKTFTNI | AWTGLANSDYT | 0.5513493 |
| VVAAAILL | AWTGLANSDYT | 0.65331876 |
| YAALDVEL | AWTGLANSDYT | 0.5399327 |
| LASVALLL | AWTGLANSDYT | 0.6405663 |
| LDAVHRV | AWTGLANSDYT | 0.6081232 |
| QGLSSSIF | AWTGLANSDYT | 0.6679906 |
| RGTFANIRL | AWTGLANSDYT | 0.6060978 |
| RQLWWGHRI | AWTGLANSDYT | 0.6337796 |
| YGPPGTGKTLL | AWTGLANSDYT | 0.59410465 |
| AALVGAPANVA | AWTGLANSDYT | 0.65886533 |
| AATFRRTE | AWTGLANSDYT | 0.5256444 |
| AGALGGGI | AWTGLANSDYT | 0.6063807 |
| AGGAGGAGGIGAG | AWTGLANSDYT | 0.5020455 |

| | | |
|---|---|---|
| AGSGALGL | AWTGLANSDYT | 0.63339096 |
| AQIADQI | AWTGLANSDYT | 0.63460875 |
| ASAVAVTYL | AWTGLANSDYT | 0.50923747 |
| ASTVLGGTGG | AWTGLANSDYT | 0.6494329 |
| AVGGEPAK | AWTGLANSDYT | 0.60886896 |
| CEQGTFGGTC | AWTGLANSDYT | 0.58785665 |
| DEGKAPAG | AWTGLANSDYT | 0.6508997 |
| DTVVFPAL | AWTGLANSDYT | 0.61858517 |
| FEGITMAG | AWTGLANSDYT | 0.64883983 |
| FPLVVQD | AWTGLANSDYT | 0.6057092 |
| GAFTTPQ | AWTGLANSDYT | 0.60093045 |
| GFGGFGTAGGVGGA | AWTGLANSDYT | 0.6370271 |
| GFLAQFL | AWTGLANSDYT | 0.5845667 |
| GGAGGAGGNAGVA | AWTGLANSDYT | 0.5673962 |
| GGVGTAGAGGAGGAGG | AWTGLANSDYT | 0.6498449 |
| GNGGGGDAGNAGSGGN | AWTGLANSDYT | 0.7153511 |
| GTGGAGGAGNDAGS | AWTGLANSDYT | 0.8192878 |
| GTTNTGWFNS | AWTGLANSDYT | 0.5686207 |
| IFMGPLFTL | AWTGLANSDYT | 0.4880321 |
| ILAHAIEAI | AWTGLANSDYT | 0.62282836 |
| IRSAIREAL | AWTGLANSDYT | 0.8184556 |
| ITFAGGG | AWTGLANSDYT | 0.78541195 |
| ITGMSGHQEV | AWTGLANSDYT | 0.7599251 |
| IVAGLACTL | AWTGLANSDYT | 0.5057979 |
| IVGGGLAGA | AWTGLANSDYT | 0.622588 |
| IVTGVLNI | AWTGLANSDYT | 0.6047864 |
| LAQQGCI | AWTGLANSDYT | 0.642782 |
| LCLHVEN | AWTGLANSDYT | 0.7573621 |
| LDAISDGS | AWTGLANSDYT | 0.7334231 |
| LGGFAAML | AWTGLANSDYT | 0.6577574 |
| LGGKNPMI | AWTGLANSDYT | 0.5663845 |
| LGPKGVAAIL | AWTGLANSDYT | 0.66881186 |
| LMVGVSI | AWTGLANSDYT | 0.76002234 |
| LVGAAFVL | AWTGLANSDYT | 0.6873412 |
| LVLAVAPL | AWTGLANSDYT | 0.84619 |
| MAAIAVIDAA | AWTGLANSDYT | 0.79290414 |
| NNGINAELQ | AWTGLANSDYT | 0.7329831 |
| RGEPLVLVHG | AWTGLANSDYT | 0.13423647 |
| RGGAGGAAGLI | AWTGLANSDYT | 0.41135678 |
| RSYRMHPAV | AWTGLANSDYT | 0.70179725 |
| SAAHLLAGQLA | AWTGLANSDYT | 0.7158042 |
| SAAVGSY | AWTGLANSDYT | 0.34336734 |
| SPHAVLHRI | AWTGLANSDYT | 0.77626663 |
| SQIAKVL | AWTGLANSDYT | 0.6377086 |
| TALGHLVP | AWTGLANSDYT | 0.6165397 |
| TAPTTSGAA | AWTGLANSDYT | 0.8011224 |
| TDGDDGNGGSPGT | AWTGLANSDYT | 0.63906825 |
| TDGLGEN | AWTGLANSDYT | 0.60837626 |
| TGGPPLPGQA | AWTGLANSDYT | 0.624071 |
| TIPAITI | AWTGLANSDYT | 0.82496387 |
| TIPGFTI | AWTGLANSDYT | 0.4165223 |

| | | |
|---|---|---|
| TSIASGALA | AWTGLANSDYT | 0.76729417 |
| TVTLPGAAI | AWTGLANSDYT | 0.6914201 |
| VASGLSAALP | AWTGLANSDYT | 0.8146498 |
| VGAGVIGI | AWTGLANSDYT | 0.38909054 |
| VGPKRFGV | AWTGLANSDYT | 0.76093614 |
| VNVAFPD | AWTGLANSDYT | 0.6317352 |
| VSMTNSGVS | AWTGLANSDYT | 0.8696711 |
| VTDVLGDLSV | AWTGLANSDYT | 0.5834457 |
| VVVVGAGFAGL | AWTGLANSDYT | 0.7171172 |
| VYTVGTVV | AWTGLANSDYT | 0.5947381 |
| WTTFSGP | AWTGLANSDYT | 0.7427457 |
| YDELVEAF | AWTGLANSDYT | 0.43693805 |
| YGTGGAGGNGG | AWTGLANSDYT | 0.58320284 |
| YTNVMAV | AWTGLANSDYT | 0.43418738 |
| FLEQVHQL | AWTGLANSDYT | 0.69375145 |
| GLSLLAVAV | AWTGLANSDYT | 0.74655175 |
| LEQVHQL | AWTGLANSDYT | 0.43552312 |
| LNAAGAI | AWTGLANSDYT | 0.75735736 |
| PAVHASL | AWTGLANSDYT | 0.71434665 |
| TELEDDA | AWTGLANSDYT | 0.502953 |
| TPPGPPN | AWTGLANSDYT | 0.8075055 |
| TPPPPPPP | AWTGLANSDYT | 0.52768356 |
| TQSPALI | ASSLAWSSYEQY | 0.7726277 |
| EFAAAFPQ | ASSLAWSSYEQY | 0.6321442 |
| LLSGNLDVL | ASSLAWSSYEQY | 0.6648367 |
| GPANLGNYNI | ASSLAWSSYEQY | 0.7739785 |
| TLGPTLP | ASSLAWSSYEQY | 0.60160315 |
| VYAYSSCCK | ASSLAWSSYEQY | 0.5760133 |
| APAPAPAPNI | ASSLAWSSYEQY | 0.7964422 |
| PGGPVPIP | ASSLAWSSYEQY | 0.51056683 |
| FLTVALEP | ASSLAWSSYEQY | 0.65510935 |
| SQYFGANL | ASSLAWSSYEQY | 0.40532517 |
| GGVKPAA | ASSLAWSSYEQY | 0.43996978 |
| GGTGGAGGAGGW | ASSLAWSSYEQY | 0.5002721 |
| GVGGVAGLGGAGG | ASSLAWSSYEQY | 0.5820531 |
| VLLVLATTI | ASSLAWSSYEQY | 0.64991844 |
| IGIFAAML | ASSLAWSSYEQY | 0.5902528 |
| AVGDLID | ASSLAWSSYEQY | 0.45935476 |
| NISPLPL | ASSLAWSSYEQY | 0.53064144 |
| VTAVFVAI | ASSLAWSSYEQY | 0.50427514 |
| SGDDNDAAL | ASSLAWSSYEQY | 0.5107022 |
| AVAALAHS | ASSLAWSSYEQY | 0.52710724 |
| GGVTVMATP | ASSLAWSSYEQY | 0.50328326 |
| AGLAANVRD | ASSLAWSSYEQY | 0.4616976 |
| TNRPDLIDPALLRPG | ASSLAWSSYEQY | 0.41481227 |
| LANLGVTAL | ASSLAWSSYEQY | 0.51247513 |
| TGERIEL | ASSLAWSSYEQY | 0.58730197 |
| NGEELAA | ASSLAWSSYEQY | 0.571322 |
| VLSLVWG | ASSLAWSSYEQY | 0.6205109 |
| IGYALWL | ASSLAWSSYEQY | 0.5769794 |
| LSLLLAPL | ASSLAWSSYEQY | 0.46552357 |

| | | |
|---|---|---|
| TLDEALK | ASSLAWSSYEQY | 0.32783544 |
| TDGAGTAEGG | ASSLAWSSYEQY | 0.39353052 |
| NAPGGAGGNGG | ASSLAWSSYEQY | 0.4863725 |
| VSVFMGMGGNF | ASSLAWSSYEQY | 0.49774063 |
| LASYLAAL | ASSLAWSSYEQY | 0.33436182 |
| LVGGTPAL | ASSLAWSSYEQY | 0.5472826 |
| VRNADVL | ASSLAWSSYEQY | 0.5194148 |
| TCSSPFLVSAM | ASSLAWSSYEQY | 0.49147707 |
| VHITDAE | ASSLAWSSYEQY | 0.5854776 |
| EYAAPSF | ASSLAWSSYEQY | 0.48733813 |
| VVDAFHGGL | ASSLAWSSYEQY | 0.4178416 |
| ASVVDPQ | ASSLAWSSYEQY | 0.4059168 |
| SYNEFGGLS | ASSLAWSSYEQY | 0.59003484 |
| DPDGSVY | ASSLAWSSYEQY | 0.36572194 |
| MVSTIHATE | ASSLAWSSYEQY | 0.5755153 |
| LAAGVADPV | ASSLAWSSYEQY | 0.46719292 |
| TKEQIAA | ASSLAWSSYEQY | 0.5945352 |
| SLASPAAL | ASSLAWSSYEQY | 0.4063732 |
| IGLVFYGV | ASSLAWSSYEQY | 0.54032236 |
| FLLDASNL | ASSLAWSSYEQY | 0.50674987 |
| YQVGNFP | ASSLAWSSYEQY | 0.64346236 |
| ESLLSQST | ASSLAWSSYEQY | 0.48397467 |
| LFTVGFAAVG | ASSLAWSSYEQY | 0.5684173 |
| ATPPTPT | ASSLAWSSYEQY | 0.43825454 |
| TDLATLG | ASSLAWSSYEQY | 0.6429176 |
| SAGGTGR | ASSLAWSSYEQY | 0.45951265 |
| GFSNSGS | ASSLAWSSYEQY | 0.46684256 |
| GGNGGSGGTSV | ASSLAWSSYEQY | 0.39117897 |
| GTSVATGGAGN | ASSLAWSSYEQY | 0.49117514 |
| NGGNGGSGGTS | ASSLAWSSYEQY | 0.57735944 |
| AVVKFPH | ASSLAWSSYEQY | 0.5604414 |
| TPPPPPP | ASSLAWSSYEQY | 0.6106737 |
| AAPAAAAAPA | ASSLAWSSYEQY | 0.6125615 |
| IALNEDL | ASSLAWSSYEQY | 0.42612106 |
| IVNVSSM | ASSLAWSSYEQY | 0.57532716 |
| TGIPPAPR | ASSLAWSSYEQY | 0.48639008 |
| FDLGGGTF | ASSLAWSSYEQY | 0.52442104 |
| FDLGGGTFDVS | ASSLAWSSYEQY | 0.38629013 |
| NEPTAAA | ASSLAWSSYEQY | 0.55277133 |
| NEPTAAALA | ASSLAWSSYEQY | 0.58474517 |
| GSIINISSI | ASSLAWSSYEQY | 0.5524043 |
| VAPGNAGTA | ASSLAWSSYEQY | 0.40757927 |
| TGGGASLE | ASSLAWSSYEQY | 0.5845547 |
| VAVYNHYKRI | ASSLAWSSYEQY | 0.5684633 |
| TDPAPATT | ASSLAWSSYEQY | 0.82320064 |
| TNRPDILDPALLRPG | ASSLAWSSYEQY | 0.5597653 |
| LTTTTTS | ASSLAWSSYEQY | 0.29402953 |
| PTGSGKTL | ASSLAWSSYEQY | 0.4487123 |
| TGSGKTL | ASSLAWSSYEQY | 0.6187712 |
| TIGHVDHGK | ASSLAWSSYEQY | 0.83280647 |
| TIGHVDHGKTT | ASSLAWSSYEQY | 0.67655647 |

| | | |
|---|---|---|
| GHVDHGKTT | ASSLAWSSYEQY | 0.2270153 |
| AQFEHTLL | ASSLAWSSYEQY | 0.6166449 |
| IVNQDEL | ASSLAWSSYEQY | 0.452048 |
| TNGSQFF | ASSLAWSSYEQY | 0.47358382 |
| TGKGIVE | ASSLAWSSYEQY | 0.5919844 |
| LISNASDA | ASSLAWSSYEQY | 0.45148116 |
| AELRTLGELTKIAKAHGAQ | ASSLAWSSYEQY | 0.58420515 |
| APNAVAAAEPSW | ASSLAWSSYEQY | 0.36650532 |
| APQVAGAATAVL | ASSLAWSSYEQY | 0.64039534 |
| ARRDPAAAAAILPTDA | ASSLAWSSYEQY | 0.647884 |
| DAAARDVLAEAGLAEYF | ASSLAWSSYEQY | 0.5361674 |
| DDQIVLSAKANLAKVEQ | ASSLAWSSYEQY | 0.7186324 |
| DELLPDTGEDSDEDGH | ASSLAWSSYEQY | 0.71983325 |
| ELPEAPANRATVPQD | ASSLAWSSYEQY | 0.6866723 |
| FGQSKIERVTGIG | ASSLAWSSYEQY | 0.13093793 |
| GAGGTGSTAGGAGGAGGAGGLYA | ASSLAWSSYEQY | 0.20616098 |
| GHGGTGAAGGNGGTGGAGGSGIDG | ASSLAWSSYEQY | 0.48410526 |
| GNGGNGGAGGEGGPGGAGGAGA | ASSLAWSSYEQY | 0.49545094 |
| GSTAGNGGNGGNATNGTIAGQPAG | ASSLAWSSYEQY | 0.17226277 |
| GTGGDGGNGGNGGTGG | ASSLAWSSYEQY | 0.4649912 |
| GVRRPAGIPADAPV | ASSLAWSSYEQY | 0.51842093 |
| GYRQVTDAHLLTLARR | ASSLAWSSYEQY | 0.41250357 |
| IAKMGQDGHDRG | ASSLAWSSYEQY | 0.75993156 |
| KAEPSIEIQYT | ASSLAWSSYEQY | 0.5917839 |
| KPVRIRMAADEDTVAVT | ASSLAWSSYEQY | 0.28984278 |
| MPPRIAGMRLLVIKPEP | ASSLAWSSYEQY | 0.29504186 |
| MKFYAVALFRSRGVPPTR | ASSLAWSSYEQY | 0.67265666 |
| NAGGFGILGAVAHSP | ASSLAWSSYEQY | 0.18551373 |
| NEGREITLAMAEM | ASSLAWSSYEQY | 0.74119085 |
| NELVEWSHPLGHRWRWE | ASSLAWSSYEQY | 0.31895176 |
| NLGAGGTGGNGGNPG | ASSLAWSSYEQY | 0.59720767 |
| NTEWPKWETVMAADPVD | ASSLAWSSYEQY | 0.18403062 |
| PASAAPAETQP | ASSLAWSSYEQY | 0.690004 |
| QQSSDGKNGVRAAEITGE | ASSLAWSSYEQY | 0.5511093 |
| RAPDAVTTSWWKE | ASSLAWSSYEQY | 0.8413597 |
| SGFGNVGAHTSGWFNQ | ASSLAWSSYEQY | 0.36071187 |
| SGVAGAVGQAASVGGLKV | ASSLAWSSYEQY | 0.4911904 |
| SNGGAGTGSAGGNGGT | ASSLAWSSYEQY | 0.324456 |
| TGGAGGEGGAAG | ASSLAWSSYEQY | 0.83640677 |
| TGGDGGAAGAGGAPGHG | ASSLAWSSYEQY | 0.25862196 |
| TMPPLLAAIANAGRHGV | ASSLAWSSYEQY | 0.41612792 |
| TSSTAGNSGGA | ASSLAWSSYEQY | 0.3198938 |
| VPATVPVGILC | ASSLAWSSYEQY | 0.60111237 |
| VQRPDSGGVVLTGRIS | ASSLAWSSYEQY | 0.68116355 |
| WTPNPVPATKGVPAGLVQAT | ASSLAWSSYEQY | 0.23851335 |
| YALKRALAKFSDRFDVVII | ASSLAWSSYEQY | 0.7677524 |
| YLIGSVIV | ASSLAWSSYEQY | 0.51821285 |
| YGQLVAVID | ASSLAWSSYEQY | 0.26244342 |
| VTVGGMTLPA | ASSLAWSSYEQY | 0.75375104 |
| VLALPAV | ASSLAWSSYEQY | 0.3658156 |
| VIPPTLNYET | ASSLAWSSYEQY | 0.7380785 |

| | | |
|---|---|---|
| VGMGGMHPSSG | ASSLAWSSYEQY | 0.3863157 |
| VGLAGGAATA | ASSLAWSSYEQY | 0.53252316 |
| VGGAGGAGGSGP | ASSLAWSSYEQY | 0.5688877 |
| VFAAFMV | ASSLAWSSYEQY | 0.5174403 |
| TLYDKLR | ASSLAWSSYEQY | 0.26669064 |
| TLAVFTVL | ASSLAWSSYEQY | 0.7562145 |
| TGSVQPARV | ASSLAWSSYEQY | 0.7288964 |
| TGGGAGSGGT | ASSLAWSSYEQY | 0.7298422 |
| SHLETSL | ASSLAWSSYEQY | 0.6895195 |
| SFTTGMT | ASSLAWSSYEQY | 0.68672776 |
| QVEVAKRMGVRQPTVS | ASSLAWSSYEQY | 0.67762136 |
| QGSGGPALAM | ASSLAWSSYEQY | 0.728658 |
| MLAPSNP | ASSLAWSSYEQY | 0.7341196 |
| LVNNAAI | ASSLAWSSYEQY | 0.73189676 |
| LTGEINTLA | ASSLAWSSYEQY | 0.6964954 |
| LSIFGPV | ASSLAWSSYEQY | 0.7270989 |
| LLALLANL | ASSLAWSSYEQY | 0.714504 |
| LISNASDALDKLR | ASSLAWSSYEQY | 0.70808136 |
| LINEAVP | ASSLAWSSYEQY | 0.7271996 |
| LGYAFSGL | ASSLAWSSYEQY | 0.7233583 |
| LDFVAVAL | ASSLAWSSYEQY | 0.68650687 |
| LAQPGAE | ASSLAWSSYEQY | 0.6976161 |
| ISNLGAVTGF | ASSLAWSSYEQY | 0.7154444 |
| HQEDPGRI | ASSLAWSSYEQY | 0.7317366 |
| GTPGEPTP | ASSLAWSSYEQY | 0.7275542 |
| GFFNSGS | ASSLAWSSYEQY | 0.7381611 |
| GAGGGGLTAGAN | ASSLAWSSYEQY | 0.7300699 |
| FVDALVNM | ASSLAWSSYEQY | 0.7333859 |
| FLTVAPEL | ASSLAWSSYEQY | 0.53471994 |
| FETLTAL | ASSLAWSSYEQY | 0.64704657 |
| DVINPIEL | ASSLAWSSYEQY | 0.6970035 |
| APVSGAPV | ASSLAWSSYEQY | 0.708332 |
| ALVGGLAW | ASSLAWSSYEQY | 0.48185027 |
| ALAAAAATAI | ASSLAWSSYEQY | 0.7269089 |
| AFGLGLLL | ASSLAWSSYEQY | 0.7159048 |
| AFFAAASL | ASSLAWSSYEQY | 0.6868182 |
| AAAALPAFSPP | ASSLAWSSYEQY | 0.72702986 |
| VVLAPAL | ASSLAWSSYEQY | 0.70239645 |
| TIGHVDHGKTTL | ASSLAWSSYEQY | 0.6807339 |
| LISNASDALDK | ASSLAWSSYEQY | 0.6999911 |
| LISNASDAL | ASSLAWSSYEQY | 0.73228014 |
| ISNASDALDK | ASSLAWSSYEQY | 0.6567292 |
| NQAETLVYQTE | ASSLAWSSYEQY | 0.7322124 |
| TGGTGGTAGAG | ASSLAWSSYEQY | 0.6558931 |
| TGSTAGGAGGA | ASSLAWSSYEQY | 0.7327005 |
| DPLAVPVAAGPV | ASSLAWSSYEQY | 0.67329264 |
| GGTGGTPGNGGN | ASSLAWSSYEQY | 0.7331534 |
| PAGLAGVDPATL | ASSLAWSSYEQY | 0.7160789 |
| GNNLVVEAPHAGASV | ASSLAWSSYEQY | 0.73849344 |
| KVTRIMSASTLEQPA | ASSLAWSSYEQY | 0.7117641 |
| LGFALPIALPRRTVV | ASSLAWSSYEQY | 0.7322742 |

| | | |
|---|---|---|
| TRLGNLTARKLVVLV | ASSLAWSSYEQY | 0.7289337 |
| WSQQRRLIYFPSAGP | ASSLAWSSYEQY | 0.7267894 |
| GPNGNPDPMAAAVDIR | ASSLAWSSYEQY | 0.6555357 |
| TQRCVSVADLRAAASR | ASSLAWSSYEQY | 0.71096504 |
| QFVLDVAEHPVEVLTETV | ASSLAWSSYEQY | 0.6619773 |
| GGAGGTGGAGGNGGTQAGDG | ASSLAWSSYEQY | 0.72089636 |
| PLLATLRAIAVDVDAKPAQV | ASSLAWSSYEQY | 0.7323744 |
| APKFRTGGYMIDFWGVGYQVAKRM | ASSLAWSSYEQY | 0.49295083 |
| GAGGAGGAGGAAGGPGGTGGTGGN | ASSLAWSSYEQY | 0.73326135 |
| IGALGLLATDVAPAGSVAFGVALA | ASSLAWSSYEQY | 0.68527246 |
| AGGAGGIGGNGGAGGAGGVLMGNG | ASSLAWSSYEQY | 0.7025001 |
| ADELIGLL | ASSLAWSSYEQY | 0.73310435 |
| ADRALKA | ASSLAWSSYEQY | 0.7180615 |
| AGAVGQVGGAGGA | ASSLAWSSYEQY | 0.728621 |
| AGGGGGLGGAP | ASSLAWSSYEQY | 0.70390314 |
| AGGLLYGNGGAG | ASSLAWSSYEQY | 0.70673203 |
| AGLESIGL | ASSLAWSSYEQY | 0.72135997 |
| AGLKTGII | ASSLAWSSYEQY | 0.5346817 |
| ASLAALLL | ASSLAWSSYEQY | 0.6546334 |
| ASVINISSI | ASSLAWSSYEQY | 0.73745394 |
| ATPPGPA | ASSLAWSSYEQY | 0.88369286 |
| ATVASALV | ASSLAWSSYEQY | 0.88523054 |
| ATVASLIL | ASSLAWSSYEQY | 0.83853674 |
| AVNGPVLIRH | ASSLAWSSYEQY | 0.82546663 |
| DGAAGAVGI | ASSLAWSSYEQY | 0.84867704 |
| DPLLHGI | ASSLAWSSYEQY | 0.89383364 |
| ELGAALGAL | ASSLAWSSYEQY | 0.8833417 |
| ELGRTQAV | ASSLAWSSYEQY | 0.8801952 |
| FVAGGYLS | ASSLAWSSYEQY | 0.7699003 |
| GASGGAGGAGGW | ASSLAWSSYEQY | 0.8876643 |
| GGAGAGGDGGAGGI | ASSLAWSSYEQY | 0.85331506 |
| GTGHNGSQPGMGGNG | ASSLAWSSYEQY | 0.85396826 |
| GYAAASASAS | ASSLAWSSYEQY | 0.87822956 |
| HQVDVAAVQQG | ASSLAWSSYEQY | 0.8562399 |
| IGAAIAHT | ASSLAWSSYEQY | 0.8694221 |
| IGMIEVAQL | ASSLAWSSYEQY | 0.86077964 |
| IRLAPEHTSAMD | ASSLAWSSYEQY | 0.87234616 |
| IYNVSVL | ASSLAWSSYEQY | 0.88324285 |
| KQGFLGM | ASSLAWSSYEQY | 0.8745226 |
| LESVITQL | ASSLAWSSYEQY | 0.9025887 |
| LGLGALVVV | ASSLAWSSYEQY | 0.8945358 |
| LIGNGGNGGSGGI | ASSLAWSSYEQY | 0.88807434 |
| LNAINAPTL | ASSLAWSSYEQY | 0.3295281 |
| LQLLNTGV | ASSLAWSSYEQY | 0.81432694 |
| LQTSIEF | ASSLAWSSYEQY | 0.8483112 |
| MSFAVSEL | ASSLAWSSYEQY | 0.8716795 |
| NTGALGL | ASSLAWSSYEQY | 0.60725105 |
| PPGAGGAGGAGTT | ASSLAWSSYEQY | 0.88099694 |
| QLVGLGI | ASSLAWSSYEQY | 0.87152267 |
| RFLEQVHQL | ASSLAWSSYEQY | 0.8368997 |
| RGVGYVLR | ASSLAWSSYEQY | 0.8957218 |

| | | |
|---|---|---|
| SATGVFAGVI | ASSLAWSSYEQY | 0.85647345 |
| SLPAITLP | ASSLAWSSYEQY | 0.8164696 |
| STALAFTV | ASSLAWSSYEQY | 0.8267305 |
| SVLTVGLGK | ASSLAWSSYEQY | 0.8963372 |
| TALAGAGSGI | ASSLAWSSYEQY | 0.7292186 |
| TGLPFTGI | ASSLAWSSYEQY | 0.8906002 |
| VAEHVTV | ASSLAWSSYEQY | 0.85388255 |
| VDIVLNSL | ASSLAWSSYEQY | 0.89943963 |
| VLTGGGAVL | ASSLAWSSYEQY | 0.71189076 |
| VNVGLAYGI | ASSLAWSSYEQY | 0.88671845 |
| VSSAAVGAV | ASSLAWSSYEQY | 0.8519617 |
| VTSVYEAGV | ASSLAWSSYEQY | 0.90326416 |
| WVALDRGA | ASSLAWSSYEQY | 0.85929835 |
| YLVGSVVI | ASSLAWSSYEQY | 0.87719446 |
| GPPGTGKTLL | ASSLAWSSYEQY | 0.8329215 |
| GVGMTKF | ASSLAWSSYEQY | 0.8878738 |
| IGIGGSDL | ASSLAWSSYEQY | 0.77141654 |
| IGTLLAEL | ASSLAWSSYEQY | 0.857128 |
| IYTSGTT | ASSLAWSSYEQY | 0.8350768 |
| LSFPAGV | ASSLAWSSYEQY | 0.87475085 |
| LYTSGTT | ASSLAWSSYEQY | 0.8837607 |
| VGAFPFGF | ASSLAWSSYEQY | 0.66499245 |
| VGAGTVEF | ASSLAWSSYEQY | 0.895771 |
| VGKSTLLL | ASSLAWSSYEQY | 0.8228829 |
| AALAHAGGGIV | ASSLAWSSYEQY | 0.8174937 |
| AALVAVSLL | ASSLAWSSYEQY | 0.8912505 |
| AAVSRAVAAL | ASSLAWSSYEQY | 0.8607359 |
| AGGSSGIGGF | ASSLAWSSYEQY | 0.8936919 |
| AGLTVANANG | ASSLAWSSYEQY | 0.82003593 |
| AGLVGAVL | ASSLAWSSYEQY | 0.87123466 |
| ALSQAFTV | ASSLAWSSYEQY | 0.882342 |
| ALTFSVSL | ASSLAWSSYEQY | 0.77623713 |
| ALTGAALVAL | ASSLAWSSYEQY | 0.7904416 |
| ASFSIGAGVL | ASSLAWSSYEQY | 0.8955539 |
| DGSVYKI | ASSLAWSSYEQY | 0.6460098 |
| EQLAEIV | ASSLAWSSYEQY | 0.65607417 |
| FAVSELTVM | ASSLAWSSYEQY | 0.6057929 |
| GAAPLAAGL | ASSLAWSSYEQY | 0.5629045 |
| GADGGDGGSGGAGG | ASSLAWSSYEQY | 0.61023664 |
| GFGLDTGGAGG | ASSLAWSSYEQY | 0.65099394 |
| GLASGGLL | ASSLAWSSYEQY | 0.65921557 |
| GRIAGAAVLL | ASSLAWSSYEQY | 0.6446609 |
| GSAGSLNTSF | ASSLAWSSYEQY | 0.53558517 |
| GTGQIGI | ASSLAWSSYEQY | 0.6541511 |
| GTLIALIL | ASSLAWSSYEQY | 0.638602 |
| IAATMGFVVG | ASSLAWSSYEQY | 0.6244178 |
| IFAQALL | ASSLAWSSYEQY | 0.64173037 |
| IGAAVALGF | ASSLAWSSYEQY | 0.64291954 |
| IGLTPLAL | ASSLAWSSYEQY | 0.5993856 |
| ILGSNTSTL | ASSLAWSSYEQY | 0.59861344 |
| IVFSLTVL | ASSLAWSSYEQY | 0.64288586 |

| | | |
|---|---|---|
| LASKATGF | ASSLAWSSYEQY | 0.656067 |
| LASVALIL | ASSLAWSSYEQY | 0.6574832 |
| LFAVGLTL | ASSLAWSSYEQY | 0.6665272 |
| LFNGGIGTYI | ASSLAWSSYEQY | 0.65932155 |
| LTLGAGII | ASSLAWSSYEQY | 0.64842695 |
| LVAAALGDI | ASSLAWSSYEQY | 0.39668506 |
| NTAFVQLGI | ASSLAWSSYEQY | 0.5128844 |
| QLFSGAEL | ASSLAWSSYEQY | 0.5885493 |
| SAATFITL | ASSLAWSSYEQY | 0.62427187 |
| SNYGLANLGV | ASSLAWSSYEQY | 0.4205253 |
| SYGGAYAVM | ASSLAWSSYEQY | 0.6426494 |
| TTGVGGLTLG | ASSLAWSSYEQY | 0.6079018 |
| TTQTIYI | ASSLAWSSYEQY | 0.59315515 |
| VAAGAGLGAVI | ASSLAWSSYEQY | 0.65595526 |
| VAAVVTLL | ASSLAWSSYEQY | 0.6081693 |
| VGFLGGLI | ASSLAWSSYEQY | 0.5655893 |
| VGVIPILL | ASSLAWSSYEQY | 0.5944881 |
| VIFASSLI | ASSLAWSSYEQY | 0.65208787 |
| VKGGDGGAAAG | ASSLAWSSYEQY | 0.5491497 |
| VTKTFTNI | ASSLAWSSYEQY | 0.65707195 |
| VVAAAILL | ASSLAWSSYEQY | 0.56687576 |
| YAALDVEL | ASSLAWSSYEQY | 0.6622816 |
| LASVALLL | ASSLAWSSYEQY | 0.5116151 |
| LDAVHRV | ASSLAWSSYEQY | 0.6571553 |
| QGLSSSIF | ASSLAWSSYEQY | 0.6307434 |
| RGTFANIRL | ASSLAWSSYEQY | 0.6705267 |
| RQLWWGHRI | ASSLAWSSYEQY | 0.6271507 |
| YGPPGTGKTLL | ASSLAWSSYEQY | 0.6512803 |
| AALVGAPANVA | ASSLAWSSYEQY | 0.6273678 |
| AATFRRTE | ASSLAWSSYEQY | 0.6592985 |
| AGALGGGI | ASSLAWSSYEQY | 0.52802753 |
| AGGAGGAGGIGAG | ASSLAWSSYEQY | 0.63067317 |
| AGSGALGL | ASSLAWSSYEQY | 0.5185791 |
| AQIADQI | ASSLAWSSYEQY | 0.63719285 |
| ASAVAVTYL | ASSLAWSSYEQY | 0.6499424 |
| ASTVLGGTGG | ASSLAWSSYEQY | 0.4278103 |
| AVGGEPAK | ASSLAWSSYEQY | 0.6575792 |
| CEQGTFGGTC | ASSLAWSSYEQY | 0.61131877 |
| DEGKAPAG | ASSLAWSSYEQY | 0.57763946 |
| DTVVFPAL | ASSLAWSSYEQY | 0.65784717 |
| FEGITMAG | ASSLAWSSYEQY | 0.6493577 |
| FPLVVQD | ASSLAWSSYEQY | 0.65033257 |
| GAFTTPQ | ASSLAWSSYEQY | 0.60133225 |
| GFGGFGTAGGVGGA | ASSLAWSSYEQY | 0.6174212 |
| GFLAQFL | ASSLAWSSYEQY | 0.63581157 |
| GGAGGAGGNAGVA | ASSLAWSSYEQY | 0.5929023 |
| GGVGTAGAGGAGGAGG | ASSLAWSSYEQY | 0.5696949 |
| GNGGGGDAGNAGSGGN | ASSLAWSSYEQY | 0.6620533 |
| GTGGAGGAGNDAGS | ASSLAWSSYEQY | 0.7966843 |
| GTTNTGWFNS | ASSLAWSSYEQY | 0.8378922 |
| IFMGPLFTL | ASSLAWSSYEQY | 0.64192474 |

| | | |
|---|---|---|
| ILAHAIEAI | ASSLAWSSYEQY | 0.62279403 |
| IRSAIREAL | ASSLAWSSYEQY | 0.7199834 |
| ITFAGGG | ASSLAWSSYEQY | 0.84558964 |
| ITGMSGHQEV | ASSLAWSSYEQY | 0.8588037 |
| IVAGLACTL | ASSLAWSSYEQY | 0.79940593 |
| IVGGGLAGA | ASSLAWSSYEQY | 0.67348254 |
| IVTGVLNI | ASSLAWSSYEQY | 0.7701889 |
| LAQQGCI | ASSLAWSSYEQY | 0.711612 |
| LCLHVEN | ASSLAWSSYEQY | 0.71142155 |
| LDAISDGS | ASSLAWSSYEQY | 0.8138058 |
| LGGFAAML | ASSLAWSSYEQY | 0.7850008 |
| LGGKNPMI | ASSLAWSSYEQY | 0.7094511 |
| LGPKGVAAIL | ASSLAWSSYEQY | 0.6939211 |
| LMVGVSI | ASSLAWSSYEQY | 0.76735723 |
| LVGAAFVL | ASSLAWSSYEQY | 0.8326062 |
| LVLAVAPL | ASSLAWSSYEQY | 0.81232333 |
| MAAIAVIDAA | ASSLAWSSYEQY | 0.8651092 |
| NNGINAELQ | ASSLAWSSYEQY | 0.82880104 |
| RGEPLVLVHG | ASSLAWSSYEQY | 0.81582713 |
| RGGAGGAAGLI | ASSLAWSSYEQY | 0.21950023 |
| RSYRMHPAV | ASSLAWSSYEQY | 0.492628 |
| SAAHLLAGQLA | ASSLAWSSYEQY | 0.69569373 |
| SAAVGSY | ASSLAWSSYEQY | 0.74375457 |
| SPHAVLHRI | ASSLAWSSYEQY | 0.34774813 |
| SQIAKVL | ASSLAWSSYEQY | 0.805464 |
| TALGHLVP | ASSLAWSSYEQY | 0.7144543 |
| TAPTTSGAA | ASSLAWSSYEQY | 0.6357999 |
| TDGDDGNGGSPGT | ASSLAWSSYEQY | 0.8316081 |
| TDGLGEN | ASSLAWSSYEQY | 0.7036903 |
| TGGPPLPGQA | ASSLAWSSYEQY | 0.68429387 |
| TIPAITI | ASSLAWSSYEQY | 0.7126007 |
| TIPGFTI | ASSLAWSSYEQY | 0.8461617 |
| TSIASGALA | ASSLAWSSYEQY | 0.5862947 |
| TVTLPGAAI | ASSLAWSSYEQY | 0.8102544 |
| VASGLSAALP | ASSLAWSSYEQY | 0.7686927 |
| VGAGVIGI | ASSLAWSSYEQY | 0.85159504 |
| VGPKRFGV | ASSLAWSSYEQY | 0.5626016 |
| VNVAFPD | ASSLAWSSYEQY | 0.8126645 |
| VSMTNSGVS | ASSLAWSSYEQY | 0.72359383 |
| VTDVLGDLSV | ASSLAWSSYEQY | 0.87467355 |
| VVVVGAGFAGL | ASSLAWSSYEQY | 0.71694815 |
| VYTVGTVV | ASSLAWSSYEQY | 0.80791426 |
| WTTFSGP | ASSLAWSSYEQY | 0.7722876 |
| YDELVEAF | ASSLAWSSYEQY | 0.8256272 |
| YGTGGAGGNGG | ASSLAWSSYEQY | 0.48911488 |
| YTNVMAV | ASSLAWSSYEQY | 0.7301135 |
| FLEQVHQL | ASSLAWSSYEQY | 0.62151754 |
| GLSLLAVAV | ASSLAWSSYEQY | 0.7686131 |
| LEQVHQL | ASSLAWSSYEQY | 0.7954054 |
| LNAAGAI | ASSLAWSSYEQY | 0.3484013 |
| PAVHASL | ASSLAWSSYEQY | 0.8207047 |

| | | |
|---|---|---|
| TELEDDA | ASSLAWSSYEQY | 0.6846601 |
| TPPGPPN | ASSLAWSSYEQY | 0.6528958 |
| TPPPPPPP | ASSLAWSSYEQY | 0.83090484 |
| TQSPALI | ASSWDLDSQNTLY | 0.7106352 |
| EFAAAFPQ | ASSWDLDSQNTLY | 0.8259825 |
| LLSGNLDVL | ASSWDLDSQNTLY | 0.7112768 |
| GPANLGNYNI | ASSWDLDSQNTLY | 0.763464 |
| TLGPTLP | ASSWDLDSQNTLY | 0.8128702 |
| VYAYSSCCK | ASSWDLDSQNTLY | 0.47139058 |
| APAPAPAPNI | ASSWDLDSQNTLY | 0.62022585 |
| PGGPVPIP | ASSWDLDSQNTLY | 0.8404244 |
| FLTVALEP | ASSWDLDSQNTLY | 0.6423429 |
| SQYFGANL | ASSWDLDSQNTLY | 0.65508974 |
| GGVKPAA | ASSWDLDSQNTLY | 0.58442646 |
| GGTGGAGGAGGW | ASSWDLDSQNTLY | 0.5609204 |
| GVGGVAGLGGAGG | ASSWDLDSQNTLY | 0.6233121 |
| VLLVLATTI | ASSWDLDSQNTLY | 0.65313005 |
| IGIFAAML | ASSWDLDSQNTLY | 0.6593433 |
| AVGDLID | ASSWDLDSQNTLY | 0.6384462 |
| NISPLPL | ASSWDLDSQNTLY | 0.5549129 |
| VTAVFVAI | ASSWDLDSQNTLY | 0.6335271 |
| SGDDNDAAL | ASSWDLDSQNTLY | 0.6242159 |
| AVAALAHS | ASSWDLDSQNTLY | 0.6111847 |
| GGVTVMATP | ASSWDLDSQNTLY | 0.64476174 |
| AGLAANVRD | ASSWDLDSQNTLY | 0.6229615 |
| TNRPDLIDPALLRPG | ASSWDLDSQNTLY | 0.57897294 |
| LANLGVTAL | ASSWDLDSQNTLY | 0.5929769 |
| TGERIEL | ASSWDLDSQNTLY | 0.6443359 |
| NGEELAA | ASSWDLDSQNTLY | 0.65775156 |
| VLSLVWG | ASSWDLDSQNTLY | 0.65730965 |
| IGYALWL | ASSWDLDSQNTLY | 0.66918266 |
| LSLLLAPL | ASSWDLDSQNTLY | 0.66017807 |
| TLDEALK | ASSWDLDSQNTLY | 0.6401484 |
| TDGAGTAEGG | ASSWDLDSQNTLY | 0.37934563 |
| NAPGGAGGNGG | ASSWDLDSQNTLY | 0.5334421 |
| VSVFMGMGGNF | ASSWDLDSQNTLY | 0.554208 |
| LASYLAAL | ASSWDLDSQNTLY | 0.63395584 |
| LVGGTPAL | ASSWDLDSQNTLY | 0.38824916 |
| VRNADVL | ASSWDLDSQNTLY | 0.6304708 |
| TCSSPFLVSAM | ASSWDLDSQNTLY | 0.6107993 |
| VHITDAE | ASSWDLDSQNTLY | 0.56756216 |
| EYAAPSF | ASSWDLDSQNTLY | 0.6669949 |
| VVDAFHGGL | ASSWDLDSQNTLY | 0.5890322 |
| ASVVDPQ | ASSWDLDSQNTLY | 0.54887044 |
| SYNEFGGLS | ASSWDLDSQNTLY | 0.56064504 |
| DPDGSVY | ASSWDLDSQNTLY | 0.66744804 |
| MVSTIHATE | ASSWDLDSQNTLY | 0.5250848 |
| LAAGVADPV | ASSWDLDSQNTLY | 0.657743 |
| TKEQIAA | ASSWDLDSQNTLY | 0.6026376 |
| SLASPAAL | ASSWDLDSQNTLY | 0.66521776 |
| IGLVFYGV | ASSWDLDSQNTLY | 0.52427435 |

| | | |
|---|---|---|
| FLLDASNL | ASSWDLDSQNTLY | 0.65530384 |
| YQVGNFP | ASSWDLDSQNTLY | 0.6170127 |
| ESLLSQST | ASSWDLDSQNTLY | 0.67521685 |
| LFTVGFAAVG | ASSWDLDSQNTLY | 0.62941414 |
| ATPPTPT | ASSWDLDSQNTLY | 0.65191406 |
| TDLATLG | ASSWDLDSQNTLY | 0.598773 |
| SAGGTGR | ASSWDLDSQNTLY | 0.6540607 |
| GFSNSGS | ASSWDLDSQNTLY | 0.52220017 |
| GGNGGSGGTSV | ASSWDLDSQNTLY | 0.62256986 |
| GTSVATGGAGN | ASSWDLDSQNTLY | 0.52362865 |
| NGGNGGSGGTS | ASSWDLDSQNTLY | 0.63331527 |
| AVVKFPH | ASSWDLDSQNTLY | 0.64631325 |
| TPPPPPP | ASSWDLDSQNTLY | 0.47251746 |
| AAPAAAAAPA | ASSWDLDSQNTLY | 0.6644861 |
| IALNEDL | ASSWDLDSQNTLY | 0.5971018 |
| IVNVSSM | ASSWDLDSQNTLY | 0.5873787 |
| TGIPPAPR | ASSWDLDSQNTLY | 0.6502843 |
| FDLGGGTF | ASSWDLDSQNTLY | 0.6123159 |
| FDLGGGTFDVS | ASSWDLDSQNTLY | 0.64755416 |
| NEPTAAA | ASSWDLDSQNTLY | 0.5346291 |
| NEPTAAALA | ASSWDLDSQNTLY | 0.63335764 |
| GSIINISSI | ASSWDLDSQNTLY | 0.64468133 |
| VAPGNAGTA | ASSWDLDSQNTLY | 0.5155418 |
| TGGGASLE | ASSWDLDSQNTLY | 0.5269084 |
| VAVYNHYKRI | ASSWDLDSQNTLY | 0.6618342 |
| TDPAPATT | ASSWDLDSQNTLY | 0.6428803 |
| TNRPDILDPALLRPG | ASSWDLDSQNTLY | 0.7014208 |
| LTTTTTS | ASSWDLDSQNTLY | 0.51033103 |
| PTGSGKTL | ASSWDLDSQNTLY | 0.47813445 |
| TGSGKTL | ASSWDLDSQNTLY | 0.5747032 |
| TIGHVDHGK | ASSWDLDSQNTLY | 0.6738625 |
| TIGHVDHGKTT | ASSWDLDSQNTLY | 0.69842815 |
| GHVDHGKTT | ASSWDLDSQNTLY | 0.65044206 |
| AQFEHTLL | ASSWDLDSQNTLY | 0.48570174 |
| IVNQDEL | ASSWDLDSQNTLY | 0.5720658 |
| TNGSQFF | ASSWDLDSQNTLY | 0.53714466 |
| TGKGIVE | ASSWDLDSQNTLY | 0.5667917 |
| LISNASDA | ASSWDLDSQNTLY | 0.6453116 |
| AELRTLGELTKIAKAHGAQ | ASSWDLDSQNTLY | 0.59749067 |
| APNAVAAAEPSW | ASSWDLDSQNTLY | 0.54177237 |
| APQVAGAATAVL | ASSWDLDSQNTLY | 0.5301045 |
| ARRDPAAAAAILPTDA | ASSWDLDSQNTLY | 0.6401231 |
| DAAARDVLAEAGLAEYF | ASSWDLDSQNTLY | 0.65377486 |
| DDQIVLSAKANLAKVEQ | ASSWDLDSQNTLY | 0.61630183 |
| DELLPDTGEDSDEDGH | ASSWDLDSQNTLY | 0.6954911 |
| ELPEAPANRATVPQD | ASSWDLDSQNTLY | 0.6614725 |
| FGQSKIERVTGIG | ASSWDLDSQNTLY | 0.6698871 |
| GAGGTGSTAGGAGGAGGAGGLYA | ASSWDLDSQNTLY | 0.29178062 |
| GHGGTGAAGGNGGTGGAGGSGIDG | ASSWDLDSQNTLY | 0.37675297 |
| GNGGNGGAGGEGGPGGAGGAGA | ASSWDLDSQNTLY | 0.5863321 |
| GSTAGNGGNGGNATNGTIAGQPAG | ASSWDLDSQNTLY | 0.5459096 |

| | | |
|---|---|---|
| GTGGDGGNGGNGGTGG | ASSWDLDSQNTLY | 0.39601964 |
| GVRRPAGIPADAPV | ASSWDLDSQNTLY | 0.6243701 |
| GYRQVTDAHLLTLARR | ASSWDLDSQNTLY | 0.51482964 |
| IAKMGQDGHDRG | ASSWDLDSQNTLY | 0.5024864 |
| KAEPSIEIQYT | ASSWDLDSQNTLY | 0.65787977 |
| KPVRIRMAADEDTVAVT | ASSWDLDSQNTLY | 0.56255436 |
| MPPRIAGMRLLVIKPEP | ASSWDLDSQNTLY | 0.5085779 |
| MKFYAVALFRSRGVPPTR | ASSWDLDSQNTLY | 0.4913597 |
| NAGGFGILGAVAHSP | ASSWDLDSQNTLY | 0.6665372 |
| NEGREITLAMAEM | ASSWDLDSQNTLY | 0.3962221 |
| NELVEWSHPLGHRWRWE | ASSWDLDSQNTLY | 0.62869936 |
| NLGAGGTGGNGGNPG | ASSWDLDSQNTLY | 0.58755827 |
| NTEWPKWETVMAADPVD | ASSWDLDSQNTLY | 0.66405386 |
| PASAAPAETQP | ASSWDLDSQNTLY | 0.40806633 |
| QQSSDGKNGVRAAEITGE | ASSWDLDSQNTLY | 0.6434132 |
| RAPDAVTTSWWKE | ASSWDLDSQNTLY | 0.57816637 |
| SGFGNVGAHTSGWFNQ | ASSWDLDSQNTLY | 0.70796895 |
| SGVAGAVGQAASVGGLKV | ASSWDLDSQNTLY | 0.52343 |
| SNGGAGTGSAGGNGGT | ASSWDLDSQNTLY | 0.59412086 |
| TGGAGGEGGAAG | ASSWDLDSQNTLY | 0.58199376 |
| TGGDGGAAGAGGAPGHG | ASSWDLDSQNTLY | 0.6940706 |
| TMPPLLAAIANAGRHGV | ASSWDLDSQNTLY | 0.4250447 |
| TSSTAGNSGGA | ASSWDLDSQNTLY | 0.5551848 |
| VPATVPVGILC | ASSWDLDSQNTLY | 0.41010103 |
| VQRPDSGGVVLTGRIS | ASSWDLDSQNTLY | 0.60955846 |
| WTPNPVPATKGVPAGLVQAT | ASSWDLDSQNTLY | 0.64326644 |
| YALKRALAKFSDRFDVVII | ASSWDLDSQNTLY | 0.41274002 |
| YLIGSVIV | ASSWDLDSQNTLY | 0.6778799 |
| YGQLVAVID | ASSWDLDSQNTLY | 0.5946484 |
| VTVGGMTLPA | ASSWDLDSQNTLY | 0.5172259 |
| VLALPAV | ASSWDLDSQNTLY | 0.680495 |
| VIPPTLNYET | ASSWDLDSQNTLY | 0.5309961 |
| VGMGGMHPSSG | ASSWDLDSQNTLY | 0.6928959 |
| VGLAGGAATA | ASSWDLDSQNTLY | 0.6100333 |
| VGGAGGAGGSGP | ASSWDLDSQNTLY | 0.6085475 |
| VFAAFMV | ASSWDLDSQNTLY | 0.64075804 |
| TLYDKLR | ASSWDLDSQNTLY | 0.5757647 |
| TLAVFTVL | ASSWDLDSQNTLY | 0.5032423 |
| TGSVQPARV | ASSWDLDSQNTLY | 0.6773718 |
| TGGGAGSGGT | ASSWDLDSQNTLY | 0.60954845 |
| SHLETSL | ASSWDLDSQNTLY | 0.632885 |
| SFTTGMT | ASSWDLDSQNTLY | 0.56430763 |
| QVEVAKRMGVRQPTVS | ASSWDLDSQNTLY | 0.54169285 |
| QGSGGPALAM | ASSWDLDSQNTLY | 0.58348763 |
| MLAPSNP | ASSWDLDSQNTLY | 0.63258386 |
| LVNNAAI | ASSWDLDSQNTLY | 0.6425009 |
| LTGEINTLA | ASSWDLDSQNTLY | 0.6126163 |
| LSIFGPV | ASSWDLDSQNTLY | 0.5492991 |
| LLALLANL | ASSWDLDSQNTLY | 0.6354697 |
| LISNASDALDKLR | ASSWDLDSQNTLY | 0.5990522 |
| LINEAVP | ASSWDLDSQNTLY | 0.5801577 |

| | | |
|---|---|---|
| LGYAFSGL | ASSWDLDSQNTLY | 0.6195533 |
| LDFVAVAL | ASSWDLDSQNTLY | 0.6233144 |
| LAQPGAE | ASSWDLDSQNTLY | 0.5911113 |
| ISNLGAVTGF | ASSWDLDSQNTLY | 0.569676 |
| HQEDPGRI | ASSWDLDSQNTLY | 0.61976993 |
| GTPGEPTP | ASSWDLDSQNTLY | 0.63375396 |
| GFFNSGS | ASSWDLDSQNTLY | 0.62707096 |
| GAGGGGLTAGAN | ASSWDLDSQNTLY | 0.6485606 |
| FVDALVNM | ASSWDLDSQNTLY | 0.62415195 |
| FLTVAPEL | ASSWDLDSQNTLY | 0.6314736 |
| FETLTAL | ASSWDLDSQNTLY | 0.39215392 |
| DVINPIEL | ASSWDLDSQNTLY | 0.44410607 |
| APVSGAPV | ASSWDLDSQNTLY | 0.5930765 |
| ALVGGLAW | ASSWDLDSQNTLY | 0.59316087 |
| ALAAAAATAI | ASSWDLDSQNTLY | 0.44074216 |
| AFGLGLLL | ASSWDLDSQNTLY | 0.63531744 |
| AFFAAASL | ASSWDLDSQNTLY | 0.6061082 |
| AAAALPAFSPP | ASSWDLDSQNTLY | 0.5804687 |
| VVLAPAL | ASSWDLDSQNTLY | 0.6338433 |
| TIGHVDHGKTTL | ASSWDLDSQNTLY | 0.5793701 |
| LISNASDALDK | ASSWDLDSQNTLY | 0.5701699 |
| LISNASDAL | ASSWDLDSQNTLY | 0.58546495 |
| ISNASDALDK | ASSWDLDSQNTLY | 0.6434178 |
| NQAETLVYQTE | ASSWDLDSQNTLY | 0.4954214 |
| TGGTGGTAGAG | ASSWDLDSQNTLY | 0.6325889 |
| TGSTAGGAGGA | ASSWDLDSQNTLY | 0.5354996 |
| DPLAVPVAAGPV | ASSWDLDSQNTLY | 0.6400162 |
| GGTGGTPGNGGN | ASSWDLDSQNTLY | 0.50155944 |
| PAGLAGVDPATL | ASSWDLDSQNTLY | 0.6182649 |
| GNNLVVEAPHAGASV | ASSWDLDSQNTLY | 0.57506186 |
| KVTRIMSASTLEQPA | ASSWDLDSQNTLY | 0.65356284 |
| LGFALPIALPRRTVV | ASSWDLDSQNTLY | 0.57926255 |
| TRLGNLTARKLVVLV | ASSWDLDSQNTLY | 0.6289151 |
| WSQQRRLIYFPSAGP | ASSWDLDSQNTLY | 0.6122951 |
| GPNGNPDPMAAAVDIR | ASSWDLDSQNTLY | 0.63672847 |
| TQRCVSVADLRAAASR | ASSWDLDSQNTLY | 0.48682576 |
| QFVLDVAEHPVEVLTETV | ASSWDLDSQNTLY | 0.59892356 |
| GGAGGTGGAGGNGGTQAGDG | ASSWDLDSQNTLY | 0.49860582 |
| PLLATLRAIAVDVDAKPAQV | ASSWDLDSQNTLY | 0.61648303 |
| APKFRTGGYMIDFWGVGYQVAKRM | ASSWDLDSQNTLY | 0.6095544 |
| GAGGAGGAGGAAGGPGGTGGTGGN | ASSWDLDSQNTLY | 0.4478856 |
| IGALGLLATDVAPAGSVAFGVALA | ASSWDLDSQNTLY | 0.6365004 |
| AGGAGGIGGNGGAGGAGGVLMGNG | ASSWDLDSQNTLY | 0.5824057 |
| ADELIGLL | ASSWDLDSQNTLY | 0.56361324 |
| ADRALKA | ASSWDLDSQNTLY | 0.6306725 |
| AGAVGQVGGAGGA | ASSWDLDSQNTLY | 0.6196143 |
| AGGGGGLGGAP | ASSWDLDSQNTLY | 0.6249155 |
| AGGLLYGNGGAG | ASSWDLDSQNTLY | 0.59727305 |
| AGLESIGL | ASSWDLDSQNTLY | 0.58433014 |
| AGLKTGII | ASSWDLDSQNTLY | 0.6263368 |
| ASLAALLL | ASSWDLDSQNTLY | 0.533877 |

| | | |
|---|---|---|
| ASVINISSI | ASSWDLDSQNTLY | 0.5625619 |
| ATPPGPA | ASSWDLDSQNTLY | 0.6217154 |
| ATVASALV | ASSWDLDSQNTLY | 0.6178915 |
| ATVASLIL | ASSWDLDSQNTLY | 0.7016816 |
| AVNGPVLIRH | ASSWDLDSQNTLY | 0.40877375 |
| DGAAGAVGI | ASSWDLDSQNTLY | 0.26882172 |
| DPLLHGI | ASSWDLDSQNTLY | 0.42921075 |
| ELGAALGAL | ASSWDLDSQNTLY | 0.6463525 |
| ELGRTQAV | ASSWDLDSQNTLY | 0.76060987 |
| FVAGGYLS | ASSWDLDSQNTLY | 0.6057182 |
| GASGGAGGAGGW | ASSWDLDSQNTLY | 0.2607872 |
| GGAGAGGDGGAGGI | ASSWDLDSQNTLY | 0.7042307 |
| GTGHNGSQPGMGGNG | ASSWDLDSQNTLY | 0.43067488 |
| GYAAASASAS | ASSWDLDSQNTLY | 0.4227211 |
| HQVDVAAVQQG | ASSWDLDSQNTLY | 0.5272596 |
| IGAAIAHT | ASSWDLDSQNTLY | 0.5300465 |
| IGMIEVAQL | ASSWDLDSQNTLY | 0.4400937 |
| IRLAPEHTSAMD | ASSWDLDSQNTLY | 0.32998583 |
| IYNVSVL | ASSWDLDSQNTLY | 0.5898049 |
| KQGFLGM | ASSWDLDSQNTLY | 0.71543765 |
| LESVITQL | ASSWDLDSQNTLY | 0.6848583 |
| LGLGALVVV | ASSWDLDSQNTLY | 0.72514796 |
| LIGNGGNGGSGGI | ASSWDLDSQNTLY | 0.7381511 |
| LNAINAPTL | ASSWDLDSQNTLY | 0.55533755 |
| LQLLNTGV | ASSWDLDSQNTLY | 0.11676362 |
| LQTSIEF | ASSWDLDSQNTLY | 0.21422212 |
| MSFAVSEL | ASSWDLDSQNTLY | 0.29655057 |
| NTGALGL | ASSWDLDSQNTLY | 0.414621 |
| PPGAGGAGGAGTT | ASSWDLDSQNTLY | 0.09819613 |
| QLVGLGI | ASSWDLDSQNTLY | 0.46326292 |
| RFLEQVHQL | ASSWDLDSQNTLY | 0.46230483 |
| RGVGYVLR | ASSWDLDSQNTLY | 0.29153067 |
| SATGVFAGVI | ASSWDLDSQNTLY | 0.746413 |
| SLPAITLP | ASSWDLDSQNTLY | 0.34652233 |
| STALAFTV | ASSWDLDSQNTLY | 0.32205415 |
| SVLTVGLGK | ASSWDLDSQNTLY | 0.32932937 |
| TALAGAGSGI | ASSWDLDSQNTLY | 0.7220594 |
| TGLPFTGI | ASSWDLDSQNTLY | 0.23345639 |
| VAEHVTV | ASSWDLDSQNTLY | 0.720613 |
| VDIVLNSL | ASSWDLDSQNTLY | 0.28026158 |
| VLTGGGAVL | ASSWDLDSQNTLY | 0.66853774 |
| VNVGLAYGI | ASSWDLDSQNTLY | 0.20814013 |
| VSSAAVGAV | ASSWDLDSQNTLY | 0.6854409 |
| VTSVYEAGV | ASSWDLDSQNTLY | 0.51614106 |
| WVALDRGA | ASSWDLDSQNTLY | 0.83355343 |
| YLVGSVVI | ASSWDLDSQNTLY | 0.4324515 |
| GPPGTGKTLL | ASSWDLDSQNTLY | 0.5539788 |
| GVGMTKF | ASSWDLDSQNTLY | 0.41199988 |
| IGIGGSDL | ASSWDLDSQNTLY | 0.7291313 |
| IGTLLAEL | ASSWDLDSQNTLY | 0.18868643 |
| IYTSGTT | ASSWDLDSQNTLY | 0.50694114 |

| | | |
|---|---|---|
| LSFPAGV | ASSWDLDSQNTLY | 0.2610261 |
| LYTSGTT | ASSWDLDSQNTLY | 0.59064937 |
| VGAFPFGF | ASSWDLDSQNTLY | 0.6265079 |
| VGAGTVEF | ASSWDLDSQNTLY | 0.10526252 |
| VGKSTLLL | ASSWDLDSQNTLY | 0.6991612 |
| AALAHAGGGIV | ASSWDLDSQNTLY | 0.24008983 |
| AALVAVSLL | ASSWDLDSQNTLY | 0.22685157 |
| AAVSRAVAAL | ASSWDLDSQNTLY | 0.6121305 |
| AGGSSGIGGF | ASSWDLDSQNTLY | 0.44775134 |
| AGLTVANANG | ASSWDLDSQNTLY | 0.75449383 |
| AGLVGAVL | ASSWDLDSQNTLY | 0.27963504 |
| ALSQAFTV | ASSWDLDSQNTLY | 0.5355685 |
| ALTFSVSL | ASSWDLDSQNTLY | 0.60603774 |
| ALTGAALVAL | ASSWDLDSQNTLY | 0.15677308 |
| ASFSIGAGVL | ASSWDLDSQNTLY | 0.25667858 |
| DGSVYKI | ASSWDLDSQNTLY | 0.75735295 |
| EQLAEIV | ASSWDLDSQNTLY | 0.8012141 |
| FAVSELTVM | ASSWDLDSQNTLY | 0.84949934 |
| GAAPLAAGL | ASSWDLDSQNTLY | 0.6980008 |
| GADGGDGGSGGAGG | ASSWDLDSQNTLY | 0.63508964 |
| GFGLDTGGAGG | ASSWDLDSQNTLY | 0.7258724 |
| GLASGGLL | ASSWDLDSQNTLY | 0.8471987 |
| GRIAGAAVLL | ASSWDLDSQNTLY | 0.8482716 |
| GSAGSLNTSF | ASSWDLDSQNTLY | 0.8041189 |
| GTGQIGI | ASSWDLDSQNTLY | 0.6166171 |
| GTLIALIL | ASSWDLDSQNTLY | 0.77855235 |
| IAATMGFVVG | ASSWDLDSQNTLY | 0.7111007 |
| IFAQALL | ASSWDLDSQNTLY | 0.7336458 |
| IGAAVALGF | ASSWDLDSQNTLY | 0.8222619 |
| IGLTPLAL | ASSWDLDSQNTLY | 0.7793181 |
| ILGSNTSTL | ASSWDLDSQNTLY | 0.69884837 |
| IVFSLTVL | ASSWDLDSQNTLY | 0.7356042 |
| LASKATGF | ASSWDLDSQNTLY | 0.7840488 |
| LASVALIL | ASSWDLDSQNTLY | 0.7973565 |
| LFAVGLTL | ASSWDLDSQNTLY | 0.79409754 |
| LFNGGIGTYI | ASSWDLDSQNTLY | 0.8688344 |
| LTLGAGII | ASSWDLDSQNTLY | 0.8318039 |
| LVAAALGDI | ASSWDLDSQNTLY | 0.8414707 |
| NTAFVQLGI | ASSWDLDSQNTLY | 0.2030027 |
| QLFSGAEL | ASSWDLDSQNTLY | 0.43435395 |
| SAATFITL | ASSWDLDSQNTLY | 0.7396255 |
| SNYGLANLGV | ASSWDLDSQNTLY | 0.71661425 |
| SYGGAYAVM | ASSWDLDSQNTLY | 0.5100758 |
| TTGVGGLTLG | ASSWDLDSQNTLY | 0.81454444 |
| TTQTIYI | ASSWDLDSQNTLY | 0.6907904 |
| VAAGAGLGAVI | ASSWDLDSQNTLY | 0.6474911 |
| VAAVVTLL | ASSWDLDSQNTLY | 0.8206476 |
| VGFLGGLI | ASSWDLDSQNTLY | 0.7123752 |
| VGVIPILL | ASSWDLDSQNTLY | 0.68619645 |
| VIFASSLI | ASSWDLDSQNTLY | 0.7047782 |
| VKGGDGGAAAG | ASSWDLDSQNTLY | 0.82790405 |

| | | |
|---|---|---|
| VTKTFTNI | ASSWDLDSQNTLY | 0.5102354 |
| VVAAAILL | ASSWDLDSQNTLY | 0.8106475 |
| YAALDVEL | ASSWDLDSQNTLY | 0.6701739 |
| LASVALLL | ASSWDLDSQNTLY | 0.84328747 |
| LDAVHRV | ASSWDLDSQNTLY | 0.538198 |
| QGLSSSIF | ASSWDLDSQNTLY | 0.81348956 |
| RGTFANIRL | ASSWDLDSQNTLY | 0.7443253 |
| RQLWWGHRI | ASSWDLDSQNTLY | 0.8728874 |
| YGPPGTGKTLL | ASSWDLDSQNTLY | 0.73236924 |
| AALVGAPANVA | ASSWDLDSQNTLY | 0.76902914 |
| AATFRRTE | ASSWDLDSQNTLY | 0.64517385 |
| AGALGGGI | ASSWDLDSQNTLY | 0.84390056 |
| AGGAGGAGGIGAG | ASSWDLDSQNTLY | 0.45227543 |
| AGSGALGL | ASSWDLDSQNTLY | 0.74125206 |
| AQIADQI | ASSWDLDSQNTLY | 0.44192356 |
| ASAVAVTYL | ASSWDLDSQNTLY | 0.803183 |
| ASTVLGGTGG | ASSWDLDSQNTLY | 0.79491854 |
| AVGGEPAK | ASSWDLDSQNTLY | 0.50512856 |
| CEQGTFGGTC | ASSWDLDSQNTLY | 0.8334174 |
| DEGKAPAG | ASSWDLDSQNTLY | 0.7614896 |
| DTVVFPAL | ASSWDLDSQNTLY | 0.6953628 |
| FEGITMAG | ASSWDLDSQNTLY | 0.8520639 |
| FPLVVQD | ASSWDLDSQNTLY | 0.7510553 |
| GAFTTPQ | ASSWDLDSQNTLY | 0.8455761 |
| GFGGFGTAGGVGGA | ASSWDLDSQNTLY | 0.7730007 |
| GFLAQFL | ASSWDLDSQNTLY | 0.6857425 |
| GGAGGAGGNAGVA | ASSWDLDSQNTLY | 0.81367284 |
| GGVGTAGAGGAGGAGG | ASSWDLDSQNTLY | 0.4993325 |
| GNGGGGDAGNAGSGGN | ASSWDLDSQNTLY | 0.6663895 |
| GTGGAGGAGNDAGS | ASSWDLDSQNTLY | 0.8412938 |
| GTTNTGWFNS | ASSWDLDSQNTLY | 0.68760324 |
| IFMGPLFTL | ASSWDLDSQNTLY | 0.6888435 |
| ILAHAIEAI | ASSWDLDSQNTLY | 0.53961027 |
| IRSAIREAL | ASSWDLDSQNTLY | 0.5775305 |
| ITFAGGG | ASSWDLDSQNTLY | 0.60521257 |
| ITGMSGHQEV | ASSWDLDSQNTLY | 0.67485714 |
| IVAGLACTL | ASSWDLDSQNTLY | 0.7150515 |
| IVGGGLAGA | ASSWDLDSQNTLY | 0.69055 |
| IVTGVLNI | ASSWDLDSQNTLY | 0.6215496 |
| LAQQGCI | ASSWDLDSQNTLY | 0.66760457 |
| LCLHVEN | ASSWDLDSQNTLY | 0.6134075 |
| LDAISDGS | ASSWDLDSQNTLY | 0.62443423 |
| LGGFAAML | ASSWDLDSQNTLY | 0.66676164 |
| LGGKNPMI | ASSWDLDSQNTLY | 0.65248525 |
| LGPKGVAAIL | ASSWDLDSQNTLY | 0.6063611 |
| LMVGVSI | ASSWDLDSQNTLY | 0.58494586 |
| LVGAAFVL | ASSWDLDSQNTLY | 0.6681461 |
| LVLAVAPL | ASSWDLDSQNTLY | 0.70522296 |
| MAAIAVIDAA | ASSWDLDSQNTLY | 0.6980792 |
| NNGINAELQ | ASSWDLDSQNTLY | 0.70955193 |
| RGEPLVLVHG | ASSWDLDSQNTLY | 0.6818226 |

| | | |
|---|---|---|
| RGGAGGAAGLI | ASSWDLDSQNTLY | 0.6910908 |
| RSYRMHPAV | ASSWDLDSQNTLY | 0.33493337 |
| SAAHLLAGQLA | ASSWDLDSQNTLY | 0.49721366 |
| SAAVGSY | ASSWDLDSQNTLY | 0.63819593 |
| SPHAVLHRI | ASSWDLDSQNTLY | 0.6265687 |
| SQIAKVL | ASSWDLDSQNTLY | 0.39354432 |
| TALGHLVP | ASSWDLDSQNTLY | 0.6722692 |
| TAPTTSGAA | ASSWDLDSQNTLY | 0.63019174 |
| TDGDDGNGGSPGT | ASSWDLDSQNTLY | 0.58264565 |
| TDGLGEN | ASSWDLDSQNTLY | 0.6874333 |
| TGGPPLPGQA | ASSWDLDSQNTLY | 0.6058403 |
| TIPAITI | ASSWDLDSQNTLY | 0.55960035 |
| TIPGFTI | ASSWDLDSQNTLY | 0.6242227 |
| TSIASGALA | ASSWDLDSQNTLY | 0.7085538 |
| TVTLPGAAI | ASSWDLDSQNTLY | 0.5531816 |
| VASGLSAALP | ASSWDLDSQNTLY | 0.6868482 |
| VGAGVIGI | ASSWDLDSQNTLY | 0.6158818 |
| VGPKRFGV | ASSWDLDSQNTLY | 0.6938858 |
| VNVAFPD | ASSWDLDSQNTLY | 0.54497683 |
| VSMTNSGVS | ASSWDLDSQNTLY | 0.6811133 |
| VTDVLGDLSV | ASSWDLDSQNTLY | 0.61591595 |
| VVVVGAGFAGL | ASSWDLDSQNTLY | 0.7218786 |
| VYTVGTVV | ASSWDLDSQNTLY | 0.6140786 |
| WTTFSGP | ASSWDLDSQNTLY | 0.6750751 |
| YDELVEAF | ASSWDLDSQNTLY | 0.6463376 |
| YGTGGAGGNGG | ASSWDLDSQNTLY | 0.704439 |
| YTNVMAV | ASSWDLDSQNTLY | 0.4870592 |
| FLEQVHQL | ASSWDLDSQNTLY | 0.6188436 |
| GLSLLAVAV | ASSWDLDSQNTLY | 0.53343874 |
| LEQVHQL | ASSWDLDSQNTLY | 0.6429105 |
| LNAAGAI | ASSWDLDSQNTLY | 0.6876873 |
| PAVHASL | ASSWDLDSQNTLY | 0.3755885 |
| TELEDDA | ASSWDLDSQNTLY | 0.6932363 |
| TPPGPPN | ASSWDLDSQNTLY | 0.5188095 |
| TPPPPPPP | ASSWDLDSQNTLY | 0.5944971 |
| TQSPALI | ASSDREVF | 0.68526363 |
| EFAAAFPQ | ASSDREVF | 0.6286382 |
| LLSGNLDVL | ASSDREVF | 0.69541466 |
| GPANLGNYNI | ASSDREVF | 0.6153713 |
| TLGPTLP | ASSDREVF | 0.64412796 |
| VYAYSSCCK | ASSDREVF | 0.68018997 |
| APAPAPAPNI | ASSDREVF | 0.50150645 |
| PGGPVPIP | ASSDREVF | 0.5780221 |
| FLTVALEP | ASSDREVF | 0.6982555 |
| SQYFGANL | ASSDREVF | 0.8151918 |
| GGVKPAA | ASSDREVF | 0.8266199 |
| GGTGGAGGAGGW | ASSDREVF | 0.7592749 |
| GVGGVAGLGGAGG | ASSDREVF | 0.6948126 |
| VLLVLATTI | ASSDREVF | 0.7635226 |
| IGIFAAML | ASSDREVF | 0.82618105 |
| AVGDLID | ASSDREVF | 0.81917524 |

| | | |
|---|---|---|
| NISPLPL | ASSDREVF | 0.8169698 |
| VTAVFVAI | ASSDREVF | 0.6880081 |
| SGDDNDAAL | ASSDREVF | 0.7947588 |
| AVAALAHS | ASSDREVF | 0.7818757 |
| GGVTVMATP | ASSDREVF | 0.7658719 |
| AGLAANVRD | ASSDREVF | 0.81795096 |
| TNRPDLIDPALLRPG | ASSDREVF | 0.7924695 |
| LANLGVTAL | ASSDREVF | 0.78291595 |
| TGERIEL | ASSDREVF | 0.7823208 |
| NGEELAA | ASSDREVF | 0.79303706 |
| VLSLVWG | ASSDREVF | 0.80759 |
| IGYALWL | ASSDREVF | 0.80205965 |
| LSLLLAPL | ASSDREVF | 0.8341925 |
| TLDEALK | ASSDREVF | 0.82249546 |
| TDGAGTAEGG | ASSDREVF | 0.8253857 |
| NAPGGAGGNGG | ASSDREVF | 0.27274248 |
| VSVFMGMGGNF | ASSDREVF | 0.57355756 |
| LASYLAAL | ASSDREVF | 0.8037982 |
| LVGGTPAL | ASSDREVF | 0.7655694 |
| VRNADVL | ASSDREVF | 0.7054247 |
| TCSSPFLVSAM | ASSDREVF | 0.81802803 |
| VHITDAE | ASSDREVF | 0.79237425 |
| EYAAPSF | ASSDREVF | 0.7320895 |
| VVDAFHGGL | ASSDREVF | 0.8061467 |
| ASVVDPQ | ASSDREVF | 0.76324195 |
| SYNEFGGLS | ASSDREVF | 0.75375676 |
| DPDGSVY | ASSDREVF | 0.778864 |
| MVSTIHATE | ASSDREVF | 0.82771635 |
| LAAGVADPV | ASSDREVF | 0.6970322 |
| TKEQIAA | ASSDREVF | 0.8049456 |
| SLASPAAL | ASSDREVF | 0.7970622 |
| IGLVFYGV | ASSDREVF | 0.8207824 |
| FLLDASNL | ASSDREVF | 0.60687435 |
| YQVGNFP | ASSDREVF | 0.8187009 |
| ESLLSQST | ASSDREVF | 0.78272486 |
| LFTVGFAAVG | ASSDREVF | 0.83181036 |
| ATPPTPT | ASSDREVF | 0.75528115 |
| TDLATLG | ASSDREVF | 0.77989054 |
| SAGGTGR | ASSDREVF | 0.74893224 |
| GFSNSGS | ASSDREVF | 0.81361794 |
| GGNGGSGGTSV | ASSDREVF | 0.51358014 |
| GTSVATGGAGN | ASSDREVF | 0.7848339 |
| NGGNGGSGGTS | ASSDREVF | 0.7169145 |
| AVVKFPH | ASSDREVF | 0.80042714 |
| TPPPPPP | ASSDREVF | 0.8149962 |
| AAPAAAAAPA | ASSDREVF | 0.6710596 |
| IALNEDL | ASSDREVF | 0.82073164 |
| IVNVSSM | ASSDREVF | 0.81124854 |
| TGIPPAPR | ASSDREVF | 0.7636448 |
| FDLGGGTF | ASSDREVF | 0.8264809 |
| FDLGGGTFDVS | ASSDREVF | 0.78868115 |

| | | |
|---|---|---|
| NEPTAAA | ASSDREVF | 0.82802534 |
| NEPTAAALA | ASSDREVF | 0.8009099 |
| GSIINISSI | ASSDREVF | 0.79634386 |
| VAPGNAGTA | ASSDREVF | 0.8120515 |
| TGGGASLE | ASSDREVF | 0.6430518 |
| VAVYNHYKRI | ASSDREVF | 0.76606613 |
| TDPAPATT | ASSDREVF | 0.82465184 |
| TNRPDILDPALLRPG | ASSDREVF | 0.75158584 |
| LTTTTTS | ASSDREVF | 0.81531507 |
| PTGSGKTL | ASSDREVF | 0.6563661 |
| TGSGKTL | ASSDREVF | 0.58337724 |
| TIGHVDHGK | ASSDREVF | 0.72387594 |
| TIGHVDHGKTT | ASSDREVF | 0.8069294 |
| GHVDHGKTT | ASSDREVF | 0.80649036 |
| AQFEHTLL | ASSDREVF | 0.78610885 |
| IVNQDEL | ASSDREVF | 0.58794326 |
| TNGSQFF | ASSDREVF | 0.7168572 |
| TGKGIVE | ASSDREVF | 0.72007805 |
| LISNASDA | ASSDREVF | 0.6685731 |
| AELRTLGELTKIAKAHGAQ | ASSDREVF | 0.7687465 |
| APNAVAAAEPSW | ASSDREVF | 0.72198087 |
| APQVAGAATAVL | ASSDREVF | 0.6093154 |
| ARRDPAAAAAILPTDA | ASSDREVF | 0.6303485 |
| DAAARDVLAEAGLAEYF | ASSDREVF | 0.75680786 |
| DDQIVLSAKANLAKVEQ | ASSDREVF | 0.785854 |
| DELLPDTGEDSDEDGH | ASSDREVF | 0.77115643 |
| ELPEAPANRATVPQD | ASSDREVF | 0.81807667 |
| FGQSKIERVTGIG | ASSDREVF | 0.7911675 |
| GAGGTGSTAGGAGGAGGAGGLYA | ASSDREVF | 0.75317943 |
| GHGGTGAAGGNGGTGGAGGSGIDG | ASSDREVF | 0.2270671 |
| GNGGNGGAGGEGGPGGAGGAGA | ASSDREVF | 0.4232739 |
| GSTAGNGGNGGNATNGTIAGQPAG | ASSDREVF | 0.70756984 |
| GTGGDGGNGGNGGTGG | ASSDREVF | 0.72414994 |
| GVRRPAGIPADAPV | ASSDREVF | 0.53178984 |
| GYRQVTDAHLLTLARR | ASSDREVF | 0.78459823 |
| IAKMGQDGHDRG | ASSDREVF | 0.71045655 |
| KAEPSIEIQYT | ASSDREVF | 0.6474412 |
| KPVRIRMAADEDTVAVT | ASSDREVF | 0.78744674 |
| MPPRIAGMRLLVIKPEP | ASSDREVF | 0.61655223 |
| MKFYAVALFRSRGVPPTR | ASSDREVF | 0.61838526 |
| NAGGFGILGAVAHSP | ASSDREVF | 0.65964264 |
| NEGREITLAMAEM | ASSDREVF | 0.79944634 |
| NELVEWSHPLGHRWRWE | ASSDREVF | 0.46675915 |
| NLGAGGTGGNGGNPG | ASSDREVF | 0.7598578 |
| NTEWPKWETVMAADPVD | ASSDREVF | 0.6851473 |
| PASAAPAETQP | ASSDREVF | 0.796976 |
| QQSSDGKNGVRAAEITGE | ASSDREVF | 0.50211394 |
| RAPDAVTTSWWKE | ASSDREVF | 0.77794796 |
| SGFGNVGAHTSGWFNQ | ASSDREVF | 0.7160825 |
| SGVAGAVGQAASVGGLKV | ASSDREVF | 0.8205546 |
| SNGGAGTGSAGGNGGT | ASSDREVF | 0.68913656 |

| | | |
|---|---|---|
| TGGAGGEGGAAG | ASSDREVF | 0.7475828 |
| TGGDGGAAGAGGAPGHG | ASSDREVF | 0.62222314 |
| TMPPLLAAIANAGRHGV | ASSDREVF | 0.7963604 |
| TSSTAGNSGGA | ASSDREVF | 0.61064005 |
| VPATVPVGILC | ASSDREVF | 0.71108925 |
| VQRPDSGGVVLTGRIS | ASSDREVF | 0.50351244 |
| WTPNPVPATKGVPAGLVQAT | ASSDREVF | 0.75207996 |
| YALKRALAKFSDRFDVVII | ASSDREVF | 0.76609504 |
| YLIGSVIV | ASSDREVF | 0.70944214 |
| YGQLVAVID | ASSDREVF | 0.7768034 |
| VTVGGMTLPA | ASSDREVF | 0.7979323 |
| VLALPAV | ASSDREVF | 0.6427548 |
| VIPPTLNYET | ASSDREVF | 0.80439514 |
| VGMGGMHPSSG | ASSDREVF | 0.72144985 |
| VGLAGGAATA | ASSDREVF | 0.7949414 |
| VGGAGGAGGSGP | ASSDREVF | 0.6812235 |
| VFAAFMV | ASSDREVF | 0.7182349 |
| TLYDKLR | ASSDREVF | 0.78551424 |
| TLAVFTVL | ASSDREVF | 0.75422513 |
| TGSVQPARV | ASSDREVF | 0.68674004 |
| TGGGAGSGGT | ASSDREVF | 0.7851906 |
| SHLETSL | ASSDREVF | 0.39806667 |
| SFTTGMT | ASSDREVF | 0.7068958 |
| QVEVAKRMGVRQPTVS | ASSDREVF | 0.25013718 |
| QGSGGPALAM | ASSDREVF | 0.24810834 |
| MLAPSNP | ASSDREVF | 0.3274048 |
| LVNNAAI | ASSDREVF | 0.62757176 |
| LTGEINTLA | ASSDREVF | 0.6581973 |
| LSIFGPV | ASSDREVF | 0.5533401 |
| LLALLANL | ASSDREVF | 0.25983036 |
| LISNASDALDKLR | ASSDREVF | 0.28152585 |
| LINEAVP | ASSDREVF | 0.29029813 |
| LGYAFSGL | ASSDREVF | 0.30392334 |
| LDFVAVAL | ASSDREVF | 0.49012455 |
| LAQPGAE | ASSDREVF | 0.37982792 |
| ISNLGAVTGF | ASSDREVF | 0.25441 |
| HQEDPGRI | ASSDREVF | 0.33503035 |
| GTPGEPTP | ASSDREVF | 0.33255747 |
| GFFNSGS | ASSDREVF | 0.58285964 |
| GAGGGGLTAGAN | ASSDREVF | 0.5044477 |
| FVDALVNM | ASSDREVF | 0.6930821 |
| FLTVAPEL | ASSDREVF | 0.50714576 |
| FETLTAL | ASSDREVF | 0.5713617 |
| DVINPIEL | ASSDREVF | 0.12542386 |
| APVSGAPV | ASSDREVF | 0.1885332 |
| ALVGGLAW | ASSDREVF | 0.3563211 |
| ALAAAAATAI | ASSDREVF | 0.37172896 |
| AFGLGLLL | ASSDREVF | 0.20935836 |
| AFFAAASL | ASSDREVF | 0.51498896 |
| AAAALPAFSPP | ASSDREVF | 0.34834418 |
| VVLAPAL | ASSDREVF | 0.23141015 |

| | | |
|---|---|---|
| TIGHVDHGKTTL | ASSDREVF | 0.5475795 |
| LISNASDALDK | ASSDREVF | 0.23882611 |
| LISNASDAL | ASSDREVF | 0.3017301 |
| ISNASDALDK | ASSDREVF | 0.35996595 |
| NQAETLVYQTE | ASSDREVF | 0.49473283 |
| TGGTGGTAGAG | ASSDREVF | 0.2145767 |
| TGSTAGGAGGA | ASSDREVF | 0.44832137 |
| DPLAVPVAAGPV | ASSDREVF | 0.3728107 |
| GGTGGTPGNGGN | ASSDREVF | 0.5223038 |
| PAGLAGVDPATL | ASSDREVF | 0.23109344 |
| GNNLVVEAPHAGASV | ASSDREVF | 0.5110684 |
| KVTRIMSASTLEQPA | ASSDREVF | 0.29668015 |
| LGFALPIALPRRTVV | ASSDREVF | 0.73472273 |
| TRLGNLTARKLVVLV | ASSDREVF | 0.3201171 |
| WSQQRRLIYFPSAGP | ASSDREVF | 0.38107768 |
| GPNGNPDPMAAAVDIR | ASSDREVF | 0.34081632 |
| TQRCVSVADLRAAASR | ASSDREVF | 0.5970094 |
| QFVLDVAEHPVEVLTETV | ASSDREVF | 0.188696 |
| GGAGGTGGAGGNGGTQAGDG | ASSDREVF | 0.2777985 |
| PLLATLRAIAVDVDAKPAQV | ASSDREVF | 0.19050325 |
| APKFRTGGYMIDFWGVGYQVAKRM | ASSDREVF | 0.38396332 |
| GAGGAGGAGGAAGGPGGTGGTGGN | ASSDREVF | 0.49603018 |
| IGALGLLATDVAPAGSVAFGVALA | ASSDREVF | 0.29590037 |
| AGGAGGIGGNGGAGGAGGVLMGNG | ASSDREVF | 0.3982492 |
| ADELIGLL | ASSDREVF | 0.53205717 |
| ADRALKA | ASSDREVF | 0.31580964 |
| AGAVGQVGGAGGA | ASSDREVF | 0.60894394 |
| AGGGGGLGGAP | ASSDREVF | 0.25258803 |
| AGGLLYGNGGAG | ASSDREVF | 0.566542 |
| AGLESIGL | ASSDREVF | 0.31989527 |
| AGLKTGII | ASSDREVF | 0.27132782 |
| ASLAALLL | ASSDREVF | 0.46864498 |
| ASVINISSI | ASSDREVF | 0.3823201 |
| ATPPGPA | ASSDREVF | 0.28966352 |
| ATVASALV | ASSDREVF | 0.56920326 |
| ATVASLIL | ASSDREVF | 0.5505693 |
| AVNGPVLIRH | ASSDREVF | 0.81152594 |
| DGAAGAVGI | ASSDREVF | 0.28130347 |
| DPLLHGI | ASSDREVF | 0.2216151 |
| ELGAALGAL | ASSDREVF | 0.3670604 |
| ELGRTQAV | ASSDREVF | 0.68031925 |
| FVAGGYLS | ASSDREVF | 0.8268315 |
| GASGGAGGAGGW | ASSDREVF | 0.61601657 |
| GGAGAGGDGGAGGI | ASSDREVF | 0.21669991 |
| GTGHNGSQPGMGGNG | ASSDREVF | 0.5311009 |
| GYAAASASAS | ASSDREVF | 0.37205487 |
| HQVDVAAVQQG | ASSDREVF | 0.36656132 |
| IGAAIAHT | ASSDREVF | 0.43492728 |
| IGMIEVAQL | ASSDREVF | 0.5473344 |
| IRLAPEHTSAMD | ASSDREVF | 0.35151166 |
| IYNVSVL | ASSDREVF | 0.21827567 |

| | | |
|---|---|---|
| KQGFLGM | ASSDREVF | 0.5651838 |
| LESVITQL | ASSDREVF | 0.7228971 |
| LGLGALVVV | ASSDREVF | 0.676024 |
| LIGNGGNGGSGGI | ASSDREVF | 0.76563865 |
| LNAINAPTL | ASSDREVF | 0.7224028 |
| LQLLNTGV | ASSDREVF | 0.46647862 |
| LQTSIEF | ASSDREVF | 0.10505917 |
| MSFAVSEL | ASSDREVF | 0.16490468 |
| NTGALGL | ASSDREVF | 0.2682963 |
| PPGAGGAGGAGTT | ASSDREVF | 0.40908313 |
| QLVGLGI | ASSDREVF | 0.11401781 |
| RFLEQVHQL | ASSDREVF | 0.60225195 |
| RGVGYVLR | ASSDREVF | 0.29715037 |
| SATGVFAGVI | ASSDREVF | 0.29263505 |
| SLPAITLP | ASSDREVF | 0.7499331 |
| STALAFTV | ASSDREVF | 0.27733794 |
| SVLTVGLGK | ASSDREVF | 0.23690464 |
| TALAGAGSGI | ASSDREVF | 0.26990193 |
| TGLPFTGI | ASSDREVF | 0.720197 |
| VAEHVTV | ASSDREVF | 0.17801517 |
| VDIVLNSL | ASSDREVF | 0.60640013 |
| VLTGGGAVL | ASSDREVF | 0.36839274 |
| VNVGLAYGI | ASSDREVF | 0.7358516 |
| VSSAAVGAV | ASSDREVF | 0.1873872 |
| VTSVYEAGV | ASSDREVF | 0.65136063 |
| WVALDRGA | ASSDREVF | 0.41774178 |
| YLVGSVVI | ASSDREVF | 0.8615608 |
| GPPGTGKTLL | ASSDREVF | 0.31873062 |
| GVGMTKF | ASSDREVF | 0.49931538 |
| IGIGGSDL | ASSDREVF | 0.35163748 |
| IGTLLAEL | ASSDREVF | 0.79311323 |
| IYTSGTT | ASSDREVF | 0.18263158 |
| LSFPAGV | ASSDREVF | 0.3172989 |
| LYTSGTT | ASSDREVF | 0.18682301 |
| VGAFPFGF | ASSDREVF | 0.35773385 |
| VGAGTVEF | ASSDREVF | 0.62775326 |
| VGKSTLLL | ASSDREVF | 0.11291925 |
| AALAHAGGGIV | ASSDREVF | 0.510494 |
| AALVAVSLL | ASSDREVF | 0.3216581 |
| AAVSRAVAAL | ASSDREVF | 0.20783879 |
| AGGSSGIGGF | ASSDREVF | 0.5705362 |
| AGLTVANANG | ASSDREVF | 0.42917907 |
| AGLVGAVL | ASSDREVF | 0.74780095 |
| ALSQAFTV | ASSDREVF | 0.25734687 |
| ALTFSVSL | ASSDREVF | 0.4378428 |
| ALTGAALVAL | ASSDREVF | 0.4177174 |
| ASFSIGAGVL | ASSDREVF | 0.21531859 |
| DGSVYKI | ASSDREVF | 0.25244364 |
| EQLAEIV | ASSDREVF | 0.7189289 |
| FAVSELTVM | ASSDREVF | 0.40215433 |
| GAAPLAAGL | ASSDREVF | 0.8094425 |

| | | |
|---|---|---|
| GADGGDGGSGGAGG | ASSDREVF | 0.2680907 |
| GFGLDTGGAGG | ASSDREVF | 0.21214233 |
| GLASGGLL | ASSDREVF | 0.4041388 |
| GRIAGAAVLL | ASSDREVF | 0.71876323 |
| GSAGSLNTSF | ASSDREVF | 0.78572917 |
| GTGQIGI | ASSDREVF | 0.6127272 |
| GTLIALIL | ASSDREVF | 0.20961386 |
| IAATMGFVVG | ASSDREVF | 0.28559017 |
| IFAQALL | ASSDREVF | 0.26885176 |
| IGAAVALGF | ASSDREVF | 0.34416085 |
| IGLTPLAL | ASSDREVF | 0.51200104 |
| ILGSNTSTL | ASSDREVF | 0.2816908 |
| IVFSLTVL | ASSDREVF | 0.21628371 |
| LASKATGF | ASSDREVF | 0.27775276 |
| LASVALIL | ASSDREVF | 0.52025247 |
| LFAVGLTL | ASSDREVF | 0.6223257 |
| LFNGGIGTYI | ASSDREVF | 0.5655235 |
| LTLGAGII | ASSDREVF | 0.77550066 |
| LVAAALGDI | ASSDREVF | 0.5848147 |
| NTAFVQLGI | ASSDREVF | 0.6365129 |
| QLFSGAEL | ASSDREVF | 0.1257108 |
| SAATFITL | ASSDREVF | 0.15871692 |
| SNYGLANLGV | ASSDREVF | 0.29179704 |
| SYGGAYAVM | ASSDREVF | 0.34300235 |
| TTGVGGLTLG | ASSDREVF | 0.19265851 |
| TTQTIYI | ASSDREVF | 0.57355976 |
| VAAGAGLGAVI | ASSDREVF | 0.2919609 |
| VAAVVTLL | ASSDREVF | 0.20552419 |
| VGFLGGLI | ASSDREVF | 0.6574425 |
| VGVIPILL | ASSDREVF | 0.22686973 |
| VIFASSLI | ASSDREVF | 0.23540014 |
| VKGGDGGAAAG | ASSDREVF | 0.32747373 |
| VTKTFTNI | ASSDREVF | 0.55581737 |
| VVAAAILL | ASSDREVF | 0.1853045 |
| YAALDVEL | ASSDREVF | 0.4797773 |
| LASVALLL | ASSDREVF | 0.375929 |
| LDAVHRV | ASSDREVF | 0.57122934 |
| QGLSSSIF | ASSDREVF | 0.18378668 |
| RGTFANIRL | ASSDREVF | 0.61979085 |
| RQLWWGHRI | ASSDREVF | 0.38073158 |
| YGPPGTGKTLL | ASSDREVF | 0.81567746 |
| AALVGAPANVA | ASSDREVF | 0.26974353 |
| AATFRRTE | ASSDREVF | 0.30811515 |
| AGALGGGI | ASSDREVF | 0.27774164 |
| AGGAGGAGGIGAG | ASSDREVF | 0.76494795 |
| AGSGALGL | ASSDREVF | 0.17825705 |
| AQIADQI | ASSDREVF | 0.2906321 |
| ASAVAVTYL | ASSDREVF | 0.17189644 |
| ASTVLGGTGG | ASSDREVF | 0.44860968 |
| AVGGEPAK | ASSDREVF | 0.50996435 |
| CEQGTFGGTC | ASSDREVF | 0.24076301 |

| | | |
|---|---|---|
| DEGKAPAG | ASSDREVF | 0.46791565 |
| DTVVFPAL | ASSDREVF | 0.6782432 |
| FEGITMAG | ASSDREVF | 0.2702028 |
| FPLVVQD | ASSDREVF | 0.70765036 |
| GAFTTPQ | ASSDREVF | 0.2885976 |
| GFGGFGTAGGVGGA | ASSDREVF | 0.68147326 |
| GFLAQFL | ASSDREVF | 0.36996213 |
| GGAGGAGGNAGVA | ASSDREVF | 0.2717046 |
| GGVGTAGAGGAGGAGG | ASSDREVF | 0.56035465 |
| GNGGGGDAGNAGSGGN | ASSDREVF | 0.27880174 |
| GTGGAGGAGNDAGS | ASSDREVF | 0.30920708 |
| GTTNTGWFNS | ASSDREVF | 0.6387914 |
| IFMGPLFTL | ASSDREVF | 0.8243636 |
| ILAHAIEAI | ASSDREVF | 0.8431231 |
| IRSAIREAL | ASSDREVF | 0.7833861 |
| ITFAGGG | ASSDREVF | 0.7520683 |
| ITGMSGHQEV | ASSDREVF | 0.7962921 |
| IVAGLACTL | ASSDREVF | 0.8358687 |
| IVGGGLAGA | ASSDREVF | 0.83803535 |
| IVTGVLNI | ASSDREVF | 0.823612 |
| LAQQGCI | ASSDREVF | 0.73163456 |
| LCLHVEN | ASSDREVF | 0.8177979 |
| LDAISDGS | ASSDREVF | 0.796682 |
| LGGFAAML | ASSDREVF | 0.78809875 |
| LGGKNPMI | ASSDREVF | 0.82770944 |
| LGPKGVAAIL | ASSDREVF | 0.8069252 |
| LMVGVSI | ASSDREVF | 0.78638154 |
| LVGAAFVL | ASSDREVF | 0.8108361 |
| LVLAVAPL | ASSDREVF | 0.79991376 |
| MAAIAVIDAA | ASSDREVF | 0.81818545 |
| NNGINAELQ | ASSDREVF | 0.8107066 |
| RGEPLVLVHG | ASSDREVF | 0.84295547 |
| RGGAGGAAGLI | ASSDREVF | 0.83364433 |
| RSYRMHPAV | ASSDREVF | 0.8248955 |
| SAAHLLAGQLA | ASSDREVF | 0.44099653 |
| SAAVGSY | ASSDREVF | 0.7168714 |
| SPHAVLHRI | ASSDREVF | 0.80137444 |
| SQIAKVL | ASSDREVF | 0.8001789 |
| TALGHLVP | ASSDREVF | 0.5966538 |
| TAPTTSGAA | ASSDREVF | 0.82263184 |
| TDGDDGNGGSPGT | ASSDREVF | 0.79325235 |
| TDGLGEN | ASSDREVF | 0.74576545 |
| TGGPPLPGQA | ASSDREVF | 0.8249396 |
| TIPAITI | ASSDREVF | 0.78606844 |
| TIPGFTI | ASSDREVF | 0.77839315 |
| TSIASGALA | ASSDREVF | 0.7899913 |
| TVTLPGAAI | ASSDREVF | 0.82376766 |
| VASGLSAALP | ASSDREVF | 0.7414291 |
| VGAGVIGI | ASSDREVF | 0.823057 |
| VGPKRFGV | ASSDREVF | 0.77706265 |
| VNVAFPD | ASSDREVF | 0.8340243 |

| | | |
|---|---|---|
| VSMTNSGVS | ASSDREVF | 0.70269156 |
| VTDVLGDLSV | ASSDREVF | 0.828332 |
| VVVVGAGFAGL | ASSDREVF | 0.8037572 |
| VYTVGTVV | ASSDREVF | 0.84420645 |
| WTTFSGP | ASSDREVF | 0.7990974 |
| YDELVEAF | ASSDREVF | 0.8183197 |
| YGTGGAGGNGG | ASSDREVF | 0.787233 |
| YTNVMAV | ASSDREVF | 0.83924854 |
| FLEQVHQL | ASSDREVF | 0.69749016 |
| GLSLLAVAV | ASSDREVF | 0.81290305 |
| LEQVHQL | ASSDREVF | 0.72990793 |
| LNAAGAI | ASSDREVF | 0.8261212 |
| PAVHASL | ASSDREVF | 0.82377553 |
| TELEDDA | ASSDREVF | 0.7136997 |
| TPPGPPN | ASSDREVF | 0.8329265 |
| TPPPPPPP | ASSDREVF | 0.8138585 |
| TQSPALI | ASSPPTGGQEDTQY | 0.76791143 |
| EFAAAFPQ | ASSPPTGGQEDTQY | 0.8373014 |
| LLSGNLDVL | ASSPPTGGQEDTQY | 0.7892182 |
| GPANLGNYNI | ASSPPTGGQEDTQY | 0.83519757 |
| TLGPTLP | ASSPPTGGQEDTQY | 0.8051115 |
| VYAYSSCCK | ASSPPTGGQEDTQY | 0.81101114 |
| APAPAPAPNI | ASSPPTGGQEDTQY | 0.8220101 |
| PGGPVPIP | ASSPPTGGQEDTQY | 0.698753 |
| FLTVALEP | ASSPPTGGQEDTQY | 0.7370318 |
| SQYFGANL | ASSPPTGGQEDTQY | 0.8374647 |
| GGVKPAA | ASSPPTGGQEDTQY | 0.57681686 |
| GGTGGAGGAGGW | ASSPPTGGQEDTQY | 0.6855359 |
| GVGGVAGLGGAGG | ASSPPTGGQEDTQY | 0.4711763 |
| VLLVLATTI | ASSPPTGGQEDTQY | 0.4485498 |
| IGIFAAML | ASSPPTGGQEDTQY | 0.5664409 |
| AVGDLID | ASSPPTGGQEDTQY | 0.66843426 |
| NISPLPL | ASSPPTGGQEDTQY | 0.6748221 |
| VTAVFVAI | ASSPPTGGQEDTQY | 0.6161623 |
| SGDDNDAAL | ASSPPTGGQEDTQY | 0.46563822 |
| AVAALAHS | ASSPPTGGQEDTQY | 0.5417482 |
| GGVTVMATP | ASSPPTGGQEDTQY | 0.5024498 |
| AGLAANVRD | ASSPPTGGQEDTQY | 0.5519798 |
| TNRPDLIDPALLRPG | ASSPPTGGQEDTQY | 0.619918 |
| LANLGVTAL | ASSPPTGGQEDTQY | 0.54199433 |
| TGERIEL | ASSPPTGGQEDTQY | 0.4742432 |
| NGEELAA | ASSPPTGGQEDTQY | 0.5173409 |
| VLSLVWG | ASSPPTGGQEDTQY | 0.61790466 |
| IGYALWL | ASSPPTGGQEDTQY | 0.586208 |
| LSLLLAPL | ASSPPTGGQEDTQY | 0.5720712 |
| TLDEALK | ASSPPTGGQEDTQY | 0.68523633 |
| TDGAGTAEGG | ASSPPTGGQEDTQY | 0.62811697 |
| NAPGGAGGNGG | ASSPPTGGQEDTQY | 0.63089263 |
| VSVFMGMGGNF | ASSPPTGGQEDTQY | 0.30009004 |
| LASYLAAL | ASSPPTGGQEDTQY | 0.35151002 |
| LVGGTPAL | ASSPPTGGQEDTQY | 0.53735334 |

| | | |
|---|---|---|
| VRNADVL | ASSPPTGGQEDTQY | 0.5537927 |
| TCSSPFLVSAM | ASSPPTGGQEDTQY | 0.36666054 |
| VHITDAE | ASSPPTGGQEDTQY | 0.6357423 |
| EYAAPSF | ASSPPTGGQEDTQY | 0.49643183 |
| VVDAFHGGL | ASSPPTGGQEDTQY | 0.4710367 |
| ASVVDPQ | ASSPPTGGQEDTQY | 0.635674 |
| SYNEFGGLS | ASSPPTGGQEDTQY | 0.49771637 |
| DPDGSVY | ASSPPTGGQEDTQY | 0.48447943 |
| MVSTIHATE | ASSPPTGGQEDTQY | 0.5237168 |
| LAAGVADPV | ASSPPTGGQEDTQY | 0.60557616 |
| TKEQIAA | ASSPPTGGQEDTQY | 0.3763649 |
| SLASPAAL | ASSPPTGGQEDTQY | 0.6026575 |
| IGLVFYGV | ASSPPTGGQEDTQY | 0.5236872 |
| FLLDASNL | ASSPPTGGQEDTQY | 0.65583557 |
| YQVGNFP | ASSPPTGGQEDTQY | 0.40394375 |
| ESLLSQST | ASSPPTGGQEDTQY | 0.6128708 |
| LFTVGFAAVG | ASSPPTGGQEDTQY | 0.56556356 |
| ATPPTPT | ASSPPTGGQEDTQY | 0.69062346 |
| TDLATLG | ASSPPTGGQEDTQY | 0.50706416 |
| SAGGTGR | ASSPPTGGQEDTQY | 0.5735069 |
| GFSNSGS | ASSPPTGGQEDTQY | 0.55758005 |
| GGNGGSGGTSV | ASSPPTGGQEDTQY | 0.6765787 |
| GTSVATGGAGN | ASSPPTGGQEDTQY | 0.43854606 |
| NGGNGGSGGTS | ASSPPTGGQEDTQY | 0.50936264 |
| AVVKFPH | ASSPPTGGQEDTQY | 0.3863931 |
| TPPPPPP | ASSPPTGGQEDTQY | 0.6082996 |
| AAPAAAAAPA | ASSPPTGGQEDTQY | 0.5888025 |
| IALNEDL | ASSPPTGGQEDTQY | 0.4528386 |
| IVNVSSM | ASSPPTGGQEDTQY | 0.6322665 |
| TGIPPAPR | ASSPPTGGQEDTQY | 0.6177931 |
| FDLGGGTF | ASSPPTGGQEDTQY | 0.48452115 |
| FDLGGGTFDVS | ASSPPTGGQEDTQY | 0.65864694 |
| NEPTAAA | ASSPPTGGQEDTQY | 0.54306674 |
| NEPTAAALA | ASSPPTGGQEDTQY | 0.6442701 |
| GSIINISSI | ASSPPTGGQEDTQY | 0.58378434 |
| VAPGNAGTA | ASSPPTGGQEDTQY | 0.47903013 |
| TGGGASLE | ASSPPTGGQEDTQY | 0.62837744 |
| VAVYNHYKRI | ASSPPTGGQEDTQY | 0.55359876 |
| TDPAPATT | ASSPPTGGQEDTQY | 0.5101408 |
| TNRPDILDPALLRPG | ASSPPTGGQEDTQY | 0.6443224 |
| LTTTTTS | ASSPPTGGQEDTQY | 0.83768755 |
| PTGSGKTL | ASSPPTGGQEDTQY | 0.8144189 |
| TGSGKTL | ASSPPTGGQEDTQY | 0.6750434 |
| TIGHVDHGK | ASSPPTGGQEDTQY | 0.6729506 |
| TIGHVDHGKTT | ASSPPTGGQEDTQY | 0.71993107 |
| GHVDHGKTT | ASSPPTGGQEDTQY | 0.83328587 |
| AQFEHTLL | ASSPPTGGQEDTQY | 0.863611 |
| IVNQDEL | ASSPPTGGQEDTQY | 0.8361223 |
| TNGSQFF | ASSPPTGGQEDTQY | 0.588098 |
| TGKGIVE | ASSPPTGGQEDTQY | 0.860126 |
| LISNASDA | ASSPPTGGQEDTQY | 0.82299256 |

| | | |
|---|---|---|
| AELRTLGELTKIAKAHGAQ | ASSPPTGGQEDTQY | 0.8314125 |
| APNAVAAAEPSW | ASSPPTGGQEDTQY | 0.79579353 |
| APQVAGAATAVL | ASSPPTGGQEDTQY | 0.77321625 |
| ARRDPAAAAAILPTDA | ASSPPTGGQEDTQY | 0.698871 |
| DAAARDVLAEAGLAEYF | ASSPPTGGQEDTQY | 0.6137102 |
| DDQIVLSAKANLAKVEQ | ASSPPTGGQEDTQY | 0.7904241 |
| DELLPDTGEDSDEDGH | ASSPPTGGQEDTQY | 0.83296126 |
| ELPEAPANRATVPQD | ASSPPTGGQEDTQY | 0.82557404 |
| FGQSKIERVTGIG | ASSPPTGGQEDTQY | 0.8733572 |
| GAGGTGSTAGGAGGAGGAGGLYA | ASSPPTGGQEDTQY | 0.8579521 |
| GHGGTGAAGGNGGTGGAGGSGIDG | ASSPPTGGQEDTQY | 0.8002444 |
| GNGGNGGAGGEGGPGGAGGAGA | ASSPPTGGQEDTQY | 0.28055093 |
| GSTAGNGGNGGNATNGTIAGQPAG | ASSPPTGGQEDTQY | 0.4770129 |
| GTGGDGGNGGNGGTGG | ASSPPTGGQEDTQY | 0.5808064 |
| GVRRPAGIPADAPV | ASSPPTGGQEDTQY | 0.7219707 |
| GYRQVTDAHLLTLARR | ASSPPTGGQEDTQY | 0.24431595 |
| IAKMGQDGHDRG | ASSPPTGGQEDTQY | 0.8028562 |
| KAEPSIEIQYT | ASSPPTGGQEDTQY | 0.75934595 |
| KPVRIRMAADEDTVAVT | ASSPPTGGQEDTQY | 0.72179115 |
| MPPRIAGMRLLVIKPEP | ASSPPTGGQEDTQY | 0.84343976 |
| MKFYAVALFRSRGVPPTR | ASSPPTGGQEDTQY | 0.7705598 |
| NAGGFGILGAVAHSP | ASSPPTGGQEDTQY | 0.55768365 |
| NEGREITLAMAEM | ASSPPTGGQEDTQY | 0.56562674 |
| NELVEWSHPLGHRWRWE | ASSPPTGGQEDTQY | 0.8642714 |
| NLGAGGTGGNGGNPG | ASSPPTGGQEDTQY | 0.6224908 |
| NTEWPKWETVMAADPVD | ASSPPTGGQEDTQY | 0.8752079 |
| PASAAPAETQP | ASSPPTGGQEDTQY | 0.5971439 |
| QQSSDGKNGVRAAEITGE | ASSPPTGGQEDTQY | 0.87393 |
| RAPDAVTTSWWKE | ASSPPTGGQEDTQY | 0.56857455 |
| SGFGNVGAHTSGWFNQ | ASSPPTGGQEDTQY | 0.86356723 |
| SGVAGAVGQAASVGGLKV | ASSPPTGGQEDTQY | 0.81656766 |
| SNGGAGTGSAGGNGGT | ASSPPTGGQEDTQY | 0.87718093 |
| TGGAGGEGGAAG | ASSPPTGGQEDTQY | 0.76485556 |
| TGGDGGAAGAGGAPGHG | ASSPPTGGQEDTQY | 0.840977 |
| TMPPLLAAIANAGRHGV | ASSPPTGGQEDTQY | 0.73657966 |
| TSSTAGNSGGA | ASSPPTGGQEDTQY | 0.8527988 |
| VPATVPVGILC | ASSPPTGGQEDTQY | 0.55394614 |
| VQRPDSGGVVLTGRIS | ASSPPTGGQEDTQY | 0.73156583 |
| WTPNPVPATKGVPAGLVQAT | ASSPPTGGQEDTQY | 0.5106665 |
| YALKRALAKFSDRFDVVII | ASSPPTGGQEDTQY | 0.7691641 |
| YLIGSVIV | ASSPPTGGQEDTQY | 0.8617468 |
| YGQLVAVID | ASSPPTGGQEDTQY | 0.1773041 |
| VTVGGMTLPA | ASSPPTGGQEDTQY | 0.8539883 |
| VLALPAV | ASSPPTGGQEDTQY | 0.56573784 |
| VIPPTLNYET | ASSPPTGGQEDTQY | 0.60224855 |
| VGMGGMHPSSG | ASSPPTGGQEDTQY | 0.83050245 |
| VGLAGGAATA | ASSPPTGGQEDTQY | 0.80293274 |
| VGGAGGAGGSGP | ASSPPTGGQEDTQY | 0.8306179 |
| VFAAFMV | ASSPPTGGQEDTQY | 0.61082 |
| TLYDKLR | ASSPPTGGQEDTQY | 0.7665731 |
| TLAVFTVL | ASSPPTGGQEDTQY | 0.7987565 |

| | | |
|---|---|---|
| TGSVQPARV | ASSPPTGGQEDTQY | 0.28473172 |
| TGGGAGSGGT | ASSPPTGGQEDTQY | 0.46718314 |
| SHLETSL | ASSPPTGGQEDTQY | 0.8789737 |
| SFTTGMT | ASSPPTGGQEDTQY | 0.79180765 |
| QVEVAKRMGVRQPTVS | ASSPPTGGQEDTQY | 0.77723897 |
| QGSGGPALAM | ASSPPTGGQEDTQY | 0.7103937 |
| MLAPSNP | ASSPPTGGQEDTQY | 0.69191974 |
| LVNNAAI | ASSPPTGGQEDTQY | 0.7459367 |
| LTGEINTLA | ASSPPTGGQEDTQY | 0.7981549 |
| LSIFGPV | ASSPPTGGQEDTQY | 0.7856244 |
| LLALLANL | ASSPPTGGQEDTQY | 0.781131 |
| LISNASDALDKLR | ASSPPTGGQEDTQY | 0.72627676 |
| LINEAVP | ASSPPTGGQEDTQY | 0.784004 |
| LGYAFSGL | ASSPPTGGQEDTQY | 0.7707494 |
| LDFVAVAL | ASSPPTGGQEDTQY | 0.73281497 |
| LAQPGAE | ASSPPTGGQEDTQY | 0.7938236 |
| ISNLGAVTGF | ASSPPTGGQEDTQY | 0.78254926 |
| HQEDPGRI | ASSPPTGGQEDTQY | 0.7355832 |
| GTPGEPTP | ASSPPTGGQEDTQY | 0.77102137 |
| GFFNSGS | ASSPPTGGQEDTQY | 0.7625747 |
| GAGGGGLTAGAN | ASSPPTGGQEDTQY | 0.7892247 |
| FVDALVNM | ASSPPTGGQEDTQY | 0.78610593 |
| FLTVAPEL | ASSPPTGGQEDTQY | 0.81708544 |
| FETLTAL | ASSPPTGGQEDTQY | 0.80093515 |
| DVINPIEL | ASSPPTGGQEDTQY | 0.7926072 |
| APVSGAPV | ASSPPTGGQEDTQY | 0.43759924 |
| ALVGGLAW | ASSPPTGGQEDTQY | 0.69415736 |
| ALAAAAATAI | ASSPPTGGQEDTQY | 0.76722324 |
| AFGLGLLL | ASSPPTGGQEDTQY | 0.76480883 |
| AFFAAASL | ASSPPTGGQEDTQY | 0.5752875 |
| AAAALPAFSPP | ASSPPTGGQEDTQY | 0.77537686 |
| VVLAPAL | ASSPPTGGQEDTQY | 0.75926054 |
| TIGHVDHGKTTL | ASSPPTGGQEDTQY | 0.6935812 |
| LISNASDALDK | ASSPPTGGQEDTQY | 0.8019883 |
| LISNASDAL | ASSPPTGGQEDTQY | 0.73685366 |
| ISNASDALDK | ASSPPTGGQEDTQY | 0.75273764 |
| NQAETLVYQTE | ASSPPTGGQEDTQY | 0.7721983 |
| TGGTGGTAGAG | ASSPPTGGQEDTQY | 0.8027824 |
| TGSTAGGAGGA | ASSPPTGGQEDTQY | 0.7183182 |
| DPLAVPVAAGPV | ASSPPTGGQEDTQY | 0.79423606 |
| GGTGGTPGNGGN | ASSPPTGGQEDTQY | 0.7420934 |
| PAGLAGVDPATL | ASSPPTGGQEDTQY | 0.8113487 |
| GNNLVVEAPHAGASV | ASSPPTGGQEDTQY | 0.6870307 |
| KVTRIMSASTLEQPA | ASSPPTGGQEDTQY | 0.79141223 |
| LGFALPIALPRRTVV | ASSPPTGGQEDTQY | 0.7538598 |
| TRLGNLTARKLVVLV | ASSPPTGGQEDTQY | 0.8164338 |
| WSQQRRLIYFPSAGP | ASSPPTGGQEDTQY | 0.7633405 |
| GPNGNPDPMAAAVDIR | ASSPPTGGQEDTQY | 0.80086136 |
| TQRCVSVADLRAAASR | ASSPPTGGQEDTQY | 0.7777573 |
| QFVLDVAEHPVEVLTETV | ASSPPTGGQEDTQY | 0.7855612 |
| GGAGGTGGAGGNGGTQAGDG | ASSPPTGGQEDTQY | 0.65192294 |

| | | |
|---|---|---|
| PLLATLRAIAVDVDAKPAQV | ASSPPTGGQEDTQY | 0.7746726 |
| APKFRTGGYMIDFWGVGYQVAKRM | ASSPPTGGQEDTQY | 0.68763274 |
| GAGGAGGAGGAAGGPGGTGGTGGN | ASSPPTGGQEDTQY | 0.7874342 |
| IGALGLLATDVAPAGSVAFGVALA | ASSPPTGGQEDTQY | 0.78432053 |
| AGGAGGIGGNGGAGGAGGVLMGNG | ASSPPTGGQEDTQY | 0.53965247 |
| ADELIGLL | ASSPPTGGQEDTQY | 0.79898536 |
| ADRALKA | ASSPPTGGQEDTQY | 0.69157517 |
| AGAVGQVGGAGGA | ASSPPTGGQEDTQY | 0.74754 |
| AGGGGGLGGAP | ASSPPTGGQEDTQY | 0.80579305 |
| AGGLLYGNGGAG | ASSPPTGGQEDTQY | 0.76130956 |
| AGLESIGL | ASSPPTGGQEDTQY | 0.8047513 |
| AGLKTGII | ASSPPTGGQEDTQY | 0.75917983 |
| ASLAALLL | ASSPPTGGQEDTQY | 0.7683448 |
| ASVINISSI | ASSPPTGGQEDTQY | 0.77599573 |
| ATPPGPA | ASSPPTGGQEDTQY | 0.64132315 |
| ATVASALV | ASSPPTGGQEDTQY | 0.7315736 |
| ATVASLIL | ASSPPTGGQEDTQY | 0.80528235 |
| AVNGPVLIRH | ASSPPTGGQEDTQY | 0.6459313 |
| DGAAGAVGI | ASSPPTGGQEDTQY | 0.7347442 |
| DPLLHGI | ASSPPTGGQEDTQY | 0.56669515 |
| ELGAALGAL | ASSPPTGGQEDTQY | 0.503768 |
| ELGRTQAV | ASSPPTGGQEDTQY | 0.6287564 |
| FVAGGYLS | ASSPPTGGQEDTQY | 0.7455999 |
| GASGGAGGAGGW | ASSPPTGGQEDTQY | 0.7224065 |
| GGAGAGGDGGAGGI | ASSPPTGGQEDTQY | 0.6855754 |
| GTGHNGSQPGMGGNG | ASSPPTGGQEDTQY | 0.54532915 |
| GYAAASASAS | ASSPPTGGQEDTQY | 0.7007196 |
| HQVDVAAVQQG | ASSPPTGGQEDTQY | 0.6162262 |
| IGAAIAHT | ASSPPTGGQEDTQY | 0.595928 |
| IGMIEVAQL | ASSPPTGGQEDTQY | 0.71759164 |
| IRLAPEHTSAMD | ASSPPTGGQEDTQY | 0.685348 |
| IYNVSVL | ASSPPTGGQEDTQY | 0.62281394 |
| KQGFLGM | ASSPPTGGQEDTQY | 0.62360257 |
| LESVITQL | ASSPPTGGQEDTQY | 0.64786947 |
| LGLGALVVV | ASSPPTGGQEDTQY | 0.66797066 |
| LIGNGGNGGSGGI | ASSPPTGGQEDTQY | 0.64817613 |
| LNAINAPTL | ASSPPTGGQEDTQY | 0.78558224 |
| LQLLNTGV | ASSPPTGGQEDTQY | 0.721173 |
| LQTSIEF | ASSPPTGGQEDTQY | 0.6977879 |
| MSFAVSEL | ASSPPTGGQEDTQY | 0.2650817 |
| NTGALGL | ASSPPTGGQEDTQY | 0.41089645 |
| PPGAGGAGGAGTT | ASSPPTGGQEDTQY | 0.61302966 |
| QLVGLGI | ASSPPTGGQEDTQY | 0.65240693 |
| RFLEQVHQL | ASSPPTGGQEDTQY | 0.35405365 |
| RGVGYVLR | ASSPPTGGQEDTQY | 0.7306988 |
| SATGVFAGVI | ASSPPTGGQEDTQY | 0.62916183 |
| SLPAITLP | ASSPPTGGQEDTQY | 0.5668365 |
| STALAFTV | ASSPPTGGQEDTQY | 0.74674916 |
| SVLTVGLGK | ASSPPTGGQEDTQY | 0.6018153 |
| TALAGAGSGI | ASSPPTGGQEDTQY | 0.6155956 |
| TGLPFTGI | ASSPPTGGQEDTQY | 0.6472118 |

| | | |
|---|---|---|
| VAEHVTV | ASSPPTGGQEDTQY | 0.7558956 |
| VDIVLNSL | ASSPPTGGQEDTQY | 0.49411762 |
| VLTGGGAVL | ASSPPTGGQEDTQY | 0.7140397 |
| VNVGLAYGI | ASSPPTGGQEDTQY | 0.58664834 |
| VSSAAVGAV | ASSPPTGGQEDTQY | 0.7477969 |
| VTSVYEAGV | ASSPPTGGQEDTQY | 0.47690743 |
| WVALDRGA | ASSPPTGGQEDTQY | 0.6994288 |
| YLVGSVVI | ASSPPTGGQEDTQY | 0.57534504 |
| GPPGTGKTLL | ASSPPTGGQEDTQY | 0.7915932 |
| GVGMTKF | ASSPPTGGQEDTQY | 0.61631835 |
| IGIGGSDL | ASSPPTGGQEDTQY | 0.6985635 |
| IGTLLAEL | ASSPPTGGQEDTQY | 0.6037808 |
| IYTSGTT | ASSPPTGGQEDTQY | 0.6999991 |
| LSFPAGV | ASSPPTGGQEDTQY | 0.43731785 |
| LYTSGTT | ASSPPTGGQEDTQY | 0.62647116 |
| VGAFPFGF | ASSPPTGGQEDTQY | 0.5215244 |
| VGAGTVEF | ASSPPTGGQEDTQY | 0.70596546 |
| VGKSTLLL | ASSPPTGGQEDTQY | 0.6672639 |
| AALAHAGGGIV | ASSPPTGGQEDTQY | 0.5021366 |
| AALVAVSLL | ASSPPTGGQEDTQY | 0.73998445 |
| AAVSRAVAAL | ASSPPTGGQEDTQY | 0.6492319 |
| AGGSSGIGGF | ASSPPTGGQEDTQY | 0.5668103 |
| AGLTVANANG | ASSPPTGGQEDTQY | 0.74945045 |
| AGLVGAVL | ASSPPTGGQEDTQY | 0.6075535 |
| ALSQAFTV | ASSPPTGGQEDTQY | 0.7049939 |
| ALTFSVSL | ASSPPTGGQEDTQY | 0.6279868 |
| ALTGAALVAL | ASSPPTGGQEDTQY | 0.6419657 |
| ASFSIGAGVL | ASSPPTGGQEDTQY | 0.72475797 |
| DGSVYKI | ASSPPTGGQEDTQY | 0.5269143 |
| EQLAEIV | ASSPPTGGQEDTQY | 0.5882136 |
| FAVSELTVM | ASSPPTGGQEDTQY | 0.7274939 |
| GAAPLAAGL | ASSPPTGGQEDTQY | 0.610113 |
| GADGGDGGSGGAGG | ASSPPTGGQEDTQY | 0.679588 |
| GFGLDTGGAGG | ASSPPTGGQEDTQY | 0.47081017 |
| GLASGGLL | ASSPPTGGQEDTQY | 0.42149597 |
| GRIAGAAVLL | ASSPPTGGQEDTQY | 0.53825796 |
| GSAGSLNTSF | ASSPPTGGQEDTQY | 0.679858 |
| GTGQIGI | ASSPPTGGQEDTQY | 0.6631521 |
| GTLIALIL | ASSPPTGGQEDTQY | 0.63464165 |
| IAATMGFVVG | ASSPPTGGQEDTQY | 0.4066108 |
| IFAQALL | ASSPPTGGQEDTQY | 0.5301194 |
| IGAAVALGF | ASSPPTGGQEDTQY | 0.51494455 |
| IGLTPLAL | ASSPPTGGQEDTQY | 0.5225499 |
| ILGSNTSTL | ASSPPTGGQEDTQY | 0.5711142 |
| IVFSLTVL | ASSPPTGGQEDTQY | 0.5233301 |
| LASKATGF | ASSPPTGGQEDTQY | 0.47327012 |
| LASVALIL | ASSPPTGGQEDTQY | 0.46595275 |
| LFAVGLTL | ASSPPTGGQEDTQY | 0.5796371 |
| LFNGGIGTYI | ASSPPTGGQEDTQY | 0.5615705 |
| LTLGAGII | ASSPPTGGQEDTQY | 0.52048314 |
| LVAAALGDI | ASSPPTGGQEDTQY | 0.69604254 |

| | | |
|---|---|---|
| NTAFVQLGI | ASSPPTGGQEDTQY | 0.65451854 |
| QLFSGAEL | ASSPPTGGQEDTQY | 0.6388509 |
| SAATFITL | ASSPPTGGQEDTQY | 0.28884155 |
| SNYGLANLGV | ASSPPTGGQEDTQY | 0.35660803 |
| SYGGAYAVM | ASSPPTGGQEDTQY | 0.43818414 |
| TTGVGGLTLG | ASSPPTGGQEDTQY | 0.49990982 |
| TTQTIYI | ASSPPTGGQEDTQY | 0.31426919 |
| VAAGAGLGAVI | ASSPPTGGQEDTQY | 0.6037523 |
| VAAVVTLL | ASSPPTGGQEDTQY | 0.54189306 |
| VGFLGGLI | ASSPPTGGQEDTQY | 0.43137017 |
| VGVIPILL | ASSPPTGGQEDTQY | 0.62544477 |
| VIFASSLI | ASSPPTGGQEDTQY | 0.43061537 |
| VKGGDGGAAAG | ASSPPTGGQEDTQY | 0.44051963 |
| VTKTFTNI | ASSPPTGGQEDTQY | 0.45057398 |
| VVAAAILL | ASSPPTGGQEDTQY | 0.641637 |
| YAALDVEL | ASSPPTGGQEDTQY | 0.37268606 |
| LASVALLL | ASSPPTGGQEDTQY | 0.60921407 |
| LDAVHRV | ASSPPTGGQEDTQY | 0.48721296 |
| QGLSSSIF | ASSPPTGGQEDTQY | 0.6765926 |
| RGTFANIRL | ASSPPTGGQEDTQY | 0.36780173 |
| RQLWWGHRI | ASSPPTGGQEDTQY | 0.6417462 |
| YGPPGTGKTLL | ASSPPTGGQEDTQY | 0.5543962 |
| AALVGAPANVA | ASSPPTGGQEDTQY | 0.6925106 |
| AATFRRTE | ASSPPTGGQEDTQY | 0.5027882 |
| AGALGGGI | ASSPPTGGQEDTQY | 0.5788854 |
| AGGAGGAGGIGAG | ASSPPTGGQEDTQY | 0.49463603 |
| AGSGALGL | ASSPPTGGQEDTQY | 0.6765693 |
| AQIADQI | ASSPPTGGQEDTQY | 0.40422946 |
| ASAVAVTYL | ASSPPTGGQEDTQY | 0.5656322 |
| ASTVLGGTGG | ASSPPTGGQEDTQY | 0.3992761 |
| AVGGEPAK | ASSPPTGGQEDTQY | 0.61865795 |
| CEQGTFGGTC | ASSPPTGGQEDTQY | 0.63084465 |
| DEGKAPAG | ASSPPTGGQEDTQY | 0.3252212 |
| DTVVFPAL | ASSPPTGGQEDTQY | 0.57419896 |
| FEGITMAG | ASSPPTGGQEDTQY | 0.5893949 |
| FPLVVQD | ASSPPTGGQEDTQY | 0.39792907 |
| GAFTTPQ | ASSPPTGGQEDTQY | 0.6410346 |
| GFGGFGTAGGVGGA | ASSPPTGGQEDTQY | 0.4682079 |
| GFLAQFL | ASSPPTGGQEDTQY | 0.67069364 |
| GGAGGAGGNAGVA | ASSPPTGGQEDTQY | 0.47696173 |
| GGVGTAGAGGAGGAGG | ASSPPTGGQEDTQY | 0.47681725 |
| GNGGGGDAGNAGSGGN | ASSPPTGGQEDTQY | 0.65040994 |
| GTGGAGGAGNDAGS | ASSPPTGGQEDTQY | 0.394634 |
| GTTNTGWFNS | ASSPPTGGQEDTQY | 0.40602478 |
| IFMGPLFTL | ASSPPTGGQEDTQY | 0.671087 |
| ILAHAIEAI | ASSPPTGGQEDTQY | 0.7938088 |
| IRSAIREAL | ASSPPTGGQEDTQY | 0.8176354 |
| ITFAGGG | ASSPPTGGQEDTQY | 0.6688396 |
| ITGMSGHQEV | ASSPPTGGQEDTQY | 0.60273296 |
| IVAGLACTL | ASSPPTGGQEDTQY | 0.7210402 |
| IVGGGLAGA | ASSPPTGGQEDTQY | 0.79981667 |

| | | |
|---|---|---|
| IVTGVLNI | ASSPPTGGQEDTQY | 0.82108176 |
| LAQQGCI | ASSPPTGGQEDTQY | 0.8038686 |
| LCLHVEN | ASSPPTGGQEDTQY | 0.577371 |
| LDAISDGS | ASSPPTGGQEDTQY | 0.76120776 |
| LGGFAAML | ASSPPTGGQEDTQY | 0.6839601 |
| LGGKNPMI | ASSPPTGGQEDTQY | 0.7104767 |
| LGPKGVAAIL | ASSPPTGGQEDTQY | 0.7982645 |
| LMVGVSI | ASSPPTGGQEDTQY | 0.75299156 |
| LVGAAFVL | ASSPPTGGQEDTQY | 0.7062577 |
| LVLAVAPL | ASSPPTGGQEDTQY | 0.6294873 |
| MAAIAVIDAA | ASSPPTGGQEDTQY | 0.80423045 |
| NNGINAELQ | ASSPPTGGQEDTQY | 0.8097619 |
| RGEPLVLVHG | ASSPPTGGQEDTQY | 0.8104306 |
| RGGAGGAAGLI | ASSPPTGGQEDTQY | 0.8140211 |
| RSYRMHPAV | ASSPPTGGQEDTQY | 0.7987559 |
| SAAHLLAGQLA | ASSPPTGGQEDTQY | 0.78448963 |
| SAAVGSY | ASSPPTGGQEDTQY | 0.24683179 |
| SPHAVLHRI | ASSPPTGGQEDTQY | 0.390513 |
| SQIAKVL | ASSPPTGGQEDTQY | 0.7679963 |
| TALGHLVP | ASSPPTGGQEDTQY | 0.6958958 |
| TAPTTSGAA | ASSPPTGGQEDTQY | 0.35547015 |
| TDGDDGNGGSPGT | ASSPPTGGQEDTQY | 0.7656942 |
| TDGLGEN | ASSPPTGGQEDTQY | 0.674646 |
| TGGPPLPGQA | ASSPPTGGQEDTQY | 0.66511923 |
| TIPAITI | ASSPPTGGQEDTQY | 0.8065875 |
| TIPGFTI | ASSPPTGGQEDTQY | 0.69626546 |
| TSIASGALA | ASSPPTGGQEDTQY | 0.5866667 |
| TVTLPGAAI | ASSPPTGGQEDTQY | 0.67940044 |
| VASGLSAALP | ASSPPTGGQEDTQY | 0.80512863 |
| VGAGVIGI | ASSPPTGGQEDTQY | 0.5566593 |
| VGPKRFGV | ASSPPTGGQEDTQY | 0.78353286 |
| VNVAFPD | ASSPPTGGQEDTQY | 0.7226227 |
| VSMTNSGVS | ASSPPTGGQEDTQY | 0.79856956 |
| VTDVLGDLSV | ASSPPTGGQEDTQY | 0.4810955 |
| VVVVGAGFAGL | ASSPPTGGQEDTQY | 0.7853683 |
| VYTVGTVV | ASSPPTGGQEDTQY | 0.73043346 |
| WTTFSGP | ASSPPTGGQEDTQY | 0.82860965 |
| YDELVEAF | ASSPPTGGQEDTQY | 0.6733457 |
| YGTGGAGGNGG | ASSPPTGGQEDTQY | 0.7393007 |
| YTNVMAV | ASSPPTGGQEDTQY | 0.72063243 |
| FLEQVHQL | ASSPPTGGQEDTQY | 0.81388545 |
| GLSLLAVAV | ASSPPTGGQEDTQY | 0.4730556 |
| LEQVHQL | ASSPPTGGQEDTQY | 0.7044843 |
| LNAAGAI | ASSPPTGGQEDTQY | 0.4660887 |
| PAVHASL | ASSPPTGGQEDTQY | 0.7442945 |
| TELEDDA | ASSPPTGGQEDTQY | 0.7832868 |
| TPPGPPN | ASSPPTGGQEDTQY | 0.43261343 |
| TPPPPPPP | ASSPPTGGQEDTQY | 0.8024184 |
| TQSPALI | ASSLSDRVNSDYT | 0.70939267 |
| EFAAAFPQ | ASSLSDRVNSDYT | 0.6498073 |
| LLSGNLDVL | ASSLSDRVNSDYT | 0.81602895 |

| | | |
|---|---|---|
| GPANLGNYNI | ASSLSDRVNSDYT | 0.75623596 |
| TLGPTLP | ASSLSDRVNSDYT | 0.8130195 |
| VYAYSSCCK | ASSLSDRVNSDYT | 0.75973374 |
| APAPAPAPNI | ASSLSDRVNSDYT | 0.7609923 |
| PGGPVPIP | ASSLSDRVNSDYT | 0.7737309 |
| FLTVALEP | ASSLSDRVNSDYT | 0.7296092 |
| SQYFGANL | ASSLSDRVNSDYT | 0.7116035 |
| GGVKPAA | ASSLSDRVNSDYT | 0.80110973 |
| GGTGGAGGAGGW | ASSLSDRVNSDYT | 0.34709483 |
| GVGGVAGLGGAGG | ASSLSDRVNSDYT | 0.73249614 |
| VLLVLATTI | ASSLSDRVNSDYT | 0.19597733 |
| IGIFAAML | ASSLSDRVNSDYT | 0.19753298 |
| AVGDLID | ASSLSDRVNSDYT | 0.2174945 |
| NISPLPL | ASSLSDRVNSDYT | 0.34425902 |
| VTAVFVAI | ASSLSDRVNSDYT | 0.7603595 |
| SGDDNDAAL | ASSLSDRVNSDYT | 0.40899783 |
| AVAALAHS | ASSLSDRVNSDYT | 0.21590191 |
| GGVTVMATP | ASSLSDRVNSDYT | 0.28832996 |
| AGLAANVRD | ASSLSDRVNSDYT | 0.22972107 |
| TNRPDLIDPALLRPG | ASSLSDRVNSDYT | 0.20968288 |
| LANLGVTAL | ASSLSDRVNSDYT | 0.3208393 |
| TGERIEL | ASSLSDRVNSDYT | 0.27940956 |
| NGEELAA | ASSLSDRVNSDYT | 0.22082907 |
| VLSLVWG | ASSLSDRVNSDYT | 0.22773348 |
| IGYALWL | ASSLSDRVNSDYT | 0.32369572 |
| LSLLLAPL | ASSLSDRVNSDYT | 0.37647313 |
| TLDEALK | ASSLSDRVNSDYT | 0.36584422 |
| TDGAGTAEGG | ASSLSDRVNSDYT | 0.41924036 |
| NAPGGAGGNGG | ASSLSDRVNSDYT | 0.36082977 |
| VSVFMGMGGNF | ASSLSDRVNSDYT | 0.3192637 |
| LASYLAAL | ASSLSDRVNSDYT | 0.118824914 |
| LVGGTPAL | ASSLSDRVNSDYT | 0.15494084 |
| VRNADVL | ASSLSDRVNSDYT | 0.3020109 |
| TCSSPFLVSAM | ASSLSDRVNSDYT | 0.22791451 |
| VHITDAE | ASSLSDRVNSDYT | 0.14259695 |
| EYAAPSF | ASSLSDRVNSDYT | 0.28948677 |
| VVDAFHGGL | ASSLSDRVNSDYT | 0.25968343 |
| ASVVDPQ | ASSLSDRVNSDYT | 0.21338116 |
| SYNEFGGLS | ASSLSDRVNSDYT | 0.42986596 |
| DPDGSVY | ASSLSDRVNSDYT | 0.2108353 |
| MVSTIHATE | ASSLSDRVNSDYT | 0.19518316 |
| LAAGVADPV | ASSLSDRVNSDYT | 0.22615461 |
| TKEQIAA | ASSLSDRVNSDYT | 0.3928237 |
| SLASPAAL | ASSLSDRVNSDYT | 0.18120113 |
| IGLVFYGV | ASSLSDRVNSDYT | 0.31626293 |
| FLLDASNL | ASSLSDRVNSDYT | 0.23900001 |
| YQVGNFP | ASSLSDRVNSDYT | 0.36558026 |
| ESLLSQST | ASSLSDRVNSDYT | 0.1747141 |
| LFTVGFAAVG | ASSLSDRVNSDYT | 0.30077448 |
| ATPPTPT | ASSLSDRVNSDYT | 0.22253008 |
| TDLATLG | ASSLSDRVNSDYT | 0.69510126 |

| | | |
|---|---|---|
| SAGGTGR | ASSLSDRVNSDYT | 0.22326237 |
| GFSNSGS | ASSLSDRVNSDYT | 0.29846993 |
| GGNGGSGGTSV | ASSLSDRVNSDYT | 0.25444362 |
| GTSVATGGAGN | ASSLSDRVNSDYT | 0.71812856 |
| NGGNGGSGGTS | ASSLSDRVNSDYT | 0.15174705 |
| AVVKFPH | ASSLSDRVNSDYT | 0.22220758 |
| TPPPPPP | ASSLSDRVNSDYT | 0.18744327 |
| AAPAAAAAPA | ASSLSDRVNSDYT | 0.25516295 |
| IALNEDL | ASSLSDRVNSDYT | 0.3534137 |
| IVNVSSM | ASSLSDRVNSDYT | 0.19315225 |
| TGIPPAPR | ASSLSDRVNSDYT | 0.40387183 |
| FDLGGGTF | ASSLSDRVNSDYT | 0.26573822 |
| FDLGGGTFDVS | ASSLSDRVNSDYT | 0.21453653 |
| NEPTAAA | ASSLSDRVNSDYT | 0.42264938 |
| NEPTAAALA | ASSLSDRVNSDYT | 0.26260582 |
| GSIINISSI | ASSLSDRVNSDYT | 0.37040204 |
| VAPGNAGTA | ASSLSDRVNSDYT | 0.22908708 |
| TGGGASLE | ASSLSDRVNSDYT | 0.29476145 |
| VAVYNHYKRI | ASSLSDRVNSDYT | 0.32815507 |
| TDPAPATT | ASSLSDRVNSDYT | 0.2869567 |
| TNRPDILDPALLRPG | ASSLSDRVNSDYT | 0.2142074 |
| LTTTTTS | ASSLSDRVNSDYT | 0.36144805 |
| PTGSGKTL | ASSLSDRVNSDYT | 0.8898705 |
| TGSGKTL | ASSLSDRVNSDYT | 0.8959831 |
| TIGHVDHGK | ASSLSDRVNSDYT | 0.84674734 |
| TIGHVDHGKTT | ASSLSDRVNSDYT | 0.82942027 |
| GHVDHGKTT | ASSLSDRVNSDYT | 0.85356647 |
| AQFEHTLL | ASSLSDRVNSDYT | 0.8972993 |
| IVNQDEL | ASSLSDRVNSDYT | 0.89743805 |
| TNGSQFF | ASSLSDRVNSDYT | 0.8943424 |
| TGKGIVE | ASSLSDRVNSDYT | 0.8105907 |
| LISNASDA | ASSLSDRVNSDYT | 0.8887106 |
| AELRTLGELTKIAKAHGAQ | ASSLSDRVNSDYT | 0.86140966 |
| APNAVAAAEPSW | ASSLSDRVNSDYT | 0.8739686 |
| APQVAGAATAVL | ASSLSDRVNSDYT | 0.88594353 |
| ARRDPAAAAAILPTDA | ASSLSDRVNSDYT | 0.8824413 |
| DAAARDVLAEAGLAEYF | ASSLSDRVNSDYT | 0.8662357 |
| DDQIVLSAKANLAKVEQ | ASSLSDRVNSDYT | 0.86796534 |
| DELLPDTGEDSDEDGH | ASSLSDRVNSDYT | 0.8790237 |
| ELPEAPANRATVPQD | ASSLSDRVNSDYT | 0.89335334 |
| FGQSKIERVTGIG | ASSLSDRVNSDYT | 0.8827517 |
| GAGGTGSTAGGAGGAGGAGGLYA | ASSLSDRVNSDYT | 0.908067 |
| GHGGTGAAGGNGGTGGAGGSGIDG | ASSLSDRVNSDYT | 0.89970493 |
| GNGGNGGAGGEGGPGGAGGAGA | ASSLSDRVNSDYT | 0.8944129 |
| GSTAGNGGNGGNATNGTIAGQPAG | ASSLSDRVNSDYT | 0.48957524 |
| GTGGDGGNGGNGGTGG | ASSLSDRVNSDYT | 0.7793104 |
| GVRRPAGIPADAPV | ASSLSDRVNSDYT | 0.8293688 |
| GYRQVTDAHLLTLARR | ASSLSDRVNSDYT | 0.8718701 |
| IAKMGQDGHDRG | ASSLSDRVNSDYT | 0.53481287 |
| KAEPSIEIQYT | ASSLSDRVNSDYT | 0.8910432 |
| KPVRIRMAADEDTVAVT | ASSLSDRVNSDYT | 0.868245 |

| | | |
|---|---|---|
| MPPRIAGMRLLVIKPEP | ASSLSDRVNSDYT | 0.8333917 |
| MKFYAVALFRSRGVPPTR | ASSLSDRVNSDYT | 0.8962724 |
| NAGGFGILGAVAHSP | ASSLSDRVNSDYT | 0.86937064 |
| NEGREITLAMAEM | ASSLSDRVNSDYT | 0.83753264 |
| NELVEWSHPLGHRWRWE | ASSLSDRVNSDYT | 0.8226851 |
| NLGAGGTGGNGGNPG | ASSLSDRVNSDYT | 0.9047045 |
| NTEWPKWETVMAADPVD | ASSLSDRVNSDYT | 0.77931124 |
| PASAAPAETQP | ASSLSDRVNSDYT | 0.90073013 |
| QQSSDGKNGVRAAEITGE | ASSLSDRVNSDYT | 0.8331933 |
| RAPDAVTTSWWKE | ASSLSDRVNSDYT | 0.9026617 |
| SGFGNVGAHTSGWFNQ | ASSLSDRVNSDYT | 0.73061854 |
| SGVAGAVGQAASVGGLKV | ASSLSDRVNSDYT | 0.8969742 |
| SNGGAGTGSAGGNGGT | ASSLSDRVNSDYT | 0.87098616 |
| TGGAGGEGGAAG | ASSLSDRVNSDYT | 0.9099167 |
| TGGDGGAAGAGGAPGHG | ASSLSDRVNSDYT | 0.8669722 |
| TMPPLLAAIANAGRHGV | ASSLSDRVNSDYT | 0.8925111 |
| TSSTAGNSGGA | ASSLSDRVNSDYT | 0.8616634 |
| VPATVPVGILC | ASSLSDRVNSDYT | 0.90093255 |
| VQRPDSGGVVLTGRIS | ASSLSDRVNSDYT | 0.7643975 |
| WTPNPVPATKGVPAGLVQAT | ASSLSDRVNSDYT | 0.8591422 |
| YALKRALAKFSDRFDVVII | ASSLSDRVNSDYT | 0.73547965 |
| YLIGSVIV | ASSLSDRVNSDYT | 0.88060933 |
| YGQLVAVID | ASSLSDRVNSDYT | 0.8946553 |
| VTVGGMTLPA | ASSLSDRVNSDYT | 0.6548304 |
| VLALPAV | ASSLSDRVNSDYT | 0.9000635 |
| VIPPTLNYET | ASSLSDRVNSDYT | 0.84778345 |
| VGMGGMHPSSG | ASSLSDRVNSDYT | 0.8193381 |
| VGLAGGAATA | ASSLSDRVNSDYT | 0.8998878 |
| VGGAGGAGGSGP | ASSLSDRVNSDYT | 0.86781615 |
| VFAAFMV | ASSLSDRVNSDYT | 0.89987457 |
| TLYDKLR | ASSLSDRVNSDYT | 0.85542643 |
| TLAVFTVL | ASSLSDRVNSDYT | 0.8593245 |
| TGSVQPARV | ASSLSDRVNSDYT | 0.8893639 |
| TGGGAGSGGT | ASSLSDRVNSDYT | 0.6620028 |
| SHLETSL | ASSLSDRVNSDYT | 0.76311374 |
| SFTTGMT | ASSLSDRVNSDYT | 0.9051664 |
| QVEVAKRMGVRQPTVS | ASSLSDRVNSDYT | 0.88519955 |
| QGSGGPALAM | ASSLSDRVNSDYT | 0.88136333 |
| MLAPSNP | ASSLSDRVNSDYT | 0.8094177 |
| LVNNAAI | ASSLSDRVNSDYT | 0.7963191 |
| LTGEINTLA | ASSLSDRVNSDYT | 0.8219559 |
| LSIFGPV | ASSLSDRVNSDYT | 0.8886266 |
| LLALLANL | ASSLSDRVNSDYT | 0.89695525 |
| LISNASDALDKLR | ASSLSDRVNSDYT | 0.8911029 |
| LINEAVP | ASSLSDRVNSDYT | 0.7267517 |
| LGYAFSGL | ASSLSDRVNSDYT | 0.89198923 |
| LDFVAVAL | ASSLSDRVNSDYT | 0.8535853 |
| LAQPGAE | ASSLSDRVNSDYT | 0.8741766 |
| ISNLGAVTGF | ASSLSDRVNSDYT | 0.86086583 |
| HQEDPGRI | ASSLSDRVNSDYT | 0.84379524 |
| GTPGEPTP | ASSLSDRVNSDYT | 0.82992256 |

| | | |
|---|---|---|
| GFFNSGS | ASSLSDRVNSDYT | 0.79955363 |
| GAGGGGLTAGAN | ASSLSDRVNSDYT | 0.8696765 |
| FVDALVNM | ASSLSDRVNSDYT | 0.89083767 |
| FLTVAPEL | ASSLSDRVNSDYT | 0.8753753 |
| FETLTAL | ASSLSDRVNSDYT | 0.90933853 |
| DVINPIEL | ASSLSDRVNSDYT | 0.8997631 |
| APVSGAPV | ASSLSDRVNSDYT | 0.88304734 |
| ALVGGLAW | ASSLSDRVNSDYT | 0.45716733 |
| ALAAAAATAI | ASSLSDRVNSDYT | 0.7456612 |
| AFGLGLLL | ASSLSDRVNSDYT | 0.6935738 |
| AFFAAASL | ASSLSDRVNSDYT | 0.8526744 |
| AAAALPAFSPP | ASSLSDRVNSDYT | 0.29356956 |
| VVLAPAL | ASSLSDRVNSDYT | 0.87811613 |
| TIGHVDHGKTTL | ASSLSDRVNSDYT | 0.843682 |
| LISNASDALDK | ASSLSDRVNSDYT | 0.80669004 |
| LISNASDAL | ASSLSDRVNSDYT | 0.8928113 |
| ISNASDALDK | ASSLSDRVNSDYT | 0.8487021 |
| NQAETLVYQTE | ASSLSDRVNSDYT | 0.7319288 |
| TGGTGGTAGAG | ASSLSDRVNSDYT | 0.72631073 |
| TGSTAGGAGGA | ASSLSDRVNSDYT | 0.90579057 |
| DPLAVPVAAGPV | ASSLSDRVNSDYT | 0.7375692 |
| GGTGGTPGNGGN | ASSLSDRVNSDYT | 0.9061388 |
| PAGLAGVDPATL | ASSLSDRVNSDYT | 0.74697155 |
| GNNLVVEAPHAGASV | ASSLSDRVNSDYT | 0.9052192 |
| KVTRIMSASTLEQPA | ASSLSDRVNSDYT | 0.6133348 |
| LGFALPIALPRRTVV | ASSLSDRVNSDYT | 0.89747596 |
| TRLGNLTARKLVVLV | ASSLSDRVNSDYT | 0.86503726 |
| WSQQRRLIYFPSAGP | ASSLSDRVNSDYT | 0.9124634 |
| GPNGNPDPMAAAVDIR | ASSLSDRVNSDYT | 0.859645 |
| TQRCVSVADLRAAASR | ASSLSDRVNSDYT | 0.89198637 |
| QFVLDVAEHPVEVLTETV | ASSLSDRVNSDYT | 0.8221717 |
| GGAGGTGGAGGNGGTQAGDG | ASSLSDRVNSDYT | 0.89183867 |
| PLLATLRAIAVDVDAKPAQV | ASSLSDRVNSDYT | 0.7086366 |
| APKFRTGGYMIDFWGVGYQVAKRM | ASSLSDRVNSDYT | 0.82870764 |
| GAGGAGGAGGAAGGPGGTGGTGGN | ASSLSDRVNSDYT | 0.69063044 |
| IGALGLLATDVAPAGSVAFGVALA | ASSLSDRVNSDYT | 0.8582829 |
| AGGAGGIGGNGGAGGAGGVLMGNG | ASSLSDRVNSDYT | 0.89423436 |
| ADELIGLL | ASSLSDRVNSDYT | 0.33191425 |
| ADRALKA | ASSLSDRVNSDYT | 0.89444304 |
| AGAVGQVGGAGGA | ASSLSDRVNSDYT | 0.75986457 |
| AGGGGGLGGAP | ASSLSDRVNSDYT | 0.70390755 |
| AGGLLYGNGGAG | ASSLSDRVNSDYT | 0.89051497 |
| AGLESIGL | ASSLSDRVNSDYT | 0.84868735 |
| AGLKTGII | ASSLSDRVNSDYT | 0.89161015 |
| ASLAALLL | ASSLSDRVNSDYT | 0.7676336 |
| ASVINISSI | ASSLSDRVNSDYT | 0.83287966 |
| ATPPGPA | ASSLSDRVNSDYT | 0.87501657 |
| ATVASALV | ASSLSDRVNSDYT | 0.44340372 |
| ATVASLIL | ASSLSDRVNSDYT | 0.61649036 |
| AVNGPVLIRH | ASSLSDRVNSDYT | 0.90781844 |
| DGAAGAVGI | ASSLSDRVNSDYT | 0.49378783 |

| | | |
|---|---|---|
| DPLLHGI | ASSLSDRVNSDYT | 0.6025562 |
| ELGAALGAL | ASSLSDRVNSDYT | 0.46938625 |
| ELGRTQAV | ASSLSDRVNSDYT | 0.4341663 |
| FVAGGYLS | ASSLSDRVNSDYT | 0.47463167 |
| GASGGAGGAGGW | ASSLSDRVNSDYT | 0.55275655 |
| GGAGAGGDGGAGGI | ASSLSDRVNSDYT | 0.6083844 |
| GTGHNGSQPGMGGNG | ASSLSDRVNSDYT | 0.50031865 |
| GYAAASASAS | ASSLSDRVNSDYT | 0.44767642 |
| HQVDVAAVQQG | ASSLSDRVNSDYT | 0.5305575 |
| IGAAIAHT | ASSLSDRVNSDYT | 0.45612264 |
| IGMIEVAQL | ASSLSDRVNSDYT | 0.4971642 |
| IRLAPEHTSAMD | ASSLSDRVNSDYT | 0.47823632 |
| IYNVSVL | ASSLSDRVNSDYT | 0.5268792 |
| KQGFLGM | ASSLSDRVNSDYT | 0.51329756 |
| LESVITQL | ASSLSDRVNSDYT | 0.4256399 |
| LGLGALVVV | ASSLSDRVNSDYT | 0.4541624 |
| LIGNGGNGGSGGI | ASSLSDRVNSDYT | 0.5245837 |
| LNAINAPTL | ASSLSDRVNSDYT | 0.45324612 |
| LQLLNTGV | ASSLSDRVNSDYT | 0.55353 |
| LQTSIEF | ASSLSDRVNSDYT | 0.5235066 |
| MSFAVSEL | ASSLSDRVNSDYT | 0.47195092 |
| NTGALGL | ASSLSDRVNSDYT | 0.32079646 |
| PPGAGGAGGAGTT | ASSLSDRVNSDYT | 0.34128907 |
| QLVGLGI | ASSLSDRVNSDYT | 0.45927694 |
| RFLEQVHQL | ASSLSDRVNSDYT | 0.43134305 |
| RGVGYVLR | ASSLSDRVNSDYT | 0.3322681 |
| SATGVFAGVI | ASSLSDRVNSDYT | 0.5779641 |
| SLPAITLP | ASSLSDRVNSDYT | 0.4897083 |
| STALAFTV | ASSLSDRVNSDYT | 0.49377185 |
| SVLTVGLGK | ASSLSDRVNSDYT | 0.5356087 |
| TALAGAGSGI | ASSLSDRVNSDYT | 0.5017379 |
| TGLPFTGI | ASSLSDRVNSDYT | 0.46285367 |
| VAEHVTV | ASSLSDRVNSDYT | 0.42447722 |
| VDIVLNSL | ASSLSDRVNSDYT | 0.55134284 |
| VLTGGGAVL | ASSLSDRVNSDYT | 0.35371047 |
| VNVGLAYGI | ASSLSDRVNSDYT | 0.5417174 |
| VSSAAVGAV | ASSLSDRVNSDYT | 0.37049207 |
| VTSVYEAGV | ASSLSDRVNSDYT | 0.53220755 |
| WVALDRGA | ASSLSDRVNSDYT | 0.41825402 |
| YLVGSVVI | ASSLSDRVNSDYT | 0.51528716 |
| GPPGTGKTLL | ASSLSDRVNSDYT | 0.4894286 |
| GVGMTKF | ASSLSDRVNSDYT | 0.62477446 |
| IGIGGSDL | ASSLSDRVNSDYT | 0.4577343 |
| IGTLLAEL | ASSLSDRVNSDYT | 0.5246477 |
| IYTSGTT | ASSLSDRVNSDYT | 0.42849112 |
| LSFPAGV | ASSLSDRVNSDYT | 0.6056256 |
| LYTSGTT | ASSLSDRVNSDYT | 0.39433807 |
| VGAFPFGF | ASSLSDRVNSDYT | 0.45307106 |
| VGAGTVEF | ASSLSDRVNSDYT | 0.38689685 |
| VGKSTLLL | ASSLSDRVNSDYT | 0.53017414 |
| AALAHAGGGIV | ASSLSDRVNSDYT | 0.50026095 |

| | | |
|---|---|---|
| AALVAVSLL | ASSLSDRVNSDYT | 0.4088307 |
| AAVSRAVAAL | ASSLSDRVNSDYT | 0.5010222 |
| AGGSSGIGGF | ASSLSDRVNSDYT | 0.51005113 |
| AGLTVANANG | ASSLSDRVNSDYT | 0.45278907 |
| AGLVGAVL | ASSLSDRVNSDYT | 0.55970913 |
| ALSQAFTV | ASSLSDRVNSDYT | 0.5048694 |
| ALTFSVSL | ASSLSDRVNSDYT | 0.54288876 |
| ALTGAALVAL | ASSLSDRVNSDYT | 0.4609315 |
| ASFSIGAGVL | ASSLSDRVNSDYT | 0.4192053 |
| DGSVYKI | ASSLSDRVNSDYT | 0.56045485 |
| EQLAEIV | ASSLSDRVNSDYT | 0.45808175 |
| FAVSELTVM | ASSLSDRVNSDYT | 0.40773976 |
| GAAPLAAGL | ASSLSDRVNSDYT | 0.54386 |
| GADGGDGGSGGAGG | ASSLSDRVNSDYT | 0.85320437 |
| GFGLDTGGAGG | ASSLSDRVNSDYT | 0.8586074 |
| GLASGGLL | ASSLSDRVNSDYT | 0.7331877 |
| GRIAGAAVLL | ASSLSDRVNSDYT | 0.69759274 |
| GSAGSLNTSF | ASSLSDRVNSDYT | 0.78975976 |
| GTGQIGI | ASSLSDRVNSDYT | 0.8657962 |
| GTLIALIL | ASSLSDRVNSDYT | 0.8792751 |
| IAATMGFVVG | ASSLSDRVNSDYT | 0.8531477 |
| IFAQALL | ASSLSDRVNSDYT | 0.6876631 |
| IGAAVALGF | ASSLSDRVNSDYT | 0.8567729 |
| IGLTPLAL | ASSLSDRVNSDYT | 0.7956254 |
| ILGSNTSTL | ASSLSDRVNSDYT | 0.8086922 |
| IVFSLTVL | ASSLSDRVNSDYT | 0.84603703 |
| LASKATGF | ASSLSDRVNSDYT | 0.82111824 |
| LASVALIL | ASSLSDRVNSDYT | 0.7782121 |
| LFAVGLTL | ASSLSDRVNSDYT | 0.6774466 |
| LFNGGIGTYI | ASSLSDRVNSDYT | 0.8622121 |
| LTLGAGII | ASSLSDRVNSDYT | 0.8760379 |
| LVAAALGDI | ASSLSDRVNSDYT | 0.86871135 |
| NTAFVQLGI | ASSLSDRVNSDYT | 0.8918922 |
| QLFSGAEL | ASSLSDRVNSDYT | 0.86961985 |
| SAATFITL | ASSLSDRVNSDYT | 0.86126924 |
| SNYGLANLGV | ASSLSDRVNSDYT | 0.27156037 |
| SYGGAYAVM | ASSLSDRVNSDYT | 0.5900458 |
| TTGVGGLTLG | ASSLSDRVNSDYT | 0.7164465 |
| TTQTIYI | ASSLSDRVNSDYT | 0.79775476 |
| VAAGAGLGAVI | ASSLSDRVNSDYT | 0.33725703 |
| VAAVVTLL | ASSLSDRVNSDYT | 0.8530356 |
| VGFLGGLI | ASSLSDRVNSDYT | 0.7368995 |
| VGVIPILL | ASSLSDRVNSDYT | 0.7315607 |
| VIFASSLI | ASSLSDRVNSDYT | 0.86974317 |
| VKGGDGGAAAG | ASSLSDRVNSDYT | 0.7664187 |
| VTKTFTNI | ASSLSDRVNSDYT | 0.6623813 |
| VVAAAILL | ASSLSDRVNSDYT | 0.7369833 |
| YAALDVEL | ASSLSDRVNSDYT | 0.8807597 |
| LASVALLL | ASSLSDRVNSDYT | 0.633222 |
| LDAVHRV | ASSLSDRVNSDYT | 0.86579645 |
| QGLSSSIF | ASSLSDRVNSDYT | 0.8068354 |

| | | |
|---|---|---|
| RGTFANIRL | ASSLSDRVNSDYT | 0.87891775 |
| RQLWWGHRI | ASSLSDRVNSDYT | 0.5534563 |
| YGPPGTGKTLL | ASSLSDRVNSDYT | 0.86615694 |
| AALVGAPANVA | ASSLSDRVNSDYT | 0.8101449 |
| AATFRRTE | ASSLSDRVNSDYT | 0.895844 |
| AGALGGGI | ASSLSDRVNSDYT | 0.798615 |
| AGGAGGAGGIGAG | ASSLSDRVNSDYT | 0.845332 |
| AGSGALGL | ASSLSDRVNSDYT | 0.82843494 |
| AQIADQI | ASSLSDRVNSDYT | 0.87767756 |
| ASAVAVTYL | ASSLSDRVNSDYT | 0.5445092 |
| ASTVLGGTGG | ASSLSDRVNSDYT | 0.7801037 |
| AVGGEPAK | ASSLSDRVNSDYT | 0.69505155 |
| CEQGTFGGTC | ASSLSDRVNSDYT | 0.8130335 |
| DEGKAPAG | ASSLSDRVNSDYT | 0.85505235 |
| DTVVFPAL | ASSLSDRVNSDYT | 0.23092684 |
| FEGITMAG | ASSLSDRVNSDYT | 0.85318595 |
| FPLVVQD | ASSLSDRVNSDYT | 0.70760983 |
| GAFTTPQ | ASSLSDRVNSDYT | 0.6804432 |
| GFGGFGTAGGVGGA | ASSLSDRVNSDYT | 0.86884093 |
| GFLAQFL | ASSLSDRVNSDYT | 0.8367765 |
| GGAGGAGGNAGVA | ASSLSDRVNSDYT | 0.87299746 |
| GGVGTAGAGGAGGAGG | ASSLSDRVNSDYT | 0.79117036 |
| GNGGGGDAGNAGSGGN | ASSLSDRVNSDYT | 0.8099386 |
| GTGGAGGAGNDAGS | ASSLSDRVNSDYT | 0.83047307 |
| GTTNTGWFNS | ASSLSDRVNSDYT | 0.5248393 |
| IFMGPLFTL | ASSLSDRVNSDYT | 0.72280896 |
| ILAHAIEAI | ASSLSDRVNSDYT | 0.8792938 |
| IRSAIREAL | ASSLSDRVNSDYT | 0.78917694 |
| ITFAGGG | ASSLSDRVNSDYT | 0.84060264 |
| ITGMSGHQEV | ASSLSDRVNSDYT | 0.62987345 |
| IVAGLACTL | ASSLSDRVNSDYT | 0.56770086 |
| IVGGGLAGA | ASSLSDRVNSDYT | 0.67543036 |
| IVTGVLNI | ASSLSDRVNSDYT | 0.85191584 |
| LAQQGCI | ASSLSDRVNSDYT | 0.8046305 |
| LCLHVEN | ASSLSDRVNSDYT | 0.7886249 |
| LDAISDGS | ASSLSDRVNSDYT | 0.54865634 |
| LGGFAAML | ASSLSDRVNSDYT | 0.7761178 |
| LGGKNPMI | ASSLSDRVNSDYT | 0.6968991 |
| LGPKGVAAIL | ASSLSDRVNSDYT | 0.7138284 |
| LMVGVSI | ASSLSDRVNSDYT | 0.8115309 |
| LVGAAFVL | ASSLSDRVNSDYT | 0.8015189 |
| LVLAVAPL | ASSLSDRVNSDYT | 0.6962371 |
| MAAIAVIDAA | ASSLSDRVNSDYT | 0.69827944 |
| NNGINAELQ | ASSLSDRVNSDYT | 0.73384225 |
| RGEPLVLVHG | ASSLSDRVNSDYT | 0.8005223 |
| RGGAGGAAGLI | ASSLSDRVNSDYT | 0.79122895 |
| RSYRMHPAV | ASSLSDRVNSDYT | 0.8797688 |
| SAAHLLAGQLA | ASSLSDRVNSDYT | 0.8321323 |
| SAAVGSY | ASSLSDRVNSDYT | 0.8511647 |
| SPHAVLHRI | ASSLSDRVNSDYT | 0.23577864 |
| SQIAKVL | ASSLSDRVNSDYT | 0.3945107 |

| | | |
|---|---|---|
| TALGHLVP | ASSLSDRVNSDYT | 0.7548842 |
| TAPTTSGAA | ASSLSDRVNSDYT | 0.71816033 |
| TDGDDGNGGSPGT | ASSLSDRVNSDYT | 0.5213794 |
| TDGLGEN | ASSLSDRVNSDYT | 0.8311156 |
| TGGPPLPGQA | ASSLSDRVNSDYT | 0.67856973 |
| TIPAITI | ASSLSDRVNSDYT | 0.6275735 |
| TIPGFTI | ASSLSDRVNSDYT | 0.82259405 |
| TSIASGALA | ASSLSDRVNSDYT | 0.6636284 |
| TVTLPGAAI | ASSLSDRVNSDYT | 0.69256306 |
| VASGLSAALP | ASSLSDRVNSDYT | 0.6808804 |
| VGAGVIGI | ASSLSDRVNSDYT | 0.8427936 |
| VGPKRFGV | ASSLSDRVNSDYT | 0.5410002 |
| VNVAFPD | ASSLSDRVNSDYT | 0.80958974 |
| VSMTNSGVS | ASSLSDRVNSDYT | 0.6991747 |
| VTDVLGDLSV | ASSLSDRVNSDYT | 0.8613452 |
| VVVVGAGFAGL | ASSLSDRVNSDYT | 0.46749154 |
| VYTVGTVV | ASSLSDRVNSDYT | 0.8254267 |
| WTTFSGP | ASSLSDRVNSDYT | 0.71947074 |
| YDELVEAF | ASSLSDRVNSDYT | 0.8860905 |
| YGTGGAGGNGG | ASSLSDRVNSDYT | 0.7097694 |
| YTNVMAV | ASSLSDRVNSDYT | 0.76029146 |
| FLEQVHQL | ASSLSDRVNSDYT | 0.6962978 |
| GLSLLAVAV | ASSLSDRVNSDYT | 0.80817443 |
| LEQVHQL | ASSLSDRVNSDYT | 0.4805519 |
| LNAAGAI | ASSLSDRVNSDYT | 0.6941844 |
| PAVHASL | ASSLSDRVNSDYT | 0.42184862 |
| TELEDDA | ASSLSDRVNSDYT | 0.77439415 |
| TPPGPPN | ASSLSDRVNSDYT | 0.796304 |
| TPPPPPPP | ASSLSDRVNSDYT | 0.5302065 |
| TQSPALI | ASSGTGENYAEQF | 0.7695389 |
| EFAAAFPQ | ASSGTGENYAEQF | 0.75859356 |
| LLSGNLDVL | ASSGTGENYAEQF | 0.61146784 |
| GPANLGNYNI | ASSGTGENYAEQF | 0.85574716 |
| TLGPTLP | ASSGTGENYAEQF | 0.7342994 |
| VYAYSSCCK | ASSGTGENYAEQF | 0.8414606 |
| APAPAPAPNI | ASSGTGENYAEQF | 0.74763 |
| PGGPVPIP | ASSGTGENYAEQF | 0.6560263 |
| FLTVALEP | ASSGTGENYAEQF | 0.81025076 |
| SQYFGANL | ASSGTGENYAEQF | 0.59347904 |
| GGVKPAA | ASSGTGENYAEQF | 0.6554926 |
| GGTGGAGGAGGW | ASSGTGENYAEQF | 0.85176206 |
| GVGGVAGLGGAGG | ASSGTGENYAEQF | 0.6597893 |
| VLLVLATTI | ASSGTGENYAEQF | 0.7085186 |
| IGIFAAML | ASSGTGENYAEQF | 0.5022931 |
| AVGDLID | ASSGTGENYAEQF | 0.51024663 |
| NISPLPL | ASSGTGENYAEQF | 0.5790013 |
| VTAVFVAI | ASSGTGENYAEQF | 0.6749903 |
| SGDDNDAAL | ASSGTGENYAEQF | 0.70855606 |
| AVAALAHS | ASSGTGENYAEQF | 0.6617442 |
| GGVTVMATP | ASSGTGENYAEQF | 0.45981854 |
| AGLAANVRD | ASSGTGENYAEQF | 0.6388833 |

| | | |
|---|---|---|
| TNRPDLIDPALLRPG | ASSGTGENYAEQF | 0.49793965 |
| LANLGVTAL | ASSGTGENYAEQF | 0.52889484 |
| TGERIEL | ASSGTGENYAEQF | 0.61437666 |
| NGEELAA | ASSGTGENYAEQF | 0.5297713 |
| VLSLVWG | ASSGTGENYAEQF | 0.48327214 |
| IGYALWL | ASSGTGENYAEQF | 0.5227229 |
| LSLLLAPL | ASSGTGENYAEQF | 0.5872165 |
| TLDEALK | ASSGTGENYAEQF | 0.67933416 |
| TDGAGTAEGG | ASSGTGENYAEQF | 0.6679724 |
| NAPGGAGGNGG | ASSGTGENYAEQF | 0.7128472 |
| VSVFMGMGGNF | ASSGTGENYAEQF | 0.69110286 |
| LASYLAAL | ASSGTGENYAEQF | 0.6571344 |
| LVGGTPAL | ASSGTGENYAEQF | 0.26252422 |
| VRNADVL | ASSGTGENYAEQF | 0.4084531 |
| TCSSPFLVSAM | ASSGTGENYAEQF | 0.3895061 |
| VHITDAE | ASSGTGENYAEQF | 0.5834087 |
| EYAAPSF | ASSGTGENYAEQF | 0.27054912 |
| VVDAFHGGL | ASSGTGENYAEQF | 0.6262245 |
| ASVVDPQ | ASSGTGENYAEQF | 0.505025 |
| SYNEFGGLS | ASSGTGENYAEQF | 0.4407314 |
| DPDGSVY | ASSGTGENYAEQF | 0.67452466 |
| MVSTIHATE | ASSGTGENYAEQF | 0.4799846 |
| LAAGVADPV | ASSGTGENYAEQF | 0.44303218 |
| TKEQIAA | ASSGTGENYAEQF | 0.42219457 |
| SLASPAAL | ASSGTGENYAEQF | 0.7091843 |
| IGLVFYGV | ASSGTGENYAEQF | 0.33636293 |
| FLLDASNL | ASSGTGENYAEQF | 0.66977787 |
| YQVGNFP | ASSGTGENYAEQF | 0.609115 |
| ESLLSQST | ASSGTGENYAEQF | 0.69741803 |
| LFTVGFAAVG | ASSGTGENYAEQF | 0.4027631 |
| ATPPTPT | ASSGTGENYAEQF | 0.6757504 |
| TDLATLG | ASSGTGENYAEQF | 0.5689939 |
| SAGGTGR | ASSGTGENYAEQF | 0.72474325 |
| GFSNSGS | ASSGTGENYAEQF | 0.5822951 |
| GGNGGSGGTSV | ASSGTGENYAEQF | 0.6030104 |
| GTSVATGGAGN | ASSGTGENYAEQF | 0.51518095 |
| NGGNGGSGGTS | ASSGTGENYAEQF | 0.7170322 |
| AVVKFPH | ASSGTGENYAEQF | 0.34763035 |
| TPPPPPP | ASSGTGENYAEQF | 0.60920954 |
| AAPAAAAAPA | ASSGTGENYAEQF | 0.43547624 |
| IALNEDL | ASSGTGENYAEQF | 0.63651246 |
| IVNVSSM | ASSGTGENYAEQF | 0.66843176 |
| TGIPPAPR | ASSGTGENYAEQF | 0.28257293 |
| FDLGGGTF | ASSGTGENYAEQF | 0.70704293 |
| FDLGGGTFDVS | ASSGTGENYAEQF | 0.53554285 |
| NEPTAAA | ASSGTGENYAEQF | 0.41967225 |
| NEPTAAALA | ASSGTGENYAEQF | 0.6660546 |
| GSIINISSI | ASSGTGENYAEQF | 0.5327988 |
| VAPGNAGTA | ASSGTGENYAEQF | 0.70789105 |
| TGGGASLE | ASSGTGENYAEQF | 0.40451202 |
| VAVYNHYKRI | ASSGTGENYAEQF | 0.6608926 |

| | | |
|---|---|---|
| TDPAPATT | ASSGTGENYAEQF | 0.65880144 |
| TNRPDILDPALLRPG | ASSGTGENYAEQF | 0.29670978 |
| LTTTTTS | ASSGTGENYAEQF | 0.37980756 |
| PTGSGKTL | ASSGTGENYAEQF | 0.7003566 |
| TGSGKTL | ASSGTGENYAEQF | 0.6580359 |
| TIGHVDHGK | ASSGTGENYAEQF | 0.6772131 |
| TIGHVDHGKTT | ASSGTGENYAEQF | 0.48013276 |
| GHVDHGKTT | ASSGTGENYAEQF | 0.46460158 |
| AQFEHTLL | ASSGTGENYAEQF | 0.58822453 |
| IVNQDEL | ASSGTGENYAEQF | 0.63966405 |
| TNGSQFF | ASSGTGENYAEQF | 0.68618023 |
| TGKGIVE | ASSGTGENYAEQF | 0.6478268 |
| LISNASDA | ASSGTGENYAEQF | 0.4495245 |
| AELRTLGELTKIAKAHGAQ | ASSGTGENYAEQF | 0.62814957 |
| APNAVAAAEPSW | ASSGTGENYAEQF | 0.5546803 |
| APQVAGAATAVL | ASSGTGENYAEQF | 0.5809212 |
| ARRDPAAAAAILPTDA | ASSGTGENYAEQF | 0.61514425 |
| DAAARDVLAEAGLAEYF | ASSGTGENYAEQF | 0.5957104 |
| DDQIVLSAKANLAKVEQ | ASSGTGENYAEQF | 0.5492641 |
| DELLPDTGEDSDEDGH | ASSGTGENYAEQF | 0.44821066 |
| ELPEAPANRATVPQD | ASSGTGENYAEQF | 0.6477155 |
| FGQSKIERVTGIG | ASSGTGENYAEQF | 0.6736994 |
| GAGGTGSTAGGAGGAGGAGGLYA | ASSGTGENYAEQF | 0.6721954 |
| GHGGTGAAGGNGGTGGAGGSGIDG | ASSGTGENYAEQF | 0.65925837 |
| GNGGNGGAGGEGGPGGAGGAGA | ASSGTGENYAEQF | 0.65806735 |
| GSTAGNGGNGGNATNGTIAGQPAG | ASSGTGENYAEQF | 0.6039862 |
| GTGGDGGNGGNGGTGG | ASSGTGENYAEQF | 0.2906827 |
| GVRRPAGIPADAPV | ASSGTGENYAEQF | 0.33718726 |
| GYRQVTDAHLLTLARR | ASSGTGENYAEQF | 0.54539174 |
| IAKMGQDGHDRG | ASSGTGENYAEQF | 0.54414463 |
| KAEPSIEIQYT | ASSGTGENYAEQF | 0.3147377 |
| KPVRIRMAADEDTVAVT | ASSGTGENYAEQF | 0.5959731 |
| MPPRIAGMRLLVIKPEP | ASSGTGENYAEQF | 0.5257727 |
| MKFYAVALFRSRGVPPTR | ASSGTGENYAEQF | 0.53684825 |
| NAGGFGILGAVAHSP | ASSGTGENYAEQF | 0.66985565 |
| NEGREITLAMAEM | ASSGTGENYAEQF | 0.55415 |
| NELVEWSHPLGHRWRWE | ASSGTGENYAEQF | 0.44224888 |
| NLGAGGTGGNGGNPG | ASSGTGENYAEQF | 0.48473835 |
| NTEWPKWETVMAADPVD | ASSGTGENYAEQF | 0.662369 |
| PASAAPAETQP | ASSGTGENYAEQF | 0.39839447 |
| QQSSDGKNGVRAAEITGE | ASSGTGENYAEQF | 0.65024006 |
| RAPDAVTTSWWKE | ASSGTGENYAEQF | 0.42112976 |
| SGFGNVGAHTSGWFNQ | ASSGTGENYAEQF | 0.66662276 |
| SGVAGAVGQAASVGGLKV | ASSGTGENYAEQF | 0.4069676 |
| SNGGAGTGSAGGNGGT | ASSGTGENYAEQF | 0.6371063 |
| TGGAGGEGGAAG | ASSGTGENYAEQF | 0.5903884 |
| TGGDGGAAGAGGAPGHG | ASSGTGENYAEQF | 0.69865394 |
| TMPPLLAAIANAGRHGV | ASSGTGENYAEQF | 0.52141076 |
| TSSTAGNSGGA | ASSGTGENYAEQF | 0.60305876 |
| VPATVPVGILC | ASSGTGENYAEQF | 0.5374585 |
| VQRPDSGGVVLTGRIS | ASSGTGENYAEQF | 0.68710065 |

| | | |
|---|---|---|
| WTPNPVPATKGVPAGLVQAT | ASSGTGENYAEQF | 0.40676284 |
| YALKRALAKFSDRFDVVII | ASSGTGENYAEQF | 0.5454864 |
| YLIGSVIV | ASSGTGENYAEQF | 0.3461473 |
| YGQLVAVID | ASSGTGENYAEQF | 0.5382954 |
| VTVGGMTLPA | ASSGTGENYAEQF | 0.6410035 |
| VLALPAV | ASSGTGENYAEQF | 0.32885328 |
| VIPPTLNYET | ASSGTGENYAEQF | 0.6543234 |
| VGMGGMHPSSG | ASSGTGENYAEQF | 0.5030334 |
| VGLAGGAATA | ASSGTGENYAEQF | 0.489293 |
| VGGAGGAGGSGP | ASSGTGENYAEQF | 0.6529262 |
| VFAAFMV | ASSGTGENYAEQF | 0.5898833 |
| TLYDKLR | ASSGTGENYAEQF | 0.6659835 |
| TLAVFTVL | ASSGTGENYAEQF | 0.51135457 |
| TGSVQPARV | ASSGTGENYAEQF | 0.5144085 |
| TGGGAGSGGT | ASSGTGENYAEQF | 0.5804625 |
| SHLETSL | ASSGTGENYAEQF | 0.47765446 |
| SFTTGMT | ASSGTGENYAEQF | 0.5030135 |
| QVEVAKRMGVRQPTVS | ASSGTGENYAEQF | 0.6653285 |
| QGSGGPALAM | ASSGTGENYAEQF | 0.12626095 |
| MLAPSNP | ASSGTGENYAEQF | 0.12583214 |
| LVNNAAI | ASSGTGENYAEQF | 0.12314896 |
| LTGEINTLA | ASSGTGENYAEQF | 0.12597592 |
| LSIFGPV | ASSGTGENYAEQF | 0.12437014 |
| LLALLANL | ASSGTGENYAEQF | 0.13018417 |
| LISNASDALDKLR | ASSGTGENYAEQF | 0.12635522 |
| LINEAVP | ASSGTGENYAEQF | 0.12559313 |
| LGYAFSGL | ASSGTGENYAEQF | 0.12689763 |
| LDFVAVAL | ASSGTGENYAEQF | 0.12597069 |
| LAQPGAE | ASSGTGENYAEQF | 0.12387099 |
| ISNLGAVTGF | ASSGTGENYAEQF | 0.12588887 |
| HQEDPGRI | ASSGTGENYAEQF | 0.12523852 |
| GTPGEPTP | ASSGTGENYAEQF | 0.1270457 |
| GFFNSGS | ASSGTGENYAEQF | 0.122943185 |
| GAGGGGLTAGAN | ASSGTGENYAEQF | 0.12556607 |
| FVDALVNM | ASSGTGENYAEQF | 0.1257232 |
| FLTVAPEL | ASSGTGENYAEQF | 0.12669548 |
| FETLTAL | ASSGTGENYAEQF | 0.12559035 |
| DVINPIEL | ASSGTGENYAEQF | 0.13221663 |
| APVSGAPV | ASSGTGENYAEQF | 0.12797086 |
| ALVGGLAW | ASSGTGENYAEQF | 0.12398604 |
| ALAAAAATAI | ASSGTGENYAEQF | 0.122462526 |
| AFGLGLLL | ASSGTGENYAEQF | 0.12390473 |
| AFFAAASL | ASSGTGENYAEQF | 0.124349214 |
| AAAALPAFSPP | ASSGTGENYAEQF | 0.12657097 |
| VVLAPAL | ASSGTGENYAEQF | 0.12167575 |
| TIGHVDHGKTTL | ASSGTGENYAEQF | 0.12883146 |
| LISNASDALDK | ASSGTGENYAEQF | 0.12635711 |
| LISNASDAL | ASSGTGENYAEQF | 0.124906495 |
| ISNASDALDK | ASSGTGENYAEQF | 0.12925209 |
| NQAETLVYQTE | ASSGTGENYAEQF | 0.12361402 |
| TGGTGGTAGAG | ASSGTGENYAEQF | 0.12513496 |

| | | |
|---|---|---|
| TGSTAGGAGGA | ASSGTGENYAEQF | 0.124282256 |
| DPLAVPVAAGPV | ASSGTGENYAEQF | 0.12979138 |
| GGTGGTPGNGGN | ASSGTGENYAEQF | 0.123470336 |
| PAGLAGVDPATL | ASSGTGENYAEQF | 0.13270721 |
| GNNLVVEAPHAGASV | ASSGTGENYAEQF | 0.12357129 |
| KVTRIMSASTLEQPA | ASSGTGENYAEQF | 0.13628602 |
| LGFALPIALPRRTVV | ASSGTGENYAEQF | 0.12576187 |
| TRLGNLTARKLVVLV | ASSGTGENYAEQF | 0.12627499 |
| WSQQRRLIYFPSAGP | ASSGTGENYAEQF | 0.12560147 |
| GPNGNPDPMAAAVDIR | ASSGTGENYAEQF | 0.13505399 |
| TQRCVSVADLRAAASR | ASSGTGENYAEQF | 0.12847555 |
| QFVLDVAEHPVEVLTETV | ASSGTGENYAEQF | 0.13081309 |
| GGAGGTGGAGGNGGTQAGDG | ASSGTGENYAEQF | 0.12492477 |
| PLLATLRAIAVDVDAKPAQV | ASSGTGENYAEQF | 0.12647092 |
| APKFRTGGYMIDFWGVGYQVAKRM | ASSGTGENYAEQF | 0.123595215 |
| GAGGAGGAGGAAGGPGGTGGTGGN | ASSGTGENYAEQF | 0.1262417 |
| IGALGLLATDVAPAGSVAFGVALA | ASSGTGENYAEQF | 0.12318099 |
| AGGAGGIGGNGGAGGAGGVLMGNG | ASSGTGENYAEQF | 0.12657706 |
| ADELIGLL | ASSGTGENYAEQF | 0.12612022 |
| ADRALKA | ASSGTGENYAEQF | 0.122361206 |
| AGAVGQVGGAGGA | ASSGTGENYAEQF | 0.124807134 |
| AGGGGGLGGAP | ASSGTGENYAEQF | 0.12255801 |
| AGGLLYGNGGAG | ASSGTGENYAEQF | 0.12438707 |
| AGLESIGL | ASSGTGENYAEQF | 0.12547049 |
| AGLKTGII | ASSGTGENYAEQF | 0.12499771 |
| ASLAALLL | ASSGTGENYAEQF | 0.12453287 |
| ASVINISSI | ASSGTGENYAEQF | 0.122547105 |
| ATPPGPA | ASSGTGENYAEQF | 0.12357445 |
| ATVASALV | ASSGTGENYAEQF | 0.12923746 |
| ATVASLIL | ASSGTGENYAEQF | 0.12311717 |
| AVNGPVLIRH | ASSGTGENYAEQF | 0.123594396 |
| DGAAGAVGI | ASSGTGENYAEQF | 0.12881228 |
| DPLLHGI | ASSGTGENYAEQF | 0.8244365 |
| ELGAALGAL | ASSGTGENYAEQF | 0.8542269 |
| ELGRTQAV | ASSGTGENYAEQF | 0.7559473 |
| FVAGGYLS | ASSGTGENYAEQF | 0.6600452 |
| GASGGAGGAGGW | ASSGTGENYAEQF | 0.748924 |
| GGAGAGGDGGAGGI | ASSGTGENYAEQF | 0.8700275 |
| GTGHNGSQPGMGGNG | ASSGTGENYAEQF | 0.8487354 |
| GYAAASASAS | ASSGTGENYAEQF | 0.8361734 |
| HQVDVAAVQQG | ASSGTGENYAEQF | 0.6321513 |
| IGAAIAHT | ASSGTGENYAEQF | 0.81921285 |
| IGMIEVAQL | ASSGTGENYAEQF | 0.75833035 |
| IRLAPEHTSAMD | ASSGTGENYAEQF | 0.77109975 |
| IYNVSVL | ASSGTGENYAEQF | 0.84735525 |
| KQGFLGM | ASSGTGENYAEQF | 0.7770721 |
| LESVITQL | ASSGTGENYAEQF | 0.77032995 |
| LGLGALVVV | ASSGTGENYAEQF | 0.7691039 |
| LIGNGGNGGSGGI | ASSGTGENYAEQF | 0.77944016 |
| LNAINAPTL | ASSGTGENYAEQF | 0.7840578 |
| LQLLNTGV | ASSGTGENYAEQF | 0.7844468 |

| | | |
|---|---|---|
| LQTSIEF | ASSGTGENYAEQF | 0.8819126 |
| MSFAVSEL | ASSGTGENYAEQF | 0.8612796 |
| NTGALGL | ASSGTGENYAEQF | 0.8598444 |
| PPGAGGAGGAGTT | ASSGTGENYAEQF | 0.20530835 |
| QLVGLGI | ASSGTGENYAEQF | 0.40972114 |
| RFLEQVHQL | ASSGTGENYAEQF | 0.8025297 |
| RGVGYVLR | ASSGTGENYAEQF | 0.731867 |
| SATGVFAGVI | ASSGTGENYAEQF | 0.5062564 |
| SLPAITLP | ASSGTGENYAEQF | 0.7875943 |
| STALAFTV | ASSGTGENYAEQF | 0.7469237 |
| SVLTVGLGK | ASSGTGENYAEQF | 0.6990836 |
| TALAGAGSGI | ASSGTGENYAEQF | 0.8433347 |
| TGLPFTGI | ASSGTGENYAEQF | 0.76655895 |
| VAEHVTV | ASSGTGENYAEQF | 0.69652736 |
| VDIVLNSL | ASSGTGENYAEQF | 0.72723615 |
| VLTGGGAVL | ASSGTGENYAEQF | 0.82452834 |
| VNVGLAYGI | ASSGTGENYAEQF | 0.5670424 |
| VSSAAVGAV | ASSGTGENYAEQF | 0.83680046 |
| VTSVYEAGV | ASSGTGENYAEQF | 0.6305089 |
| WVALDRGA | ASSGTGENYAEQF | 0.85978514 |
| YLVGSVVI | ASSGTGENYAEQF | 0.5513504 |
| GPPGTGKTLL | ASSGTGENYAEQF | 0.85053897 |
| GVGMTKF | ASSGTGENYAEQF | 0.790895 |
| IGIGGSDL | ASSGTGENYAEQF | 0.88844824 |
| IGTLLAEL | ASSGTGENYAEQF | 0.7554543 |
| IYTSGTT | ASSGTGENYAEQF | 0.7822225 |
| LSFPAGV | ASSGTGENYAEQF | 0.55464387 |
| LYTSGTT | ASSGTGENYAEQF | 0.84696656 |
| VGAFPFGF | ASSGTGENYAEQF | 0.5062959 |
| VGAGTVEF | ASSGTGENYAEQF | 0.76889277 |
| VGKSTLLL | ASSGTGENYAEQF | 0.47628927 |
| AALAHAGGGIV | ASSGTGENYAEQF | 0.83222586 |
| AALVAVSLL | ASSGTGENYAEQF | 0.8368424 |
| AAVSRAVAAL | ASSGTGENYAEQF | 0.60077894 |
| AGGSSGIGGF | ASSGTGENYAEQF | 0.85260797 |
| AGLTVANANG | ASSGTGENYAEQF | 0.8167865 |
| AGLVGAVL | ASSGTGENYAEQF | 0.7216438 |
| ALSQAFTV | ASSGTGENYAEQF | 0.877219 |
| ALTFSVSL | ASSGTGENYAEQF | 0.7673809 |
| ALTGAALVAL | ASSGTGENYAEQF | 0.8558809 |
| ASFSIGAGVL | ASSGTGENYAEQF | 0.8118212 |
| DGSVYKI | ASSGTGENYAEQF | 0.72217596 |
| EQLAEIV | ASSGTGENYAEQF | 0.8286688 |
| FAVSELTVM | ASSGTGENYAEQF | 0.42272806 |
| GAAPLAAGL | ASSGTGENYAEQF | 0.67765355 |
| GADGGDGGSGGAGG | ASSGTGENYAEQF | 0.8678147 |
| GFGLDTGGAGG | ASSGTGENYAEQF | 0.71063215 |
| GLASGGLL | ASSGTGENYAEQF | 0.79631805 |
| GRIAGAAVLL | ASSGTGENYAEQF | 0.56396365 |
| GSAGSLNTSF | ASSGTGENYAEQF | 0.5043216 |
| GTGQIGI | ASSGTGENYAEQF | 0.6340301 |

| | | |
|---|---|---|
| GTLIALIL | ASSGTGENYAEQF | 0.77612233 |
| IAATMGFVVG | ASSGTGENYAEQF | 0.77734566 |
| IFAQALL | ASSGTGENYAEQF | 0.724393 |
| IGAAVALGF | ASSGTGENYAEQF | 0.4949997 |
| IGLTPLAL | ASSGTGENYAEQF | 0.6507914 |
| ILGSNTSTL | ASSGTGENYAEQF | 0.58393455 |
| IVFSLTVL | ASSGTGENYAEQF | 0.6233173 |
| LASKATGF | ASSGTGENYAEQF | 0.75915766 |
| LASVALIL | ASSGTGENYAEQF | 0.6730431 |
| LFAVGLTL | ASSGTGENYAEQF | 0.57285595 |
| LFNGGIGTYI | ASSGTGENYAEQF | 0.62424296 |
| LTLGAGII | ASSGTGENYAEQF | 0.7007854 |
| LVAAALGDI | ASSGTGENYAEQF | 0.75585026 |
| NTAFVQLGI | ASSGTGENYAEQF | 0.76116794 |
| QLFSGAEL | ASSGTGENYAEQF | 0.81575024 |
| SAATFITL | ASSGTGENYAEQF | 0.75010145 |
| SNYGLANLGV | ASSGTGENYAEQF | 0.7830628 |
| SYGGAYAVM | ASSGTGENYAEQF | 0.24250321 |
| TTGVGGLTLG | ASSGTGENYAEQF | 0.38042825 |
| TTQTIYI | ASSGTGENYAEQF | 0.6757017 |
| VAAGAGLGAVI | ASSGTGENYAEQF | 0.6561314 |
| VAAVVTLL | ASSGTGENYAEQF | 0.4960161 |
| VGFLGGLI | ASSGTGENYAEQF | 0.7487867 |
| VGVIPILL | ASSGTGENYAEQF | 0.5834329 |
| VIFASSLI | ASSGTGENYAEQF | 0.5267846 |
| VKGGDGGAAAG | ASSGTGENYAEQF | 0.758777 |
| VTKTFTNI | ASSGTGENYAEQF | 0.5634259 |
| VVAAAILL | ASSGTGENYAEQF | 0.6146987 |
| YAALDVEL | ASSGTGENYAEQF | 0.6364372 |
| LASVALLL | ASSGTGENYAEQF | 0.7615855 |
| LDAVHRV | ASSGTGENYAEQF | 0.4584395 |
| QGLSSSIF | ASSGTGENYAEQF | 0.72211516 |
| RGTFANIRL | ASSGTGENYAEQF | 0.63420314 |
| RQLWWGHRI | ASSGTGENYAEQF | 0.7668991 |
| YGPPGTGKTLL | ASSGTGENYAEQF | 0.43470353 |
| AALVGAPANVA | ASSGTGENYAEQF | 0.7396348 |
| AATFRRTE | ASSGTGENYAEQF | 0.6123937 |
| AGALGGGI | ASSGTGENYAEQF | 0.81807625 |
| AGGAGGAGGIGAG | ASSGTGENYAEQF | 0.6351403 |
| AGSGALGL | ASSGTGENYAEQF | 0.660648 |
| AQIADQI | ASSGTGENYAEQF | 0.6284798 |
| ASAVAVTYL | ASSGTGENYAEQF | 0.7571598 |
| ASTVLGGTGG | ASSGTGENYAEQF | 0.3989888 |
| AVGGEPAK | ASSGTGENYAEQF | 0.6375909 |
| CEQGTFGGTC | ASSGTGENYAEQF | 0.40503064 |
| DEGKAPAG | ASSGTGENYAEQF | 0.7016749 |
| DTVVFPAL | ASSGTGENYAEQF | 0.6937542 |
| FEGITMAG | ASSGTGENYAEQF | 0.46393043 |
| FPLVVQD | ASSGTGENYAEQF | 0.74673647 |
| GAFTTPQ | ASSGTGENYAEQF | 0.6871749 |
| GFGGFGTAGGVGGA | ASSGTGENYAEQF | 0.58240604 |

| | | |
|---|---|---|
| GFLAQFL | ASSGTGENYAEQF | 0.80316365 |
| GGAGGAGGNAGVA | ASSGTGENYAEQF | 0.6760527 |
| GGVGTAGAGGAGGAGG | ASSGTGENYAEQF | 0.7704 |
| GNGGGGDAGNAGSGGN | ASSGTGENYAEQF | 0.69010866 |
| GTGGAGGAGNDAGS | ASSGTGENYAEQF | 0.58514047 |
| GTTNTGWFNS | ASSGTGENYAEQF | 0.7183403 |
| IFMGPLFTL | ASSGTGENYAEQF | 0.66514206 |
| ILAHAIEAI | ASSGTGENYAEQF | 0.62659556 |
| IRSAIREAL | ASSGTGENYAEQF | 0.7654147 |
| ITFAGGG | ASSGTGENYAEQF | 0.7851962 |
| ITGMSGHQEV | ASSGTGENYAEQF | 0.86352575 |
| IVAGLACTL | ASSGTGENYAEQF | 0.6902461 |
| IVGGGLAGA | ASSGTGENYAEQF | 0.6035217 |
| IVTGVLNI | ASSGTGENYAEQF | 0.7544544 |
| LAQQGCI | ASSGTGENYAEQF | 0.85287786 |
| LCLHVEN | ASSGTGENYAEQF | 0.8539977 |
| LDAISDGS | ASSGTGENYAEQF | 0.8197465 |
| LGGFAAML | ASSGTGENYAEQF | 0.54739785 |
| LGGKNPMI | ASSGTGENYAEQF | 0.76778316 |
| LGPKGVAAIL | ASSGTGENYAEQF | 0.7188345 |
| LMVGVSI | ASSGTGENYAEQF | 0.6825947 |
| LVGAAFVL | ASSGTGENYAEQF | 0.82013977 |
| LVLAVAPL | ASSGTGENYAEQF | 0.7801678 |
| MAAIAVIDAA | ASSGTGENYAEQF | 0.70218796 |
| NNGINAELQ | ASSGTGENYAEQF | 0.6163584 |
| RGEPLVLVHG | ASSGTGENYAEQF | 0.8049131 |
| RGGAGGAAGLI | ASSGTGENYAEQF | 0.82877606 |
| RSYRMHPAV | ASSGTGENYAEQF | 0.8183296 |
| SAAHLLAGQLA | ASSGTGENYAEQF | 0.86750484 |
| SAAVGSY | ASSGTGENYAEQF | 0.8345394 |
| SPHAVLHRI | ASSGTGENYAEQF | 0.821289 |
| SQIAKVL | ASSGTGENYAEQF | 0.17323586 |
| TALGHLVP | ASSGTGENYAEQF | 0.35553244 |
| TAPTTSGAA | ASSGTGENYAEQF | 0.7434983 |
| TDGDDGNGGSPGT | ASSGTGENYAEQF | 0.71202135 |
| TDGLGEN | ASSGTGENYAEQF | 0.49119663 |
| TGGPPLPGQA | ASSGTGENYAEQF | 0.8161018 |
| TIPAITI | ASSGTGENYAEQF | 0.71473575 |
| TIPGFTI | ASSGTGENYAEQF | 0.65593934 |
| TSIASGALA | ASSGTGENYAEQF | 0.8402871 |
| TVTLPGAAI | ASSGTGENYAEQF | 0.66295993 |
| VASGLSAALP | ASSGTGENYAEQF | 0.6027329 |
| VGAGVIGI | ASSGTGENYAEQF | 0.63981664 |
| VGPKRFGV | ASSGTGENYAEQF | 0.8250363 |
| VNVAFPD | ASSGTGENYAEQF | 0.44638854 |
| VSMTNSGVS | ASSGTGENYAEQF | 0.7934922 |
| VTDVLGDLSV | ASSGTGENYAEQF | 0.6259201 |
| VVVVGAGFAGL | ASSGTGENYAEQF | 0.85375273 |
| VYTVGTVV | ASSGTGENYAEQF | 0.4639702 |
| WTTFSGP | ASSGTGENYAEQF | 0.8190588 |
| YDELVEAF | ASSGTGENYAEQF | 0.7406721 |

| | | |
|---|---|---|
| YGTGGAGGNGG | ASSGTGENYAEQF | 0.87936676 |
| YTNVMAV | ASSGTGENYAEQF | 0.70703554 |
| FLEQVHQL | ASSGTGENYAEQF | 0.75426763 |
| GLSLLAVAV | ASSGTGENYAEQF | 0.5815222 |
| LEQVHQL | ASSGTGENYAEQF | 0.85064805 |
| LNAAGAI | ASSGTGENYAEQF | 0.54383856 |
| PAVHASL | ASSGTGENYAEQF | 0.7472036 |
| TELEDDA | ASSGTGENYAEQF | 0.42277905 |
| TPPGPPN | ASSGTGENYAEQF | 0.8045471 |
| TPPPPPPP | ASSGTGENYAEQF | 0.8012456 |
| TQSPALI | ASTDGGYEQY | 0.5146178 |
| EFAAAFPQ | ASTDGGYEQY | 0.79100764 |
| LLSGNLDVL | ASTDGGYEQY | 0.79438454 |
| GPANLGNYNI | ASTDGGYEQY | 0.62344223 |
| TLGPTLP | ASTDGGYEQY | 0.8647331 |
| VYAYSSCCK | ASTDGGYEQY | 0.7807915 |
| APAPAPAPNI | ASTDGGYEQY | 0.8503485 |
| PGGPVPIP | ASTDGGYEQY | 0.77468866 |
| FLTVALEP | ASTDGGYEQY | 0.63590103 |
| SQYFGANL | ASTDGGYEQY | 0.81170946 |
| GGVKPAA | ASTDGGYEQY | 0.7510661 |
| GGTGGAGGAGGW | ASTDGGYEQY | 0.6948333 |
| GVGGVAGLGGAGG | ASTDGGYEQY | 0.8346652 |
| VLLVLATTI | ASTDGGYEQY | 0.4345762 |
| IGIFAAML | ASTDGGYEQY | 0.8541141 |
| AVGDLID | ASTDGGYEQY | 0.24187744 |
| NISPLPL | ASTDGGYEQY | 0.2797653 |
| VTAVFVAI | ASTDGGYEQY | 0.46859467 |
| SGDDNDAAL | ASTDGGYEQY | 0.64755774 |
| AVAALAHS | ASTDGGYEQY | 0.8216857 |
| GGVTVMATP | ASTDGGYEQY | 0.57741344 |
| AGLAANVRD | ASTDGGYEQY | 0.25472796 |
| TNRPDLIDPALLRPG | ASTDGGYEQY | 0.37130517 |
| LANLGVTAL | ASTDGGYEQY | 0.23591945 |
| TGERIEL | ASTDGGYEQY | 0.36861098 |
| NGEELAA | ASTDGGYEQY | 0.5473231 |
| VLSLVWG | ASTDGGYEQY | 0.36924124 |
| IGYALWL | ASTDGGYEQY | 0.24085307 |
| LSLLLAPL | ASTDGGYEQY | 0.34890184 |
| TLDEALK | ASTDGGYEQY | 0.56582606 |
| TDGAGTAEGG | ASTDGGYEQY | 0.53686106 |
| NAPGGAGGNGG | ASTDGGYEQY | 0.40779546 |
| VSVFMGMGGNF | ASTDGGYEQY | 0.7464041 |
| LASYLAAL | ASTDGGYEQY | 0.6190555 |
| LVGGTPAL | ASTDGGYEQY | 0.59481835 |
| VRNADVL | ASTDGGYEQY | 0.11096532 |
| TCSSPFLVSAM | ASTDGGYEQY | 0.2183286 |
| VHITDAE | ASTDGGYEQY | 0.26914677 |
| EYAAPSF | ASTDGGYEQY | 0.44503438 |
| VVDAFHGGL | ASTDGGYEQY | 0.15135093 |
| ASVVDPQ | ASTDGGYEQY | 0.6311934 |

| | | |
|---|---|---|
| SYNEFGGLS | ASTDGGYEQY | 0.29530072 |
| DPDGSVY | ASTDGGYEQY | 0.30175444 |
| MVSTIHATE | ASTDGGYEQY | 0.675103 |
| LAAGVADPV | ASTDGGYEQY | 0.25545138 |
| TKEQIAA | ASTDGGYEQY | 0.30758262 |
| SLASPAAL | ASTDGGYEQY | 0.25807565 |
| IGLVFYGV | ASTDGGYEQY | 0.7059512 |
| FLLDASNL | ASTDGGYEQY | 0.15748262 |
| YQVGNFP | ASTDGGYEQY | 0.50994027 |
| ESLLSQST | ASTDGGYEQY | 0.26675683 |
| LFTVGFAAVG | ASTDGGYEQY | 0.58801615 |
| ATPPTPT | ASTDGGYEQY | 0.20612895 |
| TDLATLG | ASTDGGYEQY | 0.5997239 |
| SAGGTGR | ASTDGGYEQY | 0.35372984 |
| GFSNSGS | ASTDGGYEQY | 0.8632156 |
| GGNGGSGGTSV | ASTDGGYEQY | 0.38706774 |
| GTSVATGGAGN | ASTDGGYEQY | 0.51046586 |
| NGGNGGSGGTS | ASTDGGYEQY | 0.303494 |
| AVVKFPH | ASTDGGYEQY | 0.8269444 |
| TPPPPPP | ASTDGGYEQY | 0.18660758 |
| AAPAAAAAPA | ASTDGGYEQY | 0.4042661 |
| IALNEDL | ASTDGGYEQY | 0.19007084 |
| IVNVSSM | ASTDGGYEQY | 0.5148015 |
| TGIPPAPR | ASTDGGYEQY | 0.5612588 |
| FDLGGGTF | ASTDGGYEQY | 0.2426385 |
| FDLGGGTFDVS | ASTDGGYEQY | 0.5192975 |
| NEPTAAA | ASTDGGYEQY | 0.5128811 |
| NEPTAAALA | ASTDGGYEQY | 0.24980655 |
| GSIINISSI | ASTDGGYEQY | 0.7074621 |
| VAPGNAGTA | ASTDGGYEQY | 0.2672929 |
| TGGGASLE | ASTDGGYEQY | 0.77364725 |
| VAVYNHYKRI | ASTDGGYEQY | 0.36224118 |
| TDPAPATT | ASTDGGYEQY | 0.31743318 |
| TNRPDILDPALLRPG | ASTDGGYEQY | 0.54623044 |
| LTTTTTS | ASTDGGYEQY | 0.28126317 |
| PTGSGKTL | ASTDGGYEQY | 0.23835926 |
| TGSGKTL | ASTDGGYEQY | 0.6873375 |
| TIGHVDHGK | ASTDGGYEQY | 0.2784964 |
| TIGHVDHGKTT | ASTDGGYEQY | 0.8351393 |
| GHVDHGKTT | ASTDGGYEQY | 0.17038822 |
| AQFEHTLL | ASTDGGYEQY | 0.21010658 |
| IVNQDEL | ASTDGGYEQY | 0.31691742 |
| TNGSQFF | ASTDGGYEQY | 0.39974946 |
| TGKGIVE | ASTDGGYEQY | 0.81385386 |
| LISNASDA | ASTDGGYEQY | 0.3224656 |
| AELRTLGELTKIAKAHGAQ | ASTDGGYEQY | 0.19075857 |
| APNAVAAAEPSW | ASTDGGYEQY | 0.2400843 |
| APQVAGAATAVL | ASTDGGYEQY | 0.16881528 |
| ARRDPAAAAAILPTDA | ASTDGGYEQY | 0.26140442 |
| DAAARDVLAEAGLAEYF | ASTDGGYEQY | 0.3317143 |
| DDQIVLSAKANLAKVEQ | ASTDGGYEQY | 0.26819286 |

| | | |
|---|---|---|
| DELLPDTGEDSDEDGH | ASTDGGYEQY | 0.18007268 |
| ELPEAPANRATVPQD | ASTDGGYEQY | 0.21934305 |
| FGQSKIERVTGIG | ASTDGGYEQY | 0.33720067 |
| GAGGTGSTAGGAGGAGGAGGLYA | ASTDGGYEQY | 0.38082325 |
| GHGGTGAAGGNGGTGGAGGSGIDG | ASTDGGYEQY | 0.30500773 |
| GNGGNGGAGGEGGPGGAGGAGA | ASTDGGYEQY | 0.5312732 |
| GSTAGNGGNGGNATNGTIAGQPAG | ASTDGGYEQY | 0.45116228 |
| GTGGDGGNGGNGGTGG | ASTDGGYEQY | 0.35926905 |
| GVRRPAGIPADAPV | ASTDGGYEQY | 0.09793778 |
| GYRQVTDAHLLTLARR | ASTDGGYEQY | 0.15907693 |
| IAKMGQDGHDRG | ASTDGGYEQY | 0.17134981 |
| KAEPSIEIQYT | ASTDGGYEQY | 0.2942505 |
| KPVRIRMAADEDTVAVT | ASTDGGYEQY | 0.10741605 |
| MPPRIAGMRLLVIKPEP | ASTDGGYEQY | 0.42558238 |
| MKFYAVALFRSRGVPPTR | ASTDGGYEQY | 0.20313281 |
| NAGGFGILGAVAHSP | ASTDGGYEQY | 0.20414592 |
| NEGREITLAMAEM | ASTDGGYEQY | 0.50747526 |
| NELVEWSHPLGHRWRWE | ASTDGGYEQY | 0.18555966 |
| NLGAGGTGGNGGNPG | ASTDGGYEQY | 0.22507189 |
| NTEWPKWETVMAADPVD | ASTDGGYEQY | 0.16819884 |
| PASAAPAETQP | ASTDGGYEQY | 0.5119649 |
| QQSSDGKNGVRAAEITGE | ASTDGGYEQY | 0.11982693 |
| RAPDAVTTSWWKE | ASTDGGYEQY | 0.38926184 |
| SGFGNVGAHTSGWFNQ | ASTDGGYEQY | 0.16410397 |
| SGVAGAVGQAASVGGLKV | ASTDGGYEQY | 0.37391168 |
| SNGGAGTGSAGGNGGT | ASTDGGYEQY | 0.15856989 |
| TGGAGGEGGAAG | ASTDGGYEQY | 0.4124849 |
| TGGDGGAAGAGGAPGHG | ASTDGGYEQY | 0.22772112 |
| TMPPLLAAIANAGRHGV | ASTDGGYEQY | 0.84152746 |
| TSSTAGNSGGA | ASTDGGYEQY | 0.31180888 |
| VPATVPVGILC | ASTDGGYEQY | 0.33988452 |
| VQRPDSGGVVLTGRIS | ASTDGGYEQY | 0.20934376 |
| WTPNPVPATKGVPAGLVQAT | ASTDGGYEQY | 0.82200193 |
| YALKRALAKFSDRFDVVII | ASTDGGYEQY | 0.12721878 |
| YLIGSVIV | ASTDGGYEQY | 0.27314034 |
| YGQLVAVID | ASTDGGYEQY | 0.13751012 |
| VTVGGMTLPA | ASTDGGYEQY | 0.3231556 |
| VLALPAV | ASTDGGYEQY | 0.36107266 |
| VIPPTLNYET | ASTDGGYEQY | 0.15714101 |
| VGMGGMHPSSG | ASTDGGYEQY | 0.3101855 |
| VGLAGGAATA | ASTDGGYEQY | 0.28316885 |
| VGGAGGAGGSGP | ASTDGGYEQY | 0.17240176 |
| VFAAFMV | ASTDGGYEQY | 0.4984989 |
| TLYDKLR | ASTDGGYEQY | 0.21293639 |
| TLAVFTVL | ASTDGGYEQY | 0.68536633 |
| TGSVQPARV | ASTDGGYEQY | 0.20674407 |
| TGGGAGSGGT | ASTDGGYEQY | 0.19559154 |
| SHLETSL | ASTDGGYEQY | 0.33621305 |
| SFTTGMT | ASTDGGYEQY | 0.17820156 |
| QVEVAKRMGVRQPTVS | ASTDGGYEQY | 0.15955392 |
| QGSGGPALAM | ASTDGGYEQY | 0.5384028 |

| | | |
|---|---|---|
| MLAPSNP | ASTDGGYEQY | 0.71279377 |
| LVNNAAI | ASTDGGYEQY | 0.73011655 |
| LTGEINTLA | ASTDGGYEQY | 0.6545268 |
| LSIFGPV | ASTDGGYEQY | 0.60826766 |
| LLALLANL | ASTDGGYEQY | 0.6907218 |
| LISNASDALDKLR | ASTDGGYEQY | 0.7266564 |
| LINEAVP | ASTDGGYEQY | 0.7234268 |
| LGYAFSGL | ASTDGGYEQY | 0.7162024 |
| LDFVAVAL | ASTDGGYEQY | 0.6106024 |
| LAQPGAE | ASTDGGYEQY | 0.68501675 |
| ISNLGAVTGF | ASTDGGYEQY | 0.66974664 |
| HQEDPGRI | ASTDGGYEQY | 0.64801496 |
| GTPGEPTP | ASTDGGYEQY | 0.7199166 |
| GFFNSGS | ASTDGGYEQY | 0.6677519 |
| GAGGGGLTAGAN | ASTDGGYEQY | 0.6599002 |
| FVDALVNM | ASTDGGYEQY | 0.6903466 |
| FLTVAPEL | ASTDGGYEQY | 0.69286764 |
| FETLTAL | ASTDGGYEQY | 0.716842 |
| DVINPIEL | ASTDGGYEQY | 0.7093734 |
| APVSGAPV | ASTDGGYEQY | 0.73383284 |
| ALVGGLAW | ASTDGGYEQY | 0.716661 |
| ALAAAAATAI | ASTDGGYEQY | 0.7185403 |
| AFGLGLLL | ASTDGGYEQY | 0.31618863 |
| AFFAAASL | ASTDGGYEQY | 0.5731528 |
| AAAALPAFSPP | ASTDGGYEQY | 0.6859693 |
| VVLAPAL | ASTDGGYEQY | 0.68784463 |
| TIGHVDHGKTTL | ASTDGGYEQY | 0.4795126 |
| LISNASDALDK | ASTDGGYEQY | 0.7084943 |
| LISNASDAL | ASTDGGYEQY | 0.69765556 |
| ISNASDALDK | ASTDGGYEQY | 0.59894717 |
| NQAETLVYQTE | ASTDGGYEQY | 0.7155766 |
| TGGTGGTAGAG | ASTDGGYEQY | 0.62182534 |
| TGSTAGGAGGA | ASTDGGYEQY | 0.6043289 |
| DPLAVPVAAGPV | ASTDGGYEQY | 0.6676345 |
| GGTGGTPGNGGN | ASTDGGYEQY | 0.72945136 |
| PAGLAGVDPATL | ASTDGGYEQY | 0.590425 |
| GNNLVVEAPHAGASV | ASTDGGYEQY | 0.70346564 |
| KVTRIMSASTLEQPA | ASTDGGYEQY | 0.70351124 |
| LGFALPIALPRRTVV | ASTDGGYEQY | 0.72062314 |
| TRLGNLTARKLVVLV | ASTDGGYEQY | 0.5439416 |
| WSQQRRLIYFPSAGP | ASTDGGYEQY | 0.714517 |
| GPNGNPDPMAAAVDIR | ASTDGGYEQY | 0.6838876 |
| TQRCVSVADLRAAASR | ASTDGGYEQY | 0.73235786 |
| QFVLDVAEHPVEVLTETV | ASTDGGYEQY | 0.6691754 |
| GGAGGTGGAGGNGGTQAGDG | ASTDGGYEQY | 0.69165987 |
| PLLATLRAIAVDVDAKPAQV | ASTDGGYEQY | 0.6413032 |
| APKFRTGGYMIDFWGVGYQVAKRM | ASTDGGYEQY | 0.7211778 |
| GAGGAGGAGGAAGGPGGTGGTGGN | ASTDGGYEQY | 0.56246233 |
| IGALGLLATDVAPAGSVAFGVALA | ASTDGGYEQY | 0.68928915 |
| AGGAGGIGGNGGAGGAGGVLMGNG | ASTDGGYEQY | 0.62034214 |
| ADELIGLL | ASTDGGYEQY | 0.701686 |

| | | |
|---|---|---|
| ADRALKA | ASTDGGYEQY | 0.7149173 |
| AGAVGQVGGAGGA | ASTDGGYEQY | 0.6417601 |
| AGGGGGLGGAP | ASTDGGYEQY | 0.7240215 |
| AGGLLYGNGGAG | ASTDGGYEQY | 0.69954 |
| AGLESIGL | ASTDGGYEQY | 0.6553854 |
| AGLKTGII | ASTDGGYEQY | 0.7234243 |
| ASLAALLL | ASTDGGYEQY | 0.62854946 |
| ASVINISSI | ASTDGGYEQY | 0.7243834 |
| ATPPGPA | ASTDGGYEQY | 0.6633666 |
| ATVASALV | ASTDGGYEQY | 0.7056588 |
| ATVASLIL | ASTDGGYEQY | 0.71873564 |
| AVNGPVLIRH | ASTDGGYEQY | 0.57178 |
| DGAAGAVGI | ASTDGGYEQY | 0.6214185 |
| DPLLHGI | ASTDGGYEQY | 0.7223245 |
| ELGAALGAL | ASTDGGYEQY | 0.7731395 |
| ELGRTQAV | ASTDGGYEQY | 0.78875476 |
| FVAGGYLS | ASTDGGYEQY | 0.7350943 |
| GASGGAGGAGGW | ASTDGGYEQY | 0.588466 |
| GGAGAGGDGGAGGI | ASTDGGYEQY | 0.72166014 |
| GTGHNGSQPGMGGNG | ASTDGGYEQY | 0.78382325 |
| GYAAASASAS | ASTDGGYEQY | 0.78818053 |
| HQVDVAAVQQG | ASTDGGYEQY | 0.77179193 |
| IGAAIAHT | ASTDGGYEQY | 0.3860013 |
| IGMIEVAQL | ASTDGGYEQY | 0.7566035 |
| IRLAPEHTSAMD | ASTDGGYEQY | 0.70520175 |
| IYNVSVL | ASTDGGYEQY | 0.7361056 |
| KQGFLGM | ASTDGGYEQY | 0.7694627 |
| LESVITQL | ASTDGGYEQY | 0.72468406 |
| LGLGALVVV | ASTDGGYEQY | 0.7436047 |
| LIGNGGNGGSGGI | ASTDGGYEQY | 0.68293434 |
| LNAINAPTL | ASTDGGYEQY | 0.73366106 |
| LQLLNTGV | ASTDGGYEQY | 0.76262987 |
| LQTSIEF | ASTDGGYEQY | 0.7633101 |
| MSFAVSEL | ASTDGGYEQY | 0.78601575 |
| NTGALGL | ASTDGGYEQY | 0.78704906 |
| PPGAGGAGGAGTT | ASTDGGYEQY | 0.77700675 |
| QLVGLGI | ASTDGGYEQY | 0.20306914 |
| RFLEQVHQL | ASTDGGYEQY | 0.49871907 |
| RGVGYVLR | ASTDGGYEQY | 0.739208 |
| SATGVFAGVI | ASTDGGYEQY | 0.7363529 |
| SLPAITLP | ASTDGGYEQY | 0.4355882 |
| STALAFTV | ASTDGGYEQY | 0.7365632 |
| SVLTVGLGK | ASTDGGYEQY | 0.6781696 |
| TALAGAGSGI | ASTDGGYEQY | 0.7088754 |
| TGLPFTGI | ASTDGGYEQY | 0.78761584 |
| VAEHVTV | ASTDGGYEQY | 0.74376655 |
| VDIVLNSL | ASTDGGYEQY | 0.6487001 |
| VLTGGGAVL | ASTDGGYEQY | 0.6498462 |
| VNVGLAYGI | ASTDGGYEQY | 0.7645462 |
| VSSAAVGAV | ASTDGGYEQY | 0.59715796 |
| VTSVYEAGV | ASTDGGYEQY | 0.7759787 |

| | | |
|---|---|---|
| WVALDRGA | ASTDGGYEQY | 0.6173166 |
| YLVGSVVI | ASTDGGYEQY | 0.7821553 |
| GPPGTGKTLL | ASTDGGYEQY | 0.38638672 |
| GVGMTKF | ASTDGGYEQY | 0.77791715 |
| IGIGGSDL | ASTDGGYEQY | 0.7461493 |
| IGTLLAEL | ASTDGGYEQY | 0.8021999 |
| IYTSGTT | ASTDGGYEQY | 0.689619 |
| LSFPAGV | ASTDGGYEQY | 0.6847898 |
| LYTSGTT | ASTDGGYEQY | 0.5378964 |
| VGAFPFGF | ASTDGGYEQY | 0.78891504 |
| VGAGTVEF | ASTDGGYEQY | 0.5047318 |
| VGKSTLLL | ASTDGGYEQY | 0.72609353 |
| AALAHAGGGIV | ASTDGGYEQY | 0.41749012 |
| AALVAVSLL | ASTDGGYEQY | 0.76127815 |
| AAVSRAVAAL | ASTDGGYEQY | 0.7705921 |
| AGGSSGIGGF | ASTDGGYEQY | 0.38177302 |
| AGLTVANANG | ASTDGGYEQY | 0.78500384 |
| AGLVGAVL | ASTDGGYEQY | 0.73063815 |
| ALSQAFTV | ASTDGGYEQY | 0.5330603 |
| ALTFSVSL | ASTDGGYEQY | 0.79094166 |
| ALTGAALVAL | ASTDGGYEQY | 0.6394986 |
| ASFSIGAGVL | ASTDGGYEQY | 0.7910349 |
| DGSVYKI | ASTDGGYEQY | 0.7418699 |
| EQLAEIV | ASTDGGYEQY | 0.7405906 |
| FAVSELTVM | ASTDGGYEQY | 0.7460147 |
| GAAPLAAGL | ASTDGGYEQY | 0.7093896 |
| GADGGDGGSGGAGG | ASTDGGYEQY | 0.65952086 |
| GFGLDTGGAGG | ASTDGGYEQY | 0.78834295 |
| GLASGGLL | ASTDGGYEQY | 0.7937851 |
| GRIAGAAVLL | ASTDGGYEQY | 0.80254596 |
| GSAGSLNTSF | ASTDGGYEQY | 0.742207 |
| GTGQIGI | ASTDGGYEQY | 0.676242 |
| GTLIALIL | ASTDGGYEQY | 0.7647121 |
| IAATMGFVVG | ASTDGGYEQY | 0.8025919 |
| IFAQALL | ASTDGGYEQY | 0.8019359 |
| IGAAVALGF | ASTDGGYEQY | 0.78744996 |
| IGLTPLAL | ASTDGGYEQY | 0.5302096 |
| ILGSNTSTL | ASTDGGYEQY | 0.75691193 |
| IVFSLTVL | ASTDGGYEQY | 0.74423087 |
| LASKATGF | ASTDGGYEQY | 0.7094978 |
| LASVALIL | ASTDGGYEQY | 0.7961802 |
| LFAVGLTL | ASTDGGYEQY | 0.7278786 |
| LFNGGIGTYI | ASTDGGYEQY | 0.72846323 |
| LTLGAGII | ASTDGGYEQY | 0.7543584 |
| LVAAALGDI | ASTDGGYEQY | 0.76661646 |
| NTAFVQLGI | ASTDGGYEQY | 0.7993382 |
| QLFSGAEL | ASTDGGYEQY | 0.799942 |
| SAATFITL | ASTDGGYEQY | 0.8133594 |
| SNYGLANLGV | ASTDGGYEQY | 0.80526936 |
| SYGGAYAVM | ASTDGGYEQY | 0.8061463 |
| TTGVGGLTLG | ASTDGGYEQY | 0.24487448 |

| | | |
|---|---|---|
| TTQTIYI | ASTDGGYEQY | 0.5136744 |
| VAAGAGLGAVI | ASTDGGYEQY | 0.6929156 |
| VAAVVTLL | ASTDGGYEQY | 0.74017 |
| VGFLGGLI | ASTDGGYEQY | 0.5621846 |
| VGVIPILL | ASTDGGYEQY | 0.7282731 |
| VIFASSLI | ASTDGGYEQY | 0.7212615 |
| VKGGDGGAAAG | ASTDGGYEQY | 0.6385342 |
| VTKTFTNI | ASTDGGYEQY | 0.80221593 |
| VVAAAILL | ASTDGGYEQY | 0.70639515 |
| YAALDVEL | ASTDGGYEQY | 0.64553684 |
| LASVALLL | ASTDGGYEQY | 0.6286648 |
| LDAVHRV | ASTDGGYEQY | 0.79254216 |
| QGLSSSIF | ASTDGGYEQY | 0.5799719 |
| RGTFANIRL | ASTDGGYEQY | 0.78574383 |
| RQLWWGHRI | ASTDGGYEQY | 0.6225219 |
| YGPPGTGKTLL | ASTDGGYEQY | 0.7957047 |
| AALVGAPANVA | ASTDGGYEQY | 0.46995306 |
| AATFRRTE | ASTDGGYEQY | 0.79559064 |
| AGALGGGI | ASTDGGYEQY | 0.7633786 |
| AGGAGGAGGIGAG | ASTDGGYEQY | 0.8198547 |
| AGSGALGL | ASTDGGYEQY | 0.7449273 |
| AQIADQI | ASTDGGYEQY | 0.72265995 |
| ASAVAVTYL | ASTDGGYEQY | 0.5604128 |
| ASTVLGGTGG | ASTDGGYEQY | 0.79824454 |
| AVGGEPAK | ASTDGGYEQY | 0.52658373 |
| CEQGTFGGTC | ASTDGGYEQY | 0.7690546 |
| DEGKAPAG | ASTDGGYEQY | 0.50445604 |
| DTVVFPAL | ASTDGGYEQY | 0.78956467 |
| FEGITMAG | ASTDGGYEQY | 0.7864841 |
| FPLVVQD | ASTDGGYEQY | 0.59351456 |
| GAFTTPQ | ASTDGGYEQY | 0.8105767 |
| GFGGFGTAGGVGGA | ASTDGGYEQY | 0.75276613 |
| GFLAQFL | ASTDGGYEQY | 0.6562764 |
| GGAGGAGGNAGVA | ASTDGGYEQY | 0.81232864 |
| GGVGTAGAGGAGGAGG | ASTDGGYEQY | 0.6860145 |
| GNGGGGDAGNAGSGGN | ASTDGGYEQY | 0.8113161 |
| GTGGAGGAGNDAGS | ASTDGGYEQY | 0.7320516 |
| GTTNTGWFNS | ASTDGGYEQY | 0.7406052 |
| IFMGPLFTL | ASTDGGYEQY | 0.7710527 |
| ILAHAIEAI | ASTDGGYEQY | 0.5896025 |
| IRSAIREAL | ASTDGGYEQY | 0.5995984 |
| ITFAGGG | ASTDGGYEQY | 0.80491686 |
| ITGMSGHQEV | ASTDGGYEQY | 0.7796992 |
| IVAGLACTL | ASTDGGYEQY | 0.8192233 |
| IVGGGLAGA | ASTDGGYEQY | 0.62236613 |
| IVTGVLNI | ASTDGGYEQY | 0.60243505 |
| LAQQGCI | ASTDGGYEQY | 0.6848172 |
| LCLHVEN | ASTDGGYEQY | 0.8383903 |
| LDAISDGS | ASTDGGYEQY | 0.80332613 |
| LGGFAAML | ASTDGGYEQY | 0.7674022 |
| LGGKNPMI | ASTDGGYEQY | 0.5775912 |

| | | |
|---|---|---|
| LGPKGVAAIL | ASTDGGYEQY | 0.72065115 |
| LMVGVSI | ASTDGGYEQY | 0.7019299 |
| LVGAAFVL | ASTDGGYEQY | 0.648399 |
| LVLAVAPL | ASTDGGYEQY | 0.7949328 |
| MAAIAVIDAA | ASTDGGYEQY | 0.7757958 |
| NNGINAELQ | ASTDGGYEQY | 0.65416074 |
| RGEPLVLVHG | ASTDGGYEQY | 0.70787686 |
| RGGAGGAAGLI | ASTDGGYEQY | 0.714645 |
| RSYRMHPAV | ASTDGGYEQY | 0.79882485 |
| SAAHLLAGQLA | ASTDGGYEQY | 0.7879433 |
| SAAVGSY | ASTDGGYEQY | 0.85469955 |
| SPHAVLHRI | ASTDGGYEQY | 0.807135 |
| SQIAKVL | ASTDGGYEQY | 0.80315495 |
| TALGHLVP | ASTDGGYEQY | 0.23272263 |
| TAPTTSGAA | ASTDGGYEQY | 0.39830935 |
| TDGDDGNGGSPGT | ASTDGGYEQY | 0.72305715 |
| TDGLGEN | ASTDGGYEQY | 0.6921803 |
| TGGPPLPGQA | ASTDGGYEQY | 0.46819466 |
| TIPAITI | ASTDGGYEQY | 0.7899718 |
| TIPGFTI | ASTDGGYEQY | 0.70361865 |
| TSIASGALA | ASTDGGYEQY | 0.63107824 |
| TVTLPGAAI | ASTDGGYEQY | 0.7935761 |
| VASGLSAALP | ASTDGGYEQY | 0.63769007 |
| VGAGVIGI | ASTDGGYEQY | 0.69392025 |
| VGPKRFGV | ASTDGGYEQY | 0.642501 |
| VNVAFPD | ASTDGGYEQY | 0.79331875 |
| VSMTNSGVS | ASTDGGYEQY | 0.5697361 |
| VTDVLGDLSV | ASTDGGYEQY | 0.78269374 |
| VVVVGAGFAGL | ASTDGGYEQY | 0.6448096 |
| VYTVGTVV | ASTDGGYEQY | 0.8329564 |
| WTTFSGP | ASTDGGYEQY | 0.5527666 |
| YDELVEAF | ASTDGGYEQY | 0.77741265 |
| YGTGGAGGNGG | ASTDGGYEQY | 0.69230914 |
| YTNVMAV | ASTDGGYEQY | 0.85509694 |
| FLEQVHQL | ASTDGGYEQY | 0.7388253 |
| GLSLLAVAV | ASTDGGYEQY | 0.7712155 |
| LEQVHQL | ASTDGGYEQY | 0.5997584 |
| LNAAGAI | ASTDGGYEQY | 0.7660086 |
| PAVHASL | ASTDGGYEQY | 0.53628623 |
| TELEDDA | ASTDGGYEQY | 0.7202879 |
| TPPGPPN | ASTDGGYEQY | 0.4964617 |
| TPPPPPPP | ASTDGGYEQY | 0.76862 |
| TQSPALI | ASGEQRANSDYT | 0.7439829 |
| EFAAAFPQ | ASGEQRANSDYT | 0.5686975 |
| LLSGNLDVL | ASGEQRANSDYT | 0.8061243 |
| GPANLGNYNI | ASGEQRANSDYT | 0.75121653 |
| TLGPTLP | ASGEQRANSDYT | 0.66938055 |
| VYAYSSCCK | ASGEQRANSDYT | 0.8359865 |
| APAPAPAPNI | ASGEQRANSDYT | 0.72886765 |
| PGGPVPIP | ASGEQRANSDYT | 0.8148099 |
| FLTVALEP | ASGEQRANSDYT | 0.6944223 |

| | | |
|---|---|---|
| SQYFGANL | ASGEQRANSDYT | 0.66331226 |
| GGVKPAA | ASGEQRANSDYT | 0.7794553 |
| GGTGGAGGAGGW | ASGEQRANSDYT | 0.6889918 |
| GVGGVAGLGGAGG | ASGEQRANSDYT | 0.65400755 |
| VLLVLATTI | ASGEQRANSDYT | 0.8151432 |
| IGIFAAML | ASGEQRANSDYT | 0.80825824 |
| AVGDLID | ASGEQRANSDYT | 0.8540358 |
| NISPLPL | ASGEQRANSDYT | 0.6924657 |
| VTAVFVAI | ASGEQRANSDYT | 0.6830181 |
| SGDDNDAAL | ASGEQRANSDYT | 0.7489203 |
| AVAALAHS | ASGEQRANSDYT | 0.8482323 |
| GGVTVMATP | ASGEQRANSDYT | 0.81801116 |
| AGLAANVRD | ASGEQRANSDYT | 0.8071352 |
| TNRPDLIDPALLRPG | ASGEQRANSDYT | 0.58880055 |
| LANLGVTAL | ASGEQRANSDYT | 0.76065254 |
| TGERIEL | ASGEQRANSDYT | 0.67437696 |
| NGEELAA | ASGEQRANSDYT | 0.7283846 |
| VLSLVWG | ASGEQRANSDYT | 0.80742466 |
| IGYALWL | ASGEQRANSDYT | 0.752999 |
| LSLLLAPL | ASGEQRANSDYT | 0.7325177 |
| TLDEALK | ASGEQRANSDYT | 0.7728723 |
| TDGAGTAEGG | ASGEQRANSDYT | 0.77482575 |
| NAPGGAGGNGG | ASGEQRANSDYT | 0.7876494 |
| VSVFMGMGGNF | ASGEQRANSDYT | 0.7506847 |
| LASYLAAL | ASGEQRANSDYT | 0.86331177 |
| LVGGTPAL | ASGEQRANSDYT | 0.8316077 |
| VRNADVL | ASGEQRANSDYT | 0.842355 |
| TCSSPFLVSAM | ASGEQRANSDYT | 0.24371973 |
| VHITDAE | ASGEQRANSDYT | 0.50658673 |
| EYAAPSF | ASGEQRANSDYT | 0.7490147 |
| VVDAFHGGL | ASGEQRANSDYT | 0.7204051 |
| ASVVDPQ | ASGEQRANSDYT | 0.5263754 |
| SYNEFGGLS | ASGEQRANSDYT | 0.81770384 |
| DPDGSVY | ASGEQRANSDYT | 0.69344354 |
| MVSTIHATE | ASGEQRANSDYT | 0.64360166 |
| LAAGVADPV | ASGEQRANSDYT | 0.8198768 |
| TKEQIAA | ASGEQRANSDYT | 0.7468773 |
| SLASPAAL | ASGEQRANSDYT | 0.6706418 |
| IGLVFYGV | ASGEQRANSDYT | 0.59379053 |
| FLLDASNL | ASGEQRANSDYT | 0.81981397 |
| YQVGNFP | ASGEQRANSDYT | 0.4161266 |
| ESLLSQST | ASGEQRANSDYT | 0.77600396 |
| LFTVGFAAVG | ASGEQRANSDYT | 0.6511065 |
| ATPPTPT | ASGEQRANSDYT | 0.843852 |
| TDLATLG | ASGEQRANSDYT | 0.5168821 |
| SAGGTGR | ASGEQRANSDYT | 0.8155116 |
| GFSNSGS | ASGEQRANSDYT | 0.7635031 |
| GGNGGSGGTSV | ASGEQRANSDYT | 0.87202644 |
| GTSVATGGAGN | ASGEQRANSDYT | 0.7496728 |
| NGGNGGSGGTS | ASGEQRANSDYT | 0.75493085 |
| AVVKFPH | ASGEQRANSDYT | 0.6420542 |

| | | |
|---|---|---|
| TPPPPPP | ASGEQRANSDYT | 0.8280047 |
| AAPAAAAAPA | ASGEQRANSDYT | 0.53209406 |
| IALNEDL | ASGEQRANSDYT | 0.75309724 |
| IVNVSSM | ASGEQRANSDYT | 0.4561572 |
| TGIPPAPR | ASGEQRANSDYT | 0.79662234 |
| FDLGGGTF | ASGEQRANSDYT | 0.80505204 |
| FDLGGGTFDVS | ASGEQRANSDYT | 0.6561227 |
| NEPTAAA | ASGEQRANSDYT | 0.7931898 |
| NEPTAAALA | ASGEQRANSDYT | 0.7654671 |
| GSIINISSI | ASGEQRANSDYT | 0.7312958 |
| VAPGNAGTA | ASGEQRANSDYT | 0.8527845 |
| TGGGASLE | ASGEQRANSDYT | 0.7111987 |
| VAVYNHYKRI | ASGEQRANSDYT | 0.8608593 |
| TDPAPATT | ASGEQRANSDYT | 0.74647427 |
| TNRPDILDPALLRPG | ASGEQRANSDYT | 0.690658 |
| LTTTTTS | ASGEQRANSDYT | 0.80067444 |
| PTGSGKTL | ASGEQRANSDYT | 0.6123408 |
| TGSGKTL | ASGEQRANSDYT | 0.5705795 |
| TIGHVDHGK | ASGEQRANSDYT | 0.8550689 |
| TIGHVDHGKTT | ASGEQRANSDYT | 0.7455906 |
| GHVDHGKTT | ASGEQRANSDYT | 0.77456784 |
| AQFEHTLL | ASGEQRANSDYT | 0.6700896 |
| IVNQDEL | ASGEQRANSDYT | 0.59006697 |
| TNGSQFF | ASGEQRANSDYT | 0.6883118 |
| TGKGIVE | ASGEQRANSDYT | 0.7637352 |
| LISNASDA | ASGEQRANSDYT | 0.775805 |
| AELRTLGELTKIAKAHGAQ | ASGEQRANSDYT | 0.7459773 |
| APNAVAAAEPSW | ASGEQRANSDYT | 0.546883 |
| APQVAGAATAVL | ASGEQRANSDYT | 0.7475184 |
| ARRDPAAAAAILPTDA | ASGEQRANSDYT | 0.70902383 |
| DAAARDVLAEAGLAEYF | ASGEQRANSDYT | 0.6786695 |
| DDQIVLSAKANLAKVEQ | ASGEQRANSDYT | 0.71061504 |
| DELLPDTGEDSDEDGH | ASGEQRANSDYT | 0.7292514 |
| ELPEAPANRATVPQD | ASGEQRANSDYT | 0.6454543 |
| FGQSKIERVTGIG | ASGEQRANSDYT | 0.64674157 |
| GAGGTGSTAGGAGGAGGAGGLYA | ASGEQRANSDYT | 0.70552504 |
| GHGGTGAAGGNGGTGGAGGSGIDG | ASGEQRANSDYT | 0.743822 |
| GNGGNGGAGGEGGPGGAGGAGA | ASGEQRANSDYT | 0.74280775 |
| GSTAGNGGNGGNATNGTIAGGQPAG | ASGEQRANSDYT | 0.7779286 |
| GTGGDGGNGGNGGTGG | ASGEQRANSDYT | 0.7751962 |
| GVRRPAGIPADAPV | ASGEQRANSDYT | 0.73713344 |
| GYRQVTDAHLLTLARR | ASGEQRANSDYT | 0.25714538 |
| IAKMGQDGHDRG | ASGEQRANSDYT | 0.38781315 |
| KAEPSIEIQYT | ASGEQRANSDYT | 0.59772074 |
| KPVRIRMAADEDTVAVT | ASGEQRANSDYT | 0.6853361 |
| MPPRIAGMRLLVIKPEP | ASGEQRANSDYT | 0.32270548 |
| MKFYAVALFRSRGVPPTR | ASGEQRANSDYT | 0.7193601 |
| NAGGFGILGAVAHSP | ASGEQRANSDYT | 0.66722393 |
| NEGREITLAMAEM | ASGEQRANSDYT | 0.62457913 |
| NELVEWSHPLGHRWRWE | ASGEQRANSDYT | 0.76303434 |
| NLGAGGTGGNGGNPG | ASGEQRANSDYT | 0.666466 |

| | | |
|---|---|---|
| NTEWPKWETVMAADPVD | ASGEQRANSDYT | 0.5857275 |
| PASAAPAETQP | ASGEQRANSDYT | 0.57205755 |
| QQSSDGKNGVRAAEITGE | ASGEQRANSDYT | 0.75446296 |
| RAPDAVTTSWWKE | ASGEQRANSDYT | 0.4554386 |
| SGFGNVGAHTSGWFNQ | ASGEQRANSDYT | 0.76156616 |
| SGVAGAVGQAASVGGLKV | ASGEQRANSDYT | 0.44789296 |
| SNGGAGTGSAGGNGGT | ASGEQRANSDYT | 0.7731103 |
| TGGAGGEGGAAG | ASGEQRANSDYT | 0.4899338 |
| TGGDGGAAGAGGAPGHG | ASGEQRANSDYT | 0.7566782 |
| TMPPLLAAIANAGRHGV | ASGEQRANSDYT | 0.70097214 |
| TSSTAGNSGGA | ASGEQRANSDYT | 0.79660404 |
| VPATVPVGILC | ASGEQRANSDYT | 0.69240624 |
| VQRPDSGGVVLTGRIS | ASGEQRANSDYT | 0.74613845 |
| WTPNPVPATKGVPAGLVQAT | ASGEQRANSDYT | 0.57413584 |
| YALKRALAKFSDRFDVVII | ASGEQRANSDYT | 0.76934785 |
| YLIGSVIV | ASGEQRANSDYT | 0.5105353 |
| YGQLVAVID | ASGEQRANSDYT | 0.7006483 |
| VTVGGMTLPA | ASGEQRANSDYT | 0.36477578 |
| VLALPAV | ASGEQRANSDYT | 0.73620325 |
| VIPPTLNYET | ASGEQRANSDYT | 0.74983 |
| VGMGGMHPSSG | ASGEQRANSDYT | 0.49875605 |
| VGLAGGAATA | ASGEQRANSDYT | 0.7729603 |
| VGGAGGAGGSGP | ASGEQRANSDYT | 0.6985594 |
| VFAAFMV | ASGEQRANSDYT | 0.62131107 |
| TLYDKLR | ASGEQRANSDYT | 0.7691664 |
| TLAVFTVL | ASGEQRANSDYT | 0.69158506 |
| TGSVQPARV | ASGEQRANSDYT | 0.77167845 |
| TGGGAGSGGT | ASGEQRANSDYT | 0.6085603 |
| SHLETSL | ASGEQRANSDYT | 0.66259736 |
| SFTTGMT | ASGEQRANSDYT | 0.72269964 |
| QVEVAKRMGVRQPTVS | ASGEQRANSDYT | 0.6332469 |
| QGSGGPALAM | ASGEQRANSDYT | 0.5443468 |
| MLAPSNP | ASGEQRANSDYT | 0.7709713 |
| LVNNAAI | ASGEQRANSDYT | 0.0633583 |
| LTGEINTLA | ASGEQRANSDYT | 0.063407995 |
| LSIFGPV | ASGEQRANSDYT | 0.06300327 |
| LLALLANL | ASGEQRANSDYT | 0.06473357 |
| LISNASDALDKLR | ASGEQRANSDYT | 0.06334048 |
| LINEAVP | ASGEQRANSDYT | 0.06340712 |
| LGYAFSGL | ASGEQRANSDYT | 0.066350214 |
| LDFVAVAL | ASGEQRANSDYT | 0.063128024 |
| LAQPGAE | ASGEQRANSDYT | 0.06626932 |
| ISNLGAVTGF | ASGEQRANSDYT | 0.06971822 |
| HQEDPGRI | ASGEQRANSDYT | 0.0629121 |
| GTPGEPTP | ASGEQRANSDYT | 0.06536027 |
| GFFNSGS | ASGEQRANSDYT | 0.06336694 |
| GAGGGGLTAGAN | ASGEQRANSDYT | 0.06528536 |
| FVDALVNM | ASGEQRANSDYT | 0.062895045 |
| FLTVAPEL | ASGEQRANSDYT | 0.06353871 |
| FETLTAL | ASGEQRANSDYT | 0.06397965 |
| DVINPIEL | ASGEQRANSDYT | 0.0666474 |

| | | |
|---|---|---|
| APVSGAPV | ASGEQRANSDYT | 0.06533386 |
| ALVGGLAW | ASGEQRANSDYT | 0.0647971 |
| ALAAAAATAI | ASGEQRANSDYT | 0.06360256 |
| AFGLGLLL | ASGEQRANSDYT | 0.06331509 |
| AFFAAASL | ASGEQRANSDYT | 0.06292236 |
| AAAALPAFSPP | ASGEQRANSDYT | 0.06383026 |
| VVLAPAL | ASGEQRANSDYT | 0.06357202 |
| TIGHVDHGKTTL | ASGEQRANSDYT | 0.06512163 |
| LISNASDALDK | ASGEQRANSDYT | 0.06208238 |
| LISNASDAL | ASGEQRANSDYT | 0.0681125 |
| ISNASDALDK | ASGEQRANSDYT | 0.06315678 |
| NQAETLVYQTE | ASGEQRANSDYT | 0.06543457 |
| TGGTGGTAGAG | ASGEQRANSDYT | 0.06581018 |
| TGSTAGGAGGA | ASGEQRANSDYT | 0.06432378 |
| DPLAVPVAAGPV | ASGEQRANSDYT | 0.06467468 |
| GGTGGTPGNGGN | ASGEQRANSDYT | 0.063800484 |
| PAGLAGVDPATL | ASGEQRANSDYT | 0.06631427 |
| GNNLVVEAPHAGASV | ASGEQRANSDYT | 0.06316365 |
| KVTRIMSASTLEQPA | ASGEQRANSDYT | 0.07021062 |
| LGFALPIALPRRTVV | ASGEQRANSDYT | 0.06270528 |
| TRLGNLTARKLVVLV | ASGEQRANSDYT | 0.06640513 |
| WSQQRRLIYFPSAGP | ASGEQRANSDYT | 0.064651795 |
| GPNGNPDPMAAAVDIR | ASGEQRANSDYT | 0.06328105 |
| TQRCVSVADLRAAASR | ASGEQRANSDYT | 0.0635178 |
| QFVLDVAEHPVEVLTETV | ASGEQRANSDYT | 0.06907647 |
| GGAGGTGGAGGNGGTQAGDG | ASGEQRANSDYT | 0.06603172 |
| PLLATLRAIAVDVDAKPAQV | ASGEQRANSDYT | 0.067653105 |
| APKFRTGGYMIDFWGVGYQVAKRM | ASGEQRANSDYT | 0.06516248 |
| GAGGAGGAGGAAGGPGGTGGTGGN | ASGEQRANSDYT | 0.06497233 |
| IGALGLLATDVAPAGSVAFGVALA | ASGEQRANSDYT | 0.06404616 |
| AGGAGGIGGNGGAGGAGGVLMGNG | ASGEQRANSDYT | 0.063353024 |
| ADELIGLL | ASGEQRANSDYT | 0.063099466 |
| ADRALKA | ASGEQRANSDYT | 0.06421669 |
| AGAVGQVGGAGGA | ASGEQRANSDYT | 0.0632912 |
| AGGGGGLGGAP | ASGEQRANSDYT | 0.07001294 |
| AGGLLYGNGGAG | ASGEQRANSDYT | 0.063194305 |
| AGLESIGL | ASGEQRANSDYT | 0.066965416 |
| AGLKTGII | ASGEQRANSDYT | 0.06354538 |
| ASLAALLL | ASGEQRANSDYT | 0.06360804 |
| ASVINISSI | ASGEQRANSDYT | 0.06789014 |
| ATPPGPA | ASGEQRANSDYT | 0.0627615 |
| ATVASALV | ASGEQRANSDYT | 0.06300521 |
| ATVASLIL | ASGEQRANSDYT | 0.062725134 |
| AVNGPVLIRH | ASGEQRANSDYT | 0.06516883 |
| DGAAGAVGI | ASGEQRANSDYT | 0.067849174 |
| DPLLHGI | ASGEQRANSDYT | 0.064050935 |
| ELGAALGAL | ASGEQRANSDYT | 0.06388358 |
| ELGRTQAV | ASGEQRANSDYT | 0.7670536 |
| FVAGGYLS | ASGEQRANSDYT | 0.78049815 |
| GASGGAGGAGGW | ASGEQRANSDYT | 0.566237 |
| GGAGAGGDGGAGGI | ASGEQRANSDYT | 0.47659755 |

| | | |
|---|---|---|
| GTGHNGSQPGMGGNG | ASGEQRANSDYT | 0.51854247 |
| GYAAASASAS | ASGEQRANSDYT | 0.7576303 |
| HQVDVAAVQQG | ASGEQRANSDYT | 0.802873 |
| IGAAIAHT | ASGEQRANSDYT | 0.7673055 |
| IGMIEVAQL | ASGEQRANSDYT | 0.38090318 |
| IRLAPEHTSAMD | ASGEQRANSDYT | 0.75067145 |
| IYNVSVL | ASGEQRANSDYT | 0.63901895 |
| KQGFLGM | ASGEQRANSDYT | 0.63487077 |
| LESVITQL | ASGEQRANSDYT | 0.6869676 |
| LGLGALVVV | ASGEQRANSDYT | 0.6018671 |
| LIGNGGNGGSGGI | ASGEQRANSDYT | 0.51098734 |
| LNAINAPTL | ASGEQRANSDYT | 0.5085046 |
| LQLLNTGV | ASGEQRANSDYT | 0.56033486 |
| LQTSIEF | ASGEQRANSDYT | 0.7366962 |
| MSFAVSEL | ASGEQRANSDYT | 0.6639123 |
| NTGALGL | ASGEQRANSDYT | 0.8059376 |
| PPGAGGAGGAGTT | ASGEQRANSDYT | 0.78628755 |
| QLVGLGI | ASGEQRANSDYT | 0.7639368 |
| RFLEQVHQL | ASGEQRANSDYT | 0.19609371 |
| RGVGYVLR | ASGEQRANSDYT | 0.3226022 |
| SATGVFAGVI | ASGEQRANSDYT | 0.40147984 |
| SLPAITLP | ASGEQRANSDYT | 0.6083466 |
| STALAFTV | ASGEQRANSDYT | 0.18844639 |
| SVLTVGLGK | ASGEQRANSDYT | 0.6473168 |
| TALAGAGSGI | ASGEQRANSDYT | 0.60214543 |
| TGLPFTGI | ASGEQRANSDYT | 0.4777793 |
| VAEHVTV | ASGEQRANSDYT | 0.7612331 |
| VDIVLNSL | ASGEQRANSDYT | 0.55432713 |
| VLTGGGAVL | ASGEQRANSDYT | 0.403561 |
| VNVGLAYGI | ASGEQRANSDYT | 0.49461752 |
| VSSAAVGAV | ASGEQRANSDYT | 0.7952728 |
| VTSVYEAGV | ASGEQRANSDYT | 0.35958332 |
| WVALDRGA | ASGEQRANSDYT | 0.79289573 |
| YLVGSVVI | ASGEQRANSDYT | 0.49308622 |
| GPPGTGKTLL | ASGEQRANSDYT | 0.77893525 |
| GVGMTKF | ASGEQRANSDYT | 0.32158604 |
| IGIGGSDL | ASGEQRANSDYT | 0.79234123 |
| IGTLLAEL | ASGEQRANSDYT | 0.7027613 |
| IYTSGTT | ASGEQRANSDYT | 0.8172472 |
| LSFPAGV | ASGEQRANSDYT | 0.6373967 |
| LYTSGTT | ASGEQRANSDYT | 0.715788 |
| VGAFPFGF | ASGEQRANSDYT | 0.5259644 |
| VGAGTVEF | ASGEQRANSDYT | 0.80281913 |
| VGKSTLLL | ASGEQRANSDYT | 0.30836207 |
| AALAHAGGGIV | ASGEQRANSDYT | 0.56748456 |
| AALVAVSLL | ASGEQRANSDYT | 0.35617155 |
| AAVSRAVAAL | ASGEQRANSDYT | 0.68102163 |
| AGGSSGIGGF | ASGEQRANSDYT | 0.78882396 |
| AGLTVANANG | ASGEQRANSDYT | 0.21107769 |
| AGLVGAVL | ASGEQRANSDYT | 0.8049469 |
| ALSQAFTV | ASGEQRANSDYT | 0.5424117 |

| | | |
|---|---|---|
| ALTFSVSL | ASGEQRANSDYT | 0.3972466 |
| ALTGAALVAL | ASGEQRANSDYT | 0.7481626 |
| ASFSIGAGVL | ASGEQRANSDYT | 0.56269175 |
| DGSVYKI | ASGEQRANSDYT | 0.7493329 |
| EQLAEIV | ASGEQRANSDYT | 0.4582436 |
| FAVSELTVM | ASGEQRANSDYT | 0.68178713 |
| GAAPLAAGL | ASGEQRANSDYT | 0.7242787 |
| GADGGDGGSGGAGG | ASGEQRANSDYT | 0.24081568 |
| GFGLDTGGAGG | ASGEQRANSDYT | 0.3589113 |
| GLASGGLL | ASGEQRANSDYT | 0.81943893 |
| GRIAGAAVLL | ASGEQRANSDYT | 0.6055401 |
| GSAGSLNTSF | ASGEQRANSDYT | 0.7333741 |
| GTGQIGI | ASGEQRANSDYT | 0.40995786 |
| GTLIALIL | ASGEQRANSDYT | 0.38519254 |
| IAATMGFVVG | ASGEQRANSDYT | 0.42532796 |
| IFAQALL | ASGEQRANSDYT | 0.54338115 |
| IGAAVALGF | ASGEQRANSDYT | 0.7382213 |
| IGLTPLAL | ASGEQRANSDYT | 0.5463805 |
| ILGSNTSTL | ASGEQRANSDYT | 0.29757342 |
| IVFSLTVL | ASGEQRANSDYT | 0.5610882 |
| LASKATGF | ASGEQRANSDYT | 0.49232805 |
| LASVALIL | ASGEQRANSDYT | 0.59753 |
| LFAVGLTL | ASGEQRANSDYT | 0.37330917 |
| LFNGGIGTYI | ASGEQRANSDYT | 0.37704274 |
| LTLGAGII | ASGEQRANSDYT | 0.34385216 |
| LVAAALGDI | ASGEQRANSDYT | 0.2924825 |
| NTAFVQLGI | ASGEQRANSDYT | 0.43853223 |
| QLFSGAEL | ASGEQRANSDYT | 0.6012568 |
| SAATFITL | ASGEQRANSDYT | 0.56212866 |
| SNYGLANLGV | ASGEQRANSDYT | 0.50621957 |
| SYGGAYAVM | ASGEQRANSDYT | 0.6849183 |
| TTGVGGLTLG | ASGEQRANSDYT | 0.3642048 |
| TTQTIYI | ASGEQRANSDYT | 0.22146384 |
| VAAGAGLGAVI | ASGEQRANSDYT | 0.27826005 |
| VAAVVTLL | ASGEQRANSDYT | 0.29926124 |
| VGFLGGLI | ASGEQRANSDYT | 0.43656695 |
| VGVIPILL | ASGEQRANSDYT | 0.21752004 |
| VIFASSLI | ASGEQRANSDYT | 0.3848671 |
| VKGGDGGAAAG | ASGEQRANSDYT | 0.3689537 |
| VTKTFTNI | ASGEQRANSDYT | 0.4244104 |
| VVAAAILL | ASGEQRANSDYT | 0.6364305 |
| YAALDVEL | ASGEQRANSDYT | 0.43547228 |
| LASVALLL | ASGEQRANSDYT | 0.28940684 |
| LDAVHRV | ASGEQRANSDYT | 0.28413817 |
| QGLSSSIF | ASGEQRANSDYT | 0.66535795 |
| RGTFANIRL | ASGEQRANSDYT | 0.26450193 |
| RQLWWGHRI | ASGEQRANSDYT | 0.63914764 |
| YGPPGTGKTLL | ASGEQRANSDYT | 0.3153182 |
| AALVGAPANVA | ASGEQRANSDYT | 0.6670594 |
| AATFRRTE | ASGEQRANSDYT | 0.29008853 |
| AGALGGGI | ASGEQRANSDYT | 0.5879922 |

| | | |
|---|---|---|
| AGGAGGAGGIGAG | ASGEQRANSDYT | 0.602919 |
| AGSGALGL | ASGEQRANSDYT | 0.7003834 |
| AQIADQI | ASGEQRANSDYT | 0.4222685 |
| ASAVAVTYL | ASGEQRANSDYT | 0.4299684 |
| ASTVLGGTGG | ASGEQRANSDYT | 0.29175815 |
| AVGGEPAK | ASGEQRANSDYT | 0.74383473 |
| CEQGTFGGTC | ASGEQRANSDYT | 0.28169772 |
| DEGKAPAG | ASGEQRANSDYT | 0.3962185 |
| DTVVFPAL | ASGEQRANSDYT | 0.2798415 |
| FEGITMAG | ASGEQRANSDYT | 0.37969935 |
| FPLVVQD | ASGEQRANSDYT | 0.6510755 |
| GAFTTPQ | ASGEQRANSDYT | 0.2244406 |
| GFGGFGTAGGVGGA | ASGEQRANSDYT | 0.5226487 |
| GFLAQFL | ASGEQRANSDYT | 0.33593118 |
| GGAGGAGGNAGVA | ASGEQRANSDYT | 0.30050293 |
| GGVGTAGAGGAGGAGG | ASGEQRANSDYT | 0.4441694 |
| GNGGGGDAGNAGSGGN | ASGEQRANSDYT | 0.37664545 |
| GTGGAGGAGNDAGS | ASGEQRANSDYT | 0.5028996 |
| GTTNTGWFNS | ASGEQRANSDYT | 0.2802631 |
| IFMGPLFTL | ASGEQRANSDYT | 0.52042335 |
| ILAHAIEAI | ASGEQRANSDYT | 0.4517377 |
| IRSAIREAL | ASGEQRANSDYT | 0.26914245 |
| ITFAGGG | ASGEQRANSDYT | 0.26512402 |
| ITGMSGHQEV | ASGEQRANSDYT | 0.66253185 |
| IVAGLACTL | ASGEQRANSDYT | 0.20897733 |
| IVGGGLAGA | ASGEQRANSDYT | 0.26244476 |
| IVTGVLNI | ASGEQRANSDYT | 0.11240978 |
| LAQQGCI | ASGEQRANSDYT | 0.12821306 |
| LCLHVEN | ASGEQRANSDYT | 0.118346736 |
| LDAISDGS | ASGEQRANSDYT | 0.15859483 |
| LGGFAAML | ASGEQRANSDYT | 0.17833194 |
| LGGKNPMI | ASGEQRANSDYT | 0.20360635 |
| LGPKGVAAIL | ASGEQRANSDYT | 0.12533869 |
| LMVGVSI | ASGEQRANSDYT | 0.13395071 |
| LVGAAFVL | ASGEQRANSDYT | 0.13337189 |
| LVLAVAPL | ASGEQRANSDYT | 0.14151351 |
| MAAIAVIDAA | ASGEQRANSDYT | 0.13697837 |
| NNGINAELQ | ASGEQRANSDYT | 0.115887545 |
| RGEPLVLVHG | ASGEQRANSDYT | 0.11150469 |
| RGGAGGAAGLI | ASGEQRANSDYT | 0.120954454 |
| RSYRMHPAV | ASGEQRANSDYT | 0.12025981 |
| SAAHLLAGQLA | ASGEQRANSDYT | 0.14943928 |
| SAAVGSY | ASGEQRANSDYT | 0.14153445 |
| SPHAVLHRI | ASGEQRANSDYT | 0.18499336 |
| SQIAKVL | ASGEQRANSDYT | 0.1808448 |
| TALGHLVP | ASGEQRANSDYT | 0.12745327 |
| TAPTTSGAA | ASGEQRANSDYT | 0.09896023 |
| TDGDDGNGGSPGT | ASGEQRANSDYT | 0.10821253 |
| TDGLGEN | ASGEQRANSDYT | 0.12247266 |
| TGGPPLPGQA | ASGEQRANSDYT | 0.117474854 |
| TIPAITI | ASGEQRANSDYT | 0.09659071 |

| | | |
|---|---|---|
| TIPGFTI | ASGEQRANSDYT | 0.13271718 |
| TSIASGALA | ASGEQRANSDYT | 0.16363564 |
| TVTLPGAAI | ASGEQRANSDYT | 0.11558879 |
| VASGLSAALP | ASGEQRANSDYT | 0.15706895 |
| VGAGVIGI | ASGEQRANSDYT | 0.11825252 |
| VGPKRFGV | ASGEQRANSDYT | 0.110496216 |
| VNVAFPD | ASGEQRANSDYT | 0.115005955 |
| VSMTNSGVS | ASGEQRANSDYT | 0.16927978 |
| VTDVLGDLSV | ASGEQRANSDYT | 0.110313915 |
| VVVVGAGFAGL | ASGEQRANSDYT | 0.17789242 |
| VYTVGTVV | ASGEQRANSDYT | 0.12576419 |
| WTTFSGP | ASGEQRANSDYT | 0.203366 |
| YDELVEAF | ASGEQRANSDYT | 0.11679621 |
| YGTGGAGGNGG | ASGEQRANSDYT | 0.16773386 |
| YTNVMAV | ASGEQRANSDYT | 0.13739996 |
| FLEQVHQL | ASGEQRANSDYT | 0.17142302 |
| GLSLLAVAV | ASGEQRANSDYT | 0.1318566 |
| LEQVHQL | ASGEQRANSDYT | 0.14811249 |
| LNAAGAI | ASGEQRANSDYT | 0.1129981 |
| PAVHASL | ASGEQRANSDYT | 0.15425253 |
| TELEDDA | ASGEQRANSDYT | 0.10809849 |
| TPPGPPN | ASGEQRANSDYT | 0.13077638 |
| TPPPPPPP | ASGEQRANSDYT | 0.1174875 |
| TQSPALI | ASSLEEGAGNTLY | 0.13112962 |
| EFAAAFPQ | ASSLEEGAGNTLY | 0.24583907 |
| LLSGNLDVL | ASSLEEGAGNTLY | 0.099447474 |
| GPANLGNYNI | ASSLEEGAGNTLY | 0.16320495 |
| TLGPTLP | ASSLEEGAGNTLY | 0.10483209 |
| VYAYSSCCK | ASSLEEGAGNTLY | 0.12392986 |
| APAPAPAPNI | ASSLEEGAGNTLY | 0.13753147 |
| PGGPVPIP | ASSLEEGAGNTLY | 0.11719578 |
| FLTVALEP | ASSLEEGAGNTLY | 0.1292534 |
| SQYFGANL | ASSLEEGAGNTLY | 0.10124514 |
| GGVKPAA | ASSLEEGAGNTLY | 0.13938174 |
| GGTGGAGGAGGW | ASSLEEGAGNTLY | 0.17032908 |
| GVGGVAGLGGAGG | ASSLEEGAGNTLY | 0.09930264 |
| VLLVLATTI | ASSLEEGAGNTLY | 0.105895236 |
| IGIFAAML | ASSLEEGAGNTLY | 0.20312251 |
| AVGDLID | ASSLEEGAGNTLY | 0.2958901 |
| NISPLPL | ASSLEEGAGNTLY | 0.288382 |
| VTAVFVAI | ASSLEEGAGNTLY | 0.2897724 |
| SGDDNDAAL | ASSLEEGAGNTLY | 0.2918704 |
| AVAALAHS | ASSLEEGAGNTLY | 0.29105508 |
| GGVTVMATP | ASSLEEGAGNTLY | 0.29419908 |
| AGLAANVRD | ASSLEEGAGNTLY | 0.29216594 |
| TNRPDLIDPALLRPG | ASSLEEGAGNTLY | 0.29917875 |
| LANLGVTAL | ASSLEEGAGNTLY | 0.2950649 |
| TGERIEL | ASSLEEGAGNTLY | 0.29915917 |
| NGEELAA | ASSLEEGAGNTLY | 0.29682738 |
| VLSLVWG | ASSLEEGAGNTLY | 0.2990711 |
| IGYALWL | ASSLEEGAGNTLY | 0.2926225 |

| | | |
|---|---|---|
| LSLLLAPL | ASSLEEGAGNTLY | 0.29133874 |
| TLDEALK | ASSLEEGAGNTLY | 0.2895813 |
| TDGAGTAEGG | ASSLEEGAGNTLY | 0.28997386 |
| NAPGGAGGNGG | ASSLEEGAGNTLY | 0.29068303 |
| VSVFMGMGGNF | ASSLEEGAGNTLY | 0.29271746 |
| LASYLAAL | ASSLEEGAGNTLY | 0.29206118 |
| LVGGTPAL | ASSLEEGAGNTLY | 0.3047265 |
| VRNADVL | ASSLEEGAGNTLY | 0.29859126 |
| TCSSPFLVSAM | ASSLEEGAGNTLY | 0.29287916 |
| VHITDAE | ASSLEEGAGNTLY | 0.28869715 |
| EYAAPSF | ASSLEEGAGNTLY | 0.2889785 |
| VVDAFHGGL | ASSLEEGAGNTLY | 0.2938054 |
| ASVVDPQ | ASSLEEGAGNTLY | 0.2907115 |
| SYNEFGGLS | ASSLEEGAGNTLY | 0.28646913 |
| DPDGSVY | ASSLEEGAGNTLY | 0.2936867 |
| MVSTIHATE | ASSLEEGAGNTLY | 0.29390076 |
| LAAGVADPV | ASSLEEGAGNTLY | 0.2931667 |
| TKEQIAA | ASSLEEGAGNTLY | 0.2988339 |
| SLASPAAL | ASSLEEGAGNTLY | 0.2923075 |
| IGLVFYGV | ASSLEEGAGNTLY | 0.2898129 |
| FLLDASNL | ASSLEEGAGNTLY | 0.29327065 |
| YQVGNFP | ASSLEEGAGNTLY | 0.30809098 |
| ESLLSQST | ASSLEEGAGNTLY | 0.29300764 |
| LFTVGFAAVG | ASSLEEGAGNTLY | 0.3111993 |
| ATPPTPT | ASSLEEGAGNTLY | 0.2903543 |
| TDLATLG | ASSLEEGAGNTLY | 0.30694824 |
| SAGGTGR | ASSLEEGAGNTLY | 0.29265332 |
| GFSNSGS | ASSLEEGAGNTLY | 0.29951432 |
| GGNGGSGGTSV | ASSLEEGAGNTLY | 0.29406434 |
| GTSVATGGAGN | ASSLEEGAGNTLY | 0.3009589 |
| NGGNGGSGGTS | ASSLEEGAGNTLY | 0.29421687 |
| AVVKFPH | ASSLEEGAGNTLY | 0.3050598 |
| TPPPPPP | ASSLEEGAGNTLY | 0.2907692 |
| AAPAAAAAPA | ASSLEEGAGNTLY | 0.29011756 |
| IALNEDL | ASSLEEGAGNTLY | 0.29044673 |
| IVNVSSM | ASSLEEGAGNTLY | 0.29081303 |
| TGIPPAPR | ASSLEEGAGNTLY | 0.2919121 |
| FDLGGGTF | ASSLEEGAGNTLY | 0.29308102 |
| FDLGGGTFDVS | ASSLEEGAGNTLY | 0.30062157 |
| NEPTAAA | ASSLEEGAGNTLY | 0.28757164 |
| NEPTAAALA | ASSLEEGAGNTLY | 0.29853255 |
| GSIINISSI | ASSLEEGAGNTLY | 0.2888288 |
| VAPGNAGTA | ASSLEEGAGNTLY | 0.29101798 |
| TGGGASLE | ASSLEEGAGNTLY | 0.29321375 |
| VAVYNHYKRI | ASSLEEGAGNTLY | 0.29462013 |
| TDPAPATT | ASSLEEGAGNTLY | 0.28981438 |
| TNRPDILDPALLRPG | ASSLEEGAGNTLY | 0.28861555 |
| LTTTTTS | ASSLEEGAGNTLY | 0.29471633 |
| PTGSGKTL | ASSLEEGAGNTLY | 0.2945977 |
| TGSGKTL | ASSLEEGAGNTLY | 0.28783122 |
| TIGHVDHGK | ASSLEEGAGNTLY | 0.29053387 |

| | | |
|---|---|---|
| TIGHVDHGKTT | ASSLEEGAGNTLY | 0.30335233 |
| GHVDHGKTT | ASSLEEGAGNTLY | 0.21955983 |
| AQFEHTLL | ASSLEEGAGNTLY | 0.21937476 |
| IVNQDEL | ASSLEEGAGNTLY | 0.21932712 |
| TNGSQFF | ASSLEEGAGNTLY | 0.21981928 |
| TGKGIVE | ASSLEEGAGNTLY | 0.21934389 |
| LISNASDA | ASSLEEGAGNTLY | 0.2194191 |
| AELRTLGELTKIAKAHGAQ | ASSLEEGAGNTLY | 0.21975894 |
| APNAVAAAEPSW | ASSLEEGAGNTLY | 0.21956944 |
| APQVAGAATAVL | ASSLEEGAGNTLY | 0.21981202 |
| ARRDPAAAAAILPTDA | ASSLEEGAGNTLY | 0.22003463 |
| DAAARDVLAEAGLAEYF | ASSLEEGAGNTLY | 0.2194775 |
| DDQIVLSAKANLAKVEQ | ASSLEEGAGNTLY | 0.21957237 |
| DELLPDTGEDSDEDGH | ASSLEEGAGNTLY | 0.21953534 |
| ELPEAPANRATVPQD | ASSLEEGAGNTLY | 0.22030044 |
| FGQSKIERVTGIG | ASSLEEGAGNTLY | 0.21941926 |
| GAGGTGSTAGGAGGAGGAGGLYA | ASSLEEGAGNTLY | 0.21932936 |
| GHGGTGAAGGNGGTGGAGGSGIDG | ASSLEEGAGNTLY | 0.21985479 |
| GNGGNGGAGGEGGPGGAGGAGA | ASSLEEGAGNTLY | 0.21995685 |
| GSTAGNGGNGGNATNGTIAGQPAG | ASSLEEGAGNTLY | 0.21991089 |
| GTGGDGGNGGNGGTGG | ASSLEEGAGNTLY | 0.21946634 |
| GVRRPAGIPADAPV | ASSLEEGAGNTLY | 0.21957919 |
| GYRQVTDAHLLTLARR | ASSLEEGAGNTLY | 0.21966787 |
| IAKMGQDGHDRG | ASSLEEGAGNTLY | 0.21942177 |
| KAEPSIEIQYT | ASSLEEGAGNTLY | 0.21959785 |
| KPVRIRMAADEDTVAVT | ASSLEEGAGNTLY | 0.21957368 |
| MPPRIAGMRLLVIKPEP | ASSLEEGAGNTLY | 0.21967354 |
| MKFYAVALFRSRGVPPTR | ASSLEEGAGNTLY | 0.21892472 |
| NAGGFGILGAVAHSP | ASSLEEGAGNTLY | 0.22020803 |
| NEGREITLAMAEM | ASSLEEGAGNTLY | 0.22001374 |
| NELVEWSHPLGHRWRWE | ASSLEEGAGNTLY | 0.21973868 |
| NLGAGGTGGNGGNPG | ASSLEEGAGNTLY | 0.22002825 |
| NTEWPKWETVMAADPVD | ASSLEEGAGNTLY | 0.21951738 |
| PASAAPAETQP | ASSLEEGAGNTLY | 0.21978587 |
| QQSSDGKNGVRAAEITGE | ASSLEEGAGNTLY | 0.21947414 |
| RAPDAVTTSWWKE | ASSLEEGAGNTLY | 0.21968645 |
| SGFGNVGAHTSGWFNQ | ASSLEEGAGNTLY | 0.2196034 |
| SGVAGAVGQAASVGGLKV | ASSLEEGAGNTLY | 0.22027342 |
| SNGGAGTGSAGGNGGT | ASSLEEGAGNTLY | 0.21926092 |
| TGGAGGEGGAAG | ASSLEEGAGNTLY | 0.21998999 |
| TGGDGGAAGAGGAPGHG | ASSLEEGAGNTLY | 0.21970642 |
| TMPPLLAAIANAGRHGV | ASSLEEGAGNTLY | 0.21957025 |
| TSSTAGNSGGA | ASSLEEGAGNTLY | 0.21949011 |
| VPATVPVGILC | ASSLEEGAGNTLY | 0.2201705 |
| VQRPDSGGVVLTGRIS | ASSLEEGAGNTLY | 0.2198878 |
| WTPNPVPATKGVPAGLVQAT | ASSLEEGAGNTLY | 0.22028309 |
| YALKRALAKFSDRFDVVII | ASSLEEGAGNTLY | 0.21987471 |
| YLIGSVIV | ASSLEEGAGNTLY | 0.219788 |
| YGQLVAVID | ASSLEEGAGNTLY | 0.21954325 |
| VTVGGMTLPA | ASSLEEGAGNTLY | 0.21959767 |
| VLALPAV | ASSLEEGAGNTLY | 0.21957578 |

| | | |
|---|---|---|
| VIPPTLNYET | ASSLEEGAGNTLY | 0.21952012 |
| VGMGGMHPSSG | ASSLEEGAGNTLY | 0.21957424 |
| VGLAGGAATA | ASSLEEGAGNTLY | 0.21912025 |
| VGGAGGAGGSGP | ASSLEEGAGNTLY | 0.21933073 |
| VFAAFMV | ASSLEEGAGNTLY | 0.21909422 |
| TLYDKLR | ASSLEEGAGNTLY | 0.21965607 |
| TLAVFTVL | ASSLEEGAGNTLY | 0.21951443 |
| TGSVQPARV | ASSLEEGAGNTLY | 0.22029674 |
| TGGGAGSGGT | ASSLEEGAGNTLY | 0.2193037 |
| SHLETSL | ASSLEEGAGNTLY | 0.21941333 |
| SFTTGMT | ASSLEEGAGNTLY | 0.2193709 |
| QVEVAKRMGVRQPTVS | ASSLEEGAGNTLY | 0.21960871 |
| QGSGGPALAM | ASSLEEGAGNTLY | 0.21961051 |
| MLAPSNP | ASSLEEGAGNTLY | 0.21956381 |
| LVNNAAI | ASSLEEGAGNTLY | 0.21959147 |
| LTGEINTLA | ASSLEEGAGNTLY | 0.29641894 |
| LSIFGPV | ASSLEEGAGNTLY | 0.2939346 |
| LLALLANL | ASSLEEGAGNTLY | 0.26432735 |
| LISNASDALDKLR | ASSLEEGAGNTLY | 0.27705356 |
| LINEAVP | ASSLEEGAGNTLY | 0.2731553 |
| LGYAFSGL | ASSLEEGAGNTLY | 0.29347724 |
| LDFVAVAL | ASSLEEGAGNTLY | 0.29121342 |
| LAQPGAE | ASSLEEGAGNTLY | 0.29384893 |
| ISNLGAVTGF | ASSLEEGAGNTLY | 0.28695947 |
| HQEDPGRI | ASSLEEGAGNTLY | 0.27894443 |
| GTPGEPTP | ASSLEEGAGNTLY | 0.28354654 |
| GFFNSGS | ASSLEEGAGNTLY | 0.271675 |
| GAGGGGLTAGAN | ASSLEEGAGNTLY | 0.29097825 |
| FVDALVNM | ASSLEEGAGNTLY | 0.27446005 |
| FLTVAPEL | ASSLEEGAGNTLY | 0.2636755 |
| FETLTAL | ASSLEEGAGNTLY | 0.28060928 |
| DVINPIEL | ASSLEEGAGNTLY | 0.2683297 |
| APVSGAPV | ASSLEEGAGNTLY | 0.2954332 |
| ALVGGLAW | ASSLEEGAGNTLY | 0.29451814 |
| ALAAAAATAI | ASSLEEGAGNTLY | 0.33276817 |
| AFGLGLLL | ASSLEEGAGNTLY | 0.30191424 |
| AFFAAASL | ASSLEEGAGNTLY | 0.27662474 |
| AAAALPAFSPP | ASSLEEGAGNTLY | 0.2557976 |
| VVLAPAL | ASSLEEGAGNTLY | 0.2682376 |
| TIGHVDHGKTTL | ASSLEEGAGNTLY | 0.27613062 |
| LISNASDALDK | ASSLEEGAGNTLY | 0.29146618 |
| LISNASDAL | ASSLEEGAGNTLY | 0.25387058 |
| ISNASDALDK | ASSLEEGAGNTLY | 0.2838782 |
| NQAETLVYQTE | ASSLEEGAGNTLY | 0.2851179 |
| TGGTGGTAGAG | ASSLEEGAGNTLY | 0.2657761 |
| TGSTAGGAGGA | ASSLEEGAGNTLY | 0.31897452 |
| DPLAVPVAAGPV | ASSLEEGAGNTLY | 0.2659053 |
| GGTGGTPGNGGN | ASSLEEGAGNTLY | 0.2743426 |
| PAGLAGVDPATL | ASSLEEGAGNTLY | 0.27565414 |
| GNNLVVEAPHAGASV | ASSLEEGAGNTLY | 0.33714017 |
| KVTRIMSASTLEQPA | ASSLEEGAGNTLY | 0.27088594 |

| | | |
|---|---|---|
| LGFALPIALPRRTVV | ASSLEEGAGNTLY | 0.31488404 |
| TRLGNLTARKLVVLV | ASSLEEGAGNTLY | 0.26614776 |
| WSQQRRLIYFPSAGP | ASSLEEGAGNTLY | 0.3109058 |
| GPNGNPDPMAAAVDIR | ASSLEEGAGNTLY | 0.27491316 |
| TQRCVSVADLRAAASR | ASSLEEGAGNTLY | 0.29032224 |
| QFVLDVAEHPVEVLTETV | ASSLEEGAGNTLY | 0.28172067 |
| GGAGGTGGAGGNGGTQAGDG | ASSLEEGAGNTLY | 0.31177995 |
| PLLATLRAIAVDVDAKPAQV | ASSLEEGAGNTLY | 0.2914961 |
| APKFRTGGYMIDFWGVGYQVAKRM | ASSLEEGAGNTLY | 0.314437 |
| GAGGAGGAGGAAGGPGGTGGTGGN | ASSLEEGAGNTLY | 0.27173495 |
| IGALGLLATDVAPAGSVAFGVALA | ASSLEEGAGNTLY | 0.27516788 |
| AGGAGGIGGNGGAGGAGGVLMGNG | ASSLEEGAGNTLY | 0.26439127 |
| ADELIGLL | ASSLEEGAGNTLY | 0.27965334 |
| ADRALKA | ASSLEEGAGNTLY | 0.2661926 |
| AGAVGQVGGAGGA | ASSLEEGAGNTLY | 0.27819797 |
| AGGGGGLGGAP | ASSLEEGAGNTLY | 0.2922882 |
| AGGLLYGNGGAG | ASSLEEGAGNTLY | 0.26197448 |
| AGLESIGL | ASSLEEGAGNTLY | 0.28779835 |
| AGLKTGII | ASSLEEGAGNTLY | 0.26212505 |
| ASLAALLL | ASSLEEGAGNTLY | 0.28037894 |
| ASVINISSI | ASSLEEGAGNTLY | 0.2906064 |
| ATPPGPA | ASSLEEGAGNTLY | 0.26770186 |
| ATVASALV | ASSLEEGAGNTLY | 0.28313738 |
| ATVASLIL | ASSLEEGAGNTLY | 0.260534 |
| AVNGPVLIRH | ASSLEEGAGNTLY | 0.27275544 |
| DGAAGAVGI | ASSLEEGAGNTLY | 0.29866922 |
| DPLLHGI | ASSLEEGAGNTLY | 0.25707352 |
| ELGAALGAL | ASSLEEGAGNTLY | 0.26757324 |
| ELGRTQAV | ASSLEEGAGNTLY | 0.3100786 |
| FVAGGYLS | ASSLEEGAGNTLY | 0.19119501 |
| GASGGAGGAGGW | ASSLEEGAGNTLY | 0.26928958 |
| GGAGAGGDGGAGGI | ASSLEEGAGNTLY | 0.13686633 |
| GTGHNGSQPGMGGNG | ASSLEEGAGNTLY | 0.1500416 |
| GYAAASASAS | ASSLEEGAGNTLY | 0.14442515 |
| HQVDVAAVQQG | ASSLEEGAGNTLY | 0.16034506 |
| IGAAIAHT | ASSLEEGAGNTLY | 0.2645093 |
| IGMIEVAQL | ASSLEEGAGNTLY | 0.2028234 |
| IRLAPEHTSAMD | ASSLEEGAGNTLY | 0.14969866 |
| IYNVSVL | ASSLEEGAGNTLY | 0.154504 |
| KQGFLGM | ASSLEEGAGNTLY | 0.13952185 |
| LESVITQL | ASSLEEGAGNTLY | 0.16207942 |
| LGLGALVVV | ASSLEEGAGNTLY | 0.15656549 |
| LIGNGGNGGSGGI | ASSLEEGAGNTLY | 0.14112759 |
| LNAINAPTL | ASSLEEGAGNTLY | 0.13884948 |
| LQLLNTGV | ASSLEEGAGNTLY | 0.14674015 |
| LQTSIEF | ASSLEEGAGNTLY | 0.14604275 |
| MSFAVSEL | ASSLEEGAGNTLY | 0.18750995 |
| NTGALGL | ASSLEEGAGNTLY | 0.17475115 |
| PPGAGGAGGAGTT | ASSLEEGAGNTLY | 0.178284 |
| QLVGLGI | ASSLEEGAGNTLY | 0.17863975 |
| RFLEQVHQL | ASSLEEGAGNTLY | 0.15393658 |

| | | |
|---|---|---|
| RGVGYVLR | ASSLEEGAGNTLY | 0.12869044 |
| SATGVFAGVI | ASSLEEGAGNTLY | 0.136214 |
| SLPAITLP | ASSLEEGAGNTLY | 0.13991876 |
| STALAFTV | ASSLEEGAGNTLY | 0.14599492 |
| SVLTVGLGK | ASSLEEGAGNTLY | 0.12787075 |
| TALAGAGSGI | ASSLEEGAGNTLY | 0.15526572 |
| TGLPFTGI | ASSLEEGAGNTLY | 0.1587728 |
| VAEHVTV | ASSLEEGAGNTLY | 0.13928826 |
| VDIVLNSL | ASSLEEGAGNTLY | 0.18082567 |
| VLTGGGAVL | ASSLEEGAGNTLY | 0.14479995 |
| VNVGLAYGI | ASSLEEGAGNTLY | 0.13934028 |
| VSSAAVGAV | ASSLEEGAGNTLY | 0.14357077 |
| VTSVYEAGV | ASSLEEGAGNTLY | 0.18232827 |
| WVALDRGA | ASSLEEGAGNTLY | 0.1363805 |
| YLVGSVVI | ASSLEEGAGNTLY | 0.19884728 |
| GPPGTGKTLL | ASSLEEGAGNTLY | 0.14464088 |
| GVGMTKF | ASSLEEGAGNTLY | 0.18218699 |
| IGIGGSDL | ASSLEEGAGNTLY | 0.13900824 |
| IGTLLAEL | ASSLEEGAGNTLY | 0.17503448 |
| IYTSGTT | ASSLEEGAGNTLY | 0.15198845 |
| LSFPAGV | ASSLEEGAGNTLY | 0.20041019 |
| LYTSGTT | ASSLEEGAGNTLY | 0.14838685 |
| VGAFPFGF | ASSLEEGAGNTLY | 0.15732366 |
| VGAGTVEF | ASSLEEGAGNTLY | 0.14375198 |
| VGKSTLLL | ASSLEEGAGNTLY | 0.25323942 |
| AALAHAGGGIV | ASSLEEGAGNTLY | 0.13139434 |
| AALVAVSLL | ASSLEEGAGNTLY | 0.14721358 |
| AAVSRAVAAL | ASSLEEGAGNTLY | 0.14187431 |
| AGGSSGIGGF | ASSLEEGAGNTLY | 0.15165645 |
| AGLTVANANG | ASSLEEGAGNTLY | 0.22886282 |
| AGLVGAVL | ASSLEEGAGNTLY | 0.13043642 |
| ALSQAFTV | ASSLEEGAGNTLY | 0.17880659 |
| ALTFSVSL | ASSLEEGAGNTLY | 0.13261622 |
| ALTGAALVAL | ASSLEEGAGNTLY | 0.14399074 |
| ASFSIGAGVL | ASSLEEGAGNTLY | 0.1623769 |
| DGSVYKI | ASSLEEGAGNTLY | 0.14098898 |
| EQLAEIV | ASSLEEGAGNTLY | 0.15749641 |
| FAVSELTVM | ASSLEEGAGNTLY | 0.13451894 |
| GAAPLAAGL | ASSLEEGAGNTLY | 0.14946488 |
| GADGGDGGSGGAGG | ASSLEEGAGNTLY | 0.17644894 |
| GFGLDTGGAGG | ASSLEEGAGNTLY | 0.13076131 |
| GLASGGLL | ASSLEEGAGNTLY | 0.13556804 |
| GRIAGAAVLL | ASSLEEGAGNTLY | 0.19540794 |
| GSAGSLNTSF | ASSLEEGAGNTLY | 0.18838994 |
| GTGQIGI | ASSLEEGAGNTLY | 0.3566116 |
| GTLIALIL | ASSLEEGAGNTLY | 0.16716956 |
| IAATMGFVVG | ASSLEEGAGNTLY | 0.17653541 |
| IFAQALL | ASSLEEGAGNTLY | 0.18077765 |
| IGAAVALGF | ASSLEEGAGNTLY | 0.19781294 |
| IGLTPLAL | ASSLEEGAGNTLY | 0.4471318 |
| ILGSNTSTL | ASSLEEGAGNTLY | 0.1898914 |

| | | |
|---|---|---|
| IVFSLTVL | ASSLEEGAGNTLY | 0.17783491 |
| LASKATGF | ASSLEEGAGNTLY | 0.17840324 |
| LASVALIL | ASSLEEGAGNTLY | 0.17010407 |
| LFAVGLTL | ASSLEEGAGNTLY | 0.1713194 |
| LFNGGIGTYI | ASSLEEGAGNTLY | 0.18192074 |
| LTLGAGII | ASSLEEGAGNTLY | 0.1783262 |
| LVAAALGDI | ASSLEEGAGNTLY | 0.17027576 |
| NTAFVQLGI | ASSLEEGAGNTLY | 0.17440942 |
| QLFSGAEL | ASSLEEGAGNTLY | 0.17497306 |
| SAATFITL | ASSLEEGAGNTLY | 0.20828357 |
| SNYGLANLGV | ASSLEEGAGNTLY | 0.19313297 |
| SYGGAYAVM | ASSLEEGAGNTLY | 0.21103387 |
| TTGVGGLTLG | ASSLEEGAGNTLY | 0.19100459 |
| TTQTIYI | ASSLEEGAGNTLY | 0.18055154 |
| VAAGAGLGAVI | ASSLEEGAGNTLY | 0.16169527 |
| VAAVVTLL | ASSLEEGAGNTLY | 0.1682747 |
| VGFLGGLI | ASSLEEGAGNTLY | 0.16934209 |
| VGVIPILL | ASSLEEGAGNTLY | 0.17999397 |
| VIFASSLI | ASSLEEGAGNTLY | 0.16192 |
| VKGGDGGAAAG | ASSLEEGAGNTLY | 0.18753411 |
| VTKTFTNI | ASSLEEGAGNTLY | 0.18336284 |
| VVAAAILL | ASSLEEGAGNTLY | 0.16926177 |
| YAALDVEL | ASSLEEGAGNTLY | 0.19872142 |
| LASVALLL | ASSLEEGAGNTLY | 0.16878274 |
| LDAVHRV | ASSLEEGAGNTLY | 0.17100853 |
| QGLSSSIF | ASSLEEGAGNTLY | 0.16698849 |
| RGTFANIRL | ASSLEEGAGNTLY | 0.19840066 |
| RQLWWGHRI | ASSLEEGAGNTLY | 0.16565175 |
| YGPPGTGKTLL | ASSLEEGAGNTLY | 0.20382494 |
| AALVGAPANVA | ASSLEEGAGNTLY | 0.17405364 |
| AATFRRTE | ASSLEEGAGNTLY | 0.19430801 |
| AGALGGGI | ASSLEEGAGNTLY | 0.16973546 |
| AGGAGGAGGIGAG | ASSLEEGAGNTLY | 0.18025826 |
| AGSGALGL | ASSLEEGAGNTLY | 0.17247003 |
| AQIADQI | ASSLEEGAGNTLY | 0.24082789 |
| ASAVAVTYL | ASSLEEGAGNTLY | 0.18520132 |
| ASTVLGGTGG | ASSLEEGAGNTLY | 0.19031724 |
| AVGGEPAK | ASSLEEGAGNTLY | 0.17180055 |
| CEQGTFGGTC | ASSLEEGAGNTLY | 0.35118166 |
| DEGKAPAG | ASSLEEGAGNTLY | 0.16602178 |
| DTVVFPAL | ASSLEEGAGNTLY | 0.17451301 |
| FEGITMAG | ASSLEEGAGNTLY | 0.17151088 |
| FPLVVQD | ASSLEEGAGNTLY | 0.17824003 |
| GAFTTPQ | ASSLEEGAGNTLY | 0.18630487 |
| GFGGFGTAGGVGGA | ASSLEEGAGNTLY | 0.16525802 |
| GFLAQFL | ASSLEEGAGNTLY | 0.18679647 |
| GGAGGAGGNAGVA | ASSLEEGAGNTLY | 0.17007257 |
| GGVGTAGAGGAGGAGG | ASSLEEGAGNTLY | 0.17113116 |
| GNGGGGDAGNAGSGGN | ASSLEEGAGNTLY | 0.18949117 |
| GTGGAGGAGNDAGS | ASSLEEGAGNTLY | 0.1710147 |
| GTTNTGWFNS | ASSLEEGAGNTLY | 0.18992133 |

| | | |
|---|---|---|
| IFMGPLFTL | ASSLEEGAGNTLY | 0.16497907 |
| ILAHAIEAI | ASSLEEGAGNTLY | 0.1760151 |
| IRSAIREAL | ASSLEEGAGNTLY | 0.1995275 |
| ITFAGGG | ASSLEEGAGNTLY | 0.16562584 |
| ITGMSGHQEV | ASSLEEGAGNTLY | 0.16566133 |
| IVAGLACTL | ASSLEEGAGNTLY | 0.19459097 |
| IVGGGLAGA | ASSLEEGAGNTLY | 0.10048373 |
| IVTGVLNI | ASSLEEGAGNTLY | 0.096643746 |
| LAQQGCI | ASSLEEGAGNTLY | 0.09178256 |
| LCLHVEN | ASSLEEGAGNTLY | 0.0992494 |
| LDAISDGS | ASSLEEGAGNTLY | 0.094942845 |
| LGGFAAML | ASSLEEGAGNTLY | 0.10055935 |
| LGGKNPMI | ASSLEEGAGNTLY | 0.101455204 |
| LGPKGVAAIL | ASSLEEGAGNTLY | 0.09878415 |
| LMVGVSI | ASSLEEGAGNTLY | 0.10484906 |
| LVGAAFVL | ASSLEEGAGNTLY | 0.10234936 |
| LVLAVAPL | ASSLEEGAGNTLY | 0.098950386 |
| MAAIAVIDAA | ASSLEEGAGNTLY | 0.10010351 |
| NNGINAELQ | ASSLEEGAGNTLY | 0.09707153 |
| RGEPLVLVHG | ASSLEEGAGNTLY | 0.09890477 |
| RGGAGGAAGLI | ASSLEEGAGNTLY | 0.09204482 |
| RSYRMHPAV | ASSLEEGAGNTLY | 0.092344455 |
| SAAHLLAGQLA | ASSLEEGAGNTLY | 0.100016415 |
| SAAVGSY | ASSLEEGAGNTLY | 0.10727678 |
| SPHAVLHRI | ASSLEEGAGNTLY | 0.11294752 |
| SQIAKVL | ASSLEEGAGNTLY | 0.105686314 |
| TALGHLVP | ASSLEEGAGNTLY | 0.106643304 |
| TAPTTSGAA | ASSLEEGAGNTLY | 0.09647399 |
| TDGDDGNGGSPGT | ASSLEEGAGNTLY | 0.09146428 |
| TDGLGEN | ASSLEEGAGNTLY | 0.09263008 |
| TGGPPLPGQA | ASSLEEGAGNTLY | 0.095455684 |
| TIPAITI | ASSLEEGAGNTLY | 0.09969045 |
| TIPGFTI | ASSLEEGAGNTLY | 0.08863141 |
| TSIASGALA | ASSLEEGAGNTLY | 0.100640945 |
| TVTLPGAAI | ASSLEEGAGNTLY | 0.099174626 |
| VASGLSAALP | ASSLEEGAGNTLY | 0.10057118 |
| VGAGVIGI | ASSLEEGAGNTLY | 0.10640677 |
| VGPKRFGV | ASSLEEGAGNTLY | 0.09405855 |
| VNVAFPD | ASSLEEGAGNTLY | 0.095724456 |
| VSMTNSGVS | ASSLEEGAGNTLY | 0.09668402 |
| VTDVLGDLSV | ASSLEEGAGNTLY | 0.099809654 |
| VVVVGAGFAGL | ASSLEEGAGNTLY | 0.094110675 |
| VYTVGTVV | ASSLEEGAGNTLY | 0.11735034 |
| WTTFSGP | ASSLEEGAGNTLY | 0.09362405 |
| YDELVEAF | ASSLEEGAGNTLY | 0.11792295 |
| YGTGGAGGNGG | ASSLEEGAGNTLY | 0.10192832 |
| YTNVMAV | ASSLEEGAGNTLY | 0.099872574 |
| FLEQVHQL | ASSLEEGAGNTLY | 0.09582558 |
| GLSLLAVAV | ASSLEEGAGNTLY | 0.1248452 |
| LEQVHQL | ASSLEEGAGNTLY | 0.10468687 |
| LNAAGAI | ASSLEEGAGNTLY | 0.11351326 |

| | | |
|---|---|---|
| PAVHASL | ASSLEEGAGNTLY | 0.096884124 |
| TELEDDA | ASSLEEGAGNTLY | 0.09833988 |
| TPPGPPN | ASSLEEGAGNTLY | 0.0966545 |
| TPPPPPPP | ASSLEEGAGNTLY | 0.0957119 |
| TQSPALI | ASRSRDKNTLY | 0.09275267 |
| EFAAAFPQ | ASRSRDKNTLY | 0.0955764 |
| LLSGNLDVL | ASRSRDKNTLY | 0.101743124 |
| GPANLGNYNI | ASRSRDKNTLY | 0.1157603 |
| TLGPTLP | ASRSRDKNTLY | 0.092705555 |
| VYAYSSCCK | ASRSRDKNTLY | 0.1435794 |
| APAPAPAPNI | ASRSRDKNTLY | 0.09348392 |
| PGGPVPIP | ASRSRDKNTLY | 0.09661172 |
| FLTVALEP | ASRSRDKNTLY | 0.10134276 |
| SQYFGANL | ASRSRDKNTLY | 0.09393698 |
| GGVKPAA | ASRSRDKNTLY | 0.0903493 |
| GGTGGAGGAGGW | ASRSRDKNTLY | 0.09289107 |
| GVGGVAGLGGAGG | ASRSRDKNTLY | 0.10548007 |
| VLLVLATTI | ASRSRDKNTLY | 0.10771416 |
| IGIFAAML | ASRSRDKNTLY | 0.09645071 |
| AVGDLID | ASRSRDKNTLY | 0.10280901 |
| NISPLPL | ASRSRDKNTLY | 0.64598626 |
| VTAVFVAI | ASRSRDKNTLY | 0.7155018 |
| SGDDNDAAL | ASRSRDKNTLY | 0.16675542 |
| AVAALAHS | ASRSRDKNTLY | 0.15921614 |
| GGVTVMATP | ASRSRDKNTLY | 0.4403436 |
| AGLAANVRD | ASRSRDKNTLY | 0.700684 |
| TNRPDLIDPALLRPG | ASRSRDKNTLY | 0.3432061 |
| LANLGVTAL | ASRSRDKNTLY | 0.65671676 |
| TGERIEL | ASRSRDKNTLY | 0.15239215 |
| NGEELAA | ASRSRDKNTLY | 0.1751139 |
| VLSLVWG | ASRSRDKNTLY | 0.17601162 |
| IGYALWL | ASRSRDKNTLY | 0.2120516 |
| LSLLLAPL | ASRSRDKNTLY | 0.51753247 |
| TLDEALK | ASRSRDKNTLY | 0.13086662 |
| TDGAGTAEGG | ASRSRDKNTLY | 0.12731308 |
| NAPGGAGGNGG | ASRSRDKNTLY | 0.28920412 |
| VSVFMGMGGNF | ASRSRDKNTLY | 0.47535563 |
| LASYLAAL | ASRSRDKNTLY | 0.727774 |
| LVGGTPAL | ASRSRDKNTLY | 0.68078387 |
| VRNADVL | ASRSRDKNTLY | 0.7575963 |
| TCSSPFLVSAM | ASRSRDKNTLY | 0.6678235 |
| VHITDAE | ASRSRDKNTLY | 0.56704473 |
| EYAAPSF | ASRSRDKNTLY | 0.09618607 |
| VVDAFHGGL | ASRSRDKNTLY | 0.15713915 |
| ASVVDPQ | ASRSRDKNTLY | 0.13772254 |
| SYNEFGGLS | ASRSRDKNTLY | 0.37315154 |
| DPDGSVY | ASRSRDKNTLY | 0.091506325 |
| MVSTIHATE | ASRSRDKNTLY | 0.25511062 |
| LAAGVADPV | ASRSRDKNTLY | 0.3134653 |
| TKEQIAA | ASRSRDKNTLY | 0.10648443 |
| SLASPAAL | ASRSRDKNTLY | 0.62469465 |

| | | |
|---|---|---|
| IGLVFYGV | ASRSRDKNTLY | 0.11999102 |
| FLLDASNL | ASRSRDKNTLY | 0.13340735 |
| YQVGNFP | ASRSRDKNTLY | 0.1680474 |
| ESLLSQST | ASRSRDKNTLY | 0.7241167 |
| LFTVGFAAVG | ASRSRDKNTLY | 0.14907277 |
| ATPPTPT | ASRSRDKNTLY | 0.51995146 |
| TDLATLG | ASRSRDKNTLY | 0.5039963 |
| SAGGTGR | ASRSRDKNTLY | 0.4352376 |
| GFSNSGS | ASRSRDKNTLY | 0.12221113 |
| GGNGGSGGTSV | ASRSRDKNTLY | 0.6110229 |
| GTSVATGGAGN | ASRSRDKNTLY | 0.3122452 |
| NGGNGGSGGTS | ASRSRDKNTLY | 0.76743305 |
| AVVKFPH | ASRSRDKNTLY | 0.30456945 |
| TPPPPPP | ASRSRDKNTLY | 0.23592257 |
| AAPAAAAAPA | ASRSRDKNTLY | 0.1738297 |
| IALNEDL | ASRSRDKNTLY | 0.15246175 |
| IVNVSSM | ASRSRDKNTLY | 0.10183822 |
| TGIPPAPR | ASRSRDKNTLY | 0.39501047 |
| FDLGGGTF | ASRSRDKNTLY | 0.17272861 |
| FDLGGGTFDVS | ASRSRDKNTLY | 0.31766832 |
| NEPTAAA | ASRSRDKNTLY | 0.61452365 |
| NEPTAAALA | ASRSRDKNTLY | 0.10966022 |
| GSIINISSI | ASRSRDKNTLY | 0.67433804 |
| VAPGNAGTA | ASRSRDKNTLY | 0.16347523 |
| TGGGASLE | ASRSRDKNTLY | 0.14746787 |
| VAVYNHYKRI | ASRSRDKNTLY | 0.65500605 |
| TDPAPATT | ASRSRDKNTLY | 0.117486246 |
| TNRPDILDPALLRPG | ASRSRDKNTLY | 0.72771704 |
| LTTTTTS | ASRSRDKNTLY | 0.10229373 |
| PTGSGKTL | ASRSRDKNTLY | 0.5251785 |
| TGSGKTL | ASRSRDKNTLY | 0.55438393 |
| TIGHVDHGK | ASRSRDKNTLY | 0.09057316 |
| TIGHVDHGKTT | ASRSRDKNTLY | 0.12150376 |
| GHVDHGKTT | ASRSRDKNTLY | 0.6751397 |
| AQFEHTLL | ASRSRDKNTLY | 0.21380739 |
| IVNQDEL | ASRSRDKNTLY | 0.5388138 |
| TNGSQFF | ASRSRDKNTLY | 0.106469065 |
| TGKGIVE | ASRSRDKNTLY | 0.116736084 |
| LISNASDA | ASRSRDKNTLY | 0.15552479 |
| AELRTLGELTKIAKAHGAQ | ASRSRDKNTLY | 0.2654817 |
| APNAVAAAEPSW | ASRSRDKNTLY | 0.14208715 |
| APQVAGAATAVL | ASRSRDKNTLY | 0.29750693 |
| ARRDPAAAAAILPTDA | ASRSRDKNTLY | 0.120270886 |
| DAAARDVLAEAGLAEYF | ASRSRDKNTLY | 0.11012417 |
| DDQIVLSAKANLAKVEQ | ASRSRDKNTLY | 0.11584417 |
| DELLPDTGEDSDEDGH | ASRSRDKNTLY | 0.13355829 |
| ELPEAPANRATVPQD | ASRSRDKNTLY | 0.183788 |
| FGQSKIERVTGIG | ASRSRDKNTLY | 0.11083106 |
| GAGGTGSTAGGAGGAGGAGGLYA | ASRSRDKNTLY | 0.103048064 |
| GHGGTGAAGGNGGTGGAGGSGIDG | ASRSRDKNTLY | 0.13310437 |
| GNGGNGGAGGEGGPGGAGGAGA | ASRSRDKNTLY | 0.14731343 |

| | | |
|---|---|---|
| GSTAGNGGNGGNATNGTIAGQPAG | ASRSRDKNTLY | 0.34501058 |
| GTGGDGGNGGNGGTGG | ASRSRDKNTLY | 0.33840215 |
| GVRRPAGIPADAPV | ASRSRDKNTLY | 0.3701688 |
| GYRQVTDAHLLTLARR | ASRSRDKNTLY | 0.28094065 |
| IAKMGQDGHDRG | ASRSRDKNTLY | 0.14533025 |
| KAEPSIEIQYT | ASRSRDKNTLY | 0.09302949 |
| KPVRIRMAADEDTVAVT | ASRSRDKNTLY | 0.11909805 |
| MPPRIAGMRLLVIKPEP | ASRSRDKNTLY | 0.109993674 |
| MKFYAVALFRSRGVPPTR | ASRSRDKNTLY | 0.19073918 |
| NAGGFGILGAVAHSP | ASRSRDKNTLY | 0.09060663 |
| NEGREITLAMAEM | ASRSRDKNTLY | 0.13612792 |
| NELVEWSHPLGHRWRWE | ASRSRDKNTLY | 0.15419696 |
| NLGAGGTGGNGGNPG | ASRSRDKNTLY | 0.098488525 |
| NTEWPKWETVMAADPVD | ASRSRDKNTLY | 0.3951534 |
| PASAAPAETQP | ASRSRDKNTLY | 0.10327485 |
| QQSSDGKNGVRAAEITGE | ASRSRDKNTLY | 0.11082547 |
| RAPDAVTTSWWKE | ASRSRDKNTLY | 0.11811496 |
| SGFGNVGAHTSGWFNQ | ASRSRDKNTLY | 0.2924067 |
| SGVAGAVGQAASVGGLKV | ASRSRDKNTLY | 0.11402987 |
| SNGGAGTGSAGGNGGT | ASRSRDKNTLY | 0.23969029 |
| TGGAGGEGGAAG | ASRSRDKNTLY | 0.16363695 |
| TGGDGGAAGAGGAPGHG | ASRSRDKNTLY | 0.17548099 |
| TMPPLLAAIANAGRHGV | ASRSRDKNTLY | 0.107774556 |
| TSSTAGNSGGA | ASRSRDKNTLY | 0.21837196 |
| VPATVPVGILC | ASRSRDKNTLY | 0.13750057 |
| VQRPDSGGVVLTGRIS | ASRSRDKNTLY | 0.69906545 |
| WTPNPVPATKGVPAGLVQAT | ASRSRDKNTLY | 0.15699959 |
| YALKRALAKFSDRFDVVII | ASRSRDKNTLY | 0.14779438 |
| YLIGSVIV | ASRSRDKNTLY | 0.11744326 |
| YGQLVAVID | ASRSRDKNTLY | 0.12695715 |
| VTVGGMTLPA | ASRSRDKNTLY | 0.098082565 |
| VLALPAV | ASRSRDKNTLY | 0.14691605 |
| VIPPTLNYET | ASRSRDKNTLY | 0.11017726 |
| VGMGGMHPSSG | ASRSRDKNTLY | 0.14064595 |
| VGLAGGAATA | ASRSRDKNTLY | 0.20946665 |
| VGGAGGAGGSGP | ASRSRDKNTLY | 0.10200342 |
| VFAAFMV | ASRSRDKNTLY | 0.2332821 |
| TLYDKLR | ASRSRDKNTLY | 0.10264733 |
| TLAVFTVL | ASRSRDKNTLY | 0.11568335 |
| TGSVQPARV | ASRSRDKNTLY | 0.23232368 |
| TGGGAGSGGT | ASRSRDKNTLY | 0.098869234 |
| SHLETSL | ASRSRDKNTLY | 0.53898925 |
| SFTTGMT | ASRSRDKNTLY | 0.09539784 |
| QVEVAKRMGVRQPTVS | ASRSRDKNTLY | 0.18028264 |
| QGSGGPALAM | ASRSRDKNTLY | 0.20251502 |
| MLAPSNP | ASRSRDKNTLY | 0.094396815 |
| LVNNAAI | ASRSRDKNTLY | 0.103233576 |
| LTGEINTLA | ASRSRDKNTLY | 0.29204857 |
| LSIFGPV | ASRSRDKNTLY | 0.17068176 |
| LLALLANL | ASRSRDKNTLY | 0.3139353 |
| LISNASDALDKLR | ASRSRDKNTLY | 0.118164 |

| | | |
|---|---|---|
| LINEAVP | ASRSRDKNTLY | 0.13705204 |
| LGYAFSGL | ASRSRDKNTLY | 0.16507417 |
| LDFVAVAL | ASRSRDKNTLY | 0.17085832 |
| LAQPGAE | ASRSRDKNTLY | 0.27429003 |
| ISNLGAVTGF | ASRSRDKNTLY | 0.16452268 |
| HQEDPGRI | ASRSRDKNTLY | 0.13609849 |
| GTPGEPTP | ASRSRDKNTLY | 0.1456303 |
| GFFNSGS | ASRSRDKNTLY | 0.1213408 |
| GAGGGGLTAGAN | ASRSRDKNTLY | 0.16574337 |
| FVDALVNM | ASRSRDKNTLY | 0.14793727 |
| FLTVAPEL | ASRSRDKNTLY | 0.15321769 |
| FETLTAL | ASRSRDKNTLY | 0.11216969 |
| DVINPIEL | ASRSRDKNTLY | 0.11568723 |
| APVSGAPV | ASRSRDKNTLY | 0.17035389 |
| ALVGGLAW | ASRSRDKNTLY | 0.38292944 |
| ALAAAAATAI | ASRSRDKNTLY | 0.3452711 |
| AFGLGLLL | ASRSRDKNTLY | 0.17370479 |
| AFFAAASL | ASRSRDKNTLY | 0.1943922 |
| AAAALPAFSPP | ASRSRDKNTLY | 0.117308654 |
| VVLAPAL | ASRSRDKNTLY | 0.10144714 |
| TIGHVDHGKTTL | ASRSRDKNTLY | 0.10828369 |
| LISNASDALDK | ASRSRDKNTLY | 0.13794184 |
| LISNASDAL | ASRSRDKNTLY | 0.15225047 |
| ISNASDALDK | ASRSRDKNTLY | 0.09744672 |
| NQAETLVYQTE | ASRSRDKNTLY | 0.16460034 |
| TGGTGGTAGAG | ASRSRDKNTLY | 0.13491859 |
| TGSTAGGAGGA | ASRSRDKNTLY | 0.124909714 |
| DPLAVPVAAGPV | ASRSRDKNTLY | 0.33509827 |
| GGTGGTPGNGGN | ASRSRDKNTLY | 0.11850537 |
| PAGLAGVDPATL | ASRSRDKNTLY | 0.12892953 |
| GNNLVVEAPHAGASV | ASRSRDKNTLY | 0.14463766 |
| KVTRIMSASTLEQPA | ASRSRDKNTLY | 0.13354026 |
| LGFALPIALPRRTVV | ASRSRDKNTLY | 0.12630196 |
| TRLGNLTARKLVVLV | ASRSRDKNTLY | 0.33683512 |
| WSQQRRLIYFPSAGP | ASRSRDKNTLY | 0.10933912 |
| GPNGNPDPMAAAVDIR | ASRSRDKNTLY | 0.19015065 |
| TQRCVSVADLRAAASR | ASRSRDKNTLY | 0.13026139 |
| QFVLDVAEHPVEVLTETV | ASRSRDKNTLY | 0.15940645 |
| GGAGGTGGAGGNGGTQAGDG | ASRSRDKNTLY | 0.195734 |
| PLLATLRAIAVDVDAKPAQV | ASRSRDKNTLY | 0.62619746 |
| APKFRTGGYMIDFWGVGYQVAKRM | ASRSRDKNTLY | 0.16788083 |
| GAGGAGGAGGAAGGPGGTGGTGGN | ASRSRDKNTLY | 0.16639256 |
| IGALGLLATDVAPAGSVAFGVALA | ASRSRDKNTLY | 0.1230894 |
| AGGAGGIGGNGGAGGAGGVLMGNG | ASRSRDKNTLY | 0.20089357 |
| ADELIGLL | ASRSRDKNTLY | 0.112845495 |
| ADRALKA | ASRSRDKNTLY | 0.15113907 |
| AGAVGQVGGAGGA | ASRSRDKNTLY | 0.101701036 |
| AGGGGGLGGAP | ASRSRDKNTLY | 0.14816166 |
| AGGLLYGNGGAG | ASRSRDKNTLY | 0.15245706 |
| AGLESIGL | ASRSRDKNTLY | 0.11139767 |
| AGLKTGII | ASRSRDKNTLY | 0.11553242 |

| | | |
|---|---|---|
| ASLAALLL | ASRSRDKNTLY | 0.123638585 |
| ASVINISSI | ASRSRDKNTLY | 0.12951818 |
| ATPPGPA | ASRSRDKNTLY | 0.19716391 |
| ATVASALV | ASRSRDKNTLY | 0.1439199 |
| ATVASLIL | ASRSRDKNTLY | 0.10274558 |
| AVNGPVLIRH | ASRSRDKNTLY | 0.112567306 |
| DGAAGAVGI | ASRSRDKNTLY | 0.11202614 |
| DPLLHGI | ASRSRDKNTLY | 0.21870604 |
| ELGAALGAL | ASRSRDKNTLY | 0.10345838 |
| ELGRTQAV | ASRSRDKNTLY | 0.13150921 |
| FVAGGYLS | ASRSRDKNTLY | 0.17511262 |
| GASGGAGGAGGW | ASRSRDKNTLY | 0.46180895 |
| GGAGAGGDGGAGGI | ASRSRDKNTLY | 0.67177653 |
| GTGHNGSQPGMGGNG | ASRSRDKNTLY | 0.12467122 |
| GYAAASASAS | ASRSRDKNTLY | 0.14623368 |
| HQVDVAAVQQG | ASRSRDKNTLY | 0.3470293 |
| IGAAIAHT | ASRSRDKNTLY | 0.5916104 |
| IGMIEVAQL | ASRSRDKNTLY | 0.2664821 |
| IRLAPEHTSAMD | ASRSRDKNTLY | 0.56357884 |
| IYNVSVL | ASRSRDKNTLY | 0.14350303 |
| KQGFLGM | ASRSRDKNTLY | 0.13107781 |
| LESVITQL | ASRSRDKNTLY | 0.14124103 |
| LGLGALVVV | ASRSRDKNTLY | 0.17506601 |
| LIGNGGNGGSGGI | ASRSRDKNTLY | 0.42037654 |
| LNAINAPTL | ASRSRDKNTLY | 0.11940331 |
| LQLLNTGV | ASRSRDKNTLY | 0.109189466 |
| LQTSIEF | ASRSRDKNTLY | 0.22931574 |
| MSFAVSEL | ASRSRDKNTLY | 0.35949206 |
| NTGALGL | ASRSRDKNTLY | 0.6331213 |
| PPGAGGAGGAGTT | ASRSRDKNTLY | 0.61840934 |
| QLVGLGI | ASRSRDKNTLY | 0.6964078 |
| RFLEQVHQL | ASRSRDKNTLY | 0.4860309 |
| RGVGYVLR | ASRSRDKNTLY | 0.42770407 |
| SATGVFAGVI | ASRSRDKNTLY | 0.097119756 |
| SLPAITLP | ASRSRDKNTLY | 0.14524925 |
| STALAFTV | ASRSRDKNTLY | 0.123830855 |
| SVLTVGLGK | ASRSRDKNTLY | 0.29060206 |
| TALAGAGSGI | ASRSRDKNTLY | 0.094488755 |
| TGLPFTGI | ASRSRDKNTLY | 0.19539946 |
| VAEHVTV | ASRSRDKNTLY | 0.23135108 |
| VDIVLNSL | ASRSRDKNTLY | 0.10384313 |
| VLTGGGAVL | ASRSRDKNTLY | 0.552717 |
| VNVGLAYGI | ASRSRDKNTLY | 0.1074761 |
| VSSAAVGAV | ASRSRDKNTLY | 0.12575953 |
| VTSVYEAGV | ASRSRDKNTLY | 0.14213 |
| WVALDRGA | ASRSRDKNTLY | 0.54796207 |
| YLVGSVVI | ASRSRDKNTLY | 0.13834108 |
| GPPGTGKTLL | ASRSRDKNTLY | 0.37105668 |
| GVGMTKF | ASRSRDKNTLY | 0.37557715 |
| IGIGGSDL | ASRSRDKNTLY | 0.302166 |
| IGTLLAEL | ASRSRDKNTLY | 0.12085868 |

| | | |
|---|---|---|
| IYTSGTT | ASRSRDKNTLY | 0.40286058 |
| LSFPAGV | ASRSRDKNTLY | 0.20767623 |
| LYTSGTT | ASRSRDKNTLY | 0.71516794 |
| VGAFPFGF | ASRSRDKNTLY | 0.24840496 |
| VGAGTVEF | ASRSRDKNTLY | 0.1921508 |
| VGKSTLLL | ASRSRDKNTLY | 0.14468256 |
| AALAHAGGGIV | ASRSRDKNTLY | 0.14797224 |
| AALVAVSLL | ASRSRDKNTLY | 0.10186925 |
| AAVSRAVAAL | ASRSRDKNTLY | 0.22732173 |
| AGGSSGIGGF | ASRSRDKNTLY | 0.13533089 |
| AGLTVANANG | ASRSRDKNTLY | 0.22372097 |
| AGLVGAVL | ASRSRDKNTLY | 0.4210655 |
| ALSQAFTV | ASRSRDKNTLY | 0.11857872 |
| ALTFSVSL | ASRSRDKNTLY | 0.45596385 |
| ALTGAALVAL | ASRSRDKNTLY | 0.14361775 |
| ASFSIGAGVL | ASRSRDKNTLY | 0.14530542 |
| DGSVYKI | ASRSRDKNTLY | 0.54769015 |
| EQLAEIV | ASRSRDKNTLY | 0.112725824 |
| FAVSELTVM | ASRSRDKNTLY | 0.6693806 |
| GAAPLAAGL | ASRSRDKNTLY | 0.10112134 |
| GADGGDGGSGGAGG | ASRSRDKNTLY | 0.2764158 |
| GFGLDTGGAGG | ASRSRDKNTLY | 0.3961482 |
| GLASGGLL | ASRSRDKNTLY | 0.0916551 |
| GRIAGAAVLL | ASRSRDKNTLY | 0.11330758 |
| GSAGSLNTSF | ASRSRDKNTLY | 0.5230771 |
| GTGQIGI | ASRSRDKNTLY | 0.7362139 |
| GTLIALIL | ASRSRDKNTLY | 0.69605607 |
| IAATMGFVVG | ASRSRDKNTLY | 0.36224535 |
| IFAQALL | ASRSRDKNTLY | 0.3761687 |
| IGAAVALGF | ASRSRDKNTLY | 0.49668834 |
| IGLTPLAL | ASRSRDKNTLY | 0.7130529 |
| ILGSNTSTL | ASRSRDKNTLY | 0.7375077 |
| IVFSLTVL | ASRSRDKNTLY | 0.7218815 |
| LASKATGF | ASRSRDKNTLY | 0.36801282 |
| LASVALIL | ASRSRDKNTLY | 0.57638115 |
| LFAVGLTL | ASRSRDKNTLY | 0.49967575 |
| LFNGGIGTYI | ASRSRDKNTLY | 0.54743147 |
| LTLGAGII | ASRSRDKNTLY | 0.6064114 |
| LVAAALGDI | ASRSRDKNTLY | 0.3748774 |
| NTAFVQLGI | ASRSRDKNTLY | 0.37894103 |
| QLFSGAEL | ASRSRDKNTLY | 0.40613976 |
| SAATFITL | ASRSRDKNTLY | 0.5597061 |
| SNYGLANLGV | ASRSRDKNTLY | 0.7124596 |
| SYGGAYAVM | ASRSRDKNTLY | 0.68402874 |
| TTGVGGLTLG | ASRSRDKNTLY | 0.7898314 |
| TTQTIYI | ASRSRDKNTLY | 0.7614096 |
| VAAGAGLGAVI | ASRSRDKNTLY | 0.61378497 |
| VAAVVTLL | ASRSRDKNTLY | 0.15282373 |
| VGFLGGLI | ASRSRDKNTLY | 0.25741673 |
| VGVIPILL | ASRSRDKNTLY | 0.3529833 |
| VIFASSLI | ASRSRDKNTLY | 0.46830583 |

| | | |
|---|---|---|
| VKGGDGGAAAG | ASRSRDKNTLY | 0.18476957 |
| VTKTFTNI | ASRSRDKNTLY | 0.5482969 |
| VVAAAILL | ASRSRDKNTLY | 0.5590109 |
| YAALDVEL | ASRSRDKNTLY | 0.30810803 |
| LASVALLL | ASRSRDKNTLY | 0.72308254 |
| LDAVHRV | ASRSRDKNTLY | 0.33256415 |
| QGLSSSIF | ASRSRDKNTLY | 0.33946693 |
| RGTFANIRL | ASRSRDKNTLY | 0.3933589 |
| RQLWWGHRI | ASRSRDKNTLY | 0.74294317 |
| YGPPGTGKTLL | ASRSRDKNTLY | 0.38084972 |
| AALVGAPANVA | ASRSRDKNTLY | 0.70779914 |
| AATFRRTE | ASRSRDKNTLY | 0.4928192 |
| AGALGGGI | ASRSRDKNTLY | 0.77378786 |
| AGGAGGAGGIGAG | ASRSRDKNTLY | 0.30500397 |
| AGSGALGL | ASRSRDKNTLY | 0.7461934 |
| AQIADQI | ASRSRDKNTLY | 0.6164676 |
| ASAVAVTYL | ASRSRDKNTLY | 0.81726885 |
| ASTVLGGTGG | ASRSRDKNTLY | 0.48481315 |
| AVGGEPAK | ASRSRDKNTLY | 0.5268997 |
| CEQGTFGGTC | ASRSRDKNTLY | 0.33608723 |
| DEGKAPAG | ASRSRDKNTLY | 0.30369207 |
| DTVVFPAL | ASRSRDKNTLY | 0.23830177 |
| FEGITMAG | ASRSRDKNTLY | 0.55208254 |
| FPLVVQD | ASRSRDKNTLY | 0.29861322 |
| GAFTTPQ | ASRSRDKNTLY | 0.6011554 |
| GFGGFGTAGGVGGA | ASRSRDKNTLY | 0.74670744 |
| GFLAQFL | ASRSRDKNTLY | 0.19877973 |
| GGAGGAGGNAGVA | ASRSRDKNTLY | 0.69721127 |
| GGVGTAGAGGAGGAGG | ASRSRDKNTLY | 0.30468163 |
| GNGGGGDAGNAGSGGN | ASRSRDKNTLY | 0.36542943 |
| GTGGAGGAGNDAGS | ASRSRDKNTLY | 0.6522134 |
| GTTNTGWFNS | ASRSRDKNTLY | 0.36657774 |
| IFMGPLFTL | ASRSRDKNTLY | 0.7458481 |
| ILAHAIEAI | ASRSRDKNTLY | 0.3304994 |
| IRSAIREAL | ASRSRDKNTLY | 0.60441923 |
| ITFAGGG | ASRSRDKNTLY | 0.6985954 |
| ITGMSGHQEV | ASRSRDKNTLY | 0.15468949 |
| IVAGLACTL | ASRSRDKNTLY | 0.27728835 |
| IVGGGLAGA | ASRSRDKNTLY | 0.793347 |
| IVTGVLNI | ASRSRDKNTLY | 0.097359486 |
| LAQQGCI | ASRSRDKNTLY | 0.095927015 |
| LCLHVEN | ASRSRDKNTLY | 0.09600718 |
| LDAISDGS | ASRSRDKNTLY | 0.10242301 |
| LGGFAAML | ASRSRDKNTLY | 0.09681923 |
| LGGKNPMI | ASRSRDKNTLY | 0.096768 |
| LGPKGVAAIL | ASRSRDKNTLY | 0.097788185 |
| LMVGVSI | ASRSRDKNTLY | 0.09758431 |
| LVGAAFVL | ASRSRDKNTLY | 0.1049133 |
| LVLAVAPL | ASRSRDKNTLY | 0.10108026 |
| MAAIAVIDAA | ASRSRDKNTLY | 0.09652398 |
| NNGINAELQ | ASRSRDKNTLY | 0.10152556 |

| | | |
|---|---|---|
| RGEPLVLVHG | ASRSRDKNTLY | 0.09629015 |
| RGGAGGAAGLI | ASRSRDKNTLY | 0.10085558 |
| RSYRMHPAV | ASRSRDKNTLY | 0.095885985 |
| SAAHLLAGQLA | ASRSRDKNTLY | 0.09585853 |
| SAAVGSY | ASRSRDKNTLY | 0.09741974 |
| SPHAVLHRI | ASRSRDKNTLY | 0.10081574 |
| SQIAKVL | ASRSRDKNTLY | 0.09919756 |
| TALGHLVP | ASRSRDKNTLY | 0.096721955 |
| TAPTTSGAA | ASRSRDKNTLY | 0.09719895 |
| TDGDDGNGGSPGT | ASRSRDKNTLY | 0.095783696 |
| TDGLGEN | ASRSRDKNTLY | 0.09684462 |
| TGGPPLPGQA | ASRSRDKNTLY | 0.09727346 |
| TIPAITI | ASRSRDKNTLY | 0.0974345 |
| TIPGFTI | ASRSRDKNTLY | 0.09837051 |
| TSIASGALA | ASRSRDKNTLY | 0.09500214 |
| TVTLPGAAI | ASRSRDKNTLY | 0.09982203 |
| VASGLSAALP | ASRSRDKNTLY | 0.09797387 |
| VGAGVIGI | ASRSRDKNTLY | 0.105626956 |
| VGPKRFGV | ASRSRDKNTLY | 0.101229645 |
| VNVAFPD | ASRSRDKNTLY | 0.09760451 |
| VSMTNSGVS | ASRSRDKNTLY | 0.09857084 |
| VTDVLGDLSV | ASRSRDKNTLY | 0.096821986 |
| VVVVGAGFAGL | ASRSRDKNTLY | 0.09735276 |
| VYTVGTVV | ASRSRDKNTLY | 0.09621099 |
| WTTFSGP | ASRSRDKNTLY | 0.10349455 |
| YDELVEAF | ASRSRDKNTLY | 0.09548491 |
| YGTGGAGGNGG | ASRSRDKNTLY | 0.10389538 |
| YTNVMAV | ASRSRDKNTLY | 0.10402376 |
| FLEQVHQL | ASRSRDKNTLY | 0.096932 |
| GLSLLAVAV | ASRSRDKNTLY | 0.09737784 |
| LEQVHQL | ASRSRDKNTLY | 0.10300187 |
| LNAAGAI | ASRSRDKNTLY | 0.10114515 |
| PAVHASL | ASRSRDKNTLY | 0.10913055 |
| TELEDDA | ASRSRDKNTLY | 0.09657811 |
| TPPGPPN | ASRSRDKNTLY | 0.09676823 |
| TPPPPPPP | ASRSRDKNTLY | 0.10123264 |
| TQSPALI | ASGGGESQNTLY | 0.09682982 |
| EFAAAFPQ | ASGGGESQNTLY | 0.0965349 |
| LLSGNLDVL | ASGGGESQNTLY | 0.09694377 |
| GPANLGNYNI | ASGGGESQNTLY | 0.09833757 |
| TLGPTLP | ASGGGESQNTLY | 0.10190959 |
| VYAYSSCCK | ASGGGESQNTLY | 0.0958623 |
| APAPAPAPNI | ASGGGESQNTLY | 0.10289339 |
| PGGPVPIP | ASGGGESQNTLY | 0.09736547 |
| FLTVALEP | ASGGGESQNTLY | 0.09614646 |
| SQYFGANL | ASGGGESQNTLY | 0.10056302 |
| GGVKPAA | ASGGGESQNTLY | 0.095429264 |
| GGTGGAGGAGGW | ASGGGESQNTLY | 0.09540208 |
| GVGGVAGLGGAGG | ASGGGESQNTLY | 0.095876664 |
| VLLVLATTI | ASGGGESQNTLY | 0.10101723 |
| IGIFAAML | ASGGGESQNTLY | 0.10646643 |

| | | |
|---|---|---|
| AVGDLID | ASGGGESQNTLY | 0.097117886 |
| NISPLPL | ASGGGESQNTLY | 0.097369865 |
| VTAVFVAI | ASGGGESQNTLY | 0.76944983 |
| SGDDNDAAL | ASGGGESQNTLY | 0.78918165 |
| AVAALAHS | ASGGGESQNTLY | 0.4891146 |
| GGVTVMATP | ASGGGESQNTLY | 0.25240576 |
| AGLAANVRD | ASGGGESQNTLY | 0.6663124 |
| TNRPDLIDPALLRPG | ASGGGESQNTLY | 0.76277435 |
| LANLGVTAL | ASGGGESQNTLY | 0.8064282 |
| TGERIEL | ASGGGESQNTLY | 0.77546084 |
| NGEELAA | ASGGGESQNTLY | 0.2663224 |
| VLSLVWG | ASGGGESQNTLY | 0.52115846 |
| IGYALWL | ASGGGESQNTLY | 0.68099445 |
| LSLLLAPL | ASGGGESQNTLY | 0.42073926 |
| TLDEALK | ASGGGESQNTLY | 0.71201795 |
| TDGAGTAEGG | ASGGGESQNTLY | 0.24405685 |
| NAPGGAGGNGG | ASGGGESQNTLY | 0.6221057 |
| VSVFMGMGGNF | ASGGGESQNTLY | 0.4018857 |
| LASYLAAL | ASGGGESQNTLY | 0.5892919 |
| LVGGTPAL | ASGGGESQNTLY | 0.743486 |
| VRNADVL | ASGGGESQNTLY | 0.69969577 |
| TCSSPFLVSAM | ASGGGESQNTLY | 0.82859635 |
| VHITDAE | ASGGGESQNTLY | 0.79426193 |
| EYAAPSF | ASGGGESQNTLY | 0.64446914 |
| VVDAFHGGL | ASGGGESQNTLY | 0.15810418 |
| ASVVDPQ | ASGGGESQNTLY | 0.20720199 |
| SYNEFGGLS | ASGGGESQNTLY | 0.43534595 |
| DPDGSVY | ASGGGESQNTLY | 0.43017793 |
| MVSTIHATE | ASGGGESQNTLY | 0.18052892 |
| LAAGVADPV | ASGGGESQNTLY | 0.5693029 |
| TKEQIAA | ASGGGESQNTLY | 0.74055755 |
| SLASPAAL | ASGGGESQNTLY | 0.22829154 |
| IGLVFYGV | ASGGGESQNTLY | 0.70968604 |
| FLLDASNL | ASGGGESQNTLY | 0.38805905 |
| YQVGNFP | ASGGGESQNTLY | 0.24139817 |
| ESLLSQST | ASGGGESQNTLY | 0.3511552 |
| LFTVGFAAVG | ASGGGESQNTLY | 0.815985 |
| ATPPTPT | ASGGGESQNTLY | 0.27153894 |
| TDLATLG | ASGGGESQNTLY | 0.626843 |
| SAGGTGR | ASGGGESQNTLY | 0.698065 |
| GFSNSGS | ASGGGESQNTLY | 0.76196855 |
| GGNGGSGGTSV | ASGGGESQNTLY | 0.1913384 |
| GTSVATGGAGN | ASGGGESQNTLY | 0.7348963 |
| NGGNGGSGGTS | ASGGGESQNTLY | 0.5385004 |
| AVVKFPH | ASGGGESQNTLY | 0.80001444 |
| TPPPPPP | ASGGGESQNTLY | 0.31731886 |
| AAPAAAAAPA | ASGGGESQNTLY | 0.42407155 |
| IALNEDL | ASGGGESQNTLY | 0.3731457 |
| IVNVSSM | ASGGGESQNTLY | 0.7903051 |
| TGIPPAPR | ASGGGESQNTLY | 0.18909882 |
| FDLGGGTF | ASGGGESQNTLY | 0.68925905 |

| | | |
|---|---|---|
| FDLGGGTFDVS | ASGGGESQNTLY | 0.3652189 |
| NEPTAAA | ASGGGESQNTLY | 0.73595613 |
| NEPTAAALA | ASGGGESQNTLY | 0.76443547 |
| GSIINISSI | ASGGGESQNTLY | 0.22656342 |
| VAPGNAGTA | ASGGGESQNTLY | 0.80890036 |
| TGGGASLE | ASGGGESQNTLY | 0.4878326 |
| VAVYNHYKRI | ASGGGESQNTLY | 0.3149414 |
| TDPAPATT | ASGGGESQNTLY | 0.7013258 |
| TNRPDILDPALLRPG | ASGGGESQNTLY | 0.33775216 |
| LTTTTTS | ASGGGESQNTLY | 0.8007848 |
| PTGSGKTL | ASGGGESQNTLY | 0.2855752 |
| TGSGKTL | ASGGGESQNTLY | 0.7850669 |
| TIGHVDHGK | ASGGGESQNTLY | 0.78302205 |
| TIGHVDHGKTT | ASGGGESQNTLY | 0.20518492 |
| GHVDHGKTT | ASGGGESQNTLY | 0.3352607 |
| AQFEHTLL | ASGGGESQNTLY | 0.7794646 |
| IVNQDEL | ASGGGESQNTLY | 0.66984904 |
| TNGSQFF | ASGGGESQNTLY | 0.72998494 |
| TGKGIVE | ASGGGESQNTLY | 0.56525105 |
| LISNASDA | ASGGGESQNTLY | 0.47873425 |
| AELRTLGELTKIAKAHGAQ | ASGGGESQNTLY | 0.48700327 |
| APNAVAAAEPSW | ASGGGESQNTLY | 0.65037715 |
| APQVAGAATAVL | ASGGGESQNTLY | 0.7346356 |
| ARRDPAAAAAILPTDA | ASGGGESQNTLY | 0.66203034 |
| DAAARDVLAEAGLAEYF | ASGGGESQNTLY | 0.31235248 |
| DDQIVLSAKANLAKVEQ | ASGGGESQNTLY | 0.55454195 |
| DELLPDTGEDSDEDGH | ASGGGESQNTLY | 0.5502907 |
| ELPEAPANRATVPQD | ASGGGESQNTLY | 0.50092655 |
| FGQSKIERVTGIG | ASGGGESQNTLY | 0.6475247 |
| GAGGTGSTAGGAGGAGGAGGLYA | ASGGGESQNTLY | 0.5401084 |
| GHGGTGAAGGNGGTGGAGGSGIDG | ASGGGESQNTLY | 0.558158 |
| GNGGNGGAGGEGGPGGAGGAGA | ASGGGESQNTLY | 0.4721923 |
| GSTAGNGGNGGNATNGTIAGQPAG | ASGGGESQNTLY | 0.45253572 |
| GTGGDGGNGGNGGTGG | ASGGGESQNTLY | 0.6315354 |
| GVRRPAGIPADAPV | ASGGGESQNTLY | 0.6625509 |
| GYRQVTDAHLLTLARR | ASGGGESQNTLY | 0.6807482 |
| IAKMGQDGHDRG | ASGGGESQNTLY | 0.7144505 |
| KAEPSIEIQYT | ASGGGESQNTLY | 0.5694152 |
| KPVRIRMAADEDTVAVT | ASGGGESQNTLY | 0.22149988 |
| MPPRIAGMRLLVIKPEP | ASGGGESQNTLY | 0.33615416 |
| MKFYAVALFRSRGVPPTR | ASGGGESQNTLY | 0.5197607 |
| NAGGFGILGAVAHSP | ASGGGESQNTLY | 0.5460726 |
| NEGREITLAMAEM | ASGGGESQNTLY | 0.25790924 |
| NELVEWSHPLGHRWRWE | ASGGGESQNTLY | 0.5394257 |
| NLGAGGTGGNGGNPG | ASGGGESQNTLY | 0.5784446 |
| NTEWPKWETVMAADPVD | ASGGGESQNTLY | 0.44237956 |
| PASAAPAETQP | ASGGGESQNTLY | 0.68904626 |
| QQSSDGKNGVRAAEITGE | ASGGGESQNTLY | 0.5636208 |
| RAPDAVTTSWWKE | ASGGGESQNTLY | 0.40467638 |
| SGFGNVGAHTSGWFNQ | ASGGGESQNTLY | 0.35048276 |
| SGVAGAVGQAASVGGLKV | ASGGGESQNTLY | 0.52636015 |

| | | |
|---|---|---|
| SNGGAGTGSAGGNGGT | ASGGGESQNTLY | 0.3478608 |
| TGGAGGEGGAAG | ASGGGESQNTLY | 0.71306205 |
| TGGDGGAAGAGGAPGHG | ASGGGESQNTLY | 0.3396602 |
| TMPPLLAAIANAGRHGV | ASGGGESQNTLY | 0.62753046 |
| TSSTAGNSGGA | ASGGGESQNTLY | 0.31275648 |
| VPATVPVGILC | ASGGGESQNTLY | 0.6769369 |
| VQRPDSGGVVLTGRIS | ASGGGESQNTLY | 0.6585382 |
| WTPNPVPATKGVPAGLVQAT | ASGGGESQNTLY | 0.7547952 |
| YALKRALAKFSDRFDVVII | ASGGGESQNTLY | 0.5934878 |
| YLIGSVIV | ASGGGESQNTLY | 0.5184586 |
| YGQLVAVID | ASGGGESQNTLY | 0.3138153 |
| VTVGGMTLPA | ASGGGESQNTLY | 0.7246554 |
| VLALPAV | ASGGGESQNTLY | 0.33689272 |
| VIPPTLNYET | ASGGGESQNTLY | 0.51122797 |
| VGMGGMHPSSG | ASGGGESQNTLY | 0.3000052 |
| VGLAGGAATA | ASGGGESQNTLY | 0.5615942 |
| VGGAGGAGGSGP | ASGGGESQNTLY | 0.67441726 |
| VFAAFMV | ASGGGESQNTLY | 0.41656673 |
| TLYDKLR | ASGGGESQNTLY | 0.6810818 |
| TLAVFTVL | ASGGGESQNTLY | 0.59942365 |
| TGSVQPARV | ASGGGESQNTLY | 0.4029992 |
| TGGGAGSGGT | ASGGGESQNTLY | 0.72038805 |
| SHLETSL | ASGGGESQNTLY | 0.4047262 |
| SFTTGMT | ASGGGESQNTLY | 0.73168623 |
| QVEVAKRMGVRQPTVS | ASGGGESQNTLY | 0.39616865 |
| QGSGGPALAM | ASGGGESQNTLY | 0.4562897 |
| MLAPSNP | ASGGGESQNTLY | 0.5011817 |
| LVNNAAI | ASGGGESQNTLY | 0.41655815 |
| LTGEINTLA | ASGGGESQNTLY | 0.3570184 |
| LSIFGPV | ASGGGESQNTLY | 0.7274527 |
| LLALLANL | ASGGGESQNTLY | 0.1652766 |
| LISNASDALDKLR | ASGGGESQNTLY | 0.13607465 |
| LINEAVP | ASGGGESQNTLY | 0.13778742 |
| LGYAFSGL | ASGGGESQNTLY | 0.16461287 |
| LDFVAVAL | ASGGGESQNTLY | 0.13940682 |
| LAQPGAE | ASGGGESQNTLY | 0.1509749 |
| ISNLGAVTGF | ASGGGESQNTLY | 0.14381734 |
| HQEDPGRI | ASGGGESQNTLY | 0.15749444 |
| GTPGEPTP | ASGGGESQNTLY | 0.17752817 |
| GFFNSGS | ASGGGESQNTLY | 0.15959239 |
| GAGGGGLTAGAN | ASGGGESQNTLY | 0.15605074 |
| FVDALVNM | ASGGGESQNTLY | 0.16096441 |
| FLTVAPEL | ASGGGESQNTLY | 0.14504321 |
| FETLTAL | ASGGGESQNTLY | 0.13863905 |
| DVINPIEL | ASGGGESQNTLY | 0.13620955 |
| APVSGAPV | ASGGGESQNTLY | 0.13659549 |
| ALVGGLAW | ASGGGESQNTLY | 0.13687919 |
| ALAAAAATAI | ASGGGESQNTLY | 0.1648161 |
| AFGLGLLL | ASGGGESQNTLY | 0.17027041 |
| AFFAAASL | ASGGGESQNTLY | 0.15994063 |
| AAAALPAFSPP | ASGGGESQNTLY | 0.16788696 |

| | | |
|---|---|---|
| VVLAPAL | ASGGGESQNTLY | 0.13613777 |
| TIGHVDHGKTTL | ASGGGESQNTLY | 0.13257852 |
| LISNASDALDK | ASGGGESQNTLY | 0.13844587 |
| LISNASDAL | ASGGGESQNTLY | 0.14700647 |
| ISNASDALDK | ASGGGESQNTLY | 0.15274917 |
| NQAETLVYQTE | ASGGGESQNTLY | 0.12640795 |
| TGGTGGTAGAG | ASGGGESQNTLY | 0.14780308 |
| TGSTAGGAGGA | ASGGGESQNTLY | 0.16168435 |
| DPLAVPVAAGPV | ASGGGESQNTLY | 0.16823801 |
| GGTGGTPGNGGN | ASGGGESQNTLY | 0.16888216 |
| PAGLAGVDPATL | ASGGGESQNTLY | 0.1420775 |
| GNNLVVEAPHAGASV | ASGGGESQNTLY | 0.14333303 |
| KVTRIMSASTLEQPA | ASGGGESQNTLY | 0.14215545 |
| LGFALPIALPRRTVV | ASGGGESQNTLY | 0.15206856 |
| TRLGNLTARKLVVLV | ASGGGESQNTLY | 0.13866131 |
| WSQQRRLIYFPSAGP | ASGGGESQNTLY | 0.1782976 |
| GPNGNPDPMAAAVDIR | ASGGGESQNTLY | 0.13261518 |
| TQRCVSVADLRAAASR | ASGGGESQNTLY | 0.20189212 |
| QFVLDVAEHPVEVLTETV | ASGGGESQNTLY | 0.16958708 |
| GGAGGTGGAGGNGGTQAGDG | ASGGGESQNTLY | 0.15109527 |
| PLLATLRAIAVDVDAKPAQV | ASGGGESQNTLY | 0.14878155 |
| APKFRTGGYMIDFWGVGYQVAKRM | ASGGGESQNTLY | 0.17814699 |
| GAGGAGGAGGAAGGPGGTGGTGGN | ASGGGESQNTLY | 0.16849469 |
| IGALGLLATDVAPAGSVAFGVALA | ASGGGESQNTLY | 0.1849541 |
| AGGAGGIGGNGGAGGAGGVLMGNG | ASGGGESQNTLY | 0.1345299 |
| ADELIGLL | ASGGGESQNTLY | 0.13446417 |
| ADRALKA | ASGGGESQNTLY | 0.1518798 |
| AGAVGQVGGAGGA | ASGGGESQNTLY | 0.14816177 |
| AGGGGGLGGAP | ASGGGESQNTLY | 0.14154829 |
| AGGLLYGNGGAG | ASGGGESQNTLY | 0.14390914 |
| AGLESIGL | ASGGGESQNTLY | 0.16918543 |
| AGLKTGII | ASGGGESQNTLY | 0.22219953 |
| ASLAALLL | ASGGGESQNTLY | 0.14677882 |
| ASVINISSI | ASGGGESQNTLY | 0.2480611 |
| ATPPGPA | ASGGGESQNTLY | 0.14526607 |
| ATVASALV | ASGGGESQNTLY | 0.14487423 |
| ATVASLIL | ASGGGESQNTLY | 0.14534616 |
| AVNGPVLIRH | ASGGGESQNTLY | 0.1383513 |
| DGAAGAVGI | ASGGGESQNTLY | 0.12913862 |
| DPLLHGI | ASGGGESQNTLY | 0.14222412 |
| ELGAALGAL | ASGGGESQNTLY | 0.17714833 |
| ELGRTQAV | ASGGGESQNTLY | 0.16681004 |
| FVAGGYLS | ASGGGESQNTLY | 0.14061268 |
| GASGGAGGAGGW | ASGGGESQNTLY | 0.16544293 |
| GGAGAGGDGGAGGI | ASGGGESQNTLY | 0.11287062 |
| GTGHNGSQPGMGGNG | ASGGGESQNTLY | 0.11282811 |
| GYAAASASAS | ASGGGESQNTLY | 0.11280303 |
| HQVDVAAVQQG | ASGGGESQNTLY | 0.11322645 |
| IGAAIAHT | ASGGGESQNTLY | 0.11279924 |
| IGMIEVAQL | ASGGGESQNTLY | 0.11285537 |
| IRLAPEHTSAMD | ASGGGESQNTLY | 0.11346253 |

| | | |
|---|---|---|
| IYNVSVL | ASGGGESQNTLY | 0.112892665 |
| KQGFLGM | ASGGGESQNTLY | 0.11348482 |
| LESVITQL | ASGGGESQNTLY | 0.113483 |
| LGLGALVVV | ASGGGESQNTLY | 0.11290392 |
| LIGNGGNGGSGGI | ASGGGESQNTLY | 0.113102846 |
| LNAINAPTL | ASGGGESQNTLY | 0.11283529 |
| LQLLNTGV | ASGGGESQNTLY | 0.11376583 |
| LQTSIEF | ASGGGESQNTLY | 0.112799406 |
| MSFAVSEL | ASGGGESQNTLY | 0.11272233 |
| NTGALGL | ASGGGESQNTLY | 0.113122225 |
| PPGAGGAGGAGTT | ASGGGESQNTLY | 0.11316225 |
| QLVGLGI | ASGGGESQNTLY | 0.11313109 |
| RFLEQVHQL | ASGGGESQNTLY | 0.11289565 |
| RGVGYVLR | ASGGGESQNTLY | 0.11289997 |
| SATGVFAGVI | ASGGGESQNTLY | 0.11288799 |
| SLPAITLP | ASGGGESQNTLY | 0.1129338 |
| STALAFTV | ASGGGESQNTLY | 0.11292282 |
| SVLTVGLGK | ASGGGESQNTLY | 0.112928174 |
| TALAGAGSGI | ASGGGESQNTLY | 0.113091156 |
| TGLPFTGI | ASGGGESQNTLY | 0.11249877 |
| VAEHVTV | ASGGGESQNTLY | 0.11396624 |
| VDIVLNSL | ASGGGESQNTLY | 0.113103874 |
| VLTGGGAVL | ASGGGESQNTLY | 0.113780506 |
| VNVGLAYGI | ASGGGESQNTLY | 0.11334044 |
| VSSAAVGAV | ASGGGESQNTLY | 0.11305399 |
| VTSVYEAGV | ASGGGESQNTLY | 0.113236636 |
| WVALDRGA | ASGGGESQNTLY | 0.112897545 |
| YLVGSVVI | ASGGGESQNTLY | 0.11295234 |
| GPPGTGKTLL | ASGGGESQNTLY | 0.112914726 |
| GVGMTKF | ASGGGESQNTLY | 0.11352775 |
| IGIGGSDL | ASGGGESQNTLY | 0.11271978 |
| IGTLLAEL | ASGGGESQNTLY | 0.1133317 |
| IYTSGTT | ASGGGESQNTLY | 0.113267995 |
| LSFPAGV | ASGGGESQNTLY | 0.11288102 |
| LYTSGTT | ASGGGESQNTLY | 0.11284671 |
| VGAFPFGF | ASGGGESQNTLY | 0.11367 |
| VGAGTVEF | ASGGGESQNTLY | 0.113205925 |
| VGKSTLLL | ASGGGESQNTLY | 0.11377073 |
| AALAHAGGGIV | ASGGGESQNTLY | 0.11315102 |
| AALVAVSLL | ASGGGESQNTLY | 0.113259755 |
| AAVSRAVAAL | ASGGGESQNTLY | 0.1130974 |
| AGGSSGIGGF | ASGGGESQNTLY | 0.11285086 |
| AGLTVANANG | ASGGGESQNTLY | 0.112922944 |
| AGLVGAVL | ASGGGESQNTLY | 0.11288573 |
| ALSQAFTV | ASGGGESQNTLY | 0.11290426 |
| ALTFSVSL | ASGGGESQNTLY | 0.11262851 |
| ALTGAALVAL | ASGGGESQNTLY | 0.11273829 |
| ASFSIGAGVL | ASGGGESQNTLY | 0.11262131 |
| DGSVYKI | ASGGGESQNTLY | 0.113008425 |
| EQLAEIV | ASGGGESQNTLY | 0.11282774 |
| FAVSELTVM | ASGGGESQNTLY | 0.11389737 |

| | | |
|---|---|---|
| GAAPLAAGL | ASGGGESQNTLY | 0.11266615 |
| GADGGDGGSGGAGG | ASGGGESQNTLY | 0.11274795 |
| GFGLDTGGAGG | ASGGGESQNTLY | 0.11276868 |
| GLASGGLL | ASGGGESQNTLY | 0.11295935 |
| GRIAGAAVLL | ASGGGESQNTLY | 0.11293016 |
| GSAGSLNTSF | ASGGGESQNTLY | 0.11308809 |
| GTGQIGI | ASGGGESQNTLY | 0.11288394 |
| GTLIALIL | ASGGGESQNTLY | 0.70587015 |
| IAATMGFVVG | ASGGGESQNTLY | 0.5983963 |
| IFAQALL | ASGGGESQNTLY | 0.31683856 |
| IGAAVALGF | ASGGGESQNTLY | 0.25124124 |
| IGLTPLAL | ASGGGESQNTLY | 0.6162202 |
| ILGSNTSTL | ASGGGESQNTLY | 0.49144834 |
| IVFSLTVL | ASGGGESQNTLY | 0.23009439 |
| LASKATGF | ASGGGESQNTLY | 0.68741757 |
| LASVALIL | ASGGGESQNTLY | 0.18167645 |
| LFAVGLTL | ASGGGESQNTLY | 0.29969576 |
| LFNGGIGTYI | ASGGGESQNTLY | 0.26501435 |
| LTLGAGII | ASGGGESQNTLY | 0.2635424 |
| LVAAALGDI | ASGGGESQNTLY | 0.5535732 |
| NTAFVQLGI | ASGGGESQNTLY | 0.1596412 |
| QLFSGAEL | ASGGGESQNTLY | 0.25761995 |
| SAATFITL | ASGGGESQNTLY | 0.35337994 |
| SNYGLANLGV | ASGGGESQNTLY | 0.43700606 |
| SYGGAYAVM | ASGGGESQNTLY | 0.7345654 |
| TTGVGGLTLG | ASGGGESQNTLY | 0.699986 |
| TTQTIYI | ASGGGESQNTLY | 0.69570875 |
| VAAGAGLGAVI | ASGGGESQNTLY | 0.7142971 |
| VAAVVTLL | ASGGGESQNTLY | 0.71369946 |
| VGFLGGLI | ASGGGESQNTLY | 0.09783926 |
| VGVIPILL | ASGGGESQNTLY | 0.15401296 |
| VIFASSLI | ASGGGESQNTLY | 0.15988815 |
| VKGGDGGAAAG | ASGGGESQNTLY | 0.30951023 |
| VTKTFTNI | ASGGGESQNTLY | 0.12302018 |
| VVAAAILL | ASGGGESQNTLY | 0.2668375 |
| YAALDVEL | ASGGGESQNTLY | 0.494112 |
| LASVALLL | ASGGGESQNTLY | 0.1220109 |
| LDAVHRV | ASGGGESQNTLY | 0.68422043 |
| QGLSSSIF | ASGGGESQNTLY | 0.18103579 |
| RGTFANIRL | ASGGGESQNTLY | 0.17047411 |
| RQLWWGHRI | ASGGGESQNTLY | 0.23593442 |
| YGPPGTGKTLL | ASGGGESQNTLY | 0.7593061 |
| AALVGAPANVA | ASGGGESQNTLY | 0.19556293 |
| AATFRRTE | ASGGGESQNTLY | 0.5067548 |
| AGALGGGI | ASGGGESQNTLY | 0.34208244 |
| AGGAGGAGGIGAG | ASGGGESQNTLY | 0.5448696 |
| AGSGALGL | ASGGGESQNTLY | 0.13642095 |
| AQIADQI | ASGGGESQNTLY | 0.64345986 |
| ASAVAVTYL | ASGGGESQNTLY | 0.40691262 |
| ASTVLGGTGG | ASGGGESQNTLY | 0.77327424 |
| AVGGEPAK | ASGGGESQNTLY | 0.3863137 |

| | | |
|---|---|---|
| CEQGTFGGTC | ASGGGESQNTLY | 0.3052137 |
| DEGKAPAG | ASGGGESQNTLY | 0.19021155 |
| DTVVFPAL | ASGGGESQNTLY | 0.11874102 |
| FEGITMAG | ASGGGESQNTLY | 0.1114361 |
| FPLVVQD | ASGGGESQNTLY | 0.5947877 |
| GAFTTPQ | ASGGGESQNTLY | 0.21241874 |
| GFGGFGTAGGVGGA | ASGGGESQNTLY | 0.48438525 |
| GFLAQFL | ASGGGESQNTLY | 0.6895479 |
| GGAGGAGGNAGVA | ASGGGESQNTLY | 0.11263455 |
| GGVGTAGAGGAGGAGG | ASGGGESQNTLY | 0.76716125 |
| GNGGGGDAGNAGSGGN | ASGGGESQNTLY | 0.1455858 |
| GTGGAGGAGNDAGS | ASGGGESQNTLY | 0.19910508 |
| GTTNTGWFNS | ASGGGESQNTLY | 0.67812467 |
| IFMGPLFTL | ASGGGESQNTLY | 0.13043226 |
| ILAHAIEAI | ASGGGESQNTLY | 0.77733314 |
| IRSAIREAL | ASGGGESQNTLY | 0.16951765 |
| ITFAGGG | ASGGGESQNTLY | 0.63678414 |
| ITGMSGHQEV | ASGGGESQNTLY | 0.6773294 |
| IVAGLACTL | ASGGGESQNTLY | 0.09481536 |
| IVGGGLAGA | ASGGGESQNTLY | 0.13968939 |
| IVTGVLNI | ASGGGESQNTLY | 0.725965 |
| LAQQGCI | ASGGGESQNTLY | 0.43602538 |
| LCLHVEN | ASGGGESQNTLY | 0.5687683 |
| LDAISDGS | ASGGGESQNTLY | 0.15905055 |
| LGGFAAML | ASGGGESQNTLY | 0.18904714 |
| LGGKNPMI | ASGGGESQNTLY | 0.19839388 |
| LGPKGVAAIL | ASGGGESQNTLY | 0.40097553 |
| LMVGVSI | ASGGGESQNTLY | 0.59370375 |
| LVGAAFVL | ASGGGESQNTLY | 0.372638 |
| LVLAVAPL | ASGGGESQNTLY | 0.17835176 |
| MAAIAVIDAA | ASGGGESQNTLY | 0.32830405 |
| NNGINAELQ | ASGGGESQNTLY | 0.30835986 |
| RGEPLVLVHG | ASGGGESQNTLY | 0.29103187 |
| RGGAGGAAGLI | ASGGGESQNTLY | 0.26547217 |
| RSYRMHPAV | ASGGGESQNTLY | 0.32749325 |
| SAAHLLAGQLA | ASGGGESQNTLY | 0.2023226 |
| SAAVGSY | ASGGGESQNTLY | 0.1383137 |
| SPHAVLHRI | ASGGGESQNTLY | 0.26523113 |
| SQIAKVL | ASGGGESQNTLY | 0.6180482 |
| TALGHLVP | ASGGGESQNTLY | 0.4882905 |
| TAPTTSGAA | ASGGGESQNTLY | 0.4518023 |
| TDGDDGNGGSPGT | ASGGGESQNTLY | 0.5775901 |
| TDGLGEN | ASGGGESQNTLY | 0.24683136 |
| TGGPPLPGQA | ASGGGESQNTLY | 0.10460063 |
| TIPAITI | ASGGGESQNTLY | 0.11658567 |
| TIPGFTI | ASGGGESQNTLY | 0.17090636 |
| TSIASGALA | ASGGGESQNTLY | 0.22453932 |
| TVTLPGAAI | ASGGGESQNTLY | 0.09937545 |
| VASGLSAALP | ASGGGESQNTLY | 0.34707886 |
| VGAGVIGI | ASGGGESQNTLY | 0.23762041 |
| VGPKRFGV | ASGGGESQNTLY | 0.21503605 |

| | | |
|---|---|---|
| VNVAFPD | ASGGGESQNTLY | 0.5819396 |
| VSMTNSGVS | ASGGGESQNTLY | 0.20438787 |
| VTDVLGDLSV | ASGGGESQNTLY | 0.17543131 |
| VVVVGAGFAGL | ASGGGESQNTLY | 0.1823799 |
| VYTVGTVV | ASGGGESQNTLY | 0.39735457 |
| WTTFSGP | ASGGGESQNTLY | 0.14523354 |
| YDELVEAF | ASGGGESQNTLY | 0.5084545 |
| YGTGGAGGNGG | ASGGGESQNTLY | 0.15459053 |
| YTNVMAV | ASGGGESQNTLY | 0.50516546 |
| FLEQVHQL | ASGGGESQNTLY | 0.16381223 |
| GLSLLAVAV | ASGGGESQNTLY | 0.5255909 |
| LEQVHQL | ASGGGESQNTLY | 0.27913478 |
| LNAAGAI | ASGGGESQNTLY | 0.7082639 |
| PAVHASL | ASGGGESQNTLY | 0.27135795 |
| TELEDDA | ASGGGESQNTLY | 0.38360572 |
| TPPGPPN | ASGGGESQNTLY | 0.18937227 |
| TPPPPPPP | ASGGGESQNTLY | 0.41913858 |
| TQSPALI | ASGGGHSGNTLY | 0.13144676 |
| EFAAAFPQ | ASGGGHSGNTLY | 0.17757861 |
| LLSGNLDVL | ASGGGHSGNTLY | 0.119098976 |
| GPANLGNYNI | ASGGGHSGNTLY | 0.21496665 |
| TLGPTLP | ASGGGHSGNTLY | 0.3840697 |
| VYAYSSCCK | ASGGGHSGNTLY | 0.100956626 |
| APAPAPAPNI | ASGGGHSGNTLY | 0.27459064 |
| PGGPVPIP | ASGGGHSGNTLY | 0.13650015 |
| FLTVALEP | ASGGGHSGNTLY | 0.18006414 |
| SQYFGANL | ASGGGHSGNTLY | 0.34669703 |
| GGVKPAA | ASGGGHSGNTLY | 0.3174002 |
| GGTGGAGGAGGW | ASGGGHSGNTLY | 0.3190528 |
| GVGGVAGLGGAGG | ASGGGHSGNTLY | 0.1396148 |
| VLLVLATTI | ASGGGHSGNTLY | 0.20367527 |
| IGIFAAML | ASGGGHSGNTLY | 0.25626683 |
| AVGDLID | ASGGGHSGNTLY | 0.115903206 |
| NISPLPL | ASGGGHSGNTLY | 0.1600358 |
| VTAVFVAI | ASGGGHSGNTLY | 0.60534286 |
| SGDDNDAAL | ASGGGHSGNTLY | 0.74033785 |
| AVAALAHS | ASGGGHSGNTLY | 0.56797796 |
| GGVTVMATP | ASGGGHSGNTLY | 0.40650135 |
| AGLAANVRD | ASGGGHSGNTLY | 0.3783574 |
| TNRPDLIDPALLRPG | ASGGGHSGNTLY | 0.3162963 |
| LANLGVTAL | ASGGGHSGNTLY | 0.57874924 |
| TGERIEL | ASGGGHSGNTLY | 0.5617902 |
| NGEELAA | ASGGGHSGNTLY | 0.5506958 |
| VLSLVWG | ASGGGHSGNTLY | 0.27900693 |
| IGYALWL | ASGGGHSGNTLY | 0.35217083 |
| LSLLLAPL | ASGGGHSGNTLY | 0.5013586 |
| TLDEALK | ASGGGHSGNTLY | 0.48153913 |
| TDGAGTAEGG | ASGGGHSGNTLY | 0.3960191 |
| NAPGGAGGNGG | ASGGGHSGNTLY | 0.46267268 |
| VSVFMGMGGNF | ASGGGHSGNTLY | 0.3184271 |
| LASYLAAL | ASGGGHSGNTLY | 0.39837882 |

| | | |
|---|---|---|
| LVGGTPAL | ASGGGHSGNTLY | 0.30657023 |
| VRNADVL | ASGGGHSGNTLY | 0.61608934 |
| TCSSPFLVSAM | ASGGGHSGNTLY | 0.56449044 |
| VHITDAE | ASGGGHSGNTLY | 0.4933675 |
| EYAAPSF | ASGGGHSGNTLY | 0.7376599 |
| VVDAFHGGL | ASGGGHSGNTLY | 0.4091702 |
| ASVVDPQ | ASGGGHSGNTLY | 0.16763285 |
| SYNEFGGLS | ASGGGHSGNTLY | 0.25516933 |
| DPDGSVY | ASGGGHSGNTLY | 0.34785643 |
| MVSTIHATE | ASGGGHSGNTLY | 0.3957705 |
| LAAGVADPV | ASGGGHSGNTLY | 0.17509855 |
| TKEQIAA | ASGGGHSGNTLY | 0.294496 |
| SLASPAAL | ASGGGHSGNTLY | 0.27746457 |
| IGLVFYGV | ASGGGHSGNTLY | 0.27794713 |
| FLLDASNL | ASGGGHSGNTLY | 0.5693754 |
| YQVGNFP | ASGGGHSGNTLY | 0.40850258 |
| ESLLSQST | ASGGGHSGNTLY | 0.33909476 |
| LFTVGFAAVG | ASGGGHSGNTLY | 0.26724693 |
| ATPPTPT | ASGGGHSGNTLY | 0.3130122 |
| TDLATLG | ASGGGHSGNTLY | 0.3617588 |
| SAGGTGR | ASGGGHSGNTLY | 0.5948465 |
| GFSNSGS | ASGGGHSGNTLY | 0.21484219 |
| GGNGGSGGTSV | ASGGGHSGNTLY | 0.663028 |
| GTSVATGGAGN | ASGGGHSGNTLY | 0.30790925 |
| NGGNGGSGGTS | ASGGGHSGNTLY | 0.6467211 |
| AVVKFPH | ASGGGHSGNTLY | 0.627772 |
| TPPPPPP | ASGGGHSGNTLY | 0.7563015 |
| AAPAAAAAPA | ASGGGHSGNTLY | 0.62001735 |
| IALNEDL | ASGGGHSGNTLY | 0.52665925 |
| IVNVSSM | ASGGGHSGNTLY | 0.27806777 |
| TGIPPAPR | ASGGGHSGNTLY | 0.5595806 |
| FDLGGGTF | ASGGGHSGNTLY | 0.32826006 |
| FDLGGGTFDVS | ASGGGHSGNTLY | 0.40348727 |
| NEPTAAA | ASGGGHSGNTLY | 0.16472249 |
| NEPTAAALA | ASGGGHSGNTLY | 0.35541564 |
| GSIINISSI | ASGGGHSGNTLY | 0.6161094 |
| VAPGNAGTA | ASGGGHSGNTLY | 0.15958008 |
| TGGGASLE | ASGGGHSGNTLY | 0.33786476 |
| VAVYNHYKRI | ASGGGHSGNTLY | 0.26320392 |
| TDPAPATT | ASGGGHSGNTLY | 0.3300303 |
| TNRPDILDPALLRPG | ASGGGHSGNTLY | 0.5768569 |
| LTTTTTS | ASGGGHSGNTLY | 0.3032645 |
| PTGSGKTL | ASGGGHSGNTLY | 0.7591922 |
| TGSGKTL | ASGGGHSGNTLY | 0.3019417 |
| TIGHVDHGK | ASGGGHSGNTLY | 0.27676332 |
| TIGHVDHGKTT | ASGGGHSGNTLY | 0.28930432 |
| GHVDHGKTT | ASGGGHSGNTLY | 0.21039087 |
| AQFEHTLL | ASGGGHSGNTLY | 0.2511007 |
| IVNQDEL | ASGGGHSGNTLY | 0.7398397 |
| TNGSQFF | ASGGGHSGNTLY | 0.08321911 |
| TGKGIVE | ASGGGHSGNTLY | 0.082786836 |

| | | |
|---|---|---|
| LISNASDA | ASGGGHSGNTLY | 0.082916476 |
| AELRTLGELTKIAKAHGAQ | ASGGGHSGNTLY | 0.083999 |
| APNAVAAAEPSW | ASGGGHSGNTLY | 0.083043456 |
| APQVAGAATAVL | ASGGGHSGNTLY | 0.083152436 |
| ARRDPAAAAAILPTDA | ASGGGHSGNTLY | 0.083380364 |
| DAAARDVLAEAGLAEYF | ASGGGHSGNTLY | 0.083183356 |
| DDQIVLSAKANLAKVEQ | ASGGGHSGNTLY | 0.08521973 |
| DELLPDTGEDSDEDGH | ASGGGHSGNTLY | 0.08409613 |
| ELPEAPANRATVPQD | ASGGGHSGNTLY | 0.0832275 |
| FGQSKIERVTGIG | ASGGGHSGNTLY | 0.083777204 |
| GAGGTGSTAGGAGGAGGAGGLYA | ASGGGHSGNTLY | 0.08291894 |
| GHGGTGAAGGNGGTGGAGGSGIDG | ASGGGHSGNTLY | 0.083809085 |
| GNGGNGGAGGEGGPGGAGGAGA | ASGGGHSGNTLY | 0.082935795 |
| GSTAGNGGNGGNATNGTIAGQPAG | ASGGGHSGNTLY | 0.08279351 |
| GTGGDGGNGGNGGTGG | ASGGGHSGNTLY | 0.08318172 |
| GVRRPAGIPADAPV | ASGGGHSGNTLY | 0.08361797 |
| GYRQVTDAHLLTLARR | ASGGGHSGNTLY | 0.083411865 |
| IAKMGQDGHDRG | ASGGGHSGNTLY | 0.08317973 |
| KAEPSIEIQYT | ASGGGHSGNTLY | 0.083282925 |
| KPVRIRMAADEDTVAVT | ASGGGHSGNTLY | 0.08288862 |
| MPPRIAGMRLLVIKPEP | ASGGGHSGNTLY | 0.0831127 |
| MKFYAVALFRSRGVPPTR | ASGGGHSGNTLY | 0.083040744 |
| NAGGFGILGAVAHSP | ASGGGHSGNTLY | 0.08326301 |
| NEGREITLAMAEM | ASGGGHSGNTLY | 0.08330351 |
| NELVEWSHPLGHRWRWE | ASGGGHSGNTLY | 0.08256797 |
| NLGAGGTGGNGGNPG | ASGGGHSGNTLY | 0.08371824 |
| NTEWPKWETVMAADPVD | ASGGGHSGNTLY | 0.08362071 |
| PASAAPAETQP | ASGGGHSGNTLY | 0.084512144 |
| QQSSDGKNGVRAAEITGE | ASGGGHSGNTLY | 0.08403128 |
| RAPDAVTTSWWKE | ASGGGHSGNTLY | 0.08323998 |
| SGFGNVGAHTSGWFNQ | ASGGGHSGNTLY | 0.08333699 |
| SGVAGAVGQAASVGGLKV | ASGGGHSGNTLY | 0.08322045 |
| SNGGAGTGSAGGNGGT | ASGGGHSGNTLY | 0.08338577 |
| TGGAGGEGGAAG | ASGGGHSGNTLY | 0.08301067 |
| TGGDGGAAGAGGAPGHG | ASGGGHSGNTLY | 0.08505653 |
| TMPPLLAAIANAGRHGV | ASGGGHSGNTLY | 0.08283122 |
| TSSTAGNSGGA | ASGGGHSGNTLY | 0.0847399 |
| VPATVPVGILC | ASGGGHSGNTLY | 0.08414497 |
| VQRPDSGGVVLTGRIS | ASGGGHSGNTLY | 0.083117515 |
| WTPNPVPATKGVPAGLVQAT | ASGGGHSGNTLY | 0.08312976 |
| YALKRALAKFSDRFDVVII | ASGGGHSGNTLY | 0.08400261 |
| YLIGSVIV | ASGGGHSGNTLY | 0.08368567 |
| YGQLVAVID | ASGGGHSGNTLY | 0.0860356 |
| VTVGGMTLPA | ASGGGHSGNTLY | 0.08325389 |
| VLALPAV | ASGGGHSGNTLY | 0.08316041 |
| VIPPTLNYET | ASGGGHSGNTLY | 0.08380972 |
| VGMGGMHPSSG | ASGGGHSGNTLY | 0.08307179 |
| VGLAGGAATA | ASGGGHSGNTLY | 0.0832408 |
| VGGAGGAGGSGP | ASGGGHSGNTLY | 0.08323519 |
| VFAAFMV | ASGGGHSGNTLY | 0.083494194 |
| TLYDKLR | ASGGGHSGNTLY | 0.083489664 |

| | | |
|---|---|---|
| TLAVFTVL | ASGGGHSGNTLY | 0.08296795 |
| TGSVQPARV | ASGGGHSGNTLY | 0.083672754 |
| TGGGAGSGGT | ASGGGHSGNTLY | 0.083143294 |
| SHLETSL | ASGGGHSGNTLY | 0.08286296 |
| SFTTGMT | ASGGGHSGNTLY | 0.08391226 |
| QVEVAKRMGVRQPTVS | ASGGGHSGNTLY | 0.08273284 |
| QGSGGPALAM | ASGGGHSGNTLY | 0.08274702 |
| MLAPSNP | ASGGGHSGNTLY | 0.08308341 |
| LVNNAAI | ASGGGHSGNTLY | 0.08438783 |
| LTGEINTLA | ASGGGHSGNTLY | 0.084339984 |
| LSIFGPV | ASGGGHSGNTLY | 0.08323383 |
| LLALLANL | ASGGGHSGNTLY | 0.08318845 |
| LISNASDALDKLR | ASGGGHSGNTLY | 0.5320349 |
| LINEAVP | ASGGGHSGNTLY | 0.7112256 |
| LGYAFSGL | ASGGGHSGNTLY | 0.12687996 |
| LDFVAVAL | ASGGGHSGNTLY | 0.15287986 |
| LAQPGAE | ASGGGHSGNTLY | 0.39336616 |
| ISNLGAVTGF | ASGGGHSGNTLY | 0.45712656 |
| HQEDPGRI | ASGGGHSGNTLY | 0.472106 |
| GTPGEPTP | ASGGGHSGNTLY | 0.50892836 |
| GFFNSGS | ASGGGHSGNTLY | 0.15525703 |
| GAGGGGLTAGAN | ASGGGHSGNTLY | 0.15322912 |
| FVDALVNM | ASGGGHSGNTLY | 0.15462402 |
| FLTVAPEL | ASGGGHSGNTLY | 0.2543281 |
| FETLTAL | ASGGGHSGNTLY | 0.4450921 |
| DVINPIEL | ASGGGHSGNTLY | 0.12670912 |
| APVSGAPV | ASGGGHSGNTLY | 0.1176547 |
| ALVGGLAW | ASGGGHSGNTLY | 0.17963794 |
| ALAAAAATAI | ASGGGHSGNTLY | 0.4780761 |
| AFGLGLLL | ASGGGHSGNTLY | 0.70360005 |
| AFFAAASL | ASGGGHSGNTLY | 0.69224066 |
| AAAALPAFSPP | ASGGGHSGNTLY | 0.6007905 |
| VVLAPAL | ASGGGHSGNTLY | 0.54382527 |
| TIGHVDHGKTTL | ASGGGHSGNTLY | 0.24454121 |
| LISNASDALDK | ASGGGHSGNTLY | 0.11161546 |
| LISNASDAL | ASGGGHSGNTLY | 0.13243343 |
| ISNASDALDK | ASGGGHSGNTLY | 0.12795694 |
| NQAETLVYQTE | ASGGGHSGNTLY | 0.33367354 |
| TGGTGGTAGAG | ASGGGHSGNTLY | 0.10708692 |
| TGSTAGGAGGA | ASGGGHSGNTLY | 0.19840337 |
| DPLAVPVAAGPV | ASGGGHSGNTLY | 0.18256131 |
| GGTGGTPGNGGN | ASGGGHSGNTLY | 0.11915395 |
| PAGLAGVDPATL | ASGGGHSGNTLY | 0.668437 |
| GNNLVVEAPHAGASV | ASGGGHSGNTLY | 0.12398388 |
| KVTRIMSASTLEQPA | ASGGGHSGNTLY | 0.12928966 |
| LGFALPIALPRRTVV | ASGGGHSGNTLY | 0.15189862 |
| TRLGNLTARKLVVLV | ASGGGHSGNTLY | 0.6550763 |
| WSQQRRLIYFPSAGP | ASGGGHSGNTLY | 0.14313677 |
| GPNGNPDPMAAAVDIR | ASGGGHSGNTLY | 0.62809575 |
| TQRCVSVADLRAAASR | ASGGGHSGNTLY | 0.2521192 |
| QFVLDVAEHPVEVLTETV | ASGGGHSGNTLY | 0.33752584 |

| | | |
|---|---|---|
| GGAGGTGGAGGNGGTQAGDG | ASGGGHSGNTLY | 0.13211824 |
| PLLATLRAIAVDVDAKPAQV | ASGGGHSGNTLY | 0.44810736 |
| APKFRTGGYMIDFWGVGYQVAKRM | ASGGGHSGNTLY | 0.30468482 |
| GAGGAGGAGGAAGGPGGTGGTGGN | ASGGGHSGNTLY | 0.7617966 |
| IGALGLLATDVAPAGSVAFGVALA | ASGGGHSGNTLY | 0.30669555 |
| AGGAGGIGGNGGAGGAGGVLMGNG | ASGGGHSGNTLY | 0.2547641 |
| ADELIGLL | ASGGGHSGNTLY | 0.14876509 |
| ADRALKA | ASGGGHSGNTLY | 0.26788312 |
| AGAVGQVGGAGGA | ASGGGHSGNTLY | 0.117395006 |
| AGGGGGLGGAP | ASGGGHSGNTLY | 0.2712766 |
| AGGLLYGNGGAG | ASGGGHSGNTLY | 0.12852181 |
| AGLESIGL | ASGGGHSGNTLY | 0.18621084 |
| AGLKTGII | ASGGGHSGNTLY | 0.3960244 |
| ASLAALLL | ASGGGHSGNTLY | 0.1181978 |
| ASVINISSI | ASGGGHSGNTLY | 0.45578447 |
| ATPPGPA | ASGGGHSGNTLY | 0.14158583 |
| ATVASALV | ASGGGHSGNTLY | 0.14076826 |
| ATVASLIL | ASGGGHSGNTLY | 0.53025794 |
| AVNGPVLIRH | ASGGGHSGNTLY | 0.119279936 |
| DGAAGAVGI | ASGGGHSGNTLY | 0.44313908 |
| DPLLHGI | ASGGGHSGNTLY | 0.115182266 |
| ELGAALGAL | ASGGGHSGNTLY | 0.2971362 |
| ELGRTQAV | ASGGGHSGNTLY | 0.35094216 |
| FVAGGYLS | ASGGGHSGNTLY | 0.11111487 |
| GASGGAGGAGGW | ASGGGHSGNTLY | 0.12538476 |
| GGAGAGGDGGAGGI | ASGGGHSGNTLY | 0.5249674 |
| GTGHNGSQPGMGGNG | ASGGGHSGNTLY | 0.31662378 |
| GYAAASASAS | ASGGGHSGNTLY | 0.41747808 |
| HQVDVAAVQQG | ASGGGHSGNTLY | 0.18030821 |
| IGAAIAHT | ASGGGHSGNTLY | 0.18021923 |
| IGMIEVAQL | ASGGGHSGNTLY | 0.21230806 |
| IRLAPEHTSAMD | ASGGGHSGNTLY | 0.33755487 |
| IYNVSVL | ASGGGHSGNTLY | 0.50333714 |
| KQGFLGM | ASGGGHSGNTLY | 0.44482857 |
| LESVITQL | ASGGGHSGNTLY | 0.1751097 |
| LGLGALVVV | ASGGGHSGNTLY | 0.33072966 |
| LIGNGGNGGSGGI | ASGGGHSGNTLY | 0.20022878 |
| LNAINAPTL | ASGGGHSGNTLY | 0.2297241 |
| LQLLNTGV | ASGGGHSGNTLY | 0.25888577 |
| LQTSIEF | ASGGGHSGNTLY | 0.18629172 |
| MSFAVSEL | ASGGGHSGNTLY | 0.22620203 |
| NTGALGL | ASGGGHSGNTLY | 0.17081425 |
| PPGAGGAGGAGTT | ASGGGHSGNTLY | 0.20536076 |
| QLVGLGI | ASGGGHSGNTLY | 0.3947295 |
| RFLEQVHQL | ASGGGHSGNTLY | 0.26536506 |
| RGVGYVLR | ASGGGHSGNTLY | 0.55428684 |
| SATGVFAGVI | ASGGGHSGNTLY | 0.46015713 |
| SLPAITLP | ASGGGHSGNTLY | 0.1938126 |
| STALAFTV | ASGGGHSGNTLY | 0.14613354 |
| SVLTVGLGK | ASGGGHSGNTLY | 0.16841327 |
| TALAGAGSGI | ASGGGHSGNTLY | 0.17265135 |

| | | |
|---|---|---|
| TGLPFTGI | ASGGGHSGNTLY | 0.20858116 |
| VAEHVTV | ASGGGHSGNTLY | 0.1448317 |
| VDIVLNSL | ASGGGHSGNTLY | 0.23289213 |
| VLTGGGAVL | ASGGGHSGNTLY | 0.2403468 |
| VNVGLAYGI | ASGGGHSGNTLY | 0.20995343 |
| VSSAAVGAV | ASGGGHSGNTLY | 0.38928702 |
| VTSVYEAGV | ASGGGHSGNTLY | 0.21072929 |
| WVALDRGA | ASGGGHSGNTLY | 0.17806147 |
| YLVGSVVI | ASGGGHSGNTLY | 0.16483626 |
| GPPGTGKTLL | ASGGGHSGNTLY | 0.71527773 |
| GVGMTKF | ASGGGHSGNTLY | 0.16104029 |
| IGIGGSDL | ASGGGHSGNTLY | 0.5785427 |
| IGTLLAEL | ASGGGHSGNTLY | 0.31300423 |
| IYTSGTT | ASGGGHSGNTLY | 0.42576125 |
| LSFPAGV | ASGGGHSGNTLY | 0.16731818 |
| LYTSGTT | ASGGGHSGNTLY | 0.3381436 |
| VGAFPFGF | ASGGGHSGNTLY | 0.25193834 |
| VGAGTVEF | ASGGGHSGNTLY | 0.6123757 |
| VGKSTLLL | ASGGGHSGNTLY | 0.20035763 |
| AALAHAGGGIV | ASGGGHSGNTLY | 0.21442923 |
| AALVAVSLL | ASGGGHSGNTLY | 0.17504822 |
| AAVSRAVAAL | ASGGGHSGNTLY | 0.5031861 |
| AGGSSGIGGF | ASGGGHSGNTLY | 0.15435782 |
| AGLTVANANG | ASGGGHSGNTLY | 0.2184594 |
| AGLVGAVL | ASGGGHSGNTLY | 0.28603774 |
| ALSQAFTV | ASGGGHSGNTLY | 0.26359785 |
| ALTFSVSL | ASGGGHSGNTLY | 0.52611804 |
| ALTGAALVAL | ASGGGHSGNTLY | 0.14709553 |
| ASFSIGAGVL | ASGGGHSGNTLY | 0.59707606 |
| DGSVYKI | ASGGGHSGNTLY | 0.1547012 |
| EQLAEIV | ASGGGHSGNTLY | 0.164228 |
| FAVSELTVM | ASGGGHSGNTLY | 0.2880171 |
| GAAPLAAGL | ASGGGHSGNTLY | 0.1643562 |
| GADGGDGGSGGAGG | ASGGGHSGNTLY | 0.31248558 |
| GFGLDTGGAGG | ASGGGHSGNTLY | 0.16354372 |
| GLASGGLL | ASGGGHSGNTLY | 0.5267381 |
| GRIAGAAVLL | ASGGGHSGNTLY | 0.45558006 |
| GSAGSLNTSF | ASGGGHSGNTLY | 0.14921986 |
| GTGQIGI | ASGGGHSGNTLY | 0.15659405 |
| GTLIALIL | ASGGGHSGNTLY | 0.47409582 |
| IAATMGFVVG | ASGGGHSGNTLY | 0.67286 |
| IFAQALL | ASGGGHSGNTLY | 0.80068886 |
| IGAAVALGF | ASGGGHSGNTLY | 0.45967016 |
| IGLTPLAL | ASGGGHSGNTLY | 0.31003103 |
| ILGSNTSTL | ASGGGHSGNTLY | 0.50546867 |
| IVFSLTVL | ASGGGHSGNTLY | 0.6381349 |
| LASKATGF | ASGGGHSGNTLY | 0.8232639 |
| LASVALIL | ASGGGHSGNTLY | 0.70050746 |
| LFAVGLTL | ASGGGHSGNTLY | 0.2571268 |
| LFNGGIGTYI | ASGGGHSGNTLY | 0.6883671 |
| LTLGAGII | ASGGGHSGNTLY | 0.5454561 |

| | | |
|---|---|---|
| LVAAALGDI | ASGGGHSGNTLY | 0.6028364 |
| NTAFVQLGI | ASGGGHSGNTLY | 0.4029217 |
| QLFSGAEL | ASGGGHSGNTLY | 0.49045923 |
| SAATFITL | ASGGGHSGNTLY | 0.38438606 |
| SNYGLANLGV | ASGGGHSGNTLY | 0.25509962 |
| SYGGAYAVM | ASGGGHSGNTLY | 0.6143889 |
| TTGVGGLTLG | ASGGGHSGNTLY | 0.77180445 |
| TTQTIYI | ASGGGHSGNTLY | 0.76877487 |
| VAAGAGLGAVI | ASGGGHSGNTLY | 0.71342045 |
| VAAVVTLL | ASGGGHSGNTLY | 0.76299644 |
| VGFLGGLI | ASGGGHSGNTLY | 0.6459459 |
| VGVIPILL | ASGGGHSGNTLY | 0.157809 |
| VIFASSLI | ASGGGHSGNTLY | 0.20425598 |
| VKGGDGGAAAG | ASGGGHSGNTLY | 0.31540906 |
| VTKTFTNI | ASGGGHSGNTLY | 0.50891477 |
| VVAAAILL | ASGGGHSGNTLY | 0.14937785 |
| YAALDVEL | ASGGGHSGNTLY | 0.4784491 |
| LASVALLL | ASGGGHSGNTLY | 0.43485838 |
| LDAVHRV | ASGGGHSGNTLY | 0.37581134 |
| QGLSSSIF | ASGGGHSGNTLY | 0.7774179 |
| RGTFANIRL | ASGGGHSGNTLY | 0.4747457 |
| RQLWWGHRI | ASGGGHSGNTLY | 0.24722737 |
| YGPPGTGKTLL | ASGGGHSGNTLY | 0.2992149 |
| AALVGAPANVA | ASGGGHSGNTLY | 0.6638551 |
| AATFRRTE | ASGGGHSGNTLY | 0.21807514 |
| AGALGGGI | ASGGGHSGNTLY | 0.75726956 |
| AGGAGGAGGIGAG | ASGGGHSGNTLY | 0.28096953 |
| AGSGALGL | ASGGGHSGNTLY | 0.74490225 |
| AQIADQI | ASGGGHSGNTLY | 0.20550482 |
| ASAVAVTYL | ASGGGHSGNTLY | 0.7302325 |
| ASTVLGGTGG | ASGGGHSGNTLY | 0.5774416 |
| AVGGEPAK | ASGGGHSGNTLY | 0.81761056 |
| CEQGTFGGTC | ASGGGHSGNTLY | 0.43229258 |
| DEGKAPAG | ASGGGHSGNTLY | 0.56560636 |
| DTVVFPAL | ASGGGHSGNTLY | 0.38631397 |
| FEGITMAG | ASGGGHSGNTLY | 0.82262814 |
| FPLVVQD | ASGGGHSGNTLY | 0.28602767 |
| GAFTTPQ | ASGGGHSGNTLY | 0.3849955 |
| GFGGFGTAGGVGGA | ASGGGHSGNTLY | 0.19173977 |
| GFLAQFL | ASGGGHSGNTLY | 0.44724205 |
| GGAGGAGGNAGVA | ASGGGHSGNTLY | 0.71573615 |
| GGVGTAGAGGAGGAGG | ASGGGHSGNTLY | 0.16240147 |
| GNGGGGDAGNAGSGGN | ASGGGHSGNTLY | 0.6433733 |
| GTGGAGGAGNDAGS | ASGGGHSGNTLY | 0.3018825 |
| GTTNTGWFNS | ASGGGHSGNTLY | 0.24935189 |
| IFMGPLFTL | ASGGGHSGNTLY | 0.59214973 |
| ILAHAIEAI | ASGGGHSGNTLY | 0.5442395 |
| IRSAIREAL | ASGGGHSGNTLY | 0.65996575 |
| ITFAGGG | ASGGGHSGNTLY | 0.24768418 |
| ITGMSGHQEV | ASGGGHSGNTLY | 0.44500577 |
| IVAGLACTL | ASGGGHSGNTLY | 0.4749875 |

| | | |
|---|---|---|
| IVGGGLAGA | ASGGGHSGNTLY | 0.28984877 |
| IVTGVLNI | ASGGGHSGNTLY | 0.2643078 |
| LAQQGCI | ASGGGHSGNTLY | 0.77362657 |
| LCLHVEN | ASGGGHSGNTLY | 0.65561295 |
| LDAISDGS | ASGGGHSGNTLY | 0.62155515 |
| LGGFAAML | ASGGGHSGNTLY | 0.36836493 |
| LGGKNPMI | ASGGGHSGNTLY | 0.28751883 |
| LGPKGVAAIL | ASGGGHSGNTLY | 0.2928376 |
| LMVGVSI | ASGGGHSGNTLY | 0.60954297 |
| LVGAAFVL | ASGGGHSGNTLY | 0.6342263 |
| LVLAVAPL | ASGGGHSGNTLY | 0.5835346 |
| MAAIAVIDAA | ASGGGHSGNTLY | 0.2485406 |
| NNGINAELQ | ASGGGHSGNTLY | 0.27665856 |
| RGEPLVLVHG | ASGGGHSGNTLY | 0.4960382 |
| RGGAGGAAGLI | ASGGGHSGNTLY | 0.2624665 |
| RSYRMHPAV | ASGGGHSGNTLY | 0.32469374 |
| SAAHLLAGQLA | ASGGGHSGNTLY | 0.32059434 |
| SAAVGSY | ASGGGHSGNTLY | 0.34329504 |
| SPHAVLHRI | ASGGGHSGNTLY | 0.31761187 |
| SQIAKVL | ASGGGHSGNTLY | 0.23058678 |
| TALGHLVP | ASGGGHSGNTLY | 0.46990323 |
| TAPTTSGAA | ASGGGHSGNTLY | 0.42772526 |
| TDGDDGNGGSPGT | ASGGGHSGNTLY | 0.47257742 |
| TDGLGEN | ASGGGHSGNTLY | 0.6610224 |
| TGGPPLPGQA | ASGGGHSGNTLY | 0.3396782 |
| TIPAITI | ASGGGHSGNTLY | 0.21100216 |
| TIPGFTI | ASGGGHSGNTLY | 0.22640285 |
| TSIASGALA | ASGGGHSGNTLY | 0.28023553 |
| TVTLPGAAI | ASGGGHSGNTLY | 0.27183342 |
| VASGLSAALP | ASGGGHSGNTLY | 0.21305571 |
| VGAGVIGI | ASGGGHSGNTLY | 0.27285552 |
| VGPKRFGV | ASGGGHSGNTLY | 0.42871675 |
| VNVAFPD | ASGGGHSGNTLY | 0.2517056 |
| VSMTNSGVS | ASGGGHSGNTLY | 0.35536566 |
| VTDVLGDLSV | ASGGGHSGNTLY | 0.25426626 |
| VVVVGAGFAGL | ASGGGHSGNTLY | 0.27850422 |
| VYTVGTVV | ASGGGHSGNTLY | 0.2424458 |
| WTTFSGP | ASGGGHSGNTLY | 0.47160053 |
| YDELVEAF | ASGGGHSGNTLY | 0.2685159 |
| YGTGGAGGNGG | ASGGGHSGNTLY | 0.40918875 |
| YTNVMAV | ASGGGHSGNTLY | 0.2424337 |
| FLEQVHQL | ASGGGHSGNTLY | 0.5211998 |
| GLSLLAVAV | ASGGGHSGNTLY | 0.24180958 |
| LEQVHQL | ASGGGHSGNTLY | 0.55811673 |
| LNAAGAI | ASGGGHSGNTLY | 0.4783611 |
| PAVHASL | ASGGGHSGNTLY | 0.5795357 |
| TELEDDA | ASGGGHSGNTLY | 0.35176748 |
| TPPGPPN | ASGGGHSGNTLY | 0.3252135 |
| TPPPPPPP | ASGGGHSGNTLY | 0.22619733 |
| TQSPALI | ASSRDWVYAEQF | 0.6306178 |
| EFAAAFPQ | ASSRDWVYAEQF | 0.24540843 |

| | | |
|---|---|---|
| LLSGNLDVL | ASSRDWVYAEQF | 0.5095359 |
| GPANLGNYNI | ASSRDWVYAEQF | 0.21859382 |
| TLGPTLP | ASSRDWVYAEQF | 0.4836241 |
| VYAYSSCCK | ASSRDWVYAEQF | 0.6068663 |
| APAPAPAPNI | ASSRDWVYAEQF | 0.26351708 |
| PGGPVPIP | ASSRDWVYAEQF | 0.5512097 |
| FLTVALEP | ASSRDWVYAEQF | 0.504064 |
| SQYFGANL | ASSRDWVYAEQF | 0.24732211 |
| GGVKPAA | ASSRDWVYAEQF | 0.44915605 |
| GGTGGAGGAGGW | ASSRDWVYAEQF | 0.23773076 |
| GVGGVAGLGGAGG | ASSRDWVYAEQF | 0.666371 |
| VLLVLATTI | ASSRDWVYAEQF | 0.23488374 |
| IGIFAAML | ASSRDWVYAEQF | 0.479577 |
| AVGDLID | ASSRDWVYAEQF | 0.48985302 |
| NISPLPL | ASSRDWVYAEQF | 0.2572214 |
| VTAVFVAI | ASSRDWVYAEQF | 0.23165727 |
| SGDDNDAAL | ASSRDWVYAEQF | 0.6403047 |
| AVAALAHS | ASSRDWVYAEQF | 0.7117133 |
| GGVTVMATP | ASSRDWVYAEQF | 0.6974315 |
| AGLAANVRD | ASSRDWVYAEQF | 0.6379175 |
| TNRPDLIDPALLRPG | ASSRDWVYAEQF | 0.5675894 |
| LANLGVTAL | ASSRDWVYAEQF | 0.6494167 |
| TGERIEL | ASSRDWVYAEQF | 0.7000592 |
| NGEELAA | ASSRDWVYAEQF | 0.69514966 |
| VLSLVWG | ASSRDWVYAEQF | 0.6899392 |
| IGYALWL | ASSRDWVYAEQF | 0.4721626 |
| LSLLLAPL | ASSRDWVYAEQF | 0.6223984 |
| TLDEALK | ASSRDWVYAEQF | 0.6629272 |
| TDGAGTAEGG | ASSRDWVYAEQF | 0.5649258 |
| NAPGGAGGNGG | ASSRDWVYAEQF | 0.67610025 |
| VSVFMGMGGNF | ASSRDWVYAEQF | 0.65753555 |
| LASYLAAL | ASSRDWVYAEQF | 0.6184701 |
| LVGGTPAL | ASSRDWVYAEQF | 0.64427257 |
| VRNADVL | ASSRDWVYAEQF | 0.60706204 |
| TCSSPFLVSAM | ASSRDWVYAEQF | 0.69241446 |
| VHITDAE | ASSRDWVYAEQF | 0.685444 |
| EYAAPSF | ASSRDWVYAEQF | 0.70925033 |
| VVDAFHGGL | ASSRDWVYAEQF | 0.70095396 |
| ASVVDPQ | ASSRDWVYAEQF | 0.70017684 |
| SYNEFGGLS | ASSRDWVYAEQF | 0.29560545 |
| DPDGSVY | ASSRDWVYAEQF | 0.37397543 |
| MVSTIHATE | ASSRDWVYAEQF | 0.64932036 |
| LAAGVADPV | ASSRDWVYAEQF | 0.5959321 |
| TKEQIAA | ASSRDWVYAEQF | 0.39028704 |
| SLASPAAL | ASSRDWVYAEQF | 0.5808098 |
| IGLVFYGV | ASSRDWVYAEQF | 0.6449356 |
| FLLDASNL | ASSRDWVYAEQF | 0.5001385 |
| YQVGNFP | ASSRDWVYAEQF | 0.6789592 |
| ESLLSQST | ASSRDWVYAEQF | 0.6027254 |
| LFTVGFAAVG | ASSRDWVYAEQF | 0.5187839 |
| ATPPTPT | ASSRDWVYAEQF | 0.529647 |

| | | |
|---|---|---|
| TDLATLG | ASSRDWVYAEQF | 0.6822326 |
| SAGGTGR | ASSRDWVYAEQF | 0.50058645 |
| GFSNSGS | ASSRDWVYAEQF | 0.64910674 |
| GGNGGSGGTSV | ASSRDWVYAEQF | 0.50268376 |
| GTSVATGGAGN | ASSRDWVYAEQF | 0.69389373 |
| NGGNGGSGGTS | ASSRDWVYAEQF | 0.47251576 |
| AVVKFPH | ASSRDWVYAEQF | 0.6839427 |
| TPPPPPP | ASSRDWVYAEQF | 0.6662134 |
| AAPAAAAAPA | ASSRDWVYAEQF | 0.7135105 |
| IALNEDL | ASSRDWVYAEQF | 0.6326104 |
| IVNVSSM | ASSRDWVYAEQF | 0.66279936 |
| TGIPPAPR | ASSRDWVYAEQF | 0.54927385 |
| FDLGGGTF | ASSRDWVYAEQF | 0.68429387 |
| FDLGGGTFDVS | ASSRDWVYAEQF | 0.50299954 |
| NEPTAAA | ASSRDWVYAEQF | 0.67171645 |
| NEPTAAALA | ASSRDWVYAEQF | 0.3241223 |
| GSIINISSI | ASSRDWVYAEQF | 0.66274333 |
| VAPGNAGTA | ASSRDWVYAEQF | 0.69570553 |
| TGGGASLE | ASSRDWVYAEQF | 0.5039279 |
| VAVYNHYKRI | ASSRDWVYAEQF | 0.6948135 |
| TDPAPATT | ASSRDWVYAEQF | 0.6521756 |
| TNRPDILDPALLRPG | ASSRDWVYAEQF | 0.61013794 |
| LTTTTTS | ASSRDWVYAEQF | 0.70364094 |
| PTGSGKTL | ASSRDWVYAEQF | 0.5431751 |
| TGSGKTL | ASSRDWVYAEQF | 0.7182628 |
| TIGHVDHGK | ASSRDWVYAEQF | 0.5609138 |
| TIGHVDHGKTT | ASSRDWVYAEQF | 0.6425591 |
| GHVDHGKTT | ASSRDWVYAEQF | 0.6576616 |
| AQFEHTLL | ASSRDWVYAEQF | 0.5818536 |
| IVNQDEL | ASSRDWVYAEQF | 0.4718892 |
| TNGSQFF | ASSRDWVYAEQF | 0.7085104 |
| TGKGIVE | ASSRDWVYAEQF | 0.16217148 |
| LISNASDA | ASSRDWVYAEQF | 0.18253273 |
| AELRTLGELTKIAKAHGAQ | ASSRDWVYAEQF | 0.15515584 |
| APNAVAAAEPSW | ASSRDWVYAEQF | 0.16307095 |
| APQVAGAATAVL | ASSRDWVYAEQF | 0.1642482 |
| ARRDPAAAAAILPTDA | ASSRDWVYAEQF | 0.16579312 |
| DAAARDVLAEAGLAEYF | ASSRDWVYAEQF | 0.20361757 |
| DDQIVLSAKANLAKVEQ | ASSRDWVYAEQF | 0.16211386 |
| DELLPDTGEDSDEDGH | ASSRDWVYAEQF | 0.16989456 |
| ELPEAPANRATVPQD | ASSRDWVYAEQF | 0.16161685 |
| FGQSKIERVTGIG | ASSRDWVYAEQF | 0.15818694 |
| GAGGTGSTAGGAGGAGGAGGLYA | ASSRDWVYAEQF | 0.15949859 |
| GHGGTGAAGGNGGTGGAGGSGIDG | ASSRDWVYAEQF | 0.17104907 |
| GNGGNGGAGGEGGPGGAGGAGA | ASSRDWVYAEQF | 0.1795029 |
| GSTAGNGGNGGNATNGTIAGQPAG | ASSRDWVYAEQF | 0.15228897 |
| GTGGDGGNGGNGGTGG | ASSRDWVYAEQF | 0.15984222 |
| GVRRPAGIPADAPV | ASSRDWVYAEQF | 0.1677389 |
| GYRQVTDAHLLTLARR | ASSRDWVYAEQF | 0.21279237 |
| IAKMGQDGHDRG | ASSRDWVYAEQF | 0.21066928 |
| KAEPSIEIQYT | ASSRDWVYAEQF | 0.17771623 |

| | | |
|---|---|---|
| KPVRIRMAADEDTVAVT | ASSRDWVYAEQF | 0.16993828 |
| MPPRIAGMRLLVIKPEP | ASSRDWVYAEQF | 0.1660222 |
| MKFYAVALFRSRGVPPTR | ASSRDWVYAEQF | 0.14996797 |
| NAGGFGILGAVAHSP | ASSRDWVYAEQF | 0.15636162 |
| NEGREITLAMAEM | ASSRDWVYAEQF | 0.16168523 |
| NELVEWSHPLGHRWRWE | ASSRDWVYAEQF | 0.17241924 |
| NLGAGGTGGNGGNPG | ASSRDWVYAEQF | 0.14652166 |
| NTEWPKWETVMAADPVD | ASSRDWVYAEQF | 0.16921724 |
| PASAAPAETQP | ASSRDWVYAEQF | 0.16499937 |
| QQSSDGKNGVRAAEITGE | ASSRDWVYAEQF | 0.15553351 |
| RAPDAVTTSWWKE | ASSRDWVYAEQF | 0.19615012 |
| SGFGNVGAHTSGWFNQ | ASSRDWVYAEQF | 0.15516955 |
| SGVAGAVGQAASVGGLKV | ASSRDWVYAEQF | 0.16534634 |
| SNGGAGTGSAGGNGGT | ASSRDWVYAEQF | 0.15981533 |
| TGGAGGEGGAAG | ASSRDWVYAEQF | 0.16660714 |
| TGGDGGAAGAGGAPGHG | ASSRDWVYAEQF | 0.1557406 |
| TMPPLLAAIANAGRHGV | ASSRDWVYAEQF | 0.18823676 |
| TSSTAGNSGGA | ASSRDWVYAEQF | 0.1547164 |
| VPATVPVGILC | ASSRDWVYAEQF | 0.1729494 |
| VQRPDSGGVVLTGRIS | ASSRDWVYAEQF | 0.15982947 |
| WTPNPVPATKGVPAGLVQAT | ASSRDWVYAEQF | 0.16373974 |
| YALKRALAKFSDRFDVVII | ASSRDWVYAEQF | 0.15649207 |
| YLIGSVIV | ASSRDWVYAEQF | 0.26311615 |
| YGQLVAVID | ASSRDWVYAEQF | 0.18793166 |
| VTVGGMTLPA | ASSRDWVYAEQF | 0.18207006 |
| VLALPAV | ASSRDWVYAEQF | 0.1676096 |
| VIPPTLNYET | ASSRDWVYAEQF | 0.16810395 |
| VGMGGMHPSSG | ASSRDWVYAEQF | 0.1528224 |
| VGLAGGAATA | ASSRDWVYAEQF | 0.16741644 |
| VGGAGGAGGSGP | ASSRDWVYAEQF | 0.15110087 |
| VFAAFMV | ASSRDWVYAEQF | 0.16474806 |
| TLYDKLR | ASSRDWVYAEQF | 0.15908743 |
| TLAVFTVL | ASSRDWVYAEQF | 0.1642093 |
| TGSVQPARV | ASSRDWVYAEQF | 0.15787672 |
| TGGGAGSGGT | ASSRDWVYAEQF | 0.1717907 |
| SHLETSL | ASSRDWVYAEQF | 0.16326837 |
| SFTTGMT | ASSRDWVYAEQF | 0.1746621 |
| QVEVAKRMGVRQPTVS | ASSRDWVYAEQF | 0.16946551 |
| QGSGGPALAM | ASSRDWVYAEQF | 0.15475462 |
| MLAPSNP | ASSRDWVYAEQF | 0.15154725 |
| LVNNAAI | ASSRDWVYAEQF | 0.15472077 |
| LTGEINTLA | ASSRDWVYAEQF | 0.18145052 |
| LSIFGPV | ASSRDWVYAEQF | 0.15262131 |
| LLALLANL | ASSRDWVYAEQF | 0.1581168 |
| LISNASDALDKLR | ASSRDWVYAEQF | 0.1673735 |
| LINEAVP | ASSRDWVYAEQF | 0.3503386 |
| LGYAFSGL | ASSRDWVYAEQF | 0.47036228 |
| LDFVAVAL | ASSRDWVYAEQF | 0.16657655 |
| LAQPGAE | ASSRDWVYAEQF | 0.19743522 |
| ISNLGAVTGF | ASSRDWVYAEQF | 0.2722066 |
| HQEDPGRI | ASSRDWVYAEQF | 0.3698298 |

| | | |
|---|---|---|
| GTPGEPTP | ASSRDWVYAEQF | 0.33021712 |
| GFFNSGS | ASSRDWVYAEQF | 0.4456815 |
| GAGGGGLTAGAN | ASSRDWVYAEQF | 0.2058211 |
| FVDALVNM | ASSRDWVYAEQF | 0.27110308 |
| FLTVAPEL | ASSRDWVYAEQF | 0.23808777 |
| FETLTAL | ASSRDWVYAEQF | 0.31838885 |
| DVINPIEL | ASSRDWVYAEQF | 0.2646581 |
| APVSGAPV | ASSRDWVYAEQF | 0.19217424 |
| ALVGGLAW | ASSRDWVYAEQF | 0.1570413 |
| ALAAAAATAI | ASSRDWVYAEQF | 0.19948374 |
| AFGLGLLL | ASSRDWVYAEQF | 0.2816117 |
| AFFAAASL | ASSRDWVYAEQF | 0.3474658 |
| AAAALPAFSPP | ASSRDWVYAEQF | 0.27312115 |
| VVLAPAL | ASSRDWVYAEQF | 0.399883 |
| TIGHVDHGKTTL | ASSRDWVYAEQF | 0.4586845 |
| LISNASDALDK | ASSRDWVYAEQF | 0.18184984 |
| LISNASDAL | ASSRDWVYAEQF | 0.12821084 |
| ISNASDALDK | ASSRDWVYAEQF | 0.15824564 |
| NQAETLVYQTE | ASSRDWVYAEQF | 0.19539091 |
| TGGTGGTAGAG | ASSRDWVYAEQF | 0.26122063 |
| TGSTAGGAGGA | ASSRDWVYAEQF | 0.11255159 |
| DPLAVPVAAGPV | ASSRDWVYAEQF | 0.28199774 |
| GGTGGTPGNGGN | ASSRDWVYAEQF | 0.2622906 |
| PAGLAGVDPATL | ASSRDWVYAEQF | 0.17028853 |
| GNNLVVEAPHAGASV | ASSRDWVYAEQF | 0.51282966 |
| KVTRIMSASTLEQPA | ASSRDWVYAEQF | 0.19383283 |
| LGFALPIALPRRTVV | ASSRDWVYAEQF | 0.18600342 |
| TRLGNLTARKLVVLV | ASSRDWVYAEQF | 0.22571944 |
| WSQQRRLIYFPSAGP | ASSRDWVYAEQF | 0.5339713 |
| GPNGNPDPMAAAVDIR | ASSRDWVYAEQF | 0.18859594 |
| TQRCVSVADLRAAASR | ASSRDWVYAEQF | 0.51972383 |
| QFVLDVAEHPVEVLTETV | ASSRDWVYAEQF | 0.19219005 |
| GGAGGTGGAGGNGGTQAGDG | ASSRDWVYAEQF | 0.40078604 |
| PLLATLRAIAVDVDAKPAQV | ASSRDWVYAEQF | 0.1749338 |
| APKFRTGGYMIDFWGVGYQVAKRM | ASSRDWVYAEQF | 0.44068414 |
| GAGGAGGAGGAAGGPGGTGGTGGN | ASSRDWVYAEQF | 0.31667256 |
| IGALGLLATDVAPAGSVAFGVALA | ASSRDWVYAEQF | 0.5858443 |
| AGGAGGIGGNGGAGGAGGVLMGNG | ASSRDWVYAEQF | 0.2590354 |
| ADELIGLL | ASSRDWVYAEQF | 0.34199497 |
| ADRALKA | ASSRDWVYAEQF | 0.1896876 |
| AGAVGQVGGAGGA | ASSRDWVYAEQF | 0.20939076 |
| AGGGGGLGGAP | ASSRDWVYAEQF | 0.15288593 |
| AGGLLYGNGGAG | ASSRDWVYAEQF | 0.21941255 |
| AGLESIGL | ASSRDWVYAEQF | 0.16091558 |
| AGLKTGII | ASSRDWVYAEQF | 0.22655414 |
| ASLAALLL | ASSRDWVYAEQF | 0.4521738 |
| ASVINISSI | ASSRDWVYAEQF | 0.14478917 |
| ATPPGPA | ASSRDWVYAEQF | 0.3050847 |
| ATVASALV | ASSRDWVYAEQF | 0.16488267 |
| ATVASLIL | ASSRDWVYAEQF | 0.18560576 |
| AVNGPVLIRH | ASSRDWVYAEQF | 0.26844758 |

| | | |
|---|---|---|
| DGAAGAVGI | ASSRDWVYAEQF | 0.17301394 |
| DPLLHGI | ASSRDWVYAEQF | 0.23108438 |
| ELGAALGAL | ASSRDWVYAEQF | 0.13813628 |
| ELGRTQAV | ASSRDWVYAEQF | 0.24256046 |
| FVAGGYLS | ASSRDWVYAEQF | 0.302211 |
| GASGGAGGAGGW | ASSRDWVYAEQF | 0.120721795 |
| GGAGAGGDGGAGGI | ASSRDWVYAEQF | 0.169106 |
| GTGHNGSQPGMGGNG | ASSRDWVYAEQF | 0.5121622 |
| GYAAASASAS | ASSRDWVYAEQF | 0.47910732 |
| HQVDVAAVQQG | ASSRDWVYAEQF | 0.71227264 |
| IGAAIAHT | ASSRDWVYAEQF | 0.1283029 |
| IGMIEVAQL | ASSRDWVYAEQF | 0.1330807 |
| IRLAPEHTSAMD | ASSRDWVYAEQF | 0.2670841 |
| IYNVSVL | ASSRDWVYAEQF | 0.465946 |
| KQGFLGM | ASSRDWVYAEQF | 0.45029694 |
| LESVITQL | ASSRDWVYAEQF | 0.43407062 |
| LGLGALVVV | ASSRDWVYAEQF | 0.1361916 |
| LIGNGGNGGSGGI | ASSRDWVYAEQF | 0.17828068 |
| LNAINAPTL | ASSRDWVYAEQF | 0.17096357 |
| LQLLNTGV | ASSRDWVYAEQF | 0.2249687 |
| LQTSIEF | ASSRDWVYAEQF | 0.3359571 |
| MSFAVSEL | ASSRDWVYAEQF | 0.1793961 |
| NTGALGL | ASSRDWVYAEQF | 0.11915751 |
| PPGAGGAGGAGTT | ASSRDWVYAEQF | 0.17227796 |
| QLVGLGI | ASSRDWVYAEQF | 0.47715446 |
| RFLEQVHQL | ASSRDWVYAEQF | 0.66710186 |
| RGVGYVLR | ASSRDWVYAEQF | 0.6504518 |
| SATGVFAGVI | ASSRDWVYAEQF | 0.63443005 |
| SLPAITLP | ASSRDWVYAEQF | 0.5816658 |
| STALAFTV | ASSRDWVYAEQF | 0.39597672 |
| SVLTVGLGK | ASSRDWVYAEQF | 0.08970509 |
| TALAGAGSGI | ASSRDWVYAEQF | 0.119015835 |
| TGLPFTGI | ASSRDWVYAEQF | 0.14867364 |
| VAEHVTV | ASSRDWVYAEQF | 0.27644446 |
| VDIVLNSL | ASSRDWVYAEQF | 0.107456885 |
| VLTGGGAVL | ASSRDWVYAEQF | 0.25101995 |
| VNVGLAYGI | ASSRDWVYAEQF | 0.19809914 |
| VSSAAVGAV | ASSRDWVYAEQF | 0.11513618 |
| VTSVYEAGV | ASSRDWVYAEQF | 0.63971615 |
| WVALDRGA | ASSRDWVYAEQF | 0.116937324 |
| YLVGSVVI | ASSRDWVYAEQF | 0.13319296 |
| GPPGTGKTLL | ASSRDWVYAEQF | 0.16394004 |
| GVGMTKF | ASSRDWVYAEQF | 0.47670943 |
| IGIGGSDL | ASSRDWVYAEQF | 0.14072739 |
| IGTLLAEL | ASSRDWVYAEQF | 0.47248197 |
| IYTSGTT | ASSRDWVYAEQF | 0.2580996 |
| LSFPAGV | ASSRDWVYAEQF | 0.41299713 |
| LYTSGTT | ASSRDWVYAEQF | 0.12026488 |
| VGAFPFGF | ASSRDWVYAEQF | 0.5083115 |
| VGAGTVEF | ASSRDWVYAEQF | 0.27387553 |
| VGKSTLLL | ASSRDWVYAEQF | 0.7887707 |

| | | |
|---|---|---|
| AALAHAGGGIV | ASSRDWVYAEQF | 0.24977049 |
| AALVAVSLL | ASSRDWVYAEQF | 0.2596445 |
| AAVSRAVAAL | ASSRDWVYAEQF | 0.17106825 |
| AGGSSGIGGF | ASSRDWVYAEQF | 0.44685447 |
| AGLTVANANG | ASSRDWVYAEQF | 0.10746607 |
| AGLVGAVL | ASSRDWVYAEQF | 0.23035252 |
| ALSQAFTV | ASSRDWVYAEQF | 0.11326025 |
| ALTFSVSL | ASSRDWVYAEQF | 0.1838793 |
| ALTGAALVAL | ASSRDWVYAEQF | 0.42777747 |
| ASFSIGAGVL | ASSRDWVYAEQF | 0.12064058 |
| DGSVYKI | ASSRDWVYAEQF | 0.3595503 |
| EQLAEIV | ASSRDWVYAEQF | 0.18263066 |
| FAVSELTVM | ASSRDWVYAEQF | 0.13873112 |
| GAAPLAAGL | ASSRDWVYAEQF | 0.47077128 |
| GADGGDGGSGGAGG | ASSRDWVYAEQF | 0.14146332 |
| GFGLDTGGAGG | ASSRDWVYAEQF | 0.6464555 |
| GLASGGLL | ASSRDWVYAEQF | 0.13413146 |
| GRIAGAAVLL | ASSRDWVYAEQF | 0.24723385 |
| GSAGSLNTSF | ASSRDWVYAEQF | 0.2592913 |
| GTGQIGI | ASSRDWVYAEQF | 0.09981308 |
| GTLIALIL | ASSRDWVYAEQF | 0.13522904 |
| IAATMGFVVG | ASSRDWVYAEQF | 0.54235137 |
| IFAQALL | ASSRDWVYAEQF | 0.72827154 |
| IGAAVALGF | ASSRDWVYAEQF | 0.73714006 |
| IGLTPLAL | ASSRDWVYAEQF | 0.21451262 |
| ILGSNTSTL | ASSRDWVYAEQF | 0.27742118 |
| IVFSLTVL | ASSRDWVYAEQF | 0.4707011 |
| LASKATGF | ASSRDWVYAEQF | 0.6207867 |
| LASVALIL | ASSRDWVYAEQF | 0.8126042 |
| LFAVGLTL | ASSRDWVYAEQF | 0.7277734 |
| LFNGGIGTYI | ASSRDWVYAEQF | 0.24980699 |
| LTLGAGII | ASSRDWVYAEQF | 0.5881609 |
| LVAAALGDI | ASSRDWVYAEQF | 0.39976 |
| NTAFVQLGI | ASSRDWVYAEQF | 0.54676795 |
| QLFSGAEL | ASSRDWVYAEQF | 0.49105787 |
| SAATFITL | ASSRDWVYAEQF | 0.40757513 |
| SNYGLANLGV | ASSRDWVYAEQF | 0.33118445 |
| SYGGAYAVM | ASSRDWVYAEQF | 0.22331762 |
| TTGVGGLTLG | ASSRDWVYAEQF | 0.5903684 |
| TTQTIYI | ASSRDWVYAEQF | 0.70295537 |
| VAAGAGLGAVI | ASSRDWVYAEQF | 0.5747677 |
| VAAVVTLL | ASSRDWVYAEQF | 0.7429973 |
| VGFLGGLI | ASSRDWVYAEQF | 0.7317631 |
| VGVIPILL | ASSRDWVYAEQF | 0.40599638 |
| VIFASSLI | ASSRDWVYAEQF | 0.14037041 |
| VKGGDGGAAAG | ASSRDWVYAEQF | 0.16425478 |
| VTKTFTNI | ASSRDWVYAEQF | 0.24422807 |
| VVAAAILL | ASSRDWVYAEQF | 0.32876557 |
| YAALDVEL | ASSRDWVYAEQF | 0.13612482 |
| LASVALLL | ASSRDWVYAEQF | 0.561695 |
| LDAVHRV | ASSRDWVYAEQF | 0.3216415 |

| | | |
|---|---|---|
| QGLSSSIF | ASSRDWVYAEQF | 0.3178167 |
| RGTFANIRL | ASSRDWVYAEQF | 0.75062126 |
| RQLWWGHRI | ASSRDWVYAEQF | 0.31209826 |
| YGPPGTGKTLL | ASSRDWVYAEQF | 0.21769716 |
| AALVGAPANVA | ASSRDWVYAEQF | 0.27500764 |
| AATFRRTE | ASSRDWVYAEQF | 0.79767454 |
| AGALGGGI | ASSRDWVYAEQF | 0.24448827 |
| AGGAGGAGGIGAG | ASSRDWVYAEQF | 0.7544691 |
| AGSGALGL | ASSRDWVYAEQF | 0.37588298 |
| AQIADQI | ASSRDWVYAEQF | 0.74538934 |
| ASAVAVTYL | ASSRDWVYAEQF | 0.20016214 |
| ASTVLGGTGG | ASSRDWVYAEQF | 0.7001389 |
| AVGGEPAK | ASSRDWVYAEQF | 0.54429114 |
| CEQGTFGGTC | ASSRDWVYAEQF | 0.8084566 |
| DEGKAPAG | ASSRDWVYAEQF | 0.32875103 |
| DTVVFPAL | ASSRDWVYAEQF | 0.50058717 |
| FEGITMAG | ASSRDWVYAEQF | 0.36991918 |
| FPLVVQD | ASSRDWVYAEQF | 0.77405345 |
| GAFTTPQ | ASSRDWVYAEQF | 0.1990447 |
| GFGGFGTAGGVGGA | ASSRDWVYAEQF | 0.2879199 |
| GFLAQFL | ASSRDWVYAEQF | 0.2255683 |
| GGAGGAGGNAGVA | ASSRDWVYAEQF | 0.31126902 |
| GGVGTAGAGGAGGAGG | ASSRDWVYAEQF | 0.70884836 |
| GNGGGGDAGNAGSGGN | ASSRDWVYAEQF | 0.1339018 |
| GTGGAGGAGNDAGS | ASSRDWVYAEQF | 0.62985754 |
| GTTNTGWFNS | ASSRDWVYAEQF | 0.2333588 |
| IFMGPLFTL | ASSRDWVYAEQF | 0.21403658 |
| ILAHAIEAI | ASSRDWVYAEQF | 0.5190064 |
| IRSAIREAL | ASSRDWVYAEQF | 0.436305 |
| ITFAGGG | ASSRDWVYAEQF | 0.56787926 |
| ITGMSGHQEV | ASSRDWVYAEQF | 0.2456546 |
| IVAGLACTL | ASSRDWVYAEQF | 0.5245776 |
| IVGGGLAGA | ASSRDWVYAEQF | 0.47666457 |
| IVTGVLNI | ASSRDWVYAEQF | 0.13780074 |
| LAQQGCI | ASSRDWVYAEQF | 0.23178455 |
| LCLHVEN | ASSRDWVYAEQF | 0.74837434 |
| LDAISDGS | ASSRDWVYAEQF | 0.12427747 |
| LGGFAAML | ASSRDWVYAEQF | 0.15933439 |
| LGGKNPMI | ASSRDWVYAEQF | 0.10306741 |
| LGPKGVAAIL | ASSRDWVYAEQF | 0.10852855 |
| LMVGVSI | ASSRDWVYAEQF | 0.110435456 |
| LVGAAFVL | ASSRDWVYAEQF | 0.13067669 |
| LVLAVAPL | ASSRDWVYAEQF | 0.15517406 |
| MAAIAVIDAA | ASSRDWVYAEQF | 0.13130553 |
| NNGINAELQ | ASSRDWVYAEQF | 0.113274895 |
| RGEPLVLVHG | ASSRDWVYAEQF | 0.11123294 |
| RGGAGGAAGLI | ASSRDWVYAEQF | 0.11022494 |
| RSYRMHPAV | ASSRDWVYAEQF | 0.114702016 |
| SAAHLLAGQLA | ASSRDWVYAEQF | 0.118385494 |
| SAAVGSY | ASSRDWVYAEQF | 0.10704511 |
| SPHAVLHRI | ASSRDWVYAEQF | 0.10210954 |

| | | |
|---|---|---|
| SQIAKVL | ASSRDWVYAEQF | 0.10805421 |
| TALGHLVP | ASSRDWVYAEQF | 0.11164736 |
| TAPTTSGAA | ASSRDWVYAEQF | 0.12164227 |
| TDGDDGNGGSPGT | ASSRDWVYAEQF | 0.117900915 |
| TDGLGEN | ASSRDWVYAEQF | 0.1432266 |
| TGGPPLPGQA | ASSRDWVYAEQF | 0.1377111 |
| TIPAITI | ASSRDWVYAEQF | 0.10870596 |
| TIPGFTI | ASSRDWVYAEQF | 0.10009786 |
| TSIASGALA | ASSRDWVYAEQF | 0.103648886 |
| TVTLPGAAI | ASSRDWVYAEQF | 0.10969583 |
| VASGLSAALP | ASSRDWVYAEQF | 0.11318024 |
| VGAGVIGI | ASSRDWVYAEQF | 0.09818919 |
| VGPKRFGV | ASSRDWVYAEQF | 0.11470934 |
| VNVAFPD | ASSRDWVYAEQF | 0.11776623 |
| VSMTNSGVS | ASSRDWVYAEQF | 0.10470542 |
| VTDVLGDLSV | ASSRDWVYAEQF | 0.15256548 |
| VVVVGAGFAGL | ASSRDWVYAEQF | 0.103752784 |
| VYTVGTVV | ASSRDWVYAEQF | 0.104641065 |
| WTTFSGP | ASSRDWVYAEQF | 0.11101243 |
| YDELVEAF | ASSRDWVYAEQF | 0.13325967 |
| YGTGGAGGNGG | ASSRDWVYAEQF | 0.10928616 |
| YTNVMAV | ASSRDWVYAEQF | 0.13886614 |
| FLEQVHQL | ASSRDWVYAEQF | 0.111258164 |
| GLSLLAVAV | ASSRDWVYAEQF | 0.14946783 |
| LEQVHQL | ASSRDWVYAEQF | 0.10828415 |
| LNAAGAI | ASSRDWVYAEQF | 0.1256072 |
| PAVHASL | ASSRDWVYAEQF | 0.11354824 |
| TELEDDA | ASSRDWVYAEQF | 0.16490142 |
| TPPGPPN | ASSRDWVYAEQF | 0.11656981 |
| TPPPPPPP | ASSRDWVYAEQF | 0.12923971 |
| TQSPALI | ASSPGLGGYEQY | 0.10751216 |
| EFAAAFPQ | ASSPGLGGYEQY | 0.16123787 |
| LLSGNLDVL | ASSPGLGGYEQY | 0.10293241 |
| GPANLGNYNI | ASSPGLGGYEQY | 0.11213447 |
| TLGPTLP | ASSPGLGGYEQY | 0.10487768 |
| VYAYSSCCK | ASSPGLGGYEQY | 0.11293571 |
| APAPAPAPNI | ASSPGLGGYEQY | 0.13122182 |
| PGGPVPIP | ASSPGLGGYEQY | 0.10010602 |
| FLTVALEP | ASSPGLGGYEQY | 0.11840395 |
| SQYFGANL | ASSPGLGGYEQY | 0.10200823 |
| GGVKPAA | ASSPGLGGYEQY | 0.1079784 |
| GGTGGAGGAGGW | ASSPGLGGYEQY | 0.11586899 |
| GVGGVAGLGGAGG | ASSPGLGGYEQY | 0.10489576 |
| VLLVLATTI | ASSPGLGGYEQY | 0.10746064 |
| IGIFAAML | ASSPGLGGYEQY | 0.10079894 |
| AVGDLID | ASSPGLGGYEQY | 0.11483192 |
| NISPLPL | ASSPGLGGYEQY | 0.13047536 |
| VTAVFVAI | ASSPGLGGYEQY | 0.10038789 |
| SGDDNDAAL | ASSPGLGGYEQY | 0.10445466 |
| AVAALAHS | ASSPGLGGYEQY | 0.13629621 |
| GGVTVMATP | ASSPGLGGYEQY | 0.09084903 |

| | | |
|---|---|---|
| AGLAANVRD | ASSPGLGGYEQY | 0.090659484 |
| TNRPDLIDPALLRPG | ASSPGLGGYEQY | 0.09075812 |
| LANLGVTAL | ASSPGLGGYEQY | 0.09123977 |
| TGERIEL | ASSPGLGGYEQY | 0.09069403 |
| NGEELAA | ASSPGLGGYEQY | 0.09085042 |
| VLSLVWG | ASSPGLGGYEQY | 0.0909281 |
| IGYALWL | ASSPGLGGYEQY | 0.090842664 |
| LSLLLAPL | ASSPGLGGYEQY | 0.091460906 |
| TLDEALK | ASSPGLGGYEQY | 0.09127198 |
| TDGAGTAEGG | ASSPGLGGYEQY | 0.0909842 |
| NAPGGAGGNGG | ASSPGLGGYEQY | 0.09113577 |
| VSVFMGMGGNF | ASSPGLGGYEQY | 0.090712756 |
| LASYLAAL | ASSPGLGGYEQY | 0.09146167 |
| LVGGTPAL | ASSPGLGGYEQY | 0.09073785 |
| VRNADVL | ASSPGLGGYEQY | 0.09060975 |
| TCSSPFLVSAM | ASSPGLGGYEQY | 0.09097377 |
| VHITDAE | ASSPGLGGYEQY | 0.090963826 |
| EYAAPSF | ASSPGLGGYEQY | 0.090938784 |
| VVDAFHGGL | ASSPGLGGYEQY | 0.09084185 |
| ASVVDPQ | ASSPGLGGYEQY | 0.090888225 |
| SYNEFGGLS | ASSPGLGGYEQY | 0.090756595 |
| DPDGSVY | ASSPGLGGYEQY | 0.09097964 |
| MVSTIHATE | ASSPGLGGYEQY | 0.09075947 |
| LAAGVADPV | ASSPGLGGYEQY | 0.090992756 |
| TKEQIAA | ASSPGLGGYEQY | 0.0908243 |
| SLASPAAL | ASSPGLGGYEQY | 0.09041618 |
| IGLVFYGV | ASSPGLGGYEQY | 0.09132078 |
| FLLDASNL | ASSPGLGGYEQY | 0.09108768 |
| YQVGNFP | ASSPGLGGYEQY | 0.092517056 |
| ESLLSQST | ASSPGLGGYEQY | 0.09111562 |
| LFTVGFAAVG | ASSPGLGGYEQY | 0.09094477 |
| ATPPTPT | ASSPGLGGYEQY | 0.09097636 |
| TDLATLG | ASSPGLGGYEQY | 0.09089388 |
| SAGGTGR | ASSPGLGGYEQY | 0.09081728 |
| GFSNSGS | ASSPGLGGYEQY | 0.09086587 |
| GGNGGSGGTSV | ASSPGLGGYEQY | 0.09114261 |
| GTSVATGGAGN | ASSPGLGGYEQY | 0.09065384 |
| NGGNGGSGGTS | ASSPGLGGYEQY | 0.092459224 |
| AVVKFPH | ASSPGLGGYEQY | 0.09162126 |
| TPPPPPP | ASSPGLGGYEQY | 0.09083551 |
| AAPAAAAAPA | ASSPGLGGYEQY | 0.090848595 |
| IALNEDL | ASSPGLGGYEQY | 0.091411784 |
| IVNVSSM | ASSPGLGGYEQY | 0.091019355 |
| TGIPPAPR | ASSPGLGGYEQY | 0.091662236 |
| FDLGGGTF | ASSPGLGGYEQY | 0.09089807 |
| FDLGGGTFDVS | ASSPGLGGYEQY | 0.09091442 |
| NEPTAAA | ASSPGLGGYEQY | 0.0916359 |
| NEPTAAALA | ASSPGLGGYEQY | 0.09077433 |
| GSIINISSI | ASSPGLGGYEQY | 0.09087088 |
| VAPGNAGTA | ASSPGLGGYEQY | 0.09084406 |
| TGGGASLE | ASSPGLGGYEQY | 0.090995476 |

| | | |
|---|---|---|
| VAVYNHYKRI | ASSPGLGGYEQY | 0.091084436 |
| TDPAPATT | ASSPGLGGYEQY | 0.09063828 |
| TNRPDILDPALLRPG | ASSPGLGGYEQY | 0.091530986 |
| LTTTTTS | ASSPGLGGYEQY | 0.0908712 |
| PTGSGKTL | ASSPGLGGYEQY | 0.090688616 |
| TGSGKTL | ASSPGLGGYEQY | 0.09155907 |
| TIGHVDHGK | ASSPGLGGYEQY | 0.09055883 |
| TIGHVDHGKTT | ASSPGLGGYEQY | 0.09063359 |
| GHVDHGKTT | ASSPGLGGYEQY | 0.09068357 |
| AQFEHTLL | ASSPGLGGYEQY | 0.09118248 |
| IVNQDEL | ASSPGLGGYEQY | 0.09328786 |
| TNGSQFF | ASSPGLGGYEQY | 0.09100846 |
| TGKGIVE | ASSPGLGGYEQY | 0.09082863 |
| LISNASDA | ASSPGLGGYEQY | 0.5586102 |
| AELRTLGELTKIAKAHGAQ | ASSPGLGGYEQY | 0.68513083 |
| APNAVAAAEPSW | ASSPGLGGYEQY | 0.4341022 |
| APQVAGAATAVL | ASSPGLGGYEQY | 0.37569684 |
| ARRDPAAAAAILPTDA | ASSPGLGGYEQY | 0.5221379 |
| DAAARDVLAEAGLAEYF | ASSPGLGGYEQY | 0.6487373 |
| DDQIVLSAKANLAKVEQ | ASSPGLGGYEQY | 0.6988623 |
| DELLPDTGEDSDEDGH | ASSPGLGGYEQY | 0.6172309 |
| ELPEAPANRATVPQD | ASSPGLGGYEQY | 0.4273787 |
| FGQSKIERVTGIG | ASSPGLGGYEQY | 0.5323255 |
| GAGGTGSTAGGAGGAGGAGGLYA | ASSPGLGGYEQY | 0.4860593 |
| GHGGTGAAGGNGGTGGAGGSGIDG | ASSPGLGGYEQY | 0.46734065 |
| GNGGNGGAGGEGGPGGAGGAGA | ASSPGLGGYEQY | 0.50870687 |
| GSTAGNGGNGGNATNGTIAGQPAG | ASSPGLGGYEQY | 0.57708883 |
| GTGGDGGNGGNGGTGG | ASSPGLGGYEQY | 0.45514643 |
| GVRRPAGIPADAPV | ASSPGLGGYEQY | 0.40973777 |
| GYRQVTDAHLLTLARR | ASSPGLGGYEQY | 0.51812565 |
| IAKMGQDGHDRG | ASSPGLGGYEQY | 0.62165403 |
| KAEPSIEIQYT | ASSPGLGGYEQY | 0.54434943 |
| KPVRIRMAADEDTVAVT | ASSPGLGGYEQY | 0.69028395 |
| MPPRIAGMRLLVIKPEP | ASSPGLGGYEQY | 0.63449913 |
| MKFYAVALFRSRGVPPTR | ASSPGLGGYEQY | 0.512627 |
| NAGGFGILGAVAHSP | ASSPGLGGYEQY | 0.20293927 |
| NEGREITLAMAEM | ASSPGLGGYEQY | 0.24238089 |
| NELVEWSHPLGHRWRWE | ASSPGLGGYEQY | 0.46665084 |
| NLGAGGTGGNGGNPG | ASSPGLGGYEQY | 0.49494505 |
| NTEWPKWETVMAADPVD | ASSPGLGGYEQY | 0.21706486 |
| PASAAPAETQP | ASSPGLGGYEQY | 0.5865054 |
| QQSSDGKNGVRAAEITGE | ASSPGLGGYEQY | 0.51383305 |
| RAPDAVTTSWWKE | ASSPGLGGYEQY | 0.4201688 |
| SGFGNVGAHTSGWFNQ | ASSPGLGGYEQY | 0.6404985 |
| SGVAGAVGQAASVGGLKV | ASSPGLGGYEQY | 0.4700215 |
| SNGGAGTGSAGGNGGT | ASSPGLGGYEQY | 0.36783636 |
| TGGAGGEGGAAG | ASSPGLGGYEQY | 0.42146277 |
| TGGDGGAAGAGGAPGHG | ASSPGLGGYEQY | 0.7171272 |
| TMPPLLAAIANAGRHGV | ASSPGLGGYEQY | 0.28556994 |
| TSSTAGNSGGA | ASSPGLGGYEQY | 0.63929045 |
| VPATVPVGILC | ASSPGLGGYEQY | 0.41456077 |

| | | |
|---|---|---|
| VQRPDSGGVVLTGRIS | ASSPGLGGYEQY | 0.6495333 |
| WTPNPVPATKGVPAGLVQAT | ASSPGLGGYEQY | 0.32856065 |
| YALKRALAKFSDRFDVVII | ASSPGLGGYEQY | 0.6177937 |
| YLIGSVIV | ASSPGLGGYEQY | 0.48814178 |
| YGQLVAVID | ASSPGLGGYEQY | 0.72187877 |
| VTVGGMTLPA | ASSPGLGGYEQY | 0.41603035 |
| VLALPAV | ASSPGLGGYEQY | 0.5931521 |
| VIPPTLNYET | ASSPGLGGYEQY | 0.4486286 |
| VGMGGMHPSSG | ASSPGLGGYEQY | 0.67081094 |
| VGLAGGAATA | ASSPGLGGYEQY | 0.30403528 |
| VGGAGGAGGSGP | ASSPGLGGYEQY | 0.4695018 |
| VFAAFMV | ASSPGLGGYEQY | 0.27960292 |
| TLYDKLR | ASSPGLGGYEQY | 0.53656745 |
| TLAVFTVL | ASSPGLGGYEQY | 0.6015042 |
| TGSVQPARV | ASSPGLGGYEQY | 0.272562 |
| TGGGAGSGGT | ASSPGLGGYEQY | 0.66095865 |
| SHLETSL | ASSPGLGGYEQY | 0.465846 |
| SFTTGMT | ASSPGLGGYEQY | 0.41183487 |
| QVEVAKRMGVRQPTVS | ASSPGLGGYEQY | 0.6200654 |
| QGSGGPALAM | ASSPGLGGYEQY | 0.46423656 |
| MLAPSNP | ASSPGLGGYEQY | 0.65288997 |
| LVNNAAI | ASSPGLGGYEQY | 0.37093425 |
| LTGEINTLA | ASSPGLGGYEQY | 0.54862165 |
| LSIFGPV | ASSPGLGGYEQY | 0.6025979 |
| LLALLANL | ASSPGLGGYEQY | 0.3219422 |
| LISNASDALDKLR | ASSPGLGGYEQY | 0.37054554 |
| LINEAVP | ASSPGLGGYEQY | 0.64533806 |
| LGYAFSGL | ASSPGLGGYEQY | 0.76534337 |
| LDFVAVAL | ASSPGLGGYEQY | 0.7648773 |
| LAQPGAE | ASSPGLGGYEQY | 0.69940233 |
| ISNLGAVTGF | ASSPGLGGYEQY | 0.6939366 |
| HQEDPGRI | ASSPGLGGYEQY | 0.69319284 |
| GTPGEPTP | ASSPGLGGYEQY | 0.7664094 |
| GFFNSGS | ASSPGLGGYEQY | 0.77302057 |
| GAGGGGLTAGAN | ASSPGLGGYEQY | 0.76303864 |
| FVDALVNM | ASSPGLGGYEQY | 0.6891217 |
| FLTVAPEL | ASSPGLGGYEQY | 0.75341505 |
| FETLTAL | ASSPGLGGYEQY | 0.7474902 |
| DVINPIEL | ASSPGLGGYEQY | 0.74096715 |
| APVSGAPV | ASSPGLGGYEQY | 0.7578969 |
| ALVGGLAW | ASSPGLGGYEQY | 0.7510879 |
| ALAAAAATAI | ASSPGLGGYEQY | 0.70286596 |
| AFGLGLLL | ASSPGLGGYEQY | 0.6781591 |
| AFFAAASL | ASSPGLGGYEQY | 0.7383522 |
| AAAALPAFSPP | ASSPGLGGYEQY | 0.76707053 |
| VVLAPAL | ASSPGLGGYEQY | 0.7606884 |
| TIGHVDHGKTTL | ASSPGLGGYEQY | 0.7733321 |
| LISNASDALDK | ASSPGLGGYEQY | 0.76669335 |
| LISNASDAL | ASSPGLGGYEQY | 0.76158744 |
| ISNASDALDK | ASSPGLGGYEQY | 0.37432662 |
| NQAETLVYQTE | ASSPGLGGYEQY | 0.58548087 |

| | | |
|---|---|---|
| TGGTGGTAGAG | ASSPGLGGYEQY | 0.70699584 |
| TGSTAGGAGGA | ASSPGLGGYEQY | 0.7282032 |
| DPLAVPVAAGPV | ASSPGLGGYEQY | 0.42913705 |
| GGTGGTPGNGGN | ASSPGLGGYEQY | 0.7500049 |
| PAGLAGVDPATL | ASSPGLGGYEQY | 0.7130656 |
| GNNLVVEAPHAGASV | ASSPGLGGYEQY | 0.7045611 |
| KVTRIMSASTLEQPA | ASSPGLGGYEQY | 0.7597846 |
| LGFALPIALPRRTVV | ASSPGLGGYEQY | 0.7300948 |
| TRLGNLTARKLVVLV | ASSPGLGGYEQY | 0.66906154 |
| WSQQRRLIYFPSAGP | ASSPGLGGYEQY | 0.66386294 |
| GPNGNPDPMAAAVDIR | ASSPGLGGYEQY | 0.767269 |
| TQRCVSVADLRAAASR | ASSPGLGGYEQY | 0.6764835 |
| QFVLDVAEHPVEVLTETV | ASSPGLGGYEQY | 0.76659584 |
| GGAGGTGGAGGNGGTQAGDG | ASSPGLGGYEQY | 0.6362076 |
| PLLATLRAIAVDVDAKPAQV | ASSPGLGGYEQY | 0.7702969 |
| APKFRTGGYMIDFWGVGYQVAKRM | ASSPGLGGYEQY | 0.6739768 |
| GAGGAGGAGGAAGGPGGTGGTGGN | ASSPGLGGYEQY | 0.7675574 |
| IGALGLLATDVAPAGSVAFGVALA | ASSPGLGGYEQY | 0.7490436 |
| AGGAGGIGGNGGAGGAGGVLMGNG | ASSPGLGGYEQY | 0.7767129 |
| ADELIGLL | ASSPGLGGYEQY | 0.739572 |
| ADRALKA | ASSPGLGGYEQY | 0.76531804 |
| AGAVGQVGGAGGA | ASSPGLGGYEQY | 0.7372717 |
| AGGGGGLGGAP | ASSPGLGGYEQY | 0.75847054 |
| AGGLLYGNGGAG | ASSPGLGGYEQY | 0.6470283 |
| AGLESIGL | ASSPGLGGYEQY | 0.73185086 |
| AGLKTGII | ASSPGLGGYEQY | 0.6460613 |
| ASLAALLL | ASSPGLGGYEQY | 0.7427349 |
| ASVINISSI | ASSPGLGGYEQY | 0.766985 |
| ATPPGPA | ASSPGLGGYEQY | 0.3948384 |
| ATVASALV | ASSPGLGGYEQY | 0.7487697 |
| ATVASLIL | ASSPGLGGYEQY | 0.7124333 |
| AVNGPVLIRH | ASSPGLGGYEQY | 0.71518743 |
| DGAAGAVGI | ASSPGLGGYEQY | 0.76738274 |
| DPLLHGI | ASSPGLGGYEQY | 0.73358727 |
| ELGAALGAL | ASSPGLGGYEQY | 0.76148593 |
| ELGRTQAV | ASSPGLGGYEQY | 0.69202566 |
| FVAGGYLS | ASSPGLGGYEQY | 0.71650815 |
| GASGGAGGAGGW | ASSPGLGGYEQY | 0.74138206 |
| GGAGAGGDGGAGGI | ASSPGLGGYEQY | 0.39029488 |
| GTGHNGSQPGMGGNG | ASSPGLGGYEQY | 0.5988865 |
| GYAAASASAS | ASSPGLGGYEQY | 0.7744854 |
| HQVDVAAVQQG | ASSPGLGGYEQY | 0.20906162 |
| IGAAIAHT | ASSPGLGGYEQY | 0.20913279 |
| IGMIEVAQL | ASSPGLGGYEQY | 0.20896195 |
| IRLAPEHTSAMD | ASSPGLGGYEQY | 0.21017657 |
| IYNVSVL | ASSPGLGGYEQY | 0.20908621 |
| KQGFLGM | ASSPGLGGYEQY | 0.20920104 |
| LESVITQL | ASSPGLGGYEQY | 0.21394353 |
| LGLGALVVV | ASSPGLGGYEQY | 0.20909472 |
| LIGNGGNGGSGGI | ASSPGLGGYEQY | 0.21089593 |
| LNAINAPTL | ASSPGLGGYEQY | 0.21091773 |

| | | |
|---|---|---|
| LQLLNTGV | ASSPGLGGYEQY | 0.20907277 |
| LQTSIEF | ASSPGLGGYEQY | 0.20925468 |
| MSFAVSEL | ASSPGLGGYEQY | 0.20910068 |
| NTGALGL | ASSPGLGGYEQY | 0.21196932 |
| PPGAGGAGGAGTT | ASSPGLGGYEQY | 0.20898962 |
| QLVGLGI | ASSPGLGGYEQY | 0.20894475 |
| RFLEQVHQL | ASSPGLGGYEQY | 0.20939887 |
| RGVGYVLR | ASSPGLGGYEQY | 0.20952085 |
| SATGVFAGVI | ASSPGLGGYEQY | 0.20941602 |
| SLPAITLP | ASSPGLGGYEQY | 0.20933147 |
| STALAFTV | ASSPGLGGYEQY | 0.20922086 |
| SVLTVGLGK | ASSPGLGGYEQY | 0.20910521 |
| TALAGAGSGI | ASSPGLGGYEQY | 0.20925748 |
| TGLPFTGI | ASSPGLGGYEQY | 0.20949763 |
| VAEHVTV | ASSPGLGGYEQY | 0.20921102 |
| VDIVLNSL | ASSPGLGGYEQY | 0.21000299 |
| VLTGGGAVL | ASSPGLGGYEQY | 0.20841904 |
| VNVGLAYGI | ASSPGLGGYEQY | 0.21152778 |
| VSSAAVGAV | ASSPGLGGYEQY | 0.20940028 |
| VTSVYEAGV | ASSPGLGGYEQY | 0.21098287 |
| WVALDRGA | ASSPGLGGYEQY | 0.20972349 |
| YLVGSVVI | ASSPGLGGYEQY | 0.20915554 |
| GPPGTGKTLL | ASSPGLGGYEQY | 0.21022311 |
| GVGMTKF | ASSPGLGGYEQY | 0.20920295 |
| IGIGGSDL | ASSPGLGGYEQY | 0.20953844 |
| IGTLLAEL | ASSPGLGGYEQY | 0.20917663 |
| IYTSGTT | ASSPGLGGYEQY | 0.21086317 |
| LSFPAGV | ASSPGLGGYEQY | 0.20890963 |
| LYTSGTT | ASSPGLGGYEQY | 0.21023484 |
| VGAFPFGF | ASSPGLGGYEQY | 0.21069282 |
| VGAGTVEF | ASSPGLGGYEQY | 0.20911376 |
| VGKSTLLL | ASSPGLGGYEQY | 0.20908612 |
| AALAHAGGGIV | ASSPGLGGYEQY | 0.21004781 |
| AALVAVSLL | ASSPGLGGYEQY | 0.2100743 |
| AAVSRAVAAL | ASSPGLGGYEQY | 0.21151759 |
| AGGSSGIGGF | ASSPGLGGYEQY | 0.20975366 |
| AGLTVANANG | ASSPGLGGYEQY | 0.2149207 |
| AGLVGAVL | ASSPGLGGYEQY | 0.21003571 |
| ALSQAFTV | ASSPGLGGYEQY | 0.20917101 |
| ALTFSVSL | ASSPGLGGYEQY | 0.20920216 |
| ALTGAALVAL | ASSPGLGGYEQY | 0.20933709 |
| ASFSIGAGVL | ASSPGLGGYEQY | 0.20906541 |
| DGSVYKI | ASSPGLGGYEQY | 0.20884402 |
| EQLAEIV | ASSPGLGGYEQY | 0.20891184 |
| FAVSELTVM | ASSPGLGGYEQY | 0.20881316 |
| GAAPLAAGL | ASSPGLGGYEQY | 0.20915467 |
| GADGGDGGSGGAGG | ASSPGLGGYEQY | 0.20901835 |
| GFGLDTGGAGG | ASSPGLGGYEQY | 0.21097422 |
| GLASGGLL | ASSPGLGGYEQY | 0.2087616 |
| GRIAGAAVLL | ASSPGLGGYEQY | 0.20889574 |
| GSAGSLNTSF | ASSPGLGGYEQY | 0.20898041 |

| | | |
|---|---|---|
| GTGQIGI | ASSPGLGGYEQY | 0.20944801 |
| GTLIALIL | ASSPGLGGYEQY | 0.20939513 |
| IAATMGFVVG | ASSPGLGGYEQY | 0.20944536 |
| IFAQALL | ASSPGLGGYEQY | 0.20916235 |
| IGAAVALGF | ASSPGLGGYEQY | 0.08976493 |
| IGLTPLAL | ASSPGLGGYEQY | 0.08746554 |
| ILGSNTSTL | ASSPGLGGYEQY | 0.08344513 |
| IVFSLTVL | ASSPGLGGYEQY | 0.08734004 |
| LASKATGF | ASSPGLGGYEQY | 0.08565955 |
| LASVALIL | ASSPGLGGYEQY | 0.08869664 |
| LFAVGLTL | ASSPGLGGYEQY | 0.09080137 |
| LFNGGIGTYI | ASSPGLGGYEQY | 0.08830339 |
| LTLGAGII | ASSPGLGGYEQY | 0.090175115 |
| LVAAALGDI | ASSPGLGGYEQY | 0.08777653 |
| NTAFVQLGI | ASSPGLGGYEQY | 0.08572643 |
| QLFSGAEL | ASSPGLGGYEQY | 0.08835532 |
| SAATFITL | ASSPGLGGYEQY | 0.08615793 |
| SNYGLANLGV | ASSPGLGGYEQY | 0.08692683 |
| SYGGAYAVM | ASSPGLGGYEQY | 0.084468566 |
| TTGVGGLTLG | ASSPGLGGYEQY | 0.084189184 |
| TTQTIYI | ASSPGLGGYEQY | 0.08790796 |
| VAAGAGLGAVI | ASSPGLGGYEQY | 0.09548396 |
| VAAVVTLL | ASSPGLGGYEQY | 0.09326639 |
| VGFLGGLI | ASSPGLGGYEQY | 0.093252204 |
| VGVIPILL | ASSPGLGGYEQY | 0.093464546 |
| VIFASSLI | ASSPGLGGYEQY | 0.085503004 |
| VKGGDGGAAAG | ASSPGLGGYEQY | 0.08324056 |
| VTKTFTNI | ASSPGLGGYEQY | 0.08518783 |
| VVAAAILL | ASSPGLGGYEQY | 0.08489868 |
| YAALDVEL | ASSPGLGGYEQY | 0.087492615 |
| LASVALLL | ASSPGLGGYEQY | 0.08121122 |
| LDAVHRV | ASSPGLGGYEQY | 0.08749962 |
| QGLSSSIF | ASSPGLGGYEQY | 0.089794524 |
| RGTFANIRL | ASSPGLGGYEQY | 0.08438225 |
| RQLWWGHRI | ASSPGLGGYEQY | 0.092639536 |
| YGPPGTGKTLL | ASSPGLGGYEQY | 0.08406665 |
| AALVGAPANVA | ASSPGLGGYEQY | 0.08565849 |
| AATFRRTE | ASSPGLGGYEQY | 0.08528597 |
| AGALGGGI | ASSPGLGGYEQY | 0.08952398 |
| AGGAGGAGGIGAG | ASSPGLGGYEQY | 0.08380038 |
| AGSGALGL | ASSPGLGGYEQY | 0.09353793 |
| AQIADQI | ASSPGLGGYEQY | 0.08542385 |
| ASAVAVTYL | ASSPGLGGYEQY | 0.09913174 |
| ASTVLGGTGG | ASSPGLGGYEQY | 0.086000025 |
| AVGGEPAK | ASSPGLGGYEQY | 0.088149905 |
| CEQGTFGGTC | ASSPGLGGYEQY | 0.08487431 |
| DEGKAPAG | ASSPGLGGYEQY | 0.097799905 |
| DTVVFPAL | ASSPGLGGYEQY | 0.09110693 |
| FEGITMAG | ASSPGLGGYEQY | 0.095786065 |
| FPLVVQD | ASSPGLGGYEQY | 0.08738009 |
| GAFTTPQ | ASSPGLGGYEQY | 0.09340123 |

| | | |
|---|---|---|
| GFGGFGTAGGVGGA | ASSPGLGGYEQY | 0.08368642 |
| GFLAQFL | ASSPGLGGYEQY | 0.087361455 |
| GGAGGAGGNAGVA | ASSPGLGGYEQY | 0.08527246 |
| GGVGTAGAGGAGGAGG | ASSPGLGGYEQY | 0.087537944 |
| GNGGGGDAGNAGSGGN | ASSPGLGGYEQY | 0.08968675 |
| GTGGAGGAGNDAGS | ASSPGLGGYEQY | 0.08205488 |
| GTTNTGWFNS | ASSPGLGGYEQY | 0.087250344 |
| IFMGPLFTL | ASSPGLGGYEQY | 0.08212726 |
| ILAHAIEAI | ASSPGLGGYEQY | 0.084795356 |
| IRSAIREAL | ASSPGLGGYEQY | 0.08612691 |
| ITFAGGG | ASSPGLGGYEQY | 0.08600894 |
| ITGMSGHQEV | ASSPGLGGYEQY | 0.08683551 |
| IVAGLACTL | ASSPGLGGYEQY | 0.082560815 |
| IVGGGLAGA | ASSPGLGGYEQY | 0.08749307 |
| IVTGVLNI | ASSPGLGGYEQY | 0.09551452 |
| LAQQGCI | ASSPGLGGYEQY | 0.08307394 |
| LCLHVEN | ASSPGLGGYEQY | 0.08392016 |
| LDAISDGS | ASSPGLGGYEQY | 0.0914424 |
| LGGFAAML | ASSPGLGGYEQY | 0.21572353 |
| LGGKNPMI | ASSPGLGGYEQY | 0.22497278 |
| LGPKGVAAIL | ASSPGLGGYEQY | 0.21278486 |
| LMVGVSI | ASSPGLGGYEQY | 0.22101341 |
| LVGAAFVL | ASSPGLGGYEQY | 0.21445082 |
| LVLAVAPL | ASSPGLGGYEQY | 0.21629529 |
| MAAIAVIDAA | ASSPGLGGYEQY | 0.2582958 |
| NNGINAELQ | ASSPGLGGYEQY | 0.21822456 |
| RGEPLVLVHG | ASSPGLGGYEQY | 0.22641 |
| RGGAGGAAGLI | ASSPGLGGYEQY | 0.22102408 |
| RSYRMHPAV | ASSPGLGGYEQY | 0.21298982 |
| SAAHLLAGQLA | ASSPGLGGYEQY | 0.21718918 |
| SAAVGSY | ASSPGLGGYEQY | 0.21473974 |
| SPHAVLHRI | ASSPGLGGYEQY | 0.22122148 |
| SQIAKVL | ASSPGLGGYEQY | 0.21224444 |
| TALGHLVP | ASSPGLGGYEQY | 0.2154519 |
| TAPTTSGAA | ASSPGLGGYEQY | 0.21843088 |
| TDGDDGNGGSPGT | ASSPGLGGYEQY | 0.22941004 |
| TDGLGEN | ASSPGLGGYEQY | 0.23253554 |
| TGGPPLPGQA | ASSPGLGGYEQY | 0.22148824 |
| TIPAITI | ASSPGLGGYEQY | 0.21708398 |
| TIPGFTI | ASSPGLGGYEQY | 0.21783912 |
| TSIASGALA | ASSPGLGGYEQY | 0.2128978 |
| TVTLPGAAI | ASSPGLGGYEQY | 0.21651657 |
| VASGLSAALP | ASSPGLGGYEQY | 0.21479462 |
| VGAGVIGI | ASSPGLGGYEQY | 0.22074838 |
| VGPKRFGV | ASSPGLGGYEQY | 0.21073942 |
| VNVAFPD | ASSPGLGGYEQY | 0.22190498 |
| VSMTNSGVS | ASSPGLGGYEQY | 0.2149654 |
| VTDVLGDLSV | ASSPGLGGYEQY | 0.2173555 |
| VVVVGAGFAGL | ASSPGLGGYEQY | 0.22329701 |
| VYTVGTVV | ASSPGLGGYEQY | 0.21429417 |
| WTTFSGP | ASSPGLGGYEQY | 0.21789397 |

| | | |
|---|---|---|
| YDELVEAF | ASSPGLGGYEQY | 0.21495092 |
| YGTGGAGGNGG | ASSPGLGGYEQY | 0.22424449 |
| YTNVMAV | ASSPGLGGYEQY | 0.21347894 |
| FLEQVHQL | ASSPGLGGYEQY | 0.22952583 |
| GLSLLAVAV | ASSPGLGGYEQY | 0.21327469 |
| LEQVHQL | ASSPGLGGYEQY | 0.22496617 |
| LNAAGAI | ASSPGLGGYEQY | 0.2191466 |
| PAVHASL | ASSPGLGGYEQY | 0.2162478 |
| TELEDDA | ASSPGLGGYEQY | 0.21456248 |
| TPPGPPN | ASSPGLGGYEQY | 0.23929016 |
| TPPPPPPP | ASSPGLGGYEQY | 0.22273012 |
| TQSPALI | ASGEPGHQAPL | 0.23000042 |
| EFAAAFPQ | ASGEPGHQAPL | 0.21922387 |
| LLSGNLDVL | ASGEPGHQAPL | 0.23263176 |
| GPANLGNYNI | ASGEPGHQAPL | 0.21426108 |
| TLGPTLP | ASGEPGHQAPL | 0.2153624 |
| VYAYSSCCK | ASGEPGHQAPL | 0.21431348 |
| APAPAPAPNI | ASGEPGHQAPL | 0.21592417 |
| PGGPVPIP | ASGEPGHQAPL | 0.2175894 |
| FLTVALEP | ASGEPGHQAPL | 0.21178868 |
| SQYFGANL | ASGEPGHQAPL | 0.2184128 |
| GGVKPAA | ASGEPGHQAPL | 0.21160027 |
| GGTGGAGGAGGW | ASGEPGHQAPL | 0.21552123 |
| GVGGVAGLGGAGG | ASGEPGHQAPL | 0.21662869 |
| VLLVLATTI | ASGEPGHQAPL | 0.22081852 |
| IGIFAAML | ASGEPGHQAPL | 0.21424025 |
| AVGDLID | ASGEPGHQAPL | 0.21224657 |
| NISPLPL | ASGEPGHQAPL | 0.21371053 |
| VTAVFVAI | ASGEPGHQAPL | 0.2202901 |
| SGDDNDAAL | ASGEPGHQAPL | 0.21392459 |
| AVAALAHS | ASGEPGHQAPL | 0.21496381 |
| GGVTVMATP | ASGEPGHQAPL | 0.21874811 |
| AGLAANVRD | ASGEPGHQAPL | 0.72601664 |
| TNRPDLIDPALLRPG | ASGEPGHQAPL | 0.79135096 |
| LANLGVTAL | ASGEPGHQAPL | 0.64013064 |
| TGERIEL | ASGEPGHQAPL | 0.5276003 |
| NGEELAA | ASGEPGHQAPL | 0.6801814 |
| VLSLVWG | ASGEPGHQAPL | 0.77422184 |
| IGYALWL | ASGEPGHQAPL | 0.783016 |
| LSLLLAPL | ASGEPGHQAPL | 0.7472155 |
| TLDEALK | ASGEPGHQAPL | 0.52742106 |
| TDGAGTAEGG | ASGEPGHQAPL | 0.6948797 |
| NAPGGAGGNGG | ASGEPGHQAPL | 0.6723532 |
| VSVFMGMGGNF | ASGEPGHQAPL | 0.6691985 |
| LASYLAAL | ASGEPGHQAPL | 0.7595727 |
| LVGGTPAL | ASGEPGHQAPL | 0.7242404 |
| VRNADVL | ASGEPGHQAPL | 0.6567657 |
| TCSSPFLVSAM | ASGEPGHQAPL | 0.6403699 |
| VHITDAE | ASGEPGHQAPL | 0.7336767 |
| EYAAPSF | ASGEPGHQAPL | 0.7594627 |
| VVDAFHGGL | ASGEPGHQAPL | 0.74012065 |

| | | |
|---|---|---|
| ASVVDPQ | ASGEPGHQAPL | 0.8058381 |
| SYNEFGGLS | ASGEPGHQAPL | 0.7708598 |
| DPDGSVY | ASGEPGHQAPL | 0.7721589 |
| MVSTIHATE | ASGEPGHQAPL | 0.28309354 |
| LAAGVADPV | ASGEPGHQAPL | 0.40877128 |
| TKEQIAA | ASGEPGHQAPL | 0.6403048 |
| SLASPAAL | ASGEPGHQAPL | 0.64282066 |
| IGLVFYGV | ASGEPGHQAPL | 0.43071637 |
| FLLDASNL | ASGEPGHQAPL | 0.7132609 |
| YQVGNFP | ASGEPGHQAPL | 0.6462208 |
| ESLLSQST | ASGEPGHQAPL | 0.55715126 |
| LFTVGFAAVG | ASGEPGHQAPL | 0.77374315 |
| ATPPTPT | ASGEPGHQAPL | 0.6488277 |
| TDLATLG | ASGEPGHQAPL | 0.6152353 |
| SAGGTGR | ASGEPGHQAPL | 0.6281519 |
| GFSNSGS | ASGEPGHQAPL | 0.7897457 |
| GGNGGSGGTSV | ASGEPGHQAPL | 0.4959703 |
| GTSVATGGAGN | ASGEPGHQAPL | 0.72778285 |
| NGGNGGSGGTS | ASGEPGHQAPL | 0.71163464 |
| AVVKFPH | ASGEPGHQAPL | 0.77723557 |
| TPPPPPP | ASGEPGHQAPL | 0.47181156 |
| AAPAAAAAPA | ASGEPGHQAPL | 0.75455576 |
| IALNEDL | ASGEPGHQAPL | 0.6667428 |
| IVNVSSM | ASGEPGHQAPL | 0.8141651 |
| TGIPPAPR | ASGEPGHQAPL | 0.65425295 |
| FDLGGGTF | ASGEPGHQAPL | 0.72589266 |
| FDLGGGTFDVS | ASGEPGHQAPL | 0.6866211 |
| NEPTAAA | ASGEPGHQAPL | 0.7691666 |
| NEPTAAALA | ASGEPGHQAPL | 0.4797328 |
| GSIINISSI | ASGEPGHQAPL | 0.67577994 |
| VAPGNAGTA | ASGEPGHQAPL | 0.544893 |
| TGGGASLE | ASGEPGHQAPL | 0.72060627 |
| VAVYNHYKRI | ASGEPGHQAPL | 0.74450827 |
| TDPAPATT | ASGEPGHQAPL | 0.48741814 |
| TNRPDILDPALLRPG | ASGEPGHQAPL | 0.79102373 |
| LTTTTTS | ASGEPGHQAPL | 0.708872 |
| PTGSGKTL | ASGEPGHQAPL | 0.6071199 |
| TGSGKTL | ASGEPGHQAPL | 0.7874594 |
| TIGHVDHGK | ASGEPGHQAPL | 0.65844965 |
| TIGHVDHGKTT | ASGEPGHQAPL | 0.7809899 |
| GHVDHGKTT | ASGEPGHQAPL | 0.6986332 |
| AQFEHTLL | ASGEPGHQAPL | 0.7112937 |
| IVNQDEL | ASGEPGHQAPL | 0.7550063 |
| TNGSQFF | ASGEPGHQAPL | 0.60791165 |
| TGKGIVE | ASGEPGHQAPL | 0.56872195 |
| LISNASDA | ASGEPGHQAPL | 0.77774644 |
| AELRTLGELTKIAKAHGAQ | ASGEPGHQAPL | 0.63612556 |
| APNAVAAAEPSW | ASGEPGHQAPL | 0.84444475 |
| APQVAGAATAVL | ASGEPGHQAPL | 0.41759342 |
| ARRDPAAAAAILPTDA | ASGEPGHQAPL | 0.46584624 |
| DAAARDVLAEAGLAEYF | ASGEPGHQAPL | 0.51042706 |

| | | |
|---|---|---|
| DDQIVLSAKANLAKVEQ | ASGEPGHQAPL | 0.6917253 |
| DELLPDTGEDSDEDGH | ASGEPGHQAPL | 0.8364042 |
| ELPEAPANRATVPQD | ASGEPGHQAPL | 0.7681781 |
| FGQSKIERVTGIG | ASGEPGHQAPL | 0.4715892 |
| GAGGTGSTAGGAGGAGGAGGLYA | ASGEPGHQAPL | 0.6129892 |
| GHGGTGAAGGNGGTGGAGGSGIDG | ASGEPGHQAPL | 0.5137621 |
| GNGGNGGAGGEGGPGGAGGAGA | ASGEPGHQAPL | 0.5384046 |
| GSTAGNGGNGGNATNGTIAGGQPAG | ASGEPGHQAPL | 0.5885927 |
| GTGGDGGNGGNGGTGG | ASGEPGHQAPL | 0.5539274 |
| GVRRPAGIPADAPV | ASGEPGHQAPL | 0.52817947 |
| GYRQVTDAHLLTLARR | ASGEPGHQAPL | 0.37988573 |
| IAKMGQDGHDRG | ASGEPGHQAPL | 0.49992576 |
| KAEPSIEIQYT | ASGEPGHQAPL | 0.7127602 |
| KPVRIRMAADEDTVAVT | ASGEPGHQAPL | 0.6196394 |
| MPPRIAGMRLLVIKPEP | ASGEPGHQAPL | 0.76724195 |
| MKFYAVALFRSRGVPPTR | ASGEPGHQAPL | 0.694507 |
| NAGGFGILGAVAHSP | ASGEPGHQAPL | 0.50090736 |
| NEGREITLAMAEM | ASGEPGHQAPL | 0.15452503 |
| NELVEWSHPLGHRWRWE | ASGEPGHQAPL | 0.27909908 |
| NLGAGGTGGNGGNPG | ASGEPGHQAPL | 0.43625838 |
| NTEWPKWETVMAADPVD | ASGEPGHQAPL | 0.5605171 |
| PASAAPAETQP | ASGEPGHQAPL | 0.16332853 |
| QQSSDGKNGVRAAEITGE | ASGEPGHQAPL | 0.6271636 |
| RAPDAVTTSWWKE | ASGEPGHQAPL | 0.55336535 |
| SGFGNVGAHTSGWFNQ | ASGEPGHQAPL | 0.5257058 |
| SGVAGAVGQAASVGGLKV | ASGEPGHQAPL | 0.72271526 |
| SNGGAGTGSAGGNGGT | ASGEPGHQAPL | 0.58891624 |
| TGGAGGEGGAAG | ASGEPGHQAPL | 0.37084413 |
| TGGDGGAAGAGGAPGHG | ASGEPGHQAPL | 0.37127596 |
| TMPPLLAAIANAGRHGV | ASGEPGHQAPL | 0.7564964 |
| TSSTAGNSGGA | ASGEPGHQAPL | 0.23235063 |
| VPATVPVGILC | ASGEPGHQAPL | 0.67244947 |
| VQRPDSGGVVLTGRIS | ASGEPGHQAPL | 0.42088297 |
| WTPNPVPATKGVPAGLVQAT | ASGEPGHQAPL | 0.71410966 |
| YALKRALAKFSDRFDVVII | ASGEPGHQAPL | 0.35435435 |
| YLIGSVIV | ASGEPGHQAPL | 0.6994395 |
| YGQLVAVID | ASGEPGHQAPL | 0.5767812 |
| VTVGGMTLPA | ASGEPGHQAPL | 0.8079356 |
| VLALPAV | ASGEPGHQAPL | 0.46610776 |
| VIPPTLNYET | ASGEPGHQAPL | 0.66098547 |
| VGMGGMHPSSG | ASGEPGHQAPL | 0.4003726 |
| VGLAGGAATA | ASGEPGHQAPL | 0.82887244 |
| VGGAGGAGGSGP | ASGEPGHQAPL | 0.38499796 |
| VFAAFMV | ASGEPGHQAPL | 0.42404726 |
| TLYDKLR | ASGEPGHQAPL | 0.33685884 |
| TLAVFTVL | ASGEPGHQAPL | 0.52868783 |
| TGSVQPARV | ASGEPGHQAPL | 0.753232 |
| TGGGAGSGGT | ASGEPGHQAPL | 0.4765268 |
| SHLETSL | ASGEPGHQAPL | 0.7641401 |
| SFTTGMT | ASGEPGHQAPL | 0.62749517 |
| QVEVAKRMGVRQPTVS | ASGEPGHQAPL | 0.45483637 |

| | | |
|---|---|---|
| QGSGGPALAM | ASGEPGHQAPL | 0.6519576 |
| MLAPSNP | ASGEPGHQAPL | 0.46243387 |
| LVNNAAI | ASGEPGHQAPL | 0.6304015 |
| LTGEINTLA | ASGEPGHQAPL | 0.34079164 |
| LSIFGPV | ASGEPGHQAPL | 0.59617054 |
| LLALLANL | ASGEPGHQAPL | 0.678685 |
| LISNASDALDKLR | ASGEPGHQAPL | 0.44051793 |
| LINEAVP | ASGEPGHQAPL | 0.30522013 |
| LGYAFSGL | ASGEPGHQAPL | 0.76319164 |
| LDFVAVAL | ASGEPGHQAPL | 0.7683828 |
| LAQPGAE | ASGEPGHQAPL | 0.7220106 |
| ISNLGAVTGF | ASGEPGHQAPL | 0.54787606 |
| HQEDPGRI | ASGEPGHQAPL | 0.46637365 |
| GTPGEPTP | ASGEPGHQAPL | 0.6211793 |
| GFFNSGS | ASGEPGHQAPL | 0.7892545 |
| GAGGGGLTAGAN | ASGEPGHQAPL | 0.7891704 |
| FVDALVNM | ASGEPGHQAPL | 0.7207992 |
| FLTVAPEL | ASGEPGHQAPL | 0.4313956 |
| FETLTAL | ASGEPGHQAPL | 0.7551557 |
| DVINPIEL | ASGEPGHQAPL | 0.68191004 |
| APVSGAPV | ASGEPGHQAPL | 0.6577992 |
| ALVGGLAW | ASGEPGHQAPL | 0.75961435 |
| ALAAAAATAI | ASGEPGHQAPL | 0.7082001 |
| AFGLGLLL | ASGEPGHQAPL | 0.6099115 |
| AFFAAASL | ASGEPGHQAPL | 0.5025393 |
| AAAALPAFSPP | ASGEPGHQAPL | 0.7474998 |
| VVLAPAL | ASGEPGHQAPL | 0.77819675 |
| TIGHVDHGKTTL | ASGEPGHQAPL | 0.7704936 |
| LISNASDALDK | ASGEPGHQAPL | 0.8274871 |
| LISNASDAL | ASGEPGHQAPL | 0.8039962 |
| ISNASDALDK | ASGEPGHQAPL | 0.7092058 |
| NQAETLVYQTE | ASGEPGHQAPL | 0.1941903 |
| TGGTGGTAGAG | ASGEPGHQAPL | 0.33607164 |
| TGSTAGGAGGA | ASGEPGHQAPL | 0.5981327 |
| DPLAVPVAAGPV | ASGEPGHQAPL | 0.6233594 |
| GGTGGTPGNGGN | ASGEPGHQAPL | 0.24913727 |
| PAGLAGVDPATL | ASGEPGHQAPL | 0.7219402 |
| GNNLVVEAPHAGASV | ASGEPGHQAPL | 0.59610385 |
| KVTRIMSASTLEQPA | ASGEPGHQAPL | 0.5365108 |
| LGFALPIALPRRTVV | ASGEPGHQAPL | 0.80301964 |
| TRLGNLTARKLVVLV | ASGEPGHQAPL | 0.5863643 |
| WSQQRRLIYFPSAGP | ASGEPGHQAPL | 0.54227775 |
| GPNGNPDPMAAAVDIR | ASGEPGHQAPL | 0.59513557 |
| TQRCVSVADLRAAASR | ASGEPGHQAPL | 0.7868546 |
| QFVLDVAEHPVEVLTETV | ASGEPGHQAPL | 0.4901568 |
| GGAGGTGGAGGNGGTQAGDG | ASGEPGHQAPL | 0.78440875 |
| PLLATLRAIAVDVDAKPAQV | ASGEPGHQAPL | 0.5482248 |
| APKFRTGGYMIDFWGVGYQVAKRM | ASGEPGHQAPL | 0.83363223 |
| GAGGAGGAGGAAGGPGGTGGTGGN | ASGEPGHQAPL | 0.39160466 |
| IGALGLLATDVAPAGSVAFGVALA | ASGEPGHQAPL | 0.7814807 |
| AGGAGGIGGNGGAGGAGGVLMGNG | ASGEPGHQAPL | 0.684871 |

| | | |
|---|---|---|
| ADELIGLL | ASGEPGHQAPL | 0.86677605 |
| ADRALKA | ASGEPGHQAPL | 0.6389865 |
| AGAVGQVGGAGGA | ASGEPGHQAPL | 0.7242651 |
| AGGGGGLGGAP | ASGEPGHQAPL | 0.60948265 |
| AGGLLYGNGGAG | ASGEPGHQAPL | 0.7672098 |
| AGLESIGL | ASGEPGHQAPL | 0.36058593 |
| AGLKTGII | ASGEPGHQAPL | 0.65485185 |
| ASLAALLL | ASGEPGHQAPL | 0.32555893 |
| ASVINISSI | ASGEPGHQAPL | 0.6942206 |
| ATPPGPA | ASGEPGHQAPL | 0.7420863 |
| ATVASALV | ASGEPGHQAPL | 0.19961345 |
| ATVASLIL | ASGEPGHQAPL | 0.75813353 |
| AVNGPVLIRH | ASGEPGHQAPL | 0.47409835 |
| DGAAGAVGI | ASGEPGHQAPL | 0.4868011 |
| DPLLHGI | ASGEPGHQAPL | 0.79669726 |
| ELGAALGAL | ASGEPGHQAPL | 0.70214486 |
| ELGRTQAV | ASGEPGHQAPL | 0.76809895 |
| FVAGGYLS | ASGEPGHQAPL | 0.61072946 |
| GASGGAGGAGGW | ASGEPGHQAPL | 0.6508914 |
| GGAGAGGDGGAGGI | ASGEPGHQAPL | 0.6916313 |
| GTGHNGSQPGMGGNG | ASGEPGHQAPL | 0.35482168 |
| GYAAASASAS | ASGEPGHQAPL | 0.55096793 |
| HQVDVAAVQQG | ASGEPGHQAPL | 0.8177536 |
| IGAAIAHT | ASGEPGHQAPL | 0.66448855 |
| IGMIEVAQL | ASGEPGHQAPL | 0.6687147 |
| IRLAPEHTSAMD | ASGEPGHQAPL | 0.6354489 |
| IYNVSVL | ASGEPGHQAPL | 0.6343331 |
| KQGFLGM | ASGEPGHQAPL | 0.652681 |
| LESVITQL | ASGEPGHQAPL | 0.66745853 |
| LGLGALVVV | ASGEPGHQAPL | 0.6753318 |
| LIGNGGNGGSGGI | ASGEPGHQAPL | 0.6699095 |
| LNAINAPTL | ASGEPGHQAPL | 0.64053774 |
| LQLLNTGV | ASGEPGHQAPL | 0.6739737 |
| LQTSIEF | ASGEPGHQAPL | 0.661789 |
| MSFAVSEL | ASGEPGHQAPL | 0.6607357 |
| NTGALGL | ASGEPGHQAPL | 0.66874605 |
| PPGAGGAGGAGTT | ASGEPGHQAPL | 0.66218984 |
| QLVGLGI | ASGEPGHQAPL | 0.6489133 |
| RFLEQVHQL | ASGEPGHQAPL | 0.6470775 |
| RGVGYVLR | ASGEPGHQAPL | 0.6701729 |
| SATGVFAGVI | ASGEPGHQAPL | 0.66964746 |
| SLPAITLP | ASGEPGHQAPL | 0.66930073 |
| STALAFTV | ASGEPGHQAPL | 0.6774758 |
| SVLTVGLGK | ASGEPGHQAPL | 0.6704359 |
| TALAGAGSGI | ASGEPGHQAPL | 0.66828376 |
| TGLPFTGI | ASGEPGHQAPL | 0.5269045 |
| VAEHVTV | ASGEPGHQAPL | 0.6203187 |
| VDIVLNSL | ASGEPGHQAPL | 0.6412426 |
| VLTGGGAVL | ASGEPGHQAPL | 0.6565564 |
| VNVGLAYGI | ASGEPGHQAPL | 0.49554676 |
| VSSAAVGAV | ASGEPGHQAPL | 0.66657835 |

| | | |
|---|---|---|
| VTSVYEAGV | ASGEPGHQAPL | 0.6606804 |
| WVALDRGA | ASGEPGHQAPL | 0.64374995 |
| YLVGSVVI | ASGEPGHQAPL | 0.6732558 |
| GPPGTGKTLL | ASGEPGHQAPL | 0.6536375 |
| GVGMTKF | ASGEPGHQAPL | 0.6370537 |
| IGIGGSDL | ASGEPGHQAPL | 0.6534637 |
| IGTLLAEL | ASGEPGHQAPL | 0.6753884 |
| IYTSGTT | ASGEPGHQAPL | 0.642816 |
| LSFPAGV | ASGEPGHQAPL | 0.6741693 |
| LYTSGTT | ASGEPGHQAPL | 0.64954364 |
| VGAFPFGF | ASGEPGHQAPL | 0.6744313 |
| VGAGTVEF | ASGEPGHQAPL | 0.61576545 |
| VGKSTLLL | ASGEPGHQAPL | 0.67196935 |
| AALAHAGGGIV | ASGEPGHQAPL | 0.6575401 |
| AALVAVSLL | ASGEPGHQAPL | 0.67832 |
| AAVSRAVAAL | ASGEPGHQAPL | 0.6560021 |
| AGGSSGIGGF | ASGEPGHQAPL | 0.6737319 |
| AGLTVANANG | ASGEPGHQAPL | 0.66422623 |
| AGLVGAVL | ASGEPGHQAPL | 0.6720996 |
| ALSQAFTV | ASGEPGHQAPL | 0.60579526 |
| ALTFSVSL | ASGEPGHQAPL | 0.6553543 |
| ALTGAALVAL | ASGEPGHQAPL | 0.64164114 |
| ASFSIGAGVL | ASGEPGHQAPL | 0.66618645 |
| DGSVYKI | ASGEPGHQAPL | 0.6694038 |
| EQLAEIV | ASGEPGHQAPL | 0.4677014 |
| FAVSELTVM | ASGEPGHQAPL | 0.6754456 |
| GAAPLAAGL | ASGEPGHQAPL | 0.6256297 |
| GADGGDGGSGGAGG | ASGEPGHQAPL | 0.63761103 |
| GFGLDTGGAGG | ASGEPGHQAPL | 0.67256 |
| GLASGGLL | ASGEPGHQAPL | 0.6635829 |
| GRIAGAAVLL | ASGEPGHQAPL | 0.6678705 |
| GSAGSLNTSF | ASGEPGHQAPL | 0.65326846 |
| GTGQIGI | ASGEPGHQAPL | 0.66206586 |
| GTLIALIL | ASGEPGHQAPL | 0.6667777 |
| IAATMGFVVG | ASGEPGHQAPL | 0.5692543 |
| IFAQALL | ASGEPGHQAPL | 0.62901187 |
| IGAAVALGF | ASGEPGHQAPL | 0.67588276 |
| IGLTPLAL | ASGEPGHQAPL | 0.45834306 |
| ILGSNTSTL | ASGEPGHQAPL | 0.8072641 |
| IVFSLTVL | ASGEPGHQAPL | 0.31071663 |
| LASKATGF | ASGEPGHQAPL | 0.2047937 |
| LASVALIL | ASGEPGHQAPL | 0.256368 |
| LFAVGLTL | ASGEPGHQAPL | 0.61239034 |
| LFNGGIGTYI | ASGEPGHQAPL | 0.8228689 |
| LTLGAGII | ASGEPGHQAPL | 0.60951966 |
| LVAAALGDI | ASGEPGHQAPL | 0.20080829 |
| NTAFVQLGI | ASGEPGHQAPL | 0.3528263 |
| QLFSGAEL | ASGEPGHQAPL | 0.25947398 |
| SAATFITL | ASGEPGHQAPL | 0.31808463 |
| SNYGLANLGV | ASGEPGHQAPL | 0.45962298 |
| SYGGAYAVM | ASGEPGHQAPL | 0.345443 |

| | | |
|---|---|---|
| TTGVGGLTLG | ASGEPGHQAPL | 0.29826653 |
| TTQTIYI | ASGEPGHQAPL | 0.25996792 |
| VAAGAGLGAVI | ASGEPGHQAPL | 0.37117338 |
| VAAVVTLL | ASGEPGHQAPL | 0.54140073 |
| VGFLGGLI | ASGEPGHQAPL | 0.35982373 |
| VGVIPILL | ASGEPGHQAPL | 0.6555648 |
| VIFASSLI | ASGEPGHQAPL | 0.65519094 |
| VKGGDGGAAAG | ASGEPGHQAPL | 0.64427054 |
| VTKTFTNI | ASGEPGHQAPL | 0.11196446 |
| VVAAAILL | ASGEPGHQAPL | 0.19413587 |
| YAALDVEL | ASGEPGHQAPL | 0.2423331 |
| LASVALLL | ASGEPGHQAPL | 0.384581 |
| LDAVHRV | ASGEPGHQAPL | 0.13914509 |
| QGLSSSIF | ASGEPGHQAPL | 0.43115553 |
| RGTFANIRL | ASGEPGHQAPL | 0.27548602 |
| RQLWWGHRI | ASGEPGHQAPL | 0.242762 |
| YGPPGTGKTLL | ASGEPGHQAPL | 0.62232584 |
| AALVGAPANVA | ASGEPGHQAPL | 0.31736934 |
| AATFRRTE | ASGEPGHQAPL | 0.21425834 |
| AGALGGGI | ASGEPGHQAPL | 0.20324242 |
| AGGAGGAGGIGAG | ASGEPGHQAPL | 0.7101972 |
| AGSGALGL | ASGEPGHQAPL | 0.16840382 |
| AQIADQI | ASGEPGHQAPL | 0.66564554 |
| ASAVAVTYL | ASGEPGHQAPL | 0.2506013 |
| ASTVLGGTGG | ASGEPGHQAPL | 0.56589043 |
| AVGGEPAK | ASGEPGHQAPL | 0.16678308 |
| CEQGTFGGTC | ASGEPGHQAPL | 0.65201 |
| DEGKAPAG | ASGEPGHQAPL | 0.4642517 |
| DTVVFPAL | ASGEPGHQAPL | 0.8484546 |
| FEGITMAG | ASGEPGHQAPL | 0.27826446 |
| FPLVVQD | ASGEPGHQAPL | 0.35061327 |
| GAFTTPQ | ASGEPGHQAPL | 0.24559535 |
| GFGGFGTAGGVGGA | ASGEPGHQAPL | 0.8107679 |
| GFLAQFL | ASGEPGHQAPL | 0.15354456 |
| GGAGGAGGNAGVA | ASGEPGHQAPL | 0.22301763 |
| GGVGTAGAGGAGGAGG | ASGEPGHQAPL | 0.17526394 |
| GNGGGGDAGNAGSGGN | ASGEPGHQAPL | 0.32482493 |
| GTGGAGGAGNDAGS | ASGEPGHQAPL | 0.6055753 |
| GTTNTGWFNS | ASGEPGHQAPL | 0.14279653 |
| IFMGPLFTL | ASGEPGHQAPL | 0.6388012 |
| ILAHAIEAI | ASGEPGHQAPL | 0.32897532 |
| IRSAIREAL | ASGEPGHQAPL | 0.19876762 |
| ITFAGGG | ASGEPGHQAPL | 0.69431925 |
| ITGMSGHQEV | ASGEPGHQAPL | 0.2320078 |
| IVAGLACTL | ASGEPGHQAPL | 0.72306186 |
| IVGGGLAGA | ASGEPGHQAPL | 0.2469278 |
| IVTGVLNI | ASGEPGHQAPL | 0.29755884 |
| LAQQGCI | ASGEPGHQAPL | 0.4328117 |
| LCLHVEN | ASGEPGHQAPL | 0.1808921 |
| LDAISDGS | ASGEPGHQAPL | 0.18359144 |
| LGGFAAML | ASGEPGHQAPL | 0.7451863 |

| | | |
|---|---|---|
| LGGKNPMI | ASGEPGHQAPL | 0.32866332 |
| LGPKGVAAIL | ASGEPGHQAPL | 0.72254324 |
| LMVGVSI | ASGEPGHQAPL | 0.192102 |
| LVGAAFVL | ASGEPGHQAPL | 0.16769919 |
| LVLAVAPL | ASGEPGHQAPL | 0.24069847 |
| MAAIAVIDAA | ASGEPGHQAPL | 0.5310337 |
| NNGINAELQ | ASGEPGHQAPL | 0.63721186 |
| RGEPLVLVHG | ASGEPGHQAPL | 0.3572514 |
| RGGAGGAAGLI | ASGEPGHQAPL | 0.19146861 |
| RSYRMHPAV | ASGEPGHQAPL | 0.27814353 |
| SAAHLLAGQLA | ASGEPGHQAPL | 0.23676822 |
| SAAVGSY | ASGEPGHQAPL | 0.20274921 |
| SPHAVLHRI | ASGEPGHQAPL | 0.36319578 |
| SQIAKVL | ASGEPGHQAPL | 0.33674166 |
| TALGHLVP | ASGEPGHQAPL | 0.20778675 |
| TAPTTSGAA | ASGEPGHQAPL | 0.24415123 |
| TDGDDGNGGSPGT | ASGEPGHQAPL | 0.3002922 |
| TDGLGEN | ASGEPGHQAPL | 0.42907575 |
| TGGPPLPGQA | ASGEPGHQAPL | 0.36417246 |
| TIPAITI | ASGEPGHQAPL | 0.5855933 |
| TIPGFTI | ASGEPGHQAPL | 0.42323002 |
| TSIASGALA | ASGEPGHQAPL | 0.38500455 |
| TVTLPGAAI | ASGEPGHQAPL | 0.10677173 |
| VASGLSAALP | ASGEPGHQAPL | 0.13369742 |
| VGAGVIGI | ASGEPGHQAPL | 0.29882252 |
| VGPKRFGV | ASGEPGHQAPL | 0.23103787 |
| VNVAFPD | ASGEPGHQAPL | 0.1505366 |
| VSMTNSGVS | ASGEPGHQAPL | 0.43096796 |
| VTDVLGDLSV | ASGEPGHQAPL | 0.2538471 |
| VVVVGAGFAGL | ASGEPGHQAPL | 0.1948863 |
| VYTVGTVV | ASGEPGHQAPL | 0.41845232 |
| WTTFSGP | ASGEPGHQAPL | 0.18768919 |
| YDELVEAF | ASGEPGHQAPL | 0.21520565 |
| YGTGGAGGNGG | ASGEPGHQAPL | 0.23557709 |
| YTNVMAV | ASGEPGHQAPL | 0.38279432 |
| FLEQVHQL | ASGEPGHQAPL | 0.17528637 |
| GLSLLAVAV | ASGEPGHQAPL | 0.32090908 |
| LEQVHQL | ASGEPGHQAPL | 0.25704032 |
| LNAAGAI | ASGEPGHQAPL | 0.4491511 |
| PAVHASL | ASGEPGHQAPL | 0.16363741 |
| TELEDDA | ASGEPGHQAPL | 0.34772113 |
| TPPGPPN | ASGEPGHQAPL | 0.21271145 |
| TPPPPPPP | ASGEPGHQAPL | 0.71679455 |
| TQSPALI | ASSQVPRDKNSDYT | 0.23462918 |
| EFAAAFPQ | ASSQVPRDKNSDYT | 0.32876948 |
| LLSGNLDVL | ASSQVPRDKNSDYT | 0.29729882 |
| GPANLGNYNI | ASSQVPRDKNSDYT | 0.642367 |
| TLGPTLP | ASSQVPRDKNSDYT | 0.14168142 |
| VYAYSSCCK | ASSQVPRDKNSDYT | 0.24479958 |
| APAPAPAPNI | ASSQVPRDKNSDYT | 0.1546956 |
| PGGPVPIP | ASSQVPRDKNSDYT | 0.32752842 |

| | | |
|---|---|---|
| FLTVALEP | ASSQVPRDKNSDYT | 0.3215693 |
| SQYFGANL | ASSQVPRDKNSDYT | 0.17963782 |
| GGVKPAA | ASSQVPRDKNSDYT | 0.36044735 |
| GGTGGAGGAGGW | ASSQVPRDKNSDYT | 0.38889694 |
| GVGGVAGLGGAGG | ASSQVPRDKNSDYT | 0.19249798 |
| VLLVLATTI | ASSQVPRDKNSDYT | 0.53916985 |
| IGIFAAML | ASSQVPRDKNSDYT | 0.3015038 |
| AVGDLID | ASSQVPRDKNSDYT | 0.5346406 |
| NISPLPL | ASSQVPRDKNSDYT | 0.28177965 |
| VTAVFVAI | ASSQVPRDKNSDYT | 0.22623557 |
| SGDDNDAAL | ASSQVPRDKNSDYT | 0.35714877 |
| AVAALAHS | ASSQVPRDKNSDYT | 0.30820903 |
| GGVTVMATP | ASSQVPRDKNSDYT | 0.21390805 |
| AGLAANVRD | ASSQVPRDKNSDYT | 0.4205402 |
| TNRPDLIDPALLRPG | ASSQVPRDKNSDYT | 0.8328234 |
| LANLGVTAL | ASSQVPRDKNSDYT | 0.8205913 |
| TGERIEL | ASSQVPRDKNSDYT | 0.6322391 |
| NGEELAA | ASSQVPRDKNSDYT | 0.6682781 |
| VLSLVWG | ASSQVPRDKNSDYT | 0.74733937 |
| IGYALWL | ASSQVPRDKNSDYT | 0.82700074 |
| LSLLLAPL | ASSQVPRDKNSDYT | 0.8513248 |
| TLDEALK | ASSQVPRDKNSDYT | 0.8303665 |
| TDGAGTAEGG | ASSQVPRDKNSDYT | 0.6187122 |
| NAPGGAGGNGG | ASSQVPRDKNSDYT | 0.8344149 |
| VSVFMGMGGNF | ASSQVPRDKNSDYT | 0.79016423 |
| LASYLAAL | ASSQVPRDKNSDYT | 0.81630504 |
| LVGGTPAL | ASSQVPRDKNSDYT | 0.7906556 |
| VRNADVL | ASSQVPRDKNSDYT | 0.73726296 |
| TCSSPFLVSAM | ASSQVPRDKNSDYT | 0.6806142 |
| VHITDAE | ASSQVPRDKNSDYT | 0.48549446 |
| EYAAPSF | ASSQVPRDKNSDYT | 0.8296759 |
| VVDAFHGGL | ASSQVPRDKNSDYT | 0.83119625 |
| ASVVDPQ | ASSQVPRDKNSDYT | 0.81483227 |
| SYNEFGGLS | ASSQVPRDKNSDYT | 0.8532541 |
| DPDGSVY | ASSQVPRDKNSDYT | 0.8448127 |
| MVSTIHATE | ASSQVPRDKNSDYT | 0.7844186 |
| LAAGVADPV | ASSQVPRDKNSDYT | 0.23707528 |
| TKEQIAA | ASSQVPRDKNSDYT | 0.38458622 |
| SLASPAAL | ASSQVPRDKNSDYT | 0.56889975 |
| IGLVFYGV | ASSQVPRDKNSDYT | 0.70670885 |
| FLLDASNL | ASSQVPRDKNSDYT | 0.2121729 |
| YQVGNFP | ASSQVPRDKNSDYT | 0.78627473 |
| ESLLSQST | ASSQVPRDKNSDYT | 0.6839144 |
| LFTVGFAAVG | ASSQVPRDKNSDYT | 0.7336097 |
| ATPPTPT | ASSQVPRDKNSDYT | 0.84588593 |
| TDLATLG | ASSQVPRDKNSDYT | 0.7145176 |
| SAGGTGR | ASSQVPRDKNSDYT | 0.4635339 |
| GFSNSGS | ASSQVPRDKNSDYT | 0.5881252 |
| GGNGGSGGTSV | ASSQVPRDKNSDYT | 0.8556005 |
| GTSVATGGAGN | ASSQVPRDKNSDYT | 0.6280991 |
| NGGNGGSGGTS | ASSQVPRDKNSDYT | 0.850331 |

| | | |
|---|---|---|
| AVVKFPH | ASSQVPRDKNSDYT | 0.7048979 |
| TPPPPPP | ASSQVPRDKNSDYT | 0.85656375 |
| AAPAAAAAPA | ASSQVPRDKNSDYT | 0.53990316 |
| IALNEDL | ASSQVPRDKNSDYT | 0.8441758 |
| IVNVSSM | ASSQVPRDKNSDYT | 0.81434476 |
| TGIPPAPR | ASSQVPRDKNSDYT | 0.8624784 |
| FDLGGGTF | ASSQVPRDKNSDYT | 0.7343161 |
| FDLGGGTFDVS | ASSQVPRDKNSDYT | 0.80597997 |
| NEPTAAA | ASSQVPRDKNSDYT | 0.6570418 |
| NEPTAAALA | ASSQVPRDKNSDYT | 0.8326225 |
| GSIINISSI | ASSQVPRDKNSDYT | 0.5873032 |
| VAPGNAGTA | ASSQVPRDKNSDYT | 0.6886854 |
| TGGGASLE | ASSQVPRDKNSDYT | 0.5639427 |
| VAVYNHYKRI | ASSQVPRDKNSDYT | 0.71940684 |
| TDPAPATT | ASSQVPRDKNSDYT | 0.8389956 |
| TNRPDILDPALLRPG | ASSQVPRDKNSDYT | 0.16741852 |
| LTTTTTS | ASSQVPRDKNSDYT | 0.8245373 |
| PTGSGKTL | ASSQVPRDKNSDYT | 0.54303634 |
| TGSGKTL | ASSQVPRDKNSDYT | 0.508229 |
| TIGHVDHGK | ASSQVPRDKNSDYT | 0.8144536 |
| TIGHVDHGKTT | ASSQVPRDKNSDYT | 0.7857305 |
| GHVDHGKTT | ASSQVPRDKNSDYT | 0.80575573 |
| AQFEHTLL | ASSQVPRDKNSDYT | 0.63805836 |
| IVNQDEL | ASSQVPRDKNSDYT | 0.78926456 |
| TNGSQFF | ASSQVPRDKNSDYT | 0.79481196 |
| TGKGIVE | ASSQVPRDKNSDYT | 0.25968647 |
| LISNASDA | ASSQVPRDKNSDYT | 0.55787146 |
| AELRTLGELTKIAKAHGAQ | ASSQVPRDKNSDYT | 0.85423565 |
| APNAVAAAEPSW | ASSQVPRDKNSDYT | 0.508184 |
| APQVAGAATAVL | ASSQVPRDKNSDYT | 0.7890878 |
| ARRDPAAAAAILPTDA | ASSQVPRDKNSDYT | 0.17116539 |
| DAAARDVLAEAGLAEYF | ASSQVPRDKNSDYT | 0.16721441 |
| DDQIVLSAKANLAKVEQ | ASSQVPRDKNSDYT | 0.3144819 |
| DELLPDTGEDSDEDGH | ASSQVPRDKNSDYT | 0.557057 |
| ELPEAPANRATVPQD | ASSQVPRDKNSDYT | 0.81142366 |
| FGQSKIERVTGIG | ASSQVPRDKNSDYT | 0.5579696 |
| GAGGTGSTAGGAGGAGGAGGLYA | ASSQVPRDKNSDYT | 0.16989037 |
| GHGGTGAAGGNGGTGGAGGSGIDG | ASSQVPRDKNSDYT | 0.3844866 |
| GNGGNGGAGGEGGPGGAGGAGA | ASSQVPRDKNSDYT | 0.25421345 |
| GSTAGNGGNGGNATNGTIAGQPAG | ASSQVPRDKNSDYT | 0.27810973 |
| GTGGDGGNGGNGGTGG | ASSQVPRDKNSDYT | 0.40242866 |
| GVRRPAGIPADAPV | ASSQVPRDKNSDYT | 0.3569422 |
| GYRQVTDAHLLTLARR | ASSQVPRDKNSDYT | 0.1917029 |
| IAKMGQDGHDRG | ASSQVPRDKNSDYT | 0.17661333 |
| KAEPSIEIQYT | ASSQVPRDKNSDYT | 0.60945696 |
| KPVRIRMAADEDTVAVT | ASSQVPRDKNSDYT | 0.7312291 |
| MPPRIAGMRLLVIKPEP | ASSQVPRDKNSDYT | 0.6725136 |
| MKFYAVALFRSRGVPPTR | ASSQVPRDKNSDYT | 0.7517729 |
| NAGGFGILGAVAHSP | ASSQVPRDKNSDYT | 0.55950224 |
| NEGREITLAMAEM | ASSQVPRDKNSDYT | 0.56091195 |
| NELVEWSHPLGHRWRWE | ASSQVPRDKNSDYT | 0.09609616 |

| | | |
|---|---|---|
| NLGAGGTGGNGGNPG | ASSQVPRDKNSDYT | 0.11330595 |
| NTEWPKWETVMAADPVD | ASSQVPRDKNSDYT | 0.21065046 |
| PASAAPAETQP | ASSQVPRDKNSDYT | 0.2983306 |
| QQSSDGKNGVRAAEITGE | ASSQVPRDKNSDYT | 0.12953447 |
| RAPDAVTTSWWKE | ASSQVPRDKNSDYT | 0.5253136 |
| SGFGNVGAHTSGWFNQ | ASSQVPRDKNSDYT | 0.22376868 |
| SGVAGAVGQAASVGGLKV | ASSQVPRDKNSDYT | 0.19073895 |
| SNGGAGTGSAGGNGGT | ASSQVPRDKNSDYT | 0.6833879 |
| TGGAGGEGGAAG | ASSQVPRDKNSDYT | 0.17579071 |
| TGGDGGAAGAGGAPGHG | ASSQVPRDKNSDYT | 0.17102863 |
| TMPPLLAAIANAGRHGV | ASSQVPRDKNSDYT | 0.23467281 |
| TSSTAGNSGGA | ASSQVPRDKNSDYT | 0.6570407 |
| VPATVPVGILC | ASSQVPRDKNSDYT | 0.15808551 |
| VQRPDSGGVVLTGRIS | ASSQVPRDKNSDYT | 0.5061941 |
| WTPNPVPATKGVPAGLVQAT | ASSQVPRDKNSDYT | 0.26833254 |
| YALKRALAKFSDRFDVVII | ASSQVPRDKNSDYT | 0.5964358 |
| YLIGSVIV | ASSQVPRDKNSDYT | 0.13930121 |
| YGQLVAVID | ASSQVPRDKNSDYT | 0.5250353 |
| VTVGGMTLPA | ASSQVPRDKNSDYT | 0.29493326 |
| VLALPAV | ASSQVPRDKNSDYT | 0.8450303 |
| VIPPTLNYET | ASSQVPRDKNSDYT | 0.2082892 |
| VGMGGMHPSSG | ASSQVPRDKNSDYT | 0.34417927 |
| VGLAGGAATA | ASSQVPRDKNSDYT | 0.24012226 |
| VGGAGGAGGSGP | ASSQVPRDKNSDYT | 0.8000002 |
| VFAAFMV | ASSQVPRDKNSDYT | 0.12803045 |
| TLYDKLR | ASSQVPRDKNSDYT | 0.22753188 |
| TLAVFTVL | ASSQVPRDKNSDYT | 0.12369781 |
| TGSVQPARV | ASSQVPRDKNSDYT | 0.2806363 |
| TGGGAGSGGT | ASSQVPRDKNSDYT | 0.49822348 |
| SHLETSL | ASSQVPRDKNSDYT | 0.10882071 |
| SFTTGMT | ASSQVPRDKNSDYT | 0.44761252 |
| QVEVAKRMGVRQPTVS | ASSQVPRDKNSDYT | 0.26608163 |
| QGSGGPALAM | ASSQVPRDKNSDYT | 0.16361131 |
| MLAPSNP | ASSQVPRDKNSDYT | 0.5425071 |
| LVNNAAI | ASSQVPRDKNSDYT | 0.305058 |
| LTGEINTLA | ASSQVPRDKNSDYT | 0.7245499 |
| LSIFGPV | ASSQVPRDKNSDYT | 0.31472874 |
| LLALLANL | ASSQVPRDKNSDYT | 0.27418303 |
| LISNASDALDKLR | ASSQVPRDKNSDYT | 0.36092603 |
| LINEAVP | ASSQVPRDKNSDYT | 0.24834919 |
| LGYAFSGL | ASSQVPRDKNSDYT | 0.21540244 |
| LDFVAVAL | ASSQVPRDKNSDYT | 0.5834728 |
| LAQPGAE | ASSQVPRDKNSDYT | 0.90386796 |
| ISNLGAVTGF | ASSQVPRDKNSDYT | 0.88475835 |
| HQEDPGRI | ASSQVPRDKNSDYT | 0.8441825 |
| GTPGEPTP | ASSQVPRDKNSDYT | 0.8333093 |
| GFFNSGS | ASSQVPRDKNSDYT | 0.8710356 |
| GAGGGGLTAGAN | ASSQVPRDKNSDYT | 0.9047679 |
| FVDALVNM | ASSQVPRDKNSDYT | 0.9132674 |
| FLTVAPEL | ASSQVPRDKNSDYT | 0.9028518 |
| FETLTAL | ASSQVPRDKNSDYT | 0.8391089 |

| | | |
|---|---|---|
| DVINPIEL | ASSQVPRDKNSDYT | 0.9070462 |
| APVSGAPV | ASSQVPRDKNSDYT | 0.8952085 |
| ALVGGLAW | ASSQVPRDKNSDYT | 0.87513375 |
| ALAAAAATAI | ASSQVPRDKNSDYT | 0.9052949 |
| AFGLGLLL | ASSQVPRDKNSDYT | 0.8997728 |
| AFFAAASL | ASSQVPRDKNSDYT | 0.85586494 |
| AAAALPAFSPP | ASSQVPRDKNSDYT | 0.86100245 |
| VVLAPAL | ASSQVPRDKNSDYT | 0.8926157 |
| TIGHVDHGKTTL | ASSQVPRDKNSDYT | 0.902913 |
| LISNASDALDK | ASSQVPRDKNSDYT | 0.9032513 |
| LISNASDAL | ASSQVPRDKNSDYT | 0.9137963 |
| ISNASDALDK | ASSQVPRDKNSDYT | 0.91084254 |
| NQAETLVYQTE | ASSQVPRDKNSDYT | 0.9022935 |
| TGGTGGTAGAG | ASSQVPRDKNSDYT | 0.47202185 |
| TGSTAGGAGGA | ASSQVPRDKNSDYT | 0.74113697 |
| DPLAVPVAAGPV | ASSQVPRDKNSDYT | 0.86863875 |
| GGTGGTPGNGGN | ASSQVPRDKNSDYT | 0.85957146 |
| PAGLAGVDPATL | ASSQVPRDKNSDYT | 0.48079818 |
| GNNLVVEAPHAGASV | ASSQVPRDKNSDYT | 0.8618407 |
| KVTRIMSASTLEQPA | ASSQVPRDKNSDYT | 0.8794167 |
| LGFALPIALPRRTVV | ASSQVPRDKNSDYT | 0.840619 |
| TRLGNLTARKLVVLV | ASSQVPRDKNSDYT | 0.90647256 |
| WSQQRRLIYFPSAGP | ASSQVPRDKNSDYT | 0.87113744 |
| GPNGNPDPMAAAVDIR | ASSQVPRDKNSDYT | 0.8283565 |
| TQRCVSVADLRAAASR | ASSQVPRDKNSDYT | 0.8739947 |
| QFVLDVAEHPVEVLTETV | ASSQVPRDKNSDYT | 0.9099835 |
| GGAGGTGGAGGNGGTQAGDG | ASSQVPRDKNSDYT | 0.85534704 |
| PLLATLRAIAVDVDAKPAQV | ASSQVPRDKNSDYT | 0.91130143 |
| APKFRTGGYMIDFWGVGYQVAKRM | ASSQVPRDKNSDYT | 0.77604043 |
| GAGGAGGAGGAAGGPGGTGGTGGN | ASSQVPRDKNSDYT | 0.914163 |
| IGALGLLATDVAPAGSVAFGVALA | ASSQVPRDKNSDYT | 0.81275874 |
| AGGAGGIGGNGGAGGAGGVLMGNG | ASSQVPRDKNSDYT | 0.9092585 |
| ADELIGLL | ASSQVPRDKNSDYT | 0.8904078 |
| ADRALKA | ASSQVPRDKNSDYT | 0.91946006 |
| AGAVGQVGGAGGA | ASSQVPRDKNSDYT | 0.88786995 |
| AGGGGGLGGAP | ASSQVPRDKNSDYT | 0.91269183 |
| AGGLLYGNGGAG | ASSQVPRDKNSDYT | 0.88780004 |
| AGLESIGL | ASSQVPRDKNSDYT | 0.9001361 |
| AGLKTGII | ASSQVPRDKNSDYT | 0.8148105 |
| ASLAALLL | ASSQVPRDKNSDYT | 0.88361865 |
| ASVINISSI | ASSQVPRDKNSDYT | 0.6995574 |
| ATPPGPA | ASSQVPRDKNSDYT | 0.8933343 |
| ATVASALV | ASSQVPRDKNSDYT | 0.9043556 |
| ATVASLIL | ASSQVPRDKNSDYT | 0.35178676 |
| AVNGPVLIRH | ASSQVPRDKNSDYT | 0.9086434 |
| DGAAGAVGI | ASSQVPRDKNSDYT | 0.8071586 |
| DPLLHGI | ASSQVPRDKNSDYT | 0.84521776 |
| ELGAALGAL | ASSQVPRDKNSDYT | 0.9128983 |
| ELGRTQAV | ASSQVPRDKNSDYT | 0.8897544 |
| FVAGGYLS | ASSQVPRDKNSDYT | 0.910043 |
| GASGGAGGAGGW | ASSQVPRDKNSDYT | 0.866852 |

| | | |
|---|---|---|
| GGAGAGGDGGAGGI | ASSQVPRDKNSDYT | 0.8857378 |
| GTGHNGSQPGMGGNG | ASSQVPRDKNSDYT | 0.88526165 |
| GYAAASASAS | ASSQVPRDKNSDYT | 0.6408527 |
| HQVDVAAVQQG | ASSQVPRDKNSDYT | 0.8335744 |
| IGAAIAHT | ASSQVPRDKNSDYT | 0.91719574 |
| IGMIEVAQL | ASSQVPRDKNSDYT | 0.6365719 |
| IRLAPEHTSAMD | ASSQVPRDKNSDYT | 0.6252891 |
| IYNVSVL | ASSQVPRDKNSDYT | 0.53952384 |
| KQGFLGM | ASSQVPRDKNSDYT | 0.52779746 |
| LESVITQL | ASSQVPRDKNSDYT | 0.57502407 |
| LGLGALVVV | ASSQVPRDKNSDYT | 0.6475499 |
| LIGNGGNGGSGGI | ASSQVPRDKNSDYT | 0.62159395 |
| LNAINAPTL | ASSQVPRDKNSDYT | 0.6313691 |
| LQLLNTGV | ASSQVPRDKNSDYT | 0.54089725 |
| LQTSIEF | ASSQVPRDKNSDYT | 0.60221726 |
| MSFAVSEL | ASSQVPRDKNSDYT | 0.59240675 |
| NTGALGL | ASSQVPRDKNSDYT | 0.5685209 |
| PPGAGGAGGAGTT | ASSQVPRDKNSDYT | 0.6276256 |
| QLVGLGI | ASSQVPRDKNSDYT | 0.58745885 |
| RFLEQVHQL | ASSQVPRDKNSDYT | 0.5745838 |
| RGVGYVLR | ASSQVPRDKNSDYT | 0.5725895 |
| SATGVFAGVI | ASSQVPRDKNSDYT | 0.59941787 |
| SLPAITLP | ASSQVPRDKNSDYT | 0.63061035 |
| STALAFTV | ASSQVPRDKNSDYT | 0.6225828 |
| SVLTVGLGK | ASSQVPRDKNSDYT | 0.65818256 |
| TALAGAGSGI | ASSQVPRDKNSDYT | 0.64442945 |
| TGLPFTGI | ASSQVPRDKNSDYT | 0.64312166 |
| VAEHVTV | ASSQVPRDKNSDYT | 0.35992384 |
| VDIVLNSL | ASSQVPRDKNSDYT | 0.48022354 |
| VLTGGGAVL | ASSQVPRDKNSDYT | 0.56718826 |
| VNVGLAYGI | ASSQVPRDKNSDYT | 0.5878806 |
| VSSAAVGAV | ASSQVPRDKNSDYT | 0.4338918 |
| VTSVYEAGV | ASSQVPRDKNSDYT | 0.6107856 |
| WVALDRGA | ASSQVPRDKNSDYT | 0.61784667 |
| YLVGSVVI | ASSQVPRDKNSDYT | 0.53078663 |
| GPPGTGKTLL | ASSQVPRDKNSDYT | 0.62754357 |
| GVGMTKF | ASSQVPRDKNSDYT | 0.54649824 |
| IGIGGSDL | ASSQVPRDKNSDYT | 0.5262515 |
| IGTLLAEL | ASSQVPRDKNSDYT | 0.538394 |
| IYTSGTT | ASSQVPRDKNSDYT | 0.64919084 |
| LSFPAGV | ASSQVPRDKNSDYT | 0.476947 |
| LYTSGTT | ASSQVPRDKNSDYT | 0.62248576 |
| VGAFPFGF | ASSQVPRDKNSDYT | 0.5778891 |
| VGAGTVEF | ASSQVPRDKNSDYT | 0.6442222 |
| VGKSTLLL | ASSQVPRDKNSDYT | 0.4939665 |
| AALAHAGGGIV | ASSQVPRDKNSDYT | 0.63421595 |
| AALVAVSLL | ASSQVPRDKNSDYT | 0.57251734 |
| AAVSRAVAAL | ASSQVPRDKNSDYT | 0.6516863 |
| AGGSSGIGGF | ASSQVPRDKNSDYT | 0.58735096 |
| AGLTVANANG | ASSQVPRDKNSDYT | 0.610786 |
| AGLVGAVL | ASSQVPRDKNSDYT | 0.5634247 |

| | | |
|---|---|---|
| ALSQAFTV | ASSQVPRDKNSDYT | 0.61122966 |
| ALTFSVSL | ASSQVPRDKNSDYT | 0.46092257 |
| ALTGAALVAL | ASSQVPRDKNSDYT | 0.5888804 |
| ASFSIGAGVL | ASSQVPRDKNSDYT | 0.491786 |
| DGSVYKI | ASSQVPRDKNSDYT | 0.6058928 |
| EQLAEIV | ASSQVPRDKNSDYT | 0.63119453 |
| FAVSELTVM | ASSQVPRDKNSDYT | 0.4848364 |
| GAAPLAAGL | ASSQVPRDKNSDYT | 0.6312188 |
| GADGGDGGSGGAGG | ASSQVPRDKNSDYT | 0.5880988 |
| GFGLDTGGAGG | ASSQVPRDKNSDYT | 0.5695321 |
| GLASGGLL | ASSQVPRDKNSDYT | 0.6363077 |
| GRIAGAAVLL | ASSQVPRDKNSDYT | 0.57793975 |
| GSAGSLNTSF | ASSQVPRDKNSDYT | 0.6482345 |
| GTGQIGI | ASSQVPRDKNSDYT | 0.5277737 |
| GTLIALIL | ASSQVPRDKNSDYT | 0.5961972 |
| IAATMGFVVG | ASSQVPRDKNSDYT | 0.6338161 |
| IFAQALL | ASSQVPRDKNSDYT | 0.45360267 |
| IGAAVALGF | ASSQVPRDKNSDYT | 0.5280438 |
| IGLTPLAL | ASSQVPRDKNSDYT | 0.64136624 |
| ILGSNTSTL | ASSQVPRDKNSDYT | 0.8249103 |
| IVFSLTVL | ASSQVPRDKNSDYT | 0.81599903 |
| LASKATGF | ASSQVPRDKNSDYT | 0.7681354 |
| LASVALIL | ASSQVPRDKNSDYT | 0.74771416 |
| LFAVGLTL | ASSQVPRDKNSDYT | 0.7787399 |
| LFNGGIGTYI | ASSQVPRDKNSDYT | 0.8272662 |
| LTLGAGII | ASSQVPRDKNSDYT | 0.8323919 |
| LVAAALGDI | ASSQVPRDKNSDYT | 0.82163405 |
| NTAFVQLGI | ASSQVPRDKNSDYT | 0.7417598 |
| QLFSGAEL | ASSQVPRDKNSDYT | 0.8273691 |
| SAATFITL | ASSQVPRDKNSDYT | 0.80855465 |
| SNYGLANLGV | ASSQVPRDKNSDYT | 0.79297227 |
| SYGGAYAVM | ASSQVPRDKNSDYT | 0.8288356 |
| TTGVGGLTLG | ASSQVPRDKNSDYT | 0.8196088 |
| TTQTIYI | ASSQVPRDKNSDYT | 0.77764887 |
| VAAGAGLGAVI | ASSQVPRDKNSDYT | 0.7736896 |
| VAAVVTLL | ASSQVPRDKNSDYT | 0.81808317 |
| VGFLGGLI | ASSQVPRDKNSDYT | 0.8310906 |
| VGVIPILL | ASSQVPRDKNSDYT | 0.8294818 |
| VIFASSLI | ASSQVPRDKNSDYT | 0.8409846 |
| VKGGDGGAAAG | ASSQVPRDKNSDYT | 0.8357301 |
| VTKTFTNI | ASSQVPRDKNSDYT | 0.8245708 |
| VVAAAILL | ASSQVPRDKNSDYT | 0.4484933 |
| YAALDVEL | ASSQVPRDKNSDYT | 0.6938134 |
| LASVALLL | ASSQVPRDKNSDYT | 0.7726544 |
| LDAVHRV | ASSQVPRDKNSDYT | 0.79637104 |
| QGLSSSIF | ASSQVPRDKNSDYT | 0.46123695 |
| RGTFANIRL | ASSQVPRDKNSDYT | 0.7948595 |
| RQLWWGHRI | ASSQVPRDKNSDYT | 0.79738855 |
| YGPPGTGKTLL | ASSQVPRDKNSDYT | 0.75429946 |
| AALVGAPANVA | ASSQVPRDKNSDYT | 0.8349565 |
| AATFRRTE | ASSQVPRDKNSDYT | 0.7843766 |

| | | |
|---|---|---|
| AGALGGGI | ASSQVPRDKNSDYT | 0.754914 |
| AGGAGGAGGIGAG | ASSQVPRDKNSDYT | 0.791193 |
| AGSGALGL | ASSQVPRDKNSDYT | 0.83790195 |
| AQIADQI | ASSQVPRDKNSDYT | 0.7422253 |
| ASAVAVTYL | ASSQVPRDKNSDYT | 0.8334527 |
| ASTVLGGTGG | ASSQVPRDKNSDYT | 0.7058042 |
| AVGGEPAK | ASSQVPRDKNSDYT | 0.84030217 |
| CEQGTFGGTC | ASSQVPRDKNSDYT | 0.6971104 |
| DEGKAPAG | ASSQVPRDKNSDYT | 0.8307353 |
| DTVVFPAL | ASSQVPRDKNSDYT | 0.8019784 |
| FEGITMAG | ASSQVPRDKNSDYT | 0.8477894 |
| FPLVVQD | ASSQVPRDKNSDYT | 0.8088053 |
| GAFTTPQ | ASSQVPRDKNSDYT | 0.8355826 |
| GFGGFGTAGGVGGA | ASSQVPRDKNSDYT | 0.811525 |
| GFLAQFL | ASSQVPRDKNSDYT | 0.81397176 |
| GGAGGAGGNAGVA | ASSQVPRDKNSDYT | 0.6746887 |
| GGVGTAGAGGAGGAGG | ASSQVPRDKNSDYT | 0.81030554 |
| GNGGGGDAGNAGSGGN | ASSQVPRDKNSDYT | 0.67905843 |
| GTGGAGGAGNDAGS | ASSQVPRDKNSDYT | 0.8177356 |
| GTTNTGWFNS | ASSQVPRDKNSDYT | 0.8208655 |
| IFMGPLFTL | ASSQVPRDKNSDYT | 0.40107888 |
| ILAHAIEAI | ASSQVPRDKNSDYT | 0.8313112 |
| IRSAIREAL | ASSQVPRDKNSDYT | 0.73718536 |
| ITFAGGG | ASSQVPRDKNSDYT | 0.7561924 |
| ITGMSGHQEV | ASSQVPRDKNSDYT | 0.8374129 |
| IVAGLACTL | ASSQVPRDKNSDYT | 0.81068915 |
| IVGGGLAGA | ASSQVPRDKNSDYT | 0.8291844 |
| IVTGVLNI | ASSQVPRDKNSDYT | 0.78941274 |
| LAQQGCI | ASSQVPRDKNSDYT | 0.7982839 |
| LCLHVEN | ASSQVPRDKNSDYT | 0.81143856 |
| LDAISDGS | ASSQVPRDKNSDYT | 0.48866326 |
| LGGFAAML | ASSQVPRDKNSDYT | 0.74747276 |
| LGGKNPMI | ASSQVPRDKNSDYT | 0.8397677 |
| LGPKGVAAIL | ASSQVPRDKNSDYT | 0.58867884 |
| LMVGVSI | ASSQVPRDKNSDYT | 0.64038086 |
| LVGAAFVL | ASSQVPRDKNSDYT | 0.5017922 |
| LVLAVAPL | ASSQVPRDKNSDYT | 0.47105822 |
| MAAIAVIDAA | ASSQVPRDKNSDYT | 0.52430624 |
| NNGINAELQ | ASSQVPRDKNSDYT | 0.6337943 |
| RGEPLVLVHG | ASSQVPRDKNSDYT | 0.63462555 |
| RGGAGGAAGLI | ASSQVPRDKNSDYT | 0.6060767 |
| RSYRMHPAV | ASSQVPRDKNSDYT | 0.4516042 |
| SAAHLLAGQLA | ASSQVPRDKNSDYT | 0.54245615 |
| SAAVGSY | ASSQVPRDKNSDYT | 0.5243488 |
| SPHAVLHRI | ASSQVPRDKNSDYT | 0.49647865 |
| SQIAKVL | ASSQVPRDKNSDYT | 0.53729343 |
| TALGHLVP | ASSQVPRDKNSDYT | 0.56834006 |
| TAPTTSGAA | ASSQVPRDKNSDYT | 0.48778915 |
| TDGDDGNGGSPGT | ASSQVPRDKNSDYT | 0.4654743 |
| TDGLGEN | ASSQVPRDKNSDYT | 0.5259519 |
| TGGPPLPGQA | ASSQVPRDKNSDYT | 0.6065302 |

| | | |
|---|---|---|
| TIPAITI | ASSQVPRDKNSDYT | 0.5897811 |
| TIPGFTI | ASSQVPRDKNSDYT | 0.6492791 |
| TSIASGALA | ASSQVPRDKNSDYT | 0.6228831 |
| TVTLPGAAI | ASSQVPRDKNSDYT | 0.5971241 |
| VASGLSAALP | ASSQVPRDKNSDYT | 0.32406592 |
| VGAGVIGI | ASSQVPRDKNSDYT | 0.38876235 |
| VGPKRFGV | ASSQVPRDKNSDYT | 0.46053436 |
| VNVAFPD | ASSQVPRDKNSDYT | 0.53392196 |
| VSMTNSGVS | ASSQVPRDKNSDYT | 0.3467606 |
| VTDVLGDLSV | ASSQVPRDKNSDYT | 0.59472424 |
| VVVVGAGFAGL | ASSQVPRDKNSDYT | 0.546826 |
| VYTVGTVV | ASSQVPRDKNSDYT | 0.48546478 |
| WTTFSGP | ASSQVPRDKNSDYT | 0.6161158 |
| YDELVEAF | ASSQVPRDKNSDYT | 0.4667135 |
| YGTGGAGGNGG | ASSQVPRDKNSDYT | 0.47816914 |
| YTNVMAV | ASSQVPRDKNSDYT | 0.47126603 |
| FLEQVHQL | ASSQVPRDKNSDYT | 0.61736643 |
| GLSLLAVAV | ASSQVPRDKNSDYT | 0.40295208 |
| LEQVHQL | ASSQVPRDKNSDYT | 0.6014581 |
| LNAAGAI | ASSQVPRDKNSDYT | 0.44820714 |
| PAVHASL | ASSQVPRDKNSDYT | 0.6355751 |
| TELEDDA | ASSQVPRDKNSDYT | 0.42351308 |
| TPPGPPN | ASSQVPRDKNSDYT | 0.6123442 |
| TPPPPPPP | ASSQVPRDKNSDYT | 0.5297767 |
| TQSPALI | ASSFYRENTGQLY | 0.6558275 |
| EFAAAFPQ | ASSFYRENTGQLY | 0.52695334 |
| LLSGNLDVL | ASSFYRENTGQLY | 0.55833745 |
| GPANLGNYNI | ASSFYRENTGQLY | 0.46036276 |
| TLGPTLP | ASSFYRENTGQLY | 0.62184054 |
| VYAYSSCCK | ASSFYRENTGQLY | 0.43436113 |
| APAPAPAPNI | ASSFYRENTGQLY | 0.5401916 |
| PGGPVPIP | ASSFYRENTGQLY | 0.39093783 |
| FLTVALEP | ASSFYRENTGQLY | 0.5910973 |
| SQYFGANL | ASSFYRENTGQLY | 0.60648465 |
| GGVKPAA | ASSFYRENTGQLY | 0.3662529 |
| GGTGGAGGAGGW | ASSFYRENTGQLY | 0.56239265 |
| GVGGVAGLGGAGG | ASSFYRENTGQLY | 0.55939686 |
| VLLVLATTI | ASSFYRENTGQLY | 0.44381747 |
| IGIFAAML | ASSFYRENTGQLY | 0.599332 |
| AVGDLID | ASSFYRENTGQLY | 0.496972 |
| NISPLPL | ASSFYRENTGQLY | 0.63012373 |
| VTAVFVAI | ASSFYRENTGQLY | 0.44665867 |
| SGDDNDAAL | ASSFYRENTGQLY | 0.47423527 |
| AVAALAHS | ASSFYRENTGQLY | 0.61018467 |
| GGVTVMATP | ASSFYRENTGQLY | 0.46394897 |
| AGLAANVRD | ASSFYRENTGQLY | 0.46300247 |
| TNRPDLIDPALLRPG | ASSFYRENTGQLY | 0.6255457 |
| LANLGVTAL | ASSFYRENTGQLY | 0.67566544 |
| TGERIEL | ASSFYRENTGQLY | 0.8465973 |
| NGEELAA | ASSFYRENTGQLY | 0.5187333 |
| VLSLVWG | ASSFYRENTGQLY | 0.44029933 |

| | | |
|---|---|---|
| IGYALWL | ASSFYRENTGQLY | 0.6612565 |
| LSLLLAPL | ASSFYRENTGQLY | 0.8073539 |
| TLDEALK | ASSFYRENTGQLY | 0.82590324 |
| TDGAGTAEGG | ASSFYRENTGQLY | 0.74035954 |
| NAPGGAGGNGG | ASSFYRENTGQLY | 0.38596204 |
| VSVFMGMGGNF | ASSFYRENTGQLY | 0.6182493 |
| LASYLAAL | ASSFYRENTGQLY | 0.508867 |
| LVGGTPAL | ASSFYRENTGQLY | 0.6184959 |
| VRNADVL | ASSFYRENTGQLY | 0.7293216 |
| TCSSPFLVSAM | ASSFYRENTGQLY | 0.65428233 |
| VHITDAE | ASSFYRENTGQLY | 0.42523974 |
| EYAAPSF | ASSFYRENTGQLY | 0.45854312 |
| VVDAFHGGL | ASSFYRENTGQLY | 0.73099816 |
| ASVVDPQ | ASSFYRENTGQLY | 0.77308214 |
| SYNEFGGLS | ASSFYRENTGQLY | 0.76831543 |
| DPDGSVY | ASSFYRENTGQLY | 0.83896434 |
| MVSTIHATE | ASSFYRENTGQLY | 0.79359686 |
| LAAGVADPV | ASSFYRENTGQLY | 0.76324874 |
| TKEQIAA | ASSFYRENTGQLY | 0.1405038 |
| SLASPAAL | ASSFYRENTGQLY | 0.2276568 |
| IGLVFYGV | ASSFYRENTGQLY | 0.44673103 |
| FLLDASNL | ASSFYRENTGQLY | 0.63964903 |
| YQVGNFP | ASSFYRENTGQLY | 0.21249884 |
| ESLLSQST | ASSFYRENTGQLY | 0.75474834 |
| LFTVGFAAVG | ASSFYRENTGQLY | 0.49941558 |
| ATPPTPT | ASSFYRENTGQLY | 0.41795665 |
| TDLATLG | ASSFYRENTGQLY | 0.80377704 |
| SAGGTGR | ASSFYRENTGQLY | 0.40197057 |
| GFSNSGS | ASSFYRENTGQLY | 0.44483584 |
| GGNGGSGGTSV | ASSFYRENTGQLY | 0.48380184 |
| GTSVATGGAGN | ASSFYRENTGQLY | 0.777449 |
| NGGNGGSGGTS | ASSFYRENTGQLY | 0.28282148 |
| AVVKFPH | ASSFYRENTGQLY | 0.76590544 |
| TPPPPPP | ASSFYRENTGQLY | 0.5790833 |
| AAPAAAAAPA | ASSFYRENTGQLY | 0.79933655 |
| IALNEDL | ASSFYRENTGQLY | 0.30986613 |
| IVNVSSM | ASSFYRENTGQLY | 0.79069346 |
| TGIPPAPR | ASSFYRENTGQLY | 0.67409605 |
| FDLGGGTF | ASSFYRENTGQLY | 0.8583217 |
| FDLGGGTFDVS | ASSFYRENTGQLY | 0.6053251 |
| NEPTAAA | ASSFYRENTGQLY | 0.651219 |
| NEPTAAALA | ASSFYRENTGQLY | 0.46749696 |
| GSIINISSI | ASSFYRENTGQLY | 0.82368374 |
| VAPGNAGTA | ASSFYRENTGQLY | 0.30610037 |
| TGGGASLE | ASSFYRENTGQLY | 0.58865607 |
| VAVYNHYKRI | ASSFYRENTGQLY | 0.2303389 |
| TDPAPATT | ASSFYRENTGQLY | 0.6546293 |
| TNRPDILDPALLRPG | ASSFYRENTGQLY | 0.73419535 |
| LTTTTTS | ASSFYRENTGQLY | 0.26158205 |
| PTGSGKTL | ASSFYRENTGQLY | 0.7112897 |
| TGSGKTL | ASSFYRENTGQLY | 0.7676581 |

| | | |
|---|---|---|
| TIGHVDHGK | ASSFYRENTGQLY | 0.42361033 |
| TIGHVDHGKTT | ASSFYRENTGQLY | 0.80836517 |
| GHVDHGKTT | ASSFYRENTGQLY | 0.56944364 |
| AQFEHTLL | ASSFYRENTGQLY | 0.8125135 |
| IVNQDEL | ASSFYRENTGQLY | 0.5925264 |
| TNGSQFF | ASSFYRENTGQLY | 0.57688797 |
| TGKGIVE | ASSFYRENTGQLY | 0.7365687 |
| LISNASDA | ASSFYRENTGQLY | 0.36387622 |
| AELRTLGELTKIAKAHGAQ | ASSFYRENTGQLY | 0.4734825 |
| APNAVAAAEPSW | ASSFYRENTGQLY | 0.8109169 |
| APQVAGAATAVL | ASSFYRENTGQLY | 0.68847203 |
| ARRDPAAAAAILPTDA | ASSFYRENTGQLY | 0.71409684 |
| DAAARDVLAEAGLAEYF | ASSFYRENTGQLY | 0.6383213 |
| DDQIVLSAKANLAKVEQ | ASSFYRENTGQLY | 0.5931009 |
| DELLPDTGEDSDEDGH | ASSFYRENTGQLY | 0.6455022 |
| ELPEAPANRATVPQD | ASSFYRENTGQLY | 0.709102 |
| FGQSKIERVTGIG | ASSFYRENTGQLY | 0.70718575 |
| GAGGTGSTAGGAGGAGGAGGLYA | ASSFYRENTGQLY | 0.6883161 |
| GHGGTGAAGGNGGTGGAGGSGIDG | ASSFYRENTGQLY | 0.5689345 |
| GNGGNGGAGGEGGPGGAGGAGA | ASSFYRENTGQLY | 0.70160896 |
| GSTAGNGGNGGNATNGTIAGQPAG | ASSFYRENTGQLY | 0.66679144 |
| GTGGDGGNGGNGGTGG | ASSFYRENTGQLY | 0.65085566 |
| GVRRPAGIPADAPV | ASSFYRENTGQLY | 0.6967385 |
| GYRQVTDAHLLTLARR | ASSFYRENTGQLY | 0.6847089 |
| IAKMGQDGHDRG | ASSFYRENTGQLY | 0.6669854 |
| KAEPSIEIQYT | ASSFYRENTGQLY | 0.6566862 |
| KPVRIRMAADEDTVAVT | ASSFYRENTGQLY | 0.64724284 |
| MPPRIAGMRLLVIKPEP | ASSFYRENTGQLY | 0.6868458 |
| MKFYAVALFRSRGVPPTR | ASSFYRENTGQLY | 0.6688565 |
| NAGGFGILGAVAHSP | ASSFYRENTGQLY | 0.718662 |
| NEGREITLAMAEM | ASSFYRENTGQLY | 0.7055701 |
| NELVEWSHPLGHRWRWE | ASSFYRENTGQLY | 0.69105697 |
| NLGAGGTGGNGGNPG | ASSFYRENTGQLY | 0.30050528 |
| NTEWPKWETVMAADPVD | ASSFYRENTGQLY | 0.57919353 |
| PASAAPAETQP | ASSFYRENTGQLY | 0.6816454 |
| QQSSDGKNGVRAAEITGE | ASSFYRENTGQLY | 0.67396736 |
| RAPDAVTTSWWKE | ASSFYRENTGQLY | 0.47128502 |
| SGFGNVGAHTSGWFNQ | ASSFYRENTGQLY | 0.7109808 |
| SGVAGAVGQAASVGGLKV | ASSFYRENTGQLY | 0.66106164 |
| SNGGAGTGSAGGNGGT | ASSFYRENTGQLY | 0.6665572 |
| TGGAGGEGGAAG | ASSFYRENTGQLY | 0.7088126 |
| TGGDGGAAGAGGAPGHG | ASSFYRENTGQLY | 0.66450524 |
| TMPPLLAAIANAGRHGV | ASSFYRENTGQLY | 0.65950036 |
| TSSTAGNSGGA | ASSFYRENTGQLY | 0.6710334 |
| VPATVPVGILC | ASSFYRENTGQLY | 0.70986235 |
| VQRPDSGGVVLTGRIS | ASSFYRENTGQLY | 0.6100904 |
| WTPNPVPATKGVPAGLVQAT | ASSFYRENTGQLY | 0.6996299 |
| YALKRALAKFSDRFDVVII | ASSFYRENTGQLY | 0.63897985 |
| YLIGSVIV | ASSFYRENTGQLY | 0.71508765 |
| YGQLVAVID | ASSFYRENTGQLY | 0.5354142 |
| VTVGGMTLPA | ASSFYRENTGQLY | 0.69873095 |

| | | |
|---|---|---|
| VLALPAV | ASSFYRENTGQLY | 0.65714157 |
| VIPPTLNYET | ASSFYRENTGQLY | 0.7238651 |
| VGMGGMHPSSG | ASSFYRENTGQLY | 0.64639974 |
| VGLAGGAATA | ASSFYRENTGQLY | 0.6920691 |
| VGGAGGAGGSGP | ASSFYRENTGQLY | 0.6511118 |
| VFAAFMV | ASSFYRENTGQLY | 0.71610034 |
| TLYDKLR | ASSFYRENTGQLY | 0.56329185 |
| TLAVFTVL | ASSFYRENTGQLY | 0.65739584 |
| TGSVQPARV | ASSFYRENTGQLY | 0.6091161 |
| TGGGAGSGGT | ASSFYRENTGQLY | 0.68420726 |
| SHLETSL | ASSFYRENTGQLY | 0.69353354 |
| SFTTGMT | ASSFYRENTGQLY | 0.5051547 |
| QVEVAKRMGVRQPTVS | ASSFYRENTGQLY | 0.7085969 |
| QGSGGPALAM | ASSFYRENTGQLY | 0.6703083 |
| MLAPSNP | ASSFYRENTGQLY | 0.61794186 |
| LVNNAAI | ASSFYRENTGQLY | 0.7075496 |
| LTGEINTLA | ASSFYRENTGQLY | 0.6671064 |
| LSIFGPV | ASSFYRENTGQLY | 0.68819773 |
| LLALLANL | ASSFYRENTGQLY | 0.6582526 |
| LISNASDALDKLR | ASSFYRENTGQLY | 0.6734047 |
| LINEAVP | ASSFYRENTGQLY | 0.695532 |
| LGYAFSGL | ASSFYRENTGQLY | 0.5963532 |
| LDFVAVAL | ASSFYRENTGQLY | 0.6371942 |
| LAQPGAE | ASSFYRENTGQLY | 0.70658743 |
| ISNLGAVTGF | ASSFYRENTGQLY | 0.7784643 |
| HQEDPGRI | ASSFYRENTGQLY | 0.82858825 |
| GTPGEPTP | ASSFYRENTGQLY | 0.6742182 |
| GFFNSGS | ASSFYRENTGQLY | 0.54870623 |
| GAGGGGLTAGAN | ASSFYRENTGQLY | 0.7123692 |
| FVDALVNM | ASSFYRENTGQLY | 0.8444578 |
| FLTVAPEL | ASSFYRENTGQLY | 0.82514226 |
| FETLTAL | ASSFYRENTGQLY | 0.7911922 |
| DVINPIEL | ASSFYRENTGQLY | 0.5752426 |
| APVSGAPV | ASSFYRENTGQLY | 0.7453858 |
| ALVGGLAW | ASSFYRENTGQLY | 0.6883304 |
| ALAAAAATAI | ASSFYRENTGQLY | 0.64379764 |
| AFGLGLLL | ASSFYRENTGQLY | 0.8172373 |
| AFFAAASL | ASSFYRENTGQLY | 0.7952299 |
| AAAALPAFSPP | ASSFYRENTGQLY | 0.6963498 |
| VVLAPAL | ASSFYRENTGQLY | 0.7037376 |
| TIGHVDHGKTTL | ASSFYRENTGQLY | 0.7723182 |
| LISNASDALDK | ASSFYRENTGQLY | 0.72734433 |
| LISNASDAL | ASSFYRENTGQLY | 0.7069032 |
| ISNASDALDK | ASSFYRENTGQLY | 0.8647214 |
| NQAETLVYQTE | ASSFYRENTGQLY | 0.82424086 |
| TGGTGGTAGAG | ASSFYRENTGQLY | 0.83245116 |
| TGSTAGGAGGA | ASSFYRENTGQLY | 0.19706915 |
| DPLAVPVAAGPV | ASSFYRENTGQLY | 0.3524658 |
| GGTGGTPGNGGN | ASSFYRENTGQLY | 0.75960606 |
| PAGLAGVDPATL | ASSFYRENTGQLY | 0.69342583 |
| GNNLVVEAPHAGASV | ASSFYRENTGQLY | 0.46561423 |

| | | |
|---|---|---|
| KVTRIMSASTLEQPA | ASSFYRENTGQLY | 0.7490734 |
| LGFALPIALPRRTVV | ASSFYRENTGQLY | 0.67283356 |
| TRLGNLTARKLVVLV | ASSFYRENTGQLY | 0.6193712 |
| WSQQRRLIYFPSAGP | ASSFYRENTGQLY | 0.80789244 |
| GPNGNPDPMAAAVDIR | ASSFYRENTGQLY | 0.6578739 |
| TQRCVSVADLRAAASR | ASSFYRENTGQLY | 0.6555998 |
| QFVLDVAEHPVEVLTETV | ASSFYRENTGQLY | 0.6785562 |
| GGAGGTGGAGGNGGTQAGDG | ASSFYRENTGQLY | 0.82070124 |
| PLLATLRAIAVDVDAKPAQV | ASSFYRENTGQLY | 0.48382038 |
| APKFRTGGYMIDFWGVGYQVAKRM | ASSFYRENTGQLY | 0.7665905 |
| GAGGAGGAGGAAGGPGGTGGTGGN | ASSFYRENTGQLY | 0.6480133 |
| IGALGLLATDVAPAGSVAFGVALA | ASSFYRENTGQLY | 0.82018626 |
| AGGAGGIGGNGGAGGAGGVLMGNG | ASSFYRENTGQLY | 0.5050044 |
| ADELIGLL | ASSFYRENTGQLY | 0.8102572 |
| ADRALKA | ASSFYRENTGQLY | 0.7125919 |
| AGAVGQVGGAGGA | ASSFYRENTGQLY | 0.8674048 |
| AGGGGGLGGAP | ASSFYRENTGQLY | 0.694507 |
| AGGLLYGNGGAG | ASSFYRENTGQLY | 0.76182675 |
| AGLESIGL | ASSFYRENTGQLY | 0.719833 |
| AGLKTGII | ASSFYRENTGQLY | 0.8140408 |
| ASLAALLL | ASSFYRENTGQLY | 0.49395007 |
| ASVINISSI | ASSFYRENTGQLY | 0.6971809 |
| ATPPGPA | ASSFYRENTGQLY | 0.44640034 |
| ATVASALV | ASSFYRENTGQLY | 0.7821188 |
| ATVASLIL | ASSFYRENTGQLY | 0.7758883 |
| AVNGPVLIRH | ASSFYRENTGQLY | 0.48260304 |
| DGAAGAVGI | ASSFYRENTGQLY | 0.82261086 |
| DPLLHGI | ASSFYRENTGQLY | 0.79813015 |
| ELGAALGAL | ASSFYRENTGQLY | 0.65257764 |
| ELGRTQAV | ASSFYRENTGQLY | 0.8529817 |
| FVAGGYLS | ASSFYRENTGQLY | 0.7456503 |
| GASGGAGGAGGW | ASSFYRENTGQLY | 0.83809984 |
| GGAGAGGDGGAGGI | ASSFYRENTGQLY | 0.78361696 |
| GTGHNGSQPGMGGNG | ASSFYRENTGQLY | 0.7213114 |
| GYAAASASAS | ASSFYRENTGQLY | 0.78549373 |
| HQVDVAAVQQG | ASSFYRENTGQLY | 0.6281831 |
| IGAAIAHT | ASSFYRENTGQLY | 0.65214145 |
| IGMIEVAQL | ASSFYRENTGQLY | 0.8327202 |
| IRLAPEHTSAMD | ASSFYRENTGQLY | 0.7882079 |
| IYNVSVL | ASSFYRENTGQLY | 0.6903665 |
| KQGFLGM | ASSFYRENTGQLY | 0.5064126 |
| LESVITQL | ASSFYRENTGQLY | 0.35280263 |
| LGLGALVVV | ASSFYRENTGQLY | 0.53193694 |
| LIGNGGNGGSGGI | ASSFYRENTGQLY | 0.73508453 |
| LNAINAPTL | ASSFYRENTGQLY | 0.64800555 |
| LQLLNTGV | ASSFYRENTGQLY | 0.7374679 |
| LQTSIEF | ASSFYRENTGQLY | 0.239481 |
| MSFAVSEL | ASSFYRENTGQLY | 0.5419426 |
| NTGALGL | ASSFYRENTGQLY | 0.5173855 |
| PPGAGGAGGAGTT | ASSFYRENTGQLY | 0.5036912 |
| QLVGLGI | ASSFYRENTGQLY | 0.64370775 |

| | | |
|---|---|---|
| RFLEQVHQL | ASSFYRENTGQLY | 0.23746787 |
| RGVGYVLR | ASSFYRENTGQLY | 0.4909425 |
| SATGVFAGVI | ASSFYRENTGQLY | 0.4712872 |
| SLPAITLP | ASSFYRENTGQLY | 0.5213142 |
| STALAFTV | ASSFYRENTGQLY | 0.7343876 |
| SVLTVGLGK | ASSFYRENTGQLY | 0.7142843 |
| TALAGAGSGI | ASSFYRENTGQLY | 0.77703744 |
| TGLPFTGI | ASSFYRENTGQLY | 0.77444804 |
| VAEHVTV | ASSFYRENTGQLY | 0.7419205 |
| VDIVLNSL | ASSFYRENTGQLY | 0.117276266 |
| VLTGGGAVL | ASSFYRENTGQLY | 0.2806483 |
| VNVGLAYGI | ASSFYRENTGQLY | 0.21467537 |
| VSSAAVGAV | ASSFYRENTGQLY | 0.5896192 |
| VTSVYEAGV | ASSFYRENTGQLY | 0.13975799 |
| WVALDRGA | ASSFYRENTGQLY | 0.5719429 |
| YLVGSVVI | ASSFYRENTGQLY | 0.68972725 |
| GPPGTGKTLL | ASSFYRENTGQLY | 0.23573153 |
| GVGMTKF | ASSFYRENTGQLY | 0.7383987 |
| IGIGGSDL | ASSFYRENTGQLY | 0.3951106 |
| IGTLLAEL | ASSFYRENTGQLY | 0.24268812 |
| IYTSGTT | ASSFYRENTGQLY | 0.2566663 |
| LSFPAGV | ASSFYRENTGQLY | 0.78158927 |
| LYTSGTT | ASSFYRENTGQLY | 0.3228954 |
| VGAFPFGF | ASSFYRENTGQLY | 0.7666528 |
| VGAGTVEF | ASSFYRENTGQLY | 0.40468496 |
| VGKSTLLL | ASSFYRENTGQLY | 0.7081085 |
| AALAHAGGGIV | ASSFYRENTGQLY | 0.17277397 |
| AALVAVSLL | ASSFYRENTGQLY | 0.74259937 |
| AAVSRAVAAL | ASSFYRENTGQLY | 0.64157695 |
| AGGSSGIGGF | ASSFYRENTGQLY | 0.77794635 |
| AGLTVANANG | ASSFYRENTGQLY | 0.5627495 |
| AGLVGAVL | ASSFYRENTGQLY | 0.49740085 |
| ALSQAFTV | ASSFYRENTGQLY | 0.30345565 |
| ALTFSVSL | ASSFYRENTGQLY | 0.4780392 |
| ALTGAALVAL | ASSFYRENTGQLY | 0.17806576 |
| ASFSIGAGVL | ASSFYRENTGQLY | 0.59452957 |
| DGSVYKI | ASSFYRENTGQLY | 0.24300359 |
| EQLAEIV | ASSFYRENTGQLY | 0.5926051 |
| FAVSELTVM | ASSFYRENTGQLY | 0.76161766 |
| GAAPLAAGL | ASSFYRENTGQLY | 0.1402745 |
| GADGGDGGSGGAGG | ASSFYRENTGQLY | 0.70639217 |
| GFGLDTGGAGG | ASSFYRENTGQLY | 0.42552805 |
| GLASGGLL | ASSFYRENTGQLY | 0.21309818 |
| GRIAGAAVLL | ASSFYRENTGQLY | 0.7080564 |
| GSAGSLNTSF | ASSFYRENTGQLY | 0.20300694 |
| GTGQIGI | ASSFYRENTGQLY | 0.770465 |
| GTLIALIL | ASSFYRENTGQLY | 0.2130693 |
| IAATMGFVVG | ASSFYRENTGQLY | 0.64162123 |
| IFAQALL | ASSFYRENTGQLY | 0.71346617 |
| IGAAVALGF | ASSFYRENTGQLY | 0.11110756 |
| IGLTPLAL | ASSFYRENTGQLY | 0.19331278 |

| | | |
|---|---|---|
| ILGSNTSTL | ASSFYRENTGQLY | 0.7763268 |
| IVFSLTVL | ASSFYRENTGQLY | 0.60342604 |
| LASKATGF | ASSFYRENTGQLY | 0.70951307 |
| LASVALIL | ASSFYRENTGQLY | 0.49986488 |
| LFAVGLTL | ASSFYRENTGQLY | 0.4411685 |
| LFNGGIGTYI | ASSFYRENTGQLY | 0.51990044 |
| LTLGAGII | ASSFYRENTGQLY | 0.6651159 |
| LVAAALGDI | ASSFYRENTGQLY | 0.704748 |
| NTAFVQLGI | ASSFYRENTGQLY | 0.648046 |
| QLFSGAEL | ASSFYRENTGQLY | 0.42340094 |
| SAATFITL | ASSFYRENTGQLY | 0.6359798 |
| SNYGLANLGV | ASSFYRENTGQLY | 0.53782326 |
| SYGGAYAVM | ASSFYRENTGQLY | 0.5924134 |
| TTGVGGLTLG | ASSFYRENTGQLY | 0.59656155 |
| TTQTIYI | ASSFYRENTGQLY | 0.6106931 |
| VAAGAGLGAVI | ASSFYRENTGQLY | 0.47394246 |
| VAAVVTLL | ASSFYRENTGQLY | 0.41948834 |
| VGFLGGLI | ASSFYRENTGQLY | 0.6793401 |
| VGVIPILL | ASSFYRENTGQLY | 0.68029374 |
| VIFASSLI | ASSFYRENTGQLY | 0.6805159 |
| VKGGDGGAAAG | ASSFYRENTGQLY | 0.68719417 |
| VTKTFTNI | ASSFYRENTGQLY | 0.6776942 |
| VVAAAILL | ASSFYRENTGQLY | 0.61509997 |
| YAALDVEL | ASSFYRENTGQLY | 0.2998041 |
| LASVALLL | ASSFYRENTGQLY | 0.3601993 |
| LDAVHRV | ASSFYRENTGQLY | 0.487575 |
| QGLSSSIF | ASSFYRENTGQLY | 0.5699919 |
| RGTFANIRL | ASSFYRENTGQLY | 0.3156778 |
| RQLWWGHRI | ASSFYRENTGQLY | 0.63445735 |
| YGPPGTGKTLL | ASSFYRENTGQLY | 0.44439057 |
| AALVGAPANVA | ASSFYRENTGQLY | 0.4892969 |
| AATFRRTE | ASSFYRENTGQLY | 0.6835743 |
| AGALGGGI | ASSFYRENTGQLY | 0.48384613 |
| AGGAGGAGGIGAG | ASSFYRENTGQLY | 0.43135503 |
| AGSGALGL | ASSFYRENTGQLY | 0.41821003 |
| AQIADQI | ASSFYRENTGQLY | 0.5880758 |
| ASAVAVTYL | ASSFYRENTGQLY | 0.37142828 |
| ASTVLGGTGG | ASSFYRENTGQLY | 0.6411906 |
| AVGGEPAK | ASSFYRENTGQLY | 0.42692614 |
| CEQGTFGGTC | ASSFYRENTGQLY | 0.6856309 |
| DEGKAPAG | ASSFYRENTGQLY | 0.3855947 |
| DTVVFPAL | ASSFYRENTGQLY | 0.6637711 |
| FEGITMAG | ASSFYRENTGQLY | 0.5907662 |
| FPLVVQD | ASSFYRENTGQLY | 0.72008336 |
| GAFTTPQ | ASSFYRENTGQLY | 0.536595 |
| GFGGFGTAGGVGGA | ASSFYRENTGQLY | 0.60040367 |
| GFLAQFL | ASSFYRENTGQLY | 0.5324752 |
| GGAGGAGGNAGVA | ASSFYRENTGQLY | 0.7025075 |
| GGVGTAGAGGAGGAGG | ASSFYRENTGQLY | 0.38360202 |
| GNGGGGDAGNAGSGGN | ASSFYRENTGQLY | 0.51692665 |
| GTGGAGGAGNDAGS | ASSFYRENTGQLY | 0.3476948 |

| | | |
|---|---|---|
| GTTNTGWFNS | ASSFYRENTGQLY | 0.55993485 |
| IFMGPLFTL | ASSFYRENTGQLY | 0.6364841 |
| ILAHAIEAI | ASSFYRENTGQLY | 0.31780913 |
| IRSAIREAL | ASSFYRENTGQLY | 0.5791849 |
| ITFAGGG | ASSFYRENTGQLY | 0.52316284 |
| ITGMSGHQEV | ASSFYRENTGQLY | 0.4250316 |
| IVAGLACTL | ASSFYRENTGQLY | 0.67653984 |
| IVGGGLAGA | ASSFYRENTGQLY | 0.624933 |
| IVTGVLNI | ASSFYRENTGQLY | 0.67575634 |
| LAQQGCI | ASSFYRENTGQLY | 0.5246297 |
| LCLHVEN | ASSFYRENTGQLY | 0.48610526 |
| LDAISDGS | ASSFYRENTGQLY | 0.6040883 |
| LGGFAAML | ASSFYRENTGQLY | 0.60099465 |
| LGGKNPMI | ASSFYRENTGQLY | 0.4622013 |
| LGPKGVAAIL | ASSFYRENTGQLY | 0.6793232 |
| LMVGVSI | ASSFYRENTGQLY | 0.7096214 |
| LVGAAFVL | ASSFYRENTGQLY | 0.7236608 |
| LVLAVAPL | ASSFYRENTGQLY | 0.6458995 |
| MAAIAVIDAA | ASSFYRENTGQLY | 0.6240716 |
| NNGINAELQ | ASSFYRENTGQLY | 0.68638605 |
| RGEPLVLVHG | ASSFYRENTGQLY | 0.71864 |
| RGGAGGAAGLI | ASSFYRENTGQLY | 0.7294352 |
| RSYRMHPAV | ASSFYRENTGQLY | 0.71026397 |
| SAAHLLAGQLA | ASSFYRENTGQLY | 0.5877889 |
| SAAVGSY | ASSFYRENTGQLY | 0.71381515 |
| SPHAVLHRI | ASSFYRENTGQLY | 0.6882496 |
| SQIAKVL | ASSFYRENTGQLY | 0.6843306 |
| TALGHLVP | ASSFYRENTGQLY | 0.7149606 |
| TAPTTSGAA | ASSFYRENTGQLY | 0.69078565 |
| TDGDDGNGGSPGT | ASSFYRENTGQLY | 0.6578765 |
| TDGLGEN | ASSFYRENTGQLY | 0.62058794 |
| TGGPPLPGQA | ASSFYRENTGQLY | 0.7175218 |
| TIPAITI | ASSFYRENTGQLY | 0.7214596 |
| TIPGFTI | ASSFYRENTGQLY | 0.719413 |
| TSIASGALA | ASSFYRENTGQLY | 0.7308049 |
| TVTLPGAAI | ASSFYRENTGQLY | 0.71988106 |
| VASGLSAALP | ASSFYRENTGQLY | 0.7040734 |
| VGAGVIGI | ASSFYRENTGQLY | 0.3392108 |
| VGPKRFGV | ASSFYRENTGQLY | 0.5030504 |
| VNVAFPD | ASSFYRENTGQLY | 0.5974144 |
| VSMTNSGVS | ASSFYRENTGQLY | 0.6825675 |
| VTDVLGDLSV | ASSFYRENTGQLY | 0.3509218 |
| VVVVGAGFAGL | ASSFYRENTGQLY | 0.7113538 |
| VYTVGTVV | ASSFYRENTGQLY | 0.6670018 |
| WTTFSGP | ASSFYRENTGQLY | 0.6350208 |
| YDELVEAF | ASSFYRENTGQLY | 0.72290754 |
| YGTGGAGGNGG | ASSFYRENTGQLY | 0.651314 |
| YTNVMAV | ASSFYRENTGQLY | 0.58034617 |
| FLEQVHQL | ASSFYRENTGQLY | 0.6525385 |
| GLSLLAVAV | ASSFYRENTGQLY | 0.7289463 |
| LEQVHQL | ASSFYRENTGQLY | 0.54258394 |

| | | |
|---|---|---|
| LNAAGAI | ASSFYRENTGQLY | 0.7214774 |
| PAVHASL | ASSFYRENTGQLY | 0.6875005 |
| TELEDDA | ASSFYRENTGQLY | 0.7263755 |
| TPPGPPN | ASSFYRENTGQLY | 0.5062287 |
| TPPPPPPP | ASSFYRENTGQLY | 0.7168945 |
| TQSPALI | ASSQDWGGDAEQF | 0.6914587 |
| EFAAAFPQ | ASSQDWGGDAEQF | 0.7343125 |
| LLSGNLDVL | ASSQDWGGDAEQF | 0.6763225 |
| GPANLGNYNI | ASSQDWGGDAEQF | 0.710562 |
| TLGPTLP | ASSQDWGGDAEQF | 0.7024591 |
| VYAYSSCCK | ASSQDWGGDAEQF | 0.7252232 |
| APAPAPAPNI | ASSQDWGGDAEQF | 0.5491657 |
| PGGPVPIP | ASSQDWGGDAEQF | 0.684358 |
| FLTVALEP | ASSQDWGGDAEQF | 0.6426929 |
| SQYFGANL | ASSQDWGGDAEQF | 0.7008153 |
| GGVKPAA | ASSQDWGGDAEQF | 0.7082449 |
| GGTGGAGGAGGW | ASSQDWGGDAEQF | 0.3871123 |
| GVGGVAGLGGAGG | ASSQDWGGDAEQF | 0.72533834 |
| VLLVLATTI | ASSQDWGGDAEQF | 0.66144264 |
| IGIFAAML | ASSQDWGGDAEQF | 0.6029136 |
| AVGDLID | ASSQDWGGDAEQF | 0.71880984 |
| NISPLPL | ASSQDWGGDAEQF | 0.6965061 |
| VTAVFVAI | ASSQDWGGDAEQF | 0.7108933 |
| SGDDNDAAL | ASSQDWGGDAEQF | 0.6401477 |
| AVAALAHS | ASSQDWGGDAEQF | 0.6951689 |
| GGVTVMATP | ASSQDWGGDAEQF | 0.7135755 |
| AGLAANVRD | ASSQDWGGDAEQF | 0.5576404 |
| TNRPDLIDPALLRPG | ASSQDWGGDAEQF | 0.6163434 |
| LANLGVTAL | ASSQDWGGDAEQF | 0.7227732 |
| TGERIEL | ASSQDWGGDAEQF | 0.8850968 |
| NGEELAA | ASSQDWGGDAEQF | 0.8869593 |
| VLSLVWG | ASSQDWGGDAEQF | 0.82932687 |
| IGYALWL | ASSQDWGGDAEQF | 0.7738749 |
| LSLLLAPL | ASSQDWGGDAEQF | 0.8408111 |
| TLDEALK | ASSQDWGGDAEQF | 0.8928704 |
| TDGAGTAEGG | ASSQDWGGDAEQF | 0.88935554 |
| NAPGGAGGNGG | ASSQDWGGDAEQF | 0.88133657 |
| VSVFMGMGGNF | ASSQDWGGDAEQF | 0.75575256 |
| LASYLAAL | ASSQDWGGDAEQF | 0.8772957 |
| LVGGTPAL | ASSQDWGGDAEQF | 0.85245126 |
| VRNADVL | ASSQDWGGDAEQF | 0.84757376 |
| TCSSPFLVSAM | ASSQDWGGDAEQF | 0.8834747 |
| VHITDAE | ASSQDWGGDAEQF | 0.87008023 |
| EYAAPSF | ASSQDWGGDAEQF | 0.8520795 |
| VVDAFHGGL | ASSQDWGGDAEQF | 0.8566781 |
| ASVVDPQ | ASSQDWGGDAEQF | 0.8627155 |
| SYNEFGGLS | ASSQDWGGDAEQF | 0.88080657 |
| DPDGSVY | ASSQDWGGDAEQF | 0.8815242 |
| MVSTIHATE | ASSQDWGGDAEQF | 0.90179294 |
| LAAGVADPV | ASSQDWGGDAEQF | 0.8966008 |
| TKEQIAA | ASSQDWGGDAEQF | 0.8879125 |

| | | |
|---|---|---|
| SLASPAAL | ASSQDWGGDAEQF | 0.2822454 |
| IGLVFYGV | ASSQDWGGDAEQF | 0.6141459 |
| FLLDASNL | ASSQDWGGDAEQF | 0.8120856 |
| YQVGNFP | ASSQDWGGDAEQF | 0.8382994 |
| ESLLSQST | ASSQDWGGDAEQF | 0.6486273 |
| LFTVGFAAVG | ASSQDWGGDAEQF | 0.8619763 |
| ATPPTPT | ASSQDWGGDAEQF | 0.8441584 |
| TDLATLG | ASSQDWGGDAEQF | 0.7811817 |
| SAGGTGR | ASSQDWGGDAEQF | 0.8895037 |
| GFSNSGS | ASSQDWGGDAEQF | 0.84884423 |
| GGNGGSGGTSV | ASSQDWGGDAEQF | 0.8067285 |
| GTSVATGGAGN | ASSQDWGGDAEQF | 0.8262471 |
| NGGNGGSGGTS | ASSQDWGGDAEQF | 0.8919685 |
| AVVKFPH | ASSQDWGGDAEQF | 0.7716963 |
| TPPPPPP | ASSQDWGGDAEQF | 0.88662934 |
| AAPAAAAAPA | ASSQDWGGDAEQF | 0.80237377 |
| IALNEDL | ASSQDWGGDAEQF | 0.89738077 |
| IVNVSSM | ASSQDWGGDAEQF | 0.6816463 |
| TGIPPAPR | ASSQDWGGDAEQF | 0.88997793 |
| FDLGGGTF | ASSQDWGGDAEQF | 0.8602352 |
| FDLGGGTFDVS | ASSQDWGGDAEQF | 0.907742 |
| NEPTAAA | ASSQDWGGDAEQF | 0.848441 |
| NEPTAAALA | ASSQDWGGDAEQF | 0.86880845 |
| GSIINISSI | ASSQDWGGDAEQF | 0.7942451 |
| VAPGNAGTA | ASSQDWGGDAEQF | 0.88433874 |
| TGGGASLE | ASSQDWGGDAEQF | 0.6660143 |
| VAVYNHYKRI | ASSQDWGGDAEQF | 0.8649322 |
| TDPAPATT | ASSQDWGGDAEQF | 0.6704071 |
| TNRPDILDPALLRPG | ASSQDWGGDAEQF | 0.8757483 |
| LTTTTTS | ASSQDWGGDAEQF | 0.87977934 |
| PTGSGKTL | ASSQDWGGDAEQF | 0.62886184 |
| TGSGKTL | ASSQDWGGDAEQF | 0.8984678 |
| TIGHVDHGK | ASSQDWGGDAEQF | 0.8542702 |
| TIGHVDHGKTT | ASSQDWGGDAEQF | 0.8030288 |
| GHVDHGKTT | ASSQDWGGDAEQF | 0.8987242 |
| AQFEHTLL | ASSQDWGGDAEQF | 0.8423952 |
| IVNQDEL | ASSQDWGGDAEQF | 0.90144813 |
| TNGSQFF | ASSQDWGGDAEQF | 0.8546988 |
| TGKGIVE | ASSQDWGGDAEQF | 0.85187215 |
| LISNASDA | ASSQDWGGDAEQF | 0.87283665 |
| AELRTLGELTKIAKAHGAQ | ASSQDWGGDAEQF | 0.60247374 |
| APNAVAAAEPSW | ASSQDWGGDAEQF | 0.7702341 |
| APQVAGAATAVL | ASSQDWGGDAEQF | 0.9000548 |
| ARRDPAAAAAILPTDA | ASSQDWGGDAEQF | 0.83144397 |
| DAAARDVLAEAGLAEYF | ASSQDWGGDAEQF | 0.8188144 |
| DDQIVLSAKANLAKVEQ | ASSQDWGGDAEQF | 0.68899953 |
| DELLPDTGEDSDEDGH | ASSQDWGGDAEQF | 0.5719845 |
| ELPEAPANRATVPQD | ASSQDWGGDAEQF | 0.694401 |
| FGQSKIERVTGIG | ASSQDWGGDAEQF | 0.8546617 |
| GAGGTGSTAGGAGGAGGAGGLYA | ASSQDWGGDAEQF | 0.8433417 |
| GHGGTGAAGGNGGTGGAGGSGIDG | ASSQDWGGDAEQF | 0.81131697 |

| | | |
|---|---|---|
| GNGGNGGAGGEGGPGGAGGAGA | ASSQDWGGDAEQF | 0.47394195 |
| GSTAGNGGNGGNATNGTIAGQPAG | ASSQDWGGDAEQF | 0.8303608 |
| GTGGDGGNGGNGGTGG | ASSQDWGGDAEQF | 0.75273913 |
| GVRRPAGIPADAPV | ASSQDWGGDAEQF | 0.7717047 |
| GYRQVTDAHLLTLARR | ASSQDWGGDAEQF | 0.81051594 |
| IAKMGQDGHDRG | ASSQDWGGDAEQF | 0.74511236 |
| KAEPSIEIQYT | ASSQDWGGDAEQF | 0.7567662 |
| KPVRIRMAADEDTVAVT | ASSQDWGGDAEQF | 0.5813118 |
| MPPRIAGMRLLVIKPEP | ASSQDWGGDAEQF | 0.7819754 |
| MKFYAVALFRSRGVPPTR | ASSQDWGGDAEQF | 0.82608044 |
| NAGGFGILGAVAHSP | ASSQDWGGDAEQF | 0.8038925 |
| NEGREITLAMAEM | ASSQDWGGDAEQF | 0.8755344 |
| NELVEWSHPLGHRWRWE | ASSQDWGGDAEQF | 0.8677516 |
| NLGAGGTGGNGGNPG | ASSQDWGGDAEQF | 0.82089424 |
| NTEWPKWETVMAADPVD | ASSQDWGGDAEQF | 0.17499492 |
| PASAAPAETQP | ASSQDWGGDAEQF | 0.47385317 |
| QQSSDGKNGVRAAEITGE | ASSQDWGGDAEQF | 0.63031256 |
| RAPDAVTTSWWKE | ASSQDWGGDAEQF | 0.7627354 |
| SGFGNVGAHTSGWFNQ | ASSQDWGGDAEQF | 0.25167328 |
| SGVAGAVGQAASVGGLKV | ASSQDWGGDAEQF | 0.7906655 |
| SNGGAGTGSAGGNGGT | ASSQDWGGDAEQF | 0.68190265 |
| TGGAGGEGGAAG | ASSQDWGGDAEQF | 0.69897026 |
| TGGDGGAAGAGGAPGHG | ASSQDWGGDAEQF | 0.87356174 |
| TMPPLLAAIANAGRHGV | ASSQDWGGDAEQF | 0.7447785 |
| TSSTAGNSGGA | ASSQDWGGDAEQF | 0.6296363 |
| VPATVPVGILC | ASSQDWGGDAEQF | 0.6523504 |
| VQRPDSGGVVLTGRIS | ASSQDWGGDAEQF | 0.8669435 |
| WTPNPVPATKGVPAGLVQAT | ASSQDWGGDAEQF | 0.4627136 |
| YALKRALAKFSDRFDVVII | ASSQDWGGDAEQF | 0.8607271 |
| YLIGSVIV | ASSQDWGGDAEQF | 0.6175271 |
| YGQLVAVID | ASSQDWGGDAEQF | 0.8838934 |
| VTVGGMTLPA | ASSQDWGGDAEQF | 0.41710535 |
| VLALPAV | ASSQDWGGDAEQF | 0.847942 |
| VIPPTLNYET | ASSQDWGGDAEQF | 0.77923644 |
| VGMGGMHPSSG | ASSQDWGGDAEQF | 0.89645016 |
| VGLAGGAATA | ASSQDWGGDAEQF | 0.7321819 |
| VGGAGGAGGSGP | ASSQDWGGDAEQF | 0.79920363 |
| VFAAFMV | ASSQDWGGDAEQF | 0.6154618 |
| TLYDKLR | ASSQDWGGDAEQF | 0.83662045 |
| TLAVFTVL | ASSQDWGGDAEQF | 0.48261845 |
| TGSVQPARV | ASSQDWGGDAEQF | 0.702151 |
| TGGGAGSGGT | ASSQDWGGDAEQF | 0.5165793 |
| SHLETSL | ASSQDWGGDAEQF | 0.78242284 |
| SFTTGMT | ASSQDWGGDAEQF | 0.8271514 |
| QVEVAKRMGVRQPTVS | ASSQDWGGDAEQF | 0.20486774 |
| QGSGGPALAM | ASSQDWGGDAEQF | 0.8290626 |
| MLAPSNP | ASSQDWGGDAEQF | 0.58858585 |
| LVNNAAI | ASSQDWGGDAEQF | 0.5123261 |
| LTGEINTLA | ASSQDWGGDAEQF | 0.8477446 |
| LSIFGPV | ASSQDWGGDAEQF | 0.70874965 |
| LLALLANL | ASSQDWGGDAEQF | 0.8394878 |

| | | |
|---|---|---|
| LISNASDALDKLR | ASSQDWGGDAEQF | 0.6651174 |
| LINEAVP | ASSQDWGGDAEQF | 0.74297726 |
| LGYAFSGL | ASSQDWGGDAEQF | 0.7988929 |
| LDFVAVAL | ASSQDWGGDAEQF | 0.3607319 |
| LAQPGAE | ASSQDWGGDAEQF | 0.58204085 |
| ISNLGAVTGF | ASSQDWGGDAEQF | 0.87276995 |
| HQEDPGRI | ASSQDWGGDAEQF | 0.6159184 |
| GTPGEPTP | ASSQDWGGDAEQF | 0.69778645 |
| GFFNSGS | ASSQDWGGDAEQF | 0.5982221 |
| GAGGGGLTAGAN | ASSQDWGGDAEQF | 0.53270483 |
| FVDALVNM | ASSQDWGGDAEQF | 0.6069864 |
| FLTVAPEL | ASSQDWGGDAEQF | 0.6868813 |
| FETLTAL | ASSQDWGGDAEQF | 0.6919879 |
| DVINPIEL | ASSQDWGGDAEQF | 0.66229475 |
| APVSGAPV | ASSQDWGGDAEQF | 0.51044554 |
| ALVGGLAW | ASSQDWGGDAEQF | 0.65538156 |
| ALAAAAATAI | ASSQDWGGDAEQF | 0.5801457 |
| AFGLGLLL | ASSQDWGGDAEQF | 0.62864697 |
| AFFAAASL | ASSQDWGGDAEQF | 0.65358126 |
| AAAALPAFSPP | ASSQDWGGDAEQF | 0.63019276 |
| VVLAPAL | ASSQDWGGDAEQF | 0.6107397 |
| TIGHVDHGKTTL | ASSQDWGGDAEQF | 0.5622582 |
| LISNASDALDK | ASSQDWGGDAEQF | 0.651285 |
| LISNASDAL | ASSQDWGGDAEQF | 0.6240181 |
| ISNASDALDK | ASSQDWGGDAEQF | 0.58566797 |
| NQAETLVYQTE | ASSQDWGGDAEQF | 0.69448006 |
| TGGTGGTAGAG | ASSQDWGGDAEQF | 0.6711705 |
| TGSTAGGAGGA | ASSQDWGGDAEQF | 0.6569737 |
| DPLAVPVAAGPV | ASSQDWGGDAEQF | 0.31387812 |
| GGTGGTPGNGGN | ASSQDWGGDAEQF | 0.39750692 |
| PAGLAGVDPATL | ASSQDWGGDAEQF | 0.606987 |
| GNNLVVEAPHAGASV | ASSQDWGGDAEQF | 0.58605146 |
| KVTRIMSASTLEQPA | ASSQDWGGDAEQF | 0.39640006 |
| LGFALPIALPRRTVV | ASSQDWGGDAEQF | 0.66674936 |
| TRLGNLTARKLVVLV | ASSQDWGGDAEQF | 0.5991474 |
| WSQQRRLIYFPSAGP | ASSQDWGGDAEQF | 0.59585994 |
| GPNGNPDPMAAAVDIR | ASSQDWGGDAEQF | 0.6720736 |
| TQRCVSVADLRAAASR | ASSQDWGGDAEQF | 0.6119386 |
| QFVLDVAEHPVEVLTETV | ASSQDWGGDAEQF | 0.575801 |
| GGAGGTGGAGGNGGTQAGDG | ASSQDWGGDAEQF | 0.5533545 |
| PLLATLRAIAVDVDAKPAQV | ASSQDWGGDAEQF | 0.67673945 |
| APKFRTGGYMIDFWGVGYQVAKRM | ASSQDWGGDAEQF | 0.45322287 |
| GAGGAGGAGGAAGGPGGTGGTGGN | ASSQDWGGDAEQF | 0.6634927 |
| IGALGLLATDVAPAGSVAFGVALA | ASSQDWGGDAEQF | 0.53205985 |
| AGGAGGIGGNGGAGGAGGVLMGNG | ASSQDWGGDAEQF | 0.6761796 |
| ADELIGLL | ASSQDWGGDAEQF | 0.46777117 |
| ADRALKA | ASSQDWGGDAEQF | 0.66660315 |
| AGAVGQVGGAGGA | ASSQDWGGDAEQF | 0.622186 |
| AGGGGGLGGAP | ASSQDWGGDAEQF | 0.7092668 |
| AGGLLYGNGGAG | ASSQDWGGDAEQF | 0.5691693 |
| AGLESIGL | ASSQDWGGDAEQF | 0.6142029 |

| | | |
|---|---|---|
| AGLKTGII | ASSQDWGGDAEQF | 0.5753541 |
| ASLAALLL | ASSQDWGGDAEQF | 0.6888971 |
| ASVINISSI | ASSQDWGGDAEQF | 0.462427 |
| ATPPGPA | ASSQDWGGDAEQF | 0.5893919 |
| ATVASALV | ASSQDWGGDAEQF | 0.51344544 |
| ATVASLIL | ASSQDWGGDAEQF | 0.664577 |
| AVNGPVLIRH | ASSQDWGGDAEQF | 0.6512657 |
| DGAAGAVGI | ASSQDWGGDAEQF | 0.4485189 |
| DPLLHGI | ASSQDWGGDAEQF | 0.6748232 |
| ELGAALGAL | ASSQDWGGDAEQF | 0.6380235 |
| ELGRTQAV | ASSQDWGGDAEQF | 0.55388176 |
| FVAGGYLS | ASSQDWGGDAEQF | 0.6873622 |
| GASGGAGGAGGW | ASSQDWGGDAEQF | 0.62480414 |
| GGAGAGGDGGAGGI | ASSQDWGGDAEQF | 0.6686701 |
| GTGHNGSQPGMGGNG | ASSQDWGGDAEQF | 0.62999064 |
| GYAAASASAS | ASSQDWGGDAEQF | 0.5991941 |
| HQVDVAAVQQG | ASSQDWGGDAEQF | 0.6672595 |
| IGAAIAHT | ASSQDWGGDAEQF | 0.61540484 |
| IGMIEVAQL | ASSQDWGGDAEQF | 0.54082 |
| IRLAPEHTSAMD | ASSQDWGGDAEQF | 0.6824316 |
| IYNVSVL | ASSQDWGGDAEQF | 0.7406752 |
| KQGFLGM | ASSQDWGGDAEQF | 0.7431843 |
| LESVITQL | ASSQDWGGDAEQF | 0.6625484 |
| LGLGALVVV | ASSQDWGGDAEQF | 0.6531613 |
| LIGNGGNGGSGGI | ASSQDWGGDAEQF | 0.69479966 |
| LNAINAPTL | ASSQDWGGDAEQF | 0.7448077 |
| LQLLNTGV | ASSQDWGGDAEQF | 0.7673827 |
| LQTSIEF | ASSQDWGGDAEQF | 0.74955535 |
| MSFAVSEL | ASSQDWGGDAEQF | 0.68903655 |
| NTGALGL | ASSQDWGGDAEQF | 0.75604993 |
| PPGAGGAGGAGTT | ASSQDWGGDAEQF | 0.73123294 |
| QLVGLGI | ASSQDWGGDAEQF | 0.72857714 |
| RFLEQVHQL | ASSQDWGGDAEQF | 0.74609536 |
| RGVGYVLR | ASSQDWGGDAEQF | 0.73423773 |
| SATGVFAGVI | ASSQDWGGDAEQF | 0.6893362 |
| SLPAITLP | ASSQDWGGDAEQF | 0.69475424 |
| STALAFTV | ASSQDWGGDAEQF | 0.743945 |
| SVLTVGLGK | ASSQDWGGDAEQF | 0.7524338 |
| TALAGAGSGI | ASSQDWGGDAEQF | 0.74636054 |
| TGLPFTGI | ASSQDWGGDAEQF | 0.77409583 |
| VAEHVTV | ASSQDWGGDAEQF | 0.75569975 |
| VDIVLNSL | ASSQDWGGDAEQF | 0.7448987 |
| VLTGGGAVL | ASSQDWGGDAEQF | 0.4949079 |
| VNVGLAYGI | ASSQDWGGDAEQF | 0.64222395 |
| VSSAAVGAV | ASSQDWGGDAEQF | 0.697323 |
| VTSVYEAGV | ASSQDWGGDAEQF | 0.711596 |
| WVALDRGA | ASSQDWGGDAEQF | 0.44323885 |
| YLVGSVVI | ASSQDWGGDAEQF | 0.7476826 |
| GPPGTGKTLL | ASSQDWGGDAEQF | 0.7327773 |
| GVGMTKF | ASSQDWGGDAEQF | 0.66898465 |
| IGIGGSDL | ASSQDWGGDAEQF | 0.75731003 |

| | | |
|---|---|---|
| IGTLLAEL | ASSQDWGGDAEQF | 0.70301944 |
| IYTSGTT | ASSQDWGGDAEQF | 0.6821174 |
| LSFPAGV | ASSQDWGGDAEQF | 0.7175421 |
| LYTSGTT | ASSQDWGGDAEQF | 0.77117586 |
| VGAFPFGF | ASSQDWGGDAEQF | 0.66871727 |
| VGAGTVEF | ASSQDWGGDAEQF | 0.76198655 |
| VGKSTLLL | ASSQDWGGDAEQF | 0.7250344 |
| AALAHAGGGIV | ASSQDWGGDAEQF | 0.7651269 |
| AALVAVSLL | ASSQDWGGDAEQF | 0.63974047 |
| AAVSRAVAAL | ASSQDWGGDAEQF | 0.7597969 |
| AGGSSGIGGF | ASSQDWGGDAEQF | 0.7193315 |
| AGLTVANANG | ASSQDWGGDAEQF | 0.7737818 |
| AGLVGAVL | ASSQDWGGDAEQF | 0.71886563 |
| ALSQAFTV | ASSQDWGGDAEQF | 0.76317215 |
| ALTFSVSL | ASSQDWGGDAEQF | 0.73998547 |
| ALTGAALVAL | ASSQDWGGDAEQF | 0.75517714 |
| ASFSIGAGVL | ASSQDWGGDAEQF | 0.61202025 |
| DGSVYKI | ASSQDWGGDAEQF | 0.7096624 |
| EQLAEIV | ASSQDWGGDAEQF | 0.6851551 |
| FAVSELTVM | ASSQDWGGDAEQF | 0.7359307 |
| GAAPLAAGL | ASSQDWGGDAEQF | 0.75410867 |
| GADGGDGGSGGAGG | ASSQDWGGDAEQF | 0.3767089 |
| GFGLDTGGAGG | ASSQDWGGDAEQF | 0.76610774 |
| GLASGGLL | ASSQDWGGDAEQF | 0.6125766 |
| GRIAGAAVLL | ASSQDWGGDAEQF | 0.67227405 |
| GSAGSLNTSF | ASSQDWGGDAEQF | 0.7562154 |
| GTGQIGI | ASSQDWGGDAEQF | 0.7275351 |
| GTLIALIL | ASSQDWGGDAEQF | 0.7451066 |
| IAATMGFVVG | ASSQDWGGDAEQF | 0.706902 |
| IFAQALL | ASSQDWGGDAEQF | 0.72846127 |
| IGAAVALGF | ASSQDWGGDAEQF | 0.74354553 |
| IGLTPLAL | ASSQDWGGDAEQF | 0.4677795 |
| ILGSNTSTL | ASSQDWGGDAEQF | 0.64319146 |
| IVFSLTVL | ASSQDWGGDAEQF | 0.7696336 |
| LASKATGF | ASSQDWGGDAEQF | 0.25932705 |
| LASVALIL | ASSQDWGGDAEQF | 0.40798962 |
| LFAVGLTL | ASSQDWGGDAEQF | 0.25260288 |
| LFNGGIGTYI | ASSQDWGGDAEQF | 0.24816576 |
| LTLGAGII | ASSQDWGGDAEQF | 0.26213646 |
| LVAAALGDI | ASSQDWGGDAEQF | 0.26550692 |
| NTAFVQLGI | ASSQDWGGDAEQF | 0.53042793 |
| QLFSGAEL | ASSQDWGGDAEQF | 0.25299543 |
| SAATFITL | ASSQDWGGDAEQF | 0.24894655 |
| SNYGLANLGV | ASSQDWGGDAEQF | 0.34671307 |
| SYGGAYAVM | ASSQDWGGDAEQF | 0.23946068 |
| TTGVGGLTLG | ASSQDWGGDAEQF | 0.24331231 |
| TTQTIYI | ASSQDWGGDAEQF | 0.24415958 |
| VAAGAGLGAVI | ASSQDWGGDAEQF | 0.2581008 |
| VAAVVTLL | ASSQDWGGDAEQF | 0.22860658 |
| VGFLGGLI | ASSQDWGGDAEQF | 0.2479089 |
| VGVIPILL | ASSQDWGGDAEQF | 0.23828813 |

| | | |
|---|---|---|
| VIFASSLI | ASSQDWGGDAEQF | 0.31310433 |
| VKGGDGGAAAG | ASSQDWGGDAEQF | 0.27400354 |
| VTKTFTNI | ASSQDWGGDAEQF | 0.2829488 |
| VVAAAILL | ASSQDWGGDAEQF | 0.27589008 |
| YAALDVEL | ASSQDWGGDAEQF | 0.23897888 |
| LASVALLL | ASSQDWGGDAEQF | 0.21956383 |
| LDAVHRV | ASSQDWGGDAEQF | 0.23252739 |
| QGLSSSIF | ASSQDWGGDAEQF | 0.23453963 |
| RGTFANIRL | ASSQDWGGDAEQF | 0.26221848 |
| RQLWWGHRI | ASSQDWGGDAEQF | 0.21257424 |
| YGPPGTGKTLL | ASSQDWGGDAEQF | 0.28920224 |
| AALVGAPANVA | ASSQDWGGDAEQF | 0.24357569 |
| AATFRRTE | ASSQDWGGDAEQF | 0.24269791 |
| AGALGGGI | ASSQDWGGDAEQF | 0.2784728 |
| AGGAGGAGGIGAG | ASSQDWGGDAEQF | 0.23291993 |
| AGSGALGL | ASSQDWGGDAEQF | 0.23718157 |
| AQIADQI | ASSQDWGGDAEQF | 0.24566272 |
| ASAVAVTYL | ASSQDWGGDAEQF | 0.31780788 |
| ASTVLGGTGG | ASSQDWGGDAEQF | 0.23270436 |
| AVGGEPAK | ASSQDWGGDAEQF | 0.38801846 |
| CEQGTFGGTC | ASSQDWGGDAEQF | 0.22320601 |
| DEGKAPAG | ASSQDWGGDAEQF | 0.305086 |
| DTVVFPAL | ASSQDWGGDAEQF | 0.23641373 |
| FEGITMAG | ASSQDWGGDAEQF | 0.25557396 |
| FPLVVQD | ASSQDWGGDAEQF | 0.29265445 |
| GAFTTPQ | ASSQDWGGDAEQF | 0.3171633 |
| GFGGFGTAGGVGGA | ASSQDWGGDAEQF | 0.2562645 |
| GFLAQFL | ASSQDWGGDAEQF | 0.29597867 |
| GGAGGAGGNAGVA | ASSQDWGGDAEQF | 0.24819562 |
| GGVGTAGAGGAGGAGG | ASSQDWGGDAEQF | 0.5443058 |
| GNGGGGDAGNAGSGGN | ASSQDWGGDAEQF | 0.24604535 |
| GTGGAGGAGNDAGS | ASSQDWGGDAEQF | 0.27753648 |
| GTTNTGWFNS | ASSQDWGGDAEQF | 0.22584133 |
| IFMGPLFTL | ASSQDWGGDAEQF | 0.31706256 |
| ILAHAIEAI | ASSQDWGGDAEQF | 0.2756744 |
| IRSAIREAL | ASSQDWGGDAEQF | 0.21537514 |
| ITFAGGG | ASSQDWGGDAEQF | 0.2602666 |
| ITGMSGHQEV | ASSQDWGGDAEQF | 0.22046724 |
| IVAGLACTL | ASSQDWGGDAEQF | 0.22702263 |
| IVGGGLAGA | ASSQDWGGDAEQF | 0.24964964 |
| IVTGVLNI | ASSQDWGGDAEQF | 0.25520244 |
| LAQQGCI | ASSQDWGGDAEQF | 0.23966138 |
| LCLHVEN | ASSQDWGGDAEQF | 0.22474863 |
| LDAISDGS | ASSQDWGGDAEQF | 0.23194501 |
| LGGFAAML | ASSQDWGGDAEQF | 0.34618783 |
| LGGKNPMI | ASSQDWGGDAEQF | 0.2261756 |
| LGPKGVAAIL | ASSQDWGGDAEQF | 0.2287011 |
| LMVGVSI | ASSQDWGGDAEQF | 0.2805368 |
| LVGAAFVL | ASSQDWGGDAEQF | 0.8431548 |
| LVLAVAPL | ASSQDWGGDAEQF | 0.87345904 |
| MAAIAVIDAA | ASSQDWGGDAEQF | 0.754871 |

| | | |
|---|---|---|
| NNGINAELQ | ASSQDWGGDAEQF | 0.6151357 |
| RGEPLVLVHG | ASSQDWGGDAEQF | 0.76107633 |
| RGGAGGAAGLI | ASSQDWGGDAEQF | 0.86446726 |
| RSYRMHPAV | ASSQDWGGDAEQF | 0.8779261 |
| SAAHLLAGQLA | ASSQDWGGDAEQF | 0.84814894 |
| SAAVGSY | ASSQDWGGDAEQF | 0.4908029 |
| SPHAVLHRI | ASSQDWGGDAEQF | 0.83830285 |
| SQIAKVL | ASSQDWGGDAEQF | 0.7619724 |
| TALGHLVP | ASSQDWGGDAEQF | 0.8190006 |
| TAPTTSGAA | ASSQDWGGDAEQF | 0.8408486 |
| TDGDDGNGGSPGT | ASSQDWGGDAEQF | 0.79068315 |
| TDGLGEN | ASSQDWGGDAEQF | 0.7909896 |
| TGGPPLPGQA | ASSQDWGGDAEQF | 0.66845363 |
| TIPAITI | ASSQDWGGDAEQF | 0.7951543 |
| TIPGFTI | ASSQDWGGDAEQF | 0.836195 |
| TSIASGALA | ASSQDWGGDAEQF | 0.78157794 |
| TVTLPGAAI | ASSQDWGGDAEQF | 0.8813574 |
| VASGLSAALP | ASSQDWGGDAEQF | 0.875327 |
| VGAGVIGI | ASSQDWGGDAEQF | 0.8468901 |
| VGPKRFGV | ASSQDWGGDAEQF | 0.1476251 |
| VNVAFPD | ASSQDWGGDAEQF | 0.47401303 |
| VSMTNSGVS | ASSQDWGGDAEQF | 0.7106314 |
| VTDVLGDLSV | ASSQDWGGDAEQF | 0.7916452 |
| VVVVGAGFAGL | ASSQDWGGDAEQF | 0.23375778 |
| VYTVGTVV | ASSQDWGGDAEQF | 0.83988476 |
| WTTFSGP | ASSQDWGGDAEQF | 0.74054694 |
| YDELVEAF | ASSQDWGGDAEQF | 0.73977184 |
| YGTGGAGGNGG | ASSQDWGGDAEQF | 0.8759749 |
| YTNVMAV | ASSQDWGGDAEQF | 0.80322504 |
| FLEQVHQL | ASSQDWGGDAEQF | 0.6412506 |
| GLSLLAVAV | ASSQDWGGDAEQF | 0.6139994 |
| LEQVHQL | ASSQDWGGDAEQF | 0.87558484 |
| LNAAGAI | ASSQDWGGDAEQF | 0.51800615 |
| PAVHASL | ASSQDWGGDAEQF | 0.8696373 |
| TELEDDA | ASSQDWGGDAEQF | 0.6768339 |
| TPPGPPN | ASSQDWGGDAEQF | 0.88035405 |
| TPPPPPPP | ASSQDWGGDAEQF | 0.397713 |
| TQSPALI | ASSRGAINTGQLY | 0.8627932 |
| EFAAAFPQ | ASSRGAINTGQLY | 0.80798846 |
| LLSGNLDVL | ASSRGAINTGQLY | 0.8996291 |
| GPANLGNYNI | ASSRGAINTGQLY | 0.69242084 |
| TLGPTLP | ASSRGAINTGQLY | 0.785167 |
| VYAYSSCCK | ASSRGAINTGQLY | 0.6564686 |
| APAPAPAPNI | ASSRGAINTGQLY | 0.87278676 |
| PGGPVPIP | ASSRGAINTGQLY | 0.48465583 |
| FLTVALEP | ASSRGAINTGQLY | 0.7165189 |
| SQYFGANL | ASSRGAINTGQLY | 0.5445781 |
| GGVKPAA | ASSRGAINTGQLY | 0.81119204 |
| GGTGGAGGAGGW | ASSRGAINTGQLY | 0.8562032 |
| GVGGVAGLGGAGG | ASSRGAINTGQLY | 0.27565905 |
| VLLVLATTI | ASSRGAINTGQLY | 0.8682285 |

| | | |
|---|---|---|
| IGIFAAML | ASSRGAINTGQLY | 0.7392535 |
| AVGDLID | ASSRGAINTGQLY | 0.5588523 |
| NISPLPL | ASSRGAINTGQLY | 0.87259334 |
| VTAVFVAI | ASSRGAINTGQLY | 0.700853 |
| SGDDNDAAL | ASSRGAINTGQLY | 0.86320317 |
| AVAALAHS | ASSRGAINTGQLY | 0.68429947 |
| GGVTVMATP | ASSRGAINTGQLY | 0.7491404 |
| AGLAANVRD | ASSRGAINTGQLY | 0.8451032 |
| TNRPDLIDPALLRPG | ASSRGAINTGQLY | 0.39584303 |
| LANLGVTAL | ASSRGAINTGQLY | 0.480688 |
| TGERIEL | ASSRGAINTGQLY | 0.8864334 |
| NGEELAA | ASSRGAINTGQLY | 0.8226683 |
| VLSLVWG | ASSRGAINTGQLY | 0.8495092 |
| IGYALWL | ASSRGAINTGQLY | 0.6710056 |
| LSLLLAPL | ASSRGAINTGQLY | 0.6584012 |
| TLDEALK | ASSRGAINTGQLY | 0.7691241 |
| TDGAGTAEGG | ASSRGAINTGQLY | 0.85579705 |
| NAPGGAGGNGG | ASSRGAINTGQLY | 0.8566751 |
| VSVFMGMGGNF | ASSRGAINTGQLY | 0.8300174 |
| LASYLAAL | ASSRGAINTGQLY | 0.73278314 |
| LVGGTPAL | ASSRGAINTGQLY | 0.8075385 |
| VRNADVL | ASSRGAINTGQLY | 0.78511786 |
| TCSSPFLVSAM | ASSRGAINTGQLY | 0.75503886 |
| VHITDAE | ASSRGAINTGQLY | 0.8481855 |
| EYAAPSF | ASSRGAINTGQLY | 0.8201352 |
| VVDAFHGGL | ASSRGAINTGQLY | 0.7403467 |
| ASVVDPQ | ASSRGAINTGQLY | 0.79184365 |
| SYNEFGGLS | ASSRGAINTGQLY | 0.8067976 |
| DPDGSVY | ASSRGAINTGQLY | 0.84151185 |
| MVSTIHATE | ASSRGAINTGQLY | 0.82979614 |
| LAAGVADPV | ASSRGAINTGQLY | 0.88217866 |
| TKEQIAA | ASSRGAINTGQLY | 0.85014004 |
| SLASPAAL | ASSRGAINTGQLY | 0.82937324 |
| IGLVFYGV | ASSRGAINTGQLY | 0.33046794 |
| FLLDASNL | ASSRGAINTGQLY | 0.59862703 |
| YQVGNFP | ASSRGAINTGQLY | 0.7851713 |
| ESLLSQST | ASSRGAINTGQLY | 0.7926674 |
| LFTVGFAAVG | ASSRGAINTGQLY | 0.5005194 |
| ATPPTPT | ASSRGAINTGQLY | 0.8440013 |
| TDLATLG | ASSRGAINTGQLY | 0.77390695 |
| SAGGTGR | ASSRGAINTGQLY | 0.6856588 |
| GFSNSGS | ASSRGAINTGQLY | 0.84750056 |
| GGNGGSGGTSV | ASSRGAINTGQLY | 0.7364359 |
| GTSVATGGAGN | ASSRGAINTGQLY | 0.75383675 |
| NGGNGGSGGTS | ASSRGAINTGQLY | 0.8009334 |
| AVVKFPH | ASSRGAINTGQLY | 0.8774014 |
| TPPPPPP | ASSRGAINTGQLY | 0.68977815 |
| AAPAAAAAPA | ASSRGAINTGQLY | 0.8473596 |
| IALNEDL | ASSRGAINTGQLY | 0.8018497 |
| IVNVSSM | ASSRGAINTGQLY | 0.86252046 |
| TGIPPAPR | ASSRGAINTGQLY | 0.62595874 |

| | | |
|---|---|---|
| FDLGGGTF | ASSRGAINTGQLY | 0.8428493 |
| FDLGGGTFDVS | ASSRGAINTGQLY | 0.77910435 |
| NEPTAAA | ASSRGAINTGQLY | 0.88225746 |
| NEPTAAALA | ASSRGAINTGQLY | 0.7871913 |
| GSIINISSI | ASSRGAINTGQLY | 0.84579176 |
| VAPGNAGTA | ASSRGAINTGQLY | 0.8037914 |
| TGGGASLE | ASSRGAINTGQLY | 0.8245969 |
| VAVYNHYKRI | ASSRGAINTGQLY | 0.58573264 |
| TDPAPATT | ASSRGAINTGQLY | 0.78947014 |
| TNRPDILDPALLRPG | ASSRGAINTGQLY | 0.61806846 |
| LTTTTTS | ASSRGAINTGQLY | 0.81763303 |
| PTGSGKTL | ASSRGAINTGQLY | 0.8283209 |
| TGSGKTL | ASSRGAINTGQLY | 0.58954793 |
| TIGHVDHGK | ASSRGAINTGQLY | 0.8429936 |
| TIGHVDHGKTT | ASSRGAINTGQLY | 0.75404537 |
| GHVDHGKTT | ASSRGAINTGQLY | 0.73904 |
| AQFEHTLL | ASSRGAINTGQLY | 0.8564247 |
| IVNQDEL | ASSRGAINTGQLY | 0.74900794 |
| TNGSQFF | ASSRGAINTGQLY | 0.8443177 |
| TGKGIVE | ASSRGAINTGQLY | 0.7585426 |
| LISNASDA | ASSRGAINTGQLY | 0.7692782 |
| AELRTLGELTKIAKAHGAQ | ASSRGAINTGQLY | 0.83285594 |
| APNAVAAAEPSW | ASSRGAINTGQLY | 0.5375984 |
| APQVAGAATAVL | ASSRGAINTGQLY | 0.72166383 |
| ARRDPAAAAAILPTDA | ASSRGAINTGQLY | 0.8617036 |
| DAAARDVLAEAGLAEYF | ASSRGAINTGQLY | 0.6203607 |
| DDQIVLSAKANLAKVEQ | ASSRGAINTGQLY | 0.69547987 |
| DELLPDTGEDSDEDGH | ASSRGAINTGQLY | 0.53664684 |
| ELPEAPANRATVPQD | ASSRGAINTGQLY | 0.4570393 |
| FGQSKIERVTGIG | ASSRGAINTGQLY | 0.54543734 |
| GAGGTGSTAGGAGGAGGAGGLYA | ASSRGAINTGQLY | 0.57248664 |
| GHGGTGAAGGNGGTGGAGGSGIDG | ASSRGAINTGQLY | 0.70151114 |
| GNGGNGGAGGEGGPGGAGGAGA | ASSRGAINTGQLY | 0.62014 |
| GSTAGNGGNGGNATNGTIAGGQPAG | ASSRGAINTGQLY | 0.38416177 |
| GTGGDGGNGGNGGTGG | ASSRGAINTGQLY | 0.64286315 |
| GVRRPAGIPADAPV | ASSRGAINTGQLY | 0.5648594 |
| GYRQVTDAHLLTLARR | ASSRGAINTGQLY | 0.5672697 |
| IAKMGQDGHDRG | ASSRGAINTGQLY | 0.49952158 |
| KAEPSIEIQYT | ASSRGAINTGQLY | 0.59109116 |
| KPVRIRMAADEDTVAVT | ASSRGAINTGQLY | 0.56988156 |
| MPPRIAGMRLLVIKPEP | ASSRGAINTGQLY | 0.4096276 |
| MKFYAVALFRSRGVPPTR | ASSRGAINTGQLY | 0.5422611 |
| NAGGFGILGAVAHSP | ASSRGAINTGQLY | 0.6048736 |
| NEGREITLAMAEM | ASSRGAINTGQLY | 0.5779394 |
| NELVEWSHPLGHRWRWE | ASSRGAINTGQLY | 0.5939951 |
| NLGAGGTGGNGGNPG | ASSRGAINTGQLY | 0.6647936 |
| NTEWPKWETVMAADPVD | ASSRGAINTGQLY | 0.59127665 |
| PASAAPAETQP | ASSRGAINTGQLY | 0.27556202 |
| QQSSDGKNGVRAAEITGE | ASSRGAINTGQLY | 0.323915 |
| RAPDAVTTSWWKE | ASSRGAINTGQLY | 0.46511376 |
| SGFGNVGAHTSGWFNQ | ASSRGAINTGQLY | 0.48654264 |

| | | |
|---|---|---|
| SGVAGAVGQAASVGGLKV | ASSRGAINTGQLY | 0.2834879 |
| SNGGAGTGSAGGNGGT | ASSRGAINTGQLY | 0.51793385 |
| TGGAGGEGGAAG | ASSRGAINTGQLY | 0.5244149 |
| TGGDGGAAGAGGAPGHG | ASSRGAINTGQLY | 0.5217835 |
| TMPPLLAAIANAGRHGV | ASSRGAINTGQLY | 0.6488581 |
| TSSTAGNSGGA | ASSRGAINTGQLY | 0.56223214 |
| VPATVPVGILC | ASSRGAINTGQLY | 0.39219743 |
| VQRPDSGGVVLTGRIS | ASSRGAINTGQLY | 0.3602021 |
| WTPNPVPATKGVPAGLVQAT | ASSRGAINTGQLY | 0.5606248 |
| YALKRALAKFSDRFDVVII | ASSRGAINTGQLY | 0.35398245 |
| YLIGSVIV | ASSRGAINTGQLY | 0.64076763 |
| YGQLVAVID | ASSRGAINTGQLY | 0.32380283 |
| VTVGGMTLPA | ASSRGAINTGQLY | 0.6119253 |
| VLALPAV | ASSRGAINTGQLY | 0.39026973 |
| VIPPTLNYET | ASSRGAINTGQLY | 0.61389846 |
| VGMGGMHPSSG | ASSRGAINTGQLY | 0.5933385 |
| VGLAGGAATA | ASSRGAINTGQLY | 0.688991 |
| VGGAGGAGGSGP | ASSRGAINTGQLY | 0.51794434 |
| VFAAFMV | ASSRGAINTGQLY | 0.55356336 |
| TLYDKLR | ASSRGAINTGQLY | 0.38766682 |
| TLAVFTVL | ASSRGAINTGQLY | 0.6960993 |
| TGSVQPARV | ASSRGAINTGQLY | 0.39885792 |
| TGGGAGSGGT | ASSRGAINTGQLY | 0.52233964 |
| SHLETSL | ASSRGAINTGQLY | 0.329869 |
| SFTTGMT | ASSRGAINTGQLY | 0.530061 |
| QVEVAKRMGVRQPTVS | ASSRGAINTGQLY | 0.6427331 |
| QGSGGPALAM | ASSRGAINTGQLY | 0.31394553 |
| MLAPSNP | ASSRGAINTGQLY | 0.61021006 |
| LVNNAAI | ASSRGAINTGQLY | 0.46274874 |
| LTGEINTLA | ASSRGAINTGQLY | 0.42809194 |
| LSIFGPV | ASSRGAINTGQLY | 0.5912926 |
| LLALLANL | ASSRGAINTGQLY | 0.57622284 |
| LISNASDALDKLR | ASSRGAINTGQLY | 0.6745705 |
| LINEAVP | ASSRGAINTGQLY | 0.40111458 |
| LGYAFSGL | ASSRGAINTGQLY | 0.46720064 |
| LDFVAVAL | ASSRGAINTGQLY | 0.56282794 |
| LAQPGAE | ASSRGAINTGQLY | 0.46492654 |
| ISNLGAVTGF | ASSRGAINTGQLY | 0.3855139 |
| HQEDPGRI | ASSRGAINTGQLY | 0.66985816 |
| GTPGEPTP | ASSRGAINTGQLY | 0.7521541 |
| GFFNSGS | ASSRGAINTGQLY | 0.7649729 |
| GAGGGGLTAGAN | ASSRGAINTGQLY | 0.7128737 |
| FVDALVNM | ASSRGAINTGQLY | 0.71223044 |
| FLTVAPEL | ASSRGAINTGQLY | 0.7178465 |
| FETLTAL | ASSRGAINTGQLY | 0.76058036 |
| DVINPIEL | ASSRGAINTGQLY | 0.7617612 |
| APVSGAPV | ASSRGAINTGQLY | 0.75223625 |
| ALVGGLAW | ASSRGAINTGQLY | 0.63999 |
| ALAAAAATAI | ASSRGAINTGQLY | 0.75072706 |
| AFGLGLLL | ASSRGAINTGQLY | 0.7225337 |
| AFFAAASL | ASSRGAINTGQLY | 0.7223333 |

| | | |
|---|---|---|
| AAAALPAFSPP | ASSRGAINTGQLY | 0.74831134 |
| VVLAPAL | ASSRGAINTGQLY | 0.739371 |
| TIGHVDHGKTTL | ASSRGAINTGQLY | 0.709082 |
| LISNASDALDK | ASSRGAINTGQLY | 0.7148914 |
| LISNASDAL | ASSRGAINTGQLY | 0.7428588 |
| ISNASDALDK | ASSRGAINTGQLY | 0.7564918 |
| NQAETLVYQTE | ASSRGAINTGQLY | 0.7538806 |
| TGGTGGTAGAG | ASSRGAINTGQLY | 0.76646984 |
| TGSTAGGAGGA | ASSRGAINTGQLY | 0.7602192 |
| DPLAVPVAAGPV | ASSRGAINTGQLY | 0.75248194 |
| GGTGGTPGNGGN | ASSRGAINTGQLY | 0.3882687 |
| PAGLAGVDPATL | ASSRGAINTGQLY | 0.64900696 |
| GNNLVVEAPHAGASV | ASSRGAINTGQLY | 0.7198076 |
| KVTRIMSASTLEQPA | ASSRGAINTGQLY | 0.7295177 |
| LGFALPIALPRRTVV | ASSRGAINTGQLY | 0.38555574 |
| TRLGNLTARKLVVLV | ASSRGAINTGQLY | 0.74592125 |
| WSQQRRLIYFPSAGP | ASSRGAINTGQLY | 0.7286517 |
| GPNGNPDPMAAAVDIR | ASSRGAINTGQLY | 0.70273834 |
| TQRCVSVADLRAAASR | ASSRGAINTGQLY | 0.75608444 |
| QFVLDVAEHPVEVLTETV | ASSRGAINTGQLY | 0.71630096 |
| GGAGGTGGAGGNGGTQAGDG | ASSRGAINTGQLY | 0.6818322 |
| PLLATLRAIAVDVDAKPAQV | ASSRGAINTGQLY | 0.6321955 |
| APKFRTGGYMIDFWGVGYQVAKRM | ASSRGAINTGQLY | 0.76005995 |
| GAGGAGGAGGAAGGPGGTGGTGGN | ASSRGAINTGQLY | 0.6237596 |
| IGALGLLATDVAPAGSVAFGVALA | ASSRGAINTGQLY | 0.75627947 |
| AGGAGGIGGNGGAGGAGGVLMGNG | ASSRGAINTGQLY | 0.6786973 |
| ADELIGLL | ASSRGAINTGQLY | 0.7624098 |
| ADRALKA | ASSRGAINTGQLY | 0.6046219 |
| AGAVGQVGGAGGA | ASSRGAINTGQLY | 0.75524724 |
| AGGGGGLGGAP | ASSRGAINTGQLY | 0.73776853 |
| AGGLLYGNGGAG | ASSRGAINTGQLY | 0.77187175 |
| AGLESIGL | ASSRGAINTGQLY | 0.7348131 |
| AGLKTGII | ASSRGAINTGQLY | 0.74548805 |
| ASLAALLL | ASSRGAINTGQLY | 0.67176825 |
| ASVINISSI | ASSRGAINTGQLY | 0.7620268 |
| ATPPGPA | ASSRGAINTGQLY | 0.62410045 |
| ATVASALV | ASSRGAINTGQLY | 0.7294402 |
| ATVASLIL | ASSRGAINTGQLY | 0.5878793 |
| AVNGPVLIRH | ASSRGAINTGQLY | 0.7436956 |
| DGAAGAVGI | ASSRGAINTGQLY | 0.7543597 |
| DPLLHGI | ASSRGAINTGQLY | 0.65933454 |
| ELGAALGAL | ASSRGAINTGQLY | 0.75816476 |
| ELGRTQAV | ASSRGAINTGQLY | 0.72554815 |
| FVAGGYLS | ASSRGAINTGQLY | 0.68248576 |
| GASGGAGGAGGW | ASSRGAINTGQLY | 0.7616614 |
| GGAGAGGDGGAGGI | ASSRGAINTGQLY | 0.73446804 |
| GTGHNGSQPGMGGNG | ASSRGAINTGQLY | 0.7610389 |
| GYAAASASAS | ASSRGAINTGQLY | 0.6451423 |
| HQVDVAAVQQG | ASSRGAINTGQLY | 0.7416415 |
| IGAAIAHT | ASSRGAINTGQLY | 0.747347 |
| IGMIEVAQL | ASSRGAINTGQLY | 0.6970328 |

| | | |
|---|---|---|
| IRLAPEHTSAMD | ASSRGAINTGQLY | 0.6556901 |
| IYNVSVL | ASSRGAINTGQLY | 0.76438546 |
| KQGFLGM | ASSRGAINTGQLY | 0.72635436 |
| LESVITQL | ASSRGAINTGQLY | 0.75403637 |
| LGLGALVVV | ASSRGAINTGQLY | 0.6585809 |
| LIGNGGNGGSGGI | ASSRGAINTGQLY | 0.53398144 |
| LNAINAPTL | ASSRGAINTGQLY | 0.6567863 |
| LQLLNTGV | ASSRGAINTGQLY | 0.75316834 |
| LQTSIEF | ASSRGAINTGQLY | 0.7557317 |
| MSFAVSEL | ASSRGAINTGQLY | 0.7371081 |
| NTGALGL | ASSRGAINTGQLY | 0.5185759 |
| PPGAGGAGGAGTT | ASSRGAINTGQLY | 0.6896812 |
| QLVGLGI | ASSRGAINTGQLY | 0.6635215 |
| RFLEQVHQL | ASSRGAINTGQLY | 0.64574856 |
| RGVGYVLR | ASSRGAINTGQLY | 0.72555053 |
| SATGVFAGVI | ASSRGAINTGQLY | 0.7242619 |
| SLPAITLP | ASSRGAINTGQLY | 0.6773429 |
| STALAFTV | ASSRGAINTGQLY | 0.62203366 |
| SVLTVGLGK | ASSRGAINTGQLY | 0.70520926 |
| TALAGAGSGI | ASSRGAINTGQLY | 0.74205726 |
| TGLPFTGI | ASSRGAINTGQLY | 0.7311096 |
| VAEHVTV | ASSRGAINTGQLY | 0.76880753 |
| VDIVLNSL | ASSRGAINTGQLY | 0.74577093 |
| VLTGGGAVL | ASSRGAINTGQLY | 0.7373991 |
| VNVGLAYGI | ASSRGAINTGQLY | 0.22361931 |
| VSSAAVGAV | ASSRGAINTGQLY | 0.28713083 |
| VTSVYEAGV | ASSRGAINTGQLY | 0.66586405 |
| WVALDRGA | ASSRGAINTGQLY | 0.59551895 |
| YLVGSVVI | ASSRGAINTGQLY | 0.3063954 |
| GPPGTGKTLL | ASSRGAINTGQLY | 0.70370543 |
| GVGMTKF | ASSRGAINTGQLY | 0.66615486 |
| IGIGGSDL | ASSRGAINTGQLY | 0.5641361 |
| IGTLLAEL | ASSRGAINTGQLY | 0.74321467 |
| IYTSGTT | ASSRGAINTGQLY | 0.64460623 |
| LSFPAGV | ASSRGAINTGQLY | 0.55538726 |
| LYTSGTT | ASSRGAINTGQLY | 0.5699103 |
| VGAFPFGF | ASSRGAINTGQLY | 0.7434951 |
| VGAGTVEF | ASSRGAINTGQLY | 0.514452 |
| VGKSTLLL | ASSRGAINTGQLY | 0.7326612 |
| AALAHAGGGIV | ASSRGAINTGQLY | 0.50934786 |
| AALVAVSLL | ASSRGAINTGQLY | 0.7503226 |
| AAVSRAVAAL | ASSRGAINTGQLY | 0.45193478 |
| AGGSSGIGGF | ASSRGAINTGQLY | 0.73790014 |
| AGLTVANANG | ASSRGAINTGQLY | 0.6961626 |
| AGLVGAVL | ASSRGAINTGQLY | 0.78147423 |
| ALSQAFTV | ASSRGAINTGQLY | 0.62051314 |
| ALTFSVSL | ASSRGAINTGQLY | 0.68183184 |
| ALTGAALVAL | ASSRGAINTGQLY | 0.60884446 |
| ASFSIGAGVL | ASSRGAINTGQLY | 0.73942053 |
| DGSVYKI | ASSRGAINTGQLY | 0.4057293 |
| EQLAEIV | ASSRGAINTGQLY | 0.67877924 |

| | | |
|---|---|---|
| FAVSELTVM | ASSRGAINTGQLY | 0.32817784 |
| GAAPLAAGL | ASSRGAINTGQLY | 0.71665704 |
| GADGGDGGSGGAGG | ASSRGAINTGQLY | 0.7247523 |
| GFGLDTGGAGG | ASSRGAINTGQLY | 0.30633044 |
| GLASGGLL | ASSRGAINTGQLY | 0.74656415 |
| GRIAGAAVLL | ASSRGAINTGQLY | 0.6198524 |
| GSAGSLNTSF | ASSRGAINTGQLY | 0.5780339 |
| GTGQIGI | ASSRGAINTGQLY | 0.7603397 |
| GTLIALIL | ASSRGAINTGQLY | 0.6874611 |
| IAATMGFVVG | ASSRGAINTGQLY | 0.75011045 |
| IFAQALL | ASSRGAINTGQLY | 0.69262373 |
| IGAAVALGF | ASSRGAINTGQLY | 0.6562574 |
| IGLTPLAL | ASSRGAINTGQLY | 0.7132778 |
| ILGSNTSTL | ASSRGAINTGQLY | 0.49646562 |
| IVFSLTVL | ASSRGAINTGQLY | 0.54283607 |
| LASKATGF | ASSRGAINTGQLY | 0.75456774 |
| LASVALIL | ASSRGAINTGQLY | 0.71260434 |
| LFAVGLTL | ASSRGAINTGQLY | 0.7449527 |
| LFNGGIGTYI | ASSRGAINTGQLY | 0.61548346 |
| LTLGAGII | ASSRGAINTGQLY | 0.50959593 |
| LVAAALGDI | ASSRGAINTGQLY | 0.66091764 |
| NTAFVQLGI | ASSRGAINTGQLY | 0.7284862 |
| QLFSGAEL | ASSRGAINTGQLY | 0.7373576 |
| SAATFITL | ASSRGAINTGQLY | 0.72241855 |
| SNYGLANLGV | ASSRGAINTGQLY | 0.40086502 |
| SYGGAYAVM | ASSRGAINTGQLY | 0.6868712 |
| TTGVGGLTLG | ASSRGAINTGQLY | 0.61340296 |
| TTQTIYI | ASSRGAINTGQLY | 0.6471479 |
| VAAGAGLGAVI | ASSRGAINTGQLY | 0.7238858 |
| VAAVVTLL | ASSRGAINTGQLY | 0.5863191 |
| VGFLGGLI | ASSRGAINTGQLY | 0.6404812 |
| VGVIPILL | ASSRGAINTGQLY | 0.62133926 |
| VIFASSLI | ASSRGAINTGQLY | 0.6899814 |
| VKGGDGGAAAG | ASSRGAINTGQLY | 0.71613777 |
| VTKTFTNI | ASSRGAINTGQLY | 0.7029294 |
| VVAAAILL | ASSRGAINTGQLY | 0.739499 |
| YAALDVEL | ASSRGAINTGQLY | 0.7220073 |
| LASVALLL | ASSRGAINTGQLY | 0.70902884 |
| LDAVHRV | ASSRGAINTGQLY | 0.24107769 |
| QGLSSSIF | ASSRGAINTGQLY | 0.5062002 |
| RGTFANIRL | ASSRGAINTGQLY | 0.66031194 |
| RQLWWGHRI | ASSRGAINTGQLY | 0.69583666 |
| YGPPGTGKTLL | ASSRGAINTGQLY | 0.39639384 |
| AALVGAPANVA | ASSRGAINTGQLY | 0.7216961 |
| AATFRRTE | ASSRGAINTGQLY | 0.63168967 |
| AGALGGGI | ASSRGAINTGQLY | 0.6005386 |
| AGGAGGAGGIGAG | ASSRGAINTGQLY | 0.7325816 |
| AGSGALGL | ASSRGAINTGQLY | 0.6270896 |
| AQIADQI | ASSRGAINTGQLY | 0.5023535 |
| ASAVAVTYL | ASSRGAINTGQLY | 0.58576685 |
| ASTVLGGTGG | ASSRGAINTGQLY | 0.7313783 |

| | | |
|---|---|---|
| AVGGEPAK | ASSRGAINTGQLY | 0.4762236 |
| CEQGTFGGTC | ASSRGAINTGQLY | 0.71161956 |
| DEGKAPAG | ASSRGAINTGQLY | 0.6823392 |
| DTVVFPAL | ASSRGAINTGQLY | 0.7176068 |
| FEGITMAG | ASSRGAINTGQLY | 0.35096022 |
| FPLVVQD | ASSRGAINTGQLY | 0.7174807 |
| GAFTTPQ | ASSRGAINTGQLY | 0.6760824 |
| GFGGFGTAGGVGGA | ASSRGAINTGQLY | 0.74518985 |
| GFLAQFL | ASSRGAINTGQLY | 0.5750004 |
| GGAGGAGGNAGVA | ASSRGAINTGQLY | 0.6366271 |
| GGVGTAGAGGAGGAGG | ASSRGAINTGQLY | 0.46323872 |
| GNGGGGDAGNAGSGGN | ASSRGAINTGQLY | 0.7357478 |
| GTGGAGGAGNDAGS | ASSRGAINTGQLY | 0.49580717 |
| GTTNTGWFNS | ASSRGAINTGQLY | 0.59647316 |
| IFMGPLFTL | ASSRGAINTGQLY | 0.53572786 |
| ILAHAIEAI | ASSRGAINTGQLY | 0.67511255 |
| IRSAIREAL | ASSRGAINTGQLY | 0.7151582 |
| ITFAGGG | ASSRGAINTGQLY | 0.57971066 |
| ITGMSGHQEV | ASSRGAINTGQLY | 0.7237205 |
| IVAGLACTL | ASSRGAINTGQLY | 0.7083554 |
| IVGGGLAGA | ASSRGAINTGQLY | 0.4640824 |
| IVTGVLNI | ASSRGAINTGQLY | 0.7323691 |
| LAQQGCI | ASSRGAINTGQLY | 0.54655886 |
| LCLHVEN | ASSRGAINTGQLY | 0.72813666 |
| LDAISDGS | ASSRGAINTGQLY | 0.6541373 |
| LGGFAAML | ASSRGAINTGQLY | 0.6925993 |
| LGGKNPMI | ASSRGAINTGQLY | 0.71121 |
| LGPKGVAAIL | ASSRGAINTGQLY | 0.66653407 |
| LMVGVSI | ASSRGAINTGQLY | 0.58828896 |
| LVGAAFVL | ASSRGAINTGQLY | 0.72721195 |
| LVLAVAPL | ASSRGAINTGQLY | 0.7314851 |
| MAAIAVIDAA | ASSRGAINTGQLY | 0.84630144 |
| NNGINAELQ | ASSRGAINTGQLY | 0.5455905 |
| RGEPLVLVHG | ASSRGAINTGQLY | 0.4930207 |
| RGGAGGAAGLI | ASSRGAINTGQLY | 0.6139846 |
| RSYRMHPAV | ASSRGAINTGQLY | 0.824034 |
| SAAHLLAGQLA | ASSRGAINTGQLY | 0.82064366 |
| SAAVGSY | ASSRGAINTGQLY | 0.75707304 |
| SPHAVLHRI | ASSRGAINTGQLY | 0.53619695 |
| SQIAKVL | ASSRGAINTGQLY | 0.6096264 |
| TALGHLVP | ASSRGAINTGQLY | 0.5709045 |
| TAPTTSGAA | ASSRGAINTGQLY | 0.61318856 |
| TDGDDGNGGSPGT | ASSRGAINTGQLY | 0.78722334 |
| TDGLGEN | ASSRGAINTGQLY | 0.71406054 |
| TGGPPLPGQA | ASSRGAINTGQLY | 0.59647644 |
| TIPAITI | ASSRGAINTGQLY | 0.6361521 |
| TIPGFTI | ASSRGAINTGQLY | 0.65969235 |
| TSIASGALA | ASSRGAINTGQLY | 0.7442499 |
| TVTLPGAAI | ASSRGAINTGQLY | 0.7023603 |
| VASGLSAALP | ASSRGAINTGQLY | 0.8659171 |
| VGAGVIGI | ASSRGAINTGQLY | 0.7781353 |

| | | |
|---|---|---|
| VGPKRFGV | ASSRGAINTGQLY | 0.80477786 |
| VNVAFPD | ASSRGAINTGQLY | 0.14420514 |
| VSMTNSGVS | ASSRGAINTGQLY | 0.35955626 |
| VTDVLGDLSV | ASSRGAINTGQLY | 0.70905775 |
| VVVVGAGFAGL | ASSRGAINTGQLY | 0.6815995 |
| VYTVGTVV | ASSRGAINTGQLY | 0.43884975 |
| WTTFSGP | ASSRGAINTGQLY | 0.7873925 |
| YDELVEAF | ASSRGAINTGQLY | 0.6119042 |
| YGTGGAGGNGG | ASSRGAINTGQLY | 0.5195394 |
| YTNVMAV | ASSRGAINTGQLY | 0.7819849 |
| FLEQVHQL | ASSRGAINTGQLY | 0.58159614 |
| GLSLLAVAV | ASSRGAINTGQLY | 0.5928855 |
| LEQVHQL | ASSRGAINTGQLY | 0.63595825 |
| LNAAGAI | ASSRGAINTGQLY | 0.8064344 |
| PAVHASL | ASSRGAINTGQLY | 0.41233248 |
| TELEDDA | ASSRGAINTGQLY | 0.7353856 |
| TPPGPPN | ASSRGAINTGQLY | 0.68071866 |
| TPPPPPPP | ASSRGAINTGQLY | 0.8061884 |
| TQSPALI | ASGERTGGAANTGQLY | 0.43004057 |
| EFAAAFPQ | ASGERTGGAANTGQLY | 0.7597553 |
| LLSGNLDVL | ASGERTGGAANTGQLY | 0.64135706 |
| GPANLGNYNI | ASGERTGGAANTGQLY | 0.8606769 |
| TLGPTLP | ASGERTGGAANTGQLY | 0.5904613 |
| VYAYSSCCK | ASGERTGGAANTGQLY | 0.70017356 |
| APAPAPAPNI | ASGERTGGAANTGQLY | 0.66260993 |
| PGGPVPIP | ASGERTGGAANTGQLY | 0.7872783 |
| FLTVALEP | ASGERTGGAANTGQLY | 0.42116582 |
| SQYFGANL | ASGERTGGAANTGQLY | 0.5948241 |
| GGVKPAA | ASGERTGGAANTGQLY | 0.38970163 |
| GGTGGAGGAGGW | ASGERTGGAANTGQLY | 0.7134354 |
| GVGGVAGLGGAGG | ASGERTGGAANTGQLY | 0.7313732 |
| VLLVLATTI | ASGERTGGAANTGQLY | 0.4352048 |
| IGIFAAML | ASGERTGGAANTGQLY | 0.7472054 |
| AVGDLID | ASGERTGGAANTGQLY | 0.70068777 |
| NISPLPL | ASGERTGGAANTGQLY | 0.5874085 |
| VTAVFVAI | ASGERTGGAANTGQLY | 0.82774985 |
| SGDDNDAAL | ASGERTGGAANTGQLY | 0.5067011 |
| AVAALAHS | ASGERTGGAANTGQLY | 0.8109057 |
| GGVTVMATP | ASGERTGGAANTGQLY | 0.70186156 |
| AGLAANVRD | ASGERTGGAANTGQLY | 0.59715056 |
| TNRPDLIDPALLRPG | ASGERTGGAANTGQLY | 0.7690387 |
| LANLGVTAL | ASGERTGGAANTGQLY | 0.5290592 |
| TGERIEL | ASGERTGGAANTGQLY | 0.5701187 |
| NGEELAA | ASGERTGGAANTGQLY | 0.8146448 |
| VLSLVWG | ASGERTGGAANTGQLY | 0.82137096 |
| IGYALWL | ASGERTGGAANTGQLY | 0.85386705 |
| LSLLLAPL | ASGERTGGAANTGQLY | 0.7717583 |
| TLDEALK | ASGERTGGAANTGQLY | 0.71541977 |
| TDGAGTAEGG | ASGERTGGAANTGQLY | 0.8064978 |
| NAPGGAGGNGG | ASGERTGGAANTGQLY | 0.85262567 |
| VSVFMGMGGNF | ASGERTGGAANTGQLY | 0.8443624 |

| | | |
|---|---|---|
| LASYLAAL | ASGERTGGAANTGQLY | 0.83677936 |
| LVGGTPAL | ASGERTGGAANTGQLY | 0.7034793 |
| VRNADVL | ASGERTGGAANTGQLY | 0.79524916 |
| TCSSPFLVSAM | ASGERTGGAANTGQLY | 0.79644406 |
| VHITDAE | ASGERTGGAANTGQLY | 0.7737024 |
| EYAAPSF | ASGERTGGAANTGQLY | 0.8423901 |
| VVDAFHGGL | ASGERTGGAANTGQLY | 0.8008162 |
| ASVVDPQ | ASGERTGGAANTGQLY | 0.74503213 |
| SYNEFGGLS | ASGERTGGAANTGQLY | 0.7976966 |
| DPDGSVY | ASGERTGGAANTGQLY | 0.81715804 |
| MVSTIHATE | ASGERTGGAANTGQLY | 0.8204486 |
| LAAGVADPV | ASGERTGGAANTGQLY | 0.81565094 |
| TKEQIAA | ASGERTGGAANTGQLY | 0.86073065 |
| SLASPAAL | ASGERTGGAANTGQLY | 0.8438524 |
| IGLVFYGV | ASGERTGGAANTGQLY | 0.83381677 |
| FLLDASNL | ASGERTGGAANTGQLY | 0.24459454 |
| YQVGNFP | ASGERTGGAANTGQLY | 0.61318475 |
| ESLLSQST | ASGERTGGAANTGQLY | 0.7805058 |
| LFTVGFAAVG | ASGERTGGAANTGQLY | 0.80852294 |
| ATPPTPT | ASGERTGGAANTGQLY | 0.67694104 |
| TDLATLG | ASGERTGGAANTGQLY | 0.84090245 |
| SAGGTGR | ASGERTGGAANTGQLY | 0.78180796 |
| GFSNSGS | ASGERTGGAANTGQLY | 0.72767115 |
| GGNGGSGGTSV | ASGERTGGAANTGQLY | 0.8380284 |
| GTSVATGGAGN | ASGERTGGAANTGQLY | 0.73165166 |
| NGGNGGSGGTS | ASGERTGGAANTGQLY | 0.75987226 |
| AVVKFPH | ASGERTGGAANTGQLY | 0.78099346 |
| TPPPPPP | ASGERTGGAANTGQLY | 0.8499695 |
| AAPAAAAAPA | ASGERTGGAANTGQLY | 0.66801775 |
| IALNEDL | ASGERTGGAANTGQLY | 0.827391 |
| IVNVSSM | ASGERTGGAANTGQLY | 0.7996919 |
| TGIPPAPR | ASGERTGGAANTGQLY | 0.84790325 |
| FDLGGGTF | ASGERTGGAANTGQLY | 0.6255245 |
| FDLGGGTFDVS | ASGERTGGAANTGQLY | 0.8389855 |
| NEPTAAA | ASGERTGGAANTGQLY | 0.81009287 |
| NEPTAAALA | ASGERTGGAANTGQLY | 0.8593672 |
| GSIINISSI | ASGERTGGAANTGQLY | 0.7955215 |
| VAPGNAGTA | ASGERTGGAANTGQLY | 0.81412804 |
| TGGGASLE | ASGERTGGAANTGQLY | 0.7435664 |
| VAVYNHYKRI | ASGERTGGAANTGQLY | 0.83945626 |
| TDPAPATT | ASGERTGGAANTGQLY | 0.70407206 |
| TNRPDILDPALLRPG | ASGERTGGAANTGQLY | 0.80086464 |
| LTTTTTS | ASGERTGGAANTGQLY | 0.68421364 |
| PTGSGKTL | ASGERTGGAANTGQLY | 0.82369685 |
| TGSGKTL | ASGERTGGAANTGQLY | 0.8283404 |
| TIGHVDHGK | ASGERTGGAANTGQLY | 0.77402914 |
| TIGHVDHGKTT | ASGERTGGAANTGQLY | 0.8415256 |
| GHVDHGKTT | ASGERTGGAANTGQLY | 0.84746754 |
| AQFEHTLL | ASGERTGGAANTGQLY | 0.7493289 |
| IVNQDEL | ASGERTGGAANTGQLY | 0.8539975 |
| TNGSQFF | ASGERTGGAANTGQLY | 0.77916473 |

| | | |
|---|---|---|
| TGKGIVE | ASGERTGGAANTGQLY | 0.8426222 |
| LISNASDA | ASGERTGGAANTGQLY | 0.8089814 |
| AELRTLGELTKIAKAHGAQ | ASGERTGGAANTGQLY | 0.79378736 |
| APNAVAAAEPSW | ASGERTGGAANTGQLY | 0.8401381 |
| APQVAGAATAVL | ASGERTGGAANTGQLY | 0.76715547 |
| ARRDPAAAAAILPTDA | ASGERTGGAANTGQLY | 0.7768857 |
| DAAARDVLAEAGLAEYF | ASGERTGGAANTGQLY | 0.8458519 |
| DDQIVLSAKANLAKVEQ | ASGERTGGAANTGQLY | 0.7322502 |
| DELLPDTGEDSDEDGH | ASGERTGGAANTGQLY | 0.7817036 |
| ELPEAPANRATVPQD | ASGERTGGAANTGQLY | 0.3337406 |
| FGQSKIERVTGIG | ASGERTGGAANTGQLY | 0.31687576 |
| GAGGTGSTAGGAGGAGGAGGLYA | ASGERTGGAANTGQLY | 0.53756356 |
| GHGGTGAAGGNGGTGGAGGSGIDG | ASGERTGGAANTGQLY | 0.67181116 |
| GNGGNGGAGGEGGPGGAGGAGA | ASGERTGGAANTGQLY | 0.83651805 |
| GSTAGNGGNGGNATNGTIAGQPAG | ASGERTGGAANTGQLY | 0.76663095 |
| GTGGDGGNGGNGGTGG | ASGERTGGAANTGQLY | 0.32376742 |
| GVRRPAGIPADAPV | ASGERTGGAANTGQLY | 0.6474843 |
| GYRQVTDAHLLTLARR | ASGERTGGAANTGQLY | 0.4583309 |
| IAKMGQDGHDRG | ASGERTGGAANTGQLY | 0.5351371 |
| KAEPSIEIQYT | ASGERTGGAANTGQLY | 0.57323444 |
| KPVRIRMAADEDTVAVT | ASGERTGGAANTGQLY | 0.43644637 |
| MPPRIAGMRLLVIKPEP | ASGERTGGAANTGQLY | 0.39222303 |
| MKFYAVALFRSRGVPPTR | ASGERTGGAANTGQLY | 0.23943368 |
| NAGGFGILGAVAHSP | ASGERTGGAANTGQLY | 0.6868752 |
| NEGREITLAMAEM | ASGERTGGAANTGQLY | 0.7985358 |
| NELVEWSHPLGHRWRWE | ASGERTGGAANTGQLY | 0.74344766 |
| NLGAGGTGGNGGNPG | ASGERTGGAANTGQLY | 0.81138146 |
| NTEWPKWETVMAADPVD | ASGERTGGAANTGQLY | 0.7773945 |
| PASAAPAETQP | ASGERTGGAANTGQLY | 0.64138234 |
| QQSSDGKNGVRAAEITGE | ASGERTGGAANTGQLY | 0.13074492 |
| RAPDAVTTSWWKE | ASGERTGGAANTGQLY | 0.19912477 |
| SGFGNVGAHTSGWFNQ | ASGERTGGAANTGQLY | 0.26428303 |
| SGVAGAVGQAASVGGLKV | ASGERTGGAANTGQLY | 0.5029506 |
| SNGGAGTGSAGGNGGT | ASGERTGGAANTGQLY | 0.13657922 |
| TGGAGGEGGAAG | ASGERTGGAANTGQLY | 0.6157776 |
| TGGDGGAAGAGGAPGHG | ASGERTGGAANTGQLY | 0.4036641 |
| TMPPLLAAIANAGRHGV | ASGERTGGAANTGQLY | 0.3391145 |
| TSSTAGNSGGA | ASGERTGGAANTGQLY | 0.7863888 |
| VPATVPVGILC | ASGERTGGAANTGQLY | 0.36081666 |
| VQRPDSGGVVLTGRIS | ASGERTGGAANTGQLY | 0.22666423 |
| WTPNPVPATKGVPAGLVQAT | ASGERTGGAANTGQLY | 0.34306598 |
| YALKRALAKFSDRFDVVII | ASGERTGGAANTGQLY | 0.8250667 |
| YLIGSVIV | ASGERTGGAANTGQLY | 0.24578938 |
| YGQLVAVID | ASGERTGGAANTGQLY | 0.77903533 |
| VTVGGMTLPA | ASGERTGGAANTGQLY | 0.48809502 |
| VLALPAV | ASGERTGGAANTGQLY | 0.7794628 |
| VIPPTLNYET | ASGERTGGAANTGQLY | 0.207032 |
| VGMGGMHPSSG | ASGERTGGAANTGQLY | 0.7502301 |
| VGLAGGAATA | ASGERTGGAANTGQLY | 0.59314495 |
| VGGAGGAGGSGP | ASGERTGGAANTGQLY | 0.84132326 |
| VFAAFMV | ASGERTGGAANTGQLY | 0.4051421 |

| | | |
|---|---|---|
| TLYDKLR | ASGERTGGAANTGQLY | 0.55554104 |
| TLAVFTVL | ASGERTGGAANTGQLY | 0.45222333 |
| TGSVQPARV | ASGERTGGAANTGQLY | 0.80524886 |
| TGGGAGSGGT | ASGERTGGAANTGQLY | 0.20753819 |
| SHLETSL | ASGERTGGAANTGQLY | 0.39444876 |
| SFTTGMT | ASGERTGGAANTGQLY | 0.28178447 |
| QVEVAKRMGVRQPTVS | ASGERTGGAANTGQLY | 0.4476561 |
| QGSGGPALAM | ASGERTGGAANTGQLY | 0.7294877 |
| MLAPSNP | ASGERTGGAANTGQLY | 0.12435269 |
| LVNNAAI | ASGERTGGAANTGQLY | 0.71091735 |
| LTGEINTLA | ASGERTGGAANTGQLY | 0.24890846 |
| LSIFGPV | ASGERTGGAANTGQLY | 0.25000468 |
| LLALLANL | ASGERTGGAANTGQLY | 0.6425928 |
| LISNASDALDKLR | ASGERTGGAANTGQLY | 0.49480587 |
| LINEAVP | ASGERTGGAANTGQLY | 0.73639643 |
| LGYAFSGL | ASGERTGGAANTGQLY | 0.27450854 |
| LDFVAVAL | ASGERTGGAANTGQLY | 0.64882755 |
| LAQPGAE | ASGERTGGAANTGQLY | 0.57207274 |
| ISNLGAVTGF | ASGERTGGAANTGQLY | 0.14585589 |
| HQEDPGRI | ASGERTGGAANTGQLY | 0.32863343 |
| GTPGEPTP | ASGERTGGAANTGQLY | 0.7833193 |
| GFFNSGS | ASGERTGGAANTGQLY | 0.7985352 |
| GAGGGGLTAGAN | ASGERTGGAANTGQLY | 0.84196365 |
| FVDALVNM | ASGERTGGAANTGQLY | 0.66811275 |
| FLTVAPEL | ASGERTGGAANTGQLY | 0.61692524 |
| FETLTAL | ASGERTGGAANTGQLY | 0.71218526 |
| DVINPIEL | ASGERTGGAANTGQLY | 0.7784606 |
| APVSGAPV | ASGERTGGAANTGQLY | 0.84772277 |
| ALVGGLAW | ASGERTGGAANTGQLY | 0.8238622 |
| ALAAAAATAI | ASGERTGGAANTGQLY | 0.5494672 |
| AFGLGLLL | ASGERTGGAANTGQLY | 0.79219717 |
| AFFAAASL | ASGERTGGAANTGQLY | 0.74593806 |
| AAAALPAFSPP | ASGERTGGAANTGQLY | 0.7144115 |
| VVLAPAL | ASGERTGGAANTGQLY | 0.6831256 |
| TIGHVDHGKTTL | ASGERTGGAANTGQLY | 0.7148647 |
| LISNASDALDK | ASGERTGGAANTGQLY | 0.6368791 |
| LISNASDAL | ASGERTGGAANTGQLY | 0.4305424 |
| ISNASDALDK | ASGERTGGAANTGQLY | 0.6520928 |
| NQAETLVYQTE | ASGERTGGAANTGQLY | 0.78708655 |
| TGGTGGTAGAG | ASGERTGGAANTGQLY | 0.77337503 |
| TGSTAGGAGGA | ASGERTGGAANTGQLY | 0.7880646 |
| DPLAVPVAAGPV | ASGERTGGAANTGQLY | 0.8171464 |
| GGTGGTPGNGGN | ASGERTGGAANTGQLY | 0.593989 |
| PAGLAGVDPATL | ASGERTGGAANTGQLY | 0.18547048 |
| GNNLVVEAPHAGASV | ASGERTGGAANTGQLY | 0.307646 |
| KVTRIMSASTLEQPA | ASGERTGGAANTGQLY | 0.6226057 |
| LGFALPIALPRRTVV | ASGERTGGAANTGQLY | 0.665962 |
| TRLGNLTARKLVVLV | ASGERTGGAANTGQLY | 0.18001956 |
| WSQQRRLIYFPSAGP | ASGERTGGAANTGQLY | 0.66929024 |
| GPNGNPDPMAAAVDIR | ASGERTGGAANTGQLY | 0.7005599 |
| TQRCVSVADLRAAASR | ASGERTGGAANTGQLY | 0.6997118 |

| | | |
|---|---|---|
| QFVLDVAEHPVEVLTETV | ASGERTGGAANTGQLY | 0.80830437 |
| GGAGGTGGAGGNGGTQAGDG | ASGERTGGAANTGQLY | 0.7523979 |
| PLLATLRAIAVDVDAKPAQV | ASGERTGGAANTGQLY | 0.38763118 |
| APKFRTGGYMIDFWGVGYQVAKRM | ASGERTGGAANTGQLY | 0.44347033 |
| GAGGAGGAGGAAGGPGGTGGTGGN | ASGERTGGAANTGQLY | 0.81393623 |
| IGALGLLATDVAPAGSVAFGVALA | ASGERTGGAANTGQLY | 0.37342185 |
| AGGAGGIGGNGGAGGAGGVLMGNG | ASGERTGGAANTGQLY | 0.812354 |
| ADELIGLL | ASGERTGGAANTGQLY | 0.3845557 |
| ADRALKA | ASGERTGGAANTGQLY | 0.8122263 |
| AGAVGQVGGAGGA | ASGERTGGAANTGQLY | 0.50600004 |
| AGGGGGLGGAP | ASGERTGGAANTGQLY | 0.79253757 |
| AGGLLYGNGGAG | ASGERTGGAANTGQLY | 0.7795631 |
| AGLESIGL | ASGERTGGAANTGQLY | 0.84148926 |
| AGLKTGII | ASGERTGGAANTGQLY | 0.6496308 |
| ASLAALLL | ASGERTGGAANTGQLY | 0.7921929 |
| ASVINISSI | ASGERTGGAANTGQLY | 0.46502328 |
| ATPPGPA | ASGERTGGAANTGQLY | 0.8415718 |
| ATVASALV | ASGERTGGAANTGQLY | 0.6277091 |
| ATVASLIL | ASGERTGGAANTGQLY | 0.6116816 |
| AVNGPVLIRH | ASGERTGGAANTGQLY | 0.34449092 |
| DGAAGAVGI | ASGERTGGAANTGQLY | 0.66297054 |
| DPLLHGI | ASGERTGGAANTGQLY | 0.82497567 |
| ELGAALGAL | ASGERTGGAANTGQLY | 0.52030045 |
| ELGRTQAV | ASGERTGGAANTGQLY | 0.8113585 |
| FVAGGYLS | ASGERTGGAANTGQLY | 0.6932359 |
| GASGGAGGAGGW | ASGERTGGAANTGQLY | 0.58702666 |
| GGAGAGGDGGAGGI | ASGERTGGAANTGQLY | 0.7621919 |
| GTGHNGSQPGMGGNG | ASGERTGGAANTGQLY | 0.6527092 |
| GYAAASASAS | ASGERTGGAANTGQLY | 0.7924843 |
| HQVDVAAVQQG | ASGERTGGAANTGQLY | 0.47740844 |
| IGAAIAHT | ASGERTGGAANTGQLY | 0.68779457 |
| IGMIEVAQL | ASGERTGGAANTGQLY | 0.7458254 |
| IRLAPEHTSAMD | ASGERTGGAANTGQLY | 0.5510196 |
| IYNVSVL | ASGERTGGAANTGQLY | 0.4057005 |
| KQGFLGM | ASGERTGGAANTGQLY | 0.82776606 |
| LESVITQL | ASGERTGGAANTGQLY | 0.77267516 |
| LGLGALVVV | ASGERTGGAANTGQLY | 0.79282063 |
| LIGNGGNGGSGGI | ASGERTGGAANTGQLY | 0.6893804 |
| LNAINAPTL | ASGERTGGAANTGQLY | 0.5759932 |
| LQLLNTGV | ASGERTGGAANTGQLY | 0.7311725 |
| LQTSIEF | ASGERTGGAANTGQLY | 0.7826228 |
| MSFAVSEL | ASGERTGGAANTGQLY | 0.8064275 |
| NTGALGL | ASGERTGGAANTGQLY | 0.75558853 |
| PPGAGGAGGAGTT | ASGERTGGAANTGQLY | 0.48368293 |
| QLVGLGI | ASGERTGGAANTGQLY | 0.7901501 |
| RFLEQVHQL | ASGERTGGAANTGQLY | 0.7122191 |
| RGVGYVLR | ASGERTGGAANTGQLY | 0.7376646 |
| SATGVFAGVI | ASGERTGGAANTGQLY | 0.7617611 |
| SLPAITLP | ASGERTGGAANTGQLY | 0.7441962 |
| STALAFTV | ASGERTGGAANTGQLY | 0.7101075 |
| SVLTVGLGK | ASGERTGGAANTGQLY | 0.620743 |

| | | |
|---|---|---|
| TALAGAGSGI | ASGERTGGAANTGQLY | 0.77762973 |
| TGLPFTGI | ASGERTGGAANTGQLY | 0.8009842 |
| VAEHVTV | ASGERTGGAANTGQLY | 0.7957999 |
| VDIVLNSL | ASGERTGGAANTGQLY | 0.8187336 |
| VLTGGGAVL | ASGERTGGAANTGQLY | 0.8107281 |
| VNVGLAYGI | ASGERTGGAANTGQLY | 0.7918032 |
| VSSAAVGAV | ASGERTGGAANTGQLY | 0.2298812 |
| VTSVYEAGV | ASGERTGGAANTGQLY | 0.43024832 |
| WVALDRGA | ASGERTGGAANTGQLY | 0.514804 |
| YLVGSVVI | ASGERTGGAANTGQLY | 0.7202009 |
| GPPGTGKTLL | ASGERTGGAANTGQLY | 0.26981753 |
| GVGMTKF | ASGERTGGAANTGQLY | 0.7025173 |
| IGIGGSDL | ASGERTGGAANTGQLY | 0.6670283 |
| IGTLLAEL | ASGERTGGAANTGQLY | 0.5985666 |
| IYTSGTT | ASGERTGGAANTGQLY | 0.8199284 |
| LSFPAGV | ASGERTGGAANTGQLY | 0.68099666 |
| LYTSGTT | ASGERTGGAANTGQLY | 0.55496013 |
| VGAFPFGF | ASGERTGGAANTGQLY | 0.57687014 |
| VGAGTVEF | ASGERTGGAANTGQLY | 0.8131021 |
| VGKSTLLL | ASGERTGGAANTGQLY | 0.4566792 |
| AALAHAGGGIV | ASGERTGGAANTGQLY | 0.80496305 |
| AALVAVSLL | ASGERTGGAANTGQLY | 0.6442253 |
| AAVSRAVAAL | ASGERTGGAANTGQLY | 0.8133358 |
| AGGSSGIGGF | ASGERTGGAANTGQLY | 0.3909107 |
| AGLTVANANG | ASGERTGGAANTGQLY | 0.7949593 |
| AGLVGAVL | ASGERTGGAANTGQLY | 0.7221551 |
| ALSQAFTV | ASGERTGGAANTGQLY | 0.83957946 |
| ALTFSVSL | ASGERTGGAANTGQLY | 0.7067441 |
| ALTGAALVAL | ASGERTGGAANTGQLY | 0.74613905 |
| ASFSIGAGVL | ASGERTGGAANTGQLY | 0.7039757 |
| DGSVYKI | ASGERTGGAANTGQLY | 0.8049786 |
| EQLAEIV | ASGERTGGAANTGQLY | 0.39960837 |
| FAVSELTVM | ASGERTGGAANTGQLY | 0.73353785 |
| GAAPLAAGL | ASGERTGGAANTGQLY | 0.44654018 |
| GADGGDGGSGGAGG | ASGERTGGAANTGQLY | 0.7640406 |
| GFGLDTGGAGG | ASGERTGGAANTGQLY | 0.77256286 |
| GLASGGLL | ASGERTGGAANTGQLY | 0.26999465 |
| GRIAGAAVLL | ASGERTGGAANTGQLY | 0.8059125 |
| GSAGSLNTSF | ASGERTGGAANTGQLY | 0.6331943 |
| GTGQIGI | ASGERTGGAANTGQLY | 0.5032264 |
| GTLIALIL | ASGERTGGAANTGQLY | 0.7956905 |
| IAATMGFVVG | ASGERTGGAANTGQLY | 0.71347123 |
| IFAQALL | ASGERTGGAANTGQLY | 0.8073815 |
| IGAAVALGF | ASGERTGGAANTGQLY | 0.632339 |
| IGLTPLAL | ASGERTGGAANTGQLY | 0.73647714 |
| ILGSNTSTL | ASGERTGGAANTGQLY | 0.78426635 |
| IVFSLTVL | ASGERTGGAANTGQLY | 0.3796726 |
| LASKATGF | ASGERTGGAANTGQLY | 0.49954414 |
| LASVALIL | ASGERTGGAANTGQLY | 0.8114679 |
| LFAVGLTL | ASGERTGGAANTGQLY | 0.87837 |
| LFNGGIGTYI | ASGERTGGAANTGQLY | 0.8753225 |

| | | |
|---|---|---|
| LTLGAGII | ASGERTGGAANTGQLY | 0.8113562 |
| LVAAALGDI | ASGERTGGAANTGQLY | 0.7534952 |
| NTAFVQLGI | ASGERTGGAANTGQLY | 0.8277576 |
| QLFSGAEL | ASGERTGGAANTGQLY | 0.8863958 |
| SAATFITL | ASGERTGGAANTGQLY | 0.88723695 |
| SNYGLANLGV | ASGERTGGAANTGQLY | 0.88183033 |
| SYGGAYAVM | ASGERTGGAANTGQLY | 0.65050846 |
| TTGVGGLTLG | ASGERTGGAANTGQLY | 0.88885295 |
| TTQTIYI | ASGERTGGAANTGQLY | 0.84829646 |
| VAAGAGLGAVI | ASGERTGGAANTGQLY | 0.839235 |
| VAAVVTLL | ASGERTGGAANTGQLY | 0.87835383 |
| VGFLGGLI | ASGERTGGAANTGQLY | 0.8506974 |
| VGVIPILL | ASGERTGGAANTGQLY | 0.8486885 |
| VIFASSLI | ASGERTGGAANTGQLY | 0.8279867 |
| VKGGDGGAAAG | ASGERTGGAANTGQLY | 0.86470336 |
| VTKTFTNI | ASGERTGGAANTGQLY | 0.8849289 |
| VVAAAILL | ASGERTGGAANTGQLY | 0.87855196 |
| YAALDVEL | ASGERTGGAANTGQLY | 0.90743744 |
| LASVALLL | ASGERTGGAANTGQLY | 0.8960991 |
| LDAVHRV | ASGERTGGAANTGQLY | 0.8900602 |
| QGLSSSIF | ASGERTGGAANTGQLY | 0.32043236 |
| RGTFANIRL | ASGERTGGAANTGQLY | 0.5923625 |
| RQLWWGHRI | ASGERTGGAANTGQLY | 0.791617 |
| YGPPGTGKTLL | ASGERTGGAANTGQLY | 0.83301055 |
| AALVGAPANVA | ASGERTGGAANTGQLY | 0.40754732 |
| AATFRRTE | ASGERTGGAANTGQLY | 0.835339 |
| AGALGGGI | ASGERTGGAANTGQLY | 0.842453 |
| AGGAGGAGGIGAG | ASGERTGGAANTGQLY | 0.77491295 |
| AGSGALGL | ASGERTGGAANTGQLY | 0.89571416 |
| AQIADQI | ASGERTGGAANTGQLY | 0.83461106 |
| ASAVAVTYL | ASGERTGGAANTGQLY | 0.7592181 |
| ASTVLGGTGG | ASGERTGGAANTGQLY | 0.79481685 |
| AVGGEPAK | ASGERTGGAANTGQLY | 0.9021274 |
| CEQGTFGGTC | ASGERTGGAANTGQLY | 0.73398936 |
| DEGKAPAG | ASGERTGGAANTGQLY | 0.89412093 |
| DTVVFPAL | ASGERTGGAANTGQLY | 0.7995922 |
| FEGITMAG | ASGERTGGAANTGQLY | 0.9016146 |
| FPLVVQD | ASGERTGGAANTGQLY | 0.55427724 |
| GAFTTPQ | ASGERTGGAANTGQLY | 0.8895304 |
| GFGGFGTAGGVGGA | ASGERTGGAANTGQLY | 0.84355724 |
| GFLAQFL | ASGERTGGAANTGQLY | 0.91214454 |
| GGAGGAGGNAGVA | ASGERTGGAANTGQLY | 0.8335166 |
| GGVGTAGAGGAGGAGG | ASGERTGGAANTGQLY | 0.8698429 |
| GNGGGGDAGNAGSGGN | ASGERTGGAANTGQLY | 0.8160208 |
| GTGGAGGAGNDAGS | ASGERTGGAANTGQLY | 0.89119595 |
| GTTNTGWFNS | ASGERTGGAANTGQLY | 0.61067927 |
| IFMGPLFTL | ASGERTGGAANTGQLY | 0.8315792 |
| ILAHAIEAI | ASGERTGGAANTGQLY | 0.69568324 |
| IRSAIREAL | ASGERTGGAANTGQLY | 0.8697566 |
| ITFAGGG | ASGERTGGAANTGQLY | 0.88273025 |
| ITGMSGHQEV | ASGERTGGAANTGQLY | 0.43121338 |

| | | |
|---|---|---|
| IVAGLACTL | ASGERTGGAANTGQLY | 0.89884317 |
| IVGGGLAGA | ASGERTGGAANTGQLY | 0.77414715 |
| IVTGVLNI | ASGERTGGAANTGQLY | 0.699445 |
| LAQQGCI | ASGERTGGAANTGQLY | 0.89602065 |
| LCLHVEN | ASGERTGGAANTGQLY | 0.8465635 |
| LDAISDGS | ASGERTGGAANTGQLY | 0.89632547 |
| LGGFAAML | ASGERTGGAANTGQLY | 0.8285246 |
| LGGKNPMI | ASGERTGGAANTGQLY | 0.8547653 |
| LGPKGVAAIL | ASGERTGGAANTGQLY | 0.87390137 |
| LMVGVSI | ASGERTGGAANTGQLY | 0.5187585 |
| LVGAAFVL | ASGERTGGAANTGQLY | 0.7438246 |
| LVLAVAPL | ASGERTGGAANTGQLY | 0.9017532 |
| MAAIAVIDAA | ASGERTGGAANTGQLY | 0.6967344 |
| NNGINAELQ | ASGERTGGAANTGQLY | 0.7109275 |
| RGEPLVLVHG | ASGERTGGAANTGQLY | 0.6519848 |
| RGGAGGAAGLI | ASGERTGGAANTGQLY | 0.61982137 |
| RSYRMHPAV | ASGERTGGAANTGQLY | 0.6549074 |
| SAAHLLAGQLA | ASGERTGGAANTGQLY | 0.7107619 |
| SAAVGSY | ASGERTGGAANTGQLY | 0.7074574 |
| SPHAVLHRI | ASGERTGGAANTGQLY | 0.70007765 |
| SQIAKVL | ASGERTGGAANTGQLY | 0.533369 |
| TALGHLVP | ASGERTGGAANTGQLY | 0.6894938 |
| TAPTTSGAA | ASGERTGGAANTGQLY | 0.66111076 |
| TDGDDGNGGSPGT | ASGERTGGAANTGQLY | 0.6486411 |
| TDGLGEN | ASGERTGGAANTGQLY | 0.6979734 |
| TGGPPLPGQA | ASGERTGGAANTGQLY | 0.6844143 |
| TIPAITI | ASGERTGGAANTGQLY | 0.6530905 |
| TIPGFTI | ASGERTGGAANTGQLY | 0.67570436 |
| TSIASGALA | ASGERTGGAANTGQLY | 0.67803067 |
| TVTLPGAAI | ASGERTGGAANTGQLY | 0.67944866 |
| VASGLSAALP | ASGERTGGAANTGQLY | 0.68057394 |
| VGAGVIGI | ASGERTGGAANTGQLY | 0.7148018 |
| VGPKRFGV | ASGERTGGAANTGQLY | 0.70792365 |
| VNVAFPD | ASGERTGGAANTGQLY | 0.705446 |
| VSMTNSGVS | ASGERTGGAANTGQLY | 0.3333112 |
| VTDVLGDLSV | ASGERTGGAANTGQLY | 0.45687598 |
| VVVVGAGFAGL | ASGERTGGAANTGQLY | 0.6463312 |
| VYTVGTVV | ASGERTGGAANTGQLY | 0.63300735 |
| WTTFSGP | ASGERTGGAANTGQLY | 0.455923 |
| YDELVEAF | ASGERTGGAANTGQLY | 0.6761099 |
| YGTGGAGGNGG | ASGERTGGAANTGQLY | 0.6621082 |
| YTNVMAV | ASGERTGGAANTGQLY | 0.6227503 |
| FLEQVHQL | ASGERTGGAANTGQLY | 0.6990725 |
| GLSLLAVAV | ASGERTGGAANTGQLY | 0.65722954 |
| LEQVHQL | ASGERTGGAANTGQLY | 0.6436284 |
| LNAAGAI | ASGERTGGAANTGQLY | 0.59670126 |
| PAVHASL | ASGERTGGAANTGQLY | 0.67059565 |
| TELEDDA | ASGERTGGAANTGQLY | 0.5148684 |
| TPPGPPN | ASGERTGGAANTGQLY | 0.70284057 |
| TPPPPPPP | ASGERTGGAANTGQLY | 0.52137 |
| TQSPALI | ASSDNAETLY | 0.704461 |

| | | |
|---|---|---|
| EFAAAFPQ | ASSDNAETLY | 0.5443265 |
| LLSGNLDVL | ASSDNAETLY | 0.7005426 |
| GPANLGNYNI | ASSDNAETLY | 0.68089867 |
| TLGPTLP | ASSDNAETLY | 0.7260579 |
| VYAYSSCCK | ASSDNAETLY | 0.6730181 |
| APAPAPAPNI | ASSDNAETLY | 0.6819825 |
| PGGPVPIP | ASSDNAETLY | 0.59739804 |
| FLTVALEP | ASSDNAETLY | 0.7101628 |
| SQYFGANL | ASSDNAETLY | 0.5314698 |
| GGVKPAA | ASSDNAETLY | 0.67337203 |
| GGTGGAGGAGGW | ASSDNAETLY | 0.46357474 |
| GVGGVAGLGGAGG | ASSDNAETLY | 0.7010739 |
| VLLVLATTI | ASSDNAETLY | 0.7004268 |
| IGIFAAML | ASSDNAETLY | 0.54778856 |
| AVGDLID | ASSDNAETLY | 0.710266 |
| NISPLPL | ASSDNAETLY | 0.67935073 |
| VTAVFVAI | ASSDNAETLY | 0.63208055 |
| SGDDNDAAL | ASSDNAETLY | 0.7174506 |
| AVAALAHS | ASSDNAETLY | 0.6581236 |
| GGVTVMATP | ASSDNAETLY | 0.71600366 |
| AGLAANVRD | ASSDNAETLY | 0.6618277 |
| TNRPDLIDPALLRPG | ASSDNAETLY | 0.6236214 |
| LANLGVTAL | ASSDNAETLY | 0.68104875 |
| TGERIEL | ASSDNAETLY | 0.5192974 |
| NGEELAA | ASSDNAETLY | 0.5569233 |
| VLSLVWG | ASSDNAETLY | 0.71542853 |
| IGYALWL | ASSDNAETLY | 0.6553277 |
| LSLLLAPL | ASSDNAETLY | 0.68756366 |
| TLDEALK | ASSDNAETLY | 0.6179413 |
| TDGAGTAEGG | ASSDNAETLY | 0.56834626 |
| NAPGGAGGNGG | ASSDNAETLY | 0.6250619 |
| VSVFMGMGGNF | ASSDNAETLY | 0.6736698 |
| LASYLAAL | ASSDNAETLY | 0.68257666 |
| LVGGTPAL | ASSDNAETLY | 0.66415644 |
| VRNADVL | ASSDNAETLY | 0.5438884 |
| TCSSPFLVSAM | ASSDNAETLY | 0.64130867 |
| VHITDAE | ASSDNAETLY | 0.6193471 |
| EYAAPSF | ASSDNAETLY | 0.63025707 |
| VVDAFHGGL | ASSDNAETLY | 0.6604821 |
| ASVVDPQ | ASSDNAETLY | 0.64576095 |
| SYNEFGGLS | ASSDNAETLY | 0.59422725 |
| DPDGSVY | ASSDNAETLY | 0.617519 |
| MVSTIHATE | ASSDNAETLY | 0.65262127 |
| LAAGVADPV | ASSDNAETLY | 0.6681719 |
| TKEQIAA | ASSDNAETLY | 0.6657611 |
| SLASPAAL | ASSDNAETLY | 0.68576926 |
| IGLVFYGV | ASSDNAETLY | 0.67393374 |
| FLLDASNL | ASSDNAETLY | 0.6756952 |
| YQVGNFP | ASSDNAETLY | 0.3683279 |
| ESLLSQST | ASSDNAETLY | 0.47924116 |
| LFTVGFAAVG | ASSDNAETLY | 0.6070286 |

| | | |
|---|---|---|
| ATPPTPT | ASSDNAETLY | 0.6335532 |
| TDLATLG | ASSDNAETLY | 0.49597356 |
| SAGGTGR | ASSDNAETLY | 0.6569706 |
| GFSNSGS | ASSDNAETLY | 0.5901502 |
| GGNGGSGGTSV | ASSDNAETLY | 0.56422263 |
| GTSVATGGAGN | ASSDNAETLY | 0.6714796 |
| NGGNGGSGGTS | ASSDNAETLY | 0.6173048 |
| AVVKFPH | ASSDNAETLY | 0.58998734 |
| TPPPPPP | ASSDNAETLY | 0.59770167 |
| AAPAAAAAPA | ASSDNAETLY | 0.67155325 |
| IALNEDL | ASSDNAETLY | 0.49504802 |
| IVNVSSM | ASSDNAETLY | 0.66504663 |
| TGIPPAPR | ASSDNAETLY | 0.5912142 |
| FDLGGGTF | ASSDNAETLY | 0.6769586 |
| FDLGGGTFDVS | ASSDNAETLY | 0.4934997 |
| NEPTAAA | ASSDNAETLY | 0.6709173 |
| NEPTAAALA | ASSDNAETLY | 0.6395546 |
| GSIINISSI | ASSDNAETLY | 0.6897933 |
| VAPGNAGTA | ASSDNAETLY | 0.6247324 |
| TGGGASLE | ASSDNAETLY | 0.6438289 |
| VAVYNHYKRI | ASSDNAETLY | 0.60486925 |
| TDPAPATT | ASSDNAETLY | 0.68124837 |
| TNRPDILDPALLRPG | ASSDNAETLY | 0.4971617 |
| LTTTTTS | ASSDNAETLY | 0.6339171 |
| PTGSGKTL | ASSDNAETLY | 0.5123471 |
| TGSGKTL | ASSDNAETLY | 0.6540006 |
| TIGHVDHGK | ASSDNAETLY | 0.6621593 |
| TIGHVDHGKTT | ASSDNAETLY | 0.511656 |
| GHVDHGKTT | ASSDNAETLY | 0.6755209 |
| AQFEHTLL | ASSDNAETLY | 0.6468827 |
| IVNQDEL | ASSDNAETLY | 0.5957302 |
| TNGSQFF | ASSDNAETLY | 0.68214095 |
| TGKGIVE | ASSDNAETLY | 0.6216796 |
| LISNASDA | ASSDNAETLY | 0.6727862 |
| AELRTLGELTKIAKAHGAQ | ASSDNAETLY | 0.6383802 |
| APNAVAAAEPSW | ASSDNAETLY | 0.6300009 |
| APQVAGAATAVL | ASSDNAETLY | 0.6578462 |
| ARRDPAAAAAILPTDA | ASSDNAETLY | 0.6346841 |
| DAAARDVLAEAGLAEYF | ASSDNAETLY | 0.56578034 |
| DDQIVLSAKANLAKVEQ | ASSDNAETLY | 0.68007326 |
| DELLPDTGEDSDEDGH | ASSDNAETLY | 0.7705086 |
| ELPEAPANRATVPQD | ASSDNAETLY | 0.79017913 |
| FGQSKIERVTGIG | ASSDNAETLY | 0.72745264 |
| GAGGTGSTAGGAGGAGGAGGLYA | ASSDNAETLY | 0.6803163 |
| GHGGTGAAGGNGGTGGAGGSGIDG | ASSDNAETLY | 0.7281528 |
| GNGGNGGAGGEGGPGGAGGAGA | ASSDNAETLY | 0.7809356 |
| GSTAGNGGNGGNATNGTIAGQPAG | ASSDNAETLY | 0.78926814 |
| GTGGDGGNGGNGGTGG | ASSDNAETLY | 0.772733 |
| GVRRPAGIPADAPV | ASSDNAETLY | 0.66180456 |
| GYRQVTDAHLLTLARR | ASSDNAETLY | 0.76289475 |
| IAKMGQDGHDRG | ASSDNAETLY | 0.746405 |

| | | |
|---|---|---|
| KAEPSIEIQYT | ASSDNAETLY | 0.7462768 |
| KPVRIRMAADEDTVAVT | ASSDNAETLY | 0.77496105 |
| MPPRIAGMRLLVIKPEP | ASSDNAETLY | 0.76207674 |
| MKFYAVALFRSRGVPPTR | ASSDNAETLY | 0.72352517 |
| NAGGFGILGAVAHSP | ASSDNAETLY | 0.7306819 |
| NEGREITLAMAEM | ASSDNAETLY | 0.7574605 |
| NELVEWSHPLGHRWRWE | ASSDNAETLY | 0.77119565 |
| NLGAGGTGGNGGNPG | ASSDNAETLY | 0.76592404 |
| NTEWPKWETVMAADPVD | ASSDNAETLY | 0.79111934 |
| PASAAPAETQP | ASSDNAETLY | 0.7822021 |
| QQSSDGKNGVRAAEITGE | ASSDNAETLY | 0.7790361 |
| RAPDAVTTSWWKE | ASSDNAETLY | 0.38503513 |
| SGFGNVGAHTSGWFNQ | ASSDNAETLY | 0.65218073 |
| SGVAGAVGQAASVGGLKV | ASSDNAETLY | 0.74026537 |
| SNGGAGTGSAGGNGGT | ASSDNAETLY | 0.74239707 |
| TGGAGGEGGAAG | ASSDNAETLY | 0.5610684 |
| TGGDGGAAGAGGAPGHG | ASSDNAETLY | 0.7595529 |
| TMPPLLAAIANAGRHGV | ASSDNAETLY | 0.7316693 |
| TSSTAGNSGGA | ASSDNAETLY | 0.68052435 |
| VPATVPVGILC | ASSDNAETLY | 0.77487445 |
| VQRPDSGGVVLTGRIS | ASSDNAETLY | 0.727141 |
| WTPNPVPATKGVPAGLVQAT | ASSDNAETLY | 0.7164687 |
| YALKRALAKFSDRFDVVII | ASSDNAETLY | 0.7243418 |
| YLIGSVIV | ASSDNAETLY | 0.78403366 |
| YGQLVAVID | ASSDNAETLY | 0.6591885 |
| VTVGGMTLPA | ASSDNAETLY | 0.77288866 |
| VLALPAV | ASSDNAETLY | 0.74945074 |
| VIPPTLNYET | ASSDNAETLY | 0.7839439 |
| VGMGGMHPSSG | ASSDNAETLY | 0.61520123 |
| VGLAGGAATA | ASSDNAETLY | 0.7784791 |
| VGGAGGAGGSGP | ASSDNAETLY | 0.7493335 |
| VFAAFMV | ASSDNAETLY | 0.7930189 |
| TLYDKLR | ASSDNAETLY | 0.73912394 |
| TLAVFTVL | ASSDNAETLY | 0.7649159 |
| TGSVQPARV | ASSDNAETLY | 0.7578399 |
| TGGGAGSGGT | ASSDNAETLY | 0.78823596 |
| SHLETSL | ASSDNAETLY | 0.5982895 |
| SFTTGMT | ASSDNAETLY | 0.75641733 |
| QVEVAKRMGVRQPTVS | ASSDNAETLY | 0.6795467 |
| QGSGGPALAM | ASSDNAETLY | 0.7693895 |
| MLAPSNP | ASSDNAETLY | 0.7743148 |
| LVNNAAI | ASSDNAETLY | 0.5101038 |
| LTGEINTLA | ASSDNAETLY | 0.78340733 |
| LSIFGPV | ASSDNAETLY | 0.7379336 |
| LLALLANL | ASSDNAETLY | 0.7068851 |
| LISNASDALDKLR | ASSDNAETLY | 0.7842276 |
| LINEAVP | ASSDNAETLY | 0.7405056 |
| LGYAFSGL | ASSDNAETLY | 0.7827681 |
| LDFVAVAL | ASSDNAETLY | 0.75430655 |
| LAQPGAE | ASSDNAETLY | 0.7659366 |
| ISNLGAVTGF | ASSDNAETLY | 0.77279496 |

| | | |
|---|---|---|
| HQEDPGRI | ASSDNAETLY | 0.6201763 |
| GTPGEPTP | ASSDNAETLY | 0.6999433 |
| GFFNSGS | ASSDNAETLY | 0.7870252 |
| GAGGGGLTAGAN | ASSDNAETLY | 0.7806827 |
| FVDALVNM | ASSDNAETLY | 0.7635346 |
| FLTVAPEL | ASSDNAETLY | 0.7131655 |
| FETLTAL | ASSDNAETLY | 0.6958398 |
| DVINPIEL | ASSDNAETLY | 0.72296786 |
| APVSGAPV | ASSDNAETLY | 0.7741674 |
| ALVGGLAW | ASSDNAETLY | 0.7838799 |
| ALAAAAATAI | ASSDNAETLY | 0.7827981 |
| AFGLGLLL | ASSDNAETLY | 0.59210014 |
| AFFAAASL | ASSDNAETLY | 0.78669155 |
| AAAALPAFSPP | ASSDNAETLY | 0.76887685 |
| VVLAPAL | ASSDNAETLY | 0.77584976 |
| TIGHVDHGKTTL | ASSDNAETLY | 0.7508725 |
| LISNASDALDK | ASSDNAETLY | 0.7537154 |
| LISNASDAL | ASSDNAETLY | 0.7246587 |
| ISNASDALDK | ASSDNAETLY | 0.6563297 |
| NQAETLVYQTE | ASSDNAETLY | 0.7687334 |
| TGGTGGTAGAG | ASSDNAETLY | 0.77681744 |
| TGSTAGGAGGA | ASSDNAETLY | 0.77520716 |
| DPLAVPVAAGPV | ASSDNAETLY | 0.7890511 |
| GGTGGTPGNGGN | ASSDNAETLY | 0.7878952 |
| PAGLAGVDPATL | ASSDNAETLY | 0.77702343 |
| GNNLVVEAPHAGASV | ASSDNAETLY | 0.3745508 |
| KVTRIMSASTLEQPA | ASSDNAETLY | 0.5670281 |
| LGFALPIALPRRTVV | ASSDNAETLY | 0.69718975 |
| TRLGNLTARKLVVLV | ASSDNAETLY | 0.71987474 |
| WSQQRRLIYFPSAGP | ASSDNAETLY | 0.31676218 |
| GPNGNPDPMAAAVDIR | ASSDNAETLY | 0.70105606 |
| TQRCVSVADLRAAASR | ASSDNAETLY | 0.70692134 |
| QFVLDVAEHPVEVLTETV | ASSDNAETLY | 0.73021436 |
| GGAGGTGGAGGNGGTQAGDG | ASSDNAETLY | 0.7815542 |
| PLLATLRAIAVDVDAKPAQV | ASSDNAETLY | 0.7535839 |
| APKFRTGGYMIDFWGVGYQVAKRM | ASSDNAETLY | 0.575783 |
| GAGGAGGAGGAAGGPGGTGGTGGN | ASSDNAETLY | 0.6533764 |
| IGALGLLATDVAPAGSVAFGVALA | ASSDNAETLY | 0.7869724 |
| AGGAGGIGGNGGAGGAGGVLMGNG | ASSDNAETLY | 0.67484456 |
| ADELIGLL | ASSDNAETLY | 0.7900008 |
| ADRALKA | ASSDNAETLY | 0.59816444 |
| AGAVGQVGGAGGA | ASSDNAETLY | 0.78847605 |
| AGGGGGLGGAP | ASSDNAETLY | 0.6100174 |
| AGGLLYGNGGAG | ASSDNAETLY | 0.78914595 |
| AGLESIGL | ASSDNAETLY | 0.77542365 |
| AGLKTGII | ASSDNAETLY | 0.79172444 |
| ASLAALLL | ASSDNAETLY | 0.7423856 |
| ASVINISSI | ASSDNAETLY | 0.7766516 |
| ATPPGPA | ASSDNAETLY | 0.72493154 |
| ATVASALV | ASSDNAETLY | 0.76948726 |
| ATVASLIL | ASSDNAETLY | 0.68241835 |

| | | |
|---|---|---|
| AVNGPVLIRH | ASSDNAETLY | 0.72840273 |
| DGAAGAVGI | ASSDNAETLY | 0.5896797 |
| DPLLHGI | ASSDNAETLY | 0.7489133 |
| ELGAALGAL | ASSDNAETLY | 0.78643614 |
| ELGRTQAV | ASSDNAETLY | 0.24161342 |
| FVAGGYLS | ASSDNAETLY | 0.7811875 |
| GASGGAGGAGGW | ASSDNAETLY | 0.67275214 |
| GGAGAGGDGGAGGI | ASSDNAETLY | 0.60893536 |
| GTGHNGSQPGMGGNG | ASSDNAETLY | 0.781392 |
| GYAAASASAS | ASSDNAETLY | 0.7639065 |
| HQVDVAAVQQG | ASSDNAETLY | 0.77471435 |
| IGAAIAHT | ASSDNAETLY | 0.7047124 |
| IGMIEVAQL | ASSDNAETLY | 0.73632526 |
| IRLAPEHTSAMD | ASSDNAETLY | 0.7464268 |
| IYNVSVL | ASSDNAETLY | 0.38310054 |
| KQGFLGM | ASSDNAETLY | 0.56948197 |
| LESVITQL | ASSDNAETLY | 0.79275846 |
| LGLGALVVV | ASSDNAETLY | 0.82281613 |
| LIGNGGNGGSGGI | ASSDNAETLY | 0.82955086 |
| LNAINAPTL | ASSDNAETLY | 0.69636804 |
| LQLLNTGV | ASSDNAETLY | 0.6357162 |
| LQTSIEF | ASSDNAETLY | 0.7302134 |
| MSFAVSEL | ASSDNAETLY | 0.8464637 |
| NTGALGL | ASSDNAETLY | 0.8592987 |
| PPGAGGAGGAGTT | ASSDNAETLY | 0.8331909 |
| QLVGLGI | ASSDNAETLY | 0.6620181 |
| RFLEQVHQL | ASSDNAETLY | 0.8369359 |
| RGVGYVLR | ASSDNAETLY | 0.7772602 |
| SATGVFAGVI | ASSDNAETLY | 0.75135016 |
| SLPAITLP | ASSDNAETLY | 0.8314943 |
| STALAFTV | ASSDNAETLY | 0.81533486 |
| SVLTVGLGK | ASSDNAETLY | 0.7396306 |
| TALAGAGSGI | ASSDNAETLY | 0.726312 |
| TGLPFTGI | ASSDNAETLY | 0.8249864 |
| VAEHVTV | ASSDNAETLY | 0.8400667 |
| VDIVLNSL | ASSDNAETLY | 0.83231056 |
| VLTGGGAVL | ASSDNAETLY | 0.8892005 |
| VNVGLAYGI | ASSDNAETLY | 0.8506604 |
| VSSAAVGAV | ASSDNAETLY | 0.8524314 |
| VTSVYEAGV | ASSDNAETLY | 0.25649345 |
| WVALDRGA | ASSDNAETLY | 0.5024384 |
| YLVGSVVI | ASSDNAETLY | 0.72312856 |
| GPPGTGKTLL | ASSDNAETLY | 0.76426923 |
| GVGMTKF | ASSDNAETLY | 0.40072757 |
| IGIGGSDL | ASSDNAETLY | 0.8265337 |
| IGTLLAEL | ASSDNAETLY | 0.7684821 |
| IYTSGTT | ASSDNAETLY | 0.67349553 |
| LSFPAGV | ASSDNAETLY | 0.8480034 |
| LYTSGTT | ASSDNAETLY | 0.72149086 |
| VGAFPFGF | ASSDNAETLY | 0.69799614 |
| VGAGTVEF | ASSDNAETLY | 0.74175763 |

| | | |
|---|---|---|
| VGKSTLLL | ASSDNAETLY | 0.8712295 |
| AALAHAGGGIV | ASSDNAETLY | 0.6221439 |
| AALVAVSLL | ASSDNAETLY | 0.8446294 |
| AAVSRAVAAL | ASSDNAETLY | 0.7200496 |
| AGGSSGIGGF | ASSDNAETLY | 0.86616635 |
| AGLTVANANG | ASSDNAETLY | 0.56620246 |
| AGLVGAVL | ASSDNAETLY | 0.8453102 |
| ALSQAFTV | ASSDNAETLY | 0.7652929 |
| ALTFSVSL | ASSDNAETLY | 0.8920814 |
| ALTGAALVAL | ASSDNAETLY | 0.7493057 |
| ASFSIGAGVL | ASSDNAETLY | 0.8332523 |
| DGSVYKI | ASSDNAETLY | 0.7903498 |
| EQLAEIV | ASSDNAETLY | 0.8366104 |
| FAVSELTVM | ASSDNAETLY | 0.48985526 |
| GAAPLAAGL | ASSDNAETLY | 0.77938086 |
| GADGGDGGSGGAGG | ASSDNAETLY | 0.5656887 |
| GFGLDTGGAGG | ASSDNAETLY | 0.8133775 |
| GLASGGLL | ASSDNAETLY | 0.826749 |
| GRIAGAAVLL | ASSDNAETLY | 0.31905252 |
| GSAGSLNTSF | ASSDNAETLY | 0.8609717 |
| GTGQIGI | ASSDNAETLY | 0.6922296 |
| GTLIALIL | ASSDNAETLY | 0.680385 |
| IAATMGFVVG | ASSDNAETLY | 0.86055815 |
| IFAQALL | ASSDNAETLY | 0.7942519 |
| IGAAVALGF | ASSDNAETLY | 0.8558184 |
| IGLTPLAL | ASSDNAETLY | 0.7740373 |
| ILGSNTSTL | ASSDNAETLY | 0.7814173 |
| IVFSLTVL | ASSDNAETLY | 0.8275834 |
| LASKATGF | ASSDNAETLY | 0.5197904 |
| LASVALIL | ASSDNAETLY | 0.70034945 |
| LFAVGLTL | ASSDNAETLY | 0.8686148 |
| LFNGGIGTYI | ASSDNAETLY | 0.8819182 |
| LTLGAGII | ASSDNAETLY | 0.88018346 |
| LVAAALGDI | ASSDNAETLY | 0.7923728 |
| NTAFVQLGI | ASSDNAETLY | 0.7588358 |
| QLFSGAEL | ASSDNAETLY | 0.82018965 |
| SAATFITL | ASSDNAETLY | 0.88706434 |
| SNYGLANLGV | ASSDNAETLY | 0.903713 |
| SYGGAYAVM | ASSDNAETLY | 0.8921672 |
| TTGVGGLTLG | ASSDNAETLY | 0.8015455 |
| TTQTIYI | ASSDNAETLY | 0.8969798 |
| VAAGAGLGAVI | ASSDNAETLY | 0.87491226 |
| VAAVVTLL | ASSDNAETLY | 0.8507749 |
| VGFLGGLI | ASSDNAETLY | 0.8841146 |
| VGVIPILL | ASSDNAETLY | 0.8894803 |
| VIFASSLI | ASSDNAETLY | 0.8219351 |
| VKGGDGGAAAG | ASSDNAETLY | 0.8426266 |
| VTKTFTNI | ASSDNAETLY | 0.87042785 |
| VVAAAILL | ASSDNAETLY | 0.88465166 |
| YAALDVEL | ASSDNAETLY | 0.8844489 |
| LASVALLL | ASSDNAETLY | 0.9149115 |

| | | |
|---|---|---|
| LDAVHRV | ASSDNAETLY | 0.89648503 |
| QGLSSSIF | ASSDNAETLY | 0.88317436 |
| RGTFANIRL | ASSDNAETLY | 0.40555748 |
| RQLWWGHRI | ASSDNAETLY | 0.73407006 |
| YGPPGTGKTLL | ASSDNAETLY | 0.842292 |
| AALVGAPANVA | ASSDNAETLY | 0.8531152 |
| AATFRRTE | ASSDNAETLY | 0.47438633 |
| AGALGGGI | ASSDNAETLY | 0.87659 |
| AGGAGGAGGIGAG | ASSDNAETLY | 0.8489592 |
| AGSGALGL | ASSDNAETLY | 0.81418514 |
| AQIADQI | ASSDNAETLY | 0.89808416 |
| ASAVAVTYL | ASSDNAETLY | 0.84802043 |
| ASTVLGGTGG | ASSDNAETLY | 0.8243295 |
| AVGGEPAK | ASSDNAETLY | 0.8494524 |
| CEQGTFGGTC | ASSDNAETLY | 0.9060188 |
| DEGKAPAG | ASSDNAETLY | 0.792572 |
| DTVVFPAL | ASSDNAETLY | 0.9047278 |
| FEGITMAG | ASSDNAETLY | 0.7734524 |
| FPLVVQD | ASSDNAETLY | 0.9024608 |
| GAFTTPQ | ASSDNAETLY | 0.76353586 |
| GFGGFGTAGGVGGA | ASSDNAETLY | 0.90053594 |
| GFLAQFL | ASSDNAETLY | 0.86212444 |
| GGAGGAGGNAGVA | ASSDNAETLY | 0.9179178 |
| GGVGTAGAGGAGGAGG | ASSDNAETLY | 0.8467869 |
| GNGGGDAGNAGSGGN | ASSDNAETLY | 0.9053477 |
| GTGGAGGAGNDAGS | ASSDNAETLY | 0.86034834 |
| GTTNTGWFNS | ASSDNAETLY | 0.90091574 |
| IFMGPLFTL | ASSDNAETLY | 0.7583711 |
| ILAHAIEAI | ASSDNAETLY | 0.85503775 |
| IRSAIREAL | ASSDNAETLY | 0.67858857 |
| ITFAGGG | ASSDNAETLY | 0.8726038 |
| ITGMSGHQEV | ASSDNAETLY | 0.89206886 |
| IVAGLACTL | ASSDNAETLY | 0.5880377 |
| IVGGGLAGA | ASSDNAETLY | 0.9069263 |
| IVTGVLNI | ASSDNAETLY | 0.79763275 |
| LAQQGCI | ASSDNAETLY | 0.8117422 |
| LCLHVEN | ASSDNAETLY | 0.9032272 |
| LDAISDGS | ASSDNAETLY | 0.87323964 |
| LGGFAAML | ASSDNAETLY | 0.89044315 |
| LGGKNPMI | ASSDNAETLY | 0.844967 |
| LGPKGVAAIL | ASSDNAETLY | 0.85532427 |
| LMVGVSI | ASSDNAETLY | 0.8687498 |
| LVGAAFVL | ASSDNAETLY | 0.61973196 |
| LVLAVAPL | ASSDNAETLY | 0.8027109 |
| MAAIAVIDAA | ASSDNAETLY | 0.91337466 |
| NNGINAELQ | ASSDNAETLY | 0.78687435 |
| RGEPLVLVHG | ASSDNAETLY | 0.77068615 |
| RGGAGGAAGLI | ASSDNAETLY | 0.59266424 |
| RSYRMHPAV | ASSDNAETLY | 0.51745427 |
| SAAHLLAGQLA | ASSDNAETLY | 0.62874013 |
| SAAVGSY | ASSDNAETLY | 0.7844429 |

| | | |
|---|---|---|
| SPHAVLHRI | ASSDNAETLY | 0.8237687 |
| SQIAKVL | ASSDNAETLY | 0.7761165 |
| TALGHLVP | ASSDNAETLY | 0.5105365 |
| TAPTTSGAA | ASSDNAETLY | 0.79424345 |
| TDGDDGNGGSPGT | ASSDNAETLY | 0.74624777 |
| TDGLGEN | ASSDNAETLY | 0.69187766 |
| TGGPPLPGQA | ASSDNAETLY | 0.7399232 |
| TIPAITI | ASSDNAETLY | 0.7342997 |
| TIPGFTI | ASSDNAETLY | 0.5654894 |
| TSIASGALA | ASSDNAETLY | 0.57097864 |
| TVTLPGAAI | ASSDNAETLY | 0.7385186 |
| VASGLSAALP | ASSDNAETLY | 0.7951589 |
| VGAGVIGI | ASSDNAETLY | 0.79514503 |
| VGPKRFGV | ASSDNAETLY | 0.8443732 |
| VNVAFPD | ASSDNAETLY | 0.8054512 |
| VSMTNSGVS | ASSDNAETLY | 0.784714 |
| VTDVLGDLSV | ASSDNAETLY | 0.25527886 |
| VVVVGAGFAGL | ASSDNAETLY | 0.35028362 |
| VYTVGTVV | ASSDNAETLY | 0.6892165 |
| WTTFSGP | ASSDNAETLY | 0.5983083 |
| YDELVEAF | ASSDNAETLY | 0.2614277 |
| YGTGGAGGNGG | ASSDNAETLY | 0.7145189 |
| YTNVMAV | ASSDNAETLY | 0.6695485 |
| FLEQVHQL | ASSDNAETLY | 0.57455826 |
| GLSLLAVAV | ASSDNAETLY | 0.79074144 |
| LEQVHQL | ASSDNAETLY | 0.63923895 |
| LNAAGAI | ASSDNAETLY | 0.48633057 |
| PAVHASL | ASSDNAETLY | 0.6750959 |
| TELEDDA | ASSDNAETLY | 0.8016054 |
| TPPGPPN | ASSDNAETLY | 0.5868779 |
| TPPPPPPP | ASSDNAETLY | 0.8117688 |
| TQSPALI | ASSETNSDYT | 0.5880806 |
| EFAAAFPQ | ASSETNSDYT | 0.8333153 |
| LLSGNLDVL | ASSETNSDYT | 0.45330745 |
| GPANLGNYNI | ASSETNSDYT | 0.8091304 |
| TLGPTLP | ASSETNSDYT | 0.71837544 |
| VYAYSSCCK | ASSETNSDYT | 0.86327773 |
| APAPAPAPNI | ASSETNSDYT | 0.6598215 |
| PGGPVPIP | ASSETNSDYT | 0.7992902 |
| FLTVALEP | ASSETNSDYT | 0.76072305 |
| SQYFGANL | ASSETNSDYT | 0.8212291 |
| GGVKPAA | ASSETNSDYT | 0.5127115 |
| GGTGGAGGAGGW | ASSETNSDYT | 0.6894626 |
| GVGGVAGLGGAGG | ASSETNSDYT | 0.41641438 |
| VLLVLATTI | ASSETNSDYT | 0.73383296 |
| IGIFAAML | ASSETNSDYT | 0.8049656 |
| AVGDLID | ASSETNSDYT | 0.19782963 |
| NISPLPL | ASSETNSDYT | 0.82154197 |
| VTAVFVAI | ASSETNSDYT | 0.4464711 |
| SGDDNDAAL | ASSETNSDYT | 0.56011784 |
| AVAALAHS | ASSETNSDYT | 0.80050236 |

| | | |
|---|---|---|
| GGVTVMATP | ASSETNSDYT | 0.73499966 |
| AGLAANVRD | ASSETNSDYT | 0.8065933 |
| TNRPDLIDPALLRPG | ASSETNSDYT | 0.6403287 |
| LANLGVTAL | ASSETNSDYT | 0.7230592 |
| TGERIEL | ASSETNSDYT | 0.73601687 |
| NGEELAA | ASSETNSDYT | 0.4093616 |
| VLSLVWG | ASSETNSDYT | 0.56195146 |
| IGYALWL | ASSETNSDYT | 0.8375016 |
| LSLLLAPL | ASSETNSDYT | 0.7162702 |
| TLDEALK | ASSETNSDYT | 0.7186885 |
| TDGAGTAEGG | ASSETNSDYT | 0.68987805 |
| NAPGGAGGNGG | ASSETNSDYT | 0.6785298 |
| VSVFMGMGGNF | ASSETNSDYT | 0.70159054 |
| LASYLAAL | ASSETNSDYT | 0.7177342 |
| LVGGTPAL | ASSETNSDYT | 0.7218553 |
| VRNADVL | ASSETNSDYT | 0.71796024 |
| TCSSPFLVSAM | ASSETNSDYT | 0.67117816 |
| VHITDAE | ASSETNSDYT | 0.72062874 |
| EYAAPSF | ASSETNSDYT | 0.7111363 |
| VVDAFHGGL | ASSETNSDYT | 0.7088547 |
| ASVVDPQ | ASSETNSDYT | 0.7150098 |
| SYNEFGGLS | ASSETNSDYT | 0.70864725 |
| DPDGSVY | ASSETNSDYT | 0.6928467 |
| MVSTIHATE | ASSETNSDYT | 0.6871267 |
| LAAGVADPV | ASSETNSDYT | 0.7146232 |
| TKEQIAA | ASSETNSDYT | 0.7170025 |
| SLASPAAL | ASSETNSDYT | 0.7182723 |
| IGLVFYGV | ASSETNSDYT | 0.7242626 |
| FLLDASNL | ASSETNSDYT | 0.7202146 |
| YQVGNFP | ASSETNSDYT | 0.7162408 |
| ESLLSQST | ASSETNSDYT | 0.4789902 |
| LFTVGFAAVG | ASSETNSDYT | 0.6467613 |
| ATPPTPT | ASSETNSDYT | 0.6811478 |
| TDLATLG | ASSETNSDYT | 0.70385176 |
| SAGGTGR | ASSETNSDYT | 0.468994 |
| GFSNSGS | ASSETNSDYT | 0.7069551 |
| GGNGGSGGTSV | ASSETNSDYT | 0.70225406 |
| GTSVATGGAGN | ASSETNSDYT | 0.69113463 |
| NGGNGGSGGTS | ASSETNSDYT | 0.72056365 |
| AVVKFPH | ASSETNSDYT | 0.70286655 |
| TPPPPPP | ASSETNSDYT | 0.66804034 |
| AAPAAAAAPA | ASSETNSDYT | 0.6961681 |
| IALNEDL | ASSETNSDYT | 0.72209656 |
| IVNVSSM | ASSETNSDYT | 0.69010293 |
| TGIPPAPR | ASSETNSDYT | 0.72161376 |
| FDLGGGTF | ASSETNSDYT | 0.6710106 |
| FDLGGGTFDVS | ASSETNSDYT | 0.722731 |
| NEPTAAA | ASSETNSDYT | 0.65113926 |
| NEPTAAALA | ASSETNSDYT | 0.72104025 |
| GSIINISSI | ASSETNSDYT | 0.71087295 |
| VAPGNAGTA | ASSETNSDYT | 0.7261211 |

| | | |
|---|---|---|
| TGGGASLE | ASSETNSDYT | 0.70512104 |
| VAVYNHYKRI | ASSETNSDYT | 0.72000855 |
| TDPAPATT | ASSETNSDYT | 0.7016523 |
| TNRPDILDPALLRPG | ASSETNSDYT | 0.7210783 |
| LTTTTTS | ASSETNSDYT | 0.6610085 |
| PTGSGKTL | ASSETNSDYT | 0.70568085 |
| TGSGKTL | ASSETNSDYT | 0.6421454 |
| TIGHVDHGK | ASSETNSDYT | 0.7138531 |
| TIGHVDHGKTT | ASSETNSDYT | 0.7193161 |
| GHVDHGKTT | ASSETNSDYT | 0.45949122 |
| AQFEHTLL | ASSETNSDYT | 0.7221847 |
| IVNQDEL | ASSETNSDYT | 0.68414503 |
| TNGSQFF | ASSETNSDYT | 0.68277776 |
| TGKGIVE | ASSETNSDYT | 0.72107005 |
| LISNASDA | ASSETNSDYT | 0.7096151 |
| AELRTLGELTKIAKAHGAQ | ASSETNSDYT | 0.71760714 |
| APNAVAAAEPSW | ASSETNSDYT | 0.691105 |
| APQVAGAATAVL | ASSETNSDYT | 0.7083818 |
| ARRDPAAAAAILPTDA | ASSETNSDYT | 0.7137836 |
| DAAARDVLAEAGLAEYF | ASSETNSDYT | 0.62130815 |
| DDQIVLSAKANLAKVEQ | ASSETNSDYT | 0.6767906 |
| DELLPDTGEDSDEDGH | ASSETNSDYT | 0.72427213 |
| ELPEAPANRATVPQD | ASSETNSDYT | 0.28193805 |
| FGQSKIERVTGIG | ASSETNSDYT | 0.82775944 |
| GAGGTGSTAGGAGGAGGAGGLYA | ASSETNSDYT | 0.185258 |
| GHGGTGAAGGNGGTGGAGGSGIDG | ASSETNSDYT | 0.22171402 |
| GNGGNGGAGGEGGPGGAGGAGA | ASSETNSDYT | 0.30210277 |
| GSTAGNGGNGGNATNGTIAGQPAG | ASSETNSDYT | 0.38289618 |
| GTGGDGGNGGNGGTGG | ASSETNSDYT | 0.8043747 |
| GVRRPAGIPADAPV | ASSETNSDYT | 0.3279071 |
| GYRQVTDAHLLTLARR | ASSETNSDYT | 0.18572472 |
| IAKMGQDGHDRG | ASSETNSDYT | 0.2380941 |
| KAEPSIEIQYT | ASSETNSDYT | 0.17068824 |
| KPVRIRMAADEDTVAVT | ASSETNSDYT | 0.26729074 |
| MPPRIAGMRLLVIKPEP | ASSETNSDYT | 0.3259269 |
| MKFYAVALFRSRGVPPTR | ASSETNSDYT | 0.26535505 |
| NAGGFGILGAVAHSP | ASSETNSDYT | 0.17680553 |
| NEGREITLAMAEM | ASSETNSDYT | 0.21865778 |
| NELVEWSHPLGHRWRWE | ASSETNSDYT | 0.3365026 |
| NLGAGGTGGNGGNPG | ASSETNSDYT | 0.36884022 |
| NTEWPKWETVMAADPVD | ASSETNSDYT | 0.29878134 |
| PASAAPAETQP | ASSETNSDYT | 0.53015065 |
| QQSSDGKNGVRAAEITGE | ASSETNSDYT | 0.446697 |
| RAPDAVTTSWWKE | ASSETNSDYT | 0.36747938 |
| SGFGNVGAHTSGWFNQ | ASSETNSDYT | 0.11231359 |
| SGVAGAVGQAASVGGLKV | ASSETNSDYT | 0.15591212 |
| SNGGAGTGSAGGNGGT | ASSETNSDYT | 0.1692062 |
| TGGAGGEGGAAG | ASSETNSDYT | 0.27375424 |
| TGGDGGAAGAGGAPGHG | ASSETNSDYT | 0.11735942 |
| TMPPLLAAIANAGRHGV | ASSETNSDYT | 0.3753379 |
| TSSTAGNSGGA | ASSETNSDYT | 0.19767037 |

| | | |
|---|---|---|
| VPATVPVGILC | ASSETNSDYT | 0.20754966 |
| VQRPDSGGVVLTGRIS | ASSETNSDYT | 0.50725263 |
| WTPNPVPATKGVPAGLVQAT | ASSETNSDYT | 0.19805947 |
| YALKRALAKFSDRFDVVII | ASSETNSDYT | 0.22169492 |
| YLIGSVIV | ASSETNSDYT | 0.16896641 |
| YGQLVAVID | ASSETNSDYT | 0.48949897 |
| VTVGGMTLPA | ASSETNSDYT | 0.12820907 |
| VLALPAV | ASSETNSDYT | 0.37589824 |
| VIPPTLNYET | ASSETNSDYT | 0.16135766 |
| VGMGGMHPSSG | ASSETNSDYT | 0.36546692 |
| VGLAGGAATA | ASSETNSDYT | 0.16445954 |
| VGGAGGAGGSGP | ASSETNSDYT | 0.4279968 |
| VFAAFMV | ASSETNSDYT | 0.24504869 |
| TLYDKLR | ASSETNSDYT | 0.8307725 |
| TLAVFTVL | ASSETNSDYT | 0.30746862 |
| TGSVQPARV | ASSETNSDYT | 0.3329857 |
| TGGGAGSGGT | ASSETNSDYT | 0.20788026 |
| SHLETSL | ASSETNSDYT | 0.8148204 |
| SFTTGMT | ASSETNSDYT | 0.13684069 |
| QVEVAKRMGVRQPTVS | ASSETNSDYT | 0.28390896 |
| QGSGGPALAM | ASSETNSDYT | 0.14223237 |
| MLAPSNP | ASSETNSDYT | 0.3280301 |
| LVNNAAI | ASSETNSDYT | 0.37857494 |
| LTGEINTLA | ASSETNSDYT | 0.15796256 |
| LSIFGPV | ASSETNSDYT | 0.31637788 |
| LLALLANL | ASSETNSDYT | 0.28710634 |
| LISNASDALDKLR | ASSETNSDYT | 0.17705403 |
| LINEAVP | ASSETNSDYT | 0.51330477 |
| LGYAFSGL | ASSETNSDYT | 0.21735385 |
| LDFVAVAL | ASSETNSDYT | 0.67515934 |
| LAQPGAE | ASSETNSDYT | 0.2099445 |
| ISNLGAVTGF | ASSETNSDYT | 0.19919536 |
| HQEDPGRI | ASSETNSDYT | 0.30792844 |
| GTPGEPTP | ASSETNSDYT | 0.16818455 |
| GFFNSGS | ASSETNSDYT | 0.16148868 |
| GAGGGGLTAGAN | ASSETNSDYT | 0.5537092 |
| FVDALVNM | ASSETNSDYT | 0.8071692 |
| FLTVAPEL | ASSETNSDYT | 0.7545202 |
| FETLTAL | ASSETNSDYT | 0.71693593 |
| DVINPIEL | ASSETNSDYT | 0.6083436 |
| APVSGAPV | ASSETNSDYT | 0.71530735 |
| ALVGGLAW | ASSETNSDYT | 0.7939217 |
| ALAAAAATAI | ASSETNSDYT | 0.7768028 |
| AFGLGLLL | ASSETNSDYT | 0.7771853 |
| AFFAAASL | ASSETNSDYT | 0.52452284 |
| AAAALPAFSPP | ASSETNSDYT | 0.7804935 |
| VVLAPAL | ASSETNSDYT | 0.75461787 |
| TIGHVDHGKTTL | ASSETNSDYT | 0.61211276 |
| LISNASDALDK | ASSETNSDYT | 0.7768855 |
| LISNASDAL | ASSETNSDYT | 0.67784977 |
| ISNASDALDK | ASSETNSDYT | 0.67381203 |

| | | |
|---|---|---|
| NQAETLVYQTE | ASSETNSDYT | 0.7270324 |
| TGGTGGTAGAG | ASSETNSDYT | 0.6242664 |
| TGSTAGGAGGA | ASSETNSDYT | 0.77327275 |
| DPLAVPVAAGPV | ASSETNSDYT | 0.74683 |
| GGTGGTPGNGGN | ASSETNSDYT | 0.8199513 |
| PAGLAGVDPATL | ASSETNSDYT | 0.8086106 |
| GNNLVVEAPHAGASV | ASSETNSDYT | 0.7644292 |
| KVTRIMSASTLEQPA | ASSETNSDYT | 0.27715564 |
| LGFALPIALPRRTVV | ASSETNSDYT | 0.45567894 |
| TRLGNLTARKLVVLV | ASSETNSDYT | 0.7250886 |
| WSQQRRLIYFPSAGP | ASSETNSDYT | 0.6665901 |
| GPNGNPDPMAAAVDIR | ASSETNSDYT | 0.28390345 |
| TQRCVSVADLRAAASR | ASSETNSDYT | 0.6942078 |
| QFVLDVAEHPVEVLTETV | ASSETNSDYT | 0.7636634 |
| GGAGGTGGAGGNGGTQAGDG | ASSETNSDYT | 0.5714227 |
| PLLATLRAIAVDVDAKPAQV | ASSETNSDYT | 0.7485189 |
| APKFRTGGYMIDFWGVGYQVAKRM | ASSETNSDYT | 0.5973662 |
| GAGGAGGAGGAAGGPGGTGGTGGN | ASSETNSDYT | 0.49741334 |
| IGALGLLATDVAPAGSVAFGVALA | ASSETNSDYT | 0.6596327 |
| AGGAGGIGGNGGAGGAGGVLMGNG | ASSETNSDYT | 0.8032606 |
| ADELIGLL | ASSETNSDYT | 0.6184815 |
| ADRALKA | ASSETNSDYT | 0.79808706 |
| AGAVGQVGGAGGA | ASSETNSDYT | 0.59882766 |
| AGGGGGLGGAP | ASSETNSDYT | 0.7967051 |
| AGGLLYGNGGAG | ASSETNSDYT | 0.4831375 |
| AGLESIGL | ASSETNSDYT | 0.7862759 |
| AGLKTGII | ASSETNSDYT | 0.750366 |
| ASLAALLL | ASSETNSDYT | 0.80519485 |
| ASVINISSI | ASSETNSDYT | 0.67803836 |
| ATPPGPA | ASSETNSDYT | 0.75723875 |
| ATVASALV | ASSETNSDYT | 0.5887269 |
| ATVASLIL | ASSETNSDYT | 0.77908033 |
| AVNGPVLIRH | ASSETNSDYT | 0.60682875 |
| DGAAGAVGI | ASSETNSDYT | 0.7824712 |
| DPLLHGI | ASSETNSDYT | 0.44284105 |
| ELGAALGAL | ASSETNSDYT | 0.79125655 |
| ELGRTQAV | ASSETNSDYT | 0.8050587 |
| FVAGGYLS | ASSETNSDYT | 0.417598 |
| GASGGAGGAGGW | ASSETNSDYT | 0.8238621 |
| GGAGAGGDGGAGGI | ASSETNSDYT | 0.681346 |
| GTGHNGSQPGMGGNG | ASSETNSDYT | 0.5811111 |
| GYAAASASAS | ASSETNSDYT | 0.80229616 |
| HQVDVAAVQQG | ASSETNSDYT | 0.6441554 |
| IGAAIAHT | ASSETNSDYT | 0.8092116 |
| IGMIEVAQL | ASSETNSDYT | 0.6110709 |
| IRLAPEHTSAMD | ASSETNSDYT | 0.7591196 |
| IYNVSVL | ASSETNSDYT | 0.80532527 |
| KQGFLGM | ASSETNSDYT | 0.5605575 |
| LESVITQL | ASSETNSDYT | 0.47036242 |
| LGLGALVVV | ASSETNSDYT | 0.8036371 |
| LIGNGGNGGSGGI | ASSETNSDYT | 0.73426104 |

| | | |
|---|---|---|
| LNAINAPTL | ASSETNSDYT | 0.7590821 |
| LQLLNTGV | ASSETNSDYT | 0.6521671 |
| LQTSIEF | ASSETNSDYT | 0.57596076 |
| MSFAVSEL | ASSETNSDYT | 0.6608937 |
| NTGALGL | ASSETNSDYT | 0.74826837 |
| PPGAGGAGGAGTT | ASSETNSDYT | 0.7639844 |
| QLVGLGI | ASSETNSDYT | 0.73004746 |
| RFLEQVHQL | ASSETNSDYT | 0.52036035 |
| RGVGYVLR | ASSETNSDYT | 0.7452328 |
| SATGVFAGVI | ASSETNSDYT | 0.70789236 |
| SLPAITLP | ASSETNSDYT | 0.6737167 |
| STALAFTV | ASSETNSDYT | 0.6952267 |
| SVLTVGLGK | ASSETNSDYT | 0.72010225 |
| TALAGAGSGI | ASSETNSDYT | 0.6273794 |
| TGLPFTGI | ASSETNSDYT | 0.6133511 |
| VAEHVTV | ASSETNSDYT | 0.7002338 |
| VDIVLNSL | ASSETNSDYT | 0.7377145 |
| VLTGGGAVL | ASSETNSDYT | 0.7436739 |
| VNVGLAYGI | ASSETNSDYT | 0.7670692 |
| VSSAAVGAV | ASSETNSDYT | 0.767455 |
| VTSVYEAGV | ASSETNSDYT | 0.72306156 |
| WVALDRGA | ASSETNSDYT | 0.2909665 |
| YLVGSVVI | ASSETNSDYT | 0.42804232 |
| GPPGTGKTLL | ASSETNSDYT | 0.5869127 |
| GVGMTKF | ASSETNSDYT | 0.67296684 |
| IGIGGSDL | ASSETNSDYT | 0.3133512 |
| IGTLLAEL | ASSETNSDYT | 0.70466995 |
| IYTSGTT | ASSETNSDYT | 0.6545714 |
| LSFPAGV | ASSETNSDYT | 0.6197815 |
| LYTSGTT | ASSETNSDYT | 0.75273293 |
| VGAFPFGF | ASSETNSDYT | 0.6533067 |
| VGAGTVEF | ASSETNSDYT | 0.57697254 |
| VGKSTLLL | ASSETNSDYT | 0.57446104 |
| AALAHAGGGIV | ASSETNSDYT | 0.7410512 |
| AALVAVSLL | ASSETNSDYT | 0.4828782 |
| AAVSRAVAAL | ASSETNSDYT | 0.7568493 |
| AGGSSGIGGF | ASSETNSDYT | 0.4543152 |
| AGLTVANANG | ASSETNSDYT | 0.76961076 |
| AGLVGAVL | ASSETNSDYT | 0.48588073 |
| ALSQAFTV | ASSETNSDYT | 0.74912554 |
| ALTFSVSL | ASSETNSDYT | 0.6947282 |
| ALTGAALVAL | ASSETNSDYT | 0.7884835 |
| ASFSIGAGVL | ASSETNSDYT | 0.68154144 |
| DGSVYKI | ASSETNSDYT | 0.7408378 |
| EQLAEIV | ASSETNSDYT | 0.5858224 |
| FAVSELTVM | ASSETNSDYT | 0.757145 |
| GAAPLAAGL | ASSETNSDYT | 0.51978767 |
| GADGGDGGSGGAGG | ASSETNSDYT | 0.69291687 |
| GFGLDTGGAGG | ASSETNSDYT | 0.39929295 |
| GLASGGLL | ASSETNSDYT | 0.7167174 |
| GRIAGAAVLL | ASSETNSDYT | 0.7377207 |

| | | |
|---|---|---|
| GSAGSLNTSF | ASSETNSDYT | 0.41933084 |
| GTGQIGI | ASSETNSDYT | 0.7588787 |
| GTLIALIL | ASSETNSDYT | 0.65621185 |
| IAATMGFVVG | ASSETNSDYT | 0.59014255 |
| IFAQALL | ASSETNSDYT | 0.75719506 |
| IGAAVALGF | ASSETNSDYT | 0.7018416 |
| IGLTPLAL | ASSETNSDYT | 0.7597647 |
| ILGSNTSTL | ASSETNSDYT | 0.5851555 |
| IVFSLTVL | ASSETNSDYT | 0.6514809 |
| LASKATGF | ASSETNSDYT | 0.7093418 |
| LASVALIL | ASSETNSDYT | 0.61531377 |
| LFAVGLTL | ASSETNSDYT | 0.5449941 |
| LFNGGIGTYI | ASSETNSDYT | 0.76389235 |
| LTLGAGII | ASSETNSDYT | 0.5105047 |
| LVAAALGDI | ASSETNSDYT | 0.60779804 |
| NTAFVQLGI | ASSETNSDYT | 0.32946712 |
| QLFSGAEL | ASSETNSDYT | 0.33755383 |
| SAATFITL | ASSETNSDYT | 0.39187512 |
| SNYGLANLGV | ASSETNSDYT | 0.5302627 |
| SYGGAYAVM | ASSETNSDYT | 0.57535183 |
| TTGVGGLTLG | ASSETNSDYT | 0.5078817 |
| TTQTIYI | ASSETNSDYT | 0.34830537 |
| VAAGAGLGAVI | ASSETNSDYT | 0.49537316 |
| VAAVVTLL | ASSETNSDYT | 0.39988288 |
| VGFLGGLI | ASSETNSDYT | 0.3412215 |
| VGVIPILL | ASSETNSDYT | 0.45757315 |
| VIFASSLI | ASSETNSDYT | 0.4433841 |
| VKGGDGGAAAG | ASSETNSDYT | 0.35632032 |
| VTKTFTNI | ASSETNSDYT | 0.4370508 |
| VVAAAILL | ASSETNSDYT | 0.41069266 |
| YAALDVEL | ASSETNSDYT | 0.49702525 |
| LASVALLL | ASSETNSDYT | 0.45299533 |
| LDAVHRV | ASSETNSDYT | 0.5966674 |
| QGLSSSIF | ASSETNSDYT | 0.54951614 |
| RGTFANIRL | ASSETNSDYT | 0.4699468 |
| RQLWWGHRI | ASSETNSDYT | 0.22380014 |
| YGPPGTGKTLL | ASSETNSDYT | 0.29658636 |
| AALVGAPANVA | ASSETNSDYT | 0.41866323 |
| AATFRRTE | ASSETNSDYT | 0.37809426 |
| AGALGGGI | ASSETNSDYT | 0.25942862 |
| AGGAGGAGGIGAG | ASSETNSDYT | 0.47533837 |
| AGSGALGL | ASSETNSDYT | 0.45036 |
| AQIADQI | ASSETNSDYT | 0.32703912 |
| ASAVAVTYL | ASSETNSDYT | 0.47369325 |
| ASTVLGGTGG | ASSETNSDYT | 0.30811745 |
| AVGGEPAK | ASSETNSDYT | 0.33579454 |
| CEQGTFGGTC | ASSETNSDYT | 0.37617293 |
| DEGKAPAG | ASSETNSDYT | 0.5299654 |
| DTVVFPAL | ASSETNSDYT | 0.3389202 |
| FEGITMAG | ASSETNSDYT | 0.4863589 |
| FPLVVQD | ASSETNSDYT | 0.39906198 |

| | | |
|---|---|---|
| GAFTTPQ | ASSETNSDYT | 0.57149935 |
| GFGGFGTAGGVGGA | ASSETNSDYT | 0.3116407 |
| GFLAQFL | ASSETNSDYT | 0.49050066 |
| GGAGGAGGNAGVA | ASSETNSDYT | 0.39513615 |
| GGVGTAGAGGAGGAGG | ASSETNSDYT | 0.5998132 |
| GNGGGGDAGNAGSGGN | ASSETNSDYT | 0.38363913 |
| GTGGAGGAGNDAGS | ASSETNSDYT | 0.47366565 |
| GTTNTGWFNS | ASSETNSDYT | 0.40760833 |
| IFMGPLFTL | ASSETNSDYT | 0.6080845 |
| ILAHAIEAI | ASSETNSDYT | 0.27618477 |
| IRSAIREAL | ASSETNSDYT | 0.418577 |
| ITFAGGG | ASSETNSDYT | 0.31920594 |
| ITGMSGHQEV | ASSETNSDYT | 0.45482635 |
| IVAGLACTL | ASSETNSDYT | 0.5303326 |
| IVGGGLAGA | ASSETNSDYT | 0.2750701 |
| IVTGVLNI | ASSETNSDYT | 0.57697797 |
| LAQQGCI | ASSETNSDYT | 0.35701454 |
| LCLHVEN | ASSETNSDYT | 0.34834644 |
| LDAISDGS | ASSETNSDYT | 0.50153095 |
| LGGFAAML | ASSETNSDYT | 0.38051343 |
| LGGKNPMI | ASSETNSDYT | 0.57932985 |
| LGPKGVAAIL | ASSETNSDYT | 0.3727733 |
| LMVGVSI | ASSETNSDYT | 0.43425813 |
| LVGAAFVL | ASSETNSDYT | 0.49103028 |
| LVLAVAPL | ASSETNSDYT | 0.31065434 |
| MAAIAVIDAA | ASSETNSDYT | 0.315539 |
| NNGINAELQ | ASSETNSDYT | 0.5604008 |
| RGEPLVLVHG | ASSETNSDYT | 0.45261112 |
| RGGAGGAAGLI | ASSETNSDYT | 0.65393466 |
| RSYRMHPAV | ASSETNSDYT | 0.31240627 |
| SAAHLLAGQLA | ASSETNSDYT | 0.30764455 |
| SAAVGSY | ASSETNSDYT | 0.3159543 |
| SPHAVLHRI | ASSETNSDYT | 0.40234607 |
| SQIAKVL | ASSETNSDYT | 0.65808254 |
| TALGHLVP | ASSETNSDYT | 0.51494324 |
| TAPTTSGAA | ASSETNSDYT | 0.3163631 |
| TDGDDGNGGSPGT | ASSETNSDYT | 0.3751402 |
| TDGLGEN | ASSETNSDYT | 0.32111174 |
| TGGPPLPGQA | ASSETNSDYT | 0.4566291 |
| TIPAITI | ASSETNSDYT | 0.36008316 |
| TIPGFTI | ASSETNSDYT | 0.33705127 |
| TSIASGALA | ASSETNSDYT | 0.36865497 |
| TVTLPGAAI | ASSETNSDYT | 0.30154672 |
| VASGLSAALP | ASSETNSDYT | 0.3286053 |
| VGAGVIGI | ASSETNSDYT | 0.5506991 |
| VGPKRFGV | ASSETNSDYT | 0.49063304 |
| VNVAFPD | ASSETNSDYT | 0.4141951 |
| VSMTNSGVS | ASSETNSDYT | 0.5707581 |
| VTDVLGDLSV | ASSETNSDYT | 0.3438839 |
| VVVVGAGFAGL | ASSETNSDYT | 0.2544088 |
| VYTVGTVV | ASSETNSDYT | 0.30376038 |

| | | |
|---|---|---|
| WTTFSGP | ASSETNSDYT | 0.31025514 |
| YDELVEAF | ASSETNSDYT | 0.3647511 |
| YGTGGAGGNGG | ASSETNSDYT | 0.25913057 |
| YTNVMAV | ASSETNSDYT | 0.35754722 |
| FLEQVHQL | ASSETNSDYT | 0.32927638 |
| GLSLLAVAV | ASSETNSDYT | 0.35741666 |
| LEQVHQL | ASSETNSDYT | 0.50954616 |
| LNAAGAI | ASSETNSDYT | 0.366286 |
| PAVHASL | ASSETNSDYT | 0.30746606 |
| TELEDDA | ASSETNSDYT | 0.3014319 |
| TPPGPPN | ASSETNSDYT | 0.5489613 |
| TPPPPPPP | ASSETNSDYT | 0.28587788 |
| TQSPALI | ASSDPGANSDYT | 0.5139159 |
| EFAAAFPQ | ASSDPGANSDYT | 0.31348073 |
| LLSGNLDVL | ASSDPGANSDYT | 0.4669928 |
| GPANLGNYNI | ASSDPGANSDYT | 0.2920704 |
| TLGPTLP | ASSDPGANSDYT | 0.52192265 |
| VYAYSSCCK | ASSDPGANSDYT | 0.4078501 |
| APAPAPAPNI | ASSDPGANSDYT | 0.6135888 |
| PGGPVPIP | ASSDPGANSDYT | 0.3300827 |
| FLTVALEP | ASSDPGANSDYT | 0.371778 |
| SQYFGANL | ASSDPGANSDYT | 0.3077086 |
| GGVKPAA | ASSDPGANSDYT | 0.66309655 |
| GGTGGAGGAGGW | ASSDPGANSDYT | 0.2843745 |
| GVGGVAGLGGAGG | ASSDPGANSDYT | 0.31369147 |
| VLLVLATTI | ASSDPGANSDYT | 0.29861537 |
| IGIFAAML | ASSDPGANSDYT | 0.31998274 |
| AVGDLID | ASSDPGANSDYT | 0.53366095 |
| NISPLPL | ASSDPGANSDYT | 0.2604514 |
| VTAVFVAI | ASSDPGANSDYT | 0.47578037 |
| SGDDNDAAL | ASSDPGANSDYT | 0.30031642 |
| AVAALAHS | ASSDPGANSDYT | 0.30048594 |
| GGVTVMATP | ASSDPGANSDYT | 0.39292735 |
| AGLAANVRD | ASSDPGANSDYT | 0.31210417 |
| TNRPDLIDPALLRPG | ASSDPGANSDYT | 0.47230715 |
| LANLGVTAL | ASSDPGANSDYT | 0.3000131 |
| TGERIEL | ASSDPGANSDYT | 0.37293175 |
| NGEELAA | ASSDPGANSDYT | 0.36460105 |
| VLSLVWG | ASSDPGANSDYT | 0.28517303 |
| IGYALWL | ASSDPGANSDYT | 0.29334095 |
| LSLLLAPL | ASSDPGANSDYT | 0.6095306 |
| TLDEALK | ASSDPGANSDYT | 0.63571227 |
| TDGAGTAEGG | ASSDPGANSDYT | 0.7569201 |
| NAPGGAGGNGG | ASSDPGANSDYT | 0.23619357 |
| VSVFMGMGGNF | ASSDPGANSDYT | 0.22484793 |
| LASYLAAL | ASSDPGANSDYT | 0.36100549 |
| LVGGTPAL | ASSDPGANSDYT | 0.6844816 |
| VRNADVL | ASSDPGANSDYT | 0.8048206 |
| TCSSPFLVSAM | ASSDPGANSDYT | 0.6449256 |
| VHITDAE | ASSDPGANSDYT | 0.27038342 |
| EYAAPSF | ASSDPGANSDYT | 0.5680988 |

| | | |
|---|---|---|
| VVDAFHGGL | ASSDPGANSDYT | 0.3750863 |
| ASVVDPQ | ASSDPGANSDYT | 0.44281152 |
| SYNEFGGLS | ASSDPGANSDYT | 0.5911813 |
| DPDGSVY | ASSDPGANSDYT | 0.54789996 |
| MVSTIHATE | ASSDPGANSDYT | 0.3691621 |
| LAAGVADPV | ASSDPGANSDYT | 0.26874045 |
| TKEQIAA | ASSDPGANSDYT | 0.67632747 |
| SLASPAAL | ASSDPGANSDYT | 0.75154406 |
| IGLVFYGV | ASSDPGANSDYT | 0.7286701 |
| FLLDASNL | ASSDPGANSDYT | 0.8060165 |
| YQVGNFP | ASSDPGANSDYT | 0.7131854 |
| ESLLSQST | ASSDPGANSDYT | 0.7077347 |
| LFTVGFAAVG | ASSDPGANSDYT | 0.118864276 |
| ATPPTPT | ASSDPGANSDYT | 0.16569884 |
| TDLATLG | ASSDPGANSDYT | 0.38463104 |
| SAGGTGR | ASSDPGANSDYT | 0.3776747 |
| GFSNSGS | ASSDPGANSDYT | 0.17087081 |
| GGNGGSGGTSV | ASSDPGANSDYT | 0.6279226 |
| GTSVATGGAGN | ASSDPGANSDYT | 0.32353875 |
| NGGNGGSGGTS | ASSDPGANSDYT | 0.3072342 |
| AVVKFPH | ASSDPGANSDYT | 0.73824215 |
| TPPPPPP | ASSDPGANSDYT | 0.29087242 |
| AAPAAAAAPA | ASSDPGANSDYT | 0.28586242 |
| IALNEDL | ASSDPGANSDYT | 0.3783764 |
| IVNVSSM | ASSDPGANSDYT | 0.7045127 |
| TGIPPAPR | ASSDPGANSDYT | 0.273813 |
| FDLGGGTF | ASSDPGANSDYT | 0.64902544 |
| FDLGGGTFDVS | ASSDPGANSDYT | 0.41942286 |
| NEPTAAA | ASSDPGANSDYT | 0.75335175 |
| NEPTAAALA | ASSDPGANSDYT | 0.22353256 |
| GSIINISSI | ASSDPGANSDYT | 0.6971365 |
| VAPGNAGTA | ASSDPGANSDYT | 0.45309988 |
| TGGGASLE | ASSDPGANSDYT | 0.84596777 |
| VAVYNHYKRI | ASSDPGANSDYT | 0.32292178 |
| TDPAPATT | ASSDPGANSDYT | 0.49981034 |
| TNRPDILDPALLRPG | ASSDPGANSDYT | 0.4423477 |
| LTTTTTS | ASSDPGANSDYT | 0.79699576 |
| PTGSGKTL | ASSDPGANSDYT | 0.19466601 |
| TGSGKTL | ASSDPGANSDYT | 0.31121323 |
| TIGHVDHGK | ASSDPGANSDYT | 0.19528793 |
| TIGHVDHGKTT | ASSDPGANSDYT | 0.40023634 |
| GHVDHGKTT | ASSDPGANSDYT | 0.60549664 |
| AQFEHTLL | ASSDPGANSDYT | 0.12190325 |
| IVNQDEL | ASSDPGANSDYT | 0.57699096 |
| TNGSQFF | ASSDPGANSDYT | 0.36280274 |
| TGKGIVE | ASSDPGANSDYT | 0.2502443 |
| LISNASDA | ASSDPGANSDYT | 0.7257298 |
| AELRTLGELTKIAKAHGAQ | ASSDPGANSDYT | 0.5617279 |
| APNAVAAAEPSW | ASSDPGANSDYT | 0.74612373 |
| APQVAGAATAVL | ASSDPGANSDYT | 0.528021 |
| ARRDPAAAAAILPTDA | ASSDPGANSDYT | 0.3855236 |

| | | |
|---|---|---|
| DAAARDVLAEAGLAEYF | ASSDPGANSDYT | 0.452458 |
| DDQIVLSAKANLAKVEQ | ASSDPGANSDYT | 0.22264889 |
| DELLPDTGEDSDEDGH | ASSDPGANSDYT | 0.36192554 |
| ELPEAPANRATVPQD | ASSDPGANSDYT | 0.752712 |
| FGQSKIERVTGIG | ASSDPGANSDYT | 0.56734395 |
| GAGGTGSTAGGAGGAGGAGGLYA | ASSDPGANSDYT | 0.7688517 |
| GHGGTGAAGGNGGTGGAGGSGIDG | ASSDPGANSDYT | 0.35022178 |
| GNGGNGGAGGEGGPGGAGGAGA | ASSDPGANSDYT | 0.29685712 |
| GSTAGNGGNGGNATNGTIAGGQPAG | ASSDPGANSDYT | 0.48447347 |
| GTGGDGGNGGNGGTGG | ASSDPGANSDYT | 0.7011262 |
| GVRRPAGIPADAPV | ASSDPGANSDYT | 0.7934268 |
| GYRQVTDAHLLTLARR | ASSDPGANSDYT | 0.69651055 |
| IAKMGQDGHDRG | ASSDPGANSDYT | 0.32676807 |
| KAEPSIEIQYT | ASSDPGANSDYT | 0.5569627 |
| KPVRIRMAADEDTVAVT | ASSDPGANSDYT | 0.4360842 |
| MPPRIAGMRLLVIKPEP | ASSDPGANSDYT | 0.44268844 |
| MKFYAVALFRSRGVPPTR | ASSDPGANSDYT | 0.5141505 |
| NAGGFGILGAVAHSP | ASSDPGANSDYT | 0.5700786 |
| NEGREITLAMAEM | ASSDPGANSDYT | 0.40486157 |
| NELVEWSHPLGHRWRWE | ASSDPGANSDYT | 0.3213334 |
| NLGAGGTGGNGGNPG | ASSDPGANSDYT | 0.5679467 |
| NTEWPKWETVMAADPVD | ASSDPGANSDYT | 0.6937648 |
| PASAAPAETQP | ASSDPGANSDYT | 0.633638 |
| QQSSDGKNGVRAAEITGE | ASSDPGANSDYT | 0.7972022 |
| RAPDAVTTSWWKE | ASSDPGANSDYT | 0.691764 |
| SGFGNVGAHTSGWFNQ | ASSDPGANSDYT | 0.53560764 |
| SGVAGAVGQAASVGGLKV | ASSDPGANSDYT | 0.12802319 |
| SNGGAGTGSAGGNGGT | ASSDPGANSDYT | 0.19357637 |
| TGGAGGEGGAAG | ASSDPGANSDYT | 0.38167095 |
| TGGDGGAAGAGGAPGHG | ASSDPGANSDYT | 0.4767515 |
| TMPPLLAAIANAGRHGV | ASSDPGANSDYT | 0.14408118 |
| TSSTAGNSGGA | ASSDPGANSDYT | 0.6447021 |
| VPATVPVGILC | ASSDPGANSDYT | 0.42372805 |
| VQRPDSGGVVLTGRIS | ASSDPGANSDYT | 0.36244744 |
| WTPNPVPATKGVPAGLVQAT | ASSDPGANSDYT | 0.7245115 |
| YALKRALAKFSDRFDVVII | ASSDPGANSDYT | 0.39083475 |
| YLIGSVIV | ASSDPGANSDYT | 0.33372498 |
| YGQLVAVID | ASSDPGANSDYT | 0.40834674 |
| VTVGGMTLPA | ASSDPGANSDYT | 0.78871596 |
| VLALPAV | ASSDPGANSDYT | 0.2516461 |
| VIPPTLNYET | ASSDPGANSDYT | 0.6831629 |
| VGMGGMHPSSG | ASSDPGANSDYT | 0.43833694 |
| VGLAGGAATA | ASSDPGANSDYT | 0.723348 |
| VGGAGGAGGSGP | ASSDPGANSDYT | 0.24944918 |
| VFAAFMV | ASSDPGANSDYT | 0.66801274 |
| TLYDKLR | ASSDPGANSDYT | 0.45632517 |
| TLAVFTVL | ASSDPGANSDYT | 0.8386569 |
| TGSVQPARV | ASSDPGANSDYT | 0.35932767 |
| TGGGAGSGGT | ASSDPGANSDYT | 0.5694697 |
| SHLETSL | ASSDPGANSDYT | 0.38759 |
| SFTTGMT | ASSDPGANSDYT | 0.768687 |

| | | |
|---|---|---|
| QVEVAKRMGVRQPTVS | ASSDPGANSDYT | 0.22781 |
| QGSGGPALAM | ASSDPGANSDYT | 0.39883038 |
| MLAPSNP | ASSDPGANSDYT | 0.22825962 |
| LVNNAAI | ASSDPGANSDYT | 0.4840831 |
| LTGEINTLA | ASSDPGANSDYT | 0.66375387 |
| LSIFGPV | ASSDPGANSDYT | 0.15484838 |
| LLALLANL | ASSDPGANSDYT | 0.69979703 |
| LISNASDALDKLR | ASSDPGANSDYT | 0.38694155 |
| LINEAVP | ASSDPGANSDYT | 0.31202948 |
| LGYAFSGL | ASSDPGANSDYT | 0.66151494 |
| LDFVAVAL | ASSDPGANSDYT | 0.4502911 |
| LAQPGAE | ASSDPGANSDYT | 0.7477187 |
| ISNLGAVTGF | ASSDPGANSDYT | 0.35132974 |
| HQEDPGRI | ASSDPGANSDYT | 0.5445037 |
| GTPGEPTP | ASSDPGANSDYT | 0.5884874 |
| GFFNSGS | ASSDPGANSDYT | 0.2671815 |
| GAGGGGLTAGAN | ASSDPGANSDYT | 0.3624279 |
| FVDALVNM | ASSDPGANSDYT | 0.7261378 |
| FLTVAPEL | ASSDPGANSDYT | 0.6393907 |
| FETLTAL | ASSDPGANSDYT | 0.64479256 |
| DVINPIEL | ASSDPGANSDYT | 0.47110778 |
| APVSGAPV | ASSDPGANSDYT | 0.51818675 |
| ALVGGLAW | ASSDPGANSDYT | 0.44426215 |
| ALAAAAATAI | ASSDPGANSDYT | 0.6135242 |
| AFGLGLLL | ASSDPGANSDYT | 0.66362095 |
| AFFAAASL | ASSDPGANSDYT | 0.63797957 |
| AAAALPAFSPP | ASSDPGANSDYT | 0.4990192 |
| VVLAPAL | ASSDPGANSDYT | 0.6336505 |
| TIGHVDHGKTTL | ASSDPGANSDYT | 0.57857037 |
| LISNASDALDK | ASSDPGANSDYT | 0.5864245 |
| LISNASDAL | ASSDPGANSDYT | 0.5681373 |
| ISNASDALDK | ASSDPGANSDYT | 0.5822459 |
| NQAETLVYQTE | ASSDPGANSDYT | 0.4447552 |
| TGGTGGTAGAG | ASSDPGANSDYT | 0.43143728 |
| TGSTAGGAGGA | ASSDPGANSDYT | 0.5355525 |
| DPLAVPVAAGPV | ASSDPGANSDYT | 0.6387723 |
| GGTGGTPGNGGN | ASSDPGANSDYT | 0.6318641 |
| PAGLAGVDPATL | ASSDPGANSDYT | 0.64777523 |
| GNNLVVEAPHAGASV | ASSDPGANSDYT | 0.64367485 |
| KVTRIMSASTLEQPA | ASSDPGANSDYT | 0.6170456 |
| LGFALPIALPRRTVV | ASSDPGANSDYT | 0.326252 |
| TRLGNLTARKLVVLV | ASSDPGANSDYT | 0.3734552 |
| WSQQRRLIYFPSAGP | ASSDPGANSDYT | 0.43745536 |
| GPNGNPDPMAAAVDIR | ASSDPGANSDYT | 0.5211402 |
| TQRCVSVADLRAAASR | ASSDPGANSDYT | 0.30396134 |
| QFVLDVAEHPVEVLTETV | ASSDPGANSDYT | 0.59134984 |
| GGAGGTGGAGGNGGTQAGDG | ASSDPGANSDYT | 0.5113482 |
| PLLATLRAIAVDVDAKPAQV | ASSDPGANSDYT | 0.53629243 |
| APKFRTGGYMIDFWGVGYQVAKRM | ASSDPGANSDYT | 0.61805254 |
| GAGGAGGAGGAAGGPGGTGGTGGN | ASSDPGANSDYT | 0.53879535 |
| IGALGLLATDVAPAGSVAFGVALA | ASSDPGANSDYT | 0.42070147 |

| | | |
|---|---|---|
| AGGAGGIGGNGGAGGAGGVLMGNG | ASSDPGANSDYT | 0.46408966 |
| ADELIGLL | ASSDPGANSDYT | 0.64317065 |
| ADRALKA | ASSDPGANSDYT | 0.43715292 |
| AGAVGQVGGAGGA | ASSDPGANSDYT | 0.6566618 |
| AGGGGGLGGAP | ASSDPGANSDYT | 0.37724456 |
| AGGLLYGNGGAG | ASSDPGANSDYT | 0.645045 |
| AGLESIGL | ASSDPGANSDYT | 0.47160292 |
| AGLKTGII | ASSDPGANSDYT | 0.6479907 |
| ASLAALLL | ASSDPGANSDYT | 0.613831 |
| ASVINISSI | ASSDPGANSDYT | 0.66885936 |
| ATPPGPA | ASSDPGANSDYT | 0.547035 |
| ATVASALV | ASSDPGANSDYT | 0.6457032 |
| ATVASLIL | ASSDPGANSDYT | 0.55671084 |
| AVNGPVLIRH | ASSDPGANSDYT | 0.63773245 |
| DGAAGAVGI | ASSDPGANSDYT | 0.45146137 |
| DPLLHGI | ASSDPGANSDYT | 0.5093715 |
| ELGAALGAL | ASSDPGANSDYT | 0.37948292 |
| ELGRTQAV | ASSDPGANSDYT | 0.5592989 |
| FVAGGYLS | ASSDPGANSDYT | 0.6477424 |
| GASGGAGGAGGW | ASSDPGANSDYT | 0.30585998 |
| GGAGAGGDGGAGGI | ASSDPGANSDYT | 0.61690456 |
| GTGHNGSQPGMGGNG | ASSDPGANSDYT | 0.40287948 |
| GYAAASASAS | ASSDPGANSDYT | 0.4675422 |
| HQVDVAAVQQG | ASSDPGANSDYT | 0.6121452 |
| IGAAIAHT | ASSDPGANSDYT | 0.5889696 |
| IGMIEVAQL | ASSDPGANSDYT | 0.6026721 |
| IRLAPEHTSAMD | ASSDPGANSDYT | 0.4320809 |
| IYNVSVL | ASSDPGANSDYT | 0.5041275 |
| KQGFLGM | ASSDPGANSDYT | 0.5915509 |
| LESVITQL | ASSDPGANSDYT | 0.31581303 |
| LGLGALVVV | ASSDPGANSDYT | 0.41108853 |
| LIGNGGNGGSGGI | ASSDPGANSDYT | 0.66454655 |
| LNAINAPTL | ASSDPGANSDYT | 0.25711894 |
| LQLLNTGV | ASSDPGANSDYT | 0.80580145 |
| LQTSIEF | ASSDPGANSDYT | 0.17761964 |
| MSFAVSEL | ASSDPGANSDYT | 0.15926471 |
| NTGALGL | ASSDPGANSDYT | 0.30041122 |
| PPGAGGAGGAGTT | ASSDPGANSDYT | 0.54589194 |
| QLVGLGI | ASSDPGANSDYT | 0.81028825 |
| RFLEQVHQL | ASSDPGANSDYT | 0.48113188 |
| RGVGYVLR | ASSDPGANSDYT | 0.15933013 |
| SATGVFAGVI | ASSDPGANSDYT | 0.20709525 |
| SLPAITLP | ASSDPGANSDYT | 0.16438028 |
| STALAFTV | ASSDPGANSDYT | 0.23958355 |
| SVLTVGLGK | ASSDPGANSDYT | 0.45744467 |
| TALAGAGSGI | ASSDPGANSDYT | 0.21043412 |
| TGLPFTGI | ASSDPGANSDYT | 0.17674954 |
| VAEHVTV | ASSDPGANSDYT | 0.21138744 |
| VDIVLNSL | ASSDPGANSDYT | 0.48309782 |
| VLTGGGAVL | ASSDPGANSDYT | 0.44823673 |
| VNVGLAYGI | ASSDPGANSDYT | 0.33621126 |

| | | |
|---|---|---|
| VSSAAVGAV | ASSDPGANSDYT | 0.6878265 |
| VTSVYEAGV | ASSDPGANSDYT | 0.3339789 |
| WVALDRGA | ASSDPGANSDYT | 0.5897088 |
| YLVGSVVI | ASSDPGANSDYT | 0.1072717 |
| GPPGTGKTLL | ASSDPGANSDYT | 0.1277392 |
| GVGMTKF | ASSDPGANSDYT | 0.2562593 |
| IGIGGSDL | ASSDPGANSDYT | 0.23154944 |
| IGTLLAEL | ASSDPGANSDYT | 0.15578382 |
| IYTSGTT | ASSDPGANSDYT | 0.43663546 |
| LSFPAGV | ASSDPGANSDYT | 0.18947971 |
| LYTSGTT | ASSDPGANSDYT | 0.1473891 |
| VGAFPFGF | ASSDPGANSDYT | 0.59976447 |
| VGAGTVEF | ASSDPGANSDYT | 0.17296718 |
| VGKSTLLL | ASSDPGANSDYT | 0.17463747 |
| AALAHAGGGIV | ASSDPGANSDYT | 0.21491474 |
| AALVAVSLL | ASSDPGANSDYT | 0.29921874 |
| AAVSRAVAAL | ASSDPGANSDYT | 0.1464798 |
| AGGSSGIGGF | ASSDPGANSDYT | 0.33210346 |
| AGLTVANANG | ASSDPGANSDYT | 0.26451853 |
| AGLVGAVL | ASSDPGANSDYT | 0.36675388 |
| ALSQAFTV | ASSDPGANSDYT | 0.14075276 |
| ALTFSVSL | ASSDPGANSDYT | 0.39846155 |
| ALTGAALVAL | ASSDPGANSDYT | 0.2687237 |
| ASFSIGAGVL | ASSDPGANSDYT | 0.8265874 |
| DGSVYKI | ASSDPGANSDYT | 0.18359311 |
| EQLAEIV | ASSDPGANSDYT | 0.21414013 |
| FAVSELTVM | ASSDPGANSDYT | 0.23957601 |
| GAAPLAAGL | ASSDPGANSDYT | 0.79728913 |
| GADGGDGGSGGAGG | ASSDPGANSDYT | 0.13033023 |
| GFGLDTGGAGG | ASSDPGANSDYT | 0.183849 |
| GLASGGLL | ASSDPGANSDYT | 0.12835714 |
| GRIAGAAVLL | ASSDPGANSDYT | 0.30030802 |
| GSAGSLNTSF | ASSDPGANSDYT | 0.3398151 |
| GTGQIGI | ASSDPGANSDYT | 0.14402738 |
| GTLIALIL | ASSDPGANSDYT | 0.26916745 |
| IAATMGFVVG | ASSDPGANSDYT | 0.31369984 |
| IFAQALL | ASSDPGANSDYT | 0.19468081 |
| IGAAVALGF | ASSDPGANSDYT | 0.67546594 |
| IGLTPLAL | ASSDPGANSDYT | 0.21020412 |
| ILGSNTSTL | ASSDPGANSDYT | 0.5123846 |
| IVFSLTVL | ASSDPGANSDYT | 0.36318618 |
| LASKATGF | ASSDPGANSDYT | 0.1498452 |
| LASVALIL | ASSDPGANSDYT | 0.38245195 |
| LFAVGLTL | ASSDPGANSDYT | 0.17158256 |
| LFNGGIGTYI | ASSDPGANSDYT | 0.18610747 |
| LTLGAGII | ASSDPGANSDYT | 0.5006949 |
| LVAAALGDI | ASSDPGANSDYT | 0.81908166 |
| NTAFVQLGI | ASSDPGANSDYT | 0.8146767 |
| QLFSGAEL | ASSDPGANSDYT | 0.61026 |
| SAATFITL | ASSDPGANSDYT | 0.63168335 |
| SNYGLANLGV | ASSDPGANSDYT | 0.6914201 |

| | | |
|---|---|---|
| SYGGAYAVM | ASSDPGANSDYT | 0.80704236 |
| TTGVGGLTLG | ASSDPGANSDYT | 0.84674585 |
| TTQTIYI | ASSDPGANSDYT | 0.8040555 |
| VAAGAGLGAVI | ASSDPGANSDYT | 0.6216736 |
| VAAVVTLL | ASSDPGANSDYT | 0.78784513 |
| VGFLGGLI | ASSDPGANSDYT | 0.7545498 |
| VGVIPILL | ASSDPGANSDYT | 0.75689685 |
| VIFASSLI | ASSDPGANSDYT | 0.7732617 |
| VKGGDGGAAAG | ASSDPGANSDYT | 0.7819164 |
| VTKTFTNI | ASSDPGANSDYT | 0.6580212 |
| VVAAAILL | ASSDPGANSDYT | 0.6028991 |
| YAALDVEL | ASSDPGANSDYT | 0.78133744 |
| LASVALLL | ASSDPGANSDYT | 0.81953007 |
| LDAVHRV | ASSDPGANSDYT | 0.8095101 |
| QGLSSSIF | ASSDPGANSDYT | 0.849829 |
| RGTFANIRL | ASSDPGANSDYT | 0.83369875 |
| RQLWWGHRI | ASSDPGANSDYT | 0.77396065 |
| YGPPGTGKTLL | ASSDPGANSDYT | 0.23263505 |
| AALVGAPANVA | ASSDPGANSDYT | 0.48222226 |
| AATFRRTE | ASSDPGANSDYT | 0.6437 |
| AGALGGGI | ASSDPGANSDYT | 0.7245882 |
| AGGAGGAGGIGAG | ASSDPGANSDYT | 0.22123766 |
| AGSGALGL | ASSDPGANSDYT | 0.7924035 |
| AQIADQI | ASSDPGANSDYT | 0.71865344 |
| ASAVAVTYL | ASSDPGANSDYT | 0.67637515 |
| ASTVLGGTGG | ASSDPGANSDYT | 0.82044613 |
| AVGGEPAK | ASSDPGANSDYT | 0.707746 |
| CEQGTFGGTC | ASSDPGANSDYT | 0.57214886 |
| DEGKAPAG | ASSDPGANSDYT | 0.60801893 |
| DTVVFPAL | ASSDPGANSDYT | 0.8527322 |
| FEGITMAG | ASSDPGANSDYT | 0.5791068 |
| FPLVVQD | ASSDPGANSDYT | 0.8404932 |
| GAFTTPQ | ASSDPGANSDYT | 0.66592425 |
| GFGGFGTAGGVGGA | ASSDPGANSDYT | 0.84690535 |
| GFLAQFL | ASSDPGANSDYT | 0.5375723 |
| GGAGGAGGNAGVA | ASSDPGANSDYT | 0.8270706 |
| GGVGTAGAGGAGGAGG | ASSDPGANSDYT | 0.7584703 |
| GNGGGGDAGNAGSGGN | ASSDPGANSDYT | 0.86263525 |
| GTGGAGGAGNDAGS | ASSDPGANSDYT | 0.74042964 |
| GTTNTGWFNS | ASSDPGANSDYT | 0.81694824 |
| IFMGPLFTL | ASSDPGANSDYT | 0.7425551 |
| ILAHAIEAI | ASSDPGANSDYT | 0.81542903 |
| IRSAIREAL | ASSDPGANSDYT | 0.5361464 |
| ITFAGGG | ASSDPGANSDYT | 0.73487574 |
| ITGMSGHQEV | ASSDPGANSDYT | 0.47817028 |
| IVAGLACTL | ASSDPGANSDYT | 0.73215544 |
| IVGGGLAGA | ASSDPGANSDYT | 0.8214531 |
| IVTGVLNI | ASSDPGANSDYT | 0.19396399 |
| LAQQGCI | ASSDPGANSDYT | 0.82627606 |
| LCLHVEN | ASSDPGANSDYT | 0.5420685 |
| LDAISDGS | ASSDPGANSDYT | 0.6202237 |

| | | |
|---|---|---|
| LGGFAAML | ASSDPGANSDYT | 0.807682 |
| LGGKNPMI | ASSDPGANSDYT | 0.74920744 |
| LGPKGVAAIL | ASSDPGANSDYT | 0.82648766 |
| LMVGVSI | ASSDPGANSDYT | 0.62583905 |
| LVGAAFVL | ASSDPGANSDYT | 0.7667405 |
| LVLAVAPL | ASSDPGANSDYT | 0.7594565 |
| MAAIAVIDAA | ASSDPGANSDYT | 0.32294714 |
| NNGINAELQ | ASSDPGANSDYT | 0.513669 |
| RGEPLVLVHG | ASSDPGANSDYT | 0.8532007 |
| RGGAGGAAGLI | ASSDPGANSDYT | 0.24134031 |
| RSYRMHPAV | ASSDPGANSDYT | 0.372374 |
| SAAHLLAGQLA | ASSDPGANSDYT | 0.13277659 |
| SAAVGSY | ASSDPGANSDYT | 0.15274628 |
| SPHAVLHRI | ASSDPGANSDYT | 0.17318937 |
| SQIAKVL | ASSDPGANSDYT | 0.22337839 |
| TALGHLVP | ASSDPGANSDYT | 0.373631 |
| TAPTTSGAA | ASSDPGANSDYT | 0.3131187 |
| TDGDDGNGGSPGT | ASSDPGANSDYT | 0.14869148 |
| TDGLGEN | ASSDPGANSDYT | 0.18637629 |
| TGGPPLPGQA | ASSDPGANSDYT | 0.15919891 |
| TIPAITI | ASSDPGANSDYT | 0.17797421 |
| TIPGFTI | ASSDPGANSDYT | 0.15785542 |
| TSIASGALA | ASSDPGANSDYT | 0.17258993 |
| TVTLPGAAI | ASSDPGANSDYT | 0.15377815 |
| VASGLSAALP | ASSDPGANSDYT | 0.13863467 |
| VGAGVIGI | ASSDPGANSDYT | 0.15753739 |
| VGPKRFGV | ASSDPGANSDYT | 0.39694244 |
| VNVAFPD | ASSDPGANSDYT | 0.23858446 |
| VSMTNSGVS | ASSDPGANSDYT | 0.29929802 |
| VTDVLGDLSV | ASSDPGANSDYT | 0.28044876 |
| VVVVGAGFAGL | ASSDPGANSDYT | 0.15115555 |
| VYTVGTVV | ASSDPGANSDYT | 0.11923188 |
| WTTFSGP | ASSDPGANSDYT | 0.12612638 |
| YDELVEAF | ASSDPGANSDYT | 0.13816653 |
| YGTGGAGGNGG | ASSDPGANSDYT | 0.15589367 |
| YTNVMAV | ASSDPGANSDYT | 0.11786224 |
| FLEQVHQL | ASSDPGANSDYT | 0.17324491 |
| GLSLLAVAV | ASSDPGANSDYT | 0.16109174 |
| LEQVHQL | ASSDPGANSDYT | 0.13908629 |
| LNAAGAI | ASSDPGANSDYT | 0.2558692 |
| PAVHASL | ASSDPGANSDYT | 0.14503628 |
| TELEDDA | ASSDPGANSDYT | 0.14372516 |
| TPPGPPN | ASSDPGANSDYT | 0.13416032 |
| TPPPPPPP | ASSDPGANSDYT | 0.46829778 |
| TQSPALI | ASSRGQKNTLY | 0.13008352 |
| EFAAAFPQ | ASSRGQKNTLY | 0.2779851 |
| LLSGNLDVL | ASSRGQKNTLY | 0.13657945 |
| GPANLGNYNI | ASSRGQKNTLY | 0.22209175 |
| TLGPTLP | ASSRGQKNTLY | 0.13784155 |
| VYAYSSCCK | ASSRGQKNTLY | 0.24307156 |
| APAPAPAPNI | ASSRGQKNTLY | 0.19888881 |

| | | |
|---|---|---|
| PGGPVPIP | ASSRGQKNTLY | 0.5327859 |
| FLTVALEP | ASSRGQKNTLY | 0.15835641 |
| SQYFGANL | ASSRGQKNTLY | 0.17548089 |
| GGVKPAA | ASSRGQKNTLY | 0.1317876 |
| GGTGGAGGAGGW | ASSRGQKNTLY | 0.53230447 |
| GVGGVAGLGGAGG | ASSRGQKNTLY | 0.12645988 |
| VLLVLATTI | ASSRGQKNTLY | 0.15861775 |
| IGIFAAML | ASSRGQKNTLY | 0.12968622 |
| AVGDLID | ASSRGQKNTLY | 0.16229671 |
| NISPLPL | ASSRGQKNTLY | 0.27919102 |
| VTAVFVAI | ASSRGQKNTLY | 0.12240176 |
| SGDDNDAAL | ASSRGQKNTLY | 0.25177303 |
| AVAALAHS | ASSRGQKNTLY | 0.12798235 |
| GGVTVMATP | ASSRGQKNTLY | 0.13830015 |
| AGLAANVRD | ASSRGQKNTLY | 0.18014537 |
| TNRPDLIDPALLRPG | ASSRGQKNTLY | 0.14584288 |
| LANLGVTAL | ASSRGQKNTLY | 0.3228861 |
| TGERIEL | ASSRGQKNTLY | 0.12814276 |
| NGEELAA | ASSRGQKNTLY | 0.19564281 |
| VLSLVWG | ASSRGQKNTLY | 0.21650112 |
| IGYALWL | ASSRGQKNTLY | 0.120551206 |
| LSLLLAPL | ASSRGQKNTLY | 0.12863341 |
| TLDEALK | ASSRGQKNTLY | 0.4028054 |
| TDGAGTAEGG | ASSRGQKNTLY | 0.15114798 |
| NAPGGAGGNGG | ASSRGQKNTLY | 0.0772749 |
| VSVFMGMGGNF | ASSRGQKNTLY | 0.09344259 |
| LASYLAAL | ASSRGQKNTLY | 0.11345403 |
| LVGGTPAL | ASSRGQKNTLY | 0.08616181 |
| VRNADVL | ASSRGQKNTLY | 0.10548389 |
| TCSSPFLVSAM | ASSRGQKNTLY | 0.11593512 |
| VHITDAE | ASSRGQKNTLY | 0.12567626 |
| EYAAPSF | ASSRGQKNTLY | 0.13712588 |
| VVDAFHGGL | ASSRGQKNTLY | 0.1506448 |
| ASVVDPQ | ASSRGQKNTLY | 0.12554875 |
| SYNEFGGLS | ASSRGQKNTLY | 0.12075166 |
| DPDGSVY | ASSRGQKNTLY | 0.08963964 |
| MVSTIHATE | ASSRGQKNTLY | 0.10191171 |
| LAAGVADPV | ASSRGQKNTLY | 0.09632639 |
| TKEQIAA | ASSRGQKNTLY | 0.078549854 |
| SLASPAAL | ASSRGQKNTLY | 0.08040389 |
| IGLVFYGV | ASSRGQKNTLY | 0.12932989 |
| FLLDASNL | ASSRGQKNTLY | 0.12631445 |
| YQVGNFP | ASSRGQKNTLY | 0.11023108 |
| ESLLSQST | ASSRGQKNTLY | 0.16025409 |
| LFTVGFAAVG | ASSRGQKNTLY | 0.07737995 |
| ATPPTPT | ASSRGQKNTLY | 0.07266308 |
| TDLATLG | ASSRGQKNTLY | 0.08205918 |
| SAGGTGR | ASSRGQKNTLY | 0.10014918 |
| GFSNSGS | ASSRGQKNTLY | 0.09618706 |
| GGNGGSGGTSV | ASSRGQKNTLY | 0.063903466 |
| GTSVATGGAGN | ASSRGQKNTLY | 0.10244371 |

| | | |
|---|---|---|
| NGGNGGSGGTS | ASSRGQKNTLY | 0.15218565 |
| AVVKFPH | ASSRGQKNTLY | 0.106714234 |
| TPPPPPP | ASSRGQKNTLY | 0.11287677 |
| AAPAAAAAPA | ASSRGQKNTLY | 0.10035377 |
| IALNEDL | ASSRGQKNTLY | 0.10122358 |
| IVNVSSM | ASSRGQKNTLY | 0.095935516 |
| TGIPPAPR | ASSRGQKNTLY | 0.10475988 |
| FDLGGGTF | ASSRGQKNTLY | 0.09059294 |
| FDLGGGTFDVS | ASSRGQKNTLY | 0.20273125 |
| NEPTAAA | ASSRGQKNTLY | 0.07282372 |
| NEPTAAALA | ASSRGQKNTLY | 0.18199416 |
| GSIINISSI | ASSRGQKNTLY | 0.109263845 |
| VAPGNAGTA | ASSRGQKNTLY | 0.11323129 |
| TGGGASLE | ASSRGQKNTLY | 0.10952329 |
| VAVYNHYKRI | ASSRGQKNTLY | 0.13128883 |
| TDPAPATT | ASSRGQKNTLY | 0.13732186 |
| TNRPDILDPALLRPG | ASSRGQKNTLY | 0.16610485 |
| LTTTTTS | ASSRGQKNTLY | 0.08039068 |
| PTGSGKTL | ASSRGQKNTLY | 0.09893153 |
| TGSGKTL | ASSRGQKNTLY | 0.08926733 |
| TIGHVDHGK | ASSRGQKNTLY | 0.12987685 |
| TIGHVDHGKTT | ASSRGQKNTLY | 0.09800995 |
| GHVDHGKTT | ASSRGQKNTLY | 0.11179644 |
| AQFEHTLL | ASSRGQKNTLY | 0.13680638 |
| IVNQDEL | ASSRGQKNTLY | 0.14141057 |
| TNGSQFF | ASSRGQKNTLY | 0.10960481 |
| TGKGIVE | ASSRGQKNTLY | 0.20205773 |
| LISNASDA | ASSRGQKNTLY | 0.10280636 |
| AELRTLGELTKIAKAHGAQ | ASSRGQKNTLY | 0.0934499 |
| APNAVAAAEPSW | ASSRGQKNTLY | 0.10449009 |
| APQVAGAATAVL | ASSRGQKNTLY | 0.083790034 |
| ARRDPAAAAAILPTDA | ASSRGQKNTLY | 0.06979422 |
| DAAARDVLAEAGLAEYF | ASSRGQKNTLY | 0.10183517 |
| DDQIVLSAKANLAKVEQ | ASSRGQKNTLY | 0.17381139 |
| DELLPDTGEDSDEDGH | ASSRGQKNTLY | 0.17562087 |
| ELPEAPANRATVPQD | ASSRGQKNTLY | 0.09102441 |
| FGQSKIERVTGIG | ASSRGQKNTLY | 0.13541569 |
| GAGGTGSTAGGAGGAGGAGGLYA | ASSRGQKNTLY | 0.8381317 |
| GHGGTGAAGGNGGTGGAGGSGIDG | ASSRGQKNTLY | 0.70140904 |
| GNGGNGGAGGEGGPGGAGGAGA | ASSRGQKNTLY | 0.51274073 |
| GSTAGNGGNGGNATNGTIAGQPAG | ASSRGQKNTLY | 0.406613 |
| GTGGDGGNGGNGGTGG | ASSRGQKNTLY | 0.5432927 |
| GVRRPAGIPADAPV | ASSRGQKNTLY | 0.8010143 |
| GYRQVTDAHLLTLARR | ASSRGQKNTLY | 0.70754135 |
| IAKMGQDGHDRG | ASSRGQKNTLY | 0.8131527 |
| KAEPSIEIQYT | ASSRGQKNTLY | 0.2954916 |
| KPVRIRMAADEDTVAVT | ASSRGQKNTLY | 0.6804323 |
| MPPRIAGMRLLVIKPEP | ASSRGQKNTLY | 0.63112015 |
| MKFYAVALFRSRGVPPTR | ASSRGQKNTLY | 0.6086476 |
| NAGGFGILGAVAHSP | ASSRGQKNTLY | 0.7027892 |
| NEGREITLAMAEM | ASSRGQKNTLY | 0.33210492 |

| | | |
|---|---|---|
| NELVEWSHPLGHRWRWE | ASSRGQKNTLY | 0.44277728 |
| NLGAGGTGGNGGNPG | ASSRGQKNTLY | 0.39043272 |
| NTEWPKWETVMAADPVD | ASSRGQKNTLY | 0.54687524 |
| PASAAPAETQP | ASSRGQKNTLY | 0.76004803 |
| QQSSDGKNGVRAAEITGE | ASSRGQKNTLY | 0.7697079 |
| RAPDAVTTSWWKE | ASSRGQKNTLY | 0.80857664 |
| SGFGNVGAHTSGWFNQ | ASSRGQKNTLY | 0.81475455 |
| SGVAGAVGQAASVGGLKV | ASSRGQKNTLY | 0.71562684 |
| SNGGAGTGSAGGNGGT | ASSRGQKNTLY | 0.090432696 |
| TGGAGGEGGAAG | ASSRGQKNTLY | 0.17746209 |
| TGGDGGAAGAGGAPGHG | ASSRGQKNTLY | 0.3529977 |
| TMPPLLAAIANAGRHGV | ASSRGQKNTLY | 0.4567324 |
| TSSTAGNSGGA | ASSRGQKNTLY | 0.10339891 |
| VPATVPVGILC | ASSRGQKNTLY | 0.5473201 |
| VQRPDSGGVVLTGRIS | ASSRGQKNTLY | 0.69129467 |
| WTPNPVPATKGVPAGLVQAT | ASSRGQKNTLY | 0.35333937 |
| YALKRALAKFSDRFDVVII | ASSRGQKNTLY | 0.7290957 |
| YLIGSVIV | ASSRGQKNTLY | 0.43053892 |
| YGQLVAVID | ASSRGQKNTLY | 0.29248872 |
| VTVGGMTLPA | ASSRGQKNTLY | 0.41959512 |
| VLALPAV | ASSRGQKNTLY | 0.7932712 |
| VIPPTLNYET | ASSRGQKNTLY | 0.5893189 |
| VGMGGMHPSSG | ASSRGQKNTLY | 0.78337944 |
| VGLAGGAATA | ASSRGQKNTLY | 0.34482932 |
| VGGAGGAGGSGP | ASSRGQKNTLY | 0.8094975 |
| VFAAFMV | ASSRGQKNTLY | 0.26136184 |
| TLYDKLR | ASSRGQKNTLY | 0.7919642 |
| TLAVFTVL | ASSRGQKNTLY | 0.7400803 |
| TGSVQPARV | ASSRGQKNTLY | 0.8455491 |
| TGGGAGSGGT | ASSRGQKNTLY | 0.5931361 |
| SHLETSL | ASSRGQKNTLY | 0.58077234 |
| SFTTGMT | ASSRGQKNTLY | 0.21886498 |
| QVEVAKRMGVRQPTVS | ASSRGQKNTLY | 0.5221638 |
| QGSGGPALAM | ASSRGQKNTLY | 0.23591922 |
| MLAPSNP | ASSRGQKNTLY | 0.68429023 |
| LVNNAAI | ASSRGQKNTLY | 0.23221686 |
| LTGEINTLA | ASSRGQKNTLY | 0.6733655 |
| LSIFGPV | ASSRGQKNTLY | 0.8308334 |
| LLALLANL | ASSRGQKNTLY | 0.098832436 |
| LISNASDALDKLR | ASSRGQKNTLY | 0.7980939 |
| LINEAVP | ASSRGQKNTLY | 0.32921174 |
| LGYAFSGL | ASSRGQKNTLY | 0.30970627 |
| LDFVAVAL | ASSRGQKNTLY | 0.7835457 |
| LAQPGAE | ASSRGQKNTLY | 0.29190505 |
| ISNLGAVTGF | ASSRGQKNTLY | 0.78772 |
| HQEDPGRI | ASSRGQKNTLY | 0.33593217 |
| GTPGEPTP | ASSRGQKNTLY | 0.645678 |
| GFFNSGS | ASSRGQKNTLY | 0.79603434 |
| GAGGGGLTAGAN | ASSRGQKNTLY | 0.08432314 |
| FVDALVNM | ASSRGQKNTLY | 0.29522517 |
| FLTVAPEL | ASSRGQKNTLY | 0.83214295 |

| | | |
|---|---|---|
| FETLTAL | ASSRGQKNTLY | 0.24714822 |
| DVINPIEL | ASSRGQKNTLY | 0.16423175 |
| APVSGAPV | ASSRGQKNTLY | 0.15358692 |
| ALVGGLAW | ASSRGQKNTLY | 0.16648561 |
| ALAAAAATAI | ASSRGQKNTLY | 0.16465782 |
| AFGLGLLL | ASSRGQKNTLY | 0.18419327 |
| AFFAAASL | ASSRGQKNTLY | 0.19404513 |
| AAAALPAFSPP | ASSRGQKNTLY | 0.27830926 |
| VVLAPAL | ASSRGQKNTLY | 0.16719808 |
| TIGHVDHGKTTL | ASSRGQKNTLY | 0.17557222 |
| LISNASDALDK | ASSRGQKNTLY | 0.15933391 |
| LISNASDAL | ASSRGQKNTLY | 0.15721719 |
| ISNASDALDK | ASSRGQKNTLY | 0.17846629 |
| NQAETLVYQTE | ASSRGQKNTLY | 0.16472659 |
| TGGTGGTAGAG | ASSRGQKNTLY | 0.1581322 |
| TGSTAGGAGGA | ASSRGQKNTLY | 0.16235521 |
| DPLAVPVAAGPV | ASSRGQKNTLY | 0.16411741 |
| GGTGGTPGNGGN | ASSRGQKNTLY | 0.2802997 |
| PAGLAGVDPATL | ASSRGQKNTLY | 0.21731757 |
| GNNLVVEAPHAGASV | ASSRGQKNTLY | 0.23560058 |
| KVTRIMSASTLEQPA | ASSRGQKNTLY | 0.21444826 |
| LGFALPIALPRRTVV | ASSRGQKNTLY | 0.20152944 |
| TRLGNLTARKLVVLV | ASSRGQKNTLY | 0.13975552 |
| WSQQRRLIYFPSAGP | ASSRGQKNTLY | 0.15207456 |
| GPNGNPDPMAAAVDIR | ASSRGQKNTLY | 0.15859993 |
| TQRCVSVADLRAAASR | ASSRGQKNTLY | 0.1589813 |
| QFVLDVAEHPVEVLTETV | ASSRGQKNTLY | 0.13923566 |
| GGAGGTGGAGGNGGTQAGDG | ASSRGQKNTLY | 0.17010324 |
| PLLATLRAIAVDVDAKPAQV | ASSRGQKNTLY | 0.2065317 |
| APKFRTGGYMIDFWGVGYQVAKRM | ASSRGQKNTLY | 0.15351976 |
| GAGGAGGAGGAAGGPGGTGGTGGN | ASSRGQKNTLY | 0.20293032 |
| IGALGLLATDVAPAGSVAFGVALA | ASSRGQKNTLY | 0.15160464 |
| AGGAGGIGGNGGAGGAGGVLMGNG | ASSRGQKNTLY | 0.15396744 |
| ADELIGLL | ASSRGQKNTLY | 0.1596646 |
| ADRALKA | ASSRGQKNTLY | 0.27252042 |
| AGAVGQVGGAGGA | ASSRGQKNTLY | 0.15411516 |
| AGGGGGLGGAP | ASSRGQKNTLY | 0.18907 |
| AGGLLYGNGGAG | ASSRGQKNTLY | 0.15939124 |
| AGLESIGL | ASSRGQKNTLY | 0.2322523 |
| AGLKTGII | ASSRGQKNTLY | 0.1572981 |
| ASLAALLL | ASSRGQKNTLY | 0.19573775 |
| ASVINISSI | ASSRGQKNTLY | 0.16480269 |
| ATPPGPA | ASSRGQKNTLY | 0.2728145 |
| ATVASALV | ASSRGQKNTLY | 0.16917382 |
| ATVASLIL | ASSRGQKNTLY | 0.18596525 |
| AVNGPVLIRH | ASSRGQKNTLY | 0.1611382 |
| DGAAGAVGI | ASSRGQKNTLY | 0.16622116 |
| DPLLHGI | ASSRGQKNTLY | 0.14522037 |
| ELGAALGAL | ASSRGQKNTLY | 0.17175193 |
| ELGRTQAV | ASSRGQKNTLY | 0.15996951 |
| FVAGGYLS | ASSRGQKNTLY | 0.17162311 |

| | | |
|---|---|---|
| GASGGAGGAGGW | ASSRGQKNTLY | 0.2742671 |
| GGAGAGGDGGAGGI | ASSRGQKNTLY | 0.1383678 |
| GTGHNGSQPGMGGNG | ASSRGQKNTLY | 0.23038486 |
| GYAAASASAS | ASSRGQKNTLY | 0.13950363 |
| HQVDVAAVQQG | ASSRGQKNTLY | 0.15983076 |
| IGAAIAHT | ASSRGQKNTLY | 0.18892387 |
| IGMIEVAQL | ASSRGQKNTLY | 0.16192086 |
| IRLAPEHTSAMD | ASSRGQKNTLY | 0.236704 |
| IYNVSVL | ASSRGQKNTLY | 0.15212338 |
| KQGFLGM | ASSRGQKNTLY | 0.17425266 |
| LESVITQL | ASSRGQKNTLY | 0.20596454 |
| LGLGALVVV | ASSRGQKNTLY | 0.14063996 |
| LIGNGGNGGSGGI | ASSRGQKNTLY | 0.14773571 |
| LNAINAPTL | ASSRGQKNTLY | 0.22084072 |
| LQLLNTGV | ASSRGQKNTLY | 0.24298117 |
| LQTSIEF | ASSRGQKNTLY | 0.4499309 |
| MSFAVSEL | ASSRGQKNTLY | 0.21305144 |
| NTGALGL | ASSRGQKNTLY | 0.20971577 |
| PPGAGGAGGAGTT | ASSRGQKNTLY | 0.23693532 |
| QLVGLGI | ASSRGQKNTLY | 0.26265034 |
| RFLEQVHQL | ASSRGQKNTLY | 0.43532372 |
| RGVGYVLR | ASSRGQKNTLY | 0.2634863 |
| SATGVFAGVI | ASSRGQKNTLY | 0.2137309 |
| SLPAITLP | ASSRGQKNTLY | 0.21807542 |
| STALAFTV | ASSRGQKNTLY | 0.21947247 |
| SVLTVGLGK | ASSRGQKNTLY | 0.22006148 |
| TALAGAGSGI | ASSRGQKNTLY | 0.24350381 |
| TGLPFTGI | ASSRGQKNTLY | 0.21271352 |
| VAEHVTV | ASSRGQKNTLY | 0.20822266 |
| VDIVLNSL | ASSRGQKNTLY | 0.21381488 |
| VLTGGGAVL | ASSRGQKNTLY | 0.23617738 |
| VNVGLAYGI | ASSRGQKNTLY | 0.30378476 |
| VSSAAVGAV | ASSRGQKNTLY | 0.28404066 |
| VTSVYEAGV | ASSRGQKNTLY | 0.30215836 |
| WVALDRGA | ASSRGQKNTLY | 0.26456764 |
| YLVGSVVI | ASSRGQKNTLY | 0.23583438 |
| GPPGTGKTLL | ASSRGQKNTLY | 0.19510959 |
| GVGMTKF | ASSRGQKNTLY | 0.20472851 |
| IGIGGSDL | ASSRGQKNTLY | 0.21610332 |
| IGTLLAEL | ASSRGQKNTLY | 0.22462013 |
| IYTSGTT | ASSRGQKNTLY | 0.19423321 |
| LSFPAGV | ASSRGQKNTLY | 0.22844526 |
| LYTSGTT | ASSRGQKNTLY | 0.24262354 |
| VGAFPFGF | ASSRGQKNTLY | 0.20043948 |
| VGAGTVEF | ASSRGQKNTLY | 0.3059256 |
| VGKSTLLL | ASSRGQKNTLY | 0.20722161 |
| AALAHAGGGIV | ASSRGQKNTLY | 0.20867883 |
| AALVAVSLL | ASSRGQKNTLY | 0.21209608 |
| AAVSRAVAAL | ASSRGQKNTLY | 0.27663344 |
| AGGSSGIGGF | ASSRGQKNTLY | 0.20415668 |
| AGLTVANANG | ASSRGQKNTLY | 0.2583346 |

| | | |
|---|---|---|
| AGLVGAVL | ASSRGQKNTLY | 0.22122921 |
| ALSQAFTV | ASSRGQKNTLY | 0.25515172 |
| ALTFSVSL | ASSRGQKNTLY | 0.2026539 |
| ALTGAALVAL | ASSRGQKNTLY | 0.251498 |
| ASFSIGAGVL | ASSRGQKNTLY | 0.22700918 |
| DGSVYKI | ASSRGQKNTLY | 0.43597135 |
| EQLAEIV | ASSRGQKNTLY | 0.22340512 |
| FAVSELTVM | ASSRGQKNTLY | 0.24184965 |
| GAAPLAAGL | ASSRGQKNTLY | 0.22562735 |
| GADGGDGGSGGAGG | ASSRGQKNTLY | 0.30185214 |
| GFGLDTGGAGG | ASSRGQKNTLY | 0.19959751 |
| GLASGGLL | ASSRGQKNTLY | 0.24082363 |
| GRIAGAAVLL | ASSRGQKNTLY | 0.20896813 |
| GSAGSLNTSF | ASSRGQKNTLY | 0.25184163 |
| GTGQIGI | ASSRGQKNTLY | 0.2695303 |
| GTLIALIL | ASSRGQKNTLY | 0.20513497 |
| IAATMGFVVG | ASSRGQKNTLY | 0.27306768 |
| IFAQALL | ASSRGQKNTLY | 0.21341938 |
| IGAAVALGF | ASSRGQKNTLY | 0.2098933 |
| IGLTPLAL | ASSRGQKNTLY | 0.25847688 |
| ILGSNTSTL | ASSRGQKNTLY | 0.20427163 |
| IVFSLTVL | ASSRGQKNTLY | 0.24607684 |
| LASKATGF | ASSRGQKNTLY | 0.20349501 |
| LASVALIL | ASSRGQKNTLY | 0.23697375 |
| LFAVGLTL | ASSRGQKNTLY | 0.31474698 |
| LFNGGIGTYI | ASSRGQKNTLY | 0.19908848 |
| LTLGAGII | ASSRGQKNTLY | 0.20448487 |
| LVAAALGDI | ASSRGQKNTLY | 0.26958507 |
| NTAFVQLGI | ASSRGQKNTLY | 0.5922461 |
| QLFSGAEL | ASSRGQKNTLY | 0.5670562 |
| SAATFITL | ASSRGQKNTLY | 0.23100127 |
| SNYGLANLGV | ASSRGQKNTLY | 0.23994046 |
| SYGGAYAVM | ASSRGQKNTLY | 0.27292898 |
| TTGVGGLTLG | ASSRGQKNTLY | 0.40005055 |
| TTQTIYI | ASSRGQKNTLY | 0.6563253 |
| VAAGAGLGAVI | ASSRGQKNTLY | 0.52635056 |
| VAAVVTLL | ASSRGQKNTLY | 0.23111537 |
| VGFLGGLI | ASSRGQKNTLY | 0.32153583 |
| VGVIPILL | ASSRGQKNTLY | 0.32661533 |
| VIFASSLI | ASSRGQKNTLY | 0.33423674 |
| VKGGDGGAAAG | ASSRGQKNTLY | 0.29023048 |
| VTKTFTNI | ASSRGQKNTLY | 0.28991035 |
| VVAAAILL | ASSRGQKNTLY | 0.25304368 |
| YAALDVEL | ASSRGQKNTLY | 0.22832729 |
| LASVALLL | ASSRGQKNTLY | 0.2821899 |
| LDAVHRV | ASSRGQKNTLY | 0.60266685 |
| QGLSSSIF | ASSRGQKNTLY | 0.6028131 |
| RGTFANIRL | ASSRGQKNTLY | 0.3664043 |
| RQLWWGHRI | ASSRGQKNTLY | 0.5811616 |
| YGPPGTGKTLL | ASSRGQKNTLY | 0.38235372 |
| AALVGAPANVA | ASSRGQKNTLY | 0.19439048 |

| | | |
|---|---|---|
| AATFRRTE | ASSRGQKNTLY | 0.20217495 |
| AGALGGGI | ASSRGQKNTLY | 0.22033443 |
| AGGAGGAGGIGAG | ASSRGQKNTLY | 0.24060364 |
| AGSGALGL | ASSRGQKNTLY | 0.19785334 |
| AQIADQI | ASSRGQKNTLY | 0.29533565 |
| ASAVAVTYL | ASSRGQKNTLY | 0.24783775 |
| ASTVLGGTGG | ASSRGQKNTLY | 0.23717493 |
| AVGGEPAK | ASSRGQKNTLY | 0.41727 |
| CEQGTFGGTC | ASSRGQKNTLY | 0.27056208 |
| DEGKAPAG | ASSRGQKNTLY | 0.22505306 |
| DTVVFPAL | ASSRGQKNTLY | 0.22509769 |
| FEGITMAG | ASSRGQKNTLY | 0.50202584 |
| FPLVVQD | ASSRGQKNTLY | 0.21932733 |
| GAFTTPQ | ASSRGQKNTLY | 0.50791335 |
| GFGGFGTAGGVGGA | ASSRGQKNTLY | 0.21607295 |
| GFLAQFL | ASSRGQKNTLY | 0.4700709 |
| GGAGGAGGNAGVA | ASSRGQKNTLY | 0.21764472 |
| GGVGTAGAGGAGGAGG | ASSRGQKNTLY | 0.50087 |
| GNGGGGDAGNAGSGGN | ASSRGQKNTLY | 0.41142997 |
| GTGGAGGAGNDAGS | ASSRGQKNTLY | 0.6928675 |
| GTTNTGWFNS | ASSRGQKNTLY | 0.27775818 |
| IFMGPLFTL | ASSRGQKNTLY | 0.3460297 |
| ILAHAIEAI | ASSRGQKNTLY | 0.22893302 |
| IRSAIREAL | ASSRGQKNTLY | 0.7005782 |
| ITFAGGG | ASSRGQKNTLY | 0.20423517 |
| ITGMSGHQEV | ASSRGQKNTLY | 0.2889744 |
| IVAGLACTL | ASSRGQKNTLY | 0.198811 |
| IVGGGLAGA | ASSRGQKNTLY | 0.26392394 |
| IVTGVLNI | ASSRGQKNTLY | 0.5372302 |
| LAQQGCI | ASSRGQKNTLY | 0.20535854 |
| LCLHVEN | ASSRGQKNTLY | 0.35728619 |
| LDAISDGS | ASSRGQKNTLY | 0.2542708 |
| LGGFAAML | ASSRGQKNTLY | 0.23490007 |
| LGGKNPMI | ASSRGQKNTLY | 0.36611515 |
| LGPKGVAAIL | ASSRGQKNTLY | 0.25871375 |
| LMVGVSI | ASSRGQKNTLY | 0.61272496 |
| LVGAAFVL | ASSRGQKNTLY | 0.22036809 |
| LVLAVAPL | ASSRGQKNTLY | 0.28271547 |
| MAAIAVIDAA | ASSRGQKNTLY | 0.28735903 |
| NNGINAELQ | ASSRGQKNTLY | 0.2086653 |
| RGEPLVLVHG | ASSRGQKNTLY | 0.21550724 |
| RGGAGGAAGLI | ASSRGQKNTLY | 0.62426996 |
| RSYRMHPAV | ASSRGQKNTLY | 0.24666111 |
| SAAHLLAGQLA | ASSRGQKNTLY | 0.19670318 |
| SAAVGSY | ASSRGQKNTLY | 0.15678251 |
| SPHAVLHRI | ASSRGQKNTLY | 0.16115154 |
| SQIAKVL | ASSRGQKNTLY | 0.14390118 |
| TALGHLVP | ASSRGQKNTLY | 0.1911897 |
| TAPTTSGAA | ASSRGQKNTLY | 0.22461744 |
| TDGDDGNGGSPGT | ASSRGQKNTLY | 0.20875731 |
| TDGLGEN | ASSRGQKNTLY | 0.15790632 |

| | | |
|---|---|---|
| TGGPPLPGQA | ASSRGQKNTLY | 0.17800795 |
| TIPAITI | ASSRGQKNTLY | 0.17459334 |
| TIPGFTI | ASSRGQKNTLY | 0.15605088 |
| TSIASGALA | ASSRGQKNTLY | 0.165882 |
| TVTLPGAAI | ASSRGQKNTLY | 0.16521382 |
| VASGLSAALP | ASSRGQKNTLY | 0.15543625 |
| VGAGVIGI | ASSRGQKNTLY | 0.16074935 |
| VGPKRFGV | ASSRGQKNTLY | 0.14475563 |
| VNVAFPD | ASSRGQKNTLY | 0.1885019 |
| VSMTNSGVS | ASSRGQKNTLY | 0.17498916 |
| VTDVLGDLSV | ASSRGQKNTLY | 0.20601729 |
| VVVVGAGFAGL | ASSRGQKNTLY | 0.24058163 |
| VYTVGTVV | ASSRGQKNTLY | 0.17320432 |
| WTTFSGP | ASSRGQKNTLY | 0.13407314 |
| YDELVEAF | ASSRGQKNTLY | 0.14420648 |
| YGTGGAGGNGG | ASSRGQKNTLY | 0.17170754 |
| YTNVMAV | ASSRGQKNTLY | 0.15473306 |
| FLEQVHQL | ASSRGQKNTLY | 0.1291632 |
| GLSLLAVAV | ASSRGQKNTLY | 0.16565214 |
| LEQVHQL | ASSRGQKNTLY | 0.1882875 |
| LNAAGAI | ASSRGQKNTLY | 0.15724646 |
| PAVHASL | ASSRGQKNTLY | 0.16271658 |
| TELEDDA | ASSRGQKNTLY | 0.14657007 |
| TPPGPPN | ASSRGQKNTLY | 0.15077698 |
| TPPPPPPP | ASSRGQKNTLY | 0.15999454 |
| TQSPALI | ASSLARGNSDYT | 0.176677 |
| EFAAAFPQ | ASSLARGNSDYT | 0.15659352 |
| LLSGNLDVL | ASSLARGNSDYT | 0.18450393 |
| GPANLGNYNI | ASSLARGNSDYT | 0.14289963 |
| TLGPTLP | ASSLARGNSDYT | 0.22783531 |
| VYAYSSCCK | ASSLARGNSDYT | 0.1552018 |
| APAPAPAPNI | ASSLARGNSDYT | 0.19653687 |
| PGGPVPIP | ASSLARGNSDYT | 0.1714059 |
| FLTVALEP | ASSLARGNSDYT | 0.19493318 |
| SQYFGANL | ASSLARGNSDYT | 0.16703518 |
| GGVKPAA | ASSLARGNSDYT | 0.18637426 |
| GGTGGAGGAGGW | ASSLARGNSDYT | 0.15651178 |
| GVGGVAGLGGAGG | ASSLARGNSDYT | 0.24411407 |
| VLLVLATTI | ASSLARGNSDYT | 0.14373626 |
| IGIFAAML | ASSLARGNSDYT | 0.18969879 |
| AVGDLID | ASSLARGNSDYT | 0.14535874 |
| NISPLPL | ASSLARGNSDYT | 0.18824038 |
| VTAVFVAI | ASSLARGNSDYT | 0.2531343 |
| SGDDNDAAL | ASSLARGNSDYT | 0.13344571 |
| AVAALAHS | ASSLARGNSDYT | 0.20364764 |
| GGVTVMATP | ASSLARGNSDYT | 0.14163885 |
| AGLAANVRD | ASSLARGNSDYT | 0.14944147 |
| TNRPDLIDPALLRPG | ASSLARGNSDYT | 0.18755208 |
| LANLGVTAL | ASSLARGNSDYT | 0.16269438 |
| TGERIEL | ASSLARGNSDYT | 0.21123421 |
| NGEELAA | ASSLARGNSDYT | 0.14606582 |

| | | |
|---|---|---|
| VLSLVWG | ASSLARGNSDYT | 0.17766964 |
| IGYALWL | ASSLARGNSDYT | 0.23515035 |
| LSLLLAPL | ASSLARGNSDYT | 0.13847783 |
| TLDEALK | ASSLARGNSDYT | 0.14794123 |
| TDGAGTAEGG | ASSLARGNSDYT | 0.23095618 |
| NAPGGAGGNGG | ASSLARGNSDYT | 0.15413983 |
| VSVFMGMGGNF | ASSLARGNSDYT | 0.14963296 |
| LASYLAAL | ASSLARGNSDYT | 0.09459994 |
| LVGGTPAL | ASSLARGNSDYT | 0.09876027 |
| VRNADVL | ASSLARGNSDYT | 0.09282075 |
| TCSSPFLVSAM | ASSLARGNSDYT | 0.108965434 |
| VHITDAE | ASSLARGNSDYT | 0.16983631 |
| EYAAPSF | ASSLARGNSDYT | 0.14025863 |
| VVDAFHGGL | ASSLARGNSDYT | 0.09542643 |
| ASVVDPQ | ASSLARGNSDYT | 0.14736451 |
| SYNEFGGLS | ASSLARGNSDYT | 0.09692602 |
| DPDGSVY | ASSLARGNSDYT | 0.10748243 |
| MVSTIHATE | ASSLARGNSDYT | 0.10021877 |
| LAAGVADPV | ASSLARGNSDYT | 0.09846401 |
| TKEQIAA | ASSLARGNSDYT | 0.08845548 |
| SLASPAAL | ASSLARGNSDYT | 0.09230242 |
| IGLVFYGV | ASSLARGNSDYT | 0.08645801 |
| FLLDASNL | ASSLARGNSDYT | 0.29341394 |
| YQVGNFP | ASSLARGNSDYT | 0.20815982 |
| ESLLSQST | ASSLARGNSDYT | 0.10923402 |
| LFTVGFAAVG | ASSLARGNSDYT | 0.14779472 |
| ATPPTPT | ASSLARGNSDYT | 0.114158876 |
| TDLATLG | ASSLARGNSDYT | 0.07925427 |
| SAGGTGR | ASSLARGNSDYT | 0.085385494 |
| GFSNSGS | ASSLARGNSDYT | 0.0877208 |
| GGNGGSGGTSV | ASSLARGNSDYT | 0.096586846 |
| GTSVATGGAGN | ASSLARGNSDYT | 0.07761235 |
| NGGNGGSGGTS | ASSLARGNSDYT | 0.099754445 |
| AVVKFPH | ASSLARGNSDYT | 0.10637251 |
| TPPPPPP | ASSLARGNSDYT | 0.09013712 |
| AAPAAAAAPA | ASSLARGNSDYT | 0.14133492 |
| IALNEDL | ASSLARGNSDYT | 0.09443466 |
| IVNVSSM | ASSLARGNSDYT | 0.09456706 |
| TGIPPAPR | ASSLARGNSDYT | 0.08982818 |
| FDLGGGTF | ASSLARGNSDYT | 0.11495706 |
| FDLGGGTFDVS | ASSLARGNSDYT | 0.08526177 |
| NEPTAAA | ASSLARGNSDYT | 0.16231507 |
| NEPTAAALA | ASSLARGNSDYT | 0.08190635 |
| GSIINISSI | ASSLARGNSDYT | 0.12010866 |
| VAPGNAGTA | ASSLARGNSDYT | 0.089975685 |
| TGGGASLE | ASSLARGNSDYT | 0.13468067 |
| VAVYNHYKRI | ASSLARGNSDYT | 0.12331584 |
| TDPAPATT | ASSLARGNSDYT | 0.28547025 |
| TNRPDILDPALLRPG | ASSLARGNSDYT | 0.11501089 |
| LTTTTTS | ASSLARGNSDYT | 0.120342486 |
| PTGSGKTL | ASSLARGNSDYT | 0.08684588 |

| | | |
|---|---|---|
| TGSGKTL | ASSLARGNSDYT | 0.11979697 |
| TIGHVDHGK | ASSLARGNSDYT | 0.08348636 |
| TIGHVDHGKTT | ASSLARGNSDYT | 0.11134779 |
| GHVDHGKTT | ASSLARGNSDYT | 0.08375798 |
| AQFEHTLL | ASSLARGNSDYT | 0.1173724 |
| IVNQDEL | ASSLARGNSDYT | 0.15918694 |
| TNGSQFF | ASSLARGNSDYT | 0.08643293 |
| TGKGIVE | ASSLARGNSDYT | 0.12622981 |
| LISNASDA | ASSLARGNSDYT | 0.09091025 |
| AELRTLGELTKIAKAHGAQ | ASSLARGNSDYT | 0.0917131 |
| APNAVAAAEPSW | ASSLARGNSDYT | 0.1231894 |
| APQVAGAATAVL | ASSLARGNSDYT | 0.09545769 |
| ARRDPAAAAAILPTDA | ASSLARGNSDYT | 0.1282628 |
| DAAARDVLAEAGLAEYF | ASSLARGNSDYT | 0.08289527 |
| DDQIVLSAKANLAKVEQ | ASSLARGNSDYT | 0.100030124 |
| DELLPDTGEDSDEDGH | ASSLARGNSDYT | 0.15774949 |
| ELPEAPANRATVPQD | ASSLARGNSDYT | 0.08262493 |
| FGQSKIERVTGIG | ASSLARGNSDYT | 0.08634465 |
| GAGGTGSTAGGAGGAGGAGGLYA | ASSLARGNSDYT | 0.1762743 |
| GHGGTGAAGGNGGTGGAGGSGIDG | ASSLARGNSDYT | 0.28224438 |
| GNGGNGGAGGEGGPGGAGGAGA | ASSLARGNSDYT | 0.18429509 |
| GSTAGNGGNGGNATNGTIAGQPAG | ASSLARGNSDYT | 0.090914614 |
| GTGGDGGNGGNGGTGG | ASSLARGNSDYT | 0.112717874 |
| GVRRPAGIPADAPV | ASSLARGNSDYT | 0.13340504 |
| GYRQVTDAHLLTLARR | ASSLARGNSDYT | 0.17838974 |
| IAKMGQDGHDRG | ASSLARGNSDYT | 0.40283552 |
| KAEPSIEIQYT | ASSLARGNSDYT | 0.29960966 |
| KPVRIRMAADEDTVAVT | ASSLARGNSDYT | 0.11780619 |
| MPPRIAGMRLLVIKPEP | ASSLARGNSDYT | 0.15837926 |
| MKFYAVALFRSRGVPPTR | ASSLARGNSDYT | 0.10679744 |
| NAGGFGILGAVAHSP | ASSLARGNSDYT | 0.16324233 |
| NEGREITLAMAEM | ASSLARGNSDYT | 0.15869604 |
| NELVEWSHPLGHRWRWE | ASSLARGNSDYT | 0.106647946 |
| NLGAGGTGGNGGNPG | ASSLARGNSDYT | 0.09173756 |
| NTEWPKWETVMAADPVD | ASSLARGNSDYT | 0.092789516 |
| PASAAPAETQP | ASSLARGNSDYT | 0.18371505 |
| QQSSDGKNGVRAAEITGE | ASSLARGNSDYT | 0.3473579 |
| RAPDAVTTSWWKE | ASSLARGNSDYT | 0.18094017 |
| SGFGNVGAHTSGWFNQ | ASSLARGNSDYT | 0.21611422 |
| SGVAGAVGQAASVGGLKV | ASSLARGNSDYT | 0.28667176 |
| SNGGAGTGSAGGNGGT | ASSLARGNSDYT | 0.10493038 |
| TGGAGGEGGAAG | ASSLARGNSDYT | 0.08249729 |
| TGGDGGAAGAGGAPGHG | ASSLARGNSDYT | 0.08755323 |
| TMPPLLAAIANAGRHGV | ASSLARGNSDYT | 0.10041963 |
| TSSTAGNSGGA | ASSLARGNSDYT | 0.11828075 |
| VPATVPVGILC | ASSLARGNSDYT | 0.077116005 |
| VQRPDSGGVVLTGRIS | ASSLARGNSDYT | 0.16495891 |
| WTPNPVPATKGVPAGLVQAT | ASSLARGNSDYT | 0.11562412 |
| YALKRALAKFSDRFDVVII | ASSLARGNSDYT | 0.09962285 |
| YLIGSVIV | ASSLARGNSDYT | 0.31879854 |
| YGQLVAVID | ASSLARGNSDYT | 0.09715763 |

| | | |
|---|---|---|
| VTVGGMTLPA | ASSLARGNSDYT | 0.09709155 |
| VLALPAV | ASSLARGNSDYT | 0.11432247 |
| VIPPTLNYET | ASSLARGNSDYT | 0.18629491 |
| VGMGGMHPSSG | ASSLARGNSDYT | 0.10485962 |
| VGLAGGAATA | ASSLARGNSDYT | 0.38036907 |
| VGGAGGAGGSGP | ASSLARGNSDYT | 0.10737911 |
| VFAAFMV | ASSLARGNSDYT | 0.2357132 |
| TLYDKLR | ASSLARGNSDYT | 0.105124265 |
| TLAVFTVL | ASSLARGNSDYT | 0.22852299 |
| TGSVQPARV | ASSLARGNSDYT | 0.1640946 |
| TGGGAGSGGT | ASSLARGNSDYT | 0.61090946 |
| SHLETSL | ASSLARGNSDYT | 0.14208269 |
| SFTTGMT | ASSLARGNSDYT | 0.18508838 |
| QVEVAKRMGVRQPTVS | ASSLARGNSDYT | 0.11062644 |
| QGSGGPALAM | ASSLARGNSDYT | 0.10586913 |
| MLAPSNP | ASSLARGNSDYT | 0.09011519 |
| LVNNAAI | ASSLARGNSDYT | 0.1001269 |
| LTGEINTLA | ASSLARGNSDYT | 0.09147369 |
| LSIFGPV | ASSLARGNSDYT | 0.10427941 |
| LLALLANL | ASSLARGNSDYT | 0.23289631 |
| LISNASDALDKLR | ASSLARGNSDYT | 0.0881836 |
| LINEAVP | ASSLARGNSDYT | 0.10414998 |
| LGYAFSGL | ASSLARGNSDYT | 0.103117876 |
| LDFVAVAL | ASSLARGNSDYT | 0.10116395 |
| LAQPGAE | ASSLARGNSDYT | 0.17815845 |
| ISNLGAVTGF | ASSLARGNSDYT | 0.118368156 |
| HQEDPGRI | ASSLARGNSDYT | 0.086115085 |
| GTPGEPTP | ASSLARGNSDYT | 0.08625935 |
| GFFNSGS | ASSLARGNSDYT | 0.10559935 |
| GAGGGGLTAGAN | ASSLARGNSDYT | 0.1449177 |
| FVDALVNM | ASSLARGNSDYT | 0.081664324 |
| FLTVAPEL | ASSLARGNSDYT | 0.09894023 |
| FETLTAL | ASSLARGNSDYT | 0.27437207 |
| DVINPIEL | ASSLARGNSDYT | 0.28710476 |
| APVSGAPV | ASSLARGNSDYT | 0.28484973 |
| ALVGGLAW | ASSLARGNSDYT | 0.2853654 |
| ALAAAAATAI | ASSLARGNSDYT | 0.28694823 |
| AFGLGLLL | ASSLARGNSDYT | 0.2851384 |
| AFFAAASL | ASSLARGNSDYT | 0.28666845 |
| AAAALPAFSPP | ASSLARGNSDYT | 0.2875327 |
| VVLAPAL | ASSLARGNSDYT | 0.2864607 |
| TIGHVDHGKTTL | ASSLARGNSDYT | 0.28742468 |
| LISNASDALDK | ASSLARGNSDYT | 0.28936103 |
| LISNASDAL | ASSLARGNSDYT | 0.2884549 |
| ISNASDALDK | ASSLARGNSDYT | 0.29025427 |
| NQAETLVYQTE | ASSLARGNSDYT | 0.28589413 |
| TGGTGGTAGAG | ASSLARGNSDYT | 0.2873801 |
| TGSTAGGAGGA | ASSLARGNSDYT | 0.28495398 |
| DPLAVPVAAGPV | ASSLARGNSDYT | 0.28476045 |
| GGTGGTPGNGGN | ASSLARGNSDYT | 0.28795275 |
| PAGLAGVDPATL | ASSLARGNSDYT | 0.28903168 |

| | | |
|---|---|---|
| GNNLVVEAPHAGASV | ASSLARGNSDYT | 0.2902372 |
| KVTRIMSASTLEQPA | ASSLARGNSDYT | 0.289615 |
| LGFALPIALPRRTVV | ASSLARGNSDYT | 0.28981248 |
| TRLGNLTARKLVVLV | ASSLARGNSDYT | 0.28666422 |
| WSQQRRLIYFPSAGP | ASSLARGNSDYT | 0.28511274 |
| GPNGNPDPMAAAVDIR | ASSLARGNSDYT | 0.28511605 |
| TQRCVSVADLRAAASR | ASSLARGNSDYT | 0.28584808 |
| QFVLDVAEHPVEVLTETV | ASSLARGNSDYT | 0.28661984 |
| GGAGGTGGAGGNGGTQAGDG | ASSLARGNSDYT | 0.28328508 |
| PLLATLRAIAVDVDAKPAQV | ASSLARGNSDYT | 0.28882897 |
| APKFRTGGYMIDFWGVGYQVAKRM | ASSLARGNSDYT | 0.28768787 |
| GAGGAGGAGGAAGGPGGTGGTGGN | ASSLARGNSDYT | 0.2874288 |
| IGALGLLATDVAPAGSVAFGVALA | ASSLARGNSDYT | 0.29217988 |
| AGGAGGIGGNGGAGGAGGVLMGNG | ASSLARGNSDYT | 0.28594586 |
| ADELIGLL | ASSLARGNSDYT | 0.28572112 |
| ADRALKA | ASSLARGNSDYT | 0.28631145 |
| AGAVGQVGGAGGA | ASSLARGNSDYT | 0.28738308 |
| AGGGGGLGGAP | ASSLARGNSDYT | 0.28592545 |
| AGGLLYGNGGAG | ASSLARGNSDYT | 0.29228356 |
| AGLESIGL | ASSLARGNSDYT | 0.28474343 |
| AGLKTGII | ASSLARGNSDYT | 0.2956404 |
| ASLAALLL | ASSLARGNSDYT | 0.28733194 |
| ASVINISSI | ASSLARGNSDYT | 0.28827012 |
| ATPPGPA | ASSLARGNSDYT | 0.28651834 |
| ATVASALV | ASSLARGNSDYT | 0.29104885 |
| ATVASLIL | ASSLARGNSDYT | 0.290388 |
| AVNGPVLIRH | ASSLARGNSDYT | 0.2934488 |
| DGAAGAVGI | ASSLARGNSDYT | 0.28766203 |
| DPLLHGI | ASSLARGNSDYT | 0.2865127 |
| ELGAALGAL | ASSLARGNSDYT | 0.28671587 |
| ELGRTQAV | ASSLARGNSDYT | 0.28607166 |
| FVAGGYLS | ASSLARGNSDYT | 0.2855183 |
| GASGGAGGAGGW | ASSLARGNSDYT | 0.2865431 |
| GGAGAGGDGGAGGI | ASSLARGNSDYT | 0.2879535 |
| GTGHNGSQPGMGGNG | ASSLARGNSDYT | 0.29029682 |
| GYAAASASAS | ASSLARGNSDYT | 0.28544158 |
| HQVDVAAVQQG | ASSLARGNSDYT | 0.30191058 |
| IGAAIAHT | ASSLARGNSDYT | 0.2858142 |
| IGMIEVAQL | ASSLARGNSDYT | 0.28583127 |
| IRLAPEHTSAMD | ASSLARGNSDYT | 0.289468 |
| IYNVSVL | ASSLARGNSDYT | 0.28538123 |
| KQGFLGM | ASSLARGNSDYT | 0.28455696 |
| LESVITQL | ASSLARGNSDYT | 0.28536642 |
| LGLGALVVV | ASSLARGNSDYT | 0.28771058 |
| LIGNGGNGGSGGI | ASSLARGNSDYT | 0.2934272 |
| LNAINAPTL | ASSLARGNSDYT | 0.28581128 |
| LQLLNTGV | ASSLARGNSDYT | 0.28952217 |
| LQTSIEF | ASSLARGNSDYT | 0.25966582 |
| MSFAVSEL | ASSLARGNSDYT | 0.360862 |
| NTGALGL | ASSLARGNSDYT | 0.2081496 |
| PPGAGGAGGAGTT | ASSLARGNSDYT | 0.21678941 |

| | | |
|---|---|---|
| QLVGLGI | ASSLARGNSDYT | 0.23338862 |
| RFLEQVHQL | ASSLARGNSDYT | 0.26566413 |
| RGVGYVLR | ASSLARGNSDYT | 0.24491194 |
| SATGVFAGVI | ASSLARGNSDYT | 0.2500484 |
| SLPAITLP | ASSLARGNSDYT | 0.22556467 |
| STALAFTV | ASSLARGNSDYT | 0.21421437 |
| SVLTVGLGK | ASSLARGNSDYT | 0.21289186 |
| TALAGAGSGI | ASSLARGNSDYT | 0.22797091 |
| TGLPFTGI | ASSLARGNSDYT | 0.24670753 |
| VAEHVTV | ASSLARGNSDYT | 0.21241719 |
| VDIVLNSL | ASSLARGNSDYT | 0.20870756 |
| VLTGGGAVL | ASSLARGNSDYT | 0.22763783 |
| VNVGLAYGI | ASSLARGNSDYT | 0.22746864 |
| VSSAAVGAV | ASSLARGNSDYT | 0.29146582 |
| VTSVYEAGV | ASSLARGNSDYT | 0.26605275 |
| WVALDRGA | ASSLARGNSDYT | 0.2744922 |
| YLVGSVVI | ASSLARGNSDYT | 0.26694614 |
| GPPGTGKTLL | ASSLARGNSDYT | 0.23048548 |
| GVGMTKF | ASSLARGNSDYT | 0.199825 |
| IGIGGSDL | ASSLARGNSDYT | 0.21898934 |
| IGTLLAEL | ASSLARGNSDYT | 0.21120107 |
| IYTSGTT | ASSLARGNSDYT | 0.23747678 |
| LSFPAGV | ASSLARGNSDYT | 0.19634804 |
| LYTSGTT | ASSLARGNSDYT | 0.22678585 |
| VGAFPFGF | ASSLARGNSDYT | 0.23773496 |
| VGAGTVEF | ASSLARGNSDYT | 0.20451686 |
| VGKSTLLL | ASSLARGNSDYT | 0.2973421 |
| AALAHAGGGIV | ASSLARGNSDYT | 0.21135512 |
| AALVAVSLL | ASSLARGNSDYT | 0.21645069 |
| AAVSRAVAAL | ASSLARGNSDYT | 0.21349166 |
| AGGSSGIGGF | ASSLARGNSDYT | 0.27293095 |
| AGLTVANANG | ASSLARGNSDYT | 0.20807919 |
| AGLVGAVL | ASSLARGNSDYT | 0.25865877 |
| ALSQAFTV | ASSLARGNSDYT | 0.21732716 |
| ALTFSVSL | ASSLARGNSDYT | 0.25181705 |
| ALTGAALVAL | ASSLARGNSDYT | 0.21341372 |
| ASFSIGAGVL | ASSLARGNSDYT | 0.24369147 |
| DGSVYKI | ASSLARGNSDYT | 0.22146928 |
| EQLAEIV | ASSLARGNSDYT | 0.46820474 |
| FAVSELTVM | ASSLARGNSDYT | 0.24055289 |
| GAAPLAAGL | ASSLARGNSDYT | 0.25234753 |
| GADGGDGGSGGAGG | ASSLARGNSDYT | 0.22708094 |
| GFGLDTGGAGG | ASSLARGNSDYT | 0.20780793 |
| GLASGGLL | ASSLARGNSDYT | 0.2032982 |
| GRIAGAAVLL | ASSLARGNSDYT | 0.2365795 |
| GSAGSLNTSF | ASSLARGNSDYT | 0.21169093 |
| GTGQIGI | ASSLARGNSDYT | 0.23263298 |
| GTLIALIL | ASSLARGNSDYT | 0.24824616 |
| IAATMGFVVG | ASSLARGNSDYT | 0.2053833 |
| IFAQALL | ASSLARGNSDYT | 0.24947216 |
| IGAAVALGF | ASSLARGNSDYT | 0.20493627 |

| | | |
|---|---|---|
| IGLTPLAL | ASSLARGNSDYT | 0.22268786 |
| ILGSNTSTL | ASSLARGNSDYT | 0.2570778 |
| IVFSLTVL | ASSLARGNSDYT | 0.2040321 |
| LASKATGF | ASSLARGNSDYT | 0.27925497 |
| LASVALIL | ASSLARGNSDYT | 0.20306614 |
| LFAVGLTL | ASSLARGNSDYT | 0.22909105 |
| LFNGGIGTYI | ASSLARGNSDYT | 0.2771275 |
| LTLGAGII | ASSLARGNSDYT | 0.19802056 |
| LVAAALGDI | ASSLARGNSDYT | 0.20539212 |
| NTAFVQLGI | ASSLARGNSDYT | 0.2850615 |
| QLFSGAEL | ASSLARGNSDYT | 0.6984951 |
| SAATFITL | ASSLARGNSDYT | 0.7073464 |
| SNYGLANLGV | ASSLARGNSDYT | 0.19449154 |
| SYGGAYAVM | ASSLARGNSDYT | 0.19607157 |
| TTGVGGLTLG | ASSLARGNSDYT | 0.5178281 |
| TTQTIYI | ASSLARGNSDYT | 0.6621803 |
| VAAGAGLGAVI | ASSLARGNSDYT | 0.41027737 |
| VAAVVTLL | ASSLARGNSDYT | 0.68550086 |
| VGFLGGLI | ASSLARGNSDYT | 0.1870303 |
| VGVIPILL | ASSLARGNSDYT | 0.31709555 |
| VIFASSLI | ASSLARGNSDYT | 0.27526623 |
| VKGGDGGAAAG | ASSLARGNSDYT | 0.373844 |
| VTKTFTNI | ASSLARGNSDYT | 0.47614086 |
| VVAAAILL | ASSLARGNSDYT | 0.1606183 |
| YAALDVEL | ASSLARGNSDYT | 0.15846132 |
| LASVALLL | ASSLARGNSDYT | 0.28356463 |
| LDAVHRV | ASSLARGNSDYT | 0.53297347 |
| QGLSSSIF | ASSLARGNSDYT | 0.7153716 |
| RGTFANIRL | ASSLARGNSDYT | 0.70374656 |
| RQLWWGHRI | ASSLARGNSDYT | 0.7479937 |
| YGPPGTGKTLL | ASSLARGNSDYT | 0.72904265 |
| AALVGAPANVA | ASSLARGNSDYT | 0.57780445 |
| AATFRRTE | ASSLARGNSDYT | 0.10331896 |
| AGALGGGI | ASSLARGNSDYT | 0.19704148 |
| AGGAGGAGGIGAG | ASSLARGNSDYT | 0.16371217 |
| AGSGALGL | ASSLARGNSDYT | 0.50440323 |
| AQIADQI | ASSLARGNSDYT | 0.11241645 |
| ASAVAVTYL | ASSLARGNSDYT | 0.36742854 |
| ASTVLGGTGG | ASSLARGNSDYT | 0.3519835 |
| AVGGEPAK | ASSLARGNSDYT | 0.13914472 |
| CEQGTFGGTC | ASSLARGNSDYT | 0.71308506 |
| DEGKAPAG | ASSLARGNSDYT | 0.18002628 |
| DTVVFPAL | ASSLARGNSDYT | 0.17106721 |
| FEGITMAG | ASSLARGNSDYT | 0.22389399 |
| FPLVVQD | ASSLARGNSDYT | 0.75389624 |
| GAFTTPQ | ASSLARGNSDYT | 0.18112536 |
| GFGGFGTAGGVGGA | ASSLARGNSDYT | 0.71048164 |
| GFLAQFL | ASSLARGNSDYT | 0.48835897 |
| GGAGGAGGNAGVA | ASSLARGNSDYT | 0.6266811 |
| GGVGTAGAGGAGGAGG | ASSLARGNSDYT | 0.14450352 |
| GNGGGGDAGNAGSGGN | ASSLARGNSDYT | 0.68028104 |

| | | |
|---|---|---|
| GTGGAGGAGNDAGS | ASSLARGNSDYT | 0.4604581 |
| GTTNTGWFNS | ASSLARGNSDYT | 0.78750306 |
| IFMGPLFTL | ASSLARGNSDYT | 0.40265876 |
| ILAHAIEAI | ASSLARGNSDYT | 0.40118507 |
| IRSAIREAL | ASSLARGNSDYT | 0.20875838 |
| ITFAGGG | ASSLARGNSDYT | 0.19149548 |
| ITGMSGHQEV | ASSLARGNSDYT | 0.124949984 |
| IVAGLACTL | ASSLARGNSDYT | 0.41891706 |
| IVGGGLAGA | ASSLARGNSDYT | 0.18722671 |
| IVTGVLNI | ASSLARGNSDYT | 0.3287047 |
| LAQQGCI | ASSLARGNSDYT | 0.67124844 |
| LCLHVEN | ASSLARGNSDYT | 0.10434757 |
| LDAISDGS | ASSLARGNSDYT | 0.64308214 |
| LGGFAAML | ASSLARGNSDYT | 0.16947909 |
| LGGKNPMI | ASSLARGNSDYT | 0.17709036 |
| LGPKGVAAIL | ASSLARGNSDYT | 0.51370466 |
| LMVGVSI | ASSLARGNSDYT | 0.13737835 |
| LVGAAFVL | ASSLARGNSDYT | 0.7359224 |
| LVLAVAPL | ASSLARGNSDYT | 0.114088066 |
| MAAIAVIDAA | ASSLARGNSDYT | 0.6002546 |
| NNGINAELQ | ASSLARGNSDYT | 0.48793584 |
| RGEPLVLVHG | ASSLARGNSDYT | 0.089660786 |
| RGGAGGAAGLI | ASSLARGNSDYT | 0.15230067 |
| RSYRMHPAV | ASSLARGNSDYT | 0.74492705 |
| SAAHLLAGQLA | ASSLARGNSDYT | 0.098434664 |
| SAAVGSY | ASSLARGNSDYT | 0.09319131 |
| SPHAVLHRI | ASSLARGNSDYT | 0.08740299 |
| SQIAKVL | ASSLARGNSDYT | 0.09457206 |
| TALGHLVP | ASSLARGNSDYT | 0.088889904 |
| TAPTTSGAA | ASSLARGNSDYT | 0.097569056 |
| TDGDDGNGGSPGT | ASSLARGNSDYT | 0.099893205 |
| TDGLGEN | ASSLARGNSDYT | 0.10525514 |
| TGGPPLPGQA | ASSLARGNSDYT | 0.10172528 |
| TIPAITI | ASSLARGNSDYT | 0.096500374 |
| TIPGFTI | ASSLARGNSDYT | 0.09264553 |
| TSIASGALA | ASSLARGNSDYT | 0.09881852 |
| TVTLPGAAI | ASSLARGNSDYT | 0.093646675 |
| VASGLSAALP | ASSLARGNSDYT | 0.08999593 |
| VGAGVIGI | ASSLARGNSDYT | 0.086665414 |
| VGPKRFGV | ASSLARGNSDYT | 0.08819676 |
| VNVAFPD | ASSLARGNSDYT | 0.09385758 |
| VSMTNSGVS | ASSLARGNSDYT | 0.10161339 |
| VTDVLGDLSV | ASSLARGNSDYT | 0.09818582 |
| VVVVGAGFAGL | ASSLARGNSDYT | 0.10868882 |
| VYTVGTVV | ASSLARGNSDYT | 0.100361645 |
| WTTFSGP | ASSLARGNSDYT | 0.09034267 |
| YDELVEAF | ASSLARGNSDYT | 0.086035214 |
| YGTGGAGGNGG | ASSLARGNSDYT | 0.08790776 |
| YTNVMAV | ASSLARGNSDYT | 0.09058775 |
| FLEQVHQL | ASSLARGNSDYT | 0.09442581 |
| GLSLLAVAV | ASSLARGNSDYT | 0.08342212 |

| | | |
|---|---|---|
| LEQVHQL | ASSLARGNSDYT | 0.093920276 |
| LNAAGAI | ASSLARGNSDYT | 0.09640878 |
| PAVHASL | ASSLARGNSDYT | 0.09224815 |
| TELEDDA | ASSLARGNSDYT | 0.105437 |
| TPPGPPN | ASSLARGNSDYT | 0.08892627 |
| TPPPPPPP | ASSLARGNSDYT | 0.08890128 |
| TQSPALI | ASSDGTGWNTLY | 0.09325921 |
| EFAAAFPQ | ASSDGTGWNTLY | 0.103606746 |
| LLSGNLDVL | ASSDGTGWNTLY | 0.08897972 |
| GPANLGNYNI | ASSDGTGWNTLY | 0.10945842 |
| TLGPTLP | ASSDGTGWNTLY | 0.08870882 |
| VYAYSSCCK | ASSDGTGWNTLY | 0.11138498 |
| APAPAPAPNI | ASSDGTGWNTLY | 0.09541096 |
| PGGPVPIP | ASSDGTGWNTLY | 0.101080514 |
| FLTVALEP | ASSDGTGWNTLY | 0.09280402 |
| SQYFGANL | ASSDGTGWNTLY | 0.109305985 |
| GGVKPAA | ASSDGTGWNTLY | 0.09692706 |
| GGTGGAGGAGGW | ASSDGTGWNTLY | 0.10775339 |
| GVGGVAGLGGAGG | ASSDGTGWNTLY | 0.08964827 |
| VLLVLATTI | ASSDGTGWNTLY | 0.09113029 |
| IGIFAAML | ASSDGTGWNTLY | 0.088075586 |
| AVGDLID | ASSDGTGWNTLY | 0.09184424 |
| NISPLPL | ASSDGTGWNTLY | 0.09197643 |
| VTAVFVAI | ASSDGTGWNTLY | 0.09409986 |
| SGDDNDAAL | ASSDGTGWNTLY | 0.1076579 |
| AVAALAHS | ASSDGTGWNTLY | 0.0846262 |
| GGVTVMATP | ASSDGTGWNTLY | 0.09419365 |
| AGLAANVRD | ASSDGTGWNTLY | 0.085370734 |
| TNRPDLIDPALLRPG | ASSDGTGWNTLY | 0.09127324 |
| LANLGVTAL | ASSDGTGWNTLY | 0.09348778 |
| TGERIEL | ASSDGTGWNTLY | 0.090213746 |
| NGEELAA | ASSDGTGWNTLY | 0.08764315 |
| VLSLVWG | ASSDGTGWNTLY | 0.08529923 |
| IGYALWL | ASSDGTGWNTLY | 0.09149445 |
| LSLLLAPL | ASSDGTGWNTLY | 0.11541373 |
| TLDEALK | ASSDGTGWNTLY | 0.08600291 |
| TDGAGTAEGG | ASSDGTGWNTLY | 0.088185124 |
| NAPGGAGGNGG | ASSDGTGWNTLY | 0.10604029 |
| VSVFMGMGGNF | ASSDGTGWNTLY | 0.21745215 |
| LASYLAAL | ASSDGTGWNTLY | 0.32281548 |
| LVGGTPAL | ASSDGTGWNTLY | 0.08895821 |
| VRNADVL | ASSDGTGWNTLY | 0.10775094 |
| TCSSPFLVSAM | ASSDGTGWNTLY | 0.14632092 |
| VHITDAE | ASSDGTGWNTLY | 0.2152614 |
| EYAAPSF | ASSDGTGWNTLY | 0.17131618 |
| VVDAFHGGL | ASSDGTGWNTLY | 0.23980379 |
| ASVVDPQ | ASSDGTGWNTLY | 0.11091534 |
| SYNEFGGLS | ASSDGTGWNTLY | 0.11240429 |
| DPDGSVY | ASSDGTGWNTLY | 0.10482277 |
| MVSTIHATE | ASSDGTGWNTLY | 0.13826683 |
| LAAGVADPV | ASSDGTGWNTLY | 0.15109923 |

| | | |
|---|---|---|
| TKEQIAA | ASSDGTGWNTLY | 0.096404135 |
| SLASPAAL | ASSDGTGWNTLY | 0.084428415 |
| IGLVFYGV | ASSDGTGWNTLY | 0.09882845 |
| FLLDASNL | ASSDGTGWNTLY | 0.19481292 |
| YQVGNFP | ASSDGTGWNTLY | 0.42762262 |
| ESLLSQST | ASSDGTGWNTLY | 0.40963218 |
| LFTVGFAAVG | ASSDGTGWNTLY | 0.2519307 |
| ATPPTPT | ASSDGTGWNTLY | 0.24682248 |
| TDLATLG | ASSDGTGWNTLY | 0.09820209 |
| SAGGTGR | ASSDGTGWNTLY | 0.07868137 |
| GFSNSGS | ASSDGTGWNTLY | 0.08675622 |
| GGNGGSGGTSV | ASSDGTGWNTLY | 0.09598992 |
| GTSVATGGAGN | ASSDGTGWNTLY | 0.13501194 |
| NGGNGGSGGTS | ASSDGTGWNTLY | 0.07401819 |
| AVVKFPH | ASSDGTGWNTLY | 0.13955043 |
| TPPPPPP | ASSDGTGWNTLY | 0.12702693 |
| AAPAAAAAPA | ASSDGTGWNTLY | 0.08663405 |
| IALNEDL | ASSDGTGWNTLY | 0.35762924 |
| IVNVSSM | ASSDGTGWNTLY | 0.088851124 |
| TGIPPAPR | ASSDGTGWNTLY | 0.092675135 |
| FDLGGGTF | ASSDGTGWNTLY | 0.10676422 |
| FDLGGGTFDVS | ASSDGTGWNTLY | 0.25688076 |
| NEPTAAA | ASSDGTGWNTLY | 0.101277635 |
| NEPTAAALA | ASSDGTGWNTLY | 0.2724836 |
| GSIINISSI | ASSDGTGWNTLY | 0.10605761 |
| VAPGNAGTA | ASSDGTGWNTLY | 0.20684284 |
| TGGGASLE | ASSDGTGWNTLY | 0.099186465 |
| VAVYNHYKRI | ASSDGTGWNTLY | 0.19008146 |
| TDPAPATT | ASSDGTGWNTLY | 0.13810632 |
| TNRPDILDPALLRPG | ASSDGTGWNTLY | 0.67134273 |
| LTTTTTS | ASSDGTGWNTLY | 0.13681063 |
| PTGSGKTL | ASSDGTGWNTLY | 0.15867884 |
| TGSGKTL | ASSDGTGWNTLY | 0.10641371 |
| TIGHVDHGK | ASSDGTGWNTLY | 0.09561555 |
| TIGHVDHGKTT | ASSDGTGWNTLY | 0.08381426 |
| GHVDHGKTT | ASSDGTGWNTLY | 0.12173483 |
| AQFEHTLL | ASSDGTGWNTLY | 0.08790661 |
| IVNQDEL | ASSDGTGWNTLY | 0.119095534 |
| TNGSQFF | ASSDGTGWNTLY | 0.21375895 |
| TGKGIVE | ASSDGTGWNTLY | 0.08230339 |
| LISNASDA | ASSDGTGWNTLY | 0.13789949 |
| AELRTLGELTKIAKAHGAQ | ASSDGTGWNTLY | 0.08298928 |
| APNAVAAAEPSW | ASSDGTGWNTLY | 0.100966565 |
| APQVAGAATAVL | ASSDGTGWNTLY | 0.17085102 |
| ARRDPAAAAAILPTDA | ASSDGTGWNTLY | 0.08981809 |
| DAAARDVLAEAGLAEYF | ASSDGTGWNTLY | 0.14689696 |
| DDQIVLSAKANLAKVEQ | ASSDGTGWNTLY | 0.082209006 |
| DELLPDTGEDSDEDGH | ASSDGTGWNTLY | 0.11853793 |
| ELPEAPANRATVPQD | ASSDGTGWNTLY | 0.18992108 |
| FGQSKIERVTGIG | ASSDGTGWNTLY | 0.07600553 |
| GAGGTGSTAGGAGGAGGAGGLYA | ASSDGTGWNTLY | 0.089916624 |

| | | |
|---|---|---|
| GHGGTGAAGGNGGTGGAGGSGIDG | ASSDGTGWNTLY | 0.22925028 |
| GNGGNGGAGGEGGPGGAGGAGA | ASSDGTGWNTLY | 0.8086355 |
| GSTAGNGGNGGNATNGTIAGQPAG | ASSDGTGWNTLY | 0.82198346 |
| GTGGDGGNGGNGGTGG | ASSDGTGWNTLY | 0.7419383 |
| GVRRPAGIPADAPV | ASSDGTGWNTLY | 0.6685958 |
| GYRQVTDAHLLTLARR | ASSDGTGWNTLY | 0.74287164 |
| IAKMGQDGHDRG | ASSDGTGWNTLY | 0.7983905 |
| KAEPSIEIQYT | ASSDGTGWNTLY | 0.8243041 |
| KPVRIRMAADEDTVAVT | ASSDGTGWNTLY | 0.81087077 |
| MPPRIAGMRLLVIKPEP | ASSDGTGWNTLY | 0.46981388 |
| MKFYAVALFRSRGVPPTR | ASSDGTGWNTLY | 0.80692077 |
| NAGGFGILGAVAHSP | ASSDGTGWNTLY | 0.7410727 |
| NEGREITLAMAEM | ASSDGTGWNTLY | 0.76314676 |
| NELVEWSHPLGHRWRWE | ASSDGTGWNTLY | 0.7622167 |
| NLGAGGTGGNGGNPG | ASSDGTGWNTLY | 0.6964826 |
| NTEWPKWETVMAADPVD | ASSDGTGWNTLY | 0.7682648 |
| PASAAPAETQP | ASSDGTGWNTLY | 0.61994195 |
| QQSSDGKNGVRAAEITGE | ASSDGTGWNTLY | 0.76104486 |
| RAPDAVTTSWWKE | ASSDGTGWNTLY | 0.80602753 |
| SGFGNVGAHTSGWFNQ | ASSDGTGWNTLY | 0.7931895 |
| SGVAGAVGQAASVGGLKV | ASSDGTGWNTLY | 0.80654144 |
| SNGGAGTGSAGGNGGT | ASSDGTGWNTLY | 0.81597245 |
| TGGAGGEGGAAG | ASSDGTGWNTLY | 0.78767943 |
| TGGDGGAAGAGGAPGHG | ASSDGTGWNTLY | 0.21099882 |
| TMPPLLAAIANAGRHGV | ASSDGTGWNTLY | 0.45906174 |
| TSSTAGNSGGA | ASSDGTGWNTLY | 0.7173236 |
| VPATVPVGILC | ASSDGTGWNTLY | 0.7243123 |
| VQRPDSGGVVLTGRIS | ASSDGTGWNTLY | 0.22706547 |
| WTPNPVPATKGVPAGLVQAT | ASSDGTGWNTLY | 0.7677917 |
| YALKRALAKFSDRFDVVII | ASSDGTGWNTLY | 0.72331643 |
| YLIGSVIV | ASSDGTGWNTLY | 0.75123405 |
| YGQLVAVID | ASSDGTGWNTLY | 0.81379235 |
| VTVGGMTLPA | ASSDGTGWNTLY | 0.77952254 |
| VLALPAV | ASSDGTGWNTLY | 0.5831201 |
| VIPPTLNYET | ASSDGTGWNTLY | 0.5654725 |
| VGMGGMHPSSG | ASSDGTGWNTLY | 0.8224231 |
| VGLAGGAATA | ASSDGTGWNTLY | 0.46339697 |
| VGGAGGAGGSGP | ASSDGTGWNTLY | 0.8150313 |
| VFAAFMV | ASSDGTGWNTLY | 0.6696775 |
| TLYDKLR | ASSDGTGWNTLY | 0.80660236 |
| TLAVFTVL | ASSDGTGWNTLY | 0.40574878 |
| TGSVQPARV | ASSDGTGWNTLY | 0.8066 |
| TGGGAGSGGT | ASSDGTGWNTLY | 0.780629 |
| SHLETSL | ASSDGTGWNTLY | 0.82984 |
| SFTTGMT | ASSDGTGWNTLY | 0.7006986 |
| QVEVAKRMGVRQPTVS | ASSDGTGWNTLY | 0.739423 |
| QGSGGPALAM | ASSDGTGWNTLY | 0.5616439 |
| MLAPSNP | ASSDGTGWNTLY | 0.8239558 |
| LVNNAAI | ASSDGTGWNTLY | 0.5824496 |
| LTGEINTLA | ASSDGTGWNTLY | 0.7055697 |
| LSIFGPV | ASSDGTGWNTLY | 0.58472973 |

| | | |
|---|---|---|
| LLALLANL | ASSDGTGWNTLY | 0.7606272 |
| LISNASDALDKLR | ASSDGTGWNTLY | 0.812281 |
| LINEAVP | ASSDGTGWNTLY | 0.37230018 |
| LGYAFSGL | ASSDGTGWNTLY | 0.8145087 |
| LDFVAVAL | ASSDGTGWNTLY | 0.71253574 |
| LAQPGAE | ASSDGTGWNTLY | 0.5351728 |
| ISNLGAVTGF | ASSDGTGWNTLY | 0.8048783 |
| HQEDPGRI | ASSDGTGWNTLY | 0.73207086 |
| GTPGEPTP | ASSDGTGWNTLY | 0.8083751 |
| GFFNSGS | ASSDGTGWNTLY | 0.64000165 |
| GAGGGGLTAGAN | ASSDGTGWNTLY | 0.79639125 |
| FVDALVNM | ASSDGTGWNTLY | 0.7741983 |
| FLTVAPEL | ASSDGTGWNTLY | 0.66009605 |
| FETLTAL | ASSDGTGWNTLY | 0.589785 |
| DVINPIEL | ASSDGTGWNTLY | 0.8223142 |
| APVSGAPV | ASSDGTGWNTLY | 0.8053011 |
| ALVGGLAW | ASSDGTGWNTLY | 0.801969 |
| ALAAAAATAI | ASSDGTGWNTLY | 0.7331611 |
| AFGLGLLL | ASSDGTGWNTLY | 0.73486435 |
| AFFAAASL | ASSDGTGWNTLY | 0.76859796 |
| AAAALPAFSPP | ASSDGTGWNTLY | 0.7949637 |
| VVLAPAL | ASSDGTGWNTLY | 0.80721486 |
| TIGHVDHGKTTL | ASSDGTGWNTLY | 0.80462563 |
| LISNASDALDK | ASSDGTGWNTLY | 0.6959962 |
| LISNASDAL | ASSDGTGWNTLY | 0.7910963 |
| ISNASDALDK | ASSDGTGWNTLY | 0.75084925 |
| NQAETLVYQTE | ASSDGTGWNTLY | 0.7652377 |
| TGGTGGTAGAG | ASSDGTGWNTLY | 0.78829104 |
| TGSTAGGAGGA | ASSDGTGWNTLY | 0.7520722 |
| DPLAVPVAAGPV | ASSDGTGWNTLY | 0.75299555 |
| GGTGGTPGNGGN | ASSDGTGWNTLY | 0.7470161 |
| PAGLAGVDPATL | ASSDGTGWNTLY | 0.788496 |
| GNNLVVEAPHAGASV | ASSDGTGWNTLY | 0.80566 |
| KVTRIMSASTLEQPA | ASSDGTGWNTLY | 0.8007414 |
| LGFALPIALPRRTVV | ASSDGTGWNTLY | 0.8096071 |
| TRLGNLTARKLVVLV | ASSDGTGWNTLY | 0.8032217 |
| WSQQRRLIYFPSAGP | ASSDGTGWNTLY | 0.80406046 |
| GPNGNPDPMAAAVDIR | ASSDGTGWNTLY | 0.3397009 |
| TQRCVSVADLRAAASR | ASSDGTGWNTLY | 0.6309471 |
| QFVLDVAEHPVEVLTETV | ASSDGTGWNTLY | 0.7615359 |
| GGAGGTGGAGGNGGTQAGDG | ASSDGTGWNTLY | 0.7611981 |
| PLLATLRAIAVDVDAKPAQV | ASSDGTGWNTLY | 0.39957473 |
| APKFRTGGYMIDFWGVGYQVAKRM | ASSDGTGWNTLY | 0.7839274 |
| GAGGAGGAGGAAGGPGGTGGTGGN | ASSDGTGWNTLY | 0.78398985 |
| IGALGLLATDVAPAGSVAFGVALA | ASSDGTGWNTLY | 0.72747993 |
| AGGAGGIGGNGGAGGAGGVLMGNG | ASSDGTGWNTLY | 0.8023583 |
| ADELIGLL | ASSDGTGWNTLY | 0.7612017 |
| ADRALKA | ASSDGTGWNTLY | 0.69515014 |
| AGAVGQVGGAGGA | ASSDGTGWNTLY | 0.71608174 |
| AGGGGGLGGAP | ASSDGTGWNTLY | 0.8017403 |
| AGGLLYGNGGAG | ASSDGTGWNTLY | 0.68132144 |

| | | |
|---|---|---|
| AGLESIGL | ASSDGTGWNTLY | 0.8045639 |
| AGLKTGII | ASSDGTGWNTLY | 0.6350993 |
| ASLAALLL | ASSDGTGWNTLY | 0.8057349 |
| ASVINISSI | ASSDGTGWNTLY | 0.68049526 |
| ATPPGPA | ASSDGTGWNTLY | 0.803643 |
| ATVASALV | ASSDGTGWNTLY | 0.77591944 |
| ATVASLIL | ASSDGTGWNTLY | 0.8151387 |
| AVNGPVLIRH | ASSDGTGWNTLY | 0.75873077 |
| DGAAGAVGI | ASSDGTGWNTLY | 0.78582966 |
| DPLLHGI | ASSDGTGWNTLY | 0.7410626 |
| ELGAALGAL | ASSDGTGWNTLY | 0.80805045 |
| ELGRTQAV | ASSDGTGWNTLY | 0.64417464 |
| FVAGGYLS | ASSDGTGWNTLY | 0.77703285 |
| GASGGAGGAGGW | ASSDGTGWNTLY | 0.5235721 |
| GGAGAGGDGGAGGI | ASSDGTGWNTLY | 0.77812034 |
| GTGHNGSQPGMGGNG | ASSDGTGWNTLY | 0.80618775 |
| GYAAASASAS | ASSDGTGWNTLY | 0.4739198 |
| HQVDVAAVQQG | ASSDGTGWNTLY | 0.8041577 |
| IGAAIAHT | ASSDGTGWNTLY | 0.7043688 |
| IGMIEVAQL | ASSDGTGWNTLY | 0.7514723 |
| IRLAPEHTSAMD | ASSDGTGWNTLY | 0.8037988 |
| IYNVSVL | ASSDGTGWNTLY | 0.76518965 |
| KQGFLGM | ASSDGTGWNTLY | 0.80794775 |
| LESVITQL | ASSDGTGWNTLY | 0.73697174 |
| LGLGALVVV | ASSDGTGWNTLY | 0.76497674 |
| LIGNGGNGGSGGI | ASSDGTGWNTLY | 0.7771849 |
| LNAINAPTL | ASSDGTGWNTLY | 0.65501857 |
| LQLLNTGV | ASSDGTGWNTLY | 0.7073167 |
| LQTSIEF | ASSDGTGWNTLY | 0.81174135 |
| MSFAVSEL | ASSDGTGWNTLY | 0.5073705 |
| NTGALGL | ASSDGTGWNTLY | 0.5091988 |
| PPGAGGAGGAGTT | ASSDGTGWNTLY | 0.168221 |
| QLVGLGI | ASSDGTGWNTLY | 0.18200684 |
| RFLEQVHQL | ASSDGTGWNTLY | 0.20163268 |
| RGVGYVLR | ASSDGTGWNTLY | 0.29586905 |
| SATGVFAGVI | ASSDGTGWNTLY | 0.7223499 |
| SLPAITLP | ASSDGTGWNTLY | 0.48858804 |
| STALAFTV | ASSDGTGWNTLY | 0.18813346 |
| SVLTVGLGK | ASSDGTGWNTLY | 0.32585785 |
| TALAGAGSGI | ASSDGTGWNTLY | 0.23951837 |
| TGLPFTGI | ASSDGTGWNTLY | 0.2984853 |
| VAEHVTV | ASSDGTGWNTLY | 0.2230251 |
| VDIVLNSL | ASSDGTGWNTLY | 0.20433617 |
| VLTGGGAVL | ASSDGTGWNTLY | 0.17419292 |
| VNVGLAYGI | ASSDGTGWNTLY | 0.16067332 |
| VSSAAVGAV | ASSDGTGWNTLY | 0.32349956 |
| VTSVYEAGV | ASSDGTGWNTLY | 0.4448997 |
| WVALDRGA | ASSDGTGWNTLY | 0.31877816 |
| YLVGSVVI | ASSDGTGWNTLY | 0.39168245 |
| GPPGTGKTLL | ASSDGTGWNTLY | 0.5296291 |
| GVGMTKF | ASSDGTGWNTLY | 0.2424323 |

| | | |
|---|---|---|
| IGIGGSDL | ASSDGTGWNTLY | 0.14670485 |
| IGTLLAEL | ASSDGTGWNTLY | 0.15084231 |
| IYTSGTT | ASSDGTGWNTLY | 0.183447 |
| LSFPAGV | ASSDGTGWNTLY | 0.18058285 |
| LYTSGTT | ASSDGTGWNTLY | 0.14426693 |
| VGAFPFGF | ASSDGTGWNTLY | 0.25130075 |
| VGAGTVEF | ASSDGTGWNTLY | 0.19679311 |
| VGKSTLLL | ASSDGTGWNTLY | 0.18526152 |
| AALAHAGGGIV | ASSDGTGWNTLY | 0.5001341 |
| AALVAVSLL | ASSDGTGWNTLY | 0.1841715 |
| AAVSRAVAAL | ASSDGTGWNTLY | 0.1629563 |
| AGGSSGIGGF | ASSDGTGWNTLY | 0.18816647 |
| AGLTVANANG | ASSDGTGWNTLY | 0.43062103 |
| AGLVGAVL | ASSDGTGWNTLY | 0.17915067 |
| ALSQAFTV | ASSDGTGWNTLY | 0.43048415 |
| ALTFSVSL | ASSDGTGWNTLY | 0.18500052 |
| ALTGAALVAL | ASSDGTGWNTLY | 0.48400992 |
| ASFSIGAGVL | ASSDGTGWNTLY | 0.16988775 |
| DGSVYKI | ASSDGTGWNTLY | 0.44603392 |
| EQLAEIV | ASSDGTGWNTLY | 0.30580282 |
| FAVSELTVM | ASSDGTGWNTLY | 0.7112589 |
| GAAPLAAGL | ASSDGTGWNTLY | 0.20198318 |
| GADGGDGGSGGAGG | ASSDGTGWNTLY | 0.30465326 |
| GFGLDTGGAGG | ASSDGTGWNTLY | 0.200452 |
| GLASGGLL | ASSDGTGWNTLY | 0.64427054 |
| GRIAGAAVLL | ASSDGTGWNTLY | 0.16252393 |
| GSAGSLNTSF | ASSDGTGWNTLY | 0.18259108 |
| GTGQIGI | ASSDGTGWNTLY | 0.16034743 |
| GTLIALIL | ASSDGTGWNTLY | 0.19072036 |
| IAATMGFVVG | ASSDGTGWNTLY | 0.5312884 |
| IFAQALL | ASSDGTGWNTLY | 0.1440071 |
| IGAAVALGF | ASSDGTGWNTLY | 0.23018585 |
| IGLTPLAL | ASSDGTGWNTLY | 0.16225488 |
| ILGSNTSTL | ASSDGTGWNTLY | 0.166374 |
| IVFSLTVL | ASSDGTGWNTLY | 0.2617087 |
| LASKATGF | ASSDGTGWNTLY | 0.24479362 |
| LASVALIL | ASSDGTGWNTLY | 0.23161763 |
| LFAVGLTL | ASSDGTGWNTLY | 0.16931662 |
| LFNGGIGTYI | ASSDGTGWNTLY | 0.22657435 |
| LTLGAGII | ASSDGTGWNTLY | 0.2417578 |
| LVAAALGDI | ASSDGTGWNTLY | 0.1507854 |
| NTAFVQLGI | ASSDGTGWNTLY | 0.17019558 |
| QLFSGAEL | ASSDGTGWNTLY | 0.50066566 |
| SAATFITL | ASSDGTGWNTLY | 0.75842273 |
| SNYGLANLGV | ASSDGTGWNTLY | 0.82425874 |
| SYGGAYAVM | ASSDGTGWNTLY | 0.36372763 |
| TTGVGGLTLG | ASSDGTGWNTLY | 0.3097506 |
| TTQTIYI | ASSDGTGWNTLY | 0.5526321 |
| VAAGAGLGAVI | ASSDGTGWNTLY | 0.78939444 |
| VAAVVTLL | ASSDGTGWNTLY | 0.8612047 |
| VGFLGGLI | ASSDGTGWNTLY | 0.79534996 |

| | | |
|---|---|---|
| VGVIPILL | ASSDGTGWNTLY | 0.3025236 |
| VIFASSLI | ASSDGTGWNTLY | 0.6706388 |
| VKGGDGGAAAG | ASSDGTGWNTLY | 0.46995524 |
| VTKTFTNI | ASSDGTGWNTLY | 0.62881637 |
| VVAAAILL | ASSDGTGWNTLY | 0.71914834 |
| YAALDVEL | ASSDGTGWNTLY | 0.665587 |
| LASVALLL | ASSDGTGWNTLY | 0.3995666 |
| LDAVHRV | ASSDGTGWNTLY | 0.33555868 |
| QGLSSSIF | ASSDGTGWNTLY | 0.7635522 |
| RGTFANIRL | ASSDGTGWNTLY | 0.8165127 |
| RQLWWGHRI | ASSDGTGWNTLY | 0.7859191 |
| YGPPGTGKTLL | ASSDGTGWNTLY | 0.85097307 |
| AALVGAPANVA | ASSDGTGWNTLY | 0.7917741 |
| AATFRRTE | ASSDGTGWNTLY | 0.76921856 |
| AGALGGGI | ASSDGTGWNTLY | 0.12165257 |
| AGGAGGAGGIGAG | ASSDGTGWNTLY | 0.18255568 |
| AGSGALGL | ASSDGTGWNTLY | 0.4186835 |
| AQIADQI | ASSDGTGWNTLY | 0.56101865 |
| ASAVAVTYL | ASSDGTGWNTLY | 0.16225925 |
| ASTVLGGTGG | ASSDGTGWNTLY | 0.7822888 |
| AVGGEPAK | ASSDGTGWNTLY | 0.43434802 |
| CEQGTFGGTC | ASSDGTGWNTLY | 0.3339457 |
| DEGKAPAG | ASSDGTGWNTLY | 0.81255335 |
| DTVVFPAL | ASSDGTGWNTLY | 0.36800754 |
| FEGITMAG | ASSDGTGWNTLY | 0.29681495 |
| FPLVVQD | ASSDGTGWNTLY | 0.45244288 |
| GAFTTPQ | ASSDGTGWNTLY | 0.78172046 |
| GFGGFGTAGGVGGA | ASSDGTGWNTLY | 0.27535647 |
| GFLAQFL | ASSDGTGWNTLY | 0.76737297 |
| GGAGGAGGNAGVA | ASSDGTGWNTLY | 0.5614425 |
| GGVGTAGAGGAGGAGG | ASSDGTGWNTLY | 0.8026116 |
| GNGGGGDAGNAGSGGN | ASSDGTGWNTLY | 0.23601727 |
| GTGGAGGAGNDAGS | ASSDGTGWNTLY | 0.80140877 |
| GTTNTGWFNS | ASSDGTGWNTLY | 0.6464129 |
| IFMGPLFTL | ASSDGTGWNTLY | 0.8715776 |
| ILAHAIEAI | ASSDGTGWNTLY | 0.44712028 |
| IRSAIREAL | ASSDGTGWNTLY | 0.6183802 |
| ITFAGGG | ASSDGTGWNTLY | 0.5761305 |
| ITGMSGHQEV | ASSDGTGWNTLY | 0.83952236 |
| IVAGLACTL | ASSDGTGWNTLY | 0.22092211 |
| IVGGGLAGA | ASSDGTGWNTLY | 0.43176752 |
| IVTGVLNI | ASSDGTGWNTLY | 0.19312444 |
| LAQQGCI | ASSDGTGWNTLY | 0.50545174 |
| LCLHVEN | ASSDGTGWNTLY | 0.77320194 |
| LDAISDGS | ASSDGTGWNTLY | 0.14103538 |
| LGGFAAML | ASSDGTGWNTLY | 0.68723726 |
| LGGKNPMI | ASSDGTGWNTLY | 0.542789 |
| LGPKGVAAIL | ASSDGTGWNTLY | 0.33767393 |
| LMVGVSI | ASSDGTGWNTLY | 0.79137135 |
| LVGAAFVL | ASSDGTGWNTLY | 0.6805428 |
| LVLAVAPL | ASSDGTGWNTLY | 0.76990867 |

| | | |
|---|---|---|
| MAAIAVIDAA | ASSDGTGWNTLY | 0.577086 |
| NNGINAELQ | ASSDGTGWNTLY | 0.5267717 |
| RGEPLVLVHG | ASSDGTGWNTLY | 0.5983143 |
| RGGAGGAAGLI | ASSDGTGWNTLY | 0.22223383 |
| RSYRMHPAV | ASSDGTGWNTLY | 0.47529685 |
| SAAHLLAGQLA | ASSDGTGWNTLY | 0.82825756 |
| SAAVGSY | ASSDGTGWNTLY | 0.67944074 |
| SPHAVLHRI | ASSDGTGWNTLY | 0.73914427 |
| SQIAKVL | ASSDGTGWNTLY | 0.20931278 |
| TALGHLVP | ASSDGTGWNTLY | 0.23416436 |
| TAPTTSGAA | ASSDGTGWNTLY | 0.49493092 |
| TDGDDGNGGSPGT | ASSDGTGWNTLY | 0.6801435 |
| TDGLGEN | ASSDGTGWNTLY | 0.817569 |
| TGGPPLPGQA | ASSDGTGWNTLY | 0.7083862 |
| TIPAITI | ASSDGTGWNTLY | 0.26837212 |
| TIPGFTI | ASSDGTGWNTLY | 0.45688623 |
| TSIASGALA | ASSDGTGWNTLY | 0.37250108 |
| TVTLPGAAI | ASSDGTGWNTLY | 0.37662596 |
| VASGLSAALP | ASSDGTGWNTLY | 0.49639133 |
| VGAGVIGI | ASSDGTGWNTLY | 0.5006186 |
| VGPKRFGV | ASSDGTGWNTLY | 0.3629994 |
| VNVAFPD | ASSDGTGWNTLY | 0.25157872 |
| VSMTNSGVS | ASSDGTGWNTLY | 0.5001256 |
| VTDVLGDLSV | ASSDGTGWNTLY | 0.7469111 |
| VVVVGAGFAGL | ASSDGTGWNTLY | 0.6213665 |
| VYTVGTVV | ASSDGTGWNTLY | 0.78551894 |
| WTTFSGP | ASSDGTGWNTLY | 0.68244064 |
| YDELVEAF | ASSDGTGWNTLY | 0.46000925 |
| YGTGGAGGNGG | ASSDGTGWNTLY | 0.10880606 |
| YTNVMAV | ASSDGTGWNTLY | 0.15593769 |
| FLEQVHQL | ASSDGTGWNTLY | 0.22354867 |
| GLSLLAVAV | ASSDGTGWNTLY | 0.48164693 |
| LEQVHQL | ASSDGTGWNTLY | 0.11350567 |
| LNAAGAI | ASSDGTGWNTLY | 0.5617151 |
| PAVHASL | ASSDGTGWNTLY | 0.36293375 |
| TELEDDA | ASSDGTGWNTLY | 0.24191512 |
| TPPGPPN | ASSDGTGWNTLY | 0.7173116 |
| TPPPPPPP | ASSDGTGWNTLY | 0.30103123 |
| TQSPALI | ASSQEGQNNQAPL | 0.21601236 |
| EFAAAFPQ | ASSQEGQNNQAPL | 0.28881162 |
| LLSGNLDVL | ASSQEGQNNQAPL | 0.7880068 |
| GPANLGNYNI | ASSQEGQNNQAPL | 0.19752884 |
| TLGPTLP | ASSQEGQNNQAPL | 0.70201087 |
| VYAYSSCCK | ASSQEGQNNQAPL | 0.4471822 |
| APAPAPAPNI | ASSQEGQNNQAPL | 0.6757195 |
| PGGPVPIP | ASSQEGQNNQAPL | 0.17880307 |
| FLTVALEP | ASSQEGQNNQAPL | 0.6491276 |
| SQYFGANL | ASSQEGQNNQAPL | 0.41629064 |
| GGVKPAA | ASSQEGQNNQAPL | 0.80145335 |
| GGTGGAGGAGGW | ASSQEGQNNQAPL | 0.28658217 |
| GVGGVAGLGGAGG | ASSQEGQNNQAPL | 0.51918656 |

| | | |
|---|---|---|
| VLLVLATTI | ASSQEGQNNQAPL | 0.40694183 |
| IGIFAAML | ASSQEGQNNQAPL | 0.7529936 |
| AVGDLID | ASSQEGQNNQAPL | 0.14699174 |
| NISPLPL | ASSQEGQNNQAPL | 0.3611713 |
| VTAVFVAI | ASSQEGQNNQAPL | 0.19032063 |
| SGDDNDAAL | ASSQEGQNNQAPL | 0.41662902 |
| AVAALAHS | ASSQEGQNNQAPL | 0.6272579 |
| GGVTVMATP | ASSQEGQNNQAPL | 0.112996206 |
| AGLAANVRD | ASSQEGQNNQAPL | 0.71619606 |
| TNRPDLIDPALLRPG | ASSQEGQNNQAPL | 0.3126017 |
| LANLGVTAL | ASSQEGQNNQAPL | 0.25325182 |
| TGERIEL | ASSQEGQNNQAPL | 0.558485 |
| NGEELAA | ASSQEGQNNQAPL | 0.38258728 |
| VLSLVWG | ASSQEGQNNQAPL | 0.7035702 |
| IGYALWL | ASSQEGQNNQAPL | 0.24338366 |
| LSLLLAPL | ASSQEGQNNQAPL | 0.5743238 |
| TLDEALK | ASSQEGQNNQAPL | 0.56526077 |
| TDGAGTAEGG | ASSQEGQNNQAPL | 0.11513954 |
| NAPGGAGGNGG | ASSQEGQNNQAPL | 0.24258381 |
| VSVFMGMGGNF | ASSQEGQNNQAPL | 0.7033738 |
| LASYLAAL | ASSQEGQNNQAPL | 0.76439583 |
| LVGGTPAL | ASSQEGQNNQAPL | 0.81169677 |
| VRNADVL | ASSQEGQNNQAPL | 0.6154644 |
| TCSSPFLVSAM | ASSQEGQNNQAPL | 0.520699 |
| VHITDAE | ASSQEGQNNQAPL | 0.6891286 |
| EYAAPSF | ASSQEGQNNQAPL | 0.8019416 |
| VVDAFHGGL | ASSQEGQNNQAPL | 0.81821704 |
| ASVVDPQ | ASSQEGQNNQAPL | 0.77042913 |
| SYNEFGGLS | ASSQEGQNNQAPL | 0.5075708 |
| DPDGSVY | ASSQEGQNNQAPL | 0.75176 |
| MVSTIHATE | ASSQEGQNNQAPL | 0.6865943 |
| LAAGVADPV | ASSQEGQNNQAPL | 0.6969403 |
| TKEQIAA | ASSQEGQNNQAPL | 0.7959819 |
| SLASPAAL | ASSQEGQNNQAPL | 0.72510344 |
| IGLVFYGV | ASSQEGQNNQAPL | 0.685761 |
| FLLDASNL | ASSQEGQNNQAPL | 0.6189638 |
| YQVGNFP | ASSQEGQNNQAPL | 0.7272008 |
| ESLLSQST | ASSQEGQNNQAPL | 0.7872044 |
| LFTVGFAAVG | ASSQEGQNNQAPL | 0.74444485 |
| ATPPTPT | ASSQEGQNNQAPL | 0.839559 |
| TDLATLG | ASSQEGQNNQAPL | 0.80558515 |
| SAGGTGR | ASSQEGQNNQAPL | 0.776422 |
| GFSNSGS | ASSQEGQNNQAPL | 0.1740664 |
| GGNGGSGGTSV | ASSQEGQNNQAPL | 0.49804 |
| GTSVATGGAGN | ASSQEGQNNQAPL | 0.6607716 |
| NGGNGGSGGTS | ASSQEGQNNQAPL | 0.73935056 |
| AVVKFPH | ASSQEGQNNQAPL | 0.3028759 |
| TPPPPPP | ASSQEGQNNQAPL | 0.7931385 |
| AAPAAAAAPA | ASSQEGQNNQAPL | 0.6907555 |
| IALNEDL | ASSQEGQNNQAPL | 0.6209384 |
| IVNVSSM | ASSQEGQNNQAPL | 0.82982874 |

| | | |
|---|---|---|
| TGIPPAPR | ASSQEGQNNQAPL | 0.6683373 |
| FDLGGGTF | ASSQEGQNNQAPL | 0.63682455 |
| FDLGGGTFDVS | ASSQEGQNNQAPL | 0.6598667 |
| NEPTAAA | ASSQEGQNNQAPL | 0.8399559 |
| NEPTAAALA | ASSQEGQNNQAPL | 0.50573176 |
| GSIINISSI | ASSQEGQNNQAPL | 0.8013976 |
| VAPGNAGTA | ASSQEGQNNQAPL | 0.7144031 |
| TGGGASLE | ASSQEGQNNQAPL | 0.8245537 |
| VAVYNHYKRI | ASSQEGQNNQAPL | 0.397245 |
| TDPAPATT | ASSQEGQNNQAPL | 0.7921263 |
| TNRPDILDPALLRPG | ASSQEGQNNQAPL | 0.68367875 |
| LTTTTTS | ASSQEGQNNQAPL | 0.8544105 |
| PTGSGKTL | ASSQEGQNNQAPL | 0.673364 |
| TGSGKTL | ASSQEGQNNQAPL | 0.7529582 |
| TIGHVDHGK | ASSQEGQNNQAPL | 0.65738434 |
| TIGHVDHGKTT | ASSQEGQNNQAPL | 0.79857856 |
| GHVDHGKTT | ASSQEGQNNQAPL | 0.42792314 |
| AQFEHTLL | ASSQEGQNNQAPL | 0.6632006 |
| IVNQDEL | ASSQEGQNNQAPL | 0.59282005 |
| TNGSQFF | ASSQEGQNNQAPL | 0.724316 |
| TGKGIVE | ASSQEGQNNQAPL | 0.7728714 |
| LISNASDA | ASSQEGQNNQAPL | 0.31720942 |
| AELRTLGELTKIAKAHGAQ | ASSQEGQNNQAPL | 0.7821378 |
| APNAVAAAEPSW | ASSQEGQNNQAPL | 0.6329056 |
| APQVAGAATAVL | ASSQEGQNNQAPL | 0.51323235 |
| ARRDPAAAAAILPTDA | ASSQEGQNNQAPL | 0.8089293 |
| DAAARDVLAEAGLAEYF | ASSQEGQNNQAPL | 0.66297704 |
| DDQIVLSAKANLAKVEQ | ASSQEGQNNQAPL | 0.78542644 |
| DELLPDTGEDSDEDGH | ASSQEGQNNQAPL | 0.6799278 |
| ELPEAPANRATVPQD | ASSQEGQNNQAPL | 0.71169776 |
| FGQSKIERVTGIG | ASSQEGQNNQAPL | 0.7808934 |
| GAGGTGSTAGGAGGAGGAGGLYA | ASSQEGQNNQAPL | 0.5094272 |
| GHGGTGAAGGNGGTGGAGGSGIDG | ASSQEGQNNQAPL | 0.56345594 |
| GNGGNGGAGGEGGPGGAGGAGA | ASSQEGQNNQAPL | 0.81612754 |
| GSTAGNGGNGGNATNGTIAGQPAG | ASSQEGQNNQAPL | 0.8961103 |
| GTGGDGGNGGNGGTGG | ASSQEGQNNQAPL | 0.8922885 |
| GVRRPAGIPADAPV | ASSQEGQNNQAPL | 0.83259577 |
| GYRQVTDAHLLTLARR | ASSQEGQNNQAPL | 0.8114618 |
| IAKMGQDGHDRG | ASSQEGQNNQAPL | 0.85400224 |
| KAEPSIEIQYT | ASSQEGQNNQAPL | 0.9006929 |
| KPVRIRMAADEDTVAVT | ASSQEGQNNQAPL | 0.899453 |
| MPPRIAGMRLLVIKPEP | ASSQEGQNNQAPL | 0.8884702 |
| MKFYAVALFRSRGVPPTR | ASSQEGQNNQAPL | 0.81849 |
| NAGGFGILGAVAHSP | ASSQEGQNNQAPL | 0.8849178 |
| NEGREITLAMAEM | ASSQEGQNNQAPL | 0.85408515 |
| NELVEWSHPLGHRWRWE | ASSQEGQNNQAPL | 0.85321134 |
| NLGAGGTGGNGGNPG | ASSQEGQNNQAPL | 0.8990192 |
| NTEWPKWETVMAADPVD | ASSQEGQNNQAPL | 0.87091935 |
| PASAAPAETQP | ASSQEGQNNQAPL | 0.861524 |
| QQSSDGKNGVRAAEITGE | ASSQEGQNNQAPL | 0.8656218 |
| RAPDAVTTSWWKE | ASSQEGQNNQAPL | 0.88347423 |

| | | |
|---|---|---|
| SGFGNVGAHTSGWFNQ | ASSQEGQNNQAPL | 0.89907897 |
| SGVAGAVGQAASVGGLKV | ASSQEGQNNQAPL | 0.8928354 |
| SNGGAGTGSAGGNGGT | ASSQEGQNNQAPL | 0.9108742 |
| TGGAGGEGGAAG | ASSQEGQNNQAPL | 0.89873075 |
| TGGDGGAAGAGGAPGHG | ASSQEGQNNQAPL | 0.8947561 |
| TMPPLLAAIANAGRHGV | ASSQEGQNNQAPL | 0.41413885 |
| TSSTAGNSGGA | ASSQEGQNNQAPL | 0.7415917 |
| VPATVPVGILC | ASSQEGQNNQAPL | 0.8618859 |
| VQRPDSGGVVLTGRIS | ASSQEGQNNQAPL | 0.8550743 |
| WTPNPVPATKGVPAGLVQAT | ASSQEGQNNQAPL | 0.61288154 |
| YALKRALAKFSDRFDVVII | ASSQEGQNNQAPL | 0.89005625 |
| YLIGSVIV | ASSQEGQNNQAPL | 0.8746965 |
| YGQLVAVID | ASSQEGQNNQAPL | 0.80249286 |
| VTVGGMTLPA | ASSQEGQNNQAPL | 0.8960264 |
| VLALPAV | ASSQEGQNNQAPL | 0.8360694 |
| VIPPTLNYET | ASSQEGQNNQAPL | 0.8233483 |
| VGMGGMHPSSG | ASSQEGQNNQAPL | 0.86314476 |
| VGLAGGAATA | ASSQEGQNNQAPL | 0.90599924 |
| VGGAGGAGGSGP | ASSQEGQNNQAPL | 0.8323252 |
| VFAAFMV | ASSQEGQNNQAPL | 0.894122 |
| TLYDKLR | ASSQEGQNNQAPL | 0.87150455 |
| TLAVFTVL | ASSQEGQNNQAPL | 0.9034567 |
| TGSVQPARV | ASSQEGQNNQAPL | 0.7494848 |
| TGGGAGSGGT | ASSQEGQNNQAPL | 0.8929764 |
| SHLETSL | ASSQEGQNNQAPL | 0.864071 |
| SFTTGMT | ASSQEGQNNQAPL | 0.91266143 |
| QVEVAKRMGVRQPTVS | ASSQEGQNNQAPL | 0.8621848 |
| QGSGGPALAM | ASSQEGQNNQAPL | 0.8833546 |
| MLAPSNP | ASSQEGQNNQAPL | 0.8325701 |
| LVNNAAI | ASSQEGQNNQAPL | 0.8879424 |
| LTGEINTLA | ASSQEGQNNQAPL | 0.6797211 |
| LSIFGPV | ASSQEGQNNQAPL | 0.8790709 |
| LLALLANL | ASSQEGQNNQAPL | 0.8111496 |
| LISNASDALDKLR | ASSQEGQNNQAPL | 0.88572836 |
| LINEAVP | ASSQEGQNNQAPL | 0.8867937 |
| LGYAFSGL | ASSQEGQNNQAPL | 0.5473764 |
| LDFVAVAL | ASSQEGQNNQAPL | 0.9012088 |
| LAQPGAE | ASSQEGQNNQAPL | 0.82221514 |
| ISNLGAVTGF | ASSQEGQNNQAPL | 0.81505656 |
| HQEDPGRI | ASSQEGQNNQAPL | 0.9053288 |
| GTPGEPTP | ASSQEGQNNQAPL | 0.8692751 |
| GFFNSGS | ASSQEGQNNQAPL | 0.9013729 |
| GAGGGGLTAGAN | ASSQEGQNNQAPL | 0.87946844 |
| FVDALVNM | ASSQEGQNNQAPL | 0.8750316 |
| FLTVAPEL | ASSQEGQNNQAPL | 0.8974601 |
| FETLTAL | ASSQEGQNNQAPL | 0.64191025 |
| DVINPIEL | ASSQEGQNNQAPL | 0.8086719 |
| APVSGAPV | ASSQEGQNNQAPL | 0.9025544 |
| ALVGGLAW | ASSQEGQNNQAPL | 0.6549635 |
| ALAAAAATAI | ASSQEGQNNQAPL | 0.6561897 |
| AFGLGLLL | ASSQEGQNNQAPL | 0.6177372 |

| | | |
|---|---|---|
| AFFAAASL | ASSQEGQNNQAPL | 0.5939647 |
| AAAALPAFSPP | ASSQEGQNNQAPL | 0.6183606 |
| VVLAPAL | ASSQEGQNNQAPL | 0.65247715 |
| TIGHVDHGKTTL | ASSQEGQNNQAPL | 0.66472393 |
| LISNASDALDK | ASSQEGQNNQAPL | 0.6555257 |
| LISNASDAL | ASSQEGQNNQAPL | 0.60330915 |
| ISNASDALDK | ASSQEGQNNQAPL | 0.6619066 |
| NQAETLVYQTE | ASSQEGQNNQAPL | 0.647895 |
| TGGTGGTAGAG | ASSQEGQNNQAPL | 0.6402973 |
| TGSTAGGAGGA | ASSQEGQNNQAPL | 0.6457334 |
| DPLAVPVAAGPV | ASSQEGQNNQAPL | 0.6574079 |
| GGTGGTPGNGGN | ASSQEGQNNQAPL | 0.62913436 |
| PAGLAGVDPATL | ASSQEGQNNQAPL | 0.61302453 |
| GNNLVVEAPHAGASV | ASSQEGQNNQAPL | 0.6483181 |
| KVTRIMSASTLEQPA | ASSQEGQNNQAPL | 0.6556591 |
| LGFALPIALPRRTVV | ASSQEGQNNQAPL | 0.65478975 |
| TRLGNLTARKLVVLV | ASSQEGQNNQAPL | 0.66645575 |
| WSQQRRLIYFPSAGP | ASSQEGQNNQAPL | 0.66008365 |
| GPNGNPDPMAAAVDIR | ASSQEGQNNQAPL | 0.6567222 |
| TQRCVSVADLRAAASR | ASSQEGQNNQAPL | 0.44113848 |
| QFVLDVAEHPVEVLTETV | ASSQEGQNNQAPL | 0.5712104 |
| GGAGGTGGAGGNGGTQAGDG | ASSQEGQNNQAPL | 0.62915653 |
| PLLATLRAIAVDVDAKPAQV | ASSQEGQNNQAPL | 0.63427097 |
| APKFRTGGYMIDFWGVGYQVAKRM | ASSQEGQNNQAPL | 0.43138215 |
| GAGGAGGAGGAAGGPGGTGGTGGN | ASSQEGQNNQAPL | 0.65473455 |
| IGALGLLATDVAPAGSVAFGVALA | ASSQEGQNNQAPL | 0.64028835 |
| AGGAGGIGGNGGAGGAGGVLMGNG | ASSQEGQNNQAPL | 0.6262135 |
| ADELIGLL | ASSQEGQNNQAPL | 0.65841955 |
| ADRALKA | ASSQEGQNNQAPL | 0.6352563 |
| AGAVGQVGGAGGA | ASSQEGQNNQAPL | 0.60196614 |
| AGGGGGLGGAP | ASSQEGQNNQAPL | 0.6281465 |
| AGGLLYGNGGAG | ASSQEGQNNQAPL | 0.6626615 |
| AGLESIGL | ASSQEGQNNQAPL | 0.60039055 |
| AGLKTGII | ASSQEGQNNQAPL | 0.6629239 |
| ASLAALLL | ASSQEGQNNQAPL | 0.5936548 |
| ASVINISSI | ASSQEGQNNQAPL | 0.6639925 |
| ATPPGPA | ASSQEGQNNQAPL | 0.57687294 |
| ATVASALV | ASSQEGQNNQAPL | 0.6607851 |
| ATVASLIL | ASSQEGQNNQAPL | 0.64112496 |
| AVNGPVLIRH | ASSQEGQNNQAPL | 0.66903275 |
| DGAAGAVGI | ASSQEGQNNQAPL | 0.6303136 |
| DPLLHGI | ASSQEGQNNQAPL | 0.6605148 |
| ELGAALGAL | ASSQEGQNNQAPL | 0.6480975 |
| ELGRTQAV | ASSQEGQNNQAPL | 0.6637445 |
| FVAGGYLS | ASSQEGQNNQAPL | 0.591748 |
| GASGGAGGAGGW | ASSQEGQNNQAPL | 0.6377914 |
| GGAGAGGDGGAGGI | ASSQEGQNNQAPL | 0.56273335 |
| GTGHNGSQPGMGGNG | ASSQEGQNNQAPL | 0.64729244 |
| GYAAASASAS | ASSQEGQNNQAPL | 0.6577225 |
| HQVDVAAVQQG | ASSQEGQNNQAPL | 0.38703135 |
| IGAAIAHT | ASSQEGQNNQAPL | 0.66257066 |

| | | |
|---|---|---|
| IGMIEVAQL | ASSQEGQNNQAPL | 0.57690185 |
| IRLAPEHTSAMD | ASSQEGQNNQAPL | 0.6045441 |
| IYNVSVL | ASSQEGQNNQAPL | 0.658371 |
| KQGFLGM | ASSQEGQNNQAPL | 0.65652764 |
| LESVITQL | ASSQEGQNNQAPL | 0.65720487 |
| LGLGALVVV | ASSQEGQNNQAPL | 0.63523877 |
| LIGNGGNGGSGGI | ASSQEGQNNQAPL | 0.64061356 |
| LNAINAPTL | ASSQEGQNNQAPL | 0.64785427 |
| LQLLNTGV | ASSQEGQNNQAPL | 0.5567763 |
| LQTSIEF | ASSQEGQNNQAPL | 0.6097768 |
| MSFAVSEL | ASSQEGQNNQAPL | 0.66529036 |
| NTGALGL | ASSQEGQNNQAPL | 0.50199133 |
| PPGAGGAGGAGTT | ASSQEGQNNQAPL | 0.62211823 |
| QLVGLGI | ASSQEGQNNQAPL | 0.49982074 |
| RFLEQVHQL | ASSQEGQNNQAPL | 0.4277776 |
| RGVGYVLR | ASSQEGQNNQAPL | 0.4848321 |
| SATGVFAGVI | ASSQEGQNNQAPL | 0.5483941 |
| SLPAITLP | ASSQEGQNNQAPL | 0.623692 |
| STALAFTV | ASSQEGQNNQAPL | 0.51251316 |
| SVLTVGLGK | ASSQEGQNNQAPL | 0.41867635 |
| TALAGAGSGI | ASSQEGQNNQAPL | 0.5636178 |
| TGLPFTGI | ASSQEGQNNQAPL | 0.4762559 |
| VAEHVTV | ASSQEGQNNQAPL | 0.52830833 |
| VDIVLNSL | ASSQEGQNNQAPL | 0.49379367 |
| VLTGGGAVL | ASSQEGQNNQAPL | 0.5558358 |
| VNVGLAYGI | ASSQEGQNNQAPL | 0.54484105 |
| VSSAAVGAV | ASSQEGQNNQAPL | 0.44058996 |
| VTSVYEAGV | ASSQEGQNNQAPL | 0.48579204 |
| WVALDRGA | ASSQEGQNNQAPL | 0.45101917 |
| YLVGSVVI | ASSQEGQNNQAPL | 0.44542047 |
| GPPGTGKTLL | ASSQEGQNNQAPL | 0.5374174 |
| GVGMTKF | ASSQEGQNNQAPL | 0.5553671 |
| IGIGGSDL | ASSQEGQNNQAPL | 0.4772211 |
| IGTLLAEL | ASSQEGQNNQAPL | 0.31854293 |
| IYTSGTT | ASSQEGQNNQAPL | 0.33607095 |
| LSFPAGV | ASSQEGQNNQAPL | 0.4687208 |
| LYTSGTT | ASSQEGQNNQAPL | 0.4247163 |
| VGAFPFGF | ASSQEGQNNQAPL | 0.331051 |
| VGAGTVEF | ASSQEGQNNQAPL | 0.57079244 |
| VGKSTLLL | ASSQEGQNNQAPL | 0.5082775 |
| AALAHAGGGIV | ASSQEGQNNQAPL | 0.50472504 |
| AALVAVSLL | ASSQEGQNNQAPL | 0.55251795 |
| AAVSRAVAAL | ASSQEGQNNQAPL | 0.52612495 |
| AGGSSGIGGF | ASSQEGQNNQAPL | 0.44334194 |
| AGLTVANANG | ASSQEGQNNQAPL | 0.4257726 |
| AGLVGAVL | ASSQEGQNNQAPL | 0.43139955 |
| ALSQAFTV | ASSQEGQNNQAPL | 0.35129368 |
| ALTFSVSL | ASSQEGQNNQAPL | 0.58374834 |
| ALTGAALVAL | ASSQEGQNNQAPL | 0.3534319 |
| ASFSIGAGVL | ASSQEGQNNQAPL | 0.5327685 |
| DGSVYKI | ASSQEGQNNQAPL | 0.41077 |

| | | |
|---|---|---|
| EQLAEIV | ASSQEGQNNQAPL | 0.54965615 |
| FAVSELTVM | ASSQEGQNNQAPL | 0.5213333 |
| GAAPLAAGL | ASSQEGQNNQAPL | 0.63931715 |
| GADGGDGGSGGAGG | ASSQEGQNNQAPL | 0.47153914 |
| GFGLDTGGAGG | ASSQEGQNNQAPL | 0.5337795 |
| GLASGGLL | ASSQEGQNNQAPL | 0.42012778 |
| GRIAGAAVLL | ASSQEGQNNQAPL | 0.62872326 |
| GSAGSLNTSF | ASSQEGQNNQAPL | 0.37727284 |
| GTGQIGI | ASSQEGQNNQAPL | 0.46986324 |
| GTLIALIL | ASSQEGQNNQAPL | 0.37205485 |
| IAATMGFVVG | ASSQEGQNNQAPL | 0.5479181 |
| IFAQALL | ASSQEGQNNQAPL | 0.52149296 |
| IGAAVALGF | ASSQEGQNNQAPL | 0.37605035 |
| IGLTPLAL | ASSQEGQNNQAPL | 0.5518663 |
| ILGSNTSTL | ASSQEGQNNQAPL | 0.5022108 |
| IVFSLTVL | ASSQEGQNNQAPL | 0.4379828 |
| LASKATGF | ASSQEGQNNQAPL | 0.58929545 |
| LASVALIL | ASSQEGQNNQAPL | 0.5423622 |
| LFAVGLTL | ASSQEGQNNQAPL | 0.58212936 |
| LFNGGIGTYI | ASSQEGQNNQAPL | 0.4933375 |
| LTLGAGII | ASSQEGQNNQAPL | 0.4242112 |
| LVAAALGDI | ASSQEGQNNQAPL | 0.47666356 |
| NTAFVQLGI | ASSQEGQNNQAPL | 0.46086136 |
| QLFSGAEL | ASSQEGQNNQAPL | 0.41262722 |
| SAATFITL | ASSQEGQNNQAPL | 0.58140504 |
| SNYGLANLGV | ASSQEGQNNQAPL | 0.7455771 |
| SYGGAYAVM | ASSQEGQNNQAPL | 0.82449853 |
| TTGVGGLTLG | ASSQEGQNNQAPL | 0.68895733 |
| TTQTIYI | ASSQEGQNNQAPL | 0.6040516 |
| VAAGAGLGAVI | ASSQEGQNNQAPL | 0.72580326 |
| VAAVVTLL | ASSQEGQNNQAPL | 0.8141297 |
| VGFLGGLI | ASSQEGQNNQAPL | 0.8187985 |
| VGVIPILL | ASSQEGQNNQAPL | 0.7856928 |
| VIFASSLI | ASSQEGQNNQAPL | 0.52686834 |
| VKGGDGGAAAG | ASSQEGQNNQAPL | 0.7633811 |
| VTKTFTNI | ASSQEGQNNQAPL | 0.6685916 |
| VVAAAILL | ASSQEGQNNQAPL | 0.733431 |
| YAALDVEL | ASSQEGQNNQAPL | 0.7870371 |
| LASVALLL | ASSQEGQNNQAPL | 0.71721935 |
| LDAVHRV | ASSQEGQNNQAPL | 0.6514547 |
| QGLSSSIF | ASSQEGQNNQAPL | 0.6705409 |
| RGTFANIRL | ASSQEGQNNQAPL | 0.798133 |
| RQLWWGHRI | ASSQEGQNNQAPL | 0.75559694 |
| YGPPGTGKTLL | ASSQEGQNNQAPL | 0.762805 |
| AALVGAPANVA | ASSQEGQNNQAPL | 0.83077437 |
| AATFRRTE | ASSQEGQNNQAPL | 0.79921883 |
| AGALGGGI | ASSQEGQNNQAPL | 0.80726045 |
| AGGAGGAGGIGAG | ASSQEGQNNQAPL | 0.2471476 |
| AGSGALGL | ASSQEGQNNQAPL | 0.43491116 |
| AQIADQI | ASSQEGQNNQAPL | 0.67390466 |
| ASAVAVTYL | ASSQEGQNNQAPL | 0.70845556 |

| | | |
|---|---|---|
| ASTVLGGTGG | ASSQEGQNNQAPL | 0.50514954 |
| AVGGEPAK | ASSQEGQNNQAPL | 0.7789583 |
| CEQGTFGGTC | ASSQEGQNNQAPL | 0.62885153 |
| DEGKAPAG | ASSQEGQNNQAPL | 0.6061137 |
| DTVVFPAL | ASSQEGQNNQAPL | 0.8050592 |
| FEGITMAG | ASSQEGQNNQAPL | 0.66975474 |
| FPLVVQD | ASSQEGQNNQAPL | 0.61783427 |
| GAFTTPQ | ASSQEGQNNQAPL | 0.6585967 |
| GFGGFGTAGGVGGA | ASSQEGQNNQAPL | 0.78851867 |
| GFLAQFL | ASSQEGQNNQAPL | 0.45968145 |
| GGAGGAGGNAGVA | ASSQEGQNNQAPL | 0.78151983 |
| GGVGTAGAGGAGGAGG | ASSQEGQNNQAPL | 0.6905291 |
| GNGGGGDAGNAGSGGN | ASSQEGQNNQAPL | 0.81478846 |
| GTGGAGGAGNDAGS | ASSQEGQNNQAPL | 0.47987175 |
| GTTNTGWFNS | ASSQEGQNNQAPL | 0.7943087 |
| IFMGPLFTL | ASSQEGQNNQAPL | 0.7448075 |
| ILAHAIEAI | ASSQEGQNNQAPL | 0.8374138 |
| IRSAIREAL | ASSQEGQNNQAPL | 0.67479694 |
| ITFAGGG | ASSQEGQNNQAPL | 0.7014823 |
| ITGMSGHQEV | ASSQEGQNNQAPL | 0.6581354 |
| IVAGLACTL | ASSQEGQNNQAPL | 0.8214572 |
| IVGGGLAGA | ASSQEGQNNQAPL | 0.46748698 |
| IVTGVLNI | ASSQEGQNNQAPL | 0.69760686 |
| LAQQGCI | ASSQEGQNNQAPL | 0.4877437 |
| LCLHVEN | ASSQEGQNNQAPL | 0.7722814 |
| LDAISDGS | ASSQEGQNNQAPL | 0.77496296 |
| LGGFAAML | ASSQEGQNNQAPL | 0.4305076 |
| LGGKNPMI | ASSQEGQNNQAPL | 0.7925204 |
| LGPKGVAAIL | ASSQEGQNNQAPL | 0.75159365 |
| LMVGVSI | ASSQEGQNNQAPL | 0.61851716 |
| LVGAAFVL | ASSQEGQNNQAPL | 0.818433 |
| LVLAVAPL | ASSQEGQNNQAPL | 0.7366768 |
| MAAIAVIDAA | ASSQEGQNNQAPL | 0.81222206 |
| NNGINAELQ | ASSQEGQNNQAPL | 0.760638 |
| RGEPLVLVHG | ASSQEGQNNQAPL | 0.7019702 |
| RGGAGGAAGLI | ASSQEGQNNQAPL | 0.7832135 |
| RSYRMHPAV | ASSQEGQNNQAPL | 0.65129775 |
| SAAHLLAGQLA | ASSQEGQNNQAPL | 0.6658436 |
| SAAVGSY | ASSQEGQNNQAPL | 0.8120474 |
| SPHAVLHRI | ASSQEGQNNQAPL | 0.87334776 |
| SQIAKVL | ASSQEGQNNQAPL | 0.8747381 |
| TALGHLVP | ASSQEGQNNQAPL | 0.8044629 |
| TAPTTSGAA | ASSQEGQNNQAPL | 0.7308177 |
| TDGDDGNGGSPGT | ASSQEGQNNQAPL | 0.8253859 |
| TDGLGEN | ASSQEGQNNQAPL | 0.87802976 |
| TGGPPLPGQA | ASSQEGQNNQAPL | 0.89338624 |
| TIPAITI | ASSQEGQNNQAPL | 0.87075317 |
| TIPGFTI | ASSQEGQNNQAPL | 0.6523923 |
| TSIASGALA | ASSQEGQNNQAPL | 0.87963474 |
| TVTLPGAAI | ASSQEGQNNQAPL | 0.86141264 |
| VASGLSAALP | ASSQEGQNNQAPL | 0.8416452 |

| | | |
|---|---|---|
| VGAGVIGI | ASSQEGQNNQAPL | 0.87863934 |
| VGPKRFGV | ASSQEGQNNQAPL | 0.85927445 |
| VNVAFPD | ASSQEGQNNQAPL | 0.8048593 |
| VSMTNSGVS | ASSQEGQNNQAPL | 0.8161115 |
| VTDVLGDLSV | ASSQEGQNNQAPL | 0.8701831 |
| VVVVGAGFAGL | ASSQEGQNNQAPL | 0.8806952 |
| VYTVGTVV | ASSQEGQNNQAPL | 0.8860028 |
| WTTFSGP | ASSQEGQNNQAPL | 0.90958476 |
| YDELVEAF | ASSQEGQNNQAPL | 0.8958434 |
| YGTGGAGGNGG | ASSQEGQNNQAPL | 0.88516676 |
| YTNVMAV | ASSQEGQNNQAPL | 0.34484375 |
| FLEQVHQL | ASSQEGQNNQAPL | 0.65394676 |
| GLSLLAVAV | ASSQEGQNNQAPL | 0.79881346 |
| LEQVHQL | ASSQEGQNNQAPL | 0.8446926 |
| LNAAGAI | ASSQEGQNNQAPL | 0.487949 |
| PAVHASL | ASSQEGQNNQAPL | 0.8376697 |
| TELEDDA | ASSQEGQNNQAPL | 0.8010013 |
| TPPGPPN | ASSQEGQNNQAPL | 0.7723984 |
| TPPPPPPP | ASSQEGQNNQAPL | 0.8965782 |
| TQSPALI | ASSTIANTEVF | 0.8248821 |
| EFAAAFPQ | ASSTIANTEVF | 0.7722717 |
| LLSGNLDVL | ASSTIANTEVF | 0.8279915 |
| GPANLGNYNI | ASSTIANTEVF | 0.8999644 |
| TLGPTLP | ASSTIANTEVF | 0.76636577 |
| VYAYSSCCK | ASSTIANTEVF | 0.88962203 |
| APAPAPAPNI | ASSTIANTEVF | 0.7383288 |
| PGGPVPIP | ASSTIANTEVF | 0.90110403 |
| FLTVALEP | ASSTIANTEVF | 0.63698953 |
| SQYFGANL | ASSTIANTEVF | 0.89618397 |
| GGVKPAA | ASSTIANTEVF | 0.8562787 |
| GGTGGAGGAGGW | ASSTIANTEVF | 0.9166807 |
| GVGGVAGLGGAGG | ASSTIANTEVF | 0.84240806 |
| VLLVLATTI | ASSTIANTEVF | 0.87596184 |
| IGIFAAML | ASSTIANTEVF | 0.7910674 |
| AVGDLID | ASSTIANTEVF | 0.8910432 |
| NISPLPL | ASSTIANTEVF | 0.6690939 |
| VTAVFVAI | ASSTIANTEVF | 0.8553791 |
| SGDDNDAAL | ASSTIANTEVF | 0.5824327 |
| AVAALAHS | ASSTIANTEVF | 0.86913013 |
| GGVTVMATP | ASSTIANTEVF | 0.8775531 |
| AGLAANVRD | ASSTIANTEVF | 0.45180434 |
| TNRPDLIDPALLRPG | ASSTIANTEVF | 0.9011178 |
| LANLGVTAL | ASSTIANTEVF | 0.780774 |
| TGERIEL | ASSTIANTEVF | 0.7431823 |
| NGEELAA | ASSTIANTEVF | 0.9026673 |
| VLSLVWG | ASSTIANTEVF | 0.8540987 |
| IGYALWL | ASSTIANTEVF | 0.88912 |
| LSLLLAPL | ASSTIANTEVF | 0.83677113 |
| TLDEALK | ASSTIANTEVF | 0.83026445 |
| TDGAGTAEGG | ASSTIANTEVF | 0.8572606 |
| NAPGGAGGNGG | ASSTIANTEVF | 0.7035261 |

| | | |
|---|---|---|
| VSVFMGMGGNF | ASSTIANTEVF | 0.7847651 |
| LASYLAAL | ASSTIANTEVF | 0.90681386 |
| LVGGTPAL | ASSTIANTEVF | 0.82849836 |
| VRNADVL | ASSTIANTEVF | 0.82262343 |
| TCSSPFLVSAM | ASSTIANTEVF | 0.7240554 |
| VHITDAE | ASSTIANTEVF | 0.66618186 |
| EYAAPSF | ASSTIANTEVF | 0.760116 |
| VVDAFHGGL | ASSTIANTEVF | 0.836787 |
| ASVVDPQ | ASSTIANTEVF | 0.8536272 |
| SYNEFGGLS | ASSTIANTEVF | 0.8256807 |
| DPDGSVY | ASSTIANTEVF | 0.66785765 |
| MVSTIHATE | ASSTIANTEVF | 0.83703685 |
| LAAGVADPV | ASSTIANTEVF | 0.8156891 |
| TKEQIAA | ASSTIANTEVF | 0.7687515 |
| SLASPAAL | ASSTIANTEVF | 0.8491781 |
| IGLVFYGV | ASSTIANTEVF | 0.8268108 |
| FLLDASNL | ASSTIANTEVF | 0.7487327 |
| YQVGNFP | ASSTIANTEVF | 0.790752 |
| ESLLSQST | ASSTIANTEVF | 0.81948674 |
| LFTVGFAAVG | ASSTIANTEVF | 0.8387991 |
| ATPPTPT | ASSTIANTEVF | 0.84531415 |
| TDLATLG | ASSTIANTEVF | 0.8769562 |
| SAGGTGR | ASSTIANTEVF | 0.8552889 |
| GFSNSGS | ASSTIANTEVF | 0.8339629 |
| GGNGGSGGTSV | ASSTIANTEVF | 0.36659318 |
| GTSVATGGAGN | ASSTIANTEVF | 0.63843143 |
| NGGNGGSGGTS | ASSTIANTEVF | 0.78642064 |
| AVVKFPH | ASSTIANTEVF | 0.79957235 |
| TPPPPPP | ASSTIANTEVF | 0.47505808 |
| AAPAAAAAPA | ASSTIANTEVF | 0.81146526 |
| IALNEDL | ASSTIANTEVF | 0.7777326 |
| IVNVSSM | ASSTIANTEVF | 0.71003854 |
| TGIPPAPR | ASSTIANTEVF | 0.85929614 |
| FDLGGGTF | ASSTIANTEVF | 0.76197195 |
| FDLGGGTFDVS | ASSTIANTEVF | 0.7735772 |
| NEPTAAA | ASSTIANTEVF | 0.81291693 |
| NEPTAAALA | ASSTIANTEVF | 0.8677169 |
| GSIINISSI | ASSTIANTEVF | 0.733706 |
| VAPGNAGTA | ASSTIANTEVF | 0.85174924 |
| TGGGASLE | ASSTIANTEVF | 0.7315839 |
| VAVYNHYKRI | ASSTIANTEVF | 0.8655969 |
| TDPAPATT | ASSTIANTEVF | 0.639608 |
| TNRPDILDPALLRPG | ASSTIANTEVF | 0.8485076 |
| LTTTTTS | ASSTIANTEVF | 0.7865153 |
| PTGSGKTL | ASSTIANTEVF | 0.88442147 |
| TGSGKTL | ASSTIANTEVF | 0.80403554 |
| TIGHVDHGK | ASSTIANTEVF | 0.85266906 |
| TIGHVDHGKTT | ASSTIANTEVF | 0.77639914 |
| GHVDHGKTT | ASSTIANTEVF | 0.82254416 |
| AQFEHTLL | ASSTIANTEVF | 0.60954094 |
| IVNQDEL | ASSTIANTEVF | 0.8177467 |

| | | |
|---|---|---|
| TNGSQFF | ASSTIANTEVF | 0.6084024 |
| TGKGIVE | ASSTIANTEVF | 0.8300764 |
| LISNASDA | ASSTIANTEVF | 0.8226617 |
| AELRTLGELTKIAKAHGAQ | ASSTIANTEVF | 0.46350515 |
| APNAVAAAEPSW | ASSTIANTEVF | 0.863532 |
| APQVAGAATAVL | ASSTIANTEVF | 0.6910434 |
| ARRDPAAAAAILPTDA | ASSTIANTEVF | 0.73625654 |
| DAAARDVLAEAGLAEYF | ASSTIANTEVF | 0.8666211 |
| DDQIVLSAKANLAKVEQ | ASSTIANTEVF | 0.8049509 |
| DELLPDTGEDSDEDGH | ASSTIANTEVF | 0.8507746 |
| ELPEAPANRATVPQD | ASSTIANTEVF | 0.8013958 |
| FGQSKIERVTGIG | ASSTIANTEVF | 0.7932848 |
| GAGGTGSTAGGAGGAGGAGGLYA | ASSTIANTEVF | 0.81367195 |
| GHGGTGAAGGNGGTGGAGGSGIDG | ASSTIANTEVF | 0.6045797 |
| GNGGNGGAGGEGGPGGAGGAGA | ASSTIANTEVF | 0.766499 |
| GSTAGNGGNGGNATNGTIAGQPAG | ASSTIANTEVF | 0.86558 |
| GTGGDGGNGGNGGTGG | ASSTIANTEVF | 0.3822161 |
| GVRRPAGIPADAPV | ASSTIANTEVF | 0.77196634 |
| GYRQVTDAHLLTLARR | ASSTIANTEVF | 0.275167 |
| IAKMGQDGHDRG | ASSTIANTEVF | 0.3523206 |
| KAEPSIEIQYT | ASSTIANTEVF | 0.33771157 |
| KPVRIRMAADEDTVAVT | ASSTIANTEVF | 0.5021174 |
| MPPRIAGMRLLVIKPEP | ASSTIANTEVF | 0.74446476 |
| MKFYAVALFRSRGVPPTR | ASSTIANTEVF | 0.5480075 |
| NAGGFGILGAVAHSP | ASSTIANTEVF | 0.2713869 |
| NEGREITLAMAEM | ASSTIANTEVF | 0.48455447 |
| NELVEWSHPLGHRWRWE | ASSTIANTEVF | 0.26816982 |
| NLGAGGTGGNGGNPG | ASSTIANTEVF | 0.32126504 |
| NTEWPKWETVMAADPVD | ASSTIANTEVF | 0.47599065 |
| PASAAPAETQP | ASSTIANTEVF | 0.36245885 |
| QQSSDGKNGVRAAEITGE | ASSTIANTEVF | 0.27495408 |
| RAPDAVTTSWWKE | ASSTIANTEVF | 0.30735597 |
| SGFGNVGAHTSGWFNQ | ASSTIANTEVF | 0.37449124 |
| SGVAGAVGQAASVGGLKV | ASSTIANTEVF | 0.43576306 |
| SNGGAGTGSAGGNGGT | ASSTIANTEVF | 0.43044823 |
| TGGAGGEGGAAG | ASSTIANTEVF | 0.6755514 |
| TGGDGGAAGAGGAPGHG | ASSTIANTEVF | 0.5626007 |
| TMPPLLAAIANAGRHGV | ASSTIANTEVF | 0.59734666 |
| TSSTAGNSGGA | ASSTIANTEVF | 0.1772949 |
| VPATVPVGILC | ASSTIANTEVF | 0.2353794 |
| VQRPDSGGVVLTGRIS | ASSTIANTEVF | 0.25877634 |
| WTPNPVPATKGVPAGLVQAT | ASSTIANTEVF | 0.3458785 |
| YALKRALAKFSDRFDVVII | ASSTIANTEVF | 0.19925688 |
| YLIGSVIV | ASSTIANTEVF | 0.4678107 |
| YGQLVAVID | ASSTIANTEVF | 0.29337612 |
| VTVGGMTLPA | ASSTIANTEVF | 0.30980843 |
| VLALPAV | ASSTIANTEVF | 0.5764293 |
| VIPPTLNYET | ASSTIANTEVF | 0.2689417 |
| VGMGGMHPSSG | ASSTIANTEVF | 0.26118132 |
| VGLAGGAATA | ASSTIANTEVF | 0.23098381 |
| VGGAGGAGGSGP | ASSTIANTEVF | 0.41709387 |

| | | |
|---|---|---|
| VFAAFMV | ASSTIANTEVF | 0.21324039 |
| TLYDKLR | ASSTIANTEVF | 0.51880705 |
| TLAVFTVL | ASSTIANTEVF | 0.25743636 |
| TGSVQPARV | ASSTIANTEVF | 0.5464333 |
| TGGGAGSGGT | ASSTIANTEVF | 0.259194 |
| SHLETSL | ASSTIANTEVF | 0.562533 |
| SFTTGMT | ASSTIANTEVF | 0.36435986 |
| QVEVAKRMGVRQPTVS | ASSTIANTEVF | 0.7970088 |
| QGSGGPALAM | ASSTIANTEVF | 0.41922957 |
| MLAPSNP | ASSTIANTEVF | 0.33329156 |
| LVNNAAI | ASSTIANTEVF | 0.23240806 |
| LTGEINTLA | ASSTIANTEVF | 0.76751643 |
| LSIFGPV | ASSTIANTEVF | 0.21381323 |
| LLALLANL | ASSTIANTEVF | 0.3105998 |
| LISNASDALDKLR | ASSTIANTEVF | 0.23095287 |
| LINEAVP | ASSTIANTEVF | 0.43773302 |
| LGYAFSGL | ASSTIANTEVF | 0.5144259 |
| LDFVAVAL | ASSTIANTEVF | 0.18939124 |
| LAQPGAE | ASSTIANTEVF | 0.49739432 |
| ISNLGAVTGF | ASSTIANTEVF | 0.432267 |
| HQEDPGRI | ASSTIANTEVF | 0.2378872 |
| GTPGEPTP | ASSTIANTEVF | 0.681013 |
| GFFNSGS | ASSTIANTEVF | 0.38246867 |
| GAGGGGLTAGAN | ASSTIANTEVF | 0.69675744 |
| FVDALVNM | ASSTIANTEVF | 0.29940233 |
| FLTVAPEL | ASSTIANTEVF | 0.27219003 |
| FETLTAL | ASSTIANTEVF | 0.37419242 |
| DVINPIEL | ASSTIANTEVF | 0.21740565 |
| APVSGAPV | ASSTIANTEVF | 0.2166947 |
| ALVGGLAW | ASSTIANTEVF | 0.6503484 |
| ALAAAAATAI | ASSTIANTEVF | 0.74660957 |
| AFGLGLLL | ASSTIANTEVF | 0.7463409 |
| AFFAAASL | ASSTIANTEVF | 0.6887418 |
| AAAALPAFSPP | ASSTIANTEVF | 0.6965883 |
| VVLAPAL | ASSTIANTEVF | 0.7265148 |
| TIGHVDHGKTTL | ASSTIANTEVF | 0.74969256 |
| LISNASDALDK | ASSTIANTEVF | 0.75719756 |
| LISNASDAL | ASSTIANTEVF | 0.75224257 |
| ISNASDALDK | ASSTIANTEVF | 0.71595675 |
| NQAETLVYQTE | ASSTIANTEVF | 0.7500782 |
| TGGTGGTAGAG | ASSTIANTEVF | 0.7332319 |
| TGSTAGGAGGA | ASSTIANTEVF | 0.7358624 |
| DPLAVPVAAGPV | ASSTIANTEVF | 0.745201 |
| GGTGGTPGNGGN | ASSTIANTEVF | 0.73592526 |
| PAGLAGVDPATL | ASSTIANTEVF | 0.7143845 |
| GNNLVVEAPHAGASV | ASSTIANTEVF | 0.6873481 |
| KVTRIMSASTLEQPA | ASSTIANTEVF | 0.7455596 |
| LGFALPIALPRRTVV | ASSTIANTEVF | 0.7543703 |
| TRLGNLTARKLVVLV | ASSTIANTEVF | 0.7507322 |
| WSQQRRLIYFPSAGP | ASSTIANTEVF | 0.7595822 |
| GPNGNPDPMAAAVDIR | ASSTIANTEVF | 0.75032 |

| | | |
|---|---|---|
| TQRCVSVADLRAAASR | ASSTIANTEVF | 0.74956465 |
| QFVLDVAEHPVEVLTETV | ASSTIANTEVF | 0.407682 |
| GGAGGTGGAGGNGGTQAGDG | ASSTIANTEVF | 0.60031927 |
| PLLATLRAIAVDVDAKPAQV | ASSTIANTEVF | 0.7108824 |
| APKFRTGGYMIDFWGVGYQVAKRM | ASSTIANTEVF | 0.7210764 |
| GAGGAGGAGGAAGGPGGTGGTGGN | ASSTIANTEVF | 0.39556134 |
| IGALGLLATDVAPAGSVAFGVALA | ASSTIANTEVF | 0.7503885 |
| AGGAGGIGGNGGAGGAGGVLMGNG | ASSTIANTEVF | 0.72928977 |
| ADELIGLL | ASSTIANTEVF | 0.7101649 |
| ADRALKA | ASSTIANTEVF | 0.75251806 |
| AGAVGQVGGAGGA | ASSTIANTEVF | 0.7254181 |
| AGGGGGLGGAP | ASSTIANTEVF | 0.6385442 |
| AGGLLYGNGGAG | ASSTIANTEVF | 0.7120238 |
| AGLESIGL | ASSTIANTEVF | 0.7576982 |
| AGLKTGII | ASSTIANTEVF | 0.7049326 |
| ASLAALLL | ASSTIANTEVF | 0.7552258 |
| ASVINISSI | ASSTIANTEVF | 0.70112765 |
| ATPPGPA | ASSTIANTEVF | 0.75508034 |
| ATVASALV | ASSTIANTEVF | 0.66963506 |
| ATVASLIL | ASSTIANTEVF | 0.75310075 |
| AVNGPVLIRH | ASSTIANTEVF | 0.7363437 |
| DGAAGAVGI | ASSTIANTEVF | 0.7609941 |
| DPLLHGI | ASSTIANTEVF | 0.7272823 |
| ELGAALGAL | ASSTIANTEVF | 0.7540735 |
| ELGRTQAV | ASSTIANTEVF | 0.742416 |
| FVAGGYLS | ASSTIANTEVF | 0.75066996 |
| GASGGAGGAGGW | ASSTIANTEVF | 0.6735739 |
| GGAGAGGDGGAGGI | ASSTIANTEVF | 0.7201698 |
| GTGHNGSQPGMGGNG | ASSTIANTEVF | 0.6554604 |
| GYAAASASAS | ASSTIANTEVF | 0.7306175 |
| HQVDVAAVQQG | ASSTIANTEVF | 0.7528108 |
| IGAAIAHT | ASSTIANTEVF | 0.31763506 |
| IGMIEVAQL | ASSTIANTEVF | 0.74626315 |
| IRLAPEHTSAMD | ASSTIANTEVF | 0.657228 |
| IYNVSVL | ASSTIANTEVF | 0.7051512 |
| KQGFLGM | ASSTIANTEVF | 0.75048745 |
| LESVITQL | ASSTIANTEVF | 0.7343826 |
| LGLGALVVV | ASSTIANTEVF | 0.74670374 |
| LIGNGGNGGSGGI | ASSTIANTEVF | 0.7028083 |
| LNAINAPTL | ASSTIANTEVF | 0.7256992 |
| LQLLNTGV | ASSTIANTEVF | 0.7411357 |
| LQTSIEF | ASSTIANTEVF | 0.40694577 |
| MSFAVSEL | ASSTIANTEVF | 0.67862463 |
| NTGALGL | ASSTIANTEVF | 0.7579284 |
| PPGAGGAGGAGTT | ASSTIANTEVF | 0.6672772 |
| QLVGLGI | ASSTIANTEVF | 0.67985743 |
| RFLEQVHQL | ASSTIANTEVF | 0.6222793 |
| RGVGYVLR | ASSTIANTEVF | 0.620692 |
| SATGVFAGVI | ASSTIANTEVF | 0.6492667 |
| SLPAITLP | ASSTIANTEVF | 0.6782114 |
| STALAFTV | ASSTIANTEVF | 0.68460953 |

| | | |
|---|---|---|
| SVLTVGLGK | ASSTIANTEVF | 0.67181337 |
| TALAGAGSGI | ASSTIANTEVF | 0.6371008 |
| TGLPFTGI | ASSTIANTEVF | 0.6725698 |
| VAEHVTV | ASSTIANTEVF | 0.6544353 |
| VDIVLNSL | ASSTIANTEVF | 0.6506624 |
| VLTGGGAVL | ASSTIANTEVF | 0.67323166 |
| VNVGLAYGI | ASSTIANTEVF | 0.6672802 |
| VSSAAVGAV | ASSTIANTEVF | 0.64563906 |
| VTSVYEAGV | ASSTIANTEVF | 0.649548 |
| WVALDRGA | ASSTIANTEVF | 0.6717973 |
| YLVGSVVI | ASSTIANTEVF | 0.67607415 |
| GPPGTGKTLL | ASSTIANTEVF | 0.67322046 |
| GVGMTKF | ASSTIANTEVF | 0.6864633 |
| IGIGGSDL | ASSTIANTEVF | 0.67404723 |
| IGTLLAEL | ASSTIANTEVF | 0.67209625 |
| IYTSGTT | ASSTIANTEVF | 0.45734295 |
| LSFPAGV | ASSTIANTEVF | 0.57467353 |
| LYTSGTT | ASSTIANTEVF | 0.65591943 |
| VGAFPFGF | ASSTIANTEVF | 0.655619 |
| VGAGTVEF | ASSTIANTEVF | 0.5140811 |
| VGKSTLLL | ASSTIANTEVF | 0.67806077 |
| AALAHAGGGIV | ASSTIANTEVF | 0.65304905 |
| AALVAVSLL | ASSTIANTEVF | 0.63615847 |
| AAVSRAVAAL | ASSTIANTEVF | 0.67517316 |
| AGGSSGIGGF | ASSTIANTEVF | 0.6461792 |
| AGLTVANANG | ASSTIANTEVF | 0.6407046 |
| AGLVGAVL | ASSTIANTEVF | 0.6558019 |
| ALSQAFTV | ASSTIANTEVF | 0.6839869 |
| ALTFSVSL | ASSTIANTEVF | 0.62556785 |
| ALTGAALVAL | ASSTIANTEVF | 0.67646646 |
| ASFSIGAGVL | ASSTIANTEVF | 0.65667003 |
| DGSVYKI | ASSTIANTEVF | 0.6812333 |
| EQLAEIV | ASSTIANTEVF | 0.60088503 |
| FAVSELTVM | ASSTIANTEVF | 0.67105985 |
| GAAPLAAGL | ASSTIANTEVF | 0.64810866 |
| GADGGDGGSGGAGG | ASSTIANTEVF | 0.68876183 |
| GFGLDTGGAGG | ASSTIANTEVF | 0.65072393 |
| GLASGGLL | ASSTIANTEVF | 0.67635685 |
| GRIAGAAVLL | ASSTIANTEVF | 0.6686554 |
| GSAGSLNTSF | ASSTIANTEVF | 0.68165517 |
| GTGQIGI | ASSTIANTEVF | 0.587806 |
| GTLIALIL | ASSTIANTEVF | 0.6557 |
| IAATMGFVVG | ASSTIANTEVF | 0.6297242 |
| IFAQALL | ASSTIANTEVF | 0.66797644 |
| IGAAVALGF | ASSTIANTEVF | 0.66690195 |
| IGLTPLAL | ASSTIANTEVF | 0.48865214 |
| ILGSNTSTL | ASSTIANTEVF | 0.68010604 |
| IVFSLTVL | ASSTIANTEVF | 0.6229404 |
| LASKATGF | ASSTIANTEVF | 0.6366501 |
| LASVALIL | ASSTIANTEVF | 0.67805564 |
| LFAVGLTL | ASSTIANTEVF | 0.6575233 |

| | | |
|---|---|---|
| LFNGGIGTYI | ASSTIANTEVF | 0.6744503 |
| LTLGAGII | ASSTIANTEVF | 0.65768325 |
| LVAAALGDI | ASSTIANTEVF | 0.66426873 |
| NTAFVQLGI | ASSTIANTEVF | 0.6733747 |
| QLFSGAEL | ASSTIANTEVF | 0.5791364 |
| SAATFITL | ASSTIANTEVF | 0.637828 |
| SNYGLANLGV | ASSTIANTEVF | 0.67907107 |
| SYGGAYAVM | ASSTIANTEVF | 0.79131913 |
| TTGVGGLTLG | ASSTIANTEVF | 0.7720578 |
| TTQTIYI | ASSTIANTEVF | 0.6204008 |
| VAAGAGLGAVI | ASSTIANTEVF | 0.5574992 |
| VAAVVTLL | ASSTIANTEVF | 0.7254506 |
| VGFLGGLI | ASSTIANTEVF | 0.78055096 |
| VGVIPILL | ASSTIANTEVF | 0.825023 |
| VIFASSLI | ASSTIANTEVF | 0.772128 |
| VKGGDGGAAAG | ASSTIANTEVF | 0.5716733 |
| VTKTFTNI | ASSTIANTEVF | 0.80122334 |
| VVAAAILL | ASSTIANTEVF | 0.7346301 |
| YAALDVEL | ASSTIANTEVF | 0.75747335 |
| LASVALLL | ASSTIANTEVF | 0.74863625 |
| LDAVHRV | ASSTIANTEVF | 0.7173334 |
| QGLSSSIF | ASSTIANTEVF | 0.66643786 |
| RGTFANIRL | ASSTIANTEVF | 0.5761295 |
| RQLWWGHRI | ASSTIANTEVF | 0.7933316 |
| YGPPGTGKTLL | ASSTIANTEVF | 0.8093959 |
| AALVGAPANVA | ASSTIANTEVF | 0.8008929 |
| AATFRRTE | ASSTIANTEVF | 0.8439351 |
| AGALGGGI | ASSTIANTEVF | 0.82207465 |
| AGGAGGAGGIGAG | ASSTIANTEVF | 0.7902186 |
| AGSGALGL | ASSTIANTEVF | 0.32243693 |
| AQIADQI | ASSTIANTEVF | 0.4772978 |
| ASAVAVTYL | ASSTIANTEVF | 0.6124774 |
| ASTVLGGTGG | ASSTIANTEVF | 0.70240366 |
| AVGGEPAK | ASSTIANTEVF | 0.3263825 |
| CEQGTFGGTC | ASSTIANTEVF | 0.7710079 |
| DEGKAPAG | ASSTIANTEVF | 0.67513835 |
| DTVVFPAL | ASSTIANTEVF | 0.6310589 |
| FEGITMAG | ASSTIANTEVF | 0.82420355 |
| FPLVVQD | ASSTIANTEVF | 0.6744733 |
| GAFTTPQ | ASSTIANTEVF | 0.5625277 |
| GFGGFGTAGGVGGA | ASSTIANTEVF | 0.6989149 |
| GFLAQFL | ASSTIANTEVF | 0.842948 |
| GGAGGAGGNAGVA | ASSTIANTEVF | 0.57524765 |
| GGVGTAGAGGAGGAGG | ASSTIANTEVF | 0.8194361 |
| GNGGGGDAGNAGSGGN | ASSTIANTEVF | 0.72286665 |
| GTGGAGGAGNDAGS | ASSTIANTEVF | 0.83979744 |
| GTTNTGWFNS | ASSTIANTEVF | 0.46639508 |
| IFMGPLFTL | ASSTIANTEVF | 0.8222829 |
| ILAHAIEAI | ASSTIANTEVF | 0.7377281 |
| IRSAIREAL | ASSTIANTEVF | 0.85573536 |
| ITFAGGG | ASSTIANTEVF | 0.70422214 |

| | | |
|---|---|---|
| ITGMSGHQEV | ASSTIANTEVF | 0.778423 |
| IVAGLACTL | ASSTIANTEVF | 0.7281841 |
| IVGGGLAGA | ASSTIANTEVF | 0.8088708 |
| IVTGVLNI | ASSTIANTEVF | 0.47316542 |
| LAQQGCI | ASSTIANTEVF | 0.6934896 |
| LCLHVEN | ASSTIANTEVF | 0.5753092 |
| LDAISDGS | ASSTIANTEVF | 0.74197996 |
| LGGFAAML | ASSTIANTEVF | 0.79816014 |
| LGGKNPMI | ASSTIANTEVF | 0.23578708 |
| LGPKGVAAIL | ASSTIANTEVF | 0.80676347 |
| LMVGVSI | ASSTIANTEVF | 0.4653826 |
| LVGAAFVL | ASSTIANTEVF | 0.5077422 |
| LVLAVAPL | ASSTIANTEVF | 0.7910328 |
| MAAIAVIDAA | ASSTIANTEVF | 0.7484534 |
| NNGINAELQ | ASSTIANTEVF | 0.81036466 |
| RGEPLVLVHG | ASSTIANTEVF | 0.65339106 |
| RGGAGGAAGLI | ASSTIANTEVF | 0.74894667 |
| RSYRMHPAV | ASSTIANTEVF | 0.77438986 |
| SAAHLLAGQLA | ASSTIANTEVF | 0.4077735 |
| SAAVGSY | ASSTIANTEVF | 0.61478025 |
| SPHAVLHRI | ASSTIANTEVF | 0.8353093 |
| SQIAKVL | ASSTIANTEVF | 0.7961829 |
| TALGHLVP | ASSTIANTEVF | 0.79502964 |
| TAPTTSGAA | ASSTIANTEVF | 0.6823118 |
| TDGDDGNGGSPGT | ASSTIANTEVF | 0.5476958 |
| TDGLGEN | ASSTIANTEVF | 0.7187421 |
| TGGPPLPGQA | ASSTIANTEVF | 0.8250203 |
| TIPAITI | ASSTIANTEVF | 0.82301867 |
| TIPGFTI | ASSTIANTEVF | 0.7758182 |
| TSIASGALA | ASSTIANTEVF | 0.49985576 |
| TVTLPGAAI | ASSTIANTEVF | 0.7959473 |
| VASGLSAALP | ASSTIANTEVF | 0.71569777 |
| VGAGVIGI | ASSTIANTEVF | 0.6987404 |
| VGPKRFGV | ASSTIANTEVF | 0.80845964 |
| VNVAFPD | ASSTIANTEVF | 0.7236665 |
| VSMTNSGVS | ASSTIANTEVF | 0.72758555 |
| VTDVLGDLSV | ASSTIANTEVF | 0.6888564 |
| VVVVGAGFAGL | ASSTIANTEVF | 0.7648817 |
| VYTVGTVV | ASSTIANTEVF | 0.809034 |
| WTTFSGP | ASSTIANTEVF | 0.80913436 |
| YDELVEAF | ASSTIANTEVF | 0.87148786 |
| YGTGGAGGNGG | ASSTIANTEVF | 0.8421235 |
| YTNVMAV | ASSTIANTEVF | 0.86255467 |
| FLEQVHQL | ASSTIANTEVF | 0.26591858 |
| GLSLLAVAV | ASSTIANTEVF | 0.4690276 |
| LEQVHQL | ASSTIANTEVF | 0.6890181 |
| LNAAGAI | ASSTIANTEVF | 0.69480324 |
| PAVHASL | ASSTIANTEVF | 0.42428568 |
| TELEDDA | ASSTIANTEVF | 0.746971 |
| TPPGPPN | ASSTIANTEVF | 0.73378307 |
| TPPPPPPP | ASSTIANTEVF | 0.5431036 |

| | | |
|---|---|---|
| TQSPALI | ASSQASTGGHEQY | 0.8199889 |
| EFAAAFPQ | ASSQASTGGHEQY | 0.6602068 |
| LLSGNLDVL | ASSQASTGGHEQY | 0.59618974 |
| GPANLGNYNI | ASSQASTGGHEQY | 0.6891992 |
| TLGPTLP | ASSQASTGGHEQY | 0.82423806 |
| VYAYSSCCK | ASSQASTGGHEQY | 0.52927816 |
| APAPAPAPNI | ASSQASTGGHEQY | 0.79112554 |
| PGGPVPIP | ASSQASTGGHEQY | 0.67108583 |
| FLTVALEP | ASSQASTGGHEQY | 0.84701955 |
| SQYFGANL | ASSQASTGGHEQY | 0.41305476 |
| GGVKPAA | ASSQASTGGHEQY | 0.8225516 |
| GGTGGAGGAGGW | ASSQASTGGHEQY | 0.6966144 |
| GVGGVAGLGGAGG | ASSQASTGGHEQY | 0.87956715 |
| VLLVLATTI | ASSQASTGGHEQY | 0.70528734 |
| IGIFAAML | ASSQASTGGHEQY | 0.73463845 |
| AVGDLID | ASSQASTGGHEQY | 0.6769006 |
| NISPLPL | ASSQASTGGHEQY | 0.82904565 |
| VTAVFVAI | ASSQASTGGHEQY | 0.35187823 |
| SGDDNDAAL | ASSQASTGGHEQY | 0.7886294 |
| AVAALAHS | ASSQASTGGHEQY | 0.5175043 |
| GGVTVMATP | ASSQASTGGHEQY | 0.8086839 |
| AGLAANVRD | ASSQASTGGHEQY | 0.7916343 |
| TNRPDLIDPALLRPG | ASSQASTGGHEQY | 0.34256607 |
| LANLGVTAL | ASSQASTGGHEQY | 0.8329185 |
| TGERIEL | ASSQASTGGHEQY | 0.6567317 |
| NGEELAA | ASSQASTGGHEQY | 0.5683829 |
| VLSLVWG | ASSQASTGGHEQY | 0.85484254 |
| IGYALWL | ASSQASTGGHEQY | 0.7275549 |
| LSLLLAPL | ASSQASTGGHEQY | 0.8654715 |
| TLDEALK | ASSQASTGGHEQY | 0.7569254 |
| TDGAGTAEGG | ASSQASTGGHEQY | 0.71214557 |
| NAPGGAGGNGG | ASSQASTGGHEQY | 0.81001437 |
| VSVFMGMGGNF | ASSQASTGGHEQY | 0.54174435 |
| LASYLAAL | ASSQASTGGHEQY | 0.6335303 |
| LVGGTPAL | ASSQASTGGHEQY | 0.8402709 |
| VRNADVL | ASSQASTGGHEQY | 0.86025536 |
| TCSSPFLVSAM | ASSQASTGGHEQY | 0.8655181 |
| VHITDAE | ASSQASTGGHEQY | 0.7756722 |
| EYAAPSF | ASSQASTGGHEQY | 0.7150774 |
| VVDAFHGGL | ASSQASTGGHEQY | 0.8079818 |
| ASVVDPQ | ASSQASTGGHEQY | 0.86822397 |
| SYNEFGGLS | ASSQASTGGHEQY | 0.8778027 |
| DPDGSVY | ASSQASTGGHEQY | 0.85976505 |
| MVSTIHATE | ASSQASTGGHEQY | 0.71767336 |
| LAAGVADPV | ASSQASTGGHEQY | 0.8443054 |
| TKEQIAA | ASSQASTGGHEQY | 0.82183266 |
| SLASPAAL | ASSQASTGGHEQY | 0.811091 |
| IGLVFYGV | ASSQASTGGHEQY | 0.865373 |
| FLLDASNL | ASSQASTGGHEQY | 0.86158895 |
| YQVGNFP | ASSQASTGGHEQY | 0.8122317 |
| ESLLSQST | ASSQASTGGHEQY | 0.7633674 |

| | | |
|---|---|---|
| LFTVGFAAVG | ASSQASTGGHEQY | 0.8643469 |
| ATPPTPT | ASSQASTGGHEQY | 0.8704823 |
| TDLATLG | ASSQASTGGHEQY | 0.86748713 |
| SAGGTGR | ASSQASTGGHEQY | 0.8804605 |
| GFSNSGS | ASSQASTGGHEQY | 0.8672186 |
| GGNGGSGGTSV | ASSQASTGGHEQY | 0.86121583 |
| GTSVATGGAGN | ASSQASTGGHEQY | 0.30748367 |
| NGGNGGSGGTS | ASSQASTGGHEQY | 0.56629723 |
| AVVKFPH | ASSQASTGGHEQY | 0.8270975 |
| TPPPPPP | ASSQASTGGHEQY | 0.8078559 |
| AAPAAAAAPA | ASSQASTGGHEQY | 0.561967 |
| IALNEDL | ASSQASTGGHEQY | 0.84749377 |
| IVNVSSM | ASSQASTGGHEQY | 0.80967325 |
| TGIPPAPR | ASSQASTGGHEQY | 0.75184405 |
| FDLGGGTF | ASSQASTGGHEQY | 0.8671929 |
| FDLGGGTFDVS | ASSQASTGGHEQY | 0.7927839 |
| NEPTAAA | ASSQASTGGHEQY | 0.736253 |
| NEPTAAALA | ASSQASTGGHEQY | 0.80298173 |
| GSIINISSI | ASSQASTGGHEQY | 0.87555826 |
| VAPGNAGTA | ASSQASTGGHEQY | 0.7674278 |
| TGGGASLE | ASSQASTGGHEQY | 0.8631962 |
| VAVYNHYKRI | ASSQASTGGHEQY | 0.84678876 |
| TDPAPATT | ASSQASTGGHEQY | 0.8716924 |
| TNRPDILDPALLRPG | ASSQASTGGHEQY | 0.62729186 |
| LTTTTTS | ASSQASTGGHEQY | 0.86378217 |
| PTGSGKTL | ASSQASTGGHEQY | 0.82021844 |
| TGSGKTL | ASSQASTGGHEQY | 0.8855477 |
| TIGHVDHGK | ASSQASTGGHEQY | 0.80036485 |
| TIGHVDHGKTT | ASSQASTGGHEQY | 0.8546009 |
| GHVDHGKTT | ASSQASTGGHEQY | 0.85807276 |
| AQFEHTLL | ASSQASTGGHEQY | 0.8687732 |
| IVNQDEL | ASSQASTGGHEQY | 0.6165578 |
| TNGSQFF | ASSQASTGGHEQY | 0.8135 |
| TGKGIVE | ASSQASTGGHEQY | 0.6397934 |
| LISNASDA | ASSQASTGGHEQY | 0.830586 |
| AELRTLGELTKIAKAHGAQ | ASSQASTGGHEQY | 0.8546495 |
| APNAVAAAEPSW | ASSQASTGGHEQY | 0.38216347 |
| APQVAGAATAVL | ASSQASTGGHEQY | 0.87507886 |
| ARRDPAAAAAILPTDA | ASSQASTGGHEQY | 0.7740599 |
| DAAARDVLAEAGLAEYF | ASSQASTGGHEQY | 0.77041376 |
| DDQIVLSAKANLAKVEQ | ASSQASTGGHEQY | 0.8724914 |
| DELLPDTGEDSDEDGH | ASSQASTGGHEQY | 0.85133016 |
| ELPEAPANRATVPQD | ASSQASTGGHEQY | 0.8642989 |
| FGQSKIERVTGIG | ASSQASTGGHEQY | 0.8445953 |
| GAGGTGSTAGGAGGAGGAGGLYA | ASSQASTGGHEQY | 0.84991145 |
| GHGGTGAAGGNGGTGGAGGSGIDG | ASSQASTGGHEQY | 0.84785247 |
| GNGGNGGAGGEGGPGGAGGAGA | ASSQASTGGHEQY | 0.7565439 |
| GSTAGNGGNGGNATNGTIAGQPAG | ASSQASTGGHEQY | 0.7891512 |
| GTGGDGGNGGNGGTGG | ASSQASTGGHEQY | 0.86852014 |
| GVRRPAGIPADAPV | ASSQASTGGHEQY | 0.6439897 |
| GYRQVTDAHLLTLARR | ASSQASTGGHEQY | 0.83388543 |

| | | |
|---|---|---|
| IAKMGQDGHDRG | ASSQASTGGHEQY | 0.38124406 |
| KAEPSIEIQYT | ASSQASTGGHEQY | 0.28280818 |
| KPVRIRMAADEDTVAVT | ASSQASTGGHEQY | 0.49394906 |
| MPPRIAGMRLLVIKPEP | ASSQASTGGHEQY | 0.70678926 |
| MKFYAVALFRSRGVPPTR | ASSQASTGGHEQY | 0.85520124 |
| NAGGFGILGAVAHSP | ASSQASTGGHEQY | 0.7041836 |
| NEGREITLAMAEM | ASSQASTGGHEQY | 0.2883935 |
| NELVEWSHPLGHRWRWE | ASSQASTGGHEQY | 0.6672289 |
| NLGAGGTGGNGGNPG | ASSQASTGGHEQY | 0.51461804 |
| NTEWPKWETVMAADPVD | ASSQASTGGHEQY | 0.49179864 |
| PASAAPAETQP | ASSQASTGGHEQY | 0.6322382 |
| QQSSDGKNGVRAAEITGE | ASSQASTGGHEQY | 0.6229628 |
| RAPDAVTTSWWKE | ASSQASTGGHEQY | 0.41115755 |
| SGFGNVGAHTSGWFNQ | ASSQASTGGHEQY | 0.26748493 |
| SGVAGAVGQAASVGGLKV | ASSQASTGGHEQY | 0.7478827 |
| SNGGAGTGSAGGNGGT | ASSQASTGGHEQY | 0.7851982 |
| TGGAGGEGGAAG | ASSQASTGGHEQY | 0.7782149 |
| TGGDGGAAGAGGAPGHG | ASSQASTGGHEQY | 0.8091588 |
| TMPPLLAAIANAGRHGV | ASSQASTGGHEQY | 0.7609391 |
| TSSTAGNSGGA | ASSQASTGGHEQY | 0.6122097 |
| VPATVPVGILC | ASSQASTGGHEQY | 0.14335588 |
| VQRPDSGGVVLTGRIS | ASSQASTGGHEQY | 0.21550936 |
| WTPNPVPATKGVPAGLVQAT | ASSQASTGGHEQY | 0.44457132 |
| YALKRALAKFSDRFDVVII | ASSQASTGGHEQY | 0.52566457 |
| YLIGSVIV | ASSQASTGGHEQY | 0.17782156 |
| YGQLVAVID | ASSQASTGGHEQY | 0.696612 |
| VTVGGMTLPA | ASSQASTGGHEQY | 0.36021554 |
| VLALPAV | ASSQASTGGHEQY | 0.36397552 |
| VIPPTLNYET | ASSQASTGGHEQY | 0.7848809 |
| VGMGGMHPSSG | ASSQASTGGHEQY | 0.38781664 |
| VGLAGGAATA | ASSQASTGGHEQY | 0.2843398 |
| VGGAGGAGGSGP | ASSQASTGGHEQY | 0.4053073 |
| VFAAFMV | ASSQASTGGHEQY | 0.7541501 |
| TLYDKLR | ASSQASTGGHEQY | 0.23775701 |
| TLAVFTVL | ASSQASTGGHEQY | 0.7336119 |
| TGSVQPARV | ASSQASTGGHEQY | 0.5711044 |
| TGGGAGSGGT | ASSQASTGGHEQY | 0.76477504 |
| SHLETSL | ASSQASTGGHEQY | 0.22431727 |
| SFTTGMT | ASSQASTGGHEQY | 0.72308385 |
| QVEVAKRMGVRQPTVS | ASSQASTGGHEQY | 0.54779017 |
| QGSGGPALAM | ASSQASTGGHEQY | 0.8521786 |
| MLAPSNP | ASSQASTGGHEQY | 0.42163807 |
| LVNNAAI | ASSQASTGGHEQY | 0.63073736 |
| LTGEINTLA | ASSQASTGGHEQY | 0.6086462 |
| LSIFGPV | ASSQASTGGHEQY | 0.825752 |
| LLALLANL | ASSQASTGGHEQY | 0.2325927 |
| LISNASDALDKLR | ASSQASTGGHEQY | 0.42259508 |
| LINEAVP | ASSQASTGGHEQY | 0.27166742 |
| LGYAFSGL | ASSQASTGGHEQY | 0.46832573 |
| LDFVAVAL | ASSQASTGGHEQY | 0.6861027 |
| LAQPGAE | ASSQASTGGHEQY | 0.14835861 |

| | | |
|---|---|---|
| ISNLGAVTGF | ASSQASTGGHEQY | 0.6705941 |
| HQEDPGRI | ASSQASTGGHEQY | 0.41595477 |
| GTPGEPTP | ASSQASTGGHEQY | 0.2826699 |
| GFFNSGS | ASSQASTGGHEQY | 0.7230646 |
| GAGGGGLTAGAN | ASSQASTGGHEQY | 0.6369895 |
| FVDALVNM | ASSQASTGGHEQY | 0.74940157 |
| FLTVAPEL | ASSQASTGGHEQY | 0.46703953 |
| FETLTAL | ASSQASTGGHEQY | 0.6011752 |
| DVINPIEL | ASSQASTGGHEQY | 0.51313007 |
| APVSGAPV | ASSQASTGGHEQY | 0.4313265 |
| ALVGGLAW | ASSQASTGGHEQY | 0.45846397 |
| ALAAAAATAI | ASSQASTGGHEQY | 0.758286 |
| AFGLGLLL | ASSQASTGGHEQY | 0.66578066 |
| AFFAAASL | ASSQASTGGHEQY | 0.65710616 |
| AAAALPAFSPP | ASSQASTGGHEQY | 0.16396576 |
| VVLAPAL | ASSQASTGGHEQY | 0.19744183 |
| TIGHVDHGKTTL | ASSQASTGGHEQY | 0.2776993 |
| LISNASDALDK | ASSQASTGGHEQY | 0.48125952 |
| LISNASDAL | ASSQASTGGHEQY | 0.6748444 |
| ISNASDALDK | ASSQASTGGHEQY | 0.672265 |
| NQAETLVYQTE | ASSQASTGGHEQY | 0.21141098 |
| TGGTGGTAGAG | ASSQASTGGHEQY | 0.3171699 |
| TGSTAGGAGGA | ASSQASTGGHEQY | 0.27483445 |
| DPLAVPVAAGPV | ASSQASTGGHEQY | 0.30912167 |
| GGTGGTPGNGGN | ASSQASTGGHEQY | 0.38414556 |
| PAGLAGVDPATL | ASSQASTGGHEQY | 0.19594803 |
| GNNLVVEAPHAGASV | ASSQASTGGHEQY | 0.1849798 |
| KVTRIMSASTLEQPA | ASSQASTGGHEQY | 0.20920768 |
| LGFALPIALPRRTVV | ASSQASTGGHEQY | 0.24150158 |
| TRLGNLTARKLVVLV | ASSQASTGGHEQY | 0.64746296 |
| WSQQRRLIYFPSAGP | ASSQASTGGHEQY | 0.65497124 |
| GPNGNPDPMAAAVDIR | ASSQASTGGHEQY | 0.51493955 |
| TQRCVSVADLRAAASR | ASSQASTGGHEQY | 0.6407759 |
| QFVLDVAEHPVEVLTETV | ASSQASTGGHEQY | 0.40982282 |
| GGAGGTGGAGGNGGTQAGDG | ASSQASTGGHEQY | 0.11772815 |
| PLLATLRAIAVDVDAKPAQV | ASSQASTGGHEQY | 0.15444078 |
| APKFRTGGYMIDFWGVGYQVAKRM | ASSQASTGGHEQY | 0.20274214 |
| GAGGAGGAGGAAGGPGGTGGTGGN | ASSQASTGGHEQY | 0.2517476 |
| IGALGLLATDVAPAGSVAFGVALA | ASSQASTGGHEQY | 0.12372327 |
| AGGAGGIGGNGGAGGAGGVLMGNG | ASSQASTGGHEQY | 0.36076042 |
| ADELIGLL | ASSQASTGGHEQY | 0.30795577 |
| ADRALKA | ASSQASTGGHEQY | 0.17883228 |
| AGAVGQVGGAGGA | ASSQASTGGHEQY | 0.5763649 |
| AGGGGGLGGAP | ASSQASTGGHEQY | 0.21582244 |
| AGGLLYGNGGAG | ASSQASTGGHEQY | 0.17405844 |
| AGLESIGL | ASSQASTGGHEQY | 0.2140066 |
| AGLKTGII | ASSQASTGGHEQY | 0.65591645 |
| ASLAALLL | ASSQASTGGHEQY | 0.19678013 |
| ASVINISSI | ASSQASTGGHEQY | 0.6134513 |
| ATPPGPA | ASSQASTGGHEQY | 0.26170066 |
| ATVASALV | ASSQASTGGHEQY | 0.5288796 |

| | | |
|---|---|---|
| ATVASLIL | ASSQASTGGHEQY | 0.16541703 |
| AVNGPVLIRH | ASSQASTGGHEQY | 0.5762807 |
| DGAAGAVGI | ASSQASTGGHEQY | 0.3735397 |
| DPLLHGI | ASSQASTGGHEQY | 0.74778545 |
| ELGAALGAL | ASSQASTGGHEQY | 0.28441155 |
| ELGRTQAV | ASSQASTGGHEQY | 0.41432285 |
| FVAGGYLS | ASSQASTGGHEQY | 0.17575073 |
| GASGGAGGAGGW | ASSQASTGGHEQY | 0.4058059 |
| GGAGAGGDGGAGGI | ASSQASTGGHEQY | 0.1470831 |
| GTGHNGSQPGMGGNG | ASSQASTGGHEQY | 0.28213367 |
| GYAAASASAS | ASSQASTGGHEQY | 0.15748182 |
| HQVDVAAVQQG | ASSQASTGGHEQY | 0.22886272 |
| IGAAIAHT | ASSQASTGGHEQY | 0.6340769 |
| IGMIEVAQL | ASSQASTGGHEQY | 0.12898263 |
| IRLAPEHTSAMD | ASSQASTGGHEQY | 0.5194086 |
| IYNVSVL | ASSQASTGGHEQY | 0.17451751 |
| KQGFLGM | ASSQASTGGHEQY | 0.20113397 |
| LESVITQL | ASSQASTGGHEQY | 0.4548526 |
| LGLGALVVV | ASSQASTGGHEQY | 0.18074651 |
| LIGNGGNGGSGGI | ASSQASTGGHEQY | 0.55920786 |
| LNAINAPTL | ASSQASTGGHEQY | 0.15201479 |
| LQLLNTGV | ASSQASTGGHEQY | 0.43103653 |
| LQTSIEF | ASSQASTGGHEQY | 0.33838862 |
| MSFAVSEL | ASSQASTGGHEQY | 0.1187674 |
| NTGALGL | ASSQASTGGHEQY | 0.18220648 |
| PPGAGGAGGAGTT | ASSQASTGGHEQY | 0.6595128 |
| QLVGLGI | ASSQASTGGHEQY | 0.2006039 |
| RFLEQVHQL | ASSQASTGGHEQY | 0.51453066 |
| RGVGYVLR | ASSQASTGGHEQY | 0.10030307 |
| SATGVFAGVI | ASSQASTGGHEQY | 0.11235358 |
| SLPAITLP | ASSQASTGGHEQY | 0.16913639 |
| STALAFTV | ASSQASTGGHEQY | 0.2589967 |
| SVLTVGLGK | ASSQASTGGHEQY | 0.14524943 |
| TALAGAGSGI | ASSQASTGGHEQY | 0.25721923 |
| TGLPFTGI | ASSQASTGGHEQY | 0.113047436 |
| VAEHVTV | ASSQASTGGHEQY | 0.10898444 |
| VDIVLNSL | ASSQASTGGHEQY | 0.10498274 |
| VLTGGGAVL | ASSQASTGGHEQY | 0.117401585 |
| VNVGLAYGI | ASSQASTGGHEQY | 0.17359164 |
| VSSAAVGAV | ASSQASTGGHEQY | 0.10261629 |
| VTSVYEAGV | ASSQASTGGHEQY | 0.10022671 |
| WVALDRGA | ASSQASTGGHEQY | 0.14226718 |
| YLVGSVVI | ASSQASTGGHEQY | 0.2105616 |
| GPPGTGKTLL | ASSQASTGGHEQY | 0.27259797 |
| GVGMTKF | ASSQASTGGHEQY | 0.23569444 |
| IGIGGSDL | ASSQASTGGHEQY | 0.42512003 |
| IGTLLAEL | ASSQASTGGHEQY | 0.20810404 |
| IYTSGTT | ASSQASTGGHEQY | 0.1458513 |
| LSFPAGV | ASSQASTGGHEQY | 0.08846388 |
| LYTSGTT | ASSQASTGGHEQY | 0.113387555 |
| VGAFPFGF | ASSQASTGGHEQY | 0.10206158 |

| | | |
|---|---|---|
| VGAGTVEF | ASSQASTGGHEQY | 0.14828254 |
| VGKSTLLL | ASSQASTGGHEQY | 0.085004486 |
| AALAHAGGGIV | ASSQASTGGHEQY | 0.13138106 |
| AALVAVSLL | ASSQASTGGHEQY | 0.14931352 |
| AAVSRAVAAL | ASSQASTGGHEQY | 0.093646586 |
| AGGSSGIGGF | ASSQASTGGHEQY | 0.243315 |
| AGLTVANANG | ASSQASTGGHEQY | 0.09548701 |
| AGLVGAVL | ASSQASTGGHEQY | 0.10706208 |
| ALSQAFTV | ASSQASTGGHEQY | 0.112718195 |
| ALTFSVSL | ASSQASTGGHEQY | 0.48062697 |
| ALTGAALVAL | ASSQASTGGHEQY | 0.10694203 |
| ASFSIGAGVL | ASSQASTGGHEQY | 0.21592495 |
| DGSVYKI | ASSQASTGGHEQY | 0.18882158 |
| EQLAEIV | ASSQASTGGHEQY | 0.15345523 |
| FAVSELTVM | ASSQASTGGHEQY | 0.0989684 |
| GAAPLAAGL | ASSQASTGGHEQY | 0.18127058 |
| GADGGDGGSGGAGG | ASSQASTGGHEQY | 0.12519008 |
| GFGLDTGGAGG | ASSQASTGGHEQY | 0.661234 |
| GLASGGLL | ASSQASTGGHEQY | 0.15343708 |
| GRIAGAAVLL | ASSQASTGGHEQY | 0.12759574 |
| GSAGSLNTSF | ASSQASTGGHEQY | 0.108194456 |
| GTGQIGI | ASSQASTGGHEQY | 0.105282374 |
| GTLIALIL | ASSQASTGGHEQY | 0.09019643 |
| IAATMGFVVG | ASSQASTGGHEQY | 0.15734129 |
| IFAQALL | ASSQASTGGHEQY | 0.13188353 |
| IGAAVALGF | ASSQASTGGHEQY | 0.1418171 |
| IGLTPLAL | ASSQASTGGHEQY | 0.19175819 |
| ILGSNTSTL | ASSQASTGGHEQY | 0.10587545 |
| IVFSLTVL | ASSQASTGGHEQY | 0.31900257 |
| LASKATGF | ASSQASTGGHEQY | 0.10997834 |
| LASVALIL | ASSQASTGGHEQY | 0.10629217 |
| LFAVGLTL | ASSQASTGGHEQY | 0.23884925 |
| LFNGGIGTYI | ASSQASTGGHEQY | 0.098727286 |
| LTLGAGII | ASSQASTGGHEQY | 0.4097014 |
| LVAAALGDI | ASSQASTGGHEQY | 0.09075022 |
| NTAFVQLGI | ASSQASTGGHEQY | 0.20186435 |
| QLFSGAEL | ASSQASTGGHEQY | 0.24644321 |
| SAATFITL | ASSQASTGGHEQY | 0.08705834 |
| SNYGLANLGV | ASSQASTGGHEQY | 0.098607734 |
| SYGGAYAVM | ASSQASTGGHEQY | 0.20374526 |
| TTGVGGLTLG | ASSQASTGGHEQY | 0.7056521 |
| TTQTIYI | ASSQASTGGHEQY | 0.7462895 |
| VAAGAGLGAVI | ASSQASTGGHEQY | 0.43909982 |
| VAAVVTLL | ASSQASTGGHEQY | 0.3883875 |
| VGFLGGLI | ASSQASTGGHEQY | 0.5444604 |
| VGVIPILL | ASSQASTGGHEQY | 0.71846217 |
| VIFASSLI | ASSQASTGGHEQY | 0.79344356 |
| VKGGDGGAAAG | ASSQASTGGHEQY | 0.71239823 |
| VTKTFTNI | ASSQASTGGHEQY | 0.45171466 |
| VVAAAILL | ASSQASTGGHEQY | 0.70281714 |
| YAALDVEL | ASSQASTGGHEQY | 0.6599009 |

| | | |
|---|---|---|
| LASVALLL | ASSQASTGGHEQY | 0.6399629 |
| LDAVHRV | ASSQASTGGHEQY | 0.642132 |
| QGLSSSIF | ASSQASTGGHEQY | 0.6794232 |
| RGTFANIRL | ASSQASTGGHEQY | 0.49714932 |
| RQLWWGHRI | ASSQASTGGHEQY | 0.43544734 |
| YGPPGTGKTLL | ASSQASTGGHEQY | 0.7098121 |
| AALVGAPANVA | ASSQASTGGHEQY | 0.7609026 |
| AATFRRTE | ASSQASTGGHEQY | 0.749568 |
| AGALGGGI | ASSQASTGGHEQY | 0.8065412 |
| AGGAGGAGGIGAG | ASSQASTGGHEQY | 0.76243174 |
| AGSGALGL | ASSQASTGGHEQY | 0.6514126 |
| AQIADQI | ASSQASTGGHEQY | 0.2277494 |
| ASAVAVTYL | ASSQASTGGHEQY | 0.31032544 |
| ASTVLGGTGG | ASSQASTGGHEQY | 0.54716384 |
| AVGGEPAK | ASSQASTGGHEQY | 0.5623941 |
| CEQGTFGGTC | ASSQASTGGHEQY | 0.20847987 |
| DEGKAPAG | ASSQASTGGHEQY | 0.6662041 |
| DTVVFPAL | ASSQASTGGHEQY | 0.5616425 |
| FEGITMAG | ASSQASTGGHEQY | 0.52477396 |
| FPLVVQD | ASSQASTGGHEQY | 0.77188045 |
| GAFTTPQ | ASSQASTGGHEQY | 0.535745 |
| GFGGFGTAGGVGGA | ASSQASTGGHEQY | 0.45632273 |
| GFLAQFL | ASSQASTGGHEQY | 0.56681633 |
| GGAGGAGGNAGVA | ASSQASTGGHEQY | 0.79244035 |
| GGVGTAGAGGAGGAGG | ASSQASTGGHEQY | 0.4328633 |
| GNGGGGDAGNAGSGGN | ASSQASTGGHEQY | 0.7869278 |
| GTGGAGGAGNDAGS | ASSQASTGGHEQY | 0.5547415 |
| GTTNTGWFNS | ASSQASTGGHEQY | 0.79886746 |
| IFMGPLFTL | ASSQASTGGHEQY | 0.3634207 |
| ILAHAIEAI | ASSQASTGGHEQY | 0.77227056 |
| IRSAIREAL | ASSQASTGGHEQY | 0.65472865 |
| ITFAGGG | ASSQASTGGHEQY | 0.84235054 |
| ITGMSGHQEV | ASSQASTGGHEQY | 0.5942682 |
| IVAGLACTL | ASSQASTGGHEQY | 0.7590094 |
| IVGGGLAGA | ASSQASTGGHEQY | 0.6213447 |
| IVTGVLNI | ASSQASTGGHEQY | 0.7639066 |
| LAQQGCI | ASSQASTGGHEQY | 0.45307815 |
| LCLHVEN | ASSQASTGGHEQY | 0.52759075 |
| LDAISDGS | ASSQASTGGHEQY | 0.3801447 |
| LGGFAAML | ASSQASTGGHEQY | 0.5491907 |
| LGGKNPMI | ASSQASTGGHEQY | 0.747188 |
| LGPKGVAAIL | ASSQASTGGHEQY | 0.23040915 |
| LMVGVSI | ASSQASTGGHEQY | 0.6913629 |
| LVGAAFVL | ASSQASTGGHEQY | 0.4416951 |
| LVLAVAPL | ASSQASTGGHEQY | 0.41521996 |
| MAAIAVIDAA | ASSQASTGGHEQY | 0.7015583 |
| NNGINAELQ | ASSQASTGGHEQY | 0.66599494 |
| RGEPLVLVHG | ASSQASTGGHEQY | 0.6751404 |
| RGGAGGAAGLI | ASSQASTGGHEQY | 0.46830073 |
| RSYRMHPAV | ASSQASTGGHEQY | 0.5737929 |
| SAAHLLAGQLA | ASSQASTGGHEQY | 0.62245566 |

| | | |
|---|---|---|
| SAAVGSY | ASSQASTGGHEQY | 0.33769774 |
| SPHAVLHRI | ASSQASTGGHEQY | 0.4739622 |
| SQIAKVL | ASSQASTGGHEQY | 0.7982183 |
| TALGHLVP | ASSQASTGGHEQY | 0.32161546 |
| TAPTTSGAA | ASSQASTGGHEQY | 0.793194 |
| TDGDDGNGGSPGT | ASSQASTGGHEQY | 0.20342363 |
| TDGLGEN | ASSQASTGGHEQY | 0.19581994 |
| TGGPPLPGQA | ASSQASTGGHEQY | 0.18326041 |
| TIPAITI | ASSQASTGGHEQY | 0.34419805 |
| TIPGFTI | ASSQASTGGHEQY | 0.7986528 |
| TSIASGALA | ASSQASTGGHEQY | 0.45351064 |
| TVTLPGAAI | ASSQASTGGHEQY | 0.1815479 |
| VASGLSAALP | ASSQASTGGHEQY | 0.30894 |
| VGAGVIGI | ASSQASTGGHEQY | 0.20549738 |
| VGPKRFGV | ASSQASTGGHEQY | 0.22051635 |
| VNVAFPD | ASSQASTGGHEQY | 0.26828197 |
| VSMTNSGVS | ASSQASTGGHEQY | 0.22941606 |
| VTDVLGDLSV | ASSQASTGGHEQY | 0.20533988 |
| VVVVGAGFAGL | ASSQASTGGHEQY | 0.19104369 |
| VYTVGTVV | ASSQASTGGHEQY | 0.19288331 |
| WTTFSGP | ASSQASTGGHEQY | 0.40322164 |
| YDELVEAF | ASSQASTGGHEQY | 0.3484763 |
| YGTGGAGGNGG | ASSQASTGGHEQY | 0.40278417 |
| YTNVMAV | ASSQASTGGHEQY | 0.3958213 |
| FLEQVHQL | ASSQASTGGHEQY | 0.25865388 |
| GLSLLAVAV | ASSQASTGGHEQY | 0.1406133 |
| LEQVHQL | ASSQASTGGHEQY | 0.16475259 |
| LNAAGAI | ASSQASTGGHEQY | 0.17923619 |
| PAVHASL | ASSQASTGGHEQY | 0.25378636 |
| TELEDDA | ASSQASTGGHEQY | 0.14139697 |
| TPPGPPN | ASSQASTGGHEQY | 0.33446613 |
| TPPPPPPP | ASSQASTGGHEQY | 0.2242401 |
| TQSPALI | ASSIQNTLY | 0.21304946 |
| EFAAAFPQ | ASSIQNTLY | 0.36438265 |
| LLSGNLDVL | ASSIQNTLY | 0.23159693 |
| GPANLGNYNI | ASSIQNTLY | 0.17653768 |
| TLGPTLP | ASSIQNTLY | 0.17619452 |
| VYAYSSCCK | ASSIQNTLY | 0.29967406 |
| APAPAPAPNI | ASSIQNTLY | 0.15191375 |
| PGGPVPIP | ASSIQNTLY | 0.36936432 |
| FLTVALEP | ASSIQNTLY | 0.1587043 |
| SQYFGANL | ASSIQNTLY | 0.25541347 |
| GGVKPAA | ASSIQNTLY | 0.16606028 |
| GGTGGAGGAGGW | ASSIQNTLY | 0.41842324 |
| GVGGVAGLGGAGG | ASSIQNTLY | 0.2448657 |
| VLLVLATTI | ASSIQNTLY | 0.5460245 |
| IGIFAAML | ASSIQNTLY | 0.21630608 |
| AVGDLID | ASSIQNTLY | 0.26480645 |
| NISPLPL | ASSIQNTLY | 0.194506 |
| VTAVFVAI | ASSIQNTLY | 0.7821009 |
| SGDDNDAAL | ASSIQNTLY | 0.1560242 |

| | | |
|---|---|---|
| AVAALAHS | ASSIQNTLY | 0.19696511 |
| GGVTVMATP | ASSIQNTLY | 0.1617488 |
| AGLAANVRD | ASSIQNTLY | 0.23675671 |
| TNRPDLIDPALLRPG | ASSIQNTLY | 0.4315827 |
| LANLGVTAL | ASSIQNTLY | 0.18320438 |
| TGERIEL | ASSIQNTLY | 0.3136345 |
| NGEELAA | ASSIQNTLY | 0.28105462 |
| VLSLVWG | ASSIQNTLY | 0.1871022 |
| IGYALWL | ASSIQNTLY | 0.3406274 |
| LSLLLAPL | ASSIQNTLY | 0.2458147 |
| TLDEALK | ASSIQNTLY | 0.3445724 |
| TDGAGTAEGG | ASSIQNTLY | 0.17349896 |
| NAPGGAGGNGG | ASSIQNTLY | 0.21912381 |
| VSVFMGMGGNF | ASSIQNTLY | 0.24391231 |
| LASYLAAL | ASSIQNTLY | 0.2052235 |
| LVGGTPAL | ASSIQNTLY | 0.16825789 |
| VRNADVL | ASSIQNTLY | 0.5229678 |
| TCSSPFLVSAM | ASSIQNTLY | 0.8741358 |
| VHITDAE | ASSIQNTLY | 0.8785157 |
| EYAAPSF | ASSIQNTLY | 0.7613068 |
| VVDAFHGGL | ASSIQNTLY | 0.77042705 |
| ASVVDPQ | ASSIQNTLY | 0.8260385 |
| SYNEFGGLS | ASSIQNTLY | 0.88193595 |
| DPDGSVY | ASSIQNTLY | 0.89953065 |
| MVSTIHATE | ASSIQNTLY | 0.8759469 |
| LAAGVADPV | ASSIQNTLY | 0.81353974 |
| TKEQIAA | ASSIQNTLY | 0.88387465 |
| SLASPAAL | ASSIQNTLY | 0.85143733 |
| IGLVFYGV | ASSIQNTLY | 0.83888036 |
| FLLDASNL | ASSIQNTLY | 0.8746865 |
| YQVGNFP | ASSIQNTLY | 0.8673719 |
| ESLLSQST | ASSIQNTLY | 0.82177496 |
| LFTVGFAAVG | ASSIQNTLY | 0.8052095 |
| ATPPTPT | ASSIQNTLY | 0.872971 |
| TDLATLG | ASSIQNTLY | 0.8857272 |
| SAGGTGR | ASSIQNTLY | 0.87474597 |
| GFSNSGS | ASSIQNTLY | 0.902039 |
| GGNGGSGGTSV | ASSIQNTLY | 0.8833715 |
| GTSVATGGAGN | ASSIQNTLY | 0.87055385 |
| NGGNGGSGGTS | ASSIQNTLY | 0.47801515 |
| AVVKFPH | ASSIQNTLY | 0.6546848 |
| TPPPPPP | ASSIQNTLY | 0.8300097 |
| AAPAAAAAPA | ASSIQNTLY | 0.8238032 |
| IALNEDL | ASSIQNTLY | 0.40384024 |
| IVNVSSM | ASSIQNTLY | 0.8798643 |
| TGIPPAPR | ASSIQNTLY | 0.84542644 |
| FDLGGGTF | ASSIQNTLY | 0.7997391 |
| FDLGGGTFDVS | ASSIQNTLY | 0.8833835 |
| NEPTAAA | ASSIQNTLY | 0.8310185 |
| NEPTAAALA | ASSIQNTLY | 0.7751691 |
| GSIINISSI | ASSIQNTLY | 0.82837105 |

| | | |
|---|---|---|
| VAPGNAGTA | ASSIQNTLY | 0.89464784 |
| TGGGASLE | ASSIQNTLY | 0.7635571 |
| VAVYNHYKRI | ASSIQNTLY | 0.8870477 |
| TDPAPATT | ASSIQNTLY | 0.80505216 |
| TNRPDILDPALLRPG | ASSIQNTLY | 0.8920795 |
| LTTTTTS | ASSIQNTLY | 0.7602298 |
| PTGSGKTL | ASSIQNTLY | 0.8839022 |
| TGSGKTL | ASSIQNTLY | 0.84330845 |
| TIGHVDHGK | ASSIQNTLY | 0.9015944 |
| TIGHVDHGKTT | ASSIQNTLY | 0.8402114 |
| GHVDHGKTT | ASSIQNTLY | 0.8903055 |
| AQFEHTLL | ASSIQNTLY | 0.86666703 |
| IVNQDEL | ASSIQNTLY | 0.89023864 |
| TNGSQFF | ASSIQNTLY | 0.7448729 |
| TGKGIVE | ASSIQNTLY | 0.8297859 |
| LISNASDA | ASSIQNTLY | 0.7609676 |
| AELRTLGELTKIAKAHGAQ | ASSIQNTLY | 0.8549905 |
| APNAVAAAEPSW | ASSIQNTLY | 0.87947476 |
| APQVAGAATAVL | ASSIQNTLY | 0.3676181 |
| ARRDPAAAAAILPTDA | ASSIQNTLY | 0.8824762 |
| DAAARDVLAEAGLAEYF | ASSIQNTLY | 0.759396 |
| DDQIVLSAKANLAKVEQ | ASSIQNTLY | 0.80252653 |
| DELLPDTGEDSDEDGH | ASSIQNTLY | 0.8823235 |
| ELPEAPANRATVPQD | ASSIQNTLY | 0.85547984 |
| FGQSKIERVTGIG | ASSIQNTLY | 0.87701917 |
| GAGGTGSTAGGAGGAGGAGGLYA | ASSIQNTLY | 0.8306266 |
| GHGGTGAAGGNGGTGGAGGSGIDG | ASSIQNTLY | 0.8384495 |
| GNGGNGGAGGEGGPGGAGGAGA | ASSIQNTLY | 0.86471266 |
| GSTAGNGGNGGNATNGTIAGQPAG | ASSIQNTLY | 0.45194376 |
| GTGGDGGNGGNGGTGG | ASSIQNTLY | 0.75055796 |
| GVRRPAGIPADAPV | ASSIQNTLY | 0.8967161 |
| GYRQVTDAHLLTLARR | ASSIQNTLY | 0.7268688 |
| IAKMGQDGHDRG | ASSIQNTLY | 0.7793846 |
| KAEPSIEIQYT | ASSIQNTLY | 0.5829827 |
| KPVRIRMAADEDTVAVT | ASSIQNTLY | 0.52745247 |
| MPPRIAGMRLLVIKPEP | ASSIQNTLY | 0.58993626 |
| MKFYAVALFRSRGVPPTR | ASSIQNTLY | 0.76058125 |
| NAGGFGILGAVAHSP | ASSIQNTLY | 0.7718819 |
| NEGREITLAMAEM | ASSIQNTLY | 0.7234616 |
| NELVEWSHPLGHRWRWE | ASSIQNTLY | 0.51461184 |
| NLGAGGTGGNGGNPG | ASSIQNTLY | 0.70431966 |
| NTEWPKWETVMAADPVD | ASSIQNTLY | 0.60913926 |
| PASAAPAETQP | ASSIQNTLY | 0.6278391 |
| QQSSDGKNGVRAAEITGE | ASSIQNTLY | 0.693637 |
| RAPDAVTTSWWKE | ASSIQNTLY | 0.7153457 |
| SGFGNVGAHTSGWFNQ | ASSIQNTLY | 0.6476097 |
| SGVAGAVGQAASVGGLKV | ASSIQNTLY | 0.566279 |
| SNGGAGTGSAGGNGGT | ASSIQNTLY | 0.67507374 |
| TGGAGGEGGAAG | ASSIQNTLY | 0.7568587 |
| TGGDGGAAGAGGAPGHG | ASSIQNTLY | 0.7366141 |
| TMPPLLAAIANAGRHGV | ASSIQNTLY | 0.819468 |

| | | |
|---|---|---|
| TSSTAGNSGGA | ASSIQNTLY | 0.7682513 |
| VPATVPVGILC | ASSIQNTLY | 0.74058676 |
| VQRPDSGGVVLTGRIS | ASSIQNTLY | 0.17435671 |
| WTPNPVPATKGVPAGLVQAT | ASSIQNTLY | 0.3992331 |
| YALKRALAKFSDRFDVVII | ASSIQNTLY | 0.56947356 |
| YLIGSVIV | ASSIQNTLY | 0.64159036 |
| YGQLVAVID | ASSIQNTLY | 0.23958756 |
| VTVGGMTLPA | ASSIQNTLY | 0.7149543 |
| VLALPAV | ASSIQNTLY | 0.62132514 |
| VIPPTLNYET | ASSIQNTLY | 0.54380673 |
| VGMGGMHPSSG | ASSIQNTLY | 0.7716714 |
| VGLAGGAATA | ASSIQNTLY | 0.63255274 |
| VGGAGGAGGSGP | ASSIQNTLY | 0.5561019 |
| VFAAFMV | ASSIQNTLY | 0.5406816 |
| TLYDKLR | ASSIQNTLY | 0.7799361 |
| TLAVFTVL | ASSIQNTLY | 0.41158006 |
| TGSVQPARV | ASSIQNTLY | 0.7544342 |
| TGGGAGSGGT | ASSIQNTLY | 0.45251286 |
| SHLETSL | ASSIQNTLY | 0.7797026 |
| SFTTGMT | ASSIQNTLY | 0.4097754 |
| QVEVAKRMGVRQPTVS | ASSIQNTLY | 0.74550736 |
| QGSGGPALAM | ASSIQNTLY | 0.65277857 |
| MLAPSNP | ASSIQNTLY | 0.8531529 |
| LVNNAAI | ASSIQNTLY | 0.61415994 |
| LTGEINTLA | ASSIQNTLY | 0.70680726 |
| LSIFGPV | ASSIQNTLY | 0.5158629 |
| LLALLANL | ASSIQNTLY | 0.76019 |
| LISNASDALDKLR | ASSIQNTLY | 0.39665228 |
| LINEAVP | ASSIQNTLY | 0.6461696 |
| LGYAFSGL | ASSIQNTLY | 0.43591332 |
| LDFVAVAL | ASSIQNTLY | 0.7077106 |
| LAQPGAE | ASSIQNTLY | 0.72841 |
| ISNLGAVTGF | ASSIQNTLY | 0.2634146 |
| HQEDPGRI | ASSIQNTLY | 0.7483362 |
| GTPGEPTP | ASSIQNTLY | 0.5179297 |
| GFFNSGS | ASSIQNTLY | 0.4682864 |
| GAGGGGLTAGAN | ASSIQNTLY | 0.7859641 |
| FVDALVNM | ASSIQNTLY | 0.6068796 |
| FLTVAPEL | ASSIQNTLY | 0.80981934 |
| FETLTAL | ASSIQNTLY | 0.59261197 |
| DVINPIEL | ASSIQNTLY | 0.5957442 |
| APVSGAPV | ASSIQNTLY | 0.73300207 |
| ALVGGLAW | ASSIQNTLY | 0.4607775 |
| ALAAAAATAI | ASSIQNTLY | 0.48229393 |
| AFGLGLLL | ASSIQNTLY | 0.802834 |
| AFFAAASL | ASSIQNTLY | 0.7681709 |
| AAAALPAFSPP | ASSIQNTLY | 0.7750802 |
| VVLAPAL | ASSIQNTLY | 0.57466775 |
| TIGHVDHGKTTL | ASSIQNTLY | 0.57325464 |
| LISNASDALDK | ASSIQNTLY | 0.6994059 |
| LISNASDAL | ASSIQNTLY | 0.78260416 |

| | | |
|---|---|---|
| ISNASDALDK | ASSIQNTLY | 0.8172345 |
| NQAETLVYQTE | ASSIQNTLY | 0.740675 |
| TGGTGGTAGAG | ASSIQNTLY | 0.59235144 |
| TGSTAGGAGGA | ASSIQNTLY | 0.790807 |
| DPLAVPVAAGPV | ASSIQNTLY | 0.7340725 |
| GGTGGTPGNGGN | ASSIQNTLY | 0.66121924 |
| PAGLAGVDPATL | ASSIQNTLY | 0.77930796 |
| GNNLVVEAPHAGASV | ASSIQNTLY | 0.7525934 |
| KVTRIMSASTLEQPA | ASSIQNTLY | 0.6870193 |
| LGFALPIALPRRTVV | ASSIQNTLY | 0.66459113 |
| TRLGNLTARKLVVLV | ASSIQNTLY | 0.75555694 |
| WSQQRRLIYFPSAGP | ASSIQNTLY | 0.7738278 |
| GPNGNPDPMAAAVDIR | ASSIQNTLY | 0.76946867 |
| TQRCVSVADLRAAASR | ASSIQNTLY | 0.84683746 |
| QFVLDVAEHPVEVLTETV | ASSIQNTLY | 0.7925342 |
| GGAGGTGGAGGNGGTQAGDG | ASSIQNTLY | 0.7341102 |
| PLLATLRAIAVDVDAKPAQV | ASSIQNTLY | 0.28882882 |
| APKFRTGGYMIDFWGVGYQVAKRM | ASSIQNTLY | 0.48912624 |
| GAGGAGGAGGAAGGPGGTGGTGGN | ASSIQNTLY | 0.72685623 |
| IGALGLLATDVAPAGSVAFGVALA | ASSIQNTLY | 0.6898506 |
| AGGAGGIGGNGGAGGAGGVLMGNG | ASSIQNTLY | 0.3239429 |
| ADELIGLL | ASSIQNTLY | 0.7858083 |
| ADRALKA | ASSIQNTLY | 0.7453991 |
| AGAVGQVGGAGGA | ASSIQNTLY | 0.64420164 |
| AGGGGGLGGAP | ASSIQNTLY | 0.79745334 |
| AGGLLYGNGGAG | ASSIQNTLY | 0.6684559 |
| AGLESIGL | ASSIQNTLY | 0.6951793 |
| AGLKTGII | ASSIQNTLY | 0.6789948 |
| ASLAALLL | ASSIQNTLY | 0.8390211 |
| ASVINISSI | ASSIQNTLY | 0.5932371 |
| ATPPGPA | ASSIQNTLY | 0.8008574 |
| ATVASALV | ASSIQNTLY | 0.7365403 |
| ATVASLIL | ASSIQNTLY | 0.8114154 |
| AVNGPVLIRH | ASSIQNTLY | 0.5512303 |
| DGAAGAVGI | ASSIQNTLY | 0.7779983 |
| DPLLHGI | ASSIQNTLY | 0.7056645 |
| ELGAALGAL | ASSIQNTLY | 0.8334791 |
| ELGRTQAV | ASSIQNTLY | 0.7178018 |
| FVAGGYLS | ASSIQNTLY | 0.79661006 |
| GASGGAGGAGGW | ASSIQNTLY | 0.69994247 |
| GGAGAGGDGGAGGI | ASSIQNTLY | 0.8220867 |
| GTGHNGSQPGMGGNG | ASSIQNTLY | 0.56236964 |
| GYAAASASAS | ASSIQNTLY | 0.73009974 |
| HQVDVAAVQQG | ASSIQNTLY | 0.672233 |
| IGAAIAHT | ASSIQNTLY | 0.7566663 |
| IGMIEVAQL | ASSIQNTLY | 0.7475418 |
| IRLAPEHTSAMD | ASSIQNTLY | 0.34144366 |
| IYNVSVL | ASSIQNTLY | 0.8349478 |
| KQGFLGM | ASSIQNTLY | 0.5702364 |
| LESVITQL | ASSIQNTLY | 0.63749367 |
| LGLGALVVV | ASSIQNTLY | 0.7881867 |

| | | |
|---|---|---|
| LIGNGGNGGSGGI | ASSIQNTLY | 0.71723044 |
| LNAINAPTL | ASSIQNTLY | 0.79969376 |
| LQLLNTGV | ASSIQNTLY | 0.69425166 |
| LQTSIEF | ASSIQNTLY | 0.768633 |
| MSFAVSEL | ASSIQNTLY | 0.77021885 |
| NTGALGL | ASSIQNTLY | 0.5041417 |
| PPGAGGAGGAGTT | ASSIQNTLY | 0.61011 |
| QLVGLGI | ASSIQNTLY | 0.8110774 |
| RFLEQVHQL | ASSIQNTLY | 0.7877562 |
| RGVGYVLR | ASSIQNTLY | 0.8685679 |
| SATGVFAGVI | ASSIQNTLY | 0.69230413 |
| SLPAITLP | ASSIQNTLY | 0.566386 |
| STALAFTV | ASSIQNTLY | 0.7433208 |
| SVLTVGLGK | ASSIQNTLY | 0.8502805 |
| TALAGAGSGI | ASSIQNTLY | 0.85549575 |
| TGLPFTGI | ASSIQNTLY | 0.8279853 |
| VAEHVTV | ASSIQNTLY | 0.5109592 |
| VDIVLNSL | ASSIQNTLY | 0.7383479 |
| VLTGGGAVL | ASSIQNTLY | 0.6746142 |
| VNVGLAYGI | ASSIQNTLY | 0.7528195 |
| VSSAAVGAV | ASSIQNTLY | 0.80594057 |
| VTSVYEAGV | ASSIQNTLY | 0.76051754 |
| WVALDRGA | ASSIQNTLY | 0.72650754 |
| YLVGSVVI | ASSIQNTLY | 0.63694656 |
| GPPGTGKTLL | ASSIQNTLY | 0.77136225 |
| GVGMTKF | ASSIQNTLY | 0.84054077 |
| IGIGGSDL | ASSIQNTLY | 0.81415546 |
| IGTLLAEL | ASSIQNTLY | 0.88193244 |
| IYTSGTT | ASSIQNTLY | 0.8441221 |
| LSFPAGV | ASSIQNTLY | 0.84748757 |
| LYTSGTT | ASSIQNTLY | 0.13978735 |
| VGAFPFGF | ASSIQNTLY | 0.30350035 |
| VGAGTVEF | ASSIQNTLY | 0.5427772 |
| VGKSTLLL | ASSIQNTLY | 0.70382357 |
| AALAHAGGGIV | ASSIQNTLY | 0.30607072 |
| AALVAVSLL | ASSIQNTLY | 0.8040174 |
| AAVSRAVAAL | ASSIQNTLY | 0.7029296 |
| AGGSSGIGGF | ASSIQNTLY | 0.59761006 |
| AGLTVANANG | ASSIQNTLY | 0.86348367 |
| AGLVGAVL | ASSIQNTLY | 0.6979214 |
| ALSQAFTV | ASSIQNTLY | 0.5906601 |
| ALTFSVSL | ASSIQNTLY | 0.59534013 |
| ALTGAALVAL | ASSIQNTLY | 0.86517656 |
| ASFSIGAGVL | ASSIQNTLY | 0.3955372 |
| DGSVYKI | ASSIQNTLY | 0.8262126 |
| EQLAEIV | ASSIQNTLY | 0.5812793 |
| FAVSELTVM | ASSIQNTLY | 0.8424485 |
| GAAPLAAGL | ASSIQNTLY | 0.41302383 |
| GADGGDGGSGGAGG | ASSIQNTLY | 0.8362671 |
| GFGLDTGGAGG | ASSIQNTLY | 0.7397441 |
| GLASGGLL | ASSIQNTLY | 0.90199566 |

| | | |
|---|---|---|
| GRIAGAAVLL | ASSIQNTLY | 0.6933809 |
| GSAGSLNTSF | ASSIQNTLY | 0.75250506 |
| GTGQIGI | ASSIQNTLY | 0.54425085 |
| GTLIALIL | ASSIQNTLY | 0.84379226 |
| IAATMGFVVG | ASSIQNTLY | 0.42977366 |
| IFAQALL | ASSIQNTLY | 0.7075467 |
| IGAAVALGF | ASSIQNTLY | 0.44235402 |
| IGLTPLAL | ASSIQNTLY | 0.80068237 |
| ILGSNTSTL | ASSIQNTLY | 0.8225772 |
| IVFSLTVL | ASSIQNTLY | 0.47816044 |
| LASKATGF | ASSIQNTLY | 0.86596346 |
| LASVALIL | ASSIQNTLY | 0.7839047 |
| LFAVGLTL | ASSIQNTLY | 0.61799735 |
| LFNGGIGTYI | ASSIQNTLY | 0.85863286 |
| LTLGAGII | ASSIQNTLY | 0.58965194 |
| LVAAALGDI | ASSIQNTLY | 0.85389066 |
| NTAFVQLGI | ASSIQNTLY | 0.6940665 |
| QLFSGAEL | ASSIQNTLY | 0.7190522 |
| SAATFITL | ASSIQNTLY | 0.85283095 |
| SNYGLANLGV | ASSIQNTLY | 0.479962 |
| SYGGAYAVM | ASSIQNTLY | 0.48990336 |
| TTGVGGLTLG | ASSIQNTLY | 0.85927665 |
| TTQTIYI | ASSIQNTLY | 0.86338294 |
| VAAGAGLGAVI | ASSIQNTLY | 0.87830055 |
| VAAVVTLL | ASSIQNTLY | 0.76395375 |
| VGFLGGLI | ASSIQNTLY | 0.6803691 |
| VGVIPILL | ASSIQNTLY | 0.7833709 |
| VIFASSLI | ASSIQNTLY | 0.87834495 |
| VKGGDGGAAAG | ASSIQNTLY | 0.86150014 |
| VTKTFTNI | ASSIQNTLY | 0.8580954 |
| VVAAAILL | ASSIQNTLY | 0.6952508 |
| YAALDVEL | ASSIQNTLY | 0.79559225 |
| LASVALLL | ASSIQNTLY | 0.773931 |
| LDAVHRV | ASSIQNTLY | 0.76721203 |
| QGLSSSIF | ASSIQNTLY | 0.8600633 |
| RGTFANIRL | ASSIQNTLY | 0.7860979 |
| RQLWWGHRI | ASSIQNTLY | 0.820786 |
| YGPPGTGKTLL | ASSIQNTLY | 0.8249449 |
| AALVGAPANVA | ASSIQNTLY | 0.8384863 |
| AATFRRTE | ASSIQNTLY | 0.8531424 |
| AGALGGGI | ASSIQNTLY | 0.8316461 |
| AGGAGGAGGIGAG | ASSIQNTLY | 0.8921292 |
| AGSGALGL | ASSIQNTLY | 0.8673326 |
| AQIADQI | ASSIQNTLY | 0.8729142 |
| ASAVAVTYL | ASSIQNTLY | 0.20763057 |
| ASTVLGGTGG | ASSIQNTLY | 0.5070532 |
| AVGGEPAK | ASSIQNTLY | 0.81970906 |
| CEQGTFGGTC | ASSIQNTLY | 0.76478213 |
| DEGKAPAG | ASSIQNTLY | 0.68730134 |
| DTVVFPAL | ASSIQNTLY | 0.8460231 |
| FEGITMAG | ASSIQNTLY | 0.8275902 |

| | | |
|---|---|---|
| FPLVVQD | ASSIQNTLY | 0.70410514 |
| GAFTTPQ | ASSIQNTLY | 0.8574977 |
| GFGGFGTAGGVGGA | ASSIQNTLY | 0.7739365 |
| GFLAQFL | ASSIQNTLY | 0.7335804 |
| GGAGGAGGNAGVA | ASSIQNTLY | 0.7956476 |
| GGVGTAGAGGAGGAGG | ASSIQNTLY | 0.87968683 |
| GNGGGGDAGNAGSGGN | ASSIQNTLY | 0.69680834 |
| GTGGAGGAGNDAGS | ASSIQNTLY | 0.81690466 |
| GTTNTGWFNS | ASSIQNTLY | 0.84118986 |
| IFMGPLFTL | ASSIQNTLY | 0.8672664 |
| ILAHAIEAI | ASSIQNTLY | 0.5511651 |
| IRSAIREAL | ASSIQNTLY | 0.85590005 |
| ITFAGGG | ASSIQNTLY | 0.7752731 |
| ITGMSGHQEV | ASSIQNTLY | 0.89376366 |
| IVAGLACTL | ASSIQNTLY | 0.7397084 |
| IVGGGLAGA | ASSIQNTLY | 0.7921653 |
| IVTGVLNI | ASSIQNTLY | 0.78011554 |
| LAQQGCI | ASSIQNTLY | 0.8493974 |
| LCLHVEN | ASSIQNTLY | 0.5128554 |
| LDAISDGS | ASSIQNTLY | 0.81068987 |
| LGGFAAML | ASSIQNTLY | 0.73394966 |
| LGGKNPMI | ASSIQNTLY | 0.83903927 |
| LGPKGVAAIL | ASSIQNTLY | 0.85204005 |
| LMVGVSI | ASSIQNTLY | 0.6884835 |
| LVGAAFVL | ASSIQNTLY | 0.88427484 |
| LVLAVAPL | ASSIQNTLY | 0.80240774 |
| MAAIAVIDAA | ASSIQNTLY | 0.7843723 |
| NNGINAELQ | ASSIQNTLY | 0.87718195 |
| RGEPLVLVHG | ASSIQNTLY | 0.7386524 |
| RGGAGGAAGLI | ASSIQNTLY | 0.88676566 |
| RSYRMHPAV | ASSIQNTLY | 0.8282317 |
| SAAHLLAGQLA | ASSIQNTLY | 0.84332097 |
| SAAVGSY | ASSIQNTLY | 0.86240214 |
| SPHAVLHRI | ASSIQNTLY | 0.7491573 |
| SQIAKVL | ASSIQNTLY | 0.7287278 |
| TALGHLVP | ASSIQNTLY | 0.8705636 |
| TAPTTSGAA | ASSIQNTLY | 0.8071138 |
| TDGDDGNGGSPGT | ASSIQNTLY | 0.84234476 |
| TDGLGEN | ASSIQNTLY | 0.7693294 |
| TGGPPLPGQA | ASSIQNTLY | 0.71532637 |
| TIPAITI | ASSIQNTLY | 0.7385875 |
| TIPGFTI | ASSIQNTLY | 0.82396746 |
| TSIASGALA | ASSIQNTLY | 0.83970606 |
| TVTLPGAAI | ASSIQNTLY | 0.81727934 |
| VASGLSAALP | ASSIQNTLY | 0.5165471 |
| VGAGVIGI | ASSIQNTLY | 0.8192401 |
| VGPKRFGV | ASSIQNTLY | 0.76884466 |
| VNVAFPD | ASSIQNTLY | 0.75838184 |
| VSMTNSGVS | ASSIQNTLY | 0.80094177 |
| VTDVLGDLSV | ASSIQNTLY | 0.79031706 |
| VVVVGAGFAGL | ASSIQNTLY | 0.7611197 |

| | | |
|---|---|---|
| VYTVGTVV | ASSIQNTLY | 0.73946947 |
| WTTFSGP | ASSIQNTLY | 0.74772656 |
| YDELVEAF | ASSIQNTLY | 0.8166218 |
| YGTGGAGGNGG | ASSIQNTLY | 0.8062469 |
| YTNVMAV | ASSIQNTLY | 0.8381344 |
| FLEQVHQL | ASSIQNTLY | 0.8373547 |
| GLSLLAVAV | ASSIQNTLY | 0.7875771 |
| LEQVHQL | ASSIQNTLY | 0.25153744 |
| LNAAGAI | ASSIQNTLY | 0.4590704 |
| PAVHASL | ASSIQNTLY | 0.70845485 |
| TELEDDA | ASSIQNTLY | 0.76409674 |
| TPPGPPN | ASSIQNTLY | 0.28728428 |
| TPPPPPPP | ASSIQNTLY | 0.78521186 |
| TQSPALI | ASSPRERTAEQF | 0.77423155 |
| EFAAAFPQ | ASSPRERTAEQF | 0.7371932 |
| LLSGNLDVL | ASSPRERTAEQF | 0.8313815 |
| GPANLGNYNI | ASSPRERTAEQF | 0.78270245 |
| TLGPTLP | ASSPRERTAEQF | 0.69358075 |
| VYAYSSCCK | ASSPRERTAEQF | 0.61444575 |
| APAPAPAPNI | ASSPRERTAEQF | 0.78945374 |
| PGGPVPIP | ASSPRERTAEQF | 0.42593545 |
| FLTVALEP | ASSPRERTAEQF | 0.8358432 |
| SQYFGANL | ASSPRERTAEQF | 0.42806077 |
| GGVKPAA | ASSPRERTAEQF | 0.8179455 |
| GGTGGAGGAGGW | ASSPRERTAEQF | 0.50324833 |
| GVGGVAGLGGAGG | ASSPRERTAEQF | 0.82530713 |
| VLLVLATTI | ASSPRERTAEQF | 0.78904927 |
| IGIFAAML | ASSPRERTAEQF | 0.86227834 |
| AVGDLID | ASSPRERTAEQF | 0.77287227 |
| NISPLPL | ASSPRERTAEQF | 0.79167825 |
| VTAVFVAI | ASSPRERTAEQF | 0.5923431 |
| SGDDNDAAL | ASSPRERTAEQF | 0.8336908 |
| AVAALAHS | ASSPRERTAEQF | 0.6318768 |
| GGVTVMATP | ASSPRERTAEQF | 0.7671016 |
| AGLAANVRD | ASSPRERTAEQF | 0.4389553 |
| TNRPDLIDPALLRPG | ASSPRERTAEQF | 0.81004524 |
| LANLGVTAL | ASSPRERTAEQF | 0.82165825 |
| TGERIEL | ASSPRERTAEQF | 0.5997814 |
| NGEELAA | ASSPRERTAEQF | 0.82312685 |
| VLSLVWG | ASSPRERTAEQF | 0.77988267 |
| IGYALWL | ASSPRERTAEQF | 0.6582083 |
| LSLLLAPL | ASSPRERTAEQF | 0.8422953 |
| TLDEALK | ASSPRERTAEQF | 0.7087392 |
| TDGAGTAEGG | ASSPRERTAEQF | 0.83826 |
| NAPGGAGGNGG | ASSPRERTAEQF | 0.7033758 |
| VSVFMGMGGNF | ASSPRERTAEQF | 0.72820675 |
| LASYLAAL | ASSPRERTAEQF | 0.77873546 |
| LVGGTPAL | ASSPRERTAEQF | 0.63913465 |
| VRNADVL | ASSPRERTAEQF | 0.5654825 |
| TCSSPFLVSAM | ASSPRERTAEQF | 0.84092784 |
| VHITDAE | ASSPRERTAEQF | 0.80721676 |

| | | |
|---|---|---|
| EYAAPSF | ASSPRERTAEQF | 0.8236487 |
| VVDAFHGGL | ASSPRERTAEQF | 0.6648462 |
| ASVVDPQ | ASSPRERTAEQF | 0.58298737 |
| SYNEFGGLS | ASSPRERTAEQF | 0.6722972 |
| DPDGSVY | ASSPRERTAEQF | 0.8130511 |
| MVSTIHATE | ASSPRERTAEQF | 0.8385378 |
| LAAGVADPV | ASSPRERTAEQF | 0.800501 |
| TKEQIAA | ASSPRERTAEQF | 0.5910754 |
| SLASPAAL | ASSPRERTAEQF | 0.7815531 |
| IGLVFYGV | ASSPRERTAEQF | 0.7321483 |
| FLLDASNL | ASSPRERTAEQF | 0.73735857 |
| YQVGNFP | ASSPRERTAEQF | 0.7848255 |
| ESLLSQST | ASSPRERTAEQF | 0.7784054 |
| LFTVGFAAVG | ASSPRERTAEQF | 0.69214916 |
| ATPPTPT | ASSPRERTAEQF | 0.6544169 |
| TDLATLG | ASSPRERTAEQF | 0.7696872 |
| SAGGTGR | ASSPRERTAEQF | 0.82474434 |
| GFSNSGS | ASSPRERTAEQF | 0.8147707 |
| GGNGGSGGTSV | ASSPRERTAEQF | 0.8501551 |
| GTSVATGGAGN | ASSPRERTAEQF | 0.8255787 |
| NGGNGGSGGTS | ASSPRERTAEQF | 0.82943225 |
| AVVKFPH | ASSPRERTAEQF | 0.29100198 |
| TPPPPPP | ASSPRERTAEQF | 0.47961846 |
| AAPAAAAAPA | ASSPRERTAEQF | 0.6551676 |
| IALNEDL | ASSPRERTAEQF | 0.7275099 |
| IVNVSSM | ASSPRERTAEQF | 0.35658723 |
| TGIPPAPR | ASSPRERTAEQF | 0.7921363 |
| FDLGGGTF | ASSPRERTAEQF | 0.71327657 |
| FDLGGGTFDVS | ASSPRERTAEQF | 0.61043674 |
| NEPTAAA | ASSPRERTAEQF | 0.8155407 |
| NEPTAAALA | ASSPRERTAEQF | 0.69019455 |
| GSIINISSI | ASSPRERTAEQF | 0.60897213 |
| VAPGNAGTA | ASSPRERTAEQF | 0.65737426 |
| TGGGASLE | ASSPRERTAEQF | 0.836313 |
| VAVYNHYKRI | ASSPRERTAEQF | 0.5907043 |
| TDPAPATT | ASSPRERTAEQF | 0.80926573 |
| TNRPDILDPALLRPG | ASSPRERTAEQF | 0.62827295 |
| LTTTTTS | ASSPRERTAEQF | 0.8352434 |
| PTGSGKTL | ASSPRERTAEQF | 0.50150275 |
| TGSGKTL | ASSPRERTAEQF | 0.8262581 |
| TIGHVDHGK | ASSPRERTAEQF | 0.75093514 |
| TIGHVDHGKTT | ASSPRERTAEQF | 0.8636539 |
| GHVDHGKTT | ASSPRERTAEQF | 0.7024999 |
| AQFEHTLL | ASSPRERTAEQF | 0.78296185 |
| IVNQDEL | ASSPRERTAEQF | 0.73302555 |
| TNGSQFF | ASSPRERTAEQF | 0.81968564 |
| TGKGIVE | ASSPRERTAEQF | 0.47996804 |
| LISNASDA | ASSPRERTAEQF | 0.70450914 |
| AELRTLGELTKIAKAHGAQ | ASSPRERTAEQF | 0.5058109 |
| APNAVAAAEPSW | ASSPRERTAEQF | 0.76534176 |
| APQVAGAATAVL | ASSPRERTAEQF | 0.8094629 |

| | | |
|---|---|---|
| ARRDPAAAAAILPTDA | ASSPRERTAEQF | 0.33589014 |
| DAAARDVLAEAGLAEYF | ASSPRERTAEQF | 0.8039416 |
| DDQIVLSAKANLAKVEQ | ASSPRERTAEQF | 0.676878 |
| DELLPDTGEDSDEDGH | ASSPRERTAEQF | 0.6195265 |
| ELPEAPANRATVPQD | ASSPRERTAEQF | 0.8299041 |
| FGQSKIERVTGIG | ASSPRERTAEQF | 0.76458013 |
| GAGGTGSTAGGAGGAGGAGGLYA | ASSPRERTAEQF | 0.82377326 |
| GHGGTGAAGGNGGTGGAGGSGIDG | ASSPRERTAEQF | 0.692686 |
| GNGGNGGAGGEGGPGGAGGAGA | ASSPRERTAEQF | 0.7102201 |
| GSTAGNGGNGGNATNGTIAGQPAG | ASSPRERTAEQF | 0.77637035 |
| GTGGDGGNGGNGGTGG | ASSPRERTAEQF | 0.49884665 |
| GVRRPAGIPADAPV | ASSPRERTAEQF | 0.5934404 |
| GYRQVTDAHLLTLARR | ASSPRERTAEQF | 0.8469155 |
| IAKMGQDGHDRG | ASSPRERTAEQF | 0.35279697 |
| KAEPSIEIQYT | ASSPRERTAEQF | 0.75167966 |
| KPVRIRMAADEDTVAVT | ASSPRERTAEQF | 0.15168166 |
| MPPRIAGMRLLVIKPEP | ASSPRERTAEQF | 0.1459253 |
| MKFYAVALFRSRGVPPTR | ASSPRERTAEQF | 0.2530283 |
| NAGGFGILGAVAHSP | ASSPRERTAEQF | 0.5738739 |
| NEGREITLAMAEM | ASSPRERTAEQF | 0.7597558 |
| NELVEWSHPLGHRWRWE | ASSPRERTAEQF | 0.49217275 |
| NLGAGGTGGNGGNPG | ASSPRERTAEQF | 0.15872198 |
| NTEWPKWETVMAADPVD | ASSPRERTAEQF | 0.1958057 |
| PASAAPAETQP | ASSPRERTAEQF | 0.19583982 |
| QQSSDGKNGVRAAEITGE | ASSPRERTAEQF | 0.18047956 |
| RAPDAVTTSWWKE | ASSPRERTAEQF | 0.32284325 |
| SGFGNVGAHTSGWFNQ | ASSPRERTAEQF | 0.22118604 |
| SGVAGAVGQAASVGGLKV | ASSPRERTAEQF | 0.17106831 |
| SNGGAGTGSAGGNGGT | ASSPRERTAEQF | 0.16454193 |
| TGGAGGEGGAAG | ASSPRERTAEQF | 0.30609006 |
| TGGDGGAAGAGGAPGHG | ASSPRERTAEQF | 0.55872667 |
| TMPPLLAAIANAGRHGV | ASSPRERTAEQF | 0.45106742 |
| TSSTAGNSGGA | ASSPRERTAEQF | 0.59960824 |
| VPATVPVGILC | ASSPRERTAEQF | 0.41083318 |
| VQRPDSGGVVLTGRIS | ASSPRERTAEQF | 0.3692571 |
| WTPNPVPATKGVPAGLVQAT | ASSPRERTAEQF | 0.10522713 |
| YALKRALAKFSDRFDVVII | ASSPRERTAEQF | 0.117935576 |
| YLIGSVIV | ASSPRERTAEQF | 0.21985278 |
| YGQLVAVID | ASSPRERTAEQF | 0.17188297 |
| VTVGGMTLPA | ASSPRERTAEQF | 0.13238588 |
| VLALPAV | ASSPRERTAEQF | 0.39201447 |
| VIPPTLNYET | ASSPRERTAEQF | 0.20839033 |
| VGMGGMHPSSG | ASSPRERTAEQF | 0.15625067 |
| VGLAGGAATA | ASSPRERTAEQF | 0.4578601 |
| VGGAGGAGGSGP | ASSPRERTAEQF | 0.15944588 |
| VFAAFMV | ASSPRERTAEQF | 0.1565742 |
| TLYDKLR | ASSPRERTAEQF | 0.16402902 |
| TLAVFTVL | ASSPRERTAEQF | 0.4443804 |
| TGSVQPARV | ASSPRERTAEQF | 0.1401367 |
| TGGGAGSGGT | ASSPRERTAEQF | 0.278018 |
| SHLETSL | ASSPRERTAEQF | 0.24057212 |

| | | |
|---|---|---|
| SFTTGMT | ASSPRERTAEQF | 0.37939674 |
| QVEVAKRMGVRQPTVS | ASSPRERTAEQF | 0.13622063 |
| QGSGGPALAM | ASSPRERTAEQF | 0.32580602 |
| MLAPSNP | ASSPRERTAEQF | 0.1838226 |
| LVNNAAI | ASSPRERTAEQF | 0.79660344 |
| LTGEINTLA | ASSPRERTAEQF | 0.17081507 |
| LSIFGPV | ASSPRERTAEQF | 0.2309428 |
| LLALLANL | ASSPRERTAEQF | 0.16958642 |
| LISNASDALDKLR | ASSPRERTAEQF | 0.71332026 |
| LINEAVP | ASSPRERTAEQF | 0.12858711 |
| LGYAFSGL | ASSPRERTAEQF | 0.20406587 |
| LDFVAVAL | ASSPRERTAEQF | 0.13274458 |
| LAQPGAE | ASSPRERTAEQF | 0.29235962 |
| ISNLGAVTGF | ASSPRERTAEQF | 0.36815354 |
| HQEDPGRI | ASSPRERTAEQF | 0.1220419 |
| GTPGEPTP | ASSPRERTAEQF | 0.39401335 |
| GFFNSGS | ASSPRERTAEQF | 0.27367634 |
| GAGGGGLTAGAN | ASSPRERTAEQF | 0.17335947 |
| FVDALVNM | ASSPRERTAEQF | 0.5010315 |
| FLTVAPEL | ASSPRERTAEQF | 0.19090745 |
| FETLTAL | ASSPRERTAEQF | 0.5998368 |
| DVINPIEL | ASSPRERTAEQF | 0.20489794 |
| APVSGAPV | ASSPRERTAEQF | 0.22407681 |
| ALVGGLAW | ASSPRERTAEQF | 0.4272016 |
| ALAAAAATAI | ASSPRERTAEQF | 0.23832679 |
| AFGLGLLL | ASSPRERTAEQF | 0.15657729 |
| AFFAAASL | ASSPRERTAEQF | 0.39690995 |
| AAAALPAFSPP | ASSPRERTAEQF | 0.7443172 |
| VVLAPAL | ASSPRERTAEQF | 0.7424415 |
| TIGHVDHGKTTL | ASSPRERTAEQF | 0.56733185 |
| LISNASDALDK | ASSPRERTAEQF | 0.5712809 |
| LISNASDAL | ASSPRERTAEQF | 0.62747717 |
| ISNASDALDK | ASSPRERTAEQF | 0.76071024 |
| NQAETLVYQTE | ASSPRERTAEQF | 0.78212297 |
| TGGTGGTAGAG | ASSPRERTAEQF | 0.74857426 |
| TGSTAGGAGGA | ASSPRERTAEQF | 0.6543036 |
| DPLAVPVAAGPV | ASSPRERTAEQF | 0.74980026 |
| GGTGGTPGNGGN | ASSPRERTAEQF | 0.6881769 |
| PAGLAGVDPATL | ASSPRERTAEQF | 0.6943326 |
| GNNLVVEAPHAGASV | ASSPRERTAEQF | 0.7340497 |
| KVTRIMSASTLEQPA | ASSPRERTAEQF | 0.7228313 |
| LGFALPIALPRRTVV | ASSPRERTAEQF | 0.63902426 |
| TRLGNLTARKLVVLV | ASSPRERTAEQF | 0.60596514 |
| WSQQRRLIYFPSAGP | ASSPRERTAEQF | 0.7521112 |
| GPNGNPDPMAAAVDIR | ASSPRERTAEQF | 0.77586913 |
| TQRCVSVADLRAAASR | ASSPRERTAEQF | 0.76293635 |
| QFVLDVAEHPVEVLTETV | ASSPRERTAEQF | 0.80703354 |
| GGAGGTGGAGGNGGTQAGDG | ASSPRERTAEQF | 0.7629433 |
| PLLATLRAIAVDVDAKPAQV | ASSPRERTAEQF | 0.7526389 |
| APKFRTGGYMIDFWGVGYQVAKRM | ASSPRERTAEQF | 0.34899303 |
| GAGGAGGAGGAAGGPGGTGGTGGN | ASSPRERTAEQF | 0.49418992 |

| | | |
|---|---|---|
| IGALGLLATDVAPAGSVAFGVALA | ASSPRERTAEQF | 0.6404592 |
| AGGAGGIGGNGGAGGAGGVLMGNG | ASSPRERTAEQF | 0.66151005 |
| ADELIGLL | ASSPRERTAEQF | 0.33138403 |
| ADRALKA | ASSPRERTAEQF | 0.7480441 |
| AGAVGQVGGAGGA | ASSPRERTAEQF | 0.7043556 |
| AGGGGGLGGAP | ASSPRERTAEQF | 0.6071527 |
| AGGLLYGNGGAG | ASSPRERTAEQF | 0.7659582 |
| AGLESIGL | ASSPRERTAEQF | 0.64870363 |
| AGLKTGII | ASSPRERTAEQF | 0.5993577 |
| ASLAALLL | ASSPRERTAEQF | 0.64066905 |
| ASVINISSI | ASSPRERTAEQF | 0.7899536 |
| ATPPGPA | ASSPRERTAEQF | 0.5116351 |
| ATVASALV | ASSPRERTAEQF | 0.7790803 |
| ATVASLIL | ASSPRERTAEQF | 0.6497807 |
| AVNGPVLIRH | ASSPRERTAEQF | 0.78614414 |
| DGAAGAVGI | ASSPRERTAEQF | 0.54676974 |
| DPLLHGI | ASSPRERTAEQF | 0.7619213 |
| ELGAALGAL | ASSPRERTAEQF | 0.6757367 |
| ELGRTQAV | ASSPRERTAEQF | 0.8086823 |
| FVAGGYLS | ASSPRERTAEQF | 0.68075776 |
| GASGGAGGAGGW | ASSPRERTAEQF | 0.77368855 |
| GGAGAGGDGGAGGI | ASSPRERTAEQF | 0.73035216 |
| GTGHNGSQPGMGGNG | ASSPRERTAEQF | 0.77183473 |
| GYAAASASAS | ASSPRERTAEQF | 0.486637 |
| HQVDVAAVQQG | ASSPRERTAEQF | 0.6583618 |
| IGAAIAHT | ASSPRERTAEQF | 0.62246084 |
| IGMIEVAQL | ASSPRERTAEQF | 0.70678455 |
| IRLAPEHTSAMD | ASSPRERTAEQF | 0.752913 |
| IYNVSVL | ASSPRERTAEQF | 0.2876968 |
| KQGFLGM | ASSPRERTAEQF | 0.7686057 |
| LESVITQL | ASSPRERTAEQF | 0.518245 |
| LGLGALVVV | ASSPRERTAEQF | 0.6078896 |
| LIGNGGNGGSGGI | ASSPRERTAEQF | 0.7598121 |
| LNAINAPTL | ASSPRERTAEQF | 0.701804 |
| LQLLNTGV | ASSPRERTAEQF | 0.75456154 |
| LQTSIEF | ASSPRERTAEQF | 0.6234094 |
| MSFAVSEL | ASSPRERTAEQF | 0.6558703 |
| NTGALGL | ASSPRERTAEQF | 0.7479171 |
| PPGAGGAGGAGTT | ASSPRERTAEQF | 0.43198916 |
| QLVGLGI | ASSPRERTAEQF | 0.5290822 |
| RFLEQVHQL | ASSPRERTAEQF | 0.79112005 |
| RGVGYVLR | ASSPRERTAEQF | 0.6348158 |
| SATGVFAGVI | ASSPRERTAEQF | 0.64469147 |
| SLPAITLP | ASSPRERTAEQF | 0.56604326 |
| STALAFTV | ASSPRERTAEQF | 0.57190955 |
| SVLTVGLGK | ASSPRERTAEQF | 0.59923226 |
| TALAGAGSGI | ASSPRERTAEQF | 0.6445512 |
| TGLPFTGI | ASSPRERTAEQF | 0.65651846 |
| VAEHVTV | ASSPRERTAEQF | 0.63563305 |
| VDIVLNSL | ASSPRERTAEQF | 0.5965724 |
| VLTGGGAVL | ASSPRERTAEQF | 0.632705 |

| | | |
|---|---|---|
| VNVGLAYGI | ASSPRERTAEQF | 0.6012613 |
| VSSAAVGAV | ASSPRERTAEQF | 0.6074505 |
| VTSVYEAGV | ASSPRERTAEQF | 0.63769054 |
| WVALDRGA | ASSPRERTAEQF | 0.61959386 |
| YLVGSVVI | ASSPRERTAEQF | 0.5963706 |
| GPPGTGKTLL | ASSPRERTAEQF | 0.59246176 |
| GVGMTKF | ASSPRERTAEQF | 0.63478065 |
| IGIGGSDL | ASSPRERTAEQF | 0.6471343 |
| IGTLLAEL | ASSPRERTAEQF | 0.6425255 |
| IYTSGTT | ASSPRERTAEQF | 0.66234386 |
| LSFPAGV | ASSPRERTAEQF | 0.64316916 |
| LYTSGTT | ASSPRERTAEQF | 0.64218897 |
| VGAFPFGF | ASSPRERTAEQF | 0.38597327 |
| VGAGTVEF | ASSPRERTAEQF | 0.52863705 |
| VGKSTLLL | ASSPRERTAEQF | 0.60473686 |
| AALAHAGGGIV | ASSPRERTAEQF | 0.6128519 |
| AALVAVSLL | ASSPRERTAEQF | 0.4318249 |
| AAVSRAVAAL | ASSPRERTAEQF | 0.6381425 |
| AGGSSGIGGF | ASSPRERTAEQF | 0.6090456 |
| AGLTVANANG | ASSPRERTAEQF | 0.5750129 |
| AGLVGAVL | ASSPRERTAEQF | 0.6447984 |
| ALSQAFTV | ASSPRERTAEQF | 0.5896946 |
| ALTFSVSL | ASSPRERTAEQF | 0.5779877 |
| ALTGAALVAL | ASSPRERTAEQF | 0.6031072 |
| ASFSIGAGVL | ASSPRERTAEQF | 0.6582854 |
| DGSVYKI | ASSPRERTAEQF | 0.5599818 |
| EQLAEIV | ASSPRERTAEQF | 0.6428937 |
| FAVSELTVM | ASSPRERTAEQF | 0.61281 |
| GAAPLAAGL | ASSPRERTAEQF | 0.6530819 |
| GADGGDGGSGGAGG | ASSPRERTAEQF | 0.53775764 |
| GFGLDTGGAGG | ASSPRERTAEQF | 0.6389391 |
| GLASGGLL | ASSPRERTAEQF | 0.6009395 |
| GRIAGAAVLL | ASSPRERTAEQF | 0.66774136 |
| GSAGSLNTSF | ASSPRERTAEQF | 0.6018501 |
| GTGQIGI | ASSPRERTAEQF | 0.63496625 |
| GTLIALIL | ASSPRERTAEQF | 0.61342657 |
| IAATMGFVVG | ASSPRERTAEQF | 0.64485747 |
| IFAQALL | ASSPRERTAEQF | 0.48960906 |
| IGAAVALGF | ASSPRERTAEQF | 0.6103722 |
| IGLTPLAL | ASSPRERTAEQF | 0.5679758 |
| ILGSNTSTL | ASSPRERTAEQF | 0.6284555 |
| IVFSLTVL | ASSPRERTAEQF | 0.6323446 |
| LASKATGF | ASSPRERTAEQF | 0.4213323 |
| LASVALIL | ASSPRERTAEQF | 0.6498376 |
| LFAVGLTL | ASSPRERTAEQF | 0.5484733 |
| LFNGGIGTYI | ASSPRERTAEQF | 0.5812812 |
| LTLGAGII | ASSPRERTAEQF | 0.64869213 |
| LVAAALGDI | ASSPRERTAEQF | 0.6117517 |
| NTAFVQLGI | ASSPRERTAEQF | 0.6487751 |
| QLFSGAEL | ASSPRERTAEQF | 0.60494924 |
| SAATFITL | ASSPRERTAEQF | 0.61496156 |

| | | |
|---|---|---|
| SNYGLANLGV | ASSPRERTAEQF | 0.6432136 |
| SYGGAYAVM | ASSPRERTAEQF | 0.5285848 |
| TTGVGGLTLG | ASSPRERTAEQF | 0.5768625 |
| TTQTIYI | ASSPRERTAEQF | 0.6506953 |
| VAAGAGLGAVI | ASSPRERTAEQF | 0.79659206 |
| VAAVVTLL | ASSPRERTAEQF | 0.8237735 |
| VGFLGGLI | ASSPRERTAEQF | 0.7303037 |
| VGVIPILL | ASSPRERTAEQF | 0.6481035 |
| VIFASSLI | ASSPRERTAEQF | 0.73972386 |
| VKGGDGGAAAG | ASSPRERTAEQF | 0.79708076 |
| VTKTFTNI | ASSPRERTAEQF | 0.82001 |
| VVAAAILL | ASSPRERTAEQF | 0.7454113 |
| YAALDVEL | ASSPRERTAEQF | 0.45073166 |
| LASVALLL | ASSPRERTAEQF | 0.72921485 |
| LDAVHRV | ASSPRERTAEQF | 0.7332376 |
| QGLSSSIF | ASSPRERTAEQF | 0.71528924 |
| RGTFANIRL | ASSPRERTAEQF | 0.7567748 |
| RQLWWGHRI | ASSPRERTAEQF | 0.74227947 |
| YGPPGTGKTLL | ASSPRERTAEQF | 0.71164167 |
| AALVGAPANVA | ASSPRERTAEQF | 0.73271245 |
| AATFRRTE | ASSPRERTAEQF | 0.6672172 |
| AGALGGGI | ASSPRERTAEQF | 0.7923025 |
| AGGAGGAGGIGAG | ASSPRERTAEQF | 0.8055235 |
| AGSGALGL | ASSPRERTAEQF | 0.7952574 |
| AQIADQI | ASSPRERTAEQF | 0.8126342 |
| ASAVAVTYL | ASSPRERTAEQF | 0.78417206 |
| ASTVLGGTGG | ASSPRERTAEQF | 0.20004463 |
| AVGGEPAK | ASSPRERTAEQF | 0.37156093 |
| CEQGTFGGTC | ASSPRERTAEQF | 0.6979921 |
| DEGKAPAG | ASSPRERTAEQF | 0.661899 |
| DTVVFPAL | ASSPRERTAEQF | 0.35154122 |
| FEGITMAG | ASSPRERTAEQF | 0.63756895 |
| FPLVVQD | ASSPRERTAEQF | 0.6704619 |
| GAFTTPQ | ASSPRERTAEQF | 0.63623416 |
| GFGGFGTAGGVGGA | ASSPRERTAEQF | 0.787611 |
| GFLAQFL | ASSPRERTAEQF | 0.725037 |
| GGAGGAGGNAGVA | ASSPRERTAEQF | 0.67904276 |
| GGVGTAGAGGAGGAGG | ASSPRERTAEQF | 0.55179787 |
| GNGGGGDAGNAGSGGN | ASSPRERTAEQF | 0.6701519 |
| GTGGAGGAGNDAGS | ASSPRERTAEQF | 0.37633955 |
| GTTNTGWFNS | ASSPRERTAEQF | 0.790437 |
| IFMGPLFTL | ASSPRERTAEQF | 0.38172552 |
| ILAHAIEAI | ASSPRERTAEQF | 0.78709173 |
| IRSAIREAL | ASSPRERTAEQF | 0.50145584 |
| ITFAGGG | ASSPRERTAEQF | 0.78961754 |
| ITGMSGHQEV | ASSPRERTAEQF | 0.77134967 |
| IVAGLACTL | ASSPRERTAEQF | 0.8404765 |
| IVGGGLAGA | ASSPRERTAEQF | 0.7640733 |
| IVTGVLNI | ASSPRERTAEQF | 0.691877 |
| LAQQGCI | ASSPRERTAEQF | 0.42476904 |
| LCLHVEN | ASSPRERTAEQF | 0.81314826 |

| | | |
|---|---|---|
| LDAISDGS | ASSPRERTAEQF | 0.63743824 |
| LGGFAAML | ASSPRERTAEQF | 0.76249737 |
| LGGKNPMI | ASSPRERTAEQF | 0.32702765 |
| LGPKGVAAIL | ASSPRERTAEQF | 0.79424554 |
| LMVGVSI | ASSPRERTAEQF | 0.7569945 |
| LVGAAFVL | ASSPRERTAEQF | 0.52935994 |
| LVLAVAPL | ASSPRERTAEQF | 0.7973007 |
| MAAIAVIDAA | ASSPRERTAEQF | 0.7752192 |
| NNGINAELQ | ASSPRERTAEQF | 0.6266187 |
| RGEPLVLVHG | ASSPRERTAEQF | 0.8220271 |
| RGGAGGAAGLI | ASSPRERTAEQF | 0.64564204 |
| RSYRMHPAV | ASSPRERTAEQF | 0.8299544 |
| SAAHLLAGQLA | ASSPRERTAEQF | 0.68668914 |
| SAAVGSY | ASSPRERTAEQF | 0.66935617 |
| SPHAVLHRI | ASSPRERTAEQF | 0.70809555 |
| SQIAKVL | ASSPRERTAEQF | 0.73104036 |
| TALGHLVP | ASSPRERTAEQF | 0.5740593 |
| TAPTTSGAA | ASSPRERTAEQF | 0.82250273 |
| TDGDDGNGGSPGT | ASSPRERTAEQF | 0.6917815 |
| TDGLGEN | ASSPRERTAEQF | 0.69786155 |
| TGGPPLPGQA | ASSPRERTAEQF | 0.62238896 |
| TIPAITI | ASSPRERTAEQF | 0.5948111 |
| TIPGFTI | ASSPRERTAEQF | 0.6266783 |
| TSIASGALA | ASSPRERTAEQF | 0.710282 |
| TVTLPGAAI | ASSPRERTAEQF | 0.71385366 |
| VASGLSAALP | ASSPRERTAEQF | 0.69225407 |
| VGAGVIGI | ASSPRERTAEQF | 0.61492455 |
| VGPKRFGV | ASSPRERTAEQF | 0.7171665 |
| VNVAFPD | ASSPRERTAEQF | 0.6717925 |
| VSMTNSGVS | ASSPRERTAEQF | 0.63908046 |
| VTDVLGDLSV | ASSPRERTAEQF | 0.69360614 |
| VVVVGAGFAGL | ASSPRERTAEQF | 0.7081341 |
| VYTVGTVV | ASSPRERTAEQF | 0.64800817 |
| WTTFSGP | ASSPRERTAEQF | 0.6497853 |
| YDELVEAF | ASSPRERTAEQF | 0.6698588 |
| YGTGGAGGNGG | ASSPRERTAEQF | 0.70373523 |
| YTNVMAV | ASSPRERTAEQF | 0.69139993 |
| FLEQVHQL | ASSPRERTAEQF | 0.7280427 |
| GLSLLAVAV | ASSPRERTAEQF | 0.70831794 |
| LEQVHQL | ASSPRERTAEQF | 0.70281243 |
| LNAAGAI | ASSPRERTAEQF | 0.36374497 |
| PAVHASL | ASSPRERTAEQF | 0.4816036 |
| TELEDDA | ASSPRERTAEQF | 0.68593425 |
| TPPGPPN | ASSPRERTAEQF | 0.65501195 |
| TPPPPPPP | ASSPRERTAEQF | 0.41347688 |
| TQSPALI | ASSSGTGGEQY | 0.7034779 |
| EFAAAFPQ | ASSSGTGGEQY | 0.6607249 |
| LLSGNLDVL | ASSSGTGGEQY | 0.66104364 |
| GPANLGNYNI | ASSSGTGGEQY | 0.71009403 |
| TLGPTLP | ASSSGTGGEQY | 0.6623079 |
| VYAYSSCCK | ASSSGTGGEQY | 0.6461688 |

| | | |
|---|---|---|
| APAPAPAPNI | ASSSGTGGEQY | 0.66071635 |
| PGGPVPIP | ASSSGTGGEQY | 0.72967124 |
| FLTVALEP | ASSSGTGGEQY | 0.5506974 |
| SQYFGANL | ASSSGTGGEQY | 0.70778966 |
| GGVKPAA | ASSSGTGGEQY | 0.57884854 |
| GGTGGAGGAGGW | ASSSGTGGEQY | 0.722291 |
| GVGGVAGLGGAGG | ASSSGTGGEQY | 0.56102306 |
| VLLVLATTI | ASSSGTGGEQY | 0.7014216 |
| IGIFAAML | ASSSGTGGEQY | 0.65084046 |
| AVGDLID | ASSSGTGGEQY | 0.74005735 |
| NISPLPL | ASSSGTGGEQY | 0.6525351 |
| VTAVFVAI | ASSSGTGGEQY | 0.71266377 |
| SGDDNDAAL | ASSSGTGGEQY | 0.68468887 |
| AVAALAHS | ASSSGTGGEQY | 0.7050591 |
| GGVTVMATP | ASSSGTGGEQY | 0.56865454 |
| AGLAANVRD | ASSSGTGGEQY | 0.6747066 |
| TNRPDLIDPALLRPG | ASSSGTGGEQY | 0.5451222 |
| LANLGVTAL | ASSSGTGGEQY | 0.6962842 |
| TGERIEL | ASSSGTGGEQY | 0.6935351 |
| NGEELAA | ASSSGTGGEQY | 0.47091335 |
| VLSLVWG | ASSSGTGGEQY | 0.7270281 |
| IGYALWL | ASSSGTGGEQY | 0.62991184 |
| LSLLLAPL | ASSSGTGGEQY | 0.63078636 |
| TLDEALK | ASSSGTGGEQY | 0.7127496 |
| TDGAGTAEGG | ASSSGTGGEQY | 0.69352454 |
| NAPGGAGGNGG | ASSSGTGGEQY | 0.70532846 |
| VSVFMGMGGNF | ASSSGTGGEQY | 0.66250503 |
| LASYLAAL | ASSSGTGGEQY | 0.66854376 |
| LVGGTPAL | ASSSGTGGEQY | 0.70443577 |
| VRNADVL | ASSSGTGGEQY | 0.5873144 |
| TCSSPFLVSAM | ASSSGTGGEQY | 0.6263318 |
| VHITDAE | ASSSGTGGEQY | 0.7163672 |
| EYAAPSF | ASSSGTGGEQY | 0.88858235 |
| VVDAFHGGL | ASSSGTGGEQY | 0.8938867 |
| ASVVDPQ | ASSSGTGGEQY | 0.7971779 |
| SYNEFGGLS | ASSSGTGGEQY | 0.7775067 |
| DPDGSVY | ASSSGTGGEQY | 0.8419514 |
| MVSTIHATE | ASSSGTGGEQY | 0.8992063 |
| LAAGVADPV | ASSSGTGGEQY | 0.89901245 |
| TKEQIAA | ASSSGTGGEQY | 0.88888943 |
| SLASPAAL | ASSSGTGGEQY | 0.80080116 |
| IGLVFYGV | ASSSGTGGEQY | 0.874948 |
| FLLDASNL | ASSSGTGGEQY | 0.8554095 |
| YQVGNFP | ASSSGTGGEQY | 0.85467947 |
| ESLLSQST | ASSSGTGGEQY | 0.89040947 |
| LFTVGFAAVG | ASSSGTGGEQY | 0.8769429 |
| ATPPTPT | ASSSGTGGEQY | 0.8306756 |
| TDLATLG | ASSSGTGGEQY | 0.8361063 |
| SAGGTGR | ASSSGTGGEQY | 0.87871444 |
| GFSNSGS | ASSSGTGGEQY | 0.89453447 |
| GGNGGSGGTSV | ASSSGTGGEQY | 0.8869965 |

| | | |
|---|---|---|
| GTSVATGGAGN | ASSSGTGGEQY | 0.9089113 |
| NGGNGGSGGTS | ASSSGTGGEQY | 0.894917 |
| AVVKFPH | ASSSGTGGEQY | 0.8893714 |
| TPPPPPP | ASSSGTGGEQY | 0.29436868 |
| AAPAAAAAPA | ASSSGTGGEQY | 0.6637019 |
| IALNEDL | ASSSGTGGEQY | 0.8367661 |
| IVNVSSM | ASSSGTGGEQY | 0.8488767 |
| TGIPPAPR | ASSSGTGGEQY | 0.603721 |
| FDLGGGTF | ASSSGTGGEQY | 0.8883605 |
| FDLGGGTFDVS | ASSSGTGGEQY | 0.83803946 |
| NEPTAAA | ASSSGTGGEQY | 0.80542815 |
| NEPTAAALA | ASSSGTGGEQY | 0.8937001 |
| GSIINISSI | ASSSGTGGEQY | 0.8326098 |
| VAPGNAGTA | ASSSGTGGEQY | 0.8086856 |
| TGGGASLE | ASSSGTGGEQY | 0.8583605 |
| VAVYNHYKRI | ASSSGTGGEQY | 0.90503824 |
| TDPAPATT | ASSSGTGGEQY | 0.7759763 |
| TNRPDILDPALLRPG | ASSSGTGGEQY | 0.89397424 |
| LTTTTTS | ASSSGTGGEQY | 0.863576 |
| PTGSGKTL | ASSSGTGGEQY | 0.9020073 |
| TGSGKTL | ASSSGTGGEQY | 0.7258091 |
| TIGHVDHGK | ASSSGTGGEQY | 0.8954127 |
| TIGHVDHGKTT | ASSSGTGGEQY | 0.8596572 |
| GHVDHGKTT | ASSSGTGGEQY | 0.91184765 |
| AQFEHTLL | ASSSGTGGEQY | 0.8493639 |
| IVNQDEL | ASSSGTGGEQY | 0.8852561 |
| TNGSQFF | ASSSGTGGEQY | 0.8667681 |
| TGKGIVE | ASSSGTGGEQY | 0.89832866 |
| LISNASDA | ASSSGTGGEQY | 0.70580435 |
| AELRTLGELTKIAKAHGAQ | ASSSGTGGEQY | 0.85152364 |
| APNAVAAAEPSW | ASSSGTGGEQY | 0.69431657 |
| APQVAGAATAVL | ASSSGTGGEQY | 0.87327284 |
| ARRDPAAAAAILPTDA | ASSSGTGGEQY | 0.8898997 |
| DAAARDVLAEAGLAEYF | ASSSGTGGEQY | 0.45170468 |
| DDQIVLSAKANLAKVEQ | ASSSGTGGEQY | 0.8958918 |
| DELLPDTGEDSDEDGH | ASSSGTGGEQY | 0.8116082 |
| ELPEAPANRATVPQD | ASSSGTGGEQY | 0.8127245 |
| FGQSKIERVTGIG | ASSSGTGGEQY | 0.8993346 |
| GAGGTGSTAGGAGGAGGAGGLYA | ASSSGTGGEQY | 0.8532776 |
| GHGGTGAAGGNGGTGGAGGSGIDG | ASSSGTGGEQY | 0.8945087 |
| GNGGNGGAGGEGGPGGAGGAGA | ASSSGTGGEQY | 0.85459447 |
| GSTAGNGGNGGNATNGTIAGGQPAG | ASSSGTGGEQY | 0.86260664 |
| GTGGDGGNGGNGGTGG | ASSSGTGGEQY | 0.8811146 |
| GVRRPAGIPADAPV | ASSSGTGGEQY | 0.66074944 |
| GYRQVTDAHLLTLARR | ASSSGTGGEQY | 0.82509553 |
| IAKMGQDGHDRG | ASSSGTGGEQY | 0.90405196 |
| KAEPSIEIQYT | ASSSGTGGEQY | 0.72940505 |
| KPVRIRMAADEDTVAVT | ASSSGTGGEQY | 0.7470779 |
| MPPRIAGMRLLVIKPEP | ASSSGTGGEQY | 0.28729138 |
| MKFYAVALFRSRGVPPTR | ASSSGTGGEQY | 0.305837 |
| NAGGFGILGAVAHSP | ASSSGTGGEQY | 0.5408467 |

| | | |
|---|---|---|
| NEGREITLAMAEM | ASSSGTGGEQY | 0.69202 |
| NELVEWSHPLGHRWRWE | ASSSGTGGEQY | 0.76582456 |
| NLGAGGTGGNGGNPG | ASSSGTGGEQY | 0.7266269 |
| NTEWPKWETVMAADPVD | ASSSGTGGEQY | 0.27652854 |
| PASAAPAETQP | ASSSGTGGEQY | 0.65783155 |
| QQSSDGKNGVRAAEITGE | ASSSGTGGEQY | 0.4550232 |
| RAPDAVTTSWWKE | ASSSGTGGEQY | 0.58039165 |
| SGFGNVGAHTSGWFNQ | ASSSGTGGEQY | 0.65929747 |
| SGVAGAVGQAASVGGLKV | ASSSGTGGEQY | 0.51092285 |
| SNGGAGTGSAGGNGGT | ASSSGTGGEQY | 0.43971384 |
| TGGAGGEGGAAG | ASSSGTGGEQY | 0.2748511 |
| TGGDGGAAGAGGAPGHG | ASSSGTGGEQY | 0.68945855 |
| TMPPLLAAIANAGRHGV | ASSSGTGGEQY | 0.73129916 |
| TSSTAGNSGGA | ASSSGTGGEQY | 0.7094954 |
| VPATVPVGILC | ASSSGTGGEQY | 0.7590928 |
| VQRPDSGGVVLTGRIS | ASSSGTGGEQY | 0.7357152 |
| WTPNPVPATKGVPAGLVQAT | ASSSGTGGEQY | 0.59927994 |
| YALKRALAKFSDRFDVVII | ASSSGTGGEQY | 0.16910735 |
| YLIGSVIV | ASSSGTGGEQY | 0.2029571 |
| YGQLVAVID | ASSSGTGGEQY | 0.3050011 |
| VTVGGMTLPA | ASSSGTGGEQY | 0.4446857 |
| VLALPAV | ASSSGTGGEQY | 0.18262158 |
| VIPPTLNYET | ASSSGTGGEQY | 0.67143196 |
| VGMGGMHPSSG | ASSSGTGGEQY | 0.3856305 |
| VGLAGGAATA | ASSSGTGGEQY | 0.3903076 |
| VGGAGGAGGSGP | ASSSGTGGEQY | 0.7386888 |
| VFAAFMV | ASSSGTGGEQY | 0.39357156 |
| TLYDKLR | ASSSGTGGEQY | 0.2604142 |
| TLAVFTVL | ASSSGTGGEQY | 0.3800085 |
| TGSVQPARV | ASSSGTGGEQY | 0.74051416 |
| TGGGAGSGGT | ASSSGTGGEQY | 0.33401465 |
| SHLETSL | ASSSGTGGEQY | 0.7332194 |
| SFTTGMT | ASSSGTGGEQY | 0.48525518 |
| QVEVAKRMGVRQPTVS | ASSSGTGGEQY | 0.7322074 |
| QGSGGPALAM | ASSSGTGGEQY | 0.22310048 |
| MLAPSNP | ASSSGTGGEQY | 0.7180271 |
| LVNNAAI | ASSSGTGGEQY | 0.6378289 |
| LTGEINTLA | ASSSGTGGEQY | 0.7873006 |
| LSIFGPV | ASSSGTGGEQY | 0.4473703 |
| LLALLANL | ASSSGTGGEQY | 0.6201908 |
| LISNASDALDKLR | ASSSGTGGEQY | 0.424825 |
| LINEAVP | ASSSGTGGEQY | 0.7248242 |
| LGYAFSGL | ASSSGTGGEQY | 0.24605192 |
| LDFVAVAL | ASSSGTGGEQY | 0.4399238 |
| LAQPGAE | ASSSGTGGEQY | 0.24308565 |
| ISNLGAVTGF | ASSSGTGGEQY | 0.40522128 |
| HQEDPGRI | ASSSGTGGEQY | 0.70774686 |
| GTPGEPTP | ASSSGTGGEQY | 0.17410506 |
| GFFNSGS | ASSSGTGGEQY | 0.6633033 |
| GAGGGGLTAGAN | ASSSGTGGEQY | 0.4067924 |
| FVDALVNM | ASSSGTGGEQY | 0.26966316 |

| | | |
|---|---|---|
| FLTVAPEL | ASSSGTGGEQY | 0.720436 |
| FETLTAL | ASSSGTGGEQY | 0.5371296 |
| DVINPIEL | ASSSGTGGEQY | 0.6958046 |
| APVSGAPV | ASSSGTGGEQY | 0.55286634 |
| ALVGGLAW | ASSSGTGGEQY | 0.54570687 |
| ALAAAAATAI | ASSSGTGGEQY | 0.52453333 |
| AFGLGLLL | ASSSGTGGEQY | 0.17431386 |
| AFFAAASL | ASSSGTGGEQY | 0.32224685 |
| AAAALPAFSPP | ASSSGTGGEQY | 0.7500216 |
| VVLAPAL | ASSSGTGGEQY | 0.82832247 |
| TIGHVDHGKTTL | ASSSGTGGEQY | 0.83997667 |
| LISNASDALDK | ASSSGTGGEQY | 0.78429246 |
| LISNASDAL | ASSSGTGGEQY | 0.7297349 |
| ISNASDALDK | ASSSGTGGEQY | 0.7813482 |
| NQAETLVYQTE | ASSSGTGGEQY | 0.8368359 |
| TGGTGGTAGAG | ASSSGTGGEQY | 0.83863544 |
| TGSTAGGAGGA | ASSSGTGGEQY | 0.82769287 |
| DPLAVPVAAGPV | ASSSGTGGEQY | 0.72159755 |
| GGTGGTPGNGGN | ASSSGTGGEQY | 0.81750286 |
| PAGLAGVDPATL | ASSSGTGGEQY | 0.80014133 |
| GNNLVVEAPHAGASV | ASSSGTGGEQY | 0.77904254 |
| KVTRIMSASTLEQPA | ASSSGTGGEQY | 0.82722116 |
| LGFALPIALPRRTVV | ASSSGTGGEQY | 0.8213159 |
| TRLGNLTARKLVVLV | ASSSGTGGEQY | 0.7823912 |
| WSQQRRLIYFPSAGP | ASSSGTGGEQY | 0.7844323 |
| GPNGNPDPMAAAVDIR | ASSSGTGGEQY | 0.8112342 |
| TQRCVSVADLRAAASR | ASSSGTGGEQY | 0.8275659 |
| QFVLDVAEHPVEVLTETV | ASSSGTGGEQY | 0.8235832 |
| GGAGGTGGAGGNGGTQAGDG | ASSSGTGGEQY | 0.8413274 |
| PLLATLRAIAVDVDAKPAQV | ASSSGTGGEQY | 0.83551514 |
| APKFRTGGYMIDFWGVGYQVAKRM | ASSSGTGGEQY | 0.8335756 |
| GAGGAGGAGGAAGGPGGTGGTGGN | ASSSGTGGEQY | 0.30685213 |
| IGALGLLATDVAPAGSVAFGVALA | ASSSGTGGEQY | 0.5592093 |
| AGGAGGIGGNGGAGGAGGVLMGNG | ASSSGTGGEQY | 0.8083002 |
| ADELIGLL | ASSSGTGGEQY | 0.7805036 |
| ADRALKA | ASSSGTGGEQY | 0.60348064 |
| AGAVGQVGGAGGA | ASSSGTGGEQY | 0.8186104 |
| AGGGGGLGGAP | ASSSGTGGEQY | 0.7793782 |
| AGGLLYGNGGAG | ASSSGTGGEQY | 0.7602884 |
| AGLESIGL | ASSSGTGGEQY | 0.83017254 |
| AGLKTGII | ASSSGTGGEQY | 0.7884915 |
| ASLAALLL | ASSSGTGGEQY | 0.776901 |
| ASVINISSI | ASSSGTGGEQY | 0.7858926 |
| ATPPGPA | ASSSGTGGEQY | 0.8367556 |
| ATVASALV | ASSSGTGGEQY | 0.72396654 |
| ATVASLIL | ASSSGTGGEQY | 0.82271075 |
| AVNGPVLIRH | ASSSGTGGEQY | 0.7653983 |
| DGAAGAVGI | ASSSGTGGEQY | 0.83704656 |
| DPLLHGI | ASSSGTGGEQY | 0.6757543 |
| ELGAALGAL | ASSSGTGGEQY | 0.8299098 |
| ELGRTQAV | ASSSGTGGEQY | 0.80339694 |

| | | |
|---|---|---|
| FVAGGYLS | ASSSGTGGEQY | 0.84742856 |
| GASGGAGGAGGW | ASSSGTGGEQY | 0.79808724 |
| GGAGAGGDGGAGGI | ASSSGTGGEQY | 0.8113116 |
| GTGHNGSQPGMGGNG | ASSSGTGGEQY | 0.77997255 |
| GYAAASASAS | ASSSGTGGEQY | 0.8332104 |
| HQVDVAAVQQG | ASSSGTGGEQY | 0.6718842 |
| IGAAIAHT | ASSSGTGGEQY | 0.80582386 |
| IGMIEVAQL | ASSSGTGGEQY | 0.63961166 |
| IRLAPEHTSAMD | ASSSGTGGEQY | 0.81899405 |
| IYNVSVL | ASSSGTGGEQY | 0.82381225 |
| KQGFLGM | ASSSGTGGEQY | 0.56625515 |
| LESVITQL | ASSSGTGGEQY | 0.8346268 |
| LGLGALVVV | ASSSGTGGEQY | 0.7981511 |
| LIGNGGNGGSGGI | ASSSGTGGEQY | 0.77563995 |
| LNAINAPTL | ASSSGTGGEQY | 0.8400668 |
| LQLLNTGV | ASSSGTGGEQY | 0.804883 |
| LQTSIEF | ASSSGTGGEQY | 0.83906025 |
| MSFAVSEL | ASSSGTGGEQY | 0.8096035 |
| NTGALGL | ASSSGTGGEQY | 0.81209856 |
| PPGAGGAGGAGTT | ASSSGTGGEQY | 0.8227915 |
| QLVGLGI | ASSSGTGGEQY | 0.65147007 |
| RFLEQVHQL | ASSSGTGGEQY | 0.7741449 |
| RGVGYVLR | ASSSGTGGEQY | 0.837363 |
| SATGVFAGVI | ASSSGTGGEQY | 0.807462 |
| SLPAITLP | ASSSGTGGEQY | 0.8334614 |
| STALAFTV | ASSSGTGGEQY | 0.6906574 |
| SVLTVGLGK | ASSSGTGGEQY | 0.6654241 |
| TALAGAGSGI | ASSSGTGGEQY | 0.69316053 |
| TGLPFTGI | ASSSGTGGEQY | 0.8195762 |
| VAEHVTV | ASSSGTGGEQY | 0.8255014 |
| VDIVLNSL | ASSSGTGGEQY | 0.8023997 |
| VLTGGGAVL | ASSSGTGGEQY | 0.66128576 |
| VNVGLAYGI | ASSSGTGGEQY | 0.77547187 |
| VSSAAVGAV | ASSSGTGGEQY | 0.7460666 |
| VTSVYEAGV | ASSSGTGGEQY | 0.73754644 |
| WVALDRGA | ASSSGTGGEQY | 0.80593693 |
| YLVGSVVI | ASSSGTGGEQY | 0.78025806 |
| GPPGTGKTLL | ASSSGTGGEQY | 0.7384695 |
| GVGMTKF | ASSSGTGGEQY | 0.7535642 |
| IGIGGSDL | ASSSGTGGEQY | 0.7434331 |
| IGTLLAEL | ASSSGTGGEQY | 0.8081974 |
| IYTSGTT | ASSSGTGGEQY | 0.7920519 |
| LSFPAGV | ASSSGTGGEQY | 0.837558 |
| LYTSGTT | ASSSGTGGEQY | 0.8181323 |
| VGAFPFGF | ASSSGTGGEQY | 0.80550164 |
| VGAGTVEF | ASSSGTGGEQY | 0.28985146 |
| VGKSTLLL | ASSSGTGGEQY | 0.66659594 |
| AALAHAGGGIV | ASSSGTGGEQY | 0.7449617 |
| AALVAVSLL | ASSSGTGGEQY | 0.7695246 |
| AAVSRAVAAL | ASSSGTGGEQY | 0.5042335 |
| AGGSSGIGGF | ASSSGTGGEQY | 0.80562294 |

| | | |
|---|---|---|
| AGLTVANANG | ASSSGTGGEQY | 0.76871645 |
| AGLVGAVL | ASSSGTGGEQY | 0.6842714 |
| ALSQAFTV | ASSSGTGGEQY | 0.8147702 |
| ALTFSVSL | ASSSGTGGEQY | 0.7215925 |
| ALTGAALVAL | ASSSGTGGEQY | 0.74599874 |
| ASFSIGAGVL | ASSSGTGGEQY | 0.72950524 |
| DGSVYKI | ASSSGTGGEQY | 0.81331503 |
| EQLAEIV | ASSSGTGGEQY | 0.67511874 |
| FAVSELTVM | ASSSGTGGEQY | 0.80427057 |
| GAAPLAAGL | ASSSGTGGEQY | 0.70324117 |
| GADGGDGGSGGAGG | ASSSGTGGEQY | 0.8243873 |
| GFGLDTGGAGG | ASSSGTGGEQY | 0.61822295 |
| GLASGGLL | ASSSGTGGEQY | 0.8185635 |
| GRIAGAAVLL | ASSSGTGGEQY | 0.7579005 |
| GSAGSLNTSF | ASSSGTGGEQY | 0.84672964 |
| GTGQIGI | ASSSGTGGEQY | 0.76064765 |
| GTLIALIL | ASSSGTGGEQY | 0.8032795 |
| IAATMGFVVG | ASSSGTGGEQY | 0.725748 |
| IFAQALL | ASSSGTGGEQY | 0.82084167 |
| IGAAVALGF | ASSSGTGGEQY | 0.5830674 |
| IGLTPLAL | ASSSGTGGEQY | 0.74393165 |
| ILGSNTSTL | ASSSGTGGEQY | 0.6427955 |
| IVFSLTVL | ASSSGTGGEQY | 0.7727848 |
| LASKATGF | ASSSGTGGEQY | 0.8079446 |
| LASVALIL | ASSSGTGGEQY | 0.54634595 |
| LFAVGLTL | ASSSGTGGEQY | 0.8149766 |
| LFNGGIGTYI | ASSSGTGGEQY | 0.7210352 |
| LTLGAGII | ASSSGTGGEQY | 0.7204198 |
| LVAAALGDI | ASSSGTGGEQY | 0.82116866 |
| NTAFVQLGI | ASSSGTGGEQY | 0.74158424 |
| QLFSGAEL | ASSSGTGGEQY | 0.82298946 |
| SAATFITL | ASSSGTGGEQY | 0.7509526 |
| SNYGLANLGV | ASSSGTGGEQY | 0.74531674 |
| SYGGAYAVM | ASSSGTGGEQY | 0.7929573 |
| TTGVGGLTLG | ASSSGTGGEQY | 0.6386124 |
| TTQTIYI | ASSSGTGGEQY | 0.67456913 |
| VAAGAGLGAVI | ASSSGTGGEQY | 0.833445 |
| VAAVVTLL | ASSSGTGGEQY | 0.8079841 |
| VGFLGGLI | ASSSGTGGEQY | 0.8027962 |
| VGVIPILL | ASSSGTGGEQY | 0.7192091 |
| VIFASSLI | ASSSGTGGEQY | 0.57480276 |
| VKGGDGGAAAG | ASSSGTGGEQY | 0.7459679 |
| VTKTFTNI | ASSSGTGGEQY | 0.8098614 |
| VVAAAILL | ASSSGTGGEQY | 0.7939953 |
| YAALDVEL | ASSSGTGGEQY | 0.7985301 |
| LASVALLL | ASSSGTGGEQY | 0.44422308 |
| LDAVHRV | ASSSGTGGEQY | 0.6557488 |
| QGLSSSIF | ASSSGTGGEQY | 0.7518665 |
| RGTFANIRL | ASSSGTGGEQY | 0.67054343 |
| RQLWWGHRI | ASSSGTGGEQY | 0.7899031 |
| YGPPGTGKTLL | ASSSGTGGEQY | 0.65042305 |

| | | |
|---|---|---|
| AALVGAPANVA | ASSSGTGGEQY | 0.7164203 |
| AATFRRTE | ASSSGTGGEQY | 0.7307918 |
| AGALGGGI | ASSSGTGGEQY | 0.74669373 |
| AGGAGGAGGIGAG | ASSSGTGGEQY | 0.77994764 |
| AGSGALGL | ASSSGTGGEQY | 0.76513386 |
| AQIADQI | ASSSGTGGEQY | 0.8182651 |
| ASAVAVTYL | ASSSGTGGEQY | 0.8095342 |
| ASTVLGGTGG | ASSSGTGGEQY | 0.80650365 |
| AVGGEPAK | ASSSGTGGEQY | 0.21634041 |
| CEQGTFGGTC | ASSSGTGGEQY | 0.41242242 |
| DEGKAPAG | ASSSGTGGEQY | 0.6767848 |
| DTVVFPAL | ASSSGTGGEQY | 0.62429947 |
| FEGITMAG | ASSSGTGGEQY | 0.6055068 |
| FPLVVQD | ASSSGTGGEQY | 0.7361488 |
| GAFTTPQ | ASSSGTGGEQY | 0.7925495 |
| GFGGFGTAGGVGGA | ASSSGTGGEQY | 0.4828919 |
| GFLAQFL | ASSSGTGGEQY | 0.7874215 |
| GGAGGAGGNAGVA | ASSSGTGGEQY | 0.6331993 |
| GGVGTAGAGGAGGAGG | ASSSGTGGEQY | 0.5781998 |
| GNGGGGDAGNAGSGGN | ASSSGTGGEQY | 0.6498646 |
| GTGGAGGAGNDAGS | ASSSGTGGEQY | 0.8031769 |
| GTTNTGWFNS | ASSSGTGGEQY | 0.5609817 |
| IFMGPLFTL | ASSSGTGGEQY | 0.7452465 |
| ILAHAIEAI | ASSSGTGGEQY | 0.7392317 |
| IRSAIREAL | ASSSGTGGEQY | 0.8029921 |
| ITFAGGG | ASSSGTGGEQY | 0.41435874 |
| ITGMSGHQEV | ASSSGTGGEQY | 0.80074215 |
| IVAGLACTL | ASSSGTGGEQY | 0.7502845 |
| IVGGGLAGA | ASSSGTGGEQY | 0.81341183 |
| IVTGVLNI | ASSSGTGGEQY | 0.6738285 |
| LAQQGCI | ASSSGTGGEQY | 0.67001164 |
| LCLHVEN | ASSSGTGGEQY | 0.6000898 |
| LDAISDGS | ASSSGTGGEQY | 0.78527004 |
| LGGFAAML | ASSSGTGGEQY | 0.41944218 |
| LGGKNPMI | ASSSGTGGEQY | 0.76723945 |
| LGPKGVAAIL | ASSSGTGGEQY | 0.61364996 |
| LMVGVSI | ASSSGTGGEQY | 0.77350086 |
| LVGAAFVL | ASSSGTGGEQY | 0.79795873 |
| LVLAVAPL | ASSSGTGGEQY | 0.5134306 |
| MAAIAVIDAA | ASSSGTGGEQY | 0.793684 |
| NNGINAELQ | ASSSGTGGEQY | 0.7741605 |
| RGEPLVLVHG | ASSSGTGGEQY | 0.68184555 |
| RGGAGGAAGLI | ASSSGTGGEQY | 0.8040184 |
| RSYRMHPAV | ASSSGTGGEQY | 0.55533314 |
| SAAHLLAGQLA | ASSSGTGGEQY | 0.81648463 |
| SAAVGSY | ASSSGTGGEQY | 0.7167396 |
| SPHAVLHRI | ASSSGTGGEQY | 0.7373233 |
| SQIAKVL | ASSSGTGGEQY | 0.7790657 |
| TALGHLVP | ASSSGTGGEQY | 0.3796503 |
| TAPTTSGAA | ASSSGTGGEQY | 0.53019893 |
| TDGDDGNGGSPGT | ASSSGTGGEQY | 0.8101023 |

| | | |
|---|---|---|
| TDGLGEN | ASSSGTGGEQY | 0.7642282 |
| TGGPPLPGQA | ASSSGTGGEQY | 0.7697899 |
| TIPAITI | ASSSGTGGEQY | 0.71424866 |
| TIPGFTI | ASSSGTGGEQY | 0.67357206 |
| TSIASGALA | ASSSGTGGEQY | 0.7004223 |
| TVTLPGAAI | ASSSGTGGEQY | 0.7659281 |
| VASGLSAALP | ASSSGTGGEQY | 0.7699125 |
| VGAGVIGI | ASSSGTGGEQY | 0.7533245 |
| VGPKRFGV | ASSSGTGGEQY | 0.6551904 |
| VNVAFPD | ASSSGTGGEQY | 0.7407783 |
| VSMTNSGVS | ASSSGTGGEQY | 0.73567593 |
| VTDVLGDLSV | ASSSGTGGEQY | 0.7174735 |
| VVVVGAGFAGL | ASSSGTGGEQY | 0.7525909 |
| VYTVGTVV | ASSSGTGGEQY | 0.7663727 |
| WTTFSGP | ASSSGTGGEQY | 0.7220433 |
| YDELVEAF | ASSSGTGGEQY | 0.7065878 |
| YGTGGAGGNGG | ASSSGTGGEQY | 0.7180434 |
| YTNVMAV | ASSSGTGGEQY | 0.7662427 |
| FLEQVHQL | ASSSGTGGEQY | 0.75437087 |
| GLSLLAVAV | ASSSGTGGEQY | 0.7785791 |
| LEQVHQL | ASSSGTGGEQY | 0.77002 |
| LNAAGAI | ASSSGTGGEQY | 0.7675115 |
| PAVHASL | ASSSGTGGEQY | 0.3562955 |
| TELEDDA | ASSSGTGGEQY | 0.49263018 |
| TPPGPPN | ASSSGTGGEQY | 0.73600346 |
| TPPPPPPP | ASSSGTGGEQY | 0.7079483 |
| TQSPALI | ASSQSGYSQNTLY | 0.49811286 |
| EFAAAFPQ | ASSQSGYSQNTLY | 0.74591553 |
| LLSGNLDVL | ASSQSGYSQNTLY | 0.7135526 |
| GPANLGNYNI | ASSQSGYSQNTLY | 0.6880866 |
| TLGPTLP | ASSQSGYSQNTLY | 0.7583878 |
| VYAYSSCCK | ASSQSGYSQNTLY | 0.72499025 |
| APAPAPAPNI | ASSQSGYSQNTLY | 0.68886304 |
| PGGPVPIP | ASSQSGYSQNTLY | 0.653878 |
| FLTVALEP | ASSQSGYSQNTLY | 0.7643089 |
| SQYFGANL | ASSQSGYSQNTLY | 0.6084603 |
| GGVKPAA | ASSQSGYSQNTLY | 0.75632495 |
| GGTGGAGGAGGW | ASSQSGYSQNTLY | 0.5991243 |
| GVGGVAGLGGAGG | ASSQSGYSQNTLY | 0.7627151 |
| VLLVLATTI | ASSQSGYSQNTLY | 0.6144717 |
| IGIFAAML | ASSQSGYSQNTLY | 0.76753473 |
| AVGDLID | ASSQSGYSQNTLY | 0.73697567 |
| NISPLPL | ASSQSGYSQNTLY | 0.78368497 |
| VTAVFVAI | ASSQSGYSQNTLY | 0.7229974 |
| SGDDNDAAL | ASSQSGYSQNTLY | 0.75548035 |
| AVAALAHS | ASSQSGYSQNTLY | 0.7219363 |
| GGVTVMATP | ASSQSGYSQNTLY | 0.7575705 |
| AGLAANVRD | ASSQSGYSQNTLY | 0.61498773 |
| TNRPDLIDPALLRPG | ASSQSGYSQNTLY | 0.726645 |
| LANLGVTAL | ASSQSGYSQNTLY | 0.506038 |
| TGERIEL | ASSQSGYSQNTLY | 0.7469208 |

| | | |
|---|---|---|
| NGEELAA | ASSQSGYSQNTLY | 0.76084363 |
| VLSLVWG | ASSQSGYSQNTLY | 0.48054683 |
| IGYALWL | ASSQSGYSQNTLY | 0.7513652 |
| LSLLLAPL | ASSQSGYSQNTLY | 0.73827463 |
| TLDEALK | ASSQSGYSQNTLY | 0.70747435 |
| TDGAGTAEGG | ASSQSGYSQNTLY | 0.7751447 |
| NAPGGAGGNGG | ASSQSGYSQNTLY | 0.73733056 |
| VSVFMGMGGNF | ASSQSGYSQNTLY | 0.77692676 |
| LASYLAAL | ASSQSGYSQNTLY | 0.7247454 |
| LVGGTPAL | ASSQSGYSQNTLY | 0.7083351 |
| VRNADVL | ASSQSGYSQNTLY | 0.73405397 |
| TCSSPFLVSAM | ASSQSGYSQNTLY | 0.45729858 |
| VHITDAE | ASSQSGYSQNTLY | 0.5961325 |
| EYAAPSF | ASSQSGYSQNTLY | 0.7755549 |
| VVDAFHGGL | ASSQSGYSQNTLY | 0.82973444 |
| ASVVDPQ | ASSQSGYSQNTLY | 0.83603 |
| SYNEFGGLS | ASSQSGYSQNTLY | 0.78309494 |
| DPDGSVY | ASSQSGYSQNTLY | 0.75892186 |
| MVSTIHATE | ASSQSGYSQNTLY | 0.7963279 |
| LAAGVADPV | ASSQSGYSQNTLY | 0.8362615 |
| TKEQIAA | ASSQSGYSQNTLY | 0.8403534 |
| SLASPAAL | ASSQSGYSQNTLY | 0.82412314 |
| IGLVFYGV | ASSQSGYSQNTLY | 0.730902 |
| FLLDASNL | ASSQSGYSQNTLY | 0.82564986 |
| YQVGNFP | ASSQSGYSQNTLY | 0.8089216 |
| ESLLSQST | ASSQSGYSQNTLY | 0.8029028 |
| LFTVGFAAVG | ASSQSGYSQNTLY | 0.8238244 |
| ATPPTPT | ASSQSGYSQNTLY | 0.82955414 |
| TDLATLG | ASSQSGYSQNTLY | 0.79204494 |
| SAGGTGR | ASSQSGYSQNTLY | 0.7932852 |
| GFSNSGS | ASSQSGYSQNTLY | 0.8140791 |
| GGNGGSGGTSV | ASSQSGYSQNTLY | 0.83243406 |
| GTSVATGGAGN | ASSQSGYSQNTLY | 0.8289908 |
| NGGNGGSGGTS | ASSQSGYSQNTLY | 0.8411256 |
| AVVKFPH | ASSQSGYSQNTLY | 0.8355609 |
| TPPPPPP | ASSQSGYSQNTLY | 0.82739866 |
| AAPAAAAAPA | ASSQSGYSQNTLY | 0.3303368 |
| IALNEDL | ASSQSGYSQNTLY | 0.52515304 |
| IVNVSSM | ASSQSGYSQNTLY | 0.7931641 |
| TGIPPAPR | ASSQSGYSQNTLY | 0.7977141 |
| FDLGGGTF | ASSQSGYSQNTLY | 0.45448875 |
| FDLGGGTFDVS | ASSQSGYSQNTLY | 0.8281276 |
| NEPTAAA | ASSQSGYSQNTLY | 0.78849554 |
| NEPTAAALA | ASSQSGYSQNTLY | 0.7731371 |
| GSIINISSI | ASSQSGYSQNTLY | 0.831419 |
| VAPGNAGTA | ASSQSGYSQNTLY | 0.80535233 |
| TGGGASLE | ASSQSGYSQNTLY | 0.7475585 |
| VAVYNHYKRI | ASSQSGYSQNTLY | 0.7810308 |
| TDPAPATT | ASSQSGYSQNTLY | 0.8245958 |
| TNRPDILDPALLRPG | ASSQSGYSQNTLY | 0.6507071 |
| LTTTTTS | ASSQSGYSQNTLY | 0.8339952 |

| | | |
|---|---|---|
| PTGSGKTL | ASSQSGYSQNTLY | 0.69890237 |
| TGSGKTL | ASSQSGYSQNTLY | 0.8376148 |
| TIGHVDHGK | ASSQSGYSQNTLY | 0.7006374 |
| TIGHVDHGKTT | ASSQSGYSQNTLY | 0.8305865 |
| GHVDHGKTT | ASSQSGYSQNTLY | 0.8123088 |
| AQFEHTLL | ASSQSGYSQNTLY | 0.848988 |
| IVNQDEL | ASSQSGYSQNTLY | 0.8021155 |
| TNGSQFF | ASSQSGYSQNTLY | 0.8281315 |
| TGKGIVE | ASSQSGYSQNTLY | 0.8135873 |
| LISNASDA | ASSQSGYSQNTLY | 0.83937085 |
| AELRTLGELTKIAKAHGAQ | ASSQSGYSQNTLY | 0.7039353 |
| APNAVAAAEPSW | ASSQSGYSQNTLY | 0.81094694 |
| APQVAGAATAVL | ASSQSGYSQNTLY | 0.52542615 |
| ARRDPAAAAAILPTDA | ASSQSGYSQNTLY | 0.82045627 |
| DAAARDVLAEAGLAEYF | ASSQSGYSQNTLY | 0.8282886 |
| DDQIVLSAKANLAKVEQ | ASSQSGYSQNTLY | 0.47990012 |
| DELLPDTGEDSDEDGH | ASSQSGYSQNTLY | 0.8322411 |
| ELPEAPANRATVPQD | ASSQSGYSQNTLY | 0.79540825 |
| FGQSKIERVTGIG | ASSQSGYSQNTLY | 0.78371465 |
| GAGGTGSTAGGAGGAGGAGGLYA | ASSQSGYSQNTLY | 0.8356573 |
| GHGGTGAAGGNGGTGGAGGSGIDG | ASSQSGYSQNTLY | 0.81870526 |
| GNGGNGGAGGEGGPGGAGGAGA | ASSQSGYSQNTLY | 0.83326334 |
| GSTAGNGGNGGNATNGTIAGQPAG | ASSQSGYSQNTLY | 0.8002053 |
| GTGGDGGNGGNGGTGG | ASSQSGYSQNTLY | 0.7917495 |
| GVRRPAGIPADAPV | ASSQSGYSQNTLY | 0.80791247 |
| GYRQVTDAHLLTLARR | ASSQSGYSQNTLY | 0.42437202 |
| IAKMGQDGHDRG | ASSQSGYSQNTLY | 0.719163 |
| KAEPSIEIQYT | ASSQSGYSQNTLY | 0.84066516 |
| KPVRIRMAADEDTVAVT | ASSQSGYSQNTLY | 0.769474 |
| MPPRIAGMRLLVIKPEP | ASSQSGYSQNTLY | 0.75607693 |
| MKFYAVALFRSRGVPPTR | ASSQSGYSQNTLY | 0.6695523 |
| NAGGFGILGAVAHSP | ASSQSGYSQNTLY | 0.6151923 |
| NEGREITLAMAEM | ASSQSGYSQNTLY | 0.722034 |
| NELVEWSHPLGHRWRWE | ASSQSGYSQNTLY | 0.7653239 |
| NLGAGGTGGNGGNPG | ASSQSGYSQNTLY | 0.79036343 |
| NTEWPKWETVMAADPVD | ASSQSGYSQNTLY | 0.76178974 |
| PASAAPAETQP | ASSQSGYSQNTLY | 0.6281718 |
| QQSSDGKNGVRAAEITGE | ASSQSGYSQNTLY | 0.783941 |
| RAPDAVTTSWWKE | ASSQSGYSQNTLY | 0.75044507 |
| SGFGNVGAHTSGWFNQ | ASSQSGYSQNTLY | 0.7241222 |
| SGVAGAVGQAASVGGLKV | ASSQSGYSQNTLY | 0.76758987 |
| SNGGAGTGSAGGNGGT | ASSQSGYSQNTLY | 0.7685679 |
| TGGAGGEGGAAG | ASSQSGYSQNTLY | 0.72263587 |
| TGGDGGAAGAGGAPGHG | ASSQSGYSQNTLY | 0.6199826 |
| TMPPLLAAIANAGRHGV | ASSQSGYSQNTLY | 0.7716956 |
| TSSTAGNSGGA | ASSQSGYSQNTLY | 0.7832451 |
| VPATVPVGILC | ASSQSGYSQNTLY | 0.7810971 |
| VQRPDSGGVVLTGRIS | ASSQSGYSQNTLY | 0.7927317 |
| WTPNPVPATKGVPAGLVQAT | ASSQSGYSQNTLY | 0.7813346 |
| YALKRALAKFSDRFDVVII | ASSQSGYSQNTLY | 0.77210486 |
| YLIGSVIV | ASSQSGYSQNTLY | 0.31055123 |

| | | |
|---|---|---|
| YGQLVAVID | ASSQSGYSQNTLY | 0.5133668 |
| VTVGGMTLPA | ASSQSGYSQNTLY | 0.67787534 |
| VLALPAV | ASSQSGYSQNTLY | 0.71837795 |
| VIPPTLNYET | ASSQSGYSQNTLY | 0.33724537 |
| VGMGGMHPSSG | ASSQSGYSQNTLY | 0.74572533 |
| VGLAGGAATA | ASSQSGYSQNTLY | 0.72470176 |
| VGGAGGAGGSGP | ASSQSGYSQNTLY | 0.6943903 |
| VFAAFMV | ASSQSGYSQNTLY | 0.78865457 |
| TLYDKLR | ASSQSGYSQNTLY | 0.71667755 |
| TLAVFTVL | ASSQSGYSQNTLY | 0.63565946 |
| TGSVQPARV | ASSQSGYSQNTLY | 0.70992184 |
| TGGGAGSGGT | ASSQSGYSQNTLY | 0.79058635 |
| SHLETSL | ASSQSGYSQNTLY | 0.63279086 |
| SFTTGMT | ASSQSGYSQNTLY | 0.78645587 |
| QVEVAKRMGVRQPTVS | ASSQSGYSQNTLY | 0.6807536 |
| QGSGGPALAM | ASSQSGYSQNTLY | 0.7891623 |
| MLAPSNP | ASSQSGYSQNTLY | 0.5507134 |
| LVNNAAI | ASSQSGYSQNTLY | 0.7780675 |
| LTGEINTLA | ASSQSGYSQNTLY | 0.7326857 |
| LSIFGPV | ASSQSGYSQNTLY | 0.80429274 |
| LLALLANL | ASSQSGYSQNTLY | 0.7225373 |
| LISNASDALDKLR | ASSQSGYSQNTLY | 0.7796917 |
| LINEAVP | ASSQSGYSQNTLY | 0.74929416 |
| LGYAFSGL | ASSQSGYSQNTLY | 0.78220445 |
| LDFVAVAL | ASSQSGYSQNTLY | 0.58272433 |
| LAQPGAE | ASSQSGYSQNTLY | 0.723365 |
| ISNLGAVTGF | ASSQSGYSQNTLY | 0.6134008 |
| HQEDPGRI | ASSQSGYSQNTLY | 0.7386348 |
| GTPGEPTP | ASSQSGYSQNTLY | 0.7609197 |
| GFFNSGS | ASSQSGYSQNTLY | 0.2917299 |
| GAGGGGLTAGAN | ASSQSGYSQNTLY | 0.7858021 |
| FVDALVNM | ASSQSGYSQNTLY | 0.6286169 |
| FLTVAPEL | ASSQSGYSQNTLY | 0.65118706 |
| FETLTAL | ASSQSGYSQNTLY | 0.7823186 |
| DVINPIEL | ASSQSGYSQNTLY | 0.7687696 |
| APVSGAPV | ASSQSGYSQNTLY | 0.7684742 |
| ALVGGLAW | ASSQSGYSQNTLY | 0.71978307 |
| ALAAAAATAI | ASSQSGYSQNTLY | 0.74089414 |
| AFGLGLLL | ASSQSGYSQNTLY | 0.7527664 |
| AFFAAASL | ASSQSGYSQNTLY | 0.440528 |
| AAAALPAFSPP | ASSQSGYSQNTLY | 0.64670247 |
| VVLAPAL | ASSQSGYSQNTLY | 0.7874639 |
| TIGHVDHGKTTL | ASSQSGYSQNTLY | 0.71777457 |
| LISNASDALDK | ASSQSGYSQNTLY | 0.78525513 |
| LISNASDAL | ASSQSGYSQNTLY | 0.5781001 |
| ISNASDALDK | ASSQSGYSQNTLY | 0.6294471 |
| NQAETLVYQTE | ASSQSGYSQNTLY | 0.68846476 |
| TGGTGGTAGAG | ASSQSGYSQNTLY | 0.74497235 |
| TGSTAGGAGGA | ASSQSGYSQNTLY | 0.78017056 |
| DPLAVPVAAGPV | ASSQSGYSQNTLY | 0.75429595 |
| GGTGGTPGNGGN | ASSQSGYSQNTLY | 0.5884869 |

| | | |
|---|---|---|
| PAGLAGVDPATL | ASSQSGYSQNTLY | 0.73576367 |
| GNNLVVEAPHAGASV | ASSQSGYSQNTLY | 0.55046535 |
| KVTRIMSASTLEQPA | ASSQSGYSQNTLY | 0.6983955 |
| LGFALPIALPRRTVV | ASSQSGYSQNTLY | 0.7301216 |
| TRLGNLTARKLVVLV | ASSQSGYSQNTLY | 0.6909367 |
| WSQQRRLIYFPSAGP | ASSQSGYSQNTLY | 0.57968897 |
| GPNGNPDPMAAAVDIR | ASSQSGYSQNTLY | 0.6060821 |
| TQRCVSVADLRAAASR | ASSQSGYSQNTLY | 0.7490927 |
| QFVLDVAEHPVEVLTETV | ASSQSGYSQNTLY | 0.77063036 |
| GGAGGTGGAGGNGGTQAGDG | ASSQSGYSQNTLY | 0.7513854 |
| PLLATLRAIAVDVDAKPAQV | ASSQSGYSQNTLY | 0.7819774 |
| APKFRTGGYMIDFWGVGYQVAKRM | ASSQSGYSQNTLY | 0.75946057 |
| GAGGAGGAGGAAGGPGGTGGTGGN | ASSQSGYSQNTLY | 0.7408282 |
| IGALGLLATDVAPAGSVAFGVALA | ASSQSGYSQNTLY | 0.24259277 |
| AGGAGGIGGNGGAGGAGGVLMGNG | ASSQSGYSQNTLY | 0.5030941 |
| ADELIGLL | ASSQSGYSQNTLY | 0.55178237 |
| ADRALKA | ASSQSGYSQNTLY | 0.7191343 |
| AGAVGQVGGAGGA | ASSQSGYSQNTLY | 0.23748615 |
| AGGGGGLGGAP | ASSQSGYSQNTLY | 0.7561051 |
| AGGLLYGNGGAG | ASSQSGYSQNTLY | 0.61558217 |
| AGLESIGL | ASSQSGYSQNTLY | 0.61504245 |
| AGLKTGII | ASSQSGYSQNTLY | 0.7745763 |
| ASLAALLL | ASSQSGYSQNTLY | 0.6452037 |
| ASVINISSI | ASSQSGYSQNTLY | 0.56681705 |
| ATPPGPA | ASSQSGYSQNTLY | 0.54758537 |
| ATVASALV | ASSQSGYSQNTLY | 0.7820169 |
| ATVASLIL | ASSQSGYSQNTLY | 0.33246115 |
| AVNGPVLIRH | ASSQSGYSQNTLY | 0.7563372 |
| DGAAGAVGI | ASSQSGYSQNTLY | 0.5710857 |
| DPLLHGI | ASSQSGYSQNTLY | 0.7570942 |
| ELGAALGAL | ASSQSGYSQNTLY | 0.47659826 |
| ELGRTQAV | ASSQSGYSQNTLY | 0.76057196 |
| FVAGGYLS | ASSQSGYSQNTLY | 0.6923216 |
| GASGGAGGAGGW | ASSQSGYSQNTLY | 0.79585004 |
| GGAGAGGDGGAGGI | ASSQSGYSQNTLY | 0.67188823 |
| GTGHNGSQPGMGGNG | ASSQSGYSQNTLY | 0.72300535 |
| GYAAASASAS | ASSQSGYSQNTLY | 0.5962502 |
| HQVDVAAVQQG | ASSQSGYSQNTLY | 0.772999 |
| IGAAIAHT | ASSQSGYSQNTLY | 0.39739037 |
| IGMIEVAQL | ASSQSGYSQNTLY | 0.6506717 |
| IRLAPEHTSAMD | ASSQSGYSQNTLY | 0.51603377 |
| IYNVSVL | ASSQSGYSQNTLY | 0.6901804 |
| KQGFLGM | ASSQSGYSQNTLY | 0.74680185 |
| LESVITQL | ASSQSGYSQNTLY | 0.36393777 |
| LGLGALVVV | ASSQSGYSQNTLY | 0.7595924 |
| LIGNGGNGGSGGI | ASSQSGYSQNTLY | 0.6543725 |
| LNAINAPTL | ASSQSGYSQNTLY | 0.5446657 |
| LQLLNTGV | ASSQSGYSQNTLY | 0.76669747 |
| LQTSIEF | ASSQSGYSQNTLY | 0.70373297 |
| MSFAVSEL | ASSQSGYSQNTLY | 0.7673074 |
| NTGALGL | ASSQSGYSQNTLY | 0.5914355 |

| | | |
|---|---|---|
| PPGAGGAGGAGTT | ASSQSGYSQNTLY | 0.6949954 |
| QLVGLGI | ASSQSGYSQNTLY | 0.7069634 |
| RFLEQVHQL | ASSQSGYSQNTLY | 0.389901 |
| RGVGYVLR | ASSQSGYSQNTLY | 0.5266017 |
| SATGVFAGVI | ASSQSGYSQNTLY | 0.7730408 |
| SLPAITLP | ASSQSGYSQNTLY | 0.76993483 |
| STALAFTV | ASSQSGYSQNTLY | 0.77265674 |
| SVLTVGLGK | ASSQSGYSQNTLY | 0.6096229 |
| TALAGAGSGI | ASSQSGYSQNTLY | 0.6138727 |
| TGLPFTGI | ASSQSGYSQNTLY | 0.6754899 |
| VAEHVTV | ASSQSGYSQNTLY | 0.7930038 |
| VDIVLNSL | ASSQSGYSQNTLY | 0.8139577 |
| VLTGGGAVL | ASSQSGYSQNTLY | 0.78460354 |
| VNVGLAYGI | ASSQSGYSQNTLY | 0.71019876 |
| VSSAAVGAV | ASSQSGYSQNTLY | 0.7753551 |
| VTSVYEAGV | ASSQSGYSQNTLY | 0.73582643 |
| WVALDRGA | ASSQSGYSQNTLY | 0.7157923 |
| YLVGSVVI | ASSQSGYSQNTLY | 0.7729347 |
| GPPGTGKTLL | ASSQSGYSQNTLY | 0.78489125 |
| GVGMTKF | ASSQSGYSQNTLY | 0.6860312 |
| IGIGGSDL | ASSQSGYSQNTLY | 0.69997805 |
| IGTLLAEL | ASSQSGYSQNTLY | 0.7512305 |
| IYTSGTT | ASSQSGYSQNTLY | 0.78696084 |
| LSFPAGV | ASSQSGYSQNTLY | 0.7684211 |
| LYTSGTT | ASSQSGYSQNTLY | 0.83121514 |
| VGAFPFGF | ASSQSGYSQNTLY | 0.7833331 |
| VGAGTVEF | ASSQSGYSQNTLY | 0.7664054 |
| VGKSTLLL | ASSQSGYSQNTLY | 0.39213264 |
| AALAHAGGGIV | ASSQSGYSQNTLY | 0.5622989 |
| AALVAVSLL | ASSQSGYSQNTLY | 0.77606666 |
| AAVSRAVAAL | ASSQSGYSQNTLY | 0.707018 |
| AGGSSGIGGF | ASSQSGYSQNTLY | 0.30836195 |
| AGLTVANANG | ASSQSGYSQNTLY | 0.79488146 |
| AGLVGAVL | ASSQSGYSQNTLY | 0.72679347 |
| ALSQAFTV | ASSQSGYSQNTLY | 0.69071424 |
| ALTFSVSL | ASSQSGYSQNTLY | 0.78152895 |
| ALTGAALVAL | ASSQSGYSQNTLY | 0.7052027 |
| ASFSIGAGVL | ASSQSGYSQNTLY | 0.6951894 |
| DGSVYKI | ASSQSGYSQNTLY | 0.7514384 |
| EQLAEIV | ASSQSGYSQNTLY | 0.81835985 |
| FAVSELTVM | ASSQSGYSQNTLY | 0.6522026 |
| GAAPLAAGL | ASSQSGYSQNTLY | 0.7981367 |
| GADGGDGGSGGAGG | ASSQSGYSQNTLY | 0.7237425 |
| GFGLDTGGAGG | ASSQSGYSQNTLY | 0.8103025 |
| GLASGGLL | ASSQSGYSQNTLY | 0.64482063 |
| GRIAGAAVLL | ASSQSGYSQNTLY | 0.7911358 |
| GSAGSLNTSF | ASSQSGYSQNTLY | 0.71260476 |
| GTGQIGI | ASSQSGYSQNTLY | 0.8361261 |
| GTLIALIL | ASSQSGYSQNTLY | 0.70321816 |
| IAATMGFVVG | ASSQSGYSQNTLY | 0.8047778 |
| IFAQALL | ASSQSGYSQNTLY | 0.7737772 |

| | | |
|---|---|---|
| IGAAVALGF | ASSQSGYSQNTLY | 0.7867569 |
| IGLTPLAL | ASSQSGYSQNTLY | 0.61822486 |
| ILGSNTSTL | ASSQSGYSQNTLY | 0.7024996 |
| IVFSLTVL | ASSQSGYSQNTLY | 0.6195839 |
| LASKATGF | ASSQSGYSQNTLY | 0.7467538 |
| LASVALIL | ASSQSGYSQNTLY | 0.7844997 |
| LFAVGLTL | ASSQSGYSQNTLY | 0.34829855 |
| LFNGGIGTYI | ASSQSGYSQNTLY | 0.78770113 |
| LTLGAGII | ASSQSGYSQNTLY | 0.5898271 |
| LVAAALGDI | ASSQSGYSQNTLY | 0.68070936 |
| NTAFVQLGI | ASSQSGYSQNTLY | 0.7952225 |
| QLFSGAEL | ASSQSGYSQNTLY | 0.7544336 |
| SAATFITL | ASSQSGYSQNTLY | 0.76991236 |
| SNYGLANLGV | ASSQSGYSQNTLY | 0.70504135 |
| SYGGAYAVM | ASSQSGYSQNTLY | 0.7164858 |
| TTGVGGLTLG | ASSQSGYSQNTLY | 0.7697579 |
| TTQTIYI | ASSQSGYSQNTLY | 0.48172146 |
| VAAGAGLGAVI | ASSQSGYSQNTLY | 0.69795233 |
| VAAVVTLL | ASSQSGYSQNTLY | 0.8121021 |
| VGFLGGLI | ASSQSGYSQNTLY | 0.8192137 |
| VGVIPILL | ASSQSGYSQNTLY | 0.83321536 |
| VIFASSLI | ASSQSGYSQNTLY | 0.7293093 |
| VKGGDGGAAAG | ASSQSGYSQNTLY | 0.7186222 |
| VTKTFTNI | ASSQSGYSQNTLY | 0.77654624 |
| VVAAAILL | ASSQSGYSQNTLY | 0.8448665 |
| YAALDVEL | ASSQSGYSQNTLY | 0.83748144 |
| LASVALLL | ASSQSGYSQNTLY | 0.8172542 |
| LDAVHRV | ASSQSGYSQNTLY | 0.69910586 |
| QGLSSSIF | ASSQSGYSQNTLY | 0.8098668 |
| RGTFANIRL | ASSQSGYSQNTLY | 0.7628461 |
| RQLWWGHRI | ASSQSGYSQNTLY | 0.7762266 |
| YGPPGTGKTLL | ASSQSGYSQNTLY | 0.83226776 |
| AALVGAPANVA | ASSQSGYSQNTLY | 0.81438076 |
| AATFRRTE | ASSQSGYSQNTLY | 0.77257174 |
| AGALGGGI | ASSQSGYSQNTLY | 0.75167835 |
| AGGAGGAGGIGAG | ASSQSGYSQNTLY | 0.8091355 |
| AGSGALGL | ASSQSGYSQNTLY | 0.83812696 |
| AQIADQI | ASSQSGYSQNTLY | 0.82909596 |
| ASAVAVTYL | ASSQSGYSQNTLY | 0.8568926 |
| ASTVLGGTGG | ASSQSGYSQNTLY | 0.83504057 |
| AVGGEPAK | ASSQSGYSQNTLY | 0.8306722 |
| CEQGTFGGTC | ASSQSGYSQNTLY | 0.32809997 |
| DEGKAPAG | ASSQSGYSQNTLY | 0.56370896 |
| DTVVFPAL | ASSQSGYSQNTLY | 0.762486 |
| FEGITMAG | ASSQSGYSQNTLY | 0.7803023 |
| FPLVVQD | ASSQSGYSQNTLY | 0.51504433 |
| GAFTTPQ | ASSQSGYSQNTLY | 0.83390474 |
| GFGGFGTAGGVGGA | ASSQSGYSQNTLY | 0.7587143 |
| GFLAQFL | ASSQSGYSQNTLY | 0.7197473 |
| GGAGGAGGNAGVA | ASSQSGYSQNTLY | 0.8354681 |
| GGVGTAGAGGAGGAGG | ASSQSGYSQNTLY | 0.75368893 |

| | | |
|---|---|---|
| GNGGGGDAGNAGSGGN | ASSQSGYSQNTLY | 0.75206375 |
| GTGGAGGAGNDAGS | ASSQSGYSQNTLY | 0.7503115 |
| GTTNTGWFNS | ASSQSGYSQNTLY | 0.84466314 |
| IFMGPLFTL | ASSQSGYSQNTLY | 0.6442447 |
| ILAHAIEAI | ASSQSGYSQNTLY | 0.8244165 |
| IRSAIREAL | ASSQSGYSQNTLY | 0.78703046 |
| ITFAGGG | ASSQSGYSQNTLY | 0.8459556 |
| ITGMSGHQEV | ASSQSGYSQNTLY | 0.59830743 |
| IVAGLACTL | ASSQSGYSQNTLY | 0.8279413 |
| IVGGGLAGA | ASSQSGYSQNTLY | 0.7843425 |
| IVTGVLNI | ASSQSGYSQNTLY | 0.85999775 |
| LAQQGCI | ASSQSGYSQNTLY | 0.7814558 |
| LCLHVEN | ASSQSGYSQNTLY | 0.8143296 |
| LDAISDGS | ASSQSGYSQNTLY | 0.78071463 |
| LGGFAAML | ASSQSGYSQNTLY | 0.8271249 |
| LGGKNPMI | ASSQSGYSQNTLY | 0.5749177 |
| LGPKGVAAIL | ASSQSGYSQNTLY | 0.78618395 |
| LMVGVSI | ASSQSGYSQNTLY | 0.7016804 |
| LVGAAFVL | ASSQSGYSQNTLY | 0.80368435 |
| LVLAVAPL | ASSQSGYSQNTLY | 0.8131288 |
| MAAIAVIDAA | ASSQSGYSQNTLY | 0.508506 |
| NNGINAELQ | ASSQSGYSQNTLY | 0.8176392 |
| RGEPLVLVHG | ASSQSGYSQNTLY | 0.75594723 |
| RGGAGGAAGLI | ASSQSGYSQNTLY | 0.7240078 |
| RSYRMHPAV | ASSQSGYSQNTLY | 0.8427793 |
| SAAHLLAGQLA | ASSQSGYSQNTLY | 0.79583347 |
| SAAVGSY | ASSQSGYSQNTLY | 0.83223903 |
| SPHAVLHRI | ASSQSGYSQNTLY | 0.7874822 |
| SQIAKVL | ASSQSGYSQNTLY | 0.76630986 |
| TALGHLVP | ASSQSGYSQNTLY | 0.83019805 |
| TAPTTSGAA | ASSQSGYSQNTLY | 0.59681404 |
| TDGDDGNGGSPGT | ASSQSGYSQNTLY | 0.7177378 |
| TDGLGEN | ASSQSGYSQNTLY | 0.843611 |
| TGGPPLPGQA | ASSQSGYSQNTLY | 0.86783683 |
| TIPAITI | ASSQSGYSQNTLY | 0.8682888 |
| TIPGFTI | ASSQSGYSQNTLY | 0.84244484 |
| TSIASGALA | ASSQSGYSQNTLY | 0.82516456 |
| TVTLPGAAI | ASSQSGYSQNTLY | 0.82600296 |
| VASGLSAALP | ASSQSGYSQNTLY | 0.8733197 |
| VGAGVIGI | ASSQSGYSQNTLY | 0.86834997 |
| VGPKRFGV | ASSQSGYSQNTLY | 0.8687725 |
| VNVAFPD | ASSQSGYSQNTLY | 0.77610695 |
| VSMTNSGVS | ASSQSGYSQNTLY | 0.8696611 |
| VTDVLGDLSV | ASSQSGYSQNTLY | 0.8530184 |
| VVVVGAGFAGL | ASSQSGYSQNTLY | 0.8466617 |
| VYTVGTVV | ASSQSGYSQNTLY | 0.8687783 |
| WTTFSGP | ASSQSGYSQNTLY | 0.8572507 |
| YDELVEAF | ASSQSGYSQNTLY | 0.8495322 |
| YGTGGAGGNGG | ASSQSGYSQNTLY | 0.8542159 |
| YTNVMAV | ASSQSGYSQNTLY | 0.85203433 |
| FLEQVHQL | ASSQSGYSQNTLY | 0.85770065 |

| | | |
|---|---|---|
| GLSLLAVAV | ASSQSGYSQNTLY | 0.8465495 |
| LEQVHQL | ASSQSGYSQNTLY | 0.87785685 |
| LNAAGAI | ASSQSGYSQNTLY | 0.872025 |
| PAVHASL | ASSQSGYSQNTLY | 0.86981326 |
| TELEDDA | ASSQSGYSQNTLY | 0.38900572 |
| TPPGPPN | ASSQSGYSQNTLY | 0.80699265 |
| TPPPPPPP | ASSQSGYSQNTLY | 0.85625464 |
| TQSPALI | ASSLPGQNYEQY | 0.8539888 |
| EFAAAFPQ | ASSLPGQNYEQY | 0.6694205 |
| LLSGNLDVL | ASSLPGQNYEQY | 0.8580933 |
| GPANLGNYNI | ASSLPGQNYEQY | 0.8475196 |
| TLGPTLP | ASSLPGQNYEQY | 0.83735955 |
| VYAYSSCCK | ASSLPGQNYEQY | 0.8711449 |
| APAPAPAPNI | ASSLPGQNYEQY | 0.8528031 |
| PGGPVPIP | ASSLPGQNYEQY | 0.8380804 |
| FLTVALEP | ASSLPGQNYEQY | 0.84895 |
| SQYFGANL | ASSLPGQNYEQY | 0.8747542 |
| GGVKPAA | ASSLPGQNYEQY | 0.8261142 |
| GGTGGAGGAGGW | ASSLPGQNYEQY | 0.8703747 |
| GVGGVAGLGGAGG | ASSLPGQNYEQY | 0.8106763 |
| VLLVLATTI | ASSLPGQNYEQY | 0.8753408 |
| IGIFAAML | ASSLPGQNYEQY | 0.7466782 |
| AVGDLID | ASSLPGQNYEQY | 0.8706232 |
| NISPLPL | ASSLPGQNYEQY | 0.85463566 |
| VTAVFVAI | ASSLPGQNYEQY | 0.8799714 |
| SGDDNDAAL | ASSLPGQNYEQY | 0.8503076 |
| AVAALAHS | ASSLPGQNYEQY | 0.8676711 |
| GGVTVMATP | ASSLPGQNYEQY | 0.8374431 |
| AGLAANVRD | ASSLPGQNYEQY | 0.86374533 |
| TNRPDLIDPALLRPG | ASSLPGQNYEQY | 0.79362327 |
| LANLGVTAL | ASSLPGQNYEQY | 0.8434427 |
| TGERIEL | ASSLPGQNYEQY | 0.81953776 |
| NGEELAA | ASSLPGQNYEQY | 0.86219823 |
| VLSLVWG | ASSLPGQNYEQY | 0.86923754 |
| IGYALWL | ASSLPGQNYEQY | 0.74667835 |
| LSLLLAPL | ASSLPGQNYEQY | 0.87339824 |
| TLDEALK | ASSLPGQNYEQY | 0.83350325 |
| TDGAGTAEGG | ASSLPGQNYEQY | 0.8337598 |
| NAPGGAGGNGG | ASSLPGQNYEQY | 0.8755543 |
| VSVFMGMGGNF | ASSLPGQNYEQY | 0.8465221 |
| LASYLAAL | ASSLPGQNYEQY | 0.8701042 |
| LVGGTPAL | ASSLPGQNYEQY | 0.8498442 |
| VRNADVL | ASSLPGQNYEQY | 0.84818405 |
| TCSSPFLVSAM | ASSLPGQNYEQY | 0.8639038 |
| VHITDAE | ASSLPGQNYEQY | 0.78634804 |
| EYAAPSF | ASSLPGQNYEQY | 0.83412755 |
| VVDAFHGGL | ASSLPGQNYEQY | 0.87649405 |
| ASVVDPQ | ASSLPGQNYEQY | 0.64863485 |
| SYNEFGGLS | ASSLPGQNYEQY | 0.70279956 |
| DPDGSVY | ASSLPGQNYEQY | 0.5763879 |
| MVSTIHATE | ASSLPGQNYEQY | 0.51651156 |

| | | |
|---|---|---|
| LAAGVADPV | ASSLPGQNYEQY | 0.5814961 |
| TKEQIAA | ASSLPGQNYEQY | 0.65755534 |
| SLASPAAL | ASSLPGQNYEQY | 0.6962905 |
| IGLVFYGV | ASSLPGQNYEQY | 0.6487963 |
| FLLDASNL | ASSLPGQNYEQY | 0.45513755 |
| YQVGNFP | ASSLPGQNYEQY | 0.6193014 |
| ESLLSQST | ASSLPGQNYEQY | 0.57299757 |
| LFTVGFAAVG | ASSLPGQNYEQY | 0.6007874 |
| ATPPTPT | ASSLPGQNYEQY | 0.58967394 |
| TDLATLG | ASSLPGQNYEQY | 0.61048776 |
| SAGGTGR | ASSLPGQNYEQY | 0.5798594 |
| GFSNSGS | ASSLPGQNYEQY | 0.49281022 |
| GGNGGSGGTSV | ASSLPGQNYEQY | 0.6068709 |
| GTSVATGGAGN | ASSLPGQNYEQY | 0.61032283 |
| NGGNGGSGGTS | ASSLPGQNYEQY | 0.5863682 |
| AVVKFPH | ASSLPGQNYEQY | 0.66067344 |
| TPPPPPP | ASSLPGQNYEQY | 0.6820579 |
| AAPAAAAAPA | ASSLPGQNYEQY | 0.6478331 |
| IALNEDL | ASSLPGQNYEQY | 0.28517535 |
| IVNVSSM | ASSLPGQNYEQY | 0.3530892 |
| TGIPPAPR | ASSLPGQNYEQY | 0.5462495 |
| FDLGGGTF | ASSLPGQNYEQY | 0.575364 |
| FDLGGGTFDVS | ASSLPGQNYEQY | 0.30088916 |
| NEPTAAA | ASSLPGQNYEQY | 0.60418844 |
| NEPTAAALA | ASSLPGQNYEQY | 0.5158369 |
| GSIINISSI | ASSLPGQNYEQY | 0.55695534 |
| VAPGNAGTA | ASSLPGQNYEQY | 0.6722222 |
| TGGGASLE | ASSLPGQNYEQY | 0.60351455 |
| VAVYNHYKRI | ASSLPGQNYEQY | 0.4822948 |
| TDPAPATT | ASSLPGQNYEQY | 0.4516772 |
| TNRPDILDPALLRPG | ASSLPGQNYEQY | 0.6280236 |
| LTTTTTS | ASSLPGQNYEQY | 0.3742176 |
| PTGSGKTL | ASSLPGQNYEQY | 0.647628 |
| TGSGKTL | ASSLPGQNYEQY | 0.38787302 |
| TIGHVDHGK | ASSLPGQNYEQY | 0.65694964 |
| TIGHVDHGKTT | ASSLPGQNYEQY | 0.4232168 |
| GHVDHGKTT | ASSLPGQNYEQY | 0.6576207 |
| AQFEHTLL | ASSLPGQNYEQY | 0.6239679 |
| IVNQDEL | ASSLPGQNYEQY | 0.70691156 |
| TNGSQFF | ASSLPGQNYEQY | 0.56999874 |
| TGKGIVE | ASSLPGQNYEQY | 0.5839622 |
| LISNASDA | ASSLPGQNYEQY | 0.46988684 |
| AELRTLGELTKIAKAHGAQ | ASSLPGQNYEQY | 0.70075196 |
| APNAVAAAEPSW | ASSLPGQNYEQY | 0.46308053 |
| APQVAGAATAVL | ASSLPGQNYEQY | 0.5749091 |
| ARRDPAAAAAILPTDA | ASSLPGQNYEQY | 0.35560805 |
| DAAARDVLAEAGLAEYF | ASSLPGQNYEQY | 0.6022544 |
| DDQIVLSAKANLAKVEQ | ASSLPGQNYEQY | 0.65742815 |
| DELLPDTGEDSDEDGH | ASSLPGQNYEQY | 0.47076094 |
| ELPEAPANRATVPQD | ASSLPGQNYEQY | 0.65506256 |
| FGQSKIERVTGIG | ASSLPGQNYEQY | 0.6209067 |

| | | |
|---|---|---|
| GAGGTGSTAGGAGGAGGAGGLYA | ASSLPGQNYEQY | 0.51761615 |
| GHGGTGAAGGNGGTGGAGGSGIDG | ASSLPGQNYEQY | 0.6721453 |
| GNGGNGGAGGEGGPGGAGGAGA | ASSLPGQNYEQY | 0.58744216 |
| GSTAGNGGNGGNATNGTIAGQPAG | ASSLPGQNYEQY | 0.68452513 |
| GTGGDGGNGGNGGTGG | ASSLPGQNYEQY | 0.5170044 |
| GVRRPAGIPADAPV | ASSLPGQNYEQY | 0.53048915 |
| GYRQVTDAHLLTLARR | ASSLPGQNYEQY | 0.60174567 |
| IAKMGQDGHDRG | ASSLPGQNYEQY | 0.55008894 |
| KAEPSIEIQYT | ASSLPGQNYEQY | 0.47644383 |
| KPVRIRMAADEDTVAVT | ASSLPGQNYEQY | 0.68508327 |
| MPPRIAGMRLLVIKPEP | ASSLPGQNYEQY | 0.8548862 |
| MKFYAVALFRSRGVPPTR | ASSLPGQNYEQY | 0.87760943 |
| NAGGFGILGAVAHSP | ASSLPGQNYEQY | 0.79273814 |
| NEGREITLAMAEM | ASSLPGQNYEQY | 0.6787941 |
| NELVEWSHPLGHRWRWE | ASSLPGQNYEQY | 0.8055661 |
| NLGAGGTGGNGGNPG | ASSLPGQNYEQY | 0.874665 |
| NTEWPKWETVMAADPVD | ASSLPGQNYEQY | 0.8808887 |
| PASAAPAETQP | ASSLPGQNYEQY | 0.8655908 |
| QQSSDGKNGVRAAEITGE | ASSLPGQNYEQY | 0.5770446 |
| RAPDAVTTSWWKE | ASSLPGQNYEQY | 0.8535869 |
| SGFGNVGAHTSGWFNQ | ASSLPGQNYEQY | 0.79831064 |
| SGVAGAVGQAASVGGLKV | ASSLPGQNYEQY | 0.81539613 |
| SNGGAGTGSAGGNGGT | ASSLPGQNYEQY | 0.8659394 |
| TGGAGGEGGAAG | ASSLPGQNYEQY | 0.8026801 |
| TGGDGGAAGAGGAPGHG | ASSLPGQNYEQY | 0.7676302 |
| TMPPLLAAIANAGRHGV | ASSLPGQNYEQY | 0.72544575 |
| TSSTAGNSGGA | ASSLPGQNYEQY | 0.86783236 |
| VPATVPVGILC | ASSLPGQNYEQY | 0.87316 |
| VQRPDSGGVVLTGRIS | ASSLPGQNYEQY | 0.872455 |
| WTPNPVPATKGVPAGLVQAT | ASSLPGQNYEQY | 0.888319 |
| YALKRALAKFSDRFDVVII | ASSLPGQNYEQY | 0.87901294 |
| YLIGSVIV | ASSLPGQNYEQY | 0.8770791 |
| YGQLVAVID | ASSLPGQNYEQY | 0.28363836 |
| VTVGGMTLPA | ASSLPGQNYEQY | 0.528621 |
| VLALPAV | ASSLPGQNYEQY | 0.7511785 |
| VIPPTLNYET | ASSLPGQNYEQY | 0.79184586 |
| VGMGGMHPSSG | ASSLPGQNYEQY | 0.48171064 |
| VGLAGGAATA | ASSLPGQNYEQY | 0.8195354 |
| VGGAGGAGGSGP | ASSLPGQNYEQY | 0.7296062 |
| VFAAFMV | ASSLPGQNYEQY | 0.69760114 |
| TLYDKLR | ASSLPGQNYEQY | 0.8745292 |
| TLAVFTVL | ASSLPGQNYEQY | 0.7411786 |
| TGSVQPARV | ASSLPGQNYEQY | 0.64596266 |
| TGGGAGSGGT | ASSLPGQNYEQY | 0.720065 |
| SHLETSL | ASSLPGQNYEQY | 0.8807161 |
| SFTTGMT | ASSLPGQNYEQY | 0.658356 |
| QVEVAKRMGVRQPTVS | ASSLPGQNYEQY | 0.8629396 |
| QGSGGPALAM | ASSLPGQNYEQY | 0.81401885 |
| MLAPSNP | ASSLPGQNYEQY | 0.8779997 |
| LVNNAAI | ASSLPGQNYEQY | 0.49120462 |
| LTGEINTLA | ASSLPGQNYEQY | 0.87528825 |

| | | |
|---|---|---|
| LSIFGPV | ASSLPGQNYEQY | 0.84014285 |
| LLALLANL | ASSLPGQNYEQY | 0.893311 |
| LISNASDALDKLR | ASSLPGQNYEQY | 0.7832681 |
| LINEAVP | ASSLPGQNYEQY | 0.7970224 |
| LGYAFSGL | ASSLPGQNYEQY | 0.72755015 |
| LDFVAVAL | ASSLPGQNYEQY | 0.8755454 |
| LAQPGAE | ASSLPGQNYEQY | 0.47005135 |
| ISNLGAVTGF | ASSLPGQNYEQY | 0.81762815 |
| HQEDPGRI | ASSLPGQNYEQY | 0.68955046 |
| GTPGEPTP | ASSLPGQNYEQY | 0.83887416 |
| GFFNSGS | ASSLPGQNYEQY | 0.8614943 |
| GAGGGGLTAGAN | ASSLPGQNYEQY | 0.32187396 |
| FVDALVNM | ASSLPGQNYEQY | 0.87559974 |
| FLTVAPEL | ASSLPGQNYEQY | 0.8137484 |
| FETLTAL | ASSLPGQNYEQY | 0.61718595 |
| DVINPIEL | ASSLPGQNYEQY | 0.8837461 |
| APVSGAPV | ASSLPGQNYEQY | 0.82435733 |
| ALVGGLAW | ASSLPGQNYEQY | 0.8821453 |
| ALAAAAATAI | ASSLPGQNYEQY | 0.8290746 |
| AFGLGLLL | ASSLPGQNYEQY | 0.85495394 |
| AFFAAASL | ASSLPGQNYEQY | 0.8570185 |
| AAAALPAFSPP | ASSLPGQNYEQY | 0.66596925 |
| VVLAPAL | ASSLPGQNYEQY | 0.7390077 |
| TIGHVDHGKTTL | ASSLPGQNYEQY | 0.8834102 |
| LISNASDALDK | ASSLPGQNYEQY | 0.77791154 |
| LISNASDAL | ASSLPGQNYEQY | 0.83402824 |
| ISNASDALDK | ASSLPGQNYEQY | 0.5920626 |
| NQAETLVYQTE | ASSLPGQNYEQY | 0.5047146 |
| TGGTGGTAGAG | ASSLPGQNYEQY | 0.6652746 |
| TGSTAGGAGGA | ASSLPGQNYEQY | 0.82032746 |
| DPLAVPVAAGPV | ASSLPGQNYEQY | 0.83171546 |
| GGTGGTPGNGGN | ASSLPGQNYEQY | 0.7960962 |
| PAGLAGVDPATL | ASSLPGQNYEQY | 0.5555742 |
| GNNLVVEAPHAGASV | ASSLPGQNYEQY | 0.7000238 |
| KVTRIMSASTLEQPA | ASSLPGQNYEQY | 0.6613935 |
| LGFALPIALPRRTVV | ASSLPGQNYEQY | 0.6804852 |
| TRLGNLTARKLVVLV | ASSLPGQNYEQY | 0.76733667 |
| WSQQRRLIYFPSAGP | ASSLPGQNYEQY | 0.758629 |
| GPNGNPDPMAAAVDIR | ASSLPGQNYEQY | 0.6584399 |
| TQRCVSVADLRAAASR | ASSLPGQNYEQY | 0.60371023 |
| QFVLDVAEHPVEVLTETV | ASSLPGQNYEQY | 0.73380446 |
| GGAGGTGGAGGNGGTQAGDG | ASSLPGQNYEQY | 0.8065729 |
| PLLATLRAIAVDVDAKPAQV | ASSLPGQNYEQY | 0.7871957 |
| APKFRTGGYMIDFWGVGYQVAKRM | ASSLPGQNYEQY | 0.8477782 |
| GAGGAGGAGGAAGGPGGTGGTGGN | ASSLPGQNYEQY | 0.8126292 |
| IGALGLLATDVAPAGSVAFGVALA | ASSLPGQNYEQY | 0.81880975 |
| AGGAGGIGGNGGAGGAGGVLMGNG | ASSLPGQNYEQY | 0.1929799 |
| ADELIGLL | ASSLPGQNYEQY | 0.36561704 |
| ADRALKA | ASSLPGQNYEQY | 0.6219213 |
| AGAVGQVGGAGGA | ASSLPGQNYEQY | 0.68322676 |
| AGGGGGLGGAP | ASSLPGQNYEQY | 0.33671996 |

| | | |
|---|---|---|
| AGGLLYGNGGAG | ASSLPGQNYEQY | 0.76484823 |
| AGLESIGL | ASSLPGQNYEQY | 0.60098195 |
| AGLKTGII | ASSLPGQNYEQY | 0.55985373 |
| ASLAALLL | ASSLPGQNYEQY | 0.80754256 |
| ASVINISSI | ASSLPGQNYEQY | 0.670367 |
| ATPPGPA | ASSLPGQNYEQY | 0.58771145 |
| ATVASALV | ASSLPGQNYEQY | 0.5834059 |
| ATVASLIL | ASSLPGQNYEQY | 0.8142408 |
| AVNGPVLIRH | ASSLPGQNYEQY | 0.41851372 |
| DGAAGAVGI | ASSLPGQNYEQY | 0.78101593 |
| DPLLHGI | ASSLPGQNYEQY | 0.7227495 |
| ELGAALGAL | ASSLPGQNYEQY | 0.8147427 |
| ELGRTQAV | ASSLPGQNYEQY | 0.434406 |
| FVAGGYLS | ASSLPGQNYEQY | 0.8064782 |
| GASGGAGGAGGW | ASSLPGQNYEQY | 0.7013848 |
| GGAGAGGDGGAGGI | ASSLPGQNYEQY | 0.8565371 |
| GTGHNGSQPGMGGNG | ASSLPGQNYEQY | 0.61766005 |
| GYAAASASAS | ASSLPGQNYEQY | 0.74344003 |
| HQVDVAAVQQG | ASSLPGQNYEQY | 0.70776737 |
| IGAAIAHT | ASSLPGQNYEQY | 0.8265196 |
| IGMIEVAQL | ASSLPGQNYEQY | 0.39194053 |
| IRLAPEHTSAMD | ASSLPGQNYEQY | 0.6310991 |
| IYNVSVL | ASSLPGQNYEQY | 0.4685663 |
| KQGFLGM | ASSLPGQNYEQY | 0.7171906 |
| LESVITQL | ASSLPGQNYEQY | 0.7966962 |
| LGLGALVVV | ASSLPGQNYEQY | 0.2953386 |
| LIGNGGNGGSGGI | ASSLPGQNYEQY | 0.80394197 |
| LNAINAPTL | ASSLPGQNYEQY | 0.72878706 |
| LQLLNTGV | ASSLPGQNYEQY | 0.57080996 |
| LQTSIEF | ASSLPGQNYEQY | 0.8210057 |
| MSFAVSEL | ASSLPGQNYEQY | 0.6398239 |
| NTGALGL | ASSLPGQNYEQY | 0.8273668 |
| PPGAGGAGGAGTT | ASSLPGQNYEQY | 0.6517692 |
| QLVGLGI | ASSLPGQNYEQY | 0.7012651 |
| RFLEQVHQL | ASSLPGQNYEQY | 0.767605 |
| RGVGYVLR | ASSLPGQNYEQY | 0.33750924 |
| SATGVFAGVI | ASSLPGQNYEQY | 0.50860703 |
| SLPAITLP | ASSLPGQNYEQY | 0.83407485 |
| STALAFTV | ASSLPGQNYEQY | 0.62255603 |
| SVLTVGLGK | ASSLPGQNYEQY | 0.6478248 |
| TALAGAGSGI | ASSLPGQNYEQY | 0.5390429 |
| TGLPFTGI | ASSLPGQNYEQY | 0.4945155 |
| VAEHVTV | ASSLPGQNYEQY | 0.53685904 |
| VDIVLNSL | ASSLPGQNYEQY | 0.62360257 |
| VLTGGGAVL | ASSLPGQNYEQY | 0.66075367 |
| VNVGLAYGI | ASSLPGQNYEQY | 0.6224741 |
| VSSAAVGAV | ASSLPGQNYEQY | 0.50166506 |
| VTSVYEAGV | ASSLPGQNYEQY | 0.62506604 |
| WVALDRGA | ASSLPGQNYEQY | 0.59036577 |
| YLVGSVVI | ASSLPGQNYEQY | 0.5679831 |
| GPPGTGKTLL | ASSLPGQNYEQY | 0.5788387 |

| | | |
|---|---|---|
| GVGMTKF | ASSLPGQNYEQY | 0.6341329 |
| IGIGGSDL | ASSLPGQNYEQY | 0.55818284 |
| IGTLLAEL | ASSLPGQNYEQY | 0.4945029 |
| IYTSGTT | ASSLPGQNYEQY | 0.580731 |
| LSFPAGV | ASSLPGQNYEQY | 0.64068425 |
| LYTSGTT | ASSLPGQNYEQY | 0.6248931 |
| VGAFPFGF | ASSLPGQNYEQY | 0.6587771 |
| VGAGTVEF | ASSLPGQNYEQY | 0.6444229 |
| VGKSTLLL | ASSLPGQNYEQY | 0.60924053 |
| AALAHAGGGIV | ASSLPGQNYEQY | 0.33671522 |
| AALVAVSLL | ASSLPGQNYEQY | 0.40036795 |
| AAVSRAVAAL | ASSLPGQNYEQY | 0.56517744 |
| AGGSSGIGGF | ASSLPGQNYEQY | 0.5615662 |
| AGLTVANANG | ASSLPGQNYEQY | 0.33751488 |
| AGLVGAVL | ASSLPGQNYEQY | 0.61786664 |
| ALSQAFTV | ASSLPGQNYEQY | 0.5520415 |
| ALTFSVSL | ASSLPGQNYEQY | 0.5514038 |
| ALTGAALVAL | ASSLPGQNYEQY | 0.6345632 |
| ASFSIGAGVL | ASSLPGQNYEQY | 0.5634073 |
| DGSVYKI | ASSLPGQNYEQY | 0.4990095 |
| EQLAEIV | ASSLPGQNYEQY | 0.53155404 |
| FAVSELTVM | ASSLPGQNYEQY | 0.65831494 |
| GAAPLAAGL | ASSLPGQNYEQY | 0.45856377 |
| GADGGDGGSGGAGG | ASSLPGQNYEQY | 0.63211644 |
| GFGLDTGGAGG | ASSLPGQNYEQY | 0.5041348 |
| GLASGGLL | ASSLPGQNYEQY | 0.651328 |
| GRIAGAAVLL | ASSLPGQNYEQY | 0.45075333 |
| GSAGSLNTSF | ASSLPGQNYEQY | 0.6342992 |
| GTGQIGI | ASSLPGQNYEQY | 0.5817656 |
| GTLIALIL | ASSLPGQNYEQY | 0.6738961 |
| IAATMGFVVG | ASSLPGQNYEQY | 0.544062 |
| IFAQALL | ASSLPGQNYEQY | 0.6230024 |
| IGAAVALGF | ASSLPGQNYEQY | 0.52946454 |
| IGLTPLAL | ASSLPGQNYEQY | 0.64125794 |
| ILGSNTSTL | ASSLPGQNYEQY | 0.4732751 |
| IVFSLTVL | ASSLPGQNYEQY | 0.5555464 |
| LASKATGF | ASSLPGQNYEQY | 0.43003017 |
| LASVALIL | ASSLPGQNYEQY | 0.58181363 |
| LFAVGLTL | ASSLPGQNYEQY | 0.6313237 |
| LFNGGIGTYI | ASSLPGQNYEQY | 0.34461534 |
| LTLGAGII | ASSLPGQNYEQY | 0.639504 |
| LVAAALGDI | ASSLPGQNYEQY | 0.4864811 |
| NTAFVQLGI | ASSLPGQNYEQY | 0.48981762 |
| QLFSGAEL | ASSLPGQNYEQY | 0.63097304 |
| SAATFITL | ASSLPGQNYEQY | 0.61118996 |
| SNYGLANLGV | ASSLPGQNYEQY | 0.6412397 |
| SYGGAYAVM | ASSLPGQNYEQY | 0.53959316 |
| TTGVGGLTLG | ASSLPGQNYEQY | 0.5827333 |
| TTQTIYI | ASSLPGQNYEQY | 0.6083605 |
| VAAGAGLGAVI | ASSLPGQNYEQY | 0.439083 |
| VAAVVTLL | ASSLPGQNYEQY | 0.5108472 |

| | | |
|---|---|---|
| VGFLGGLI | ASSLPGQNYEQY | 0.6495633 |
| VGVIPILL | ASSLPGQNYEQY | 0.5903413 |
| VIFASSLI | ASSLPGQNYEQY | 0.82256913 |
| VKGGDGGAAAG | ASSLPGQNYEQY | 0.46988085 |
| VTKTFTNI | ASSLPGQNYEQY | 0.3463663 |
| VVAAAILL | ASSLPGQNYEQY | 0.49772412 |
| YAALDVEL | ASSLPGQNYEQY | 0.6692985 |
| LASVALLL | ASSLPGQNYEQY | 0.8182197 |
| LDAVHRV | ASSLPGQNYEQY | 0.6440402 |
| QGLSSSIF | ASSLPGQNYEQY | 0.27098194 |
| RGTFANIRL | ASSLPGQNYEQY | 0.534414 |
| RQLWWGHRI | ASSLPGQNYEQY | 0.40287983 |
| YGPPGTGKTLL | ASSLPGQNYEQY | 0.45403206 |
| AALVGAPANVA | ASSLPGQNYEQY | 0.54007506 |
| AATFRRTE | ASSLPGQNYEQY | 0.57621706 |
| AGALGGGI | ASSLPGQNYEQY | 0.5023989 |
| AGGAGGAGGIGAG | ASSLPGQNYEQY | 0.32602125 |
| AGSGALGL | ASSLPGQNYEQY | 0.6118237 |
| AQIADQI | ASSLPGQNYEQY | 0.7090875 |
| ASAVAVTYL | ASSLPGQNYEQY | 0.638395 |
| ASTVLGGTGG | ASSLPGQNYEQY | 0.72887087 |
| AVGGEPAK | ASSLPGQNYEQY | 0.70647657 |
| CEQGTFGGTC | ASSLPGQNYEQY | 0.558462 |
| DEGKAPAG | ASSLPGQNYEQY | 0.14804703 |
| DTVVFPAL | ASSLPGQNYEQY | 0.20536782 |
| FEGITMAG | ASSLPGQNYEQY | 0.41290274 |
| FPLVVQD | ASSLPGQNYEQY | 0.44552726 |
| GAFTTPQ | ASSLPGQNYEQY | 0.17984442 |
| GFGGFGTAGGVGGA | ASSLPGQNYEQY | 0.53653234 |
| GFLAQFL | ASSLPGQNYEQY | 0.418499 |
| GGAGGAGGNAGVA | ASSLPGQNYEQY | 0.40775698 |
| GGVGTAGAGGAGGAGG | ASSLPGQNYEQY | 0.7661288 |
| GNGGGGDAGNAGSGGN | ASSLPGQNYEQY | 0.46661496 |
| GTGGAGGAGNDAGS | ASSLPGQNYEQY | 0.37395287 |
| GTTNTGWFNS | ASSLPGQNYEQY | 0.30645698 |
| IFMGPLFTL | ASSLPGQNYEQY | 0.6928618 |
| ILAHAIEAI | ASSLPGQNYEQY | 0.23008063 |
| IRSAIREAL | ASSLPGQNYEQY | 0.6143204 |
| ITFAGGG | ASSLPGQNYEQY | 0.29541004 |
| ITGMSGHQEV | ASSLPGQNYEQY | 0.6561856 |
| IVAGLACTL | ASSLPGQNYEQY | 0.23027432 |
| IVGGGLAGA | ASSLPGQNYEQY | 0.65825665 |
| IVTGVLNI | ASSLPGQNYEQY | 0.5453342 |
| LAQQGCI | ASSLPGQNYEQY | 0.84023863 |
| LCLHVEN | ASSLPGQNYEQY | 0.42126906 |
| LDAISDGS | ASSLPGQNYEQY | 0.49192038 |
| LGGFAAML | ASSLPGQNYEQY | 0.2966033 |
| LGGKNPMI | ASSLPGQNYEQY | 0.8039876 |
| LGPKGVAAIL | ASSLPGQNYEQY | 0.27805862 |
| LMVGVSI | ASSLPGQNYEQY | 0.45127618 |
| LVGAAFVL | ASSLPGQNYEQY | 0.23071085 |

| | | |
|---|---|---|
| LVLAVAPL | ASSLPGQNYEQY | 0.5869109 |
| MAAIAVIDAA | ASSLPGQNYEQY | 0.623925 |
| NNGINAELQ | ASSLPGQNYEQY | 0.21939151 |
| RGEPLVLVHG | ASSLPGQNYEQY | 0.6298608 |
| RGGAGGAAGLI | ASSLPGQNYEQY | 0.5568047 |
| RSYRMHPAV | ASSLPGQNYEQY | 0.27913165 |
| SAAHLLAGQLA | ASSLPGQNYEQY | 0.7176839 |
| SAAVGSY | ASSLPGQNYEQY | 0.3393683 |
| SPHAVLHRI | ASSLPGQNYEQY | 0.75437653 |
| SQIAKVL | ASSLPGQNYEQY | 0.39673787 |
| TALGHLVP | ASSLPGQNYEQY | 0.3988364 |
| TAPTTSGAA | ASSLPGQNYEQY | 0.6176634 |
| TDGDDGNGGSPGT | ASSLPGQNYEQY | 0.4341058 |
| TDGLGEN | ASSLPGQNYEQY | 0.29648328 |
| TGGPPLPGQA | ASSLPGQNYEQY | 0.72480583 |
| TIPAITI | ASSLPGQNYEQY | 0.76422215 |
| TIPGFTI | ASSLPGQNYEQY | 0.7900232 |
| TSIASGALA | ASSLPGQNYEQY | 0.6838526 |
| TVTLPGAAI | ASSLPGQNYEQY | 0.69546485 |
| VASGLSAALP | ASSLPGQNYEQY | 0.75254405 |
| VGAGVIGI | ASSLPGQNYEQY | 0.7776358 |
| VGPKRFGV | ASSLPGQNYEQY | 0.8035506 |
| VNVAFPD | ASSLPGQNYEQY | 0.7720388 |
| VSMTNSGVS | ASSLPGQNYEQY | 0.72953284 |
| VTDVLGDLSV | ASSLPGQNYEQY | 0.76980805 |
| VVVVGAGFAGL | ASSLPGQNYEQY | 0.7452383 |
| VYTVGTVV | ASSLPGQNYEQY | 0.74672735 |
| WTTFSGP | ASSLPGQNYEQY | 0.7866634 |
| YDELVEAF | ASSLPGQNYEQY | 0.7462762 |
| YGTGGAGGNGG | ASSLPGQNYEQY | 0.6985204 |
| YTNVMAV | ASSLPGQNYEQY | 0.7134302 |
| FLEQVHQL | ASSLPGQNYEQY | 0.783154 |
| GLSLLAVAV | ASSLPGQNYEQY | 0.7839906 |
| LEQVHQL | ASSLPGQNYEQY | 0.78116924 |
| LNAAGAI | ASSLPGQNYEQY | 0.8046493 |
| PAVHASL | ASSLPGQNYEQY | 0.78065056 |
| TELEDDA | ASSLPGQNYEQY | 0.7659244 |
| TPPGPPN | ASSLPGQNYEQY | 0.46799773 |
| TPPPPPPP | ASSLPGQNYEQY | 0.66070104 |
| TQSPALI | ASSLEGWGGSYEQY | 0.7065722 |
| EFAAAFPQ | ASSLEGWGGSYEQY | 0.7562039 |
| LLSGNLDVL | ASSLEGWGGSYEQY | 0.4367991 |
| GPANLGNYNI | ASSLEGWGGSYEQY | 0.76750016 |
| TLGPTLP | ASSLEGWGGSYEQY | 0.74945766 |
| VYAYSSCCK | ASSLEGWGGSYEQY | 0.68604434 |
| APAPAPAPNI | ASSLEGWGGSYEQY | 0.7904002 |
| PGGPVPIP | ASSLEGWGGSYEQY | 0.70025754 |
| FLTVALEP | ASSLEGWGGSYEQY | 0.69963485 |
| SQYFGANL | ASSLEGWGGSYEQY | 0.7409246 |
| GGVKPAA | ASSLEGWGGSYEQY | 0.8033032 |
| GGTGGAGGAGGW | ASSLEGWGGSYEQY | 0.6848504 |

| | | |
|---|---|---|
| GVGGVAGLGGAGG | ASSLEGWGGSYEQY | 0.7913029 |
| VLLVLATTI | ASSLEGWGGSYEQY | 0.7714249 |
| IGIFAAML | ASSLEGWGGSYEQY | 0.7930279 |
| AVGDLID | ASSLEGWGGSYEQY | 0.6586453 |
| NISPLPL | ASSLEGWGGSYEQY | 0.78401595 |
| VTAVFVAI | ASSLEGWGGSYEQY | 0.73927176 |
| SGDDNDAAL | ASSLEGWGGSYEQY | 0.804137 |
| AVAALAHS | ASSLEGWGGSYEQY | 0.7556399 |
| GGVTVMATP | ASSLEGWGGSYEQY | 0.7869462 |
| AGLAANVRD | ASSLEGWGGSYEQY | 0.7703667 |
| TNRPDLIDPALLRPG | ASSLEGWGGSYEQY | 0.7962251 |
| LANLGVTAL | ASSLEGWGGSYEQY | 0.62191296 |
| TGERIEL | ASSLEGWGGSYEQY | 0.7440963 |
| NGEELAA | ASSLEGWGGSYEQY | 0.7314024 |
| VLSLVWG | ASSLEGWGGSYEQY | 0.7639285 |
| IGYALWL | ASSLEGWGGSYEQY | 0.7748683 |
| LSLLLAPL | ASSLEGWGGSYEQY | 0.40455696 |
| TLDEALK | ASSLEGWGGSYEQY | 0.79016197 |
| TDGAGTAEGG | ASSLEGWGGSYEQY | 0.64279413 |
| NAPGGAGGNGG | ASSLEGWGGSYEQY | 0.7061882 |
| VSVFMGMGGNF | ASSLEGWGGSYEQY | 0.7844388 |
| LASYLAAL | ASSLEGWGGSYEQY | 0.73097646 |
| LVGGTPAL | ASSLEGWGGSYEQY | 0.7763717 |
| VRNADVL | ASSLEGWGGSYEQY | 0.71804 |
| TCSSPFLVSAM | ASSLEGWGGSYEQY | 0.7637228 |
| VHITDAE | ASSLEGWGGSYEQY | 0.77807164 |
| EYAAPSF | ASSLEGWGGSYEQY | 0.46154767 |
| VVDAFHGGL | ASSLEGWGGSYEQY | 0.67322075 |
| ASVVDPQ | ASSLEGWGGSYEQY | 0.7939248 |
| SYNEFGGLS | ASSLEGWGGSYEQY | 0.7240579 |
| DPDGSVY | ASSLEGWGGSYEQY | 0.75124323 |
| MVSTIHATE | ASSLEGWGGSYEQY | 0.6833058 |
| LAAGVADPV | ASSLEGWGGSYEQY | 0.59798586 |
| TKEQIAA | ASSLEGWGGSYEQY | 0.69856364 |
| SLASPAAL | ASSLEGWGGSYEQY | 0.74072754 |
| IGLVFYGV | ASSLEGWGGSYEQY | 0.74958026 |
| FLLDASNL | ASSLEGWGGSYEQY | 0.7320871 |
| YQVGNFP | ASSLEGWGGSYEQY | 0.57584167 |
| ESLLSQST | ASSLEGWGGSYEQY | 0.70636827 |
| LFTVGFAAVG | ASSLEGWGGSYEQY | 0.6845957 |
| ATPPTPT | ASSLEGWGGSYEQY | 0.695336 |
| TDLATLG | ASSLEGWGGSYEQY | 0.7335075 |
| SAGGTGR | ASSLEGWGGSYEQY | 0.71832776 |
| GFSNSGS | ASSLEGWGGSYEQY | 0.67485976 |
| GGNGGSGGTSV | ASSLEGWGGSYEQY | 0.6494095 |
| GTSVATGGAGN | ASSLEGWGGSYEQY | 0.7164078 |
| NGGNGGSGGTS | ASSLEGWGGSYEQY | 0.7414663 |
| AVVKFPH | ASSLEGWGGSYEQY | 0.7336248 |
| TPPPPPP | ASSLEGWGGSYEQY | 0.75302243 |
| AAPAAAAAPA | ASSLEGWGGSYEQY | 0.743669 |
| IALNEDL | ASSLEGWGGSYEQY | 0.7429516 |

| | | |
|---|---|---|
| IVNVSSM | ASSLEGWGGSYEQY | 0.3091372 |
| TGIPPAPR | ASSLEGWGGSYEQY | 0.47876203 |
| FDLGGGTF | ASSLEGWGGSYEQY | 0.6424962 |
| FDLGGGTFDVS | ASSLEGWGGSYEQY | 0.6820208 |
| NEPTAAA | ASSLEGWGGSYEQY | 0.4721133 |
| NEPTAAALA | ASSLEGWGGSYEQY | 0.708112 |
| GSIINISSI | ASSLEGWGGSYEQY | 0.6567234 |
| VAPGNAGTA | ASSLEGWGGSYEQY | 0.6202407 |
| TGGGASLE | ASSLEGWGGSYEQY | 0.7438549 |
| VAVYNHYKRI | ASSLEGWGGSYEQY | 0.6780631 |
| TDPAPATT | ASSLEGWGGSYEQY | 0.6388664 |
| TNRPDILDPALLRPG | ASSLEGWGGSYEQY | 0.643033 |
| LTTTTTS | ASSLEGWGGSYEQY | 0.75067735 |
| PTGSGKTL | ASSLEGWGGSYEQY | 0.5144386 |
| TGSGKTL | ASSLEGWGGSYEQY | 0.7258377 |
| TIGHVDHGK | ASSLEGWGGSYEQY | 0.69899035 |
| TIGHVDHGKTT | ASSLEGWGGSYEQY | 0.7413599 |
| GHVDHGKTT | ASSLEGWGGSYEQY | 0.51267624 |
| AQFEHTLL | ASSLEGWGGSYEQY | 0.7390623 |
| IVNQDEL | ASSLEGWGGSYEQY | 0.70387936 |
| TNGSQFF | ASSLEGWGGSYEQY | 0.7582219 |
| TGKGIVE | ASSLEGWGGSYEQY | 0.68323094 |
| LISNASDA | ASSLEGWGGSYEQY | 0.7149599 |
| AELRTLGELTKIAKAHGAQ | ASSLEGWGGSYEQY | 0.6819924 |
| APNAVAAAEPSW | ASSLEGWGGSYEQY | 0.74524033 |
| APQVAGAATAVL | ASSLEGWGGSYEQY | 0.53045607 |
| ARRDPAAAAAILPTDA | ASSLEGWGGSYEQY | 0.6928277 |
| DAAARDVLAEAGLAEYF | ASSLEGWGGSYEQY | 0.57015276 |
| DDQIVLSAKANLAKVEQ | ASSLEGWGGSYEQY | 0.71837455 |
| DELLPDTGEDSDEDGH | ASSLEGWGGSYEQY | 0.7318072 |
| ELPEAPANRATVPQD | ASSLEGWGGSYEQY | 0.5281389 |
| FGQSKIERVTGIG | ASSLEGWGGSYEQY | 0.748922 |
| GAGGTGSTAGGAGGAGGAGGLYA | ASSLEGWGGSYEQY | 0.7199062 |
| GHGGTGAAGGNGGTGGAGGSGIDG | ASSLEGWGGSYEQY | 0.63610375 |
| GNGGNGGAGGEGGPGGAGGAGA | ASSLEGWGGSYEQY | 0.7484413 |
| GSTAGNGGNGGNATNGTIAGGQPAG | ASSLEGWGGSYEQY | 0.67673916 |
| GTGGDGGNGGNGGTGG | ASSLEGWGGSYEQY | 0.74737227 |
| GVRRPAGIPADAPV | ASSLEGWGGSYEQY | 0.69350725 |
| GYRQVTDAHLLTLARR | ASSLEGWGGSYEQY | 0.71484375 |
| IAKMGQDGHDRG | ASSLEGWGGSYEQY | 0.7333519 |
| KAEPSIEIQYT | ASSLEGWGGSYEQY | 0.57914186 |
| KPVRIRMAADEDTVAVT | ASSLEGWGGSYEQY | 0.6071218 |
| MPPRIAGMRLLVIKPEP | ASSLEGWGGSYEQY | 0.7467748 |
| MKFYAVALFRSRGVPPTR | ASSLEGWGGSYEQY | 0.8108503 |
| NAGGFGILGAVAHSP | ASSLEGWGGSYEQY | 0.82570976 |
| NEGREITLAMAEM | ASSLEGWGGSYEQY | 0.6796074 |
| NELVEWSHPLGHRWRWE | ASSLEGWGGSYEQY | 0.59299016 |
| NLGAGGTGGNGGNPG | ASSLEGWGGSYEQY | 0.7393143 |
| NTEWPKWETVMAADPVD | ASSLEGWGGSYEQY | 0.8226238 |
| PASAAPAETQP | ASSLEGWGGSYEQY | 0.8221078 |
| QQSSDGKNGVRAAEITGE | ASSLEGWGGSYEQY | 0.8130109 |

| | | |
|---|---|---|
| RAPDAVTTSWWKE | ASSLEGWGGSYEQY | 0.51406676 |
| SGFGNVGAHTSGWFNQ | ASSLEGWGGSYEQY | 0.7722828 |
| SGVAGAVGQAASVGGLKV | ASSLEGWGGSYEQY | 0.72673535 |
| SNGGAGTGSAGGNGGT | ASSLEGWGGSYEQY | 0.7571169 |
| TGGAGGEGGAAG | ASSLEGWGGSYEQY | 0.80509126 |
| TGGDGGAAGAGGAPGHG | ASSLEGWGGSYEQY | 0.75669175 |
| TMPPLLAAIANAGRHGV | ASSLEGWGGSYEQY | 0.70634955 |
| TSSTAGNSGGA | ASSLEGWGGSYEQY | 0.68245023 |
| VPATVPVGILC | ASSLEGWGGSYEQY | 0.80922127 |
| VQRPDSGGVVLTGRIS | ASSLEGWGGSYEQY | 0.8180202 |
| WTPNPVPATKGVPAGLVQAT | ASSLEGWGGSYEQY | 0.8137852 |
| YALKRALAKFSDRFDVVII | ASSLEGWGGSYEQY | 0.83185637 |
| YLIGSVIV | ASSLEGWGGSYEQY | 0.81521094 |
| YGQLVAVID | ASSLEGWGGSYEQY | 0.80874765 |
| VTVGGMTLPA | ASSLEGWGGSYEQY | 0.21617323 |
| VLALPAV | ASSLEGWGGSYEQY | 0.4456926 |
| VIPPTLNYET | ASSLEGWGGSYEQY | 0.6826452 |
| VGMGGMHPSSG | ASSLEGWGGSYEQY | 0.733794 |
| VGLAGGAATA | ASSLEGWGGSYEQY | 0.35850054 |
| VGGAGGAGGSGP | ASSLEGWGGSYEQY | 0.8021184 |
| VFAAFMV | ASSLEGWGGSYEQY | 0.7158743 |
| TLYDKLR | ASSLEGWGGSYEQY | 0.59042865 |
| TLAVFTVL | ASSLEGWGGSYEQY | 0.8131739 |
| TGSVQPARV | ASSLEGWGGSYEQY | 0.66542304 |
| TGGGAGSGGT | ASSLEGWGGSYEQY | 0.60939324 |
| SHLETSL | ASSLEGWGGSYEQY | 0.69850314 |
| SFTTGMT | ASSLEGWGGSYEQY | 0.81415176 |
| QVEVAKRMGVRQPTVS | ASSLEGWGGSYEQY | 0.57092965 |
| QGSGGPALAM | ASSLEGWGGSYEQY | 0.8073551 |
| MLAPSNP | ASSLEGWGGSYEQY | 0.748253 |
| LVNNAAI | ASSLEGWGGSYEQY | 0.8204309 |
| LTGEINTLA | ASSLEGWGGSYEQY | 0.45930895 |
| LSIFGPV | ASSLEGWGGSYEQY | 0.81557673 |
| LLALLANL | ASSLEGWGGSYEQY | 0.778213 |
| LISNASDALDKLR | ASSLEGWGGSYEQY | 0.8376475 |
| LINEAVP | ASSLEGWGGSYEQY | 0.72856534 |
| LGYAFSGL | ASSLEGWGGSYEQY | 0.7553425 |
| LDFVAVAL | ASSLEGWGGSYEQY | 0.64343643 |
| LAQPGAE | ASSLEGWGGSYEQY | 0.82285845 |
| ISNLGAVTGF | ASSLEGWGGSYEQY | 0.39152712 |
| HQEDPGRI | ASSLEGWGGSYEQY | 0.7496722 |
| GTPGEPTP | ASSLEGWGGSYEQY | 0.4874946 |
| GFFNSGS | ASSLEGWGGSYEQY | 0.7585776 |
| GAGGGGLTAGAN | ASSLEGWGGSYEQY | 0.8108032 |
| FVDALVNM | ASSLEGWGGSYEQY | 0.32201385 |
| FLTVAPEL | ASSLEGWGGSYEQY | 0.8073838 |
| FETLTAL | ASSLEGWGGSYEQY | 0.7433772 |
| DVINPIEL | ASSLEGWGGSYEQY | 0.62946635 |
| APVSGAPV | ASSLEGWGGSYEQY | 0.8218368 |
| ALVGGLAW | ASSLEGWGGSYEQY | 0.7487309 |
| ALAAAAATAI | ASSLEGWGGSYEQY | 0.8266442 |

| | | |
|---|---|---|
| AFGLGLLL | ASSLEGWGGSYEQY | 0.76131225 |
| AFFAAASL | ASSLEGWGGSYEQY | 0.76587653 |
| AAAALPAFSPP | ASSLEGWGGSYEQY | 0.7892812 |
| VVLAPAL | ASSLEGWGGSYEQY | 0.55059856 |
| TIGHVDHGKTTL | ASSLEGWGGSYEQY | 0.64959586 |
| LISNASDALDK | ASSLEGWGGSYEQY | 0.8268808 |
| LISNASDAL | ASSLEGWGGSYEQY | 0.7660732 |
| ISNASDALDK | ASSLEGWGGSYEQY | 0.8427002 |
| NQAETLVYQTE | ASSLEGWGGSYEQY | 0.6340623 |
| TGGTGGTAGAG | ASSLEGWGGSYEQY | 0.60001576 |
| TGSTAGGAGGA | ASSLEGWGGSYEQY | 0.6835052 |
| DPLAVPVAAGPV | ASSLEGWGGSYEQY | 0.8563969 |
| GGTGGTPGNGGN | ASSLEGWGGSYEQY | 0.83146477 |
| PAGLAGVDPATL | ASSLEGWGGSYEQY | 0.7838264 |
| GNNLVVEAPHAGASV | ASSLEGWGGSYEQY | 0.61823314 |
| KVTRIMSASTLEQPA | ASSLEGWGGSYEQY | 0.7849112 |
| LGFALPIALPRRTVV | ASSLEGWGGSYEQY | 0.67383075 |
| TRLGNLTARKLVVLV | ASSLEGWGGSYEQY | 0.6842206 |
| WSQQRRLIYFPSAGP | ASSLEGWGGSYEQY | 0.80350286 |
| GPNGNPDPMAAAVDIR | ASSLEGWGGSYEQY | 0.77912045 |
| TQRCVSVADLRAAASR | ASSLEGWGGSYEQY | 0.7127427 |
| QFVLDVAEHPVEVLTETV | ASSLEGWGGSYEQY | 0.68855053 |
| GGAGGTGGAGGNGGTQAGDG | ASSLEGWGGSYEQY | 0.76340485 |
| PLLATLRAIAVDVDAKPAQV | ASSLEGWGGSYEQY | 0.7912745 |
| APKFRTGGYMIDFWGVGYQVAKRM | ASSLEGWGGSYEQY | 0.73499703 |
| GAGGAGGAGGAAGGPGGTGGTGGN | ASSLEGWGGSYEQY | 0.88406885 |
| IGALGLLATDVAPAGSVAFGVALA | ASSLEGWGGSYEQY | 0.8200668 |
| AGGAGGIGGNGGAGGAGGVLMGNG | ASSLEGWGGSYEQY | 0.813853 |
| ADELIGLL | ASSLEGWGGSYEQY | 0.21704996 |
| ADRALKA | ASSLEGWGGSYEQY | 0.44659352 |
| AGAVGQVGGAGGA | ASSLEGWGGSYEQY | 0.7202971 |
| AGGGGGLGGAP | ASSLEGWGGSYEQY | 0.6957304 |
| AGGLLYGNGGAG | ASSLEGWGGSYEQY | 0.41825622 |
| AGLESIGL | ASSLEGWGGSYEQY | 0.8306222 |
| AGLKTGII | ASSLEGWGGSYEQY | 0.7115924 |
| ASLAALLL | ASSLEGWGGSYEQY | 0.64084756 |
| ASVINISSI | ASSLEGWGGSYEQY | 0.80107135 |
| ATPPGPA | ASSLEGWGGSYEQY | 0.678449 |
| ATVASALV | ASSLEGWGGSYEQY | 0.70000136 |
| ATVASLIL | ASSLEGWGGSYEQY | 0.6830119 |
| AVNGPVLIRH | ASSLEGWGGSYEQY | 0.86124265 |
| DGAAGAVGI | ASSLEGWGGSYEQY | 0.5271617 |
| DPLLHGI | ASSLEGWGGSYEQY | 0.7948236 |
| ELGAALGAL | ASSLEGWGGSYEQY | 0.7226882 |
| ELGRTQAV | ASSLEGWGGSYEQY | 0.85110956 |
| FVAGGYLS | ASSLEGWGGSYEQY | 0.51137066 |
| GASGGAGGAGGW | ASSLEGWGGSYEQY | 0.7886516 |
| GGAGAGGDGGAGGI | ASSLEGWGGSYEQY | 0.68989146 |
| GTGHNGSQPGMGGNG | ASSLEGWGGSYEQY | 0.8812079 |
| GYAAASASAS | ASSLEGWGGSYEQY | 0.6887784 |
| HQVDVAAVQQG | ASSLEGWGGSYEQY | 0.76932895 |

| | | |
|---|---|---|
| IGAAIAHT | ASSLEGWGGSYEQY | 0.66988796 |
| IGMIEVAQL | ASSLEGWGGSYEQY | 0.8337654 |
| IRLAPEHTSAMD | ASSLEGWGGSYEQY | 0.49414814 |
| IYNVSVL | ASSLEGWGGSYEQY | 0.71481454 |
| KQGFLGM | ASSLEGWGGSYEQY | 0.61810434 |
| LESVITQL | ASSLEGWGGSYEQY | 0.79980254 |
| LGLGALVVV | ASSLEGWGGSYEQY | 0.78034836 |
| LIGNGGNGGSGGI | ASSLEGWGGSYEQY | 0.46241698 |
| LNAINAPTL | ASSLEGWGGSYEQY | 0.822893 |
| LQLLNTGV | ASSLEGWGGSYEQY | 0.72656953 |
| LQTSIEF | ASSLEGWGGSYEQY | 0.64081573 |
| MSFAVSEL | ASSLEGWGGSYEQY | 0.83910435 |
| NTGALGL | ASSLEGWGGSYEQY | 0.71173906 |
| PPGAGGAGGAGTT | ASSLEGWGGSYEQY | 0.8456746 |
| QLVGLGI | ASSLEGWGGSYEQY | 0.76136005 |
| RFLEQVHQL | ASSLEGWGGSYEQY | 0.70818055 |
| RGVGYVLR | ASSLEGWGGSYEQY | 0.83962023 |
| SATGVFAGVI | ASSLEGWGGSYEQY | 0.6022332 |
| SLPAITLP | ASSLEGWGGSYEQY | 0.63323057 |
| STALAFTV | ASSLEGWGGSYEQY | 0.8326764 |
| SVLTVGLGK | ASSLEGWGGSYEQY | 0.7110336 |
| TALAGAGSGI | ASSLEGWGGSYEQY | 0.77219653 |
| TGLPFTGI | ASSLEGWGGSYEQY | 0.5882097 |
| VAEHVTV | ASSLEGWGGSYEQY | 0.5124856 |
| VDIVLNSL | ASSLEGWGGSYEQY | 0.65538645 |
| VLTGGGAVL | ASSLEGWGGSYEQY | 0.78504264 |
| VNVGLAYGI | ASSLEGWGGSYEQY | 0.7580941 |
| VSSAAVGAV | ASSLEGWGGSYEQY | 0.73599315 |
| VTSVYEAGV | ASSLEGWGGSYEQY | 0.52858156 |
| WVALDRGA | ASSLEGWGGSYEQY | 0.6875638 |
| YLVGSVVI | ASSLEGWGGSYEQY | 0.63349026 |
| GPPGTGKTLL | ASSLEGWGGSYEQY | 0.6281729 |
| GVGMTKF | ASSLEGWGGSYEQY | 0.75794053 |
| IGIGGSDL | ASSLEGWGGSYEQY | 0.69349265 |
| IGTLLAEL | ASSLEGWGGSYEQY | 0.60610497 |
| IYTSGTT | ASSLEGWGGSYEQY | 0.6534637 |
| LSFPAGV | ASSLEGWGGSYEQY | 0.73115367 |
| LYTSGTT | ASSLEGWGGSYEQY | 0.72075236 |
| VGAFPFGF | ASSLEGWGGSYEQY | 0.70384914 |
| VGAGTVEF | ASSLEGWGGSYEQY | 0.8170425 |
| VGKSTLLL | ASSLEGWGGSYEQY | 0.75933194 |
| AALAHAGGGIV | ASSLEGWGGSYEQY | 0.7598978 |
| AALVAVSLL | ASSLEGWGGSYEQY | 0.28910854 |
| AAVSRAVAAL | ASSLEGWGGSYEQY | 0.40162376 |
| AGGSSGIGGF | ASSLEGWGGSYEQY | 0.67121637 |
| AGLTVANANG | ASSLEGWGGSYEQY | 0.6471828 |
| AGLVGAVL | ASSLEGWGGSYEQY | 0.43464255 |
| ALSQAFTV | ASSLEGWGGSYEQY | 0.76577485 |
| ALTFSVSL | ASSLEGWGGSYEQY | 0.62210906 |
| ALTGAALVAL | ASSLEGWGGSYEQY | 0.5643612 |
| ASFSIGAGVL | ASSLEGWGGSYEQY | 0.7593861 |

| | | |
|---|---|---|
| DGSVYKI | ASSLEGWGGSYEQY | 0.6048638 |
| EQLAEIV | ASSLEGWGGSYEQY | 0.6114863 |
| FAVSELTVM | ASSLEGWGGSYEQY | 0.63396466 |
| GAAPLAAGL | ASSLEGWGGSYEQY | 0.7751081 |
| GADGGDGGSGGAGG | ASSLEGWGGSYEQY | 0.51576096 |
| GFGLDTGGAGG | ASSLEGWGGSYEQY | 0.7179975 |
| GLASGGLL | ASSLEGWGGSYEQY | 0.62500614 |
| GRIAGAAVLL | ASSLEGWGGSYEQY | 0.77803206 |
| GSAGSLNTSF | ASSLEGWGGSYEQY | 0.46229702 |
| GTGQIGI | ASSLEGWGGSYEQY | 0.7460079 |
| GTLIALIL | ASSLEGWGGSYEQY | 0.64664286 |
| IAATMGFVVG | ASSLEGWGGSYEQY | 0.8193252 |
| IFAQALL | ASSLEGWGGSYEQY | 0.6451191 |
| IGAAVALGF | ASSLEGWGGSYEQY | 0.68799883 |
| IGLTPLAL | ASSLEGWGGSYEQY | 0.6337468 |
| ILGSNTSTL | ASSLEGWGGSYEQY | 0.7493211 |
| IVFSLTVL | ASSLEGWGGSYEQY | 0.43838894 |
| LASKATGF | ASSLEGWGGSYEQY | 0.6400566 |
| LASVALIL | ASSLEGWGGSYEQY | 0.48683783 |
| LFAVGLTL | ASSLEGWGGSYEQY | 0.7117278 |
| LFNGGIGTYI | ASSLEGWGGSYEQY | 0.7207475 |
| LTLGAGII | ASSLEGWGGSYEQY | 0.5137577 |
| LVAAALGDI | ASSLEGWGGSYEQY | 0.7555234 |
| NTAFVQLGI | ASSLEGWGGSYEQY | 0.7035999 |
| QLFSGAEL | ASSLEGWGGSYEQY | 0.5940075 |
| SAATFITL | ASSLEGWGGSYEQY | 0.7930502 |
| SNYGLANLGV | ASSLEGWGGSYEQY | 0.66332704 |
| SYGGAYAVM | ASSLEGWGGSYEQY | 0.779971 |
| TTGVGGLTLG | ASSLEGWGGSYEQY | 0.69173086 |
| TTQTIYI | ASSLEGWGGSYEQY | 0.6100942 |
| VAAGAGLGAVI | ASSLEGWGGSYEQY | 0.7352129 |
| VAAVVTLL | ASSLEGWGGSYEQY | 0.6403317 |
| VGFLGGLI | ASSLEGWGGSYEQY | 0.60498583 |
| VGVIPILL | ASSLEGWGGSYEQY | 0.7735719 |
| VIFASSLI | ASSLEGWGGSYEQY | 0.8078283 |
| VKGGDGGAAAG | ASSLEGWGGSYEQY | 0.78815126 |
| VTKTFTNI | ASSLEGWGGSYEQY | 0.68525606 |
| VVAAAILL | ASSLEGWGGSYEQY | 0.633817 |
| YAALDVEL | ASSLEGWGGSYEQY | 0.7297261 |
| LASVALLL | ASSLEGWGGSYEQY | 0.82308096 |
| LDAVHRV | ASSLEGWGGSYEQY | 0.82132655 |
| QGLSSSIF | ASSLEGWGGSYEQY | 0.7896832 |
| RGTFANIRL | ASSLEGWGGSYEQY | 0.59688324 |
| RQLWWGHRI | ASSLEGWGGSYEQY | 0.8150863 |
| YGPPGTGKTLL | ASSLEGWGGSYEQY | 0.76799816 |
| AALVGAPANVA | ASSLEGWGGSYEQY | 0.7565291 |
| AATFRRTE | ASSLEGWGGSYEQY | 0.8017106 |
| AGALGGGI | ASSLEGWGGSYEQY | 0.7969477 |
| AGGAGGAGGIGAG | ASSLEGWGGSYEQY | 0.75207204 |
| AGSGALGL | ASSLEGWGGSYEQY | 0.71761143 |
| AQIADQI | ASSLEGWGGSYEQY | 0.8062544 |

| | | |
|---|---|---|
| ASAVAVTYL | ASSLEGWGGSYEQY | 0.830817 |
| ASTVLGGTGG | ASSLEGWGGSYEQY | 0.82041264 |
| AVGGEPAK | ASSLEGWGGSYEQY | 0.8531635 |
| CEQGTFGGTC | ASSLEGWGGSYEQY | 0.82949555 |
| DEGKAPAG | ASSLEGWGGSYEQY | 0.8211926 |
| DTVVFPAL | ASSLEGWGGSYEQY | 0.2930462 |
| FEGITMAG | ASSLEGWGGSYEQY | 0.5108007 |
| FPLVVQD | ASSLEGWGGSYEQY | 0.7105148 |
| GAFTTPQ | ASSLEGWGGSYEQY | 0.7455335 |
| GFGGFGTAGGVGGA | ASSLEGWGGSYEQY | 0.36630324 |
| GFLAQFL | ASSLEGWGGSYEQY | 0.7987313 |
| GGAGGAGGNAGVA | ASSLEGWGGSYEQY | 0.7474277 |
| GGVGTAGAGGAGGAGG | ASSLEGWGGSYEQY | 0.6681361 |
| GNGGGGDAGNAGSGGN | ASSLEGWGGSYEQY | 0.82684577 |
| GTGGAGGAGNDAGS | ASSLEGWGGSYEQY | 0.7288569 |
| GTTNTGWFNS | ASSLEGWGGSYEQY | 0.70081115 |
| IFMGPLFTL | ASSLEGWGGSYEQY | 0.68360883 |
| ILAHAIEAI | ASSLEGWGGSYEQY | 0.84460926 |
| IRSAIREAL | ASSLEGWGGSYEQY | 0.6167837 |
| ITFAGGG | ASSLEGWGGSYEQY | 0.8249757 |
| ITGMSGHQEV | ASSLEGWGGSYEQY | 0.7031009 |
| IVAGLACTL | ASSLEGWGGSYEQY | 0.8495946 |
| IVGGGLAGA | ASSLEGWGGSYEQY | 0.50532055 |
| IVTGVLNI | ASSLEGWGGSYEQY | 0.8200763 |
| LAQQGCI | ASSLEGWGGSYEQY | 0.75719124 |
| LCLHVEN | ASSLEGWGGSYEQY | 0.86138844 |
| LDAISDGS | ASSLEGWGGSYEQY | 0.7547761 |
| LGGFAAML | ASSLEGWGGSYEQY | 0.801314 |
| LGGKNPMI | ASSLEGWGGSYEQY | 0.72845745 |
| LGPKGVAAIL | ASSLEGWGGSYEQY | 0.8100916 |
| LMVGVSI | ASSLEGWGGSYEQY | 0.50616884 |
| LVGAAFVL | ASSLEGWGGSYEQY | 0.7575513 |
| LVLAVAPL | ASSLEGWGGSYEQY | 0.5950465 |
| MAAIAVIDAA | ASSLEGWGGSYEQY | 0.7944058 |
| NNGINAELQ | ASSLEGWGGSYEQY | 0.8031553 |
| RGEPLVLVHG | ASSLEGWGGSYEQY | 0.33228385 |
| RGGAGGAAGLI | ASSLEGWGGSYEQY | 0.8167648 |
| RSYRMHPAV | ASSLEGWGGSYEQY | 0.6625477 |
| SAAHLLAGQLA | ASSLEGWGGSYEQY | 0.63164276 |
| SAAVGSY | ASSLEGWGGSYEQY | 0.8304186 |
| SPHAVLHRI | ASSLEGWGGSYEQY | 0.77959985 |
| SQIAKVL | ASSLEGWGGSYEQY | 0.8338125 |
| TALGHLVP | ASSLEGWGGSYEQY | 0.73016655 |
| TAPTTSGAA | ASSLEGWGGSYEQY | 0.7406033 |
| TDGDDGNGGSPGT | ASSLEGWGGSYEQY | 0.80950195 |
| TDGLGEN | ASSLEGWGGSYEQY | 0.48678234 |
| TGGPPLPGQA | ASSLEGWGGSYEQY | 0.6147719 |
| TIPAITI | ASSLEGWGGSYEQY | 0.84248304 |
| TIPGFTI | ASSLEGWGGSYEQY | 0.81605613 |
| TSIASGALA | ASSLEGWGGSYEQY | 0.86072934 |
| TVTLPGAAI | ASSLEGWGGSYEQY | 0.6942942 |

| | | |
|---|---|---|
| VASGLSAALP | ASSLEGWGGSYEQY | 0.60471094 |
| VGAGVIGI | ASSLEGWGGSYEQY | 0.76441324 |
| VGPKRFGV | ASSLEGWGGSYEQY | 0.87376666 |
| VNVAFPD | ASSLEGWGGSYEQY | 0.8538001 |
| VSMTNSGVS | ASSLEGWGGSYEQY | 0.8287346 |
| VTDVLGDLSV | ASSLEGWGGSYEQY | 0.5602973 |
| VVVVGAGFAGL | ASSLEGWGGSYEQY | 0.7631265 |
| VYTVGTVV | ASSLEGWGGSYEQY | 0.71879673 |
| WTTFSGP | ASSLEGWGGSYEQY | 0.7301804 |
| YDELVEAF | ASSLEGWGGSYEQY | 0.84839237 |
| YGTGGAGGNGG | ASSLEGWGGSYEQY | 0.7918881 |
| YTNVMAV | ASSLEGWGGSYEQY | 0.7158745 |
| FLEQVHQL | ASSLEGWGGSYEQY | 0.692916 |
| GLSLLAVAV | ASSLEGWGGSYEQY | 0.8051808 |
| LEQVHQL | ASSLEGWGGSYEQY | 0.863377 |
| LNAAGAI | ASSLEGWGGSYEQY | 0.84974706 |
| PAVHASL | ASSLEGWGGSYEQY | 0.8922652 |
| TELEDDA | ASSLEGWGGSYEQY | 0.85352933 |
| TPPGPPN | ASSLEGWGGSYEQY | 0.8638672 |
| TPPPPPPP | ASSLEGWGGSYEQY | 0.18238789 |
| TQSPALI | ASLSYSGNTLY | 0.3707407 |
| EFAAAFPQ | ASLSYSGNTLY | 0.7573265 |
| LLSGNLDVL | ASLSYSGNTLY | 0.7129737 |
| GPANLGNYNI | ASLSYSGNTLY | 0.5262456 |
| TLGPTLP | ASLSYSGNTLY | 0.80195475 |
| VYAYSSCCK | ASLSYSGNTLY | 0.6988411 |
| APAPAPAPNI | ASLSYSGNTLY | 0.64038885 |
| PGGPVPIP | ASLSYSGNTLY | 0.85567594 |
| FLTVALEP | ASLSYSGNTLY | 0.6973125 |
| SQYFGANL | ASLSYSGNTLY | 0.6761123 |
| GGVKPAA | ASLSYSGNTLY | 0.6701687 |
| GGTGGAGGAGGW | ASLSYSGNTLY | 0.8630835 |
| GVGGVAGLGGAGG | ASLSYSGNTLY | 0.5388725 |
| VLLVLATTI | ASLSYSGNTLY | 0.81839085 |
| IGIFAAML | ASLSYSGNTLY | 0.73536193 |
| AVGDLID | ASLSYSGNTLY | 0.874272 |
| NISPLPL | ASLSYSGNTLY | 0.477127 |
| VTAVFVAI | ASLSYSGNTLY | 0.83661085 |
| SGDDNDAAL | ASLSYSGNTLY | 0.7378016 |
| AVAALAHS | ASLSYSGNTLY | 0.89775467 |
| GGVTVMATP | ASLSYSGNTLY | 0.7353758 |
| AGLAANVRD | ASLSYSGNTLY | 0.79010487 |
| TNRPDLIDPALLRPG | ASLSYSGNTLY | 0.6691943 |
| LANLGVTAL | ASLSYSGNTLY | 0.83797944 |
| TGERIEL | ASLSYSGNTLY | 0.5357293 |
| NGEELAA | ASLSYSGNTLY | 0.74581206 |
| VLSLVWG | ASLSYSGNTLY | 0.514968 |
| IGYALWL | ASLSYSGNTLY | 0.8118762 |
| LSLLLAPL | ASLSYSGNTLY | 0.814013 |
| TLDEALK | ASLSYSGNTLY | 0.47053713 |
| TDGAGTAEGG | ASLSYSGNTLY | 0.86116016 |

| | | |
|---|---|---|
| NAPGGAGGNGG | ASLSYSGNTLY | 0.77658975 |
| VSVFMGMGGNF | ASLSYSGNTLY | 0.6902235 |
| LASYLAAL | ASLSYSGNTLY | 0.88191265 |
| LVGGTPAL | ASLSYSGNTLY | 0.7618797 |
| VRNADVL | ASLSYSGNTLY | 0.85921323 |
| TCSSPFLVSAM | ASLSYSGNTLY | 0.79343945 |
| VHITDAE | ASLSYSGNTLY | 0.7262641 |
| EYAAPSF | ASLSYSGNTLY | 0.85103667 |
| VVDAFHGGL | ASLSYSGNTLY | 0.7482039 |
| ASVVDPQ | ASLSYSGNTLY | 0.67769337 |
| SYNEFGGLS | ASLSYSGNTLY | 0.85939205 |
| DPDGSVY | ASLSYSGNTLY | 0.73000526 |
| MVSTIHATE | ASLSYSGNTLY | 0.7606522 |
| LAAGVADPV | ASLSYSGNTLY | 0.7015122 |
| TKEQIAA | ASLSYSGNTLY | 0.63376564 |
| SLASPAAL | ASLSYSGNTLY | 0.70263827 |
| IGLVFYGV | ASLSYSGNTLY | 0.7540598 |
| FLLDASNL | ASLSYSGNTLY | 0.7577981 |
| YQVGNFP | ASLSYSGNTLY | 0.7354691 |
| ESLLSQST | ASLSYSGNTLY | 0.61259735 |
| LFTVGFAAVG | ASLSYSGNTLY | 0.72490776 |
| ATPPTPT | ASLSYSGNTLY | 0.70381534 |
| TDLATLG | ASLSYSGNTLY | 0.70342696 |
| SAGGTGR | ASLSYSGNTLY | 0.7216072 |
| GFSNSGS | ASLSYSGNTLY | 0.7409178 |
| GGNGGSGGTSV | ASLSYSGNTLY | 0.6958247 |
| GTSVATGGAGN | ASLSYSGNTLY | 0.6694861 |
| NGGNGGSGGTS | ASLSYSGNTLY | 0.72978586 |
| AVVKFPH | ASLSYSGNTLY | 0.7192942 |
| TPPPPPP | ASLSYSGNTLY | 0.7027317 |
| AAPAAAAAPA | ASLSYSGNTLY | 0.7601072 |
| IALNEDL | ASLSYSGNTLY | 0.7523836 |
| IVNVSSM | ASLSYSGNTLY | 0.7429684 |
| TGIPPAPR | ASLSYSGNTLY | 0.28326085 |
| FDLGGGTF | ASLSYSGNTLY | 0.40280914 |
| FDLGGGTFDVS | ASLSYSGNTLY | 0.70207644 |
| NEPTAAA | ASLSYSGNTLY | 0.692144 |
| NEPTAAALA | ASLSYSGNTLY | 0.4375451 |
| GSIINISSI | ASLSYSGNTLY | 0.7310958 |
| VAPGNAGTA | ASLSYSGNTLY | 0.6589631 |
| TGGGASLE | ASLSYSGNTLY | 0.6798246 |
| VAVYNHYKRI | ASLSYSGNTLY | 0.7480401 |
| TDPAPATT | ASLSYSGNTLY | 0.7026115 |
| TNRPDILDPALLRPG | ASLSYSGNTLY | 0.6574186 |
| LTTTTTS | ASLSYSGNTLY | 0.6670843 |
| PTGSGKTL | ASLSYSGNTLY | 0.75124955 |
| TGSGKTL | ASLSYSGNTLY | 0.5835899 |
| TIGHVDHGK | ASLSYSGNTLY | 0.7353991 |
| TIGHVDHGKTT | ASLSYSGNTLY | 0.6349775 |
| GHVDHGKTT | ASLSYSGNTLY | 0.7549797 |
| AQFEHTLL | ASLSYSGNTLY | 0.5773848 |

| | | |
|---|---|---|
| IVNQDEL | ASLSYSGNTLY | 0.7462766 |
| TNGSQFF | ASLSYSGNTLY | 0.71866727 |
| TGKGIVE | ASLSYSGNTLY | 0.7705089 |
| LISNASDA | ASLSYSGNTLY | 0.6945158 |
| AELRTLGELTKIAKAHGAQ | ASLSYSGNTLY | 0.7203884 |
| APNAVAAAEPSW | ASLSYSGNTLY | 0.65370625 |
| APQVAGAATAVL | ASLSYSGNTLY | 0.75665855 |
| ARRDPAAAAAILPTDA | ASLSYSGNTLY | 0.587419 |
| DAAARDVLAEAGLAEYF | ASLSYSGNTLY | 0.69372976 |
| DDQIVLSAKANLAKVEQ | ASLSYSGNTLY | 0.4604475 |
| DELLPDTGEDSDEDGH | ASLSYSGNTLY | 0.7301938 |
| ELPEAPANRATVPQD | ASLSYSGNTLY | 0.73206776 |
| FGQSKIERVTGIG | ASLSYSGNTLY | 0.463876 |
| GAGGTGSTAGGAGGAGGAGGLYA | ASLSYSGNTLY | 0.74912256 |
| GHGGTGAAGGNGGTGGAGGSGIDG | ASLSYSGNTLY | 0.70532745 |
| GNGGNGGAGGEGGPGGAGGAGA | ASLSYSGNTLY | 0.67128706 |
| GSTAGNGGNGGNATNGTIAGQPAG | ASLSYSGNTLY | 0.7541213 |
| GTGGDGGNGGNGGTGG | ASLSYSGNTLY | 0.71984434 |
| GVRRPAGIPADAPV | ASLSYSGNTLY | 0.7527781 |
| GYRQVTDAHLLTLARR | ASLSYSGNTLY | 0.7203342 |
| IAKMGQDGHDRG | ASLSYSGNTLY | 0.71327245 |
| KAEPSIEIQYT | ASLSYSGNTLY | 0.73345184 |
| KPVRIRMAADEDTVAVT | ASLSYSGNTLY | 0.69083375 |
| MPPRIAGMRLLVIKPEP | ASLSYSGNTLY | 0.6833258 |
| MKFYAVALFRSRGVPPTR | ASLSYSGNTLY | 0.75382316 |
| NAGGFGILGAVAHSP | ASLSYSGNTLY | 0.83232224 |
| NEGREITLAMAEM | ASLSYSGNTLY | 0.8725646 |
| NELVEWSHPLGHRWRWE | ASLSYSGNTLY | 0.73268116 |
| NLGAGGTGGNGGNPG | ASLSYSGNTLY | 0.62804544 |
| NTEWPKWETVMAADPVD | ASLSYSGNTLY | 0.78675306 |
| PASAAPAETQP | ASLSYSGNTLY | 0.8810846 |
| QQSSDGKNGVRAAEITGE | ASLSYSGNTLY | 0.86729205 |
| RAPDAVTTSWWKE | ASLSYSGNTLY | 0.8423817 |
| SGFGNVGAHTSGWFNQ | ASLSYSGNTLY | 0.5842799 |
| SGVAGAVGQAASVGGLKV | ASLSYSGNTLY | 0.7853371 |
| SNGGAGTGSAGGNGGT | ASLSYSGNTLY | 0.75056064 |
| TGGAGGEGGAAG | ASLSYSGNTLY | 0.7592816 |
| TGGDGGAAGAGGAPGHG | ASLSYSGNTLY | 0.85615486 |
| TMPPLLAAIANAGRHGV | ASLSYSGNTLY | 0.8106003 |
| TSSTAGNSGGA | ASLSYSGNTLY | 0.73266757 |
| VPATVPVGILC | ASLSYSGNTLY | 0.7201004 |
| VQRPDSGGVVLTGRIS | ASLSYSGNTLY | 0.8289284 |
| WTPNPVPATKGVPAGLVQAT | ASLSYSGNTLY | 0.87376153 |
| YALKRALAKFSDRFDVVII | ASLSYSGNTLY | 0.8650766 |
| YLIGSVIV | ASLSYSGNTLY | 0.89640534 |
| YGQLVAVID | ASLSYSGNTLY | 0.8663657 |
| VTVGGMTLPA | ASLSYSGNTLY | 0.8754271 |
| VLALPAV | ASLSYSGNTLY | 0.19330056 |
| VIPPTLNYET | ASLSYSGNTLY | 0.4280934 |
| VGMGGMHPSSG | ASLSYSGNTLY | 0.76499635 |
| VGLAGGAATA | ASLSYSGNTLY | 0.758029 |

| | | |
|---|---|---|
| VGGAGGAGGSGP | ASLSYSGNTLY | 0.5653057 |
| VFAAFMV | ASLSYSGNTLY | 0.81624293 |
| TLYDKLR | ASLSYSGNTLY | 0.7153386 |
| TLAVFTVL | ASLSYSGNTLY | 0.6668186 |
| TGSVQPARV | ASLSYSGNTLY | 0.8692694 |
| TGGGAGSGGT | ASLSYSGNTLY | 0.717582 |
| SHLETSL | ASLSYSGNTLY | 0.69877344 |
| SFTTGMT | ASLSYSGNTLY | 0.70100856 |
| QVEVAKRMGVRQPTVS | ASLSYSGNTLY | 0.8762952 |
| QGSGGPALAM | ASLSYSGNTLY | 0.537079 |
| MLAPSNP | ASLSYSGNTLY | 0.8376981 |
| LVNNAAI | ASLSYSGNTLY | 0.754846 |
| LTGEINTLA | ASLSYSGNTLY | 0.8832059 |
| LSIFGPV | ASLSYSGNTLY | 0.4967381 |
| LLALLANL | ASLSYSGNTLY | 0.85228616 |
| LISNASDALDKLR | ASLSYSGNTLY | 0.7709388 |
| LINEAVP | ASLSYSGNTLY | 0.90202284 |
| LGYAFSGL | ASLSYSGNTLY | 0.76256764 |
| LDFVAVAL | ASLSYSGNTLY | 0.8006739 |
| LAQPGAE | ASLSYSGNTLY | 0.6852807 |
| ISNLGAVTGF | ASLSYSGNTLY | 0.8557011 |
| HQEDPGRI | ASLSYSGNTLY | 0.5714998 |
| GTPGEPTP | ASLSYSGNTLY | 0.77717924 |
| GFFNSGS | ASLSYSGNTLY | 0.55309236 |
| GAGGGGLTAGAN | ASLSYSGNTLY | 0.8252671 |
| FVDALVNM | ASLSYSGNTLY | 0.8338514 |
| FLTVAPEL | ASLSYSGNTLY | 0.50865024 |
| FETLTAL | ASLSYSGNTLY | 0.8700095 |
| DVINPIEL | ASLSYSGNTLY | 0.8049493 |
| APVSGAPV | ASLSYSGNTLY | 0.71269107 |
| ALVGGLAW | ASLSYSGNTLY | 0.8889208 |
| ALAAAAATAI | ASLSYSGNTLY | 0.7959161 |
| AFGLGLLL | ASLSYSGNTLY | 0.86926115 |
| AFFAAASL | ASLSYSGNTLY | 0.8092867 |
| AAAALPAFSPP | ASLSYSGNTLY | 0.7491385 |
| VVLAPAL | ASLSYSGNTLY | 0.8602463 |
| TIGHVDHGKTTL | ASLSYSGNTLY | 0.7749514 |
| LISNASDALDK | ASLSYSGNTLY | 0.7090825 |
| LISNASDAL | ASLSYSGNTLY | 0.87046695 |
| ISNASDALDK | ASLSYSGNTLY | 0.7466692 |
| NQAETLVYQTE | ASLSYSGNTLY | 0.78002304 |
| TGGTGGTAGAG | ASLSYSGNTLY | 0.54379636 |
| TGSTAGGAGGA | ASLSYSGNTLY | 0.5753027 |
| DPLAVPVAAGPV | ASLSYSGNTLY | 0.6700016 |
| GGTGGTPGNGGN | ASLSYSGNTLY | 0.8120929 |
| PAGLAGVDPATL | ASLSYSGNTLY | 0.8230274 |
| GNNLVVEAPHAGASV | ASLSYSGNTLY | 0.77206993 |
| KVTRIMSASTLEQPA | ASLSYSGNTLY | 0.6544363 |
| LGFALPIALPRRTVV | ASLSYSGNTLY | 0.77031016 |
| TRLGNLTARKLVVLV | ASLSYSGNTLY | 0.6668431 |
| WSQQRRLIYFPSAGP | ASLSYSGNTLY | 0.693275 |

| | | |
|---|---|---|
| GPNGNPDPMAAAVDIR | ASLSYSGNTLY | 0.7822745 |
| TQRCVSVADLRAAASR | ASLSYSGNTLY | 0.74130917 |
| QFVLDVAEHPVEVLTETV | ASLSYSGNTLY | 0.6891848 |
| GGAGGTGGAGGNGGTQAGDG | ASLSYSGNTLY | 0.62497675 |
| PLLATLRAIAVDVDAKPAQV | ASLSYSGNTLY | 0.77586555 |
| APKFRTGGYMIDFWGVGYQVAKRM | ASLSYSGNTLY | 0.8069384 |
| GAGGAGGAGGAAGGPGGTGGTGGN | ASLSYSGNTLY | 0.7772835 |
| IGALGLLATDVAPAGSVAFGVALA | ASLSYSGNTLY | 0.8502234 |
| AGGAGGIGGNGGAGGAGGVLMGNG | ASLSYSGNTLY | 0.77600324 |
| ADELIGLL | ASLSYSGNTLY | 0.7830546 |
| ADRALKA | ASLSYSGNTLY | 0.26522204 |
| AGAVGQVGGAGGA | ASLSYSGNTLY | 0.44954917 |
| AGGGGGLGGAP | ASLSYSGNTLY | 0.69750655 |
| AGGLLYGNGGAG | ASLSYSGNTLY | 0.69931185 |
| AGLESIGL | ASLSYSGNTLY | 0.33014894 |
| AGLKTGII | ASLSYSGNTLY | 0.7974175 |
| ASLAALLL | ASLSYSGNTLY | 0.6900737 |
| ASVINISSI | ASLSYSGNTLY | 0.62876713 |
| ATPPGPA | ASLSYSGNTLY | 0.7955122 |
| ATVASALV | ASLSYSGNTLY | 0.65853995 |
| ATVASLIL | ASLSYSGNTLY | 0.6370257 |
| AVNGPVLIRH | ASLSYSGNTLY | 0.6819626 |
| DGAAGAVGI | ASLSYSGNTLY | 0.8302088 |
| DPLLHGI | ASLSYSGNTLY | 0.58699346 |
| ELGAALGAL | ASLSYSGNTLY | 0.78103125 |
| ELGRTQAV | ASLSYSGNTLY | 0.7383407 |
| FVAGGYLS | ASLSYSGNTLY | 0.81802404 |
| GASGGAGGAGGW | ASLSYSGNTLY | 0.5194495 |
| GGAGAGGDGGAGGI | ASLSYSGNTLY | 0.7728783 |
| GTGHNGSQPGMGGNG | ASLSYSGNTLY | 0.6592144 |
| GYAAASASAS | ASLSYSGNTLY | 0.8498448 |
| HQVDVAAVQQG | ASLSYSGNTLY | 0.65977573 |
| IGAAIAHT | ASLSYSGNTLY | 0.77965117 |
| IGMIEVAQL | ASLSYSGNTLY | 0.76627004 |
| IRLAPEHTSAMD | ASLSYSGNTLY | 0.7756188 |
| IYNVSVL | ASLSYSGNTLY | 0.47216648 |
| KQGFLGM | ASLSYSGNTLY | 0.6487036 |
| LESVITQL | ASLSYSGNTLY | 0.679129 |
| LGLGALVVV | ASLSYSGNTLY | 0.74005705 |
| LIGNGGNGGSGGI | ASLSYSGNTLY | 0.7608912 |
| LNAINAPTL | ASLSYSGNTLY | 0.31899703 |
| LQLLNTGV | ASLSYSGNTLY | 0.7818762 |
| LQTSIEF | ASLSYSGNTLY | 0.57914764 |
| MSFAVSEL | ASLSYSGNTLY | 0.62821585 |
| NTGALGL | ASLSYSGNTLY | 0.79587084 |
| PPGAGGAGGAGTT | ASLSYSGNTLY | 0.71773565 |
| QLVGLGI | ASLSYSGNTLY | 0.7668326 |
| RFLEQVHQL | ASLSYSGNTLY | 0.6969608 |
| RGVGYVLR | ASLSYSGNTLY | 0.69357896 |
| SATGVFAGVI | ASLSYSGNTLY | 0.7869023 |
| SLPAITLP | ASLSYSGNTLY | 0.4588181 |

| | | |
|---|---|---|
| STALAFTV | ASLSYSGNTLY | 0.5950651 |
| SVLTVGLGK | ASLSYSGNTLY | 0.80996007 |
| TALAGAGSGI | ASLSYSGNTLY | 0.84394485 |
| TGLPFTGI | ASLSYSGNTLY | 0.846887 |
| VAEHVTV | ASLSYSGNTLY | 0.7769255 |
| VDIVLNSL | ASLSYSGNTLY | 0.7341244 |
| VLTGGGAVL | ASLSYSGNTLY | 0.79474384 |
| VNVGLAYGI | ASLSYSGNTLY | 0.8495743 |
| VSSAAVGAV | ASLSYSGNTLY | 0.8511207 |
| VTSVYEAGV | ASLSYSGNTLY | 0.8384596 |
| WVALDRGA | ASLSYSGNTLY | 0.63019943 |
| YLVGSVVI | ASLSYSGNTLY | 0.837951 |
| GPPGTGKTLL | ASLSYSGNTLY | 0.80407 |
| GVGMTKF | ASLSYSGNTLY | 0.80664515 |
| IGIGGSDL | ASLSYSGNTLY | 0.8408482 |
| IGTLLAEL | ASLSYSGNTLY | 0.82539845 |
| IYTSGTT | ASLSYSGNTLY | 0.80154604 |
| LSFPAGV | ASLSYSGNTLY | 0.7464863 |
| LYTSGTT | ASLSYSGNTLY | 0.8368881 |
| VGAFPFGF | ASLSYSGNTLY | 0.8517713 |
| VGAGTVEF | ASLSYSGNTLY | 0.8490716 |
| VGKSTLLL | ASLSYSGNTLY | 0.854639 |
| AALAHAGGGIV | ASLSYSGNTLY | 0.8515056 |
| AALVAVSLL | ASLSYSGNTLY | 0.849586 |
| AAVSRAVAAL | ASLSYSGNTLY | 0.27762017 |
| AGGSSGIGGF | ASLSYSGNTLY | 0.5136151 |
| AGLTVANANG | ASLSYSGNTLY | 0.7805384 |
| AGLVGAVL | ASLSYSGNTLY | 0.7749748 |
| ALSQAFTV | ASLSYSGNTLY | 0.44060338 |
| ALTFSVSL | ASLSYSGNTLY | 0.8301096 |
| ALTGAALVAL | ASLSYSGNTLY | 0.7757712 |
| ASFSIGAGVL | ASLSYSGNTLY | 0.75433254 |
| DGSVYKI | ASLSYSGNTLY | 0.8449449 |
| EQLAEIV | ASLSYSGNTLY | 0.79668385 |
| FAVSELTVM | ASLSYSGNTLY | 0.7460287 |
| GAAPLAAGL | ASLSYSGNTLY | 0.7412132 |
| GADGGDGGSGGAGG | ASLSYSGNTLY | 0.85239565 |
| GFGLDTGGAGG | ASLSYSGNTLY | 0.7043835 |
| GLASGGLL | ASLSYSGNTLY | 0.8461292 |
| GRIAGAAVLL | ASLSYSGNTLY | 0.7015732 |
| GSAGSLNTSF | ASLSYSGNTLY | 0.8533099 |
| GTGQIGI | ASLSYSGNTLY | 0.5564853 |
| GTLIALIL | ASLSYSGNTLY | 0.84965503 |
| IAATMGFVVG | ASLSYSGNTLY | 0.83254725 |
| IFAQALL | ASLSYSGNTLY | 0.8622308 |
| IGAAVALGF | ASLSYSGNTLY | 0.8140015 |
| IGLTPLAL | ASLSYSGNTLY | 0.8282155 |
| ILGSNTSTL | ASLSYSGNTLY | 0.7832625 |
| IVFSLTVL | ASLSYSGNTLY | 0.8443432 |
| LASKATGF | ASLSYSGNTLY | 0.6059692 |
| LASVALIL | ASLSYSGNTLY | 0.81186676 |

| | | |
|---|---|---|
| LFAVGLTL | ASLSYSGNTLY | 0.5799125 |
| LFNGGIGTYI | ASLSYSGNTLY | 0.829813 |
| LTLGAGII | ASLSYSGNTLY | 0.8389893 |
| LVAAALGDI | ASLSYSGNTLY | 0.40564242 |
| NTAFVQLGI | ASLSYSGNTLY | 0.83897924 |
| QLFSGAEL | ASLSYSGNTLY | 0.808337 |
| SAATFITL | ASLSYSGNTLY | 0.7115915 |
| SNYGLANLGV | ASLSYSGNTLY | 0.85510373 |
| SYGGAYAVM | ASLSYSGNTLY | 0.81750876 |
| TTGVGGLTLG | ASLSYSGNTLY | 0.8517722 |
| TTQTIYI | ASLSYSGNTLY | 0.7984836 |
| VAAGAGLGAVI | ASLSYSGNTLY | 0.78734756 |
| VAAVVTLL | ASLSYSGNTLY | 0.82484114 |
| VGFLGGLI | ASLSYSGNTLY | 0.53664494 |
| VGVIPILL | ASLSYSGNTLY | 0.7046311 |
| VIFASSLI | ASLSYSGNTLY | 0.8586493 |
| VKGGDGGAAAG | ASLSYSGNTLY | 0.7053169 |
| VTKTFTNI | ASLSYSGNTLY | 0.7867321 |
| VVAAAILL | ASLSYSGNTLY | 0.57678354 |
| YAALDVEL | ASLSYSGNTLY | 0.5413074 |
| LASVALLL | ASLSYSGNTLY | 0.6583668 |
| LDAVHRV | ASLSYSGNTLY | 0.73304945 |
| QGLSSSIF | ASLSYSGNTLY | 0.78289837 |
| RGTFANIRL | ASLSYSGNTLY | 0.7417509 |
| RQLWWGHRI | ASLSYSGNTLY | 0.4814608 |
| YGPPGTGKTLL | ASLSYSGNTLY | 0.6887078 |
| AALVGAPANVA | ASLSYSGNTLY | 0.53316313 |
| AATFRRTE | ASLSYSGNTLY | 0.62286747 |
| AGALGGGI | ASLSYSGNTLY | 0.6979842 |
| AGGAGGAGGIGAG | ASLSYSGNTLY | 0.5703161 |
| AGSGALGL | ASLSYSGNTLY | 0.5559625 |
| AQIADQI | ASLSYSGNTLY | 0.5447256 |
| ASAVAVTYL | ASLSYSGNTLY | 0.67667747 |
| ASTVLGGTGG | ASLSYSGNTLY | 0.7771367 |
| AVGGEPAK | ASLSYSGNTLY | 0.7567909 |
| CEQGTFGGTC | ASLSYSGNTLY | 0.7852025 |
| DEGKAPAG | ASLSYSGNTLY | 0.76291555 |
| DTVVFPAL | ASLSYSGNTLY | 0.76679134 |
| FEGITMAG | ASLSYSGNTLY | 0.22251126 |
| FPLVVQD | ASLSYSGNTLY | 0.38771528 |
| GAFTTPQ | ASLSYSGNTLY | 0.4764651 |
| GFGGFGTAGGVGGA | ASLSYSGNTLY | 0.6257987 |
| GFLAQFL | ASLSYSGNTLY | 0.27183297 |
| GGAGGAGGNAGVA | ASLSYSGNTLY | 0.7026311 |
| GGVGTAGAGGAGGAGG | ASLSYSGNTLY | 0.6000829 |
| GNGGGGDAGNAGSGGN | ASLSYSGNTLY | 0.48556072 |
| GTGGAGGAGNDAGS | ASLSYSGNTLY | 0.7603085 |
| GTTNTGWFNS | ASLSYSGNTLY | 0.52686703 |
| IFMGPLFTL | ASLSYSGNTLY | 0.4302796 |
| ILAHAIEAI | ASLSYSGNTLY | 0.48486644 |
| IRSAIREAL | ASLSYSGNTLY | 0.7541121 |

| | | |
|---|---|---|
| ITFAGGG | ASLSYSGNTLY | 0.33327413 |
| ITGMSGHQEV | ASLSYSGNTLY | 0.7201604 |
| IVAGLACTL | ASLSYSGNTLY | 0.40012938 |
| IVGGGLAGA | ASLSYSGNTLY | 0.7602246 |
| IVTGVLNI | ASLSYSGNTLY | 0.378579 |
| LAQQGCI | ASLSYSGNTLY | 0.74843335 |
| LCLHVEN | ASLSYSGNTLY | 0.61396337 |
| LDAISDGS | ASLSYSGNTLY | 0.81041014 |
| LGGFAAML | ASLSYSGNTLY | 0.61561537 |
| LGGKNPMI | ASLSYSGNTLY | 0.6538148 |
| LGPKGVAAIL | ASLSYSGNTLY | 0.41132963 |
| LMVGVSI | ASLSYSGNTLY | 0.77638817 |
| LVGAAFVL | ASLSYSGNTLY | 0.32953057 |
| LVLAVAPL | ASLSYSGNTLY | 0.6612861 |
| MAAIAVIDAA | ASLSYSGNTLY | 0.34945786 |
| NNGINAELQ | ASLSYSGNTLY | 0.7042471 |
| RGEPLVLVHG | ASLSYSGNTLY | 0.73720634 |
| RGGAGGAAGLI | ASLSYSGNTLY | 0.32478112 |
| RSYRMHPAV | ASLSYSGNTLY | 0.6875071 |
| SAAHLLAGQLA | ASLSYSGNTLY | 0.5531538 |
| SAAVGSY | ASLSYSGNTLY | 0.4950401 |
| SPHAVLHRI | ASLSYSGNTLY | 0.7719363 |
| SQIAKVL | ASLSYSGNTLY | 0.64007306 |
| TALGHLVP | ASLSYSGNTLY | 0.7725946 |
| TAPTTSGAA | ASLSYSGNTLY | 0.5198142 |
| TDGDDGNGGSPGT | ASLSYSGNTLY | 0.5024605 |
| TDGLGEN | ASLSYSGNTLY | 0.72097605 |
| TGGPPLPGQA | ASLSYSGNTLY | 0.5063852 |
| TIPAITI | ASLSYSGNTLY | 0.42063892 |
| TIPGFTI | ASLSYSGNTLY | 0.7687309 |
| TSIASGALA | ASLSYSGNTLY | 0.68965614 |
| TVTLPGAAI | ASLSYSGNTLY | 0.7038357 |
| VASGLSAALP | ASLSYSGNTLY | 0.63055074 |
| VGAGVIGI | ASLSYSGNTLY | 0.63475084 |
| VGPKRFGV | ASLSYSGNTLY | 0.64586747 |
| VNVAFPD | ASLSYSGNTLY | 0.6996262 |
| VSMTNSGVS | ASLSYSGNTLY | 0.71048295 |
| VTDVLGDLSV | ASLSYSGNTLY | 0.6999519 |
| VVVVGAGFAGL | ASLSYSGNTLY | 0.65555274 |
| VYTVGTVV | ASLSYSGNTLY | 0.6959441 |
| WTTFSGP | ASLSYSGNTLY | 0.6696617 |
| YDELVEAF | ASLSYSGNTLY | 0.67234504 |
| YGTGGAGGNGG | ASLSYSGNTLY | 0.69149524 |
| YTNVMAV | ASLSYSGNTLY | 0.6914693 |
| FLEQVHQL | ASLSYSGNTLY | 0.64219666 |
| GLSLLAVAV | ASLSYSGNTLY | 0.64047873 |
| LEQVHQL | ASLSYSGNTLY | 0.67940223 |
| LNAAGAI | ASLSYSGNTLY | 0.69828296 |
| PAVHASL | ASLSYSGNTLY | 0.6854899 |
| TELEDDA | ASLSYSGNTLY | 0.7135769 |
| TPPGPPN | ASLSYSGNTLY | 0.6960253 |

| | | |
|---|---|---|
| TPPPPPPP | ASLSYSGNTLY | 0.6907909 |
| TQSPALI | ASSPPGGTGNTLY | 0.40889794 |
| EFAAAFPQ | ASSPPGGTGNTLY | 0.5855614 |
| LLSGNLDVL | ASSPPGGTGNTLY | 0.6458594 |
| GPANLGNYNI | ASSPPGGTGNTLY | 0.67397624 |
| TLGPTLP | ASSPPGGTGNTLY | 0.3313627 |
| VYAYSSCCK | ASSPPGGTGNTLY | 0.6985253 |
| APAPAPAPNI | ASSPPGGTGNTLY | 0.672687 |
| PGGPVPIP | ASSPPGGTGNTLY | 0.64677095 |
| FLTVALEP | ASSPPGGTGNTLY | 0.69768983 |
| SQYFGANL | ASSPPGGTGNTLY | 0.6600529 |
| GGVKPAA | ASSPPGGTGNTLY | 0.6409649 |
| GGTGGAGGAGGW | ASSPPGGTGNTLY | 0.6650092 |
| GVGGVAGLGGAGG | ASSPPGGTGNTLY | 0.70882726 |
| VLLVLATTI | ASSPPGGTGNTLY | 0.59545434 |
| IGIFAAML | ASSPPGGTGNTLY | 0.7032026 |
| AVGDLID | ASSPPGGTGNTLY | 0.6538987 |
| NISPLPL | ASSPPGGTGNTLY | 0.70373106 |
| VTAVFVAI | ASSPPGGTGNTLY | 0.6238822 |
| SGDDNDAAL | ASSPPGGTGNTLY | 0.70291317 |
| AVAALAHS | ASSPPGGTGNTLY | 0.67135215 |
| GGVTVMATP | ASSPPGGTGNTLY | 0.71399456 |
| AGLAANVRD | ASSPPGGTGNTLY | 0.6680263 |
| TNRPDLIDPALLRPG | ASSPPGGTGNTLY | 0.701779 |
| LANLGVTAL | ASSPPGGTGNTLY | 0.6804953 |
| TGERIEL | ASSPPGGTGNTLY | 0.6988889 |
| NGEELAA | ASSPPGGTGNTLY | 0.59578764 |
| VLSLVWG | ASSPPGGTGNTLY | 0.66053784 |
| IGYALWL | ASSPPGGTGNTLY | 0.6198679 |
| LSLLLAPL | ASSPPGGTGNTLY | 0.6824154 |
| TLDEALK | ASSPPGGTGNTLY | 0.70028377 |
| TDGAGTAEGG | ASSPPGGTGNTLY | 0.43053862 |
| NAPGGAGGNGG | ASSPPGGTGNTLY | 0.69287133 |
| VSVFMGMGGNF | ASSPPGGTGNTLY | 0.63311404 |
| LASYLAAL | ASSPPGGTGNTLY | 0.6442698 |
| LVGGTPAL | ASSPPGGTGNTLY | 0.69910896 |
| VRNADVL | ASSPPGGTGNTLY | 0.67524624 |
| TCSSPFLVSAM | ASSPPGGTGNTLY | 0.6902895 |
| VHITDAE | ASSPPGGTGNTLY | 0.6489423 |
| EYAAPSF | ASSPPGGTGNTLY | 0.6663712 |
| VVDAFHGGL | ASSPPGGTGNTLY | 0.69089675 |
| ASVVDPQ | ASSPPGGTGNTLY | 0.42390785 |
| SYNEFGGLS | ASSPPGGTGNTLY | 0.6014148 |
| DPDGSVY | ASSPPGGTGNTLY | 0.70976937 |
| MVSTIHATE | ASSPPGGTGNTLY | 0.6864321 |
| LAAGVADPV | ASSPPGGTGNTLY | 0.6963798 |
| TKEQIAA | ASSPPGGTGNTLY | 0.6320643 |
| SLASPAAL | ASSPPGGTGNTLY | 0.6030957 |
| IGLVFYGV | ASSPPGGTGNTLY | 0.6237032 |
| FLLDASNL | ASSPPGGTGNTLY | 0.6813648 |
| YQVGNFP | ASSPPGGTGNTLY | 0.7060694 |

| | | |
|---|---|---|
| ESLLSQST | ASSPPGGTGNTLY | 0.6891247 |
| LFTVGFAAVG | ASSPPGGTGNTLY | 0.5574498 |
| ATPPTPT | ASSPPGGTGNTLY | 0.7083995 |
| TDLATLG | ASSPPGGTGNTLY | 0.67677575 |
| SAGGTGR | ASSPPGGTGNTLY | 0.6685795 |
| GFSNSGS | ASSPPGGTGNTLY | 0.67076445 |
| GGNGGSGGTSV | ASSPPGGTGNTLY | 0.6735225 |
| GTSVATGGAGN | ASSPPGGTGNTLY | 0.6356356 |
| NGGNGGSGGTS | ASSPPGGTGNTLY | 0.57132345 |
| AVVKFPH | ASSPPGGTGNTLY | 0.664885 |
| TPPPPPP | ASSPPGGTGNTLY | 0.69421685 |
| AAPAAAAAPA | ASSPPGGTGNTLY | 0.6891557 |
| IALNEDL | ASSPPGGTGNTLY | 0.70890707 |
| IVNVSSM | ASSPPGGTGNTLY | 0.70221406 |
| TGIPPAPR | ASSPPGGTGNTLY | 0.67957103 |
| FDLGGGTF | ASSPPGGTGNTLY | 0.37871873 |
| FDLGGGTFDVS | ASSPPGGTGNTLY | 0.567718 |
| NEPTAAA | ASSPPGGTGNTLY | 0.54320794 |
| NEPTAAALA | ASSPPGGTGNTLY | 0.6659939 |
| GSIINISSI | ASSPPGGTGNTLY | 0.33603144 |
| VAPGNAGTA | ASSPPGGTGNTLY | 0.66898376 |
| TGGGASLE | ASSPPGGTGNTLY | 0.6608859 |
| VAVYNHYKRI | ASSPPGGTGNTLY | 0.63142085 |
| TDPAPATT | ASSPPGGTGNTLY | 0.7017748 |
| TNRPDILDPALLRPG | ASSPPGGTGNTLY | 0.64720756 |
| LTTTTTS | ASSPPGGTGNTLY | 0.57068074 |
| PTGSGKTL | ASSPPGGTGNTLY | 0.57775813 |
| TGSGKTL | ASSPPGGTGNTLY | 0.7028699 |
| TIGHVDHGK | ASSPPGGTGNTLY | 0.56987715 |
| TIGHVDHGKTT | ASSPPGGTGNTLY | 0.709118 |
| GHVDHGKTT | ASSPPGGTGNTLY | 0.51024336 |
| AQFEHTLL | ASSPPGGTGNTLY | 0.7064242 |
| IVNQDEL | ASSPPGGTGNTLY | 0.5384741 |
| TNGSQFF | ASSPPGGTGNTLY | 0.7015927 |
| TGKGIVE | ASSPPGGTGNTLY | 0.6705228 |
| LISNASDA | ASSPPGGTGNTLY | 0.715512 |
| AELRTLGELTKIAKAHGAQ | ASSPPGGTGNTLY | 0.65831393 |
| APNAVAAAEPSW | ASSPPGGTGNTLY | 0.69695145 |
| APQVAGAATAVL | ASSPPGGTGNTLY | 0.611897 |
| ARRDPAAAAAILPTDA | ASSPPGGTGNTLY | 0.7034731 |
| DAAARDVLAEAGLAEYF | ASSPPGGTGNTLY | 0.55163366 |
| DDQIVLSAKANLAKVEQ | ASSPPGGTGNTLY | 0.6626179 |
| DELLPDTGEDSDEDGH | ASSPPGGTGNTLY | 0.5549928 |
| ELPEAPANRATVPQD | ASSPPGGTGNTLY | 0.6735828 |
| FGQSKIERVTGIG | ASSPPGGTGNTLY | 0.6988446 |
| GAGGTGSTAGGAGGAGGAGGLYA | ASSPPGGTGNTLY | 0.36172935 |
| GHGGTGAAGGNGGTGGAGGSGIDG | ASSPPGGTGNTLY | 0.69800234 |
| GNGGNGGAGGEGGPGGAGGAGA | ASSPPGGTGNTLY | 0.5864806 |
| GSTAGNGGNGGNATNGTIAGQPAG | ASSPPGGTGNTLY | 0.5780807 |
| GTGGDGGNGGNGGTGG | ASSPPGGTGNTLY | 0.6942197 |
| GVRRPAGIPADAPV | ASSPPGGTGNTLY | 0.6847378 |

| | | |
|---|---|---|
| GYRQVTDAHLLTLARR | ASSPPGGTGNTLY | 0.687986 |
| IAKMGQDGHDRG | ASSPPGGTGNTLY | 0.5693698 |
| KAEPSIEIQYT | ASSPPGGTGNTLY | 0.6636125 |
| KPVRIRMAADEDTVAVT | ASSPPGGTGNTLY | 0.6836895 |
| MPPRIAGMRLLVIKPEP | ASSPPGGTGNTLY | 0.40883982 |
| MKFYAVALFRSRGVPPTR | ASSPPGGTGNTLY | 0.51881826 |
| NAGGFGILGAVAHSP | ASSPPGGTGNTLY | 0.71075815 |
| NEGREITLAMAEM | ASSPPGGTGNTLY | 0.6808887 |
| NELVEWSHPLGHRWRWE | ASSPPGGTGNTLY | 0.6807864 |
| NLGAGGTGGNGGNPG | ASSPPGGTGNTLY | 0.6399187 |
| NTEWPKWETVMAADPVD | ASSPPGGTGNTLY | 0.6168888 |
| PASAAPAETQP | ASSPPGGTGNTLY | 0.61441416 |
| QQSSDGKNGVRAAEITGE | ASSPPGGTGNTLY | 0.6783133 |
| RAPDAVTTSWWKE | ASSPPGGTGNTLY | 0.68703663 |
| SGFGNVGAHTSGWFNQ | ASSPPGGTGNTLY | 0.6823983 |
| SGVAGAVGQAASVGGLKV | ASSPPGGTGNTLY | 0.5295108 |
| SNGGAGTGSAGGNGGT | ASSPPGGTGNTLY | 0.6826575 |
| TGGAGGEGGAAG | ASSPPGGTGNTLY | 0.66756594 |
| TGGDGGAAGAGGAPGHG | ASSPPGGTGNTLY | 0.6678679 |
| TMPPLLAAIANAGRHGV | ASSPPGGTGNTLY | 0.6665077 |
| TSSTAGNSGGA | ASSPPGGTGNTLY | 0.6564237 |
| VPATVPVGILC | ASSPPGGTGNTLY | 0.6004938 |
| VQRPDSGGVVLTGRIS | ASSPPGGTGNTLY | 0.59710103 |
| WTPNPVPATKGVPAGLVQAT | ASSPPGGTGNTLY | 0.6550491 |
| YALKRALAKFSDRFDVVII | ASSPPGGTGNTLY | 0.68692046 |
| YLIGSVIV | ASSPPGGTGNTLY | 0.6867837 |
| YGQLVAVID | ASSPPGGTGNTLY | 0.6904751 |
| VTVGGMTLPA | ASSPPGGTGNTLY | 0.6897433 |
| VLALPAV | ASSPPGGTGNTLY | 0.68158746 |
| VIPPTLNYET | ASSPPGGTGNTLY | 0.38680312 |
| VGMGGMHPSSG | ASSPPGGTGNTLY | 0.55888206 |
| VGLAGGAATA | ASSPPGGTGNTLY | 0.58697486 |
| VGGAGGAGGSGP | ASSPPGGTGNTLY | 0.6530935 |
| VFAAFMV | ASSPPGGTGNTLY | 0.40901145 |
| TLYDKLR | ASSPPGGTGNTLY | 0.6104648 |
| TLAVFTVL | ASSPPGGTGNTLY | 0.6380918 |
| TGSVQPARV | ASSPPGGTGNTLY | 0.624529 |
| TGGGAGSGGT | ASSPPGGTGNTLY | 0.6843306 |
| SHLETSL | ASSPPGGTGNTLY | 0.64746964 |
| SFTTGMT | ASSPPGGTGNTLY | 0.5782487 |
| QVEVAKRMGVRQPTVS | ASSPPGGTGNTLY | 0.558609 |
| QGSGGPALAM | ASSPPGGTGNTLY | 0.68072695 |
| MLAPSNP | ASSPPGGTGNTLY | 0.5972194 |
| LVNNAAI | ASSPPGGTGNTLY | 0.6902494 |
| LTGEINTLA | ASSPPGGTGNTLY | 0.49532312 |
| LSIFGPV | ASSPPGGTGNTLY | 0.69314647 |
| LLALLANL | ASSPPGGTGNTLY | 0.55231166 |
| LISNASDALDKLR | ASSPPGGTGNTLY | 0.69024765 |
| LINEAVP | ASSPPGGTGNTLY | 0.6705762 |
| LGYAFSGL | ASSPPGGTGNTLY | 0.69899726 |
| LDFVAVAL | ASSPPGGTGNTLY | 0.6637258 |

| | | |
|---|---|---|
| LAQPGAE | ASSPPGGTGNTLY | 0.67613006 |
| ISNLGAVTGF | ASSPPGGTGNTLY | 0.5663971 |
| HQEDPGRI | ASSPPGGTGNTLY | 0.68229073 |
| GTPGEPTP | ASSPPGGTGNTLY | 0.55282307 |
| GFFNSGS | ASSPPGGTGNTLY | 0.65290165 |
| GAGGGGLTAGAN | ASSPPGGTGNTLY | 0.53718233 |
| FVDALVNM | ASSPPGGTGNTLY | 0.66020566 |
| FLTVAPEL | ASSPPGGTGNTLY | 0.6855241 |
| FETLTAL | ASSPPGGTGNTLY | 0.3905962 |
| DVINPIEL | ASSPPGGTGNTLY | 0.6825502 |
| APVSGAPV | ASSPPGGTGNTLY | 0.63429177 |
| ALVGGLAW | ASSPPGGTGNTLY | 0.6037326 |
| ALAAAAATAI | ASSPPGGTGNTLY | 0.6893513 |
| AFGLGLLL | ASSPPGGTGNTLY | 0.6646494 |
| AFFAAASL | ASSPPGGTGNTLY | 0.68161714 |
| AAAALPAFSPP | ASSPPGGTGNTLY | 0.57409614 |
| VVLAPAL | ASSPPGGTGNTLY | 0.6335316 |
| TIGHVDHGKTTL | ASSPPGGTGNTLY | 0.6621448 |
| LISNASDALDK | ASSPPGGTGNTLY | 0.6087922 |
| LISNASDAL | ASSPPGGTGNTLY | 0.5404495 |
| ISNASDALDK | ASSPPGGTGNTLY | 0.69469273 |
| NQAETLVYQTE | ASSPPGGTGNTLY | 0.6982275 |
| TGGTGGTAGAG | ASSPPGGTGNTLY | 0.7771275 |
| TGSTAGGAGGA | ASSPPGGTGNTLY | 0.49612415 |
| DPLAVPVAAGPV | ASSPPGGTGNTLY | 0.43727678 |
| GGTGGTPGNGGN | ASSPPGGTGNTLY | 0.6290843 |
| PAGLAGVDPATL | ASSPPGGTGNTLY | 0.7773807 |
| GNNLVVEAPHAGASV | ASSPPGGTGNTLY | 0.7754373 |
| KVTRIMSASTLEQPA | ASSPPGGTGNTLY | 0.7274918 |
| LGFALPIALPRRTVV | ASSPPGGTGNTLY | 0.4090879 |
| TRLGNLTARKLVVLV | ASSPPGGTGNTLY | 0.6085856 |
| WSQQRRLIYFPSAGP | ASSPPGGTGNTLY | 0.5738784 |
| GPNGNPDPMAAAVDIR | ASSPPGGTGNTLY | 0.5648127 |
| TQRCVSVADLRAAASR | ASSPPGGTGNTLY | 0.70601475 |
| QFVLDVAEHPVEVLTETV | ASSPPGGTGNTLY | 0.6122515 |
| GGAGGTGGAGGNGGTQAGDG | ASSPPGGTGNTLY | 0.47819257 |
| PLLATLRAIAVDVDAKPAQV | ASSPPGGTGNTLY | 0.5094434 |
| APKFRTGGYMIDFWGVGYQVAKRM | ASSPPGGTGNTLY | 0.6261629 |
| GAGGAGGAGGAAGGPGGTGGTGGN | ASSPPGGTGNTLY | 0.75820136 |
| IGALGLLATDVAPAGSVAFGVALA | ASSPPGGTGNTLY | 0.73227227 |
| AGGAGGIGGNGGAGGAGGVLMGNG | ASSPPGGTGNTLY | 0.7948258 |
| ADELIGLL | ASSPPGGTGNTLY | 0.74362123 |
| ADRALKA | ASSPPGGTGNTLY | 0.7387997 |
| AGAVGQVGGAGGA | ASSPPGGTGNTLY | 0.18468039 |
| AGGGGGLGGAP | ASSPPGGTGNTLY | 0.2654702 |
| AGGLLYGNGGAG | ASSPPGGTGNTLY | 0.5554941 |
| AGLESIGL | ASSPPGGTGNTLY | 0.60121363 |
| AGLKTGII | ASSPPGGTGNTLY | 0.35431093 |
| ASLAALLL | ASSPPGGTGNTLY | 0.7382946 |
| ASVINISSI | ASSPPGGTGNTLY | 0.5593434 |
| ATPPGPA | ASSPPGGTGNTLY | 0.46785718 |

| | | |
|---|---|---|
| ATVASALV | ASSPPGGTGNTLY | 0.75149345 |
| ATVASLIL | ASSPPGGTGNTLY | 0.48605168 |
| AVNGPVLIRH | ASSPPGGTGNTLY | 0.46903458 |
| DGAAGAVGI | ASSPPGGTGNTLY | 0.49084878 |
| DPLLHGI | ASSPPGGTGNTLY | 0.7571917 |
| ELGAALGAL | ASSPPGGTGNTLY | 0.34972197 |
| ELGRTQAV | ASSPPGGTGNTLY | 0.691479 |
| FVAGGYLS | ASSPPGGTGNTLY | 0.53416246 |
| GASGGAGGAGGW | ASSPPGGTGNTLY | 0.744313 |
| GGAGAGGDGGAGGI | ASSPPGGTGNTLY | 0.3570828 |
| GTGHNGSQPGMGGNG | ASSPPGGTGNTLY | 0.7184001 |
| GYAAASASAS | ASSPPGGTGNTLY | 0.59047705 |
| HQVDVAAVQQG | ASSPPGGTGNTLY | 0.8000536 |
| IGAAIAHT | ASSPPGGTGNTLY | 0.5706029 |
| IGMIEVAQL | ASSPPGGTGNTLY | 0.6567751 |
| IRLAPEHTSAMD | ASSPPGGTGNTLY | 0.46347594 |
| IYNVSVL | ASSPPGGTGNTLY | 0.7368882 |
| KQGFLGM | ASSPPGGTGNTLY | 0.42560565 |
| LESVITQL | ASSPPGGTGNTLY | 0.597702 |
| LGLGALVVV | ASSPPGGTGNTLY | 0.33935246 |
| LIGNGGNGGSGGI | ASSPPGGTGNTLY | 0.67619574 |
| LNAINAPTL | ASSPPGGTGNTLY | 0.6938801 |
| LQLLNTGV | ASSPPGGTGNTLY | 0.39027685 |
| LQTSIEF | ASSPPGGTGNTLY | 0.6675509 |
| MSFAVSEL | ASSPPGGTGNTLY | 0.69727457 |
| NTGALGL | ASSPPGGTGNTLY | 0.5217482 |
| PPGAGGAGGAGTT | ASSPPGGTGNTLY | 0.76555586 |
| QLVGLGI | ASSPPGGTGNTLY | 0.5823768 |
| RFLEQVHQL | ASSPPGGTGNTLY | 0.7469362 |
| RGVGYVLR | ASSPPGGTGNTLY | 0.53577435 |
| SATGVFAGVI | ASSPPGGTGNTLY | 0.52891016 |
| SLPAITLP | ASSPPGGTGNTLY | 0.7335538 |
| STALAFTV | ASSPPGGTGNTLY | 0.55652785 |
| SVLTVGLGK | ASSPPGGTGNTLY | 0.48503712 |
| TALAGAGSGI | ASSPPGGTGNTLY | 0.7336965 |
| TGLPFTGI | ASSPPGGTGNTLY | 0.6566634 |
| VAEHVTV | ASSPPGGTGNTLY | 0.7059827 |
| VDIVLNSL | ASSPPGGTGNTLY | 0.39428818 |
| VLTGGGAVL | ASSPPGGTGNTLY | 0.35263902 |
| VNVGLAYGI | ASSPPGGTGNTLY | 0.39419731 |
| VSSAAVGAV | ASSPPGGTGNTLY | 0.7120633 |
| VTSVYEAGV | ASSPPGGTGNTLY | 0.73969626 |
| WVALDRGA | ASSPPGGTGNTLY | 0.6683389 |
| YLVGSVVI | ASSPPGGTGNTLY | 0.36858177 |
| GPPGTGKTLL | ASSPPGGTGNTLY | 0.5927409 |
| GVGMTKF | ASSPPGGTGNTLY | 0.43931046 |
| IGIGGSDL | ASSPPGGTGNTLY | 0.51130265 |
| IGTLLAEL | ASSPPGGTGNTLY | 0.52259403 |
| IYTSGTT | ASSPPGGTGNTLY | 0.6205448 |
| LSFPAGV | ASSPPGGTGNTLY | 0.43492717 |
| LYTSGTT | ASSPPGGTGNTLY | 0.38821915 |

| | | |
|---|---|---|
| VGAFPFGF | ASSPPGGTGNTLY | 0.56431293 |
| VGAGTVEF | ASSPPGGTGNTLY | 0.7204174 |
| VGKSTLLL | ASSPPGGTGNTLY | 0.65218794 |
| AALAHAGGGIV | ASSPPGGTGNTLY | 0.75847214 |
| AALVAVSLL | ASSPPGGTGNTLY | 0.70210147 |
| AAVSRAVAAL | ASSPPGGTGNTLY | 0.63010347 |
| AGGSSGIGGF | ASSPPGGTGNTLY | 0.17497063 |
| AGLTVANANG | ASSPPGGTGNTLY | 0.27692595 |
| AGLVGAVL | ASSPPGGTGNTLY | 0.3958197 |
| ALSQAFTV | ASSPPGGTGNTLY | 0.4896242 |
| ALTFSVSL | ASSPPGGTGNTLY | 0.16823155 |
| ALTGAALVAL | ASSPPGGTGNTLY | 0.61920846 |
| ASFSIGAGVL | ASSPPGGTGNTLY | 0.46681568 |
| DGSVYKI | ASSPPGGTGNTLY | 0.3802337 |
| EQLAEIV | ASSPPGGTGNTLY | 0.6605854 |
| FAVSELTVM | ASSPPGGTGNTLY | 0.44409198 |
| GAAPLAAGL | ASSPPGGTGNTLY | 0.4208318 |
| GADGGDGGSGGAGG | ASSPPGGTGNTLY | 0.41682407 |
| GFGLDTGGAGG | ASSPPGGTGNTLY | 0.7127309 |
| GLASGGLL | ASSPPGGTGNTLY | 0.27320728 |
| GRIAGAAVLL | ASSPPGGTGNTLY | 0.6834508 |
| GSAGSLNTSF | ASSPPGGTGNTLY | 0.33003044 |
| GTGQIGI | ASSPPGGTGNTLY | 0.7121761 |
| GTLIALIL | ASSPPGGTGNTLY | 0.31937382 |
| IAATMGFVVG | ASSPPGGTGNTLY | 0.7068533 |
| IFAQALL | ASSPPGGTGNTLY | 0.56184435 |
| IGAAVALGF | ASSPPGGTGNTLY | 0.80571735 |
| IGLTPLAL | ASSPPGGTGNTLY | 0.46124822 |
| ILGSNTSTL | ASSPPGGTGNTLY | 0.587439 |
| IVFSLTVL | ASSPPGGTGNTLY | 0.504511 |
| LASKATGF | ASSPPGGTGNTLY | 0.7224369 |
| LASVALIL | ASSPPGGTGNTLY | 0.25467902 |
| LFAVGLTL | ASSPPGGTGNTLY | 0.4807561 |
| LFNGGIGTYI | ASSPPGGTGNTLY | 0.26864737 |
| LTLGAGII | ASSPPGGTGNTLY | 0.5714999 |
| LVAAALGDI | ASSPPGGTGNTLY | 0.69047356 |
| NTAFVQLGI | ASSPPGGTGNTLY | 0.1728208 |
| QLFSGAEL | ASSPPGGTGNTLY | 0.5343037 |
| SAATFITL | ASSPPGGTGNTLY | 0.3765684 |
| SNYGLANLGV | ASSPPGGTGNTLY | 0.3519846 |
| SYGGAYAVM | ASSPPGGTGNTLY | 0.6360818 |
| TTGVGGLTLG | ASSPPGGTGNTLY | 0.54338735 |
| TTQTIYI | ASSPPGGTGNTLY | 0.7003857 |
| VAAGAGLGAVI | ASSPPGGTGNTLY | 0.40336582 |
| VAAVVTLL | ASSPPGGTGNTLY | 0.38813084 |
| VGFLGGLI | ASSPPGGTGNTLY | 0.62397826 |
| VGVIPILL | ASSPPGGTGNTLY | 0.2527951 |
| VIFASSLI | ASSPPGGTGNTLY | 0.32996288 |
| VKGGDGGAAAG | ASSPPGGTGNTLY | 0.7623122 |
| VTKTFTNI | ASSPPGGTGNTLY | 0.80085444 |
| VVAAAILL | ASSPPGGTGNTLY | 0.83826387 |

| | | |
|---|---|---|
| YAALDVEL | ASSPPGGTGNTLY | 0.6606102 |
| LASVALLL | ASSPPGGTGNTLY | 0.60722816 |
| LDAVHRV | ASSPPGGTGNTLY | 0.7069081 |
| QGLSSSIF | ASSPPGGTGNTLY | 0.8255177 |
| RGTFANIRL | ASSPPGGTGNTLY | 0.85003674 |
| RQLWWGHRI | ASSPPGGTGNTLY | 0.8089099 |
| YGPPGTGKTLL | ASSPPGGTGNTLY | 0.6353762 |
| AALVGAPANVA | ASSPPGGTGNTLY | 0.80459166 |
| AATFRRTE | ASSPPGGTGNTLY | 0.74944466 |
| AGALGGGI | ASSPPGGTGNTLY | 0.73997027 |
| AGGAGGAGGIGAG | ASSPPGGTGNTLY | 0.8048745 |
| AGSGALGL | ASSPPGGTGNTLY | 0.78664887 |
| AQIADQI | ASSPPGGTGNTLY | 0.70752376 |
| ASAVAVTYL | ASSPPGGTGNTLY | 0.6972053 |
| ASTVLGGTGG | ASSPPGGTGNTLY | 0.7875532 |
| AVGGEPAK | ASSPPGGTGNTLY | 0.8358016 |
| CEQGTFGGTC | ASSPPGGTGNTLY | 0.82360333 |
| DEGKAPAG | ASSPPGGTGNTLY | 0.8709428 |
| DTVVFPAL | ASSPPGGTGNTLY | 0.833648 |
| FEGITMAG | ASSPPGGTGNTLY | 0.83819824 |
| FPLVVQD | ASSPPGGTGNTLY | 0.28734314 |
| GAFTTPQ | ASSPPGGTGNTLY | 0.5597978 |
| GFGGFGTAGGVGGA | ASSPPGGTGNTLY | 0.68622017 |
| GFLAQFL | ASSPPGGTGNTLY | 0.74823886 |
| GGAGGAGGNAGVA | ASSPPGGTGNTLY | 0.37664348 |
| GGVGTAGAGGAGGAGG | ASSPPGGTGNTLY | 0.82258475 |
| GNGGGGDAGNAGSGGN | ASSPPGGTGNTLY | 0.7408105 |
| GTGGAGGAGNDAGS | ASSPPGGTGNTLY | 0.6292292 |
| GTTNTGWFNS | ASSPPGGTGNTLY | 0.8369411 |
| IFMGPLFTL | ASSPPGGTGNTLY | 0.7051998 |
| ILAHAIEAI | ASSPPGGTGNTLY | 0.67196923 |
| IRSAIREAL | ASSPPGGTGNTLY | 0.716079 |
| ITFAGGG | ASSPPGGTGNTLY | 0.85414326 |
| ITGMSGHQEV | ASSPPGGTGNTLY | 0.60258996 |
| IVAGLACTL | ASSPPGGTGNTLY | 0.8263064 |
| IVGGGLAGA | ASSPPGGTGNTLY | 0.70169234 |
| IVTGVLNI | ASSPPGGTGNTLY | 0.85515416 |
| LAQQGCI | ASSPPGGTGNTLY | 0.5310998 |
| LCLHVEN | ASSPPGGTGNTLY | 0.83354706 |
| LDAISDGS | ASSPPGGTGNTLY | 0.74442697 |
| LGGFAAML | ASSPPGGTGNTLY | 0.8884933 |
| LGGKNPMI | ASSPPGGTGNTLY | 0.7354602 |
| LGPKGVAAIL | ASSPPGGTGNTLY | 0.8064052 |
| LMVGVSI | ASSPPGGTGNTLY | 0.7467818 |
| LVGAAFVL | ASSPPGGTGNTLY | 0.84110296 |
| LVLAVAPL | ASSPPGGTGNTLY | 0.4719211 |
| MAAIAVIDAA | ASSPPGGTGNTLY | 0.7522724 |
| NNGINAELQ | ASSPPGGTGNTLY | 0.61590934 |
| RGEPLVLVHG | ASSPPGGTGNTLY | 0.7866812 |
| RGGAGGAAGLI | ASSPPGGTGNTLY | 0.81276655 |
| RSYRMHPAV | ASSPPGGTGNTLY | 0.33145532 |

| | | |
|---|---|---|
| SAAHLLAGQLA | ASSPPGGTGNTLY | 0.8313956 |
| SAAVGSY | ASSPPGGTGNTLY | 0.670461 |
| SPHAVLHRI | ASSPPGGTGNTLY | 0.6328299 |
| SQIAKVL | ASSPPGGTGNTLY | 0.84895205 |
| TALGHLVP | ASSPPGGTGNTLY | 0.7709651 |
| TAPTTSGAA | ASSPPGGTGNTLY | 0.8367017 |
| TDGDDGNGGSPGT | ASSPPGGTGNTLY | 0.7518332 |
| TDGLGEN | ASSPPGGTGNTLY | 0.7589407 |
| TGGPPLPGQA | ASSPPGGTGNTLY | 0.81115806 |
| TIPAITI | ASSPPGGTGNTLY | 0.58329415 |
| TIPGFTI | ASSPPGGTGNTLY | 0.649343 |
| TSIASGALA | ASSPPGGTGNTLY | 0.8613116 |
| TVTLPGAAI | ASSPPGGTGNTLY | 0.72449124 |
| VASGLSAALP | ASSPPGGTGNTLY | 0.8086207 |
| VGAGVIGI | ASSPPGGTGNTLY | 0.62381434 |
| VGPKRFGV | ASSPPGGTGNTLY | 0.49324393 |
| VNVAFPD | ASSPPGGTGNTLY | 0.72204447 |
| VSMTNSGVS | ASSPPGGTGNTLY | 0.71559834 |
| VTDVLGDLSV | ASSPPGGTGNTLY | 0.81970143 |
| VVVVGAGFAGL | ASSPPGGTGNTLY | 0.7463875 |
| VYTVGTVV | ASSPPGGTGNTLY | 0.39038154 |
| WTTFSGP | ASSPPGGTGNTLY | 0.766663 |
| YDELVEAF | ASSPPGGTGNTLY | 0.67163396 |
| YGTGGAGGNGG | ASSPPGGTGNTLY | 0.61554545 |
| YTNVMAV | ASSPPGGTGNTLY | 0.66534996 |
| FLEQVHQL | ASSPPGGTGNTLY | 0.6939564 |
| GLSLLAVAV | ASSPPGGTGNTLY | 0.63291055 |
| LEQVHQL | ASSPPGGTGNTLY | 0.4566459 |
| LNAAGAI | ASSPPGGTGNTLY | 0.73723304 |
| PAVHASL | ASSPPGGTGNTLY | 0.78202957 |
| TELEDDA | ASSPPGGTGNTLY | 0.7862562 |
| TPPGPPN | ASSPPGGTGNTLY | 0.7595595 |
| TPPPPPPP | ASSPPGGTGNTLY | 0.7698528 |
| TQSPALI | ASSPPVYEQY | 0.72698283 |
| EFAAAFPQ | ASSPPVYEQY | 0.19032271 |
| LLSGNLDVL | ASSPPVYEQY | 0.2559687 |
| GPANLGNYNI | ASSPPVYEQY | 0.48171538 |
| TLGPTLP | ASSPPVYEQY | 0.56116915 |
| VYAYSSCCK | ASSPPVYEQY | 0.23870316 |
| APAPAPAPNI | ASSPPVYEQY | 0.59826523 |
| PGGPVPIP | ASSPPVYEQY | 0.6155391 |
| FLTVALEP | ASSPPVYEQY | 0.5028228 |
| SQYFGANL | ASSPPVYEQY | 0.8029889 |
| GGVKPAA | ASSPPVYEQY | 0.6034353 |
| GGTGGAGGAGGW | ASSPPVYEQY | 0.4121909 |
| GVGGVAGLGGAGG | ASSPPVYEQY | 0.45390898 |
| VLLVLATTI | ASSPPVYEQY | 0.72299266 |
| IGIFAAML | ASSPPVYEQY | 0.331874 |
| AVGDLID | ASSPPVYEQY | 0.76443756 |
| NISPLPL | ASSPPVYEQY | 0.42014468 |
| VTAVFVAI | ASSPPVYEQY | 0.755088 |

| | | |
|---|---|---|
| SGDDNDAAL | ASSPPVYEQY | 0.35083005 |
| AVAALAHS | ASSPPVYEQY | 0.7343366 |
| GGVTVMATP | ASSPPVYEQY | 0.6625501 |
| AGLAANVRD | ASSPPVYEQY | 0.83261085 |
| TNRPDLIDPALLRPG | ASSPPVYEQY | 0.59720206 |
| LANLGVTAL | ASSPPVYEQY | 0.68196636 |
| TGERIEL | ASSPPVYEQY | 0.44730186 |
| NGEELAA | ASSPPVYEQY | 0.8076408 |
| VLSLVWG | ASSPPVYEQY | 0.36680442 |
| IGYALWL | ASSPPVYEQY | 0.6397881 |
| LSLLLAPL | ASSPPVYEQY | 0.29683864 |
| TLDEALK | ASSPPVYEQY | 0.67668223 |
| TDGAGTAEGG | ASSPPVYEQY | 0.71479475 |
| NAPGGAGGNGG | ASSPPVYEQY | 0.2857421 |
| VSVFMGMGGNF | ASSPPVYEQY | 0.79364777 |
| LASYLAAL | ASSPPVYEQY | 0.5793926 |
| LVGGTPAL | ASSPPVYEQY | 0.49794504 |
| VRNADVL | ASSPPVYEQY | 0.77253896 |
| TCSSPFLVSAM | ASSPPVYEQY | 0.65633404 |
| VHITDAE | ASSPPVYEQY | 0.79042244 |
| EYAAPSF | ASSPPVYEQY | 0.5093461 |
| VVDAFHGGL | ASSPPVYEQY | 0.6312035 |
| ASVVDPQ | ASSPPVYEQY | 0.68641675 |
| SYNEFGGLS | ASSPPVYEQY | 0.5359895 |
| DPDGSVY | ASSPPVYEQY | 0.45309776 |
| MVSTIHATE | ASSPPVYEQY | 0.7644124 |
| LAAGVADPV | ASSPPVYEQY | 0.6399102 |
| TKEQIAA | ASSPPVYEQY | 0.6859475 |
| SLASPAAL | ASSPPVYEQY | 0.6052374 |
| IGLVFYGV | ASSPPVYEQY | 0.54873663 |
| FLLDASNL | ASSPPVYEQY | 0.62548816 |
| YQVGNFP | ASSPPVYEQY | 0.67315257 |
| ESLLSQST | ASSPPVYEQY | 0.68805915 |
| LFTVGFAAVG | ASSPPVYEQY | 0.6659177 |
| ATPPTPT | ASSPPVYEQY | 0.51268363 |
| TDLATLG | ASSPPVYEQY | 0.68140924 |
| SAGGTGR | ASSPPVYEQY | 0.61105406 |
| GFSNSGS | ASSPPVYEQY | 0.62684107 |
| GGNGGSGGTSV | ASSPPVYEQY | 0.66371554 |
| GTSVATGGAGN | ASSPPVYEQY | 0.6414934 |
| NGGNGGSGGTS | ASSPPVYEQY | 0.62151337 |
| AVVKFPH | ASSPPVYEQY | 0.5821637 |
| TPPPPPP | ASSPPVYEQY | 0.65864325 |
| AAPAAAAAPA | ASSPPVYEQY | 0.6711874 |
| IALNEDL | ASSPPVYEQY | 0.6650293 |
| IVNVSSM | ASSPPVYEQY | 0.7012781 |
| TGIPPAPR | ASSPPVYEQY | 0.6770625 |
| FDLGGGTF | ASSPPVYEQY | 0.6779543 |
| FDLGGGTFDVS | ASSPPVYEQY | 0.34646857 |
| NEPTAAA | ASSPPVYEQY | 0.4405673 |
| NEPTAAALA | ASSPPVYEQY | 0.56021076 |

| | | |
|---|---|---|
| GSIINISSI | ASSPPVYEQY | 0.61838657 |
| VAPGNAGTA | ASSPPVYEQY | 0.37350106 |
| TGGGASLE | ASSPPVYEQY | 0.6548409 |
| VAVYNHYKRI | ASSPPVYEQY | 0.6271919 |
| TDPAPATT | ASSPPVYEQY | 0.5641892 |
| TNRPDILDPALLRPG | ASSPPVYEQY | 0.6843027 |
| LTTTTTS | ASSPPVYEQY | 0.6070509 |
| PTGSGKTL | ASSPPVYEQY | 0.5639137 |
| TGSGKTL | ASSPPVYEQY | 0.5627834 |
| TIGHVDHGK | ASSPPVYEQY | 0.6809126 |
| TIGHVDHGKTT | ASSPPVYEQY | 0.47010148 |
| GHVDHGKTT | ASSPPVYEQY | 0.6789773 |
| AQFEHTLL | ASSPPVYEQY | 0.53038895 |
| IVNQDEL | ASSPPVYEQY | 0.67296326 |
| TNGSQFF | ASSPPVYEQY | 0.46939886 |
| TGKGIVE | ASSPPVYEQY | 0.67584 |
| LISNASDA | ASSPPVYEQY | 0.61681 |
| AELRTLGELTKIAKAHGAQ | ASSPPVYEQY | 0.7076917 |
| APNAVAAAEPSW | ASSPPVYEQY | 0.613196 |
| APQVAGAATAVL | ASSPPVYEQY | 0.6487403 |
| ARRDPAAAAAILPTDA | ASSPPVYEQY | 0.53806925 |
| DAAARDVLAEAGLAEYF | ASSPPVYEQY | 0.68753934 |
| DDQIVLSAKANLAKVEQ | ASSPPVYEQY | 0.42984393 |
| DELLPDTGEDSDEDGH | ASSPPVYEQY | 0.6263168 |
| ELPEAPANRATVPQD | ASSPPVYEQY | 0.5279294 |
| FGQSKIERVTGIG | ASSPPVYEQY | 0.66527784 |
| GAGGTGSTAGGAGGAGGAGGLYA | ASSPPVYEQY | 0.66236407 |
| GHGGTGAAGGNGGTGGAGGSGIDG | ASSPPVYEQY | 0.4706412 |
| GNGGNGGAGGEGGPGGAGGAGA | ASSPPVYEQY | 0.69172096 |
| GSTAGNGGNGGNATNGTIAGQPAG | ASSPPVYEQY | 0.6370813 |
| GTGGDGGNGGNGGTGG | ASSPPVYEQY | 0.557545 |
| GVRRPAGIPADAPV | ASSPPVYEQY | 0.68783057 |
| GYRQVTDAHLLTLARR | ASSPPVYEQY | 0.6558173 |
| IAKMGQDGHDRG | ASSPPVYEQY | 0.66968024 |
| KAEPSIEIQYT | ASSPPVYEQY | 0.62181085 |
| KPVRIRMAADEDTVAVT | ASSPPVYEQY | 0.62510026 |
| MPPRIAGMRLLVIKPEP | ASSPPVYEQY | 0.6706731 |
| MKFYAVALFRSRGVPPTR | ASSPPVYEQY | 0.51891375 |
| NAGGFGILGAVAHSP | ASSPPVYEQY | 0.5087594 |
| NEGREITLAMAEM | ASSPPVYEQY | 0.68549156 |
| NELVEWSHPLGHRWRWE | ASSPPVYEQY | 0.8240542 |
| NLGAGGTGGNGGNPG | ASSPPVYEQY | 0.8123746 |
| NTEWPKWETVMAADPVD | ASSPPVYEQY | 0.6836258 |
| PASAAPAETQP | ASSPPVYEQY | 0.60374767 |
| QQSSDGKNGVRAAEITGE | ASSPPVYEQY | 0.6700609 |
| RAPDAVTTSWWKE | ASSPPVYEQY | 0.80766046 |
| SGFGNVGAHTSGWFNQ | ASSPPVYEQY | 0.8310634 |
| SGVAGAVGQAASVGGLKV | ASSPPVYEQY | 0.8131359 |
| SNGGAGTGSAGGNGGT | ASSPPVYEQY | 0.4853835 |
| TGGAGGEGGAAG | ASSPPVYEQY | 0.82213664 |
| TGGDGGAAGAGGAPGHG | ASSPPVYEQY | 0.7833061 |

| | | |
|---|---|---|
| TMPPLLAAIANAGRHGV | ASSPPVYEQY | 0.7940182 |
| TSSTAGNSGGA | ASSPPVYEQY | 0.74134207 |
| VPATVPVGILC | ASSPPVYEQY | 0.7253995 |
| VQRPDSGGVVLTGRIS | ASSPPVYEQY | 0.65433466 |
| WTPNPVPATKGVPAGLVQAT | ASSPPVYEQY | 0.5368632 |
| YALKRALAKFSDRFDVVII | ASSPPVYEQY | 0.775398 |
| YLIGSVIV | ASSPPVYEQY | 0.8182506 |
| YGQLVAVID | ASSPPVYEQY | 0.82050633 |
| VTVGGMTLPA | ASSPPVYEQY | 0.84861887 |
| VLALPAV | ASSPPVYEQY | 0.8463613 |
| VIPPTLNYET | ASSPPVYEQY | 0.80867773 |
| VGMGGMHPSSG | ASSPPVYEQY | 0.26758277 |
| VGLAGGAATA | ASSPPVYEQY | 0.48169023 |
| VGGAGGAGGSGP | ASSPPVYEQY | 0.56237113 |
| VFAAFMV | ASSPPVYEQY | 0.71586436 |
| TLYDKLR | ASSPPVYEQY | 0.2868298 |
| TLAVFTVL | ASSPPVYEQY | 0.62844896 |
| TGSVQPARV | ASSPPVYEQY | 0.659631 |
| TGGGAGSGGT | ASSPPVYEQY | 0.6513599 |
| SHLETSL | ASSPPVYEQY | 0.826306 |
| SFTTGMT | ASSPPVYEQY | 0.7131239 |
| QVEVAKRMGVRQPTVS | ASSPPVYEQY | 0.5002342 |
| QGSGGPALAM | ASSPPVYEQY | 0.5141657 |
| MLAPSNP | ASSPPVYEQY | 0.82593596 |
| LVNNAAI | ASSPPVYEQY | 0.58023006 |
| LTGEINTLA | ASSPPVYEQY | 0.84444547 |
| LSIFGPV | ASSPPVYEQY | 0.4980754 |
| LLALLANL | ASSPPVYEQY | 0.8503773 |
| LISNASDALDKLR | ASSPPVYEQY | 0.46711725 |
| LINEAVP | ASSPPVYEQY | 0.8433403 |
| LGYAFSGL | ASSPPVYEQY | 0.79305553 |
| LDFVAVAL | ASSPPVYEQY | 0.8659695 |
| LAQPGAE | ASSPPVYEQY | 0.739226 |
| ISNLGAVTGF | ASSPPVYEQY | 0.78282696 |
| HQEDPGRI | ASSPPVYEQY | 0.6289658 |
| GTPGEPTP | ASSPPVYEQY | 0.8082176 |
| GFFNSGS | ASSPPVYEQY | 0.47979563 |
| GAGGGGLTAGAN | ASSPPVYEQY | 0.7100328 |
| FVDALVNM | ASSPPVYEQY | 0.48330265 |
| FLTVAPEL | ASSPPVYEQY | 0.74210715 |
| FETLTAL | ASSPPVYEQY | 0.8302803 |
| DVINPIEL | ASSPPVYEQY | 0.20872785 |
| APVSGAPV | ASSPPVYEQY | 0.8015147 |
| ALVGGLAW | ASSPPVYEQY | 0.6463256 |
| ALAAAAATAI | ASSPPVYEQY | 0.5146315 |
| AFGLGLLL | ASSPPVYEQY | 0.8284125 |
| AFFAAASL | ASSPPVYEQY | 0.7675277 |
| AAAALPAFSPP | ASSPPVYEQY | 0.8285369 |
| VVLAPAL | ASSPPVYEQY | 0.6050191 |
| TIGHVDHGKTTL | ASSPPVYEQY | 0.7046876 |
| LISNASDALDK | ASSPPVYEQY | 0.75281197 |

| | | |
|---|---|---|
| LISNASDAL | ASSPPVYEQY | 0.35357457 |
| ISNASDALDK | ASSPPVYEQY | 0.45831177 |
| NQAETLVYQTE | ASSPPVYEQY | 0.85849 |
| TGGTGGTAGAG | ASSPPVYEQY | 0.72384983 |
| TGSTAGGAGGA | ASSPPVYEQY | 0.75135034 |
| DPLAVPVAAGPV | ASSPPVYEQY | 0.6316393 |
| GGTGGTPGNGGN | ASSPPVYEQY | 0.5669028 |
| PAGLAGVDPATL | ASSPPVYEQY | 0.60725546 |
| GNNLVVEAPHAGASV | ASSPPVYEQY | 0.7392103 |
| KVTRIMSASTLEQPA | ASSPPVYEQY | 0.7479011 |
| LGFALPIALPRRTVV | ASSPPVYEQY | 0.7044464 |
| TRLGNLTARKLVVLV | ASSPPVYEQY | 0.52678806 |
| WSQQRRLIYFPSAGP | ASSPPVYEQY | 0.7020075 |
| GPNGNPDPMAAAVDIR | ASSPPVYEQY | 0.66238475 |
| TQRCVSVADLRAAASR | ASSPPVYEQY | 0.66222596 |
| QFVLDVAEHPVEVLTETV | ASSPPVYEQY | 0.6569675 |
| GGAGGTGGAGGNGGTQAGDG | ASSPPVYEQY | 0.7276271 |
| PLLATLRAIAVDVDAKPAQV | ASSPPVYEQY | 0.6407167 |
| APKFRTGGYMIDFWGVGYQVAKRM | ASSPPVYEQY | 0.5503197 |
| GAGGAGGAGGAAGGPGGTGGTGGN | ASSPPVYEQY | 0.6802695 |
| IGALGLLATDVAPAGSVAFGVALA | ASSPPVYEQY | 0.7402125 |
| AGGAGGIGGNGGAGGAGGVLMGNG | ASSPPVYEQY | 0.7182497 |
| ADELIGLL | ASSPPVYEQY | 0.7472931 |
| ADRALKA | ASSPPVYEQY | 0.74189466 |
| AGAVGQVGGAGGA | ASSPPVYEQY | 0.71061677 |
| AGGGGGLGGAP | ASSPPVYEQY | 0.2699734 |
| AGGLLYGNGGAG | ASSPPVYEQY | 0.33120692 |
| AGLESIGL | ASSPPVYEQY | 0.5873642 |
| AGLKTGII | ASSPPVYEQY | 0.6070058 |
| ASLAALLL | ASSPPVYEQY | 0.3082769 |
| ASVINISSI | ASSPPVYEQY | 0.6981876 |
| ATPPGPA | ASSPPVYEQY | 0.5941292 |
| ATVASALV | ASSPPVYEQY | 0.6146878 |
| ATVASLIL | ASSPPVYEQY | 0.7318317 |
| AVNGPVLIRH | ASSPPVYEQY | 0.6280519 |
| DGAAGAVGI | ASSPPVYEQY | 0.5730315 |
| DPLLHGI | ASSPPVYEQY | 0.49570185 |
| ELGAALGAL | ASSPPVYEQY | 0.73384774 |
| ELGRTQAV | ASSPPVYEQY | 0.37289977 |
| FVAGGYLS | ASSPPVYEQY | 0.7233329 |
| GASGGAGGAGGW | ASSPPVYEQY | 0.46692318 |
| GGAGAGGDGGAGGI | ASSPPVYEQY | 0.7388717 |
| GTGHNGSQPGMGGNG | ASSPPVYEQY | 0.51025045 |
| GYAAASASAS | ASSPPVYEQY | 0.73181397 |
| HQVDVAAVQQG | ASSPPVYEQY | 0.6800772 |
| IGAAIAHT | ASSPPVYEQY | 0.76735306 |
| IGMIEVAQL | ASSPPVYEQY | 0.64487165 |
| IRLAPEHTSAMD | ASSPPVYEQY | 0.68598807 |
| IYNVSVL | ASSPPVYEQY | 0.5594491 |
| KQGFLGM | ASSPPVYEQY | 0.7361541 |
| LESVITQL | ASSPPVYEQY | 0.48204035 |

| | | |
|---|---|---|
| LGLGALVVV | ASSPPVYEQY | 0.6505843 |
| LIGNGGNGGSGGI | ASSPPVYEQY | 0.42849767 |
| LNAINAPTL | ASSPPVYEQY | 0.68224347 |
| LQLLNTGV | ASSPPVYEQY | 0.7188262 |
| LQTSIEF | ASSPPVYEQY | 0.31330305 |
| MSFAVSEL | ASSPPVYEQY | 0.7036028 |
| NTGALGL | ASSPPVYEQY | 0.6266874 |
| PPGAGGAGGAGTT | ASSPPVYEQY | 0.5632696 |
| QLVGLGI | ASSPPVYEQY | 0.7323668 |
| RFLEQVHQL | ASSPPVYEQY | 0.70413715 |
| RGVGYVLR | ASSPPVYEQY | 0.7419578 |
| SATGVFAGVI | ASSPPVYEQY | 0.6263846 |
| SLPAITLP | ASSPPVYEQY | 0.64345014 |
| STALAFTV | ASSPPVYEQY | 0.7062103 |
| SVLTVGLGK | ASSPPVYEQY | 0.5777303 |
| TALAGAGSGI | ASSPPVYEQY | 0.4670992 |
| TGLPFTGI | ASSPPVYEQY | 0.7435944 |
| VAEHVTV | ASSPPVYEQY | 0.63857645 |
| VDIVLNSL | ASSPPVYEQY | 0.72939545 |
| VLTGGGAVL | ASSPPVYEQY | 0.5130689 |
| VNVGLAYGI | ASSPPVYEQY | 0.4868098 |
| VSSAAVGAV | ASSPPVYEQY | 0.5691519 |
| VTSVYEAGV | ASSPPVYEQY | 0.72391284 |
| WVALDRGA | ASSPPVYEQY | 0.7218535 |
| YLVGSVVI | ASSPPVYEQY | 0.65418863 |
| GPPGTGKTLL | ASSPPVYEQY | 0.50510395 |
| GVGMTKF | ASSPPVYEQY | 0.59871376 |
| IGIGGSDL | ASSPPVYEQY | 0.53730595 |
| IGTLLAEL | ASSPPVYEQY | 0.57115376 |
| IYTSGTT | ASSPPVYEQY | 0.6758828 |
| LSFPAGV | ASSPPVYEQY | 0.6628011 |
| LYTSGTT | ASSPPVYEQY | 0.57463026 |
| VGAFPFGF | ASSPPVYEQY | 0.57312787 |
| VGAGTVEF | ASSPPVYEQY | 0.5925486 |
| VGKSTLLL | ASSPPVYEQY | 0.6964342 |
| AALAHAGGGIV | ASSPPVYEQY | 0.66492456 |
| AALVAVSLL | ASSPPVYEQY | 0.73572874 |
| AAVSRAVAAL | ASSPPVYEQY | 0.68356866 |
| AGGSSGIGGF | ASSPPVYEQY | 0.64411676 |
| AGLTVANANG | ASSPPVYEQY | 0.23854436 |
| AGLVGAVL | ASSPPVYEQY | 0.3420984 |
| ALSQAFTV | ASSPPVYEQY | 0.6041597 |
| ALTFSVSL | ASSPPVYEQY | 0.583831 |
| ALTGAALVAL | ASSPPVYEQY | 0.33590677 |
| ASFSIGAGVL | ASSPPVYEQY | 0.70981044 |
| DGSVYKI | ASSPPVYEQY | 0.56399506 |
| EQLAEIV | ASSPPVYEQY | 0.55641514 |
| FAVSELTVM | ASSPPVYEQY | 0.70716476 |
| GAAPLAAGL | ASSPPVYEQY | 0.59391785 |
| GADGGDGGSGGAGG | ASSPPVYEQY | 0.576016 |
| GFGLDTGGAGG | ASSPPVYEQY | 0.5415188 |

| | | |
|---|---|---|
| GLASGGLL | ASSPPVYEQY | 0.69030243 |
| GRIAGAAVLL | ASSPPVYEQY | 0.42834195 |
| GSAGSLNTSF | ASSPPVYEQY | 0.6632147 |
| GTGQIGI | ASSPPVYEQY | 0.46899223 |
| GTLIALIL | ASSPPVYEQY | 0.70590746 |
| IAATMGFVVG | ASSPPVYEQY | 0.43968835 |
| IFAQALL | ASSPPVYEQY | 0.65006006 |
| IGAAVALGF | ASSPPVYEQY | 0.5592993 |
| IGLTPLAL | ASSPPVYEQY | 0.77967215 |
| ILGSNTSTL | ASSPPVYEQY | 0.58934915 |
| IVFSLTVL | ASSPPVYEQY | 0.64100266 |
| LASKATGF | ASSPPVYEQY | 0.52088815 |
| LASVALIL | ASSPPVYEQY | 0.69639015 |
| LFAVGLTL | ASSPPVYEQY | 0.4091601 |
| LFNGGIGTYI | ASSPPVYEQY | 0.5656066 |
| LTLGAGII | ASSPPVYEQY | 0.39244452 |
| LVAAALGDI | ASSPPVYEQY | 0.64597344 |
| NTAFVQLGI | ASSPPVYEQY | 0.6211038 |
| QLFSGAEL | ASSPPVYEQY | 0.44053987 |
| SAATFITL | ASSPPVYEQY | 0.69341934 |
| SNYGLANLGV | ASSPPVYEQY | 0.61647546 |
| SYGGAYAVM | ASSPPVYEQY | 0.55856586 |
| TTGVGGLTLG | ASSPPVYEQY | 0.7274307 |
| TTQTIYI | ASSPPVYEQY | 0.540954 |
| VAAGAGLGAVI | ASSPPVYEQY | 0.7223413 |
| VAAVVTLL | ASSPPVYEQY | 0.60260254 |
| VGFLGGLI | ASSPPVYEQY | 0.49955764 |
| VGVIPILL | ASSPPVYEQY | 0.65606534 |
| VIFASSLI | ASSPPVYEQY | 0.5565613 |
| VKGGDGGAAAG | ASSPPVYEQY | 0.5522249 |
| VTKTFTNI | ASSPPVYEQY | 0.6948892 |
| VVAAAILL | ASSPPVYEQY | 0.7663597 |
| YAALDVEL | ASSPPVYEQY | 0.75793767 |
| LASVALLL | ASSPPVYEQY | 0.7138401 |
| LDAVHRV | ASSPPVYEQY | 0.69446754 |
| QGLSSSIF | ASSPPVYEQY | 0.74409264 |
| RGTFANIRL | ASSPPVYEQY | 0.7555703 |
| RQLWWGHRI | ASSPPVYEQY | 0.77077746 |
| YGPPGTGKTLL | ASSPPVYEQY | 0.76129436 |
| AALVGAPANVA | ASSPPVYEQY | 0.63370645 |
| AATFRRTE | ASSPPVYEQY | 0.7673156 |
| AGALGGGI | ASSPPVYEQY | 0.74497664 |
| AGGAGGAGGIGAG | ASSPPVYEQY | 0.7457898 |
| AGSGALGL | ASSPPVYEQY | 0.752519 |
| AQIADQI | ASSPPVYEQY | 0.74632055 |
| ASAVAVTYL | ASSPPVYEQY | 0.7366606 |
| ASTVLGGTGG | ASSPPVYEQY | 0.71147203 |
| AVGGEPAK | ASSPPVYEQY | 0.75534534 |
| CEQGTFGGTC | ASSPPVYEQY | 0.7684597 |
| DEGKAPAG | ASSPPVYEQY | 0.7671788 |
| DTVVFPAL | ASSPPVYEQY | 0.77023697 |

| | | |
|---|---|---|
| FEGITMAG | ASSPPVYEQY | 0.7701453 |
| FPLVVQD | ASSPPVYEQY | 0.7655766 |
| GAFTTPQ | ASSPPVYEQY | 0.4406567 |
| GFGGFGTAGGVGGA | ASSPPVYEQY | 0.61224353 |
| GFLAQFL | ASSPPVYEQY | 0.7244942 |
| GGAGGAGGNAGVA | ASSPPVYEQY | 0.7285783 |
| GGVGTAGAGGAGGAGG | ASSPPVYEQY | 0.41083154 |
| GNGGGGDAGNAGSGGN | ASSPPVYEQY | 0.7346519 |
| GTGGAGGAGNDAGS | ASSPPVYEQY | 0.7410753 |
| GTTNTGWFNS | ASSPPVYEQY | 0.70099527 |
| IFMGPLFTL | ASSPPVYEQY | 0.7698648 |
| ILAHAIEAI | ASSPPVYEQY | 0.7392447 |
| IRSAIREAL | ASSPPVYEQY | 0.64555734 |
| ITFAGGG | ASSPPVYEQY | 0.71349144 |
| ITGMSGHQEV | ASSPPVYEQY | 0.7590722 |
| IVAGLACTL | ASSPPVYEQY | 0.6833006 |
| IVGGGLAGA | ASSPPVYEQY | 0.77110934 |
| IVTGVLNI | ASSPPVYEQY | 0.6619925 |
| LAQQGCI | ASSPPVYEQY | 0.7714788 |
| LCLHVEN | ASSPPVYEQY | 0.57367945 |
| LDAISDGS | ASSPPVYEQY | 0.7640078 |
| LGGFAAML | ASSPPVYEQY | 0.7462521 |
| LGGKNPMI | ASSPPVYEQY | 0.781924 |
| LGPKGVAAIL | ASSPPVYEQY | 0.7336806 |
| LMVGVSI | ASSPPVYEQY | 0.76372707 |
| LVGAAFVL | ASSPPVYEQY | 0.7023001 |
| LVLAVAPL | ASSPPVYEQY | 0.7701566 |
| MAAIAVIDAA | ASSPPVYEQY | 0.6677331 |
| NNGINAELQ | ASSPPVYEQY | 0.7425953 |
| RGEPLVLVHG | ASSPPVYEQY | 0.5772141 |
| RGGAGGAAGLI | ASSPPVYEQY | 0.74768865 |
| RSYRMHPAV | ASSPPVYEQY | 0.7625295 |
| SAAHLLAGQLA | ASSPPVYEQY | 0.3410398 |
| SAAVGSY | ASSPPVYEQY | 0.77169776 |
| SPHAVLHRI | ASSPPVYEQY | 0.63434 |
| SQIAKVL | ASSPPVYEQY | 0.6619583 |
| TALGHLVP | ASSPPVYEQY | 0.7707369 |
| TAPTTSGAA | ASSPPVYEQY | 0.73285466 |
| TDGDDGNGGSPGT | ASSPPVYEQY | 0.77567035 |
| TDGLGEN | ASSPPVYEQY | 0.71718997 |
| TGGPPLPGQA | ASSPPVYEQY | 0.74760497 |
| TIPAITI | ASSPPVYEQY | 0.7414406 |
| TIPGFTI | ASSPPVYEQY | 0.6767672 |
| TSIASGALA | ASSPPVYEQY | 0.68155676 |
| TVTLPGAAI | ASSPPVYEQY | 0.7752363 |
| VASGLSAALP | ASSPPVYEQY | 0.74047947 |
| VGAGVIGI | ASSPPVYEQY | 0.7272793 |
| VGPKRFGV | ASSPPVYEQY | 0.66792893 |
| VNVAFPD | ASSPPVYEQY | 0.5600354 |
| VSMTNSGVS | ASSPPVYEQY | 0.649297 |
| VTDVLGDLSV | ASSPPVYEQY | 0.7372305 |

| | | |
|---|---|---|
| VVVVGAGFAGL | ASSPPVYEQY | 0.7451096 |
| VYTVGTVV | ASSPPVYEQY | 0.73202646 |
| WTTFSGP | ASSPPVYEQY | 0.5098226 |
| YDELVEAF | ASSPPVYEQY | 0.71509385 |
| YGTGGAGGNGG | ASSPPVYEQY | 0.67236745 |
| YTNVMAV | ASSPPVYEQY | 0.6578369 |
| FLEQVHQL | ASSPPVYEQY | 0.7202619 |
| GLSLLAVAV | ASSPPVYEQY | 0.71685815 |
| LEQVHQL | ASSPPVYEQY | 0.652742 |
| LNAAGAI | ASSPPVYEQY | 0.60925066 |
| PAVHASL | ASSPPVYEQY | 0.7207625 |
| TELEDDA | ASSPPVYEQY | 0.7567526 |
| TPPGPPN | ASSPPVYEQY | 0.7524147 |
| TPPPPPPP | ASSPPVYEQY | 0.7662001 |
| TQSPALI | ASRVQGAEQY | 0.7613473 |
| EFAAAFPQ | ASRVQGAEQY | 0.7568029 |
| LLSGNLDVL | ASRVQGAEQY | 0.23933397 |
| GPANLGNYNI | ASRVQGAEQY | 0.46808603 |
| TLGPTLP | ASRVQGAEQY | 0.6050813 |
| VYAYSSCCK | ASRVQGAEQY | 0.69974804 |
| APAPAPAPNI | ASRVQGAEQY | 0.32774767 |
| PGGPVPIP | ASRVQGAEQY | 0.62030077 |
| FLTVALEP | ASRVQGAEQY | 0.6605984 |
| SQYFGANL | ASRVQGAEQY | 0.53722525 |
| GGVKPAA | ASRVQGAEQY | 0.7737623 |
| GGTGGAGGAGGW | ASRVQGAEQY | 0.6406605 |
| GVGGVAGLGGAGG | ASRVQGAEQY | 0.5354241 |
| VLLVLATTI | ASRVQGAEQY | 0.6193241 |
| IGIFAAML | ASRVQGAEQY | 0.74490964 |
| AVGDLID | ASRVQGAEQY | 0.52394664 |
| NISPLPL | ASRVQGAEQY | 0.7354197 |
| VTAVFVAI | ASRVQGAEQY | 0.5434852 |
| SGDDNDAAL | ASRVQGAEQY | 0.7536831 |
| AVAALAHS | ASRVQGAEQY | 0.43671975 |
| GGVTVMATP | ASRVQGAEQY | 0.7479671 |
| AGLAANVRD | ASRVQGAEQY | 0.6764411 |
| TNRPDLIDPALLRPG | ASRVQGAEQY | 0.7844727 |
| LANLGVTAL | ASRVQGAEQY | 0.6603422 |
| TGERIEL | ASRVQGAEQY | 0.7333073 |
| NGEELAA | ASRVQGAEQY | 0.62816757 |
| VLSLVWG | ASRVQGAEQY | 0.7379359 |
| IGYALWL | ASRVQGAEQY | 0.47361663 |
| LSLLLAPL | ASRVQGAEQY | 0.6631394 |
| TLDEALK | ASRVQGAEQY | 0.38198888 |
| TDGAGTAEGG | ASRVQGAEQY | 0.7046392 |
| NAPGGAGGNGG | ASRVQGAEQY | 0.73415804 |
| VSVFMGMGGNF | ASRVQGAEQY | 0.31585118 |
| LASYLAAL | ASRVQGAEQY | 0.75014985 |
| LVGGTPAL | ASRVQGAEQY | 0.6069936 |
| VRNADVL | ASRVQGAEQY | 0.52341586 |
| TCSSPFLVSAM | ASRVQGAEQY | 0.7650168 |

| | | |
|---|---|---|
| VHITDAE | ASRVQGAEQY | 0.6377209 |
| EYAAPSF | ASRVQGAEQY | 0.7621766 |
| VVDAFHGGL | ASRVQGAEQY | 0.6219053 |
| ASVVDPQ | ASRVQGAEQY | 0.6567447 |
| SYNEFGGLS | ASRVQGAEQY | 0.7134609 |
| DPDGSVY | ASRVQGAEQY | 0.6059226 |
| MVSTIHATE | ASRVQGAEQY | 0.5809558 |
| LAAGVADPV | ASRVQGAEQY | 0.7583158 |
| TKEQIAA | ASRVQGAEQY | 0.7309264 |
| SLASPAAL | ASRVQGAEQY | 0.7340982 |
| IGLVFYGV | ASRVQGAEQY | 0.6654199 |
| FLLDASNL | ASRVQGAEQY | 0.6650933 |
| YQVGNFP | ASRVQGAEQY | 0.69597846 |
| ESLLSQST | ASRVQGAEQY | 0.73031116 |
| LFTVGFAAVG | ASRVQGAEQY | 0.74238294 |
| ATPPTPT | ASRVQGAEQY | 0.73378277 |
| TDLATLG | ASRVQGAEQY | 0.6700072 |
| SAGGTGR | ASRVQGAEQY | 0.73231494 |
| GFSNSGS | ASRVQGAEQY | 0.7127214 |
| GGNGGSGGTSV | ASRVQGAEQY | 0.7163782 |
| GTSVATGGAGN | ASRVQGAEQY | 0.7138356 |
| NGGNGGSGGTS | ASRVQGAEQY | 0.7024669 |
| AVVKFPH | ASRVQGAEQY | 0.67382175 |
| TPPPPPP | ASRVQGAEQY | 0.595886 |
| AAPAAAAAPA | ASRVQGAEQY | 0.7273 |
| IALNEDL | ASRVQGAEQY | 0.73478127 |
| IVNVSSM | ASRVQGAEQY | 0.72856057 |
| TGIPPAPR | ASRVQGAEQY | 0.7448033 |
| FDLGGGTF | ASRVQGAEQY | 0.73522466 |
| FDLGGGTFDVS | ASRVQGAEQY | 0.7238021 |
| NEPTAAA | ASRVQGAEQY | 0.42703182 |
| NEPTAAALA | ASRVQGAEQY | 0.5713005 |
| GSIINISSI | ASRVQGAEQY | 0.6209115 |
| VAPGNAGTA | ASRVQGAEQY | 0.6943139 |
| TGGGASLE | ASRVQGAEQY | 0.332024 |
| VAVYNHYKRI | ASRVQGAEQY | 0.72518766 |
| TDPAPATT | ASRVQGAEQY | 0.6964376 |
| TNRPDILDPALLRPG | ASRVQGAEQY | 0.6770508 |
| LTTTTTS | ASRVQGAEQY | 0.7350215 |
| PTGSGKTL | ASRVQGAEQY | 0.6899955 |
| TGSGKTL | ASRVQGAEQY | 0.5706551 |
| TIGHVDHGK | ASRVQGAEQY | 0.66308594 |
| TIGHVDHGKTT | ASRVQGAEQY | 0.73872876 |
| GHVDHGKTT | ASRVQGAEQY | 0.62470794 |
| AQFEHTLL | ASRVQGAEQY | 0.7388521 |
| IVNQDEL | ASRVQGAEQY | 0.69654244 |
| TNGSQFF | ASRVQGAEQY | 0.74040794 |
| TGKGIVE | ASRVQGAEQY | 0.61003447 |
| LISNASDA | ASRVQGAEQY | 0.73769975 |
| AELRTLGELTKIAKAHGAQ | ASRVQGAEQY | 0.715346 |
| APNAVAAAEPSW | ASRVQGAEQY | 0.7460982 |

| | | |
|---|---|---|
| APQVAGAATAVL | ASRVQGAEQY | 0.7004709 |
| ARRDPAAAAAILPTDA | ASRVQGAEQY | 0.7331207 |
| DAAARDVLAEAGLAEYF | ASRVQGAEQY | 0.716463 |
| DDQIVLSAKANLAKVEQ | ASRVQGAEQY | 0.7362893 |
| DELLPDTGEDSDEDGH | ASRVQGAEQY | 0.6204609 |
| ELPEAPANRATVPQD | ASRVQGAEQY | 0.6789518 |
| FGQSKIERVTGIG | ASRVQGAEQY | 0.6694559 |
| GAGGTGSTAGGAGGAGGAGGLYA | ASRVQGAEQY | 0.69870144 |
| GHGGTGAAGGNGGTGGAGGSGIDG | ASRVQGAEQY | 0.7365804 |
| GNGGNGGAGGEGGPGGAGGAGA | ASRVQGAEQY | 0.30176967 |
| GSTAGNGGNGGNATNGTIAGGQPAG | ASRVQGAEQY | 0.717859 |
| GTGGDGGNGGNGGTGG | ASRVQGAEQY | 0.64751697 |
| GVRRPAGIPADAPV | ASRVQGAEQY | 0.60835034 |
| GYRQVTDAHLLTLARR | ASRVQGAEQY | 0.7278013 |
| IAKMGQDGHDRG | ASRVQGAEQY | 0.7136471 |
| KAEPSIEIQYT | ASRVQGAEQY | 0.72829205 |
| KPVRIRMAADEDTVAVT | ASRVQGAEQY | 0.633463 |
| MPPRIAGMRLLVIKPEP | ASRVQGAEQY | 0.70291877 |
| MKFYAVALFRSRGVPPTR | ASRVQGAEQY | 0.718626 |
| NAGGFGILGAVAHSP | ASRVQGAEQY | 0.45256957 |
| NEGREITLAMAEM | ASRVQGAEQY | 0.6283988 |
| NELVEWSHPLGHRWRWE | ASRVQGAEQY | 0.7429888 |
| NLGAGGTGGNGGNPG | ASRVQGAEQY | 0.17724496 |
| NTEWPKWETVMAADPVD | ASRVQGAEQY | 0.403885 |
| PASAAPAETQP | ASRVQGAEQY | 0.13197376 |
| QQSSDGKNGVRAAEITGE | ASRVQGAEQY | 0.14308424 |
| RAPDAVTTSWWKE | ASRVQGAEQY | 0.14740156 |
| SGFGNVGAHTSGWFNQ | ASRVQGAEQY | 0.19837004 |
| SGVAGAVGQAASVGGLKV | ASRVQGAEQY | 0.3009539 |
| SNGGAGTGSAGGNGGT | ASRVQGAEQY | 0.22159664 |
| TGGAGGEGGAAG | ASRVQGAEQY | 0.1458876 |
| TGGDGGAAGAGGAPGHG | ASRVQGAEQY | 0.14261672 |
| TMPPLLAAIANAGRHGV | ASRVQGAEQY | 0.13787907 |
| TSSTAGNSGGA | ASRVQGAEQY | 0.14559962 |
| VPATVPVGILC | ASRVQGAEQY | 0.16338438 |
| VQRPDSGGVVLTGRIS | ASRVQGAEQY | 0.1391631 |
| WTPNPVPATKGVPAGLVQAT | ASRVQGAEQY | 0.13390908 |
| YALKRALAKFSDRFDVVII | ASRVQGAEQY | 0.14988047 |
| YLIGSVIV | ASRVQGAEQY | 0.14767358 |
| YGQLVAVID | ASRVQGAEQY | 0.18109551 |
| VTVGGMTLPA | ASRVQGAEQY | 0.1755303 |
| VLALPAV | ASRVQGAEQY | 0.22441444 |
| VIPPTLNYET | ASRVQGAEQY | 0.17463148 |
| VGMGGMHPSSG | ASRVQGAEQY | 0.15895844 |
| VGLAGGAATA | ASRVQGAEQY | 0.12484053 |
| VGGAGGAGGSGP | ASRVQGAEQY | 0.13229564 |
| VFAAFMV | ASRVQGAEQY | 0.14833236 |
| TLYDKLR | ASRVQGAEQY | 0.14980835 |
| TLAVFTVL | ASRVQGAEQY | 0.1262833 |
| TGSVQPARV | ASRVQGAEQY | 0.15950197 |
| TGGGAGSGGT | ASRVQGAEQY | 0.16358736 |

| | | |
|---|---|---|
| SHLETSL | ASRVQGAEQY | 0.13187784 |
| SFTTGMT | ASRVQGAEQY | 0.17497371 |
| QVEVAKRMGVRQPTVS | ASRVQGAEQY | 0.13472012 |
| QGSGGPALAM | ASRVQGAEQY | 0.1348049 |
| MLAPSNP | ASRVQGAEQY | 0.14257677 |
| LVNNAAI | ASRVQGAEQY | 0.18540004 |
| LTGEINTLA | ASRVQGAEQY | 0.13719061 |
| LSIFGPV | ASRVQGAEQY | 0.1739271 |
| LLALLANL | ASRVQGAEQY | 0.16281681 |
| LISNASDALDKLR | ASRVQGAEQY | 0.1808471 |
| LINEAVP | ASRVQGAEQY | 0.13613501 |
| LGYAFSGL | ASRVQGAEQY | 0.16292754 |
| LDFVAVAL | ASRVQGAEQY | 0.13947731 |
| LAQPGAE | ASRVQGAEQY | 0.23161857 |
| ISNLGAVTGF | ASRVQGAEQY | 0.15002814 |
| HQEDPGRI | ASRVQGAEQY | 0.16647881 |
| GTPGEPTP | ASRVQGAEQY | 0.14507504 |
| GFFNSGS | ASRVQGAEQY | 0.2053203 |
| GAGGGGLTAGAN | ASRVQGAEQY | 0.12821974 |
| FVDALVNM | ASRVQGAEQY | 0.14021158 |
| FLTVAPEL | ASRVQGAEQY | 0.14107452 |
| FETLTAL | ASRVQGAEQY | 0.14512852 |
| DVINPIEL | ASRVQGAEQY | 0.20223169 |
| APVSGAPV | ASRVQGAEQY | 0.13951068 |
| ALVGGLAW | ASRVQGAEQY | 0.16454679 |
| ALAAAAATAI | ASRVQGAEQY | 0.14908186 |
| AFGLGLLL | ASRVQGAEQY | 0.1476243 |
| AFFAAASL | ASRVQGAEQY | 0.16629265 |
| AAAALPAFSPP | ASRVQGAEQY | 0.13223225 |
| VVLAPAL | ASRVQGAEQY | 0.16091281 |
| TIGHVDHGKTTL | ASRVQGAEQY | 0.12822658 |
| LISNASDALDK | ASRVQGAEQY | 0.15065357 |
| LISNASDAL | ASRVQGAEQY | 0.18407156 |
| ISNASDALDK | ASRVQGAEQY | 0.13076854 |
| NQAETLVYQTE | ASRVQGAEQY | 0.13170932 |
| TGGTGGTAGAG | ASRVQGAEQY | 0.18388967 |
| TGSTAGGAGGA | ASRVQGAEQY | 0.69704336 |
| DPLAVPVAAGPV | ASRVQGAEQY | 0.7537371 |
| GGTGGTPGNGGN | ASRVQGAEQY | 0.3158864 |
| PAGLAGVDPATL | ASRVQGAEQY | 0.38192046 |
| GNNLVVEAPHAGASV | ASRVQGAEQY | 0.5329218 |
| KVTRIMSASTLEQPA | ASRVQGAEQY | 0.7534667 |
| LGFALPIALPRRTVV | ASRVQGAEQY | 0.8041611 |
| TRLGNLTARKLVVLV | ASRVQGAEQY | 0.73067796 |
| WSQQRRLIYFPSAGP | ASRVQGAEQY | 0.39346805 |
| GPNGNPDPMAAAVDIR | ASRVQGAEQY | 0.66069126 |
| TQRCVSVADLRAAASR | ASRVQGAEQY | 0.5407789 |
| QFVLDVAEHPVEVLTETV | ASRVQGAEQY | 0.54342324 |
| GGAGGTGGAGGNGGTQAGDG | ASRVQGAEQY | 0.6550198 |
| PLLATLRAIAVDVDAKPAQV | ASRVQGAEQY | 0.61912674 |
| APKFRTGGYMIDFWGVGYQVAKRM | ASRVQGAEQY | 0.41525888 |

| | | |
|---|---|---|
| GAGGAGGAGGAAGGPGGTGGTGGN | ASRVQGAEQY | 0.34652367 |
| IGALGLLATDVAPAGSVAFGVALA | ASRVQGAEQY | 0.68009657 |
| AGGAGGIGGNGGAGGAGGVLMGNG | ASRVQGAEQY | 0.75923645 |
| ADELIGLL | ASRVQGAEQY | 0.73387474 |
| ADRALKA | ASRVQGAEQY | 0.8365363 |
| AGAVGQVGGAGGA | ASRVQGAEQY | 0.74540627 |
| AGGGGGLGGAP | ASRVQGAEQY | 0.6594515 |
| AGGLLYGNGGAG | ASRVQGAEQY | 0.1363601 |
| AGLESIGL | ASRVQGAEQY | 0.26470083 |
| AGLKTGII | ASRVQGAEQY | 0.3993513 |
| ASLAALLL | ASRVQGAEQY | 0.6103987 |
| ASVINISSI | ASRVQGAEQY | 0.16860941 |
| ATPPGPA | ASRVQGAEQY | 0.71044934 |
| ATVASALV | ASRVQGAEQY | 0.4372394 |
| ATVASLIL | ASRVQGAEQY | 0.41791397 |
| AVNGPVLIRH | ASRVQGAEQY | 0.758111 |
| DGAAGAVGI | ASRVQGAEQY | 0.36593965 |
| DPLLHGI | ASRVQGAEQY | 0.39491528 |
| ELGAALGAL | ASRVQGAEQY | 0.4177376 |
| ELGRTQAV | ASRVQGAEQY | 0.7742648 |
| FVAGGYLS | ASRVQGAEQY | 0.32160872 |
| GASGGAGGAGGW | ASRVQGAEQY | 0.7557324 |
| GGAGAGGDGGAGGI | ASRVQGAEQY | 0.4781521 |
| GTGHNGSQPGMGGNG | ASRVQGAEQY | 0.7729663 |
| GYAAASASAS | ASRVQGAEQY | 0.3190722 |
| HQVDVAAVQQG | ASRVQGAEQY | 0.7233387 |
| IGAAIAHT | ASRVQGAEQY | 0.58568037 |
| IGMIEVAQL | ASRVQGAEQY | 0.83359766 |
| IRLAPEHTSAMD | ASRVQGAEQY | 0.56530035 |
| IYNVSVL | ASRVQGAEQY | 0.733873 |
| KQGFLGM | ASRVQGAEQY | 0.5395723 |
| LESVITQL | ASRVQGAEQY | 0.76052123 |
| LGLGALVVV | ASRVQGAEQY | 0.27954942 |
| LIGNGGNGGSGGI | ASRVQGAEQY | 0.44267708 |
| LNAINAPTL | ASRVQGAEQY | 0.2764622 |
| LQLLNTGV | ASRVQGAEQY | 0.5166168 |
| LQTSIEF | ASRVQGAEQY | 0.68382585 |
| MSFAVSEL | ASRVQGAEQY | 0.14099514 |
| NTGALGL | ASRVQGAEQY | 0.54499507 |
| PPGAGGAGGAGTT | ASRVQGAEQY | 0.41520828 |
| QLVGLGI | ASRVQGAEQY | 0.3578717 |
| RFLEQVHQL | ASRVQGAEQY | 0.71729565 |
| RGVGYVLR | ASRVQGAEQY | 0.5883963 |
| SATGVFAGVI | ASRVQGAEQY | 0.7470813 |
| SLPAITLP | ASRVQGAEQY | 0.44096762 |
| STALAFTV | ASRVQGAEQY | 0.4651043 |
| SVLTVGLGK | ASRVQGAEQY | 0.5794187 |
| TALAGAGSGI | ASRVQGAEQY | 0.20498759 |
| TGLPFTGI | ASRVQGAEQY | 0.38553274 |
| VAEHVTV | ASRVQGAEQY | 0.7885637 |
| VDIVLNSL | ASRVQGAEQY | 0.7506167 |

| | | |
|---|---|---|
| VLTGGGAVL | ASRVQGAEQY | 0.7607467 |
| VNVGLAYGI | ASRVQGAEQY | 0.6996937 |
| VSSAAVGAV | ASRVQGAEQY | 0.6669005 |
| VTSVYEAGV | ASRVQGAEQY | 0.73027205 |
| WVALDRGA | ASRVQGAEQY | 0.75654817 |
| YLVGSVVI | ASRVQGAEQY | 0.764769 |
| GPPGTGKTLL | ASRVQGAEQY | 0.7554903 |
| GVGMTKF | ASRVQGAEQY | 0.6668549 |
| IGIGGSDL | ASRVQGAEQY | 0.74788904 |
| IGTLLAEL | ASRVQGAEQY | 0.73311543 |
| IYTSGTT | ASRVQGAEQY | 0.7283835 |
| LSFPAGV | ASRVQGAEQY | 0.7554516 |
| LYTSGTT | ASRVQGAEQY | 0.73207414 |
| VGAFPFGF | ASRVQGAEQY | 0.6913922 |
| VGAGTVEF | ASRVQGAEQY | 0.6918837 |
| VGKSTLLL | ASRVQGAEQY | 0.75339794 |
| AALAHAGGGIV | ASRVQGAEQY | 0.76078546 |
| AALVAVSLL | ASRVQGAEQY | 0.75948817 |
| AAVSRAVAAL | ASRVQGAEQY | 0.76765597 |
| AGGSSGIGGF | ASRVQGAEQY | 0.75913465 |
| AGLTVANANG | ASRVQGAEQY | 0.7531878 |
| AGLVGAVL | ASRVQGAEQY | 0.3490082 |
| ALSQAFTV | ASRVQGAEQY | 0.5961502 |
| ALTFSVSL | ASRVQGAEQY | 0.6982256 |
| ALTGAALVAL | ASRVQGAEQY | 0.7346056 |
| ASFSIGAGVL | ASRVQGAEQY | 0.49898595 |
| DGSVYKI | ASRVQGAEQY | 0.75098103 |
| EQLAEIV | ASRVQGAEQY | 0.7002466 |
| FAVSELTVM | ASRVQGAEQY | 0.67373013 |
| GAAPLAAGL | ASRVQGAEQY | 0.7604668 |
| GADGGDGGSGGAGG | ASRVQGAEQY | 0.6880137 |
| GFGLDTGGAGG | ASRVQGAEQY | 0.6311445 |
| GLASGGLL | ASRVQGAEQY | 0.71896124 |
| GRIAGAAVLL | ASRVQGAEQY | 0.7653239 |
| GSAGSLNTSF | ASRVQGAEQY | 0.64228106 |
| GTGQIGI | ASRVQGAEQY | 0.7580211 |
| GTLIALIL | ASRVQGAEQY | 0.73896575 |
| IAATMGFVVG | ASRVQGAEQY | 0.76300395 |
| IFAQALL | ASRVQGAEQY | 0.5793042 |
| IGAAVALGF | ASRVQGAEQY | 0.75770193 |
| IGLTPLAL | ASRVQGAEQY | 0.7321171 |
| ILGSNTSTL | ASRVQGAEQY | 0.76887363 |
| IVFSLTVL | ASRVQGAEQY | 0.72923446 |
| LASKATGF | ASRVQGAEQY | 0.74821997 |
| LASVALIL | ASRVQGAEQY | 0.73004943 |
| LFAVGLTL | ASRVQGAEQY | 0.7599387 |
| LFNGGIGTYI | ASRVQGAEQY | 0.59910333 |
| LTLGAGII | ASRVQGAEQY | 0.7249166 |
| LVAAALGDI | ASRVQGAEQY | 0.6687514 |
| NTAFVQLGI | ASRVQGAEQY | 0.73455894 |
| QLFSGAEL | ASRVQGAEQY | 0.75222933 |

| | | |
|---|---|---|
| SAATFITL | ASRVQGAEQY | 0.4175507 |
| SNYGLANLGV | ASRVQGAEQY | 0.7584412 |
| SYGGAYAVM | ASRVQGAEQY | 0.7095041 |
| TTGVGGLTLG | ASRVQGAEQY | 0.67202985 |
| TTQTIYI | ASRVQGAEQY | 0.7605312 |
| VAAGAGLGAVI | ASRVQGAEQY | 0.739638 |
| VAAVVTLL | ASRVQGAEQY | 0.75445235 |
| VGFLGGLI | ASRVQGAEQY | 0.71601605 |
| VGVIPILL | ASRVQGAEQY | 0.7426014 |
| VIFASSLI | ASRVQGAEQY | 0.7455508 |
| VKGGDGGAAAG | ASRVQGAEQY | 0.69846547 |
| VTKTFTNI | ASRVQGAEQY | 0.7038306 |
| VVAAAILL | ASRVQGAEQY | 0.7611935 |
| YAALDVEL | ASRVQGAEQY | 0.8631104 |
| LASVALLL | ASRVQGAEQY | 0.8545021 |
| LDAVHRV | ASRVQGAEQY | 0.7424189 |
| QGLSSSIF | ASRVQGAEQY | 0.7573533 |
| RGTFANIRL | ASRVQGAEQY | 0.7956196 |
| RQLWWGHRI | ASRVQGAEQY | 0.8666177 |
| YGPPGTGKTLL | ASRVQGAEQY | 0.8700917 |
| AALVGAPANVA | ASRVQGAEQY | 0.8663807 |
| AATFRRTE | ASRVQGAEQY | 0.7856051 |
| AGALGGGI | ASRVQGAEQY | 0.8486651 |
| AGGAGGAGGIGAG | ASRVQGAEQY | 0.8157471 |
| AGSGALGL | ASRVQGAEQY | 0.8350063 |
| AQIADQI | ASRVQGAEQY | 0.85436666 |
| ASAVAVTYL | ASRVQGAEQY | 0.8318474 |
| ASTVLGGTGG | ASRVQGAEQY | 0.7587979 |
| AVGGEPAK | ASRVQGAEQY | 0.7851418 |
| CEQGTFGGTC | ASRVQGAEQY | 0.8464323 |
| DEGKAPAG | ASRVQGAEQY | 0.86154604 |
| DTVVFPAL | ASRVQGAEQY | 0.8603006 |
| FEGITMAG | ASRVQGAEQY | 0.8919674 |
| FPLVVQD | ASRVQGAEQY | 0.8708335 |
| GAFTTPQ | ASRVQGAEQY | 0.8520761 |
| GFGGFGTAGGVGGA | ASRVQGAEQY | 0.449856 |
| GFLAQFL | ASRVQGAEQY | 0.6846186 |
| GGAGGAGGNAGVA | ASRVQGAEQY | 0.79157186 |
| GGVGTAGAGGAGGAGG | ASRVQGAEQY | 0.8016174 |
| GNGGGGDAGNAGSGGN | ASRVQGAEQY | 0.40506506 |
| GTGGAGGAGNDAGS | ASRVQGAEQY | 0.854323 |
| GTTNTGWFNS | ASRVQGAEQY | 0.8254664 |
| IFMGPLFTL | ASRVQGAEQY | 0.7502639 |
| ILAHAIEAI | ASRVQGAEQY | 0.8709449 |
| IRSAIREAL | ASRVQGAEQY | 0.7902397 |
| ITFAGGG | ASRVQGAEQY | 0.75461906 |
| ITGMSGHQEV | ASRVQGAEQY | 0.80548173 |
| IVAGLACTL | ASRVQGAEQY | 0.8768512 |
| IVGGGLAGA | ASRVQGAEQY | 0.7950933 |
| IVTGVLNI | ASRVQGAEQY | 0.8754411 |
| LAQQGCI | ASRVQGAEQY | 0.7878588 |

| | | |
|---|---|---|
| LCLHVEN | ASRVQGAEQY | 0.8839302 |
| LDAISDGS | ASRVQGAEQY | 0.7421801 |
| LGGFAAML | ASRVQGAEQY | 0.8789816 |
| LGGKNPMI | ASRVQGAEQY | 0.83655083 |
| LGPKGVAAIL | ASRVQGAEQY | 0.8995493 |
| LMVGVSI | ASRVQGAEQY | 0.83560777 |
| LVGAAFVL | ASRVQGAEQY | 0.8714634 |
| LVLAVAPL | ASRVQGAEQY | 0.78578025 |
| MAAIAVIDAA | ASRVQGAEQY | 0.84776735 |
| NNGINAELQ | ASRVQGAEQY | 0.69525075 |
| RGEPLVLVHG | ASRVQGAEQY | 0.8011405 |
| RGGAGGAAGLI | ASRVQGAEQY | 0.6912154 |
| RSYRMHPAV | ASRVQGAEQY | 0.82892513 |
| SAAHLLAGQLA | ASRVQGAEQY | 0.8727213 |
| SAAVGSY | ASRVQGAEQY | 0.35615245 |
| SPHAVLHRI | ASRVQGAEQY | 0.8583391 |
| SQIAKVL | ASRVQGAEQY | 0.6762444 |
| TALGHLVP | ASRVQGAEQY | 0.7708323 |
| TAPTTSGAA | ASRVQGAEQY | 0.8721012 |
| TDGDDGNGGSPGT | ASRVQGAEQY | 0.8229933 |
| TDGLGEN | ASRVQGAEQY | 0.8653111 |
| TGGPPLPGQA | ASRVQGAEQY | 0.7679058 |
| TIPAITI | ASRVQGAEQY | 0.80752134 |
| TIPGFTI | ASRVQGAEQY | 0.8459475 |
| TSIASGALA | ASRVQGAEQY | 0.5494733 |
| TVTLPGAAI | ASRVQGAEQY | 0.735957 |
| VASGLSAALP | ASRVQGAEQY | 0.8920015 |
| VGAGVIGI | ASRVQGAEQY | 0.6035302 |
| VGPKRFGV | ASRVQGAEQY | 0.64350027 |
| VNVAFPD | ASRVQGAEQY | 0.5133322 |
| VSMTNSGVS | ASRVQGAEQY | 0.49615827 |
| VTDVLGDLSV | ASRVQGAEQY | 0.5305543 |
| VVVVGAGFAGL | ASRVQGAEQY | 0.63722605 |
| VYTVGTVV | ASRVQGAEQY | 0.6373571 |
| WTTFSGP | ASRVQGAEQY | 0.6177756 |
| YDELVEAF | ASRVQGAEQY | 0.52817047 |
| YGTGGAGGNGG | ASRVQGAEQY | 0.59896445 |
| YTNVMAV | ASRVQGAEQY | 0.5595378 |
| FLEQVHQL | ASRVQGAEQY | 0.55689406 |
| GLSLLAVAV | ASRVQGAEQY | 0.61671865 |
| LEQVHQL | ASRVQGAEQY | 0.62797946 |
| LNAAGAI | ASRVQGAEQY | 0.5632502 |
| PAVHASL | ASRVQGAEQY | 0.5305393 |
| TELEDDA | ASRVQGAEQY | 0.6031493 |
| TPPGPPN | ASRVQGAEQY | 0.6336779 |
| TPPPPPPP | ASRVQGAEQY | 0.6190819 |
| TQSPALI | AWGDNERLF | 0.6607329 |
| EFAAAFPQ | AWGDNERLF | 0.62398386 |
| LLSGNLDVL | AWGDNERLF | 0.62497044 |
| GPANLGNYNI | AWGDNERLF | 0.34892914 |
| TLGPTLP | AWGDNERLF | 0.41217563 |

| | | |
|---|---|---|
| VYAYSSCCK | AWGDNERLF | 0.60494524 |
| APAPAPAPNI | AWGDNERLF | 0.5679616 |
| PGGPVPIP | AWGDNERLF | 0.44227505 |
| FLTVALEP | AWGDNERLF | 0.6186236 |
| SQYFGANL | AWGDNERLF | 0.55638194 |
| GGVKPAA | AWGDNERLF | 0.5559349 |
| GGTGGAGGAGGW | AWGDNERLF | 0.6199318 |
| GVGGVAGLGGAGG | AWGDNERLF | 0.54037654 |
| VLLVLATTI | AWGDNERLF | 0.5316157 |
| IGIFAAML | AWGDNERLF | 0.54921407 |
| AVGDLID | AWGDNERLF | 0.641281 |
| NISPLPL | AWGDNERLF | 0.45324692 |
| VTAVFVAI | AWGDNERLF | 0.61099863 |
| SGDDNDAAL | AWGDNERLF | 0.5611012 |
| AVAALAHS | AWGDNERLF | 0.6410101 |
| GGVTVMATP | AWGDNERLF | 0.4850783 |
| AGLAANVRD | AWGDNERLF | 0.6236454 |
| TNRPDLIDPALLRPG | AWGDNERLF | 0.54699725 |
| LANLGVTAL | AWGDNERLF | 0.6641288 |
| TGERIEL | AWGDNERLF | 0.5527817 |
| NGEELAA | AWGDNERLF | 0.61612236 |
| VLSLVWG | AWGDNERLF | 0.60826695 |
| IGYALWL | AWGDNERLF | 0.61275315 |
| LSLLLAPL | AWGDNERLF | 0.44051957 |
| TLDEALK | AWGDNERLF | 0.5554902 |
| TDGAGTAEGG | AWGDNERLF | 0.46356764 |
| NAPGGAGGNGG | AWGDNERLF | 0.5863971 |
| VSVFMGMGGNF | AWGDNERLF | 0.616469 |
| LASYLAAL | AWGDNERLF | 0.4247301 |
| LVGGTPAL | AWGDNERLF | 0.6135075 |
| VRNADVL | AWGDNERLF | 0.56658643 |
| TCSSPFLVSAM | AWGDNERLF | 0.54246795 |
| VHITDAE | AWGDNERLF | 0.6416095 |
| EYAAPSF | AWGDNERLF | 0.5973924 |
| VVDAFHGGL | AWGDNERLF | 0.630427 |
| ASVVDPQ | AWGDNERLF | 0.5870303 |
| SYNEFGGLS | AWGDNERLF | 0.5456931 |
| DPDGSVY | AWGDNERLF | 0.6133441 |
| MVSTIHATE | AWGDNERLF | 0.5416766 |
| LAAGVADPV | AWGDNERLF | 0.5318643 |
| TKEQIAA | AWGDNERLF | 0.63149315 |
| SLASPAAL | AWGDNERLF | 0.83787334 |
| IGLVFYGV | AWGDNERLF | 0.8853466 |
| FLLDASNL | AWGDNERLF | 0.7266039 |
| YQVGNFP | AWGDNERLF | 0.70863813 |
| ESLLSQST | AWGDNERLF | 0.78696096 |
| LFTVGFAAVG | AWGDNERLF | 0.8824435 |
| ATPPTPT | AWGDNERLF | 0.873338 |
| TDLATLG | AWGDNERLF | 0.84775615 |
| SAGGTGR | AWGDNERLF | 0.7122784 |
| GFSNSGS | AWGDNERLF | 0.8124205 |

| | | |
|---|---|---|
| GGNGGSGGTSV | AWGDNERLF | 0.7448185 |
| GTSVATGGAGN | AWGDNERLF | 0.7810229 |
| NGGNGGSGGTS | AWGDNERLF | 0.85912955 |
| AVVKFPH | AWGDNERLF | 0.8002881 |
| TPPPPPP | AWGDNERLF | 0.76157755 |
| AAPAAAAAPA | AWGDNERLF | 0.787822 |
| IALNEDL | AWGDNERLF | 0.82431674 |
| IVNVSSM | AWGDNERLF | 0.86548316 |
| TGIPPAPR | AWGDNERLF | 0.83923674 |
| FDLGGGTF | AWGDNERLF | 0.90008366 |
| FDLGGGTFDVS | AWGDNERLF | 0.85757315 |
| NEPTAAA | AWGDNERLF | 0.87052906 |
| NEPTAAALA | AWGDNERLF | 0.22161528 |
| GSIINISSI | AWGDNERLF | 0.54331267 |
| VAPGNAGTA | AWGDNERLF | 0.7758118 |
| TGGGASLE | AWGDNERLF | 0.79800874 |
| VAVYNHYKRI | AWGDNERLF | 0.5591163 |
| TDPAPATT | AWGDNERLF | 0.8615457 |
| TNRPDILDPALLRPG | AWGDNERLF | 0.77423733 |
| LTTTTTS | AWGDNERLF | 0.7186084 |
| PTGSGKTL | AWGDNERLF | 0.85968506 |
| TGSGKTL | AWGDNERLF | 0.7627029 |
| TIGHVDHGK | AWGDNERLF | 0.7428822 |
| TIGHVDHGKTT | AWGDNERLF | 0.73643154 |
| GHVDHGKTT | AWGDNERLF | 0.8809961 |
| AQFEHTLL | AWGDNERLF | 0.5173832 |
| IVNQDEL | AWGDNERLF | 0.83995354 |
| TNGSQFF | AWGDNERLF | 0.81434685 |
| TGKGIVE | AWGDNERLF | 0.87223464 |
| LISNASDA | AWGDNERLF | 0.58907664 |
| AELRTLGELTKIAKAHGAQ | AWGDNERLF | 0.8440306 |
| APNAVAAAEPSW | AWGDNERLF | 0.7737844 |
| APQVAGAATAVL | AWGDNERLF | 0.9003626 |
| ARRDPAAAAAILPTDA | AWGDNERLF | 0.7727257 |
| DAAARDVLAEAGLAEYF | AWGDNERLF | 0.81828 |
| DDQIVLSAKANLAKVEQ | AWGDNERLF | 0.73324394 |
| DELLPDTGEDSDEDGH | AWGDNERLF | 0.85444975 |
| ELPEAPANRATVPQD | AWGDNERLF | 0.60824066 |
| FGQSKIERVTGIG | AWGDNERLF | 0.787194 |
| GAGGTGSTAGGAGGAGGAGGLYA | AWGDNERLF | 0.7213267 |
| GHGGTGAAGGNGGTGGAGGSGIDG | AWGDNERLF | 0.8378858 |
| GNGGNGGAGGEGGPGGAGGAGA | AWGDNERLF | 0.8389201 |
| GSTAGNGGNGGNATNGTIAGQPAG | AWGDNERLF | 0.6725862 |
| GTGGDGGNGGNGGTGG | AWGDNERLF | 0.86432624 |
| GVRRPAGIPADAPV | AWGDNERLF | 0.80398834 |
| GYRQVTDAHLLTLARR | AWGDNERLF | 0.7523798 |
| IAKMGQDGHDRG | AWGDNERLF | 0.879946 |
| KAEPSIEIQYT | AWGDNERLF | 0.76407886 |
| KPVRIRMAADEDTVAVT | AWGDNERLF | 0.8680863 |
| MPPRIAGMRLLVIKPEP | AWGDNERLF | 0.789069 |
| MKFYAVALFRSRGVPPTR | AWGDNERLF | 0.8073316 |

| | | |
|---|---|---|
| NAGGFGILGAVAHSP | AWGDNERLF | 0.8702364 |
| NEGREITLAMAEM | AWGDNERLF | 0.6719084 |
| NELVEWSHPLGHRWRWE | AWGDNERLF | 0.6698421 |
| NLGAGGTGGNGGNPG | AWGDNERLF | 0.8711555 |
| NTEWPKWETVMAADPVD | AWGDNERLF | 0.811105 |
| PASAAPAETQP | AWGDNERLF | 0.8280312 |
| QQSSDGKNGVRAAEITGE | AWGDNERLF | 0.67250586 |
| RAPDAVTTSWWKE | AWGDNERLF | 0.61981344 |
| SGFGNVGAHTSGWFNQ | AWGDNERLF | 0.70846343 |
| SGVAGAVGQAASVGGLKV | AWGDNERLF | 0.8253773 |
| SNGGAGTGSAGGNGGT | AWGDNERLF | 0.8523207 |
| TGGAGGEGGAAG | AWGDNERLF | 0.7955735 |
| TGGDGGAAGAGGAPGHG | AWGDNERLF | 0.66797674 |
| TMPPLLAAIANAGRHGV | AWGDNERLF | 0.80161285 |
| TSSTAGNSGGA | AWGDNERLF | 0.76039106 |
| VPATVPVGILC | AWGDNERLF | 0.74688995 |
| VQRPDSGGVVLTGRIS | AWGDNERLF | 0.80567706 |
| WTPNPVPATKGVPAGLVQAT | AWGDNERLF | 0.798611 |
| YALKRALAKFSDRFDVVII | AWGDNERLF | 0.72982526 |
| YLIGSVIV | AWGDNERLF | 0.69249284 |
| YGQLVAVID | AWGDNERLF | 0.79326 |
| VTVGGMTLPA | AWGDNERLF | 0.83566 |
| VLALPAV | AWGDNERLF | 0.8294561 |
| VIPPTLNYET | AWGDNERLF | 0.86337644 |
| VGMGGMHPSSG | AWGDNERLF | 0.8396311 |
| VGLAGGAATA | AWGDNERLF | 0.81502545 |
| VGGAGGAGGSGP | AWGDNERLF | 0.26812288 |
| VFAAFMV | AWGDNERLF | 0.55392194 |
| TLYDKLR | AWGDNERLF | 0.65220356 |
| TLAVFTVL | AWGDNERLF | 0.7593178 |
| TGSVQPARV | AWGDNERLF | 0.32734516 |
| TGGGAGSGGT | AWGDNERLF | 0.80149496 |
| SHLETSL | AWGDNERLF | 0.73819816 |
| SFTTGMT | AWGDNERLF | 0.6564407 |
| QVEVAKRMGVRQPTVS | AWGDNERLF | 0.8383957 |
| QGSGGPALAM | AWGDNERLF | 0.721617 |
| MLAPSNP | AWGDNERLF | 0.69512594 |
| LVNNAAI | AWGDNERLF | 0.70587486 |
| LTGEINTLA | AWGDNERLF | 0.8622483 |
| LSIFGPV | AWGDNERLF | 0.56644475 |
| LLALLANL | AWGDNERLF | 0.83108294 |
| LISNASDALDKLR | AWGDNERLF | 0.71238536 |
| LINEAVP | AWGDNERLF | 0.85480535 |
| LGYAFSGL | AWGDNERLF | 0.5466235 |
| LDFVAVAL | AWGDNERLF | 0.82598114 |
| LAQPGAE | AWGDNERLF | 0.7524444 |
| ISNLGAVTGF | AWGDNERLF | 0.8773296 |
| HQEDPGRI | AWGDNERLF | 0.76012254 |
| GTPGEPTP | AWGDNERLF | 0.81604743 |
| GFFNSGS | AWGDNERLF | 0.7248019 |
| GAGGGGLTAGAN | AWGDNERLF | 0.8292012 |

| | | |
|---|---|---|
| FVDALVNM | AWGDNERLF | 0.5166856 |
| FLTVAPEL | AWGDNERLF | 0.768972 |
| FETLTAL | AWGDNERLF | 0.64600456 |
| DVINPIEL | AWGDNERLF | 0.7877177 |
| APVSGAPV | AWGDNERLF | 0.80278575 |
| ALVGGLAW | AWGDNERLF | 0.33414525 |
| ALAAAAATAI | AWGDNERLF | 0.83024514 |
| AFGLGLLL | AWGDNERLF | 0.66315174 |
| AFFAAASL | AWGDNERLF | 0.6504177 |
| AAAALPAFSPP | AWGDNERLF | 0.8383935 |
| VVLAPAL | AWGDNERLF | 0.7627124 |
| TIGHVDHGKTTL | AWGDNERLF | 0.8365042 |
| LISNASDALDK | AWGDNERLF | 0.7104378 |
| LISNASDAL | AWGDNERLF | 0.74981505 |
| ISNASDALDK | AWGDNERLF | 0.81367123 |
| NQAETLVYQTE | AWGDNERLF | 0.4694814 |
| TGGTGGTAGAG | AWGDNERLF | 0.6223936 |
| TGSTAGGAGGA | AWGDNERLF | 0.85285556 |
| DPLAVPVAAGPV | AWGDNERLF | 0.8023646 |
| GGTGGTPGNGGN | AWGDNERLF | 0.7374368 |
| PAGLAGVDPATL | AWGDNERLF | 0.6175033 |
| GNNLVVEAPHAGASV | AWGDNERLF | 0.57017654 |
| KVTRIMSASTLEQPA | AWGDNERLF | 0.6968051 |
| LGFALPIALPRRTVV | AWGDNERLF | 0.80364645 |
| TRLGNLTARKLVVLV | AWGDNERLF | 0.80941033 |
| WSQQRRLIYFPSAGP | AWGDNERLF | 0.7938384 |
| GPNGNPDPMAAAVDIR | AWGDNERLF | 0.601475 |
| TQRCVSVADLRAAASR | AWGDNERLF | 0.798736 |
| QFVLDVAEHPVEVLTETV | AWGDNERLF | 0.71756244 |
| GGAGGTGGAGGNGGTQAGDG | AWGDNERLF | 0.7095967 |
| PLLATLRAIAVDVDAKPAQV | AWGDNERLF | 0.787048 |
| APKFRTGGYMIDFWGVGYQVAKRM | AWGDNERLF | 0.7258125 |
| GAGGAGGAGGAAGGPGGTGGTGGN | AWGDNERLF | 0.65565765 |
| IGALGLLATDVAPAGSVAFGVALA | AWGDNERLF | 0.5715036 |
| AGGAGGIGGNGGAGGAGGVLMGNG | AWGDNERLF | 0.80372417 |
| ADELIGLL | AWGDNERLF | 0.8120315 |
| ADRALKA | AWGDNERLF | 0.783425 |
| AGAVGQVGGAGGA | AWGDNERLF | 0.8509238 |
| AGGGGGLGGAP | AWGDNERLF | 0.8156738 |
| AGGLLYGNGGAG | AWGDNERLF | 0.7781347 |
| AGLESIGL | AWGDNERLF | 0.29160744 |
| AGLKTGII | AWGDNERLF | 0.41202202 |
| ASLAALLL | AWGDNERLF | 0.6485174 |
| ASVINISSI | AWGDNERLF | 0.6681099 |
| ATPPGPA | AWGDNERLF | 0.19398206 |
| ATVASALV | AWGDNERLF | 0.691155 |
| ATVASLIL | AWGDNERLF | 0.72054017 |
| AVNGPVLIRH | AWGDNERLF | 0.5746068 |
| DGAAGAVGI | AWGDNERLF | 0.82407403 |
| DPLLHGI | AWGDNERLF | 0.65274113 |
| ELGAALGAL | AWGDNERLF | 0.5316381 |

| | | |
|---|---|---|
| ELGRTQAV | AWGDNERLF | 0.7062504 |
| FVAGGYLS | AWGDNERLF | 0.84592736 |
| GASGGAGGAGGW | AWGDNERLF | 0.6336872 |
| GGAGAGGDGGAGGI | AWGDNERLF | 0.806113 |
| GTGHNGSQPGMGGNG | AWGDNERLF | 0.68971 |
| GYAAASASAS | AWGDNERLF | 0.837188 |
| HQVDVAAVQQG | AWGDNERLF | 0.47059274 |
| IGAAIAHT | AWGDNERLF | 0.8058748 |
| IGMIEVAQL | AWGDNERLF | 0.70280874 |
| IRLAPEHTSAMD | AWGDNERLF | 0.85466903 |
| IYNVSVL | AWGDNERLF | 0.6871073 |
| KQGFLGM | AWGDNERLF | 0.812901 |
| LESVITQL | AWGDNERLF | 0.78078234 |
| LGLGALVVV | AWGDNERLF | 0.78827137 |
| LIGNGGNGGSGGI | AWGDNERLF | 0.43350762 |
| LNAINAPTL | AWGDNERLF | 0.7003914 |
| LQLLNTGV | AWGDNERLF | 0.6111219 |
| LQTSIEF | AWGDNERLF | 0.75349385 |
| MSFAVSEL | AWGDNERLF | 0.79662067 |
| NTGALGL | AWGDNERLF | 0.14707105 |
| PPGAGGAGGAGTT | AWGDNERLF | 0.8209061 |
| QLVGLGI | AWGDNERLF | 0.40480968 |
| RFLEQVHQL | AWGDNERLF | 0.5616684 |
| RGVGYVLR | AWGDNERLF | 0.81151843 |
| SATGVFAGVI | AWGDNERLF | 0.7358825 |
| SLPAITLP | AWGDNERLF | 0.79110646 |
| STALAFTV | AWGDNERLF | 0.63428295 |
| SVLTVGLGK | AWGDNERLF | 0.7584423 |
| TALAGAGSGI | AWGDNERLF | 0.7839277 |
| TGLPFTGI | AWGDNERLF | 0.28357342 |
| VAEHVTV | AWGDNERLF | 0.5784867 |
| VDIVLNSL | AWGDNERLF | 0.8288878 |
| VLTGGGAVL | AWGDNERLF | 0.88763016 |
| VNVGLAYGI | AWGDNERLF | 0.8884444 |
| VSSAAVGAV | AWGDNERLF | 0.83541906 |
| VTSVYEAGV | AWGDNERLF | 0.8065988 |
| WVALDRGA | AWGDNERLF | 0.84996957 |
| YLVGSVVI | AWGDNERLF | 0.8887982 |
| GPPGTGKTLL | AWGDNERLF | 0.8968053 |
| GVGMTKF | AWGDNERLF | 0.8853563 |
| IGIGGSDL | AWGDNERLF | 0.7948253 |
| IGTLLAEL | AWGDNERLF | 0.8924609 |
| IYTSGTT | AWGDNERLF | 0.8712228 |
| LSFPAGV | AWGDNERLF | 0.86172295 |
| LYTSGTT | AWGDNERLF | 0.8842392 |
| VGAFPFGF | AWGDNERLF | 0.87502205 |
| VGAGTVEF | AWGDNERLF | 0.8420211 |
| VGKSTLLL | AWGDNERLF | 0.81937784 |
| AALAHAGGGIV | AWGDNERLF | 0.8803349 |
| AALVAVSLL | AWGDNERLF | 0.89273745 |
| AAVSRAVAAL | AWGDNERLF | 0.89236015 |

| | | |
|---|---|---|
| AGGSSGIGGF | AWGDNERLF | 0.9009731 |
| AGLTVANANG | AWGDNERLF | 0.89521277 |
| AGLVGAVL | AWGDNERLF | 0.8812518 |
| ALSQAFTV | AWGDNERLF | 0.3454075 |
| ALTFSVSL | AWGDNERLF | 0.76067746 |
| ALTGAALVAL | AWGDNERLF | 0.7914964 |
| ASFSIGAGVL | AWGDNERLF | 0.86438924 |
| DGSVYKI | AWGDNERLF | 0.35868412 |
| EQLAEIV | AWGDNERLF | 0.853804 |
| FAVSELTVM | AWGDNERLF | 0.86434907 |
| GAAPLAAGL | AWGDNERLF | 0.8268039 |
| GADGGDGGSGGAGG | AWGDNERLF | 0.8938595 |
| GFGLDTGGAGG | AWGDNERLF | 0.84189624 |
| GLASGGLL | AWGDNERLF | 0.76988864 |
| GRIAGAAVLL | AWGDNERLF | 0.8029394 |
| GSAGSLNTSF | AWGDNERLF | 0.8993434 |
| GTGQIGI | AWGDNERLF | 0.7957418 |
| GTLIALIL | AWGDNERLF | 0.8954608 |
| IAATMGFVVG | AWGDNERLF | 0.7928321 |
| IFAQALL | AWGDNERLF | 0.8990758 |
| IGAAVALGF | AWGDNERLF | 0.7435688 |
| IGLTPLAL | AWGDNERLF | 0.8913127 |
| ILGSNTSTL | AWGDNERLF | 0.870912 |
| IVFSLTVL | AWGDNERLF | 0.9048422 |
| LASKATGF | AWGDNERLF | 0.8616455 |
| LASVALIL | AWGDNERLF | 0.8923459 |
| LFAVGLTL | AWGDNERLF | 0.8503678 |
| LFNGGIGTYI | AWGDNERLF | 0.89022315 |
| LTLGAGII | AWGDNERLF | 0.76012814 |
| LVAAALGDI | AWGDNERLF | 0.8720082 |
| NTAFVQLGI | AWGDNERLF | 0.7653395 |
| QLFSGAEL | AWGDNERLF | 0.88063264 |
| SAATFITL | AWGDNERLF | 0.88803905 |
| SNYGLANLGV | AWGDNERLF | 0.405586 |
| SYGGAYAVM | AWGDNERLF | 0.89706165 |
| TTGVGGLTLG | AWGDNERLF | 0.81458044 |
| TTQTIYI | AWGDNERLF | 0.81087005 |
| VAAGAGLGAVI | AWGDNERLF | 0.8930244 |
| VAAVVTLL | AWGDNERLF | 0.8657515 |
| VGFLGGLI | AWGDNERLF | 0.8919221 |
| VGVIPILL | AWGDNERLF | 0.79784834 |
| VIFASSLI | AWGDNERLF | 0.8737204 |
| VKGGDGGAAAG | AWGDNERLF | 0.8818852 |
| VTKTFTNI | AWGDNERLF | 0.6086717 |
| VVAAAILL | AWGDNERLF | 0.7062478 |
| YAALDVEL | AWGDNERLF | 0.8989881 |
| LASVALLL | AWGDNERLF | 0.8390775 |
| LDAVHRV | AWGDNERLF | 0.8268855 |
| QGLSSSIF | AWGDNERLF | 0.7274985 |
| RGTFANIRL | AWGDNERLF | 0.7480564 |
| RQLWWGHRI | AWGDNERLF | 0.7796418 |

| | | |
|---|---|---|
| YGPPGTGKTLL | AWGDNERLF | 0.8415675 |
| AALVGAPANVA | AWGDNERLF | 0.8516067 |
| AATFRRTE | AWGDNERLF | 0.84298086 |
| AGALGGGI | AWGDNERLF | 0.7730721 |
| AGGAGGAGGIGAG | AWGDNERLF | 0.8370563 |
| AGSGALGL | AWGDNERLF | 0.80347717 |
| AQIADQI | AWGDNERLF | 0.80271983 |
| ASAVAVTYL | AWGDNERLF | 0.82835364 |
| ASTVLGGTGG | AWGDNERLF | 0.8249465 |
| AVGGEPAK | AWGDNERLF | 0.76296544 |
| CEQGTFGGTC | AWGDNERLF | 0.696752 |
| DEGKAPAG | AWGDNERLF | 0.8202548 |
| DTVVFPAL | AWGDNERLF | 0.8463888 |
| FEGITMAG | AWGDNERLF | 0.8306309 |
| FPLVVQD | AWGDNERLF | 0.8567627 |
| GAFTTPQ | AWGDNERLF | 0.8417938 |
| GFGGFGTAGGVGGA | AWGDNERLF | 0.8360613 |
| GFLAQFL | AWGDNERLF | 0.30338287 |
| GGAGGAGGNAGVA | AWGDNERLF | 0.54728484 |
| GGVGTAGAGGAGGAGG | AWGDNERLF | 0.73902977 |
| GNGGGGDAGNAGSGGN | AWGDNERLF | 0.77738106 |
| GTGGAGGAGNDAGS | AWGDNERLF | 0.27934116 |
| GTTNTGWFNS | AWGDNERLF | 0.8253733 |
| IFMGPLFTL | AWGDNERLF | 0.7840142 |
| ILAHAIEAI | AWGDNERLF | 0.77301955 |
| IRSAIREAL | AWGDNERLF | 0.8413749 |
| ITFAGGG | AWGDNERLF | 0.78741443 |
| ITGMSGHQEV | AWGDNERLF | 0.661945 |
| IVAGLACTL | AWGDNERLF | 0.7597825 |
| IVGGGLAGA | AWGDNERLF | 0.8512481 |
| IVTGVLNI | AWGDNERLF | 0.69953376 |
| LAQQGCI | AWGDNERLF | 0.8455628 |
| LCLHVEN | AWGDNERLF | 0.6886308 |
| LDAISDGS | AWGDNERLF | 0.8507979 |
| LGGFAAML | AWGDNERLF | 0.7291305 |
| LGGKNPMI | AWGDNERLF | 0.84663695 |
| LGPKGVAAIL | AWGDNERLF | 0.81404114 |
| LMVGVSI | AWGDNERLF | 0.8593309 |
| LVGAAFVL | AWGDNERLF | 0.7970549 |
| LVLAVAPL | AWGDNERLF | 0.84727037 |
| MAAIAVIDAA | AWGDNERLF | 0.81689525 |
| NNGINAELQ | AWGDNERLF | 0.8351998 |
| RGEPLVLVHG | AWGDNERLF | 0.6947468 |
| RGGAGGAAGLI | AWGDNERLF | 0.78004235 |
| RSYRMHPAV | AWGDNERLF | 0.66083646 |
| SAAHLLAGQLA | AWGDNERLF | 0.80615723 |
| SAAVGSY | AWGDNERLF | 0.84635633 |
| SPHAVLHRI | AWGDNERLF | 0.24316922 |
| SQIAKVL | AWGDNERLF | 0.82203925 |
| TALGHLVP | AWGDNERLF | 0.70810056 |
| TAPTTSGAA | AWGDNERLF | 0.7456893 |

| | | |
|---|---|---|
| TDGDDGNGGSPGT | AWGDNERLF | 0.8402976 |
| TDGLGEN | AWGDNERLF | 0.8089472 |
| TGGPPLPGQA | AWGDNERLF | 0.8349547 |
| TIPAITI | AWGDNERLF | 0.72859526 |
| TIPGFTI | AWGDNERLF | 0.77500427 |
| TSIASGALA | AWGDNERLF | 0.8215667 |
| TVTLPGAAI | AWGDNERLF | 0.33513427 |
| VASGLSAALP | AWGDNERLF | 0.6391802 |
| VGAGVIGI | AWGDNERLF | 0.8567246 |
| VGPKRFGV | AWGDNERLF | 0.8022922 |
| VNVAFPD | AWGDNERLF | 0.81382537 |
| VSMTNSGVS | AWGDNERLF | 0.67606694 |
| VTDVLGDLSV | AWGDNERLF | 0.64398843 |
| VVVVGAGFAGL | AWGDNERLF | 0.70165217 |
| VYTVGTVV | AWGDNERLF | 0.8290642 |
| WTTFSGP | AWGDNERLF | 0.82578087 |
| YDELVEAF | AWGDNERLF | 0.8041004 |
| YGTGGAGGNGG | AWGDNERLF | 0.6577074 |
| YTNVMAV | AWGDNERLF | 0.8098055 |
| FLEQVHQL | AWGDNERLF | 0.74915445 |
| GLSLLAVAV | AWGDNERLF | 0.7387856 |
| LEQVHQL | AWGDNERLF | 0.8152314 |
| LNAAGAI | AWGDNERLF | 0.79189575 |
| PAVHASL | AWGDNERLF | 0.72502005 |
| TELEDDA | AWGDNERLF | 0.73231137 |
| TPPGPPN | AWGDNERLF | 0.80003595 |
| TPPPPPPP | AWGDNERLF | 0.8189042 |
| TQSPALI | ASSRDRGWDSYEQY | 0.8120136 |
| EFAAAFPQ | ASSRDRGWDSYEQY | 0.8576797 |
| LLSGNLDVL | ASSRDRGWDSYEQY | 0.8235564 |
| GPANLGNYNI | ASSRDRGWDSYEQY | 0.81186163 |
| TLGPTLP | ASSRDRGWDSYEQY | 0.3527778 |
| VYAYSSCCK | ASSRDRGWDSYEQY | 0.5608741 |
| APAPAPAPNI | ASSRDRGWDSYEQY | 0.72589153 |
| PGGPVPIP | ASSRDRGWDSYEQY | 0.7470012 |
| FLTVALEP | ASSRDRGWDSYEQY | 0.3863725 |
| SQYFGANL | ASSRDRGWDSYEQY | 0.8156187 |
| GGVKPAA | ASSRDRGWDSYEQY | 0.75440085 |
| GGTGGAGGAGGW | ASSRDRGWDSYEQY | 0.6644412 |
| GVGGVAGLGGAGG | ASSRDRGWDSYEQY | 0.81933004 |
| VLLVLATTI | ASSRDRGWDSYEQY | 0.71342444 |
| IGIFAAML | ASSRDRGWDSYEQY | 0.71346724 |
| AVGDLID | ASSRDRGWDSYEQY | 0.7438303 |
| NISPLPL | ASSRDRGWDSYEQY | 0.84210825 |
| VTAVFVAI | ASSRDRGWDSYEQY | 0.648313 |
| SGDDNDAAL | ASSRDRGWDSYEQY | 0.81906915 |
| AVAALAHS | ASSRDRGWDSYEQY | 0.7154132 |
| GGVTVMATP | ASSRDRGWDSYEQY | 0.8419478 |
| AGLAANVRD | ASSRDRGWDSYEQY | 0.585252 |
| TNRPDLIDPALLRPG | ASSRDRGWDSYEQY | 0.8190256 |
| LANLGVTAL | ASSRDRGWDSYEQY | 0.7496959 |

| | | |
|---|---|---|
| TGERIEL | ASSRDRGWDSYEQY | 0.86298877 |
| NGEELAA | ASSRDRGWDSYEQY | 0.7397311 |
| VLSLVWG | ASSRDRGWDSYEQY | 0.8066055 |
| IGYALWL | ASSRDRGWDSYEQY | 0.7745303 |
| LSLLLAPL | ASSRDRGWDSYEQY | 0.80142945 |
| TLDEALK | ASSRDRGWDSYEQY | 0.5022861 |
| TDGAGTAEGG | ASSRDRGWDSYEQY | 0.7650907 |
| NAPGGAGGNGG | ASSRDRGWDSYEQY | 0.6609948 |
| VSVFMGMGGNF | ASSRDRGWDSYEQY | 0.79629743 |
| LASYLAAL | ASSRDRGWDSYEQY | 0.8097545 |
| LVGGTPAL | ASSRDRGWDSYEQY | 0.3994943 |
| VRNADVL | ASSRDRGWDSYEQY | 0.8255805 |
| TCSSPFLVSAM | ASSRDRGWDSYEQY | 0.69150543 |
| VHITDAE | ASSRDRGWDSYEQY | 0.6929041 |
| EYAAPSF | ASSRDRGWDSYEQY | 0.8373194 |
| VVDAFHGGL | ASSRDRGWDSYEQY | 0.77492213 |
| ASVVDPQ | ASSRDRGWDSYEQY | 0.8240151 |
| SYNEFGGLS | ASSRDRGWDSYEQY | 0.75116813 |
| DPDGSVY | ASSRDRGWDSYEQY | 0.73313236 |
| MVSTIHATE | ASSRDRGWDSYEQY | 0.8149264 |
| LAAGVADPV | ASSRDRGWDSYEQY | 0.63451916 |
| TKEQIAA | ASSRDRGWDSYEQY | 0.6723323 |
| SLASPAAL | ASSRDRGWDSYEQY | 0.8402616 |
| IGLVFYGV | ASSRDRGWDSYEQY | 0.7609097 |
| FLLDASNL | ASSRDRGWDSYEQY | 0.7722386 |
| YQVGNFP | ASSRDRGWDSYEQY | 0.64934015 |
| ESLLSQST | ASSRDRGWDSYEQY | 0.5812464 |
| LFTVGFAAVG | ASSRDRGWDSYEQY | 0.7127515 |
| ATPPTPT | ASSRDRGWDSYEQY | 0.7771622 |
| TDLATLG | ASSRDRGWDSYEQY | 0.785242 |
| SAGGTGR | ASSRDRGWDSYEQY | 0.74710923 |
| GFSNSGS | ASSRDRGWDSYEQY | 0.5658523 |
| GGNGGSGGTSV | ASSRDRGWDSYEQY | 0.75423586 |
| GTSVATGGAGN | ASSRDRGWDSYEQY | 0.7099925 |
| NGGNGGSGGTS | ASSRDRGWDSYEQY | 0.6899864 |
| AVVKFPH | ASSRDRGWDSYEQY | 0.74980336 |
| TPPPPPP | ASSRDRGWDSYEQY | 0.73089933 |
| AAPAAAAAPA | ASSRDRGWDSYEQY | 0.67655057 |
| IALNEDL | ASSRDRGWDSYEQY | 0.6086482 |
| IVNVSSM | ASSRDRGWDSYEQY | 0.74944615 |
| TGIPPAPR | ASSRDRGWDSYEQY | 0.77806026 |
| FDLGGGTF | ASSRDRGWDSYEQY | 0.7757176 |
| FDLGGGTFDVS | ASSRDRGWDSYEQY | 0.79207146 |
| NEPTAAA | ASSRDRGWDSYEQY | 0.77947927 |
| NEPTAAALA | ASSRDRGWDSYEQY | 0.76852477 |
| GSIINISSI | ASSRDRGWDSYEQY | 0.2665566 |
| VAPGNAGTA | ASSRDRGWDSYEQY | 0.35329613 |
| TGGGASLE | ASSRDRGWDSYEQY | 0.68682975 |
| VAVYNHYKRI | ASSRDRGWDSYEQY | 0.6783583 |
| TDPAPATT | ASSRDRGWDSYEQY | 0.35943875 |
| TNRPDILDPALLRPG | ASSRDRGWDSYEQY | 0.740391 |

| | | |
|---|---|---|
| LTTTTTS | ASSRDRGWDSYEQY | 0.65658456 |
| PTGSGKTL | ASSRDRGWDSYEQY | 0.6238008 |
| TGSGKTL | ASSRDRGWDSYEQY | 0.7793846 |
| TIGHVDHGK | ASSRDRGWDSYEQY | 0.66389394 |
| TIGHVDHGKTT | ASSRDRGWDSYEQY | 0.61457825 |
| GHVDHGKTT | ASSRDRGWDSYEQY | 0.6389048 |
| AQFEHTLL | ASSRDRGWDSYEQY | 0.7607441 |
| IVNQDEL | ASSRDRGWDSYEQY | 0.5387223 |
| TNGSQFF | ASSRDRGWDSYEQY | 0.7660752 |
| TGKGIVE | ASSRDRGWDSYEQY | 0.5353904 |
| LISNASDA | ASSRDRGWDSYEQY | 0.7794079 |
| AELRTLGELTKIAKAHGAQ | ASSRDRGWDSYEQY | 0.4948343 |
| APNAVAAAEPSW | ASSRDRGWDSYEQY | 0.7601899 |
| APQVAGAATAVL | ASSRDRGWDSYEQY | 0.70292056 |
| ARRDPAAAAAILPTDA | ASSRDRGWDSYEQY | 0.80117714 |
| DAAARDVLAEAGLAEYF | ASSRDRGWDSYEQY | 0.7014191 |
| DDQIVLSAKANLAKVEQ | ASSRDRGWDSYEQY | 0.739563 |
| DELLPDTGEDSDEDGH | ASSRDRGWDSYEQY | 0.6153693 |
| ELPEAPANRATVPQD | ASSRDRGWDSYEQY | 0.7706076 |
| FGQSKIERVTGIG | ASSRDRGWDSYEQY | 0.44525892 |
| GAGGTGSTAGGAGGAGGAGGLYA | ASSRDRGWDSYEQY | 0.723786 |
| GHGGTGAAGGNGGTGGAGGSGIDG | ASSRDRGWDSYEQY | 0.4194451 |
| GNGGNGGAGGEGGPGGAGGAGA | ASSRDRGWDSYEQY | 0.73290366 |
| GSTAGNGGNGGNATNGTIAGQPAG | ASSRDRGWDSYEQY | 0.73757607 |
| GTGGDGGNGGNGGTGG | ASSRDRGWDSYEQY | 0.42548627 |
| GVRRPAGIPADAPV | ASSRDRGWDSYEQY | 0.7564603 |
| GYRQVTDAHLLTLARR | ASSRDRGWDSYEQY | 0.6869836 |
| IAKMGQDGHDRG | ASSRDRGWDSYEQY | 0.62972546 |
| KAEPSIEIQYT | ASSRDRGWDSYEQY | 0.7851195 |
| KPVRIRMAADEDTVAVT | ASSRDRGWDSYEQY | 0.7517273 |
| MPPRIAGMRLLVIKPEP | ASSRDRGWDSYEQY | 0.7759659 |
| MKFYAVALFRSRGVPPTR | ASSRDRGWDSYEQY | 0.68465126 |
| NAGGFGILGAVAHSP | ASSRDRGWDSYEQY | 0.6262115 |
| NEGREITLAMAEM | ASSRDRGWDSYEQY | 0.74077916 |
| NELVEWSHPLGHRWRWE | ASSRDRGWDSYEQY | 0.61246693 |
| NLGAGGTGGNGGNPG | ASSRDRGWDSYEQY | 0.64869964 |
| NTEWPKWETVMAADPVD | ASSRDRGWDSYEQY | 0.772261 |
| PASAAPAETQP | ASSRDRGWDSYEQY | 0.8030705 |
| QQSSDGKNGVRAAEITGE | ASSRDRGWDSYEQY | 0.819315 |
| RAPDAVTTSWWKE | ASSRDRGWDSYEQY | 0.7489351 |
| SGFGNVGAHTSGWFNQ | ASSRDRGWDSYEQY | 0.65531313 |
| SGVAGAVGQAASVGGLKV | ASSRDRGWDSYEQY | 0.7519624 |
| SNGGAGTGSAGGNGGT | ASSRDRGWDSYEQY | 0.81544024 |
| TGGAGGEGGAAG | ASSRDRGWDSYEQY | 0.81990457 |
| TGGDGGAAGAGGAPGHG | ASSRDRGWDSYEQY | 0.7887044 |
| TMPPLLAAIANAGRHGV | ASSRDRGWDSYEQY | 0.521513 |
| TSSTAGNSGGA | ASSRDRGWDSYEQY | 0.79016346 |
| VPATVPVGILC | ASSRDRGWDSYEQY | 0.7589958 |
| VQRPDSGGVVLTGRIS | ASSRDRGWDSYEQY | 0.71236956 |
| WTPNPVPATKGVPAGLVQAT | ASSRDRGWDSYEQY | 0.79549724 |
| YALKRALAKFSDRFDVVII | ASSRDRGWDSYEQY | 0.7739067 |

| | | |
|---|---|---|
| YLIGSVIV | ASSRDRGWDSYEQY | 0.7012541 |
| YGQLVAVID | ASSRDRGWDSYEQY | 0.7000041 |
| VTVGGMTLPA | ASSRDRGWDSYEQY | 0.7707535 |
| VLALPAV | ASSRDRGWDSYEQY | 0.8120324 |
| VIPPTLNYET | ASSRDRGWDSYEQY | 0.8165318 |
| VGMGGMHPSSG | ASSRDRGWDSYEQY | 0.82694274 |
| VGLAGGAATA | ASSRDRGWDSYEQY | 0.8304497 |
| VGGAGGAGGSGP | ASSRDRGWDSYEQY | 0.814834 |
| VFAAFMV | ASSRDRGWDSYEQY | 0.248002 |
| TLYDKLR | ASSRDRGWDSYEQY | 0.35057884 |
| TLAVFTVL | ASSRDRGWDSYEQY | 0.7126531 |
| TGSVQPARV | ASSRDRGWDSYEQY | 0.69642043 |
| TGGGAGSGGT | ASSRDRGWDSYEQY | 0.4134899 |
| SHLETSL | ASSRDRGWDSYEQY | 0.63107955 |
| SFTTGMT | ASSRDRGWDSYEQY | 0.7017901 |
| QVEVAKRMGVRQPTVS | ASSRDRGWDSYEQY | 0.62104666 |
| QGSGGPALAM | ASSRDRGWDSYEQY | 0.8221223 |
| MLAPSNP | ASSRDRGWDSYEQY | 0.7175813 |
| LVNNAAI | ASSRDRGWDSYEQY | 0.6478412 |
| LTGEINTLA | ASSRDRGWDSYEQY | 0.67302644 |
| LSIFGPV | ASSRDRGWDSYEQY | 0.7755446 |
| LLALLANL | ASSRDRGWDSYEQY | 0.58175474 |
| LISNASDALDKLR | ASSRDRGWDSYEQY | 0.80390584 |
| LINEAVP | ASSRDRGWDSYEQY | 0.49528745 |
| LGYAFSGL | ASSRDRGWDSYEQY | 0.8246388 |
| LDFVAVAL | ASSRDRGWDSYEQY | 0.52621776 |
| LAQPGAE | ASSRDRGWDSYEQY | 0.8116622 |
| ISNLGAVTGF | ASSRDRGWDSYEQY | 0.7691449 |
| HQEDPGRI | ASSRDRGWDSYEQY | 0.85053796 |
| GTPGEPTP | ASSRDRGWDSYEQY | 0.77200127 |
| GFFNSGS | ASSRDRGWDSYEQY | 0.7779788 |
| GAGGGGLTAGAN | ASSRDRGWDSYEQY | 0.59758615 |
| FVDALVNM | ASSRDRGWDSYEQY | 0.8044125 |
| FLTVAPEL | ASSRDRGWDSYEQY | 0.55714417 |
| FETLTAL | ASSRDRGWDSYEQY | 0.7776356 |
| DVINPIEL | ASSRDRGWDSYEQY | 0.3922315 |
| APVSGAPV | ASSRDRGWDSYEQY | 0.7996991 |
| ALVGGLAW | ASSRDRGWDSYEQY | 0.79233503 |
| ALAAAAATAI | ASSRDRGWDSYEQY | 0.4454275 |
| AFGLGLLL | ASSRDRGWDSYEQY | 0.8114325 |
| AFFAAASL | ASSRDRGWDSYEQY | 0.7571164 |
| AAAALPAFSPP | ASSRDRGWDSYEQY | 0.66602373 |
| VVLAPAL | ASSRDRGWDSYEQY | 0.83686066 |
| TIGHVDHGKTTL | ASSRDRGWDSYEQY | 0.7449399 |
| LISNASDALDK | ASSRDRGWDSYEQY | 0.8330229 |
| LISNASDAL | ASSRDRGWDSYEQY | 0.7323001 |
| ISNASDALDK | ASSRDRGWDSYEQY | 0.70216954 |
| NQAETLVYQTE | ASSRDRGWDSYEQY | 0.76840895 |
| TGGTGGTAGAG | ASSRDRGWDSYEQY | 0.7019884 |
| TGSTAGGAGGA | ASSRDRGWDSYEQY | 0.6676258 |
| DPLAVPVAAGPV | ASSRDRGWDSYEQY | 0.82636 |

| | | |
|---|---|---|
| GGTGGTPGNGGN | ASSRDRGWDSYEQY | 0.8876395 |
| PAGLAGVDPATL | ASSRDRGWDSYEQY | 0.8890732 |
| GNNLVVEAPHAGASV | ASSRDRGWDSYEQY | 0.8491287 |
| KVTRIMSASTLEQPA | ASSRDRGWDSYEQY | 0.8231973 |
| LGFALPIALPRRTVV | ASSRDRGWDSYEQY | 0.86152315 |
| TRLGNLTARKLVVLV | ASSRDRGWDSYEQY | 0.89161146 |
| WSQQRRLIYFPSAGP | ASSRDRGWDSYEQY | 0.8920661 |
| GPNGNPDPMAAAVDIR | ASSRDRGWDSYEQY | 0.8819215 |
| TQRCVSVADLRAAASR | ASSRDRGWDSYEQY | 0.82712036 |
| QFVLDVAEHPVEVLTETV | ASSRDRGWDSYEQY | 0.878559 |
| GGAGGTGGAGGNGGTQAGDG | ASSRDRGWDSYEQY | 0.86401254 |
| PLLATLRAIAVDVDAKPAQV | ASSRDRGWDSYEQY | 0.8582964 |
| APKFRTGGYMIDFWGVGYQVAKRM | ASSRDRGWDSYEQY | 0.8853505 |
| GAGGAGGAGGAAGGPGGTGGTGGN | ASSRDRGWDSYEQY | 0.878599 |
| IGALGLLATDVAPAGSVAFGVALA | ASSRDRGWDSYEQY | 0.860078 |
| AGGAGGIGGNGGAGGAGGVLMGNG | ASSRDRGWDSYEQY | 0.84742653 |
| ADELIGLL | ASSRDRGWDSYEQY | 0.873448 |
| ADRALKA | ASSRDRGWDSYEQY | 0.8904858 |
| AGAVGQVGGAGGA | ASSRDRGWDSYEQY | 0.8849456 |
| AGGGGGLGGAP | ASSRDRGWDSYEQY | 0.90169823 |
| AGGLLYGNGGAG | ASSRDRGWDSYEQY | 0.89311147 |
| AGLESIGL | ASSRDRGWDSYEQY | 0.88807714 |
| AGLKTGII | ASSRDRGWDSYEQY | 0.38609773 |
| ASLAALLL | ASSRDRGWDSYEQY | 0.7410623 |
| ASVINISSI | ASSRDRGWDSYEQY | 0.8717608 |
| ATPPGPA | ASSRDRGWDSYEQY | 0.8661494 |
| ATVASALV | ASSRDRGWDSYEQY | 0.5955576 |
| ATVASLIL | ASSRDRGWDSYEQY | 0.87934625 |
| AVNGPVLIRH | ASSRDRGWDSYEQY | 0.86786526 |
| DGAAGAVGI | ASSRDRGWDSYEQY | 0.8318322 |
| DPLLHGI | ASSRDRGWDSYEQY | 0.89266443 |
| ELGAALGAL | ASSRDRGWDSYEQY | 0.852669 |
| ELGRTQAV | ASSRDRGWDSYEQY | 0.8236588 |
| FVAGGYLS | ASSRDRGWDSYEQY | 0.85804564 |
| GASGGAGGAGGW | ASSRDRGWDSYEQY | 0.89773 |
| GGAGAGGDGGAGGI | ASSRDRGWDSYEQY | 0.8162148 |
| GTGHNGSQPGMGGNG | ASSRDRGWDSYEQY | 0.8878511 |
| GYAAASASAS | ASSRDRGWDSYEQY | 0.8716843 |
| HQVDVAAVQQG | ASSRDRGWDSYEQY | 0.8972868 |
| IGAAIAHT | ASSRDRGWDSYEQY | 0.772506 |
| IGMIEVAQL | ASSRDRGWDSYEQY | 0.8866161 |
| IRLAPEHTSAMD | ASSRDRGWDSYEQY | 0.8595937 |
| IYNVSVL | ASSRDRGWDSYEQY | 0.90287244 |
| KQGFLGM | ASSRDRGWDSYEQY | 0.8610946 |
| LESVITQL | ASSRDRGWDSYEQY | 0.8848027 |
| LGLGALVVV | ASSRDRGWDSYEQY | 0.8505573 |
| LIGNGGNGGSGGI | ASSRDRGWDSYEQY | 0.88058406 |
| LNAINAPTL | ASSRDRGWDSYEQY | 0.7914572 |
| LQLLNTGV | ASSRDRGWDSYEQY | 0.87290883 |
| LQTSIEF | ASSRDRGWDSYEQY | 0.7915009 |
| MSFAVSEL | ASSRDRGWDSYEQY | 0.88004243 |

| | | |
|---|---|---|
| NTGALGL | ASSRDRGWDSYEQY | 0.88428533 |
| PPGAGGAGGAGTT | ASSRDRGWDSYEQY | 0.6627829 |
| QLVGLGI | ASSRDRGWDSYEQY | 0.89345413 |
| RFLEQVHQL | ASSRDRGWDSYEQY | 0.837305 |
| RGVGYVLR | ASSRDRGWDSYEQY | 0.8445279 |
| SATGVFAGVI | ASSRDRGWDSYEQY | 0.89496917 |
| SLPAITLP | ASSRDRGWDSYEQY | 0.8690249 |
| STALAFTV | ASSRDRGWDSYEQY | 0.89253885 |
| SVLTVGLGK | ASSRDRGWDSYEQY | 0.85584694 |
| TALAGAGSGI | ASSRDRGWDSYEQY | 0.87431437 |
| TGLPFTGI | ASSRDRGWDSYEQY | 0.88885516 |
| VAEHVTV | ASSRDRGWDSYEQY | 0.7897564 |
| VDIVLNSL | ASSRDRGWDSYEQY | 0.84229565 |
| VLTGGGAVL | ASSRDRGWDSYEQY | 0.89276123 |
| VNVGLAYGI | ASSRDRGWDSYEQY | 0.82176566 |
| VSSAAVGAV | ASSRDRGWDSYEQY | 0.86208665 |
| VTSVYEAGV | ASSRDRGWDSYEQY | 0.7362114 |
| WVALDRGA | ASSRDRGWDSYEQY | 0.64275455 |
| YLVGSVVI | ASSRDRGWDSYEQY | 0.73253953 |
| GPPGTGKTLL | ASSRDRGWDSYEQY | 0.82926905 |
| GVGMTKF | ASSRDRGWDSYEQY | 0.8708527 |
| IGIGGSDL | ASSRDRGWDSYEQY | 0.8359331 |
| IGTLLAEL | ASSRDRGWDSYEQY | 0.6298584 |
| IYTSGTT | ASSRDRGWDSYEQY | 0.85396874 |
| LSFPAGV | ASSRDRGWDSYEQY | 0.7911105 |
| LYTSGTT | ASSRDRGWDSYEQY | 0.7665721 |
| VGAFPFGF | ASSRDRGWDSYEQY | 0.80807745 |
| VGAGTVEF | ASSRDRGWDSYEQY | 0.83556795 |
| VGKSTLLL | ASSRDRGWDSYEQY | 0.7650143 |
| AALAHAGGGIV | ASSRDRGWDSYEQY | 0.6698077 |
| AALVAVSLL | ASSRDRGWDSYEQY | 0.7819307 |
| AAVSRAVAAL | ASSRDRGWDSYEQY | 0.8416033 |
| AGGSSGIGGF | ASSRDRGWDSYEQY | 0.8324003 |
| AGLTVANANG | ASSRDRGWDSYEQY | 0.8829396 |
| AGLVGAVL | ASSRDRGWDSYEQY | 0.86372936 |
| ALSQAFTV | ASSRDRGWDSYEQY | 0.8062264 |
| ALTFSVSL | ASSRDRGWDSYEQY | 0.2283912 |
| ALTGAALVAL | ASSRDRGWDSYEQY | 0.49648076 |
| ASFSIGAGVL | ASSRDRGWDSYEQY | 0.68452215 |
| DGSVYKI | ASSRDRGWDSYEQY | 0.76593804 |
| EQLAEIV | ASSRDRGWDSYEQY | 0.27745977 |
| FAVSELTVM | ASSRDRGWDSYEQY | 0.7892848 |
| GAAPLAAGL | ASSRDRGWDSYEQY | 0.79017985 |
| GADGGDGGSGGAGG | ASSRDRGWDSYEQY | 0.69688267 |
| GFGLDTGGAGG | ASSRDRGWDSYEQY | 0.86026484 |
| GLASGGLL | ASSRDRGWDSYEQY | 0.7445456 |
| GRIAGAAVLL | ASSRDRGWDSYEQY | 0.65027773 |
| GSAGSLNTSF | ASSRDRGWDSYEQY | 0.6810279 |
| GTGQIGI | ASSRDRGWDSYEQY | 0.8640008 |
| GTLIALIL | ASSRDRGWDSYEQY | 0.5455451 |
| IAATMGFVVG | ASSRDRGWDSYEQY | 0.85737085 |

| | | |
|---|---|---|
| IFAQALL | ASSRDRGWDSYEQY | 0.60525095 |
| IGAAVALGF | ASSRDRGWDSYEQY | 0.86770236 |
| IGLTPLAL | ASSRDRGWDSYEQY | 0.5322778 |
| ILGSNTSTL | ASSRDRGWDSYEQY | 0.8478397 |
| IVFSLTVL | ASSRDRGWDSYEQY | 0.76430726 |
| LASKATGF | ASSRDRGWDSYEQY | 0.90107083 |
| LASVALIL | ASSRDRGWDSYEQY | 0.7489635 |
| LFAVGLTL | ASSRDRGWDSYEQY | 0.8469678 |
| LFNGGIGTYI | ASSRDRGWDSYEQY | 0.6887572 |
| LTLGAGII | ASSRDRGWDSYEQY | 0.84831774 |
| LVAAALGDI | ASSRDRGWDSYEQY | 0.5348826 |
| NTAFVQLGI | ASSRDRGWDSYEQY | 0.77693343 |
| QLFSGAEL | ASSRDRGWDSYEQY | 0.51509166 |
| SAATFITL | ASSRDRGWDSYEQY | 0.8107736 |
| SNYGLANLGV | ASSRDRGWDSYEQY | 0.8368144 |
| SYGGAYAVM | ASSRDRGWDSYEQY | 0.35410586 |
| TTGVGGLTLG | ASSRDRGWDSYEQY | 0.87097406 |
| TTQTIYI | ASSRDRGWDSYEQY | 0.6945216 |
| VAAGAGLGAVI | ASSRDRGWDSYEQY | 0.64018893 |
| VAAVVTLL | ASSRDRGWDSYEQY | 0.86191684 |
| VGFLGGLI | ASSRDRGWDSYEQY | 0.7975726 |
| VGVIPILL | ASSRDRGWDSYEQY | 0.86710095 |
| VIFASSLI | ASSRDRGWDSYEQY | 0.70654714 |
| VKGGDGGAAAG | ASSRDRGWDSYEQY | 0.7890098 |
| VTKTFTNI | ASSRDRGWDSYEQY | 0.8185315 |
| VVAAAILL | ASSRDRGWDSYEQY | 0.58098984 |
| YAALDVEL | ASSRDRGWDSYEQY | 0.61081785 |
| LASVALLL | ASSRDRGWDSYEQY | 0.8712092 |
| LDAVHRV | ASSRDRGWDSYEQY | 0.7959257 |
| QGLSSSIF | ASSRDRGWDSYEQY | 0.7885774 |
| RGTFANIRL | ASSRDRGWDSYEQY | 0.630787 |
| RQLWWGHRI | ASSRDRGWDSYEQY | 0.618227 |
| YGPPGTGKTLL | ASSRDRGWDSYEQY | 0.6920613 |
| AALVGAPANVA | ASSRDRGWDSYEQY | 0.8181279 |
| AATFRRTE | ASSRDRGWDSYEQY | 0.83932966 |
| AGALGGGI | ASSRDRGWDSYEQY | 0.80561435 |
| AGGAGGAGGIGAG | ASSRDRGWDSYEQY | 0.6905183 |
| AGSGALGL | ASSRDRGWDSYEQY | 0.82408625 |
| AQIADQI | ASSRDRGWDSYEQY | 0.76089233 |
| ASAVAVTYL | ASSRDRGWDSYEQY | 0.74866253 |
| ASTVLGGTGG | ASSRDRGWDSYEQY | 0.81909406 |
| AVGGEPAK | ASSRDRGWDSYEQY | 0.7967746 |
| CEQGTFGGTC | ASSRDRGWDSYEQY | 0.7116247 |
| DEGKAPAG | ASSRDRGWDSYEQY | 0.69271183 |
| DTVVFPAL | ASSRDRGWDSYEQY | 0.7975714 |
| FEGITMAG | ASSRDRGWDSYEQY | 0.8244489 |
| FPLVVQD | ASSRDRGWDSYEQY | 0.81135464 |
| GAFTTPQ | ASSRDRGWDSYEQY | 0.86792445 |
| GFGGFGTAGGVGGA | ASSRDRGWDSYEQY | 0.8254779 |
| GFLAQFL | ASSRDRGWDSYEQY | 0.79496145 |
| GGAGGAGGNAGVA | ASSRDRGWDSYEQY | 0.32222062 |

| | | |
|---|---|---|
| GGVGTAGAGGAGGAGG | ASSRDRGWDSYEQY | 0.5999819 |
| GNGGGGDAGNAGSGGN | ASSRDRGWDSYEQY | 0.70876956 |
| GTGGAGGAGNDAGS | ASSRDRGWDSYEQY | 0.75615036 |
| GTTNTGWFNS | ASSRDRGWDSYEQY | 0.32222468 |
| IFMGPLFTL | ASSRDRGWDSYEQY | 0.7999417 |
| ILAHAIEAI | ASSRDRGWDSYEQY | 0.76086235 |
| IRSAIREAL | ASSRDRGWDSYEQY | 0.6649225 |
| ITFAGGG | ASSRDRGWDSYEQY | 0.8301542 |
| ITGMSGHQEV | ASSRDRGWDSYEQY | 0.69058037 |
| IVAGLACTL | ASSRDRGWDSYEQY | 0.7047493 |
| IVGGGLAGA | ASSRDRGWDSYEQY | 0.7476633 |
| IVTGVLNI | ASSRDRGWDSYEQY | 0.85705787 |
| LAQQGCI | ASSRDRGWDSYEQY | 0.644497 |
| LCLHVEN | ASSRDRGWDSYEQY | 0.8353037 |
| LDAISDGS | ASSRDRGWDSYEQY | 0.71171486 |
| LGGFAAML | ASSRDRGWDSYEQY | 0.84930515 |
| LGGKNPMI | ASSRDRGWDSYEQY | 0.597996 |
| LGPKGVAAIL | ASSRDRGWDSYEQY | 0.82287735 |
| LMVGVSI | ASSRDRGWDSYEQY | 0.73478544 |
| LVGAAFVL | ASSRDRGWDSYEQY | 0.87163115 |
| LVLAVAPL | ASSRDRGWDSYEQY | 0.7454888 |
| MAAIAVIDAA | ASSRDRGWDSYEQY | 0.83985126 |
| NNGINAELQ | ASSRDRGWDSYEQY | 0.78858614 |
| RGEPLVLVHG | ASSRDRGWDSYEQY | 0.7910052 |
| RGGAGGAAGLI | ASSRDRGWDSYEQY | 0.5229714 |
| RSYRMHPAV | ASSRDRGWDSYEQY | 0.7439842 |
| SAAHLLAGQLA | ASSRDRGWDSYEQY | 0.64878 |
| SAAVGSY | ASSRDRGWDSYEQY | 0.79168165 |
| SPHAVLHRI | ASSRDRGWDSYEQY | 0.8025833 |
| SQIAKVL | ASSRDRGWDSYEQY | 0.29958 |
| TALGHLVP | ASSRDRGWDSYEQY | 0.8309397 |
| TAPTTSGAA | ASSRDRGWDSYEQY | 0.588029 |
| TDGDDGNGGSPGT | ASSRDRGWDSYEQY | 0.6590823 |
| TDGLGEN | ASSRDRGWDSYEQY | 0.83331805 |
| TGGPPLPGQA | ASSRDRGWDSYEQY | 0.7694929 |
| TIPAITI | ASSRDRGWDSYEQY | 0.8058634 |
| TIPGFTI | ASSRDRGWDSYEQY | 0.73622453 |
| TSIASGALA | ASSRDRGWDSYEQY | 0.7652168 |
| TVTLPGAAI | ASSRDRGWDSYEQY | 0.8057839 |
| VASGLSAALP | ASSRDRGWDSYEQY | 0.47022563 |
| VGAGVIGI | ASSRDRGWDSYEQY | 0.6576367 |
| VGPKRFGV | ASSRDRGWDSYEQY | 0.8468223 |
| VNVAFPD | ASSRDRGWDSYEQY | 0.79594177 |
| VSMTNSGVS | ASSRDRGWDSYEQY | 0.84466696 |
| VTDVLGDLSV | ASSRDRGWDSYEQY | 0.64530957 |
| VVVVGAGFAGL | ASSRDRGWDSYEQY | 0.5409898 |
| VYTVGTVV | ASSRDRGWDSYEQY | 0.73561996 |
| WTTFSGP | ASSRDRGWDSYEQY | 0.84058094 |
| YDELVEAF | ASSRDRGWDSYEQY | 0.845034 |
| YGTGGAGGNGG | ASSRDRGWDSYEQY | 0.81297743 |
| YTNVMAV | ASSRDRGWDSYEQY | 0.56062317 |

| | | |
|---|---|---|
| FLEQVHQL | ASSRDRGWDSYEQY | 0.6574276 |
| GLSLLAVAV | ASSRDRGWDSYEQY | 0.6576971 |
| LEQVHQL | ASSRDRGWDSYEQY | 0.6581849 |
| LNAAGAI | ASSRDRGWDSYEQY | 0.809676 |
| PAVHASL | ASSRDRGWDSYEQY | 0.7724196 |
| TELEDDA | ASSRDRGWDSYEQY | 0.6690042 |
| TPPGPPN | ASSRDRGWDSYEQY | 0.6607671 |
| TPPPPPPP | ASSRDRGWDSYEQY | 0.79217416 |
| TQSPALI | ASSFGGGGNTEVF | 0.8188647 |
| EFAAAFPQ | ASSFGGGGNTEVF | 0.7582655 |
| LLSGNLDVL | ASSFGGGGNTEVF | 0.8679771 |
| GPANLGNYNI | ASSFGGGGNTEVF | 0.82004845 |
| TLGPTLP | ASSFGGGGNTEVF | 0.843874 |
| VYAYSSCCK | ASSFGGGGNTEVF | 0.14580274 |
| APAPAPAPNI | ASSFGGGGNTEVF | 0.2748559 |
| PGGPVPIP | ASSFGGGGNTEVF | 0.65391237 |
| FLTVALEP | ASSFGGGGNTEVF | 0.63870066 |
| SQYFGANL | ASSFGGGGNTEVF | 0.37851068 |
| GGVKPAA | ASSFGGGGNTEVF | 0.7618283 |
| GGTGGAGGAGGW | ASSFGGGGNTEVF | 0.6068623 |
| GVGGVAGLGGAGG | ASSFGGGGNTEVF | 0.54481214 |
| VLLVLATTI | ASSFGGGGNTEVF | 0.8320391 |
| IGIFAAML | ASSFGGGGNTEVF | 0.66517407 |
| AVGDLID | ASSFGGGGNTEVF | 0.5884641 |
| NISPLPL | ASSFGGGGNTEVF | 0.5160493 |
| VTAVFVAI | ASSFGGGGNTEVF | 0.836508 |
| SGDDNDAAL | ASSFGGGGNTEVF | 0.29962912 |
| AVAALAHS | ASSFGGGGNTEVF | 0.77566826 |
| GGVTVMATP | ASSFGGGGNTEVF | 0.63081014 |
| AGLAANVRD | ASSFGGGGNTEVF | 0.8189858 |
| TNRPDLIDPALLRPG | ASSFGGGGNTEVF | 0.45696968 |
| LANLGVTAL | ASSFGGGGNTEVF | 0.81239253 |
| TGERIEL | ASSFGGGGNTEVF | 0.7389034 |
| NGEELAA | ASSFGGGGNTEVF | 0.87895596 |
| VLSLVWG | ASSFGGGGNTEVF | 0.67323434 |
| IGYALWL | ASSFGGGGNTEVF | 0.7593671 |
| LSLLLAPL | ASSFGGGGNTEVF | 0.6411346 |
| TLDEALK | ASSFGGGGNTEVF | 0.8270628 |
| TDGAGTAEGG | ASSFGGGGNTEVF | 0.48622978 |
| NAPGGAGGNGG | ASSFGGGGNTEVF | 0.67738396 |
| VSVFMGMGGNF | ASSFGGGGNTEVF | 0.3634798 |
| LASYLAAL | ASSFGGGGNTEVF | 0.7658727 |
| LVGGTPAL | ASSFGGGGNTEVF | 0.79384184 |
| VRNADVL | ASSFGGGGNTEVF | 0.36564207 |
| TCSSPFLVSAM | ASSFGGGGNTEVF | 0.8290771 |
| VHITDAE | ASSFGGGGNTEVF | 0.76818883 |
| EYAAPSF | ASSFGGGGNTEVF | 0.65660477 |
| VVDAFHGGL | ASSFGGGGNTEVF | 0.85633767 |
| ASVVDPQ | ASSFGGGGNTEVF | 0.6265865 |
| SYNEFGGLS | ASSFGGGGNTEVF | 0.8632554 |
| DPDGSVY | ASSFGGGGNTEVF | 0.71580595 |

| | | |
|---|---|---|
| MVSTIHATE | ASSFGGGGNTEVF | 0.66954684 |
| LAAGVADPV | ASSFGGGGNTEVF | 0.80878633 |
| TKEQIAA | ASSFGGGGNTEVF | 0.38334072 |
| SLASPAAL | ASSFGGGGNTEVF | 0.4737602 |
| IGLVFYGV | ASSFGGGGNTEVF | 0.84555733 |
| FLLDASNL | ASSFGGGGNTEVF | 0.8801351 |
| YQVGNFP | ASSFGGGGNTEVF | 0.87886316 |
| ESLLSQST | ASSFGGGGNTEVF | 0.7990379 |
| LFTVGFAAVG | ASSFGGGGNTEVF | 0.7032191 |
| ATPPTPT | ASSFGGGGNTEVF | 0.8324062 |
| TDLATLG | ASSFGGGGNTEVF | 0.8830007 |
| SAGGTGR | ASSFGGGGNTEVF | 0.88845956 |
| GFSNSGS | ASSFGGGGNTEVF | 0.87065566 |
| GGNGGSGGTSV | ASSFGGGGNTEVF | 0.6837654 |
| GTSVATGGAGN | ASSFGGGGNTEVF | 0.8650815 |
| NGGNGGSGGTS | ASSFGGGGNTEVF | 0.8329467 |
| AVVKFPH | ASSFGGGGNTEVF | 0.8251706 |
| TPPPPPP | ASSFGGGGNTEVF | 0.88096255 |
| AAPAAAAAPA | ASSFGGGGNTEVF | 0.8636128 |
| IALNEDL | ASSFGGGGNTEVF | 0.82243514 |
| IVNVSSM | ASSFGGGGNTEVF | 0.7793476 |
| TGIPPAPR | ASSFGGGGNTEVF | 0.8714489 |
| FDLGGGTF | ASSFGGGGNTEVF | 0.88801277 |
| FDLGGGTFDVS | ASSFGGGGNTEVF | 0.8869709 |
| NEPTAAA | ASSFGGGGNTEVF | 0.8968379 |
| NEPTAAALA | ASSFGGGGNTEVF | 0.8890219 |
| GSIINISSI | ASSFGGGGNTEVF | 0.8804276 |
| VAPGNAGTA | ASSFGGGGNTEVF | 0.24586023 |
| TGGGASLE | ASSFGGGGNTEVF | 0.5481368 |
| VAVYNHYKRI | ASSFGGGGNTEVF | 0.7942602 |
| TDPAPATT | ASSFGGGGNTEVF | 0.8263534 |
| TNRPDILDPALLRPG | ASSFGGGGNTEVF | 0.49464947 |
| LTTTTTS | ASSFGGGGNTEVF | 0.8354475 |
| PTGSGKTL | ASSFGGGGNTEVF | 0.8073109 |
| TGSGKTL | ASSFGGGGNTEVF | 0.7422097 |
| TIGHVDHGK | ASSFGGGGNTEVF | 0.8895809 |
| TIGHVDHGKTT | ASSFGGGGNTEVF | 0.8056767 |
| GHVDHGKTT | ASSFGGGGNTEVF | 0.747406 |
| AQFEHTLL | ASSFGGGGNTEVF | 0.78976357 |
| IVNQDEL | ASSFGGGGNTEVF | 0.89529324 |
| TNGSQFF | ASSFGGGGNTEVF | 0.7223049 |
| TGKGIVE | ASSFGGGGNTEVF | 0.87868404 |
| LISNASDA | ASSFGGGGNTEVF | 0.80665004 |
| AELRTLGELTKIAKAHGAQ | ASSFGGGGNTEVF | 0.89154136 |
| APNAVAAAEPSW | ASSFGGGGNTEVF | 0.57950634 |
| APQVAGAATAVL | ASSFGGGGNTEVF | 0.8820301 |
| ARRDPAAAAAILPTDA | ASSFGGGGNTEVF | 0.84195006 |
| DAAARDVLAEAGLAEYF | ASSFGGGGNTEVF | 0.90580714 |
| DDQIVLSAKANLAKVEQ | ASSFGGGGNTEVF | 0.825567 |
| DELLPDTGEDSDEDGH | ASSFGGGGNTEVF | 0.8622948 |
| ELPEAPANRATVPQD | ASSFGGGGNTEVF | 0.8324347 |

| | | |
|---|---|---|
| FGQSKIERVTGIG | ASSFGGGGNTEVF | 0.87955666 |
| GAGGTGSTAGGAGGAGGAGGLYA | ASSFGGGGNTEVF | 0.6220447 |
| GHGGTGAAGGNGGTGGAGGSGIDG | ASSFGGGGNTEVF | 0.844832 |
| GNGGNGGAGGEGGPGGAGGAGA | ASSFGGGGNTEVF | 0.6058756 |
| GSTAGNGGNGGNATNGTIAGQPAG | ASSFGGGGNTEVF | 0.85336876 |
| GTGGDGGNGGNGGTGG | ASSFGGGGNTEVF | 0.8679158 |
| GVRRPAGIPADAPV | ASSFGGGGNTEVF | 0.35596693 |
| GYRQVTDAHLLTLARR | ASSFGGGGNTEVF | 0.88594615 |
| IAKMGQDGHDRG | ASSFGGGGNTEVF | 0.79514754 |
| KAEPSIEIQYT | ASSFGGGGNTEVF | 0.75769377 |
| KPVRIRMAADEDTVAVT | ASSFGGGGNTEVF | 0.89368105 |
| MPPRIAGMRLLVIKPEP | ASSFGGGGNTEVF | 0.84833944 |
| MKFYAVALFRSRGVPPTR | ASSFGGGGNTEVF | 0.89066476 |
| NAGGFGILGAVAHSP | ASSFGGGGNTEVF | 0.84239566 |
| NEGREITLAMAEM | ASSFGGGGNTEVF | 0.85173124 |
| NELVEWSHPLGHRWRWE | ASSFGGGGNTEVF | 0.86131436 |
| NLGAGGTGGNGGNPG | ASSFGGGGNTEVF | 0.6037434 |
| NTEWPKWETVMAADPVD | ASSFGGGGNTEVF | 0.7634481 |
| PASAAPAETQP | ASSFGGGGNTEVF | 0.8920232 |
| QQSSDGKNGVRAAEITGE | ASSFGGGGNTEVF | 0.82265824 |
| RAPDAVTTSWWKE | ASSFGGGGNTEVF | 0.81023824 |
| SGFGNVGAHTSGWFNQ | ASSFGGGGNTEVF | 0.681461 |
| SGVAGAVGQAASVGGLKV | ASSFGGGGNTEVF | 0.6923649 |
| SNGGAGTGSAGGNGGT | ASSFGGGGNTEVF | 0.753959 |
| TGGAGGEGGAAG | ASSFGGGGNTEVF | 0.80660653 |
| TGGDGGAAGAGGAPGHG | ASSFGGGGNTEVF | 0.845391 |
| TMPPLLAAIANAGRHGV | ASSFGGGGNTEVF | 0.81925523 |
| TSSTAGNSGGA | ASSFGGGGNTEVF | 0.735947 |
| VPATVPVGILC | ASSFGGGGNTEVF | 0.81221604 |
| VQRPDSGGVVLTGRIS | ASSFGGGGNTEVF | 0.78625274 |
| WTPNPVPATKGVPAGLVQAT | ASSFGGGGNTEVF | 0.7782563 |
| YALKRALAKFSDRFDVVII | ASSFGGGGNTEVF | 0.80149716 |
| YLIGSVIV | ASSFGGGGNTEVF | 0.78917205 |
| YGQLVAVID | ASSFGGGGNTEVF | 0.7028337 |
| VTVGGMTLPA | ASSFGGGGNTEVF | 0.69249296 |
| VLALPAV | ASSFGGGGNTEVF | 0.81307834 |
| VIPPTLNYET | ASSFGGGGNTEVF | 0.8301492 |
| VGMGGMHPSSG | ASSFGGGGNTEVF | 0.82840335 |
| VGLAGGAATA | ASSFGGGGNTEVF | 0.84585536 |
| VGGAGGAGGSGP | ASSFGGGGNTEVF | 0.83486676 |
| VFAAFMV | ASSFGGGGNTEVF | 0.8156667 |
| TLYDKLR | ASSFGGGGNTEVF | 0.40023333 |
| TLAVFTVL | ASSFGGGGNTEVF | 0.60115784 |
| TGSVQPARV | ASSFGGGGNTEVF | 0.74091494 |
| TGGGAGSGGT | ASSFGGGGNTEVF | 0.75299203 |
| SHLETSL | ASSFGGGGNTEVF | 0.39354667 |
| SFTTGMT | ASSFGGGGNTEVF | 0.8014803 |
| QVEVAKRMGVRQPTVS | ASSFGGGGNTEVF | 0.7651403 |
| QGSGGPALAM | ASSFGGGGNTEVF | 0.7022776 |
| MLAPSNP | ASSFGGGGNTEVF | 0.82993925 |
| LVNNAAI | ASSFGGGGNTEVF | 0.7438519 |

| | | |
|---|---|---|
| LTGEINTLA | ASSFGGGGNTEVF | 0.6454469 |
| LSIFGPV | ASSFGGGGNTEVF | 0.73606753 |
| LLALLANL | ASSFGGGGNTEVF | 0.8470882 |
| LISNASDALDKLR | ASSFGGGGNTEVF | 0.71176314 |
| LINEAVP | ASSFGGGGNTEVF | 0.83718723 |
| LGYAFSGL | ASSFGGGGNTEVF | 0.74967897 |
| LDFVAVAL | ASSFGGGGNTEVF | 0.83800054 |
| LAQPGAE | ASSFGGGGNTEVF | 0.66428703 |
| ISNLGAVTGF | ASSFGGGGNTEVF | 0.83684057 |
| HQEDPGRI | ASSFGGGGNTEVF | 0.79243934 |
| GTPGEPTP | ASSFGGGGNTEVF | 0.85323155 |
| GFFNSGS | ASSFGGGGNTEVF | 0.7753986 |
| GAGGGGLTAGAN | ASSFGGGGNTEVF | 0.8246399 |
| FVDALVNM | ASSFGGGGNTEVF | 0.7811485 |
| FLTVAPEL | ASSFGGGGNTEVF | 0.83046657 |
| FETLTAL | ASSFGGGGNTEVF | 0.6473143 |
| DVINPIEL | ASSFGGGGNTEVF | 0.7584748 |
| APVSGAPV | ASSFGGGGNTEVF | 0.6616224 |
| ALVGGLAW | ASSFGGGGNTEVF | 0.76548016 |
| ALAAAAATAI | ASSFGGGGNTEVF | 0.8279277 |
| AFGLGLLL | ASSFGGGGNTEVF | 0.3074366 |
| AFFAAASL | ASSFGGGGNTEVF | 0.8307981 |
| AAAALPAFSPP | ASSFGGGGNTEVF | 0.5906359 |
| VVLAPAL | ASSFGGGGNTEVF | 0.6930384 |
| TIGHVDHGKTTL | ASSFGGGGNTEVF | 0.8270521 |
| LISNASDALDK | ASSFGGGGNTEVF | 0.7873206 |
| LISNASDAL | ASSFGGGGNTEVF | 0.8304179 |
| ISNASDALDK | ASSFGGGGNTEVF | 0.7286551 |
| NQAETLVYQTE | ASSFGGGGNTEVF | 0.7869556 |
| TGGTGGTAGAG | ASSFGGGGNTEVF | 0.79050195 |
| TGSTAGGAGGA | ASSFGGGGNTEVF | 0.54328644 |
| DPLAVPVAAGPV | ASSFGGGGNTEVF | 0.67636853 |
| GGTGGTPGNGGN | ASSFGGGGNTEVF | 0.84880614 |
| PAGLAGVDPATL | ASSFGGGGNTEVF | 0.7614557 |
| GNNLVVEAPHAGASV | ASSFGGGGNTEVF | 0.76860344 |
| KVTRIMSASTLEQPA | ASSFGGGGNTEVF | 0.64882743 |
| LGFALPIALPRRTVV | ASSFGGGGNTEVF | 0.64601487 |
| TRLGNLTARKLVVLV | ASSFGGGGNTEVF | 0.6824434 |
| WSQQRRLIYFPSAGP | ASSFGGGGNTEVF | 0.7716928 |
| GPNGNPDPMAAAVDIR | ASSFGGGGNTEVF | 0.7817229 |
| TQRCVSVADLRAAASR | ASSFGGGGNTEVF | 0.7635981 |
| QFVLDVAEHPVEVLTETV | ASSFGGGGNTEVF | 0.62437797 |
| GGAGGTGGAGGNGGTQAGDG | ASSFGGGGNTEVF | 0.7390926 |
| PLLATLRAIAVDVDAKPAQV | ASSFGGGGNTEVF | 0.69567347 |
| APKFRTGGYMIDFWGVGYQVAKRM | ASSFGGGGNTEVF | 0.70613796 |
| GAGGAGGAGGAAGGPGGTGGTGGN | ASSFGGGGNTEVF | 0.74918437 |
| IGALGLLATDVAPAGSVAFGVALA | ASSFGGGGNTEVF | 0.73149955 |
| AGGAGGIGGNGGAGGAGGVLMGNG | ASSFGGGGNTEVF | 0.7007828 |
| ADELIGLL | ASSFGGGGNTEVF | 0.5830108 |
| ADRALKA | ASSFGGGGNTEVF | 0.74822307 |
| AGAVGQVGGAGGA | ASSFGGGGNTEVF | 0.78611267 |

| | | |
|---|---|---|
| AGGGGGLGGAP | ASSFGGGGNTEVF | 0.772164 |
| AGGLLYGNGGAG | ASSFGGGGNTEVF | 0.7877131 |
| AGLESIGL | ASSFGGGGNTEVF | 0.7713301 |
| AGLKTGII | ASSFGGGGNTEVF | 0.7679243 |
| ASLAALLL | ASSFGGGGNTEVF | 0.26347774 |
| ASVINISSI | ASSFGGGGNTEVF | 0.43641132 |
| ATPPGPA | ASSFGGGGNTEVF | 0.55785286 |
| ATVASALV | ASSFGGGGNTEVF | 0.69660723 |
| ATVASLIL | ASSFGGGGNTEVF | 0.30449274 |
| AVNGPVLIRH | ASSFGGGGNTEVF | 0.7472564 |
| DGAAGAVGI | ASSFGGGGNTEVF | 0.6907202 |
| DPLLHGI | ASSFGGGGNTEVF | 0.6607335 |
| ELGAALGAL | ASSFGGGGNTEVF | 0.77378035 |
| ELGRTQAV | ASSFGGGGNTEVF | 0.6905016 |
| FVAGGYLS | ASSFGGGGNTEVF | 0.5951497 |
| GASGGAGGAGGW | ASSFGGGGNTEVF | 0.59996414 |
| GGAGAGGDGGAGGI | ASSFGGGGNTEVF | 0.7877831 |
| GTGHNGSQPGMGGNG | ASSFGGGGNTEVF | 0.44310308 |
| GYAAASASAS | ASSFGGGGNTEVF | 0.7671089 |
| HQVDVAAVQQG | ASSFGGGGNTEVF | 0.59472203 |
| IGAAIAHT | ASSFGGGGNTEVF | 0.7780735 |
| IGMIEVAQL | ASSFGGGGNTEVF | 0.55655193 |
| IRLAPEHTSAMD | ASSFGGGGNTEVF | 0.772691 |
| IYNVSVL | ASSFGGGGNTEVF | 0.7176153 |
| KQGFLGM | ASSFGGGGNTEVF | 0.7986499 |
| LESVITQL | ASSFGGGGNTEVF | 0.70054865 |
| LGLGALVVV | ASSFGGGGNTEVF | 0.7533779 |
| LIGNGGNGGSGGI | ASSFGGGGNTEVF | 0.7167245 |
| LNAINAPTL | ASSFGGGGNTEVF | 0.7669697 |
| LQLLNTGV | ASSFGGGGNTEVF | 0.46672332 |
| LQTSIEF | ASSFGGGGNTEVF | 0.6933658 |
| MSFAVSEL | ASSFGGGGNTEVF | 0.59367895 |
| NTGALGL | ASSFGGGGNTEVF | 0.72254884 |
| PPGAGGAGGAGTT | ASSFGGGGNTEVF | 0.7675371 |
| QLVGLGI | ASSFGGGGNTEVF | 0.30004507 |
| RFLEQVHQL | ASSFGGGGNTEVF | 0.74810183 |
| RGVGYVLR | ASSFGGGGNTEVF | 0.65192246 |
| SATGVFAGVI | ASSFGGGGNTEVF | 0.65072227 |
| SLPAITLP | ASSFGGGGNTEVF | 0.76606023 |
| STALAFTV | ASSFGGGGNTEVF | 0.7182095 |
| SVLTVGLGK | ASSFGGGGNTEVF | 0.7619152 |
| TALAGAGSGI | ASSFGGGGNTEVF | 0.61384153 |
| TGLPFTGI | ASSFGGGGNTEVF | 0.6638422 |
| VAEHVTV | ASSFGGGGNTEVF | 0.7517582 |
| VDIVLNSL | ASSFGGGGNTEVF | 0.37444696 |
| VLTGGGAVL | ASSFGGGGNTEVF | 0.5043089 |
| VNVGLAYGI | ASSFGGGGNTEVF | 0.7845073 |
| VSSAAVGAV | ASSFGGGGNTEVF | 0.6707293 |
| VTSVYEAGV | ASSFGGGGNTEVF | 0.6800804 |
| WVALDRGA | ASSFGGGGNTEVF | 0.63691556 |
| YLVGSVVI | ASSFGGGGNTEVF | 0.6030546 |

| | | |
|---|---|---|
| GPPGTGKTLL | ASSFGGGGNTEVF | 0.6343422 |
| GVGMTKF | ASSFGGGGNTEVF | 0.6824764 |
| IGIGGSDL | ASSFGGGGNTEVF | 0.67636573 |
| IGTLLAEL | ASSFGGGGNTEVF | 0.66458666 |
| IYTSGTT | ASSFGGGGNTEVF | 0.5166646 |
| LSFPAGV | ASSFGGGGNTEVF | 0.66196847 |
| LYTSGTT | ASSFGGGGNTEVF | 0.6494815 |
| VGAFPFGF | ASSFGGGGNTEVF | 0.6326117 |
| VGAGTVEF | ASSFGGGGNTEVF | 0.6614986 |
| VGKSTLLL | ASSFGGGGNTEVF | 0.6670786 |
| AALAHAGGGIV | ASSFGGGGNTEVF | 0.6169703 |
| AALVAVSLL | ASSFGGGGNTEVF | 0.640151 |
| AAVSRAVAAL | ASSFGGGGNTEVF | 0.65706915 |
| AGGSSGIGGF | ASSFGGGGNTEVF | 0.67602843 |
| AGLTVANANG | ASSFGGGGNTEVF | 0.6761329 |
| AGLVGAVL | ASSFGGGGNTEVF | 0.68776256 |
| ALSQAFTV | ASSFGGGGNTEVF | 0.68359375 |
| ALTFSVSL | ASSFGGGGNTEVF | 0.6815003 |
| ALTGAALVAL | ASSFGGGGNTEVF | 0.37495598 |
| ASFSIGAGVL | ASSFGGGGNTEVF | 0.49249464 |
| DGSVYKI | ASSFGGGGNTEVF | 0.6327317 |
| EQLAEIV | ASSFGGGGNTEVF | 0.6355009 |
| FAVSELTVM | ASSFGGGGNTEVF | 0.507735 |
| GAAPLAAGL | ASSFGGGGNTEVF | 0.64794326 |
| GADGGDGGSGGAGG | ASSFGGGGNTEVF | 0.6192772 |
| GFGLDTGGAGG | ASSFGGGGNTEVF | 0.5962961 |
| GLASGGLL | ASSFGGGGNTEVF | 0.677199 |
| GRIAGAAVLL | ASSFGGGGNTEVF | 0.6282279 |
| GSAGSLNTSF | ASSFGGGGNTEVF | 0.6203972 |
| GTGQIGI | ASSFGGGGNTEVF | 0.58518195 |
| GTLIALIL | ASSFGGGGNTEVF | 0.67556465 |
| IAATMGFVVG | ASSFGGGGNTEVF | 0.54276943 |
| IFAQALL | ASSFGGGGNTEVF | 0.67074394 |
| IGAAVALGF | ASSFGGGGNTEVF | 0.53706545 |
| IGLTPLAL | ASSFGGGGNTEVF | 0.68531835 |
| ILGSNTSTL | ASSFGGGGNTEVF | 0.5147404 |
| IVFSLTVL | ASSFGGGGNTEVF | 0.67574656 |
| LASKATGF | ASSFGGGGNTEVF | 0.64888805 |
| LASVALIL | ASSFGGGGNTEVF | 0.6955316 |
| LFAVGLTL | ASSFGGGGNTEVF | 0.6568489 |
| LFNGGIGTYI | ASSFGGGGNTEVF | 0.6568847 |
| LTLGAGII | ASSFGGGGNTEVF | 0.58683074 |
| LVAAALGDI | ASSFGGGGNTEVF | 0.6763808 |
| NTAFVQLGI | ASSFGGGGNTEVF | 0.5228194 |
| QLFSGAEL | ASSFGGGGNTEVF | 0.65637964 |
| SAATFITL | ASSFGGGGNTEVF | 0.47861353 |
| SNYGLANLGV | ASSFGGGGNTEVF | 0.6660656 |
| SYGGAYAVM | ASSFGGGGNTEVF | 0.6677685 |
| TTGVGGLTLG | ASSFGGGGNTEVF | 0.5201019 |
| TTQTIYI | ASSFGGGGNTEVF | 0.6746088 |
| VAAGAGLGAVI | ASSFGGGGNTEVF | 0.6465232 |

| | | |
|---|---|---|
| VAAVVTLL | ASSFGGGGNTEVF | 0.606468 |
| VGFLGGLI | ASSFGGGGNTEVF | 0.687456 |
| VGVIPILL | ASSFGGGGNTEVF | 0.65453637 |
| VIFASSLI | ASSFGGGGNTEVF | 0.6828472 |
| VKGGDGGAAAG | ASSFGGGGNTEVF | 0.6407182 |
| VTKTFTNI | ASSFGGGGNTEVF | 0.6380374 |
| VVAAAILL | ASSFGGGGNTEVF | 0.6632572 |
| YAALDVEL | ASSFGGGGNTEVF | 0.6433865 |
| LASVALLL | ASSFGGGGNTEVF | 0.5970509 |
| LDAVHRV | ASSFGGGGNTEVF | 0.68385017 |
| QGLSSSIF | ASSFGGGGNTEVF | 0.73855054 |
| RGTFANIRL | ASSFGGGGNTEVF | 0.727961 |
| RQLWWGHRI | ASSFGGGGNTEVF | 0.69838446 |
| YGPPGTGKTLL | ASSFGGGGNTEVF | 0.6775757 |
| AALVGAPANVA | ASSFGGGGNTEVF | 0.7061318 |
| AATFRRTE | ASSFGGGGNTEVF | 0.7370863 |
| AGALGGGI | ASSFGGGGNTEVF | 0.73978937 |
| AGGAGGAGGIGAG | ASSFGGGGNTEVF | 0.73173076 |
| AGSGALGL | ASSFGGGGNTEVF | 0.68784773 |
| AQIADQI | ASSFGGGGNTEVF | 0.7332895 |
| ASAVAVTYL | ASSFGGGGNTEVF | 0.7289 |
| ASTVLGGTGG | ASSFGGGGNTEVF | 0.700601 |
| AVGGEPAK | ASSFGGGGNTEVF | 0.73735905 |
| CEQGTFGGTC | ASSFGGGGNTEVF | 0.7375266 |
| DEGKAPAG | ASSFGGGGNTEVF | 0.7081604 |
| DTVVFPAL | ASSFGGGGNTEVF | 0.7170004 |
| FEGITMAG | ASSFGGGGNTEVF | 0.7292826 |
| FPLVVQD | ASSFGGGGNTEVF | 0.73833966 |
| GAFTTPQ | ASSFGGGGNTEVF | 0.7370552 |
| GFGGFGTAGGVGGA | ASSFGGGGNTEVF | 0.7442024 |
| GFLAQFL | ASSFGGGGNTEVF | 0.7413875 |
| GGAGGAGGNAGVA | ASSFGGGGNTEVF | 0.7361435 |
| GGVGTAGAGGAGGAGG | ASSFGGGGNTEVF | 0.5830363 |
| GNGGGGDAGNAGSGGN | ASSFGGGGNTEVF | 0.6313139 |
| GTGGAGGAGNDAGS | ASSFGGGGNTEVF | 0.73138326 |
| GTTNTGWFNS | ASSFGGGGNTEVF | 0.7110082 |
| IFMGPLFTL | ASSFGGGGNTEVF | 0.5877509 |
| ILAHAIEAI | ASSFGGGGNTEVF | 0.72439003 |
| IRSAIREAL | ASSFGGGGNTEVF | 0.7174382 |
| ITFAGGG | ASSFGGGGNTEVF | 0.6906378 |
| ITGMSGHQEV | ASSFGGGGNTEVF | 0.7365813 |
| IVAGLACTL | ASSFGGGGNTEVF | 0.71321297 |
| IVGGGLAGA | ASSFGGGGNTEVF | 0.7032317 |
| IVTGVLNI | ASSFGGGGNTEVF | 0.7196647 |
| LAQQGCI | ASSFGGGGNTEVF | 0.73803115 |
| LCLHVEN | ASSFGGGGNTEVF | 0.6929378 |
| LDAISDGS | ASSFGGGGNTEVF | 0.7365327 |
| LGGFAAML | ASSFGGGGNTEVF | 0.6903069 |
| LGGKNPMI | ASSFGGGGNTEVF | 0.74311274 |
| LGPKGVAAIL | ASSFGGGGNTEVF | 0.6602991 |
| LMVGVSI | ASSFGGGGNTEVF | 0.73612505 |

| | | |
|---|---|---|
| LVGAAFVL | ASSFGGGGNTEVF | 0.7176029 |
| LVLAVAPL | ASSFGGGGNTEVF | 0.74867904 |
| MAAIAVIDAA | ASSFGGGGNTEVF | 0.7219423 |
| NNGINAELQ | ASSFGGGGNTEVF | 0.7400578 |
| RGEPLVLVHG | ASSFGGGGNTEVF | 0.7302996 |
| RGGAGGAAGLI | ASSFGGGGNTEVF | 0.7312084 |
| RSYRMHPAV | ASSFGGGGNTEVF | 0.66894454 |
| SAAHLLAGQLA | ASSFGGGGNTEVF | 0.72946405 |
| SAAVGSY | ASSFGGGGNTEVF | 0.6664772 |
| SPHAVLHRI | ASSFGGGGNTEVF | 0.7301698 |
| SQIAKVL | ASSFGGGGNTEVF | 0.7334143 |
| TALGHLVP | ASSFGGGGNTEVF | 0.5106019 |
| TAPTTSGAA | ASSFGGGGNTEVF | 0.7375958 |
| TDGDDGNGGSPGT | ASSFGGGGNTEVF | 0.6917077 |
| TDGLGEN | ASSFGGGGNTEVF | 0.7079031 |
| TGGPPLPGQA | ASSFGGGGNTEVF | 0.744582 |
| TIPAITI | ASSFGGGGNTEVF | 0.7346847 |
| TIPGFTI | ASSFGGGGNTEVF | 0.7423638 |
| TSIASGALA | ASSFGGGGNTEVF | 0.72410834 |
| TVTLPGAAI | ASSFGGGGNTEVF | 0.71913636 |
| VASGLSAALP | ASSFGGGGNTEVF | 0.7230583 |
| VGAGVIGI | ASSFGGGGNTEVF | 0.6042977 |
| VGPKRFGV | ASSFGGGGNTEVF | 0.7035001 |
| VNVAFPD | ASSFGGGGNTEVF | 0.7430242 |
| VSMTNSGVS | ASSFGGGGNTEVF | 0.6875708 |
| VTDVLGDLSV | ASSFGGGGNTEVF | 0.75526965 |
| VVVVGAGFAGL | ASSFGGGGNTEVF | 0.5028872 |
| VYTVGTVV | ASSFGGGGNTEVF | 0.4215807 |
| WTTFSGP | ASSFGGGGNTEVF | 0.47513002 |
| YDELVEAF | ASSFGGGGNTEVF | 0.7714149 |
| YGTGGAGGNGG | ASSFGGGGNTEVF | 0.7138519 |
| YTNVMAV | ASSFGGGGNTEVF | 0.7261342 |
| FLEQVHQL | ASSFGGGGNTEVF | 0.4388564 |
| GLSLLAVAV | ASSFGGGGNTEVF | 0.5791461 |
| LEQVHQL | ASSFGGGGNTEVF | 0.5075742 |
| LNAAGAI | ASSFGGGGNTEVF | 0.5888474 |
| PAVHASL | ASSFGGGGNTEVF | 0.66446537 |
| TELEDDA | ASSFGGGGNTEVF | 0.7239369 |
| TPPGPPN | ASSFGGGGNTEVF | 0.5432215 |
| TPPPPPPP | ASSFGGGGNTEVF | 0.48522562 |
| TQSPALI | ASSSRISNERLF | 0.57939583 |
| EFAAAFPQ | ASSSRISNERLF | 0.6935901 |
| LLSGNLDVL | ASSSRISNERLF | 0.64342666 |
| GPANLGNYNI | ASSSRISNERLF | 0.7980677 |
| TLGPTLP | ASSSRISNERLF | 0.7324655 |
| VYAYSSCCK | ASSSRISNERLF | 0.724317 |
| APAPAPAPNI | ASSSRISNERLF | 0.18464863 |
| PGGPVPIP | ASSSRISNERLF | 0.2853673 |
| FLTVALEP | ASSSRISNERLF | 0.5063839 |
| SQYFGANL | ASSSRISNERLF | 0.5804137 |
| GGVKPAA | ASSSRISNERLF | 0.2476841 |

| | | |
|---|---|---|
| GGTGGAGGAGGW | ASSSRISNERLF | 0.6893182 |
| GVGGVAGLGGAGG | ASSSRISNERLF | 0.53856635 |
| VLLVLATTI | ASSSRISNERLF | 0.50984204 |
| IGIFAAML | ASSSRISNERLF | 0.7126704 |
| AVGDLID | ASSSRISNERLF | 0.54699826 |
| NISPLPL | ASSSRISNERLF | 0.52778363 |
| VTAVFVAI | ASSSRISNERLF | 0.509606 |
| SGDDNDAAL | ASSSRISNERLF | 0.71933436 |
| AVAALAHS | ASSSRISNERLF | 0.3039616 |
| GGVTVMATP | ASSSRISNERLF | 0.6977759 |
| AGLAANVRD | ASSSRISNERLF | 0.46572632 |
| TNRPDLIDPALLRPG | ASSSRISNERLF | 0.7488506 |
| LANLGVTAL | ASSSRISNERLF | 0.39139116 |
| TGERIEL | ASSSRISNERLF | 0.74399364 |
| NGEELAA | ASSSRISNERLF | 0.61132264 |
| VLSLVWG | ASSSRISNERLF | 0.80812025 |
| IGYALWL | ASSSRISNERLF | 0.5203875 |
| LSLLLAPL | ASSSRISNERLF | 0.6302289 |
| TLDEALK | ASSSRISNERLF | 0.56457967 |
| TDGAGTAEGG | ASSSRISNERLF | 0.71702635 |
| NAPGGAGGNGG | ASSSRISNERLF | 0.321457 |
| VSVFMGMGGNF | ASSSRISNERLF | 0.5403531 |
| LASYLAAL | ASSSRISNERLF | 0.28998932 |
| LVGGTPAL | ASSSRISNERLF | 0.65057915 |
| VRNADVL | ASSSRISNERLF | 0.7385122 |
| TCSSPFLVSAM | ASSSRISNERLF | 0.3004903 |
| VHITDAE | ASSSRISNERLF | 0.67122436 |
| EYAAPSF | ASSSRISNERLF | 0.65707856 |
| VVDAFHGGL | ASSSRISNERLF | 0.4935724 |
| ASVVDPQ | ASSSRISNERLF | 0.74467444 |
| SYNEFGGLS | ASSSRISNERLF | 0.5661391 |
| DPDGSVY | ASSSRISNERLF | 0.7559959 |
| MVSTIHATE | ASSSRISNERLF | 0.6030171 |
| LAAGVADPV | ASSSRISNERLF | 0.47783473 |
| TKEQIAA | ASSSRISNERLF | 0.7028737 |
| SLASPAAL | ASSSRISNERLF | 0.28446257 |
| IGLVFYGV | ASSSRISNERLF | 0.428036 |
| FLLDASNL | ASSSRISNERLF | 0.77123004 |
| YQVGNFP | ASSSRISNERLF | 0.7800793 |
| ESLLSQST | ASSSRISNERLF | 0.8601353 |
| LFTVGFAAVG | ASSSRISNERLF | 0.6078423 |
| ATPPTPT | ASSSRISNERLF | 0.4777396 |
| TDLATLG | ASSSRISNERLF | 0.6666405 |
| SAGGTGR | ASSSRISNERLF | 0.84513414 |
| GFSNSGS | ASSSRISNERLF | 0.8522328 |
| GGNGGSGGTSV | ASSSRISNERLF | 0.8041494 |
| GTSVATGGAGN | ASSSRISNERLF | 0.44037914 |
| NGGNGGSGGTS | ASSSRISNERLF | 0.7334945 |
| AVVKFPH | ASSSRISNERLF | 0.6485585 |
| TPPPPPP | ASSSRISNERLF | 0.6969279 |
| AAPAAAAAPA | ASSSRISNERLF | 0.76535076 |

| | | |
|---|---|---|
| IALNEDL | ASSSRISNERLF | 0.78512424 |
| IVNVSSM | ASSSRISNERLF | 0.65194786 |
| TGIPPAPR | ASSSRISNERLF | 0.5245749 |
| FDLGGGTF | ASSSRISNERLF | 0.78897536 |
| FDLGGGTFDVS | ASSSRISNERLF | 0.7956822 |
| NEPTAAA | ASSSRISNERLF | 0.8008537 |
| NEPTAAALA | ASSSRISNERLF | 0.8574185 |
| GSIINISSI | ASSSRISNERLF | 0.8406116 |
| VAPGNAGTA | ASSSRISNERLF | 0.8265466 |
| TGGGASLE | ASSSRISNERLF | 0.14436096 |
| VAVYNHYKRI | ASSSRISNERLF | 0.25010014 |
| TDPAPATT | ASSSRISNERLF | 0.55952626 |
| TNRPDILDPALLRPG | ASSSRISNERLF | 0.6826279 |
| LTTTTS | ASSSRISNERLF | 0.33946553 |
| PTGSGKTL | ASSSRISNERLF | 0.7544155 |
| TGSGKTL | ASSSRISNERLF | 0.5458276 |
| TIGHVDHGK | ASSSRISNERLF | 0.564796 |
| TIGHVDHGKTT | ASSSRISNERLF | 0.83149797 |
| GHVDHGKTT | ASSSRISNERLF | 0.6254846 |
| AQFEHTLL | ASSSRISNERLF | 0.54733944 |
| IVNQDEL | ASSSRISNERLF | 0.575474 |
| TNGSQFF | ASSSRISNERLF | 0.7887654 |
| TGKGIVE | ASSSRISNERLF | 0.47946516 |
| LISNASDA | ASSSRISNERLF | 0.7863668 |
| AELRTLGELTKIAKAHGAQ | ASSSRISNERLF | 0.65770316 |
| APNAVAAAEPSW | ASSSRISNERLF | 0.8418365 |
| APQVAGAATAVL | ASSSRISNERLF | 0.37768513 |
| ARRDPAAAAAILPTDA | ASSSRISNERLF | 0.8295011 |
| DAAARDVLAEAGLAEYF | ASSSRISNERLF | 0.73896074 |
| DDQIVLSAKANLAKVEQ | ASSSRISNERLF | 0.87788653 |
| DELLPDTGEDSDEDGH | ASSSRISNERLF | 0.647812 |
| ELPEAPANRATVPQD | ASSSRISNERLF | 0.73615557 |
| FGQSKIERVTGIG | ASSSRISNERLF | 0.65342104 |
| GAGGTGSTAGGAGGAGGAGGLYA | ASSSRISNERLF | 0.85107803 |
| GHGGTGAAGGNGGTGGAGGSGIDG | ASSSRISNERLF | 0.4080158 |
| GNGGNGGAGGEGGPGGAGGAGA | ASSSRISNERLF | 0.5939894 |
| GSTAGNGGNGGNATNGTIAGQPAG | ASSSRISNERLF | 0.22402641 |
| GTGGDGGNGGNGGTGG | ASSSRISNERLF | 0.71970534 |
| GVRRPAGIPADAPV | ASSSRISNERLF | 0.7942321 |
| GYRQVTDAHLLTLARR | ASSSRISNERLF | 0.27050024 |
| IAKMGQDGHDRG | ASSSRISNERLF | 0.8337923 |
| KAEPSIEIQYT | ASSSRISNERLF | 0.8052981 |
| KPVRIRMAADEDTVAVT | ASSSRISNERLF | 0.5324062 |
| MPPRIAGMRLLVIKPEP | ASSSRISNERLF | 0.8409415 |
| MKFYAVALFRSRGVPPTR | ASSSRISNERLF | 0.68792117 |
| NAGGFGILGAVAHSP | ASSSRISNERLF | 0.85376346 |
| NEGREITLAMAEM | ASSSRISNERLF | 0.7542951 |
| NELVEWSHPLGHRWRWE | ASSSRISNERLF | 0.7105894 |
| NLGAGGTGGNGGNPG | ASSSRISNERLF | 0.77362067 |
| NTEWPKWETVMAADPVD | ASSSRISNERLF | 0.4779705 |
| PASAAPAETQP | ASSSRISNERLF | 0.5665345 |

| | | |
|---|---|---|
| QQSSDGKNGVRAAEITGE | ASSSRISNERLF | 0.84668064 |
| RAPDAVTTSWWKE | ASSSRISNERLF | 0.48520896 |
| SGFGNVGAHTSGWFNQ | ASSSRISNERLF | 0.7271036 |
| SGVAGAVGQAASVGGLKV | ASSSRISNERLF | 0.24215722 |
| SNGGAGTGSAGGNGGT | ASSSRISNERLF | 0.18704256 |
| TGGAGGEGGAAG | ASSSRISNERLF | 0.29000807 |
| TGGDGGAAGAGGAPGHG | ASSSRISNERLF | 0.61533195 |
| TMPPLLAAIANAGRHGV | ASSSRISNERLF | 0.78753626 |
| TSSTAGNSGGA | ASSSRISNERLF | 0.5368476 |
| VPATVPVGILC | ASSSRISNERLF | 0.20286983 |
| VQRPDSGGVVLTGRIS | ASSSRISNERLF | 0.4106164 |
| WTPNPVPATKGVPAGLVQAT | ASSSRISNERLF | 0.26132348 |
| YALKRALAKFSDRFDVVII | ASSSRISNERLF | 0.3186865 |
| YLIGSVIV | ASSSRISNERLF | 0.43311167 |
| YGQLVAVID | ASSSRISNERLF | 0.38026989 |
| VTVGGMTLPA | ASSSRISNERLF | 0.3067427 |
| VLALPAV | ASSSRISNERLF | 0.2091034 |
| VIPPTLNYET | ASSSRISNERLF | 0.63644314 |
| VGMGGMHPSSG | ASSSRISNERLF | 0.7025545 |
| VGLAGGAATA | ASSSRISNERLF | 0.661314 |
| VGGAGGAGGSGP | ASSSRISNERLF | 0.72656745 |
| VFAAFMV | ASSSRISNERLF | 0.6281769 |
| TLYDKLR | ASSSRISNERLF | 0.52613056 |
| TLAVFTVL | ASSSRISNERLF | 0.09452147 |
| TGSVQPARV | ASSSRISNERLF | 0.14347261 |
| TGGGAGSGGT | ASSSRISNERLF | 0.26439437 |
| SHLETSL | ASSSRISNERLF | 0.32774782 |
| SFTTGMT | ASSSRISNERLF | 0.103265315 |
| QVEVAKRMGVRQPTVS | ASSSRISNERLF | 0.44739014 |
| QGSGGPALAM | ASSSRISNERLF | 0.2644514 |
| MLAPSNP | ASSSRISNERLF | 0.21353742 |
| LVNNAAI | ASSSRISNERLF | 0.71078396 |
| LTGEINTLA | ASSSRISNERLF | 0.23004654 |
| LSIFGPV | ASSSRISNERLF | 0.22492144 |
| LLALLANL | ASSSRISNERLF | 0.2583034 |
| LISNASDALDKLR | ASSSRISNERLF | 0.66113114 |
| LINEAVP | ASSSRISNERLF | 0.17009057 |
| LGYAFSGL | ASSSRISNERLF | 0.5224962 |
| LDFVAVAL | ASSSRISNERLF | 0.30218366 |
| LAQPGAE | ASSSRISNERLF | 0.6466288 |
| ISNLGAVTGF | ASSSRISNERLF | 0.15583588 |
| HQEDPGRI | ASSSRISNERLF | 0.58108974 |
| GTPGEPTP | ASSSRISNERLF | 0.35296062 |
| GFFNSGS | ASSSRISNERLF | 0.8400731 |
| GAGGGGLTAGAN | ASSSRISNERLF | 0.2445027 |
| FVDALVNM | ASSSRISNERLF | 0.3115735 |
| FLTVAPEL | ASSSRISNERLF | 0.2754047 |
| FETLTAL | ASSSRISNERLF | 0.754325 |
| DVINPIEL | ASSSRISNERLF | 0.13461053 |
| APVSGAPV | ASSSRISNERLF | 0.29787198 |
| ALVGGLAW | ASSSRISNERLF | 0.17128311 |

| | | |
|---|---|---|
| ALAAAAATAI | ASSSRISNERLF | 0.37312186 |
| AFGLGLLL | ASSSRISNERLF | 0.47931513 |
| AFFAAASL | ASSSRISNERLF | 0.096700475 |
| AAAALPAFSPP | ASSSRISNERLF | 0.5570848 |
| VVLAPAL | ASSSRISNERLF | 0.1836583 |
| TIGHVDHGKTTL | ASSSRISNERLF | 0.17870046 |
| LISNASDALDK | ASSSRISNERLF | 0.62541807 |
| LISNASDAL | ASSSRISNERLF | 0.31532001 |
| ISNASDALDK | ASSSRISNERLF | 0.71840554 |
| NQAETLVYQTE | ASSSRISNERLF | 0.2715667 |
| TGGTGGTAGAG | ASSSRISNERLF | 0.4082821 |
| TGSTAGGAGGA | ASSSRISNERLF | 0.4712289 |
| DPLAVPVAAGPV | ASSSRISNERLF | 0.13477412 |
| GGTGGTPGNGGN | ASSSRISNERLF | 0.2572508 |
| PAGLAGVDPATL | ASSSRISNERLF | 0.6399452 |
| GNNLVVEAPHAGASV | ASSSRISNERLF | 0.8364189 |
| KVTRIMSASTLEQPA | ASSSRISNERLF | 0.8861209 |
| LGFALPIALPRRTVV | ASSSRISNERLF | 0.746783 |
| TRLGNLTARKLVVLV | ASSSRISNERLF | 0.7039113 |
| WSQQRRLIYFPSAGP | ASSSRISNERLF | 0.7807615 |
| GPNGNPDPMAAAVDIR | ASSSRISNERLF | 0.8748285 |
| TQRCVSVADLRAAASR | ASSSRISNERLF | 0.8810869 |
| QFVLDVAEHPVEVLTETV | ASSSRISNERLF | 0.84806764 |
| GGAGGTGGAGGNGGTQAGDG | ASSSRISNERLF | 0.6006308 |
| PLLATLRAIAVDVDAKPAQV | ASSSRISNERLF | 0.7818201 |
| APKFRTGGYMIDFWGVGYQVAKRM | ASSSRISNERLF | 0.7603608 |
| GAGGAGGAGGAAGGPGGTGGTGGN | ASSSRISNERLF | 0.7435876 |
| IGALGLLATDVAPAGSVAFGVALA | ASSSRISNERLF | 0.8654419 |
| AGGAGGIGGNGGAGGAGGVLMGNG | ASSSRISNERLF | 0.8069917 |
| ADELIGLL | ASSSRISNERLF | 0.7398952 |
| ADRALKA | ASSSRISNERLF | 0.78870916 |
| AGAVGQVGGAGGA | ASSSRISNERLF | 0.7942022 |
| AGGGGGLGGAP | ASSSRISNERLF | 0.8262322 |
| AGGLLYGNGGAG | ASSSRISNERLF | 0.8389945 |
| AGLESIGL | ASSSRISNERLF | 0.8803865 |
| AGLKTGII | ASSSRISNERLF | 0.86604893 |
| ASLAALLL | ASSSRISNERLF | 0.8232904 |
| ASVINISSI | ASSSRISNERLF | 0.1868131 |
| ATPPGPA | ASSSRISNERLF | 0.4962911 |
| ATVASALV | ASSSRISNERLF | 0.81259847 |
| ATVASLIL | ASSSRISNERLF | 0.7427511 |
| AVNGPVLIRH | ASSSRISNERLF | 0.3371398 |
| DGAAGAVGI | ASSSRISNERLF | 0.7527462 |
| DPLLHGI | ASSSRISNERLF | 0.75205743 |
| ELGAALGAL | ASSSRISNERLF | 0.6681946 |
| ELGRTQAV | ASSSRISNERLF | 0.8600286 |
| FVAGGYLS | ASSSRISNERLF | 0.7856136 |
| GASGGAGGAGGW | ASSSRISNERLF | 0.65951645 |
| GGAGAGGDGGAGGI | ASSSRISNERLF | 0.6825719 |
| GTGHNGSQPGMGGNG | ASSSRISNERLF | 0.86814356 |
| GYAAASASAS | ASSSRISNERLF | 0.47823673 |

| | | |
|---|---|---|
| HQVDVAAVQQG | ASSSRISNERLF | 0.8370261 |
| IGAAIAHT | ASSSRISNERLF | 0.7742605 |
| IGMIEVAQL | ASSSRISNERLF | 0.8697318 |
| IRLAPEHTSAMD | ASSSRISNERLF | 0.5645278 |
| IYNVSVL | ASSSRISNERLF | 0.843343 |
| KQGFLGM | ASSSRISNERLF | 0.7988236 |
| LESVITQL | ASSSRISNERLF | 0.9025738 |
| LGLGALVVV | ASSSRISNERLF | 0.789044 |
| LIGNGGNGGSGGI | ASSSRISNERLF | 0.79391754 |
| LNAINAPTL | ASSSRISNERLF | 0.7123216 |
| LQLLNTGV | ASSSRISNERLF | 0.8756899 |
| LQTSIEF | ASSSRISNERLF | 0.61469233 |
| MSFAVSEL | ASSSRISNERLF | 0.8032304 |
| NTGALGL | ASSSRISNERLF | 0.46103242 |
| PPGAGGAGGAGTT | ASSSRISNERLF | 0.83179367 |
| QLVGLGI | ASSSRISNERLF | 0.8468995 |
| RFLEQVHQL | ASSSRISNERLF | 0.7317204 |
| RGVGYVLR | ASSSRISNERLF | 0.8856612 |
| SATGVFAGVI | ASSSRISNERLF | 0.81493133 |
| SLPAITLP | ASSSRISNERLF | 0.75771683 |
| STALAFTV | ASSSRISNERLF | 0.89194655 |
| SVLTVGLGK | ASSSRISNERLF | 0.7064717 |
| TALAGAGSGI | ASSSRISNERLF | 0.88966495 |
| TGLPFTGI | ASSSRISNERLF | 0.75823015 |
| VAEHVTV | ASSSRISNERLF | 0.8361681 |
| VDIVLNSL | ASSSRISNERLF | 0.85054976 |
| VLTGGGAVL | ASSSRISNERLF | 0.70877326 |
| VNVGLAYGI | ASSSRISNERLF | 0.66550314 |
| VSSAAVGAV | ASSSRISNERLF | 0.876521 |
| VTSVYEAGV | ASSSRISNERLF | 0.82240176 |
| WVALDRGA | ASSSRISNERLF | 0.8185445 |
| YLVGSVVI | ASSSRISNERLF | 0.74366623 |
| GPPGTGKTLL | ASSSRISNERLF | 0.7513376 |
| GVGMTKF | ASSSRISNERLF | 0.74992466 |
| IGIGGSDL | ASSSRISNERLF | 0.81495714 |
| IGTLLAEL | ASSSRISNERLF | 0.82912695 |
| IYTSGTT | ASSSRISNERLF | 0.82295954 |
| LSFPAGV | ASSSRISNERLF | 0.69775975 |
| LYTSGTT | ASSSRISNERLF | 0.8225522 |
| VGAFPFGF | ASSSRISNERLF | 0.78369343 |
| VGAGTVEF | ASSSRISNERLF | 0.8045338 |
| VGKSTLLL | ASSSRISNERLF | 0.79610014 |
| AALAHAGGGIV | ASSSRISNERLF | 0.79497534 |
| AALVAVSLL | ASSSRISNERLF | 0.7484195 |
| AAVSRAVAAL | ASSSRISNERLF | 0.6117284 |
| AGGSSGIGGF | ASSSRISNERLF | 0.81126153 |
| AGLTVANANG | ASSSRISNERLF | 0.8231367 |
| AGLVGAVL | ASSSRISNERLF | 0.8124094 |
| ALSQAFTV | ASSSRISNERLF | 0.8295922 |
| ALTFSVSL | ASSSRISNERLF | 0.8265372 |
| ALTGAALVAL | ASSSRISNERLF | 0.81334436 |

| | | |
|---|---|---|
| ASFSIGAGVL | ASSSRISNERLF | 0.32694858 |
| DGSVYKI | ASSSRISNERLF | 0.5583631 |
| EQLAEIV | ASSSRISNERLF | 0.6064211 |
| FAVSELTVM | ASSSRISNERLF | 0.7681693 |
| GAAPLAAGL | ASSSRISNERLF | 0.25044727 |
| GADGGDGGSGGAGG | ASSSRISNERLF | 0.80121595 |
| GFGLDTGGAGG | ASSSRISNERLF | 0.7360031 |
| GLASGGLL | ASSSRISNERLF | 0.76521707 |
| GRIAGAAVLL | ASSSRISNERLF | 0.82421106 |
| GSAGSLNTSF | ASSSRISNERLF | 0.7779795 |
| GTGQIGI | ASSSRISNERLF | 0.6059648 |
| GTLIALIL | ASSSRISNERLF | 0.6771656 |
| IAATMGFVVG | ASSSRISNERLF | 0.830743 |
| IFAQALL | ASSSRISNERLF | 0.5794879 |
| IGAAVALGF | ASSSRISNERLF | 0.8271397 |
| IGLTPLAL | ASSSRISNERLF | 0.7202391 |
| ILGSNTSTL | ASSSRISNERLF | 0.8267734 |
| IVFSLTVL | ASSSRISNERLF | 0.65487814 |
| LASKATGF | ASSSRISNERLF | 0.82745385 |
| LASVALIL | ASSSRISNERLF | 0.8064943 |
| LFAVGLTL | ASSSRISNERLF | 0.83423555 |
| LFNGGIGTYI | ASSSRISNERLF | 0.78225446 |
| LTLGAGII | ASSSRISNERLF | 0.8129083 |
| LVAAALGDI | ASSSRISNERLF | 0.76414084 |
| NTAFVQLGI | ASSSRISNERLF | 0.81672674 |
| QLFSGAEL | ASSSRISNERLF | 0.62164515 |
| SAATFITL | ASSSRISNERLF | 0.7633679 |
| SNYGLANLGV | ASSSRISNERLF | 0.72437614 |
| SYGGAYAVM | ASSSRISNERLF | 0.7801487 |
| TTGVGGLTLG | ASSSRISNERLF | 0.82608116 |
| TTQTIYI | ASSSRISNERLF | 0.2244038 |
| VAAGAGLGAVI | ASSSRISNERLF | 0.81526566 |
| VAAVVTLL | ASSSRISNERLF | 0.7046515 |
| VGFLGGLI | ASSSRISNERLF | 0.63076794 |
| VGVIPILL | ASSSRISNERLF | 0.81907654 |
| VIFASSLI | ASSSRISNERLF | 0.7978141 |
| VKGGDGGAAAG | ASSSRISNERLF | 0.81632286 |
| VTKTFTNI | ASSSRISNERLF | 0.70768005 |
| VVAAAILL | ASSSRISNERLF | 0.79451275 |
| YAALDVEL | ASSSRISNERLF | 0.7904042 |
| LASVALLL | ASSSRISNERLF | 0.3027192 |
| LDAVHRV | ASSSRISNERLF | 0.5851068 |
| QGLSSSIF | ASSSRISNERLF | 0.83220387 |
| RGTFANIRL | ASSSRISNERLF | 0.846033 |
| RQLWWGHRI | ASSSRISNERLF | 0.8377698 |
| YGPPGTGKTLL | ASSSRISNERLF | 0.80878556 |
| AALVGAPANVA | ASSSRISNERLF | 0.803731 |
| AATFRRTE | ASSSRISNERLF | 0.8084642 |
| AGALGGGI | ASSSRISNERLF | 0.84445065 |
| AGGAGGAGGIGAG | ASSSRISNERLF | 0.8519807 |
| AGSGALGL | ASSSRISNERLF | 0.8495599 |

| | | |
|---|---|---|
| AQIADQI | ASSSRISNERLF | 0.80469126 |
| ASAVAVTYL | ASSSRISNERLF | 0.85367775 |
| ASTVLGGTGG | ASSSRISNERLF | 0.8343334 |
| AVGGEPAK | ASSSRISNERLF | 0.8307294 |
| CEQGTFGGTC | ASSSRISNERLF | 0.8463975 |
| DEGKAPAG | ASSSRISNERLF | 0.84043634 |
| DTVVFPAL | ASSSRISNERLF | 0.81847084 |
| FEGITMAG | ASSSRISNERLF | 0.8057213 |
| FPLVVQD | ASSSRISNERLF | 0.8377733 |
| GAFTTPQ | ASSSRISNERLF | 0.84981155 |
| GFGGFGTAGGVGGA | ASSSRISNERLF | 0.8449788 |
| GFLAQFL | ASSSRISNERLF | 0.85710645 |
| GGAGGAGGNAGVA | ASSSRISNERLF | 0.85027087 |
| GGVGTAGAGGAGGAGG | ASSSRISNERLF | 0.84467065 |
| GNGGGGDAGNAGSGGN | ASSSRISNERLF | 0.6300626 |
| GTGGAGGAGNDAGS | ASSSRISNERLF | 0.7803677 |
| GTTNTGWFNS | ASSSRISNERLF | 0.804046 |
| IFMGPLFTL | ASSSRISNERLF | 0.829558 |
| ILAHAIEAI | ASSSRISNERLF | 0.44308564 |
| IRSAIREAL | ASSSRISNERLF | 0.83511996 |
| ITFAGGG | ASSSRISNERLF | 0.83743954 |
| ITGMSGHQEV | ASSSRISNERLF | 0.8121513 |
| IVAGLACTL | ASSSRISNERLF | 0.85119957 |
| IVGGGLAGA | ASSSRISNERLF | 0.8232708 |
| IVTGVLNI | ASSSRISNERLF | 0.7935711 |
| LAQQGCI | ASSSRISNERLF | 0.8238127 |
| LCLHVEN | ASSSRISNERLF | 0.85553837 |
| LDAISDGS | ASSSRISNERLF | 0.7908066 |
| LGGFAAML | ASSSRISNERLF | 0.85387254 |
| LGGKNPMI | ASSSRISNERLF | 0.7453203 |
| LGPKGVAAIL | ASSSRISNERLF | 0.8536251 |
| LMVGVSI | ASSSRISNERLF | 0.7749433 |
| LVGAAFVL | ASSSRISNERLF | 0.8515769 |
| LVLAVAPL | ASSSRISNERLF | 0.831725 |
| MAAIAVIDAA | ASSSRISNERLF | 0.8589149 |
| NNGINAELQ | ASSSRISNERLF | 0.8304238 |
| RGEPLVLVHG | ASSSRISNERLF | 0.85330236 |
| RGGAGGAAGLI | ASSSRISNERLF | 0.83236 |
| RSYRMHPAV | ASSSRISNERLF | 0.831502 |
| SAAHLLAGQLA | ASSSRISNERLF | 0.76207066 |
| SAAVGSY | ASSSRISNERLF | 0.8319342 |
| SPHAVLHRI | ASSSRISNERLF | 0.78258216 |
| SQIAKVL | ASSSRISNERLF | 0.8407732 |
| TALGHLVP | ASSSRISNERLF | 0.84978104 |
| TAPTTSGAA | ASSSRISNERLF | 0.44635108 |
| TDGDDGNGGSPGT | ASSSRISNERLF | 0.8526044 |
| TDGLGEN | ASSSRISNERLF | 0.7796011 |
| TGGPPLPGQA | ASSSRISNERLF | 0.7955137 |
| TIPAITI | ASSSRISNERLF | 0.852121 |
| TIPGFTI | ASSSRISNERLF | 0.8366449 |
| TSIASGALA | ASSSRISNERLF | 0.8463588 |

| | | |
|---|---|---|
| TVTLPGAAI | ASSSRISNERLF | 0.80788875 |
| VASGLSAALP | ASSSRISNERLF | 0.8282543 |
| VGAGVIGI | ASSSRISNERLF | 0.8432463 |
| VGPKRFGV | ASSSRISNERLF | 0.4406905 |
| VNVAFPD | ASSSRISNERLF | 0.77903414 |
| VSMTNSGVS | ASSSRISNERLF | 0.85644865 |
| VTDVLGDLSV | ASSSRISNERLF | 0.85209703 |
| VVVVGAGFAGL | ASSSRISNERLF | 0.8730028 |
| VYTVGTVV | ASSSRISNERLF | 0.78655064 |
| WTTFSGP | ASSSRISNERLF | 0.74235505 |
| YDELVEAF | ASSSRISNERLF | 0.8240534 |
| YGTGGAGGNGG | ASSSRISNERLF | 0.8742226 |
| YTNVMAV | ASSSRISNERLF | 0.87643015 |
| FLEQVHQL | ASSSRISNERLF | 0.86164117 |
| GLSLLAVAV | ASSSRISNERLF | 0.72839403 |
| LEQVHQL | ASSSRISNERLF | 0.84985083 |
| LNAAGAI | ASSSRISNERLF | 0.78807926 |
| PAVHASL | ASSSRISNERLF | 0.82688034 |
| TELEDDA | ASSSRISNERLF | 0.86475813 |
| TPPGPPN | ASSSRISNERLF | 0.8421918 |
| TPPPPPPP | ASSSRISNERLF | 0.8112409 |
| TQSPALI | ASSQGQGANTEVF | 0.7740438 |
| EFAAAFPQ | ASSQGQGANTEVF | 0.8581001 |
| LLSGNLDVL | ASSQGQGANTEVF | 0.8698724 |
| GPANLGNYNI | ASSQGQGANTEVF | 0.8674022 |
| TLGPTLP | ASSQGQGANTEVF | 0.884231 |
| VYAYSSCCK | ASSQGQGANTEVF | 0.8692306 |
| APAPAPAPNI | ASSQGQGANTEVF | 0.8641177 |
| PGGPVPIP | ASSQGQGANTEVF | 0.2412515 |
| FLTVALEP | ASSQGQGANTEVF | 0.49747765 |
| SQYFGANL | ASSQGQGANTEVF | 0.7896916 |
| GGVKPAA | ASSQGQGANTEVF | 0.8237528 |
| GGTGGAGGAGGW | ASSQGQGANTEVF | 0.52759904 |
| GVGGVAGLGGAGG | ASSQGQGANTEVF | 0.8622909 |
| VLLVLATTI | ASSQGQGANTEVF | 0.7778443 |
| IGIFAAML | ASSQGQGANTEVF | 0.7670852 |
| AVGDLID | ASSQGQGANTEVF | 0.8715182 |
| NISPLPL | ASSQGQGANTEVF | 0.7986602 |
| VTAVFVAI | ASSQGQGANTEVF | 0.76736736 |
| SGDDNDAAL | ASSQGQGANTEVF | 0.78851914 |
| AVAALAHS | ASSQGQGANTEVF | 0.88200444 |
| GGVTVMATP | ASSQGQGANTEVF | 0.695531 |
| AGLAANVRD | ASSQGQGANTEVF | 0.86332476 |
| TNRPDLIDPALLRPG | ASSQGQGANTEVF | 0.8154316 |
| LANLGVTAL | ASSQGQGANTEVF | 0.8749472 |
| TGERIEL | ASSQGQGANTEVF | 0.6058923 |
| NGEELAA | ASSQGQGANTEVF | 0.86422443 |
| VLSLVWG | ASSQGQGANTEVF | 0.8335254 |
| IGYALWL | ASSQGQGANTEVF | 0.88819104 |
| LSLLLAPL | ASSQGQGANTEVF | 0.7986135 |
| TLDEALK | ASSQGQGANTEVF | 0.8287306 |

| | | |
|---|---|---|
| TDGAGTAEGG | ASSQGQGANTEVF | 0.76555026 |
| NAPGGAGGNGG | ASSQGQGANTEVF | 0.8668362 |
| VSVFMGMGGNF | ASSQGQGANTEVF | 0.5739981 |
| LASYLAAL | ASSQGQGANTEVF | 0.8193289 |
| LVGGTPAL | ASSQGQGANTEVF | 0.6969171 |
| VRNADVL | ASSQGQGANTEVF | 0.84444857 |
| TCSSPFLVSAM | ASSQGQGANTEVF | 0.8516941 |
| VHITDAE | ASSQGQGANTEVF | 0.43608987 |
| EYAAPSF | ASSQGQGANTEVF | 0.8697586 |
| VVDAFHGGL | ASSQGQGANTEVF | 0.8079701 |
| ASVVDPQ | ASSQGQGANTEVF | 0.72917485 |
| SYNEFGGLS | ASSQGQGANTEVF | 0.8779964 |
| DPDGSVY | ASSQGQGANTEVF | 0.8161286 |
| MVSTIHATE | ASSQGQGANTEVF | 0.87227666 |
| LAAGVADPV | ASSQGQGANTEVF | 0.8384725 |
| TKEQIAA | ASSQGQGANTEVF | 0.8302026 |
| SLASPAAL | ASSQGQGANTEVF | 0.86585534 |
| IGLVFYGV | ASSQGQGANTEVF | 0.63373613 |
| FLLDASNL | ASSQGQGANTEVF | 0.7728726 |
| YQVGNFP | ASSQGQGANTEVF | 0.87457615 |
| ESLLSQST | ASSQGQGANTEVF | 0.777974 |
| LFTVGFAAVG | ASSQGQGANTEVF | 0.8687608 |
| ATPPTPT | ASSQGQGANTEVF | 0.7236886 |
| TDLATLG | ASSQGQGANTEVF | 0.603614 |
| SAGGTGR | ASSQGQGANTEVF | 0.7371705 |
| GFSNSGS | ASSQGQGANTEVF | 0.8192587 |
| GGNGGSGGTSV | ASSQGQGANTEVF | 0.86474144 |
| GTSVATGGAGN | ASSQGQGANTEVF | 0.79416984 |
| NGGNGGSGGTS | ASSQGQGANTEVF | 0.48930496 |
| AVVKFPH | ASSQGQGANTEVF | 0.77983344 |
| TPPPPPP | ASSQGQGANTEVF | 0.6834872 |
| AAPAAAAAPA | ASSQGQGANTEVF | 0.7154751 |
| IALNEDL | ASSQGQGANTEVF | 0.79416275 |
| IVNVSSM | ASSQGQGANTEVF | 0.7593492 |
| TGIPPAPR | ASSQGQGANTEVF | 0.7114803 |
| FDLGGGTF | ASSQGQGANTEVF | 0.6271465 |
| FDLGGGTFDVS | ASSQGQGANTEVF | 0.78907853 |
| NEPTAAA | ASSQGQGANTEVF | 0.8155348 |
| NEPTAAALA | ASSQGQGANTEVF | 0.80002296 |
| GSIINISSI | ASSQGQGANTEVF | 0.85334975 |
| VAPGNAGTA | ASSQGQGANTEVF | 0.8353587 |
| TGGGASLE | ASSQGQGANTEVF | 0.82901424 |
| VAVYNHYKRI | ASSQGQGANTEVF | 0.1757617 |
| TDPAPATT | ASSQGQGANTEVF | 0.29698986 |
| TNRPDILDPALLRPG | ASSQGQGANTEVF | 0.58001506 |
| LTTTTTS | ASSQGQGANTEVF | 0.65344286 |
| PTGSGKTL | ASSQGQGANTEVF | 0.25502768 |
| TGSGKTL | ASSQGQGANTEVF | 0.78603536 |
| TIGHVDHGK | ASSQGQGANTEVF | 0.665122 |
| TIGHVDHGKTT | ASSQGQGANTEVF | 0.6269494 |
| GHVDHGKTT | ASSQGQGANTEVF | 0.83955634 |

| | | |
|---|---|---|
| AQFEHTLL | ASSQGQGANTEVF | 0.7263112 |
| IVNQDEL | ASSQGQGANTEVF | 0.6126517 |
| TNGSQFF | ASSQGQGANTEVF | 0.5795368 |
| TGKGIVE | ASSQGQGANTEVF | 0.8265334 |
| LISNASDA | ASSQGQGANTEVF | 0.3418274 |
| AELRTLGELTKIAKAHGAQ | ASSQGQGANTEVF | 0.8120917 |
| APNAVAAAEPSW | ASSQGQGANTEVF | 0.45024496 |
| APQVAGAATAVL | ASSQGQGANTEVF | 0.8212471 |
| ARRDPAAAAAILPTDA | ASSQGQGANTEVF | 0.41145492 |
| DAAARDVLAEAGLAEYF | ASSQGQGANTEVF | 0.8210776 |
| DDQIVLSAKANLAKVEQ | ASSQGQGANTEVF | 0.7725072 |
| DELLPDTGEDSDEDGH | ASSQGQGANTEVF | 0.8898517 |
| ELPEAPANRATVPQD | ASSQGQGANTEVF | 0.70496917 |
| FGQSKIERVTGIG | ASSQGQGANTEVF | 0.7132479 |
| GAGGTGSTAGGAGGAGGAGGLYA | ASSQGQGANTEVF | 0.54381216 |
| GHGGTGAAGGNGGTGGAGGSGIDG | ASSQGQGANTEVF | 0.85962886 |
| GNGGNGGAGGEGGPGGAGGAGA | ASSQGQGANTEVF | 0.3685267 |
| GSTAGNGGNGGNATNGTIAGQPAG | ASSQGQGANTEVF | 0.73658514 |
| GTGGDGGNGGNGGTGG | ASSQGQGANTEVF | 0.4739328 |
| GVRRPAGIPADAPV | ASSQGQGANTEVF | 0.80750334 |
| GYRQVTDAHLLTLARR | ASSQGQGANTEVF | 0.79354334 |
| IAKMGQDGHDRG | ASSQGQGANTEVF | 0.3133566 |
| KAEPSIEIQYT | ASSQGQGANTEVF | 0.8264088 |
| KPVRIRMAADEDTVAVT | ASSQGQGANTEVF | 0.75384784 |
| MPPRIAGMRLLVIKPEP | ASSQGQGANTEVF | 0.57138574 |
| MKFYAVALFRSRGVPPTR | ASSQGQGANTEVF | 0.85666025 |
| NAGGFGILGAVAHSP | ASSQGQGANTEVF | 0.70662564 |
| NEGREITLAMAEM | ASSQGQGANTEVF | 0.86361647 |
| NELVEWSHPLGHRWRWE | ASSQGQGANTEVF | 0.677752 |
| NLGAGGTGGNGGNPG | ASSQGQGANTEVF | 0.5981033 |
| NTEWPKWETVMAADPVD | ASSQGQGANTEVF | 0.7916217 |
| PASAAPAETQP | ASSQGQGANTEVF | 0.41516024 |
| QQSSDGKNGVRAAEITGE | ASSQGQGANTEVF | 0.50094426 |
| RAPDAVTTSWWKE | ASSQGQGANTEVF | 0.85685873 |
| SGFGNVGAHTSGWFNQ | ASSQGQGANTEVF | 0.7815529 |
| SGVAGAVGQAASVGGLKV | ASSQGQGANTEVF | 0.7896231 |
| SNGGAGTGSAGGNGGT | ASSQGQGANTEVF | 0.7342276 |
| TGGAGGEGGAAG | ASSQGQGANTEVF | 0.69857204 |
| TGGDGGAAGAGGAPGHG | ASSQGQGANTEVF | 0.7619347 |
| TMPPLLAAIANAGRHGV | ASSQGQGANTEVF | 0.79617083 |
| TSSTAGNSGGA | ASSQGQGANTEVF | 0.79939365 |
| VPATVPVGILC | ASSQGQGANTEVF | 0.7886948 |
| VQRPDSGGVVLTGRIS | ASSQGQGANTEVF | 0.7018588 |
| WTPNPVPATKGVPAGLVQAT | ASSQGQGANTEVF | 0.7936914 |
| YALKRALAKFSDRFDVVII | ASSQGQGANTEVF | 0.7722195 |
| YLIGSVIV | ASSQGQGANTEVF | 0.7578467 |
| YGQLVAVID | ASSQGQGANTEVF | 0.79738843 |
| VTVGGMTLPA | ASSQGQGANTEVF | 0.7830376 |
| VLALPAV | ASSQGQGANTEVF | 0.7457919 |
| VIPPTLNYET | ASSQGQGANTEVF | 0.74837184 |
| VGMGGMHPSSG | ASSQGQGANTEVF | 0.7805214 |

| | | |
|---|---|---|
| VGLAGGAATA | ASSQGQGANTEVF | 0.7910551 |
| VGGAGGAGGSGP | ASSQGQGANTEVF | 0.7942983 |
| VFAAFMV | ASSQGQGANTEVF | 0.8110354 |
| TLYDKLR | ASSQGQGANTEVF | 0.79817545 |
| TLAVFTVL | ASSQGQGANTEVF | 0.7870759 |
| TGSVQPARV | ASSQGQGANTEVF | 0.3334057 |
| TGGGAGSGGT | ASSQGQGANTEVF | 0.5473958 |
| SHLETSL | ASSQGQGANTEVF | 0.75473595 |
| SFTTGMT | ASSQGQGANTEVF | 0.76782995 |
| QVEVAKRMGVRQPTVS | ASSQGQGANTEVF | 0.47447997 |
| QGSGGPALAM | ASSQGQGANTEVF | 0.7746735 |
| MLAPSNP | ASSQGQGANTEVF | 0.7542713 |
| LVNNAAI | ASSQGQGANTEVF | 0.72925234 |
| LTGEINTLA | ASSQGQGANTEVF | 0.8032302 |
| LSIFGPV | ASSQGQGANTEVF | 0.75506544 |
| LLALLANL | ASSQGQGANTEVF | 0.7222134 |
| LISNASDALDKLR | ASSQGQGANTEVF | 0.75984144 |
| LINEAVP | ASSQGQGANTEVF | 0.80506563 |
| LGYAFSGL | ASSQGQGANTEVF | 0.71425307 |
| LDFVAVAL | ASSQGQGANTEVF | 0.79849845 |
| LAQPGAE | ASSQGQGANTEVF | 0.67120314 |
| ISNLGAVTGF | ASSQGQGANTEVF | 0.80167764 |
| HQEDPGRI | ASSQGQGANTEVF | 0.65596026 |
| GTPGEPTP | ASSQGQGANTEVF | 0.7953743 |
| GFFNSGS | ASSQGQGANTEVF | 0.7653731 |
| GAGGGGLTAGAN | ASSQGQGANTEVF | 0.8146949 |
| FVDALVNM | ASSQGQGANTEVF | 0.7620336 |
| FLTVAPEL | ASSQGQGANTEVF | 0.7968762 |
| FETLTAL | ASSQGQGANTEVF | 0.7460341 |
| DVINPIEL | ASSQGQGANTEVF | 0.7926984 |
| APVSGAPV | ASSQGQGANTEVF | 0.644328 |
| ALVGGLAW | ASSQGQGANTEVF | 0.7748352 |
| ALAAAAATAI | ASSQGQGANTEVF | 0.5340202 |
| AFGLGLLL | ASSQGQGANTEVF | 0.78279424 |
| AFFAAASL | ASSQGQGANTEVF | 0.7849359 |
| AAAALPAFSPP | ASSQGQGANTEVF | 0.62763315 |
| VVLAPAL | ASSQGQGANTEVF | 0.8013825 |
| TIGHVDHGKTTL | ASSQGQGANTEVF | 0.76220596 |
| LISNASDALDK | ASSQGQGANTEVF | 0.72689444 |
| LISNASDAL | ASSQGQGANTEVF | 0.8056929 |
| ISNASDALDK | ASSQGQGANTEVF | 0.7665893 |
| NQAETLVYQTE | ASSQGQGANTEVF | 0.7929241 |
| TGGTGGTAGAG | ASSQGQGANTEVF | 0.7587341 |
| TGSTAGGAGGA | ASSQGQGANTEVF | 0.7461031 |
| DPLAVPVAAGPV | ASSQGQGANTEVF | 0.7835093 |
| GGTGGTPGNGGN | ASSQGQGANTEVF | 0.6477242 |
| PAGLAGVDPATL | ASSQGQGANTEVF | 0.7334838 |
| GNNLVVEAPHAGASV | ASSQGQGANTEVF | 0.8039199 |
| KVTRIMSASTLEQPA | ASSQGQGANTEVF | 0.83501387 |
| LGFALPIALPRRTVV | ASSQGQGANTEVF | 0.8424077 |
| TRLGNLTARKLVVLV | ASSQGQGANTEVF | 0.63942885 |

| | | |
|---|---|---|
| WSQQRRLIYFPSAGP | ASSQGQGANTEVF | 0.6554326 |
| GPNGNPDPMAAAVDIR | ASSQGQGANTEVF | 0.7345764 |
| TQRCVSVADLRAAASR | ASSQGQGANTEVF | 0.8423105 |
| QFVLDVAEHPVEVLTETV | ASSQGQGANTEVF | 0.8613353 |
| GGAGGTGGAGGNGGTQAGDG | ASSQGQGANTEVF | 0.8410311 |
| PLLATLRAIAVDVDAKPAQV | ASSQGQGANTEVF | 0.683336 |
| APKFRTGGYMIDFWGVGYQVAKRM | ASSQGQGANTEVF | 0.818035 |
| GAGGAGGAGGAAGGPGGTGGTGGN | ASSQGQGANTEVF | 0.75427294 |
| IGALGLLATDVAPAGSVAFGVALA | ASSQGQGANTEVF | 0.75253224 |
| AGGAGGIGGNGGAGGAGGVLMGNG | ASSQGQGANTEVF | 0.8468558 |
| ADELIGLL | ASSQGQGANTEVF | 0.78748 |
| ADRALKA | ASSQGQGANTEVF | 0.6838498 |
| AGAVGQVGGAGGA | ASSQGQGANTEVF | 0.68891037 |
| AGGGGGLGGAP | ASSQGQGANTEVF | 0.80198205 |
| AGGLLYGNGGAG | ASSQGQGANTEVF | 0.8322988 |
| AGLESIGL | ASSQGQGANTEVF | 0.82497984 |
| AGLKTGII | ASSQGQGANTEVF | 0.89272153 |
| ASLAALLL | ASSQGQGANTEVF | 0.85197634 |
| ASVINISSI | ASSQGQGANTEVF | 0.8413956 |
| ATPPGPA | ASSQGQGANTEVF | 0.25369602 |
| ATVASALV | ASSQGQGANTEVF | 0.5027964 |
| ATVASLIL | ASSQGQGANTEVF | 0.74445295 |
| AVNGPVLIRH | ASSQGQGANTEVF | 0.74919486 |
| DGAAGAVGI | ASSQGQGANTEVF | 0.38284636 |
| DPLLHGI | ASSQGQGANTEVF | 0.8240682 |
| ELGAALGAL | ASSQGQGANTEVF | 0.75363505 |
| ELGRTQAV | ASSQGQGANTEVF | 0.6463332 |
| FVAGGYLS | ASSQGQGANTEVF | 0.8397206 |
| GASGGAGGAGGW | ASSQGQGANTEVF | 0.6276411 |
| GGAGAGGDGGAGGI | ASSQGQGANTEVF | 0.5825874 |
| GTGHNGSQPGMGGNG | ASSQGQGANTEVF | 0.7538248 |
| GYAAASASAS | ASSQGQGANTEVF | 0.8708639 |
| HQVDVAAVQQG | ASSQGQGANTEVF | 0.68234503 |
| IGAAIAHT | ASSQGQGANTEVF | 0.8476131 |
| IGMIEVAQL | ASSQGQGANTEVF | 0.79519254 |
| IRLAPEHTSAMD | ASSQGQGANTEVF | 0.86180484 |
| IYNVSVL | ASSQGQGANTEVF | 0.5564997 |
| KQGFLGM | ASSQGQGANTEVF | 0.8521369 |
| LESVITQL | ASSQGQGANTEVF | 0.75671804 |
| LGLGALVVV | ASSQGQGANTEVF | 0.88142425 |
| LIGNGGNGGSGGI | ASSQGQGANTEVF | 0.7687032 |
| LNAINAPTL | ASSQGQGANTEVF | 0.8421147 |
| LQLLNTGV | ASSQGQGANTEVF | 0.7821053 |
| LQTSIEF | ASSQGQGANTEVF | 0.858544 |
| MSFAVSEL | ASSQGQGANTEVF | 0.5297564 |
| NTGALGL | ASSQGQGANTEVF | 0.74699533 |
| PPGAGGAGGAGTT | ASSQGQGANTEVF | 0.5430554 |
| QLVGLGI | ASSQGQGANTEVF | 0.77833223 |
| RFLEQVHQL | ASSQGQGANTEVF | 0.8339212 |
| RGVGYVLR | ASSQGQGANTEVF | 0.2158774 |
| SATGVFAGVI | ASSQGQGANTEVF | 0.8493932 |

| | | |
|---|---|---|
| SLPAITLP | ASSQGQGANTEVF | 0.59684336 |
| STALAFTV | ASSQGQGANTEVF | 0.631421 |
| SVLTVGLGK | ASSQGQGANTEVF | 0.8565305 |
| TALAGAGSGI | ASSQGQGANTEVF | 0.7546563 |
| TGLPFTGI | ASSQGQGANTEVF | 0.86619973 |
| VAEHVTV | ASSQGQGANTEVF | 0.74842167 |
| VDIVLNSL | ASSQGQGANTEVF | 0.78697526 |
| VLTGGGAVL | ASSQGQGANTEVF | 0.7965555 |
| VNVGLAYGI | ASSQGQGANTEVF | 0.37624094 |
| VSSAAVGAV | ASSQGQGANTEVF | 0.6550975 |
| VTSVYEAGV | ASSQGQGANTEVF | 0.8712437 |
| WVALDRGA | ASSQGQGANTEVF | 0.85033 |
| YLVGSVVI | ASSQGQGANTEVF | 0.8708602 |
| GPPGTGKTLL | ASSQGQGANTEVF | 0.77049637 |
| GVGMTKF | ASSQGQGANTEVF | 0.74370444 |
| IGIGGSDL | ASSQGQGANTEVF | 0.7926408 |
| IGTLLAEL | ASSQGQGANTEVF | 0.8740273 |
| IYTSGTT | ASSQGQGANTEVF | 0.8550163 |
| LSFPAGV | ASSQGQGANTEVF | 0.8445 |
| LYTSGTT | ASSQGQGANTEVF | 0.6927849 |
| VGAFPFGF | ASSQGQGANTEVF | 0.78303623 |
| VGAGTVEF | ASSQGQGANTEVF | 0.7568228 |
| VGKSTLLL | ASSQGQGANTEVF | 0.77288544 |
| AALAHAGGGIV | ASSQGQGANTEVF | 0.8628739 |
| AALVAVSLL | ASSQGQGANTEVF | 0.77617764 |
| AAVSRAVAAL | ASSQGQGANTEVF | 0.7678627 |
| AGGSSGIGGF | ASSQGQGANTEVF | 0.8208624 |
| AGLTVANANG | ASSQGQGANTEVF | 0.7728629 |
| AGLVGAVL | ASSQGQGANTEVF | 0.8123256 |
| ALSQAFTV | ASSQGQGANTEVF | 0.79594505 |
| ALTFSVSL | ASSQGQGANTEVF | 0.8877015 |
| ALTGAALVAL | ASSQGQGANTEVF | 0.8663299 |
| ASFSIGAGVL | ASSQGQGANTEVF | 0.8628695 |
| DGSVYKI | ASSQGQGANTEVF | 0.20122328 |
| EQLAEIV | ASSQGQGANTEVF | 0.57091445 |
| FAVSELTVM | ASSQGQGANTEVF | 0.82335734 |
| GAAPLAAGL | ASSQGQGANTEVF | 0.77561116 |
| GADGGDGGSGGAGG | ASSQGQGANTEVF | 0.51296926 |
| GFGLDTGGAGG | ASSQGQGANTEVF | 0.8236039 |
| GLASGGLL | ASSQGQGANTEVF | 0.790177 |
| GRIAGAAVLL | ASSQGQGANTEVF | 0.6765158 |
| GSAGSLNTSF | ASSQGQGANTEVF | 0.854252 |
| GTGQIGI | ASSQGQGANTEVF | 0.7815216 |
| GTLIALIL | ASSQGQGANTEVF | 0.67588556 |
| IAATMGFVVG | ASSQGQGANTEVF | 0.76221377 |
| IFAQALL | ASSQGQGANTEVF | 0.8534378 |
| IGAAVALGF | ASSQGQGANTEVF | 0.6187563 |
| IGLTPLAL | ASSQGQGANTEVF | 0.8350195 |
| ILGSNTSTL | ASSQGQGANTEVF | 0.7243892 |
| IVFSLTVL | ASSQGQGANTEVF | 0.8764777 |
| LASKATGF | ASSQGQGANTEVF | 0.634432 |

| | | |
|---|---|---|
| LASVALIL | ASSQGQGANTEVF | 0.85326046 |
| LFAVGLTL | ASSQGQGANTEVF | 0.80540866 |
| LFNGGIGTYI | ASSQGQGANTEVF | 0.89595973 |
| LTLGAGII | ASSQGQGANTEVF | 0.7984614 |
| LVAAALGDI | ASSQGQGANTEVF | 0.8251352 |
| NTAFVQLGI | ASSQGQGANTEVF | 0.6988735 |
| QLFSGAEL | ASSQGQGANTEVF | 0.82382905 |
| SAATFITL | ASSQGQGANTEVF | 0.5783073 |
| SNYGLANLGV | ASSQGQGANTEVF | 0.81715953 |
| SYGGAYAVM | ASSQGQGANTEVF | 0.56661797 |
| TTGVGGLTLG | ASSQGQGANTEVF | 0.8473693 |
| TTQTIYI | ASSQGQGANTEVF | 0.84033245 |
| VAAGAGLGAVI | ASSQGQGANTEVF | 0.60192955 |
| VAAVVTLL | ASSQGQGANTEVF | 0.8656379 |
| VGFLGGLI | ASSQGQGANTEVF | 0.77269316 |
| VGVIPILL | ASSQGQGANTEVF | 0.7730485 |
| VIFASSLI | ASSQGQGANTEVF | 0.8869681 |
| VKGGDGGAAAG | ASSQGQGANTEVF | 0.74531335 |
| VTKTFTNI | ASSQGQGANTEVF | 0.8819843 |
| VVAAAILL | ASSQGQGANTEVF | 0.7831853 |
| YAALDVEL | ASSQGQGANTEVF | 0.7814408 |
| LASVALLL | ASSQGQGANTEVF | 0.840158 |
| LDAVHRV | ASSQGQGANTEVF | 0.3287851 |
| QGLSSSIF | ASSQGQGANTEVF | 0.68286765 |
| RGTFANIRL | ASSQGQGANTEVF | 0.8773202 |
| RQLWWGHRI | ASSQGQGANTEVF | 0.8363265 |
| YGPPGTGKTLL | ASSQGQGANTEVF | 0.8242358 |
| AALVGAPANVA | ASSQGQGANTEVF | 0.7904419 |
| AATFRRTE | ASSQGQGANTEVF | 0.7716768 |
| AGALGGGI | ASSQGQGANTEVF | 0.8124334 |
| AGGAGGAGGIGAG | ASSQGQGANTEVF | 0.819973 |
| AGSGALGL | ASSQGQGANTEVF | 0.8460573 |
| AQIADQI | ASSQGQGANTEVF | 0.8363167 |
| ASAVAVTYL | ASSQGQGANTEVF | 0.73380494 |
| ASTVLGGTGG | ASSQGQGANTEVF | 0.8444343 |
| AVGGEPAK | ASSQGQGANTEVF | 0.8273438 |
| CEQGTFGGTC | ASSQGQGANTEVF | 0.8229453 |
| DEGKAPAG | ASSQGQGANTEVF | 0.82109725 |
| DTVVFPAL | ASSQGQGANTEVF | 0.80725217 |
| FEGITMAG | ASSQGQGANTEVF | 0.79951715 |
| FPLVVQD | ASSQGQGANTEVF | 0.6947533 |
| GAFTTPQ | ASSQGQGANTEVF | 0.8290493 |
| GFGGFGTAGGVGGA | ASSQGQGANTEVF | 0.8383851 |
| GFLAQFL | ASSQGQGANTEVF | 0.83749974 |
| GGAGGAGGNAGVA | ASSQGQGANTEVF | 0.8406671 |
| GGVGTAGAGGAGGAGG | ASSQGQGANTEVF | 0.8426676 |
| GNGGGGDAGNAGSGGN | ASSQGQGANTEVF | 0.8293768 |
| GTGGAGGAGNDAGS | ASSQGQGANTEVF | 0.44142112 |
| GTTNTGWFNS | ASSQGQGANTEVF | 0.611576 |
| IFMGPLFTL | ASSQGQGANTEVF | 0.71327704 |
| ILAHAIEAI | ASSQGQGANTEVF | 0.79432654 |

| | | |
|---|---|---|
| IRSAIREAL | ASSQGQGANTEVF | 0.2994244 |
| ITFAGGG | ASSQGQGANTEVF | 0.7723057 |
| ITGMSGHQEV | ASSQGQGANTEVF | 0.810522 |
| IVAGLACTL | ASSQGQGANTEVF | 0.7853631 |
| IVGGGLAGA | ASSQGQGANTEVF | 0.84178567 |
| IVTGVLNI | ASSQGQGANTEVF | 0.8095275 |
| LAQQGCI | ASSQGQGANTEVF | 0.5995157 |
| LCLHVEN | ASSQGQGANTEVF | 0.77000344 |
| LDAISDGS | ASSQGQGANTEVF | 0.83649665 |
| LGGFAAML | ASSQGQGANTEVF | 0.75573224 |
| LGGKNPMI | ASSQGQGANTEVF | 0.84551793 |
| LGPKGVAAIL | ASSQGQGANTEVF | 0.6839272 |
| LMVGVSI | ASSQGQGANTEVF | 0.8431561 |
| LVGAAFVL | ASSQGQGANTEVF | 0.686543 |
| LVLAVAPL | ASSQGQGANTEVF | 0.84210706 |
| MAAIAVIDAA | ASSQGQGANTEVF | 0.8267784 |
| NNGINAELQ | ASSQGQGANTEVF | 0.8480808 |
| RGEPLVLVHG | ASSQGQGANTEVF | 0.8054214 |
| RGGAGGAAGLI | ASSQGQGANTEVF | 0.83781826 |
| RSYRMHPAV | ASSQGQGANTEVF | 0.8025862 |
| SAAHLLAGQLA | ASSQGQGANTEVF | 0.8308674 |
| SAAVGSY | ASSQGQGANTEVF | 0.72546947 |
| SPHAVLHRI | ASSQGQGANTEVF | 0.80340695 |
| SQIAKVL | ASSQGQGANTEVF | 0.6921153 |
| TALGHLVP | ASSQGQGANTEVF | 0.8093563 |
| TAPTTSGAA | ASSQGQGANTEVF | 0.83972746 |
| TDGDDGNGGSPGT | ASSQGQGANTEVF | 0.22246356 |
| TDGLGEN | ASSQGQGANTEVF | 0.84050626 |
| TGGPPLPGQA | ASSQGQGANTEVF | 0.68173957 |
| TIPAITI | ASSQGQGANTEVF | 0.7002871 |
| TIPGFTI | ASSQGQGANTEVF | 0.8390084 |
| TSIASGALA | ASSQGQGANTEVF | 0.82589 |
| TVTLPGAAI | ASSQGQGANTEVF | 0.8328053 |
| VASGLSAALP | ASSQGQGANTEVF | 0.7379397 |
| VGAGVIGI | ASSQGQGANTEVF | 0.8068572 |
| VGPKRFGV | ASSQGQGANTEVF | 0.81776327 |
| VNVAFPD | ASSQGQGANTEVF | 0.3376701 |
| VSMTNSGVS | ASSQGQGANTEVF | 0.6692543 |
| VTDVLGDLSV | ASSQGQGANTEVF | 0.8469643 |
| VVVVGAGFAGL | ASSQGQGANTEVF | 0.7147647 |
| VYTVGTVV | ASSQGQGANTEVF | 0.7183598 |
| WTTFSGP | ASSQGQGANTEVF | 0.66543174 |
| YDELVEAF | ASSQGQGANTEVF | 0.66618055 |
| YGTGGAGGNGG | ASSQGQGANTEVF | 0.69791734 |
| YTNVMAV | ASSQGQGANTEVF | 0.717305 |
| FLEQVHQL | ASSQGQGANTEVF | 0.72960544 |
| GLSLLAVAV | ASSQGQGANTEVF | 0.7160327 |
| LEQVHQL | ASSQGQGANTEVF | 0.69003975 |
| LNAAGAI | ASSQGQGANTEVF | 0.71650064 |
| PAVHASL | ASSQGQGANTEVF | 0.7081834 |
| TELEDDA | ASSQGQGANTEVF | 0.6963301 |

| | | |
|---|---|---|
| TPPGPPN | ASSQGQGANTEVF | 0.72228837 |
| TPPPPPPP | ASSQGQGANTEVF | 0.7095514 |
| TQSPALI | ASSLEGSSYEQY | 0.67779905 |
| EFAAAFPQ | ASSLEGSSYEQY | 0.6970644 |
| LLSGNLDVL | ASSLEGSSYEQY | 0.71722347 |
| GPANLGNYNI | ASSLEGSSYEQY | 0.7219424 |
| TLGPTLP | ASSLEGSSYEQY | 0.7183008 |
| VYAYSSCCK | ASSLEGSSYEQY | 0.7329986 |
| APAPAPAPNI | ASSLEGSSYEQY | 0.72289586 |
| PGGPVPIP | ASSLEGSSYEQY | 0.7174016 |
| FLTVALEP | ASSLEGSSYEQY | 0.5738892 |
| SQYFGANL | ASSLEGSSYEQY | 0.66190606 |
| GGVKPAA | ASSLEGSSYEQY | 0.6911775 |
| GGTGGAGGAGGW | ASSLEGSSYEQY | 0.70077443 |
| GVGGVAGLGGAGG | ASSLEGSSYEQY | 0.50932884 |
| VLLVLATTI | ASSLEGSSYEQY | 0.71116555 |
| IGIFAAML | ASSLEGSSYEQY | 0.7091317 |
| AVGDLID | ASSLEGSSYEQY | 0.6691677 |
| NISPLPL | ASSLEGSSYEQY | 0.7235092 |
| VTAVFVAI | ASSLEGSSYEQY | 0.6855692 |
| SGDDNDAAL | ASSLEGSSYEQY | 0.6908212 |
| AVAALAHS | ASSLEGSSYEQY | 0.69415754 |
| GGVTVMATP | ASSLEGSSYEQY | 0.7328424 |
| AGLAANVRD | ASSLEGSSYEQY | 0.6745868 |
| TNRPDLIDPALLRPG | ASSLEGSSYEQY | 0.72614235 |
| LANLGVTAL | ASSLEGSSYEQY | 0.7149196 |
| TGERIEL | ASSLEGSSYEQY | 0.7276059 |
| NGEELAA | ASSLEGSSYEQY | 0.65847737 |
| VLSLVWG | ASSLEGSSYEQY | 0.72295445 |
| IGYALWL | ASSLEGSSYEQY | 0.7033106 |
| LSLLLAPL | ASSLEGSSYEQY | 0.73207426 |
| TLDEALK | ASSLEGSSYEQY | 0.7091561 |
| TDGAGTAEGG | ASSLEGSSYEQY | 0.72773015 |
| NAPGGAGGNGG | ASSLEGSSYEQY | 0.71113783 |
| VSVFMGMGGNF | ASSLEGSSYEQY | 0.7200997 |
| LASYLAAL | ASSLEGSSYEQY | 0.64549965 |
| LVGGTPAL | ASSLEGSSYEQY | 0.7039357 |
| VRNADVL | ASSLEGSSYEQY | 0.68804324 |
| TCSSPFLVSAM | ASSLEGSSYEQY | 0.71196187 |
| VHITDAE | ASSLEGSSYEQY | 0.71688825 |
| EYAAPSF | ASSLEGSSYEQY | 0.496024 |
| VVDAFHGGL | ASSLEGSSYEQY | 0.7299098 |
| ASVVDPQ | ASSLEGSSYEQY | 0.6424004 |
| SYNEFGGLS | ASSLEGSSYEQY | 0.6897303 |
| DPDGSVY | ASSLEGSSYEQY | 0.7242596 |
| MVSTIHATE | ASSLEGSSYEQY | 0.69814616 |
| LAAGVADPV | ASSLEGSSYEQY | 0.7190942 |
| TKEQIAA | ASSLEGSSYEQY | 0.68644893 |
| SLASPAAL | ASSLEGSSYEQY | 0.71320367 |
| IGLVFYGV | ASSLEGSSYEQY | 0.71526355 |
| FLLDASNL | ASSLEGSSYEQY | 0.5259188 |

| | | |
|---|---|---|
| YQVGNFP | ASSLEGSSYEQY | 0.6512214 |
| ESLLSQST | ASSLEGSSYEQY | 0.7296735 |
| LFTVGFAAVG | ASSLEGSSYEQY | 0.7511704 |
| ATPPTPT | ASSLEGSSYEQY | 0.78017664 |
| TDLATLG | ASSLEGSSYEQY | 0.7173872 |
| SAGGTGR | ASSLEGSSYEQY | 0.63716936 |
| GFSNSGS | ASSLEGSSYEQY | 0.70350593 |
| GGNGGSGGTSV | ASSLEGSSYEQY | 0.7528637 |
| GTSVATGGAGN | ASSLEGSSYEQY | 0.78094953 |
| NGGNGGSGGTS | ASSLEGSSYEQY | 0.76728046 |
| AVVKFPH | ASSLEGSSYEQY | 0.57305574 |
| TPPPPPP | ASSLEGSSYEQY | 0.77429324 |
| AAPAAAAAPA | ASSLEGSSYEQY | 0.72576326 |
| IALNEDL | ASSLEGSSYEQY | 0.731839 |
| IVNVSSM | ASSLEGSSYEQY | 0.7158908 |
| TGIPPAPR | ASSLEGSSYEQY | 0.748476 |
| FDLGGGTF | ASSLEGSSYEQY | 0.7204011 |
| FDLGGGTFDVS | ASSLEGSSYEQY | 0.594874 |
| NEPTAAA | ASSLEGSSYEQY | 0.7264607 |
| NEPTAAALA | ASSLEGSSYEQY | 0.7637473 |
| GSIINISSI | ASSLEGSSYEQY | 0.7577132 |
| VAPGNAGTA | ASSLEGSSYEQY | 0.78362054 |
| TGGGASLE | ASSLEGSSYEQY | 0.77559936 |
| VAVYNHYKRI | ASSLEGSSYEQY | 0.7543124 |
| TDPAPATT | ASSLEGSSYEQY | 0.27388868 |
| TNRPDILDPALLRPG | ASSLEGSSYEQY | 0.4885625 |
| LTTTTTS | ASSLEGSSYEQY | 0.6052111 |
| PTGSGKTL | ASSLEGSSYEQY | 0.72243035 |
| TGSGKTL | ASSLEGSSYEQY | 0.2958534 |
| TIGHVDHGK | ASSLEGSSYEQY | 0.71815574 |
| TIGHVDHGKTT | ASSLEGSSYEQY | 0.721298 |
| GHVDHGKTT | ASSLEGSSYEQY | 0.6725994 |
| AQFEHTLL | ASSLEGSSYEQY | 0.7733947 |
| IVNQDEL | ASSLEGSSYEQY | 0.7126993 |
| TNGSQFF | ASSLEGSSYEQY | 0.5757497 |
| TGKGIVE | ASSLEGSSYEQY | 0.5519657 |
| LISNASDA | ASSLEGSSYEQY | 0.7713419 |
| AELRTLGELTKIAKAHGAQ | ASSLEGSSYEQY | 0.45542008 |
| APNAVAAAEPSW | ASSLEGSSYEQY | 0.7728789 |
| APQVAGAATAVL | ASSLEGSSYEQY | 0.53220105 |
| ARRDPAAAAAILPTDA | ASSLEGSSYEQY | 0.76969856 |
| DAAARDVLAEAGLAEYF | ASSLEGSSYEQY | 0.50500995 |
| DDQIVLSAKANLAKVEQ | ASSLEGSSYEQY | 0.7717092 |
| DELLPDTGEDSDEDGH | ASSLEGSSYEQY | 0.73293424 |
| ELPEAPANRATVPQD | ASSLEGSSYEQY | 0.78994864 |
| FGQSKIERVTGIG | ASSLEGSSYEQY | 0.67988527 |
| GAGGTGSTAGGAGGAGGAGGLYA | ASSLEGSSYEQY | 0.7473499 |
| GHGGTGAAGGNGGTGGAGGSGIDG | ASSLEGSSYEQY | 0.60034513 |
| GNGGNGGAGGEGGPGGAGGAGA | ASSLEGSSYEQY | 0.7761222 |
| GSTAGNGGNGGNATNGTIAGQPAG | ASSLEGSSYEQY | 0.5319872 |
| GTGGDGGNGGNGGTGG | ASSLEGSSYEQY | 0.70346653 |

| | | |
|---|---|---|
| GVRRPAGIPADAPV | ASSLEGSSYEQY | 0.46919844 |
| GYRQVTDAHLLTLARR | ASSLEGSSYEQY | 0.7385167 |
| IAKMGQDGHDRG | ASSLEGSSYEQY | 0.7689327 |
| KAEPSIEIQYT | ASSLEGSSYEQY | 0.37614238 |
| KPVRIRMAADEDTVAVT | ASSLEGSSYEQY | 0.77600646 |
| MPPRIAGMRLLVIKPEP | ASSLEGSSYEQY | 0.6776658 |
| MKFYAVALFRSRGVPPTR | ASSLEGSSYEQY | 0.5963482 |
| NAGGFGILGAVAHSP | ASSLEGSSYEQY | 0.76479626 |
| NEGREITLAMAEM | ASSLEGSSYEQY | 0.73277795 |
| NELVEWSHPLGHRWRWE | ASSLEGSSYEQY | 0.7741613 |
| NLGAGGTGGNGGNPG | ASSLEGSSYEQY | 0.59988296 |
| NTEWPKWETVMAADPVD | ASSLEGSSYEQY | 0.71725965 |
| PASAAPAETQP | ASSLEGSSYEQY | 0.74707365 |
| QQSSDGKNGVRAAEITGE | ASSLEGSSYEQY | 0.596856 |
| RAPDAVTTSWWKE | ASSLEGSSYEQY | 0.5310316 |
| SGFGNVGAHTSGWFNQ | ASSLEGSSYEQY | 0.78072464 |
| SGVAGAVGQAASVGGLKV | ASSLEGSSYEQY | 0.66801906 |
| SNGGAGTGSAGGNGGT | ASSLEGSSYEQY | 0.67681825 |
| TGGAGGEGGAAG | ASSLEGSSYEQY | 0.6002606 |
| TGGDGGAAGAGGAPGHG | ASSLEGSSYEQY | 0.61594635 |
| TMPPLLAAIANAGRHGV | ASSLEGSSYEQY | 0.6146263 |
| TSSTAGNSGGA | ASSLEGSSYEQY | 0.66494787 |
| VPATVPVGILC | ASSLEGSSYEQY | 0.6799613 |
| VQRPDSGGVVLTGRIS | ASSLEGSSYEQY | 0.6698197 |
| WTPNPVPATKGVPAGLVQAT | ASSLEGSSYEQY | 0.592726 |
| YALKRALAKFSDRFDVVII | ASSLEGSSYEQY | 0.6637076 |
| YLIGSVIV | ASSLEGSSYEQY | 0.630503 |
| YGQLVAVID | ASSLEGSSYEQY | 0.63884115 |
| VTVGGMTLPA | ASSLEGSSYEQY | 0.65065837 |
| VLALPAV | ASSLEGSSYEQY | 0.64648265 |
| VIPPTLNYET | ASSLEGSSYEQY | 0.6234951 |
| VGMGGMHPSSG | ASSLEGSSYEQY | 0.57037973 |
| VGLAGGAATA | ASSLEGSSYEQY | 0.6561919 |
| VGGAGGAGGSGP | ASSLEGSSYEQY | 0.67439926 |
| VFAAFMV | ASSLEGSSYEQY | 0.66867 |
| TLYDKLR | ASSLEGSSYEQY | 0.6730373 |
| TLAVFTVL | ASSLEGSSYEQY | 0.66914827 |
| TGSVQPARV | ASSLEGSSYEQY | 0.65916723 |
| TGGGAGSGGT | ASSLEGSSYEQY | 0.38838136 |
| SHLETSL | ASSLEGSSYEQY | 0.52947676 |
| SFTTGMT | ASSLEGSSYEQY | 0.5976205 |
| QVEVAKRMGVRQPTVS | ASSLEGSSYEQY | 0.64051926 |
| QGSGGPALAM | ASSLEGSSYEQY | 0.3607441 |
| MLAPSNP | ASSLEGSSYEQY | 0.6629832 |
| LVNNAAI | ASSLEGSSYEQY | 0.6352879 |
| LTGEINTLA | ASSLEGSSYEQY | 0.6244232 |
| LSIFGPV | ASSLEGSSYEQY | 0.66897 |
| LLALLANL | ASSLEGSSYEQY | 0.62921053 |
| LISNASDALDKLR | ASSLEGSSYEQY | 0.5801146 |
| LINEAVP | ASSLEGSSYEQY | 0.54920334 |
| LGYAFSGL | ASSLEGSSYEQY | 0.6729721 |

| | | |
|---|---|---|
| LDFVAVAL | ASSLEGSSYEQY | 0.52486 |
| LAQPGAE | ASSLEGSSYEQY | 0.6712211 |
| ISNLGAVTGF | ASSLEGSSYEQY | 0.4918077 |
| HQEDPGRI | ASSLEGSSYEQY | 0.67057806 |
| GTPGEPTP | ASSLEGSSYEQY | 0.5682971 |
| GFFNSGS | ASSLEGSSYEQY | 0.6684933 |
| GAGGGGLTAGAN | ASSLEGSSYEQY | 0.63964033 |
| FVDALVNM | ASSLEGSSYEQY | 0.68404347 |
| FLTVAPEL | ASSLEGSSYEQY | 0.6318979 |
| FETLTAL | ASSLEGSSYEQY | 0.6603422 |
| DVINPIEL | ASSLEGSSYEQY | 0.6225199 |
| APVSGAPV | ASSLEGSSYEQY | 0.6785193 |
| ALVGGLAW | ASSLEGSSYEQY | 0.5327505 |
| ALAAAAATAI | ASSLEGSSYEQY | 0.634591 |
| AFGLGLLL | ASSLEGSSYEQY | 0.51213974 |
| AFFAAASL | ASSLEGSSYEQY | 0.64256185 |
| AAAALPAFSPP | ASSLEGSSYEQY | 0.66904914 |
| VVLAPAL | ASSLEGSSYEQY | 0.4416244 |
| TIGHVDHGKTTL | ASSLEGSSYEQY | 0.65791595 |
| LISNASDALDK | ASSLEGSSYEQY | 0.6029277 |
| LISNASDAL | ASSLEGSSYEQY | 0.6062282 |
| ISNASDALDK | ASSLEGSSYEQY | 0.66427237 |
| NQAETLVYQTE | ASSLEGSSYEQY | 0.64933074 |
| TGGTGGTAGAG | ASSLEGSSYEQY | 0.6663817 |
| TGSTAGGAGGA | ASSLEGSSYEQY | 0.57490146 |
| DPLAVPVAAGPV | ASSLEGSSYEQY | 0.6094 |
| GGTGGTPGNGGN | ASSLEGSSYEQY | 0.65010357 |
| PAGLAGVDPATL | ASSLEGSSYEQY | 0.57787675 |
| GNNLVVEAPHAGASV | ASSLEGSSYEQY | 0.57358325 |
| KVTRIMSASTLEQPA | ASSLEGSSYEQY | 0.67601293 |
| LGFALPIALPRRTVV | ASSLEGSSYEQY | 0.6240363 |
| TRLGNLTARKLVVLV | ASSLEGSSYEQY | 0.64047796 |
| WSQQRRLIYFPSAGP | ASSLEGSSYEQY | 0.56705713 |
| GPNGNPDPMAAAVDIR | ASSLEGSSYEQY | 0.554867 |
| TQRCVSVADLRAAASR | ASSLEGSSYEQY | 0.5884847 |
| QFVLDVAEHPVEVLTETV | ASSLEGSSYEQY | 0.6451005 |
| GGAGGTGGAGGNGGTQAGDG | ASSLEGSSYEQY | 0.64110297 |
| PLLATLRAIAVDVDAKPAQV | ASSLEGSSYEQY | 0.61787826 |
| APKFRTGGYMIDFWGVGYQVAKRM | ASSLEGSSYEQY | 0.5454469 |
| GAGGAGGAGGAAGGPGGTGGTGGN | ASSLEGSSYEQY | 0.61962694 |
| IGALGLLATDVAPAGSVAFGVALA | ASSLEGSSYEQY | 0.6106169 |
| AGGAGGIGGNGGAGGAGGVLMGNG | ASSLEGSSYEQY | 0.5857735 |
| ADELIGLL | ASSLEGSSYEQY | 0.63353753 |
| ADRALKA | ASSLEGSSYEQY | 0.62269235 |
| AGAVGQVGGAGGA | ASSLEGSSYEQY | 0.5801691 |
| AGGGGGLGGAP | ASSLEGSSYEQY | 0.59041476 |
| AGGLLYGNGGAG | ASSLEGSSYEQY | 0.61095655 |
| AGLESIGL | ASSLEGSSYEQY | 0.63715464 |
| AGLKTGII | ASSLEGSSYEQY | 0.6356119 |
| ASLAALLL | ASSLEGSSYEQY | 0.65622646 |
| ASVINISSI | ASSLEGSSYEQY | 0.6402879 |

| | | |
|---|---|---|
| ATPPGPA | ASSLEGSSYEQY | 0.62623703 |
| ATVASALV | ASSLEGSSYEQY | 0.38962904 |
| ATVASLIL | ASSLEGSSYEQY | 0.44593096 |
| AVNGPVLIRH | ASSLEGSSYEQY | 0.5909265 |
| DGAAGAVGI | ASSLEGSSYEQY | 0.60457945 |
| DPLLHGI | ASSLEGSSYEQY | 0.47563475 |
| ELGAALGAL | ASSLEGSSYEQY | 0.6366458 |
| ELGRTQAV | ASSLEGSSYEQY | 0.59338087 |
| FVAGGYLS | ASSLEGSSYEQY | 0.57316005 |
| GASGGAGGAGGW | ASSLEGSSYEQY | 0.6375945 |
| GGAGAGGDGGAGGI | ASSLEGSSYEQY | 0.5762016 |
| GTGHNGSQPGMGGNG | ASSLEGSSYEQY | 0.58887815 |
| GYAAASASAS | ASSLEGSSYEQY | 0.5846813 |
| HQVDVAAVQQG | ASSLEGSSYEQY | 0.6430709 |
| IGAAIAHT | ASSLEGSSYEQY | 0.54364485 |
| IGMIEVAQL | ASSLEGSSYEQY | 0.6324011 |
| IRLAPEHTSAMD | ASSLEGSSYEQY | 0.54501545 |
| IYNVSVL | ASSLEGSSYEQY | 0.6484166 |
| KQGFLGM | ASSLEGSSYEQY | 0.52015483 |
| LESVITQL | ASSLEGSSYEQY | 0.62848324 |
| LGLGALVVV | ASSLEGSSYEQY | 0.589695 |
| LIGNGGNGGSGGI | ASSLEGSSYEQY | 0.6571611 |
| LNAINAPTL | ASSLEGSSYEQY | 0.6119645 |
| LQLLNTGV | ASSLEGSSYEQY | 0.63225406 |
| LQTSIEF | ASSLEGSSYEQY | 0.5894487 |
| MSFAVSEL | ASSLEGSSYEQY | 0.6274494 |
| NTGALGL | ASSLEGSSYEQY | 0.5188415 |
| PPGAGGAGGAGTT | ASSLEGSSYEQY | 0.6115277 |
| QLVGLGI | ASSLEGSSYEQY | 0.51794785 |
| RFLEQVHQL | ASSLEGSSYEQY | 0.6208925 |
| RGVGYVLR | ASSLEGSSYEQY | 0.61611444 |
| SATGVFAGVI | ASSLEGSSYEQY | 0.54005307 |
| SLPAITLP | ASSLEGSSYEQY | 0.6307268 |
| STALAFTV | ASSLEGSSYEQY | 0.6127994 |
| SVLTVGLGK | ASSLEGSSYEQY | 0.5762979 |
| TALAGAGSGI | ASSLEGSSYEQY | 0.64464384 |
| TGLPFTGI | ASSLEGSSYEQY | 0.6129761 |
| VAEHVTV | ASSLEGSSYEQY | 0.63354486 |
| VDIVLNSL | ASSLEGSSYEQY | 0.5887003 |
| VLTGGGAVL | ASSLEGSSYEQY | 0.5603953 |
| VNVGLAYGI | ASSLEGSSYEQY | 0.62746036 |
| VSSAAVGAV | ASSLEGSSYEQY | 0.58801484 |
| VTSVYEAGV | ASSLEGSSYEQY | 0.58275753 |
| WVALDRGA | ASSLEGSSYEQY | 0.6360308 |
| YLVGSVVI | ASSLEGSSYEQY | 0.8259618 |
| GPPGTGKTLL | ASSLEGSSYEQY | 0.8161638 |
| GVGMTKF | ASSLEGSSYEQY | 0.6490574 |
| IGIGGSDL | ASSLEGSSYEQY | 0.605734 |
| IGTLLAEL | ASSLEGSSYEQY | 0.6998174 |
| IYTSGTT | ASSLEGSSYEQY | 0.8369626 |
| LSFPAGV | ASSLEGSSYEQY | 0.7974782 |

| | | |
|---|---|---|
| LYTSGTT | ASSLEGSSYEQY | 0.80749446 |
| VGAFPFGF | ASSLEGSSYEQY | 0.54690534 |
| VGAGTVEF | ASSLEGSSYEQY | 0.73929703 |
| VGKSTLLL | ASSLEGSSYEQY | 0.724452 |
| AALAHAGGGIV | ASSLEGSSYEQY | 0.725253 |
| AALVAVSLL | ASSLEGSSYEQY | 0.80782604 |
| AAVSRAVAAL | ASSLEGSSYEQY | 0.7115915 |
| AGGSSGIGGF | ASSLEGSSYEQY | 0.7260536 |
| AGLTVANANG | ASSLEGSSYEQY | 0.6875653 |
| AGLVGAVL | ASSLEGSSYEQY | 0.7488916 |
| ALSQAFTV | ASSLEGSSYEQY | 0.8077034 |
| ALTFSVSL | ASSLEGSSYEQY | 0.79180884 |
| ALTGAALVAL | ASSLEGSSYEQY | 0.86336696 |
| ASFSIGAGVL | ASSLEGSSYEQY | 0.82598567 |
| DGSVYKI | ASSLEGSSYEQY | 0.81571585 |
| EQLAEIV | ASSLEGSSYEQY | 0.20188843 |
| FAVSELTVM | ASSLEGSSYEQY | 0.43791318 |
| GAAPLAAGL | ASSLEGSSYEQY | 0.6698458 |
| GADGGDGGSGGAGG | ASSLEGSSYEQY | 0.6678585 |
| GFGLDTGGAGG | ASSLEGSSYEQY | 0.3365206 |
| GLASGGLL | ASSLEGSSYEQY | 0.7895744 |
| GRIAGAAVLL | ASSLEGSSYEQY | 0.7920893 |
| GSAGSLNTSF | ASSLEGSSYEQY | 0.56763995 |
| GTGQIGI | ASSLEGSSYEQY | 0.8178123 |
| GTLIALIL | ASSLEGSSYEQY | 0.6630906 |
| IAATMGFVVG | ASSLEGSSYEQY | 0.6333694 |
| IFAQALL | ASSLEGSSYEQY | 0.65242386 |
| IGAAVALGF | ASSLEGSSYEQY | 0.8303557 |
| IGLTPLAL | ASSLEGSSYEQY | 0.5960307 |
| ILGSNTSTL | ASSLEGSSYEQY | 0.8106732 |
| IVFSLTVL | ASSLEGSSYEQY | 0.6721468 |
| LASKATGF | ASSLEGSSYEQY | 0.8449408 |
| LASVALIL | ASSLEGSSYEQY | 0.5133722 |
| LFAVGLTL | ASSLEGSSYEQY | 0.81320745 |
| LFNGGIGTYI | ASSLEGSSYEQY | 0.7329895 |
| LTLGAGII | ASSLEGSSYEQY | 0.8717099 |
| LVAAALGDI | ASSLEGSSYEQY | 0.7117355 |
| NTAFVQLGI | ASSLEGSSYEQY | 0.71861404 |
| QLFSGAEL | ASSLEGSSYEQY | 0.5810812 |
| SAATFITL | ASSLEGSSYEQY | 0.76385266 |
| SNYGLANLGV | ASSLEGSSYEQY | 0.39646286 |
| SYGGAYAVM | ASSLEGSSYEQY | 0.7304627 |
| TTGVGGLTLG | ASSLEGSSYEQY | 0.54600084 |
| TTQTIYI | ASSLEGSSYEQY | 0.7559173 |
| VAAGAGLGAVI | ASSLEGSSYEQY | 0.8065322 |
| VAAVVTLL | ASSLEGSSYEQY | 0.3638129 |
| VGFLGGLI | ASSLEGSSYEQY | 0.7873777 |
| VGVIPILL | ASSLEGSSYEQY | 0.668756 |
| VIFASSLI | ASSLEGSSYEQY | 0.7031888 |
| VKGGDGGAAAG | ASSLEGSSYEQY | 0.8275673 |
| VTKTFTNI | ASSLEGSSYEQY | 0.66015255 |

| | | |
|---|---|---|
| VVAAAILL | ASSLEGSSYEQY | 0.83475363 |
| YAALDVEL | ASSLEGSSYEQY | 0.6623126 |
| LASVALLL | ASSLEGSSYEQY | 0.6954243 |
| LDAVHRV | ASSLEGSSYEQY | 0.8063022 |
| QGLSSSIF | ASSLEGSSYEQY | 0.3311783 |
| RGTFANIRL | ASSLEGSSYEQY | 0.5540809 |
| RQLWWGHRI | ASSLEGSSYEQY | 0.8419863 |
| YGPPGTGKTLL | ASSLEGSSYEQY | 0.70931506 |
| AALVGAPANVA | ASSLEGSSYEQY | 0.7222361 |
| AATFRRTE | ASSLEGSSYEQY | 0.60108775 |
| AGALGGGI | ASSLEGSSYEQY | 0.5566684 |
| AGGAGGAGGIGAG | ASSLEGSSYEQY | 0.65806735 |
| AGSGALGL | ASSLEGSSYEQY | 0.71948266 |
| AQIADQI | ASSLEGSSYEQY | 0.71695274 |
| ASAVAVTYL | ASSLEGSSYEQY | 0.7104775 |
| ASTVLGGTGG | ASSLEGSSYEQY | 0.5427189 |
| AVGGEPAK | ASSLEGSSYEQY | 0.6681348 |
| CEQGTFGGTC | ASSLEGSSYEQY | 0.65393806 |
| DEGKAPAG | ASSLEGSSYEQY | 0.6462669 |
| DTVVFPAL | ASSLEGSSYEQY | 0.7006246 |
| FEGITMAG | ASSLEGSSYEQY | 0.59395623 |
| FPLVVQD | ASSLEGSSYEQY | 0.65610677 |
| GAFTTPQ | ASSLEGSSYEQY | 0.6301261 |
| GFGGFGTAGGVGGA | ASSLEGSSYEQY | 0.6935338 |
| GFLAQFL | ASSLEGSSYEQY | 0.7036106 |
| GGAGGAGGNAGVA | ASSLEGSSYEQY | 0.695153 |
| GGVGTAGAGGAGGAGG | ASSLEGSSYEQY | 0.72526807 |
| GNGGGGDAGNAGSGGN | ASSLEGSSYEQY | 0.7128497 |
| GTGGAGGAGNDAGS | ASSLEGSSYEQY | 0.70904267 |
| GTTNTGWFNS | ASSLEGSSYEQY | 0.29939473 |
| IFMGPLFTL | ASSLEGSSYEQY | 0.44461426 |
| ILAHAIEAI | ASSLEGSSYEQY | 0.6400287 |
| IRSAIREAL | ASSLEGSSYEQY | 0.6212431 |
| ITFAGGG | ASSLEGSSYEQY | 0.49150127 |
| ITGMSGHQEV | ASSLEGSSYEQY | 0.6847235 |
| IVAGLACTL | ASSLEGSSYEQY | 0.69195735 |
| IVGGGLAGA | ASSLEGSSYEQY | 0.5703325 |
| IVTGVLNI | ASSLEGSSYEQY | 0.69564867 |
| LAQQGCI | ASSLEGSSYEQY | 0.5990815 |
| LCLHVEN | ASSLEGSSYEQY | 0.5604688 |
| LDAISDGS | ASSLEGSSYEQY | 0.5816257 |
| LGGFAAML | ASSLEGSSYEQY | 0.71484685 |
| LGGKNPMI | ASSLEGSSYEQY | 0.5015965 |
| LGPKGVAAIL | ASSLEGSSYEQY | 0.67934835 |
| LMVGVSI | ASSLEGSSYEQY | 0.68901885 |
| LVGAAFVL | ASSLEGSSYEQY | 0.71183425 |
| LVLAVAPL | ASSLEGSSYEQY | 0.4354691 |
| MAAIAVIDAA | ASSLEGSSYEQY | 0.7053871 |
| NNGINAELQ | ASSLEGSSYEQY | 0.6440845 |
| RGEPLVLVHG | ASSLEGSSYEQY | 0.7215553 |
| RGGAGGAAGLI | ASSLEGSSYEQY | 0.5978613 |

| | | |
|---|---|---|
| RSYRMHPAV | ASSLEGSSYEQY | 0.6342364 |
| SAAHLLAGQLA | ASSLEGSSYEQY | 0.60799944 |
| SAAVGSY | ASSLEGSSYEQY | 0.70996594 |
| SPHAVLHRI | ASSLEGSSYEQY | 0.40164274 |
| SQIAKVL | ASSLEGSSYEQY | 0.6710683 |
| TALGHLVP | ASSLEGSSYEQY | 0.6389636 |
| TAPTTSGAA | ASSLEGSSYEQY | 0.68925524 |
| TDGDDGNGGSPGT | ASSLEGSSYEQY | 0.70730776 |
| TDGLGEN | ASSLEGSSYEQY | 0.50580597 |
| TGGPPLPGQA | ASSLEGSSYEQY | 0.7158979 |
| TIPAITI | ASSLEGSSYEQY | 0.66597366 |
| TIPGFTI | ASSLEGSSYEQY | 0.6040354 |
| TSIASGALA | ASSLEGSSYEQY | 0.70910543 |
| TVTLPGAAI | ASSLEGSSYEQY | 0.62810135 |
| VASGLSAALP | ASSLEGSSYEQY | 0.7194401 |
| VGAGVIGI | ASSLEGSSYEQY | 0.6613872 |
| VGPKRFGV | ASSLEGSSYEQY | 0.6954044 |
| VNVAFPD | ASSLEGSSYEQY | 0.7040782 |
| VSMTNSGVS | ASSLEGSSYEQY | 0.5954162 |
| VTDVLGDLSV | ASSLEGSSYEQY | 0.5517366 |
| VVVVGAGFAGL | ASSLEGSSYEQY | 0.7131957 |
| VYTVGTVV | ASSLEGSSYEQY | 0.75791377 |
| WTTFSGP | ASSLEGSSYEQY | 0.8301445 |
| YDELVEAF | ASSLEGSSYEQY | 0.4605295 |
| YGTGGAGGNGG | ASSLEGSSYEQY | 0.4390933 |
| YTNVMAV | ASSLEGSSYEQY | 0.6604688 |
| FLEQVHQL | ASSLEGSSYEQY | 0.8110375 |
| GLSLLAVAV | ASSLEGSSYEQY | 0.8283931 |
| LEQVHQL | ASSLEGSSYEQY | 0.7859501 |
| LNAAGAI | ASSLEGSSYEQY | 0.38144535 |
| PAVHASL | ASSLEGSSYEQY | 0.6713631 |
| TELEDDA | ASSLEGSSYEQY | 0.52789223 |
| TPPGPPN | ASSLEGSSYEQY | 0.6740843 |
| TPPPPPPP | ASSLEGSSYEQY | 0.7525892 |
| TQSPALI | TCSALGISGNTLY | 0.67448825 |
| EFAAAFPQ | TCSALGISGNTLY | 0.5692072 |
| LLSGNLDVL | TCSALGISGNTLY | 0.5004093 |
| GPANLGNYNI | TCSALGISGNTLY | 0.78376025 |
| TLGPTLP | TCSALGISGNTLY | 0.8073205 |
| VYAYSSCCK | TCSALGISGNTLY | 0.7828656 |
| APAPAPAPNI | TCSALGISGNTLY | 0.8413899 |
| PGGPVPIP | TCSALGISGNTLY | 0.7889291 |
| FLTVALEP | TCSALGISGNTLY | 0.7632724 |
| SQYFGANL | TCSALGISGNTLY | 0.1655879 |
| GGVKPAA | TCSALGISGNTLY | 0.2831906 |
| GGTGGAGGAGGW | TCSALGISGNTLY | 0.4867212 |
| GVGGVAGLGGAGG | TCSALGISGNTLY | 0.6202153 |
| VLLVLATTI | TCSALGISGNTLY | 0.22469765 |
| IGIFAAML | TCSALGISGNTLY | 0.74649024 |
| AVGDLID | TCSALGISGNTLY | 0.5080895 |
| NISPLPL | TCSALGISGNTLY | 0.45805776 |

| | | |
|---|---|---|
| VTAVFVAI | TCSALGISGNTLY | 0.7972959 |
| SGDDNDAAL | TCSALGISGNTLY | 0.49169248 |
| AVAALAHS | TCSALGISGNTLY | 0.46663415 |
| GGVTVMATP | TCSALGISGNTLY | 0.54094565 |
| AGLAANVRD | TCSALGISGNTLY | 0.7852779 |
| TNRPDLIDPALLRPG | TCSALGISGNTLY | 0.36128134 |
| LANLGVTAL | TCSALGISGNTLY | 0.75962734 |
| TGERIEL | TCSALGISGNTLY | 0.6743243 |
| NGEELAA | TCSALGISGNTLY | 0.818434 |
| VLSLVWG | TCSALGISGNTLY | 0.3343578 |
| IGYALWL | TCSALGISGNTLY | 0.7811265 |
| LSLLLAPL | TCSALGISGNTLY | 0.6808964 |
| TLDEALK | TCSALGISGNTLY | 0.8523028 |
| TDGAGTAEGG | TCSALGISGNTLY | 0.5403815 |
| NAPGGAGGNGG | TCSALGISGNTLY | 0.64374137 |
| VSVFMGMGGNF | TCSALGISGNTLY | 0.5240553 |
| LASYLAAL | TCSALGISGNTLY | 0.81879115 |
| LVGGTPAL | TCSALGISGNTLY | 0.28114837 |
| VRNADVL | TCSALGISGNTLY | 0.59921956 |
| TCSSPFLVSAM | TCSALGISGNTLY | 0.35768017 |
| VHITDAE | TCSALGISGNTLY | 0.6463824 |
| EYAAPSF | TCSALGISGNTLY | 0.772606 |
| VVDAFHGGL | TCSALGISGNTLY | 0.21074504 |
| ASVVDPQ | TCSALGISGNTLY | 0.77013505 |
| SYNEFGGLS | TCSALGISGNTLY | 0.57625437 |
| DPDGSVY | TCSALGISGNTLY | 0.43369496 |
| MVSTIHATE | TCSALGISGNTLY | 0.8070208 |
| LAAGVADPV | TCSALGISGNTLY | 0.59391433 |
| TKEQIAA | TCSALGISGNTLY | 0.81667113 |
| SLASPAAL | TCSALGISGNTLY | 0.6720547 |
| IGLVFYGV | TCSALGISGNTLY | 0.64582825 |
| FLLDASNL | TCSALGISGNTLY | 0.738799 |
| YQVGNFP | TCSALGISGNTLY | 0.29542583 |
| ESLLSQST | TCSALGISGNTLY | 0.49130565 |
| LFTVGFAAVG | TCSALGISGNTLY | 0.8178297 |
| ATPPTPT | TCSALGISGNTLY | 0.6890776 |
| TDLATLG | TCSALGISGNTLY | 0.6990124 |
| SAGGTGR | TCSALGISGNTLY | 0.62850416 |
| GFSNSGS | TCSALGISGNTLY | 0.6184245 |
| GGNGGSGGTSV | TCSALGISGNTLY | 0.648644 |
| GTSVATGGAGN | TCSALGISGNTLY | 0.68651915 |
| NGGNGGSGGTS | TCSALGISGNTLY | 0.70993984 |
| AVVKFPH | TCSALGISGNTLY | 0.6960666 |
| TPPPPPP | TCSALGISGNTLY | 0.5908926 |
| AAPAAAAAPA | TCSALGISGNTLY | 0.6942397 |
| IALNEDL | TCSALGISGNTLY | 0.66762495 |
| IVNVSSM | TCSALGISGNTLY | 0.6847892 |
| TGIPPAPR | TCSALGISGNTLY | 0.6731077 |
| FDLGGGTF | TCSALGISGNTLY | 0.6378038 |
| FDLGGGTFDVS | TCSALGISGNTLY | 0.6217258 |
| NEPTAAA | TCSALGISGNTLY | 0.57216483 |

| | | |
|---|---|---|
| NEPTAAALA | TCSALGISGNTLY | 0.66300046 |
| GSIINISSI | TCSALGISGNTLY | 0.6954836 |
| VAPGNAGTA | TCSALGISGNTLY | 0.6857538 |
| TGGGASLE | TCSALGISGNTLY | 0.7082125 |
| VAVYNHYKRI | TCSALGISGNTLY | 0.69857883 |
| TDPAPATT | TCSALGISGNTLY | 0.6733433 |
| TNRPDILDPALLRPG | TCSALGISGNTLY | 0.33793217 |
| LTTTTTS | TCSALGISGNTLY | 0.5573358 |
| PTGSGKTL | TCSALGISGNTLY | 0.51907754 |
| TGSGKTL | TCSALGISGNTLY | 0.6646896 |
| TIGHVDHGK | TCSALGISGNTLY | 0.2980296 |
| TIGHVDHGKTT | TCSALGISGNTLY | 0.6693046 |
| GHVDHGKTT | TCSALGISGNTLY | 0.66272247 |
| AQFEHTLL | TCSALGISGNTLY | 0.6212652 |
| IVNQDEL | TCSALGISGNTLY | 0.6977482 |
| TNGSQFF | TCSALGISGNTLY | 0.656523 |
| TGKGIVE | TCSALGISGNTLY | 0.5318545 |
| LISNASDA | TCSALGISGNTLY | 0.5362233 |
| AELRTLGELTKIAKAHGAQ | TCSALGISGNTLY | 0.71319926 |
| APNAVAAAEPSW | TCSALGISGNTLY | 0.53091407 |
| APQVAGAATAVL | TCSALGISGNTLY | 0.7097037 |
| ARRDPAAAAAILPTDA | TCSALGISGNTLY | 0.57870483 |
| DAAARDVLAEAGLAEYF | TCSALGISGNTLY | 0.70448756 |
| DDQIVLSAKANLAKVEQ | TCSALGISGNTLY | 0.5193489 |
| DELLPDTGEDSDEDGH | TCSALGISGNTLY | 0.698614 |
| ELPEAPANRATVPQD | TCSALGISGNTLY | 0.6771953 |
| FGQSKIERVTGIG | TCSALGISGNTLY | 0.7124501 |
| GAGGTGSTAGGAGGAGGAGGLYA | TCSALGISGNTLY | 0.6463524 |
| GHGGTGAAGGNGGTGGAGGSGIDG | TCSALGISGNTLY | 0.69202447 |
| GNGGNGGAGGEGGPGGAGGAGA | TCSALGISGNTLY | 0.5748162 |
| GSTAGNGGNGGNATNGTIAGQPAG | TCSALGISGNTLY | 0.71020895 |
| GTGGDGGNGGNGGTGG | TCSALGISGNTLY | 0.5300108 |
| GVRRPAGIPADAPV | TCSALGISGNTLY | 0.65668535 |
| GYRQVTDAHLLTLARR | TCSALGISGNTLY | 0.5955026 |
| IAKMGQDGHDRG | TCSALGISGNTLY | 0.6707736 |
| KAEPSIEIQYT | TCSALGISGNTLY | 0.6981081 |
| KPVRIRMAADEDTVAVT | TCSALGISGNTLY | 0.3380031 |
| MPPRIAGMRLLVIKPEP | TCSALGISGNTLY | 0.7048032 |
| MKFYAVALFRSRGVPPTR | TCSALGISGNTLY | 0.61233264 |
| NAGGFGILGAVAHSP | TCSALGISGNTLY | 0.57467467 |
| NEGREITLAMAEM | TCSALGISGNTLY | 0.69321144 |
| NELVEWSHPLGHRWRWE | TCSALGISGNTLY | 0.62245506 |
| NLGAGGTGGNGGNPG | TCSALGISGNTLY | 0.68871886 |
| NTEWPKWETVMAADPVD | TCSALGISGNTLY | 0.53481483 |
| PASAAPAETQP | TCSALGISGNTLY | 0.68000805 |
| QQSSDGKNGVRAAEITGE | TCSALGISGNTLY | 0.6932076 |
| RAPDAVTTSWWKE | TCSALGISGNTLY | 0.38641226 |
| SGFGNVGAHTSGWFNQ | TCSALGISGNTLY | 0.42237556 |
| SGVAGAVGQAASVGGLKV | TCSALGISGNTLY | 0.7086135 |
| SNGGAGTGSAGGNGGT | TCSALGISGNTLY | 0.8490422 |
| TGGAGGEGGAAG | TCSALGISGNTLY | 0.8441926 |

| | | |
|---|---|---|
| TGGDGGAAGAGGAPGHG | TCSALGISGNTLY | 0.7507941 |
| TMPPLLAAIANAGRHGV | TCSALGISGNTLY | 0.71996117 |
| TSSTAGNSGGA | TCSALGISGNTLY | 0.79173064 |
| VPATVPVGILC | TCSALGISGNTLY | 0.8727806 |
| VQRPDSGGVVLTGRIS | TCSALGISGNTLY | 0.8504261 |
| WTPNPVPATKGVPAGLVQAT | TCSALGISGNTLY | 0.8447359 |
| YALKRALAKFSDRFDVVII | TCSALGISGNTLY | 0.71859425 |
| YLIGSVIV | TCSALGISGNTLY | 0.86763954 |
| YGQLVAVID | TCSALGISGNTLY | 0.78579855 |
| VTVGGMTLPA | TCSALGISGNTLY | 0.8034773 |
| VLALPAV | TCSALGISGNTLY | 0.8688279 |
| VIPPTLNYET | TCSALGISGNTLY | 0.8339709 |
| VGMGGMHPSSG | TCSALGISGNTLY | 0.81885874 |
| VGLAGGAATA | TCSALGISGNTLY | 0.8149046 |
| VGGAGGAGGSGP | TCSALGISGNTLY | 0.8488732 |
| VFAAFMV | TCSALGISGNTLY | 0.8637816 |
| TLYDKLR | TCSALGISGNTLY | 0.8410573 |
| TLAVFTVL | TCSALGISGNTLY | 0.9027633 |
| TGSVQPARV | TCSALGISGNTLY | 0.86029905 |
| TGGGAGSGGT | TCSALGISGNTLY | 0.88549924 |
| SHLETSL | TCSALGISGNTLY | 0.38190365 |
| SFTTGMT | TCSALGISGNTLY | 0.6744721 |
| QVEVAKRMGVRQPTVS | TCSALGISGNTLY | 0.84771717 |
| QGSGGPALAM | TCSALGISGNTLY | 0.8052671 |
| MLAPSNP | TCSALGISGNTLY | 0.55096745 |
| LVNNAAI | TCSALGISGNTLY | 0.8492981 |
| LTGEINTLA | TCSALGISGNTLY | 0.8119275 |
| LSIFGPV | TCSALGISGNTLY | 0.7312244 |
| LLALLANL | TCSALGISGNTLY | 0.87241495 |
| LISNASDALDKLR | TCSALGISGNTLY | 0.78893137 |
| LINEAVP | TCSALGISGNTLY | 0.8162115 |
| LGYAFSGL | TCSALGISGNTLY | 0.8061305 |
| LDFVAVAL | TCSALGISGNTLY | 0.88822424 |
| LAQPGAE | TCSALGISGNTLY | 0.7326386 |
| ISNLGAVTGF | TCSALGISGNTLY | 0.85583246 |
| HQEDPGRI | TCSALGISGNTLY | 0.8236504 |
| GTPGEPTP | TCSALGISGNTLY | 0.8816247 |
| GFFNSGS | TCSALGISGNTLY | 0.6380888 |
| GAGGGGLTAGAN | TCSALGISGNTLY | 0.86009604 |
| FVDALVNM | TCSALGISGNTLY | 0.78960073 |
| FLTVAPEL | TCSALGISGNTLY | 0.90257275 |
| FETLTAL | TCSALGISGNTLY | 0.80023646 |
| DVINPIEL | TCSALGISGNTLY | 0.8372762 |
| APVSGAPV | TCSALGISGNTLY | 0.84334266 |
| ALVGGLAW | TCSALGISGNTLY | 0.8458404 |
| ALAAAAATAI | TCSALGISGNTLY | 0.5928575 |
| AFGLGLLL | TCSALGISGNTLY | 0.8024607 |
| AFFAAASL | TCSALGISGNTLY | 0.7654747 |
| AAAALPAFSPP | TCSALGISGNTLY | 0.8470913 |
| VVLAPAL | TCSALGISGNTLY | 0.8431599 |
| TIGHVDHGKTTL | TCSALGISGNTLY | 0.4788466 |

| | | |
|---|---|---|
| LISNASDALDK | TCSALGISGNTLY | 0.87406313 |
| LISNASDAL | TCSALGISGNTLY | 0.7494582 |
| ISNASDALDK | TCSALGISGNTLY | 0.74660003 |
| NQAETLVYQTE | TCSALGISGNTLY | 0.8889121 |
| TGGTGGTAGAG | TCSALGISGNTLY | 0.84819347 |
| TGSTAGGAGGA | TCSALGISGNTLY | 0.8780869 |
| DPLAVPVAAGPV | TCSALGISGNTLY | 0.83238333 |
| GGTGGTPGNGGN | TCSALGISGNTLY | 0.8002378 |
| PAGLAGVDPATL | TCSALGISGNTLY | 0.86725855 |
| GNNLVVEAPHAGASV | TCSALGISGNTLY | 0.625387 |
| KVTRIMSASTLEQPA | TCSALGISGNTLY | 0.76099706 |
| LGFALPIALPRRTVV | TCSALGISGNTLY | 0.88320094 |
| TRLGNLTARKLVVLV | TCSALGISGNTLY | 0.88794136 |
| WSQQRRLIYFPSAGP | TCSALGISGNTLY | 0.9040675 |
| GPNGNPDPMAAAVDIR | TCSALGISGNTLY | 0.7951285 |
| TQRCVSVADLRAAASR | TCSALGISGNTLY | 0.7957024 |
| QFVLDVAEHPVEVLTETV | TCSALGISGNTLY | 0.8603012 |
| GGAGGTGGAGGNGGTQAGDG | TCSALGISGNTLY | 0.90310293 |
| PLLATLRAIAVDVDAKPAQV | TCSALGISGNTLY | 0.91451794 |
| APKFRTGGYMIDFWGVGYQVAKRM | TCSALGISGNTLY | 0.8978603 |
| GAGGAGGAGGAAGGPGGTGGTGGN | TCSALGISGNTLY | 0.8231739 |
| IGALGLLATDVAPAGSVAFGVALA | TCSALGISGNTLY | 0.89107627 |
| AGGAGGIGGNGGAGGAGGVLMGNG | TCSALGISGNTLY | 0.86457956 |
| ADELIGLL | TCSALGISGNTLY | 0.86506927 |
| ADRALKA | TCSALGISGNTLY | 0.9008415 |
| AGAVGQVGGAGGA | TCSALGISGNTLY | 0.8731791 |
| AGGGGGLGGAP | TCSALGISGNTLY | 0.8353637 |
| AGGLLYGNGGAG | TCSALGISGNTLY | 0.8111174 |
| AGLESIGL | TCSALGISGNTLY | 0.891436 |
| AGLKTGII | TCSALGISGNTLY | 0.90100056 |
| ASLAALLL | TCSALGISGNTLY | 0.8967466 |
| ASVINISSI | TCSALGISGNTLY | 0.9165994 |
| ATPPGPA | TCSALGISGNTLY | 0.89857256 |
| ATVASALV | TCSALGISGNTLY | 0.893388 |
| ATVASLIL | TCSALGISGNTLY | 0.36865452 |
| AVNGPVLIRH | TCSALGISGNTLY | 0.74473023 |
| DGAAGAVGI | TCSALGISGNTLY | 0.8319268 |
| DPLLHGI | TCSALGISGNTLY | 0.8663162 |
| ELGAALGAL | TCSALGISGNTLY | 0.58571965 |
| ELGRTQAV | TCSALGISGNTLY | 0.8963752 |
| FVAGGYLS | TCSALGISGNTLY | 0.8562724 |
| GASGGAGGAGGW | TCSALGISGNTLY | 0.8107064 |
| GGAGAGGDGGAGGI | TCSALGISGNTLY | 0.90146214 |
| GTGHNGSQPGMGGNG | TCSALGISGNTLY | 0.8339 |
| GYAAASASAS | TCSALGISGNTLY | 0.7856175 |
| HQVDVAAVQQG | TCSALGISGNTLY | 0.85918033 |
| IGAAIAHT | TCSALGISGNTLY | 0.9129161 |
| IGMIEVAQL | TCSALGISGNTLY | 0.8171679 |
| IRLAPEHTSAMD | TCSALGISGNTLY | 0.90388346 |
| IYNVSVL | TCSALGISGNTLY | 0.8837478 |
| KQGFLGM | TCSALGISGNTLY | 0.90808266 |

| | | |
|---|---|---|
| LESVITQL | TCSALGISGNTLY | 0.70248413 |
| LGLGALVVV | TCSALGISGNTLY | 0.9002384 |
| LIGNGGNGGSGGI | TCSALGISGNTLY | 0.8640212 |
| LNAINAPTL | TCSALGISGNTLY | 0.9183649 |
| LQLLNTGV | TCSALGISGNTLY | 0.8619567 |
| LQTSIEF | TCSALGISGNTLY | 0.8974955 |
| MSFAVSEL | TCSALGISGNTLY | 0.88233876 |
| NTGALGL | TCSALGISGNTLY | 0.90523154 |
| PPGAGGAGGAGTT | TCSALGISGNTLY | 0.7137959 |
| QLVGLGI | TCSALGISGNTLY | 0.8513645 |
| RFLEQVHQL | TCSALGISGNTLY | 0.84110564 |
| RGVGYVLR | TCSALGISGNTLY | 0.87218654 |
| SATGVFAGVI | TCSALGISGNTLY | 0.8922887 |
| SLPAITLP | TCSALGISGNTLY | 0.3819977 |
| STALAFTV | TCSALGISGNTLY | 0.8992049 |
| SVLTVGLGK | TCSALGISGNTLY | 0.80259633 |
| TALAGAGSGI | TCSALGISGNTLY | 0.80172896 |
| TGLPFTGI | TCSALGISGNTLY | 0.90479434 |
| VAEHVTV | TCSALGISGNTLY | 0.86754024 |
| VDIVLNSL | TCSALGISGNTLY | 0.8933565 |
| VLTGGGAVL | TCSALGISGNTLY | 0.8703286 |
| VNVGLAYGI | TCSALGISGNTLY | 0.8723695 |
| VSSAAVGAV | TCSALGISGNTLY | 0.890916 |
| VTSVYEAGV | TCSALGISGNTLY | 0.61861706 |
| WVALDRGA | TCSALGISGNTLY | 0.821612 |
| YLVGSVVI | TCSALGISGNTLY | 0.90850544 |
| GPPGTGKTLL | TCSALGISGNTLY | 0.71306634 |
| GVGMTKF | TCSALGISGNTLY | 0.80200016 |
| IGIGGSDL | TCSALGISGNTLY | 0.26760522 |
| IGTLLAEL | TCSALGISGNTLY | 0.24173059 |
| IYTSGTT | TCSALGISGNTLY | 0.5272599 |
| LSFPAGV | TCSALGISGNTLY | 0.7245579 |
| LYTSGTT | TCSALGISGNTLY | 0.78445715 |
| VGAFPFGF | TCSALGISGNTLY | 0.7318877 |
| VGAGTVEF | TCSALGISGNTLY | 0.22019969 |
| VGKSTLLL | TCSALGISGNTLY | 0.5291601 |
| AALAHAGGGIV | TCSALGISGNTLY | 0.308136 |
| AALVAVSLL | TCSALGISGNTLY | 0.50466335 |
| AAVSRAVAAL | TCSALGISGNTLY | 0.5923339 |
| AGGSSGIGGF | TCSALGISGNTLY | 0.33754975 |
| AGLTVANANG | TCSALGISGNTLY | 0.306256 |
| AGLVGAVL | TCSALGISGNTLY | 0.24700849 |
| ALSQAFTV | TCSALGISGNTLY | 0.75431865 |
| ALTFSVSL | TCSALGISGNTLY | 0.78144175 |
| ALTGAALVAL | TCSALGISGNTLY | 0.753131 |
| ASFSIGAGVL | TCSALGISGNTLY | 0.80063343 |
| DGSVYKI | TCSALGISGNTLY | 0.7383926 |
| EQLAEIV | TCSALGISGNTLY | 0.7155815 |
| FAVSELTVM | TCSALGISGNTLY | 0.12308827 |
| GAAPLAAGL | TCSALGISGNTLY | 0.16184108 |
| GADGGDGGSGGAGG | TCSALGISGNTLY | 0.21603149 |

| | | |
|---|---|---|
| GFGLDTGGAGG | TCSALGISGNTLY | 0.48639333 |
| GLASGGLL | TCSALGISGNTLY | 0.1526405 |
| GRIAGAAVLL | TCSALGISGNTLY | 0.60046875 |
| GSAGSLNTSF | TCSALGISGNTLY | 0.3066104 |
| GTGQIGI | TCSALGISGNTLY | 0.22319527 |
| GTLIALIL | TCSALGISGNTLY | 0.7761216 |
| IAATMGFVVG | TCSALGISGNTLY | 0.27684003 |
| IFAQALL | TCSALGISGNTLY | 0.19708039 |
| IGAAVALGF | TCSALGISGNTLY | 0.25710085 |
| IGLTPLAL | TCSALGISGNTLY | 0.768471 |
| ILGSNTSTL | TCSALGISGNTLY | 0.17545608 |
| IVFSLTVL | TCSALGISGNTLY | 0.7084673 |
| LASKATGF | TCSALGISGNTLY | 0.48093098 |
| LASVALIL | TCSALGISGNTLY | 0.6739395 |
| LFAVGLTL | TCSALGISGNTLY | 0.16432202 |
| LFNGGIGTYI | TCSALGISGNTLY | 0.73649514 |
| LTLGAGII | TCSALGISGNTLY | 0.5651394 |
| LVAAALGDI | TCSALGISGNTLY | 0.83807135 |
| NTAFVQLGI | TCSALGISGNTLY | 0.3220495 |
| QLFSGAEL | TCSALGISGNTLY | 0.44006965 |
| SAATFITL | TCSALGISGNTLY | 0.3302714 |
| SNYGLANLGV | TCSALGISGNTLY | 0.74344885 |
| SYGGAYAVM | TCSALGISGNTLY | 0.15965937 |
| TTGVGGLTLG | TCSALGISGNTLY | 0.36754665 |
| TTQTIYI | TCSALGISGNTLY | 0.19423433 |
| VAAGAGLGAVI | TCSALGISGNTLY | 0.39625508 |
| VAAVVTLL | TCSALGISGNTLY | 0.720459 |
| VGFLGGLI | TCSALGISGNTLY | 0.13339864 |
| VGVIPILL | TCSALGISGNTLY | 0.66570556 |
| VIFASSLI | TCSALGISGNTLY | 0.5472287 |
| VKGGDGGAAAG | TCSALGISGNTLY | 0.21262442 |
| VTKTFTNI | TCSALGISGNTLY | 0.7195045 |
| VVAAAILL | TCSALGISGNTLY | 0.34403366 |
| YAALDVEL | TCSALGISGNTLY | 0.78225374 |
| LASVALLL | TCSALGISGNTLY | 0.41742045 |
| LDAVHRV | TCSALGISGNTLY | 0.5488627 |
| QGLSSSIF | TCSALGISGNTLY | 0.54753125 |
| RGTFANIRL | TCSALGISGNTLY | 0.12686251 |
| RQLWWGHRI | TCSALGISGNTLY | 0.21416438 |
| YGPPGTGKTLL | TCSALGISGNTLY | 0.78103876 |
| AALVGAPANVA | TCSALGISGNTLY | 0.70953065 |
| AATFRRTE | TCSALGISGNTLY | 0.8163595 |
| AGALGGGI | TCSALGISGNTLY | 0.5398284 |
| AGGAGGAGGIGAG | TCSALGISGNTLY | 0.45262164 |
| AGSGALGL | TCSALGISGNTLY | 0.60017174 |
| AQIADQI | TCSALGISGNTLY | 0.76828325 |
| ASAVAVTYL | TCSALGISGNTLY | 0.8146244 |
| ASTVLGGTGG | TCSALGISGNTLY | 0.73755795 |
| AVGGEPAK | TCSALGISGNTLY | 0.48902723 |
| CEQGTFGGTC | TCSALGISGNTLY | 0.65565896 |
| DEGKAPAG | TCSALGISGNTLY | 0.5968723 |

| | | |
|---|---|---|
| DTVVFPAL | TCSALGISGNTLY | 0.58763146 |
| FEGITMAG | TCSALGISGNTLY | 0.6864337 |
| FPLVVQD | TCSALGISGNTLY | 0.71298856 |
| GAFTTPQ | TCSALGISGNTLY | 0.5419451 |
| GFGGFGTAGGVGGA | TCSALGISGNTLY | 0.5362865 |
| GFLAQFL | TCSALGISGNTLY | 0.6789552 |
| GGAGGAGGNAGVA | TCSALGISGNTLY | 0.76622653 |
| GGVGTAGAGGAGGAGG | TCSALGISGNTLY | 0.7315036 |
| GNGGGGDAGNAGSGGN | TCSALGISGNTLY | 0.82899153 |
| GTGGAGGAGNDAGS | TCSALGISGNTLY | 0.77364135 |
| GTTNTGWFNS | TCSALGISGNTLY | 0.7558356 |
| IFMGPLFTL | TCSALGISGNTLY | 0.20173942 |
| ILAHAIEAI | TCSALGISGNTLY | 0.33948785 |
| IRSAIREAL | TCSALGISGNTLY | 0.5959834 |
| ITFAGGG | TCSALGISGNTLY | 0.62396944 |
| ITGMSGHQEV | TCSALGISGNTLY | 0.32230723 |
| IVAGLACTL | TCSALGISGNTLY | 0.72933435 |
| IVGGGLAGA | TCSALGISGNTLY | 0.6065748 |
| IVTGVLNI | TCSALGISGNTLY | 0.48378283 |
| LAQQGCI | TCSALGISGNTLY | 0.7685585 |
| LCLHVEN | TCSALGISGNTLY | 0.5281735 |
| LDAISDGS | TCSALGISGNTLY | 0.5100039 |
| LGGFAAML | TCSALGISGNTLY | 0.5651975 |
| LGGKNPMI | TCSALGISGNTLY | 0.7862369 |
| LGPKGVAAIL | TCSALGISGNTLY | 0.39555672 |
| LMVGVSI | TCSALGISGNTLY | 0.7352345 |
| LVGAAFVL | TCSALGISGNTLY | 0.6020781 |
| LVLAVAPL | TCSALGISGNTLY | 0.7951856 |
| MAAIAVIDAA | TCSALGISGNTLY | 0.3950452 |
| NNGINAELQ | TCSALGISGNTLY | 0.7605905 |
| RGEPLVLVHG | TCSALGISGNTLY | 0.60071826 |
| RGGAGGAAGLI | TCSALGISGNTLY | 0.857135 |
| RSYRMHPAV | TCSALGISGNTLY | 0.59603107 |
| SAAHLLAGQLA | TCSALGISGNTLY | 0.7010923 |
| SAAVGSY | TCSALGISGNTLY | 0.6177857 |
| SPHAVLHRI | TCSALGISGNTLY | 0.7731418 |
| SQIAKVL | TCSALGISGNTLY | 0.3582633 |
| TALGHLVP | TCSALGISGNTLY | 0.5844609 |
| TAPTTSGAA | TCSALGISGNTLY | 0.35963216 |
| TDGDDGNGGSPGT | TCSALGISGNTLY | 0.64392126 |
| TDGLGEN | TCSALGISGNTLY | 0.7384778 |
| TGGPPLPGQA | TCSALGISGNTLY | 0.31762195 |
| TIPAITI | TCSALGISGNTLY | 0.7300074 |
| TIPGFTI | TCSALGISGNTLY | 0.613108 |
| TSIASGALA | TCSALGISGNTLY | 0.5008322 |
| TVTLPGAAI | TCSALGISGNTLY | 0.789433 |
| VASGLSAALP | TCSALGISGNTLY | 0.63799894 |
| VGAGVIGI | TCSALGISGNTLY | 0.7864896 |
| VGPKRFGV | TCSALGISGNTLY | 0.6034565 |
| VNVAFPD | TCSALGISGNTLY | 0.60658956 |
| VSMTNSGVS | TCSALGISGNTLY | 0.7175772 |

| | | |
|---|---|---|
| VTDVLGDLSV | TCSALGISGNTLY | 0.53252286 |
| VVVVGAGFAGL | TCSALGISGNTLY | 0.5189885 |
| VYTVGTVV | TCSALGISGNTLY | 0.7942712 |
| WTTFSGP | TCSALGISGNTLY | 0.7544784 |
| YDELVEAF | TCSALGISGNTLY | 0.7627583 |
| YGTGGAGGNGG | TCSALGISGNTLY | 0.6538918 |
| YTNVMAV | TCSALGISGNTLY | 0.6521198 |
| FLEQVHQL | TCSALGISGNTLY | 0.68374074 |
| GLSLLAVAV | TCSALGISGNTLY | 0.764602 |
| LEQVHQL | TCSALGISGNTLY | 0.7741111 |
| LNAAGAI | TCSALGISGNTLY | 0.7559281 |
| PAVHASL | TCSALGISGNTLY | 0.68692416 |
| TELEDDA | TCSALGISGNTLY | 0.7476851 |
| TPPGPPN | TCSALGISGNTLY | 0.7067569 |
| TPPPPPPP | TCSALGISGNTLY | 0.69987124 |
| TQSPALI | ASSLEHPNTEVF | 0.7546431 |
| EFAAAFPQ | ASSLEHPNTEVF | 0.73822343 |
| LLSGNLDVL | ASSLEHPNTEVF | 0.6695824 |
| GPANLGNYNI | ASSLEHPNTEVF | 0.72953403 |
| TLGPTLP | ASSLEHPNTEVF | 0.7273512 |
| VYAYSSCCK | ASSLEHPNTEVF | 0.75659823 |
| APAPAPAPNI | ASSLEHPNTEVF | 0.7573931 |
| PGGPVPIP | ASSLEHPNTEVF | 0.78112864 |
| FLTVALEP | ASSLEHPNTEVF | 0.7652966 |
| SQYFGANL | ASSLEHPNTEVF | 0.75345075 |
| GGVKPAA | ASSLEHPNTEVF | 0.4067863 |
| GGTGGAGGAGGW | ASSLEHPNTEVF | 0.59851205 |
| GVGGVAGLGGAGG | ASSLEHPNTEVF | 0.72023225 |
| VLLVLATTI | ASSLEHPNTEVF | 0.7214023 |
| IGIFAAML | ASSLEHPNTEVF | 0.51361966 |
| AVGDLID | ASSLEHPNTEVF | 0.7475867 |
| NISPLPL | ASSLEHPNTEVF | 0.72457933 |
| VTAVFVAI | ASSLEHPNTEVF | 0.66317064 |
| SGDDNDAAL | ASSLEHPNTEVF | 0.7622093 |
| AVAALAHS | ASSLEHPNTEVF | 0.678647 |
| GGVTVMATP | ASSLEHPNTEVF | 0.6974895 |
| AGLAANVRD | ASSLEHPNTEVF | 0.7101941 |
| TNRPDLIDPALLRPG | ASSLEHPNTEVF | 0.7702579 |
| LANLGVTAL | ASSLEHPNTEVF | 0.6687498 |
| TGERIEL | ASSLEHPNTEVF | 0.76072097 |
| NGEELAA | ASSLEHPNTEVF | 0.6614878 |
| VLSLVWG | ASSLEHPNTEVF | 0.7709408 |
| IGYALWL | ASSLEHPNTEVF | 0.64882267 |
| LSLLLAPL | ASSLEHPNTEVF | 0.75522816 |
| TLDEALK | ASSLEHPNTEVF | 0.71169734 |
| TDGAGTAEGG | ASSLEHPNTEVF | 0.79219973 |
| NAPGGAGGNGG | ASSLEHPNTEVF | 0.7248858 |
| VSVFMGMGGNF | ASSLEHPNTEVF | 0.75802565 |
| LASYLAAL | ASSLEHPNTEVF | 0.7224198 |
| LVGGTPAL | ASSLEHPNTEVF | 0.76795137 |
| VRNADVL | ASSLEHPNTEVF | 0.5726261 |

| | | |
|---|---|---|
| TCSSPFLVSAM | ASSLEHPNTEVF | 0.728371 |
| VHITDAE | ASSLEHPNTEVF | 0.57747734 |
| EYAAPSF | ASSLEHPNTEVF | 0.74439585 |
| VVDAFHGGL | ASSLEHPNTEVF | 0.75367093 |
| ASVVDPQ | ASSLEHPNTEVF | 0.5319505 |
| SYNEFGGLS | ASSLEHPNTEVF | 0.7767266 |
| DPDGSVY | ASSLEHPNTEVF | 0.654986 |
| MVSTIHATE | ASSLEHPNTEVF | 0.7067758 |
| LAAGVADPV | ASSLEHPNTEVF | 0.77252495 |
| TKEQIAA | ASSLEHPNTEVF | 0.71001256 |
| SLASPAAL | ASSLEHPNTEVF | 0.77743435 |
| IGLVFYGV | ASSLEHPNTEVF | 0.7077459 |
| FLLDASNL | ASSLEHPNTEVF | 0.7254018 |
| YQVGNFP | ASSLEHPNTEVF | 0.75273496 |
| ESLLSQST | ASSLEHPNTEVF | 0.60308313 |
| LFTVGFAAVG | ASSLEHPNTEVF | 0.6714279 |
| ATPPTPT | ASSLEHPNTEVF | 0.77548075 |
| TDLATLG | ASSLEHPNTEVF | 0.6501505 |
| SAGGTGR | ASSLEHPNTEVF | 0.6487778 |
| GFSNSGS | ASSLEHPNTEVF | 0.5908096 |
| GGNGGSGGTSV | ASSLEHPNTEVF | 0.5436729 |
| GTSVATGGAGN | ASSLEHPNTEVF | 0.5804347 |
| NGGNGGSGGTS | ASSLEHPNTEVF | 0.6450441 |
| AVVKFPH | ASSLEHPNTEVF | 0.6588094 |
| TPPPPPP | ASSLEHPNTEVF | 0.643333 |
| AAPAAAAAPA | ASSLEHPNTEVF | 0.5305302 |
| IALNEDL | ASSLEHPNTEVF | 0.6460884 |
| IVNVSSM | ASSLEHPNTEVF | 0.62947917 |
| TGIPPAPR | ASSLEHPNTEVF | 0.6225965 |
| FDLGGGTF | ASSLEHPNTEVF | 0.6250595 |
| FDLGGGTFDVS | ASSLEHPNTEVF | 0.6529967 |
| NEPTAAA | ASSLEHPNTEVF | 0.6165066 |
| NEPTAAALA | ASSLEHPNTEVF | 0.50646454 |
| GSIINISSI | ASSLEHPNTEVF | 0.63687867 |
| VAPGNAGTA | ASSLEHPNTEVF | 0.65774554 |
| TGGGASLE | ASSLEHPNTEVF | 0.651071 |
| VAVYNHYKRI | ASSLEHPNTEVF | 0.6596842 |
| TDPAPATT | ASSLEHPNTEVF | 0.6569009 |
| TNRPDILDPALLRPG | ASSLEHPNTEVF | 0.6344882 |
| LTTTTTS | ASSLEHPNTEVF | 0.3659472 |
| PTGSGKTL | ASSLEHPNTEVF | 0.49204192 |
| TGSGKTL | ASSLEHPNTEVF | 0.55693734 |
| TIGHVDHGK | ASSLEHPNTEVF | 0.6076618 |
| TIGHVDHGKTT | ASSLEHPNTEVF | 0.36219287 |
| GHVDHGKTT | ASSLEHPNTEVF | 0.6332324 |
| AQFEHTLL | ASSLEHPNTEVF | 0.5994806 |
| IVNQDEL | ASSLEHPNTEVF | 0.6045153 |
| TNGSQFF | ASSLEHPNTEVF | 0.65435135 |
| TGKGIVE | ASSLEHPNTEVF | 0.6132136 |
| LISNASDA | ASSLEHPNTEVF | 0.55068934 |
| AELRTLGELTKIAKAHGAQ | ASSLEHPNTEVF | 0.5672715 |

| | | |
|---|---|---|
| APNAVAAAEPSW | ASSLEHPNTEVF | 0.6615084 |
| APQVAGAATAVL | ASSLEHPNTEVF | 0.5122048 |
| ARRDPAAAAAILPTDA | ASSLEHPNTEVF | 0.6564586 |
| DAAARDVLAEAGLAEYF | ASSLEHPNTEVF | 0.5702296 |
| DDQIVLSAKANLAKVEQ | ASSLEHPNTEVF | 0.6609505 |
| DELLPDTGEDSDEDGH | ASSLEHPNTEVF | 0.5090166 |
| ELPEAPANRATVPQD | ASSLEHPNTEVF | 0.65500236 |
| FGQSKIERVTGIG | ASSLEHPNTEVF | 0.6279779 |
| GAGGTGSTAGGAGGAGGAGGLYA | ASSLEHPNTEVF | 0.66882986 |
| GHGGTGAAGGNGGTGGAGGSGIDG | ASSLEHPNTEVF | 0.6054972 |
| GNGGNGGAGGEGGPGGAGGAGA | ASSLEHPNTEVF | 0.65098774 |
| GSTAGNGGNGGNATNGTIAGGQPAG | ASSLEHPNTEVF | 0.62080884 |
| GTGGDGGNGGNGGTGG | ASSLEHPNTEVF | 0.64876753 |
| GVRRPAGIPADAPV | ASSLEHPNTEVF | 0.5380174 |
| GYRQVTDAHLLTLARR | ASSLEHPNTEVF | 0.6052338 |
| IAKMGQDGHDRG | ASSLEHPNTEVF | 0.51324046 |
| KAEPSIEIQYT | ASSLEHPNTEVF | 0.6162042 |
| KPVRIRMAADEDTVAVT | ASSLEHPNTEVF | 0.6507776 |
| MPPRIAGMRLLVIKPEP | ASSLEHPNTEVF | 0.341928 |
| MKFYAVALFRSRGVPPTR | ASSLEHPNTEVF | 0.64875317 |
| NAGGFGILGAVAHSP | ASSLEHPNTEVF | 0.51124656 |
| NEGREITLAMAEM | ASSLEHPNTEVF | 0.5392126 |
| NELVEWSHPLGHRWRWE | ASSLEHPNTEVF | 0.6466259 |
| NLGAGGTGGNGGNPG | ASSLEHPNTEVF | 0.64451504 |
| NTEWPKWETVMAADPVD | ASSLEHPNTEVF | 0.6480605 |
| PASAAPAETQP | ASSLEHPNTEVF | 0.59329295 |
| QQSSDGKNGVRAAEITGE | ASSLEHPNTEVF | 0.6150607 |
| RAPDAVTTSWWKE | ASSLEHPNTEVF | 0.6327965 |
| SGFGNVGAHTSGWFNQ | ASSLEHPNTEVF | 0.4930222 |
| SGVAGAVGQAASVGGLKV | ASSLEHPNTEVF | 0.53103286 |
| SNGGAGTGSAGGNGGT | ASSLEHPNTEVF | 0.66007674 |
| TGGAGGEGGAAG | ASSLEHPNTEVF | 0.61778176 |
| TGGDGGAAGAGGAPGHG | ASSLEHPNTEVF | 0.7477107 |
| TMPPLLAAIANAGRHGV | ASSLEHPNTEVF | 0.4758304 |
| TSSTAGNSGGA | ASSLEHPNTEVF | 0.34969223 |
| VPATVPVGILC | ASSLEHPNTEVF | 0.4917567 |
| VQRPDSGGVVLTGRIS | ASSLEHPNTEVF | 0.7197052 |
| WTPNPVPATKGVPAGLVQAT | ASSLEHPNTEVF | 0.7517665 |
| YALKRALAKFSDRFDVVII | ASSLEHPNTEVF | 0.6697637 |
| YLIGSVIV | ASSLEHPNTEVF | 0.3779037 |
| YGQLVAVID | ASSLEHPNTEVF | 0.54157597 |
| VTVGGMTLPA | ASSLEHPNTEVF | 0.4656504 |
| VLALPAV | ASSLEHPNTEVF | 0.50786686 |
| VIPPTLNYET | ASSLEHPNTEVF | 0.591089 |
| VGMGGMHPSSG | ASSLEHPNTEVF | 0.592631 |
| VGLAGGAATA | ASSLEHPNTEVF | 0.51012266 |
| VGGAGGAGGSGP | ASSLEHPNTEVF | 0.407588 |
| VFAAFMV | ASSLEHPNTEVF | 0.54514647 |
| TLYDKLR | ASSLEHPNTEVF | 0.6592275 |
| TLAVFTVL | ASSLEHPNTEVF | 0.55909324 |
| TGSVQPARV | ASSLEHPNTEVF | 0.7547602 |

| | | |
|---|---|---|
| TGGGAGSGGT | ASSLEHPNTEVF | 0.6968855 |
| SHLETSL | ASSLEHPNTEVF | 0.63443017 |
| SFTTGMT | ASSLEHPNTEVF | 0.1951896 |
| QVEVAKRMGVRQPTVS | ASSLEHPNTEVF | 0.25425565 |
| QGSGGPALAM | ASSLEHPNTEVF | 0.47767514 |
| MLAPSNP | ASSLEHPNTEVF | 0.5006506 |
| LVNNAAI | ASSLEHPNTEVF | 0.2109731 |
| LTGEINTLA | ASSLEHPNTEVF | 0.64153564 |
| LSIFGPV | ASSLEHPNTEVF | 0.4676407 |
| LLALLANL | ASSLEHPNTEVF | 0.40423527 |
| LISNASDALDKLR | ASSLEHPNTEVF | 0.70004535 |
| LINEAVP | ASSLEHPNTEVF | 0.438967 |
| LGYAFSGL | ASSLEHPNTEVF | 0.41651422 |
| LDFVAVAL | ASSLEHPNTEVF | 0.44230926 |
| LAQPGAE | ASSLEHPNTEVF | 0.7025311 |
| ISNLGAVTGF | ASSLEHPNTEVF | 0.27863052 |
| HQEDPGRI | ASSLEHPNTEVF | 0.6439231 |
| GTPGEPTP | ASSLEHPNTEVF | 0.4978604 |
| GFFNSGS | ASSLEHPNTEVF | 0.6898191 |
| GAGGGGLTAGAN | ASSLEHPNTEVF | 0.299971 |
| FVDALVNM | ASSLEHPNTEVF | 0.696027 |
| FLTVAPEL | ASSLEHPNTEVF | 0.55485183 |
| FETLTAL | ASSLEHPNTEVF | 0.7701951 |
| DVINPIEL | ASSLEHPNTEVF | 0.4356236 |
| APVSGAPV | ASSLEHPNTEVF | 0.50491107 |
| ALVGGLAW | ASSLEHPNTEVF | 0.45843506 |
| ALAAAAATAI | ASSLEHPNTEVF | 0.73249614 |
| AFGLGLLL | ASSLEHPNTEVF | 0.268851 |
| AFFAAASL | ASSLEHPNTEVF | 0.4964801 |
| AAAALPAFSPP | ASSLEHPNTEVF | 0.27964053 |
| VVLAPAL | ASSLEHPNTEVF | 0.5964848 |
| TIGHVDHGKTTL | ASSLEHPNTEVF | 0.65461487 |
| LISNASDALDK | ASSLEHPNTEVF | 0.23114724 |
| LISNASDAL | ASSLEHPNTEVF | 0.6707559 |
| ISNASDALDK | ASSLEHPNTEVF | 0.469397 |
| NQAETLVYQTE | ASSLEHPNTEVF | 0.3841862 |
| TGGTGGTAGAG | ASSLEHPNTEVF | 0.70351464 |
| TGSTAGGAGGA | ASSLEHPNTEVF | 0.48693436 |
| DPLAVPVAAGPV | ASSLEHPNTEVF | 0.7276345 |
| GGTGGTPGNGGN | ASSLEHPNTEVF | 0.5029408 |
| PAGLAGVDPATL | ASSLEHPNTEVF | 0.56980914 |
| GNNLVVEAPHAGASV | ASSLEHPNTEVF | 0.61370814 |
| KVTRIMSASTLEQPA | ASSLEHPNTEVF | 0.310778 |
| LGFALPIALPRRTVV | ASSLEHPNTEVF | 0.43845314 |
| TRLGNLTARKLVVLV | ASSLEHPNTEVF | 0.71028787 |
| WSQQRRLIYFPSAGP | ASSLEHPNTEVF | 0.8633489 |
| GPNGNPDPMAAAVDIR | ASSLEHPNTEVF | 0.88851756 |
| TQRCVSVADLRAAASR | ASSLEHPNTEVF | 0.7944646 |
| QFVLDVAEHPVEVLTETV | ASSLEHPNTEVF | 0.74332577 |
| GGAGGTGGAGGNGGTQAGDG | ASSLEHPNTEVF | 0.80522066 |
| PLLATLRAIAVDVDAKPAQV | ASSLEHPNTEVF | 0.8835522 |

| | | |
|---|---|---|
| APKFRTGGYMIDFWGVGYQVAKRM | ASSLEHPNTEVF | 0.87434304 |
| GAGGAGGAGGAAGGPGGTGGTGGN | ASSLEHPNTEVF | 0.8611075 |
| IGALGLLATDVAPAGSVAFGVALA | ASSLEHPNTEVF | 0.60450804 |
| AGGAGGIGGNGGAGGAGGVLMGNG | ASSLEHPNTEVF | 0.8103168 |
| ADELIGLL | ASSLEHPNTEVF | 0.77967864 |
| ADRALKA | ASSLEHPNTEVF | 0.7873806 |
| AGAVGQVGGAGGA | ASSLEHPNTEVF | 0.8615856 |
| AGGGGGLGGAP | ASSLEHPNTEVF | 0.83058214 |
| AGGLLYGNGGAG | ASSLEHPNTEVF | 0.769764 |
| AGLESIGL | ASSLEHPNTEVF | 0.7766438 |
| AGLKTGII | ASSLEHPNTEVF | 0.82475597 |
| ASLAALLL | ASSLEHPNTEVF | 0.8506037 |
| ASVINISSI | ASSLEHPNTEVF | 0.84454143 |
| ATPPGPA | ASSLEHPNTEVF | 0.9001321 |
| ATVASALV | ASSLEHPNTEVF | 0.8798444 |
| ATVASLIL | ASSLEHPNTEVF | 0.8824086 |
| AVNGPVLIRH | ASSLEHPNTEVF | 0.15482679 |
| DGAAGAVGI | ASSLEHPNTEVF | 0.60333514 |
| DPLLHGI | ASSLEHPNTEVF | 0.7641026 |
| ELGAALGAL | ASSLEHPNTEVF | 0.8317281 |
| ELGRTQAV | ASSLEHPNTEVF | 0.56125116 |
| FVAGGYLS | ASSLEHPNTEVF | 0.85129964 |
| GASGGAGGAGGW | ASSLEHPNTEVF | 0.79902905 |
| GGAGAGGDGGAGGI | ASSLEHPNTEVF | 0.70887876 |
| GTGHNGSQPGMGGNG | ASSLEHPNTEVF | 0.8769468 |
| GYAAASASAS | ASSLEHPNTEVF | 0.76012826 |
| HQVDVAAVQQG | ASSLEHPNTEVF | 0.6996776 |
| IGAAIAHT | ASSLEHPNTEVF | 0.68004847 |
| IGMIEVAQL | ASSLEHPNTEVF | 0.87394875 |
| IRLAPEHTSAMD | ASSLEHPNTEVF | 0.597816 |
| IYNVSVL | ASSLEHPNTEVF | 0.8664721 |
| KQGFLGM | ASSLEHPNTEVF | 0.7226876 |
| LESVITQL | ASSLEHPNTEVF | 0.88745373 |
| LGLGALVVV | ASSLEHPNTEVF | 0.55545616 |
| LIGNGGNGGSGGI | ASSLEHPNTEVF | 0.8756435 |
| LNAINAPTL | ASSLEHPNTEVF | 0.83252645 |
| LQLLNTGV | ASSLEHPNTEVF | 0.9002955 |
| LQTSIEF | ASSLEHPNTEVF | 0.7927234 |
| MSFAVSEL | ASSLEHPNTEVF | 0.80991244 |
| NTGALGL | ASSLEHPNTEVF | 0.6347537 |
| PPGAGGAGGAGTT | ASSLEHPNTEVF | 0.87968475 |
| QLVGLGI | ASSLEHPNTEVF | 0.61433387 |
| RFLEQVHQL | ASSLEHPNTEVF | 0.82997835 |
| RGVGYVLR | ASSLEHPNTEVF | 0.52253115 |
| SATGVFAGVI | ASSLEHPNTEVF | 0.8608328 |
| SLPAITLP | ASSLEHPNTEVF | 0.86519074 |
| STALAFTV | ASSLEHPNTEVF | 0.32713598 |
| SVLTVGLGK | ASSLEHPNTEVF | 0.8647747 |
| TALAGAGSGI | ASSLEHPNTEVF | 0.8145534 |
| TGLPFTGI | ASSLEHPNTEVF | 0.5957204 |
| VAEHVTV | ASSLEHPNTEVF | 0.8867749 |

| | | |
|---|---|---|
| VDIVLNSL | ASSLEHPNTEVF | 0.7028059 |
| VLTGGGAVL | ASSLEHPNTEVF | 0.89337516 |
| VNVGLAYGI | ASSLEHPNTEVF | 0.7967899 |
| VSSAAVGAV | ASSLEHPNTEVF | 0.7957515 |
| VTSVYEAGV | ASSLEHPNTEVF | 0.85136354 |
| WVALDRGA | ASSLEHPNTEVF | 0.6053947 |
| YLVGSVVI | ASSLEHPNTEVF | 0.64059913 |
| GPPGTGKTLL | ASSLEHPNTEVF | 0.8889667 |
| GVGMTKF | ASSLEHPNTEVF | 0.7190069 |
| IGIGGSDL | ASSLEHPNTEVF | 0.85912824 |
| IGTLLAEL | ASSLEHPNTEVF | 0.5534112 |
| IYTSGTT | ASSLEHPNTEVF | 0.5313479 |
| LSFPAGV | ASSLEHPNTEVF | 0.64601445 |
| LYTSGTT | ASSLEHPNTEVF | 0.7984232 |
| VGAFPFGF | ASSLEHPNTEVF | 0.86149216 |
| VGAGTVEF | ASSLEHPNTEVF | 0.79815495 |
| VGKSTLLL | ASSLEHPNTEVF | 0.42030287 |
| AALAHAGGGIV | ASSLEHPNTEVF | 0.6803256 |
| AALVAVSLL | ASSLEHPNTEVF | 0.5917839 |
| AAVSRAVAAL | ASSLEHPNTEVF | 0.6623401 |
| AGGSSGIGGF | ASSLEHPNTEVF | 0.77643144 |
| AGLTVANANG | ASSLEHPNTEVF | 0.6176204 |
| AGLVGAVL | ASSLEHPNTEVF | 0.58103687 |
| ALSQAFTV | ASSLEHPNTEVF | 0.46526685 |
| ALTFSVSL | ASSLEHPNTEVF | 0.7605833 |
| ALTGAALVAL | ASSLEHPNTEVF | 0.79622316 |
| ASFSIGAGVL | ASSLEHPNTEVF | 0.7686393 |
| DGSVYKI | ASSLEHPNTEVF | 0.83505416 |
| EQLAEIV | ASSLEHPNTEVF | 0.80527437 |
| FAVSELTVM | ASSLEHPNTEVF | 0.665298 |
| GAAPLAAGL | ASSLEHPNTEVF | 0.15617591 |
| GADGGDGGSGGAGG | ASSLEHPNTEVF | 0.361418 |
| GFGLDTGGAGG | ASSLEHPNTEVF | 0.6650317 |
| GLASGGLL | ASSLEHPNTEVF | 0.68208265 |
| GRIAGAAVLL | ASSLEHPNTEVF | 0.19105504 |
| GSAGSLNTSF | ASSLEHPNTEVF | 0.78183746 |
| GTGQIGI | ASSLEHPNTEVF | 0.57768035 |
| GTLIALIL | ASSLEHPNTEVF | 0.59328157 |
| IAATMGFVVG | ASSLEHPNTEVF | 0.82216996 |
| IFAQALL | ASSLEHPNTEVF | 0.64306945 |
| IGAAVALGF | ASSLEHPNTEVF | 0.4488621 |
| IGLTPLAL | ASSLEHPNTEVF | 0.4013995 |
| ILGSNTSTL | ASSLEHPNTEVF | 0.78506285 |
| IVFSLTVL | ASSLEHPNTEVF | 0.33530694 |
| LASKATGF | ASSLEHPNTEVF | 0.7727372 |
| LASVALIL | ASSLEHPNTEVF | 0.5598272 |
| LFAVGLTL | ASSLEHPNTEVF | 0.78085816 |
| LFNGGIGTYI | ASSLEHPNTEVF | 0.30509835 |
| LTLGAGII | ASSLEHPNTEVF | 0.7763925 |
| LVAAALGDI | ASSLEHPNTEVF | 0.7036991 |
| NTAFVQLGI | ASSLEHPNTEVF | 0.859452 |

| | | |
|---|---|---|
| QLFSGAEL | ASSLEHPNTEVF | 0.5992515 |
| SAATFITL | ASSLEHPNTEVF | 0.69372284 |
| SNYGLANLGV | ASSLEHPNTEVF | 0.5387001 |
| SYGGAYAVM | ASSLEHPNTEVF | 0.8539568 |
| TTGVGGLTLG | ASSLEHPNTEVF | 0.48356804 |
| TTQTIYI | ASSLEHPNTEVF | 0.49601737 |
| VAAGAGLGAVI | ASSLEHPNTEVF | 0.33013967 |
| VAAVVTLL | ASSLEHPNTEVF | 0.661738 |
| VGFLGGLI | ASSLEHPNTEVF | 0.77565813 |
| VGVIPILL | ASSLEHPNTEVF | 0.6219449 |
| VIFASSLI | ASSLEHPNTEVF | 0.7378197 |
| VKGGDGGAAAG | ASSLEHPNTEVF | 0.72490704 |
| VTKTFTNI | ASSLEHPNTEVF | 0.44550294 |
| VVAAAILL | ASSLEHPNTEVF | 0.8220508 |
| YAALDVEL | ASSLEHPNTEVF | 0.53945017 |
| LASVALLL | ASSLEHPNTEVF | 0.7949089 |
| LDAVHRV | ASSLEHPNTEVF | 0.5184027 |
| QGLSSSIF | ASSLEHPNTEVF | 0.6519936 |
| RGTFANIRL | ASSLEHPNTEVF | 0.7102572 |
| RQLWWGHRI | ASSLEHPNTEVF | 0.6752311 |
| YGPPGTGKTLL | ASSLEHPNTEVF | 0.45156917 |
| AALVGAPANVA | ASSLEHPNTEVF | 0.8189601 |
| AATFRRTE | ASSLEHPNTEVF | 0.8441154 |
| AGALGGGI | ASSLEHPNTEVF | 0.85242367 |
| AGGAGGAGGIGAG | ASSLEHPNTEVF | 0.7551836 |
| AGSGALGL | ASSLEHPNTEVF | 0.70404506 |
| AQIADQI | ASSLEHPNTEVF | 0.7908044 |
| ASAVAVTYL | ASSLEHPNTEVF | 0.84496087 |
| ASTVLGGTGG | ASSLEHPNTEVF | 0.85388625 |
| AVGGEPAK | ASSLEHPNTEVF | 0.8369232 |
| CEQGTFGGTC | ASSLEHPNTEVF | 0.64698744 |
| DEGKAPAG | ASSLEHPNTEVF | 0.81245947 |
| DTVVFPAL | ASSLEHPNTEVF | 0.79515326 |
| FEGITMAG | ASSLEHPNTEVF | 0.7869054 |
| FPLVVQD | ASSLEHPNTEVF | 0.81899774 |
| GAFTTPQ | ASSLEHPNTEVF | 0.7981158 |
| GFGGFGTAGGVGGA | ASSLEHPNTEVF | 0.7654202 |
| GFLAQFL | ASSLEHPNTEVF | 0.7532056 |
| GGAGGAGGNAGVA | ASSLEHPNTEVF | 0.81468296 |
| GGVGTAGAGGAGGAGG | ASSLEHPNTEVF | 0.8435006 |
| GNGGGGDAGNAGSGGN | ASSLEHPNTEVF | 0.8338404 |
| GTGGAGGAGNDAGS | ASSLEHPNTEVF | 0.86181426 |
| GTTNTGWFNS | ASSLEHPNTEVF | 0.8519009 |
| IFMGPLFTL | ASSLEHPNTEVF | 0.8369653 |
| ILAHAIEAI | ASSLEHPNTEVF | 0.259707 |
| IRSAIREAL | ASSLEHPNTEVF | 0.56986856 |
| ITFAGGG | ASSLEHPNTEVF | 0.72367 |
| ITGMSGHQEV | ASSLEHPNTEVF | 0.80879915 |
| IVAGLACTL | ASSLEHPNTEVF | 0.3609478 |
| IVGGGLAGA | ASSLEHPNTEVF | 0.8226466 |
| IVTGVLNI | ASSLEHPNTEVF | 0.7851611 |

| | | |
|---|---|---|
| LAQQGCI | ASSLEHPNTEVF | 0.73584735 |
| LCLHVEN | ASSLEHPNTEVF | 0.84808546 |
| LDAISDGS | ASSLEHPNTEVF | 0.77397406 |
| LGGFAAML | ASSLEHPNTEVF | 0.6765748 |
| LGGKNPMI | ASSLEHPNTEVF | 0.6308551 |
| LGPKGVAAIL | ASSLEHPNTEVF | 0.8476801 |
| LMVGVSI | ASSLEHPNTEVF | 0.5854148 |
| LVGAAFVL | ASSLEHPNTEVF | 0.8480641 |
| LVLAVAPL | ASSLEHPNTEVF | 0.69232345 |
| MAAIAVIDAA | ASSLEHPNTEVF | 0.85109365 |
| NNGINAELQ | ASSLEHPNTEVF | 0.54616296 |
| RGEPLVLVHG | ASSLEHPNTEVF | 0.84789705 |
| RGGAGGAAGLI | ASSLEHPNTEVF | 0.8185625 |
| RSYRMHPAV | ASSLEHPNTEVF | 0.8644775 |
| SAAHLLAGQLA | ASSLEHPNTEVF | 0.790367 |
| SAAVGSY | ASSLEHPNTEVF | 0.81985885 |
| SPHAVLHRI | ASSLEHPNTEVF | 0.7884527 |
| SQIAKVL | ASSLEHPNTEVF | 0.8503449 |
| TALGHLVP | ASSLEHPNTEVF | 0.70867085 |
| TAPTTSGAA | ASSLEHPNTEVF | 0.7822065 |
| TDGDDGNGGSPGT | ASSLEHPNTEVF | 0.41395172 |
| TDGLGEN | ASSLEHPNTEVF | 0.7998364 |
| TGGPPLPGQA | ASSLEHPNTEVF | 0.8419467 |
| TIPAITI | ASSLEHPNTEVF | 0.3921064 |
| TIPGFTI | ASSLEHPNTEVF | 0.8077023 |
| TSIASGALA | ASSLEHPNTEVF | 0.76858294 |
| TVTLPGAAI | ASSLEHPNTEVF | 0.6706296 |
| VASGLSAALP | ASSLEHPNTEVF | 0.84709775 |
| VGAGVIGI | ASSLEHPNTEVF | 0.7763233 |
| VGPKRFGV | ASSLEHPNTEVF | 0.8577679 |
| VNVAFPD | ASSLEHPNTEVF | 0.72385246 |
| VSMTNSGVS | ASSLEHPNTEVF | 0.76940495 |
| VTDVLGDLSV | ASSLEHPNTEVF | 0.8090148 |
| VVVVGAGFAGL | ASSLEHPNTEVF | 0.55838907 |
| VYTVGTVV | ASSLEHPNTEVF | 0.66854745 |
| WTTFSGP | ASSLEHPNTEVF | 0.8561341 |
| YDELVEAF | ASSLEHPNTEVF | 0.6847094 |
| YGTGGAGGNGG | ASSLEHPNTEVF | 0.710145 |
| YTNVMAV | ASSLEHPNTEVF | 0.65027297 |
| FLEQVHQL | ASSLEHPNTEVF | 0.6021583 |
| GLSLLAVAV | ASSLEHPNTEVF | 0.65831316 |
| LEQVHQL | ASSLEHPNTEVF | 0.70034885 |
| LNAAGAI | ASSLEHPNTEVF | 0.7072078 |
| PAVHASL | ASSLEHPNTEVF | 0.69022185 |
| TELEDDA | ASSLEHPNTEVF | 0.5488726 |
| TPPGPPN | ASSLEHPNTEVF | 0.677323 |
| TPPPPPPP | ASSLEHPNTEVF | 0.6572546 |
| TQSPALI | TCSAGTGLYEQY | 0.6586392 |
| EFAAAFPQ | TCSAGTGLYEQY | 0.68465984 |
| LLSGNLDVL | TCSAGTGLYEQY | 0.6783078 |
| GPANLGNYNI | TCSAGTGLYEQY | 0.6378776 |

| | | |
|---|---|---|
| TLGPTLP | TCSAGTGLYEQY | 0.61619973 |
| VYAYSSCCK | TCSAGTGLYEQY | 0.6760801 |
| APAPAPAPNI | TCSAGTGLYEQY | 0.68364096 |
| PGGPVPIP | TCSAGTGLYEQY | 0.6820973 |
| FLTVALEP | TCSAGTGLYEQY | 0.7110306 |
| SQYFGANL | TCSAGTGLYEQY | 0.70696783 |
| GGVKPAA | TCSAGTGLYEQY | 0.69526565 |
| GGTGGAGGAGGW | TCSAGTGLYEQY | 0.3230405 |
| GVGGVAGLGGAGG | TCSAGTGLYEQY | 0.4876435 |
| VLLVLATTI | TCSAGTGLYEQY | 0.6206869 |
| IGIFAAML | TCSAGTGLYEQY | 0.6575248 |
| AVGDLID | TCSAGTGLYEQY | 0.3737976 |
| NISPLPL | TCSAGTGLYEQY | 0.6618103 |
| VTAVFVAI | TCSAGTGLYEQY | 0.61327577 |
| SGDDNDAAL | TCSAGTGLYEQY | 0.62027085 |
| AVAALAHS | TCSAGTGLYEQY | 0.7028153 |
| GGVTVMATP | TCSAGTGLYEQY | 0.65565056 |
| AGLAANVRD | TCSAGTGLYEQY | 0.6134778 |
| TNRPDLIDPALLRPG | TCSAGTGLYEQY | 0.5851147 |
| LANLGVTAL | TCSAGTGLYEQY | 0.6859602 |
| TGERIEL | TCSAGTGLYEQY | 0.48641312 |
| NGEELAA | TCSAGTGLYEQY | 0.69877386 |
| VLSLVWG | TCSAGTGLYEQY | 0.51977175 |
| IGYALWL | TCSAGTGLYEQY | 0.70423406 |
| LSLLLAPL | TCSAGTGLYEQY | 0.521975 |
| TLDEALK | TCSAGTGLYEQY | 0.69992733 |
| TDGAGTAEGG | TCSAGTGLYEQY | 0.67657536 |
| NAPGGAGGNGG | TCSAGTGLYEQY | 0.72139347 |
| VSVFMGMGGNF | TCSAGTGLYEQY | 0.65412176 |
| LASYLAAL | TCSAGTGLYEQY | 0.6753119 |
| LVGGTPAL | TCSAGTGLYEQY | 0.5868416 |
| VRNADVL | TCSAGTGLYEQY | 0.7106994 |
| TCSSPFLVSAM | TCSAGTGLYEQY | 0.52274704 |
| VHITDAE | TCSAGTGLYEQY | 0.66576886 |
| EYAAPSF | TCSAGTGLYEQY | 0.45518357 |
| VVDAFHGGL | TCSAGTGLYEQY | 0.68786365 |
| ASVVDPQ | TCSAGTGLYEQY | 0.6919962 |
| SYNEFGGLS | TCSAGTGLYEQY | 0.39688513 |
| DPDGSVY | TCSAGTGLYEQY | 0.69905865 |
| MVSTIHATE | TCSAGTGLYEQY | 0.64895475 |
| LAAGVADPV | TCSAGTGLYEQY | 0.59623677 |
| TKEQIAA | TCSAGTGLYEQY | 0.7085035 |
| SLASPAAL | TCSAGTGLYEQY | 0.6412219 |
| IGLVFYGV | TCSAGTGLYEQY | 0.70887434 |
| FLLDASNL | TCSAGTGLYEQY | 0.6333097 |
| YQVGNFP | TCSAGTGLYEQY | 0.6526258 |
| ESLLSQST | TCSAGTGLYEQY | 0.67392206 |
| LFTVGFAAVG | TCSAGTGLYEQY | 0.5237349 |
| ATPPTPT | TCSAGTGLYEQY | 0.55863094 |
| TDLATLG | TCSAGTGLYEQY | 0.71060723 |
| SAGGTGR | TCSAGTGLYEQY | 0.71852934 |

| | | |
|---|---|---|
| GFSNSGS | TCSAGTGLYEQY | 0.82399225 |
| GGNGGSGGTSV | TCSAGTGLYEQY | 0.5269391 |
| GTSVATGGAGN | TCSAGTGLYEQY | 0.38978127 |
| NGGNGGSGGTS | TCSAGTGLYEQY | 0.6026206 |
| AVVKFPH | TCSAGTGLYEQY | 0.7589744 |
| TPPPPPP | TCSAGTGLYEQY | 0.85851586 |
| AAPAAAAAPA | TCSAGTGLYEQY | 0.721267 |
| IALNEDL | TCSAGTGLYEQY | 0.31289423 |
| IVNVSSM | TCSAGTGLYEQY | 0.69761944 |
| TGIPPAPR | TCSAGTGLYEQY | 0.49394706 |
| FDLGGGTF | TCSAGTGLYEQY | 0.54297864 |
| FDLGGGTFDVS | TCSAGTGLYEQY | 0.65256596 |
| NEPTAAA | TCSAGTGLYEQY | 0.60788035 |
| NEPTAAALA | TCSAGTGLYEQY | 0.62462187 |
| GSIINISSI | TCSAGTGLYEQY | 0.44919485 |
| VAPGNAGTA | TCSAGTGLYEQY | 0.6690722 |
| TGGGASLE | TCSAGTGLYEQY | 0.76252294 |
| VAVYNHYKRI | TCSAGTGLYEQY | 0.57427853 |
| TDPAPATT | TCSAGTGLYEQY | 0.8282787 |
| TNRPDILDPALLRPG | TCSAGTGLYEQY | 0.7648009 |
| LTTTTTS | TCSAGTGLYEQY | 0.76669943 |
| PTGSGKTL | TCSAGTGLYEQY | 0.11856085 |
| TGSGKTL | TCSAGTGLYEQY | 0.21361238 |
| TIGHVDHGK | TCSAGTGLYEQY | 0.42498112 |
| TIGHVDHGKTT | TCSAGTGLYEQY | 0.52801436 |
| GHVDHGKTT | TCSAGTGLYEQY | 0.16291992 |
| AQFEHTLL | TCSAGTGLYEQY | 0.7027022 |
| IVNQDEL | TCSAGTGLYEQY | 0.5084363 |
| TNGSQFF | TCSAGTGLYEQY | 0.45532188 |
| TGKGIVE | TCSAGTGLYEQY | 0.80072296 |
| LISNASDA | TCSAGTGLYEQY | 0.5788586 |
| AELRTLGELTKIAKAHGAQ | TCSAGTGLYEQY | 0.37027705 |
| APNAVAAAEPSW | TCSAGTGLYEQY | 0.32109624 |
| APQVAGAATAVL | TCSAGTGLYEQY | 0.7768881 |
| ARRDPAAAAAILPTDA | TCSAGTGLYEQY | 0.22210926 |
| DAAARDVLAEAGLAEYF | TCSAGTGLYEQY | 0.75643957 |
| DDQIVLSAKANLAKVEQ | TCSAGTGLYEQY | 0.40008032 |
| DELLPDTGEDSDEDGH | TCSAGTGLYEQY | 0.7187774 |
| ELPEAPANRATVPQD | TCSAGTGLYEQY | 0.22732124 |
| FGQSKIERVTGIG | TCSAGTGLYEQY | 0.7362454 |
| GAGGTGSTAGGAGGAGGAGGLYA | TCSAGTGLYEQY | 0.61668146 |
| GHGGTGAAGGNGGTGGAGGSGIDG | TCSAGTGLYEQY | 0.8721311 |
| GNGGNGGAGGEGGPGGAGGAGA | TCSAGTGLYEQY | 0.4652368 |
| GSTAGNGGNGGNATNGTIAGQPAG | TCSAGTGLYEQY | 0.5788396 |
| GTGGDGGNGGNGGTGG | TCSAGTGLYEQY | 0.58219266 |
| GVRRPAGIPADAPV | TCSAGTGLYEQY | 0.8444998 |
| GYRQVTDAHLLTLARR | TCSAGTGLYEQY | 0.3011777 |
| IAKMGQDGHDRG | TCSAGTGLYEQY | 0.40784806 |
| KAEPSIEIQYT | TCSAGTGLYEQY | 0.29281956 |
| KPVRIRMAADEDTVAVT | TCSAGTGLYEQY | 0.62326366 |
| MPPRIAGMRLLVIKPEP | TCSAGTGLYEQY | 0.7215551 |

| | | |
|---|---|---|
| MKFYAVALFRSRGVPPTR | TCSAGTGLYEQY | 0.16714925 |
| NAGGFGILGAVAHSP | TCSAGTGLYEQY | 0.68131673 |
| NEGREITLAMAEM | TCSAGTGLYEQY | 0.53053916 |
| NELVEWSHPLGHRWRWE | TCSAGTGLYEQY | 0.328121 |
| NLGAGGTGGNGGNPG | TCSAGTGLYEQY | 0.78872097 |
| NTEWPKWETVMAADPVD | TCSAGTGLYEQY | 0.5074887 |
| PASAAPAETQP | TCSAGTGLYEQY | 0.84874964 |
| QQSSDGKNGVRAAEITGE | TCSAGTGLYEQY | 0.4459006 |
| RAPDAVTTSWWKE | TCSAGTGLYEQY | 0.41196698 |
| SGFGNVGAHTSGWFNQ | TCSAGTGLYEQY | 0.6599388 |
| SGVAGAVGQAASVGGLKV | TCSAGTGLYEQY | 0.3072953 |
| SNGGAGTGSAGGNGGT | TCSAGTGLYEQY | 0.29409635 |
| TGGAGGEGGAAG | TCSAGTGLYEQY | 0.80247533 |
| TGGDGGAAGAGGAPGHG | TCSAGTGLYEQY | 0.63342965 |
| TMPPLLAAIANAGRHGV | TCSAGTGLYEQY | 0.65546656 |
| TSSTAGNSGGA | TCSAGTGLYEQY | 0.58796346 |
| VPATVPVGILC | TCSAGTGLYEQY | 0.57549965 |
| VQRPDSGGVVLTGRIS | TCSAGTGLYEQY | 0.58848274 |
| WTPNPVPATKGVPAGLVQAT | TCSAGTGLYEQY | 0.6486712 |
| YALKRALAKFSDRFDVVII | TCSAGTGLYEQY | 0.6542046 |
| YLIGSVIV | TCSAGTGLYEQY | 0.6398804 |
| YGQLVAVID | TCSAGTGLYEQY | 0.58818376 |
| VTVGGMTLPA | TCSAGTGLYEQY | 0.6322051 |
| VLALPAV | TCSAGTGLYEQY | 0.60800254 |
| VIPPTLNYET | TCSAGTGLYEQY | 0.6018986 |
| VGMGGMHPSSG | TCSAGTGLYEQY | 0.6345078 |
| VGLAGGAATA | TCSAGTGLYEQY | 0.6351009 |
| VGGAGGAGGSGP | TCSAGTGLYEQY | 0.5904804 |
| VFAAFMV | TCSAGTGLYEQY | 0.59814537 |
| TLYDKLR | TCSAGTGLYEQY | 0.621657 |
| TLAVFTVL | TCSAGTGLYEQY | 0.64352185 |
| TGSVQPARV | TCSAGTGLYEQY | 0.63419366 |
| TGGGAGSGGT | TCSAGTGLYEQY | 0.6583384 |
| SHLETSL | TCSAGTGLYEQY | 0.6437617 |
| SFTTGMT | TCSAGTGLYEQY | 0.63897634 |
| QVEVAKRMGVRQPTVS | TCSAGTGLYEQY | 0.3753829 |
| QGSGGPALAM | TCSAGTGLYEQY | 0.49647307 |
| MLAPSNP | TCSAGTGLYEQY | 0.56362194 |
| LVNNAAI | TCSAGTGLYEQY | 0.6171691 |
| LTGEINTLA | TCSAGTGLYEQY | 0.4137642 |
| LSIFGPV | TCSAGTGLYEQY | 0.644338 |
| LLALLANL | TCSAGTGLYEQY | 0.6093781 |
| LISNASDALDKLR | TCSAGTGLYEQY | 0.5811843 |
| LINEAVP | TCSAGTGLYEQY | 0.64538497 |
| LGYAFSGL | TCSAGTGLYEQY | 0.6030697 |
| LDFVAVAL | TCSAGTGLYEQY | 0.5885581 |
| LAQPGAE | TCSAGTGLYEQY | 0.5811091 |
| ISNLGAVTGF | TCSAGTGLYEQY | 0.6476972 |
| HQEDPGRI | TCSAGTGLYEQY | 0.49013105 |
| GTPGEPTP | TCSAGTGLYEQY | 0.6449483 |
| GFFNSGS | TCSAGTGLYEQY | 0.5286585 |

| | | |
|---|---|---|
| GAGGGGLTAGAN | TCSAGTGLYEQY | 0.649752 |
| FVDALVNM | TCSAGTGLYEQY | 0.5507798 |
| FLTVAPEL | TCSAGTGLYEQY | 0.6420227 |
| FETLTAL | TCSAGTGLYEQY | 0.6120818 |
| DVINPIEL | TCSAGTGLYEQY | 0.6629648 |
| APVSGAPV | TCSAGTGLYEQY | 0.6079853 |
| ALVGGLAW | TCSAGTGLYEQY | 0.6427035 |
| ALAAAAATAI | TCSAGTGLYEQY | 0.6234361 |
| AFGLGLLL | TCSAGTGLYEQY | 0.653399 |
| AFFAAASL | TCSAGTGLYEQY | 0.5076215 |
| AAAALPAFSPP | TCSAGTGLYEQY | 0.61039925 |
| VVLAPAL | TCSAGTGLYEQY | 0.49574423 |
| TIGHVDHGKTTL | TCSAGTGLYEQY | 0.6320534 |
| LISNASDALDK | TCSAGTGLYEQY | 0.63968885 |
| LISNASDAL | TCSAGTGLYEQY | 0.4673751 |
| ISNASDALDK | TCSAGTGLYEQY | 0.65218014 |
| NQAETLVYQTE | TCSAGTGLYEQY | 0.60132945 |
| TGGTGGTAGAG | TCSAGTGLYEQY | 0.5966657 |
| TGSTAGGAGGA | TCSAGTGLYEQY | 0.6447755 |
| DPLAVPVAAGPV | TCSAGTGLYEQY | 0.6116998 |
| GGTGGTPGNGGN | TCSAGTGLYEQY | 0.64361596 |
| PAGLAGVDPATL | TCSAGTGLYEQY | 0.58780867 |
| GNNLVVEAPHAGASV | TCSAGTGLYEQY | 0.6021018 |
| KVTRIMSASTLEQPA | TCSAGTGLYEQY | 0.6400184 |
| LGFALPIALPRRTVV | TCSAGTGLYEQY | 0.5172744 |
| TRLGNLTARKLVVLV | TCSAGTGLYEQY | 0.53490806 |
| WSQQRRLIYFPSAGP | TCSAGTGLYEQY | 0.65120465 |
| GPNGNPDPMAAAVDIR | TCSAGTGLYEQY | 0.78297746 |
| TQRCVSVADLRAAASR | TCSAGTGLYEQY | 0.80281687 |
| QFVLDVAEHPVEVLTETV | TCSAGTGLYEQY | 0.727704 |
| GGAGGTGGAGGNGGTQAGDG | TCSAGTGLYEQY | 0.71108645 |
| PLLATLRAIAVDVDAKPAQV | TCSAGTGLYEQY | 0.7399055 |
| APKFRTGGYMIDFWGVGYQVAKRM | TCSAGTGLYEQY | 0.80037665 |
| GAGGAGGAGGAAGGPGGTGGTGGN | TCSAGTGLYEQY | 0.799337 |
| IGALGLLATDVAPAGSVAFGVALA | TCSAGTGLYEQY | 0.7833119 |
| AGGAGGIGGNGGAGGAGGVLMGNG | TCSAGTGLYEQY | 0.73451614 |
| ADELIGLL | TCSAGTGLYEQY | 0.7776682 |
| ADRALKA | TCSAGTGLYEQY | 0.7578634 |
| AGAVGQVGGAGGA | TCSAGTGLYEQY | 0.7483005 |
| AGGGGGLGGAP | TCSAGTGLYEQY | 0.78918326 |
| AGGLLYGNGGAG | TCSAGTGLYEQY | 0.78031486 |
| AGLESIGL | TCSAGTGLYEQY | 0.7450376 |
| AGLKTGII | TCSAGTGLYEQY | 0.75783825 |
| ASLAALLL | TCSAGTGLYEQY | 0.7698647 |
| ASVINISSI | TCSAGTGLYEQY | 0.7856402 |
| ATPPGPA | TCSAGTGLYEQY | 0.7792214 |
| ATVASALV | TCSAGTGLYEQY | 0.81256795 |
| ATVASLIL | TCSAGTGLYEQY | 0.794155 |
| AVNGPVLIRH | TCSAGTGLYEQY | 0.79352826 |
| DGAAGAVGI | TCSAGTGLYEQY | 0.4295174 |
| DPLLHGI | TCSAGTGLYEQY | 0.6668085 |

| | | |
|---|---|---|
| ELGAALGAL | TCSAGTGLYEQY | 0.770038 |
| ELGRTQAV | TCSAGTGLYEQY | 0.759569 |
| FVAGGYLS | TCSAGTGLYEQY | 0.5897239 |
| GASGGAGGAGGW | TCSAGTGLYEQY | 0.7900049 |
| GGAGAGGDGGAGGI | TCSAGTGLYEQY | 0.746004 |
| GTGHNGSQPGMGGNG | TCSAGTGLYEQY | 0.7189245 |
| GYAAASASAS | TCSAGTGLYEQY | 0.78675747 |
| HQVDVAAVQQG | TCSAGTGLYEQY | 0.74679255 |
| IGAAIAHT | TCSAGTGLYEQY | 0.74841714 |
| IGMIEVAQL | TCSAGTGLYEQY | 0.7556692 |
| IRLAPEHTSAMD | TCSAGTGLYEQY | 0.80188316 |
| IYNVSVL | TCSAGTGLYEQY | 0.6894028 |
| KQGFLGM | TCSAGTGLYEQY | 0.7858592 |
| LESVITQL | TCSAGTGLYEQY | 0.7612418 |
| LGLGALVVV | TCSAGTGLYEQY | 0.8019837 |
| LIGNGGNGGSGGI | TCSAGTGLYEQY | 0.68938565 |
| LNAINAPTL | TCSAGTGLYEQY | 0.788412 |
| LQLLNTGV | TCSAGTGLYEQY | 0.75469744 |
| LQTSIEF | TCSAGTGLYEQY | 0.81177074 |
| MSFAVSEL | TCSAGTGLYEQY | 0.758553 |
| NTGALGL | TCSAGTGLYEQY | 0.78627133 |
| PPGAGGAGGAGTT | TCSAGTGLYEQY | 0.7795246 |
| QLVGLGI | TCSAGTGLYEQY | 0.79756093 |
| RFLEQVHQL | TCSAGTGLYEQY | 0.6285013 |
| RGVGYVLR | TCSAGTGLYEQY | 0.76103985 |
| SATGVFAGVI | TCSAGTGLYEQY | 0.70949435 |
| SLPAITLP | TCSAGTGLYEQY | 0.77732503 |
| STALAFTV | TCSAGTGLYEQY | 0.7843773 |
| SVLTVGLGK | TCSAGTGLYEQY | 0.6479969 |
| TALAGAGSGI | TCSAGTGLYEQY | 0.7879152 |
| TGLPFTGI | TCSAGTGLYEQY | 0.7506459 |
| VAEHVTV | TCSAGTGLYEQY | 0.7462305 |
| VDIVLNSL | TCSAGTGLYEQY | 0.80196565 |
| VLTGGGAVL | TCSAGTGLYEQY | 0.7614904 |
| VNVGLAYGI | TCSAGTGLYEQY | 0.7953944 |
| VSSAAVGAV | TCSAGTGLYEQY | 0.75971824 |
| VTSVYEAGV | TCSAGTGLYEQY | 0.7531828 |
| WVALDRGA | TCSAGTGLYEQY | 0.78992087 |
| YLVGSVVI | TCSAGTGLYEQY | 0.68762535 |
| GPPGTGKTLL | TCSAGTGLYEQY | 0.7265228 |
| GVGMTKF | TCSAGTGLYEQY | 0.8002924 |
| IGIGGSDL | TCSAGTGLYEQY | 0.12994418 |
| IGTLLAEL | TCSAGTGLYEQY | 0.12967339 |
| IYTSGTT | TCSAGTGLYEQY | 0.10440951 |
| LSFPAGV | TCSAGTGLYEQY | 0.113226086 |
| LYTSGTT | TCSAGTGLYEQY | 0.10850064 |
| VGAFPFGF | TCSAGTGLYEQY | 0.13352638 |
| VGAGTVEF | TCSAGTGLYEQY | 0.2385433 |
| VGKSTLLL | TCSAGTGLYEQY | 0.1303725 |
| AALAHAGGGIV | TCSAGTGLYEQY | 0.1185917 |
| AALVAVSLL | TCSAGTGLYEQY | 0.11740026 |

| | | |
|---|---|---|
| AAVSRAVAAL | TCSAGTGLYEQY | 0.11692645 |
| AGGSSGIGGF | TCSAGTGLYEQY | 0.11399704 |
| AGLTVANANG | TCSAGTGLYEQY | 0.114180796 |
| AGLVGAVL | TCSAGTGLYEQY | 0.11815121 |
| ALSQAFTV | TCSAGTGLYEQY | 0.10950926 |
| ALTFSVSL | TCSAGTGLYEQY | 0.104347 |
| ALTGAALVAL | TCSAGTGLYEQY | 0.11144094 |
| ASFSIGAGVL | TCSAGTGLYEQY | 0.14051504 |
| DGSVYKI | TCSAGTGLYEQY | 0.13374528 |
| EQLAEIV | TCSAGTGLYEQY | 0.1399588 |
| FAVSELTVM | TCSAGTGLYEQY | 0.1551428 |
| GAAPLAAGL | TCSAGTGLYEQY | 0.107107416 |
| GADGGDGGSGGAGG | TCSAGTGLYEQY | 0.098407164 |
| GFGLDTGGAGG | TCSAGTGLYEQY | 0.10446049 |
| GLASGGLL | TCSAGTGLYEQY | 0.110075094 |
| GRIAGAAVLL | TCSAGTGLYEQY | 0.114507645 |
| GSAGSLNTSF | TCSAGTGLYEQY | 0.09568254 |
| GTGQIGI | TCSAGTGLYEQY | 0.1249313 |
| GTLIALIL | TCSAGTGLYEQY | 0.11978366 |
| IAATMGFVVG | TCSAGTGLYEQY | 0.10859871 |
| IFAQALL | TCSAGTGLYEQY | 0.14007986 |
| IGAAVALGF | TCSAGTGLYEQY | 0.1057263 |
| IGLTPLAL | TCSAGTGLYEQY | 0.11389127 |
| ILGSNTSTL | TCSAGTGLYEQY | 0.11251818 |
| IVFSLTVL | TCSAGTGLYEQY | 0.13297425 |
| LASKATGF | TCSAGTGLYEQY | 0.10684852 |
| LASVALIL | TCSAGTGLYEQY | 0.13527742 |
| LFAVGLTL | TCSAGTGLYEQY | 0.108865775 |
| LFNGGIGTYI | TCSAGTGLYEQY | 0.15788382 |
| LTLGAGII | TCSAGTGLYEQY | 0.11025465 |
| LVAAALGDI | TCSAGTGLYEQY | 0.12762824 |
| NTAFVQLGI | TCSAGTGLYEQY | 0.11125557 |
| QLFSGAEL | TCSAGTGLYEQY | 0.26010785 |
| SAATFITL | TCSAGTGLYEQY | 0.12313135 |
| SNYGLANLGV | TCSAGTGLYEQY | 0.13674082 |
| SYGGAYAVM | TCSAGTGLYEQY | 0.11152033 |
| TTGVGGLTLG | TCSAGTGLYEQY | 0.14801645 |
| TTQTIYI | TCSAGTGLYEQY | 0.10296678 |
| VAAGAGLGAVI | TCSAGTGLYEQY | 0.11598107 |
| VAAVVTLL | TCSAGTGLYEQY | 0.10471068 |
| VGFLGGLI | TCSAGTGLYEQY | 0.116492316 |
| VGVIPILL | TCSAGTGLYEQY | 0.1279048 |
| VIFASSLI | TCSAGTGLYEQY | 0.098304905 |
| VKGGDGGAAAG | TCSAGTGLYEQY | 0.114501394 |
| VTKTFTNI | TCSAGTGLYEQY | 0.10032659 |
| VVAAAILL | TCSAGTGLYEQY | 0.106467515 |
| YAALDVEL | TCSAGTGLYEQY | 0.11489135 |
| LASVALLL | TCSAGTGLYEQY | 0.10881627 |
| LDAVHRV | TCSAGTGLYEQY | 0.12431959 |
| QGLSSSIF | TCSAGTGLYEQY | 0.10200246 |
| RGTFANIRL | TCSAGTGLYEQY | 0.11593861 |

| | | |
|---|---|---|
| RQLWWGHRI | TCSAGTGLYEQY | 0.13804501 |
| YGPPGTGKTLL | TCSAGTGLYEQY | 0.10013273 |
| AALVGAPANVA | TCSAGTGLYEQY | 0.10694495 |
| AATFRRTE | TCSAGTGLYEQY | 0.13963471 |
| AGALGGGI | TCSAGTGLYEQY | 0.7987964 |
| AGGAGGAGGIGAG | TCSAGTGLYEQY | 0.8001354 |
| AGSGALGL | TCSAGTGLYEQY | 0.76674974 |
| AQIADQI | TCSAGTGLYEQY | 0.76027477 |
| ASAVAVTYL | TCSAGTGLYEQY | 0.78354895 |
| ASTVLGGTGG | TCSAGTGLYEQY | 0.79997534 |
| AVGGEPAK | TCSAGTGLYEQY | 0.80483174 |
| CEQGTFGGTC | TCSAGTGLYEQY | 0.7977426 |
| DEGKAPAG | TCSAGTGLYEQY | 0.7663908 |
| DTVVFPAL | TCSAGTGLYEQY | 0.79417366 |
| FEGITMAG | TCSAGTGLYEQY | 0.7931117 |
| FPLVVQD | TCSAGTGLYEQY | 0.7814084 |
| GAFTTPQ | TCSAGTGLYEQY | 0.79895294 |
| GFGGFGTAGGVGGA | TCSAGTGLYEQY | 0.79438925 |
| GFLAQFL | TCSAGTGLYEQY | 0.77294135 |
| GGAGGAGGNAGVA | TCSAGTGLYEQY | 0.77546775 |
| GGVGTAGAGGAGGAGG | TCSAGTGLYEQY | 0.7956592 |
| GNGGGGDAGNAGSGGN | TCSAGTGLYEQY | 0.8000118 |
| GTGGAGGAGNDAGS | TCSAGTGLYEQY | 0.7999367 |
| GTTNTGWFNS | TCSAGTGLYEQY | 0.8053037 |
| IFMGPLFTL | TCSAGTGLYEQY | 0.80172527 |
| ILAHAIEAI | TCSAGTGLYEQY | 0.79746807 |
| IRSAIREAL | TCSAGTGLYEQY | 0.49445567 |
| ITFAGGG | TCSAGTGLYEQY | 0.72898614 |
| ITGMSGHQEV | TCSAGTGLYEQY | 0.7735021 |
| IVAGLACTL | TCSAGTGLYEQY | 0.78793985 |
| IVGGGLAGA | TCSAGTGLYEQY | 0.5862602 |
| IVTGVLNI | TCSAGTGLYEQY | 0.7945709 |
| LAQQGCI | TCSAGTGLYEQY | 0.78292084 |
| LCLHVEN | TCSAGTGLYEQY | 0.76147956 |
| LDAISDGS | TCSAGTGLYEQY | 0.80086315 |
| LGGFAAML | TCSAGTGLYEQY | 0.7776679 |
| LGGKNPMI | TCSAGTGLYEQY | 0.76742303 |
| LGPKGVAAIL | TCSAGTGLYEQY | 0.7788121 |
| LMVGVSI | TCSAGTGLYEQY | 0.80432343 |
| LVGAAFVL | TCSAGTGLYEQY | 0.77285314 |
| LVLAVAPL | TCSAGTGLYEQY | 0.80085194 |
| MAAIAVIDAA | TCSAGTGLYEQY | 0.7752714 |
| NNGINAELQ | TCSAGTGLYEQY | 0.80362296 |
| RGEPLVLVHG | TCSAGTGLYEQY | 0.7441441 |
| RGGAGGAAGLI | TCSAGTGLYEQY | 0.80060524 |
| RSYRMHPAV | TCSAGTGLYEQY | 0.7910204 |
| SAAHLLAGQLA | TCSAGTGLYEQY | 0.8074851 |
| SAAVGSY | TCSAGTGLYEQY | 0.7914963 |
| SPHAVLHRI | TCSAGTGLYEQY | 0.8012785 |
| SQIAKVL | TCSAGTGLYEQY | 0.77716637 |
| TALGHLVP | TCSAGTGLYEQY | 0.8017299 |

| | | |
|---|---|---|
| TAPTTSGAA | TCSAGTGLYEQY | 0.7440871 |
| TDGDDGNGGSPGT | TCSAGTGLYEQY | 0.78865504 |
| TDGLGEN | TCSAGTGLYEQY | 0.7147159 |
| TGGPPLPGQA | TCSAGTGLYEQY | 0.790621 |
| TIPAITI | TCSAGTGLYEQY | 0.7967975 |
| TIPGFTI | TCSAGTGLYEQY | 0.57281554 |
| TSIASGALA | TCSAGTGLYEQY | 0.8012675 |
| TVTLPGAAI | TCSAGTGLYEQY | 0.76555425 |
| VASGLSAALP | TCSAGTGLYEQY | 0.7746362 |
| VGAGVIGI | TCSAGTGLYEQY | 0.80269325 |
| VGPKRFGV | TCSAGTGLYEQY | 0.7841176 |
| VNVAFPD | TCSAGTGLYEQY | 0.8018173 |
| VSMTNSGVS | TCSAGTGLYEQY | 0.7784698 |
| VTDVLGDLSV | TCSAGTGLYEQY | 0.7860725 |
| VVVVGAGFAGL | TCSAGTGLYEQY | 0.7930168 |
| VYTVGTVV | TCSAGTGLYEQY | 0.6896783 |
| WTTFSGP | TCSAGTGLYEQY | 0.7680054 |
| YDELVEAF | TCSAGTGLYEQY | 0.8038172 |
| YGTGGAGGNGG | TCSAGTGLYEQY | 0.59041363 |
| YTNVMAV | TCSAGTGLYEQY | 0.52295953 |
| FLEQVHQL | TCSAGTGLYEQY | 0.16588236 |
| GLSLLAVAV | TCSAGTGLYEQY | 0.20533898 |
| LEQVHQL | TCSAGTGLYEQY | 0.24381138 |
| LNAAGAI | TCSAGTGLYEQY | 0.31525058 |
| PAVHASL | TCSAGTGLYEQY | 0.7428265 |
| TELEDDA | TCSAGTGLYEQY | 0.5881265 |
| TPPGPPN | TCSAGTGLYEQY | 0.20436586 |
| TPPPPPPP | TCSAGTGLYEQY | 0.39099473 |
| TQSPALI | ASSATGGSQNTLY | 0.24525094 |
| EFAAAFPQ | ASSATGGSQNTLY | 0.33317578 |
| LLSGNLDVL | ASSATGGSQNTLY | 0.25756246 |
| GPANLGNYNI | ASSATGGSQNTLY | 0.2828998 |
| TLGPTLP | ASSATGGSQNTLY | 0.1923244 |
| VYAYSSCCK | ASSATGGSQNTLY | 0.1627945 |
| APAPAPAPNI | ASSATGGSQNTLY | 0.44207007 |
| PGGPVPIP | ASSATGGSQNTLY | 0.6485646 |
| FLTVALEP | ASSATGGSQNTLY | 0.48939887 |
| SQYFGANL | ASSATGGSQNTLY | 0.4208616 |
| GGVKPAA | ASSATGGSQNTLY | 0.56821746 |
| GGTGGAGGAGGW | ASSATGGSQNTLY | 0.3271485 |
| GVGGVAGLGGAGG | ASSATGGSQNTLY | 0.13736993 |
| VLLVLATTI | ASSATGGSQNTLY | 0.14491232 |
| IGIFAAML | ASSATGGSQNTLY | 0.19401602 |
| AVGDLID | ASSATGGSQNTLY | 0.19793877 |
| NISPLPL | ASSATGGSQNTLY | 0.13524099 |
| VTAVFVAI | ASSATGGSQNTLY | 0.33071956 |
| SGDDNDAAL | ASSATGGSQNTLY | 0.23021641 |
| AVAALAHS | ASSATGGSQNTLY | 0.20857856 |
| GGVTVMATP | ASSATGGSQNTLY | 0.5884636 |
| AGLAANVRD | ASSATGGSQNTLY | 0.20372915 |
| TNRPDLIDPALLRPG | ASSATGGSQNTLY | 0.16887677 |

| | | |
|---|---|---|
| LANLGVTAL | ASSATGGSQNTLY | 0.20188612 |
| TGERIEL | ASSATGGSQNTLY | 0.40331966 |
| NGEELAA | ASSATGGSQNTLY | 0.17574027 |
| VLSLVWG | ASSATGGSQNTLY | 0.54417896 |
| IGYALWL | ASSATGGSQNTLY | 0.19943677 |
| LSLLLAPL | ASSATGGSQNTLY | 0.49513623 |
| TLDEALK | ASSATGGSQNTLY | 0.17811081 |
| TDGAGTAEGG | ASSATGGSQNTLY | 0.52289176 |
| NAPGGAGGNGG | ASSATGGSQNTLY | 0.32988358 |
| VSVFMGMGGNF | ASSATGGSQNTLY | 0.75368947 |
| LASYLAAL | ASSATGGSQNTLY | 0.24596903 |
| LVGGTPAL | ASSATGGSQNTLY | 0.3989371 |
| VRNADVL | ASSATGGSQNTLY | 0.25553802 |
| TCSSPFLVSAM | ASSATGGSQNTLY | 0.6779671 |
| VHITDAE | ASSATGGSQNTLY | 0.16171359 |
| EYAAPSF | ASSATGGSQNTLY | 0.18358462 |
| VVDAFHGGL | ASSATGGSQNTLY | 0.15044932 |
| ASVVDPQ | ASSATGGSQNTLY | 0.19282678 |
| SYNEFGGLS | ASSATGGSQNTLY | 0.5964363 |
| DPDGSVY | ASSATGGSQNTLY | 0.1328952 |
| MVSTIHATE | ASSATGGSQNTLY | 0.2225259 |
| LAAGVADPV | ASSATGGSQNTLY | 0.15696216 |
| TKEQIAA | ASSATGGSQNTLY | 0.18504196 |
| SLASPAAL | ASSATGGSQNTLY | 0.31882104 |
| IGLVFYGV | ASSATGGSQNTLY | 0.35344684 |
| FLLDASNL | ASSATGGSQNTLY | 0.35310137 |
| YQVGNFP | ASSATGGSQNTLY | 0.19176883 |
| ESLLSQST | ASSATGGSQNTLY | 0.21282238 |
| LFTVGFAAVG | ASSATGGSQNTLY | 0.23901743 |
| ATPPTPT | ASSATGGSQNTLY | 0.14635755 |
| TDLATLG | ASSATGGSQNTLY | 0.17888014 |
| SAGGTGR | ASSATGGSQNTLY | 0.59662664 |
| GFSNSGS | ASSATGGSQNTLY | 0.122973904 |
| GGNGGSGGTSV | ASSATGGSQNTLY | 0.11205649 |
| GTSVATGGAGN | ASSATGGSQNTLY | 0.110982865 |
| NGGNGGSGGTS | ASSATGGSQNTLY | 0.11749422 |
| AVVKFPH | ASSATGGSQNTLY | 0.11314094 |
| TPPPPPP | ASSATGGSQNTLY | 0.1335657 |
| AAPAAAAAPA | ASSATGGSQNTLY | 0.12553093 |
| IALNEDL | ASSATGGSQNTLY | 0.12695819 |
| IVNVSSM | ASSATGGSQNTLY | 0.11994866 |
| TGIPPAPR | ASSATGGSQNTLY | 0.12294388 |
| FDLGGGTF | ASSATGGSQNTLY | 0.119730815 |
| FDLGGGTFDVS | ASSATGGSQNTLY | 0.12134001 |
| NEPTAAA | ASSATGGSQNTLY | 0.114657186 |
| NEPTAAALA | ASSATGGSQNTLY | 0.122287154 |
| GSIINISSI | ASSATGGSQNTLY | 0.1131874 |
| VAPGNAGTA | ASSATGGSQNTLY | 0.11005595 |
| TGGGASLE | ASSATGGSQNTLY | 0.114533365 |
| VAVYNHYKRI | ASSATGGSQNTLY | 0.12427865 |
| TDPAPATT | ASSATGGSQNTLY | 0.12159832 |

| | | |
|---|---|---|
| TNRPDILDPALLRPG | ASSATGGSQNTLY | 0.13367014 |
| LTTTTTS | ASSATGGSQNTLY | 0.13425748 |
| PTGSGKTL | ASSATGGSQNTLY | 0.11071362 |
| TGSGKTL | ASSATGGSQNTLY | 0.107068375 |
| TIGHVDHGK | ASSATGGSQNTLY | 0.1099197 |
| TIGHVDHGKTT | ASSATGGSQNTLY | 0.11599819 |
| GHVDHGKTT | ASSATGGSQNTLY | 0.118131414 |
| AQFEHTLL | ASSATGGSQNTLY | 0.10367084 |
| IVNQDEL | ASSATGGSQNTLY | 0.12656173 |
| TNGSQFF | ASSATGGSQNTLY | 0.12099999 |
| TGKGIVE | ASSATGGSQNTLY | 0.115433514 |
| LISNASDA | ASSATGGSQNTLY | 0.12705183 |
| AELRTLGELTKIAKAHGAQ | ASSATGGSQNTLY | 0.11196284 |
| APNAVAAAEPSW | ASSATGGSQNTLY | 0.11746017 |
| APQVAGAATAVL | ASSATGGSQNTLY | 0.12323956 |
| ARRDPAAAAAILPTDA | ASSATGGSQNTLY | 0.12612174 |
| DAAARDVLAEAGLAEYF | ASSATGGSQNTLY | 0.115440786 |
| DDQIVLSAKANLAKVEQ | ASSATGGSQNTLY | 0.13738327 |
| DELLPDTGEDSDEDGH | ASSATGGSQNTLY | 0.112909585 |
| ELPEAPANRATVPQD | ASSATGGSQNTLY | 0.1561137 |
| FGQSKIERVTGIG | ASSATGGSQNTLY | 0.11722618 |
| GAGGTGSTAGGAGGAGGAGGLYA | ASSATGGSQNTLY | 0.12428282 |
| GHGGTGAAGGNGGTGGAGGSGIDG | ASSATGGSQNTLY | 0.11620755 |
| GNGGNGGAGGEGGPGGAGGAGA | ASSATGGSQNTLY | 0.17926641 |
| GSTAGNGGNGGNATNGTIAGQPAG | ASSATGGSQNTLY | 0.12183114 |
| GTGGDGGNGGNGGTGG | ASSATGGSQNTLY | 0.13356288 |
| GVRRPAGIPADAPV | ASSATGGSQNTLY | 0.11453234 |
| GYRQVTDAHLLTLARR | ASSATGGSQNTLY | 0.120740056 |
| IAKMGQDGHDRG | ASSATGGSQNTLY | 0.110053204 |
| KAEPSIEIQYT | ASSATGGSQNTLY | 0.11494644 |
| KPVRIRMAADEDTVAVT | ASSATGGSQNTLY | 0.11017052 |
| MPPRIAGMRLLVIKPEP | ASSATGGSQNTLY | 0.120449394 |
| MKFYAVALFRSRGVPPTR | ASSATGGSQNTLY | 0.12719235 |
| NAGGFGILGAVAHSP | ASSATGGSQNTLY | 0.10531631 |
| NEGREITLAMAEM | ASSATGGSQNTLY | 0.113801114 |
| NELVEWSHPLGHRWRWE | ASSATGGSQNTLY | 0.105247445 |
| NLGAGGTGGNGGNPG | ASSATGGSQNTLY | 0.11173576 |
| NTEWPKWETVMAADPVD | ASSATGGSQNTLY | 0.11356909 |
| PASAAPAETQP | ASSATGGSQNTLY | 0.118399076 |
| QQSSDGKNGVRAAEITGE | ASSATGGSQNTLY | 0.11520922 |
| RAPDAVTTSWWKE | ASSATGGSQNTLY | 0.1091454 |
| SGFGNVGAHTSGWFNQ | ASSATGGSQNTLY | 0.11387752 |
| SGVAGAVGQAASVGGLKV | ASSATGGSQNTLY | 0.13306536 |
| SNGGAGTGSAGGNGGT | ASSATGGSQNTLY | 0.10690231 |
| TGGAGGEGGAAG | ASSATGGSQNTLY | 0.116035484 |
| TGGDGGAAGAGGAPGHG | ASSATGGSQNTLY | 0.1341117 |
| TMPPLLAAIANAGRHGV | ASSATGGSQNTLY | 0.2519747 |
| TSSTAGNSGGA | ASSATGGSQNTLY | 0.6727566 |
| VPATVPVGILC | ASSATGGSQNTLY | 0.107106395 |
| VQRPDSGGVVLTGRIS | ASSATGGSQNTLY | 0.13598981 |
| WTPNPVPATKGVPAGLVQAT | ASSATGGSQNTLY | 0.3045619 |

| | | |
|---|---|---|
| YALKRALAKFSDRFDVVII | ASSATGGSQNTLY | 0.33633327 |
| YLIGSVIV | ASSATGGSQNTLY | 0.12896307 |
| YGQLVAVID | ASSATGGSQNTLY | 0.35938746 |
| VTVGGMTLPA | ASSATGGSQNTLY | 0.12767985 |
| VLALPAV | ASSATGGSQNTLY | 0.1246252 |
| VIPPTLNYET | ASSATGGSQNTLY | 0.12219638 |
| VGMGGMHPSSG | ASSATGGSQNTLY | 0.18938848 |
| VGLAGGAATA | ASSATGGSQNTLY | 0.27388847 |
| VGGAGGAGGSGP | ASSATGGSQNTLY | 0.1121241 |
| VFAAFMV | ASSATGGSQNTLY | 0.09407067 |
| TLYDKLR | ASSATGGSQNTLY | 0.17220421 |
| TLAVFTVL | ASSATGGSQNTLY | 0.41001832 |
| TGSVQPARV | ASSATGGSQNTLY | 0.64858496 |
| TGGGAGSGGT | ASSATGGSQNTLY | 0.65009904 |
| SHLETSL | ASSATGGSQNTLY | 0.5058579 |
| SFTTGMT | ASSATGGSQNTLY | 0.3588603 |
| QVEVAKRMGVRQPTVS | ASSATGGSQNTLY | 0.25776938 |
| QGSGGPALAM | ASSATGGSQNTLY | 0.084451854 |
| MLAPSNP | ASSATGGSQNTLY | 0.12350085 |
| LVNNAAI | ASSATGGSQNTLY | 0.11012101 |
| LTGEINTLA | ASSATGGSQNTLY | 0.27136916 |
| LSIFGPV | ASSATGGSQNTLY | 0.07623825 |
| LLALLANL | ASSATGGSQNTLY | 0.16912092 |
| LISNASDALDKLR | ASSATGGSQNTLY | 0.18053086 |
| LINEAVP | ASSATGGSQNTLY | 0.09177969 |
| LGYAFSGL | ASSATGGSQNTLY | 0.55217147 |
| LDFVAVAL | ASSATGGSQNTLY | 0.098530374 |
| LAQPGAE | ASSATGGSQNTLY | 0.11598812 |
| ISNLGAVTGF | ASSATGGSQNTLY | 0.13416547 |
| HQEDPGRI | ASSATGGSQNTLY | 0.53972954 |
| GTPGEPTP | ASSATGGSQNTLY | 0.1257482 |
| GFFNSGS | ASSATGGSQNTLY | 0.41167596 |
| GAGGGGLTAGAN | ASSATGGSQNTLY | 0.18482153 |
| FVDALVNM | ASSATGGSQNTLY | 0.2580961 |
| FLTVAPEL | ASSATGGSQNTLY | 0.106025316 |
| FETLTAL | ASSATGGSQNTLY | 0.3330261 |
| DVINPIEL | ASSATGGSQNTLY | 0.1969957 |
| APVSGAPV | ASSATGGSQNTLY | 0.76985884 |
| ALVGGLAW | ASSATGGSQNTLY | 0.24119045 |
| ALAAAAATAI | ASSATGGSQNTLY | 0.18304041 |
| AFGLGLLL | ASSATGGSQNTLY | 0.11451744 |
| AFFAAASL | ASSATGGSQNTLY | 0.098504305 |
| AAAALPAFSPP | ASSATGGSQNTLY | 0.09235587 |
| VVLAPAL | ASSATGGSQNTLY | 0.20708062 |
| TIGHVDHGKTTL | ASSATGGSQNTLY | 0.116166234 |
| LISNASDALDK | ASSATGGSQNTLY | 0.16885552 |
| LISNASDAL | ASSATGGSQNTLY | 0.27781814 |
| ISNASDALDK | ASSATGGSQNTLY | 0.109619096 |
| NQAETLVYQTE | ASSATGGSQNTLY | 0.3172078 |
| TGGTGGTAGAG | ASSATGGSQNTLY | 0.12407233 |
| TGSTAGGAGGA | ASSATGGSQNTLY | 0.12162883 |

| | | |
|---|---|---|
| DPLAVPVAAGPV | ASSATGGSQNTLY | 0.3572644 |
| GGTGGTPGNGGN | ASSATGGSQNTLY | 0.097069286 |
| PAGLAGVDPATL | ASSATGGSQNTLY | 0.62030476 |
| GNNLVVEAPHAGASV | ASSATGGSQNTLY | 0.085258074 |
| KVTRIMSASTLEQPA | ASSATGGSQNTLY | 0.20591417 |
| LGFALPIALPRRTVV | ASSATGGSQNTLY | 0.31264794 |
| TRLGNLTARKLVVLV | ASSATGGSQNTLY | 0.07899773 |
| WSQQRRLIYFPSAGP | ASSATGGSQNTLY | 0.10416396 |
| GPNGNPDPMAAAVDIR | ASSATGGSQNTLY | 0.38996166 |
| TQRCVSVADLRAAASR | ASSATGGSQNTLY | 0.38860548 |
| QFVLDVAEHPVEVLTETV | ASSATGGSQNTLY | 0.7535172 |
| GGAGGTGGAGGNGGTQAGDG | ASSATGGSQNTLY | 0.17346638 |
| PLLATLRAIAVDVDAKPAQV | ASSATGGSQNTLY | 0.16987267 |
| APKFRTGGYMIDFWGVGYQVAKRM | ASSATGGSQNTLY | 0.3258764 |
| GAGGAGGAGGAAGGPGGTGGTGGN | ASSATGGSQNTLY | 0.4799708 |
| IGALGLLATDVAPAGSVAFGVALA | ASSATGGSQNTLY | 0.46372163 |
| AGGAGGIGGNGGAGGAGGVLMGNG | ASSATGGSQNTLY | 0.44614312 |
| ADELIGLL | ASSATGGSQNTLY | 0.15929933 |
| ADRALKA | ASSATGGSQNTLY | 0.26415038 |
| AGAVGQVGGAGGA | ASSATGGSQNTLY | 0.18660755 |
| AGGGGGLGGAP | ASSATGGSQNTLY | 0.24450547 |
| AGGLLYGNGGAG | ASSATGGSQNTLY | 0.35151786 |
| AGLESIGL | ASSATGGSQNTLY | 0.22351262 |
| AGLKTGII | ASSATGGSQNTLY | 0.16183923 |
| ASLAALLL | ASSATGGSQNTLY | 0.21012454 |
| ASVINISSI | ASSATGGSQNTLY | 0.5685452 |
| ATPPGPA | ASSATGGSQNTLY | 0.6197078 |
| ATVASALV | ASSATGGSQNTLY | 0.55991733 |
| ATVASLIL | ASSATGGSQNTLY | 0.6620221 |
| AVNGPVLIRH | ASSATGGSQNTLY | 0.5503885 |
| DGAAGAVGI | ASSATGGSQNTLY | 0.48018092 |
| DPLLHGI | ASSATGGSQNTLY | 0.111095496 |
| ELGAALGAL | ASSATGGSQNTLY | 0.14667591 |
| ELGRTQAV | ASSATGGSQNTLY | 0.17580783 |
| FVAGGYLS | ASSATGGSQNTLY | 0.30701917 |
| GASGGAGGAGGW | ASSATGGSQNTLY | 0.12493417 |
| GGAGAGGDGGAGGI | ASSATGGSQNTLY | 0.3627755 |
| GTGHNGSQPGMGGNG | ASSATGGSQNTLY | 0.20913967 |
| GYAAASASAS | ASSATGGSQNTLY | 0.14723802 |
| HQVDVAAVQQG | ASSATGGSQNTLY | 0.66498166 |
| IGAAIAHT | ASSATGGSQNTLY | 0.14936274 |
| IGMIEVAQL | ASSATGGSQNTLY | 0.17360708 |
| IRLAPEHTSAMD | ASSATGGSQNTLY | 0.19824527 |
| IYNVSVL | ASSATGGSQNTLY | 0.5033996 |
| KQGFLGM | ASSATGGSQNTLY | 0.15313442 |
| LESVITQL | ASSATGGSQNTLY | 0.48403555 |
| LGLGALVVV | ASSATGGSQNTLY | 0.32917476 |
| LIGNGGNGGSGGI | ASSATGGSQNTLY | 0.5281047 |
| LNAINAPTL | ASSATGGSQNTLY | 0.14118282 |
| LQLLNTGV | ASSATGGSQNTLY | 0.5214486 |
| LQTSIEF | ASSATGGSQNTLY | 0.31080082 |

| | | |
|---|---|---|
| MSFAVSEL | ASSATGGSQNTLY | 0.8172618 |
| NTGALGL | ASSATGGSQNTLY | 0.26664048 |
| PPGAGGAGGAGTT | ASSATGGSQNTLY | 0.2747066 |
| QLVGLGI | ASSATGGSQNTLY | 0.20407808 |
| RFLEQVHQL | ASSATGGSQNTLY | 0.51966345 |
| RGVGYVLR | ASSATGGSQNTLY | 0.13099308 |
| SATGVFAGVI | ASSATGGSQNTLY | 0.2364704 |
| SLPAITLP | ASSATGGSQNTLY | 0.14175102 |
| STALAFTV | ASSATGGSQNTLY | 0.24513716 |
| SVLTVGLGK | ASSATGGSQNTLY | 0.42735967 |
| TALAGAGSGI | ASSATGGSQNTLY | 0.1332037 |
| TGLPFTGI | ASSATGGSQNTLY | 0.41187495 |
| VAEHVTV | ASSATGGSQNTLY | 0.20917521 |
| VDIVLNSL | ASSATGGSQNTLY | 0.16399945 |
| VLTGGGAVL | ASSATGGSQNTLY | 0.4694205 |
| VNVGLAYGI | ASSATGGSQNTLY | 0.18978323 |
| VSSAAVGAV | ASSATGGSQNTLY | 0.68873143 |
| VTSVYEAGV | ASSATGGSQNTLY | 0.19044296 |
| WVALDRGA | ASSATGGSQNTLY | 0.32929185 |
| YLVGSVVI | ASSATGGSQNTLY | 0.36001366 |
| GPPGTGKTLL | ASSATGGSQNTLY | 0.12113218 |
| GVGMTKF | ASSATGGSQNTLY | 0.17314483 |
| IGIGGSDL | ASSATGGSQNTLY | 0.5858325 |
| IGTLLAEL | ASSATGGSQNTLY | 0.45791382 |
| IYTSGTT | ASSATGGSQNTLY | 0.80719006 |
| LSFPAGV | ASSATGGSQNTLY | 0.20632583 |
| LYTSGTT | ASSATGGSQNTLY | 0.18144795 |
| VGAFPFGF | ASSATGGSQNTLY | 0.26251122 |
| VGAGTVEF | ASSATGGSQNTLY | 0.61027265 |
| VGKSTLLL | ASSATGGSQNTLY | 0.79699826 |
| AALAHAGGGIV | ASSATGGSQNTLY | 0.55415785 |
| AALVAVSLL | ASSATGGSQNTLY | 0.20448588 |
| AAVSRAVAAL | ASSATGGSQNTLY | 0.24351679 |
| AGGSSGIGGF | ASSATGGSQNTLY | 0.24639452 |
| AGLTVANANG | ASSATGGSQNTLY | 0.21726802 |
| AGLVGAVL | ASSATGGSQNTLY | 0.5152718 |
| ALSQAFTV | ASSATGGSQNTLY | 0.28736874 |
| ALTFSVSL | ASSATGGSQNTLY | 0.19626534 |
| ALTGAALVAL | ASSATGGSQNTLY | 0.26064748 |
| ASFSIGAGVL | ASSATGGSQNTLY | 0.36983716 |
| DGSVYKI | ASSATGGSQNTLY | 0.6751071 |
| EQLAEIV | ASSATGGSQNTLY | 0.6248282 |
| FAVSELTVM | ASSATGGSQNTLY | 0.7045187 |
| GAAPLAAGL | ASSATGGSQNTLY | 0.54331464 |
| GADGGDGGSGGAGG | ASSATGGSQNTLY | 0.62482744 |
| GFGLDTGGAGG | ASSATGGSQNTLY | 0.09857003 |
| GLASGGLL | ASSATGGSQNTLY | 0.14449081 |
| GRIAGAAVLL | ASSATGGSQNTLY | 0.2810398 |
| GSAGSLNTSF | ASSATGGSQNTLY | 0.24799676 |
| GTGQIGI | ASSATGGSQNTLY | 0.13943726 |
| GTLIALIL | ASSATGGSQNTLY | 0.38101885 |

| | | |
|---|---|---|
| IAATMGFVVG | ASSATGGSQNTLY | 0.264104 |
| IFAQALL | ASSATGGSQNTLY | 0.15593758 |
| IGAAVALGF | ASSATGGSQNTLY | 0.52750075 |
| IGLTPLAL | ASSATGGSQNTLY | 0.17980559 |
| ILGSNTSTL | ASSATGGSQNTLY | 0.219012 |
| IVFSLTVL | ASSATGGSQNTLY | 0.22670484 |
| LASKATGF | ASSATGGSQNTLY | 0.59491247 |
| LASVALIL | ASSATGGSQNTLY | 0.16977498 |
| LFAVGLTL | ASSATGGSQNTLY | 0.38664663 |
| LFNGGIGTYI | ASSATGGSQNTLY | 0.31553283 |
| LTLGAGII | ASSATGGSQNTLY | 0.52930826 |
| LVAAALGDI | ASSATGGSQNTLY | 0.14981638 |
| NTAFVQLGI | ASSATGGSQNTLY | 0.4993226 |
| QLFSGAEL | ASSATGGSQNTLY | 0.256149 |
| SAATFITL | ASSATGGSQNTLY | 0.807376 |
| SNYGLANLGV | ASSATGGSQNTLY | 0.24870068 |
| SYGGAYAVM | ASSATGGSQNTLY | 0.2888589 |
| TTGVGGLTLG | ASSATGGSQNTLY | 0.24002561 |
| TTQTIYI | ASSATGGSQNTLY | 0.75998867 |
| VAAGAGLGAVI | ASSATGGSQNTLY | 0.1211395 |
| VAAVVTLL | ASSATGGSQNTLY | 0.2777785 |
| VGFLGGLI | ASSATGGSQNTLY | 0.17940155 |
| VGVIPILL | ASSATGGSQNTLY | 0.40282845 |
| VIFASSLI | ASSATGGSQNTLY | 0.48266336 |
| VKGGDGGAAAG | ASSATGGSQNTLY | 0.15874083 |
| VTKTFTNI | ASSATGGSQNTLY | 0.5765507 |
| VVAAAILL | ASSATGGSQNTLY | 0.39655408 |
| YAALDVEL | ASSATGGSQNTLY | 0.217009 |
| LASVALLL | ASSATGGSQNTLY | 0.6893213 |
| LDAVHRV | ASSATGGSQNTLY | 0.23109199 |
| QGLSSSIF | ASSATGGSQNTLY | 0.65819865 |
| RGTFANIRL | ASSATGGSQNTLY | 0.28808594 |
| RQLWWGHRI | ASSATGGSQNTLY | 0.32469216 |
| YGPPGTGKTLL | ASSATGGSQNTLY | 0.53326935 |
| AALVGAPANVA | ASSATGGSQNTLY | 0.22338073 |
| AATFRRTE | ASSATGGSQNTLY | 0.18984914 |
| AGALGGGI | ASSATGGSQNTLY | 0.60979724 |
| AGGAGGAGGIGAG | ASSATGGSQNTLY | 0.8250977 |
| AGSGALGL | ASSATGGSQNTLY | 0.8203572 |
| AQIADQI | ASSATGGSQNTLY | 0.67665464 |
| ASAVAVTYL | ASSATGGSQNTLY | 0.58630943 |
| ASTVLGGTGG | ASSATGGSQNTLY | 0.68041164 |
| AVGGEPAK | ASSATGGSQNTLY | 0.8592303 |
| CEQGTFGGTC | ASSATGGSQNTLY | 0.835126 |
| DEGKAPAG | ASSATGGSQNTLY | 0.8153113 |
| DTVVFPAL | ASSATGGSQNTLY | 0.5520238 |
| FEGITMAG | ASSATGGSQNTLY | 0.78456646 |
| FPLVVQD | ASSATGGSQNTLY | 0.7626519 |
| GAFTTPQ | ASSATGGSQNTLY | 0.68783444 |
| GFGGFGTAGGVGGA | ASSATGGSQNTLY | 0.84609437 |
| GFLAQFL | ASSATGGSQNTLY | 0.82278347 |

| | | |
|---|---|---|
| GGAGGAGGNAGVA | ASSATGGSQNTLY | 0.70890385 |
| GGVGTAGAGGAGGAGG | ASSATGGSQNTLY | 0.7399124 |
| GNGGGGDAGNAGSGGN | ASSATGGSQNTLY | 0.80751383 |
| GTGGAGGAGNDAGS | ASSATGGSQNTLY | 0.8285167 |
| GTTNTGWFNS | ASSATGGSQNTLY | 0.82784 |
| IFMGPLFTL | ASSATGGSQNTLY | 0.8822603 |
| ILAHAIEAI | ASSATGGSQNTLY | 0.8513589 |
| IRSAIREAL | ASSATGGSQNTLY | 0.8354436 |
| ITFAGGG | ASSATGGSQNTLY | 0.23759931 |
| ITGMSGHQEV | ASSATGGSQNTLY | 0.44554186 |
| IVAGLACTL | ASSATGGSQNTLY | 0.76241153 |
| IVGGGLAGA | ASSATGGSQNTLY | 0.70258224 |
| IVTGVLNI | ASSATGGSQNTLY | 0.40592283 |
| LAQQGCI | ASSATGGSQNTLY | 0.74448586 |
| LCLHVEN | ASSATGGSQNTLY | 0.7080457 |
| LDAISDGS | ASSATGGSQNTLY | 0.6073267 |
| LGGFAAML | ASSATGGSQNTLY | 0.8412074 |
| LGGKNPMI | ASSATGGSQNTLY | 0.70250016 |
| LGPKGVAAIL | ASSATGGSQNTLY | 0.7266027 |
| LMVGVSI | ASSATGGSQNTLY | 0.686599 |
| LVGAAFVL | ASSATGGSQNTLY | 0.83715296 |
| LVLAVAPL | ASSATGGSQNTLY | 0.62084097 |
| MAAIAVIDAA | ASSATGGSQNTLY | 0.8127471 |
| NNGINAELQ | ASSATGGSQNTLY | 0.66172355 |
| RGEPLVLVHG | ASSATGGSQNTLY | 0.8577502 |
| RGGAGGAAGLI | ASSATGGSQNTLY | 0.5412791 |
| RSYRMHPAV | ASSATGGSQNTLY | 0.82925606 |
| SAAHLLAGQLA | ASSATGGSQNTLY | 0.7516556 |
| SAAVGSY | ASSATGGSQNTLY | 0.8892622 |
| SPHAVLHRI | ASSATGGSQNTLY | 0.75007796 |
| SQIAKVL | ASSATGGSQNTLY | 0.8130994 |
| TALGHLVP | ASSATGGSQNTLY | 0.7586576 |
| TAPTTSGAA | ASSATGGSQNTLY | 0.793771 |
| TDGDDGNGGSPGT | ASSATGGSQNTLY | 0.4695073 |
| TDGLGEN | ASSATGGSQNTLY | 0.76855534 |
| TGGPPLPGQA | ASSATGGSQNTLY | 0.4642906 |
| TIPAITI | ASSATGGSQNTLY | 0.7921728 |
| TIPGFTI | ASSATGGSQNTLY | 0.79821616 |
| TSIASGALA | ASSATGGSQNTLY | 0.43044272 |
| TVTLPGAAI | ASSATGGSQNTLY | 0.845899 |
| VASGLSAALP | ASSATGGSQNTLY | 0.7238596 |
| VGAGVIGI | ASSATGGSQNTLY | 0.69563377 |
| VGPKRFGV | ASSATGGSQNTLY | 0.87025756 |
| VNVAFPD | ASSATGGSQNTLY | 0.76070917 |
| VSMTNSGVS | ASSATGGSQNTLY | 0.8629291 |
| VTDVLGDLSV | ASSATGGSQNTLY | 0.78689665 |
| VVVVGAGFAGL | ASSATGGSQNTLY | 0.7215258 |
| VYTVGTVV | ASSATGGSQNTLY | 0.7786221 |
| WTTFSGP | ASSATGGSQNTLY | 0.55999994 |
| YDELVEAF | ASSATGGSQNTLY | 0.6880086 |
| YGTGGAGGNGG | ASSATGGSQNTLY | 0.8591637 |

| | | |
|---|---|---|
| YTNVMAV | ASSATGGSQNTLY | 0.8105607 |
| FLEQVHQL | ASSATGGSQNTLY | 0.7961374 |
| GLSLLAVAV | ASSATGGSQNTLY | 0.7665565 |
| LEQVHQL | ASSATGGSQNTLY | 0.74596953 |
| LNAAGAI | ASSATGGSQNTLY | 0.7746484 |
| PAVHASL | ASSATGGSQNTLY | 0.8067184 |
| TELEDDA | ASSATGGSQNTLY | 0.81245005 |
| TPPGPPN | ASSATGGSQNTLY | 0.8008877 |
| TPPPPPPP | ASSATGGSQNTLY | 0.6288229 |
| TQSPALI | ASSLGWGNYAEQF | 0.8132657 |
| EFAAAFPQ | ASSLGWGNYAEQF | 0.7978046 |
| LLSGNLDVL | ASSLGWGNYAEQF | 0.7933433 |
| GPANLGNYNI | ASSLGWGNYAEQF | 0.80456024 |
| TLGPTLP | ASSLGWGNYAEQF | 0.80044186 |
| VYAYSSCCK | ASSLGWGNYAEQF | 0.7742446 |
| APAPAPAPNI | ASSLGWGNYAEQF | 0.76494074 |
| PGGPVPIP | ASSLGWGNYAEQF | 0.7950822 |
| FLTVALEP | ASSLGWGNYAEQF | 0.81208694 |
| SQYFGANL | ASSLGWGNYAEQF | 0.80988204 |
| GGVKPAA | ASSLGWGNYAEQF | 0.8132752 |
| GGTGGAGGAGGW | ASSLGWGNYAEQF | 0.81542504 |
| GVGGVAGLGGAGG | ASSLGWGNYAEQF | 0.8037834 |
| VLLVLATTI | ASSLGWGNYAEQF | 0.50762326 |
| IGIFAAML | ASSLGWGNYAEQF | 0.6513268 |
| AVGDLID | ASSLGWGNYAEQF | 0.74439776 |
| NISPLPL | ASSLGWGNYAEQF | 0.77690285 |
| VTAVFVAI | ASSLGWGNYAEQF | 0.43413228 |
| SGDDNDAAL | ASSLGWGNYAEQF | 0.73947597 |
| AVAALAHS | ASSLGWGNYAEQF | 0.7864746 |
| GGVTVMATP | ASSLGWGNYAEQF | 0.7525974 |
| AGLAANVRD | ASSLGWGNYAEQF | 0.8115325 |
| TNRPDLIDPALLRPG | ASSLGWGNYAEQF | 0.7771976 |
| LANLGVTAL | ASSLGWGNYAEQF | 0.75158536 |
| TGERIEL | ASSLGWGNYAEQF | 0.73339427 |
| NGEELAA | ASSLGWGNYAEQF | 0.8071641 |
| VLSLVWG | ASSLGWGNYAEQF | 0.731714 |
| IGYALWL | ASSLGWGNYAEQF | 0.8155237 |
| LSLLLAPL | ASSLGWGNYAEQF | 0.7040742 |
| TLDEALK | ASSLGWGNYAEQF | 0.81754124 |
| TDGAGTAEGG | ASSLGWGNYAEQF | 0.6381029 |
| NAPGGAGGNGG | ASSLGWGNYAEQF | 0.8121043 |
| VSVFMGMGGNF | ASSLGWGNYAEQF | 0.7935792 |
| LASYLAAL | ASSLGWGNYAEQF | 0.82245034 |
| LVGGTPAL | ASSLGWGNYAEQF | 0.7974397 |
| VRNADVL | ASSLGWGNYAEQF | 0.8102423 |
| TCSSPFLVSAM | ASSLGWGNYAEQF | 0.79047024 |
| VHITDAE | ASSLGWGNYAEQF | 0.80429626 |
| EYAAPSF | ASSLGWGNYAEQF | 0.68080306 |
| VVDAFHGGL | ASSLGWGNYAEQF | 0.7958826 |
| ASVVDPQ | ASSLGWGNYAEQF | 0.6326044 |
| SYNEFGGLS | ASSLGWGNYAEQF | 0.80171615 |

| | | |
|---|---|---|
| DPDGSVY | ASSLGWGNYAEQF | 0.80661815 |
| MVSTIHATE | ASSLGWGNYAEQF | 0.39408407 |
| LAAGVADPV | ASSLGWGNYAEQF | 0.8091849 |
| TKEQIAA | ASSLGWGNYAEQF | 0.73123294 |
| SLASPAAL | ASSLGWGNYAEQF | 0.7178244 |
| IGLVFYGV | ASSLGWGNYAEQF | 0.81557286 |
| FLLDASNL | ASSLGWGNYAEQF | 0.7939894 |
| YQVGNFP | ASSLGWGNYAEQF | 0.8135522 |
| ESLLSQST | ASSLGWGNYAEQF | 0.7634571 |
| LFTVGFAAVG | ASSLGWGNYAEQF | 0.7881739 |
| ATPPTPT | ASSLGWGNYAEQF | 0.7940712 |
| TDLATLG | ASSLGWGNYAEQF | 0.4739694 |
| SAGGTGR | ASSLGWGNYAEQF | 0.67657495 |
| GFSNSGS | ASSLGWGNYAEQF | 0.81920457 |
| GGNGGSGGTSV | ASSLGWGNYAEQF | 0.8538384 |
| GTSVATGGAGN | ASSLGWGNYAEQF | 0.8306439 |
| NGGNGGSGGTS | ASSLGWGNYAEQF | 0.7191482 |
| AVVKFPH | ASSLGWGNYAEQF | 0.7207781 |
| TPPPPPP | ASSLGWGNYAEQF | 0.76516545 |
| AAPAAAAAPA | ASSLGWGNYAEQF | 0.85899985 |
| IALNEDL | ASSLGWGNYAEQF | 0.84389067 |
| IVNVSSM | ASSLGWGNYAEQF | 0.84162927 |
| TGIPPAPR | ASSLGWGNYAEQF | 0.68841565 |
| FDLGGGTF | ASSLGWGNYAEQF | 0.8451743 |
| FDLGGGTFDVS | ASSLGWGNYAEQF | 0.8023238 |
| NEPTAAA | ASSLGWGNYAEQF | 0.7699896 |
| NEPTAAALA | ASSLGWGNYAEQF | 0.82092464 |
| GSIINISSI | ASSLGWGNYAEQF | 0.83262837 |
| VAPGNAGTA | ASSLGWGNYAEQF | 0.79754674 |
| TGGGASLE | ASSLGWGNYAEQF | 0.75127906 |
| VAVYNHYKRI | ASSLGWGNYAEQF | 0.80985963 |
| TDPAPATT | ASSLGWGNYAEQF | 0.8601848 |
| TNRPDILDPALLRPG | ASSLGWGNYAEQF | 0.8411794 |
| LTTTTTS | ASSLGWGNYAEQF | 0.8822058 |
| PTGSGKTL | ASSLGWGNYAEQF | 0.86236537 |
| TGSGKTL | ASSLGWGNYAEQF | 0.83204454 |
| TIGHVDHGK | ASSLGWGNYAEQF | 0.35959095 |
| TIGHVDHGKTT | ASSLGWGNYAEQF | 0.5115325 |
| GHVDHGKTT | ASSLGWGNYAEQF | 0.7421512 |
| AQFEHTLL | ASSLGWGNYAEQF | 0.7666228 |
| IVNQDEL | ASSLGWGNYAEQF | 0.32765895 |
| TNGSQFF | ASSLGWGNYAEQF | 0.83675504 |
| TGKGIVE | ASSLGWGNYAEQF | 0.83733994 |
| LISNASDA | ASSLGWGNYAEQF | 0.7321196 |
| AELRTLGELTKIAKAHGAQ | ASSLGWGNYAEQF | 0.8563607 |
| APNAVAAAEPSW | ASSLGWGNYAEQF | 0.7560564 |
| APQVAGAATAVL | ASSLGWGNYAEQF | 0.7332086 |
| ARRDPAAAAAILPTDA | ASSLGWGNYAEQF | 0.6846639 |
| DAAARDVLAEAGLAEYF | ASSLGWGNYAEQF | 0.8542544 |
| DDQIVLSAKANLAKVEQ | ASSLGWGNYAEQF | 0.5851179 |
| DELLPDTGEDSDEDGH | ASSLGWGNYAEQF | 0.8583467 |

| | | |
|---|---|---|
| ELPEAPANRATVPQD | ASSLGWGNYAEQF | 0.635137 |
| FGQSKIERVTGIG | ASSLGWGNYAEQF | 0.8796226 |
| GAGGTGSTAGGAGGAGGAGGLYA | ASSLGWGNYAEQF | 0.6385883 |
| GHGGTGAAGGNGGTGGAGGSGIDG | ASSLGWGNYAEQF | 0.84769535 |
| GNGGNGGAGGEGGPGGAGGAGA | ASSLGWGNYAEQF | 0.7743906 |
| GSTAGNGGNGGNATNGTIAGQPAG | ASSLGWGNYAEQF | 0.88770425 |
| GTGGDGGNGGNGGTGG | ASSLGWGNYAEQF | 0.80214745 |
| GVRRPAGIPADAPV | ASSLGWGNYAEQF | 0.85424745 |
| GYRQVTDAHLLTLARR | ASSLGWGNYAEQF | 0.7315787 |
| IAKMGQDGHDRG | ASSLGWGNYAEQF | 0.832281 |
| KAEPSIEIQYT | ASSLGWGNYAEQF | 0.63344234 |
| KPVRIRMAADEDTVAVT | ASSLGWGNYAEQF | 0.81535006 |
| MPPRIAGMRLLVIKPEP | ASSLGWGNYAEQF | 0.51248527 |
| MKFYAVALFRSRGVPPTR | ASSLGWGNYAEQF | 0.8203063 |
| NAGGFGILGAVAHSP | ASSLGWGNYAEQF | 0.84769326 |
| NEGREITLAMAEM | ASSLGWGNYAEQF | 0.3429807 |
| NELVEWSHPLGHRWRWE | ASSLGWGNYAEQF | 0.8545382 |
| NLGAGGTGGNGGNPG | ASSLGWGNYAEQF | 0.69219923 |
| NTEWPKWETVMAADPVD | ASSLGWGNYAEQF | 0.72465384 |
| PASAAPAETQP | ASSLGWGNYAEQF | 0.8471501 |
| QQSSDGKNGVRAAEITGE | ASSLGWGNYAEQF | 0.820959 |
| RAPDAVTTSWWKE | ASSLGWGNYAEQF | 0.87473506 |
| SGFGNVGAHTSGWFNQ | ASSLGWGNYAEQF | 0.70112467 |
| SGVAGAVGQAASVGGLKV | ASSLGWGNYAEQF | 0.7613215 |
| SNGGAGTGSAGGNGGT | ASSLGWGNYAEQF | 0.8194107 |
| TGGAGGEGGAAG | ASSLGWGNYAEQF | 0.51289946 |
| TGGDGGAAGAGGAPGHG | ASSLGWGNYAEQF | 0.61959565 |
| TMPPLLAAIANAGRHGV | ASSLGWGNYAEQF | 0.87502944 |
| TSSTAGNSGGA | ASSLGWGNYAEQF | 0.87154865 |
| VPATVPVGILC | ASSLGWGNYAEQF | 0.8924351 |
| VQRPDSGGVVLTGRIS | ASSLGWGNYAEQF | 0.7526959 |
| WTPNPVPATKGVPAGLVQAT | ASSLGWGNYAEQF | 0.756711 |
| YALKRALAKFSDRFDVVII | ASSLGWGNYAEQF | 0.8281102 |
| YLIGSVIV | ASSLGWGNYAEQF | 0.8994713 |
| YGQLVAVID | ASSLGWGNYAEQF | 0.9042145 |
| VTVGGMTLPA | ASSLGWGNYAEQF | 0.8845097 |
| VLALPAV | ASSLGWGNYAEQF | 0.80953455 |
| VIPPTLNYET | ASSLGWGNYAEQF | 0.8648995 |
| VGMGGMHPSSG | ASSLGWGNYAEQF | 0.82737863 |
| VGLAGGAATA | ASSLGWGNYAEQF | 0.83310276 |
| VGGAGGAGGSGP | ASSLGWGNYAEQF | 0.8897145 |
| VFAAFMV | ASSLGWGNYAEQF | 0.86549914 |
| TLYDKLR | ASSLGWGNYAEQF | 0.8077528 |
| TLAVFTVL | ASSLGWGNYAEQF | 0.81955326 |
| TGSVQPARV | ASSLGWGNYAEQF | 0.86554146 |
| TGGGAGSGGT | ASSLGWGNYAEQF | 0.88583994 |
| SHLETSL | ASSLGWGNYAEQF | 0.8800423 |
| SFTTGMT | ASSLGWGNYAEQF | 0.9142599 |
| QVEVAKRMGVRQPTVS | ASSLGWGNYAEQF | 0.88521767 |
| QGSGGPALAM | ASSLGWGNYAEQF | 0.88291514 |
| MLAPSNP | ASSLGWGNYAEQF | 0.29113045 |

| | | |
|---|---|---|
| LVNNAAI | ASSLGWGNYAEQF | 0.6270751 |
| LTGEINTLA | ASSLGWGNYAEQF | 0.85814595 |
| LSIFGPV | ASSLGWGNYAEQF | 0.83781624 |
| LLALLANL | ASSLGWGNYAEQF | 0.5578368 |
| LISNASDALDKLR | ASSLGWGNYAEQF | 0.88365006 |
| LINEAVP | ASSLGWGNYAEQF | 0.8205267 |
| LGYAFSGL | ASSLGWGNYAEQF | 0.78415966 |
| LDFVAVAL | ASSLGWGNYAEQF | 0.8860509 |
| LAQPGAE | ASSLGWGNYAEQF | 0.8129351 |
| ISNLGAVTGF | ASSLGWGNYAEQF | 0.7985765 |
| HQEDPGRI | ASSLGWGNYAEQF | 0.8561974 |
| GTPGEPTP | ASSLGWGNYAEQF | 0.9093428 |
| GFFNSGS | ASSLGWGNYAEQF | 0.8019879 |
| GAGGGGLTAGAN | ASSLGWGNYAEQF | 0.88862616 |
| FVDALVNM | ASSLGWGNYAEQF | 0.856632 |
| FLTVAPEL | ASSLGWGNYAEQF | 0.90023446 |
| FETLTAL | ASSLGWGNYAEQF | 0.7272397 |
| DVINPIEL | ASSLGWGNYAEQF | 0.88099277 |
| APVSGAPV | ASSLGWGNYAEQF | 0.8286856 |
| ALVGGLAW | ASSLGWGNYAEQF | 0.9154111 |
| ALAAAAATAI | ASSLGWGNYAEQF | 0.82110614 |
| AFGLGLLL | ASSLGWGNYAEQF | 0.880949 |
| AFFAAASL | ASSLGWGNYAEQF | 0.84989566 |
| AAAALPAFSPP | ASSLGWGNYAEQF | 0.8940848 |
| VVLAPAL | ASSLGWGNYAEQF | 0.68633306 |
| TIGHVDHGKTTL | ASSLGWGNYAEQF | 0.8221278 |
| LISNASDALDK | ASSLGWGNYAEQF | 0.7251538 |
| LISNASDAL | ASSLGWGNYAEQF | 0.86458755 |
| ISNASDALDK | ASSLGWGNYAEQF | 0.87233984 |
| NQAETLVYQTE | ASSLGWGNYAEQF | 0.46413484 |
| TGGTGGTAGAG | ASSLGWGNYAEQF | 0.8906612 |
| TGSTAGGAGGA | ASSLGWGNYAEQF | 0.7756007 |
| DPLAVPVAAGPV | ASSLGWGNYAEQF | 0.7918302 |
| GGTGGTPGNGGN | ASSLGWGNYAEQF | 0.8994719 |
| PAGLAGVDPATL | ASSLGWGNYAEQF | 0.8265035 |
| GNNLVVEAPHAGASV | ASSLGWGNYAEQF | 0.88475245 |
| KVTRIMSASTLEQPA | ASSLGWGNYAEQF | 0.8594958 |
| LGFALPIALPRRTVV | ASSLGWGNYAEQF | 0.84166706 |
| TRLGNLTARKLVVLV | ASSLGWGNYAEQF | 0.88586587 |
| WSQQRRLIYFPSAGP | ASSLGWGNYAEQF | 0.68484604 |
| GPNGNPDPMAAAVDIR | ASSLGWGNYAEQF | 0.81474304 |
| TQRCVSVADLRAAASR | ASSLGWGNYAEQF | 0.89840984 |
| QFVLDVAEHPVEVLTETV | ASSLGWGNYAEQF | 0.6187817 |
| GGAGGTGGAGGNGGTQAGDG | ASSLGWGNYAEQF | 0.8409827 |
| PLLATLRAIAVDVDAKPAQV | ASSLGWGNYAEQF | 0.38357294 |
| APKFRTGGYMIDFWGVGYQVAKRM | ASSLGWGNYAEQF | 0.31512538 |
| GAGGAGGAGGAAGGPGGTGGTGGN | ASSLGWGNYAEQF | 0.34028193 |
| IGALGLLATDVAPAGSVAFGVALA | ASSLGWGNYAEQF | 0.7798412 |
| AGGAGGIGGNGGAGGAGGVLMGNG | ASSLGWGNYAEQF | 0.8044403 |
| ADELIGLL | ASSLGWGNYAEQF | 0.7406864 |
| ADRALKA | ASSLGWGNYAEQF | 0.32340008 |

| | | |
|---|---|---|
| AGAVGQVGGAGGA | ASSLGWGNYAEQF | 0.52247804 |
| AGGGGGLGGAP | ASSLGWGNYAEQF | 0.31599218 |
| AGGLLYGNGGAG | ASSLGWGNYAEQF | 0.44751227 |
| AGLESIGL | ASSLGWGNYAEQF | 0.7110387 |
| AGLKTGII | ASSLGWGNYAEQF | 0.46098155 |
| ASLAALLL | ASSLGWGNYAEQF | 0.4149118 |
| ASVINISSI | ASSLGWGNYAEQF | 0.42324835 |
| ATPPGPA | ASSLGWGNYAEQF | 0.62351286 |
| ATVASALV | ASSLGWGNYAEQF | 0.64838636 |
| ATVASLIL | ASSLGWGNYAEQF | 0.48207378 |
| AVNGPVLIRH | ASSLGWGNYAEQF | 0.83509004 |
| DGAAGAVGI | ASSLGWGNYAEQF | 0.69568086 |
| DPLLHGI | ASSLGWGNYAEQF | 0.76356065 |
| ELGAALGAL | ASSLGWGNYAEQF | 0.11199025 |
| ELGRTQAV | ASSLGWGNYAEQF | 0.25435287 |
| FVAGGYLS | ASSLGWGNYAEQF | 0.42423174 |
| GASGGAGGAGGW | ASSLGWGNYAEQF | 0.5674814 |
| GGAGAGGDGGAGGI | ASSLGWGNYAEQF | 0.21643981 |
| GTGHNGSQPGMGGNG | ASSLGWGNYAEQF | 0.6398731 |
| GYAAASASAS | ASSLGWGNYAEQF | 0.45002395 |
| HQVDVAAVQQG | ASSLGWGNYAEQF | 0.3804611 |
| IGAAIAHT | ASSLGWGNYAEQF | 0.744074 |
| IGMIEVAQL | ASSLGWGNYAEQF | 0.38805014 |
| IRLAPEHTSAMD | ASSLGWGNYAEQF | 0.3983999 |
| IYNVSVL | ASSLGWGNYAEQF | 0.42188585 |
| KQGFLGM | ASSLGWGNYAEQF | 0.8114121 |
| LESVITQL | ASSLGWGNYAEQF | 0.25898433 |
| LGLGALVVV | ASSLGWGNYAEQF | 0.69279695 |
| LIGNGGNGGSGGI | ASSLGWGNYAEQF | 0.43615907 |
| LNAINAPTL | ASSLGWGNYAEQF | 0.64305955 |
| LQLLNTGV | ASSLGWGNYAEQF | 0.25669548 |
| LQTSIEF | ASSLGWGNYAEQF | 0.72888124 |
| MSFAVSEL | ASSLGWGNYAEQF | 0.46604818 |
| NTGALGL | ASSLGWGNYAEQF | 0.8650935 |
| PPGAGGAGGAGTT | ASSLGWGNYAEQF | 0.36179748 |
| QLVGLGI | ASSLGWGNYAEQF | 0.47755018 |
| RFLEQVHQL | ASSLGWGNYAEQF | 0.37103134 |
| RGVGYVLR | ASSLGWGNYAEQF | 0.8042048 |
| SATGVFAGVI | ASSLGWGNYAEQF | 0.19109191 |
| SLPAITLP | ASSLGWGNYAEQF | 0.37243053 |
| STALAFTV | ASSLGWGNYAEQF | 0.37173194 |
| SVLTVGLGK | ASSLGWGNYAEQF | 0.5807352 |
| TALAGAGSGI | ASSLGWGNYAEQF | 0.7017719 |
| TGLPFTGI | ASSLGWGNYAEQF | 0.32463536 |
| VAEHVTV | ASSLGWGNYAEQF | 0.77853703 |
| VDIVLNSL | ASSLGWGNYAEQF | 0.6433346 |
| VLTGGGAVL | ASSLGWGNYAEQF | 0.31608328 |
| VNVGLAYGI | ASSLGWGNYAEQF | 0.79893035 |
| VSSAAVGAV | ASSLGWGNYAEQF | 0.31472883 |
| VTSVYEAGV | ASSLGWGNYAEQF | 0.77577925 |
| WVALDRGA | ASSLGWGNYAEQF | 0.58688384 |

| | | |
|---|---|---|
| YLVGSVVI | ASSLGWGNYAEQF | 0.52969193 |
| GPPGTGKTLL | ASSLGWGNYAEQF | 0.632698 |
| GVGMTKF | ASSLGWGNYAEQF | 0.43435085 |
| IGIGGSDL | ASSLGWGNYAEQF | 0.3334152 |
| IGTLLAEL | ASSLGWGNYAEQF | 0.7553269 |
| IYTSGTT | ASSLGWGNYAEQF | 0.86980045 |
| LSFPAGV | ASSLGWGNYAEQF | 0.87844336 |
| LYTSGTT | ASSLGWGNYAEQF | 0.7845111 |
| VGAFPFGF | ASSLGWGNYAEQF | 0.7221703 |
| VGAGTVEF | ASSLGWGNYAEQF | 0.8089578 |
| VGKSTLLL | ASSLGWGNYAEQF | 0.86234915 |
| AALAHAGGGIV | ASSLGWGNYAEQF | 0.8886625 |
| AALVAVSLL | ASSLGWGNYAEQF | 0.8654456 |
| AAVSRAVAAL | ASSLGWGNYAEQF | 0.6941442 |
| AGGSSGIGGF | ASSLGWGNYAEQF | 0.8632282 |
| AGLTVANANG | ASSLGWGNYAEQF | 0.8399246 |
| AGLVGAVL | ASSLGWGNYAEQF | 0.8309152 |
| ALSQAFTV | ASSLGWGNYAEQF | 0.86123276 |
| ALTFSVSL | ASSLGWGNYAEQF | 0.8370966 |
| ALTGAALVAL | ASSLGWGNYAEQF | 0.7824009 |
| ASFSIGAGVL | ASSLGWGNYAEQF | 0.7168283 |
| DGSVYKI | ASSLGWGNYAEQF | 0.8643828 |
| EQLAEIV | ASSLGWGNYAEQF | 0.8745333 |
| FAVSELTVM | ASSLGWGNYAEQF | 0.87241983 |
| GAAPLAAGL | ASSLGWGNYAEQF | 0.8841858 |
| GADGGDGGSGGAGG | ASSLGWGNYAEQF | 0.8768113 |
| GFGLDTGGAGG | ASSLGWGNYAEQF | 0.86978793 |
| GLASGGLL | ASSLGWGNYAEQF | 0.38830906 |
| GRIAGAAVLL | ASSLGWGNYAEQF | 0.6072742 |
| GSAGSLNTSF | ASSLGWGNYAEQF | 0.78537536 |
| GTGQIGI | ASSLGWGNYAEQF | 0.8018073 |
| GTLIALIL | ASSLGWGNYAEQF | 0.43746847 |
| IAATMGFVVG | ASSLGWGNYAEQF | 0.8126353 |
| IFAQALL | ASSLGWGNYAEQF | 0.80217886 |
| IGAAVALGF | ASSLGWGNYAEQF | 0.75326914 |
| IGLTPLAL | ASSLGWGNYAEQF | 0.873793 |
| ILGSNTSTL | ASSLGWGNYAEQF | 0.80346954 |
| IVFSLTVL | ASSLGWGNYAEQF | 0.67697495 |
| LASKATGF | ASSLGWGNYAEQF | 0.79432124 |
| LASVALIL | ASSLGWGNYAEQF | 0.8741459 |
| LFAVGLTL | ASSLGWGNYAEQF | 0.77046907 |
| LFNGGIGTYI | ASSLGWGNYAEQF | 0.87090766 |
| LTLGAGII | ASSLGWGNYAEQF | 0.802399 |
| LVAAALGDI | ASSLGWGNYAEQF | 0.87856394 |
| NTAFVQLGI | ASSLGWGNYAEQF | 0.63393724 |
| QLFSGAEL | ASSLGWGNYAEQF | 0.875072 |
| SAATFITL | ASSLGWGNYAEQF | 0.83988494 |
| SNYGLANLGV | ASSLGWGNYAEQF | 0.88834906 |
| SYGGAYAVM | ASSLGWGNYAEQF | 0.8297926 |
| TTGVGGLTLG | ASSLGWGNYAEQF | 0.8646843 |
| TTQTIYI | ASSLGWGNYAEQF | 0.8358389 |

| | | |
|---|---|---|
| VAAGAGLGAVI | ASSLGWGNYAEQF | 0.88310087 |
| VAAVVTLL | ASSLGWGNYAEQF | 0.6793351 |
| VGFLGGLI | ASSLGWGNYAEQF | 0.8198403 |
| VGVIPILL | ASSLGWGNYAEQF | 0.7536031 |
| VIFASSLI | ASSLGWGNYAEQF | 0.83740646 |
| VKGGDGGAAAG | ASSLGWGNYAEQF | 0.8658399 |
| VTKTFTNI | ASSLGWGNYAEQF | 0.29308578 |
| VVAAAILL | ASSLGWGNYAEQF | 0.87077415 |
| YAALDVEL | ASSLGWGNYAEQF | 0.7400522 |
| LASVALLL | ASSLGWGNYAEQF | 0.7314253 |
| LDAVHRV | ASSLGWGNYAEQF | 0.87838936 |
| QGLSSSIF | ASSLGWGNYAEQF | 0.84417886 |
| RGTFANIRL | ASSLGWGNYAEQF | 0.87418014 |
| RQLWWGHRI | ASSLGWGNYAEQF | 0.82298076 |
| YGPPGTGKTLL | ASSLGWGNYAEQF | 0.8502071 |
| AALVGAPANVA | ASSLGWGNYAEQF | 0.8401519 |
| AATFRRTE | ASSLGWGNYAEQF | 0.50683594 |
| AGALGGGI | ASSLGWGNYAEQF | 0.73675966 |
| AGGAGGAGGIGAG | ASSLGWGNYAEQF | 0.8833583 |
| AGSGALGL | ASSLGWGNYAEQF | 0.7852913 |
| AQIADQI | ASSLGWGNYAEQF | 0.8457414 |
| ASAVAVTYL | ASSLGWGNYAEQF | 0.61327606 |
| ASTVLGGTGG | ASSLGWGNYAEQF | 0.53880703 |
| AVGGEPAK | ASSLGWGNYAEQF | 0.67399275 |
| CEQGTFGGTC | ASSLGWGNYAEQF | 0.8257041 |
| DEGKAPAG | ASSLGWGNYAEQF | 0.853263 |
| DTVVFPAL | ASSLGWGNYAEQF | 0.80999553 |
| FEGITMAG | ASSLGWGNYAEQF | 0.4662999 |
| FPLVVQD | ASSLGWGNYAEQF | 0.81576526 |
| GAFTTPQ | ASSLGWGNYAEQF | 0.628963 |
| GFGGFGTAGGVGGA | ASSLGWGNYAEQF | 0.73068726 |
| GFLAQFL | ASSLGWGNYAEQF | 0.7848562 |
| GGAGGAGGNAGVA | ASSLGWGNYAEQF | 0.74048066 |
| GGVGTAGAGGAGGAGG | ASSLGWGNYAEQF | 0.6725377 |
| GNGGGGDAGNAGSGGN | ASSLGWGNYAEQF | 0.46401352 |
| GTGGAGGAGNDAGS | ASSLGWGNYAEQF | 0.84146625 |
| GTTNTGWFNS | ASSLGWGNYAEQF | 0.847314 |
| IFMGPLFTL | ASSLGWGNYAEQF | 0.83741343 |
| ILAHAIEAI | ASSLGWGNYAEQF | 0.8506318 |
| IRSAIREAL | ASSLGWGNYAEQF | 0.8347877 |
| ITFAGGG | ASSLGWGNYAEQF | 0.81838155 |
| ITGMSGHQEV | ASSLGWGNYAEQF | 0.1952425 |
| IVAGLACTL | ASSLGWGNYAEQF | 0.3564548 |
| IVGGGLAGA | ASSLGWGNYAEQF | 0.49017122 |
| IVTGVLNI | ASSLGWGNYAEQF | 0.7027442 |
| LAQQGCI | ASSLGWGNYAEQF | 0.2565359 |
| LCLHVEN | ASSLGWGNYAEQF | 0.81346154 |
| LDAISDGS | ASSLGWGNYAEQF | 0.57674176 |
| LGGFAAML | ASSLGWGNYAEQF | 0.624823 |
| LGGKNPMI | ASSLGWGNYAEQF | 0.840047 |
| LGPKGVAAIL | ASSLGWGNYAEQF | 0.61569124 |

| | | |
|---|---|---|
| LMVGVSI | ASSLGWGNYAEQF | 0.49689156 |
| LVGAAFVL | ASSLGWGNYAEQF | 0.55993474 |
| LVLAVAPL | ASSLGWGNYAEQF | 0.8391862 |
| MAAIAVIDAA | ASSLGWGNYAEQF | 0.43621755 |
| NNGINAELQ | ASSLGWGNYAEQF | 0.8242334 |
| RGEPLVLVHG | ASSLGWGNYAEQF | 0.5695261 |
| RGGAGGAAGLI | ASSLGWGNYAEQF | 0.83483255 |
| RSYRMHPAV | ASSLGWGNYAEQF | 0.3840583 |
| SAAHLLAGQLA | ASSLGWGNYAEQF | 0.8297463 |
| SAAVGSY | ASSLGWGNYAEQF | 0.7499393 |
| SPHAVLHRI | ASSLGWGNYAEQF | 0.86727035 |
| SQIAKVL | ASSLGWGNYAEQF | 0.6687133 |
| TALGHLVP | ASSLGWGNYAEQF | 0.7327135 |
| TAPTTSGAA | ASSLGWGNYAEQF | 0.6614797 |
| TDGDDGNGGSPGT | ASSLGWGNYAEQF | 0.8469339 |
| TDGLGEN | ASSLGWGNYAEQF | 0.33736894 |
| TGGPPLPGQA | ASSLGWGNYAEQF | 0.664479 |
| TIPAITI | ASSLGWGNYAEQF | 0.47453764 |
| TIPGFTI | ASSLGWGNYAEQF | 0.72698855 |
| TSIASGALA | ASSLGWGNYAEQF | 0.79952496 |
| TVTLPGAAI | ASSLGWGNYAEQF | 0.18387339 |
| VASGLSAALP | ASSLGWGNYAEQF | 0.7736442 |
| VGAGVIGI | ASSLGWGNYAEQF | 0.6436531 |
| VGPKRFGV | ASSLGWGNYAEQF | 0.4476014 |
| VNVAFPD | ASSLGWGNYAEQF | 0.83109117 |
| VSMTNSGVS | ASSLGWGNYAEQF | 0.7835763 |
| VTDVLGDLSV | ASSLGWGNYAEQF | 0.8407215 |
| VVVVGAGFAGL | ASSLGWGNYAEQF | 0.7135738 |
| VYTVGTVV | ASSLGWGNYAEQF | 0.6673323 |
| WTTFSGP | ASSLGWGNYAEQF | 0.76005375 |
| YDELVEAF | ASSLGWGNYAEQF | 0.42053658 |
| YGTGGAGGNGG | ASSLGWGNYAEQF | 0.5720812 |
| YTNVMAV | ASSLGWGNYAEQF | 0.8437594 |
| FLEQVHQL | ASSLGWGNYAEQF | 0.8252498 |
| GLSLLAVAV | ASSLGWGNYAEQF | 0.8617344 |
| LEQVHQL | ASSLGWGNYAEQF | 0.7470842 |
| LNAAGAI | ASSLGWGNYAEQF | 0.60950214 |
| PAVHASL | ASSLGWGNYAEQF | 0.73854244 |
| TELEDDA | ASSLGWGNYAEQF | 0.85718226 |
| TPPGPPN | ASSLGWGNYAEQF | 0.8650634 |
| TPPPPPPP | ASSLGWGNYAEQF | 0.84333825 |
| TQSPALI | ASGDRGDTQY | 0.5780281 |
| EFAAAFPQ | ASGDRGDTQY | 0.74977636 |
| LLSGNLDVL | ASGDRGDTQY | 0.70510244 |
| GPANLGNYNI | ASGDRGDTQY | 0.7253902 |
| TLGPTLP | ASGDRGDTQY | 0.85155916 |
| VYAYSSCCK | ASGDRGDTQY | 0.79903656 |
| APAPAPAPNI | ASGDRGDTQY | 0.7378111 |
| PGGPVPIP | ASGDRGDTQY | 0.7511823 |
| FLTVALEP | ASGDRGDTQY | 0.75496995 |
| SQYFGANL | ASGDRGDTQY | 0.820485 |

| | | |
|---|---|---|
| GGVKPAA | ASGDRGDTQY | 0.80025125 |
| GGTGGAGGAGGW | ASGDRGDTQY | 0.88270056 |
| GVGGVAGLGGAGG | ASGDRGDTQY | 0.85190296 |
| VLLVLATTI | ASGDRGDTQY | 0.8662224 |
| IGIFAAML | ASGDRGDTQY | 0.18580958 |
| AVGDLID | ASGDRGDTQY | 0.37197524 |
| NISPLPL | ASGDRGDTQY | 0.7530412 |
| VTAVFVAI | ASGDRGDTQY | 0.7021842 |
| SGDDNDAAL | ASGDRGDTQY | 0.6035569 |
| AVAALAHS | ASGDRGDTQY | 0.78660023 |
| GGVTVMATP | ASGDRGDTQY | 0.6719513 |
| AGLAANVRD | ASGDRGDTQY | 0.6327081 |
| TNRPDLIDPALLRPG | ASGDRGDTQY | 0.84890616 |
| LANLGVTAL | ASGDRGDTQY | 0.7516142 |
| TGERIEL | ASGDRGDTQY | 0.6786825 |
| NGEELAA | ASGDRGDTQY | 0.6992842 |
| VLSLVWG | ASGDRGDTQY | 0.8358947 |
| IGYALWL | ASGDRGDTQY | 0.48214895 |
| LSLLLAPL | ASGDRGDTQY | 0.80011594 |
| TLDEALK | ASGDRGDTQY | 0.64059037 |
| TDGAGTAEGG | ASGDRGDTQY | 0.8345079 |
| NAPGGAGGNGG | ASGDRGDTQY | 0.4871431 |
| VSVFMGMGGNF | ASGDRGDTQY | 0.8513674 |
| LASYLAAL | ASGDRGDTQY | 0.7823827 |
| LVGGTPAL | ASGDRGDTQY | 0.89230794 |
| VRNADVL | ASGDRGDTQY | 0.71073794 |
| TCSSPFLVSAM | ASGDRGDTQY | 0.742398 |
| VHITDAE | ASGDRGDTQY | 0.64023465 |
| EYAAPSF | ASGDRGDTQY | 0.84358615 |
| VVDAFHGGL | ASGDRGDTQY | 0.49580175 |
| ASVVDPQ | ASGDRGDTQY | 0.73759544 |
| SYNEFGGLS | ASGDRGDTQY | 0.42264968 |
| DPDGSVY | ASGDRGDTQY | 0.8154207 |
| MVSTIHATE | ASGDRGDTQY | 0.83294773 |
| LAAGVADPV | ASGDRGDTQY | 0.50515467 |
| TKEQIAA | ASGDRGDTQY | 0.86274314 |
| SLASPAAL | ASGDRGDTQY | 0.80332613 |
| IGLVFYGV | ASGDRGDTQY | 0.6823282 |
| FLLDASNL | ASGDRGDTQY | 0.8808644 |
| YQVGNFP | ASGDRGDTQY | 0.6790432 |
| ESLLSQST | ASGDRGDTQY | 0.8785099 |
| LFTVGFAAVG | ASGDRGDTQY | 0.8176054 |
| ATPPTPT | ASGDRGDTQY | 0.68288267 |
| TDLATLG | ASGDRGDTQY | 0.8104633 |
| SAGGTGR | ASGDRGDTQY | 0.46448773 |
| GFSNSGS | ASGDRGDTQY | 0.61812705 |
| GGNGGSGGTSV | ASGDRGDTQY | 0.8734304 |
| GTSVATGGAGN | ASGDRGDTQY | 0.78955704 |
| NGGNGGSGGTS | ASGDRGDTQY | 0.8139707 |
| AVVKFPH | ASGDRGDTQY | 0.7390105 |
| TPPPPPP | ASGDRGDTQY | 0.7148831 |

| | | |
|---|---|---|
| AAPAAAAAPA | ASGDRGDTQY | 0.7743425 |
| IALNEDL | ASGDRGDTQY | 0.79817295 |
| IVNVSSM | ASGDRGDTQY | 0.81463367 |
| TGIPPAPR | ASGDRGDTQY | 0.79274976 |
| FDLGGGTF | ASGDRGDTQY | 0.69563115 |
| FDLGGGTFDVS | ASGDRGDTQY | 0.7862893 |
| NEPTAAA | ASGDRGDTQY | 0.752854 |
| NEPTAAALA | ASGDRGDTQY | 0.74772406 |
| GSIINISSI | ASGDRGDTQY | 0.7981447 |
| VAPGNAGTA | ASGDRGDTQY | 0.74170196 |
| TGGGASLE | ASGDRGDTQY | 0.74957204 |
| VAVYNHYKRI | ASGDRGDTQY | 0.75816524 |
| TDPAPATT | ASGDRGDTQY | 0.7854121 |
| TNRPDILDPALLRPG | ASGDRGDTQY | 0.80309296 |
| LTTTTTS | ASGDRGDTQY | 0.80109864 |
| PTGSGKTL | ASGDRGDTQY | 0.8114569 |
| TGSGKTL | ASGDRGDTQY | 0.79551196 |
| TIGHVDHGK | ASGDRGDTQY | 0.7811295 |
| TIGHVDHGKTT | ASGDRGDTQY | 0.38444942 |
| GHVDHGKTT | ASGDRGDTQY | 0.6657226 |
| AQFEHTLL | ASGDRGDTQY | 0.76634926 |
| IVNQDEL | ASGDRGDTQY | 0.75758916 |
| TNGSQFF | ASGDRGDTQY | 0.3958097 |
| TGKGIVE | ASGDRGDTQY | 0.7800034 |
| LISNASDA | ASGDRGDTQY | 0.77256715 |
| AELRTLGELTKIAKAHGAQ | ASGDRGDTQY | 0.70935214 |
| APNAVAAAEPSW | ASGDRGDTQY | 0.7961962 |
| APQVAGAATAVL | ASGDRGDTQY | 0.74225104 |
| ARRDPAAAAAILPTDA | ASGDRGDTQY | 0.6640738 |
| DAAARDVLAEAGLAEYF | ASGDRGDTQY | 0.74473006 |
| DDQIVLSAKANLAKVEQ | ASGDRGDTQY | 0.80782676 |
| DELLPDTGEDSDEDGH | ASGDRGDTQY | 0.694381 |
| ELPEAPANRATVPQD | ASGDRGDTQY | 0.7922307 |
| FGQSKIERVTGIG | ASGDRGDTQY | 0.7723954 |
| GAGGTGSTAGGAGGAGGAGGLYA | ASGDRGDTQY | 0.800298 |
| GHGGTGAAGGNGGTGGAGGSGIDG | ASGDRGDTQY | 0.6147362 |
| GNGGNGGAGGEGGPGGAGGAGA | ASGDRGDTQY | 0.78356063 |
| GSTAGNGGNGGNATNGTIAGQPAG | ASGDRGDTQY | 0.75803155 |
| GTGGDGGNGGNGGTGG | ASGDRGDTQY | 0.81597614 |
| GVRRPAGIPADAPV | ASGDRGDTQY | 0.746794 |
| GYRQVTDAHLLTLARR | ASGDRGDTQY | 0.78349805 |
| IAKMGQDGHDRG | ASGDRGDTQY | 0.7560066 |
| KAEPSIEIQYT | ASGDRGDTQY | 0.8127527 |
| KPVRIRMAADEDTVAVT | ASGDRGDTQY | 0.6333168 |
| MPPRIAGMRLLVIKPEP | ASGDRGDTQY | 0.7809384 |
| MKFYAVALFRSRGVPPTR | ASGDRGDTQY | 0.73326766 |
| NAGGFGILGAVAHSP | ASGDRGDTQY | 0.78977966 |
| NEGREITLAMAEM | ASGDRGDTQY | 0.79040265 |
| NELVEWSHPLGHRWRWE | ASGDRGDTQY | 0.69587725 |
| NLGAGGTGGNGGNPG | ASGDRGDTQY | 0.80877614 |
| NTEWPKWETVMAADPVD | ASGDRGDTQY | 0.7371642 |

| | | |
|---|---|---|
| PASAAPAETQP | ASGDRGDTQY | 0.7277211 |
| QQSSDGKNGVRAAEITGE | ASGDRGDTQY | 0.80287635 |
| RAPDAVTTSWWKE | ASGDRGDTQY | 0.75659287 |
| SGFGNVGAHTSGWFNQ | ASGDRGDTQY | 0.79780185 |
| SGVAGAVGQAASVGGLKV | ASGDRGDTQY | 0.73243594 |
| SNGGAGTGSAGGNGGT | ASGDRGDTQY | 0.7989535 |
| TGGAGGEGGAAG | ASGDRGDTQY | 0.8002257 |
| TGGDGGAAGAGGAPGHG | ASGDRGDTQY | 0.7265545 |
| TMPPLLAAIANAGRHGV | ASGDRGDTQY | 0.7022066 |
| TSSTAGNSGGA | ASGDRGDTQY | 0.796308 |
| VPATVPVGILC | ASGDRGDTQY | 0.77645195 |
| VQRPDSGGVVLTGRIS | ASGDRGDTQY | 0.8002258 |
| WTPNPVPATKGVPAGLVQAT | ASGDRGDTQY | 0.69554555 |
| YALKRALAKFSDRFDVVII | ASGDRGDTQY | 0.6709542 |
| YLIGSVIV | ASGDRGDTQY | 0.7073859 |
| YGQLVAVID | ASGDRGDTQY | 0.79148966 |
| VTVGGMTLPA | ASGDRGDTQY | 0.80433637 |
| VLALPAV | ASGDRGDTQY | 0.7680533 |
| VIPPTLNYET | ASGDRGDTQY | 0.6996249 |
| VGMGGMHPSSG | ASGDRGDTQY | 0.7575666 |
| VGLAGGAATA | ASGDRGDTQY | 0.74050474 |
| VGGAGGAGGSGP | ASGDRGDTQY | 0.7220503 |
| VFAAFMV | ASGDRGDTQY | 0.7738745 |
| TLYDKLR | ASGDRGDTQY | 0.77063197 |
| TLAVFTVL | ASGDRGDTQY | 0.7160337 |
| TGSVQPARV | ASGDRGDTQY | 0.7330657 |
| TGGGAGSGGT | ASGDRGDTQY | 0.7439339 |
| SHLETSL | ASGDRGDTQY | 0.7871131 |
| SFTTGMT | ASGDRGDTQY | 0.77141565 |
| QVEVAKRMGVRQPTVS | ASGDRGDTQY | 0.80789435 |
| QGSGGPALAM | ASGDRGDTQY | 0.7871049 |
| MLAPSNP | ASGDRGDTQY | 0.78685427 |
| LVNNAAI | ASGDRGDTQY | 0.39579776 |
| LTGEINTLA | ASGDRGDTQY | 0.5964816 |
| LSIFGPV | ASGDRGDTQY | 0.72410834 |
| LLALLANL | ASGDRGDTQY | 0.73894155 |
| LISNASDALDKLR | ASGDRGDTQY | 0.48514563 |
| LINEAVP | ASGDRGDTQY | 0.77363765 |
| LGYAFSGL | ASGDRGDTQY | 0.73752517 |
| LDFVAVAL | ASGDRGDTQY | 0.67155993 |
| LAQPGAE | ASGDRGDTQY | 0.78345 |
| ISNLGAVTGF | ASGDRGDTQY | 0.7185155 |
| HQEDPGRI | ASGDRGDTQY | 0.7182706 |
| GTPGEPTP | ASGDRGDTQY | 0.7259124 |
| GFFNSGS | ASGDRGDTQY | 0.800825 |
| GAGGGGLTAGAN | ASGDRGDTQY | 0.6004987 |
| FVDALVNM | ASGDRGDTQY | 0.7806907 |
| FLTVAPEL | ASGDRGDTQY | 0.7137533 |
| FETLTAL | ASGDRGDTQY | 0.7950603 |
| DVINPIEL | ASGDRGDTQY | 0.6247492 |
| APVSGAPV | ASGDRGDTQY | 0.7808442 |

| | | |
|---|---|---|
| ALVGGLAW | ASGDRGDTQY | 0.73537135 |
| ALAAAAATAI | ASGDRGDTQY | 0.8123573 |
| AFGLGLLL | ASGDRGDTQY | 0.7326223 |
| AFFAAASL | ASGDRGDTQY | 0.7835455 |
| AAAALPAFSPP | ASGDRGDTQY | 0.7636366 |
| VVLAPAL | ASGDRGDTQY | 0.79289556 |
| TIGHVDHGKTTL | ASGDRGDTQY | 0.61582744 |
| LISNASDALDK | ASGDRGDTQY | 0.74897456 |
| LISNASDAL | ASGDRGDTQY | 0.6756879 |
| ISNASDALDK | ASGDRGDTQY | 0.77167714 |
| NQAETLVYQTE | ASGDRGDTQY | 0.77432406 |
| TGGTGGTAGAG | ASGDRGDTQY | 0.53083223 |
| TGSTAGGAGGA | ASGDRGDTQY | 0.796818 |
| DPLAVPVAAGPV | ASGDRGDTQY | 0.69624823 |
| GGTGGTPGNGGN | ASGDRGDTQY | 0.70250916 |
| PAGLAGVDPATL | ASGDRGDTQY | 0.7903372 |
| GNNLVVEAPHAGASV | ASGDRGDTQY | 0.7292117 |
| KVTRIMSASTLEQPA | ASGDRGDTQY | 0.78850603 |
| LGFALPIALPRRTVV | ASGDRGDTQY | 0.7224145 |
| TRLGNLTARKLVVLV | ASGDRGDTQY | 0.74898565 |
| WSQQRRLIYFPSAGP | ASGDRGDTQY | 0.77944255 |
| GPNGNPDPMAAAVDIR | ASGDRGDTQY | 0.6280527 |
| TQRCVSVADLRAAASR | ASGDRGDTQY | 0.6585612 |
| QFVLDVAEHPVEVLTETV | ASGDRGDTQY | 0.79644144 |
| GGAGGTGGAGGNGGTQAGDG | ASGDRGDTQY | 0.6270125 |
| PLLATLRAIAVDVDAKPAQV | ASGDRGDTQY | 0.6849451 |
| APKFRTGGYMIDFWGVGYQVAKRM | ASGDRGDTQY | 0.48458982 |
| GAGGAGGAGGAAGGPGGTGGTGGN | ASGDRGDTQY | 0.49384376 |
| IGALGLLATDVAPAGSVAFGVALA | ASGDRGDTQY | 0.55222356 |
| AGGAGGIGGNGGAGGAGGVLMGNG | ASGDRGDTQY | 0.65566325 |
| ADELIGLL | ASGDRGDTQY | 0.69561434 |
| ADRALKA | ASGDRGDTQY | 0.64495295 |
| AGAVGQVGGAGGA | ASGDRGDTQY | 0.5363006 |
| AGGGGGLGGAP | ASGDRGDTQY | 0.63627774 |
| AGGLLYGNGGAG | ASGDRGDTQY | 0.55826336 |
| AGLESIGL | ASGDRGDTQY | 0.57532775 |
| AGLKTGII | ASGDRGDTQY | 0.6168478 |
| ASLAALLL | ASGDRGDTQY | 0.6127356 |
| ASVINISSI | ASGDRGDTQY | 0.5423067 |
| ATPPGPA | ASGDRGDTQY | 0.5388055 |
| ATVASALV | ASGDRGDTQY | 0.64180636 |
| ATVASLIL | ASGDRGDTQY | 0.67187494 |
| AVNGPVLIRH | ASGDRGDTQY | 0.6527229 |
| DGAAGAVGI | ASGDRGDTQY | 0.702964 |
| DPLLHGI | ASGDRGDTQY | 0.6489314 |
| ELGAALGAL | ASGDRGDTQY | 0.649546 |
| ELGRTQAV | ASGDRGDTQY | 0.319731 |
| FVAGGYLS | ASGDRGDTQY | 0.4159998 |
| GASGGAGGAGGW | ASGDRGDTQY | 0.56183106 |
| GGAGAGGDGGAGGI | ASGDRGDTQY | 0.59011877 |
| GTGHNGSQPGMGGNG | ASGDRGDTQY | 0.35518208 |

| | | |
|---|---|---|
| GYAAASASAS | ASGDRGDTQY | 0.660603 |
| HQVDVAAVQQG | ASGDRGDTQY | 0.576964 |
| IGAAIAHT | ASGDRGDTQY | 0.52899134 |
| IGMIEVAQL | ASGDRGDTQY | 0.65713155 |
| IRLAPEHTSAMD | ASGDRGDTQY | 0.54156476 |
| IYNVSVL | ASGDRGDTQY | 0.51141506 |
| KQGFLGM | ASGDRGDTQY | 0.54051614 |
| LESVITQL | ASGDRGDTQY | 0.69999444 |
| LGLGALVVV | ASGDRGDTQY | 0.42338622 |
| LIGNGGNGGSGGI | ASGDRGDTQY | 0.66043323 |
| LNAINAPTL | ASGDRGDTQY | 0.55364835 |
| LQLLNTGV | ASGDRGDTQY | 0.6677295 |
| LQTSIEF | ASGDRGDTQY | 0.45861676 |
| MSFAVSEL | ASGDRGDTQY | 0.644182 |
| NTGALGL | ASGDRGDTQY | 0.5446702 |
| PPGAGGAGGAGTT | ASGDRGDTQY | 0.71081406 |
| QLVGLGI | ASGDRGDTQY | 0.54997593 |
| RFLEQVHQL | ASGDRGDTQY | 0.63290715 |
| RGVGYVLR | ASGDRGDTQY | 0.63133395 |
| SATGVFAGVI | ASGDRGDTQY | 0.6663486 |
| SLPAITLP | ASGDRGDTQY | 0.38803285 |
| STALAFTV | ASGDRGDTQY | 0.54669887 |
| SVLTVGLGK | ASGDRGDTQY | 0.5160012 |
| TALAGAGSGI | ASGDRGDTQY | 0.5958889 |
| TGLPFTGI | ASGDRGDTQY | 0.64245677 |
| VAEHVTV | ASGDRGDTQY | 0.436558 |
| VDIVLNSL | ASGDRGDTQY | 0.65961593 |
| VLTGGGAVL | ASGDRGDTQY | 0.550362 |
| VNVGLAYGI | ASGDRGDTQY | 0.5344929 |
| VSSAAVGAV | ASGDRGDTQY | 0.6505194 |
| VTSVYEAGV | ASGDRGDTQY | 0.600963 |
| WVALDRGA | ASGDRGDTQY | 0.65990543 |
| YLVGSVVI | ASGDRGDTQY | 0.5394145 |
| GPPGTGKTLL | ASGDRGDTQY | 0.57833683 |
| GVGMTKF | ASGDRGDTQY | 0.64200747 |
| IGIGGSDL | ASGDRGDTQY | 0.53918886 |
| IGTLLAEL | ASGDRGDTQY | 0.5003303 |
| IYTSGTT | ASGDRGDTQY | 0.66781175 |
| LSFPAGV | ASGDRGDTQY | 0.7873436 |
| LYTSGTT | ASGDRGDTQY | 0.8004557 |
| VGAFPFGF | ASGDRGDTQY | 0.64724636 |
| VGAGTVEF | ASGDRGDTQY | 0.5710213 |
| VGKSTLLL | ASGDRGDTQY | 0.6927567 |
| AALAHAGGGIV | ASGDRGDTQY | 0.8310144 |
| AALVAVSLL | ASGDRGDTQY | 0.8375889 |
| AAVSRAVAAL | ASGDRGDTQY | 0.7761969 |
| AGGSSGIGGF | ASGDRGDTQY | 0.5871892 |
| AGLTVANANG | ASGDRGDTQY | 0.78079045 |
| AGLVGAVL | ASGDRGDTQY | 0.7198213 |
| ALSQAFTV | ASGDRGDTQY | 0.71426785 |
| ALTFSVSL | ASGDRGDTQY | 0.78742635 |

| | | |
|---|---|---|
| ALTGAALVAL | ASGDRGDTQY | 0.78528714 |
| ASFSIGAGVL | ASGDRGDTQY | 0.713184 |
| DGSVYKI | ASGDRGDTQY | 0.66132367 |
| EQLAEIV | ASGDRGDTQY | 0.75964355 |
| FAVSELTVM | ASGDRGDTQY | 0.8209617 |
| GAAPLAAGL | ASGDRGDTQY | 0.7943028 |
| GADGGDGGSGGAGG | ASGDRGDTQY | 0.86249006 |
| GFGLDTGGAGG | ASGDRGDTQY | 0.8256166 |
| GLASGGLL | ASGDRGDTQY | 0.8063048 |
| GRIAGAAVLL | ASGDRGDTQY | 0.2206478 |
| GSAGSLNTSF | ASGDRGDTQY | 0.43266577 |
| GTGQIGI | ASGDRGDTQY | 0.6311871 |
| GTLIALIL | ASGDRGDTQY | 0.7260565 |
| IAATMGFVVG | ASGDRGDTQY | 0.3181336 |
| IFAQALL | ASGDRGDTQY | 0.8057218 |
| IGAAVALGF | ASGDRGDTQY | 0.7108621 |
| IGLTPLAL | ASGDRGDTQY | 0.6289643 |
| ILGSNTSTL | ASGDRGDTQY | 0.81340724 |
| IVFSLTVL | ASGDRGDTQY | 0.69362986 |
| LASKATGF | ASGDRGDTQY | 0.6746984 |
| LASVALIL | ASGDRGDTQY | 0.6341305 |
| LFAVGLTL | ASGDRGDTQY | 0.8523103 |
| LFNGGIGTYI | ASGDRGDTQY | 0.48431626 |
| LTLGAGII | ASGDRGDTQY | 0.81605613 |
| LVAAALGDI | ASGDRGDTQY | 0.656355 |
| NTAFVQLGI | ASGDRGDTQY | 0.84848344 |
| QLFSGAEL | ASGDRGDTQY | 0.4932671 |
| SAATFITL | ASGDRGDTQY | 0.8124567 |
| SNYGLANLGV | ASGDRGDTQY | 0.7151463 |
| SYGGAYAVM | ASGDRGDTQY | 0.8721342 |
| TTGVGGLTLG | ASGDRGDTQY | 0.71433604 |
| TTQTIYI | ASGDRGDTQY | 0.7800243 |
| VAAGAGLGAVI | ASGDRGDTQY | 0.7244544 |
| VAAVVTLL | ASGDRGDTQY | 0.79953444 |
| VGFLGGLI | ASGDRGDTQY | 0.42193738 |
| VGVIPILL | ASGDRGDTQY | 0.7439182 |
| VIFASSLI | ASGDRGDTQY | 0.5276252 |
| VKGGDGGAAAG | ASGDRGDTQY | 0.7784806 |
| VTKTFTNI | ASGDRGDTQY | 0.78880185 |
| VVAAAILL | ASGDRGDTQY | 0.3464521 |
| YAALDVEL | ASGDRGDTQY | 0.82172847 |
| LASVALLL | ASGDRGDTQY | 0.6822235 |
| LDAVHRV | ASGDRGDTQY | 0.627591 |
| QGLSSSIF | ASGDRGDTQY | 0.8219962 |
| RGTFANIRL | ASGDRGDTQY | 0.72858256 |
| RQLWWGHRI | ASGDRGDTQY | 0.8314686 |
| YGPPGTGKTLL | ASGDRGDTQY | 0.69282246 |
| AALVGAPANVA | ASGDRGDTQY | 0.71905434 |
| AATFRRTE | ASGDRGDTQY | 0.8048762 |
| AGALGGGI | ASGDRGDTQY | 0.44532675 |
| AGGAGGAGGIGAG | ASGDRGDTQY | 0.553171 |

| | | |
|---|---|---|
| AGSGALGL | ASGDRGDTQY | 0.84351134 |
| AQIADQI | ASGDRGDTQY | 0.7719759 |
| ASAVAVTYL | ASGDRGDTQY | 0.8357891 |
| ASTVLGGTGG | ASGDRGDTQY | 0.5866165 |
| AVGGEPAK | ASGDRGDTQY | 0.47987026 |
| CEQGTFGGTC | ASGDRGDTQY | 0.6750095 |
| DEGKAPAG | ASGDRGDTQY | 0.8411747 |
| DTVVFPAL | ASGDRGDTQY | 0.84630954 |
| FEGITMAG | ASGDRGDTQY | 0.78061116 |
| FPLVVQD | ASGDRGDTQY | 0.49258685 |
| GAFTTPQ | ASGDRGDTQY | 0.72262144 |
| GFGGFGTAGGVGGA | ASGDRGDTQY | 0.61869615 |
| GFLAQFL | ASGDRGDTQY | 0.6446907 |
| GGAGGAGGNAGVA | ASGDRGDTQY | 0.817258 |
| GGVGTAGAGGAGGAGG | ASGDRGDTQY | 0.6355474 |
| GNGGGGDAGNAGSGGN | ASGDRGDTQY | 0.6484158 |
| GTGGAGGAGNDAGS | ASGDRGDTQY | 0.639772 |
| GTTNTGWFNS | ASGDRGDTQY | 0.73517025 |
| IFMGPLFTL | ASGDRGDTQY | 0.813383 |
| ILAHAIEAI | ASGDRGDTQY | 0.7739377 |
| IRSAIREAL | ASGDRGDTQY | 0.889498 |
| ITFAGGG | ASGDRGDTQY | 0.8214948 |
| ITGMSGHQEV | ASGDRGDTQY | 0.83890474 |
| IVAGLACTL | ASGDRGDTQY | 0.17361233 |
| IVGGGLAGA | ASGDRGDTQY | 0.3760076 |
| IVTGVLNI | ASGDRGDTQY | 0.6705269 |
| LAQQGCI | ASGDRGDTQY | 0.64570427 |
| LCLHVEN | ASGDRGDTQY | 0.39764434 |
| LDAISDGS | ASGDRGDTQY | 0.80003035 |
| LGGFAAML | ASGDRGDTQY | 0.67150784 |
| LGGKNPMI | ASGDRGDTQY | 0.45625514 |
| LGPKGVAAIL | ASGDRGDTQY | 0.79250646 |
| LMVGVSI | ASGDRGDTQY | 0.5742228 |
| LVGAAFVL | ASGDRGDTQY | 0.53617436 |
| LVLAVAPL | ASGDRGDTQY | 0.63303244 |
| MAAIAVIDAA | ASGDRGDTQY | 0.82641554 |
| NNGINAELQ | ASGDRGDTQY | 0.46165788 |
| RGEPLVLVHG | ASGDRGDTQY | 0.7835187 |
| RGGAGGAAGLI | ASGDRGDTQY | 0.6820024 |
| RSYRMHPAV | ASGDRGDTQY | 0.8365935 |
| SAAHLLAGQLA | ASGDRGDTQY | 0.35079548 |
| SAAVGSY | ASGDRGDTQY | 0.80378413 |
| SPHAVLHRI | ASGDRGDTQY | 0.67833066 |
| SQIAKVL | ASGDRGDTQY | 0.8844639 |
| TALGHLVP | ASGDRGDTQY | 0.6182222 |
| TAPTTSGAA | ASGDRGDTQY | 0.66951436 |
| TDGDDGNGGSPGT | ASGDRGDTQY | 0.59257805 |
| TDGLGEN | ASGDRGDTQY | 0.8250805 |
| TGGPPLPGQA | ASGDRGDTQY | 0.3553948 |
| TIPAITI | ASGDRGDTQY | 0.69741154 |
| TIPGFTI | ASGDRGDTQY | 0.45550022 |

| | | |
|---|---|---|
| TSIASGALA | ASGDRGDTQY | 0.77232563 |
| TVTLPGAAI | ASGDRGDTQY | 0.75200367 |
| VASGLSAALP | ASGDRGDTQY | 0.42328307 |
| VGAGVIGI | ASGDRGDTQY | 0.80523324 |
| VGPKRFGV | ASGDRGDTQY | 0.7139919 |
| VNVAFPD | ASGDRGDTQY | 0.55113876 |
| VSMTNSGVS | ASGDRGDTQY | 0.85684574 |
| VTDVLGDLSV | ASGDRGDTQY | 0.6298269 |
| VVVVGAGFAGL | ASGDRGDTQY | 0.8564377 |
| VYTVGTVV | ASGDRGDTQY | 0.75160605 |
| WTTFSGP | ASGDRGDTQY | 0.62083673 |
| YDELVEAF | ASGDRGDTQY | 0.8193251 |
| YGTGGAGGNGG | ASGDRGDTQY | 0.38491762 |
| YTNVMAV | ASGDRGDTQY | 0.5878608 |
| FLEQVHQL | ASGDRGDTQY | 0.83722126 |
| GLSLLAVAV | ASGDRGDTQY | 0.86041427 |
| LEQVHQL | ASGDRGDTQY | 0.8496752 |
| LNAAGAI | ASGDRGDTQY | 0.7326335 |
| PAVHASL | ASGDRGDTQY | 0.64443815 |
| TELEDDA | ASGDRGDTQY | 0.71591216 |
| TPPGPPN | ASGDRGDTQY | 0.85701376 |
| TPPPPPPP | ASGDRGDTQY | 0.8603426 |
| TQSPALI | ASGEDWGGTETLY | 0.84197134 |
| EFAAAFPQ | ASGEDWGGTETLY | 0.4657923 |
| LLSGNLDVL | ASGEDWGGTETLY | 0.8141912 |
| GPANLGNYNI | ASGEDWGGTETLY | 0.749465 |
| TLGPTLP | ASGEDWGGTETLY | 0.7929244 |
| VYAYSSCCK | ASGEDWGGTETLY | 0.7943555 |
| APAPAPAPNI | ASGEDWGGTETLY | 0.7189107 |
| PGGPVPIP | ASGEDWGGTETLY | 0.69629294 |
| FLTVALEP | ASGEDWGGTETLY | 0.5901838 |
| SQYFGANL | ASGEDWGGTETLY | 0.8405249 |
| GGVKPAA | ASGEDWGGTETLY | 0.87089646 |
| GGTGGAGGAGGW | ASGEDWGGTETLY | 0.8615444 |
| GVGGVAGLGGAGG | ASGEDWGGTETLY | 0.8650869 |
| VLLVLATTI | ASGEDWGGTETLY | 0.86988294 |
| IGIFAAML | ASGEDWGGTETLY | 0.84776527 |
| AVGDLID | ASGEDWGGTETLY | 0.20777354 |
| NISPLPL | ASGEDWGGTETLY | 0.37032002 |
| VTAVFVAI | ASGEDWGGTETLY | 0.579438 |
| SGDDNDAAL | ASGEDWGGTETLY | 0.7206539 |
| AVAALAHS | ASGEDWGGTETLY | 0.22195321 |
| GGVTVMATP | ASGEDWGGTETLY | 0.769935 |
| AGLAANVRD | ASGEDWGGTETLY | 0.69445443 |
| TNRPDLIDPALLRPG | ASGEDWGGTETLY | 0.59892434 |
| LANLGVTAL | ASGEDWGGTETLY | 0.8574964 |
| TGERIEL | ASGEDWGGTETLY | 0.70442355 |
| NGEELAA | ASGEDWGGTETLY | 0.559134 |
| VLSLVWG | ASGEDWGGTETLY | 0.50328493 |
| IGYALWL | ASGEDWGGTETLY | 0.83930916 |
| LSLLLAPL | ASGEDWGGTETLY | 0.5479396 |

| | | |
|---|---|---|
| TLDEALK | ASGEDWGGTETLY | 0.84851575 |
| TDGAGTAEGG | ASGEDWGGTETLY | 0.4634732 |
| NAPGGAGGNGG | ASGEDWGGTETLY | 0.86798424 |
| VSVFMGMGGNF | ASGEDWGGTETLY | 0.48671123 |
| LASYLAAL | ASGEDWGGTETLY | 0.8648196 |
| LVGGTPAL | ASGEDWGGTETLY | 0.8144553 |
| VRNADVL | ASGEDWGGTETLY | 0.8937436 |
| TCSSPFLVSAM | ASGEDWGGTETLY | 0.7548558 |
| VHITDAE | ASGEDWGGTETLY | 0.74485576 |
| EYAAPSF | ASGEDWGGTETLY | 0.54203737 |
| VVDAFHGGL | ASGEDWGGTETLY | 0.84478474 |
| ASVVDPQ | ASGEDWGGTETLY | 0.42115635 |
| SYNEFGGLS | ASGEDWGGTETLY | 0.7726065 |
| DPDGSVY | ASGEDWGGTETLY | 0.4120681 |
| MVSTIHATE | ASGEDWGGTETLY | 0.787811 |
| LAAGVADPV | ASGEDWGGTETLY | 0.8568934 |
| TKEQIAA | ASGEDWGGTETLY | 0.20236538 |
| SLASPAAL | ASGEDWGGTETLY | 0.7942474 |
| IGLVFYGV | ASGEDWGGTETLY | 0.63958776 |
| FLLDASNL | ASGEDWGGTETLY | 0.58320427 |
| YQVGNFP | ASGEDWGGTETLY | 0.8539773 |
| ESLLSQST | ASGEDWGGTETLY | 0.7848512 |
| LFTVGFAAVG | ASGEDWGGTETLY | 0.8713908 |
| ATPPTPT | ASGEDWGGTETLY | 0.6352715 |
| TDLATLG | ASGEDWGGTETLY | 0.71578944 |
| SAGGTGR | ASGEDWGGTETLY | 0.81244075 |
| GFSNSGS | ASGEDWGGTETLY | 0.33376187 |
| GGNGGSGGTSV | ASGEDWGGTETLY | 0.4973167 |
| GTSVATGGAGN | ASGEDWGGTETLY | 0.8794471 |
| NGGNGGSGGTS | ASGEDWGGTETLY | 0.8360085 |
| AVVKFPH | ASGEDWGGTETLY | 0.85494167 |
| TPPPPPP | ASGEDWGGTETLY | 0.74530214 |
| AAPAAAAAPA | ASGEDWGGTETLY | 0.68775785 |
| IALNEDL | ASGEDWGGTETLY | 0.74565494 |
| IVNVSSM | ASGEDWGGTETLY | 0.8704632 |
| TGIPPAPR | ASGEDWGGTETLY | 0.8433532 |
| FDLGGGTF | ASGEDWGGTETLY | 0.83807755 |
| FDLGGGTFDVS | ASGEDWGGTETLY | 0.7237435 |
| NEPTAAA | ASGEDWGGTETLY | 0.82515633 |
| NEPTAAALA | ASGEDWGGTETLY | 0.81175596 |
| GSIINISSI | ASGEDWGGTETLY | 0.75870466 |
| VAPGNAGTA | ASGEDWGGTETLY | 0.8579532 |
| TGGGASLE | ASGEDWGGTETLY | 0.8436005 |
| VAVYNHYKRI | ASGEDWGGTETLY | 0.7817321 |
| TDPAPATT | ASGEDWGGTETLY | 0.81198823 |
| TNRPDILDPALLRPG | ASGEDWGGTETLY | 0.7750211 |
| LTTTTTS | ASGEDWGGTETLY | 0.8107695 |
| PTGSGKTL | ASGEDWGGTETLY | 0.78597933 |
| TGSGKTL | ASGEDWGGTETLY | 0.89050305 |
| TIGHVDHGK | ASGEDWGGTETLY | 0.8656077 |
| TIGHVDHGKTT | ASGEDWGGTETLY | 0.83841765 |

| | | |
|---|---|---|
| GHVDHGKTT | ASGEDWGGTETLY | 0.3872267 |
| AQFEHTLL | ASGEDWGGTETLY | 0.6285857 |
| IVNQDEL | ASGEDWGGTETLY | 0.8189353 |
| TNGSQFF | ASGEDWGGTETLY | 0.7970921 |
| TGKGIVE | ASGEDWGGTETLY | 0.55982095 |
| LISNASDA | ASGEDWGGTETLY | 0.8278402 |
| AELRTLGELTKIAKAHGAQ | ASGEDWGGTETLY | 0.7893214 |
| APNAVAAAEPSW | ASGEDWGGTETLY | 0.7455634 |
| APQVAGAATAVL | ASGEDWGGTETLY | 0.85193706 |
| ARRDPAAAAAILPTDA | ASGEDWGGTETLY | 0.79208755 |
| DAAARDVLAEAGLAEYF | ASGEDWGGTETLY | 0.80086684 |
| DDQIVLSAKANLAKVEQ | ASGEDWGGTETLY | 0.8043164 |
| DELLPDTGEDSDEDGH | ASGEDWGGTETLY | 0.87264776 |
| ELPEAPANRATVPQD | ASGEDWGGTETLY | 0.7000543 |
| FGQSKIERVTGIG | ASGEDWGGTETLY | 0.8330873 |
| GAGGTGSTAGGAGGAGGAGGLYA | ASGEDWGGTETLY | 0.7378347 |
| GHGGTGAAGGNGGTGGAGGSGIDG | ASGEDWGGTETLY | 0.8759754 |
| GNGGNGGAGGEGGPGGAGGAGA | ASGEDWGGTETLY | 0.682943 |
| GSTAGNGGNGGNATNGTIAGQPAG | ASGEDWGGTETLY | 0.8520843 |
| GTGGDGGNGGNGGTGG | ASGEDWGGTETLY | 0.77361166 |
| GVRRPAGIPADAPV | ASGEDWGGTETLY | 0.89643025 |
| GYRQVTDAHLLTLARR | ASGEDWGGTETLY | 0.7905222 |
| IAKMGQDGHDRG | ASGEDWGGTETLY | 0.8512598 |
| KAEPSIEIQYT | ASGEDWGGTETLY | 0.81775117 |
| KPVRIRMAADEDTVAVT | ASGEDWGGTETLY | 0.83442503 |
| MPPRIAGMRLLVIKPEP | ASGEDWGGTETLY | 0.6637906 |
| MKFYAVALFRSRGVPPTR | ASGEDWGGTETLY | 0.804765 |
| NAGGFGILGAVAHSP | ASGEDWGGTETLY | 0.7082094 |
| NEGREITLAMAEM | ASGEDWGGTETLY | 0.84142876 |
| NELVEWSHPLGHRWRWE | ASGEDWGGTETLY | 0.8347382 |
| NLGAGGTGGNGGNPG | ASGEDWGGTETLY | 0.6767186 |
| NTEWPKWETVMAADPVD | ASGEDWGGTETLY | 0.8664181 |
| PASAAPAETQP | ASGEDWGGTETLY | 0.79801667 |
| QQSSDGKNGVRAAEITGE | ASGEDWGGTETLY | 0.78026843 |
| RAPDAVTTSWWKE | ASGEDWGGTETLY | 0.87635016 |
| SGFGNVGAHTSGWFNQ | ASGEDWGGTETLY | 0.7898191 |
| SGVAGAVGQAASVGGLKV | ASGEDWGGTETLY | 0.8576536 |
| SNGGAGTGSAGGNGGT | ASGEDWGGTETLY | 0.8081959 |
| TGGAGGEGGAAG | ASGEDWGGTETLY | 0.81043005 |
| TGGDGGAAGAGGAPGHG | ASGEDWGGTETLY | 0.83447033 |
| TMPPLLAAIANAGRHGV | ASGEDWGGTETLY | 0.6902997 |
| TSSTAGNSGGA | ASGEDWGGTETLY | 0.75289893 |
| VPATVPVGILC | ASGEDWGGTETLY | 0.87049353 |
| VQRPDSGGVVLTGRIS | ASGEDWGGTETLY | 0.8687553 |
| WTPNPVPATKGVPAGLVQAT | ASGEDWGGTETLY | 0.859249 |
| YALKRALAKFSDRFDVVII | ASGEDWGGTETLY | 0.756312 |
| YLIGSVIV | ASGEDWGGTETLY | 0.63161755 |
| YGQLVAVID | ASGEDWGGTETLY | 0.8154468 |
| VTVGGMTLPA | ASGEDWGGTETLY | 0.8502668 |
| VLALPAV | ASGEDWGGTETLY | 0.88975877 |
| VIPPTLNYET | ASGEDWGGTETLY | 0.8583374 |

| | | |
|---|---|---|
| VGMGGMHPSSG | ASGEDWGGTETLY | 0.5840185 |
| VGLAGGAATA | ASGEDWGGTETLY | 0.87943614 |
| VGGAGGAGGSGP | ASGEDWGGTETLY | 0.81898403 |
| VFAAFMV | ASGEDWGGTETLY | 0.8118776 |
| TLYDKLR | ASGEDWGGTETLY | 0.8299327 |
| TLAVFTVL | ASGEDWGGTETLY | 0.80649036 |
| TGSVQPARV | ASGEDWGGTETLY | 0.7889045 |
| TGGGAGSGGT | ASGEDWGGTETLY | 0.6242516 |
| SHLETSL | ASGEDWGGTETLY | 0.8631247 |
| SFTTGMT | ASGEDWGGTETLY | 0.881294 |
| QVEVAKRMGVRQPTVS | ASGEDWGGTETLY | 0.87685907 |
| QGSGGPALAM | ASGEDWGGTETLY | 0.87700343 |
| MLAPSNP | ASGEDWGGTETLY | 0.881654 |
| LVNNAAI | ASGEDWGGTETLY | 0.8682864 |
| LTGEINTLA | ASGEDWGGTETLY | 0.20261855 |
| LSIFGPV | ASGEDWGGTETLY | 0.4167775 |
| LLALLANL | ASGEDWGGTETLY | 0.5662115 |
| LISNASDALDKLR | ASGEDWGGTETLY | 0.7800591 |
| LINEAVP | ASGEDWGGTETLY | 0.2697125 |
| LGYAFSGL | ASGEDWGGTETLY | 0.7961843 |
| LDFVAVAL | ASGEDWGGTETLY | 0.7576499 |
| LAQPGAE | ASGEDWGGTETLY | 0.70025766 |
| ISNLGAVTGF | ASGEDWGGTETLY | 0.8857914 |
| HQEDPGRI | ASGEDWGGTETLY | 0.7864177 |
| GTPGEPTP | ASGEDWGGTETLY | 0.554029 |
| GFFNSGS | ASGEDWGGTETLY | 0.7128449 |
| GAGGGGLTAGAN | ASGEDWGGTETLY | 0.8797524 |
| FVDALVNM | ASGEDWGGTETLY | 0.6154507 |
| FLTVAPEL | ASGEDWGGTETLY | 0.8785995 |
| FETLTAL | ASGEDWGGTETLY | 0.5992545 |
| DVINPIEL | ASGEDWGGTETLY | 0.8817849 |
| APVSGAPV | ASGEDWGGTETLY | 0.4854777 |
| ALVGGLAW | ASGEDWGGTETLY | 0.8769231 |
| ALAAAAATAI | ASGEDWGGTETLY | 0.8309696 |
| AFGLGLLL | ASGEDWGGTETLY | 0.90369564 |
| AFFAAASL | ASGEDWGGTETLY | 0.7669256 |
| AAAALPAFSPP | ASGEDWGGTETLY | 0.84807336 |
| VVLAPAL | ASGEDWGGTETLY | 0.741964 |
| TIGHVDHGKTTL | ASGEDWGGTETLY | 0.8807639 |
| LISNASDALDK | ASGEDWGGTETLY | 0.51858693 |
| LISNASDAL | ASGEDWGGTETLY | 0.7908825 |
| ISNASDALDK | ASGEDWGGTETLY | 0.43835598 |
| NQAETLVYQTE | ASGEDWGGTETLY | 0.8160826 |
| TGGTGGTAGAG | ASGEDWGGTETLY | 0.86010873 |
| TGSTAGGAGGA | ASGEDWGGTETLY | 0.20699398 |
| DPLAVPVAAGPV | ASGEDWGGTETLY | 0.883664 |
| GGTGGTPGNGGN | ASGEDWGGTETLY | 0.6659149 |
| PAGLAGVDPATL | ASGEDWGGTETLY | 0.6036308 |
| GNNLVVEAPHAGASV | ASGEDWGGTETLY | 0.8720191 |
| KVTRIMSASTLEQPA | ASGEDWGGTETLY | 0.82347316 |
| LGFALPIALPRRTVV | ASGEDWGGTETLY | 0.8818247 |

| | | |
|---|---|---|
| TRLGNLTARKLVVLV | ASGEDWGGTETLY | 0.71060467 |
| WSQQRRLIYFPSAGP | ASGEDWGGTETLY | 0.7943403 |
| GPNGNPDPMAAAVDIR | ASGEDWGGTETLY | 0.8256396 |
| TQRCVSVADLRAAASR | ASGEDWGGTETLY | 0.39414844 |
| QFVLDVAEHPVEVLTETV | ASGEDWGGTETLY | 0.61714345 |
| GGAGGTGGAGGNGGTQAGDG | ASGEDWGGTETLY | 0.8903084 |
| PLLATLRAIAVDVDAKPAQV | ASGEDWGGTETLY | 0.76646924 |
| APKFRTGGYMIDFWGVGYQVAKRM | ASGEDWGGTETLY | 0.7774519 |
| GAGGAGGAGGAAGGPGGTGGTGGN | ASGEDWGGTETLY | 0.70223844 |
| IGALGLLATDVAPAGSVAFGVALA | ASGEDWGGTETLY | 0.6970403 |
| AGGAGGIGGNGGAGGAGGVLMGNG | ASGEDWGGTETLY | 0.70138067 |
| ADELIGLL | ASGEDWGGTETLY | 0.7706433 |
| ADRALKA | ASGEDWGGTETLY | 0.78792214 |
| AGAVGQVGGAGGA | ASGEDWGGTETLY | 0.7750896 |
| AGGGGGLGGAP | ASGEDWGGTETLY | 0.7310225 |
| AGGLLYGNGGAG | ASGEDWGGTETLY | 0.77362025 |
| AGLESIGL | ASGEDWGGTETLY | 0.75033367 |
| AGLKTGII | ASGEDWGGTETLY | 0.745905 |
| ASLAALLL | ASGEDWGGTETLY | 0.7689573 |
| ASVINISSI | ASGEDWGGTETLY | 0.76011515 |
| ATPPGPA | ASGEDWGGTETLY | 0.6924111 |
| ATVASALV | ASGEDWGGTETLY | 0.7239227 |
| ATVASLIL | ASGEDWGGTETLY | 0.7501893 |
| AVNGPVLIRH | ASGEDWGGTETLY | 0.7727023 |
| DGAAGAVGI | ASGEDWGGTETLY | 0.7667633 |
| DPLLHGI | ASGEDWGGTETLY | 0.7895912 |
| ELGAALGAL | ASGEDWGGTETLY | 0.7737036 |
| ELGRTQAV | ASGEDWGGTETLY | 0.7753244 |
| FVAGGYLS | ASGEDWGGTETLY | 0.4689388 |
| GASGGAGGAGGW | ASGEDWGGTETLY | 0.6707204 |
| GGAGAGGDGGAGGI | ASGEDWGGTETLY | 0.7219599 |
| GTGHNGSQPGMGGNG | ASGEDWGGTETLY | 0.7444084 |
| GYAAASASAS | ASGEDWGGTETLY | 0.41423285 |
| HQVDVAAVQQG | ASGEDWGGTETLY | 0.7701401 |
| IGAAIAHT | ASGEDWGGTETLY | 0.7505243 |
| IGMIEVAQL | ASGEDWGGTETLY | 0.7067046 |
| IRLAPEHTSAMD | ASGEDWGGTETLY | 0.7753067 |
| IYNVSVL | ASGEDWGGTETLY | 0.7264618 |
| KQGFLGM | ASGEDWGGTETLY | 0.71093655 |
| LESVITQL | ASGEDWGGTETLY | 0.75124615 |
| LGLGALVVV | ASGEDWGGTETLY | 0.78297234 |
| LIGNGGNGGSGGI | ASGEDWGGTETLY | 0.6798325 |
| LNAINAPTL | ASGEDWGGTETLY | 0.7822703 |
| LQLLNTGV | ASGEDWGGTETLY | 0.672271 |
| LQTSIEF | ASGEDWGGTETLY | 0.7795245 |
| MSFAVSEL | ASGEDWGGTETLY | 0.6906135 |
| NTGALGL | ASGEDWGGTETLY | 0.7806124 |
| PPGAGGAGGAGTT | ASGEDWGGTETLY | 0.7507367 |
| QLVGLGI | ASGEDWGGTETLY | 0.7888243 |
| RFLEQVHQL | ASGEDWGGTETLY | 0.7457669 |
| RGVGYVLR | ASGEDWGGTETLY | 0.7827232 |

| | | |
|---|---|---|
| SATGVFAGVI | ASGEDWGGTETLY | 0.7648177 |
| SLPAITLP | ASGEDWGGTETLY | 0.7808155 |
| STALAFTV | ASGEDWGGTETLY | 0.65941954 |
| SVLTVGLGK | ASGEDWGGTETLY | 0.7419475 |
| TALAGAGSGI | ASGEDWGGTETLY | 0.6565198 |
| TGLPFTGI | ASGEDWGGTETLY | 0.76372975 |
| VAEHVTV | ASGEDWGGTETLY | 0.77694774 |
| VDIVLNSL | ASGEDWGGTETLY | 0.4578293 |
| VLTGGGAVL | ASGEDWGGTETLY | 0.78388417 |
| VNVGLAYGI | ASGEDWGGTETLY | 0.67773616 |
| VSSAAVGAV | ASGEDWGGTETLY | 0.71727717 |
| VTSVYEAGV | ASGEDWGGTETLY | 0.7793986 |
| WVALDRGA | ASGEDWGGTETLY | 0.7480525 |
| YLVGSVVI | ASGEDWGGTETLY | 0.7681495 |
| GPPGTGKTLL | ASGEDWGGTETLY | 0.73217356 |
| GVGMTKF | ASGEDWGGTETLY | 0.74451005 |
| IGIGGSDL | ASGEDWGGTETLY | 0.76517284 |
| IGTLLAEL | ASGEDWGGTETLY | 0.4904893 |
| IYTSGTT | ASGEDWGGTETLY | 0.68050325 |
| LSFPAGV | ASGEDWGGTETLY | 0.7879208 |
| LYTSGTT | ASGEDWGGTETLY | 0.72819054 |
| VGAFPFGF | ASGEDWGGTETLY | 0.6732563 |
| VGAGTVEF | ASGEDWGGTETLY | 0.37985247 |
| VGKSTLLL | ASGEDWGGTETLY | 0.4249626 |
| AALAHAGGGIV | ASGEDWGGTETLY | 0.48291785 |
| AALVAVSLL | ASGEDWGGTETLY | 0.57354146 |
| AAVSRAVAAL | ASGEDWGGTETLY | 0.81551147 |
| AGGSSGIGGF | ASGEDWGGTETLY | 0.68787634 |
| AGLTVANANG | ASGEDWGGTETLY | 0.41442788 |
| AGLVGAVL | ASGEDWGGTETLY | 0.81997514 |
| ALSQAFTV | ASGEDWGGTETLY | 0.64313245 |
| ALTFSVSL | ASGEDWGGTETLY | 0.6822609 |
| ALTGAALVAL | ASGEDWGGTETLY | 0.58524215 |
| ASFSIGAGVL | ASGEDWGGTETLY | 0.5627235 |
| DGSVYKI | ASGEDWGGTETLY | 0.5241303 |
| EQLAEIV | ASGEDWGGTETLY | 0.4213599 |
| FAVSELTVM | ASGEDWGGTETLY | 0.65460825 |
| GAAPLAAGL | ASGEDWGGTETLY | 0.75568056 |
| GADGGDGGSGGAGG | ASGEDWGGTETLY | 0.7429395 |
| GFGLDTGGAGG | ASGEDWGGTETLY | 0.6856131 |
| GLASGGLL | ASGEDWGGTETLY | 0.7686871 |
| GRIAGAAVLL | ASGEDWGGTETLY | 0.5401311 |
| GSAGSLNTSF | ASGEDWGGTETLY | 0.21316099 |
| GTGQIGI | ASGEDWGGTETLY | 0.3400591 |
| GTLIALIL | ASGEDWGGTETLY | 0.44070137 |
| IAATMGFVVG | ASGEDWGGTETLY | 0.46046874 |
| IFAQALL | ASGEDWGGTETLY | 0.18900214 |
| IGAAVALGF | ASGEDWGGTETLY | 0.53966933 |
| IGLTPLAL | ASGEDWGGTETLY | 0.5991864 |
| ILGSNTSTL | ASGEDWGGTETLY | 0.5353185 |
| IVFSLTVL | ASGEDWGGTETLY | 0.73384404 |

| | | |
|---|---|---|
| LASKATGF | ASGEDWGGTETLY | 0.53923595 |
| LASVALIL | ASGEDWGGTETLY | 0.41257346 |
| LFAVGLTL | ASGEDWGGTETLY | 0.4457096 |
| LFNGGIGTYI | ASGEDWGGTETLY | 0.7911866 |
| LTLGAGII | ASGEDWGGTETLY | 0.40857744 |
| LVAAALGDI | ASGEDWGGTETLY | 0.83664626 |
| NTAFVQLGI | ASGEDWGGTETLY | 0.44618887 |
| QLFSGAEL | ASGEDWGGTETLY | 0.757759 |
| SAATFITL | ASGEDWGGTETLY | 0.3530868 |
| SNYGLANLGV | ASGEDWGGTETLY | 0.71541095 |
| SYGGAYAVM | ASGEDWGGTETLY | 0.6406297 |
| TTGVGGLTLG | ASGEDWGGTETLY | 0.7616019 |
| TTQTIYI | ASGEDWGGTETLY | 0.5101678 |
| VAAGAGLGAVI | ASGEDWGGTETLY | 0.6959438 |
| VAAVVTLL | ASGEDWGGTETLY | 0.53767985 |
| VGFLGGLI | ASGEDWGGTETLY | 0.74316776 |
| VGVIPILL | ASGEDWGGTETLY | 0.3403512 |
| VIFASSLI | ASGEDWGGTETLY | 0.5340775 |
| VKGGDGGAAAG | ASGEDWGGTETLY | 0.45089373 |
| VTKTFTNI | ASGEDWGGTETLY | 0.5352726 |
| VVAAAILL | ASGEDWGGTETLY | 0.7692157 |
| YAALDVEL | ASGEDWGGTETLY | 0.1346237 |
| LASVALLL | ASGEDWGGTETLY | 0.7290245 |
| LDAVHRV | ASGEDWGGTETLY | 0.25129598 |
| QGLSSSIF | ASGEDWGGTETLY | 0.39078987 |
| RGTFANIRL | ASGEDWGGTETLY | 0.6223817 |
| RQLWWGHRI | ASGEDWGGTETLY | 0.6380504 |
| YGPPGTGKTLL | ASGEDWGGTETLY | 0.6302625 |
| AALVGAPANVA | ASGEDWGGTETLY | 0.42183945 |
| AATFRRTE | ASGEDWGGTETLY | 0.5786 |
| AGALGGGI | ASGEDWGGTETLY | 0.6152878 |
| AGGAGGAGGIGAG | ASGEDWGGTETLY | 0.21718344 |
| AGSGALGL | ASGEDWGGTETLY | 0.35962898 |
| AQIADQI | ASGEDWGGTETLY | 0.7882467 |
| ASAVAVTYL | ASGEDWGGTETLY | 0.6221355 |
| ASTVLGGTGG | ASGEDWGGTETLY | 0.77515584 |
| AVGGEPAK | ASGEDWGGTETLY | 0.4417462 |
| CEQGTFGGTC | ASGEDWGGTETLY | 0.47304946 |
| DEGKAPAG | ASGEDWGGTETLY | 0.51893336 |
| DTVVFPAL | ASGEDWGGTETLY | 0.7154637 |
| FEGITMAG | ASGEDWGGTETLY | 0.72306174 |
| FPLVVQD | ASGEDWGGTETLY | 0.63782954 |
| GAFTTPQ | ASGEDWGGTETLY | 0.4489585 |
| GFGGFGTAGGVGGA | ASGEDWGGTETLY | 0.5365796 |
| GFLAQFL | ASGEDWGGTETLY | 0.45643672 |
| GGAGGAGGNAGVA | ASGEDWGGTETLY | 0.5175576 |
| GGVGTAGAGGAGGAGG | ASGEDWGGTETLY | 0.59972245 |
| GNGGGGDAGNAGSGGN | ASGEDWGGTETLY | 0.62664247 |
| GTGGAGGAGNDAGS | ASGEDWGGTETLY | 0.47213995 |
| GTTNTGWFNS | ASGEDWGGTETLY | 0.52270204 |
| IFMGPLFTL | ASGEDWGGTETLY | 0.60617864 |

| | | |
|---|---|---|
| ILAHAIEAI | ASGEDWGGTETLY | 0.7011797 |
| IRSAIREAL | ASGEDWGGTETLY | 0.6752967 |
| ITFAGGG | ASGEDWGGTETLY | 0.75737226 |
| ITGMSGHQEV | ASGEDWGGTETLY | 0.7192904 |
| IVAGLACTL | ASGEDWGGTETLY | 0.6732652 |
| IVGGGLAGA | ASGEDWGGTETLY | 0.19340031 |
| IVTGVLNI | ASGEDWGGTETLY | 0.3825662 |
| LAQQGCI | ASGEDWGGTETLY | 0.45409122 |
| LCLHVEN | ASGEDWGGTETLY | 0.5517191 |
| LDAISDGS | ASGEDWGGTETLY | 0.2792623 |
| LGGFAAML | ASGEDWGGTETLY | 0.64207065 |
| LGGKNPMI | ASGEDWGGTETLY | 0.5392824 |
| LGPKGVAAIL | ASGEDWGGTETLY | 0.43590888 |
| LMVGVSI | ASGEDWGGTETLY | 0.7035987 |
| LVGAAFVL | ASGEDWGGTETLY | 0.48460665 |
| LVLAVAPL | ASGEDWGGTETLY | 0.4950199 |
| MAAIAVIDAA | ASGEDWGGTETLY | 0.38512635 |
| NNGINAELQ | ASGEDWGGTETLY | 0.674194 |
| RGEPLVLVHG | ASGEDWGGTETLY | 0.3165618 |
| RGGAGGAAGLI | ASGEDWGGTETLY | 0.68052226 |
| RSYRMHPAV | ASGEDWGGTETLY | 0.44581822 |
| SAAHLLAGQLA | ASGEDWGGTETLY | 0.7171625 |
| SAAVGSY | ASGEDWGGTETLY | 0.38148388 |
| SPHAVLHRI | ASGEDWGGTETLY | 0.6781986 |
| SQIAKVL | ASGEDWGGTETLY | 0.5548179 |
| TALGHLVP | ASGEDWGGTETLY | 0.82067776 |
| TAPTTSGAA | ASGEDWGGTETLY | 0.58551264 |
| TDGDDGNGGSPGT | ASGEDWGGTETLY | 0.5858667 |
| TDGLGEN | ASGEDWGGTETLY | 0.49587044 |
| TGGPPLPGQA | ASGEDWGGTETLY | 0.73565555 |
| TIPAITI | ASGEDWGGTETLY | 0.30783182 |
| TIPGFTI | ASGEDWGGTETLY | 0.5169508 |
| TSIASGALA | ASGEDWGGTETLY | 0.334083 |
| TVTLPGAAI | ASGEDWGGTETLY | 0.5592233 |
| VASGLSAALP | ASGEDWGGTETLY | 0.66273 |
| VGAGVIGI | ASGEDWGGTETLY | 0.34990925 |
| VGPKRFGV | ASGEDWGGTETLY | 0.6119807 |
| VNVAFPD | ASGEDWGGTETLY | 0.5588639 |
| VSMTNSGVS | ASGEDWGGTETLY | 0.47730038 |
| VTDVLGDLSV | ASGEDWGGTETLY | 0.71334887 |
| VVVVGAGFAGL | ASGEDWGGTETLY | 0.51395845 |
| VYTVGTVV | ASGEDWGGTETLY | 0.7646094 |
| WTTFSGP | ASGEDWGGTETLY | 0.44440538 |
| YDELVEAF | ASGEDWGGTETLY | 0.45383054 |
| YGTGGAGGNGG | ASGEDWGGTETLY | 0.6403949 |
| YTNVMAV | ASGEDWGGTETLY | 0.4681041 |
| FLEQVHQL | ASGEDWGGTETLY | 0.3426197 |
| GLSLLAVAV | ASGEDWGGTETLY | 0.7681476 |
| LEQVHQL | ASGEDWGGTETLY | 0.6043469 |
| LNAAGAI | ASGEDWGGTETLY | 0.78195137 |
| PAVHASL | ASGEDWGGTETLY | 0.3342084 |

| | | |
|---|---|---|
| TELEDDA | ASGEDWGGTETLY | 0.25545377 |
| TPPGPPN | ASGEDWGGTETLY | 0.3874013 |
| TPPPPPPP | ASGEDWGGTETLY | 0.67671263 |
| TQSPALI | ASGDRGENYAEQF | 0.77385736 |
| EFAAAFPQ | ASGDRGENYAEQF | 0.625722 |
| LLSGNLDVL | ASGDRGENYAEQF | 0.20951346 |
| GPANLGNYNI | ASGDRGENYAEQF | 0.39787152 |
| TLGPTLP | ASGDRGENYAEQF | 0.3681296 |
| VYAYSSCCK | ASGDRGENYAEQF | 0.38088673 |
| APAPAPAPNI | ASGDRGENYAEQF | 0.48360187 |
| PGGPVPIP | ASGDRGENYAEQF | 0.39925563 |
| FLTVALEP | ASGDRGENYAEQF | 0.3552224 |
| SQYFGANL | ASGDRGENYAEQF | 0.26959413 |
| GGVKPAA | ASGDRGENYAEQF | 0.52254695 |
| GGTGGAGGAGGW | ASGDRGENYAEQF | 0.5760521 |
| GVGGVAGLGGAGG | ASGDRGENYAEQF | 0.41099188 |
| VLLVLATTI | ASGDRGENYAEQF | 0.7166269 |
| IGIFAAML | ASGDRGENYAEQF | 0.73739773 |
| AVGDLID | ASGDRGENYAEQF | 0.6142617 |
| NISPLPL | ASGDRGENYAEQF | 0.13764158 |
| VTAVFVAI | ASGDRGENYAEQF | 0.21701764 |
| SGDDNDAAL | ASGDRGENYAEQF | 0.27013934 |
| AVAALAHS | ASGDRGENYAEQF | 0.46692917 |
| GGVTVMATP | ASGDRGENYAEQF | 0.15780494 |
| AGLAANVRD | ASGDRGENYAEQF | 0.54206705 |
| TNRPDLIDPALLRPG | ASGDRGENYAEQF | 0.3959725 |
| LANLGVTAL | ASGDRGENYAEQF | 0.25387406 |
| TGERIEL | ASGDRGENYAEQF | 0.72017527 |
| NGEELAA | ASGDRGENYAEQF | 0.34960198 |
| VLSLVWG | ASGDRGENYAEQF | 0.23744464 |
| IGYALWL | ASGDRGENYAEQF | 0.23727441 |
| LSLLLAPL | ASGDRGENYAEQF | 0.6130477 |
| TLDEALK | ASGDRGENYAEQF | 0.21278007 |
| TDGAGTAEGG | ASGDRGENYAEQF | 0.6391295 |
| NAPGGAGGNGG | ASGDRGENYAEQF | 0.23498824 |
| VSVFMGMGGNF | ASGDRGENYAEQF | 0.64205545 |
| LASYLAAL | ASGDRGENYAEQF | 0.19116847 |
| LVGGTPAL | ASGDRGENYAEQF | 0.70016646 |
| VRNADVL | ASGDRGENYAEQF | 0.48664948 |
| TCSSPFLVSAM | ASGDRGENYAEQF | 0.82045186 |
| VHITDAE | ASGDRGENYAEQF | 0.37135017 |
| EYAAPSF | ASGDRGENYAEQF | 0.3903317 |
| VVDAFHGGL | ASGDRGENYAEQF | 0.30698797 |
| ASVVDPQ | ASGDRGENYAEQF | 0.77714825 |
| SYNEFGGLS | ASGDRGENYAEQF | 0.18063822 |
| DPDGSVY | ASGDRGENYAEQF | 0.25795007 |
| MVSTIHATE | ASGDRGENYAEQF | 0.18859029 |
| LAAGVADPV | ASGDRGENYAEQF | 0.35291815 |
| TKEQIAA | ASGDRGENYAEQF | 0.62531 |
| SLASPAAL | ASGDRGENYAEQF | 0.16633862 |
| IGLVFYGV | ASGDRGENYAEQF | 0.30249807 |

| | | |
|---|---|---|
| FLLDASNL | ASGDRGENYAEQF | 0.4692588 |
| YQVGNFP | ASGDRGENYAEQF | 0.22469589 |
| ESLLSQST | ASGDRGENYAEQF | 0.64168453 |
| LFTVGFAAVG | ASGDRGENYAEQF | 0.31077588 |
| ATPPTPT | ASGDRGENYAEQF | 0.7629839 |
| TDLATLG | ASGDRGENYAEQF | 0.3044616 |
| SAGGTGR | ASGDRGENYAEQF | 0.30005318 |
| GFSNSGS | ASGDRGENYAEQF | 0.34315068 |
| GGNGGSGGTSV | ASGDRGENYAEQF | 0.2636733 |
| GTSVATGGAGN | ASGDRGENYAEQF | 0.22833748 |
| NGGNGGSGGTS | ASGDRGENYAEQF | 0.7597989 |
| AVVKFPH | ASGDRGENYAEQF | 0.87240463 |
| TPPPPPP | ASGDRGENYAEQF | 0.86438555 |
| AAPAAAAAPA | ASGDRGENYAEQF | 0.7718772 |
| IALNEDL | ASGDRGENYAEQF | 0.7867143 |
| IVNVSSM | ASGDRGENYAEQF | 0.8302649 |
| TGIPPAPR | ASGDRGENYAEQF | 0.8809817 |
| FDLGGGTF | ASGDRGENYAEQF | 0.88369185 |
| FDLGGGTFDVS | ASGDRGENYAEQF | 0.87696993 |
| NEPTAAA | ASGDRGENYAEQF | 0.7746179 |
| NEPTAAALA | ASGDRGENYAEQF | 0.8614074 |
| GSIINISSI | ASGDRGENYAEQF | 0.8350614 |
| VAPGNAGTA | ASGDRGENYAEQF | 0.8503576 |
| TGGGASLE | ASGDRGENYAEQF | 0.85943747 |
| VAVYNHYKRI | ASGDRGENYAEQF | 0.85156894 |
| TDPAPATT | ASGDRGENYAEQF | 0.80879176 |
| TNRPDILDPALLRPG | ASGDRGENYAEQF | 0.76579 |
| LTTTTTS | ASGDRGENYAEQF | 0.8616305 |
| PTGSGKTL | ASGDRGENYAEQF | 0.8759196 |
| TGSGKTL | ASGDRGENYAEQF | 0.8654338 |
| TIGHVDHGK | ASGDRGENYAEQF | 0.8902953 |
| TIGHVDHGKTT | ASGDRGENYAEQF | 0.8747358 |
| GHVDHGKTT | ASGDRGENYAEQF | 0.86547446 |
| AQFEHTLL | ASGDRGENYAEQF | 0.34864557 |
| IVNQDEL | ASGDRGENYAEQF | 0.65334857 |
| TNGSQFF | ASGDRGENYAEQF | 0.7772148 |
| TGKGIVE | ASGDRGENYAEQF | 0.83302855 |
| LISNASDA | ASGDRGENYAEQF | 0.3863088 |
| AELRTLGELTKIAKAHGAQ | ASGDRGENYAEQF | 0.87120146 |
| APNAVAAAEPSW | ASGDRGENYAEQF | 0.84253025 |
| APQVAGAATAVL | ASGDRGENYAEQF | 0.7842018 |
| ARRDPAAAAAILPTDA | ASGDRGENYAEQF | 0.87699175 |
| DAAARDVLAEAGLAEYF | ASGDRGENYAEQF | 0.8134732 |
| DDQIVLSAKANLAKVEQ | ASGDRGENYAEQF | 0.7553576 |
| DELLPDTGEDSDEDGH | ASGDRGENYAEQF | 0.8046337 |
| ELPEAPANRATVPQD | ASGDRGENYAEQF | 0.8837173 |
| FGQSKIERVTGIG | ASGDRGENYAEQF | 0.7100202 |
| GAGGTGSTAGGAGGAGGAGGLYA | ASGDRGENYAEQF | 0.8808093 |
| GHGGTGAAGGNGGTGGAGGSGIDG | ASGDRGENYAEQF | 0.79627895 |
| GNGGNGGAGGEGGPGGAGGAGA | ASGDRGENYAEQF | 0.8834672 |
| GSTAGNGGNGGNATNGTIAGQPAG | ASGDRGENYAEQF | 0.74149877 |

| | | |
|---|---|---|
| GTGGDGGNGGNGGTGG | ASGDRGENYAEQF | 0.877718 |
| GVRRPAGIPADAPV | ASGDRGENYAEQF | 0.8459863 |
| GYRQVTDAHLLTLARR | ASGDRGENYAEQF | 0.8933672 |
| IAKMGQDGHDRG | ASGDRGENYAEQF | 0.84013855 |
| KAEPSIEIQYT | ASGDRGENYAEQF | 0.86965656 |
| KPVRIRMAADEDTVAVT | ASGDRGENYAEQF | 0.8340034 |
| MPPRIAGMRLLVIKPEP | ASGDRGENYAEQF | 0.86839163 |
| MKFYAVALFRSRGVPPTR | ASGDRGENYAEQF | 0.7080076 |
| NAGGFGILGAVAHSP | ASGDRGENYAEQF | 0.82866144 |
| NEGREITLAMAEM | ASGDRGENYAEQF | 0.68489516 |
| NELVEWSHPLGHRWRWE | ASGDRGENYAEQF | 0.8424885 |
| NLGAGGTGGNGGNPG | ASGDRGENYAEQF | 0.8782885 |
| NTEWPKWETVMAADPVD | ASGDRGENYAEQF | 0.3206706 |
| PASAAPAETQP | ASGDRGENYAEQF | 0.86230177 |
| QQSSDGKNGVRAAEITGE | ASGDRGENYAEQF | 0.7707865 |
| RAPDAVTTSWWKE | ASGDRGENYAEQF | 0.79541916 |
| SGFGNVGAHTSGWFNQ | ASGDRGENYAEQF | 0.8728123 |
| SGVAGAVGQAASVGGLKV | ASGDRGENYAEQF | 0.84545964 |
| SNGGAGTGSAGGNGGT | ASGDRGENYAEQF | 0.86945534 |
| TGGAGGEGGAAG | ASGDRGENYAEQF | 0.7980022 |
| TGGDGGAAGAGGAPGHG | ASGDRGENYAEQF | 0.8395363 |
| TMPPLLAAIANAGRHGV | ASGDRGENYAEQF | 0.8583025 |
| TSSTAGNSGGA | ASGDRGENYAEQF | 0.5382629 |
| VPATVPVGILC | ASGDRGENYAEQF | 0.7387661 |
| VQRPDSGGVVLTGRIS | ASGDRGENYAEQF | 0.88757837 |
| WTPNPVPATKGVPAGLVQAT | ASGDRGENYAEQF | 0.744383 |
| YALKRALAKFSDRFDVVII | ASGDRGENYAEQF | 0.7602763 |
| YLIGSVIV | ASGDRGENYAEQF | 0.57584924 |
| YGQLVAVID | ASGDRGENYAEQF | 0.48446816 |
| VTVGGMTLPA | ASGDRGENYAEQF | 0.5854204 |
| VLALPAV | ASGDRGENYAEQF | 0.7555516 |
| VIPPTLNYET | ASGDRGENYAEQF | 0.73814493 |
| VGMGGMHPSSG | ASGDRGENYAEQF | 0.7273103 |
| VGLAGGAATA | ASGDRGENYAEQF | 0.5145704 |
| VGGAGGAGGSGP | ASGDRGENYAEQF | 0.6483034 |
| VFAAFMV | ASGDRGENYAEQF | 0.66356426 |
| TLYDKLR | ASGDRGENYAEQF | 0.57711476 |
| TLAVFTVL | ASGDRGENYAEQF | 0.7137725 |
| TGSVQPARV | ASGDRGENYAEQF | 0.66826844 |
| TGGGAGSGGT | ASGDRGENYAEQF | 0.6569127 |
| SHLETSL | ASGDRGENYAEQF | 0.6354946 |
| SFTTGMT | ASGDRGENYAEQF | 0.6670479 |
| QVEVAKRMGVRQPTVS | ASGDRGENYAEQF | 0.70731914 |
| QGSGGPALAM | ASGDRGENYAEQF | 0.673465 |
| MLAPSNP | ASGDRGENYAEQF | 0.7793486 |
| LVNNAAI | ASGDRGENYAEQF | 0.7515174 |
| LTGEINTLA | ASGDRGENYAEQF | 0.73893535 |
| LSIFGPV | ASGDRGENYAEQF | 0.31554383 |
| LLALLANL | ASGDRGENYAEQF | 0.399116 |
| LISNASDALDKLR | ASGDRGENYAEQF | 0.70107126 |
| LINEAVP | ASGDRGENYAEQF | 0.5628422 |

| | | |
|---|---|---|
| LGYAFSGL | ASGDRGENYAEQF | 0.41501158 |
| LDFVAVAL | ASGDRGENYAEQF | 0.691113 |
| LAQPGAE | ASGDRGENYAEQF | 0.71430445 |
| ISNLGAVTGF | ASGDRGENYAEQF | 0.5356767 |
| HQEDPGRI | ASGDRGENYAEQF | 0.70306367 |
| GTPGEPTP | ASGDRGENYAEQF | 0.5541584 |
| GFFNSGS | ASGDRGENYAEQF | 0.5650388 |
| GAGGGGLTAGAN | ASGDRGENYAEQF | 0.6145202 |
| FVDALVNM | ASGDRGENYAEQF | 0.7394811 |
| FLTVAPEL | ASGDRGENYAEQF | 0.51440537 |
| FETLTAL | ASGDRGENYAEQF | 0.6987613 |
| DVINPIEL | ASGDRGENYAEQF | 0.6474038 |
| APVSGAPV | ASGDRGENYAEQF | 0.7487362 |
| ALVGGLAW | ASGDRGENYAEQF | 0.4605391 |
| ALAAAAATAI | ASGDRGENYAEQF | 0.73198366 |
| AFGLGLLL | ASGDRGENYAEQF | 0.6337882 |
| AFFAAASL | ASGDRGENYAEQF | 0.76694393 |
| AAAALPAFSPP | ASGDRGENYAEQF | 0.5819577 |
| VVLAPAL | ASGDRGENYAEQF | 0.6534227 |
| TIGHVDHGKTTL | ASGDRGENYAEQF | 0.6617465 |
| LISNASDALDK | ASGDRGENYAEQF | 0.7183431 |
| LISNASDAL | ASGDRGENYAEQF | 0.44985297 |
| ISNASDALDK | ASGDRGENYAEQF | 0.64857197 |
| NQAETLVYQTE | ASGDRGENYAEQF | 0.55783165 |
| TGGTGGTAGAG | ASGDRGENYAEQF | 0.68593514 |
| TGSTAGGAGGA | ASGDRGENYAEQF | 0.7265899 |
| DPLAVPVAAGPV | ASGDRGENYAEQF | 0.43331626 |
| GGTGGTPGNGGN | ASGDRGENYAEQF | 0.73920643 |
| PAGLAGVDPATL | ASGDRGENYAEQF | 0.6724774 |
| GNNLVVEAPHAGASV | ASGDRGENYAEQF | 0.5925613 |
| KVTRIMSASTLEQPA | ASGDRGENYAEQF | 0.74865824 |
| LGFALPIALPRRTVV | ASGDRGENYAEQF | 0.64422035 |
| TRLGNLTARKLVVLV | ASGDRGENYAEQF | 0.76247674 |
| WSQQRRLIYFPSAGP | ASGDRGENYAEQF | 0.6472742 |
| GPNGNPDPMAAAVDIR | ASGDRGENYAEQF | 0.6685403 |
| TQRCVSVADLRAAASR | ASGDRGENYAEQF | 0.72619903 |
| QFVLDVAEHPVEVLTETV | ASGDRGENYAEQF | 0.54000956 |
| GGAGGTGGAGGNGGTQAGDG | ASGDRGENYAEQF | 0.53970647 |
| PLLATLRAIAVDVDAKPAQV | ASGDRGENYAEQF | 0.75098675 |
| APKFRTGGYMIDFWGVGYQVAKRM | ASGDRGENYAEQF | 0.71112275 |
| GAGGAGGAGGAAGGPGGTGGTGGN | ASGDRGENYAEQF | 0.7098652 |
| IGALGLLATDVAPAGSVAFGVALA | ASGDRGENYAEQF | 0.67246014 |
| AGGAGGIGGNGGAGGAGGVLMGNG | ASGDRGENYAEQF | 0.67217803 |
| ADELIGLL | ASGDRGENYAEQF | 0.6771481 |
| ADRALKA | ASGDRGENYAEQF | 0.7063551 |
| AGAVGQVGGAGGA | ASGDRGENYAEQF | 0.717612 |
| AGGGGGLGGAP | ASGDRGENYAEQF | 0.7105721 |
| AGGLLYGNGGAG | ASGDRGENYAEQF | 0.67246825 |
| AGLESIGL | ASGDRGENYAEQF | 0.71232665 |
| AGLKTGII | ASGDRGENYAEQF | 0.7050434 |
| ASLAALLL | ASGDRGENYAEQF | 0.69368994 |

| | | |
|---|---|---|
| ASVINISSI | ASGDRGENYAEQF | 0.7000456 |
| ATPPGPA | ASGDRGENYAEQF | 0.69420063 |
| ATVASALV | ASGDRGENYAEQF | 0.6791363 |
| ATVASLIL | ASGDRGENYAEQF | 0.63382494 |
| AVNGPVLIRH | ASGDRGENYAEQF | 0.6951997 |
| DGAAGAVGI | ASGDRGENYAEQF | 0.71276784 |
| DPLLHGI | ASGDRGENYAEQF | 0.7085391 |
| ELGAALGAL | ASGDRGENYAEQF | 0.72011757 |
| ELGRTQAV | ASGDRGENYAEQF | 0.7128782 |
| FVAGGYLS | ASGDRGENYAEQF | 0.70258343 |
| GASGGAGGAGGW | ASGDRGENYAEQF | 0.49157834 |
| GGAGAGGDGGAGGI | ASGDRGENYAEQF | 0.61923593 |
| GTGHNGSQPGMGGNG | ASGDRGENYAEQF | 0.6569618 |
| GYAAASASAS | ASGDRGENYAEQF | 0.6908114 |
| HQVDVAAVQQG | ASGDRGENYAEQF | 0.38847816 |
| IGAAIAHT | ASGDRGENYAEQF | 0.70416653 |
| IGMIEVAQL | ASGDRGENYAEQF | 0.6992378 |
| IRLAPEHTSAMD | ASGDRGENYAEQF | 0.67961323 |
| IYNVSVL | ASGDRGENYAEQF | 0.71004957 |
| KQGFLGM | ASGDRGENYAEQF | 0.69153154 |
| LESVITQL | ASGDRGENYAEQF | 0.6410854 |
| LGLGALVVV | ASGDRGENYAEQF | 0.6730521 |
| LIGNGGNGGSGGI | ASGDRGENYAEQF | 0.7208203 |
| LNAINAPTL | ASGDRGENYAEQF | 0.6304686 |
| LQLLNTGV | ASGDRGENYAEQF | 0.718239 |
| LQTSIEF | ASGDRGENYAEQF | 0.6270757 |
| MSFAVSEL | ASGDRGENYAEQF | 0.7175357 |
| NTGALGL | ASGDRGENYAEQF | 0.65009093 |
| PPGAGGAGGAGTT | ASGDRGENYAEQF | 0.7149204 |
| QLVGLGI | ASGDRGENYAEQF | 0.6979307 |
| RFLEQVHQL | ASGDRGENYAEQF | 0.722353 |
| RGVGYVLR | ASGDRGENYAEQF | 0.68826675 |
| SATGVFAGVI | ASGDRGENYAEQF | 0.71397877 |
| SLPAITLP | ASGDRGENYAEQF | 0.68412024 |
| STALAFTV | ASGDRGENYAEQF | 0.71086884 |
| SVLTVGLGK | ASGDRGENYAEQF | 0.64002514 |
| TALAGAGSGI | ASGDRGENYAEQF | 0.68754506 |
| TGLPFTGI | ASGDRGENYAEQF | 0.6530646 |
| VAEHVTV | ASGDRGENYAEQF | 0.6951789 |
| VDIVLNSL | ASGDRGENYAEQF | 0.7152791 |
| VLTGGGAVL | ASGDRGENYAEQF | 0.5273316 |
| VNVGLAYGI | ASGDRGENYAEQF | 0.71321714 |
| VSSAAVGAV | ASGDRGENYAEQF | 0.6444056 |
| VTSVYEAGV | ASGDRGENYAEQF | 0.6672225 |
| WVALDRGA | ASGDRGENYAEQF | 0.70808595 |
| YLVGSVVI | ASGDRGENYAEQF | 0.7001618 |
| GPPGTGKTLL | ASGDRGENYAEQF | 0.71076 |
| GVGMTKF | ASGDRGENYAEQF | 0.6468985 |
| IGIGGSDL | ASGDRGENYAEQF | 0.69651496 |
| IGTLLAEL | ASGDRGENYAEQF | 0.7032124 |
| IYTSGTT | ASGDRGENYAEQF | 0.5115578 |

| | | |
|---|---|---|
| LSFPAGV | ASGDRGENYAEQF | 0.6443173 |
| LYTSGTT | ASGDRGENYAEQF | 0.7186033 |
| VGAFPFGF | ASGDRGENYAEQF | 0.67977405 |
| VGAGTVEF | ASGDRGENYAEQF | 0.6988323 |
| VGKSTLLL | ASGDRGENYAEQF | 0.6047004 |
| AALAHAGGGIV | ASGDRGENYAEQF | 0.5891649 |
| AALVAVSLL | ASGDRGENYAEQF | 0.6401571 |
| AAVSRAVAAL | ASGDRGENYAEQF | 0.6925545 |
| AGGSSGIGGF | ASGDRGENYAEQF | 0.70948774 |
| AGLTVANANG | ASGDRGENYAEQF | 0.68488455 |
| AGLVGAVL | ASGDRGENYAEQF | 0.6307212 |
| ALSQAFTV | ASGDRGENYAEQF | 0.6743438 |
| ALTFSVSL | ASGDRGENYAEQF | 0.66419315 |
| ALTGAALVAL | ASGDRGENYAEQF | 0.64717436 |
| ASFSIGAGVL | ASGDRGENYAEQF | 0.69319165 |
| DGSVYKI | ASGDRGENYAEQF | 0.6817407 |
| EQLAEIV | ASGDRGENYAEQF | 0.6323949 |
| FAVSELTVM | ASGDRGENYAEQF | 0.64620745 |
| GAAPLAAGL | ASGDRGENYAEQF | 0.67936695 |
| GADGGDGGSGGAGG | ASGDRGENYAEQF | 0.69300765 |
| GFGLDTGGAGG | ASGDRGENYAEQF | 0.6867801 |
| GLASGGLL | ASGDRGENYAEQF | 0.7139555 |
| GRIAGAAVLL | ASGDRGENYAEQF | 0.6930665 |
| GSAGSLNTSF | ASGDRGENYAEQF | 0.6800732 |
| GTGQIGI | ASGDRGENYAEQF | 0.440384 |
| GTLIALIL | ASGDRGENYAEQF | 0.5766453 |
| IAATMGFVVG | ASGDRGENYAEQF | 0.6755901 |
| IFAQALL | ASGDRGENYAEQF | 0.658005 |
| IGAAVALGF | ASGDRGENYAEQF | 0.44897583 |
| IGLTPLAL | ASGDRGENYAEQF | 0.6834724 |
| ILGSNTSTL | ASGDRGENYAEQF | 0.6526722 |
| IVFSLTVL | ASGDRGENYAEQF | 0.62432295 |
| LASKATGF | ASGDRGENYAEQF | 0.69638395 |
| LASVALIL | ASGDRGENYAEQF | 0.63866544 |
| LFAVGLTL | ASGDRGENYAEQF | 0.6414491 |
| LFNGGIGTYI | ASGDRGENYAEQF | 0.6714055 |
| LTLGAGII | ASGDRGENYAEQF | 0.7093487 |
| LVAAALGDI | ASGDRGENYAEQF | 0.62159795 |
| NTAFVQLGI | ASGDRGENYAEQF | 0.69749284 |
| QLFSGAEL | ASGDRGENYAEQF | 0.6835888 |
| SAATFITL | ASGDRGENYAEQF | 0.7023939 |
| SNYGLANLGV | ASGDRGENYAEQF | 0.587747 |
| SYGGAYAVM | ASGDRGENYAEQF | 0.691893 |
| TTGVGGLTLG | ASGDRGENYAEQF | 0.64792573 |
| TTQTIYI | ASGDRGENYAEQF | 0.71592253 |
| VAAGAGLGAVI | ASGDRGENYAEQF | 0.65098685 |
| VAAVVTLL | ASGDRGENYAEQF | 0.69930756 |
| VGFLGGLI | ASGDRGENYAEQF | 0.6820286 |
| VGVIPILL | ASGDRGENYAEQF | 0.6957588 |
| VIFASSLI | ASGDRGENYAEQF | 0.58883023 |
| VKGGDGGAAAG | ASGDRGENYAEQF | 0.6530942 |

| | | |
|---|---|---|
| VTKTFTNI | ASGDRGENYAEQF | 0.62049735 |
| VVAAAILL | ASGDRGENYAEQF | 0.67020667 |
| YAALDVEL | ASGDRGENYAEQF | 0.6874991 |
| LASVALLL | ASGDRGENYAEQF | 0.48248845 |
| LDAVHRV | ASGDRGENYAEQF | 0.70262665 |
| QGLSSSIF | ASGDRGENYAEQF | 0.6491705 |
| RGTFANIRL | ASGDRGENYAEQF | 0.63025117 |
| RQLWWGHRI | ASGDRGENYAEQF | 0.6972454 |
| YGPPGTGKTLL | ASGDRGENYAEQF | 0.6560751 |
| AALVGAPANVA | ASGDRGENYAEQF | 0.67667896 |
| AATFRRTE | ASGDRGENYAEQF | 0.64843994 |
| AGALGGGI | ASGDRGENYAEQF | 0.67261684 |
| AGGAGGAGGIGAG | ASGDRGENYAEQF | 0.6810634 |
| AGSGALGL | ASGDRGENYAEQF | 0.5414211 |
| AQIADQI | ASGDRGENYAEQF | 0.6362073 |
| ASAVAVTYL | ASGDRGENYAEQF | 0.7003849 |
| ASTVLGGTGG | ASGDRGENYAEQF | 0.6866073 |
| AVGGEPAK | ASGDRGENYAEQF | 0.7046331 |
| CEQGTFGGTC | ASGDRGENYAEQF | 0.5831138 |
| DEGKAPAG | ASGDRGENYAEQF | 0.56687677 |
| DTVVFPAL | ASGDRGENYAEQF | 0.6333773 |
| FEGITMAG | ASGDRGENYAEQF | 0.69715166 |
| FPLVVQD | ASGDRGENYAEQF | 0.70013505 |
| GAFTTPQ | ASGDRGENYAEQF | 0.6876793 |
| GFGGFGTAGGVGGA | ASGDRGENYAEQF | 0.5587013 |
| GFLAQFL | ASGDRGENYAEQF | 0.6610376 |
| GGAGGAGGNAGVA | ASGDRGENYAEQF | 0.6378875 |
| GGVGTAGAGGAGGAGG | ASGDRGENYAEQF | 0.62985647 |
| GNGGGGDAGNAGSGGN | ASGDRGENYAEQF | 0.66601104 |
| GTGGAGGAGNDAGS | ASGDRGENYAEQF | 0.66823685 |
| GTTNTGWFNS | ASGDRGENYAEQF | 0.612273 |
| IFMGPLFTL | ASGDRGENYAEQF | 0.61136854 |
| ILAHAIEAI | ASGDRGENYAEQF | 0.63089395 |
| IRSAIREAL | ASGDRGENYAEQF | 0.68314844 |
| ITFAGGG | ASGDRGENYAEQF | 0.6664124 |
| ITGMSGHQEV | ASGDRGENYAEQF | 0.7058152 |
| IVAGLACTL | ASGDRGENYAEQF | 0.6917934 |
| IVGGGLAGA | ASGDRGENYAEQF | 0.682366 |
| IVTGVLNI | ASGDRGENYAEQF | 0.30738622 |
| LAQQGCI | ASGDRGENYAEQF | 0.37413728 |
| LCLHVEN | ASGDRGENYAEQF | 0.5953397 |
| LDAISDGS | ASGDRGENYAEQF | 0.62933075 |
| LGGFAAML | ASGDRGENYAEQF | 0.3399885 |
| LGGKNPMI | ASGDRGENYAEQF | 0.66827846 |
| LGPKGVAAIL | ASGDRGENYAEQF | 0.6435405 |
| LMVGVSI | ASGDRGENYAEQF | 0.58683705 |
| LVGAAFVL | ASGDRGENYAEQF | 0.6725851 |
| LVLAVAPL | ASGDRGENYAEQF | 0.6162895 |
| MAAIAVIDAA | ASGDRGENYAEQF | 0.5720689 |
| NNGINAELQ | ASGDRGENYAEQF | 0.5554486 |
| RGEPLVLVHG | ASGDRGENYAEQF | 0.680781 |

| | | |
|---|---|---|
| RGGAGGAAGLI | ASGDRGENYAEQF | 0.46178287 |
| RSYRMHPAV | ASGDRGENYAEQF | 0.679917 |
| SAAHLLAGQLA | ASGDRGENYAEQF | 0.49973893 |
| SAAVGSY | ASGDRGENYAEQF | 0.6870768 |
| SPHAVLHRI | ASGDRGENYAEQF | 0.5453683 |
| SQIAKVL | ASGDRGENYAEQF | 0.6917978 |
| TALGHLVP | ASGDRGENYAEQF | 0.651227 |
| TAPTTSGAA | ASGDRGENYAEQF | 0.7064502 |
| TDGDDGNGGSPGT | ASGDRGENYAEQF | 0.6102319 |
| TDGLGEN | ASGDRGENYAEQF | 0.668821 |
| TGGPPLPGQA | ASGDRGENYAEQF | 0.6247522 |
| TIPAITI | ASGDRGENYAEQF | 0.70426977 |
| TIPGFTI | ASGDRGENYAEQF | 0.47042045 |
| TSIASGALA | ASGDRGENYAEQF | 0.64646137 |
| TVTLPGAAI | ASGDRGENYAEQF | 0.39729828 |
| VASGLSAALP | ASGDRGENYAEQF | 0.6681588 |
| VGAGVIGI | ASGDRGENYAEQF | 0.69195783 |
| VGPKRFGV | ASGDRGENYAEQF | 0.39031908 |
| VNVAFPD | ASGDRGENYAEQF | 0.6809194 |
| VSMTNSGVS | ASGDRGENYAEQF | 0.66264427 |
| VTDVLGDLSV | ASGDRGENYAEQF | 0.60744375 |
| VVVVGAGFAGL | ASGDRGENYAEQF | 0.687806 |
| VYTVGTVV | ASGDRGENYAEQF | 0.63078415 |
| WTTFSGP | ASGDRGENYAEQF | 0.69532585 |
| YDELVEAF | ASGDRGENYAEQF | 0.5979988 |
| YGTGGAGGNGG | ASGDRGENYAEQF | 0.6164659 |
| YTNVMAV | ASGDRGENYAEQF | 0.66564715 |
| FLEQVHQL | ASGDRGENYAEQF | 0.39852 |
| GLSLLAVAV | ASGDRGENYAEQF | 0.47981137 |
| LEQVHQL | ASGDRGENYAEQF | 0.7023237 |
| LNAAGAI | ASGDRGENYAEQF | 0.672751 |
| PAVHASL | ASGDRGENYAEQF | 0.68003416 |
| TELEDDA | ASGDRGENYAEQF | 0.6287337 |
| TPPGPPN | ASGDRGENYAEQF | 0.6158809 |
| TPPPPPPP | ASGDRGENYAEQF | 0.6350242 |
| TQSPALI | ASSHHREGTEVF | 0.6736811 |
| EFAAAFPQ | ASSHHREGTEVF | 0.6802085 |
| LLSGNLDVL | ASSHHREGTEVF | 0.67364806 |
| GPANLGNYNI | ASSHHREGTEVF | 0.6149526 |
| TLGPTLP | ASSHHREGTEVF | 0.6730459 |
| VYAYSSCCK | ASSHHREGTEVF | 0.6479692 |
| APAPAPAPNI | ASSHHREGTEVF | 0.64784646 |
| PGGPVPIP | ASSHHREGTEVF | 0.66554165 |
| FLTVALEP | ASSHHREGTEVF | 0.6713237 |
| SQYFGANL | ASSHHREGTEVF | 0.64256424 |
| GGVKPAA | ASSHHREGTEVF | 0.6424652 |
| GGTGGAGGAGGW | ASSHHREGTEVF | 0.66514283 |
| GVGGVAGLGGAGG | ASSHHREGTEVF | 0.6726284 |
| VLLVLATTI | ASSHHREGTEVF | 0.6698396 |
| IGIFAAML | ASSHHREGTEVF | 0.68463063 |
| AVGDLID | ASSHHREGTEVF | 0.6758796 |

| | | |
|---|---|---|
| NISPLPL | ASSHHREGTEVF | 0.6759187 |
| VTAVFVAI | ASSHHREGTEVF | 0.40748197 |
| SGDDNDAAL | ASSHHREGTEVF | 0.524082 |
| AVAALAHS | ASSHHREGTEVF | 0.6536822 |
| GGVTVMATP | ASSHHREGTEVF | 0.6494779 |
| AGLAANVRD | ASSHHREGTEVF | 0.46363306 |
| TNRPDLIDPALLRPG | ASSHHREGTEVF | 0.67594594 |
| LANLGVTAL | ASSHHREGTEVF | 0.6453759 |
| TGERIEL | ASSHHREGTEVF | 0.6379559 |
| NGEELAA | ASSHHREGTEVF | 0.6759218 |
| VLSLVWG | ASSHHREGTEVF | 0.65174115 |
| IGYALWL | ASSHHREGTEVF | 0.6272358 |
| LSLLLAPL | ASSHHREGTEVF | 0.640148 |
| TLDEALK | ASSHHREGTEVF | 0.6685965 |
| TDGAGTAEGG | ASSHHREGTEVF | 0.60327375 |
| NAPGGAGGNGG | ASSHHREGTEVF | 0.6745554 |
| VSVFMGMGGNF | ASSHHREGTEVF | 0.55608124 |
| LASYLAAL | ASSHHREGTEVF | 0.67817605 |
| LVGGTPAL | ASSHHREGTEVF | 0.5945107 |
| VRNADVL | ASSHHREGTEVF | 0.6744217 |
| TCSSPFLVSAM | ASSHHREGTEVF | 0.6523257 |
| VHITDAE | ASSHHREGTEVF | 0.691373 |
| EYAAPSF | ASSHHREGTEVF | 0.6458194 |
| VVDAFHGGL | ASSHHREGTEVF | 0.6706402 |
| ASVVDPQ | ASSHHREGTEVF | 0.6535892 |
| SYNEFGGLS | ASSHHREGTEVF | 0.68148136 |
| DPDGSVY | ASSHHREGTEVF | 0.57112217 |
| MVSTIHATE | ASSHHREGTEVF | 0.65144193 |
| LAAGVADPV | ASSHHREGTEVF | 0.47240072 |
| TKEQIAA | ASSHHREGTEVF | 0.66399276 |
| SLASPAAL | ASSHHREGTEVF | 0.6739383 |
| IGLVFYGV | ASSHHREGTEVF | 0.53997767 |
| FLLDASNL | ASSHHREGTEVF | 0.6792928 |
| YQVGNFP | ASSHHREGTEVF | 0.63534534 |
| ESLLSQST | ASSHHREGTEVF | 0.63214314 |
| LFTVGFAAVG | ASSHHREGTEVF | 0.6800208 |
| ATPPTPT | ASSHHREGTEVF | 0.6680866 |
| TDLATLG | ASSHHREGTEVF | 0.67969966 |
| SAGGTGR | ASSHHREGTEVF | 0.65412927 |
| GFSNSGS | ASSHHREGTEVF | 0.6406921 |
| GGNGGSGGTSV | ASSHHREGTEVF | 0.66533804 |
| GTSVATGGAGN | ASSHHREGTEVF | 0.6193322 |
| NGGNGGSGGTS | ASSHHREGTEVF | 0.63141483 |
| AVVKFPH | ASSHHREGTEVF | 0.68286926 |
| TPPPPPP | ASSHHREGTEVF | 0.6514127 |
| AAPAAAAAPA | ASSHHREGTEVF | 0.6650094 |
| IALNEDL | ASSHHREGTEVF | 0.5704985 |
| IVNVSSM | ASSHHREGTEVF | 0.5445176 |
| TGIPPAPR | ASSHHREGTEVF | 0.5945717 |
| FDLGGGTF | ASSHHREGTEVF | 0.66134423 |
| FDLGGGTFDVS | ASSHHREGTEVF | 0.6776346 |

| | | |
|---|---|---|
| NEPTAAA | ASSHHREGTEVF | 0.64784086 |
| NEPTAAALA | ASSHHREGTEVF | 0.5707849 |
| GSIINISSI | ASSHHREGTEVF | 0.643504 |
| VAPGNAGTA | ASSHHREGTEVF | 0.6182276 |
| TGGGASLE | ASSHHREGTEVF | 0.6078464 |
| VAVYNHYKRI | ASSHHREGTEVF | 0.65750796 |
| TDPAPATT | ASSHHREGTEVF | 0.6485435 |
| TNRPDILDPALLRPG | ASSHHREGTEVF | 0.6010527 |
| LTTTTTS | ASSHHREGTEVF | 0.5985832 |
| PTGSGKTL | ASSHHREGTEVF | 0.6523748 |
| TGSGKTL | ASSHHREGTEVF | 0.66572034 |
| TIGHVDHGK | ASSHHREGTEVF | 0.6571696 |
| TIGHVDHGKTT | ASSHHREGTEVF | 0.68671465 |
| GHVDHGKTT | ASSHHREGTEVF | 0.660479 |
| AQFEHTLL | ASSHHREGTEVF | 0.67338085 |
| IVNQDEL | ASSHHREGTEVF | 0.40314204 |
| TNGSQFF | ASSHHREGTEVF | 0.5029149 |
| TGKGIVE | ASSHHREGTEVF | 0.62136376 |
| LISNASDA | ASSHHREGTEVF | 0.6115974 |
| AELRTLGELTKIAKAHGAQ | ASSHHREGTEVF | 0.47050506 |
| APNAVAAAEPSW | ASSHHREGTEVF | 0.6540618 |
| APQVAGAATAVL | ASSHHREGTEVF | 0.61822534 |
| ARRDPAAAAAILPTDA | ASSHHREGTEVF | 0.55356944 |
| DAAARDVLAEAGLAEYF | ASSHHREGTEVF | 0.656614 |
| DDQIVLSAKANLAKVEQ | ASSHHREGTEVF | 0.5882364 |
| DELLPDTGEDSDEDGH | ASSHHREGTEVF | 0.56960857 |
| ELPEAPANRATVPQD | ASSHHREGTEVF | 0.6054876 |
| FGQSKIERVTGIG | ASSHHREGTEVF | 0.6705893 |
| GAGGTGSTAGGAGGAGGAGGLYA | ASSHHREGTEVF | 0.5473568 |
| GHGGTGAAGGNGGTGGAGGSGIDG | ASSHHREGTEVF | 0.6537672 |
| GNGGNGGAGGEGGPGGAGGAGA | ASSHHREGTEVF | 0.6166469 |
| GSTAGNGGNGGNATNGTIAGQPAG | ASSHHREGTEVF | 0.6702582 |
| GTGGDGGNGGNGGTGG | ASSHHREGTEVF | 0.51689327 |
| GVRRPAGIPADAPV | ASSHHREGTEVF | 0.6586145 |
| GYRQVTDAHLLTLARR | ASSHHREGTEVF | 0.60763466 |
| IAKMGQDGHDRG | ASSHHREGTEVF | 0.689327 |
| KAEPSIEIQYT | ASSHHREGTEVF | 0.6046802 |
| KPVRIRMAADEDTVAVT | ASSHHREGTEVF | 0.6563736 |
| MPPRIAGMRLLVIKPEP | ASSHHREGTEVF | 0.65906787 |
| MKFYAVALFRSRGVPPTR | ASSHHREGTEVF | 0.66830736 |
| NAGGFGILGAVAHSP | ASSHHREGTEVF | 0.48360416 |
| NEGREITLAMAEM | ASSHHREGTEVF | 0.6178435 |
| NELVEWSHPLGHRWRWE | ASSHHREGTEVF | 0.5552081 |
| NLGAGGTGGNGGNPG | ASSHHREGTEVF | 0.63489425 |
| NTEWPKWETVMAADPVD | ASSHHREGTEVF | 0.64722204 |
| PASAAPAETQP | ASSHHREGTEVF | 0.41736814 |
| QQSSDGKNGVRAAEITGE | ASSHHREGTEVF | 0.66737217 |
| RAPDAVTTSWWKE | ASSHHREGTEVF | 0.5758759 |
| SGFGNVGAHTSGWFNQ | ASSHHREGTEVF | 0.5872484 |
| SGVAGAVGQAASVGGLKV | ASSHHREGTEVF | 0.6716988 |
| SNGGAGTGSAGGNGGT | ASSHHREGTEVF | 0.64009887 |

| | | |
|---|---|---|
| TGGAGGEGGAAG | ASSHHREGTEVF | 0.6672723 |
| TGGDGGAAGAGGAPGHG | ASSHHREGTEVF | 0.63424736 |
| TMPPLLAAIANAGRHGV | ASSHHREGTEVF | 0.6175639 |
| TSSTAGNSGGA | ASSHHREGTEVF | 0.64224887 |
| VPATVPVGILC | ASSHHREGTEVF | 0.51189834 |
| VQRPDSGGVVLTGRIS | ASSHHREGTEVF | 0.56903243 |
| WTPNPVPATKGVPAGLVQAT | ASSHHREGTEVF | 0.67324394 |
| YALKRALAKFSDRFDVVII | ASSHHREGTEVF | 0.80679435 |
| YLIGSVIV | ASSHHREGTEVF | 0.8135743 |
| YGQLVAVID | ASSHHREGTEVF | 0.6857181 |
| VTVGGMTLPA | ASSHHREGTEVF | 0.67835283 |
| VLALPAV | ASSHHREGTEVF | 0.7330575 |
| VIPPTLNYET | ASSHHREGTEVF | 0.8332647 |
| VGMGGMHPSSG | ASSHHREGTEVF | 0.83705705 |
| VGLAGGAATA | ASSHHREGTEVF | 0.7917341 |
| VGGAGGAGGSGP | ASSHHREGTEVF | 0.730999 |
| VFAAFMV | ASSHHREGTEVF | 0.8313016 |
| TLYDKLR | ASSHHREGTEVF | 0.7932043 |
| TLAVFTVL | ASSHHREGTEVF | 0.7637096 |
| TGSVQPARV | ASSHHREGTEVF | 0.82589424 |
| TGGGAGSGGT | ASSHHREGTEVF | 0.8168389 |
| SHLETSL | ASSHHREGTEVF | 0.7513244 |
| SFTTGMT | ASSHHREGTEVF | 0.7310306 |
| QVEVAKRMGVRQPTVS | ASSHHREGTEVF | 0.807632 |
| QGSGGPALAM | ASSHHREGTEVF | 0.8262649 |
| MLAPSNP | ASSHHREGTEVF | 0.821087 |
| LVNNAAI | ASSHHREGTEVF | 0.8713072 |
| LTGEINTLA | ASSHHREGTEVF | 0.8371308 |
| LSIFGPV | ASSHHREGTEVF | 0.8084323 |
| LLALLANL | ASSHHREGTEVF | 0.43758503 |
| LISNASDALDKLR | ASSHHREGTEVF | 0.64944845 |
| LINEAVP | ASSHHREGTEVF | 0.77204573 |
| LGYAFSGL | ASSHHREGTEVF | 0.7824459 |
| LDFVAVAL | ASSHHREGTEVF | 0.3895123 |
| LAQPGAE | ASSHHREGTEVF | 0.8329415 |
| ISNLGAVTGF | ASSHHREGTEVF | 0.75656533 |
| HQEDPGRI | ASSHHREGTEVF | 0.7064004 |
| GTPGEPTP | ASSHHREGTEVF | 0.83388424 |
| GFFNSGS | ASSHHREGTEVF | 0.72593844 |
| GAGGGGLTAGAN | ASSHHREGTEVF | 0.74976134 |
| FVDALVNM | ASSHHREGTEVF | 0.78246266 |
| FLTVAPEL | ASSHHREGTEVF | 0.8556305 |
| FETLTAL | ASSHHREGTEVF | 0.69495463 |
| DVINPIEL | ASSHHREGTEVF | 0.83985853 |
| APVSGAPV | ASSHHREGTEVF | 0.78622866 |
| ALVGGLAW | ASSHHREGTEVF | 0.8596865 |
| ALAAAAATAI | ASSHHREGTEVF | 0.6616149 |
| AFGLGLLL | ASSHHREGTEVF | 0.82654023 |
| AFFAAASL | ASSHHREGTEVF | 0.76150453 |
| AAAALPAFSPP | ASSHHREGTEVF | 0.8685305 |
| VVLAPAL | ASSHHREGTEVF | 0.7832981 |

| | | |
|---|---|---|
| TIGHVDHGKTTL | ASSHHREGTEVF | 0.8421099 |
| LISNASDALDK | ASSHHREGTEVF | 0.8115844 |
| LISNASDAL | ASSHHREGTEVF | 0.8389766 |
| ISNASDALDK | ASSHHREGTEVF | 0.6023352 |
| NQAETLVYQTE | ASSHHREGTEVF | 0.7723847 |
| TGGTGGTAGAG | ASSHHREGTEVF | 0.72251856 |
| TGSTAGGAGGA | ASSHHREGTEVF | 0.80603784 |
| DPLAVPVAAGPV | ASSHHREGTEVF | 0.8107908 |
| GGTGGTPGNGGN | ASSHHREGTEVF | 0.4208482 |
| PAGLAGVDPATL | ASSHHREGTEVF | 0.82265806 |
| GNNLVVEAPHAGASV | ASSHHREGTEVF | 0.6723529 |
| KVTRIMSASTLEQPA | ASSHHREGTEVF | 0.70140326 |
| LGFALPIALPRRTVV | ASSHHREGTEVF | 0.8337306 |
| TRLGNLTARKLVVLV | ASSHHREGTEVF | 0.80950934 |
| WSQQRRLIYFPSAGP | ASSHHREGTEVF | 0.8295408 |
| GPNGNPDPMAAAVDIR | ASSHHREGTEVF | 0.76184994 |
| TQRCVSVADLRAAASR | ASSHHREGTEVF | 0.7674371 |
| QFVLDVAEHPVEVLTETV | ASSHHREGTEVF | 0.81784594 |
| GGAGGTGGAGGNGGTQAGDG | ASSHHREGTEVF | 0.6228603 |
| PLLATLRAIAVDVDAKPAQV | ASSHHREGTEVF | 0.72508264 |
| APKFRTGGYMIDFWGVGYQVAKRM | ASSHHREGTEVF | 0.8466505 |
| GAGGAGGAGGAAGGPGGTGGTGGN | ASSHHREGTEVF | 0.43418995 |
| IGALGLLATDVAPAGSVAFGVALA | ASSHHREGTEVF | 0.69170064 |
| AGGAGGIGGNGGAGGAGGVLMGNG | ASSHHREGTEVF | 0.23379469 |
| ADELIGLL | ASSHHREGTEVF | 0.24561217 |
| ADRALKA | ASSHHREGTEVF | 0.34160116 |
| AGAVGQVGGAGGA | ASSHHREGTEVF | 0.5159604 |
| AGGGGGLGGAP | ASSHHREGTEVF | 0.6905513 |
| AGGLLYGNGGAG | ASSHHREGTEVF | 0.508533 |
| AGLESIGL | ASSHHREGTEVF | 0.25233358 |
| AGLKTGII | ASSHHREGTEVF | 0.33886912 |
| ASLAALLL | ASSHHREGTEVF | 0.29223314 |
| ASVINISSI | ASSHHREGTEVF | 0.2876674 |
| ATPPGPA | ASSHHREGTEVF | 0.43616745 |
| ATVASALV | ASSHHREGTEVF | 0.2780222 |
| ATVASLIL | ASSHHREGTEVF | 0.2567578 |
| AVNGPVLIRH | ASSHHREGTEVF | 0.26440367 |
| DGAAGAVGI | ASSHHREGTEVF | 0.3395388 |
| DPLLHGI | ASSHHREGTEVF | 0.53736395 |
| ELGAALGAL | ASSHHREGTEVF | 0.55301946 |
| ELGRTQAV | ASSHHREGTEVF | 0.59860754 |
| FVAGGYLS | ASSHHREGTEVF | 0.44596624 |
| GASGGAGGAGGW | ASSHHREGTEVF | 0.32914668 |
| GGAGAGGDGGAGGI | ASSHHREGTEVF | 0.2008322 |
| GTGHNGSQPGMGGNG | ASSHHREGTEVF | 0.22370481 |
| GYAAASASAS | ASSHHREGTEVF | 0.2957955 |
| HQVDVAAVQQG | ASSHHREGTEVF | 0.30760795 |
| IGAAIAHT | ASSHHREGTEVF | 0.20632686 |
| IGMIEVAQL | ASSHHREGTEVF | 0.4613921 |
| IRLAPEHTSAMD | ASSHHREGTEVF | 0.3378417 |
| IYNVSVL | ASSHHREGTEVF | 0.28337952 |

| | | |
|---|---|---|
| KQGFLGM | ASSHHREGTEVF | 0.5255813 |
| LESVITQL | ASSHHREGTEVF | 0.26106 |
| LGLGALVVV | ASSHHREGTEVF | 0.24397284 |
| LIGNGGNGGSGGI | ASSHHREGTEVF | 0.24900475 |
| LNAINAPTL | ASSHHREGTEVF | 0.50313526 |
| LQLLNTGV | ASSHHREGTEVF | 0.22494216 |
| LQTSIEF | ASSHHREGTEVF | 0.45885372 |
| MSFAVSEL | ASSHHREGTEVF | 0.31748465 |
| NTGALGL | ASSHHREGTEVF | 0.48732018 |
| PPGAGGAGGAGTT | ASSHHREGTEVF | 0.23213054 |
| QLVGLGI | ASSHHREGTEVF | 0.37344247 |
| RFLEQVHQL | ASSHHREGTEVF | 0.29314482 |
| RGVGYVLR | ASSHHREGTEVF | 0.6754411 |
| SATGVFAGVI | ASSHHREGTEVF | 0.30031186 |
| SLPAITLP | ASSHHREGTEVF | 0.3670531 |
| STALAFTV | ASSHHREGTEVF | 0.23658876 |
| SVLTVGLGK | ASSHHREGTEVF | 0.64845717 |
| TALAGAGSGI | ASSHHREGTEVF | 0.2410501 |
| TGLPFTGI | ASSHHREGTEVF | 0.32305527 |
| VAEHVTV | ASSHHREGTEVF | 0.22379658 |
| VDIVLNSL | ASSHHREGTEVF | 0.3396014 |
| VLTGGGAVL | ASSHHREGTEVF | 0.41159898 |
| VNVGLAYGI | ASSHHREGTEVF | 0.26428583 |
| VSSAAVGAV | ASSHHREGTEVF | 0.4202162 |
| VTSVYEAGV | ASSHHREGTEVF | 0.3836758 |
| WVALDRGA | ASSHHREGTEVF | 0.26709378 |
| YLVGSVVI | ASSHHREGTEVF | 0.5324369 |
| GPPGTGKTLL | ASSHHREGTEVF | 0.31399983 |
| GVGMTKF | ASSHHREGTEVF | 0.43330285 |
| IGIGGSDL | ASSHHREGTEVF | 0.2648803 |
| IGTLLAEL | ASSHHREGTEVF | 0.3431154 |
| IYTSGTT | ASSHHREGTEVF | 0.4643743 |
| LSFPAGV | ASSHHREGTEVF | 0.25134444 |
| LYTSGTT | ASSHHREGTEVF | 0.26417518 |
| VGAFPFGF | ASSHHREGTEVF | 0.42092663 |
| VGAGTVEF | ASSHHREGTEVF | 0.62333727 |
| VGKSTLLL | ASSHHREGTEVF | 0.6643058 |
| AALAHAGGGIV | ASSHHREGTEVF | 0.43032813 |
| AALVAVSLL | ASSHHREGTEVF | 0.453192 |
| AAVSRAVAAL | ASSHHREGTEVF | 0.49983016 |
| AGGSSGIGGF | ASSHHREGTEVF | 0.6456525 |
| AGLTVANANG | ASSHHREGTEVF | 0.6715802 |
| AGLVGAVL | ASSHHREGTEVF | 0.62601495 |
| ALSQAFTV | ASSHHREGTEVF | 0.47549534 |
| ALTFSVSL | ASSHHREGTEVF | 0.55712306 |
| ALTGAALVAL | ASSHHREGTEVF | 0.47354364 |
| ASFSIGAGVL | ASSHHREGTEVF | 0.5174502 |
| DGSVYKI | ASSHHREGTEVF | 0.5991117 |
| EQLAEIV | ASSHHREGTEVF | 0.5853789 |
| FAVSELTVM | ASSHHREGTEVF | 0.5213883 |
| GAAPLAAGL | ASSHHREGTEVF | 0.49688715 |

| | | |
|---|---|---|
| GADGGDGGSGGAGG | ASSHHREGTEVF | 0.6015849 |
| GFGLDTGGAGG | ASSHHREGTEVF | 0.65382487 |
| GLASGGLL | ASSHHREGTEVF | 0.630633 |
| GRIAGAAVLL | ASSHHREGTEVF | 0.6766019 |
| GSAGSLNTSF | ASSHHREGTEVF | 0.6289061 |
| GTGQIGI | ASSHHREGTEVF | 0.6257608 |
| GTLIALIL | ASSHHREGTEVF | 0.27827793 |
| IAATMGFVVG | ASSHHREGTEVF | 0.3653158 |
| IFAQALL | ASSHHREGTEVF | 0.55971044 |
| IGAAVALGF | ASSHHREGTEVF | 0.51608986 |
| IGLTPLAL | ASSHHREGTEVF | 0.34969938 |
| ILGSNTSTL | ASSHHREGTEVF | 0.61717665 |
| IVFSLTVL | ASSHHREGTEVF | 0.5234199 |
| LASKATGF | ASSHHREGTEVF | 0.47796947 |
| LASVALIL | ASSHHREGTEVF | 0.63546735 |
| LFAVGLTL | ASSHHREGTEVF | 0.488761 |
| LFNGGIGTYI | ASSHHREGTEVF | 0.47321087 |
| LTLGAGII | ASSHHREGTEVF | 0.47260904 |
| LVAAALGDI | ASSHHREGTEVF | 0.66091275 |
| NTAFVQLGI | ASSHHREGTEVF | 0.4008259 |
| QLFSGAEL | ASSHHREGTEVF | 0.61540806 |
| SAATFITL | ASSHHREGTEVF | 0.49618486 |
| SNYGLANLGV | ASSHHREGTEVF | 0.65155756 |
| SYGGAYAVM | ASSHHREGTEVF | 0.41232097 |
| TTGVGGLTLG | ASSHHREGTEVF | 0.61004007 |
| TTQTIYI | ASSHHREGTEVF | 0.50676763 |
| VAAGAGLGAVI | ASSHHREGTEVF | 0.71001583 |
| VAAVVTLL | ASSHHREGTEVF | 0.50005776 |
| VGFLGGLI | ASSHHREGTEVF | 0.5589318 |
| VGVIPILL | ASSHHREGTEVF | 0.53170854 |
| VIFASSLI | ASSHHREGTEVF | 0.66782445 |
| VKGGDGGAAAG | ASSHHREGTEVF | 0.36343938 |
| VTKTFTNI | ASSHHREGTEVF | 0.5134194 |
| VVAAAILL | ASSHHREGTEVF | 0.40800568 |
| YAALDVEL | ASSHHREGTEVF | 0.54925805 |
| LASVALLL | ASSHHREGTEVF | 0.61359847 |
| LDAVHRV | ASSHHREGTEVF | 0.34055513 |
| QGLSSSIF | ASSHHREGTEVF | 0.6304697 |
| RGTFANIRL | ASSHHREGTEVF | 0.49049 |
| RQLWWGHRI | ASSHHREGTEVF | 0.47354034 |
| YGPPGTGKTLL | ASSHHREGTEVF | 0.64374477 |
| AALVGAPANVA | ASSHHREGTEVF | 0.51398224 |
| AATFRRTE | ASSHHREGTEVF | 0.67139584 |
| AGALGGGI | ASSHHREGTEVF | 0.5181526 |
| AGGAGGAGGIGAG | ASSHHREGTEVF | 0.5145892 |
| AGSGALGL | ASSHHREGTEVF | 0.61606675 |
| AQIADQI | ASSHHREGTEVF | 0.47240418 |
| ASAVAVTYL | ASSHHREGTEVF | 0.45243225 |
| ASTVLGGTGG | ASSHHREGTEVF | 0.6442578 |
| AVGGEPAK | ASSHHREGTEVF | 0.47823057 |
| CEQGTFGGTC | ASSHHREGTEVF | 0.6953125 |

| | | |
|---|---|---|
| DEGKAPAG | ASSHHREGTEVF | 0.2597333 |
| DTVVFPAL | ASSHHREGTEVF | 0.31259876 |
| FEGITMAG | ASSHHREGTEVF | 0.36920616 |
| FPLVVQD | ASSHHREGTEVF | 0.57872564 |
| GAFTTPQ | ASSHHREGTEVF | 0.71622646 |
| GFGGFGTAGGVGGA | ASSHHREGTEVF | 0.5000724 |
| GFLAQFL | ASSHHREGTEVF | 0.32651195 |
| GGAGGAGGNAGVA | ASSHHREGTEVF | 0.4179151 |
| GGVGTAGAGGAGGAGG | ASSHHREGTEVF | 0.37896088 |
| GNGGGGDAGNAGSGGN | ASSHHREGTEVF | 0.43694887 |
| GTGGAGGAGNDAGS | ASSHHREGTEVF | 0.45647234 |
| GTTNTGWFNS | ASSHHREGTEVF | 0.4465106 |
| IFMGPLFTL | ASSHHREGTEVF | 0.32053185 |
| ILAHAIEAI | ASSHHREGTEVF | 0.33636186 |
| IRSAIREAL | ASSHHREGTEVF | 0.5353488 |
| ITFAGGG | ASSHHREGTEVF | 0.6223177 |
| ITGMSGHQEV | ASSHHREGTEVF | 0.60419226 |
| IVAGLACTL | ASSHHREGTEVF | 0.63368285 |
| IVGGGLAGA | ASSHHREGTEVF | 0.57653564 |
| IVTGVLNI | ASSHHREGTEVF | 0.41743883 |
| LAQQGCI | ASSHHREGTEVF | 0.18457812 |
| LCLHVEN | ASSHHREGTEVF | 0.24643975 |
| LDAISDGS | ASSHHREGTEVF | 0.37559018 |
| LGGFAAML | ASSHHREGTEVF | 0.4059991 |
| LGGKNPMI | ASSHHREGTEVF | 0.22969915 |
| LGPKGVAAIL | ASSHHREGTEVF | 0.5079232 |
| LMVGVSI | ASSHHREGTEVF | 0.3978151 |
| LVGAAFVL | ASSHHREGTEVF | 0.33762744 |
| LVLAVAPL | ASSHHREGTEVF | 0.6277224 |
| MAAIAVIDAA | ASSHHREGTEVF | 0.29877424 |
| NNGINAELQ | ASSHHREGTEVF | 0.31280494 |
| RGEPLVLVHG | ASSHHREGTEVF | 0.33144602 |
| RGGAGGAAGLI | ASSHHREGTEVF | 0.50353944 |
| RSYRMHPAV | ASSHHREGTEVF | 0.28774744 |
| SAAHLLAGQLA | ASSHHREGTEVF | 0.58855075 |
| SAAVGSY | ASSHHREGTEVF | 0.36009222 |
| SPHAVLHRI | ASSHHREGTEVF | 0.6389139 |
| SQIAKVL | ASSHHREGTEVF | 0.29355487 |
| TALGHLVP | ASSHHREGTEVF | 0.53989756 |
| TAPTTSGAA | ASSHHREGTEVF | 0.38945535 |
| TDGDDGNGGSPGT | ASSHHREGTEVF | 0.7457244 |
| TDGLGEN | ASSHHREGTEVF | 0.39460486 |
| TGGPPLPGQA | ASSHHREGTEVF | 0.48630962 |
| TIPAITI | ASSHHREGTEVF | 0.39405861 |
| TIPGFTI | ASSHHREGTEVF | 0.68421173 |
| TSIASGALA | ASSHHREGTEVF | 0.24549638 |
| TVTLPGAAI | ASSHHREGTEVF | 0.34234294 |
| VASGLSAALP | ASSHHREGTEVF | 0.23625898 |
| VGAGVIGI | ASSHHREGTEVF | 0.37645268 |
| VGPKRFGV | ASSHHREGTEVF | 0.50939965 |
| VNVAFPD | ASSHHREGTEVF | 0.2445142 |

| | | |
|---|---|---|
| VSMTNSGVS | ASSHHREGTEVF | 0.383221 |
| VTDVLGDLSV | ASSHHREGTEVF | 0.34997487 |
| VVVVGAGFAGL | ASSHHREGTEVF | 0.3216653 |
| VYTVGTVV | ASSHHREGTEVF | 0.48588544 |
| WTTFSGP | ASSHHREGTEVF | 0.41580302 |
| YDELVEAF | ASSHHREGTEVF | 0.56353587 |
| YGTGGAGGNGG | ASSHHREGTEVF | 0.3029511 |
| YTNVMAV | ASSHHREGTEVF | 0.30979136 |
| FLEQVHQL | ASSHHREGTEVF | 0.44263956 |
| GLSLLAVAV | ASSHHREGTEVF | 0.34181064 |
| LEQVHQL | ASSHHREGTEVF | 0.28103906 |
| LNAAGAI | ASSHHREGTEVF | 0.6161623 |
| PAVHASL | ASSHHREGTEVF | 0.8847209 |
| TELEDDA | ASSHHREGTEVF | 0.8715813 |
| TPPGPPN | ASSHHREGTEVF | 0.7960794 |
| TPPPPPPP | ASSHHREGTEVF | 0.77561754 |
| TQSPALI | ASSSGQSANTEVF | 0.8235667 |
| EFAAAFPQ | ASSSGQSANTEVF | 0.90043056 |
| LLSGNLDVL | ASSSGQSANTEVF | 0.8940417 |
| GPANLGNYNI | ASSSGQSANTEVF | 0.8758688 |
| TLGPTLP | ASSSGQSANTEVF | 0.7753447 |
| VYAYSSCCK | ASSSGQSANTEVF | 0.87710714 |
| APAPAPAPNI | ASSSGQSANTEVF | 0.8553628 |
| PGGPVPIP | ASSSGQSANTEVF | 0.83843327 |
| FLTVALEP | ASSSGQSANTEVF | 0.88953817 |
| SQYFGANL | ASSSGQSANTEVF | 0.8819343 |
| GGVKPAA | ASSSGQSANTEVF | 0.83725536 |
| GGTGGAGGAGGW | ASSSGQSANTEVF | 0.84787947 |
| GVGGVAGLGGAGG | ASSSGQSANTEVF | 0.86479115 |
| VLLVLATTI | ASSSGQSANTEVF | 0.8853267 |
| IGIFAAML | ASSSGQSANTEVF | 0.88574016 |
| AVGDLID | ASSSGQSANTEVF | 0.90273786 |
| NISPLPL | ASSSGQSANTEVF | 0.8960791 |
| VTAVFVAI | ASSSGQSANTEVF | 0.88038874 |
| SGDDNDAAL | ASSSGQSANTEVF | 0.31335545 |
| AVAALAHS | ASSSGQSANTEVF | 0.5715643 |
| GGVTVMATP | ASSSGQSANTEVF | 0.84726906 |
| AGLAANVRD | ASSSGQSANTEVF | 0.83013046 |
| TNRPDLIDPALLRPG | ASSSGQSANTEVF | 0.5296365 |
| LANLGVTAL | ASSSGQSANTEVF | 0.8650506 |
| TGERIEL | ASSSGQSANTEVF | 0.8377262 |
| NGEELAA | ASSSGQSANTEVF | 0.79653907 |
| VLSLVWG | ASSSGQSANTEVF | 0.88922244 |
| IGYALWL | ASSSGQSANTEVF | 0.84032094 |
| LSLLLAPL | ASSSGQSANTEVF | 0.8174465 |
| TLDEALK | ASSSGQSANTEVF | 0.830428 |
| TDGAGTAEGG | ASSSGQSANTEVF | 0.8976821 |
| NAPGGAGGNGG | ASSSGQSANTEVF | 0.7563901 |
| VSVFMGMGGNF | ASSSGQSANTEVF | 0.88973886 |
| LASYLAAL | ASSSGQSANTEVF | 0.7667242 |
| LVGGTPAL | ASSSGQSANTEVF | 0.9052114 |

| | | |
|---|---|---|
| VRNADVL | ASSSGQSANTEVF | 0.739028 |
| TCSSPFLVSAM | ASSSGQSANTEVF | 0.8860073 |
| VHITDAE | ASSSGQSANTEVF | 0.84452784 |
| EYAAPSF | ASSSGQSANTEVF | 0.9164956 |
| VVDAFHGGL | ASSSGQSANTEVF | 0.8527192 |
| ASVVDPQ | ASSSGQSANTEVF | 0.8913827 |
| SYNEFGGLS | ASSSGQSANTEVF | 0.8517605 |
| DPDGSVY | ASSSGQSANTEVF | 0.8821175 |
| MVSTIHATE | ASSSGQSANTEVF | 0.7097447 |
| LAAGVADPV | ASSSGQSANTEVF | 0.8662604 |
| TKEQIAA | ASSSGQSANTEVF | 0.5760914 |
| SLASPAAL | ASSSGQSANTEVF | 0.8757647 |
| IGLVFYGV | ASSSGQSANTEVF | 0.8760824 |
| FLLDASNL | ASSSGQSANTEVF | 0.5039102 |
| YQVGNFP | ASSSGQSANTEVF | 0.8922622 |
| ESLLSQST | ASSSGQSANTEVF | 0.7965944 |
| LFTVGFAAVG | ASSSGQSANTEVF | 0.82685244 |
| ATPPTPT | ASSSGQSANTEVF | 0.9014653 |
| TDLATLG | ASSSGQSANTEVF | 0.852788 |
| SAGGTGR | ASSSGQSANTEVF | 0.8971822 |
| GFSNSGS | ASSSGQSANTEVF | 0.85624796 |
| GGNGGSGGTSV | ASSSGQSANTEVF | 0.8488833 |
| GTSVATGGAGN | ASSSGQSANTEVF | 0.8698678 |
| NGGNGGSGGTS | ASSSGQSANTEVF | 0.5345924 |
| AVVKFPH | ASSSGQSANTEVF | 0.8004726 |
| TPPPPPP | ASSSGQSANTEVF | 0.904638 |
| AAPAAAAAPA | ASSSGQSANTEVF | 0.8270389 |
| IALNEDL | ASSSGQSANTEVF | 0.86478305 |
| IVNVSSM | ASSSGQSANTEVF | 0.7324175 |
| TGIPPAPR | ASSSGQSANTEVF | 0.66532356 |
| FDLGGGTF | ASSSGQSANTEVF | 0.75699174 |
| FDLGGGTFDVS | ASSSGQSANTEVF | 0.88129807 |
| NEPTAAA | ASSSGQSANTEVF | 0.8590038 |
| NEPTAAALA | ASSSGQSANTEVF | 0.82981217 |
| GSIINISSI | ASSSGQSANTEVF | 0.6089027 |
| VAPGNAGTA | ASSSGQSANTEVF | 0.78864795 |
| TGGGASLE | ASSSGQSANTEVF | 0.7487521 |
| VAVYNHYKRI | ASSSGQSANTEVF | 0.7297262 |
| TDPAPATT | ASSSGQSANTEVF | 0.86234295 |
| TNRPDILDPALLRPG | ASSSGQSANTEVF | 0.8222798 |
| LTTTTTS | ASSSGQSANTEVF | 0.7404567 |
| PTGSGKTL | ASSSGQSANTEVF | 0.7873821 |
| TGSGKTL | ASSSGQSANTEVF | 0.80663526 |
| TIGHVDHGK | ASSSGQSANTEVF | 0.7850243 |
| TIGHVDHGKTT | ASSSGQSANTEVF | 0.7754822 |
| GHVDHGKTT | ASSSGQSANTEVF | 0.8839151 |
| AQFEHTLL | ASSSGQSANTEVF | 0.8538171 |
| IVNQDEL | ASSSGQSANTEVF | 0.86188555 |
| TNGSQFF | ASSSGQSANTEVF | 0.19623071 |
| TGKGIVE | ASSSGQSANTEVF | 0.38300318 |
| LISNASDA | ASSSGQSANTEVF | 0.8145001 |

| | | |
|---|---|---|
| AELRTLGELTKIAKAHGAQ | ASSSGQSANTEVF | 0.6966783 |
| APNAVAAAEPSW | ASSSGQSANTEVF | 0.5778968 |
| APQVAGAATAVL | ASSSGQSANTEVF | 0.8084458 |
| ARRDPAAAAAILPTDA | ASSSGQSANTEVF | 0.73105407 |
| DAAARDVLAEAGLAEYF | ASSSGQSANTEVF | 0.65953815 |
| DDQIVLSAKANLAKVEQ | ASSSGQSANTEVF | 0.8418531 |
| DELLPDTGEDSDEDGH | ASSSGQSANTEVF | 0.7573241 |
| ELPEAPANRATVPQD | ASSSGQSANTEVF | 0.712311 |
| FGQSKIERVTGIG | ASSSGQSANTEVF | 0.7128581 |
| GAGGTGSTAGGAGGAGGAGGLYA | ASSSGQSANTEVF | 0.82610106 |
| GHGGTGAAGGNGGTGGAGGSGIDG | ASSSGQSANTEVF | 0.58700943 |
| GNGGNGGAGGEGGPGGAGGAGA | ASSSGQSANTEVF | 0.8160259 |
| GSTAGNGGNGGNATNGTIAGQPAG | ASSSGQSANTEVF | 0.6646991 |
| GTGGDGGNGGNGGTGG | ASSSGQSANTEVF | 0.864059 |
| GVRRPAGIPADAPV | ASSSGQSANTEVF | 0.5706485 |
| GYRQVTDAHLLTLARR | ASSSGQSANTEVF | 0.8328479 |
| IAKMGQDGHDRG | ASSSGQSANTEVF | 0.76714903 |
| KAEPSIEIQYT | ASSSGQSANTEVF | 0.90543675 |
| KPVRIRMAADEDTVAVT | ASSSGQSANTEVF | 0.76543677 |
| MPPRIAGMRLLVIKPEP | ASSSGQSANTEVF | 0.79898834 |
| MKFYAVALFRSRGVPPTR | ASSSGQSANTEVF | 0.70840824 |
| NAGGFGILGAVAHSP | ASSSGQSANTEVF | 0.8440887 |
| NEGREITLAMAEM | ASSSGQSANTEVF | 0.5893122 |
| NELVEWSHPLGHRWRWE | ASSSGQSANTEVF | 0.7718576 |
| NLGAGGTGGNGGNPG | ASSSGQSANTEVF | 0.44928578 |
| NTEWPKWETVMAADPVD | ASSSGQSANTEVF | 0.82952106 |
| PASAAPAETQP | ASSSGQSANTEVF | 0.8050004 |
| QQSSDGKNGVRAAEITGE | ASSSGQSANTEVF | 0.5983362 |
| RAPDAVTTSWWKE | ASSSGQSANTEVF | 0.8505837 |
| SGFGNVGAHTSGWFNQ | ASSSGQSANTEVF | 0.8071341 |
| SGVAGAVGQAASVGGLKV | ASSSGQSANTEVF | 0.7597625 |
| SNGGAGTGSAGGNGGT | ASSSGQSANTEVF | 0.8915774 |
| TGGAGGEGGAAG | ASSSGQSANTEVF | 0.7725147 |
| TGGDGGAAGAGGAPGHG | ASSSGQSANTEVF | 0.87977016 |
| TMPPLLAAIANAGRHGV | ASSSGQSANTEVF | 0.83286786 |
| TSSTAGNSGGA | ASSSGQSANTEVF | 0.71862555 |
| VPATVPVGILC | ASSSGQSANTEVF | 0.81836426 |
| VQRPDSGGVVLTGRIS | ASSSGQSANTEVF | 0.64692396 |
| WTPNPVPATKGVPAGLVQAT | ASSSGQSANTEVF | 0.72380006 |
| YALKRALAKFSDRFDVVII | ASSSGQSANTEVF | 0.86710525 |
| YLIGSVIV | ASSSGQSANTEVF | 0.68145454 |
| YGQLVAVID | ASSSGQSANTEVF | 0.69667614 |
| VTVGGMTLPA | ASSSGQSANTEVF | 0.6194936 |
| VLALPAV | ASSSGQSANTEVF | 0.5636716 |
| VIPPTLNYET | ASSSGQSANTEVF | 0.63315254 |
| VGMGGMHPSSG | ASSSGQSANTEVF | 0.6946279 |
| VGLAGGAATA | ASSSGQSANTEVF | 0.7046442 |
| VGGAGGAGGSGP | ASSSGQSANTEVF | 0.68385863 |
| VFAAFMV | ASSSGQSANTEVF | 0.5544163 |
| TLYDKLR | ASSSGQSANTEVF | 0.6942713 |
| TLAVFTVL | ASSSGQSANTEVF | 0.6631888 |

| | | |
|---|---|---|
| TGSVQPARV | ASSSGQSANTEVF | 0.66448236 |
| TGGGAGSGGT | ASSSGQSANTEVF | 0.6777804 |
| SHLETSL | ASSSGQSANTEVF | 0.6839311 |
| SFTTGMT | ASSSGQSANTEVF | 0.63998246 |
| QVEVAKRMGVRQPTVS | ASSSGQSANTEVF | 0.5766572 |
| QGSGGPALAM | ASSSGQSANTEVF | 0.6849796 |
| MLAPSNP | ASSSGQSANTEVF | 0.6941627 |
| LVNNAAI | ASSSGQSANTEVF | 0.69225276 |
| LTGEINTLA | ASSSGQSANTEVF | 0.70871204 |
| LSIFGPV | ASSSGQSANTEVF | 0.6989732 |
| LLALLANL | ASSSGQSANTEVF | 0.6735513 |
| LISNASDALDKLR | ASSSGQSANTEVF | 0.34599227 |
| LINEAVP | ASSSGQSANTEVF | 0.4811231 |
| LGYAFSGL | ASSSGQSANTEVF | 0.61581206 |
| LDFVAVAL | ASSSGQSANTEVF | 0.6512788 |
| LAQPGAE | ASSSGQSANTEVF | 0.34994665 |
| ISNLGAVTGF | ASSSGQSANTEVF | 0.6812781 |
| HQEDPGRI | ASSSGQSANTEVF | 0.61526585 |
| GTPGEPTP | ASSSGQSANTEVF | 0.6368742 |
| GFFNSGS | ASSSGQSANTEVF | 0.69684386 |
| GAGGGGLTAGAN | ASSSGQSANTEVF | 0.63115346 |
| FVDALVNM | ASSSGQSANTEVF | 0.573477 |
| FLTVAPEL | ASSSGQSANTEVF | 0.6331726 |
| FETLTAL | ASSSGQSANTEVF | 0.70746195 |
| DVINPIEL | ASSSGQSANTEVF | 0.5654259 |
| APVSGAPV | ASSSGQSANTEVF | 0.70035064 |
| ALVGGLAW | ASSSGQSANTEVF | 0.62224287 |
| ALAAAAATAI | ASSSGQSANTEVF | 0.7063011 |
| AFGLGLLL | ASSSGQSANTEVF | 0.5187254 |
| AFFAAASL | ASSSGQSANTEVF | 0.696323 |
| AAAALPAFSPP | ASSSGQSANTEVF | 0.66793 |
| VVLAPAL | ASSSGQSANTEVF | 0.7161263 |
| TIGHVDHGKTTL | ASSSGQSANTEVF | 0.6362622 |
| LISNASDALDK | ASSSGQSANTEVF | 0.68437976 |
| LISNASDAL | ASSSGQSANTEVF | 0.64849347 |
| ISNASDALDK | ASSSGQSANTEVF | 0.700451 |
| NQAETLVYQTE | ASSSGQSANTEVF | 0.5424567 |
| TGGTGGTAGAG | ASSSGQSANTEVF | 0.6398706 |
| TGSTAGGAGGA | ASSSGQSANTEVF | 0.52844 |
| DPLAVPVAAGPV | ASSSGQSANTEVF | 0.6646433 |
| GGTGGTPGNGGN | ASSSGQSANTEVF | 0.6825969 |
| PAGLAGVDPATL | ASSSGQSANTEVF | 0.34921488 |
| GNNLVVEAPHAGASV | ASSSGQSANTEVF | 0.69843984 |
| KVTRIMSASTLEQPA | ASSSGQSANTEVF | 0.55443853 |
| LGFALPIALPRRTVV | ASSSGQSANTEVF | 0.55181944 |
| TRLGNLTARKLVVLV | ASSSGQSANTEVF | 0.69583267 |
| WSQQRRLIYFPSAGP | ASSSGQSANTEVF | 0.68097186 |
| GPNGNPDPMAAAVDIR | ASSSGQSANTEVF | 0.6891516 |
| TQRCVSVADLRAAASR | ASSSGQSANTEVF | 0.6452322 |
| QFVLDVAEHPVEVLTETV | ASSSGQSANTEVF | 0.66889125 |
| GGAGGTGGAGGNGGTQAGDG | ASSSGQSANTEVF | 0.6783948 |

| | | |
|---|---|---|
| PLLATLRAIAVDVDAKPAQV | ASSSGQSANTEVF | 0.5578664 |
| APKFRTGGYMIDFWGVGYQVAKRM | ASSSGQSANTEVF | 0.6163697 |
| GAGGAGGAGGAAGGPGGTGGTGGN | ASSSGQSANTEVF | 0.70265126 |
| IGALGLLATDVAPAGSVAFGVALA | ASSSGQSANTEVF | 0.78956383 |
| AGGAGGIGGNGGAGGAGGVLMGNG | ASSSGQSANTEVF | 0.8069701 |
| ADELIGLL | ASSSGQSANTEVF | 0.73615456 |
| ADRALKA | ASSSGQSANTEVF | 0.6436909 |
| AGAVGQVGGAGGA | ASSSGQSANTEVF | 0.7595349 |
| AGGGGGLGGAP | ASSSGQSANTEVF | 0.8105406 |
| AGGLLYGNGGAG | ASSSGQSANTEVF | 0.8184098 |
| AGLESIGL | ASSSGQSANTEVF | 0.8015004 |
| AGLKTGII | ASSSGQSANTEVF | 0.5635207 |
| ASLAALLL | ASSSGQSANTEVF | 0.80314755 |
| ASVINISSI | ASSSGQSANTEVF | 0.747193 |
| ATPPGPA | ASSSGQSANTEVF | 0.7612785 |
| ATVASALV | ASSSGQSANTEVF | 0.7963044 |
| ATVASLIL | ASSSGQSANTEVF | 0.75163954 |
| AVNGPVLIRH | ASSSGQSANTEVF | 0.74835044 |
| DGAAGAVGI | ASSSGQSANTEVF | 0.66702855 |
| DPLLHGI | ASSSGQSANTEVF | 0.80957735 |
| ELGAALGAL | ASSSGQSANTEVF | 0.81334096 |
| ELGRTQAV | ASSSGQSANTEVF | 0.810264 |
| FVAGGYLS | ASSSGQSANTEVF | 0.8270384 |
| GASGGAGGAGGW | ASSSGQSANTEVF | 0.8128664 |
| GGAGAGGDGGAGGI | ASSSGQSANTEVF | 0.79217595 |
| GTGHNGSQPGMGGNG | ASSSGQSANTEVF | 0.26069376 |
| GYAAASASAS | ASSSGQSANTEVF | 0.4892745 |
| HQVDVAAVQQG | ASSSGQSANTEVF | 0.662607 |
| IGAAIAHT | ASSSGQSANTEVF | 0.74563324 |
| IGMIEVAQL | ASSSGQSANTEVF | 0.26724294 |
| IRLAPEHTSAMD | ASSSGQSANTEVF | 0.75812185 |
| IYNVSVL | ASSSGQSANTEVF | 0.7242665 |
| KQGFLGM | ASSSGQSANTEVF | 0.66262555 |
| LESVITQL | ASSSGQSANTEVF | 0.8169848 |
| LGLGALVVV | ASSSGQSANTEVF | 0.70673555 |
| LIGNGGNGGSGGI | ASSSGQSANTEVF | 0.64931107 |
| LNAINAPTL | ASSSGQSANTEVF | 0.7108785 |
| LQLLNTGV | ASSSGQSANTEVF | 0.8250281 |
| LQTSIEF | ASSSGQSANTEVF | 0.6045431 |
| MSFAVSEL | ASSSGQSANTEVF | 0.8103266 |
| NTGALGL | ASSSGQSANTEVF | 0.74870574 |
| PPGAGGAGGAGTT | ASSSGQSANTEVF | 0.81942284 |
| QLVGLGI | ASSSGQSANTEVF | 0.46064615 |
| RFLEQVHQL | ASSSGQSANTEVF | 0.8081353 |
| RGVGYVLR | ASSSGQSANTEVF | 0.7758826 |
| SATGVFAGVI | ASSSGQSANTEVF | 0.8325274 |
| SLPAITLP | ASSSGQSANTEVF | 0.733134 |
| STALAFTV | ASSSGQSANTEVF | 0.7620063 |
| SVLTVGLGK | ASSSGQSANTEVF | 0.64644754 |
| TALAGAGSGI | ASSSGQSANTEVF | 0.8131939 |
| TGLPFTGI | ASSSGQSANTEVF | 0.48025835 |

| | | |
|---|---|---|
| VAEHVTV | ASSSGQSANTEVF | 0.76190054 |
| VDIVLNSL | ASSSGQSANTEVF | 0.61797786 |
| VLTGGGAVL | ASSSGQSANTEVF | 0.7892322 |
| VNVGLAYGI | ASSSGQSANTEVF | 0.7976952 |
| VSSAAVGAV | ASSSGQSANTEVF | 0.28082263 |
| VTSVYEAGV | ASSSGQSANTEVF | 0.8188132 |
| WVALDRGA | ASSSGQSANTEVF | 0.6629578 |
| YLVGSVVI | ASSSGQSANTEVF | 0.5953784 |
| GPPGTGKTLL | ASSSGQSANTEVF | 0.8126638 |
| GVGMTKF | ASSSGQSANTEVF | 0.7504606 |
| IGIGGSDL | ASSSGQSANTEVF | 0.8100307 |
| IGTLLAEL | ASSSGQSANTEVF | 0.71411467 |
| IYTSGTT | ASSSGQSANTEVF | 0.7896172 |
| LSFPAGV | ASSSGQSANTEVF | 0.8068723 |
| LYTSGTT | ASSSGQSANTEVF | 0.4422556 |
| VGAFPFGF | ASSSGQSANTEVF | 0.675468 |
| VGAGTVEF | ASSSGQSANTEVF | 0.81912315 |
| VGKSTLLL | ASSSGQSANTEVF | 0.6523516 |
| AALAHAGGGIV | ASSSGQSANTEVF | 0.6815292 |
| AALVAVSLL | ASSSGQSANTEVF | 0.5521404 |
| AAVSRAVAAL | ASSSGQSANTEVF | 0.42828318 |
| AGGSSGIGGF | ASSSGQSANTEVF | 0.5215076 |
| AGLTVANANG | ASSSGQSANTEVF | 0.6724617 |
| AGLVGAVL | ASSSGQSANTEVF | 0.69912744 |
| ALSQAFTV | ASSSGQSANTEVF | 0.6338416 |
| ALTFSVSL | ASSSGQSANTEVF | 0.40361288 |
| ALTGAALVAL | ASSSGQSANTEVF | 0.66928554 |
| ASFSIGAGVL | ASSSGQSANTEVF | 0.62395686 |
| DGSVYKI | ASSSGQSANTEVF | 0.5392238 |
| EQLAEIV | ASSSGQSANTEVF | 0.6020764 |
| FAVSELTVM | ASSSGQSANTEVF | 0.64085656 |
| GAAPLAAGL | ASSSGQSANTEVF | 0.5063051 |
| GADGGDGGSGGAGG | ASSSGQSANTEVF | 0.4874832 |
| GFGLDTGGAGG | ASSSGQSANTEVF | 0.633838 |
| GLASGGLL | ASSSGQSANTEVF | 0.6756834 |
| GRIAGAAVLL | ASSSGQSANTEVF | 0.6771222 |
| GSAGSLNTSF | ASSSGQSANTEVF | 0.73279405 |
| GTGQIGI | ASSSGQSANTEVF | 0.70837027 |
| GTLIALIL | ASSSGQSANTEVF | 0.68055975 |
| IAATMGFVVG | ASSSGQSANTEVF | 0.2864595 |
| IFAQALL | ASSSGQSANTEVF | 0.35863757 |
| IGAAVALGF | ASSSGQSANTEVF | 0.5652228 |
| IGLTPLAL | ASSSGQSANTEVF | 0.5381856 |
| ILGSNTSTL | ASSSGQSANTEVF | 0.30064562 |
| IVFSLTVL | ASSSGQSANTEVF | 0.6047143 |
| LASKATGF | ASSSGQSANTEVF | 0.5387881 |
| LASVALIL | ASSSGQSANTEVF | 0.46611142 |
| LFAVGLTL | ASSSGQSANTEVF | 0.68816626 |
| LFNGGIGTYI | ASSSGQSANTEVF | 0.53048146 |
| LTLGAGII | ASSSGQSANTEVF | 0.46259922 |
| LVAAALGDI | ASSSGQSANTEVF | 0.51290596 |

| | | |
|---|---|---|
| NTAFVQLGI | ASSSGQSANTEVF | 0.6549436 |
| QLFSGAEL | ASSSGQSANTEVF | 0.41869578 |
| SAATFITL | ASSSGQSANTEVF | 0.6724365 |
| SNYGLANLGV | ASSSGQSANTEVF | 0.4455424 |
| SYGGAYAVM | ASSSGQSANTEVF | 0.72196305 |
| TTGVGGLTLG | ASSSGQSANTEVF | 0.37636703 |
| TTQTIYI | ASSSGQSANTEVF | 0.68877435 |
| VAAGAGLGAVI | ASSSGQSANTEVF | 0.5862928 |
| VAAVVTLL | ASSSGQSANTEVF | 0.7615862 |
| VGFLGGLI | ASSSGQSANTEVF | 0.56960773 |
| VGVIPILL | ASSSGQSANTEVF | 0.6508961 |
| VIFASSLI | ASSSGQSANTEVF | 0.57193744 |
| VKGGDGGAAAG | ASSSGQSANTEVF | 0.6804067 |
| VTKTFTNI | ASSSGQSANTEVF | 0.3770104 |
| VVAAAILL | ASSSGQSANTEVF | 0.5943209 |
| YAALDVEL | ASSSGQSANTEVF | 0.37454838 |
| LASVALLL | ASSSGQSANTEVF | 0.6153087 |
| LDAVHRV | ASSSGQSANTEVF | 0.66233075 |
| QGLSSSIF | ASSSGQSANTEVF | 0.2757411 |
| RGTFANIRL | ASSSGQSANTEVF | 0.667461 |
| RQLWWGHRI | ASSSGQSANTEVF | 0.46399426 |
| YGPPGTGKTLL | ASSSGQSANTEVF | 0.47453505 |
| AALVGAPANVA | ASSSGQSANTEVF | 0.70595145 |
| AATFRRTE | ASSSGQSANTEVF | 0.65594137 |
| AGALGGGI | ASSSGQSANTEVF | 0.6955248 |
| AGGAGGAGGIGAG | ASSSGQSANTEVF | 0.52371895 |
| AGSGALGL | ASSSGQSANTEVF | 0.551407 |
| AQIADQI | ASSSGQSANTEVF | 0.5846334 |
| ASAVAVTYL | ASSSGQSANTEVF | 0.49314165 |
| ASTVLGGTGG | ASSSGQSANTEVF | 0.49726453 |
| AVGGEPAK | ASSSGQSANTEVF | 0.7164818 |
| CEQGTFGGTC | ASSSGQSANTEVF | 0.8622898 |
| DEGKAPAG | ASSSGQSANTEVF | 0.8672913 |
| DTVVFPAL | ASSSGQSANTEVF | 0.7576283 |
| FEGITMAG | ASSSGQSANTEVF | 0.7240299 |
| FPLVVQD | ASSSGQSANTEVF | 0.8177974 |
| GAFTTPQ | ASSSGQSANTEVF | 0.86564827 |
| GFGGFGTAGGVGGA | ASSSGQSANTEVF | 0.8792687 |
| GFLAQFL | ASSSGQSANTEVF | 0.85692585 |
| GGAGGAGGNAGVA | ASSSGQSANTEVF | 0.7371692 |
| GGVGTAGAGGAGGAGG | ASSSGQSANTEVF | 0.8464432 |
| GNGGGGDAGNAGSGGN | ASSSGQSANTEVF | 0.8182548 |
| GTGGAGGAGNDAGS | ASSSGQSANTEVF | 0.8083863 |
| GTTNTGWFNS | ASSSGQSANTEVF | 0.8620286 |
| IFMGPLFTL | ASSSGQSANTEVF | 0.84411204 |
| ILAHAIEAI | ASSSGQSANTEVF | 0.8136685 |
| IRSAIREAL | ASSSGQSANTEVF | 0.79000926 |
| ITFAGGG | ASSSGQSANTEVF | 0.84751356 |
| ITGMSGHQEV | ASSSGQSANTEVF | 0.875147 |
| IVAGLACTL | ASSSGQSANTEVF | 0.86584646 |
| IVGGGLAGA | ASSSGQSANTEVF | 0.8827202 |

| | | |
|---|---|---|
| IVTGVLNI | ASSSGQSANTEVF | 0.8685711 |
| LAQQGCI | ASSSGQSANTEVF | 0.8646339 |
| LCLHVEN | ASSSGQSANTEVF | 0.35889408 |
| LDAISDGS | ASSSGQSANTEVF | 0.6073711 |
| LGGFAAML | ASSSGQSANTEVF | 0.78910005 |
| LGGKNPMI | ASSSGQSANTEVF | 0.81081116 |
| LGPKGVAAIL | ASSSGQSANTEVF | 0.5813632 |
| LMVGVSI | ASSSGQSANTEVF | 0.8593956 |
| LVGAAFVL | ASSSGQSANTEVF | 0.8083576 |
| LVLAVAPL | ASSSGQSANTEVF | 0.73201233 |
| MAAIAVIDAA | ASSSGQSANTEVF | 0.8679831 |
| NNGINAELQ | ASSSGQSANTEVF | 0.80791557 |
| RGEPLVLVHG | ASSSGQSANTEVF | 0.7414142 |
| RGGAGGAAGLI | ASSSGQSANTEVF | 0.77164364 |
| RSYRMHPAV | ASSSGQSANTEVF | 0.8775022 |
| SAAHLLAGQLA | ASSSGQSANTEVF | 0.6900008 |
| SAAVGSY | ASSSGQSANTEVF | 0.8591502 |
| SPHAVLHRI | ASSSGQSANTEVF | 0.8141775 |
| SQIAKVL | ASSSGQSANTEVF | 0.8700688 |
| TALGHLVP | ASSSGQSANTEVF | 0.59356666 |
| TAPTTSGAA | ASSSGQSANTEVF | 0.8646716 |
| TDGDDGNGGSPGT | ASSSGQSANTEVF | 0.8240964 |
| TDGLGEN | ASSSGQSANTEVF | 0.8868344 |
| TGGPPLPGQA | ASSSGQSANTEVF | 0.8138244 |
| TIPAITI | ASSSGQSANTEVF | 0.85605776 |
| TIPGFTI | ASSSGQSANTEVF | 0.83865446 |
| TSIASGALA | ASSSGQSANTEVF | 0.86007017 |
| TVTLPGAAI | ASSSGQSANTEVF | 0.63307846 |
| VASGLSAALP | ASSSGQSANTEVF | 0.8168434 |
| VGAGVIGI | ASSSGQSANTEVF | 0.72870266 |
| VGPKRFGV | ASSSGQSANTEVF | 0.8344865 |
| VNVAFPD | ASSSGQSANTEVF | 0.85565734 |
| VSMTNSGVS | ASSSGQSANTEVF | 0.4290874 |
| VTDVLGDLSV | ASSSGQSANTEVF | 0.86050063 |
| VVVVGAGFAGL | ASSSGQSANTEVF | 0.77896965 |
| VYTVGTVV | ASSSGQSANTEVF | 0.7647516 |
| WTTFSGP | ASSSGQSANTEVF | 0.8729643 |
| YDELVEAF | ASSSGQSANTEVF | 0.82774687 |
| YGTGGAGGNGG | ASSSGQSANTEVF | 0.8719442 |
| YTNVMAV | ASSSGQSANTEVF | 0.823015 |
| FLEQVHQL | ASSSGQSANTEVF | 0.8196659 |
| GLSLLAVAV | ASSSGQSANTEVF | 0.8347033 |
| LEQVHQL | ASSSGQSANTEVF | 0.6081101 |
| LNAAGAI | ASSSGQSANTEVF | 0.7195597 |
| PAVHASL | ASSSGQSANTEVF | 0.8783474 |
| TELEDDA | ASSSGQSANTEVF | 0.7345206 |
| TPPGPPN | ASSSGQSANTEVF | 0.8510379 |
| TPPPPPPP | ASSSGQSANTEVF | 0.54099166 |
| TQSPALI | GGRDSSGNTLY | 0.4662999 |
| EFAAAFPQ | GGRDSSGNTLY | 0.69473225 |
| LLSGNLDVL | GGRDSSGNTLY | 0.7997631 |

| | | |
|---|---|---|
| GPANLGNYNI | GGRDSSGNTLY | 0.8502567 |
| TLGPTLP | GGRDSSGNTLY | 0.773456 |
| VYAYSSCCK | GGRDSSGNTLY | 0.4269557 |
| APAPAPAPNI | GGRDSSGNTLY | 0.6896876 |
| PGGPVPIP | GGRDSSGNTLY | 0.5312536 |
| FLTVALEP | GGRDSSGNTLY | 0.66007864 |
| SQYFGANL | GGRDSSGNTLY | 0.78102016 |
| GGVKPAA | GGRDSSGNTLY | 0.5832628 |
| GGTGGAGGAGGW | GGRDSSGNTLY | 0.61298275 |
| GVGGVAGLGGAGG | GGRDSSGNTLY | 0.56591815 |
| VLLVLATTI | GGRDSSGNTLY | 0.8133169 |
| IGIFAAML | GGRDSSGNTLY | 0.81061274 |
| AVGDLID | GGRDSSGNTLY | 0.7704153 |
| NISPLPL | GGRDSSGNTLY | 0.8736528 |
| VTAVFVAI | GGRDSSGNTLY | 0.79344845 |
| SGDDNDAAL | GGRDSSGNTLY | 0.8325628 |
| AVAALAHS | GGRDSSGNTLY | 0.17572784 |
| GGVTVMATP | GGRDSSGNTLY | 0.31751162 |
| AGLAANVRD | GGRDSSGNTLY | 0.60381305 |
| TNRPDLIDPALLRPG | GGRDSSGNTLY | 0.62452286 |
| LANLGVTAL | GGRDSSGNTLY | 0.33447623 |
| TGERIEL | GGRDSSGNTLY | 0.79158247 |
| NGEELAA | GGRDSSGNTLY | 0.6212628 |
| VLSLVWG | GGRDSSGNTLY | 0.42000538 |
| IGYALWL | GGRDSSGNTLY | 0.8208426 |
| LSLLLAPL | GGRDSSGNTLY | 0.6219301 |
| TLDEALK | GGRDSSGNTLY | 0.5500271 |
| TDGAGTAEGG | GGRDSSGNTLY | 0.6010778 |
| NAPGGAGGNGG | GGRDSSGNTLY | 0.8087754 |
| VSVFMGMGGNF | GGRDSSGNTLY | 0.38386166 |
| LASYLAAL | GGRDSSGNTLY | 0.7292118 |
| LVGGTPAL | GGRDSSGNTLY | 0.6408323 |
| VRNADVL | GGRDSSGNTLY | 0.7953379 |
| TCSSPFLVSAM | GGRDSSGNTLY | 0.30938116 |
| VHITDAE | GGRDSSGNTLY | 0.7679592 |
| EYAAPSF | GGRDSSGNTLY | 0.6199069 |
| VVDAFHGGL | GGRDSSGNTLY | 0.8893311 |
| ASVVDPQ | GGRDSSGNTLY | 0.5603938 |
| SYNEFGGLS | GGRDSSGNTLY | 0.6114676 |
| DPDGSVY | GGRDSSGNTLY | 0.632564 |
| MVSTIHATE | GGRDSSGNTLY | 0.8295618 |
| LAAGVADPV | GGRDSSGNTLY | 0.28852487 |
| TKEQIAA | GGRDSSGNTLY | 0.63345885 |
| SLASPAAL | GGRDSSGNTLY | 0.4152615 |
| IGLVFYGV | GGRDSSGNTLY | 0.73295987 |
| FLLDASNL | GGRDSSGNTLY | 0.7315481 |
| YQVGNFP | GGRDSSGNTLY | 0.37857017 |
| ESLLSQST | GGRDSSGNTLY | 0.7755904 |
| LFTVGFAAVG | GGRDSSGNTLY | 0.61362267 |
| ATPPTPT | GGRDSSGNTLY | 0.6049717 |
| TDLATLG | GGRDSSGNTLY | 0.8396074 |

| | | |
|---|---|---|
| SAGGTGR | GGRDSSGNTLY | 0.6440314 |
| GFSNSGS | GGRDSSGNTLY | 0.861371 |
| GGNGGSGGTSV | GGRDSSGNTLY | 0.7300496 |
| GTSVATGGAGN | GGRDSSGNTLY | 0.640712 |
| NGGNGGSGGTS | GGRDSSGNTLY | 0.7921032 |
| AVVKFPH | GGRDSSGNTLY | 0.5325293 |
| TPPPPPP | GGRDSSGNTLY | 0.61424613 |
| AAPAAAAAPA | GGRDSSGNTLY | 0.8052368 |
| IALNEDL | GGRDSSGNTLY | 0.80836105 |
| IVNVSSM | GGRDSSGNTLY | 0.81785 |
| TGIPPAPR | GGRDSSGNTLY | 0.7537738 |
| FDLGGGTF | GGRDSSGNTLY | 0.74773914 |
| FDLGGGTFDVS | GGRDSSGNTLY | 0.7762395 |
| NEPTAAA | GGRDSSGNTLY | 0.81944776 |
| NEPTAAALA | GGRDSSGNTLY | 0.818506 |
| GSIINISSI | GGRDSSGNTLY | 0.81127733 |
| VAPGNAGTA | GGRDSSGNTLY | 0.7556443 |
| TGGGASLE | GGRDSSGNTLY | 0.80359364 |
| VAVYNHYKRI | GGRDSSGNTLY | 0.7943574 |
| TDPAPATT | GGRDSSGNTLY | 0.7816359 |
| TNRPDILDPALLRPG | GGRDSSGNTLY | 0.81140393 |
| LTTTTTS | GGRDSSGNTLY | 0.8056673 |
| PTGSGKTL | GGRDSSGNTLY | 0.75950426 |
| TGSGKTL | GGRDSSGNTLY | 0.7917903 |
| TIGHVDHGK | GGRDSSGNTLY | 0.8013803 |
| TIGHVDHGKTT | GGRDSSGNTLY | 0.80832756 |
| GHVDHGKTT | GGRDSSGNTLY | 0.8077114 |
| AQFEHTLL | GGRDSSGNTLY | 0.82694924 |
| IVNQDEL | GGRDSSGNTLY | 0.81693625 |
| TNGSQFF | GGRDSSGNTLY | 0.80980945 |
| TGKGIVE | GGRDSSGNTLY | 0.46248794 |
| LISNASDA | GGRDSSGNTLY | 0.6758868 |
| AELRTLGELTKIAKAHGAQ | GGRDSSGNTLY | 0.7843691 |
| APNAVAAAEPSW | GGRDSSGNTLY | 0.7875279 |
| APQVAGAATAVL | GGRDSSGNTLY | 0.5744494 |
| ARRDPAAAAAILPTDA | GGRDSSGNTLY | 0.810233 |
| DAAARDVLAEAGLAEYF | GGRDSSGNTLY | 0.7690987 |
| DDQIVLSAKANLAKVEQ | GGRDSSGNTLY | 0.7593578 |
| DELLPDTGEDSDEDGH | GGRDSSGNTLY | 0.8150569 |
| ELPEAPANRATVPQD | GGRDSSGNTLY | 0.78574 |
| FGQSKIERVTGIG | GGRDSSGNTLY | 0.77843255 |
| GAGGTGSTAGGAGGAGGAGGLYA | GGRDSSGNTLY | 0.78988487 |
| GHGGTGAAGGNGGTGGAGGSGIDG | GGRDSSGNTLY | 0.8166915 |
| GNGGNGGAGGEGGPGGAGGAGA | GGRDSSGNTLY | 0.74158424 |
| GSTAGNGGNGGNATNGTIAGQPAG | GGRDSSGNTLY | 0.8163246 |
| GTGGDGGNGGNGGTGG | GGRDSSGNTLY | 0.7376387 |
| GVRRPAGIPADAPV | GGRDSSGNTLY | 0.82310075 |
| GYRQVTDAHLLTLARR | GGRDSSGNTLY | 0.72918236 |
| IAKMGQDGHDRG | GGRDSSGNTLY | 0.81487054 |
| KAEPSIEIQYT | GGRDSSGNTLY | 0.79489744 |
| KPVRIRMAADEDTVAVT | GGRDSSGNTLY | 0.8304879 |

| | | |
|---|---|---|
| MPPRIAGMRLLVIKPEP | GGRDSSGNTLY | 0.789098 |
| MKFYAVALFRSRGVPPTR | GGRDSSGNTLY | 0.81497586 |
| NAGGFGILGAVAHSP | GGRDSSGNTLY | 0.78567827 |
| NEGREITLAMAEM | GGRDSSGNTLY | 0.8196875 |
| NELVEWSHPLGHRWRWE | GGRDSSGNTLY | 0.71608305 |
| NLGAGGTGGNGGNPG | GGRDSSGNTLY | 0.7943652 |
| NTEWPKWETVMAADPVD | GGRDSSGNTLY | 0.6435678 |
| PASAAPAETQP | GGRDSSGNTLY | 0.8081175 |
| QQSSDGKNGVRAAEITGE | GGRDSSGNTLY | 0.8107865 |
| RAPDAVTTSWWKE | GGRDSSGNTLY | 0.57254696 |
| SGFGNVGAHTSGWFNQ | GGRDSSGNTLY | 0.8199941 |
| SGVAGAVGQAASVGGLKV | GGRDSSGNTLY | 0.7569953 |
| SNGGAGTGSAGGNGGT | GGRDSSGNTLY | 0.77291274 |
| TGGAGGEGGAAG | GGRDSSGNTLY | 0.8208202 |
| TGGDGGAAGAGGAPGHG | GGRDSSGNTLY | 0.7844745 |
| TMPPLLAAIANAGRHGV | GGRDSSGNTLY | 0.81416166 |
| TSSTAGNSGGA | GGRDSSGNTLY | 0.77324754 |
| VPATVPVGILC | GGRDSSGNTLY | 0.7809706 |
| VQRPDSGGVVLTGRIS | GGRDSSGNTLY | 0.80249554 |
| WTPNPVPATKGVPAGLVQAT | GGRDSSGNTLY | 0.66557956 |
| YALKRALAKFSDRFDVVII | GGRDSSGNTLY | 0.76323366 |
| YLIGSVIV | GGRDSSGNTLY | 0.82472134 |
| YGQLVAVID | GGRDSSGNTLY | 0.63859373 |
| VTVGGMTLPA | GGRDSSGNTLY | 0.7066731 |
| VLALPAV | GGRDSSGNTLY | 0.53934824 |
| VIPPTLNYET | GGRDSSGNTLY | 0.51135314 |
| VGMGGMHPSSG | GGRDSSGNTLY | 0.5369369 |
| VGLAGGAATA | GGRDSSGNTLY | 0.6994611 |
| VGGAGGAGGSGP | GGRDSSGNTLY | 0.6882129 |
| VFAAFMV | GGRDSSGNTLY | 0.6405535 |
| TLYDKLR | GGRDSSGNTLY | 0.54893225 |
| TLAVFTVL | GGRDSSGNTLY | 0.62103623 |
| TGSVQPARV | GGRDSSGNTLY | 0.5919423 |
| TGGGAGSGGT | GGRDSSGNTLY | 0.540781 |
| SHLETSL | GGRDSSGNTLY | 0.6419468 |
| SFTTGMT | GGRDSSGNTLY | 0.66011745 |
| QVEVAKRMGVRQPTVS | GGRDSSGNTLY | 0.5675555 |
| QGSGGPALAM | GGRDSSGNTLY | 0.5846289 |
| MLAPSNP | GGRDSSGNTLY | 0.58386725 |
| LVNNAAI | GGRDSSGNTLY | 0.66563904 |
| LTGEINTLA | GGRDSSGNTLY | 0.63416475 |
| LSIFGPV | GGRDSSGNTLY | 0.73027396 |
| LLALLANL | GGRDSSGNTLY | 0.67424786 |
| LISNASDALDKLR | GGRDSSGNTLY | 0.65567034 |
| LINEAVP | GGRDSSGNTLY | 0.30877885 |
| LGYAFSGL | GGRDSSGNTLY | 0.4141206 |
| LDFVAVAL | GGRDSSGNTLY | 0.6587751 |
| LAQPGAE | GGRDSSGNTLY | 0.57634544 |
| ISNLGAVTGF | GGRDSSGNTLY | 0.37913546 |
| HQEDPGRI | GGRDSSGNTLY | 0.6815744 |
| GTPGEPTP | GGRDSSGNTLY | 0.5951924 |

| | | |
|---|---|---|
| GFFNSGS | GGRDSSGNTLY | 0.56959337 |
| GAGGGGLTAGAN | GGRDSSGNTLY | 0.6535737 |
| FVDALVNM | GGRDSSGNTLY | 0.5660152 |
| FLTVAPEL | GGRDSSGNTLY | 0.5709308 |
| FETLTAL | GGRDSSGNTLY | 0.57901245 |
| DVINPIEL | GGRDSSGNTLY | 0.7008119 |
| APVSGAPV | GGRDSSGNTLY | 0.4817023 |
| ALVGGLAW | GGRDSSGNTLY | 0.6454488 |
| ALAAAAATAI | GGRDSSGNTLY | 0.5545499 |
| AFGLGLLL | GGRDSSGNTLY | 0.7055867 |
| AFFAAASL | GGRDSSGNTLY | 0.48547798 |
| AAAALPAFSPP | GGRDSSGNTLY | 0.64134467 |
| VVLAPAL | GGRDSSGNTLY | 0.5592492 |
| TIGHVDHGKTTL | GGRDSSGNTLY | 0.74263686 |
| LISNASDALDK | GGRDSSGNTLY | 0.58738995 |
| LISNASDAL | GGRDSSGNTLY | 0.6537807 |
| ISNASDALDK | GGRDSSGNTLY | 0.59330255 |
| NQAETLVYQTE | GGRDSSGNTLY | 0.6563734 |
| TGGTGGTAGAG | GGRDSSGNTLY | 0.4501627 |
| TGSTAGGAGGA | GGRDSSGNTLY | 0.57533824 |
| DPLAVPVAAGPV | GGRDSSGNTLY | 0.4700703 |
| GGTGGTPGNGGN | GGRDSSGNTLY | 0.6368904 |
| PAGLAGVDPATL | GGRDSSGNTLY | 0.63820606 |
| GNNLVVEAPHAGASV | GGRDSSGNTLY | 0.52560747 |
| KVTRIMSASTLEQPA | GGRDSSGNTLY | 0.663756 |
| LGFALPIALPRRTVV | GGRDSSGNTLY | 0.63317525 |
| TRLGNLTARKLVVLV | GGRDSSGNTLY | 0.5643101 |
| WSQQRRLIYFPSAGP | GGRDSSGNTLY | 0.7045861 |
| GPNGNPDPMAAAVDIR | GGRDSSGNTLY | 0.6112889 |
| TQRCVSVADLRAAASR | GGRDSSGNTLY | 0.6759122 |
| QFVLDVAEHPVEVLTETV | GGRDSSGNTLY | 0.5692768 |
| GGAGGTGGAGGNGGTQAGDG | GGRDSSGNTLY | 0.5564949 |
| PLLATLRAIAVDVDAKPAQV | GGRDSSGNTLY | 0.6841655 |
| APKFRTGGYMIDFWGVGYQVAKRM | GGRDSSGNTLY | 0.59401524 |
| GAGGAGGAGGAAGGPGGTGGTGGN | GGRDSSGNTLY | 0.5691834 |
| IGALGLLATDVAPAGSVAFGVALA | GGRDSSGNTLY | 0.6658397 |
| AGGAGGIGGNGGAGGAGGVLMGNG | GGRDSSGNTLY | 0.5968363 |
| ADELIGLL | GGRDSSGNTLY | 0.8670577 |
| ADRALKA | GGRDSSGNTLY | 0.48196697 |
| AGAVGQVGGAGGA | GGRDSSGNTLY | 0.49566874 |
| AGGGGGLGGAP | GGRDSSGNTLY | 0.39343694 |
| AGGLLYGNGGAG | GGRDSSGNTLY | 0.7034321 |
| AGLESIGL | GGRDSSGNTLY | 0.8609435 |
| AGLKTGII | GGRDSSGNTLY | 0.7753258 |
| ASLAALLL | GGRDSSGNTLY | 0.4060226 |
| ASVINISSI | GGRDSSGNTLY | 0.6320378 |
| ATPPGPA | GGRDSSGNTLY | 0.4943774 |
| ATVASALV | GGRDSSGNTLY | 0.49883163 |
| ATVASLIL | GGRDSSGNTLY | 0.59702015 |
| AVNGPVLIRH | GGRDSSGNTLY | 0.5834893 |
| DGAAGAVGI | GGRDSSGNTLY | 0.4065319 |

| | | |
|---|---|---|
| DPLLHGI | GGRDSSGNTLY | 0.31464317 |
| ELGAALGAL | GGRDSSGNTLY | 0.3875007 |
| ELGRTQAV | GGRDSSGNTLY | 0.6191589 |
| FVAGGYLS | GGRDSSGNTLY | 0.5302321 |
| GASGGAGGAGGW | GGRDSSGNTLY | 0.76767254 |
| GGAGAGGDGGAGGI | GGRDSSGNTLY | 0.71651465 |
| GTGHNGSQPGMGGNG | GGRDSSGNTLY | 0.41026348 |
| GYAAASASAS | GGRDSSGNTLY | 0.11351741 |
| HQVDVAAVQQG | GGRDSSGNTLY | 0.23013648 |
| IGAAIAHT | GGRDSSGNTLY | 0.42349184 |
| IGMIEVAQL | GGRDSSGNTLY | 0.59853053 |
| IRLAPEHTSAMD | GGRDSSGNTLY | 0.10873074 |
| IYNVSVL | GGRDSSGNTLY | 0.57885665 |
| KQGFLGM | GGRDSSGNTLY | 0.5250913 |
| LESVITQL | GGRDSSGNTLY | 0.58310026 |
| LGLGALVVV | GGRDSSGNTLY | 0.7609468 |
| LIGNGGNGGSGGI | GGRDSSGNTLY | 0.55110186 |
| LNAINAPTL | GGRDSSGNTLY | 0.3418971 |
| LQLLNTGV | GGRDSSGNTLY | 0.3001687 |
| LQTSIEF | GGRDSSGNTLY | 0.77901757 |
| MSFAVSEL | GGRDSSGNTLY | 0.18834415 |
| NTGALGL | GGRDSSGNTLY | 0.7170092 |
| PPGAGGAGGAGTT | GGRDSSGNTLY | 0.27006966 |
| QLVGLGI | GGRDSSGNTLY | 0.67446065 |
| RFLEQVHQL | GGRDSSGNTLY | 0.34155852 |
| RGVGYVLR | GGRDSSGNTLY | 0.715095 |
| SATGVFAGVI | GGRDSSGNTLY | 0.6224187 |
| SLPAITLP | GGRDSSGNTLY | 0.8481328 |
| STALAFTV | GGRDSSGNTLY | 0.5055952 |
| SVLTVGLGK | GGRDSSGNTLY | 0.65869415 |
| TALAGAGSGI | GGRDSSGNTLY | 0.2642865 |
| TGLPFTGI | GGRDSSGNTLY | 0.8482654 |
| VAEHVTV | GGRDSSGNTLY | 0.40684468 |
| VDIVLNSL | GGRDSSGNTLY | 0.4145398 |
| VLTGGGAVL | GGRDSSGNTLY | 0.2130522 |
| VNVGLAYGI | GGRDSSGNTLY | 0.5357413 |
| VSSAAVGAV | GGRDSSGNTLY | 0.74895835 |
| VTSVYEAGV | GGRDSSGNTLY | 0.49142253 |
| WVALDRGA | GGRDSSGNTLY | 0.7548742 |
| YLVGSVVI | GGRDSSGNTLY | 0.6987387 |
| GPPGTGKTLL | GGRDSSGNTLY | 0.3934164 |
| GVGMTKF | GGRDSSGNTLY | 0.7174196 |
| IGIGGSDL | GGRDSSGNTLY | 0.43710038 |
| IGTLLAEL | GGRDSSGNTLY | 0.58049905 |
| IYTSGTT | GGRDSSGNTLY | 0.24108396 |
| LSFPAGV | GGRDSSGNTLY | 0.5974168 |
| LYTSGTT | GGRDSSGNTLY | 0.6165242 |
| VGAFPFGF | GGRDSSGNTLY | 0.4259809 |
| VGAGTVEF | GGRDSSGNTLY | 0.3124134 |
| VGKSTLLL | GGRDSSGNTLY | 0.77310944 |
| AALAHAGGGIV | GGRDSSGNTLY | 0.5002352 |

| | | |
|---|---|---|
| AALVAVSLL | GGRDSSGNTLY | 0.5939226 |
| AAVSRAVAAL | GGRDSSGNTLY | 0.3550333 |
| AGGSSGIGGF | GGRDSSGNTLY | 0.31877512 |
| AGLTVANANG | GGRDSSGNTLY | 0.39286754 |
| AGLVGAVL | GGRDSSGNTLY | 0.52021885 |
| ALSQAFTV | GGRDSSGNTLY | 0.6485386 |
| ALTFSVSL | GGRDSSGNTLY | 0.5150421 |
| ALTGAALVAL | GGRDSSGNTLY | 0.3415407 |
| ASFSIGAGVL | GGRDSSGNTLY | 0.5443119 |
| DGSVYKI | GGRDSSGNTLY | 0.4184314 |
| EQLAEIV | GGRDSSGNTLY | 0.44310513 |
| FAVSELTVM | GGRDSSGNTLY | 0.40766487 |
| GAAPLAAGL | GGRDSSGNTLY | 0.4277507 |
| GADGGDGGSGGAGG | GGRDSSGNTLY | 0.3852361 |
| GFGLDTGGAGG | GGRDSSGNTLY | 0.325133 |
| GLASGGLL | GGRDSSGNTLY | 0.48473784 |
| GRIAGAAVLL | GGRDSSGNTLY | 0.6017959 |
| GSAGSLNTSF | GGRDSSGNTLY | 0.54342544 |
| GTGQIGI | GGRDSSGNTLY | 0.59792936 |
| GTLIALIL | GGRDSSGNTLY | 0.59371805 |
| IAATMGFVVG | GGRDSSGNTLY | 0.49719983 |
| IFAQALL | GGRDSSGNTLY | 0.25930157 |
| IGAAVALGF | GGRDSSGNTLY | 0.30266896 |
| IGLTPLAL | GGRDSSGNTLY | 0.34557033 |
| ILGSNTSTL | GGRDSSGNTLY | 0.42041823 |
| IVFSLTVL | GGRDSSGNTLY | 0.25800055 |
| LASKATGF | GGRDSSGNTLY | 0.43752187 |
| LASVALIL | GGRDSSGNTLY | 0.36805683 |
| LFAVGLTL | GGRDSSGNTLY | 0.36020577 |
| LFNGGIGTYI | GGRDSSGNTLY | 0.5989622 |
| LTLGAGII | GGRDSSGNTLY | 0.38142097 |
| LVAAALGDI | GGRDSSGNTLY | 0.33103767 |
| NTAFVQLGI | GGRDSSGNTLY | 0.36442646 |
| QLFSGAEL | GGRDSSGNTLY | 0.6389533 |
| SAATFITL | GGRDSSGNTLY | 0.3167229 |
| SNYGLANLGV | GGRDSSGNTLY | 0.57134247 |
| SYGGAYAVM | GGRDSSGNTLY | 0.3890827 |
| TTGVGGLTLG | GGRDSSGNTLY | 0.6025926 |
| TTQTIYI | GGRDSSGNTLY | 0.3008021 |
| VAAGAGLGAVI | GGRDSSGNTLY | 0.5546494 |
| VAAVVTLL | GGRDSSGNTLY | 0.4456821 |
| VGFLGGLI | GGRDSSGNTLY | 0.6681363 |
| VGVIPILL | GGRDSSGNTLY | 0.3699721 |
| VIFASSLI | GGRDSSGNTLY | 0.48337498 |
| VKGGDGGAAAG | GGRDSSGNTLY | 0.391968 |
| VTKTFTNI | GGRDSSGNTLY | 0.61732405 |
| VVAAAILL | GGRDSSGNTLY | 0.29558325 |
| YAALDVEL | GGRDSSGNTLY | 0.36988795 |
| LASVALLL | GGRDSSGNTLY | 0.353422 |
| LDAVHRV | GGRDSSGNTLY | 0.43595755 |
| QGLSSSIF | GGRDSSGNTLY | 0.52820396 |

| | | |
|---|---|---|
| RGTFANIRL | GGRDSSGNTLY | 0.25175565 |
| RQLWWGHRI | GGRDSSGNTLY | 0.52420324 |
| YGPPGTGKTLL | GGRDSSGNTLY | 0.30342746 |
| AALVGAPANVA | GGRDSSGNTLY | 0.30668807 |
| AATFRRTE | GGRDSSGNTLY | 0.46598205 |
| AGALGGGI | GGRDSSGNTLY | 0.4129975 |
| AGGAGGAGGIGAG | GGRDSSGNTLY | 0.52104646 |
| AGSGALGL | GGRDSSGNTLY | 0.3433593 |
| AQIADQI | GGRDSSGNTLY | 0.49637604 |
| ASAVAVTYL | GGRDSSGNTLY | 0.47992745 |
| ASTVLGGTGG | GGRDSSGNTLY | 0.2772623 |
| AVGGEPAK | GGRDSSGNTLY | 0.32447398 |
| CEQGTFGGTC | GGRDSSGNTLY | 0.6005623 |
| DEGKAPAG | GGRDSSGNTLY | 0.7687031 |
| DTVVFPAL | GGRDSSGNTLY | 0.86254275 |
| FEGITMAG | GGRDSSGNTLY | 0.7138238 |
| FPLVVQD | GGRDSSGNTLY | 0.5148189 |
| GAFTTPQ | GGRDSSGNTLY | 0.7290443 |
| GFGGFGTAGGVGGA | GGRDSSGNTLY | 0.8569086 |
| GFLAQFL | GGRDSSGNTLY | 0.8586076 |
| GGAGGAGGNAGVA | GGRDSSGNTLY | 0.826719 |
| GGVGTAGAGGAGGAGG | GGRDSSGNTLY | 0.4686929 |
| GNGGGGDAGNAGSGGN | GGRDSSGNTLY | 0.7269993 |
| GTGGAGGAGNDAGS | GGRDSSGNTLY | 0.6215966 |
| GTTNTGWFNS | GGRDSSGNTLY | 0.75122327 |
| IFMGPLFTL | GGRDSSGNTLY | 0.8254081 |
| ILAHAIEAI | GGRDSSGNTLY | 0.7270398 |
| IRSAIREAL | GGRDSSGNTLY | 0.72269344 |
| ITFAGGG | GGRDSSGNTLY | 0.6290529 |
| ITGMSGHQEV | GGRDSSGNTLY | 0.7947868 |
| IVAGLACTL | GGRDSSGNTLY | 0.8125603 |
| IVGGGLAGA | GGRDSSGNTLY | 0.7774875 |
| IVTGVLNI | GGRDSSGNTLY | 0.88592005 |
| LAQQGCI | GGRDSSGNTLY | 0.8502103 |
| LCLHVEN | GGRDSSGNTLY | 0.8440911 |
| LDAISDGS | GGRDSSGNTLY | 0.14480293 |
| LGGFAAML | GGRDSSGNTLY | 0.31923667 |
| LGGKNPMI | GGRDSSGNTLY | 0.66639054 |
| LGPKGVAAIL | GGRDSSGNTLY | 0.6815178 |
| LMVGVSI | GGRDSSGNTLY | 0.30582726 |
| LVGAAFVL | GGRDSSGNTLY | 0.7750217 |
| LVLAVAPL | GGRDSSGNTLY | 0.61787295 |
| MAAIAVIDAA | GGRDSSGNTLY | 0.55533105 |
| NNGINAELQ | GGRDSSGNTLY | 0.86364293 |
| RGEPLVLVHG | GGRDSSGNTLY | 0.7023604 |
| RGGAGGAAGLI | GGRDSSGNTLY | 0.5864179 |
| RSYRMHPAV | GGRDSSGNTLY | 0.65116596 |
| SAAHLLAGQLA | GGRDSSGNTLY | 0.86713284 |
| SAAVGSY | GGRDSSGNTLY | 0.4324367 |
| SPHAVLHRI | GGRDSSGNTLY | 0.810211 |
| SQIAKVL | GGRDSSGNTLY | 0.69167215 |

| | | |
|---|---|---|
| TALGHLVP | GGRDSSGNTLY | 0.85148954 |
| TAPTTSGAA | GGRDSSGNTLY | 0.34955847 |
| TDGDDGNGGSPGT | GGRDSSGNTLY | 0.8422146 |
| TDGLGEN | GGRDSSGNTLY | 0.765625 |
| TGGPPLPGQA | GGRDSSGNTLY | 0.9041168 |
| TIPAITI | GGRDSSGNTLY | 0.6248155 |
| TIPGFTI | GGRDSSGNTLY | 0.686668 |
| TSIASGALA | GGRDSSGNTLY | 0.51515526 |
| TVTLPGAAI | GGRDSSGNTLY | 0.85652274 |
| VASGLSAALP | GGRDSSGNTLY | 0.37532556 |
| VGAGVIGI | GGRDSSGNTLY | 0.66839075 |
| VGPKRFGV | GGRDSSGNTLY | 0.434385 |
| VNVAFPD | GGRDSSGNTLY | 0.79079676 |
| VSMTNSGVS | GGRDSSGNTLY | 0.81702745 |
| VTDVLGDLSV | GGRDSSGNTLY | 0.32311347 |
| VVVVGAGFAGL | GGRDSSGNTLY | 0.863858 |
| VYTVGTVV | GGRDSSGNTLY | 0.7388717 |
| WTTFSGP | GGRDSSGNTLY | 0.5252397 |
| YDELVEAF | GGRDSSGNTLY | 0.87497926 |
| YGTGGAGGNGG | GGRDSSGNTLY | 0.5413351 |
| YTNVMAV | GGRDSSGNTLY | 0.8634163 |
| FLEQVHQL | GGRDSSGNTLY | 0.75603366 |
| GLSLLAVAV | GGRDSSGNTLY | 0.72623837 |
| LEQVHQL | GGRDSSGNTLY | 0.8339721 |
| LNAAGAI | GGRDSSGNTLY | 0.391879 |
| PAVHASL | GGRDSSGNTLY | 0.5673782 |
| TELEDDA | GGRDSSGNTLY | 0.8723125 |
| TPPGPPN | GGRDSSGNTLY | 0.7509851 |
| TPPPPPPP | GGRDSSGNTLY | 0.8818009 |
| TQSPALI | ASSDGQGNNQAPL | 0.6147772 |
| EFAAAFPQ | ASSDGQGNNQAPL | 0.47455564 |
| LLSGNLDVL | ASSDGQGNNQAPL | 0.7348915 |
| GPANLGNYNI | ASSDGQGNNQAPL | 0.85108507 |
| TLGPTLP | ASSDGQGNNQAPL | 0.87524986 |
| VYAYSSCCK | ASSDGQGNNQAPL | 0.81001335 |
| APAPAPAPNI | ASSDGQGNNQAPL | 0.48841318 |
| PGGPVPIP | ASSDGQGNNQAPL | 0.6263893 |
| FLTVALEP | ASSDGQGNNQAPL | 0.61619395 |
| SQYFGANL | ASSDGQGNNQAPL | 0.67804325 |
| GGVKPAA | ASSDGQGNNQAPL | 0.83371353 |
| GGTGGAGGAGGW | ASSDGQGNNQAPL | 0.59313315 |
| GVGGVAGLGGAGG | ASSDGQGNNQAPL | 0.55060923 |
| VLLVLATTI | ASSDGQGNNQAPL | 0.59996516 |
| IGIFAAML | ASSDGQGNNQAPL | 0.79815125 |
| AVGDLID | ASSDGQGNNQAPL | 0.81634057 |
| NISPLPL | ASSDGQGNNQAPL | 0.7935512 |
| VTAVFVAI | ASSDGQGNNQAPL | 0.8835287 |
| SGDDNDAAL | ASSDGQGNNQAPL | 0.8253979 |
| AVAALAHS | ASSDGQGNNQAPL | 0.837796 |
| GGVTVMATP | ASSDGQGNNQAPL | 0.15955406 |
| AGLAANVRD | ASSDGQGNNQAPL | 0.31695879 |

| | | |
|---|---|---|
| TNRPDLIDPALLRPG | ASSDGQGNNQAPL | 0.6372213 |
| LANLGVTAL | ASSDGQGNNQAPL | 0.6815464 |
| TGERIEL | ASSDGQGNNQAPL | 0.32134277 |
| NGEELAA | ASSDGQGNNQAPL | 0.7495215 |
| VLSLVWG | ASSDGQGNNQAPL | 0.56894445 |
| IGYALWL | ASSDGQGNNQAPL | 0.4295327 |
| LSLLLAPL | ASSDGQGNNQAPL | 0.84126574 |
| TLDEALK | ASSDGQGNNQAPL | 0.5485612 |
| TDGAGTAEGG | ASSDGQGNNQAPL | 0.51098853 |
| NAPGGAGGNGG | ASSDGQGNNQAPL | 0.554209 |
| VSVFMGMGGNF | ASSDGQGNNQAPL | 0.8575967 |
| LASYLAAL | ASSDGQGNNQAPL | 0.31071126 |
| LVGGTPAL | ASSDGQGNNQAPL | 0.7784552 |
| VRNADVL | ASSDGQGNNQAPL | 0.76817065 |
| TCSSPFLVSAM | ASSDGQGNNQAPL | 0.8255147 |
| VHITDAE | ASSDGQGNNQAPL | 0.32148346 |
| EYAAPSF | ASSDGQGNNQAPL | 0.81637776 |
| VVDAFHGGL | ASSDGQGNNQAPL | 0.6917279 |
| ASVVDPQ | ASSDGQGNNQAPL | 0.88649595 |
| SYNEFGGLS | ASSDGQGNNQAPL | 0.6199291 |
| DPDGSVY | ASSDGQGNNQAPL | 0.67798615 |
| MVSTIHATE | ASSDGQGNNQAPL | 0.59477 |
| LAAGVADPV | ASSDGQGNNQAPL | 0.8591861 |
| TKEQIAA | ASSDGQGNNQAPL | 0.3797533 |
| SLASPAAL | ASSDGQGNNQAPL | 0.6464299 |
| IGLVFYGV | ASSDGQGNNQAPL | 0.49350435 |
| FLLDASNL | ASSDGQGNNQAPL | 0.76993215 |
| YQVGNFP | ASSDGQGNNQAPL | 0.7792338 |
| ESLLSQST | ASSDGQGNNQAPL | 0.512184 |
| LFTVGFAAVG | ASSDGQGNNQAPL | 0.8485577 |
| ATPPTPT | ASSDGQGNNQAPL | 0.7958476 |
| TDLATLG | ASSDGQGNNQAPL | 0.56548214 |
| SAGGTGR | ASSDGQGNNQAPL | 0.86258197 |
| GFSNSGS | ASSDGQGNNQAPL | 0.52273256 |
| GGNGGSGGTSV | ASSDGQGNNQAPL | 0.83331245 |
| GTSVATGGAGN | ASSDGQGNNQAPL | 0.7316958 |
| NGGNGGSGGTS | ASSDGQGNNQAPL | 0.75770015 |
| AVVKFPH | ASSDGQGNNQAPL | 0.8333044 |
| TPPPPPP | ASSDGQGNNQAPL | 0.44231254 |
| AAPAAAAAPA | ASSDGQGNNQAPL | 0.49547687 |
| IALNEDL | ASSDGQGNNQAPL | 0.84952366 |
| IVNVSSM | ASSDGQGNNQAPL | 0.7651234 |
| TGIPPAPR | ASSDGQGNNQAPL | 0.7810062 |
| FDLGGGTF | ASSDGQGNNQAPL | 0.67768717 |
| FDLGGGTFDVS | ASSDGQGNNQAPL | 0.6021056 |
| NEPTAAA | ASSDGQGNNQAPL | 0.6653386 |
| NEPTAAALA | ASSDGQGNNQAPL | 0.7586569 |
| GSIINISSI | ASSDGQGNNQAPL | 0.7861588 |
| VAPGNAGTA | ASSDGQGNNQAPL | 0.76881754 |
| TGGGASLE | ASSDGQGNNQAPL | 0.4508772 |
| VAVYNHYKRI | ASSDGQGNNQAPL | 0.75678 |

| | | |
|---|---|---|
| TDPAPATT | ASSDGQGNNQAPL | 0.67875373 |
| TNRPDILDPALLRPG | ASSDGQGNNQAPL | 0.7020153 |
| LTTTTTS | ASSDGQGNNQAPL | 0.6970652 |
| PTGSGKTL | ASSDGQGNNQAPL | 0.67322755 |
| TGSGKTL | ASSDGQGNNQAPL | 0.72683585 |
| TIGHVDHGK | ASSDGQGNNQAPL | 0.53896725 |
| TIGHVDHGKTT | ASSDGQGNNQAPL | 0.67346394 |
| GHVDHGKTT | ASSDGQGNNQAPL | 0.77090144 |
| AQFEHTLL | ASSDGQGNNQAPL | 0.7455431 |
| IVNQDEL | ASSDGQGNNQAPL | 0.7762754 |
| TNGSQFF | ASSDGQGNNQAPL | 0.77283585 |
| TGKGIVE | ASSDGQGNNQAPL | 0.7469046 |
| LISNASDA | ASSDGQGNNQAPL | 0.22440062 |
| AELRTLGELTKIAKAHGAQ | ASSDGQGNNQAPL | 0.4556299 |
| APNAVAAAEPSW | ASSDGQGNNQAPL | 0.6403841 |
| APQVAGAATAVL | ASSDGQGNNQAPL | 0.6969607 |
| ARRDPAAAAAILPTDA | ASSDGQGNNQAPL | 0.24479088 |
| DAAARDVLAEAGLAEYF | ASSDGQGNNQAPL | 0.7049563 |
| DDQIVLSAKANLAKVEQ | ASSDGQGNNQAPL | 0.65113753 |
| DELLPDTGEDSDEDGH | ASSDGQGNNQAPL | 0.7015467 |
| ELPEAPANRATVPQD | ASSDGQGNNQAPL | 0.77214086 |
| FGQSKIERVTGIG | ASSDGQGNNQAPL | 0.7286267 |
| GAGGTGSTAGGAGGAGGAGGLYA | ASSDGQGNNQAPL | 0.5340988 |
| GHGGTGAAGGNGGTGGAGGSGIDG | ASSDGQGNNQAPL | 0.5141465 |
| GNGGNGGAGGEGGPGGAGGAGA | ASSDGQGNNQAPL | 0.78876287 |
| GSTAGNGGNGGNATNGTIAGQPAG | ASSDGQGNNQAPL | 0.4106396 |
| GTGGDGGNGGNGGTGG | ASSDGQGNNQAPL | 0.7718686 |
| GVRRPAGIPADAPV | ASSDGQGNNQAPL | 0.6634919 |
| GYRQVTDAHLLTLARR | ASSDGQGNNQAPL | 0.7699286 |
| IAKMGQDGHDRG | ASSDGQGNNQAPL | 0.37230182 |
| KAEPSIEIQYT | ASSDGQGNNQAPL | 0.76564413 |
| KPVRIRMAADEDTVAVT | ASSDGQGNNQAPL | 0.7238636 |
| MPPRIAGMRLLVIKPEP | ASSDGQGNNQAPL | 0.7895576 |
| MKFYAVALFRSRGVPPTR | ASSDGQGNNQAPL | 0.63759595 |
| NAGGFGILGAVAHSP | ASSDGQGNNQAPL | 0.7160995 |
| NEGREITLAMAEM | ASSDGQGNNQAPL | 0.5794906 |
| NELVEWSHPLGHRWRWE | ASSDGQGNNQAPL | 0.78511155 |
| NLGAGGTGGNGGNPG | ASSDGQGNNQAPL | 0.538211 |
| NTEWPKWETVMAADPVD | ASSDGQGNNQAPL | 0.61643934 |
| PASAAPAETQP | ASSDGQGNNQAPL | 0.5713454 |
| QQSSDGKNGVRAAEITGE | ASSDGQGNNQAPL | 0.7049202 |
| RAPDAVTTSWWKE | ASSDGQGNNQAPL | 0.7703009 |
| SGFGNVGAHTSGWFNQ | ASSDGQGNNQAPL | 0.34938812 |
| SGVAGAVGQAASVGGLKV | ASSDGQGNNQAPL | 0.785364 |
| SNGGAGTGSAGGNGGT | ASSDGQGNNQAPL | 0.67403424 |
| TGGAGGEGGAAG | ASSDGQGNNQAPL | 0.506886 |
| TGGDGGAAGAGGAPGHG | ASSDGQGNNQAPL | 0.7459332 |
| TMPPLLAAIANAGRHGV | ASSDGQGNNQAPL | 0.64184284 |
| TSSTAGNSGGA | ASSDGQGNNQAPL | 0.7674097 |
| VPATVPVGILC | ASSDGQGNNQAPL | 0.52270067 |
| VQRPDSGGVVLTGRIS | ASSDGQGNNQAPL | 0.74981046 |

| | | |
|---|---|---|
| WTPNPVPATKGVPAGLVQAT | ASSDGQGNNQAPL | 0.71153724 |
| YALKRALAKFSDRFDVVII | ASSDGQGNNQAPL | 0.584773 |
| YLIGSVIV | ASSDGQGNNQAPL | 0.52781093 |
| YGQLVAVID | ASSDGQGNNQAPL | 0.78074133 |
| VTVGGMTLPA | ASSDGQGNNQAPL | 0.36308455 |
| VLALPAV | ASSDGQGNNQAPL | 0.7834575 |
| VIPPTLNYET | ASSDGQGNNQAPL | 0.2114245 |
| VGMGGMHPSSG | ASSDGQGNNQAPL | 0.19586726 |
| VGLAGGAATA | ASSDGQGNNQAPL | 0.36418593 |
| VGGAGGAGGSGP | ASSDGQGNNQAPL | 0.64572334 |
| VFAAFMV | ASSDGQGNNQAPL | 0.8056152 |
| TLYDKLR | ASSDGQGNNQAPL | 0.5958442 |
| TLAVFTVL | ASSDGQGNNQAPL | 0.21029794 |
| TGSVQPARV | ASSDGQGNNQAPL | 0.3123396 |
| TGGGAGSGGT | ASSDGQGNNQAPL | 0.2257632 |
| SHLETSL | ASSDGQGNNQAPL | 0.32335627 |
| SFTTGMT | ASSDGQGNNQAPL | 0.53315485 |
| QVEVAKRMGVRQPTVS | ASSDGQGNNQAPL | 0.4011984 |
| QGSGGPALAM | ASSDGQGNNQAPL | 0.27363133 |
| MLAPSNP | ASSDGQGNNQAPL | 0.28909114 |
| LVNNAAI | ASSDGQGNNQAPL | 0.5781073 |
| LTGEINTLA | ASSDGQGNNQAPL | 0.57574165 |
| LSIFGPV | ASSDGQGNNQAPL | 0.47814885 |
| LLALLANL | ASSDGQGNNQAPL | 0.76933086 |
| LISNASDALDKLR | ASSDGQGNNQAPL | 0.489067 |
| LINEAVP | ASSDGQGNNQAPL | 0.64721596 |
| LGYAFSGL | ASSDGQGNNQAPL | 0.11039659 |
| LDFVAVAL | ASSDGQGNNQAPL | 0.1485112 |
| LAQPGAE | ASSDGQGNNQAPL | 0.34687567 |
| ISNLGAVTGF | ASSDGQGNNQAPL | 0.3252458 |
| HQEDPGRI | ASSDGQGNNQAPL | 0.17509143 |
| GTPGEPTP | ASSDGQGNNQAPL | 0.57483715 |
| GFFNSGS | ASSDGQGNNQAPL | 0.24410126 |
| GAGGGGLTAGAN | ASSDGQGNNQAPL | 0.20184195 |
| FVDALVNM | ASSDGQGNNQAPL | 0.65436184 |
| FLTVAPEL | ASSDGQGNNQAPL | 0.25789735 |
| FETLTAL | ASSDGQGNNQAPL | 0.25302762 |
| DVINPIEL | ASSDGQGNNQAPL | 0.30901837 |
| APVSGAPV | ASSDGQGNNQAPL | 0.4105459 |
| ALVGGLAW | ASSDGQGNNQAPL | 0.18381815 |
| ALAAAAATAI | ASSDGQGNNQAPL | 0.47061613 |
| AFGLGLLL | ASSDGQGNNQAPL | 0.40516296 |
| AFFAAASL | ASSDGQGNNQAPL | 0.51498806 |
| AAAALPAFSPP | ASSDGQGNNQAPL | 0.18110593 |
| VVLAPAL | ASSDGQGNNQAPL | 0.54377323 |
| TIGHVDHGKTTL | ASSDGQGNNQAPL | 0.34033725 |
| LISNASDALDK | ASSDGQGNNQAPL | 0.8219107 |
| LISNASDAL | ASSDGQGNNQAPL | 0.24532814 |
| ISNASDALDK | ASSDGQGNNQAPL | 0.32686958 |
| NQAETLVYQTE | ASSDGQGNNQAPL | 0.3908757 |
| TGGTGGTAGAG | ASSDGQGNNQAPL | 0.76705235 |

| | | |
|---|---|---|
| TGSTAGGAGGA | ASSDGQGNNQAPL | 0.14768983 |
| DPLAVPVAAGPV | ASSDGQGNNQAPL | 0.21913671 |
| GGTGGTPGNGGN | ASSDGQGNNQAPL | 0.15477154 |
| PAGLAGVDPATL | ASSDGQGNNQAPL | 0.36111048 |
| GNNLVVEAPHAGASV | ASSDGQGNNQAPL | 0.4929309 |
| KVTRIMSASTLEQPA | ASSDGQGNNQAPL | 0.15152258 |
| LGFALPIALPRRTVV | ASSDGQGNNQAPL | 0.42935604 |
| TRLGNLTARKLVVLV | ASSDGQGNNQAPL | 0.41177407 |
| WSQQRRLIYFPSAGP | ASSDGQGNNQAPL | 0.27359602 |
| GPNGNPDPMAAAVDIR | ASSDGQGNNQAPL | 0.69278735 |
| TQRCVSVADLRAAASR | ASSDGQGNNQAPL | 0.30960307 |
| QFVLDVAEHPVEVLTETV | ASSDGQGNNQAPL | 0.66008174 |
| GGAGGTGGAGGNGGTQAGDG | ASSDGQGNNQAPL | 0.50056213 |
| PLLATLRAIAVDVDAKPAQV | ASSDGQGNNQAPL | 0.22706398 |
| APKFRTGGYMIDFWGVGYQVAKRM | ASSDGQGNNQAPL | 0.43996587 |
| GAGGAGGAGGAAGGPGGTGGTGGN | ASSDGQGNNQAPL | 0.23106316 |
| IGALGLLATDVAPAGSVAFGVALA | ASSDGQGNNQAPL | 0.2906385 |
| AGGAGGIGGNGGAGGAGGVLMGNG | ASSDGQGNNQAPL | 0.64259905 |
| ADELIGLL | ASSDGQGNNQAPL | 0.7959014 |
| ADRALKA | ASSDGQGNNQAPL | 0.86511314 |
| AGAVGQVGGAGGA | ASSDGQGNNQAPL | 0.69282097 |
| AGGGGGLGGAP | ASSDGQGNNQAPL | 0.58977926 |
| AGGLLYGNGGAG | ASSDGQGNNQAPL | 0.73762524 |
| AGLESIGL | ASSDGQGNNQAPL | 0.85208803 |
| AGLKTGII | ASSDGQGNNQAPL | 0.8599804 |
| ASLAALLL | ASSDGQGNNQAPL | 0.8165174 |
| ASVINISSI | ASSDGQGNNQAPL | 0.61443007 |
| ATPPGPA | ASSDGQGNNQAPL | 0.7698362 |
| ATVASALV | ASSDGQGNNQAPL | 0.6990297 |
| ATVASLIL | ASSDGQGNNQAPL | 0.7621386 |
| AVNGPVLIRH | ASSDGQGNNQAPL | 0.8133398 |
| DGAAGAVGI | ASSDGQGNNQAPL | 0.7762289 |
| DPLLHGI | ASSDGQGNNQAPL | 0.7475008 |
| ELGAALGAL | ASSDGQGNNQAPL | 0.699247 |
| ELGRTQAV | ASSDGQGNNQAPL | 0.78966665 |
| FVAGGYLS | ASSDGQGNNQAPL | 0.82636 |
| GASGGAGGAGGW | ASSDGQGNNQAPL | 0.7908653 |
| GGAGAGGDGGAGGI | ASSDGQGNNQAPL | 0.8807472 |
| GTGHNGSQPGMGGNG | ASSDGQGNNQAPL | 0.84437245 |
| GYAAASASAS | ASSDGQGNNQAPL | 0.8389014 |
| HQVDVAAVQQG | ASSDGQGNNQAPL | 0.18779337 |
| IGAAIAHT | ASSDGQGNNQAPL | 0.45717573 |
| IGMIEVAQL | ASSDGQGNNQAPL | 0.68234026 |
| IRLAPEHTSAMD | ASSDGQGNNQAPL | 0.75031555 |
| IYNVSVL | ASSDGQGNNQAPL | 0.31759837 |
| KQGFLGM | ASSDGQGNNQAPL | 0.80890584 |
| LESVITQL | ASSDGQGNNQAPL | 0.698274 |
| LGLGALVVV | ASSDGQGNNQAPL | 0.64451635 |
| LIGNGGNGGSGGI | ASSDGQGNNQAPL | 0.8494513 |
| LNAINAPTL | ASSDGQGNNQAPL | 0.7485274 |
| LQLLNTGV | ASSDGQGNNQAPL | 0.6712485 |

| | | |
|---|---|---|
| LQTSIEF | ASSDGQGNNQAPL | 0.66189814 |
| MSFAVSEL | ASSDGQGNNQAPL | 0.8698656 |
| NTGALGL | ASSDGQGNNQAPL | 0.4446714 |
| PPGAGGAGGAGTT | ASSDGQGNNQAPL | 0.8299943 |
| QLVGLGI | ASSDGQGNNQAPL | 0.77099055 |
| RFLEQVHQL | ASSDGQGNNQAPL | 0.85418713 |
| RGVGYVLR | ASSDGQGNNQAPL | 0.48664123 |
| SATGVFAGVI | ASSDGQGNNQAPL | 0.835102 |
| SLPAITLP | ASSDGQGNNQAPL | 0.7509128 |
| STALAFTV | ASSDGQGNNQAPL | 0.8896071 |
| SVLTVGLGK | ASSDGQGNNQAPL | 0.7002239 |
| TALAGAGSGI | ASSDGQGNNQAPL | 0.77221775 |
| TGLPFTGI | ASSDGQGNNQAPL | 0.7171755 |
| VAEHVTV | ASSDGQGNNQAPL | 0.85320604 |
| VDIVLNSL | ASSDGQGNNQAPL | 0.50080585 |
| VLTGGGAVL | ASSDGQGNNQAPL | 0.7117144 |
| VNVGLAYGI | ASSDGQGNNQAPL | 0.5575244 |
| VSSAAVGAV | ASSDGQGNNQAPL | 0.7921951 |
| VTSVYEAGV | ASSDGQGNNQAPL | 0.8184364 |
| WVALDRGA | ASSDGQGNNQAPL | 0.41211742 |
| YLVGSVVI | ASSDGQGNNQAPL | 0.8589593 |
| GPPGTGKTLL | ASSDGQGNNQAPL | 0.7582171 |
| GVGMTKF | ASSDGQGNNQAPL | 0.6528851 |
| IGIGGSDL | ASSDGQGNNQAPL | 0.8543893 |
| IGTLLAEL | ASSDGQGNNQAPL | 0.6674595 |
| IYTSGTT | ASSDGQGNNQAPL | 0.84911776 |
| LSFPAGV | ASSDGQGNNQAPL | 0.72187793 |
| LYTSGTT | ASSDGQGNNQAPL | 0.757761 |
| VGAFPFGF | ASSDGQGNNQAPL | 0.8381802 |
| VGAGTVEF | ASSDGQGNNQAPL | 0.5208273 |
| VGKSTLLL | ASSDGQGNNQAPL | 0.5635699 |
| AALAHAGGGIV | ASSDGQGNNQAPL | 0.8624867 |
| AALVAVSLL | ASSDGQGNNQAPL | 0.75085247 |
| AAVSRAVAAL | ASSDGQGNNQAPL | 0.772689 |
| AGGSSGIGGF | ASSDGQGNNQAPL | 0.7068511 |
| AGLTVANANG | ASSDGQGNNQAPL | 0.639616 |
| AGLVGAVL | ASSDGQGNNQAPL | 0.6846918 |
| ALSQAFTV | ASSDGQGNNQAPL | 0.7639737 |
| ALTFSVSL | ASSDGQGNNQAPL | 0.77323985 |
| ALTGAALVAL | ASSDGQGNNQAPL | 0.75447875 |
| ASFSIGAGVL | ASSDGQGNNQAPL | 0.508932 |
| DGSVYKI | ASSDGQGNNQAPL | 0.7550856 |
| EQLAEIV | ASSDGQGNNQAPL | 0.6806916 |
| FAVSELTVM | ASSDGQGNNQAPL | 0.6917294 |
| GAAPLAAGL | ASSDGQGNNQAPL | 0.7447926 |
| GADGGDGGSGGAGG | ASSDGQGNNQAPL | 0.666558 |
| GFGLDTGGAGG | ASSDGQGNNQAPL | 0.6953546 |
| GLASGGLL | ASSDGQGNNQAPL | 0.6827419 |
| GRIAGAAVLL | ASSDGQGNNQAPL | 0.7288052 |
| GSAGSLNTSF | ASSDGQGNNQAPL | 0.7593364 |
| GTGQIGI | ASSDGQGNNQAPL | 0.74521226 |

| | | |
|---|---|---|
| GTLIALIL | ASSDGQGNNQAPL | 0.77118266 |
| IAATMGFVVG | ASSDGQGNNQAPL | 0.76281786 |
| IFAQALL | ASSDGQGNNQAPL | 0.7609161 |
| IGAAVALGF | ASSDGQGNNQAPL | 0.23339729 |
| IGLTPLAL | ASSDGQGNNQAPL | 0.5314555 |
| ILGSNTSTL | ASSDGQGNNQAPL | 0.6073122 |
| IVFSLTVL | ASSDGQGNNQAPL | 0.7320056 |
| LASKATGF | ASSDGQGNNQAPL | 0.32672995 |
| LASVALIL | ASSDGQGNNQAPL | 0.74126345 |
| LFAVGLTL | ASSDGQGNNQAPL | 0.7116256 |
| LFNGGIGTYI | ASSDGQGNNQAPL | 0.66436446 |
| LTLGAGII | ASSDGQGNNQAPL | 0.769773 |
| LVAAALGDI | ASSDGQGNNQAPL | 0.6977955 |
| NTAFVQLGI | ASSDGQGNNQAPL | 0.5628216 |
| QLFSGAEL | ASSDGQGNNQAPL | 0.66155255 |
| SAATFITL | ASSDGQGNNQAPL | 0.7680719 |
| SNYGLANLGV | ASSDGQGNNQAPL | 0.49089193 |
| SYGGAYAVM | ASSDGQGNNQAPL | 0.7620525 |
| TTGVGGLTLG | ASSDGQGNNQAPL | 0.6401764 |
| TTQTIYI | ASSDGQGNNQAPL | 0.7526778 |
| VAAGAGLGAVI | ASSDGQGNNQAPL | 0.38679472 |
| VAAVVTLL | ASSDGQGNNQAPL | 0.7594081 |
| VGFLGGLI | ASSDGQGNNQAPL | 0.71924245 |
| VGVIPILL | ASSDGQGNNQAPL | 0.7798502 |
| VIFASSLI | ASSDGQGNNQAPL | 0.68453765 |
| VKGGDGGAAAG | ASSDGQGNNQAPL | 0.72809696 |
| VTKTFTNI | ASSDGQGNNQAPL | 0.6616129 |
| VVAAAILL | ASSDGQGNNQAPL | 0.7712376 |
| YAALDVEL | ASSDGQGNNQAPL | 0.47718197 |
| LASVALLL | ASSDGQGNNQAPL | 0.66080904 |
| LDAVHRV | ASSDGQGNNQAPL | 0.6051638 |
| QGLSSSIF | ASSDGQGNNQAPL | 0.7359276 |
| RGTFANIRL | ASSDGQGNNQAPL | 0.7560259 |
| RQLWWGHRI | ASSDGQGNNQAPL | 0.5213349 |
| YGPPGTGKTLL | ASSDGQGNNQAPL | 0.7710884 |
| AALVGAPANVA | ASSDGQGNNQAPL | 0.70996106 |
| AATFRRTE | ASSDGQGNNQAPL | 0.5624713 |
| AGALGGGI | ASSDGQGNNQAPL | 0.7596133 |
| AGGAGGAGGIGAG | ASSDGQGNNQAPL | 0.67494357 |
| AGSGALGL | ASSDGQGNNQAPL | 0.7619244 |
| AQIADQI | ASSDGQGNNQAPL | 0.6417955 |
| ASAVAVTYL | ASSDGQGNNQAPL | 0.72950625 |
| ASTVLGGTGG | ASSDGQGNNQAPL | 0.7523885 |
| AVGGEPAK | ASSDGQGNNQAPL | 0.5545542 |
| CEQGTFGGTC | ASSDGQGNNQAPL | 0.58543754 |
| DEGKAPAG | ASSDGQGNNQAPL | 0.766796 |
| DTVVFPAL | ASSDGQGNNQAPL | 0.72220004 |
| FEGITMAG | ASSDGQGNNQAPL | 0.7529223 |
| FPLVVQD | ASSDGQGNNQAPL | 0.6420103 |
| GAFTTPQ | ASSDGQGNNQAPL | 0.57314 |
| GFGGFGTAGGVGGA | ASSDGQGNNQAPL | 0.6375551 |

| | | |
|---|---|---|
| GFLAQFL | ASSDGQGNNQAPL | 0.7384808 |
| GGAGGAGGNAGVA | ASSDGQGNNQAPL | 0.7638352 |
| GGVGTAGAGGAGGAGG | ASSDGQGNNQAPL | 0.73810863 |
| GNGGGGDAGNAGSGGN | ASSDGQGNNQAPL | 0.5858998 |
| GTGGAGGAGNDAGS | ASSDGQGNNQAPL | 0.7281915 |
| GTTNTGWFNS | ASSDGQGNNQAPL | 0.6937406 |
| IFMGPLFTL | ASSDGQGNNQAPL | 0.69267803 |
| ILAHAIEAI | ASSDGQGNNQAPL | 0.696538 |
| IRSAIREAL | ASSDGQGNNQAPL | 0.70522 |
| ITFAGGG | ASSDGQGNNQAPL | 0.6455718 |
| ITGMSGHQEV | ASSDGQGNNQAPL | 0.58713126 |
| IVAGLACTL | ASSDGQGNNQAPL | 0.69541943 |
| IVGGGLAGA | ASSDGQGNNQAPL | 0.75137484 |
| IVTGVLNI | ASSDGQGNNQAPL | 0.7433883 |
| LAQQGCI | ASSDGQGNNQAPL | 0.772017 |
| LCLHVEN | ASSDGQGNNQAPL | 0.754112 |
| LDAISDGS | ASSDGQGNNQAPL | 0.7269002 |
| LGGFAAML | ASSDGQGNNQAPL | 0.31276008 |
| LGGKNPMI | ASSDGQGNNQAPL | 0.45981556 |
| LGPKGVAAIL | ASSDGQGNNQAPL | 0.60682327 |
| LMVGVSI | ASSDGQGNNQAPL | 0.6769933 |
| LVGAAFVL | ASSDGQGNNQAPL | 0.31066677 |
| LVLAVAPL | ASSDGQGNNQAPL | 0.71079624 |
| MAAIAVIDAA | ASSDGQGNNQAPL | 0.6687508 |
| NNGINAELQ | ASSDGQGNNQAPL | 0.594658 |
| RGEPLVLVHG | ASSDGQGNNQAPL | 0.748471 |
| RGGAGGAAGLI | ASSDGQGNNQAPL | 0.65775746 |
| RSYRMHPAV | ASSDGQGNNQAPL | 0.56201756 |
| SAAHLLAGQLA | ASSDGQGNNQAPL | 0.59523976 |
| SAAVGSY | ASSDGQGNNQAPL | 0.76680744 |
| SPHAVLHRI | ASSDGQGNNQAPL | 0.53608954 |
| SQIAKVL | ASSDGQGNNQAPL | 0.74361414 |
| TALGHLVP | ASSDGQGNNQAPL | 0.6228597 |
| TAPTTSGAA | ASSDGQGNNQAPL | 0.7600846 |
| TDGDDGNGGSPGT | ASSDGQGNNQAPL | 0.47916168 |
| TDGLGEN | ASSDGQGNNQAPL | 0.74930894 |
| TGGPPLPGQA | ASSDGQGNNQAPL | 0.6894096 |
| TIPAITI | ASSDGQGNNQAPL | 0.78237057 |
| TIPGFTI | ASSDGQGNNQAPL | 0.644651 |
| TSIASGALA | ASSDGQGNNQAPL | 0.7235102 |
| TVTLPGAAI | ASSDGQGNNQAPL | 0.6011426 |
| VASGLSAALP | ASSDGQGNNQAPL | 0.7419224 |
| VGAGVIGI | ASSDGQGNNQAPL | 0.4835814 |
| VGPKRFGV | ASSDGQGNNQAPL | 0.6587108 |
| VNVAFPD | ASSDGQGNNQAPL | 0.57804775 |
| VSMTNSGVS | ASSDGQGNNQAPL | 0.70353425 |
| VTDVLGDLSV | ASSDGQGNNQAPL | 0.7422267 |
| VVVVGAGFAGL | ASSDGQGNNQAPL | 0.35169432 |
| VYTVGTVV | ASSDGQGNNQAPL | 0.74976295 |
| WTTFSGP | ASSDGQGNNQAPL | 0.6203856 |
| YDELVEAF | ASSDGQGNNQAPL | 0.5538183 |

| | | |
|---|---|---|
| YGTGGAGGNGG | ASSDGQGNNQAPL | 0.7480775 |
| YTNVMAV | ASSDGQGNNQAPL | 0.67647576 |
| FLEQVHQL | ASSDGQGNNQAPL | 0.7477108 |
| GLSLLAVAV | ASSDGQGNNQAPL | 0.57776535 |
| LEQVHQL | ASSDGQGNNQAPL | 0.70013106 |
| LNAAGAI | ASSDGQGNNQAPL | 0.7343972 |
| PAVHASL | ASSDGQGNNQAPL | 0.53204423 |
| TELEDDA | ASSDGQGNNQAPL | 0.5470457 |
| TPPGPPN | ASSDGQGNNQAPL | 0.7655481 |
| TPPPPPPP | ASSDGQGNNQAPL | 0.6945363 |
| TQSPALI | ASSLWDWGYEQY | 0.7310566 |
| EFAAAFPQ | ASSLWDWGYEQY | 0.6280527 |
| LLSGNLDVL | ASSLWDWGYEQY | 0.6386279 |
| GPANLGNYNI | ASSLWDWGYEQY | 0.6308051 |
| TLGPTLP | ASSLWDWGYEQY | 0.718351 |
| VYAYSSCCK | ASSLWDWGYEQY | 0.7267865 |
| APAPAPAPNI | ASSLWDWGYEQY | 0.71153414 |
| PGGPVPIP | ASSLWDWGYEQY | 0.53644615 |
| FLTVALEP | ASSLWDWGYEQY | 0.6974236 |
| SQYFGANL | ASSLWDWGYEQY | 0.6392648 |
| GGVKPAA | ASSLWDWGYEQY | 0.66564566 |
| GGTGGAGGAGGW | ASSLWDWGYEQY | 0.69296384 |
| GVGGVAGLGGAGG | ASSLWDWGYEQY | 0.64490515 |
| VLLVLATTI | ASSLWDWGYEQY | 0.6423137 |
| IGIFAAML | ASSLWDWGYEQY | 0.61670834 |
| AVGDLID | ASSLWDWGYEQY | 0.66401374 |
| NISPLPL | ASSLWDWGYEQY | 0.70770645 |
| VTAVFVAI | ASSLWDWGYEQY | 0.69847125 |
| SGDDNDAAL | ASSLWDWGYEQY | 0.7316128 |
| AVAALAHS | ASSLWDWGYEQY | 0.7194495 |
| GGVTVMATP | ASSLWDWGYEQY | 0.68448406 |
| AGLAANVRD | ASSLWDWGYEQY | 0.30949056 |
| TNRPDLIDPALLRPG | ASSLWDWGYEQY | 0.5538776 |
| LANLGVTAL | ASSLWDWGYEQY | 0.591636 |
| TGERIEL | ASSLWDWGYEQY | 0.6930741 |
| NGEELAA | ASSLWDWGYEQY | 0.33072057 |
| VLSLVWG | ASSLWDWGYEQY | 0.69445074 |
| IGYALWL | ASSLWDWGYEQY | 0.66769457 |
| LSLLLAPL | ASSLWDWGYEQY | 0.6398132 |
| TLDEALK | ASSLWDWGYEQY | 0.720209 |
| TDGAGTAEGG | ASSLWDWGYEQY | 0.6603973 |
| NAPGGAGGNGG | ASSLWDWGYEQY | 0.5732454 |
| VSVFMGMGGNF | ASSLWDWGYEQY | 0.4901518 |
| LASYLAAL | ASSLWDWGYEQY | 0.707001 |
| LVGGTPAL | ASSLWDWGYEQY | 0.46871096 |
| VRNADVL | ASSLWDWGYEQY | 0.715017 |
| TCSSPFLVSAM | ASSLWDWGYEQY | 0.49469894 |
| VHITDAE | ASSLWDWGYEQY | 0.7167212 |
| EYAAPSF | ASSLWDWGYEQY | 0.47851276 |
| VVDAFHGGL | ASSLWDWGYEQY | 0.7122867 |
| ASVVDPQ | ASSLWDWGYEQY | 0.66419846 |

| | | |
|---|---|---|
| SYNEFGGLS | ASSLWDWGYEQY | 0.7427433 |
| DPDGSVY | ASSLWDWGYEQY | 0.6588596 |
| MVSTIHATE | ASSLWDWGYEQY | 0.67073834 |
| LAAGVADPV | ASSLWDWGYEQY | 0.5390314 |
| TKEQIAA | ASSLWDWGYEQY | 0.72442055 |
| SLASPAAL | ASSLWDWGYEQY | 0.5260877 |
| IGLVFYGV | ASSLWDWGYEQY | 0.6215647 |
| FLLDASNL | ASSLWDWGYEQY | 0.49651557 |
| YQVGNFP | ASSLWDWGYEQY | 0.6848678 |
| ESLLSQST | ASSLWDWGYEQY | 0.7114836 |
| LFTVGFAAVG | ASSLWDWGYEQY | 0.62555444 |
| ATPPTPT | ASSLWDWGYEQY | 0.71301246 |
| TDLATLG | ASSLWDWGYEQY | 0.66253847 |
| SAGGTGR | ASSLWDWGYEQY | 0.5740439 |
| GFSNSGS | ASSLWDWGYEQY | 0.71832114 |
| GGNGGSGGTSV | ASSLWDWGYEQY | 0.6060953 |
| GTSVATGGAGN | ASSLWDWGYEQY | 0.71393186 |
| NGGNGGSGGTS | ASSLWDWGYEQY | 0.53432536 |
| AVVKFPH | ASSLWDWGYEQY | 0.6555902 |
| TPPPPPP | ASSLWDWGYEQY | 0.6959622 |
| AAPAAAAAPA | ASSLWDWGYEQY | 0.63286734 |
| IALNEDL | ASSLWDWGYEQY | 0.5327657 |
| IVNVSSM | ASSLWDWGYEQY | 0.722902 |
| TGIPPAPR | ASSLWDWGYEQY | 0.8904679 |
| FDLGGGTF | ASSLWDWGYEQY | 0.88278544 |
| FDLGGGTFDVS | ASSLWDWGYEQY | 0.79729724 |
| NEPTAAA | ASSLWDWGYEQY | 0.8044902 |
| NEPTAAALA | ASSLWDWGYEQY | 0.84560204 |
| GSIINISSI | ASSLWDWGYEQY | 0.8977016 |
| VAPGNAGTA | ASSLWDWGYEQY | 0.90679336 |
| TGGGASLE | ASSLWDWGYEQY | 0.8957406 |
| VAVYNHYKRI | ASSLWDWGYEQY | 0.8411892 |
| TDPAPATT | ASSLWDWGYEQY | 0.8999605 |
| TNRPDILDPALLRPG | ASSLWDWGYEQY | 0.87569433 |
| LTTTTTS | ASSLWDWGYEQY | 0.86683136 |
| PTGSGKTL | ASSLWDWGYEQY | 0.8988507 |
| TGSGKTL | ASSLWDWGYEQY | 0.8859799 |
| TIGHVDHGK | ASSLWDWGYEQY | 0.83954203 |
| TIGHVDHGKTT | ASSLWDWGYEQY | 0.85493875 |
| GHVDHGKTT | ASSLWDWGYEQY | 0.8869153 |
| AQFEHTLL | ASSLWDWGYEQY | 0.89639175 |
| IVNQDEL | ASSLWDWGYEQY | 0.89129555 |
| TNGSQFF | ASSLWDWGYEQY | 0.9138197 |
| TGKGIVE | ASSLWDWGYEQY | 0.90026534 |
| LISNASDA | ASSLWDWGYEQY | 0.8855715 |
| AELRTLGELTKIAKAHGAQ | ASSLWDWGYEQY | 0.5614169 |
| APNAVAAAEPSW | ASSLWDWGYEQY | 0.7437563 |
| APQVAGAATAVL | ASSLWDWGYEQY | 0.8735 |
| ARRDPAAAAAILPTDA | ASSLWDWGYEQY | 0.84623873 |
| DAAARDVLAEAGLAEYF | ASSLWDWGYEQY | 0.54093105 |
| DDQIVLSAKANLAKVEQ | ASSLWDWGYEQY | 0.896844 |

| | | |
|---|---|---|
| DELLPDTGEDSDEDGH | ASSLWDWGYEQY | 0.87226033 |
| ELPEAPANRATVPQD | ASSLWDWGYEQY | 0.8323202 |
| FGQSKIERVTGIG | ASSLWDWGYEQY | 0.8997642 |
| GAGGTGSTAGGAGGAGGAGGLYA | ASSLWDWGYEQY | 0.8575971 |
| GHGGTGAAGGNGGTGGAGGSGIDG | ASSLWDWGYEQY | 0.8426982 |
| GNGGNGGAGGEGGPGGAGGAGA | ASSLWDWGYEQY | 0.8798366 |
| GSTAGNGGNGGNATNGTIAGQPAG | ASSLWDWGYEQY | 0.9109698 |
| GTGGDGGNGGNGGTGG | ASSLWDWGYEQY | 0.8478985 |
| GVRRPAGIPADAPV | ASSLWDWGYEQY | 0.9050166 |
| GYRQVTDAHLLTLARR | ASSLWDWGYEQY | 0.8626131 |
| IAKMGQDGHDRG | ASSLWDWGYEQY | 0.90925264 |
| KAEPSIEIQYT | ASSLWDWGYEQY | 0.80037904 |
| KPVRIRMAADEDTVAVT | ASSLWDWGYEQY | 0.90138805 |
| MPPRIAGMRLLVIKPEP | ASSLWDWGYEQY | 0.8709369 |
| MKFYAVALFRSRGVPPTR | ASSLWDWGYEQY | 0.9164759 |
| NAGGFGILGAVAHSP | ASSLWDWGYEQY | 0.8692533 |
| NEGREITLAMAEM | ASSLWDWGYEQY | 0.9062683 |
| NELVEWSHPLGHRWRWE | ASSLWDWGYEQY | 0.88073224 |
| NLGAGGTGGNGGNPG | ASSLWDWGYEQY | 0.88904065 |
| NTEWPKWETVMAADPVD | ASSLWDWGYEQY | 0.77484167 |
| PASAAPAETQP | ASSLWDWGYEQY | 0.85977846 |
| QQSSDGKNGVRAAEITGE | ASSLWDWGYEQY | 0.8040397 |
| RAPDAVTTSWWKE | ASSLWDWGYEQY | 0.88035816 |
| SGFGNVGAHTSGWFNQ | ASSLWDWGYEQY | 0.8952243 |
| SGVAGAVGQAASVGGLKV | ASSLWDWGYEQY | 0.42044163 |
| SNGGAGTGSAGGNGGT | ASSLWDWGYEQY | 0.9068824 |
| TGGAGGEGGAAG | ASSLWDWGYEQY | 0.7583503 |
| TGGDGGAAGAGGAPGHG | ASSLWDWGYEQY | 0.83397675 |
| TMPPLLAAIANAGRHGV | ASSLWDWGYEQY | 0.9049634 |
| TSSTAGNSGGA | ASSLWDWGYEQY | 0.87406594 |
| VPATVPVGILC | ASSLWDWGYEQY | 0.8970165 |
| VQRPDSGGVVLTGRIS | ASSLWDWGYEQY | 0.86615056 |
| WTPNPVPATKGVPAGLVQAT | ASSLWDWGYEQY | 0.8722886 |
| YALKRALAKFSDRFDVVII | ASSLWDWGYEQY | 0.88647515 |
| YLIGSVIV | ASSLWDWGYEQY | 0.58935547 |
| YGQLVAVID | ASSLWDWGYEQY | 0.83630776 |
| VTVGGMTLPA | ASSLWDWGYEQY | 0.91145825 |
| VLALPAV | ASSLWDWGYEQY | 0.6760603 |
| VIPPTLNYET | ASSLWDWGYEQY | 0.6860228 |
| VGMGGMHPSSG | ASSLWDWGYEQY | 0.65104425 |
| VGLAGGAATA | ASSLWDWGYEQY | 0.6308166 |
| VGGAGGAGGSGP | ASSLWDWGYEQY | 0.65179163 |
| VFAAFMV | ASSLWDWGYEQY | 0.6793834 |
| TLYDKLR | ASSLWDWGYEQY | 0.68568116 |
| TLAVFTVL | ASSLWDWGYEQY | 0.674351 |
| TGSVQPARV | ASSLWDWGYEQY | 0.6407708 |
| TGGGAGSGGT | ASSLWDWGYEQY | 0.67229795 |
| SHLETSL | ASSLWDWGYEQY | 0.66658044 |
| SFTTGMT | ASSLWDWGYEQY | 0.65684354 |
| QVEVAKRMGVRQPTVS | ASSLWDWGYEQY | 0.675866 |
| QGSGGPALAM | ASSLWDWGYEQY | 0.67157346 |

| | | |
|---|---|---|
| MLAPSNP | ASSLWDWGYEQY | 0.64439535 |
| LVNNAAI | ASSLWDWGYEQY | 0.6615309 |
| LTGEINTLA | ASSLWDWGYEQY | 0.66296023 |
| LSIFGPV | ASSLWDWGYEQY | 0.6753479 |
| LLALLANL | ASSLWDWGYEQY | 0.6729543 |
| LISNASDALDKLR | ASSLWDWGYEQY | 0.68780404 |
| LINEAVP | ASSLWDWGYEQY | 0.6811111 |
| LGYAFSGL | ASSLWDWGYEQY | 0.67880964 |
| LDFVAVAL | ASSLWDWGYEQY | 0.47735396 |
| LAQPGAE | ASSLWDWGYEQY | 0.608721 |
| ISNLGAVTGF | ASSLWDWGYEQY | 0.6596132 |
| HQEDPGRI | ASSLWDWGYEQY | 0.6636882 |
| GTPGEPTP | ASSLWDWGYEQY | 0.54118216 |
| GFFNSGS | ASSLWDWGYEQY | 0.6771153 |
| GAGGGGLTAGAN | ASSLWDWGYEQY | 0.65883374 |
| FVDALVNM | ASSLWDWGYEQY | 0.6359434 |
| FLTVAPEL | ASSLWDWGYEQY | 0.6782901 |
| FETLTAL | ASSLWDWGYEQY | 0.65453815 |
| DVINPIEL | ASSLWDWGYEQY | 0.65130514 |
| APVSGAPV | ASSLWDWGYEQY | 0.66330457 |
| ALVGGLAW | ASSLWDWGYEQY | 0.6831871 |
| ALAAAAATAI | ASSLWDWGYEQY | 0.6277636 |
| AFGLGLLL | ASSLWDWGYEQY | 0.6804807 |
| AFFAAASL | ASSLWDWGYEQY | 0.646799 |
| AAAALPAFSPP | ASSLWDWGYEQY | 0.68446904 |
| VVLAPAL | ASSLWDWGYEQY | 0.61669093 |
| TIGHVDHGKTTL | ASSLWDWGYEQY | 0.6785097 |
| LISNASDALDK | ASSLWDWGYEQY | 0.665742 |
| LISNASDAL | ASSLWDWGYEQY | 0.6896681 |
| ISNASDALDK | ASSLWDWGYEQY | 0.66209376 |
| NQAETLVYQTE | ASSLWDWGYEQY | 0.68011975 |
| TGGTGGTAGAG | ASSLWDWGYEQY | 0.672983 |
| TGSTAGGAGGA | ASSLWDWGYEQY | 0.68473107 |
| DPLAVPVAAGPV | ASSLWDWGYEQY | 0.61807895 |
| GGTGGTPGNGGN | ASSLWDWGYEQY | 0.6686282 |
| PAGLAGVDPATL | ASSLWDWGYEQY | 0.60366476 |
| GNNLVVEAPHAGASV | ASSLWDWGYEQY | 0.67381346 |
| KVTRIMSASTLEQPA | ASSLWDWGYEQY | 0.67661625 |
| LGFALPIALPRRTVV | ASSLWDWGYEQY | 0.5987184 |
| TRLGNLTARKLVVLV | ASSLWDWGYEQY | 0.6848823 |
| WSQQRRLIYFPSAGP | ASSLWDWGYEQY | 0.6574013 |
| GPNGNPDPMAAAVDIR | ASSLWDWGYEQY | 0.64974564 |
| TQRCVSVADLRAAASR | ASSLWDWGYEQY | 0.6832071 |
| QFVLDVAEHPVEVLTETV | ASSLWDWGYEQY | 0.66075736 |
| GGAGGTGGAGGNGGTQAGDG | ASSLWDWGYEQY | 0.67878485 |
| PLLATLRAIAVDVDAKPAQV | ASSLWDWGYEQY | 0.65694964 |
| APKFRTGGYMIDFWGVGYQVAKRM | ASSLWDWGYEQY | 0.666664 |
| GAGGAGGAGGAAGGPGGTGGTGGN | ASSLWDWGYEQY | 0.67387676 |
| IGALGLLATDVAPAGSVAFGVALA | ASSLWDWGYEQY | 0.611378 |
| AGGAGGIGGNGGAGGAGGVLMGNG | ASSLWDWGYEQY | 0.63245976 |
| ADELIGLL | ASSLWDWGYEQY | 0.68431526 |

| | | |
|---|---|---|
| ADRALKA | ASSLWDWGYEQY | 0.7291608 |
| AGAVGQVGGAGGA | ASSLWDWGYEQY | 0.7251965 |
| AGGGGGLGGAP | ASSLWDWGYEQY | 0.6829567 |
| AGGLLYGNGGAG | ASSLWDWGYEQY | 0.668116 |
| AGLESIGL | ASSLWDWGYEQY | 0.69904035 |
| AGLKTGII | ASSLWDWGYEQY | 0.74269366 |
| ASLAALLL | ASSLWDWGYEQY | 0.7320564 |
| ASVINISSI | ASSLWDWGYEQY | 0.7327266 |
| ATPPGPA | ASSLWDWGYEQY | 0.6549307 |
| ATVASALV | ASSLWDWGYEQY | 0.7425878 |
| ATVASLIL | ASSLWDWGYEQY | 0.71875536 |
| AVNGPVLIRH | ASSLWDWGYEQY | 0.71864265 |
| DGAAGAVGI | ASSLWDWGYEQY | 0.7424201 |
| DPLLHGI | ASSLWDWGYEQY | 0.7133093 |
| ELGAALGAL | ASSLWDWGYEQY | 0.70459574 |
| ELGRTQAV | ASSLWDWGYEQY | 0.713603 |
| FVAGGYLS | ASSLWDWGYEQY | 0.71057034 |
| GASGGAGGAGGW | ASSLWDWGYEQY | 0.72629076 |
| GGAGAGGDGGAGGI | ASSLWDWGYEQY | 0.7188105 |
| GTGHNGSQPGMGGNG | ASSLWDWGYEQY | 0.7549001 |
| GYAAASASAS | ASSLWDWGYEQY | 0.74354863 |
| HQVDVAAVQQG | ASSLWDWGYEQY | 0.7309449 |
| IGAAIAHT | ASSLWDWGYEQY | 0.36251527 |
| IGMIEVAQL | ASSLWDWGYEQY | 0.67261547 |
| IRLAPEHTSAMD | ASSLWDWGYEQY | 0.7115122 |
| IYNVSVL | ASSLWDWGYEQY | 0.7256297 |
| KQGFLGM | ASSLWDWGYEQY | 0.48487407 |
| LESVITQL | ASSLWDWGYEQY | 0.7348596 |
| LGLGALVVV | ASSLWDWGYEQY | 0.71107507 |
| LIGNGGNGGSGGI | ASSLWDWGYEQY | 0.6960273 |
| LNAINAPTL | ASSLWDWGYEQY | 0.74917233 |
| LQLLNTGV | ASSLWDWGYEQY | 0.69949305 |
| LQTSIEF | ASSLWDWGYEQY | 0.7080468 |
| MSFAVSEL | ASSLWDWGYEQY | 0.70044976 |
| NTGALGL | ASSLWDWGYEQY | 0.75589323 |
| PPGAGGAGGAGTT | ASSLWDWGYEQY | 0.68547714 |
| QLVGLGI | ASSLWDWGYEQY | 0.74818283 |
| RFLEQVHQL | ASSLWDWGYEQY | 0.71178967 |
| RGVGYVLR | ASSLWDWGYEQY | 0.7514921 |
| SATGVFAGVI | ASSLWDWGYEQY | 0.6209769 |
| SLPAITLP | ASSLWDWGYEQY | 0.7414462 |
| STALAFTV | ASSLWDWGYEQY | 0.7112774 |
| SVLTVGLGK | ASSLWDWGYEQY | 0.7553241 |
| TALAGAGSGI | ASSLWDWGYEQY | 0.7183751 |
| TGLPFTGI | ASSLWDWGYEQY | 0.73825234 |
| VAEHVTV | ASSLWDWGYEQY | 0.70349395 |
| VDIVLNSL | ASSLWDWGYEQY | 0.73537564 |
| VLTGGGAVL | ASSLWDWGYEQY | 0.6330972 |
| VNVGLAYGI | ASSLWDWGYEQY | 0.70377606 |
| VSSAAVGAV | ASSLWDWGYEQY | 0.6623784 |
| VTSVYEAGV | ASSLWDWGYEQY | 0.72873783 |

| | | |
|---|---|---|
| WVALDRGA | ASSLWDWGYEQY | 0.73502004 |
| YLVGSVVI | ASSLWDWGYEQY | 0.5489893 |
| GPPGTGKTLL | ASSLWDWGYEQY | 0.74673915 |
| GVGMTKF | ASSLWDWGYEQY | 0.65681684 |
| IGIGGSDL | ASSLWDWGYEQY | 0.6842575 |
| IGTLLAEL | ASSLWDWGYEQY | 0.74362385 |
| IYTSGTT | ASSLWDWGYEQY | 0.71040493 |
| LSFPAGV | ASSLWDWGYEQY | 0.7384902 |
| LYTSGTT | ASSLWDWGYEQY | 0.69652414 |
| VGAFPFGF | ASSLWDWGYEQY | 0.7146784 |
| VGAGTVEF | ASSLWDWGYEQY | 0.7348946 |
| VGKSTLLL | ASSLWDWGYEQY | 0.56551546 |
| AALAHAGGGIV | ASSLWDWGYEQY | 0.6748334 |
| AALVAVSLL | ASSLWDWGYEQY | 0.7486707 |
| AAVSRAVAAL | ASSLWDWGYEQY | 0.58272266 |
| AGGSSGIGGF | ASSLWDWGYEQY | 0.67921287 |
| AGLTVANANG | ASSLWDWGYEQY | 0.4681733 |
| AGLVGAVL | ASSLWDWGYEQY | 0.44979873 |
| ALSQAFTV | ASSLWDWGYEQY | 0.5465036 |
| ALTFSVSL | ASSLWDWGYEQY | 0.6647162 |
| ALTGAALVAL | ASSLWDWGYEQY | 0.6560343 |
| ASFSIGAGVL | ASSLWDWGYEQY | 0.6175797 |
| DGSVYKI | ASSLWDWGYEQY | 0.4775219 |
| EQLAEIV | ASSLWDWGYEQY | 0.53317386 |
| FAVSELTVM | ASSLWDWGYEQY | 0.49987906 |
| GAAPLAAGL | ASSLWDWGYEQY | 0.5424735 |
| GADGGDGGSGGAGG | ASSLWDWGYEQY | 0.6271751 |
| GFGLDTGGAGG | ASSLWDWGYEQY | 0.5447812 |
| GLASGGLL | ASSLWDWGYEQY | 0.49100322 |
| GRIAGAAVLL | ASSLWDWGYEQY | 0.52802414 |
| GSAGSLNTSF | ASSLWDWGYEQY | 0.60243213 |
| GTGQIGI | ASSLWDWGYEQY | 0.5891412 |
| GTLIALIL | ASSLWDWGYEQY | 0.57910395 |
| IAATMGFVVG | ASSLWDWGYEQY | 0.6840191 |
| IFAQALL | ASSLWDWGYEQY | 0.62216103 |
| IGAAVALGF | ASSLWDWGYEQY | 0.63763857 |
| IGLTPLAL | ASSLWDWGYEQY | 0.31805456 |
| ILGSNTSTL | ASSLWDWGYEQY | 0.37198442 |
| IVFSLTVL | ASSLWDWGYEQY | 0.5684751 |
| LASKATGF | ASSLWDWGYEQY | 0.54276955 |
| LASVALIL | ASSLWDWGYEQY | 0.42379686 |
| LFAVGLTL | ASSLWDWGYEQY | 0.6295377 |
| LFNGGIGTYI | ASSLWDWGYEQY | 0.5229564 |
| LTLGAGII | ASSLWDWGYEQY | 0.47206196 |
| LVAAALGDI | ASSLWDWGYEQY | 0.6209562 |
| NTAFVQLGI | ASSLWDWGYEQY | 0.49253973 |
| QLFSGAEL | ASSLWDWGYEQY | 0.4952452 |
| SAATFITL | ASSLWDWGYEQY | 0.5393616 |
| SNYGLANLGV | ASSLWDWGYEQY | 0.62381184 |
| SYGGAYAVM | ASSLWDWGYEQY | 0.41857514 |
| TTGVGGLTLG | ASSLWDWGYEQY | 0.593746 |

| | | |
|---|---|---|
| TTQTIYI | ASSLWDWGYEQY | 0.5561899 |
| VAAGAGLGAVI | ASSLWDWGYEQY | 0.6476304 |
| VAAVVTLL | ASSLWDWGYEQY | 0.426403 |
| VGFLGGLI | ASSLWDWGYEQY | 0.6142088 |
| VGVIPILL | ASSLWDWGYEQY | 0.54622436 |
| VIFASSLI | ASSLWDWGYEQY | 0.68623304 |
| VKGGDGGAAAG | ASSLWDWGYEQY | 0.50614005 |
| VTKTFTNI | ASSLWDWGYEQY | 0.56857294 |
| VVAAAILL | ASSLWDWGYEQY | 0.56708306 |
| YAALDVEL | ASSLWDWGYEQY | 0.656091 |
| LASVALLL | ASSLWDWGYEQY | 0.42950636 |
| LDAVHRV | ASSLWDWGYEQY | 0.50859934 |
| QGLSSSIF | ASSLWDWGYEQY | 0.41117287 |
| RGTFANIRL | ASSLWDWGYEQY | 0.59643596 |
| RQLWWGHRI | ASSLWDWGYEQY | 0.59528995 |
| YGPPGTGKTLL | ASSLWDWGYEQY | 0.45441422 |
| AALVGAPANVA | ASSLWDWGYEQY | 0.6260751 |
| AATFRRTE | ASSLWDWGYEQY | 0.6168192 |
| AGALGGGI | ASSLWDWGYEQY | 0.5000126 |
| AGGAGGAGGIGAG | ASSLWDWGYEQY | 0.65920097 |
| AGSGALGL | ASSLWDWGYEQY | 0.5411442 |
| AQIADQI | ASSLWDWGYEQY | 0.64308715 |
| ASAVAVTYL | ASSLWDWGYEQY | 0.57786494 |
| ASTVLGGTGG | ASSLWDWGYEQY | 0.49497557 |
| AVGGEPAK | ASSLWDWGYEQY | 0.63173723 |
| CEQGTFGGTC | ASSLWDWGYEQY | 0.5620845 |
| DEGKAPAG | ASSLWDWGYEQY | 0.5103554 |
| DTVVFPAL | ASSLWDWGYEQY | 0.64537245 |
| FEGITMAG | ASSLWDWGYEQY | 0.53448975 |
| FPLVVQD | ASSLWDWGYEQY | 0.6650639 |
| GAFTTPQ | ASSLWDWGYEQY | 0.4131938 |
| GFGGFGTAGGVGGA | ASSLWDWGYEQY | 0.40112102 |
| GFLAQFL | ASSLWDWGYEQY | 0.52647775 |
| GGAGGAGGNAGVA | ASSLWDWGYEQY | 0.64220107 |
| GGVGTAGAGGAGGAGG | ASSLWDWGYEQY | 0.656551 |
| GNGGGGDAGNAGSGGN | ASSLWDWGYEQY | 0.58351296 |
| GTGGAGGAGNDAGS | ASSLWDWGYEQY | 0.4221254 |
| GTTNTGWFNS | ASSLWDWGYEQY | 0.4780531 |
| IFMGPLFTL | ASSLWDWGYEQY | 0.44903195 |
| ILAHAIEAI | ASSLWDWGYEQY | 0.49677497 |
| IRSAIREAL | ASSLWDWGYEQY | 0.5792501 |
| ITFAGGG | ASSLWDWGYEQY | 0.49485344 |
| ITGMSGHQEV | ASSLWDWGYEQY | 0.41697067 |
| IVAGLACTL | ASSLWDWGYEQY | 0.4662541 |
| IVGGGLAGA | ASSLWDWGYEQY | 0.5881052 |
| IVTGVLNI | ASSLWDWGYEQY | 0.55588764 |
| LAQQGCI | ASSLWDWGYEQY | 0.53446484 |
| LCLHVEN | ASSLWDWGYEQY | 0.6651536 |
| LDAISDGS | ASSLWDWGYEQY | 0.5907676 |
| LGGFAAML | ASSLWDWGYEQY | 0.60561717 |
| LGGKNPMI | ASSLWDWGYEQY | 0.29875827 |

| | | |
|---|---|---|
| LGPKGVAAIL | ASSLWDWGYEQY | 0.32920322 |
| LMVGVSI | ASSLWDWGYEQY | 0.47605464 |
| LVGAAFVL | ASSLWDWGYEQY | 0.5029959 |
| LVLAVAPL | ASSLWDWGYEQY | 0.3527145 |
| MAAIAVIDAA | ASSLWDWGYEQY | 0.59957653 |
| NNGINAELQ | ASSLWDWGYEQY | 0.45555276 |
| RGEPLVLVHG | ASSLWDWGYEQY | 0.4154493 |
| RGGAGGAAGLI | ASSLWDWGYEQY | 0.6071199 |
| RSYRMHPAV | ASSLWDWGYEQY | 0.43948567 |
| SAAHLLAGQLA | ASSLWDWGYEQY | 0.42860353 |
| SAAVGSY | ASSLWDWGYEQY | 0.45972323 |
| SPHAVLHRI | ASSLWDWGYEQY | 0.5518865 |
| SQIAKVL | ASSLWDWGYEQY | 0.34767795 |
| TALGHLVP | ASSLWDWGYEQY | 0.55883145 |
| TAPTTSGAA | ASSLWDWGYEQY | 0.46701264 |
| TDGDDGNGGSPGT | ASSLWDWGYEQY | 0.6228748 |
| TDGLGEN | ASSLWDWGYEQY | 0.37116805 |
| TGGPPLPGQA | ASSLWDWGYEQY | 0.5735376 |
| TIPAITI | ASSLWDWGYEQY | 0.5225595 |
| TIPGFTI | ASSLWDWGYEQY | 0.6728622 |
| TSIASGALA | ASSLWDWGYEQY | 0.45333278 |
| TVTLPGAAI | ASSLWDWGYEQY | 0.5316546 |
| VASGLSAALP | ASSLWDWGYEQY | 0.5236849 |
| VGAGVIGI | ASSLWDWGYEQY | 0.6550946 |
| VGPKRFGV | ASSLWDWGYEQY | 0.40088618 |
| VNVAFPD | ASSLWDWGYEQY | 0.4566579 |
| VSMTNSGVS | ASSLWDWGYEQY | 0.3501779 |
| VTDVLGDLSV | ASSLWDWGYEQY | 0.5615404 |
| VVVVGAGFAGL | ASSLWDWGYEQY | 0.5482038 |
| VYTVGTVV | ASSLWDWGYEQY | 0.4104722 |
| WTTFSGP | ASSLWDWGYEQY | 0.5911664 |
| YDELVEAF | ASSLWDWGYEQY | 0.58371353 |
| YGTGGAGGNGG | ASSLWDWGYEQY | 0.4402955 |
| YTNVMAV | ASSLWDWGYEQY | 0.6309664 |
| FLEQVHQL | ASSLWDWGYEQY | 0.49257398 |
| GLSLLAVAV | ASSLWDWGYEQY | 0.6177246 |
| LEQVHQL | ASSLWDWGYEQY | 0.54205316 |
| LNAAGAI | ASSLWDWGYEQY | 0.41719776 |
| PAVHASL | ASSLWDWGYEQY | 0.5853611 |
| TELEDDA | ASSLWDWGYEQY | 0.48673436 |
| TPPGPPN | ASSLWDWGYEQY | 0.45007092 |
| TPPPPPPP | ASSLWDWGYEQY | 0.60737586 |

PanPep_cluster3

| Peptide | CDR3 | Score | |
|---|---|---|---|
| TQSPALI | ASSQEGQPGERLF | | 0.36801848 |
| EFAAAFPQ | ASSQEGQPGERLF | | 0.6998399 |
| LLSGNLDVL | ASSQEGQPGERLF | | 0.6533816 |
| GPANLGNYNI | ASSQEGQPGERLF | | 0.58931303 |
| TLGPTLP | ASSQEGQPGERLF | | 0.5274655 |
| VYAYSSCCK | ASSQEGQPGERLF | | 0.3413203 |
| APAPAPAPNI | ASSQEGQPGERLF | | 0.5047952 |
| PGGPVPIP | ASSQEGQPGERLF | | 0.4688177 |
| FLTVALEP | ASSQEGQPGERLF | | 0.70737255 |
| SQYFGANL | ASSQEGQPGERLF | | 0.67770255 |
| GGVKPAA | ASSQEGQPGERLF | | 0.51718235 |
| GGTGGAGGAGGW | ASSQEGQPGERLF | | 0.6882988 |
| GVGGVAGLGGAGG | ASSQEGQPGERLF | | 0.62368655 |
| VLLVLATTI | ASSQEGQPGERLF | | 0.38211074 |
| IGIFAAML | ASSQEGQPGERLF | | 0.58023334 |
| AVGDLID | ASSQEGQPGERLF | | 0.38194776 |
| NISPLPL | ASSQEGQPGERLF | | 0.54783547 |
| VTAVFVAI | ASSQEGQPGERLF | | 0.30297083 |
| SGDDNDAAL | ASSQEGQPGERLF | | 0.6412419 |
| AVAALAHS | ASSQEGQPGERLF | | 0.5335578 |
| GGVTVMATP | ASSQEGQPGERLF | | 0.29026744 |
| AGLAANVRD | ASSQEGQPGERLF | | 0.65759254 |
| TNRPDLIDPALLRPG | ASSQEGQPGERLF | | 0.5258607 |
| LANLGVTAL | ASSQEGQPGERLF | | 0.52123845 |
| TGERIEL | ASSQEGQPGERLF | | 0.6761222 |
| NGEELAA | ASSQEGQPGERLF | | 0.565118 |
| VLSLVWG | ASSQEGQPGERLF | | 0.4562728 |
| IGYALWL | ASSQEGQPGERLF | | 0.6835643 |
| LSLLLAPL | ASSQEGQPGERLF | | 0.7082344 |
| TLDEALK | ASSQEGQPGERLF | | 0.397135 |
| TDGAGTAEGG | ASSQEGQPGERLF | | 0.6374999 |
| NAPGGAGGNGG | ASSQEGQPGERLF | | 0.706486 |
| VSVFMGMGGNF | ASSQEGQPGERLF | | 0.6485741 |
| LASYLAAL | ASSQEGQPGERLF | | 0.6719018 |
| LVGGTPAL | ASSQEGQPGERLF | | 0.6907734 |
| VRNADVL | ASSQEGQPGERLF | | 0.52230924 |
| TCSSPFLVSAM | ASSQEGQPGERLF | | 0.7003191 |
| VHITDAE | ASSQEGQPGERLF | | 0.7052721 |
| EYAAPSF | ASSQEGQPGERLF | | 0.5549834 |
| VVDAFHGGL | ASSQEGQPGERLF | | 0.56530714 |
| ASVVDPQ | ASSQEGQPGERLF | | 0.3869705 |
| SYNEFGGLS | ASSQEGQPGERLF | | 0.5824926 |
| DPDGSVY | ASSQEGQPGERLF | | 0.46540502 |
| MVSTIHATE | ASSQEGQPGERLF | | 0.66755664 |
| LAAGVADPV | ASSQEGQPGERLF | | 0.7014655 |
| TKEQIAA | ASSQEGQPGERLF | | 0.5797781 |
| SLASPAAL | ASSQEGQPGERLF | | 0.39959255 |
| IGLVFYGV | ASSQEGQPGERLF | | 0.4383317 |
| FLLDASNL | ASSQEGQPGERLF | | 0.65781033 |

| | | |
|---|---|---|
| YQVGNFP | ASSQEGQPGERLF | 0.37700978 |
| ESLLSQST | ASSQEGQPGERLF | 0.34594575 |
| LFTVGFAAVG | ASSQEGQPGERLF | 0.47300613 |
| ATPPTPT | ASSQEGQPGERLF | 0.68530285 |
| TDLATLG | ASSQEGQPGERLF | 0.682755 |
| SAGGTGR | ASSQEGQPGERLF | 0.5180192 |
| GFSNSGS | ASSQEGQPGERLF | 0.70528066 |
| GGNGGSGGTSV | ASSQEGQPGERLF | 0.69547427 |
| GTSVATGGAGN | ASSQEGQPGERLF | 0.6683823 |
| NGGNGGSGGTS | ASSQEGQPGERLF | 0.5162626 |
| AVVKFPH | ASSQEGQPGERLF | 0.39247483 |
| TPPPPPP | ASSQEGQPGERLF | 0.7078661 |
| AAPAAAAAPA | ASSQEGQPGERLF | 0.6339294 |
| IALNEDL | ASSQEGQPGERLF | 0.62853193 |
| IVNVSSM | ASSQEGQPGERLF | 0.6632315 |
| TGIPPAPR | ASSQEGQPGERLF | 0.575953 |
| FDLGGGTF | ASSQEGQPGERLF | 0.48847598 |
| FDLGGGTFDVS | ASSQEGQPGERLF | 0.6565378 |
| NEPTAAA | ASSQEGQPGERLF | 0.6945641 |
| NEPTAAALA | ASSQEGQPGERLF | 0.48238945 |
| GSIINISSI | ASSQEGQPGERLF | 0.39224893 |
| VAPGNAGTA | ASSQEGQPGERLF | 0.56150687 |
| TGGGASLE | ASSQEGQPGERLF | 0.5981225 |
| VAVYNHYKRI | ASSQEGQPGERLF | 0.70293003 |
| TDPAPATT | ASSQEGQPGERLF | 0.5916232 |
| TNRPDILDPALLRPG | ASSQEGQPGERLF | 0.5692742 |
| LTTTTTS | ASSQEGQPGERLF | 0.6640603 |
| PTGSGKTL | ASSQEGQPGERLF | 0.6242362 |
| TGSGKTL | ASSQEGQPGERLF | 0.60354286 |
| TIGHVDHGK | ASSQEGQPGERLF | 0.6409936 |
| TIGHVDHGKTT | ASSQEGQPGERLF | 0.5299619 |
| GHVDHGKTT | ASSQEGQPGERLF | 0.68393993 |
| AQFEHTLL | ASSQEGQPGERLF | 0.6511091 |
| IVNQDEL | ASSQEGQPGERLF | 0.6016384 |
| TNGSQFF | ASSQEGQPGERLF | 0.6817695 |
| TGKGIVE | ASSQEGQPGERLF | 0.6386126 |
| LISNASDA | ASSQEGQPGERLF | 0.6118958 |
| AELRTLGELTKIAKAHGAQ | ASSQEGQPGERLF | 0.452213 |
| APNAVAAAEPSW | ASSQEGQPGERLF | 0.51542836 |
| APQVAGAATAVL | ASSQEGQPGERLF | 0.6346823 |
| ARRDPAAAAAILPTDA | ASSQEGQPGERLF | 0.63856363 |
| DAAARDVLAEAGLAEYF | ASSQEGQPGERLF | 0.69033635 |
| DDQIVLSAKANLAKVEQ | ASSQEGQPGERLF | 0.5829494 |
| DELLPDTGEDSDEDGH | ASSQEGQPGERLF | 0.5630158 |
| ELPEAPANRATVPQD | ASSQEGQPGERLF | 0.687245 |
| FGQSKIERVTGIG | ASSQEGQPGERLF | 0.6963241 |
| GAGGTGSTAGGAGGAGGAGGLY | ASSQEGQPGERLF | 0.3930762 |
| GHGGTGAAGGNGGTGGAGGSGI | ASSQEGQPGERLF | 0.5996045 |
| GNGGNGGAGGEGGPGGAGGAGA | ASSQEGQPGERLF | 0.7125956 |
| GSTAGNGGNGGNATNGTIAGQP | ASSQEGQPGERLF | 0.6720227 |
| GTGGDGGNGGNGGTGG | ASSQEGQPGERLF | 0.58511245 |

| | | |
|---|---|---|
| GVRRPAGIPADAPV | ASSQEGQPGERLF | 0.5250454 |
| GYRQVTDAHLLTLARR | ASSQEGQPGERLF | 0.71439826 |
| IAKMGQDGHDRG | ASSQEGQPGERLF | 0.5768678 |
| KAEPSIEIQYT | ASSQEGQPGERLF | 0.7134315 |
| KPVRIRMAADEDTVAVT | ASSQEGQPGERLF | 0.3054887 |
| MPPRIAGMRLLVIKPEP | ASSQEGQPGERLF | 0.47856092 |
| MKFYAVALFRSRGVPPTR | ASSQEGQPGERLF | 0.4571911 |
| NAGGFGILGAVAHSP | ASSQEGQPGERLF | 0.28190464 |
| NEGREITLAMAEM | ASSQEGQPGERLF | 0.6919645 |
| NELVEWSHPLGHRWRWE | ASSQEGQPGERLF | 0.44134843 |
| NLGAGGTGGNGGNPG | ASSQEGQPGERLF | 0.6350687 |
| NTEWPKWETVMAADPVD | ASSQEGQPGERLF | 0.5192311 |
| PASAAPAETQP | ASSQEGQPGERLF | 0.69097936 |
| QQSSDGKNGVRAAEITGE | ASSQEGQPGERLF | 0.6311145 |
| RAPDAVTTSWWKE | ASSQEGQPGERLF | 0.57117134 |
| SGFGNVGAHTSGWFNQ | ASSQEGQPGERLF | 0.43195555 |
| SGVAGAVGQAASVGGLKV | ASSQEGQPGERLF | 0.6618829 |
| SNGGAGTGSAGGNGGT | ASSQEGQPGERLF | 0.6995279 |
| TGGAGGEGGAAG | ASSQEGQPGERLF | 0.65426636 |
| TGGDGGAAGAGGAPGHG | ASSQEGQPGERLF | 0.33938816 |
| TMPPLLAAIANAGRHGV | ASSQEGQPGERLF | 0.66949785 |
| TSSTAGNSGGA | ASSQEGQPGERLF | 0.6080996 |
| VPATVPVGILC | ASSQEGQPGERLF | 0.4051842 |
| VQRPDSGGVVLTGRIS | ASSQEGQPGERLF | 0.31317106 |
| WTPNPVPATKGVPAGLVQAT | ASSQEGQPGERLF | 0.55134505 |
| YALKRALAKFSDRFDVVII | ASSQEGQPGERLF | 0.6191261 |
| YLIGSVIV | ASSQEGQPGERLF | 0.6811147 |
| YGQLVAVID | ASSQEGQPGERLF | 0.6867906 |
| VTVGGMTLPA | ASSQEGQPGERLF | 0.50866866 |
| VLALPAV | ASSQEGQPGERLF | 0.677652 |
| VIPPTLNYET | ASSQEGQPGERLF | 0.66032344 |
| VGMGGMHPSSG | ASSQEGQPGERLF | 0.24357346 |
| VGLAGGAATA | ASSQEGQPGERLF | 0.47525722 |
| VGGAGGAGGSGP | ASSQEGQPGERLF | 0.3832236 |
| VFAAFMV | ASSQEGQPGERLF | 0.66786885 |
| TLYDKLR | ASSQEGQPGERLF | 0.4961981 |
| TLAVFTVL | ASSQEGQPGERLF | 0.5522306 |
| TGSVQPARV | ASSQEGQPGERLF | 0.5898421 |
| TGGGAGSGGT | ASSQEGQPGERLF | 0.6824131 |
| SHLETSL | ASSQEGQPGERLF | 0.6232513 |
| SFTTGMT | ASSQEGQPGERLF | 0.46764067 |
| QVEVAKRMGVRQPTVS | ASSQEGQPGERLF | 0.41623282 |
| QGSGGPALAM | ASSQEGQPGERLF | 0.6591029 |
| MLAPSNP | ASSQEGQPGERLF | 0.54681873 |
| LVNNAAI | ASSQEGQPGERLF | 0.46986908 |
| LTGEINTLA | ASSQEGQPGERLF | 0.42003733 |
| LSIFGPV | ASSQEGQPGERLF | 0.6892871 |
| LLALLANL | ASSQEGQPGERLF | 0.30155054 |
| LISNASDALDKLR | ASSQEGQPGERLF | 0.6379461 |
| LINEAVP | ASSQEGQPGERLF | 0.4733684 |
| LGYAFSGL | ASSQEGQPGERLF | 0.5924275 |

| | | |
|---|---|---|
| LDFVAVAL | ASSQEGQPGERLF | 0.71375334 |
| LAQPGAE | ASSQEGQPGERLF | 0.67510015 |
| ISNLGAVTGF | ASSQEGQPGERLF | 0.5632546 |
| HQEDPGRI | ASSQEGQPGERLF | 0.48825184 |
| GTPGEPTP | ASSQEGQPGERLF | 0.42634138 |
| GFFNSGS | ASSQEGQPGERLF | 0.5025011 |
| GAGGGGLTAGAN | ASSQEGQPGERLF | 0.46710438 |
| FVDALVNM | ASSQEGQPGERLF | 0.69010377 |
| FLTVAPEL | ASSQEGQPGERLF | 0.55771375 |
| FETLTAL | ASSQEGQPGERLF | 0.3412733 |
| DVINPIEL | ASSQEGQPGERLF | 0.56417257 |
| APVSGAPV | ASSQEGQPGERLF | 0.70131886 |
| ALVGGLAW | ASSQEGQPGERLF | 0.7201725 |
| ALAAAAATAI | ASSQEGQPGERLF | 0.5306906 |
| AFGLGLLL | ASSQEGQPGERLF | 0.42991996 |
| AFFAAASL | ASSQEGQPGERLF | 0.4999335 |
| AAAALPAFSPP | ASSQEGQPGERLF | 0.6144576 |
| VVLAPAL | ASSQEGQPGERLF | 0.41882464 |
| TIGHVDHGKTTL | ASSQEGQPGERLF | 0.41412702 |
| LISNASDALDK | ASSQEGQPGERLF | 0.6707486 |
| LISNASDAL | ASSQEGQPGERLF | 0.7234844 |
| ISNASDALDK | ASSQEGQPGERLF | 0.5725747 |
| NQAETLVYQTE | ASSQEGQPGERLF | 0.2878357 |
| TGGTGGTAGAG | ASSQEGQPGERLF | 0.6123252 |
| TGSTAGGAGGA | ASSQEGQPGERLF | 0.46552017 |
| DPLAVPVAAGPV | ASSQEGQPGERLF | 0.5038209 |
| GGTGGTPGNGGN | ASSQEGQPGERLF | 0.5432448 |
| PAGLAGVDPATL | ASSQEGQPGERLF | 0.34505707 |
| GNNLVVEAPHAGASV | ASSQEGQPGERLF | 0.5770669 |
| KVTRIMSASTLEQPA | ASSQEGQPGERLF | 0.7146822 |
| LGFALPIALPRRTVV | ASSQEGQPGERLF | 0.6267171 |
| TRLGNLTARKLVVLV | ASSQEGQPGERLF | 0.6004653 |
| WSQQRRLIYFPSAGP | ASSQEGQPGERLF | 0.7062495 |
| GPNGNPDPMAAAVDIR | ASSQEGQPGERLF | 0.66932243 |
| TQRCVSVADLRAAASR | ASSQEGQPGERLF | 0.70465505 |
| QFVLDVAEHPVEVLTETV | ASSQEGQPGERLF | 0.70163316 |
| GGAGGTGGAGGNGGTQAGDG | ASSQEGQPGERLF | 0.54352915 |
| PLLATLRAIAVDVDAKPAQV | ASSQEGQPGERLF | 0.5935723 |
| APKFRTGGYMIDFWGVGYQVAK | ASSQEGQPGERLF | 0.7087983 |
| GAGGAGGAGGAAGGPGGTGGTG | ASSQEGQPGERLF | 0.6063092 |
| IGALGLLATDVAPAGSVAFGVA | ASSQEGQPGERLF | 0.62403095 |
| AGGAGGIGGNGGAGGAGGVLMG | ASSQEGQPGERLF | 0.53039706 |
| ADELIGLL | ASSQEGQPGERLF | 0.684861 |
| ADRALKA | ASSQEGQPGERLF | 0.547795 |
| AGAVGQVGGAGGA | ASSQEGQPGERLF | 0.7097571 |
| AGGGGGLGGAP | ASSQEGQPGERLF | 0.28889313 |
| AGGLLYGNGGAG | ASSQEGQPGERLF | 0.39559895 |
| AGLESIGL | ASSQEGQPGERLF | 0.6958323 |
| AGLKTGII | ASSQEGQPGERLF | 0.2660176 |
| ASLAALLL | ASSQEGQPGERLF | 0.31166363 |
| ASVINISSI | ASSQEGQPGERLF | 0.6065325 |

| | | |
|---|---|---|
| ATPPGPA | ASSQEGQPGERLF | 0.69769984 |
| ATVASALV | ASSQEGQPGERLF | 0.27832395 |
| ATVASLIL | ASSQEGQPGERLF | 0.49703336 |
| AVNGPVLIRH | ASSQEGQPGERLF | 0.35074937 |
| DGAAGAVGI | ASSQEGQPGERLF | 0.6111329 |
| DPLLHGI | ASSQEGQPGERLF | 0.5771483 |
| ELGAALGAL | ASSQEGQPGERLF | 0.34139732 |
| ELGRTQAV | ASSQEGQPGERLF | 0.64803356 |
| FVAGGYLS | ASSQEGQPGERLF | 0.29519516 |
| GASGGAGGAGGW | ASSQEGQPGERLF | 0.66707206 |
| GGAGAGGDGGAGGI | ASSQEGQPGERLF | 0.6831863 |
| GTGHNGSQPGMGGNG | ASSQEGQPGERLF | 0.7194903 |
| GYAAASASAS | ASSQEGQPGERLF | 0.46941012 |
| HQVDVAAVQQG | ASSQEGQPGERLF | 0.6627336 |
| IGAAIAHT | ASSQEGQPGERLF | 0.5104702 |
| IGMIEVAQL | ASSQEGQPGERLF | 0.7256521 |
| IRLAPEHTSAMD | ASSQEGQPGERLF | 0.55928266 |
| IYNVSVL | ASSQEGQPGERLF | 0.33542067 |
| KQGFLGM | ASSQEGQPGERLF | 0.6787624 |
| LESVITQL | ASSQEGQPGERLF | 0.63855994 |
| LGLGALVVV | ASSQEGQPGERLF | 0.5715339 |
| LIGNGGNGGSGGI | ASSQEGQPGERLF | 0.6951802 |
| LNAINAPTL | ASSQEGQPGERLF | 0.49738398 |
| LQLLNTGV | ASSQEGQPGERLF | 0.57580113 |
| LQTSIEF | ASSQEGQPGERLF | 0.6811625 |
| MSFAVSEL | ASSQEGQPGERLF | 0.67761564 |
| NTGALGL | ASSQEGQPGERLF | 0.4609847 |
| PPGAGGAGGAGTT | ASSQEGQPGERLF | 0.6886375 |
| QLVGLGI | ASSQEGQPGERLF | 0.62459445 |
| RFLEQVHQL | ASSQEGQPGERLF | 0.42220306 |
| RGVGYVLR | ASSQEGQPGERLF | 0.7206832 |
| SATGVFAGVI | ASSQEGQPGERLF | 0.49642658 |
| SLPAITLP | ASSQEGQPGERLF | 0.67614615 |
| STALAFTV | ASSQEGQPGERLF | 0.6408247 |
| SVLTVGLGK | ASSQEGQPGERLF | 0.7149558 |
| TALAGAGSGI | ASSQEGQPGERLF | 0.40992194 |
| TGLPFTGI | ASSQEGQPGERLF | 0.63243484 |
| VAEHVTV | ASSQEGQPGERLF | 0.63107884 |
| VDIVLNSL | ASSQEGQPGERLF | 0.65384686 |
| VLTGGGAVL | ASSQEGQPGERLF | 0.52634084 |
| VNVGLAYGI | ASSQEGQPGERLF | 0.6453755 |
| VSSAAVGAV | ASSQEGQPGERLF | 0.65475047 |
| VTSVYEAGV | ASSQEGQPGERLF | 0.6934203 |
| WVALDRGA | ASSQEGQPGERLF | 0.58487386 |
| YLVGSVVI | ASSQEGQPGERLF | 0.63079464 |
| GPPGTGKTLL | ASSQEGQPGERLF | 0.71481955 |
| GVGMTKF | ASSQEGQPGERLF | 0.4409433 |
| IGIGGSDL | ASSQEGQPGERLF | 0.5850691 |
| IGTLLAEL | ASSQEGQPGERLF | 0.6774306 |
| IYTSGTT | ASSQEGQPGERLF | 0.6900842 |
| LSFPAGV | ASSQEGQPGERLF | 0.50690466 |

| | | |
|---|---|---|
| LYTSGTT | ASSQEGQPGERLF | 0.7047571 |
| VGAFPFGF | ASSQEGQPGERLF | 0.56746024 |
| VGAGTVEF | ASSQEGQPGERLF | 0.67427695 |
| VGKSTLLL | ASSQEGQPGERLF | 0.5998776 |
| AALAHAGGGIV | ASSQEGQPGERLF | 0.5156255 |
| AALVAVSLL | ASSQEGQPGERLF | 0.50360745 |
| AAVSRAVAAL | ASSQEGQPGERLF | 0.66772056 |
| AGGSSGIGGF | ASSQEGQPGERLF | 0.6845652 |
| AGLTVANANG | ASSQEGQPGERLF | 0.6614848 |
| AGLVGAVL | ASSQEGQPGERLF | 0.2902823 |
| ALSQAFTV | ASSQEGQPGERLF | 0.66956556 |
| ALTFSVSL | ASSQEGQPGERLF | 0.6720449 |
| ALTGAALVAL | ASSQEGQPGERLF | 0.72008693 |
| ASFSIGAGVL | ASSQEGQPGERLF | 0.5721296 |
| DGSVYKI | ASSQEGQPGERLF | 0.65310824 |
| EQLAEIV | ASSQEGQPGERLF | 0.5896408 |
| FAVSELTVM | ASSQEGQPGERLF | 0.5700511 |
| GAAPLAAGL | ASSQEGQPGERLF | 0.36986735 |
| GADGGDGGSGGAGG | ASSQEGQPGERLF | 0.5003384 |
| GFGLDTGGAGG | ASSQEGQPGERLF | 0.59729457 |
| GLASGGLL | ASSQEGQPGERLF | 0.51272345 |
| GRIAGAAVLL | ASSQEGQPGERLF | 0.7026725 |
| GSAGSLNTSF | ASSQEGQPGERLF | 0.69924057 |
| GTGQIGI | ASSQEGQPGERLF | 0.6903737 |
| GTLIALIL | ASSQEGQPGERLF | 0.7008333 |
| IAATMGFVVG | ASSQEGQPGERLF | 0.7105723 |
| IFAQALL | ASSQEGQPGERLF | 0.6332943 |
| IGAAVALGF | ASSQEGQPGERLF | 0.5517385 |
| IGLTPLAL | ASSQEGQPGERLF | 0.57811075 |
| ILGSNTSTL | ASSQEGQPGERLF | 0.36508703 |
| IVFSLTVL | ASSQEGQPGERLF | 0.5448985 |
| LASKATGF | ASSQEGQPGERLF | 0.5358758 |
| LASVALIL | ASSQEGQPGERLF | 0.60918105 |
| LFAVGLTL | ASSQEGQPGERLF | 0.2613704 |
| LFNGGIGTYI | ASSQEGQPGERLF | 0.6826117 |
| LTLGAGII | ASSQEGQPGERLF | 0.69672436 |
| LVAAALGDI | ASSQEGQPGERLF | 0.6555368 |
| NTAFVQLGI | ASSQEGQPGERLF | 0.65232307 |
| QLFSGAEL | ASSQEGQPGERLF | 0.67246604 |
| SAATFITL | ASSQEGQPGERLF | 0.4913748 |
| SNYGLANLGV | ASSQEGQPGERLF | 0.67393875 |
| SYGGAYAVM | ASSQEGQPGERLF | 0.48547477 |
| TTGVGGLTLG | ASSQEGQPGERLF | 0.65699077 |
| TTQTIYI | ASSQEGQPGERLF | 0.7021624 |
| VAAGAGLGAVI | ASSQEGQPGERLF | 0.6517787 |
| VAAVVTLL | ASSQEGQPGERLF | 0.6131925 |
| VGFLGGLI | ASSQEGQPGERLF | 0.56306326 |
| VGVIPILL | ASSQEGQPGERLF | 0.34937137 |
| VIFASSLI | ASSQEGQPGERLF | 0.6403786 |
| VKGGDGGAAAG | ASSQEGQPGERLF | 0.62030107 |
| VTKTFTNI | ASSQEGQPGERLF | 0.6395195 |

| | | |
|---|---|---|
| VVAAAILL | ASSQEGQPGERLF | 0.4839471 |
| YAALDVEL | ASSQEGQPGERLF | 0.67556953 |
| LASVALLL | ASSQEGQPGERLF | 0.6348299 |
| LDAVHRV | ASSQEGQPGERLF | 0.6414019 |
| QGLSSSIF | ASSQEGQPGERLF | 0.4844942 |
| RGTFANIRL | ASSQEGQPGERLF | 0.6680863 |
| RQLWWGHRI | ASSQEGQPGERLF | 0.6468489 |
| YGPPGTGKTLL | ASSQEGQPGERLF | 0.3904664 |
| AALVGAPANVA | ASSQEGQPGERLF | 0.65601146 |
| AATFRRTE | ASSQEGQPGERLF | 0.5411254 |
| AGALGGGI | ASSQEGQPGERLF | 0.6118773 |
| AGGAGGAGGIGAG | ASSQEGQPGERLF | 0.6896527 |
| AGSGALGL | ASSQEGQPGERLF | 0.6059411 |
| AQIADQI | ASSQEGQPGERLF | 0.30782032 |
| ASAVAVTYL | ASSQEGQPGERLF | 0.5896597 |
| ASTVLGGTGG | ASSQEGQPGERLF | 0.6458938 |
| AVGGEPAK | ASSQEGQPGERLF | 0.59782696 |
| CEQGTFGGTC | ASSQEGQPGERLF | 0.27634028 |
| DEGKAPAG | ASSQEGQPGERLF | 0.6656744 |
| DTVVFPAL | ASSQEGQPGERLF | 0.41930175 |
| FEGITMAG | ASSQEGQPGERLF | 0.5023527 |
| FPLVVQD | ASSQEGQPGERLF | 0.5352998 |
| GAFTTPQ | ASSQEGQPGERLF | 0.6860998 |
| GFGGFGTAGGVGGA | ASSQEGQPGERLF | 0.69012403 |
| GFLAQFL | ASSQEGQPGERLF | 0.5067111 |
| GGAGGAGGNAGVA | ASSQEGQPGERLF | 0.66597307 |
| GGVGTAGAGGAGGAGG | ASSQEGQPGERLF | 0.64496124 |
| GNGGGGDAGNAGSGGN | ASSQEGQPGERLF | 0.6979035 |
| GTGGAGGAGNDAGS | ASSQEGQPGERLF | 0.64060134 |
| GTTNTGWFNS | ASSQEGQPGERLF | 0.5705857 |
| IFMGPLFTL | ASSQEGQPGERLF | 0.5894072 |
| ILAHAIEAI | ASSQEGQPGERLF | 0.44676095 |
| IRSAIREAL | ASSQEGQPGERLF | 0.49352843 |
| ITFAGGG | ASSQEGQPGERLF | 0.39728743 |
| ITGMSGHQEV | ASSQEGQPGERLF | 0.45116034 |
| IVAGLACTL | ASSQEGQPGERLF | 0.42936307 |
| IVGGGLAGA | ASSQEGQPGERLF | 0.65665424 |
| IVTGVLNI | ASSQEGQPGERLF | 0.6632953 |
| LAQQGCI | ASSQEGQPGERLF | 0.556496 |
| LCLHVEN | ASSQEGQPGERLF | 0.35375434 |
| LDAISDGS | ASSQEGQPGERLF | 0.565556 |
| LGGFAAML | ASSQEGQPGERLF | 0.49932867 |
| LGGKNPMI | ASSQEGQPGERLF | 0.67647046 |
| LGPKGVAAIL | ASSQEGQPGERLF | 0.6690117 |
| LMVGVSI | ASSQEGQPGERLF | 0.7184905 |
| LVGAAFVL | ASSQEGQPGERLF | 0.6945565 |
| LVLAVAPL | ASSQEGQPGERLF | 0.5548836 |
| MAAIAVIDAA | ASSQEGQPGERLF | 0.6843926 |
| NNGINAELQ | ASSQEGQPGERLF | 0.4920455 |
| RGEPLVLVHG | ASSQEGQPGERLF | 0.552177 |
| RGGAGGAAGLI | ASSQEGQPGERLF | 0.60112906 |

| | | |
|---|---|---|
| RSYRMHPAV | ASSQEGQPGERLF | 0.64512396 |
| SAAHLLAGQLA | ASSQEGQPGERLF | 0.5277902 |
| SAAVGSY | ASSQEGQPGERLF | 0.54993016 |
| SPHAVLHRI | ASSQEGQPGERLF | 0.32107067 |
| SQIAKVL | ASSQEGQPGERLF | 0.6849792 |
| TALGHLVP | ASSQEGQPGERLF | 0.42236036 |
| TAPTTSGAA | ASSQEGQPGERLF | 0.36002913 |
| TDGDDGNGGSPGT | ASSQEGQPGERLF | 0.55094725 |
| TDGLGEN | ASSQEGQPGERLF | 0.72145444 |
| TGGPPLPGQA | ASSQEGQPGERLF | 0.710182 |
| TIPAITI | ASSQEGQPGERLF | 0.55656517 |
| TIPGFTI | ASSQEGQPGERLF | 0.50230163 |
| TSIASGALA | ASSQEGQPGERLF | 0.691677 |
| TVTLPGAAI | ASSQEGQPGERLF | 0.523288 |
| VASGLSAALP | ASSQEGQPGERLF | 0.62783957 |
| VGAGVIGI | ASSQEGQPGERLF | 0.6363498 |
| VGPKRFGV | ASSQEGQPGERLF | 0.616232 |
| VNVAFPD | ASSQEGQPGERLF | 0.43074608 |
| VSMTNSGVS | ASSQEGQPGERLF | 0.6627047 |
| VTDVLGDLSV | ASSQEGQPGERLF | 0.6009442 |
| VVVVGAGFAGL | ASSQEGQPGERLF | 0.5729468 |
| VYTVGTVV | ASSQEGQPGERLF | 0.36358133 |
| WTTFSGP | ASSQEGQPGERLF | 0.6895795 |
| YDELVEAF | ASSQEGQPGERLF | 0.48870486 |
| YGTGGAGGNGG | ASSQEGQPGERLF | 0.43248963 |
| YTNVMAV | ASSQEGQPGERLF | 0.28497824 |
| FLEQVHQL | ASSQEGQPGERLF | 0.4946363 |
| GLSLLAVAV | ASSQEGQPGERLF | 0.671999 |
| LEQVHQL | ASSQEGQPGERLF | 0.70667815 |
| LNAAGAI | ASSQEGQPGERLF | 0.62458456 |
| PAVHASL | ASSQEGQPGERLF | 0.66561043 |
| TELEDDA | ASSQEGQPGERLF | 0.6982999 |
| TPPGPPN | ASSQEGQPGERLF | 0.5452276 |
| TPPPPPPP | ASSQEGQPGERLF | 0.32050818 |
| TQSPALI | ASGDADTQY | 0.5761594 |
| EFAAAFPQ | ASGDADTQY | 0.6281597 |
| LLSGNLDVL | ASGDADTQY | 0.6849623 |
| GPANLGNYNI | ASGDADTQY | 0.6954533 |
| TLGPTLP | ASGDADTQY | 0.39011943 |
| VYAYSSCCK | ASGDADTQY | 0.6074604 |
| APAPAPAPNI | ASGDADTQY | 0.65162855 |
| PGGPVPIP | ASGDADTQY | 0.3948269 |
| FLTVALEP | ASGDADTQY | 0.4491825 |
| SQYFGANL | ASGDADTQY | 0.40871668 |
| GGVKPAA | ASGDADTQY | 0.40950853 |
| GGTGGAGGAGGW | ASGDADTQY | 0.6491517 |
| GVGGVAGLGGAGG | ASGDADTQY | 0.5605161 |
| VLLVLATTI | ASGDADTQY | 0.5272682 |
| IGIFAAML | ASGDADTQY | 0.5372982 |
| AVGDLID | ASGDADTQY | 0.46478003 |
| NISPLPL | ASGDADTQY | 0.45779073 |

| | | |
|---|---|---|
| VTAVFVAI | ASGDADTQY | 0.5264502 |
| SGDDNDAAL | ASGDADTQY | 0.62059593 |
| AVAALAHS | ASGDADTQY | 0.6819798 |
| GGVTVMATP | ASGDADTQY | 0.53652084 |
| AGLAANVRD | ASGDADTQY | 0.5111555 |
| TNRPDLIDPALLRPG | ASGDADTQY | 0.70724034 |
| LANLGVTAL | ASGDADTQY | 0.3691079 |
| TGERIEL | ASGDADTQY | 0.4632213 |
| NGEELAA | ASGDADTQY | 0.64625883 |
| VLSLVWG | ASGDADTQY | 0.5110397 |
| IGYALWL | ASGDADTQY | 0.6902294 |
| LSLLLAPL | ASGDADTQY | 0.6554733 |
| TLDEALK | ASGDADTQY | 0.6390846 |
| TDGAGTAEGG | ASGDADTQY | 0.62350994 |
| NAPGGAGGNGG | ASGDADTQY | 0.72006035 |
| VSVFMGMGGNF | ASGDADTQY | 0.6432098 |
| LASYLAAL | ASGDADTQY | 0.4679008 |
| LVGGTPAL | ASGDADTQY | 0.69819593 |
| VRNADVL | ASGDADTQY | 0.35431325 |
| TCSSPFLVSAM | ASGDADTQY | 0.6685344 |
| VHITDAE | ASGDADTQY | 0.6520949 |
| EYAAPSF | ASGDADTQY | 0.48029765 |
| VVDAFHGGL | ASGDADTQY | 0.58890843 |
| ASVVDPQ | ASGDADTQY | 0.70534277 |
| SYNEFGGLS | ASGDADTQY | 0.69381404 |
| DPDGSVY | ASGDADTQY | 0.47061354 |
| MVSTIHATE | ASGDADTQY | 0.49425933 |
| LAAGVADPV | ASGDADTQY | 0.47019082 |
| TKEQIAA | ASGDADTQY | 0.47391015 |
| SLASPAAL | ASGDADTQY | 0.3361009 |
| IGLVFYGV | ASGDADTQY | 0.66373175 |
| FLLDASNL | ASGDADTQY | 0.5662492 |
| YQVGNFP | ASGDADTQY | 0.6998787 |
| ESLLSQST | ASGDADTQY | 0.65721786 |
| LFTVGFAAVG | ASGDADTQY | 0.6937955 |
| ATPPTPT | ASGDADTQY | 0.5520353 |
| TDLATLG | ASGDADTQY | 0.59964055 |
| SAGGTGR | ASGDADTQY | 0.32773656 |
| GFSNSGS | ASGDADTQY | 0.5871636 |
| GGNGGSGGTSV | ASGDADTQY | 0.4228393 |
| GTSVATGGAGN | ASGDADTQY | 0.56454307 |
| NGGNGGSGGTS | ASGDADTQY | 0.39048308 |
| AVVKFPH | ASGDADTQY | 0.7015898 |
| TPPPPPP | ASGDADTQY | 0.5573562 |
| AAPAAAAAPA | ASGDADTQY | 0.659629 |
| IALNEDL | ASGDADTQY | 0.50568575 |
| IVNVSSM | ASGDADTQY | 0.40321937 |
| TGIPPAPR | ASGDADTQY | 0.54002523 |
| FDLGGGTF | ASGDADTQY | 0.686803 |
| FDLGGGTFDVS | ASGDADTQY | 0.44217592 |
| NEPTAAA | ASGDADTQY | 0.27608487 |

| | | |
|---|---|---|
| NEPTAAALA | ASGDADTQY | 0.41685325 |
| GSIINISSI | ASGDADTQY | 0.34397793 |
| VAPGNAGTA | ASGDADTQY | 0.7182872 |
| TGGGASLE | ASGDADTQY | 0.6965966 |
| VAVYNHYKRI | ASGDADTQY | 0.63077426 |
| TDPAPATT | ASGDADTQY | 0.7166892 |
| TNRPDILDPALLRPG | ASGDADTQY | 0.2619575 |
| LTTTTTS | ASGDADTQY | 0.5557082 |
| PTGSGKTL | ASGDADTQY | 0.7031745 |
| TGSGKTL | ASGDADTQY | 0.65745074 |
| TIGHVDHGK | ASGDADTQY | 0.5950953 |
| TIGHVDHGKTT | ASGDADTQY | 0.6813395 |
| GHVDHGKTT | ASGDADTQY | 0.509923 |
| AQFEHTLL | ASGDADTQY | 0.5997894 |
| IVNQDEL | ASGDADTQY | 0.5946135 |
| TNGSQFF | ASGDADTQY | 0.6611115 |
| TGKGIVE | ASGDADTQY | 0.60992783 |
| LISNASDA | ASGDADTQY | 0.36245924 |
| AELRTLGELTKIAKAHGAQ | ASGDADTQY | 0.61371034 |
| APNAVAAAEPSW | ASGDADTQY | 0.7276503 |
| APQVAGAATAVL | ASGDADTQY | 0.59756505 |
| ARRDPAAAAAILPTDA | ASGDADTQY | 0.6524798 |
| DAAARDVLAEAGLAEYF | ASGDADTQY | 0.5966509 |
| DDQIVLSAKANLAKVEQ | ASGDADTQY | 0.6495003 |
| DELLPDTGEDSDEDGH | ASGDADTQY | 0.5150691 |
| ELPEAPANRATVPQD | ASGDADTQY | 0.45284247 |
| FGQSKIERVTGIG | ASGDADTQY | 0.66492105 |
| GAGGTGSTAGGAGGAGGAGGLY | ASGDADTQY | 0.44069648 |
| GHGGTGAAGGNGGTGGAGGSGI | ASGDADTQY | 0.5477231 |
| GNGGNGGAGGEGGPGGAGGAGA | ASGDADTQY | 0.509125 |
| GSTAGNGGNGGNATNGTIAGQP | ASGDADTQY | 0.729292 |
| GTGGDGGNGGNGGTGG | ASGDADTQY | 0.6568769 |
| GVRRPAGIPADAPV | ASGDADTQY | 0.6465033 |
| GYRQVTDAHLLTLARR | ASGDADTQY | 0.5210962 |
| IAKMGQDGHDRG | ASGDADTQY | 0.62119436 |
| KAEPSIEIQYT | ASGDADTQY | 0.6739249 |
| KPVRIRMAADEDTVAVT | ASGDADTQY | 0.32721305 |
| MPPRIAGMRLLVIKPEP | ASGDADTQY | 0.6981633 |
| MKFYAVALFRSRGVPPTR | ASGDADTQY | 0.4105116 |
| NAGGFGILGAVAHSP | ASGDADTQY | 0.62677234 |
| NEGREITLAMAEM | ASGDADTQY | 0.5775572 |
| NELVEWSHPLGHRWRWE | ASGDADTQY | 0.7113165 |
| NLGAGGTGGNGGNPG | ASGDADTQY | 0.5878013 |
| NTEWPKWETVMAADPVD | ASGDADTQY | 0.6530279 |
| PASAAPAETQP | ASGDADTQY | 0.45037365 |
| QQSSDGKNGVRAAEITGE | ASGDADTQY | 0.6656087 |
| RAPDAVTTSWWKE | ASGDADTQY | 0.7096425 |
| SGFGNVGAHTSGWFNQ | ASGDADTQY | 0.6577605 |
| SGVAGAVGQAASVGGLKV | ASGDADTQY | 0.72379863 |
| SNGGAGTGSAGGNGGT | ASGDADTQY | 0.5765232 |
| TGGAGGEGGAAG | ASGDADTQY | 0.31103346 |

| | | |
|---|---|---|
| TGGDGGAAGAGGAPGHG | ASGDADTQY | 0.6019087 |
| TMPPLLAAIANAGRHGV | ASGDADTQY | 0.70001936 |
| TSSTAGNSGGA | ASGDADTQY | 0.7077939 |
| VPATVPVGILC | ASGDADTQY | 0.6586623 |
| VQRPDSGGVVLTGRIS | ASGDADTQY | 0.6817349 |
| WTPNPVPATKGVPAGLVQAT | ASGDADTQY | 0.6195632 |
| YALKRALAKFSDRFDVVII | ASGDADTQY | 0.70499086 |
| YLIGSVIV | ASGDADTQY | 0.72246766 |
| YGQLVAVID | ASGDADTQY | 0.64082134 |
| VTVGGMTLPA | ASGDADTQY | 0.44245195 |
| VLALPAV | ASGDADTQY | 0.3949326 |
| VIPPTLNYET | ASGDADTQY | 0.6577725 |
| VGMGGMHPSSG | ASGDADTQY | 0.69721794 |
| VGLAGGAATA | ASGDADTQY | 0.34948915 |
| VGGAGGAGGSGP | ASGDADTQY | 0.6134385 |
| VFAAFMV | ASGDADTQY | 0.6245692 |
| TLYDKLR | ASGDADTQY | 0.30926612 |
| TLAVFTVL | ASGDADTQY | 0.65077186 |
| TGSVQPARV | ASGDADTQY | 0.70917845 |
| TGGGAGSGGT | ASGDADTQY | 0.5797943 |
| SHLETSL | ASGDADTQY | 0.38058984 |
| SFTTGMT | ASGDADTQY | 0.7051203 |
| QVEVAKRMGVRQPTVS | ASGDADTQY | 0.7218752 |
| QGSGGPALAM | ASGDADTQY | 0.6976906 |
| MLAPSNP | ASGDADTQY | 0.3180929 |
| LVNNAAI | ASGDADTQY | 0.26338777 |
| LTGEINTLA | ASGDADTQY | 0.66547024 |
| LSIFGPV | ASGDADTQY | 0.68799067 |
| LLALLANL | ASGDADTQY | 0.5177437 |
| LISNASDALDKLR | ASGDADTQY | 0.4787071 |
| LINEAVP | ASGDADTQY | 0.62445056 |
| LGYAFSGL | ASGDADTQY | 0.6842048 |
| LDFVAVAL | ASGDADTQY | 0.41576895 |
| LAQPGAE | ASGDADTQY | 0.38455927 |
| ISNLGAVTGF | ASGDADTQY | 0.60848254 |
| HQEDPGRI | ASGDADTQY | 0.67455447 |
| GTPGEPTP | ASGDADTQY | 0.4996361 |
| GFFNSGS | ASGDADTQY | 0.59172934 |
| GAGGGGLTAGAN | ASGDADTQY | 0.67439723 |
| FVDALVNM | ASGDADTQY | 0.6688651 |
| FLTVAPEL | ASGDADTQY | 0.64471114 |
| FETLTAL | ASGDADTQY | 0.42107266 |
| DVINPIEL | ASGDADTQY | 0.70034444 |
| APVSGAPV | ASGDADTQY | 0.36106902 |
| ALVGGLAW | ASGDADTQY | 0.6839148 |
| ALAAAAATAI | ASGDADTQY | 0.6267885 |
| AFGLGLLL | ASGDADTQY | 0.6528491 |
| AFFAAASL | ASGDADTQY | 0.5869175 |
| AAAALPAFSPP | ASGDADTQY | 0.42935842 |
| VVLAPAL | ASGDADTQY | 0.42433128 |
| TIGHVDHGKTTL | ASGDADTQY | 0.3970837 |

| | | |
|---|---|---|
| LISNASDALDK | ASGDADTQY | 0.69801956 |
| LISNASDAL | ASGDADTQY | 0.6693481 |
| ISNASDALDK | ASGDADTQY | 0.6796281 |
| NQAETLVYQTE | ASGDADTQY | 0.61825573 |
| TGGTGGTAGAG | ASGDADTQY | 0.48075294 |
| TGSTAGGAGGA | ASGDADTQY | 0.6417124 |
| DPLAVPVAAGPV | ASGDADTQY | 0.4103759 |
| GGTGGTPGNGGN | ASGDADTQY | 0.6139118 |
| PAGLAGVDPATL | ASGDADTQY | 0.56519496 |
| GNNLVVEAPHAGASV | ASGDADTQY | 0.60425353 |
| KVTRIMSASTLEQPA | ASGDADTQY | 0.49799514 |
| LGFALPIALPRRTVV | ASGDADTQY | 0.60018176 |
| TRLGNLTARKLVVLV | ASGDADTQY | 0.6610121 |
| WSQQRRLIYFPSAGP | ASGDADTQY | 0.40185714 |
| GPNGNPDPMAAAVDIR | ASGDADTQY | 0.57552433 |
| TQRCVSVADLRAAASR | ASGDADTQY | 0.6906312 |
| QFVLDVAEHPVEVLTETV | ASGDADTQY | 0.5916116 |
| GGAGGTGGAGGNGGTQAGDG | ASGDADTQY | 0.37410247 |
| PLLATLRAIAVDVDAKPAQV | ASGDADTQY | 0.6419656 |
| APKFRTGGYMIDFWGVGYQVAK | ASGDADTQY | 0.5213449 |
| GAGGAGGAGGAAGGPGGTGGTG | ASGDADTQY | 0.6596738 |
| IGALGLLATDVAPAGSVAFGVA | ASGDADTQY | 0.6212934 |
| AGGAGGIGGNGGAGGAGGVLMG | ASGDADTQY | 0.48363924 |
| ADELIGLL | ASGDADTQY | 0.65578115 |
| ADRALKA | ASGDADTQY | 0.33732152 |
| AGAVGQVGGAGGA | ASGDADTQY | 0.5547924 |
| AGGGGGLGGAP | ASGDADTQY | 0.58958435 |
| AGGLLYGNGGAG | ASGDADTQY | 0.64992535 |
| AGLESIGL | ASGDADTQY | 0.6415177 |
| AGLKTGII | ASGDADTQY | 0.41932505 |
| ASLAALLL | ASGDADTQY | 0.6482121 |
| ASVINISSI | ASGDADTQY | 0.54989856 |
| ATPPGPA | ASGDADTQY | 0.46470922 |
| ATVASALV | ASGDADTQY | 0.50516784 |
| ATVASLIL | ASGDADTQY | 0.48847347 |
| AVNGPVLIRH | ASGDADTQY | 0.36136115 |
| DGAAGAVGI | ASGDADTQY | 0.66421646 |
| DPLLHGI | ASGDADTQY | 0.4413807 |
| ELGAALGAL | ASGDADTQY | 0.6401878 |
| ELGRTQAV | ASGDADTQY | 0.65271425 |
| FVAGGYLS | ASGDADTQY | 0.36577296 |
| GASGGAGGAGGW | ASGDADTQY | 0.6650562 |
| GGAGAGGDGGAGGI | ASGDADTQY | 0.62729824 |
| GTGHNGSQPGMGGNG | ASGDADTQY | 0.64568377 |
| GYAAASASAS | ASGDADTQY | 0.5685406 |
| HQVDVAAVQQG | ASGDADTQY | 0.44792196 |
| IGAAIAHT | ASGDADTQY | 0.6961479 |
| IGMIEVAQL | ASGDADTQY | 0.65955126 |
| IRLAPEHTSAMD | ASGDADTQY | 0.5922749 |
| IYNVSVL | ASGDADTQY | 0.64147544 |
| KQGFLGM | ASGDADTQY | 0.6824337 |

| | | |
|---|---|---|
| LESVITQL | ASGDADTQY | 0.675419 |
| LGLGALVVV | ASGDADTQY | 0.5169797 |
| LIGNGGNGGSGGI | ASGDADTQY | 0.32774252 |
| LNAINAPTL | ASGDADTQY | 0.5839437 |
| LQLLNTGV | ASGDADTQY | 0.59440696 |
| LQTSIEF | ASGDADTQY | 0.41295666 |
| MSFAVSEL | ASGDADTQY | 0.30891514 |
| NTGALGL | ASGDADTQY | 0.6737032 |
| PPGAGGAGGAGTT | ASGDADTQY | 0.7270309 |
| QLVGLGI | ASGDADTQY | 0.52831686 |
| RFLEQVHQL | ASGDADTQY | 0.7254516 |
| RGVGYVLR | ASGDADTQY | 0.3536076 |
| SATGVFAGVI | ASGDADTQY | 0.6811318 |
| SLPAITLP | ASGDADTQY | 0.68380266 |
| STALAFTV | ASGDADTQY | 0.43292853 |
| SVLTVGLGK | ASGDADTQY | 0.63176066 |
| TALAGAGSGI | ASGDADTQY | 0.40590537 |
| TGLPFTGI | ASGDADTQY | 0.49012136 |
| VAEHVTV | ASGDADTQY | 0.69392556 |
| VDIVLNSL | ASGDADTQY | 0.52014136 |
| VLTGGGAVL | ASGDADTQY | 0.61484563 |
| VNVGLAYGI | ASGDADTQY | 0.5081174 |
| VSSAAVGAV | ASGDADTQY | 0.4625684 |
| VTSVYEAGV | ASGDADTQY | 0.6294109 |
| WVALDRGA | ASGDADTQY | 0.6434273 |
| YLVGSVVI | ASGDADTQY | 0.5870962 |
| GPPGTGKTLL | ASGDADTQY | 0.7138385 |
| GVGMTKF | ASGDADTQY | 0.52799374 |
| IGIGGSDL | ASGDADTQY | 0.70988345 |
| IGTLLAEL | ASGDADTQY | 0.7129199 |
| IYTSGTT | ASGDADTQY | 0.6427168 |
| LSFPAGV | ASGDADTQY | 0.51477003 |
| LYTSGTT | ASGDADTQY | 0.6972966 |
| VGAFPFGF | ASGDADTQY | 0.5804297 |
| VGAGTVEF | ASGDADTQY | 0.65382516 |
| VGKSTLLL | ASGDADTQY | 0.67471385 |
| AALAHAGGGIV | ASGDADTQY | 0.70729446 |
| AALVAVSLL | ASGDADTQY | 0.8680105 |
| AAVSRAVAAL | ASGDADTQY | 0.8399972 |
| AGGSSGIGGF | ASGDADTQY | 0.76178765 |
| AGLTVANANG | ASGDADTQY | 0.695315 |
| AGLVGAVL | ASGDADTQY | 0.40652668 |
| ALSQAFTV | ASGDADTQY | 0.77377313 |
| ALTFSVSL | ASGDADTQY | 0.614286 |
| ALTGAALVAL | ASGDADTQY | 0.83129513 |
| ASFSIGAGVL | ASGDADTQY | 0.85077673 |
| DGSVYKI | ASGDADTQY | 0.68476963 |
| EQLAEIV | ASGDADTQY | 0.8900142 |
| FAVSELTVM | ASGDADTQY | 0.8583963 |
| GAAPLAAGL | ASGDADTQY | 0.5271536 |
| GADGGDGGSGGAGG | ASGDADTQY | 0.72097325 |

| | | |
|---|---|---|
| GFGLDTGGAGG | ASGDADTQY | 0.6827157 |
| GLASGGLL | ASGDADTQY | 0.7689355 |
| GRIAGAAVLL | ASGDADTQY | 0.38409388 |
| GSAGSLNTSF | ASGDADTQY | 0.8569615 |
| GTGQIGI | ASGDADTQY | 0.68197984 |
| GTLIALIL | ASGDADTQY | 0.14665642 |
| IAATMGFVVG | ASGDADTQY | 0.80159295 |
| IFAQALL | ASGDADTQY | 0.762079 |
| IGAAVALGF | ASGDADTQY | 0.76198447 |
| IGLTPLAL | ASGDADTQY | 0.86771667 |
| ILGSNTSTL | ASGDADTQY | 0.39456755 |
| IVFSLTVL | ASGDADTQY | 0.64736545 |
| LASKATGF | ASGDADTQY | 0.8530152 |
| LASVALIL | ASGDADTQY | 0.8812132 |
| LFAVGLTL | ASGDADTQY | 0.5674636 |
| LFNGGIGTYI | ASGDADTQY | 0.82433474 |
| LTLGAGII | ASGDADTQY | 0.86697674 |
| LVAAALGDI | ASGDADTQY | 0.8430622 |
| NTAFVQLGI | ASGDADTQY | 0.8503156 |
| QLFSGAEL | ASGDADTQY | 0.8778098 |
| SAATFITL | ASGDADTQY | 0.6399947 |
| SNYGLANLGV | ASGDADTQY | 0.8702834 |
| SYGGAYAVM | ASGDADTQY | 0.8974478 |
| TTGVGGLTLG | ASGDADTQY | 0.7020104 |
| TTQTIYI | ASGDADTQY | 0.7980373 |
| VAAGAGLGAVI | ASGDADTQY | 0.6620913 |
| VAAVVTLL | ASGDADTQY | 0.7487507 |
| VGFLGGLI | ASGDADTQY | 0.2808498 |
| VGVIPILL | ASGDADTQY | 0.82290566 |
| VIFASSLI | ASGDADTQY | 0.8732899 |
| VKGGDGGAAAG | ASGDADTQY | 0.74442714 |
| VTKTFTNI | ASGDADTQY | 0.6064094 |
| VVAAAILL | ASGDADTQY | 0.6379331 |
| YAALDVEL | ASGDADTQY | 0.6986726 |
| LASVALLL | ASGDADTQY | 0.77681005 |
| LDAVHRV | ASGDADTQY | 0.26141542 |
| QGLSSSIF | ASGDADTQY | 0.683663 |
| RGTFANIRL | ASGDADTQY | 0.8773718 |
| RQLWWGHRI | ASGDADTQY | 0.86937547 |
| YGPPGTGKTLL | ASGDADTQY | 0.35607064 |
| AALVGAPANVA | ASGDADTQY | 0.88695276 |
| AATFRRTE | ASGDADTQY | 0.8530487 |
| AGALGGGI | ASGDADTQY | 0.8814392 |
| AGGAGGAGGIGAG | ASGDADTQY | 0.72588056 |
| AGSGALGL | ASGDADTQY | 0.6651057 |
| AQIADQI | ASGDADTQY | 0.8400469 |
| ASAVAVTYL | ASGDADTQY | 0.74212027 |
| ASTVLGGTGG | ASGDADTQY | 0.52981716 |
| AVGGEPAK | ASGDADTQY | 0.86468107 |
| CEQGTFGGTC | ASGDADTQY | 0.76656437 |
| DEGKAPAG | ASGDADTQY | 0.6759964 |

| | | |
|---|---|---|
| DTVVFPAL | ASGDADTQY | 0.6555953 |
| FEGITMAG | ASGDADTQY | 0.8706462 |
| FPLVVQD | ASGDADTQY | 0.560611 |
| GAFTTPQ | ASGDADTQY | 0.65758723 |
| GFGGFGTAGGVGGA | ASGDADTQY | 0.50043374 |
| GFLAQFL | ASGDADTQY | 0.7992666 |
| GGAGGAGGNAGVA | ASGDADTQY | 0.86788565 |
| GGVGTAGAGGAGGAGG | ASGDADTQY | 0.8237027 |
| GNGGGGDAGNAGSGGN | ASGDADTQY | 0.68902296 |
| GTGGAGGAGNDAGS | ASGDADTQY | 0.83588934 |
| GTTNTGWFNS | ASGDADTQY | 0.8003582 |
| IFMGPLFTL | ASGDADTQY | 0.79214424 |
| ILAHAIEAI | ASGDADTQY | 0.8593427 |
| IRSAIREAL | ASGDADTQY | 0.62905645 |
| ITFAGGG | ASGDADTQY | 0.8637314 |
| ITGMSGHQEV | ASGDADTQY | 0.81725776 |
| IVAGLACTL | ASGDADTQY | 0.84615266 |
| IVGGGLAGA | ASGDADTQY | 0.8516337 |
| IVTGVLNI | ASGDADTQY | 0.8063309 |
| LAQQGCI | ASGDADTQY | 0.7727675 |
| LCLHVEN | ASGDADTQY | 0.28848344 |
| LDAISDGS | ASGDADTQY | 0.6631267 |
| LGGFAAML | ASGDADTQY | 0.8494179 |
| LGGKNPMI | ASGDADTQY | 0.7990386 |
| LGPKGVAAIL | ASGDADTQY | 0.86121583 |
| LMVGVSI | ASGDADTQY | 0.80089414 |
| LVGAAFVL | ASGDADTQY | 0.69904315 |
| LVLAVAPL | ASGDADTQY | 0.8607315 |
| MAAIAVIDAA | ASGDADTQY | 0.85239863 |
| NNGINAELQ | ASGDADTQY | 0.429354 |
| RGEPLVLVHG | ASGDADTQY | 0.7303466 |
| RGGAGGAAGLI | ASGDADTQY | 0.8928168 |
| RSYRMHPAV | ASGDADTQY | 0.8147517 |
| SAAHLLAGQLA | ASGDADTQY | 0.77208745 |
| SAAVGSY | ASGDADTQY | 0.67930776 |
| SPHAVLHRI | ASGDADTQY | 0.8893337 |
| SQIAKVL | ASGDADTQY | 0.58693826 |
| TALGHLVP | ASGDADTQY | 0.88932407 |
| TAPTTSGAA | ASGDADTQY | 0.18512848 |
| TDGDDGNGGSPGT | ASGDADTQY | 0.63806677 |
| TDGLGEN | ASGDADTQY | 0.56257355 |
| TGGPPLPGQA | ASGDADTQY | 0.5070665 |
| TIPAITI | ASGDADTQY | 0.8678634 |
| TIPGFTI | ASGDADTQY | 0.6877805 |
| TSIASGALA | ASGDADTQY | 0.85347414 |
| TVTLPGAAI | ASGDADTQY | 0.7906501 |
| VASGLSAALP | ASGDADTQY | 0.85550165 |
| VGAGVIGI | ASGDADTQY | 0.8016869 |
| VGPKRFGV | ASGDADTQY | 0.80744374 |
| VNVAFPD | ASGDADTQY | 0.60140985 |
| VSMTNSGVS | ASGDADTQY | 0.8630028 |

| | | |
|---|---|---|
| VTDVLGDLSV | ASGDADTQY | 0.87611663 |
| VVVVGAGFAGL | ASGDADTQY | 0.80815744 |
| VYTVGTVV | ASGDADTQY | 0.5054439 |
| WTTFSGP | ASGDADTQY | 0.62581825 |
| YDELVEAF | ASGDADTQY | 0.81471616 |
| YGTGGAGGNGG | ASGDADTQY | 0.28829613 |
| YTNVMAV | ASGDADTQY | 0.21587095 |
| FLEQVHQL | ASGDADTQY | 0.698601 |
| GLSLLAVAV | ASGDADTQY | 0.6002114 |
| LEQVHQL | ASGDADTQY | 0.88035774 |
| LNAAGAI | ASGDADTQY | 0.85355633 |
| PAVHASL | ASGDADTQY | 0.7925352 |
| TELEDDA | ASGDADTQY | 0.85239977 |
| TPPGPPN | ASGDADTQY | 0.8044133 |
| TPPPPPPP | ASGDADTQY | 0.10867839 |
| TQSPALI | ASRRTGGDSAETLY | 0.64443815 |
| EFAAAFPQ | ASRRTGGDSAETLY | 0.54866624 |
| LLSGNLDVL | ASRRTGGDSAETLY | 0.8656305 |
| GPANLGNYNI | ASRRTGGDSAETLY | 0.6808212 |
| TLGPTLP | ASRRTGGDSAETLY | 0.6205268 |
| VYAYSSCCK | ASRRTGGDSAETLY | 0.8020755 |
| APAPAPAPNI | ASRRTGGDSAETLY | 0.78332335 |
| PGGPVPIP | ASRRTGGDSAETLY | 0.75073195 |
| FLTVALEP | ASRRTGGDSAETLY | 0.5070643 |
| SQYFGANL | ASRRTGGDSAETLY | 0.15146428 |
| GGVKPAA | ASRRTGGDSAETLY | 0.8689492 |
| GGTGGAGGAGGW | ASRRTGGDSAETLY | 0.34299287 |
| GVGGVAGLGGAGG | ASRRTGGDSAETLY | 0.6241152 |
| VLLVLATTI | ASRRTGGDSAETLY | 0.24145992 |
| IGIFAAML | ASRRTGGDSAETLY | 0.4672122 |
| AVGDLID | ASRRTGGDSAETLY | 0.3904034 |
| NISPLPL | ASRRTGGDSAETLY | 0.8469219 |
| VTAVFVAI | ASRRTGGDSAETLY | 0.7369262 |
| SGDDNDAAL | ASRRTGGDSAETLY | 0.70603037 |
| AVAALAHS | ASRRTGGDSAETLY | 0.88964325 |
| GGVTVMATP | ASRRTGGDSAETLY | 0.7747547 |
| AGLAANVRD | ASRRTGGDSAETLY | 0.73342144 |
| TNRPDLIDPALLRPG | ASRRTGGDSAETLY | 0.64533263 |
| LANLGVTAL | ASRRTGGDSAETLY | 0.65880716 |
| TGERIEL | ASRRTGGDSAETLY | 0.64823455 |
| NGEELAA | ASRRTGGDSAETLY | 0.62901103 |
| VLSLVWG | ASRRTGGDSAETLY | 0.8888149 |
| IGYALWL | ASRRTGGDSAETLY | 0.65374804 |
| LSLLLAPL | ASRRTGGDSAETLY | 0.49143744 |
| TLDEALK | ASRRTGGDSAETLY | 0.64861846 |
| TDGAGTAEGG | ASRRTGGDSAETLY | 0.88034153 |
| NAPGGAGGNGG | ASRRTGGDSAETLY | 0.90121365 |
| VSVFMGMGGNF | ASRRTGGDSAETLY | 0.71372116 |
| LASYLAAL | ASRRTGGDSAETLY | 0.6576068 |
| LVGGTPAL | ASRRTGGDSAETLY | 0.68140084 |
| VRNADVL | ASRRTGGDSAETLY | 0.78668165 |

| | | |
|---|---|---|
| TCSSPFLVSAM | ASRRTGGDSAETLY | 0.6987998 |
| VHITDAE | ASRRTGGDSAETLY | 0.48193997 |
| EYAAPSF | ASRRTGGDSAETLY | 0.8405893 |
| VVDAFHGGL | ASRRTGGDSAETLY | 0.8974161 |
| ASVVDPQ | ASRRTGGDSAETLY | 0.69947493 |
| SYNEFGGLS | ASRRTGGDSAETLY | 0.503361 |
| DPDGSVY | ASRRTGGDSAETLY | 0.78018963 |
| MVSTIHATE | ASRRTGGDSAETLY | 0.6460612 |
| LAAGVADPV | ASRRTGGDSAETLY | 0.75384176 |
| TKEQIAA | ASRRTGGDSAETLY | 0.817966 |
| SLASPAAL | ASRRTGGDSAETLY | 0.41952252 |
| IGLVFYGV | ASRRTGGDSAETLY | 0.68490154 |
| FLLDASNL | ASRRTGGDSAETLY | 0.898789 |
| YQVGNFP | ASRRTGGDSAETLY | 0.7261231 |
| ESLLSQST | ASRRTGGDSAETLY | 0.811591 |
| LFTVGFAAVG | ASRRTGGDSAETLY | 0.87393606 |
| ATPPTPT | ASRRTGGDSAETLY | 0.84745467 |
| TDLATLG | ASRRTGGDSAETLY | 0.8806992 |
| SAGGTGR | ASRRTGGDSAETLY | 0.85109425 |
| GFSNSGS | ASRRTGGDSAETLY | 0.7289692 |
| GGNGGSGGTSV | ASRRTGGDSAETLY | 0.782307 |
| GTSVATGGAGN | ASRRTGGDSAETLY | 0.80535936 |
| NGGNGGSGGTS | ASRRTGGDSAETLY | 0.80570793 |
| AVVKFPH | ASRRTGGDSAETLY | 0.764899 |
| TPPPPPP | ASRRTGGDSAETLY | 0.5569667 |
| AAPAAAAAPA | ASRRTGGDSAETLY | 0.8724319 |
| IALNEDL | ASRRTGGDSAETLY | 0.7355896 |
| IVNVSSM | ASRRTGGDSAETLY | 0.8664168 |
| TGIPPAPR | ASRRTGGDSAETLY | 0.5797446 |
| FDLGGGTF | ASRRTGGDSAETLY | 0.5966417 |
| FDLGGGTFDVS | ASRRTGGDSAETLY | 0.85055393 |
| NEPTAAA | ASRRTGGDSAETLY | 0.13543907 |
| NEPTAAALA | ASRRTGGDSAETLY | 0.19468136 |
| GSIINISSI | ASRRTGGDSAETLY | 0.7907388 |
| VAPGNAGTA | ASRRTGGDSAETLY | 0.8853533 |
| TGGGASLE | ASRRTGGDSAETLY | 0.16769804 |
| VAVYNHYKRI | ASRRTGGDSAETLY | 0.594733 |
| TDPAPATT | ASRRTGGDSAETLY | 0.4762797 |
| TNRPDILDPALLRPG | ASRRTGGDSAETLY | 0.7494591 |
| LTTTTTS | ASRRTGGDSAETLY | 0.808164 |
| PTGSGKTL | ASRRTGGDSAETLY | 0.4205338 |
| TGSGKTL | ASRRTGGDSAETLY | 0.838055 |
| TIGHVDHGK | ASRRTGGDSAETLY | 0.39298207 |
| TIGHVDHGKTT | ASRRTGGDSAETLY | 0.4691317 |
| GHVDHGKTT | ASRRTGGDSAETLY | 0.85706866 |
| AQFEHTLL | ASRRTGGDSAETLY | 0.8733939 |
| IVNQDEL | ASRRTGGDSAETLY | 0.6329694 |
| TNGSQFF | ASRRTGGDSAETLY | 0.86362517 |
| TGKGIVE | ASRRTGGDSAETLY | 0.5903604 |
| LISNASDA | ASRRTGGDSAETLY | 0.9020665 |
| AELRTLGELTKIAKAHGAQ | ASRRTGGDSAETLY | 0.73316956 |

| | | |
|---|---|---|
| APNAVAAAEPSW | ASRRTGGDSAETLY | 0.41941574 |
| APQVAGAATAVL | ASRRTGGDSAETLY | 0.85674196 |
| ARRDPAAAAAILPTDA | ASRRTGGDSAETLY | 0.8341597 |
| DAAARDVLAEAGLAEYF | ASRRTGGDSAETLY | 0.8201517 |
| DDQIVLSAKANLAKVEQ | ASRRTGGDSAETLY | 0.8571332 |
| DELLPDTGEDSDEDGH | ASRRTGGDSAETLY | 0.5787828 |
| ELPEAPANRATVPQD | ASRRTGGDSAETLY | 0.7505084 |
| FGQSKIERVTGIG | ASRRTGGDSAETLY | 0.87411845 |
| GAGGTGSTAGGAGGAGGAGGLY | ASRRTGGDSAETLY | 0.8646661 |
| GHGGTGAAGGNGGTGGAGGSGI | ASRRTGGDSAETLY | 0.53300345 |
| GNGGNGGAGGEGGPGGAGGAGA | ASRRTGGDSAETLY | 0.8623191 |
| GSTAGNGGNGGNATNGTIAGQP | ASRRTGGDSAETLY | 0.7669684 |
| GTGGDGGNGGNGGTGG | ASRRTGGDSAETLY | 0.7077518 |
| GVRRPAGIPADAPV | ASRRTGGDSAETLY | 0.89118934 |
| GYRQVTDAHLLTLARR | ASRRTGGDSAETLY | 0.7518902 |
| IAKMGQDGHDRG | ASRRTGGDSAETLY | 0.86513954 |
| KAEPSIEIQYT | ASRRTGGDSAETLY | 0.86031437 |
| KPVRIRMAADEDTVAVT | ASRRTGGDSAETLY | 0.8750883 |
| MPPRIAGMRLLVIKPEP | ASRRTGGDSAETLY | 0.5244006 |
| MKFYAVALFRSRGVPPTR | ASRRTGGDSAETLY | 0.8454395 |
| NAGGFGILGAVAHSP | ASRRTGGDSAETLY | 0.7897094 |
| NEGREITLAMAEM | ASRRTGGDSAETLY | 0.82937014 |
| NELVEWSHPLGHRWRWE | ASRRTGGDSAETLY | 0.7959441 |
| NLGAGGTGGNGGNPG | ASRRTGGDSAETLY | 0.5394178 |
| NTEWPKWETVMAADPVD | ASRRTGGDSAETLY | 0.81822944 |
| PASAAPAETQP | ASRRTGGDSAETLY | 0.8789276 |
| QQSSDGKNGVRAAEITGE | ASRRTGGDSAETLY | 0.79988253 |
| RAPDAVTTSWWKE | ASRRTGGDSAETLY | 0.80714834 |
| SGFGNVGAHTSGWFNQ | ASRRTGGDSAETLY | 0.8883141 |
| SGVAGAVGQAASVGGLKV | ASRRTGGDSAETLY | 0.72727406 |
| SNGGAGTGSAGGNGGT | ASRRTGGDSAETLY | 0.7470037 |
| TGGAGGEGGAAG | ASRRTGGDSAETLY | 0.85061526 |
| TGGDGGAAGAGGAPGHG | ASRRTGGDSAETLY | 0.86503583 |
| TMPPLLAAIANAGRHGV | ASRRTGGDSAETLY | 0.7035989 |
| TSSTAGNSGGA | ASRRTGGDSAETLY | 0.8817295 |
| VPATVPVGILC | ASRRTGGDSAETLY | 0.63269293 |
| VQRPDSGGVVLTGRIS | ASRRTGGDSAETLY | 0.8651406 |
| WTPNPVPATKGVPAGLVQAT | ASRRTGGDSAETLY | 0.6971972 |
| YALKRALAKFSDRFDVVII | ASRRTGGDSAETLY | 0.8326198 |
| YLIGSVIV | ASRRTGGDSAETLY | 0.62858605 |
| YGQLVAVID | ASRRTGGDSAETLY | 0.8658059 |
| VTVGGMTLPA | ASRRTGGDSAETLY | 0.865574 |
| VLALPAV | ASRRTGGDSAETLY | 0.8745555 |
| VIPPTLNYET | ASRRTGGDSAETLY | 0.18957348 |
| VGMGGMHPSSG | ASRRTGGDSAETLY | 0.86409247 |
| VGLAGGAATA | ASRRTGGDSAETLY | 0.85744697 |
| VGGAGGAGGSGP | ASRRTGGDSAETLY | 0.899837 |
| VFAAFMV | ASRRTGGDSAETLY | 0.7722026 |
| TLYDKLR | ASRRTGGDSAETLY | 0.8075372 |
| TLAVFTVL | ASRRTGGDSAETLY | 0.7699882 |
| TGSVQPARV | ASRRTGGDSAETLY | 0.72465336 |

| | | |
|---|---|---|
| TGGGAGSGGT | ASRRTGGDSAETLY | 0.58779573 |
| SHLETSL | ASRRTGGDSAETLY | 0.77345014 |
| SFTTGMT | ASRRTGGDSAETLY | 0.73270833 |
| QVEVAKRMGVRQPTVS | ASRRTGGDSAETLY | 0.6885417 |
| QGSGGPALAM | ASRRTGGDSAETLY | 0.85545164 |
| MLAPSNP | ASRRTGGDSAETLY | 0.8761688 |
| LVNNAAI | ASRRTGGDSAETLY | 0.8720697 |
| LTGEINTLA | ASRRTGGDSAETLY | 0.8629501 |
| LSIFGPV | ASRRTGGDSAETLY | 0.88284314 |
| LLALLANL | ASRRTGGDSAETLY | 0.8591223 |
| LISNASDALDKLR | ASRRTGGDSAETLY | 0.6934455 |
| LINEAVP | ASRRTGGDSAETLY | 0.74679625 |
| LGYAFSGL | ASRRTGGDSAETLY | 0.59738 |
| LDFVAVAL | ASRRTGGDSAETLY | 0.7019347 |
| LAQPGAE | ASRRTGGDSAETLY | 0.6888385 |
| ISNLGAVTGF | ASRRTGGDSAETLY | 0.81432307 |
| HQEDPGRI | ASRRTGGDSAETLY | 0.12843437 |
| GTPGEPTP | ASRRTGGDSAETLY | 0.7734786 |
| GFFNSGS | ASRRTGGDSAETLY | 0.8881917 |
| GAGGGGLTAGAN | ASRRTGGDSAETLY | 0.62608397 |
| FVDALVNM | ASRRTGGDSAETLY | 0.82437944 |
| FLTVAPEL | ASRRTGGDSAETLY | 0.76606715 |
| FETLTAL | ASRRTGGDSAETLY | 0.6492356 |
| DVINPIEL | ASRRTGGDSAETLY | 0.8499713 |
| APVSGAPV | ASRRTGGDSAETLY | 0.577859 |
| ALVGGLAW | ASRRTGGDSAETLY | 0.8179332 |
| ALAAAAATAI | ASRRTGGDSAETLY | 0.89108485 |
| AFGLGLLL | ASRRTGGDSAETLY | 0.79302436 |
| AFFAAASL | ASRRTGGDSAETLY | 0.80836904 |
| AAAALPAFSPP | ASRRTGGDSAETLY | 0.7433917 |
| VVLAPAL | ASRRTGGDSAETLY | 0.43398368 |
| TIGHVDHGKTTL | ASRRTGGDSAETLY | 0.8268631 |
| LISNASDALDK | ASRRTGGDSAETLY | 0.2800881 |
| LISNASDAL | ASRRTGGDSAETLY | 0.7523899 |
| ISNASDALDK | ASRRTGGDSAETLY | 0.60811245 |
| NQAETLVYQTE | ASRRTGGDSAETLY | 0.8215425 |
| TGGTGGTAGAG | ASRRTGGDSAETLY | 0.7980893 |
| TGSTAGGAGGA | ASRRTGGDSAETLY | 0.8325491 |
| DPLAVPVAAGPV | ASRRTGGDSAETLY | 0.6373131 |
| GGTGGTPGNGGN | ASRRTGGDSAETLY | 0.6993699 |
| PAGLAGVDPATL | ASRRTGGDSAETLY | 0.59806776 |
| GNNLVVEAPHAGASV | ASRRTGGDSAETLY | 0.5156573 |
| KVTRIMSASTLEQPA | ASRRTGGDSAETLY | 0.8656762 |
| LGFALPIALPRRTVV | ASRRTGGDSAETLY | 0.74825144 |
| TRLGNLTARKLVVLV | ASRRTGGDSAETLY | 0.72432315 |
| WSQQRRLIYFPSAGP | ASRRTGGDSAETLY | 0.88037145 |
| GPNGNPDPMAAAVDIR | ASRRTGGDSAETLY | 0.81998086 |
| TQRCVSVADLRAAASR | ASRRTGGDSAETLY | 0.43759495 |
| QFVLDVAEHPVEVLTETV | ASRRTGGDSAETLY | 0.78903985 |
| GGAGGTGGAGGNGGTQAGDG | ASRRTGGDSAETLY | 0.7769825 |
| PLLATLRAIAVDVDAKPAQV | ASRRTGGDSAETLY | 0.76890707 |

| | | |
|---|---|---|
| APKFRTGGYMIDFWGVGYQVAK | ASRRTGGDSAETLY | 0.16142498 |
| GAGGAGGAGGAAGGPGGTGGTG | ASRRTGGDSAETLY | 0.81304383 |
| IGALGLLATDVAPAGSVAFGVA | ASRRTGGDSAETLY | 0.7153359 |
| AGGAGGIGGNGGAGGAGGVLMG | ASRRTGGDSAETLY | 0.58792365 |
| ADELIGLL | ASRRTGGDSAETLY | 0.66017985 |
| ADRALKA | ASRRTGGDSAETLY | 0.85429835 |
| AGAVGQVGGAGGA | ASRRTGGDSAETLY | 0.7081975 |
| AGGGGGLGGAP | ASRRTGGDSAETLY | 0.81447804 |
| AGGLLYGNGGAG | ASRRTGGDSAETLY | 0.85621864 |
| AGLESIGL | ASRRTGGDSAETLY | 0.8655059 |
| AGLKTGII | ASRRTGGDSAETLY | 0.88797283 |
| ASLAALLL | ASRRTGGDSAETLY | 0.8205825 |
| ASVINISSI | ASRRTGGDSAETLY | 0.71276176 |
| ATPPGPA | ASRRTGGDSAETLY | 0.81009907 |
| ATVASALV | ASRRTGGDSAETLY | 0.73452985 |
| ATVASLIL | ASRRTGGDSAETLY | 0.29592335 |
| AVNGPVLIRH | ASRRTGGDSAETLY | 0.22526973 |
| DGAAGAVGI | ASRRTGGDSAETLY | 0.83024156 |
| DPLLHGI | ASRRTGGDSAETLY | 0.26212093 |
| ELGAALGAL | ASRRTGGDSAETLY | 0.78282183 |
| ELGRTQAV | ASRRTGGDSAETLY | 0.85134363 |
| FVAGGYLS | ASRRTGGDSAETLY | 0.75413215 |
| GASGGAGGAGGW | ASRRTGGDSAETLY | 0.5841502 |
| GGAGAGGDGGAGGI | ASRRTGGDSAETLY | 0.72019017 |
| GTGHNGSQPGMGGNG | ASRRTGGDSAETLY | 0.72131896 |
| GYAAASASAS | ASRRTGGDSAETLY | 0.8622524 |
| HQVDVAAVQQG | ASRRTGGDSAETLY | 0.8618694 |
| IGAAIAHT | ASRRTGGDSAETLY | 0.89446187 |
| IGMIEVAQL | ASRRTGGDSAETLY | 0.8640962 |
| IRLAPEHTSAMD | ASRRTGGDSAETLY | 0.73011637 |
| IYNVSVL | ASRRTGGDSAETLY | 0.8453972 |
| KQGFLGM | ASRRTGGDSAETLY | 0.6933141 |
| LESVITQL | ASRRTGGDSAETLY | 0.7694783 |
| LGLGALVVV | ASRRTGGDSAETLY | 0.76354647 |
| LIGNGGNGGSGGI | ASRRTGGDSAETLY | 0.8068691 |
| LNAINAPTL | ASRRTGGDSAETLY | 0.7664584 |
| LQLLNTGV | ASRRTGGDSAETLY | 0.7329097 |
| LQTSIEF | ASRRTGGDSAETLY | 0.39505395 |
| MSFAVSEL | ASRRTGGDSAETLY | 0.84920776 |
| NTGALGL | ASRRTGGDSAETLY | 0.84635025 |
| PPGAGGAGGAGTT | ASRRTGGDSAETLY | 0.31432915 |
| QLVGLGI | ASRRTGGDSAETLY | 0.66298974 |
| RFLEQVHQL | ASRRTGGDSAETLY | 0.8967155 |
| RGVGYVLR | ASRRTGGDSAETLY | 0.8796925 |
| SATGVFAGVI | ASRRTGGDSAETLY | 0.6040759 |
| SLPAITLP | ASRRTGGDSAETLY | 0.6115025 |
| STALAFTV | ASRRTGGDSAETLY | 0.7584837 |
| SVLTVGLGK | ASRRTGGDSAETLY | 0.30126983 |
| TALAGAGSGI | ASRRTGGDSAETLY | 0.7994518 |
| TGLPFTGI | ASRRTGGDSAETLY | 0.7555152 |
| VAEHVTV | ASRRTGGDSAETLY | 0.7638713 |

| | | |
|---|---|---|
| VDIVLNSL | ASRRTGGDSAETLY | 0.57069385 |
| VLTGGGAVL | ASRRTGGDSAETLY | 0.7679217 |
| VNVGLAYGI | ASRRTGGDSAETLY | 0.70305824 |
| VSSAAVGAV | ASRRTGGDSAETLY | 0.83511776 |
| VTSVYEAGV | ASRRTGGDSAETLY | 0.17443196 |
| WVALDRGA | ASRRTGGDSAETLY | 0.8124529 |
| YLVGSVVI | ASRRTGGDSAETLY | 0.6217389 |
| GPPGTGKTLL | ASRRTGGDSAETLY | 0.630991 |
| GVGMTKF | ASRRTGGDSAETLY | 0.43641478 |
| IGIGGSDL | ASRRTGGDSAETLY | 0.7667626 |
| IGTLLAEL | ASRRTGGDSAETLY | 0.87113523 |
| IYTSGTT | ASRRTGGDSAETLY | 0.8539103 |
| LSFPAGV | ASRRTGGDSAETLY | 0.42522845 |
| LYTSGTT | ASRRTGGDSAETLY | 0.8686784 |
| VGAFPFGF | ASRRTGGDSAETLY | 0.9002136 |
| VGAGTVEF | ASRRTGGDSAETLY | 0.68950236 |
| VGKSTLLL | ASRRTGGDSAETLY | 0.4426508 |
| AALAHAGGGIV | ASRRTGGDSAETLY | 0.832777 |
| AALVAVSLL | ASRRTGGDSAETLY | 0.75136924 |
| AAVSRAVAAL | ASRRTGGDSAETLY | 0.8189179 |
| AGGSSGIGGF | ASRRTGGDSAETLY | 0.8431767 |
| AGLTVANANG | ASRRTGGDSAETLY | 0.5763343 |
| AGLVGAVL | ASRRTGGDSAETLY | 0.8358563 |
| ALSQAFTV | ASRRTGGDSAETLY | 0.8665023 |
| ALTFSVSL | ASRRTGGDSAETLY | 0.6744493 |
| ALTGAALVAL | ASRRTGGDSAETLY | 0.67180145 |
| ASFSIGAGVL | ASRRTGGDSAETLY | 0.5911661 |
| DGSVYKI | ASRRTGGDSAETLY | 0.295366 |
| EQLAEIV | ASRRTGGDSAETLY | 0.85849 |
| FAVSELTVM | ASRRTGGDSAETLY | 0.6601763 |
| GAAPLAAGL | ASRRTGGDSAETLY | 0.6915215 |
| GADGGDGGSGGAGG | ASRRTGGDSAETLY | 0.8087932 |
| GFGLDTGGAGG | ASRRTGGDSAETLY | 0.66652805 |
| GLASGGLL | ASRRTGGDSAETLY | 0.7151839 |
| GRIAGAAVLL | ASRRTGGDSAETLY | 0.72538865 |
| GSAGSLNTSF | ASRRTGGDSAETLY | 0.78565735 |
| GTGQIGI | ASRRTGGDSAETLY | 0.7971772 |
| GTLIALIL | ASRRTGGDSAETLY | 0.7837237 |
| IAATMGFVVG | ASRRTGGDSAETLY | 0.7696965 |
| IFAQALL | ASRRTGGDSAETLY | 0.88301414 |
| IGAAVALGF | ASRRTGGDSAETLY | 0.48571354 |
| IGLTPLAL | ASRRTGGDSAETLY | 0.60088265 |
| ILGSNTSTL | ASRRTGGDSAETLY | 0.81212175 |
| IVFSLTVL | ASRRTGGDSAETLY | 0.37069714 |
| LASKATGF | ASRRTGGDSAETLY | 0.8362043 |
| LASVALIL | ASRRTGGDSAETLY | 0.7911967 |
| LFAVGLTL | ASRRTGGDSAETLY | 0.818323 |
| LFNGGIGTYI | ASRRTGGDSAETLY | 0.7524483 |
| LTLGAGII | ASRRTGGDSAETLY | 0.896332 |
| LVAAALGDI | ASRRTGGDSAETLY | 0.8685554 |
| NTAFVQLGI | ASRRTGGDSAETLY | 0.7378834 |

| | | |
|---|---|---|
| QLFSGAEL | ASRRTGGDSAETLY | 0.85104 |
| SAATFITL | ASRRTGGDSAETLY | 0.5158503 |
| SNYGLANLGV | ASRRTGGDSAETLY | 0.86423755 |
| SYGGAYAVM | ASRRTGGDSAETLY | 0.82221055 |
| TTGVGGLTLG | ASRRTGGDSAETLY | 0.5691581 |
| TTQTIYI | ASRRTGGDSAETLY | 0.7546345 |
| VAAGAGLGAVI | ASRRTGGDSAETLY | 0.8809334 |
| VAAVVTLL | ASRRTGGDSAETLY | 0.8737192 |
| VGFLGGLI | ASRRTGGDSAETLY | 0.30142775 |
| VGVIPILL | ASRRTGGDSAETLY | 0.7463299 |
| VIFASSLI | ASRRTGGDSAETLY | 0.30650592 |
| VKGGDGGAAAG | ASRRTGGDSAETLY | 0.6227875 |
| VTKTFTNI | ASRRTGGDSAETLY | 0.57899207 |
| VVAAAILL | ASRRTGGDSAETLY | 0.83968014 |
| YAALDVEL | ASRRTGGDSAETLY | 0.74532986 |
| LASVALLL | ASRRTGGDSAETLY | 0.8332579 |
| LDAVHRV | ASRRTGGDSAETLY | 0.7979543 |
| QGLSSSIF | ASRRTGGDSAETLY | 0.8847048 |
| RGTFANIRL | ASRRTGGDSAETLY | 0.68138564 |
| RQLWWGHRI | ASRRTGGDSAETLY | 0.7745226 |
| YGPPGTGKTLL | ASRRTGGDSAETLY | 0.4753446 |
| AALVGAPANVA | ASRRTGGDSAETLY | 0.4613949 |
| AATFRRTE | ASRRTGGDSAETLY | 0.66362613 |
| AGALGGGI | ASRRTGGDSAETLY | 0.72079885 |
| AGGAGGAGGIGAG | ASRRTGGDSAETLY | 0.22982968 |
| AGSGALGL | ASRRTGGDSAETLY | 0.86821187 |
| AQIADQI | ASRRTGGDSAETLY | 0.81691515 |
| ASAVAVTYL | ASRRTGGDSAETLY | 0.86302733 |
| ASTVLGGTGG | ASRRTGGDSAETLY | 0.73301053 |
| AVGGEPAK | ASRRTGGDSAETLY | 0.75838655 |
| CEQGTFGGTC | ASRRTGGDSAETLY | 0.6829413 |
| DEGKAPAG | ASRRTGGDSAETLY | 0.8360442 |
| DTVVFPAL | ASRRTGGDSAETLY | 0.67308843 |
| FEGITMAG | ASRRTGGDSAETLY | 0.42292878 |
| FPLVVQD | ASRRTGGDSAETLY | 0.68337655 |
| GAFTTPQ | ASRRTGGDSAETLY | 0.5239562 |
| GFGGFGTAGGVGGA | ASRRTGGDSAETLY | 0.8876382 |
| GFLAQFL | ASRRTGGDSAETLY | 0.86580074 |
| GGAGGAGGNAGVA | ASRRTGGDSAETLY | 0.82011616 |
| GGVGTAGAGGAGGAGG | ASRRTGGDSAETLY | 0.8649315 |
| GNGGGGDAGNAGSGGN | ASRRTGGDSAETLY | 0.26869643 |
| GTGGAGGAGNDAGS | ASRRTGGDSAETLY | 0.7253871 |
| GTTNTGWFNS | ASRRTGGDSAETLY | 0.8847305 |
| IFMGPLFTL | ASRRTGGDSAETLY | 0.82511884 |
| ILAHAIEAI | ASRRTGGDSAETLY | 0.7647371 |
| IRSAIREAL | ASRRTGGDSAETLY | 0.8829833 |
| ITFAGGG | ASRRTGGDSAETLY | 0.70312154 |
| ITGMSGHQEV | ASRRTGGDSAETLY | 0.76546466 |
| IVAGLACTL | ASRRTGGDSAETLY | 0.78640544 |
| IVGGGLAGA | ASRRTGGDSAETLY | 0.82184017 |
| IVTGVLNI | ASRRTGGDSAETLY | 0.8279296 |

| | | |
|---|---|---|
| LAQQGCI | ASRRTGGDSAETLY | 0.6526618 |
| LCLHVEN | ASRRTGGDSAETLY | 0.7947526 |
| LDAISDGS | ASRRTGGDSAETLY | 0.9023349 |
| LGGFAAML | ASRRTGGDSAETLY | 0.7618131 |
| LGGKNPMI | ASRRTGGDSAETLY | 0.8745 |
| LGPKGVAAIL | ASRRTGGDSAETLY | 0.8459152 |
| LMVGVSI | ASRRTGGDSAETLY | 0.8483402 |
| LVGAAFVL | ASRRTGGDSAETLY | 0.65628946 |
| LVLAVAPL | ASRRTGGDSAETLY | 0.73992705 |
| MAAIAVIDAA | ASRRTGGDSAETLY | 0.704285 |
| NNGINAELQ | ASRRTGGDSAETLY | 0.66084576 |
| RGEPLVLVHG | ASRRTGGDSAETLY | 0.72342956 |
| RGGAGGAAGLI | ASRRTGGDSAETLY | 0.6552004 |
| RSYRMHPAV | ASRRTGGDSAETLY | 0.90805393 |
| SAAHLLAGQLA | ASRRTGGDSAETLY | 0.84887975 |
| SAAVGSY | ASRRTGGDSAETLY | 0.8220371 |
| SPHAVLHRI | ASRRTGGDSAETLY | 0.64714396 |
| SQIAKVL | ASRRTGGDSAETLY | 0.81618476 |
| TALGHLVP | ASRRTGGDSAETLY | 0.82717663 |
| TAPTTSGAA | ASRRTGGDSAETLY | 0.4606167 |
| TDGDDGNGGSPGT | ASRRTGGDSAETLY | 0.88375103 |
| TDGLGEN | ASRRTGGDSAETLY | 0.6894628 |
| TGGPPLPGQA | ASRRTGGDSAETLY | 0.66392785 |
| TIPAITI | ASRRTGGDSAETLY | 0.81288695 |
| TIPGFTI | ASRRTGGDSAETLY | 0.8857579 |
| TSIASGALA | ASRRTGGDSAETLY | 0.7139405 |
| TVTLPGAAI | ASRRTGGDSAETLY | 0.6522782 |
| VASGLSAALP | ASRRTGGDSAETLY | 0.66102743 |
| VGAGVIGI | ASRRTGGDSAETLY | 0.85520864 |
| VGPKRFGV | ASRRTGGDSAETLY | 0.874575 |
| VNVAFPD | ASRRTGGDSAETLY | 0.79575634 |
| VSMTNSGVS | ASRRTGGDSAETLY | 0.89223945 |
| VTDVLGDLSV | ASRRTGGDSAETLY | 0.7282279 |
| VVVVGAGFAGL | ASRRTGGDSAETLY | 0.43893188 |
| VYTVGTVV | ASRRTGGDSAETLY | 0.795352 |
| WTTFSGP | ASRRTGGDSAETLY | 0.86175036 |
| YDELVEAF | ASRRTGGDSAETLY | 0.8643087 |
| YGTGGAGGNGG | ASRRTGGDSAETLY | 0.9025316 |
| YTNVMAV | ASRRTGGDSAETLY | 0.857486 |
| FLEQVHQL | ASRRTGGDSAETLY | 0.7080542 |
| GLSLLAVAV | ASRRTGGDSAETLY | 0.8744911 |
| LEQVHQL | ASRRTGGDSAETLY | 0.889572 |
| LNAAGAI | ASRRTGGDSAETLY | 0.8566129 |
| PAVHASL | ASRRTGGDSAETLY | 0.7112494 |
| TELEDDA | ASRRTGGDSAETLY | 0.56884015 |
| TPPGPPN | ASRRTGGDSAETLY | 0.7831948 |
| TPPPPPPP | ASRRTGGDSAETLY | 0.8673271 |
| TQSPALI | ASSSTGDAETLY | 0.5510276 |
| EFAAAFPQ | ASSSTGDAETLY | 0.7868607 |
| LLSGNLDVL | ASSSTGDAETLY | 0.8270274 |
| GPANLGNYNI | ASSSTGDAETLY | 0.54717183 |

| | | |
|---|---|---|
| TLGPTLP | ASSSTGDAETLY | 0.82811105 |
| VYAYSSCCK | ASSSTGDAETLY | 0.88089824 |
| APAPAPAPNI | ASSSTGDAETLY | 0.7065096 |
| PGGPVPIP | ASSSTGDAETLY | 0.6859865 |
| FLTVALEP | ASSSTGDAETLY | 0.87203884 |
| SQYFGANL | ASSSTGDAETLY | 0.90846205 |
| GGVKPAA | ASSSTGDAETLY | 0.86669815 |
| GGTGGAGGAGGW | ASSSTGDAETLY | 0.47166288 |
| GVGGVAGLGGAGG | ASSSTGDAETLY | 0.26104107 |
| VLLVLATTI | ASSSTGDAETLY | 0.6573515 |
| IGIFAAML | ASSSTGDAETLY | 0.853726 |
| AVGDLID | ASSSTGDAETLY | 0.72182405 |
| NISPLPL | ASSSTGDAETLY | 0.70902276 |
| VTAVFVAI | ASSSTGDAETLY | 0.84200203 |
| SGDDNDAAL | ASSSTGDAETLY | 0.8641621 |
| AVAALAHS | ASSSTGDAETLY | 0.70146316 |
| GGVTVMATP | ASSSTGDAETLY | 0.47382072 |
| AGLAANVRD | ASSSTGDAETLY | 0.8319713 |
| TNRPDLIDPALLRPG | ASSSTGDAETLY | 0.8276582 |
| LANLGVTAL | ASSSTGDAETLY | 0.674935 |
| TGERIEL | ASSSTGDAETLY | 0.77718866 |
| NGEELAA | ASSSTGDAETLY | 0.62116134 |
| VLSLVWG | ASSSTGDAETLY | 0.4965772 |
| IGYALWL | ASSSTGDAETLY | 0.80404484 |
| LSLLLAPL | ASSSTGDAETLY | 0.600242 |
| TLDEALK | ASSSTGDAETLY | 0.8833668 |
| TDGAGTAEGG | ASSSTGDAETLY | 0.66382277 |
| NAPGGAGGNGG | ASSSTGDAETLY | 0.8567041 |
| VSVFMGMGGNF | ASSSTGDAETLY | 0.84615 |
| LASYLAAL | ASSSTGDAETLY | 0.8356041 |
| LVGGTPAL | ASSSTGDAETLY | 0.7486128 |
| VRNADVL | ASSSTGDAETLY | 0.6700818 |
| TCSSPFLVSAM | ASSSTGDAETLY | 0.54915977 |
| VHITDAE | ASSSTGDAETLY | 0.25190422 |
| EYAAPSF | ASSSTGDAETLY | 0.8766903 |
| VVDAFHGGL | ASSSTGDAETLY | 0.6002275 |
| ASVVDPQ | ASSSTGDAETLY | 0.87515426 |
| SYNEFGGLS | ASSSTGDAETLY | 0.8586644 |
| DPDGSVY | ASSSTGDAETLY | 0.6269991 |
| MVSTIHATE | ASSSTGDAETLY | 0.8300053 |
| LAAGVADPV | ASSSTGDAETLY | 0.6195859 |
| TKEQIAA | ASSSTGDAETLY | 0.5285511 |
| SLASPAAL | ASSSTGDAETLY | 0.8680726 |
| IGLVFYGV | ASSSTGDAETLY | 0.8166208 |
| FLLDASNL | ASSSTGDAETLY | 0.6551875 |
| YQVGNFP | ASSSTGDAETLY | 0.80461437 |
| ESLLSQST | ASSSTGDAETLY | 0.79454756 |
| LFTVGFAAVG | ASSSTGDAETLY | 0.63825244 |
| ATPPTPT | ASSSTGDAETLY | 0.72553587 |
| TDLATLG | ASSSTGDAETLY | 0.8630054 |
| SAGGTGR | ASSSTGDAETLY | 0.77799124 |

| | | |
|---|---|---|
| GFSNSGS | ASSSTGDAETLY | 0.7246388 |
| GGNGGSGGTSV | ASSSTGDAETLY | 0.85008585 |
| GTSVATGGAGN | ASSSTGDAETLY | 0.56034917 |
| NGGNGGSGGTS | ASSSTGDAETLY | 0.82945096 |
| AVVKFPH | ASSSTGDAETLY | 0.76503277 |
| TPPPPPP | ASSSTGDAETLY | 0.7418801 |
| AAPAAAAAPA | ASSSTGDAETLY | 0.8592684 |
| IALNEDL | ASSSTGDAETLY | 0.4829792 |
| IVNVSSM | ASSSTGDAETLY | 0.6504886 |
| TGIPPAPR | ASSSTGDAETLY | 0.74035454 |
| FDLGGGTF | ASSSTGDAETLY | 0.8219526 |
| FDLGGGTFDVS | ASSSTGDAETLY | 0.85692286 |
| NEPTAAA | ASSSTGDAETLY | 0.63937116 |
| NEPTAAALA | ASSSTGDAETLY | 0.7885979 |
| GSIINISSI | ASSSTGDAETLY | 0.6772258 |
| VAPGNAGTA | ASSSTGDAETLY | 0.60275567 |
| TGGGASLE | ASSSTGDAETLY | 0.7039101 |
| VAVYNHYKRI | ASSSTGDAETLY | 0.7376983 |
| TDPAPATT | ASSSTGDAETLY | 0.6841914 |
| TNRPDILDPALLRPG | ASSSTGDAETLY | 0.8234399 |
| LTTTTTS | ASSSTGDAETLY | 0.6365054 |
| PTGSGKTL | ASSSTGDAETLY | 0.82071584 |
| TGSGKTL | ASSSTGDAETLY | 0.86878574 |
| TIGHVDHGK | ASSSTGDAETLY | 0.7500119 |
| TIGHVDHGKTT | ASSSTGDAETLY | 0.8449365 |
| GHVDHGKTT | ASSSTGDAETLY | 0.8258002 |
| AQFEHTLL | ASSSTGDAETLY | 0.8470806 |
| IVNQDEL | ASSSTGDAETLY | 0.7393197 |
| TNGSQFF | ASSSTGDAETLY | 0.6880877 |
| TGKGIVE | ASSSTGDAETLY | 0.87666523 |
| LISNASDA | ASSSTGDAETLY | 0.8544704 |
| AELRTLGELTKIAKAHGAQ | ASSSTGDAETLY | 0.84969985 |
| APNAVAAAEPSW | ASSSTGDAETLY | 0.8338111 |
| APQVAGAATAVL | ASSSTGDAETLY | 0.8318198 |
| ARRDPAAAAAILPTDA | ASSSTGDAETLY | 0.83415985 |
| DAAARDVLAEAGLAEYF | ASSSTGDAETLY | 0.7234371 |
| DDQIVLSAKANLAKVEQ | ASSSTGDAETLY | 0.46554527 |
| DELLPDTGEDSDEDGH | ASSSTGDAETLY | 0.7442221 |
| ELPEAPANRATVPQD | ASSSTGDAETLY | 0.74569124 |
| FGQSKIERVTGIG | ASSSTGDAETLY | 0.72235316 |
| GAGGTGSTAGGAGGAGGAGGLY | ASSSTGDAETLY | 0.34570894 |
| GHGGTGAAGGNGGTGGAGGSGI | ASSSTGDAETLY | 0.87143755 |
| GNGGNGGAGGEGGPGGAGGAGA | ASSSTGDAETLY | 0.90393054 |
| GSTAGNGGNGGNATNGTIAGQP | ASSSTGDAETLY | 0.65834785 |
| GTGGDGGNGGNGGTGG | ASSSTGDAETLY | 0.8973843 |
| GVRRPAGIPADAPV | ASSSTGDAETLY | 0.46020633 |
| GYRQVTDAHLLTLARR | ASSSTGDAETLY | 0.8522625 |
| IAKMGQDGHDRG | ASSSTGDAETLY | 0.87624395 |
| KAEPSIEIQYT | ASSSTGDAETLY | 0.61979395 |
| KPVRIRMAADEDTVAVT | ASSSTGDAETLY | 0.82774174 |
| MPPRIAGMRLLVIKPEP | ASSSTGDAETLY | 0.6890524 |

| | | |
|---|---|---|
| MKFYAVALFRSRGVPPTR | ASSSTGDAETLY | 0.7410534 |
| NAGGFGILGAVAHSP | ASSSTGDAETLY | 0.8853376 |
| NEGREITLAMAEM | ASSSTGDAETLY | 0.728198 |
| NELVEWSHPLGHRWRWE | ASSSTGDAETLY | 0.8077491 |
| NLGAGGTGGNGGNPG | ASSSTGDAETLY | 0.7746409 |
| NTEWPKWETVMAADPVD | ASSSTGDAETLY | 0.6538412 |
| PASAAPAETQP | ASSSTGDAETLY | 0.80344605 |
| QQSSDGKNGVRAAEITGE | ASSSTGDAETLY | 0.8688845 |
| RAPDAVTTSWWKE | ASSSTGDAETLY | 0.5053845 |
| SGFGNVGAHTSGWFNQ | ASSSTGDAETLY | 0.88914084 |
| SGVAGAVGQAASVGGLKV | ASSSTGDAETLY | 0.6925689 |
| SNGGAGTGSAGGNGGT | ASSSTGDAETLY | 0.89632046 |
| TGGAGGEGGAAG | ASSSTGDAETLY | 0.8409692 |
| TGGDGGAAGAGGAPGHG | ASSSTGDAETLY | 0.787562 |
| TMPPLLAAIANAGRHGV | ASSSTGDAETLY | 0.7114879 |
| TSSTAGNSGGA | ASSSTGDAETLY | 0.87701166 |
| VPATVPVGILC | ASSSTGDAETLY | 0.76783 |
| VQRPDSGGVVLTGRIS | ASSSTGDAETLY | 0.84939337 |
| WTPNPVPATKGVPAGLVQAT | ASSSTGDAETLY | 0.8711927 |
| YALKRALAKFSDRFDVVII | ASSSTGDAETLY | 0.43402112 |
| YLIGSVIV | ASSSTGDAETLY | 0.6984767 |
| YGQLVAVID | ASSSTGDAETLY | 0.6308206 |
| VTVGGMTLPA | ASSSTGDAETLY | 0.586117 |
| VLALPAV | ASSSTGDAETLY | 0.516503 |
| VIPPTLNYET | ASSSTGDAETLY | 0.40303424 |
| VGMGGMHPSSG | ASSSTGDAETLY | 0.56124246 |
| VGLAGGAATA | ASSSTGDAETLY | 0.46462867 |
| VGGAGGAGGSGP | ASSSTGDAETLY | 0.6842426 |
| VFAAFMV | ASSSTGDAETLY | 0.6695516 |
| TLYDKLR | ASSSTGDAETLY | 0.5035795 |
| TLAVFTVL | ASSSTGDAETLY | 0.68820226 |
| TGSVQPARV | ASSSTGDAETLY | 0.64425415 |
| TGGGAGSGGT | ASSSTGDAETLY | 0.40938288 |
| SHLETSL | ASSSTGDAETLY | 0.5734626 |
| SFTTGMT | ASSSTGDAETLY | 0.5691613 |
| QVEVAKRMGVRQPTVS | ASSSTGDAETLY | 0.5462333 |
| QGSGGPALAM | ASSSTGDAETLY | 0.39793256 |
| MLAPSNP | ASSSTGDAETLY | 0.6446675 |
| LVNNAAI | ASSSTGDAETLY | 0.52356625 |
| LTGEINTLA | ASSSTGDAETLY | 0.2927122 |
| LSIFGPV | ASSSTGDAETLY | 0.6335708 |
| LLALLANL | ASSSTGDAETLY | 0.59008193 |
| LISNASDALDKLR | ASSSTGDAETLY | 0.5298842 |
| LINEAVP | ASSSTGDAETLY | 0.6758195 |
| LGYAFSGL | ASSSTGDAETLY | 0.53484464 |
| LDFVAVAL | ASSSTGDAETLY | 0.502563 |
| LAQPGAE | ASSSTGDAETLY | 0.675477 |
| ISNLGAVTGF | ASSSTGDAETLY | 0.6882436 |
| HQEDPGRI | ASSSTGDAETLY | 0.42089444 |
| GTPGEPTP | ASSSTGDAETLY | 0.6234009 |
| GFFNSGS | ASSSTGDAETLY | 0.68424314 |

| | | |
|---|---|---|
| GAGGGGLTAGAN | ASSSTGDAETLY | 0.6153206 |
| FVDALVNM | ASSSTGDAETLY | 0.6925558 |
| FLTVAPEL | ASSSTGDAETLY | 0.6941598 |
| FETLTAL | ASSSTGDAETLY | 0.4932692 |
| DVINPIEL | ASSSTGDAETLY | 0.6924082 |
| APVSGAPV | ASSSTGDAETLY | 0.7060702 |
| ALVGGLAW | ASSSTGDAETLY | 0.57298577 |
| ALAAAAATAI | ASSSTGDAETLY | 0.5924469 |
| AFGLGLLL | ASSSTGDAETLY | 0.4319323 |
| AFFAAASL | ASSSTGDAETLY | 0.5741272 |
| AAAALPAFSPP | ASSSTGDAETLY | 0.41215682 |
| VVLAPAL | ASSSTGDAETLY | 0.6554091 |
| TIGHVDHGKTTL | ASSSTGDAETLY | 0.6960995 |
| LISNASDALDK | ASSSTGDAETLY | 0.5798958 |
| LISNASDAL | ASSSTGDAETLY | 0.4147519 |
| ISNASDALDK | ASSSTGDAETLY | 0.4518265 |
| NQAETLVYQTE | ASSSTGDAETLY | 0.62011904 |
| TGGTGGTAGAG | ASSSTGDAETLY | 0.41679314 |
| TGSTAGGAGGA | ASSSTGDAETLY | 0.39031976 |
| DPLAVPVAAGPV | ASSSTGDAETLY | 0.5474244 |
| GGTGGTPGNGGN | ASSSTGDAETLY | 0.6798469 |
| PAGLAGVDPATL | ASSSTGDAETLY | 0.67959404 |
| GNNLVVEAPHAGASV | ASSSTGDAETLY | 0.44772702 |
| KVTRIMSASTLEQPA | ASSSTGDAETLY | 0.7010428 |
| LGFALPIALPRRTVV | ASSSTGDAETLY | 0.6655674 |
| TRLGNLTARKLVVLV | ASSSTGDAETLY | 0.6871742 |
| WSQQRRLIYFPSAGP | ASSSTGDAETLY | 0.60391855 |
| GPNGNPDPMAAAVDIR | ASSSTGDAETLY | 0.45128325 |
| TQRCVSVADLRAAASR | ASSSTGDAETLY | 0.6834917 |
| QFVLDVAEHPVEVLTETV | ASSSTGDAETLY | 0.56996846 |
| GGAGGTGGAGGNGGTQAGDG | ASSSTGDAETLY | 0.6054261 |
| PLLATLRAIAVDVDAKPAQV | ASSSTGDAETLY | 0.66714805 |
| APKFRTGGYMIDFWGVGYQVAK | ASSSTGDAETLY | 0.5839317 |
| GAGGAGGAGGAAGGPGGTGGTG | ASSSTGDAETLY | 0.53089774 |
| IGALGLLATDVAPAGSVAFGVA | ASSSTGDAETLY | 0.59467006 |
| AGGAGGIGGNGGAGGAGGVLMG | ASSSTGDAETLY | 0.6734538 |
| ADELIGLL | ASSSTGDAETLY | 0.4928004 |
| ADRALKA | ASSSTGDAETLY | 0.42300904 |
| AGAVGQVGGAGGA | ASSSTGDAETLY | 0.55610627 |
| AGGGGGLGGAP | ASSSTGDAETLY | 0.59540546 |
| AGGLLYGNGGAG | ASSSTGDAETLY | 0.6915194 |
| AGLESIGL | ASSSTGDAETLY | 0.6262806 |
| AGLKTGII | ASSSTGDAETLY | 0.5827823 |
| ASLAALLL | ASSSTGDAETLY | 0.65384233 |
| ASVINISSI | ASSSTGDAETLY | 0.65411437 |
| ATPPGPA | ASSSTGDAETLY | 0.6106426 |
| ATVASALV | ASSSTGDAETLY | 0.61257017 |
| ATVASLIL | ASSSTGDAETLY | 0.5400202 |
| AVNGPVLIRH | ASSSTGDAETLY | 0.6755921 |
| DGAAGAVGI | ASSSTGDAETLY | 0.65999806 |
| DPLLHGI | ASSSTGDAETLY | 0.6475526 |

| | | |
|---|---|---|
| ELGAALGAL | ASSSTGDAETLY | 0.68877596 |
| ELGRTQAV | ASSSTGDAETLY | 0.64248335 |
| FVAGGYLS | ASSSTGDAETLY | 0.60817397 |
| GASGGAGGAGGW | ASSSTGDAETLY | 0.38368666 |
| GGAGAGGDGGAGGI | ASSSTGDAETLY | 0.5354344 |
| GTGHNGSQPGMGGNG | ASSSTGDAETLY | 0.61739445 |
| GYAAASASAS | ASSSTGDAETLY | 0.64671546 |
| HQVDVAAVQQG | ASSSTGDAETLY | 0.66880655 |
| IGAAIAHT | ASSSTGDAETLY | 0.62295973 |
| IGMIEVAQL | ASSSTGDAETLY | 0.5289575 |
| IRLAPEHTSAMD | ASSSTGDAETLY | 0.6618497 |
| IYNVSVL | ASSSTGDAETLY | 0.6845924 |
| KQGFLGM | ASSSTGDAETLY | 0.41509688 |
| LESVITQL | ASSSTGDAETLY | 0.5640881 |
| LGLGALVVV | ASSSTGDAETLY | 0.7092742 |
| LIGNGGNGGSGGI | ASSSTGDAETLY | 0.6367206 |
| LNAINAPTL | ASSSTGDAETLY | 0.57482266 |
| LQLLNTGV | ASSSTGDAETLY | 0.516165 |
| LQTSIEF | ASSSTGDAETLY | 0.6939992 |
| MSFAVSEL | ASSSTGDAETLY | 0.53812206 |
| NTGALGL | ASSSTGDAETLY | 0.6971563 |
| PPGAGGAGGAGTT | ASSSTGDAETLY | 0.3188417 |
| QLVGLGI | ASSSTGDAETLY | 0.5060446 |
| RFLEQVHQL | ASSSTGDAETLY | 0.4705678 |
| RGVGYVLR | ASSSTGDAETLY | 0.306292 |
| SATGVFAGVI | ASSSTGDAETLY | 0.6788527 |
| SLPAITLP | ASSSTGDAETLY | 0.46767846 |
| STALAFTV | ASSSTGDAETLY | 0.6343863 |
| SVLTVGLGK | ASSSTGDAETLY | 0.6074534 |
| TALAGAGSGI | ASSSTGDAETLY | 0.6860322 |
| TGLPFTGI | ASSSTGDAETLY | 0.6383489 |
| VAEHVTV | ASSSTGDAETLY | 0.57024324 |
| VDIVLNSL | ASSSTGDAETLY | 0.49431795 |
| VLTGGGAVL | ASSSTGDAETLY | 0.65612316 |
| VNVGLAYGI | ASSSTGDAETLY | 0.6774248 |
| VSSAAVGAV | ASSSTGDAETLY | 0.660578 |
| VTSVYEAGV | ASSSTGDAETLY | 0.36179823 |
| WVALDRGA | ASSSTGDAETLY | 0.61562717 |
| YLVGSVVI | ASSSTGDAETLY | 0.5838051 |
| GPPGTGKTLL | ASSSTGDAETLY | 0.40881738 |
| GVGMTKF | ASSSTGDAETLY | 0.32388014 |
| IGIGGSDL | ASSSTGDAETLY | 0.51850116 |
| IGTLLAEL | ASSSTGDAETLY | 0.5906924 |
| IYTSGTT | ASSSTGDAETLY | 0.6732031 |
| LSFPAGV | ASSSTGDAETLY | 0.68469095 |
| LYTSGTT | ASSSTGDAETLY | 0.53895974 |
| VGAFPFGF | ASSSTGDAETLY | 0.66660595 |
| VGAGTVEF | ASSSTGDAETLY | 0.5985793 |
| VGKSTLLL | ASSSTGDAETLY | 0.2549637 |
| AALAHAGGGIV | ASSSTGDAETLY | 0.5469276 |
| AALVAVSLL | ASSSTGDAETLY | 0.42110968 |

| | | |
|---|---|---|
| AAVSRAVAAL | ASSSTGDAETLY | 0.6899245 |
| AGGSSGIGGF | ASSSTGDAETLY | 0.46846268 |
| AGLTVANANG | ASSSTGDAETLY | 0.5665896 |
| AGLVGAVL | ASSSTGDAETLY | 0.5983287 |
| ALSQAFTV | ASSSTGDAETLY | 0.64101464 |
| ALTFSVSL | ASSSTGDAETLY | 0.6113002 |
| ALTGAALVAL | ASSSTGDAETLY | 0.48819414 |
| ASFSIGAGVL | ASSSTGDAETLY | 0.36043763 |
| DGSVYKI | ASSSTGDAETLY | 0.6599103 |
| EQLAEIV | ASSSTGDAETLY | 0.5685288 |
| FAVSELTVM | ASSSTGDAETLY | 0.4345502 |
| GAAPLAAGL | ASSSTGDAETLY | 0.41796005 |
| GADGGDGGSGGAGG | ASSSTGDAETLY | 0.6614282 |
| GFGLDTGGAGG | ASSSTGDAETLY | 0.38098198 |
| GLASGGLL | ASSSTGDAETLY | 0.66728735 |
| GRIAGAAVLL | ASSSTGDAETLY | 0.5359048 |
| GSAGSLNTSF | ASSSTGDAETLY | 0.59200394 |
| GTGQIGI | ASSSTGDAETLY | 0.7107477 |
| GTLIALIL | ASSSTGDAETLY | 0.6322061 |
| IAATMGFVVG | ASSSTGDAETLY | 0.5827633 |
| IFAQALL | ASSSTGDAETLY | 0.5009659 |
| IGAAVALGF | ASSSTGDAETLY | 0.47015983 |
| IGLTPLAL | ASSSTGDAETLY | 0.48320577 |
| ILGSNTSTL | ASSSTGDAETLY | 0.5001796 |
| IVFSLTVL | ASSSTGDAETLY | 0.6951291 |
| LASKATGF | ASSSTGDAETLY | 0.5500076 |
| LASVALIL | ASSSTGDAETLY | 0.43829003 |
| LFAVGLTL | ASSSTGDAETLY | 0.5808288 |
| LFNGGIGTYI | ASSSTGDAETLY | 0.69900113 |
| LTLGAGII | ASSSTGDAETLY | 0.7079003 |
| LVAAALGDI | ASSSTGDAETLY | 0.49938893 |
| NTAFVQLGI | ASSSTGDAETLY | 0.5298239 |
| QLFSGAEL | ASSSTGDAETLY | 0.50704104 |
| SAATFITL | ASSSTGDAETLY | 0.6001273 |
| SNYGLANLGV | ASSSTGDAETLY | 0.4668786 |
| SYGGAYAVM | ASSSTGDAETLY | 0.4462774 |
| TTGVGGLTLG | ASSSTGDAETLY | 0.65606236 |
| TTQTIYI | ASSSTGDAETLY | 0.7055435 |
| VAAGAGLGAVI | ASSSTGDAETLY | 0.5613797 |
| VAAVVTLL | ASSSTGDAETLY | 0.46979332 |
| VGFLGGLI | ASSSTGDAETLY | 0.5920015 |
| VGVIPILL | ASSSTGDAETLY | 0.51958686 |
| VIFASSLI | ASSSTGDAETLY | 0.50758934 |
| VKGGDGGAAAG | ASSSTGDAETLY | 0.5734716 |
| VTKTFTNI | ASSSTGDAETLY | 0.37173036 |
| VVAAAILL | ASSSTGDAETLY | 0.58825535 |
| YAALDVEL | ASSSTGDAETLY | 0.6999352 |
| LASVALLL | ASSSTGDAETLY | 0.603953 |
| LDAVHRV | ASSSTGDAETLY | 0.61866724 |
| QGLSSSIF | ASSSTGDAETLY | 0.69695824 |
| RGTFANIRL | ASSSTGDAETLY | 0.65701103 |

| | | |
|---|---|---|
| RQLWWGHRI | ASSSTGDAETLY | 0.688324 |
| YGPPGTGKTLL | ASSSTGDAETLY | 0.68278074 |
| AALVGAPANVA | ASSSTGDAETLY | 0.5833863 |
| AATFRRTE | ASSSTGDAETLY | 0.59799194 |
| AGALGGGI | ASSSTGDAETLY | 0.6920136 |
| AGGAGGAGGIGAG | ASSSTGDAETLY | 0.59136546 |
| AGSGALGL | ASSSTGDAETLY | 0.6029302 |
| AQIADQI | ASSSTGDAETLY | 0.54674345 |
| ASAVAVTYL | ASSSTGDAETLY | 0.6722429 |
| ASTVLGGTGG | ASSSTGDAETLY | 0.5837962 |
| AVGGEPAK | ASSSTGDAETLY | 0.6831126 |
| CEQGTFGGTC | ASSSTGDAETLY | 0.33422655 |
| DEGKAPAG | ASSSTGDAETLY | 0.42533368 |
| DTVVFPAL | ASSSTGDAETLY | 0.6826155 |
| FEGITMAG | ASSSTGDAETLY | 0.27920285 |
| FPLVVQD | ASSSTGDAETLY | 0.32839283 |
| GAFTTPQ | ASSSTGDAETLY | 0.59636843 |
| GFGGFGTAGGVGGA | ASSSTGDAETLY | 0.683671 |
| GFLAQFL | ASSSTGDAETLY | 0.28955013 |
| GGAGGAGGNAGVA | ASSSTGDAETLY | 0.50893545 |
| GGVGTAGAGGAGGAGG | ASSSTGDAETLY | 0.40240595 |
| GNGGGDAGNAGSGGN | ASSSTGDAETLY | 0.6409813 |
| GTGGAGGAGNDAGS | ASSSTGDAETLY | 0.5989236 |
| GTTNTGWFNS | ASSSTGDAETLY | 0.37815517 |
| IFMGPLFTL | ASSSTGDAETLY | 0.6513669 |
| ILAHAIEAI | ASSSTGDAETLY | 0.41082814 |
| IRSAIREAL | ASSSTGDAETLY | 0.6301968 |
| ITFAGGG | ASSSTGDAETLY | 0.68618023 |
| ITGMSGHQEV | ASSSTGDAETLY | 0.70245755 |
| IVAGLACTL | ASSSTGDAETLY | 0.43851626 |
| IVGGGLAGA | ASSSTGDAETLY | 0.6636877 |
| IVTGVLNI | ASSSTGDAETLY | 0.53325635 |
| LAQQGCI | ASSSTGDAETLY | 0.7152991 |
| LCLHVEN | ASSSTGDAETLY | 0.56672037 |
| LDAISDGS | ASSSTGDAETLY | 0.38059807 |
| LGGFAAML | ASSSTGDAETLY | 0.66937554 |
| LGGKNPMI | ASSSTGDAETLY | 0.62680066 |
| LGPKGVAAIL | ASSSTGDAETLY | 0.5295476 |
| LMVGVSI | ASSSTGDAETLY | 0.66903704 |
| LVGAAFVL | ASSSTGDAETLY | 0.51300985 |
| LVLAVAPL | ASSSTGDAETLY | 0.5833514 |
| MAAIAVIDAA | ASSSTGDAETLY | 0.6862359 |
| NNGINAELQ | ASSSTGDAETLY | 0.66848207 |
| RGEPLVLVHG | ASSSTGDAETLY | 0.47697636 |
| RGGAGGAAGLI | ASSSTGDAETLY | 0.6703686 |
| RSYRMHPAV | ASSSTGDAETLY | 0.597805 |
| SAAHLLAGQLA | ASSSTGDAETLY | 0.4553638 |
| SAAVGSY | ASSSTGDAETLY | 0.7054307 |
| SPHAVLHRI | ASSSTGDAETLY | 0.57030714 |
| SQIAKVL | ASSSTGDAETLY | 0.6698065 |
| TALGHLVP | ASSSTGDAETLY | 0.6379992 |

| | | |
|---|---|---|
| TAPTTSGAA | ASSSTGDAETLY | 0.6873021 |
| TDGDDGNGGSPGT | ASSSTGDAETLY | 0.43582237 |
| TDGLGEN | ASSSTGDAETLY | 0.5943097 |
| TGGPPLPGQA | ASSSTGDAETLY | 0.59935653 |
| TIPAITI | ASSSTGDAETLY | 0.6758499 |
| TIPGFTI | ASSSTGDAETLY | 0.563435 |
| TSIASGALA | ASSSTGDAETLY | 0.64418185 |
| TVTLPGAAI | ASSSTGDAETLY | 0.63894165 |
| VASGLSAALP | ASSSTGDAETLY | 0.6930876 |
| VGAGVIGI | ASSSTGDAETLY | 0.58678085 |
| VGPKRFGV | ASSSTGDAETLY | 0.61674666 |
| VNVAFPD | ASSSTGDAETLY | 0.6987432 |
| VSMTNSGVS | ASSSTGDAETLY | 0.45574516 |
| VTDVLGDLSV | ASSSTGDAETLY | 0.5809455 |
| VVVVGAGFAGL | ASSSTGDAETLY | 0.65492594 |
| VYTVGTVV | ASSSTGDAETLY | 0.67733634 |
| WTTFSGP | ASSSTGDAETLY | 0.56178206 |
| YDELVEAF | ASSSTGDAETLY | 0.68763816 |
| YGTGGAGGNGG | ASSSTGDAETLY | 0.54898953 |
| YTNVMAV | ASSSTGDAETLY | 0.6685683 |
| FLEQVHQL | ASSSTGDAETLY | 0.59772444 |
| GLSLLAVAV | ASSSTGDAETLY | 0.6030817 |
| LEQVHQL | ASSSTGDAETLY | 0.4730218 |
| LNAAGAI | ASSSTGDAETLY | 0.6581972 |
| PAVHASL | ASSSTGDAETLY | 0.6857337 |
| TELEDDA | ASSSTGDAETLY | 0.6683835 |
| TPPGPPN | ASSSTGDAETLY | 0.33033082 |
| TPPPPPPP | ASSSTGDAETLY | 0.6712891 |
| TQSPALI | ASSQDWGSQNTLY | 0.6726662 |
| EFAAAFPQ | ASSQDWGSQNTLY | 0.6997886 |
| LLSGNLDVL | ASSQDWGSQNTLY | 0.57735443 |
| GPANLGNYNI | ASSQDWGSQNTLY | 0.6216261 |
| TLGPTLP | ASSQDWGSQNTLY | 0.55361015 |
| VYAYSSCCK | ASSQDWGSQNTLY | 0.5885251 |
| APAPAPAPNI | ASSQDWGSQNTLY | 0.41156703 |
| PGGPVPIP | ASSQDWGSQNTLY | 0.5245601 |
| FLTVALEP | ASSQDWGSQNTLY | 0.5831932 |
| SQYFGANL | ASSQDWGSQNTLY | 0.5081111 |
| GGVKPAA | ASSQDWGSQNTLY | 0.6678081 |
| GGTGGAGGAGGW | ASSQDWGSQNTLY | 0.68273485 |
| GVGGVAGLGGAGG | ASSQDWGSQNTLY | 0.68121034 |
| VLLVLATTI | ASSQDWGSQNTLY | 0.6971252 |
| IGIFAAML | ASSQDWGSQNTLY | 0.6965672 |
| AVGDLID | ASSQDWGSQNTLY | 0.6572181 |
| NISPLPL | ASSQDWGSQNTLY | 0.5223838 |
| VTAVFVAI | ASSQDWGSQNTLY | 0.60966825 |
| SGDDNDAAL | ASSQDWGSQNTLY | 0.40490383 |
| AVAALAHS | ASSQDWGSQNTLY | 0.5898402 |
| GGVTVMATP | ASSQDWGSQNTLY | 0.5361842 |
| AGLAANVRD | ASSQDWGSQNTLY | 0.63721704 |
| TNRPDLIDPALLRPG | ASSQDWGSQNTLY | 0.2782439 |

| | | |
|---|---|---|
| LANLGVTAL | ASSQDWGSQNTLY | 0.640628 |
| TGERIEL | ASSQDWGSQNTLY | 0.7007148 |
| NGEELAA | ASSQDWGSQNTLY | 0.5702282 |
| VLSLVWG | ASSQDWGSQNTLY | 0.6320441 |
| IGYALWL | ASSQDWGSQNTLY | 0.6609514 |
| LSLLLAPL | ASSQDWGSQNTLY | 0.5185007 |
| TLDEALK | ASSQDWGSQNTLY | 0.66866875 |
| TDGAGTAEGG | ASSQDWGSQNTLY | 0.5112984 |
| NAPGGAGGNGG | ASSQDWGSQNTLY | 0.66006076 |
| VSVFMGMGGNF | ASSQDWGSQNTLY | 0.6955929 |
| LASYLAAL | ASSQDWGSQNTLY | 0.6334282 |
| LVGGTPAL | ASSQDWGSQNTLY | 0.5320115 |
| VRNADVL | ASSQDWGSQNTLY | 0.5671072 |
| TCSSPFLVSAM | ASSQDWGSQNTLY | 0.4064038 |
| VHITDAE | ASSQDWGSQNTLY | 0.58799785 |
| EYAAPSF | ASSQDWGSQNTLY | 0.50211424 |
| VVDAFHGGL | ASSQDWGSQNTLY | 0.6224193 |
| ASVVDPQ | ASSQDWGSQNTLY | 0.47326082 |
| SYNEFGGLS | ASSQDWGSQNTLY | 0.6380913 |
| DPDGSVY | ASSQDWGSQNTLY | 0.62134784 |
| MVSTIHATE | ASSQDWGSQNTLY | 0.61770606 |
| LAAGVADPV | ASSQDWGSQNTLY | 0.4955427 |
| TKEQIAA | ASSQDWGSQNTLY | 0.6342914 |
| SLASPAAL | ASSQDWGSQNTLY | 0.5965283 |
| IGLVFYGV | ASSQDWGSQNTLY | 0.41672134 |
| FLLDASNL | ASSQDWGSQNTLY | 0.669721 |
| YQVGNFP | ASSQDWGSQNTLY | 0.54510343 |
| ESLLSQST | ASSQDWGSQNTLY | 0.59237766 |
| LFTVGFAAVG | ASSQDWGSQNTLY | 0.68243325 |
| ATPPTPT | ASSQDWGSQNTLY | 0.600399 |
| TDLATLG | ASSQDWGSQNTLY | 0.4138788 |
| SAGGTGR | ASSQDWGSQNTLY | 0.57848924 |
| GFSNSGS | ASSQDWGSQNTLY | 0.6186576 |
| GGNGGSGGTSV | ASSQDWGSQNTLY | 0.56621885 |
| GTSVATGGAGN | ASSQDWGSQNTLY | 0.29788136 |
| NGGNGGSGGTS | ASSQDWGSQNTLY | 0.6596783 |
| AVVKFPH | ASSQDWGSQNTLY | 0.45579588 |
| TPPPPPP | ASSQDWGSQNTLY | 0.52370894 |
| AAPAAAAAPA | ASSQDWGSQNTLY | 0.51771796 |
| IALNEDL | ASSQDWGSQNTLY | 0.674937 |
| IVNVSSM | ASSQDWGSQNTLY | 0.60498446 |
| TGIPPAPR | ASSQDWGSQNTLY | 0.5078285 |
| FDLGGGTF | ASSQDWGSQNTLY | 0.65279686 |
| FDLGGGTFDVS | ASSQDWGSQNTLY | 0.6324962 |
| NEPTAAA | ASSQDWGSQNTLY | 0.6782079 |
| NEPTAAALA | ASSQDWGSQNTLY | 0.61512184 |
| GSIINISSI | ASSQDWGSQNTLY | 0.5886313 |
| VAPGNAGTA | ASSQDWGSQNTLY | 0.5843099 |
| TGGGASLE | ASSQDWGSQNTLY | 0.5438304 |
| VAVYNHYKRI | ASSQDWGSQNTLY | 0.44055083 |
| TDPAPATT | ASSQDWGSQNTLY | 0.37221026 |

| | | |
|---|---|---|
| TNRPDILDPALLRPG | ASSQDWGSQNTLY | 0.5838142 |
| LTTTTTS | ASSQDWGSQNTLY | 0.39650658 |
| PTGSGKTL | ASSQDWGSQNTLY | 0.6273964 |
| TGSGKTL | ASSQDWGSQNTLY | 0.68091094 |
| TIGHVDHGK | ASSQDWGSQNTLY | 0.583231 |
| TIGHVDHGKTT | ASSQDWGSQNTLY | 0.3927766 |
| GHVDHGKTT | ASSQDWGSQNTLY | 0.5662981 |
| AQFEHTLL | ASSQDWGSQNTLY | 0.5670006 |
| IVNQDEL | ASSQDWGSQNTLY | 0.6680155 |
| TNGSQFF | ASSQDWGSQNTLY | 0.64657795 |
| TGKGIVE | ASSQDWGSQNTLY | 0.70412815 |
| LISNASDA | ASSQDWGSQNTLY | 0.6768968 |
| AELRTLGELTKIAKAHGAQ | ASSQDWGSQNTLY | 0.5363654 |
| APNAVAAAEPSW | ASSQDWGSQNTLY | 0.66333663 |
| APQVAGAATAVL | ASSQDWGSQNTLY | 0.53677404 |
| ARRDPAAAAAILPTDA | ASSQDWGSQNTLY | 0.61518824 |
| DAAARDVLAEAGLAEYF | ASSQDWGSQNTLY | 0.5988966 |
| DDQIVLSAKANLAKVEQ | ASSQDWGSQNTLY | 0.6262034 |
| DELLPDTGEDSDEDGH | ASSQDWGSQNTLY | 0.53370535 |
| ELPEAPANRATVPQD | ASSQDWGSQNTLY | 0.54565614 |
| FGQSKIERVTGIG | ASSQDWGSQNTLY | 0.41453055 |
| GAGGTGSTAGGAGGAGGAGGLY | ASSQDWGSQNTLY | 0.63736403 |
| GHGGTGAAGGNGGTGGAGGSGI | ASSQDWGSQNTLY | 0.46700308 |
| GNGGNGGAGGEGGPGGAGGAGA | ASSQDWGSQNTLY | 0.39501762 |
| GSTAGNGGNGGNATNGTIAGQP | ASSQDWGSQNTLY | 0.53399175 |
| GTGGDGGNGGNGGTGG | ASSQDWGSQNTLY | 0.70469856 |
| GVRRPAGIPADAPV | ASSQDWGSQNTLY | 0.69947296 |
| GYRQVTDAHLLTLARR | ASSQDWGSQNTLY | 0.5516771 |
| IAKMGQDGHDRG | ASSQDWGSQNTLY | 0.50353986 |
| KAEPSIEIQYT | ASSQDWGSQNTLY | 0.6538111 |
| KPVRIRMAADEDTVAVT | ASSQDWGSQNTLY | 0.43232733 |
| MPPRIAGMRLLVIKPEP | ASSQDWGSQNTLY | 0.6273459 |
| MKFYAVALFRSRGVPPTR | ASSQDWGSQNTLY | 0.6225642 |
| NAGGFGILGAVAHSP | ASSQDWGSQNTLY | 0.5988877 |
| NEGREITLAMAEM | ASSQDWGSQNTLY | 0.4342061 |
| NELVEWSHPLGHRWRWE | ASSQDWGSQNTLY | 0.6563804 |
| NLGAGGTGGNGGNPG | ASSQDWGSQNTLY | 0.58767503 |
| NTEWPKWETVMAADPVD | ASSQDWGSQNTLY | 0.6224675 |
| PASAAPAETQP | ASSQDWGSQNTLY | 0.39550555 |
| QQSSDGKNGVRAAEITGE | ASSQDWGSQNTLY | 0.6701814 |
| RAPDAVTTSWWKE | ASSQDWGSQNTLY | 0.47353324 |
| SGFGNVGAHTSGWFNQ | ASSQDWGSQNTLY | 0.4811352 |
| SGVAGAVGQAASVGGLKV | ASSQDWGSQNTLY | 0.32404166 |
| SNGGAGTGSAGGNGGT | ASSQDWGSQNTLY | 0.5773492 |
| TGGAGGEGGAAG | ASSQDWGSQNTLY | 0.67919683 |
| TGGDGGAAGAGGAPGHG | ASSQDWGSQNTLY | 0.6914961 |
| TMPPLLAAIANAGRHGV | ASSQDWGSQNTLY | 0.5660945 |
| TSSTAGNSGGA | ASSQDWGSQNTLY | 0.6739296 |
| VPATVPVGILC | ASSQDWGSQNTLY | 0.7012544 |
| VQRPDSGGVVLTGRIS | ASSQDWGSQNTLY | 0.5806229 |
| WTPNPVPATKGVPAGLVQAT | ASSQDWGSQNTLY | 0.36316192 |

| | | |
|---|---|---|
| YALKRALAKFSDRFDVVII | ASSQDWGSQNTLY | 0.61639184 |
| YLIGSVIV | ASSQDWGSQNTLY | 0.6179805 |
| YGQLVAVID | ASSQDWGSQNTLY | 0.689005 |
| VTVGGMTLPA | ASSQDWGSQNTLY | 0.6519486 |
| VLALPAV | ASSQDWGSQNTLY | 0.42778093 |
| VIPPTLNYET | ASSQDWGSQNTLY | 0.58713675 |
| VGMGGMHPSSG | ASSQDWGSQNTLY | 0.66981506 |
| VGLAGGAATA | ASSQDWGSQNTLY | 0.41683298 |
| VGGAGGAGGSGP | ASSQDWGSQNTLY | 0.4866959 |
| VFAAFMV | ASSQDWGSQNTLY | 0.5069348 |
| TLYDKLR | ASSQDWGSQNTLY | 0.38685665 |
| TLAVFTVL | ASSQDWGSQNTLY | 0.6366202 |
| TGSVQPARV | ASSQDWGSQNTLY | 0.57044685 |
| TGGGAGSGGT | ASSQDWGSQNTLY | 0.5416938 |
| SHLETSL | ASSQDWGSQNTLY | 0.574211 |
| SFTTGMT | ASSQDWGSQNTLY | 0.48425254 |
| QVEVAKRMGVRQPTVS | ASSQDWGSQNTLY | 0.5094912 |
| QGSGGPALAM | ASSQDWGSQNTLY | 0.54613817 |
| MLAPSNP | ASSQDWGSQNTLY | 0.65223 |
| LVNNAAI | ASSQDWGSQNTLY | 0.67057943 |
| LTGEINTLA | ASSQDWGSQNTLY | 0.60809934 |
| LSIFGPV | ASSQDWGSQNTLY | 0.58102524 |
| LLALLANL | ASSQDWGSQNTLY | 0.70034873 |
| LISNASDALDKLR | ASSQDWGSQNTLY | 0.42721295 |
| LINEAVP | ASSQDWGSQNTLY | 0.4716484 |
| LGYAFSGL | ASSQDWGSQNTLY | 0.6484276 |
| LDFVAVAL | ASSQDWGSQNTLY | 0.47961187 |
| LAQPGAE | ASSQDWGSQNTLY | 0.67916596 |
| ISNLGAVTGF | ASSQDWGSQNTLY | 0.6243031 |
| HQEDPGRI | ASSQDWGSQNTLY | 0.64753455 |
| GTPGEPTP | ASSQDWGSQNTLY | 0.6267429 |
| GFFNSGS | ASSQDWGSQNTLY | 0.70907253 |
| GAGGGGLTAGAN | ASSQDWGSQNTLY | 0.6557049 |
| FVDALVNM | ASSQDWGSQNTLY | 0.5385878 |
| FLTVAPEL | ASSQDWGSQNTLY | 0.66207325 |
| FETLTAL | ASSQDWGSQNTLY | 0.41230953 |
| DVINPIEL | ASSQDWGSQNTLY | 0.6748712 |
| APVSGAPV | ASSQDWGSQNTLY | 0.6442605 |
| ALVGGLAW | ASSQDWGSQNTLY | 0.46951646 |
| ALAAAAATAI | ASSQDWGSQNTLY | 0.5933262 |
| AFGLGLLL | ASSQDWGSQNTLY | 0.68974125 |
| AFFAAASL | ASSQDWGSQNTLY | 0.6863314 |
| AAAALPAFSPP | ASSQDWGSQNTLY | 0.43386546 |
| VVLAPAL | ASSQDWGSQNTLY | 0.5342912 |
| TIGHVDHGKTTL | ASSQDWGSQNTLY | 0.43388265 |
| LISNASDALDK | ASSQDWGSQNTLY | 0.45805377 |
| LISNASDAL | ASSQDWGSQNTLY | 0.3924746 |
| ISNASDALDK | ASSQDWGSQNTLY | 0.6580742 |
| NQAETLVYQTE | ASSQDWGSQNTLY | 0.55567116 |
| TGGTGGTAGAG | ASSQDWGSQNTLY | 0.66105556 |
| TGSTAGGAGGA | ASSQDWGSQNTLY | 0.63368535 |

| | | |
|---|---|---|
| DPLAVPVAAGPV | ASSQDWGSQNTLY | 0.68625796 |
| GGTGGTPGNGGN | ASSQDWGSQNTLY | 0.52186286 |
| PAGLAGVDPATL | ASSQDWGSQNTLY | 0.5706806 |
| GNNLVVEAPHAGASV | ASSQDWGSQNTLY | 0.35247025 |
| KVTRIMSASTLEQPA | ASSQDWGSQNTLY | 0.50497067 |
| LGFALPIALPRRTVV | ASSQDWGSQNTLY | 0.48019287 |
| TRLGNLTARKLVVLV | ASSQDWGSQNTLY | 0.5614448 |
| WSQQRRLIYFPSAGP | ASSQDWGSQNTLY | 0.41834694 |
| GPNGNPDPMAAAVDIR | ASSQDWGSQNTLY | 0.6984502 |
| TQRCVSVADLRAAASR | ASSQDWGSQNTLY | 0.5720894 |
| QFVLDVAEHPVEVLTETV | ASSQDWGSQNTLY | 0.65627205 |
| GGAGGTGGAGGNGGTQAGDG | ASSQDWGSQNTLY | 0.5256289 |
| PLLATLRAIAVDVDAKPAQV | ASSQDWGSQNTLY | 0.4639596 |
| APKFRTGGYMIDFWGVGYQVAK | ASSQDWGSQNTLY | 0.51985323 |
| GAGGAGGAGGAAGGPGGTGGTG | ASSQDWGSQNTLY | 0.6557001 |
| IGALGLLATDVAPAGSVAFGVA | ASSQDWGSQNTLY | 0.4467676 |
| AGGAGGIGGNGGAGGAGGVLMG | ASSQDWGSQNTLY | 0.37434357 |
| ADELIGLL | ASSQDWGSQNTLY | 0.5071748 |
| ADRALKA | ASSQDWGSQNTLY | 0.5695758 |
| AGAVGQVGGAGGA | ASSQDWGSQNTLY | 0.698111 |
| AGGGGGLGGAP | ASSQDWGSQNTLY | 0.69199014 |
| AGGLLYGNGGAG | ASSQDWGSQNTLY | 0.62547183 |
| AGLESIGL | ASSQDWGSQNTLY | 0.6925678 |
| AGLKTGII | ASSQDWGSQNTLY | 0.30016965 |
| ASLAALLL | ASSQDWGSQNTLY | 0.5682968 |
| ASVINISSI | ASSQDWGSQNTLY | 0.69324404 |
| ATPPGPA | ASSQDWGSQNTLY | 0.6553359 |
| ATVASALV | ASSQDWGSQNTLY | 0.589733 |
| ATVASLIL | ASSQDWGSQNTLY | 0.68922853 |
| AVNGPVLIRH | ASSQDWGSQNTLY | 0.4779249 |
| DGAAGAVGI | ASSQDWGSQNTLY | 0.56385183 |
| DPLLHGI | ASSQDWGSQNTLY | 0.6002866 |
| ELGAALGAL | ASSQDWGSQNTLY | 0.64658606 |
| ELGRTQAV | ASSQDWGSQNTLY | 0.6273818 |
| FVAGGYLS | ASSQDWGSQNTLY | 0.41097024 |
| GASGGAGGAGGW | ASSQDWGSQNTLY | 0.603238 |
| GGAGAGGDGGAGGI | ASSQDWGSQNTLY | 0.7172464 |
| GTGHNGSQPGMGGNG | ASSQDWGSQNTLY | 0.562258 |
| GYAAASASAS | ASSQDWGSQNTLY | 0.6851938 |
| HQVDVAAVQQG | ASSQDWGSQNTLY | 0.61360526 |
| IGAAIAHT | ASSQDWGSQNTLY | 0.63682026 |
| IGMIEVAQL | ASSQDWGSQNTLY | 0.49431962 |
| IRLAPEHTSAMD | ASSQDWGSQNTLY | 0.48151696 |
| IYNVSVL | ASSQDWGSQNTLY | 0.63294226 |
| KQGFLGM | ASSQDWGSQNTLY | 0.46867698 |
| LESVITQL | ASSQDWGSQNTLY | 0.5657164 |
| LGLGALVVV | ASSQDWGSQNTLY | 0.56951296 |
| LIGNGGNGGSGGI | ASSQDWGSQNTLY | 0.71507436 |
| LNAINAPTL | ASSQDWGSQNTLY | 0.6429736 |
| LQLLNTGV | ASSQDWGSQNTLY | 0.6504477 |
| LQTSIEF | ASSQDWGSQNTLY | 0.47252494 |

| | | |
|---|---|---|
| MSFAVSEL | ASSQDWGSQNTLY | 0.61060333 |
| NTGALGL | ASSQDWGSQNTLY | 0.66020894 |
| PPGAGGAGGAGTT | ASSQDWGSQNTLY | 0.43997154 |
| QLVGLGI | ASSQDWGSQNTLY | 0.6890224 |
| RFLEQVHQL | ASSQDWGSQNTLY | 0.47472742 |
| RGVGYVLR | ASSQDWGSQNTLY | 0.6166893 |
| SATGVFAGVI | ASSQDWGSQNTLY | 0.64256394 |
| SLPAITLP | ASSQDWGSQNTLY | 0.68733865 |
| STALAFTV | ASSQDWGSQNTLY | 0.5614275 |
| SVLTVGLGK | ASSQDWGSQNTLY | 0.62034225 |
| TALAGAGSGI | ASSQDWGSQNTLY | 0.44985062 |
| TGLPFTGI | ASSQDWGSQNTLY | 0.66360277 |
| VAEHVTV | ASSQDWGSQNTLY | 0.68670833 |
| VDIVLNSL | ASSQDWGSQNTLY | 0.6195905 |
| VLTGGGAVL | ASSQDWGSQNTLY | 0.70549643 |
| VNVGLAYGI | ASSQDWGSQNTLY | 0.6221087 |
| VSSAAVGAV | ASSQDWGSQNTLY | 0.35763484 |
| VTSVYEAGV | ASSQDWGSQNTLY | 0.57879287 |
| WVALDRGA | ASSQDWGSQNTLY | 0.6893395 |
| YLVGSVVI | ASSQDWGSQNTLY | 0.6980712 |
| GPPGTGKTLL | ASSQDWGSQNTLY | 0.69180083 |
| GVGMTKF | ASSQDWGSQNTLY | 0.6460301 |
| IGIGGSDL | ASSQDWGSQNTLY | 0.5680389 |
| IGTLLAEL | ASSQDWGSQNTLY | 0.69967306 |
| IYTSGTT | ASSQDWGSQNTLY | 0.71440923 |
| LSFPAGV | ASSQDWGSQNTLY | 0.6517496 |
| LYTSGTT | ASSQDWGSQNTLY | 0.48966384 |
| VGAFPFGF | ASSQDWGSQNTLY | 0.46487033 |
| VGAGTVEF | ASSQDWGSQNTLY | 0.63690627 |
| VGKSTLLL | ASSQDWGSQNTLY | 0.67068756 |
| AALAHAGGGIV | ASSQDWGSQNTLY | 0.40046683 |
| AALVAVSLL | ASSQDWGSQNTLY | 0.6133167 |
| AAVSRAVAAL | ASSQDWGSQNTLY | 0.6270902 |
| AGGSSGIGGF | ASSQDWGSQNTLY | 0.34752184 |
| AGLTVANANG | ASSQDWGSQNTLY | 0.66943157 |
| AGLVGAVL | ASSQDWGSQNTLY | 0.6963095 |
| ALSQAFTV | ASSQDWGSQNTLY | 0.59253925 |
| ALTFSVSL | ASSQDWGSQNTLY | 0.44436204 |
| ALTGAALVAL | ASSQDWGSQNTLY | 0.69145024 |
| ASFSIGAGVL | ASSQDWGSQNTLY | 0.7110612 |
| DGSVYKI | ASSQDWGSQNTLY | 0.6909092 |
| EQLAEIV | ASSQDWGSQNTLY | 0.36928636 |
| FAVSELTVM | ASSQDWGSQNTLY | 0.29462212 |
| GAAPLAAGL | ASSQDWGSQNTLY | 0.59155244 |
| GADGGDGGSGGAGG | ASSQDWGSQNTLY | 0.665457 |
| GFGLDTGGAGG | ASSQDWGSQNTLY | 0.53516877 |
| GLASGGLL | ASSQDWGSQNTLY | 0.46808404 |
| GRIAGAAVLL | ASSQDWGSQNTLY | 0.6308863 |
| GSAGSLNTSF | ASSQDWGSQNTLY | 0.6858379 |
| GTGQIGI | ASSQDWGSQNTLY | 0.43209815 |
| GTLIALIL | ASSQDWGSQNTLY | 0.41351494 |

| | | |
|---|---|---|
| IAATMGFVVG | ASSQDWGSQNTLY | 0.67349553 |
| IFAQALL | ASSQDWGSQNTLY | 0.6632052 |
| IGAAVALGF | ASSQDWGSQNTLY | 0.48887944 |
| IGLTPLAL | ASSQDWGSQNTLY | 0.56051505 |
| ILGSNTSTL | ASSQDWGSQNTLY | 0.6277633 |
| IVFSLTVL | ASSQDWGSQNTLY | 0.6377501 |
| LASKATGF | ASSQDWGSQNTLY | 0.6467117 |
| LASVALIL | ASSQDWGSQNTLY | 0.43044463 |
| LFAVGLTL | ASSQDWGSQNTLY | 0.69802487 |
| LFNGGIGTYI | ASSQDWGSQNTLY | 0.43214524 |
| LTLGAGII | ASSQDWGSQNTLY | 0.65691423 |
| LVAAALGDI | ASSQDWGSQNTLY | 0.61406547 |
| NTAFVQLGI | ASSQDWGSQNTLY | 0.62491006 |
| QLFSGAEL | ASSQDWGSQNTLY | 0.55299103 |
| SAATFITL | ASSQDWGSQNTLY | 0.5137882 |
| SNYGLANLGV | ASSQDWGSQNTLY | 0.43372536 |
| SYGGAYAVM | ASSQDWGSQNTLY | 0.37271434 |
| TTGVGGLTLG | ASSQDWGSQNTLY | 0.6866385 |
| TTQTIYI | ASSQDWGSQNTLY | 0.6393988 |
| VAAGAGLGAVI | ASSQDWGSQNTLY | 0.6740361 |
| VAAVVTLL | ASSQDWGSQNTLY | 0.63040805 |
| VGFLGGLI | ASSQDWGSQNTLY | 0.4992128 |
| VGVIPILL | ASSQDWGSQNTLY | 0.64231974 |
| VIFASSLI | ASSQDWGSQNTLY | 0.41467565 |
| VKGGDGGAAAG | ASSQDWGSQNTLY | 0.5598508 |
| VTKTFTNI | ASSQDWGSQNTLY | 0.56932217 |
| VVAAAILL | ASSQDWGSQNTLY | 0.625898 |
| YAALDVEL | ASSQDWGSQNTLY | 0.510129 |
| LASVALLL | ASSQDWGSQNTLY | 0.6259723 |
| LDAVHRV | ASSQDWGSQNTLY | 0.64865524 |
| QGLSSSIF | ASSQDWGSQNTLY | 0.47940916 |
| RGTFANIRL | ASSQDWGSQNTLY | 0.57118475 |
| RQLWWGHRI | ASSQDWGSQNTLY | 0.69050974 |
| YGPPGTGKTLL | ASSQDWGSQNTLY | 0.6013445 |
| AALVGAPANVA | ASSQDWGSQNTLY | 0.43043852 |
| AATFRRTE | ASSQDWGSQNTLY | 0.6524389 |
| AGALGGGI | ASSQDWGSQNTLY | 0.49703538 |
| AGGAGGAGGIGAG | ASSQDWGSQNTLY | 0.64669853 |
| AGSGALGL | ASSQDWGSQNTLY | 0.6230161 |
| AQIADQI | ASSQDWGSQNTLY | 0.48847374 |
| ASAVAVTYL | ASSQDWGSQNTLY | 0.6466082 |
| ASTVLGGTGG | ASSQDWGSQNTLY | 0.43148428 |
| AVGGEPAK | ASSQDWGSQNTLY | 0.5201407 |
| CEQGTFGGTC | ASSQDWGSQNTLY | 0.52591854 |
| DEGKAPAG | ASSQDWGSQNTLY | 0.6051501 |
| DTVVFPAL | ASSQDWGSQNTLY | 0.66289973 |
| FEGITMAG | ASSQDWGSQNTLY | 0.4464626 |
| FPLVVQD | ASSQDWGSQNTLY | 0.6364793 |
| GAFTTPQ | ASSQDWGSQNTLY | 0.5539766 |
| GFGGFGTAGGVGGA | ASSQDWGSQNTLY | 0.47039673 |
| GFLAQFL | ASSQDWGSQNTLY | 0.55104816 |

| | | |
|---|---|---|
| GGAGGAGGNAGVA | ASSQDWGSQNTLY | 0.5468013 |
| GGVGTAGAGGAGGAGG | ASSQDWGSQNTLY | 0.42662317 |
| GNGGGGDAGNAGSGGN | ASSQDWGSQNTLY | 0.6533939 |
| GTGGAGGAGNDAGS | ASSQDWGSQNTLY | 0.4647727 |
| GTTNTGWFNS | ASSQDWGSQNTLY | 0.6178201 |
| IFMGPLFTL | ASSQDWGSQNTLY | 0.70011425 |
| ILAHAIEAI | ASSQDWGSQNTLY | 0.40805784 |
| IRSAIREAL | ASSQDWGSQNTLY | 0.6505168 |
| ITFAGGG | ASSQDWGSQNTLY | 0.63293153 |
| ITGMSGHQEV | ASSQDWGSQNTLY | 0.63969135 |
| IVAGLACTL | ASSQDWGSQNTLY | 0.56993246 |
| IVGGGLAGA | ASSQDWGSQNTLY | 0.5127057 |
| IVTGVLNI | ASSQDWGSQNTLY | 0.6917012 |
| LAQQGCI | ASSQDWGSQNTLY | 0.6622883 |
| LCLHVEN | ASSQDWGSQNTLY | 0.5900971 |
| LDAISDGS | ASSQDWGSQNTLY | 0.6437453 |
| LGGFAAML | ASSQDWGSQNTLY | 0.6645156 |
| LGGKNPMI | ASSQDWGSQNTLY | 0.66614914 |
| LGPKGVAAIL | ASSQDWGSQNTLY | 0.5411571 |
| LMVGVSI | ASSQDWGSQNTLY | 0.3548591 |
| LVGAAFVL | ASSQDWGSQNTLY | 0.57611513 |
| LVLAVAPL | ASSQDWGSQNTLY | 0.5658958 |
| MAAIAVIDAA | ASSQDWGSQNTLY | 0.5072564 |
| NNGINAELQ | ASSQDWGSQNTLY | 0.3733685 |
| RGEPLVLVHG | ASSQDWGSQNTLY | 0.68146574 |
| RGGAGGAAGLI | ASSQDWGSQNTLY | 0.7136779 |
| RSYRMHPAV | ASSQDWGSQNTLY | 0.5301299 |
| SAAHLLAGQLA | ASSQDWGSQNTLY | 0.7112713 |
| SAAVGSY | ASSQDWGSQNTLY | 0.39012006 |
| SPHAVLHRI | ASSQDWGSQNTLY | 0.65658367 |
| SQIAKVL | ASSQDWGSQNTLY | 0.68015206 |
| TALGHLVP | ASSQDWGSQNTLY | 0.47905996 |
| TAPTTSGAA | ASSQDWGSQNTLY | 0.6495108 |
| TDGDDGNGGSPGT | ASSQDWGSQNTLY | 0.5247758 |
| TDGLGEN | ASSQDWGSQNTLY | 0.5164684 |
| TGGPPLPGQA | ASSQDWGSQNTLY | 0.68511754 |
| TIPAITI | ASSQDWGSQNTLY | 0.4894897 |
| TIPGFTI | ASSQDWGSQNTLY | 0.6198964 |
| TSIASGALA | ASSQDWGSQNTLY | 0.5132224 |
| TVTLPGAAI | ASSQDWGSQNTLY | 0.46729037 |
| VASGLSAALP | ASSQDWGSQNTLY | 0.62768793 |
| VGAGVIGI | ASSQDWGSQNTLY | 0.6544405 |
| VGPKRFGV | ASSQDWGSQNTLY | 0.56139547 |
| VNVAFPD | ASSQDWGSQNTLY | 0.7117006 |
| VSMTNSGVS | ASSQDWGSQNTLY | 0.5113859 |
| VTDVLGDLSV | ASSQDWGSQNTLY | 0.7139073 |
| VVVVGAGFAGL | ASSQDWGSQNTLY | 0.6852882 |
| VYTVGTVV | ASSQDWGSQNTLY | 0.6291506 |
| WTTFSGP | ASSQDWGSQNTLY | 0.5449448 |
| YDELVEAF | ASSQDWGSQNTLY | 0.6861992 |
| YGTGGAGGNGG | ASSQDWGSQNTLY | 0.5071267 |

| | | |
|---|---|---|
| YTNVMAV | ASSQDWGSQNTLY | 0.6249048 |
| FLEQVHQL | ASSQDWGSQNTLY | 0.69001496 |
| GLSLLAVAV | ASSQDWGSQNTLY | 0.7172048 |
| LEQVHQL | ASSQDWGSQNTLY | 0.7959183 |
| LNAAGAI | ASSQDWGSQNTLY | 0.7736545 |
| PAVHASL | ASSQDWGSQNTLY | 0.7873131 |
| TELEDDA | ASSQDWGSQNTLY | 0.74218094 |
| TPPGPPN | ASSQDWGSQNTLY | 0.52669305 |
| TPPPPPPP | ASSQDWGSQNTLY | 0.7776542 |
| TQSPALI | ASSLGGAKDTQY | 0.6933096 |
| EFAAAFPQ | ASSLGGAKDTQY | 0.78035444 |
| LLSGNLDVL | ASSLGGAKDTQY | 0.79203796 |
| GPANLGNYNI | ASSLGGAKDTQY | 0.7464678 |
| TLGPTLP | ASSLGGAKDTQY | 0.7931312 |
| VYAYSSCCK | ASSLGGAKDTQY | 0.79756224 |
| APAPAPAPNI | ASSLGGAKDTQY | 0.6106839 |
| PGGPVPIP | ASSLGGAKDTQY | 0.75731874 |
| FLTVALEP | ASSLGGAKDTQY | 0.7362895 |
| SQYFGANL | ASSLGGAKDTQY | 0.7683606 |
| GGVKPAA | ASSLGGAKDTQY | 0.41793656 |
| GGTGGAGGAGGW | ASSLGGAKDTQY | 0.7956476 |
| GVGGVAGLGGAGG | ASSLGGAKDTQY | 0.7357919 |
| VLLVLATTI | ASSLGGAKDTQY | 0.24633878 |
| IGIFAAML | ASSLGGAKDTQY | 0.7713 |
| AVGDLID | ASSLGGAKDTQY | 0.75434196 |
| NISPLPL | ASSLGGAKDTQY | 0.76361847 |
| VTAVFVAI | ASSLGGAKDTQY | 0.7901271 |
| SGDDNDAAL | ASSLGGAKDTQY | 0.520039 |
| AVAALAHS | ASSLGGAKDTQY | 0.7140008 |
| GGVTVMATP | ASSLGGAKDTQY | 0.7783866 |
| AGLAANVRD | ASSLGGAKDTQY | 0.79803765 |
| TNRPDLIDPALLRPG | ASSLGGAKDTQY | 0.68421626 |
| LANLGVTAL | ASSLGGAKDTQY | 0.7743836 |
| TGERIEL | ASSLGGAKDTQY | 0.80348986 |
| NGEELAA | ASSLGGAKDTQY | 0.7783376 |
| VLSLVWG | ASSLGGAKDTQY | 0.79621035 |
| IGYALWL | ASSLGGAKDTQY | 0.80056167 |
| LSLLLAPL | ASSLGGAKDTQY | 0.73358744 |
| TLDEALK | ASSLGGAKDTQY | 0.7932459 |
| TDGAGTAEGG | ASSLGGAKDTQY | 0.7934112 |
| NAPGGAGGNGG | ASSLGGAKDTQY | 0.7585665 |
| VSVFMGMGGNF | ASSLGGAKDTQY | 0.76930463 |
| LASYLAAL | ASSLGGAKDTQY | 0.7281909 |
| LVGGTPAL | ASSLGGAKDTQY | 0.7592406 |
| VRNADVL | ASSLGGAKDTQY | 0.49761012 |
| TCSSPFLVSAM | ASSLGGAKDTQY | 0.77284133 |
| VHITDAE | ASSLGGAKDTQY | 0.79635054 |
| EYAAPSF | ASSLGGAKDTQY | 0.7638729 |
| VVDAFHGGL | ASSLGGAKDTQY | 0.6756706 |
| ASVVDPQ | ASSLGGAKDTQY | 0.70034575 |
| SYNEFGGLS | ASSLGGAKDTQY | 0.7515484 |

| | | |
|---|---|---|
| DPDGSVY | ASSLGGAKDTQY | 0.76370525 |
| MVSTIHATE | ASSLGGAKDTQY | 0.4317065 |
| LAAGVADPV | ASSLGGAKDTQY | 0.70635957 |
| TKEQIAA | ASSLGGAKDTQY | 0.7972034 |
| SLASPAAL | ASSLGGAKDTQY | 0.79972094 |
| IGLVFYGV | ASSLGGAKDTQY | 0.59957653 |
| FLLDASNL | ASSLGGAKDTQY | 0.7949271 |
| YQVGNFP | ASSLGGAKDTQY | 0.78944165 |
| ESLLSQST | ASSLGGAKDTQY | 0.8011476 |
| LFTVGFAAVG | ASSLGGAKDTQY | 0.77650356 |
| ATPPTPT | ASSLGGAKDTQY | 0.70251155 |
| TDLATLG | ASSLGGAKDTQY | 0.79177314 |
| SAGGTGR | ASSLGGAKDTQY | 0.7461234 |
| GFSNSGS | ASSLGGAKDTQY | 0.72028214 |
| GGNGGSGGTSV | ASSLGGAKDTQY | 0.79306436 |
| GTSVATGGAGN | ASSLGGAKDTQY | 0.7463 |
| NGGNGGSGGTS | ASSLGGAKDTQY | 0.7216182 |
| AVVKFPH | ASSLGGAKDTQY | 0.7587037 |
| TPPPPPP | ASSLGGAKDTQY | 0.79695785 |
| AAPAAAAAPA | ASSLGGAKDTQY | 0.687935 |
| IALNEDL | ASSLGGAKDTQY | 0.69474506 |
| IVNVSSM | ASSLGGAKDTQY | 0.7261831 |
| TGIPPAPR | ASSLGGAKDTQY | 0.77143145 |
| FDLGGGTF | ASSLGGAKDTQY | 0.7976773 |
| FDLGGGTFDVS | ASSLGGAKDTQY | 0.7874733 |
| NEPTAAA | ASSLGGAKDTQY | 0.7467048 |
| NEPTAAALA | ASSLGGAKDTQY | 0.7843656 |
| GSIINISSI | ASSLGGAKDTQY | 0.79349613 |
| VAPGNAGTA | ASSLGGAKDTQY | 0.7765054 |
| TGGGASLE | ASSLGGAKDTQY | 0.7871783 |
| VAVYNHYKRI | ASSLGGAKDTQY | 0.707713 |
| TDPAPATT | ASSLGGAKDTQY | 0.79158765 |
| TNRPDILDPALLRPG | ASSLGGAKDTQY | 0.7917666 |
| LTTTTTS | ASSLGGAKDTQY | 0.7957866 |
| PTGSGKTL | ASSLGGAKDTQY | 0.79186773 |
| TGSGKTL | ASSLGGAKDTQY | 0.7727978 |
| TIGHVDHGK | ASSLGGAKDTQY | 0.7817678 |
| TIGHVDHGKTT | ASSLGGAKDTQY | 0.58467054 |
| GHVDHGKTT | ASSLGGAKDTQY | 0.7470075 |
| AQFEHTLL | ASSLGGAKDTQY | 0.7715852 |
| IVNQDEL | ASSLGGAKDTQY | 0.775462 |
| TNGSQFF | ASSLGGAKDTQY | 0.7834469 |
| TGKGIVE | ASSLGGAKDTQY | 0.7966918 |
| LISNASDA | ASSLGGAKDTQY | 0.72595847 |
| AELRTLGELTKIAKAHGAQ | ASSLGGAKDTQY | 0.796696 |
| APNAVAAAEPSW | ASSLGGAKDTQY | 0.79435766 |
| APQVAGAATAVL | ASSLGGAKDTQY | 0.6333251 |
| ARRDPAAAAAILPTDA | ASSLGGAKDTQY | 0.753258 |
| DAAARDVLAEAGLAEYF | ASSLGGAKDTQY | 0.7992927 |
| DDQIVLSAKANLAKVEQ | ASSLGGAKDTQY | 0.78242856 |
| DELLPDTGEDSDEDGH | ASSLGGAKDTQY | 0.75759906 |

| | | |
|---|---|---|
| ELPEAPANRATVPQD | ASSLGGAKDTQY | 0.76417106 |
| FGQSKIERVTGIG | ASSLGGAKDTQY | 0.79651904 |
| GAGGTGSTAGGAGGAGGAGGLY | ASSLGGAKDTQY | 0.7318279 |
| GHGGTGAAGGNGGTGGAGGSGI | ASSLGGAKDTQY | 0.7999 |
| GNGGNGGAGGEGGPGGAGGAGA | ASSLGGAKDTQY | 0.47150165 |
| GSTAGNGGNGGNATNGTIAGQP | ASSLGGAKDTQY | 0.7629755 |
| GTGGDGGNGGNGGTGG | ASSLGGAKDTQY | 0.72361106 |
| GVRRPAGIPADAPV | ASSLGGAKDTQY | 0.4408455 |
| GYRQVTDAHLLTLARR | ASSLGGAKDTQY | 0.7922275 |
| IAKMGQDGHDRG | ASSLGGAKDTQY | 0.7480153 |
| KAEPSIEIQYT | ASSLGGAKDTQY | 0.79233 |
| KPVRIRMAADEDTVAVT | ASSLGGAKDTQY | 0.7614391 |
| MPPRIAGMRLLVIKPEP | ASSLGGAKDTQY | 0.79413223 |
| MKFYAVALFRSRGVPPTR | ASSLGGAKDTQY | 0.76838225 |
| NAGGFGILGAVAHSP | ASSLGGAKDTQY | 0.76938474 |
| NEGREITLAMAEM | ASSLGGAKDTQY | 0.7000673 |
| NELVEWSHPLGHRWRWE | ASSLGGAKDTQY | 0.79207337 |
| NLGAGGTGGNGGNPG | ASSLGGAKDTQY | 0.7937051 |
| NTEWPKWETVMAADPVD | ASSLGGAKDTQY | 0.7906345 |
| PASAAPAETQP | ASSLGGAKDTQY | 0.6128338 |
| QQSSDGKNGVRAAEITGE | ASSLGGAKDTQY | 0.7409488 |
| RAPDAVTTSWWKE | ASSLGGAKDTQY | 0.7798144 |
| SGFGNVGAHTSGWFNQ | ASSLGGAKDTQY | 0.31764567 |
| SGVAGAVGQAASVGGLKV | ASSLGGAKDTQY | 0.27702385 |
| SNGGAGTGSAGGNGGT | ASSLGGAKDTQY | 0.68934953 |
| TGGAGGEGGAAG | ASSLGGAKDTQY | 0.6976491 |
| TGGDGGAAGAGGAPGHG | ASSLGGAKDTQY | 0.7903899 |
| TMPPLLAAIANAGRHGV | ASSLGGAKDTQY | 0.7827537 |
| TSSTAGNSGGA | ASSLGGAKDTQY | 0.7686675 |
| VPATVPVGILC | ASSLGGAKDTQY | 0.78200567 |
| VQRPDSGGVVLTGRIS | ASSLGGAKDTQY | 0.76815015 |
| WTPNPVPATKGVPAGLVQAT | ASSLGGAKDTQY | 0.18048461 |
| YALKRALAKFSDRFDVVII | ASSLGGAKDTQY | 0.74954236 |
| YLIGSVIV | ASSLGGAKDTQY | 0.6687244 |
| YGQLVAVID | ASSLGGAKDTQY | 0.80091155 |
| VTVGGMTLPA | ASSLGGAKDTQY | 0.67187953 |
| VLALPAV | ASSLGGAKDTQY | 0.71039784 |
| VIPPTLNYET | ASSLGGAKDTQY | 0.7740524 |
| VGMGGMHPSSG | ASSLGGAKDTQY | 0.7707406 |
| VGLAGGAATA | ASSLGGAKDTQY | 0.7722673 |
| VGGAGGAGGSGP | ASSLGGAKDTQY | 0.6850341 |
| VFAAFMV | ASSLGGAKDTQY | 0.25381756 |
| TLYDKLR | ASSLGGAKDTQY | 0.7956201 |
| TLAVFTVL | ASSLGGAKDTQY | 0.6304209 |
| TGSVQPARV | ASSLGGAKDTQY | 0.7118229 |
| TGGGAGSGGT | ASSLGGAKDTQY | 0.4324214 |
| SHLETSL | ASSLGGAKDTQY | 0.30684355 |
| SFTTGMT | ASSLGGAKDTQY | 0.62537014 |
| QVEVAKRMGVRQPTVS | ASSLGGAKDTQY | 0.795768 |
| QGSGGPALAM | ASSLGGAKDTQY | 0.7335613 |
| MLAPSNP | ASSLGGAKDTQY | 0.7410418 |

| | | |
|---|---|---|
| LVNNAAI | ASSLGGAKDTQY | 0.7938335 |
| LTGEINTLA | ASSLGGAKDTQY | 0.7708203 |
| LSIFGPV | ASSLGGAKDTQY | 0.75225997 |
| LLALLANL | ASSLGGAKDTQY | 0.7284304 |
| LISNASDALDKLR | ASSLGGAKDTQY | 0.71091866 |
| LINEAVP | ASSLGGAKDTQY | 0.7122439 |
| LGYAFSGL | ASSLGGAKDTQY | 0.7133616 |
| LDFVAVAL | ASSLGGAKDTQY | 0.79790974 |
| LAQPGAE | ASSLGGAKDTQY | 0.75457585 |
| ISNLGAVTGF | ASSLGGAKDTQY | 0.66271937 |
| HQEDPGRI | ASSLGGAKDTQY | 0.78672194 |
| GTPGEPTP | ASSLGGAKDTQY | 0.799644 |
| GFFNSGS | ASSLGGAKDTQY | 0.80180717 |
| GAGGGGLTAGAN | ASSLGGAKDTQY | 0.68986905 |
| FVDALVNM | ASSLGGAKDTQY | 0.7803842 |
| FLTVAPEL | ASSLGGAKDTQY | 0.7639942 |
| FETLTAL | ASSLGGAKDTQY | 0.7604002 |
| DVINPIEL | ASSLGGAKDTQY | 0.74579024 |
| APVSGAPV | ASSLGGAKDTQY | 0.59891975 |
| ALVGGLAW | ASSLGGAKDTQY | 0.7919541 |
| ALAAAAATAI | ASSLGGAKDTQY | 0.80555284 |
| AFGLGLLL | ASSLGGAKDTQY | 0.7582247 |
| AFFAAASL | ASSLGGAKDTQY | 0.68750525 |
| AAAALPAFSPP | ASSLGGAKDTQY | 0.7623912 |
| VVLAPAL | ASSLGGAKDTQY | 0.7279286 |
| TIGHVDHGKTTL | ASSLGGAKDTQY | 0.7377971 |
| LISNASDALDK | ASSLGGAKDTQY | 0.776773 |
| LISNASDAL | ASSLGGAKDTQY | 0.6053152 |
| ISNASDALDK | ASSLGGAKDTQY | 0.7554041 |
| NQAETLVYQTE | ASSLGGAKDTQY | 0.802967 |
| TGGTGGTAGAG | ASSLGGAKDTQY | 0.75499636 |
| TGSTAGGAGGA | ASSLGGAKDTQY | 0.77206904 |
| DPLAVPVAAGPV | ASSLGGAKDTQY | 0.7931612 |
| GGTGGTPGNGGN | ASSLGGAKDTQY | 0.7803717 |
| PAGLAGVDPATL | ASSLGGAKDTQY | 0.79600513 |
| GNNLVVEAPHAGASV | ASSLGGAKDTQY | 0.7900497 |
| KVTRIMSASTLEQPA | ASSLGGAKDTQY | 0.7610137 |
| LGFALPIALPRRTVV | ASSLGGAKDTQY | 0.76961243 |
| TRLGNLTARKLVVLV | ASSLGGAKDTQY | 0.76854134 |
| WSQQRRLIYFPSAGP | ASSLGGAKDTQY | 0.76762456 |
| GPNGNPDPMAAAVDIR | ASSLGGAKDTQY | 0.7715081 |
| TQRCVSVADLRAAASR | ASSLGGAKDTQY | 0.7451954 |
| QFVLDVAEHPVEVLTETV | ASSLGGAKDTQY | 0.79477507 |
| GGAGGTGGAGGNGGTQAGDG | ASSLGGAKDTQY | 0.7713701 |
| PLLATLRAIAVDVDAKPAQV | ASSLGGAKDTQY | 0.79917204 |
| APKFRTGGYMIDFWGVGYQVAK | ASSLGGAKDTQY | 0.6484086 |
| GAGGAGGAGGAAGGPGGTGGTG | ASSLGGAKDTQY | 0.67928994 |
| IGALGLLATDVAPAGSVAFGVA | ASSLGGAKDTQY | 0.7918888 |
| AGGAGGIGGNGGAGGAGGVLMG | ASSLGGAKDTQY | 0.28095496 |
| ADELIGLL | ASSLGGAKDTQY | 0.34997374 |
| ADRALKA | ASSLGGAKDTQY | 0.7721933 |

| | | |
|---|---|---|
| AGAVGQVGGAGGA | ASSLGGAKDTQY | 0.79822326 |
| AGGGGGLGGAP | ASSLGGAKDTQY | 0.21697916 |
| AGGLLYGNGGAG | ASSLGGAKDTQY | 0.69193804 |
| AGLESIGL | ASSLGGAKDTQY | 0.6587171 |
| AGLKTGII | ASSLGGAKDTQY | 0.7935313 |
| ASLAALLL | ASSLGGAKDTQY | 0.7748294 |
| ASVINISSI | ASSLGGAKDTQY | 0.5358995 |
| ATPPGPA | ASSLGGAKDTQY | 0.7961247 |
| ATVASALV | ASSLGGAKDTQY | 0.61503536 |
| ATVASLIL | ASSLGGAKDTQY | 0.5408969 |
| AVNGPVLIRH | ASSLGGAKDTQY | 0.79212046 |
| DGAAGAVGI | ASSLGGAKDTQY | 0.7976332 |
| DPLLHGI | ASSLGGAKDTQY | 0.65655607 |
| ELGAALGAL | ASSLGGAKDTQY | 0.7946955 |
| ELGRTQAV | ASSLGGAKDTQY | 0.75056285 |
| FVAGGYLS | ASSLGGAKDTQY | 0.8019891 |
| GASGGAGGAGGW | ASSLGGAKDTQY | 0.7543112 |
| GGAGAGGDGGAGGI | ASSLGGAKDTQY | 0.5742303 |
| GTGHNGSQPGMGGNG | ASSLGGAKDTQY | 0.79231185 |
| GYAAASASAS | ASSLGGAKDTQY | 0.7812083 |
| HQVDVAAVQQG | ASSLGGAKDTQY | 0.78986996 |
| IGAAIAHT | ASSLGGAKDTQY | 0.78728527 |
| IGMIEVAQL | ASSLGGAKDTQY | 0.7397343 |
| IRLAPEHTSAMD | ASSLGGAKDTQY | 0.75811964 |
| IYNVSVL | ASSLGGAKDTQY | 0.79268676 |
| KQGFLGM | ASSLGGAKDTQY | 0.7887714 |
| LESVITQL | ASSLGGAKDTQY | 0.5677031 |
| LGLGALVVV | ASSLGGAKDTQY | 0.79435784 |
| LIGNGGNGGSGGI | ASSLGGAKDTQY | 0.7683039 |
| LNAINAPTL | ASSLGGAKDTQY | 0.7358617 |
| LQLLNTGV | ASSLGGAKDTQY | 0.8008483 |
| LQTSIEF | ASSLGGAKDTQY | 0.7781116 |
| MSFAVSEL | ASSLGGAKDTQY | 0.7919377 |
| NTGALGL | ASSLGGAKDTQY | 0.7958915 |
| PPGAGGAGGAGTT | ASSLGGAKDTQY | 0.8037144 |
| QLVGLGI | ASSLGGAKDTQY | 0.5513802 |
| RFLEQVHQL | ASSLGGAKDTQY | 0.7772428 |
| RGVGYVLR | ASSLGGAKDTQY | 0.76961374 |
| SATGVFAGVI | ASSLGGAKDTQY | 0.7919426 |
| SLPAITLP | ASSLGGAKDTQY | 0.76568216 |
| STALAFTV | ASSLGGAKDTQY | 0.65905786 |
| SVLTVGLGK | ASSLGGAKDTQY | 0.7886048 |
| TALAGAGSGI | ASSLGGAKDTQY | 0.7966758 |
| TGLPFTGI | ASSLGGAKDTQY | 0.76938623 |
| VAEHVTV | ASSLGGAKDTQY | 0.78298825 |
| VDIVLNSL | ASSLGGAKDTQY | 0.80146646 |
| VLTGGGAVL | ASSLGGAKDTQY | 0.71579075 |
| VNVGLAYGI | ASSLGGAKDTQY | 0.7678884 |
| VSSAAVGAV | ASSLGGAKDTQY | 0.7927934 |
| VTSVYEAGV | ASSLGGAKDTQY | 0.79243386 |
| WVALDRGA | ASSLGGAKDTQY | 0.7692443 |

| | | |
|---|---|---|
| YLVGSVVI | ASSLGGAKDTQY | 0.79495996 |
| GPPGTGKTLL | ASSLGGAKDTQY | 0.6856376 |
| GVGMTKF | ASSLGGAKDTQY | 0.78158617 |
| IGIGGSDL | ASSLGGAKDTQY | 0.70700747 |
| IGTLLAEL | ASSLGGAKDTQY | 0.77674866 |
| IYTSGTT | ASSLGGAKDTQY | 0.63565457 |
| LSFPAGV | ASSLGGAKDTQY | 0.7790654 |
| LYTSGTT | ASSLGGAKDTQY | 0.78691936 |
| VGAFPFGF | ASSLGGAKDTQY | 0.7847099 |
| VGAGTVEF | ASSLGGAKDTQY | 0.3687238 |
| VGKSTLLL | ASSLGGAKDTQY | 0.7827777 |
| AALAHAGGGIV | ASSLGGAKDTQY | 0.7896783 |
| AALVAVSLL | ASSLGGAKDTQY | 0.80383337 |
| AAVSRAVAAL | ASSLGGAKDTQY | 0.77243173 |
| AGGSSGIGGF | ASSLGGAKDTQY | 0.7955729 |
| AGLTVANANG | ASSLGGAKDTQY | 0.74698746 |
| AGLVGAVL | ASSLGGAKDTQY | 0.7698721 |
| ALSQAFTV | ASSLGGAKDTQY | 0.7056681 |
| ALTFSVSL | ASSLGGAKDTQY | 0.75113773 |
| ALTGAALVAL | ASSLGGAKDTQY | 0.7330577 |
| ASFSIGAGVL | ASSLGGAKDTQY | 0.71540046 |
| DGSVYKI | ASSLGGAKDTQY | 0.78633904 |
| EQLAEIV | ASSLGGAKDTQY | 0.7906716 |
| FAVSELTVM | ASSLGGAKDTQY | 0.7920801 |
| GAAPLAAGL | ASSLGGAKDTQY | 0.7947308 |
| GADGGDGGSGGAGG | ASSLGGAKDTQY | 0.7943746 |
| GFGLDTGGAGG | ASSLGGAKDTQY | 0.7964827 |
| GLASGGLL | ASSLGGAKDTQY | 0.7491113 |
| GRIAGAAVLL | ASSLGGAKDTQY | 0.75456184 |
| GSAGSLNTSF | ASSLGGAKDTQY | 0.6660327 |
| GTGQIGI | ASSLGGAKDTQY | 0.74625385 |
| GTLIALIL | ASSLGGAKDTQY | 0.7481928 |
| IAATMGFVVG | ASSLGGAKDTQY | 0.77424824 |
| IFAQALL | ASSLGGAKDTQY | 0.2041173 |
| IGAAVALGF | ASSLGGAKDTQY | 0.7651467 |
| IGLTPLAL | ASSLGGAKDTQY | 0.7963106 |
| ILGSNTSTL | ASSLGGAKDTQY | 0.7206573 |
| IVFSLTVL | ASSLGGAKDTQY | 0.79051125 |
| LASKATGF | ASSLGGAKDTQY | 0.76050544 |
| LASVALIL | ASSLGGAKDTQY | 0.72057027 |
| LFAVGLTL | ASSLGGAKDTQY | 0.7882527 |
| LFNGGIGTYI | ASSLGGAKDTQY | 0.73780084 |
| LTLGAGII | ASSLGGAKDTQY | 0.79508287 |
| LVAAALGDI | ASSLGGAKDTQY | 0.80107826 |
| NTAFVQLGI | ASSLGGAKDTQY | 0.77540195 |
| QLFSGAEL | ASSLGGAKDTQY | 0.7692126 |
| SAATFITL | ASSLGGAKDTQY | 0.781734 |
| SNYGLANLGV | ASSLGGAKDTQY | 0.5242348 |
| SYGGAYAVM | ASSLGGAKDTQY | 0.772225 |
| TTGVGGLTLG | ASSLGGAKDTQY | 0.190917 |
| TTQTIYI | ASSLGGAKDTQY | 0.7639138 |

| | | |
|---|---|---|
| VAAGAGLGAVI | ASSLGGAKDTQY | 0.7219125 |
| VAAVVTLL | ASSLGGAKDTQY | 0.7772719 |
| VGFLGGLI | ASSLGGAKDTQY | 0.7615633 |
| VGVIPILL | ASSLGGAKDTQY | 0.78601897 |
| VIFASSLI | ASSLGGAKDTQY | 0.7248673 |
| VKGGDGGAAAG | ASSLGGAKDTQY | 0.7601044 |
| VTKTFTNI | ASSLGGAKDTQY | 0.7352313 |
| VVAAAILL | ASSLGGAKDTQY | 0.65149987 |
| YAALDVEL | ASSLGGAKDTQY | 0.7855243 |
| LASVALLL | ASSLGGAKDTQY | 0.76896 |
| LDAVHRV | ASSLGGAKDTQY | 0.7427522 |
| QGLSSSIF | ASSLGGAKDTQY | 0.7959951 |
| RGTFANIRL | ASSLGGAKDTQY | 0.7832513 |
| RQLWWGHRI | ASSLGGAKDTQY | 0.5648423 |
| YGPPGTGKTLL | ASSLGGAKDTQY | 0.75360894 |
| AALVGAPANVA | ASSLGGAKDTQY | 0.7733178 |
| AATFRRTE | ASSLGGAKDTQY | 0.76637554 |
| AGALGGGI | ASSLGGAKDTQY | 0.19356404 |
| AGGAGGAGGIGAG | ASSLGGAKDTQY | 0.7767713 |
| AGSGALGL | ASSLGGAKDTQY | 0.73468256 |
| AQIADQI | ASSLGGAKDTQY | 0.6879732 |
| ASAVAVTYL | ASSLGGAKDTQY | 0.7417976 |
| ASTVLGGTGG | ASSLGGAKDTQY | 0.79327214 |
| AVGGEPAK | ASSLGGAKDTQY | 0.6196563 |
| CEQGTFGGTC | ASSLGGAKDTQY | 0.7792667 |
| DEGKAPAG | ASSLGGAKDTQY | 0.78718877 |
| DTVVFPAL | ASSLGGAKDTQY | 0.7891519 |
| FEGITMAG | ASSLGGAKDTQY | 0.7986102 |
| FPLVVQD | ASSLGGAKDTQY | 0.77987325 |
| GAFTTPQ | ASSLGGAKDTQY | 0.76495177 |
| GFGGFGTAGGVGGA | ASSLGGAKDTQY | 0.7823808 |
| GFLAQFL | ASSLGGAKDTQY | 0.78068453 |
| GGAGGAGGNAGVA | ASSLGGAKDTQY | 0.47590154 |
| GGVGTAGAGGAGGAGG | ASSLGGAKDTQY | 0.52480936 |
| GNGGGGDAGNAGSGGN | ASSLGGAKDTQY | 0.7760798 |
| GTGGAGGAGNDAGS | ASSLGGAKDTQY | 0.4915173 |
| GTTNTGWFNS | ASSLGGAKDTQY | 0.7794976 |
| IFMGPLFTL | ASSLGGAKDTQY | 0.80100834 |
| ILAHAIEAI | ASSLGGAKDTQY | 0.77015173 |
| IRSAIREAL | ASSLGGAKDTQY | 0.6481708 |
| ITFAGGG | ASSLGGAKDTQY | 0.75290877 |
| ITGMSGHQEV | ASSLGGAKDTQY | 0.7162028 |
| IVAGLACTL | ASSLGGAKDTQY | 0.7872853 |
| IVGGGLAGA | ASSLGGAKDTQY | 0.79252887 |
| IVTGVLNI | ASSLGGAKDTQY | 0.7994032 |
| LAQQGCI | ASSLGGAKDTQY | 0.7914388 |
| LCLHVEN | ASSLGGAKDTQY | 0.74411654 |
| LDAISDGS | ASSLGGAKDTQY | 0.7917114 |
| LGGFAAML | ASSLGGAKDTQY | 0.7607987 |
| LGGKNPMI | ASSLGGAKDTQY | 0.73297226 |
| LGPKGVAAIL | ASSLGGAKDTQY | 0.7699647 |

| | | |
|---|---|---|
| LMVGVSI | ASSLGGAKDTQY | 0.77497333 |
| LVGAAFVL | ASSLGGAKDTQY | 0.7644199 |
| LVLAVAPL | ASSLGGAKDTQY | 0.73175997 |
| MAAIAVIDAA | ASSLGGAKDTQY | 0.6382041 |
| NNGINAELQ | ASSLGGAKDTQY | 0.7833837 |
| RGEPLVLVHG | ASSLGGAKDTQY | 0.7783672 |
| RGGAGGAAGLI | ASSLGGAKDTQY | 0.5862703 |
| RSYRMHPAV | ASSLGGAKDTQY | 0.74408925 |
| SAAHLLAGQLA | ASSLGGAKDTQY | 0.79943764 |
| SAAVGSY | ASSLGGAKDTQY | 0.7995622 |
| SPHAVLHRI | ASSLGGAKDTQY | 0.73137534 |
| SQIAKVL | ASSLGGAKDTQY | 0.7258744 |
| TALGHLVP | ASSLGGAKDTQY | 0.76716524 |
| TAPTTSGAA | ASSLGGAKDTQY | 0.617581 |
| TDGDDGNGGSPGT | ASSLGGAKDTQY | 0.78293777 |
| TDGLGEN | ASSLGGAKDTQY | 0.7582139 |
| TGGPPLPGQA | ASSLGGAKDTQY | 0.75413823 |
| TIPAITI | ASSLGGAKDTQY | 0.73389137 |
| TIPGFTI | ASSLGGAKDTQY | 0.7730957 |
| TSIASGALA | ASSLGGAKDTQY | 0.7338488 |
| TVTLPGAAI | ASSLGGAKDTQY | 0.7946366 |
| VASGLSAALP | ASSLGGAKDTQY | 0.2398398 |
| VGAGVIGI | ASSLGGAKDTQY | 0.780727 |
| VGPKRFGV | ASSLGGAKDTQY | 0.69675654 |
| VNVAFPD | ASSLGGAKDTQY | 0.7221523 |
| VSMTNSGVS | ASSLGGAKDTQY | 0.47952878 |
| VTDVLGDLSV | ASSLGGAKDTQY | 0.77090144 |
| VVVVGAGFAGL | ASSLGGAKDTQY | 0.787248 |
| VYTVGTVV | ASSLGGAKDTQY | 0.7929922 |
| WTTFSGP | ASSLGGAKDTQY | 0.68171966 |
| YDELVEAF | ASSLGGAKDTQY | 0.786206 |
| YGTGGAGGNGG | ASSLGGAKDTQY | 0.79858875 |
| YTNVMAV | ASSLGGAKDTQY | 0.7323058 |
| FLEQVHQL | ASSLGGAKDTQY | 0.43077862 |
| GLSLLAVAV | ASSLGGAKDTQY | 0.78841966 |
| LEQVHQL | ASSLGGAKDTQY | 0.75172603 |
| LNAAGAI | ASSLGGAKDTQY | 0.79104275 |
| PAVHASL | ASSLGGAKDTQY | 0.78947663 |
| TELEDDA | ASSLGGAKDTQY | 0.658285 |
| TPPGPPN | ASSLGGAKDTQY | 0.8009323 |
| TPPPPPPP | ASSLGGAKDTQY | 0.7881808 |
| TQSPALI | ASSPQGSNERLF | 0.7086247 |
| EFAAAFPQ | ASSPQGSNERLF | 0.7352548 |
| LLSGNLDVL | ASSPQGSNERLF | 0.7064186 |
| GPANLGNYNI | ASSPQGSNERLF | 0.38479176 |
| TLGPTLP | ASSPQGSNERLF | 0.78862464 |
| VYAYSSCCK | ASSPQGSNERLF | 0.7281542 |
| APAPAPAPNI | ASSPQGSNERLF | 0.7502485 |
| PGGPVPIP | ASSPQGSNERLF | 0.7773093 |
| FLTVALEP | ASSPQGSNERLF | 0.7380249 |
| SQYFGANL | ASSPQGSNERLF | 0.7569572 |

| | | |
|---|---|---|
| GGVKPAA | ASSPQGSNERLF | 0.7443198 |
| GGTGGAGGAGGW | ASSPQGSNERLF | 0.7775737 |
| GVGGVAGLGGAGG | ASSPQGSNERLF | 0.7786509 |
| VLLVLATTI | ASSPQGSNERLF | 0.7687253 |
| IGIFAAML | ASSPQGSNERLF | 0.78186893 |
| AVGDLID | ASSPQGSNERLF | 0.7960056 |
| NISPLPL | ASSPQGSNERLF | 0.55196154 |
| VTAVFVAI | ASSPQGSNERLF | 0.7014995 |
| SGDDNDAAL | ASSPQGSNERLF | 0.78802776 |
| AVAALAHS | ASSPQGSNERLF | 0.65140116 |
| GGVTVMATP | ASSPQGSNERLF | 0.78880155 |
| AGLAANVRD | ASSPQGSNERLF | 0.7667413 |
| TNRPDLIDPALLRPG | ASSPQGSNERLF | 0.78009856 |
| LANLGVTAL | ASSPQGSNERLF | 0.763765 |
| TGERIEL | ASSPQGSNERLF | 0.79819226 |
| NGEELAA | ASSPQGSNERLF | 0.7910882 |
| VLSLVWG | ASSPQGSNERLF | 0.76093477 |
| IGYALWL | ASSPQGSNERLF | 0.73285484 |
| LSLLLAPL | ASSPQGSNERLF | 0.66202164 |
| TLDEALK | ASSPQGSNERLF | 0.7864717 |
| TDGAGTAEGG | ASSPQGSNERLF | 0.7932854 |
| NAPGGAGGNGG | ASSPQGSNERLF | 0.6837835 |
| VSVFMGMGGNF | ASSPQGSNERLF | 0.7592721 |
| LASYLAAL | ASSPQGSNERLF | 0.7987429 |
| LVGGTPAL | ASSPQGSNERLF | 0.7902546 |
| VRNADVL | ASSPQGSNERLF | 0.60193074 |
| TCSSPFLVSAM | ASSPQGSNERLF | 0.7463448 |
| VHITDAE | ASSPQGSNERLF | 0.57708484 |
| EYAAPSF | ASSPQGSNERLF | 0.75618714 |
| VVDAFHGGL | ASSPQGSNERLF | 0.6057615 |
| ASVVDPQ | ASSPQGSNERLF | 0.78418565 |
| SYNEFGGLS | ASSPQGSNERLF | 0.7525692 |
| DPDGSVY | ASSPQGSNERLF | 0.79354537 |
| MVSTIHATE | ASSPQGSNERLF | 0.77931356 |
| LAAGVADPV | ASSPQGSNERLF | 0.8002565 |
| TKEQIAA | ASSPQGSNERLF | 0.75028574 |
| SLASPAAL | ASSPQGSNERLF | 0.7474148 |
| IGLVFYGV | ASSPQGSNERLF | 0.57583165 |
| FLLDASNL | ASSPQGSNERLF | 0.64042574 |
| YQVGNFP | ASSPQGSNERLF | 0.7097734 |
| ESLLSQST | ASSPQGSNERLF | 0.7546797 |
| LFTVGFAAVG | ASSPQGSNERLF | 0.304701 |
| ATPPTPT | ASSPQGSNERLF | 0.79839367 |
| TDLATLG | ASSPQGSNERLF | 0.77620786 |
| SAGGTGR | ASSPQGSNERLF | 0.79504037 |
| GFSNSGS | ASSPQGSNERLF | 0.7557792 |
| GGNGGSGGTSV | ASSPQGSNERLF | 0.7489399 |
| GTSVATGGAGN | ASSPQGSNERLF | 0.7294472 |
| NGGNGGSGGTS | ASSPQGSNERLF | 0.7916467 |
| AVVKFPH | ASSPQGSNERLF | 0.64947724 |
| TPPPPPP | ASSPQGSNERLF | 0.5948919 |

| | | |
|---|---|---|
| AAPAAAAAPA | ASSPQGSNERLF | 0.7643769 |
| IALNEDL | ASSPQGSNERLF | 0.64038503 |
| IVNVSSM | ASSPQGSNERLF | 0.800503 |
| TGIPPAPR | ASSPQGSNERLF | 0.79296 |
| FDLGGGTF | ASSPQGSNERLF | 0.7762124 |
| FDLGGGTFDVS | ASSPQGSNERLF | 0.8001157 |
| NEPTAAA | ASSPQGSNERLF | 0.31914797 |
| NEPTAAALA | ASSPQGSNERLF | 0.7300397 |
| GSIINISSI | ASSPQGSNERLF | 0.80251896 |
| VAPGNAGTA | ASSPQGSNERLF | 0.7771156 |
| TGGGASLE | ASSPQGSNERLF | 0.7604674 |
| VAVYNHYKRI | ASSPQGSNERLF | 0.79686403 |
| TDPAPATT | ASSPQGSNERLF | 0.72794485 |
| TNRPDILDPALLRPG | ASSPQGSNERLF | 0.7694503 |
| LTTTTTS | ASSPQGSNERLF | 0.76458657 |
| PTGSGKTL | ASSPQGSNERLF | 0.7834013 |
| TGSGKTL | ASSPQGSNERLF | 0.76917076 |
| TIGHVDHGK | ASSPQGSNERLF | 0.69277275 |
| TIGHVDHGKTT | ASSPQGSNERLF | 0.79424304 |
| GHVDHGKTT | ASSPQGSNERLF | 0.7995225 |
| AQFEHTLL | ASSPQGSNERLF | 0.7515522 |
| IVNQDEL | ASSPQGSNERLF | 0.7933135 |
| TNGSQFF | ASSPQGSNERLF | 0.7868655 |
| TGKGIVE | ASSPQGSNERLF | 0.7849301 |
| LISNASDA | ASSPQGSNERLF | 0.7321037 |
| AELRTLGELTKIAKAHGAQ | ASSPQGSNERLF | 0.72102284 |
| APNAVAAAEPSW | ASSPQGSNERLF | 0.7622186 |
| APQVAGAATAVL | ASSPQGSNERLF | 0.6870614 |
| ARRDPAAAAAILPTDA | ASSPQGSNERLF | 0.7750713 |
| DAAARDVLAEAGLAEYF | ASSPQGSNERLF | 0.75726694 |
| DDQIVLSAKANLAKVEQ | ASSPQGSNERLF | 0.8040781 |
| DELLPDTGEDSDEDGH | ASSPQGSNERLF | 0.78817207 |
| ELPEAPANRATVPQD | ASSPQGSNERLF | 0.7815374 |
| FGQSKIERVTGIG | ASSPQGSNERLF | 0.6839291 |
| GAGGTGSTAGGAGGAGGAGGLY | ASSPQGSNERLF | 0.77441466 |
| GHGGTGAAGGNGGTGGAGGSGI | ASSPQGSNERLF | 0.7807622 |
| GNGGNGGAGGEGGPGGAGGAGA | ASSPQGSNERLF | 0.6127077 |
| GSTAGNGGNGGNATNGTIAGQP | ASSPQGSNERLF | 0.7975197 |
| GTGGDGGNGGNGGTGG | ASSPQGSNERLF | 0.73552155 |
| GVRRPAGIPADAPV | ASSPQGSNERLF | 0.75588393 |
| GYRQVTDAHLLTLARR | ASSPQGSNERLF | 0.79350245 |
| IAKMGQDGHDRG | ASSPQGSNERLF | 0.7994594 |
| KAEPSIEIQYT | ASSPQGSNERLF | 0.7584866 |
| KPVRIRMAADEDTVAVT | ASSPQGSNERLF | 0.75646615 |
| MPPRIAGMRLLVIKPEP | ASSPQGSNERLF | 0.6894007 |
| MKFYAVALFRSRGVPPTR | ASSPQGSNERLF | 0.7938665 |
| NAGGFGILGAVAHSP | ASSPQGSNERLF | 0.7942713 |
| NEGREITLAMAEM | ASSPQGSNERLF | 0.7604652 |
| NELVEWSHPLGHRWRWE | ASSPQGSNERLF | 0.8006643 |
| NLGAGGTGGNGGNPG | ASSPQGSNERLF | 0.78990185 |
| NTEWPKWETVMAADPVD | ASSPQGSNERLF | 0.53844315 |

| | | |
|---|---|---|
| PASAAPAETQP | ASSPQGSNERLF | 0.7756521 |
| QQSSDGKNGVRAAEITGE | ASSPQGSNERLF | 0.792426 |
| RAPDAVTTSWWKE | ASSPQGSNERLF | 0.78959775 |
| SGFGNVGAHTSGWFNQ | ASSPQGSNERLF | 0.80085635 |
| SGVAGAVGQAASVGGLKV | ASSPQGSNERLF | 0.76433563 |
| SNGGAGTGSAGGNGGT | ASSPQGSNERLF | 0.73463386 |
| TGGAGGEGGAAG | ASSPQGSNERLF | 0.79833627 |
| TGGDGGAAGAGGAPGHG | ASSPQGSNERLF | 0.80487645 |
| TMPPLLAAIANAGRHGV | ASSPQGSNERLF | 0.79564774 |
| TSSTAGNSGGA | ASSPQGSNERLF | 0.76453435 |
| VPATVPVGILC | ASSPQGSNERLF | 0.7480153 |
| VQRPDSGGVVLTGRIS | ASSPQGSNERLF | 0.7805958 |
| WTPNPVPATKGVPAGLVQAT | ASSPQGSNERLF | 0.78693706 |
| YALKRALAKFSDRFDVVII | ASSPQGSNERLF | 0.7094241 |
| YLIGSVIV | ASSPQGSNERLF | 0.77105695 |
| YGQLVAVID | ASSPQGSNERLF | 0.7800504 |
| VTVGGMTLPA | ASSPQGSNERLF | 0.5823742 |
| VLALPAV | ASSPQGSNERLF | 0.77923477 |
| VIPPTLNYET | ASSPQGSNERLF | 0.7875426 |
| VGMGGMHPSSG | ASSPQGSNERLF | 0.7693157 |
| VGLAGGAATA | ASSPQGSNERLF | 0.7402711 |
| VGGAGGAGGSGP | ASSPQGSNERLF | 0.7918842 |
| VFAAFMV | ASSPQGSNERLF | 0.80075014 |
| TLYDKLR | ASSPQGSNERLF | 0.80047774 |
| TLAVFTVL | ASSPQGSNERLF | 0.6077901 |
| TGSVQPARV | ASSPQGSNERLF | 0.46238863 |
| TGGGAGSGGT | ASSPQGSNERLF | 0.73671496 |
| SHLETSL | ASSPQGSNERLF | 0.7877791 |
| SFTTGMT | ASSPQGSNERLF | 0.7507718 |
| QVEVAKRMGVRQPTVS | ASSPQGSNERLF | 0.746277 |
| QGSGGPALAM | ASSPQGSNERLF | 0.7632329 |
| MLAPSNP | ASSPQGSNERLF | 0.79582524 |
| LVNNAAI | ASSPQGSNERLF | 0.7085697 |
| LTGEINTLA | ASSPQGSNERLF | 0.5871483 |
| LSIFGPV | ASSPQGSNERLF | 0.7757259 |
| LLALLANL | ASSPQGSNERLF | 0.7981545 |
| LISNASDALDKLR | ASSPQGSNERLF | 0.7087538 |
| LINEAVP | ASSPQGSNERLF | 0.7473675 |
| LGYAFSGL | ASSPQGSNERLF | 0.7457216 |
| LDFVAVAL | ASSPQGSNERLF | 0.69339585 |
| LAQPGAE | ASSPQGSNERLF | 0.77217174 |
| ISNLGAVTGF | ASSPQGSNERLF | 0.69498754 |
| HQEDPGRI | ASSPQGSNERLF | 0.7896249 |
| GTPGEPTP | ASSPQGSNERLF | 0.68127775 |
| GFFNSGS | ASSPQGSNERLF | 0.78808767 |
| GAGGGGLTAGAN | ASSPQGSNERLF | 0.77489257 |
| FVDALVNM | ASSPQGSNERLF | 0.7850605 |
| FLTVAPEL | ASSPQGSNERLF | 0.7525574 |
| FETLTAL | ASSPQGSNERLF | 0.7332474 |
| DVINPIEL | ASSPQGSNERLF | 0.65821713 |
| APVSGAPV | ASSPQGSNERLF | 0.5418903 |

| | | |
|---|---|---|
| ALVGGLAW | ASSPQGSNERLF | 0.79012537 |
| ALAAAAATAI | ASSPQGSNERLF | 0.25571346 |
| AFGLGLLL | ASSPQGSNERLF | 0.79432744 |
| AFFAAASL | ASSPQGSNERLF | 0.7893114 |
| AAAALPAFSPP | ASSPQGSNERLF | 0.68723667 |
| VVLAPAL | ASSPQGSNERLF | 0.7824998 |
| TIGHVDHGKTTL | ASSPQGSNERLF | 0.63134086 |
| LISNASDALDK | ASSPQGSNERLF | 0.7196542 |
| LISNASDAL | ASSPQGSNERLF | 0.789106 |
| ISNASDALDK | ASSPQGSNERLF | 0.79346657 |
| NQAETLVYQTE | ASSPQGSNERLF | 0.72523344 |
| TGGTGGTAGAG | ASSPQGSNERLF | 0.7850182 |
| TGSTAGGAGGA | ASSPQGSNERLF | 0.7772117 |
| DPLAVPVAAGPV | ASSPQGSNERLF | 0.7410437 |
| GGTGGTPGNGGN | ASSPQGSNERLF | 0.77150744 |
| PAGLAGVDPATL | ASSPQGSNERLF | 0.79303944 |
| GNNLVVEAPHAGASV | ASSPQGSNERLF | 0.78462094 |
| KVTRIMSASTLEQPA | ASSPQGSNERLF | 0.71679485 |
| LGFALPIALPRRTVV | ASSPQGSNERLF | 0.78492206 |
| TRLGNLTARKLVVLV | ASSPQGSNERLF | 0.7042229 |
| WSQQRRLIYFPSAGP | ASSPQGSNERLF | 0.7793385 |
| GPNGNPDPMAAAVDIR | ASSPQGSNERLF | 0.7646259 |
| TQRCVSVADLRAAASR | ASSPQGSNERLF | 0.7680895 |
| QFVLDVAEHPVEVLTETV | ASSPQGSNERLF | 0.78934544 |
| GGAGGTGGAGGNGGTQAGDG | ASSPQGSNERLF | 0.7004609 |
| PLLATLRAIAVDVDAKPAQV | ASSPQGSNERLF | 0.71798897 |
| APKFRTGGYMIDFWGVGYQVAK | ASSPQGSNERLF | 0.7624373 |
| GAGGAGGAGGAAGGPGGTGGTG | ASSPQGSNERLF | 0.7749326 |
| IGALGLLATDVAPAGSVAFGVA | ASSPQGSNERLF | 0.79180264 |
| AGGAGGIGGNGGAGGAGGVLMG | ASSPQGSNERLF | 0.704798 |
| ADELIGLL | ASSPQGSNERLF | 0.7783228 |
| ADRALKA | ASSPQGSNERLF | 0.7629964 |
| AGAVGQVGGAGGA | ASSPQGSNERLF | 0.7279935 |
| AGGGGGLGGAP | ASSPQGSNERLF | 0.77913797 |
| AGGLLYGNGGAG | ASSPQGSNERLF | 0.76159567 |
| AGLESIGL | ASSPQGSNERLF | 0.70923007 |
| AGLKTGII | ASSPQGSNERLF | 0.78619355 |
| ASLAALLL | ASSPQGSNERLF | 0.67724967 |
| ASVINISSI | ASSPQGSNERLF | 0.7896166 |
| ATPPGPA | ASSPQGSNERLF | 0.7906117 |
| ATVASALV | ASSPQGSNERLF | 0.7553781 |
| ATVASLIL | ASSPQGSNERLF | 0.7856668 |
| AVNGPVLIRH | ASSPQGSNERLF | 0.77651334 |
| DGAAGAVGI | ASSPQGSNERLF | 0.7706672 |
| DPLLHGI | ASSPQGSNERLF | 0.7506175 |
| ELGAALGAL | ASSPQGSNERLF | 0.7204882 |
| ELGRTQAV | ASSPQGSNERLF | 0.7955272 |
| FVAGGYLS | ASSPQGSNERLF | 0.7912569 |
| GASGGAGGAGGW | ASSPQGSNERLF | 0.7866576 |
| GGAGAGGDGGAGGI | ASSPQGSNERLF | 0.7792258 |
| GTGHNGSQPGMGGNG | ASSPQGSNERLF | 0.785233 |

| | | |
|---|---|---|
| GYAAASASAS | ASSPQGSNERLF | 0.793386 |
| HQVDVAAVQQG | ASSPQGSNERLF | 0.7375382 |
| IGAAIAHT | ASSPQGSNERLF | 0.6591667 |
| IGMIEVAQL | ASSPQGSNERLF | 0.75006783 |
| IRLAPEHTSAMD | ASSPQGSNERLF | 0.7153607 |
| IYNVSVL | ASSPQGSNERLF | 0.75734955 |
| KQGFLGM | ASSPQGSNERLF | 0.549279 |
| LESVITQL | ASSPQGSNERLF | 0.7940335 |
| LGLGALVVV | ASSPQGSNERLF | 0.80309194 |
| LIGNGGNGGSGGI | ASSPQGSNERLF | 0.7153883 |
| LNAINAPTL | ASSPQGSNERLF | 0.8012566 |
| LQLLNTGV | ASSPQGSNERLF | 0.6707449 |
| LQTSIEF | ASSPQGSNERLF | 0.7924367 |
| MSFAVSEL | ASSPQGSNERLF | 0.7957738 |
| NTGALGL | ASSPQGSNERLF | 0.70522213 |
| PPGAGGAGGAGTT | ASSPQGSNERLF | 0.7875619 |
| QLVGLGI | ASSPQGSNERLF | 0.72440994 |
| RFLEQVHQL | ASSPQGSNERLF | 0.74586976 |
| RGVGYVLR | ASSPQGSNERLF | 0.79328054 |
| SATGVFAGVI | ASSPQGSNERLF | 0.7597215 |
| SLPAITLP | ASSPQGSNERLF | 0.7758305 |
| STALAFTV | ASSPQGSNERLF | 0.77452934 |
| SVLTVGLGK | ASSPQGSNERLF | 0.7256113 |
| TALAGAGSGI | ASSPQGSNERLF | 0.7804328 |
| TGLPFTGI | ASSPQGSNERLF | 0.7934303 |
| VAEHVTV | ASSPQGSNERLF | 0.72874314 |
| VDIVLNSL | ASSPQGSNERLF | 0.80503416 |
| VLTGGGAVL | ASSPQGSNERLF | 0.7090956 |
| VNVGLAYGI | ASSPQGSNERLF | 0.79985845 |
| VSSAAVGAV | ASSPQGSNERLF | 0.79246867 |
| VTSVYEAGV | ASSPQGSNERLF | 0.7668176 |
| WVALDRGA | ASSPQGSNERLF | 0.76380515 |
| YLVGSVVI | ASSPQGSNERLF | 0.79339695 |
| GPPGTGKTLL | ASSPQGSNERLF | 0.7686756 |
| GVGMTKF | ASSPQGSNERLF | 0.7853744 |
| IGIGGSDL | ASSPQGSNERLF | 0.79926455 |
| IGTLLAEL | ASSPQGSNERLF | 0.3785288 |
| IYTSGTT | ASSPQGSNERLF | 0.73055035 |
| LSFPAGV | ASSPQGSNERLF | 0.64250016 |
| LYTSGTT | ASSPQGSNERLF | 0.681201 |
| VGAFPFGF | ASSPQGSNERLF | 0.55257404 |
| VGAGTVEF | ASSPQGSNERLF | 0.40672073 |
| VGKSTLLL | ASSPQGSNERLF | 0.6092941 |
| AALAHAGGGIV | ASSPQGSNERLF | 0.45991012 |
| AALVAVSLL | ASSPQGSNERLF | 0.72103405 |
| AAVSRAVAAL | ASSPQGSNERLF | 0.7192239 |
| AGGSSGIGGF | ASSPQGSNERLF | 0.50231075 |
| AGLTVANANG | ASSPQGSNERLF | 0.71393406 |
| AGLVGAVL | ASSPQGSNERLF | 0.7159705 |
| ALSQAFTV | ASSPQGSNERLF | 0.3752835 |
| ALTFSVSL | ASSPQGSNERLF | 0.6275706 |

| | | |
|---|---|---|
| ALTGAALVAL | ASSPQGSNERLF | 0.5914081 |
| ASFSIGAGVL | ASSPQGSNERLF | 0.5880771 |
| DGSVYKI | ASSPQGSNERLF | 0.30787194 |
| EQLAEIV | ASSPQGSNERLF | 0.6919214 |
| FAVSELTVM | ASSPQGSNERLF | 0.5137009 |
| GAAPLAAGL | ASSPQGSNERLF | 0.2312513 |
| GADGGDGGSGGAGG | ASSPQGSNERLF | 0.67503995 |
| GFGLDTGGAGG | ASSPQGSNERLF | 0.5893496 |
| GLASGGLL | ASSPQGSNERLF | 0.5480878 |
| GRIAGAAVLL | ASSPQGSNERLF | 0.70270544 |
| GSAGSLNTSF | ASSPQGSNERLF | 0.53690916 |
| GTGQIGI | ASSPQGSNERLF | 0.5408884 |
| GTLIALIL | ASSPQGSNERLF | 0.6896943 |
| IAATMGFVVG | ASSPQGSNERLF | 0.72617215 |
| IFAQALL | ASSPQGSNERLF | 0.4634217 |
| IGAAVALGF | ASSPQGSNERLF | 0.6431825 |
| IGLTPLAL | ASSPQGSNERLF | 0.73749375 |
| ILGSNTSTL | ASSPQGSNERLF | 0.6249817 |
| IVFSLTVL | ASSPQGSNERLF | 0.73710024 |
| LASKATGF | ASSPQGSNERLF | 0.7430701 |
| LASVALIL | ASSPQGSNERLF | 0.55425 |
| LFAVGLTL | ASSPQGSNERLF | 0.72288954 |
| LFNGGIGTYI | ASSPQGSNERLF | 0.7548795 |
| LTLGAGII | ASSPQGSNERLF | 0.60372305 |
| LVAAALGDI | ASSPQGSNERLF | 0.62285924 |
| NTAFVQLGI | ASSPQGSNERLF | 0.36811876 |
| QLFSGAEL | ASSPQGSNERLF | 0.62454116 |
| SAATFITL | ASSPQGSNERLF | 0.36859652 |
| SNYGLANLGV | ASSPQGSNERLF | 0.6562008 |
| SYGGAYAVM | ASSPQGSNERLF | 0.7393911 |
| TTGVGGLTLG | ASSPQGSNERLF | 0.60470974 |
| TTQTIYI | ASSPQGSNERLF | 0.34070864 |
| VAAGAGLGAVI | ASSPQGSNERLF | 0.43888783 |
| VAAVVTLL | ASSPQGSNERLF | 0.6422292 |
| VGFLGGLI | ASSPQGSNERLF | 0.42638105 |
| VGVIPILL | ASSPQGSNERLF | 0.36605036 |
| VIFASSLI | ASSPQGSNERLF | 0.4743107 |
| VKGGDGGAAAG | ASSPQGSNERLF | 0.72355783 |
| VTKTFTNI | ASSPQGSNERLF | 0.7267928 |
| VVAAAILL | ASSPQGSNERLF | 0.45114887 |
| YAALDVEL | ASSPQGSNERLF | 0.7404022 |
| LASVALLL | ASSPQGSNERLF | 0.702797 |
| LDAVHRV | ASSPQGSNERLF | 0.74561983 |
| QGLSSSIF | ASSPQGSNERLF | 0.6563653 |
| RGTFANIRL | ASSPQGSNERLF | 0.3745688 |
| RQLWWGHRI | ASSPQGSNERLF | 0.7274006 |
| YGPPGTGKTLL | ASSPQGSNERLF | 0.5474746 |
| AALVGAPANVA | ASSPQGSNERLF | 0.6486318 |
| AATFRRTE | ASSPQGSNERLF | 0.7046486 |
| AGALGGGI | ASSPQGSNERLF | 0.61538905 |
| AGGAGGAGGIGAG | ASSPQGSNERLF | 0.51376355 |

| | | |
|---|---|---|
| AGSGALGL | ASSPQGSNERLF | 0.6428585 |
| AQIADQI | ASSPQGSNERLF | 0.73540163 |
| ASAVAVTYL | ASSPQGSNERLF | 0.49189374 |
| ASTVLGGTGG | ASSPQGSNERLF | 0.40378585 |
| AVGGEPAK | ASSPQGSNERLF | 0.5683286 |
| CEQGTFGGTC | ASSPQGSNERLF | 0.6013216 |
| DEGKAPAG | ASSPQGSNERLF | 0.73005235 |
| DTVVFPAL | ASSPQGSNERLF | 0.68936557 |
| FEGITMAG | ASSPQGSNERLF | 0.61190045 |
| FPLVVQD | ASSPQGSNERLF | 0.6756519 |
| GAFTTPQ | ASSPQGSNERLF | 0.6992923 |
| GFGGFGTAGGVGGA | ASSPQGSNERLF | 0.6483609 |
| GFLAQFL | ASSPQGSNERLF | 0.6335432 |
| GGAGGAGGNAGVA | ASSPQGSNERLF | 0.5113302 |
| GGVGTAGAGGAGGAGG | ASSPQGSNERLF | 0.72812665 |
| GNGGGDAGNAGSGGN | ASSPQGSNERLF | 0.7057446 |
| GTGGAGGAGNDAGS | ASSPQGSNERLF | 0.71186644 |
| GTTNTGWFNS | ASSPQGSNERLF | 0.7309338 |
| IFMGPLFTL | ASSPQGSNERLF | 0.662279 |
| ILAHAIEAI | ASSPQGSNERLF | 0.66492105 |
| IRSAIREAL | ASSPQGSNERLF | 0.38563117 |
| ITFAGGG | ASSPQGSNERLF | 0.5396347 |
| ITGMSGHQEV | ASSPQGSNERLF | 0.6196302 |
| IVAGLACTL | ASSPQGSNERLF | 0.66094875 |
| IVGGGLAGA | ASSPQGSNERLF | 0.6925077 |
| IVTGVLNI | ASSPQGSNERLF | 0.71476114 |
| LAQQGCI | ASSPQGSNERLF | 0.5667396 |
| LCLHVEN | ASSPQGSNERLF | 0.72297287 |
| LDAISDGS | ASSPQGSNERLF | 0.7272496 |
| LGGFAAML | ASSPQGSNERLF | 0.43362185 |
| LGGKNPMI | ASSPQGSNERLF | 0.55275226 |
| LGPKGVAAIL | ASSPQGSNERLF | 0.7381836 |
| LMVGVSI | ASSPQGSNERLF | 0.6960404 |
| LVGAAFVL | ASSPQGSNERLF | 0.58359265 |
| LVLAVAPL | ASSPQGSNERLF | 0.6274227 |
| MAAIAVIDAA | ASSPQGSNERLF | 0.7573236 |
| NNGINAELQ | ASSPQGSNERLF | 0.5975964 |
| RGEPLVLVHG | ASSPQGSNERLF | 0.73227024 |
| RGGAGGAAGLI | ASSPQGSNERLF | 0.32994962 |
| RSYRMHPAV | ASSPQGSNERLF | 0.6179072 |
| SAAHLLAGQLA | ASSPQGSNERLF | 0.56571376 |
| SAAVGSY | ASSPQGSNERLF | 0.24500561 |
| SPHAVLHRI | ASSPQGSNERLF | 0.71944416 |
| SQIAKVL | ASSPQGSNERLF | 0.5418487 |
| TALGHLVP | ASSPQGSNERLF | 0.6982888 |
| TAPTTSGAA | ASSPQGSNERLF | 0.5495067 |
| TDGDDGNGGSPGT | ASSPQGSNERLF | 0.7281977 |
| TDGLGEN | ASSPQGSNERLF | 0.65954554 |
| TGGPPLPGQA | ASSPQGSNERLF | 0.5990623 |
| TIPAITI | ASSPQGSNERLF | 0.47968864 |
| TIPGFTI | ASSPQGSNERLF | 0.69571793 |

| | | |
|---|---|---|
| TSIASGALA | ASSPQGSNERLF | 0.73220754 |
| TVTLPGAAI | ASSPQGSNERLF | 0.7368829 |
| VASGLSAALP | ASSPQGSNERLF | 0.36719233 |
| VGAGVIGI | ASSPQGSNERLF | 0.63724077 |
| VGPKRFGV | ASSPQGSNERLF | 0.641561 |
| VNVAFPD | ASSPQGSNERLF | 0.2938876 |
| VSMTNSGVS | ASSPQGSNERLF | 0.25610518 |
| VTDVLGDLSV | ASSPQGSNERLF | 0.6006235 |
| VVVVGAGFAGL | ASSPQGSNERLF | 0.5674598 |
| VYTVGTVV | ASSPQGSNERLF | 0.6916101 |
| WTTFSGP | ASSPQGSNERLF | 0.7035134 |
| YDELVEAF | ASSPQGSNERLF | 0.5519357 |
| YGTGGAGGNGG | ASSPQGSNERLF | 0.6855931 |
| YTNVMAV | ASSPQGSNERLF | 0.60483813 |
| FLEQVHQL | ASSPQGSNERLF | 0.20027958 |
| GLSLLAVAV | ASSPQGSNERLF | 0.5243983 |
| LEQVHQL | ASSPQGSNERLF | 0.38254753 |
| LNAAGAI | ASSPQGSNERLF | 0.7452773 |
| PAVHASL | ASSPQGSNERLF | 0.40717977 |
| TELEDDA | ASSPQGSNERLF | 0.5815286 |
| TPPGPPN | ASSPQGSNERLF | 0.6142477 |
| TPPPPPPP | ASSPQGSNERLF | 0.69800967 |
| TQSPALI | ASSQDSGQQDTQY | 0.6515677 |
| EFAAAFPQ | ASSQDSGQQDTQY | 0.5152044 |
| LLSGNLDVL | ASSQDSGQQDTQY | 0.2771725 |
| GPANLGNYNI | ASSQDSGQQDTQY | 0.7040838 |
| TLGPTLP | ASSQDSGQQDTQY | 0.59496653 |
| VYAYSSCCK | ASSQDSGQQDTQY | 0.4964375 |
| APAPAPAPNI | ASSQDSGQQDTQY | 0.39609897 |
| PGGPVPIP | ASSQDSGQQDTQY | 0.4778977 |
| FLTVALEP | ASSQDSGQQDTQY | 0.3451506 |
| SQYFGANL | ASSQDSGQQDTQY | 0.71603286 |
| GGVKPAA | ASSQDSGQQDTQY | 0.5339875 |
| GGTGGAGGAGGW | ASSQDSGQQDTQY | 0.59811294 |
| GVGGVAGLGGAGG | ASSQDSGQQDTQY | 0.75130004 |
| VLLVLATTI | ASSQDSGQQDTQY | 0.669504 |
| IGIFAAML | ASSQDSGQQDTQY | 0.59254414 |
| AVGDLID | ASSQDSGQQDTQY | 0.5289035 |
| NISPLPL | ASSQDSGQQDTQY | 0.4319217 |
| VTAVFVAI | ASSQDSGQQDTQY | 0.4993739 |
| SGDDNDAAL | ASSQDSGQQDTQY | 0.5508316 |
| AVAALAHS | ASSQDSGQQDTQY | 0.73703885 |
| GGVTVMATP | ASSQDSGQQDTQY | 0.64918417 |
| AGLAANVRD | ASSQDSGQQDTQY | 0.40891302 |
| TNRPDLIDPALLRPG | ASSQDSGQQDTQY | 0.695306 |
| LANLGVTAL | ASSQDSGQQDTQY | 0.72910887 |
| TGERIEL | ASSQDSGQQDTQY | 0.74332285 |
| NGEELAA | ASSQDSGQQDTQY | 0.45388263 |
| VLSLVWG | ASSQDSGQQDTQY | 0.6166848 |
| IGYALWL | ASSQDSGQQDTQY | 0.61113113 |
| LSLLLAPL | ASSQDSGQQDTQY | 0.5996471 |

| | | |
|---|---|---|
| TLDEALK | ASSQDSGQQDTQY | 0.4547916 |
| TDGAGTAEGG | ASSQDSGQQDTQY | 0.4779059 |
| NAPGGAGGNGG | ASSQDSGQQDTQY | 0.7170421 |
| VSVFMGMGGNF | ASSQDSGQQDTQY | 0.7575725 |
| LASYLAAL | ASSQDSGQQDTQY | 0.59294915 |
| LVGGTPAL | ASSQDSGQQDTQY | 0.35397294 |
| VRNADVL | ASSQDSGQQDTQY | 0.5988983 |
| TCSSPFLVSAM | ASSQDSGQQDTQY | 0.51806706 |
| VHITDAE | ASSQDSGQQDTQY | 0.5021217 |
| EYAAPSF | ASSQDSGQQDTQY | 0.5950695 |
| VVDAFHGGL | ASSQDSGQQDTQY | 0.35342744 |
| ASVVDPQ | ASSQDSGQQDTQY | 0.634429 |
| SYNEFGGLS | ASSQDSGQQDTQY | 0.74395585 |
| DPDGSVY | ASSQDSGQQDTQY | 0.6369169 |
| MVSTIHATE | ASSQDSGQQDTQY | 0.7012804 |
| LAAGVADPV | ASSQDSGQQDTQY | 0.7259826 |
| TKEQIAA | ASSQDSGQQDTQY | 0.683589 |
| SLASPAAL | ASSQDSGQQDTQY | 0.7176829 |
| IGLVFYGV | ASSQDSGQQDTQY | 0.71569645 |
| FLLDASNL | ASSQDSGQQDTQY | 0.6427331 |
| YQVGNFP | ASSQDSGQQDTQY | 0.6493398 |
| ESLLSQST | ASSQDSGQQDTQY | 0.6843562 |
| LFTVGFAAVG | ASSQDSGQQDTQY | 0.5881568 |
| ATPPTPT | ASSQDSGQQDTQY | 0.62198246 |
| TDLATLG | ASSQDSGQQDTQY | 0.6141192 |
| SAGGTGR | ASSQDSGQQDTQY | 0.702423 |
| GFSNSGS | ASSQDSGQQDTQY | 0.6090293 |
| GGNGGSGGTSV | ASSQDSGQQDTQY | 0.7380309 |
| GTSVATGGAGN | ASSQDSGQQDTQY | 0.2799173 |
| NGGNGGSGGTS | ASSQDSGQQDTQY | 0.45240748 |
| AVVKFPH | ASSQDSGQQDTQY | 0.714207 |
| TPPPPPP | ASSQDSGQQDTQY | 0.23109838 |
| AAPAAAAAPA | ASSQDSGQQDTQY | 0.31039175 |
| IALNEDL | ASSQDSGQQDTQY | 0.6513074 |
| IVNVSSM | ASSQDSGQQDTQY | 0.72249854 |
| TGIPPAPR | ASSQDSGQQDTQY | 0.23553532 |
| FDLGGGTF | ASSQDSGQQDTQY | 0.60269344 |
| FDLGGGTFDVS | ASSQDSGQQDTQY | 0.42484778 |
| NEPTAAA | ASSQDSGQQDTQY | 0.71334195 |
| NEPTAAALA | ASSQDSGQQDTQY | 0.61124057 |
| GSIINISSI | ASSQDSGQQDTQY | 0.35338145 |
| VAPGNAGTA | ASSQDSGQQDTQY | 0.70005274 |
| TGGGASLE | ASSQDSGQQDTQY | 0.38052553 |
| VAVYNHYKRI | ASSQDSGQQDTQY | 0.5522603 |
| TDPAPATT | ASSQDSGQQDTQY | 0.72488034 |
| TNRPDILDPALLRPG | ASSQDSGQQDTQY | 0.7425251 |
| LTTTTTS | ASSQDSGQQDTQY | 0.38178867 |
| PTGSGKTL | ASSQDSGQQDTQY | 0.70424104 |
| TGSGKTL | ASSQDSGQQDTQY | 0.5647322 |
| TIGHVDHGK | ASSQDSGQQDTQY | 0.7501657 |
| TIGHVDHGKTT | ASSQDSGQQDTQY | 0.6047689 |

| | | |
|---|---|---|
| GHVDHGKTT | ASSQDSGQQDTQY | 0.33059877 |
| AQFEHTLL | ASSQDSGQQDTQY | 0.71692944 |
| IVNQDEL | ASSQDSGQQDTQY | 0.6947138 |
| TNGSQFF | ASSQDSGQQDTQY | 0.6107472 |
| TGKGIVE | ASSQDSGQQDTQY | 0.7012063 |
| LISNASDA | ASSQDSGQQDTQY | 0.59163356 |
| AELRTLGELTKIAKAHGAQ | ASSQDSGQQDTQY | 0.59078276 |
| APNAVAAAEPSW | ASSQDSGQQDTQY | 0.7104361 |
| APQVAGAATAVL | ASSQDSGQQDTQY | 0.680326 |
| ARRDPAAAAAILPTDA | ASSQDSGQQDTQY | 0.4880161 |
| DAAARDVLAEAGLAEYF | ASSQDSGQQDTQY | 0.73510396 |
| DDQIVLSAKANLAKVEQ | ASSQDSGQQDTQY | 0.62008166 |
| DELLPDTGEDSDEDGH | ASSQDSGQQDTQY | 0.44503176 |
| ELPEAPANRATVPQD | ASSQDSGQQDTQY | 0.7509943 |
| FGQSKIERVTGIG | ASSQDSGQQDTQY | 0.62306917 |
| GAGGTGSTAGGAGGAGGAGGLY | ASSQDSGQQDTQY | 0.69306564 |
| GHGGTGAAGGNGGTGGAGGSGI | ASSQDSGQQDTQY | 0.71017206 |
| GNGGNGGAGGEGGPGGAGGAGA | ASSQDSGQQDTQY | 0.7452295 |
| GSTAGNGGNGGNATNGTIAGQP | ASSQDSGQQDTQY | 0.44371402 |
| GTGGDGGNGGNGGTGG | ASSQDSGQQDTQY | 0.6305615 |
| GVRRPAGIPADAPV | ASSQDSGQQDTQY | 0.61520374 |
| GYRQVTDAHLLTLARR | ASSQDSGQQDTQY | 0.71856 |
| IAKMGQDGHDRG | ASSQDSGQQDTQY | 0.6014092 |
| KAEPSIEIQYT | ASSQDSGQQDTQY | 0.64745045 |
| KPVRIRMAADEDTVAVT | ASSQDSGQQDTQY | 0.69454503 |
| MPPRIAGMRLLVIKPEP | ASSQDSGQQDTQY | 0.7298937 |
| MKFYAVALFRSRGVPPTR | ASSQDSGQQDTQY | 0.6302793 |
| NAGGFGILGAVAHSP | ASSQDSGQQDTQY | 0.6359161 |
| NEGREITLAMAEM | ASSQDSGQQDTQY | 0.73717475 |
| NELVEWSHPLGHRWRWE | ASSQDSGQQDTQY | 0.4227745 |
| NLGAGGTGGNGGNPG | ASSQDSGQQDTQY | 0.6305305 |
| NTEWPKWETVMAADPVD | ASSQDSGQQDTQY | 0.7154082 |
| PASAAPAETQP | ASSQDSGQQDTQY | 0.7288723 |
| QQSSDGKNGVRAAEITGE | ASSQDSGQQDTQY | 0.62246567 |
| RAPDAVTTSWWKE | ASSQDSGQQDTQY | 0.7257397 |
| SGFGNVGAHTSGWFNQ | ASSQDSGQQDTQY | 0.5714382 |
| SGVAGAVGQAASVGGLKV | ASSQDSGQQDTQY | 0.68622303 |
| SNGGAGTGSAGGNGGT | ASSQDSGQQDTQY | 0.54826766 |
| TGGAGGEGGAAG | ASSQDSGQQDTQY | 0.6174576 |
| TGGDGGAAGAGGAPGHG | ASSQDSGQQDTQY | 0.43931466 |
| TMPPLLAAIANAGRHGV | ASSQDSGQQDTQY | 0.669093 |
| TSSTAGNSGGA | ASSQDSGQQDTQY | 0.6949166 |
| VPATVPVGILC | ASSQDSGQQDTQY | 0.6896701 |
| VQRPDSGGVVLTGRIS | ASSQDSGQQDTQY | 0.2863321 |
| WTPNPVPATKGVPAGLVQAT | ASSQDSGQQDTQY | 0.6752628 |
| YALKRALAKFSDRFDVVII | ASSQDSGQQDTQY | 0.7000093 |
| YLIGSVIV | ASSQDSGQQDTQY | 0.7526232 |
| YGQLVAVID | ASSQDSGQQDTQY | 0.6034211 |
| VTVGGMTLPA | ASSQDSGQQDTQY | 0.71342576 |
| VLALPAV | ASSQDSGQQDTQY | 0.5760485 |
| VIPPTLNYET | ASSQDSGQQDTQY | 0.6459474 |

| | | |
|---|---|---|
| VGMGGMHPSSG | ASSQDSGQQDTQY | 0.37250593 |
| VGLAGGAATA | ASSQDSGQQDTQY | 0.47119087 |
| VGGAGGAGGSGP | ASSQDSGQQDTQY | 0.57675934 |
| VFAAFMV | ASSQDSGQQDTQY | 0.4769916 |
| TLYDKLR | ASSQDSGQQDTQY | 0.7109651 |
| TLAVFTVL | ASSQDSGQQDTQY | 0.71162415 |
| TGSVQPARV | ASSQDSGQQDTQY | 0.72533596 |
| TGGGAGSGGT | ASSQDSGQQDTQY | 0.72115266 |
| SHLETSL | ASSQDSGQQDTQY | 0.72757363 |
| SFTTGMT | ASSQDSGQQDTQY | 0.72870785 |
| QVEVAKRMGVRQPTVS | ASSQDSGQQDTQY | 0.5674198 |
| QGSGGPALAM | ASSQDSGQQDTQY | 0.59245896 |
| MLAPSNP | ASSQDSGQQDTQY | 0.31304502 |
| LVNNAAI | ASSQDSGQQDTQY | 0.5808176 |
| LTGEINTLA | ASSQDSGQQDTQY | 0.5587647 |
| LSIFGPV | ASSQDSGQQDTQY | 0.63935244 |
| LLALLANL | ASSQDSGQQDTQY | 0.23303029 |
| LISNASDALDKLR | ASSQDSGQQDTQY | 0.65925634 |
| LINEAVP | ASSQDSGQQDTQY | 0.74028987 |
| LGYAFSGL | ASSQDSGQQDTQY | 0.547802 |
| LDFVAVAL | ASSQDSGQQDTQY | 0.6859187 |
| LAQPGAE | ASSQDSGQQDTQY | 0.68728065 |
| ISNLGAVTGF | ASSQDSGQQDTQY | 0.52891636 |
| HQEDPGRI | ASSQDSGQQDTQY | 0.68939126 |
| GTPGEPTP | ASSQDSGQQDTQY | 0.5606959 |
| GFFNSGS | ASSQDSGQQDTQY | 0.7093066 |
| GAGGGGLTAGAN | ASSQDSGQQDTQY | 0.7525972 |
| FVDALVNM | ASSQDSGQQDTQY | 0.677426 |
| FLTVAPEL | ASSQDSGQQDTQY | 0.58296305 |
| FETLTAL | ASSQDSGQQDTQY | 0.67695004 |
| DVINPIEL | ASSQDSGQQDTQY | 0.43616575 |
| APVSGAPV | ASSQDSGQQDTQY | 0.62934554 |
| ALVGGLAW | ASSQDSGQQDTQY | 0.27381524 |
| ALAAAAATAI | ASSQDSGQQDTQY | 0.64940894 |
| AFGLGLLL | ASSQDSGQQDTQY | 0.5372859 |
| AFFAAASL | ASSQDSGQQDTQY | 0.679653 |
| AAAALPAFSPP | ASSQDSGQQDTQY | 0.6283549 |
| VVLAPAL | ASSQDSGQQDTQY | 0.67810893 |
| TIGHVDHGKTTL | ASSQDSGQQDTQY | 0.5057619 |
| LISNASDALDK | ASSQDSGQQDTQY | 0.6652777 |
| LISNASDAL | ASSQDSGQQDTQY | 0.6505493 |
| ISNASDALDK | ASSQDSGQQDTQY | 0.43105948 |
| NQAETLVYQTE | ASSQDSGQQDTQY | 0.70629233 |
| TGGTGGTAGAG | ASSQDSGQQDTQY | 0.6281469 |
| TGSTAGGAGGA | ASSQDSGQQDTQY | 0.6027341 |
| DPLAVPVAAGPV | ASSQDSGQQDTQY | 0.72807765 |
| GGTGGTPGNGGN | ASSQDSGQQDTQY | 0.6444739 |
| PAGLAGVDPATL | ASSQDSGQQDTQY | 0.4094346 |
| GNNLVVEAPHAGASV | ASSQDSGQQDTQY | 0.56071603 |
| KVTRIMSASTLEQPA | ASSQDSGQQDTQY | 0.6462412 |
| LGFALPIALPRRTVV | ASSQDSGQQDTQY | 0.6028831 |

| | | |
|---|---|---|
| TRLGNLTARKLVVLV | ASSQDSGQQDTQY | 0.24697709 |
| WSQQRRLIYFPSAGP | ASSQDSGQQDTQY | 0.68430734 |
| GPNGNPDPMAAAVDIR | ASSQDSGQQDTQY | 0.44904006 |
| TQRCVSVADLRAAASR | ASSQDSGQQDTQY | 0.5404054 |
| QFVLDVAEHPVEVLTETV | ASSQDSGQQDTQY | 0.5358533 |
| GGAGGTGGAGGNGGTQAGDG | ASSQDSGQQDTQY | 0.7066753 |
| PLLATLRAIAVDVDAKPAQV | ASSQDSGQQDTQY | 0.597116 |
| APKFRTGGYMIDFWGVGYQVAK | ASSQDSGQQDTQY | 0.6264064 |
| GAGGAGGAGGAAGGPGGTGGTG | ASSQDSGQQDTQY | 0.66771126 |
| IGALGLLATDVAPAGSVAFGVA | ASSQDSGQQDTQY | 0.6731607 |
| AGGAGGIGGNGGAGGAGGVLMG | ASSQDSGQQDTQY | 0.7208854 |
| ADELIGLL | ASSQDSGQQDTQY | 0.62675375 |
| ADRALKA | ASSQDSGQQDTQY | 0.6647072 |
| AGAVGQVGGAGGA | ASSQDSGQQDTQY | 0.6870043 |
| AGGGGGLGGAP | ASSQDSGQQDTQY | 0.5278814 |
| AGGLLYGNGGAG | ASSQDSGQQDTQY | 0.4101469 |
| AGLESIGL | ASSQDSGQQDTQY | 0.46887296 |
| AGLKTGII | ASSQDSGQQDTQY | 0.56313026 |
| ASLAALLL | ASSQDSGQQDTQY | 0.38017514 |
| ASVINISSI | ASSQDSGQQDTQY | 0.6883793 |
| ATPPGPA | ASSQDSGQQDTQY | 0.74567986 |
| ATVASALV | ASSQDSGQQDTQY | 0.6237277 |
| ATVASLIL | ASSQDSGQQDTQY | 0.3396 |
| AVNGPVLIRH | ASSQDSGQQDTQY | 0.5928819 |
| DGAAGAVGI | ASSQDSGQQDTQY | 0.58360046 |
| DPLLHGI | ASSQDSGQQDTQY | 0.68076587 |
| ELGAALGAL | ASSQDSGQQDTQY | 0.71169317 |
| ELGRTQAV | ASSQDSGQQDTQY | 0.75041425 |
| FVAGGYLS | ASSQDSGQQDTQY | 0.7075808 |
| GASGGAGGAGGW | ASSQDSGQQDTQY | 0.54760504 |
| GGAGAGGDGGAGGI | ASSQDSGQQDTQY | 0.69803435 |
| GTGHNGSQPGMGGNG | ASSQDSGQQDTQY | 0.59914935 |
| GYAAASASAS | ASSQDSGQQDTQY | 0.6615876 |
| HQVDVAAVQQG | ASSQDSGQQDTQY | 0.6520219 |
| IGAAIAHT | ASSQDSGQQDTQY | 0.67839944 |
| IGMIEVAQL | ASSQDSGQQDTQY | 0.5352208 |
| IRLAPEHTSAMD | ASSQDSGQQDTQY | 0.5981028 |
| IYNVSVL | ASSQDSGQQDTQY | 0.3997211 |
| KQGFLGM | ASSQDSGQQDTQY | 0.7087152 |
| LESVITQL | ASSQDSGQQDTQY | 0.5769367 |
| LGLGALVVV | ASSQDSGQQDTQY | 0.40846264 |
| LIGNGGNGGSGGI | ASSQDSGQQDTQY | 0.6195761 |
| LNAINAPTL | ASSQDSGQQDTQY | 0.73648596 |
| LQLLNTGV | ASSQDSGQQDTQY | 0.7370422 |
| LQTSIEF | ASSQDSGQQDTQY | 0.5583027 |
| MSFAVSEL | ASSQDSGQQDTQY | 0.5502645 |
| NTGALGL | ASSQDSGQQDTQY | 0.68016946 |
| PPGAGGAGGAGTT | ASSQDSGQQDTQY | 0.48384136 |
| QLVGLGI | ASSQDSGQQDTQY | 0.69967306 |
| RFLEQVHQL | ASSQDSGQQDTQY | 0.62788194 |
| RGVGYVLR | ASSQDSGQQDTQY | 0.5798348 |

| | | |
|---|---|---|
| SATGVFAGVI | ASSQDSGQQDTQY | 0.57529587 |
| SLPAITLP | ASSQDSGQQDTQY | 0.6835437 |
| STALAFTV | ASSQDSGQQDTQY | 0.56646127 |
| SVLTVGLGK | ASSQDSGQQDTQY | 0.7011802 |
| TALAGAGSGI | ASSQDSGQQDTQY | 0.25447002 |
| TGLPFTGI | ASSQDSGQQDTQY | 0.70844734 |
| VAEHVTV | ASSQDSGQQDTQY | 0.4755804 |
| VDIVLNSL | ASSQDSGQQDTQY | 0.4937478 |
| VLTGGGAVL | ASSQDSGQQDTQY | 0.26071766 |
| VNVGLAYGI | ASSQDSGQQDTQY | 0.6164849 |
| VSSAAVGAV | ASSQDSGQQDTQY | 0.67380935 |
| VTSVYEAGV | ASSQDSGQQDTQY | 0.723158 |
| WVALDRGA | ASSQDSGQQDTQY | 0.58678085 |
| YLVGSVVI | ASSQDSGQQDTQY | 0.6952309 |
| GPPGTGKTLL | ASSQDSGQQDTQY | 0.7313503 |
| GVGMTKF | ASSQDSGQQDTQY | 0.5584939 |
| IGIGGSDL | ASSQDSGQQDTQY | 0.31943557 |
| IGTLLAEL | ASSQDSGQQDTQY | 0.65081465 |
| IYTSGTT | ASSQDSGQQDTQY | 0.6360017 |
| LSFPAGV | ASSQDSGQQDTQY | 0.72254926 |
| LYTSGTT | ASSQDSGQQDTQY | 0.68625045 |
| VGAFPFGF | ASSQDSGQQDTQY | 0.34847033 |
| VGAGTVEF | ASSQDSGQQDTQY | 0.7065507 |
| VGKSTLLL | ASSQDSGQQDTQY | 0.72760403 |
| AALAHAGGGIV | ASSQDSGQQDTQY | 0.43581772 |
| AALVAVSLL | ASSQDSGQQDTQY | 0.46524543 |
| AAVSRAVAAL | ASSQDSGQQDTQY | 0.4913697 |
| AGGSSGIGGF | ASSQDSGQQDTQY | 0.3530981 |
| AGLTVANANG | ASSQDSGQQDTQY | 0.6968142 |
| AGLVGAVL | ASSQDSGQQDTQY | 0.57288754 |
| ALSQAFTV | ASSQDSGQQDTQY | 0.60881186 |
| ALTFSVSL | ASSQDSGQQDTQY | 0.60481095 |
| ALTGAALVAL | ASSQDSGQQDTQY | 0.5697143 |
| ASFSIGAGVL | ASSQDSGQQDTQY | 0.57979083 |
| DGSVYKI | ASSQDSGQQDTQY | 0.59354806 |
| EQLAEIV | ASSQDSGQQDTQY | 0.679884 |
| FAVSELTVM | ASSQDSGQQDTQY | 0.69643664 |
| GAAPLAAGL | ASSQDSGQQDTQY | 0.6011816 |
| GADGGDGGSGGAGG | ASSQDSGQQDTQY | 0.6560073 |
| GFGLDTGGAGG | ASSQDSGQQDTQY | 0.72884345 |
| GLASGGLL | ASSQDSGQQDTQY | 0.40653542 |
| GRIAGAAVLL | ASSQDSGQQDTQY | 0.51579255 |
| GSAGSLNTSF | ASSQDSGQQDTQY | 0.6854594 |
| GTGQIGI | ASSQDSGQQDTQY | 0.53223586 |
| GTLIALIL | ASSQDSGQQDTQY | 0.7196394 |
| IAATMGFVVG | ASSQDSGQQDTQY | 0.6519884 |
| IFAQALL | ASSQDSGQQDTQY | 0.657367 |
| IGAAVALGF | ASSQDSGQQDTQY | 0.6459809 |
| IGLTPLAL | ASSQDSGQQDTQY | 0.74733865 |
| ILGSNTSTL | ASSQDSGQQDTQY | 0.6859587 |
| IVFSLTVL | ASSQDSGQQDTQY | 0.5860144 |

| | | |
|---|---|---|
| LASKATGF | ASSQDSGQQDTQY | 0.71564054 |
| LASVALIL | ASSQDSGQQDTQY | 0.39484888 |
| LFAVGLTL | ASSQDSGQQDTQY | 0.70323783 |
| LFNGGIGTYI | ASSQDSGQQDTQY | 0.7030598 |
| LTLGAGII | ASSQDSGQQDTQY | 0.4552478 |
| LVAAALGDI | ASSQDSGQQDTQY | 0.5978173 |
| NTAFVQLGI | ASSQDSGQQDTQY | 0.732971 |
| QLFSGAEL | ASSQDSGQQDTQY | 0.7264221 |
| SAATFITL | ASSQDSGQQDTQY | 0.5229145 |
| SNYGLANLGV | ASSQDSGQQDTQY | 0.5193298 |
| SYGGAYAVM | ASSQDSGQQDTQY | 0.44794348 |
| TTGVGGLTLG | ASSQDSGQQDTQY | 0.578678 |
| TTQTIYI | ASSQDSGQQDTQY | 0.357837 |
| VAAGAGLGAVI | ASSQDSGQQDTQY | 0.6850016 |
| VAAVVTLL | ASSQDSGQQDTQY | 0.5753963 |
| VGFLGGLI | ASSQDSGQQDTQY | 0.70857716 |
| VGVIPILL | ASSQDSGQQDTQY | 0.673329 |
| VIFASSLI | ASSQDSGQQDTQY | 0.7315061 |
| VKGGDGGAAAG | ASSQDSGQQDTQY | 0.5690547 |
| VTKTFTNI | ASSQDSGQQDTQY | 0.59816575 |
| VVAAAILL | ASSQDSGQQDTQY | 0.33123156 |
| YAALDVEL | ASSQDSGQQDTQY | 0.5322296 |
| LASVALLL | ASSQDSGQQDTQY | 0.48668957 |
| LDAVHRV | ASSQDSGQQDTQY | 0.6275264 |
| QGLSSSIF | ASSQDSGQQDTQY | 0.30121773 |
| RGTFANIRL | ASSQDSGQQDTQY | 0.7375575 |
| RQLWWGHRI | ASSQDSGQQDTQY | 0.5732981 |
| YGPPGTGKTLL | ASSQDSGQQDTQY | 0.70016557 |
| AALVGAPANVA | ASSQDSGQQDTQY | 0.5994099 |
| AATFRRTE | ASSQDSGQQDTQY | 0.41043088 |
| AGALGGGI | ASSQDSGQQDTQY | 0.5108119 |
| AGGAGGAGGIGAG | ASSQDSGQQDTQY | 0.69538146 |
| AGSGALGL | ASSQDSGQQDTQY | 0.37187865 |
| AQIADQI | ASSQDSGQQDTQY | 0.3513408 |
| ASAVAVTYL | ASSQDSGQQDTQY | 0.4664876 |
| ASTVLGGTGG | ASSQDSGQQDTQY | 0.5660187 |
| AVGGEPAK | ASSQDSGQQDTQY | 0.7383864 |
| CEQGTFGGTC | ASSQDSGQQDTQY | 0.7261343 |
| DEGKAPAG | ASSQDSGQQDTQY | 0.6510937 |
| DTVVFPAL | ASSQDSGQQDTQY | 0.73636365 |
| FEGITMAG | ASSQDSGQQDTQY | 0.23730403 |
| FPLVVQD | ASSQDSGQQDTQY | 0.614668 |
| GAFTTPQ | ASSQDSGQQDTQY | 0.7376701 |
| GFGGFGTAGGVGGA | ASSQDSGQQDTQY | 0.6849302 |
| GFLAQFL | ASSQDSGQQDTQY | 0.61854637 |
| GGAGGAGGNAGVA | ASSQDSGQQDTQY | 0.7162819 |
| GGVGTAGAGGAGGAGG | ASSQDSGQQDTQY | 0.5163302 |
| GNGGGGDAGNAGSGGN | ASSQDSGQQDTQY | 0.6182349 |
| GTGGAGGAGNDAGS | ASSQDSGQQDTQY | 0.5892942 |
| GTTNTGWFNS | ASSQDSGQQDTQY | 0.7083199 |
| IFMGPLFTL | ASSQDSGQQDTQY | 0.63330984 |

| | | |
|---|---|---|
| ILAHAIEAI | ASSQDSGQQDTQY | 0.34564367 |
| IRSAIREAL | ASSQDSGQQDTQY | 0.70810324 |
| ITFAGGG | ASSQDSGQQDTQY | 0.7515824 |
| ITGMSGHQEV | ASSQDSGQQDTQY | 0.5587046 |
| IVAGLACTL | ASSQDSGQQDTQY | 0.70983684 |
| IVGGGLAGA | ASSQDSGQQDTQY | 0.6495006 |
| IVTGVLNI | ASSQDSGQQDTQY | 0.68944705 |
| LAQQGCI | ASSQDSGQQDTQY | 0.49659002 |
| LCLHVEN | ASSQDSGQQDTQY | 0.42049497 |
| LDAISDGS | ASSQDSGQQDTQY | 0.6696853 |
| LGGFAAML | ASSQDSGQQDTQY | 0.44181728 |
| LGGKNPMI | ASSQDSGQQDTQY | 0.62841326 |
| LGPKGVAAIL | ASSQDSGQQDTQY | 0.61940825 |
| LMVGVSI | ASSQDSGQQDTQY | 0.7576976 |
| LVGAAFVL | ASSQDSGQQDTQY | 0.6727264 |
| LVLAVAPL | ASSQDSGQQDTQY | 0.6620114 |
| MAAIAVIDAA | ASSQDSGQQDTQY | 0.50091326 |
| NNGINAELQ | ASSQDSGQQDTQY | 0.67205936 |
| RGEPLVLVHG | ASSQDSGQQDTQY | 0.6795324 |
| RGGAGGAAGLI | ASSQDSGQQDTQY | 0.40960497 |
| RSYRMHPAV | ASSQDSGQQDTQY | 0.71842 |
| SAAHLLAGQLA | ASSQDSGQQDTQY | 0.46697712 |
| SAAVGSY | ASSQDSGQQDTQY | 0.6440651 |
| SPHAVLHRI | ASSQDSGQQDTQY | 0.6917052 |
| SQIAKVL | ASSQDSGQQDTQY | 0.7278036 |
| TALGHLVP | ASSQDSGQQDTQY | 0.59408677 |
| TAPTTSGAA | ASSQDSGQQDTQY | 0.6833497 |
| TDGDDGNGGSPGT | ASSQDSGQQDTQY | 0.43055856 |
| TDGLGEN | ASSQDSGQQDTQY | 0.70451844 |
| TGGPPLPGQA | ASSQDSGQQDTQY | 0.71926653 |
| TIPAITI | ASSQDSGQQDTQY | 0.6933908 |
| TIPGFTI | ASSQDSGQQDTQY | 0.7513863 |
| TSIASGALA | ASSQDSGQQDTQY | 0.6687181 |
| TVTLPGAAI | ASSQDSGQQDTQY | 0.3177676 |
| VASGLSAALP | ASSQDSGQQDTQY | 0.6319175 |
| VGAGVIGI | ASSQDSGQQDTQY | 0.72209823 |
| VGPKRFGV | ASSQDSGQQDTQY | 0.7217023 |
| VNVAFPD | ASSQDSGQQDTQY | 0.75047266 |
| VSMTNSGVS | ASSQDSGQQDTQY | 0.6928146 |
| VTDVLGDLSV | ASSQDSGQQDTQY | 0.5977639 |
| VVVVGAGFAGL | ASSQDSGQQDTQY | 0.73459554 |
| VYTVGTVV | ASSQDSGQQDTQY | 0.76483035 |
| WTTFSGP | ASSQDSGQQDTQY | 0.6927116 |
| YDELVEAF | ASSQDSGQQDTQY | 0.49743354 |
| YGTGGAGGNGG | ASSQDSGQQDTQY | 0.47944254 |
| YTNVMAV | ASSQDSGQQDTQY | 0.6839044 |
| FLEQVHQL | ASSQDSGQQDTQY | 0.6949831 |
| GLSLLAVAV | ASSQDSGQQDTQY | 0.41889748 |
| LEQVHQL | ASSQDSGQQDTQY | 0.61296964 |
| LNAAGAI | ASSQDSGQQDTQY | 0.64499426 |
| PAVHASL | ASSQDSGQQDTQY | 0.28826264 |

| | | |
|---|---|---|
| TELEDDA | ASSQDSGQQDTQY | 0.69089186 |
| TPPGPPN | ASSQDSGQQDTQY | 0.73154813 |
| TPPPPPPP | ASSQDSGQQDTQY | 0.6799934 |
| TQSPALI | AWRQGGAETLY | 0.41639978 |
| EFAAAFPQ | AWRQGGAETLY | 0.7155334 |
| LLSGNLDVL | AWRQGGAETLY | 0.73813224 |
| GPANLGNYNI | AWRQGGAETLY | 0.7369587 |
| TLGPTLP | AWRQGGAETLY | 0.40250587 |
| VYAYSSCCK | AWRQGGAETLY | 0.24041449 |
| APAPAPAPNI | AWRQGGAETLY | 0.6184293 |
| PGGPVPIP | AWRQGGAETLY | 0.72417104 |
| FLTVALEP | AWRQGGAETLY | 0.60348976 |
| SQYFGANL | AWRQGGAETLY | 0.50713027 |
| GGVKPAA | AWRQGGAETLY | 0.6249653 |
| GGTGGAGGAGGW | AWRQGGAETLY | 0.72084516 |
| GVGGVAGLGGAGG | AWRQGGAETLY | 0.40655142 |
| VLLVLATTI | AWRQGGAETLY | 0.4163797 |
| IGIFAAML | AWRQGGAETLY | 0.6970687 |
| AVGDLID | AWRQGGAETLY | 0.7147621 |
| NISPLPL | AWRQGGAETLY | 0.505327 |
| VTAVFVAI | AWRQGGAETLY | 0.5588826 |
| SGDDNDAAL | AWRQGGAETLY | 0.6663273 |
| AVAALAHS | AWRQGGAETLY | 0.6499101 |
| GGVTVMATP | AWRQGGAETLY | 0.6612727 |
| AGLAANVRD | AWRQGGAETLY | 0.4166848 |
| TNRPDLIDPALLRPG | AWRQGGAETLY | 0.7097906 |
| LANLGVTAL | AWRQGGAETLY | 0.39785028 |
| TGERIEL | AWRQGGAETLY | 0.68602693 |
| NGEELAA | AWRQGGAETLY | 0.64847845 |
| VLSLVWG | AWRQGGAETLY | 0.6886502 |
| IGYALWL | AWRQGGAETLY | 0.5452607 |
| LSLLLAPL | AWRQGGAETLY | 0.49019504 |
| TLDEALK | AWRQGGAETLY | 0.40607613 |
| TDGAGTAEGG | AWRQGGAETLY | 0.42139944 |
| NAPGGAGGNGG | AWRQGGAETLY | 0.7198973 |
| VSVFMGMGGNF | AWRQGGAETLY | 0.3338181 |
| LASYLAAL | AWRQGGAETLY | 0.7115867 |
| LVGGTPAL | AWRQGGAETLY | 0.6668842 |
| VRNADVL | AWRQGGAETLY | 0.46470165 |
| TCSSPFLVSAM | AWRQGGAETLY | 0.6795806 |
| VHITDAE | AWRQGGAETLY | 0.34810215 |
| EYAAPSF | AWRQGGAETLY | 0.6390214 |
| VVDAFHGGL | AWRQGGAETLY | 0.6432065 |
| ASVVDPQ | AWRQGGAETLY | 0.6945814 |
| SYNEFGGLS | AWRQGGAETLY | 0.5436357 |
| DPDGSVY | AWRQGGAETLY | 0.6883851 |
| MVSTIHATE | AWRQGGAETLY | 0.682011 |
| LAAGVADPV | AWRQGGAETLY | 0.47678378 |
| TKEQIAA | AWRQGGAETLY | 0.68556476 |
| SLASPAAL | AWRQGGAETLY | 0.71841097 |
| IGLVFYGV | AWRQGGAETLY | 0.69075096 |

| | | |
|---|---|---|
| FLLDASNL | AWRQGGAETLY | 0.32044268 |
| YQVGNFP | AWRQGGAETLY | 0.67829895 |
| ESLLSQST | AWRQGGAETLY | 0.4928807 |
| LFTVGFAAVG | AWRQGGAETLY | 0.65764856 |
| ATPPTPT | AWRQGGAETLY | 0.6144824 |
| TDLATLG | AWRQGGAETLY | 0.5831699 |
| SAGGTGR | AWRQGGAETLY | 0.7024065 |
| GFSNSGS | AWRQGGAETLY | 0.36970142 |
| GGNGGSGGTSV | AWRQGGAETLY | 0.54984444 |
| GTSVATGGAGN | AWRQGGAETLY | 0.55540717 |
| NGGNGGSGGTS | AWRQGGAETLY | 0.66910934 |
| AVVKFPH | AWRQGGAETLY | 0.6936543 |
| TPPPPPP | AWRQGGAETLY | 0.4412496 |
| AAPAAAAAPA | AWRQGGAETLY | 0.6648966 |
| IALNEDL | AWRQGGAETLY | 0.6180786 |
| IVNVSSM | AWRQGGAETLY | 0.5204989 |
| TGIPPAPR | AWRQGGAETLY | 0.66902006 |
| FDLGGGTF | AWRQGGAETLY | 0.5358429 |
| FDLGGGTFDVS | AWRQGGAETLY | 0.3650084 |
| NEPTAAA | AWRQGGAETLY | 0.7012025 |
| NEPTAAALA | AWRQGGAETLY | 0.48478308 |
| GSIINISSI | AWRQGGAETLY | 0.694119 |
| VAPGNAGTA | AWRQGGAETLY | 0.75392354 |
| TGGGASLE | AWRQGGAETLY | 0.33787513 |
| VAVYNHYKRI | AWRQGGAETLY | 0.6766399 |
| TDPAPATT | AWRQGGAETLY | 0.63944584 |
| TNRPDILDPALLRPG | AWRQGGAETLY | 0.6414795 |
| LTTTTTS | AWRQGGAETLY | 0.6200781 |
| PTGSGKTL | AWRQGGAETLY | 0.4839116 |
| TGSGKTL | AWRQGGAETLY | 0.7275952 |
| TIGHVDHGK | AWRQGGAETLY | 0.6955426 |
| TIGHVDHGKTT | AWRQGGAETLY | 0.67195725 |
| GHVDHGKTT | AWRQGGAETLY | 0.65815103 |
| AQFEHTLL | AWRQGGAETLY | 0.7065351 |
| IVNQDEL | AWRQGGAETLY | 0.70803267 |
| TNGSQFF | AWRQGGAETLY | 0.5290848 |
| TGKGIVE | AWRQGGAETLY | 0.34668502 |
| LISNASDA | AWRQGGAETLY | 0.602131 |
| AELRTLGELTKIAKAHGAQ | AWRQGGAETLY | 0.6018617 |
| APNAVAAAEPSW | AWRQGGAETLY | 0.5125797 |
| APQVAGAATAVL | AWRQGGAETLY | 0.368169 |
| ARRDPAAAAAILPTDA | AWRQGGAETLY | 0.71017545 |
| DAAARDVLAEAGLAEYF | AWRQGGAETLY | 0.75671124 |
| DDQIVLSAKANLAKVEQ | AWRQGGAETLY | 0.57528365 |
| DELLPDTGEDSDEDGH | AWRQGGAETLY | 0.74835217 |
| ELPEAPANRATVPQD | AWRQGGAETLY | 0.3626113 |
| FGQSKIERVTGIG | AWRQGGAETLY | 0.69922745 |
| GAGGTGSTAGGAGGAGGAGGLY | AWRQGGAETLY | 0.72595656 |
| GHGGTGAAGGNGGTGGAGGSGI | AWRQGGAETLY | 0.45195305 |
| GNGGNGGAGGEGGPGGAGGAGA | AWRQGGAETLY | 0.69461846 |
| GSTAGNGGNGGNATNGTIAGQP | AWRQGGAETLY | 0.48019683 |

| | | |
|---|---|---|
| GTGGDGGNGGNGGTGG | AWRQGGAETLY | 0.45756763 |
| GVRRPAGIPADAPV | AWRQGGAETLY | 0.71675265 |
| GYRQVTDAHLLTLARR | AWRQGGAETLY | 0.5819792 |
| IAKMGQDGHDRG | AWRQGGAETLY | 0.69347125 |
| KAEPSIEIQYT | AWRQGGAETLY | 0.5733435 |
| KPVRIRMAADEDTVAVT | AWRQGGAETLY | 0.4559434 |
| MPPRIAGMRLLVIKPEP | AWRQGGAETLY | 0.6733029 |
| MKFYAVALFRSRGVPPTR | AWRQGGAETLY | 0.71028954 |
| NAGGFGILGAVAHSP | AWRQGGAETLY | 0.5988107 |
| NEGREITLAMAEM | AWRQGGAETLY | 0.7640953 |
| NELVEWSHPLGHRWRWE | AWRQGGAETLY | 0.50900555 |
| NLGAGGTGGNGGNPG | AWRQGGAETLY | 0.74765366 |
| NTEWPKWETVMAADPVD | AWRQGGAETLY | 0.7266896 |
| PASAAPAETQP | AWRQGGAETLY | 0.63688993 |
| QQSSDGKNGVRAAEITGE | AWRQGGAETLY | 0.5755765 |
| RAPDAVTTSWWKE | AWRQGGAETLY | 0.7138473 |
| SGFGNVGAHTSGWFNQ | AWRQGGAETLY | 0.6039766 |
| SGVAGAVGQAASVGGLKV | AWRQGGAETLY | 0.63770306 |
| SNGGAGTGSAGGNGGT | AWRQGGAETLY | 0.74098146 |
| TGGAGGEGGAAG | AWRQGGAETLY | 0.38130513 |
| TGGDGGAAGAGGAPGHG | AWRQGGAETLY | 0.7203855 |
| TMPPLLAAIANAGRHGV | AWRQGGAETLY | 0.57300675 |
| TSSTAGNSGGA | AWRQGGAETLY | 0.6366216 |
| VPATVPVGILC | AWRQGGAETLY | 0.43934017 |
| VQRPDSGGVVLTGRIS | AWRQGGAETLY | 0.3338083 |
| WTPNPVPATKGVPAGLVQAT | AWRQGGAETLY | 0.6427664 |
| YALKRALAKFSDRFDVVII | AWRQGGAETLY | 0.3708125 |
| YLIGSVIV | AWRQGGAETLY | 0.76999867 |
| YGQLVAVID | AWRQGGAETLY | 0.7251507 |
| VTVGGMTLPA | AWRQGGAETLY | 0.46669647 |
| VLALPAV | AWRQGGAETLY | 0.7588325 |
| VIPPTLNYET | AWRQGGAETLY | 0.76955485 |
| VGMGGMHPSSG | AWRQGGAETLY | 0.25532532 |
| VGLAGGAATA | AWRQGGAETLY | 0.41763604 |
| VGGAGGAGGSGP | AWRQGGAETLY | 0.46590254 |
| VFAAFMV | AWRQGGAETLY | 0.5162232 |
| TLYDKLR | AWRQGGAETLY | 0.2565242 |
| TLAVFTVL | AWRQGGAETLY | 0.6770546 |
| TGSVQPARV | AWRQGGAETLY | 0.41297424 |
| TGGGAGSGGT | AWRQGGAETLY | 0.20668103 |
| SHLETSL | AWRQGGAETLY | 0.62223625 |
| SFTTGMT | AWRQGGAETLY | 0.54753095 |
| QVEVAKRMGVRQPTVS | AWRQGGAETLY | 0.4482828 |
| QGSGGPALAM | AWRQGGAETLY | 0.7024038 |
| MLAPSNP | AWRQGGAETLY | 0.4746951 |
| LVNNAAI | AWRQGGAETLY | 0.42671302 |
| LTGEINTLA | AWRQGGAETLY | 0.693789 |
| LSIFGPV | AWRQGGAETLY | 0.78109455 |
| LLALLANL | AWRQGGAETLY | 0.3802017 |
| LISNASDALDKLR | AWRQGGAETLY | 0.6446497 |
| LINEAVP | AWRQGGAETLY | 0.7984258 |

| | | |
|---|---|---|
| LGYAFSGL | AWRQGGAETLY | 0.69616526 |
| LDFVAVAL | AWRQGGAETLY | 0.767398 |
| LAQPGAE | AWRQGGAETLY | 0.78808635 |
| ISNLGAVTGF | AWRQGGAETLY | 0.5242172 |
| HQEDPGRI | AWRQGGAETLY | 0.7471565 |
| GTPGEPTP | AWRQGGAETLY | 0.786288 |
| GFFNSGS | AWRQGGAETLY | 0.50842494 |
| GAGGGGLTAGAN | AWRQGGAETLY | 0.6532651 |
| FVDALVNM | AWRQGGAETLY | 0.4286866 |
| FLTVAPEL | AWRQGGAETLY | 0.589165 |
| FETLTAL | AWRQGGAETLY | 0.24960434 |
| DVINPIEL | AWRQGGAETLY | 0.64098346 |
| APVSGAPV | AWRQGGAETLY | 0.76173604 |
| ALVGGLAW | AWRQGGAETLY | 0.54559284 |
| ALAAAAATAI | AWRQGGAETLY | 0.31249002 |
| AFGLGLLL | AWRQGGAETLY | 0.39427656 |
| AFFAAASL | AWRQGGAETLY | 0.43567112 |
| AAAALPAFSPP | AWRQGGAETLY | 0.44172645 |
| VVLAPAL | AWRQGGAETLY | 0.39341566 |
| TIGHVDHGKTTL | AWRQGGAETLY | 0.35641694 |
| LISNASDALDK | AWRQGGAETLY | 0.7598933 |
| LISNASDAL | AWRQGGAETLY | 0.776073 |
| ISNASDALDK | AWRQGGAETLY | 0.26842815 |
| NQAETLVYQTE | AWRQGGAETLY | 0.7511188 |
| TGGTGGTAGAG | AWRQGGAETLY | 0.7521658 |
| TGSTAGGAGGA | AWRQGGAETLY | 0.79025686 |
| DPLAVPVAAGPV | AWRQGGAETLY | 0.627188 |
| GGTGGTPGNGGN | AWRQGGAETLY | 0.33420753 |
| PAGLAGVDPATL | AWRQGGAETLY | 0.7478416 |
| GNNLVVEAPHAGASV | AWRQGGAETLY | 0.55037266 |
| KVTRIMSASTLEQPA | AWRQGGAETLY | 0.7231164 |
| LGFALPIALPRRTVV | AWRQGGAETLY | 0.70236754 |
| TRLGNLTARKLVVLV | AWRQGGAETLY | 0.44669193 |
| WSQQRRLIYFPSAGP | AWRQGGAETLY | 0.45361114 |
| GPNGNPDPMAAAVDIR | AWRQGGAETLY | 0.6516204 |
| TQRCVSVADLRAAASR | AWRQGGAETLY | 0.76615095 |
| QFVLDVAEHPVEVLTETV | AWRQGGAETLY | 0.3144636 |
| GGAGGTGGAGGNGGTQAGDG | AWRQGGAETLY | 0.36707428 |
| PLLATLRAIAVDVDAKPAQV | AWRQGGAETLY | 0.5089616 |
| APKFRTGGYMIDFWGVGYQVAK | AWRQGGAETLY | 0.61533266 |
| GAGGAGGAGGAAGGPGGTGGTG | AWRQGGAETLY | 0.75361097 |
| IGALGLLATDVAPAGSVAFGVA | AWRQGGAETLY | 0.642973 |
| AGGAGGIGGNGGAGGAGGVLMG | AWRQGGAETLY | 0.5386045 |
| ADELIGLL | AWRQGGAETLY | 0.6954546 |
| ADRALKA | AWRQGGAETLY | 0.686394 |
| AGAVGQVGGAGGA | AWRQGGAETLY | 0.6437 |
| AGGGGGLGGAP | AWRQGGAETLY | 0.7450975 |
| AGGLLYGNGGAG | AWRQGGAETLY | 0.41524518 |
| AGLESIGL | AWRQGGAETLY | 0.7458558 |
| AGLKTGII | AWRQGGAETLY | 0.68211097 |
| ASLAALLL | AWRQGGAETLY | 0.7210837 |

| | | |
|---|---|---|
| ASVINISSI | AWRQGGAETLY | 0.74219626 |
| ATPPGPA | AWRQGGAETLY | 0.61340797 |
| ATVASALV | AWRQGGAETLY | 0.6294506 |
| ATVASLIL | AWRQGGAETLY | 0.29195288 |
| AVNGPVLIRH | AWRQGGAETLY | 0.6115622 |
| DGAAGAVGI | AWRQGGAETLY | 0.680179 |
| DPLLHGI | AWRQGGAETLY | 0.5363631 |
| ELGAALGAL | AWRQGGAETLY | 0.7280587 |
| ELGRTQAV | AWRQGGAETLY | 0.70798737 |
| FVAGGYLS | AWRQGGAETLY | 0.47043175 |
| GASGGAGGAGGW | AWRQGGAETLY | 0.79326767 |
| GGAGAGGDGGAGGI | AWRQGGAETLY | 0.77281797 |
| GTGHNGSQPGMGGNG | AWRQGGAETLY | 0.30457723 |
| GYAAASASAS | AWRQGGAETLY | 0.48479918 |
| HQVDVAAVQQG | AWRQGGAETLY | 0.7486044 |
| IGAAIAHT | AWRQGGAETLY | 0.74371135 |
| IGMIEVAQL | AWRQGGAETLY | 0.5312172 |
| IRLAPEHTSAMD | AWRQGGAETLY | 0.6147662 |
| IYNVSVL | AWRQGGAETLY | 0.81233245 |
| KQGFLGM | AWRQGGAETLY | 0.71618545 |
| LESVITQL | AWRQGGAETLY | 0.78598577 |
| LGLGALVVV | AWRQGGAETLY | 0.21495204 |
| LIGNGGNGGSGGI | AWRQGGAETLY | 0.49388838 |
| LNAINAPTL | AWRQGGAETLY | 0.40962538 |
| LQLLNTGV | AWRQGGAETLY | 0.2317611 |
| LQTSIEF | AWRQGGAETLY | 0.74943435 |
| MSFAVSEL | AWRQGGAETLY | 0.5680446 |
| NTGALGL | AWRQGGAETLY | 0.7119969 |
| PPGAGGAGGAGTT | AWRQGGAETLY | 0.5979781 |
| QLVGLGI | AWRQGGAETLY | 0.74159014 |
| RFLEQVHQL | AWRQGGAETLY | 0.6367583 |
| RGVGYVLR | AWRQGGAETLY | 0.59114 |
| SATGVFAGVI | AWRQGGAETLY | 0.40514287 |
| SLPAITLP | AWRQGGAETLY | 0.6941996 |
| STALAFTV | AWRQGGAETLY | 0.7967994 |
| SVLTVGLGK | AWRQGGAETLY | 0.7146307 |
| TALAGAGSGI | AWRQGGAETLY | 0.3283702 |
| TGLPFTGI | AWRQGGAETLY | 0.7373065 |
| VAEHVTV | AWRQGGAETLY | 0.65373796 |
| VDIVLNSL | AWRQGGAETLY | 0.2948425 |
| VLTGGGAVL | AWRQGGAETLY | 0.32120386 |
| VNVGLAYGI | AWRQGGAETLY | 0.32412842 |
| VSSAAVGAV | AWRQGGAETLY | 0.39136863 |
| VTSVYEAGV | AWRQGGAETLY | 0.7253133 |
| WVALDRGA | AWRQGGAETLY | 0.7089129 |
| YLVGSVVI | AWRQGGAETLY | 0.53068256 |
| GPPGTGKTLL | AWRQGGAETLY | 0.73780787 |
| GVGMTKF | AWRQGGAETLY | 0.6643672 |
| IGIGGSDL | AWRQGGAETLY | 0.14348023 |
| IGTLLAEL | AWRQGGAETLY | 0.4762704 |
| IYTSGTT | AWRQGGAETLY | 0.2650255 |

| | | |
|---|---|---|
| LSFPAGV | AWRQGGAETLY | 0.7744277 |
| LYTSGTT | AWRQGGAETLY | 0.36388928 |
| VGAFPFGF | AWRQGGAETLY | 0.3983528 |
| VGAGTVEF | AWRQGGAETLY | 0.6005834 |
| VGKSTLLL | AWRQGGAETLY | 0.68975383 |
| AALAHAGGGIV | AWRQGGAETLY | 0.6334108 |
| AALVAVSLL | AWRQGGAETLY | 0.40424606 |
| AAVSRAVAAL | AWRQGGAETLY | 0.20863046 |
| AGGSSGIGGF | AWRQGGAETLY | 0.72215617 |
| AGLTVANANG | AWRQGGAETLY | 0.44705456 |
| AGLVGAVL | AWRQGGAETLY | 0.37198648 |
| ALSQAFTV | AWRQGGAETLY | 0.32827982 |
| ALTFSVSL | AWRQGGAETLY | 0.57883424 |
| ALTGAALVAL | AWRQGGAETLY | 0.2991333 |
| ASFSIGAGVL | AWRQGGAETLY | 0.76836354 |
| DGSVYKI | AWRQGGAETLY | 0.38576412 |
| EQLAEIV | AWRQGGAETLY | 0.4720887 |
| FAVSELTVM | AWRQGGAETLY | 0.76823425 |
| GAAPLAAGL | AWRQGGAETLY | 0.67177796 |
| GADGGDGGSGGAGG | AWRQGGAETLY | 0.5343038 |
| GFGLDTGGAGG | AWRQGGAETLY | 0.3879513 |
| GLASGGLL | AWRQGGAETLY | 0.34378287 |
| GRIAGAAVLL | AWRQGGAETLY | 0.42677066 |
| GSAGSLNTSF | AWRQGGAETLY | 0.40572536 |
| GTGQIGI | AWRQGGAETLY | 0.7845216 |
| GTLIALIL | AWRQGGAETLY | 0.52330047 |
| IAATMGFVVG | AWRQGGAETLY | 0.36333388 |
| IFAQALL | AWRQGGAETLY | 0.5824609 |
| IGAAVALGF | AWRQGGAETLY | 0.7749758 |
| IGLTPLAL | AWRQGGAETLY | 0.8104733 |
| ILGSNTSTL | AWRQGGAETLY | 0.33930975 |
| IVFSLTVL | AWRQGGAETLY | 0.57647246 |
| LASKATGF | AWRQGGAETLY | 0.60533524 |
| LASVALIL | AWRQGGAETLY | 0.60017926 |
| LFAVGLTL | AWRQGGAETLY | 0.5135166 |
| LFNGGIGTYI | AWRQGGAETLY | 0.39646208 |
| LTLGAGII | AWRQGGAETLY | 0.7264313 |
| LVAAALGDI | AWRQGGAETLY | 0.8280157 |
| NTAFVQLGI | AWRQGGAETLY | 0.51771104 |
| QLFSGAEL | AWRQGGAETLY | 0.3625678 |
| SAATFITL | AWRQGGAETLY | 0.5987689 |
| SNYGLANLGV | AWRQGGAETLY | 0.3968387 |
| SYGGAYAVM | AWRQGGAETLY | 0.5128683 |
| TTGVGGLTLG | AWRQGGAETLY | 0.6612633 |
| TTQTIYI | AWRQGGAETLY | 0.25252342 |
| VAAGAGLGAVI | AWRQGGAETLY | 0.5390525 |
| VAAVVTLL | AWRQGGAETLY | 0.80980235 |
| VGFLGGLI | AWRQGGAETLY | 0.5120903 |
| VGVIPILL | AWRQGGAETLY | 0.6363827 |
| VIFASSLI | AWRQGGAETLY | 0.750064 |
| VKGGDGGAAAG | AWRQGGAETLY | 0.6401027 |

| | | |
|---|---|---|
| VTKTFTNI | AWRQGGAETLY | 0.7620264 |
| VVAAAILL | AWRQGGAETLY | 0.7576647 |
| YAALDVEL | AWRQGGAETLY | 0.5368211 |
| LASVALLL | AWRQGGAETLY | 0.5811672 |
| LDAVHRV | AWRQGGAETLY | 0.6355115 |
| QGLSSSIF | AWRQGGAETLY | 0.654834 |
| RGTFANIRL | AWRQGGAETLY | 0.56819856 |
| RQLWWGHRI | AWRQGGAETLY | 0.5727563 |
| YGPPGTGKTLL | AWRQGGAETLY | 0.73850965 |
| AALVGAPANVA | AWRQGGAETLY | 0.54572606 |
| AATFRRTE | AWRQGGAETLY | 0.7807852 |
| AGALGGGI | AWRQGGAETLY | 0.28333825 |
| AGGAGGAGGIGAG | AWRQGGAETLY | 0.3746363 |
| AGSGALGL | AWRQGGAETLY | 0.69779134 |
| AQIADQI | AWRQGGAETLY | 0.19432607 |
| ASAVAVTYL | AWRQGGAETLY | 0.36014393 |
| ASTVLGGTGG | AWRQGGAETLY | 0.589889 |
| AVGGEPAK | AWRQGGAETLY | 0.7683257 |
| CEQGTFGGTC | AWRQGGAETLY | 0.19547014 |
| DEGKAPAG | AWRQGGAETLY | 0.42083028 |
| DTVVFPAL | AWRQGGAETLY | 0.27168095 |
| FEGITMAG | AWRQGGAETLY | 0.64989233 |
| FPLVVQD | AWRQGGAETLY | 0.6328161 |
| GAFTTPQ | AWRQGGAETLY | 0.28234154 |
| GFGGFGTAGGVGGA | AWRQGGAETLY | 0.68348885 |
| GFLAQFL | AWRQGGAETLY | 0.27155557 |
| GGAGGAGGNAGVA | AWRQGGAETLY | 0.47665653 |
| GGVGTAGAGGAGGAGG | AWRQGGAETLY | 0.7264105 |
| GNGGGGDAGNAGSGGN | AWRQGGAETLY | 0.7769369 |
| GTGGAGGAGNDAGS | AWRQGGAETLY | 0.34298548 |
| GTTNTGWFNS | AWRQGGAETLY | 0.73658586 |
| IFMGPLFTL | AWRQGGAETLY | 0.47949326 |
| ILAHAIEAI | AWRQGGAETLY | 0.8161106 |
| IRSAIREAL | AWRQGGAETLY | 0.48542517 |
| ITFAGGG | AWRQGGAETLY | 0.3008503 |
| ITGMSGHQEV | AWRQGGAETLY | 0.70938027 |
| IVAGLACTL | AWRQGGAETLY | 0.62958527 |
| IVGGGLAGA | AWRQGGAETLY | 0.6300726 |
| IVTGVLNI | AWRQGGAETLY | 0.7310183 |
| LAQQGCI | AWRQGGAETLY | 0.4294641 |
| LCLHVEN | AWRQGGAETLY | 0.555228 |
| LDAISDGS | AWRQGGAETLY | 0.7461562 |
| LGGFAAML | AWRQGGAETLY | 0.71931654 |
| LGGKNPMI | AWRQGGAETLY | 0.36431155 |
| LGPKGVAAIL | AWRQGGAETLY | 0.7639352 |
| LMVGVSI | AWRQGGAETLY | 0.5695401 |
| LVGAAFVL | AWRQGGAETLY | 0.40564084 |
| LVLAVAPL | AWRQGGAETLY | 0.795668 |
| MAAIAVIDAA | AWRQGGAETLY | 0.62600243 |
| NNGINAELQ | AWRQGGAETLY | 0.7326979 |
| RGEPLVLVHG | AWRQGGAETLY | 0.73058856 |

| | | |
|---|---|---|
| RGGAGGAAGLI | AWRQGGAETLY | 0.8154267 |
| RSYRMHPAV | AWRQGGAETLY | 0.34696746 |
| SAAHLLAGQLA | AWRQGGAETLY | 0.7033249 |
| SAAVGSY | AWRQGGAETLY | 0.65779805 |
| SPHAVLHRI | AWRQGGAETLY | 0.73130584 |
| SQIAKVL | AWRQGGAETLY | 0.57576096 |
| TALGHLVP | AWRQGGAETLY | 0.47423723 |
| TAPTTSGAA | AWRQGGAETLY | 0.67021686 |
| TDGDDGNGGSPGT | AWRQGGAETLY | 0.75743055 |
| TDGLGEN | AWRQGGAETLY | 0.5472721 |
| TGGPPLPGQA | AWRQGGAETLY | 0.70524466 |
| TIPAITI | AWRQGGAETLY | 0.7657452 |
| TIPGFTI | AWRQGGAETLY | 0.41142884 |
| TSIASGALA | AWRQGGAETLY | 0.58446926 |
| TVTLPGAAI | AWRQGGAETLY | 0.70556295 |
| VASGLSAALP | AWRQGGAETLY | 0.7740673 |
| VGAGVIGI | AWRQGGAETLY | 0.53005713 |
| VGPKRFGV | AWRQGGAETLY | 0.7776588 |
| VNVAFPD | AWRQGGAETLY | 0.4336208 |
| VSMTNSGVS | AWRQGGAETLY | 0.71702397 |
| VTDVLGDLSV | AWRQGGAETLY | 0.44943303 |
| VVVVGAGFAGL | AWRQGGAETLY | 0.66814566 |
| VYTVGTVV | AWRQGGAETLY | 0.2602027 |
| WTTFSGP | AWRQGGAETLY | 0.69445956 |
| YDELVEAF | AWRQGGAETLY | 0.7255156 |
| YGTGGAGGNGG | AWRQGGAETLY | 0.7243105 |
| YTNVMAV | AWRQGGAETLY | 0.35028356 |
| FLEQVHQL | AWRQGGAETLY | 0.71007705 |
| GLSLLAVAV | AWRQGGAETLY | 0.73838073 |
| LEQVHQL | AWRQGGAETLY | 0.8044555 |
| LNAAGAI | AWRQGGAETLY | 0.5941431 |
| PAVHASL | AWRQGGAETLY | 0.74275297 |
| TELEDDA | AWRQGGAETLY | 0.50701183 |
| TPPGPPN | AWRQGGAETLY | 0.55907357 |
| TPPPPPPP | AWRQGGAETLY | 0.3168799 |
| TQSPALI | ASSLERAYEQY | 0.492441 |
| EFAAAFPQ | ASSLERAYEQY | 0.58559674 |
| LLSGNLDVL | ASSLERAYEQY | 0.36863112 |
| GPANLGNYNI | ASSLERAYEQY | 0.7522268 |
| TLGPTLP | ASSLERAYEQY | 0.78698325 |
| VYAYSSCCK | ASSLERAYEQY | 0.7521229 |
| APAPAPAPNI | ASSLERAYEQY | 0.73335314 |
| PGGPVPIP | ASSLERAYEQY | 0.75958663 |
| FLTVALEP | ASSLERAYEQY | 0.735194 |
| SQYFGANL | ASSLERAYEQY | 0.4796276 |
| GGVKPAA | ASSLERAYEQY | 0.5827469 |
| GGTGGAGGAGGW | ASSLERAYEQY | 0.2814201 |
| GVGGVAGLGGAGG | ASSLERAYEQY | 0.510795 |
| VLLVLATTI | ASSLERAYEQY | 0.4155678 |
| IGIFAAML | ASSLERAYEQY | 0.6610224 |
| AVGDLID | ASSLERAYEQY | 0.18174098 |

| | | |
|---|---|---|
| NISPLPL | ASSLERAYEQY | 0.64530814 |
| VTAVFVAI | ASSLERAYEQY | 0.7767209 |
| SGDDNDAAL | ASSLERAYEQY | 0.5199851 |
| AVAALAHS | ASSLERAYEQY | 0.7198459 |
| GGVTVMATP | ASSLERAYEQY | 0.61230344 |
| AGLAANVRD | ASSLERAYEQY | 0.3579791 |
| TNRPDLIDPALLRPG | ASSLERAYEQY | 0.6972512 |
| LANLGVTAL | ASSLERAYEQY | 0.40428573 |
| TGERIEL | ASSLERAYEQY | 0.7291624 |
| NGEELAA | ASSLERAYEQY | 0.7941437 |
| VLSLVWG | ASSLERAYEQY | 0.65921026 |
| IGYALWL | ASSLERAYEQY | 0.68915296 |
| LSLLLAPL | ASSLERAYEQY | 0.67050296 |
| TLDEALK | ASSLERAYEQY | 0.31908575 |
| TDGAGTAEGG | ASSLERAYEQY | 0.7090883 |
| NAPGGAGGNGG | ASSLERAYEQY | 0.28855324 |
| VSVFMGMGGNF | ASSLERAYEQY | 0.59082234 |
| LASYLAAL | ASSLERAYEQY | 0.3813281 |
| LVGGTPAL | ASSLERAYEQY | 0.62057626 |
| VRNADVL | ASSLERAYEQY | 0.5800919 |
| TCSSPFLVSAM | ASSLERAYEQY | 0.6614595 |
| VHITDAE | ASSLERAYEQY | 0.41623166 |
| EYAAPSF | ASSLERAYEQY | 0.7382406 |
| VVDAFHGGL | ASSLERAYEQY | 0.66386 |
| ASVVDPQ | ASSLERAYEQY | 0.33266652 |
| SYNEFGGLS | ASSLERAYEQY | 0.7341599 |
| DPDGSVY | ASSLERAYEQY | 0.53991413 |
| MVSTIHATE | ASSLERAYEQY | 0.4955058 |
| LAAGVADPV | ASSLERAYEQY | 0.75797886 |
| TKEQIAA | ASSLERAYEQY | 0.6166198 |
| SLASPAAL | ASSLERAYEQY | 0.31108302 |
| IGLVFYGV | ASSLERAYEQY | 0.5507455 |
| FLLDASNL | ASSLERAYEQY | 0.5735327 |
| YQVGNFP | ASSLERAYEQY | 0.52498955 |
| ESLLSQST | ASSLERAYEQY | 0.20675972 |
| LFTVGFAAVG | ASSLERAYEQY | 0.60602796 |
| ATPPTPT | ASSLERAYEQY | 0.3916195 |
| TDLATLG | ASSLERAYEQY | 0.3466846 |
| SAGGTGR | ASSLERAYEQY | 0.41829145 |
| GFSNSGS | ASSLERAYEQY | 0.7009853 |
| GGNGGSGGTSV | ASSLERAYEQY | 0.6748085 |
| GTSVATGGAGN | ASSLERAYEQY | 0.5698665 |
| NGGNGGSGGTS | ASSLERAYEQY | 0.71762264 |
| AVVKFPH | ASSLERAYEQY | 0.74103963 |
| TPPPPPP | ASSLERAYEQY | 0.78560627 |
| AAPAAAAAPA | ASSLERAYEQY | 0.5633737 |
| IALNEDL | ASSLERAYEQY | 0.5349802 |
| IVNVSSM | ASSLERAYEQY | 0.6708422 |
| TGIPPAPR | ASSLERAYEQY | 0.6489333 |
| FDLGGGTF | ASSLERAYEQY | 0.38449013 |
| FDLGGGTFDVS | ASSLERAYEQY | 0.3863812 |

| | | |
|---|---|---|
| NEPTAAA | ASSLERAYEQY | 0.6330328 |
| NEPTAAALA | ASSLERAYEQY | 0.36780366 |
| GSIINISSI | ASSLERAYEQY | 0.6327914 |
| VAPGNAGTA | ASSLERAYEQY | 0.7711325 |
| TGGGASLE | ASSLERAYEQY | 0.49802798 |
| VAVYNHYKRI | ASSLERAYEQY | 0.34079522 |
| TDPAPATT | ASSLERAYEQY | 0.41236088 |
| TNRPDILDPALLRPG | ASSLERAYEQY | 0.4239277 |
| LTTTTTS | ASSLERAYEQY | 0.70948684 |
| PTGSGKTL | ASSLERAYEQY | 0.6562897 |
| TGSGKTL | ASSLERAYEQY | 0.80384874 |
| TIGHVDHGK | ASSLERAYEQY | 0.7409931 |
| TIGHVDHGKTT | ASSLERAYEQY | 0.48871398 |
| GHVDHGKTT | ASSLERAYEQY | 0.67296517 |
| AQFEHTLL | ASSLERAYEQY | 0.546038 |
| IVNQDEL | ASSLERAYEQY | 0.56842834 |
| TNGSQFF | ASSLERAYEQY | 0.57299685 |
| TGKGIVE | ASSLERAYEQY | 0.59375954 |
| LISNASDA | ASSLERAYEQY | 0.55508333 |
| AELRTLGELTKIAKAHGAQ | ASSLERAYEQY | 0.4997615 |
| APNAVAAAEPSW | ASSLERAYEQY | 0.3343497 |
| APQVAGAATAVL | ASSLERAYEQY | 0.6891248 |
| ARRDPAAAAAILPTDA | ASSLERAYEQY | 0.6035637 |
| DAAARDVLAEAGLAEYF | ASSLERAYEQY | 0.5140748 |
| DDQIVLSAKANLAKVEQ | ASSLERAYEQY | 0.5143075 |
| DELLPDTGEDSDEDGH | ASSLERAYEQY | 0.78678584 |
| ELPEAPANRATVPQD | ASSLERAYEQY | 0.78152734 |
| FGQSKIERVTGIG | ASSLERAYEQY | 0.4709444 |
| GAGGTGSTAGGAGGAGGAGGLY | ASSLERAYEQY | 0.39643678 |
| GHGGTGAAGGNGGTGGAGGSGI | ASSLERAYEQY | 0.6302515 |
| GNGGNGGAGGEGGPGGAGGAGA | ASSLERAYEQY | 0.29705533 |
| GSTAGNGGNGGNATNGTIAGQP | ASSLERAYEQY | 0.63787246 |
| GTGGDGGNGGNGGTGG | ASSLERAYEQY | 0.6033531 |
| GVRRPAGIPADAPV | ASSLERAYEQY | 0.5429465 |
| GYRQVTDAHLLTLARR | ASSLERAYEQY | 0.5479379 |
| IAKMGQDGHDRG | ASSLERAYEQY | 0.5777435 |
| KAEPSIEIQYT | ASSLERAYEQY | 0.42997453 |
| KPVRIRMAADEDTVAVT | ASSLERAYEQY | 0.74522305 |
| MPPRIAGMRLLVIKPEP | ASSLERAYEQY | 0.19094816 |
| MKFYAVALFRSRGVPPTR | ASSLERAYEQY | 0.66236174 |
| NAGGFGILGAVAHSP | ASSLERAYEQY | 0.3554042 |
| NEGREITLAMAEM | ASSLERAYEQY | 0.41751602 |
| NELVEWSHPLGHRWRWE | ASSLERAYEQY | 0.26794803 |
| NLGAGGTGGNGGNPG | ASSLERAYEQY | 0.61273926 |
| NTEWPKWETVMAADPVD | ASSLERAYEQY | 0.7249714 |
| PASAAPAETQP | ASSLERAYEQY | 0.6947933 |
| QQSSDGKNGVRAAEITGE | ASSLERAYEQY | 0.6208862 |
| RAPDAVTTSWWKE | ASSLERAYEQY | 0.73259366 |
| SGFGNVGAHTSGWFNQ | ASSLERAYEQY | 0.7979172 |
| SGVAGAVGQAASVGGLKV | ASSLERAYEQY | 0.4887154 |
| SNGGAGTGSAGGNGGT | ASSLERAYEQY | 0.27405068 |

| | | |
|---|---|---|
| TGGAGGEGGAAG | ASSLERAYEQY | 0.6688556 |
| TGGDGGAAGAGGAPGHG | ASSLERAYEQY | 0.65240204 |
| TMPPLLAAIANAGRHGV | ASSLERAYEQY | 0.68410206 |
| TSSTAGNSGGA | ASSLERAYEQY | 0.74740636 |
| VPATVPVGILC | ASSLERAYEQY | 0.27077773 |
| VQRPDSGGVVLTGRIS | ASSLERAYEQY | 0.7737217 |
| WTPNPVPATKGVPAGLVQAT | ASSLERAYEQY | 0.7047622 |
| YALKRALAKFSDRFDVVII | ASSLERAYEQY | 0.3686406 |
| YLIGSVIV | ASSLERAYEQY | 0.52908784 |
| YGQLVAVID | ASSLERAYEQY | 0.47781247 |
| VTVGGMTLPA | ASSLERAYEQY | 0.2770144 |
| VLALPAV | ASSLERAYEQY | 0.7335093 |
| VIPPTLNYET | ASSLERAYEQY | 0.37983468 |
| VGMGGMHPSSG | ASSLERAYEQY | 0.50973845 |
| VGLAGGAATA | ASSLERAYEQY | 0.5053975 |
| VGGAGGAGGSGP | ASSLERAYEQY | 0.45996255 |
| VFAAFMV | ASSLERAYEQY | 0.49357063 |
| TLYDKLR | ASSLERAYEQY | 0.4588008 |
| TLAVFTVL | ASSLERAYEQY | 0.6483525 |
| TGSVQPARV | ASSLERAYEQY | 0.613008 |
| TGGGAGSGGT | ASSLERAYEQY | 0.58241904 |
| SHLETSL | ASSLERAYEQY | 0.6695861 |
| SFTTGMT | ASSLERAYEQY | 0.7554794 |
| QVEVAKRMGVRQPTVS | ASSLERAYEQY | 0.31398076 |
| QGSGGPALAM | ASSLERAYEQY | 0.4577381 |
| MLAPSNP | ASSLERAYEQY | 0.64574313 |
| LVNNAAI | ASSLERAYEQY | 0.38521218 |
| LTGEINTLA | ASSLERAYEQY | 0.7238431 |
| LSIFGPV | ASSLERAYEQY | 0.6062757 |
| LLALLANL | ASSLERAYEQY | 0.64390296 |
| LISNASDALDKLR | ASSLERAYEQY | 0.60657364 |
| LINEAVP | ASSLERAYEQY | 0.79809225 |
| LGYAFSGL | ASSLERAYEQY | 0.6954273 |
| LDFVAVAL | ASSLERAYEQY | 0.58967364 |
| LAQPGAE | ASSLERAYEQY | 0.68168676 |
| ISNLGAVTGF | ASSLERAYEQY | 0.41124195 |
| HQEDPGRI | ASSLERAYEQY | 0.7501289 |
| GTPGEPTP | ASSLERAYEQY | 0.7006787 |
| GFFNSGS | ASSLERAYEQY | 0.33745155 |
| GAGGGGLTAGAN | ASSLERAYEQY | 0.5646349 |
| FVDALVNM | ASSLERAYEQY | 0.7802208 |
| FLTVAPEL | ASSLERAYEQY | 0.7308774 |
| FETLTAL | ASSLERAYEQY | 0.4096211 |
| DVINPIEL | ASSLERAYEQY | 0.4080218 |
| APVSGAPV | ASSLERAYEQY | 0.30381986 |
| ALVGGLAW | ASSLERAYEQY | 0.4170495 |
| ALAAAAATAI | ASSLERAYEQY | 0.30923662 |
| AFGLGLLL | ASSLERAYEQY | 0.6990862 |
| AFFAAASL | ASSLERAYEQY | 0.5533907 |
| AAAALPAFSPP | ASSLERAYEQY | 0.7737112 |
| VVLAPAL | ASSLERAYEQY | 0.6171385 |

| | | |
|---|---|---|
| TIGHVDHGKTTL | ASSLERAYEQY | 0.7701497 |
| LISNASDALDK | ASSLERAYEQY | 0.4737476 |
| LISNASDAL | ASSLERAYEQY | 0.4858563 |
| ISNASDALDK | ASSLERAYEQY | 0.30395868 |
| NQAETLVYQTE | ASSLERAYEQY | 0.54425085 |
| TGGTGGTAGAG | ASSLERAYEQY | 0.42759672 |
| TGSTAGGAGGA | ASSLERAYEQY | 0.4764979 |
| DPLAVPVAAGPV | ASSLERAYEQY | 0.24625345 |
| GGTGGTPGNGGN | ASSLERAYEQY | 0.7586123 |
| PAGLAGVDPATL | ASSLERAYEQY | 0.63397294 |
| GNNLVVEAPHAGASV | ASSLERAYEQY | 0.70984113 |
| KVTRIMSASTLEQPA | ASSLERAYEQY | 0.5227166 |
| LGFALPIALPRRTVV | ASSLERAYEQY | 0.5427518 |
| TRLGNLTARKLVVLV | ASSLERAYEQY | 0.42928368 |
| WSQQRRLIYFPSAGP | ASSLERAYEQY | 0.71717584 |
| GPNGNPDPMAAAVDIR | ASSLERAYEQY | 0.3130789 |
| TQRCVSVADLRAAASR | ASSLERAYEQY | 0.29928333 |
| QFVLDVAEHPVEVLTETV | ASSLERAYEQY | 0.5804488 |
| GGAGGTGGAGGNGGTQAGDG | ASSLERAYEQY | 0.32533544 |
| PLLATLRAIAVDVDAKPAQV | ASSLERAYEQY | 0.77356684 |
| APKFRTGGYMIDFWGVGYQVAK | ASSLERAYEQY | 0.7434223 |
| GAGGAGGAGGAAGGPGGTGGTG | ASSLERAYEQY | 0.5993229 |
| IGALGLLATDVAPAGSVAFGVA | ASSLERAYEQY | 0.75891316 |
| AGGAGGIGGNGGAGGAGGVLMG | ASSLERAYEQY | 0.21802908 |
| ADELIGLL | ASSLERAYEQY | 0.52926755 |
| ADRALKA | ASSLERAYEQY | 0.79573894 |
| AGAVGQVGGAGGA | ASSLERAYEQY | 0.6816901 |
| AGGGGGLGGAP | ASSLERAYEQY | 0.56606555 |
| AGGLLYGNGGAG | ASSLERAYEQY | 0.7660295 |
| AGLESIGL | ASSLERAYEQY | 0.43684548 |
| AGLKTGII | ASSLERAYEQY | 0.55199933 |
| ASLAALLL | ASSLERAYEQY | 0.5789116 |
| ASVINISSI | ASSLERAYEQY | 0.6965064 |
| ATPPGPA | ASSLERAYEQY | 0.6559987 |
| ATVASALV | ASSLERAYEQY | 0.37701535 |
| ATVASLIL | ASSLERAYEQY | 0.7148869 |
| AVNGPVLIRH | ASSLERAYEQY | 0.78499436 |
| DGAAGAVGI | ASSLERAYEQY | 0.4587116 |
| DPLLHGI | ASSLERAYEQY | 0.731469 |
| ELGAALGAL | ASSLERAYEQY | 0.67828596 |
| ELGRTQAV | ASSLERAYEQY | 0.6747843 |
| FVAGGYLS | ASSLERAYEQY | 0.42189032 |
| GASGGAGGAGGW | ASSLERAYEQY | 0.34669158 |
| GGAGAGGDGGAGGI | ASSLERAYEQY | 0.7490097 |
| GTGHNGSQPGMGGNG | ASSLERAYEQY | 0.4522556 |
| GYAAASASAS | ASSLERAYEQY | 0.58275235 |
| HQVDVAAVQQG | ASSLERAYEQY | 0.52952296 |
| IGAAIAHT | ASSLERAYEQY | 0.817165 |
| IGMIEVAQL | ASSLERAYEQY | 0.7230144 |
| IRLAPEHTSAMD | ASSLERAYEQY | 0.6609815 |
| IYNVSVL | ASSLERAYEQY | 0.3857374 |

| | | |
|---|---|---|
| KQGFLGM | ASSLERAYEQY | 0.63625216 |
| LESVITQL | ASSLERAYEQY | 0.65549004 |
| LGLGALVVV | ASSLERAYEQY | 0.3442547 |
| LIGNGGNGGSGGI | ASSLERAYEQY | 0.76575387 |
| LNAINAPTL | ASSLERAYEQY | 0.42154825 |
| LQLLNTGV | ASSLERAYEQY | 0.5308109 |
| LQTSIEF | ASSLERAYEQY | 0.6878756 |
| MSFAVSEL | ASSLERAYEQY | 0.80009186 |
| NTGALGL | ASSLERAYEQY | 0.6123383 |
| PPGAGGAGGAGTT | ASSLERAYEQY | 0.74424005 |
| QLVGLGI | ASSLERAYEQY | 0.31094918 |
| RFLEQVHQL | ASSLERAYEQY | 0.72970295 |
| RGVGYVLR | ASSLERAYEQY | 0.7184415 |
| SATGVFAGVI | ASSLERAYEQY | 0.7234402 |
| SLPAITLP | ASSLERAYEQY | 0.7857021 |
| STALAFTV | ASSLERAYEQY | 0.64570796 |
| SVLTVGLGK | ASSLERAYEQY | 0.2780247 |
| TALAGAGSGI | ASSLERAYEQY | 0.56291187 |
| TGLPFTGI | ASSLERAYEQY | 0.7324973 |
| VAEHVTV | ASSLERAYEQY | 0.68384945 |
| VDIVLNSL | ASSLERAYEQY | 0.7922939 |
| VLTGGGAVL | ASSLERAYEQY | 0.72083056 |
| VNVGLAYGI | ASSLERAYEQY | 0.5464921 |
| VSSAAVGAV | ASSLERAYEQY | 0.77372617 |
| VTSVYEAGV | ASSLERAYEQY | 0.8105712 |
| WVALDRGA | ASSLERAYEQY | 0.7250076 |
| YLVGSVVI | ASSLERAYEQY | 0.51948273 |
| GPPGTGKTLL | ASSLERAYEQY | 0.40959805 |
| GVGMTKF | ASSLERAYEQY | 0.6774344 |
| IGIGGSDL | ASSLERAYEQY | 0.7083614 |
| IGTLLAEL | ASSLERAYEQY | 0.38632658 |
| IYTSGTT | ASSLERAYEQY | 0.65892243 |
| LSFPAGV | ASSLERAYEQY | 0.59582394 |
| LYTSGTT | ASSLERAYEQY | 0.2654954 |
| VGAFPFGF | ASSLERAYEQY | 0.6782317 |
| VGAGTVEF | ASSLERAYEQY | 0.74817544 |
| VGKSTLLL | ASSLERAYEQY | 0.6115447 |
| AALAHAGGGIV | ASSLERAYEQY | 0.39583004 |
| AALVAVSLL | ASSLERAYEQY | 0.7635962 |
| AAVSRAVAAL | ASSLERAYEQY | 0.7854674 |
| AGGSSGIGGF | ASSLERAYEQY | 0.7537786 |
| AGLTVANANG | ASSLERAYEQY | 0.33319244 |
| AGLVGAVL | ASSLERAYEQY | 0.20536068 |
| ALSQAFTV | ASSLERAYEQY | 0.5562757 |
| ALTFSVSL | ASSLERAYEQY | 0.7587905 |
| ALTGAALVAL | ASSLERAYEQY | 0.47774947 |
| ASFSIGAGVL | ASSLERAYEQY | 0.40051574 |
| DGSVYKI | ASSLERAYEQY | 0.6078036 |
| EQLAEIV | ASSLERAYEQY | 0.74615216 |
| FAVSELTVM | ASSLERAYEQY | 0.40417624 |
| GAAPLAAGL | ASSLERAYEQY | 0.3480868 |

| | | |
|---|---|---|
| GADGGDGGSGGAGG | ASSLERAYEQY | 0.6348934 |
| GFGLDTGGAGG | ASSLERAYEQY | 0.7022443 |
| GLASGGLL | ASSLERAYEQY | 0.3857648 |
| GRIAGAAVLL | ASSLERAYEQY | 0.47854984 |
| GSAGSLNTSF | ASSLERAYEQY | 0.53474027 |
| GTGQIGI | ASSLERAYEQY | 0.6050165 |
| GTLIALIL | ASSLERAYEQY | 0.6081287 |
| IAATMGFVVG | ASSLERAYEQY | 0.36022416 |
| IFAQALL | ASSLERAYEQY | 0.72557926 |
| IGAAVALGF | ASSLERAYEQY | 0.34174615 |
| IGLTPLAL | ASSLERAYEQY | 0.7339118 |
| ILGSNTSTL | ASSLERAYEQY | 0.64238524 |
| IVFSLTVL | ASSLERAYEQY | 0.6628233 |
| LASKATGF | ASSLERAYEQY | 0.5477712 |
| LASVALIL | ASSLERAYEQY | 0.51653576 |
| LFAVGLTL | ASSLERAYEQY | 0.31328875 |
| LFNGGIGTYI | ASSLERAYEQY | 0.3168877 |
| LTLGAGII | ASSLERAYEQY | 0.7784848 |
| LVAAALGDI | ASSLERAYEQY | 0.59954554 |
| NTAFVQLGI | ASSLERAYEQY | 0.7394608 |
| QLFSGAEL | ASSLERAYEQY | 0.72056544 |
| SAATFITL | ASSLERAYEQY | 0.31944108 |
| SNYGLANLGV | ASSLERAYEQY | 0.6654793 |
| SYGGAYAVM | ASSLERAYEQY | 0.28143236 |
| TTGVGGLTLG | ASSLERAYEQY | 0.6129866 |
| TTQTIYI | ASSLERAYEQY | 0.6994121 |
| VAAGAGLGAVI | ASSLERAYEQY | 0.67802435 |
| VAAVVTLL | ASSLERAYEQY | 0.49462453 |
| VGFLGGLI | ASSLERAYEQY | 0.6263305 |
| VGVIPILL | ASSLERAYEQY | 0.6775869 |
| VIFASSLI | ASSLERAYEQY | 0.47304907 |
| VKGGDGGAAAG | ASSLERAYEQY | 0.6470281 |
| VTKTFTNI | ASSLERAYEQY | 0.74027234 |
| VVAAAILL | ASSLERAYEQY | 0.6984622 |
| YAALDVEL | ASSLERAYEQY | 0.31897026 |
| LASVALLL | ASSLERAYEQY | 0.7079889 |
| LDAVHRV | ASSLERAYEQY | 0.3710739 |
| QGLSSSIF | ASSLERAYEQY | 0.6551552 |
| RGTFANIRL | ASSLERAYEQY | 0.5178478 |
| RQLWWGHRI | ASSLERAYEQY | 0.6270749 |
| YGPPGTGKTLL | ASSLERAYEQY | 0.7042063 |
| AALVGAPANVA | ASSLERAYEQY | 0.44559428 |
| AATFRRTE | ASSLERAYEQY | 0.4545447 |
| AGALGGGI | ASSLERAYEQY | 0.6070948 |
| AGGAGGAGGIGAG | ASSLERAYEQY | 0.6041953 |
| AGSGALGL | ASSLERAYEQY | 0.7200161 |
| AQIADQI | ASSLERAYEQY | 0.38447267 |
| ASAVAVTYL | ASSLERAYEQY | 0.6711988 |
| ASTVLGGTGG | ASSLERAYEQY | 0.4484825 |
| AVGGEPAK | ASSLERAYEQY | 0.40425402 |
| CEQGTFGGTC | ASSLERAYEQY | 0.56761456 |

| | | |
|---|---|---|
| DEGKAPAG | ASSLERAYEQY | 0.61098766 |
| DTVVFPAL | ASSLERAYEQY | 0.40833715 |
| FEGITMAG | ASSLERAYEQY | 0.67787313 |
| FPLVVQD | ASSLERAYEQY | 0.36047885 |
| GAFTTPQ | ASSLERAYEQY | 0.69085705 |
| GFGGFGTAGGVGGA | ASSLERAYEQY | 0.78373766 |
| GFLAQFL | ASSLERAYEQY | 0.49183318 |
| GGAGGAGGNAGVA | ASSLERAYEQY | 0.72631407 |
| GGVGTAGAGGAGGAGG | ASSLERAYEQY | 0.6569998 |
| GNGGGGDAGNAGSGGN | ASSLERAYEQY | 0.6854104 |
| GTGGAGGAGNDAGS | ASSLERAYEQY | 0.44491658 |
| GTTNTGWFNS | ASSLERAYEQY | 0.4016883 |
| IFMGPLFTL | ASSLERAYEQY | 0.73076415 |
| ILAHAIEAI | ASSLERAYEQY | 0.7372296 |
| IRSAIREAL | ASSLERAYEQY | 0.7671524 |
| ITFAGGG | ASSLERAYEQY | 0.64398557 |
| ITGMSGHQEV | ASSLERAYEQY | 0.76422894 |
| IVAGLACTL | ASSLERAYEQY | 0.6958532 |
| IVGGGLAGA | ASSLERAYEQY | 0.51754785 |
| IVTGVLNI | ASSLERAYEQY | 0.2569708 |
| LAQQGCI | ASSLERAYEQY | 0.49202722 |
| LCLHVEN | ASSLERAYEQY | 0.38980928 |
| LDAISDGS | ASSLERAYEQY | 0.52895135 |
| LGGFAAML | ASSLERAYEQY | 0.33437335 |
| LGGKNPMI | ASSLERAYEQY | 0.74788904 |
| LGPKGVAAIL | ASSLERAYEQY | 0.80683494 |
| LMVGVSI | ASSLERAYEQY | 0.46694008 |
| LVGAAFVL | ASSLERAYEQY | 0.7754216 |
| LVLAVAPL | ASSLERAYEQY | 0.35255384 |
| MAAIAVIDAA | ASSLERAYEQY | 0.76127803 |
| NNGINAELQ | ASSLERAYEQY | 0.77439296 |
| RGEPLVLVHG | ASSLERAYEQY | 0.36540872 |
| RGGAGGAAGLI | ASSLERAYEQY | 0.72634566 |
| RSYRMHPAV | ASSLERAYEQY | 0.41038156 |
| SAAHLLAGQLA | ASSLERAYEQY | 0.47962472 |
| SAAVGSY | ASSLERAYEQY | 0.77921563 |
| SPHAVLHRI | ASSLERAYEQY | 0.5086798 |
| SQIAKVL | ASSLERAYEQY | 0.6727165 |
| TALGHLVP | ASSLERAYEQY | 0.5072171 |
| TAPTTSGAA | ASSLERAYEQY | 0.35112572 |
| TDGDDGNGGSPGT | ASSLERAYEQY | 0.59066045 |
| TDGLGEN | ASSLERAYEQY | 0.7645196 |
| TGGPPLPGQA | ASSLERAYEQY | 0.6504768 |
| TIPAITI | ASSLERAYEQY | 0.8040617 |
| TIPGFTI | ASSLERAYEQY | 0.45221466 |
| TSIASGALA | ASSLERAYEQY | 0.75974786 |
| TVTLPGAAI | ASSLERAYEQY | 0.733139 |
| VASGLSAALP | ASSLERAYEQY | 0.57927495 |
| VGAGVIGI | ASSLERAYEQY | 0.49298304 |
| VGPKRFGV | ASSLERAYEQY | 0.75598836 |
| VNVAFPD | ASSLERAYEQY | 0.5260131 |

| | | |
|---|---|---|
| VSMTNSGVS | ASSLERAYEQY | 0.7629031 |
| VTDVLGDLSV | ASSLERAYEQY | 0.75042546 |
| VVVVGAGFAGL | ASSLERAYEQY | 0.41916528 |
| VYTVGTVV | ASSLERAYEQY | 0.7270504 |
| WTTFSGP | ASSLERAYEQY | 0.68197215 |
| YDELVEAF | ASSLERAYEQY | 0.5969989 |
| YGTGGAGGNGG | ASSLERAYEQY | 0.5757983 |
| YTNVMAV | ASSLERAYEQY | 0.3223288 |
| FLEQVHQL | ASSLERAYEQY | 0.57301855 |
| GLSLLAVAV | ASSLERAYEQY | 0.47312587 |
| LEQVHQL | ASSLERAYEQY | 0.7217779 |
| LNAAGAI | ASSLERAYEQY | 0.6632179 |
| PAVHASL | ASSLERAYEQY | 0.5185671 |
| TELEDDA | ASSLERAYEQY | 0.7292588 |
| TPPGPPN | ASSLERAYEQY | 0.6885601 |
| TPPPPPPP | ASSLERAYEQY | 0.4131574 |
| TQSPALI | ASSDWDNSGNTLY | 0.5837282 |
| EFAAAFPQ | ASSDWDNSGNTLY | 0.6041868 |
| LLSGNLDVL | ASSDWDNSGNTLY | 0.6106849 |
| GPANLGNYNI | ASSDWDNSGNTLY | 0.3031751 |
| TLGPTLP | ASSDWDNSGNTLY | 0.7125541 |
| VYAYSSCCK | ASSDWDNSGNTLY | 0.5655956 |
| APAPAPAPNI | ASSDWDNSGNTLY | 0.24291416 |
| PGGPVPIP | ASSDWDNSGNTLY | 0.6216957 |
| FLTVALEP | ASSDWDNSGNTLY | 0.62963796 |
| SQYFGANL | ASSDWDNSGNTLY | 0.5870631 |
| GGVKPAA | ASSDWDNSGNTLY | 0.7275601 |
| GGTGGAGGAGGW | ASSDWDNSGNTLY | 0.3098377 |
| GVGGVAGLGGAGG | ASSDWDNSGNTLY | 0.47203 |
| VLLVLATTI | ASSDWDNSGNTLY | 0.7197789 |
| IGIFAAML | ASSDWDNSGNTLY | 0.73730445 |
| AVGDLID | ASSDWDNSGNTLY | 0.39870888 |
| NISPLPL | ASSDWDNSGNTLY | 0.6947101 |
| VTAVFVAI | ASSDWDNSGNTLY | 0.69909436 |
| SGDDNDAAL | ASSDWDNSGNTLY | 0.709224 |
| AVAALAHS | ASSDWDNSGNTLY | 0.69460696 |
| GGVTVMATP | ASSDWDNSGNTLY | 0.7106994 |
| AGLAANVRD | ASSDWDNSGNTLY | 0.4121319 |
| TNRPDLIDPALLRPG | ASSDWDNSGNTLY | 0.7315085 |
| LANLGVTAL | ASSDWDNSGNTLY | 0.73561645 |
| TGERIEL | ASSDWDNSGNTLY | 0.50553435 |
| NGEELAA | ASSDWDNSGNTLY | 0.65839064 |
| VLSLVWG | ASSDWDNSGNTLY | 0.38717026 |
| IGYALWL | ASSDWDNSGNTLY | 0.49464965 |
| LSLLLAPL | ASSDWDNSGNTLY | 0.33631176 |
| TLDEALK | ASSDWDNSGNTLY | 0.71287984 |
| TDGAGTAEGG | ASSDWDNSGNTLY | 0.7285428 |
| NAPGGAGGNGG | ASSDWDNSGNTLY | 0.67184734 |
| VSVFMGMGGNF | ASSDWDNSGNTLY | 0.45634863 |
| LASYLAAL | ASSDWDNSGNTLY | 0.53207546 |
| LVGGTPAL | ASSDWDNSGNTLY | 0.422027 |

| | | |
|---|---|---|
| VRNADVL | ASSDWDNSGNTLY | 0.55642384 |
| TCSSPFLVSAM | ASSDWDNSGNTLY | 0.2764538 |
| VHITDAE | ASSDWDNSGNTLY | 0.43033022 |
| EYAAPSF | ASSDWDNSGNTLY | 0.73465455 |
| VVDAFHGGL | ASSDWDNSGNTLY | 0.72763205 |
| ASVVDPQ | ASSDWDNSGNTLY | 0.38590804 |
| SYNEFGGLS | ASSDWDNSGNTLY | 0.7313466 |
| DPDGSVY | ASSDWDNSGNTLY | 0.7177763 |
| MVSTIHATE | ASSDWDNSGNTLY | 0.7289388 |
| LAAGVADPV | ASSDWDNSGNTLY | 0.58033836 |
| TKEQIAA | ASSDWDNSGNTLY | 0.38469434 |
| SLASPAAL | ASSDWDNSGNTLY | 0.7307036 |
| IGLVFYGV | ASSDWDNSGNTLY | 0.65733886 |
| FLLDASNL | ASSDWDNSGNTLY | 0.46810102 |
| YQVGNFP | ASSDWDNSGNTLY | 0.7078959 |
| ESLLSQST | ASSDWDNSGNTLY | 0.55250984 |
| LFTVGFAAVG | ASSDWDNSGNTLY | 0.5861269 |
| ATPPTPT | ASSDWDNSGNTLY | 0.68252957 |
| TDLATLG | ASSDWDNSGNTLY | 0.6544341 |
| SAGGTGR | ASSDWDNSGNTLY | 0.5377998 |
| GFSNSGS | ASSDWDNSGNTLY | 0.4340542 |
| GGNGGSGGTSV | ASSDWDNSGNTLY | 0.43795913 |
| GTSVATGGAGN | ASSDWDNSGNTLY | 0.6850208 |
| NGGNGGSGGTS | ASSDWDNSGNTLY | 0.73025334 |
| AVVKFPH | ASSDWDNSGNTLY | 0.62214446 |
| TPPPPPP | ASSDWDNSGNTLY | 0.5002139 |
| AAPAAAAAPA | ASSDWDNSGNTLY | 0.70088845 |
| IALNEDL | ASSDWDNSGNTLY | 0.6590312 |
| IVNVSSM | ASSDWDNSGNTLY | 0.64321285 |
| TGIPPAPR | ASSDWDNSGNTLY | 0.5604511 |
| FDLGGGTF | ASSDWDNSGNTLY | 0.5712805 |
| FDLGGGTFDVS | ASSDWDNSGNTLY | 0.65439355 |
| NEPTAAA | ASSDWDNSGNTLY | 0.6967899 |
| NEPTAAALA | ASSDWDNSGNTLY | 0.66793096 |
| GSIINISSI | ASSDWDNSGNTLY | 0.68793 |
| VAPGNAGTA | ASSDWDNSGNTLY | 0.5864527 |
| TGGGASLE | ASSDWDNSGNTLY | 0.620289 |
| VAVYNHYKRI | ASSDWDNSGNTLY | 0.32388276 |
| TDPAPATT | ASSDWDNSGNTLY | 0.47364813 |
| TNRPDILDPALLRPG | ASSDWDNSGNTLY | 0.6538918 |
| LTTTTTS | ASSDWDNSGNTLY | 0.6857072 |
| PTGSGKTL | ASSDWDNSGNTLY | 0.68664 |
| TGSGKTL | ASSDWDNSGNTLY | 0.5517672 |
| TIGHVDHGK | ASSDWDNSGNTLY | 0.5503927 |
| TIGHVDHGKTT | ASSDWDNSGNTLY | 0.5607856 |
| GHVDHGKTT | ASSDWDNSGNTLY | 0.7020903 |
| AQFEHTLL | ASSDWDNSGNTLY | 0.41606084 |
| IVNQDEL | ASSDWDNSGNTLY | 0.53477895 |
| TNGSQFF | ASSDWDNSGNTLY | 0.73909837 |
| TGKGIVE | ASSDWDNSGNTLY | 0.69801843 |
| LISNASDA | ASSDWDNSGNTLY | 0.63014793 |

| | | |
|---|---|---|
| AELRTLGELTKIAKAHGAQ | ASSDWDNSGNTLY | 0.4582883 |
| APNAVAAAEPSW | ASSDWDNSGNTLY | 0.74072754 |
| APQVAGAATAVL | ASSDWDNSGNTLY | 0.49407366 |
| ARRDPAAAAAILPTDA | ASSDWDNSGNTLY | 0.71575844 |
| DAAARDVLAEAGLAEYF | ASSDWDNSGNTLY | 0.27410844 |
| DDQIVLSAKANLAKVEQ | ASSDWDNSGNTLY | 0.41659006 |
| DELLPDTGEDSDEDGH | ASSDWDNSGNTLY | 0.34841537 |
| ELPEAPANRATVPQD | ASSDWDNSGNTLY | 0.2888096 |
| FGQSKIERVTGIG | ASSDWDNSGNTLY | 0.7286123 |
| GAGGTGSTAGGAGGAGGAGGLY | ASSDWDNSGNTLY | 0.46663827 |
| GHGGTGAAGGNGGTGGAGGSGI | ASSDWDNSGNTLY | 0.68084157 |
| GNGGNGGAGGEGGPGGAGGAGA | ASSDWDNSGNTLY | 0.65905166 |
| GSTAGNGGNGGNATNGTIAGQP | ASSDWDNSGNTLY | 0.70943195 |
| GTGGDGGNGGNGGTGG | ASSDWDNSGNTLY | 0.69955647 |
| GVRRPAGIPADAPV | ASSDWDNSGNTLY | 0.6046387 |
| GYRQVTDAHLLTLARR | ASSDWDNSGNTLY | 0.4091892 |
| IAKMGQDGHDRG | ASSDWDNSGNTLY | 0.7182468 |
| KAEPSIEIQYT | ASSDWDNSGNTLY | 0.6166568 |
| KPVRIRMAADEDTVAVT | ASSDWDNSGNTLY | 0.5749428 |
| MPPRIAGMRLLVIKPEP | ASSDWDNSGNTLY | 0.33909184 |
| MKFYAVALFRSRGVPPTR | ASSDWDNSGNTLY | 0.54569626 |
| NAGGFGILGAVAHSP | ASSDWDNSGNTLY | 0.6754174 |
| NEGREITLAMAEM | ASSDWDNSGNTLY | 0.293953 |
| NELVEWSHPLGHRWRWE | ASSDWDNSGNTLY | 0.24164878 |
| NLGAGGTGGNGGNPG | ASSDWDNSGNTLY | 0.47035205 |
| NTEWPKWETVMAADPVD | ASSDWDNSGNTLY | 0.3972271 |
| PASAAPAETQP | ASSDWDNSGNTLY | 0.73181784 |
| QQSSDGKNGVRAAEITGE | ASSDWDNSGNTLY | 0.7216319 |
| RAPDAVTTSWWKE | ASSDWDNSGNTLY | 0.670424 |
| SGFGNVGAHTSGWFNQ | ASSDWDNSGNTLY | 0.6891931 |
| SGVAGAVGQAASVGGLKV | ASSDWDNSGNTLY | 0.6440114 |
| SNGGAGTGSAGGNGGT | ASSDWDNSGNTLY | 0.2239646 |
| TGGAGGEGGAAG | ASSDWDNSGNTLY | 0.4712863 |
| TGGDGGAAGAGGAPGHG | ASSDWDNSGNTLY | 0.44387895 |
| TMPPLLAAIANAGRHGV | ASSDWDNSGNTLY | 0.7039238 |
| TSSTAGNSGGA | ASSDWDNSGNTLY | 0.43484738 |
| VPATVPVGILC | ASSDWDNSGNTLY | 0.58116794 |
| VQRPDSGGVVLTGRIS | ASSDWDNSGNTLY | 0.6965337 |
| WTPNPVPATKGVPAGLVQAT | ASSDWDNSGNTLY | 0.7131425 |
| YALKRALAKFSDRFDVVII | ASSDWDNSGNTLY | 0.6543565 |
| YLIGSVIV | ASSDWDNSGNTLY | 0.37545466 |
| YGQLVAVID | ASSDWDNSGNTLY | 0.23777711 |
| VTVGGMTLPA | ASSDWDNSGNTLY | 0.719507 |
| VLALPAV | ASSDWDNSGNTLY | 0.35328162 |
| VIPPTLNYET | ASSDWDNSGNTLY | 0.37963474 |
| VGMGGMHPSSG | ASSDWDNSGNTLY | 0.25937918 |
| VGLAGGAATA | ASSDWDNSGNTLY | 0.24667104 |
| VGGAGGAGGSGP | ASSDWDNSGNTLY | 0.30691588 |
| VFAAFMV | ASSDWDNSGNTLY | 0.64828825 |
| TLYDKLR | ASSDWDNSGNTLY | 0.5610019 |
| TLAVFTVL | ASSDWDNSGNTLY | 0.6468942 |

| | | |
|---|---|---|
| TGSVQPARV | ASSDWDNSGNTLY | 0.74283457 |
| TGGGAGSGGT | ASSDWDNSGNTLY | 0.715522 |
| SHLETSL | ASSDWDNSGNTLY | 0.63744503 |
| SFTTGMT | ASSDWDNSGNTLY | 0.45764664 |
| QVEVAKRMGVRQPTVS | ASSDWDNSGNTLY | 0.505518 |
| QGSGGPALAM | ASSDWDNSGNTLY | 0.48176405 |
| MLAPSNP | ASSDWDNSGNTLY | 0.45263356 |
| LVNNAAI | ASSDWDNSGNTLY | 0.72835964 |
| LTGEINTLA | ASSDWDNSGNTLY | 0.42649493 |
| LSIFGPV | ASSDWDNSGNTLY | 0.37569395 |
| LLALLANL | ASSDWDNSGNTLY | 0.4602239 |
| LISNASDALDKLR | ASSDWDNSGNTLY | 0.72905636 |
| LINEAVP | ASSDWDNSGNTLY | 0.74641657 |
| LGYAFSGL | ASSDWDNSGNTLY | 0.530146 |
| LDFVAVAL | ASSDWDNSGNTLY | 0.42835972 |
| LAQPGAE | ASSDWDNSGNTLY | 0.46881735 |
| ISNLGAVTGF | ASSDWDNSGNTLY | 0.687186 |
| HQEDPGRI | ASSDWDNSGNTLY | 0.39200383 |
| GTPGEPTP | ASSDWDNSGNTLY | 0.3601901 |
| GFFNSGS | ASSDWDNSGNTLY | 0.641016 |
| GAGGGGLTAGAN | ASSDWDNSGNTLY | 0.7525112 |
| FVDALVNM | ASSDWDNSGNTLY | 0.5266229 |
| FLTVAPEL | ASSDWDNSGNTLY | 0.4773773 |
| FETLTAL | ASSDWDNSGNTLY | 0.6695114 |
| DVINPIEL | ASSDWDNSGNTLY | 0.57957286 |
| APVSGAPV | ASSDWDNSGNTLY | 0.6382861 |
| ALVGGLAW | ASSDWDNSGNTLY | 0.66450477 |
| ALAAAAATAI | ASSDWDNSGNTLY | 0.36723843 |
| AFGLGLLL | ASSDWDNSGNTLY | 0.58253974 |
| AFFAAASL | ASSDWDNSGNTLY | 0.7434889 |
| AAAALPAFSPP | ASSDWDNSGNTLY | 0.6838813 |
| VVLAPAL | ASSDWDNSGNTLY | 0.6510755 |
| TIGHVDHGKTTL | ASSDWDNSGNTLY | 0.7353468 |
| LISNASDALDK | ASSDWDNSGNTLY | 0.7161524 |
| LISNASDAL | ASSDWDNSGNTLY | 0.7383842 |
| ISNASDALDK | ASSDWDNSGNTLY | 0.72210175 |
| NQAETLVYQTE | ASSDWDNSGNTLY | 0.5008845 |
| TGGTGGTAGAG | ASSDWDNSGNTLY | 0.5089989 |
| TGSTAGGAGGA | ASSDWDNSGNTLY | 0.68306035 |
| DPLAVPVAAGPV | ASSDWDNSGNTLY | 0.6728107 |
| GGTGGTPGNGGN | ASSDWDNSGNTLY | 0.6621748 |
| PAGLAGVDPATL | ASSDWDNSGNTLY | 0.43954447 |
| GNNLVVEAPHAGASV | ASSDWDNSGNTLY | 0.73231196 |
| KVTRIMSASTLEQPA | ASSDWDNSGNTLY | 0.53186476 |
| LGFALPIALPRRTVV | ASSDWDNSGNTLY | 0.73376614 |
| TRLGNLTARKLVVLV | ASSDWDNSGNTLY | 0.32482392 |
| WSQQRRLIYFPSAGP | ASSDWDNSGNTLY | 0.42317608 |
| GPNGNPDPMAAAVDIR | ASSDWDNSGNTLY | 0.71894145 |
| TQRCVSVADLRAAASR | ASSDWDNSGNTLY | 0.22693919 |
| QFVLDVAEHPVEVLTETV | ASSDWDNSGNTLY | 0.24765743 |
| GGAGGTGGAGGNGGTQAGDG | ASSDWDNSGNTLY | 0.56899416 |

| | | |
|---|---|---|
| PLLATLRAIAVDVDAKPAQV | ASSDWDNSGNTLY | 0.73356277 |
| APKFRTGGYMIDFWGVGYQVAK | ASSDWDNSGNTLY | 0.23273936 |
| GAGGAGGAGGAAGGPGGTGGTG | ASSDWDNSGNTLY | 0.39910966 |
| IGALGLLATDVAPAGSVAFGVA | ASSDWDNSGNTLY | 0.34832093 |
| AGGAGGIGGNGGAGGAGGVLMG | ASSDWDNSGNTLY | 0.5306374 |
| ADELIGLL | ASSDWDNSGNTLY | 0.6613617 |
| ADRALKA | ASSDWDNSGNTLY | 0.35395178 |
| AGAVGQVGGAGGA | ASSDWDNSGNTLY | 0.69065726 |
| AGGGGGLGGAP | ASSDWDNSGNTLY | 0.37371787 |
| AGGLLYGNGGAG | ASSDWDNSGNTLY | 0.5099244 |
| AGLESIGL | ASSDWDNSGNTLY | 0.6948161 |
| AGLKTGII | ASSDWDNSGNTLY | 0.7361606 |
| ASLAALLL | ASSDWDNSGNTLY | 0.39718017 |
| ASVINISSI | ASSDWDNSGNTLY | 0.7104467 |
| ATPPGPA | ASSDWDNSGNTLY | 0.49728996 |
| ATVASALV | ASSDWDNSGNTLY | 0.7474289 |
| ATVASLIL | ASSDWDNSGNTLY | 0.6070966 |
| AVNGPVLIRH | ASSDWDNSGNTLY | 0.32434866 |
| DGAAGAVGI | ASSDWDNSGNTLY | 0.71671164 |
| DPLLHGI | ASSDWDNSGNTLY | 0.63924634 |
| ELGAALGAL | ASSDWDNSGNTLY | 0.5060542 |
| ELGRTQAV | ASSDWDNSGNTLY | 0.72142345 |
| FVAGGYLS | ASSDWDNSGNTLY | 0.36764222 |
| GASGGAGGAGGW | ASSDWDNSGNTLY | 0.6845239 |
| GGAGAGGDGGAGGI | ASSDWDNSGNTLY | 0.73380435 |
| GTGHNGSQPGMGGNG | ASSDWDNSGNTLY | 0.7116258 |
| GYAAASASAS | ASSDWDNSGNTLY | 0.3707893 |
| HQVDVAAVQQG | ASSDWDNSGNTLY | 0.6532483 |
| IGAAIAHT | ASSDWDNSGNTLY | 0.6674304 |
| IGMIEVAQL | ASSDWDNSGNTLY | 0.41109246 |
| IRLAPEHTSAMD | ASSDWDNSGNTLY | 0.7398708 |
| IYNVSVL | ASSDWDNSGNTLY | 0.56643724 |
| KQGFLGM | ASSDWDNSGNTLY | 0.71491265 |
| LESVITQL | ASSDWDNSGNTLY | 0.71302104 |
| LGLGALVVV | ASSDWDNSGNTLY | 0.7450247 |
| LIGNGGNGGSGGI | ASSDWDNSGNTLY | 0.36234552 |
| LNAINAPTL | ASSDWDNSGNTLY | 0.5649023 |
| LQLLNTGV | ASSDWDNSGNTLY | 0.6506867 |
| LQTSIEF | ASSDWDNSGNTLY | 0.69434917 |
| MSFAVSEL | ASSDWDNSGNTLY | 0.5501881 |
| NTGALGL | ASSDWDNSGNTLY | 0.4726552 |
| PPGAGGAGGAGTT | ASSDWDNSGNTLY | 0.6585442 |
| QLVGLGI | ASSDWDNSGNTLY | 0.72551864 |
| RFLEQVHQL | ASSDWDNSGNTLY | 0.5212236 |
| RGVGYVLR | ASSDWDNSGNTLY | 0.53698015 |
| SATGVFAGVI | ASSDWDNSGNTLY | 0.7454954 |
| SLPAITLP | ASSDWDNSGNTLY | 0.4076896 |
| STALAFTV | ASSDWDNSGNTLY | 0.6308898 |
| SVLTVGLGK | ASSDWDNSGNTLY | 0.65348876 |
| TALAGAGSGI | ASSDWDNSGNTLY | 0.6915901 |
| TGLPFTGI | ASSDWDNSGNTLY | 0.51280856 |

| | | |
|---|---|---|
| VAEHVTV | ASSDWDNSGNTLY | 0.70763683 |
| VDIVLNSL | ASSDWDNSGNTLY | 0.44298843 |
| VLTGGGAVL | ASSDWDNSGNTLY | 0.7260357 |
| VNVGLAYGI | ASSDWDNSGNTLY | 0.5610262 |
| VSSAAVGAV | ASSDWDNSGNTLY | 0.68172485 |
| VTSVYEAGV | ASSDWDNSGNTLY | 0.43127978 |
| WVALDRGA | ASSDWDNSGNTLY | 0.71158683 |
| YLVGSVVI | ASSDWDNSGNTLY | 0.7266716 |
| GPPGTGKTLL | ASSDWDNSGNTLY | 0.72275424 |
| GVGMTKF | ASSDWDNSGNTLY | 0.24559529 |
| IGIGGSDL | ASSDWDNSGNTLY | 0.72950304 |
| IGTLLAEL | ASSDWDNSGNTLY | 0.71071494 |
| IYTSGTT | ASSDWDNSGNTLY | 0.74881923 |
| LSFPAGV | ASSDWDNSGNTLY | 0.55838585 |
| LYTSGTT | ASSDWDNSGNTLY | 0.62961227 |
| VGAFPFGF | ASSDWDNSGNTLY | 0.6048614 |
| VGAGTVEF | ASSDWDNSGNTLY | 0.57896113 |
| VGKSTLLL | ASSDWDNSGNTLY | 0.46216607 |
| AALAHAGGGIV | ASSDWDNSGNTLY | 0.41909105 |
| AALVAVSLL | ASSDWDNSGNTLY | 0.6635915 |
| AAVSRAVAAL | ASSDWDNSGNTLY | 0.6106184 |
| AGGSSGIGGF | ASSDWDNSGNTLY | 0.724625 |
| AGLTVANANG | ASSDWDNSGNTLY | 0.7049662 |
| AGLVGAVL | ASSDWDNSGNTLY | 0.71278226 |
| ALSQAFTV | ASSDWDNSGNTLY | 0.7245697 |
| ALTFSVSL | ASSDWDNSGNTLY | 0.72751486 |
| ALTGAALVAL | ASSDWDNSGNTLY | 0.65326476 |
| ASFSIGAGVL | ASSDWDNSGNTLY | 0.57171166 |
| DGSVYKI | ASSDWDNSGNTLY | 0.61606604 |
| EQLAEIV | ASSDWDNSGNTLY | 0.4919408 |
| FAVSELTVM | ASSDWDNSGNTLY | 0.5894977 |
| GAAPLAAGL | ASSDWDNSGNTLY | 0.56733704 |
| GADGGDGGSGGAGG | ASSDWDNSGNTLY | 0.6942474 |
| GFGLDTGGAGG | ASSDWDNSGNTLY | 0.22833085 |
| GLASGGLL | ASSDWDNSGNTLY | 0.7029765 |
| GRIAGAAVLL | ASSDWDNSGNTLY | 0.7319753 |
| GSAGSLNTSF | ASSDWDNSGNTLY | 0.6087725 |
| GTGQIGI | ASSDWDNSGNTLY | 0.68807495 |
| GTLIALIL | ASSDWDNSGNTLY | 0.62564373 |
| IAATMGFVVG | ASSDWDNSGNTLY | 0.45272928 |
| IFAQALL | ASSDWDNSGNTLY | 0.6984478 |
| IGAAVALGF | ASSDWDNSGNTLY | 0.4998061 |
| IGLTPLAL | ASSDWDNSGNTLY | 0.6737982 |
| ILGSNTSTL | ASSDWDNSGNTLY | 0.73973334 |
| IVFSLTVL | ASSDWDNSGNTLY | 0.67981195 |
| LASKATGF | ASSDWDNSGNTLY | 0.440853 |
| LASVALIL | ASSDWDNSGNTLY | 0.5586038 |
| LFAVGLTL | ASSDWDNSGNTLY | 0.32871997 |
| LFNGGIGTYI | ASSDWDNSGNTLY | 0.54252195 |
| LTLGAGII | ASSDWDNSGNTLY | 0.2234493 |
| LVAAALGDI | ASSDWDNSGNTLY | 0.67453676 |

| | | |
|---|---|---|
| NTAFVQLGI | ASSDWDNSGNTLY | 0.5067936 |
| QLFSGAEL | ASSDWDNSGNTLY | 0.68723094 |
| SAATFITL | ASSDWDNSGNTLY | 0.6601372 |
| SNYGLANLGV | ASSDWDNSGNTLY | 0.6014037 |
| SYGGAYAVM | ASSDWDNSGNTLY | 0.5649477 |
| TTGVGGLTLG | ASSDWDNSGNTLY | 0.66495395 |
| TTQTIYI | ASSDWDNSGNTLY | 0.59875786 |
| VAAGAGLGAVI | ASSDWDNSGNTLY | 0.3630069 |
| VAAVVTLL | ASSDWDNSGNTLY | 0.69038665 |
| VGFLGGLI | ASSDWDNSGNTLY | 0.46894956 |
| VGVIPILL | ASSDWDNSGNTLY | 0.5827395 |
| VIFASSLI | ASSDWDNSGNTLY | 0.7333391 |
| VKGGDGGAAAG | ASSDWDNSGNTLY | 0.6405693 |
| VTKTFTNI | ASSDWDNSGNTLY | 0.34691432 |
| VVAAAILL | ASSDWDNSGNTLY | 0.6854408 |
| YAALDVEL | ASSDWDNSGNTLY | 0.6418231 |
| LASVALLL | ASSDWDNSGNTLY | 0.6641824 |
| LDAVHRV | ASSDWDNSGNTLY | 0.22636132 |
| QGLSSSIF | ASSDWDNSGNTLY | 0.5983349 |
| RGTFANIRL | ASSDWDNSGNTLY | 0.4050488 |
| RQLWWGHRI | ASSDWDNSGNTLY | 0.52801514 |
| YGPPGTGKTLL | ASSDWDNSGNTLY | 0.46984977 |
| AALVGAPANVA | ASSDWDNSGNTLY | 0.7286422 |
| AATFRRTE | ASSDWDNSGNTLY | 0.4168319 |
| AGALGGGI | ASSDWDNSGNTLY | 0.5737523 |
| AGGAGGAGGIGAG | ASSDWDNSGNTLY | 0.6946931 |
| AGSGALGL | ASSDWDNSGNTLY | 0.63842154 |
| AQIADQI | ASSDWDNSGNTLY | 0.73694086 |
| ASAVAVTYL | ASSDWDNSGNTLY | 0.56838304 |
| ASTVLGGTGG | ASSDWDNSGNTLY | 0.44448894 |
| AVGGEPAK | ASSDWDNSGNTLY | 0.5296889 |
| CEQGTFGGTC | ASSDWDNSGNTLY | 0.4812101 |
| DEGKAPAG | ASSDWDNSGNTLY | 0.29416737 |
| DTVVFPAL | ASSDWDNSGNTLY | 0.32083747 |
| FEGITMAG | ASSDWDNSGNTLY | 0.7095866 |
| FPLVVQD | ASSDWDNSGNTLY | 0.27909246 |
| GAFTTPQ | ASSDWDNSGNTLY | 0.668512 |
| GFGGFGTAGGVGGA | ASSDWDNSGNTLY | 0.65577316 |
| GFLAQFL | ASSDWDNSGNTLY | 0.46301123 |
| GGAGGAGGNAGVA | ASSDWDNSGNTLY | 0.30358922 |
| GGVGTAGAGGAGGAGG | ASSDWDNSGNTLY | 0.535002 |
| GNGGGGDAGNAGSGGN | ASSDWDNSGNTLY | 0.60180414 |
| GTGGAGGAGNDAGS | ASSDWDNSGNTLY | 0.71762586 |
| GTTNTGWFNS | ASSDWDNSGNTLY | 0.6451682 |
| IFMGPLFTL | ASSDWDNSGNTLY | 0.74698293 |
| ILAHAIEAI | ASSDWDNSGNTLY | 0.71886384 |
| IRSAIREAL | ASSDWDNSGNTLY | 0.6200185 |
| ITFAGGG | ASSDWDNSGNTLY | 0.7132703 |
| ITGMSGHQEV | ASSDWDNSGNTLY | 0.5894018 |
| IVAGLACTL | ASSDWDNSGNTLY | 0.43852347 |
| IVGGGLAGA | ASSDWDNSGNTLY | 0.55375695 |

| | | |
|---|---|---|
| IVTGVLNI | ASSDWDNSGNTLY | 0.5347191 |
| LAQQGCI | ASSDWDNSGNTLY | 0.4618786 |
| LCLHVEN | ASSDWDNSGNTLY | 0.44973284 |
| LDAISDGS | ASSDWDNSGNTLY | 0.36848497 |
| LGGFAAML | ASSDWDNSGNTLY | 0.63602304 |
| LGGKNPMI | ASSDWDNSGNTLY | 0.68750376 |
| LGPKGVAAIL | ASSDWDNSGNTLY | 0.31327057 |
| LMVGVSI | ASSDWDNSGNTLY | 0.5226989 |
| LVGAAFVL | ASSDWDNSGNTLY | 0.7457658 |
| LVLAVAPL | ASSDWDNSGNTLY | 0.73111105 |
| MAAIAVIDAA | ASSDWDNSGNTLY | 0.57181156 |
| NNGINAELQ | ASSDWDNSGNTLY | 0.49053696 |
| RGEPLVLVHG | ASSDWDNSGNTLY | 0.69506735 |
| RGGAGGAAGLI | ASSDWDNSGNTLY | 0.33979303 |
| RSYRMHPAV | ASSDWDNSGNTLY | 0.5690535 |
| SAAHLLAGQLA | ASSDWDNSGNTLY | 0.696519 |
| SAAVGSY | ASSDWDNSGNTLY | 0.6630516 |
| SPHAVLHRI | ASSDWDNSGNTLY | 0.35360175 |
| SQIAKVL | ASSDWDNSGNTLY | 0.6500723 |
| TALGHLVP | ASSDWDNSGNTLY | 0.57579374 |
| TAPTTSGAA | ASSDWDNSGNTLY | 0.6081921 |
| TDGDDGNGGSPGT | ASSDWDNSGNTLY | 0.24972329 |
| TDGLGEN | ASSDWDNSGNTLY | 0.6112311 |
| TGGPPLPGQA | ASSDWDNSGNTLY | 0.55469596 |
| TIPAITI | ASSDWDNSGNTLY | 0.38648397 |
| TIPGFTI | ASSDWDNSGNTLY | 0.2811507 |
| TSIASGALA | ASSDWDNSGNTLY | 0.55195874 |
| TVTLPGAAI | ASSDWDNSGNTLY | 0.72547174 |
| VASGLSAALP | ASSDWDNSGNTLY | 0.7207713 |
| VGAGVIGI | ASSDWDNSGNTLY | 0.38738507 |
| VGPKRFGV | ASSDWDNSGNTLY | 0.71514606 |
| VNVAFPD | ASSDWDNSGNTLY | 0.7363964 |
| VSMTNSGVS | ASSDWDNSGNTLY | 0.6208493 |
| VTDVLGDLSV | ASSDWDNSGNTLY | 0.28384972 |
| VVVVGAGFAGL | ASSDWDNSGNTLY | 0.6976714 |
| VYTVGTVV | ASSDWDNSGNTLY | 0.5966823 |
| WTTFSGP | ASSDWDNSGNTLY | 0.66292137 |
| YDELVEAF | ASSDWDNSGNTLY | 0.7009446 |
| YGTGGAGGNGG | ASSDWDNSGNTLY | 0.4762417 |
| YTNVMAV | ASSDWDNSGNTLY | 0.65287477 |
| FLEQVHQL | ASSDWDNSGNTLY | 0.72625613 |
| GLSLLAVAV | ASSDWDNSGNTLY | 0.49189347 |
| LEQVHQL | ASSDWDNSGNTLY | 0.37741065 |
| LNAAGAI | ASSDWDNSGNTLY | 0.36745244 |
| PAVHASL | ASSDWDNSGNTLY | 0.28311217 |
| TELEDDA | ASSDWDNSGNTLY | 0.6339934 |
| TPPGPPN | ASSDWDNSGNTLY | 0.6207707 |
| TPPPPPPP | ASSDWDNSGNTLY | 0.54545164 |
| TQSPALI | ASSPPTEVF | 0.68248737 |
| EFAAAFPQ | ASSPPTEVF | 0.4618248 |
| LLSGNLDVL | ASSPPTEVF | 0.56031597 |

| | | |
|---|---|---|
| GPANLGNYNI | ASSPPTEVF | 0.55947125 |
| TLGPTLP | ASSPPTEVF | 0.64251184 |
| VYAYSSCCK | ASSPPTEVF | 0.682183 |
| APAPAPAPNI | ASSPPTEVF | 0.65902615 |
| PGGPVPIP | ASSPPTEVF | 0.56739384 |
| FLTVALEP | ASSPPTEVF | 0.74073964 |
| SQYFGANL | ASSPPTEVF | 0.32420087 |
| GGVKPAA | ASSPPTEVF | 0.4348866 |
| GGTGGAGGAGGW | ASSPPTEVF | 0.67958033 |
| GVGGVAGLGGAGG | ASSPPTEVF | 0.38608718 |
| VLLVLATTI | ASSPPTEVF | 0.71465755 |
| IGIFAAML | ASSPPTEVF | 0.63809216 |
| AVGDLID | ASSPPTEVF | 0.70629054 |
| NISPLPL | ASSPPTEVF | 0.57239115 |
| VTAVFVAI | ASSPPTEVF | 0.735861 |
| SGDDNDAAL | ASSPPTEVF | 0.72348976 |
| AVAALAHS | ASSPPTEVF | 0.5298418 |
| GGVTVMATP | ASSPPTEVF | 0.61756384 |
| AGLAANVRD | ASSPPTEVF | 0.29999056 |
| TNRPDLIDPALLRPG | ASSPPTEVF | 0.6933471 |
| LANLGVTAL | ASSPPTEVF | 0.6516372 |
| TGERIEL | ASSPPTEVF | 0.5292797 |
| NGEELAA | ASSPPTEVF | 0.68684447 |
| VLSLVWG | ASSPPTEVF | 0.70327693 |
| IGYALWL | ASSPPTEVF | 0.7252492 |
| LSLLLAPL | ASSPPTEVF | 0.3415228 |
| TLDEALK | ASSPPTEVF | 0.64420915 |
| TDGAGTAEGG | ASSPPTEVF | 0.27265385 |
| NAPGGAGGNGG | ASSPPTEVF | 0.317464 |
| VSVFMGMGGNF | ASSPPTEVF | 0.31530705 |
| LASYLAAL | ASSPPTEVF | 0.702392 |
| LVGGTPAL | ASSPPTEVF | 0.6193069 |
| VRNADVL | ASSPPTEVF | 0.71758604 |
| TCSSPFLVSAM | ASSPPTEVF | 0.6850508 |
| VHITDAE | ASSPPTEVF | 0.73939526 |
| EYAAPSF | ASSPPTEVF | 0.5146766 |
| VVDAFHGGL | ASSPPTEVF | 0.55749565 |
| ASVVDPQ | ASSPPTEVF | 0.29924065 |
| SYNEFGGLS | ASSPPTEVF | 0.4145041 |
| DPDGSVY | ASSPPTEVF | 0.38782507 |
| MVSTIHATE | ASSPPTEVF | 0.54213804 |
| LAAGVADPV | ASSPPTEVF | 0.26444086 |
| TKEQIAA | ASSPPTEVF | 0.7177894 |
| SLASPAAL | ASSPPTEVF | 0.67893755 |
| IGLVFYGV | ASSPPTEVF | 0.72403514 |
| FLLDASNL | ASSPPTEVF | 0.48610651 |
| YQVGNFP | ASSPPTEVF | 0.4783049 |
| ESLLSQST | ASSPPTEVF | 0.56272745 |
| LFTVGFAAVG | ASSPPTEVF | 0.70376086 |
| ATPPTPT | ASSPPTEVF | 0.46186197 |
| TDLATLG | ASSPPTEVF | 0.3468178 |

| | | |
|---|---|---|
| SAGGTGR | ASSPPTEVF | 0.45291638 |
| GFSNSGS | ASSPPTEVF | 0.43812603 |
| GGNGGSGGTSV | ASSPPTEVF | 0.7360635 |
| GTSVATGGAGN | ASSPPTEVF | 0.7302063 |
| NGGNGGSGGTS | ASSPPTEVF | 0.69981146 |
| AVVKFPH | ASSPPTEVF | 0.7264271 |
| TPPPPPP | ASSPPTEVF | 0.24278748 |
| AAPAAAAAPA | ASSPPTEVF | 0.47609115 |
| IALNEDL | ASSPPTEVF | 0.7367898 |
| IVNVSSM | ASSPPTEVF | 0.6891365 |
| TGIPPAPR | ASSPPTEVF | 0.63344 |
| FDLGGGTF | ASSPPTEVF | 0.7326521 |
| FDLGGGTFDVS | ASSPPTEVF | 0.53582364 |
| NEPTAAA | ASSPPTEVF | 0.6518947 |
| NEPTAAALA | ASSPPTEVF | 0.6482675 |
| GSIINISSI | ASSPPTEVF | 0.60330516 |
| VAPGNAGTA | ASSPPTEVF | 0.5389115 |
| TGGGASLE | ASSPPTEVF | 0.4328503 |
| VAVYNHYKRI | ASSPPTEVF | 0.6470802 |
| TDPAPATT | ASSPPTEVF | 0.74287945 |
| TNRPDILDPALLRPG | ASSPPTEVF | 0.6664275 |
| LTTTTTS | ASSPPTEVF | 0.73670924 |
| PTGSGKTL | ASSPPTEVF | 0.6885712 |
| TGSGKTL | ASSPPTEVF | 0.7282762 |
| TIGHVDHGK | ASSPPTEVF | 0.56213605 |
| TIGHVDHGKTT | ASSPPTEVF | 0.55884635 |
| GHVDHGKTT | ASSPPTEVF | 0.64222884 |
| AQFEHTLL | ASSPPTEVF | 0.5885148 |
| IVNQDEL | ASSPPTEVF | 0.528979 |
| TNGSQFF | ASSPPTEVF | 0.57943463 |
| TGKGIVE | ASSPPTEVF | 0.7486007 |
| LISNASDA | ASSPPTEVF | 0.6865612 |
| AELRTLGELTKIAKAHGAQ | ASSPPTEVF | 0.68447614 |
| APNAVAAAEPSW | ASSPPTEVF | 0.5239773 |
| APQVAGAATAVL | ASSPPTEVF | 0.5426822 |
| ARRDPAAAAAILPTDA | ASSPPTEVF | 0.6965304 |
| DAAARDVLAEAGLAEYF | ASSPPTEVF | 0.3832718 |
| DDQIVLSAKANLAKVEQ | ASSPPTEVF | 0.72477734 |
| DELLPDTGEDSDEDGH | ASSPPTEVF | 0.55083764 |
| ELPEAPANRATVPQD | ASSPPTEVF | 0.52321476 |
| FGQSKIERVTGIG | ASSPPTEVF | 0.6500207 |
| GAGGTGSTAGGAGGAGGAGGLY | ASSPPTEVF | 0.74458015 |
| GHGGTGAAGGNGGTGGAGGSGI | ASSPPTEVF | 0.5994889 |
| GNGGNGGAGGEGGPGGAGGAGA | ASSPPTEVF | 0.64838976 |
| GSTAGNGGNGGNATNGTIAGQP | ASSPPTEVF | 0.49736914 |
| GTGGDGGNGGNGGTGG | ASSPPTEVF | 0.69176304 |
| GVRRPAGIPADAPV | ASSPPTEVF | 0.73657 |
| GYRQVTDAHLLTLARR | ASSPPTEVF | 0.62530684 |
| IAKMGQDGHDRG | ASSPPTEVF | 0.7452439 |
| KAEPSIEIQYT | ASSPPTEVF | 0.55890274 |
| KPVRIRMAADEDTVAVT | ASSPPTEVF | 0.281306 |

| | | |
|---|---|---|
| MPPRIAGMRLLVIKPEP | ASSPPTEVF | 0.60334575 |
| MKFYAVALFRSRGVPPTR | ASSPPTEVF | 0.7066177 |
| NAGGFGILGAVAHSP | ASSPPTEVF | 0.72392094 |
| NEGREITLAMAEM | ASSPPTEVF | 0.74080724 |
| NELVEWSHPLGHRWRWE | ASSPPTEVF | 0.50252235 |
| NLGAGGTGGNGGNPG | ASSPPTEVF | 0.5877567 |
| NTEWPKWETVMAADPVD | ASSPPTEVF | 0.73726594 |
| PASAAPAETQP | ASSPPTEVF | 0.7281964 |
| QQSSDGKNGVRAAEITGE | ASSPPTEVF | 0.72155404 |
| RAPDAVTTSWWKE | ASSPPTEVF | 0.5155177 |
| SGFGNVGAHTSGWFNQ | ASSPPTEVF | 0.3567294 |
| SGVAGAVGQAASVGGLKV | ASSPPTEVF | 0.6380352 |
| SNGGAGTGSAGGNGGT | ASSPPTEVF | 0.71592104 |
| TGGAGGEGGAAG | ASSPPTEVF | 0.34144577 |
| TGGDGGAAGAGGAPGHG | ASSPPTEVF | 0.68469894 |
| TMPPLLAAIANAGRHGV | ASSPPTEVF | 0.69832915 |
| TSSTAGNSGGA | ASSPPTEVF | 0.28787503 |
| VPATVPVGILC | ASSPPTEVF | 0.654794 |
| VQRPDSGGVVLTGRIS | ASSPPTEVF | 0.7140721 |
| WTPNPVPATKGVPAGLVQAT | ASSPPTEVF | 0.53523445 |
| YALKRALAKFSDRFDVVII | ASSPPTEVF | 0.5076716 |
| YLIGSVIV | ASSPPTEVF | 0.72289413 |
| YGQLVAVID | ASSPPTEVF | 0.74861467 |
| VTVGGMTLPA | ASSPPTEVF | 0.7204734 |
| VLALPAV | ASSPPTEVF | 0.32535613 |
| VIPPTLNYET | ASSPPTEVF | 0.27372256 |
| VGMGGMHPSSG | ASSPPTEVF | 0.54744315 |
| VGLAGGAATA | ASSPPTEVF | 0.70618486 |
| VGGAGGAGGSGP | ASSPPTEVF | 0.53846747 |
| VFAAFMV | ASSPPTEVF | 0.51212645 |
| TLYDKLR | ASSPPTEVF | 0.6858262 |
| TLAVFTVL | ASSPPTEVF | 0.7327584 |
| TGSVQPARV | ASSPPTEVF | 0.3717292 |
| TGGGAGSGGT | ASSPPTEVF | 0.30292577 |
| SHLETSL | ASSPPTEVF | 0.63330805 |
| SFTTGMT | ASSPPTEVF | 0.6833041 |
| QVEVAKRMGVRQPTVS | ASSPPTEVF | 0.5479589 |
| QGSGGPALAM | ASSPPTEVF | 0.6723611 |
| MLAPSNP | ASSPPTEVF | 0.6008139 |
| LVNNAAI | ASSPPTEVF | 0.43760753 |
| LTGEINTLA | ASSPPTEVF | 0.6905096 |
| LSIFGPV | ASSPPTEVF | 0.4461066 |
| LLALLANL | ASSPPTEVF | 0.71871686 |
| LISNASDALDKLR | ASSPPTEVF | 0.53649366 |
| LINEAVP | ASSPPTEVF | 0.6641593 |
| LGYAFSGL | ASSPPTEVF | 0.6721827 |
| LDFVAVAL | ASSPPTEVF | 0.6566471 |
| LAQPGAE | ASSPPTEVF | 0.6466764 |
| ISNLGAVTGF | ASSPPTEVF | 0.58579695 |
| HQEDPGRI | ASSPPTEVF | 0.43472275 |
| GTPGEPTP | ASSPPTEVF | 0.29491907 |

| | | |
|---|---|---|
| GFFNSGS | ASSPPTEVF | 0.7150471 |
| GAGGGGLTAGAN | ASSPPTEVF | 0.24392048 |
| FVDALVNM | ASSPPTEVF | 0.70881927 |
| FLTVAPEL | ASSPPTEVF | 0.7184 |
| FETLTAL | ASSPPTEVF | 0.44740406 |
| DVINPIEL | ASSPPTEVF | 0.64377296 |
| APVSGAPV | ASSPPTEVF | 0.40523398 |
| ALVGGLAW | ASSPPTEVF | 0.47450054 |
| ALAAAAATAI | ASSPPTEVF | 0.6999209 |
| AFGLGLLL | ASSPPTEVF | 0.6549775 |
| AFFAAASL | ASSPPTEVF | 0.50529003 |
| AAAALPAFSPP | ASSPPTEVF | 0.5850732 |
| VVLAPAL | ASSPPTEVF | 0.69128865 |
| TIGHVDHGKTTL | ASSPPTEVF | 0.45389628 |
| LISNASDALDK | ASSPPTEVF | 0.5162307 |
| LISNASDAL | ASSPPTEVF | 0.7248627 |
| ISNASDALDK | ASSPPTEVF | 0.5832797 |
| NQAETLVYQTE | ASSPPTEVF | 0.5454741 |
| TGGTGGTAGAG | ASSPPTEVF | 0.6981054 |
| TGSTAGGAGGA | ASSPPTEVF | 0.5153291 |
| DPLAVPVAAGPV | ASSPPTEVF | 0.71036935 |
| GGTGGTPGNGGN | ASSPPTEVF | 0.5874615 |
| PAGLAGVDPATL | ASSPPTEVF | 0.4875672 |
| GNNLVVEAPHAGASV | ASSPPTEVF | 0.6776105 |
| KVTRIMSASTLEQPA | ASSPPTEVF | 0.31809342 |
| LGFALPIALPRRTVV | ASSPPTEVF | 0.4587651 |
| TRLGNLTARKLVVLV | ASSPPTEVF | 0.593662 |
| WSQQRRLIYFPSAGP | ASSPPTEVF | 0.6410365 |
| GPNGNPDPMAAAVDIR | ASSPPTEVF | 0.7237154 |
| TQRCVSVADLRAAASR | ASSPPTEVF | 0.5469415 |
| QFVLDVAEHPVEVLTETV | ASSPPTEVF | 0.67604744 |
| GGAGGTGGAGGNGGTQAGDG | ASSPPTEVF | 0.59077287 |
| PLLATLRAIAVDVDAKPAQV | ASSPPTEVF | 0.5209193 |
| APKFRTGGYMIDFWGVGYQVAK | ASSPPTEVF | 0.49365988 |
| GAGGAGGAGGAAGGPGGTGGTG | ASSPPTEVF | 0.49249247 |
| IGALGLLATDVAPAGSVAFGVA | ASSPPTEVF | 0.423091 |
| AGGAGGIGGNGGAGGAGGVLMG | ASSPPTEVF | 0.6423459 |
| ADELIGLL | ASSPPTEVF | 0.36108023 |
| ADRALKA | ASSPPTEVF | 0.56815094 |
| AGAVGQVGGAGGA | ASSPPTEVF | 0.7199031 |
| AGGGGGLGGAP | ASSPPTEVF | 0.4589628 |
| AGGLLYGNGGAG | ASSPPTEVF | 0.65393984 |
| AGLESIGL | ASSPPTEVF | 0.6670511 |
| AGLKTGII | ASSPPTEVF | 0.67796075 |
| ASLAALLL | ASSPPTEVF | 0.58047485 |
| ASVINISSI | ASSPPTEVF | 0.56006163 |
| ATPPGPA | ASSPPTEVF | 0.7333636 |
| ATVASALV | ASSPPTEVF | 0.7087624 |
| ATVASLIL | ASSPPTEVF | 0.618353 |
| AVNGPVLIRH | ASSPPTEVF | 0.6143391 |
| DGAAGAVGI | ASSPPTEVF | 0.70126015 |

| | | |
|---|---|---|
| DPLLHGI | ASSPPTEVF | 0.70631456 |
| ELGAALGAL | ASSPPTEVF | 0.6419045 |
| ELGRTQAV | ASSPPTEVF | 0.3279217 |
| FVAGGYLS | ASSPPTEVF | 0.5689299 |
| GASGGAGGAGGW | ASSPPTEVF | 0.59492993 |
| GGAGAGGDGGAGGI | ASSPPTEVF | 0.53351325 |
| GTGHNGSQPGMGGNG | ASSPPTEVF | 0.28801164 |
| GYAAASASAS | ASSPPTEVF | 0.730039 |
| HQVDVAAVQQG | ASSPPTEVF | 0.75014275 |
| IGAAIAHT | ASSPPTEVF | 0.44235313 |
| IGMIEVAQL | ASSPPTEVF | 0.74175894 |
| IRLAPEHTSAMD | ASSPPTEVF | 0.38942808 |
| IYNVSVL | ASSPPTEVF | 0.6992465 |
| KQGFLGM | ASSPPTEVF | 0.7253097 |
| LESVITQL | ASSPPTEVF | 0.53084624 |
| LGLGALVVV | ASSPPTEVF | 0.6566081 |
| LIGNGGNGGSGGI | ASSPPTEVF | 0.5553738 |
| LNAINAPTL | ASSPPTEVF | 0.54149544 |
| LQLLNTGV | ASSPPTEVF | 0.728173 |
| LQTSIEF | ASSPPTEVF | 0.5214032 |
| MSFAVSEL | ASSPPTEVF | 0.5439082 |
| NTGALGL | ASSPPTEVF | 0.5872324 |
| PPGAGGAGGAGTT | ASSPPTEVF | 0.46714783 |
| QLVGLGI | ASSPPTEVF | 0.62988436 |
| RFLEQVHQL | ASSPPTEVF | 0.66435003 |
| RGVGYVLR | ASSPPTEVF | 0.42475832 |
| SATGVFAGVI | ASSPPTEVF | 0.7265729 |
| SLPAITLP | ASSPPTEVF | 0.46545324 |
| STALAFTV | ASSPPTEVF | 0.74778545 |
| SVLTVGLGK | ASSPPTEVF | 0.7086336 |
| TALAGAGSGI | ASSPPTEVF | 0.7047906 |
| TGLPFTGI | ASSPPTEVF | 0.534554 |
| VAEHVTV | ASSPPTEVF | 0.73525274 |
| VDIVLNSL | ASSPPTEVF | 0.59493613 |
| VLTGGGAVL | ASSPPTEVF | 0.5700439 |
| VNVGLAYGI | ASSPPTEVF | 0.69550776 |
| VSSAAVGAV | ASSPPTEVF | 0.2536749 |
| VTSVYEAGV | ASSPPTEVF | 0.72667587 |
| WVALDRGA | ASSPPTEVF | 0.6168817 |
| YLVGSVVI | ASSPPTEVF | 0.6800069 |
| GPPGTGKTLL | ASSPPTEVF | 0.45293123 |
| GVGMTKF | ASSPPTEVF | 0.2820317 |
| IGIGGSDL | ASSPPTEVF | 0.48406345 |
| IGTLLAEL | ASSPPTEVF | 0.3963907 |
| IYTSGTT | ASSPPTEVF | 0.7571183 |
| LSFPAGV | ASSPPTEVF | 0.68865466 |
| LYTSGTT | ASSPPTEVF | 0.42435163 |
| VGAFPFGF | ASSPPTEVF | 0.63812006 |
| VGAGTVEF | ASSPPTEVF | 0.6854454 |
| VGKSTLLL | ASSPPTEVF | 0.28689426 |
| AALAHAGGGIV | ASSPPTEVF | 0.51681095 |

| | | |
|---|---|---|
| AALVAVSLL | ASSPPTEVF | 0.38085112 |
| AAVSRAVAAL | ASSPPTEVF | 0.543588 |
| AGGSSGIGGF | ASSPPTEVF | 0.1820254 |
| AGLTVANANG | ASSPPTEVF | 0.66652906 |
| AGLVGAVL | ASSPPTEVF | 0.39311862 |
| ALSQAFTV | ASSPPTEVF | 0.1793944 |
| ALTFSVSL | ASSPPTEVF | 0.6183282 |
| ALTGAALVAL | ASSPPTEVF | 0.4248049 |
| ASFSIGAGVL | ASSPPTEVF | 0.4909647 |
| DGSVYKI | ASSPPTEVF | 0.66448593 |
| EQLAEIV | ASSPPTEVF | 0.50107574 |
| FAVSELTVM | ASSPPTEVF | 0.36880434 |
| GAAPLAAGL | ASSPPTEVF | 0.60954857 |
| GADGGDGGSGGAGG | ASSPPTEVF | 0.7166069 |
| GFGLDTGGAGG | ASSPPTEVF | 0.35379863 |
| GLASGGLL | ASSPPTEVF | 0.5826 |
| GRIAGAAVLL | ASSPPTEVF | 0.7672745 |
| GSAGSLNTSF | ASSPPTEVF | 0.6045393 |
| GTGQIGI | ASSPPTEVF | 0.7417877 |
| GTLIALIL | ASSPPTEVF | 0.7611663 |
| IAATMGFVVG | ASSPPTEVF | 0.46875918 |
| IFAQALL | ASSPPTEVF | 0.7053114 |
| IGAAVALGF | ASSPPTEVF | 0.67004764 |
| IGLTPLAL | ASSPPTEVF | 0.5209756 |
| ILGSNTSTL | ASSPPTEVF | 0.5383651 |
| IVFSLTVL | ASSPPTEVF | 0.25647396 |
| LASKATGF | ASSPPTEVF | 0.4894241 |
| LASVALIL | ASSPPTEVF | 0.35815942 |
| LFAVGLTL | ASSPPTEVF | 0.59336627 |
| LFNGGIGTYI | ASSPPTEVF | 0.75170946 |
| LTLGAGII | ASSPPTEVF | 0.5513038 |
| LVAAALGDI | ASSPPTEVF | 0.2133908 |
| NTAFVQLGI | ASSPPTEVF | 0.26087826 |
| QLFSGAEL | ASSPPTEVF | 0.6016067 |
| SAATFITL | ASSPPTEVF | 0.31259072 |
| SNYGLANLGV | ASSPPTEVF | 0.32889622 |
| SYGGAYAVM | ASSPPTEVF | 0.40193173 |
| TTGVGGLTLG | ASSPPTEVF | 0.72173345 |
| TTQTIYI | ASSPPTEVF | 0.72985613 |
| VAAGAGLGAVI | ASSPPTEVF | 0.45395803 |
| VAAVVTLL | ASSPPTEVF | 0.72882605 |
| VGFLGGLI | ASSPPTEVF | 0.72426 |
| VGVIPILL | ASSPPTEVF | 0.7775775 |
| VIFASSLI | ASSPPTEVF | 0.5688107 |
| VKGGDGGAAAG | ASSPPTEVF | 0.27107918 |
| VTKTFTNI | ASSPPTEVF | 0.72126555 |
| VVAAAILL | ASSPPTEVF | 0.47054702 |
| YAALDVEL | ASSPPTEVF | 0.68843734 |
| LASVALLL | ASSPPTEVF | 0.6629876 |
| LDAVHRV | ASSPPTEVF | 0.49684384 |
| QGLSSSIF | ASSPPTEVF | 0.38099867 |

| | | |
|---|---|---|
| RGTFANIRL | ASSPPTEVF | 0.65918064 |
| RQLWWGHRI | ASSPPTEVF | 0.7409148 |
| YGPPGTGKTLL | ASSPPTEVF | 0.34235814 |
| AALVGAPANVA | ASSPPTEVF | 0.3473751 |
| AATFRRTE | ASSPPTEVF | 0.5662677 |
| AGALGGGI | ASSPPTEVF | 0.54204464 |
| AGGAGGAGGIGAG | ASSPPTEVF | 0.7283313 |
| AGSGALGL | ASSPPTEVF | 0.66305363 |
| AQIADQI | ASSPPTEVF | 0.5000236 |
| ASAVAVTYL | ASSPPTEVF | 0.63017225 |
| ASTVLGGTGG | ASSPPTEVF | 0.69052607 |
| AVGGEPAK | ASSPPTEVF | 0.6063074 |
| CEQGTFGGTC | ASSPPTEVF | 0.5224674 |
| DEGKAPAG | ASSPPTEVF | 0.3877626 |
| DTVVFPAL | ASSPPTEVF | 0.71336156 |
| FEGITMAG | ASSPPTEVF | 0.7123829 |
| FPLVVQD | ASSPPTEVF | 0.66741335 |
| GAFTTPQ | ASSPPTEVF | 0.7294344 |
| GFGGFGTAGGVGGA | ASSPPTEVF | 0.60007936 |
| GFLAQFL | ASSPPTEVF | 0.60769135 |
| GGAGGAGGNAGVA | ASSPPTEVF | 0.34868944 |
| GGVGTAGAGGAGGAGG | ASSPPTEVF | 0.4855519 |
| GNGGGGDAGNAGSGGN | ASSPPTEVF | 0.41320714 |
| GTGGAGGAGNDAGS | ASSPPTEVF | 0.59886634 |
| GTTNTGWFNS | ASSPPTEVF | 0.6395368 |
| IFMGPLFTL | ASSPPTEVF | 0.67990553 |
| ILAHAIEAI | ASSPPTEVF | 0.43556148 |
| IRSAIREAL | ASSPPTEVF | 0.6845318 |
| ITFAGGG | ASSPPTEVF | 0.7264462 |
| ITGMSGHQEV | ASSPPTEVF | 0.38284126 |
| IVAGLACTL | ASSPPTEVF | 0.48847204 |
| IVGGGLAGA | ASSPPTEVF | 0.7367306 |
| IVTGVLNI | ASSPPTEVF | 0.6822581 |
| LAQQGCI | ASSPPTEVF | 0.46765047 |
| LCLHVEN | ASSPPTEVF | 0.5982943 |
| LDAISDGS | ASSPPTEVF | 0.78829455 |
| LGGFAAML | ASSPPTEVF | 0.64400876 |
| LGGKNPMI | ASSPPTEVF | 0.726694 |
| LGPKGVAAIL | ASSPPTEVF | 0.29256338 |
| LMVGVSI | ASSPPTEVF | 0.5571786 |
| LVGAAFVL | ASSPPTEVF | 0.43110102 |
| LVLAVAPL | ASSPPTEVF | 0.16227363 |
| MAAIAVIDAA | ASSPPTEVF | 0.7123443 |
| NNGINAELQ | ASSPPTEVF | 0.4739635 |
| RGEPLVLVHG | ASSPPTEVF | 0.644484 |
| RGGAGGAAGLI | ASSPPTEVF | 0.39512086 |
| RSYRMHPAV | ASSPPTEVF | 0.73860097 |
| SAAHLLAGQLA | ASSPPTEVF | 0.60467553 |
| SAAVGSY | ASSPPTEVF | 0.5328612 |
| SPHAVLHRI | ASSPPTEVF | 0.41973072 |
| SQIAKVL | ASSPPTEVF | 0.6480434 |

| | | |
|---|---|---|
| TALGHLVP | ASSPPTEVF | 0.6714346 |
| TAPTTSGAA | ASSPPTEVF | 0.75628805 |
| TDGDDGNGGSPGT | ASSPPTEVF | 0.25269955 |
| TDGLGEN | ASSPPTEVF | 0.66259176 |
| TGGPPLPGQA | ASSPPTEVF | 0.5868677 |
| TIPAITI | ASSPPTEVF | 0.23100391 |
| TIPGFTI | ASSPPTEVF | 0.18948276 |
| TSIASGALA | ASSPPTEVF | 0.39146408 |
| TVTLPGAAI | ASSPPTEVF | 0.47515282 |
| VASGLSAALP | ASSPPTEVF | 0.5929193 |
| VGAGVIGI | ASSPPTEVF | 0.62012905 |
| VGPKRFGV | ASSPPTEVF | 0.44681215 |
| VNVAFPD | ASSPPTEVF | 0.58715004 |
| VSMTNSGVS | ASSPPTEVF | 0.58301866 |
| VTDVLGDLSV | ASSPPTEVF | 0.11901439 |
| VVVVGAGFAGL | ASSPPTEVF | 0.40041137 |
| VYTVGTVV | ASSPPTEVF | 0.26736194 |
| WTTFSGP | ASSPPTEVF | 0.76592255 |
| YDELVEAF | ASSPPTEVF | 0.32677352 |
| YGTGGAGGNGG | ASSPPTEVF | 0.46778256 |
| YTNVMAV | ASSPPTEVF | 0.5422419 |
| FLEQVHQL | ASSPPTEVF | 0.7060691 |
| GLSLLAVAV | ASSPPTEVF | 0.5658942 |
| LEQVHQL | ASSPPTEVF | 0.40817922 |
| LNAAGAI | ASSPPTEVF | 0.23647557 |
| PAVHASL | ASSPPTEVF | 0.67765796 |
| TELEDDA | ASSPPTEVF | 0.5872809 |
| TPPGPPN | ASSPPTEVF | 0.39864966 |
| TPPPPPPP | ASSPPTEVF | 0.3638165 |
| TQSPALI | ASSQDTDTEVF | 0.48254377 |
| EFAAAFPQ | ASSQDTDTEVF | 0.22984593 |
| LLSGNLDVL | ASSQDTDTEVF | 0.7073188 |
| GPANLGNYNI | ASSQDTDTEVF | 0.32270354 |
| TLGPTLP | ASSQDTDTEVF | 0.45828432 |
| VYAYSSCCK | ASSQDTDTEVF | 0.7546042 |
| APAPAPAPNI | ASSQDTDTEVF | 0.6826724 |
| PGGPVPIP | ASSQDTDTEVF | 0.4887127 |
| FLTVALEP | ASSQDTDTEVF | 0.44470742 |
| SQYFGANL | ASSQDTDTEVF | 0.32958668 |
| GGVKPAA | ASSQDTDTEVF | 0.39744636 |
| GGTGGAGGAGGW | ASSQDTDTEVF | 0.4238935 |
| GVGGVAGLGGAGG | ASSQDTDTEVF | 0.69613874 |
| VLLVLATTI | ASSQDTDTEVF | 0.63336515 |
| IGIFAAML | ASSQDTDTEVF | 0.34018058 |
| AVGDLID | ASSQDTDTEVF | 0.73695505 |
| NISPLPL | ASSQDTDTEVF | 0.72282326 |
| VTAVFVAI | ASSQDTDTEVF | 0.77064514 |
| SGDDNDAAL | ASSQDTDTEVF | 0.36189076 |
| AVAALAHS | ASSQDTDTEVF | 0.54536605 |
| GGVTVMATP | ASSQDTDTEVF | 0.52380943 |
| AGLAANVRD | ASSQDTDTEVF | 0.52002734 |

| | | |
|---|---|---|
| TNRPDLIDPALLRPG | ASSQDTDTEVF | 0.38161463 |
| LANLGVTAL | ASSQDTDTEVF | 0.3996904 |
| TGERIEL | ASSQDTDTEVF | 0.7080921 |
| NGEELAA | ASSQDTDTEVF | 0.82444656 |
| VLSLVWG | ASSQDTDTEVF | 0.5529897 |
| IGYALWL | ASSQDTDTEVF | 0.18871184 |
| LSLLLAPL | ASSQDTDTEVF | 0.49372146 |
| TLDEALK | ASSQDTDTEVF | 0.35589758 |
| TDGAGTAEGG | ASSQDTDTEVF | 0.37960282 |
| NAPGGAGGNGG | ASSQDTDTEVF | 0.48447043 |
| VSVFMGMGGNF | ASSQDTDTEVF | 0.26586518 |
| LASYLAAL | ASSQDTDTEVF | 0.52228475 |
| LVGGTPAL | ASSQDTDTEVF | 0.7718847 |
| VRNADVL | ASSQDTDTEVF | 0.5252263 |
| TCSSPFLVSAM | ASSQDTDTEVF | 0.6213834 |
| VHITDAE | ASSQDTDTEVF | 0.70664465 |
| EYAAPSF | ASSQDTDTEVF | 0.6192381 |
| VVDAFHGGL | ASSQDTDTEVF | 0.7294031 |
| ASVVDPQ | ASSQDTDTEVF | 0.7404976 |
| SYNEFGGLS | ASSQDTDTEVF | 0.5844946 |
| DPDGSVY | ASSQDTDTEVF | 0.5491667 |
| MVSTIHATE | ASSQDTDTEVF | 0.66340274 |
| LAAGVADPV | ASSQDTDTEVF | 0.48784488 |
| TKEQIAA | ASSQDTDTEVF | 0.5811295 |
| SLASPAAL | ASSQDTDTEVF | 0.5132389 |
| IGLVFYGV | ASSQDTDTEVF | 0.6684149 |
| FLLDASNL | ASSQDTDTEVF | 0.48668164 |
| YQVGNFP | ASSQDTDTEVF | 0.7865422 |
| ESLLSQST | ASSQDTDTEVF | 0.17785297 |
| LFTVGFAAVG | ASSQDTDTEVF | 0.36903405 |
| ATPPTPT | ASSQDTDTEVF | 0.7147196 |
| TDLATLG | ASSQDTDTEVF | 0.16409636 |
| SAGGTGR | ASSQDTDTEVF | 0.28738654 |
| GFSNSGS | ASSQDTDTEVF | 0.53092855 |
| GGNGGSGGTSV | ASSQDTDTEVF | 0.73300827 |
| GTSVATGGAGN | ASSQDTDTEVF | 0.16862346 |
| NGGNGGSGGTS | ASSQDTDTEVF | 0.5281283 |
| AVVKFPH | ASSQDTDTEVF | 0.25821152 |
| TPPPPPP | ASSQDTDTEVF | 0.73833174 |
| AAPAAAAAPA | ASSQDTDTEVF | 0.5290384 |
| IALNEDL | ASSQDTDTEVF | 0.28910977 |
| IVNVSSM | ASSQDTDTEVF | 0.7113881 |
| TGIPPAPR | ASSQDTDTEVF | 0.2240041 |
| FDLGGGTF | ASSQDTDTEVF | 0.56665313 |
| FDLGGGTFDVS | ASSQDTDTEVF | 0.7022542 |
| NEPTAAA | ASSQDTDTEVF | 0.7842525 |
| NEPTAAALA | ASSQDTDTEVF | 0.26528022 |
| GSIINISSI | ASSQDTDTEVF | 0.6204679 |
| VAPGNAGTA | ASSQDTDTEVF | 0.45292282 |
| TGGGASLE | ASSQDTDTEVF | 0.78309685 |
| VAVYNHYKRI | ASSQDTDTEVF | 0.46739256 |

| | | |
|---|---|---|
| TDPAPATT | ASSQDTDTEVF | 0.24626015 |
| TNRPDILDPALLRPG | ASSQDTDTEVF | 0.6970955 |
| LTTTTTS | ASSQDTDTEVF | 0.686872 |
| PTGSGKTL | ASSQDTDTEVF | 0.53687644 |
| TGSGKTL | ASSQDTDTEVF | 0.7069373 |
| TIGHVDHGK | ASSQDTDTEVF | 0.60948896 |
| TIGHVDHGKTT | ASSQDTDTEVF | 0.47954854 |
| GHVDHGKTT | ASSQDTDTEVF | 0.6697515 |
| AQFEHTLL | ASSQDTDTEVF | 0.60674566 |
| IVNQDEL | ASSQDTDTEVF | 0.45202458 |
| TNGSQFF | ASSQDTDTEVF | 0.74472463 |
| TGKGIVE | ASSQDTDTEVF | 0.5912769 |
| LISNASDA | ASSQDTDTEVF | 0.3214328 |
| AELRTLGELTKIAKAHGAQ | ASSQDTDTEVF | 0.78159463 |
| APNAVAAAEPSW | ASSQDTDTEVF | 0.5430309 |
| APQVAGAATAVL | ASSQDTDTEVF | 0.6474614 |
| ARRDPAAAAAILPTDA | ASSQDTDTEVF | 0.69875497 |
| DAAARDVLAEAGLAEYF | ASSQDTDTEVF | 0.8198426 |
| DDQIVLSAKANLAKVEQ | ASSQDTDTEVF | 0.41130704 |
| DELLPDTGEDSDEDGH | ASSQDTDTEVF | 0.6102245 |
| ELPEAPANRATVPQD | ASSQDTDTEVF | 0.5593506 |
| FGQSKIERVTGIG | ASSQDTDTEVF | 0.6871095 |
| GAGGTGSTAGGAGGAGGAGGLY | ASSQDTDTEVF | 0.56751204 |
| GHGGTGAAGGNGGTGGAGGSGI | ASSQDTDTEVF | 0.6065863 |
| GNGGNGGAGGEGGPGGAGGAGA | ASSQDTDTEVF | 0.6687353 |
| GSTAGNGGNGGNATNGTIAGQP | ASSQDTDTEVF | 0.7198484 |
| GTGGDGGNGGNGGTGG | ASSQDTDTEVF | 0.5194259 |
| GVRRPAGIPADAPV | ASSQDTDTEVF | 0.47660446 |
| GYRQVTDAHLLTLARR | ASSQDTDTEVF | 0.76658285 |
| IAKMGQDGHDRG | ASSQDTDTEVF | 0.2992432 |
| KAEPSIEIQYT | ASSQDTDTEVF | 0.6168933 |
| KPVRIRMAADEDTVAVT | ASSQDTDTEVF | 0.7013831 |
| MPPRIAGMRLLVIKPEP | ASSQDTDTEVF | 0.64764714 |
| MKFYAVALFRSRGVPPTR | ASSQDTDTEVF | 0.5450524 |
| NAGGFGILGAVAHSP | ASSQDTDTEVF | 0.72083557 |
| NEGREITLAMAEM | ASSQDTDTEVF | 0.48413724 |
| NELVEWSHPLGHRWRWE | ASSQDTDTEVF | 0.58727425 |
| NLGAGGTGGNGGNPG | ASSQDTDTEVF | 0.50585884 |
| NTEWPKWETVMAADPVD | ASSQDTDTEVF | 0.40620595 |
| PASAAPAETQP | ASSQDTDTEVF | 0.31664374 |
| QQSSDGKNGVRAAEITGE | ASSQDTDTEVF | 0.48737037 |
| RAPDAVTTSWWKE | ASSQDTDTEVF | 0.6393812 |
| SGFGNVGAHTSGWFNQ | ASSQDTDTEVF | 0.5798358 |
| SGVAGAVGQAASVGGLKV | ASSQDTDTEVF | 0.24169694 |
| SNGGAGTGSAGGNGGT | ASSQDTDTEVF | 0.6359012 |
| TGGAGGEGGAAG | ASSQDTDTEVF | 0.6299783 |
| TGGDGGAAGAGGAPGHG | ASSQDTDTEVF | 0.7817621 |
| TMPPLLAAIANAGRHGV | ASSQDTDTEVF | 0.5226695 |
| TSSTAGNSGGA | ASSQDTDTEVF | 0.7155677 |
| VPATVPVGILC | ASSQDTDTEVF | 0.4450392 |
| VQRPDSGGVVLTGRIS | ASSQDTDTEVF | 0.63615286 |

| | | |
|---|---|---|
| WTPNPVPATKGVPAGLVQAT | ASSQDTDTEVF | 0.24584746 |
| YALKRALAKFSDRFDVVII | ASSQDTDTEVF | 0.35081235 |
| YLIGSVIV | ASSQDTDTEVF | 0.5171578 |
| YGQLVAVID | ASSQDTDTEVF | 0.3625026 |
| VTVGGMTLPA | ASSQDTDTEVF | 0.7185878 |
| VLALPAV | ASSQDTDTEVF | 0.65208215 |
| VIPPTLNYET | ASSQDTDTEVF | 0.6892076 |
| VGMGGMHPSSG | ASSQDTDTEVF | 0.708483 |
| VGLAGGAATA | ASSQDTDTEVF | 0.7207905 |
| VGGAGGAGGSGP | ASSQDTDTEVF | 0.7248924 |
| VFAAFMV | ASSQDTDTEVF | 0.46911252 |
| TLYDKLR | ASSQDTDTEVF | 0.48248515 |
| TLAVFTVL | ASSQDTDTEVF | 0.23282383 |
| TGSVQPARV | ASSQDTDTEVF | 0.43170297 |
| TGGGAGSGGT | ASSQDTDTEVF | 0.44744238 |
| SHLETSL | ASSQDTDTEVF | 0.5230578 |
| SFTTGMT | ASSQDTDTEVF | 0.16053885 |
| QVEVAKRMGVRQPTVS | ASSQDTDTEVF | 0.6454054 |
| QGSGGPALAM | ASSQDTDTEVF | 0.69686866 |
| MLAPSNP | ASSQDTDTEVF | 0.5466604 |
| LVNNAAI | ASSQDTDTEVF | 0.67065513 |
| LTGEINTLA | ASSQDTDTEVF | 0.6215912 |
| LSIFGPV | ASSQDTDTEVF | 0.3870048 |
| LLALLANL | ASSQDTDTEVF | 0.63072646 |
| LISNASDALDKLR | ASSQDTDTEVF | 0.4631829 |
| LINEAVP | ASSQDTDTEVF | 0.72839534 |
| LGYAFSGL | ASSQDTDTEVF | 0.74726325 |
| LDFVAVAL | ASSQDTDTEVF | 0.628187 |
| LAQPGAE | ASSQDTDTEVF | 0.37203735 |
| ISNLGAVTGF | ASSQDTDTEVF | 0.64709723 |
| HQEDPGRI | ASSQDTDTEVF | 0.37400407 |
| GTPGEPTP | ASSQDTDTEVF | 0.40899456 |
| GFFNSGS | ASSQDTDTEVF | 0.20603047 |
| GAGGGGLTAGAN | ASSQDTDTEVF | 0.59503245 |
| FVDALVNM | ASSQDTDTEVF | 0.4094552 |
| FLTVAPEL | ASSQDTDTEVF | 0.6440822 |
| FETLTAL | ASSQDTDTEVF | 0.55136096 |
| DVINPIEL | ASSQDTDTEVF | 0.59239197 |
| APVSGAPV | ASSQDTDTEVF | 0.34727836 |
| ALVGGLAW | ASSQDTDTEVF | 0.714275 |
| ALAAAAATAI | ASSQDTDTEVF | 0.6823276 |
| AFGLGLLL | ASSQDTDTEVF | 0.3054742 |
| AFFAAASL | ASSQDTDTEVF | 0.68209445 |
| AAAALPAFSPP | ASSQDTDTEVF | 0.5196039 |
| VVLAPAL | ASSQDTDTEVF | 0.52832234 |
| TIGHVDHGKTTL | ASSQDTDTEVF | 0.72107184 |
| LISNASDALDK | ASSQDTDTEVF | 0.5350141 |
| LISNASDAL | ASSQDTDTEVF | 0.28531665 |
| ISNASDALDK | ASSQDTDTEVF | 0.43267322 |
| NQAETLVYQTE | ASSQDTDTEVF | 0.58528316 |
| TGGTGGTAGAG | ASSQDTDTEVF | 0.52304775 |

| | | |
|---|---|---|
| TGSTAGGAGGA | ASSQDTDTEVF | 0.17491135 |
| DPLAVPVAAGPV | ASSQDTDTEVF | 0.6571075 |
| GGTGGTPGNGGN | ASSQDTDTEVF | 0.32746905 |
| PAGLAGVDPATL | ASSQDTDTEVF | 0.44308895 |
| GNNLVVEAPHAGASV | ASSQDTDTEVF | 0.47523037 |
| KVTRIMSASTLEQPA | ASSQDTDTEVF | 0.7100464 |
| LGFALPIALPRRTVV | ASSQDTDTEVF | 0.6203711 |
| TRLGNLTARKLVVLV | ASSQDTDTEVF | 0.5465465 |
| WSQQRRLIYFPSAGP | ASSQDTDTEVF | 0.5609082 |
| GPNGNPDPMAAAVDIR | ASSQDTDTEVF | 0.5897167 |
| TQRCVSVADLRAAASR | ASSQDTDTEVF | 0.7030819 |
| QFVLDVAEHPVEVLTETV | ASSQDTDTEVF | 0.6079388 |
| GGAGGTGGAGGNGGTQAGDG | ASSQDTDTEVF | 0.5509886 |
| PLLATLRAIAVDVDAKPAQV | ASSQDTDTEVF | 0.63936454 |
| APKFRTGGYMIDFWGVGYQVAK | ASSQDTDTEVF | 0.47089124 |
| GAGGAGGAGGAAGGPGGTGGTG | ASSQDTDTEVF | 0.3729647 |
| IGALGLLATDVAPAGSVAFGVA | ASSQDTDTEVF | 0.4934489 |
| AGGAGGIGGNGGAGGAGGVLMG | ASSQDTDTEVF | 0.33171073 |
| ADELIGLL | ASSQDTDTEVF | 0.35601017 |
| ADRALKA | ASSQDTDTEVF | 0.65024984 |
| AGAVGQVGGAGGA | ASSQDTDTEVF | 0.7784157 |
| AGGGGGLGGAP | ASSQDTDTEVF | 0.6037415 |
| AGGLLYGNGGAG | ASSQDTDTEVF | 0.21387379 |
| AGLESIGL | ASSQDTDTEVF | 0.4262207 |
| AGLKTGII | ASSQDTDTEVF | 0.51938975 |
| ASLAALLL | ASSQDTDTEVF | 0.6317669 |
| ASVINISSI | ASSQDTDTEVF | 0.70604396 |
| ATPPGPA | ASSQDTDTEVF | 0.78752416 |
| ATVASALV | ASSQDTDTEVF | 0.69283164 |
| ATVASLIL | ASSQDTDTEVF | 0.45708108 |
| AVNGPVLIRH | ASSQDTDTEVF | 0.6875938 |
| DGAAGAVGI | ASSQDTDTEVF | 0.5154548 |
| DPLLHGI | ASSQDTDTEVF | 0.50908166 |
| ELGAALGAL | ASSQDTDTEVF | 0.5654495 |
| ELGRTQAV | ASSQDTDTEVF | 0.65074396 |
| FVAGGYLS | ASSQDTDTEVF | 0.36887172 |
| GASGGAGGAGGW | ASSQDTDTEVF | 0.46215332 |
| GGAGAGGDGGAGGI | ASSQDTDTEVF | 0.25731242 |
| GTGHNGSQPGMGGNG | ASSQDTDTEVF | 0.68898773 |
| GYAAASASAS | ASSQDTDTEVF | 0.29859275 |
| HQVDVAAVQQG | ASSQDTDTEVF | 0.4073858 |
| IGAAIAHT | ASSQDTDTEVF | 0.50949144 |
| IGMIEVAQL | ASSQDTDTEVF | 0.7603395 |
| IRLAPEHTSAMD | ASSQDTDTEVF | 0.74458075 |
| IYNVSVL | ASSQDTDTEVF | 0.46049142 |
| KQGFLGM | ASSQDTDTEVF | 0.450799 |
| LESVITQL | ASSQDTDTEVF | 0.63940495 |
| LGLGALVVV | ASSQDTDTEVF | 0.49662098 |
| LIGNGGNGGSGGI | ASSQDTDTEVF | 0.695603 |
| LNAINAPTL | ASSQDTDTEVF | 0.5708736 |
| LQLLNTGV | ASSQDTDTEVF | 0.5110513 |

| | | |
|---|---|---|
| LQTSIEF | ASSQDTDTEVF | 0.4759693 |
| MSFAVSEL | ASSQDTDTEVF | 0.6483495 |
| NTGALGL | ASSQDTDTEVF | 0.44625622 |
| PPGAGGAGGAGTT | ASSQDTDTEVF | 0.6487005 |
| QLVGLGI | ASSQDTDTEVF | 0.19789802 |
| RFLEQVHQL | ASSQDTDTEVF | 0.7337525 |
| RGVGYVLR | ASSQDTDTEVF | 0.31460014 |
| SATGVFAGVI | ASSQDTDTEVF | 0.38884878 |
| SLPAITLP | ASSQDTDTEVF | 0.17100573 |
| STALAFTV | ASSQDTDTEVF | 0.52460927 |
| SVLTVGLGK | ASSQDTDTEVF | 0.61262894 |
| TALAGAGSGI | ASSQDTDTEVF | 0.7176261 |
| TGLPFTGI | ASSQDTDTEVF | 0.6038983 |
| VAEHVTV | ASSQDTDTEVF | 0.59841603 |
| VDIVLNSL | ASSQDTDTEVF | 0.68136466 |
| VLTGGGAVL | ASSQDTDTEVF | 0.36771458 |
| VNVGLAYGI | ASSQDTDTEVF | 0.22329621 |
| VSSAAVGAV | ASSQDTDTEVF | 0.6305196 |
| VTSVYEAGV | ASSQDTDTEVF | 0.56463313 |
| WVALDRGA | ASSQDTDTEVF | 0.73815167 |
| YLVGSVVI | ASSQDTDTEVF | 0.67614114 |
| GPPGTGKTLL | ASSQDTDTEVF | 0.22608872 |
| GVGMTKF | ASSQDTDTEVF | 0.67660964 |
| IGIGGSDL | ASSQDTDTEVF | 0.6844405 |
| IGTLLAEL | ASSQDTDTEVF | 0.24993172 |
| IYTSGTT | ASSQDTDTEVF | 0.33575588 |
| LSFPAGV | ASSQDTDTEVF | 0.39604777 |
| LYTSGTT | ASSQDTDTEVF | 0.29940307 |
| VGAFPFGF | ASSQDTDTEVF | 0.61721206 |
| VGAGTVEF | ASSQDTDTEVF | 0.44317302 |
| VGKSTLLL | ASSQDTDTEVF | 0.5520133 |
| AALAHAGGGIV | ASSQDTDTEVF | 0.5159391 |
| AALVAVSLL | ASSQDTDTEVF | 0.4231271 |
| AAVSRAVAAL | ASSQDTDTEVF | 0.55716735 |
| AGGSSGIGGF | ASSQDTDTEVF | 0.5100424 |
| AGLTVANANG | ASSQDTDTEVF | 0.66894066 |
| AGLVGAVL | ASSQDTDTEVF | 0.6641145 |
| ALSQAFTV | ASSQDTDTEVF | 0.5313573 |
| ALTFSVSL | ASSQDTDTEVF | 0.6497241 |
| ALTGAALVAL | ASSQDTDTEVF | 0.6904645 |
| ASFSIGAGVL | ASSQDTDTEVF | 0.28244624 |
| DGSVYKI | ASSQDTDTEVF | 0.4107263 |
| EQLAEIV | ASSQDTDTEVF | 0.68075275 |
| FAVSELTVM | ASSQDTDTEVF | 0.55677634 |
| GAAPLAAGL | ASSQDTDTEVF | 0.73882073 |
| GADGGDGGSGGAGG | ASSQDTDTEVF | 0.6058227 |
| GFGLDTGGAGG | ASSQDTDTEVF | 0.58544934 |
| GLASGGLL | ASSQDTDTEVF | 0.5497871 |
| GRIAGAAVLL | ASSQDTDTEVF | 0.75303066 |
| GSAGSLNTSF | ASSQDTDTEVF | 0.6375234 |
| GTGQIGI | ASSQDTDTEVF | 0.5014208 |

| | | |
|---|---|---|
| GTLIALIL | ASSQDTDTEVF | 0.6837612 |
| IAATMGFVVG | ASSQDTDTEVF | 0.29936033 |
| IFAQALL | ASSQDTDTEVF | 0.6486932 |
| IGAAVALGF | ASSQDTDTEVF | 0.6859478 |
| IGLTPLAL | ASSQDTDTEVF | 0.3479759 |
| ILGSNTSTL | ASSQDTDTEVF | 0.4879626 |
| IVFSLTVL | ASSQDTDTEVF | 0.73001206 |
| LASKATGF | ASSQDTDTEVF | 0.69224036 |
| LASVALIL | ASSQDTDTEVF | 0.51269406 |
| LFAVGLTL | ASSQDTDTEVF | 0.4480738 |
| LFNGGIGTYI | ASSQDTDTEVF | 0.3788796 |
| LTLGAGII | ASSQDTDTEVF | 0.4180742 |
| LVAAALGDI | ASSQDTDTEVF | 0.26258075 |
| NTAFVQLGI | ASSQDTDTEVF | 0.6229413 |
| QLFSGAEL | ASSQDTDTEVF | 0.45118678 |
| SAATFITL | ASSQDTDTEVF | 0.7540883 |
| SNYGLANLGV | ASSQDTDTEVF | 0.6366853 |
| SYGGAYAVM | ASSQDTDTEVF | 0.73521495 |
| TTGVGGLTLG | ASSQDTDTEVF | 0.45056075 |
| TTQTIYI | ASSQDTDTEVF | 0.54734707 |
| VAAGAGLGAVI | ASSQDTDTEVF | 0.24651212 |
| VAAVVTLL | ASSQDTDTEVF | 0.39114743 |
| VGFLGGLI | ASSQDTDTEVF | 0.34207705 |
| VGVIPILL | ASSQDTDTEVF | 0.57524425 |
| VIFASSLI | ASSQDTDTEVF | 0.24037346 |
| VKGGDGGAAAG | ASSQDTDTEVF | 0.7639601 |
| VTKTFTNI | ASSQDTDTEVF | 0.4455685 |
| VVAAAILL | ASSQDTDTEVF | 0.69158864 |
| YAALDVEL | ASSQDTDTEVF | 0.46482688 |
| LASVALLL | ASSQDTDTEVF | 0.2715927 |
| LDAVHRV | ASSQDTDTEVF | 0.46381557 |
| QGLSSSIF | ASSQDTDTEVF | 0.68255955 |
| RGTFANIRL | ASSQDTDTEVF | 0.28759527 |
| RQLWWGHRI | ASSQDTDTEVF | 0.17357744 |
| YGPPGTGKTLL | ASSQDTDTEVF | 0.36605853 |
| AALVGAPANVA | ASSQDTDTEVF | 0.30954596 |
| AATFRRTE | ASSQDTDTEVF | 0.7736175 |
| AGALGGGI | ASSQDTDTEVF | 0.7203346 |
| AGGAGGAGGIGAG | ASSQDTDTEVF | 0.54324913 |
| AGSGALGL | ASSQDTDTEVF | 0.78128314 |
| AQIADQI | ASSQDTDTEVF | 0.14400238 |
| ASAVAVTYL | ASSQDTDTEVF | 0.524205 |
| ASTVLGGTGG | ASSQDTDTEVF | 0.7572496 |
| AVGGEPAK | ASSQDTDTEVF | 0.6424475 |
| CEQGTFGGTC | ASSQDTDTEVF | 0.49267453 |
| DEGKAPAG | ASSQDTDTEVF | 0.69315493 |
| DTVVFPAL | ASSQDTDTEVF | 0.42563674 |
| FEGITMAG | ASSQDTDTEVF | 0.5363388 |
| FPLVVQD | ASSQDTDTEVF | 0.559283 |
| GAFTTPQ | ASSQDTDTEVF | 0.6673572 |
| GFGGFGTAGGVGGA | ASSQDTDTEVF | 0.6021692 |

| | | |
|---|---|---|
| GFLAQFL | ASSQDTDTEVF | 0.244742 |
| GGAGGAGGNAGVA | ASSQDTDTEVF | 0.7009231 |
| GGVGTAGAGGAGGAGG | ASSQDTDTEVF | 0.7434763 |
| GNGGGGDAGNAGSGGN | ASSQDTDTEVF | 0.44556618 |
| GTGGAGGAGNDAGS | ASSQDTDTEVF | 0.6480465 |
| GTTNTGWFNS | ASSQDTDTEVF | 0.5592145 |
| IFMGPLFTL | ASSQDTDTEVF | 0.50215554 |
| ILAHAIEAI | ASSQDTDTEVF | 0.37982216 |
| IRSAIREAL | ASSQDTDTEVF | 0.2579757 |
| ITFAGGG | ASSQDTDTEVF | 0.7170032 |
| ITGMSGHQEV | ASSQDTDTEVF | 0.3457737 |
| IVAGLACTL | ASSQDTDTEVF | 0.56868196 |
| IVGGGLAGA | ASSQDTDTEVF | 0.60687286 |
| IVTGVLNI | ASSQDTDTEVF | 0.78290915 |
| LAQQGCI | ASSQDTDTEVF | 0.61505497 |
| LCLHVEN | ASSQDTDTEVF | 0.6015862 |
| LDAISDGS | ASSQDTDTEVF | 0.44382656 |
| LGGFAAML | ASSQDTDTEVF | 0.573635 |
| LGGKNPMI | ASSQDTDTEVF | 0.6473898 |
| LGPKGVAAIL | ASSQDTDTEVF | 0.25658858 |
| LMVGVSI | ASSQDTDTEVF | 0.6981597 |
| LVGAAFVL | ASSQDTDTEVF | 0.324847 |
| LVLAVAPL | ASSQDTDTEVF | 0.5957227 |
| MAAIAVIDAA | ASSQDTDTEVF | 0.64604694 |
| NNGINAELQ | ASSQDTDTEVF | 0.77558684 |
| RGEPLVLVHG | ASSQDTDTEVF | 0.5510498 |
| RGGAGGAAGLI | ASSQDTDTEVF | 0.730381 |
| RSYRMHPAV | ASSQDTDTEVF | 0.38154274 |
| SAAHLLAGQLA | ASSQDTDTEVF | 0.6863649 |
| SAAVGSY | ASSQDTDTEVF | 0.7154778 |
| SPHAVLHRI | ASSQDTDTEVF | 0.67547 |
| SQIAKVL | ASSQDTDTEVF | 0.77575195 |
| TALGHLVP | ASSQDTDTEVF | 0.709133 |
| TAPTTSGAA | ASSQDTDTEVF | 0.22463505 |
| TDGDDGNGGSPGT | ASSQDTDTEVF | 0.5607915 |
| TDGLGEN | ASSQDTDTEVF | 0.74374795 |
| TGGPPLPGQA | ASSQDTDTEVF | 0.7068426 |
| TIPAITI | ASSQDTDTEVF | 0.69546974 |
| TIPGFTI | ASSQDTDTEVF | 0.57000065 |
| TSIASGALA | ASSQDTDTEVF | 0.5018108 |
| TVTLPGAAI | ASSQDTDTEVF | 0.7363863 |
| VASGLSAALP | ASSQDTDTEVF | 0.83418435 |
| VGAGVIGI | ASSQDTDTEVF | 0.67959595 |
| VGPKRFGV | ASSQDTDTEVF | 0.38685685 |
| VNVAFPD | ASSQDTDTEVF | 0.3702583 |
| VSMTNSGVS | ASSQDTDTEVF | 0.6596786 |
| VTDVLGDLSV | ASSQDTDTEVF | 0.70305634 |
| VVVVGAGFAGL | ASSQDTDTEVF | 0.30791655 |
| VYTVGTVV | ASSQDTDTEVF | 0.5144454 |
| WTTFSGP | ASSQDTDTEVF | 0.579775 |
| YDELVEAF | ASSQDTDTEVF | 0.17492321 |

| | | |
|---|---|---|
| YGTGGAGGNGG | ASSQDTDTEVF | 0.67501783 |
| YTNVMAV | ASSQDTDTEVF | 0.71475065 |
| FLEQVHQL | ASSQDTDTEVF | 0.6581172 |
| GLSLLAVAV | ASSQDTDTEVF | 0.27911395 |
| LEQVHQL | ASSQDTDTEVF | 0.68436384 |
| LNAAGAI | ASSQDTDTEVF | 0.68375564 |
| PAVHASL | ASSQDTDTEVF | 0.7742518 |
| TELEDDA | ASSQDTDTEVF | 0.29168588 |
| TPPGPPN | ASSQDTDTEVF | 0.15511721 |
| TPPPPPPP | ASSQDTDTEVF | 0.62608516 |
| TQSPALI | ASSQDRGQSNERLF | 0.7388438 |
| EFAAAFPQ | ASSQDRGQSNERLF | 0.48771077 |
| LLSGNLDVL | ASSQDRGQSNERLF | 0.40475154 |
| GPANLGNYNI | ASSQDRGQSNERLF | 0.5245732 |
| TLGPTLP | ASSQDRGQSNERLF | 0.7200477 |
| VYAYSSCCK | ASSQDRGQSNERLF | 0.28139958 |
| APAPAPAPNI | ASSQDRGQSNERLF | 0.29241157 |
| PGGPVPIP | ASSQDRGQSNERLF | 0.61142755 |
| FLTVALEP | ASSQDRGQSNERLF | 0.73015606 |
| SQYFGANL | ASSQDRGQSNERLF | 0.33778572 |
| GGVKPAA | ASSQDRGQSNERLF | 0.46505105 |
| GGTGGAGGAGGW | ASSQDRGQSNERLF | 0.65580213 |
| GVGGVAGLGGAGG | ASSQDRGQSNERLF | 0.69840646 |
| VLLVLATTI | ASSQDRGQSNERLF | 0.5743889 |
| IGIFAAML | ASSQDRGQSNERLF | 0.3048707 |
| AVGDLID | ASSQDRGQSNERLF | 0.62200487 |
| NISPLPL | ASSQDRGQSNERLF | 0.27636382 |
| VTAVFVAI | ASSQDRGQSNERLF | 0.66923934 |
| SGDDNDAAL | ASSQDRGQSNERLF | 0.49932146 |
| AVAALAHS | ASSQDRGQSNERLF | 0.633687 |
| GGVTVMATP | ASSQDRGQSNERLF | 0.48064595 |
| AGLAANVRD | ASSQDRGQSNERLF | 0.27806163 |
| TNRPDLIDPALLRPG | ASSQDRGQSNERLF | 0.34862193 |
| LANLGVTAL | ASSQDRGQSNERLF | 0.3562256 |
| TGERIEL | ASSQDRGQSNERLF | 0.65209126 |
| NGEELAA | ASSQDRGQSNERLF | 0.36474776 |
| VLSLVWG | ASSQDRGQSNERLF | 0.6178174 |
| IGYALWL | ASSQDRGQSNERLF | 0.597749 |
| LSLLLAPL | ASSQDRGQSNERLF | 0.35854873 |
| TLDEALK | ASSQDRGQSNERLF | 0.6337943 |
| TDGAGTAEGG | ASSQDRGQSNERLF | 0.27903026 |
| NAPGGAGGNGG | ASSQDRGQSNERLF | 0.6446527 |
| VSVFMGMGGNF | ASSQDRGQSNERLF | 0.5595858 |
| LASYLAAL | ASSQDRGQSNERLF | 0.7163464 |
| LVGGTPAL | ASSQDRGQSNERLF | 0.3948145 |
| VRNADVL | ASSQDRGQSNERLF | 0.65009034 |
| TCSSPFLVSAM | ASSQDRGQSNERLF | 0.63324124 |
| VHITDAE | ASSQDRGQSNERLF | 0.3468384 |
| EYAAPSF | ASSQDRGQSNERLF | 0.65624285 |
| VVDAFHGGL | ASSQDRGQSNERLF | 0.6980872 |
| ASVVDPQ | ASSQDRGQSNERLF | 0.6602381 |

| | | |
|---|---|---|
| SYNEFGGLS | ASSQDRGQSNERLF | 0.21278036 |
| DPDGSVY | ASSQDRGQSNERLF | 0.57758594 |
| MVSTIHATE | ASSQDRGQSNERLF | 0.397706 |
| LAAGVADPV | ASSQDRGQSNERLF | 0.61001456 |
| TKEQIAA | ASSQDRGQSNERLF | 0.55071336 |
| SLASPAAL | ASSQDRGQSNERLF | 0.4603356 |
| IGLVFYGV | ASSQDRGQSNERLF | 0.6262269 |
| FLLDASNL | ASSQDRGQSNERLF | 0.347704 |
| YQVGNFP | ASSQDRGQSNERLF | 0.41403416 |
| ESLLSQST | ASSQDRGQSNERLF | 0.5005617 |
| LFTVGFAAVG | ASSQDRGQSNERLF | 0.6245692 |
| ATPPTPT | ASSQDRGQSNERLF | 0.65459806 |
| TDLATLG | ASSQDRGQSNERLF | 0.3400737 |
| SAGGTGR | ASSQDRGQSNERLF | 0.58802944 |
| GFSNSGS | ASSQDRGQSNERLF | 0.540699 |
| GGNGGSGGTSV | ASSQDRGQSNERLF | 0.39686227 |
| GTSVATGGAGN | ASSQDRGQSNERLF | 0.6078873 |
| NGGNGGSGGTS | ASSQDRGQSNERLF | 0.4793346 |
| AVVKFPH | ASSQDRGQSNERLF | 0.22862229 |
| TPPPPPP | ASSQDRGQSNERLF | 0.6726261 |
| AAPAAAAAPA | ASSQDRGQSNERLF | 0.3525042 |
| IALNEDL | ASSQDRGQSNERLF | 0.62484086 |
| IVNVSSM | ASSQDRGQSNERLF | 0.759194 |
| TGIPPAPR | ASSQDRGQSNERLF | 0.22598836 |
| FDLGGGTF | ASSQDRGQSNERLF | 0.65100074 |
| FDLGGGTFDVS | ASSQDRGQSNERLF | 0.5487732 |
| NEPTAAA | ASSQDRGQSNERLF | 0.46119255 |
| NEPTAAALA | ASSQDRGQSNERLF | 0.56506383 |
| GSIINISSI | ASSQDRGQSNERLF | 0.42555547 |
| VAPGNAGTA | ASSQDRGQSNERLF | 0.73154116 |
| TGGGASLE | ASSQDRGQSNERLF | 0.65039194 |
| VAVYNHYKRI | ASSQDRGQSNERLF | 0.5950884 |
| TDPAPATT | ASSQDRGQSNERLF | 0.5822092 |
| TNRPDILDPALLRPG | ASSQDRGQSNERLF | 0.6923163 |
| LTTTTTS | ASSQDRGQSNERLF | 0.71696913 |
| PTGSGKTL | ASSQDRGQSNERLF | 0.375459 |
| TGSGKTL | ASSQDRGQSNERLF | 0.23491225 |
| TIGHVDHGK | ASSQDRGQSNERLF | 0.54318404 |
| TIGHVDHGKTT | ASSQDRGQSNERLF | 0.51405704 |
| GHVDHGKTT | ASSQDRGQSNERLF | 0.3806115 |
| AQFEHTLL | ASSQDRGQSNERLF | 0.27070725 |
| IVNQDEL | ASSQDRGQSNERLF | 0.6769948 |
| TNGSQFF | ASSQDRGQSNERLF | 0.7903725 |
| TGKGIVE | ASSQDRGQSNERLF | 0.43520314 |
| LISNASDA | ASSQDRGQSNERLF | 0.7979659 |
| AELRTLGELTKIAKAHGAQ | ASSQDRGQSNERLF | 0.30767426 |
| APNAVAAAEPSW | ASSQDRGQSNERLF | 0.6789784 |
| APQVAGAATAVL | ASSQDRGQSNERLF | 0.7014735 |
| ARRDPAAAAAILPTDA | ASSQDRGQSNERLF | 0.30562958 |
| DAAARDVLAEAGLAEYF | ASSQDRGQSNERLF | 0.65039295 |
| DDQIVLSAKANLAKVEQ | ASSQDRGQSNERLF | 0.3436193 |

| | | |
|---|---|---|
| DELLPDTGEDSDEDGH | ASSQDRGQSNERLF | 0.3752034 |
| ELPEAPANRATVPQD | ASSQDRGQSNERLF | 0.66185546 |
| FGQSKIERVTGIG | ASSQDRGQSNERLF | 0.5569724 |
| GAGGTGSTAGGAGGAGGAGGLY | ASSQDRGQSNERLF | 0.6169643 |
| GHGGTGAAGGNGGTGGAGGSGI | ASSQDRGQSNERLF | 0.54431015 |
| GNGGNGGAGGEGGPGGAGGAGA | ASSQDRGQSNERLF | 0.40936446 |
| GSTAGNGGNGGNATNGTIAGQP | ASSQDRGQSNERLF | 0.6339297 |
| GTGGDGGNGGNGGTGG | ASSQDRGQSNERLF | 0.6253948 |
| GVRRPAGIPADAPV | ASSQDRGQSNERLF | 0.6175271 |
| GYRQVTDAHLLTLARR | ASSQDRGQSNERLF | 0.8276104 |
| IAKMGQDGHDRG | ASSQDRGQSNERLF | 0.28894606 |
| KAEPSIEIQYT | ASSQDRGQSNERLF | 0.72227967 |
| KPVRIRMAADEDTVAVT | ASSQDRGQSNERLF | 0.72483367 |
| MPPRIAGMRLLVIKPEP | ASSQDRGQSNERLF | 0.5969218 |
| MKFYAVALFRSRGVPPTR | ASSQDRGQSNERLF | 0.52344877 |
| NAGGFGILGAVAHSP | ASSQDRGQSNERLF | 0.69291866 |
| NEGREITLAMAEM | ASSQDRGQSNERLF | 0.5628003 |
| NELVEWSHPLGHRWRWE | ASSQDRGQSNERLF | 0.49037153 |
| NLGAGGTGGNGGNPG | ASSQDRGQSNERLF | 0.7495077 |
| NTEWPKWETVMAADPVD | ASSQDRGQSNERLF | 0.47667745 |
| PASAAPAETQP | ASSQDRGQSNERLF | 0.82165194 |
| QQSSDGKNGVRAAEITGE | ASSQDRGQSNERLF | 0.738197 |
| RAPDAVTTSWWKE | ASSQDRGQSNERLF | 0.6859498 |
| SGFGNVGAHTSGWFNQ | ASSQDRGQSNERLF | 0.50466794 |
| SGVAGAVGQAASVGGLKV | ASSQDRGQSNERLF | 0.3561487 |
| SNGGAGTGSAGGNGGT | ASSQDRGQSNERLF | 0.5954853 |
| TGGAGGEGGAAG | ASSQDRGQSNERLF | 0.42636234 |
| TGGDGGAAGAGGAPGHG | ASSQDRGQSNERLF | 0.78120065 |
| TMPPLLAAIANAGRHGV | ASSQDRGQSNERLF | 0.78406537 |
| TSSTAGNSGGA | ASSQDRGQSNERLF | 0.50736797 |
| VPATVPVGILC | ASSQDRGQSNERLF | 0.81636435 |
| VQRPDSGGVVLTGRIS | ASSQDRGQSNERLF | 0.79683995 |
| WTPNPVPATKGVPAGLVQAT | ASSQDRGQSNERLF | 0.39940393 |
| YALKRALAKFSDRFDVVII | ASSQDRGQSNERLF | 0.6200409 |
| YLIGSVIV | ASSQDRGQSNERLF | 0.53724396 |
| YGQLVAVID | ASSQDRGQSNERLF | 0.6563383 |
| VTVGGMTLPA | ASSQDRGQSNERLF | 0.26747465 |
| VLALPAV | ASSQDRGQSNERLF | 0.7818847 |
| VIPPTLNYET | ASSQDRGQSNERLF | 0.54693836 |
| VGMGGMHPSSG | ASSQDRGQSNERLF | 0.17049533 |
| VGLAGGAATA | ASSQDRGQSNERLF | 0.72411966 |
| VGGAGGAGGSGP | ASSQDRGQSNERLF | 0.65532136 |
| VFAAFMV | ASSQDRGQSNERLF | 0.59314054 |
| TLYDKLR | ASSQDRGQSNERLF | 0.8077426 |
| TLAVFTVL | ASSQDRGQSNERLF | 0.48365664 |
| TGSVQPARV | ASSQDRGQSNERLF | 0.46441558 |
| TGGGAGSGGT | ASSQDRGQSNERLF | 0.7777833 |
| SHLETSL | ASSQDRGQSNERLF | 0.79695845 |
| SFTTGMT | ASSQDRGQSNERLF | 0.4443764 |
| QVEVAKRMGVRQPTVS | ASSQDRGQSNERLF | 0.74853086 |
| QGSGGPALAM | ASSQDRGQSNERLF | 0.7975966 |

| | | |
|---|---|---|
| MLAPSNP | ASSQDRGQSNERLF | 0.74434173 |
| LVNNAAI | ASSQDRGQSNERLF | 0.7936462 |
| LTGEINTLA | ASSQDRGQSNERLF | 0.8128767 |
| LSIFGPV | ASSQDRGQSNERLF | 0.6146275 |
| LLALLANL | ASSQDRGQSNERLF | 0.7988547 |
| LISNASDALDKLR | ASSQDRGQSNERLF | 0.83703005 |
| LINEAVP | ASSQDRGQSNERLF | 0.5568852 |
| LGYAFSGL | ASSQDRGQSNERLF | 0.712268 |
| LDFVAVAL | ASSQDRGQSNERLF | 0.4261143 |
| LAQPGAE | ASSQDRGQSNERLF | 0.65254366 |
| ISNLGAVTGF | ASSQDRGQSNERLF | 0.33541355 |
| HQEDPGRI | ASSQDRGQSNERLF | 0.75113887 |
| GTPGEPTP | ASSQDRGQSNERLF | 0.80962265 |
| GFFNSGS | ASSQDRGQSNERLF | 0.66077745 |
| GAGGGGLTAGAN | ASSQDRGQSNERLF | 0.43903422 |
| FVDALVNM | ASSQDRGQSNERLF | 0.35803527 |
| FLTVAPEL | ASSQDRGQSNERLF | 0.69831574 |
| FETLTAL | ASSQDRGQSNERLF | 0.5433438 |
| DVINPIEL | ASSQDRGQSNERLF | 0.2964704 |
| APVSGAPV | ASSQDRGQSNERLF | 0.66864824 |
| ALVGGLAW | ASSQDRGQSNERLF | 0.7952428 |
| ALAAAAATAI | ASSQDRGQSNERLF | 0.7848058 |
| AFGLGLLL | ASSQDRGQSNERLF | 0.4049753 |
| AFFAAASL | ASSQDRGQSNERLF | 0.8348373 |
| AAAALPAFSPP | ASSQDRGQSNERLF | 0.76945937 |
| VVLAPAL | ASSQDRGQSNERLF | 0.817363 |
| TIGHVDHGKTTL | ASSQDRGQSNERLF | 0.5977831 |
| LISNASDALDK | ASSQDRGQSNERLF | 0.46375293 |
| LISNASDAL | ASSQDRGQSNERLF | 0.80178815 |
| ISNASDALDK | ASSQDRGQSNERLF | 0.637865 |
| NQAETLVYQTE | ASSQDRGQSNERLF | 0.5759719 |
| TGGTGGTAGAG | ASSQDRGQSNERLF | 0.7964246 |
| TGSTAGGAGGA | ASSQDRGQSNERLF | 0.6784584 |
| DPLAVPVAAGPV | ASSQDRGQSNERLF | 0.5114658 |
| GGTGGTPGNGGN | ASSQDRGQSNERLF | 0.6587883 |
| PAGLAGVDPATL | ASSQDRGQSNERLF | 0.79461217 |
| GNNLVVEAPHAGASV | ASSQDRGQSNERLF | 0.49712428 |
| KVTRIMSASTLEQPA | ASSQDRGQSNERLF | 0.46063876 |
| LGFALPIALPRRTVV | ASSQDRGQSNERLF | 0.49613887 |
| TRLGNLTARKLVVLV | ASSQDRGQSNERLF | 0.66743165 |
| WSQQRRLIYFPSAGP | ASSQDRGQSNERLF | 0.7934468 |
| GPNGNPDPMAAAVDIR | ASSQDRGQSNERLF | 0.73640287 |
| TQRCVSVADLRAAASR | ASSQDRGQSNERLF | 0.5854419 |
| QFVLDVAEHPVEVLTETV | ASSQDRGQSNERLF | 0.75203395 |
| GGAGGTGGAGGNGGTQAGDG | ASSQDRGQSNERLF | 0.7283031 |
| PLLATLRAIAVDVDAKPAQV | ASSQDRGQSNERLF | 0.70237434 |
| APKFRTGGYMIDFWGVGYQVAK | ASSQDRGQSNERLF | 0.7228164 |
| GAGGAGGAGGAAGGPGGTGGTG | ASSQDRGQSNERLF | 0.5579218 |
| IGALGLLATDVAPAGSVAFGVA | ASSQDRGQSNERLF | 0.7811751 |
| AGGAGGIGGNGGAGGAGGVLMG | ASSQDRGQSNERLF | 0.76085746 |
| ADELIGLL | ASSQDRGQSNERLF | 0.7545527 |

| | | |
|---|---|---|
| ADRALKA | ASSQDRGQSNERLF | 0.79557294 |
| AGAVGQVGGAGGA | ASSQDRGQSNERLF | 0.73790514 |
| AGGGGGLGGAP | ASSQDRGQSNERLF | 0.699527 |
| AGGLLYGNGGAG | ASSQDRGQSNERLF | 0.23791897 |
| AGLESIGL | ASSQDRGQSNERLF | 0.62966096 |
| AGLKTGII | ASSQDRGQSNERLF | 0.73118556 |
| ASLAALLL | ASSQDRGQSNERLF | 0.7373121 |
| ASVINISSI | ASSQDRGQSNERLF | 0.7789545 |
| ATPPGPA | ASSQDRGQSNERLF | 0.741446 |
| ATVASALV | ASSQDRGQSNERLF | 0.58353007 |
| ATVASLIL | ASSQDRGQSNERLF | 0.7816334 |
| AVNGPVLIRH | ASSQDRGQSNERLF | 0.8026339 |
| DGAAGAVGI | ASSQDRGQSNERLF | 0.3588606 |
| DPLLHGI | ASSQDRGQSNERLF | 0.6128363 |
| ELGAALGAL | ASSQDRGQSNERLF | 0.83833003 |
| ELGRTQAV | ASSQDRGQSNERLF | 0.74471146 |
| FVAGGYLS | ASSQDRGQSNERLF | 0.6349523 |
| GASGGAGGAGGW | ASSQDRGQSNERLF | 0.5988068 |
| GGAGAGGDGGAGGI | ASSQDRGQSNERLF | 0.829046 |
| GTGHNGSQPGMGGNG | ASSQDRGQSNERLF | 0.5732342 |
| GYAAASASAS | ASSQDRGQSNERLF | 0.8224939 |
| HQVDVAAVQQG | ASSQDRGQSNERLF | 0.22493967 |
| IGAAIAHT | ASSQDRGQSNERLF | 0.5603541 |
| IGMIEVAQL | ASSQDRGQSNERLF | 0.41009954 |
| IRLAPEHTSAMD | ASSQDRGQSNERLF | 0.24220283 |
| IYNVSVL | ASSQDRGQSNERLF | 0.78610146 |
| KQGFLGM | ASSQDRGQSNERLF | 0.5922718 |
| LESVITQL | ASSQDRGQSNERLF | 0.7838369 |
| LGLGALVVV | ASSQDRGQSNERLF | 0.67819864 |
| LIGNGGNGGSGGI | ASSQDRGQSNERLF | 0.7981366 |
| LNAINAPTL | ASSQDRGQSNERLF | 0.737316 |
| LQLLNTGV | ASSQDRGQSNERLF | 0.67058146 |
| LQTSIEF | ASSQDRGQSNERLF | 0.4837598 |
| MSFAVSEL | ASSQDRGQSNERLF | 0.75712943 |
| NTGALGL | ASSQDRGQSNERLF | 0.8060951 |
| PPGAGGAGGAGTT | ASSQDRGQSNERLF | 0.7406616 |
| QLVGLGI | ASSQDRGQSNERLF | 0.36239502 |
| RFLEQVHQL | ASSQDRGQSNERLF | 0.6690628 |
| RGVGYVLR | ASSQDRGQSNERLF | 0.62266713 |
| SATGVFAGVI | ASSQDRGQSNERLF | 0.23652449 |
| SLPAITLP | ASSQDRGQSNERLF | 0.20690376 |
| STALAFTV | ASSQDRGQSNERLF | 0.6289008 |
| SVLTVGLGK | ASSQDRGQSNERLF | 0.6354245 |
| TALAGAGSGI | ASSQDRGQSNERLF | 0.79354906 |
| TGLPFTGI | ASSQDRGQSNERLF | 0.7745787 |
| VAEHVTV | ASSQDRGQSNERLF | 0.6555319 |
| VDIVLNSL | ASSQDRGQSNERLF | 0.7666905 |
| VLTGGGAVL | ASSQDRGQSNERLF | 0.724466 |
| VNVGLAYGI | ASSQDRGQSNERLF | 0.13336675 |
| VSSAAVGAV | ASSQDRGQSNERLF | 0.5328086 |
| VTSVYEAGV | ASSQDRGQSNERLF | 0.39153498 |

| | | |
|---|---|---|
| WVALDRGA | ASSQDRGQSNERLF | 0.8094014 |
| YLVGSVVI | ASSQDRGQSNERLF | 0.5320662 |
| GPPGTGKTLL | ASSQDRGQSNERLF | 0.5802814 |
| GVGMTKF | ASSQDRGQSNERLF | 0.6997894 |
| IGIGGSDL | ASSQDRGQSNERLF | 0.75142056 |
| IGTLLAEL | ASSQDRGQSNERLF | 0.6976194 |
| IYTSGTT | ASSQDRGQSNERLF | 0.4382469 |
| LSFPAGV | ASSQDRGQSNERLF | 0.20095003 |
| LYTSGTT | ASSQDRGQSNERLF | 0.7659195 |
| VGAFPFGF | ASSQDRGQSNERLF | 0.4085446 |
| VGAGTVEF | ASSQDRGQSNERLF | 0.45557827 |
| VGKSTLLL | ASSQDRGQSNERLF | 0.23566882 |
| AALAHAGGGIV | ASSQDRGQSNERLF | 0.24839996 |
| AALVAVSLL | ASSQDRGQSNERLF | 0.32112208 |
| AAVSRAVAAL | ASSQDRGQSNERLF | 0.7842772 |
| AGGSSGIGGF | ASSQDRGQSNERLF | 0.6022066 |
| AGLTVANANG | ASSQDRGQSNERLF | 0.6642339 |
| AGLVGAVL | ASSQDRGQSNERLF | 0.84157455 |
| ALSQAFTV | ASSQDRGQSNERLF | 0.7243781 |
| ALTFSVSL | ASSQDRGQSNERLF | 0.665178 |
| ALTGAALVAL | ASSQDRGQSNERLF | 0.51353115 |
| ASFSIGAGVL | ASSQDRGQSNERLF | 0.4699729 |
| DGSVYKI | ASSQDRGQSNERLF | 0.5053245 |
| EQLAEIV | ASSQDRGQSNERLF | 0.5109965 |
| FAVSELTVM | ASSQDRGQSNERLF | 0.8200172 |
| GAAPLAAGL | ASSQDRGQSNERLF | 0.5688075 |
| GADGGDGGSGGAGG | ASSQDRGQSNERLF | 0.42866623 |
| GFGLDTGGAGG | ASSQDRGQSNERLF | 0.6323378 |
| GLASGGLL | ASSQDRGQSNERLF | 0.8238909 |
| GRIAGAAVLL | ASSQDRGQSNERLF | 0.83402026 |
| GSAGSLNTSF | ASSQDRGQSNERLF | 0.5364019 |
| GTGQIGI | ASSQDRGQSNERLF | 0.52632517 |
| GTLIALIL | ASSQDRGQSNERLF | 0.5397936 |
| IAATMGFVVG | ASSQDRGQSNERLF | 0.6826544 |
| IFAQALL | ASSQDRGQSNERLF | 0.5244923 |
| IGAAVALGF | ASSQDRGQSNERLF | 0.40391576 |
| IGLTPLAL | ASSQDRGQSNERLF | 0.7609912 |
| ILGSNTSTL | ASSQDRGQSNERLF | 0.8318971 |
| IVFSLTVL | ASSQDRGQSNERLF | 0.6314571 |
| LASKATGF | ASSQDRGQSNERLF | 0.26883787 |
| LASVALIL | ASSQDRGQSNERLF | 0.6508099 |
| LFAVGLTL | ASSQDRGQSNERLF | 0.478545 |
| LFNGGIGTYI | ASSQDRGQSNERLF | 0.5870465 |
| LTLGAGII | ASSQDRGQSNERLF | 0.6208848 |
| LVAAALGDI | ASSQDRGQSNERLF | 0.38672417 |
| NTAFVQLGI | ASSQDRGQSNERLF | 0.702265 |
| QLFSGAEL | ASSQDRGQSNERLF | 0.84087 |
| SAATFITL | ASSQDRGQSNERLF | 0.6307081 |
| SNYGLANLGV | ASSQDRGQSNERLF | 0.7669568 |
| SYGGAYAVM | ASSQDRGQSNERLF | 0.8215206 |
| TTGVGGLTLG | ASSQDRGQSNERLF | 0.7842039 |

| | | |
|---|---|---|
| TTQTIYI | ASSQDRGQSNERLF | 0.81607234 |
| VAAGAGLGAVI | ASSQDRGQSNERLF | 0.77731204 |
| VAAVVTLL | ASSQDRGQSNERLF | 0.6965989 |
| VGFLGGLI | ASSQDRGQSNERLF | 0.64789903 |
| VGVIPILL | ASSQDRGQSNERLF | 0.7811265 |
| VIFASSLI | ASSQDRGQSNERLF | 0.66806376 |
| VKGGDGGAAAG | ASSQDRGQSNERLF | 0.6928776 |
| VTKTFTNI | ASSQDRGQSNERLF | 0.52542984 |
| VVAAAILL | ASSQDRGQSNERLF | 0.8009255 |
| YAALDVEL | ASSQDRGQSNERLF | 0.5527165 |
| LASVALLL | ASSQDRGQSNERLF | 0.8061463 |
| LDAVHRV | ASSQDRGQSNERLF | 0.3053592 |
| QGLSSSIF | ASSQDRGQSNERLF | 0.41502324 |
| RGTFANIRL | ASSQDRGQSNERLF | 0.7969613 |
| RQLWWGHRI | ASSQDRGQSNERLF | 0.14913528 |
| YGPPGTGKTLL | ASSQDRGQSNERLF | 0.19607648 |
| AALVGAPANVA | ASSQDRGQSNERLF | 0.6719378 |
| AATFRRTE | ASSQDRGQSNERLF | 0.8202249 |
| AGALGGGI | ASSQDRGQSNERLF | 0.16691856 |
| AGGAGGAGGIGAG | ASSQDRGQSNERLF | 0.45168224 |
| AGSGALGL | ASSQDRGQSNERLF | 0.34368205 |
| AQIADQI | ASSQDRGQSNERLF | 0.7537534 |
| ASAVAVTYL | ASSQDRGQSNERLF | 0.700418 |
| ASTVLGGTGG | ASSQDRGQSNERLF | 0.3344644 |
| AVGGEPAK | ASSQDRGQSNERLF | 0.7664975 |
| CEQGTFGGTC | ASSQDRGQSNERLF | 0.28675193 |
| DEGKAPAG | ASSQDRGQSNERLF | 0.437652 |
| DTVVFPAL | ASSQDRGQSNERLF | 0.77701914 |
| FEGITMAG | ASSQDRGQSNERLF | 0.8099022 |
| FPLVVQD | ASSQDRGQSNERLF | 0.43952587 |
| GAFTTPQ | ASSQDRGQSNERLF | 0.7547935 |
| GFGGFGTAGGVGGA | ASSQDRGQSNERLF | 0.57207286 |
| GFLAQFL | ASSQDRGQSNERLF | 0.84177804 |
| GGAGGAGGNAGVA | ASSQDRGQSNERLF | 0.5798627 |
| GGVGTAGAGGAGGAGG | ASSQDRGQSNERLF | 0.36407593 |
| GNGGGGDAGNAGSGGN | ASSQDRGQSNERLF | 0.79791903 |
| GTGGAGGAGNDAGS | ASSQDRGQSNERLF | 0.7877391 |
| GTTNTGWFNS | ASSQDRGQSNERLF | 0.66458356 |
| IFMGPLFTL | ASSQDRGQSNERLF | 0.77893543 |
| ILAHAIEAI | ASSQDRGQSNERLF | 0.5402553 |
| IRSAIREAL | ASSQDRGQSNERLF | 0.6636139 |
| ITFAGGG | ASSQDRGQSNERLF | 0.8057002 |
| ITGMSGHQEV | ASSQDRGQSNERLF | 0.7875528 |
| IVAGLACTL | ASSQDRGQSNERLF | 0.47677976 |
| IVGGGLAGA | ASSQDRGQSNERLF | 0.80681646 |
| IVTGVLNI | ASSQDRGQSNERLF | 0.7013501 |
| LAQQGCI | ASSQDRGQSNERLF | 0.49160463 |
| LCLHVEN | ASSQDRGQSNERLF | 0.8354387 |
| LDAISDGS | ASSQDRGQSNERLF | 0.66676855 |
| LGGFAAML | ASSQDRGQSNERLF | 0.77989244 |
| LGGKNPMI | ASSQDRGQSNERLF | 0.77013886 |

| | | |
|---|---|---|
| LGPKGVAAIL | ASSQDRGQSNERLF | 0.78931475 |
| LMVGVSI | ASSQDRGQSNERLF | 0.43526345 |
| LVGAAFVL | ASSQDRGQSNERLF | 0.7237358 |
| LVLAVAPL | ASSQDRGQSNERLF | 0.6961766 |
| MAAIAVIDAA | ASSQDRGQSNERLF | 0.752914 |
| NNGINAELQ | ASSQDRGQSNERLF | 0.67616117 |
| RGEPLVLVHG | ASSQDRGQSNERLF | 0.5432945 |
| RGGAGGAAGLI | ASSQDRGQSNERLF | 0.76233256 |
| RSYRMHPAV | ASSQDRGQSNERLF | 0.8224724 |
| SAAHLLAGQLA | ASSQDRGQSNERLF | 0.66925097 |
| SAAVGSY | ASSQDRGQSNERLF | 0.7247704 |
| SPHAVLHRI | ASSQDRGQSNERLF | 0.82614493 |
| SQIAKVL | ASSQDRGQSNERLF | 0.4896958 |
| TALGHLVP | ASSQDRGQSNERLF | 0.6754229 |
| TAPTTSGAA | ASSQDRGQSNERLF | 0.7625103 |
| TDGDDGNGGSPGT | ASSQDRGQSNERLF | 0.7970604 |
| TDGLGEN | ASSQDRGQSNERLF | 0.6220554 |
| TGGPPLPGQA | ASSQDRGQSNERLF | 0.8179904 |
| TIPAITI | ASSQDRGQSNERLF | 0.6074059 |
| TIPGFTI | ASSQDRGQSNERLF | 0.78373766 |
| TSIASGALA | ASSQDRGQSNERLF | 0.646768 |
| TVTLPGAAI | ASSQDRGQSNERLF | 0.7209561 |
| VASGLSAALP | ASSQDRGQSNERLF | 0.4975257 |
| VGAGVIGI | ASSQDRGQSNERLF | 0.77250576 |
| VGPKRFGV | ASSQDRGQSNERLF | 0.79415876 |
| VNVAFPD | ASSQDRGQSNERLF | 0.7974601 |
| VSMTNSGVS | ASSQDRGQSNERLF | 0.20227318 |
| VTDVLGDLSV | ASSQDRGQSNERLF | 0.78642225 |
| VVVVGAGFAGL | ASSQDRGQSNERLF | 0.7693627 |
| VYTVGTVV | ASSQDRGQSNERLF | 0.83749914 |
| WTTFSGP | ASSQDRGQSNERLF | 0.58668905 |
| YDELVEAF | ASSQDRGQSNERLF | 0.72290957 |
| YGTGGAGGNGG | ASSQDRGQSNERLF | 0.58025664 |
| YTNVMAV | ASSQDRGQSNERLF | 0.6383605 |
| FLEQVHQL | ASSQDRGQSNERLF | 0.37365386 |
| GLSLLAVAV | ASSQDRGQSNERLF | 0.5769046 |
| LEQVHQL | ASSQDRGQSNERLF | 0.62884915 |
| LNAAGAI | ASSQDRGQSNERLF | 0.5246179 |
| PAVHASL | ASSQDRGQSNERLF | 0.7942525 |
| TELEDDA | ASSQDRGQSNERLF | 0.8048778 |
| TPPGPPN | ASSQDRGQSNERLF | 0.8175052 |
| TPPPPPPP | ASSQDRGQSNERLF | 0.7982632 |
| TQSPALI | ASSQVGASSAETLY | 0.82449514 |
| EFAAAFPQ | ASSQVGASSAETLY | 0.77817297 |
| LLSGNLDVL | ASSQVGASSAETLY | 0.5486022 |
| GPANLGNYNI | ASSQVGASSAETLY | 0.65822315 |
| TLGPTLP | ASSQVGASSAETLY | 0.34807715 |
| VYAYSSCCK | ASSQVGASSAETLY | 0.65987146 |
| APAPAPAPNI | ASSQVGASSAETLY | 0.5878504 |
| PGGPVPIP | ASSQVGASSAETLY | 0.72263914 |
| FLTVALEP | ASSQVGASSAETLY | 0.15195125 |

| | | |
|---|---|---|
| SQYFGANL | ASSQVGASSAETLY | 0.73882574 |
| GGVKPAA | ASSQVGASSAETLY | 0.8301446 |
| GGTGGAGGAGGW | ASSQVGASSAETLY | 0.56058514 |
| GVGGVAGLGGAGG | ASSQVGASSAETLY | 0.7420024 |
| VLLVLATTI | ASSQVGASSAETLY | 0.77742904 |
| IGIFAAML | ASSQVGASSAETLY | 0.5693945 |
| AVGDLID | ASSQVGASSAETLY | 0.759017 |
| NISPLPL | ASSQVGASSAETLY | 0.5022913 |
| VTAVFVAI | ASSQVGASSAETLY | 0.7640544 |
| SGDDNDAAL | ASSQVGASSAETLY | 0.8306223 |
| AVAALAHS | ASSQVGASSAETLY | 0.74298596 |
| GGVTVMATP | ASSQVGASSAETLY | 0.70412195 |
| AGLAANVRD | ASSQVGASSAETLY | 0.67741346 |
| TNRPDLIDPALLRPG | ASSQVGASSAETLY | 0.33931231 |
| LANLGVTAL | ASSQVGASSAETLY | 0.70756257 |
| TGERIEL | ASSQVGASSAETLY | 0.15033904 |
| NGEELAA | ASSQVGASSAETLY | 0.6984924 |
| VLSLVWG | ASSQVGASSAETLY | 0.4894544 |
| IGYALWL | ASSQVGASSAETLY | 0.7227386 |
| LSLLLAPL | ASSQVGASSAETLY | 0.71471506 |
| TLDEALK | ASSQVGASSAETLY | 0.6978185 |
| TDGAGTAEGG | ASSQVGASSAETLY | 0.48897007 |
| NAPGGAGGNGG | ASSQVGASSAETLY | 0.71449554 |
| VSVFMGMGGNF | ASSQVGASSAETLY | 0.6316433 |
| LASYLAAL | ASSQVGASSAETLY | 0.35988343 |
| LVGGTPAL | ASSQVGASSAETLY | 0.80865884 |
| VRNADVL | ASSQVGASSAETLY | 0.5888528 |
| TCSSPFLVSAM | ASSQVGASSAETLY | 0.6966846 |
| VHITDAE | ASSQVGASSAETLY | 0.8076273 |
| EYAAPSF | ASSQVGASSAETLY | 0.70434964 |
| VVDAFHGGL | ASSQVGASSAETLY | 0.28898448 |
| ASVVDPQ | ASSQVGASSAETLY | 0.6837492 |
| SYNEFGGLS | ASSQVGASSAETLY | 0.6957873 |
| DPDGSVY | ASSQVGASSAETLY | 0.61546504 |
| MVSTIHATE | ASSQVGASSAETLY | 0.15901566 |
| LAAGVADPV | ASSQVGASSAETLY | 0.77253366 |
| TKEQIAA | ASSQVGASSAETLY | 0.50543666 |
| SLASPAAL | ASSQVGASSAETLY | 0.5422195 |
| IGLVFYGV | ASSQVGASSAETLY | 0.5813447 |
| FLLDASNL | ASSQVGASSAETLY | 0.7979604 |
| YQVGNFP | ASSQVGASSAETLY | 0.6047971 |
| ESLLSQST | ASSQVGASSAETLY | 0.6740997 |
| LFTVGFAAVG | ASSQVGASSAETLY | 0.7753366 |
| ATPPTPT | ASSQVGASSAETLY | 0.76265574 |
| TDLATLG | ASSQVGASSAETLY | 0.8205199 |
| SAGGTGR | ASSQVGASSAETLY | 0.6903921 |
| GFSNSGS | ASSQVGASSAETLY | 0.5532945 |
| GGNGGSGGTSV | ASSQVGASSAETLY | 0.72633356 |
| GTSVATGGAGN | ASSQVGASSAETLY | 0.5430988 |
| NGGNGGSGGTS | ASSQVGASSAETLY | 0.33463326 |
| AVVKFPH | ASSQVGASSAETLY | 0.28592175 |

| | | |
|---|---|---|
| TPPPPPP | ASSQVGASSAETLY | 0.6018034 |
| AAPAAAAAPA | ASSQVGASSAETLY | 0.294886 |
| IALNEDL | ASSQVGASSAETLY | 0.68258977 |
| IVNVSSM | ASSQVGASSAETLY | 0.79358625 |
| TGIPPAPR | ASSQVGASSAETLY | 0.68551105 |
| FDLGGGTF | ASSQVGASSAETLY | 0.33830824 |
| FDLGGGTFDVS | ASSQVGASSAETLY | 0.56664383 |
| NEPTAAA | ASSQVGASSAETLY | 0.6300662 |
| NEPTAAALA | ASSQVGASSAETLY | 0.76950717 |
| GSIINISSI | ASSQVGASSAETLY | 0.7647999 |
| VAPGNAGTA | ASSQVGASSAETLY | 0.8305446 |
| TGGGASLE | ASSQVGASSAETLY | 0.7772728 |
| VAVYNHYKRI | ASSQVGASSAETLY | 0.5817231 |
| TDPAPATT | ASSQVGASSAETLY | 0.78590244 |
| TNRPDILDPALLRPG | ASSQVGASSAETLY | 0.6285107 |
| LTTTTTS | ASSQVGASSAETLY | 0.7226254 |
| PTGSGKTL | ASSQVGASSAETLY | 0.70754695 |
| TGSGKTL | ASSQVGASSAETLY | 0.72541 |
| TIGHVDHGK | ASSQVGASSAETLY | 0.64401984 |
| TIGHVDHGKTT | ASSQVGASSAETLY | 0.6288419 |
| GHVDHGKTT | ASSQVGASSAETLY | 0.27382457 |
| AQFEHTLL | ASSQVGASSAETLY | 0.75270855 |
| IVNQDEL | ASSQVGASSAETLY | 0.7266917 |
| TNGSQFF | ASSQVGASSAETLY | 0.30966455 |
| TGKGIVE | ASSQVGASSAETLY | 0.615202 |
| LISNASDA | ASSQVGASSAETLY | 0.82852674 |
| AELRTLGELTKIAKAHGAQ | ASSQVGASSAETLY | 0.80449784 |
| APNAVAAAEPSW | ASSQVGASSAETLY | 0.53626704 |
| APQVAGAATAVL | ASSQVGASSAETLY | 0.4617437 |
| ARRDPAAAAAILPTDA | ASSQVGASSAETLY | 0.7344061 |
| DAAARDVLAEAGLAEYF | ASSQVGASSAETLY | 0.33218294 |
| DDQIVLSAKANLAKVEQ | ASSQVGASSAETLY | 0.7399814 |
| DELLPDTGEDSDEDGH | ASSQVGASSAETLY | 0.6892921 |
| ELPEAPANRATVPQD | ASSQVGASSAETLY | 0.72754973 |
| FGQSKIERVTGIG | ASSQVGASSAETLY | 0.5002863 |
| GAGGTGSTAGGAGGAGGAGGLY | ASSQVGASSAETLY | 0.7393545 |
| GHGGTGAAGGNGGTGGAGGSGI | ASSQVGASSAETLY | 0.59426534 |
| GNGGNGGAGGEGGPGGAGGAGA | ASSQVGASSAETLY | 0.7446109 |
| GSTAGNGGNGGNATNGTIAGQP | ASSQVGASSAETLY | 0.15922385 |
| GTGGDGGNGGNGGTGG | ASSQVGASSAETLY | 0.7668443 |
| GVRRPAGIPADAPV | ASSQVGASSAETLY | 0.5140226 |
| GYRQVTDAHLLTLARR | ASSQVGASSAETLY | 0.46227062 |
| IAKMGQDGHDRG | ASSQVGASSAETLY | 0.2571603 |
| KAEPSIEIQYT | ASSQVGASSAETLY | 0.63775384 |
| KPVRIRMAADEDTVAVT | ASSQVGASSAETLY | 0.78034925 |
| MPPRIAGMRLLVIKPEP | ASSQVGASSAETLY | 0.7917372 |
| MKFYAVALFRSRGVPPTR | ASSQVGASSAETLY | 0.52648723 |
| NAGGFGILGAVAHSP | ASSQVGASSAETLY | 0.7828594 |
| NEGREITLAMAEM | ASSQVGASSAETLY | 0.80895746 |
| NELVEWSHPLGHRWRWE | ASSQVGASSAETLY | 0.64546466 |
| NLGAGGTGGNGGNPG | ASSQVGASSAETLY | 0.29833713 |

| | | |
|---|---|---|
| NTEWPKWETVMAADPVD | ASSQVGASSAETLY | 0.7193794 |
| PASAAPAETQP | ASSQVGASSAETLY | 0.64791787 |
| QQSSDGKNGVRAAEITGE | ASSQVGASSAETLY | 0.78194904 |
| RAPDAVTTSWWKE | ASSQVGASSAETLY | 0.76791155 |
| SGFGNVGAHTSGWFNQ | ASSQVGASSAETLY | 0.38660508 |
| SGVAGAVGQAASVGGLKV | ASSQVGASSAETLY | 0.75821304 |
| SNGGAGTGSAGGNGGT | ASSQVGASSAETLY | 0.79787487 |
| TGGAGGEGGAAG | ASSQVGASSAETLY | 0.49322617 |
| TGGDGGAAGAGGAPGHG | ASSQVGASSAETLY | 0.45053554 |
| TMPPLLAAIANAGRHGV | ASSQVGASSAETLY | 0.48274344 |
| TSSTAGNSGGA | ASSQVGASSAETLY | 0.28246188 |
| VPATVPVGILC | ASSQVGASSAETLY | 0.7896104 |
| VQRPDSGGVVLTGRIS | ASSQVGASSAETLY | 0.6257051 |
| WTPNPVPATKGVPAGLVQAT | ASSQVGASSAETLY | 0.5816371 |
| YALKRALAKFSDRFDVVII | ASSQVGASSAETLY | 0.71761113 |
| YLIGSVIV | ASSQVGASSAETLY | 0.5884825 |
| YGQLVAVID | ASSQVGASSAETLY | 0.557225 |
| VTVGGMTLPA | ASSQVGASSAETLY | 0.62551504 |
| VLALPAV | ASSQVGASSAETLY | 0.69490004 |
| VIPPTLNYET | ASSQVGASSAETLY | 0.7577256 |
| VGMGGMHPSSG | ASSQVGASSAETLY | 0.7214438 |
| VGLAGGAATA | ASSQVGASSAETLY | 0.72124076 |
| VGGAGGAGGSGP | ASSQVGASSAETLY | 0.82699007 |
| VFAAFMV | ASSQVGASSAETLY | 0.39136618 |
| TLYDKLR | ASSQVGASSAETLY | 0.57544845 |
| TLAVFTVL | ASSQVGASSAETLY | 0.7671115 |
| TGSVQPARV | ASSQVGASSAETLY | 0.40225196 |
| TGGGAGSGGT | ASSQVGASSAETLY | 0.79712176 |
| SHLETSL | ASSQVGASSAETLY | 0.7145883 |
| SFTTGMT | ASSQVGASSAETLY | 0.7507049 |
| QVEVAKRMGVRQPTVS | ASSQVGASSAETLY | 0.6487928 |
| QGSGGPALAM | ASSQVGASSAETLY | 0.8379059 |
| MLAPSNP | ASSQVGASSAETLY | 0.7842149 |
| LVNNAAI | ASSQVGASSAETLY | 0.6715581 |
| LTGEINTLA | ASSQVGASSAETLY | 0.7924584 |
| LSIFGPV | ASSQVGASSAETLY | 0.34836587 |
| LLALLANL | ASSQVGASSAETLY | 0.79173875 |
| LISNASDALDKLR | ASSQVGASSAETLY | 0.7365949 |
| LINEAVP | ASSQVGASSAETLY | 0.46706358 |
| LGYAFSGL | ASSQVGASSAETLY | 0.66955966 |
| LDFVAVAL | ASSQVGASSAETLY | 0.80890524 |
| LAQPGAE | ASSQVGASSAETLY | 0.8144397 |
| ISNLGAVTGF | ASSQVGASSAETLY | 0.32334968 |
| HQEDPGRI | ASSQVGASSAETLY | 0.64238465 |
| GTPGEPTP | ASSQVGASSAETLY | 0.28550214 |
| GFFNSGS | ASSQVGASSAETLY | 0.39512303 |
| GAGGGGLTAGAN | ASSQVGASSAETLY | 0.39277354 |
| FVDALVNM | ASSQVGASSAETLY | 0.76914865 |
| FLTVAPEL | ASSQVGASSAETLY | 0.6210617 |
| FETLTAL | ASSQVGASSAETLY | 0.75497967 |
| DVINPIEL | ASSQVGASSAETLY | 0.6882967 |

| | | |
|---|---|---|
| APVSGAPV | ASSQVGASSAETLY | 0.80585253 |
| ALVGGLAW | ASSQVGASSAETLY | 0.5239856 |
| ALAAAAATAI | ASSQVGASSAETLY | 0.6344758 |
| AFGLGLLL | ASSQVGASSAETLY | 0.36569422 |
| AFFAAASL | ASSQVGASSAETLY | 0.44618005 |
| AAAALPAFSPP | ASSQVGASSAETLY | 0.48456645 |
| VVLAPAL | ASSQVGASSAETLY | 0.647596 |
| TIGHVDHGKTTL | ASSQVGASSAETLY | 0.21095866 |
| LISNASDALDK | ASSQVGASSAETLY | 0.8208957 |
| LISNASDAL | ASSQVGASSAETLY | 0.6681831 |
| ISNASDALDK | ASSQVGASSAETLY | 0.7722963 |
| NQAETLVYQTE | ASSQVGASSAETLY | 0.6078214 |
| TGGTGGTAGAG | ASSQVGASSAETLY | 0.5502341 |
| TGSTAGGAGGA | ASSQVGASSAETLY | 0.5372916 |
| DPLAVPVAAGPV | ASSQVGASSAETLY | 0.7497752 |
| GGTGGTPGNGGN | ASSQVGASSAETLY | 0.4683107 |
| PAGLAGVDPATL | ASSQVGASSAETLY | 0.24093896 |
| GNNLVVEAPHAGASV | ASSQVGASSAETLY | 0.4884059 |
| KVTRIMSASTLEQPA | ASSQVGASSAETLY | 0.35221133 |
| LGFALPIALPRRTVV | ASSQVGASSAETLY | 0.8306538 |
| TRLGNLTARKLVVLV | ASSQVGASSAETLY | 0.8032918 |
| WSQQRRLIYFPSAGP | ASSQVGASSAETLY | 0.71813023 |
| GPNGNPDPMAAAVDIR | ASSQVGASSAETLY | 0.81004024 |
| TQRCVSVADLRAAASR | ASSQVGASSAETLY | 0.19511084 |
| QFVLDVAEHPVEVLTETV | ASSQVGASSAETLY | 0.6166853 |
| GGAGGTGGAGGNGGTQAGDG | ASSQVGASSAETLY | 0.82151544 |
| PLLATLRAIAVDVDAKPAQV | ASSQVGASSAETLY | 0.74315244 |
| APKFRTGGYMIDFWGVGYQVAK | ASSQVGASSAETLY | 0.6790389 |
| GAGGAGGAGGAAGGPGGTGGTG | ASSQVGASSAETLY | 0.8064791 |
| IGALGLLATDVAPAGSVAFGVA | ASSQVGASSAETLY | 0.57003736 |
| AGGAGGIGGNGGAGGAGGVLMG | ASSQVGASSAETLY | 0.6047877 |
| ADELIGLL | ASSQVGASSAETLY | 0.67804646 |
| ADRALKA | ASSQVGASSAETLY | 0.7695004 |
| AGAVGQVGGAGGA | ASSQVGASSAETLY | 0.7182915 |
| AGGGGGLGGAP | ASSQVGASSAETLY | 0.39359763 |
| AGGLLYGNGGAG | ASSQVGASSAETLY | 0.72330564 |
| AGLESIGL | ASSQVGASSAETLY | 0.8445066 |
| AGLKTGII | ASSQVGASSAETLY | 0.6036078 |
| ASLAALLL | ASSQVGASSAETLY | 0.7760224 |
| ASVINISSI | ASSQVGASSAETLY | 0.71646047 |
| ATPPGPA | ASSQVGASSAETLY | 0.76213664 |
| ATVASALV | ASSQVGASSAETLY | 0.5090716 |
| ATVASLIL | ASSQVGASSAETLY | 0.5412353 |
| AVNGPVLIRH | ASSQVGASSAETLY | 0.70356405 |
| DGAAGAVGI | ASSQVGASSAETLY | 0.48629588 |
| DPLLHGI | ASSQVGASSAETLY | 0.6252745 |
| ELGAALGAL | ASSQVGASSAETLY | 0.63881624 |
| ELGRTQAV | ASSQVGASSAETLY | 0.8482419 |
| FVAGGYLS | ASSQVGASSAETLY | 0.7515104 |
| GASGGAGGAGGW | ASSQVGASSAETLY | 0.7553209 |
| GGAGAGGDGGAGGI | ASSQVGASSAETLY | 0.4497692 |

| | | |
|---|---|---|
| GTGHNGSQPGMGGNG | ASSQVGASSAETLY | 0.6993631 |
| GYAAASASAS | ASSQVGASSAETLY | 0.75661945 |
| HQVDVAAVQQG | ASSQVGASSAETLY | 0.33675542 |
| IGAAIAHT | ASSQVGASSAETLY | 0.81140924 |
| IGMIEVAQL | ASSQVGASSAETLY | 0.4780523 |
| IRLAPEHTSAMD | ASSQVGASSAETLY | 0.6755888 |
| IYNVSVL | ASSQVGASSAETLY | 0.7668604 |
| KQGFLGM | ASSQVGASSAETLY | 0.8075446 |
| LESVITQL | ASSQVGASSAETLY | 0.64470816 |
| LGLGALVVV | ASSQVGASSAETLY | 0.69674265 |
| LIGNGGNGGSGGI | ASSQVGASSAETLY | 0.49296135 |
| LNAINAPTL | ASSQVGASSAETLY | 0.7817826 |
| LQLLNTGV | ASSQVGASSAETLY | 0.79941785 |
| LQTSIEF | ASSQVGASSAETLY | 0.6964767 |
| MSFAVSEL | ASSQVGASSAETLY | 0.8264512 |
| NTGALGL | ASSQVGASSAETLY | 0.6844499 |
| PPGAGGAGGAGTT | ASSQVGASSAETLY | 0.2827413 |
| QLVGLGI | ASSQVGASSAETLY | 0.64287186 |
| RFLEQVHQL | ASSQVGASSAETLY | 0.81080914 |
| RGVGYVLR | ASSQVGASSAETLY | 0.8087115 |
| SATGVFAGVI | ASSQVGASSAETLY | 0.82700354 |
| SLPAITLP | ASSQVGASSAETLY | 0.7790693 |
| STALAFTV | ASSQVGASSAETLY | 0.5865136 |
| SVLTVGLGK | ASSQVGASSAETLY | 0.81331444 |
| TALAGAGSGI | ASSQVGASSAETLY | 0.83499503 |
| TGLPFTGI | ASSQVGASSAETLY | 0.7475162 |
| VAEHVTV | ASSQVGASSAETLY | 0.49719313 |
| VDIVLNSL | ASSQVGASSAETLY | 0.4226359 |
| VLTGGGAVL | ASSQVGASSAETLY | 0.7528883 |
| VNVGLAYGI | ASSQVGASSAETLY | 0.7988308 |
| VSSAAVGAV | ASSQVGASSAETLY | 0.3665369 |
| VTSVYEAGV | ASSQVGASSAETLY | 0.73495007 |
| WVALDRGA | ASSQVGASSAETLY | 0.72018963 |
| YLVGSVVI | ASSQVGASSAETLY | 0.28750715 |
| GPPGTGKTLL | ASSQVGASSAETLY | 0.77868015 |
| GVGMTKF | ASSQVGASSAETLY | 0.82338417 |
| IGIGGSDL | ASSQVGASSAETLY | 0.72449183 |
| IGTLLAEL | ASSQVGASSAETLY | 0.50891405 |
| IYTSGTT | ASSQVGASSAETLY | 0.81062245 |
| LSFPAGV | ASSQVGASSAETLY | 0.83771026 |
| LYTSGTT | ASSQVGASSAETLY | 0.8190715 |
| VGAFPFGF | ASSQVGASSAETLY | 0.33171666 |
| VGAGTVEF | ASSQVGASSAETLY | 0.19414495 |
| VGKSTLLL | ASSQVGASSAETLY | 0.65056276 |
| AALAHAGGGIV | ASSQVGASSAETLY | 0.7874063 |
| AALVAVSLL | ASSQVGASSAETLY | 0.61436003 |
| AAVSRAVAAL | ASSQVGASSAETLY | 0.5207657 |
| AGGSSGIGGF | ASSQVGASSAETLY | 0.7638706 |
| AGLTVANANG | ASSQVGASSAETLY | 0.7988119 |
| AGLVGAVL | ASSQVGASSAETLY | 0.45316613 |
| ALSQAFTV | ASSQVGASSAETLY | 0.3156659 |

| | | |
|---|---|---|
| ALTFSVSL | ASSQVGASSAETLY | 0.77980536 |
| ALTGAALVAL | ASSQVGASSAETLY | 0.74195564 |
| ASFSIGAGVL | ASSQVGASSAETLY | 0.57975006 |
| DGSVYKI | ASSQVGASSAETLY | 0.64915204 |
| EQLAEIV | ASSQVGASSAETLY | 0.66944045 |
| FAVSELTVM | ASSQVGASSAETLY | 0.62099373 |
| GAAPLAAGL | ASSQVGASSAETLY | 0.728828 |
| GADGGDGGSGGAGG | ASSQVGASSAETLY | 0.34827834 |
| GFGLDTGGAGG | ASSQVGASSAETLY | 0.8207061 |
| GLASGGLL | ASSQVGASSAETLY | 0.41389245 |
| GRIAGAAVLL | ASSQVGASSAETLY | 0.7688754 |
| GSAGSLNTSF | ASSQVGASSAETLY | 0.7165065 |
| GTGQIGI | ASSQVGASSAETLY | 0.74710673 |
| GTLIALIL | ASSQVGASSAETLY | 0.6401553 |
| IAATMGFVVG | ASSQVGASSAETLY | 0.51643527 |
| IFAQALL | ASSQVGASSAETLY | 0.4490019 |
| IGAAVALGF | ASSQVGASSAETLY | 0.26361948 |
| IGLTPLAL | ASSQVGASSAETLY | 0.809633 |
| ILGSNTSTL | ASSQVGASSAETLY | 0.20786501 |
| IVFSLTVL | ASSQVGASSAETLY | 0.8085449 |
| LASKATGF | ASSQVGASSAETLY | 0.72184294 |
| LASVALIL | ASSQVGASSAETLY | 0.5083106 |
| LFAVGLTL | ASSQVGASSAETLY | 0.72362256 |
| LFNGGIGTYI | ASSQVGASSAETLY | 0.42941886 |
| LTLGAGII | ASSQVGASSAETLY | 0.58677626 |
| LVAAALGDI | ASSQVGASSAETLY | 0.7721017 |
| NTAFVQLGI | ASSQVGASSAETLY | 0.7678404 |
| QLFSGAEL | ASSQVGASSAETLY | 0.48498172 |
| SAATFITL | ASSQVGASSAETLY | 0.73684525 |
| SNYGLANLGV | ASSQVGASSAETLY | 0.74430346 |
| SYGGAYAVM | ASSQVGASSAETLY | 0.4409907 |
| TTGVGGLTLG | ASSQVGASSAETLY | 0.7170536 |
| TTQTIYI | ASSQVGASSAETLY | 0.7923385 |
| VAAGAGLGAVI | ASSQVGASSAETLY | 0.71355355 |
| VAAVVTLL | ASSQVGASSAETLY | 0.38606662 |
| VGFLGGLI | ASSQVGASSAETLY | 0.7820996 |
| VGVIPILL | ASSQVGASSAETLY | 0.5065619 |
| VIFASSLI | ASSQVGASSAETLY | 0.7479414 |
| VKGGDGGAAAG | ASSQVGASSAETLY | 0.6971445 |
| VTKTFTNI | ASSQVGASSAETLY | 0.5724923 |
| VVAAAILL | ASSQVGASSAETLY | 0.8048789 |
| YAALDVEL | ASSQVGASSAETLY | 0.33925822 |
| LASVALLL | ASSQVGASSAETLY | 0.5945667 |
| LDAVHRV | ASSQVGASSAETLY | 0.62332046 |
| QGLSSSIF | ASSQVGASSAETLY | 0.69095844 |
| RGTFANIRL | ASSQVGASSAETLY | 0.7768365 |
| RQLWWGHRI | ASSQVGASSAETLY | 0.4656055 |
| YGPPGTGKTLL | ASSQVGASSAETLY | 0.69267106 |
| AALVGAPANVA | ASSQVGASSAETLY | 0.5967485 |
| AATFRRTE | ASSQVGASSAETLY | 0.41996253 |
| AGALGGGI | ASSQVGASSAETLY | 0.64311373 |

| | | |
|---|---|---|
| AGGAGGAGGIGAG | ASSQVGASSAETLY | 0.61098135 |
| AGSGALGL | ASSQVGASSAETLY | 0.41567373 |
| AQIADQI | ASSQVGASSAETLY | 0.7642387 |
| ASAVAVTYL | ASSQVGASSAETLY | 0.5023873 |
| ASTVLGGTGG | ASSQVGASSAETLY | 0.72944796 |
| AVGGEPAK | ASSQVGASSAETLY | 0.8415925 |
| CEQGTFGGTC | ASSQVGASSAETLY | 0.43598896 |
| DEGKAPAG | ASSQVGASSAETLY | 0.7459579 |
| DTVVFPAL | ASSQVGASSAETLY | 0.73144746 |
| FEGITMAG | ASSQVGASSAETLY | 0.75328183 |
| FPLVVQD | ASSQVGASSAETLY | 0.63300765 |
| GAFTTPQ | ASSQVGASSAETLY | 0.55056554 |
| GFGGFGTAGGVGGA | ASSQVGASSAETLY | 0.8248713 |
| GFLAQFL | ASSQVGASSAETLY | 0.7738689 |
| GGAGGAGGNAGVA | ASSQVGASSAETLY | 0.76729536 |
| GGVGTAGAGGAGGAGG | ASSQVGASSAETLY | 0.7695675 |
| GNGGGGDAGNAGSGGN | ASSQVGASSAETLY | 0.7616296 |
| GTGGAGGAGNDAGS | ASSQVGASSAETLY | 0.78310937 |
| GTTNTGWFNS | ASSQVGASSAETLY | 0.56799036 |
| IFMGPLFTL | ASSQVGASSAETLY | 0.32089508 |
| ILAHAIEAI | ASSQVGASSAETLY | 0.6139428 |
| IRSAIREAL | ASSQVGASSAETLY | 0.6861471 |
| ITFAGGG | ASSQVGASSAETLY | 0.52044606 |
| ITGMSGHQEV | ASSQVGASSAETLY | 0.25943604 |
| IVAGLACTL | ASSQVGASSAETLY | 0.80572045 |
| IVGGGLAGA | ASSQVGASSAETLY | 0.8389715 |
| IVTGVLNI | ASSQVGASSAETLY | 0.50189066 |
| LAQQGCI | ASSQVGASSAETLY | 0.84132135 |
| LCLHVEN | ASSQVGASSAETLY | 0.39580607 |
| LDAISDGS | ASSQVGASSAETLY | 0.7804346 |
| LGGFAAML | ASSQVGASSAETLY | 0.81526816 |
| LGGKNPMI | ASSQVGASSAETLY | 0.40748817 |
| LGPKGVAAIL | ASSQVGASSAETLY | 0.74307716 |
| LMVGVSI | ASSQVGASSAETLY | 0.48754674 |
| LVGAAFVL | ASSQVGASSAETLY | 0.57714176 |
| LVLAVAPL | ASSQVGASSAETLY | 0.8123937 |
| MAAIAVIDAA | ASSQVGASSAETLY | 0.61432654 |
| NNGINAELQ | ASSQVGASSAETLY | 0.7721168 |
| RGEPLVLVHG | ASSQVGASSAETLY | 0.6059294 |
| RGGAGGAAGLI | ASSQVGASSAETLY | 0.44315767 |
| RSYRMHPAV | ASSQVGASSAETLY | 0.7338914 |
| SAAHLLAGQLA | ASSQVGASSAETLY | 0.78165984 |
| SAAVGSY | ASSQVGASSAETLY | 0.53817564 |
| SPHAVLHRI | ASSQVGASSAETLY | 0.83044255 |
| SQIAKVL | ASSQVGASSAETLY | 0.4894027 |
| TALGHLVP | ASSQVGASSAETLY | 0.8409045 |
| TAPTTSGAA | ASSQVGASSAETLY | 0.79362506 |
| TDGDDGNGGSPGT | ASSQVGASSAETLY | 0.71995926 |
| TDGLGEN | ASSQVGASSAETLY | 0.59807324 |
| TGGPPLPGQA | ASSQVGASSAETLY | 0.79799557 |
| TIPAITI | ASSQVGASSAETLY | 0.5519066 |

| | | |
|---|---|---|
| TIPGFTI | ASSQVGASSAETLY | 0.74862325 |
| TSIASGALA | ASSQVGASSAETLY | 0.80592406 |
| TVTLPGAAI | ASSQVGASSAETLY | 0.41222525 |
| VASGLSAALP | ASSQVGASSAETLY | 0.62625116 |
| VGAGVIGI | ASSQVGASSAETLY | 0.5492999 |
| VGPKRFGV | ASSQVGASSAETLY | 0.5845946 |
| VNVAFPD | ASSQVGASSAETLY | 0.4672192 |
| VSMTNSGVS | ASSQVGASSAETLY | 0.42012578 |
| VTDVLGDLSV | ASSQVGASSAETLY | 0.50930965 |
| VVVVGAGFAGL | ASSQVGASSAETLY | 0.44122258 |
| VYTVGTVV | ASSQVGASSAETLY | 0.61961454 |
| WTTFSGP | ASSQVGASSAETLY | 0.6085762 |
| YDELVEAF | ASSQVGASSAETLY | 0.46888828 |
| YGTGGAGGNGG | ASSQVGASSAETLY | 0.5972038 |
| YTNVMAV | ASSQVGASSAETLY | 0.6258334 |
| FLEQVHQL | ASSQVGASSAETLY | 0.39578587 |
| GLSLLAVAV | ASSQVGASSAETLY | 0.49943998 |
| LEQVHQL | ASSQVGASSAETLY | 0.51037365 |
| LNAAGAI | ASSQVGASSAETLY | 0.4972566 |
| PAVHASL | ASSQVGASSAETLY | 0.37365392 |
| TELEDDA | ASSQVGASSAETLY | 0.5873704 |
| TPPGPPN | ASSQVGASSAETLY | 0.43148008 |
| TPPPPPPP | ASSQVGASSAETLY | 0.33217776 |
| TQSPALI | ASSPGLGGRTQY | 0.5633738 |
| EFAAAFPQ | ASSPGLGGRTQY | 0.5035646 |
| LLSGNLDVL | ASSPGLGGRTQY | 0.4966582 |
| GPANLGNYNI | ASSPGLGGRTQY | 0.59583634 |
| TLGPTLP | ASSPGLGGRTQY | 0.5025865 |
| VYAYSSCCK | ASSPGLGGRTQY | 0.45769858 |
| APAPAPAPNI | ASSPGLGGRTQY | 0.57516456 |
| PGGPVPIP | ASSPGLGGRTQY | 0.61878246 |
| FLTVALEP | ASSPGLGGRTQY | 0.43170288 |
| SQYFGANL | ASSPGLGGRTQY | 0.54462063 |
| GGVKPAA | ASSPGLGGRTQY | 0.6298901 |
| GGTGGAGGAGGW | ASSPGLGGRTQY | 0.5393653 |
| GVGGVAGLGGAGG | ASSPGLGGRTQY | 0.6221222 |
| VLLVLATTI | ASSPGLGGRTQY | 0.63366777 |
| IGIFAAML | ASSPGLGGRTQY | 0.5186677 |
| AVGDLID | ASSPGLGGRTQY | 0.615421 |
| NISPLPL | ASSPGLGGRTQY | 0.62409675 |
| VTAVFVAI | ASSPGLGGRTQY | 0.5088183 |
| SGDDNDAAL | ASSPGLGGRTQY | 0.52459687 |
| AVAALAHS | ASSPGLGGRTQY | 0.41291162 |
| GGVTVMATP | ASSPGLGGRTQY | 0.55078304 |
| AGLAANVRD | ASSPGLGGRTQY | 0.43752104 |
| TNRPDLIDPALLRPG | ASSPGLGGRTQY | 0.5475292 |
| LANLGVTAL | ASSPGLGGRTQY | 0.63202244 |
| TGERIEL | ASSPGLGGRTQY | 0.49212635 |
| NGEELAA | ASSPGLGGRTQY | 0.39413005 |
| VLSLVWG | ASSPGLGGRTQY | 0.40164846 |
| IGYALWL | ASSPGLGGRTQY | 0.5826913 |

| | | |
|---|---|---|
| LSLLLAPL | ASSPGLGGRTQY | 0.4153121 |
| TLDEALK | ASSPGLGGRTQY | 0.40737155 |
| TDGAGTAEGG | ASSPGLGGRTQY | 0.48921296 |
| NAPGGAGGNGG | ASSPGLGGRTQY | 0.61378556 |
| VSVFMGMGGNF | ASSPGLGGRTQY | 0.6089253 |
| LASYLAAL | ASSPGLGGRTQY | 0.43973547 |
| LVGGTPAL | ASSPGLGGRTQY | 0.62988657 |
| VRNADVL | ASSPGLGGRTQY | 0.59341264 |
| TCSSPFLVSAM | ASSPGLGGRTQY | 0.6376324 |
| VHITDAE | ASSPGLGGRTQY | 0.52538174 |
| EYAAPSF | ASSPGLGGRTQY | 0.41137943 |
| VVDAFHGGL | ASSPGLGGRTQY | 0.5992915 |
| ASVVDPQ | ASSPGLGGRTQY | 0.48300707 |
| SYNEFGGLS | ASSPGLGGRTQY | 0.5833313 |
| DPDGSVY | ASSPGLGGRTQY | 0.5929626 |
| MVSTIHATE | ASSPGLGGRTQY | 0.51148576 |
| LAAGVADPV | ASSPGLGGRTQY | 0.448687 |
| TKEQIAA | ASSPGLGGRTQY | 0.52257234 |
| SLASPAAL | ASSPGLGGRTQY | 0.629177 |
| IGLVFYGV | ASSPGLGGRTQY | 0.41641313 |
| FLLDASNL | ASSPGLGGRTQY | 0.45566764 |
| YQVGNFP | ASSPGLGGRTQY | 0.54412484 |
| ESLLSQST | ASSPGLGGRTQY | 0.51353925 |
| LFTVGFAAVG | ASSPGLGGRTQY | 0.6225815 |
| ATPPTPT | ASSPGLGGRTQY | 0.574846 |
| TDLATLG | ASSPGLGGRTQY | 0.5168716 |
| SAGGTGR | ASSPGLGGRTQY | 0.572451 |
| GFSNSGS | ASSPGLGGRTQY | 0.59234273 |
| GGNGGSGGTSV | ASSPGLGGRTQY | 0.56150436 |
| GTSVATGGAGN | ASSPGLGGRTQY | 0.5941097 |
| NGGNGGSGGTS | ASSPGLGGRTQY | 0.44796038 |
| AVVKFPH | ASSPGLGGRTQY | 0.62016094 |
| TPPPPPP | ASSPGLGGRTQY | 0.59072304 |
| AAPAAAAAPA | ASSPGLGGRTQY | 0.5931919 |
| IALNEDL | ASSPGLGGRTQY | 0.62213933 |
| IVNVSSM | ASSPGLGGRTQY | 0.57634044 |
| TGIPPAPR | ASSPGLGGRTQY | 0.5460894 |
| FDLGGGTF | ASSPGLGGRTQY | 0.38601202 |
| FDLGGGTFDVS | ASSPGLGGRTQY | 0.52747196 |
| NEPTAAA | ASSPGLGGRTQY | 0.54612356 |
| NEPTAAALA | ASSPGLGGRTQY | 0.5443637 |
| GSIINISSI | ASSPGLGGRTQY | 0.60264766 |
| VAPGNAGTA | ASSPGLGGRTQY | 0.59285223 |
| TGGGASLE | ASSPGLGGRTQY | 0.4672212 |
| VAVYNHYKRI | ASSPGLGGRTQY | 0.63727343 |
| TDPAPATT | ASSPGLGGRTQY | 0.6142031 |
| TNRPDILDPALLRPG | ASSPGLGGRTQY | 0.40040892 |
| LTTTTTS | ASSPGLGGRTQY | 0.50776076 |
| PTGSGKTL | ASSPGLGGRTQY | 0.63564646 |
| TGSGKTL | ASSPGLGGRTQY | 0.55845106 |
| TIGHVDHGK | ASSPGLGGRTQY | 0.4951085 |

| | | |
|---|---|---|
| TIGHVDHGKTT | ASSPGLGGRTQY | 0.5274668 |
| GHVDHGKTT | ASSPGLGGRTQY | 0.62770844 |
| AQFEHTLL | ASSPGLGGRTQY | 0.5593245 |
| IVNQDEL | ASSPGLGGRTQY | 0.6336751 |
| TNGSQFF | ASSPGLGGRTQY | 0.34273452 |
| TGKGIVE | ASSPGLGGRTQY | 0.50061166 |
| LISNASDA | ASSPGLGGRTQY | 0.4693784 |
| AELRTLGELTKIAKAHGAQ | ASSPGLGGRTQY | 0.33120686 |
| APNAVAAAEPSW | ASSPGLGGRTQY | 0.5984273 |
| APQVAGAATAVL | ASSPGLGGRTQY | 0.49678475 |
| ARRDPAAAAAILPTDA | ASSPGLGGRTQY | 0.5867614 |
| DAAARDVLAEAGLAEYF | ASSPGLGGRTQY | 0.4944715 |
| DDQIVLSAKANLAKVEQ | ASSPGLGGRTQY | 0.6098189 |
| DELLPDTGEDSDEDGH | ASSPGLGGRTQY | 0.54725015 |
| ELPEAPANRATVPQD | ASSPGLGGRTQY | 0.54853827 |
| FGQSKIERVTGIG | ASSPGLGGRTQY | 0.4715003 |
| GAGGTGSTAGGAGGAGGAGGLY | ASSPGLGGRTQY | 0.5684342 |
| GHGGTGAAGGNGGTGGAGGSGI | ASSPGLGGRTQY | 0.6427232 |
| GNGGNGGAGGEGGPGGAGGAGA | ASSPGLGGRTQY | 0.62343615 |
| GSTAGNGGNGGNATNGTIAGQP | ASSPGLGGRTQY | 0.39326158 |
| GTGGDGGNGGNGGTGG | ASSPGLGGRTQY | 0.5806606 |
| GVRRPAGIPADAPV | ASSPGLGGRTQY | 0.5175298 |
| GYRQVTDAHLLTLARR | ASSPGLGGRTQY | 0.407908 |
| IAKMGQDGHDRG | ASSPGLGGRTQY | 0.3638993 |
| KAEPSIEIQYT | ASSPGLGGRTQY | 0.48990136 |
| KPVRIRMAADEDTVAVT | ASSPGLGGRTQY | 0.51723146 |
| MPPRIAGMRLLVIKPEP | ASSPGLGGRTQY | 0.57230574 |
| MKFYAVALFRSRGVPPTR | ASSPGLGGRTQY | 0.58576524 |
| NAGGFGILGAVAHSP | ASSPGLGGRTQY | 0.47004968 |
| NEGREITLAMAEM | ASSPGLGGRTQY | 0.57787967 |
| NELVEWSHPLGHRWRWE | ASSPGLGGRTQY | 0.5678334 |
| NLGAGGTGGNGGNPG | ASSPGLGGRTQY | 0.29149097 |
| NTEWPKWETVMAADPVD | ASSPGLGGRTQY | 0.49100947 |
| PASAAPAETQP | ASSPGLGGRTQY | 0.39897257 |
| QQSSDGKNGVRAAEITGE | ASSPGLGGRTQY | 0.64382255 |
| RAPDAVTTSWWKE | ASSPGLGGRTQY | 0.4434341 |
| SGFGNVGAHTSGWFNQ | ASSPGLGGRTQY | 0.48309645 |
| SGVAGAVGQAASVGGLKV | ASSPGLGGRTQY | 0.5202975 |
| SNGGAGTGSAGGNGGT | ASSPGLGGRTQY | 0.5586145 |
| TGGAGGEGGAAG | ASSPGLGGRTQY | 0.5220047 |
| TGGDGGAAGAGGAPGHG | ASSPGLGGRTQY | 0.45285016 |
| TMPPLLAAIANAGRHGV | ASSPGLGGRTQY | 0.3744948 |
| TSSTAGNSGGA | ASSPGLGGRTQY | 0.5811051 |
| VPATVPVGILC | ASSPGLGGRTQY | 0.5385386 |
| VQRPDSGGVVLTGRIS | ASSPGLGGRTQY | 0.45883495 |
| WTPNPVPATKGVPAGLVQAT | ASSPGLGGRTQY | 0.43016207 |
| YALKRALAKFSDRFDVVII | ASSPGLGGRTQY | 0.5454869 |
| YLIGSVIV | ASSPGLGGRTQY | 0.379932 |
| YGQLVAVID | ASSPGLGGRTQY | 0.6101869 |
| VTVGGMTLPA | ASSPGLGGRTQY | 0.43466526 |
| VLALPAV | ASSPGLGGRTQY | 0.48073855 |

| | | |
|---|---|---|
| VIPPTLNYET | ASSPGLGGRTQY | 0.6379038 |
| VGMGGMHPSSG | ASSPGLGGRTQY | 0.5487496 |
| VGLAGGAATA | ASSPGLGGRTQY | 0.48898286 |
| VGGAGGAGGSGP | ASSPGLGGRTQY | 0.46374545 |
| VFAAFMV | ASSPGLGGRTQY | 0.43073818 |
| TLYDKLR | ASSPGLGGRTQY | 0.450149 |
| TLAVFTVL | ASSPGLGGRTQY | 0.46931782 |
| TGSVQPARV | ASSPGLGGRTQY | 0.62917507 |
| TGGGAGSGGT | ASSPGLGGRTQY | 0.54202414 |
| SHLETSL | ASSPGLGGRTQY | 0.43075347 |
| SFTTGMT | ASSPGLGGRTQY | 0.5946715 |
| QVEVAKRMGVRQPTVS | ASSPGLGGRTQY | 0.62829256 |
| QGSGGPALAM | ASSPGLGGRTQY | 0.6449336 |
| MLAPSNP | ASSPGLGGRTQY | 0.44877762 |
| LVNNAAI | ASSPGLGGRTQY | 0.52086926 |
| LTGEINTLA | ASSPGLGGRTQY | 0.5272361 |
| LSIFGPV | ASSPGLGGRTQY | 0.5097506 |
| LLALLANL | ASSPGLGGRTQY | 0.4608781 |
| LISNASDALDKLR | ASSPGLGGRTQY | 0.44438663 |
| LINEAVP | ASSPGLGGRTQY | 0.6037342 |
| LGYAFSGL | ASSPGLGGRTQY | 0.6551993 |
| LDFVAVAL | ASSPGLGGRTQY | 0.5249482 |
| LAQPGAE | ASSPGLGGRTQY | 0.37137452 |
| ISNLGAVTGF | ASSPGLGGRTQY | 0.51096314 |
| HQEDPGRI | ASSPGLGGRTQY | 0.43565983 |
| GTPGEPTP | ASSPGLGGRTQY | 0.44018194 |
| GFFNSGS | ASSPGLGGRTQY | 0.5423094 |
| GAGGGGLTAGAN | ASSPGLGGRTQY | 0.3653161 |
| FVDALVNM | ASSPGLGGRTQY | 0.5176679 |
| FLTVAPEL | ASSPGLGGRTQY | 0.6407801 |
| FETLTAL | ASSPGLGGRTQY | 0.48923492 |
| DVINPIEL | ASSPGLGGRTQY | 0.5630123 |
| APVSGAPV | ASSPGLGGRTQY | 0.61447304 |
| ALVGGLAW | ASSPGLGGRTQY | 0.5702304 |
| ALAAAAATAI | ASSPGLGGRTQY | 0.6141814 |
| AFGLGLLL | ASSPGLGGRTQY | 0.60684365 |
| AFFAAASL | ASSPGLGGRTQY | 0.55310345 |
| AAAALPAFSPP | ASSPGLGGRTQY | 0.53242064 |
| VVLAPAL | ASSPGLGGRTQY | 0.6081838 |
| TIGHVDHGKTTL | ASSPGLGGRTQY | 0.5167558 |
| LISNASDALDK | ASSPGLGGRTQY | 0.5303037 |
| LISNASDAL | ASSPGLGGRTQY | 0.49816477 |
| ISNASDALDK | ASSPGLGGRTQY | 0.5868518 |
| NQAETLVYQTE | ASSPGLGGRTQY | 0.5047444 |
| TGGTGGTAGAG | ASSPGLGGRTQY | 0.62901866 |
| TGSTAGGAGGA | ASSPGLGGRTQY | 0.34880447 |
| DPLAVPVAAGPV | ASSPGLGGRTQY | 0.43433565 |
| GGTGGTPGNGGN | ASSPGLGGRTQY | 0.6065624 |
| PAGLAGVDPATL | ASSPGLGGRTQY | 0.3196817 |
| GNNLVVEAPHAGASV | ASSPGLGGRTQY | 0.38293305 |
| KVTRIMSASTLEQPA | ASSPGLGGRTQY | 0.52626103 |

| | | |
|---|---|---|
| LGFALPIALPRRTVV | ASSPGLGGRTQY | 0.62079597 |
| TRLGNLTARKLVVLV | ASSPGLGGRTQY | 0.32518056 |
| WSQQRRLIYFPSAGP | ASSPGLGGRTQY | 0.50841 |
| GPNGNPDPMAAAVDIR | ASSPGLGGRTQY | 0.39829567 |
| TQRCVSVADLRAAASR | ASSPGLGGRTQY | 0.6250269 |
| QFVLDVAEHPVEVLTETV | ASSPGLGGRTQY | 0.5251234 |
| GGAGGTGGAGGNGGTQAGDG | ASSPGLGGRTQY | 0.4079448 |
| PLLATLRAIAVDVDAKPAQV | ASSPGLGGRTQY | 0.598014 |
| APKFRTGGYMIDFWGVGYQVAK | ASSPGLGGRTQY | 0.38679877 |
| GAGGAGGAGGAAGGPGGTGGTG | ASSPGLGGRTQY | 0.5687418 |
| IGALGLLATDVAPAGSVAFGVA | ASSPGLGGRTQY | 0.5999303 |
| AGGAGGIGGNGGAGGAGGVLMG | ASSPGLGGRTQY | 0.63667864 |
| ADELIGLL | ASSPGLGGRTQY | 0.40280494 |
| ADRALKA | ASSPGLGGRTQY | 0.58008826 |
| AGAVGQVGGAGGA | ASSPGLGGRTQY | 0.46062317 |
| AGGGGGLGGAP | ASSPGLGGRTQY | 0.6529047 |
| AGGLLYGNGGAG | ASSPGLGGRTQY | 0.47987235 |
| AGLESIGL | ASSPGLGGRTQY | 0.3810802 |
| AGLKTGII | ASSPGLGGRTQY | 0.6078068 |
| ASLAALLL | ASSPGLGGRTQY | 0.59754825 |
| ASVINISSI | ASSPGLGGRTQY | 0.5484765 |
| ATPPGPA | ASSPGLGGRTQY | 0.5818745 |
| ATVASALV | ASSPGLGGRTQY | 0.53310364 |
| ATVASLIL | ASSPGLGGRTQY | 0.49424773 |
| AVNGPVLIRH | ASSPGLGGRTQY | 0.5868983 |
| DGAAGAVGI | ASSPGLGGRTQY | 0.5882823 |
| DPLLHGI | ASSPGLGGRTQY | 0.46261126 |
| ELGAALGAL | ASSPGLGGRTQY | 0.6235293 |
| ELGRTQAV | ASSPGLGGRTQY | 0.5425555 |
| FVAGGYLS | ASSPGLGGRTQY | 0.43821964 |
| GASGGAGGAGGW | ASSPGLGGRTQY | 0.6442997 |
| GGAGAGGDGGAGGI | ASSPGLGGRTQY | 0.5419254 |
| GTGHNGSQPGMGGNG | ASSPGLGGRTQY | 0.5827 |
| GYAAASASAS | ASSPGLGGRTQY | 0.5819075 |
| HQVDVAAVQQG | ASSPGLGGRTQY | 0.63940644 |
| IGAAIAHT | ASSPGLGGRTQY | 0.44818363 |
| IGMIEVAQL | ASSPGLGGRTQY | 0.55792415 |
| IRLAPEHTSAMD | ASSPGLGGRTQY | 0.5259845 |
| IYNVSVL | ASSPGLGGRTQY | 0.6075838 |
| KQGFLGM | ASSPGLGGRTQY | 0.5443023 |
| LESVITQL | ASSPGLGGRTQY | 0.5521249 |
| LGLGALVVV | ASSPGLGGRTQY | 0.5865355 |
| LIGNGGNGGSGGI | ASSPGLGGRTQY | 0.6203886 |
| LNAINAPTL | ASSPGLGGRTQY | 0.52981526 |
| LQLLNTGV | ASSPGLGGRTQY | 0.58835936 |
| LQTSIEF | ASSPGLGGRTQY | 0.62762415 |
| MSFAVSEL | ASSPGLGGRTQY | 0.43828234 |
| NTGALGL | ASSPGLGGRTQY | 0.5403331 |
| PPGAGGAGGAGTT | ASSPGLGGRTQY | 0.59703934 |
| QLVGLGI | ASSPGLGGRTQY | 0.6301103 |
| RFLEQVHQL | ASSPGLGGRTQY | 0.51514804 |

| | | |
|---|---|---|
| RGVGYVLR | ASSPGLGGRTQY | 0.6258594 |
| SATGVFAGVI | ASSPGLGGRTQY | 0.5273002 |
| SLPAITLP | ASSPGLGGRTQY | 0.57853544 |
| STALAFTV | ASSPGLGGRTQY | 0.5095432 |
| SVLTVGLGK | ASSPGLGGRTQY | 0.49160728 |
| TALAGAGSGI | ASSPGLGGRTQY | 0.43517026 |
| TGLPFTGI | ASSPGLGGRTQY | 0.55464023 |
| VAEHVTV | ASSPGLGGRTQY | 0.59397763 |
| VDIVLNSL | ASSPGLGGRTQY | 0.5684657 |
| VLTGGGAVL | ASSPGLGGRTQY | 0.36272764 |
| VNVGLAYGI | ASSPGLGGRTQY | 0.5746305 |
| VSSAAVGAV | ASSPGLGGRTQY | 0.5771743 |
| VTSVYEAGV | ASSPGLGGRTQY | 0.6297295 |
| WVALDRGA | ASSPGLGGRTQY | 0.5131348 |
| YLVGSVVI | ASSPGLGGRTQY | 0.6083612 |
| GPPGTGKTLL | ASSPGLGGRTQY | 0.4966871 |
| GVGMTKF | ASSPGLGGRTQY | 0.5478082 |
| IGIGGSDL | ASSPGLGGRTQY | 0.37863696 |
| IGTLLAEL | ASSPGLGGRTQY | 0.45941517 |
| IYTSGTT | ASSPGLGGRTQY | 0.50870985 |
| LSFPAGV | ASSPGLGGRTQY | 0.42754453 |
| LYTSGTT | ASSPGLGGRTQY | 0.5906098 |
| VGAFPFGF | ASSPGLGGRTQY | 0.615915 |
| VGAGTVEF | ASSPGLGGRTQY | 0.60905796 |
| VGKSTLLL | ASSPGLGGRTQY | 0.61479044 |
| AALAHAGGGIV | ASSPGLGGRTQY | 0.6194237 |
| AALVAVSLL | ASSPGLGGRTQY | 0.61577797 |
| AAVSRAVAAL | ASSPGLGGRTQY | 0.47141558 |
| AGGSSGIGGF | ASSPGLGGRTQY | 0.5372075 |
| AGLTVANANG | ASSPGLGGRTQY | 0.37072593 |
| AGLVGAVL | ASSPGLGGRTQY | 0.50769395 |
| ALSQAFTV | ASSPGLGGRTQY | 0.48194495 |
| ALTFSVSL | ASSPGLGGRTQY | 0.5334772 |
| ALTGAALVAL | ASSPGLGGRTQY | 0.31497312 |
| ASFSIGAGVL | ASSPGLGGRTQY | 0.55118203 |
| DGSVYKI | ASSPGLGGRTQY | 0.6321994 |
| EQLAEIV | ASSPGLGGRTQY | 0.54174876 |
| FAVSELTVM | ASSPGLGGRTQY | 0.5765999 |
| GAAPLAAGL | ASSPGLGGRTQY | 0.57587796 |
| GADGGDGGSGGAGG | ASSPGLGGRTQY | 0.44458106 |
| GFGLDTGGAGG | ASSPGLGGRTQY | 0.5834414 |
| GLASGGLL | ASSPGLGGRTQY | 0.45748866 |
| GRIAGAAVLL | ASSPGLGGRTQY | 0.5928879 |
| GSAGSLNTSF | ASSPGLGGRTQY | 0.6154769 |
| GTGQIGI | ASSPGLGGRTQY | 0.55904603 |
| GTLIALIL | ASSPGLGGRTQY | 0.5543617 |
| IAATMGFVVG | ASSPGLGGRTQY | 0.5600722 |
| IFAQALL | ASSPGLGGRTQY | 0.43096158 |
| IGAAVALGF | ASSPGLGGRTQY | 0.5709008 |
| IGLTPLAL | ASSPGLGGRTQY | 0.45195732 |
| ILGSNTSTL | ASSPGLGGRTQY | 0.5270513 |

| | | |
|---|---|---|
| IVFSLTVL | ASSPGLGGRTQY | 0.44231796 |
| LASKATGF | ASSPGLGGRTQY | 0.5678705 |
| LASVALIL | ASSPGLGGRTQY | 0.549357 |
| LFAVGLTL | ASSPGLGGRTQY | 0.56504893 |
| LFNGGIGTYI | ASSPGLGGRTQY | 0.44498974 |
| LTLGAGII | ASSPGLGGRTQY | 0.5985174 |
| LVAAALGDI | ASSPGLGGRTQY | 0.5763948 |
| NTAFVQLGI | ASSPGLGGRTQY | 0.41327855 |
| QLFSGAEL | ASSPGLGGRTQY | 0.59157914 |
| SAATFITL | ASSPGLGGRTQY | 0.5069857 |
| SNYGLANLGV | ASSPGLGGRTQY | 0.511181 |
| SYGGAYAVM | ASSPGLGGRTQY | 0.6180517 |
| TTGVGGLTLG | ASSPGLGGRTQY | 0.54283017 |
| TTQTIYI | ASSPGLGGRTQY | 0.4277436 |
| VAAGAGLGAVI | ASSPGLGGRTQY | 0.48206666 |
| VAAVVTLL | ASSPGLGGRTQY | 0.54652125 |
| VGFLGGLI | ASSPGLGGRTQY | 0.48507142 |
| VGVIPILL | ASSPGLGGRTQY | 0.33461675 |
| VIFASSLI | ASSPGLGGRTQY | 0.59711486 |
| VKGGDGGAAAG | ASSPGLGGRTQY | 0.44423765 |
| VTKTFTNI | ASSPGLGGRTQY | 0.47167623 |
| VVAAAILL | ASSPGLGGRTQY | 0.49514034 |
| YAALDVEL | ASSPGLGGRTQY | 0.5942602 |
| LASVALLL | ASSPGLGGRTQY | 0.5532696 |
| LDAVHRV | ASSPGLGGRTQY | 0.5176822 |
| QGLSSSIF | ASSPGLGGRTQY | 0.5699363 |
| RGTFANIRL | ASSPGLGGRTQY | 0.589594 |
| RQLWWGHRI | ASSPGLGGRTQY | 0.60222155 |
| YGPPGTGKTLL | ASSPGLGGRTQY | 0.5742796 |
| AALVGAPANVA | ASSPGLGGRTQY | 0.5404197 |
| AATFRRTE | ASSPGLGGRTQY | 0.59520185 |
| AGALGGGI | ASSPGLGGRTQY | 0.4951151 |
| AGGAGGAGGIGAG | ASSPGLGGRTQY | 0.48601565 |
| AGSGALGL | ASSPGLGGRTQY | 0.4248866 |
| AQIADQI | ASSPGLGGRTQY | 0.47193617 |
| ASAVAVTYL | ASSPGLGGRTQY | 0.43995512 |
| ASTVLGGTGG | ASSPGLGGRTQY | 0.5522457 |
| AVGGEPAK | ASSPGLGGRTQY | 0.6364619 |
| CEQGTFGGTC | ASSPGLGGRTQY | 0.5552959 |
| DEGKAPAG | ASSPGLGGRTQY | 0.39045036 |
| DTVVFPAL | ASSPGLGGRTQY | 0.4824282 |
| FEGITMAG | ASSPGLGGRTQY | 0.5211908 |
| FPLVVQD | ASSPGLGGRTQY | 0.577234 |
| GAFTTPQ | ASSPGLGGRTQY | 0.6210172 |
| GFGGFGTAGGVGGA | ASSPGLGGRTQY | 0.64274025 |
| GFLAQFL | ASSPGLGGRTQY | 0.59787184 |
| GGAGGAGGNAGVA | ASSPGLGGRTQY | 0.48305774 |
| GGVGTAGAGGAGGAGG | ASSPGLGGRTQY | 0.5827372 |
| GNGGGGDAGNAGSGGN | ASSPGLGGRTQY | 0.4740007 |
| GTGGAGGAGNDAGS | ASSPGLGGRTQY | 0.5367987 |
| GTTNTGWFNS | ASSPGLGGRTQY | 0.5373596 |

| | | |
|---|---|---|
| IFMGPLFTL | ASSPGLGGRTQY | 0.57614684 |
| ILAHAIEAI | ASSPGLGGRTQY | 0.51775014 |
| IRSAIREAL | ASSPGLGGRTQY | 0.51894855 |
| ITFAGGG | ASSPGLGGRTQY | 0.38685086 |
| ITGMSGHQEV | ASSPGLGGRTQY | 0.5959556 |
| IVAGLACTL | ASSPGLGGRTQY | 0.46516517 |
| IVGGGLAGA | ASSPGLGGRTQY | 0.4547169 |
| IVTGVLNI | ASSPGLGGRTQY | 0.4867521 |
| LAQQGCI | ASSPGLGGRTQY | 0.6353089 |
| LCLHVEN | ASSPGLGGRTQY | 0.63330954 |
| LDAISDGS | ASSPGLGGRTQY | 0.46416593 |
| LGGFAAML | ASSPGLGGRTQY | 0.45783806 |
| LGGKNPMI | ASSPGLGGRTQY | 0.5644229 |
| LGPKGVAAIL | ASSPGLGGRTQY | 0.4324329 |
| LMVGVSI | ASSPGLGGRTQY | 0.5919639 |
| LVGAAFVL | ASSPGLGGRTQY | 0.52681917 |
| LVLAVAPL | ASSPGLGGRTQY | 0.540144 |
| MAAIAVIDAA | ASSPGLGGRTQY | 0.4881717 |
| NNGINAELQ | ASSPGLGGRTQY | 0.5682285 |
| RGEPLVLVHG | ASSPGLGGRTQY | 0.5226594 |
| RGGAGGAAGLI | ASSPGLGGRTQY | 0.58961356 |
| RSYRMHPAV | ASSPGLGGRTQY | 0.3424779 |
| SAAHLLAGQLA | ASSPGLGGRTQY | 0.61564267 |
| SAAVGSY | ASSPGLGGRTQY | 0.42082506 |
| SPHAVLHRI | ASSPGLGGRTQY | 0.44953266 |
| SQIAKVL | ASSPGLGGRTQY | 0.33522555 |
| TALGHLVP | ASSPGLGGRTQY | 0.5301848 |
| TAPTTSGAA | ASSPGLGGRTQY | 0.58202094 |
| TDGDDGNGGSPGT | ASSPGLGGRTQY | 0.61172336 |
| TDGLGEN | ASSPGLGGRTQY | 0.5524474 |
| TGGPPLPGQA | ASSPGLGGRTQY | 0.5982265 |
| TIPAITI | ASSPGLGGRTQY | 0.62306154 |
| TIPGFTI | ASSPGLGGRTQY | 0.46365085 |
| TSIASGALA | ASSPGLGGRTQY | 0.3786189 |
| TVTLPGAAI | ASSPGLGGRTQY | 0.5718716 |
| VASGLSAALP | ASSPGLGGRTQY | 0.52824086 |
| VGAGVIGI | ASSPGLGGRTQY | 0.6186448 |
| VGPKRFGV | ASSPGLGGRTQY | 0.5955213 |
| VNVAFPD | ASSPGLGGRTQY | 0.37830254 |
| VSMTNSGVS | ASSPGLGGRTQY | 0.62937844 |
| VTDVLGDLSV | ASSPGLGGRTQY | 0.5799508 |
| VVVVGAGFAGL | ASSPGLGGRTQY | 0.41059047 |
| VYTVGTVV | ASSPGLGGRTQY | 0.4532618 |
| WTTFSGP | ASSPGLGGRTQY | 0.48761564 |
| YDELVEAF | ASSPGLGGRTQY | 0.37502682 |
| YGTGGAGGNGG | ASSPGLGGRTQY | 0.611251 |
| YTNVMAV | ASSPGLGGRTQY | 0.46544093 |
| FLEQVHQL | ASSPGLGGRTQY | 0.50971895 |
| GLSLLAVAV | ASSPGLGGRTQY | 0.52314895 |
| LEQVHQL | ASSPGLGGRTQY | 0.48448932 |
| LNAAGAI | ASSPGLGGRTQY | 0.48026848 |

| | | |
|---|---|---|
| PAVHASL | ASSPGLGGRTQY | 0.5042711 |
| TELEDDA | ASSPGLGGRTQY | 0.6031636 |
| TPPGPPN | ASSPGLGGRTQY | 0.5848954 |
| TPPPPPPP | ASSPGLGGRTQY | 0.5280121 |
| TQSPALI | ASSQEPGEGTLY | 0.5644624 |
| EFAAAFPQ | ASSQEPGEGTLY | 0.60991836 |
| LLSGNLDVL | ASSQEPGEGTLY | 0.41388613 |
| GPANLGNYNI | ASSQEPGEGTLY | 0.4816644 |
| TLGPTLP | ASSQEPGEGTLY | 0.5893486 |
| VYAYSSCCK | ASSQEPGEGTLY | 0.48125833 |
| APAPAPAPNI | ASSQEPGEGTLY | 0.61467123 |
| PGGPVPIP | ASSQEPGEGTLY | 0.569841 |
| FLTVALEP | ASSQEPGEGTLY | 0.54919916 |
| SQYFGANL | ASSQEPGEGTLY | 0.5309734 |
| GGVKPAA | ASSQEPGEGTLY | 0.64469975 |
| GGTGGAGGAGGW | ASSQEPGEGTLY | 0.5676704 |
| GVGGVAGLGGAGG | ASSQEPGEGTLY | 0.5212645 |
| VLLVLATTI | ASSQEPGEGTLY | 0.58171093 |
| IGIFAAML | ASSQEPGEGTLY | 0.43105534 |
| AVGDLID | ASSQEPGEGTLY | 0.6084224 |
| NISPLPL | ASSQEPGEGTLY | 0.586396 |
| VTAVFVAI | ASSQEPGEGTLY | 0.40896428 |
| SGDDNDAAL | ASSQEPGEGTLY | 0.49952978 |
| AVAALAHS | ASSQEPGEGTLY | 0.6259943 |
| GGVTVMATP | ASSQEPGEGTLY | 0.59465545 |
| AGLAANVRD | ASSQEPGEGTLY | 0.48791766 |
| TNRPDLIDPALLRPG | ASSQEPGEGTLY | 0.46894267 |
| LANLGVTAL | ASSQEPGEGTLY | 0.44349727 |
| TGERIEL | ASSQEPGEGTLY | 0.47525814 |
| NGEELAA | ASSQEPGEGTLY | 0.40857843 |
| VLSLVWG | ASSQEPGEGTLY | 0.5765394 |
| IGYALWL | ASSQEPGEGTLY | 0.48655808 |
| LSLLLAPL | ASSQEPGEGTLY | 0.600345 |
| TLDEALK | ASSQEPGEGTLY | 0.5567392 |
| TDGAGTAEGG | ASSQEPGEGTLY | 0.60998344 |
| NAPGGAGGNGG | ASSQEPGEGTLY | 0.4738434 |
| VSVFMGMGGNF | ASSQEPGEGTLY | 0.5158315 |
| LASYLAAL | ASSQEPGEGTLY | 0.3693096 |
| LVGGTPAL | ASSQEPGEGTLY | 0.5210005 |
| VRNADVL | ASSQEPGEGTLY | 0.4506713 |
| TCSSPFLVSAM | ASSQEPGEGTLY | 0.52342606 |
| VHITDAE | ASSQEPGEGTLY | 0.37797463 |
| EYAAPSF | ASSQEPGEGTLY | 0.6397893 |
| VVDAFHGGL | ASSQEPGEGTLY | 0.49331886 |
| ASVVDPQ | ASSQEPGEGTLY | 0.57767963 |
| SYNEFGGLS | ASSQEPGEGTLY | 0.523053 |
| DPDGSVY | ASSQEPGEGTLY | 0.4453649 |
| MVSTIHATE | ASSQEPGEGTLY | 0.4747003 |
| LAAGVADPV | ASSQEPGEGTLY | 0.5880753 |
| TKEQIAA | ASSQEPGEGTLY | 0.42250073 |
| SLASPAAL | ASSQEPGEGTLY | 0.3604556 |

| | | |
|---|---|---|
| IGLVFYGV | ASSQEPGEGTLY | 0.45777905 |
| FLLDASNL | ASSQEPGEGTLY | 0.46502823 |
| YQVGNFP | ASSQEPGEGTLY | 0.6381952 |
| ESLLSQST | ASSQEPGEGTLY | 0.6169321 |
| LFTVGFAAVG | ASSQEPGEGTLY | 0.5436165 |
| ATPPTPT | ASSQEPGEGTLY | 0.6403555 |
| TDLATLG | ASSQEPGEGTLY | 0.31964067 |
| SAGGTGR | ASSQEPGEGTLY | 0.5398258 |
| GFSNSGS | ASSQEPGEGTLY | 0.6273003 |
| GGNGGSGGTSV | ASSQEPGEGTLY | 0.5746661 |
| GTSVATGGAGN | ASSQEPGEGTLY | 0.5108026 |
| NGGNGGSGGTS | ASSQEPGEGTLY | 0.6031429 |
| AVVKFPH | ASSQEPGEGTLY | 0.46999663 |
| TPPPPPP | ASSQEPGEGTLY | 0.4887814 |
| AAPAAAAAPA | ASSQEPGEGTLY | 0.52679145 |
| IALNEDL | ASSQEPGEGTLY | 0.5986819 |
| IVNVSSM | ASSQEPGEGTLY | 0.57507485 |
| TGIPPAPR | ASSQEPGEGTLY | 0.39608338 |
| FDLGGGTF | ASSQEPGEGTLY | 0.58673155 |
| FDLGGGTFDVS | ASSQEPGEGTLY | 0.6394177 |
| NEPTAAA | ASSQEPGEGTLY | 0.46993798 |
| NEPTAAALA | ASSQEPGEGTLY | 0.6025317 |
| GSIINISSI | ASSQEPGEGTLY | 0.5498385 |
| VAPGNAGTA | ASSQEPGEGTLY | 0.53903013 |
| TGGGASLE | ASSQEPGEGTLY | 0.43970567 |
| VAVYNHYKRI | ASSQEPGEGTLY | 0.40187615 |
| TDPAPATT | ASSQEPGEGTLY | 0.6050475 |
| TNRPDILDPALLRPG | ASSQEPGEGTLY | 0.43679583 |
| LTTTTTS | ASSQEPGEGTLY | 0.5237737 |
| PTGSGKTL | ASSQEPGEGTLY | 0.5393005 |
| TGSGKTL | ASSQEPGEGTLY | 0.6510896 |
| TIGHVDHGK | ASSQEPGEGTLY | 0.5692291 |
| TIGHVDHGKTT | ASSQEPGEGTLY | 0.57559 |
| GHVDHGKTT | ASSQEPGEGTLY | 0.44807002 |
| AQFEHTLL | ASSQEPGEGTLY | 0.5564271 |
| IVNQDEL | ASSQEPGEGTLY | 0.56973666 |
| TNGSQFF | ASSQEPGEGTLY | 0.41419208 |
| TGKGIVE | ASSQEPGEGTLY | 0.60804534 |
| LISNASDA | ASSQEPGEGTLY | 0.41403648 |
| AELRTLGELTKIAKAHGAQ | ASSQEPGEGTLY | 0.549239 |
| APNAVAAAEPSW | ASSQEPGEGTLY | 0.5690747 |
| APQVAGAATAVL | ASSQEPGEGTLY | 0.6225075 |
| ARRDPAAAAAILPTDA | ASSQEPGEGTLY | 0.49737775 |
| DAAARDVLAEAGLAEYF | ASSQEPGEGTLY | 0.59638083 |
| DDQIVLSAKANLAKVEQ | ASSQEPGEGTLY | 0.43350604 |
| DELLPDTGEDSDEDGH | ASSQEPGEGTLY | 0.6009445 |
| ELPEAPANRATVPQD | ASSQEPGEGTLY | 0.6102822 |
| FGQSKIERVTGIG | ASSQEPGEGTLY | 0.55147594 |
| GAGGTGSTAGGAGGAGGAGGLY | ASSQEPGEGTLY | 0.6435814 |
| GHGGTGAAGGNGGTGGAGGSGI | ASSQEPGEGTLY | 0.6131886 |
| GNGGNGGAGGEGGPGGAGGAGA | ASSQEPGEGTLY | 0.37500638 |

| | | |
|---|---|---|
| GSTAGNGGNGGNATNGTIAGQP | ASSQEPGEGTLY | 0.5399152 |
| GTGGDGGNGGNGGTGG | ASSQEPGEGTLY | 0.61324906 |
| GVRRPAGIPADAPV | ASSQEPGEGTLY | 0.6151649 |
| GYRQVTDAHLLTLARR | ASSQEPGEGTLY | 0.6236124 |
| IAKMGQDGHDRG | ASSQEPGEGTLY | 0.60751545 |
| KAEPSIEIQYT | ASSQEPGEGTLY | 0.5084797 |
| KPVRIRMAADEDTVAVT | ASSQEPGEGTLY | 0.62539935 |
| MPPRIAGMRLLVIKPEP | ASSQEPGEGTLY | 0.6659537 |
| MKFYAVALFRSRGVPPTR | ASSQEPGEGTLY | 0.58626664 |
| NAGGFGILGAVAHSP | ASSQEPGEGTLY | 0.4367777 |
| NEGREITLAMAEM | ASSQEPGEGTLY | 0.45320654 |
| NELVEWSHPLGHRWRWE | ASSQEPGEGTLY | 0.5739907 |
| NLGAGGTGGNGGNPG | ASSQEPGEGTLY | 0.6115305 |
| NTEWPKWETVMAADPVD | ASSQEPGEGTLY | 0.3987504 |
| PASAAPAETQP | ASSQEPGEGTLY | 0.5215889 |
| QQSSDGKNGVRAAEITGE | ASSQEPGEGTLY | 0.5451897 |
| RAPDAVTTSWWKE | ASSQEPGEGTLY | 0.35197115 |
| SGFGNVGAHTSGWFNQ | ASSQEPGEGTLY | 0.5950916 |
| SGVAGAVGQAASVGGLKV | ASSQEPGEGTLY | 0.62402064 |
| SNGGAGTGSAGGNGGT | ASSQEPGEGTLY | 0.56064725 |
| TGGAGGEGGAAG | ASSQEPGEGTLY | 0.4139629 |
| TGGDGGAAGAGGAPGHG | ASSQEPGEGTLY | 0.6096917 |
| TMPPLLAAIANAGRHGV | ASSQEPGEGTLY | 0.61847407 |
| TSSTAGNSGGA | ASSQEPGEGTLY | 0.63226354 |
| VPATVPVGILC | ASSQEPGEGTLY | 0.39163494 |
| VQRPDSGGVVLTGRIS | ASSQEPGEGTLY | 0.3172148 |
| WTPNPVPATKGVPAGLVQAT | ASSQEPGEGTLY | 0.5408658 |
| YALKRALAKFSDRFDVVII | ASSQEPGEGTLY | 0.5822176 |
| YLIGSVIV | ASSQEPGEGTLY | 0.50839823 |
| YGQLVAVID | ASSQEPGEGTLY | 0.45466793 |
| VTVGGMTLPA | ASSQEPGEGTLY | 0.5361271 |
| VLALPAV | ASSQEPGEGTLY | 0.61046934 |
| VIPPTLNYET | ASSQEPGEGTLY | 0.4327101 |
| VGMGGMHPSSG | ASSQEPGEGTLY | 0.4146834 |
| VGLAGGAATA | ASSQEPGEGTLY | 0.5715996 |
| VGGAGGAGGSGP | ASSQEPGEGTLY | 0.601549 |
| VFAAFMV | ASSQEPGEGTLY | 0.42258346 |
| TLYDKLR | ASSQEPGEGTLY | 0.477104 |
| TLAVFTVL | ASSQEPGEGTLY | 0.5617716 |
| TGSVQPARV | ASSQEPGEGTLY | 0.57227963 |
| TGGGAGSGGT | ASSQEPGEGTLY | 0.54518664 |
| SHLETSL | ASSQEPGEGTLY | 0.40441433 |
| SFTTGMT | ASSQEPGEGTLY | 0.6083498 |
| QVEVAKRMGVRQPTVS | ASSQEPGEGTLY | 0.39174756 |
| QGSGGPALAM | ASSQEPGEGTLY | 0.59270394 |
| MLAPSNP | ASSQEPGEGTLY | 0.55030704 |
| LVNNAAI | ASSQEPGEGTLY | 0.57880914 |
| LTGEINTLA | ASSQEPGEGTLY | 0.47697502 |
| LSIFGPV | ASSQEPGEGTLY | 0.42657587 |
| LLALLANL | ASSQEPGEGTLY | 0.42764536 |
| LISNASDALDKLR | ASSQEPGEGTLY | 0.4006894 |

| | | |
|---|---|---|
| LINEAVP | ASSQEPGEGTLY | 0.619316 |
| LGYAFSGL | ASSQEPGEGTLY | 0.5551573 |
| LDFVAVAL | ASSQEPGEGTLY | 0.6015561 |
| LAQPGAE | ASSQEPGEGTLY | 0.5340929 |
| ISNLGAVTGF | ASSQEPGEGTLY | 0.44508067 |
| HQEDPGRI | ASSQEPGEGTLY | 0.56722873 |
| GTPGEPTP | ASSQEPGEGTLY | 0.40767848 |
| GFFNSGS | ASSQEPGEGTLY | 0.55949396 |
| GAGGGGLTAGAN | ASSQEPGEGTLY | 0.5305961 |
| FVDALVNM | ASSQEPGEGTLY | 0.59353817 |
| FLTVAPEL | ASSQEPGEGTLY | 0.4638542 |
| FETLTAL | ASSQEPGEGTLY | 0.5703254 |
| DVINPIEL | ASSQEPGEGTLY | 0.56539494 |
| APVSGAPV | ASSQEPGEGTLY | 0.43655583 |
| ALVGGLAW | ASSQEPGEGTLY | 0.57211375 |
| ALAAAAATAI | ASSQEPGEGTLY | 0.60565436 |
| AFGLGLLL | ASSQEPGEGTLY | 0.5683599 |
| AFFAAASL | ASSQEPGEGTLY | 0.3783561 |
| AAAALPAFSPP | ASSQEPGEGTLY | 0.55942327 |
| VVLAPAL | ASSQEPGEGTLY | 0.45595965 |
| TIGHVDHGKTTL | ASSQEPGEGTLY | 0.56088275 |
| LISNASDALDK | ASSQEPGEGTLY | 0.53923386 |
| LISNASDAL | ASSQEPGEGTLY | 0.47914225 |
| ISNASDALDK | ASSQEPGEGTLY | 0.5932215 |
| NQAETLVYQTE | ASSQEPGEGTLY | 0.43918937 |
| TGGTGGTAGAG | ASSQEPGEGTLY | 0.46567824 |
| TGSTAGGAGGA | ASSQEPGEGTLY | 0.4965254 |
| DPLAVPVAAGPV | ASSQEPGEGTLY | 0.55705684 |
| GGTGGTPGNGGN | ASSQEPGEGTLY | 0.5879015 |
| PAGLAGVDPATL | ASSQEPGEGTLY | 0.42332 |
| GNNLVVEAPHAGASV | ASSQEPGEGTLY | 0.54873526 |
| KVTRIMSASTLEQPA | ASSQEPGEGTLY | 0.49181455 |
| LGFALPIALPRRTVV | ASSQEPGEGTLY | 0.41761518 |
| TRLGNLTARKLVVLV | ASSQEPGEGTLY | 0.53612936 |
| WSQQRRLIYFPSAGP | ASSQEPGEGTLY | 0.48434168 |
| GPNGNPDPMAAAVDIR | ASSQEPGEGTLY | 0.39756367 |
| TQRCVSVADLRAAASR | ASSQEPGEGTLY | 0.599085 |
| QFVLDVAEHPVEVLTETV | ASSQEPGEGTLY | 0.45489836 |
| GGAGGTGGAGGNGGTQAGDG | ASSQEPGEGTLY | 0.5870597 |
| PLLATLRAIAVDVDAKPAQV | ASSQEPGEGTLY | 0.6038193 |
| APKFRTGGYMIDFWGVGYQVAK | ASSQEPGEGTLY | 0.40266448 |
| GAGGAGGAGGAAGGPGGTGGTG | ASSQEPGEGTLY | 0.58073926 |
| IGALGLLATDVAPAGSVAFGVA | ASSQEPGEGTLY | 0.5627015 |
| AGGAGGIGGNGGAGGAGGVLMG | ASSQEPGEGTLY | 0.5542438 |
| ADELIGLL | ASSQEPGEGTLY | 0.5204624 |
| ADRALKA | ASSQEPGEGTLY | 0.4584557 |
| AGAVGQVGGAGGA | ASSQEPGEGTLY | 0.6262467 |
| AGGGGGLGGAP | ASSQEPGEGTLY | 0.59328806 |
| AGGLLYGNGGAG | ASSQEPGEGTLY | 0.6059988 |
| AGLESIGL | ASSQEPGEGTLY | 0.5980118 |
| AGLKTGII | ASSQEPGEGTLY | 0.57750434 |

| | | |
|---|---|---|
| ASLAALLL | ASSQEPGEGTLY | 0.59837204 |
| ASVINISSI | ASSQEPGEGTLY | 0.46585488 |
| ATPPGPA | ASSQEPGEGTLY | 0.36256537 |
| ATVASALV | ASSQEPGEGTLY | 0.518121 |
| ATVASLIL | ASSQEPGEGTLY | 0.499334 |
| AVNGPVLIRH | ASSQEPGEGTLY | 0.4634385 |
| DGAAGAVGI | ASSQEPGEGTLY | 0.3998508 |
| DPLLHGI | ASSQEPGEGTLY | 0.5932575 |
| ELGAALGAL | ASSQEPGEGTLY | 0.6557374 |
| ELGRTQAV | ASSQEPGEGTLY | 0.47393745 |
| FVAGGYLS | ASSQEPGEGTLY | 0.655818 |
| GASGGAGGAGGW | ASSQEPGEGTLY | 0.39479625 |
| GGAGAGGDGGAGGI | ASSQEPGEGTLY | 0.597062 |
| GTGHNGSQPGMGGNG | ASSQEPGEGTLY | 0.60379934 |
| GYAAASASAS | ASSQEPGEGTLY | 0.40077832 |
| HQVDVAAVQQG | ASSQEPGEGTLY | 0.5941949 |
| IGAAIAHT | ASSQEPGEGTLY | 0.4486414 |
| IGMIEVAQL | ASSQEPGEGTLY | 0.46282405 |
| IRLAPEHTSAMD | ASSQEPGEGTLY | 0.6143534 |
| IYNVSVL | ASSQEPGEGTLY | 0.51675326 |
| KQGFLGM | ASSQEPGEGTLY | 0.589756 |
| LESVITQL | ASSQEPGEGTLY | 0.5028533 |
| LGLGALVVV | ASSQEPGEGTLY | 0.4452646 |
| LIGNGGNGGSGGI | ASSQEPGEGTLY | 0.56749827 |
| LNAINAPTL | ASSQEPGEGTLY | 0.60860455 |
| LQLLNTGV | ASSQEPGEGTLY | 0.56057215 |
| LQTSIEF | ASSQEPGEGTLY | 0.6629692 |
| MSFAVSEL | ASSQEPGEGTLY | 0.45980006 |
| NTGALGL | ASSQEPGEGTLY | 0.63629216 |
| PPGAGGAGGAGTT | ASSQEPGEGTLY | 0.6175996 |
| QLVGLGI | ASSQEPGEGTLY | 0.5354529 |
| RFLEQVHQL | ASSQEPGEGTLY | 0.4967234 |
| RGVGYVLR | ASSQEPGEGTLY | 0.5918592 |
| SATGVFAGVI | ASSQEPGEGTLY | 0.49270585 |
| SLPAITLP | ASSQEPGEGTLY | 0.59675777 |
| STALAFTV | ASSQEPGEGTLY | 0.62947655 |
| SVLTVGLGK | ASSQEPGEGTLY | 0.72093457 |
| TALAGAGSGI | ASSQEPGEGTLY | 0.8763082 |
| TGLPFTGI | ASSQEPGEGTLY | 0.81952304 |
| VAEHVTV | ASSQEPGEGTLY | 0.8443122 |
| VDIVLNSL | ASSQEPGEGTLY | 0.7547817 |
| VLTGGGAVL | ASSQEPGEGTLY | 0.5334237 |
| VNVGLAYGI | ASSQEPGEGTLY | 0.82229173 |
| VSSAAVGAV | ASSQEPGEGTLY | 0.68579876 |
| VTSVYEAGV | ASSQEPGEGTLY | 0.869812 |
| WVALDRGA | ASSQEPGEGTLY | 0.87761986 |
| YLVGSVVI | ASSQEPGEGTLY | 0.7749573 |
| GPPGTGKTLL | ASSQEPGEGTLY | 0.86261714 |
| GVGMTKF | ASSQEPGEGTLY | 0.87544703 |
| IGIGGSDL | ASSQEPGEGTLY | 0.66963464 |
| IGTLLAEL | ASSQEPGEGTLY | 0.7962556 |

| | | |
|---|---|---|
| IYTSGTT | ASSQEPGEGTLY | 0.7946191 |
| LSFPAGV | ASSQEPGEGTLY | 0.83055437 |
| LYTSGTT | ASSQEPGEGTLY | 0.53179485 |
| VGAFPFGF | ASSQEPGEGTLY | 0.8699796 |
| VGAGTVEF | ASSQEPGEGTLY | 0.7634423 |
| VGKSTLLL | ASSQEPGEGTLY | 0.31730112 |
| AALAHAGGGIV | ASSQEPGEGTLY | 0.83699375 |
| AALVAVSLL | ASSQEPGEGTLY | 0.81792593 |
| AAVSRAVAAL | ASSQEPGEGTLY | 0.8061091 |
| AGGSSGIGGF | ASSQEPGEGTLY | 0.87273824 |
| AGLTVANANG | ASSQEPGEGTLY | 0.76184213 |
| AGLVGAVL | ASSQEPGEGTLY | 0.66490424 |
| ALSQAFTV | ASSQEPGEGTLY | 0.83999395 |
| ALTFSVSL | ASSQEPGEGTLY | 0.8832408 |
| ALTGAALVAL | ASSQEPGEGTLY | 0.6209382 |
| ASFSIGAGVL | ASSQEPGEGTLY | 0.83846354 |
| DGSVYKI | ASSQEPGEGTLY | 0.88681084 |
| EQLAEIV | ASSQEPGEGTLY | 0.82632804 |
| FAVSELTVM | ASSQEPGEGTLY | 0.88447905 |
| GAAPLAAGL | ASSQEPGEGTLY | 0.88812953 |
| GADGGDGGSGGAGG | ASSQEPGEGTLY | 0.768335 |
| GFGLDTGGAGG | ASSQEPGEGTLY | 0.8766902 |
| GLASGGLL | ASSQEPGEGTLY | 0.8659109 |
| GRIAGAAVLL | ASSQEPGEGTLY | 0.8118435 |
| GSAGSLNTSF | ASSQEPGEGTLY | 0.8203645 |
| GTGQIGI | ASSQEPGEGTLY | 0.72207654 |
| GTLIALIL | ASSQEPGEGTLY | 0.82355225 |
| IAATMGFVVG | ASSQEPGEGTLY | 0.54082227 |
| IFAQALL | ASSQEPGEGTLY | 0.83707076 |
| IGAAVALGF | ASSQEPGEGTLY | 0.8797715 |
| IGLTPLAL | ASSQEPGEGTLY | 0.79703385 |
| ILGSNTSTL | ASSQEPGEGTLY | 0.66736245 |
| IVFSLTVL | ASSQEPGEGTLY | 0.61536616 |
| LASKATGF | ASSQEPGEGTLY | 0.8253422 |
| LASVALIL | ASSQEPGEGTLY | 0.7418957 |
| LFAVGLTL | ASSQEPGEGTLY | 0.5027399 |
| LFNGGIGTYI | ASSQEPGEGTLY | 0.78777957 |
| LTLGAGII | ASSQEPGEGTLY | 0.87092173 |
| LVAAALGDI | ASSQEPGEGTLY | 0.87587214 |
| NTAFVQLGI | ASSQEPGEGTLY | 0.64867866 |
| QLFSGAEL | ASSQEPGEGTLY | 0.8847142 |
| SAATFITL | ASSQEPGEGTLY | 0.8678001 |
| SNYGLANLGV | ASSQEPGEGTLY | 0.8703382 |
| SYGGAYAVM | ASSQEPGEGTLY | 0.82088304 |
| TTGVGGLTLG | ASSQEPGEGTLY | 0.6970792 |
| TTQTIYI | ASSQEPGEGTLY | 0.87898254 |
| VAAGAGLGAVI | ASSQEPGEGTLY | 0.7633443 |
| VAAVVTLL | ASSQEPGEGTLY | 0.8122757 |
| VGFLGGLI | ASSQEPGEGTLY | 0.87145317 |
| VGVIPILL | ASSQEPGEGTLY | 0.7568787 |
| VIFASSLI | ASSQEPGEGTLY | 0.73460114 |

| | | |
|---|---|---|
| VKGGDGGAAAG | ASSQEPGEGTLY | 0.80672073 |
| VTKTFTNI | ASSQEPGEGTLY | 0.8843479 |
| VVAAAILL | ASSQEPGEGTLY | 0.64197254 |
| YAALDVEL | ASSQEPGEGTLY | 0.67962205 |
| LASVALLL | ASSQEPGEGTLY | 0.7775001 |
| LDAVHRV | ASSQEPGEGTLY | 0.8078556 |
| QGLSSSIF | ASSQEPGEGTLY | 0.8815465 |
| RGTFANIRL | ASSQEPGEGTLY | 0.8624593 |
| RQLWWGHRI | ASSQEPGEGTLY | 0.7882117 |
| YGPPGTGKTLL | ASSQEPGEGTLY | 0.8483759 |
| AALVGAPANVA | ASSQEPGEGTLY | 0.8658991 |
| AATFRRTE | ASSQEPGEGTLY | 0.838804 |
| AGALGGGI | ASSQEPGEGTLY | 0.8467929 |
| AGGAGGAGGIGAG | ASSQEPGEGTLY | 0.7368752 |
| AGSGALGL | ASSQEPGEGTLY | 0.88297606 |
| AQIADQI | ASSQEPGEGTLY | 0.86428213 |
| ASAVAVTYL | ASSQEPGEGTLY | 0.8775121 |
| ASTVLGGTGG | ASSQEPGEGTLY | 0.87897146 |
| AVGGEPAK | ASSQEPGEGTLY | 0.83587384 |
| CEQGTFGGTC | ASSQEPGEGTLY | 0.8328755 |
| DEGKAPAG | ASSQEPGEGTLY | 0.4926326 |
| DTVVFPAL | ASSQEPGEGTLY | 0.78335357 |
| FEGITMAG | ASSQEPGEGTLY | 0.8181889 |
| FPLVVQD | ASSQEPGEGTLY | 0.83505833 |
| GAFTTPQ | ASSQEPGEGTLY | 0.8513516 |
| GFGGFGTAGGVGGA | ASSQEPGEGTLY | 0.8689744 |
| GFLAQFL | ASSQEPGEGTLY | 0.7597156 |
| GGAGGAGGNAGVA | ASSQEPGEGTLY | 0.8728925 |
| GGVGTAGAGGAGGAGG | ASSQEPGEGTLY | 0.87673324 |
| GNGGGGDAGNAGSGGN | ASSQEPGEGTLY | 0.63884634 |
| GTGGAGGAGNDAGS | ASSQEPGEGTLY | 0.78555995 |
| GTTNTGWFNS | ASSQEPGEGTLY | 0.8914753 |
| IFMGPLFTL | ASSQEPGEGTLY | 0.84194297 |
| ILAHAIEAI | ASSQEPGEGTLY | 0.779987 |
| IRSAIREAL | ASSQEPGEGTLY | 0.7984359 |
| ITFAGGG | ASSQEPGEGTLY | 0.8899783 |
| ITGMSGHQEV | ASSQEPGEGTLY | 0.8276105 |
| IVAGLACTL | ASSQEPGEGTLY | 0.882612 |
| IVGGGLAGA | ASSQEPGEGTLY | 0.39778972 |
| IVTGVLNI | ASSQEPGEGTLY | 0.8137399 |
| LAQQGCI | ASSQEPGEGTLY | 0.68178105 |
| LCLHVEN | ASSQEPGEGTLY | 0.3493586 |
| LDAISDGS | ASSQEPGEGTLY | 0.8636372 |
| LGGFAAML | ASSQEPGEGTLY | 0.7815819 |
| LGGKNPMI | ASSQEPGEGTLY | 0.86140025 |
| LGPKGVAAIL | ASSQEPGEGTLY | 0.8030715 |
| LMVGVSI | ASSQEPGEGTLY | 0.87348944 |
| LVGAAFVL | ASSQEPGEGTLY | 0.8143586 |
| LVLAVAPL | ASSQEPGEGTLY | 0.82960135 |
| MAAIAVIDAA | ASSQEPGEGTLY | 0.7789848 |
| NNGINAELQ | ASSQEPGEGTLY | 0.84284794 |

| | | |
|---|---|---|
| RGEPLVLVHG | ASSQEPGEGTLY | 0.87361145 |
| RGGAGGAAGLI | ASSQEPGEGTLY | 0.8823151 |
| RSYRMHPAV | ASSQEPGEGTLY | 0.4127223 |
| SAAHLLAGQLA | ASSQEPGEGTLY | 0.8105087 |
| SAAVGSY | ASSQEPGEGTLY | 0.8065487 |
| SPHAVLHRI | ASSQEPGEGTLY | 0.42516762 |
| SQIAKVL | ASSQEPGEGTLY | 0.40315294 |
| TALGHLVP | ASSQEPGEGTLY | 0.7338964 |
| TAPTTSGAA | ASSQEPGEGTLY | 0.7930814 |
| TDGDDGNGGSPGT | ASSQEPGEGTLY | 0.8556725 |
| TDGLGEN | ASSQEPGEGTLY | 0.8522235 |
| TGGPPLPGQA | ASSQEPGEGTLY | 0.7850555 |
| TIPAITI | ASSQEPGEGTLY | 0.8524433 |
| TIPGFTI | ASSQEPGEGTLY | 0.8043891 |
| TSIASGALA | ASSQEPGEGTLY | 0.114496574 |
| TVTLPGAAI | ASSQEPGEGTLY | 0.80538434 |
| VASGLSAALP | ASSQEPGEGTLY | 0.6736398 |
| VGAGVIGI | ASSQEPGEGTLY | 0.8889017 |
| VGPKRFGV | ASSQEPGEGTLY | 0.7231041 |
| VNVAFPD | ASSQEPGEGTLY | 0.7597255 |
| VSMTNSGVS | ASSQEPGEGTLY | 0.8180077 |
| VTDVLGDLSV | ASSQEPGEGTLY | 0.8439925 |
| VVVVGAGFAGL | ASSQEPGEGTLY | 0.8213843 |
| VYTVGTVV | ASSQEPGEGTLY | 0.70884 |
| WTTFSGP | ASSQEPGEGTLY | 0.2577515 |
| YDELVEAF | ASSQEPGEGTLY | 0.8538529 |
| YGTGGAGGNGG | ASSQEPGEGTLY | 0.7227187 |
| YTNVMAV | ASSQEPGEGTLY | 0.6513618 |
| FLEQVHQL | ASSQEPGEGTLY | 0.6512768 |
| GLSLLAVAV | ASSQEPGEGTLY | 0.4703035 |
| LEQVHQL | ASSQEPGEGTLY | 0.56684256 |
| LNAAGAI | ASSQEPGEGTLY | 0.8786744 |
| PAVHASL | ASSQEPGEGTLY | 0.734582 |
| TELEDDA | ASSQEPGEGTLY | 0.7647065 |
| TPPGPPN | ASSQEPGEGTLY | 0.87971795 |
| TPPPPPPP | ASSQEPGEGTLY | 0.83691096 |
| TQSPALI | ASSQDVREGRNERLF | 0.7936243 |
| EFAAAFPQ | ASSQDVREGRNERLF | 0.74011964 |
| LLSGNLDVL | ASSQDVREGRNERLF | 0.7590989 |
| GPANLGNYNI | ASSQDVREGRNERLF | 0.69062006 |
| TLGPTLP | ASSQDVREGRNERLF | 0.7022954 |
| VYAYSSCCK | ASSQDVREGRNERLF | 0.8777612 |
| APAPAPAPNI | ASSQDVREGRNERLF | 0.8269832 |
| PGGPVPIP | ASSQDVREGRNERLF | 0.6825901 |
| FLTVALEP | ASSQDVREGRNERLF | 0.8672069 |
| SQYFGANL | ASSQDVREGRNERLF | 0.88168025 |
| GGVKPAA | ASSQDVREGRNERLF | 0.885705 |
| GGTGGAGGAGGW | ASSQDVREGRNERLF | 0.7398974 |
| GVGGVAGLGGAGG | ASSQDVREGRNERLF | 0.7905185 |
| VLLVLATTI | ASSQDVREGRNERLF | 0.79605997 |
| IGIFAAML | ASSQDVREGRNERLF | 0.79732114 |

| | | |
|---|---|---|
| AVGDLID | ASSQDVREGRNERLF | 0.78696537 |
| NISPLPL | ASSQDVREGRNERLF | 0.6521774 |
| VTAVFVAI | ASSQDVREGRNERLF | 0.87751013 |
| SGDDNDAAL | ASSQDVREGRNERLF | 0.89958674 |
| AVAALAHS | ASSQDVREGRNERLF | 0.80297256 |
| GGVTVMATP | ASSQDVREGRNERLF | 0.64756656 |
| AGLAANVRD | ASSQDVREGRNERLF | 0.79115295 |
| TNRPDLIDPALLRPG | ASSQDVREGRNERLF | 0.6914226 |
| LANLGVTAL | ASSQDVREGRNERLF | 0.7545427 |
| TGERIEL | ASSQDVREGRNERLF | 0.82467663 |
| NGEELAA | ASSQDVREGRNERLF | 0.6158865 |
| VLSLVWG | ASSQDVREGRNERLF | 0.7925422 |
| IGYALWL | ASSQDVREGRNERLF | 0.8851809 |
| LSLLLAPL | ASSQDVREGRNERLF | 0.7820176 |
| TLDEALK | ASSQDVREGRNERLF | 0.8317555 |
| TDGAGTAEGG | ASSQDVREGRNERLF | 0.8724543 |
| NAPGGAGGNGG | ASSQDVREGRNERLF | 0.85962534 |
| VSVFMGMGGNF | ASSQDVREGRNERLF | 0.88536215 |
| LASYLAAL | ASSQDVREGRNERLF | 0.875895 |
| LVGGTPAL | ASSQDVREGRNERLF | 0.8499565 |
| VRNADVL | ASSQDVREGRNERLF | 0.8273916 |
| TCSSPFLVSAM | ASSQDVREGRNERLF | 0.8474105 |
| VHITDAE | ASSQDVREGRNERLF | 0.81333995 |
| EYAAPSF | ASSQDVREGRNERLF | 0.8187678 |
| VVDAFHGGL | ASSQDVREGRNERLF | 0.7896286 |
| ASVVDPQ | ASSQDVREGRNERLF | 0.8672656 |
| SYNEFGGLS | ASSQDVREGRNERLF | 0.8211278 |
| DPDGSVY | ASSQDVREGRNERLF | 0.8890259 |
| MVSTIHATE | ASSQDVREGRNERLF | 0.48576576 |
| LAAGVADPV | ASSQDVREGRNERLF | 0.67603505 |
| TKEQIAA | ASSQDVREGRNERLF | 0.8732648 |
| SLASPAAL | ASSQDVREGRNERLF | 0.19243352 |
| IGLVFYGV | ASSQDVREGRNERLF | 0.46385133 |
| FLLDASNL | ASSQDVREGRNERLF | 0.8314842 |
| YQVGNFP | ASSQDVREGRNERLF | 0.8695377 |
| ESLLSQST | ASSQDVREGRNERLF | 0.19662783 |
| LFTVGFAAVG | ASSQDVREGRNERLF | 0.7507568 |
| ATPPTPT | ASSQDVREGRNERLF | 0.5886613 |
| TDLATLG | ASSQDVREGRNERLF | 0.8786098 |
| SAGGTGR | ASSQDVREGRNERLF | 0.81919175 |
| GFSNSGS | ASSQDVREGRNERLF | 0.5814809 |
| GGNGGSGGTSV | ASSQDVREGRNERLF | 0.87607443 |
| GTSVATGGAGN | ASSQDVREGRNERLF | 0.6203841 |
| NGGNGGSGGTS | ASSQDVREGRNERLF | 0.771889 |
| AVVKFPH | ASSQDVREGRNERLF | 0.86672693 |
| TPPPPPP | ASSQDVREGRNERLF | 0.89050066 |
| AAPAAAAAPA | ASSQDVREGRNERLF | 0.662103 |
| IALNEDL | ASSQDVREGRNERLF | 0.86840165 |
| IVNVSSM | ASSQDVREGRNERLF | 0.7855203 |
| TGIPPAPR | ASSQDVREGRNERLF | 0.8886383 |
| FDLGGGTF | ASSQDVREGRNERLF | 0.7607225 |

| | | |
|---|---|---|
| FDLGGGTFDVS | ASSQDVREGRNERLF | 0.617618 |
| NEPTAAA | ASSQDVREGRNERLF | 0.87010247 |
| NEPTAAALA | ASSQDVREGRNERLF | 0.86075956 |
| GSIINISSI | ASSQDVREGRNERLF | 0.83262277 |
| VAPGNAGTA | ASSQDVREGRNERLF | 0.8735151 |
| TGGGASLE | ASSQDVREGRNERLF | 0.8249038 |
| VAVYNHYKRI | ASSQDVREGRNERLF | 0.79327625 |
| TDPAPATT | ASSQDVREGRNERLF | 0.87336063 |
| TNRPDILDPALLRPG | ASSQDVREGRNERLF | 0.8518189 |
| LTTTTTS | ASSQDVREGRNERLF | 0.64375085 |
| PTGSGKTL | ASSQDVREGRNERLF | 0.8803288 |
| TGSGKTL | ASSQDVREGRNERLF | 0.8071315 |
| TIGHVDHGK | ASSQDVREGRNERLF | 0.73076355 |
| TIGHVDHGKTT | ASSQDVREGRNERLF | 0.8855798 |
| GHVDHGKTT | ASSQDVREGRNERLF | 0.85701704 |
| AQFEHTLL | ASSQDVREGRNERLF | 0.8612964 |
| IVNQDEL | ASSQDVREGRNERLF | 0.8562579 |
| TNGSQFF | ASSQDVREGRNERLF | 0.89087814 |
| TGKGIVE | ASSQDVREGRNERLF | 0.6501857 |
| LISNASDA | ASSQDVREGRNERLF | 0.83123916 |
| AELRTLGELTKIAKAHGAQ | ASSQDVREGRNERLF | 0.80906016 |
| APNAVAAAEPSW | ASSQDVREGRNERLF | 0.87579083 |
| APQVAGAATAVL | ASSQDVREGRNERLF | 0.8219155 |
| ARRDPAAAAAILPTDA | ASSQDVREGRNERLF | 0.72343016 |
| DAAARDVLAEAGLAEYF | ASSQDVREGRNERLF | 0.8659023 |
| DDQIVLSAKANLAKVEQ | ASSQDVREGRNERLF | 0.874276 |
| DELLPDTGEDSDEDGH | ASSQDVREGRNERLF | 0.8308455 |
| ELPEAPANRATVPQD | ASSQDVREGRNERLF | 0.84083205 |
| FGQSKIERVTGIG | ASSQDVREGRNERLF | 0.89004266 |
| GAGGTGSTAGGAGGAGGAGGLY | ASSQDVREGRNERLF | 0.70905614 |
| GHGGTGAAGGNGGTGGAGGSGI | ASSQDVREGRNERLF | 0.81614065 |
| GNGGNGGAGGEGGPGGAGGAGA | ASSQDVREGRNERLF | 0.8666645 |
| GSTAGNGGNGGNATNGTIAGQP | ASSQDVREGRNERLF | 0.8705152 |
| GTGGDGGNGGNGGTGG | ASSQDVREGRNERLF | 0.8384893 |
| GVRRPAGIPADAPV | ASSQDVREGRNERLF | 0.87613773 |
| GYRQVTDAHLLTLARR | ASSQDVREGRNERLF | 0.7473366 |
| IAKMGQDGHDRG | ASSQDVREGRNERLF | 0.84467506 |
| KAEPSIEIQYT | ASSQDVREGRNERLF | 0.7060257 |
| KPVRIRMAADEDTVAVT | ASSQDVREGRNERLF | 0.82276034 |
| MPPRIAGMRLLVIKPEP | ASSQDVREGRNERLF | 0.66568 |
| MKFYAVALFRSRGVPPTR | ASSQDVREGRNERLF | 0.8323358 |
| NAGGFGILGAVAHSP | ASSQDVREGRNERLF | 0.85758865 |
| NEGREITLAMAEM | ASSQDVREGRNERLF | 0.84841275 |
| NELVEWSHPLGHRWRWE | ASSQDVREGRNERLF | 0.41297203 |
| NLGAGGTGGNGGNPG | ASSQDVREGRNERLF | 0.8585338 |
| NTEWPKWETVMAADPVD | ASSQDVREGRNERLF | 0.8597926 |
| PASAAPAETQP | ASSQDVREGRNERLF | 0.89692545 |
| QQSSDGKNGVRAAEITGE | ASSQDVREGRNERLF | 0.8125326 |
| RAPDAVTTSWWKE | ASSQDVREGRNERLF | 0.8591896 |
| SGFGNVGAHTSGWFNQ | ASSQDVREGRNERLF | 0.7477832 |
| SGVAGAVGQAASVGGLKV | ASSQDVREGRNERLF | 0.8496639 |

| | | |
|---|---|---|
| SNGGAGTGSAGGNGGT | ASSQDVREGRNERLF | 0.6665741 |
| TGGAGGEGGAAG | ASSQDVREGRNERLF | 0.7468838 |
| TGGDGGAAGAGGAPGHG | ASSQDVREGRNERLF | 0.7768773 |
| TMPPLLAAIANAGRHGV | ASSQDVREGRNERLF | 0.74367666 |
| TSSTAGNSGGA | ASSQDVREGRNERLF | 0.8677024 |
| VPATVPVGILC | ASSQDVREGRNERLF | 0.86395526 |
| VQRPDSGGVVLTGRIS | ASSQDVREGRNERLF | 0.86578083 |
| WTPNPVPATKGVPAGLVQAT | ASSQDVREGRNERLF | 0.8719945 |
| YALKRALAKFSDRFDVVII | ASSQDVREGRNERLF | 0.87796605 |
| YLIGSVIV | ASSQDVREGRNERLF | 0.88257396 |
| YGQLVAVID | ASSQDVREGRNERLF | 0.75158066 |
| VTVGGMTLPA | ASSQDVREGRNERLF | 0.82071614 |
| VLALPAV | ASSQDVREGRNERLF | 0.6616527 |
| VIPPTLNYET | ASSQDVREGRNERLF | 0.8028382 |
| VGMGGMHPSSG | ASSQDVREGRNERLF | 0.7669153 |
| VGLAGGAATA | ASSQDVREGRNERLF | 0.84115744 |
| VGGAGGAGGSGP | ASSQDVREGRNERLF | 0.17417888 |
| VFAAFMV | ASSQDVREGRNERLF | 0.8342947 |
| TLYDKLR | ASSQDVREGRNERLF | 0.8782261 |
| TLAVFTVL | ASSQDVREGRNERLF | 0.6947173 |
| TGSVQPARV | ASSQDVREGRNERLF | 0.8517183 |
| TGGGAGSGGT | ASSQDVREGRNERLF | 0.8364967 |
| SHLETSL | ASSQDVREGRNERLF | 0.754111 |
| SFTTGMT | ASSQDVREGRNERLF | 0.853112 |
| QVEVAKRMGVRQPTVS | ASSQDVREGRNERLF | 0.7343602 |
| QGSGGPALAM | ASSQDVREGRNERLF | 0.86896855 |
| MLAPSNP | ASSQDVREGRNERLF | 0.87745416 |
| LVNNAAI | ASSQDVREGRNERLF | 0.8258272 |
| LTGEINTLA | ASSQDVREGRNERLF | 0.7949955 |
| LSIFGPV | ASSQDVREGRNERLF | 0.8304968 |
| LLALLANL | ASSQDVREGRNERLF | 0.57584864 |
| LISNASDALDKLR | ASSQDVREGRNERLF | 0.8135919 |
| LINEAVP | ASSQDVREGRNERLF | 0.20157415 |
| LGYAFSGL | ASSQDVREGRNERLF | 0.8188653 |
| LDFVAVAL | ASSQDVREGRNERLF | 0.6929656 |
| LAQPGAE | ASSQDVREGRNERLF | 0.8349405 |
| ISNLGAVTGF | ASSQDVREGRNERLF | 0.80962205 |
| HQEDPGRI | ASSQDVREGRNERLF | 0.8496659 |
| GTPGEPTP | ASSQDVREGRNERLF | 0.6822324 |
| GFFNSGS | ASSQDVREGRNERLF | 0.8486313 |
| GAGGGGLTAGAN | ASSQDVREGRNERLF | 0.80186164 |
| FVDALVNM | ASSQDVREGRNERLF | 0.54594404 |
| FLTVAPEL | ASSQDVREGRNERLF | 0.857018 |
| FETLTAL | ASSQDVREGRNERLF | 0.816692 |
| DVINPIEL | ASSQDVREGRNERLF | 0.8014198 |
| APVSGAPV | ASSQDVREGRNERLF | 0.87377197 |
| ALVGGLAW | ASSQDVREGRNERLF | 0.843597 |
| ALAAAAATAI | ASSQDVREGRNERLF | 0.6639352 |
| AFGLGLLL | ASSQDVREGRNERLF | 0.7648797 |
| AFFAAASL | ASSQDVREGRNERLF | 0.81283873 |
| AAAALPAFSPP | ASSQDVREGRNERLF | 0.7638085 |

| | | |
|---|---|---|
| VVLAPAL | ASSQDVREGRNERLF | 0.19725183 |
| TIGHVDHGKTTL | ASSQDVREGRNERLF | 0.8627077 |
| LISNASDALDK | ASSQDVREGRNERLF | 0.7287444 |
| LISNASDAL | ASSQDVREGRNERLF | 0.70319533 |
| ISNASDALDK | ASSQDVREGRNERLF | 0.7841754 |
| NQAETLVYQTE | ASSQDVREGRNERLF | 0.87146425 |
| TGGTGGTAGAG | ASSQDVREGRNERLF | 0.76173985 |
| TGSTAGGAGGA | ASSQDVREGRNERLF | 0.81282896 |
| DPLAVPVAAGPV | ASSQDVREGRNERLF | 0.8519039 |
| GGTGGTPGNGGN | ASSQDVREGRNERLF | 0.84243035 |
| PAGLAGVDPATL | ASSQDVREGRNERLF | 0.8741627 |
| GNNLVVEAPHAGASV | ASSQDVREGRNERLF | 0.8442781 |
| KVTRIMSASTLEQPA | ASSQDVREGRNERLF | 0.8088243 |
| LGFALPIALPRRTVV | ASSQDVREGRNERLF | 0.8644457 |
| TRLGNLTARKLVVLV | ASSQDVREGRNERLF | 0.8381463 |
| WSQQRRLIYFPSAGP | ASSQDVREGRNERLF | 0.73112 |
| GPNGNPDPMAAAVDIR | ASSQDVREGRNERLF | 0.6751629 |
| TQRCVSVADLRAAASR | ASSQDVREGRNERLF | 0.7739035 |
| QFVLDVAEHPVEVLTETV | ASSQDVREGRNERLF | 0.60828316 |
| GGAGGTGGAGGNGGTQAGDG | ASSQDVREGRNERLF | 0.82862324 |
| PLLATLRAIAVDVDAKPAQV | ASSQDVREGRNERLF | 0.89298445 |
| APKFRTGGYMIDFWGVGYQVAK | ASSQDVREGRNERLF | 0.8350574 |
| GAGGAGGAGGAAGGPGGTGGTG | ASSQDVREGRNERLF | 0.6042225 |
| IGALGLLATDVAPAGSVAFGVA | ASSQDVREGRNERLF | 0.7669556 |
| AGGAGGIGGNGGAGGAGGVLMG | ASSQDVREGRNERLF | 0.8258654 |
| ADELIGLL | ASSQDVREGRNERLF | 0.8623059 |
| ADRALKA | ASSQDVREGRNERLF | 0.8552472 |
| AGAVGQVGGAGGA | ASSQDVREGRNERLF | 0.8937427 |
| AGGGGGLGGAP | ASSQDVREGRNERLF | 0.8667621 |
| AGGLLYGNGGAG | ASSQDVREGRNERLF | 0.7501808 |
| AGLESIGL | ASSQDVREGRNERLF | 0.87015426 |
| AGLKTGII | ASSQDVREGRNERLF | 0.8064941 |
| ASLAALLL | ASSQDVREGRNERLF | 0.79857653 |
| ASVINISSI | ASSQDVREGRNERLF | 0.8342445 |
| ATPPGPA | ASSQDVREGRNERLF | 0.8455845 |
| ATVASALV | ASSQDVREGRNERLF | 0.7904904 |
| ATVASLIL | ASSQDVREGRNERLF | 0.7797917 |
| AVNGPVLIRH | ASSQDVREGRNERLF | 0.5919138 |
| DGAAGAVGI | ASSQDVREGRNERLF | 0.874033 |
| DPLLHGI | ASSQDVREGRNERLF | 0.6907249 |
| ELGAALGAL | ASSQDVREGRNERLF | 0.71744585 |
| ELGRTQAV | ASSQDVREGRNERLF | 0.78713274 |
| FVAGGYLS | ASSQDVREGRNERLF | 0.887241 |
| GASGGAGGAGGW | ASSQDVREGRNERLF | 0.8796238 |
| GGAGAGGDGGAGGI | ASSQDVREGRNERLF | 0.7350335 |
| GTGHNGSQPGMGGNG | ASSQDVREGRNERLF | 0.71902645 |
| GYAAASASAS | ASSQDVREGRNERLF | 0.83651453 |
| HQVDVAAVQQG | ASSQDVREGRNERLF | 0.6520246 |
| IGAAIAHT | ASSQDVREGRNERLF | 0.8689864 |
| IGMIEVAQL | ASSQDVREGRNERLF | 0.8089136 |
| IRLAPEHTSAMD | ASSQDVREGRNERLF | 0.7871405 |

| | | |
|---|---|---|
| IYNVSVL | ASSQDVREGRNERLF | 0.73984885 |
| KQGFLGM | ASSQDVREGRNERLF | 0.8410182 |
| LESVITQL | ASSQDVREGRNERLF | 0.7523159 |
| LGLGALVVV | ASSQDVREGRNERLF | 0.8764 |
| LIGNGGNGGSGGI | ASSQDVREGRNERLF | 0.28520405 |
| LNAINAPTL | ASSQDVREGRNERLF | 0.86084306 |
| LQLLNTGV | ASSQDVREGRNERLF | 0.68903875 |
| LQTSIEF | ASSQDVREGRNERLF | 0.78370464 |
| MSFAVSEL | ASSQDVREGRNERLF | 0.41583306 |
| NTGALGL | ASSQDVREGRNERLF | 0.831041 |
| PPGAGGAGGAGTT | ASSQDVREGRNERLF | 0.84114707 |
| QLVGLGI | ASSQDVREGRNERLF | 0.8758219 |
| RFLEQVHQL | ASSQDVREGRNERLF | 0.80126876 |
| RGVGYVLR | ASSQDVREGRNERLF | 0.84966236 |
| SATGVFAGVI | ASSQDVREGRNERLF | 0.86931556 |
| SLPAITLP | ASSQDVREGRNERLF | 0.76836383 |
| STALAFTV | ASSQDVREGRNERLF | 0.4099782 |
| SVLTVGLGK | ASSQDVREGRNERLF | 0.85424036 |
| TALAGAGSGI | ASSQDVREGRNERLF | 0.8087046 |
| TGLPFTGI | ASSQDVREGRNERLF | 0.8774168 |
| VAEHVTV | ASSQDVREGRNERLF | 0.83846045 |
| VDIVLNSL | ASSQDVREGRNERLF | 0.67578256 |
| VLTGGGAVL | ASSQDVREGRNERLF | 0.88287693 |
| VNVGLAYGI | ASSQDVREGRNERLF | 0.8629582 |
| VSSAAVGAV | ASSQDVREGRNERLF | 0.6608943 |
| VTSVYEAGV | ASSQDVREGRNERLF | 0.76358736 |
| WVALDRGA | ASSQDVREGRNERLF | 0.774439 |
| YLVGSVVI | ASSQDVREGRNERLF | 0.42607543 |
| GPPGTGKTLL | ASSQDVREGRNERLF | 0.8439995 |
| GVGMTKF | ASSQDVREGRNERLF | 0.7369582 |
| IGIGGSDL | ASSQDVREGRNERLF | 0.762821 |
| IGTLLAEL | ASSQDVREGRNERLF | 0.8344214 |
| IYTSGTT | ASSQDVREGRNERLF | 0.72948384 |
| LSFPAGV | ASSQDVREGRNERLF | 0.75572383 |
| LYTSGTT | ASSQDVREGRNERLF | 0.73853636 |
| VGAFPFGF | ASSQDVREGRNERLF | 0.84087867 |
| VGAGTVEF | ASSQDVREGRNERLF | 0.85168934 |
| VGKSTLLL | ASSQDVREGRNERLF | 0.811795 |
| AALAHAGGGIV | ASSQDVREGRNERLF | 0.853245 |
| AALVAVSLL | ASSQDVREGRNERLF | 0.8749495 |
| AAVSRAVAAL | ASSQDVREGRNERLF | 0.5658774 |
| AGGSSGIGGF | ASSQDVREGRNERLF | 0.72313386 |
| AGLTVANANG | ASSQDVREGRNERLF | 0.8654715 |
| AGLVGAVL | ASSQDVREGRNERLF | 0.6988002 |
| ALSQAFTV | ASSQDVREGRNERLF | 0.872598 |
| ALTFSVSL | ASSQDVREGRNERLF | 0.81527853 |
| ALTGAALVAL | ASSQDVREGRNERLF | 0.8488759 |
| ASFSIGAGVL | ASSQDVREGRNERLF | 0.8310176 |
| DGSVYKI | ASSQDVREGRNERLF | 0.8803563 |
| EQLAEIV | ASSQDVREGRNERLF | 0.86423343 |
| FAVSELTVM | ASSQDVREGRNERLF | 0.8145217 |

| | | |
|---|---|---|
| GAAPLAAGL | ASSQDVREGRNERLF | 0.8561976 |
| GADGGDGGSGGAGG | ASSQDVREGRNERLF | 0.6807817 |
| GFGLDTGGAGG | ASSQDVREGRNERLF | 0.8521178 |
| GLASGGLL | ASSQDVREGRNERLF | 0.865392 |
| GRIAGAAVLL | ASSQDVREGRNERLF | 0.6773044 |
| GSAGSLNTSF | ASSQDVREGRNERLF | 0.7986733 |
| GTGQIGI | ASSQDVREGRNERLF | 0.8823767 |
| GTLIALIL | ASSQDVREGRNERLF | 0.87182426 |
| IAATMGFVVG | ASSQDVREGRNERLF | 0.62688607 |
| IFAQALL | ASSQDVREGRNERLF | 0.7964389 |
| IGAAVALGF | ASSQDVREGRNERLF | 0.6283251 |
| IGLTPLAL | ASSQDVREGRNERLF | 0.75089395 |
| ILGSNTSTL | ASSQDVREGRNERLF | 0.5671115 |
| IVFSLTVL | ASSQDVREGRNERLF | 0.8573582 |
| LASKATGF | ASSQDVREGRNERLF | 0.7620189 |
| LASVALIL | ASSQDVREGRNERLF | 0.8709581 |
| LFAVGLTL | ASSQDVREGRNERLF | 0.8370733 |
| LFNGGIGTYI | ASSQDVREGRNERLF | 0.88018584 |
| LTLGAGII | ASSQDVREGRNERLF | 0.7463735 |
| LVAAALGDI | ASSQDVREGRNERLF | 0.7746637 |
| NTAFVQLGI | ASSQDVREGRNERLF | 0.52932996 |
| QLFSGAEL | ASSQDVREGRNERLF | 0.45923394 |
| SAATFITL | ASSQDVREGRNERLF | 0.7146301 |
| SNYGLANLGV | ASSQDVREGRNERLF | 0.7601558 |
| SYGGAYAVM | ASSQDVREGRNERLF | 0.35583383 |
| TTGVGGLTLG | ASSQDVREGRNERLF | 0.88529605 |
| TTQTIYI | ASSQDVREGRNERLF | 0.83398664 |
| VAAGAGLGAVI | ASSQDVREGRNERLF | 0.8585061 |
| VAAVVTLL | ASSQDVREGRNERLF | 0.7957493 |
| VGFLGGLI | ASSQDVREGRNERLF | 0.72067326 |
| VGVIPILL | ASSQDVREGRNERLF | 0.75656444 |
| VIFASSLI | ASSQDVREGRNERLF | 0.8595685 |
| VKGGDGGAAAG | ASSQDVREGRNERLF | 0.6988473 |
| VTKTFTNI | ASSQDVREGRNERLF | 0.4905333 |
| VVAAAILL | ASSQDVREGRNERLF | 0.78203183 |
| YAALDVEL | ASSQDVREGRNERLF | 0.7392656 |
| LASVALLL | ASSQDVREGRNERLF | 0.88284695 |
| LDAVHRV | ASSQDVREGRNERLF | 0.86448574 |
| QGLSSSIF | ASSQDVREGRNERLF | 0.84931636 |
| RGTFANIRL | ASSQDVREGRNERLF | 0.8879975 |
| RQLWWGHRI | ASSQDVREGRNERLF | 0.2670709 |
| YGPPGTGKTLL | ASSQDVREGRNERLF | 0.81174475 |
| AALVGAPANVA | ASSQDVREGRNERLF | 0.88464725 |
| AATFRRTE | ASSQDVREGRNERLF | 0.84392005 |
| AGALGGGI | ASSQDVREGRNERLF | 0.7881355 |
| AGGAGGAGGIGAG | ASSQDVREGRNERLF | 0.87825394 |
| AGSGALGL | ASSQDVREGRNERLF | 0.6985446 |
| AQIADQI | ASSQDVREGRNERLF | 0.7616645 |
| ASAVAVTYL | ASSQDVREGRNERLF | 0.826944 |
| ASTVLGGTGG | ASSQDVREGRNERLF | 0.8670341 |
| AVGGEPAK | ASSQDVREGRNERLF | 0.8314926 |

| | | |
|---|---|---|
| CEQGTFGGTC | ASSQDVREGRNERLF | 0.62026 |
| DEGKAPAG | ASSQDVREGRNERLF | 0.85874116 |
| DTVVFPAL | ASSQDVREGRNERLF | 0.8832491 |
| FEGITMAG | ASSQDVREGRNERLF | 0.7725495 |
| FPLVVQD | ASSQDVREGRNERLF | 0.86622125 |
| GAFTTPQ | ASSQDVREGRNERLF | 0.8531629 |
| GFGGFGTAGGVGGA | ASSQDVREGRNERLF | 0.7663503 |
| GFLAQFL | ASSQDVREGRNERLF | 0.73083234 |
| GGAGGAGGNAGVA | ASSQDVREGRNERLF | 0.7194313 |
| GGVGTAGAGGAGGAGG | ASSQDVREGRNERLF | 0.84844613 |
| GNGGGGDAGNAGSGGN | ASSQDVREGRNERLF | 0.7038438 |
| GTGGAGGAGNDAGS | ASSQDVREGRNERLF | 0.8198747 |
| GTTNTGWFNS | ASSQDVREGRNERLF | 0.8282026 |
| IFMGPLFTL | ASSQDVREGRNERLF | 0.8899781 |
| ILAHAIEAI | ASSQDVREGRNERLF | 0.85052377 |
| IRSAIREAL | ASSQDVREGRNERLF | 0.86053884 |
| ITFAGGG | ASSQDVREGRNERLF | 0.6960748 |
| ITGMSGHQEV | ASSQDVREGRNERLF | 0.84008133 |
| IVAGLACTL | ASSQDVREGRNERLF | 0.84264755 |
| IVGGGLAGA | ASSQDVREGRNERLF | 0.61430466 |
| IVTGVLNI | ASSQDVREGRNERLF | 0.8773967 |
| LAQQGCI | ASSQDVREGRNERLF | 0.6842789 |
| LCLHVEN | ASSQDVREGRNERLF | 0.8141908 |
| LDAISDGS | ASSQDVREGRNERLF | 0.8685818 |
| LGGFAAML | ASSQDVREGRNERLF | 0.8830413 |
| LGGKNPMI | ASSQDVREGRNERLF | 0.8015147 |
| LGPKGVAAIL | ASSQDVREGRNERLF | 0.84209245 |
| LMVGVSI | ASSQDVREGRNERLF | 0.7064265 |
| LVGAAFVL | ASSQDVREGRNERLF | 0.8747331 |
| LVLAVAPL | ASSQDVREGRNERLF | 0.877902 |
| MAAIAVIDAA | ASSQDVREGRNERLF | 0.8074897 |
| NNGINAELQ | ASSQDVREGRNERLF | 0.89496374 |
| RGEPLVLVHG | ASSQDVREGRNERLF | 0.87363505 |
| RGGAGGAAGLI | ASSQDVREGRNERLF | 0.4865403 |
| RSYRMHPAV | ASSQDVREGRNERLF | 0.83333135 |
| SAAHLLAGQLA | ASSQDVREGRNERLF | 0.87188697 |
| SAAVGSY | ASSQDVREGRNERLF | 0.8684257 |
| SPHAVLHRI | ASSQDVREGRNERLF | 0.86991096 |
| SQIAKVL | ASSQDVREGRNERLF | 0.8206244 |
| TALGHLVP | ASSQDVREGRNERLF | 0.748966 |
| TAPTTSGAA | ASSQDVREGRNERLF | 0.87997615 |
| TDGDDGNGGSPGT | ASSQDVREGRNERLF | 0.890447 |
| TDGLGEN | ASSQDVREGRNERLF | 0.8737608 |
| TGGPPLPGQA | ASSQDVREGRNERLF | 0.78609765 |
| TIPAITI | ASSQDVREGRNERLF | 0.7863134 |
| TIPGFTI | ASSQDVREGRNERLF | 0.8295847 |
| TSIASGALA | ASSQDVREGRNERLF | 0.86244833 |
| TVTLPGAAI | ASSQDVREGRNERLF | 0.6631645 |
| VASGLSAALP | ASSQDVREGRNERLF | 0.828295 |
| VGAGVIGI | ASSQDVREGRNERLF | 0.8274186 |
| VGPKRFGV | ASSQDVREGRNERLF | 0.4329964 |

| | | |
|---|---|---|
| VNVAFPD | ASSQDVREGRNERLF | 0.83825934 |
| VSMTNSGVS | ASSQDVREGRNERLF | 0.8659587 |
| VTDVLGDLSV | ASSQDVREGRNERLF | 0.83775616 |
| VVVVGAGFAGL | ASSQDVREGRNERLF | 0.7412401 |
| VYTVGTVV | ASSQDVREGRNERLF | 0.87196577 |
| WTTFSGP | ASSQDVREGRNERLF | 0.88260436 |
| YDELVEAF | ASSQDVREGRNERLF | 0.88905835 |
| YGTGGAGGNGG | ASSQDVREGRNERLF | 0.51577187 |
| YTNVMAV | ASSQDVREGRNERLF | 0.35408577 |
| FLEQVHQL | ASSQDVREGRNERLF | 0.76892257 |
| GLSLLAVAV | ASSQDVREGRNERLF | 0.8646091 |
| LEQVHQL | ASSQDVREGRNERLF | 0.7669481 |
| LNAAGAI | ASSQDVREGRNERLF | 0.7474951 |
| PAVHASL | ASSQDVREGRNERLF | 0.8280186 |
| TELEDDA | ASSQDVREGRNERLF | 0.8762046 |
| TPPGPPN | ASSQDVREGRNERLF | 0.67489934 |
| TPPPPPPP | ASSQDVREGRNERLF | 0.6276654 |
| TQSPALI | ASSLEGNTGQLY | 0.8387342 |
| EFAAAFPQ | ASSLEGNTGQLY | 0.8778448 |
| LLSGNLDVL | ASSLEGNTGQLY | 0.70603526 |
| GPANLGNYNI | ASSLEGNTGQLY | 0.7898025 |
| TLGPTLP | ASSLEGNTGQLY | 0.7851808 |
| VYAYSSCCK | ASSLEGNTGQLY | 0.809713 |
| APAPAPAPNI | ASSLEGNTGQLY | 0.8199695 |
| PGGPVPIP | ASSLEGNTGQLY | 0.67466104 |
| FLTVALEP | ASSLEGNTGQLY | 0.86879826 |
| SQYFGANL | ASSLEGNTGQLY | 0.6666311 |
| GGVKPAA | ASSLEGNTGQLY | 0.8652036 |
| GGTGGAGGAGGW | ASSLEGNTGQLY | 0.8135687 |
| GVGGVAGLGGAGG | ASSLEGNTGQLY | 0.85353243 |
| VLLVLATTI | ASSLEGNTGQLY | 0.77627397 |
| IGIFAAML | ASSLEGNTGQLY | 0.76456213 |
| AVGDLID | ASSLEGNTGQLY | 0.66904664 |
| NISPLPL | ASSLEGNTGQLY | 0.62126946 |
| VTAVFVAI | ASSLEGNTGQLY | 0.8647621 |
| SGDDNDAAL | ASSLEGNTGQLY | 0.40952727 |
| AVAALAHS | ASSLEGNTGQLY | 0.87246895 |
| GGVTVMATP | ASSLEGNTGQLY | 0.83695424 |
| AGLAANVRD | ASSLEGNTGQLY | 0.7407887 |
| TNRPDLIDPALLRPG | ASSLEGNTGQLY | 0.86223304 |
| LANLGVTAL | ASSLEGNTGQLY | 0.6160569 |
| TGERIEL | ASSLEGNTGQLY | 0.8426975 |
| NGEELAA | ASSLEGNTGQLY | 0.829955 |
| VLSLVWG | ASSLEGNTGQLY | 0.8726151 |
| IGYALWL | ASSLEGNTGQLY | 0.74437726 |
| LSLLLAPL | ASSLEGNTGQLY | 0.8624916 |
| TLDEALK | ASSLEGNTGQLY | 0.8420831 |
| TDGAGTAEGG | ASSLEGNTGQLY | 0.7132384 |
| NAPGGAGGNGG | ASSLEGNTGQLY | 0.8334998 |
| VSVFMGMGGNF | ASSLEGNTGQLY | 0.87577844 |
| LASYLAAL | ASSLEGNTGQLY | 0.8438522 |

| | | |
|---|---|---|
| LVGGTPAL | ASSLEGNTGQLY | 0.6497843 |
| VRNADVL | ASSLEGNTGQLY | 0.8441714 |
| TCSSPFLVSAM | ASSLEGNTGQLY | 0.76061296 |
| VHITDAE | ASSLEGNTGQLY | 0.8471892 |
| EYAAPSF | ASSLEGNTGQLY | 0.83876663 |
| VVDAFHGGL | ASSLEGNTGQLY | 0.79245377 |
| ASVVDPQ | ASSLEGNTGQLY | 0.8677868 |
| SYNEFGGLS | ASSLEGNTGQLY | 0.70880425 |
| DPDGSVY | ASSLEGNTGQLY | 0.71779287 |
| MVSTIHATE | ASSLEGNTGQLY | 0.7989801 |
| LAAGVADPV | ASSLEGNTGQLY | 0.8060417 |
| TKEQIAA | ASSLEGNTGQLY | 0.87421125 |
| SLASPAAL | ASSLEGNTGQLY | 0.69271284 |
| IGLVFYGV | ASSLEGNTGQLY | 0.8357993 |
| FLLDASNL | ASSLEGNTGQLY | 0.7572451 |
| YQVGNFP | ASSLEGNTGQLY | 0.6599349 |
| ESLLSQST | ASSLEGNTGQLY | 0.8499086 |
| LFTVGFAAVG | ASSLEGNTGQLY | 0.8034274 |
| ATPPTPT | ASSLEGNTGQLY | 0.71131396 |
| TDLATLG | ASSLEGNTGQLY | 0.86387897 |
| SAGGTGR | ASSLEGNTGQLY | 0.6714653 |
| GFSNSGS | ASSLEGNTGQLY | 0.84002495 |
| GGNGGSGGTSV | ASSLEGNTGQLY | 0.86405176 |
| GTSVATGGAGN | ASSLEGNTGQLY | 0.69131374 |
| NGGNGGSGGTS | ASSLEGNTGQLY | 0.8670388 |
| AVVKFPH | ASSLEGNTGQLY | 0.8495661 |
| TPPPPPP | ASSLEGNTGQLY | 0.8359653 |
| AAPAAAAAPA | ASSLEGNTGQLY | 0.75654083 |
| IALNEDL | ASSLEGNTGQLY | 0.74676156 |
| IVNVSSM | ASSLEGNTGQLY | 0.8861673 |
| TGIPPAPR | ASSLEGNTGQLY | 0.8591031 |
| FDLGGGTF | ASSLEGNTGQLY | 0.85539234 |
| FDLGGGTFDVS | ASSLEGNTGQLY | 0.858004 |
| NEPTAAA | ASSLEGNTGQLY | 0.86880434 |
| NEPTAAALA | ASSLEGNTGQLY | 0.8738115 |
| GSIINISSI | ASSLEGNTGQLY | 0.7548965 |
| VAPGNAGTA | ASSLEGNTGQLY | 0.5580847 |
| TGGGASLE | ASSLEGNTGQLY | 0.78012645 |
| VAVYNHYKRI | ASSLEGNTGQLY | 0.7745279 |
| TDPAPATT | ASSLEGNTGQLY | 0.79829913 |
| TNRPDILDPALLRPG | ASSLEGNTGQLY | 0.6106461 |
| LTTTTTS | ASSLEGNTGQLY | 0.8708317 |
| PTGSGKTL | ASSLEGNTGQLY | 0.8934113 |
| TGSGKTL | ASSLEGNTGQLY | 0.7162131 |
| TIGHVDHGK | ASSLEGNTGQLY | 0.8932056 |
| TIGHVDHGKTT | ASSLEGNTGQLY | 0.6548567 |
| GHVDHGKTT | ASSLEGNTGQLY | 0.8516363 |
| AQFEHTLL | ASSLEGNTGQLY | 0.8732477 |
| IVNQDEL | ASSLEGNTGQLY | 0.684083 |
| TNGSQFF | ASSLEGNTGQLY | 0.85257435 |
| TGKGIVE | ASSLEGNTGQLY | 0.7468427 |

| | | |
|---|---|---|
| LISNASDA | ASSLEGNTGQLY | 0.77369225 |
| AELRTLGELTKIAKAHGAQ | ASSLEGNTGQLY | 0.8645437 |
| APNAVAAAEPSW | ASSLEGNTGQLY | 0.74999064 |
| APQVAGAATAVL | ASSLEGNTGQLY | 0.8557233 |
| ARRDPAAAAAILPTDA | ASSLEGNTGQLY | 0.8159602 |
| DAAARDVLAEAGLAEYF | ASSLEGNTGQLY | 0.730026 |
| DDQIVLSAKANLAKVEQ | ASSLEGNTGQLY | 0.8540786 |
| DELLPDTGEDSDEDGH | ASSLEGNTGQLY | 0.8698025 |
| ELPEAPANRATVPQD | ASSLEGNTGQLY | 0.82363605 |
| FGQSKIERVTGIG | ASSLEGNTGQLY | 0.8904544 |
| GAGGTGSTAGGAGGAGGAGGLY | ASSLEGNTGQLY | 0.7020703 |
| GHGGTGAAGGNGGTGGAGGSGI | ASSLEGNTGQLY | 0.88889074 |
| GNGGNGGAGGEGGPGGAGGAGA | ASSLEGNTGQLY | 0.8809459 |
| GSTAGNGGNGGNATNGTIAGQP | ASSLEGNTGQLY | 0.825696 |
| GTGGDGGNGGNGGTGG | ASSLEGNTGQLY | 0.80518746 |
| GVRRPAGIPADAPV | ASSLEGNTGQLY | 0.8702079 |
| GYRQVTDAHLLTLARR | ASSLEGNTGQLY | 0.79019034 |
| IAKMGQDGHDRG | ASSLEGNTGQLY | 0.85068375 |
| KAEPSIEIQYT | ASSLEGNTGQLY | 0.8863953 |
| KPVRIRMAADEDTVAVT | ASSLEGNTGQLY | 0.19823527 |
| MPPRIAGMRLLVIKPEP | ASSLEGNTGQLY | 0.7989471 |
| MKFYAVALFRSRGVPPTR | ASSLEGNTGQLY | 0.54586345 |
| NAGGFGILGAVAHSP | ASSLEGNTGQLY | 0.43418753 |
| NEGREITLAMAEM | ASSLEGNTGQLY | 0.27151847 |
| NELVEWSHPLGHRWRWE | ASSLEGNTGQLY | 0.16491339 |
| NLGAGGTGGNGGNPG | ASSLEGNTGQLY | 0.38394964 |
| NTEWPKWETVMAADPVD | ASSLEGNTGQLY | 0.21607056 |
| PASAAPAETQP | ASSLEGNTGQLY | 0.77401954 |
| QQSSDGKNGVRAAEITGE | ASSLEGNTGQLY | 0.6056088 |
| RAPDAVTTSWWKE | ASSLEGNTGQLY | 0.28611517 |
| SGFGNVGAHTSGWFNQ | ASSLEGNTGQLY | 0.69726014 |
| SGVAGAVGQAASVGGLKV | ASSLEGNTGQLY | 0.69740546 |
| SNGGAGTGSAGGNGGT | ASSLEGNTGQLY | 0.15056694 |
| TGGAGGEGGAAG | ASSLEGNTGQLY | 0.3478264 |
| TGGDGGAAGAGGAPGHG | ASSLEGNTGQLY | 0.2750715 |
| TMPPLLAAIANAGRHGV | ASSLEGNTGQLY | 0.31429198 |
| TSSTAGNSGGA | ASSLEGNTGQLY | 0.11518662 |
| VPATVPVGILC | ASSLEGNTGQLY | 0.6136862 |
| VQRPDSGGVVLTGRIS | ASSLEGNTGQLY | 0.31740087 |
| WTPNPVPATKGVPAGLVQAT | ASSLEGNTGQLY | 0.08637479 |
| YALKRALAKFSDRFDVVII | ASSLEGNTGQLY | 0.35930386 |
| YLIGSVIV | ASSLEGNTGQLY | 0.31765 |
| YGQLVAVID | ASSLEGNTGQLY | 0.2811671 |
| VTVGGMTLPA | ASSLEGNTGQLY | 0.6718223 |
| VLALPAV | ASSLEGNTGQLY | 0.365914 |
| VIPPTLNYET | ASSLEGNTGQLY | 0.2441545 |
| VGMGGMHPSSG | ASSLEGNTGQLY | 0.6413522 |
| VGLAGGAATA | ASSLEGNTGQLY | 0.7390813 |
| VGGAGGAGGSGP | ASSLEGNTGQLY | 0.21263108 |
| VFAAFMV | ASSLEGNTGQLY | 0.5660115 |
| TLYDKLR | ASSLEGNTGQLY | 0.8082019 |

| | | |
|---|---|---|
| TLAVFTVL | ASSLEGNTGQLY | 0.44748765 |
| TGSVQPARV | ASSLEGNTGQLY | 0.73013103 |
| TGGGAGSGGT | ASSLEGNTGQLY | 0.7679443 |
| SHLETSL | ASSLEGNTGQLY | 0.27962908 |
| SFTTGMT | ASSLEGNTGQLY | 0.6909021 |
| QVEVAKRMGVRQPTVS | ASSLEGNTGQLY | 0.79171765 |
| QGSGGPALAM | ASSLEGNTGQLY | 0.37018397 |
| MLAPSNP | ASSLEGNTGQLY | 0.3549172 |
| LVNNAAI | ASSLEGNTGQLY | 0.22808538 |
| LTGEINTLA | ASSLEGNTGQLY | 0.329143 |
| LSIFGPV | ASSLEGNTGQLY | 0.14487942 |
| LLALLANL | ASSLEGNTGQLY | 0.5998761 |
| LISNASDALDKLR | ASSLEGNTGQLY | 0.6988162 |
| LINEAVP | ASSLEGNTGQLY | 0.32538795 |
| LGYAFSGL | ASSLEGNTGQLY | 0.15134022 |
| LDFVAVAL | ASSLEGNTGQLY | 0.17480521 |
| LAQPGAE | ASSLEGNTGQLY | 0.47463244 |
| ISNLGAVTGF | ASSLEGNTGQLY | 0.28417847 |
| HQEDPGRI | ASSLEGNTGQLY | 0.16061963 |
| GTPGEPTP | ASSLEGNTGQLY | 0.26225135 |
| GFFNSGS | ASSLEGNTGQLY | 0.6526544 |
| GAGGGGLTAGAN | ASSLEGNTGQLY | 0.6910775 |
| FVDALVNM | ASSLEGNTGQLY | 0.13883501 |
| FLTVAPEL | ASSLEGNTGQLY | 0.7657052 |
| FETLTAL | ASSLEGNTGQLY | 0.7797049 |
| DVINPIEL | ASSLEGNTGQLY | 0.81839097 |
| APVSGAPV | ASSLEGNTGQLY | 0.45197254 |
| ALVGGLAW | ASSLEGNTGQLY | 0.20547639 |
| ALAAAAATAI | ASSLEGNTGQLY | 0.7877395 |
| AFGLGLLL | ASSLEGNTGQLY | 0.33030346 |
| AFFAAASL | ASSLEGNTGQLY | 0.29955655 |
| AAAALPAFSPP | ASSLEGNTGQLY | 0.5132688 |
| VVLAPAL | ASSLEGNTGQLY | 0.40471798 |
| TIGHVDHGKTTL | ASSLEGNTGQLY | 0.25918025 |
| LISNASDALDK | ASSLEGNTGQLY | 0.65144914 |
| LISNASDAL | ASSLEGNTGQLY | 0.6997023 |
| ISNASDALDK | ASSLEGNTGQLY | 0.18471345 |
| NQAETLVYQTE | ASSLEGNTGQLY | 0.2291148 |
| TGGTGGTAGAG | ASSLEGNTGQLY | 0.23423564 |
| TGSTAGGAGGA | ASSLEGNTGQLY | 0.40367925 |
| DPLAVPVAAGPV | ASSLEGNTGQLY | 0.68379796 |
| GGTGGTPGNGGN | ASSLEGNTGQLY | 0.4777506 |
| PAGLAGVDPATL | ASSLEGNTGQLY | 0.46922198 |
| GNNLVVEAPHAGASV | ASSLEGNTGQLY | 0.6093628 |
| KVTRIMSASTLEQPA | ASSLEGNTGQLY | 0.6569464 |
| LGFALPIALPRRTVV | ASSLEGNTGQLY | 0.43128943 |
| TRLGNLTARKLVVLV | ASSLEGNTGQLY | 0.5859155 |
| WSQQRRLIYFPSAGP | ASSLEGNTGQLY | 0.25631443 |
| GPNGNPDPMAAAVDIR | ASSLEGNTGQLY | 0.69947827 |
| TQRCVSVADLRAAASR | ASSLEGNTGQLY | 0.5679816 |
| QFVLDVAEHPVEVLTETV | ASSLEGNTGQLY | 0.62364984 |

| | | |
|---|---|---|
| GGAGGTGGAGGNGGTQAGDG | ASSLEGNTGQLY | 0.718027 |
| PLLATLRAIAVDVDAKPAQV | ASSLEGNTGQLY | 0.6602913 |
| APKFRTGGYMIDFWGVGYQVAK | ASSLEGNTGQLY | 0.4714665 |
| GAGGAGGAGGAAGGPGGTGGTG | ASSLEGNTGQLY | 0.12749584 |
| IGALGLLATDVAPAGSVAFGVA | ASSLEGNTGQLY | 0.30601645 |
| AGGAGGIGGNGGAGGAGGVLMG | ASSLEGNTGQLY | 0.45123392 |
| ADELIGLL | ASSLEGNTGQLY | 0.5232456 |
| ADRALKA | ASSLEGNTGQLY | 0.5594686 |
| AGAVGQVGGAGGA | ASSLEGNTGQLY | 0.64390504 |
| AGGGGGLGGAP | ASSLEGNTGQLY | 0.27263182 |
| AGGLLYGNGGAG | ASSLEGNTGQLY | 0.7062895 |
| AGLESIGL | ASSLEGNTGQLY | 0.74790835 |
| AGLKTGII | ASSLEGNTGQLY | 0.1511453 |
| ASLAALLL | ASSLEGNTGQLY | 0.28424525 |
| ASVINISSI | ASSLEGNTGQLY | 0.7810602 |
| ATPPGPA | ASSLEGNTGQLY | 0.6942282 |
| ATVASALV | ASSLEGNTGQLY | 0.3180261 |
| ATVASLIL | ASSLEGNTGQLY | 0.3716717 |
| AVNGPVLIRH | ASSLEGNTGQLY | 0.8318764 |
| DGAAGAVGI | ASSLEGNTGQLY | 0.31613883 |
| DPLLHGI | ASSLEGNTGQLY | 0.78458065 |
| ELGAALGAL | ASSLEGNTGQLY | 0.106960125 |
| ELGRTQAV | ASSLEGNTGQLY | 0.39826775 |
| FVAGGYLS | ASSLEGNTGQLY | 0.2996716 |
| GASGGAGGAGGW | ASSLEGNTGQLY | 0.10745742 |
| GGAGAGGDGGAGGI | ASSLEGNTGQLY | 0.6337855 |
| GTGHNGSQPGMGGNG | ASSLEGNTGQLY | 0.2784262 |
| GYAAASASAS | ASSLEGNTGQLY | 0.64017236 |
| HQVDVAAVQQG | ASSLEGNTGQLY | 0.3925682 |
| IGAAIAHT | ASSLEGNTGQLY | 0.6261059 |
| IGMIEVAQL | ASSLEGNTGQLY | 0.5659795 |
| IRLAPEHTSAMD | ASSLEGNTGQLY | 0.33064815 |
| IYNVSVL | ASSLEGNTGQLY | 0.221814 |
| KQGFLGM | ASSLEGNTGQLY | 0.56254834 |
| LESVITQL | ASSLEGNTGQLY | 0.689525 |
| LGLGALVVV | ASSLEGNTGQLY | 0.63480055 |
| LIGNGGNGGSGGI | ASSLEGNTGQLY | 0.1620969 |
| LNAINAPTL | ASSLEGNTGQLY | 0.3859145 |
| LQLLNTGV | ASSLEGNTGQLY | 0.39706057 |
| LQTSIEF | ASSLEGNTGQLY | 0.13588636 |
| MSFAVSEL | ASSLEGNTGQLY | 0.09436259 |
| NTGALGL | ASSLEGNTGQLY | 0.33275467 |
| PPGAGGAGGAGTT | ASSLEGNTGQLY | 0.3089043 |
| QLVGLGI | ASSLEGNTGQLY | 0.641749 |
| RFLEQVHQL | ASSLEGNTGQLY | 0.6445374 |
| RGVGYVLR | ASSLEGNTGQLY | 0.28679323 |
| SATGVFAGVI | ASSLEGNTGQLY | 0.68789697 |
| SLPAITLP | ASSLEGNTGQLY | 0.5186504 |
| STALAFTV | ASSLEGNTGQLY | 0.070033014 |
| SVLTVGLGK | ASSLEGNTGQLY | 0.34323743 |
| TALAGAGSGI | ASSLEGNTGQLY | 0.14467344 |

| | | |
|---|---|---|
| TGLPFTGI | ASSLEGNTGQLY | 0.75360847 |
| VAEHVTV | ASSLEGNTGQLY | 0.16998702 |
| VDIVLNSL | ASSLEGNTGQLY | 0.2550524 |
| VLTGGGAVL | ASSLEGNTGQLY | 0.34149635 |
| VNVGLAYGI | ASSLEGNTGQLY | 0.71994543 |
| VSSAAVGAV | ASSLEGNTGQLY | 0.5378618 |
| VTSVYEAGV | ASSLEGNTGQLY | 0.2683777 |
| WVALDRGA | ASSLEGNTGQLY | 0.09529913 |
| YLVGSVVI | ASSLEGNTGQLY | 0.5450786 |
| GPPGTGKTLL | ASSLEGNTGQLY | 0.2939956 |
| GVGMTKF | ASSLEGNTGQLY | 0.2207794 |
| IGIGGSDL | ASSLEGNTGQLY | 0.13482079 |
| IGTLLAEL | ASSLEGNTGQLY | 0.1379361 |
| IYTSGTT | ASSLEGNTGQLY | 0.21591017 |
| LSFPAGV | ASSLEGNTGQLY | 0.69022894 |
| LYTSGTT | ASSLEGNTGQLY | 0.29054788 |
| VGAFPFGF | ASSLEGNTGQLY | 0.32666904 |
| VGAGTVEF | ASSLEGNTGQLY | 0.7849587 |
| VGKSTLLL | ASSLEGNTGQLY | 0.7169082 |
| AALAHAGGGIV | ASSLEGNTGQLY | 0.30038804 |
| AALVAVSLL | ASSLEGNTGQLY | 0.2846808 |
| AAVSRAVAAL | ASSLEGNTGQLY | 0.1899701 |
| AGGSSGIGGF | ASSLEGNTGQLY | 0.24495438 |
| AGLTVANANG | ASSLEGNTGQLY | 0.25329068 |
| AGLVGAVL | ASSLEGNTGQLY | 0.71320087 |
| ALSQAFTV | ASSLEGNTGQLY | 0.40324557 |
| ALTFSVSL | ASSLEGNTGQLY | 0.20407172 |
| ALTGAALVAL | ASSLEGNTGQLY | 0.7297907 |
| ASFSIGAGVL | ASSLEGNTGQLY | 0.75926375 |
| DGSVYKI | ASSLEGNTGQLY | 0.83895135 |
| EQLAEIV | ASSLEGNTGQLY | 0.17767383 |
| FAVSELTVM | ASSLEGNTGQLY | 0.42998445 |
| GAAPLAAGL | ASSLEGNTGQLY | 0.33628076 |
| GADGGDGGSGGAGG | ASSLEGNTGQLY | 0.40782106 |
| GFGLDTGGAGG | ASSLEGNTGQLY | 0.2961412 |
| GLASGGLL | ASSLEGNTGQLY | 0.2230756 |
| GRIAGAAVLL | ASSLEGNTGQLY | 0.6881182 |
| GSAGSLNTSF | ASSLEGNTGQLY | 0.8739886 |
| GTGQIGI | ASSLEGNTGQLY | 0.30520824 |
| GTLIALIL | ASSLEGNTGQLY | 0.20430042 |
| IAATMGFVVG | ASSLEGNTGQLY | 0.33572912 |
| IFAQALL | ASSLEGNTGQLY | 0.24105795 |
| IGAAVALGF | ASSLEGNTGQLY | 0.24066755 |
| IGLTPLAL | ASSLEGNTGQLY | 0.46875918 |
| ILGSNTSTL | ASSLEGNTGQLY | 0.13308874 |
| IVFSLTVL | ASSLEGNTGQLY | 0.32585517 |
| LASKATGF | ASSLEGNTGQLY | 0.84231365 |
| LASVALIL | ASSLEGNTGQLY | 0.41404134 |
| LFAVGLTL | ASSLEGNTGQLY | 0.57925856 |
| LFNGGIGTYI | ASSLEGNTGQLY | 0.7308539 |
| LTLGAGII | ASSLEGNTGQLY | 0.57694423 |

| | | |
|---|---|---|
| LVAAALGDI | ASSLEGNTGQLY | 0.7552979 |
| NTAFVQLGI | ASSLEGNTGQLY | 0.7949978 |
| QLFSGAEL | ASSLEGNTGQLY | 0.33046383 |
| SAATFITL | ASSLEGNTGQLY | 0.53364897 |
| SNYGLANLGV | ASSLEGNTGQLY | 0.5492985 |
| SYGGAYAVM | ASSLEGNTGQLY | 0.34759772 |
| TTGVGGLTLG | ASSLEGNTGQLY | 0.36026728 |
| TTQTIYI | ASSLEGNTGQLY | 0.34974653 |
| VAAGAGLGAVI | ASSLEGNTGQLY | 0.68156636 |
| VAAVVTLL | ASSLEGNTGQLY | 0.5793108 |
| VGFLGGLI | ASSLEGNTGQLY | 0.7464589 |
| VGVIPILL | ASSLEGNTGQLY | 0.14656281 |
| VIFASSLI | ASSLEGNTGQLY | 0.21163599 |
| VKGGDGGAAAG | ASSLEGNTGQLY | 0.72771966 |
| VTKTFTNI | ASSLEGNTGQLY | 0.09888958 |
| VVAAAILL | ASSLEGNTGQLY | 0.14672665 |
| YAALDVEL | ASSLEGNTGQLY | 0.4989267 |
| LASVALLL | ASSLEGNTGQLY | 0.79978585 |
| LDAVHRV | ASSLEGNTGQLY | 0.09772186 |
| QGLSSSIF | ASSLEGNTGQLY | 0.23678586 |
| RGTFANIRL | ASSLEGNTGQLY | 0.17263997 |
| RQLWWGHRI | ASSLEGNTGQLY | 0.7362572 |
| YGPPGTGKTLL | ASSLEGNTGQLY | 0.4343865 |
| AALVGAPANVA | ASSLEGNTGQLY | 0.13884419 |
| AATFRRTE | ASSLEGNTGQLY | 0.56061816 |
| AGALGGGI | ASSLEGNTGQLY | 0.13552018 |
| AGGAGGAGGIGAG | ASSLEGNTGQLY | 0.20468168 |
| AGSGALGL | ASSLEGNTGQLY | 0.638831 |
| AQIADQI | ASSLEGNTGQLY | 0.81181896 |
| ASAVAVTYL | ASSLEGNTGQLY | 0.1618437 |
| ASTVLGGTGG | ASSLEGNTGQLY | 0.6864537 |
| AVGGEPAK | ASSLEGNTGQLY | 0.35653085 |
| CEQGTFGGTC | ASSLEGNTGQLY | 0.8482016 |
| DEGKAPAG | ASSLEGNTGQLY | 0.28391352 |
| DTVVFPAL | ASSLEGNTGQLY | 0.15888089 |
| FEGITMAG | ASSLEGNTGQLY | 0.66027844 |
| FPLVVQD | ASSLEGNTGQLY | 0.61767817 |
| GAFTTPQ | ASSLEGNTGQLY | 0.39879966 |
| GFGGFGTAGGVGGA | ASSLEGNTGQLY | 0.7385874 |
| GFLAQFL | ASSLEGNTGQLY | 0.5011533 |
| GGAGGAGGNAGVA | ASSLEGNTGQLY | 0.34315538 |
| GGVGTAGAGGAGGAGG | ASSLEGNTGQLY | 0.68741846 |
| GNGGGGDAGNAGSGGN | ASSLEGNTGQLY | 0.5616797 |
| GTGGAGGAGNDAGS | ASSLEGNTGQLY | 0.22485964 |
| GTTNTGWFNS | ASSLEGNTGQLY | 0.61751556 |
| IFMGPLFTL | ASSLEGNTGQLY | 0.40692297 |
| ILAHAIEAI | ASSLEGNTGQLY | 0.2206388 |
| IRSAIREAL | ASSLEGNTGQLY | 0.8168812 |
| ITFAGGG | ASSLEGNTGQLY | 0.46351412 |
| ITGMSGHQEV | ASSLEGNTGQLY | 0.6707321 |
| IVAGLACTL | ASSLEGNTGQLY | 0.575258 |

| | | |
|---|---|---|
| IVGGGLAGA | ASSLEGNTGQLY | 0.86984247 |
| IVTGVLNI | ASSLEGNTGQLY | 0.16968206 |
| LAQQGCI | ASSLEGNTGQLY | 0.60923505 |
| LCLHVEN | ASSLEGNTGQLY | 0.40901196 |
| LDAISDGS | ASSLEGNTGQLY | 0.69029844 |
| LGGFAAML | ASSLEGNTGQLY | 0.55717003 |
| LGGKNPMI | ASSLEGNTGQLY | 0.2730744 |
| LGPKGVAAIL | ASSLEGNTGQLY | 0.49536353 |
| LMVGVSI | ASSLEGNTGQLY | 0.7328787 |
| LVGAAFVL | ASSLEGNTGQLY | 0.39440644 |
| LVLAVAPL | ASSLEGNTGQLY | 0.5168365 |
| MAAIAVIDAA | ASSLEGNTGQLY | 0.7862135 |
| NNGINAELQ | ASSLEGNTGQLY | 0.21147224 |
| RGEPLVLVHG | ASSLEGNTGQLY | 0.37866232 |
| RGGAGGAAGLI | ASSLEGNTGQLY | 0.63595223 |
| RSYRMHPAV | ASSLEGNTGQLY | 0.682533 |
| SAAHLLAGQLA | ASSLEGNTGQLY | 0.41103292 |
| SAAVGSY | ASSLEGNTGQLY | 0.73463976 |
| SPHAVLHRI | ASSLEGNTGQLY | 0.27707264 |
| SQIAKVL | ASSLEGNTGQLY | 0.5724803 |
| TALGHLVP | ASSLEGNTGQLY | 0.37158284 |
| TAPTTSGAA | ASSLEGNTGQLY | 0.5063306 |
| TDGDDGNGGSPGT | ASSLEGNTGQLY | 0.18733987 |
| TDGLGEN | ASSLEGNTGQLY | 0.5871418 |
| TGGPPLPGQA | ASSLEGNTGQLY | 0.70030504 |
| TIPAITI | ASSLEGNTGQLY | 0.6401578 |
| TIPGFTI | ASSLEGNTGQLY | 0.10346646 |
| TSIASGALA | ASSLEGNTGQLY | 0.63113916 |
| TVTLPGAAI | ASSLEGNTGQLY | 0.67554665 |
| VASGLSAALP | ASSLEGNTGQLY | 0.85453117 |
| VGAGVIGI | ASSLEGNTGQLY | 0.5214206 |
| VGPKRFGV | ASSLEGNTGQLY | 0.5846783 |
| VNVAFPD | ASSLEGNTGQLY | 0.3145382 |
| VSMTNSGVS | ASSLEGNTGQLY | 0.44322193 |
| VTDVLGDLSV | ASSLEGNTGQLY | 0.17383271 |
| VVVVGAGFAGL | ASSLEGNTGQLY | 0.36008653 |
| VYTVGTVV | ASSLEGNTGQLY | 0.32782084 |
| WTTFSGP | ASSLEGNTGQLY | 0.18999869 |
| YDELVEAF | ASSLEGNTGQLY | 0.7805232 |
| YGTGGAGGNGG | ASSLEGNTGQLY | 0.71007097 |
| YTNVMAV | ASSLEGNTGQLY | 0.7432597 |
| FLEQVHQL | ASSLEGNTGQLY | 0.75645185 |
| GLSLLAVAV | ASSLEGNTGQLY | 0.7584014 |
| LEQVHQL | ASSLEGNTGQLY | 0.61153996 |
| LNAAGAI | ASSLEGNTGQLY | 0.29165557 |
| PAVHASL | ASSLEGNTGQLY | 0.3328293 |
| TELEDDA | ASSLEGNTGQLY | 0.15261482 |
| TPPGPPN | ASSLEGNTGQLY | 0.33682156 |
| TPPPPPPP | ASSLEGNTGQLY | 0.3282708 |
| TQSPALI | AWSFPGGRSAETLY | 0.4206289 |
| EFAAAFPQ | AWSFPGGRSAETLY | 0.088002324 |

| | | |
|---|---|---|
| LLSGNLDVL | AWSFPGGRSAETLY | 0.6823385 |
| GPANLGNYNI | AWSFPGGRSAETLY | 0.71896625 |
| TLGPTLP | AWSFPGGRSAETLY | 0.40700608 |
| VYAYSSCCK | AWSFPGGRSAETLY | 0.5774052 |
| APAPAPAPNI | AWSFPGGRSAETLY | 0.54234064 |
| PGGPVPIP | AWSFPGGRSAETLY | 0.2591624 |
| FLTVALEP | AWSFPGGRSAETLY | 0.6728004 |
| SQYFGANL | AWSFPGGRSAETLY | 0.22814104 |
| GGVKPAA | AWSFPGGRSAETLY | 0.65615535 |
| GGTGGAGGAGGW | AWSFPGGRSAETLY | 0.8145479 |
| GVGGVAGLGGAGG | AWSFPGGRSAETLY | 0.46197078 |
| VLLVLATTI | AWSFPGGRSAETLY | 0.43076667 |
| IGIFAAML | AWSFPGGRSAETLY | 0.44827354 |
| AVGDLID | AWSFPGGRSAETLY | 0.15367961 |
| NISPLPL | AWSFPGGRSAETLY | 0.45673013 |
| VTAVFVAI | AWSFPGGRSAETLY | 0.07939959 |
| SGDDNDAAL | AWSFPGGRSAETLY | 0.52546644 |
| AVAALAHS | AWSFPGGRSAETLY | 0.21411489 |
| GGVTVMATP | AWSFPGGRSAETLY | 0.44863924 |
| AGLAANVRD | AWSFPGGRSAETLY | 0.40843278 |
| TNRPDLIDPALLRPG | AWSFPGGRSAETLY | 0.5501089 |
| LANLGVTAL | AWSFPGGRSAETLY | 0.20224768 |
| TGERIEL | AWSFPGGRSAETLY | 0.4777612 |
| NGEELAA | AWSFPGGRSAETLY | 0.39154518 |
| VLSLVWG | AWSFPGGRSAETLY | 0.18895279 |
| IGYALWL | AWSFPGGRSAETLY | 0.691128 |
| LSLLLAPL | AWSFPGGRSAETLY | 0.37126324 |
| TLDEALK | AWSFPGGRSAETLY | 0.49964908 |
| TDGAGTAEGG | AWSFPGGRSAETLY | 0.7026019 |
| NAPGGAGGNGG | AWSFPGGRSAETLY | 0.4348902 |
| VSVFMGMGGNF | AWSFPGGRSAETLY | 0.17726459 |
| LASYLAAL | AWSFPGGRSAETLY | 0.44525212 |
| LVGGTPAL | AWSFPGGRSAETLY | 0.40820518 |
| VRNADVL | AWSFPGGRSAETLY | 0.30372193 |
| TCSSPFLVSAM | AWSFPGGRSAETLY | 0.08732047 |
| VHITDAE | AWSFPGGRSAETLY | 0.5079168 |
| EYAAPSF | AWSFPGGRSAETLY | 0.22446865 |
| VVDAFHGGL | AWSFPGGRSAETLY | 0.28041434 |
| ASVVDPQ | AWSFPGGRSAETLY | 0.25235265 |
| SYNEFGGLS | AWSFPGGRSAETLY | 0.6542125 |
| DPDGSVY | AWSFPGGRSAETLY | 0.5323125 |
| MVSTIHATE | AWSFPGGRSAETLY | 0.5499555 |
| LAAGVADPV | AWSFPGGRSAETLY | 0.6128495 |
| TKEQIAA | AWSFPGGRSAETLY | 0.5483186 |
| SLASPAAL | AWSFPGGRSAETLY | 0.7455925 |
| IGLVFYGV | AWSFPGGRSAETLY | 0.56168026 |
| FLLDASNL | AWSFPGGRSAETLY | 0.45971516 |
| YQVGNFP | AWSFPGGRSAETLY | 0.46398318 |
| ESLLSQST | AWSFPGGRSAETLY | 0.4624114 |
| LFTVGFAAVG | AWSFPGGRSAETLY | 0.12448894 |
| ATPPTPT | AWSFPGGRSAETLY | 0.15146247 |

| | | |
|---|---|---|
| TDLATLG | AWSFPGGRSAETLY | 0.5056648 |
| SAGGTGR | AWSFPGGRSAETLY | 0.12680547 |
| GFSNSGS | AWSFPGGRSAETLY | 0.47535712 |
| GGNGGSGGTSV | AWSFPGGRSAETLY | 0.7585468 |
| GTSVATGGAGN | AWSFPGGRSAETLY | 0.5959509 |
| NGGNGGSGGTS | AWSFPGGRSAETLY | 0.17336304 |
| AVVKFPH | AWSFPGGRSAETLY | 0.30484095 |
| TPPPPPP | AWSFPGGRSAETLY | 0.25216103 |
| AAPAAAAAPA | AWSFPGGRSAETLY | 0.60256886 |
| IALNEDL | AWSFPGGRSAETLY | 0.65979135 |
| IVNVSSM | AWSFPGGRSAETLY | 0.82928324 |
| TGIPPAPR | AWSFPGGRSAETLY | 0.7149976 |
| FDLGGGTF | AWSFPGGRSAETLY | 0.290723 |
| FDLGGGTFDVS | AWSFPGGRSAETLY | 0.6161287 |
| NEPTAAA | AWSFPGGRSAETLY | 0.35932893 |
| NEPTAAALA | AWSFPGGRSAETLY | 0.4113114 |
| GSIINISSI | AWSFPGGRSAETLY | 0.3491035 |
| VAPGNAGTA | AWSFPGGRSAETLY | 0.60232335 |
| TGGGASLE | AWSFPGGRSAETLY | 0.32835862 |
| VAVYNHYKRI | AWSFPGGRSAETLY | 0.2551191 |
| TDPAPATT | AWSFPGGRSAETLY | 0.19340762 |
| TNRPDILDPALLRPG | AWSFPGGRSAETLY | 0.5671934 |
| LTTTTTS | AWSFPGGRSAETLY | 0.57249516 |
| PTGSGKTL | AWSFPGGRSAETLY | 0.20578417 |
| TGSGKTL | AWSFPGGRSAETLY | 0.30874115 |
| TIGHVDHGK | AWSFPGGRSAETLY | 0.8232169 |
| TIGHVDHGKTT | AWSFPGGRSAETLY | 0.8065566 |
| GHVDHGKTT | AWSFPGGRSAETLY | 0.24773154 |
| AQFEHTLL | AWSFPGGRSAETLY | 0.2643904 |
| IVNQDEL | AWSFPGGRSAETLY | 0.7058979 |
| TNGSQFF | AWSFPGGRSAETLY | 0.14502275 |
| TGKGIVE | AWSFPGGRSAETLY | 0.5789991 |
| LISNASDA | AWSFPGGRSAETLY | 0.5301923 |
| AELRTLGELTKIAKAHGAQ | AWSFPGGRSAETLY | 0.37455764 |
| APNAVAAAEPSW | AWSFPGGRSAETLY | 0.27101108 |
| APQVAGAATAVL | AWSFPGGRSAETLY | 0.5455849 |
| ARRDPAAAAAILPTDA | AWSFPGGRSAETLY | 0.3578074 |
| DAAARDVLAEAGLAEYF | AWSFPGGRSAETLY | 0.6663127 |
| DDQIVLSAKANLAKVEQ | AWSFPGGRSAETLY | 0.09409467 |
| DELLPDTGEDSDEDGH | AWSFPGGRSAETLY | 0.6140768 |
| ELPEAPANRATVPQD | AWSFPGGRSAETLY | 0.18945965 |
| FGQSKIERVTGIG | AWSFPGGRSAETLY | 0.36118585 |
| GAGGTGSTAGGAGGAGGAGGLY | AWSFPGGRSAETLY | 0.13037501 |
| GHGGTGAAGGNGGTGGAGGSGI | AWSFPGGRSAETLY | 0.34712613 |
| GNGGNGGAGGEGGPGGAGGAGA | AWSFPGGRSAETLY | 0.73504966 |
| GSTAGNGGNGGNATNGTIAGQP | AWSFPGGRSAETLY | 0.7371762 |
| GTGGDGGNGGNGGTGG | AWSFPGGRSAETLY | 0.18322167 |
| GVRRPAGIPADAPV | AWSFPGGRSAETLY | 0.62423503 |
| GYRQVTDAHLLTLARR | AWSFPGGRSAETLY | 0.83602047 |
| IAKMGQDGHDRG | AWSFPGGRSAETLY | 0.30867982 |
| KAEPSIEIQYT | AWSFPGGRSAETLY | 0.129015 |

| | | |
|---|---|---|
| KPVRIRMAADEDTVAVT | AWSFPGGRSAETLY | 0.53680557 |
| MPPRIAGMRLLVIKPEP | AWSFPGGRSAETLY | 0.63100463 |
| MKFYAVALFRSRGVPPTR | AWSFPGGRSAETLY | 0.7152989 |
| NAGGFGILGAVAHSP | AWSFPGGRSAETLY | 0.7638444 |
| NEGREITLAMAEM | AWSFPGGRSAETLY | 0.12524624 |
| NELVEWSHPLGHRWRWE | AWSFPGGRSAETLY | 0.71133745 |
| NLGAGGTGGNGGNPG | AWSFPGGRSAETLY | 0.72042036 |
| NTEWPKWETVMAADPVD | AWSFPGGRSAETLY | 0.19963554 |
| PASAAPAETQP | AWSFPGGRSAETLY | 0.25592464 |
| QQSSDGKNGVRAAEITGE | AWSFPGGRSAETLY | 0.2587461 |
| RAPDAVTTSWWKE | AWSFPGGRSAETLY | 0.12960196 |
| SGFGNVGAHTSGWFNQ | AWSFPGGRSAETLY | 0.58637536 |
| SGVAGAVGQAASVGGLKV | AWSFPGGRSAETLY | 0.2858739 |
| SNGGAGTGSAGGNGGT | AWSFPGGRSAETLY | 0.33176288 |
| TGGAGGEGGAAG | AWSFPGGRSAETLY | 0.34095338 |
| TGGDGGAAGAGGAPGHG | AWSFPGGRSAETLY | 0.28894067 |
| TMPPLLAAIANAGRHGV | AWSFPGGRSAETLY | 0.2880968 |
| TSSTAGNSGGA | AWSFPGGRSAETLY | 0.29519784 |
| VPATVPVGILC | AWSFPGGRSAETLY | 0.5321684 |
| VQRPDSGGVVLTGRIS | AWSFPGGRSAETLY | 0.6154079 |
| WTPNPVPATKGVPAGLVQAT | AWSFPGGRSAETLY | 0.57351017 |
| YALKRALAKFSDRFDVVII | AWSFPGGRSAETLY | 0.6058418 |
| YLIGSVIV | AWSFPGGRSAETLY | 0.7574323 |
| YGQLVAVID | AWSFPGGRSAETLY | 0.19807488 |
| VTVGGMTLPA | AWSFPGGRSAETLY | 0.21872884 |
| VLALPAV | AWSFPGGRSAETLY | 0.67869043 |
| VIPPTLNYET | AWSFPGGRSAETLY | 0.20378111 |
| VGMGGMHPSSG | AWSFPGGRSAETLY | 0.716768 |
| VGLAGGAATA | AWSFPGGRSAETLY | 0.45473576 |
| VGGAGGAGGSGP | AWSFPGGRSAETLY | 0.48739147 |
| VFAAFMV | AWSFPGGRSAETLY | 0.6420121 |
| TLYDKLR | AWSFPGGRSAETLY | 0.8088089 |
| TLAVFTVL | AWSFPGGRSAETLY | 0.5842069 |
| TGSVQPARV | AWSFPGGRSAETLY | 0.343694 |
| TGGGAGSGGT | AWSFPGGRSAETLY | 0.6883605 |
| SHLETSL | AWSFPGGRSAETLY | 0.24499889 |
| SFTTGMT | AWSFPGGRSAETLY | 0.6287973 |
| QVEVAKRMGVRQPTVS | AWSFPGGRSAETLY | 0.6674714 |
| QGSGGPALAM | AWSFPGGRSAETLY | 0.1786097 |
| MLAPSNP | AWSFPGGRSAETLY | 0.36102298 |
| LVNNAAI | AWSFPGGRSAETLY | 0.73490775 |
| LTGEINTLA | AWSFPGGRSAETLY | 0.6839639 |
| LSIFGPV | AWSFPGGRSAETLY | 0.16821441 |
| LLALLANL | AWSFPGGRSAETLY | 0.24076532 |
| LISNASDALDKLR | AWSFPGGRSAETLY | 0.18251777 |
| LINEAVP | AWSFPGGRSAETLY | 0.48568675 |
| LGYAFSGL | AWSFPGGRSAETLY | 0.19343363 |
| LDFVAVAL | AWSFPGGRSAETLY | 0.63209045 |
| LAQPGAE | AWSFPGGRSAETLY | 0.30294955 |
| ISNLGAVTGF | AWSFPGGRSAETLY | 0.7747985 |
| HQEDPGRI | AWSFPGGRSAETLY | 0.4555419 |

| | | |
|---|---|---|
| GTPGEPTP | AWSFPGGRSAETLY | 0.68185145 |
| GFFNSGS | AWSFPGGRSAETLY | 0.29063976 |
| GAGGGGLTAGAN | AWSFPGGRSAETLY | 0.3313238 |
| FVDALVNM | AWSFPGGRSAETLY | 0.16526507 |
| FLTVAPEL | AWSFPGGRSAETLY | 0.20268296 |
| FETLTAL | AWSFPGGRSAETLY | 0.23745108 |
| DVINPIEL | AWSFPGGRSAETLY | 0.37189412 |
| APVSGAPV | AWSFPGGRSAETLY | 0.12517814 |
| ALVGGLAW | AWSFPGGRSAETLY | 0.7881154 |
| ALAAAAATAI | AWSFPGGRSAETLY | 0.4270184 |
| AFGLGLLL | AWSFPGGRSAETLY | 0.57016134 |
| AFFAAASL | AWSFPGGRSAETLY | 0.34471852 |
| AAAALPAFSPP | AWSFPGGRSAETLY | 0.24141458 |
| VVLAPAL | AWSFPGGRSAETLY | 0.27183285 |
| TIGHVDHGKTTL | AWSFPGGRSAETLY | 0.6761513 |
| LISNASDALDK | AWSFPGGRSAETLY | 0.15227492 |
| LISNASDAL | AWSFPGGRSAETLY | 0.14312373 |
| ISNASDALDK | AWSFPGGRSAETLY | 0.35105774 |
| NQAETLVYQTE | AWSFPGGRSAETLY | 0.18296176 |
| TGGTGGTAGAG | AWSFPGGRSAETLY | 0.79286313 |
| TGSTAGGAGGA | AWSFPGGRSAETLY | 0.7173389 |
| DPLAVPVAAGPV | AWSFPGGRSAETLY | 0.4830105 |
| GGTGGTPGNGGN | AWSFPGGRSAETLY | 0.7458194 |
| PAGLAGVDPATL | AWSFPGGRSAETLY | 0.11256268 |
| GNNLVVEAPHAGASV | AWSFPGGRSAETLY | 0.3454851 |
| KVTRIMSASTLEQPA | AWSFPGGRSAETLY | 0.7868608 |
| LGFALPIALPRRTVV | AWSFPGGRSAETLY | 0.5729734 |
| TRLGNLTARKLVVLV | AWSFPGGRSAETLY | 0.33912513 |
| WSQQRRLIYFPSAGP | AWSFPGGRSAETLY | 0.70296043 |
| GPNGNPDPMAAAVDIR | AWSFPGGRSAETLY | 0.24866994 |
| TQRCVSVADLRAAASR | AWSFPGGRSAETLY | 0.31549555 |
| QFVLDVAEHPVEVLTETV | AWSFPGGRSAETLY | 0.38407835 |
| GGAGGTGGAGGNGGTQAGDG | AWSFPGGRSAETLY | 0.45264375 |
| PLLATLRAIAVDVDAKPAQV | AWSFPGGRSAETLY | 0.5790957 |
| APKFRTGGYMIDFWGVGYQVAK | AWSFPGGRSAETLY | 0.1663694 |
| GAGGAGGAGGAAGGPGGTGGTG | AWSFPGGRSAETLY | 0.55640376 |
| IGALGLLATDVAPAGSVAFGVA | AWSFPGGRSAETLY | 0.82970786 |
| AGGAGGIGGNGGAGGAGGVLMG | AWSFPGGRSAETLY | 0.2955767 |
| ADELIGLL | AWSFPGGRSAETLY | 0.717445 |
| ADRALKA | AWSFPGGRSAETLY | 0.52759135 |
| AGAVGQVGGAGGA | AWSFPGGRSAETLY | 0.76172537 |
| AGGGGGLGGAP | AWSFPGGRSAETLY | 0.23281546 |
| AGGLLYGNGGAG | AWSFPGGRSAETLY | 0.17575671 |
| AGLESIGL | AWSFPGGRSAETLY | 0.49569792 |
| AGLKTGII | AWSFPGGRSAETLY | 0.17401111 |
| ASLAALLL | AWSFPGGRSAETLY | 0.4836889 |
| ASVINISSI | AWSFPGGRSAETLY | 0.32245183 |
| ATPPGPA | AWSFPGGRSAETLY | 0.8584819 |
| ATVASALV | AWSFPGGRSAETLY | 0.6441339 |
| ATVASLIL | AWSFPGGRSAETLY | 0.5144709 |
| AVNGPVLIRH | AWSFPGGRSAETLY | 0.18503124 |

| | | |
|---|---|---|
| DGAAGAVGI | AWSFPGGRSAETLY | 0.45497224 |
| DPLLHGI | AWSFPGGRSAETLY | 0.6209566 |
| ELGAALGAL | AWSFPGGRSAETLY | 0.19025698 |
| ELGRTQAV | AWSFPGGRSAETLY | 0.7436773 |
| FVAGGYLS | AWSFPGGRSAETLY | 0.22434685 |
| GASGGAGGAGGW | AWSFPGGRSAETLY | 0.5141526 |
| GGAGAGGDGGAGGI | AWSFPGGRSAETLY | 0.6242449 |
| GTGHNGSQPGMGGNG | AWSFPGGRSAETLY | 0.8252759 |
| GYAAASASAS | AWSFPGGRSAETLY | 0.31332675 |
| HQVDVAAVQQG | AWSFPGGRSAETLY | 0.43601874 |
| IGAAIAHT | AWSFPGGRSAETLY | 0.16880159 |
| IGMIEVAQL | AWSFPGGRSAETLY | 0.71275663 |
| IRLAPEHTSAMD | AWSFPGGRSAETLY | 0.7055739 |
| IYNVSVL | AWSFPGGRSAETLY | 0.7533604 |
| KQGFLGM | AWSFPGGRSAETLY | 0.7902498 |
| LESVITQL | AWSFPGGRSAETLY | 0.6826001 |
| LGLGALVVV | AWSFPGGRSAETLY | 0.18824472 |
| LIGNGGNGGSGGI | AWSFPGGRSAETLY | 0.33883086 |
| LNAINAPTL | AWSFPGGRSAETLY | 0.6808195 |
| LQLLNTGV | AWSFPGGRSAETLY | 0.75229067 |
| LQTSIEF | AWSFPGGRSAETLY | 0.7368783 |
| MSFAVSEL | AWSFPGGRSAETLY | 0.6113894 |
| NTGALGL | AWSFPGGRSAETLY | 0.33834177 |
| PPGAGGAGGAGTT | AWSFPGGRSAETLY | 0.7784241 |
| QLVGLGI | AWSFPGGRSAETLY | 0.8740593 |
| RFLEQVHQL | AWSFPGGRSAETLY | 0.6413879 |
| RGVGYVLR | AWSFPGGRSAETLY | 0.339091 |
| SATGVFAGVI | AWSFPGGRSAETLY | 0.36002633 |
| SLPAITLP | AWSFPGGRSAETLY | 0.4446854 |
| STALAFTV | AWSFPGGRSAETLY | 0.7167171 |
| SVLTVGLGK | AWSFPGGRSAETLY | 0.24026285 |
| TALAGAGSGI | AWSFPGGRSAETLY | 0.43756843 |
| TGLPFTGI | AWSFPGGRSAETLY | 0.474022 |
| VAEHVTV | AWSFPGGRSAETLY | 0.13762026 |
| VDIVLNSL | AWSFPGGRSAETLY | 0.6626725 |
| VLTGGGAVL | AWSFPGGRSAETLY | 0.7594071 |
| VNVGLAYGI | AWSFPGGRSAETLY | 0.39107478 |
| VSSAAVGAV | AWSFPGGRSAETLY | 0.24690226 |
| VTSVYEAGV | AWSFPGGRSAETLY | 0.7357069 |
| WVALDRGA | AWSFPGGRSAETLY | 0.82444406 |
| YLVGSVVI | AWSFPGGRSAETLY | 0.7019462 |
| GPPGTGKTLL | AWSFPGGRSAETLY | 0.16828296 |
| GVGMTKF | AWSFPGGRSAETLY | 0.108629525 |
| IGIGGSDL | AWSFPGGRSAETLY | 0.22825077 |
| IGTLLAEL | AWSFPGGRSAETLY | 0.79867816 |
| IYTSGTT | AWSFPGGRSAETLY | 0.34026277 |
| LSFPAGV | AWSFPGGRSAETLY | 0.26189324 |
| LYTSGTT | AWSFPGGRSAETLY | 0.57301927 |
| VGAFPFGF | AWSFPGGRSAETLY | 0.7318262 |
| VGAGTVEF | AWSFPGGRSAETLY | 0.20860544 |
| VGKSTLLL | AWSFPGGRSAETLY | 0.20366162 |

| | | |
|---|---|---|
| AALAHAGGGIV | AWSFPGGRSAETLY | 0.63098043 |
| AALVAVSLL | AWSFPGGRSAETLY | 0.69779027 |
| AAVSRAVAAL | AWSFPGGRSAETLY | 0.17981605 |
| AGGSSGIGGF | AWSFPGGRSAETLY | 0.30702814 |
| AGLTVANANG | AWSFPGGRSAETLY | 0.31123796 |
| AGLVGAVL | AWSFPGGRSAETLY | 0.57511985 |
| ALSQAFTV | AWSFPGGRSAETLY | 0.58085835 |
| ALTFSVSL | AWSFPGGRSAETLY | 0.21379268 |
| ALTGAALVAL | AWSFPGGRSAETLY | 0.74201196 |
| ASFSIGAGVL | AWSFPGGRSAETLY | 0.16580081 |
| DGSVYKI | AWSFPGGRSAETLY | 0.67152005 |
| EQLAEIV | AWSFPGGRSAETLY | 0.38679028 |
| FAVSELTVM | AWSFPGGRSAETLY | 0.45144087 |
| GAAPLAAGL | AWSFPGGRSAETLY | 0.2900469 |
| GADGGDGGSGGAGG | AWSFPGGRSAETLY | 0.26956946 |
| GFGLDTGGAGG | AWSFPGGRSAETLY | 0.1836773 |
| GLASGGLL | AWSFPGGRSAETLY | 0.13829342 |
| GRIAGAAVLL | AWSFPGGRSAETLY | 0.6875645 |
| GSAGSLNTSF | AWSFPGGRSAETLY | 0.1213315 |
| GTGQIGI | AWSFPGGRSAETLY | 0.6107956 |
| GTLIALIL | AWSFPGGRSAETLY | 0.49865434 |
| IAATMGFVVG | AWSFPGGRSAETLY | 0.18236661 |
| IFAQALL | AWSFPGGRSAETLY | 0.5357819 |
| IGAAVALGF | AWSFPGGRSAETLY | 0.19438794 |
| IGLTPLAL | AWSFPGGRSAETLY | 0.2656806 |
| ILGSNTSTL | AWSFPGGRSAETLY | 0.7396598 |
| IVFSLTVL | AWSFPGGRSAETLY | 0.5458252 |
| LASKATGF | AWSFPGGRSAETLY | 0.23372382 |
| LASVALIL | AWSFPGGRSAETLY | 0.5371566 |
| LFAVGLTL | AWSFPGGRSAETLY | 0.52053356 |
| LFNGGIGTYI | AWSFPGGRSAETLY | 0.26449668 |
| LTLGAGII | AWSFPGGRSAETLY | 0.37375414 |
| LVAAALGDI | AWSFPGGRSAETLY | 0.7447151 |
| NTAFVQLGI | AWSFPGGRSAETLY | 0.46634686 |
| QLFSGAEL | AWSFPGGRSAETLY | 0.15425023 |
| SAATFITL | AWSFPGGRSAETLY | 0.6840521 |
| SNYGLANLGV | AWSFPGGRSAETLY | 0.19360681 |
| SYGGAYAVM | AWSFPGGRSAETLY | 0.5196308 |
| TTGVGGLTLG | AWSFPGGRSAETLY | 0.31346622 |
| TTQTIYI | AWSFPGGRSAETLY | 0.35607374 |
| VAAGAGLGAVI | AWSFPGGRSAETLY | 0.66071784 |
| VAAVVTLL | AWSFPGGRSAETLY | 0.34591585 |
| VGFLGGLI | AWSFPGGRSAETLY | 0.2576244 |
| VGVIPILL | AWSFPGGRSAETLY | 0.32316625 |
| VIFASSLI | AWSFPGGRSAETLY | 0.40122265 |
| VKGGDGGAAAG | AWSFPGGRSAETLY | 0.63769746 |
| VTKTFTNI | AWSFPGGRSAETLY | 0.2026834 |
| VVAAAILL | AWSFPGGRSAETLY | 0.5055784 |
| YAALDVEL | AWSFPGGRSAETLY | 0.29491708 |
| LASVALLL | AWSFPGGRSAETLY | 0.20715785 |
| LDAVHRV | AWSFPGGRSAETLY | 0.4375146 |

| | | |
|---|---|---|
| QGLSSSIF | AWSFPGGRSAETLY | 0.34830454 |
| RGTFANIRL | AWSFPGGRSAETLY | 0.22356918 |
| RQLWWGHRI | AWSFPGGRSAETLY | 0.48775315 |
| YGPPGTGKTLL | AWSFPGGRSAETLY | 0.2504603 |
| AALVGAPANVA | AWSFPGGRSAETLY | 0.5948128 |
| AATFRRTE | AWSFPGGRSAETLY | 0.8304638 |
| AGALGGGI | AWSFPGGRSAETLY | 0.22328287 |
| AGGAGGAGGIGAG | AWSFPGGRSAETLY | 0.62494075 |
| AGSGALGL | AWSFPGGRSAETLY | 0.5977388 |
| AQIADQI | AWSFPGGRSAETLY | 0.48129028 |
| ASAVAVTYL | AWSFPGGRSAETLY | 0.33185726 |
| ASTVLGGTGG | AWSFPGGRSAETLY | 0.23604064 |
| AVGGEPAK | AWSFPGGRSAETLY | 0.73821974 |
| CEQGTFGGTC | AWSFPGGRSAETLY | 0.59741646 |
| DEGKAPAG | AWSFPGGRSAETLY | 0.60859764 |
| DTVVFPAL | AWSFPGGRSAETLY | 0.5748025 |
| FEGITMAG | AWSFPGGRSAETLY | 0.6966989 |
| FPLVVQD | AWSFPGGRSAETLY | 0.5875293 |
| GAFTTPQ | AWSFPGGRSAETLY | 0.26026207 |
| GFGGFGTAGGVGGA | AWSFPGGRSAETLY | 0.15669389 |
| GFLAQFL | AWSFPGGRSAETLY | 0.3386808 |
| GGAGGAGGNAGVA | AWSFPGGRSAETLY | 0.36943775 |
| GGVGTAGAGGAGGAGG | AWSFPGGRSAETLY | 0.28214303 |
| GNGGGGDAGNAGSGGN | AWSFPGGRSAETLY | 0.19474515 |
| GTGGAGGAGNDAGS | AWSFPGGRSAETLY | 0.713199 |
| GTTNTGWFNS | AWSFPGGRSAETLY | 0.8428103 |
| IFMGPLFTL | AWSFPGGRSAETLY | 0.30333224 |
| ILAHAIEAI | AWSFPGGRSAETLY | 0.82851374 |
| IRSAIREAL | AWSFPGGRSAETLY | 0.17939341 |
| ITFAGGG | AWSFPGGRSAETLY | 0.6530435 |
| ITGMSGHQEV | AWSFPGGRSAETLY | 0.7341576 |
| IVAGLACTL | AWSFPGGRSAETLY | 0.21982966 |
| IVGGGLAGA | AWSFPGGRSAETLY | 0.54783463 |
| IVTGVLNI | AWSFPGGRSAETLY | 0.19572794 |
| LAQQGCI | AWSFPGGRSAETLY | 0.25713974 |
| LCLHVEN | AWSFPGGRSAETLY | 0.671139 |
| LDAISDGS | AWSFPGGRSAETLY | 0.34771448 |
| LGGFAAML | AWSFPGGRSAETLY | 0.42522892 |
| LGGKNPMI | AWSFPGGRSAETLY | 0.31282902 |
| LGPKGVAAIL | AWSFPGGRSAETLY | 0.20928992 |
| LMVGVSI | AWSFPGGRSAETLY | 0.35672203 |
| LVGAAFVL | AWSFPGGRSAETLY | 0.680626 |
| LVLAVAPL | AWSFPGGRSAETLY | 0.3254502 |
| MAAIAVIDAA | AWSFPGGRSAETLY | 0.8683563 |
| NNGINAELQ | AWSFPGGRSAETLY | 0.2069732 |
| RGEPLVLVHG | AWSFPGGRSAETLY | 0.81265044 |
| RGGAGGAAGLI | AWSFPGGRSAETLY | 0.7721847 |
| RSYRMHPAV | AWSFPGGRSAETLY | 0.46395403 |
| SAAHLLAGQLA | AWSFPGGRSAETLY | 0.33984956 |
| SAAVGSY | AWSFPGGRSAETLY | 0.7333101 |
| SPHAVLHRI | AWSFPGGRSAETLY | 0.35046816 |

| | | |
|---|---|---|
| SQIAKVL | AWSFPGGRSAETLY | 0.57658553 |
| TALGHLVP | AWSFPGGRSAETLY | 0.7214328 |
| TAPTTSGAA | AWSFPGGRSAETLY | 0.1771059 |
| TDGDDGNGGSPGT | AWSFPGGRSAETLY | 0.634429 |
| TDGLGEN | AWSFPGGRSAETLY | 0.4794278 |
| TGGPPLPGQA | AWSFPGGRSAETLY | 0.41479936 |
| TIPAITI | AWSFPGGRSAETLY | 0.20978354 |
| TIPGFTI | AWSFPGGRSAETLY | 0.15665989 |
| TSIASGALA | AWSFPGGRSAETLY | 0.23704892 |
| TVTLPGAAI | AWSFPGGRSAETLY | 0.17223129 |
| VASGLSAALP | AWSFPGGRSAETLY | 0.4018913 |
| VGAGVIGI | AWSFPGGRSAETLY | 0.38713962 |
| VGPKRFGV | AWSFPGGRSAETLY | 0.23174988 |
| VNVAFPD | AWSFPGGRSAETLY | 0.5200112 |
| VSMTNSGVS | AWSFPGGRSAETLY | 0.6234089 |
| VTDVLGDLSV | AWSFPGGRSAETLY | 0.15740262 |
| VVVVGAGFAGL | AWSFPGGRSAETLY | 0.31656423 |
| VYTVGTVV | AWSFPGGRSAETLY | 0.18394224 |
| WTTFSGP | AWSFPGGRSAETLY | 0.23434108 |
| YDELVEAF | AWSFPGGRSAETLY | 0.14100192 |
| YGTGGAGGNGG | AWSFPGGRSAETLY | 0.39881074 |
| YTNVMAV | AWSFPGGRSAETLY | 0.20222826 |
| FLEQVHQL | AWSFPGGRSAETLY | 0.13719362 |
| GLSLLAVAV | AWSFPGGRSAETLY | 0.32436797 |
| LEQVHQL | AWSFPGGRSAETLY | 0.252805 |
| LNAAGAI | AWSFPGGRSAETLY | 0.20578745 |
| PAVHASL | AWSFPGGRSAETLY | 0.53528047 |
| TELEDDA | AWSFPGGRSAETLY | 0.1681781 |
| TPPGPPN | AWSFPGGRSAETLY | 0.17382607 |
| TPPPPPPP | AWSFPGGRSAETLY | 0.43334115 |
| TQSPALI | TCSADSRDRGYEQY | 0.61764324 |
| EFAAAFPQ | TCSADSRDRGYEQY | 0.1730938 |
| LLSGNLDVL | TCSADSRDRGYEQY | 0.42045796 |
| GPANLGNYNI | TCSADSRDRGYEQY | 0.70717615 |
| TLGPTLP | TCSADSRDRGYEQY | 0.2915308 |
| VYAYSSCCK | TCSADSRDRGYEQY | 0.5888548 |
| APAPAPAPNI | TCSADSRDRGYEQY | 0.67229646 |
| PGGPVPIP | TCSADSRDRGYEQY | 0.23254113 |
| FLTVALEP | TCSADSRDRGYEQY | 0.5125857 |
| SQYFGANL | TCSADSRDRGYEQY | 0.66562134 |
| GGVKPAA | TCSADSRDRGYEQY | 0.26550335 |
| GGTGGAGGAGGW | TCSADSRDRGYEQY | 0.25557685 |
| GVGGVAGLGGAGG | TCSADSRDRGYEQY | 0.1816448 |
| VLLVLATTI | TCSADSRDRGYEQY | 0.28413042 |
| IGIFAAML | TCSADSRDRGYEQY | 0.14492913 |
| AVGDLID | TCSADSRDRGYEQY | 0.45552593 |
| NISPLPL | TCSADSRDRGYEQY | 0.46552005 |
| VTAVFVAI | TCSADSRDRGYEQY | 0.22938326 |
| SGDDNDAAL | TCSADSRDRGYEQY | 0.15928724 |
| AVAALAHS | TCSADSRDRGYEQY | 0.15808047 |
| GGVTVMATP | TCSADSRDRGYEQY | 0.24108653 |

| | | |
|---|---|---|
| AGLAANVRD | TCSADSRDRGYEQY | 0.23773564 |
| TNRPDLIDPALLRPG | TCSADSRDRGYEQY | 0.15263231 |
| LANLGVTAL | TCSADSRDRGYEQY | 0.21698612 |
| TGERIEL | TCSADSRDRGYEQY | 0.42175934 |
| NGEELAA | TCSADSRDRGYEQY | 0.48051304 |
| VLSLVWG | TCSADSRDRGYEQY | 0.16525522 |
| IGYALWL | TCSADSRDRGYEQY | 0.67963254 |
| LSLLLAPL | TCSADSRDRGYEQY | 0.5763384 |
| TLDEALK | TCSADSRDRGYEQY | 0.64804184 |
| TDGAGTAEGG | TCSADSRDRGYEQY | 0.26581717 |
| NAPGGAGGNGG | TCSADSRDRGYEQY | 0.17169753 |
| VSVFMGMGGNF | TCSADSRDRGYEQY | 0.69163656 |
| LASYLAAL | TCSADSRDRGYEQY | 0.30231112 |
| LVGGTPAL | TCSADSRDRGYEQY | 0.19200984 |
| VRNADVL | TCSADSRDRGYEQY | 0.41873467 |
| TCSSPFLVSAM | TCSADSRDRGYEQY | 0.20370163 |
| VHITDAE | TCSADSRDRGYEQY | 0.19366693 |
| EYAAPSF | TCSADSRDRGYEQY | 0.5383039 |
| VVDAFHGGL | TCSADSRDRGYEQY | 0.4552396 |
| ASVVDPQ | TCSADSRDRGYEQY | 0.17250738 |
| SYNEFGGLS | TCSADSRDRGYEQY | 0.1662357 |
| DPDGSVY | TCSADSRDRGYEQY | 0.22056273 |
| MVSTIHATE | TCSADSRDRGYEQY | 0.32758802 |
| LAAGVADPV | TCSADSRDRGYEQY | 0.5116104 |
| TKEQIAA | TCSADSRDRGYEQY | 0.3107099 |
| SLASPAAL | TCSADSRDRGYEQY | 0.27313825 |
| IGLVFYGV | TCSADSRDRGYEQY | 0.56705105 |
| FLLDASNL | TCSADSRDRGYEQY | 0.45058054 |
| YQVGNFP | TCSADSRDRGYEQY | 0.29058632 |
| ESLLSQST | TCSADSRDRGYEQY | 0.6059408 |
| LFTVGFAAVG | TCSADSRDRGYEQY | 0.19041228 |
| ATPPTPT | TCSADSRDRGYEQY | 0.45833856 |
| TDLATLG | TCSADSRDRGYEQY | 0.442097 |
| SAGGTGR | TCSADSRDRGYEQY | 0.48957565 |
| GFSNSGS | TCSADSRDRGYEQY | 0.6035743 |
| GGNGGSGGTSV | TCSADSRDRGYEQY | 0.2984057 |
| GTSVATGGAGN | TCSADSRDRGYEQY | 0.35365924 |
| NGGNGGSGGTS | TCSADSRDRGYEQY | 0.14963661 |
| AVVKFPH | TCSADSRDRGYEQY | 0.23724785 |
| TPPPPPP | TCSADSRDRGYEQY | 0.4752426 |
| AAPAAAAAPA | TCSADSRDRGYEQY | 0.38903373 |
| IALNEDL | TCSADSRDRGYEQY | 0.5716149 |
| IVNVSSM | TCSADSRDRGYEQY | 0.45428824 |
| TGIPPAPR | TCSADSRDRGYEQY | 0.21628174 |
| FDLGGGTF | TCSADSRDRGYEQY | 0.697484 |
| FDLGGGTFDVS | TCSADSRDRGYEQY | 0.65851474 |
| NEPTAAA | TCSADSRDRGYEQY | 0.15777206 |
| NEPTAAALA | TCSADSRDRGYEQY | 0.25151807 |
| GSIINISSI | TCSADSRDRGYEQY | 0.661544 |
| VAPGNAGTA | TCSADSRDRGYEQY | 0.5429418 |
| TGGGASLE | TCSADSRDRGYEQY | 0.25835034 |

| | | |
|---|---|---|
| VAVYNHYKRI | TCSADSRDRGYEQY | 0.3018558 |
| TDPAPATT | TCSADSRDRGYEQY | 0.5186547 |
| TNRPDILDPALLRPG | TCSADSRDRGYEQY | 0.19102539 |
| LTTTTTS | TCSADSRDRGYEQY | 0.69866794 |
| PTGSGKTL | TCSADSRDRGYEQY | 0.14726406 |
| TGSGKTL | TCSADSRDRGYEQY | 0.33256707 |
| TIGHVDHGK | TCSADSRDRGYEQY | 0.19472045 |
| TIGHVDHGKTT | TCSADSRDRGYEQY | 0.14328285 |
| GHVDHGKTT | TCSADSRDRGYEQY | 0.37559047 |
| AQFEHTLL | TCSADSRDRGYEQY | 0.22669114 |
| IVNQDEL | TCSADSRDRGYEQY | 0.5434204 |
| TNGSQFF | TCSADSRDRGYEQY | 0.23031518 |
| TGKGIVE | TCSADSRDRGYEQY | 0.45078295 |
| LISNASDA | TCSADSRDRGYEQY | 0.3998688 |
| AELRTLGELTKIAKAHGAQ | TCSADSRDRGYEQY | 0.3243321 |
| APNAVAAAEPSW | TCSADSRDRGYEQY | 0.18469036 |
| APQVAGAATAVL | TCSADSRDRGYEQY | 0.37517834 |
| ARRDPAAAAAILPTDA | TCSADSRDRGYEQY | 0.6904193 |
| DAAARDVLAEAGLAEYF | TCSADSRDRGYEQY | 0.39358306 |
| DDQIVLSAKANLAKVEQ | TCSADSRDRGYEQY | 0.1538948 |
| DELLPDTGEDSDEDGH | TCSADSRDRGYEQY | 0.31086606 |
| ELPEAPANRATVPQD | TCSADSRDRGYEQY | 0.23766668 |
| FGQSKIERVTGIG | TCSADSRDRGYEQY | 0.13868399 |
| GAGGTGSTAGGAGGAGGAGGLY | TCSADSRDRGYEQY | 0.1414955 |
| GHGGTGAAGGNGGTGGAGGSGI | TCSADSRDRGYEQY | 0.16628435 |
| GNGGNGGAGGEGGPGGAGGAGA | TCSADSRDRGYEQY | 0.20621175 |
| GSTAGNGGNGGNATNGTIAGQP | TCSADSRDRGYEQY | 0.4792524 |
| GTGGDGGNGGNGGTGG | TCSADSRDRGYEQY | 0.4017501 |
| GVRRPAGIPADAPV | TCSADSRDRGYEQY | 0.22158255 |
| GYRQVTDAHLLTLARR | TCSADSRDRGYEQY | 0.50982 |
| IAKMGQDGHDRG | TCSADSRDRGYEQY | 0.4861548 |
| KAEPSIEIQYT | TCSADSRDRGYEQY | 0.13391797 |
| KPVRIRMAADEDTVAVT | TCSADSRDRGYEQY | 0.22050121 |
| MPPRIAGMRLLVIKPEP | TCSADSRDRGYEQY | 0.15975922 |
| MKFYAVALFRSRGVPPTR | TCSADSRDRGYEQY | 0.6267072 |
| NAGGFGILGAVAHSP | TCSADSRDRGYEQY | 0.17406695 |
| NEGREITLAMAEM | TCSADSRDRGYEQY | 0.19524853 |
| NELVEWSHPLGHRWRWE | TCSADSRDRGYEQY | 0.2449755 |
| NLGAGGTGGNGGNPG | TCSADSRDRGYEQY | 0.5129361 |
| NTEWPKWETVMAADPVD | TCSADSRDRGYEQY | 0.3837443 |
| PASAAPAETQP | TCSADSRDRGYEQY | 0.19151397 |
| QQSSDGKNGVRAAEITGE | TCSADSRDRGYEQY | 0.14280911 |
| RAPDAVTTSWWKE | TCSADSRDRGYEQY | 0.3395984 |
| SGFGNVGAHTSGWFNQ | TCSADSRDRGYEQY | 0.14873129 |
| SGVAGAVGQAASVGGLKV | TCSADSRDRGYEQY | 0.18480973 |
| SNGGAGTGSAGGNGGT | TCSADSRDRGYEQY | 0.15346766 |
| TGGAGGEGGAAG | TCSADSRDRGYEQY | 0.13540882 |
| TGGDGGAAGAGGAPGHG | TCSADSRDRGYEQY | 0.1687914 |
| TMPPLLAAIANAGRHGV | TCSADSRDRGYEQY | 0.5568198 |
| TSSTAGNSGGA | TCSADSRDRGYEQY | 0.2565371 |
| VPATVPVGILC | TCSADSRDRGYEQY | 0.2512836 |

| | | |
|---|---|---|
| VQRPDSGGVVLTGRIS | TCSADSRDRGYEQY | 0.6036003 |
| WTPNPVPATKGVPAGLVQAT | TCSADSRDRGYEQY | 0.61437637 |
| YALKRALAKFSDRFDVVII | TCSADSRDRGYEQY | 0.23053029 |
| YLIGSVIV | TCSADSRDRGYEQY | 0.21101388 |
| YGQLVAVID | TCSADSRDRGYEQY | 0.16914636 |
| VTVGGMTLPA | TCSADSRDRGYEQY | 0.17212085 |
| VLALPAV | TCSADSRDRGYEQY | 0.1894129 |
| VIPPTLNYET | TCSADSRDRGYEQY | 0.64014226 |
| VGMGGMHPSSG | TCSADSRDRGYEQY | 0.29694983 |
| VGLAGGAATA | TCSADSRDRGYEQY | 0.16266812 |
| VGGAGGAGGSGP | TCSADSRDRGYEQY | 0.41821188 |
| VFAAFMV | TCSADSRDRGYEQY | 0.66399926 |
| TLYDKLR | TCSADSRDRGYEQY | 0.7539164 |
| TLAVFTVL | TCSADSRDRGYEQY | 0.17358507 |
| TGSVQPARV | TCSADSRDRGYEQY | 0.26666525 |
| TGGGAGSGGT | TCSADSRDRGYEQY | 0.2763783 |
| SHLETSL | TCSADSRDRGYEQY | 0.28374678 |
| SFTTGMT | TCSADSRDRGYEQY | 0.24129856 |
| QVEVAKRMGVRQPTVS | TCSADSRDRGYEQY | 0.17128488 |
| QGSGGPALAM | TCSADSRDRGYEQY | 0.43629348 |
| MLAPSNP | TCSADSRDRGYEQY | 0.77252364 |
| LVNNAAI | TCSADSRDRGYEQY | 0.23002699 |
| LTGEINTLA | TCSADSRDRGYEQY | 0.14657407 |
| LSIFGPV | TCSADSRDRGYEQY | 0.29434896 |
| LLALLANL | TCSADSRDRGYEQY | 0.18576494 |
| LISNASDALDKLR | TCSADSRDRGYEQY | 0.18530142 |
| LINEAVP | TCSADSRDRGYEQY | 0.2505849 |
| LGYAFSGL | TCSADSRDRGYEQY | 0.15595356 |
| LDFVAVAL | TCSADSRDRGYEQY | 0.25752747 |
| LAQPGAE | TCSADSRDRGYEQY | 0.7034116 |
| ISNLGAVTGF | TCSADSRDRGYEQY | 0.28087616 |
| HQEDPGRI | TCSADSRDRGYEQY | 0.56891775 |
| GTPGEPTP | TCSADSRDRGYEQY | 0.61981636 |
| GFFNSGS | TCSADSRDRGYEQY | 0.4288277 |
| GAGGGGLTAGAN | TCSADSRDRGYEQY | 0.68438697 |
| FVDALVNM | TCSADSRDRGYEQY | 0.6505286 |
| FLTVAPEL | TCSADSRDRGYEQY | 0.2845042 |
| FETLTAL | TCSADSRDRGYEQY | 0.3526178 |
| DVINPIEL | TCSADSRDRGYEQY | 0.4712386 |
| APVSGAPV | TCSADSRDRGYEQY | 0.3027719 |
| ALVGGLAW | TCSADSRDRGYEQY | 0.30261695 |
| ALAAAAATAI | TCSADSRDRGYEQY | 0.2376109 |
| AFGLGLLL | TCSADSRDRGYEQY | 0.57529575 |
| AFFAAASL | TCSADSRDRGYEQY | 0.31682473 |
| AAAALPAFSPP | TCSADSRDRGYEQY | 0.5850414 |
| VVLAPAL | TCSADSRDRGYEQY | 0.17201129 |
| TIGHVDHGKTTL | TCSADSRDRGYEQY | 0.17202805 |
| LISNASDALDK | TCSADSRDRGYEQY | 0.65890425 |
| LISNASDAL | TCSADSRDRGYEQY | 0.13892832 |
| ISNASDALDK | TCSADSRDRGYEQY | 0.15631494 |
| NQAETLVYQTE | TCSADSRDRGYEQY | 0.28044972 |

| | | |
|---|---|---|
| TGGTGGTAGAG | TCSADSRDRGYEQY | 0.70930564 |
| TGSTAGGAGGA | TCSADSRDRGYEQY | 0.13529186 |
| DPLAVPVAAGPV | TCSADSRDRGYEQY | 0.18334213 |
| GGTGGTPGNGGN | TCSADSRDRGYEQY | 0.17250083 |
| PAGLAGVDPATL | TCSADSRDRGYEQY | 0.4357319 |
| GNNLVVEAPHAGASV | TCSADSRDRGYEQY | 0.2485801 |
| KVTRIMSASTLEQPA | TCSADSRDRGYEQY | 0.15840891 |
| LGFALPIALPRRTVV | TCSADSRDRGYEQY | 0.45701343 |
| TRLGNLTARKLVVLV | TCSADSRDRGYEQY | 0.14527817 |
| WSQQRRLIYFPSAGP | TCSADSRDRGYEQY | 0.14728633 |
| GPNGNPDPMAAAVDIR | TCSADSRDRGYEQY | 0.40135986 |
| TQRCVSVADLRAAASR | TCSADSRDRGYEQY | 0.6971762 |
| QFVLDVAEHPVEVLTETV | TCSADSRDRGYEQY | 0.17033957 |
| GGAGGTGGAGGNGGTQAGDG | TCSADSRDRGYEQY | 0.40323037 |
| PLLATLRAIAVDVDAKPAQV | TCSADSRDRGYEQY | 0.2370351 |
| APKFRTGGYMIDFWGVGYQVAK | TCSADSRDRGYEQY | 0.7729683 |
| GAGGAGGAGGAAGGPGGTGGTG | TCSADSRDRGYEQY | 0.23617294 |
| IGALGLLATDVAPAGSVAFGVA | TCSADSRDRGYEQY | 0.16223949 |
| AGGAGGIGGNGGAGGAGGVLMG | TCSADSRDRGYEQY | 0.53002906 |
| ADELIGLL | TCSADSRDRGYEQY | 0.6073785 |
| ADRALKA | TCSADSRDRGYEQY | 0.27468476 |
| AGAVGQVGGAGGA | TCSADSRDRGYEQY | 0.56215465 |
| AGGGGGLGGAP | TCSADSRDRGYEQY | 0.24434672 |
| AGGLLYGNGGAG | TCSADSRDRGYEQY | 0.25026444 |
| AGLESIGL | TCSADSRDRGYEQY | 0.52586997 |
| AGLKTGII | TCSADSRDRGYEQY | 0.556564 |
| ASLAALLL | TCSADSRDRGYEQY | 0.16217186 |
| ASVINISSI | TCSADSRDRGYEQY | 0.47472125 |
| ATPPGPA | TCSADSRDRGYEQY | 0.32661343 |
| ATVASALV | TCSADSRDRGYEQY | 0.19979268 |
| ATVASLIL | TCSADSRDRGYEQY | 0.6262511 |
| AVNGPVLIRH | TCSADSRDRGYEQY | 0.30532044 |
| DGAAGAVGI | TCSADSRDRGYEQY | 0.48941067 |
| DPLLHGI | TCSADSRDRGYEQY | 0.3407438 |
| ELGAALGAL | TCSADSRDRGYEQY | 0.7208538 |
| ELGRTQAV | TCSADSRDRGYEQY | 0.1658563 |
| FVAGGYLS | TCSADSRDRGYEQY | 0.5981586 |
| GASGGAGGAGGW | TCSADSRDRGYEQY | 0.3262925 |
| GGAGAGGDGGAGGI | TCSADSRDRGYEQY | 0.54510355 |
| GTGHNGSQPGMGGNG | TCSADSRDRGYEQY | 0.38737398 |
| GYAAASASAS | TCSADSRDRGYEQY | 0.17307302 |
| HQVDVAAVQQG | TCSADSRDRGYEQY | 0.3788693 |
| IGAAIAHT | TCSADSRDRGYEQY | 0.55961466 |
| IGMIEVAQL | TCSADSRDRGYEQY | 0.24565493 |
| IRLAPEHTSAMD | TCSADSRDRGYEQY | 0.5055503 |
| IYNVSVL | TCSADSRDRGYEQY | 0.70440817 |
| KQGFLGM | TCSADSRDRGYEQY | 0.1884124 |
| LESVITQL | TCSADSRDRGYEQY | 0.266618 |
| LGLGALVVV | TCSADSRDRGYEQY | 0.42037666 |
| LIGNGGNGGSGGI | TCSADSRDRGYEQY | 0.6517063 |
| LNAINAPTL | TCSADSRDRGYEQY | 0.29452318 |

| | | |
|---|---|---|
| LQLLNTGV | TCSADSRDRGYEQY | 0.655668 |
| LQTSIEF | TCSADSRDRGYEQY | 0.22011429 |
| MSFAVSEL | TCSADSRDRGYEQY | 0.4115926 |
| NTGALGL | TCSADSRDRGYEQY | 0.23994754 |
| PPGAGGAGGAGTT | TCSADSRDRGYEQY | 0.29643428 |
| QLVGLGI | TCSADSRDRGYEQY | 0.1526057 |
| RFLEQVHQL | TCSADSRDRGYEQY | 0.3843512 |
| RGVGYVLR | TCSADSRDRGYEQY | 0.52162135 |
| SATGVFAGVI | TCSADSRDRGYEQY | 0.46926665 |
| SLPAITLP | TCSADSRDRGYEQY | 0.14234081 |
| STALAFTV | TCSADSRDRGYEQY | 0.4314525 |
| SVLTVGLGK | TCSADSRDRGYEQY | 0.5048128 |
| TALAGAGSGI | TCSADSRDRGYEQY | 0.775663 |
| TGLPFTGI | TCSADSRDRGYEQY | 0.35638487 |
| VAEHVTV | TCSADSRDRGYEQY | 0.5364528 |
| VDIVLNSL | TCSADSRDRGYEQY | 0.20159838 |
| VLTGGGAVL | TCSADSRDRGYEQY | 0.26575676 |
| VNVGLAYGI | TCSADSRDRGYEQY | 0.17232251 |
| VSSAAVGAV | TCSADSRDRGYEQY | 0.29522893 |
| VTSVYEAGV | TCSADSRDRGYEQY | 0.23278484 |
| WVALDRGA | TCSADSRDRGYEQY | 0.18407436 |
| YLVGSVVI | TCSADSRDRGYEQY | 0.6350239 |
| GPPGTGKTLL | TCSADSRDRGYEQY | 0.6772596 |
| GVGMTKF | TCSADSRDRGYEQY | 0.5210073 |
| IGIGGSDL | TCSADSRDRGYEQY | 0.68300664 |
| IGTLLAEL | TCSADSRDRGYEQY | 0.68517876 |
| IYTSGTT | TCSADSRDRGYEQY | 0.46803087 |
| LSFPAGV | TCSADSRDRGYEQY | 0.19502501 |
| LYTSGTT | TCSADSRDRGYEQY | 0.24030684 |
| VGAFPFGF | TCSADSRDRGYEQY | 0.15649131 |
| VGAGTVEF | TCSADSRDRGYEQY | 0.23808467 |
| VGKSTLLL | TCSADSRDRGYEQY | 0.19641198 |
| AALAHAGGGIV | TCSADSRDRGYEQY | 0.29372185 |
| AALVAVSLL | TCSADSRDRGYEQY | 0.13458167 |
| AAVSRAVAAL | TCSADSRDRGYEQY | 0.5244795 |
| AGGSSGIGGF | TCSADSRDRGYEQY | 0.64851743 |
| AGLTVANANG | TCSADSRDRGYEQY | 0.21900514 |
| AGLVGAVL | TCSADSRDRGYEQY | 0.46319532 |
| ALSQAFTV | TCSADSRDRGYEQY | 0.5710871 |
| ALTFSVSL | TCSADSRDRGYEQY | 0.20630325 |
| ALTGAALVAL | TCSADSRDRGYEQY | 0.5803731 |
| ASFSIGAGVL | TCSADSRDRGYEQY | 0.19863755 |
| DGSVYKI | TCSADSRDRGYEQY | 0.5014209 |
| EQLAEIV | TCSADSRDRGYEQY | 0.59224385 |
| FAVSELTVM | TCSADSRDRGYEQY | 0.35915318 |
| GAAPLAAGL | TCSADSRDRGYEQY | 0.468351 |
| GADGGDGGSGGAGG | TCSADSRDRGYEQY | 0.34743685 |
| GFGLDTGGAGG | TCSADSRDRGYEQY | 0.15797003 |
| GLASGGLL | TCSADSRDRGYEQY | 0.4938928 |
| GRIAGAAVLL | TCSADSRDRGYEQY | 0.13373782 |
| GSAGSLNTSF | TCSADSRDRGYEQY | 0.4087876 |

| | | |
|---|---|---|
| GTGQIGI | TCSADSRDRGYEQY | 0.17228761 |
| GTLIALIL | TCSADSRDRGYEQY | 0.37115347 |
| IAATMGFVVG | TCSADSRDRGYEQY | 0.29110417 |
| IFAQALL | TCSADSRDRGYEQY | 0.34704417 |
| IGAAVALGF | TCSADSRDRGYEQY | 0.20085391 |
| IGLTPLAL | TCSADSRDRGYEQY | 0.2546604 |
| ILGSNTSTL | TCSADSRDRGYEQY | 0.21289377 |
| IVFSLTVL | TCSADSRDRGYEQY | 0.15972415 |
| LASKATGF | TCSADSRDRGYEQY | 0.57152313 |
| LASVALIL | TCSADSRDRGYEQY | 0.2251215 |
| LFAVGLTL | TCSADSRDRGYEQY | 0.26028407 |
| LFNGGIGTYI | TCSADSRDRGYEQY | 0.4705225 |
| LTLGAGII | TCSADSRDRGYEQY | 0.273144 |
| LVAAALGDI | TCSADSRDRGYEQY | 0.14870974 |
| NTAFVQLGI | TCSADSRDRGYEQY | 0.34568882 |
| QLFSGAEL | TCSADSRDRGYEQY | 0.3784455 |
| SAATFITL | TCSADSRDRGYEQY | 0.25036636 |
| SNYGLANLGV | TCSADSRDRGYEQY | 0.13389595 |
| SYGGAYAVM | TCSADSRDRGYEQY | 0.38240272 |
| TTGVGGLTLG | TCSADSRDRGYEQY | 0.20425074 |
| TTQTIYI | TCSADSRDRGYEQY | 0.19652146 |
| VAAGAGLGAVI | TCSADSRDRGYEQY | 0.19872998 |
| VAAVVTLL | TCSADSRDRGYEQY | 0.57132155 |
| VGFLGGLI | TCSADSRDRGYEQY | 0.1962649 |
| VGVIPILL | TCSADSRDRGYEQY | 0.2401703 |
| VIFASSLI | TCSADSRDRGYEQY | 0.5175396 |
| VKGGDGGAAAG | TCSADSRDRGYEQY | 0.35398722 |
| VTKTFTNI | TCSADSRDRGYEQY | 0.6679107 |
| VVAAAILL | TCSADSRDRGYEQY | 0.27017835 |
| YAALDVEL | TCSADSRDRGYEQY | 0.26283994 |
| LASVALLL | TCSADSRDRGYEQY | 0.42954013 |
| LDAVHRV | TCSADSRDRGYEQY | 0.27393827 |
| QGLSSSIF | TCSADSRDRGYEQY | 0.14942876 |
| RGTFANIRL | TCSADSRDRGYEQY | 0.14646067 |
| RQLWWGHRI | TCSADSRDRGYEQY | 0.22994474 |
| YGPPGTGKTLL | TCSADSRDRGYEQY | 0.15385717 |
| AALVGAPANVA | TCSADSRDRGYEQY | 0.39156067 |
| AATFRRTE | TCSADSRDRGYEQY | 0.5892693 |
| AGALGGGI | TCSADSRDRGYEQY | 0.36083442 |
| AGGAGGAGGIGAG | TCSADSRDRGYEQY | 0.1651239 |
| AGSGALGL | TCSADSRDRGYEQY | 0.25037023 |
| AQIADQI | TCSADSRDRGYEQY | 0.20998779 |
| ASAVAVTYL | TCSADSRDRGYEQY | 0.43946904 |
| ASTVLGGTGG | TCSADSRDRGYEQY | 0.32573462 |
| AVGGEPAK | TCSADSRDRGYEQY | 0.7508843 |
| CEQGTFGGTC | TCSADSRDRGYEQY | 0.5692409 |
| DEGKAPAG | TCSADSRDRGYEQY | 0.22520086 |
| DTVVFPAL | TCSADSRDRGYEQY | 0.48843488 |
| FEGITMAG | TCSADSRDRGYEQY | 0.24828295 |
| FPLVVQD | TCSADSRDRGYEQY | 0.3317358 |
| GAFTTPQ | TCSADSRDRGYEQY | 0.27465504 |

| | | |
|---|---|---|
| GFGGFGTAGGVGGA | TCSADSRDRGYEQY | 0.2622886 |
| GFLAQFL | TCSADSRDRGYEQY | 0.3242421 |
| GGAGGAGGNAGVA | TCSADSRDRGYEQY | 0.25353628 |
| GGVGTAGAGGAGGAGG | TCSADSRDRGYEQY | 0.15907006 |
| GNGGGGDAGNAGSGGN | TCSADSRDRGYEQY | 0.33683252 |
| GTGGAGGAGNDAGS | TCSADSRDRGYEQY | 0.18320714 |
| GTTNTGWFNS | TCSADSRDRGYEQY | 0.15685582 |
| IFMGPLFTL | TCSADSRDRGYEQY | 0.31280154 |
| ILAHAIEAI | TCSADSRDRGYEQY | 0.7407628 |
| IRSAIREAL | TCSADSRDRGYEQY | 0.7167222 |
| ITFAGGG | TCSADSRDRGYEQY | 0.20704718 |
| ITGMSGHQEV | TCSADSRDRGYEQY | 0.18103687 |
| IVAGLACTL | TCSADSRDRGYEQY | 0.62871575 |
| IVGGGLAGA | TCSADSRDRGYEQY | 0.15918928 |
| IVTGVLNI | TCSADSRDRGYEQY | 0.4206249 |
| LAQQGCI | TCSADSRDRGYEQY | 0.3496068 |
| LCLHVEN | TCSADSRDRGYEQY | 0.23772013 |
| LDAISDGS | TCSADSRDRGYEQY | 0.25139797 |
| LGGFAAML | TCSADSRDRGYEQY | 0.4249797 |
| LGGKNPMI | TCSADSRDRGYEQY | 0.20475753 |
| LGPKGVAAIL | TCSADSRDRGYEQY | 0.4890902 |
| LMVGVSI | TCSADSRDRGYEQY | 0.13433655 |
| LVGAAFVL | TCSADSRDRGYEQY | 0.27585655 |
| LVLAVAPL | TCSADSRDRGYEQY | 0.18096563 |
| MAAIAVIDAA | TCSADSRDRGYEQY | 0.19085947 |
| NNGINAELQ | TCSADSRDRGYEQY | 0.1518367 |
| RGEPLVLVHG | TCSADSRDRGYEQY | 0.22937725 |
| RGGAGGAAGLI | TCSADSRDRGYEQY | 0.5864047 |
| RSYRMHPAV | TCSADSRDRGYEQY | 0.6254672 |
| SAAHLLAGQLA | TCSADSRDRGYEQY | 0.17698997 |
| SAAVGSY | TCSADSRDRGYEQY | 0.5632684 |
| SPHAVLHRI | TCSADSRDRGYEQY | 0.667058 |
| SQIAKVL | TCSADSRDRGYEQY | 0.21220884 |
| TALGHLVP | TCSADSRDRGYEQY | 0.15005033 |
| TAPTTSGAA | TCSADSRDRGYEQY | 0.313167 |
| TDGDDGNGGSPGT | TCSADSRDRGYEQY | 0.15874973 |
| TDGLGEN | TCSADSRDRGYEQY | 0.46161732 |
| TGGPPLPGQA | TCSADSRDRGYEQY | 0.7227453 |
| TIPAITI | TCSADSRDRGYEQY | 0.15167637 |
| TIPGFTI | TCSADSRDRGYEQY | 0.5067279 |
| TSIASGALA | TCSADSRDRGYEQY | 0.5733657 |
| TVTLPGAAI | TCSADSRDRGYEQY | 0.18684824 |
| VASGLSAALP | TCSADSRDRGYEQY | 0.18016115 |
| VGAGVIGI | TCSADSRDRGYEQY | 0.18888058 |
| VGPKRFGV | TCSADSRDRGYEQY | 0.1536 |
| VNVAFPD | TCSADSRDRGYEQY | 0.37814993 |
| VSMTNSGVS | TCSADSRDRGYEQY | 0.23256722 |
| VTDVLGDLSV | TCSADSRDRGYEQY | 0.20651308 |
| VVVVGAGFAGL | TCSADSRDRGYEQY | 0.24783018 |
| VYTVGTVV | TCSADSRDRGYEQY | 0.20514338 |
| WTTFSGP | TCSADSRDRGYEQY | 0.20266482 |

| | | |
|---|---|---|
| YDELVEAF | TCSADSRDRGYEQY | 0.20718321 |
| YGTGGAGGNGG | TCSADSRDRGYEQY | 0.20423198 |
| YTNVMAV | TCSADSRDRGYEQY | 0.48045582 |
| FLEQVHQL | TCSADSRDRGYEQY | 0.223896 |
| GLSLLAVAV | TCSADSRDRGYEQY | 0.4280576 |
| LEQVHQL | TCSADSRDRGYEQY | 0.6020889 |
| LNAAGAI | TCSADSRDRGYEQY | 0.14975774 |
| PAVHASL | TCSADSRDRGYEQY | 0.19953448 |
| TELEDDA | TCSADSRDRGYEQY | 0.49556944 |
| TPPGPPN | TCSADSRDRGYEQY | 0.15751177 |
| TPPPPPPP | TCSADSRDRGYEQY | 0.6084681 |
| TQSPALI | ASSRPQGASTEVF | 0.40999216 |
| EFAAAFPQ | ASSRPQGASTEVF | 0.35170674 |
| LLSGNLDVL | ASSRPQGASTEVF | 0.31632093 |
| GPANLGNYNI | ASSRPQGASTEVF | 0.7566192 |
| TLGPTLP | ASSRPQGASTEVF | 0.42134959 |
| VYAYSSCCK | ASSRPQGASTEVF | 0.2296356 |
| APAPAPAPNI | ASSRPQGASTEVF | 0.25139993 |
| PGGPVPIP | ASSRPQGASTEVF | 0.18827948 |
| FLTVALEP | ASSRPQGASTEVF | 0.57146144 |
| SQYFGANL | ASSRPQGASTEVF | 0.6309043 |
| GGVKPAA | ASSRPQGASTEVF | 0.1760395 |
| GGTGGAGGAGGW | ASSRPQGASTEVF | 0.2603181 |
| GVGGVAGLGGAGG | ASSRPQGASTEVF | 0.57556075 |
| VLLVLATTI | ASSRPQGASTEVF | 0.5774172 |
| IGIFAAML | ASSRPQGASTEVF | 0.1574065 |
| AVGDLID | ASSRPQGASTEVF | 0.18813995 |
| NISPLPL | ASSRPQGASTEVF | 0.17379475 |
| VTAVFVAI | ASSRPQGASTEVF | 0.32157376 |
| SGDDNDAAL | ASSRPQGASTEVF | 0.17828518 |
| AVAALAHS | ASSRPQGASTEVF | 0.4671509 |
| GGVTVMATP | ASSRPQGASTEVF | 0.24177721 |
| AGLAANVRD | ASSRPQGASTEVF | 0.67728233 |
| TNRPDLIDPALLRPG | ASSRPQGASTEVF | 0.34772485 |
| LANLGVTAL | ASSRPQGASTEVF | 0.46916664 |
| TGERIEL | ASSRPQGASTEVF | 0.2580223 |
| NGEELAA | ASSRPQGASTEVF | 0.23884636 |
| VLSLVWG | ASSRPQGASTEVF | 0.16376822 |
| IGYALWL | ASSRPQGASTEVF | 0.14755635 |
| LSLLLAPL | ASSRPQGASTEVF | 0.17945294 |
| TLDEALK | ASSRPQGASTEVF | 0.22012368 |
| TDGAGTAEGG | ASSRPQGASTEVF | 0.1365936 |
| NAPGGAGGNGG | ASSRPQGASTEVF | 0.6368363 |
| VSVFMGMGGNF | ASSRPQGASTEVF | 0.29949355 |
| LASYLAAL | ASSRPQGASTEVF | 0.36865127 |
| LVGGTPAL | ASSRPQGASTEVF | 0.21119812 |
| VRNADVL | ASSRPQGASTEVF | 0.22779141 |
| TCSSPFLVSAM | ASSRPQGASTEVF | 0.18123613 |
| VHITDAE | ASSRPQGASTEVF | 0.6487898 |
| EYAAPSF | ASSRPQGASTEVF | 0.16167542 |
| VVDAFHGGL | ASSRPQGASTEVF | 0.14371291 |

| | | |
|---|---|---|
| ASVVDPQ | ASSRPQGASTEVF | 0.20051618 |
| SYNEFGGLS | ASSRPQGASTEVF | 0.1506629 |
| DPDGSVY | ASSRPQGASTEVF | 0.7045262 |
| MVSTIHATE | ASSRPQGASTEVF | 0.566983 |
| LAAGVADPV | ASSRPQGASTEVF | 0.35896677 |
| TKEQIAA | ASSRPQGASTEVF | 0.5940793 |
| SLASPAAL | ASSRPQGASTEVF | 0.14244626 |
| IGLVFYGV | ASSRPQGASTEVF | 0.20115213 |
| FLLDASNL | ASSRPQGASTEVF | 0.68287766 |
| YQVGNFP | ASSRPQGASTEVF | 0.3102414 |
| ESLLSQST | ASSRPQGASTEVF | 0.28013724 |
| LFTVGFAAVG | ASSRPQGASTEVF | 0.5413828 |
| ATPPTPT | ASSRPQGASTEVF | 0.20516394 |
| TDLATLG | ASSRPQGASTEVF | 0.26664773 |
| SAGGTGR | ASSRPQGASTEVF | 0.24356261 |
| GFSNSGS | ASSRPQGASTEVF | 0.40895647 |
| GGNGGSGGTSV | ASSRPQGASTEVF | 0.26467943 |
| GTSVATGGAGN | ASSRPQGASTEVF | 0.16693145 |
| NGGNGGSGGTS | ASSRPQGASTEVF | 0.46600324 |
| AVVKFPH | ASSRPQGASTEVF | 0.7537397 |
| TPPPPPP | ASSRPQGASTEVF | 0.2385706 |
| AAPAAAAAPA | ASSRPQGASTEVF | 0.41205552 |
| IALNEDL | ASSRPQGASTEVF | 0.3258053 |
| IVNVSSM | ASSRPQGASTEVF | 0.6224555 |
| TGIPPAPR | ASSRPQGASTEVF | 0.19959147 |
| FDLGGGTF | ASSRPQGASTEVF | 0.16592203 |
| FDLGGGTFDVS | ASSRPQGASTEVF | 0.26281267 |
| NEPTAAA | ASSRPQGASTEVF | 0.16864672 |
| NEPTAAALA | ASSRPQGASTEVF | 0.41821766 |
| GSIINISSI | ASSRPQGASTEVF | 0.2247733 |
| VAPGNAGTA | ASSRPQGASTEVF | 0.78443396 |
| TGGGASLE | ASSRPQGASTEVF | 0.5126785 |
| VAVYNHYKRI | ASSRPQGASTEVF | 0.29285318 |
| TDPAPATT | ASSRPQGASTEVF | 0.1634185 |
| TNRPDILDPALLRPG | ASSRPQGASTEVF | 0.2739334 |
| LTTTTTS | ASSRPQGASTEVF | 0.4760487 |
| PTGSGKTL | ASSRPQGASTEVF | 0.15751271 |
| TGSGKTL | ASSRPQGASTEVF | 0.6318325 |
| TIGHVDHGK | ASSRPQGASTEVF | 0.18289866 |
| TIGHVDHGKTT | ASSRPQGASTEVF | 0.3072977 |
| GHVDHGKTT | ASSRPQGASTEVF | 0.48354918 |
| AQFEHTLL | ASSRPQGASTEVF | 0.7254894 |
| IVNQDEL | ASSRPQGASTEVF | 0.24755247 |
| TNGSQFF | ASSRPQGASTEVF | 0.24447042 |
| TGKGIVE | ASSRPQGASTEVF | 0.16169305 |
| LISNASDA | ASSRPQGASTEVF | 0.54207855 |
| AELRTLGELTKIAKAHGAQ | ASSRPQGASTEVF | 0.59678197 |
| APNAVAAAEPSW | ASSRPQGASTEVF | 0.19322062 |
| APQVAGAATAVL | ASSRPQGASTEVF | 0.72260284 |
| ARRDPAAAAAILPTDA | ASSRPQGASTEVF | 0.31928962 |
| DAAARDVLAEAGLAEYF | ASSRPQGASTEVF | 0.17062329 |

| | | |
|---|---|---|
| DDQIVLSAKANLAKVEQ | ASSRPQGASTEVF | 0.2752073 |
| DELLPDTGEDSDEDGH | ASSRPQGASTEVF | 0.5411029 |
| ELPEAPANRATVPQD | ASSRPQGASTEVF | 0.6833946 |
| FGQSKIERVTGIG | ASSRPQGASTEVF | 0.486747 |
| GAGGTGSTAGGAGGAGGAGGLY | ASSRPQGASTEVF | 0.63341177 |
| GHGGTGAAGGNGGTGGAGGSGI | ASSRPQGASTEVF | 0.24010761 |
| GNGGNGGAGGEGGPGGAGGAGA | ASSRPQGASTEVF | 0.6612302 |
| GSTAGNGGNGGNATNGTIAGQP | ASSRPQGASTEVF | 0.7411643 |
| GTGGDGGNGGNGGTGG | ASSRPQGASTEVF | 0.3747155 |
| GVRRPAGIPADAPV | ASSRPQGASTEVF | 0.23147449 |
| GYRQVTDAHLLTLARR | ASSRPQGASTEVF | 0.25493747 |
| IAKMGQDGHDRG | ASSRPQGASTEVF | 0.35365063 |
| KAEPSIEIQYT | ASSRPQGASTEVF | 0.5216515 |
| KPVRIRMAADEDTVAVT | ASSRPQGASTEVF | 0.20463297 |
| MPPRIAGMRLLVIKPEP | ASSRPQGASTEVF | 0.36570916 |
| MKFYAVALFRSRGVPPTR | ASSRPQGASTEVF | 0.37627846 |
| NAGGFGILGAVAHSP | ASSRPQGASTEVF | 0.15722317 |
| NEGREITLAMAEM | ASSRPQGASTEVF | 0.46938366 |
| NELVEWSHPLGHRWRWE | ASSRPQGASTEVF | 0.7259627 |
| NLGAGGTGGNGGNPG | ASSRPQGASTEVF | 0.33137578 |
| NTEWPKWETVMAADPVD | ASSRPQGASTEVF | 0.21213269 |
| PASAAPAETQP | ASSRPQGASTEVF | 0.63861716 |
| QQSSDGKNGVRAAEITGE | ASSRPQGASTEVF | 0.7547022 |
| RAPDAVTTSWWKE | ASSRPQGASTEVF | 0.6064863 |
| SGFGNVGAHTSGWFNQ | ASSRPQGASTEVF | 0.15505381 |
| SGVAGAVGQAASVGGLKV | ASSRPQGASTEVF | 0.14160778 |
| SNGGAGTGSAGGNGGT | ASSRPQGASTEVF | 0.28364593 |
| TGGAGGEGGAAG | ASSRPQGASTEVF | 0.38509712 |
| TGGDGGAAGAGGAPGHG | ASSRPQGASTEVF | 0.23030871 |
| TMPPLLAAIANAGRHGV | ASSRPQGASTEVF | 0.18450521 |
| TSSTAGNSGGA | ASSRPQGASTEVF | 0.5451696 |
| VPATVPVGILC | ASSRPQGASTEVF | 0.54767334 |
| VQRPDSGGVVLTGRIS | ASSRPQGASTEVF | 0.19149397 |
| WTPNPVPATKGVPAGLVQAT | ASSRPQGASTEVF | 0.16153689 |
| YALKRALAKFSDRFDVVII | ASSRPQGASTEVF | 0.4223355 |
| YLIGSVIV | ASSRPQGASTEVF | 0.6192986 |
| YGQLVAVID | ASSRPQGASTEVF | 0.17495796 |
| VTVGGMTLPA | ASSRPQGASTEVF | 0.23824728 |
| VLALPAV | ASSRPQGASTEVF | 0.25832346 |
| VIPPTLNYET | ASSRPQGASTEVF | 0.26308072 |
| VGMGGMHPSSG | ASSRPQGASTEVF | 0.3943681 |
| VGLAGGAATA | ASSRPQGASTEVF | 0.17994694 |
| VGGAGGAGGSGP | ASSRPQGASTEVF | 0.56543183 |
| VFAAFMV | ASSRPQGASTEVF | 0.15945242 |
| TLYDKLR | ASSRPQGASTEVF | 0.47610837 |
| TLAVFTVL | ASSRPQGASTEVF | 0.38518858 |
| TGSVQPARV | ASSRPQGASTEVF | 0.36137813 |
| TGGGAGSGGT | ASSRPQGASTEVF | 0.2760627 |
| SHLETSL | ASSRPQGASTEVF | 0.18854034 |
| SFTTGMT | ASSRPQGASTEVF | 0.17811364 |
| QVEVAKRMGVRQPTVS | ASSRPQGASTEVF | 0.14659443 |

| | | |
|---|---|---|
| QGSGGPALAM | ASSRPQGASTEVF | 0.6756637 |
| MLAPSNP | ASSRPQGASTEVF | 0.13492583 |
| LVNNAAI | ASSRPQGASTEVF | 0.5769272 |
| LTGEINTLA | ASSRPQGASTEVF | 0.27585763 |
| LSIFGPV | ASSRPQGASTEVF | 0.17216358 |
| LLALLANL | ASSRPQGASTEVF | 0.334328 |
| LISNASDALDKLR | ASSRPQGASTEVF | 0.15220226 |
| LINEAVP | ASSRPQGASTEVF | 0.19711497 |
| LGYAFSGL | ASSRPQGASTEVF | 0.6417023 |
| LDFVAVAL | ASSRPQGASTEVF | 0.4720531 |
| LAQPGAE | ASSRPQGASTEVF | 0.19576593 |
| ISNLGAVTGF | ASSRPQGASTEVF | 0.39723998 |
| HQEDPGRI | ASSRPQGASTEVF | 0.3863489 |
| GTPGEPTP | ASSRPQGASTEVF | 0.19903803 |
| GFFNSGS | ASSRPQGASTEVF | 0.35149544 |
| GAGGGGLTAGAN | ASSRPQGASTEVF | 0.5598439 |
| FVDALVNM | ASSRPQGASTEVF | 0.36663896 |
| FLTVAPEL | ASSRPQGASTEVF | 0.15835632 |
| FETLTAL | ASSRPQGASTEVF | 0.3410061 |
| DVINPIEL | ASSRPQGASTEVF | 0.18739024 |
| APVSGAPV | ASSRPQGASTEVF | 0.35842332 |
| ALVGGLAW | ASSRPQGASTEVF | 0.25276026 |
| ALAAAAATAI | ASSRPQGASTEVF | 0.30686128 |
| AFGLGLLL | ASSRPQGASTEVF | 0.3894474 |
| AFFAAASL | ASSRPQGASTEVF | 0.16643876 |
| AAAALPAFSPP | ASSRPQGASTEVF | 0.19125642 |
| VVLAPAL | ASSRPQGASTEVF | 0.28020552 |
| TIGHVDHGKTTL | ASSRPQGASTEVF | 0.26216465 |
| LISNASDALDK | ASSRPQGASTEVF | 0.40029985 |
| LISNASDAL | ASSRPQGASTEVF | 0.17556806 |
| ISNASDALDK | ASSRPQGASTEVF | 0.3338278 |
| NQAETLVYQTE | ASSRPQGASTEVF | 0.2422598 |
| TGGTGGTAGAG | ASSRPQGASTEVF | 0.19088228 |
| TGSTAGGAGGA | ASSRPQGASTEVF | 0.34630543 |
| DPLAVPVAAGPV | ASSRPQGASTEVF | 0.23994151 |
| GGTGGTPGNGGN | ASSRPQGASTEVF | 0.17919493 |
| PAGLAGVDPATL | ASSRPQGASTEVF | 0.46570432 |
| GNNLVVEAPHAGASV | ASSRPQGASTEVF | 0.18043551 |
| KVTRIMSASTLEQPA | ASSRPQGASTEVF | 0.3436556 |
| LGFALPIALPRRTVV | ASSRPQGASTEVF | 0.63943636 |
| TRLGNLTARKLVVLV | ASSRPQGASTEVF | 0.21621826 |
| WSQQRRLIYFPSAGP | ASSRPQGASTEVF | 0.37888378 |
| GPNGNPDPMAAAVDIR | ASSRPQGASTEVF | 0.25811872 |
| TQRCVSVADLRAAASR | ASSRPQGASTEVF | 0.48178178 |
| QFVLDVAEHPVEVLTETV | ASSRPQGASTEVF | 0.23127508 |
| GGAGGTGGAGGNGGTQAGDG | ASSRPQGASTEVF | 0.19258052 |
| PLLATLRAIAVDVDAKPAQV | ASSRPQGASTEVF | 0.4998516 |
| APKFRTGGYMIDFWGVGYQVAK | ASSRPQGASTEVF | 0.34490803 |
| GAGGAGGAGGAAGGPGGTGGTG | ASSRPQGASTEVF | 0.58504796 |
| IGALGLLATDVAPAGSVAFGVA | ASSRPQGASTEVF | 0.44441864 |
| AGGAGGIGGNGGAGGAGGVLMG | ASSRPQGASTEVF | 0.5520932 |

| | | |
|---|---|---|
| ADELIGLL | ASSRPQGASTEVF | 0.47640258 |
| ADRALKA | ASSRPQGASTEVF | 0.21521275 |
| AGAVGQVGGAGGA | ASSRPQGASTEVF | 0.16013877 |
| AGGGGGLGGAP | ASSRPQGASTEVF | 0.20558734 |
| AGGLLYGNGGAG | ASSRPQGASTEVF | 0.18293902 |
| AGLESIGL | ASSRPQGASTEVF | 0.19508067 |
| AGLKTGII | ASSRPQGASTEVF | 0.16488044 |
| ASLAALLL | ASSRPQGASTEVF | 0.5851494 |
| ASVINISSI | ASSRPQGASTEVF | 0.76314247 |
| ATPPGPA | ASSRPQGASTEVF | 0.18115523 |
| ATVASALV | ASSRPQGASTEVF | 0.6750184 |
| ATVASLIL | ASSRPQGASTEVF | 0.16930617 |
| AVNGPVLIRH | ASSRPQGASTEVF | 0.43711412 |
| DGAAGAVGI | ASSRPQGASTEVF | 0.6386026 |
| DPLLHGI | ASSRPQGASTEVF | 0.16796033 |
| ELGAALGAL | ASSRPQGASTEVF | 0.33976173 |
| ELGRTQAV | ASSRPQGASTEVF | 0.17351592 |
| FVAGGYLS | ASSRPQGASTEVF | 0.20270246 |
| GASGGAGGAGGW | ASSRPQGASTEVF | 0.65914106 |
| GGAGAGGDGGAGGI | ASSRPQGASTEVF | 0.24829191 |
| GTGHNGSQPGMGGNG | ASSRPQGASTEVF | 0.3976214 |
| GYAAASASAS | ASSRPQGASTEVF | 0.22166798 |
| HQVDVAAVQQG | ASSRPQGASTEVF | 0.17168921 |
| IGAAIAHT | ASSRPQGASTEVF | 0.30869257 |
| IGMIEVAQL | ASSRPQGASTEVF | 0.6415655 |
| IRLAPEHTSAMD | ASSRPQGASTEVF | 0.22736341 |
| IYNVSVL | ASSRPQGASTEVF | 0.7092352 |
| KQGFLGM | ASSRPQGASTEVF | 0.1881022 |
| LESVITQL | ASSRPQGASTEVF | 0.71157795 |
| LGLGALVVV | ASSRPQGASTEVF | 0.7093965 |
| LIGNGGNGGSGGI | ASSRPQGASTEVF | 0.33333522 |
| LNAINAPTL | ASSRPQGASTEVF | 0.23345141 |
| LQLLNTGV | ASSRPQGASTEVF | 0.58837926 |
| LQTSIEF | ASSRPQGASTEVF | 0.3608657 |
| MSFAVSEL | ASSRPQGASTEVF | 0.58700407 |
| NTGALGL | ASSRPQGASTEVF | 0.58691573 |
| PPGAGGAGGAGTT | ASSRPQGASTEVF | 0.5906669 |
| QLVGLGI | ASSRPQGASTEVF | 0.7658062 |
| RFLEQVHQL | ASSRPQGASTEVF | 0.73180914 |
| RGVGYVLR | ASSRPQGASTEVF | 0.7049461 |
| SATGVFAGVI | ASSRPQGASTEVF | 0.6418449 |
| SLPAITLP | ASSRPQGASTEVF | 0.44270396 |
| STALAFTV | ASSRPQGASTEVF | 0.69979405 |
| SVLTVGLGK | ASSRPQGASTEVF | 0.5865452 |
| TALAGAGSGI | ASSRPQGASTEVF | 0.7152183 |
| TGLPFTGI | ASSRPQGASTEVF | 0.74694 |
| VAEHVTV | ASSRPQGASTEVF | 0.6369413 |
| VDIVLNSL | ASSRPQGASTEVF | 0.7594414 |
| VLTGGGAVL | ASSRPQGASTEVF | 0.75437754 |
| VNVGLAYGI | ASSRPQGASTEVF | 0.4870873 |
| VSSAAVGAV | ASSRPQGASTEVF | 0.6943104 |

| | | |
|---|---|---|
| VTSVYEAGV | ASSRPQGASTEVF | 0.6801908 |
| WVALDRGA | ASSRPQGASTEVF | 0.6858904 |
| YLVGSVVI | ASSRPQGASTEVF | 0.44321597 |
| GPPGTGKTLL | ASSRPQGASTEVF | 0.7408641 |
| GVGMTKF | ASSRPQGASTEVF | 0.65568787 |
| IGIGGSDL | ASSRPQGASTEVF | 0.23816717 |
| IGTLLAEL | ASSRPQGASTEVF | 0.7144096 |
| IYTSGTT | ASSRPQGASTEVF | 0.70548713 |
| LSFPAGV | ASSRPQGASTEVF | 0.66910124 |
| LYTSGTT | ASSRPQGASTEVF | 0.7472271 |
| VGAFPFGF | ASSRPQGASTEVF | 0.4762132 |
| VGAGTVEF | ASSRPQGASTEVF | 0.6214348 |
| VGKSTLLL | ASSRPQGASTEVF | 0.75127053 |
| AALAHAGGGIV | ASSRPQGASTEVF | 0.7599997 |
| AALVAVSLL | ASSRPQGASTEVF | 0.5268821 |
| AAVSRAVAAL | ASSRPQGASTEVF | 0.7232174 |
| AGGSSGIGGF | ASSRPQGASTEVF | 0.75444734 |
| AGLTVANANG | ASSRPQGASTEVF | 0.7281663 |
| AGLVGAVL | ASSRPQGASTEVF | 0.75118756 |
| ALSQAFTV | ASSRPQGASTEVF | 0.7590638 |
| ALTFSVSL | ASSRPQGASTEVF | 0.6653525 |
| ALTGAALVAL | ASSRPQGASTEVF | 0.7529104 |
| ASFSIGAGVL | ASSRPQGASTEVF | 0.76390123 |
| DGSVYKI | ASSRPQGASTEVF | 0.67534614 |
| EQLAEIV | ASSRPQGASTEVF | 0.7067037 |
| FAVSELTVM | ASSRPQGASTEVF | 0.5850588 |
| GAAPLAAGL | ASSRPQGASTEVF | 0.690377 |
| GADGGDGGSGGAGG | ASSRPQGASTEVF | 0.39551476 |
| GFGLDTGGAGG | ASSRPQGASTEVF | 0.72182995 |
| GLASGGLL | ASSRPQGASTEVF | 0.7542565 |
| GRIAGAAVLL | ASSRPQGASTEVF | 0.67856383 |
| GSAGSLNTSF | ASSRPQGASTEVF | 0.61070883 |
| GTGQIGI | ASSRPQGASTEVF | 0.5494236 |
| GTLIALIL | ASSRPQGASTEVF | 0.7149756 |
| IAATMGFVVG | ASSRPQGASTEVF | 0.6370108 |
| IFAQALL | ASSRPQGASTEVF | 0.36465505 |
| IGAAVALGF | ASSRPQGASTEVF | 0.70203257 |
| IGLTPLAL | ASSRPQGASTEVF | 0.7506161 |
| ILGSNTSTL | ASSRPQGASTEVF | 0.7470325 |
| IVFSLTVL | ASSRPQGASTEVF | 0.4605164 |
| LASKATGF | ASSRPQGASTEVF | 0.7629237 |
| LASVALIL | ASSRPQGASTEVF | 0.72775096 |
| LFAVGLTL | ASSRPQGASTEVF | 0.7604274 |
| LFNGGIGTYI | ASSRPQGASTEVF | 0.7113497 |
| LTLGAGII | ASSRPQGASTEVF | 0.6139997 |
| LVAAALGDI | ASSRPQGASTEVF | 0.7336981 |
| NTAFVQLGI | ASSRPQGASTEVF | 0.6825981 |
| QLFSGAEL | ASSRPQGASTEVF | 0.56508386 |
| SAATFITL | ASSRPQGASTEVF | 0.74475324 |
| SNYGLANLGV | ASSRPQGASTEVF | 0.68513453 |
| SYGGAYAVM | ASSRPQGASTEVF | 0.6314432 |

| | | |
|---|---|---|
| TTGVGGLTLG | ASSRPQGASTEVF | 0.6450914 |
| TTQTIYI | ASSRPQGASTEVF | 0.7546381 |
| VAAGAGLGAVI | ASSRPQGASTEVF | 0.62142015 |
| VAAVVTLL | ASSRPQGASTEVF | 0.51921594 |
| VGFLGGLI | ASSRPQGASTEVF | 0.5917106 |
| VGVIPILL | ASSRPQGASTEVF | 0.7106443 |
| VIFASSLI | ASSRPQGASTEVF | 0.75479096 |
| VKGGDGGAAAG | ASSRPQGASTEVF | 0.7261781 |
| VTKTFTNI | ASSRPQGASTEVF | 0.6424085 |
| VVAAAILL | ASSRPQGASTEVF | 0.73110914 |
| YAALDVEL | ASSRPQGASTEVF | 0.7223549 |
| LASVALLL | ASSRPQGASTEVF | 0.7150273 |
| LDAVHRV | ASSRPQGASTEVF | 0.73506784 |
| QGLSSSIF | ASSRPQGASTEVF | 0.6408161 |
| RGTFANIRL | ASSRPQGASTEVF | 0.7548661 |
| RQLWWGHRI | ASSRPQGASTEVF | 0.7272447 |
| YGPPGTGKTLL | ASSRPQGASTEVF | 0.7403593 |
| AALVGAPANVA | ASSRPQGASTEVF | 0.75351846 |
| AATFRRTE | ASSRPQGASTEVF | 0.7339946 |
| AGALGGGI | ASSRPQGASTEVF | 0.7150226 |
| AGGAGGAGGIGAG | ASSRPQGASTEVF | 0.35817987 |
| AGSGALGL | ASSRPQGASTEVF | 0.70921636 |
| AQIADQI | ASSRPQGASTEVF | 0.72676563 |
| ASAVAVTYL | ASSRPQGASTEVF | 0.72260046 |
| ASTVLGGTGG | ASSRPQGASTEVF | 0.73751515 |
| AVGGEPAK | ASSRPQGASTEVF | 0.72684777 |
| CEQGTFGGTC | ASSRPQGASTEVF | 0.653504 |
| DEGKAPAG | ASSRPQGASTEVF | 0.7543378 |
| DTVVFPAL | ASSRPQGASTEVF | 0.7511932 |
| FEGITMAG | ASSRPQGASTEVF | 0.512821 |
| FPLVVQD | ASSRPQGASTEVF | 0.6937628 |
| GAFTTPQ | ASSRPQGASTEVF | 0.767191 |
| GFGGFGTAGGVGGA | ASSRPQGASTEVF | 0.709386 |
| GFLAQFL | ASSRPQGASTEVF | 0.6877142 |
| GGAGGAGGNAGVA | ASSRPQGASTEVF | 0.6213559 |
| GGVGTAGAGGAGGAGG | ASSRPQGASTEVF | 0.74271643 |
| GNGGGGDAGNAGSGGN | ASSRPQGASTEVF | 0.6136645 |
| GTGGAGGAGNDAGS | ASSRPQGASTEVF | 0.766693 |
| GTTNTGWFNS | ASSRPQGASTEVF | 0.2643438 |
| IFMGPLFTL | ASSRPQGASTEVF | 0.63278717 |
| ILAHAIEAI | ASSRPQGASTEVF | 0.60550845 |
| IRSAIREAL | ASSRPQGASTEVF | 0.31525046 |
| ITFAGGG | ASSRPQGASTEVF | 0.7534126 |
| ITGMSGHQEV | ASSRPQGASTEVF | 0.65473235 |
| IVAGLACTL | ASSRPQGASTEVF | 0.73385197 |
| IVGGGLAGA | ASSRPQGASTEVF | 0.72327125 |
| IVTGVLNI | ASSRPQGASTEVF | 0.755688 |
| LAQQGCI | ASSRPQGASTEVF | 0.7185855 |
| LCLHVEN | ASSRPQGASTEVF | 0.6916859 |
| LDAISDGS | ASSRPQGASTEVF | 0.51404566 |
| LGGFAAML | ASSRPQGASTEVF | 0.745264 |

| | | |
|---|---|---|
| LGGKNPMI | ASSRPQGASTEVF | 0.761433 |
| LGPKGVAAIL | ASSRPQGASTEVF | 0.73445725 |
| LMVGVSI | ASSRPQGASTEVF | 0.44190955 |
| LVGAAFVL | ASSRPQGASTEVF | 0.6726734 |
| LVLAVAPL | ASSRPQGASTEVF | 0.7163557 |
| MAAIAVIDAA | ASSRPQGASTEVF | 0.26191342 |
| NNGINAELQ | ASSRPQGASTEVF | 0.26568657 |
| RGEPLVLVHG | ASSRPQGASTEVF | 0.70801985 |
| RGGAGGAAGLI | ASSRPQGASTEVF | 0.66280335 |
| RSYRMHPAV | ASSRPQGASTEVF | 0.7518661 |
| SAAHLLAGQLA | ASSRPQGASTEVF | 0.75215197 |
| SAAVGSY | ASSRPQGASTEVF | 0.67352194 |
| SPHAVLHRI | ASSRPQGASTEVF | 0.74622655 |
| SQIAKVL | ASSRPQGASTEVF | 0.71583915 |
| TALGHLVP | ASSRPQGASTEVF | 0.21123037 |
| TAPTTSGAA | ASSRPQGASTEVF | 0.6755609 |
| TDGDDGNGGSPGT | ASSRPQGASTEVF | 0.5052209 |
| TDGLGEN | ASSRPQGASTEVF | 0.7580422 |
| TGGPPLPGQA | ASSRPQGASTEVF | 0.65168077 |
| TIPAITI | ASSRPQGASTEVF | 0.6533586 |
| TIPGFTI | ASSRPQGASTEVF | 0.69691133 |
| TSIASGALA | ASSRPQGASTEVF | 0.69919366 |
| TVTLPGAAI | ASSRPQGASTEVF | 0.70151204 |
| VASGLSAALP | ASSRPQGASTEVF | 0.5971768 |
| VGAGVIGI | ASSRPQGASTEVF | 0.2720787 |
| VGPKRFGV | ASSRPQGASTEVF | 0.74692523 |
| VNVAFPD | ASSRPQGASTEVF | 0.4653308 |
| VSMTNSGVS | ASSRPQGASTEVF | 0.56975746 |
| VTDVLGDLSV | ASSRPQGASTEVF | 0.32417443 |
| VVVVGAGFAGL | ASSRPQGASTEVF | 0.2634576 |
| VYTVGTVV | ASSRPQGASTEVF | 0.41198948 |
| WTTFSGP | ASSRPQGASTEVF | 0.7406042 |
| YDELVEAF | ASSRPQGASTEVF | 0.7000055 |
| YGTGGAGGNGG | ASSRPQGASTEVF | 0.69535106 |
| YTNVMAV | ASSRPQGASTEVF | 0.7696432 |
| FLEQVHQL | ASSRPQGASTEVF | 0.686443 |
| GLSLLAVAV | ASSRPQGASTEVF | 0.6778872 |
| LEQVHQL | ASSRPQGASTEVF | 0.57857287 |
| LNAAGAI | ASSRPQGASTEVF | 0.5914148 |
| PAVHASL | ASSRPQGASTEVF | 0.6193812 |
| TELEDDA | ASSRPQGASTEVF | 0.60986334 |
| TPPGPPN | ASSRPQGASTEVF | 0.7656418 |
| TPPPPPPP | ASSRPQGASTEVF | 0.6478636 |
| TQSPALI | ASSQVPVNTGQLY | 0.54031646 |
| EFAAAFPQ | ASSQVPVNTGQLY | 0.69587874 |
| LLSGNLDVL | ASSQVPVNTGQLY | 0.7605821 |
| GPANLGNYNI | ASSQVPVNTGQLY | 0.7698915 |
| TLGPTLP | ASSQVPVNTGQLY | 0.60199946 |
| VYAYSSCCK | ASSQVPVNTGQLY | 0.66771495 |
| APAPAPAPNI | ASSQVPVNTGQLY | 0.6791275 |
| PGGPVPIP | ASSQVPVNTGQLY | 0.7068005 |

| | | |
|---|---|---|
| FLTVALEP | ASSQVPVNTGQLY | 0.65013 |
| SQYFGANL | ASSQVPVNTGQLY | 0.48998687 |
| GGVKPAA | ASSQVPVNTGQLY | 0.7417835 |
| GGTGGAGGAGGW | ASSQVPVNTGQLY | 0.7481679 |
| GVGGVAGLGGAGG | ASSQVPVNTGQLY | 0.6709354 |
| VLLVLATTI | ASSQVPVNTGQLY | 0.52078176 |
| IGIFAAML | ASSQVPVNTGQLY | 0.6904632 |
| AVGDLID | ASSQVPVNTGQLY | 0.6177083 |
| NISPLPL | ASSQVPVNTGQLY | 0.6564028 |
| VTAVFVAI | ASSQVPVNTGQLY | 0.70177335 |
| SGDDNDAAL | ASSQVPVNTGQLY | 0.53530157 |
| AVAALAHS | ASSQVPVNTGQLY | 0.7046263 |
| GGVTVMATP | ASSQVPVNTGQLY | 0.7702881 |
| AGLAANVRD | ASSQVPVNTGQLY | 0.6720922 |
| TNRPDLIDPALLRPG | ASSQVPVNTGQLY | 0.7398334 |
| LANLGVTAL | ASSQVPVNTGQLY | 0.7565527 |
| TGERIEL | ASSQVPVNTGQLY | 0.73553133 |
| NGEELAA | ASSQVPVNTGQLY | 0.7570323 |
| VLSLVWG | ASSQVPVNTGQLY | 0.7318394 |
| IGYALWL | ASSQVPVNTGQLY | 0.6777558 |
| LSLLLAPL | ASSQVPVNTGQLY | 0.7144022 |
| TLDEALK | ASSQVPVNTGQLY | 0.73387283 |
| TDGAGTAEGG | ASSQVPVNTGQLY | 0.6981666 |
| NAPGGAGGNGG | ASSQVPVNTGQLY | 0.68689275 |
| VSVFMGMGGNF | ASSQVPVNTGQLY | 0.6380915 |
| LASYLAAL | ASSQVPVNTGQLY | 0.7572585 |
| LVGGTPAL | ASSQVPVNTGQLY | 0.71208715 |
| VRNADVL | ASSQVPVNTGQLY | 0.72987396 |
| TCSSPFLVSAM | ASSQVPVNTGQLY | 0.42783126 |
| VHITDAE | ASSQVPVNTGQLY | 0.4886807 |
| EYAAPSF | ASSQVPVNTGQLY | 0.7420522 |
| VVDAFHGGL | ASSQVPVNTGQLY | 0.22528513 |
| ASVVDPQ | ASSQVPVNTGQLY | 0.26271445 |
| SYNEFGGLS | ASSQVPVNTGQLY | 0.7200266 |
| DPDGSVY | ASSQVPVNTGQLY | 0.7584255 |
| MVSTIHATE | ASSQVPVNTGQLY | 0.23139887 |
| LAAGVADPV | ASSQVPVNTGQLY | 0.5985683 |
| TKEQIAA | ASSQVPVNTGQLY | 0.5182431 |
| SLASPAAL | ASSQVPVNTGQLY | 0.732031 |
| IGLVFYGV | ASSQVPVNTGQLY | 0.69987285 |
| FLLDASNL | ASSQVPVNTGQLY | 0.43678284 |
| YQVGNFP | ASSQVPVNTGQLY | 0.73777103 |
| ESLLSQST | ASSQVPVNTGQLY | 0.43740714 |
| LFTVGFAAVG | ASSQVPVNTGQLY | 0.43245792 |
| ATPPTPT | ASSQVPVNTGQLY | 0.7545793 |
| TDLATLG | ASSQVPVNTGQLY | 0.7412846 |
| SAGGTGR | ASSQVPVNTGQLY | 0.6156013 |
| GFSNSGS | ASSQVPVNTGQLY | 0.74430716 |
| GGNGGSGGTSV | ASSQVPVNTGQLY | 0.6546792 |
| GTSVATGGAGN | ASSQVPVNTGQLY | 0.77194923 |
| NGGNGGSGGTS | ASSQVPVNTGQLY | 0.6743803 |

| | | |
|---|---|---|
| AVVKFPH | ASSQVPVNTGQLY | 0.46348634 |
| TPPPPPP | ASSQVPVNTGQLY | 0.7482528 |
| AAPAAAAAPA | ASSQVPVNTGQLY | 0.74061215 |
| IALNEDL | ASSQVPVNTGQLY | 0.69519025 |
| IVNVSSM | ASSQVPVNTGQLY | 0.72810805 |
| TGIPPAPR | ASSQVPVNTGQLY | 0.6050894 |
| FDLGGGTF | ASSQVPVNTGQLY | 0.68515 |
| FDLGGGTFDVS | ASSQVPVNTGQLY | 0.75118685 |
| NEPTAAA | ASSQVPVNTGQLY | 0.75424474 |
| NEPTAAALA | ASSQVPVNTGQLY | 0.56727487 |
| GSIINISSI | ASSQVPVNTGQLY | 0.7510706 |
| VAPGNAGTA | ASSQVPVNTGQLY | 0.68104565 |
| TGGGASLE | ASSQVPVNTGQLY | 0.6145785 |
| VAVYNHYKRI | ASSQVPVNTGQLY | 0.7697221 |
| TDPAPATT | ASSQVPVNTGQLY | 0.6800096 |
| TNRPDILDPALLRPG | ASSQVPVNTGQLY | 0.7419002 |
| LTTTTTS | ASSQVPVNTGQLY | 0.7447635 |
| PTGSGKTL | ASSQVPVNTGQLY | 0.7056199 |
| TGSGKTL | ASSQVPVNTGQLY | 0.5063078 |
| TIGHVDHGK | ASSQVPVNTGQLY | 0.738245 |
| TIGHVDHGKTT | ASSQVPVNTGQLY | 0.7188902 |
| GHVDHGKTT | ASSQVPVNTGQLY | 0.74450445 |
| AQFEHTLL | ASSQVPVNTGQLY | 0.71537924 |
| IVNQDEL | ASSQVPVNTGQLY | 0.5913526 |
| TNGSQFF | ASSQVPVNTGQLY | 0.7321528 |
| TGKGIVE | ASSQVPVNTGQLY | 0.76097655 |
| LISNASDA | ASSQVPVNTGQLY | 0.7207025 |
| AELRTLGELTKIAKAHGAQ | ASSQVPVNTGQLY | 0.7307631 |
| APNAVAAAEPSW | ASSQVPVNTGQLY | 0.75950485 |
| APQVAGAATAVL | ASSQVPVNTGQLY | 0.5925092 |
| ARRDPAAAAAILPTDA | ASSQVPVNTGQLY | 0.67224073 |
| DAAARDVLAEAGLAEYF | ASSQVPVNTGQLY | 0.75236845 |
| DDQIVLSAKANLAKVEQ | ASSQVPVNTGQLY | 0.7561381 |
| DELLPDTGEDSDEDGH | ASSQVPVNTGQLY | 0.67232025 |
| ELPEAPANRATVPQD | ASSQVPVNTGQLY | 0.757335 |
| FGQSKIERVTGIG | ASSQVPVNTGQLY | 0.66852 |
| GAGGTGSTAGGAGGAGGAGGLY | ASSQVPVNTGQLY | 0.7378953 |
| GHGGTGAAGGNGGTGGAGGSGI | ASSQVPVNTGQLY | 0.690047 |
| GNGGNGGAGGEGGPGGAGGAGA | ASSQVPVNTGQLY | 0.7287743 |
| GSTAGNGGNGGNATNGTIAGQP | ASSQVPVNTGQLY | 0.65227926 |
| GTGGDGGNGGNGGTGG | ASSQVPVNTGQLY | 0.7438947 |
| GVRRPAGIPADAPV | ASSQVPVNTGQLY | 0.74814963 |
| GYRQVTDAHLLTLARR | ASSQVPVNTGQLY | 0.74572086 |
| IAKMGQDGHDRG | ASSQVPVNTGQLY | 0.27658746 |
| KAEPSIEIQYT | ASSQVPVNTGQLY | 0.7447841 |
| KPVRIRMAADEDTVAVT | ASSQVPVNTGQLY | 0.72966766 |
| MPPRIAGMRLLVIKPEP | ASSQVPVNTGQLY | 0.7595146 |
| MKFYAVALFRSRGVPPTR | ASSQVPVNTGQLY | 0.6993313 |
| NAGGFGILGAVAHSP | ASSQVPVNTGQLY | 0.7348833 |
| NEGREITLAMAEM | ASSQVPVNTGQLY | 0.67124903 |
| NELVEWSHPLGHRWRWE | ASSQVPVNTGQLY | 0.6525316 |

| | | |
|---|---|---|
| NLGAGGTGGNGGNPG | ASSQVPVNTGQLY | 0.5479549 |
| NTEWPKWETVMAADPVD | ASSQVPVNTGQLY | 0.6707952 |
| PASAAPAETQP | ASSQVPVNTGQLY | 0.67985404 |
| QQSSDGKNGVRAAEITGE | ASSQVPVNTGQLY | 0.6281947 |
| RAPDAVTTSWWKE | ASSQVPVNTGQLY | 0.7334045 |
| SGFGNVGAHTSGWFNQ | ASSQVPVNTGQLY | 0.75209725 |
| SGVAGAVGQAASVGGLKV | ASSQVPVNTGQLY | 0.75821793 |
| SNGGAGTGSAGGNGGT | ASSQVPVNTGQLY | 0.75720054 |
| TGGAGGEGGAAG | ASSQVPVNTGQLY | 0.75970185 |
| TGGDGGAAGAGGAPGHG | ASSQVPVNTGQLY | 0.7391701 |
| TMPPLLAAIANAGRHGV | ASSQVPVNTGQLY | 0.62977445 |
| TSSTAGNSGGA | ASSQVPVNTGQLY | 0.7139833 |
| VPATVPVGILC | ASSQVPVNTGQLY | 0.5347095 |
| VQRPDSGGVVLTGRIS | ASSQVPVNTGQLY | 0.7087514 |
| WTPNPVPATKGVPAGLVQAT | ASSQVPVNTGQLY | 0.65668756 |
| YALKRALAKFSDRFDVVII | ASSQVPVNTGQLY | 0.7260103 |
| YLIGSVIV | ASSQVPVNTGQLY | 0.21114591 |
| YGQLVAVID | ASSQVPVNTGQLY | 0.6975517 |
| VTVGGMTLPA | ASSQVPVNTGQLY | 0.76536167 |
| VLALPAV | ASSQVPVNTGQLY | 0.6373112 |
| VIPPTLNYET | ASSQVPVNTGQLY | 0.72536254 |
| VGMGGMHPSSG | ASSQVPVNTGQLY | 0.7332128 |
| VGLAGGAATA | ASSQVPVNTGQLY | 0.65417147 |
| VGGAGGAGGSGP | ASSQVPVNTGQLY | 0.744615 |
| VFAAFMV | ASSQVPVNTGQLY | 0.6040492 |
| TLYDKLR | ASSQVPVNTGQLY | 0.7253548 |
| TLAVFTVL | ASSQVPVNTGQLY | 0.76523376 |
| TGSVQPARV | ASSQVPVNTGQLY | 0.72482437 |
| TGGGAGSGGT | ASSQVPVNTGQLY | 0.72629493 |
| SHLETSL | ASSQVPVNTGQLY | 0.68979746 |
| SFTTGMT | ASSQVPVNTGQLY | 0.41592672 |
| QVEVAKRMGVRQPTVS | ASSQVPVNTGQLY | 0.73229337 |
| QGSGGPALAM | ASSQVPVNTGQLY | 0.20901124 |
| MLAPSNP | ASSQVPVNTGQLY | 0.7086224 |
| LVNNAAI | ASSQVPVNTGQLY | 0.58207047 |
| LTGEINTLA | ASSQVPVNTGQLY | 0.72420204 |
| LSIFGPV | ASSQVPVNTGQLY | 0.7157576 |
| LLALLANL | ASSQVPVNTGQLY | 0.7385856 |
| LISNASDALDKLR | ASSQVPVNTGQLY | 0.61649334 |
| LINEAVP | ASSQVPVNTGQLY | 0.67965066 |
| LGYAFSGL | ASSQVPVNTGQLY | 0.64460665 |
| LDFVAVAL | ASSQVPVNTGQLY | 0.5178461 |
| LAQPGAE | ASSQVPVNTGQLY | 0.75314 |
| ISNLGAVTGF | ASSQVPVNTGQLY | 0.6899743 |
| HQEDPGRI | ASSQVPVNTGQLY | 0.70130956 |
| GTPGEPTP | ASSQVPVNTGQLY | 0.7549584 |
| GFFNSGS | ASSQVPVNTGQLY | 0.7196802 |
| GAGGGGLTAGAN | ASSQVPVNTGQLY | 0.41668886 |
| FVDALVNM | ASSQVPVNTGQLY | 0.7093023 |
| FLTVAPEL | ASSQVPVNTGQLY | 0.70701265 |
| FETLTAL | ASSQVPVNTGQLY | 0.6645307 |

| | | |
|---|---|---|
| DVINPIEL | ASSQVPVNTGQLY | 0.22380352 |
| APVSGAPV | ASSQVPVNTGQLY | 0.73851085 |
| ALVGGLAW | ASSQVPVNTGQLY | 0.61062884 |
| ALAAAAATAI | ASSQVPVNTGQLY | 0.6622485 |
| AFGLGLLL | ASSQVPVNTGQLY | 0.6639354 |
| AFFAAASL | ASSQVPVNTGQLY | 0.7425536 |
| AAAALPAFSPP | ASSQVPVNTGQLY | 0.5235152 |
| VVLAPAL | ASSQVPVNTGQLY | 0.650069 |
| TIGHVDHGKTTL | ASSQVPVNTGQLY | 0.73985255 |
| LISNASDALDK | ASSQVPVNTGQLY | 0.7394962 |
| LISNASDAL | ASSQVPVNTGQLY | 0.7592366 |
| ISNASDALDK | ASSQVPVNTGQLY | 0.7208539 |
| NQAETLVYQTE | ASSQVPVNTGQLY | 0.6929748 |
| TGGTGGTAGAG | ASSQVPVNTGQLY | 0.7254753 |
| TGSTAGGAGGA | ASSQVPVNTGQLY | 0.7002927 |
| DPLAVPVAAGPV | ASSQVPVNTGQLY | 0.4442892 |
| GGTGGTPGNGGN | ASSQVPVNTGQLY | 0.3811897 |
| PAGLAGVDPATL | ASSQVPVNTGQLY | 0.70816344 |
| GNNLVVEAPHAGASV | ASSQVPVNTGQLY | 0.34670606 |
| KVTRIMSASTLEQPA | ASSQVPVNTGQLY | 0.7197976 |
| LGFALPIALPRRTVV | ASSQVPVNTGQLY | 0.75222677 |
| TRLGNLTARKLVVLV | ASSQVPVNTGQLY | 0.7164803 |
| WSQQRRLIYFPSAGP | ASSQVPVNTGQLY | 0.52781653 |
| GPNGNPDPMAAAVDIR | ASSQVPVNTGQLY | 0.697579 |
| TQRCVSVADLRAAASR | ASSQVPVNTGQLY | 0.6702213 |
| QFVLDVAEHPVEVLTETV | ASSQVPVNTGQLY | 0.7318528 |
| GGAGGTGGAGGNGGTQAGDG | ASSQVPVNTGQLY | 0.7493562 |
| PLLATLRAIAVDVDAKPAQV | ASSQVPVNTGQLY | 0.7485812 |
| APKFRTGGYMIDFWGVGYQVAK | ASSQVPVNTGQLY | 0.7511995 |
| GAGGAGGAGGAAGGPGGTGGTG | ASSQVPVNTGQLY | 0.65619075 |
| IGALGLLATDVAPAGSVAFGVA | ASSQVPVNTGQLY | 0.7387264 |
| AGGAGGIGGNGGAGGAGGVLMG | ASSQVPVNTGQLY | 0.669353 |
| ADELIGLL | ASSQVPVNTGQLY | 0.7257713 |
| ADRALKA | ASSQVPVNTGQLY | 0.7057619 |
| AGAVGQVGGAGGA | ASSQVPVNTGQLY | 0.7207513 |
| AGGGGGLGGAP | ASSQVPVNTGQLY | 0.6820388 |
| AGGLLYGNGGAG | ASSQVPVNTGQLY | 0.6556053 |
| AGLESIGL | ASSQVPVNTGQLY | 0.4280085 |
| AGLKTGII | ASSQVPVNTGQLY | 0.71409225 |
| ASLAALLL | ASSQVPVNTGQLY | 0.7343968 |
| ASVINISSI | ASSQVPVNTGQLY | 0.38135082 |
| ATPPGPA | ASSQVPVNTGQLY | 0.6610903 |
| ATVASALV | ASSQVPVNTGQLY | 0.76984787 |
| ATVASLIL | ASSQVPVNTGQLY | 0.76126003 |
| AVNGPVLIRH | ASSQVPVNTGQLY | 0.62627554 |
| DGAAGAVGI | ASSQVPVNTGQLY | 0.5743606 |
| DPLLHGI | ASSQVPVNTGQLY | 0.708777 |
| ELGAALGAL | ASSQVPVNTGQLY | 0.40072194 |
| ELGRTQAV | ASSQVPVNTGQLY | 0.71838 |
| FVAGGYLS | ASSQVPVNTGQLY | 0.7012558 |
| GASGGAGGAGGW | ASSQVPVNTGQLY | 0.7086426 |

| | | |
|---|---|---|
| GGAGAGGDGGAGGI | ASSQVPVNTGQLY | 0.6261064 |
| GTGHNGSQPGMGGNG | ASSQVPVNTGQLY | 0.7210465 |
| GYAAASASAS | ASSQVPVNTGQLY | 0.6834798 |
| HQVDVAAVQQG | ASSQVPVNTGQLY | 0.7165983 |
| IGAAIAHT | ASSQVPVNTGQLY | 0.22507955 |
| IGMIEVAQL | ASSQVPVNTGQLY | 0.7493308 |
| IRLAPEHTSAMD | ASSQVPVNTGQLY | 0.6231843 |
| IYNVSVL | ASSQVPVNTGQLY | 0.61465156 |
| KQGFLGM | ASSQVPVNTGQLY | 0.36714482 |
| LESVITQL | ASSQVPVNTGQLY | 0.71473324 |
| LGLGALVVV | ASSQVPVNTGQLY | 0.7547641 |
| LIGNGGNGGSGGI | ASSQVPVNTGQLY | 0.7489097 |
| LNAINAPTL | ASSQVPVNTGQLY | 0.57152915 |
| LQLLNTGV | ASSQVPVNTGQLY | 0.752482 |
| LQTSIEF | ASSQVPVNTGQLY | 0.7642902 |
| MSFAVSEL | ASSQVPVNTGQLY | 0.6855702 |
| NTGALGL | ASSQVPVNTGQLY | 0.3783688 |
| PPGAGGAGGAGTT | ASSQVPVNTGQLY | 0.719933 |
| QLVGLGI | ASSQVPVNTGQLY | 0.6867918 |
| RFLEQVHQL | ASSQVPVNTGQLY | 0.7419915 |
| RGVGYVLR | ASSQVPVNTGQLY | 0.73660505 |
| SATGVFAGVI | ASSQVPVNTGQLY | 0.5535402 |
| SLPAITLP | ASSQVPVNTGQLY | 0.7415452 |
| STALAFTV | ASSQVPVNTGQLY | 0.75863385 |
| SVLTVGLGK | ASSQVPVNTGQLY | 0.61889696 |
| TALAGAGSGI | ASSQVPVNTGQLY | 0.6647669 |
| TGLPFTGI | ASSQVPVNTGQLY | 0.64187896 |
| VAEHVTV | ASSQVPVNTGQLY | 0.41996711 |
| VDIVLNSL | ASSQVPVNTGQLY | 0.74421394 |
| VLTGGGAVL | ASSQVPVNTGQLY | 0.6676283 |
| VNVGLAYGI | ASSQVPVNTGQLY | 0.61688936 |
| VSSAAVGAV | ASSQVPVNTGQLY | 0.70355403 |
| VTSVYEAGV | ASSQVPVNTGQLY | 0.63206124 |
| WVALDRGA | ASSQVPVNTGQLY | 0.60573316 |
| YLVGSVVI | ASSQVPVNTGQLY | 0.6614121 |
| GPPGTGKTLL | ASSQVPVNTGQLY | 0.7340943 |
| GVGMTKF | ASSQVPVNTGQLY | 0.73089904 |
| IGIGGSDL | ASSQVPVNTGQLY | 0.7195394 |
| IGTLLAEL | ASSQVPVNTGQLY | 0.7122474 |
| IYTSGTT | ASSQVPVNTGQLY | 0.76233876 |
| LSFPAGV | ASSQVPVNTGQLY | 0.52602047 |
| LYTSGTT | ASSQVPVNTGQLY | 0.6177537 |
| VGAFPFGF | ASSQVPVNTGQLY | 0.7325697 |
| VGAGTVEF | ASSQVPVNTGQLY | 0.5042305 |
| VGKSTLLL | ASSQVPVNTGQLY | 0.73987395 |
| AALAHAGGGIV | ASSQVPVNTGQLY | 0.71803755 |
| AALVAVSLL | ASSQVPVNTGQLY | 0.7231592 |
| AAVSRAVAAL | ASSQVPVNTGQLY | 0.70181143 |
| AGGSSGIGGF | ASSQVPVNTGQLY | 0.77035695 |
| AGLTVANANG | ASSQVPVNTGQLY | 0.74432194 |
| AGLVGAVL | ASSQVPVNTGQLY | 0.7099321 |

| | | |
|---|---|---|
| ALSQAFTV | ASSQVPVNTGQLY | 0.72411925 |
| ALTFSVSL | ASSQVPVNTGQLY | 0.541358 |
| ALTGAALVAL | ASSQVPVNTGQLY | 0.7526344 |
| ASFSIGAGVL | ASSQVPVNTGQLY | 0.7271708 |
| DGSVYKI | ASSQVPVNTGQLY | 0.5989131 |
| EQLAEIV | ASSQVPVNTGQLY | 0.69153285 |
| FAVSELTVM | ASSQVPVNTGQLY | 0.7613169 |
| GAAPLAAGL | ASSQVPVNTGQLY | 0.7535124 |
| GADGGDGGSGGAGG | ASSQVPVNTGQLY | 0.37369713 |
| GFGLDTGGAGG | ASSQVPVNTGQLY | 0.66013265 |
| GLASGGLL | ASSQVPVNTGQLY | 0.39406586 |
| GRIAGAAVLL | ASSQVPVNTGQLY | 0.6426881 |
| GSAGSLNTSF | ASSQVPVNTGQLY | 0.4664408 |
| GTGQIGI | ASSQVPVNTGQLY | 0.7349869 |
| GTLIALIL | ASSQVPVNTGQLY | 0.66975605 |
| IAATMGFVVG | ASSQVPVNTGQLY | 0.6966685 |
| IFAQALL | ASSQVPVNTGQLY | 0.7150869 |
| IGAAVALGF | ASSQVPVNTGQLY | 0.75841904 |
| IGLTPLAL | ASSQVPVNTGQLY | 0.64393026 |
| ILGSNTSTL | ASSQVPVNTGQLY | 0.67923176 |
| IVFSLTVL | ASSQVPVNTGQLY | 0.38678202 |
| LASKATGF | ASSQVPVNTGQLY | 0.48085096 |
| LASVALIL | ASSQVPVNTGQLY | 0.6438039 |
| LFAVGLTL | ASSQVPVNTGQLY | 0.68418705 |
| LFNGGIGTYI | ASSQVPVNTGQLY | 0.28141943 |
| LTLGAGII | ASSQVPVNTGQLY | 0.7561629 |
| LVAAALGDI | ASSQVPVNTGQLY | 0.710609 |
| NTAFVQLGI | ASSQVPVNTGQLY | 0.7366675 |
| QLFSGAEL | ASSQVPVNTGQLY | 0.6546327 |
| SAATFITL | ASSQVPVNTGQLY | 0.6710035 |
| SNYGLANLGV | ASSQVPVNTGQLY | 0.6407951 |
| SYGGAYAVM | ASSQVPVNTGQLY | 0.74041003 |
| TTGVGGLTLG | ASSQVPVNTGQLY | 0.6125051 |
| TTQTIYI | ASSQVPVNTGQLY | 0.3971119 |
| VAAGAGLGAVI | ASSQVPVNTGQLY | 0.65258336 |
| VAAVVTLL | ASSQVPVNTGQLY | 0.58992684 |
| VGFLGGLI | ASSQVPVNTGQLY | 0.7602253 |
| VGVIPILL | ASSQVPVNTGQLY | 0.75070626 |
| VIFASSLI | ASSQVPVNTGQLY | 0.721275 |
| VKGGDGGAAAG | ASSQVPVNTGQLY | 0.74665093 |
| VTKTFTNI | ASSQVPVNTGQLY | 0.26906958 |
| VVAAAILL | ASSQVPVNTGQLY | 0.667106 |
| YAALDVEL | ASSQVPVNTGQLY | 0.7602024 |
| LASVALLL | ASSQVPVNTGQLY | 0.72571164 |
| LDAVHRV | ASSQVPVNTGQLY | 0.6940874 |
| QGLSSSIF | ASSQVPVNTGQLY | 0.75348735 |
| RGTFANIRL | ASSQVPVNTGQLY | 0.61537874 |
| RQLWWGHRI | ASSQVPVNTGQLY | 0.64729893 |
| YGPPGTGKTLL | ASSQVPVNTGQLY | 0.70059943 |
| AALVGAPANVA | ASSQVPVNTGQLY | 0.7384845 |
| AATFRRTE | ASSQVPVNTGQLY | 0.7275207 |

| | | |
|---|---|---|
| AGALGGGI | ASSQVPVNTGQLY | 0.5643252 |
| AGGAGGAGGIGAG | ASSQVPVNTGQLY | 0.72359973 |
| AGSGALGL | ASSQVPVNTGQLY | 0.7715485 |
| AQIADQI | ASSQVPVNTGQLY | 0.6807449 |
| ASAVAVTYL | ASSQVPVNTGQLY | 0.74645364 |
| ASTVLGGTGG | ASSQVPVNTGQLY | 0.7271084 |
| AVGGEPAK | ASSQVPVNTGQLY | 0.7153617 |
| CEQGTFGGTC | ASSQVPVNTGQLY | 0.61612314 |
| DEGKAPAG | ASSQVPVNTGQLY | 0.6303344 |
| DTVVFPAL | ASSQVPVNTGQLY | 0.6842701 |
| FEGITMAG | ASSQVPVNTGQLY | 0.60321885 |
| FPLVVQD | ASSQVPVNTGQLY | 0.6900647 |
| GAFTTPQ | ASSQVPVNTGQLY | 0.6811364 |
| GFGGFGTAGGVGGA | ASSQVPVNTGQLY | 0.7742747 |
| GFLAQFL | ASSQVPVNTGQLY | 0.727568 |
| GGAGGAGGNAGVA | ASSQVPVNTGQLY | 0.7275435 |
| GGVGTAGAGGAGGAGG | ASSQVPVNTGQLY | 0.5971074 |
| GNGGGGDAGNAGSGGN | ASSQVPVNTGQLY | 0.7271824 |
| GTGGAGGAGNDAGS | ASSQVPVNTGQLY | 0.7263927 |
| GTTNTGWFNS | ASSQVPVNTGQLY | 0.48167962 |
| IFMGPLFTL | ASSQVPVNTGQLY | 0.7627667 |
| ILAHAIEAI | ASSQVPVNTGQLY | 0.58700526 |
| IRSAIREAL | ASSQVPVNTGQLY | 0.692793 |
| ITFAGGG | ASSQVPVNTGQLY | 0.733737 |
| ITGMSGHQEV | ASSQVPVNTGQLY | 0.7289166 |
| IVAGLACTL | ASSQVPVNTGQLY | 0.68541026 |
| IVGGGLAGA | ASSQVPVNTGQLY | 0.6582179 |
| IVTGVLNI | ASSQVPVNTGQLY | 0.57479227 |
| LAQQGCI | ASSQVPVNTGQLY | 0.74463534 |
| LCLHVEN | ASSQVPVNTGQLY | 0.7506622 |
| LDAISDGS | ASSQVPVNTGQLY | 0.66954124 |
| LGGFAAML | ASSQVPVNTGQLY | 0.75259626 |
| LGGKNPMI | ASSQVPVNTGQLY | 0.7295587 |
| LGPKGVAAIL | ASSQVPVNTGQLY | 0.3737609 |
| LMVGVSI | ASSQVPVNTGQLY | 0.6784946 |
| LVGAAFVL | ASSQVPVNTGQLY | 0.7596799 |
| LVLAVAPL | ASSQVPVNTGQLY | 0.7505076 |
| MAAIAVIDAA | ASSQVPVNTGQLY | 0.76442695 |
| NNGINAELQ | ASSQVPVNTGQLY | 0.7486757 |
| RGEPLVLVHG | ASSQVPVNTGQLY | 0.68068075 |
| RGGAGGAAGLI | ASSQVPVNTGQLY | 0.75895834 |
| RSYRMHPAV | ASSQVPVNTGQLY | 0.77120733 |
| SAAHLLAGQLA | ASSQVPVNTGQLY | 0.7417414 |
| SAAVGSY | ASSQVPVNTGQLY | 0.61916995 |
| SPHAVLHRI | ASSQVPVNTGQLY | 0.56500936 |
| SQIAKVL | ASSQVPVNTGQLY | 0.7322229 |
| TALGHLVP | ASSQVPVNTGQLY | 0.745734 |
| TAPTTSGAA | ASSQVPVNTGQLY | 0.5287726 |
| TDGDDGNGGSPGT | ASSQVPVNTGQLY | 0.71985656 |
| TDGLGEN | ASSQVPVNTGQLY | 0.7193572 |
| TGGPPLPGQA | ASSQVPVNTGQLY | 0.42295688 |

| | | |
|---|---|---|
| TIPAITI | ASSQVPVNTGQLY | 0.75464344 |
| TIPGFTI | ASSQVPVNTGQLY | 0.76427543 |
| TSIASGALA | ASSQVPVNTGQLY | 0.7033819 |
| TVTLPGAAI | ASSQVPVNTGQLY | 0.66433144 |
| VASGLSAALP | ASSQVPVNTGQLY | 0.7608976 |
| VGAGVIGI | ASSQVPVNTGQLY | 0.7732594 |
| VGPKRFGV | ASSQVPVNTGQLY | 0.7564793 |
| VNVAFPD | ASSQVPVNTGQLY | 0.41755253 |
| VSMTNSGVS | ASSQVPVNTGQLY | 0.29465738 |
| VTDVLGDLSV | ASSQVPVNTGQLY | 0.6475516 |
| VVVVGAGFAGL | ASSQVPVNTGQLY | 0.7157482 |
| VYTVGTVV | ASSQVPVNTGQLY | 0.6732857 |
| WTTFSGP | ASSQVPVNTGQLY | 0.6216266 |
| YDELVEAF | ASSQVPVNTGQLY | 0.73296165 |
| YGTGGAGGNGG | ASSQVPVNTGQLY | 0.75290954 |
| YTNVMAV | ASSQVPVNTGQLY | 0.5597967 |
| FLEQVHQL | ASSQVPVNTGQLY | 0.44342375 |
| GLSLLAVAV | ASSQVPVNTGQLY | 0.7473609 |
| LEQVHQL | ASSQVPVNTGQLY | 0.73499596 |
| LNAAGAI | ASSQVPVNTGQLY | 0.6225661 |
| PAVHASL | ASSQVPVNTGQLY | 0.68134904 |
| TELEDDA | ASSQVPVNTGQLY | 0.66773325 |
| TPPGPPN | ASSQVPVNTGQLY | 0.47315866 |
| TPPPPPPP | ASSQVPVNTGQLY | 0.72188544 |
| TQSPALI | ASSQEGWGDEQY | 0.5407019 |
| EFAAAFPQ | ASSQEGWGDEQY | 0.76353407 |
| LLSGNLDVL | ASSQEGWGDEQY | 0.5808911 |
| GPANLGNYNI | ASSQEGWGDEQY | 0.7441294 |
| TLGPTLP | ASSQEGWGDEQY | 0.72073627 |
| VYAYSSCCK | ASSQEGWGDEQY | 0.7284001 |
| APAPAPAPNI | ASSQEGWGDEQY | 0.68492186 |
| PGGPVPIP | ASSQEGWGDEQY | 0.64877737 |
| FLTVALEP | ASSQEGWGDEQY | 0.5772567 |
| SQYFGANL | ASSQEGWGDEQY | 0.40185907 |
| GGVKPAA | ASSQEGWGDEQY | 0.7584157 |
| GGTGGAGGAGGW | ASSQEGWGDEQY | 0.32070866 |
| GVGGVAGLGGAGG | ASSQEGWGDEQY | 0.75043523 |
| VLLVLATTI | ASSQEGWGDEQY | 0.73893607 |
| IGIFAAML | ASSQEGWGDEQY | 0.57629085 |
| AVGDLID | ASSQEGWGDEQY | 0.72568405 |
| NISPLPL | ASSQEGWGDEQY | 0.58467185 |
| VTAVFVAI | ASSQEGWGDEQY | 0.5839547 |
| SGDDNDAAL | ASSQEGWGDEQY | 0.7461057 |
| AVAALAHS | ASSQEGWGDEQY | 0.73272586 |
| GGVTVMATP | ASSQEGWGDEQY | 0.63970244 |
| AGLAANVRD | ASSQEGWGDEQY | 0.7157023 |
| TNRPDLIDPALLRPG | ASSQEGWGDEQY | 0.7260463 |
| LANLGVTAL | ASSQEGWGDEQY | 0.6524801 |
| TGERIEL | ASSQEGWGDEQY | 0.71039814 |
| NGEELAA | ASSQEGWGDEQY | 0.75214696 |
| VLSLVWG | ASSQEGWGDEQY | 0.7193664 |

| | | |
|---|---|---|
| IGYALWL | ASSQEGWGDEQY | 0.6038967 |
| LSLLLAPL | ASSQEGWGDEQY | 0.74681604 |
| TLDEALK | ASSQEGWGDEQY | 0.5885119 |
| TDGAGTAEGG | ASSQEGWGDEQY | 0.723397 |
| NAPGGAGGNGG | ASSQEGWGDEQY | 0.6965457 |
| VSVFMGMGGNF | ASSQEGWGDEQY | 0.669659 |
| LASYLAAL | ASSQEGWGDEQY | 0.75281036 |
| LVGGTPAL | ASSQEGWGDEQY | 0.5506525 |
| VRNADVL | ASSQEGWGDEQY | 0.651387 |
| TCSSPFLVSAM | ASSQEGWGDEQY | 0.6529014 |
| VHITDAE | ASSQEGWGDEQY | 0.7009289 |
| EYAAPSF | ASSQEGWGDEQY | 0.74181134 |
| VVDAFHGGL | ASSQEGWGDEQY | 0.5972762 |
| ASVVDPQ | ASSQEGWGDEQY | 0.7110038 |
| SYNEFGGLS | ASSQEGWGDEQY | 0.6515349 |
| DPDGSVY | ASSQEGWGDEQY | 0.5309309 |
| MVSTIHATE | ASSQEGWGDEQY | 0.68142104 |
| LAAGVADPV | ASSQEGWGDEQY | 0.69902015 |
| TKEQIAA | ASSQEGWGDEQY | 0.5692278 |
| SLASPAAL | ASSQEGWGDEQY | 0.73463523 |
| IGLVFYGV | ASSQEGWGDEQY | 0.6029513 |
| FLLDASNL | ASSQEGWGDEQY | 0.7335494 |
| YQVGNFP | ASSQEGWGDEQY | 0.7713584 |
| ESLLSQST | ASSQEGWGDEQY | 0.6019431 |
| LFTVGFAAVG | ASSQEGWGDEQY | 0.74285555 |
| ATPPTPT | ASSQEGWGDEQY | 0.7327271 |
| TDLATLG | ASSQEGWGDEQY | 0.73558724 |
| SAGGTGR | ASSQEGWGDEQY | 0.67525774 |
| GFSNSGS | ASSQEGWGDEQY | 0.6663126 |
| GGNGGSGGTSV | ASSQEGWGDEQY | 0.7649712 |
| GTSVATGGAGN | ASSQEGWGDEQY | 0.7408098 |
| NGGNGGSGGTS | ASSQEGWGDEQY | 0.74938494 |
| AVVKFPH | ASSQEGWGDEQY | 0.7408747 |
| TPPPPPP | ASSQEGWGDEQY | 0.72875845 |
| AAPAAAAAPA | ASSQEGWGDEQY | 0.7377082 |
| IALNEDL | ASSQEGWGDEQY | 0.6394346 |
| IVNVSSM | ASSQEGWGDEQY | 0.48512796 |
| TGIPPAPR | ASSQEGWGDEQY | 0.66171336 |
| FDLGGGTF | ASSQEGWGDEQY | 0.7244581 |
| FDLGGGTFDVS | ASSQEGWGDEQY | 0.66439366 |
| NEPTAAA | ASSQEGWGDEQY | 0.3450464 |
| NEPTAAALA | ASSQEGWGDEQY | 0.749593 |
| GSIINISSI | ASSQEGWGDEQY | 0.7712304 |
| VAPGNAGTA | ASSQEGWGDEQY | 0.6248683 |
| TGGGASLE | ASSQEGWGDEQY | 0.76328444 |
| VAVYNHYKRI | ASSQEGWGDEQY | 0.5448327 |
| TDPAPATT | ASSQEGWGDEQY | 0.74078065 |
| TNRPDILDPALLRPG | ASSQEGWGDEQY | 0.7552659 |
| LTTTTTS | ASSQEGWGDEQY | 0.571211 |
| PTGSGKTL | ASSQEGWGDEQY | 0.7366984 |
| TGSGKTL | ASSQEGWGDEQY | 0.6632756 |

| | | |
|---|---|---|
| TIGHVDHGK | ASSQEGWGDEQY | 0.68740094 |
| TIGHVDHGKTT | ASSQEGWGDEQY | 0.75814414 |
| GHVDHGKTT | ASSQEGWGDEQY | 0.6506808 |
| AQFEHTLL | ASSQEGWGDEQY | 0.7251712 |
| IVNQDEL | ASSQEGWGDEQY | 0.6539106 |
| TNGSQFF | ASSQEGWGDEQY | 0.5528091 |
| TGKGIVE | ASSQEGWGDEQY | 0.7215843 |
| LISNASDA | ASSQEGWGDEQY | 0.7534718 |
| AELRTLGELTKIAKAHGAQ | ASSQEGWGDEQY | 0.6226268 |
| APNAVAAAEPSW | ASSQEGWGDEQY | 0.7679238 |
| APQVAGAATAVL | ASSQEGWGDEQY | 0.6562448 |
| ARRDPAAAAAILPTDA | ASSQEGWGDEQY | 0.773756 |
| DAAARDVLAEAGLAEYF | ASSQEGWGDEQY | 0.7416385 |
| DDQIVLSAKANLAKVEQ | ASSQEGWGDEQY | 0.70397186 |
| DELLPDTGEDSDEDGH | ASSQEGWGDEQY | 0.6457063 |
| ELPEAPANRATVPQD | ASSQEGWGDEQY | 0.7530022 |
| FGQSKIERVTGIG | ASSQEGWGDEQY | 0.6350895 |
| GAGGTGSTAGGAGGAGGAGGLY | ASSQEGWGDEQY | 0.74596184 |
| GHGGTGAAGGNGGTGGAGGSGI | ASSQEGWGDEQY | 0.75653875 |
| GNGGNGGAGGEGGPGGAGGAGA | ASSQEGWGDEQY | 0.5761 |
| GSTAGNGGNGGNATNGTIAGQP | ASSQEGWGDEQY | 0.84666437 |
| GTGGDGGNGGNGGTGG | ASSQEGWGDEQY | 0.7619483 |
| GVRRPAGIPADAPV | ASSQEGWGDEQY | 0.7650647 |
| GYRQVTDAHLLTLARR | ASSQEGWGDEQY | 0.6061585 |
| IAKMGQDGHDRG | ASSQEGWGDEQY | 0.39967042 |
| KAEPSIEIQYT | ASSQEGWGDEQY | 0.70842934 |
| KPVRIRMAADEDTVAVT | ASSQEGWGDEQY | 0.5124595 |
| MPPRIAGMRLLVIKPEP | ASSQEGWGDEQY | 0.77605 |
| MKFYAVALFRSRGVPPTR | ASSQEGWGDEQY | 0.811265 |
| NAGGFGILGAVAHSP | ASSQEGWGDEQY | 0.58759654 |
| NEGREITLAMAEM | ASSQEGWGDEQY | 0.84888935 |
| NELVEWSHPLGHRWRWE | ASSQEGWGDEQY | 0.8494519 |
| NLGAGGTGGNGGNPG | ASSQEGWGDEQY | 0.42200565 |
| NTEWPKWETVMAADPVD | ASSQEGWGDEQY | 0.6576767 |
| PASAAPAETQP | ASSQEGWGDEQY | 0.5970813 |
| QQSSDGKNGVRAAEITGE | ASSQEGWGDEQY | 0.70899177 |
| RAPDAVTTSWWKE | ASSQEGWGDEQY | 0.3137074 |
| SGFGNVGAHTSGWFNQ | ASSQEGWGDEQY | 0.81388354 |
| SGVAGAVGQAASVGGLKV | ASSQEGWGDEQY | 0.5685253 |
| SNGGAGTGSAGGNGGT | ASSQEGWGDEQY | 0.17809726 |
| TGGAGGEGGAAG | ASSQEGWGDEQY | 0.77549374 |
| TGGDGGAAGAGGAPGHG | ASSQEGWGDEQY | 0.6738602 |
| TMPPLLAAIANAGRHGV | ASSQEGWGDEQY | 0.66705465 |
| TSSTAGNSGGA | ASSQEGWGDEQY | 0.83189946 |
| VPATVPVGILC | ASSQEGWGDEQY | 0.52458775 |
| VQRPDSGGVVLTGRIS | ASSQEGWGDEQY | 0.5069224 |
| WTPNPVPATKGVPAGLVQAT | ASSQEGWGDEQY | 0.7974858 |
| YALKRALAKFSDRFDVVII | ASSQEGWGDEQY | 0.8355779 |
| YLIGSVIV | ASSQEGWGDEQY | 0.45671868 |
| YGQLVAVID | ASSQEGWGDEQY | 0.758464 |
| VTVGGMTLPA | ASSQEGWGDEQY | 0.8410129 |

| | | |
|---|---|---|
| VLALPAV | ASSQEGWGDEQY | 0.77207303 |
| VIPPTLNYET | ASSQEGWGDEQY | 0.83139634 |
| VGMGGMHPSSG | ASSQEGWGDEQY | 0.8480722 |
| VGLAGGAATA | ASSQEGWGDEQY | 0.69115126 |
| VGGAGGAGGSGP | ASSQEGWGDEQY | 0.82497275 |
| VFAAFMV | ASSQEGWGDEQY | 0.8584 |
| TLYDKLR | ASSQEGWGDEQY | 0.65366983 |
| TLAVFTVL | ASSQEGWGDEQY | 0.7494818 |
| TGSVQPARV | ASSQEGWGDEQY | 0.5366831 |
| TGGGAGSGGT | ASSQEGWGDEQY | 0.72135127 |
| SHLETSL | ASSQEGWGDEQY | 0.33432537 |
| SFTTGMT | ASSQEGWGDEQY | 0.74827474 |
| QVEVAKRMGVRQPTVS | ASSQEGWGDEQY | 0.8408321 |
| QGSGGPALAM | ASSQEGWGDEQY | 0.67883325 |
| MLAPSNP | ASSQEGWGDEQY | 0.50213987 |
| LVNNAAI | ASSQEGWGDEQY | 0.4162261 |
| LTGEINTLA | ASSQEGWGDEQY | 0.758834 |
| LSIFGPV | ASSQEGWGDEQY | 0.6236025 |
| LLALLANL | ASSQEGWGDEQY | 0.34444746 |
| LISNASDALDKLR | ASSQEGWGDEQY | 0.73278624 |
| LINEAVP | ASSQEGWGDEQY | 0.83787715 |
| LGYAFSGL | ASSQEGWGDEQY | 0.818764 |
| LDFVAVAL | ASSQEGWGDEQY | 0.49314493 |
| LAQPGAE | ASSQEGWGDEQY | 0.8582338 |
| ISNLGAVTGF | ASSQEGWGDEQY | 0.7845476 |
| HQEDPGRI | ASSQEGWGDEQY | 0.8376611 |
| GTPGEPTP | ASSQEGWGDEQY | 0.6524986 |
| GFFNSGS | ASSQEGWGDEQY | 0.55844367 |
| GAGGGGLTAGAN | ASSQEGWGDEQY | 0.8229139 |
| FVDALVNM | ASSQEGWGDEQY | 0.6881577 |
| FLTVAPEL | ASSQEGWGDEQY | 0.67028403 |
| FETLTAL | ASSQEGWGDEQY | 0.83301497 |
| DVINPIEL | ASSQEGWGDEQY | 0.6883698 |
| APVSGAPV | ASSQEGWGDEQY | 0.58762455 |
| ALVGGLAW | ASSQEGWGDEQY | 0.6187139 |
| ALAAAAATAI | ASSQEGWGDEQY | 0.84048915 |
| AFGLGLLL | ASSQEGWGDEQY | 0.53576934 |
| AFFAAASL | ASSQEGWGDEQY | 0.5286339 |
| AAAALPAFSPP | ASSQEGWGDEQY | 0.6238175 |
| VVLAPAL | ASSQEGWGDEQY | 0.7169856 |
| TIGHVDHGKTTL | ASSQEGWGDEQY | 0.83068407 |
| LISNASDALDK | ASSQEGWGDEQY | 0.77535087 |
| LISNASDAL | ASSQEGWGDEQY | 0.67208725 |
| ISNASDALDK | ASSQEGWGDEQY | 0.7699765 |
| NQAETLVYQTE | ASSQEGWGDEQY | 0.78743863 |
| TGGTGGTAGAG | ASSQEGWGDEQY | 0.7578811 |
| TGSTAGGAGGA | ASSQEGWGDEQY | 0.8200915 |
| DPLAVPVAAGPV | ASSQEGWGDEQY | 0.58658314 |
| GGTGGTPGNGGN | ASSQEGWGDEQY | 0.83888215 |
| PAGLAGVDPATL | ASSQEGWGDEQY | 0.81158054 |
| GNNLVVEAPHAGASV | ASSQEGWGDEQY | 0.81400025 |

| | | |
|---|---|---|
| KVTRIMSASTLEQPA | ASSQEGWGDEQY | 0.82947546 |
| LGFALPIALPRRTVV | ASSQEGWGDEQY | 0.7729392 |
| TRLGNLTARKLVVLV | ASSQEGWGDEQY | 0.75111115 |
| WSQQRRLIYFPSAGP | ASSQEGWGDEQY | 0.33561862 |
| GPNGNPDPMAAAVDIR | ASSQEGWGDEQY | 0.70186055 |
| TQRCVSVADLRAAASR | ASSQEGWGDEQY | 0.769501 |
| QFVLDVAEHPVEVLTETV | ASSQEGWGDEQY | 0.7574981 |
| GGAGGTGGAGGNGGTQAGDG | ASSQEGWGDEQY | 0.8073995 |
| PLLATLRAIAVDVDAKPAQV | ASSQEGWGDEQY | 0.79712224 |
| APKFRTGGYMIDFWGVGYQVAK | ASSQEGWGDEQY | 0.609569 |
| GAGGAGGAGGAAGGPGGTGGTG | ASSQEGWGDEQY | 0.84025073 |
| IGALGLLATDVAPAGSVAFGVA | ASSQEGWGDEQY | 0.83343434 |
| AGGAGGIGGNGGAGGAGGVLMG | ASSQEGWGDEQY | 0.41532052 |
| ADELIGLL | ASSQEGWGDEQY | 0.7067795 |
| ADRALKA | ASSQEGWGDEQY | 0.86697066 |
| AGAVGQVGGAGGA | ASSQEGWGDEQY | 0.7580377 |
| AGGGGGLGGAP | ASSQEGWGDEQY | 0.67967606 |
| AGGLLYGNGGAG | ASSQEGWGDEQY | 0.67428946 |
| AGLESIGL | ASSQEGWGDEQY | 0.829318 |
| AGLKTGII | ASSQEGWGDEQY | 0.657885 |
| ASLAALLL | ASSQEGWGDEQY | 0.8590202 |
| ASVINISSI | ASSQEGWGDEQY | 0.24436642 |
| ATPPGPA | ASSQEGWGDEQY | 0.6319556 |
| ATVASALV | ASSQEGWGDEQY | 0.6030803 |
| ATVASLIL | ASSQEGWGDEQY | 0.28624618 |
| AVNGPVLIRH | ASSQEGWGDEQY | 0.8209238 |
| DGAAGAVGI | ASSQEGWGDEQY | 0.6776903 |
| DPLLHGI | ASSQEGWGDEQY | 0.8230325 |
| ELGAALGAL | ASSQEGWGDEQY | 0.71516025 |
| ELGRTQAV | ASSQEGWGDEQY | 0.8285784 |
| FVAGGYLS | ASSQEGWGDEQY | 0.74779755 |
| GASGGAGGAGGW | ASSQEGWGDEQY | 0.7631551 |
| GGAGAGGDGGAGGI | ASSQEGWGDEQY | 0.5788185 |
| GTGHNGSQPGMGGNG | ASSQEGWGDEQY | 0.804268 |
| GYAAASASAS | ASSQEGWGDEQY | 0.8549461 |
| HQVDVAAVQQG | ASSQEGWGDEQY | 0.8114399 |
| IGAAIAHT | ASSQEGWGDEQY | 0.37706313 |
| IGMIEVAQL | ASSQEGWGDEQY | 0.7057884 |
| IRLAPEHTSAMD | ASSQEGWGDEQY | 0.7081604 |
| IYNVSVL | ASSQEGWGDEQY | 0.30661428 |
| KQGFLGM | ASSQEGWGDEQY | 0.26172706 |
| LESVITQL | ASSQEGWGDEQY | 0.6645556 |
| LGLGALVVV | ASSQEGWGDEQY | 0.6732173 |
| LIGNGGNGGSGGI | ASSQEGWGDEQY | 0.81235623 |
| LNAINAPTL | ASSQEGWGDEQY | 0.80129635 |
| LQLLNTGV | ASSQEGWGDEQY | 0.6822771 |
| LQTSIEF | ASSQEGWGDEQY | 0.80191946 |
| MSFAVSEL | ASSQEGWGDEQY | 0.7769822 |
| NTGALGL | ASSQEGWGDEQY | 0.11430752 |
| PPGAGGAGGAGTT | ASSQEGWGDEQY | 0.65720475 |
| QLVGLGI | ASSQEGWGDEQY | 0.40814364 |

| | | |
|---|---|---|
| RFLEQVHQL | ASSQEGWGDEQY | 0.8501184 |
| RGVGYVLR | ASSQEGWGDEQY | 0.6163843 |
| SATGVFAGVI | ASSQEGWGDEQY | 0.6087494 |
| SLPAITLP | ASSQEGWGDEQY | 0.7306223 |
| STALAFTV | ASSQEGWGDEQY | 0.7255509 |
| SVLTVGLGK | ASSQEGWGDEQY | 0.708169 |
| TALAGAGSGI | ASSQEGWGDEQY | 0.4902516 |
| TGLPFTGI | ASSQEGWGDEQY | 0.24112374 |
| VAEHVTV | ASSQEGWGDEQY | 0.827587 |
| VDIVLNSL | ASSQEGWGDEQY | 0.47453874 |
| VLTGGGAVL | ASSQEGWGDEQY | 0.5030185 |
| VNVGLAYGI | ASSQEGWGDEQY | 0.41093767 |
| VSSAAVGAV | ASSQEGWGDEQY | 0.30532748 |
| VTSVYEAGV | ASSQEGWGDEQY | 0.35287315 |
| WVALDRGA | ASSQEGWGDEQY | 0.82090354 |
| YLVGSVVI | ASSQEGWGDEQY | 0.61632156 |
| GPPGTGKTLL | ASSQEGWGDEQY | 0.6953684 |
| GVGMTKF | ASSQEGWGDEQY | 0.8642653 |
| IGIGGSDL | ASSQEGWGDEQY | 0.69616115 |
| IGTLLAEL | ASSQEGWGDEQY | 0.70969206 |
| IYTSGTT | ASSQEGWGDEQY | 0.58683383 |
| LSFPAGV | ASSQEGWGDEQY | 0.516519 |
| LYTSGTT | ASSQEGWGDEQY | 0.5559746 |
| VGAFPFGF | ASSQEGWGDEQY | 0.5704029 |
| VGAGTVEF | ASSQEGWGDEQY | 0.865077 |
| VGKSTLLL | ASSQEGWGDEQY | 0.6865433 |
| AALAHAGGGIV | ASSQEGWGDEQY | 0.50735044 |
| AALVAVSLL | ASSQEGWGDEQY | 0.7388266 |
| AAVSRAVAAL | ASSQEGWGDEQY | 0.85483575 |
| AGGSSGIGGF | ASSQEGWGDEQY | 0.87202644 |
| AGLTVANANG | ASSQEGWGDEQY | 0.5890846 |
| AGLVGAVL | ASSQEGWGDEQY | 0.64367855 |
| ALSQAFTV | ASSQEGWGDEQY | 0.65840113 |
| ALTFSVSL | ASSQEGWGDEQY | 0.702772 |
| ALTGAALVAL | ASSQEGWGDEQY | 0.6562442 |
| ASFSIGAGVL | ASSQEGWGDEQY | 0.4900831 |
| DGSVYKI | ASSQEGWGDEQY | 0.81058854 |
| EQLAEIV | ASSQEGWGDEQY | 0.8527454 |
| FAVSELTVM | ASSQEGWGDEQY | 0.70736945 |
| GAAPLAAGL | ASSQEGWGDEQY | 0.31818074 |
| GADGGDGGSGGAGG | ASSQEGWGDEQY | 0.66989076 |
| GFGLDTGGAGG | ASSQEGWGDEQY | 0.5105634 |
| GLASGGLL | ASSQEGWGDEQY | 0.58928853 |
| GRIAGAAVLL | ASSQEGWGDEQY | 0.6802135 |
| GSAGSLNTSF | ASSQEGWGDEQY | 0.40854102 |
| GTGQIGI | ASSQEGWGDEQY | 0.70698076 |
| GTLIALIL | ASSQEGWGDEQY | 0.86445963 |
| IAATMGFVVG | ASSQEGWGDEQY | 0.6132901 |
| IFAQALL | ASSQEGWGDEQY | 0.79362583 |
| IGAAVALGF | ASSQEGWGDEQY | 0.84304583 |
| IGLTPLAL | ASSQEGWGDEQY | 0.8084099 |

| | | |
|---|---|---|
| ILGSNTSTL | ASSQEGWGDEQY | 0.84439456 |
| IVFSLTVL | ASSQEGWGDEQY | 0.7841667 |
| LASKATGF | ASSQEGWGDEQY | 0.7649242 |
| LASVALIL | ASSQEGWGDEQY | 0.74841106 |
| LFAVGLTL | ASSQEGWGDEQY | 0.79748726 |
| LFNGGIGTYI | ASSQEGWGDEQY | 0.70714164 |
| LTLGAGII | ASSQEGWGDEQY | 0.70961607 |
| LVAAALGDI | ASSQEGWGDEQY | 0.64250505 |
| NTAFVQLGI | ASSQEGWGDEQY | 0.8180879 |
| QLFSGAEL | ASSQEGWGDEQY | 0.66824514 |
| SAATFITL | ASSQEGWGDEQY | 0.82589394 |
| SNYGLANLGV | ASSQEGWGDEQY | 0.37009427 |
| SYGGAYAVM | ASSQEGWGDEQY | 0.46625268 |
| TTGVGGLTLG | ASSQEGWGDEQY | 0.82480836 |
| TTQTIYI | ASSQEGWGDEQY | 0.1553488 |
| VAAGAGLGAVI | ASSQEGWGDEQY | 0.24472415 |
| VAAVVTLL | ASSQEGWGDEQY | 0.72974586 |
| VGFLGGLI | ASSQEGWGDEQY | 0.84477746 |
| VGVIPILL | ASSQEGWGDEQY | 0.16904967 |
| VIFASSLI | ASSQEGWGDEQY | 0.53705597 |
| VKGGDGGAAAG | ASSQEGWGDEQY | 0.38064522 |
| VTKTFTNI | ASSQEGWGDEQY | 0.81111866 |
| VVAAAILL | ASSQEGWGDEQY | 0.7189603 |
| YAALDVEL | ASSQEGWGDEQY | 0.39742267 |
| LASVALLL | ASSQEGWGDEQY | 0.81250787 |
| LDAVHRV | ASSQEGWGDEQY | 0.35624614 |
| QGLSSSIF | ASSQEGWGDEQY | 0.56974447 |
| RGTFANIRL | ASSQEGWGDEQY | 0.81655484 |
| RQLWWGHRI | ASSQEGWGDEQY | 0.8319541 |
| YGPPGTGKTLL | ASSQEGWGDEQY | 0.5234992 |
| AALVGAPANVA | ASSQEGWGDEQY | 0.7905541 |
| AATFRRTE | ASSQEGWGDEQY | 0.60253966 |
| AGALGGGI | ASSQEGWGDEQY | 0.8755771 |
| AGGAGGAGGIGAG | ASSQEGWGDEQY | 0.66271746 |
| AGSGALGL | ASSQEGWGDEQY | 0.39862582 |
| AQIADQI | ASSQEGWGDEQY | 0.819142 |
| ASAVAVTYL | ASSQEGWGDEQY | 0.81932044 |
| ASTVLGGTGG | ASSQEGWGDEQY | 0.75496674 |
| AVGGEPAK | ASSQEGWGDEQY | 0.78874624 |
| CEQGTFGGTC | ASSQEGWGDEQY | 0.6317083 |
| DEGKAPAG | ASSQEGWGDEQY | 0.6647897 |
| DTVVFPAL | ASSQEGWGDEQY | 0.8230507 |
| FEGITMAG | ASSQEGWGDEQY | 0.8257857 |
| FPLVVQD | ASSQEGWGDEQY | 0.55436885 |
| GAFTTPQ | ASSQEGWGDEQY | 0.84368265 |
| GFGGFGTAGGVGGA | ASSQEGWGDEQY | 0.73066807 |
| GFLAQFL | ASSQEGWGDEQY | 0.61090386 |
| GGAGGAGGNAGVA | ASSQEGWGDEQY | 0.8621367 |
| GGVGTAGAGGAGGAGG | ASSQEGWGDEQY | 0.7177776 |
| GNGGGDAGNAGSGGN | ASSQEGWGDEQY | 0.8168647 |
| GTGGAGGAGNDAGS | ASSQEGWGDEQY | 0.80353594 |

| | | |
|---|---|---|
| GTTNTGWFNS | ASSQEGWGDEQY | 0.7616122 |
| IFMGPLFTL | ASSQEGWGDEQY | 0.5196151 |
| ILAHAIEAI | ASSQEGWGDEQY | 0.81195104 |
| IRSAIREAL | ASSQEGWGDEQY | 0.73319066 |
| ITFAGGG | ASSQEGWGDEQY | 0.8023995 |
| ITGMSGHQEV | ASSQEGWGDEQY | 0.7338379 |
| IVAGLACTL | ASSQEGWGDEQY | 0.600129 |
| IVGGGLAGA | ASSQEGWGDEQY | 0.8072145 |
| IVTGVLNI | ASSQEGWGDEQY | 0.85264003 |
| LAQQGCI | ASSQEGWGDEQY | 0.7760412 |
| LCLHVEN | ASSQEGWGDEQY | 0.7905716 |
| LDAISDGS | ASSQEGWGDEQY | 0.8551828 |
| LGGFAAML | ASSQEGWGDEQY | 0.5628901 |
| LGGKNPMI | ASSQEGWGDEQY | 0.7354686 |
| LGPKGVAAIL | ASSQEGWGDEQY | 0.814762 |
| LMVGVSI | ASSQEGWGDEQY | 0.84939086 |
| LVGAAFVL | ASSQEGWGDEQY | 0.69726837 |
| LVLAVAPL | ASSQEGWGDEQY | 0.84299415 |
| MAAIAVIDAA | ASSQEGWGDEQY | 0.63280964 |
| NNGINAELQ | ASSQEGWGDEQY | 0.8038448 |
| RGEPLVLVHG | ASSQEGWGDEQY | 0.62451404 |
| RGGAGGAAGLI | ASSQEGWGDEQY | 0.7607199 |
| RSYRMHPAV | ASSQEGWGDEQY | 0.5588039 |
| SAAHLLAGQLA | ASSQEGWGDEQY | 0.7961669 |
| SAAVGSY | ASSQEGWGDEQY | 0.8130701 |
| SPHAVLHRI | ASSQEGWGDEQY | 0.82023597 |
| SQIAKVL | ASSQEGWGDEQY | 0.25127614 |
| TALGHLVP | ASSQEGWGDEQY | 0.7995281 |
| TAPTTSGAA | ASSQEGWGDEQY | 0.8076389 |
| TDGDDGNGGSPGT | ASSQEGWGDEQY | 0.8670083 |
| TDGLGEN | ASSQEGWGDEQY | 0.70887077 |
| TGGPPLPGQA | ASSQEGWGDEQY | 0.78924483 |
| TIPAITI | ASSQEGWGDEQY | 0.6363126 |
| TIPGFTI | ASSQEGWGDEQY | 0.70060533 |
| TSIASGALA | ASSQEGWGDEQY | 0.4537258 |
| TVTLPGAAI | ASSQEGWGDEQY | 0.66835326 |
| VASGLSAALP | ASSQEGWGDEQY | 0.64180255 |
| VGAGVIGI | ASSQEGWGDEQY | 0.5381839 |
| VGPKRFGV | ASSQEGWGDEQY | 0.7983586 |
| VNVAFPD | ASSQEGWGDEQY | 0.832232 |
| VSMTNSGVS | ASSQEGWGDEQY | 0.8357176 |
| VTDVLGDLSV | ASSQEGWGDEQY | 0.82198274 |
| VVVVGAGFAGL | ASSQEGWGDEQY | 0.8539721 |
| VYTVGTVV | ASSQEGWGDEQY | 0.81182975 |
| WTTFSGP | ASSQEGWGDEQY | 0.59748745 |
| YDELVEAF | ASSQEGWGDEQY | 0.7117257 |
| YGTGGAGGNGG | ASSQEGWGDEQY | 0.43505406 |
| YTNVMAV | ASSQEGWGDEQY | 0.6725372 |
| FLEQVHQL | ASSQEGWGDEQY | 0.6539911 |
| GLSLLAVAV | ASSQEGWGDEQY | 0.7532271 |
| LEQVHQL | ASSQEGWGDEQY | 0.13566558 |

| | | |
|---|---|---|
| LNAAGAI | ASSQEGWGDEQY | 0.72735834 |
| PAVHASL | ASSQEGWGDEQY | 0.8666483 |
| TELEDDA | ASSQEGWGDEQY | 0.66694796 |
| TPPGPPN | ASSQEGWGDEQY | 0.76370096 |
| TPPPPPPP | ASSQEGWGDEQY | 0.7936621 |
| TQSPALI | ASSPGTGGGEQY | 0.63786805 |
| EFAAAFPQ | ASSPGTGGGEQY | 0.7837381 |
| LLSGNLDVL | ASSPGTGGGEQY | 0.57249206 |
| GPANLGNYNI | ASSPGTGGGEQY | 0.80344296 |
| TLGPTLP | ASSPGTGGGEQY | 0.8554357 |
| VYAYSSCCK | ASSPGTGGGEQY | 0.7552843 |
| APAPAPAPNI | ASSPGTGGGEQY | 0.75863135 |
| PGGPVPIP | ASSPGTGGGEQY | 0.7307571 |
| FLTVALEP | ASSPGTGGGEQY | 0.41229925 |
| SQYFGANL | ASSPGTGGGEQY | 0.7750852 |
| GGVKPAA | ASSPGTGGGEQY | 0.14428791 |
| GGTGGAGGAGGW | ASSPGTGGGEQY | 0.69949794 |
| GVGGVAGLGGAGG | ASSPGTGGGEQY | 0.5432726 |
| VLLVLATTI | ASSPGTGGGEQY | 0.7458497 |
| IGIFAAML | ASSPGTGGGEQY | 0.76241493 |
| AVGDLID | ASSPGTGGGEQY | 0.79013216 |
| NISPLPL | ASSPGTGGGEQY | 0.52401423 |
| VTAVFVAI | ASSPGTGGGEQY | 0.7440753 |
| SGDDNDAAL | ASSPGTGGGEQY | 0.67954576 |
| AVAALAHS | ASSPGTGGGEQY | 0.43363962 |
| GGVTVMATP | ASSPGTGGGEQY | 0.83664805 |
| AGLAANVRD | ASSPGTGGGEQY | 0.7107719 |
| TNRPDLIDPALLRPG | ASSPGTGGGEQY | 0.7163769 |
| LANLGVTAL | ASSPGTGGGEQY | 0.83747166 |
| TGERIEL | ASSPGTGGGEQY | 0.76171136 |
| NGEELAA | ASSPGTGGGEQY | 0.36924133 |
| VLSLVWG | ASSPGTGGGEQY | 0.67996824 |
| IGYALWL | ASSPGTGGGEQY | 0.7191658 |
| LSLLLAPL | ASSPGTGGGEQY | 0.6699562 |
| TLDEALK | ASSPGTGGGEQY | 0.15331134 |
| TDGAGTAEGG | ASSPGTGGGEQY | 0.81957495 |
| NAPGGAGGNGG | ASSPGTGGGEQY | 0.6250453 |
| VSVFMGMGGNF | ASSPGTGGGEQY | 0.53689086 |
| LASYLAAL | ASSPGTGGGEQY | 0.6573681 |
| LVGGTPAL | ASSPGTGGGEQY | 0.8160378 |
| VRNADVL | ASSPGTGGGEQY | 0.66399527 |
| TCSSPFLVSAM | ASSPGTGGGEQY | 0.685735 |
| VHITDAE | ASSPGTGGGEQY | 0.80180085 |
| EYAAPSF | ASSPGTGGGEQY | 0.8005098 |
| VVDAFHGGL | ASSPGTGGGEQY | 0.85164785 |
| ASVVDPQ | ASSPGTGGGEQY | 0.77068967 |
| SYNEFGGLS | ASSPGTGGGEQY | 0.73472154 |
| DPDGSVY | ASSPGTGGGEQY | 0.8110497 |
| MVSTIHATE | ASSPGTGGGEQY | 0.7203482 |
| LAAGVADPV | ASSPGTGGGEQY | 0.46593857 |
| TKEQIAA | ASSPGTGGGEQY | 0.32587442 |

| | | |
|---|---|---|
| SLASPAAL | ASSPGTGGGEQY | 0.68927586 |
| IGLVFYGV | ASSPGTGGGEQY | 0.39029932 |
| FLLDASNL | ASSPGTGGGEQY | 0.74298966 |
| YQVGNFP | ASSPGTGGGEQY | 0.8370197 |
| ESLLSQST | ASSPGTGGGEQY | 0.71329236 |
| LFTVGFAAVG | ASSPGTGGGEQY | 0.47876936 |
| ATPPTPT | ASSPGTGGGEQY | 0.65532684 |
| TDLATLG | ASSPGTGGGEQY | 0.70262414 |
| SAGGTGR | ASSPGTGGGEQY | 0.7978445 |
| GFSNSGS | ASSPGTGGGEQY | 0.8175127 |
| GGNGGSGGTSV | ASSPGTGGGEQY | 0.849195 |
| GTSVATGGAGN | ASSPGTGGGEQY | 0.8138107 |
| NGGNGGSGGTS | ASSPGTGGGEQY | 0.6459956 |
| AVVKFPH | ASSPGTGGGEQY | 0.8068253 |
| TPPPPPP | ASSPGTGGGEQY | 0.67425907 |
| AAPAAAAAPA | ASSPGTGGGEQY | 0.76496553 |
| IALNEDL | ASSPGTGGGEQY | 0.752704 |
| IVNVSSM | ASSPGTGGGEQY | 0.77792525 |
| TGIPPAPR | ASSPGTGGGEQY | 0.70073825 |
| FDLGGGTF | ASSPGTGGGEQY | 0.69409406 |
| FDLGGGTFDVS | ASSPGTGGGEQY | 0.3437983 |
| NEPTAAA | ASSPGTGGGEQY | 0.77061015 |
| NEPTAAALA | ASSPGTGGGEQY | 0.7473469 |
| GSIINISSI | ASSPGTGGGEQY | 0.37801382 |
| VAPGNAGTA | ASSPGTGGGEQY | 0.6666173 |
| TGGGASLE | ASSPGTGGGEQY | 0.85822296 |
| VAVYNHYKRI | ASSPGTGGGEQY | 0.83375216 |
| TDPAPATT | ASSPGTGGGEQY | 0.57645786 |
| TNRPDILDPALLRPG | ASSPGTGGGEQY | 0.55491066 |
| LTTTTTS | ASSPGTGGGEQY | 0.74056965 |
| PTGSGKTL | ASSPGTGGGEQY | 0.4082609 |
| TGSGKTL | ASSPGTGGGEQY | 0.78406763 |
| TIGHVDHGK | ASSPGTGGGEQY | 0.69955635 |
| TIGHVDHGKTT | ASSPGTGGGEQY | 0.73475534 |
| GHVDHGKTT | ASSPGTGGGEQY | 0.6114526 |
| AQFEHTLL | ASSPGTGGGEQY | 0.7639599 |
| IVNQDEL | ASSPGTGGGEQY | 0.6885864 |
| TNGSQFF | ASSPGTGGGEQY | 0.78876793 |
| TGKGIVE | ASSPGTGGGEQY | 0.16733709 |
| LISNASDA | ASSPGTGGGEQY | 0.8166993 |
| AELRTLGELTKIAKAHGAQ | ASSPGTGGGEQY | 0.5487765 |
| APNAVAAAEPSW | ASSPGTGGGEQY | 0.552837 |
| APQVAGAATAVL | ASSPGTGGGEQY | 0.30990207 |
| ARRDPAAAAAILPTDA | ASSPGTGGGEQY | 0.76607454 |
| DAAARDVLAEAGLAEYF | ASSPGTGGGEQY | 0.8096014 |
| DDQIVLSAKANLAKVEQ | ASSPGTGGGEQY | 0.822554 |
| DELLPDTGEDSDEDGH | ASSPGTGGGEQY | 0.6482485 |
| ELPEAPANRATVPQD | ASSPGTGGGEQY | 0.818157 |
| FGQSKIERVTGIG | ASSPGTGGGEQY | 0.84657 |
| GAGGTGSTAGGAGGAGGAGGLY | ASSPGTGGGEQY | 0.67377675 |
| GHGGTGAAGGNGGTGGAGGSGI | ASSPGTGGGEQY | 0.3556208 |

| | | |
|---|---|---|
| GNGGNGGAGGEGGPGGAGGAGA | ASSPGTGGGEQY | 0.7640004 |
| GSTAGNGGNGGNATNGTIAGQP | ASSPGTGGGEQY | 0.6889318 |
| GTGGDGGNGGNGGTGG | ASSPGTGGGEQY | 0.8151622 |
| GVRRPAGIPADAPV | ASSPGTGGGEQY | 0.81952786 |
| GYRQVTDAHLLTLARR | ASSPGTGGGEQY | 0.45871186 |
| IAKMGQDGHDRG | ASSPGTGGGEQY | 0.80227697 |
| KAEPSIEIQYT | ASSPGTGGGEQY | 0.8216256 |
| KPVRIRMAADEDTVAVT | ASSPGTGGGEQY | 0.5288296 |
| MPPRIAGMRLLVIKPEP | ASSPGTGGGEQY | 0.6502136 |
| MKFYAVALFRSRGVPPTR | ASSPGTGGGEQY | 0.6956102 |
| NAGGFGILGAVAHSP | ASSPGTGGGEQY | 0.33811295 |
| NEGREITLAMAEM | ASSPGTGGGEQY | 0.8158665 |
| NELVEWSHPLGHRWRWE | ASSPGTGGGEQY | 0.64278996 |
| NLGAGGTGGNGGNPG | ASSPGTGGGEQY | 0.56513983 |
| NTEWPKWETVMAADPVD | ASSPGTGGGEQY | 0.7511579 |
| PASAAPAETQP | ASSPGTGGGEQY | 0.5927315 |
| QQSSDGKNGVRAAEITGE | ASSPGTGGGEQY | 0.60701877 |
| RAPDAVTTSWWKE | ASSPGTGGGEQY | 0.6454777 |
| SGFGNVGAHTSGWFNQ | ASSPGTGGGEQY | 0.74860793 |
| SGVAGAVGQAASVGGLKV | ASSPGTGGGEQY | 0.77928376 |
| SNGGAGTGSAGGNGGT | ASSPGTGGGEQY | 0.71888673 |
| TGGAGGEGGAAG | ASSPGTGGGEQY | 0.7613554 |
| TGGDGGAAGAGGAPGHG | ASSPGTGGGEQY | 0.85125035 |
| TMPPLLAAIANAGRHGV | ASSPGTGGGEQY | 0.41902006 |
| TSSTAGNSGGA | ASSPGTGGGEQY | 0.62642133 |
| VPATVPVGILC | ASSPGTGGGEQY | 0.7812594 |
| VQRPDSGGVVLTGRIS | ASSPGTGGGEQY | 0.5180917 |
| WTPNPVPATKGVPAGLVQAT | ASSPGTGGGEQY | 0.8186239 |
| YALKRALAKFSDRFDVVII | ASSPGTGGGEQY | 0.76777655 |
| YLIGSVIV | ASSPGTGGGEQY | 0.7764319 |
| YGQLVAVID | ASSPGTGGGEQY | 0.66988355 |
| VTVGGMTLPA | ASSPGTGGGEQY | 0.86921704 |
| VLALPAV | ASSPGTGGGEQY | 0.8061316 |
| VIPPTLNYET | ASSPGTGGGEQY | 0.7540457 |
| VGMGGMHPSSG | ASSPGTGGGEQY | 0.78938544 |
| VGLAGGAATA | ASSPGTGGGEQY | 0.5634669 |
| VGGAGGAGGSGP | ASSPGTGGGEQY | 0.82650423 |
| VFAAFMV | ASSPGTGGGEQY | 0.76990163 |
| TLYDKLR | ASSPGTGGGEQY | 0.4988038 |
| TLAVFTVL | ASSPGTGGGEQY | 0.6729996 |
| TGSVQPARV | ASSPGTGGGEQY | 0.8504927 |
| TGGGAGSGGT | ASSPGTGGGEQY | 0.83673024 |
| SHLETSL | ASSPGTGGGEQY | 0.4020143 |
| SFTTGMT | ASSPGTGGGEQY | 0.663529 |
| QVEVAKRMGVRQPTVS | ASSPGTGGGEQY | 0.43663386 |
| QGSGGPALAM | ASSPGTGGGEQY | 0.6308849 |
| MLAPSNP | ASSPGTGGGEQY | 0.515608 |
| LVNNAAI | ASSPGTGGGEQY | 0.7880331 |
| LTGEINTLA | ASSPGTGGGEQY | 0.6541276 |
| LSIFGPV | ASSPGTGGGEQY | 0.7641523 |
| LLALLANL | ASSPGTGGGEQY | 0.7418951 |

| | | |
|---|---|---|
| LISNASDALDKLR | ASSPGTGGGEQY | 0.83891135 |
| LINEAVP | ASSPGTGGGEQY | 0.5732519 |
| LGYAFSGL | ASSPGTGGGEQY | 0.7088492 |
| LDFVAVAL | ASSPGTGGGEQY | 0.39608854 |
| LAQPGAE | ASSPGTGGGEQY | 0.4858946 |
| ISNLGAVTGF | ASSPGTGGGEQY | 0.5911035 |
| HQEDPGRI | ASSPGTGGGEQY | 0.6414733 |
| GTPGEPTP | ASSPGTGGGEQY | 0.25135455 |
| GFFNSGS | ASSPGTGGGEQY | 0.85266626 |
| GAGGGGLTAGAN | ASSPGTGGGEQY | 0.71872073 |
| FVDALVNM | ASSPGTGGGEQY | 0.8075277 |
| FLTVAPEL | ASSPGTGGGEQY | 0.64057106 |
| FETLTAL | ASSPGTGGGEQY | 0.63736355 |
| DVINPIEL | ASSPGTGGGEQY | 0.62324184 |
| APVSGAPV | ASSPGTGGGEQY | 0.7757588 |
| ALVGGLAW | ASSPGTGGGEQY | 0.5598724 |
| ALAAAAATAI | ASSPGTGGGEQY | 0.29781193 |
| AFGLGLLL | ASSPGTGGGEQY | 0.60421205 |
| AFFAAASL | ASSPGTGGGEQY | 0.43928432 |
| AAAALPAFSPP | ASSPGTGGGEQY | 0.861313 |
| VVLAPAL | ASSPGTGGGEQY | 0.8229108 |
| TIGHVDHGKTTL | ASSPGTGGGEQY | 0.74648154 |
| LISNASDALDK | ASSPGTGGGEQY | 0.84842014 |
| LISNASDAL | ASSPGTGGGEQY | 0.21655063 |
| ISNASDALDK | ASSPGTGGGEQY | 0.6705861 |
| NQAETLVYQTE | ASSPGTGGGEQY | 0.85597974 |
| TGGTGGTAGAG | ASSPGTGGGEQY | 0.7720361 |
| TGSTAGGAGGA | ASSPGTGGGEQY | 0.71213317 |
| DPLAVPVAAGPV | ASSPGTGGGEQY | 0.83403337 |
| GGTGGTPGNGGN | ASSPGTGGGEQY | 0.5997915 |
| PAGLAGVDPATL | ASSPGTGGGEQY | 0.661571 |
| GNNLVVEAPHAGASV | ASSPGTGGGEQY | 0.70969963 |
| KVTRIMSASTLEQPA | ASSPGTGGGEQY | 0.8190401 |
| LGFALPIALPRRTVV | ASSPGTGGGEQY | 0.7617227 |
| TRLGNLTARKLVVLV | ASSPGTGGGEQY | 0.4935697 |
| WSQQRRLIYFPSAGP | ASSPGTGGGEQY | 0.771979 |
| GPNGNPDPMAAAVDIR | ASSPGTGGGEQY | 0.8727366 |
| TQRCVSVADLRAAASR | ASSPGTGGGEQY | 0.61412096 |
| QFVLDVAEHPVEVLTETV | ASSPGTGGGEQY | 0.8177488 |
| GGAGGTGGAGGNGGTQAGDG | ASSPGTGGGEQY | 0.76727855 |
| PLLATLRAIAVDVDAKPAQV | ASSPGTGGGEQY | 0.719975 |
| APKFRTGGYMIDFWGVGYQVAK | ASSPGTGGGEQY | 0.5346502 |
| GAGGAGGAGGAAGGPGGTGGTG | ASSPGTGGGEQY | 0.58591396 |
| IGALGLLATDVAPAGSVAFGVA | ASSPGTGGGEQY | 0.7618183 |
| AGGAGGIGGNGGAGGAGGVLMG | ASSPGTGGGEQY | 0.5087018 |
| ADELIGLL | ASSPGTGGGEQY | 0.6983674 |
| ADRALKA | ASSPGTGGGEQY | 0.6904515 |
| AGAVGQVGGAGGA | ASSPGTGGGEQY | 0.88097405 |
| AGGGGGLGGAP | ASSPGTGGGEQY | 0.794171 |
| AGGLLYGNGGAG | ASSPGTGGGEQY | 0.79563975 |
| AGLESIGL | ASSPGTGGGEQY | 0.5125488 |

| | | |
|---|---|---|
| AGLKTGII | ASSPGTGGGEQY | 0.7778387 |
| ASLAALLL | ASSPGTGGGEQY | 0.7803642 |
| ASVINISSI | ASSPGTGGGEQY | 0.4180248 |
| ATPPGPA | ASSPGTGGGEQY | 0.85141927 |
| ATVASALV | ASSPGTGGGEQY | 0.54022837 |
| ATVASLIL | ASSPGTGGGEQY | 0.70495033 |
| AVNGPVLIRH | ASSPGTGGGEQY | 0.8043449 |
| DGAAGAVGI | ASSPGTGGGEQY | 0.8184362 |
| DPLLHGI | ASSPGTGGGEQY | 0.70267916 |
| ELGAALGAL | ASSPGTGGGEQY | 0.7579371 |
| ELGRTQAV | ASSPGTGGGEQY | 0.56426805 |
| FVAGGYLS | ASSPGTGGGEQY | 0.8119432 |
| GASGGAGGAGGW | ASSPGTGGGEQY | 0.82403237 |
| GGAGAGGDGGAGGI | ASSPGTGGGEQY | 0.66196597 |
| GTGHNGSQPGMGGNG | ASSPGTGGGEQY | 0.8555803 |
| GYAAASASAS | ASSPGTGGGEQY | 0.7730627 |
| HQVDVAAVQQG | ASSPGTGGGEQY | 0.40031272 |
| IGAAIAHT | ASSPGTGGGEQY | 0.7115731 |
| IGMIEVAQL | ASSPGTGGGEQY | 0.83995783 |
| IRLAPEHTSAMD | ASSPGTGGGEQY | 0.82992345 |
| IYNVSVL | ASSPGTGGGEQY | 0.83428437 |
| KQGFLGM | ASSPGTGGGEQY | 0.81350756 |
| LESVITQL | ASSPGTGGGEQY | 0.6361958 |
| LGLGALVVV | ASSPGTGGGEQY | 0.8445629 |
| LIGNGGNGGSGGI | ASSPGTGGGEQY | 0.8683497 |
| LNAINAPTL | ASSPGTGGGEQY | 0.80060196 |
| LQLLNTGV | ASSPGTGGGEQY | 0.561593 |
| LQTSIEF | ASSPGTGGGEQY | 0.54131037 |
| MSFAVSEL | ASSPGTGGGEQY | 0.7752211 |
| NTGALGL | ASSPGTGGGEQY | 0.82361424 |
| PPGAGGAGGAGTT | ASSPGTGGGEQY | 0.4642919 |
| QLVGLGI | ASSPGTGGGEQY | 0.7586756 |
| RFLEQVHQL | ASSPGTGGGEQY | 0.75817806 |
| RGVGYVLR | ASSPGTGGGEQY | 0.36692935 |
| SATGVFAGVI | ASSPGTGGGEQY | 0.80650485 |
| SLPAITLP | ASSPGTGGGEQY | 0.85113335 |
| STALAFTV | ASSPGTGGGEQY | 0.77521765 |
| SVLTVGLGK | ASSPGTGGGEQY | 0.5500766 |
| TALAGAGSGI | ASSPGTGGGEQY | 0.8414881 |
| TGLPFTGI | ASSPGTGGGEQY | 0.8656702 |
| VAEHVTV | ASSPGTGGGEQY | 0.855646 |
| VDIVLNSL | ASSPGTGGGEQY | 0.3408187 |
| VLTGGGAVL | ASSPGTGGGEQY | 0.21420664 |
| VNVGLAYGI | ASSPGTGGGEQY | 0.6875738 |
| VSSAAVGAV | ASSPGTGGGEQY | 0.7756121 |
| VTSVYEAGV | ASSPGTGGGEQY | 0.6682237 |
| WVALDRGA | ASSPGTGGGEQY | 0.5998031 |
| YLVGSVVI | ASSPGTGGGEQY | 0.78064966 |
| GPPGTGKTLL | ASSPGTGGGEQY | 0.83297366 |
| GVGMTKF | ASSPGTGGGEQY | 0.56307846 |
| IGIGGSDL | ASSPGTGGGEQY | 0.42279425 |

| | | |
|---|---|---|
| IGTLLAEL | ASSPGTGGGEQY | 0.81239027 |
| IYTSGTT | ASSPGTGGGEQY | 0.7914081 |
| LSFPAGV | ASSPGTGGGEQY | 0.5767055 |
| LYTSGTT | ASSPGTGGGEQY | 0.6618409 |
| VGAFPFGF | ASSPGTGGGEQY | 0.72804856 |
| VGAGTVEF | ASSPGTGGGEQY | 0.6461137 |
| VGKSTLLL | ASSPGTGGGEQY | 0.7558201 |
| AALAHAGGGIV | ASSPGTGGGEQY | 0.4070617 |
| AALVAVSLL | ASSPGTGGGEQY | 0.84586835 |
| AAVSRAVAAL | ASSPGTGGGEQY | 0.44348824 |
| AGGSSGIGGF | ASSPGTGGGEQY | 0.82014894 |
| AGLTVANANG | ASSPGTGGGEQY | 0.7672348 |
| AGLVGAVL | ASSPGTGGGEQY | 0.78060013 |
| ALSQAFTV | ASSPGTGGGEQY | 0.6636739 |
| ALTFSVSL | ASSPGTGGGEQY | 0.5287068 |
| ALTGAALVAL | ASSPGTGGGEQY | 0.54072976 |
| ASFSIGAGVL | ASSPGTGGGEQY | 0.3258276 |
| DGSVYKI | ASSPGTGGGEQY | 0.8445946 |
| EQLAEIV | ASSPGTGGGEQY | 0.31015077 |
| FAVSELTVM | ASSPGTGGGEQY | 0.83192027 |
| GAAPLAAGL | ASSPGTGGGEQY | 0.77795494 |
| GADGGDGGSGGAGG | ASSPGTGGGEQY | 0.58241373 |
| GFGLDTGGAGG | ASSPGTGGGEQY | 0.7849772 |
| GLASGGLL | ASSPGTGGGEQY | 0.4635201 |
| GRIAGAAVLL | ASSPGTGGGEQY | 0.6996913 |
| GSAGSLNTSF | ASSPGTGGGEQY | 0.79457223 |
| GTGQIGI | ASSPGTGGGEQY | 0.81704926 |
| GTLIALIL | ASSPGTGGGEQY | 0.5435842 |
| IAATMGFVVG | ASSPGTGGGEQY | 0.7858956 |
| IFAQALL | ASSPGTGGGEQY | 0.76997733 |
| IGAAVALGF | ASSPGTGGGEQY | 0.54030085 |
| IGLTPLAL | ASSPGTGGGEQY | 0.76935065 |
| ILGSNTSTL | ASSPGTGGGEQY | 0.8125285 |
| IVFSLTVL | ASSPGTGGGEQY | 0.78319204 |
| LASKATGF | ASSPGTGGGEQY | 0.4376613 |
| LASVALIL | ASSPGTGGGEQY | 0.7947328 |
| LFAVGLTL | ASSPGTGGGEQY | 0.5772439 |
| LFNGGIGTYI | ASSPGTGGGEQY | 0.7695142 |
| LTLGAGII | ASSPGTGGGEQY | 0.76135564 |
| LVAAALGDI | ASSPGTGGGEQY | 0.65367126 |
| NTAFVQLGI | ASSPGTGGGEQY | 0.82733136 |
| QLFSGAEL | ASSPGTGGGEQY | 0.54863393 |
| SAATFITL | ASSPGTGGGEQY | 0.5656682 |
| SNYGLANLGV | ASSPGTGGGEQY | 0.6906209 |
| SYGGAYAVM | ASSPGTGGGEQY | 0.72708774 |
| TTGVGGLTLG | ASSPGTGGGEQY | 0.8074311 |
| TTQTIYI | ASSPGTGGGEQY | 0.5335737 |
| VAAGAGLGAVI | ASSPGTGGGEQY | 0.72484946 |
| VAAVVTLL | ASSPGTGGGEQY | 0.669579 |
| VGFLGGLI | ASSPGTGGGEQY | 0.4974463 |
| VGVIPILL | ASSPGTGGGEQY | 0.7312736 |

| | | |
|---|---|---|
| VIFASSLI | ASSPGTGGGEQY | 0.70826244 |
| VKGGDGGAAAG | ASSPGTGGGEQY | 0.4561491 |
| VTKTFTNI | ASSPGTGGGEQY | 0.8189494 |
| VVAAAILL | ASSPGTGGGEQY | 0.5779691 |
| YAALDVEL | ASSPGTGGGEQY | 0.77693236 |
| LASVALLL | ASSPGTGGGEQY | 0.86390567 |
| LDAVHRV | ASSPGTGGGEQY | 0.5383183 |
| QGLSSSIF | ASSPGTGGGEQY | 0.80782163 |
| RGTFANIRL | ASSPGTGGGEQY | 0.7707863 |
| RQLWWGHRI | ASSPGTGGGEQY | 0.7819407 |
| YGPPGTGKTLL | ASSPGTGGGEQY | 0.66423786 |
| AALVGAPANVA | ASSPGTGGGEQY | 0.6223062 |
| AATFRRTE | ASSPGTGGGEQY | 0.8498397 |
| AGALGGGI | ASSPGTGGGEQY | 0.8008739 |
| AGGAGGAGGIGAG | ASSPGTGGGEQY | 0.8293871 |
| AGSGALGL | ASSPGTGGGEQY | 0.8172069 |
| AQIADQI | ASSPGTGGGEQY | 0.79184663 |
| ASAVAVTYL | ASSPGTGGGEQY | 0.81763387 |
| ASTVLGGTGG | ASSPGTGGGEQY | 0.5902615 |
| AVGGEPAK | ASSPGTGGGEQY | 0.36775967 |
| CEQGTFGGTC | ASSPGTGGGEQY | 0.6676076 |
| DEGKAPAG | ASSPGTGGGEQY | 0.73174655 |
| DTVVFPAL | ASSPGTGGGEQY | 0.6306968 |
| FEGITMAG | ASSPGTGGGEQY | 0.3300547 |
| FPLVVQD | ASSPGTGGGEQY | 0.82660043 |
| GAFTTPQ | ASSPGTGGGEQY | 0.8697525 |
| GFGGFGTAGGVGGA | ASSPGTGGGEQY | 0.5476233 |
| GFLAQFL | ASSPGTGGGEQY | 0.8643136 |
| GGAGGAGGNAGVA | ASSPGTGGGEQY | 0.45971555 |
| GGVGTAGAGGAGGAGG | ASSPGTGGGEQY | 0.79580253 |
| GNGGGGDAGNAGSGGN | ASSPGTGGGEQY | 0.8339138 |
| GTGGAGGAGNDAGS | ASSPGTGGGEQY | 0.4867345 |
| GTTNTGWFNS | ASSPGTGGGEQY | 0.7910918 |
| IFMGPLFTL | ASSPGTGGGEQY | 0.5934387 |
| ILAHAIEAI | ASSPGTGGGEQY | 0.6868197 |
| IRSAIREAL | ASSPGTGGGEQY | 0.84457815 |
| ITFAGGG | ASSPGTGGGEQY | 0.636698 |
| ITGMSGHQEV | ASSPGTGGGEQY | 0.810027 |
| IVAGLACTL | ASSPGTGGGEQY | 0.6687948 |
| IVGGGLAGA | ASSPGTGGGEQY | 0.5525688 |
| IVTGVLNI | ASSPGTGGGEQY | 0.7826967 |
| LAQQGCI | ASSPGTGGGEQY | 0.84562486 |
| LCLHVEN | ASSPGTGGGEQY | 0.6596745 |
| LDAISDGS | ASSPGTGGGEQY | 0.86211777 |
| LGGFAAML | ASSPGTGGGEQY | 0.60602635 |
| LGGKNPMI | ASSPGTGGGEQY | 0.86358535 |
| LGPKGVAAIL | ASSPGTGGGEQY | 0.8259101 |
| LMVGVSI | ASSPGTGGGEQY | 0.71308017 |
| LVGAAFVL | ASSPGTGGGEQY | 0.6678801 |
| LVLAVAPL | ASSPGTGGGEQY | 0.8080718 |
| MAAIAVIDAA | ASSPGTGGGEQY | 0.59049803 |

| | | |
|---|---|---|
| NNGINAELQ | ASSPGTGGGEQY | 0.8121699 |
| RGEPLVLVHG | ASSPGTGGGEQY | 0.8421774 |
| RGGAGGAAGLI | ASSPGTGGGEQY | 0.41858393 |
| RSYRMHPAV | ASSPGTGGGEQY | 0.78194314 |
| SAAHLLAGQLA | ASSPGTGGGEQY | 0.7454973 |
| SAAVGSY | ASSPGTGGGEQY | 0.7006769 |
| SPHAVLHRI | ASSPGTGGGEQY | 0.5943967 |
| SQIAKVL | ASSPGTGGGEQY | 0.33614382 |
| TALGHLVP | ASSPGTGGGEQY | 0.62037396 |
| TAPTTSGAA | ASSPGTGGGEQY | 0.52304614 |
| TDGDDGNGGSPGT | ASSPGTGGGEQY | 0.7972764 |
| TDGLGEN | ASSPGTGGGEQY | 0.76680565 |
| TGGPPLPGQA | ASSPGTGGGEQY | 0.538484 |
| TIPAITI | ASSPGTGGGEQY | 0.7854172 |
| TIPGFTI | ASSPGTGGGEQY | 0.68559223 |
| TSIASGALA | ASSPGTGGGEQY | 0.47161227 |
| TVTLPGAAI | ASSPGTGGGEQY | 0.6517909 |
| VASGLSAALP | ASSPGTGGGEQY | 0.63067454 |
| VGAGVIGI | ASSPGTGGGEQY | 0.69137454 |
| VGPKRFGV | ASSPGTGGGEQY | 0.40371943 |
| VNVAFPD | ASSPGTGGGEQY | 0.7362751 |
| VSMTNSGVS | ASSPGTGGGEQY | 0.71335167 |
| VTDVLGDLSV | ASSPGTGGGEQY | 0.29131788 |
| VVVVGAGFAGL | ASSPGTGGGEQY | 0.75032985 |
| VYTVGTVV | ASSPGTGGGEQY | 0.57980496 |
| WTTFSGP | ASSPGTGGGEQY | 0.56846005 |
| YDELVEAF | ASSPGTGGGEQY | 0.76870763 |
| YGTGGAGGNGG | ASSPGTGGGEQY | 0.37177768 |
| YTNVMAV | ASSPGTGGGEQY | 0.4269898 |
| FLEQVHQL | ASSPGTGGGEQY | 0.77781284 |
| GLSLLAVAV | ASSPGTGGGEQY | 0.77999794 |
| LEQVHQL | ASSPGTGGGEQY | 0.34815973 |
| LNAAGAI | ASSPGTGGGEQY | 0.73600614 |
| PAVHASL | ASSPGTGGGEQY | 0.7811134 |
| TELEDDA | ASSPGTGGGEQY | 0.74569046 |
| TPPGPPN | ASSPGTGGGEQY | 0.77033937 |
| TPPPPPPP | ASSPGTGGGEQY | 0.77472734 |
| TQSPALI | GARVESQNTLY | 0.70210254 |
| EFAAAFPQ | GARVESQNTLY | 0.7818763 |
| LLSGNLDVL | GARVESQNTLY | 0.7979752 |
| GPANLGNYNI | GARVESQNTLY | 0.62164783 |
| TLGPTLP | GARVESQNTLY | 0.7099882 |
| VYAYSSCCK | GARVESQNTLY | 0.41470772 |
| APAPAPAPNI | GARVESQNTLY | 0.63878167 |
| PGGPVPIP | GARVESQNTLY | 0.41854694 |
| FLTVALEP | GARVESQNTLY | 0.7559172 |
| SQYFGANL | GARVESQNTLY | 0.78464013 |
| GGVKPAA | GARVESQNTLY | 0.6875857 |
| GGTGGAGGAGGW | GARVESQNTLY | 0.46628883 |
| GVGGVAGLGGAGG | GARVESQNTLY | 0.58224344 |
| VLLVLATTI | GARVESQNTLY | 0.7241429 |

| | | |
|---|---|---|
| IGIFAAML | GARVESQNTLY | 0.41495955 |
| AVGDLID | GARVESQNTLY | 0.41734284 |
| NISPLPL | GARVESQNTLY | 0.68876696 |
| VTAVFVAI | GARVESQNTLY | 0.7724073 |
| SGDDNDAAL | GARVESQNTLY | 0.757033 |
| AVAALAHS | GARVESQNTLY | 0.52259296 |
| GGVTVMATP | GARVESQNTLY | 0.7847382 |
| AGLAANVRD | GARVESQNTLY | 0.776784 |
| TNRPDLIDPALLRPG | GARVESQNTLY | 0.77076304 |
| LANLGVTAL | GARVESQNTLY | 0.5993776 |
| TGERIEL | GARVESQNTLY | 0.41748425 |
| NGEELAA | GARVESQNTLY | 0.78930193 |
| VLSLVWG | GARVESQNTLY | 0.69695795 |
| IGYALWL | GARVESQNTLY | 0.69878066 |
| LSLLLAPL | GARVESQNTLY | 0.7543057 |
| TLDEALK | GARVESQNTLY | 0.6110119 |
| TDGAGTAEGG | GARVESQNTLY | 0.52560204 |
| NAPGGAGGNGG | GARVESQNTLY | 0.7312846 |
| VSVFMGMGGNF | GARVESQNTLY | 0.7822058 |
| LASYLAAL | GARVESQNTLY | 0.55233437 |
| LVGGTPAL | GARVESQNTLY | 0.46816084 |
| VRNADVL | GARVESQNTLY | 0.572603 |
| TCSSPFLVSAM | GARVESQNTLY | 0.6244011 |
| VHITDAE | GARVESQNTLY | 0.7725544 |
| EYAAPSF | GARVESQNTLY | 0.69786346 |
| VVDAFHGGL | GARVESQNTLY | 0.6823138 |
| ASVVDPQ | GARVESQNTLY | 0.7552935 |
| SYNEFGGLS | GARVESQNTLY | 0.7196334 |
| DPDGSVY | GARVESQNTLY | 0.6867603 |
| MVSTIHATE | GARVESQNTLY | 0.6804199 |
| LAAGVADPV | GARVESQNTLY | 0.62967646 |
| TKEQIAA | GARVESQNTLY | 0.77070534 |
| SLASPAAL | GARVESQNTLY | 0.74772274 |
| IGLVFYGV | GARVESQNTLY | 0.712677 |
| FLLDASNL | GARVESQNTLY | 0.76455545 |
| YQVGNFP | GARVESQNTLY | 0.68358135 |
| ESLLSQST | GARVESQNTLY | 0.7221352 |
| LFTVGFAAVG | GARVESQNTLY | 0.47752464 |
| ATPPTPT | GARVESQNTLY | 0.59994304 |
| TDLATLG | GARVESQNTLY | 0.7356682 |
| SAGGTGR | GARVESQNTLY | 0.7280166 |
| GFSNSGS | GARVESQNTLY | 0.757895 |
| GGNGGSGGTSV | GARVESQNTLY | 0.6669222 |
| GTSVATGGAGN | GARVESQNTLY | 0.6722788 |
| NGGNGGSGGTS | GARVESQNTLY | 0.74221337 |
| AVVKFPH | GARVESQNTLY | 0.7834656 |
| TPPPPPP | GARVESQNTLY | 0.5137101 |
| AAPAAAAAPA | GARVESQNTLY | 0.71591324 |
| IALNEDL | GARVESQNTLY | 0.79189634 |
| IVNVSSM | GARVESQNTLY | 0.76008964 |
| TGIPPAPR | GARVESQNTLY | 0.67458314 |

| | | |
|---|---|---|
| FDLGGGTF | GARVESQNTLY | 0.6340032 |
| FDLGGGTFDVS | GARVESQNTLY | 0.8003978 |
| NEPTAAA | GARVESQNTLY | 0.6729815 |
| NEPTAAALA | GARVESQNTLY | 0.80155516 |
| GSIINISSI | GARVESQNTLY | 0.2974682 |
| VAPGNAGTA | GARVESQNTLY | 0.6253524 |
| TGGGASLE | GARVESQNTLY | 0.5154823 |
| VAVYNHYKRI | GARVESQNTLY | 0.24418451 |
| TDPAPATT | GARVESQNTLY | 0.7623325 |
| TNRPDILDPALLRPG | GARVESQNTLY | 0.5726229 |
| LTTTTS | GARVESQNTLY | 0.7262558 |
| PTGSGKTL | GARVESQNTLY | 0.7591133 |
| TGSGKTL | GARVESQNTLY | 0.7655996 |
| TIGHVDHGK | GARVESQNTLY | 0.7444033 |
| TIGHVDHGKTT | GARVESQNTLY | 0.6332721 |
| GHVDHGKTT | GARVESQNTLY | 0.51261854 |
| AQFEHTLL | GARVESQNTLY | 0.73140943 |
| IVNQDEL | GARVESQNTLY | 0.7690355 |
| TNGSQFF | GARVESQNTLY | 0.73915577 |
| TGKGIVE | GARVESQNTLY | 0.32156426 |
| LISNASDA | GARVESQNTLY | 0.747143 |
| AELRTLGELTKIAKAHGAQ | GARVESQNTLY | 0.66367054 |
| APNAVAAAEPSW | GARVESQNTLY | 0.2978444 |
| APQVAGAATAVL | GARVESQNTLY | 0.30624932 |
| ARRDPAAAAAILPTDA | GARVESQNTLY | 0.6341381 |
| DAAARDVLAEAGLAEYF | GARVESQNTLY | 0.6355119 |
| DDQIVLSAKANLAKVEQ | GARVESQNTLY | 0.7743374 |
| DELLPDTGEDSDEDGH | GARVESQNTLY | 0.7742646 |
| ELPEAPANRATVPQD | GARVESQNTLY | 0.64615625 |
| FGQSKIERVTGIG | GARVESQNTLY | 0.76517093 |
| GAGGTGSTAGGAGGAGGAGGLY | GARVESQNTLY | 0.72061086 |
| GHGGTGAAGGNGGTGGAGGSGI | GARVESQNTLY | 0.17478476 |
| GNGGNGGAGGEGGPGGAGGAGA | GARVESQNTLY | 0.64789987 |
| GSTAGNGGNGGNATNGTIAGQP | GARVESQNTLY | 0.4565645 |
| GTGGDGGNGGNGGTGG | GARVESQNTLY | 0.76003516 |
| GVRRPAGIPADAPV | GARVESQNTLY | 0.654333 |
| GYRQVTDAHLLTLARR | GARVESQNTLY | 0.67961735 |
| IAKMGQDGHDRG | GARVESQNTLY | 0.6873152 |
| KAEPSIEIQYT | GARVESQNTLY | 0.76654387 |
| KPVRIRMAADEDTVAVT | GARVESQNTLY | 0.6904657 |
| MPPRIAGMRLLVIKPEP | GARVESQNTLY | 0.5257574 |
| MKFYAVALFRSRGVPPTR | GARVESQNTLY | 0.3069895 |
| NAGGFGILGAVAHSP | GARVESQNTLY | 0.7390187 |
| NEGREITLAMAEM | GARVESQNTLY | 0.5780795 |
| NELVEWSHPLGHRWRWE | GARVESQNTLY | 0.42439798 |
| NLGAGGTGGNGGNPG | GARVESQNTLY | 0.52330077 |
| NTEWPKWETVMAADPVD | GARVESQNTLY | 0.7377445 |
| PASAAPAETQP | GARVESQNTLY | 0.3150782 |
| QQSSDGKNGVRAAEITGE | GARVESQNTLY | 0.74521005 |
| RAPDAVTTSWWKE | GARVESQNTLY | 0.6075001 |
| SGFGNVGAHTSGWFNQ | GARVESQNTLY | 0.64438385 |

| | | |
|---|---|---|
| SGVAGAVGQAASVGGLKV | GARVESQNTLY | 0.7958071 |
| SNGGAGTGSAGGNGGT | GARVESQNTLY | 0.7573492 |
| TGGAGGEGGAAG | GARVESQNTLY | 0.684819 |
| TGGDGGAAGAGGAPGHG | GARVESQNTLY | 0.5261109 |
| TMPPLLAAIANAGRHGV | GARVESQNTLY | 0.6008096 |
| TSSTAGNSGGA | GARVESQNTLY | 0.5654726 |
| VPATVPVGILC | GARVESQNTLY | 0.47822857 |
| VQRPDSGGVVLTGRIS | GARVESQNTLY | 0.7553862 |
| WTPNPVPATKGVPAGLVQAT | GARVESQNTLY | 0.65992963 |
| YALKRALAKFSDRFDVVII | GARVESQNTLY | 0.49042913 |
| YLIGSVIV | GARVESQNTLY | 0.60265195 |
| YGQLVAVID | GARVESQNTLY | 0.7899257 |
| VTVGGMTLPA | GARVESQNTLY | 0.80133474 |
| VLALPAV | GARVESQNTLY | 0.6401332 |
| VIPPTLNYET | GARVESQNTLY | 0.5345026 |
| VGMGGMHPSSG | GARVESQNTLY | 0.56587785 |
| VGLAGGAATA | GARVESQNTLY | 0.707835 |
| VGGAGGAGGSGP | GARVESQNTLY | 0.4621148 |
| VFAAFMV | GARVESQNTLY | 0.3886897 |
| TLYDKLR | GARVESQNTLY | 0.7705022 |
| TLAVFTVL | GARVESQNTLY | 0.80432785 |
| TGSVQPARV | GARVESQNTLY | 0.65719914 |
| TGGGAGSGGT | GARVESQNTLY | 0.47199798 |
| SHLETSL | GARVESQNTLY | 0.6717155 |
| SFTTGMT | GARVESQNTLY | 0.5456493 |
| QVEVAKRMGVRQPTVS | GARVESQNTLY | 0.5894011 |
| QGSGGPALAM | GARVESQNTLY | 0.6285611 |
| MLAPSNP | GARVESQNTLY | 0.44084698 |
| LVNNAAI | GARVESQNTLY | 0.6766226 |
| LTGEINTLA | GARVESQNTLY | 0.7923848 |
| LSIFGPV | GARVESQNTLY | 0.7418192 |
| LLALLANL | GARVESQNTLY | 0.7487577 |
| LISNASDALDKLR | GARVESQNTLY | 0.79630405 |
| LINEAVP | GARVESQNTLY | 0.7681978 |
| LGYAFSGL | GARVESQNTLY | 0.7915993 |
| LDFVAVAL | GARVESQNTLY | 0.7827081 |
| LAQPGAE | GARVESQNTLY | 0.6340859 |
| ISNLGAVTGF | GARVESQNTLY | 0.674558 |
| HQEDPGRI | GARVESQNTLY | 0.7864568 |
| GTPGEPTP | GARVESQNTLY | 0.6693474 |
| GFFNSGS | GARVESQNTLY | 0.7213966 |
| GAGGGGLTAGAN | GARVESQNTLY | 0.65261143 |
| FVDALVNM | GARVESQNTLY | 0.7817941 |
| FLTVAPEL | GARVESQNTLY | 0.61518055 |
| FETLTAL | GARVESQNTLY | 0.7950498 |
| DVINPIEL | GARVESQNTLY | 0.25962812 |
| APVSGAPV | GARVESQNTLY | 0.36598253 |
| ALVGGLAW | GARVESQNTLY | 0.78205794 |
| ALAAAAATAI | GARVESQNTLY | 0.19220224 |
| AFGLGLLL | GARVESQNTLY | 0.2942657 |
| AFFAAASL | GARVESQNTLY | 0.72942436 |

| | | |
|---|---|---|
| AAAALPAFSPP | GARVESQNTLY | 0.78349453 |
| VVLAPAL | GARVESQNTLY | 0.2051724 |
| TIGHVDHGKTTL | GARVESQNTLY | 0.5105387 |
| LISNASDALDK | GARVESQNTLY | 0.39515287 |
| LISNASDAL | GARVESQNTLY | 0.5991796 |
| ISNASDALDK | GARVESQNTLY | 0.64055157 |
| NQAETLVYQTE | GARVESQNTLY | 0.3896473 |
| TGGTGGTAGAG | GARVESQNTLY | 0.72644794 |
| TGSTAGGAGGA | GARVESQNTLY | 0.39922827 |
| DPLAVPVAAGPV | GARVESQNTLY | 0.6477922 |
| GGTGGTPGNGGN | GARVESQNTLY | 0.7671652 |
| PAGLAGVDPATL | GARVESQNTLY | 0.7969574 |
| GNNLVVEAPHAGASV | GARVESQNTLY | 0.6169712 |
| KVTRIMSASTLEQPA | GARVESQNTLY | 0.7264595 |
| LGFALPIALPRRTVV | GARVESQNTLY | 0.62482566 |
| TRLGNLTARKLVVLV | GARVESQNTLY | 0.8081134 |
| WSQQRRLIYFPSAGP | GARVESQNTLY | 0.6406622 |
| GPNGNPDPMAAAVDIR | GARVESQNTLY | 0.3921353 |
| TQRCVSVADLRAAASR | GARVESQNTLY | 0.7635714 |
| QFVLDVAEHPVEVLTETV | GARVESQNTLY | 0.73945594 |
| GGAGGTGGAGGNGGTQAGDG | GARVESQNTLY | 0.5917218 |
| PLLATLRAIAVDVDAKPAQV | GARVESQNTLY | 0.7814926 |
| APKFRTGGYMIDFWGVGYQVAK | GARVESQNTLY | 0.5151606 |
| GAGGAGGAGGAAGGPGGTGGTG | GARVESQNTLY | 0.6736299 |
| IGALGLLATDVAPAGSVAFGVA | GARVESQNTLY | 0.7726208 |
| AGGAGGIGGNGGAGGAGGVLMG | GARVESQNTLY | 0.7751524 |
| ADELIGLL | GARVESQNTLY | 0.42173308 |
| ADRALKA | GARVESQNTLY | 0.75211287 |
| AGAVGQVGGAGGA | GARVESQNTLY | 0.6872385 |
| AGGGGGLGGAP | GARVESQNTLY | 0.49015462 |
| AGGLLYGNGGAG | GARVESQNTLY | 0.7971085 |
| AGLESIGL | GARVESQNTLY | 0.6322096 |
| AGLKTGII | GARVESQNTLY | 0.76547635 |
| ASLAALLL | GARVESQNTLY | 0.7418207 |
| ASVINISSI | GARVESQNTLY | 0.7965584 |
| ATPPGPA | GARVESQNTLY | 0.53921205 |
| ATVASALV | GARVESQNTLY | 0.7404118 |
| ATVASLIL | GARVESQNTLY | 0.7229451 |
| AVNGPVLIRH | GARVESQNTLY | 0.7398106 |
| DGAAGAVGI | GARVESQNTLY | 0.61509037 |
| DPLLHGI | GARVESQNTLY | 0.7442695 |
| ELGAALGAL | GARVESQNTLY | 0.7554525 |
| ELGRTQAV | GARVESQNTLY | 0.7774662 |
| FVAGGYLS | GARVESQNTLY | 0.7181928 |
| GASGGAGGAGGW | GARVESQNTLY | 0.64702183 |
| GGAGAGGDGGAGGI | GARVESQNTLY | 0.79375887 |
| GTGHNGSQPGMGGNG | GARVESQNTLY | 0.47855288 |
| GYAAASASAS | GARVESQNTLY | 0.6664235 |
| HQVDVAAVQQG | GARVESQNTLY | 0.76297724 |
| IGAAIAHT | GARVESQNTLY | 0.74442613 |
| IGMIEVAQL | GARVESQNTLY | 0.64980274 |

| | | |
|---|---|---|
| IRLAPEHTSAMD | GARVESQNTLY | 0.7922991 |
| IYNVSVL | GARVESQNTLY | 0.62554395 |
| KQGFLGM | GARVESQNTLY | 0.77834845 |
| LESVITQL | GARVESQNTLY | 0.7337902 |
| LGLGALVVV | GARVESQNTLY | 0.67105657 |
| LIGNGGNGGSGGI | GARVESQNTLY | 0.5094217 |
| LNAINAPTL | GARVESQNTLY | 0.781263 |
| LQLLNTGV | GARVESQNTLY | 0.77759695 |
| LQTSIEF | GARVESQNTLY | 0.7724002 |
| MSFAVSEL | GARVESQNTLY | 0.3188373 |
| NTGALGL | GARVESQNTLY | 0.7592148 |
| PPGAGGAGGAGTT | GARVESQNTLY | 0.7774368 |
| QLVGLGI | GARVESQNTLY | 0.8024536 |
| RFLEQVHQL | GARVESQNTLY | 0.64835083 |
| RGVGYVLR | GARVESQNTLY | 0.71948195 |
| SATGVFAGVI | GARVESQNTLY | 0.6558152 |
| SLPAITLP | GARVESQNTLY | 0.69023514 |
| STALAFTV | GARVESQNTLY | 0.4617196 |
| SVLTVGLGK | GARVESQNTLY | 0.6232592 |
| TALAGAGSGI | GARVESQNTLY | 0.69877195 |
| TGLPFTGI | GARVESQNTLY | 0.59173405 |
| VAEHVTV | GARVESQNTLY | 0.7858162 |
| VDIVLNSL | GARVESQNTLY | 0.77055776 |
| VLTGGGAVL | GARVESQNTLY | 0.75399184 |
| VNVGLAYGI | GARVESQNTLY | 0.7814684 |
| VSSAAVGAV | GARVESQNTLY | 0.792552 |
| VTSVYEAGV | GARVESQNTLY | 0.70110726 |
| WVALDRGA | GARVESQNTLY | 0.6282916 |
| YLVGSVVI | GARVESQNTLY | 0.7362227 |
| GPPGTGKTLL | GARVESQNTLY | 0.43807817 |
| GVGMTKF | GARVESQNTLY | 0.6579697 |
| IGIGGSDL | GARVESQNTLY | 0.6221722 |
| IGTLLAEL | GARVESQNTLY | 0.6883515 |
| IYTSGTT | GARVESQNTLY | 0.18454576 |
| LSFPAGV | GARVESQNTLY | 0.7618318 |
| LYTSGTT | GARVESQNTLY | 0.7769103 |
| VGAFPFGF | GARVESQNTLY | 0.7221137 |
| VGAGTVEF | GARVESQNTLY | 0.71799195 |
| VGKSTLLL | GARVESQNTLY | 0.76628304 |
| AALAHAGGGIV | GARVESQNTLY | 0.6077599 |
| AALVAVSLL | GARVESQNTLY | 0.74577117 |
| AAVSRAVAAL | GARVESQNTLY | 0.550126 |
| AGGSSGIGGF | GARVESQNTLY | 0.76027966 |
| AGLTVANANG | GARVESQNTLY | 0.7809086 |
| AGLVGAVL | GARVESQNTLY | 0.7433387 |
| ALSQAFTV | GARVESQNTLY | 0.6128602 |
| ALTFSVSL | GARVESQNTLY | 0.665665 |
| ALTGAALVAL | GARVESQNTLY | 0.3597939 |
| ASFSIGAGVL | GARVESQNTLY | 0.6337113 |
| DGSVYKI | GARVESQNTLY | 0.4564582 |
| EQLAEIV | GARVESQNTLY | 0.73252547 |

| | | |
|---|---|---|
| FAVSELTVM | GARVESQNTLY | 0.62920094 |
| GAAPLAAGL | GARVESQNTLY | 0.7499206 |
| GADGGDGGSGGAGG | GARVESQNTLY | 0.6896884 |
| GFGLDTGGAGG | GARVESQNTLY | 0.725786 |
| GLASGGLL | GARVESQNTLY | 0.62347674 |
| GRIAGAAVLL | GARVESQNTLY | 0.74906725 |
| GSAGSLNTSF | GARVESQNTLY | 0.751506 |
| GTGQIGI | GARVESQNTLY | 0.3845864 |
| GTLIALIL | GARVESQNTLY | 0.78988296 |
| IAATMGFVVG | GARVESQNTLY | 0.6782501 |
| IFAQALL | GARVESQNTLY | 0.6912008 |
| IGAAVALGF | GARVESQNTLY | 0.7794795 |
| IGLTPLAL | GARVESQNTLY | 0.6569101 |
| ILGSNTSTL | GARVESQNTLY | 0.4474217 |
| IVFSLTVL | GARVESQNTLY | 0.69190377 |
| LASKATGF | GARVESQNTLY | 0.7404874 |
| LASVALIL | GARVESQNTLY | 0.69065595 |
| LFAVGLTL | GARVESQNTLY | 0.22163634 |
| LFNGGIGTYI | GARVESQNTLY | 0.7517939 |
| LTLGAGII | GARVESQNTLY | 0.4784516 |
| LVAAALGDI | GARVESQNTLY | 0.63162583 |
| NTAFVQLGI | GARVESQNTLY | 0.65207493 |
| QLFSGAEL | GARVESQNTLY | 0.77042496 |
| SAATFITL | GARVESQNTLY | 0.66357386 |
| SNYGLANLGV | GARVESQNTLY | 0.5375755 |
| SYGGAYAVM | GARVESQNTLY | 0.7540267 |
| TTGVGGLTLG | GARVESQNTLY | 0.6924338 |
| TTQTIYI | GARVESQNTLY | 0.7752193 |
| VAAGAGLGAVI | GARVESQNTLY | 0.687757 |
| VAAVVTLL | GARVESQNTLY | 0.6380503 |
| VGFLGGLI | GARVESQNTLY | 0.6576294 |
| VGVIPILL | GARVESQNTLY | 0.5195942 |
| VIFASSLI | GARVESQNTLY | 0.59564555 |
| VKGGDGGAAAG | GARVESQNTLY | 0.5123856 |
| VTKTFTNI | GARVESQNTLY | 0.6979693 |
| VVAAAILL | GARVESQNTLY | 0.47918057 |
| YAALDVEL | GARVESQNTLY | 0.7349793 |
| LASVALLL | GARVESQNTLY | 0.73584116 |
| LDAVHRV | GARVESQNTLY | 0.6731936 |
| QGLSSSIF | GARVESQNTLY | 0.3787019 |
| RGTFANIRL | GARVESQNTLY | 0.6498615 |
| RQLWWGHRI | GARVESQNTLY | 0.6804306 |
| YGPPGTGKTLL | GARVESQNTLY | 0.7648941 |
| AALVGAPANVA | GARVESQNTLY | 0.7363738 |
| AATFRRTE | GARVESQNTLY | 0.8007441 |
| AGALGGGI | GARVESQNTLY | 0.772647 |
| AGGAGGAGGIGAG | GARVESQNTLY | 0.61237776 |
| AGSGALGL | GARVESQNTLY | 0.7740594 |
| AQIADQI | GARVESQNTLY | 0.6068089 |
| ASAVAVTYL | GARVESQNTLY | 0.7322279 |
| ASTVLGGTGG | GARVESQNTLY | 0.7252984 |

| | | |
|---|---|---|
| AVGGEPAK | GARVESQNTLY | 0.7071357 |
| CEQGTFGGTC | GARVESQNTLY | 0.5692184 |
| DEGKAPAG | GARVESQNTLY | 0.64134 |
| DTVVFPAL | GARVESQNTLY | 0.3954491 |
| FEGITMAG | GARVESQNTLY | 0.7632683 |
| FPLVVQD | GARVESQNTLY | 0.5868407 |
| GAFTTPQ | GARVESQNTLY | 0.4408715 |
| GFGGFGTAGGVGGA | GARVESQNTLY | 0.6244796 |
| GFLAQFL | GARVESQNTLY | 0.7990943 |
| GGAGGAGGNAGVA | GARVESQNTLY | 0.7742659 |
| GGVGTAGAGGAGGAGG | GARVESQNTLY | 0.63529056 |
| GNGGGGDAGNAGSGGN | GARVESQNTLY | 0.5744101 |
| GTGGAGGAGNDAGS | GARVESQNTLY | 0.7672348 |
| GTTNTGWFNS | GARVESQNTLY | 0.5432282 |
| IFMGPLFTL | GARVESQNTLY | 0.7342031 |
| ILAHAIEAI | GARVESQNTLY | 0.70207 |
| IRSAIREAL | GARVESQNTLY | 0.7166109 |
| ITFAGGG | GARVESQNTLY | 0.53342605 |
| ITGMSGHQEV | GARVESQNTLY | 0.7514506 |
| IVAGLACTL | GARVESQNTLY | 0.6723975 |
| IVGGGLAGA | GARVESQNTLY | 0.6691485 |
| IVTGVLNI | GARVESQNTLY | 0.27141708 |
| LAQQGCI | GARVESQNTLY | 0.74757683 |
| LCLHVEN | GARVESQNTLY | 0.6125727 |
| LDAISDGS | GARVESQNTLY | 0.55707675 |
| LGGFAAML | GARVESQNTLY | 0.30565023 |
| LGGKNPMI | GARVESQNTLY | 0.6654494 |
| LGPKGVAAIL | GARVESQNTLY | 0.7824391 |
| LMVGVSI | GARVESQNTLY | 0.78058887 |
| LVGAAFVL | GARVESQNTLY | 0.6872158 |
| LVLAVAPL | GARVESQNTLY | 0.72322464 |
| MAAIAVIDAA | GARVESQNTLY | 0.79152954 |
| NNGINAELQ | GARVESQNTLY | 0.6632577 |
| RGEPLVLVHG | GARVESQNTLY | 0.36924738 |
| RGGAGGAAGLI | GARVESQNTLY | 0.6602303 |
| RSYRMHPAV | GARVESQNTLY | 0.744832 |
| SAAHLLAGQLA | GARVESQNTLY | 0.71996 |
| SAAVGSY | GARVESQNTLY | 0.7708205 |
| SPHAVLHRI | GARVESQNTLY | 0.4931675 |
| SQIAKVL | GARVESQNTLY | 0.7181556 |
| TALGHLVP | GARVESQNTLY | 0.7674613 |
| TAPTTSGAA | GARVESQNTLY | 0.43926704 |
| TDGDDGNGGSPGT | GARVESQNTLY | 0.59915286 |
| TDGLGEN | GARVESQNTLY | 0.5573014 |
| TGGPPLPGQA | GARVESQNTLY | 0.457524 |
| TIPAITI | GARVESQNTLY | 0.72318316 |
| TIPGFTI | GARVESQNTLY | 0.6372784 |
| TSIASGALA | GARVESQNTLY | 0.49515623 |
| TVTLPGAAI | GARVESQNTLY | 0.6768173 |
| VASGLSAALP | GARVESQNTLY | 0.48526508 |
| VGAGVIGI | GARVESQNTLY | 0.49088866 |

| | | |
|---|---|---|
| VGPKRFGV | GARVESQNTLY | 0.53535175 |
| VNVAFPD | GARVESQNTLY | 0.7316729 |
| VSMTNSGVS | GARVESQNTLY | 0.7627958 |
| VTDVLGDLSV | GARVESQNTLY | 0.6557008 |
| VVVVGAGFAGL | GARVESQNTLY | 0.6532787 |
| VYTVGTVV | GARVESQNTLY | 0.7963456 |
| WTTFSGP | GARVESQNTLY | 0.32807934 |
| YDELVEAF | GARVESQNTLY | 0.5223314 |
| YGTGGAGGNGG | GARVESQNTLY | 0.7122717 |
| YTNVMAV | GARVESQNTLY | 0.63132596 |
| FLEQVHQL | GARVESQNTLY | 0.7845932 |
| GLSLLAVAV | GARVESQNTLY | 0.71365863 |
| LEQVHQL | GARVESQNTLY | 0.7237926 |
| LNAAGAI | GARVESQNTLY | 0.7650027 |
| PAVHASL | GARVESQNTLY | 0.7953086 |
| TELEDDA | GARVESQNTLY | 0.7405859 |
| TPPGPPN | GARVESQNTLY | 0.63145244 |
| TPPPPPPP | GARVESQNTLY | 0.78772867 |
| TQSPALI | ASSPRLNQDTQY | 0.37894258 |
| EFAAAFPQ | ASSPRLNQDTQY | 0.74545026 |
| LLSGNLDVL | ASSPRLNQDTQY | 0.7488202 |
| GPANLGNYNI | ASSPRLNQDTQY | 0.53694785 |
| TLGPTLP | ASSPRLNQDTQY | 0.6812555 |
| VYAYSSCCK | ASSPRLNQDTQY | 0.7912663 |
| APAPAPAPNI | ASSPRLNQDTQY | 0.78208756 |
| PGGPVPIP | ASSPRLNQDTQY | 0.5265379 |
| FLTVALEP | ASSPRLNQDTQY | 0.6227555 |
| SQYFGANL | ASSPRLNQDTQY | 0.42681175 |
| GGVKPAA | ASSPRLNQDTQY | 0.5294887 |
| GGTGGAGGAGGW | ASSPRLNQDTQY | 0.33670086 |
| GVGGVAGLGGAGG | ASSPRLNQDTQY | 0.73782337 |
| VLLVLATTI | ASSPRLNQDTQY | 0.666909 |
| IGIFAAML | ASSPRLNQDTQY | 0.7815902 |
| AVGDLID | ASSPRLNQDTQY | 0.73244655 |
| NISPLPL | ASSPRLNQDTQY | 0.77179444 |
| VTAVFVAI | ASSPRLNQDTQY | 0.65248615 |
| SGDDNDAAL | ASSPRLNQDTQY | 0.6461159 |
| AVAALAHS | ASSPRLNQDTQY | 0.36032856 |
| GGVTVMATP | ASSPRLNQDTQY | 0.5814839 |
| AGLAANVRD | ASSPRLNQDTQY | 0.66219854 |
| TNRPDLIDPALLRPG | ASSPRLNQDTQY | 0.5732562 |
| LANLGVTAL | ASSPRLNQDTQY | 0.35908586 |
| TGERIEL | ASSPRLNQDTQY | 0.77589333 |
| NGEELAA | ASSPRLNQDTQY | 0.6506974 |
| VLSLVWG | ASSPRLNQDTQY | 0.74408126 |
| IGYALWL | ASSPRLNQDTQY | 0.57281697 |
| LSLLLAPL | ASSPRLNQDTQY | 0.45371655 |
| TLDEALK | ASSPRLNQDTQY | 0.63398355 |
| TDGAGTAEGG | ASSPRLNQDTQY | 0.7652452 |
| NAPGGAGGNGG | ASSPRLNQDTQY | 0.6351377 |
| VSVFMGMGGNF | ASSPRLNQDTQY | 0.30030316 |

| | | |
|---|---|---|
| LASYLAAL | ASSPRLNQDTQY | 0.48831296 |
| LVGGTPAL | ASSPRLNQDTQY | 0.57491636 |
| VRNADVL | ASSPRLNQDTQY | 0.7977065 |
| TCSSPFLVSAM | ASSPRLNQDTQY | 0.78672063 |
| VHITDAE | ASSPRLNQDTQY | 0.73758197 |
| EYAAPSF | ASSPRLNQDTQY | 0.7981668 |
| VVDAFHGGL | ASSPRLNQDTQY | 0.2037625 |
| ASVVDPQ | ASSPRLNQDTQY | 0.6005676 |
| SYNEFGGLS | ASSPRLNQDTQY | 0.77396023 |
| DPDGSVY | ASSPRLNQDTQY | 0.7620845 |
| MVSTIHATE | ASSPRLNQDTQY | 0.68985903 |
| LAAGVADPV | ASSPRLNQDTQY | 0.77813053 |
| TKEQIAA | ASSPRLNQDTQY | 0.5799475 |
| SLASPAAL | ASSPRLNQDTQY | 0.6912278 |
| IGLVFYGV | ASSPRLNQDTQY | 0.6746786 |
| FLLDASNL | ASSPRLNQDTQY | 0.7574805 |
| YQVGNFP | ASSPRLNQDTQY | 0.71885026 |
| ESLLSQST | ASSPRLNQDTQY | 0.50589097 |
| LFTVGFAAVG | ASSPRLNQDTQY | 0.70539665 |
| ATPPTPT | ASSPRLNQDTQY | 0.8077475 |
| TDLATLG | ASSPRLNQDTQY | 0.6539457 |
| SAGGTGR | ASSPRLNQDTQY | 0.7572509 |
| GFSNSGS | ASSPRLNQDTQY | 0.67169577 |
| GGNGGSGGTSV | ASSPRLNQDTQY | 0.7506661 |
| GTSVATGGAGN | ASSPRLNQDTQY | 0.5751204 |
| NGGNGGSGGTS | ASSPRLNQDTQY | 0.50273865 |
| AVVKFPH | ASSPRLNQDTQY | 0.7501323 |
| TPPPPPP | ASSPRLNQDTQY | 0.5761692 |
| AAPAAAAAPA | ASSPRLNQDTQY | 0.6966011 |
| IALNEDL | ASSPRLNQDTQY | 0.62791336 |
| IVNVSSM | ASSPRLNQDTQY | 0.80962455 |
| TGIPPAPR | ASSPRLNQDTQY | 0.7417649 |
| FDLGGGTF | ASSPRLNQDTQY | 0.7198799 |
| FDLGGGTFDVS | ASSPRLNQDTQY | 0.5900866 |
| NEPTAAA | ASSPRLNQDTQY | 0.718158 |
| NEPTAAALA | ASSPRLNQDTQY | 0.75470865 |
| GSIINISSI | ASSPRLNQDTQY | 0.41567466 |
| VAPGNAGTA | ASSPRLNQDTQY | 0.7840432 |
| TGGGASLE | ASSPRLNQDTQY | 0.42713857 |
| VAVYNHYKRI | ASSPRLNQDTQY | 0.6734025 |
| TDPAPATT | ASSPRLNQDTQY | 0.66063994 |
| TNRPDILDPALLRPG | ASSPRLNQDTQY | 0.793326 |
| LTTTTTS | ASSPRLNQDTQY | 0.7035456 |
| PTGSGKTL | ASSPRLNQDTQY | 0.73694634 |
| TGSGKTL | ASSPRLNQDTQY | 0.5188199 |
| TIGHVDHGK | ASSPRLNQDTQY | 0.74719393 |
| TIGHVDHGKTT | ASSPRLNQDTQY | 0.7851789 |
| GHVDHGKTT | ASSPRLNQDTQY | 0.7565906 |
| AQFEHTLL | ASSPRLNQDTQY | 0.80269456 |
| IVNQDEL | ASSPRLNQDTQY | 0.53997207 |
| TNGSQFF | ASSPRLNQDTQY | 0.27452222 |

| | | |
|---|---|---|
| TGKGIVE | ASSPRLNQDTQY | 0.66366917 |
| LISNASDA | ASSPRLNQDTQY | 0.7759825 |
| AELRTLGELTKIAKAHGAQ | ASSPRLNQDTQY | 0.7894559 |
| APNAVAAAEPSW | ASSPRLNQDTQY | 0.7750596 |
| APQVAGAATAVL | ASSPRLNQDTQY | 0.7151308 |
| ARRDPAAAAAILPTDA | ASSPRLNQDTQY | 0.6902901 |
| DAAARDVLAEAGLAEYF | ASSPRLNQDTQY | 0.7913339 |
| DDQIVLSAKANLAKVEQ | ASSPRLNQDTQY | 0.7961856 |
| DELLPDTGEDSDEDGH | ASSPRLNQDTQY | 0.724922 |
| ELPEAPANRATVPQD | ASSPRLNQDTQY | 0.44408122 |
| FGQSKIERVTGIG | ASSPRLNQDTQY | 0.36310506 |
| GAGGTGSTAGGAGGAGGAGGLY | ASSPRLNQDTQY | 0.75633657 |
| GHGGTGAAGGNGGTGGAGGSGI | ASSPRLNQDTQY | 0.77667475 |
| GNGGNGGAGGEGGPGGAGGAGA | ASSPRLNQDTQY | 0.42418456 |
| GSTAGNGGNGGNATNGTIAGQP | ASSPRLNQDTQY | 0.7065711 |
| GTGGDGGNGGNGGTGG | ASSPRLNQDTQY | 0.66575265 |
| GVRRPAGIPADAPV | ASSPRLNQDTQY | 0.29128397 |
| GYRQVTDAHLLTLARR | ASSPRLNQDTQY | 0.7512857 |
| IAKMGQDGHDRG | ASSPRLNQDTQY | 0.7900011 |
| KAEPSIEIQYT | ASSPRLNQDTQY | 0.7199975 |
| KPVRIRMAADEDTVAVT | ASSPRLNQDTQY | 0.53507257 |
| MPPRIAGMRLLVIKPEP | ASSPRLNQDTQY | 0.78434527 |
| MKFYAVALFRSRGVPPTR | ASSPRLNQDTQY | 0.8042209 |
| NAGGFGILGAVAHSP | ASSPRLNQDTQY | 0.78876656 |
| NEGREITLAMAEM | ASSPRLNQDTQY | 0.3048962 |
| NELVEWSHPLGHRWRWE | ASSPRLNQDTQY | 0.2156183 |
| NLGAGGTGGNGGNPG | ASSPRLNQDTQY | 0.71450436 |
| NTEWPKWETVMAADPVD | ASSPRLNQDTQY | 0.77614194 |
| PASAAPAETQP | ASSPRLNQDTQY | 0.5475084 |
| QQSSDGKNGVRAAEITGE | ASSPRLNQDTQY | 0.5262177 |
| RAPDAVTTSWWKE | ASSPRLNQDTQY | 0.7221447 |
| SGFGNVGAHTSGWFNQ | ASSPRLNQDTQY | 0.7751606 |
| SGVAGAVGQAASVGGLKV | ASSPRLNQDTQY | 0.46864235 |
| SNGGAGTGSAGGNGGT | ASSPRLNQDTQY | 0.39843974 |
| TGGAGGEGGAAG | ASSPRLNQDTQY | 0.76008666 |
| TGGDGGAAGAGGAPGHG | ASSPRLNQDTQY | 0.7634591 |
| TMPPLLAAIANAGRHGV | ASSPRLNQDTQY | 0.60083807 |
| TSSTAGNSGGA | ASSPRLNQDTQY | 0.7067187 |
| VPATVPVGILC | ASSPRLNQDTQY | 0.7431196 |
| VQRPDSGGVVLTGRIS | ASSPRLNQDTQY | 0.66240585 |
| WTPNPVPATKGVPAGLVQAT | ASSPRLNQDTQY | 0.7159312 |
| YALKRALAKFSDRFDVVII | ASSPRLNQDTQY | 0.39319715 |
| YLIGSVIV | ASSPRLNQDTQY | 0.7843932 |
| YGQLVAVID | ASSPRLNQDTQY | 0.49801433 |
| VTVGGMTLPA | ASSPRLNQDTQY | 0.7766717 |
| VLALPAV | ASSPRLNQDTQY | 0.659082 |
| VIPPTLNYET | ASSPRLNQDTQY | 0.7229791 |
| VGMGGMHPSSG | ASSPRLNQDTQY | 0.68803954 |
| VGLAGGAATA | ASSPRLNQDTQY | 0.47121856 |
| VGGAGGAGGSGP | ASSPRLNQDTQY | 0.5793724 |
| VFAAFMV | ASSPRLNQDTQY | 0.46896487 |

| | | |
|---|---|---|
| TLYDKLR | ASSPRLNQDTQY | 0.7757324 |
| TLAVFTVL | ASSPRLNQDTQY | 0.7068569 |
| TGSVQPARV | ASSPRLNQDTQY | 0.74565125 |
| TGGGAGSGGT | ASSPRLNQDTQY | 0.7084085 |
| SHLETSL | ASSPRLNQDTQY | 0.52425265 |
| SFTTGMT | ASSPRLNQDTQY | 0.74060404 |
| QVEVAKRMGVRQPTVS | ASSPRLNQDTQY | 0.3637215 |
| QGSGGPALAM | ASSPRLNQDTQY | 0.7209335 |
| MLAPSNP | ASSPRLNQDTQY | 0.69251096 |
| LVNNAAI | ASSPRLNQDTQY | 0.70403516 |
| LTGEINTLA | ASSPRLNQDTQY | 0.636958 |
| LSIFGPV | ASSPRLNQDTQY | 0.7271013 |
| LLALLANL | ASSPRLNQDTQY | 0.7438327 |
| LISNASDALDKLR | ASSPRLNQDTQY | 0.63125265 |
| LINEAVP | ASSPRLNQDTQY | 0.71507406 |
| LGYAFSGL | ASSPRLNQDTQY | 0.7856904 |
| LDFVAVAL | ASSPRLNQDTQY | 0.71334 |
| LAQPGAE | ASSPRLNQDTQY | 0.62163764 |
| ISNLGAVTGF | ASSPRLNQDTQY | 0.7041725 |
| HQEDPGRI | ASSPRLNQDTQY | 0.63184243 |
| GTPGEPTP | ASSPRLNQDTQY | 0.74866 |
| GFFNSGS | ASSPRLNQDTQY | 0.7086289 |
| GAGGGGLTAGAN | ASSPRLNQDTQY | 0.5313513 |
| FVDALVNM | ASSPRLNQDTQY | 0.7015286 |
| FLTVAPEL | ASSPRLNQDTQY | 0.30388975 |
| FETLTAL | ASSPRLNQDTQY | 0.63630843 |
| DVINPIEL | ASSPRLNQDTQY | 0.6336788 |
| APVSGAPV | ASSPRLNQDTQY | 0.6974747 |
| ALVGGLAW | ASSPRLNQDTQY | 0.7338965 |
| ALAAAAATAI | ASSPRLNQDTQY | 0.47030455 |
| AFGLGLLL | ASSPRLNQDTQY | 0.7107079 |
| AFFAAASL | ASSPRLNQDTQY | 0.6745466 |
| AAAALPAFSPP | ASSPRLNQDTQY | 0.53508294 |
| VVLAPAL | ASSPRLNQDTQY | 0.65301764 |
| TIGHVDHGKTTL | ASSPRLNQDTQY | 0.61419106 |
| LISNASDALDK | ASSPRLNQDTQY | 0.39451683 |
| LISNASDAL | ASSPRLNQDTQY | 0.751171 |
| ISNASDALDK | ASSPRLNQDTQY | 0.44697586 |
| NQAETLVYQTE | ASSPRLNQDTQY | 0.7208944 |
| TGGTGGTAGAG | ASSPRLNQDTQY | 0.77849156 |
| TGSTAGGAGGA | ASSPRLNQDTQY | 0.39910418 |
| DPLAVPVAAGPV | ASSPRLNQDTQY | 0.7715874 |
| GGTGGTPGNGGN | ASSPRLNQDTQY | 0.6925713 |
| PAGLAGVDPATL | ASSPRLNQDTQY | 0.74145424 |
| GNNLVVEAPHAGASV | ASSPRLNQDTQY | 0.59806776 |
| KVTRIMSASTLEQPA | ASSPRLNQDTQY | 0.70252866 |
| LGFALPIALPRRTVV | ASSPRLNQDTQY | 0.7773298 |
| TRLGNLTARKLVVLV | ASSPRLNQDTQY | 0.715422 |
| WSQQRRLIYFPSAGP | ASSPRLNQDTQY | 0.61667144 |
| GPNGNPDPMAAAVDIR | ASSPRLNQDTQY | 0.66053414 |
| TQRCVSVADLRAAASR | ASSPRLNQDTQY | 0.7850597 |

| | | |
|---|---|---|
| QFVLDVAEHPVEVLTETV | ASSPRLNQDTQY | 0.7527527 |
| GGAGGTGGAGGNGGTQAGDG | ASSPRLNQDTQY | 0.5573603 |
| PLLATLRAIAVDVDAKPAQV | ASSPRLNQDTQY | 0.29914284 |
| APKFRTGGYMIDFWGVGYQVAK | ASSPRLNQDTQY | 0.65445435 |
| GAGGAGGAGGAAGGPGGTGGTG | ASSPRLNQDTQY | 0.6745199 |
| IGALGLLATDVAPAGSVAFGVA | ASSPRLNQDTQY | 0.55911076 |
| AGGAGGIGGNGGAGGAGGVLMG | ASSPRLNQDTQY | 0.26593632 |
| ADELIGLL | ASSPRLNQDTQY | 0.76305985 |
| ADRALKA | ASSPRLNQDTQY | 0.80715364 |
| AGAVGQVGGAGGA | ASSPRLNQDTQY | 0.43620062 |
| AGGGGGLGGAP | ASSPRLNQDTQY | 0.80084014 |
| AGGLLYGNGGAG | ASSPRLNQDTQY | 0.43768203 |
| AGLESIGL | ASSPRLNQDTQY | 0.74213207 |
| AGLKTGII | ASSPRLNQDTQY | 0.77758545 |
| ASLAALLL | ASSPRLNQDTQY | 0.4025856 |
| ASVINISSI | ASSPRLNQDTQY | 0.6528773 |
| ATPPGPA | ASSPRLNQDTQY | 0.74812555 |
| ATVASALV | ASSPRLNQDTQY | 0.661064 |
| ATVASLIL | ASSPRLNQDTQY | 0.7823018 |
| AVNGPVLIRH | ASSPRLNQDTQY | 0.52314794 |
| DGAAGAVGI | ASSPRLNQDTQY | 0.7275939 |
| DPLLHGI | ASSPRLNQDTQY | 0.54884887 |
| ELGAALGAL | ASSPRLNQDTQY | 0.5031935 |
| ELGRTQAV | ASSPRLNQDTQY | 0.72798073 |
| FVAGGYLS | ASSPRLNQDTQY | 0.7089169 |
| GASGGAGGAGGW | ASSPRLNQDTQY | 0.67227983 |
| GGAGAGGDGGAGGI | ASSPRLNQDTQY | 0.7836347 |
| GTGHNGSQPGMGGNG | ASSPRLNQDTQY | 0.6508602 |
| GYAAASASAS | ASSPRLNQDTQY | 0.8034719 |
| HQVDVAAVQQG | ASSPRLNQDTQY | 0.78589994 |
| IGAAIAHT | ASSPRLNQDTQY | 0.7335017 |
| IGMIEVAQL | ASSPRLNQDTQY | 0.6437015 |
| IRLAPEHTSAMD | ASSPRLNQDTQY | 0.7832589 |
| IYNVSVL | ASSPRLNQDTQY | 0.6262193 |
| KQGFLGM | ASSPRLNQDTQY | 0.68864566 |
| LESVITQL | ASSPRLNQDTQY | 0.7581684 |
| LGLGALVVV | ASSPRLNQDTQY | 0.42300624 |
| LIGNGGNGGSGGI | ASSPRLNQDTQY | 0.79341424 |
| LNAINAPTL | ASSPRLNQDTQY | 0.7568414 |
| LQLLNTGV | ASSPRLNQDTQY | 0.76430583 |
| LQTSIEF | ASSPRLNQDTQY | 0.62606704 |
| MSFAVSEL | ASSPRLNQDTQY | 0.4221872 |
| NTGALGL | ASSPRLNQDTQY | 0.6792611 |
| PPGAGGAGGAGTT | ASSPRLNQDTQY | 0.55016994 |
| QLVGLGI | ASSPRLNQDTQY | 0.8116591 |
| RFLEQVHQL | ASSPRLNQDTQY | 0.7891655 |
| RGVGYVLR | ASSPRLNQDTQY | 0.5998688 |
| SATGVFAGVI | ASSPRLNQDTQY | 0.7960739 |
| SLPAITLP | ASSPRLNQDTQY | 0.7884284 |
| STALAFTV | ASSPRLNQDTQY | 0.43013 |
| SVLTVGLGK | ASSPRLNQDTQY | 0.70570976 |

| | | |
|---|---|---|
| TALAGAGSGI | ASSPRLNQDTQY | 0.61211216 |
| TGLPFTGI | ASSPRLNQDTQY | 0.6799383 |
| VAEHVTV | ASSPRLNQDTQY | 0.32134277 |
| VDIVLNSL | ASSPRLNQDTQY | 0.7781394 |
| VLTGGGAVL | ASSPRLNQDTQY | 0.66828334 |
| VNVGLAYGI | ASSPRLNQDTQY | 0.29179505 |
| VSSAAVGAV | ASSPRLNQDTQY | 0.7706789 |
| VTSVYEAGV | ASSPRLNQDTQY | 0.6650014 |
| WVALDRGA | ASSPRLNQDTQY | 0.68343705 |
| YLVGSVVI | ASSPRLNQDTQY | 0.7965246 |
| GPPGTGKTLL | ASSPRLNQDTQY | 0.7149052 |
| GVGMTKF | ASSPRLNQDTQY | 0.6134979 |
| IGIGGSDL | ASSPRLNQDTQY | 0.77971804 |
| IGTLLAEL | ASSPRLNQDTQY | 0.80334747 |
| IYTSGTT | ASSPRLNQDTQY | 0.55536675 |
| LSFPAGV | ASSPRLNQDTQY | 0.75180435 |
| LYTSGTT | ASSPRLNQDTQY | 0.8049216 |
| VGAFPFGF | ASSPRLNQDTQY | 0.75016046 |
| VGAGTVEF | ASSPRLNQDTQY | 0.8073073 |
| VGKSTLLL | ASSPRLNQDTQY | 0.8101835 |
| AALAHAGGGIV | ASSPRLNQDTQY | 0.6439694 |
| AALVAVSLL | ASSPRLNQDTQY | 0.8046019 |
| AAVSRAVAAL | ASSPRLNQDTQY | 0.81054324 |
| AGGSSGIGGF | ASSPRLNQDTQY | 0.6569425 |
| AGLTVANANG | ASSPRLNQDTQY | 0.71583384 |
| AGLVGAVL | ASSPRLNQDTQY | 0.456466 |
| ALSQAFTV | ASSPRLNQDTQY | 0.7334838 |
| ALTFSVSL | ASSPRLNQDTQY | 0.6019779 |
| ALTGAALVAL | ASSPRLNQDTQY | 0.77797866 |
| ASFSIGAGVL | ASSPRLNQDTQY | 0.81099355 |
| DGSVYKI | ASSPRLNQDTQY | 0.7389235 |
| EQLAEIV | ASSPRLNQDTQY | 0.44613463 |
| FAVSELTVM | ASSPRLNQDTQY | 0.48301414 |
| GAAPLAAGL | ASSPRLNQDTQY | 0.76233685 |
| GADGGDGGSGGAGG | ASSPRLNQDTQY | 0.475177 |
| GFGLDTGGAGG | ASSPRLNQDTQY | 0.46774966 |
| GLASGGLL | ASSPRLNQDTQY | 0.63492525 |
| GRIAGAAVLL | ASSPRLNQDTQY | 0.80287004 |
| GSAGSLNTSF | ASSPRLNQDTQY | 0.80284464 |
| GTGQIGI | ASSPRLNQDTQY | 0.63991916 |
| GTLIALIL | ASSPRLNQDTQY | 0.8101103 |
| IAATMGFVVG | ASSPRLNQDTQY | 0.8013201 |
| IFAQALL | ASSPRLNQDTQY | 0.7993621 |
| IGAAVALGF | ASSPRLNQDTQY | 0.70187426 |
| IGLTPLAL | ASSPRLNQDTQY | 0.39103544 |
| ILGSNTSTL | ASSPRLNQDTQY | 0.8109047 |
| IVFSLTVL | ASSPRLNQDTQY | 0.7330961 |
| LASKATGF | ASSPRLNQDTQY | 0.7831473 |
| LASVALIL | ASSPRLNQDTQY | 0.7899675 |
| LFAVGLTL | ASSPRLNQDTQY | 0.7576928 |
| LFNGGIGTYI | ASSPRLNQDTQY | 0.6243086 |

| | | |
|---|---|---|
| LTLGAGII | ASSPRLNQDTQY | 0.7700157 |
| LVAAALGDI | ASSPRLNQDTQY | 0.8018979 |
| NTAFVQLGI | ASSPRLNQDTQY | 0.5979846 |
| QLFSGAEL | ASSPRLNQDTQY | 0.44713247 |
| SAATFITL | ASSPRLNQDTQY | 0.74438643 |
| SNYGLANLGV | ASSPRLNQDTQY | 0.7293699 |
| SYGGAYAVM | ASSPRLNQDTQY | 0.80516064 |
| TTGVGGLTLG | ASSPRLNQDTQY | 0.7458478 |
| TTQTIYI | ASSPRLNQDTQY | 0.68693525 |
| VAAGAGLGAVI | ASSPRLNQDTQY | 0.7787955 |
| VAAVVTLL | ASSPRLNQDTQY | 0.7750778 |
| VGFLGGLI | ASSPRLNQDTQY | 0.7412156 |
| VGVIPILL | ASSPRLNQDTQY | 0.7213063 |
| VIFASSLI | ASSPRLNQDTQY | 0.6720189 |
| VKGGDGGAAAG | ASSPRLNQDTQY | 0.79116 |
| VTKTFTNI | ASSPRLNQDTQY | 0.7972035 |
| VVAAAILL | ASSPRLNQDTQY | 0.77524495 |
| YAALDVEL | ASSPRLNQDTQY | 0.80397046 |
| LASVALLL | ASSPRLNQDTQY | 0.7392359 |
| LDAVHRV | ASSPRLNQDTQY | 0.74421084 |
| QGLSSSIF | ASSPRLNQDTQY | 0.5439951 |
| RGTFANIRL | ASSPRLNQDTQY | 0.70718384 |
| RQLWWGHRI | ASSPRLNQDTQY | 0.73226845 |
| YGPPGTGKTLL | ASSPRLNQDTQY | 0.77085793 |
| AALVGAPANVA | ASSPRLNQDTQY | 0.7984127 |
| AATFRRTE | ASSPRLNQDTQY | 0.75966895 |
| AGALGGGI | ASSPRLNQDTQY | 0.68072295 |
| AGGAGGAGGIGAG | ASSPRLNQDTQY | 0.79577214 |
| AGSGALGL | ASSPRLNQDTQY | 0.8030473 |
| AQIADQI | ASSPRLNQDTQY | 0.5142354 |
| ASAVAVTYL | ASSPRLNQDTQY | 0.67418927 |
| ASTVLGGTGG | ASSPRLNQDTQY | 0.8098543 |
| AVGGEPAK | ASSPRLNQDTQY | 0.79078937 |
| CEQGTFGGTC | ASSPRLNQDTQY | 0.7126265 |
| DEGKAPAG | ASSPRLNQDTQY | 0.72281086 |
| DTVVFPAL | ASSPRLNQDTQY | 0.8174998 |
| FEGITMAG | ASSPRLNQDTQY | 0.7599547 |
| FPLVVQD | ASSPRLNQDTQY | 0.8103937 |
| GAFTTPQ | ASSPRLNQDTQY | 0.39953667 |
| GFGGFGTAGGVGGA | ASSPRLNQDTQY | 0.67684984 |
| GFLAQFL | ASSPRLNQDTQY | 0.55267537 |
| GGAGGAGGNAGVA | ASSPRLNQDTQY | 0.21352018 |
| GGVGTAGAGGAGGAGG | ASSPRLNQDTQY | 0.8010793 |
| GNGGGGDAGNAGSGGN | ASSPRLNQDTQY | 0.63509524 |
| GTGGAGGAGNDAGS | ASSPRLNQDTQY | 0.76474094 |
| GTTNTGWFNS | ASSPRLNQDTQY | 0.6778277 |
| IFMGPLFTL | ASSPRLNQDTQY | 0.7990192 |
| ILAHAIEAI | ASSPRLNQDTQY | 0.7494028 |
| IRSAIREAL | ASSPRLNQDTQY | 0.6970676 |
| ITFAGGG | ASSPRLNQDTQY | 0.5396351 |
| ITGMSGHQEV | ASSPRLNQDTQY | 0.78339744 |

| | | |
|---|---|---|
| IVAGLACTL | ASSPRLNQDTQY | 0.80511445 |
| IVGGGLAGA | ASSPRLNQDTQY | 0.7925941 |
| IVTGVLNI | ASSPRLNQDTQY | 0.37331834 |
| LAQQGCI | ASSPRLNQDTQY | 0.78591764 |
| LCLHVEN | ASSPRLNQDTQY | 0.71253824 |
| LDAISDGS | ASSPRLNQDTQY | 0.5781751 |
| LGGFAAML | ASSPRLNQDTQY | 0.3417076 |
| LGGKNPMI | ASSPRLNQDTQY | 0.6799178 |
| LGPKGVAAIL | ASSPRLNQDTQY | 0.7210417 |
| LMVGVSI | ASSPRLNQDTQY | 0.7830199 |
| LVGAAFVL | ASSPRLNQDTQY | 0.7869166 |
| LVLAVAPL | ASSPRLNQDTQY | 0.6664009 |
| MAAIAVIDAA | ASSPRLNQDTQY | 0.77892274 |
| NNGINAELQ | ASSPRLNQDTQY | 0.74556494 |
| RGEPLVLVHG | ASSPRLNQDTQY | 0.16203882 |
| RGGAGGAAGLI | ASSPRLNQDTQY | 0.58752227 |
| RSYRMHPAV | ASSPRLNQDTQY | 0.44639152 |
| SAAHLLAGQLA | ASSPRLNQDTQY | 0.8067782 |
| SAAVGSY | ASSPRLNQDTQY | 0.5394146 |
| SPHAVLHRI | ASSPRLNQDTQY | 0.6973125 |
| SQIAKVL | ASSPRLNQDTQY | 0.7159904 |
| TALGHLVP | ASSPRLNQDTQY | 0.7920638 |
| TAPTTSGAA | ASSPRLNQDTQY | 0.7620614 |
| TDGDDGNGGSPGT | ASSPRLNQDTQY | 0.531667 |
| TDGLGEN | ASSPRLNQDTQY | 0.565806 |
| TGGPPLPGQA | ASSPRLNQDTQY | 0.77575785 |
| TIPAITI | ASSPRLNQDTQY | 0.7496498 |
| TIPGFTI | ASSPRLNQDTQY | 0.5614343 |
| TSIASGALA | ASSPRLNQDTQY | 0.59932935 |
| TVTLPGAAI | ASSPRLNQDTQY | 0.8017394 |
| VASGLSAALP | ASSPRLNQDTQY | 0.31582457 |
| VGAGVIGI | ASSPRLNQDTQY | 0.79266995 |
| VGPKRFGV | ASSPRLNQDTQY | 0.57317126 |
| VNVAFPD | ASSPRLNQDTQY | 0.7190358 |
| VSMTNSGVS | ASSPRLNQDTQY | 0.81660295 |
| VTDVLGDLSV | ASSPRLNQDTQY | 0.7841909 |
| VVVVGAGFAGL | ASSPRLNQDTQY | 0.72495866 |
| VYTVGTVV | ASSPRLNQDTQY | 0.58511424 |
| WTTFSGP | ASSPRLNQDTQY | 0.5380995 |
| YDELVEAF | ASSPRLNQDTQY | 0.6526462 |
| YGTGGAGGNGG | ASSPRLNQDTQY | 0.63748837 |
| YTNVMAV | ASSPRLNQDTQY | 0.8059677 |
| FLEQVHQL | ASSPRLNQDTQY | 0.72824895 |
| GLSLLAVAV | ASSPRLNQDTQY | 0.45013446 |
| LEQVHQL | ASSPRLNQDTQY | 0.74572617 |
| LNAAGAI | ASSPRLNQDTQY | 0.81068945 |
| PAVHASL | ASSPRLNQDTQY | 0.8167759 |
| TELEDDA | ASSPRLNQDTQY | 0.5782204 |
| TPPGPPN | ASSPRLNQDTQY | 0.5863643 |
| TPPPPPPP | ASSPRLNQDTQY | 0.6693959 |
| TQSPALI | ASSLKDTQY | 0.72971624 |

| | | |
|---|---|---|
| EFAAAFPQ | ASSLKDTQY | 0.45019567 |
| LLSGNLDVL | ASSLKDTQY | 0.58722997 |
| GPANLGNYNI | ASSLKDTQY | 0.7835866 |
| TLGPTLP | ASSLKDTQY | 0.825721 |
| VYAYSSCCK | ASSLKDTQY | 0.6910258 |
| APAPAPAPNI | ASSLKDTQY | 0.32339108 |
| PGGPVPIP | ASSLKDTQY | 0.7175185 |
| FLTVALEP | ASSLKDTQY | 0.64854604 |
| SQYFGANL | ASSLKDTQY | 0.61164284 |
| GGVKPAA | ASSLKDTQY | 0.723614 |
| GGTGGAGGAGGW | ASSLKDTQY | 0.38359356 |
| GVGGVAGLGGAGG | ASSLKDTQY | 0.72503996 |
| VLLVLATTI | ASSLKDTQY | 0.8157222 |
| IGIFAAML | ASSLKDTQY | 0.73970515 |
| AVGDLID | ASSLKDTQY | 0.7629713 |
| NISPLPL | ASSLKDTQY | 0.8091663 |
| VTAVFVAI | ASSLKDTQY | 0.78338385 |
| SGDDNDAAL | ASSLKDTQY | 0.8063729 |
| AVAALAHS | ASSLKDTQY | 0.807583 |
| GGVTVMATP | ASSLKDTQY | 0.71214557 |
| AGLAANVRD | ASSLKDTQY | 0.7043989 |
| TNRPDLIDPALLRPG | ASSLKDTQY | 0.8056061 |
| LANLGVTAL | ASSLKDTQY | 0.6974874 |
| TGERIEL | ASSLKDTQY | 0.758392 |
| NGEELAA | ASSLKDTQY | 0.6976941 |
| VLSLVWG | ASSLKDTQY | 0.78956723 |
| IGYALWL | ASSLKDTQY | 0.64066184 |
| LSLLLAPL | ASSLKDTQY | 0.81812215 |
| TLDEALK | ASSLKDTQY | 0.23367962 |
| TDGAGTAEGG | ASSLKDTQY | 0.61928535 |
| NAPGGAGGNGG | ASSLKDTQY | 0.8049564 |
| VSVFMGMGGNF | ASSLKDTQY | 0.1907002 |
| LASYLAAL | ASSLKDTQY | 0.35494393 |
| LVGGTPAL | ASSLKDTQY | 0.7228869 |
| VRNADVL | ASSLKDTQY | 0.8037975 |
| TCSSPFLVSAM | ASSLKDTQY | 0.2559994 |
| VHITDAE | ASSLKDTQY | 0.6801531 |
| EYAAPSF | ASSLKDTQY | 0.39214832 |
| VVDAFHGGL | ASSLKDTQY | 0.7709625 |
| ASVVDPQ | ASSLKDTQY | 0.72131646 |
| SYNEFGGLS | ASSLKDTQY | 0.39682645 |
| DPDGSVY | ASSLKDTQY | 0.7891183 |
| MVSTIHATE | ASSLKDTQY | 0.40566832 |
| LAAGVADPV | ASSLKDTQY | 0.7826051 |
| TKEQIAA | ASSLKDTQY | 0.79825664 |
| SLASPAAL | ASSLKDTQY | 0.8200897 |
| IGLVFYGV | ASSLKDTQY | 0.4588144 |
| FLLDASNL | ASSLKDTQY | 0.7828045 |
| YQVGNFP | ASSLKDTQY | 0.6718737 |
| ESLLSQST | ASSLKDTQY | 0.8220802 |
| LFTVGFAAVG | ASSLKDTQY | 0.6847232 |

| | | |
|---|---|---|
| ATPPTPT | ASSLKDTQY | 0.33941847 |
| TDLATLG | ASSLKDTQY | 0.7976912 |
| SAGGTGR | ASSLKDTQY | 0.75964046 |
| GFSNSGS | ASSLKDTQY | 0.6659579 |
| GGNGGSGGTSV | ASSLKDTQY | 0.79949284 |
| GTSVATGGAGN | ASSLKDTQY | 0.64966786 |
| NGGNGGSGGTS | ASSLKDTQY | 0.6988878 |
| AVVKFPH | ASSLKDTQY | 0.7978575 |
| TPPPPPP | ASSLKDTQY | 0.77380204 |
| AAPAAAAAPA | ASSLKDTQY | 0.6775929 |
| IALNEDL | ASSLKDTQY | 0.8053522 |
| IVNVSSM | ASSLKDTQY | 0.750106 |
| TGIPPAPR | ASSLKDTQY | 0.48063576 |
| FDLGGGTF | ASSLKDTQY | 0.8196309 |
| FDLGGGTFDVS | ASSLKDTQY | 0.6899098 |
| NEPTAAA | ASSLKDTQY | 0.7934215 |
| NEPTAAALA | ASSLKDTQY | 0.78522325 |
| GSIINISSI | ASSLKDTQY | 0.81685424 |
| VAPGNAGTA | ASSLKDTQY | 0.6308179 |
| TGGGASLE | ASSLKDTQY | 0.7478324 |
| VAVYNHYKRI | ASSLKDTQY | 0.72992074 |
| TDPAPATT | ASSLKDTQY | 0.78726375 |
| TNRPDILDPALLRPG | ASSLKDTQY | 0.7074132 |
| LTTTTTS | ASSLKDTQY | 0.7811569 |
| PTGSGKTL | ASSLKDTQY | 0.7787578 |
| TGSGKTL | ASSLKDTQY | 0.808161 |
| TIGHVDHGK | ASSLKDTQY | 0.6701318 |
| TIGHVDHGKTT | ASSLKDTQY | 0.70461166 |
| GHVDHGKTT | ASSLKDTQY | 0.81508505 |
| AQFEHTLL | ASSLKDTQY | 0.52994806 |
| IVNQDEL | ASSLKDTQY | 0.7324046 |
| TNGSQFF | ASSLKDTQY | 0.7725318 |
| TGKGIVE | ASSLKDTQY | 0.80567193 |
| LISNASDA | ASSLKDTQY | 0.67942363 |
| AELRTLGELTKIAKAHGAQ | ASSLKDTQY | 0.8003371 |
| APNAVAAAEPSW | ASSLKDTQY | 0.708901 |
| APQVAGAATAVL | ASSLKDTQY | 0.78437555 |
| ARRDPAAAAAILPTDA | ASSLKDTQY | 0.7260721 |
| DAAARDVLAEAGLAEYF | ASSLKDTQY | 0.6831516 |
| DDQIVLSAKANLAKVEQ | ASSLKDTQY | 0.61873424 |
| DELLPDTGEDSDEDGH | ASSLKDTQY | 0.75596184 |
| ELPEAPANRATVPQD | ASSLKDTQY | 0.7978761 |
| FGQSKIERVTGIG | ASSLKDTQY | 0.7796329 |
| GAGGTGSTAGGAGGAGGAGGLY | ASSLKDTQY | 0.38791335 |
| GHGGTGAAGGNGGTGGAGGSGI | ASSLKDTQY | 0.7817725 |
| GNGGNGGAGGEGGPGGAGGAGA | ASSLKDTQY | 0.7810283 |
| GSTAGNGGNGGNATNGTIAGQP | ASSLKDTQY | 0.82141614 |
| GTGGDGGNGGNGGTGG | ASSLKDTQY | 0.68748534 |
| GVRRPAGIPADAPV | ASSLKDTQY | 0.78239405 |
| GYRQVTDAHLLTLARR | ASSLKDTQY | 0.6839296 |
| IAKMGQDGHDRG | ASSLKDTQY | 0.73282874 |

| | | |
|---|---|---|
| KAEPSIEIQYT | ASSLKDTQY | 0.4389814 |
| KPVRIRMAADEDTVAVT | ASSLKDTQY | 0.5661323 |
| MPPRIAGMRLLVIKPEP | ASSLKDTQY | 0.7255833 |
| MKFYAVALFRSRGVPPTR | ASSLKDTQY | 0.63632977 |
| NAGGFGILGAVAHSP | ASSLKDTQY | 0.80144835 |
| NEGREITLAMAEM | ASSLKDTQY | 0.7989876 |
| NELVEWSHPLGHRWRWE | ASSLKDTQY | 0.7991882 |
| NLGAGGTGGNGGNPG | ASSLKDTQY | 0.8048459 |
| NTEWPKWETVMAADPVD | ASSLKDTQY | 0.8103421 |
| PASAAPAETQP | ASSLKDTQY | 0.7922034 |
| QQSSDGKNGVRAAEITGE | ASSLKDTQY | 0.66383475 |
| RAPDAVTTSWWKE | ASSLKDTQY | 0.73285407 |
| SGFGNVGAHTSGWFNQ | ASSLKDTQY | 0.40851623 |
| SGVAGAVGQAASVGGLKV | ASSLKDTQY | 0.669906 |
| SNGGAGTGSAGGNGGT | ASSLKDTQY | 0.66821223 |
| TGGAGGEGGAAG | ASSLKDTQY | 0.75151694 |
| TGGDGGAAGAGGAPGHG | ASSLKDTQY | 0.20075622 |
| TMPPLLAAIANAGRHGV | ASSLKDTQY | 0.78549606 |
| TSSTAGNSGGA | ASSLKDTQY | 0.8112247 |
| VPATVPVGILC | ASSLKDTQY | 0.75658107 |
| VQRPDSGGVVLTGRIS | ASSLKDTQY | 0.7813407 |
| WTPNPVPATKGVPAGLVQAT | ASSLKDTQY | 0.78311616 |
| YALKRALAKFSDRFDVVII | ASSLKDTQY | 0.63327014 |
| YLIGSVIV | ASSLKDTQY | 0.78234303 |
| YGQLVAVID | ASSLKDTQY | 0.69332457 |
| VTVGGMTLPA | ASSLKDTQY | 0.7988661 |
| VLALPAV | ASSLKDTQY | 0.81088555 |
| VIPPTLNYET | ASSLKDTQY | 0.76988035 |
| VGMGGMHPSSG | ASSLKDTQY | 0.67027926 |
| VGLAGGAATA | ASSLKDTQY | 0.7605973 |
| VGGAGGAGGSGP | ASSLKDTQY | 0.5612234 |
| VFAAFMV | ASSLKDTQY | 0.71410996 |
| TLYDKLR | ASSLKDTQY | 0.69968 |
| TLAVFTVL | ASSLKDTQY | 0.76163864 |
| TGSVQPARV | ASSLKDTQY | 0.61420983 |
| TGGGAGSGGT | ASSLKDTQY | 0.7778663 |
| SHLETSL | ASSLKDTQY | 0.7418239 |
| SFTTGMT | ASSLKDTQY | 0.72842944 |
| QVEVAKRMGVRQPTVS | ASSLKDTQY | 0.59030807 |
| QGSGGPALAM | ASSLKDTQY | 0.793713 |
| MLAPSNP | ASSLKDTQY | 0.76769066 |
| LVNNAAI | ASSLKDTQY | 0.47439766 |
| LTGEINTLA | ASSLKDTQY | 0.77233493 |
| LSIFGPV | ASSLKDTQY | 0.6648278 |
| LLALLANL | ASSLKDTQY | 0.7351911 |
| LISNASDALDKLR | ASSLKDTQY | 0.8056199 |
| LINEAVP | ASSLKDTQY | 0.73895824 |
| LGYAFSGL | ASSLKDTQY | 0.4229781 |
| LDFVAVAL | ASSLKDTQY | 0.70435685 |
| LAQPGAE | ASSLKDTQY | 0.75074387 |
| ISNLGAVTGF | ASSLKDTQY | 0.7183423 |

| | | |
|---|---|---|
| HQEDPGRI | ASSLKDTQY | 0.25816613 |
| GTPGEPTP | ASSLKDTQY | 0.76218003 |
| GFFNSGS | ASSLKDTQY | 0.4817142 |
| GAGGGGLTAGAN | ASSLKDTQY | 0.59246933 |
| FVDALVNM | ASSLKDTQY | 0.6382127 |
| FLTVAPEL | ASSLKDTQY | 0.79848677 |
| FETLTAL | ASSLKDTQY | 0.7780751 |
| DVINPIEL | ASSLKDTQY | 0.6536975 |
| APVSGAPV | ASSLKDTQY | 0.7772359 |
| ALVGGLAW | ASSLKDTQY | 0.7513489 |
| ALAAAAATAI | ASSLKDTQY | 0.7986599 |
| AFGLGLLL | ASSLKDTQY | 0.73986495 |
| AFFAAASL | ASSLKDTQY | 0.6921975 |
| AAAALPAFSPP | ASSLKDTQY | 0.75983495 |
| VVLAPAL | ASSLKDTQY | 0.5641684 |
| TIGHVDHGKTTL | ASSLKDTQY | 0.6575613 |
| LISNASDALDK | ASSLKDTQY | 0.62895083 |
| LISNASDAL | ASSLKDTQY | 0.5408379 |
| ISNASDALDK | ASSLKDTQY | 0.6010546 |
| NQAETLVYQTE | ASSLKDTQY | 0.76415515 |
| TGGTGGTAGAG | ASSLKDTQY | 0.80486387 |
| TGSTAGGAGGA | ASSLKDTQY | 0.68172103 |
| DPLAVPVAAGPV | ASSLKDTQY | 0.35118425 |
| GGTGGTPGNGGN | ASSLKDTQY | 0.6604216 |
| PAGLAGVDPATL | ASSLKDTQY | 0.72509277 |
| GNNLVVEAPHAGASV | ASSLKDTQY | 0.7948771 |
| KVTRIMSASTLEQPA | ASSLKDTQY | 0.78644896 |
| LGFALPIALPRRTVV | ASSLKDTQY | 0.8224601 |
| TRLGNLTARKLVVLV | ASSLKDTQY | 0.78526574 |
| WSQQRRLIYFPSAGP | ASSLKDTQY | 0.6758746 |
| GPNGNPDPMAAAVDIR | ASSLKDTQY | 0.7898796 |
| TQRCVSVADLRAAASR | ASSLKDTQY | 0.68625534 |
| QFVLDVAEHPVEVLTETV | ASSLKDTQY | 0.72462475 |
| GGAGGTGGAGGNGGTQAGDG | ASSLKDTQY | 0.74512374 |
| PLLATLRAIAVDVDAKPAQV | ASSLKDTQY | 0.7503193 |
| APKFRTGGYMIDFWGVGYQVAK | ASSLKDTQY | 0.6151679 |
| GAGGAGGAGGAAGGPGGTGGTG | ASSLKDTQY | 0.6890197 |
| IGALGLLATDVAPAGSVAFGVA | ASSLKDTQY | 0.35173953 |
| AGGAGGIGGNGGAGGAGGVLMG | ASSLKDTQY | 0.7884774 |
| ADELIGLL | ASSLKDTQY | 0.53827465 |
| ADRALKA | ASSLKDTQY | 0.58789694 |
| AGAVGQVGGAGGA | ASSLKDTQY | 0.70308936 |
| AGGGGGLGGAP | ASSLKDTQY | 0.8157214 |
| AGGLLYGNGGAG | ASSLKDTQY | 0.81364584 |
| AGLESIGL | ASSLKDTQY | 0.6858553 |
| AGLKTGII | ASSLKDTQY | 0.61418086 |
| ASLAALLL | ASSLKDTQY | 0.79200697 |
| ASVINISSI | ASSLKDTQY | 0.6606456 |
| ATPPGPA | ASSLKDTQY | 0.764171 |
| ATVASALV | ASSLKDTQY | 0.7523215 |
| ATVASLIL | ASSLKDTQY | 0.709903 |

| | | |
|---|---|---|
| AVNGPVLIRH | ASSLKDTQY | 0.6022569 |
| DGAAGAVGI | ASSLKDTQY | 0.78503525 |
| DPLLHGI | ASSLKDTQY | 0.6619116 |
| ELGAALGAL | ASSLKDTQY | 0.76249075 |
| ELGRTQAV | ASSLKDTQY | 0.61388224 |
| FVAGGYLS | ASSLKDTQY | 0.79175955 |
| GASGGAGGAGGW | ASSLKDTQY | 0.601544 |
| GGAGAGGDGGAGGI | ASSLKDTQY | 0.49753004 |
| GTGHNGSQPGMGGNG | ASSLKDTQY | 0.24306722 |
| GYAAASASAS | ASSLKDTQY | 0.6753005 |
| HQVDVAAVQQG | ASSLKDTQY | 0.7863064 |
| IGAAIAHT | ASSLKDTQY | 0.80927974 |
| IGMIEVAQL | ASSLKDTQY | 0.7712759 |
| IRLAPEHTSAMD | ASSLKDTQY | 0.7862586 |
| IYNVSVL | ASSLKDTQY | 0.80522496 |
| KQGFLGM | ASSLKDTQY | 0.70348006 |
| LESVITQL | ASSLKDTQY | 0.32603216 |
| LGLGALVVV | ASSLKDTQY | 0.7631628 |
| LIGNGGNGGSGGI | ASSLKDTQY | 0.7274512 |
| LNAINAPTL | ASSLKDTQY | 0.8016168 |
| LQLLNTGV | ASSLKDTQY | 0.78651875 |
| LQTSIEF | ASSLKDTQY | 0.4637075 |
| MSFAVSEL | ASSLKDTQY | 0.7715335 |
| NTGALGL | ASSLKDTQY | 0.77441865 |
| PPGAGGAGGAGTT | ASSLKDTQY | 0.47009158 |
| QLVGLGI | ASSLKDTQY | 0.52577746 |
| RFLEQVHQL | ASSLKDTQY | 0.5315252 |
| RGVGYVLR | ASSLKDTQY | 0.5618621 |
| SATGVFAGVI | ASSLKDTQY | 0.7654407 |
| SLPAITLP | ASSLKDTQY | 0.7071339 |
| STALAFTV | ASSLKDTQY | 0.73496366 |
| SVLTVGLGK | ASSLKDTQY | 0.67420614 |
| TALAGAGSGI | ASSLKDTQY | 0.6231562 |
| TGLPFTGI | ASSLKDTQY | 0.6807222 |
| VAEHVTV | ASSLKDTQY | 0.7203413 |
| VDIVLNSL | ASSLKDTQY | 0.75114655 |
| VLTGGGAVL | ASSLKDTQY | 0.7944548 |
| VNVGLAYGI | ASSLKDTQY | 0.7076071 |
| VSSAAVGAV | ASSLKDTQY | 0.73565435 |
| VTSVYEAGV | ASSLKDTQY | 0.81033957 |
| WVALDRGA | ASSLKDTQY | 0.5055899 |
| YLVGSVVI | ASSLKDTQY | 0.55662704 |
| GPPGTGKTLL | ASSLKDTQY | 0.76977056 |
| GVGMTKF | ASSLKDTQY | 0.7059776 |
| IGIGGSDL | ASSLKDTQY | 0.8030788 |
| IGTLLAEL | ASSLKDTQY | 0.75516516 |
| IYTSGTT | ASSLKDTQY | 0.7712064 |
| LSFPAGV | ASSLKDTQY | 0.7459214 |
| LYTSGTT | ASSLKDTQY | 0.81834304 |
| VGAFPFGF | ASSLKDTQY | 0.7827712 |
| VGAGTVEF | ASSLKDTQY | 0.6336228 |

| | | |
|---|---|---|
| VGKSTLLL | ASSLKDTQY | 0.79158634 |
| AALAHAGGGIV | ASSLKDTQY | 0.36468416 |
| AALVAVSLL | ASSLKDTQY | 0.79171854 |
| AAVSRAVAAL | ASSLKDTQY | 0.78565 |
| AGGSSGIGGF | ASSLKDTQY | 0.60067546 |
| AGLTVANANG | ASSLKDTQY | 0.7011876 |
| AGLVGAVL | ASSLKDTQY | 0.80519235 |
| ALSQAFTV | ASSLKDTQY | 0.80167186 |
| ALTFSVSL | ASSLKDTQY | 0.64321977 |
| ALTGAALVAL | ASSLKDTQY | 0.626199 |
| ASFSIGAGVL | ASSLKDTQY | 0.5709499 |
| DGSVYKI | ASSLKDTQY | 0.4363419 |
| EQLAEIV | ASSLKDTQY | 0.29769665 |
| FAVSELTVM | ASSLKDTQY | 0.78440666 |
| GAAPLAAGL | ASSLKDTQY | 0.7000608 |
| GADGGDGGSGGAGG | ASSLKDTQY | 0.8052342 |
| GFGLDTGGAGG | ASSLKDTQY | 0.7672273 |
| GLASGGLL | ASSLKDTQY | 0.80752134 |
| GRIAGAAVLL | ASSLKDTQY | 0.66139686 |
| GSAGSLNTSF | ASSLKDTQY | 0.71181023 |
| GTGQIGI | ASSLKDTQY | 0.3109102 |
| GTLIALIL | ASSLKDTQY | 0.7154601 |
| IAATMGFVVG | ASSLKDTQY | 0.5122174 |
| IFAQALL | ASSLKDTQY | 0.7243737 |
| IGAAVALGF | ASSLKDTQY | 0.55371207 |
| IGLTPLAL | ASSLKDTQY | 0.8102838 |
| ILGSNTSTL | ASSLKDTQY | 0.69101214 |
| IVFSLTVL | ASSLKDTQY | 0.79210645 |
| LASKATGF | ASSLKDTQY | 0.6493176 |
| LASVALIL | ASSLKDTQY | 0.45455843 |
| LFAVGLTL | ASSLKDTQY | 0.665426 |
| LFNGGIGTYI | ASSLKDTQY | 0.7866523 |
| LTLGAGII | ASSLKDTQY | 0.5029032 |
| LVAAALGDI | ASSLKDTQY | 0.2502113 |
| NTAFVQLGI | ASSLKDTQY | 0.49914846 |
| QLFSGAEL | ASSLKDTQY | 0.53080714 |
| SAATFITL | ASSLKDTQY | 0.81691 |
| SNYGLANLGV | ASSLKDTQY | 0.808433 |
| SYGGAYAVM | ASSLKDTQY | 0.7472601 |
| TTGVGGLTLG | ASSLKDTQY | 0.81677574 |
| TTQTIYI | ASSLKDTQY | 0.1837191 |
| VAAGAGLGAVI | ASSLKDTQY | 0.6840745 |
| VAAVVTLL | ASSLKDTQY | 0.80846405 |
| VGFLGGLI | ASSLKDTQY | 0.77013385 |
| VGVIPILL | ASSLKDTQY | 0.72624886 |
| VIFASSLI | ASSLKDTQY | 0.803095 |
| VKGGDGGAAAG | ASSLKDTQY | 0.6302552 |
| VTKTFTNI | ASSLKDTQY | 0.71194714 |
| VVAAAILL | ASSLKDTQY | 0.7432375 |
| YAALDVEL | ASSLKDTQY | 0.7833885 |
| LASVALLL | ASSLKDTQY | 0.7685631 |

| | | |
|---|---|---|
| LDAVHRV | ASSLKDTQY | 0.35974932 |
| QGLSSSIF | ASSLKDTQY | 0.7826569 |
| RGTFANIRL | ASSLKDTQY | 0.8208978 |
| RQLWWGHRI | ASSLKDTQY | 0.7075013 |
| YGPPGTGKTLL | ASSLKDTQY | 0.7881317 |
| AALVGAPANVA | ASSLKDTQY | 0.75729454 |
| AATFRRTE | ASSLKDTQY | 0.7436727 |
| AGALGGGI | ASSLKDTQY | 0.6354991 |
| AGGAGGAGGIGAG | ASSLKDTQY | 0.5555182 |
| AGSGALGL | ASSLKDTQY | 0.79293334 |
| AQIADQI | ASSLKDTQY | 0.6162205 |
| ASAVAVTYL | ASSLKDTQY | 0.6900301 |
| ASTVLGGTGG | ASSLKDTQY | 0.7207488 |
| AVGGEPAK | ASSLKDTQY | 0.82286507 |
| CEQGTFGGTC | ASSLKDTQY | 0.77274424 |
| DEGKAPAG | ASSLKDTQY | 0.7746762 |
| DTVVFPAL | ASSLKDTQY | 0.6451494 |
| FEGITMAG | ASSLKDTQY | 0.73853457 |
| FPLVVQD | ASSLKDTQY | 0.78963286 |
| GAFTTPQ | ASSLKDTQY | 0.38774282 |
| GFGGFGTAGGVGGA | ASSLKDTQY | 0.80012465 |
| GFLAQFL | ASSLKDTQY | 0.55573523 |
| GGAGGAGGNAGVA | ASSLKDTQY | 0.74046296 |
| GGVGTAGAGGAGGAGG | ASSLKDTQY | 0.7345499 |
| GNGGGGDAGNAGSGGN | ASSLKDTQY | 0.8132682 |
| GTGGAGGAGNDAGS | ASSLKDTQY | 0.7226571 |
| GTTNTGWFNS | ASSLKDTQY | 0.7984761 |
| IFMGPLFTL | ASSLKDTQY | 0.5835778 |
| ILAHAIEAI | ASSLKDTQY | 0.783977 |
| IRSAIREAL | ASSLKDTQY | 0.80787 |
| ITFAGGG | ASSLKDTQY | 0.76781607 |
| ITGMSGHQEV | ASSLKDTQY | 0.82283753 |
| IVAGLACTL | ASSLKDTQY | 0.7678171 |
| IVGGGLAGA | ASSLKDTQY | 0.24921732 |
| IVTGVLNI | ASSLKDTQY | 0.729926 |
| LAQQGCI | ASSLKDTQY | 0.803915 |
| LCLHVEN | ASSLKDTQY | 0.81006664 |
| LDAISDGS | ASSLKDTQY | 0.7925519 |
| LGGFAAML | ASSLKDTQY | 0.7806947 |
| LGGKNPMI | ASSLKDTQY | 0.7077855 |
| LGPKGVAAIL | ASSLKDTQY | 0.812081 |
| LMVGVSI | ASSLKDTQY | 0.82412535 |
| LVGAAFVL | ASSLKDTQY | 0.79384184 |
| LVLAVAPL | ASSLKDTQY | 0.5388938 |
| MAAIAVIDAA | ASSLKDTQY | 0.47650436 |
| NNGINAELQ | ASSLKDTQY | 0.7778386 |
| RGEPLVLVHG | ASSLKDTQY | 0.79344 |
| RGGAGGAAGLI | ASSLKDTQY | 0.4353872 |
| RSYRMHPAV | ASSLKDTQY | 0.7465451 |
| SAAHLLAGQLA | ASSLKDTQY | 0.76284146 |
| SAAVGSY | ASSLKDTQY | 0.2634831 |

| | | |
|---|---|---|
| SPHAVLHRI | ASSLKDTQY | 0.75880647 |
| SQIAKVL | ASSLKDTQY | 0.80911547 |
| TALGHLVP | ASSLKDTQY | 0.7597976 |
| TAPTTSGAA | ASSLKDTQY | 0.4496184 |
| TDGDDGNGGSPGT | ASSLKDTQY | 0.804005 |
| TDGLGEN | ASSLKDTQY | 0.81755674 |
| TGGPPLPGQA | ASSLKDTQY | 0.81485355 |
| TIPAITI | ASSLKDTQY | 0.3842623 |
| TIPGFTI | ASSLKDTQY | 0.21109538 |
| TSIASGALA | ASSLKDTQY | 0.7780699 |
| TVTLPGAAI | ASSLKDTQY | 0.79387647 |
| VASGLSAALP | ASSLKDTQY | 0.6786096 |
| VGAGVIGI | ASSLKDTQY | 0.5888216 |
| VGPKRFGV | ASSLKDTQY | 0.75950295 |
| VNVAFPD | ASSLKDTQY | 0.80031466 |
| VSMTNSGVS | ASSLKDTQY | 0.49833333 |
| VTDVLGDLSV | ASSLKDTQY | 0.41504902 |
| VVVVGAGFAGL | ASSLKDTQY | 0.7719966 |
| VYTVGTVV | ASSLKDTQY | 0.7925142 |
| WTTFSGP | ASSLKDTQY | 0.58703303 |
| YDELVEAF | ASSLKDTQY | 0.69273925 |
| YGTGGAGGNGG | ASSLKDTQY | 0.78573895 |
| YTNVMAV | ASSLKDTQY | 0.7926829 |
| FLEQVHQL | ASSLKDTQY | 0.76458514 |
| GLSLLAVAV | ASSLKDTQY | 0.4199586 |
| LEQVHQL | ASSLKDTQY | 0.7816305 |
| LNAAGAI | ASSLKDTQY | 0.44643772 |
| PAVHASL | ASSLKDTQY | 0.7789739 |
| TELEDDA | ASSLKDTQY | 0.751604 |
| TPPGPPN | ASSLKDTQY | 0.75682807 |
| TPPPPPPP | ASSLKDTQY | 0.70609397 |
| TQSPALI | ASSRQSSYEQY | 0.54748607 |
| EFAAAFPQ | ASSRQSSYEQY | 0.5113257 |
| LLSGNLDVL | ASSRQSSYEQY | 0.5797267 |
| GPANLGNYNI | ASSRQSSYEQY | 0.80436367 |
| TLGPTLP | ASSRQSSYEQY | 0.78528905 |
| VYAYSSCCK | ASSRQSSYEQY | 0.7905037 |
| APAPAPAPNI | ASSRQSSYEQY | 0.7659442 |
| PGGPVPIP | ASSRQSSYEQY | 0.54756093 |
| FLTVALEP | ASSRQSSYEQY | 0.7696694 |
| SQYFGANL | ASSRQSSYEQY | 0.48838308 |
| GGVKPAA | ASSRQSSYEQY | 0.7812451 |
| GGTGGAGGAGGW | ASSRQSSYEQY | 0.6577424 |
| GVGGVAGLGGAGG | ASSRQSSYEQY | 0.7739138 |
| VLLVLATTI | ASSRQSSYEQY | 0.6064287 |
| IGIFAAML | ASSRQSSYEQY | 0.76157296 |
| AVGDLID | ASSRQSSYEQY | 0.77486783 |
| NISPLPL | ASSRQSSYEQY | 0.47598422 |
| VTAVFVAI | ASSRQSSYEQY | 0.7548411 |
| SGDDNDAAL | ASSRQSSYEQY | 0.80112934 |
| AVAALAHS | ASSRQSSYEQY | 0.7698597 |

| | | |
|---|---|---|
| GGVTVMATP | ASSRQSSYEQY | 0.41850132 |
| AGLAANVRD | ASSRQSSYEQY | 0.77300876 |
| TNRPDLIDPALLRPG | ASSRQSSYEQY | 0.6386351 |
| LANLGVTAL | ASSRQSSYEQY | 0.7633722 |
| TGERIEL | ASSRQSSYEQY | 0.7416905 |
| NGEELAA | ASSRQSSYEQY | 0.57614434 |
| VLSLVWG | ASSRQSSYEQY | 0.77503633 |
| IGYALWL | ASSRQSSYEQY | 0.36326438 |
| LSLLLAPL | ASSRQSSYEQY | 0.6521956 |
| TLDEALK | ASSRQSSYEQY | 0.69144523 |
| TDGAGTAEGG | ASSRQSSYEQY | 0.7638464 |
| NAPGGAGGNGG | ASSRQSSYEQY | 0.78890604 |
| VSVFMGMGGNF | ASSRQSSYEQY | 0.55119336 |
| LASYLAAL | ASSRQSSYEQY | 0.76368326 |
| LVGGTPAL | ASSRQSSYEQY | 0.69594455 |
| VRNADVL | ASSRQSSYEQY | 0.59817946 |
| TCSSPFLVSAM | ASSRQSSYEQY | 0.7176086 |
| VHITDAE | ASSRQSSYEQY | 0.6320923 |
| EYAAPSF | ASSRQSSYEQY | 0.41285533 |
| VVDAFHGGL | ASSRQSSYEQY | 0.7829828 |
| ASVVDPQ | ASSRQSSYEQY | 0.584284 |
| SYNEFGGLS | ASSRQSSYEQY | 0.75036776 |
| DPDGSVY | ASSRQSSYEQY | 0.8012191 |
| MVSTIHATE | ASSRQSSYEQY | 0.41273972 |
| LAAGVADPV | ASSRQSSYEQY | 0.7670848 |
| TKEQIAA | ASSRQSSYEQY | 0.7389703 |
| SLASPAAL | ASSRQSSYEQY | 0.7560679 |
| IGLVFYGV | ASSRQSSYEQY | 0.73973787 |
| FLLDASNL | ASSRQSSYEQY | 0.5579046 |
| YQVGNFP | ASSRQSSYEQY | 0.8050685 |
| ESLLSQST | ASSRQSSYEQY | 0.78671956 |
| LFTVGFAAVG | ASSRQSSYEQY | 0.7658291 |
| ATPPTPT | ASSRQSSYEQY | 0.7495066 |
| TDLATLG | ASSRQSSYEQY | 0.79607284 |
| SAGGTGR | ASSRQSSYEQY | 0.7963877 |
| GFSNSGS | ASSRQSSYEQY | 0.69153583 |
| GGNGGSGGTSV | ASSRQSSYEQY | 0.35280824 |
| GTSVATGGAGN | ASSRQSSYEQY | 0.73271835 |
| NGGNGGSGGTS | ASSRQSSYEQY | 0.73737836 |
| AVVKFPH | ASSRQSSYEQY | 0.5721567 |
| TPPPPPP | ASSRQSSYEQY | 0.3164522 |
| AAPAAAAAPA | ASSRQSSYEQY | 0.7941704 |
| IALNEDL | ASSRQSSYEQY | 0.82213175 |
| IVNVSSM | ASSRQSSYEQY | 0.70551956 |
| TGIPPAPR | ASSRQSSYEQY | 0.82396793 |
| FDLGGGTF | ASSRQSSYEQY | 0.42919168 |
| FDLGGGTFDVS | ASSRQSSYEQY | 0.7922809 |
| NEPTAAA | ASSRQSSYEQY | 0.79816705 |
| NEPTAAALA | ASSRQSSYEQY | 0.58547187 |
| GSIINISSI | ASSRQSSYEQY | 0.77033114 |
| VAPGNAGTA | ASSRQSSYEQY | 0.659309 |

| | | |
|---|---|---|
| TGGGASLE | ASSRQSSYEQY | 0.61559 |
| VAVYNHYKRI | ASSRQSSYEQY | 0.80250275 |
| TDPAPATT | ASSRQSSYEQY | 0.6933736 |
| TNRPDILDPALLRPG | ASSRQSSYEQY | 0.7672807 |
| LTTTTTS | ASSRQSSYEQY | 0.6961291 |
| PTGSGKTL | ASSRQSSYEQY | 0.5572519 |
| TGSGKTL | ASSRQSSYEQY | 0.7661976 |
| TIGHVDHGK | ASSRQSSYEQY | 0.7791335 |
| TIGHVDHGKTT | ASSRQSSYEQY | 0.76128834 |
| GHVDHGKTT | ASSRQSSYEQY | 0.8239654 |
| AQFEHTLL | ASSRQSSYEQY | 0.5985702 |
| IVNQDEL | ASSRQSSYEQY | 0.8111413 |
| TNGSQFF | ASSRQSSYEQY | 0.8106964 |
| TGKGIVE | ASSRQSSYEQY | 0.7603868 |
| LISNASDA | ASSRQSSYEQY | 0.6781979 |
| AELRTLGELTKIAKAHGAQ | ASSRQSSYEQY | 0.8036555 |
| APNAVAAAEPSW | ASSRQSSYEQY | 0.68746006 |
| APQVAGAATAVL | ASSRQSSYEQY | 0.7219306 |
| ARRDPAAAAAILPTDA | ASSRQSSYEQY | 0.80782944 |
| DAAARDVLAEAGLAEYF | ASSRQSSYEQY | 0.67812794 |
| DDQIVLSAKANLAKVEQ | ASSRQSSYEQY | 0.85346913 |
| DELLPDTGEDSDEDGH | ASSRQSSYEQY | 0.82068324 |
| ELPEAPANRATVPQD | ASSRQSSYEQY | 0.7747822 |
| FGQSKIERVTGIG | ASSRQSSYEQY | 0.7207254 |
| GAGGTGSTAGGAGGAGGAGGLY | ASSRQSSYEQY | 0.3034526 |
| GHGGTGAAGGNGGTGGAGGSGI | ASSRQSSYEQY | 0.7694779 |
| GNGGNGGAGGEGGPGGAGGAGA | ASSRQSSYEQY | 0.6687385 |
| GSTAGNGGNGGNATNGTIAGQP | ASSRQSSYEQY | 0.82212234 |
| GTGGDGGNGGNGGTGG | ASSRQSSYEQY | 0.821942 |
| GVRRPAGIPADAPV | ASSRQSSYEQY | 0.6952719 |
| GYRQVTDAHLLTLARR | ASSRQSSYEQY | 0.85147357 |
| IAKMGQDGHDRG | ASSRQSSYEQY | 0.82005763 |
| KAEPSIEIQYT | ASSRQSSYEQY | 0.511387 |
| KPVRIRMAADEDTVAVT | ASSRQSSYEQY | 0.7918211 |
| MPPRIAGMRLLVIKPEP | ASSRQSSYEQY | 0.71817553 |
| MKFYAVALFRSRGVPPTR | ASSRQSSYEQY | 0.7773708 |
| NAGGFGILGAVAHSP | ASSRQSSYEQY | 0.4596272 |
| NEGREITLAMAEM | ASSRQSSYEQY | 0.8278297 |
| NELVEWSHPLGHRWRWE | ASSRQSSYEQY | 0.75576043 |
| NLGAGGTGGNGGNPG | ASSRQSSYEQY | 0.1500416 |
| NTEWPKWETVMAADPVD | ASSRQSSYEQY | 0.80523664 |
| PASAAPAETQP | ASSRQSSYEQY | 0.7831428 |
| QQSSDGKNGVRAAEITGE | ASSRQSSYEQY | 0.7197878 |
| RAPDAVTTSWWKE | ASSRQSSYEQY | 0.8453334 |
| SGFGNVGAHTSGWFNQ | ASSRQSSYEQY | 0.24480563 |
| SGVAGAVGQAASVGGLKV | ASSRQSSYEQY | 0.637871 |
| SNGGAGTGSAGGNGGT | ASSRQSSYEQY | 0.84479505 |
| TGGAGGEGGAAG | ASSRQSSYEQY | 0.8483232 |
| TGGDGGAAGAGGAPGHG | ASSRQSSYEQY | 0.5208759 |
| TMPPLLAAIANAGRHGV | ASSRQSSYEQY | 0.82032543 |
| TSSTAGNSGGA | ASSRQSSYEQY | 0.84059805 |

| | | |
|---|---|---|
| VPATVPVGILC | ASSRQSSYEQY | 0.8171656 |
| VQRPDSGGVVLTGRIS | ASSRQSSYEQY | 0.83144003 |
| WTPNPVPATKGVPAGLVQAT | ASSRQSSYEQY | 0.83838177 |
| YALKRALAKFSDRFDVVII | ASSRQSSYEQY | 0.59645635 |
| YLIGSVIV | ASSRQSSYEQY | 0.8489313 |
| YGQLVAVID | ASSRQSSYEQY | 0.853341 |
| VTVGGMTLPA | ASSRQSSYEQY | 0.7040095 |
| VLALPAV | ASSRQSSYEQY | 0.7970875 |
| VIPPTLNYET | ASSRQSSYEQY | 0.65069896 |
| VGMGGMHPSSG | ASSRQSSYEQY | 0.7162728 |
| VGLAGGAATA | ASSRQSSYEQY | 0.23960832 |
| VGGAGGAGGSGP | ASSRQSSYEQY | 0.8309625 |
| VFAAFMV | ASSRQSSYEQY | 0.8227899 |
| TLYDKLR | ASSRQSSYEQY | 0.7809862 |
| TLAVFTVL | ASSRQSSYEQY | 0.69588256 |
| TGSVQPARV | ASSRQSSYEQY | 0.5917039 |
| TGGGAGSGGT | ASSRQSSYEQY | 0.7765401 |
| SHLETSL | ASSRQSSYEQY | 0.6867776 |
| SFTTGMT | ASSRQSSYEQY | 0.239261 |
| QVEVAKRMGVRQPTVS | ASSRQSSYEQY | 0.68530667 |
| QGSGGPALAM | ASSRQSSYEQY | 0.83880216 |
| MLAPSNP | ASSRQSSYEQY | 0.8364728 |
| LVNNAAI | ASSRQSSYEQY | 0.5282608 |
| LTGEINTLA | ASSRQSSYEQY | 0.85528934 |
| LSIFGPV | ASSRQSSYEQY | 0.8335769 |
| LLALLANL | ASSRQSSYEQY | 0.8449678 |
| LISNASDALDKLR | ASSRQSSYEQY | 0.73060966 |
| LINEAVP | ASSRQSSYEQY | 0.6590124 |
| LGYAFSGL | ASSRQSSYEQY | 0.84406036 |
| LDFVAVAL | ASSRQSSYEQY | 0.7966049 |
| LAQPGAE | ASSRQSSYEQY | 0.6018238 |
| ISNLGAVTGF | ASSRQSSYEQY | 0.8424618 |
| HQEDPGRI | ASSRQSSYEQY | 0.72981036 |
| GTPGEPTP | ASSRQSSYEQY | 0.707178 |
| GFFNSGS | ASSRQSSYEQY | 0.7711044 |
| GAGGGGLTAGAN | ASSRQSSYEQY | 0.83556634 |
| FVDALVNM | ASSRQSSYEQY | 0.7070919 |
| FLTVAPEL | ASSRQSSYEQY | 0.4575014 |
| FETLTAL | ASSRQSSYEQY | 0.5096531 |
| DVINPIEL | ASSRQSSYEQY | 0.7902148 |
| APVSGAPV | ASSRQSSYEQY | 0.84507734 |
| ALVGGLAW | ASSRQSSYEQY | 0.7895225 |
| ALAAAAATAI | ASSRQSSYEQY | 0.6350448 |
| AFGLGLLL | ASSRQSSYEQY | 0.8270873 |
| AFFAAASL | ASSRQSSYEQY | 0.7774118 |
| AAAALPAFSPP | ASSRQSSYEQY | 0.78684133 |
| VVLAPAL | ASSRQSSYEQY | 0.7814276 |
| TIGHVDHGKTTL | ASSRQSSYEQY | 0.7562397 |
| LISNASDALDK | ASSRQSSYEQY | 0.84202975 |
| LISNASDAL | ASSRQSSYEQY | 0.8204185 |
| ISNASDALDK | ASSRQSSYEQY | 0.8118564 |

| | | |
|---|---|---|
| NQAETLVYQTE | ASSRQSSYEQY | 0.8269039 |
| TGGTGGTAGAG | ASSRQSSYEQY | 0.7306553 |
| TGSTAGGAGGA | ASSRQSSYEQY | 0.7737977 |
| DPLAVPVAAGPV | ASSRQSSYEQY | 0.38748235 |
| GGTGGTPGNGGN | ASSRQSSYEQY | 0.5129241 |
| PAGLAGVDPATL | ASSRQSSYEQY | 0.8231632 |
| GNNLVVEAPHAGASV | ASSRQSSYEQY | 0.8280983 |
| KVTRIMSASTLEQPA | ASSRQSSYEQY | 0.83341694 |
| LGFALPIALPRRTVV | ASSRQSSYEQY | 0.74425304 |
| TRLGNLTARKLVVLV | ASSRQSSYEQY | 0.71525407 |
| WSQQRRLIYFPSAGP | ASSRQSSYEQY | 0.8034966 |
| GPNGNPDPMAAAVDIR | ASSRQSSYEQY | 0.8419092 |
| TQRCVSVADLRAAASR | ASSRQSSYEQY | 0.5094323 |
| QFVLDVAEHPVEVLTETV | ASSRQSSYEQY | 0.744654 |
| GGAGGTGGAGGNGGTQAGDG | ASSRQSSYEQY | 0.8595056 |
| PLLATLRAIAVDVDAKPAQV | ASSRQSSYEQY | 0.8226854 |
| APKFRTGGYMIDFWGVGYQVAK | ASSRQSSYEQY | 0.7814678 |
| GAGGAGGAGGAAGGPGGTGGTG | ASSRQSSYEQY | 0.64234835 |
| IGALGLLATDVAPAGSVAFGVA | ASSRQSSYEQY | 0.8438487 |
| AGGAGGIGGNGGAGGAGGVLMG | ASSRQSSYEQY | 0.6360038 |
| ADELIGLL | ASSRQSSYEQY | 0.8586147 |
| ADRALKA | ASSRQSSYEQY | 0.21737453 |
| AGAVGQVGGAGGA | ASSRQSSYEQY | 0.61405504 |
| AGGGGGLGGAP | ASSRQSSYEQY | 0.5538783 |
| AGGLLYGNGGAG | ASSRQSSYEQY | 0.2885394 |
| AGLESIGL | ASSRQSSYEQY | 0.84767586 |
| AGLKTGII | ASSRQSSYEQY | 0.6645384 |
| ASLAALLL | ASSRQSSYEQY | 0.8170753 |
| ASVINISSI | ASSRQSSYEQY | 0.80589235 |
| ATPPGPA | ASSRQSSYEQY | 0.8428669 |
| ATVASALV | ASSRQSSYEQY | 0.8246139 |
| ATVASLIL | ASSRQSSYEQY | 0.7332074 |
| AVNGPVLIRH | ASSRQSSYEQY | 0.45632035 |
| DGAAGAVGI | ASSRQSSYEQY | 0.84036297 |
| DPLLHGI | ASSRQSSYEQY | 0.8298055 |
| ELGAALGAL | ASSRQSSYEQY | 0.7779519 |
| ELGRTQAV | ASSRQSSYEQY | 0.40427086 |
| FVAGGYLS | ASSRQSSYEQY | 0.78298765 |
| GASGGAGGAGGW | ASSRQSSYEQY | 0.8156264 |
| GGAGAGGDGGAGGI | ASSRQSSYEQY | 0.18712616 |
| GTGHNGSQPGMGGNG | ASSRQSSYEQY | 0.20096883 |
| GYAAASASAS | ASSRQSSYEQY | 0.7534344 |
| HQVDVAAVQQG | ASSRQSSYEQY | 0.7141514 |
| IGAAIAHT | ASSRQSSYEQY | 0.8498001 |
| IGMIEVAQL | ASSRQSSYEQY | 0.84489334 |
| IRLAPEHTSAMD | ASSRQSSYEQY | 0.7870064 |
| IYNVSVL | ASSRQSSYEQY | 0.8196495 |
| KQGFLGM | ASSRQSSYEQY | 0.7997479 |
| LESVITQL | ASSRQSSYEQY | 0.12479983 |
| LGLGALVVV | ASSRQSSYEQY | 0.59255683 |
| LIGNGGNGGSGGI | ASSRQSSYEQY | 0.55725443 |

| | | |
|---|---|---|
| LNAINAPTL | ASSRQSSYEQY | 0.8314922 |
| LQLLNTGV | ASSRQSSYEQY | 0.67797333 |
| LQTSIEF | ASSRQSSYEQY | 0.73735183 |
| MSFAVSEL | ASSRQSSYEQY | 0.80494076 |
| NTGALGL | ASSRQSSYEQY | 0.8171785 |
| PPGAGGAGGAGTT | ASSRQSSYEQY | 0.80148697 |
| QLVGLGI | ASSRQSSYEQY | 0.5566913 |
| RFLEQVHQL | ASSRQSSYEQY | 0.1590675 |
| RGVGYVLR | ASSRQSSYEQY | 0.8331908 |
| SATGVFAGVI | ASSRQSSYEQY | 0.31744915 |
| SLPAITLP | ASSRQSSYEQY | 0.5963666 |
| STALAFTV | ASSRQSSYEQY | 0.25929087 |
| SVLTVGLGK | ASSRQSSYEQY | 0.19985242 |
| TALAGAGSGI | ASSRQSSYEQY | 0.34434083 |
| TGLPFTGI | ASSRQSSYEQY | 0.8136133 |
| VAEHVTV | ASSRQSSYEQY | 0.780056 |
| VDIVLNSL | ASSRQSSYEQY | 0.7949808 |
| VLTGGGAVL | ASSRQSSYEQY | 0.8581259 |
| VNVGLAYGI | ASSRQSSYEQY | 0.8104076 |
| VSSAAVGAV | ASSRQSSYEQY | 0.791204 |
| VTSVYEAGV | ASSRQSSYEQY | 0.6308033 |
| WVALDRGA | ASSRQSSYEQY | 0.6753885 |
| YLVGSVVI | ASSRQSSYEQY | 0.645523 |
| GPPGTGKTLL | ASSRQSSYEQY | 0.6116754 |
| GVGMTKF | ASSRQSSYEQY | 0.8523061 |
| IGIGGSDL | ASSRQSSYEQY | 0.66268164 |
| IGTLLAEL | ASSRQSSYEQY | 0.4862338 |
| IYTSGTT | ASSRQSSYEQY | 0.73329914 |
| LSFPAGV | ASSRQSSYEQY | 0.8534271 |
| LYTSGTT | ASSRQSSYEQY | 0.8593706 |
| VGAFPFGF | ASSRQSSYEQY | 0.69039667 |
| VGAGTVEF | ASSRQSSYEQY | 0.612982 |
| VGKSTLLL | ASSRQSSYEQY | 0.7111642 |
| AALAHAGGGIV | ASSRQSSYEQY | 0.8154534 |
| AALVAVSLL | ASSRQSSYEQY | 0.6276162 |
| AAVSRAVAAL | ASSRQSSYEQY | 0.403499 |
| AGGSSGIGGF | ASSRQSSYEQY | 0.8221336 |
| AGLTVANANG | ASSRQSSYEQY | 0.85076207 |
| AGLVGAVL | ASSRQSSYEQY | 0.633088 |
| ALSQAFTV | ASSRQSSYEQY | 0.6750172 |
| ALTFSVSL | ASSRQSSYEQY | 0.7999438 |
| ALTGAALVAL | ASSRQSSYEQY | 0.6823069 |
| ASFSIGAGVL | ASSRQSSYEQY | 0.7602245 |
| DGSVYKI | ASSRQSSYEQY | 0.7055788 |
| EQLAEIV | ASSRQSSYEQY | 0.527412 |
| FAVSELTVM | ASSRQSSYEQY | 0.76634514 |
| GAAPLAAGL | ASSRQSSYEQY | 0.85694444 |
| GADGGDGGSGGAGG | ASSRQSSYEQY | 0.78772056 |
| GFGLDTGGAGG | ASSRQSSYEQY | 0.8189049 |
| GLASGGLL | ASSRQSSYEQY | 0.8548562 |
| GRIAGAAVLL | ASSRQSSYEQY | 0.84297657 |

| | | |
|---|---|---|
| GSAGSLNTSF | ASSRQSSYEQY | 0.8534937 |
| GTGQIGI | ASSRQSSYEQY | 0.83119726 |
| GTLIALIL | ASSRQSSYEQY | 0.7121052 |
| IAATMGFVVG | ASSRQSSYEQY | 0.7716774 |
| IFAQALL | ASSRQSSYEQY | 0.8289604 |
| IGAAVALGF | ASSRQSSYEQY | 0.80530834 |
| IGLTPLAL | ASSRQSSYEQY | 0.791566 |
| ILGSNTSTL | ASSRQSSYEQY | 0.7076466 |
| IVFSLTVL | ASSRQSSYEQY | 0.8496052 |
| LASKATGF | ASSRQSSYEQY | 0.7822516 |
| LASVALIL | ASSRQSSYEQY | 0.8395407 |
| LFAVGLTL | ASSRQSSYEQY | 0.3587406 |
| LFNGGIGTYI | ASSRQSSYEQY | 0.51410127 |
| LTLGAGII | ASSRQSSYEQY | 0.8445122 |
| LVAAALGDI | ASSRQSSYEQY | 0.13723586 |
| NTAFVQLGI | ASSRQSSYEQY | 0.17876291 |
| QLFSGAEL | ASSRQSSYEQY | 0.8034856 |
| SAATFITL | ASSRQSSYEQY | 0.8472721 |
| SNYGLANLGV | ASSRQSSYEQY | 0.14286853 |
| SYGGAYAVM | ASSRQSSYEQY | 0.46841568 |
| TTGVGGLTLG | ASSRQSSYEQY | 0.4420877 |
| TTQTIYI | ASSRQSSYEQY | 0.7711612 |
| VAAGAGLGAVI | ASSRQSSYEQY | 0.79469013 |
| VAAVVTLL | ASSRQSSYEQY | 0.29590607 |
| VGFLGGLI | ASSRQSSYEQY | 0.8135056 |
| VGVIPILL | ASSRQSSYEQY | 0.52202195 |
| VIFASSLI | ASSRQSSYEQY | 0.49061087 |
| VKGGDGGAAAG | ASSRQSSYEQY | 0.8371366 |
| VTKTFTNI | ASSRQSSYEQY | 0.84788597 |
| VVAAAILL | ASSRQSSYEQY | 0.5713066 |
| YAALDVEL | ASSRQSSYEQY | 0.8363294 |
| LASVALLL | ASSRQSSYEQY | 0.70227736 |
| LDAVHRV | ASSRQSSYEQY | 0.85827255 |
| QGLSSSIF | ASSRQSSYEQY | 0.7522938 |
| RGTFANIRL | ASSRQSSYEQY | 0.37246555 |
| RQLWWGHRI | ASSRQSSYEQY | 0.83878964 |
| YGPPGTGKTLL | ASSRQSSYEQY | 0.82596815 |
| AALVGAPANVA | ASSRQSSYEQY | 0.7332267 |
| AATFRRTE | ASSRQSSYEQY | 0.84204775 |
| AGALGGGI | ASSRQSSYEQY | 0.5815352 |
| AGGAGGAGGIGAG | ASSRQSSYEQY | 0.8004298 |
| AGSGALGL | ASSRQSSYEQY | 0.848969 |
| AQIADQI | ASSRQSSYEQY | 0.8437371 |
| ASAVAVTYL | ASSRQSSYEQY | 0.3911748 |
| ASTVLGGTGG | ASSRQSSYEQY | 0.8241316 |
| AVGGEPAK | ASSRQSSYEQY | 0.78310823 |
| CEQGTFGGTC | ASSRQSSYEQY | 0.6394356 |
| DEGKAPAG | ASSRQSSYEQY | 0.8572473 |
| DTVVFPAL | ASSRQSSYEQY | 0.65204483 |
| FEGITMAG | ASSRQSSYEQY | 0.844275 |
| FPLVVQD | ASSRQSSYEQY | 0.83001065 |

| | | |
|---|---|---|
| GAFTTPQ | ASSRQSSYEQY | 0.8215572 |
| GFGGFGTAGGVGGA | ASSRQSSYEQY | 0.3360604 |
| GFLAQFL | ASSRQSSYEQY | 0.76107943 |
| GGAGGAGGNAGVA | ASSRQSSYEQY | 0.7890357 |
| GGVGTAGAGGAGGAGG | ASSRQSSYEQY | 0.82375884 |
| GNGGGGDAGNAGSGGN | ASSRQSSYEQY | 0.75503814 |
| GTGGAGGAGNDAGS | ASSRQSSYEQY | 0.57049114 |
| GTTNTGWFNS | ASSRQSSYEQY | 0.81148237 |
| IFMGPLFTL | ASSRQSSYEQY | 0.84948516 |
| ILAHAIEAI | ASSRQSSYEQY | 0.7757995 |
| IRSAIREAL | ASSRQSSYEQY | 0.7722836 |
| ITFAGGG | ASSRQSSYEQY | 0.85631645 |
| ITGMSGHQEV | ASSRQSSYEQY | 0.66365063 |
| IVAGLACTL | ASSRQSSYEQY | 0.74921626 |
| IVGGGLAGA | ASSRQSSYEQY | 0.8327482 |
| IVTGVLNI | ASSRQSSYEQY | 0.8365413 |
| LAQQGCI | ASSRQSSYEQY | 0.7230077 |
| LCLHVEN | ASSRQSSYEQY | 0.8458278 |
| LDAISDGS | ASSRQSSYEQY | 0.48944408 |
| LGGFAAML | ASSRQSSYEQY | 0.83143556 |
| LGGKNPMI | ASSRQSSYEQY | 0.72934294 |
| LGPKGVAAIL | ASSRQSSYEQY | 0.8204974 |
| LMVGVSI | ASSRQSSYEQY | 0.6636755 |
| LVGAAFVL | ASSRQSSYEQY | 0.84563136 |
| LVLAVAPL | ASSRQSSYEQY | 0.84578407 |
| MAAIAVIDAA | ASSRQSSYEQY | 0.8455146 |
| NNGINAELQ | ASSRQSSYEQY | 0.19960693 |
| RGEPLVLVHG | ASSRQSSYEQY | 0.83870625 |
| RGGAGGAAGLI | ASSRQSSYEQY | 0.8262134 |
| RSYRMHPAV | ASSRQSSYEQY | 0.8605963 |
| SAAHLLAGQLA | ASSRQSSYEQY | 0.7723822 |
| SAAVGSY | ASSRQSSYEQY | 0.80925435 |
| SPHAVLHRI | ASSRQSSYEQY | 0.7364298 |
| SQIAKVL | ASSRQSSYEQY | 0.6755874 |
| TALGHLVP | ASSRQSSYEQY | 0.6117261 |
| TAPTTSGAA | ASSRQSSYEQY | 0.7174869 |
| TDGDDGNGGSPGT | ASSRQSSYEQY | 0.7402039 |
| TDGLGEN | ASSRQSSYEQY | 0.75192773 |
| TGGPPLPGQA | ASSRQSSYEQY | 0.83952475 |
| TIPAITI | ASSRQSSYEQY | 0.840849 |
| TIPGFTI | ASSRQSSYEQY | 0.8394761 |
| TSIASGALA | ASSRQSSYEQY | 0.8475435 |
| TVTLPGAAI | ASSRQSSYEQY | 0.8535458 |
| VASGLSAALP | ASSRQSSYEQY | 0.7927481 |
| VGAGVIGI | ASSRQSSYEQY | 0.7063539 |
| VGPKRFGV | ASSRQSSYEQY | 0.6995944 |
| VNVAFPD | ASSRQSSYEQY | 0.6739053 |
| VSMTNSGVS | ASSRQSSYEQY | 0.7771412 |
| VTDVLGDLSV | ASSRQSSYEQY | 0.69352007 |
| VVVVGAGFAGL | ASSRQSSYEQY | 0.81759965 |
| VYTVGTVV | ASSRQSSYEQY | 0.12962744 |

| | | |
|---|---|---|
| WTTFSGP | ASSRQSSYEQY | 0.81863916 |
| YDELVEAF | ASSRQSSYEQY | 0.85453 |
| YGTGGAGGNGG | ASSRQSSYEQY | 0.72624695 |
| YTNVMAV | ASSRQSSYEQY | 0.8116088 |
| FLEQVHQL | ASSRQSSYEQY | 0.81066597 |
| GLSLLAVAV | ASSRQSSYEQY | 0.73468256 |
| LEQVHQL | ASSRQSSYEQY | 0.83320796 |
| LNAAGAI | ASSRQSSYEQY | 0.585215 |
| PAVHASL | ASSRQSSYEQY | 0.7990644 |
| TELEDDA | ASSRQSSYEQY | 0.8571271 |
| TPPGPPN | ASSRQSSYEQY | 0.82208276 |
| TPPPPPPP | ASSRQSSYEQY | 0.725458 |
| TQSPALI | ASSFGGAQDTQY | 0.7091597 |
| EFAAAFPQ | ASSFGGAQDTQY | 0.31933984 |
| LLSGNLDVL | ASSFGGAQDTQY | 0.7934959 |
| GPANLGNYNI | ASSFGGAQDTQY | 0.12982467 |
| TLGPTLP | ASSFGGAQDTQY | 0.8103304 |
| VYAYSSCCK | ASSFGGAQDTQY | 0.49693134 |
| APAPAPAPNI | ASSFGGAQDTQY | 0.8178719 |
| PGGPVPIP | ASSFGGAQDTQY | 0.8105493 |
| FLTVALEP | ASSFGGAQDTQY | 0.8127639 |
| SQYFGANL | ASSFGGAQDTQY | 0.7176137 |
| GGVKPAA | ASSFGGAQDTQY | 0.7652564 |
| GGTGGAGGAGGW | ASSFGGAQDTQY | 0.70253026 |
| GVGGVAGLGGAGG | ASSFGGAQDTQY | 0.47704548 |
| VLLVLATTI | ASSFGGAQDTQY | 0.84039205 |
| IGIFAAML | ASSFGGAQDTQY | 0.7101914 |
| AVGDLID | ASSFGGAQDTQY | 0.79161656 |
| NISPLPL | ASSFGGAQDTQY | 0.84250504 |
| VTAVFVAI | ASSFGGAQDTQY | 0.80356336 |
| SGDDNDAAL | ASSFGGAQDTQY | 0.40433997 |
| AVAALAHS | ASSFGGAQDTQY | 0.8023556 |
| GGVTVMATP | ASSFGGAQDTQY | 0.800562 |
| AGLAANVRD | ASSFGGAQDTQY | 0.7540865 |
| TNRPDLIDPALLRPG | ASSFGGAQDTQY | 0.13085385 |
| LANLGVTAL | ASSFGGAQDTQY | 0.79869616 |
| TGERIEL | ASSFGGAQDTQY | 0.62968624 |
| NGEELAA | ASSFGGAQDTQY | 0.7406013 |
| VLSLVWG | ASSFGGAQDTQY | 0.73581135 |
| IGYALWL | ASSFGGAQDTQY | 0.84095687 |
| LSLLLAPL | ASSFGGAQDTQY | 0.38852412 |
| TLDEALK | ASSFGGAQDTQY | 0.5797949 |
| TDGAGTAEGG | ASSFGGAQDTQY | 0.83535445 |
| NAPGGAGGNGG | ASSFGGAQDTQY | 0.810812 |
| VSVFMGMGGNF | ASSFGGAQDTQY | 0.8466616 |
| LASYLAAL | ASSFGGAQDTQY | 0.76594543 |
| LVGGTPAL | ASSFGGAQDTQY | 0.69347966 |
| VRNADVL | ASSFGGAQDTQY | 0.76682734 |
| TCSSPFLVSAM | ASSFGGAQDTQY | 0.69309366 |
| VHITDAE | ASSFGGAQDTQY | 0.34770647 |
| EYAAPSF | ASSFGGAQDTQY | 0.26948065 |

| | | |
|---|---|---|
| VVDAFHGGL | ASSFGGAQDTQY | 0.8077887 |
| ASVVDPQ | ASSFGGAQDTQY | 0.2611974 |
| SYNEFGGLS | ASSFGGAQDTQY | 0.802075 |
| DPDGSVY | ASSFGGAQDTQY | 0.81541216 |
| MVSTIHATE | ASSFGGAQDTQY | 0.67687607 |
| LAAGVADPV | ASSFGGAQDTQY | 0.47252107 |
| TKEQIAA | ASSFGGAQDTQY | 0.7797743 |
| SLASPAAL | ASSFGGAQDTQY | 0.66845816 |
| IGLVFYGV | ASSFGGAQDTQY | 0.83789086 |
| FLLDASNL | ASSFGGAQDTQY | 0.7949772 |
| YQVGNFP | ASSFGGAQDTQY | 0.8522347 |
| ESLLSQST | ASSFGGAQDTQY | 0.8434179 |
| LFTVGFAAVG | ASSFGGAQDTQY | 0.7651313 |
| ATPPTPT | ASSFGGAQDTQY | 0.8406714 |
| TDLATLG | ASSFGGAQDTQY | 0.7366578 |
| SAGGTGR | ASSFGGAQDTQY | 0.75978696 |
| GFSNSGS | ASSFGGAQDTQY | 0.7711489 |
| GGNGGSGGTSV | ASSFGGAQDTQY | 0.7953056 |
| GTSVATGGAGN | ASSFGGAQDTQY | 0.65975404 |
| NGGNGGSGGTS | ASSFGGAQDTQY | 0.7243755 |
| AVVKFPH | ASSFGGAQDTQY | 0.44921172 |
| TPPPPPP | ASSFGGAQDTQY | 0.8092334 |
| AAPAAAAAPA | ASSFGGAQDTQY | 0.77766573 |
| IALNEDL | ASSFGGAQDTQY | 0.29546568 |
| IVNVSSM | ASSFGGAQDTQY | 0.7173089 |
| TGIPPAPR | ASSFGGAQDTQY | 0.85621846 |
| FDLGGGTF | ASSFGGAQDTQY | 0.8413791 |
| FDLGGGTFDVS | ASSFGGAQDTQY | 0.69545186 |
| NEPTAAA | ASSFGGAQDTQY | 0.5893658 |
| NEPTAAALA | ASSFGGAQDTQY | 0.824159 |
| GSIINISSI | ASSFGGAQDTQY | 0.37605962 |
| VAPGNAGTA | ASSFGGAQDTQY | 0.7722538 |
| TGGGASLE | ASSFGGAQDTQY | 0.8096874 |
| VAVYNHYKRI | ASSFGGAQDTQY | 0.8067188 |
| TDPAPATT | ASSFGGAQDTQY | 0.57023406 |
| TNRPDILDPALLRPG | ASSFGGAQDTQY | 0.82555246 |
| LTTTTTS | ASSFGGAQDTQY | 0.7656615 |
| PTGSGKTL | ASSFGGAQDTQY | 0.729845 |
| TGSGKTL | ASSFGGAQDTQY | 0.15210354 |
| TIGHVDHGK | ASSFGGAQDTQY | 0.7995931 |
| TIGHVDHGKTT | ASSFGGAQDTQY | 0.7053283 |
| GHVDHGKTT | ASSFGGAQDTQY | 0.5711087 |
| AQFEHTLL | ASSFGGAQDTQY | 0.2681464 |
| IVNQDEL | ASSFGGAQDTQY | 0.78052765 |
| TNGSQFF | ASSFGGAQDTQY | 0.84335077 |
| TGKGIVE | ASSFGGAQDTQY | 0.8494645 |
| LISNASDA | ASSFGGAQDTQY | 0.60729504 |
| AELRTLGELTKIAKAHGAQ | ASSFGGAQDTQY | 0.8407056 |
| APNAVAAAEPSW | ASSFGGAQDTQY | 0.8540686 |
| APQVAGAATAVL | ASSFGGAQDTQY | 0.78857315 |
| ARRDPAAAAAILPTDA | ASSFGGAQDTQY | 0.25288284 |

| | | |
|---|---|---|
| DAAARDVLAEAGLAEYF | ASSFGGAQDTQY | 0.8005266 |
| DDQIVLSAKANLAKVEQ | ASSFGGAQDTQY | 0.7885942 |
| DELLPDTGEDSDEDGH | ASSFGGAQDTQY | 0.83016217 |
| ELPEAPANRATVPQD | ASSFGGAQDTQY | 0.81443965 |
| FGQSKIERVTGIG | ASSFGGAQDTQY | 0.67405653 |
| GAGGTGSTAGGAGGAGGAGGLY | ASSFGGAQDTQY | 0.8074787 |
| GHGGTGAAGGNGGTGGAGGSGI | ASSFGGAQDTQY | 0.8415461 |
| GNGGNGGAGGEGGPGGAGGAGA | ASSFGGAQDTQY | 0.65298414 |
| GSTAGNGGNGGNATNGTIAGQP | ASSFGGAQDTQY | 0.67246395 |
| GTGGDGGNGGNGGTGG | ASSFGGAQDTQY | 0.6124081 |
| GVRRPAGIPADAPV | ASSFGGAQDTQY | 0.33540994 |
| GYRQVTDAHLLTLARR | ASSFGGAQDTQY | 0.81115264 |
| IAKMGQDGHDRG | ASSFGGAQDTQY | 0.7776327 |
| KAEPSIEIQYT | ASSFGGAQDTQY | 0.62592787 |
| KPVRIRMAADEDTVAVT | ASSFGGAQDTQY | 0.8033484 |
| MPPRIAGMRLLVIKPEP | ASSFGGAQDTQY | 0.61723727 |
| MKFYAVALFRSRGVPPTR | ASSFGGAQDTQY | 0.6308905 |
| NAGGFGILGAVAHSP | ASSFGGAQDTQY | 0.71484774 |
| NEGREITLAMAEM | ASSFGGAQDTQY | 0.7685467 |
| NELVEWSHPLGHRWRWE | ASSFGGAQDTQY | 0.83204377 |
| NLGAGGTGGNGGNPG | ASSFGGAQDTQY | 0.7783099 |
| NTEWPKWETVMAADPVD | ASSFGGAQDTQY | 0.7675257 |
| PASAAPAETQP | ASSFGGAQDTQY | 0.8573426 |
| QQSSDGKNGVRAAEITGE | ASSFGGAQDTQY | 0.35920277 |
| RAPDAVTTSWWKE | ASSFGGAQDTQY | 0.6210862 |
| SGFGNVGAHTSGWFNQ | ASSFGGAQDTQY | 0.8015307 |
| SGVAGAVGQAASVGGLKV | ASSFGGAQDTQY | 0.45211172 |
| SNGGAGTGSAGGNGGT | ASSFGGAQDTQY | 0.81490827 |
| TGGAGGEGGAAG | ASSFGGAQDTQY | 0.8043388 |
| TGGDGGAAGAGGAPGHG | ASSFGGAQDTQY | 0.82545793 |
| TMPPLLAAIANAGRHGV | ASSFGGAQDTQY | 0.7774379 |
| TSSTAGNSGGA | ASSFGGAQDTQY | 0.86006474 |
| VPATVPVGILC | ASSFGGAQDTQY | 0.83884823 |
| VQRPDSGGVVLTGRIS | ASSFGGAQDTQY | 0.7604828 |
| WTPNPVPATKGVPAGLVQAT | ASSFGGAQDTQY | 0.8082509 |
| YALKRALAKFSDRFDVVII | ASSFGGAQDTQY | 0.38935632 |
| YLIGSVIV | ASSFGGAQDTQY | 0.83255357 |
| YGQLVAVID | ASSFGGAQDTQY | 0.8157589 |
| VTVGGMTLPA | ASSFGGAQDTQY | 0.67638683 |
| VLALPAV | ASSFGGAQDTQY | 0.8035246 |
| VIPPTLNYET | ASSFGGAQDTQY | 0.850238 |
| VGMGGMHPSSG | ASSFGGAQDTQY | 0.8511922 |
| VGLAGGAATA | ASSFGGAQDTQY | 0.2716176 |
| VGGAGGAGGSGP | ASSFGGAQDTQY | 0.7665956 |
| VFAAFMV | ASSFGGAQDTQY | 0.2873658 |
| TLYDKLR | ASSFGGAQDTQY | 0.5959495 |
| TLAVFTVL | ASSFGGAQDTQY | 0.42704076 |
| TGSVQPARV | ASSFGGAQDTQY | 0.8329565 |
| TGGGAGSGGT | ASSFGGAQDTQY | 0.7666094 |
| SHLETSL | ASSFGGAQDTQY | 0.81774706 |
| SFTTGMT | ASSFGGAQDTQY | 0.8021405 |

| | | |
|---|---|---|
| QVEVAKRMGVRQPTVS | ASSFGGAQDTQY | 0.85243726 |
| QGSGGPALAM | ASSFGGAQDTQY | 0.72112644 |
| MLAPSNP | ASSFGGAQDTQY | 0.7639272 |
| LVNNAAI | ASSFGGAQDTQY | 0.35252434 |
| LTGEINTLA | ASSFGGAQDTQY | 0.29974264 |
| LSIFGPV | ASSFGGAQDTQY | 0.6405994 |
| LLALLANL | ASSFGGAQDTQY | 0.7271158 |
| LISNASDALDKLR | ASSFGGAQDTQY | 0.17038304 |
| LINEAVP | ASSFGGAQDTQY | 0.84576833 |
| LGYAFSGL | ASSFGGAQDTQY | 0.8014009 |
| LDFVAVAL | ASSFGGAQDTQY | 0.8318281 |
| LAQPGAE | ASSFGGAQDTQY | 0.63909477 |
| ISNLGAVTGF | ASSFGGAQDTQY | 0.71872675 |
| HQEDPGRI | ASSFGGAQDTQY | 0.72161007 |
| GTPGEPTP | ASSFGGAQDTQY | 0.8263086 |
| GFFNSGS | ASSFGGAQDTQY | 0.65031725 |
| GAGGGGLTAGAN | ASSFGGAQDTQY | 0.42604437 |
| FVDALVNM | ASSFGGAQDTQY | 0.69949746 |
| FLTVAPEL | ASSFGGAQDTQY | 0.66200304 |
| FETLTAL | ASSFGGAQDTQY | 0.85498834 |
| DVINPIEL | ASSFGGAQDTQY | 0.8436538 |
| APVSGAPV | ASSFGGAQDTQY | 0.82654345 |
| ALVGGLAW | ASSFGGAQDTQY | 0.8442136 |
| ALAAAAATAI | ASSFGGAQDTQY | 0.1748863 |
| AFGLGLLL | ASSFGGAQDTQY | 0.65502584 |
| AFFAAASL | ASSFGGAQDTQY | 0.846877 |
| AAAALPAFSPP | ASSFGGAQDTQY | 0.7849788 |
| VVLAPAL | ASSFGGAQDTQY | 0.7805572 |
| TIGHVDHGKTTL | ASSFGGAQDTQY | 0.84744656 |
| LISNASDALDK | ASSFGGAQDTQY | 0.7111903 |
| LISNASDAL | ASSFGGAQDTQY | 0.73815596 |
| ISNASDALDK | ASSFGGAQDTQY | 0.7689887 |
| NQAETLVYQTE | ASSFGGAQDTQY | 0.81227785 |
| TGGTGGTAGAG | ASSFGGAQDTQY | 0.7863587 |
| TGSTAGGAGGA | ASSFGGAQDTQY | 0.6472069 |
| DPLAVPVAAGPV | ASSFGGAQDTQY | 0.7915727 |
| GGTGGTPGNGGN | ASSFGGAQDTQY | 0.86404884 |
| PAGLAGVDPATL | ASSFGGAQDTQY | 0.792994 |
| GNNLVVEAPHAGASV | ASSFGGAQDTQY | 0.8408313 |
| KVTRIMSASTLEQPA | ASSFGGAQDTQY | 0.81604075 |
| LGFALPIALPRRTVV | ASSFGGAQDTQY | 0.79377097 |
| TRLGNLTARKLVVLV | ASSFGGAQDTQY | 0.7220232 |
| WSQQRRLIYFPSAGP | ASSFGGAQDTQY | 0.7473129 |
| GPNGNPDPMAAAVDIR | ASSFGGAQDTQY | 0.77782047 |
| TQRCVSVADLRAAASR | ASSFGGAQDTQY | 0.5978618 |
| QFVLDVAEHPVEVLTETV | ASSFGGAQDTQY | 0.763669 |
| GGAGGTGGAGGNGGTQAGDG | ASSFGGAQDTQY | 0.7134472 |
| PLLATLRAIAVDVDAKPAQV | ASSFGGAQDTQY | 0.8635684 |
| APKFRTGGYMIDFWGVGYQVAK | ASSFGGAQDTQY | 0.82834065 |
| GAGGAGGAGGAAGGPGGTGGTG | ASSFGGAQDTQY | 0.80436754 |
| IGALGLLATDVAPAGSVAFGVA | ASSFGGAQDTQY | 0.67933095 |

| | | |
|---|---|---|
| AGGAGGIGGNGGAGGAGGVLMG | ASSFGGAQDTQY | 0.8015435 |
| ADELIGLL | ASSFGGAQDTQY | 0.82369214 |
| ADRALKA | ASSFGGAQDTQY | 0.5575341 |
| AGAVGQVGGAGGA | ASSFGGAQDTQY | 0.8483937 |
| AGGGGGLGGAP | ASSFGGAQDTQY | 0.65459883 |
| AGGLLYGNGGAG | ASSFGGAQDTQY | 0.76850563 |
| AGLESIGL | ASSFGGAQDTQY | 0.7727907 |
| AGLKTGII | ASSFGGAQDTQY | 0.8440141 |
| ASLAALLL | ASSFGGAQDTQY | 0.7605953 |
| ASVINISSI | ASSFGGAQDTQY | 0.73428655 |
| ATPPGPA | ASSFGGAQDTQY | 0.67138976 |
| ATVASALV | ASSFGGAQDTQY | 0.83235765 |
| ATVASLIL | ASSFGGAQDTQY | 0.8494997 |
| AVNGPVLIRH | ASSFGGAQDTQY | 0.7445582 |
| DGAAGAVGI | ASSFGGAQDTQY | 0.85637724 |
| DPLLHGI | ASSFGGAQDTQY | 0.7389422 |
| ELGAALGAL | ASSFGGAQDTQY | 0.30618146 |
| ELGRTQAV | ASSFGGAQDTQY | 0.7291941 |
| FVAGGYLS | ASSFGGAQDTQY | 0.8401505 |
| GASGGAGGAGGW | ASSFGGAQDTQY | 0.8526598 |
| GGAGAGGDGGAGGI | ASSFGGAQDTQY | 0.8458095 |
| GTGHNGSQPGMGGNG | ASSFGGAQDTQY | 0.78932154 |
| GYAAASASAS | ASSFGGAQDTQY | 0.7789128 |
| HQVDVAAVQQG | ASSFGGAQDTQY | 0.8533257 |
| IGAAIAHT | ASSFGGAQDTQY | 0.8529502 |
| IGMIEVAQL | ASSFGGAQDTQY | 0.8245115 |
| IRLAPEHTSAMD | ASSFGGAQDTQY | 0.6585266 |
| IYNVSVL | ASSFGGAQDTQY | 0.47237772 |
| KQGFLGM | ASSFGGAQDTQY | 0.80720246 |
| LESVITQL | ASSFGGAQDTQY | 0.8448547 |
| LGLGALVVV | ASSFGGAQDTQY | 0.5227438 |
| LIGNGGNGGSGGI | ASSFGGAQDTQY | 0.8127192 |
| LNAINAPTL | ASSFGGAQDTQY | 0.81496745 |
| LQLLNTGV | ASSFGGAQDTQY | 0.4162696 |
| LQTSIEF | ASSFGGAQDTQY | 0.8024422 |
| MSFAVSEL | ASSFGGAQDTQY | 0.8522339 |
| NTGALGL | ASSFGGAQDTQY | 0.7466441 |
| PPGAGGAGGAGTT | ASSFGGAQDTQY | 0.65293837 |
| QLVGLGI | ASSFGGAQDTQY | 0.8486701 |
| RFLEQVHQL | ASSFGGAQDTQY | 0.86358494 |
| RGVGYVLR | ASSFGGAQDTQY | 0.84107256 |
| SATGVFAGVI | ASSFGGAQDTQY | 0.32604814 |
| SLPAITLP | ASSFGGAQDTQY | 0.26313812 |
| STALAFTV | ASSFGGAQDTQY | 0.73383546 |
| SVLTVGLGK | ASSFGGAQDTQY | 0.8169464 |
| TALAGAGSGI | ASSFGGAQDTQY | 0.68727434 |
| TGLPFTGI | ASSFGGAQDTQY | 0.57373047 |
| VAEHVTV | ASSFGGAQDTQY | 0.8197396 |
| VDIVLNSL | ASSFGGAQDTQY | 0.8484006 |
| VLTGGGAVL | ASSFGGAQDTQY | 0.62485945 |
| VNVGLAYGI | ASSFGGAQDTQY | 0.34581453 |

| | | |
|---|---|---|
| VSSAAVGAV | ASSFGGAQDTQY | 0.8323314 |
| VTSVYEAGV | ASSFGGAQDTQY | 0.8248972 |
| WVALDRGA | ASSFGGAQDTQY | 0.70922375 |
| YLVGSVVI | ASSFGGAQDTQY | 0.79755104 |
| GPPGTGKTLL | ASSFGGAQDTQY | 0.7437805 |
| GVGMTKF | ASSFGGAQDTQY | 0.46141493 |
| IGIGGSDL | ASSFGGAQDTQY | 0.82185125 |
| IGTLLAEL | ASSFGGAQDTQY | 0.57217515 |
| IYTSGTT | ASSFGGAQDTQY | 0.85518146 |
| LSFPAGV | ASSFGGAQDTQY | 0.65224874 |
| LYTSGTT | ASSFGGAQDTQY | 0.82319033 |
| VGAFPFGF | ASSFGGAQDTQY | 0.82154673 |
| VGAGTVEF | ASSFGGAQDTQY | 0.81826043 |
| VGKSTLLL | ASSFGGAQDTQY | 0.7831835 |
| AALAHAGGGIV | ASSFGGAQDTQY | 0.7053615 |
| AALVAVSLL | ASSFGGAQDTQY | 0.58235955 |
| AAVSRAVAAL | ASSFGGAQDTQY | 0.28335828 |
| AGGSSGIGGF | ASSFGGAQDTQY | 0.84522724 |
| AGLTVANANG | ASSFGGAQDTQY | 0.15651488 |
| AGLVGAVL | ASSFGGAQDTQY | 0.84705687 |
| ALSQAFTV | ASSFGGAQDTQY | 0.8392076 |
| ALTFSVSL | ASSFGGAQDTQY | 0.66224307 |
| ALTGAALVAL | ASSFGGAQDTQY | 0.823614 |
| ASFSIGAGVL | ASSFGGAQDTQY | 0.4479299 |
| DGSVYKI | ASSFGGAQDTQY | 0.6160461 |
| EQLAEIV | ASSFGGAQDTQY | 0.82917124 |
| FAVSELTVM | ASSFGGAQDTQY | 0.80065364 |
| GAAPLAAGL | ASSFGGAQDTQY | 0.6978835 |
| GADGGDGGSGGAGG | ASSFGGAQDTQY | 0.7761327 |
| GFGLDTGGAGG | ASSFGGAQDTQY | 0.82114196 |
| GLASGGLL | ASSFGGAQDTQY | 0.604918 |
| GRIAGAAVLL | ASSFGGAQDTQY | 0.75215137 |
| GSAGSLNTSF | ASSFGGAQDTQY | 0.84636444 |
| GTGQIGI | ASSFGGAQDTQY | 0.7774745 |
| GTLIALIL | ASSFGGAQDTQY | 0.688274 |
| IAATMGFVVG | ASSFGGAQDTQY | 0.8268508 |
| IFAQALL | ASSFGGAQDTQY | 0.66342074 |
| IGAAVALGF | ASSFGGAQDTQY | 0.82755786 |
| IGLTPLAL | ASSFGGAQDTQY | 0.79203695 |
| ILGSNTSTL | ASSFGGAQDTQY | 0.7274048 |
| IVFSLTVL | ASSFGGAQDTQY | 0.82607955 |
| LASKATGF | ASSFGGAQDTQY | 0.323057 |
| LASVALIL | ASSFGGAQDTQY | 0.6988714 |
| LFAVGLTL | ASSFGGAQDTQY | 0.737371 |
| LFNGGIGTYI | ASSFGGAQDTQY | 0.7927475 |
| LTLGAGII | ASSFGGAQDTQY | 0.8327943 |
| LVAAALGDI | ASSFGGAQDTQY | 0.7102264 |
| NTAFVQLGI | ASSFGGAQDTQY | 0.7914493 |
| QLFSGAEL | ASSFGGAQDTQY | 0.70723593 |
| SAATFITL | ASSFGGAQDTQY | 0.5922893 |
| SNYGLANLGV | ASSFGGAQDTQY | 0.7136741 |

| | | |
|---|---|---|
| SYGGAYAVM | ASSFGGAQDTQY | 0.7196275 |
| TTGVGGLTLG | ASSFGGAQDTQY | 0.5148579 |
| TTQTIYI | ASSFGGAQDTQY | 0.81556976 |
| VAAGAGLGAVI | ASSFGGAQDTQY | 0.535817 |
| VAAVVTLL | ASSFGGAQDTQY | 0.7645126 |
| VGFLGGLI | ASSFGGAQDTQY | 0.8517637 |
| VGVIPILL | ASSFGGAQDTQY | 0.6389016 |
| VIFASSLI | ASSFGGAQDTQY | 0.8278577 |
| VKGGDGGAAAG | ASSFGGAQDTQY | 0.8128881 |
| VTKTFTNI | ASSFGGAQDTQY | 0.8314694 |
| VVAAAILL | ASSFGGAQDTQY | 0.75762814 |
| YAALDVEL | ASSFGGAQDTQY | 0.71727073 |
| LASVALLL | ASSFGGAQDTQY | 0.8510601 |
| LDAVHRV | ASSFGGAQDTQY | 0.82370055 |
| QGLSSSIF | ASSFGGAQDTQY | 0.78040236 |
| RGTFANIRL | ASSFGGAQDTQY | 0.81902975 |
| RQLWWGHRI | ASSFGGAQDTQY | 0.8315904 |
| YGPPGTGKTLL | ASSFGGAQDTQY | 0.8246783 |
| AALVGAPANVA | ASSFGGAQDTQY | 0.73631746 |
| AATFRRTE | ASSFGGAQDTQY | 0.3827612 |
| AGALGGGI | ASSFGGAQDTQY | 0.7562125 |
| AGGAGGAGGIGAG | ASSFGGAQDTQY | 0.7712336 |
| AGSGALGL | ASSFGGAQDTQY | 0.70153314 |
| AQIADQI | ASSFGGAQDTQY | 0.23533262 |
| ASAVAVTYL | ASSFGGAQDTQY | 0.84653664 |
| ASTVLGGTGG | ASSFGGAQDTQY | 0.8603909 |
| AVGGEPAK | ASSFGGAQDTQY | 0.61174256 |
| CEQGTFGGTC | ASSFGGAQDTQY | 0.8574916 |
| DEGKAPAG | ASSFGGAQDTQY | 0.517905 |
| DTVVFPAL | ASSFGGAQDTQY | 0.8170309 |
| FEGITMAG | ASSFGGAQDTQY | 0.84245646 |
| FPLVVQD | ASSFGGAQDTQY | 0.64102006 |
| GAFTTPQ | ASSFGGAQDTQY | 0.790788 |
| GFGGFGTAGGVGGA | ASSFGGAQDTQY | 0.6691897 |
| GFLAQFL | ASSFGGAQDTQY | 0.7312121 |
| GGAGGAGGNAGVA | ASSFGGAQDTQY | 0.8475328 |
| GGVGTAGAGGAGGAGG | ASSFGGAQDTQY | 0.6882875 |
| GNGGGGDAGNAGSGGN | ASSFGGAQDTQY | 0.78493667 |
| GTGGAGGAGNDAGS | ASSFGGAQDTQY | 0.6725982 |
| GTTNTGWFNS | ASSFGGAQDTQY | 0.59614277 |
| IFMGPLFTL | ASSFGGAQDTQY | 0.8034987 |
| ILAHAIEAI | ASSFGGAQDTQY | 0.8307445 |
| IRSAIREAL | ASSFGGAQDTQY | 0.6446885 |
| ITFAGGG | ASSFGGAQDTQY | 0.8515338 |
| ITGMSGHQEV | ASSFGGAQDTQY | 0.7240348 |
| IVAGLACTL | ASSFGGAQDTQY | 0.860027 |
| IVGGGLAGA | ASSFGGAQDTQY | 0.8449367 |
| IVTGVLNI | ASSFGGAQDTQY | 0.8148074 |
| LAQQGCI | ASSFGGAQDTQY | 0.6977519 |
| LCLHVEN | ASSFGGAQDTQY | 0.84663856 |
| LDAISDGS | ASSFGGAQDTQY | 0.73675376 |

| | | |
|---|---|---|
| LGGFAAML | ASSFGGAQDTQY | 0.80686194 |
| LGGKNPMI | ASSFGGAQDTQY | 0.8372188 |
| LGPKGVAAIL | ASSFGGAQDTQY | 0.39095184 |
| LMVGVSI | ASSFGGAQDTQY | 0.5979269 |
| LVGAAFVL | ASSFGGAQDTQY | 0.4788801 |
| LVLAVAPL | ASSFGGAQDTQY | 0.49560603 |
| MAAIAVIDAA | ASSFGGAQDTQY | 0.42160246 |
| NNGINAELQ | ASSFGGAQDTQY | 0.39505982 |
| RGEPLVLVHG | ASSFGGAQDTQY | 0.48349527 |
| RGGAGGAAGLI | ASSFGGAQDTQY | 0.41432118 |
| RSYRMHPAV | ASSFGGAQDTQY | 0.64178085 |
| SAAHLLAGQLA | ASSFGGAQDTQY | 0.5600097 |
| SAAVGSY | ASSFGGAQDTQY | 0.41386494 |
| SPHAVLHRI | ASSFGGAQDTQY | 0.621577 |
| SQIAKVL | ASSFGGAQDTQY | 0.5919894 |
| TALGHLVP | ASSFGGAQDTQY | 0.37785643 |
| TAPTTSGAA | ASSFGGAQDTQY | 0.44872642 |
| TDGDDGNGGSPGT | ASSFGGAQDTQY | 0.44442084 |
| TDGLGEN | ASSFGGAQDTQY | 0.43860552 |
| TGGPPLPGQA | ASSFGGAQDTQY | 0.35764048 |
| TIPAITI | ASSFGGAQDTQY | 0.49929997 |
| TIPGFTI | ASSFGGAQDTQY | 0.5458082 |
| TSIASGALA | ASSFGGAQDTQY | 0.378309 |
| TVTLPGAAI | ASSFGGAQDTQY | 0.5168103 |
| VASGLSAALP | ASSFGGAQDTQY | 0.4676044 |
| VGAGVIGI | ASSFGGAQDTQY | 0.4844407 |
| VGPKRFGV | ASSFGGAQDTQY | 0.5592771 |
| VNVAFPD | ASSFGGAQDTQY | 0.44055575 |
| VSMTNSGVS | ASSFGGAQDTQY | 0.45537776 |
| VTDVLGDLSV | ASSFGGAQDTQY | 0.5838793 |
| VVVVGAGFAGL | ASSFGGAQDTQY | 0.62440485 |
| VYTVGTVV | ASSFGGAQDTQY | 0.40454167 |
| WTTFSGP | ASSFGGAQDTQY | 0.47126508 |
| YDELVEAF | ASSFGGAQDTQY | 0.6099342 |
| YGTGGAGGNGG | ASSFGGAQDTQY | 0.52502495 |
| YTNVMAV | ASSFGGAQDTQY | 0.590103 |
| FLEQVHQL | ASSFGGAQDTQY | 0.5897967 |
| GLSLLAVAV | ASSFGGAQDTQY | 0.41276345 |
| LEQVHQL | ASSFGGAQDTQY | 0.6162929 |
| LNAAGAI | ASSFGGAQDTQY | 0.6317539 |
| PAVHASL | ASSFGGAQDTQY | 0.4883516 |
| TELEDDA | ASSFGGAQDTQY | 0.48889986 |
| TPPGPPN | ASSFGGAQDTQY | 0.38795874 |
| TPPPPPPP | ASSFGGAQDTQY | 0.47131193 |
| TQSPALI | ASSLRLGGYEQY | 0.3753438 |
| EFAAAFPQ | ASSLRLGGYEQY | 0.5572221 |
| LLSGNLDVL | ASSLRLGGYEQY | 0.6051295 |
| GPANLGNYNI | ASSLRLGGYEQY | 0.49199188 |
| TLGPTLP | ASSLRLGGYEQY | 0.36221194 |
| VYAYSSCCK | ASSLRLGGYEQY | 0.3829431 |
| APAPAPAPNI | ASSLRLGGYEQY | 0.41499406 |

| | | |
|---|---|---|
| PGGPVPIP | ASSLRLGGYEQY | 0.386144 |
| FLTVALEP | ASSLRLGGYEQY | 0.435704 |
| SQYFGANL | ASSLRLGGYEQY | 0.40477294 |
| GGVKPAA | ASSLRLGGYEQY | 0.6150206 |
| GGTGGAGGAGGW | ASSLRLGGYEQY | 0.6050435 |
| GVGGVAGLGGAGG | ASSLRLGGYEQY | 0.43069714 |
| VLLVLATTI | ASSLRLGGYEQY | 0.57648087 |
| IGIFAAML | ASSLRLGGYEQY | 0.60495764 |
| AVGDLID | ASSLRLGGYEQY | 0.5421481 |
| NISPLPL | ASSLRLGGYEQY | 0.49284774 |
| VTAVFVAI | ASSLRLGGYEQY | 0.3760048 |
| SGDDNDAAL | ASSLRLGGYEQY | 0.6381319 |
| AVAALAHS | ASSLRLGGYEQY | 0.47282815 |
| GGVTVMATP | ASSLRLGGYEQY | 0.6175634 |
| AGLAANVRD | ASSLRLGGYEQY | 0.54053134 |
| TNRPDLIDPALLRPG | ASSLRLGGYEQY | 0.5107772 |
| LANLGVTAL | ASSLRLGGYEQY | 0.4110679 |
| TGERIEL | ASSLRLGGYEQY | 0.56591547 |
| NGEELAA | ASSLRLGGYEQY | 0.48984465 |
| VLSLVWG | ASSLRLGGYEQY | 0.3829052 |
| IGYALWL | ASSLRLGGYEQY | 0.38198635 |
| LSLLLAPL | ASSLRLGGYEQY | 0.5786048 |
| TLDEALK | ASSLRLGGYEQY | 0.49835527 |
| TDGAGTAEGG | ASSLRLGGYEQY | 0.6141768 |
| NAPGGAGGNGG | ASSLRLGGYEQY | 0.49196598 |
| VSVFMGMGGNF | ASSLRLGGYEQY | 0.4330228 |
| LASYLAAL | ASSLRLGGYEQY | 0.58497494 |
| LVGGTPAL | ASSLRLGGYEQY | 0.5369678 |
| VRNADVL | ASSLRLGGYEQY | 0.46993265 |
| TCSSPFLVSAM | ASSLRLGGYEQY | 0.46766606 |
| VHITDAE | ASSLRLGGYEQY | 0.4549033 |
| EYAAPSF | ASSLRLGGYEQY | 0.49465382 |
| VVDAFHGGL | ASSLRLGGYEQY | 0.5359443 |
| ASVVDPQ | ASSLRLGGYEQY | 0.5040882 |
| SYNEFGGLS | ASSLRLGGYEQY | 0.5451907 |
| DPDGSVY | ASSLRLGGYEQY | 0.52653164 |
| MVSTIHATE | ASSLRLGGYEQY | 0.46674246 |
| LAAGVADPV | ASSLRLGGYEQY | 0.40133455 |
| TKEQIAA | ASSLRLGGYEQY | 0.49659324 |
| SLASPAAL | ASSLRLGGYEQY | 0.48478708 |
| IGLVFYGV | ASSLRLGGYEQY | 0.5050577 |
| FLLDASNL | ASSLRLGGYEQY | 0.5653888 |
| YQVGNFP | ASSLRLGGYEQY | 0.5038302 |
| ESLLSQST | ASSLRLGGYEQY | 0.4352532 |
| LFTVGFAAVG | ASSLRLGGYEQY | 0.48751414 |
| ATPPTPT | ASSLRLGGYEQY | 0.60651875 |
| TDLATLG | ASSLRLGGYEQY | 0.36937058 |
| SAGGTGR | ASSLRLGGYEQY | 0.4899007 |
| GFSNSGS | ASSLRLGGYEQY | 0.61177695 |
| GGNGGSGGTSV | ASSLRLGGYEQY | 0.6034905 |
| GTSVATGGAGN | ASSLRLGGYEQY | 0.4565406 |

| | | |
|---|---|---|
| NGGNGGSGGTS | ASSLRLGGYEQY | 0.46660328 |
| AVVKFPH | ASSLRLGGYEQY | 0.65023357 |
| TPPPPPP | ASSLRLGGYEQY | 0.58664894 |
| AAPAAAAAPA | ASSLRLGGYEQY | 0.6174164 |
| IALNEDL | ASSLRLGGYEQY | 0.35324928 |
| IVNVSSM | ASSLRLGGYEQY | 0.4213002 |
| TGIPPAPR | ASSLRLGGYEQY | 0.39012158 |
| FDLGGGTF | ASSLRLGGYEQY | 0.3270513 |
| FDLGGGTFDVS | ASSLRLGGYEQY | 0.5745705 |
| NEPTAAA | ASSLRLGGYEQY | 0.44523194 |
| NEPTAAALA | ASSLRLGGYEQY | 0.57380354 |
| GSIINISSI | ASSLRLGGYEQY | 0.5152699 |
| VAPGNAGTA | ASSLRLGGYEQY | 0.56434387 |
| TGGGASLE | ASSLRLGGYEQY | 0.47360772 |
| VAVYNHYKRI | ASSLRLGGYEQY | 0.47792435 |
| TDPAPATT | ASSLRLGGYEQY | 0.42145142 |
| TNRPDILDPALLRPG | ASSLRLGGYEQY | 0.5519698 |
| LTTTTTS | ASSLRLGGYEQY | 0.52321035 |
| PTGSGKTL | ASSLRLGGYEQY | 0.47911343 |
| TGSGKTL | ASSLRLGGYEQY | 0.3765504 |
| TIGHVDHGK | ASSLRLGGYEQY | 0.5867451 |
| TIGHVDHGKTT | ASSLRLGGYEQY | 0.486048 |
| GHVDHGKTT | ASSLRLGGYEQY | 0.4036875 |
| AQFEHTLL | ASSLRLGGYEQY | 0.39822695 |
| IVNQDEL | ASSLRLGGYEQY | 0.38845277 |
| TNGSQFF | ASSLRLGGYEQY | 0.40082064 |
| TGKGIVE | ASSLRLGGYEQY | 0.5772513 |
| LISNASDA | ASSLRLGGYEQY | 0.60787785 |
| AELRTLGELTKIAKAHGAQ | ASSLRLGGYEQY | 0.42251062 |
| APNAVAAAEPSW | ASSLRLGGYEQY | 0.5725666 |
| APQVAGAATAVL | ASSLRLGGYEQY | 0.5178791 |
| ARRDPAAAAAILPTDA | ASSLRLGGYEQY | 0.30668524 |
| DAAARDVLAEAGLAEYF | ASSLRLGGYEQY | 0.4812652 |
| DDQIVLSAKANLAKVEQ | ASSLRLGGYEQY | 0.3832734 |
| DELLPDTGEDSDEDGH | ASSLRLGGYEQY | 0.55980337 |
| ELPEAPANRATVPQD | ASSLRLGGYEQY | 0.4231436 |
| FGQSKIERVTGIG | ASSLRLGGYEQY | 0.4682003 |
| GAGGTGSTAGGAGGAGGAGGLY | ASSLRLGGYEQY | 0.4722012 |
| GHGGTGAAGGNGGTGGAGGSGI | ASSLRLGGYEQY | 0.60023516 |
| GNGGNGGAGGEGGPGGAGGAGA | ASSLRLGGYEQY | 0.5020696 |
| GSTAGNGGNGGNATNGTIAGQP | ASSLRLGGYEQY | 0.37720278 |
| GTGGDGGNGGNGGTGG | ASSLRLGGYEQY | 0.34482527 |
| GVRRPAGIPADAPV | ASSLRLGGYEQY | 0.5496248 |
| GYRQVTDAHLLTLARR | ASSLRLGGYEQY | 0.50507224 |
| IAKMGQDGHDRG | ASSLRLGGYEQY | 0.42309022 |
| KAEPSIEIQYT | ASSLRLGGYEQY | 0.41299874 |
| KPVRIRMAADEDTVAVT | ASSLRLGGYEQY | 0.55019 |
| MPPRIAGMRLLVIKPEP | ASSLRLGGYEQY | 0.37108016 |
| MKFYAVALFRSRGVPPTR | ASSLRLGGYEQY | 0.5460279 |
| NAGGFGILGAVAHSP | ASSLRLGGYEQY | 0.40152904 |
| NEGREITLAMAEM | ASSLRLGGYEQY | 0.4730855 |

| | | |
|---|---|---|
| NELVEWSHPLGHRWRWE | ASSLRLGGYEQY | 0.62670064 |
| NLGAGGTGGNGGNPG | ASSLRLGGYEQY | 0.6069082 |
| NTEWPKWETVMAADPVD | ASSLRLGGYEQY | 0.43737578 |
| PASAAPAETQP | ASSLRLGGYEQY | 0.47830594 |
| QQSSDGKNGVRAAEITGE | ASSLRLGGYEQY | 0.39285284 |
| RAPDAVTTSWWKE | ASSLRLGGYEQY | 0.42954516 |
| SGFGNVGAHTSGWFNQ | ASSLRLGGYEQY | 0.43907383 |
| SGVAGAVGQAASVGGLKV | ASSLRLGGYEQY | 0.5989388 |
| SNGGAGTGSAGGNGGT | ASSLRLGGYEQY | 0.41882947 |
| TGGAGGEGGAAG | ASSLRLGGYEQY | 0.39711365 |
| TGGDGGAAGAGGAPGHG | ASSLRLGGYEQY | 0.44537723 |
| TMPPLLAAIANAGRHGV | ASSLRLGGYEQY | 0.59883064 |
| TSSTAGNSGGA | ASSLRLGGYEQY | 0.6212027 |
| VPATVPVGILC | ASSLRLGGYEQY | 0.42765298 |
| VQRPDSGGVVLTGRIS | ASSLRLGGYEQY | 0.45312774 |
| WTPNPVPATKGVPAGLVQAT | ASSLRLGGYEQY | 0.44883296 |
| YALKRALAKFSDRFDVVII | ASSLRLGGYEQY | 0.49489242 |
| YLIGSVIV | ASSLRLGGYEQY | 0.41498506 |
| YGQLVAVID | ASSLRLGGYEQY | 0.4134091 |
| VTVGGMTLPA | ASSLRLGGYEQY | 0.49319035 |
| VLALPAV | ASSLRLGGYEQY | 0.6569781 |
| VIPPTLNYET | ASSLRLGGYEQY | 0.45338795 |
| VGMGGMHPSSG | ASSLRLGGYEQY | 0.4165234 |
| VGLAGGAATA | ASSLRLGGYEQY | 0.49737194 |
| VGGAGGAGGSGP | ASSLRLGGYEQY | 0.40561897 |
| VFAAFMV | ASSLRLGGYEQY | 0.4137158 |
| TLYDKLR | ASSLRLGGYEQY | 0.534348 |
| TLAVFTVL | ASSLRLGGYEQY | 0.34555963 |
| TGSVQPARV | ASSLRLGGYEQY | 0.46946335 |
| TGGGAGSGGT | ASSLRLGGYEQY | 0.6138272 |
| SHLETSL | ASSLRLGGYEQY | 0.57791007 |
| SFTTGMT | ASSLRLGGYEQY | 0.47682145 |
| QVEVAKRMGVRQPTVS | ASSLRLGGYEQY | 0.60567886 |
| QGSGGPALAM | ASSLRLGGYEQY | 0.53020006 |
| MLAPSNP | ASSLRLGGYEQY | 0.60892284 |
| LVNNAAI | ASSLRLGGYEQY | 0.6221043 |
| LTGEINTLA | ASSLRLGGYEQY | 0.47078547 |
| LSIFGPV | ASSLRLGGYEQY | 0.47014683 |
| LLALLANL | ASSLRLGGYEQY | 0.56166875 |
| LISNASDALDKLR | ASSLRLGGYEQY | 0.5036656 |
| LINEAVP | ASSLRLGGYEQY | 0.47461125 |
| LGYAFSGL | ASSLRLGGYEQY | 0.44016233 |
| LDFVAVAL | ASSLRLGGYEQY | 0.5825677 |
| LAQPGAE | ASSLRLGGYEQY | 0.48231503 |
| ISNLGAVTGF | ASSLRLGGYEQY | 0.6324645 |
| HQEDPGRI | ASSLRLGGYEQY | 0.35647538 |
| GTPGEPTP | ASSLRLGGYEQY | 0.4066111 |
| GFFNSGS | ASSLRLGGYEQY | 0.58325374 |
| GAGGGGLTAGAN | ASSLRLGGYEQY | 0.32777664 |
| FVDALVNM | ASSLRLGGYEQY | 0.40005508 |
| FLTVAPEL | ASSLRLGGYEQY | 0.4605042 |

| | | |
|---|---|---|
| FETLTAL | ASSLRLGGYEQY | 0.5388219 |
| DVINPIEL | ASSLRLGGYEQY | 0.32820788 |
| APVSGAPV | ASSLRLGGYEQY | 0.48905683 |
| ALVGGLAW | ASSLRLGGYEQY | 0.3665631 |
| ALAAAAATAI | ASSLRLGGYEQY | 0.47570473 |
| AFGLGLLL | ASSLRLGGYEQY | 0.5007317 |
| AFFAAASL | ASSLRLGGYEQY | 0.3737146 |
| AAAALPAFSPP | ASSLRLGGYEQY | 0.5245963 |
| VVLAPAL | ASSLRLGGYEQY | 0.36546174 |
| TIGHVDHGKTTL | ASSLRLGGYEQY | 0.49499005 |
| LISNASDALDK | ASSLRLGGYEQY | 0.58417165 |
| LISNASDAL | ASSLRLGGYEQY | 0.6435381 |
| ISNASDALDK | ASSLRLGGYEQY | 0.40352005 |
| NQAETLVYQTE | ASSLRLGGYEQY | 0.5856044 |
| TGGTGGTAGAG | ASSLRLGGYEQY | 0.38555562 |
| TGSTAGGAGGA | ASSLRLGGYEQY | 0.6471527 |
| DPLAVPVAAGPV | ASSLRLGGYEQY | 0.46171278 |
| GGTGGTPGNGGN | ASSLRLGGYEQY | 0.36470276 |
| PAGLAGVDPATL | ASSLRLGGYEQY | 0.5621494 |
| GNNLVVEAPHAGASV | ASSLRLGGYEQY | 0.43599498 |
| KVTRIMSASTLEQPA | ASSLRLGGYEQY | 0.4669185 |
| LGFALPIALPRRTVV | ASSLRLGGYEQY | 0.6093117 |
| TRLGNLTARKLVVLV | ASSLRLGGYEQY | 0.3973987 |
| WSQQRRLIYFPSAGP | ASSLRLGGYEQY | 0.4755851 |
| GPNGNPDPMAAAVDIR | ASSLRLGGYEQY | 0.6089602 |
| TQRCVSVADLRAAASR | ASSLRLGGYEQY | 0.5132032 |
| QFVLDVAEHPVEVLTETV | ASSLRLGGYEQY | 0.4070902 |
| GGAGGTGGAGGNGGTQAGDG | ASSLRLGGYEQY | 0.51311433 |
| PLLATLRAIAVDVDAKPAQV | ASSLRLGGYEQY | 0.52950156 |
| APKFRTGGYMIDFWGVGYQVAK | ASSLRLGGYEQY | 0.40921122 |
| GAGGAGGAGGAAGGPGGTGGTG | ASSLRLGGYEQY | 0.58996046 |
| IGALGLLATDVAPAGSVAFGVA | ASSLRLGGYEQY | 0.4505738 |
| AGGAGGIGGNGGAGGAGGVLMG | ASSLRLGGYEQY | 0.54677224 |
| ADELIGLL | ASSLRLGGYEQY | 0.5208134 |
| ADRALKA | ASSLRLGGYEQY | 0.6478561 |
| AGAVGQVGGAGGA | ASSLRLGGYEQY | 0.4292946 |
| AGGGGGLGGAP | ASSLRLGGYEQY | 0.4714247 |
| AGGLLYGNGGAG | ASSLRLGGYEQY | 0.5122672 |
| AGLESIGL | ASSLRLGGYEQY | 0.556682 |
| AGLKTGII | ASSLRLGGYEQY | 0.4106345 |
| ASLAALLL | ASSLRLGGYEQY | 0.5512745 |
| ASVINISSI | ASSLRLGGYEQY | 0.4789483 |
| ATPPGPA | ASSLRLGGYEQY | 0.60448027 |
| ATVASALV | ASSLRLGGYEQY | 0.4342031 |
| ATVASLIL | ASSLRLGGYEQY | 0.46013567 |
| AVNGPVLIRH | ASSLRLGGYEQY | 0.6232186 |
| DGAAGAVGI | ASSLRLGGYEQY | 0.40783063 |
| DPLLHGI | ASSLRLGGYEQY | 0.49096617 |
| ELGAALGAL | ASSLRLGGYEQY | 0.5239841 |
| ELGRTQAV | ASSLRLGGYEQY | 0.5450573 |
| FVAGGYLS | ASSLRLGGYEQY | 0.43240002 |

| | | |
|---|---|---|
| GASGGAGGAGGW | ASSLRLGGYEQY | 0.57332546 |
| GGAGAGGDGGAGGI | ASSLRLGGYEQY | 0.48869723 |
| GTGHNGSQPGMGGNG | ASSLRLGGYEQY | 0.5988562 |
| GYAAASASAS | ASSLRLGGYEQY | 0.3988637 |
| HQVDVAAVQQG | ASSLRLGGYEQY | 0.5440784 |
| IGAAIAHT | ASSLRLGGYEQY | 0.40984982 |
| IGMIEVAQL | ASSLRLGGYEQY | 0.5785586 |
| IRLAPEHTSAMD | ASSLRLGGYEQY | 0.58268243 |
| IYNVSVL | ASSLRLGGYEQY | 0.5920299 |
| KQGFLGM | ASSLRLGGYEQY | 0.44683617 |
| LESVITQL | ASSLRLGGYEQY | 0.5880881 |
| LGLGALVVV | ASSLRLGGYEQY | 0.601493 |
| LIGNGGNGGSGGI | ASSLRLGGYEQY | 0.65155673 |
| LNAINAPTL | ASSLRLGGYEQY | 0.47192085 |
| LQLLNTGV | ASSLRLGGYEQY | 0.5308484 |
| LQTSIEF | ASSLRLGGYEQY | 0.5395696 |
| MSFAVSEL | ASSLRLGGYEQY | 0.45202526 |
| NTGALGL | ASSLRLGGYEQY | 0.36537945 |
| PPGAGGAGGAGTT | ASSLRLGGYEQY | 0.41559055 |
| QLVGLGI | ASSLRLGGYEQY | 0.51315176 |
| RFLEQVHQL | ASSLRLGGYEQY | 0.41707623 |
| RGVGYVLR | ASSLRLGGYEQY | 0.631615 |
| SATGVFAGVI | ASSLRLGGYEQY | 0.57817423 |
| SLPAITLP | ASSLRLGGYEQY | 0.585832 |
| STALAFTV | ASSLRLGGYEQY | 0.6114485 |
| SVLTVGLGK | ASSLRLGGYEQY | 0.57122153 |
| TALAGAGSGI | ASSLRLGGYEQY | 0.4981408 |
| TGLPFTGI | ASSLRLGGYEQY | 0.4205995 |
| VAEHVTV | ASSLRLGGYEQY | 0.56033266 |
| VDIVLNSL | ASSLRLGGYEQY | 0.36879474 |
| VLTGGGAVL | ASSLRLGGYEQY | 0.4700644 |
| VNVGLAYGI | ASSLRLGGYEQY | 0.43335795 |
| VSSAAVGAV | ASSLRLGGYEQY | 0.51727915 |
| VTSVYEAGV | ASSLRLGGYEQY | 0.32266638 |
| WVALDRGA | ASSLRLGGYEQY | 0.59960175 |
| YLVGSVVI | ASSLRLGGYEQY | 0.6047789 |
| GPPGTGKTLL | ASSLRLGGYEQY | 0.5208738 |
| GVGMTKF | ASSLRLGGYEQY | 0.5283232 |
| IGIGGSDL | ASSLRLGGYEQY | 0.48456684 |
| IGTLLAEL | ASSLRLGGYEQY | 0.37866634 |
| IYTSGTT | ASSLRLGGYEQY | 0.5632317 |
| LSFPAGV | ASSLRLGGYEQY | 0.40918413 |
| LYTSGTT | ASSLRLGGYEQY | 0.5541987 |
| VGAFPFGF | ASSLRLGGYEQY | 0.57053447 |
| VGAGTVEF | ASSLRLGGYEQY | 0.49522042 |
| VGKSTLLL | ASSLRLGGYEQY | 0.4052962 |
| AALAHAGGGIV | ASSLRLGGYEQY | 0.47980508 |
| AALVAVSLL | ASSLRLGGYEQY | 0.41475585 |
| AAVSRAVAAL | ASSLRLGGYEQY | 0.4316236 |
| AGGSSGIGGF | ASSLRLGGYEQY | 0.3896786 |
| AGLTVANANG | ASSLRLGGYEQY | 0.5014344 |

| | | |
|---|---|---|
| AGLVGAVL | ASSLRLGGYEQY | 0.39450428 |
| ALSQAFTV | ASSLRLGGYEQY | 0.560239 |
| ALTFSVSL | ASSLRLGGYEQY | 0.52666575 |
| ALTGAALVAL | ASSLRLGGYEQY | 0.47852916 |
| ASFSIGAGVL | ASSLRLGGYEQY | 0.41203398 |
| DGSVYKI | ASSLRLGGYEQY | 0.6173158 |
| EQLAEIV | ASSLRLGGYEQY | 0.6062113 |
| FAVSELTVM | ASSLRLGGYEQY | 0.39867422 |
| GAAPLAAGL | ASSLRLGGYEQY | 0.47987658 |
| GADGGDGGSGGAGG | ASSLRLGGYEQY | 0.42211828 |
| GFGLDTGGAGG | ASSLRLGGYEQY | 0.41510403 |
| GLASGGLL | ASSLRLGGYEQY | 0.6136391 |
| GRIAGAAVLL | ASSLRLGGYEQY | 0.49294358 |
| GSAGSLNTSF | ASSLRLGGYEQY | 0.3892022 |
| GTGQIGI | ASSLRLGGYEQY | 0.46790513 |
| GTLIALIL | ASSLRLGGYEQY | 0.4983092 |
| IAATMGFVVG | ASSLRLGGYEQY | 0.4425162 |
| IFAQALL | ASSLRLGGYEQY | 0.33757952 |
| IGAAVALGF | ASSLRLGGYEQY | 0.48230475 |
| IGLTPLAL | ASSLRLGGYEQY | 0.41123146 |
| ILGSNTSTL | ASSLRLGGYEQY | 0.39773357 |
| IVFSLTVL | ASSLRLGGYEQY | 0.4194867 |
| LASKATGF | ASSLRLGGYEQY | 0.5841748 |
| LASVALIL | ASSLRLGGYEQY | 0.5402784 |
| LFAVGLTL | ASSLRLGGYEQY | 0.47855747 |
| LFNGGIGTYI | ASSLRLGGYEQY | 0.53530514 |
| LTLGAGII | ASSLRLGGYEQY | 0.54273593 |
| LVAAALGDI | ASSLRLGGYEQY | 0.6151565 |
| NTAFVQLGI | ASSLRLGGYEQY | 0.5469526 |
| QLFSGAEL | ASSLRLGGYEQY | 0.43005258 |
| SAATFITL | ASSLRLGGYEQY | 0.5371336 |
| SNYGLANLGV | ASSLRLGGYEQY | 0.4838388 |
| SYGGAYAVM | ASSLRLGGYEQY | 0.5415112 |
| TTGVGGLTLG | ASSLRLGGYEQY | 0.47517374 |
| TTQTIYI | ASSLRLGGYEQY | 0.47775415 |
| VAAGAGLGAVI | ASSLRLGGYEQY | 0.50341624 |
| VAAVVTLL | ASSLRLGGYEQY | 0.5310675 |
| VGFLGGLI | ASSLRLGGYEQY | 0.53208405 |
| VGVIPILL | ASSLRLGGYEQY | 0.44007295 |
| VIFASSLI | ASSLRLGGYEQY | 0.3750139 |
| VKGGDGGAAAG | ASSLRLGGYEQY | 0.44028962 |
| VTKTFTNI | ASSLRLGGYEQY | 0.49161422 |
| VVAAAILL | ASSLRLGGYEQY | 0.56611115 |
| YAALDVEL | ASSLRLGGYEQY | 0.4604149 |
| LASVALLL | ASSLRLGGYEQY | 0.6546444 |
| LDAVHRV | ASSLRLGGYEQY | 0.6068264 |
| QGLSSSIF | ASSLRLGGYEQY | 0.45051312 |
| RGTFANIRL | ASSLRLGGYEQY | 0.5367889 |
| RQLWWGHRI | ASSLRLGGYEQY | 0.4039354 |
| YGPPGTGKTLL | ASSLRLGGYEQY | 0.4270064 |
| AALVGAPANVA | ASSLRLGGYEQY | 0.44552416 |

| | | |
|---|---|---|
| AATFRRTE | ASSLRLGGYEQY | 0.43349507 |
| AGALGGGI | ASSLRLGGYEQY | 0.4548606 |
| AGGAGGAGGIGAG | ASSLRLGGYEQY | 0.44428387 |
| AGSGALGL | ASSLRLGGYEQY | 0.37827185 |
| AQIADQI | ASSLRLGGYEQY | 0.5797691 |
| ASAVAVTYL | ASSLRLGGYEQY | 0.4928268 |
| ASTVLGGTGG | ASSLRLGGYEQY | 0.44117543 |
| AVGGEPAK | ASSLRLGGYEQY | 0.4217481 |
| CEQGTFGGTC | ASSLRLGGYEQY | 0.6310557 |
| DEGKAPAG | ASSLRLGGYEQY | 0.635036 |
| DTVVFPAL | ASSLRLGGYEQY | 0.44692114 |
| FEGITMAG | ASSLRLGGYEQY | 0.4047861 |
| FPLVVQD | ASSLRLGGYEQY | 0.60459554 |
| GAFTTPQ | ASSLRLGGYEQY | 0.414764 |
| GFGGFGTAGGVGGA | ASSLRLGGYEQY | 0.44464794 |
| GFLAQFL | ASSLRLGGYEQY | 0.52745545 |
| GGAGGAGGNAGVA | ASSLRLGGYEQY | 0.46696064 |
| GGVGTAGAGGAGGAGG | ASSLRLGGYEQY | 0.4123105 |
| GNGGGGDAGNAGSGGN | ASSLRLGGYEQY | 0.5256293 |
| GTGGAGGAGNDAGS | ASSLRLGGYEQY | 0.48605853 |
| GTTNTGWFNS | ASSLRLGGYEQY | 0.4722945 |
| IFMGPLFTL | ASSLRLGGYEQY | 0.3418442 |
| ILAHAIEAI | ASSLRLGGYEQY | 0.5051498 |
| IRSAIREAL | ASSLRLGGYEQY | 0.41222525 |
| ITFAGGG | ASSLRLGGYEQY | 0.4309295 |
| ITGMSGHQEV | ASSLRLGGYEQY | 0.34569585 |
| IVAGLACTL | ASSLRLGGYEQY | 0.4609406 |
| IVGGGLAGA | ASSLRLGGYEQY | 0.59190434 |
| IVTGVLNI | ASSLRLGGYEQY | 0.6009178 |
| LAQQGCI | ASSLRLGGYEQY | 0.55527717 |
| LCLHVEN | ASSLRLGGYEQY | 0.5518817 |
| LDAISDGS | ASSLRLGGYEQY | 0.6164092 |
| LGGFAAML | ASSLRLGGYEQY | 0.44907326 |
| LGGKNPMI | ASSLRLGGYEQY | 0.3920741 |
| LGPKGVAAIL | ASSLRLGGYEQY | 0.52700716 |
| LMVGVSI | ASSLRLGGYEQY | 0.5773181 |
| LVGAAFVL | ASSLRLGGYEQY | 0.51602954 |
| LVLAVAPL | ASSLRLGGYEQY | 0.55586827 |
| MAAIAVIDAA | ASSLRLGGYEQY | 0.37464342 |
| NNGINAELQ | ASSLRLGGYEQY | 0.52857286 |
| RGEPLVLVHG | ASSLRLGGYEQY | 0.562128 |
| RGGAGGAAGLI | ASSLRLGGYEQY | 0.372073 |
| RSYRMHPAV | ASSLRLGGYEQY | 0.40751055 |
| SAAHLLAGQLA | ASSLRLGGYEQY | 0.44639155 |
| SAAVGSY | ASSLRLGGYEQY | 0.36553693 |
| SPHAVLHRI | ASSLRLGGYEQY | 0.5247191 |
| SQIAKVL | ASSLRLGGYEQY | 0.42496347 |
| TALGHLVP | ASSLRLGGYEQY | 0.5119002 |
| TAPTTSGAA | ASSLRLGGYEQY | 0.4342698 |
| TDGDDGNGGSPGT | ASSLRLGGYEQY | 0.44214663 |
| TDGLGEN | ASSLRLGGYEQY | 0.46379685 |

| | | |
|---|---|---|
| TGGPPLPGQA | ASSLRLGGYEQY | 0.506796 |
| TIPAITI | ASSLRLGGYEQY | 0.5291627 |
| TIPGFTI | ASSLRLGGYEQY | 0.52752435 |
| TSIASGALA | ASSLRLGGYEQY | 0.46201748 |
| TVTLPGAAI | ASSLRLGGYEQY | 0.454242 |
| VASGLSAALP | ASSLRLGGYEQY | 0.6296977 |
| VGAGVIGI | ASSLRLGGYEQY | 0.40630648 |
| VGPKRFGV | ASSLRLGGYEQY | 0.4151083 |
| VNVAFPD | ASSLRLGGYEQY | 0.48873904 |
| VSMTNSGVS | ASSLRLGGYEQY | 0.49262545 |
| VTDVLGDLSV | ASSLRLGGYEQY | 0.58021057 |
| VVVVGAGFAGL | ASSLRLGGYEQY | 0.52884775 |
| VYTVGTVV | ASSLRLGGYEQY | 0.5149449 |
| WTTFSGP | ASSLRLGGYEQY | 0.62054104 |
| YDELVEAF | ASSLRLGGYEQY | 0.5950005 |
| YGTGGAGGNGG | ASSLRLGGYEQY | 0.5545505 |
| YTNVMAV | ASSLRLGGYEQY | 0.43007168 |
| FLEQVHQL | ASSLRLGGYEQY | 0.6212404 |
| GLSLLAVAV | ASSLRLGGYEQY | 0.3954328 |
| LEQVHQL | ASSLRLGGYEQY | 0.5285317 |
| LNAAGAI | ASSLRLGGYEQY | 0.53324836 |
| PAVHASL | ASSLRLGGYEQY | 0.3824647 |
| TELEDDA | ASSLRLGGYEQY | 0.48519203 |
| TPPGPPN | ASSLRLGGYEQY | 0.5613272 |
| TPPPPPPP | ASSLRLGGYEQY | 0.58318704 |
| TQSPALI | ASSSDSAETLY | 0.47350618 |
| EFAAAFPQ | ASSSDSAETLY | 0.42627645 |
| LLSGNLDVL | ASSSDSAETLY | 0.43569416 |
| GPANLGNYNI | ASSSDSAETLY | 0.40765455 |
| TLGPTLP | ASSSDSAETLY | 0.3948438 |
| VYAYSSCCK | ASSSDSAETLY | 0.52255964 |
| APAPAPAPNI | ASSSDSAETLY | 0.47197527 |
| PGGPVPIP | ASSSDSAETLY | 0.6259045 |
| FLTVALEP | ASSSDSAETLY | 0.52162296 |
| SQYFGANL | ASSSDSAETLY | 0.5834235 |
| GGVKPAA | ASSSDSAETLY | 0.41398785 |
| GGTGGAGGAGGW | ASSSDSAETLY | 0.5079106 |
| GVGGVAGLGGAGG | ASSSDSAETLY | 0.354935 |
| VLLVLATTI | ASSSDSAETLY | 0.5125109 |
| IGIFAAML | ASSSDSAETLY | 0.4009095 |
| AVGDLID | ASSSDSAETLY | 0.5030485 |
| NISPLPL | ASSSDSAETLY | 0.386078 |
| VTAVFVAI | ASSSDSAETLY | 0.5677441 |
| SGDDNDAAL | ASSSDSAETLY | 0.48075867 |
| AVAALAHS | ASSSDSAETLY | 0.55367005 |
| GGVTVMATP | ASSSDSAETLY | 0.46049115 |
| AGLAANVRD | ASSSDSAETLY | 0.41153544 |
| TNRPDLIDPALLRPG | ASSSDSAETLY | 0.44536564 |
| LANLGVTAL | ASSSDSAETLY | 0.589192 |
| TGERIEL | ASSSDSAETLY | 0.41687828 |
| NGEELAA | ASSSDSAETLY | 0.3595575 |

| | | |
|---|---|---|
| VLSLVWG | ASSSDSAETLY | 0.4554032 |
| IGYALWL | ASSSDSAETLY | 0.45027536 |
| LSLLLAPL | ASSSDSAETLY | 0.57436347 |
| TLDEALK | ASSSDSAETLY | 0.5822152 |
| TDGAGTAEGG | ASSSDSAETLY | 0.5135138 |
| NAPGGAGGNGG | ASSSDSAETLY | 0.63559365 |
| VSVFMGMGGNF | ASSSDSAETLY | 0.32774758 |
| LASYLAAL | ASSSDSAETLY | 0.50427014 |
| LVGGTPAL | ASSSDSAETLY | 0.6293466 |
| VRNADVL | ASSSDSAETLY | 0.5432144 |
| TCSSPFLVSAM | ASSSDSAETLY | 0.47601083 |
| VHITDAE | ASSSDSAETLY | 0.6118911 |
| EYAAPSF | ASSSDSAETLY | 0.44051787 |
| VVDAFHGGL | ASSSDSAETLY | 0.46178558 |
| ASVVDPQ | ASSSDSAETLY | 0.52701455 |
| SYNEFGGLS | ASSSDSAETLY | 0.48471424 |
| DPDGSVY | ASSSDSAETLY | 0.42117673 |
| MVSTIHATE | ASSSDSAETLY | 0.38683188 |
| LAAGVADPV | ASSSDSAETLY | 0.5165961 |
| TKEQIAA | ASSSDSAETLY | 0.64134055 |
| SLASPAAL | ASSSDSAETLY | 0.49938828 |
| IGLVFYGV | ASSSDSAETLY | 0.5656777 |
| FLLDASNL | ASSSDSAETLY | 0.50769806 |
| YQVGNFP | ASSSDSAETLY | 0.6059087 |
| ESLLSQST | ASSSDSAETLY | 0.40676892 |
| LFTVGFAAVG | ASSSDSAETLY | 0.3780134 |
| ATPPTPT | ASSSDSAETLY | 0.62191796 |
| TDLATLG | ASSSDSAETLY | 0.4133025 |
| SAGGTGR | ASSSDSAETLY | 0.49085414 |
| GFSNSGS | ASSSDSAETLY | 0.4330337 |
| GGNGGSGGTSV | ASSSDSAETLY | 0.646508 |
| GTSVATGGAGN | ASSSDSAETLY | 0.5314505 |
| NGGNGGSGGTS | ASSSDSAETLY | 0.5414718 |
| AVVKFPH | ASSSDSAETLY | 0.45661506 |
| TPPPPPP | ASSSDSAETLY | 0.4433743 |
| AAPAAAAAPA | ASSSDSAETLY | 0.5486807 |
| IALNEDL | ASSSDSAETLY | 0.38490728 |
| IVNVSSM | ASSSDSAETLY | 0.6029052 |
| TGIPPAPR | ASSSDSAETLY | 0.4011439 |
| FDLGGGTF | ASSSDSAETLY | 0.44560418 |
| FDLGGGTFDVS | ASSSDSAETLY | 0.5663854 |
| NEPTAAA | ASSSDSAETLY | 0.64513254 |
| NEPTAAALA | ASSSDSAETLY | 0.43168086 |
| GSIINISSI | ASSSDSAETLY | 0.6179273 |
| VAPGNAGTA | ASSSDSAETLY | 0.41580144 |
| TGGGASLE | ASSSDSAETLY | 0.54276365 |
| VAVYNHYKRI | ASSSDSAETLY | 0.6183704 |
| TDPAPATT | ASSSDSAETLY | 0.57959384 |
| TNRPDILDPALLRPG | ASSSDSAETLY | 0.64898175 |
| LTTTTTS | ASSSDSAETLY | 0.46497992 |
| PTGSGKTL | ASSSDSAETLY | 0.37470996 |

| | | |
|---|---|---|
| TGSGKTL | ASSSDSAETLY | 0.4523705 |
| TIGHVDHGK | ASSSDSAETLY | 0.53277797 |
| TIGHVDHGKTT | ASSSDSAETLY | 0.5728857 |
| GHVDHGKTT | ASSSDSAETLY | 0.57963985 |
| AQFEHTLL | ASSSDSAETLY | 0.4354438 |
| IVNQDEL | ASSSDSAETLY | 0.44239873 |
| TNGSQFF | ASSSDSAETLY | 0.6035014 |
| TGKGIVE | ASSSDSAETLY | 0.5857139 |
| LISNASDA | ASSSDSAETLY | 0.5726097 |
| AELRTLGELTKIAKAHGAQ | ASSSDSAETLY | 0.39671403 |
| APNAVAAAEPSW | ASSSDSAETLY | 0.41494742 |
| APQVAGAATAVL | ASSSDSAETLY | 0.47973964 |
| ARRDPAAAAAILPTDA | ASSSDSAETLY | 0.52593625 |
| DAAARDVLAEAGLAEYF | ASSSDSAETLY | 0.39603308 |
| DDQIVLSAKANLAKVEQ | ASSSDSAETLY | 0.49130833 |
| DELLPDTGEDSDEDGH | ASSSDSAETLY | 0.52786326 |
| ELPEAPANRATVPQD | ASSSDSAETLY | 0.36812872 |
| FGQSKIERVTGIG | ASSSDSAETLY | 0.49923247 |
| GAGGTGSTAGGAGGAGGAGGLY | ASSSDSAETLY | 0.57211196 |
| GHGGTGAAGGNGGTGGAGGSGI | ASSSDSAETLY | 0.43862033 |
| GNGGNGGAGGEGGPGGAGGAGA | ASSSDSAETLY | 0.40216064 |
| GSTAGNGGNGGNATNGTIAGQP | ASSSDSAETLY | 0.6080997 |
| GTGGDGGNGGNGGTGG | ASSSDSAETLY | 0.63535434 |
| GVRRPAGIPADAPV | ASSSDSAETLY | 0.55935335 |
| GYRQVTDAHLLTLARR | ASSSDSAETLY | 0.38591245 |
| IAKMGQDGHDRG | ASSSDSAETLY | 0.322897 |
| KAEPSIEIQYT | ASSSDSAETLY | 0.54597646 |
| KPVRIRMAADEDTVAVT | ASSSDSAETLY | 0.63266397 |
| MPPRIAGMRLLVIKPEP | ASSSDSAETLY | 0.47128364 |
| MKFYAVALFRSRGVPPTR | ASSSDSAETLY | 0.43347165 |
| NAGGFGILGAVAHSP | ASSSDSAETLY | 0.44291726 |
| NEGREITLAMAEM | ASSSDSAETLY | 0.5763338 |
| NELVEWSHPLGHRWRWE | ASSSDSAETLY | 0.41261253 |
| NLGAGGTGGNGGNPG | ASSSDSAETLY | 0.3761833 |
| NTEWPKWETVMAADPVD | ASSSDSAETLY | 0.47881728 |
| PASAAPAETQP | ASSSDSAETLY | 0.5601437 |
| QQSSDGKNGVRAAEITGE | ASSSDSAETLY | 0.40294468 |
| RAPDAVTTSWWKE | ASSSDSAETLY | 0.43435478 |
| SGFGNVGAHTSGWFNQ | ASSSDSAETLY | 0.5872483 |
| SGVAGAVGQAASVGGLKV | ASSSDSAETLY | 0.4771421 |
| SNGGAGTGSAGGNGGT | ASSSDSAETLY | 0.50824404 |
| TGGAGGEGGAAG | ASSSDSAETLY | 0.3891585 |
| TGGDGGAAGAGGAPGHG | ASSSDSAETLY | 0.59936476 |
| TMPPLLAAIANAGRHGV | ASSSDSAETLY | 0.3835071 |
| TSSTAGNSGGA | ASSSDSAETLY | 0.49745968 |
| VPATVPVGILC | ASSSDSAETLY | 0.49888563 |
| VQRPDSGGVVLTGRIS | ASSSDSAETLY | 0.47044402 |
| WTPNPVPATKGVPAGLVQAT | ASSSDSAETLY | 0.44093814 |
| YALKRALAKFSDRFDVVII | ASSSDSAETLY | 0.42569005 |
| YLIGSVIV | ASSSDSAETLY | 0.39361528 |
| YGQLVAVID | ASSSDSAETLY | 0.43889856 |

| | | |
|---|---|---|
| VTVGGMTLPA | ASSSDSAETLY | 0.5704822 |
| VLALPAV | ASSSDSAETLY | 0.50651544 |
| VIPPTLNYET | ASSSDSAETLY | 0.53567445 |
| VGMGGMHPSSG | ASSSDSAETLY | 0.53909945 |
| VGLAGGAATA | ASSSDSAETLY | 0.42006183 |
| VGGAGGAGGSGP | ASSSDSAETLY | 0.4778101 |
| VFAAFMV | ASSSDSAETLY | 0.37341863 |
| TLYDKLR | ASSSDSAETLY | 0.59854203 |
| TLAVFTVL | ASSSDSAETLY | 0.44797027 |
| TGSVQPARV | ASSSDSAETLY | 0.48488554 |
| TGGGAGSGGT | ASSSDSAETLY | 0.4296407 |
| SHLETSL | ASSSDSAETLY | 0.46675777 |
| SFTTGMT | ASSSDSAETLY | 0.5108146 |
| QVEVAKRMGVRQPTVS | ASSSDSAETLY | 0.44035318 |
| QGSGGPALAM | ASSSDSAETLY | 0.44772574 |
| MLAPSNP | ASSSDSAETLY | 0.61524045 |
| LVNNAAI | ASSSDSAETLY | 0.4766621 |
| LTGEINTLA | ASSSDSAETLY | 0.4097129 |
| LSIFGPV | ASSSDSAETLY | 0.5706646 |
| LLALLANL | ASSSDSAETLY | 0.41666007 |
| LISNASDALDKLR | ASSSDSAETLY | 0.52096456 |
| LINEAVP | ASSSDSAETLY | 0.45648393 |
| LGYAFSGL | ASSSDSAETLY | 0.45051476 |
| LDFVAVAL | ASSSDSAETLY | 0.49379686 |
| LAQPGAE | ASSSDSAETLY | 0.4065189 |
| ISNLGAVTGF | ASSSDSAETLY | 0.42004436 |
| HQEDPGRI | ASSSDSAETLY | 0.43597105 |
| GTPGEPTP | ASSSDSAETLY | 0.5612814 |
| GFFNSGS | ASSSDSAETLY | 0.5877031 |
| GAGGGGLTAGAN | ASSSDSAETLY | 0.38348058 |
| FVDALVNM | ASSSDSAETLY | 0.5124076 |
| FLTVAPEL | ASSSDSAETLY | 0.4453097 |
| FETLTAL | ASSSDSAETLY | 0.40927258 |
| DVINPIEL | ASSSDSAETLY | 0.44219258 |
| APVSGAPV | ASSSDSAETLY | 0.44887367 |
| ALVGGLAW | ASSSDSAETLY | 0.39126837 |
| ALAAAAATAI | ASSSDSAETLY | 0.4829974 |
| AFGLGLLL | ASSSDSAETLY | 0.4085927 |
| AFFAAASL | ASSSDSAETLY | 0.5242601 |
| AAAALPAFSPP | ASSSDSAETLY | 0.5714081 |
| VVLAPAL | ASSSDSAETLY | 0.38864732 |
| TIGHVDHGKTTL | ASSSDSAETLY | 0.49164206 |
| LISNASDALDK | ASSSDSAETLY | 0.50965095 |
| LISNASDAL | ASSSDSAETLY | 0.50516284 |
| ISNASDALDK | ASSSDSAETLY | 0.5146289 |
| NQAETLVYQTE | ASSSDSAETLY | 0.42232877 |
| TGGTGGTAGAG | ASSSDSAETLY | 0.57543206 |
| TGSTAGGAGGA | ASSSDSAETLY | 0.58553153 |
| DPLAVPVAAGPV | ASSSDSAETLY | 0.548383 |
| GGTGGTPGNGGN | ASSSDSAETLY | 0.46790457 |
| PAGLAGVDPATL | ASSSDSAETLY | 0.6321685 |

| | | |
|---|---|---|
| GNNLVVEAPHAGASV | ASSSDSAETLY | 0.53524375 |
| KVTRIMSASTLEQPA | ASSSDSAETLY | 0.45668983 |
| LGFALPIALPRRTVV | ASSSDSAETLY | 0.34106177 |
| TRLGNLTARKLVVLV | ASSSDSAETLY | 0.5321386 |
| WSQQRRLIYFPSAGP | ASSSDSAETLY | 0.5334748 |
| GPNGNPDPMAAAVDIR | ASSSDSAETLY | 0.42735985 |
| TQRCVSVADLRAAASR | ASSSDSAETLY | 0.36782935 |
| QFVLDVAEHPVEVLTETV | ASSSDSAETLY | 0.6074664 |
| GGAGGTGGAGGNGGTQAGDG | ASSSDSAETLY | 0.6420433 |
| PLLATLRAIAVDVDAKPAQV | ASSSDSAETLY | 0.48796132 |
| APKFRTGGYMIDFWGVGYQVAK | ASSSDSAETLY | 0.5778862 |
| GAGGAGGAGGAAGGPGGTGGTG | ASSSDSAETLY | 0.36609498 |
| IGALGLLATDVAPAGSVAFGVA | ASSSDSAETLY | 0.5671913 |
| AGGAGGIGGNGGAGGAGGVLMG | ASSSDSAETLY | 0.6148782 |
| ADELIGLL | ASSSDSAETLY | 0.39534163 |
| ADRALKA | ASSSDSAETLY | 0.5663177 |
| AGAVGQVGGAGGA | ASSSDSAETLY | 0.44364223 |
| AGGGGGLGGAP | ASSSDSAETLY | 0.44991353 |
| AGGLLYGNGGAG | ASSSDSAETLY | 0.61750335 |
| AGLESIGL | ASSSDSAETLY | 0.49974996 |
| AGLKTGII | ASSSDSAETLY | 0.50143254 |
| ASLAALLL | ASSSDSAETLY | 0.47764948 |
| ASVINISSI | ASSSDSAETLY | 0.4020748 |
| ATPPGPA | ASSSDSAETLY | 0.465593 |
| ATVASALV | ASSSDSAETLY | 0.54484534 |
| ATVASLIL | ASSSDSAETLY | 0.57866263 |
| AVNGPVLIRH | ASSSDSAETLY | 0.57365763 |
| DGAAGAVGI | ASSSDSAETLY | 0.42343912 |
| DPLLHGI | ASSSDSAETLY | 0.6393375 |
| ELGAALGAL | ASSSDSAETLY | 0.63452697 |
| ELGRTQAV | ASSSDSAETLY | 0.5280009 |
| FVAGGYLS | ASSSDSAETLY | 0.41721293 |
| GASGGAGGAGGW | ASSSDSAETLY | 0.6242137 |
| GGAGAGGDGGAGGI | ASSSDSAETLY | 0.45819843 |
| GTGHNGSQPGMGGNG | ASSSDSAETLY | 0.4870314 |
| GYAAASASAS | ASSSDSAETLY | 0.52789366 |
| HQVDVAAVQQG | ASSSDSAETLY | 0.7547264 |
| IGAAIAHT | ASSSDSAETLY | 0.8420697 |
| IGMIEVAQL | ASSSDSAETLY | 0.805946 |
| IRLAPEHTSAMD | ASSSDSAETLY | 0.7828543 |
| IYNVSVL | ASSSDSAETLY | 0.75336874 |
| KQGFLGM | ASSSDSAETLY | 0.68122196 |
| LESVITQL | ASSSDSAETLY | 0.7780241 |
| LGLGALVVV | ASSSDSAETLY | 0.75227326 |
| LIGNGGNGGSGGI | ASSSDSAETLY | 0.83135235 |
| LNAINAPTL | ASSSDSAETLY | 0.83237153 |
| LQLLNTGV | ASSSDSAETLY | 0.7418197 |
| LQTSIEF | ASSSDSAETLY | 0.8440809 |
| MSFAVSEL | ASSSDSAETLY | 0.83178526 |
| NTGALGL | ASSSDSAETLY | 0.656773 |
| PPGAGGAGGAGTT | ASSSDSAETLY | 0.7550485 |

| | | |
|---|---|---|
| QLVGLGI | ASSSDSAETLY | 0.7332624 |
| RFLEQVHQL | ASSSDSAETLY | 0.7949529 |
| RGVGYVLR | ASSSDSAETLY | 0.55676943 |
| SATGVFAGVI | ASSSDSAETLY | 0.8346558 |
| SLPAITLP | ASSSDSAETLY | 0.71831685 |
| STALAFTV | ASSSDSAETLY | 0.27733904 |
| SVLTVGLGK | ASSSDSAETLY | 0.7963693 |
| TALAGAGSGI | ASSSDSAETLY | 0.78096294 |
| TGLPFTGI | ASSSDSAETLY | 0.81654644 |
| VAEHVTV | ASSSDSAETLY | 0.8235785 |
| VDIVLNSL | ASSSDSAETLY | 0.629009 |
| VLTGGGAVL | ASSSDSAETLY | 0.7638455 |
| VNVGLAYGI | ASSSDSAETLY | 0.8259367 |
| VSSAAVGAV | ASSSDSAETLY | 0.8406028 |
| VTSVYEAGV | ASSSDSAETLY | 0.73645693 |
| WVALDRGA | ASSSDSAETLY | 0.81030214 |
| YLVGSVVI | ASSSDSAETLY | 0.8290222 |
| GPPGTGKTLL | ASSSDSAETLY | 0.8142251 |
| GVGMTKF | ASSSDSAETLY | 0.8293737 |
| IGIGGSDL | ASSSDSAETLY | 0.84259176 |
| IGTLLAEL | ASSSDSAETLY | 0.69828945 |
| IYTSGTT | ASSSDSAETLY | 0.8395899 |
| LSFPAGV | ASSSDSAETLY | 0.8445319 |
| LYTSGTT | ASSSDSAETLY | 0.79760253 |
| VGAFPFGF | ASSSDSAETLY | 0.78175175 |
| VGAGTVEF | ASSSDSAETLY | 0.7094496 |
| VGKSTLLL | ASSSDSAETLY | 0.7559694 |
| AALAHAGGGIV | ASSSDSAETLY | 0.5430124 |
| AALVAVSLL | ASSSDSAETLY | 0.8007329 |
| AAVSRAVAAL | ASSSDSAETLY | 0.8378624 |
| AGGSSGIGGF | ASSSDSAETLY | 0.73514706 |
| AGLTVANANG | ASSSDSAETLY | 0.6760101 |
| AGLVGAVL | ASSSDSAETLY | 0.71575314 |
| ALSQAFTV | ASSSDSAETLY | 0.75809884 |
| ALTFSVSL | ASSSDSAETLY | 0.77273226 |
| ALTGAALVAL | ASSSDSAETLY | 0.52188843 |
| ASFSIGAGVL | ASSSDSAETLY | 0.7407514 |
| DGSVYKI | ASSSDSAETLY | 0.834838 |
| EQLAEIV | ASSSDSAETLY | 0.8348227 |
| FAVSELTVM | ASSSDSAETLY | 0.49607453 |
| GAAPLAAGL | ASSSDSAETLY | 0.8363389 |
| GADGGDGGSGGAGG | ASSSDSAETLY | 0.83226645 |
| GFGLDTGGAGG | ASSSDSAETLY | 0.8387424 |
| GLASGGLL | ASSSDSAETLY | 0.79246354 |
| GRIAGAAVLL | ASSSDSAETLY | 0.73251367 |
| GSAGSLNTSF | ASSSDSAETLY | 0.80810606 |
| GTGQIGI | ASSSDSAETLY | 0.70701927 |
| GTLIALIL | ASSSDSAETLY | 0.69079185 |
| IAATMGFVVG | ASSSDSAETLY | 0.8242824 |
| IFAQALL | ASSSDSAETLY | 0.8044793 |
| IGAAVALGF | ASSSDSAETLY | 0.71943 |

| | | |
|---|---|---|
| IGLTPLAL | ASSSDSAETLY | 0.6910819 |
| ILGSNTSTL | ASSSDSAETLY | 0.84059286 |
| IVFSLTVL | ASSSDSAETLY | 0.61048794 |
| LASKATGF | ASSSDSAETLY | 0.7794492 |
| LASVALIL | ASSSDSAETLY | 0.662707 |
| LFAVGLTL | ASSSDSAETLY | 0.7837691 |
| LFNGGIGTYI | ASSSDSAETLY | 0.834211 |
| LTLGAGII | ASSSDSAETLY | 0.82348573 |
| LVAAALGDI | ASSSDSAETLY | 0.7318395 |
| NTAFVQLGI | ASSSDSAETLY | 0.8104994 |
| QLFSGAEL | ASSSDSAETLY | 0.8218088 |
| SAATFITL | ASSSDSAETLY | 0.8063179 |
| SNYGLANLGV | ASSSDSAETLY | 0.82120436 |
| SYGGAYAVM | ASSSDSAETLY | 0.6962961 |
| TTGVGGLTLG | ASSSDSAETLY | 0.8386034 |
| TTQTIYI | ASSSDSAETLY | 0.8077049 |
| VAAGAGLGAVI | ASSSDSAETLY | 0.8355483 |
| VAAVVTLL | ASSSDSAETLY | 0.8316247 |
| VGFLGGLI | ASSSDSAETLY | 0.830425 |
| VGVIPILL | ASSSDSAETLY | 0.7981414 |
| VIFASSLI | ASSSDSAETLY | 0.46957713 |
| VKGGDGGAAAG | ASSSDSAETLY | 0.7296442 |
| VTKTFTNI | ASSSDSAETLY | 0.80097854 |
| VVAAAILL | ASSSDSAETLY | 0.78730357 |
| YAALDVEL | ASSSDSAETLY | 0.8246357 |
| LASVALLL | ASSSDSAETLY | 0.8265706 |
| LDAVHRV | ASSSDSAETLY | 0.7224277 |
| QGLSSSIF | ASSSDSAETLY | 0.82244664 |
| RGTFANIRL | ASSSDSAETLY | 0.8181701 |
| RQLWWGHRI | ASSSDSAETLY | 0.6001549 |
| YGPPGTGKTLL | ASSSDSAETLY | 0.7672864 |
| AALVGAPANVA | ASSSDSAETLY | 0.8543972 |
| AATFRRTE | ASSSDSAETLY | 0.78788793 |
| AGALGGGI | ASSSDSAETLY | 0.7515966 |
| AGGAGGAGGIGAG | ASSSDSAETLY | 0.75652987 |
| AGSGALGL | ASSSDSAETLY | 0.8517028 |
| AQIADQI | ASSSDSAETLY | 0.739388 |
| ASAVAVTYL | ASSSDSAETLY | 0.84366584 |
| ASTVLGGTGG | ASSSDSAETLY | 0.34642935 |
| AVGGEPAK | ASSSDSAETLY | 0.75941217 |
| CEQGTFGGTC | ASSSDSAETLY | 0.7260798 |
| DEGKAPAG | ASSSDSAETLY | 0.6700398 |
| DTVVFPAL | ASSSDSAETLY | 0.8343278 |
| FEGITMAG | ASSSDSAETLY | 0.77241623 |
| FPLVVQD | ASSSDSAETLY | 0.8172891 |
| GAFTTPQ | ASSSDSAETLY | 0.78156793 |
| GFGGFGTAGGVGGA | ASSSDSAETLY | 0.8377302 |
| GFLAQFL | ASSSDSAETLY | 0.77148736 |
| GGAGGAGGNAGVA | ASSSDSAETLY | 0.81042266 |
| GGVGTAGAGGAGGAGG | ASSSDSAETLY | 0.7857219 |
| GNGGGGDAGNAGSGGN | ASSSDSAETLY | 0.82659674 |

| | | |
|---|---|---|
| GTGGAGGAGNDAGS | ASSSDSAETLY | 0.83273906 |
| GTTNTGWFNS | ASSSDSAETLY | 0.82906175 |
| IFMGPLFTL | ASSSDSAETLY | 0.71339875 |
| ILAHAIEAI | ASSSDSAETLY | 0.72292304 |
| IRSAIREAL | ASSSDSAETLY | 0.81444484 |
| ITFAGGG | ASSSDSAETLY | 0.45844087 |
| ITGMSGHQEV | ASSSDSAETLY | 0.40525153 |
| IVAGLACTL | ASSSDSAETLY | 0.78880477 |
| IVGGGLAGA | ASSSDSAETLY | 0.6491558 |
| IVTGVLNI | ASSSDSAETLY | 0.8361658 |
| LAQQGCI | ASSSDSAETLY | 0.824521 |
| LCLHVEN | ASSSDSAETLY | 0.7577587 |
| LDAISDGS | ASSSDSAETLY | 0.8133883 |
| LGGFAAML | ASSSDSAETLY | 0.78727406 |
| LGGKNPMI | ASSSDSAETLY | 0.21003564 |
| LGPKGVAAIL | ASSSDSAETLY | 0.74640393 |
| LMVGVSI | ASSSDSAETLY | 0.6602372 |
| LVGAAFVL | ASSSDSAETLY | 0.8483489 |
| LVLAVAPL | ASSSDSAETLY | 0.77349734 |
| MAAIAVIDAA | ASSSDSAETLY | 0.6362775 |
| NNGINAELQ | ASSSDSAETLY | 0.7699642 |
| RGEPLVLVHG | ASSSDSAETLY | 0.7792778 |
| RGGAGGAAGLI | ASSSDSAETLY | 0.7352856 |
| RSYRMHPAV | ASSSDSAETLY | 0.69285774 |
| SAAHLLAGQLA | ASSSDSAETLY | 0.30764258 |
| SAAVGSY | ASSSDSAETLY | 0.8281318 |
| SPHAVLHRI | ASSSDSAETLY | 0.62033606 |
| SQIAKVL | ASSSDSAETLY | 0.7646469 |
| TALGHLVP | ASSSDSAETLY | 0.39080554 |
| TAPTTSGAA | ASSSDSAETLY | 0.6392349 |
| TDGDDGNGGSPGT | ASSSDSAETLY | 0.6503401 |
| TDGLGEN | ASSSDSAETLY | 0.8263439 |
| TGGPPLPGQA | ASSSDSAETLY | 0.737702 |
| TIPAITI | ASSSDSAETLY | 0.6965937 |
| TIPGFTI | ASSSDSAETLY | 0.84431595 |
| TSIASGALA | ASSSDSAETLY | 0.7882516 |
| TVTLPGAAI | ASSSDSAETLY | 0.708782 |
| VASGLSAALP | ASSSDSAETLY | 0.7834994 |
| VGAGVIGI | ASSSDSAETLY | 0.7013021 |
| VGPKRFGV | ASSSDSAETLY | 0.77217424 |
| VNVAFPD | ASSSDSAETLY | 0.7767621 |
| VSMTNSGVS | ASSSDSAETLY | 0.84477913 |
| VTDVLGDLSV | ASSSDSAETLY | 0.7732973 |
| VVVVGAGFAGL | ASSSDSAETLY | 0.7155447 |
| VYTVGTVV | ASSSDSAETLY | 0.79209286 |
| WTTFSGP | ASSSDSAETLY | 0.84196246 |
| YDELVEAF | ASSSDSAETLY | 0.849343 |
| YGTGGAGGNGG | ASSSDSAETLY | 0.7257365 |
| YTNVMAV | ASSSDSAETLY | 0.76699907 |
| FLEQVHQL | ASSSDSAETLY | 0.73966134 |
| GLSLLAVAV | ASSSDSAETLY | 0.77382183 |

| | | |
|---|---|---|
| LEQVHQL | ASSSDSAETLY | 0.7462607 |
| LNAAGAI | ASSSDSAETLY | 0.75144166 |
| PAVHASL | ASSSDSAETLY | 0.8271704 |
| TELEDDA | ASSSDSAETLY | 0.8572728 |
| TPPGPPN | ASSSDSAETLY | 0.78289324 |
| TPPPPPPP | ASSSDSAETLY | 0.6159073 |
| TQSPALI | ASSRDRGEQY | 0.75823164 |
| EFAAAFPQ | ASSRDRGEQY | 0.6880322 |
| LLSGNLDVL | ASSRDRGEQY | 0.7428004 |
| GPANLGNYNI | ASSRDRGEQY | 0.80841637 |
| TLGPTLP | ASSRDRGEQY | 0.56831014 |
| VYAYSSCCK | ASSRDRGEQY | 0.74722946 |
| APAPAPAPNI | ASSRDRGEQY | 0.8494368 |
| PGGPVPIP | ASSRDRGEQY | 0.71594274 |
| FLTVALEP | ASSRDRGEQY | 0.7890457 |
| SQYFGANL | ASSRDRGEQY | 0.8331907 |
| GGVKPAA | ASSRDRGEQY | 0.8063996 |
| GGTGGAGGAGGW | ASSRDRGEQY | 0.8376373 |
| GVGGVAGLGGAGG | ASSRDRGEQY | 0.8393108 |
| VLLVLATTI | ASSRDRGEQY | 0.813413 |
| IGIFAAML | ASSRDRGEQY | 0.77690506 |
| AVGDLID | ASSRDRGEQY | 0.7816285 |
| NISPLPL | ASSRDRGEQY | 0.77826464 |
| VTAVFVAI | ASSRDRGEQY | 0.74193597 |
| SGDDNDAAL | ASSRDRGEQY | 0.66319716 |
| AVAALAHS | ASSRDRGEQY | 0.8356489 |
| GGVTVMATP | ASSRDRGEQY | 0.77775764 |
| AGLAANVRD | ASSRDRGEQY | 0.8352603 |
| TNRPDLIDPALLRPG | ASSRDRGEQY | 0.71485895 |
| LANLGVTAL | ASSRDRGEQY | 0.75120944 |
| TGERIEL | ASSRDRGEQY | 0.8247434 |
| NGEELAA | ASSRDRGEQY | 0.23022455 |
| VLSLVWG | ASSRDRGEQY | 0.41835183 |
| IGYALWL | ASSRDRGEQY | 0.7976028 |
| LSLLLAPL | ASSRDRGEQY | 0.83693004 |
| TLDEALK | ASSRDRGEQY | 0.2990546 |
| TDGAGTAEGG | ASSRDRGEQY | 0.7468656 |
| NAPGGAGGNGG | ASSRDRGEQY | 0.59236264 |
| VSVFMGMGGNF | ASSRDRGEQY | 0.8243874 |
| LASYLAAL | ASSRDRGEQY | 0.8144919 |
| LVGGTPAL | ASSRDRGEQY | 0.6266922 |
| VRNADVL | ASSRDRGEQY | 0.82881856 |
| TCSSPFLVSAM | ASSRDRGEQY | 0.49938333 |
| VHITDAE | ASSRDRGEQY | 0.63796306 |
| EYAAPSF | ASSRDRGEQY | 0.83308154 |
| VVDAFHGGL | ASSRDRGEQY | 0.8409028 |
| ASVVDPQ | ASSRDRGEQY | 0.70213413 |
| SYNEFGGLS | ASSRDRGEQY | 0.82579035 |
| DPDGSVY | ASSRDRGEQY | 0.71711653 |
| MVSTIHATE | ASSRDRGEQY | 0.8478653 |
| LAAGVADPV | ASSRDRGEQY | 0.7413884 |

| | | |
|---|---|---|
| TKEQIAA | ASSRDRGEQY | 0.65258336 |
| SLASPAAL | ASSRDRGEQY | 0.82908463 |
| IGLVFYGV | ASSRDRGEQY | 0.8048276 |
| FLLDASNL | ASSRDRGEQY | 0.81422824 |
| YQVGNFP | ASSRDRGEQY | 0.8300646 |
| ESLLSQST | ASSRDRGEQY | 0.7456316 |
| LFTVGFAAVG | ASSRDRGEQY | 0.74867845 |
| ATPPTPT | ASSRDRGEQY | 0.8389025 |
| TDLATLG | ASSRDRGEQY | 0.83101517 |
| SAGGTGR | ASSRDRGEQY | 0.75944775 |
| GFSNSGS | ASSRDRGEQY | 0.8286949 |
| GGNGGSGGTSV | ASSRDRGEQY | 0.76181054 |
| GTSVATGGAGN | ASSRDRGEQY | 0.7177847 |
| NGGNGGSGGTS | ASSRDRGEQY | 0.8453143 |
| AVVKFPH | ASSRDRGEQY | 0.8018019 |
| TPPPPPP | ASSRDRGEQY | 0.81718844 |
| AAPAAAAAPA | ASSRDRGEQY | 0.8358215 |
| IALNEDL | ASSRDRGEQY | 0.8447945 |
| IVNVSSM | ASSRDRGEQY | 0.72136396 |
| TGIPPAPR | ASSRDRGEQY | 0.7877793 |
| FDLGGGTF | ASSRDRGEQY | 0.7632332 |
| FDLGGGTFDVS | ASSRDRGEQY | 0.8278532 |
| NEPTAAA | ASSRDRGEQY | 0.799274 |
| NEPTAAALA | ASSRDRGEQY | 0.6560179 |
| GSIINISSI | ASSRDRGEQY | 0.81659806 |
| VAPGNAGTA | ASSRDRGEQY | 0.83283633 |
| TGGGASLE | ASSRDRGEQY | 0.80419016 |
| VAVYNHYKRI | ASSRDRGEQY | 0.80184364 |
| TDPAPATT | ASSRDRGEQY | 0.84494495 |
| TNRPDILDPALLRPG | ASSRDRGEQY | 0.7660712 |
| LTTTTTS | ASSRDRGEQY | 0.76047254 |
| PTGSGKTL | ASSRDRGEQY | 0.82657826 |
| TGSGKTL | ASSRDRGEQY | 0.82169914 |
| TIGHVDHGK | ASSRDRGEQY | 0.7826208 |
| TIGHVDHGKTT | ASSRDRGEQY | 0.8388926 |
| GHVDHGKTT | ASSRDRGEQY | 0.679229 |
| AQFEHTLL | ASSRDRGEQY | 0.83182156 |
| IVNQDEL | ASSRDRGEQY | 0.6553053 |
| TNGSQFF | ASSRDRGEQY | 0.8080439 |
| TGKGIVE | ASSRDRGEQY | 0.72719467 |
| LISNASDA | ASSRDRGEQY | 0.82570404 |
| AELRTLGELTKIAKAHGAQ | ASSRDRGEQY | 0.82063514 |
| APNAVAAAEPSW | ASSRDRGEQY | 0.83209646 |
| APQVAGAATAVL | ASSRDRGEQY | 0.35578567 |
| ARRDPAAAAAILPTDA | ASSRDRGEQY | 0.83150595 |
| DAAARDVLAEAGLAEYF | ASSRDRGEQY | 0.8360989 |
| DDQIVLSAKANLAKVEQ | ASSRDRGEQY | 0.85244685 |
| DELLPDTGEDSDEDGH | ASSRDRGEQY | 0.76969016 |
| ELPEAPANRATVPQD | ASSRDRGEQY | 0.7972546 |
| FGQSKIERVTGIG | ASSRDRGEQY | 0.77094203 |
| GAGGTGSTAGGAGGAGGAGGLY | ASSRDRGEQY | 0.7790494 |

| | | |
|---|---|---|
| GHGGTGAAGGNGGTGGAGGSGI | ASSRDRGEQY | 0.6353189 |
| GNGGNGGAGGEGGPGGAGGAGA | ASSRDRGEQY | 0.7389226 |
| GSTAGNGGNGGNATNGTIAGQP | ASSRDRGEQY | 0.70246136 |
| GTGGDGGNGGNGGTGG | ASSRDRGEQY | 0.69627136 |
| GVRRPAGIPADAPV | ASSRDRGEQY | 0.81847084 |
| GYRQVTDAHLLTLARR | ASSRDRGEQY | 0.8284443 |
| IAKMGQDGHDRG | ASSRDRGEQY | 0.8285211 |
| KAEPSIEIQYT | ASSRDRGEQY | 0.837372 |
| KPVRIRMAADEDTVAVT | ASSRDRGEQY | 0.8319613 |
| MPPRIAGMRLLVIKPEP | ASSRDRGEQY | 0.8388406 |
| MKFYAVALFRSRGVPPTR | ASSRDRGEQY | 0.76153547 |
| NAGGFGILGAVAHSP | ASSRDRGEQY | 0.7714181 |
| NEGREITLAMAEM | ASSRDRGEQY | 0.6713568 |
| NELVEWSHPLGHRWRWE | ASSRDRGEQY | 0.76149005 |
| NLGAGGTGGNGGNPG | ASSRDRGEQY | 0.7770228 |
| NTEWPKWETVMAADPVD | ASSRDRGEQY | 0.79404277 |
| PASAAPAETQP | ASSRDRGEQY | 0.25023258 |
| QQSSDGKNGVRAAEITGE | ASSRDRGEQY | 0.7655939 |
| RAPDAVTTSWWKE | ASSRDRGEQY | 0.839345 |
| SGFGNVGAHTSGWFNQ | ASSRDRGEQY | 0.61891395 |
| SGVAGAVGQAASVGGLKV | ASSRDRGEQY | 0.8040091 |
| SNGGAGTGSAGGNGGT | ASSRDRGEQY | 0.7444101 |
| TGGAGGEGGAAG | ASSRDRGEQY | 0.73116064 |
| TGGDGGAAGAGGAPGHG | ASSRDRGEQY | 0.8201576 |
| TMPPLLAAIANAGRHGV | ASSRDRGEQY | 0.70509875 |
| TSSTAGNSGGA | ASSRDRGEQY | 0.80923814 |
| VPATVPVGILC | ASSRDRGEQY | 0.83975387 |
| VQRPDSGGVVLTGRIS | ASSRDRGEQY | 0.7877231 |
| WTPNPVPATKGVPAGLVQAT | ASSRDRGEQY | 0.7724765 |
| YALKRALAKFSDRFDVVII | ASSRDRGEQY | 0.7774176 |
| YLIGSVIV | ASSRDRGEQY | 0.6684253 |
| YGQLVAVID | ASSRDRGEQY | 0.7977418 |
| VTVGGMTLPA | ASSRDRGEQY | 0.4642528 |
| VLALPAV | ASSRDRGEQY | 0.75763655 |
| VIPPTLNYET | ASSRDRGEQY | 0.73441267 |
| VGMGGMHPSSG | ASSRDRGEQY | 0.8033823 |
| VGLAGGAATA | ASSRDRGEQY | 0.7781296 |
| VGGAGGAGGSGP | ASSRDRGEQY | 0.804773 |
| VFAAFMV | ASSRDRGEQY | 0.66634905 |
| TLYDKLR | ASSRDRGEQY | 0.7782662 |
| TLAVFTVL | ASSRDRGEQY | 0.6933555 |
| TGSVQPARV | ASSRDRGEQY | 0.7432188 |
| TGGGAGSGGT | ASSRDRGEQY | 0.8173921 |
| SHLETSL | ASSRDRGEQY | 0.8036983 |
| SFTTGMT | ASSRDRGEQY | 0.7286257 |
| QVEVAKRMGVRQPTVS | ASSRDRGEQY | 0.83833 |
| QGSGGPALAM | ASSRDRGEQY | 0.8057517 |
| MLAPSNP | ASSRDRGEQY | 0.6381928 |
| LVNNAAI | ASSRDRGEQY | 0.75322133 |
| LTGEINTLA | ASSRDRGEQY | 0.758732 |
| LSIFGPV | ASSRDRGEQY | 0.762867 |

| | | |
|---|---|---|
| LLALLANL | ASSRDRGEQY | 0.28871718 |
| LISNASDALDKLR | ASSRDRGEQY | 0.81715035 |
| LINEAVP | ASSRDRGEQY | 0.71848327 |
| LGYAFSGL | ASSRDRGEQY | 0.653955 |
| LDFVAVAL | ASSRDRGEQY | 0.76837945 |
| LAQPGAE | ASSRDRGEQY | 0.81867075 |
| ISNLGAVTGF | ASSRDRGEQY | 0.6979388 |
| HQEDPGRI | ASSRDRGEQY | 0.8252651 |
| GTPGEPTP | ASSRDRGEQY | 0.8113776 |
| GFFNSGS | ASSRDRGEQY | 0.8324757 |
| GAGGGGLTAGAN | ASSRDRGEQY | 0.8387418 |
| FVDALVNM | ASSRDRGEQY | 0.8353754 |
| FLTVAPEL | ASSRDRGEQY | 0.76365834 |
| FETLTAL | ASSRDRGEQY | 0.8101673 |
| DVINPIEL | ASSRDRGEQY | 0.78916603 |
| APVSGAPV | ASSRDRGEQY | 0.580371 |
| ALVGGLAW | ASSRDRGEQY | 0.48708946 |
| ALAAAAATAI | ASSRDRGEQY | 0.7901247 |
| AFGLGLLL | ASSRDRGEQY | 0.46873614 |
| AFFAAASL | ASSRDRGEQY | 0.7802795 |
| AAAALPAFSPP | ASSRDRGEQY | 0.8439598 |
| VVLAPAL | ASSRDRGEQY | 0.77853996 |
| TIGHVDHGKTTL | ASSRDRGEQY | 0.65757716 |
| LISNASDALDK | ASSRDRGEQY | 0.74343973 |
| LISNASDAL | ASSRDRGEQY | 0.7623294 |
| ISNASDALDK | ASSRDRGEQY | 0.8195181 |
| NQAETLVYQTE | ASSRDRGEQY | 0.83221585 |
| TGGTGGTAGAG | ASSRDRGEQY | 0.8525696 |
| TGSTAGGAGGA | ASSRDRGEQY | 0.84022737 |
| DPLAVPVAAGPV | ASSRDRGEQY | 0.7252836 |
| GGTGGTPGNGGN | ASSRDRGEQY | 0.80323803 |
| PAGLAGVDPATL | ASSRDRGEQY | 0.74531984 |
| GNNLVVEAPHAGASV | ASSRDRGEQY | 0.7416377 |
| KVTRIMSASTLEQPA | ASSRDRGEQY | 0.7923067 |
| LGFALPIALPRRTVV | ASSRDRGEQY | 0.804062 |
| TRLGNLTARKLVVLV | ASSRDRGEQY | 0.8147493 |
| WSQQRRLIYFPSAGP | ASSRDRGEQY | 0.7542938 |
| GPNGNPDPMAAAVDIR | ASSRDRGEQY | 0.597625 |
| TQRCVSVADLRAAASR | ASSRDRGEQY | 0.82931316 |
| QFVLDVAEHPVEVLTETV | ASSRDRGEQY | 0.8170619 |
| GGAGGTGGAGGNGGTQAGDG | ASSRDRGEQY | 0.56321216 |
| PLLATLRAIAVDVDAKPAQV | ASSRDRGEQY | 0.7128469 |
| APKFRTGGYMIDFWGVGYQVAK | ASSRDRGEQY | 0.84871876 |
| GAGGAGGAGGAAGGPGGTGGTG | ASSRDRGEQY | 0.8380531 |
| IGALGLLATDVAPAGSVAFGVA | ASSRDRGEQY | 0.6882329 |
| AGGAGGIGGNGGAGGAGGVLMG | ASSRDRGEQY | 0.7662933 |
| ADELIGLL | ASSRDRGEQY | 0.748026 |
| ADRALKA | ASSRDRGEQY | 0.48445073 |
| AGAVGQVGGAGGA | ASSRDRGEQY | 0.81494397 |
| AGGGGGLGGAP | ASSRDRGEQY | 0.76517713 |
| AGGLLYGNGGAG | ASSRDRGEQY | 0.7607679 |

| | | |
|---|---|---|
| AGLESIGL | ASSRDRGEQY | 0.7141081 |
| AGLKTGII | ASSRDRGEQY | 0.77002746 |
| ASLAALLL | ASSRDRGEQY | 0.7575574 |
| ASVINISSI | ASSRDRGEQY | 0.82962084 |
| ATPPGPA | ASSRDRGEQY | 0.25797606 |
| ATVASALV | ASSRDRGEQY | 0.8322146 |
| ATVASLIL | ASSRDRGEQY | 0.6377774 |
| AVNGPVLIRH | ASSRDRGEQY | 0.755159 |
| DGAAGAVGI | ASSRDRGEQY | 0.60666716 |
| DPLLHGI | ASSRDRGEQY | 0.76737225 |
| ELGAALGAL | ASSRDRGEQY | 0.8253217 |
| ELGRTQAV | ASSRDRGEQY | 0.82087976 |
| FVAGGYLS | ASSRDRGEQY | 0.6253482 |
| GASGGAGGAGGW | ASSRDRGEQY | 0.8323566 |
| GGAGAGGDGGAGGI | ASSRDRGEQY | 0.8465975 |
| GTGHNGSQPGMGGNG | ASSRDRGEQY | 0.69970584 |
| GYAAASASAS | ASSRDRGEQY | 0.661346 |
| HQVDVAAVQQG | ASSRDRGEQY | 0.8281063 |
| IGAAIAHT | ASSRDRGEQY | 0.77082884 |
| IGMIEVAQL | ASSRDRGEQY | 0.8226563 |
| IRLAPEHTSAMD | ASSRDRGEQY | 0.78936213 |
| IYNVSVL | ASSRDRGEQY | 0.6153088 |
| KQGFLGM | ASSRDRGEQY | 0.82772315 |
| LESVITQL | ASSRDRGEQY | 0.8325423 |
| LGLGALVVV | ASSRDRGEQY | 0.6756307 |
| LIGNGGNGGSGGI | ASSRDRGEQY | 0.70804673 |
| LNAINAPTL | ASSRDRGEQY | 0.74711955 |
| LQLLNTGV | ASSRDRGEQY | 0.4969554 |
| LQTSIEF | ASSRDRGEQY | 0.8368918 |
| MSFAVSEL | ASSRDRGEQY | 0.65946215 |
| NTGALGL | ASSRDRGEQY | 0.7740836 |
| PPGAGGAGGAGTT | ASSRDRGEQY | 0.7941835 |
| QLVGLGI | ASSRDRGEQY | 0.79156387 |
| RFLEQVHQL | ASSRDRGEQY | 0.78386563 |
| RGVGYVLR | ASSRDRGEQY | 0.7868147 |
| SATGVFAGVI | ASSRDRGEQY | 0.82602805 |
| SLPAITLP | ASSRDRGEQY | 0.78984225 |
| STALAFTV | ASSRDRGEQY | 0.78845847 |
| SVLTVGLGK | ASSRDRGEQY | 0.7779575 |
| TALAGAGSGI | ASSRDRGEQY | 0.8342351 |
| TGLPFTGI | ASSRDRGEQY | 0.73311305 |
| VAEHVTV | ASSRDRGEQY | 0.74617434 |
| VDIVLNSL | ASSRDRGEQY | 0.80895424 |
| VLTGGGAVL | ASSRDRGEQY | 0.6321738 |
| VNVGLAYGI | ASSRDRGEQY | 0.82002187 |
| VSSAAVGAV | ASSRDRGEQY | 0.77528095 |
| VTSVYEAGV | ASSRDRGEQY | 0.7864722 |
| WVALDRGA | ASSRDRGEQY | 0.77139413 |
| YLVGSVVI | ASSRDRGEQY | 0.847102 |
| GPPGTGKTLL | ASSRDRGEQY | 0.83061165 |
| GVGMTKF | ASSRDRGEQY | 0.7882688 |

| | | |
|---|---|---|
| IGIGGSDL | ASSRDRGEQY | 0.81992763 |
| IGTLLAEL | ASSRDRGEQY | 0.70635617 |
| IYTSGTT | ASSRDRGEQY | 0.83130574 |
| LSFPAGV | ASSRDRGEQY | 0.8071009 |
| LYTSGTT | ASSRDRGEQY | 0.63387233 |
| VGAFPFGF | ASSRDRGEQY | 0.76114655 |
| VGAGTVEF | ASSRDRGEQY | 0.8349497 |
| VGKSTLLL | ASSRDRGEQY | 0.83052146 |
| AALAHAGGGIV | ASSRDRGEQY | 0.56625265 |
| AALVAVSLL | ASSRDRGEQY | 0.75047815 |
| AAVSRAVAAL | ASSRDRGEQY | 0.45214498 |
| AGGSSGIGGF | ASSRDRGEQY | 0.7085716 |
| AGLTVANANG | ASSRDRGEQY | 0.71949136 |
| AGLVGAVL | ASSRDRGEQY | 0.8057224 |
| ALSQAFTV | ASSRDRGEQY | 0.7213508 |
| ALTFSVSL | ASSRDRGEQY | 0.81309366 |
| ALTGAALVAL | ASSRDRGEQY | 0.7892864 |
| ASFSIGAGVL | ASSRDRGEQY | 0.84271526 |
| DGSVYKI | ASSRDRGEQY | 0.71563536 |
| EQLAEIV | ASSRDRGEQY | 0.797907 |
| FAVSELTVM | ASSRDRGEQY | 0.6341189 |
| GAAPLAAGL | ASSRDRGEQY | 0.6785778 |
| GADGGDGGSGGAGG | ASSRDRGEQY | 0.75003654 |
| GFGLDTGGAGG | ASSRDRGEQY | 0.8034489 |
| GLASGGLL | ASSRDRGEQY | 0.41726 |
| GRIAGAAVLL | ASSRDRGEQY | 0.84042203 |
| GSAGSLNTSF | ASSRDRGEQY | 0.79598534 |
| GTGQIGI | ASSRDRGEQY | 0.8235531 |
| GTLIALIL | ASSRDRGEQY | 0.77881384 |
| IAATMGFVVG | ASSRDRGEQY | 0.778713 |
| IFAQALL | ASSRDRGEQY | 0.76829845 |
| IGAAVALGF | ASSRDRGEQY | 0.81487554 |
| IGLTPLAL | ASSRDRGEQY | 0.76051223 |
| ILGSNTSTL | ASSRDRGEQY | 0.597845 |
| IVFSLTVL | ASSRDRGEQY | 0.7497237 |
| LASKATGF | ASSRDRGEQY | 0.6168722 |
| LASVALIL | ASSRDRGEQY | 0.83679366 |
| LFAVGLTL | ASSRDRGEQY | 0.83322465 |
| LFNGGIGTYI | ASSRDRGEQY | 0.80804265 |
| LTLGAGII | ASSRDRGEQY | 0.83013624 |
| LVAAALGDI | ASSRDRGEQY | 0.4241979 |
| NTAFVQLGI | ASSRDRGEQY | 0.79192007 |
| QLFSGAEL | ASSRDRGEQY | 0.8428303 |
| SAATFITL | ASSRDRGEQY | 0.8130025 |
| SNYGLANLGV | ASSRDRGEQY | 0.73647374 |
| SYGGAYAVM | ASSRDRGEQY | 0.8353702 |
| TTGVGGLTLG | ASSRDRGEQY | 0.72480416 |
| TTQTIYI | ASSRDRGEQY | 0.7619284 |
| VAAGAGLGAVI | ASSRDRGEQY | 0.8099069 |
| VAAVVTLL | ASSRDRGEQY | 0.8223491 |
| VGFLGGLI | ASSRDRGEQY | 0.80436146 |

| | | |
|---|---|---|
| VGVIPILL | ASSRDRGEQY | 0.74520177 |
| VIFASSLI | ASSRDRGEQY | 0.8031964 |
| VKGGDGGAAAG | ASSRDRGEQY | 0.8512785 |
| VTKTFTNI | ASSRDRGEQY | 0.75766826 |
| VVAAAILL | ASSRDRGEQY | 0.8256906 |
| YAALDVEL | ASSRDRGEQY | 0.80723524 |
| LASVALLL | ASSRDRGEQY | 0.82367253 |
| LDAVHRV | ASSRDRGEQY | 0.6884727 |
| QGLSSSIF | ASSRDRGEQY | 0.73417693 |
| RGTFANIRL | ASSRDRGEQY | 0.78358245 |
| RQLWWGHRI | ASSRDRGEQY | 0.634916 |
| YGPPGTGKTLL | ASSRDRGEQY | 0.7582052 |
| AALVGAPANVA | ASSRDRGEQY | 0.77053934 |
| AATFRRTE | ASSRDRGEQY | 0.85603166 |
| AGALGGGI | ASSRDRGEQY | 0.79982674 |
| AGGAGGAGGIGAG | ASSRDRGEQY | 0.8163501 |
| AGSGALGL | ASSRDRGEQY | 0.7410229 |
| AQIADQI | ASSRDRGEQY | 0.8072822 |
| ASAVAVTYL | ASSRDRGEQY | 0.80628204 |
| ASTVLGGTGG | ASSRDRGEQY | 0.6573288 |
| AVGGEPAK | ASSRDRGEQY | 0.83537567 |
| CEQGTFGGTC | ASSRDRGEQY | 0.7029212 |
| DEGKAPAG | ASSRDRGEQY | 0.7223222 |
| DTVVFPAL | ASSRDRGEQY | 0.83110607 |
| FEGITMAG | ASSRDRGEQY | 0.8539721 |
| FPLVVQD | ASSRDRGEQY | 0.7392044 |
| GAFTTPQ | ASSRDRGEQY | 0.73043364 |
| GFGGFGTAGGVGGA | ASSRDRGEQY | 0.74002105 |
| GFLAQFL | ASSRDRGEQY | 0.82880634 |
| GGAGGAGGNAGVA | ASSRDRGEQY | 0.83627164 |
| GGVGTAGAGGAGGAGG | ASSRDRGEQY | 0.7973632 |
| GNGGGGDAGNAGSGGN | ASSRDRGEQY | 0.8456506 |
| GTGGAGGAGNDAGS | ASSRDRGEQY | 0.8277873 |
| GTTNTGWFNS | ASSRDRGEQY | 0.63438237 |
| IFMGPLFTL | ASSRDRGEQY | 0.79062593 |
| ILAHAIEAI | ASSRDRGEQY | 0.8304991 |
| IRSAIREAL | ASSRDRGEQY | 0.81999075 |
| ITFAGGG | ASSRDRGEQY | 0.8528279 |
| ITGMSGHQEV | ASSRDRGEQY | 0.8158531 |
| IVAGLACTL | ASSRDRGEQY | 0.71821123 |
| IVGGGLAGA | ASSRDRGEQY | 0.8410029 |
| IVTGVLNI | ASSRDRGEQY | 0.85152054 |
| LAQQGCI | ASSRDRGEQY | 0.8260424 |
| LCLHVEN | ASSRDRGEQY | 0.7532115 |
| LDAISDGS | ASSRDRGEQY | 0.7437997 |
| LGGFAAML | ASSRDRGEQY | 0.7895988 |
| LGGKNPMI | ASSRDRGEQY | 0.8307728 |
| LGPKGVAAIL | ASSRDRGEQY | 0.68676734 |
| LMVGVSI | ASSRDRGEQY | 0.764467 |
| LVGAAFVL | ASSRDRGEQY | 0.79720676 |
| LVLAVAPL | ASSRDRGEQY | 0.60621434 |

| | | |
|---|---|---|
| MAAIAVIDAA | ASSRDRGEQY | 0.8313115 |
| NNGINAELQ | ASSRDRGEQY | 0.8339843 |
| RGEPLVLVHG | ASSRDRGEQY | 0.7817935 |
| RGGAGGAAGLI | ASSRDRGEQY | 0.75887895 |
| RSYRMHPAV | ASSRDRGEQY | 0.8243001 |
| SAAHLLAGQLA | ASSRDRGEQY | 0.8560794 |
| SAAVGSY | ASSRDRGEQY | 0.83820707 |
| SPHAVLHRI | ASSRDRGEQY | 0.6813828 |
| SQIAKVL | ASSRDRGEQY | 0.43133143 |
| TALGHLVP | ASSRDRGEQY | 0.6927762 |
| TAPTTSGAA | ASSRDRGEQY | 0.83060205 |
| TDGDDGNGGSPGT | ASSRDRGEQY | 0.801163 |
| TDGLGEN | ASSRDRGEQY | 0.8043375 |
| TGGPPLPGQA | ASSRDRGEQY | 0.7983214 |
| TIPAITI | ASSRDRGEQY | 0.8353589 |
| TIPGFTI | ASSRDRGEQY | 0.7468432 |
| TSIASGALA | ASSRDRGEQY | 0.7232419 |
| TVTLPGAAI | ASSRDRGEQY | 0.789541 |
| VASGLSAALP | ASSRDRGEQY | 0.81487584 |
| VGAGVIGI | ASSRDRGEQY | 0.69826525 |
| VGPKRFGV | ASSRDRGEQY | 0.7649106 |
| VNVAFPD | ASSRDRGEQY | 0.6755674 |
| VSMTNSGVS | ASSRDRGEQY | 0.70788425 |
| VTDVLGDLSV | ASSRDRGEQY | 0.79263556 |
| VVVVGAGFAGL | ASSRDRGEQY | 0.6979294 |
| VYTVGTVV | ASSRDRGEQY | 0.84438354 |
| WTTFSGP | ASSRDRGEQY | 0.69126916 |
| YDELVEAF | ASSRDRGEQY | 0.8142306 |
| YGTGGAGGNGG | ASSRDRGEQY | 0.80138963 |
| YTNVMAV | ASSRDRGEQY | 0.819411 |
| FLEQVHQL | ASSRDRGEQY | 0.7279295 |
| GLSLLAVAV | ASSRDRGEQY | 0.71082807 |
| LEQVHQL | ASSRDRGEQY | 0.6744002 |
| LNAAGAI | ASSRDRGEQY | 0.4841141 |
| PAVHASL | ASSRDRGEQY | 0.82603127 |
| TELEDDA | ASSRDRGEQY | 0.6465481 |
| TPPGPPN | ASSRDRGEQY | 0.83490634 |
| TPPPPPPP | ASSRDRGEQY | 0.8240896 |
| TQSPALI | ASSRGGAYAEQF | 0.7292043 |
| EFAAAFPQ | ASSRGGAYAEQF | 0.80836934 |
| LLSGNLDVL | ASSRGGAYAEQF | 0.75405926 |
| GPANLGNYNI | ASSRGGAYAEQF | 0.69411296 |
| TLGPTLP | ASSRGGAYAEQF | 0.8171365 |
| VYAYSSCCK | ASSRGGAYAEQF | 0.82302505 |
| APAPAPAPNI | ASSRGGAYAEQF | 0.67613995 |
| PGGPVPIP | ASSRGGAYAEQF | 0.79846376 |
| FLTVALEP | ASSRGGAYAEQF | 0.7887093 |
| SQYFGANL | ASSRGGAYAEQF | 0.70964366 |
| GGVKPAA | ASSRGGAYAEQF | 0.800395 |
| GGTGGAGGAGGW | ASSRGGAYAEQF | 0.830067 |
| GVGGVAGLGGAGG | ASSRGGAYAEQF | 0.80312425 |

| | | |
|---|---|---|
| VLLVLATTI | ASSRGGAYAEQF | 0.7362176 |
| IGIFAAML | ASSRGGAYAEQF | 0.8177313 |
| AVGDLID | ASSRGGAYAEQF | 0.62438947 |
| NISPLPL | ASSRGGAYAEQF | 0.8043828 |
| VTAVFVAI | ASSRGGAYAEQF | 0.7906791 |
| SGDDNDAAL | ASSRGGAYAEQF | 0.7465295 |
| AVAALAHS | ASSRGGAYAEQF | 0.81731933 |
| GGVTVMATP | ASSRGGAYAEQF | 0.7169794 |
| AGLAANVRD | ASSRGGAYAEQF | 0.6838153 |
| TNRPDLIDPALLRPG | ASSRGGAYAEQF | 0.7468738 |
| LANLGVTAL | ASSRGGAYAEQF | 0.8087519 |
| TGERIEL | ASSRGGAYAEQF | 0.82647675 |
| NGEELAA | ASSRGGAYAEQF | 0.6971774 |
| VLSLVWG | ASSRGGAYAEQF | 0.7897231 |
| IGYALWL | ASSRGGAYAEQF | 0.7388272 |
| LSLLLAPL | ASSRGGAYAEQF | 0.6847894 |
| TLDEALK | ASSRGGAYAEQF | 0.7931701 |
| TDGAGTAEGG | ASSRGGAYAEQF | 0.78504884 |
| NAPGGAGGNGG | ASSRGGAYAEQF | 0.73132694 |
| VSVFMGMGGNF | ASSRGGAYAEQF | 0.81203794 |
| LASYLAAL | ASSRGGAYAEQF | 0.7814356 |
| LVGGTPAL | ASSRGGAYAEQF | 0.82087123 |
| VRNADVL | ASSRGGAYAEQF | 0.8407943 |
| TCSSPFLVSAM | ASSRGGAYAEQF | 0.7441937 |
| VHITDAE | ASSRGGAYAEQF | 0.8139043 |
| EYAAPSF | ASSRGGAYAEQF | 0.81168497 |
| VVDAFHGGL | ASSRGGAYAEQF | 0.80109644 |
| ASVVDPQ | ASSRGGAYAEQF | 0.7956653 |
| SYNEFGGLS | ASSRGGAYAEQF | 0.7292589 |
| DPDGSVY | ASSRGGAYAEQF | 0.8402917 |
| MVSTIHATE | ASSRGGAYAEQF | 0.8230047 |
| LAAGVADPV | ASSRGGAYAEQF | 0.80871385 |
| TKEQIAA | ASSRGGAYAEQF | 0.81350166 |
| SLASPAAL | ASSRGGAYAEQF | 0.8085487 |
| IGLVFYGV | ASSRGGAYAEQF | 0.8167476 |
| FLLDASNL | ASSRGGAYAEQF | 0.7102153 |
| YQVGNFP | ASSRGGAYAEQF | 0.6263654 |
| ESLLSQST | ASSRGGAYAEQF | 0.7922161 |
| LFTVGFAAVG | ASSRGGAYAEQF | 0.76519454 |
| ATPPTPT | ASSRGGAYAEQF | 0.7752351 |
| TDLATLG | ASSRGGAYAEQF | 0.63068235 |
| SAGGTGR | ASSRGGAYAEQF | 0.83257645 |
| GFSNSGS | ASSRGGAYAEQF | 0.85396695 |
| GGNGGSGGTSV | ASSRGGAYAEQF | 0.76580536 |
| GTSVATGGAGN | ASSRGGAYAEQF | 0.8488929 |
| NGGNGGSGGTS | ASSRGGAYAEQF | 0.63414645 |
| AVVKFPH | ASSRGGAYAEQF | 0.82275337 |
| TPPPPPP | ASSRGGAYAEQF | 0.82985944 |
| AAPAAAAAPA | ASSRGGAYAEQF | 0.655723 |
| IALNEDL | ASSRGGAYAEQF | 0.83474636 |
| IVNVSSM | ASSRGGAYAEQF | 0.7251196 |

| | | |
|---|---|---|
| TGIPPAPR | ASSRGGAYAEQF | 0.78689325 |
| FDLGGGTF | ASSRGGAYAEQF | 0.8366981 |
| FDLGGGTFDVS | ASSRGGAYAEQF | 0.7831919 |
| NEPTAAA | ASSRGGAYAEQF | 0.8048163 |
| NEPTAAALA | ASSRGGAYAEQF | 0.82295597 |
| GSIINISSI | ASSRGGAYAEQF | 0.77321637 |
| VAPGNAGTA | ASSRGGAYAEQF | 0.81147087 |
| TGGGASLE | ASSRGGAYAEQF | 0.8291503 |
| VAVYNHYKRI | ASSRGGAYAEQF | 0.70390314 |
| TDPAPATT | ASSRGGAYAEQF | 0.8496237 |
| TNRPDILDPALLRPG | ASSRGGAYAEQF | 0.6885573 |
| LTTTTTS | ASSRGGAYAEQF | 0.8549943 |
| PTGSGKTL | ASSRGGAYAEQF | 0.79659 |
| TGSGKTL | ASSRGGAYAEQF | 0.7809013 |
| TIGHVDHGK | ASSRGGAYAEQF | 0.76387787 |
| TIGHVDHGKTT | ASSRGGAYAEQF | 0.8359964 |
| GHVDHGKTT | ASSRGGAYAEQF | 0.76452136 |
| AQFEHTLL | ASSRGGAYAEQF | 0.80655384 |
| IVNQDEL | ASSRGGAYAEQF | 0.84654033 |
| TNGSQFF | ASSRGGAYAEQF | 0.37289435 |
| TGKGIVE | ASSRGGAYAEQF | 0.82608443 |
| LISNASDA | ASSRGGAYAEQF | 0.7364713 |
| AELRTLGELTKIAKAHGAQ | ASSRGGAYAEQF | 0.75257564 |
| APNAVAAAEPSW | ASSRGGAYAEQF | 0.5521331 |
| APQVAGAATAVL | ASSRGGAYAEQF | 0.29259637 |
| ARRDPAAAAAILPTDA | ASSRGGAYAEQF | 0.60626245 |
| DAAARDVLAEAGLAEYF | ASSRGGAYAEQF | 0.46208534 |
| DDQIVLSAKANLAKVEQ | ASSRGGAYAEQF | 0.81945664 |
| DELLPDTGEDSDEDGH | ASSRGGAYAEQF | 0.7789263 |
| ELPEAPANRATVPQD | ASSRGGAYAEQF | 0.5349169 |
| FGQSKIERVTGIG | ASSRGGAYAEQF | 0.78264487 |
| GAGGTGSTAGGAGGAGGAGGLY | ASSRGGAYAEQF | 0.81933683 |
| GHGGTGAAGGNGGTGGAGGSGI | ASSRGGAYAEQF | 0.34497005 |
| GNGGNGGAGGEGGPGGAGGAGA | ASSRGGAYAEQF | 0.63022757 |
| GSTAGNGGNGGNATNGTIAGQP | ASSRGGAYAEQF | 0.5559447 |
| GTGGDGGNGGNGGTGG | ASSRGGAYAEQF | 0.6512848 |
| GVRRPAGIPADAPV | ASSRGGAYAEQF | 0.24654505 |
| GYRQVTDAHLLTLARR | ASSRGGAYAEQF | 0.7745395 |
| IAKMGQDGHDRG | ASSRGGAYAEQF | 0.5531184 |
| KAEPSIEIQYT | ASSRGGAYAEQF | 0.15080932 |
| KPVRIRMAADEDTVAVT | ASSRGGAYAEQF | 0.6958816 |
| MPPRIAGMRLLVIKPEP | ASSRGGAYAEQF | 0.5934912 |
| MKFYAVALFRSRGVPPTR | ASSRGGAYAEQF | 0.60682726 |
| NAGGFGILGAVAHSP | ASSRGGAYAEQF | 0.80310136 |
| NEGREITLAMAEM | ASSRGGAYAEQF | 0.5597241 |
| NELVEWSHPLGHRWRWE | ASSRGGAYAEQF | 0.39590615 |
| NLGAGGTGGNGGNPG | ASSRGGAYAEQF | 0.73644805 |
| NTEWPKWETVMAADPVD | ASSRGGAYAEQF | 0.80401516 |
| PASAAPAETQP | ASSRGGAYAEQF | 0.35175747 |
| QQSSDGKNGVRAAEITGE | ASSRGGAYAEQF | 0.7256141 |
| RAPDAVTTSWWKE | ASSRGGAYAEQF | 0.8296236 |

| | | |
|---|---|---|
| SGFGNVGAHTSGWFNQ | ASSRGGAYAEQF | 0.7081697 |
| SGVAGAVGQAASVGGLKV | ASSRGGAYAEQF | 0.8170536 |
| SNGGAGTGSAGGNGGT | ASSRGGAYAEQF | 0.8307886 |
| TGGAGGEGGAAG | ASSRGGAYAEQF | 0.56620383 |
| TGGDGGAAGAGGAPGHG | ASSRGGAYAEQF | 0.80717623 |
| TMPPLLAAIANAGRHGV | ASSRGGAYAEQF | 0.8130511 |
| TSSTAGNSGGA | ASSRGGAYAEQF | 0.5908189 |
| VPATVPVGILC | ASSRGGAYAEQF | 0.6387607 |
| VQRPDSGGVVLTGRIS | ASSRGGAYAEQF | 0.3558267 |
| WTPNPVPATKGVPAGLVQAT | ASSRGGAYAEQF | 0.6617941 |
| YALKRALAKFSDRFDVVII | ASSRGGAYAEQF | 0.41146737 |
| YLIGSVIV | ASSRGGAYAEQF | 0.74091554 |
| YGQLVAVID | ASSRGGAYAEQF | 0.81392026 |
| VTVGGMTLPA | ASSRGGAYAEQF | 0.64764297 |
| VLALPAV | ASSRGGAYAEQF | 0.33561447 |
| VIPPTLNYET | ASSRGGAYAEQF | 0.27230263 |
| VGMGGMHPSSG | ASSRGGAYAEQF | 0.7253297 |
| VGLAGGAATA | ASSRGGAYAEQF | 0.41590607 |
| VGGAGGAGGSGP | ASSRGGAYAEQF | 0.31714073 |
| VFAAFMV | ASSRGGAYAEQF | 0.5995883 |
| TLYDKLR | ASSRGGAYAEQF | 0.78823483 |
| TLAVFTVL | ASSRGGAYAEQF | 0.8081306 |
| TGSVQPARV | ASSRGGAYAEQF | 0.4739225 |
| TGGGAGSGGT | ASSRGGAYAEQF | 0.84345996 |
| SHLETSL | ASSRGGAYAEQF | 0.7897004 |
| SFTTGMT | ASSRGGAYAEQF | 0.83373773 |
| QVEVAKRMGVRQPTVS | ASSRGGAYAEQF | 0.67076683 |
| QGSGGPALAM | ASSRGGAYAEQF | 0.36660695 |
| MLAPSNP | ASSRGGAYAEQF | 0.83572805 |
| LVNNAAI | ASSRGGAYAEQF | 0.60495394 |
| LTGEINTLA | ASSRGGAYAEQF | 0.6613461 |
| LSIFGPV | ASSRGGAYAEQF | 0.78734255 |
| LLALLANL | ASSRGGAYAEQF | 0.61363006 |
| LISNASDALDKLR | ASSRGGAYAEQF | 0.46329197 |
| LINEAVP | ASSRGGAYAEQF | 0.7127439 |
| LGYAFSGL | ASSRGGAYAEQF | 0.82173944 |
| LDFVAVAL | ASSRGGAYAEQF | 0.41204402 |
| LAQPGAE | ASSRGGAYAEQF | 0.4187206 |
| ISNLGAVTGF | ASSRGGAYAEQF | 0.6191888 |
| HQEDPGRI | ASSRGGAYAEQF | 0.66277766 |
| GTPGEPTP | ASSRGGAYAEQF | 0.80571467 |
| GFFNSGS | ASSRGGAYAEQF | 0.74984944 |
| GAGGGGLTAGAN | ASSRGGAYAEQF | 0.60467273 |
| FVDALVNM | ASSRGGAYAEQF | 0.74284375 |
| FLTVAPEL | ASSRGGAYAEQF | 0.7773622 |
| FETLTAL | ASSRGGAYAEQF | 0.70626724 |
| DVINPIEL | ASSRGGAYAEQF | 0.73525393 |
| APVSGAPV | ASSRGGAYAEQF | 0.5613035 |
| ALVGGLAW | ASSRGGAYAEQF | 0.80144656 |
| ALAAAAATAI | ASSRGGAYAEQF | 0.79975885 |
| AFGLGLLL | ASSRGGAYAEQF | 0.77181923 |

| | | |
|---|---|---|
| AFFAAASL | ASSRGGAYAEQF | 0.80715954 |
| AAAALPAFSPP | ASSRGGAYAEQF | 0.68831784 |
| VVLAPAL | ASSRGGAYAEQF | 0.7093476 |
| TIGHVDHGKTTL | ASSRGGAYAEQF | 0.29252678 |
| LISNASDALDK | ASSRGGAYAEQF | 0.6595485 |
| LISNASDAL | ASSRGGAYAEQF | 0.66322315 |
| ISNASDALDK | ASSRGGAYAEQF | 0.7337822 |
| NQAETLVYQTE | ASSRGGAYAEQF | 0.7609737 |
| TGGTGGTAGAG | ASSRGGAYAEQF | 0.72853535 |
| TGSTAGGAGGA | ASSRGGAYAEQF | 0.5087037 |
| DPLAVPVAAGPV | ASSRGGAYAEQF | 0.7909156 |
| GGTGGTPGNGGN | ASSRGGAYAEQF | 0.8212206 |
| PAGLAGVDPATL | ASSRGGAYAEQF | 0.37891722 |
| GNNLVVEAPHAGASV | ASSRGGAYAEQF | 0.58247316 |
| KVTRIMSASTLEQPA | ASSRGGAYAEQF | 0.8533827 |
| LGFALPIALPRRTVV | ASSRGGAYAEQF | 0.76112956 |
| TRLGNLTARKLVVLV | ASSRGGAYAEQF | 0.5873215 |
| WSQQRRLIYFPSAGP | ASSRGGAYAEQF | 0.6548636 |
| GPNGNPDPMAAAVDIR | ASSRGGAYAEQF | 0.8522197 |
| TQRCVSVADLRAAASR | ASSRGGAYAEQF | 0.6660763 |
| QFVLDVAEHPVEVLTETV | ASSRGGAYAEQF | 0.82825327 |
| GGAGGTGGAGGNGGTQAGDG | ASSRGGAYAEQF | 0.24429941 |
| PLLATLRAIAVDVDAKPAQV | ASSRGGAYAEQF | 0.6565938 |
| APKFRTGGYMIDFWGVGYQVAK | ASSRGGAYAEQF | 0.5099233 |
| GAGGAGGAGGAAGGPGGTGGTG | ASSRGGAYAEQF | 0.17699611 |
| IGALGLLATDVAPAGSVAFGVA | ASSRGGAYAEQF | 0.7912476 |
| AGGAGGIGGNGGAGGAGGVLMG | ASSRGGAYAEQF | 0.53310746 |
| ADELIGLL | ASSRGGAYAEQF | 0.773728 |
| ADRALKA | ASSRGGAYAEQF | 0.5977182 |
| AGAVGQVGGAGGA | ASSRGGAYAEQF | 0.8065553 |
| AGGGGGLGGAP | ASSRGGAYAEQF | 0.72091174 |
| AGGLLYGNGGAG | ASSRGGAYAEQF | 0.67190266 |
| AGLESIGL | ASSRGGAYAEQF | 0.46305627 |
| AGLKTGII | ASSRGGAYAEQF | 0.7374532 |
| ASLAALLL | ASSRGGAYAEQF | 0.78943515 |
| ASVINISSI | ASSRGGAYAEQF | 0.7975017 |
| ATPPGPA | ASSRGGAYAEQF | 0.22476555 |
| ATVASALV | ASSRGGAYAEQF | 0.7073394 |
| ATVASLIL | ASSRGGAYAEQF | 0.6276558 |
| AVNGPVLIRH | ASSRGGAYAEQF | 0.26888797 |
| DGAAGAVGI | ASSRGGAYAEQF | 0.18356392 |
| DPLLHGI | ASSRGGAYAEQF | 0.5607986 |
| ELGAALGAL | ASSRGGAYAEQF | 0.65593797 |
| ELGRTQAV | ASSRGGAYAEQF | 0.7585939 |
| FVAGGYLS | ASSRGGAYAEQF | 0.7527655 |
| GASGGAGGAGGW | ASSRGGAYAEQF | 0.5994551 |
| GGAGAGGDGGAGGI | ASSRGGAYAEQF | 0.75737584 |
| GTGHNGSQPGMGGNG | ASSRGGAYAEQF | 0.6697935 |
| GYAAASASAS | ASSRGGAYAEQF | 0.10131934 |
| HQVDVAAVQQG | ASSRGGAYAEQF | 0.57370234 |
| IGAAIAHT | ASSRGGAYAEQF | 0.32027867 |

| | | |
|---|---|---|
| IGMIEVAQL | ASSRGGAYAEQF | 0.8297728 |
| IRLAPEHTSAMD | ASSRGGAYAEQF | 0.44167373 |
| IYNVSVL | ASSRGGAYAEQF | 0.58618 |
| KQGFLGM | ASSRGGAYAEQF | 0.6559111 |
| LESVITQL | ASSRGGAYAEQF | 0.7789408 |
| LGLGALVVV | ASSRGGAYAEQF | 0.70272636 |
| LIGNGGNGGSGGI | ASSRGGAYAEQF | 0.41923648 |
| LNAINAPTL | ASSRGGAYAEQF | 0.21331117 |
| LQLLNTGV | ASSRGGAYAEQF | 0.736104 |
| LQTSIEF | ASSRGGAYAEQF | 0.5548445 |
| MSFAVSEL | ASSRGGAYAEQF | 0.38160616 |
| NTGALGL | ASSRGGAYAEQF | 0.38567412 |
| PPGAGGAGGAGTT | ASSRGGAYAEQF | 0.43570077 |
| QLVGLGI | ASSRGGAYAEQF | 0.3041746 |
| RFLEQVHQL | ASSRGGAYAEQF | 0.78875387 |
| RGVGYVLR | ASSRGGAYAEQF | 0.46546692 |
| SATGVFAGVI | ASSRGGAYAEQF | 0.5740854 |
| SLPAITLP | ASSRGGAYAEQF | 0.84162307 |
| STALAFTV | ASSRGGAYAEQF | 0.75679713 |
| SVLTVGLGK | ASSRGGAYAEQF | 0.6403892 |
| TALAGAGSGI | ASSRGGAYAEQF | 0.48709702 |
| TGLPFTGI | ASSRGGAYAEQF | 0.47477475 |
| VAEHVTV | ASSRGGAYAEQF | 0.44583774 |
| VDIVLNSL | ASSRGGAYAEQF | 0.4755282 |
| VLTGGGAVL | ASSRGGAYAEQF | 0.8237705 |
| VNVGLAYGI | ASSRGGAYAEQF | 0.70697534 |
| VSSAAVGAV | ASSRGGAYAEQF | 0.42390338 |
| VTSVYEAGV | ASSRGGAYAEQF | 0.81035197 |
| WVALDRGA | ASSRGGAYAEQF | 0.8364924 |
| YLVGSVVI | ASSRGGAYAEQF | 0.8468179 |
| GPPGTGKTLL | ASSRGGAYAEQF | 0.4342425 |
| GVGMTKF | ASSRGGAYAEQF | 0.6205646 |
| IGIGGSDL | ASSRGGAYAEQF | 0.6127291 |
| IGTLLAEL | ASSRGGAYAEQF | 0.64913106 |
| IYTSGTT | ASSRGGAYAEQF | 0.49673235 |
| LSFPAGV | ASSRGGAYAEQF | 0.41654772 |
| LYTSGTT | ASSRGGAYAEQF | 0.7907331 |
| VGAFPFGF | ASSRGGAYAEQF | 0.86842394 |
| VGAGTVEF | ASSRGGAYAEQF | 0.63203895 |
| VGKSTLLL | ASSRGGAYAEQF | 0.3411063 |
| AALAHAGGGIV | ASSRGGAYAEQF | 0.59865004 |
| AALVAVSLL | ASSRGGAYAEQF | 0.4259869 |
| AAVSRAVAAL | ASSRGGAYAEQF | 0.485627 |
| AGGSSGIGGF | ASSRGGAYAEQF | 0.6084661 |
| AGLTVANANG | ASSRGGAYAEQF | 0.33579177 |
| AGLVGAVL | ASSRGGAYAEQF | 0.6362574 |
| ALSQAFTV | ASSRGGAYAEQF | 0.8407014 |
| ALTFSVSL | ASSRGGAYAEQF | 0.66037285 |
| ALTGAALVAL | ASSRGGAYAEQF | 0.75519323 |
| ASFSIGAGVL | ASSRGGAYAEQF | 0.82918346 |
| DGSVYKI | ASSRGGAYAEQF | 0.7722131 |

| | | |
|---|---|---|
| EQLAEIV | ASSRGGAYAEQF | 0.83206445 |
| FAVSELTVM | ASSRGGAYAEQF | 0.7948485 |
| GAAPLAAGL | ASSRGGAYAEQF | 0.6691465 |
| GADGGDGGSGGAGG | ASSRGGAYAEQF | 0.66950536 |
| GFGLDTGGAGG | ASSRGGAYAEQF | 0.797139 |
| GLASGGLL | ASSRGGAYAEQF | 0.5854206 |
| GRIAGAAVLL | ASSRGGAYAEQF | 0.69382703 |
| GSAGSLNTSF | ASSRGGAYAEQF | 0.62174845 |
| GTGQIGI | ASSRGGAYAEQF | 0.7912023 |
| GTLIALIL | ASSRGGAYAEQF | 0.62376654 |
| IAATMGFVVG | ASSRGGAYAEQF | 0.8523435 |
| IFAQALL | ASSRGGAYAEQF | 0.21958911 |
| IGAAVALGF | ASSRGGAYAEQF | 0.40489894 |
| IGLTPLAL | ASSRGGAYAEQF | 0.8216249 |
| ILGSNTSTL | ASSRGGAYAEQF | 0.12823957 |
| IVFSLTVL | ASSRGGAYAEQF | 0.24528171 |
| LASKATGF | ASSRGGAYAEQF | 0.6499834 |
| LASVALIL | ASSRGGAYAEQF | 0.8213839 |
| LFAVGLTL | ASSRGGAYAEQF | 0.13919327 |
| LFNGGIGTYI | ASSRGGAYAEQF | 0.5050605 |
| LTLGAGII | ASSRGGAYAEQF | 0.2773109 |
| LVAAALGDI | ASSRGGAYAEQF | 0.7899111 |
| NTAFVQLGI | ASSRGGAYAEQF | 0.6408108 |
| QLFSGAEL | ASSRGGAYAEQF | 0.2810585 |
| SAATFITL | ASSRGGAYAEQF | 0.79980695 |
| SNYGLANLGV | ASSRGGAYAEQF | 0.34336877 |
| SYGGAYAVM | ASSRGGAYAEQF | 0.6858686 |
| TTGVGGLTLG | ASSRGGAYAEQF | 0.7829541 |
| TTQTIYI | ASSRGGAYAEQF | 0.8445037 |
| VAAGAGLGAVI | ASSRGGAYAEQF | 0.33634028 |
| VAAVVTLL | ASSRGGAYAEQF | 0.73526573 |
| VGFLGGLI | ASSRGGAYAEQF | 0.5809228 |
| VGVIPILL | ASSRGGAYAEQF | 0.86051726 |
| VIFASSLI | ASSRGGAYAEQF | 0.5466602 |
| VKGGDGGAAAG | ASSRGGAYAEQF | 0.3149784 |
| VTKTFTNI | ASSRGGAYAEQF | 0.7970179 |
| VVAAAILL | ASSRGGAYAEQF | 0.8027437 |
| YAALDVEL | ASSRGGAYAEQF | 0.6109536 |
| LASVALLL | ASSRGGAYAEQF | 0.80674404 |
| LDAVHRV | ASSRGGAYAEQF | 0.682763 |
| QGLSSSIF | ASSRGGAYAEQF | 0.63578486 |
| RGTFANIRL | ASSRGGAYAEQF | 0.8087513 |
| RQLWWGHRI | ASSRGGAYAEQF | 0.74051654 |
| YGPPGTGKTLL | ASSRGGAYAEQF | 0.4733559 |
| AALVGAPANVA | ASSRGGAYAEQF | 0.81055677 |
| AATFRRTE | ASSRGGAYAEQF | 0.6802696 |
| AGALGGGI | ASSRGGAYAEQF | 0.3941427 |
| AGGAGGAGGIGAG | ASSRGGAYAEQF | 0.85428345 |
| AGSGALGL | ASSRGGAYAEQF | 0.6436324 |
| AQIADQI | ASSRGGAYAEQF | 0.7908107 |
| ASAVAVTYL | ASSRGGAYAEQF | 0.77008533 |

| | | |
|---|---|---|
| ASTVLGGTGG | ASSRGGAYAEQF | 0.85422075 |
| AVGGEPAK | ASSRGGAYAEQF | 0.39371267 |
| CEQGTFGGTC | ASSRGGAYAEQF | 0.75623137 |
| DEGKAPAG | ASSRGGAYAEQF | 0.6825681 |
| DTVVFPAL | ASSRGGAYAEQF | 0.7734717 |
| FEGITMAG | ASSRGGAYAEQF | 0.6647039 |
| FPLVVQD | ASSRGGAYAEQF | 0.643215 |
| GAFTTPQ | ASSRGGAYAEQF | 0.77581483 |
| GFGGFGTAGGVGGA | ASSRGGAYAEQF | 0.81706285 |
| GFLAQFL | ASSRGGAYAEQF | 0.63553405 |
| GGAGGAGGNAGVA | ASSRGGAYAEQF | 0.6951704 |
| GGVGTAGAGGAGGAGG | ASSRGGAYAEQF | 0.84699607 |
| GNGGGGDAGNAGSGGN | ASSRGGAYAEQF | 0.38253155 |
| GTGGAGGAGNDAGS | ASSRGGAYAEQF | 0.6620022 |
| GTTNTGWFNS | ASSRGGAYAEQF | 0.7672609 |
| IFMGPLFTL | ASSRGGAYAEQF | 0.78458035 |
| ILAHAIEAI | ASSRGGAYAEQF | 0.69677347 |
| IRSAIREAL | ASSRGGAYAEQF | 0.81031454 |
| ITFAGGG | ASSRGGAYAEQF | 0.55338115 |
| ITGMSGHQEV | ASSRGGAYAEQF | 0.7422709 |
| IVAGLACTL | ASSRGGAYAEQF | 0.6401828 |
| IVGGGLAGA | ASSRGGAYAEQF | 0.62978244 |
| IVTGVLNI | ASSRGGAYAEQF | 0.4162343 |
| LAQQGCI | ASSRGGAYAEQF | 0.70316947 |
| LCLHVEN | ASSRGGAYAEQF | 0.7860209 |
| LDAISDGS | ASSRGGAYAEQF | 0.75228065 |
| LGGFAAML | ASSRGGAYAEQF | 0.22333688 |
| LGGKNPMI | ASSRGGAYAEQF | 0.7693448 |
| LGPKGVAAIL | ASSRGGAYAEQF | 0.75433385 |
| LMVGVSI | ASSRGGAYAEQF | 0.86334 |
| LVGAAFVL | ASSRGGAYAEQF | 0.6351658 |
| LVLAVAPL | ASSRGGAYAEQF | 0.77881706 |
| MAAIAVIDAA | ASSRGGAYAEQF | 0.52612305 |
| NNGINAELQ | ASSRGGAYAEQF | 0.71492743 |
| RGEPLVLVHG | ASSRGGAYAEQF | 0.34713432 |
| RGGAGGAAGLI | ASSRGGAYAEQF | 0.47151905 |
| RSYRMHPAV | ASSRGGAYAEQF | 0.6249345 |
| SAAHLLAGQLA | ASSRGGAYAEQF | 0.49977145 |
| SAAVGSY | ASSRGGAYAEQF | 0.8125868 |
| SPHAVLHRI | ASSRGGAYAEQF | 0.7774182 |
| SQIAKVL | ASSRGGAYAEQF | 0.78630435 |
| TALGHLVP | ASSRGGAYAEQF | 0.8164874 |
| TAPTTSGAA | ASSRGGAYAEQF | 0.83515424 |
| TDGDDGNGGSPGT | ASSRGGAYAEQF | 0.81057847 |
| TDGLGEN | ASSRGGAYAEQF | 0.5743996 |
| TGGPPLPGQA | ASSRGGAYAEQF | 0.67184997 |
| TIPAITI | ASSRGGAYAEQF | 0.35070062 |
| TIPGFTI | ASSRGGAYAEQF | 0.5828762 |
| TSIASGALA | ASSRGGAYAEQF | 0.5976902 |
| TVTLPGAAI | ASSRGGAYAEQF | 0.6781562 |
| VASGLSAALP | ASSRGGAYAEQF | 0.12787783 |

| | | |
|---|---|---|
| VGAGVIGI | ASSRGGAYAEQF | 0.7527553 |
| VGPKRFGV | ASSRGGAYAEQF | 0.8330614 |
| VNVAFPD | ASSRGGAYAEQF | 0.5799635 |
| VSMTNSGVS | ASSRGGAYAEQF | 0.76927096 |
| VTDVLGDLSV | ASSRGGAYAEQF | 0.7838749 |
| VVVVGAGFAGL | ASSRGGAYAEQF | 0.54410803 |
| VYTVGTVV | ASSRGGAYAEQF | 0.7574376 |
| WTTFSGP | ASSRGGAYAEQF | 0.5079322 |
| YDELVEAF | ASSRGGAYAEQF | 0.80203485 |
| YGTGGAGGNGG | ASSRGGAYAEQF | 0.82907665 |
| YTNVMAV | ASSRGGAYAEQF | 0.7336863 |
| FLEQVHQL | ASSRGGAYAEQF | 0.5960331 |
| GLSLLAVAV | ASSRGGAYAEQF | 0.69826233 |
| LEQVHQL | ASSRGGAYAEQF | 0.3294622 |
| LNAAGAI | ASSRGGAYAEQF | 0.62016076 |
| PAVHASL | ASSRGGAYAEQF | 0.15511796 |
| TELEDDA | ASSRGGAYAEQF | 0.707815 |
| TPPGPPN | ASSRGGAYAEQF | 0.46255416 |
| TPPPPPPP | ASSRGGAYAEQF | 0.678877 |
| TQSPALI | ASSRLGVYAEQF | 0.66166025 |
| EFAAAFPQ | ASSRLGVYAEQF | 0.6832443 |
| LLSGNLDVL | ASSRLGVYAEQF | 0.43395343 |
| GPANLGNYNI | ASSRLGVYAEQF | 0.76862144 |
| TLGPTLP | ASSRLGVYAEQF | 0.6915261 |
| VYAYSSCCK | ASSRLGVYAEQF | 0.30146605 |
| APAPAPAPNI | ASSRLGVYAEQF | 0.8039136 |
| PGGPVPIP | ASSRLGVYAEQF | 0.5790386 |
| FLTVALEP | ASSRLGVYAEQF | 0.68424 |
| SQYFGANL | ASSRLGVYAEQF | 0.79848576 |
| GGVKPAA | ASSRLGVYAEQF | 0.6695837 |
| GGTGGAGGAGGW | ASSRLGVYAEQF | 0.33866963 |
| GVGGVAGLGGAGG | ASSRLGVYAEQF | 0.6324665 |
| VLLVLATTI | ASSRLGVYAEQF | 0.6808832 |
| IGIFAAML | ASSRLGVYAEQF | 0.5903909 |
| AVGDLID | ASSRLGVYAEQF | 0.14705348 |
| NISPLPL | ASSRLGVYAEQF | 0.77689517 |
| VTAVFVAI | ASSRLGVYAEQF | 0.3982346 |
| SGDDNDAAL | ASSRLGVYAEQF | 0.5398828 |
| AVAALAHS | ASSRLGVYAEQF | 0.62973195 |
| GGVTVMATP | ASSRLGVYAEQF | 0.8069868 |
| AGLAANVRD | ASSRLGVYAEQF | 0.64321893 |
| TNRPDLIDPALLRPG | ASSRLGVYAEQF | 0.60514975 |
| LANLGVTAL | ASSRLGVYAEQF | 0.73618287 |
| TGERIEL | ASSRLGVYAEQF | 0.6474538 |
| NGEELAA | ASSRLGVYAEQF | 0.8125976 |
| VLSLVWG | ASSRLGVYAEQF | 0.7249402 |
| IGYALWL | ASSRLGVYAEQF | 0.65947545 |
| LSLLLAPL | ASSRLGVYAEQF | 0.7521386 |
| TLDEALK | ASSRLGVYAEQF | 0.5975554 |
| TDGAGTAEGG | ASSRLGVYAEQF | 0.38311628 |
| NAPGGAGGNGG | ASSRLGVYAEQF | 0.41191852 |

| | | |
|---|---|---|
| VSVFMGMGGNF | ASSRLGVYAEQF | 0.54056406 |
| LASYLAAL | ASSRLGVYAEQF | 0.31628466 |
| LVGGTPAL | ASSRLGVYAEQF | 0.7201803 |
| VRNADVL | ASSRLGVYAEQF | 0.8279165 |
| TCSSPFLVSAM | ASSRLGVYAEQF | 0.68854725 |
| VHITDAE | ASSRLGVYAEQF | 0.28304458 |
| EYAAPSF | ASSRLGVYAEQF | 0.5493536 |
| VVDAFHGGL | ASSRLGVYAEQF | 0.6369598 |
| ASVVDPQ | ASSRLGVYAEQF | 0.7777778 |
| SYNEFGGLS | ASSRLGVYAEQF | 0.80075073 |
| DPDGSVY | ASSRLGVYAEQF | 0.84759915 |
| MVSTIHATE | ASSRLGVYAEQF | 0.7789055 |
| LAAGVADPV | ASSRLGVYAEQF | 0.5624138 |
| TKEQIAA | ASSRLGVYAEQF | 0.796434 |
| SLASPAAL | ASSRLGVYAEQF | 0.6342023 |
| IGLVFYGV | ASSRLGVYAEQF | 0.7057252 |
| FLLDASNL | ASSRLGVYAEQF | 0.67594373 |
| YQVGNFP | ASSRLGVYAEQF | 0.7303829 |
| ESLLSQST | ASSRLGVYAEQF | 0.5211042 |
| LFTVGFAAVG | ASSRLGVYAEQF | 0.5489891 |
| ATPPTPT | ASSRLGVYAEQF | 0.31247687 |
| TDLATLG | ASSRLGVYAEQF | 0.7735791 |
| SAGGTGR | ASSRLGVYAEQF | 0.49863002 |
| GFSNSGS | ASSRLGVYAEQF | 0.39273745 |
| GGNGGSGGTSV | ASSRLGVYAEQF | 0.5715106 |
| GTSVATGGAGN | ASSRLGVYAEQF | 0.8472617 |
| NGGNGGSGGTS | ASSRLGVYAEQF | 0.82188237 |
| AVVKFPH | ASSRLGVYAEQF | 0.50898075 |
| TPPPPPP | ASSRLGVYAEQF | 0.47881973 |
| AAPAAAAAPA | ASSRLGVYAEQF | 0.7664552 |
| IALNEDL | ASSRLGVYAEQF | 0.46932954 |
| IVNVSSM | ASSRLGVYAEQF | 0.78858006 |
| TGIPPAPR | ASSRLGVYAEQF | 0.6895374 |
| FDLGGGTF | ASSRLGVYAEQF | 0.6592236 |
| FDLGGGTFDVS | ASSRLGVYAEQF | 0.5151327 |
| NEPTAAA | ASSRLGVYAEQF | 0.7607057 |
| NEPTAAALA | ASSRLGVYAEQF | 0.5955295 |
| GSIINISSI | ASSRLGVYAEQF | 0.74197316 |
| VAPGNAGTA | ASSRLGVYAEQF | 0.17772502 |
| TGGGASLE | ASSRLGVYAEQF | 0.7936535 |
| VAVYNHYKRI | ASSRLGVYAEQF | 0.42623547 |
| TDPAPATT | ASSRLGVYAEQF | 0.48814037 |
| TNRPDILDPALLRPG | ASSRLGVYAEQF | 0.19695896 |
| LTTTTTS | ASSRLGVYAEQF | 0.6565784 |
| PTGSGKTL | ASSRLGVYAEQF | 0.7757188 |
| TGSGKTL | ASSRLGVYAEQF | 0.82148516 |
| TIGHVDHGK | ASSRLGVYAEQF | 0.6291388 |
| TIGHVDHGKTT | ASSRLGVYAEQF | 0.76382947 |
| GHVDHGKTT | ASSRLGVYAEQF | 0.79803735 |
| AQFEHTLL | ASSRLGVYAEQF | 0.57078445 |
| IVNQDEL | ASSRLGVYAEQF | 0.21352136 |

| | | |
|---|---|---|
| TNGSQFF | ASSRLGVYAEQF | 0.73184955 |
| TGKGIVE | ASSRLGVYAEQF | 0.7047268 |
| LISNASDA | ASSRLGVYAEQF | 0.81649077 |
| AELRTLGELTKIAKAHGAQ | ASSRLGVYAEQF | 0.7630998 |
| APNAVAAAEPSW | ASSRLGVYAEQF | 0.33636898 |
| APQVAGAATAVL | ASSRLGVYAEQF | 0.7861724 |
| ARRDPAAAAAILPTDA | ASSRLGVYAEQF | 0.74626213 |
| DAAARDVLAEAGLAEYF | ASSRLGVYAEQF | 0.34287083 |
| DDQIVLSAKANLAKVEQ | ASSRLGVYAEQF | 0.5147768 |
| DELLPDTGEDSDEDGH | ASSRLGVYAEQF | 0.53685415 |
| ELPEAPANRATVPQD | ASSRLGVYAEQF | 0.3296312 |
| FGQSKIERVTGIG | ASSRLGVYAEQF | 0.68632174 |
| GAGGTGSTAGGAGGAGGAGGLY | ASSRLGVYAEQF | 0.5872302 |
| GHGGTGAAGGNGGTGGAGGSGI | ASSRLGVYAEQF | 0.52644277 |
| GNGGNGGAGGEGGPGGAGGAGA | ASSRLGVYAEQF | 0.6527872 |
| GSTAGNGGNGGNATNGTIAGQP | ASSRLGVYAEQF | 0.46772295 |
| GTGGDGGNGGNGGTGG | ASSRLGVYAEQF | 0.5193686 |
| GVRRPAGIPADAPV | ASSRLGVYAEQF | 0.58393365 |
| GYRQVTDAHLLTLARR | ASSRLGVYAEQF | 0.75583637 |
| IAKMGQDGHDRG | ASSRLGVYAEQF | 0.78017855 |
| KAEPSIEIQYT | ASSRLGVYAEQF | 0.7115091 |
| KPVRIRMAADEDTVAVT | ASSRLGVYAEQF | 0.7511002 |
| MPPRIAGMRLLVIKPEP | ASSRLGVYAEQF | 0.8266939 |
| MKFYAVALFRSRGVPPTR | ASSRLGVYAEQF | 0.314525 |
| NAGGFGILGAVAHSP | ASSRLGVYAEQF | 0.43561512 |
| NEGREITLAMAEM | ASSRLGVYAEQF | 0.77550006 |
| NELVEWSHPLGHRWRWE | ASSRLGVYAEQF | 0.58337307 |
| NLGAGGTGGNGGNPG | ASSRLGVYAEQF | 0.8061131 |
| NTEWPKWETVMAADPVD | ASSRLGVYAEQF | 0.69290495 |
| PASAAPAETQP | ASSRLGVYAEQF | 0.7399804 |
| QQSSDGKNGVRAAEITGE | ASSRLGVYAEQF | 0.6903639 |
| RAPDAVTTSWWKE | ASSRLGVYAEQF | 0.840403 |
| SGFGNVGAHTSGWFNQ | ASSRLGVYAEQF | 0.76592 |
| SGVAGAVGQAASVGGLKV | ASSRLGVYAEQF | 0.64202094 |
| SNGGAGTGSAGGNGGT | ASSRLGVYAEQF | 0.80088836 |
| TGGAGGEGGAAG | ASSRLGVYAEQF | 0.38089594 |
| TGGDGGAAGAGGAPGHG | ASSRLGVYAEQF | 0.7729054 |
| TMPPLLAAIANAGRHGV | ASSRLGVYAEQF | 0.7822633 |
| TSSTAGNSGGA | ASSRLGVYAEQF | 0.4535915 |
| VPATVPVGILC | ASSRLGVYAEQF | 0.6324245 |
| VQRPDSGGVVLTGRIS | ASSRLGVYAEQF | 0.820359 |
| WTPNPVPATKGVPAGLVQAT | ASSRLGVYAEQF | 0.8124589 |
| YALKRALAKFSDRFDVVII | ASSRLGVYAEQF | 0.394935 |
| YLIGSVIV | ASSRLGVYAEQF | 0.57111657 |
| YGQLVAVID | ASSRLGVYAEQF | 0.367362 |
| VTVGGMTLPA | ASSRLGVYAEQF | 0.49583492 |
| VLALPAV | ASSRLGVYAEQF | 0.29374647 |
| VIPPTLNYET | ASSRLGVYAEQF | 0.7631556 |
| VGMGGMHPSSG | ASSRLGVYAEQF | 0.5495863 |
| VGLAGGAATA | ASSRLGVYAEQF | 0.8169118 |
| VGGAGGAGGSGP | ASSRLGVYAEQF | 0.72883224 |

| | | |
|---|---|---|
| VFAAFMV | ASSRLGVYAEQF | 0.8165523 |
| TLYDKLR | ASSRLGVYAEQF | 0.5163468 |
| TLAVFTVL | ASSRLGVYAEQF | 0.58104265 |
| TGSVQPARV | ASSRLGVYAEQF | 0.2702226 |
| TGGGAGSGGT | ASSRLGVYAEQF | 0.34339193 |
| SHLETSL | ASSRLGVYAEQF | 0.42329544 |
| SFTTGMT | ASSRLGVYAEQF | 0.6285732 |
| QVEVAKRMGVRQPTVS | ASSRLGVYAEQF | 0.25045472 |
| QGSGGPALAM | ASSRLGVYAEQF | 0.8521261 |
| MLAPSNP | ASSRLGVYAEQF | 0.624447 |
| LVNNAAI | ASSRLGVYAEQF | 0.76734495 |
| LTGEINTLA | ASSRLGVYAEQF | 0.514505 |
| LSIFGPV | ASSRLGVYAEQF | 0.4085319 |
| LLALLANL | ASSRLGVYAEQF | 0.58399326 |
| LISNASDALDKLR | ASSRLGVYAEQF | 0.76091 |
| LINEAVP | ASSRLGVYAEQF | 0.39721107 |
| LGYAFSGL | ASSRLGVYAEQF | 0.22211719 |
| LDFVAVAL | ASSRLGVYAEQF | 0.47011226 |
| LAQPGAE | ASSRLGVYAEQF | 0.4848534 |
| ISNLGAVTGF | ASSRLGVYAEQF | 0.8520482 |
| HQEDPGRI | ASSRLGVYAEQF | 0.80807316 |
| GTPGEPTP | ASSRLGVYAEQF | 0.70301104 |
| GFFNSGS | ASSRLGVYAEQF | 0.85861045 |
| GAGGGGLTAGAN | ASSRLGVYAEQF | 0.14216712 |
| FVDALVNM | ASSRLGVYAEQF | 0.5858823 |
| FLTVAPEL | ASSRLGVYAEQF | 0.83620447 |
| FETLTAL | ASSRLGVYAEQF | 0.7309369 |
| DVINPIEL | ASSRLGVYAEQF | 0.60685384 |
| APVSGAPV | ASSRLGVYAEQF | 0.8144461 |
| ALVGGLAW | ASSRLGVYAEQF | 0.4625267 |
| ALAAAAATAI | ASSRLGVYAEQF | 0.5787195 |
| AFGLGLLL | ASSRLGVYAEQF | 0.6625874 |
| AFFAAASL | ASSRLGVYAEQF | 0.75515896 |
| AAAALPAFSPP | ASSRLGVYAEQF | 0.7020223 |
| VVLAPAL | ASSRLGVYAEQF | 0.30512747 |
| TIGHVDHGKTTL | ASSRLGVYAEQF | 0.7566848 |
| LISNASDALDK | ASSRLGVYAEQF | 0.8532797 |
| LISNASDAL | ASSRLGVYAEQF | 0.5699109 |
| ISNASDALDK | ASSRLGVYAEQF | 0.7936747 |
| NQAETLVYQTE | ASSRLGVYAEQF | 0.7230228 |
| TGGTGGTAGAG | ASSRLGVYAEQF | 0.69316614 |
| TGSTAGGAGGA | ASSRLGVYAEQF | 0.5123468 |
| DPLAVPVAAGPV | ASSRLGVYAEQF | 0.39311013 |
| GGTGGTPGNGGN | ASSRLGVYAEQF | 0.75650984 |
| PAGLAGVDPATL | ASSRLGVYAEQF | 0.4379813 |
| GNNLVVEAPHAGASV | ASSRLGVYAEQF | 0.6608827 |
| KVTRIMSASTLEQPA | ASSRLGVYAEQF | 0.6811748 |
| LGFALPIALPRRTVV | ASSRLGVYAEQF | 0.8635436 |
| TRLGNLTARKLVVLV | ASSRLGVYAEQF | 0.76081777 |
| WSQQRRLIYFPSAGP | ASSRLGVYAEQF | 0.75743604 |
| GPNGNPDPMAAAVDIR | ASSRLGVYAEQF | 0.45195168 |

| | | |
|---|---|---|
| TQRCVSVADLRAAASR | ASSRLGVYAEQF | 0.6902914 |
| QFVLDVAEHPVEVLTETV | ASSRLGVYAEQF | 0.7643357 |
| GGAGGTGGAGGNGGTQAGDG | ASSRLGVYAEQF | 0.3413059 |
| PLLATLRAIAVDVDAKPAQV | ASSRLGVYAEQF | 0.81549466 |
| APKFRTGGYMIDFWGVGYQVAK | ASSRLGVYAEQF | 0.39231235 |
| GAGGAGGAGGAAGGPGGTGGTG | ASSRLGVYAEQF | 0.72527385 |
| IGALGLLATDVAPAGSVAFGVA | ASSRLGVYAEQF | 0.7697858 |
| AGGAGGIGGNGGAGGAGGVLMG | ASSRLGVYAEQF | 0.82673264 |
| ADELIGLL | ASSRLGVYAEQF | 0.66560507 |
| ADRALKA | ASSRLGVYAEQF | 0.75577456 |
| AGAVGQVGGAGGA | ASSRLGVYAEQF | 0.4133798 |
| AGGGGGLGGAP | ASSRLGVYAEQF | 0.7966025 |
| AGGLLYGNGGAG | ASSRLGVYAEQF | 0.8127882 |
| AGLESIGL | ASSRLGVYAEQF | 0.7102454 |
| AGLKTGII | ASSRLGVYAEQF | 0.85321623 |
| ASLAALLL | ASSRLGVYAEQF | 0.74818635 |
| ASVINISSI | ASSRLGVYAEQF | 0.23990242 |
| ATPPGPA | ASSRLGVYAEQF | 0.62435925 |
| ATVASALV | ASSRLGVYAEQF | 0.8195144 |
| ATVASLIL | ASSRLGVYAEQF | 0.82265604 |
| AVNGPVLIRH | ASSRLGVYAEQF | 0.8073381 |
| DGAAGAVGI | ASSRLGVYAEQF | 0.7382995 |
| DPLLHGI | ASSRLGVYAEQF | 0.5600227 |
| ELGAALGAL | ASSRLGVYAEQF | 0.8350384 |
| ELGRTQAV | ASSRLGVYAEQF | 0.87038195 |
| FVAGGYLS | ASSRLGVYAEQF | 0.78488356 |
| GASGGAGGAGGW | ASSRLGVYAEQF | 0.47906145 |
| GGAGAGGDGGAGGI | ASSRLGVYAEQF | 0.49923372 |
| GTGHNGSQPGMGGNG | ASSRLGVYAEQF | 0.7544869 |
| GYAAASASAS | ASSRLGVYAEQF | 0.81809855 |
| HQVDVAAVQQG | ASSRLGVYAEQF | 0.37584022 |
| IGAAIAHT | ASSRLGVYAEQF | 0.6927073 |
| IGMIEVAQL | ASSRLGVYAEQF | 0.70505524 |
| IRLAPEHTSAMD | ASSRLGVYAEQF | 0.20837937 |
| IYNVSVL | ASSRLGVYAEQF | 0.7637133 |
| KQGFLGM | ASSRLGVYAEQF | 0.8127405 |
| LESVITQL | ASSRLGVYAEQF | 0.75354624 |
| LGLGALVVV | ASSRLGVYAEQF | 0.38387248 |
| LIGNGGNGGSGGI | ASSRLGVYAEQF | 0.8154073 |
| LNAINAPTL | ASSRLGVYAEQF | 0.8312396 |
| LQLLNTGV | ASSRLGVYAEQF | 0.85220134 |
| LQTSIEF | ASSRLGVYAEQF | 0.25366327 |
| MSFAVSEL | ASSRLGVYAEQF | 0.16194932 |
| NTGALGL | ASSRLGVYAEQF | 0.65379167 |
| PPGAGGAGGAGTT | ASSRLGVYAEQF | 0.80653954 |
| QLVGLGI | ASSRLGVYAEQF | 0.5553751 |
| RFLEQVHQL | ASSRLGVYAEQF | 0.46828353 |
| RGVGYVLR | ASSRLGVYAEQF | 0.7399218 |
| SATGVFAGVI | ASSRLGVYAEQF | 0.8199494 |
| SLPAITLP | ASSRLGVYAEQF | 0.35176355 |
| STALAFTV | ASSRLGVYAEQF | 0.36841837 |

| | | |
|---|---|---|
| SVLTVGLGK | ASSRLGVYAEQF | 0.7945837 |
| TALAGAGSGI | ASSRLGVYAEQF | 0.813061 |
| TGLPFTGI | ASSRLGVYAEQF | 0.41248927 |
| VAEHVTV | ASSRLGVYAEQF | 0.5785263 |
| VDIVLNSL | ASSRLGVYAEQF | 0.67864406 |
| VLTGGGAVL | ASSRLGVYAEQF | 0.7514627 |
| VNVGLAYGI | ASSRLGVYAEQF | 0.70803463 |
| VSSAAVGAV | ASSRLGVYAEQF | 0.34751123 |
| VTSVYEAGV | ASSRLGVYAEQF | 0.7973249 |
| WVALDRGA | ASSRLGVYAEQF | 0.35882407 |
| YLVGSVVI | ASSRLGVYAEQF | 0.7680522 |
| GPPGTGKTLL | ASSRLGVYAEQF | 0.64169693 |
| GVGMTKF | ASSRLGVYAEQF | 0.71746016 |
| IGIGGSDL | ASSRLGVYAEQF | 0.62423563 |
| IGTLLAEL | ASSRLGVYAEQF | 0.3960563 |
| IYTSGTT | ASSRLGVYAEQF | 0.39416468 |
| LSFPAGV | ASSRLGVYAEQF | 0.3410583 |
| LYTSGTT | ASSRLGVYAEQF | 0.7877136 |
| VGAFPFGF | ASSRLGVYAEQF | 0.42665395 |
| VGAGTVEF | ASSRLGVYAEQF | 0.7861809 |
| VGKSTLLL | ASSRLGVYAEQF | 0.68873954 |
| AALAHAGGGIV | ASSRLGVYAEQF | 0.42705554 |
| AALVAVSLL | ASSRLGVYAEQF | 0.74752986 |
| AAVSRAVAAL | ASSRLGVYAEQF | 0.3348651 |
| AGGSSGIGGF | ASSRLGVYAEQF | 0.67166 |
| AGLTVANANG | ASSRLGVYAEQF | 0.73164463 |
| AGLVGAVL | ASSRLGVYAEQF | 0.790069 |
| ALSQAFTV | ASSRLGVYAEQF | 0.46920592 |
| ALTFSVSL | ASSRLGVYAEQF | 0.7716073 |
| ALTGAALVAL | ASSRLGVYAEQF | 0.7460103 |
| ASFSIGAGVL | ASSRLGVYAEQF | 0.44224763 |
| DGSVYKI | ASSRLGVYAEQF | 0.73583746 |
| EQLAEIV | ASSRLGVYAEQF | 0.80362713 |
| FAVSELTVM | ASSRLGVYAEQF | 0.7374212 |
| GAAPLAAGL | ASSRLGVYAEQF | 0.30830812 |
| GADGGDGGSGGAGG | ASSRLGVYAEQF | 0.7137338 |
| GFGLDTGGAGG | ASSRLGVYAEQF | 0.5292526 |
| GLASGGLL | ASSRLGVYAEQF | 0.7228495 |
| GRIAGAAVLL | ASSRLGVYAEQF | 0.68711555 |
| GSAGSLNTSF | ASSRLGVYAEQF | 0.5892808 |
| GTGQIGI | ASSRLGVYAEQF | 0.7677494 |
| GTLIALIL | ASSRLGVYAEQF | 0.4329239 |
| IAATMGFVVG | ASSRLGVYAEQF | 0.49711108 |
| IFAQALL | ASSRLGVYAEQF | 0.58728313 |
| IGAAVALGF | ASSRLGVYAEQF | 0.6579135 |
| IGLTPLAL | ASSRLGVYAEQF | 0.7861507 |
| ILGSNTSTL | ASSRLGVYAEQF | 0.41217986 |
| IVFSLTVL | ASSRLGVYAEQF | 0.685413 |
| LASKATGF | ASSRLGVYAEQF | 0.59293604 |
| LASVALIL | ASSRLGVYAEQF | 0.4155122 |
| LFAVGLTL | ASSRLGVYAEQF | 0.751909 |

| | | |
|---|---|---|
| LFNGGIGTYI | ASSRLGVYAEQF | 0.596291 |
| LTLGAGII | ASSRLGVYAEQF | 0.3195327 |
| LVAAALGDI | ASSRLGVYAEQF | 0.7626474 |
| NTAFVQLGI | ASSRLGVYAEQF | 0.4349422 |
| QLFSGAEL | ASSRLGVYAEQF | 0.6373229 |
| SAATFITL | ASSRLGVYAEQF | 0.8374709 |
| SNYGLANLGV | ASSRLGVYAEQF | 0.3106784 |
| SYGGAYAVM | ASSRLGVYAEQF | 0.7523936 |
| TTGVGGLTLG | ASSRLGVYAEQF | 0.7193903 |
| TTQTIYI | ASSRLGVYAEQF | 0.6967634 |
| VAAGAGLGAVI | ASSRLGVYAEQF | 0.6356677 |
| VAAVVTLL | ASSRLGVYAEQF | 0.46521994 |
| VGFLGGLI | ASSRLGVYAEQF | 0.8442867 |
| VGVIPILL | ASSRLGVYAEQF | 0.718431 |
| VIFASSLI | ASSRLGVYAEQF | 0.7699149 |
| VKGGDGGAAAG | ASSRLGVYAEQF | 0.7720635 |
| VTKTFTNI | ASSRLGVYAEQF | 0.7933546 |
| VVAAAILL | ASSRLGVYAEQF | 0.7913652 |
| YAALDVEL | ASSRLGVYAEQF | 0.47644892 |
| LASVALLL | ASSRLGVYAEQF | 0.26708683 |
| LDAVHRV | ASSRLGVYAEQF | 0.59895504 |
| QGLSSSIF | ASSRLGVYAEQF | 0.64571905 |
| RGTFANIRL | ASSRLGVYAEQF | 0.5062983 |
| RQLWWGHRI | ASSRLGVYAEQF | 0.29050842 |
| YGPPGTGKTLL | ASSRLGVYAEQF | 0.80082595 |
| AALVGAPANVA | ASSRLGVYAEQF | 0.85654813 |
| AATFRRTE | ASSRLGVYAEQF | 0.42774874 |
| AGALGGGI | ASSRLGVYAEQF | 0.86125296 |
| AGGAGGAGGIGAG | ASSRLGVYAEQF | 0.3940953 |
| AGSGALGL | ASSRLGVYAEQF | 0.73086596 |
| AQIADQI | ASSRLGVYAEQF | 0.8122947 |
| ASAVAVTYL | ASSRLGVYAEQF | 0.36976576 |
| ASTVLGGTGG | ASSRLGVYAEQF | 0.7476647 |
| AVGGEPAK | ASSRLGVYAEQF | 0.5023573 |
| CEQGTFGGTC | ASSRLGVYAEQF | 0.47649947 |
| DEGKAPAG | ASSRLGVYAEQF | 0.80395013 |
| DTVVFPAL | ASSRLGVYAEQF | 0.5415515 |
| FEGITMAG | ASSRLGVYAEQF | 0.74199605 |
| FPLVVQD | ASSRLGVYAEQF | 0.6186451 |
| GAFTTPQ | ASSRLGVYAEQF | 0.47656476 |
| GFGGFGTAGGVGGA | ASSRLGVYAEQF | 0.7323157 |
| GFLAQFL | ASSRLGVYAEQF | 0.7909403 |
| GGAGGAGGNAGVA | ASSRLGVYAEQF | 0.673985 |
| GGVGTAGAGGAGGAGG | ASSRLGVYAEQF | 0.86897254 |
| GNGGGGDAGNAGSGGN | ASSRLGVYAEQF | 0.39878154 |
| GTGGAGGAGNDAGS | ASSRLGVYAEQF | 0.8513506 |
| GTTNTGWFNS | ASSRLGVYAEQF | 0.8383699 |
| IFMGPLFTL | ASSRLGVYAEQF | 0.709502 |
| ILAHAIEAI | ASSRLGVYAEQF | 0.58801705 |
| IRSAIREAL | ASSRLGVYAEQF | 0.79260933 |
| ITFAGGG | ASSRLGVYAEQF | 0.55939925 |

| | | |
|---|---|---|
| ITGMSGHQEV | ASSRLGVYAEQF | 0.72246575 |
| IVAGLACTL | ASSRLGVYAEQF | 0.824206 |
| IVGGGLAGA | ASSRLGVYAEQF | 0.3569863 |
| IVTGVLNI | ASSRLGVYAEQF | 0.8075866 |
| LAQQGCI | ASSRLGVYAEQF | 0.6556459 |
| LCLHVEN | ASSRLGVYAEQF | 0.61697227 |
| LDAISDGS | ASSRLGVYAEQF | 0.5324364 |
| LGGFAAML | ASSRLGVYAEQF | 0.26956925 |
| LGGKNPMI | ASSRLGVYAEQF | 0.5021823 |
| LGPKGVAAIL | ASSRLGVYAEQF | 0.4661922 |
| LMVGVSI | ASSRLGVYAEQF | 0.8049747 |
| LVGAAFVL | ASSRLGVYAEQF | 0.68917894 |
| LVLAVAPL | ASSRLGVYAEQF | 0.5157124 |
| MAAIAVIDAA | ASSRLGVYAEQF | 0.83123 |
| NNGINAELQ | ASSRLGVYAEQF | 0.80307513 |
| RGEPLVLVHG | ASSRLGVYAEQF | 0.31897113 |
| RGGAGGAAGLI | ASSRLGVYAEQF | 0.60435444 |
| RSYRMHPAV | ASSRLGVYAEQF | 0.32575127 |
| SAAHLLAGQLA | ASSRLGVYAEQF | 0.5446006 |
| SAAVGSY | ASSRLGVYAEQF | 0.13894667 |
| SPHAVLHRI | ASSRLGVYAEQF | 0.70472735 |
| SQIAKVL | ASSRLGVYAEQF | 0.61478436 |
| TALGHLVP | ASSRLGVYAEQF | 0.1387671 |
| TAPTTSGAA | ASSRLGVYAEQF | 0.5916255 |
| TDGDDGNGGSPGT | ASSRLGVYAEQF | 0.5763098 |
| TDGLGEN | ASSRLGVYAEQF | 0.65123934 |
| TGGPPLPGQA | ASSRLGVYAEQF | 0.78638995 |
| TIPAITI | ASSRLGVYAEQF | 0.31728265 |
| TIPGFTI | ASSRLGVYAEQF | 0.45650858 |
| TSIASGALA | ASSRLGVYAEQF | 0.774917 |
| TVTLPGAAI | ASSRLGVYAEQF | 0.8526889 |
| VASGLSAALP | ASSRLGVYAEQF | 0.38606554 |
| VGAGVIGI | ASSRLGVYAEQF | 0.6942169 |
| VGPKRFGV | ASSRLGVYAEQF | 0.82964635 |
| VNVAFPD | ASSRLGVYAEQF | 0.655198 |
| VSMTNSGVS | ASSRLGVYAEQF | 0.7177403 |
| VTDVLGDLSV | ASSRLGVYAEQF | 0.7676314 |
| VVVVGAGFAGL | ASSRLGVYAEQF | 0.36425 |
| VYTVGTVV | ASSRLGVYAEQF | 0.82766545 |
| WTTFSGP | ASSRLGVYAEQF | 0.81575847 |
| YDELVEAF | ASSRLGVYAEQF | 0.6064424 |
| YGTGGAGGNGG | ASSRLGVYAEQF | 0.59537977 |
| YTNVMAV | ASSRLGVYAEQF | 0.33623266 |
| FLEQVHQL | ASSRLGVYAEQF | 0.4719732 |
| GLSLLAVAV | ASSRLGVYAEQF | 0.25156304 |
| LEQVHQL | ASSRLGVYAEQF | 0.71038043 |
| LNAAGAI | ASSRLGVYAEQF | 0.7899465 |
| PAVHASL | ASSRLGVYAEQF | 0.6226382 |
| TELEDDA | ASSRLGVYAEQF | 0.28910542 |
| TPPGPPN | ASSRLGVYAEQF | 0.3614493 |
| TPPPPPPP | ASSRLGVYAEQF | 0.5987853 |

| | | |
|---|---|---|
| TQSPALI | ASSLDLDRANTEVF | 0.46015888 |
| EFAAAFPQ | ASSLDLDRANTEVF | 0.25300828 |
| LLSGNLDVL | ASSLDLDRANTEVF | 0.31257924 |
| GPANLGNYNI | ASSLDLDRANTEVF | 0.8151868 |
| TLGPTLP | ASSLDLDRANTEVF | 0.8240795 |
| VYAYSSCCK | ASSLDLDRANTEVF | 0.36654156 |
| APAPAPAPNI | ASSLDLDRANTEVF | 0.8105626 |
| PGGPVPIP | ASSLDLDRANTEVF | 0.7887756 |
| FLTVALEP | ASSLDLDRANTEVF | 0.74884355 |
| SQYFGANL | ASSLDLDRANTEVF | 0.5328476 |
| GGVKPAA | ASSLDLDRANTEVF | 0.31586727 |
| GGTGGAGGAGGW | ASSLDLDRANTEVF | 0.83227694 |
| GVGGVAGLGGAGG | ASSLDLDRANTEVF | 0.64772755 |
| VLLVLATTI | ASSLDLDRANTEVF | 0.5693386 |
| IGIFAAML | ASSLDLDRANTEVF | 0.7399119 |
| AVGDLID | ASSLDLDRANTEVF | 0.6055681 |
| NISPLPL | ASSLDLDRANTEVF | 0.50449795 |
| VTAVFVAI | ASSLDLDRANTEVF | 0.6677233 |
| SGDDNDAAL | ASSLDLDRANTEVF | 0.63949317 |
| AVAALAHS | ASSLDLDRANTEVF | 0.34944326 |
| GGVTVMATP | ASSLDLDRANTEVF | 0.39341325 |
| AGLAANVRD | ASSLDLDRANTEVF | 0.55171597 |
| TNRPDLIDPALLRPG | ASSLDLDRANTEVF | 0.68890965 |
| LANLGVTAL | ASSLDLDRANTEVF | 0.83915573 |
| TGERIEL | ASSLDLDRANTEVF | 0.5737264 |
| NGEELAA | ASSLDLDRANTEVF | 0.44757593 |
| VLSLVWG | ASSLDLDRANTEVF | 0.7979038 |
| IGYALWL | ASSLDLDRANTEVF | 0.66671187 |
| LSLLLAPL | ASSLDLDRANTEVF | 0.5717545 |
| TLDEALK | ASSLDLDRANTEVF | 0.6242573 |
| TDGAGTAEGG | ASSLDLDRANTEVF | 0.5366364 |
| NAPGGAGGNGG | ASSLDLDRANTEVF | 0.6434814 |
| VSVFMGMGGNF | ASSLDLDRANTEVF | 0.6699939 |
| LASYLAAL | ASSLDLDRANTEVF | 0.63018745 |
| LVGGTPAL | ASSLDLDRANTEVF | 0.66956925 |
| VRNADVL | ASSLDLDRANTEVF | 0.69991803 |
| TCSSPFLVSAM | ASSLDLDRANTEVF | 0.5539925 |
| VHITDAE | ASSLDLDRANTEVF | 0.24570312 |
| EYAAPSF | ASSLDLDRANTEVF | 0.38808736 |
| VVDAFHGGL | ASSLDLDRANTEVF | 0.6826198 |
| ASVVDPQ | ASSLDLDRANTEVF | 0.6807125 |
| SYNEFGGLS | ASSLDLDRANTEVF | 0.7430443 |
| DPDGSVY | ASSLDLDRANTEVF | 0.6207972 |
| MVSTIHATE | ASSLDLDRANTEVF | 0.51818573 |
| LAAGVADPV | ASSLDLDRANTEVF | 0.63987404 |
| TKEQIAA | ASSLDLDRANTEVF | 0.8191662 |
| SLASPAAL | ASSLDLDRANTEVF | 0.2599417 |
| IGLVFYGV | ASSLDLDRANTEVF | 0.55585784 |
| FLLDASNL | ASSLDLDRANTEVF | 0.84102964 |
| YQVGNFP | ASSLDLDRANTEVF | 0.7729229 |
| ESLLSQST | ASSLDLDRANTEVF | 0.62573075 |

| | | |
|---|---|---|
| LFTVGFAAVG | ASSLDLDRANTEVF | 0.4637819 |
| ATPPTPT | ASSLDLDRANTEVF | 0.86384153 |
| TDLATLG | ASSLDLDRANTEVF | 0.6388993 |
| SAGGTGR | ASSLDLDRANTEVF | 0.8251201 |
| GFSNSGS | ASSLDLDRANTEVF | 0.18761238 |
| GGNGGSGGTSV | ASSLDLDRANTEVF | 0.46760023 |
| GTSVATGGAGN | ASSLDLDRANTEVF | 0.31792337 |
| NGGNGGSGGTS | ASSLDLDRANTEVF | 0.20223215 |
| AVVKFPH | ASSLDLDRANTEVF | 0.7482526 |
| TPPPPPP | ASSLDLDRANTEVF | 0.49227512 |
| AAPAAAAAPA | ASSLDLDRANTEVF | 0.77448475 |
| IALNEDL | ASSLDLDRANTEVF | 0.53739357 |
| IVNVSSM | ASSLDLDRANTEVF | 0.7412692 |
| TGIPPAPR | ASSLDLDRANTEVF | 0.5686325 |
| FDLGGGTF | ASSLDLDRANTEVF | 0.58019507 |
| FDLGGGTFDVS | ASSLDLDRANTEVF | 0.38777232 |
| NEPTAAA | ASSLDLDRANTEVF | 0.743176 |
| NEPTAAALA | ASSLDLDRANTEVF | 0.7123414 |
| GSIINISSI | ASSLDLDRANTEVF | 0.56956476 |
| VAPGNAGTA | ASSLDLDRANTEVF | 0.288913 |
| TGGGASLE | ASSLDLDRANTEVF | 0.59832263 |
| VAVYNHYKRI | ASSLDLDRANTEVF | 0.66053504 |
| TDPAPATT | ASSLDLDRANTEVF | 0.20049345 |
| TNRPDILDPALLRPG | ASSLDLDRANTEVF | 0.19207917 |
| LTTTTTS | ASSLDLDRANTEVF | 0.5502068 |
| PTGSGKTL | ASSLDLDRANTEVF | 0.41710645 |
| TGSGKTL | ASSLDLDRANTEVF | 0.8094239 |
| TIGHVDHGK | ASSLDLDRANTEVF | 0.7967794 |
| TIGHVDHGKTT | ASSLDLDRANTEVF | 0.5992677 |
| GHVDHGKTT | ASSLDLDRANTEVF | 0.7555111 |
| AQFEHTLL | ASSLDLDRANTEVF | 0.70641345 |
| IVNQDEL | ASSLDLDRANTEVF | 0.09607774 |
| TNGSQFF | ASSLDLDRANTEVF | 0.4367785 |
| TGKGIVE | ASSLDLDRANTEVF | 0.34114516 |
| LISNASDA | ASSLDLDRANTEVF | 0.70242006 |
| AELRTLGELTKIAKAHGAQ | ASSLDLDRANTEVF | 0.39734125 |
| APNAVAAAEPSW | ASSLDLDRANTEVF | 0.48721382 |
| APQVAGAATAVL | ASSLDLDRANTEVF | 0.69241273 |
| ARRDPAAAAAILPTDA | ASSLDLDRANTEVF | 0.76384175 |
| DAAARDVLAEAGLAEYF | ASSLDLDRANTEVF | 0.60193896 |
| DDQIVLSAKANLAKVEQ | ASSLDLDRANTEVF | 0.2980094 |
| DELLPDTGEDSDEDGH | ASSLDLDRANTEVF | 0.13571863 |
| ELPEAPANRATVPQD | ASSLDLDRANTEVF | 0.7754689 |
| FGQSKIERVTGIG | ASSLDLDRANTEVF | 0.30242315 |
| GAGGTGSTAGGAGGAGGAGGLY | ASSLDLDRANTEVF | 0.44812137 |
| GHGGTGAAGGNGGTGGAGGSGI | ASSLDLDRANTEVF | 0.1980933 |
| GNGGNGGAGGEGGPGGAGGAGA | ASSLDLDRANTEVF | 0.26981395 |
| GSTAGNGGNGGNATNGTIAGQP | ASSLDLDRANTEVF | 0.25727823 |
| GTGGDGGNGGNGGTGG | ASSLDLDRANTEVF | 0.63618815 |
| GVRRPAGIPADAPV | ASSLDLDRANTEVF | 0.53016186 |
| GYRQVTDAHLLTLARR | ASSLDLDRANTEVF | 0.5801673 |

| | | |
|---|---|---|
| IAKMGQDGHDRG | ASSLDLDRANTEVF | 0.83104414 |
| KAEPSIEIQYT | ASSLDLDRANTEVF | 0.7828788 |
| KPVRIRMAADEDTVAVT | ASSLDLDRANTEVF | 0.6008665 |
| MPPRIAGMRLLVIKPEP | ASSLDLDRANTEVF | 0.56997365 |
| MKFYAVALFRSRGVPPTR | ASSLDLDRANTEVF | 0.3601269 |
| NAGGFGILGAVAHSP | ASSLDLDRANTEVF | 0.44044635 |
| NEGREITLAMAEM | ASSLDLDRANTEVF | 0.4678614 |
| NELVEWSHPLGHRWRWE | ASSLDLDRANTEVF | 0.8289109 |
| NLGAGGTGGNGGNPG | ASSLDLDRANTEVF | 0.43795165 |
| NTEWPKWETVMAADPVD | ASSLDLDRANTEVF | 0.26788875 |
| PASAAPAETQP | ASSLDLDRANTEVF | 0.5112553 |
| QQSSDGKNGVRAAEITGE | ASSLDLDRANTEVF | 0.81438243 |
| RAPDAVTTSWWKE | ASSLDLDRANTEVF | 0.8500093 |
| SGFGNVGAHTSGWFNQ | ASSLDLDRANTEVF | 0.48544762 |
| SGVAGAVGQAASVGGLKV | ASSLDLDRANTEVF | 0.42924932 |
| SNGGAGTGSAGGNGGT | ASSLDLDRANTEVF | 0.49871746 |
| TGGAGGEGGAAG | ASSLDLDRANTEVF | 0.6526574 |
| TGGDGGAAGAGGAPGHG | ASSLDLDRANTEVF | 0.3699853 |
| TMPPLLAAIANAGRHGV | ASSLDLDRANTEVF | 0.3654385 |
| TSSTAGNSGGA | ASSLDLDRANTEVF | 0.6265081 |
| VPATVPVGILC | ASSLDLDRANTEVF | 0.88605416 |
| VQRPDSGGVVLTGRIS | ASSLDLDRANTEVF | 0.48327285 |
| WTPNPVPATKGVPAGLVQAT | ASSLDLDRANTEVF | 0.18788616 |
| YALKRALAKFSDRFDVVII | ASSLDLDRANTEVF | 0.67474955 |
| YLIGSVIV | ASSLDLDRANTEVF | 0.4170076 |
| YGQLVAVID | ASSLDLDRANTEVF | 0.56825894 |
| VTVGGMTLPA | ASSLDLDRANTEVF | 0.68376935 |
| VLALPAV | ASSLDLDRANTEVF | 0.23332006 |
| VIPPTLNYET | ASSLDLDRANTEVF | 0.52359176 |
| VGMGGMHPSSG | ASSLDLDRANTEVF | 0.8392725 |
| VGLAGGAATA | ASSLDLDRANTEVF | 0.715344 |
| VGGAGGAGGSGP | ASSLDLDRANTEVF | 0.6863853 |
| VFAAFMV | ASSLDLDRANTEVF | 0.8179375 |
| TLYDKLR | ASSLDLDRANTEVF | 0.733084 |
| TLAVFTVL | ASSLDLDRANTEVF | 0.8296244 |
| TGSVQPARV | ASSLDLDRANTEVF | 0.822984 |
| TGGGAGSGGT | ASSLDLDRANTEVF | 0.4786409 |
| SHLETSL | ASSLDLDRANTEVF | 0.49748042 |
| SFTTGMT | ASSLDLDRANTEVF | 0.7521181 |
| QVEVAKRMGVRQPTVS | ASSLDLDRANTEVF | 0.7088525 |
| QGSGGPALAM | ASSLDLDRANTEVF | 0.6699252 |
| MLAPSNP | ASSLDLDRANTEVF | 0.3943552 |
| LVNNAAI | ASSLDLDRANTEVF | 0.829784 |
| LTGEINTLA | ASSLDLDRANTEVF | 0.4803649 |
| LSIFGPV | ASSLDLDRANTEVF | 0.8504101 |
| LLALLANL | ASSLDLDRANTEVF | 0.27393377 |
| LISNASDALDKLR | ASSLDLDRANTEVF | 0.40604922 |
| LINEAVP | ASSLDLDRANTEVF | 0.7766417 |
| LGYAFSGL | ASSLDLDRANTEVF | 0.12913784 |
| LDFVAVAL | ASSLDLDRANTEVF | 0.23093657 |
| LAQPGAE | ASSLDLDRANTEVF | 0.55554676 |

| | | |
|---|---|---|
| ISNLGAVTGF | ASSLDLDRANTEVF | 0.7918464 |
| HQEDPGRI | ASSLDLDRANTEVF | 0.119604625 |
| GTPGEPTP | ASSLDLDRANTEVF | 0.48800194 |
| GFFNSGS | ASSLDLDRANTEVF | 0.28365606 |
| GAGGGGLTAGAN | ASSLDLDRANTEVF | 0.6576673 |
| FVDALVNM | ASSLDLDRANTEVF | 0.67960864 |
| FLTVAPEL | ASSLDLDRANTEVF | 0.25795773 |
| FETLTAL | ASSLDLDRANTEVF | 0.7198343 |
| DVINPIEL | ASSLDLDRANTEVF | 0.19321793 |
| APVSGAPV | ASSLDLDRANTEVF | 0.40545434 |
| ALVGGLAW | ASSLDLDRANTEVF | 0.71552193 |
| ALAAAAATAI | ASSLDLDRANTEVF | 0.8515222 |
| AFGLGLLL | ASSLDLDRANTEVF | 0.35092926 |
| AFFAAASL | ASSLDLDRANTEVF | 0.7799285 |
| AAAALPAFSPP | ASSLDLDRANTEVF | 0.35512134 |
| VVLAPAL | ASSLDLDRANTEVF | 0.8599628 |
| TIGHVDHGKTTL | ASSLDLDRANTEVF | 0.5897947 |
| LISNASDALDK | ASSLDLDRANTEVF | 0.25384638 |
| LISNASDAL | ASSLDLDRANTEVF | 0.758778 |
| ISNASDALDK | ASSLDLDRANTEVF | 0.6272839 |
| NQAETLVYQTE | ASSLDLDRANTEVF | 0.619212 |
| TGGTGGTAGAG | ASSLDLDRANTEVF | 0.8078234 |
| TGSTAGGAGGA | ASSLDLDRANTEVF | 0.4334783 |
| DPLAVPVAAGPV | ASSLDLDRANTEVF | 0.6722501 |
| GGTGGTPGNGGN | ASSLDLDRANTEVF | 0.82625127 |
| PAGLAGVDPATL | ASSLDLDRANTEVF | 0.7172691 |
| GNNLVVEAPHAGASV | ASSLDLDRANTEVF | 0.42022663 |
| KVTRIMSASTLEQPA | ASSLDLDRANTEVF | 0.6671347 |
| LGFALPIALPRRTVV | ASSLDLDRANTEVF | 0.6895883 |
| TRLGNLTARKLVVLV | ASSLDLDRANTEVF | 0.40832624 |
| WSQQRRLIYFPSAGP | ASSLDLDRANTEVF | 0.79873204 |
| GPNGNPDPMAAAVDIR | ASSLDLDRANTEVF | 0.49294585 |
| TQRCVSVADLRAAASR | ASSLDLDRANTEVF | 0.7594855 |
| QFVLDVAEHPVEVLTETV | ASSLDLDRANTEVF | 0.70144284 |
| GGAGGTGGAGGNGGTQAGDG | ASSLDLDRANTEVF | 0.8654425 |
| PLLATLRAIAVDVDAKPAQV | ASSLDLDRANTEVF | 0.30151123 |
| APKFRTGGYMIDFWGVGYQVAK | ASSLDLDRANTEVF | 0.5266752 |
| GAGGAGGAGGAAGGPGGTGGTG | ASSLDLDRANTEVF | 0.6772027 |
| IGALGLLATDVAPAGSVAFGVA | ASSLDLDRANTEVF | 0.71053416 |
| AGGAGGIGGNGGAGGAGGVLMG | ASSLDLDRANTEVF | 0.40859842 |
| ADELIGLL | ASSLDLDRANTEVF | 0.3778187 |
| ADRALKA | ASSLDLDRANTEVF | 0.6143134 |
| AGAVGQVGGAGGA | ASSLDLDRANTEVF | 0.81198347 |
| AGGGGGLGGAP | ASSLDLDRANTEVF | 0.46677145 |
| AGGLLYGNGGAG | ASSLDLDRANTEVF | 0.52924407 |
| AGLESIGL | ASSLDLDRANTEVF | 0.85914886 |
| AGLKTGII | ASSLDLDRANTEVF | 0.43301848 |
| ASLAALLL | ASSLDLDRANTEVF | 0.6412792 |
| ASVINISSI | ASSLDLDRANTEVF | 0.65827435 |
| ATPPGPA | ASSLDLDRANTEVF | 0.75463396 |
| ATVASALV | ASSLDLDRANTEVF | 0.47093856 |

| | | |
|---|---|---|
| ATVASLIL | ASSLDLDRANTEVF | 0.76520675 |
| AVNGPVLIRH | ASSLDLDRANTEVF | 0.37371278 |
| DGAAGAVGI | ASSLDLDRANTEVF | 0.7966666 |
| DPLLHGI | ASSLDLDRANTEVF | 0.37532794 |
| ELGAALGAL | ASSLDLDRANTEVF | 0.71361136 |
| ELGRTQAV | ASSLDLDRANTEVF | 0.5168998 |
| FVAGGYLS | ASSLDLDRANTEVF | 0.7682625 |
| GASGGAGGAGGW | ASSLDLDRANTEVF | 0.7802469 |
| GGAGAGGDGGAGGI | ASSLDLDRANTEVF | 0.7975816 |
| GTGHNGSQPGMGGNG | ASSLDLDRANTEVF | 0.17474328 |
| GYAAASASAS | ASSLDLDRANTEVF | 0.80207264 |
| HQVDVAAVQQG | ASSLDLDRANTEVF | 0.81165355 |
| IGAAIAHT | ASSLDLDRANTEVF | 0.88160074 |
| IGMIEVAQL | ASSLDLDRANTEVF | 0.5073786 |
| IRLAPEHTSAMD | ASSLDLDRANTEVF | 0.7220497 |
| IYNVSVL | ASSLDLDRANTEVF | 0.6618526 |
| KQGFLGM | ASSLDLDRANTEVF | 0.5049524 |
| LESVITQL | ASSLDLDRANTEVF | 0.37623563 |
| LGLGALVVV | ASSLDLDRANTEVF | 0.4972486 |
| LIGNGGNGGSGGI | ASSLDLDRANTEVF | 0.6154325 |
| LNAINAPTL | ASSLDLDRANTEVF | 0.5579679 |
| LQLLNTGV | ASSLDLDRANTEVF | 0.8265223 |
| LQTSIEF | ASSLDLDRANTEVF | 0.7397865 |
| MSFAVSEL | ASSLDLDRANTEVF | 0.7823137 |
| NTGALGL | ASSLDLDRANTEVF | 0.84560156 |
| PPGAGGAGGAGTT | ASSLDLDRANTEVF | 0.8267778 |
| QLVGLGI | ASSLDLDRANTEVF | 0.6891968 |
| RFLEQVHQL | ASSLDLDRANTEVF | 0.49122328 |
| RGVGYVLR | ASSLDLDRANTEVF | 0.56513494 |
| SATGVFAGVI | ASSLDLDRANTEVF | 0.33255726 |
| SLPAITLP | ASSLDLDRANTEVF | 0.4731492 |
| STALAFTV | ASSLDLDRANTEVF | 0.54388785 |
| SVLTVGLGK | ASSLDLDRANTEVF | 0.692538 |
| TALAGAGSGI | ASSLDLDRANTEVF | 0.112330504 |
| TGLPFTGI | ASSLDLDRANTEVF | 0.7466867 |
| VAEHVTV | ASSLDLDRANTEVF | 0.8279015 |
| VDIVLNSL | ASSLDLDRANTEVF | 0.5282919 |
| VLTGGGAVL | ASSLDLDRANTEVF | 0.718675 |
| VNVGLAYGI | ASSLDLDRANTEVF | 0.6211506 |
| VSSAAVGAV | ASSLDLDRANTEVF | 0.34972858 |
| VTSVYEAGV | ASSLDLDRANTEVF | 0.8051455 |
| WVALDRGA | ASSLDLDRANTEVF | 0.38528064 |
| YLVGSVVI | ASSLDLDRANTEVF | 0.69011706 |
| GPPGTGKTLL | ASSLDLDRANTEVF | 0.80536824 |
| GVGMTKF | ASSLDLDRANTEVF | 0.6568781 |
| IGIGGSDL | ASSLDLDRANTEVF | 0.4588486 |
| IGTLLAEL | ASSLDLDRANTEVF | 0.5326946 |
| IYTSGTT | ASSLDLDRANTEVF | 0.26922482 |
| LSFPAGV | ASSLDLDRANTEVF | 0.5571321 |
| LYTSGTT | ASSLDLDRANTEVF | 0.17071828 |
| VGAFPFGF | ASSLDLDRANTEVF | 0.6685637 |

| | | |
|---|---|---|
| VGAGTVEF | ASSLDLDRANTEVF | 0.37082398 |
| VGKSTLLL | ASSLDLDRANTEVF | 0.7865873 |
| AALAHAGGGIV | ASSLDLDRANTEVF | 0.7056457 |
| AALVAVSLL | ASSLDLDRANTEVF | 0.5854834 |
| AAVSRAVAAL | ASSLDLDRANTEVF | 0.46605143 |
| AGGSSGIGGF | ASSLDLDRANTEVF | 0.7284473 |
| AGLTVANANG | ASSLDLDRANTEVF | 0.60679543 |
| AGLVGAVL | ASSLDLDRANTEVF | 0.3528025 |
| ALSQAFTV | ASSLDLDRANTEVF | 0.61939883 |
| ALTFSVSL | ASSLDLDRANTEVF | 0.46060926 |
| ALTGAALVAL | ASSLDLDRANTEVF | 0.4768568 |
| ASFSIGAGVL | ASSLDLDRANTEVF | 0.8047822 |
| DGSVYKI | ASSLDLDRANTEVF | 0.6734127 |
| EQLAEIV | ASSLDLDRANTEVF | 0.22400549 |
| FAVSELTVM | ASSLDLDRANTEVF | 0.6432079 |
| GAAPLAAGL | ASSLDLDRANTEVF | 0.7079168 |
| GADGGDGGSGGAGG | ASSLDLDRANTEVF | 0.6217451 |
| GFGLDTGGAGG | ASSLDLDRANTEVF | 0.10950459 |
| GLASGGLL | ASSLDLDRANTEVF | 0.6318173 |
| GRIAGAAVLL | ASSLDLDRANTEVF | 0.4126168 |
| GSAGSLNTSF | ASSLDLDRANTEVF | 0.37088174 |
| GTGQIGI | ASSLDLDRANTEVF | 0.41836008 |
| GTLIALIL | ASSLDLDRANTEVF | 0.8297926 |
| IAATMGFVVG | ASSLDLDRANTEVF | 0.5622481 |
| IFAQALL | ASSLDLDRANTEVF | 0.69565666 |
| IGAAVALGF | ASSLDLDRANTEVF | 0.73341674 |
| IGLTPLAL | ASSLDLDRANTEVF | 0.74956167 |
| ILGSNTSTL | ASSLDLDRANTEVF | 0.8364605 |
| IVFSLTVL | ASSLDLDRANTEVF | 0.7439587 |
| LASKATGF | ASSLDLDRANTEVF | 0.39262158 |
| LASVALIL | ASSLDLDRANTEVF | 0.6522302 |
| LFAVGLTL | ASSLDLDRANTEVF | 0.46588525 |
| LFNGGIGTYI | ASSLDLDRANTEVF | 0.3276957 |
| LTLGAGII | ASSLDLDRANTEVF | 0.28728488 |
| LVAAALGDI | ASSLDLDRANTEVF | 0.5834669 |
| NTAFVQLGI | ASSLDLDRANTEVF | 0.27929628 |
| QLFSGAEL | ASSLDLDRANTEVF | 0.7255691 |
| SAATFITL | ASSLDLDRANTEVF | 0.69975245 |
| SNYGLANLGV | ASSLDLDRANTEVF | 0.5326781 |
| SYGGAYAVM | ASSLDLDRANTEVF | 0.29255486 |
| TTGVGGLTLG | ASSLDLDRANTEVF | 0.48567516 |
| TTQTIYI | ASSLDLDRANTEVF | 0.4597498 |
| VAAGAGLGAVI | ASSLDLDRANTEVF | 0.78079677 |
| VAAVVTLL | ASSLDLDRANTEVF | 0.6232357 |
| VGFLGGLI | ASSLDLDRANTEVF | 0.87111187 |
| VGVIPILL | ASSLDLDRANTEVF | 0.8090135 |
| VIFASSLI | ASSLDLDRANTEVF | 0.6038316 |
| VKGGDGGAAAG | ASSLDLDRANTEVF | 0.78004813 |
| VTKTFTNI | ASSLDLDRANTEVF | 0.44118756 |
| VVAAAILL | ASSLDLDRANTEVF | 0.39746702 |
| YAALDVEL | ASSLDLDRANTEVF | 0.44407424 |

| | | |
|---|---|---|
| LASVALLL | ASSLDLDRANTEVF | 0.56651723 |
| LDAVHRV | ASSLDLDRANTEVF | 0.58317405 |
| QGLSSSIF | ASSLDLDRANTEVF | 0.42212412 |
| RGTFANIRL | ASSLDLDRANTEVF | 0.24800767 |
| RQLWWGHRI | ASSLDLDRANTEVF | 0.7673571 |
| YGPPGTGKTLL | ASSLDLDRANTEVF | 0.7339797 |
| AALVGAPANVA | ASSLDLDRANTEVF | 0.28552932 |
| AATFRRTE | ASSLDLDRANTEVF | 0.41545495 |
| AGALGGGI | ASSLDLDRANTEVF | 0.8612919 |
| AGGAGGAGGIGAG | ASSLDLDRANTEVF | 0.85775435 |
| AGSGALGL | ASSLDLDRANTEVF | 0.5014768 |
| AQIADQI | ASSLDLDRANTEVF | 0.44669044 |
| ASAVAVTYL | ASSLDLDRANTEVF | 0.74647444 |
| ASTVLGGTGG | ASSLDLDRANTEVF | 0.33814353 |
| AVGGEPAK | ASSLDLDRANTEVF | 0.55526996 |
| CEQGTFGGTC | ASSLDLDRANTEVF | 0.6657785 |
| DEGKAPAG | ASSLDLDRANTEVF | 0.6067248 |
| DTVVFPAL | ASSLDLDRANTEVF | 0.41846997 |
| FEGITMAG | ASSLDLDRANTEVF | 0.6558667 |
| FPLVVQD | ASSLDLDRANTEVF | 0.55952907 |
| GAFTTPQ | ASSLDLDRANTEVF | 0.590544 |
| GFGGFGTAGGVGGA | ASSLDLDRANTEVF | 0.12852338 |
| GFLAQFL | ASSLDLDRANTEVF | 0.5886282 |
| GGAGGAGGNAGVA | ASSLDLDRANTEVF | 0.47324783 |
| GGVGTAGAGGAGGAGG | ASSLDLDRANTEVF | 0.3695366 |
| GNGGGGDAGNAGSGGN | ASSLDLDRANTEVF | 0.2060784 |
| GTGGAGGAGNDAGS | ASSLDLDRANTEVF | 0.4596589 |
| GTTNTGWFNS | ASSLDLDRANTEVF | 0.7950001 |
| IFMGPLFTL | ASSLDLDRANTEVF | 0.79956645 |
| ILAHAIEAI | ASSLDLDRANTEVF | 0.4272045 |
| IRSAIREAL | ASSLDLDRANTEVF | 0.7514074 |
| ITFAGGG | ASSLDLDRANTEVF | 0.8108213 |
| ITGMSGHQEV | ASSLDLDRANTEVF | 0.46907052 |
| IVAGLACTL | ASSLDLDRANTEVF | 0.23621167 |
| IVGGGLAGA | ASSLDLDRANTEVF | 0.75269413 |
| IVTGVLNI | ASSLDLDRANTEVF | 0.60238624 |
| LAQQGCI | ASSLDLDRANTEVF | 0.6953394 |
| LCLHVEN | ASSLDLDRANTEVF | 0.7846571 |
| LDAISDGS | ASSLDLDRANTEVF | 0.36743858 |
| LGGFAAML | ASSLDLDRANTEVF | 0.76381135 |
| LGGKNPMI | ASSLDLDRANTEVF | 0.8044258 |
| LGPKGVAAIL | ASSLDLDRANTEVF | 0.3805281 |
| LMVGVSI | ASSLDLDRANTEVF | 0.29302183 |
| LVGAAFVL | ASSLDLDRANTEVF | 0.34306428 |
| LVLAVAPL | ASSLDLDRANTEVF | 0.1627275 |
| MAAIAVIDAA | ASSLDLDRANTEVF | 0.69531214 |
| NNGINAELQ | ASSLDLDRANTEVF | 0.49395275 |
| RGEPLVLVHG | ASSLDLDRANTEVF | 0.54856735 |
| RGGAGGAAGLI | ASSLDLDRANTEVF | 0.6181287 |
| RSYRMHPAV | ASSLDLDRANTEVF | 0.49214575 |
| SAAHLLAGQLA | ASSLDLDRANTEVF | 0.58220947 |

| | | |
|---|---|---|
| SAAVGSY | ASSLDLDRANTEVF | 0.5947224 |
| SPHAVLHRI | ASSLDLDRANTEVF | 0.55340135 |
| SQIAKVL | ASSLDLDRANTEVF | 0.67359585 |
| TALGHLVP | ASSLDLDRANTEVF | 0.43793836 |
| TAPTTSGAA | ASSLDLDRANTEVF | 0.47630993 |
| TDGDDGNGGSPGT | ASSLDLDRANTEVF | 0.8428712 |
| TDGLGEN | ASSLDLDRANTEVF | 0.36189005 |
| TGGPPLPGQA | ASSLDLDRANTEVF | 0.41902322 |
| TIPAITI | ASSLDLDRANTEVF | 0.69116956 |
| TIPGFTI | ASSLDLDRANTEVF | 0.42997098 |
| TSIASGALA | ASSLDLDRANTEVF | 0.7454457 |
| TVTLPGAAI | ASSLDLDRANTEVF | 0.69822204 |
| VASGLSAALP | ASSLDLDRANTEVF | 0.71501255 |
| VGAGVIGI | ASSLDLDRANTEVF | 0.7010144 |
| VGPKRFGV | ASSLDLDRANTEVF | 0.8238438 |
| VNVAFPD | ASSLDLDRANTEVF | 0.78604007 |
| VSMTNSGVS | ASSLDLDRANTEVF | 0.4220278 |
| VTDVLGDLSV | ASSLDLDRANTEVF | 0.7668319 |
| VVVVGAGFAGL | ASSLDLDRANTEVF | 0.3275293 |
| VYTVGTVV | ASSLDLDRANTEVF | 0.7026918 |
| WTTFSGP | ASSLDLDRANTEVF | 0.73780036 |
| YDELVEAF | ASSLDLDRANTEVF | 0.40800118 |
| YGTGGAGGNGG | ASSLDLDRANTEVF | 0.6754472 |
| YTNVMAV | ASSLDLDRANTEVF | 0.72181106 |
| FLEQVHQL | ASSLDLDRANTEVF | 0.8128655 |
| GLSLLAVAV | ASSLDLDRANTEVF | 0.284604 |
| LEQVHQL | ASSLDLDRANTEVF | 0.5599756 |
| LNAAGAI | ASSLDLDRANTEVF | 0.28024673 |
| PAVHASL | ASSLDLDRANTEVF | 0.36647242 |
| TELEDDA | ASSLDLDRANTEVF | 0.31414345 |
| TPPGPPN | ASSLDLDRANTEVF | 0.7228695 |
| TPPPPPPP | ASSLDLDRANTEVF | 0.62906724 |
| TQSPALI | ASSLNNSYEQY | 0.82617396 |
| EFAAAFPQ | ASSLNNSYEQY | 0.72754776 |
| LLSGNLDVL | ASSLNNSYEQY | 0.8233705 |
| GPANLGNYNI | ASSLNNSYEQY | 0.5014112 |
| TLGPTLP | ASSLNNSYEQY | 0.68258536 |
| VYAYSSCCK | ASSLNNSYEQY | 0.24731237 |
| APAPAPAPNI | ASSLNNSYEQY | 0.30963415 |
| PGGPVPIP | ASSLNNSYEQY | 0.32105213 |
| FLTVALEP | ASSLNNSYEQY | 0.60961103 |
| SQYFGANL | ASSLNNSYEQY | 0.1604715 |
| GGVKPAA | ASSLNNSYEQY | 0.8106365 |
| GGTGGAGGAGGW | ASSLNNSYEQY | 0.6882257 |
| GVGGVAGLGGAGG | ASSLNNSYEQY | 0.7706133 |
| VLLVLATTI | ASSLNNSYEQY | 0.517249 |
| IGIFAAML | ASSLNNSYEQY | 0.37520352 |
| AVGDLID | ASSLNNSYEQY | 0.54687953 |
| NISPLPL | ASSLNNSYEQY | 0.82977414 |
| VTAVFVAI | ASSLNNSYEQY | 0.38419408 |
| SGDDNDAAL | ASSLNNSYEQY | 0.19930737 |

| | | |
|---|---|---|
| AVAALAHS | ASSLNNSYEQY | 0.3899576 |
| GGVTVMATP | ASSLNNSYEQY | 0.23505138 |
| AGLAANVRD | ASSLNNSYEQY | 0.8153575 |
| TNRPDLIDPALLRPG | ASSLNNSYEQY | 0.7736834 |
| LANLGVTAL | ASSLNNSYEQY | 0.7349405 |
| TGERIEL | ASSLNNSYEQY | 0.8640163 |
| NGEELAA | ASSLNNSYEQY | 0.16228496 |
| VLSLVWG | ASSLNNSYEQY | 0.5139816 |
| IGYALWL | ASSLNNSYEQY | 0.85466605 |
| LSLLLAPL | ASSLNNSYEQY | 0.6800147 |
| TLDEALK | ASSLNNSYEQY | 0.67087674 |
| TDGAGTAEGG | ASSLNNSYEQY | 0.80820525 |
| NAPGGAGGNGG | ASSLNNSYEQY | 0.5727122 |
| VSVFMGMGGNF | ASSLNNSYEQY | 0.6438742 |
| LASYLAAL | ASSLNNSYEQY | 0.6480434 |
| LVGGTPAL | ASSLNNSYEQY | 0.5684294 |
| VRNADVL | ASSLNNSYEQY | 0.4332379 |
| TCSSPFLVSAM | ASSLNNSYEQY | 0.2982133 |
| VHITDAE | ASSLNNSYEQY | 0.6597082 |
| EYAAPSF | ASSLNNSYEQY | 0.8488047 |
| VVDAFHGGL | ASSLNNSYEQY | 0.7094793 |
| ASVVDPQ | ASSLNNSYEQY | 0.7205 |
| SYNEFGGLS | ASSLNNSYEQY | 0.7032133 |
| DPDGSVY | ASSLNNSYEQY | 0.79505444 |
| MVSTIHATE | ASSLNNSYEQY | 0.5159399 |
| LAAGVADPV | ASSLNNSYEQY | 0.40020004 |
| TKEQIAA | ASSLNNSYEQY | 0.7197552 |
| SLASPAAL | ASSLNNSYEQY | 0.41353396 |
| IGLVFYGV | ASSLNNSYEQY | 0.5577987 |
| FLLDASNL | ASSLNNSYEQY | 0.4172821 |
| YQVGNFP | ASSLNNSYEQY | 0.8771215 |
| ESLLSQST | ASSLNNSYEQY | 0.7400888 |
| LFTVGFAAVG | ASSLNNSYEQY | 0.71206856 |
| ATPPTPT | ASSLNNSYEQY | 0.50145113 |
| TDLATLG | ASSLNNSYEQY | 0.48795152 |
| SAGGTGR | ASSLNNSYEQY | 0.69014496 |
| GFSNSGS | ASSLNNSYEQY | 0.25616482 |
| GGNGGSGGTSV | ASSLNNSYEQY | 0.82593906 |
| GTSVATGGAGN | ASSLNNSYEQY | 0.47676605 |
| NGGNGGSGGTS | ASSLNNSYEQY | 0.49748084 |
| AVVKFPH | ASSLNNSYEQY | 0.74407685 |
| TPPPPPP | ASSLNNSYEQY | 0.8615689 |
| AAPAAAAAPA | ASSLNNSYEQY | 0.44964188 |
| IALNEDL | ASSLNNSYEQY | 0.6587516 |
| IVNVSSM | ASSLNNSYEQY | 0.5045444 |
| TGIPPAPR | ASSLNNSYEQY | 0.7352756 |
| FDLGGGTF | ASSLNNSYEQY | 0.8439275 |
| FDLGGGTFDVS | ASSLNNSYEQY | 0.7216113 |
| NEPTAAA | ASSLNNSYEQY | 0.8699846 |
| NEPTAAALA | ASSLNNSYEQY | 0.51964307 |
| GSIINISSI | ASSLNNSYEQY | 0.27757818 |

| | | |
|---|---|---|
| VAPGNAGTA | ASSLNNSYEQY | 0.57499087 |
| TGGGASLE | ASSLNNSYEQY | 0.67600775 |
| VAVYNHYKRI | ASSLNNSYEQY | 0.7687374 |
| TDPAPATT | ASSLNNSYEQY | 0.7830148 |
| TNRPDILDPALLRPG | ASSLNNSYEQY | 0.5285668 |
| LTTTTTS | ASSLNNSYEQY | 0.4530799 |
| PTGSGKTL | ASSLNNSYEQY | 0.817078 |
| TGSGKTL | ASSLNNSYEQY | 0.82906604 |
| TIGHVDHGK | ASSLNNSYEQY | 0.7942599 |
| TIGHVDHGKTT | ASSLNNSYEQY | 0.4972139 |
| GHVDHGKTT | ASSLNNSYEQY | 0.4178092 |
| AQFEHTLL | ASSLNNSYEQY | 0.58786315 |
| IVNQDEL | ASSLNNSYEQY | 0.7668419 |
| TNGSQFF | ASSLNNSYEQY | 0.36893 |
| TGKGIVE | ASSLNNSYEQY | 0.64233565 |
| LISNASDA | ASSLNNSYEQY | 0.72254 |
| AELRTLGELTKIAKAHGAQ | ASSLNNSYEQY | 0.26033503 |
| APNAVAAAEPSW | ASSLNNSYEQY | 0.5241102 |
| APQVAGAATAVL | ASSLNNSYEQY | 0.76857984 |
| ARRDPAAAAAILPTDA | ASSLNNSYEQY | 0.4635195 |
| DAAARDVLAEAGLAEYF | ASSLNNSYEQY | 0.42852277 |
| DDQIVLSAKANLAKVEQ | ASSLNNSYEQY | 0.8232 |
| DELLPDTGEDSDEDGH | ASSLNNSYEQY | 0.8706815 |
| ELPEAPANRATVPQD | ASSLNNSYEQY | 0.763507 |
| FGQSKIERVTGIG | ASSLNNSYEQY | 0.27649775 |
| GAGGTGSTAGGAGGAGGAGGLY | ASSLNNSYEQY | 0.15027763 |
| GHGGTGAAGGNGGTGGAGGSGI | ASSLNNSYEQY | 0.5816666 |
| GNGGNGGAGGEGGPGGAGGAGA | ASSLNNSYEQY | 0.82910967 |
| GSTAGNGGNGGNATNGTIAGQP | ASSLNNSYEQY | 0.57677877 |
| GTGGDGGNGGNGGTGG | ASSLNNSYEQY | 0.5437036 |
| GVRRPAGIPADAPV | ASSLNNSYEQY | 0.58511263 |
| GYRQVTDAHLLTLARR | ASSLNNSYEQY | 0.7781125 |
| IAKMGQDGHDRG | ASSLNNSYEQY | 0.43593463 |
| KAEPSIEIQYT | ASSLNNSYEQY | 0.28789857 |
| KPVRIRMAADEDTVAVT | ASSLNNSYEQY | 0.47585595 |
| MPPRIAGMRLLVIKPEP | ASSLNNSYEQY | 0.77625394 |
| MKFYAVALFRSRGVPPTR | ASSLNNSYEQY | 0.43002272 |
| NAGGFGILGAVAHSP | ASSLNNSYEQY | 0.654794 |
| NEGREITLAMAEM | ASSLNNSYEQY | 0.61940336 |
| NELVEWSHPLGHRWRWE | ASSLNNSYEQY | 0.37539968 |
| NLGAGGTGGNGGNPG | ASSLNNSYEQY | 0.66908 |
| NTEWPKWETVMAADPVD | ASSLNNSYEQY | 0.41498095 |
| PASAAPAETQP | ASSLNNSYEQY | 0.7859719 |
| QQSSDGKNGVRAAEITGE | ASSLNNSYEQY | 0.36228132 |
| RAPDAVTTSWWKE | ASSLNNSYEQY | 0.63407475 |
| SGFGNVGAHTSGWFNQ | ASSLNNSYEQY | 0.71635795 |
| SGVAGAVGQAASVGGLKV | ASSLNNSYEQY | 0.62657535 |
| SNGGAGTGSAGGNGGT | ASSLNNSYEQY | 0.6314267 |
| TGGAGGEGGAAG | ASSLNNSYEQY | 0.43123972 |
| TGGDGGAAGAGGAPGHG | ASSLNNSYEQY | 0.36222777 |
| TMPPLLAAIANAGRHGV | ASSLNNSYEQY | 0.310134 |

| | | |
|---|---|---|
| TSSTAGNSGGA | ASSLNNSYEQY | 0.73690176 |
| VPATVPVGILC | ASSLNNSYEQY | 0.27353054 |
| VQRPDSGGVVLTGRIS | ASSLNNSYEQY | 0.7903197 |
| WTPNPVPATKGVPAGLVQAT | ASSLNNSYEQY | 0.72208774 |
| YALKRALAKFSDRFDVVII | ASSLNNSYEQY | 0.43853986 |
| YLIGSVIV | ASSLNNSYEQY | 0.59973085 |
| YGQLVAVID | ASSLNNSYEQY | 0.46194896 |
| VTVGGMTLPA | ASSLNNSYEQY | 0.5314745 |
| VLALPAV | ASSLNNSYEQY | 0.7306905 |
| VIPPTLNYET | ASSLNNSYEQY | 0.66480964 |
| VGMGGMHPSSG | ASSLNNSYEQY | 0.475692 |
| VGLAGGAATA | ASSLNNSYEQY | 0.5747243 |
| VGGAGGAGGSGP | ASSLNNSYEQY | 0.6802772 |
| VFAAFMV | ASSLNNSYEQY | 0.41467237 |
| TLYDKLR | ASSLNNSYEQY | 0.5060188 |
| TLAVFTVL | ASSLNNSYEQY | 0.8057685 |
| TGSVQPARV | ASSLNNSYEQY | 0.52769935 |
| TGGGAGSGGT | ASSLNNSYEQY | 0.4212142 |
| SHLETSL | ASSLNNSYEQY | 0.6980417 |
| SFTTGMT | ASSLNNSYEQY | 0.40842095 |
| QVEVAKRMGVRQPTVS | ASSLNNSYEQY | 0.7478252 |
| QGSGGPALAM | ASSLNNSYEQY | 0.52356154 |
| MLAPSNP | ASSLNNSYEQY | 0.5950928 |
| LVNNAAI | ASSLNNSYEQY | 0.679052 |
| LTGEINTLA | ASSLNNSYEQY | 0.2672593 |
| LSIFGPV | ASSLNNSYEQY | 0.40309668 |
| LLALLANL | ASSLNNSYEQY | 0.5002688 |
| LISNASDALDKLR | ASSLNNSYEQY | 0.74499977 |
| LINEAVP | ASSLNNSYEQY | 0.78609014 |
| LGYAFSGL | ASSLNNSYEQY | 0.41011018 |
| LDFVAVAL | ASSLNNSYEQY | 0.6817383 |
| LAQPGAE | ASSLNNSYEQY | 0.52804023 |
| ISNLGAVTGF | ASSLNNSYEQY | 0.45402294 |
| HQEDPGRI | ASSLNNSYEQY | 0.4935255 |
| GTPGEPTP | ASSLNNSYEQY | 0.40971923 |
| GFFNSGS | ASSLNNSYEQY | 0.3803329 |
| GAGGGGLTAGAN | ASSLNNSYEQY | 0.6124239 |
| FVDALVNM | ASSLNNSYEQY | 0.4360936 |
| FLTVAPEL | ASSLNNSYEQY | 0.6493143 |
| FETLTAL | ASSLNNSYEQY | 0.6742003 |
| DVINPIEL | ASSLNNSYEQY | 0.4233591 |
| APVSGAPV | ASSLNNSYEQY | 0.596959 |
| ALVGGLAW | ASSLNNSYEQY | 0.6854514 |
| ALAAAAATAI | ASSLNNSYEQY | 0.69993216 |
| AFGLGLLL | ASSLNNSYEQY | 0.59065175 |
| AFFAAASL | ASSLNNSYEQY | 0.43298078 |
| AAAALPAFSPP | ASSLNNSYEQY | 0.71813357 |
| VVLAPAL | ASSLNNSYEQY | 0.7923062 |
| TIGHVDHGKTTL | ASSLNNSYEQY | 0.7084906 |
| LISNASDALDK | ASSLNNSYEQY | 0.7022027 |
| LISNASDAL | ASSLNNSYEQY | 0.7857261 |

| | | |
|---|---|---|
| ISNASDALDK | ASSLNNSYEQY | 0.75658387 |
| NQAETLVYQTE | ASSLNNSYEQY | 0.5969887 |
| TGGTGGTAGAG | ASSLNNSYEQY | 0.24457034 |
| TGSTAGGAGGA | ASSLNNSYEQY | 0.64478004 |
| DPLAVPVAAGPV | ASSLNNSYEQY | 0.724757 |
| GGTGGTPGNGGN | ASSLNNSYEQY | 0.39468744 |
| PAGLAGVDPATL | ASSLNNSYEQY | 0.26829386 |
| GNNLVVEAPHAGASV | ASSLNNSYEQY | 0.81969786 |
| KVTRIMSASTLEQPA | ASSLNNSYEQY | 0.874591 |
| LGFALPIALPRRTVV | ASSLNNSYEQY | 0.4697523 |
| TRLGNLTARKLVVLV | ASSLNNSYEQY | 0.8423053 |
| WSQQRRLIYFPSAGP | ASSLNNSYEQY | 0.24563356 |
| GPNGNPDPMAAAVDIR | ASSLNNSYEQY | 0.76607394 |
| TQRCVSVADLRAAASR | ASSLNNSYEQY | 0.8161217 |
| QFVLDVAEHPVEVLTETV | ASSLNNSYEQY | 0.45434272 |
| GGAGGTGGAGGNGGTQAGDG | ASSLNNSYEQY | 0.73233557 |
| PLLATLRAIAVDVDAKPAQV | ASSLNNSYEQY | 0.35697374 |
| APKFRTGGYMIDFWGVGYQVAK | ASSLNNSYEQY | 0.42681912 |
| GAGGAGGAGGAAGGPGGTGGTG | ASSLNNSYEQY | 0.8360785 |
| IGALGLLATDVAPAGSVAFGVA | ASSLNNSYEQY | 0.60662454 |
| AGGAGGIGGNGGAGGAGGVLMG | ASSLNNSYEQY | 0.5290157 |
| ADELIGLL | ASSLNNSYEQY | 0.66331935 |
| ADRALKA | ASSLNNSYEQY | 0.44280052 |
| AGAVGQVGGAGGA | ASSLNNSYEQY | 0.53002936 |
| AGGGGGLGGAP | ASSLNNSYEQY | 0.7722904 |
| AGGLLYGNGGAG | ASSLNNSYEQY | 0.5275731 |
| AGLESIGL | ASSLNNSYEQY | 0.8168479 |
| AGLKTGII | ASSLNNSYEQY | 0.3900585 |
| ASLAALLL | ASSLNNSYEQY | 0.84515905 |
| ASVINISSI | ASSLNNSYEQY | 0.8431533 |
| ATPPGPA | ASSLNNSYEQY | 0.7186942 |
| ATVASALV | ASSLNNSYEQY | 0.41906548 |
| ATVASLIL | ASSLNNSYEQY | 0.8321616 |
| AVNGPVLIRH | ASSLNNSYEQY | 0.7055825 |
| DGAAGAVGI | ASSLNNSYEQY | 0.5723958 |
| DPLLHGI | ASSLNNSYEQY | 0.6879367 |
| ELGAALGAL | ASSLNNSYEQY | 0.116424 |
| ELGRTQAV | ASSLNNSYEQY | 0.12100014 |
| FVAGGYLS | ASSLNNSYEQY | 0.117435776 |
| GASGGAGGAGGW | ASSLNNSYEQY | 0.12455371 |
| GGAGAGGDGGAGGI | ASSLNNSYEQY | 0.12158685 |
| GTGHNGSQPGMGGNG | ASSLNNSYEQY | 0.116607204 |
| GYAAASASAS | ASSLNNSYEQY | 0.118528254 |
| HQVDVAAVQQG | ASSLNNSYEQY | 0.11904987 |
| IGAAIAHT | ASSLNNSYEQY | 0.11786152 |
| IGMIEVAQL | ASSLNNSYEQY | 0.12105976 |
| IRLAPEHTSAMD | ASSLNNSYEQY | 0.121327884 |
| IYNVSVL | ASSLNNSYEQY | 0.12906669 |
| KQGFLGM | ASSLNNSYEQY | 0.1252152 |
| LESVITQL | ASSLNNSYEQY | 0.11682624 |
| LGLGALVVV | ASSLNNSYEQY | 0.11770341 |

| | | |
|---|---|---|
| LIGNGGNGGSGGI | ASSLNNSYEQY | 0.11837903 |
| LNAINAPTL | ASSLNNSYEQY | 0.11988506 |
| LQLLNTGV | ASSLNNSYEQY | 0.11535299 |
| LQTSIEF | ASSLNNSYEQY | 0.12092518 |
| MSFAVSEL | ASSLNNSYEQY | 0.115635976 |
| NTGALGL | ASSLNNSYEQY | 0.11521424 |
| PPGAGGAGGAGTT | ASSLNNSYEQY | 0.12085873 |
| QLVGLGI | ASSLNNSYEQY | 0.11998464 |
| RFLEQVHQL | ASSLNNSYEQY | 0.11824023 |
| RGVGYVLR | ASSLNNSYEQY | 0.12287048 |
| SATGVFAGVI | ASSLNNSYEQY | 0.116688535 |
| SLPAITLP | ASSLNNSYEQY | 0.11836988 |
| STALAFTV | ASSLNNSYEQY | 0.12113958 |
| SVLTVGLGK | ASSLNNSYEQY | 0.13183391 |
| TALAGAGSGI | ASSLNNSYEQY | 0.11887333 |
| TGLPFTGI | ASSLNNSYEQY | 0.11793431 |
| VAEHVTV | ASSLNNSYEQY | 0.13248022 |
| VDIVLNSL | ASSLNNSYEQY | 0.122465014 |
| VLTGGGAVL | ASSLNNSYEQY | 0.12236131 |
| VNVGLAYGI | ASSLNNSYEQY | 0.123700716 |
| VSSAAVGAV | ASSLNNSYEQY | 0.11616511 |
| VTSVYEAGV | ASSLNNSYEQY | 0.12675573 |
| WVALDRGA | ASSLNNSYEQY | 0.12092446 |
| YLVGSVVI | ASSLNNSYEQY | 0.12035143 |
| GPPGTGKTLL | ASSLNNSYEQY | 0.12399009 |
| GVGMTKF | ASSLNNSYEQY | 0.116941154 |
| IGIGGSDL | ASSLNNSYEQY | 0.11752135 |
| IGTLLAEL | ASSLNNSYEQY | 0.115440615 |
| IYTSGTT | ASSLNNSYEQY | 0.119216256 |
| LSFPAGV | ASSLNNSYEQY | 0.123664424 |
| LYTSGTT | ASSLNNSYEQY | 0.123318695 |
| VGAFPFGF | ASSLNNSYEQY | 0.11608592 |
| VGAGTVEF | ASSLNNSYEQY | 0.117228806 |
| VGKSTLLL | ASSLNNSYEQY | 0.115710065 |
| AALAHAGGGIV | ASSLNNSYEQY | 0.11747344 |
| AALVAVSLL | ASSLNNSYEQY | 0.11582664 |
| AAVSRAVAAL | ASSLNNSYEQY | 0.116205886 |
| AGGSSGIGGF | ASSLNNSYEQY | 0.13287978 |
| AGLTVANANG | ASSLNNSYEQY | 0.12922978 |
| AGLVGAVL | ASSLNNSYEQY | 0.12078962 |
| ALSQAFTV | ASSLNNSYEQY | 0.12315523 |
| ALTFSVSL | ASSLNNSYEQY | 0.11654425 |
| ALTGAALVAL | ASSLNNSYEQY | 0.121764116 |
| ASFSIGAGVL | ASSLNNSYEQY | 0.12105463 |
| DGSVYKI | ASSLNNSYEQY | 0.11631128 |
| EQLAEIV | ASSLNNSYEQY | 0.12195896 |
| FAVSELTVM | ASSLNNSYEQY | 0.120010465 |
| GAAPLAAGL | ASSLNNSYEQY | 0.11857054 |
| GADGGDGGSGGAGG | ASSLNNSYEQY | 0.1266854 |
| GFGLDTGGAGG | ASSLNNSYEQY | 0.119313814 |
| GLASGGLL | ASSLNNSYEQY | 0.119278066 |

| | | |
|---|---|---|
| GRIAGAAVLL | ASSLNNSYEQY | 0.11651531 |
| GSAGSLNTSF | ASSLNNSYEQY | 0.11975857 |
| GTGQIGI | ASSLNNSYEQY | 0.11650318 |
| GTLIALIL | ASSLNNSYEQY | 0.11804816 |
| IAATMGFVVG | ASSLNNSYEQY | 0.116506524 |
| IFAQALL | ASSLNNSYEQY | 0.12081426 |
| IGAAVALGF | ASSLNNSYEQY | 0.13550358 |
| IGLTPLAL | ASSLNNSYEQY | 0.12281229 |
| ILGSNTSTL | ASSLNNSYEQY | 0.11670646 |
| IVFSLTVL | ASSLNNSYEQY | 0.1211105 |
| LASKATGF | ASSLNNSYEQY | 0.122413054 |
| LASVALIL | ASSLNNSYEQY | 0.12079546 |
| LFAVGLTL | ASSLNNSYEQY | 0.11736924 |
| LFNGGIGTYI | ASSLNNSYEQY | 0.11745972 |
| LTLGAGII | ASSLNNSYEQY | 0.119835086 |
| LVAAALGDI | ASSLNNSYEQY | 0.1295596 |
| NTAFVQLGI | ASSLNNSYEQY | 0.12307893 |
| QLFSGAEL | ASSLNNSYEQY | 0.120188385 |
| SAATFITL | ASSLNNSYEQY | 0.11954856 |
| SNYGLANLGV | ASSLNNSYEQY | 0.11836742 |
| SYGGAYAVM | ASSLNNSYEQY | 0.119011894 |
| TTGVGGLTLG | ASSLNNSYEQY | 0.117320314 |
| TTQTIYI | ASSLNNSYEQY | 0.11787689 |
| VAAGAGLGAVI | ASSLNNSYEQY | 0.11881716 |
| VAAVVTLL | ASSLNNSYEQY | 0.11873367 |
| VGFLGGLI | ASSLNNSYEQY | 0.12120503 |
| VGVIPILL | ASSLNNSYEQY | 0.116921484 |
| VIFASSLI | ASSLNNSYEQY | 0.1187068 |
| VKGGDGGAAAG | ASSLNNSYEQY | 0.12391205 |
| VTKTFTNI | ASSLNNSYEQY | 0.11789915 |
| VVAAAILL | ASSLNNSYEQY | 0.118328355 |
| YAALDVEL | ASSLNNSYEQY | 0.1233793 |
| LASVALLL | ASSLNNSYEQY | 0.118711725 |
| LDAVHRV | ASSLNNSYEQY | 0.12140156 |
| QGLSSSIF | ASSLNNSYEQY | 0.12052567 |
| RGTFANIRL | ASSLNNSYEQY | 0.12145295 |
| RQLWWGHRI | ASSLNNSYEQY | 0.116197646 |
| YGPPGTGKTLL | ASSLNNSYEQY | 0.121637456 |
| AALVGAPANVA | ASSLNNSYEQY | 0.117956236 |
| AATFRRTE | ASSLNNSYEQY | 0.11785498 |
| AGALGGGI | ASSLNNSYEQY | 0.11606243 |
| AGGAGGAGGIGAG | ASSLNNSYEQY | 0.115445144 |
| AGSGALGL | ASSLNNSYEQY | 0.1281049 |
| AQIADQI | ASSLNNSYEQY | 0.1170343 |
| ASAVAVTYL | ASSLNNSYEQY | 0.12111357 |
| ASTVLGGTGG | ASSLNNSYEQY | 0.117913276 |
| AVGGEPAK | ASSLNNSYEQY | 0.12633638 |
| CEQGTFGGTC | ASSLNNSYEQY | 0.12034638 |
| DEGKAPAG | ASSLNNSYEQY | 0.11855145 |
| DTVVFPAL | ASSLNNSYEQY | 0.11753442 |
| FEGITMAG | ASSLNNSYEQY | 0.12498131 |

| | | |
|---|---|---|
| FPLVVQD | ASSLNNSYEQY | 0.118906304 |
| GAFTTPQ | ASSLNNSYEQY | 0.12163493 |
| GFGGFGTAGGVGGA | ASSLNNSYEQY | 0.11717004 |
| GFLAQFL | ASSLNNSYEQY | 0.11600508 |
| GGAGGAGGNAGVA | ASSLNNSYEQY | 0.1237588 |
| GGVGTAGAGGAGGAGG | ASSLNNSYEQY | 0.11576955 |
| GNGGGGDAGNAGSGGN | ASSLNNSYEQY | 0.1152156 |
| GTGGAGGAGNDAGS | ASSLNNSYEQY | 0.115422614 |
| GTTNTGWFNS | ASSLNNSYEQY | 0.11610071 |
| IFMGPLFTL | ASSLNNSYEQY | 0.12582609 |
| ILAHAIEAI | ASSLNNSYEQY | 0.12620233 |
| IRSAIREAL | ASSLNNSYEQY | 0.122357756 |
| ITFAGGG | ASSLNNSYEQY | 0.121170506 |
| ITGMSGHQEV | ASSLNNSYEQY | 0.12062055 |
| IVAGLACTL | ASSLNNSYEQY | 0.115041696 |
| IVGGGLAGA | ASSLNNSYEQY | 0.11786021 |
| IVTGVLNI | ASSLNNSYEQY | 0.11705994 |
| LAQQGCI | ASSLNNSYEQY | 0.1231334 |
| LCLHVEN | ASSLNNSYEQY | 0.116068214 |
| LDAISDGS | ASSLNNSYEQY | 0.11772622 |
| LGGFAAML | ASSLNNSYEQY | 0.12688562 |
| LGGKNPMI | ASSLNNSYEQY | 0.11793725 |
| LGPKGVAAIL | ASSLNNSYEQY | 0.11943704 |
| LMVGVSI | ASSLNNSYEQY | 0.116222374 |
| LVGAAFVL | ASSLNNSYEQY | 0.11520854 |
| LVLAVAPL | ASSLNNSYEQY | 0.13061485 |
| MAAIAVIDAA | ASSLNNSYEQY | 0.11623682 |
| NNGINAELQ | ASSLNNSYEQY | 0.1176284 |
| RGEPLVLVHG | ASSLNNSYEQY | 0.11562426 |
| RGGAGGAAGLI | ASSLNNSYEQY | 0.11653341 |
| RSYRMHPAV | ASSLNNSYEQY | 0.11619384 |
| SAAHLLAGQLA | ASSLNNSYEQY | 0.121754006 |
| SAAVGSY | ASSLNNSYEQY | 0.117101066 |
| SPHAVLHRI | ASSLNNSYEQY | 0.120517366 |
| SQIAKVL | ASSLNNSYEQY | 0.12316112 |
| TALGHLVP | ASSLNNSYEQY | 0.12036272 |
| TAPTTSGAA | ASSLNNSYEQY | 0.12147812 |
| TDGDDGNGGSPGT | ASSLNNSYEQY | 0.119056344 |
| TDGLGEN | ASSLNNSYEQY | 0.1184183 |
| TGGPPLPGQA | ASSLNNSYEQY | 0.11775728 |
| TIPAITI | ASSLNNSYEQY | 0.11888926 |
| TIPGFTI | ASSLNNSYEQY | 0.12937288 |
| TSIASGALA | ASSLNNSYEQY | 0.118485145 |
| TVTLPGAAI | ASSLNNSYEQY | 0.11637171 |
| VASGLSAALP | ASSLNNSYEQY | 0.11853273 |
| VGAGVIGI | ASSLNNSYEQY | 0.124776416 |
| VGPKRFGV | ASSLNNSYEQY | 0.12607722 |
| VNVAFPD | ASSLNNSYEQY | 0.11681081 |
| VSMTNSGVS | ASSLNNSYEQY | 0.11890485 |
| VTDVLGDLSV | ASSLNNSYEQY | 0.11891238 |
| VVVVGAGFAGL | ASSLNNSYEQY | 0.12017394 |

| | | |
|---|---|---|
| VYTVGTVV | ASSLNNSYEQY | 0.11677715 |
| WTTFSGP | ASSLNNSYEQY | 0.11973177 |
| YDELVEAF | ASSLNNSYEQY | 0.11939825 |
| YGTGGAGGNGG | ASSLNNSYEQY | 0.12624624 |
| YTNVMAV | ASSLNNSYEQY | 0.117802426 |
| FLEQVHQL | ASSLNNSYEQY | 0.11599181 |
| GLSLLAVAV | ASSLNNSYEQY | 0.121871375 |
| LEQVHQL | ASSLNNSYEQY | 0.1167544 |
| LNAAGAI | ASSLNNSYEQY | 0.12092617 |
| PAVHASL | ASSLNNSYEQY | 0.12009226 |
| TELEDDA | ASSLNNSYEQY | 0.11658387 |
| TPPGPPN | ASSLNNSYEQY | 0.11898609 |
| TPPPPPPP | ASSLNNSYEQY | 0.12739702 |
| TQSPALI | ASKTGTEVF | 0.117651924 |
| EFAAAFPQ | ASKTGTEVF | 0.118575364 |
| LLSGNLDVL | ASKTGTEVF | 0.12522066 |
| GPANLGNYNI | ASKTGTEVF | 0.12172234 |
| TLGPTLP | ASKTGTEVF | 0.124003224 |
| VYAYSSCCK | ASKTGTEVF | 0.11704133 |
| APAPAPAPNI | ASKTGTEVF | 0.120676845 |
| PGGPVPIP | ASKTGTEVF | 0.11769876 |
| FLTVALEP | ASKTGTEVF | 0.11786153 |
| SQYFGANL | ASKTGTEVF | 0.12363447 |
| GGVKPAA | ASKTGTEVF | 0.12386313 |
| GGTGGAGGAGGW | ASKTGTEVF | 0.11903952 |
| GVGGVAGLGGAGG | ASKTGTEVF | 0.1280222 |
| VLLVLATTI | ASKTGTEVF | 0.11807934 |
| IGIFAAML | ASKTGTEVF | 0.12035314 |
| AVGDLID | ASKTGTEVF | 0.11611737 |
| NISPLPL | ASKTGTEVF | 0.11856769 |
| VTAVFVAI | ASKTGTEVF | 0.11992098 |
| SGDDNDAAL | ASKTGTEVF | 0.11579746 |
| AVAALAHS | ASKTGTEVF | 0.11539968 |
| GGVTVMATP | ASKTGTEVF | 0.117501386 |
| AGLAANVRD | ASKTGTEVF | 0.121165894 |
| TNRPDLIDPALLRPG | ASKTGTEVF | 0.11506001 |
| LANLGVTAL | ASKTGTEVF | 0.11922523 |
| TGERIEL | ASKTGTEVF | 0.11661814 |
| NGEELAA | ASKTGTEVF | 0.11760471 |
| VLSLVWG | ASKTGTEVF | 0.12350931 |
| IGYALWL | ASKTGTEVF | 0.11716195 |
| LSLLLAPL | ASKTGTEVF | 0.122547604 |
| TLDEALK | ASKTGTEVF | 0.11619877 |
| TDGAGTAEGG | ASKTGTEVF | 0.11761062 |
| NAPGGAGGNGG | ASKTGTEVF | 0.123937085 |
| VSVFMGMGGNF | ASKTGTEVF | 0.120017394 |
| LASYLAAL | ASKTGTEVF | 0.116825 |
| LVGGTPAL | ASKTGTEVF | 0.1263611 |
| VRNADVL | ASKTGTEVF | 0.11606636 |
| TCSSPFLVSAM | ASKTGTEVF | 0.12525053 |
| VHITDAE | ASKTGTEVF | 0.12280574 |

| | | |
|---|---|---|
| EYAAPSF | ASKTGTEVF | 0.1159953 |
| VVDAFHGGL | ASKTGTEVF | 0.120200746 |
| ASVVDPQ | ASKTGTEVF | 0.11922246 |
| SYNEFGGLS | ASKTGTEVF | 0.12071786 |
| DPDGSVY | ASKTGTEVF | 0.11769349 |
| MVSTIHATE | ASKTGTEVF | 0.116556756 |
| LAAGVADPV | ASKTGTEVF | 0.12108514 |
| TKEQIAA | ASKTGTEVF | 0.12809978 |
| SLASPAAL | ASKTGTEVF | 0.11886147 |
| IGLVFYGV | ASKTGTEVF | 0.118122034 |
| FLLDASNL | ASKTGTEVF | 0.12528832 |
| YQVGNFP | ASKTGTEVF | 0.121368356 |
| ESLLSQST | ASKTGTEVF | 0.11750378 |
| LFTVGFAAVG | ASKTGTEVF | 0.123378456 |
| ATPPTPT | ASKTGTEVF | 0.11918667 |
| TDLATLG | ASKTGTEVF | 0.12501091 |
| SAGGTGR | ASKTGTEVF | 0.12328297 |
| GFSNSGS | ASKTGTEVF | 0.11940879 |
| GGNGGSGGTSV | ASKTGTEVF | 0.116840646 |
| GTSVATGGAGN | ASKTGTEVF | 0.11646244 |
| NGGNGGSGGTS | ASKTGTEVF | 0.11953649 |
| AVVKFPH | ASKTGTEVF | 0.1219393 |
| TPPPPPP | ASKTGTEVF | 0.116274044 |
| AAPAAAAAPA | ASKTGTEVF | 0.11680082 |
| IALNEDL | ASKTGTEVF | 0.120645724 |
| IVNVSSM | ASKTGTEVF | 0.12782162 |
| TGIPPAPR | ASKTGTEVF | 0.11734091 |
| FDLGGGTF | ASKTGTEVF | 0.117457435 |
| FDLGGGTFDVS | ASKTGTEVF | 0.1314675 |
| NEPTAAA | ASKTGTEVF | 0.11683267 |
| NEPTAAALA | ASKTGTEVF | 0.123870686 |
| GSIINISSI | ASKTGTEVF | 0.1204143 |
| VAPGNAGTA | ASKTGTEVF | 0.12096578 |
| TGGGASLE | ASKTGTEVF | 0.119171605 |
| VAVYNHYKRI | ASKTGTEVF | 0.12038626 |
| TDPAPATT | ASKTGTEVF | 0.11587574 |
| TNRPDILDPALLRPG | ASKTGTEVF | 0.12377965 |
| LTTTTTS | ASKTGTEVF | 0.11531121 |
| PTGSGKTL | ASKTGTEVF | 0.12193522 |
| TGSGKTL | ASKTGTEVF | 0.116389565 |
| TIGHVDHGK | ASKTGTEVF | 0.12049569 |
| TIGHVDHGKTT | ASKTGTEVF | 0.122488864 |
| GHVDHGKTT | ASKTGTEVF | 0.12531956 |
| AQFEHTLL | ASKTGTEVF | 0.11535816 |
| IVNQDEL | ASKTGTEVF | 0.12565145 |
| TNGSQFF | ASKTGTEVF | 0.12339082 |
| TGKGIVE | ASKTGTEVF | 0.124248326 |
| LISNASDA | ASKTGTEVF | 0.11805306 |
| AELRTLGELTKIAKAHGAQ | ASKTGTEVF | 0.12445279 |
| APNAVAAAEPSW | ASKTGTEVF | 0.11884843 |
| APQVAGAATAVL | ASKTGTEVF | 0.11958075 |

| | | |
|---|---|---|
| ARRDPAAAAAILPTDA | ASKTGTEVF | 0.11653244 |
| DAAARDVLAEAGLAEYF | ASKTGTEVF | 0.116421394 |
| DDQIVLSAKANLAKVEQ | ASKTGTEVF | 0.11919503 |
| DELLPDTGEDSDEDGH | ASKTGTEVF | 0.119420566 |
| ELPEAPANRATVPQD | ASKTGTEVF | 0.12118204 |
| FGQSKIERVTGIG | ASKTGTEVF | 0.11971314 |
| GAGGTGSTAGGAGGAGGAGGLY | ASKTGTEVF | 0.124101155 |
| GHGGTGAAGGNGGTGGAGGSGI | ASKTGTEVF | 0.124601275 |
| GNGGNGGAGGEGGPGGAGGAGA | ASKTGTEVF | 0.122981414 |
| GSTAGNGGNGGNATNGTIAGQP | ASKTGTEVF | 0.12089758 |
| GTGGDGGNGGNGGTGG | ASKTGTEVF | 0.11895691 |
| GVRRPAGIPADAPV | ASKTGTEVF | 0.11917754 |
| GYRQVTDAHLLTLARR | ASKTGTEVF | 0.117003426 |
| IAKMGQDGHDRG | ASKTGTEVF | 0.118342645 |
| KAEPSIEIQYT | ASKTGTEVF | 0.118291326 |
| KPVRIRMAADEDTVAVT | ASKTGTEVF | 0.12586498 |
| MPPRIAGMRLLVIKPEP | ASKTGTEVF | 0.11499526 |
| MKFYAVALFRSRGVPPTR | ASKTGTEVF | 0.1166617 |
| NAGGFGILGAVAHSP | ASKTGTEVF | 0.12871085 |
| NEGREITLAMAEM | ASKTGTEVF | 0.11630386 |
| NELVEWSHPLGHRWRWE | ASKTGTEVF | 0.12156725 |
| NLGAGGTGGNGGNPG | ASKTGTEVF | 0.11906333 |
| NTEWPKWETVMAADPVD | ASKTGTEVF | 0.11603725 |
| PASAAPAETQP | ASKTGTEVF | 0.124496475 |
| QQSSDGKNGVRAAEITGE | ASKTGTEVF | 0.11732398 |
| RAPDAVTTSWWKE | ASKTGTEVF | 0.12340594 |
| SGFGNVGAHTSGWFNQ | ASKTGTEVF | 0.12562123 |
| SGVAGAVGQAASVGGLKV | ASKTGTEVF | 0.12013649 |
| SNGGAGTGSAGGNGGT | ASKTGTEVF | 0.11653614 |
| TGGAGGEGGAAG | ASKTGTEVF | 0.12161432 |
| TGGDGGAAGAGGAPGHG | ASKTGTEVF | 0.11732578 |
| TMPPLLAAIANAGRHGV | ASKTGTEVF | 0.116972595 |
| TSSTAGNSGGA | ASKTGTEVF | 0.11583136 |
| VPATVPVGILC | ASKTGTEVF | 0.11979245 |
| VQRPDSGGVVLTGRIS | ASKTGTEVF | 0.11697466 |
| WTPNPVPATKGVPAGLVQAT | ASKTGTEVF | 0.12599982 |
| YALKRALAKFSDRFDVVII | ASKTGTEVF | 0.12195219 |
| YLIGSVIV | ASKTGTEVF | 0.11868457 |
| YGQLVAVID | ASKTGTEVF | 0.11797415 |
| VTVGGMTLPA | ASKTGTEVF | 0.117408365 |
| VLALPAV | ASKTGTEVF | 0.11638277 |
| VIPPTLNYET | ASKTGTEVF | 0.11706501 |
| VGMGGMHPSSG | ASKTGTEVF | 0.11720409 |
| VGLAGGAATA | ASKTGTEVF | 0.11764966 |
| VGGAGGAGGSGP | ASKTGTEVF | 0.117134795 |
| VFAAFMV | ASKTGTEVF | 0.12567404 |
| TLYDKLR | ASKTGTEVF | 0.12164413 |
| TLAVFTVL | ASKTGTEVF | 0.11649287 |
| TGSVQPARV | ASKTGTEVF | 0.1175895 |
| TGGGAGSGGT | ASKTGTEVF | 0.119076565 |
| SHLETSL | ASKTGTEVF | 0.121642545 |

| | | |
|---|---|---|
| SFTTGMT | ASKTGTEVF | 0.11503999 |
| QVEVAKRMGVRQPTVS | ASKTGTEVF | 0.11737553 |
| QGSGGPALAM | ASKTGTEVF | 0.1176213 |
| MLAPSNP | ASKTGTEVF | 0.11673549 |
| LVNNAAI | ASKTGTEVF | 0.11816285 |
| LTGEINTLA | ASKTGTEVF | 0.12769045 |
| LSIFGPV | ASKTGTEVF | 0.119151205 |
| LLALLANL | ASKTGTEVF | 0.11803595 |
| LISNASDALDKLR | ASKTGTEVF | 0.11902875 |
| LINEAVP | ASKTGTEVF | 0.12698899 |
| LGYAFSGL | ASKTGTEVF | 0.13054012 |
| LDFVAVAL | ASKTGTEVF | 0.1209533 |
| LAQPGAE | ASKTGTEVF | 0.117253445 |
| ISNLGAVTGF | ASKTGTEVF | 0.12083571 |
| HQEDPGRI | ASKTGTEVF | 0.119637065 |
| GTPGEPTP | ASKTGTEVF | 0.11571996 |
| GFFNSGS | ASKTGTEVF | 0.11535195 |
| GAGGGGLTAGAN | ASKTGTEVF | 0.1201394 |
| FVDALVNM | ASKTGTEVF | 0.11588734 |
| FLTVAPEL | ASKTGTEVF | 0.12588911 |
| FETLTAL | ASKTGTEVF | 0.12471572 |
| DVINPIEL | ASKTGTEVF | 0.11616764 |
| APVSGAPV | ASKTGTEVF | 0.115902826 |
| ALVGGLAW | ASKTGTEVF | 0.116989456 |
| ALAAAAATAI | ASKTGTEVF | 0.1190936 |
| AFGLGLLL | ASKTGTEVF | 0.12250064 |
| AFFAAASL | ASKTGTEVF | 0.11666233 |
| AAAALPAFSPP | ASKTGTEVF | 0.12223553 |
| VVLAPAL | ASKTGTEVF | 0.1249652 |
| TIGHVDHGKTTL | ASKTGTEVF | 0.11881046 |
| LISNASDALDK | ASKTGTEVF | 0.1241573 |
| LISNASDAL | ASKTGTEVF | 0.11886629 |
| ISNASDALDK | ASKTGTEVF | 0.11602349 |
| NQAETLVYQTE | ASKTGTEVF | 0.118932955 |
| TGGTGGTAGAG | ASKTGTEVF | 0.11644707 |
| TGSTAGGAGGA | ASKTGTEVF | 0.1167004 |
| DPLAVPVAAGPV | ASKTGTEVF | 0.11749049 |
| GGTGGTPGNGGN | ASKTGTEVF | 0.11698167 |
| PAGLAGVDPATL | ASKTGTEVF | 0.11663241 |
| GNNLVVEAPHAGASV | ASKTGTEVF | 0.11885566 |
| KVTRIMSASTLEQPA | ASKTGTEVF | 0.11573655 |
| LGFALPIALPRRTVV | ASKTGTEVF | 0.11912501 |
| TRLGNLTARKLVVLV | ASKTGTEVF | 0.12674369 |
| WSQQRRLIYFPSAGP | ASKTGTEVF | 0.13121185 |
| GPNGNPDPMAAAVDIR | ASKTGTEVF | 0.11805417 |
| TQRCVSVADLRAAASR | ASKTGTEVF | 0.11807031 |
| QFVLDVAEHPVEVLTETV | ASKTGTEVF | 0.11682501 |
| GGAGGTGGAGGNGGTQAGDG | ASKTGTEVF | 0.12206614 |
| PLLATLRAIAVDVDAKPAQV | ASKTGTEVF | 0.11964288 |
| APKFRTGGYMIDFWGVGYQVAK | ASKTGTEVF | 0.119535126 |
| GAGGAGGAGGAAGGPGGTGGTG | ASKTGTEVF | 0.11886117 |

| | | |
|---|---|---|
| IGALGLLATDVAPAGSVAFGVA | ASKTGTEVF | 0.11732724 |
| AGGAGGIGGNGGAGGAGGVLMG | ASKTGTEVF | 0.11790995 |
| ADELIGLL | ASKTGTEVF | 0.117469184 |
| ADRALKA | ASKTGTEVF | 0.12087203 |
| AGAVGQVGGAGGA | ASKTGTEVF | 0.115174115 |
| AGGGGGLGGAP | ASKTGTEVF | 0.1190946 |
| AGGLLYGNGGAG | ASKTGTEVF | 0.11795248 |
| AGLESIGL | ASKTGTEVF | 0.11716649 |
| AGLKTGII | ASKTGTEVF | 0.1155095 |
| ASLAALLL | ASKTGTEVF | 0.1190009 |
| ASVINISSI | ASKTGTEVF | 0.11873605 |
| ATPPGPA | ASKTGTEVF | 0.122734845 |
| ATVASALV | ASKTGTEVF | 0.1160177 |
| ATVASLIL | ASKTGTEVF | 0.12183877 |
| AVNGPVLIRH | ASKTGTEVF | 0.12387055 |
| DGAAGAVGI | ASKTGTEVF | 0.1192794 |
| DPLLHGI | ASKTGTEVF | 0.116155855 |
| ELGAALGAL | ASKTGTEVF | 0.12564696 |
| ELGRTQAV | ASKTGTEVF | 0.11556099 |
| FVAGGYLS | ASKTGTEVF | 0.11847058 |
| GASGGAGGAGGW | ASKTGTEVF | 0.11992995 |
| GGAGAGGDGGAGGI | ASKTGTEVF | 0.117023796 |
| GTGHNGSQPGMGGNG | ASKTGTEVF | 0.12385565 |
| GYAAASASAS | ASKTGTEVF | 0.12534119 |
| HQVDVAAVQQG | ASKTGTEVF | 0.11679812 |
| IGAAIAHT | ASKTGTEVF | 0.11658237 |
| IGMIEVAQL | ASKTGTEVF | 0.11692416 |
| IRLAPEHTSAMD | ASKTGTEVF | 0.11546298 |
| IYNVSVL | ASKTGTEVF | 0.12223492 |
| KQGFLGM | ASKTGTEVF | 0.11777416 |
| LESVITQL | ASKTGTEVF | 0.12044962 |
| LGLGALVVV | ASKTGTEVF | 0.12165048 |
| LIGNGGNGGSGGI | ASKTGTEVF | 0.118979014 |
| LNAINAPTL | ASKTGTEVF | 0.12415401 |
| LQLLNTGV | ASKTGTEVF | 0.122329205 |
| LQTSIEF | ASKTGTEVF | 0.11801786 |
| MSFAVSEL | ASKTGTEVF | 0.1179503 |
| NTGALGL | ASKTGTEVF | 0.1167185 |
| PPGAGGAGGAGTT | ASKTGTEVF | 0.11791124 |
| QLVGLGI | ASKTGTEVF | 0.13463765 |
| RFLEQVHQL | ASKTGTEVF | 0.11701244 |
| RGVGYVLR | ASKTGTEVF | 0.11971115 |
| SATGVFAGVI | ASKTGTEVF | 0.11845104 |
| SLPAITLP | ASKTGTEVF | 0.11630343 |
| STALAFTV | ASKTGTEVF | 0.119302176 |
| SVLTVGLGK | ASKTGTEVF | 0.1206559 |
| TALAGAGSGI | ASKTGTEVF | 0.12562585 |
| TGLPFTGI | ASKTGTEVF | 0.116034746 |
| VAEHVTV | ASKTGTEVF | 0.12283315 |
| VDIVLNSL | ASKTGTEVF | 0.123646885 |
| VLTGGGAVL | ASKTGTEVF | 0.11744158 |

| | | |
|---|---|---|
| VNVGLAYGI | ASKTGTEVF | 0.11602523 |
| VSSAAVGAV | ASKTGTEVF | 0.11620948 |
| VTSVYEAGV | ASKTGTEVF | 0.11979449 |
| WVALDRGA | ASKTGTEVF | 0.12152171 |
| YLVGSVVI | ASKTGTEVF | 0.11826616 |
| GPPGTGKTLL | ASKTGTEVF | 0.121005416 |
| GVGMTKF | ASKTGTEVF | 0.12004743 |
| IGIGGSDL | ASKTGTEVF | 0.1227542 |
| IGTLLAEL | ASKTGTEVF | 0.115956314 |
| IYTSGTT | ASKTGTEVF | 0.118902944 |
| LSFPAGV | ASKTGTEVF | 0.11796934 |
| LYTSGTT | ASKTGTEVF | 0.11655061 |
| VGAFPFGF | ASKTGTEVF | 0.116649434 |
| VGAGTVEF | ASKTGTEVF | 0.12002203 |
| VGKSTLLL | ASKTGTEVF | 0.121130474 |
| AALAHAGGGIV | ASKTGTEVF | 0.12017002 |
| AALVAVSLL | ASKTGTEVF | 0.12759472 |
| AAVSRAVAAL | ASKTGTEVF | 0.13362686 |
| AGGSSGIGGF | ASKTGTEVF | 0.11774857 |
| AGLTVANANG | ASKTGTEVF | 0.12168707 |
| AGLVGAVL | ASKTGTEVF | 0.11611986 |
| ALSQAFTV | ASKTGTEVF | 0.11707544 |
| ALTFSVSL | ASKTGTEVF | 0.11595179 |
| ALTGAALVAL | ASKTGTEVF | 0.11993346 |
| ASFSIGAGVL | ASKTGTEVF | 0.1153807 |
| DGSVYKI | ASKTGTEVF | 0.12267133 |
| EQLAEIV | ASKTGTEVF | 0.11892535 |
| FAVSELTVM | ASKTGTEVF | 0.12812518 |
| GAAPLAAGL | ASKTGTEVF | 0.11995216 |
| GADGGDGGSGGAGG | ASKTGTEVF | 0.11685017 |
| GFGLDTGGAGG | ASKTGTEVF | 0.11918734 |
| GLASGGLL | ASKTGTEVF | 0.12077608 |
| GRIAGAAVLL | ASKTGTEVF | 0.11624078 |
| GSAGSLNTSF | ASKTGTEVF | 0.11614237 |
| GTGQIGI | ASKTGTEVF | 0.117683664 |
| GTLIALIL | ASKTGTEVF | 0.116426684 |
| IAATMGFVVG | ASKTGTEVF | 0.12099653 |
| IFAQALL | ASKTGTEVF | 0.12441474 |
| IGAAVALGF | ASKTGTEVF | 0.12129805 |
| IGLTPLAL | ASKTGTEVF | 0.12101646 |
| ILGSNTSTL | ASKTGTEVF | 0.11530078 |
| IVFSLTVL | ASKTGTEVF | 0.120189585 |
| LASKATGF | ASKTGTEVF | 0.13788307 |
| LASVALIL | ASKTGTEVF | 0.11888558 |
| LFAVGLTL | ASKTGTEVF | 0.122212544 |
| LFNGGIGTYI | ASKTGTEVF | 0.12795895 |
| LTLGAGII | ASKTGTEVF | 0.11863394 |
| LVAAALGDI | ASKTGTEVF | 0.121628195 |
| NTAFVQLGI | ASKTGTEVF | 0.11845748 |
| QLFSGAEL | ASKTGTEVF | 0.12049619 |
| SAATFITL | ASKTGTEVF | 0.116500296 |

| | | |
|---|---|---|
| SNYGLANLGV | ASKTGTEVF | 0.1167255 |
| SYGGAYAVM | ASKTGTEVF | 0.12560286 |
| TTGVGGLTLG | ASKTGTEVF | 0.12436131 |
| TTQTIYI | ASKTGTEVF | 0.12028585 |
| VAAGAGLGAVI | ASKTGTEVF | 0.124170594 |
| VAAVVTLL | ASKTGTEVF | 0.12369915 |
| VGFLGGLI | ASKTGTEVF | 0.117879875 |
| VGVIPILL | ASKTGTEVF | 0.11956268 |
| VIFASSLI | ASKTGTEVF | 0.11766468 |
| VKGGDGGAAAG | ASKTGTEVF | 0.11703254 |
| VTKTFTNI | ASKTGTEVF | 0.11716832 |
| VVAAAILL | ASKTGTEVF | 0.11925492 |
| YAALDVEL | ASKTGTEVF | 0.12111835 |
| LASVALLL | ASKTGTEVF | 0.12809429 |
| LDAVHRV | ASKTGTEVF | 0.122229755 |
| QGLSSSIF | ASKTGTEVF | 0.12368883 |
| RGTFANIRL | ASKTGTEVF | 0.11817012 |
| RQLWWGHRI | ASKTGTEVF | 0.117188945 |
| YGPPGTGKTLL | ASKTGTEVF | 0.117948264 |
| AALVGAPANVA | ASKTGTEVF | 0.11743758 |
| AATFRRTE | ASKTGTEVF | 0.12912162 |
| AGALGGGI | ASKTGTEVF | 0.12015286 |
| AGGAGGAGGIGAG | ASKTGTEVF | 0.116759345 |
| AGSGALGL | ASKTGTEVF | 0.12143572 |
| AQIADQI | ASKTGTEVF | 0.12244752 |
| ASAVAVTYL | ASKTGTEVF | 0.12120415 |
| ASTVLGGTGG | ASKTGTEVF | 0.117561236 |
| AVGGEPAK | ASKTGTEVF | 0.119220845 |
| CEQGTFGGTC | ASKTGTEVF | 0.12059264 |
| DEGKAPAG | ASKTGTEVF | 0.13329585 |
| DTVVFPAL | ASKTGTEVF | 0.11563968 |
| FEGITMAG | ASKTGTEVF | 0.123416744 |
| FPLVVQD | ASKTGTEVF | 0.11755881 |
| GAFTTPQ | ASKTGTEVF | 0.11604432 |
| GFGGFGTAGGVGGA | ASKTGTEVF | 0.1199822 |
| GFLAQFL | ASKTGTEVF | 0.12199004 |
| GGAGGAGGNAGVA | ASKTGTEVF | 0.11906467 |
| GGVGTAGAGGAGGAGG | ASKTGTEVF | 0.121755466 |
| GNGGGGDAGNAGSGGN | ASKTGTEVF | 0.115631565 |
| GTGGAGGAGNDAGS | ASKTGTEVF | 0.11713731 |
| GTTNTGWFNS | ASKTGTEVF | 0.12836652 |
| IFMGPLFTL | ASKTGTEVF | 0.121967256 |
| ILAHAIEAI | ASKTGTEVF | 0.12869877 |
| IRSAIREAL | ASKTGTEVF | 0.118080966 |
| ITFAGGG | ASKTGTEVF | 0.11683933 |
| ITGMSGHQEV | ASKTGTEVF | 0.118246816 |
| IVAGLACTL | ASKTGTEVF | 0.12022562 |
| IVGGGLAGA | ASKTGTEVF | 0.11715674 |
| IVTGVLNI | ASKTGTEVF | 0.12263039 |
| LAQQGCI | ASKTGTEVF | 0.12212834 |
| LCLHVEN | ASKTGTEVF | 0.11593381 |

| | | |
|---|---|---|
| LDAISDGS | ASKTGTEVF | 0.11781036 |
| LGGFAAML | ASKTGTEVF | 0.12023014 |
| LGGKNPMI | ASKTGTEVF | 0.11983323 |
| LGPKGVAAIL | ASKTGTEVF | 0.11641184 |
| LMVGVSI | ASKTGTEVF | 0.12562236 |
| LVGAAFVL | ASKTGTEVF | 0.12814471 |
| LVLAVAPL | ASKTGTEVF | 0.12379617 |
| MAAIAVIDAA | ASKTGTEVF | 0.11721098 |
| NNGINAELQ | ASKTGTEVF | 0.11547419 |
| RGEPLVLVHG | ASKTGTEVF | 0.12097933 |
| RGGAGGAAGLI | ASKTGTEVF | 0.12065037 |
| RSYRMHPAV | ASKTGTEVF | 0.11998775 |
| SAAHLLAGQLA | ASKTGTEVF | 0.11695237 |
| SAAVGSY | ASKTGTEVF | 0.11667739 |
| SPHAVLHRI | ASKTGTEVF | 0.13096952 |
| SQIAKVL | ASKTGTEVF | 0.11656389 |
| TALGHLVP | ASKTGTEVF | 0.116078034 |
| TAPTTSGAA | ASKTGTEVF | 0.11661519 |
| TDGDDGNGGSPGT | ASKTGTEVF | 0.12600537 |
| TDGLGEN | ASKTGTEVF | 0.11720321 |
| TGGPPLPGQA | ASKTGTEVF | 0.1188646 |
| TIPAITI | ASKTGTEVF | 0.12508643 |
| TIPGFTI | ASKTGTEVF | 0.11933631 |
| TSIASGALA | ASKTGTEVF | 0.11848278 |
| TVTLPGAAI | ASKTGTEVF | 0.11794591 |
| VASGLSAALP | ASKTGTEVF | 0.11933014 |
| VGAGVIGI | ASKTGTEVF | 0.118589595 |
| VGPKRFGV | ASKTGTEVF | 0.11834059 |
| VNVAFPD | ASKTGTEVF | 0.11929005 |
| VSMTNSGVS | ASKTGTEVF | 0.120681465 |
| VTDVLGDLSV | ASKTGTEVF | 0.120391205 |
| VVVVGAGFAGL | ASKTGTEVF | 0.11878701 |
| VYTVGTVV | ASKTGTEVF | 0.11687114 |
| WTTFSGP | ASKTGTEVF | 0.11523742 |
| YDELVEAF | ASKTGTEVF | 0.12045908 |
| YGTGGAGGNGG | ASKTGTEVF | 0.11735853 |
| YTNVMAV | ASKTGTEVF | 0.122539856 |
| FLEQVHQL | ASKTGTEVF | 0.12633292 |
| GLSLLAVAV | ASKTGTEVF | 0.11793706 |
| LEQVHQL | ASKTGTEVF | 0.11920386 |
| LNAAGAI | ASKTGTEVF | 0.11591347 |
| PAVHASL | ASKTGTEVF | 0.11635673 |
| TELEDDA | ASKTGTEVF | 0.11774136 |
| TPPGPPN | ASKTGTEVF | 0.12117572 |
| TPPPPPPP | ASKTGTEVF | 0.11861877 |
| TQSPALI | ASSDANSDYT | 0.12112409 |
| EFAAAFPQ | ASSDANSDYT | 0.11954352 |
| LLSGNLDVL | ASSDANSDYT | 0.116596416 |
| GPANLGNYNI | ASSDANSDYT | 0.12014042 |
| TLGPTLP | ASSDANSDYT | 0.12650931 |
| VYAYSSCCK | ASSDANSDYT | 0.12237003 |

| | | |
|---|---|---|
| APAPAPAPNI | ASSDANSDYT | 0.11697259 |
| PGGPVPIP | ASSDANSDYT | 0.12113516 |
| FLTVALEP | ASSDANSDYT | 0.11767482 |
| SQYFGANL | ASSDANSDYT | 0.12238497 |
| GGVKPAA | ASSDANSDYT | 0.119523264 |
| GGTGGAGGAGGW | ASSDANSDYT | 0.119829826 |
| GVGGVAGLGGAGG | ASSDANSDYT | 0.12009257 |
| VLLVLATTI | ASSDANSDYT | 0.115932114 |
| IGIFAAML | ASSDANSDYT | 0.11690211 |
| AVGDLID | ASSDANSDYT | 0.119352914 |
| NISPLPL | ASSDANSDYT | 0.1253737 |
| VTAVFVAI | ASSDANSDYT | 0.12615314 |
| SGDDNDAAL | ASSDANSDYT | 0.11810258 |
| AVAALAHS | ASSDANSDYT | 0.11972446 |
| GGVTVMATP | ASSDANSDYT | 0.11873035 |
| AGLAANVRD | ASSDANSDYT | 0.11981221 |
| TNRPDLIDPALLRPG | ASSDANSDYT | 0.12029641 |
| LANLGVTAL | ASSDANSDYT | 0.117565855 |
| TGERIEL | ASSDANSDYT | 0.11674391 |
| NGEELAA | ASSDANSDYT | 0.120835975 |
| VLSLVWG | ASSDANSDYT | 0.117222644 |
| IGYALWL | ASSDANSDYT | 0.12040898 |
| LSLLLAPL | ASSDANSDYT | 0.11633157 |
| TLDEALK | ASSDANSDYT | 0.11661162 |
| TDGAGTAEGG | ASSDANSDYT | 0.11888732 |
| NAPGGAGGNGG | ASSDANSDYT | 0.11907123 |
| VSVFMGMGGNF | ASSDANSDYT | 0.11841061 |
| LASYLAAL | ASSDANSDYT | 0.121645145 |
| LVGGTPAL | ASSDANSDYT | 0.11756428 |
| VRNADVL | ASSDANSDYT | 0.12195174 |
| TCSSPFLVSAM | ASSDANSDYT | 0.13259107 |
| VHITDAE | ASSDANSDYT | 0.120908156 |
| EYAAPSF | ASSDANSDYT | 0.11737924 |
| VVDAFHGGL | ASSDANSDYT | 0.116971694 |
| ASVVDPQ | ASSDANSDYT | 0.1254241 |
| SYNEFGGLS | ASSDANSDYT | 0.11908451 |
| DPDGSVY | ASSDANSDYT | 0.115798794 |
| MVSTIHATE | ASSDANSDYT | 0.12102715 |
| LAAGVADPV | ASSDANSDYT | 0.11790199 |
| TKEQIAA | ASSDANSDYT | 0.11855359 |
| SLASPAAL | ASSDANSDYT | 0.1161857 |
| IGLVFYGV | ASSDANSDYT | 0.12777406 |
| FLLDASNL | ASSDANSDYT | 0.1320049 |
| YQVGNFP | ASSDANSDYT | 0.11891186 |
| ESLLSQST | ASSDANSDYT | 0.12039931 |
| LFTVGFAAVG | ASSDANSDYT | 0.115890115 |
| ATPPTPT | ASSDANSDYT | 0.12469036 |
| TDLATLG | ASSDANSDYT | 0.12647942 |
| SAGGTGR | ASSDANSDYT | 0.118306875 |
| GFSNSGS | ASSDANSDYT | 0.12393603 |
| GGNGGSGGTSV | ASSDANSDYT | 0.117384255 |

| | | |
|---|---|---|
| GTSVATGGAGN | ASSDANSDYT | 0.11834906 |
| NGGNGGSGGTS | ASSDANSDYT | 0.12526695 |
| AVVKFPH | ASSDANSDYT | 0.12131912 |
| TPPPPPP | ASSDANSDYT | 0.11988674 |
| AAPAAAAAPA | ASSDANSDYT | 0.11912516 |
| IALNEDL | ASSDANSDYT | 0.11901105 |
| IVNVSSM | ASSDANSDYT | 0.12093421 |
| TGIPPAPR | ASSDANSDYT | 0.12260167 |
| FDLGGGTF | ASSDANSDYT | 0.11625759 |
| FDLGGGTFDVS | ASSDANSDYT | 0.12235507 |
| NEPTAAA | ASSDANSDYT | 0.11745704 |
| NEPTAAALA | ASSDANSDYT | 0.12243048 |
| GSIINISSI | ASSDANSDYT | 0.119819775 |
| VAPGNAGTA | ASSDANSDYT | 0.12116447 |
| TGGGASLE | ASSDANSDYT | 0.11889845 |
| VAVYNHYKRI | ASSDANSDYT | 0.13083634 |
| TDPAPATT | ASSDANSDYT | 0.11880948 |
| TNRPDILDPALLRPG | ASSDANSDYT | 0.117792994 |
| LTTTTTS | ASSDANSDYT | 0.12255485 |
| PTGSGKTL | ASSDANSDYT | 0.6908699 |
| TGSGKTL | ASSDANSDYT | 0.87452614 |
| TIGHVDHGK | ASSDANSDYT | 0.78184485 |
| TIGHVDHGKTT | ASSDANSDYT | 0.8188032 |
| GHVDHGKTT | ASSDANSDYT | 0.72113264 |
| AQFEHTLL | ASSDANSDYT | 0.44950065 |
| IVNQDEL | ASSDANSDYT | 0.78874695 |
| TNGSQFF | ASSDANSDYT | 0.64806604 |
| TGKGIVE | ASSDANSDYT | 0.75578463 |
| LISNASDA | ASSDANSDYT | 0.8475181 |
| AELRTLGELTKIAKAHGAQ | ASSDANSDYT | 0.7156497 |
| APNAVAAAEPSW | ASSDANSDYT | 0.8735448 |
| APQVAGAATAVL | ASSDANSDYT | 0.87944293 |
| ARRDPAAAAAILPTDA | ASSDANSDYT | 0.56123126 |
| DAAARDVLAEAGLAEYF | ASSDANSDYT | 0.7539065 |
| DDQIVLSAKANLAKVEQ | ASSDANSDYT | 0.74770397 |
| DELLPDTGEDSDEDGH | ASSDANSDYT | 0.76368195 |
| ELPEAPANRATVPQD | ASSDANSDYT | 0.45939332 |
| FGQSKIERVTGIG | ASSDANSDYT | 0.82848907 |
| GAGGTGSTAGGAGGAGGAGGLY | ASSDANSDYT | 0.6144382 |
| GHGGTGAAGGNGGTGGAGGSGI | ASSDANSDYT | 0.13219507 |
| GNGGNGGAGGEGGPGGAGGAGA | ASSDANSDYT | 0.81319386 |
| GSTAGNGGNGGNATNGTIAGQP | ASSDANSDYT | 0.77958643 |
| GTGGDGGNGGNGGTGG | ASSDANSDYT | 0.72934216 |
| GVRRPAGIPADAPV | ASSDANSDYT | 0.85068977 |
| GYRQVTDAHLLTLARR | ASSDANSDYT | 0.3310088 |
| IAKMGQDGHDRG | ASSDANSDYT | 0.68662316 |
| KAEPSIEIQYT | ASSDANSDYT | 0.8411503 |
| KPVRIRMAADEDTVAVT | ASSDANSDYT | 0.8757403 |
| MPPRIAGMRLLVIKPEP | ASSDANSDYT | 0.5730685 |
| MKFYAVALFRSRGVPPTR | ASSDANSDYT | 0.7933911 |
| NAGGFGILGAVAHSP | ASSDANSDYT | 0.87365365 |

| | | |
|---|---|---|
| NEGREITLAMAEM | ASSDANSDYT | 0.82562757 |
| NELVEWSHPLGHRWRWE | ASSDANSDYT | 0.86335266 |
| NLGAGGTGGNGGNPG | ASSDANSDYT | 0.87473536 |
| NTEWPKWETVMAADPVD | ASSDANSDYT | 0.6302105 |
| PASAAPAETQP | ASSDANSDYT | 0.8625133 |
| QQSSDGKNGVRAAEITGE | ASSDANSDYT | 0.8627114 |
| RAPDAVTTSWWKE | ASSDANSDYT | 0.78381586 |
| SGFGNVGAHTSGWFNQ | ASSDANSDYT | 0.79418004 |
| SGVAGAVGQAASVGGLKV | ASSDANSDYT | 0.68488514 |
| SNGGAGTGSAGGNGGT | ASSDANSDYT | 0.73208773 |
| TGGAGGEGGAAG | ASSDANSDYT | 0.22881548 |
| TGGDGGAAGAGGAPGHG | ASSDANSDYT | 0.783712 |
| TMPPLLAAIANAGRHGV | ASSDANSDYT | 0.85478616 |
| TSSTAGNSGGA | ASSDANSDYT | 0.7193194 |
| VPATVPVGILC | ASSDANSDYT | 0.5833601 |
| VQRPDSGGVVLTGRIS | ASSDANSDYT | 0.6000222 |
| WTPNPVPATKGVPAGLVQAT | ASSDANSDYT | 0.72366655 |
| YALKRALAKFSDRFDVVII | ASSDANSDYT | 0.73403084 |
| YLIGSVIV | ASSDANSDYT | 0.2618007 |
| YGQLVAVID | ASSDANSDYT | 0.7333783 |
| VTVGGMTLPA | ASSDANSDYT | 0.8739425 |
| VLALPAV | ASSDANSDYT | 0.8674066 |
| VIPPTLNYET | ASSDANSDYT | 0.42429227 |
| VGMGGMHPSSG | ASSDANSDYT | 0.8746941 |
| VGLAGGAATA | ASSDANSDYT | 0.7860404 |
| VGGAGGAGGSGP | ASSDANSDYT | 0.85756075 |
| VFAAFMV | ASSDANSDYT | 0.7918265 |
| TLYDKLR | ASSDANSDYT | 0.70220065 |
| TLAVFTVL | ASSDANSDYT | 0.8081267 |
| TGSVQPARV | ASSDANSDYT | 0.72204316 |
| TGGGAGSGGT | ASSDANSDYT | 0.55430615 |
| SHLETSL | ASSDANSDYT | 0.87233615 |
| SFTTGMT | ASSDANSDYT | 0.73659706 |
| QVEVAKRMGVRQPTVS | ASSDANSDYT | 0.6954353 |
| QGSGGPALAM | ASSDANSDYT | 0.5812985 |
| MLAPSNP | ASSDANSDYT | 0.855504 |
| LVNNAAI | ASSDANSDYT | 0.5926702 |
| LTGEINTLA | ASSDANSDYT | 0.5817709 |
| LSIFGPV | ASSDANSDYT | 0.4684347 |
| LLALLANL | ASSDANSDYT | 0.7632282 |
| LISNASDALDKLR | ASSDANSDYT | 0.8715981 |
| LINEAVP | ASSDANSDYT | 0.8241462 |
| LGYAFSGL | ASSDANSDYT | 0.7018508 |
| LDFVAVAL | ASSDANSDYT | 0.8206228 |
| LAQPGAE | ASSDANSDYT | 0.82541597 |
| ISNLGAVTGF | ASSDANSDYT | 0.80228555 |
| HQEDPGRI | ASSDANSDYT | 0.86239386 |
| GTPGEPTP | ASSDANSDYT | 0.6659608 |
| GFFNSGS | ASSDANSDYT | 0.8597891 |
| GAGGGGLTAGAN | ASSDANSDYT | 0.83376116 |
| FVDALVNM | ASSDANSDYT | 0.85302633 |

| | | |
|---|---|---|
| FLTVAPEL | ASSDANSDYT | 0.8555358 |
| FETLTAL | ASSDANSDYT | 0.8143469 |
| DVINPIEL | ASSDANSDYT | 0.7772919 |
| APVSGAPV | ASSDANSDYT | 0.32606292 |
| ALVGGLAW | ASSDANSDYT | 0.7358322 |
| ALAAAAATAI | ASSDANSDYT | 0.8224373 |
| AFGLGLLL | ASSDANSDYT | 0.7892848 |
| AFFAAASL | ASSDANSDYT | 0.83445376 |
| AAAALPAFSPP | ASSDANSDYT | 0.83203185 |
| VVLAPAL | ASSDANSDYT | 0.65995014 |
| TIGHVDHGKTTL | ASSDANSDYT | 0.8609336 |
| LISNASDALDK | ASSDANSDYT | 0.8539801 |
| LISNASDAL | ASSDANSDYT | 0.49659827 |
| ISNASDALDK | ASSDANSDYT | 0.7690648 |
| NQAETLVYQTE | ASSDANSDYT | 0.8879973 |
| TGGTGGTAGAG | ASSDANSDYT | 0.7531752 |
| TGSTAGGAGGA | ASSDANSDYT | 0.75535595 |
| DPLAVPVAAGPV | ASSDANSDYT | 0.68900764 |
| GGTGGTPGNGGN | ASSDANSDYT | 0.8448255 |
| PAGLAGVDPATL | ASSDANSDYT | 0.5621302 |
| GNNLVVEAPHAGASV | ASSDANSDYT | 0.8798522 |
| KVTRIMSASTLEQPA | ASSDANSDYT | 0.23444176 |
| LGFALPIALPRRTVV | ASSDANSDYT | 0.67303973 |
| TRLGNLTARKLVVLV | ASSDANSDYT | 0.6199295 |
| WSQQRRLIYFPSAGP | ASSDANSDYT | 0.34429967 |
| GPNGNPDPMAAAVDIR | ASSDANSDYT | 0.86023366 |
| TQRCVSVADLRAAASR | ASSDANSDYT | 0.73513746 |
| QFVLDVAEHPVEVLTETV | ASSDANSDYT | 0.8511817 |
| GGAGGTGGAGGNGGTQAGDG | ASSDANSDYT | 0.7722827 |
| PLLATLRAIAVDVDAKPAQV | ASSDANSDYT | 0.8706068 |
| APKFRTGGYMIDFWGVGYQVAK | ASSDANSDYT | 0.77230537 |
| GAGGAGGAGGAAGGPGGTGGTG | ASSDANSDYT | 0.81484926 |
| IGALGLLATDVAPAGSVAFGVA | ASSDANSDYT | 0.65397406 |
| AGGAGGIGGNGGAGGAGGVLMG | ASSDANSDYT | 0.8509121 |
| ADELIGLL | ASSDANSDYT | 0.8741323 |
| ADRALKA | ASSDANSDYT | 0.823606 |
| AGAVGQVGGAGGA | ASSDANSDYT | 0.49627334 |
| AGGGGGLGGAP | ASSDANSDYT | 0.6505182 |
| AGGLLYGNGGAG | ASSDANSDYT | 0.811128 |
| AGLESIGL | ASSDANSDYT | 0.24692115 |
| AGLKTGII | ASSDANSDYT | 0.19346863 |
| ASLAALLL | ASSDANSDYT | 0.6545328 |
| ASVINISSI | ASSDANSDYT | 0.6360124 |
| ATPPGPA | ASSDANSDYT | 0.8598356 |
| ATVASALV | ASSDANSDYT | 0.8588692 |
| ATVASLIL | ASSDANSDYT | 0.7426399 |
| AVNGPVLIRH | ASSDANSDYT | 0.82702714 |
| DGAAGAVGI | ASSDANSDYT | 0.79166055 |
| DPLLHGI | ASSDANSDYT | 0.097550996 |
| ELGAALGAL | ASSDANSDYT | 0.7404864 |
| ELGRTQAV | ASSDANSDYT | 0.58917326 |

| | | |
|---|---|---|
| FVAGGYLS | ASSDANSDYT | 0.8757553 |
| GASGGAGGAGGW | ASSDANSDYT | 0.6935558 |
| GGAGAGGDGGAGGI | ASSDANSDYT | 0.6668005 |
| GTGHNGSQPGMGGNG | ASSDANSDYT | 0.7812815 |
| GYAAASASAS | ASSDANSDYT | 0.7068577 |
| HQVDVAAVQQG | ASSDANSDYT | 0.74493074 |
| IGAAIAHT | ASSDANSDYT | 0.6129505 |
| IGMIEVAQL | ASSDANSDYT | 0.13499519 |
| IRLAPEHTSAMD | ASSDANSDYT | 0.86744004 |
| IYNVSVL | ASSDANSDYT | 0.41904303 |
| KQGFLGM | ASSDANSDYT | 0.65997815 |
| LESVITQL | ASSDANSDYT | 0.3213026 |
| LGLGALVVV | ASSDANSDYT | 0.17372583 |
| LIGNGGNGGSGGI | ASSDANSDYT | 0.4498263 |
| LNAINAPTL | ASSDANSDYT | 0.8572736 |
| LQLLNTGV | ASSDANSDYT | 0.7283118 |
| LQTSIEF | ASSDANSDYT | 0.73182034 |
| MSFAVSEL | ASSDANSDYT | 0.876886 |
| NTGALGL | ASSDANSDYT | 0.7122603 |
| PPGAGGAGGAGTT | ASSDANSDYT | 0.73967636 |
| QLVGLGI | ASSDANSDYT | 0.7144255 |
| RFLEQVHQL | ASSDANSDYT | 0.6596993 |
| RGVGYVLR | ASSDANSDYT | 0.65445256 |
| SATGVFAGVI | ASSDANSDYT | 0.636789 |
| SLPAITLP | ASSDANSDYT | 0.8903565 |
| STALAFTV | ASSDANSDYT | 0.67942536 |
| SVLTVGLGK | ASSDANSDYT | 0.628987 |
| TALAGAGSGI | ASSDANSDYT | 0.75437534 |
| TGLPFTGI | ASSDANSDYT | 0.8630887 |
| VAEHVTV | ASSDANSDYT | 0.8848474 |
| VDIVLNSL | ASSDANSDYT | 0.691681 |
| VLTGGGAVL | ASSDANSDYT | 0.75074846 |
| VNVGLAYGI | ASSDANSDYT | 0.76430094 |
| VSSAAVGAV | ASSDANSDYT | 0.76450914 |
| VTSVYEAGV | ASSDANSDYT | 0.70584124 |
| WVALDRGA | ASSDANSDYT | 0.5397909 |
| YLVGSVVI | ASSDANSDYT | 0.8371457 |
| GPPGTGKTLL | ASSDANSDYT | 0.8603081 |
| GVGMTKF | ASSDANSDYT | 0.73399746 |
| IGIGGSDL | ASSDANSDYT | 0.60037625 |
| IGTLLAEL | ASSDANSDYT | 0.75761354 |
| IYTSGTT | ASSDANSDYT | 0.63025945 |
| LSFPAGV | ASSDANSDYT | 0.7000895 |
| LYTSGTT | ASSDANSDYT | 0.8132243 |
| VGAFPFGF | ASSDANSDYT | 0.5112747 |
| VGAGTVEF | ASSDANSDYT | 0.72534245 |
| VGKSTLLL | ASSDANSDYT | 0.88531244 |
| AALAHAGGGIV | ASSDANSDYT | 0.6863624 |
| AALVAVSLL | ASSDANSDYT | 0.8104188 |
| AAVSRAVAAL | ASSDANSDYT | 0.845112 |
| AGGSSGIGGF | ASSDANSDYT | 0.80614996 |

| | | |
|---|---|---|
| AGLTVANANG | ASSDANSDYT | 0.8646295 |
| AGLVGAVL | ASSDANSDYT | 0.8087407 |
| ALSQAFTV | ASSDANSDYT | 0.7965474 |
| ALTFSVSL | ASSDANSDYT | 0.77013755 |
| ALTGAALVAL | ASSDANSDYT | 0.76523757 |
| ASFSIGAGVL | ASSDANSDYT | 0.7842821 |
| DGSVYKI | ASSDANSDYT | 0.76409066 |
| EQLAEIV | ASSDANSDYT | 0.68785256 |
| FAVSELTVM | ASSDANSDYT | 0.8620112 |
| GAAPLAAGL | ASSDANSDYT | 0.7899828 |
| GADGGDGGSGGAGG | ASSDANSDYT | 0.796593 |
| GFGLDTGGAGG | ASSDANSDYT | 0.53425527 |
| GLASGGLL | ASSDANSDYT | 0.57341415 |
| GRIAGAAVLL | ASSDANSDYT | 0.8272351 |
| GSAGSLNTSF | ASSDANSDYT | 0.15023673 |
| GTGQIGI | ASSDANSDYT | 0.22701244 |
| GTLIALIL | ASSDANSDYT | 0.787043 |
| IAATMGFVVG | ASSDANSDYT | 0.85604775 |
| IFAQALL | ASSDANSDYT | 0.14616981 |
| IGAAVALGF | ASSDANSDYT | 0.6585437 |
| IGLTPLAL | ASSDANSDYT | 0.52776414 |
| ILGSNTSTL | ASSDANSDYT | 0.8209177 |
| IVFSLTVL | ASSDANSDYT | 0.802363 |
| LASKATGF | ASSDANSDYT | 0.47528425 |
| LASVALIL | ASSDANSDYT | 0.83546865 |
| LFAVGLTL | ASSDANSDYT | 0.47367388 |
| LFNGGIGTYI | ASSDANSDYT | 0.37793478 |
| LTLGAGII | ASSDANSDYT | 0.8658826 |
| LVAAALGDI | ASSDANSDYT | 0.83900917 |
| NTAFVQLGI | ASSDANSDYT | 0.67163986 |
| QLFSGAEL | ASSDANSDYT | 0.85126364 |
| SAATFITL | ASSDANSDYT | 0.64134187 |
| SNYGLANLGV | ASSDANSDYT | 0.88292974 |
| SYGGAYAVM | ASSDANSDYT | 0.74290293 |
| TTGVGGLTLG | ASSDANSDYT | 0.41301492 |
| TTQTIYI | ASSDANSDYT | 0.83184624 |
| VAAGAGLGAVI | ASSDANSDYT | 0.8188979 |
| VAAVVTLL | ASSDANSDYT | 0.81613827 |
| VGFLGGLI | ASSDANSDYT | 0.7980536 |
| VGVIPILL | ASSDANSDYT | 0.62645686 |
| VIFASSLI | ASSDANSDYT | 0.7143033 |
| VKGGDGGAAAG | ASSDANSDYT | 0.8642061 |
| VTKTFTNI | ASSDANSDYT | 0.8672112 |
| VVAAAILL | ASSDANSDYT | 0.56382746 |
| YAALDVEL | ASSDANSDYT | 0.85218376 |
| LASVALLL | ASSDANSDYT | 0.7731267 |
| LDAVHRV | ASSDANSDYT | 0.72549707 |
| QGLSSSIF | ASSDANSDYT | 0.88086843 |
| RGTFANIRL | ASSDANSDYT | 0.7892558 |
| RQLWWGHRI | ASSDANSDYT | 0.8299631 |
| YGPPGTGKTLL | ASSDANSDYT | 0.8341309 |

| | | |
|---|---|---|
| AALVGAPANVA | ASSDANSDYT | 0.8264668 |
| AATFRRTE | ASSDANSDYT | 0.5440867 |
| AGALGGGI | ASSDANSDYT | 0.80274093 |
| AGGAGGAGGIGAG | ASSDANSDYT | 0.78222024 |
| AGSGALGL | ASSDANSDYT | 0.85834765 |
| AQIADQI | ASSDANSDYT | 0.76435435 |
| ASAVAVTYL | ASSDANSDYT | 0.4642363 |
| ASTVLGGTGG | ASSDANSDYT | 0.79768884 |
| AVGGEPAK | ASSDANSDYT | 0.865774 |
| CEQGTFGGTC | ASSDANSDYT | 0.8132483 |
| DEGKAPAG | ASSDANSDYT | 0.8373102 |
| DTVVFPAL | ASSDANSDYT | 0.8742985 |
| FEGITMAG | ASSDANSDYT | 0.67555857 |
| FPLVVQD | ASSDANSDYT | 0.7732974 |
| GAFTTPQ | ASSDANSDYT | 0.8487297 |
| GFGGFGTAGGVGGA | ASSDANSDYT | 0.8658047 |
| GFLAQFL | ASSDANSDYT | 0.74734706 |
| GGAGGAGGNAGVA | ASSDANSDYT | 0.86810255 |
| GGVGTAGAGGAGGAGG | ASSDANSDYT | 0.6557083 |
| GNGGGGDAGNAGSGGN | ASSDANSDYT | 0.83025825 |
| GTGGAGGAGNDAGS | ASSDANSDYT | 0.5726459 |
| GTTNTGWFNS | ASSDANSDYT | 0.8300484 |
| IFMGPLFTL | ASSDANSDYT | 0.5911057 |
| ILAHAIEAI | ASSDANSDYT | 0.8473605 |
| IRSAIREAL | ASSDANSDYT | 0.8352766 |
| ITFAGGG | ASSDANSDYT | 0.8548831 |
| ITGMSGHQEV | ASSDANSDYT | 0.2091833 |
| IVAGLACTL | ASSDANSDYT | 0.84651005 |
| IVGGGLAGA | ASSDANSDYT | 0.8467673 |
| IVTGVLNI | ASSDANSDYT | 0.8749279 |
| LAQQGCI | ASSDANSDYT | 0.76279485 |
| LCLHVEN | ASSDANSDYT | 0.8435761 |
| LDAISDGS | ASSDANSDYT | 0.75324124 |
| LGGFAAML | ASSDANSDYT | 0.7307223 |
| LGGKNPMI | ASSDANSDYT | 0.60642266 |
| LGPKGVAAIL | ASSDANSDYT | 0.71917915 |
| LMVGVSI | ASSDANSDYT | 0.7158348 |
| LVGAAFVL | ASSDANSDYT | 0.64262015 |
| LVLAVAPL | ASSDANSDYT | 0.80466956 |
| MAAIAVIDAA | ASSDANSDYT | 0.8617377 |
| NNGINAELQ | ASSDANSDYT | 0.8568826 |
| RGEPLVLVHG | ASSDANSDYT | 0.85176766 |
| RGGAGGAAGLI | ASSDANSDYT | 0.857798 |
| RSYRMHPAV | ASSDANSDYT | 0.85451937 |
| SAAHLLAGQLA | ASSDANSDYT | 0.6889964 |
| SAAVGSY | ASSDANSDYT | 0.7775043 |
| SPHAVLHRI | ASSDANSDYT | 0.59778714 |
| SQIAKVL | ASSDANSDYT | 0.7448875 |
| TALGHLVP | ASSDANSDYT | 0.69586545 |
| TAPTTSGAA | ASSDANSDYT | 0.81556535 |
| TDGDDGNGGSPGT | ASSDANSDYT | 0.12146313 |

| | | |
|---|---|---|
| TDGLGEN | ASSDANSDYT | 0.69082665 |
| TGGPPLPGQA | ASSDANSDYT | 0.88387156 |
| TIPAITI | ASSDANSDYT | 0.5807078 |
| TIPGFTI | ASSDANSDYT | 0.8070451 |
| TSIASGALA | ASSDANSDYT | 0.7582755 |
| TVTLPGAAI | ASSDANSDYT | 0.67917264 |
| VASGLSAALP | ASSDANSDYT | 0.83471155 |
| VGAGVIGI | ASSDANSDYT | 0.5983802 |
| VGPKRFGV | ASSDANSDYT | 0.811429 |
| VNVAFPD | ASSDANSDYT | 0.87461776 |
| VSMTNSGVS | ASSDANSDYT | 0.7880692 |
| VTDVLGDLSV | ASSDANSDYT | 0.80027294 |
| VVVVGAGFAGL | ASSDANSDYT | 0.7609972 |
| VYTVGTVV | ASSDANSDYT | 0.45925817 |
| WTTFSGP | ASSDANSDYT | 0.8250568 |
| YDELVEAF | ASSDANSDYT | 0.114428945 |
| YGTGGAGGNGG | ASSDANSDYT | 0.74629784 |
| YTNVMAV | ASSDANSDYT | 0.58549774 |
| FLEQVHQL | ASSDANSDYT | 0.8153823 |
| GLSLLAVAV | ASSDANSDYT | 0.80428207 |
| LEQVHQL | ASSDANSDYT | 0.8364178 |
| LNAAGAI | ASSDANSDYT | 0.65890133 |
| PAVHASL | ASSDANSDYT | 0.66309416 |
| TELEDDA | ASSDANSDYT | 0.5591823 |
| TPPGPPN | ASSDANSDYT | 0.56138724 |
| TPPPPPPP | ASSDANSDYT | 0.8343048 |
| TQSPALI | ASSTPGQAGSDYT | 0.75503534 |
| EFAAAFPQ | ASSTPGQAGSDYT | 0.717764 |
| LLSGNLDVL | ASSTPGQAGSDYT | 0.8675703 |
| GPANLGNYNI | ASSTPGQAGSDYT | 0.81041443 |
| TLGPTLP | ASSTPGQAGSDYT | 0.500571 |
| VYAYSSCCK | ASSTPGQAGSDYT | 0.7166374 |
| APAPAPAPNI | ASSTPGQAGSDYT | 0.7683688 |
| PGGPVPIP | ASSTPGQAGSDYT | 0.7433018 |
| FLTVALEP | ASSTPGQAGSDYT | 0.11328428 |
| SQYFGANL | ASSTPGQAGSDYT | 0.82129383 |
| GGVKPAA | ASSTPGQAGSDYT | 0.735324 |
| GGTGGAGGAGGW | ASSTPGQAGSDYT | 0.6354049 |
| GVGGVAGLGGAGG | ASSTPGQAGSDYT | 0.70497847 |
| VLLVLATTI | ASSTPGQAGSDYT | 0.84131414 |
| IGIFAAML | ASSTPGQAGSDYT | 0.41342232 |
| AVGDLID | ASSTPGQAGSDYT | 0.72398984 |
| NISPLPL | ASSTPGQAGSDYT | 0.84231496 |
| VTAVFVAI | ASSTPGQAGSDYT | 0.8553474 |
| SGDDNDAAL | ASSTPGQAGSDYT | 0.8769101 |
| AVAALAHS | ASSTPGQAGSDYT | 0.8339265 |
| GGVTVMATP | ASSTPGQAGSDYT | 0.7439563 |
| AGLAANVRD | ASSTPGQAGSDYT | 0.8378495 |
| TNRPDLIDPALLRPG | ASSTPGQAGSDYT | 0.8153895 |
| LANLGVTAL | ASSTPGQAGSDYT | 0.28138676 |
| TGERIEL | ASSTPGQAGSDYT | 0.23652041 |

| | | |
|---|---|---|
| NGEELAA | ASSTPGQAGSDYT | 0.78092486 |
| VLSLVWG | ASSTPGQAGSDYT | 0.2816539 |
| IGYALWL | ASSTPGQAGSDYT | 0.79761976 |
| LSLLLAPL | ASSTPGQAGSDYT | 0.86855567 |
| TLDEALK | ASSTPGQAGSDYT | 0.742121 |
| TDGAGTAEGG | ASSTPGQAGSDYT | 0.53811383 |
| NAPGGAGGNGG | ASSTPGQAGSDYT | 0.74465823 |
| VSVFMGMGGNF | ASSTPGQAGSDYT | 0.77430296 |
| LASYLAAL | ASSTPGQAGSDYT | 0.8342855 |
| LVGGTPAL | ASSTPGQAGSDYT | 0.80886614 |
| VRNADVL | ASSTPGQAGSDYT | 0.83571523 |
| TCSSPFLVSAM | ASSTPGQAGSDYT | 0.8557489 |
| VHITDAE | ASSTPGQAGSDYT | 0.7074357 |
| EYAAPSF | ASSTPGQAGSDYT | 0.8346316 |
| VVDAFHGGL | ASSTPGQAGSDYT | 0.70528543 |
| ASVVDPQ | ASSTPGQAGSDYT | 0.7457967 |
| SYNEFGGLS | ASSTPGQAGSDYT | 0.7739829 |
| DPDGSVY | ASSTPGQAGSDYT | 0.78679633 |
| MVSTIHATE | ASSTPGQAGSDYT | 0.77583474 |
| LAAGVADPV | ASSTPGQAGSDYT | 0.7489561 |
| TKEQIAA | ASSTPGQAGSDYT | 0.52187943 |
| SLASPAAL | ASSTPGQAGSDYT | 0.7797631 |
| IGLVFYGV | ASSTPGQAGSDYT | 0.78477305 |
| FLLDASNL | ASSTPGQAGSDYT | 0.2768607 |
| YQVGNFP | ASSTPGQAGSDYT | 0.693698 |
| ESLLSQST | ASSTPGQAGSDYT | 0.8869791 |
| LFTVGFAAVG | ASSTPGQAGSDYT | 0.86902857 |
| ATPPTPT | ASSTPGQAGSDYT | 0.5963429 |
| TDLATLG | ASSTPGQAGSDYT | 0.64492977 |
| SAGGTGR | ASSTPGQAGSDYT | 0.7102163 |
| GFSNSGS | ASSTPGQAGSDYT | 0.3768524 |
| GGNGGSGGTSV | ASSTPGQAGSDYT | 0.80493814 |
| GTSVATGGAGN | ASSTPGQAGSDYT | 0.7520466 |
| NGGNGGSGGTS | ASSTPGQAGSDYT | 0.74937904 |
| AVVKFPH | ASSTPGQAGSDYT | 0.6951425 |
| TPPPPPP | ASSTPGQAGSDYT | 0.770324 |
| AAPAAAAAPA | ASSTPGQAGSDYT | 0.72055054 |
| IALNEDL | ASSTPGQAGSDYT | 0.8326836 |
| IVNVSSM | ASSTPGQAGSDYT | 0.15034403 |
| TGIPPAPR | ASSTPGQAGSDYT | 0.83634675 |
| FDLGGGTF | ASSTPGQAGSDYT | 0.6314069 |
| FDLGGGTFDVS | ASSTPGQAGSDYT | 0.6832995 |
| NEPTAAA | ASSTPGQAGSDYT | 0.37012994 |
| NEPTAAALA | ASSTPGQAGSDYT | 0.8104775 |
| GSIINISSI | ASSTPGQAGSDYT | 0.83709943 |
| VAPGNAGTA | ASSTPGQAGSDYT | 0.8447242 |
| TGGGASLE | ASSTPGQAGSDYT | 0.45181668 |
| VAVYNHYKRI | ASSTPGQAGSDYT | 0.86856174 |
| TDPAPATT | ASSTPGQAGSDYT | 0.87692654 |
| TNRPDILDPALLRPG | ASSTPGQAGSDYT | 0.71603394 |
| LTTTTTS | ASSTPGQAGSDYT | 0.44145465 |

| | | |
|---|---|---|
| PTGSGKTL | ASSTPGQAGSDYT | 0.83533937 |
| TGSGKTL | ASSTPGQAGSDYT | 0.7101864 |
| TIGHVDHGK | ASSTPGQAGSDYT | 0.84178174 |
| TIGHVDHGKTT | ASSTPGQAGSDYT | 0.80708194 |
| GHVDHGKTT | ASSTPGQAGSDYT | 0.57146734 |
| AQFEHTLL | ASSTPGQAGSDYT | 0.85224783 |
| IVNQDEL | ASSTPGQAGSDYT | 0.86250526 |
| TNGSQFF | ASSTPGQAGSDYT | 0.6393087 |
| TGKGIVE | ASSTPGQAGSDYT | 0.73114794 |
| LISNASDA | ASSTPGQAGSDYT | 0.7355605 |
| AELRTLGELTKIAKAHGAQ | ASSTPGQAGSDYT | 0.25841326 |
| APNAVAAAEPSW | ASSTPGQAGSDYT | 0.84597754 |
| APQVAGAATAVL | ASSTPGQAGSDYT | 0.662745 |
| ARRDPAAAAAILPTDA | ASSTPGQAGSDYT | 0.6718379 |
| DAAARDVLAEAGLAEYF | ASSTPGQAGSDYT | 0.772028 |
| DDQIVLSAKANLAKVEQ | ASSTPGQAGSDYT | 0.6817469 |
| DELLPDTGEDSDEDGH | ASSTPGQAGSDYT | 0.74030495 |
| ELPEAPANRATVPQD | ASSTPGQAGSDYT | 0.7409019 |
| FGQSKIERVTGIG | ASSTPGQAGSDYT | 0.79780257 |
| GAGGTGSTAGGAGGAGGAGGLY | ASSTPGQAGSDYT | 0.7837477 |
| GHGGTGAAGGNGGTGGAGGSGI | ASSTPGQAGSDYT | 0.76997626 |
| GNGGNGGAGGEGGPGGAGGAGA | ASSTPGQAGSDYT | 0.79591745 |
| GSTAGNGGNGGNATNGTIAGQP | ASSTPGQAGSDYT | 0.86868733 |
| GTGGDGGNGGNGGTGG | ASSTPGQAGSDYT | 0.541061 |
| GVRRPAGIPADAPV | ASSTPGQAGSDYT | 0.6517865 |
| GYRQVTDAHLLTLARR | ASSTPGQAGSDYT | 0.81300354 |
| IAKMGQDGHDRG | ASSTPGQAGSDYT | 0.38629332 |
| KAEPSIEIQYT | ASSTPGQAGSDYT | 0.81936264 |
| KPVRIRMAADEDTVAVT | ASSTPGQAGSDYT | 0.7982563 |
| MPPRIAGMRLLVIKPEP | ASSTPGQAGSDYT | 0.81037784 |
| MKFYAVALFRSRGVPPTR | ASSTPGQAGSDYT | 0.7039795 |
| NAGGFGILGAVAHSP | ASSTPGQAGSDYT | 0.8907231 |
| NEGREITLAMAEM | ASSTPGQAGSDYT | 0.84492433 |
| NELVEWSHPLGHRWRWE | ASSTPGQAGSDYT | 0.76551765 |
| NLGAGGTGGNGGNPG | ASSTPGQAGSDYT | 0.7060245 |
| NTEWPKWETVMAADPVD | ASSTPGQAGSDYT | 0.65555096 |
| PASAAPAETQP | ASSTPGQAGSDYT | 0.859583 |
| QQSSDGKNGVRAAEITGE | ASSTPGQAGSDYT | 0.8313899 |
| RAPDAVTTSWWKE | ASSTPGQAGSDYT | 0.5851598 |
| SGFGNVGAHTSGWFNQ | ASSTPGQAGSDYT | 0.73027265 |
| SGVAGAVGQAASVGGLKV | ASSTPGQAGSDYT | 0.867522 |
| SNGGAGTGSAGGNGGT | ASSTPGQAGSDYT | 0.8610488 |
| TGGAGGEGGAAG | ASSTPGQAGSDYT | 0.2945597 |
| TGGDGGAAGAGGAPGHG | ASSTPGQAGSDYT | 0.73261476 |
| TMPPLLAAIANAGRHGV | ASSTPGQAGSDYT | 0.37835738 |
| TSSTAGNSGGA | ASSTPGQAGSDYT | 0.70966744 |
| VPATVPVGILC | ASSTPGQAGSDYT | 0.61935997 |
| VQRPDSGGVVLTGRIS | ASSTPGQAGSDYT | 0.80221295 |
| WTPNPVPATKGVPAGLVQAT | ASSTPGQAGSDYT | 0.740572 |
| YALKRALAKFSDRFDVVII | ASSTPGQAGSDYT | 0.7552246 |
| YLIGSVIV | ASSTPGQAGSDYT | 0.8061582 |

| | | |
|---|---|---|
| YGQLVAVID | ASSTPGQAGSDYT | 0.8781055 |
| VTVGGMTLPA | ASSTPGQAGSDYT | 0.6740043 |
| VLALPAV | ASSTPGQAGSDYT | 0.7825395 |
| VIPPTLNYET | ASSTPGQAGSDYT | 0.45257095 |
| VGMGGMHPSSG | ASSTPGQAGSDYT | 0.37163773 |
| VGLAGGAATA | ASSTPGQAGSDYT | 0.630179 |
| VGGAGGAGGSGP | ASSTPGQAGSDYT | 0.74205565 |
| VFAAFMV | ASSTPGQAGSDYT | 0.21811002 |
| TLYDKLR | ASSTPGQAGSDYT | 0.86350274 |
| TLAVFTVL | ASSTPGQAGSDYT | 0.79221296 |
| TGSVQPARV | ASSTPGQAGSDYT | 0.85363245 |
| TGGGAGSGGT | ASSTPGQAGSDYT | 0.72622204 |
| SHLETSL | ASSTPGQAGSDYT | 0.7707679 |
| SFTTGMT | ASSTPGQAGSDYT | 0.6840742 |
| QVEVAKRMGVRQPTVS | ASSTPGQAGSDYT | 0.8278991 |
| QGSGGPALAM | ASSTPGQAGSDYT | 0.6804273 |
| MLAPSNP | ASSTPGQAGSDYT | 0.47851038 |
| LVNNAAI | ASSTPGQAGSDYT | 0.7493795 |
| LTGEINTLA | ASSTPGQAGSDYT | 0.63549626 |
| LSIFGPV | ASSTPGQAGSDYT | 0.84834135 |
| LLALLANL | ASSTPGQAGSDYT | 0.8407722 |
| LISNASDALDKLR | ASSTPGQAGSDYT | 0.7927991 |
| LINEAVP | ASSTPGQAGSDYT | 0.8360808 |
| LGYAFSGL | ASSTPGQAGSDYT | 0.2657737 |
| LDFVAVAL | ASSTPGQAGSDYT | 0.74001044 |
| LAQPGAE | ASSTPGQAGSDYT | 0.8850736 |
| ISNLGAVTGF | ASSTPGQAGSDYT | 0.8045768 |
| HQEDPGRI | ASSTPGQAGSDYT | 0.76376945 |
| GTPGEPTP | ASSTPGQAGSDYT | 0.8680489 |
| GFFNSGS | ASSTPGQAGSDYT | 0.6649318 |
| GAGGGGLTAGAN | ASSTPGQAGSDYT | 0.7484867 |
| FVDALVNM | ASSTPGQAGSDYT | 0.7714741 |
| FLTVAPEL | ASSTPGQAGSDYT | 0.83326244 |
| FETLTAL | ASSTPGQAGSDYT | 0.76784825 |
| DVINPIEL | ASSTPGQAGSDYT | 0.6815753 |
| APVSGAPV | ASSTPGQAGSDYT | 0.8219376 |
| ALVGGLAW | ASSTPGQAGSDYT | 0.88677245 |
| ALAAAAATAI | ASSTPGQAGSDYT | 0.7284014 |
| AFGLGLLL | ASSTPGQAGSDYT | 0.8574985 |
| AFFAAASL | ASSTPGQAGSDYT | 0.8252563 |
| AAAALPAFSPP | ASSTPGQAGSDYT | 0.72282696 |
| VVLAPAL | ASSTPGQAGSDYT | 0.648136 |
| TIGHVDHGKTTL | ASSTPGQAGSDYT | 0.6803807 |
| LISNASDALDK | ASSTPGQAGSDYT | 0.6677735 |
| LISNASDAL | ASSTPGQAGSDYT | 0.6238282 |
| ISNASDALDK | ASSTPGQAGSDYT | 0.7798088 |
| NQAETLVYQTE | ASSTPGQAGSDYT | 0.7509584 |
| TGGTGGTAGAG | ASSTPGQAGSDYT | 0.89038444 |
| TGSTAGGAGGA | ASSTPGQAGSDYT | 0.81323713 |
| DPLAVPVAAGPV | ASSTPGQAGSDYT | 0.8274492 |
| GGTGGTPGNGGN | ASSTPGQAGSDYT | 0.62760794 |

| | | |
|---|---|---|
| PAGLAGVDPATL | ASSTPGQAGSDYT | 0.8006231 |
| GNNLVVEAPHAGASV | ASSTPGQAGSDYT | 0.7914952 |
| KVTRIMSASTLEQPA | ASSTPGQAGSDYT | 0.58518445 |
| LGFALPIALPRRTVV | ASSTPGQAGSDYT | 0.87468684 |
| TRLGNLTARKLVVLV | ASSTPGQAGSDYT | 0.6908556 |
| WSQQRRLIYFPSAGP | ASSTPGQAGSDYT | 0.69025004 |
| GPNGNPDPMAAAVDIR | ASSTPGQAGSDYT | 0.84943527 |
| TQRCVSVADLRAAASR | ASSTPGQAGSDYT | 0.81406116 |
| QFVLDVAEHPVEVLTETV | ASSTPGQAGSDYT | 0.7172371 |
| GGAGGTGGAGGNGGTQAGDG | ASSTPGQAGSDYT | 0.64124674 |
| PLLATLRAIAVDVDAKPAQV | ASSTPGQAGSDYT | 0.66093445 |
| APKFRTGGYMIDFWGVGYQVAK | ASSTPGQAGSDYT | 0.8285955 |
| GAGGAGGAGGAAGGPGGTGGTG | ASSTPGQAGSDYT | 0.8569504 |
| IGALGLLATDVAPAGSVAFGVA | ASSTPGQAGSDYT | 0.6034384 |
| AGGAGGIGGNGGAGGAGGVLMG | ASSTPGQAGSDYT | 0.8563392 |
| ADELIGLL | ASSTPGQAGSDYT | 0.8066 |
| ADRALKA | ASSTPGQAGSDYT | 0.49603298 |
| AGAVGQVGGAGGA | ASSTPGQAGSDYT | 0.7725449 |
| AGGGGGLGGAP | ASSTPGQAGSDYT | 0.8647516 |
| AGGLLYGNGGAG | ASSTPGQAGSDYT | 0.8378537 |
| AGLESIGL | ASSTPGQAGSDYT | 0.8730892 |
| AGLKTGII | ASSTPGQAGSDYT | 0.81403315 |
| ASLAALLL | ASSTPGQAGSDYT | 0.6898233 |
| ASVINISSI | ASSTPGQAGSDYT | 0.8626853 |
| ATPPGPA | ASSTPGQAGSDYT | 0.8820429 |
| ATVASALV | ASSTPGQAGSDYT | 0.85912323 |
| ATVASLIL | ASSTPGQAGSDYT | 0.74000984 |
| AVNGPVLIRH | ASSTPGQAGSDYT | 0.6362873 |
| DGAAGAVGI | ASSTPGQAGSDYT | 0.77560127 |
| DPLLHGI | ASSTPGQAGSDYT | 0.83631146 |
| ELGAALGAL | ASSTPGQAGSDYT | 0.6085035 |
| ELGRTQAV | ASSTPGQAGSDYT | 0.7918075 |
| FVAGGYLS | ASSTPGQAGSDYT | 0.78810215 |
| GASGGAGGAGGW | ASSTPGQAGSDYT | 0.5258375 |
| GGAGAGGDGGAGGI | ASSTPGQAGSDYT | 0.82193065 |
| GTGHNGSQPGMGGNG | ASSTPGQAGSDYT | 0.8708637 |
| GYAAASASAS | ASSTPGQAGSDYT | 0.73591983 |
| HQVDVAAVQQG | ASSTPGQAGSDYT | 0.68002284 |
| IGAAIAHT | ASSTPGQAGSDYT | 0.8547081 |
| IGMIEVAQL | ASSTPGQAGSDYT | 0.89033586 |
| IRLAPEHTSAMD | ASSTPGQAGSDYT | 0.85829717 |
| IYNVSVL | ASSTPGQAGSDYT | 0.47696835 |
| KQGFLGM | ASSTPGQAGSDYT | 0.26405698 |
| LESVITQL | ASSTPGQAGSDYT | 0.6572678 |
| LGLGALVVV | ASSTPGQAGSDYT | 0.7789432 |
| LIGNGGNGGSGGI | ASSTPGQAGSDYT | 0.71925116 |
| LNAINAPTL | ASSTPGQAGSDYT | 0.6497839 |
| LQLLNTGV | ASSTPGQAGSDYT | 0.7821447 |
| LQTSIEF | ASSTPGQAGSDYT | 0.86010027 |
| MSFAVSEL | ASSTPGQAGSDYT | 0.6571756 |
| NTGALGL | ASSTPGQAGSDYT | 0.5329511 |

| | | |
|---|---|---|
| PPGAGGAGGAGTT | ASSTPGQAGSDYT | 0.8111551 |
| QLVGLGI | ASSTPGQAGSDYT | 0.8410164 |
| RFLEQVHQL | ASSTPGQAGSDYT | 0.6481037 |
| RGVGYVLR | ASSTPGQAGSDYT | 0.71774495 |
| SATGVFAGVI | ASSTPGQAGSDYT | 0.6708081 |
| SLPAITLP | ASSTPGQAGSDYT | 0.41546154 |
| STALAFTV | ASSTPGQAGSDYT | 0.787808 |
| SVLTVGLGK | ASSTPGQAGSDYT | 0.60173607 |
| TALAGAGSGI | ASSTPGQAGSDYT | 0.8762454 |
| TGLPFTGI | ASSTPGQAGSDYT | 0.6166252 |
| VAEHVTV | ASSTPGQAGSDYT | 0.83757836 |
| VDIVLNSL | ASSTPGQAGSDYT | 0.8295914 |
| VLTGGGAVL | ASSTPGQAGSDYT | 0.83120215 |
| VNVGLAYGI | ASSTPGQAGSDYT | 0.7301368 |
| VSSAAVGAV | ASSTPGQAGSDYT | 0.7099121 |
| VTSVYEAGV | ASSTPGQAGSDYT | 0.6207472 |
| WVALDRGA | ASSTPGQAGSDYT | 0.2731346 |
| YLVGSVVI | ASSTPGQAGSDYT | 0.86976075 |
| GPPGTGKTLL | ASSTPGQAGSDYT | 0.15011574 |
| GVGMTKF | ASSTPGQAGSDYT | 0.86678886 |
| IGIGGSDL | ASSTPGQAGSDYT | 0.8399852 |
| IGTLLAEL | ASSTPGQAGSDYT | 0.6896803 |
| IYTSGTT | ASSTPGQAGSDYT | 0.81589514 |
| LSFPAGV | ASSTPGQAGSDYT | 0.5532073 |
| LYTSGTT | ASSTPGQAGSDYT | 0.48970836 |
| VGAFPFGF | ASSTPGQAGSDYT | 0.8197493 |
| VGAGTVEF | ASSTPGQAGSDYT | 0.83673644 |
| VGKSTLLL | ASSTPGQAGSDYT | 0.6850024 |
| AALAHAGGGIV | ASSTPGQAGSDYT | 0.8024891 |
| AALVAVSLL | ASSTPGQAGSDYT | 0.78912884 |
| AAVSRAVAAL | ASSTPGQAGSDYT | 0.68223816 |
| AGGSSGIGGF | ASSTPGQAGSDYT | 0.75741726 |
| AGLTVANANG | ASSTPGQAGSDYT | 0.856992 |
| AGLVGAVL | ASSTPGQAGSDYT | 0.79749 |
| ALSQAFTV | ASSTPGQAGSDYT | 0.677089 |
| ALTFSVSL | ASSTPGQAGSDYT | 0.8333943 |
| ALTGAALVAL | ASSTPGQAGSDYT | 0.62836313 |
| ASFSIGAGVL | ASSTPGQAGSDYT | 0.7982991 |
| DGSVYKI | ASSTPGQAGSDYT | 0.7979948 |
| EQLAEIV | ASSTPGQAGSDYT | 0.7558632 |
| FAVSELTVM | ASSTPGQAGSDYT | 0.8444979 |
| GAAPLAAGL | ASSTPGQAGSDYT | 0.5831536 |
| GADGGDGGSGGAGG | ASSTPGQAGSDYT | 0.62750256 |
| GFGLDTGGAGG | ASSTPGQAGSDYT | 0.7026664 |
| GLASGGLL | ASSTPGQAGSDYT | 0.81649196 |
| GRIAGAAVLL | ASSTPGQAGSDYT | 0.8563979 |
| GSAGSLNTSF | ASSTPGQAGSDYT | 0.62098265 |
| GTGQIGI | ASSTPGQAGSDYT | 0.7785431 |
| GTLIALIL | ASSTPGQAGSDYT | 0.71729964 |
| IAATMGFVVG | ASSTPGQAGSDYT | 0.61937004 |
| IFAQALL | ASSTPGQAGSDYT | 0.7376314 |

| | | |
|---|---|---|
| IGAAVALGF | ASSTPGQAGSDYT | 0.7400903 |
| IGLTPLAL | ASSTPGQAGSDYT | 0.6642112 |
| ILGSNTSTL | ASSTPGQAGSDYT | 0.83560157 |
| IVFSLTVL | ASSTPGQAGSDYT | 0.6236189 |
| LASKATGF | ASSTPGQAGSDYT | 0.8231983 |
| LASVALIL | ASSTPGQAGSDYT | 0.8712466 |
| LFAVGLTL | ASSTPGQAGSDYT | 0.7545321 |
| LFNGGIGTYI | ASSTPGQAGSDYT | 0.8362322 |
| LTLGAGII | ASSTPGQAGSDYT | 0.8282989 |
| LVAAALGDI | ASSTPGQAGSDYT | 0.82579494 |
| NTAFVQLGI | ASSTPGQAGSDYT | 0.75050175 |
| QLFSGAEL | ASSTPGQAGSDYT | 0.69542444 |
| SAATFITL | ASSTPGQAGSDYT | 0.8724257 |
| SNYGLANLGV | ASSTPGQAGSDYT | 0.85527515 |
| SYGGAYAVM | ASSTPGQAGSDYT | 0.8622316 |
| TTGVGGLTLG | ASSTPGQAGSDYT | 0.8304273 |
| TTQTIYI | ASSTPGQAGSDYT | 0.80334044 |
| VAAGAGLGAVI | ASSTPGQAGSDYT | 0.8352933 |
| VAAVVTLL | ASSTPGQAGSDYT | 0.6741373 |
| VGFLGGLI | ASSTPGQAGSDYT | 0.47317997 |
| VGVIPILL | ASSTPGQAGSDYT | 0.7346704 |
| VIFASSLI | ASSTPGQAGSDYT | 0.76670396 |
| VKGGDGGAAAG | ASSTPGQAGSDYT | 0.7513984 |
| VTKTFTNI | ASSTPGQAGSDYT | 0.41657168 |
| VVAAAILL | ASSTPGQAGSDYT | 0.86560017 |
| YAALDVEL | ASSTPGQAGSDYT | 0.8955358 |
| LASVALLL | ASSTPGQAGSDYT | 0.66257197 |
| LDAVHRV | ASSTPGQAGSDYT | 0.8716859 |
| QGLSSSIF | ASSTPGQAGSDYT | 0.48540452 |
| RGTFANIRL | ASSTPGQAGSDYT | 0.8360319 |
| RQLWWGHRI | ASSTPGQAGSDYT | 0.8670175 |
| YGPPGTGKTLL | ASSTPGQAGSDYT | 0.62317944 |
| AALVGAPANVA | ASSTPGQAGSDYT | 0.84542143 |
| AATFRRTE | ASSTPGQAGSDYT | 0.70425 |
| AGALGGGI | ASSTPGQAGSDYT | 0.77453524 |
| AGGAGGAGGIGAG | ASSTPGQAGSDYT | 0.87026966 |
| AGSGALGL | ASSTPGQAGSDYT | 0.73462653 |
| AQIADQI | ASSTPGQAGSDYT | 0.8179718 |
| ASAVAVTYL | ASSTPGQAGSDYT | 0.73450005 |
| ASTVLGGTGG | ASSTPGQAGSDYT | 0.6489046 |
| AVGGEPAK | ASSTPGQAGSDYT | 0.8222636 |
| CEQGTFGGTC | ASSTPGQAGSDYT | 0.8722466 |
| DEGKAPAG | ASSTPGQAGSDYT | 0.5001266 |
| DTVVFPAL | ASSTPGQAGSDYT | 0.8771144 |
| FEGITMAG | ASSTPGQAGSDYT | 0.692858 |
| FPLVVQD | ASSTPGQAGSDYT | 0.8898585 |
| GAFTTPQ | ASSTPGQAGSDYT | 0.82629716 |
| GFGGFGTAGGVGGA | ASSTPGQAGSDYT | 0.7484468 |
| GFLAQFL | ASSTPGQAGSDYT | 0.7271863 |
| GGAGGAGGNAGVA | ASSTPGQAGSDYT | 0.8592062 |
| GGVGTAGAGGAGGAGG | ASSTPGQAGSDYT | 0.7575183 |

| | | |
|---|---|---|
| GNGGGGDAGNAGSGGN | ASSTPGQAGSDYT | 0.8556825 |
| GTGGAGGAGNDAGS | ASSTPGQAGSDYT | 0.87185127 |
| GTTNTGWFNS | ASSTPGQAGSDYT | 0.7385516 |
| IFMGPLFTL | ASSTPGQAGSDYT | 0.89850163 |
| ILAHAIEAI | ASSTPGQAGSDYT | 0.8468394 |
| IRSAIREAL | ASSTPGQAGSDYT | 0.83691317 |
| ITFAGGG | ASSTPGQAGSDYT | 0.6298901 |
| ITGMSGHQEV | ASSTPGQAGSDYT | 0.27768102 |
| IVAGLACTL | ASSTPGQAGSDYT | 0.8463074 |
| IVGGGLAGA | ASSTPGQAGSDYT | 0.6558541 |
| IVTGVLNI | ASSTPGQAGSDYT | 0.8614068 |
| LAQQGCI | ASSTPGQAGSDYT | 0.88234174 |
| LCLHVEN | ASSTPGQAGSDYT | 0.7277185 |
| LDAISDGS | ASSTPGQAGSDYT | 0.8806217 |
| LGGFAAML | ASSTPGQAGSDYT | 0.88141495 |
| LGGKNPMI | ASSTPGQAGSDYT | 0.60936844 |
| LGPKGVAAIL | ASSTPGQAGSDYT | 0.804159 |
| LMVGVSI | ASSTPGQAGSDYT | 0.6007598 |
| LVGAAFVL | ASSTPGQAGSDYT | 0.84476125 |
| LVLAVAPL | ASSTPGQAGSDYT | 0.62657994 |
| MAAIAVIDAA | ASSTPGQAGSDYT | 0.87472624 |
| NNGINAELQ | ASSTPGQAGSDYT | 0.75293624 |
| RGEPLVLVHG | ASSTPGQAGSDYT | 0.284034 |
| RGGAGGAAGLI | ASSTPGQAGSDYT | 0.8579099 |
| RSYRMHPAV | ASSTPGQAGSDYT | 0.81361884 |
| SAAHLLAGQLA | ASSTPGQAGSDYT | 0.7973199 |
| SAAVGSY | ASSTPGQAGSDYT | 0.885892 |
| SPHAVLHRI | ASSTPGQAGSDYT | 0.768544 |
| SQIAKVL | ASSTPGQAGSDYT | 0.6354686 |
| TALGHLVP | ASSTPGQAGSDYT | 0.8679036 |
| TAPTTSGAA | ASSTPGQAGSDYT | 0.88499653 |
| TDGDDGNGGSPGT | ASSTPGQAGSDYT | 0.539919 |
| TDGLGEN | ASSTPGQAGSDYT | 0.85205036 |
| TGGPPLPGQA | ASSTPGQAGSDYT | 0.8888441 |
| TIPAITI | ASSTPGQAGSDYT | 0.8190411 |
| TIPGFTI | ASSTPGQAGSDYT | 0.8935455 |
| TSIASGALA | ASSTPGQAGSDYT | 0.8949503 |
| TVTLPGAAI | ASSTPGQAGSDYT | 0.72075874 |
| VASGLSAALP | ASSTPGQAGSDYT | 0.88377774 |
| VGAGVIGI | ASSTPGQAGSDYT | 0.8862957 |
| VGPKRFGV | ASSTPGQAGSDYT | 0.78881705 |
| VNVAFPD | ASSTPGQAGSDYT | 0.8411269 |
| VSMTNSGVS | ASSTPGQAGSDYT | 0.6129579 |
| VTDVLGDLSV | ASSTPGQAGSDYT | 0.7750143 |
| VVVVGAGFAGL | ASSTPGQAGSDYT | 0.32641542 |
| VYTVGTVV | ASSTPGQAGSDYT | 0.8555857 |
| WTTFSGP | ASSTPGQAGSDYT | 0.8819096 |
| YDELVEAF | ASSTPGQAGSDYT | 0.8181765 |
| YGTGGAGGNGG | ASSTPGQAGSDYT | 0.752343 |
| YTNVMAV | ASSTPGQAGSDYT | 0.5298343 |
| FLEQVHQL | ASSTPGQAGSDYT | 0.8684437 |

| | | |
|---|---|---|
| GLSLLAVAV | ASSTPGQAGSDYT | 0.6672719 |
| LEQVHQL | ASSTPGQAGSDYT | 0.523148 |
| LNAAGAI | ASSTPGQAGSDYT | 0.780689 |
| PAVHASL | ASSTPGQAGSDYT | 0.86435485 |
| TELEDDA | ASSTPGQAGSDYT | 0.87043726 |
| TPPGPPN | ASSTPGQAGSDYT | 0.6773745 |
| TPPPPPPP | ASSTPGQAGSDYT | 0.89750504 |
| TQSPALI | ASSPDRGTNERLF | 0.8588874 |
| EFAAAFPQ | ASSPDRGTNERLF | 0.8828701 |
| LLSGNLDVL | ASSPDRGTNERLF | 0.82744896 |
| GPANLGNYNI | ASSPDRGTNERLF | 0.74166214 |
| TLGPTLP | ASSPDRGTNERLF | 0.8854817 |
| VYAYSSCCK | ASSPDRGTNERLF | 0.7793577 |
| APAPAPAPNI | ASSPDRGTNERLF | 0.7816527 |
| PGGPVPIP | ASSPDRGTNERLF | 0.8585551 |
| FLTVALEP | ASSPDRGTNERLF | 0.7335756 |
| SQYFGANL | ASSPDRGTNERLF | 0.74146175 |
| GGVKPAA | ASSPDRGTNERLF | 0.7952647 |
| GGTGGAGGAGGW | ASSPDRGTNERLF | 0.8929755 |
| GVGGVAGLGGAGG | ASSPDRGTNERLF | 0.65678734 |
| VLLVLATTI | ASSPDRGTNERLF | 0.5902829 |
| IGIFAAML | ASSPDRGTNERLF | 0.7402339 |
| AVGDLID | ASSPDRGTNERLF | 0.82594365 |
| NISPLPL | ASSPDRGTNERLF | 0.88669056 |
| VTAVFVAI | ASSPDRGTNERLF | 0.871619 |
| SGDDNDAAL | ASSPDRGTNERLF | 0.8226099 |
| AVAALAHS | ASSPDRGTNERLF | 0.86060524 |
| GGVTVMATP | ASSPDRGTNERLF | 0.8723755 |
| AGLAANVRD | ASSPDRGTNERLF | 0.82864285 |
| TNRPDLIDPALLRPG | ASSPDRGTNERLF | 0.84943485 |
| LANLGVTAL | ASSPDRGTNERLF | 0.7813613 |
| TGERIEL | ASSPDRGTNERLF | 0.8912952 |
| NGEELAA | ASSPDRGTNERLF | 0.86999977 |
| VLSLVWG | ASSPDRGTNERLF | 0.88188934 |
| IGYALWL | ASSPDRGTNERLF | 0.8845187 |
| LSLLLAPL | ASSPDRGTNERLF | 0.8213854 |
| TLDEALK | ASSPDRGTNERLF | 0.8102673 |
| TDGAGTAEGG | ASSPDRGTNERLF | 0.3217442 |
| NAPGGAGGNGG | ASSPDRGTNERLF | 0.7207353 |
| VSVFMGMGGNF | ASSPDRGTNERLF | 0.84019923 |
| LASYLAAL | ASSPDRGTNERLF | 0.84962034 |
| LVGGTPAL | ASSPDRGTNERLF | 0.8612485 |
| VRNADVL | ASSPDRGTNERLF | 0.86480516 |
| TCSSPFLVSAM | ASSPDRGTNERLF | 0.7716627 |
| VHITDAE | ASSPDRGTNERLF | 0.86841697 |
| EYAAPSF | ASSPDRGTNERLF | 0.88670456 |
| VVDAFHGGL | ASSPDRGTNERLF | 0.5451724 |
| ASVVDPQ | ASSPDRGTNERLF | 0.78978854 |
| SYNEFGGLS | ASSPDRGTNERLF | 0.9061857 |
| DPDGSVY | ASSPDRGTNERLF | 0.850935 |
| MVSTIHATE | ASSPDRGTNERLF | 0.7415973 |

| | | |
|---|---|---|
| LAAGVADPV | ASSPDRGTNERLF | 0.7796672 |
| TKEQIAA | ASSPDRGTNERLF | 0.8892561 |
| SLASPAAL | ASSPDRGTNERLF | 0.77910036 |
| IGLVFYGV | ASSPDRGTNERLF | 0.89489067 |
| FLLDASNL | ASSPDRGTNERLF | 0.29829344 |
| YQVGNFP | ASSPDRGTNERLF | 0.81807804 |
| ESLLSQST | ASSPDRGTNERLF | 0.6239413 |
| LFTVGFAAVG | ASSPDRGTNERLF | 0.4243487 |
| ATPPTPT | ASSPDRGTNERLF | 0.86822784 |
| TDLATLG | ASSPDRGTNERLF | 0.82779366 |
| SAGGTGR | ASSPDRGTNERLF | 0.86972797 |
| GFSNSGS | ASSPDRGTNERLF | 0.80678153 |
| GGNGGSGGTSV | ASSPDRGTNERLF | 0.879809 |
| GTSVATGGAGN | ASSPDRGTNERLF | 0.85783565 |
| NGGNGGSGGTS | ASSPDRGTNERLF | 0.8214451 |
| AVVKFPH | ASSPDRGTNERLF | 0.7726973 |
| TPPPPPP | ASSPDRGTNERLF | 0.79165465 |
| AAPAAAAAPA | ASSPDRGTNERLF | 0.8773592 |
| IALNEDL | ASSPDRGTNERLF | 0.8818938 |
| IVNVSSM | ASSPDRGTNERLF | 0.28337118 |
| TGIPPAPR | ASSPDRGTNERLF | 0.83804893 |
| FDLGGGTF | ASSPDRGTNERLF | 0.7567459 |
| FDLGGGTFDVS | ASSPDRGTNERLF | 0.4741658 |
| NEPTAAA | ASSPDRGTNERLF | 0.26436234 |
| NEPTAAALA | ASSPDRGTNERLF | 0.7240442 |
| GSIINISSI | ASSPDRGTNERLF | 0.8367114 |
| VAPGNAGTA | ASSPDRGTNERLF | 0.87210834 |
| TGGGASLE | ASSPDRGTNERLF | 0.87769 |
| VAVYNHYKRI | ASSPDRGTNERLF | 0.76042604 |
| TDPAPATT | ASSPDRGTNERLF | 0.87723446 |
| TNRPDILDPALLRPG | ASSPDRGTNERLF | 0.8161335 |
| LTTTTTS | ASSPDRGTNERLF | 0.111161575 |
| PTGSGKTL | ASSPDRGTNERLF | 0.76639295 |
| TGSGKTL | ASSPDRGTNERLF | 0.6114664 |
| TIGHVDHGK | ASSPDRGTNERLF | 0.8849097 |
| TIGHVDHGKTT | ASSPDRGTNERLF | 0.7546139 |
| GHVDHGKTT | ASSPDRGTNERLF | 0.7954154 |
| AQFEHTLL | ASSPDRGTNERLF | 0.8230876 |
| IVNQDEL | ASSPDRGTNERLF | 0.848343 |
| TNGSQFF | ASSPDRGTNERLF | 0.807425 |
| TGKGIVE | ASSPDRGTNERLF | 0.6617092 |
| LISNASDA | ASSPDRGTNERLF | 0.17748383 |
| AELRTLGELTKIAKAHGAQ | ASSPDRGTNERLF | 0.827593 |
| APNAVAAAEPSW | ASSPDRGTNERLF | 0.62528646 |
| APQVAGAATAVL | ASSPDRGTNERLF | 0.5721985 |
| ARRDPAAAAAILPTDA | ASSPDRGTNERLF | 0.66743326 |
| DAAARDVLAEAGLAEYF | ASSPDRGTNERLF | 0.5865547 |
| DDQIVLSAKANLAKVEQ | ASSPDRGTNERLF | 0.45665148 |
| DELLPDTGEDSDEDGH | ASSPDRGTNERLF | 0.8881676 |
| ELPEAPANRATVPQD | ASSPDRGTNERLF | 0.69057214 |
| FGQSKIERVTGIG | ASSPDRGTNERLF | 0.80409247 |

| | | |
|---|---|---|
| GAGGTGSTAGGAGGAGGAGGLY | ASSPDRGTNERLF | 0.8951353 |
| GHGGTGAAGGNGGTGGAGGSGI | ASSPDRGTNERLF | 0.8210387 |
| GNGGNGGAGGEGGPGGAGGAGA | ASSPDRGTNERLF | 0.81457055 |
| GSTAGNGGNGGNATNGTIAGQP | ASSPDRGTNERLF | 0.70322734 |
| GTGGDGGNGGNGGTGG | ASSPDRGTNERLF | 0.7293318 |
| GVRRPAGIPADAPV | ASSPDRGTNERLF | 0.71675384 |
| GYRQVTDAHLLTLARR | ASSPDRGTNERLF | 0.6964688 |
| IAKMGQDGHDRG | ASSPDRGTNERLF | 0.8910759 |
| KAEPSIEIQYT | ASSPDRGTNERLF | 0.828442 |
| KPVRIRMAADEDTVAVT | ASSPDRGTNERLF | 0.7343211 |
| MPPRIAGMRLLVIKPEP | ASSPDRGTNERLF | 0.88093644 |
| MKFYAVALFRSRGVPPTR | ASSPDRGTNERLF | 0.8918542 |
| NAGGFGILGAVAHSP | ASSPDRGTNERLF | 0.90100694 |
| NEGREITLAMAEM | ASSPDRGTNERLF | 0.7435222 |
| NELVEWSHPLGHRWRWE | ASSPDRGTNERLF | 0.7488971 |
| NLGAGGTGGNGGNPG | ASSPDRGTNERLF | 0.7957258 |
| NTEWPKWETVMAADPVD | ASSPDRGTNERLF | 0.8202354 |
| PASAAPAETQP | ASSPDRGTNERLF | 0.8156035 |
| QQSSDGKNGVRAAEITGE | ASSPDRGTNERLF | 0.6477728 |
| RAPDAVTTSWWKE | ASSPDRGTNERLF | 0.88558185 |
| SGFGNVGAHTSGWFNQ | ASSPDRGTNERLF | 0.9014806 |
| SGVAGAVGQAASVGGLKV | ASSPDRGTNERLF | 0.81444025 |
| SNGGAGTGSAGGNGGT | ASSPDRGTNERLF | 0.5299231 |
| TGGAGGEGGAAG | ASSPDRGTNERLF | 0.75814205 |
| TGGDGGAAGAGGAPGHG | ASSPDRGTNERLF | 0.6584091 |
| TMPPLLAAIANAGRHGV | ASSPDRGTNERLF | 0.7615988 |
| TSSTAGNSGGA | ASSPDRGTNERLF | 0.79142046 |
| VPATVPVGILC | ASSPDRGTNERLF | 0.63874686 |
| VQRPDSGGVVLTGRIS | ASSPDRGTNERLF | 0.80370545 |
| WTPNPVPATKGVPAGLVQAT | ASSPDRGTNERLF | 0.89508986 |
| YALKRALAKFSDRFDVVII | ASSPDRGTNERLF | 0.7091502 |
| YLIGSVIV | ASSPDRGTNERLF | 0.85768473 |
| YGQLVAVID | ASSPDRGTNERLF | 0.89623976 |
| VTVGGMTLPA | ASSPDRGTNERLF | 0.87559855 |
| VLALPAV | ASSPDRGTNERLF | 0.89623225 |
| VIPPTLNYET | ASSPDRGTNERLF | 0.87238926 |
| VGMGGMHPSSG | ASSPDRGTNERLF | 0.819724 |
| VGLAGGAATA | ASSPDRGTNERLF | 0.8182732 |
| VGGAGGAGGSGP | ASSPDRGTNERLF | 0.86093223 |
| VFAAFMV | ASSPDRGTNERLF | 0.79810005 |
| TLYDKLR | ASSPDRGTNERLF | 0.7988832 |
| TLAVFTVL | ASSPDRGTNERLF | 0.80408627 |
| TGSVQPARV | ASSPDRGTNERLF | 0.8802842 |
| TGGGAGSGGT | ASSPDRGTNERLF | 0.81322193 |
| SHLETSL | ASSPDRGTNERLF | 0.884127 |
| SFTTGMT | ASSPDRGTNERLF | 0.38568717 |
| QVEVAKRMGVRQPTVS | ASSPDRGTNERLF | 0.60462976 |
| QGSGGPALAM | ASSPDRGTNERLF | 0.88840777 |
| MLAPSNP | ASSPDRGTNERLF | 0.14787775 |
| LVNNAAI | ASSPDRGTNERLF | 0.33415174 |
| LTGEINTLA | ASSPDRGTNERLF | 0.8494408 |

| | | |
|---|---|---|
| LSIFGPV | ASSPDRGTNERLF | 0.8877045 |
| LLALLANL | ASSPDRGTNERLF | 0.20961063 |
| LISNASDALDKLR | ASSPDRGTNERLF | 0.62819135 |
| LINEAVP | ASSPDRGTNERLF | 0.47379002 |
| LGYAFSGL | ASSPDRGTNERLF | 0.87593186 |
| LDFVAVAL | ASSPDRGTNERLF | 0.80302685 |
| LAQPGAE | ASSPDRGTNERLF | 0.29420114 |
| ISNLGAVTGF | ASSPDRGTNERLF | 0.8783017 |
| HQEDPGRI | ASSPDRGTNERLF | 0.63267726 |
| GTPGEPTP | ASSPDRGTNERLF | 0.7665943 |
| GFFNSGS | ASSPDRGTNERLF | 0.87933207 |
| GAGGGGLTAGAN | ASSPDRGTNERLF | 0.8944197 |
| FVDALVNM | ASSPDRGTNERLF | 0.6933366 |
| FLTVAPEL | ASSPDRGTNERLF | 0.86111677 |
| FETLTAL | ASSPDRGTNERLF | 0.79589117 |
| DVINPIEL | ASSPDRGTNERLF | 0.9009252 |
| APVSGAPV | ASSPDRGTNERLF | 0.59914076 |
| ALVGGLAW | ASSPDRGTNERLF | 0.6387243 |
| ALAAAAATAI | ASSPDRGTNERLF | 0.8823246 |
| AFGLGLLL | ASSPDRGTNERLF | 0.872172 |
| AFFAAASL | ASSPDRGTNERLF | 0.7881013 |
| AAAALPAFSPP | ASSPDRGTNERLF | 0.8815204 |
| VVLAPAL | ASSPDRGTNERLF | 0.8329214 |
| TIGHVDHGKTTL | ASSPDRGTNERLF | 0.7980308 |
| LISNASDALDK | ASSPDRGTNERLF | 0.8885556 |
| LISNASDAL | ASSPDRGTNERLF | 0.8695876 |
| ISNASDALDK | ASSPDRGTNERLF | 0.66520405 |
| NQAETLVYQTE | ASSPDRGTNERLF | 0.8869008 |
| TGGTGGTAGAG | ASSPDRGTNERLF | 0.8185214 |
| TGSTAGGAGGA | ASSPDRGTNERLF | 0.57712793 |
| DPLAVPVAAGPV | ASSPDRGTNERLF | 0.89659333 |
| GGTGGTPGNGGN | ASSPDRGTNERLF | 0.85120153 |
| PAGLAGVDPATL | ASSPDRGTNERLF | 0.8789566 |
| GNNLVVEAPHAGASV | ASSPDRGTNERLF | 0.83869386 |
| KVTRIMSASTLEQPA | ASSPDRGTNERLF | 0.8823458 |
| LGFALPIALPRRTVV | ASSPDRGTNERLF | 0.31082195 |
| TRLGNLTARKLVVLV | ASSPDRGTNERLF | 0.85176367 |
| WSQQRRLIYFPSAGP | ASSPDRGTNERLF | 0.7762947 |
| GPNGNPDPMAAAVDIR | ASSPDRGTNERLF | 0.8711467 |
| TQRCVSVADLRAAASR | ASSPDRGTNERLF | 0.8380456 |
| QFVLDVAEHPVEVLTETV | ASSPDRGTNERLF | 0.65368474 |
| GGAGGTGGAGGNGGTQAGDG | ASSPDRGTNERLF | 0.84538245 |
| PLLATLRAIAVDVDAKPAQV | ASSPDRGTNERLF | 0.89158785 |
| APKFRTGGYMIDFWGVGYQVAK | ASSPDRGTNERLF | 0.8414724 |
| GAGGAGGAGGAAGGPGGTGGTG | ASSPDRGTNERLF | 0.83812034 |
| IGALGLLATDVAPAGSVAFGVA | ASSPDRGTNERLF | 0.90158117 |
| AGGAGGIGGNGGAGGAGGVLMG | ASSPDRGTNERLF | 0.64309883 |
| ADELIGLL | ASSPDRGTNERLF | 0.8281461 |
| ADRALKA | ASSPDRGTNERLF | 0.8763432 |
| AGAVGQVGGAGGA | ASSPDRGTNERLF | 0.8774561 |
| AGGGGGLGGAP | ASSPDRGTNERLF | 0.8446082 |

| | | |
|---|---|---|
| AGGLLYGNGGAG | ASSPDRGTNERLF | 0.8937638 |
| AGLESIGL | ASSPDRGTNERLF | 0.67043066 |
| AGLKTGII | ASSPDRGTNERLF | 0.8749086 |
| ASLAALLL | ASSPDRGTNERLF | 0.76652175 |
| ASVINISSI | ASSPDRGTNERLF | 0.8269545 |
| ATPPGPA | ASSPDRGTNERLF | 0.7182192 |
| ATVASALV | ASSPDRGTNERLF | 0.8607508 |
| ATVASLIL | ASSPDRGTNERLF | 0.88432854 |
| AVNGPVLIRH | ASSPDRGTNERLF | 0.8651883 |
| DGAAGAVGI | ASSPDRGTNERLF | 0.3998127 |
| DPLLHGI | ASSPDRGTNERLF | 0.88772595 |
| ELGAALGAL | ASSPDRGTNERLF | 0.8698412 |
| ELGRTQAV | ASSPDRGTNERLF | 0.9044533 |
| FVAGGYLS | ASSPDRGTNERLF | 0.7792755 |
| GASGGAGGAGGW | ASSPDRGTNERLF | 0.86863256 |
| GGAGAGGDGGAGGI | ASSPDRGTNERLF | 0.7784368 |
| GTGHNGSQPGMGGNG | ASSPDRGTNERLF | 0.85654926 |
| GYAAASASAS | ASSPDRGTNERLF | 0.68328655 |
| HQVDVAAVQQG | ASSPDRGTNERLF | 0.66485226 |
| IGAAIAHT | ASSPDRGTNERLF | 0.7959527 |
| IGMIEVAQL | ASSPDRGTNERLF | 0.7629146 |
| IRLAPEHTSAMD | ASSPDRGTNERLF | 0.8758414 |
| IYNVSVL | ASSPDRGTNERLF | 0.8728689 |
| KQGFLGM | ASSPDRGTNERLF | 0.8627306 |
| LESVITQL | ASSPDRGTNERLF | 0.88330144 |
| LGLGALVVV | ASSPDRGTNERLF | 0.8914075 |
| LIGNGGNGGSGGI | ASSPDRGTNERLF | 0.88703156 |
| LNAINAPTL | ASSPDRGTNERLF | 0.74961656 |
| LQLLNTGV | ASSPDRGTNERLF | 0.7621592 |
| LQTSIEF | ASSPDRGTNERLF | 0.72196513 |
| MSFAVSEL | ASSPDRGTNERLF | 0.80453223 |
| NTGALGL | ASSPDRGTNERLF | 0.75356126 |
| PPGAGGAGGAGTT | ASSPDRGTNERLF | 0.86014736 |
| QLVGLGI | ASSPDRGTNERLF | 0.12941019 |
| RFLEQVHQL | ASSPDRGTNERLF | 0.8528224 |
| RGVGYVLR | ASSPDRGTNERLF | 0.8928415 |
| SATGVFAGVI | ASSPDRGTNERLF | 0.66801274 |
| SLPAITLP | ASSPDRGTNERLF | 0.8361237 |
| STALAFTV | ASSPDRGTNERLF | 0.85974616 |
| SVLTVGLGK | ASSPDRGTNERLF | 0.7743556 |
| TALAGAGSGI | ASSPDRGTNERLF | 0.8604189 |
| TGLPFTGI | ASSPDRGTNERLF | 0.7247199 |
| VAEHVTV | ASSPDRGTNERLF | 0.8760114 |
| VDIVLNSL | ASSPDRGTNERLF | 0.8769749 |
| VLTGGGAVL | ASSPDRGTNERLF | 0.83052623 |
| VNVGLAYGI | ASSPDRGTNERLF | 0.77486694 |
| VSSAAVGAV | ASSPDRGTNERLF | 0.81895596 |
| VTSVYEAGV | ASSPDRGTNERLF | 0.26302558 |
| WVALDRGA | ASSPDRGTNERLF | 0.7934722 |
| YLVGSVVI | ASSPDRGTNERLF | 0.14907074 |
| GPPGTGKTLL | ASSPDRGTNERLF | 0.8180338 |

| | | |
|---|---|---|
| GVGMTKF | ASSPDRGTNERLF | 0.57316303 |
| IGIGGSDL | ASSPDRGTNERLF | 0.8395008 |
| IGTLLAEL | ASSPDRGTNERLF | 0.8283571 |
| IYTSGTT | ASSPDRGTNERLF | 0.85235715 |
| LSFPAGV | ASSPDRGTNERLF | 0.5427026 |
| LYTSGTT | ASSPDRGTNERLF | 0.8450055 |
| VGAFPFGF | ASSPDRGTNERLF | 0.8174518 |
| VGAGTVEF | ASSPDRGTNERLF | 0.47430056 |
| VGKSTLLL | ASSPDRGTNERLF | 0.88754255 |
| AALAHAGGGIV | ASSPDRGTNERLF | 0.8241676 |
| AALVAVSLL | ASSPDRGTNERLF | 0.8256192 |
| AAVSRAVAAL | ASSPDRGTNERLF | 0.8701932 |
| AGGSSGIGGF | ASSPDRGTNERLF | 0.81637776 |
| AGLTVANANG | ASSPDRGTNERLF | 0.4217837 |
| AGLVGAVL | ASSPDRGTNERLF | 0.7629845 |
| ALSQAFTV | ASSPDRGTNERLF | 0.81227565 |
| ALTFSVSL | ASSPDRGTNERLF | 0.59534395 |
| ALTGAALVAL | ASSPDRGTNERLF | 0.23829295 |
| ASFSIGAGVL | ASSPDRGTNERLF | 0.8759397 |
| DGSVYKI | ASSPDRGTNERLF | 0.5664586 |
| EQLAEIV | ASSPDRGTNERLF | 0.67099094 |
| FAVSELTVM | ASSPDRGTNERLF | 0.8038602 |
| GAAPLAAGL | ASSPDRGTNERLF | 0.88402903 |
| GADGGDGGSGGAGG | ASSPDRGTNERLF | 0.6795572 |
| GFGLDTGGAGG | ASSPDRGTNERLF | 0.740538 |
| GLASGGLL | ASSPDRGTNERLF | 0.86585724 |
| GRIAGAAVLL | ASSPDRGTNERLF | 0.8082527 |
| GSAGSLNTSF | ASSPDRGTNERLF | 0.8850795 |
| GTGQIGI | ASSPDRGTNERLF | 0.8531828 |
| GTLIALIL | ASSPDRGTNERLF | 0.77262086 |
| IAATMGFVVG | ASSPDRGTNERLF | 0.8574883 |
| IFAQALL | ASSPDRGTNERLF | 0.8497435 |
| IGAAVALGF | ASSPDRGTNERLF | 0.694243 |
| IGLTPLAL | ASSPDRGTNERLF | 0.62604046 |
| ILGSNTSTL | ASSPDRGTNERLF | 0.75618976 |
| IVFSLTVL | ASSPDRGTNERLF | 0.46468908 |
| LASKATGF | ASSPDRGTNERLF | 0.76360035 |
| LASVALIL | ASSPDRGTNERLF | 0.8957516 |
| LFAVGLTL | ASSPDRGTNERLF | 0.8142591 |
| LFNGGIGTYI | ASSPDRGTNERLF | 0.5107382 |
| LTLGAGII | ASSPDRGTNERLF | 0.8000253 |
| LVAAALGDI | ASSPDRGTNERLF | 0.5115378 |
| NTAFVQLGI | ASSPDRGTNERLF | 0.85690486 |
| QLFSGAEL | ASSPDRGTNERLF | 0.81412286 |
| SAATFITL | ASSPDRGTNERLF | 0.89979136 |
| SNYGLANLGV | ASSPDRGTNERLF | 0.8805046 |
| SYGGAYAVM | ASSPDRGTNERLF | 0.78411174 |
| TTGVGGLTLG | ASSPDRGTNERLF | 0.88900805 |
| TTQTIYI | ASSPDRGTNERLF | 0.7587911 |
| VAAGAGLGAVI | ASSPDRGTNERLF | 0.8331496 |
| VAAVVTLL | ASSPDRGTNERLF | 0.8311918 |

| | | |
|---|---|---|
| VGFLGGLI | ASSPDRGTNERLF | 0.8644663 |
| VGVIPILL | ASSPDRGTNERLF | 0.78853256 |
| VIFASSLI | ASSPDRGTNERLF | 0.76666796 |
| VKGGDGGAAAG | ASSPDRGTNERLF | 0.386343 |
| VTKTFTNI | ASSPDRGTNERLF | 0.86486757 |
| VVAAAILL | ASSPDRGTNERLF | 0.50358874 |
| YAALDVEL | ASSPDRGTNERLF | 0.5623363 |
| LASVALLL | ASSPDRGTNERLF | 0.70958906 |
| LDAVHRV | ASSPDRGTNERLF | 0.898394 |
| QGLSSSIF | ASSPDRGTNERLF | 0.8930494 |
| RGTFANIRL | ASSPDRGTNERLF | 0.7436399 |
| RQLWWGHRI | ASSPDRGTNERLF | 0.67065775 |
| YGPPGTGKTLL | ASSPDRGTNERLF | 0.85814136 |
| AALVGAPANVA | ASSPDRGTNERLF | 0.60753953 |
| AATFRRTE | ASSPDRGTNERLF | 0.8779813 |
| AGALGGGI | ASSPDRGTNERLF | 0.829428 |
| AGGAGGAGGIGAG | ASSPDRGTNERLF | 0.8379917 |
| AGSGALGL | ASSPDRGTNERLF | 0.708567 |
| AQIADQI | ASSPDRGTNERLF | 0.8580551 |
| ASAVAVTYL | ASSPDRGTNERLF | 0.7836365 |
| ASTVLGGTGG | ASSPDRGTNERLF | 0.8758306 |
| AVGGEPAK | ASSPDRGTNERLF | 0.3264181 |
| CEQGTFGGTC | ASSPDRGTNERLF | 0.87544596 |
| DEGKAPAG | ASSPDRGTNERLF | 0.7306564 |
| DTVVFPAL | ASSPDRGTNERLF | 0.72277904 |
| FEGITMAG | ASSPDRGTNERLF | 0.53643453 |
| FPLVVQD | ASSPDRGTNERLF | 0.84466076 |
| GAFTTPQ | ASSPDRGTNERLF | 0.86776114 |
| GFGGFGTAGGVGGA | ASSPDRGTNERLF | 0.8871151 |
| GFLAQFL | ASSPDRGTNERLF | 0.7331806 |
| GGAGGAGGNAGVA | ASSPDRGTNERLF | 0.87128127 |
| GGVGTAGAGGAGGAGG | ASSPDRGTNERLF | 0.8769712 |
| GNGGGGDAGNAGSGGN | ASSPDRGTNERLF | 0.81702054 |
| GTGGAGGAGNDAGS | ASSPDRGTNERLF | 0.30789638 |
| GTTNTGWFNS | ASSPDRGTNERLF | 0.8454834 |
| IFMGPLFTL | ASSPDRGTNERLF | 0.82771605 |
| ILAHAIEAI | ASSPDRGTNERLF | 0.8871182 |
| IRSAIREAL | ASSPDRGTNERLF | 0.8395759 |
| ITFAGGG | ASSPDRGTNERLF | 0.660308 |
| ITGMSGHQEV | ASSPDRGTNERLF | 0.858781 |
| IVAGLACTL | ASSPDRGTNERLF | 0.87914443 |
| IVGGGLAGA | ASSPDRGTNERLF | 0.63446665 |
| IVTGVLNI | ASSPDRGTNERLF | 0.8069178 |
| LAQQGCI | ASSPDRGTNERLF | 0.7966415 |
| LCLHVEN | ASSPDRGTNERLF | 0.54165316 |
| LDAISDGS | ASSPDRGTNERLF | 0.8242263 |
| LGGFAAML | ASSPDRGTNERLF | 0.78388035 |
| LGGKNPMI | ASSPDRGTNERLF | 0.7091751 |
| LGPKGVAAIL | ASSPDRGTNERLF | 0.8464494 |
| LMVGVSI | ASSPDRGTNERLF | 0.700378 |
| LVGAAFVL | ASSPDRGTNERLF | 0.6819325 |

| | | |
|---|---|---|
| LVLAVAPL | ASSPDRGTNERLF | 0.71822774 |
| MAAIAVIDAA | ASSPDRGTNERLF | 0.784802 |
| NNGINAELQ | ASSPDRGTNERLF | 0.8699503 |
| RGEPLVLVHG | ASSPDRGTNERLF | 0.8390774 |
| RGGAGGAAGLI | ASSPDRGTNERLF | 0.8612915 |
| RSYRMHPAV | ASSPDRGTNERLF | 0.8965579 |
| SAAHLLAGQLA | ASSPDRGTNERLF | 0.47421914 |
| SAAVGSY | ASSPDRGTNERLF | 0.6329283 |
| SPHAVLHRI | ASSPDRGTNERLF | 0.8765372 |
| SQIAKVL | ASSPDRGTNERLF | 0.6623327 |
| TALGHLVP | ASSPDRGTNERLF | 0.8897517 |
| TAPTTSGAA | ASSPDRGTNERLF | 0.81577384 |
| TDGDDGNGGSPGT | ASSPDRGTNERLF | 0.86395967 |
| TDGLGEN | ASSPDRGTNERLF | 0.8048167 |
| TGGPPLPGQA | ASSPDRGTNERLF | 0.90194273 |
| TIPAITI | ASSPDRGTNERLF | 0.8767429 |
| TIPGFTI | ASSPDRGTNERLF | 0.8581923 |
| TSIASGALA | ASSPDRGTNERLF | 0.8711798 |
| TVTLPGAAI | ASSPDRGTNERLF | 0.66640747 |
| VASGLSAALP | ASSPDRGTNERLF | 0.8713539 |
| VGAGVIGI | ASSPDRGTNERLF | 0.87127507 |
| VGPKRFGV | ASSPDRGTNERLF | 0.61154604 |
| VNVAFPD | ASSPDRGTNERLF | 0.79563737 |
| VSMTNSGVS | ASSPDRGTNERLF | 0.891472 |
| VTDVLGDLSV | ASSPDRGTNERLF | 0.89533365 |
| VVVVGAGFAGL | ASSPDRGTNERLF | 0.4070717 |
| VYTVGTVV | ASSPDRGTNERLF | 0.82501924 |
| WTTFSGP | ASSPDRGTNERLF | 0.5160425 |
| YDELVEAF | ASSPDRGTNERLF | 0.7160007 |
| YGTGGAGGNGG | ASSPDRGTNERLF | 0.41318163 |
| YTNVMAV | ASSPDRGTNERLF | 0.85057914 |
| FLEQVHQL | ASSPDRGTNERLF | 0.7415042 |
| GLSLLAVAV | ASSPDRGTNERLF | 0.86194074 |
| LEQVHQL | ASSPDRGTNERLF | 0.7891151 |
| LNAAGAI | ASSPDRGTNERLF | 0.8734709 |
| PAVHASL | ASSPDRGTNERLF | 0.73362195 |
| TELEDDA | ASSPDRGTNERLF | 0.77331793 |
| TPPGPPN | ASSPDRGTNERLF | 0.5562646 |
| TPPPPPPP | ASSPDRGTNERLF | 0.27263892 |
| TQSPALI | ASSLPRAPNTEVF | 0.7496066 |
| EFAAAFPQ | ASSLPRAPNTEVF | 0.7423281 |
| LLSGNLDVL | ASSLPRAPNTEVF | 0.28109077 |
| GPANLGNYNI | ASSLPRAPNTEVF | 0.901618 |
| TLGPTLP | ASSLPRAPNTEVF | 0.8323887 |
| VYAYSSCCK | ASSLPRAPNTEVF | 0.83399343 |
| APAPAPAPNI | ASSLPRAPNTEVF | 0.72897315 |
| PGGPVPIP | ASSLPRAPNTEVF | 0.7640259 |
| FLTVALEP | ASSLPRAPNTEVF | 0.76734364 |
| SQYFGANL | ASSLPRAPNTEVF | 0.86460274 |
| GGVKPAA | ASSLPRAPNTEVF | 0.6768875 |
| GGTGGAGGAGGW | ASSLPRAPNTEVF | 0.4496749 |

| | | |
|---|---|---|
| GVGGVAGLGGAGG | ASSLPRAPNTEVF | 0.7956767 |
| VLLVLATTI | ASSLPRAPNTEVF | 0.70356643 |
| IGIFAAML | ASSLPRAPNTEVF | 0.8921874 |
| AVGDLID | ASSLPRAPNTEVF | 0.8840941 |
| NISPLPL | ASSLPRAPNTEVF | 0.8563216 |
| VTAVFVAI | ASSLPRAPNTEVF | 0.89367515 |
| SGDDNDAAL | ASSLPRAPNTEVF | 0.23422973 |
| AVAALAHS | ASSLPRAPNTEVF | 0.78881955 |
| GGVTVMATP | ASSLPRAPNTEVF | 0.89399856 |
| AGLAANVRD | ASSLPRAPNTEVF | 0.8404522 |
| TNRPDLIDPALLRPG | ASSLPRAPNTEVF | 0.7489568 |
| LANLGVTAL | ASSLPRAPNTEVF | 0.8934871 |
| TGERIEL | ASSLPRAPNTEVF | 0.71789557 |
| NGEELAA | ASSLPRAPNTEVF | 0.58612776 |
| VLSLVWG | ASSLPRAPNTEVF | 0.84182245 |
| IGYALWL | ASSLPRAPNTEVF | 0.8686814 |
| LSLLLAPL | ASSLPRAPNTEVF | 0.86026394 |
| TLDEALK | ASSLPRAPNTEVF | 0.6316895 |
| TDGAGTAEGG | ASSLPRAPNTEVF | 0.85384965 |
| NAPGGAGGNGG | ASSLPRAPNTEVF | 0.90533423 |
| VSVFMGMGGNF | ASSLPRAPNTEVF | 0.7929372 |
| LASYLAAL | ASSLPRAPNTEVF | 0.86457914 |
| LVGGTPAL | ASSLPRAPNTEVF | 0.8465203 |
| VRNADVL | ASSLPRAPNTEVF | 0.66228193 |
| TCSSPFLVSAM | ASSLPRAPNTEVF | 0.69358414 |
| VHITDAE | ASSLPRAPNTEVF | 0.70134497 |
| EYAAPSF | ASSLPRAPNTEVF | 0.8370873 |
| VVDAFHGGL | ASSLPRAPNTEVF | 0.7141506 |
| ASVVDPQ | ASSLPRAPNTEVF | 0.8270769 |
| SYNEFGGLS | ASSLPRAPNTEVF | 0.8207372 |
| DPDGSVY | ASSLPRAPNTEVF | 0.90684855 |
| MVSTIHATE | ASSLPRAPNTEVF | 0.8633855 |
| LAAGVADPV | ASSLPRAPNTEVF | 0.8724693 |
| TKEQIAA | ASSLPRAPNTEVF | 0.7145462 |
| SLASPAAL | ASSLPRAPNTEVF | 0.8455342 |
| IGLVFYGV | ASSLPRAPNTEVF | 0.85655844 |
| FLLDASNL | ASSLPRAPNTEVF | 0.41839117 |
| YQVGNFP | ASSLPRAPNTEVF | 0.89123285 |
| ESLLSQST | ASSLPRAPNTEVF | 0.6646923 |
| LFTVGFAAVG | ASSLPRAPNTEVF | 0.8368734 |
| ATPPTPT | ASSLPRAPNTEVF | 0.8671447 |
| TDLATLG | ASSLPRAPNTEVF | 0.87943846 |
| SAGGTGR | ASSLPRAPNTEVF | 0.8358095 |
| GFSNSGS | ASSLPRAPNTEVF | 0.8485106 |
| GGNGGSGGTSV | ASSLPRAPNTEVF | 0.72673255 |
| GTSVATGGAGN | ASSLPRAPNTEVF | 0.8683572 |
| NGGNGGSGGTS | ASSLPRAPNTEVF | 0.8890709 |
| AVVKFPH | ASSLPRAPNTEVF | 0.78214025 |
| TPPPPPP | ASSLPRAPNTEVF | 0.8948774 |
| AAPAAAAAPA | ASSLPRAPNTEVF | 0.87510157 |
| IALNEDL | ASSLPRAPNTEVF | 0.349643 |

| | | |
|---|---|---|
| IVNVSSM | ASSLPRAPNTEVF | 0.75050515 |
| TGIPPAPR | ASSLPRAPNTEVF | 0.8829651 |
| FDLGGGTF | ASSLPRAPNTEVF | 0.88333917 |
| FDLGGGTFDVS | ASSLPRAPNTEVF | 0.86204857 |
| NEPTAAA | ASSLPRAPNTEVF | 0.85652757 |
| NEPTAAALA | ASSLPRAPNTEVF | 0.70464224 |
| GSIINISSI | ASSLPRAPNTEVF | 0.89524615 |
| VAPGNAGTA | ASSLPRAPNTEVF | 0.9056571 |
| TGGGASLE | ASSLPRAPNTEVF | 0.8705702 |
| VAVYNHYKRI | ASSLPRAPNTEVF | 0.6786754 |
| TDPAPATT | ASSLPRAPNTEVF | 0.5838148 |
| TNRPDILDPALLRPG | ASSLPRAPNTEVF | 0.8460053 |
| LTTTTTS | ASSLPRAPNTEVF | 0.89142346 |
| PTGSGKTL | ASSLPRAPNTEVF | 0.65888137 |
| TGSGKTL | ASSLPRAPNTEVF | 0.8447329 |
| TIGHVDHGK | ASSLPRAPNTEVF | 0.82704574 |
| TIGHVDHGKTT | ASSLPRAPNTEVF | 0.28244677 |
| GHVDHGKTT | ASSLPRAPNTEVF | 0.87818587 |
| AQFEHTLL | ASSLPRAPNTEVF | 0.89285684 |
| IVNQDEL | ASSLPRAPNTEVF | 0.85244036 |
| TNGSQFF | ASSLPRAPNTEVF | 0.6513709 |
| TGKGIVE | ASSLPRAPNTEVF | 0.8926846 |
| LISNASDA | ASSLPRAPNTEVF | 0.9008249 |
| AELRTLGELTKIAKAHGAQ | ASSLPRAPNTEVF | 0.8957498 |
| APNAVAAAEPSW | ASSLPRAPNTEVF | 0.40669107 |
| APQVAGAATAVL | ASSLPRAPNTEVF | 0.37168878 |
| ARRDPAAAAAILPTDA | ASSLPRAPNTEVF | 0.8144612 |
| DAAARDVLAEAGLAEYF | ASSLPRAPNTEVF | 0.86129266 |
| DDQIVLSAKANLAKVEQ | ASSLPRAPNTEVF | 0.7500088 |
| DELLPDTGEDSDEDGH | ASSLPRAPNTEVF | 0.7019268 |
| ELPEAPANRATVPQD | ASSLPRAPNTEVF | 0.84543896 |
| FGQSKIERVTGIG | ASSLPRAPNTEVF | 0.89212716 |
| GAGGTGSTAGGAGGAGGAGGLY | ASSLPRAPNTEVF | 0.61539805 |
| GHGGTGAAGGNGGTGGAGGSGI | ASSLPRAPNTEVF | 0.48622292 |
| GNGGNGGAGGEGGPGGAGGAGA | ASSLPRAPNTEVF | 0.8762784 |
| GSTAGNGGNGGNATNGTIAGQP | ASSLPRAPNTEVF | 0.8828527 |
| GTGGDGGNGGNGGTGG | ASSLPRAPNTEVF | 0.739105 |
| GVRRPAGIPADAPV | ASSLPRAPNTEVF | 0.8063996 |
| GYRQVTDAHLLTLARR | ASSLPRAPNTEVF | 0.767255 |
| IAKMGQDGHDRG | ASSLPRAPNTEVF | 0.8448299 |
| KAEPSIEIQYT | ASSLPRAPNTEVF | 0.8256266 |
| KPVRIRMAADEDTVAVT | ASSLPRAPNTEVF | 0.5833291 |
| MPPRIAGMRLLVIKPEP | ASSLPRAPNTEVF | 0.89252496 |
| MKFYAVALFRSRGVPPTR | ASSLPRAPNTEVF | 0.63189316 |
| NAGGFGILGAVAHSP | ASSLPRAPNTEVF | 0.8807498 |
| NEGREITLAMAEM | ASSLPRAPNTEVF | 0.8386489 |
| NELVEWSHPLGHRWRWE | ASSLPRAPNTEVF | 0.8496441 |
| NLGAGGTGGNGGNPG | ASSLPRAPNTEVF | 0.7878275 |
| NTEWPKWETVMAADPVD | ASSLPRAPNTEVF | 0.6644449 |
| PASAAPAETQP | ASSLPRAPNTEVF | 0.5372958 |
| QQSSDGKNGVRAAEITGE | ASSLPRAPNTEVF | 0.5478992 |

| | | |
|---|---|---|
| RAPDAVTTSWWKE | ASSLPRAPNTEVF | 0.87653023 |
| SGFGNVGAHTSGWFNQ | ASSLPRAPNTEVF | 0.21092415 |
| SGVAGAVGQAASVGGLKV | ASSLPRAPNTEVF | 0.88313365 |
| SNGGAGTGSAGGNGGT | ASSLPRAPNTEVF | 0.8122014 |
| TGGAGGEGGAAG | ASSLPRAPNTEVF | 0.6854073 |
| TGGDGGAAGAGGAPGHG | ASSLPRAPNTEVF | 0.87368727 |
| TMPPLLAAIANAGRHGV | ASSLPRAPNTEVF | 0.59382385 |
| TSSTAGNSGGA | ASSLPRAPNTEVF | 0.7531844 |
| VPATVPVGILC | ASSLPRAPNTEVF | 0.8031674 |
| VQRPDSGGVVLTGRIS | ASSLPRAPNTEVF | 0.8689474 |
| WTPNPVPATKGVPAGLVQAT | ASSLPRAPNTEVF | 0.6908103 |
| YALKRALAKFSDRFDVVII | ASSLPRAPNTEVF | 0.8743388 |
| YLIGSVIV | ASSLPRAPNTEVF | 0.82944477 |
| YGQLVAVID | ASSLPRAPNTEVF | 0.66577053 |
| VTVGGMTLPA | ASSLPRAPNTEVF | 0.840271 |
| VLALPAV | ASSLPRAPNTEVF | 0.88638747 |
| VIPPTLNYET | ASSLPRAPNTEVF | 0.8532815 |
| VGMGGMHPSSG | ASSLPRAPNTEVF | 0.5806904 |
| VGLAGGAATA | ASSLPRAPNTEVF | 0.855519 |
| VGGAGGAGGSGP | ASSLPRAPNTEVF | 0.7418443 |
| VFAAFMV | ASSLPRAPNTEVF | 0.85600394 |
| TLYDKLR | ASSLPRAPNTEVF | 0.8626312 |
| TLAVFTVL | ASSLPRAPNTEVF | 0.7441248 |
| TGSVQPARV | ASSLPRAPNTEVF | 0.86924887 |
| TGGGAGSGGT | ASSLPRAPNTEVF | 0.73609257 |
| SHLETSL | ASSLPRAPNTEVF | 0.7543107 |
| SFTTGMT | ASSLPRAPNTEVF | 0.79438394 |
| QVEVAKRMGVRQPTVS | ASSLPRAPNTEVF | 0.82489645 |
| QGSGGPALAM | ASSLPRAPNTEVF | 0.8782916 |
| MLAPSNP | ASSLPRAPNTEVF | 0.6535769 |
| LVNNAAI | ASSLPRAPNTEVF | 0.80456716 |
| LTGEINTLA | ASSLPRAPNTEVF | 0.6023022 |
| LSIFGPV | ASSLPRAPNTEVF | 0.46797773 |
| LLALLANL | ASSLPRAPNTEVF | 0.8641281 |
| LISNASDALDKLR | ASSLPRAPNTEVF | 0.84157515 |
| LINEAVP | ASSLPRAPNTEVF | 0.6058431 |
| LGYAFSGL | ASSLPRAPNTEVF | 0.8574405 |
| LDFVAVAL | ASSLPRAPNTEVF | 0.6725201 |
| LAQPGAE | ASSLPRAPNTEVF | 0.81745344 |
| ISNLGAVTGF | ASSLPRAPNTEVF | 0.9008653 |
| HQEDPGRI | ASSLPRAPNTEVF | 0.7005355 |
| GTPGEPTP | ASSLPRAPNTEVF | 0.87864006 |
| GFFNSGS | ASSLPRAPNTEVF | 0.85767686 |
| GAGGGGLTAGAN | ASSLPRAPNTEVF | 0.8492928 |
| FVDALVNM | ASSLPRAPNTEVF | 0.718696 |
| FLTVAPEL | ASSLPRAPNTEVF | 0.633716 |
| FETLTAL | ASSLPRAPNTEVF | 0.8985777 |
| DVINPIEL | ASSLPRAPNTEVF | 0.859004 |
| APVSGAPV | ASSLPRAPNTEVF | 0.85873896 |
| ALVGGLAW | ASSLPRAPNTEVF | 0.87531775 |
| ALAAAAATAI | ASSLPRAPNTEVF | 0.87868583 |

| | | |
|---|---|---|
| AFGLGLLL | ASSLPRAPNTEVF | 0.88252914 |
| AFFAAASL | ASSLPRAPNTEVF | 0.7833384 |
| AAAALPAFSPP | ASSLPRAPNTEVF | 0.50288624 |
| VVLAPAL | ASSLPRAPNTEVF | 0.80117166 |
| TIGHVDHGKTTL | ASSLPRAPNTEVF | 0.7684717 |
| LISNASDALDK | ASSLPRAPNTEVF | 0.8090012 |
| LISNASDAL | ASSLPRAPNTEVF | 0.46165645 |
| ISNASDALDK | ASSLPRAPNTEVF | 0.8813518 |
| NQAETLVYQTE | ASSLPRAPNTEVF | 0.9049434 |
| TGGTGGTAGAG | ASSLPRAPNTEVF | 0.6098139 |
| TGSTAGGAGGA | ASSLPRAPNTEVF | 0.9030713 |
| DPLAVPVAAGPV | ASSLPRAPNTEVF | 0.70182204 |
| GGTGGTPGNGGN | ASSLPRAPNTEVF | 0.84320164 |
| PAGLAGVDPATL | ASSLPRAPNTEVF | 0.88973975 |
| GNNLVVEAPHAGASV | ASSLPRAPNTEVF | 0.62542665 |
| KVTRIMSASTLEQPA | ASSLPRAPNTEVF | 0.8396899 |
| LGFALPIALPRRTVV | ASSLPRAPNTEVF | 0.5577134 |
| TRLGNLTARKLVVLV | ASSLPRAPNTEVF | 0.67680454 |
| WSQQRRLIYFPSAGP | ASSLPRAPNTEVF | 0.8893127 |
| GPNGNPDPMAAAVDIR | ASSLPRAPNTEVF | 0.64432347 |
| TQRCVSVADLRAAASR | ASSLPRAPNTEVF | 0.83859456 |
| QFVLDVAEHPVEVLTETV | ASSLPRAPNTEVF | 0.7843086 |
| GGAGGTGGAGGNGGTQAGDG | ASSLPRAPNTEVF | 0.69733024 |
| PLLATLRAIAVDVDAKPAQV | ASSLPRAPNTEVF | 0.85772586 |
| APKFRTGGYMIDFWGVGYQVAK | ASSLPRAPNTEVF | 0.8766173 |
| GAGGAGGAGGAAGGPGGTGGTG | ASSLPRAPNTEVF | 0.7092638 |
| IGALGLLATDVAPAGSVAFGVA | ASSLPRAPNTEVF | 0.90443474 |
| AGGAGGIGGNGGAGGAGGVLMG | ASSLPRAPNTEVF | 0.6636343 |
| ADELIGLL | ASSLPRAPNTEVF | 0.9013057 |
| ADRALKA | ASSLPRAPNTEVF | 0.8991674 |
| AGAVGQVGGAGGA | ASSLPRAPNTEVF | 0.8442924 |
| AGGGGGLGGAP | ASSLPRAPNTEVF | 0.81829035 |
| AGGLLYGNGGAG | ASSLPRAPNTEVF | 0.8821591 |
| AGLESIGL | ASSLPRAPNTEVF | 0.77969646 |
| AGLKTGII | ASSLPRAPNTEVF | 0.8495463 |
| ASLAALLL | ASSLPRAPNTEVF | 0.891412 |
| ASVINISSI | ASSLPRAPNTEVF | 0.68701744 |
| ATPPGPA | ASSLPRAPNTEVF | 0.8614758 |
| ATVASALV | ASSLPRAPNTEVF | 0.80047005 |
| ATVASLIL | ASSLPRAPNTEVF | 0.8135813 |
| AVNGPVLIRH | ASSLPRAPNTEVF | 0.46401787 |
| DGAAGAVGI | ASSLPRAPNTEVF | 0.2640419 |
| DPLLHGI | ASSLPRAPNTEVF | 0.7929822 |
| ELGAALGAL | ASSLPRAPNTEVF | 0.551249 |
| ELGRTQAV | ASSLPRAPNTEVF | 0.8111016 |
| FVAGGYLS | ASSLPRAPNTEVF | 0.84891903 |
| GASGGAGGAGGW | ASSLPRAPNTEVF | 0.64506197 |
| GGAGAGGDGGAGGI | ASSLPRAPNTEVF | 0.858161 |
| GTGHNGSQPGMGGNG | ASSLPRAPNTEVF | 0.8636304 |
| GYAAASASAS | ASSLPRAPNTEVF | 0.43008536 |
| HQVDVAAVQQG | ASSLPRAPNTEVF | 0.7840979 |

| | | |
|---|---|---|
| IGAAIAHT | ASSLPRAPNTEVF | 0.6016053 |
| IGMIEVAQL | ASSLPRAPNTEVF | 0.77755404 |
| IRLAPEHTSAMD | ASSLPRAPNTEVF | 0.52817667 |
| IYNVSVL | ASSLPRAPNTEVF | 0.8456147 |
| KQGFLGM | ASSLPRAPNTEVF | 0.71281683 |
| LESVITQL | ASSLPRAPNTEVF | 0.21021456 |
| LGLGALVVV | ASSLPRAPNTEVF | 0.78739077 |
| LIGNGGNGGSGGI | ASSLPRAPNTEVF | 0.79372114 |
| LNAINAPTL | ASSLPRAPNTEVF | 0.7851262 |
| LQLLNTGV | ASSLPRAPNTEVF | 0.85406864 |
| LQTSIEF | ASSLPRAPNTEVF | 0.65971375 |
| MSFAVSEL | ASSLPRAPNTEVF | 0.6407884 |
| NTGALGL | ASSLPRAPNTEVF | 0.8414846 |
| PPGAGGAGGAGTT | ASSLPRAPNTEVF | 0.8595037 |
| QLVGLGI | ASSLPRAPNTEVF | 0.5735476 |
| RFLEQVHQL | ASSLPRAPNTEVF | 0.81150347 |
| RGVGYVLR | ASSLPRAPNTEVF | 0.85894567 |
| SATGVFAGVI | ASSLPRAPNTEVF | 0.73677087 |
| SLPAITLP | ASSLPRAPNTEVF | 0.85232663 |
| STALAFTV | ASSLPRAPNTEVF | 0.8559638 |
| SVLTVGLGK | ASSLPRAPNTEVF | 0.6555139 |
| TALAGAGSGI | ASSLPRAPNTEVF | 0.8496973 |
| TGLPFTGI | ASSLPRAPNTEVF | 0.85829836 |
| VAEHVTV | ASSLPRAPNTEVF | 0.76572585 |
| VDIVLNSL | ASSLPRAPNTEVF | 0.8112324 |
| VLTGGGAVL | ASSLPRAPNTEVF | 0.5189446 |
| VNVGLAYGI | ASSLPRAPNTEVF | 0.6774045 |
| VSSAAVGAV | ASSLPRAPNTEVF | 0.23292635 |
| VTSVYEAGV | ASSLPRAPNTEVF | 0.8218032 |
| WVALDRGA | ASSLPRAPNTEVF | 0.847366 |
| YLVGSVVI | ASSLPRAPNTEVF | 0.7754407 |
| GPPGTGKTLL | ASSLPRAPNTEVF | 0.70493925 |
| GVGMTKF | ASSLPRAPNTEVF | 0.43405926 |
| IGIGGSDL | ASSLPRAPNTEVF | 0.81830204 |
| IGTLLAEL | ASSLPRAPNTEVF | 0.71804225 |
| IYTSGTT | ASSLPRAPNTEVF | 0.41630796 |
| LSFPAGV | ASSLPRAPNTEVF | 0.62015843 |
| LYTSGTT | ASSLPRAPNTEVF | 0.836986 |
| VGAFPFGF | ASSLPRAPNTEVF | 0.83502686 |
| VGAGTVEF | ASSLPRAPNTEVF | 0.6319149 |
| VGKSTLLL | ASSLPRAPNTEVF | 0.8611312 |
| AALAHAGGGIV | ASSLPRAPNTEVF | 0.82854205 |
| AALVAVSLL | ASSLPRAPNTEVF | 0.8418414 |
| AAVSRAVAAL | ASSLPRAPNTEVF | 0.79600227 |
| AGGSSGIGGF | ASSLPRAPNTEVF | 0.6331476 |
| AGLTVANANG | ASSLPRAPNTEVF | 0.8429996 |
| AGLVGAVL | ASSLPRAPNTEVF | 0.7199891 |
| ALSQAFTV | ASSLPRAPNTEVF | 0.6766475 |
| ALTFSVSL | ASSLPRAPNTEVF | 0.78988177 |
| ALTGAALVAL | ASSLPRAPNTEVF | 0.7387898 |
| ASFSIGAGVL | ASSLPRAPNTEVF | 0.74343705 |

| | | |
|---|---|---|
| DGSVYKI | ASSLPRAPNTEVF | 0.7668589 |
| EQLAEIV | ASSLPRAPNTEVF | 0.83933055 |
| FAVSELTVM | ASSLPRAPNTEVF | 0.65717125 |
| GAAPLAAGL | ASSLPRAPNTEVF | 0.40749174 |
| GADGGDGGSGGAGG | ASSLPRAPNTEVF | 0.7430443 |
| GFGLDTGGAGG | ASSLPRAPNTEVF | 0.80929565 |
| GLASGGLL | ASSLPRAPNTEVF | 0.8563982 |
| GRIAGAAVLL | ASSLPRAPNTEVF | 0.82303643 |
| GSAGSLNTSF | ASSLPRAPNTEVF | 0.77607733 |
| GTGQIGI | ASSLPRAPNTEVF | 0.8293067 |
| GTLIALIL | ASSLPRAPNTEVF | 0.836846 |
| IAATMGFVVG | ASSLPRAPNTEVF | 0.75247806 |
| IFAQALL | ASSLPRAPNTEVF | 0.8190911 |
| IGAAVALGF | ASSLPRAPNTEVF | 0.75513947 |
| IGLTPLAL | ASSLPRAPNTEVF | 0.8417413 |
| ILGSNTSTL | ASSLPRAPNTEVF | 0.8245207 |
| IVFSLTVL | ASSLPRAPNTEVF | 0.8421765 |
| LASKATGF | ASSLPRAPNTEVF | 0.85036105 |
| LASVALIL | ASSLPRAPNTEVF | 0.802661 |
| LFAVGLTL | ASSLPRAPNTEVF | 0.7256774 |
| LFNGGIGTYI | ASSLPRAPNTEVF | 0.21717674 |
| LTLGAGII | ASSLPRAPNTEVF | 0.66651106 |
| LVAAALGDI | ASSLPRAPNTEVF | 0.79181314 |
| NTAFVQLGI | ASSLPRAPNTEVF | 0.8194306 |
| QLFSGAEL | ASSLPRAPNTEVF | 0.8124776 |
| SAATFITL | ASSLPRAPNTEVF | 0.83296686 |
| SNYGLANLGV | ASSLPRAPNTEVF | 0.72830915 |
| SYGGAYAVM | ASSLPRAPNTEVF | 0.83335036 |
| TTGVGGLTLG | ASSLPRAPNTEVF | 0.85056233 |
| TTQTIYI | ASSLPRAPNTEVF | 0.36188138 |
| VAAGAGLGAVI | ASSLPRAPNTEVF | 0.735047 |
| VAAVVTLL | ASSLPRAPNTEVF | 0.86680645 |
| VGFLGGLI | ASSLPRAPNTEVF | 0.81483746 |
| VGVIPILL | ASSLPRAPNTEVF | 0.7017928 |
| VIFASSLI | ASSLPRAPNTEVF | 0.7541789 |
| VKGGDGGAAAG | ASSLPRAPNTEVF | 0.8567825 |
| VTKTFTNI | ASSLPRAPNTEVF | 0.764834 |
| VVAAAILL | ASSLPRAPNTEVF | 0.8614203 |
| YAALDVEL | ASSLPRAPNTEVF | 0.22118104 |
| LASVALLL | ASSLPRAPNTEVF | 0.7381628 |
| LDAVHRV | ASSLPRAPNTEVF | 0.44065 |
| QGLSSSIF | ASSLPRAPNTEVF | 0.33596832 |
| RGTFANIRL | ASSLPRAPNTEVF | 0.8155364 |
| RQLWWGHRI | ASSLPRAPNTEVF | 0.7856878 |
| YGPPGTGKTLL | ASSLPRAPNTEVF | 0.84455836 |
| AALVGAPANVA | ASSLPRAPNTEVF | 0.77329016 |
| AATFRRTE | ASSLPRAPNTEVF | 0.84811074 |
| AGALGGGI | ASSLPRAPNTEVF | 0.78877383 |
| AGGAGGAGGIGAG | ASSLPRAPNTEVF | 0.7627611 |
| AGSGALGL | ASSLPRAPNTEVF | 0.56135535 |
| AQIADQI | ASSLPRAPNTEVF | 0.79963124 |

| | | |
|---|---|---|
| ASAVAVTYL | ASSLPRAPNTEVF | 0.8401362 |
| ASTVLGGTGG | ASSLPRAPNTEVF | 0.826736 |
| AVGGEPAK | ASSLPRAPNTEVF | 0.40158278 |
| CEQGTFGGTC | ASSLPRAPNTEVF | 0.7746308 |
| DEGKAPAG | ASSLPRAPNTEVF | 0.71734834 |
| DTVVFPAL | ASSLPRAPNTEVF | 0.24971569 |
| FEGITMAG | ASSLPRAPNTEVF | 0.21038927 |
| FPLVVQD | ASSLPRAPNTEVF | 0.7269913 |
| GAFTTPQ | ASSLPRAPNTEVF | 0.7484705 |
| GFGGFGTAGGVGGA | ASSLPRAPNTEVF | 0.8488208 |
| GFLAQFL | ASSLPRAPNTEVF | 0.8457644 |
| GGAGGAGGNAGVA | ASSLPRAPNTEVF | 0.74271524 |
| GGVGTAGAGGAGGAGG | ASSLPRAPNTEVF | 0.8379475 |
| GNGGGGDAGNAGSGGN | ASSLPRAPNTEVF | 0.8190275 |
| GTGGAGGAGNDAGS | ASSLPRAPNTEVF | 0.13017069 |
| GTTNTGWFNS | ASSLPRAPNTEVF | 0.7320152 |
| IFMGPLFTL | ASSLPRAPNTEVF | 0.4595626 |
| ILAHAIEAI | ASSLPRAPNTEVF | 0.8625213 |
| IRSAIREAL | ASSLPRAPNTEVF | 0.66892153 |
| ITFAGGG | ASSLPRAPNTEVF | 0.7544147 |
| ITGMSGHQEV | ASSLPRAPNTEVF | 0.77572286 |
| IVAGLACTL | ASSLPRAPNTEVF | 0.8201946 |
| IVGGGLAGA | ASSLPRAPNTEVF | 0.7537284 |
| IVTGVLNI | ASSLPRAPNTEVF | 0.6385927 |
| LAQQGCI | ASSLPRAPNTEVF | 0.1422822 |
| LCLHVEN | ASSLPRAPNTEVF | 0.80429506 |
| LDAISDGS | ASSLPRAPNTEVF | 0.4899405 |
| LGGFAAML | ASSLPRAPNTEVF | 0.6067689 |
| LGGKNPMI | ASSLPRAPNTEVF | 0.42102075 |
| LGPKGVAAIL | ASSLPRAPNTEVF | 0.21596643 |
| LMVGVSI | ASSLPRAPNTEVF | 0.5314249 |
| LVGAAFVL | ASSLPRAPNTEVF | 0.8442482 |
| LVLAVAPL | ASSLPRAPNTEVF | 0.67940295 |
| MAAIAVIDAA | ASSLPRAPNTEVF | 0.7846439 |
| NNGINAELQ | ASSLPRAPNTEVF | 0.8579957 |
| RGEPLVLVHG | ASSLPRAPNTEVF | 0.8177794 |
| RGGAGGAAGLI | ASSLPRAPNTEVF | 0.7406882 |
| RSYRMHPAV | ASSLPRAPNTEVF | 0.70178336 |
| SAAHLLAGQLA | ASSLPRAPNTEVF | 0.5869014 |
| SAAVGSY | ASSLPRAPNTEVF | 0.6203878 |
| SPHAVLHRI | ASSLPRAPNTEVF | 0.66638696 |
| SQIAKVL | ASSLPRAPNTEVF | 0.8647418 |
| TALGHLVP | ASSLPRAPNTEVF | 0.7262825 |
| TAPTTSGAA | ASSLPRAPNTEVF | 0.60898745 |
| TDGDDGNGGSPGT | ASSLPRAPNTEVF | 0.8179083 |
| TDGLGEN | ASSLPRAPNTEVF | 0.85498995 |
| TGGPPLPGQA | ASSLPRAPNTEVF | 0.8574985 |
| TIPAITI | ASSLPRAPNTEVF | 0.6320758 |
| TIPGFTI | ASSLPRAPNTEVF | 0.7227563 |
| TSIASGALA | ASSLPRAPNTEVF | 0.7611796 |
| TVTLPGAAI | ASSLPRAPNTEVF | 0.7988498 |

| | | |
|---|---|---|
| VASGLSAALP | ASSLPRAPNTEVF | 0.7543757 |
| VGAGVIGI | ASSLPRAPNTEVF | 0.5331228 |
| VGPKRFGV | ASSLPRAPNTEVF | 0.8299791 |
| VNVAFPD | ASSLPRAPNTEVF | 0.8702165 |
| VSMTNSGVS | ASSLPRAPNTEVF | 0.7501693 |
| VTDVLGDLSV | ASSLPRAPNTEVF | 0.39918655 |
| VVVVGAGFAGL | ASSLPRAPNTEVF | 0.75655544 |
| VYTVGTVV | ASSLPRAPNTEVF | 0.66586876 |
| WTTFSGP | ASSLPRAPNTEVF | 0.6612968 |
| YDELVEAF | ASSLPRAPNTEVF | 0.7987154 |
| YGTGGAGGNGG | ASSLPRAPNTEVF | 0.58506453 |
| YTNVMAV | ASSLPRAPNTEVF | 0.75915706 |
| FLEQVHQL | ASSLPRAPNTEVF | 0.8551209 |
| GLSLLAVAV | ASSLPRAPNTEVF | 0.5353457 |
| LEQVHQL | ASSLPRAPNTEVF | 0.81779087 |
| LNAAGAI | ASSLPRAPNTEVF | 0.8539647 |
| PAVHASL | ASSLPRAPNTEVF | 0.83087754 |
| TELEDDA | ASSLPRAPNTEVF | 0.85843146 |
| TPPGPPN | ASSLPRAPNTEVF | 0.84297156 |
| TPPPPPPP | ASSLPRAPNTEVF | 0.7289146 |
| TQSPALI | GARADWGSQNTLY | 0.7783288 |
| EFAAAFPQ | GARADWGSQNTLY | 0.7828932 |
| LLSGNLDVL | GARADWGSQNTLY | 0.7855089 |
| GPANLGNYNI | GARADWGSQNTLY | 0.7433674 |
| TLGPTLP | GARADWGSQNTLY | 0.70750374 |
| VYAYSSCCK | GARADWGSQNTLY | 0.8530812 |
| APAPAPAPNI | GARADWGSQNTLY | 0.75045663 |
| PGGPVPIP | GARADWGSQNTLY | 0.843454 |
| FLTVALEP | GARADWGSQNTLY | 0.4487241 |
| SQYFGANL | GARADWGSQNTLY | 0.5502025 |
| GGVKPAA | GARADWGSQNTLY | 0.8487642 |
| GGTGGAGGAGGW | GARADWGSQNTLY | 0.14128575 |
| GVGGVAGLGGAGG | GARADWGSQNTLY | 0.2696378 |
| VLLVLATTI | GARADWGSQNTLY | 0.78489774 |
| IGIFAAML | GARADWGSQNTLY | 0.8543435 |
| AVGDLID | GARADWGSQNTLY | 0.15422018 |
| NISPLPL | GARADWGSQNTLY | 0.650862 |
| VTAVFVAI | GARADWGSQNTLY | 0.45114306 |
| SGDDNDAAL | GARADWGSQNTLY | 0.83600116 |
| AVAALAHS | GARADWGSQNTLY | 0.78976464 |
| GGVTVMATP | GARADWGSQNTLY | 0.30314523 |
| AGLAANVRD | GARADWGSQNTLY | 0.84100163 |
| TNRPDLIDPALLRPG | GARADWGSQNTLY | 0.5065198 |
| LANLGVTAL | GARADWGSQNTLY | 0.5330126 |
| TGERIEL | GARADWGSQNTLY | 0.83763814 |
| NGEELAA | GARADWGSQNTLY | 0.8556305 |
| VLSLVWG | GARADWGSQNTLY | 0.58091795 |
| IGYALWL | GARADWGSQNTLY | 0.8431988 |
| LSLLLAPL | GARADWGSQNTLY | 0.76736605 |
| TLDEALK | GARADWGSQNTLY | 0.85998416 |
| TDGAGTAEGG | GARADWGSQNTLY | 0.50376785 |

| | | |
|---|---|---|
| NAPGGAGGNGG | GARADWGSQNTLY | 0.48788434 |
| VSVFMGMGGNF | GARADWGSQNTLY | 0.84895885 |
| LASYLAAL | GARADWGSQNTLY | 0.806981 |
| LVGGTPAL | GARADWGSQNTLY | 0.76851535 |
| VRNADVL | GARADWGSQNTLY | 0.83192444 |
| TCSSPFLVSAM | GARADWGSQNTLY | 0.75507283 |
| VHITDAE | GARADWGSQNTLY | 0.7574856 |
| EYAAPSF | GARADWGSQNTLY | 0.85329705 |
| VVDAFHGGL | GARADWGSQNTLY | 0.833886 |
| ASVVDPQ | GARADWGSQNTLY | 0.53816056 |
| SYNEFGGLS | GARADWGSQNTLY | 0.8317019 |
| DPDGSVY | GARADWGSQNTLY | 0.8096512 |
| MVSTIHATE | GARADWGSQNTLY | 0.5485722 |
| LAAGVADPV | GARADWGSQNTLY | 0.84898317 |
| TKEQIAA | GARADWGSQNTLY | 0.7908081 |
| SLASPAAL | GARADWGSQNTLY | 0.8334665 |
| IGLVFYGV | GARADWGSQNTLY | 0.7963711 |
| FLLDASNL | GARADWGSQNTLY | 0.85823286 |
| YQVGNFP | GARADWGSQNTLY | 0.27821216 |
| ESLLSQST | GARADWGSQNTLY | 0.814805 |
| LFTVGFAAVG | GARADWGSQNTLY | 0.7317146 |
| ATPPTPT | GARADWGSQNTLY | 0.8406527 |
| TDLATLG | GARADWGSQNTLY | 0.7883933 |
| SAGGTGR | GARADWGSQNTLY | 0.47340018 |
| GFSNSGS | GARADWGSQNTLY | 0.7691883 |
| GGNGGSGGTSV | GARADWGSQNTLY | 0.8547375 |
| GTSVATGGAGN | GARADWGSQNTLY | 0.7871481 |
| NGGNGGSGGTS | GARADWGSQNTLY | 0.8043484 |
| AVVKFPH | GARADWGSQNTLY | 0.8638986 |
| TPPPPPP | GARADWGSQNTLY | 0.52101755 |
| AAPAAAAAPA | GARADWGSQNTLY | 0.8008838 |
| IALNEDL | GARADWGSQNTLY | 0.8339217 |
| IVNVSSM | GARADWGSQNTLY | 0.83803964 |
| TGIPPAPR | GARADWGSQNTLY | 0.77463794 |
| FDLGGGTF | GARADWGSQNTLY | 0.8582114 |
| FDLGGGTFDVS | GARADWGSQNTLY | 0.61850524 |
| NEPTAAA | GARADWGSQNTLY | 0.84489584 |
| NEPTAAALA | GARADWGSQNTLY | 0.6624485 |
| GSIINISSI | GARADWGSQNTLY | 0.81653595 |
| VAPGNAGTA | GARADWGSQNTLY | 0.6845485 |
| TGGGASLE | GARADWGSQNTLY | 0.82629746 |
| VAVYNHYKRI | GARADWGSQNTLY | 0.84476054 |
| TDPAPATT | GARADWGSQNTLY | 0.8490548 |
| TNRPDILDPALLRPG | GARADWGSQNTLY | 0.26178366 |
| LTTTTTS | GARADWGSQNTLY | 0.845562 |
| PTGSGKTL | GARADWGSQNTLY | 0.8475524 |
| TGSGKTL | GARADWGSQNTLY | 0.869194 |
| TIGHVDHGK | GARADWGSQNTLY | 0.7118218 |
| TIGHVDHGKTT | GARADWGSQNTLY | 0.83567107 |
| GHVDHGKTT | GARADWGSQNTLY | 0.74492115 |
| AQFEHTLL | GARADWGSQNTLY | 0.7626439 |

| | | |
|---|---|---|
| IVNQDEL | GARADWGSQNTLY | 0.66741 |
| TNGSQFF | GARADWGSQNTLY | 0.6486938 |
| TGKGIVE | GARADWGSQNTLY | 0.76814044 |
| LISNASDA | GARADWGSQNTLY | 0.71514106 |
| AELRTLGELTKIAKAHGAQ | GARADWGSQNTLY | 0.848518 |
| APNAVAAAEPSW | GARADWGSQNTLY | 0.8300479 |
| APQVAGAATAVL | GARADWGSQNTLY | 0.83289707 |
| ARRDPAAAAAILPTDA | GARADWGSQNTLY | 0.85509443 |
| DAAARDVLAEAGLAEYF | GARADWGSQNTLY | 0.8452412 |
| DDQIVLSAKANLAKVEQ | GARADWGSQNTLY | 0.8441216 |
| DELLPDTGEDSDEDGH | GARADWGSQNTLY | 0.65404254 |
| ELPEAPANRATVPQD | GARADWGSQNTLY | 0.75127214 |
| FGQSKIERVTGIG | GARADWGSQNTLY | 0.6307237 |
| GAGGTGSTAGGAGGAGGAGGLY | GARADWGSQNTLY | 0.7753111 |
| GHGGTGAAGGNGGTGGAGGSGI | GARADWGSQNTLY | 0.6916794 |
| GNGGNGGAGGEGGPGGAGGAGA | GARADWGSQNTLY | 0.8188453 |
| GSTAGNGGNGGNATNGTIAGQP | GARADWGSQNTLY | 0.12796351 |
| GTGGDGGNGGNGGTGG | GARADWGSQNTLY | 0.8197036 |
| GVRRPAGIPADAPV | GARADWGSQNTLY | 0.864825 |
| GYRQVTDAHLLTLARR | GARADWGSQNTLY | 0.6377406 |
| IAKMGQDGHDRG | GARADWGSQNTLY | 0.79060304 |
| KAEPSIEIQYT | GARADWGSQNTLY | 0.78753805 |
| KPVRIRMAADEDTVAVT | GARADWGSQNTLY | 0.74536186 |
| MPPRIAGMRLLVIKPEP | GARADWGSQNTLY | 0.840466 |
| MKFYAVALFRSRGVPPTR | GARADWGSQNTLY | 0.6887375 |
| NAGGFGILGAVAHSP | GARADWGSQNTLY | 0.83495545 |
| NEGREITLAMAEM | GARADWGSQNTLY | 0.8424572 |
| NELVEWSHPLGHRWRWE | GARADWGSQNTLY | 0.76245385 |
| NLGAGGTGGNGGNPG | GARADWGSQNTLY | 0.7334274 |
| NTEWPKWETVMAADPVD | GARADWGSQNTLY | 0.7879865 |
| PASAAPAETQP | GARADWGSQNTLY | 0.2228264 |
| QQSSDGKNGVRAAEITGE | GARADWGSQNTLY | 0.76540184 |
| RAPDAVTTSWWKE | GARADWGSQNTLY | 0.11994702 |
| SGFGNVGAHTSGWFNQ | GARADWGSQNTLY | 0.7386083 |
| SGVAGAVGQAASVGGLKV | GARADWGSQNTLY | 0.41591686 |
| SNGGAGTGSAGGNGGT | GARADWGSQNTLY | 0.81081593 |
| TGGAGGEGGAAG | GARADWGSQNTLY | 0.81717837 |
| TGGDGGAAGAGGAPGHG | GARADWGSQNTLY | 0.81992817 |
| TMPPLLAAIANAGRHGV | GARADWGSQNTLY | 0.46768016 |
| TSSTAGNSGGA | GARADWGSQNTLY | 0.7959899 |
| VPATVPVGILC | GARADWGSQNTLY | 0.75622356 |
| VQRPDSGGVVLTGRIS | GARADWGSQNTLY | 0.48037165 |
| WTPNPVPATKGVPAGLVQAT | GARADWGSQNTLY | 0.833329 |
| YALKRALAKFSDRFDVVII | GARADWGSQNTLY | 0.7108796 |
| YLIGSVIV | GARADWGSQNTLY | 0.7475457 |
| YGQLVAVID | GARADWGSQNTLY | 0.84163487 |
| VTVGGMTLPA | GARADWGSQNTLY | 0.7821155 |
| VLALPAV | GARADWGSQNTLY | 0.38669503 |
| VIPPTLNYET | GARADWGSQNTLY | 0.7418441 |
| VGMGGMHPSSG | GARADWGSQNTLY | 0.7886843 |
| VGLAGGAATA | GARADWGSQNTLY | 0.4537226 |

| | | |
|---|---|---|
| VGGAGGAGGSGP | GARADWGSQNTLY | 0.15937427 |
| VFAAFMV | GARADWGSQNTLY | 0.8039149 |
| TLYDKLR | GARADWGSQNTLY | 0.5588281 |
| TLAVFTVL | GARADWGSQNTLY | 0.5967847 |
| TGSVQPARV | GARADWGSQNTLY | 0.75865835 |
| TGGGAGSGGT | GARADWGSQNTLY | 0.8491213 |
| SHLETSL | GARADWGSQNTLY | 0.6015669 |
| SFTTGMT | GARADWGSQNTLY | 0.78521127 |
| QVEVAKRMGVRQPTVS | GARADWGSQNTLY | 0.8158041 |
| QGSGGPALAM | GARADWGSQNTLY | 0.8251154 |
| MLAPSNP | GARADWGSQNTLY | 0.8607083 |
| LVNNAAI | GARADWGSQNTLY | 0.8150008 |
| LTGEINTLA | GARADWGSQNTLY | 0.63326234 |
| LSIFGPV | GARADWGSQNTLY | 0.813504 |
| LLALLANL | GARADWGSQNTLY | 0.7961963 |
| LISNASDALDKLR | GARADWGSQNTLY | 0.5479535 |
| LINEAVP | GARADWGSQNTLY | 0.45120135 |
| LGYAFSGL | GARADWGSQNTLY | 0.71766645 |
| LDFVAVAL | GARADWGSQNTLY | 0.37093478 |
| LAQPGAE | GARADWGSQNTLY | 0.69669795 |
| ISNLGAVTGF | GARADWGSQNTLY | 0.85563445 |
| HQEDPGRI | GARADWGSQNTLY | 0.758114 |
| GTPGEPTP | GARADWGSQNTLY | 0.43017972 |
| GFFNSGS | GARADWGSQNTLY | 0.77914023 |
| GAGGGGLTAGAN | GARADWGSQNTLY | 0.5146681 |
| FVDALVNM | GARADWGSQNTLY | 0.8185291 |
| FLTVAPEL | GARADWGSQNTLY | 0.7041961 |
| FETLTAL | GARADWGSQNTLY | 0.86809164 |
| DVINPIEL | GARADWGSQNTLY | 0.8554378 |
| APVSGAPV | GARADWGSQNTLY | 0.76537716 |
| ALVGGLAW | GARADWGSQNTLY | 0.8417951 |
| ALAAAAATAI | GARADWGSQNTLY | 0.68736917 |
| AFGLGLLL | GARADWGSQNTLY | 0.77942646 |
| AFFAAASL | GARADWGSQNTLY | 0.72872895 |
| AAAALPAFSPP | GARADWGSQNTLY | 0.80239224 |
| VVLAPAL | GARADWGSQNTLY | 0.75035876 |
| TIGHVDHGKTTL | GARADWGSQNTLY | 0.6530022 |
| LISNASDALDK | GARADWGSQNTLY | 0.27171594 |
| LISNASDAL | GARADWGSQNTLY | 0.8297479 |
| ISNASDALDK | GARADWGSQNTLY | 0.6655127 |
| NQAETLVYQTE | GARADWGSQNTLY | 0.4933487 |
| TGGTGGTAGAG | GARADWGSQNTLY | 0.6838791 |
| TGSTAGGAGGA | GARADWGSQNTLY | 0.8632244 |
| DPLAVPVAAGPV | GARADWGSQNTLY | 0.8603074 |
| GGTGGTPGNGGN | GARADWGSQNTLY | 0.72413814 |
| PAGLAGVDPATL | GARADWGSQNTLY | 0.5829477 |
| GNNLVVEAPHAGASV | GARADWGSQNTLY | 0.8251166 |
| KVTRIMSASTLEQPA | GARADWGSQNTLY | 0.43569803 |
| LGFALPIALPRRTVV | GARADWGSQNTLY | 0.8168018 |
| TRLGNLTARKLVVLV | GARADWGSQNTLY | 0.8008886 |
| WSQQRRLIYFPSAGP | GARADWGSQNTLY | 0.7940384 |

| | | |
|---|---|---|
| GPNGNPDPMAAAVDIR | GARADWGSQNTLY | 0.6965406 |
| TQRCVSVADLRAAASR | GARADWGSQNTLY | 0.818447 |
| QFVLDVAEHPVEVLTETV | GARADWGSQNTLY | 0.76535535 |
| GGAGGTGGAGGNGGTQAGDG | GARADWGSQNTLY | 0.82308227 |
| PLLATLRAIAVDVDAKPAQV | GARADWGSQNTLY | 0.17403337 |
| APKFRTGGYMIDFWGVGYQVAK | GARADWGSQNTLY | 0.8313133 |
| GAGGAGGAGGAAGGPGGTGGTG | GARADWGSQNTLY | 0.7365228 |
| IGALGLLATDVAPAGSVAFGVA | GARADWGSQNTLY | 0.5843078 |
| AGGAGGIGGNGGAGGAGGVLMG | GARADWGSQNTLY | 0.3475777 |
| ADELIGLL | GARADWGSQNTLY | 0.79713535 |
| ADRALKA | GARADWGSQNTLY | 0.83185774 |
| AGAVGQVGGAGGA | GARADWGSQNTLY | 0.846779 |
| AGGGGGLGGAP | GARADWGSQNTLY | 0.470557 |
| AGGLLYGNGGAG | GARADWGSQNTLY | 0.8400178 |
| AGLESIGL | GARADWGSQNTLY | 0.8490249 |
| AGLKTGII | GARADWGSQNTLY | 0.73251534 |
| ASLAALLL | GARADWGSQNTLY | 0.22063547 |
| ASVINISSI | GARADWGSQNTLY | 0.8334169 |
| ATPPGPA | GARADWGSQNTLY | 0.7848766 |
| ATVASALV | GARADWGSQNTLY | 0.84615695 |
| ATVASLIL | GARADWGSQNTLY | 0.8082783 |
| AVNGPVLIRH | GARADWGSQNTLY | 0.5515214 |
| DGAAGAVGI | GARADWGSQNTLY | 0.85707545 |
| DPLLHGI | GARADWGSQNTLY | 0.8455025 |
| ELGAALGAL | GARADWGSQNTLY | 0.5903759 |
| ELGRTQAV | GARADWGSQNTLY | 0.72298324 |
| FVAGGYLS | GARADWGSQNTLY | 0.7344061 |
| GASGGAGGAGGW | GARADWGSQNTLY | 0.33903456 |
| GGAGAGGDGGAGGI | GARADWGSQNTLY | 0.8320243 |
| GTGHNGSQPGMGGNG | GARADWGSQNTLY | 0.7536279 |
| GYAAASASAS | GARADWGSQNTLY | 0.7115443 |
| HQVDVAAVQQG | GARADWGSQNTLY | 0.8137219 |
| IGAAIAHT | GARADWGSQNTLY | 0.7097447 |
| IGMIEVAQL | GARADWGSQNTLY | 0.68155456 |
| IRLAPEHTSAMD | GARADWGSQNTLY | 0.7287741 |
| IYNVSVL | GARADWGSQNTLY | 0.76720554 |
| KQGFLGM | GARADWGSQNTLY | 0.8255894 |
| LESVITQL | GARADWGSQNTLY | 0.8095355 |
| LGLGALVVV | GARADWGSQNTLY | 0.82029164 |
| LIGNGGNGGSGGI | GARADWGSQNTLY | 0.8626109 |
| LNAINAPTL | GARADWGSQNTLY | 0.4464428 |
| LQLLNTGV | GARADWGSQNTLY | 0.71597207 |
| LQTSIEF | GARADWGSQNTLY | 0.84014857 |
| MSFAVSEL | GARADWGSQNTLY | 0.48550785 |
| NTGALGL | GARADWGSQNTLY | 0.8507105 |
| PPGAGGAGGAGTT | GARADWGSQNTLY | 0.8219743 |
| QLVGLGI | GARADWGSQNTLY | 0.820505 |
| RFLEQVHQL | GARADWGSQNTLY | 0.7655037 |
| RGVGYVLR | GARADWGSQNTLY | 0.864478 |
| SATGVFAGVI | GARADWGSQNTLY | 0.84622073 |
| SLPAITLP | GARADWGSQNTLY | 0.8139102 |

| | | |
|---|---|---|
| STALAFTV | GARADWGSQNTLY | 0.83222586 |
| SVLTVGLGK | GARADWGSQNTLY | 0.6683148 |
| TALAGAGSGI | GARADWGSQNTLY | 0.84388256 |
| TGLPFTGI | GARADWGSQNTLY | 0.8291204 |
| VAEHVTV | GARADWGSQNTLY | 0.4717433 |
| VDIVLNSL | GARADWGSQNTLY | 0.7668083 |
| VLTGGGAVL | GARADWGSQNTLY | 0.8445648 |
| VNVGLAYGI | GARADWGSQNTLY | 0.85058737 |
| VSSAAVGAV | GARADWGSQNTLY | 0.2677384 |
| VTSVYEAGV | GARADWGSQNTLY | 0.78316474 |
| WVALDRGA | GARADWGSQNTLY | 0.4125458 |
| YLVGSVVI | GARADWGSQNTLY | 0.59557843 |
| GPPGTGKTLL | GARADWGSQNTLY | 0.45359737 |
| GVGMTKF | GARADWGSQNTLY | 0.79587317 |
| IGIGGSDL | GARADWGSQNTLY | 0.70275724 |
| IGTLLAEL | GARADWGSQNTLY | 0.83220404 |
| IYTSGTT | GARADWGSQNTLY | 0.6704862 |
| LSFPAGV | GARADWGSQNTLY | 0.8367349 |
| LYTSGTT | GARADWGSQNTLY | 0.6736897 |
| VGAFPFGF | GARADWGSQNTLY | 0.7815825 |
| VGAGTVEF | GARADWGSQNTLY | 0.37338966 |
| VGKSTLLL | GARADWGSQNTLY | 0.21021724 |
| AALAHAGGGIV | GARADWGSQNTLY | 0.6308528 |
| AALVAVSLL | GARADWGSQNTLY | 0.7598095 |
| AAVSRAVAAL | GARADWGSQNTLY | 0.19886772 |
| AGGSSGIGGF | GARADWGSQNTLY | 0.8624656 |
| AGLTVANANG | GARADWGSQNTLY | 0.80922365 |
| AGLVGAVL | GARADWGSQNTLY | 0.793879 |
| ALSQAFTV | GARADWGSQNTLY | 0.7229016 |
| ALTFSVSL | GARADWGSQNTLY | 0.744125 |
| ALTGAALVAL | GARADWGSQNTLY | 0.6687904 |
| ASFSIGAGVL | GARADWGSQNTLY | 0.8374427 |
| DGSVYKI | GARADWGSQNTLY | 0.53846574 |
| EQLAEIV | GARADWGSQNTLY | 0.3821634 |
| FAVSELTVM | GARADWGSQNTLY | 0.7134769 |
| GAAPLAAGL | GARADWGSQNTLY | 0.62625015 |
| GADGGDGGSGGAGG | GARADWGSQNTLY | 0.83896875 |
| GFGLDTGGAGG | GARADWGSQNTLY | 0.84177697 |
| GLASGGLL | GARADWGSQNTLY | 0.8221457 |
| GRIAGAAVLL | GARADWGSQNTLY | 0.84556544 |
| GSAGSLNTSF | GARADWGSQNTLY | 0.21171728 |
| GTGQIGI | GARADWGSQNTLY | 0.7628611 |
| GTLIALIL | GARADWGSQNTLY | 0.86581516 |
| IAATMGFVVG | GARADWGSQNTLY | 0.76722014 |
| IFAQALL | GARADWGSQNTLY | 0.72252476 |
| IGAAVALGF | GARADWGSQNTLY | 0.85396624 |
| IGLTPLAL | GARADWGSQNTLY | 0.74738914 |
| ILGSNTSTL | GARADWGSQNTLY | 0.49166423 |
| IVFSLTVL | GARADWGSQNTLY | 0.80501306 |
| LASKATGF | GARADWGSQNTLY | 0.7857163 |
| LASVALIL | GARADWGSQNTLY | 0.7808492 |

| | | |
|---|---|---|
| LFAVGLTL | GARADWGSQNTLY | 0.48369786 |
| LFNGGIGTYI | GARADWGSQNTLY | 0.82444626 |
| LTLGAGII | GARADWGSQNTLY | 0.8690744 |
| LVAAALGDI | GARADWGSQNTLY | 0.7558185 |
| NTAFVQLGI | GARADWGSQNTLY | 0.83067703 |
| QLFSGAEL | GARADWGSQNTLY | 0.8322878 |
| SAATFITL | GARADWGSQNTLY | 0.81259 |
| SNYGLANLGV | GARADWGSQNTLY | 0.6055967 |
| SYGGAYAVM | GARADWGSQNTLY | 0.7005681 |
| TTGVGGLTLG | GARADWGSQNTLY | 0.7984745 |
| TTQTIYI | GARADWGSQNTLY | 0.7127488 |
| VAAGAGLGAVI | GARADWGSQNTLY | 0.7916789 |
| VAAVVTLL | GARADWGSQNTLY | 0.7334118 |
| VGFLGGLI | GARADWGSQNTLY | 0.86796963 |
| VGVIPILL | GARADWGSQNTLY | 0.81984854 |
| VIFASSLI | GARADWGSQNTLY | 0.8301167 |
| VKGGDGGAAAG | GARADWGSQNTLY | 0.68240404 |
| VTKTFTNI | GARADWGSQNTLY | 0.78290087 |
| VVAAAILL | GARADWGSQNTLY | 0.82183504 |
| YAALDVEL | GARADWGSQNTLY | 0.27975348 |
| LASVALLL | GARADWGSQNTLY | 0.85960525 |
| LDAVHRV | GARADWGSQNTLY | 0.68953073 |
| QGLSSSIF | GARADWGSQNTLY | 0.76116455 |
| RGTFANIRL | GARADWGSQNTLY | 0.8489828 |
| RQLWWGHRI | GARADWGSQNTLY | 0.85910064 |
| YGPPGTGKTLL | GARADWGSQNTLY | 0.7151612 |
| AALVGAPANVA | GARADWGSQNTLY | 0.79140043 |
| AATFRRTE | GARADWGSQNTLY | 0.68086255 |
| AGALGGGI | GARADWGSQNTLY | 0.82153004 |
| AGGAGGAGGIGAG | GARADWGSQNTLY | 0.848214 |
| AGSGALGL | GARADWGSQNTLY | 0.69805866 |
| AQIADQI | GARADWGSQNTLY | 0.84694684 |
| ASAVAVTYL | GARADWGSQNTLY | 0.8428224 |
| ASTVLGGTGG | GARADWGSQNTLY | 0.34130022 |
| AVGGEPAK | GARADWGSQNTLY | 0.671798 |
| CEQGTFGGTC | GARADWGSQNTLY | 0.84339213 |
| DEGKAPAG | GARADWGSQNTLY | 0.8437048 |
| DTVVFPAL | GARADWGSQNTLY | 0.84374046 |
| FEGITMAG | GARADWGSQNTLY | 0.7963635 |
| FPLVVQD | GARADWGSQNTLY | 0.54572743 |
| GAFTTPQ | GARADWGSQNTLY | 0.8614442 |
| GFGGFGTAGGVGGA | GARADWGSQNTLY | 0.8633491 |
| GFLAQFL | GARADWGSQNTLY | 0.8421854 |
| GGAGGAGGNAGVA | GARADWGSQNTLY | 0.65240204 |
| GGVGTAGAGGAGGAGG | GARADWGSQNTLY | 0.59978294 |
| GNGGGGDAGNAGSGGN | GARADWGSQNTLY | 0.7731021 |
| GTGGAGGAGNDAGS | GARADWGSQNTLY | 0.843899 |
| GTTNTGWFNS | GARADWGSQNTLY | 0.5547991 |
| IFMGPLFTL | GARADWGSQNTLY | 0.79483503 |
| ILAHAIEAI | GARADWGSQNTLY | 0.8172746 |
| IRSAIREAL | GARADWGSQNTLY | 0.29534963 |

| | | |
|---|---|---|
| ITFAGGG | GARADWGSQNTLY | 0.8255942 |
| ITGMSGHQEV | GARADWGSQNTLY | 0.8518033 |
| IVAGLACTL | GARADWGSQNTLY | 0.7889955 |
| IVGGGLAGA | GARADWGSQNTLY | 0.7378596 |
| IVTGVLNI | GARADWGSQNTLY | 0.8553417 |
| LAQQGCI | GARADWGSQNTLY | 0.86514294 |
| LCLHVEN | GARADWGSQNTLY | 0.85691535 |
| LDAISDGS | GARADWGSQNTLY | 0.40551764 |
| LGGFAAML | GARADWGSQNTLY | 0.31570145 |
| LGGKNPMI | GARADWGSQNTLY | 0.7499173 |
| LGPKGVAAIL | GARADWGSQNTLY | 0.8307345 |
| LMVGVSI | GARADWGSQNTLY | 0.7648283 |
| LVGAAFVL | GARADWGSQNTLY | 0.7242679 |
| LVLAVAPL | GARADWGSQNTLY | 0.81063235 |
| MAAIAVIDAA | GARADWGSQNTLY | 0.8572695 |
| NNGINAELQ | GARADWGSQNTLY | 0.52608144 |
| RGEPLVLVHG | GARADWGSQNTLY | 0.32730615 |
| RGGAGGAAGLI | GARADWGSQNTLY | 0.82948256 |
| RSYRMHPAV | GARADWGSQNTLY | 0.8341079 |
| SAAHLLAGQLA | GARADWGSQNTLY | 0.70656437 |
| SAAVGSY | GARADWGSQNTLY | 0.7329171 |
| SPHAVLHRI | GARADWGSQNTLY | 0.6952871 |
| SQIAKVL | GARADWGSQNTLY | 0.75139916 |
| TALGHLVP | GARADWGSQNTLY | 0.80643356 |
| TAPTTSGAA | GARADWGSQNTLY | 0.60008293 |
| TDGDDGNGGSPGT | GARADWGSQNTLY | 0.85123706 |
| TDGLGEN | GARADWGSQNTLY | 0.544175 |
| TGGPPLPGQA | GARADWGSQNTLY | 0.82988846 |
| TIPAITI | GARADWGSQNTLY | 0.80560017 |
| TIPGFTI | GARADWGSQNTLY | 0.76654744 |
| TSIASGALA | GARADWGSQNTLY | 0.72251856 |
| TVTLPGAAI | GARADWGSQNTLY | 0.64729834 |
| VASGLSAALP | GARADWGSQNTLY | 0.4494513 |
| VGAGVIGI | GARADWGSQNTLY | 0.50112677 |
| VGPKRFGV | GARADWGSQNTLY | 0.8378625 |
| VNVAFPD | GARADWGSQNTLY | 0.14820868 |
| VSMTNSGVS | GARADWGSQNTLY | 0.8399017 |
| VTDVLGDLSV | GARADWGSQNTLY | 0.77112675 |
| VVVVGAGFAGL | GARADWGSQNTLY | 0.5676716 |
| VYTVGTVV | GARADWGSQNTLY | 0.81054497 |
| WTTFSGP | GARADWGSQNTLY | 0.5991727 |
| YDELVEAF | GARADWGSQNTLY | 0.57366306 |
| YGTGGAGGNGG | GARADWGSQNTLY | 0.77804977 |
| YTNVMAV | GARADWGSQNTLY | 0.8401506 |
| FLEQVHQL | GARADWGSQNTLY | 0.587142 |
| GLSLLAVAV | GARADWGSQNTLY | 0.81402624 |
| LEQVHQL | GARADWGSQNTLY | 0.7570007 |
| LNAAGAI | GARADWGSQNTLY | 0.63754594 |
| PAVHASL | GARADWGSQNTLY | 0.8002578 |
| TELEDDA | GARADWGSQNTLY | 0.85750854 |
| TPPGPPN | GARADWGSQNTLY | 0.801516 |

| | | |
|---|---|---|
| TPPPPPPP | GARADWGSQNTLY | 0.4958512 |
| TQSPALI | ASSQPGLGGLYEQY | 0.82663286 |
| EFAAAFPQ | ASSQPGLGGLYEQY | 0.62879944 |
| LLSGNLDVL | ASSQPGLGGLYEQY | 0.81448615 |
| GPANLGNYNI | ASSQPGLGGLYEQY | 0.777051 |
| TLGPTLP | ASSQPGLGGLYEQY | 0.69477844 |
| VYAYSSCCK | ASSQPGLGGLYEQY | 0.8224933 |
| APAPAPAPNI | ASSQPGLGGLYEQY | 0.61108893 |
| PGGPVPIP | ASSQPGLGGLYEQY | 0.60213065 |
| FLTVALEP | ASSQPGLGGLYEQY | 0.70697343 |
| SQYFGANL | ASSQPGLGGLYEQY | 0.79125416 |
| GGVKPAA | ASSQPGLGGLYEQY | 0.84739566 |
| GGTGGAGGAGGW | ASSQPGLGGLYEQY | 0.5966607 |
| GVGGVAGLGGAGG | ASSQPGLGGLYEQY | 0.73399174 |
| VLLVLATTI | ASSQPGLGGLYEQY | 0.5481487 |
| IGIFAAML | ASSQPGLGGLYEQY | 0.35417345 |
| AVGDLID | ASSQPGLGGLYEQY | 0.7912197 |
| NISPLPL | ASSQPGLGGLYEQY | 0.75451297 |
| VTAVFVAI | ASSQPGLGGLYEQY | 0.65075403 |
| SGDDNDAAL | ASSQPGLGGLYEQY | 0.7984298 |
| AVAALAHS | ASSQPGLGGLYEQY | 0.5356492 |
| GGVTVMATP | ASSQPGLGGLYEQY | 0.8014567 |
| AGLAANVRD | ASSQPGLGGLYEQY | 0.8622254 |
| TNRPDLIDPALLRPG | ASSQPGLGGLYEQY | 0.6840974 |
| LANLGVTAL | ASSQPGLGGLYEQY | 0.8236939 |
| TGERIEL | ASSQPGLGGLYEQY | 0.82303464 |
| NGEELAA | ASSQPGLGGLYEQY | 0.8076047 |
| VLSLVWG | ASSQPGLGGLYEQY | 0.7382063 |
| IGYALWL | ASSQPGLGGLYEQY | 0.6357833 |
| LSLLLAPL | ASSQPGLGGLYEQY | 0.860719 |
| TLDEALK | ASSQPGLGGLYEQY | 0.8323719 |
| TDGAGTAEGG | ASSQPGLGGLYEQY | 0.840629 |
| NAPGGAGGNGG | ASSQPGLGGLYEQY | 0.82200396 |
| VSVFMGMGGNF | ASSQPGLGGLYEQY | 0.8418826 |
| LASYLAAL | ASSQPGLGGLYEQY | 0.84543324 |
| LVGGTPAL | ASSQPGLGGLYEQY | 0.76534045 |
| VRNADVL | ASSQPGLGGLYEQY | 0.45088083 |
| TCSSPFLVSAM | ASSQPGLGGLYEQY | 0.75943244 |
| VHITDAE | ASSQPGLGGLYEQY | 0.7936116 |
| EYAAPSF | ASSQPGLGGLYEQY | 0.76286197 |
| VVDAFHGGL | ASSQPGLGGLYEQY | 0.47155792 |
| ASVVDPQ | ASSQPGLGGLYEQY | 0.854982 |
| SYNEFGGLS | ASSQPGLGGLYEQY | 0.8673271 |
| DPDGSVY | ASSQPGLGGLYEQY | 0.6772839 |
| MVSTIHATE | ASSQPGLGGLYEQY | 0.86011773 |
| LAAGVADPV | ASSQPGLGGLYEQY | 0.6441786 |
| TKEQIAA | ASSQPGLGGLYEQY | 0.81765634 |
| SLASPAAL | ASSQPGLGGLYEQY | 0.85812664 |
| IGLVFYGV | ASSQPGLGGLYEQY | 0.6666871 |
| FLLDASNL | ASSQPGLGGLYEQY | 0.8315807 |
| YQVGNFP | ASSQPGLGGLYEQY | 0.49568737 |

| | | |
|---|---|---|
| ESLLSQST | ASSQPGLGGLYEQY | 0.65709263 |
| LFTVGFAAVG | ASSQPGLGGLYEQY | 0.8544611 |
| ATPPTPT | ASSQPGLGGLYEQY | 0.7537929 |
| TDLATLG | ASSQPGLGGLYEQY | 0.7860577 |
| SAGGTGR | ASSQPGLGGLYEQY | 0.7873025 |
| GFSNSGS | ASSQPGLGGLYEQY | 0.59034276 |
| GGNGGSGGTSV | ASSQPGLGGLYEQY | 0.76273805 |
| GTSVATGGAGN | ASSQPGLGGLYEQY | 0.8547933 |
| NGGNGGSGGTS | ASSQPGLGGLYEQY | 0.6219481 |
| AVVKFPH | ASSQPGLGGLYEQY | 0.86414397 |
| TPPPPPP | ASSQPGLGGLYEQY | 0.56577027 |
| AAPAAAAAPA | ASSQPGLGGLYEQY | 0.8670842 |
| IALNEDL | ASSQPGLGGLYEQY | 0.85874677 |
| IVNVSSM | ASSQPGLGGLYEQY | 0.8032743 |
| TGIPPAPR | ASSQPGLGGLYEQY | 0.707808 |
| FDLGGGTF | ASSQPGLGGLYEQY | 0.84979767 |
| FDLGGGTFDVS | ASSQPGLGGLYEQY | 0.74462855 |
| NEPTAAA | ASSQPGLGGLYEQY | 0.8149284 |
| NEPTAAALA | ASSQPGLGGLYEQY | 0.85453594 |
| GSIINISSI | ASSQPGLGGLYEQY | 0.44600508 |
| VAPGNAGTA | ASSQPGLGGLYEQY | 0.75049895 |
| TGGGASLE | ASSQPGLGGLYEQY | 0.70270616 |
| VAVYNHYKRI | ASSQPGLGGLYEQY | 0.67977613 |
| TDPAPATT | ASSQPGLGGLYEQY | 0.5434259 |
| TNRPDILDPALLRPG | ASSQPGLGGLYEQY | 0.39149174 |
| LTTTTTS | ASSQPGLGGLYEQY | 0.6030607 |
| PTGSGKTL | ASSQPGLGGLYEQY | 0.48918068 |
| TGSGKTL | ASSQPGLGGLYEQY | 0.77265954 |
| TIGHVDHGK | ASSQPGLGGLYEQY | 0.71629626 |
| TIGHVDHGKTT | ASSQPGLGGLYEQY | 0.52571636 |
| GHVDHGKTT | ASSQPGLGGLYEQY | 0.7597534 |
| AQFEHTLL | ASSQPGLGGLYEQY | 0.7701956 |
| IVNQDEL | ASSQPGLGGLYEQY | 0.33702084 |
| TNGSQFF | ASSQPGLGGLYEQY | 0.6348982 |
| TGKGIVE | ASSQPGLGGLYEQY | 0.48555875 |
| LISNASDA | ASSQPGLGGLYEQY | 0.5705906 |
| AELRTLGELTKIAKAHGAQ | ASSQPGLGGLYEQY | 0.27643776 |
| APNAVAAAEPSW | ASSQPGLGGLYEQY | 0.7158826 |
| APQVAGAATAVL | ASSQPGLGGLYEQY | 0.57453597 |
| ARRDPAAAAAILPTDA | ASSQPGLGGLYEQY | 0.23576102 |
| DAAARDVLAEAGLAEYF | ASSQPGLGGLYEQY | 0.69820046 |
| DDQIVLSAKANLAKVEQ | ASSQPGLGGLYEQY | 0.633138 |
| DELLPDTGEDSDEDGH | ASSQPGLGGLYEQY | 0.60875607 |
| ELPEAPANRATVPQD | ASSQPGLGGLYEQY | 0.7412329 |
| FGQSKIERVTGIG | ASSQPGLGGLYEQY | 0.5678518 |
| GAGGTGSTAGGAGGAGGAGGLY | ASSQPGLGGLYEQY | 0.516453 |
| GHGGTGAAGGNGGTGGAGGSGI | ASSQPGLGGLYEQY | 0.72552586 |
| GNGGNGGAGGEGGPGGAGGAGA | ASSQPGLGGLYEQY | 0.75427467 |
| GSTAGNGGNGGNATNGTIAGQP | ASSQPGLGGLYEQY | 0.46718812 |
| GTGGDGGNGGNGGTGG | ASSQPGLGGLYEQY | 0.6814463 |
| GVRRPAGIPADAPV | ASSQPGLGGLYEQY | 0.76668024 |

| | | |
|---|---|---|
| GYRQVTDAHLLTLARR | ASSQPGLGGLYEQY | 0.6673845 |
| IAKMGQDGHDRG | ASSQPGLGGLYEQY | 0.7574523 |
| KAEPSIEIQYT | ASSQPGLGGLYEQY | 0.76912165 |
| KPVRIRMAADEDTVAVT | ASSQPGLGGLYEQY | 0.5530501 |
| MPPRIAGMRLLVIKPEP | ASSQPGLGGLYEQY | 0.74888074 |
| MKFYAVALFRSRGVPPTR | ASSQPGLGGLYEQY | 0.77237076 |
| NAGGFGILGAVAHSP | ASSQPGLGGLYEQY | 0.5381881 |
| NEGREITLAMAEM | ASSQPGLGGLYEQY | 0.6469479 |
| NELVEWSHPLGHRWRWE | ASSQPGLGGLYEQY | 0.4723015 |
| NLGAGGTGGNGGNPG | ASSQPGLGGLYEQY | 0.670274 |
| NTEWPKWETVMAADPVD | ASSQPGLGGLYEQY | 0.3869868 |
| PASAAPAETQP | ASSQPGLGGLYEQY | 0.7135756 |
| QQSSDGKNGVRAAEITGE | ASSQPGLGGLYEQY | 0.75168514 |
| RAPDAVTTSWWKE | ASSQPGLGGLYEQY | 0.64028764 |
| SGFGNVGAHTSGWFNQ | ASSQPGLGGLYEQY | 0.39030737 |
| SGVAGAVGQAASVGGLKV | ASSQPGLGGLYEQY | 0.37741122 |
| SNGGAGTGSAGGNGGT | ASSQPGLGGLYEQY | 0.6619954 |
| TGGAGGEGGAAG | ASSQPGLGGLYEQY | 0.46980882 |
| TGGDGGAAGAGGAPGHG | ASSQPGLGGLYEQY | 0.33144805 |
| TMPPLLAAIANAGRHGV | ASSQPGLGGLYEQY | 0.5904796 |
| TSSTAGNSGGA | ASSQPGLGGLYEQY | 0.7534642 |
| VPATVPVGILC | ASSQPGLGGLYEQY | 0.7551683 |
| VQRPDSGGVVLTGRIS | ASSQPGLGGLYEQY | 0.47859624 |
| WTPNPVPATKGVPAGLVQAT | ASSQPGLGGLYEQY | 0.76986563 |
| YALKRALAKFSDRFDVVII | ASSQPGLGGLYEQY | 0.76160693 |
| YLIGSVIV | ASSQPGLGGLYEQY | 0.7576117 |
| YGQLVAVID | ASSQPGLGGLYEQY | 0.6311362 |
| VTVGGMTLPA | ASSQPGLGGLYEQY | 0.41899848 |
| VLALPAV | ASSQPGLGGLYEQY | 0.76298445 |
| VIPPTLNYET | ASSQPGLGGLYEQY | 0.65242183 |
| VGMGGMHPSSG | ASSQPGLGGLYEQY | 0.63611495 |
| VGLAGGAATA | ASSQPGLGGLYEQY | 0.7442601 |
| VGGAGGAGGSGP | ASSQPGLGGLYEQY | 0.68229914 |
| VFAAFMV | ASSQPGLGGLYEQY | 0.53766155 |
| TLYDKLR | ASSQPGLGGLYEQY | 0.7018678 |
| TLAVFTVL | ASSQPGLGGLYEQY | 0.73576444 |
| TGSVQPARV | ASSQPGLGGLYEQY | 0.5058373 |
| TGGGAGSGGT | ASSQPGLGGLYEQY | 0.39938372 |
| SHLETSL | ASSQPGLGGLYEQY | 0.6465583 |
| SFTTGMT | ASSQPGLGGLYEQY | 0.6840351 |
| QVEVAKRMGVRQPTVS | ASSQPGLGGLYEQY | 0.74769044 |
| QGSGGPALAM | ASSQPGLGGLYEQY | 0.6836939 |
| MLAPSNP | ASSQPGLGGLYEQY | 0.56690645 |
| LVNNAAI | ASSQPGLGGLYEQY | 0.7273538 |
| LTGEINTLA | ASSQPGLGGLYEQY | 0.6984078 |
| LSIFGPV | ASSQPGLGGLYEQY | 0.6650089 |
| LLALLANL | ASSQPGLGGLYEQY | 0.7566478 |
| LISNASDALDKLR | ASSQPGLGGLYEQY | 0.5648346 |
| LINEAVP | ASSQPGLGGLYEQY | 0.7360332 |
| LGYAFSGL | ASSQPGLGGLYEQY | 0.73197424 |
| LDFVAVAL | ASSQPGLGGLYEQY | 0.71655166 |

| | | |
|---|---|---|
| LAQPGAE | ASSQPGLGGLYEQY | 0.7583902 |
| ISNLGAVTGF | ASSQPGLGGLYEQY | 0.657577 |
| HQEDPGRI | ASSQPGLGGLYEQY | 0.6393209 |
| GTPGEPTP | ASSQPGLGGLYEQY | 0.3626189 |
| GFFNSGS | ASSQPGLGGLYEQY | 0.5942092 |
| GAGGGGLTAGAN | ASSQPGLGGLYEQY | 0.7199248 |
| FVDALVNM | ASSQPGLGGLYEQY | 0.6970915 |
| FLTVAPEL | ASSQPGLGGLYEQY | 0.75678194 |
| FETLTAL | ASSQPGLGGLYEQY | 0.7205169 |
| DVINPIEL | ASSQPGLGGLYEQY | 0.56589484 |
| APVSGAPV | ASSQPGLGGLYEQY | 0.7771209 |
| ALVGGLAW | ASSQPGLGGLYEQY | 0.75605595 |
| ALAAAAATAI | ASSQPGLGGLYEQY | 0.36067134 |
| AFGLGLLL | ASSQPGLGGLYEQY | 0.6123525 |
| AFFAAASL | ASSQPGLGGLYEQY | 0.7685034 |
| AAAALPAFSPP | ASSQPGLGGLYEQY | 0.7444757 |
| VVLAPAL | ASSQPGLGGLYEQY | 0.62544817 |
| TIGHVDHGKTTL | ASSQPGLGGLYEQY | 0.6073077 |
| LISNASDALDK | ASSQPGLGGLYEQY | 0.77940655 |
| LISNASDAL | ASSQPGLGGLYEQY | 0.6314377 |
| ISNASDALDK | ASSQPGLGGLYEQY | 0.7711334 |
| NQAETLVYQTE | ASSQPGLGGLYEQY | 0.2990375 |
| TGGTGGTAGAG | ASSQPGLGGLYEQY | 0.57021815 |
| TGSTAGGAGGA | ASSQPGLGGLYEQY | 0.5120568 |
| DPLAVPVAAGPV | ASSQPGLGGLYEQY | 0.2506757 |
| GGTGGTPGNGGN | ASSQPGLGGLYEQY | 0.74205947 |
| PAGLAGVDPATL | ASSQPGLGGLYEQY | 0.5667063 |
| GNNLVVEAPHAGASV | ASSQPGLGGLYEQY | 0.73763525 |
| KVTRIMSASTLEQPA | ASSQPGLGGLYEQY | 0.6191634 |
| LGFALPIALPRRTVV | ASSQPGLGGLYEQY | 0.74224544 |
| TRLGNLTARKLVVLV | ASSQPGLGGLYEQY | 0.66658425 |
| WSQQRRLIYFPSAGP | ASSQPGLGGLYEQY | 0.67295325 |
| GPNGNPDPMAAAVDIR | ASSQPGLGGLYEQY | 0.39571977 |
| TQRCVSVADLRAAASR | ASSQPGLGGLYEQY | 0.7326852 |
| QFVLDVAEHPVEVLTETV | ASSQPGLGGLYEQY | 0.77781916 |
| GGAGGTGGAGGNGGTQAGDG | ASSQPGLGGLYEQY | 0.70363164 |
| PLLATLRAIAVDVDAKPAQV | ASSQPGLGGLYEQY | 0.346189 |
| APKFRTGGYMIDFWGVGYQVAK | ASSQPGLGGLYEQY | 0.6869581 |
| GAGGAGGAGGAAGGPGGTGGTG | ASSQPGLGGLYEQY | 0.6418929 |
| IGALGLLATDVAPAGSVAFGVA | ASSQPGLGGLYEQY | 0.30111954 |
| AGGAGGIGGNGGAGGAGGVLMG | ASSQPGLGGLYEQY | 0.30020827 |
| ADELIGLL | ASSQPGLGGLYEQY | 0.5226587 |
| ADRALKA | ASSQPGLGGLYEQY | 0.62630403 |
| AGAVGQVGGAGGA | ASSQPGLGGLYEQY | 0.74520904 |
| AGGGGGLGGAP | ASSQPGLGGLYEQY | 0.72844726 |
| AGGLLYGNGGAG | ASSQPGLGGLYEQY | 0.58600694 |
| AGLESIGL | ASSQPGLGGLYEQY | 0.7206626 |
| AGLKTGII | ASSQPGLGGLYEQY | 0.7064085 |
| ASLAALLL | ASSQPGLGGLYEQY | 0.1963363 |
| ASVINISSI | ASSQPGLGGLYEQY | 0.5114751 |
| ATPPGPA | ASSQPGLGGLYEQY | 0.37143394 |

| | | |
|---|---|---|
| ATVASALV | ASSQPGLGGLYEQY | 0.765991 |
| ATVASLIL | ASSQPGLGGLYEQY | 0.49452966 |
| AVNGPVLIRH | ASSQPGLGGLYEQY | 0.5585023 |
| DGAAGAVGI | ASSQPGLGGLYEQY | 0.6476675 |
| DPLLHGI | ASSQPGLGGLYEQY | 0.73784316 |
| ELGAALGAL | ASSQPGLGGLYEQY | 0.6977525 |
| ELGRTQAV | ASSQPGLGGLYEQY | 0.4721301 |
| FVAGGYLS | ASSQPGLGGLYEQY | 0.32806873 |
| GASGGAGGAGGW | ASSQPGLGGLYEQY | 0.72419137 |
| GGAGAGGDGGAGGI | ASSQPGLGGLYEQY | 0.523452 |
| GTGHNGSQPGMGGNG | ASSQPGLGGLYEQY | 0.45713297 |
| GYAAASASAS | ASSQPGLGGLYEQY | 0.37686044 |
| HQVDVAAVQQG | ASSQPGLGGLYEQY | 0.68971646 |
| IGAAIAHT | ASSQPGLGGLYEQY | 0.35806867 |
| IGMIEVAQL | ASSQPGLGGLYEQY | 0.74255335 |
| IRLAPEHTSAMD | ASSQPGLGGLYEQY | 0.548841 |
| IYNVSVL | ASSQPGLGGLYEQY | 0.64762163 |
| KQGFLGM | ASSQPGLGGLYEQY | 0.7768477 |
| LESVITQL | ASSQPGLGGLYEQY | 0.7307056 |
| LGLGALVVV | ASSQPGLGGLYEQY | 0.61633855 |
| LIGNGGNGGSGGI | ASSQPGLGGLYEQY | 0.51740885 |
| LNAINAPTL | ASSQPGLGGLYEQY | 0.4044702 |
| LQLLNTGV | ASSQPGLGGLYEQY | 0.4776554 |
| LQTSIEF | ASSQPGLGGLYEQY | 0.57600975 |
| MSFAVSEL | ASSQPGLGGLYEQY | 0.77290404 |
| NTGALGL | ASSQPGLGGLYEQY | 0.60170853 |
| PPGAGGAGGAGTT | ASSQPGLGGLYEQY | 0.3900575 |
| QLVGLGI | ASSQPGLGGLYEQY | 0.65702885 |
| RFLEQVHQL | ASSQPGLGGLYEQY | 0.77081877 |
| RGVGYVLR | ASSQPGLGGLYEQY | 0.77525777 |
| SATGVFAGVI | ASSQPGLGGLYEQY | 0.45431274 |
| SLPAITLP | ASSQPGLGGLYEQY | 0.63701797 |
| STALAFTV | ASSQPGLGGLYEQY | 0.5733636 |
| SVLTVGLGK | ASSQPGLGGLYEQY | 0.6629839 |
| TALAGAGSGI | ASSQPGLGGLYEQY | 0.47166264 |
| TGLPFTGI | ASSQPGLGGLYEQY | 0.4658273 |
| VAEHVTV | ASSQPGLGGLYEQY | 0.70341814 |
| VDIVLNSL | ASSQPGLGGLYEQY | 0.7878604 |
| VLTGGGAVL | ASSQPGLGGLYEQY | 0.5719906 |
| VNVGLAYGI | ASSQPGLGGLYEQY | 0.2995402 |
| VSSAAVGAV | ASSQPGLGGLYEQY | 0.65151644 |
| VTSVYEAGV | ASSQPGLGGLYEQY | 0.50268966 |
| WVALDRGA | ASSQPGLGGLYEQY | 0.51057523 |
| YLVGSVVI | ASSQPGLGGLYEQY | 0.5943538 |
| GPPGTGKTLL | ASSQPGLGGLYEQY | 0.32230613 |
| GVGMTKF | ASSQPGLGGLYEQY | 0.65334606 |
| IGIGGSDL | ASSQPGLGGLYEQY | 0.77276874 |
| IGTLLAEL | ASSQPGLGGLYEQY | 0.67368996 |
| IYTSGTT | ASSQPGLGGLYEQY | 0.70720965 |
| LSFPAGV | ASSQPGLGGLYEQY | 0.7592064 |
| LYTSGTT | ASSQPGLGGLYEQY | 0.72065127 |

| | | |
|---|---|---|
| VGAFPFGF | ASSQPGLGGLYEQY | 0.7559701 |
| VGAGTVEF | ASSQPGLGGLYEQY | 0.7660807 |
| VGKSTLLL | ASSQPGLGGLYEQY | 0.63407046 |
| AALAHAGGGIV | ASSQPGLGGLYEQY | 0.6610333 |
| AALVAVSLL | ASSQPGLGGLYEQY | 0.7543942 |
| AAVSRAVAAL | ASSQPGLGGLYEQY | 0.6646214 |
| AGGSSGIGGF | ASSQPGLGGLYEQY | 0.6616593 |
| AGLTVANANG | ASSQPGLGGLYEQY | 0.5610692 |
| AGLVGAVL | ASSQPGLGGLYEQY | 0.7600385 |
| ALSQAFTV | ASSQPGLGGLYEQY | 0.60024285 |
| ALTFSVSL | ASSQPGLGGLYEQY | 0.7679611 |
| ALTGAALVAL | ASSQPGLGGLYEQY | 0.3148849 |
| ASFSIGAGVL | ASSQPGLGGLYEQY | 0.45322454 |
| DGSVYKI | ASSQPGLGGLYEQY | 0.7456951 |
| EQLAEIV | ASSQPGLGGLYEQY | 0.22334224 |
| FAVSELTVM | ASSQPGLGGLYEQY | 0.2928571 |
| GAAPLAAGL | ASSQPGLGGLYEQY | 0.6163385 |
| GADGGDGGSGGAGG | ASSQPGLGGLYEQY | 0.7618531 |
| GFGLDTGGAGG | ASSQPGLGGLYEQY | 0.22727457 |
| GLASGGLL | ASSQPGLGGLYEQY | 0.50483835 |
| GRIAGAAVLL | ASSQPGLGGLYEQY | 0.34523824 |
| GSAGSLNTSF | ASSQPGLGGLYEQY | 0.7184119 |
| GTGQIGI | ASSQPGLGGLYEQY | 0.66473126 |
| GTLIALIL | ASSQPGLGGLYEQY | 0.32015124 |
| IAATMGFVVG | ASSQPGLGGLYEQY | 0.72129536 |
| IFAQALL | ASSQPGLGGLYEQY | 0.30084425 |
| IGAAVALGF | ASSQPGLGGLYEQY | 0.634619 |
| IGLTPLAL | ASSQPGLGGLYEQY | 0.7297884 |
| ILGSNTSTL | ASSQPGLGGLYEQY | 0.77421355 |
| IVFSLTVL | ASSQPGLGGLYEQY | 0.38119334 |
| LASKATGF | ASSQPGLGGLYEQY | 0.7128118 |
| LASVALIL | ASSQPGLGGLYEQY | 0.5790218 |
| LFAVGLTL | ASSQPGLGGLYEQY | 0.7822583 |
| LFNGGIGTYI | ASSQPGLGGLYEQY | 0.6213618 |
| LTLGAGII | ASSQPGLGGLYEQY | 0.3385925 |
| LVAAALGDI | ASSQPGLGGLYEQY | 0.7452003 |
| NTAFVQLGI | ASSQPGLGGLYEQY | 0.69828254 |
| QLFSGAEL | ASSQPGLGGLYEQY | 0.61961406 |
| SAATFITL | ASSQPGLGGLYEQY | 0.75224817 |
| SNYGLANLGV | ASSQPGLGGLYEQY | 0.5358833 |
| SYGGAYAVM | ASSQPGLGGLYEQY | 0.6165539 |
| TTGVGGLTLG | ASSQPGLGGLYEQY | 0.7507577 |
| TTQTIYI | ASSQPGLGGLYEQY | 0.7482636 |
| VAAGAGLGAVI | ASSQPGLGGLYEQY | 0.511889 |
| VAAVVTLL | ASSQPGLGGLYEQY | 0.74228823 |
| VGFLGGLI | ASSQPGLGGLYEQY | 0.6764273 |
| VGVIPILL | ASSQPGLGGLYEQY | 0.52783906 |
| VIFASSLI | ASSQPGLGGLYEQY | 0.77516246 |
| VKGGDGGAAAG | ASSQPGLGGLYEQY | 0.56342757 |
| VTKTFTNI | ASSQPGLGGLYEQY | 0.7388292 |
| VVAAAILL | ASSQPGLGGLYEQY | 0.7256416 |

| | | |
|---|---|---|
| YAALDVEL | ASSQPGLGGLYEQY | 0.7806282 |
| LASVALLL | ASSQPGLGGLYEQY | 0.42604724 |
| LDAVHRV | ASSQPGLGGLYEQY | 0.73747975 |
| QGLSSSIF | ASSQPGLGGLYEQY | 0.6835638 |
| RGTFANIRL | ASSQPGLGGLYEQY | 0.73988587 |
| RQLWWGHRI | ASSQPGLGGLYEQY | 0.6632133 |
| YGPPGTGKTLL | ASSQPGLGGLYEQY | 0.6561558 |
| AALVGAPANVA | ASSQPGLGGLYEQY | 0.69983435 |
| AATFRRTE | ASSQPGLGGLYEQY | 0.7638471 |
| AGALGGGI | ASSQPGLGGLYEQY | 0.60494155 |
| AGGAGGAGGIGAG | ASSQPGLGGLYEQY | 0.73794645 |
| AGSGALGL | ASSQPGLGGLYEQY | 0.767279 |
| AQIADQI | ASSQPGLGGLYEQY | 0.48506606 |
| ASAVAVTYL | ASSQPGLGGLYEQY | 0.658373 |
| ASTVLGGTGG | ASSQPGLGGLYEQY | 0.7107126 |
| AVGGEPAK | ASSQPGLGGLYEQY | 0.7723935 |
| CEQGTFGGTC | ASSQPGLGGLYEQY | 0.5850357 |
| DEGKAPAG | ASSQPGLGGLYEQY | 0.75687486 |
| DTVVFPAL | ASSQPGLGGLYEQY | 0.57220656 |
| FEGITMAG | ASSQPGLGGLYEQY | 0.71284354 |
| FPLVVQD | ASSQPGLGGLYEQY | 0.6251874 |
| GAFTTPQ | ASSQPGLGGLYEQY | 0.67514586 |
| GFGGFGTAGGVGGA | ASSQPGLGGLYEQY | 0.5091444 |
| GFLAQFL | ASSQPGLGGLYEQY | 0.7227557 |
| GGAGGAGGNAGVA | ASSQPGLGGLYEQY | 0.73868716 |
| GGVGTAGAGGAGGAGG | ASSQPGLGGLYEQY | 0.74149907 |
| GNGGGGDAGNAGSGGN | ASSQPGLGGLYEQY | 0.294662 |
| GTGGAGGAGNDAGS | ASSQPGLGGLYEQY | 0.71380687 |
| GTTNTGWFNS | ASSQPGLGGLYEQY | 0.7361891 |
| IFMGPLFTL | ASSQPGLGGLYEQY | 0.7807577 |
| ILAHAIEAI | ASSQPGLGGLYEQY | 0.6340547 |
| IRSAIREAL | ASSQPGLGGLYEQY | 0.72279894 |
| ITFAGGG | ASSQPGLGGLYEQY | 0.5657215 |
| ITGMSGHQEV | ASSQPGLGGLYEQY | 0.61061287 |
| IVAGLACTL | ASSQPGLGGLYEQY | 0.41719466 |
| IVGGGLAGA | ASSQPGLGGLYEQY | 0.63867605 |
| IVTGVLNI | ASSQPGLGGLYEQY | 0.59682083 |
| LAQQGCI | ASSQPGLGGLYEQY | 0.53102934 |
| LCLHVEN | ASSQPGLGGLYEQY | 0.75585115 |
| LDAISDGS | ASSQPGLGGLYEQY | 0.7669896 |
| LGGFAAML | ASSQPGLGGLYEQY | 0.7508632 |
| LGGKNPMI | ASSQPGLGGLYEQY | 0.7607011 |
| LGPKGVAAIL | ASSQPGLGGLYEQY | 0.76209974 |
| LMVGVSI | ASSQPGLGGLYEQY | 0.7311942 |
| LVGAAFVL | ASSQPGLGGLYEQY | 0.5881655 |
| LVLAVAPL | ASSQPGLGGLYEQY | 0.63904285 |
| MAAIAVIDAA | ASSQPGLGGLYEQY | 0.3689748 |
| NNGINAELQ | ASSQPGLGGLYEQY | 0.5918076 |
| RGEPLVLVHG | ASSQPGLGGLYEQY | 0.58438605 |
| RGGAGGAAGLI | ASSQPGLGGLYEQY | 0.6821923 |
| RSYRMHPAV | ASSQPGLGGLYEQY | 0.21694249 |

| | | |
|---|---|---|
| SAAHLLAGQLA | ASSQPGLGGLYEQY | 0.7215095 |
| SAAVGSY | ASSQPGLGGLYEQY | 0.7751981 |
| SPHAVLHRI | ASSQPGLGGLYEQY | 0.6472679 |
| SQIAKVL | ASSQPGLGGLYEQY | 0.7325903 |
| TALGHLVP | ASSQPGLGGLYEQY | 0.7259722 |
| TAPTTSGAA | ASSQPGLGGLYEQY | 0.540441 |
| TDGDDGNGGSPGT | ASSQPGLGGLYEQY | 0.738626 |
| TDGLGEN | ASSQPGLGGLYEQY | 0.54330856 |
| TGGPPLPGQA | ASSQPGLGGLYEQY | 0.73099005 |
| TIPAITI | ASSQPGLGGLYEQY | 0.77161247 |
| TIPGFTI | ASSQPGLGGLYEQY | 0.69218844 |
| TSIASGALA | ASSQPGLGGLYEQY | 0.7243818 |
| TVTLPGAAI | ASSQPGLGGLYEQY | 0.6524089 |
| VASGLSAALP | ASSQPGLGGLYEQY | 0.38694704 |
| VGAGVIGI | ASSQPGLGGLYEQY | 0.733012 |
| VGPKRFGV | ASSQPGLGGLYEQY | 0.47775984 |
| VNVAFPD | ASSQPGLGGLYEQY | 0.6738074 |
| VSMTNSGVS | ASSQPGLGGLYEQY | 0.44799718 |
| VTDVLGDLSV | ASSQPGLGGLYEQY | 0.71240413 |
| VVVVGAGFAGL | ASSQPGLGGLYEQY | 0.6862031 |
| VYTVGTVV | ASSQPGLGGLYEQY | 0.66096556 |
| WTTFSGP | ASSQPGLGGLYEQY | 0.53669524 |
| YDELVEAF | ASSQPGLGGLYEQY | 0.6971918 |
| YGTGGAGGNGG | ASSQPGLGGLYEQY | 0.6540916 |
| YTNVMAV | ASSQPGLGGLYEQY | 0.42348382 |
| FLEQVHQL | ASSQPGLGGLYEQY | 0.7451966 |
| GLSLLAVAV | ASSQPGLGGLYEQY | 0.5395369 |
| LEQVHQL | ASSQPGLGGLYEQY | 0.6504557 |
| LNAAGAI | ASSQPGLGGLYEQY | 0.7576806 |
| PAVHASL | ASSQPGLGGLYEQY | 0.6583892 |
| TELEDDA | ASSQPGLGGLYEQY | 0.28840023 |
| TPPGPPN | ASSQPGLGGLYEQY | 0.64855146 |
| TPPPPPPP | ASSQPGLGGLYEQY | 0.6755843 |
| TQSPALI | ASSLGETGQLY | 0.6230961 |
| EFAAAFPQ | ASSLGETGQLY | 0.2342046 |
| LLSGNLDVL | ASSLGETGQLY | 0.6908234 |
| GPANLGNYNI | ASSLGETGQLY | 0.52251625 |
| TLGPTLP | ASSLGETGQLY | 0.47153562 |
| VYAYSSCCK | ASSLGETGQLY | 0.5577878 |
| APAPAPAPNI | ASSLGETGQLY | 0.743667 |
| PGGPVPIP | ASSLGETGQLY | 0.6418771 |
| FLTVALEP | ASSLGETGQLY | 0.6345849 |
| SQYFGANL | ASSLGETGQLY | 0.7419398 |
| GGVKPAA | ASSLGETGQLY | 0.70008504 |
| GGTGGAGGAGGW | ASSLGETGQLY | 0.76601446 |
| GVGGVAGLGGAGG | ASSLGETGQLY | 0.7064056 |
| VLLVLATTI | ASSLGETGQLY | 0.6432878 |
| IGIFAAML | ASSLGETGQLY | 0.73454225 |
| AVGDLID | ASSLGETGQLY | 0.5513612 |
| NISPLPL | ASSLGETGQLY | 0.43829036 |
| VTAVFVAI | ASSLGETGQLY | 0.3519407 |

| | | |
|---|---|---|
| SGDDNDAAL | ASSLGETGQLY | 0.53496146 |
| AVAALAHS | ASSLGETGQLY | 0.38848865 |
| GGVTVMATP | ASSLGETGQLY | 0.70588696 |
| AGLAANVRD | ASSLGETGQLY | 0.75189143 |
| TNRPDLIDPALLRPG | ASSLGETGQLY | 0.63801384 |
| LANLGVTAL | ASSLGETGQLY | 0.38503772 |
| TGERIEL | ASSLGETGQLY | 0.5774579 |
| NGEELAA | ASSLGETGQLY | 0.54277456 |
| VLSLVWG | ASSLGETGQLY | 0.7317565 |
| IGYALWL | ASSLGETGQLY | 0.7291975 |
| LSLLLAPL | ASSLGETGQLY | 0.781788 |
| TLDEALK | ASSLGETGQLY | 0.7449273 |
| TDGAGTAEGG | ASSLGETGQLY | 0.60028195 |
| NAPGGAGGNGG | ASSLGETGQLY | 0.72515786 |
| VSVFMGMGGNF | ASSLGETGQLY | 0.59005797 |
| LASYLAAL | ASSLGETGQLY | 0.70358443 |
| LVGGTPAL | ASSLGETGQLY | 0.6474333 |
| VRNADVL | ASSLGETGQLY | 0.6675727 |
| TCSSPFLVSAM | ASSLGETGQLY | 0.64093864 |
| VHITDAE | ASSLGETGQLY | 0.60679126 |
| EYAAPSF | ASSLGETGQLY | 0.31482327 |
| VVDAFHGGL | ASSLGETGQLY | 0.70785034 |
| ASVVDPQ | ASSLGETGQLY | 0.50968975 |
| SYNEFGGLS | ASSLGETGQLY | 0.36678508 |
| DPDGSVY | ASSLGETGQLY | 0.5917623 |
| MVSTIHATE | ASSLGETGQLY | 0.77253854 |
| LAAGVADPV | ASSLGETGQLY | 0.76765704 |
| TKEQIAA | ASSLGETGQLY | 0.5688509 |
| SLASPAAL | ASSLGETGQLY | 0.51653653 |
| IGLVFYGV | ASSLGETGQLY | 0.7208295 |
| FLLDASNL | ASSLGETGQLY | 0.4446658 |
| YQVGNFP | ASSLGETGQLY | 0.68245244 |
| ESLLSQST | ASSLGETGQLY | 0.6745939 |
| LFTVGFAAVG | ASSLGETGQLY | 0.6240225 |
| ATPPTPT | ASSLGETGQLY | 0.5255529 |
| TDLATLG | ASSLGETGQLY | 0.7169421 |
| SAGGTGR | ASSLGETGQLY | 0.6073756 |
| GFSNSGS | ASSLGETGQLY | 0.69954276 |
| GGNGGSGGTSV | ASSLGETGQLY | 0.32582974 |
| GTSVATGGAGN | ASSLGETGQLY | 0.7294463 |
| NGGNGGSGGTS | ASSLGETGQLY | 0.48193318 |
| AVVKFPH | ASSLGETGQLY | 0.44754893 |
| TPPPPPP | ASSLGETGQLY | 0.26204458 |
| AAPAAAAAPA | ASSLGETGQLY | 0.65353477 |
| IALNEDL | ASSLGETGQLY | 0.7375604 |
| IVNVSSM | ASSLGETGQLY | 0.75450885 |
| TGIPPAPR | ASSLGETGQLY | 0.60074365 |
| FDLGGGTF | ASSLGETGQLY | 0.75012577 |
| FDLGGGTFDVS | ASSLGETGQLY | 0.7514764 |
| NEPTAAA | ASSLGETGQLY | 0.60831726 |
| NEPTAAALA | ASSLGETGQLY | 0.31875935 |

| | | |
|---|---|---|
| GSIINISSI | ASSLGETGQLY | 0.6978973 |
| VAPGNAGTA | ASSLGETGQLY | 0.68613076 |
| TGGGASLE | ASSLGETGQLY | 0.754118 |
| VAVYNHYKRI | ASSLGETGQLY | 0.74799204 |
| TDPAPATT | ASSLGETGQLY | 0.39210618 |
| TNRPDILDPALLRPG | ASSLGETGQLY | 0.730235 |
| LTTTTTS | ASSLGETGQLY | 0.69992054 |
| PTGSGKTL | ASSLGETGQLY | 0.4325472 |
| TGSGKTL | ASSLGETGQLY | 0.4742729 |
| TIGHVDHGK | ASSLGETGQLY | 0.5241196 |
| TIGHVDHGKTT | ASSLGETGQLY | 0.36745867 |
| GHVDHGKTT | ASSLGETGQLY | 0.7199954 |
| AQFEHTLL | ASSLGETGQLY | 0.6118272 |
| IVNQDEL | ASSLGETGQLY | 0.61022484 |
| TNGSQFF | ASSLGETGQLY | 0.6177193 |
| TGKGIVE | ASSLGETGQLY | 0.5900757 |
| LISNASDA | ASSLGETGQLY | 0.5516309 |
| AELRTLGELTKIAKAHGAQ | ASSLGETGQLY | 0.6090172 |
| APNAVAAAEPSW | ASSLGETGQLY | 0.68701684 |
| APQVAGAATAVL | ASSLGETGQLY | 0.7194288 |
| ARRDPAAAAAILPTDA | ASSLGETGQLY | 0.6328626 |
| DAAARDVLAEAGLAEYF | ASSLGETGQLY | 0.69106793 |
| DDQIVLSAKANLAKVEQ | ASSLGETGQLY | 0.7653713 |
| DELLPDTGEDSDEDGH | ASSLGETGQLY | 0.41488785 |
| ELPEAPANRATVPQD | ASSLGETGQLY | 0.51494175 |
| FGQSKIERVTGIG | ASSLGETGQLY | 0.7055212 |
| GAGGTGSTAGGAGGAGGAGGLY | ASSLGETGQLY | 0.5192546 |
| GHGGTGAAGGNGGTGGAGGSGI | ASSLGETGQLY | 0.7364687 |
| GNGGNGGAGGEGGPGGAGGAGA | ASSLGETGQLY | 0.69474876 |
| GSTAGNGGNGGNATNGTIAGQP | ASSLGETGQLY | 0.70572424 |
| GTGGDGGNGGNGGTGG | ASSLGETGQLY | 0.6817969 |
| GVRRPAGIPADAPV | ASSLGETGQLY | 0.7789695 |
| GYRQVTDAHLLTLARR | ASSLGETGQLY | 0.7232055 |
| IAKMGQDGHDRG | ASSLGETGQLY | 0.6223165 |
| KAEPSIEIQYT | ASSLGETGQLY | 0.7550748 |
| KPVRIRMAADEDTVAVT | ASSLGETGQLY | 0.42026824 |
| MPPRIAGMRLLVIKPEP | ASSLGETGQLY | 0.7511002 |
| MKFYAVALFRSRGVPPTR | ASSLGETGQLY | 0.73250294 |
| NAGGFGILGAVAHSP | ASSLGETGQLY | 0.47729877 |
| NEGREITLAMAEM | ASSLGETGQLY | 0.6236025 |
| NELVEWSHPLGHRWRWE | ASSLGETGQLY | 0.7576827 |
| NLGAGGTGGNGGNPG | ASSLGETGQLY | 0.7471791 |
| NTEWPKWETVMAADPVD | ASSLGETGQLY | 0.42184287 |
| PASAAPAETQP | ASSLGETGQLY | 0.5305532 |
| QQSSDGKNGVRAAEITGE | ASSLGETGQLY | 0.42168528 |
| RAPDAVTTSWWKE | ASSLGETGQLY | 0.47673613 |
| SGFGNVGAHTSGWFNQ | ASSLGETGQLY | 0.37933877 |
| SGVAGAVGQAASVGGLKV | ASSLGETGQLY | 0.71036226 |
| SNGGAGTGSAGGNGGT | ASSLGETGQLY | 0.60549855 |
| TGGAGGEGGAAG | ASSLGETGQLY | 0.75852525 |
| TGGDGGAAGAGGAPGHG | ASSLGETGQLY | 0.6919831 |

| | | |
|---|---|---|
| TMPPLLAAIANAGRHGV | ASSLGETGQLY | 0.76008475 |
| TSSTAGNSGGA | ASSLGETGQLY | 0.5594053 |
| VPATVPVGILC | ASSLGETGQLY | 0.642501 |
| VQRPDSGGVVLTGRIS | ASSLGETGQLY | 0.301776 |
| WTPNPVPATKGVPAGLVQAT | ASSLGETGQLY | 0.5647093 |
| YALKRALAKFSDRFDVVII | ASSLGETGQLY | 0.43390697 |
| YLIGSVIV | ASSLGETGQLY | 0.6569549 |
| YGQLVAVID | ASSLGETGQLY | 0.3516872 |
| VTVGGMTLPA | ASSLGETGQLY | 0.761338 |
| VLALPAV | ASSLGETGQLY | 0.62371725 |
| VIPPTLNYET | ASSLGETGQLY | 0.7253183 |
| VGMGGMHPSSG | ASSLGETGQLY | 0.48787418 |
| VGLAGGAATA | ASSLGETGQLY | 0.5561644 |
| VGGAGGAGGSGP | ASSLGETGQLY | 0.55895495 |
| VFAAFMV | ASSLGETGQLY | 0.7394756 |
| TLYDKLR | ASSLGETGQLY | 0.43689376 |
| TLAVFTVL | ASSLGETGQLY | 0.25808617 |
| TGSVQPARV | ASSLGETGQLY | 0.49426067 |
| TGGGAGSGGT | ASSLGETGQLY | 0.36815196 |
| SHLETSL | ASSLGETGQLY | 0.7710472 |
| SFTTGMT | ASSLGETGQLY | 0.7525538 |
| QVEVAKRMGVRQPTVS | ASSLGETGQLY | 0.68295455 |
| QGSGGPALAM | ASSLGETGQLY | 0.762882 |
| MLAPSNP | ASSLGETGQLY | 0.23176375 |
| LVNNAAI | ASSLGETGQLY | 0.5411152 |
| LTGEINTLA | ASSLGETGQLY | 0.77238894 |
| LSIFGPV | ASSLGETGQLY | 0.69701844 |
| LLALLANL | ASSLGETGQLY | 0.65324664 |
| LISNASDALDKLR | ASSLGETGQLY | 0.7448505 |
| LINEAVP | ASSLGETGQLY | 0.54600275 |
| LGYAFSGL | ASSLGETGQLY | 0.6493387 |
| LDFVAVAL | ASSLGETGQLY | 0.6572397 |
| LAQPGAE | ASSLGETGQLY | 0.72202826 |
| ISNLGAVTGF | ASSLGETGQLY | 0.6876329 |
| HQEDPGRI | ASSLGETGQLY | 0.4188816 |
| GTPGEPTP | ASSLGETGQLY | 0.68942976 |
| GFFNSGS | ASSLGETGQLY | 0.7806858 |
| GAGGGGLTAGAN | ASSLGETGQLY | 0.61714983 |
| FVDALVNM | ASSLGETGQLY | 0.72646093 |
| FLTVAPEL | ASSLGETGQLY | 0.68405545 |
| FETLTAL | ASSLGETGQLY | 0.7360493 |
| DVINPIEL | ASSLGETGQLY | 0.527874 |
| APVSGAPV | ASSLGETGQLY | 0.45643422 |
| ALVGGLAW | ASSLGETGQLY | 0.70977294 |
| ALAAAAATAI | ASSLGETGQLY | 0.4345717 |
| AFGLGLLL | ASSLGETGQLY | 0.6339915 |
| AFFAAASL | ASSLGETGQLY | 0.5757484 |
| AAAALPAFSPP | ASSLGETGQLY | 0.78466076 |
| VVLAPAL | ASSLGETGQLY | 0.73080873 |
| TIGHVDHGKTTL | ASSLGETGQLY | 0.67461085 |
| LISNASDALDK | ASSLGETGQLY | 0.49991333 |

| | | |
|---|---|---|
| LISNASDAL | ASSLGETGQLY | 0.6621519 |
| ISNASDALDK | ASSLGETGQLY | 0.72692263 |
| NQAETLVYQTE | ASSLGETGQLY | 0.3355179 |
| TGGTGGTAGAG | ASSLGETGQLY | 0.75809526 |
| TGSTAGGAGGA | ASSLGETGQLY | 0.47617596 |
| DPLAVPVAAGPV | ASSLGETGQLY | 0.6345539 |
| GGTGGTPGNGGN | ASSLGETGQLY | 0.705616 |
| PAGLAGVDPATL | ASSLGETGQLY | 0.77607906 |
| GNNLVVEAPHAGASV | ASSLGETGQLY | 0.604384 |
| KVTRIMSASTLEQPA | ASSLGETGQLY | 0.6900666 |
| LGFALPIALPRRTVV | ASSLGETGQLY | 0.4738102 |
| TRLGNLTARKLVVLV | ASSLGETGQLY | 0.7180308 |
| WSQQRRLIYFPSAGP | ASSLGETGQLY | 0.7564249 |
| GPNGNPDPMAAAVDIR | ASSLGETGQLY | 0.7405866 |
| TQRCVSVADLRAAASR | ASSLGETGQLY | 0.77464664 |
| QFVLDVAEHPVEVLTETV | ASSLGETGQLY | 0.7012394 |
| GGAGGTGGAGGNGGTQAGDG | ASSLGETGQLY | 0.32574293 |
| PLLATLRAIAVDVDAKPAQV | ASSLGETGQLY | 0.5789294 |
| APKFRTGGYMIDFWGVGYQVAK | ASSLGETGQLY | 0.7497036 |
| GAGGAGGAGGAAGGPGGTGGTG | ASSLGETGQLY | 0.7599783 |
| IGALGLLATDVAPAGSVAFGVA | ASSLGETGQLY | 0.75945276 |
| AGGAGGIGGNGGAGGAGGVLMG | ASSLGETGQLY | 0.75925136 |
| ADELIGLL | ASSLGETGQLY | 0.59379506 |
| ADRALKA | ASSLGETGQLY | 0.762602 |
| AGAVGQVGGAGGA | ASSLGETGQLY | 0.7822375 |
| AGGGGGLGGAP | ASSLGETGQLY | 0.72497225 |
| AGGLLYGNGGAG | ASSLGETGQLY | 0.4620577 |
| AGLESIGL | ASSLGETGQLY | 0.44326258 |
| AGLKTGII | ASSLGETGQLY | 0.70625174 |
| ASLAALLL | ASSLGETGQLY | 0.74028707 |
| ASVINISSI | ASSLGETGQLY | 0.39732873 |
| ATPPGPA | ASSLGETGQLY | 0.69312984 |
| ATVASALV | ASSLGETGQLY | 0.7135225 |
| ATVASLIL | ASSLGETGQLY | 0.331917 |
| AVNGPVLIRH | ASSLGETGQLY | 0.64793885 |
| DGAAGAVGI | ASSLGETGQLY | 0.76750225 |
| DPLLHGI | ASSLGETGQLY | 0.6567352 |
| ELGAALGAL | ASSLGETGQLY | 0.47410613 |
| ELGRTQAV | ASSLGETGQLY | 0.758703 |
| FVAGGYLS | ASSLGETGQLY | 0.7780125 |
| GASGGAGGAGGW | ASSLGETGQLY | 0.7656286 |
| GGAGAGGDGGAGGI | ASSLGETGQLY | 0.3220549 |
| GTGHNGSQPGMGGNG | ASSLGETGQLY | 0.23194343 |
| GYAAASASAS | ASSLGETGQLY | 0.6788729 |
| HQVDVAAVQQG | ASSLGETGQLY | 0.7572044 |
| IGAAIAHT | ASSLGETGQLY | 0.63022447 |
| IGMIEVAQL | ASSLGETGQLY | 0.5253968 |
| IRLAPEHTSAMD | ASSLGETGQLY | 0.7129972 |
| IYNVSVL | ASSLGETGQLY | 0.75458133 |
| KQGFLGM | ASSLGETGQLY | 0.49431098 |
| LESVITQL | ASSLGETGQLY | 0.35407296 |

| | | |
|---|---|---|
| LGLGALVVV | ASSLGETGQLY | 0.74648654 |
| LIGNGGNGGSGGI | ASSLGETGQLY | 0.7286977 |
| LNAINAPTL | ASSLGETGQLY | 0.4662354 |
| LQLLNTGV | ASSLGETGQLY | 0.60697436 |
| LQTSIEF | ASSLGETGQLY | 0.6867372 |
| MSFAVSEL | ASSLGETGQLY | 0.62783855 |
| NTGALGL | ASSLGETGQLY | 0.702816 |
| PPGAGGAGGAGTT | ASSLGETGQLY | 0.38987118 |
| QLVGLGI | ASSLGETGQLY | 0.74492425 |
| RFLEQVHQL | ASSLGETGQLY | 0.3514009 |
| RGVGYVLR | ASSLGETGQLY | 0.71418935 |
| SATGVFAGVI | ASSLGETGQLY | 0.7222794 |
| SLPAITLP | ASSLGETGQLY | 0.6974473 |
| STALAFTV | ASSLGETGQLY | 0.60852826 |
| SVLTVGLGK | ASSLGETGQLY | 0.5012259 |
| TALAGAGSGI | ASSLGETGQLY | 0.43899664 |
| TGLPFTGI | ASSLGETGQLY | 0.36381623 |
| VAEHVTV | ASSLGETGQLY | 0.7702282 |
| VDIVLNSL | ASSLGETGQLY | 0.6810429 |
| VLTGGGAVL | ASSLGETGQLY | 0.7562659 |
| VNVGLAYGI | ASSLGETGQLY | 0.70182645 |
| VSSAAVGAV | ASSLGETGQLY | 0.4368298 |
| VTSVYEAGV | ASSLGETGQLY | 0.70339704 |
| WVALDRGA | ASSLGETGQLY | 0.44908226 |
| YLVGSVVI | ASSLGETGQLY | 0.6035859 |
| GPPGTGKTLL | ASSLGETGQLY | 0.6801789 |
| GVGMTKF | ASSLGETGQLY | 0.7246704 |
| IGIGGSDL | ASSLGETGQLY | 0.48999274 |
| IGTLLAEL | ASSLGETGQLY | 0.67676777 |
| IYTSGTT | ASSLGETGQLY | 0.6895724 |
| LSFPAGV | ASSLGETGQLY | 0.47806054 |
| LYTSGTT | ASSLGETGQLY | 0.6674415 |
| VGAFPFGF | ASSLGETGQLY | 0.7519604 |
| VGAGTVEF | ASSLGETGQLY | 0.6675602 |
| VGKSTLLL | ASSLGETGQLY | 0.37724185 |
| AALAHAGGGIV | ASSLGETGQLY | 0.7014575 |
| AALVAVSLL | ASSLGETGQLY | 0.50451076 |
| AAVSRAVAAL | ASSLGETGQLY | 0.69775593 |
| AGGSSGIGGF | ASSLGETGQLY | 0.6346632 |
| AGLTVANANG | ASSLGETGQLY | 0.62103856 |
| AGLVGAVL | ASSLGETGQLY | 0.71047807 |
| ALSQAFTV | ASSLGETGQLY | 0.3613592 |
| ALTFSVSL | ASSLGETGQLY | 0.52224505 |
| ALTGAALVAL | ASSLGETGQLY | 0.5910034 |
| ASFSIGAGVL | ASSLGETGQLY | 0.6711955 |
| DGSVYKI | ASSLGETGQLY | 0.7224678 |
| EQLAEIV | ASSLGETGQLY | 0.50310653 |
| FAVSELTVM | ASSLGETGQLY | 0.65178084 |
| GAAPLAAGL | ASSLGETGQLY | 0.61716664 |
| GADGGDGGSGGAGG | ASSLGETGQLY | 0.52257013 |
| GFGLDTGGAGG | ASSLGETGQLY | 0.6359322 |

| | | |
|---|---|---|
| GLASGGLL | ASSLGETGQLY | 0.5877749 |
| GRIAGAAVLL | ASSLGETGQLY | 0.36162525 |
| GSAGSLNTSF | ASSLGETGQLY | 0.7239172 |
| GTGQIGI | ASSLGETGQLY | 0.52372956 |
| GTLIALIL | ASSLGETGQLY | 0.66948223 |
| IAATMGFVVG | ASSLGETGQLY | 0.7596814 |
| IFAQALL | ASSLGETGQLY | 0.56202525 |
| IGAAVALGF | ASSLGETGQLY | 0.6867617 |
| IGLTPLAL | ASSLGETGQLY | 0.67410654 |
| ILGSNTSTL | ASSLGETGQLY | 0.7276741 |
| IVFSLTVL | ASSLGETGQLY | 0.65084875 |
| LASKATGF | ASSLGETGQLY | 0.47956645 |
| LASVALIL | ASSLGETGQLY | 0.75711507 |
| LFAVGLTL | ASSLGETGQLY | 0.7151071 |
| LFNGGIGTYI | ASSLGETGQLY | 0.75252515 |
| LTLGAGII | ASSLGETGQLY | 0.7294331 |
| LVAAALGDI | ASSLGETGQLY | 0.7514355 |
| NTAFVQLGI | ASSLGETGQLY | 0.7314428 |
| QLFSGAEL | ASSLGETGQLY | 0.57570523 |
| SAATFITL | ASSLGETGQLY | 0.31222498 |
| SNYGLANLGV | ASSLGETGQLY | 0.6459125 |
| SYGGAYAVM | ASSLGETGQLY | 0.6357305 |
| TTGVGGLTLG | ASSLGETGQLY | 0.5090237 |
| TTQTIYI | ASSLGETGQLY | 0.3070601 |
| VAAGAGLGAVI | ASSLGETGQLY | 0.7551247 |
| VAAVVTLL | ASSLGETGQLY | 0.7807385 |
| VGFLGGLI | ASSLGETGQLY | 0.5388884 |
| VGVIPILL | ASSLGETGQLY | 0.7773166 |
| VIFASSLI | ASSLGETGQLY | 0.3729187 |
| VKGGDGGAAAG | ASSLGETGQLY | 0.7311369 |
| VTKTFTNI | ASSLGETGQLY | 0.7578795 |
| VVAAAILL | ASSLGETGQLY | 0.4536081 |
| YAALDVEL | ASSLGETGQLY | 0.7410298 |
| LASVALLL | ASSLGETGQLY | 0.4618894 |
| LDAVHRV | ASSLGETGQLY | 0.5261589 |
| QGLSSSIF | ASSLGETGQLY | 0.7638239 |
| RGTFANIRL | ASSLGETGQLY | 0.6213914 |
| RQLWWGHRI | ASSLGETGQLY | 0.72388434 |
| YGPPGTGKTLL | ASSLGETGQLY | 0.60255027 |
| AALVGAPANVA | ASSLGETGQLY | 0.49623495 |
| AATFRRTE | ASSLGETGQLY | 0.6849797 |
| AGALGGGI | ASSLGETGQLY | 0.75509334 |
| AGGAGGAGGIGAG | ASSLGETGQLY | 0.62879896 |
| AGSGALGL | ASSLGETGQLY | 0.7776303 |
| AQIADQI | ASSLGETGQLY | 0.4904118 |
| ASAVAVTYL | ASSLGETGQLY | 0.7724123 |
| ASTVLGGTGG | ASSLGETGQLY | 0.7504557 |
| AVGGEPAK | ASSLGETGQLY | 0.67661726 |
| CEQGTFGGTC | ASSLGETGQLY | 0.553954 |
| DEGKAPAG | ASSLGETGQLY | 0.75578356 |
| DTVVFPAL | ASSLGETGQLY | 0.607128 |

| | | |
|---|---|---|
| FEGITMAG | ASSLGETGQLY | 0.7505119 |
| FPLVVQD | ASSLGETGQLY | 0.7602256 |
| GAFTTPQ | ASSLGETGQLY | 0.6529419 |
| GFGGFGTAGGVGGA | ASSLGETGQLY | 0.8930057 |
| GFLAQFL | ASSLGETGQLY | 0.85987884 |
| GGAGGAGGNAGVA | ASSLGETGQLY | 0.8440143 |
| GGVGTAGAGGAGGAGG | ASSLGETGQLY | 0.71267307 |
| GNGGGGDAGNAGSGGN | ASSLGETGQLY | 0.3524812 |
| GTGGAGGAGNDAGS | ASSLGETGQLY | 0.7726036 |
| GTTNTGWFNS | ASSLGETGQLY | 0.63112843 |
| IFMGPLFTL | ASSLGETGQLY | 0.8938464 |
| ILAHAIEAI | ASSLGETGQLY | 0.8801383 |
| IRSAIREAL | ASSLGETGQLY | 0.69064236 |
| ITFAGGG | ASSLGETGQLY | 0.8881399 |
| ITGMSGHQEV | ASSLGETGQLY | 0.892248 |
| IVAGLACTL | ASSLGETGQLY | 0.39112648 |
| IVGGGLAGA | ASSLGETGQLY | 0.7753409 |
| IVTGVLNI | ASSLGETGQLY | 0.6583264 |
| LAQQGCI | ASSLGETGQLY | 0.8050436 |
| LCLHVEN | ASSLGETGQLY | 0.25731656 |
| LDAISDGS | ASSLGETGQLY | 0.8852321 |
| LGGFAAML | ASSLGETGQLY | 0.7676381 |
| LGGKNPMI | ASSLGETGQLY | 0.15003693 |
| LGPKGVAAIL | ASSLGETGQLY | 0.8282311 |
| LMVGVSI | ASSLGETGQLY | 0.7951457 |
| LVGAAFVL | ASSLGETGQLY | 0.79595613 |
| LVLAVAPL | ASSLGETGQLY | 0.88944775 |
| MAAIAVIDAA | ASSLGETGQLY | 0.73184 |
| NNGINAELQ | ASSLGETGQLY | 0.6240652 |
| RGEPLVLVHG | ASSLGETGQLY | 0.8634683 |
| RGGAGGAAGLI | ASSLGETGQLY | 0.8983525 |
| RSYRMHPAV | ASSLGETGQLY | 0.5327209 |
| SAAHLLAGQLA | ASSLGETGQLY | 0.866108 |
| SAAVGSY | ASSLGETGQLY | 0.89777094 |
| SPHAVLHRI | ASSLGETGQLY | 0.8590293 |
| SQIAKVL | ASSLGETGQLY | 0.8886683 |
| TALGHLVP | ASSLGETGQLY | 0.9026616 |
| TAPTTSGAA | ASSLGETGQLY | 0.6615567 |
| TDGDDGNGGSPGT | ASSLGETGQLY | 0.8898752 |
| TDGLGEN | ASSLGETGQLY | 0.8893931 |
| TGGPPLPGQA | ASSLGETGQLY | 0.7090641 |
| TIPAITI | ASSLGETGQLY | 0.81251156 |
| TIPGFTI | ASSLGETGQLY | 0.64997655 |
| TSIASGALA | ASSLGETGQLY | 0.7987795 |
| TVTLPGAAI | ASSLGETGQLY | 0.38541114 |
| VASGLSAALP | ASSLGETGQLY | 0.85640705 |
| VGAGVIGI | ASSLGETGQLY | 0.88896024 |
| VGPKRFGV | ASSLGETGQLY | 0.79287493 |
| VNVAFPD | ASSLGETGQLY | 0.6067959 |
| VSMTNSGVS | ASSLGETGQLY | 0.60221744 |
| VTDVLGDLSV | ASSLGETGQLY | 0.8378668 |

| | | |
|---|---|---|
| VVVVGAGFAGL | ASSLGETGQLY | 0.80306685 |
| VYTVGTVV | ASSLGETGQLY | 0.41318655 |
| WTTFSGP | ASSLGETGQLY | 0.76997244 |
| YDELVEAF | ASSLGETGQLY | 0.8860526 |
| YGTGGAGGNGG | ASSLGETGQLY | 0.88840723 |
| YTNVMAV | ASSLGETGQLY | 0.45645612 |
| FLEQVHQL | ASSLGETGQLY | 0.8986381 |
| GLSLLAVAV | ASSLGETGQLY | 0.89652634 |
| LEQVHQL | ASSLGETGQLY | 0.9040773 |
| LNAAGAI | ASSLGETGQLY | 0.8009959 |
| PAVHASL | ASSLGETGQLY | 0.6229709 |
| TELEDDA | ASSLGETGQLY | 0.89442515 |
| TPPGPPN | ASSLGETGQLY | 0.7892931 |
| TPPPPPPP | ASSLGETGQLY | 0.73395777 |
| TQSPALI | ASSPGGGASSYEQY | 0.8807807 |
| EFAAAFPQ | ASSPGGGASSYEQY | 0.79388064 |
| LLSGNLDVL | ASSPGGGASSYEQY | 0.71505463 |
| GPANLGNYNI | ASSPGGGASSYEQY | 0.8214836 |
| TLGPTLP | ASSPGGGASSYEQY | 0.9039775 |
| VYAYSSCCK | ASSPGGGASSYEQY | 0.598549 |
| APAPAPAPNI | ASSPGGGASSYEQY | 0.6654061 |
| PGGPVPIP | ASSPGGGASSYEQY | 0.74258727 |
| FLTVALEP | ASSPGGGASSYEQY | 0.8298396 |
| SQYFGANL | ASSPGGGASSYEQY | 0.888548 |
| GGVKPAA | ASSPGGGASSYEQY | 0.8493122 |
| GGTGGAGGAGGW | ASSPGGGASSYEQY | 0.7586968 |
| GVGGVAGLGGAGG | ASSPGGGASSYEQY | 0.85883284 |
| VLLVLATTI | ASSPGGGASSYEQY | 0.8727205 |
| IGIFAAML | ASSPGGGASSYEQY | 0.8426003 |
| AVGDLID | ASSPGGGASSYEQY | 0.88797677 |
| NISPLPL | ASSPGGGASSYEQY | 0.6904737 |
| VTAVFVAI | ASSPGGGASSYEQY | 0.89980865 |
| SGDDNDAAL | ASSPGGGASSYEQY | 0.8705643 |
| AVAALAHS | ASSPGGGASSYEQY | 0.88478935 |
| GGVTVMATP | ASSPGGGASSYEQY | 0.8855338 |
| AGLAANVRD | ASSPGGGASSYEQY | 0.84917307 |
| TNRPDLIDPALLRPG | ASSPGGGASSYEQY | 0.85489047 |
| LANLGVTAL | ASSPGGGASSYEQY | 0.39244685 |
| TGERIEL | ASSPGGGASSYEQY | 0.7179954 |
| NGEELAA | ASSPGGGASSYEQY | 0.8606763 |
| VLSLVWG | ASSPGGGASSYEQY | 0.85013795 |
| IGYALWL | ASSPGGGASSYEQY | 0.8773817 |
| LSLLLAPL | ASSPGGGASSYEQY | 0.8566673 |
| TLDEALK | ASSPGGGASSYEQY | 0.73634595 |
| TDGAGTAEGG | ASSPGGGASSYEQY | 0.8952371 |
| NAPGGAGGNGG | ASSPGGGASSYEQY | 0.88500607 |
| VSVFMGMGGNF | ASSPGGGASSYEQY | 0.43634814 |
| LASYLAAL | ASSPGGGASSYEQY | 0.7729765 |
| LVGGTPAL | ASSPGGGASSYEQY | 0.9027357 |
| VRNADVL | ASSPGGGASSYEQY | 0.8825072 |
| TCSSPFLVSAM | ASSPGGGASSYEQY | 0.76455784 |

| | | |
|---|---|---|
| VHITDAE | ASSPGGGASSYEQY | 0.7832184 |
| EYAAPSF | ASSPGGGASSYEQY | 0.91343427 |
| VVDAFHGGL | ASSPGGGASSYEQY | 0.8038399 |
| ASVVDPQ | ASSPGGGASSYEQY | 0.90303874 |
| SYNEFGGLS | ASSPGGGASSYEQY | 0.20631993 |
| DPDGSVY | ASSPGGGASSYEQY | 0.79598117 |
| MVSTIHATE | ASSPGGGASSYEQY | 0.65148187 |
| LAAGVADPV | ASSPGGGASSYEQY | 0.44249904 |
| TKEQIAA | ASSPGGGASSYEQY | 0.8814411 |
| SLASPAAL | ASSPGGGASSYEQY | 0.7449868 |
| IGLVFYGV | ASSPGGGASSYEQY | 0.8757445 |
| FLLDASNL | ASSPGGGASSYEQY | 0.78374475 |
| YQVGNFP | ASSPGGGASSYEQY | 0.8883723 |
| ESLLSQST | ASSPGGGASSYEQY | 0.83560205 |
| LFTVGFAAVG | ASSPGGGASSYEQY | 0.8378421 |
| ATPPTPT | ASSPGGGASSYEQY | 0.63057387 |
| TDLATLG | ASSPGGGASSYEQY | 0.8773992 |
| SAGGTGR | ASSPGGGASSYEQY | 0.899868 |
| GFSNSGS | ASSPGGGASSYEQY | 0.88524514 |
| GGNGGSGGTSV | ASSPGGGASSYEQY | 0.40269336 |
| GTSVATGGAGN | ASSPGGGASSYEQY | 0.82832307 |
| NGGNGGSGGTS | ASSPGGGASSYEQY | 0.82728374 |
| AVVKFPH | ASSPGGGASSYEQY | 0.30617577 |
| TPPPPPP | ASSPGGGASSYEQY | 0.2638104 |
| AAPAAAAAPA | ASSPGGGASSYEQY | 0.8104205 |
| IALNEDL | ASSPGGGASSYEQY | 0.7681702 |
| IVNVSSM | ASSPGGGASSYEQY | 0.8844726 |
| TGIPPAPR | ASSPGGGASSYEQY | 0.8608351 |
| FDLGGGTF | ASSPGGGASSYEQY | 0.8068147 |
| FDLGGGTFDVS | ASSPGGGASSYEQY | 0.87314934 |
| NEPTAAA | ASSPGGGASSYEQY | 0.8377985 |
| NEPTAAALA | ASSPGGGASSYEQY | 0.10408165 |
| GSIINISSI | ASSPGGGASSYEQY | 0.66638565 |
| VAPGNAGTA | ASSPGGGASSYEQY | 0.4173855 |
| TGGGASLE | ASSPGGGASSYEQY | 0.8981301 |
| VAVYNHYKRI | ASSPGGGASSYEQY | 0.6910079 |
| TDPAPATT | ASSPGGGASSYEQY | 0.6686251 |
| TNRPDILDPALLRPG | ASSPGGGASSYEQY | 0.8183882 |
| LTTTTTS | ASSPGGGASSYEQY | 0.86543405 |
| PTGSGKTL | ASSPGGGASSYEQY | 0.818261 |
| TGSGKTL | ASSPGGGASSYEQY | 0.66262805 |
| TIGHVDHGK | ASSPGGGASSYEQY | 0.20616975 |
| TIGHVDHGKTT | ASSPGGGASSYEQY | 0.8744815 |
| GHVDHGKTT | ASSPGGGASSYEQY | 0.5165006 |
| AQFEHTLL | ASSPGGGASSYEQY | 0.5849531 |
| IVNQDEL | ASSPGGGASSYEQY | 0.3053267 |
| TNGSQFF | ASSPGGGASSYEQY | 0.38293505 |
| TGKGIVE | ASSPGGGASSYEQY | 0.3733813 |
| LISNASDA | ASSPGGGASSYEQY | 0.88564664 |
| AELRTLGELTKIAKAHGAQ | ASSPGGGASSYEQY | 0.7678915 |
| APNAVAAAEPSW | ASSPGGGASSYEQY | 0.76343304 |

| | | |
|---|---|---|
| APQVAGAATAVL | ASSPGGGASSYEQY | 0.89886117 |
| ARRDPAAAAAILPTDA | ASSPGGGASSYEQY | 0.8528724 |
| DAAARDVLAEAGLAEYF | ASSPGGGASSYEQY | 0.79011965 |
| DDQIVLSAKANLAKVEQ | ASSPGGGASSYEQY | 0.64062476 |
| DELLPDTGEDSDEDGH | ASSPGGGASSYEQY | 0.6266062 |
| ELPEAPANRATVPQD | ASSPGGGASSYEQY | 0.6635603 |
| FGQSKIERVTGIG | ASSPGGGASSYEQY | 0.6488346 |
| GAGGTGSTAGGAGGAGGAGGLY | ASSPGGGASSYEQY | 0.90173477 |
| GHGGTGAAGGNGGTGGAGGSGI | ASSPGGGASSYEQY | 0.7885784 |
| GNGGNGGAGGEGGPGGAGGAGA | ASSPGGGASSYEQY | 0.42381635 |
| GSTAGNGGNGGNATNGTIAGQP | ASSPGGGASSYEQY | 0.8394306 |
| GTGGDGGNGGNGGTGG | ASSPGGGASSYEQY | 0.89937025 |
| GVRRPAGIPADAPV | ASSPGGGASSYEQY | 0.91319174 |
| GYRQVTDAHLLTLARR | ASSPGGGASSYEQY | 0.6597282 |
| IAKMGQDGHDRG | ASSPGGGASSYEQY | 0.73889387 |
| KAEPSIEIQYT | ASSPGGGASSYEQY | 0.7952752 |
| KPVRIRMAADEDTVAVT | ASSPGGGASSYEQY | 0.8234103 |
| MPPRIAGMRLLVIKPEP | ASSPGGGASSYEQY | 0.73395944 |
| MKFYAVALFRSRGVPPTR | ASSPGGGASSYEQY | 0.5159522 |
| NAGGFGILGAVAHSP | ASSPGGGASSYEQY | 0.8912579 |
| NEGREITLAMAEM | ASSPGGGASSYEQY | 0.9173568 |
| NELVEWSHPLGHRWRWE | ASSPGGGASSYEQY | 0.7913594 |
| NLGAGGTGGNGGNPG | ASSPGGGASSYEQY | 0.37792253 |
| NTEWPKWETVMAADPVD | ASSPGGGASSYEQY | 0.79900914 |
| PASAAPAETQP | ASSPGGGASSYEQY | 0.69078076 |
| QQSSDGKNGVRAAEITGE | ASSPGGGASSYEQY | 0.7616279 |
| RAPDAVTTSWWKE | ASSPGGGASSYEQY | 0.838431 |
| SGFGNVGAHTSGWFNQ | ASSPGGGASSYEQY | 0.42277572 |
| SGVAGAVGQAASVGGLKV | ASSPGGGASSYEQY | 0.780313 |
| SNGGAGTGSAGGNGGT | ASSPGGGASSYEQY | 0.9094503 |
| TGGAGGEGGAAG | ASSPGGGASSYEQY | 0.7834166 |
| TGGDGGAAGAGGAPGHG | ASSPGGGASSYEQY | 0.86052346 |
| TMPPLLAAIANAGRHGV | ASSPGGGASSYEQY | 0.8914272 |
| TSSTAGNSGGA | ASSPGGGASSYEQY | 0.87295735 |
| VPATVPVGILC | ASSPGGGASSYEQY | 0.900334 |
| VQRPDSGGVVLTGRIS | ASSPGGGASSYEQY | 0.8980309 |
| WTPNPVPATKGVPAGLVQAT | ASSPGGGASSYEQY | 0.78981924 |
| YALKRALAKFSDRFDVVII | ASSPGGGASSYEQY | 0.8324752 |
| YLIGSVIV | ASSPGGGASSYEQY | 0.8398446 |
| YGQLVAVID | ASSPGGGASSYEQY | 0.8131761 |
| VTVGGMTLPA | ASSPGGGASSYEQY | 0.80618787 |
| VLALPAV | ASSPGGGASSYEQY | 0.7206818 |
| VIPPTLNYET | ASSPGGGASSYEQY | 0.8822956 |
| VGMGGMHPSSG | ASSPGGGASSYEQY | 0.8217495 |
| VGLAGGAATA | ASSPGGGASSYEQY | 0.9058932 |
| VGGAGGAGGSGP | ASSPGGGASSYEQY | 0.47902656 |
| VFAAFMV | ASSPGGGASSYEQY | 0.579764 |
| TLYDKLR | ASSPGGGASSYEQY | 0.8878331 |
| TLAVFTVL | ASSPGGGASSYEQY | 0.1608747 |
| TGSVQPARV | ASSPGGGASSYEQY | 0.26324856 |
| TGGGAGSGGT | ASSPGGGASSYEQY | 0.8517279 |

| | | |
|---|---|---|
| SHLETSL | ASSPGGGASSYEQY | 0.9041742 |
| SFTTGMT | ASSPGGGASSYEQY | 0.21048892 |
| QVEVAKRMGVRQPTVS | ASSPGGGASSYEQY | 0.5498488 |
| QGSGGPALAM | ASSPGGGASSYEQY | 0.5521373 |
| MLAPSNP | ASSPGGGASSYEQY | 0.8733767 |
| LVNNAAI | ASSPGGGASSYEQY | 0.81271374 |
| LTGEINTLA | ASSPGGGASSYEQY | 0.37485853 |
| LSIFGPV | ASSPGGGASSYEQY | 0.88435364 |
| LLALLANL | ASSPGGGASSYEQY | 0.365103 |
| LISNASDALDKLR | ASSPGGGASSYEQY | 0.64193827 |
| LINEAVP | ASSPGGGASSYEQY | 0.8835509 |
| LGYAFSGL | ASSPGGGASSYEQY | 0.90589523 |
| LDFVAVAL | ASSPGGGASSYEQY | 0.52184206 |
| LAQPGAE | ASSPGGGASSYEQY | 0.8857941 |
| ISNLGAVTGF | ASSPGGGASSYEQY | 0.7590257 |
| HQEDPGRI | ASSPGGGASSYEQY | 0.91392803 |
| GTPGEPTP | ASSPGGGASSYEQY | 0.7604053 |
| GFFNSGS | ASSPGGGASSYEQY | 0.42137703 |
| GAGGGGLTAGAN | ASSPGGGASSYEQY | 0.8855188 |
| FVDALVNM | ASSPGGGASSYEQY | 0.87716687 |
| FLTVAPEL | ASSPGGGASSYEQY | 0.8320247 |
| FETLTAL | ASSPGGGASSYEQY | 0.89857495 |
| DVINPIEL | ASSPGGGASSYEQY | 0.7907624 |
| APVSGAPV | ASSPGGGASSYEQY | 0.7869363 |
| ALVGGLAW | ASSPGGGASSYEQY | 0.88754046 |
| ALAAAAATAI | ASSPGGGASSYEQY | 0.8848072 |
| AFGLGLLL | ASSPGGGASSYEQY | 0.5620811 |
| AFFAAASL | ASSPGGGASSYEQY | 0.8939688 |
| AAAALPAFSPP | ASSPGGGASSYEQY | 0.8111789 |
| VVLAPAL | ASSPGGGASSYEQY | 0.62867355 |
| TIGHVDHGKTTL | ASSPGGGASSYEQY | 0.9087024 |
| LISNASDALDK | ASSPGGGASSYEQY | 0.833547 |
| LISNASDAL | ASSPGGGASSYEQY | 0.8859351 |
| ISNASDALDK | ASSPGGGASSYEQY | 0.886381 |
| NQAETLVYQTE | ASSPGGGASSYEQY | 0.906497 |
| TGGTGGTAGAG | ASSPGGGASSYEQY | 0.45793608 |
| TGSTAGGAGGA | ASSPGGGASSYEQY | 0.85816026 |
| DPLAVPVAAGPV | ASSPGGGASSYEQY | 0.826807 |
| GGTGGTPGNGGN | ASSPGGGASSYEQY | 0.8740882 |
| PAGLAGVDPATL | ASSPGGGASSYEQY | 0.8262963 |
| GNNLVVEAPHAGASV | ASSPGGGASSYEQY | 0.6530906 |
| KVTRIMSASTLEQPA | ASSPGGGASSYEQY | 0.8700178 |
| LGFALPIALPRRTVV | ASSPGGGASSYEQY | 0.8937398 |
| TRLGNLTARKLVVLV | ASSPGGGASSYEQY | 0.8220289 |
| WSQQRRLIYFPSAGP | ASSPGGGASSYEQY | 0.8559425 |
| GPNGNPDPMAAAVDIR | ASSPGGGASSYEQY | 0.90451515 |
| TQRCVSVADLRAAASR | ASSPGGGASSYEQY | 0.7059167 |
| QFVLDVAEHPVEVLTETV | ASSPGGGASSYEQY | 0.7902533 |
| GGAGGTGGAGGNGGTQAGDG | ASSPGGGASSYEQY | 0.876031 |
| PLLATLRAIAVDVDAKPAQV | ASSPGGGASSYEQY | 0.8920165 |
| APKFRTGGYMIDFWGVGYQVAK | ASSPGGGASSYEQY | 0.8039094 |

| | | |
|---|---|---|
| GAGGAGGAGGAAGGPGGTGGTG | ASSPGGGASSYEQY | 0.90393984 |
| IGALGLLATDVAPAGSVAFGVA | ASSPGGGASSYEQY | 0.7031222 |
| AGGAGGIGGNGGAGGAGGVLMG | ASSPGGGASSYEQY | 0.86878276 |
| ADELIGLL | ASSPGGGASSYEQY | 0.7389424 |
| ADRALKA | ASSPGGGASSYEQY | 0.8387781 |
| AGAVGQVGGAGGA | ASSPGGGASSYEQY | 0.6906788 |
| AGGGGGLGGAP | ASSPGGGASSYEQY | 0.85841316 |
| AGGLLYGNGGAG | ASSPGGGASSYEQY | 0.8825697 |
| AGLESIGL | ASSPGGGASSYEQY | 0.86906576 |
| AGLKTGII | ASSPGGGASSYEQY | 0.21102671 |
| ASLAALLL | ASSPGGGASSYEQY | 0.87483686 |
| ASVINISSI | ASSPGGGASSYEQY | 0.86383545 |
| ATPPGPA | ASSPGGGASSYEQY | 0.91701835 |
| ATVASALV | ASSPGGGASSYEQY | 0.8340841 |
| ATVASLIL | ASSPGGGASSYEQY | 0.8787095 |
| AVNGPVLIRH | ASSPGGGASSYEQY | 0.7342132 |
| DGAAGAVGI | ASSPGGGASSYEQY | 0.827318 |
| DPLLHGI | ASSPGGGASSYEQY | 0.6027677 |
| ELGAALGAL | ASSPGGGASSYEQY | 0.76222503 |
| ELGRTQAV | ASSPGGGASSYEQY | 0.75774616 |
| FVAGGYLS | ASSPGGGASSYEQY | 0.7025708 |
| GASGGAGGAGGW | ASSPGGGASSYEQY | 0.8878808 |
| GGAGAGGDGGAGGI | ASSPGGGASSYEQY | 0.89351887 |
| GTGHNGSQPGMGGNG | ASSPGGGASSYEQY | 0.8845468 |
| GYAAASASAS | ASSPGGGASSYEQY | 0.8900667 |
| HQVDVAAVQQG | ASSPGGGASSYEQY | 0.8989531 |
| IGAAIAHT | ASSPGGGASSYEQY | 0.8983395 |
| IGMIEVAQL | ASSPGGGASSYEQY | 0.75728095 |
| IRLAPEHTSAMD | ASSPGGGASSYEQY | 0.74829745 |
| IYNVSVL | ASSPGGGASSYEQY | 0.5160624 |
| KQGFLGM | ASSPGGGASSYEQY | 0.7529767 |
| LESVITQL | ASSPGGGASSYEQY | 0.74867517 |
| LGLGALVVV | ASSPGGGASSYEQY | 0.8418913 |
| LIGNGGNGGSGGI | ASSPGGGASSYEQY | 0.14201607 |
| LNAINAPTL | ASSPGGGASSYEQY | 0.86540747 |
| LQLLNTGV | ASSPGGGASSYEQY | 0.900618 |
| LQTSIEF | ASSPGGGASSYEQY | 0.78614116 |
| MSFAVSEL | ASSPGGGASSYEQY | 0.85612345 |
| NTGALGL | ASSPGGGASSYEQY | 0.83346754 |
| PPGAGGAGGAGTT | ASSPGGGASSYEQY | 0.7543518 |
| QLVGLGI | ASSPGGGASSYEQY | 0.873958 |
| RFLEQVHQL | ASSPGGGASSYEQY | 0.6948496 |
| RGVGYVLR | ASSPGGGASSYEQY | 0.88427347 |
| SATGVFAGVI | ASSPGGGASSYEQY | 0.9035657 |
| SLPAITLP | ASSPGGGASSYEQY | 0.83248127 |
| STALAFTV | ASSPGGGASSYEQY | 0.85643256 |
| SVLTVGLGK | ASSPGGGASSYEQY | 0.8454108 |
| TALAGAGSGI | ASSPGGGASSYEQY | 0.35278943 |
| TGLPFTGI | ASSPGGGASSYEQY | 0.8620024 |
| VAEHVTV | ASSPGGGASSYEQY | 0.115882725 |
| VDIVLNSL | ASSPGGGASSYEQY | 0.8132509 |

| | | |
|---|---|---|
| VLTGGGAVL | ASSPGGGASSYEQY | 0.686285 |
| VNVGLAYGI | ASSPGGGASSYEQY | 0.8436407 |
| VSSAAVGAV | ASSPGGGASSYEQY | 0.81346095 |
| VTSVYEAGV | ASSPGGGASSYEQY | 0.86080205 |
| WVALDRGA | ASSPGGGASSYEQY | 0.6000495 |
| YLVGSVVI | ASSPGGGASSYEQY | 0.8404101 |
| GPPGTGKTLL | ASSPGGGASSYEQY | 0.76406586 |
| GVGMTKF | ASSPGGGASSYEQY | 0.49779478 |
| IGIGGSDL | ASSPGGGASSYEQY | 0.8859942 |
| IGTLLAEL | ASSPGGGASSYEQY | 0.7886443 |
| IYTSGTT | ASSPGGGASSYEQY | 0.80747557 |
| LSFPAGV | ASSPGGGASSYEQY | 0.891171 |
| LYTSGTT | ASSPGGGASSYEQY | 0.8528291 |
| VGAFPFGF | ASSPGGGASSYEQY | 0.37098694 |
| VGAGTVEF | ASSPGGGASSYEQY | 0.8095606 |
| VGKSTLLL | ASSPGGGASSYEQY | 0.81797993 |
| AALAHAGGGIV | ASSPGGGASSYEQY | 0.80062366 |
| AALVAVSLL | ASSPGGGASSYEQY | 0.14321744 |
| AAVSRAVAAL | ASSPGGGASSYEQY | 0.86184967 |
| AGGSSGIGGF | ASSPGGGASSYEQY | 0.62834513 |
| AGLTVANANG | ASSPGGGASSYEQY | 0.63387966 |
| AGLVGAVL | ASSPGGGASSYEQY | 0.7497227 |
| ALSQAFTV | ASSPGGGASSYEQY | 0.8804945 |
| ALTFSVSL | ASSPGGGASSYEQY | 0.81981456 |
| ALTGAALVAL | ASSPGGGASSYEQY | 0.83298147 |
| ASFSIGAGVL | ASSPGGGASSYEQY | 0.8897288 |
| DGSVYKI | ASSPGGGASSYEQY | 0.8576731 |
| EQLAEIV | ASSPGGGASSYEQY | 0.898008 |
| FAVSELTVM | ASSPGGGASSYEQY | 0.84741026 |
| GAAPLAAGL | ASSPGGGASSYEQY | 0.804562 |
| GADGGDGGSGGAGG | ASSPGGGASSYEQY | 0.86235774 |
| GFGLDTGGAGG | ASSPGGGASSYEQY | 0.76945764 |
| GLASGGLL | ASSPGGGASSYEQY | 0.5253612 |
| GRIAGAAVLL | ASSPGGGASSYEQY | 0.43089026 |
| GSAGSLNTSF | ASSPGGGASSYEQY | 0.8320284 |
| GTGQIGI | ASSPGGGASSYEQY | 0.36070073 |
| GTLIALIL | ASSPGGGASSYEQY | 0.8293224 |
| IAATMGFVVG | ASSPGGGASSYEQY | 0.89540726 |
| IFAQALL | ASSPGGGASSYEQY | 0.842259 |
| IGAAVALGF | ASSPGGGASSYEQY | 0.47224605 |
| IGLTPLAL | ASSPGGGASSYEQY | 0.76719797 |
| ILGSNTSTL | ASSPGGGASSYEQY | 0.69337666 |
| IVFSLTVL | ASSPGGGASSYEQY | 0.88309777 |
| LASKATGF | ASSPGGGASSYEQY | 0.881852 |
| LASVALIL | ASSPGGGASSYEQY | 0.91246796 |
| LFAVGLTL | ASSPGGGASSYEQY | 0.87542266 |
| LFNGGIGTYI | ASSPGGGASSYEQY | 0.7451916 |
| LTLGAGII | ASSPGGGASSYEQY | 0.88082075 |
| LVAAALGDI | ASSPGGGASSYEQY | 0.79720634 |
| NTAFVQLGI | ASSPGGGASSYEQY | 0.7888235 |
| QLFSGAEL | ASSPGGGASSYEQY | 0.82359105 |

| | | |
|---|---|---|
| SAATFITL | ASSPGGGASSYEQY | 0.86480826 |
| SNYGLANLGV | ASSPGGGASSYEQY | 0.8199109 |
| SYGGAYAVM | ASSPGGGASSYEQY | 0.76038945 |
| TTGVGGLTLG | ASSPGGGASSYEQY | 0.34178975 |
| TTQTIYI | ASSPGGGASSYEQY | 0.88127875 |
| VAAGAGLGAVI | ASSPGGGASSYEQY | 0.8912076 |
| VAAVVTLL | ASSPGGGASSYEQY | 0.61431366 |
| VGFLGGLI | ASSPGGGASSYEQY | 0.73439234 |
| VGVIPILL | ASSPGGGASSYEQY | 0.9069746 |
| VIFASSLI | ASSPGGGASSYEQY | 0.89919317 |
| VKGGDGGAAAG | ASSPGGGASSYEQY | 0.7329842 |
| VTKTFTNI | ASSPGGGASSYEQY | 0.6625309 |
| VVAAAILL | ASSPGGGASSYEQY | 0.85060143 |
| YAALDVEL | ASSPGGGASSYEQY | 0.4477104 |
| LASVALLL | ASSPGGGASSYEQY | 0.875698 |
| LDAVHRV | ASSPGGGASSYEQY | 0.81089747 |
| QGLSSSIF | ASSPGGGASSYEQY | 0.793105 |
| RGTFANIRL | ASSPGGGASSYEQY | 0.7381879 |
| RQLWWGHRI | ASSPGGGASSYEQY | 0.84252405 |
| YGPPGTGKTLL | ASSPGGGASSYEQY | 0.7489183 |
| AALVGAPANVA | ASSPGGGASSYEQY | 0.8909454 |
| AATFRRTE | ASSPGGGASSYEQY | 0.1728867 |
| AGALGGGI | ASSPGGGASSYEQY | 0.86500144 |
| AGGAGGAGGIGAG | ASSPGGGASSYEQY | 0.6475395 |
| AGSGALGL | ASSPGGGASSYEQY | 0.72536296 |
| AQIADQI | ASSPGGGASSYEQY | 0.37069216 |
| ASAVAVTYL | ASSPGGGASSYEQY | 0.7749511 |
| ASTVLGGTGG | ASSPGGGASSYEQY | 0.885151 |
| AVGGEPAK | ASSPGGGASSYEQY | 0.8890165 |
| CEQGTFGGTC | ASSPGGGASSYEQY | 0.7326113 |
| DEGKAPAG | ASSPGGGASSYEQY | 0.87981105 |
| DTVVFPAL | ASSPGGGASSYEQY | 0.911038 |
| FEGITMAG | ASSPGGGASSYEQY | 0.7621764 |
| FPLVVQD | ASSPGGGASSYEQY | 0.30631626 |
| GAFTTPQ | ASSPGGGASSYEQY | 0.86089027 |
| GFGGFGTAGGVGGA | ASSPGGGASSYEQY | 0.82636863 |
| GFLAQFL | ASSPGGGASSYEQY | 0.8789768 |
| GGAGGAGGNAGVA | ASSPGGGASSYEQY | 0.88609076 |
| GGVGTAGAGGAGGAGG | ASSPGGGASSYEQY | 0.54366255 |
| GNGGGGDAGNAGSGGN | ASSPGGGASSYEQY | 0.89169484 |
| GTGGAGGAGNDAGS | ASSPGGGASSYEQY | 0.88435304 |
| GTTNTGWFNS | ASSPGGGASSYEQY | 0.6929261 |
| IFMGPLFTL | ASSPGGGASSYEQY | 0.65778494 |
| ILAHAIEAI | ASSPGGGASSYEQY | 0.6130261 |
| IRSAIREAL | ASSPGGGASSYEQY | 0.44147938 |
| ITFAGGG | ASSPGGGASSYEQY | 0.87491715 |
| ITGMSGHQEV | ASSPGGGASSYEQY | 0.729018 |
| IVAGLACTL | ASSPGGGASSYEQY | 0.69620645 |
| IVGGGLAGA | ASSPGGGASSYEQY | 0.8206768 |
| IVTGVLNI | ASSPGGGASSYEQY | 0.6861789 |
| LAQQGCI | ASSPGGGASSYEQY | 0.68738484 |

| | | |
|---|---|---|
| LCLHVEN | ASSPGGGASSYEQY | 0.7310044 |
| LDAISDGS | ASSPGGGASSYEQY | 0.833246 |
| LGGFAAML | ASSPGGGASSYEQY | 0.8643045 |
| LGGKNPMI | ASSPGGGASSYEQY | 0.8101145 |
| LGPKGVAAIL | ASSPGGGASSYEQY | 0.82858133 |
| LMVGVSI | ASSPGGGASSYEQY | 0.8927294 |
| LVGAAFVL | ASSPGGGASSYEQY | 0.501172 |
| LVLAVAPL | ASSPGGGASSYEQY | 0.62807715 |
| MAAIAVIDAA | ASSPGGGASSYEQY | 0.8802598 |
| NNGINAELQ | ASSPGGGASSYEQY | 0.5580068 |
| RGEPLVLVHG | ASSPGGGASSYEQY | 0.88206935 |
| RGGAGGAAGLI | ASSPGGGASSYEQY | 0.8201548 |
| RSYRMHPAV | ASSPGGGASSYEQY | 0.855639 |
| SAAHLLAGQLA | ASSPGGGASSYEQY | 0.82802665 |
| SAAVGSY | ASSPGGGASSYEQY | 0.90901524 |
| SPHAVLHRI | ASSPGGGASSYEQY | 0.8814193 |
| SQIAKVL | ASSPGGGASSYEQY | 0.7798077 |
| TALGHLVP | ASSPGGGASSYEQY | 0.8845898 |
| TAPTTSGAA | ASSPGGGASSYEQY | 0.5640909 |
| TDGDDGNGGSPGT | ASSPGGGASSYEQY | 0.8850452 |
| TDGLGEN | ASSPGGGASSYEQY | 0.87480444 |
| TGGPPLPGQA | ASSPGGGASSYEQY | 0.61503327 |
| TIPAITI | ASSPGGGASSYEQY | 0.79612935 |
| TIPGFTI | ASSPGGGASSYEQY | 0.9024892 |
| TSIASGALA | ASSPGGGASSYEQY | 0.8936347 |
| TVTLPGAAI | ASSPGGGASSYEQY | 0.47806308 |
| VASGLSAALP | ASSPGGGASSYEQY | 0.7468848 |
| VGAGVIGI | ASSPGGGASSYEQY | 0.43054295 |
| VGPKRFGV | ASSPGGGASSYEQY | 0.7272151 |
| VNVAFPD | ASSPGGGASSYEQY | 0.44968212 |
| VSMTNSGVS | ASSPGGGASSYEQY | 0.87576354 |
| VTDVLGDLSV | ASSPGGGASSYEQY | 0.7415728 |
| VVVVGAGFAGL | ASSPGGGASSYEQY | 0.8953186 |
| VYTVGTVV | ASSPGGGASSYEQY | 0.8256357 |
| WTTFSGP | ASSPGGGASSYEQY | 0.89753664 |
| YDELVEAF | ASSPGGGASSYEQY | 0.7265262 |
| YGTGGAGGNGG | ASSPGGGASSYEQY | 0.73957705 |
| YTNVMAV | ASSPGGGASSYEQY | 0.43776023 |
| FLEQVHQL | ASSPGGGASSYEQY | 0.5601891 |
| GLSLLAVAV | ASSPGGGASSYEQY | 0.6842022 |
| LEQVHQL | ASSPGGGASSYEQY | 0.74234915 |
| LNAAGAI | ASSPGGGASSYEQY | 0.32565212 |
| PAVHASL | ASSPGGGASSYEQY | 0.89733756 |
| TELEDDA | ASSPGGGASSYEQY | 0.85201156 |
| TPPGPPN | ASSPGGGASSYEQY | 0.87767875 |
| TPPPPPPP | ASSPGGGASSYEQY | 0.7081221 |
| TQSPALI | ASSLPGLGGREDTQY | 0.7727181 |
| EFAAAFPQ | ASSLPGLGGREDTQY | 0.7272465 |
| LLSGNLDVL | ASSLPGLGGREDTQY | 0.8722764 |
| GPANLGNYNI | ASSLPGLGGREDTQY | 0.6584412 |
| TLGPTLP | ASSLPGLGGREDTQY | 0.31503376 |

| | | |
|---|---|---|
| VYAYSSCCK | ASSLPGLGGREDTQY | 0.6664256 |
| APAPAPAPNI | ASSLPGLGGREDTQY | 0.41939425 |
| PGGPVPIP | ASSLPGLGGREDTQY | 0.9084815 |
| FLTVALEP | ASSLPGLGGREDTQY | 0.8892015 |
| SQYFGANL | ASSLPGLGGREDTQY | 0.85478926 |
| GGVKPAA | ASSLPGLGGREDTQY | 0.9030064 |
| GGTGGAGGAGGW | ASSLPGLGGREDTQY | 0.2232155 |
| GVGGVAGLGGAGG | ASSLPGLGGREDTQY | 0.68352246 |
| VLLVLATTI | ASSLPGLGGREDTQY | 0.89993274 |
| IGIFAAML | ASSLPGLGGREDTQY | 0.86793005 |
| AVGDLID | ASSLPGLGGREDTQY | 0.7761275 |
| NISPLPL | ASSLPGLGGREDTQY | 0.89831674 |
| VTAVFVAI | ASSLPGLGGREDTQY | 0.6935936 |
| SGDDNDAAL | ASSLPGLGGREDTQY | 0.80055416 |
| AVAALAHS | ASSLPGLGGREDTQY | 0.82645905 |
| GGVTVMATP | ASSLPGLGGREDTQY | 0.8764509 |
| AGLAANVRD | ASSLPGLGGREDTQY | 0.8614977 |
| TNRPDLIDPALLRPG | ASSLPGLGGREDTQY | 0.62687904 |
| LANLGVTAL | ASSLPGLGGREDTQY | 0.866967 |
| TGERIEL | ASSLPGLGGREDTQY | 0.90690756 |
| NGEELAA | ASSLPGLGGREDTQY | 0.7755406 |
| VLSLVWG | ASSLPGLGGREDTQY | 0.88937557 |
| IGYALWL | ASSLPGLGGREDTQY | 0.8653599 |
| LSLLLAPL | ASSLPGLGGREDTQY | 0.87055576 |
| TLDEALK | ASSLPGLGGREDTQY | 0.67472357 |
| TDGAGTAEGG | ASSLPGLGGREDTQY | 0.74060404 |
| NAPGGAGGNGG | ASSLPGLGGREDTQY | 0.84317315 |
| VSVFMGMGGNF | ASSLPGLGGREDTQY | 0.654865 |
| LASYLAAL | ASSLPGLGGREDTQY | 0.7914646 |
| LVGGTPAL | ASSLPGLGGREDTQY | 0.73749447 |
| VRNADVL | ASSLPGLGGREDTQY | 0.9166935 |
| TCSSPFLVSAM | ASSLPGLGGREDTQY | 0.8708314 |
| VHITDAE | ASSLPGLGGREDTQY | 0.8694571 |
| EYAAPSF | ASSLPGLGGREDTQY | 0.6699803 |
| VVDAFHGGL | ASSLPGLGGREDTQY | 0.8571998 |
| ASVVDPQ | ASSLPGLGGREDTQY | 0.8652883 |
| SYNEFGGLS | ASSLPGLGGREDTQY | 0.39192078 |
| DPDGSVY | ASSLPGLGGREDTQY | 0.8965782 |
| MVSTIHATE | ASSLPGLGGREDTQY | 0.67243624 |
| LAAGVADPV | ASSLPGLGGREDTQY | 0.79462445 |
| TKEQIAA | ASSLPGLGGREDTQY | 0.8685614 |
| SLASPAAL | ASSLPGLGGREDTQY | 0.9070854 |
| IGLVFYGV | ASSLPGLGGREDTQY | 0.7904379 |
| FLLDASNL | ASSLPGLGGREDTQY | 0.83504665 |
| YQVGNFP | ASSLPGLGGREDTQY | 0.5875321 |
| ESLLSQST | ASSLPGLGGREDTQY | 0.89438134 |
| LFTVGFAAVG | ASSLPGLGGREDTQY | 0.8896413 |
| ATPPTPT | ASSLPGLGGREDTQY | 0.8785012 |
| TDLATLG | ASSLPGLGGREDTQY | 0.9109022 |
| SAGGTGR | ASSLPGLGGREDTQY | 0.85523456 |
| GFSNSGS | ASSLPGLGGREDTQY | 0.36926496 |

| | | |
|---|---|---|
| GGNGGSGGTSV | ASSLPGLGGREDTQY | 0.82582533 |
| GTSVATGGAGN | ASSLPGLGGREDTQY | 0.88667995 |
| NGGNGGSGGTS | ASSLPGLGGREDTQY | 0.89322704 |
| AVVKFPH | ASSLPGLGGREDTQY | 0.90140486 |
| TPPPPPP | ASSLPGLGGREDTQY | 0.87591916 |
| AAPAAAAAPA | ASSLPGLGGREDTQY | 0.77397615 |
| IALNEDL | ASSLPGLGGREDTQY | 0.8987859 |
| IVNVSSM | ASSLPGLGGREDTQY | 0.9140991 |
| TGIPPAPR | ASSLPGLGGREDTQY | 0.88485813 |
| FDLGGGTF | ASSLPGLGGREDTQY | 0.7719636 |
| FDLGGGTFDVS | ASSLPGLGGREDTQY | 0.69608456 |
| NEPTAAA | ASSLPGLGGREDTQY | 0.84066474 |
| NEPTAAALA | ASSLPGLGGREDTQY | 0.89238214 |
| GSIINISSI | ASSLPGLGGREDTQY | 0.62647706 |
| VAPGNAGTA | ASSLPGLGGREDTQY | 0.83888817 |
| TGGGASLE | ASSLPGLGGREDTQY | 0.84966314 |
| VAVYNHYKRI | ASSLPGLGGREDTQY | 0.3291133 |
| TDPAPATT | ASSLPGLGGREDTQY | 0.8531619 |
| TNRPDILDPALLRPG | ASSLPGLGGREDTQY | 0.89843774 |
| LTTTTTS | ASSLPGLGGREDTQY | 0.8331927 |
| PTGSGKTL | ASSLPGLGGREDTQY | 0.71572936 |
| TGSGKTL | ASSLPGLGGREDTQY | 0.88843495 |
| TIGHVDHGK | ASSLPGLGGREDTQY | 0.9134455 |
| TIGHVDHGKTT | ASSLPGLGGREDTQY | 0.9042731 |
| GHVDHGKTT | ASSLPGLGGREDTQY | 0.34445798 |
| AQFEHTLL | ASSLPGLGGREDTQY | 0.23612116 |
| IVNQDEL | ASSLPGLGGREDTQY | 0.7897396 |
| TNGSQFF | ASSLPGLGGREDTQY | 0.8945677 |
| TGKGIVE | ASSLPGLGGREDTQY | 0.7537579 |
| LISNASDA | ASSLPGLGGREDTQY | 0.75462174 |
| AELRTLGELTKIAKAHGAQ | ASSLPGLGGREDTQY | 0.86072826 |
| APNAVAAAEPSW | ASSLPGLGGREDTQY | 0.88956606 |
| APQVAGAATAVL | ASSLPGLGGREDTQY | 0.645457 |
| ARRDPAAAAAILPTDA | ASSLPGLGGREDTQY | 0.5131774 |
| DAAARDVLAEAGLAEYF | ASSLPGLGGREDTQY | 0.8516935 |
| DDQIVLSAKANLAKVEQ | ASSLPGLGGREDTQY | 0.8768991 |
| DELLPDTGEDSDEDGH | ASSLPGLGGREDTQY | 0.65096927 |
| ELPEAPANRATVPQD | ASSLPGLGGREDTQY | 0.8140781 |
| FGQSKIERVTGIG | ASSLPGLGGREDTQY | 0.7594752 |
| GAGGTGSTAGGAGGAGGAGGLY | ASSLPGLGGREDTQY | 0.79587597 |
| GHGGTGAAGGNGGTGGAGGSGI | ASSLPGLGGREDTQY | 0.8422301 |
| GNGGNGGAGGEGGPGGAGGAGA | ASSLPGLGGREDTQY | 0.5948215 |
| GSTAGNGGNGGNATNGTIAGQP | ASSLPGLGGREDTQY | 0.8873575 |
| GTGGDGGNGGNGGTGG | ASSLPGLGGREDTQY | 0.5873839 |
| GVRRPAGIPADAPV | ASSLPGLGGREDTQY | 0.8911691 |
| GYRQVTDAHLLTLARR | ASSLPGLGGREDTQY | 0.847617 |
| IAKMGQDGHDRG | ASSLPGLGGREDTQY | 0.86193955 |
| KAEPSIEIQYT | ASSLPGLGGREDTQY | 0.7715006 |
| KPVRIRMAADEDTVAVT | ASSLPGLGGREDTQY | 0.6951529 |
| MPPRIAGMRLLVIKPEP | ASSLPGLGGREDTQY | 0.5579052 |
| MKFYAVALFRSRGVPPTR | ASSLPGLGGREDTQY | 0.4775716 |

| | | |
|---|---|---|
| NAGGFGILGAVAHSP | ASSLPGLGGREDTQY | 0.8908913 |
| NEGREITLAMAEM | ASSLPGLGGREDTQY | 0.40494508 |
| NELVEWSHPLGHRWRWE | ASSLPGLGGREDTQY | 0.89737654 |
| NLGAGGTGGNGGNPG | ASSLPGLGGREDTQY | 0.86771375 |
| NTEWPKWETVMAADPVD | ASSLPGLGGREDTQY | 0.5771386 |
| PASAAPAETQP | ASSLPGLGGREDTQY | 0.868785 |
| QQSSDGKNGVRAAEITGE | ASSLPGLGGREDTQY | 0.6888176 |
| RAPDAVTTSWWKE | ASSLPGLGGREDTQY | 0.79736197 |
| SGFGNVGAHTSGWFNQ | ASSLPGLGGREDTQY | 0.88691175 |
| SGVAGAVGQAASVGGLKV | ASSLPGLGGREDTQY | 0.8755233 |
| SNGGAGTGSAGGNGGT | ASSLPGLGGREDTQY | 0.6254834 |
| TGGAGGEGGAAG | ASSLPGLGGREDTQY | 0.8648554 |
| TGGDGGAAGAGGAPGHG | ASSLPGLGGREDTQY | 0.8480642 |
| TMPPLLAAIANAGRHGV | ASSLPGLGGREDTQY | 0.6428832 |
| TSSTAGNSGGA | ASSLPGLGGREDTQY | 0.82884943 |
| VPATVPVGILC | ASSLPGLGGREDTQY | 0.8822042 |
| VQRPDSGGVVLTGRIS | ASSLPGLGGREDTQY | 0.85916454 |
| WTPNPVPATKGVPAGLVQAT | ASSLPGLGGREDTQY | 0.6183885 |
| YALKRALAKFSDRFDVVII | ASSLPGLGGREDTQY | 0.8661319 |
| YLIGSVIV | ASSLPGLGGREDTQY | 0.69807845 |
| YGQLVAVID | ASSLPGLGGREDTQY | 0.8465286 |
| VTVGGMTLPA | ASSLPGLGGREDTQY | 0.81557274 |
| VLALPAV | ASSLPGLGGREDTQY | 0.7571348 |
| VIPPTLNYET | ASSLPGLGGREDTQY | 0.8818846 |
| VGMGGMHPSSG | ASSLPGLGGREDTQY | 0.5938424 |
| VGLAGGAATA | ASSLPGLGGREDTQY | 0.71266806 |
| VGGAGGAGGSGP | ASSLPGLGGREDTQY | 0.80935955 |
| VFAAFMV | ASSLPGLGGREDTQY | 0.8039542 |
| TLYDKLR | ASSLPGLGGREDTQY | 0.8823508 |
| TLAVFTVL | ASSLPGLGGREDTQY | 0.6752912 |
| TGSVQPARV | ASSLPGLGGREDTQY | 0.84217703 |
| TGGGAGSGGT | ASSLPGLGGREDTQY | 0.7566886 |
| SHLETSL | ASSLPGLGGREDTQY | 0.64231336 |
| SFTTGMT | ASSLPGLGGREDTQY | 0.8425206 |
| QVEVAKRMGVRQPTVS | ASSLPGLGGREDTQY | 0.7689539 |
| QGSGGPALAM | ASSLPGLGGREDTQY | 0.6523477 |
| MLAPSNP | ASSLPGLGGREDTQY | 0.8644327 |
| LVNNAAI | ASSLPGLGGREDTQY | 0.62062097 |
| LTGEINTLA | ASSLPGLGGREDTQY | 0.86246943 |
| LSIFGPV | ASSLPGLGGREDTQY | 0.9058633 |
| LLALLANL | ASSLPGLGGREDTQY | 0.78702104 |
| LISNASDALDKLR | ASSLPGLGGREDTQY | 0.8847151 |
| LINEAVP | ASSLPGLGGREDTQY | 0.8624161 |
| LGYAFSGL | ASSLPGLGGREDTQY | 0.8631079 |
| LDFVAVAL | ASSLPGLGGREDTQY | 0.75109565 |
| LAQPGAE | ASSLPGLGGREDTQY | 0.6223843 |
| ISNLGAVTGF | ASSLPGLGGREDTQY | 0.89393723 |
| HQEDPGRI | ASSLPGLGGREDTQY | 0.86646795 |
| GTPGEPTP | ASSLPGLGGREDTQY | 0.8823646 |
| GFFNSGS | ASSLPGLGGREDTQY | 0.8685813 |
| GAGGGGLTAGAN | ASSLPGLGGREDTQY | 0.8783609 |

| | | |
|---|---|---|
| FVDALVNM | ASSLPGLGGREDTQY | 0.87915945 |
| FLTVAPEL | ASSLPGLGGREDTQY | 0.7531787 |
| FETLTAL | ASSLPGLGGREDTQY | 0.57609355 |
| DVINPIEL | ASSLPGLGGREDTQY | 0.7545802 |
| APVSGAPV | ASSLPGLGGREDTQY | 0.7917993 |
| ALVGGLAW | ASSLPGLGGREDTQY | 0.72631764 |
| ALAAAAATAI | ASSLPGLGGREDTQY | 0.39729625 |
| AFGLGLLL | ASSLPGLGGREDTQY | 0.8809051 |
| AFFAAASL | ASSLPGLGGREDTQY | 0.9130406 |
| AAAALPAFSPP | ASSLPGLGGREDTQY | 0.65226406 |
| VVLAPAL | ASSLPGLGGREDTQY | 0.9117575 |
| TIGHVDHGKTTL | ASSLPGLGGREDTQY | 0.5365441 |
| LISNASDALDK | ASSLPGLGGREDTQY | 0.87875044 |
| LISNASDAL | ASSLPGLGGREDTQY | 0.88705885 |
| ISNASDALDK | ASSLPGLGGREDTQY | 0.662795 |
| NQAETLVYQTE | ASSLPGLGGREDTQY | 0.85927117 |
| TGGTGGTAGAG | ASSLPGLGGREDTQY | 0.665264 |
| TGSTAGGAGGA | ASSLPGLGGREDTQY | 0.7515477 |
| DPLAVPVAAGPV | ASSLPGLGGREDTQY | 0.8964124 |
| GGTGGTPGNGGN | ASSLPGLGGREDTQY | 0.74577355 |
| PAGLAGVDPATL | ASSLPGLGGREDTQY | 0.84879124 |
| GNNLVVEAPHAGASV | ASSLPGLGGREDTQY | 0.80191255 |
| KVTRIMSASTLEQPA | ASSLPGLGGREDTQY | 0.66611993 |
| LGFALPIALPRRTVV | ASSLPGLGGREDTQY | 0.8517371 |
| TRLGNLTARKLVVLV | ASSLPGLGGREDTQY | 0.887316 |
| WSQQRRLIYFPSAGP | ASSLPGLGGREDTQY | 0.8043622 |
| GPNGNPDPMAAAVDIR | ASSLPGLGGREDTQY | 0.91162455 |
| TQRCVSVADLRAAASR | ASSLPGLGGREDTQY | 0.6520098 |
| QFVLDVAEHPVEVLTETV | ASSLPGLGGREDTQY | 0.9026928 |
| GGAGGTGGAGGNGGTQAGDG | ASSLPGLGGREDTQY | 0.8879126 |
| PLLATLRAIAVDVDAKPAQV | ASSLPGLGGREDTQY | 0.83103335 |
| APKFRTGGYMIDFWGVGYQVAK | ASSLPGLGGREDTQY | 0.7817012 |
| GAGGAGGAGGAAGGPGGTGGTG | ASSLPGLGGREDTQY | 0.8808579 |
| IGALGLLATDVAPAGSVAFGVA | ASSLPGLGGREDTQY | 0.7901228 |
| AGGAGGIGGNGGAGGAGGVLMG | ASSLPGLGGREDTQY | 0.8795775 |
| ADELIGLL | ASSLPGLGGREDTQY | 0.90007997 |
| ADRALKA | ASSLPGLGGREDTQY | 0.58373284 |
| AGAVGQVGGAGGA | ASSLPGLGGREDTQY | 0.659141 |
| AGGGGGLGGAP | ASSLPGLGGREDTQY | 0.63343227 |
| AGGLLYGNGGAG | ASSLPGLGGREDTQY | 0.62612844 |
| AGLESIGL | ASSLPGLGGREDTQY | 0.59875464 |
| AGLKTGII | ASSLPGLGGREDTQY | 0.53944755 |
| ASLAALLL | ASSLPGLGGREDTQY | 0.62782687 |
| ASVINISSI | ASSLPGLGGREDTQY | 0.59150755 |
| ATPPGPA | ASSLPGLGGREDTQY | 0.6542454 |
| ATVASALV | ASSLPGLGGREDTQY | 0.6572106 |
| ATVASLIL | ASSLPGLGGREDTQY | 0.5884677 |
| AVNGPVLIRH | ASSLPGLGGREDTQY | 0.662562 |
| DGAAGAVGI | ASSLPGLGGREDTQY | 0.6584983 |
| DPLLHGI | ASSLPGLGGREDTQY | 0.52480876 |
| ELGAALGAL | ASSLPGLGGREDTQY | 0.609311 |

| | | |
|---|---|---|
| ELGRTQAV | ASSLPGLGGREDTQY | 0.6001482 |
| FVAGGYLS | ASSLPGLGGREDTQY | 0.61733454 |
| GASGGAGGAGGW | ASSLPGLGGREDTQY | 0.4782542 |
| GGAGAGGDGGAGGI | ASSLPGLGGREDTQY | 0.6499557 |
| GTGHNGSQPGMGGNG | ASSLPGLGGREDTQY | 0.58177817 |
| GYAAASASAS | ASSLPGLGGREDTQY | 0.34456182 |
| HQVDVAAVQQG | ASSLPGLGGREDTQY | 0.63440627 |
| IGAAIAHT | ASSLPGLGGREDTQY | 0.62590975 |
| IGMIEVAQL | ASSLPGLGGREDTQY | 0.6294185 |
| IRLAPEHTSAMD | ASSLPGLGGREDTQY | 0.64950776 |
| IYNVSVL | ASSLPGLGGREDTQY | 0.54714084 |
| KQGFLGM | ASSLPGLGGREDTQY | 0.6098058 |
| LESVITQL | ASSLPGLGGREDTQY | 0.65187603 |
| LGLGALVVV | ASSLPGLGGREDTQY | 0.65909857 |
| LIGNGGNGGSGGI | ASSLPGLGGREDTQY | 0.582063 |
| LNAINAPTL | ASSLPGLGGREDTQY | 0.6303263 |
| LQLLNTGV | ASSLPGLGGREDTQY | 0.6584451 |
| LQTSIEF | ASSLPGLGGREDTQY | 0.6385884 |
| MSFAVSEL | ASSLPGLGGREDTQY | 0.6567131 |
| NTGALGL | ASSLPGLGGREDTQY | 0.66259724 |
| PPGAGGAGGAGTT | ASSLPGLGGREDTQY | 0.5933143 |
| QLVGLGI | ASSLPGLGGREDTQY | 0.6589817 |
| RFLEQVHQL | ASSLPGLGGREDTQY | 0.66244507 |
| RGVGYVLR | ASSLPGLGGREDTQY | 0.6232369 |
| SATGVFAGVI | ASSLPGLGGREDTQY | 0.6237177 |
| SLPAITLP | ASSLPGLGGREDTQY | 0.56402844 |
| STALAFTV | ASSLPGLGGREDTQY | 0.619091 |
| SVLTVGLGK | ASSLPGLGGREDTQY | 0.49320692 |
| TALAGAGSGI | ASSLPGLGGREDTQY | 0.6324971 |
| TGLPFTGI | ASSLPGLGGREDTQY | 0.6611018 |
| VAEHVTV | ASSLPGLGGREDTQY | 0.6012192 |
| VDIVLNSL | ASSLPGLGGREDTQY | 0.5440041 |
| VLTGGGAVL | ASSLPGLGGREDTQY | 0.56256133 |
| VNVGLAYGI | ASSLPGLGGREDTQY | 0.6147806 |
| VSSAAVGAV | ASSLPGLGGREDTQY | 0.5987607 |
| VTSVYEAGV | ASSLPGLGGREDTQY | 0.42158124 |
| WVALDRGA | ASSLPGLGGREDTQY | 0.6093675 |
| YLVGSVVI | ASSLPGLGGREDTQY | 0.6584945 |
| GPPGTGKTLL | ASSLPGLGGREDTQY | 0.6572586 |
| GVGMTKF | ASSLPGLGGREDTQY | 0.49419874 |
| IGIGGSDL | ASSLPGLGGREDTQY | 0.66154206 |
| IGTLLAEL | ASSLPGLGGREDTQY | 0.6529153 |
| IYTSGTT | ASSLPGLGGREDTQY | 0.66045743 |
| LSFPAGV | ASSLPGLGGREDTQY | 0.6276774 |
| LYTSGTT | ASSLPGLGGREDTQY | 0.5739751 |
| VGAFPFGF | ASSLPGLGGREDTQY | 0.65234804 |
| VGAGTVEF | ASSLPGLGGREDTQY | 0.5905045 |
| VGKSTLLL | ASSLPGLGGREDTQY | 0.56200206 |
| AALAHAGGGIV | ASSLPGLGGREDTQY | 0.65094006 |
| AALVAVSLL | ASSLPGLGGREDTQY | 0.6355647 |
| AAVSRAVAAL | ASSLPGLGGREDTQY | 0.5914384 |

| | | |
|---|---|---|
| AGGSSGIGGF | ASSLPGLGGREDTQY | 0.592753 |
| AGLTVANANG | ASSLPGLGGREDTQY | 0.6579551 |
| AGLVGAVL | ASSLPGLGGREDTQY | 0.55944073 |
| ALSQAFTV | ASSLPGLGGREDTQY | 0.6033374 |
| ALTFSVSL | ASSLPGLGGREDTQY | 0.5666749 |
| ALTGAALVAL | ASSLPGLGGREDTQY | 0.6224817 |
| ASFSIGAGVL | ASSLPGLGGREDTQY | 0.65723836 |
| DGSVYKI | ASSLPGLGGREDTQY | 0.64722097 |
| EQLAEIV | ASSLPGLGGREDTQY | 0.600991 |
| FAVSELTVM | ASSLPGLGGREDTQY | 0.635037 |
| GAAPLAAGL | ASSLPGLGGREDTQY | 0.64441466 |
| GADGGDGGSGGAGG | ASSLPGLGGREDTQY | 0.63691294 |
| GFGLDTGGAGG | ASSLPGLGGREDTQY | 0.64587045 |
| GLASGGLL | ASSLPGLGGREDTQY | 0.5760838 |
| GRIAGAAVLL | ASSLPGLGGREDTQY | 0.65758574 |
| GSAGSLNTSF | ASSLPGLGGREDTQY | 0.6411843 |
| GTGQIGI | ASSLPGLGGREDTQY | 0.6560685 |
| GTLIALIL | ASSLPGLGGREDTQY | 0.6587467 |
| IAATMGFVVG | ASSLPGLGGREDTQY | 0.6479623 |
| IFAQALL | ASSLPGLGGREDTQY | 0.62649655 |
| IGAAVALGF | ASSLPGLGGREDTQY | 0.4487623 |
| IGLTPLAL | ASSLPGLGGREDTQY | 0.6073599 |
| ILGSNTSTL | ASSLPGLGGREDTQY | 0.6325975 |
| IVFSLTVL | ASSLPGLGGREDTQY | 0.6286355 |
| LASKATGF | ASSLPGLGGREDTQY | 0.6521561 |
| LASVALIL | ASSLPGLGGREDTQY | 0.6508845 |
| LFAVGLTL | ASSLPGLGGREDTQY | 0.59277505 |
| LFNGGIGTYI | ASSLPGLGGREDTQY | 0.6570553 |
| LTLGAGII | ASSLPGLGGREDTQY | 0.6547562 |
| LVAAALGDI | ASSLPGLGGREDTQY | 0.5018443 |
| NTAFVQLGI | ASSLPGLGGREDTQY | 0.62196946 |
| QLFSGAEL | ASSLPGLGGREDTQY | 0.66634 |
| SAATFITL | ASSLPGLGGREDTQY | 0.62804806 |
| SNYGLANLGV | ASSLPGLGGREDTQY | 0.6000875 |
| SYGGAYAVM | ASSLPGLGGREDTQY | 0.5912216 |
| TTGVGGLTLG | ASSLPGLGGREDTQY | 0.66784143 |
| TTQTIYI | ASSLPGLGGREDTQY | 0.57508504 |
| VAAGAGLGAVI | ASSLPGLGGREDTQY | 0.66206384 |
| VAAVVTLL | ASSLPGLGGREDTQY | 0.40508986 |
| VGFLGGLI | ASSLPGLGGREDTQY | 0.5845337 |
| VGVIPILL | ASSLPGLGGREDTQY | 0.5583793 |
| VIFASSLI | ASSLPGLGGREDTQY | 0.44400713 |
| VKGGDGGAAAG | ASSLPGLGGREDTQY | 0.6544843 |
| VTKTFTNI | ASSLPGLGGREDTQY | 0.60768455 |
| VVAAAILL | ASSLPGLGGREDTQY | 0.64645433 |
| YAALDVEL | ASSLPGLGGREDTQY | 0.62814426 |
| LASVALLL | ASSLPGLGGREDTQY | 0.6603826 |
| LDAVHRV | ASSLPGLGGREDTQY | 0.6202406 |
| QGLSSSIF | ASSLPGLGGREDTQY | 0.63850904 |
| RGTFANIRL | ASSLPGLGGREDTQY | 0.57535404 |
| RQLWWGHRI | ASSLPGLGGREDTQY | 0.6516588 |

| | | |
|---|---|---|
| YGPPGTGKTLL | ASSLPGLGGREDTQY | 0.6621294 |
| AALVGAPANVA | ASSLPGLGGREDTQY | 0.6473705 |
| AATFRRTE | ASSLPGLGGREDTQY | 0.56453544 |
| AGALGGGI | ASSLPGLGGREDTQY | 0.5891764 |
| AGGAGGAGGIGAG | ASSLPGLGGREDTQY | 0.63839823 |
| AGSGALGL | ASSLPGLGGREDTQY | 0.42637607 |
| AQIADQI | ASSLPGLGGREDTQY | 0.37124518 |
| ASAVAVTYL | ASSLPGLGGREDTQY | 0.6192338 |
| ASTVLGGTGG | ASSLPGLGGREDTQY | 0.579026 |
| AVGGEPAK | ASSLPGLGGREDTQY | 0.6550122 |
| CEQGTFGGTC | ASSLPGLGGREDTQY | 0.65061325 |
| DEGKAPAG | ASSLPGLGGREDTQY | 0.60569257 |
| DTVVFPAL | ASSLPGLGGREDTQY | 0.64522636 |
| FEGITMAG | ASSLPGLGGREDTQY | 0.63257235 |
| FPLVVQD | ASSLPGLGGREDTQY | 0.3138726 |
| GAFTTPQ | ASSLPGLGGREDTQY | 0.6036601 |
| GFGGFGTAGGVGGA | ASSLPGLGGREDTQY | 0.541518 |
| GFLAQFL | ASSLPGLGGREDTQY | 0.6674249 |
| GGAGGAGGNAGVA | ASSLPGLGGREDTQY | 0.61022305 |
| GGVGTAGAGGAGGAGG | ASSLPGLGGREDTQY | 0.5711936 |
| GNGGGGDAGNAGSGGN | ASSLPGLGGREDTQY | 0.6212809 |
| GTGGAGGAGNDAGS | ASSLPGLGGREDTQY | 0.6269136 |
| GTTNTGWFNS | ASSLPGLGGREDTQY | 0.61225444 |
| IFMGPLFTL | ASSLPGLGGREDTQY | 0.5766896 |
| ILAHAIEAI | ASSLPGLGGREDTQY | 0.38106984 |
| IRSAIREAL | ASSLPGLGGREDTQY | 0.65552825 |
| ITFAGGG | ASSLPGLGGREDTQY | 0.5542922 |
| ITGMSGHQEV | ASSLPGLGGREDTQY | 0.599728 |
| IVAGLACTL | ASSLPGLGGREDTQY | 0.43840575 |
| IVGGGLAGA | ASSLPGLGGREDTQY | 0.57098347 |
| IVTGVLNI | ASSLPGLGGREDTQY | 0.5190562 |
| LAQQGCI | ASSLPGLGGREDTQY | 0.65127516 |
| LCLHVEN | ASSLPGLGGREDTQY | 0.60171753 |
| LDAISDGS | ASSLPGLGGREDTQY | 0.6017286 |
| LGGFAAML | ASSLPGLGGREDTQY | 0.6662276 |
| LGGKNPMI | ASSLPGLGGREDTQY | 0.6315361 |
| LGPKGVAAIL | ASSLPGLGGREDTQY | 0.5971281 |
| LMVGVSI | ASSLPGLGGREDTQY | 0.6068165 |
| LVGAAFVL | ASSLPGLGGREDTQY | 0.5589019 |
| LVLAVAPL | ASSLPGLGGREDTQY | 0.60204387 |
| MAAIAVIDAA | ASSLPGLGGREDTQY | 0.6139166 |
| NNGINAELQ | ASSLPGLGGREDTQY | 0.6646568 |
| RGEPLVLVHG | ASSLPGLGGREDTQY | 0.6052603 |
| RGGAGGAAGLI | ASSLPGLGGREDTQY | 0.57405955 |
| RSYRMHPAV | ASSLPGLGGREDTQY | 0.62464386 |
| SAAHLLAGQLA | ASSLPGLGGREDTQY | 0.6630049 |
| SAAVGSY | ASSLPGLGGREDTQY | 0.6660075 |
| SPHAVLHRI | ASSLPGLGGREDTQY | 0.5802692 |
| SQIAKVL | ASSLPGLGGREDTQY | 0.6097466 |
| TALGHLVP | ASSLPGLGGREDTQY | 0.5930451 |
| TAPTTSGAA | ASSLPGLGGREDTQY | 0.6158276 |

| | | |
|---|---|---|
| TDGDDGNGGSPGT | ASSLPGLGGREDTQY | 0.59283334 |
| TDGLGEN | ASSLPGLGGREDTQY | 0.57486176 |
| TGGPPLPGQA | ASSLPGLGGREDTQY | 0.6479757 |
| TIPAITI | ASSLPGLGGREDTQY | 0.6729049 |
| TIPGFTI | ASSLPGLGGREDTQY | 0.6142272 |
| TSIASGALA | ASSLPGLGGREDTQY | 0.5201763 |
| TVTLPGAAI | ASSLPGLGGREDTQY | 0.6114151 |
| VASGLSAALP | ASSLPGLGGREDTQY | 0.5778872 |
| VGAGVIGI | ASSLPGLGGREDTQY | 0.58805954 |
| VGPKRFGV | ASSLPGLGGREDTQY | 0.63387066 |
| VNVAFPD | ASSLPGLGGREDTQY | 0.49040005 |
| VSMTNSGVS | ASSLPGLGGREDTQY | 0.6170309 |
| VTDVLGDLSV | ASSLPGLGGREDTQY | 0.6669182 |
| VVVVGAGFAGL | ASSLPGLGGREDTQY | 0.5936371 |
| VYTVGTVV | ASSLPGLGGREDTQY | 0.6343614 |
| WTTFSGP | ASSLPGLGGREDTQY | 0.6546917 |
| YDELVEAF | ASSLPGLGGREDTQY | 0.63791525 |
| YGTGGAGGNGG | ASSLPGLGGREDTQY | 0.651734 |
| YTNVMAV | ASSLPGLGGREDTQY | 0.6601426 |
| FLEQVHQL | ASSLPGLGGREDTQY | 0.6363428 |
| GLSLLAVAV | ASSLPGLGGREDTQY | 0.6215476 |
| LEQVHQL | ASSLPGLGGREDTQY | 0.6395215 |
| LNAAGAI | ASSLPGLGGREDTQY | 0.6192084 |
| PAVHASL | ASSLPGLGGREDTQY | 0.6023773 |
| TELEDDA | ASSLPGLGGREDTQY | 0.5628455 |
| TPPGPPN | ASSLPGLGGREDTQY | 0.65607786 |
| TPPPPPPP | ASSLPGLGGREDTQY | 0.6107623 |
| TQSPALI | ASSLRDSHNSPLY | 0.65834206 |
| EFAAAFPQ | ASSLRDSHNSPLY | 0.51141244 |
| LLSGNLDVL | ASSLRDSHNSPLY | 0.58268684 |
| GPANLGNYNI | ASSLRDSHNSPLY | 0.64859915 |
| TLGPTLP | ASSLRDSHNSPLY | 0.323246 |
| VYAYSSCCK | ASSLRDSHNSPLY | 0.36914837 |
| APAPAPAPNI | ASSLRDSHNSPLY | 0.62275493 |
| PGGPVPIP | ASSLRDSHNSPLY | 0.65805274 |
| FLTVALEP | ASSLRDSHNSPLY | 0.33910394 |
| SQYFGANL | ASSLRDSHNSPLY | 0.59913003 |
| GGVKPAA | ASSLRDSHNSPLY | 0.5276824 |
| GGTGGAGGAGGW | ASSLRDSHNSPLY | 0.6492974 |
| GVGGVAGLGGAGG | ASSLRDSHNSPLY | 0.6398139 |
| VLLVLATTI | ASSLRDSHNSPLY | 0.53038603 |
| IGIFAAML | ASSLRDSHNSPLY | 0.649304 |
| AVGDLID | ASSLRDSHNSPLY | 0.47943065 |
| NISPLPL | ASSLRDSHNSPLY | 0.56827843 |
| VTAVFVAI | ASSLRDSHNSPLY | 0.6562611 |
| SGDDNDAAL | ASSLRDSHNSPLY | 0.6632432 |
| AVAALAHS | ASSLRDSHNSPLY | 0.56428325 |
| GGVTVMATP | ASSLRDSHNSPLY | 0.64934266 |
| AGLAANVRD | ASSLRDSHNSPLY | 0.5890996 |
| TNRPDLIDPALLRPG | ASSLRDSHNSPLY | 0.6673351 |
| LANLGVTAL | ASSLRDSHNSPLY | 0.60282123 |

| | | |
|---|---|---|
| TGERIEL | ASSLRDSHNSPLY | 0.49580684 |
| NGEELAA | ASSLRDSHNSPLY | 0.65269077 |
| VLSLVWG | ASSLRDSHNSPLY | 0.6407587 |
| IGYALWL | ASSLRDSHNSPLY | 0.64178383 |
| LSLLLAPL | ASSLRDSHNSPLY | 0.65097153 |
| TLDEALK | ASSLRDSHNSPLY | 0.5991012 |
| TDGAGTAEGG | ASSLRDSHNSPLY | 0.6041931 |
| NAPGGAGGNGG | ASSLRDSHNSPLY | 0.6595879 |
| VSVFMGMGGNF | ASSLRDSHNSPLY | 0.6530632 |
| LASYLAAL | ASSLRDSHNSPLY | 0.58893144 |
| LVGGTPAL | ASSLRDSHNSPLY | 0.6595996 |
| VRNADVL | ASSLRDSHNSPLY | 0.61629325 |
| TCSSPFLVSAM | ASSLRDSHNSPLY | 0.5825596 |
| VHITDAE | ASSLRDSHNSPLY | 0.66547227 |
| EYAAPSF | ASSLRDSHNSPLY | 0.6339535 |
| VVDAFHGGL | ASSLRDSHNSPLY | 0.6445018 |
| ASVVDPQ | ASSLRDSHNSPLY | 0.65451884 |
| SYNEFGGLS | ASSLRDSHNSPLY | 0.6625085 |
| DPDGSVY | ASSLRDSHNSPLY | 0.56939673 |
| MVSTIHATE | ASSLRDSHNSPLY | 0.63821566 |
| LAAGVADPV | ASSLRDSHNSPLY | 0.6128134 |
| TKEQIAA | ASSLRDSHNSPLY | 0.6557081 |
| SLASPAAL | ASSLRDSHNSPLY | 0.6294006 |
| IGLVFYGV | ASSLRDSHNSPLY | 0.59081495 |
| FLLDASNL | ASSLRDSHNSPLY | 0.64360464 |
| YQVGNFP | ASSLRDSHNSPLY | 0.6590756 |
| ESLLSQST | ASSLRDSHNSPLY | 0.63328063 |
| LFTVGFAAVG | ASSLRDSHNSPLY | 0.64297324 |
| ATPPTPT | ASSLRDSHNSPLY | 0.6599432 |
| TDLATLG | ASSLRDSHNSPLY | 0.57373905 |
| SAGGTGR | ASSLRDSHNSPLY | 0.6186842 |
| GFSNSGS | ASSLRDSHNSPLY | 0.65041316 |
| GGNGGSGGTSV | ASSLRDSHNSPLY | 0.6560607 |
| GTSVATGGAGN | ASSLRDSHNSPLY | 0.61831474 |
| NGGNGGSGGTS | ASSLRDSHNSPLY | 0.65841484 |
| AVVKFPH | ASSLRDSHNSPLY | 0.5812917 |
| TPPPPPP | ASSLRDSHNSPLY | 0.65396976 |
| AAPAAAAAPA | ASSLRDSHNSPLY | 0.57246375 |
| IALNEDL | ASSLRDSHNSPLY | 0.637589 |
| IVNVSSM | ASSLRDSHNSPLY | 0.5870112 |
| TGIPPAPR | ASSLRDSHNSPLY | 0.64881456 |
| FDLGGGTF | ASSLRDSHNSPLY | 0.6485268 |
| FDLGGGTFDVS | ASSLRDSHNSPLY | 0.6527884 |
| NEPTAAA | ASSLRDSHNSPLY | 0.3684499 |
| NEPTAAALA | ASSLRDSHNSPLY | 0.648038 |
| GSIINISSI | ASSLRDSHNSPLY | 0.65749776 |
| VAPGNAGTA | ASSLRDSHNSPLY | 0.66699517 |
| TGGGASLE | ASSLRDSHNSPLY | 0.61164284 |
| VAVYNHYKRI | ASSLRDSHNSPLY | 0.63691926 |
| TDPAPATT | ASSLRDSHNSPLY | 0.6137461 |
| TNRPDILDPALLRPG | ASSLRDSHNSPLY | 0.616071 |

| | | |
|---|---|---|
| LTTTTTS | ASSLRDSHNSPLY | 0.53780663 |
| PTGSGKTL | ASSLRDSHNSPLY | 0.59915596 |
| TGSGKTL | ASSLRDSHNSPLY | 0.58739245 |
| TIGHVDHGK | ASSLRDSHNSPLY | 0.5635677 |
| TIGHVDHGKTT | ASSLRDSHNSPLY | 0.64695984 |
| GHVDHGKTT | ASSLRDSHNSPLY | 0.65513784 |
| AQFEHTLL | ASSLRDSHNSPLY | 0.65551305 |
| IVNQDEL | ASSLRDSHNSPLY | 0.65855175 |
| TNGSQFF | ASSLRDSHNSPLY | 0.65773445 |
| TGKGIVE | ASSLRDSHNSPLY | 0.65663326 |
| LISNASDA | ASSLRDSHNSPLY | 0.6096068 |
| AELRTLGELTKIAKAHGAQ | ASSLRDSHNSPLY | 0.62645364 |
| APNAVAAAEPSW | ASSLRDSHNSPLY | 0.5480577 |
| APQVAGAATAVL | ASSLRDSHNSPLY | 0.617166 |
| ARRDPAAAAAILPTDA | ASSLRDSHNSPLY | 0.6158952 |
| DAAARDVLAEAGLAEYF | ASSLRDSHNSPLY | 0.63221353 |
| DDQIVLSAKANLAKVEQ | ASSLRDSHNSPLY | 0.33147854 |
| DELLPDTGEDSDEDGH | ASSLRDSHNSPLY | 0.626675 |
| ELPEAPANRATVPQD | ASSLRDSHNSPLY | 0.662945 |
| FGQSKIERVTGIG | ASSLRDSHNSPLY | 0.5701401 |
| GAGGTGSTAGGAGGAGGAGGLY | ASSLRDSHNSPLY | 0.6361896 |
| GHGGTGAAGGNGGTGGAGGSGI | ASSLRDSHNSPLY | 0.62790996 |
| GNGGNGGAGGEGGPGGAGGAGA | ASSLRDSHNSPLY | 0.5900798 |
| GSTAGNGGNGGNATNGTIAGQP | ASSLRDSHNSPLY | 0.6469118 |
| GTGGDGGNGGNGGTGG | ASSLRDSHNSPLY | 0.5780591 |
| GVRRPAGIPADAPV | ASSLRDSHNSPLY | 0.6384492 |
| GYRQVTDAHLLTLARR | ASSLRDSHNSPLY | 0.6610175 |
| IAKMGQDGHDRG | ASSLRDSHNSPLY | 0.6280131 |
| KAEPSIEIQYT | ASSLRDSHNSPLY | 0.62475014 |
| KPVRIRMAADEDTVAVT | ASSLRDSHNSPLY | 0.6195678 |
| MPPRIAGMRLLVIKPEP | ASSLRDSHNSPLY | 0.5205375 |
| MKFYAVALFRSRGVPPTR | ASSLRDSHNSPLY | 0.6389753 |
| NAGGFGILGAVAHSP | ASSLRDSHNSPLY | 0.48015362 |
| NEGREITLAMAEM | ASSLRDSHNSPLY | 0.6133839 |
| NELVEWSHPLGHRWRWE | ASSLRDSHNSPLY | 0.58417666 |
| NLGAGGTGGNGGNPG | ASSLRDSHNSPLY | 0.6376941 |
| NTEWPKWETVMAADPVD | ASSLRDSHNSPLY | 0.6275233 |
| PASAAPAETQP | ASSLRDSHNSPLY | 0.6374082 |
| QQSSDGKNGVRAAEITGE | ASSLRDSHNSPLY | 0.5601333 |
| RAPDAVTTSWWKE | ASSLRDSHNSPLY | 0.6140179 |
| SGFGNVGAHTSGWFNQ | ASSLRDSHNSPLY | 0.5850626 |
| SGVAGAVGQAASVGGLKV | ASSLRDSHNSPLY | 0.58338857 |
| SNGGAGTGSAGGNGGT | ASSLRDSHNSPLY | 0.64868784 |
| TGGAGGEGGAAG | ASSLRDSHNSPLY | 0.62326837 |
| TGGDGGAAGAGGAPGHG | ASSLRDSHNSPLY | 0.6105537 |
| TMPPLLAAIANAGRHGV | ASSLRDSHNSPLY | 0.66010344 |
| TSSTAGNSGGA | ASSLRDSHNSPLY | 0.6375339 |
| VPATVPVGILC | ASSLRDSHNSPLY | 0.5005067 |
| VQRPDSGGVVLTGRIS | ASSLRDSHNSPLY | 0.6058296 |
| WTPNPVPATKGVPAGLVQAT | ASSLRDSHNSPLY | 0.61252856 |
| YALKRALAKFSDRFDVVII | ASSLRDSHNSPLY | 0.6085371 |

| | | |
|---|---|---|
| YLIGSVIV | ASSLRDSHNSPLY | 0.35373116 |
| YGQLVAVID | ASSLRDSHNSPLY | 0.64422095 |
| VTVGGMTLPA | ASSLRDSHNSPLY | 0.584255 |
| VLALPAV | ASSLRDSHNSPLY | 0.56373227 |
| VIPPTLNYET | ASSLRDSHNSPLY | 0.60879505 |
| VGMGGMHPSSG | ASSLRDSHNSPLY | 0.64561343 |
| VGLAGGAATA | ASSLRDSHNSPLY | 0.5865182 |
| VGGAGGAGGSGP | ASSLRDSHNSPLY | 0.6431466 |
| VFAAFMV | ASSLRDSHNSPLY | 0.64340854 |
| TLYDKLR | ASSLRDSHNSPLY | 0.65669173 |
| TLAVFTVL | ASSLRDSHNSPLY | 0.66039824 |
| TGSVQPARV | ASSLRDSHNSPLY | 0.65290767 |
| TGGGAGSGGT | ASSLRDSHNSPLY | 0.60996944 |
| SHLETSL | ASSLRDSHNSPLY | 0.6447774 |
| SFTTGMT | ASSLRDSHNSPLY | 0.62129325 |
| QVEVAKRMGVRQPTVS | ASSLRDSHNSPLY | 0.47980607 |
| QGSGGPALAM | ASSLRDSHNSPLY | 0.40644205 |
| MLAPSNP | ASSLRDSHNSPLY | 0.6219384 |
| LVNNAAI | ASSLRDSHNSPLY | 0.42542467 |
| LTGEINTLA | ASSLRDSHNSPLY | 0.62771016 |
| LSIFGPV | ASSLRDSHNSPLY | 0.6633792 |
| LLALLANL | ASSLRDSHNSPLY | 0.6241876 |
| LISNASDALDKLR | ASSLRDSHNSPLY | 0.51452774 |
| LINEAVP | ASSLRDSHNSPLY | 0.60340893 |
| LGYAFSGL | ASSLRDSHNSPLY | 0.6221472 |
| LDFVAVAL | ASSLRDSHNSPLY | 0.6415995 |
| LAQPGAE | ASSLRDSHNSPLY | 0.6531393 |
| ISNLGAVTGF | ASSLRDSHNSPLY | 0.67042357 |
| HQEDPGRI | ASSLRDSHNSPLY | 0.65797436 |
| GTPGEPTP | ASSLRDSHNSPLY | 0.59046066 |
| GFFNSGS | ASSLRDSHNSPLY | 0.6385394 |
| GAGGGGLTAGAN | ASSLRDSHNSPLY | 0.5976808 |
| FVDALVNM | ASSLRDSHNSPLY | 0.6211735 |
| FLTVAPEL | ASSLRDSHNSPLY | 0.62615234 |
| FETLTAL | ASSLRDSHNSPLY | 0.63682723 |
| DVINPIEL | ASSLRDSHNSPLY | 0.63056433 |
| APVSGAPV | ASSLRDSHNSPLY | 0.60606045 |
| ALVGGLAW | ASSLRDSHNSPLY | 0.4925935 |
| ALAAAAATAI | ASSLRDSHNSPLY | 0.64751714 |
| AFGLGLLL | ASSLRDSHNSPLY | 0.6349803 |
| AFFAAASL | ASSLRDSHNSPLY | 0.42147964 |
| AAAALPAFSPP | ASSLRDSHNSPLY | 0.6030362 |
| VVLAPAL | ASSLRDSHNSPLY | 0.6645117 |
| TIGHVDHGKTTL | ASSLRDSHNSPLY | 0.6613301 |
| LISNASDALDK | ASSLRDSHNSPLY | 0.57646286 |
| LISNASDAL | ASSLRDSHNSPLY | 0.6076604 |
| ISNASDALDK | ASSLRDSHNSPLY | 0.6213909 |
| NQAETLVYQTE | ASSLRDSHNSPLY | 0.47820437 |
| TGGTGGTAGAG | ASSLRDSHNSPLY | 0.6397215 |
| TGSTAGGAGGA | ASSLRDSHNSPLY | 0.6209326 |
| DPLAVPVAAGPV | ASSLRDSHNSPLY | 0.6170349 |

| | | |
|---|---|---|
| GGTGGTPGNGGN | ASSLRDSHNSPLY | 0.56603116 |
| PAGLAGVDPATL | ASSLRDSHNSPLY | 0.62565684 |
| GNNLVVEAPHAGASV | ASSLRDSHNSPLY | 0.61124206 |
| KVTRIMSASTLEQPA | ASSLRDSHNSPLY | 0.65330374 |
| LGFALPIALPRRTVV | ASSLRDSHNSPLY | 0.34680927 |
| TRLGNLTARKLVVLV | ASSLRDSHNSPLY | 0.6583954 |
| WSQQRRLIYFPSAGP | ASSLRDSHNSPLY | 0.56420743 |
| GPNGNPDPMAAAVDIR | ASSLRDSHNSPLY | 0.56843334 |
| TQRCVSVADLRAAASR | ASSLRDSHNSPLY | 0.4319725 |
| QFVLDVAEHPVEVLTETV | ASSLRDSHNSPLY | 0.62804055 |
| GGAGGTGGAGGNGGTQAGDG | ASSLRDSHNSPLY | 0.6484889 |
| PLLATLRAIAVDVDAKPAQV | ASSLRDSHNSPLY | 0.649811 |
| APKFRTGGYMIDFWGVGYQVAK | ASSLRDSHNSPLY | 0.5338458 |
| GAGGAGGAGGAAGGPGGTGGTG | ASSLRDSHNSPLY | 0.6558082 |
| IGALGLLATDVAPAGSVAFGVA | ASSLRDSHNSPLY | 0.6620139 |
| AGGAGGIGGNGGAGGAGGVLMG | ASSLRDSHNSPLY | 0.5920613 |
| ADELIGLL | ASSLRDSHNSPLY | 0.50203615 |
| ADRALKA | ASSLRDSHNSPLY | 0.6500598 |
| AGAVGQVGGAGGA | ASSLRDSHNSPLY | 0.61972004 |
| AGGGGGLGGAP | ASSLRDSHNSPLY | 0.65344185 |
| AGGLLYGNGGAG | ASSLRDSHNSPLY | 0.64375407 |
| AGLESIGL | ASSLRDSHNSPLY | 0.529508 |
| AGLKTGII | ASSLRDSHNSPLY | 0.65383583 |
| ASLAALLL | ASSLRDSHNSPLY | 0.65803653 |
| ASVINISSI | ASSLRDSHNSPLY | 0.5574439 |
| ATPPGPA | ASSLRDSHNSPLY | 0.56690544 |
| ATVASALV | ASSLRDSHNSPLY | 0.5968641 |
| ATVASLIL | ASSLRDSHNSPLY | 0.47515327 |
| AVNGPVLIRH | ASSLRDSHNSPLY | 0.6616356 |
| DGAAGAVGI | ASSLRDSHNSPLY | 0.58533657 |
| DPLLHGI | ASSLRDSHNSPLY | 0.6138998 |
| ELGAALGAL | ASSLRDSHNSPLY | 0.6269371 |
| ELGRTQAV | ASSLRDSHNSPLY | 0.6214768 |
| FVAGGYLS | ASSLRDSHNSPLY | 0.62022907 |
| GASGGAGGAGGW | ASSLRDSHNSPLY | 0.6223721 |
| GGAGAGGDGGAGGI | ASSLRDSHNSPLY | 0.6499866 |
| GTGHNGSQPGMGGNG | ASSLRDSHNSPLY | 0.63324916 |
| GYAAASASAS | ASSLRDSHNSPLY | 0.6208755 |
| HQVDVAAVQQG | ASSLRDSHNSPLY | 0.6348948 |
| IGAAIAHT | ASSLRDSHNSPLY | 0.65827507 |
| IGMIEVAQL | ASSLRDSHNSPLY | 0.5765095 |
| IRLAPEHTSAMD | ASSLRDSHNSPLY | 0.59996057 |
| IYNVSVL | ASSLRDSHNSPLY | 0.6361898 |
| KQGFLGM | ASSLRDSHNSPLY | 0.50241375 |
| LESVITQL | ASSLRDSHNSPLY | 0.6467633 |
| LGLGALVVV | ASSLRDSHNSPLY | 0.6293003 |
| LIGNGGNGGSGGI | ASSLRDSHNSPLY | 0.63187754 |
| LNAINAPTL | ASSLRDSHNSPLY | 0.61691713 |
| LQLLNTGV | ASSLRDSHNSPLY | 0.66609085 |
| LQTSIEF | ASSLRDSHNSPLY | 0.64696705 |
| MSFAVSEL | ASSLRDSHNSPLY | 0.62961924 |

| | | |
|---|---|---|
| NTGALGL | ASSLRDSHNSPLY | 0.6505915 |
| PPGAGGAGGAGTT | ASSLRDSHNSPLY | 0.5565009 |
| QLVGLGI | ASSLRDSHNSPLY | 0.65564346 |
| RFLEQVHQL | ASSLRDSHNSPLY | 0.6389414 |
| RGVGYVLR | ASSLRDSHNSPLY | 0.54013956 |
| SATGVFAGVI | ASSLRDSHNSPLY | 0.61046135 |
| SLPAITLP | ASSLRDSHNSPLY | 0.6570146 |
| STALAFTV | ASSLRDSHNSPLY | 0.6524044 |
| SVLTVGLGK | ASSLRDSHNSPLY | 0.48777714 |
| TALAGAGSGI | ASSLRDSHNSPLY | 0.5995017 |
| TGLPFTGI | ASSLRDSHNSPLY | 0.44829148 |
| VAEHVTV | ASSLRDSHNSPLY | 0.54191065 |
| VDIVLNSL | ASSLRDSHNSPLY | 0.5452808 |
| VLTGGGAVL | ASSLRDSHNSPLY | 0.63790536 |
| VNVGLAYGI | ASSLRDSHNSPLY | 0.58376586 |
| VSSAAVGAV | ASSLRDSHNSPLY | 0.6461751 |
| VTSVYEAGV | ASSLRDSHNSPLY | 0.6274781 |
| WVALDRGA | ASSLRDSHNSPLY | 0.6614649 |
| YLVGSVVI | ASSLRDSHNSPLY | 0.57157046 |
| GPPGTGKTLL | ASSLRDSHNSPLY | 0.6345093 |
| GVGMTKF | ASSLRDSHNSPLY | 0.5042826 |
| IGIGGSDL | ASSLRDSHNSPLY | 0.59115696 |
| IGTLLAEL | ASSLRDSHNSPLY | 0.5815909 |
| IYTSGTT | ASSLRDSHNSPLY | 0.6354357 |
| LSFPAGV | ASSLRDSHNSPLY | 0.43108088 |
| LYTSGTT | ASSLRDSHNSPLY | 0.65963113 |
| VGAFPFGF | ASSLRDSHNSPLY | 0.6206238 |
| VGAGTVEF | ASSLRDSHNSPLY | 0.64928657 |
| VGKSTLLL | ASSLRDSHNSPLY | 0.6237048 |
| AALAHAGGGIV | ASSLRDSHNSPLY | 0.6097795 |
| AALVAVSLL | ASSLRDSHNSPLY | 0.60562396 |
| AAVSRAVAAL | ASSLRDSHNSPLY | 0.64110285 |
| AGGSSGIGGF | ASSLRDSHNSPLY | 0.59639484 |
| AGLTVANANG | ASSLRDSHNSPLY | 0.47226274 |
| AGLVGAVL | ASSLRDSHNSPLY | 0.5954993 |
| ALSQAFTV | ASSLRDSHNSPLY | 0.5394507 |
| ALTFSVSL | ASSLRDSHNSPLY | 0.65980923 |
| ALTGAALVAL | ASSLRDSHNSPLY | 0.65753776 |
| ASFSIGAGVL | ASSLRDSHNSPLY | 0.63081867 |
| DGSVYKI | ASSLRDSHNSPLY | 0.6548228 |
| EQLAEIV | ASSLRDSHNSPLY | 0.36577073 |
| FAVSELTVM | ASSLRDSHNSPLY | 0.63154125 |
| GAAPLAAGL | ASSLRDSHNSPLY | 0.6642554 |
| GADGGDGGSGGAGG | ASSLRDSHNSPLY | 0.6382259 |
| GFGLDTGGAGG | ASSLRDSHNSPLY | 0.6022655 |
| GLASGGLL | ASSLRDSHNSPLY | 0.6546276 |
| GRIAGAAVLL | ASSLRDSHNSPLY | 0.5909703 |
| GSAGSLNTSF | ASSLRDSHNSPLY | 0.611019 |
| GTGQIGI | ASSLRDSHNSPLY | 0.63190764 |
| GTLIALIL | ASSLRDSHNSPLY | 0.6504704 |
| IAATMGFVVG | ASSLRDSHNSPLY | 0.6293751 |

| | | |
|---|---|---|
| IFAQALL | ASSLRDSHNSPLY | 0.57518995 |
| IGAAVALGF | ASSLRDSHNSPLY | 0.6378801 |
| IGLTPLAL | ASSLRDSHNSPLY | 0.66808385 |
| ILGSNTSTL | ASSLRDSHNSPLY | 0.59911394 |
| IVFSLTVL | ASSLRDSHNSPLY | 0.65333456 |
| LASKATGF | ASSLRDSHNSPLY | 0.6384429 |
| LASVALIL | ASSLRDSHNSPLY | 0.6450971 |
| LFAVGLTL | ASSLRDSHNSPLY | 0.55895746 |
| LFNGGIGTYI | ASSLRDSHNSPLY | 0.581185 |
| LTLGAGII | ASSLRDSHNSPLY | 0.6200415 |
| LVAAALGDI | ASSLRDSHNSPLY | 0.5571132 |
| NTAFVQLGI | ASSLRDSHNSPLY | 0.6088685 |
| QLFSGAEL | ASSLRDSHNSPLY | 0.6131003 |
| SAATFITL | ASSLRDSHNSPLY | 0.67161024 |
| SNYGLANLGV | ASSLRDSHNSPLY | 0.6379075 |
| SYGGAYAVM | ASSLRDSHNSPLY | 0.6407542 |
| TTGVGGLTLG | ASSLRDSHNSPLY | 0.5886594 |
| TTQTIYI | ASSLRDSHNSPLY | 0.63467425 |
| VAAGAGLGAVI | ASSLRDSHNSPLY | 0.6405645 |
| VAAVVTLL | ASSLRDSHNSPLY | 0.51797783 |
| VGFLGGLI | ASSLRDSHNSPLY | 0.6608184 |
| VGVIPILL | ASSLRDSHNSPLY | 0.5806278 |
| VIFASSLI | ASSLRDSHNSPLY | 0.6039101 |
| VKGGDGGAAAG | ASSLRDSHNSPLY | 0.6502199 |
| VTKTFTNI | ASSLRDSHNSPLY | 0.6674974 |
| VVAAAILL | ASSLRDSHNSPLY | 0.59750307 |
| YAALDVEL | ASSLRDSHNSPLY | 0.5955774 |
| LASVALLL | ASSLRDSHNSPLY | 0.58128625 |
| LDAVHRV | ASSLRDSHNSPLY | 0.6494907 |
| QGLSSSIF | ASSLRDSHNSPLY | 0.6546297 |
| RGTFANIRL | ASSLRDSHNSPLY | 0.6312113 |
| RQLWWGHRI | ASSLRDSHNSPLY | 0.66706085 |
| YGPPGTGKTLL | ASSLRDSHNSPLY | 0.6534207 |
| AALVGAPANVA | ASSLRDSHNSPLY | 0.47143364 |
| AATFRRTE | ASSLRDSHNSPLY | 0.62629324 |
| AGALGGGI | ASSLRDSHNSPLY | 0.657061 |
| AGGAGGAGGIGAG | ASSLRDSHNSPLY | 0.6495492 |
| AGSGALGL | ASSLRDSHNSPLY | 0.667267 |
| AQIADQI | ASSLRDSHNSPLY | 0.64503384 |
| ASAVAVTYL | ASSLRDSHNSPLY | 0.5835508 |
| ASTVLGGTGG | ASSLRDSHNSPLY | 0.6612837 |
| AVGGEPAK | ASSLRDSHNSPLY | 0.6697117 |
| CEQGTFGGTC | ASSLRDSHNSPLY | 0.6502961 |
| DEGKAPAG | ASSLRDSHNSPLY | 0.5924692 |
| DTVVFPAL | ASSLRDSHNSPLY | 0.5664303 |
| FEGITMAG | ASSLRDSHNSPLY | 0.6284026 |
| FPLVVQD | ASSLRDSHNSPLY | 0.648274 |
| GAFTTPQ | ASSLRDSHNSPLY | 0.5281067 |
| GFGGFGTAGGVGGA | ASSLRDSHNSPLY | 0.62339723 |
| GFLAQFL | ASSLRDSHNSPLY | 0.6323067 |
| GGAGGAGGNAGVA | ASSLRDSHNSPLY | 0.48782253 |

| | | |
|---|---|---|
| GGVGTAGAGGAGGAGG | ASSLRDSHNSPLY | 0.65429556 |
| GNGGGGDAGNAGSGGN | ASSLRDSHNSPLY | 0.6590218 |
| GTGGAGGAGNDAGS | ASSLRDSHNSPLY | 0.62021744 |
| GTTNTGWFNS | ASSLRDSHNSPLY | 0.59693253 |
| IFMGPLFTL | ASSLRDSHNSPLY | 0.65377206 |
| ILAHAIEAI | ASSLRDSHNSPLY | 0.6697079 |
| IRSAIREAL | ASSLRDSHNSPLY | 0.65727377 |
| ITFAGGG | ASSLRDSHNSPLY | 0.53001 |
| ITGMSGHQEV | ASSLRDSHNSPLY | 0.38885534 |
| IVAGLACTL | ASSLRDSHNSPLY | 0.5930215 |
| IVGGGLAGA | ASSLRDSHNSPLY | 0.65004003 |
| IVTGVLNI | ASSLRDSHNSPLY | 0.63426065 |
| LAQQGCI | ASSLRDSHNSPLY | 0.6207098 |
| LCLHVEN | ASSLRDSHNSPLY | 0.6303234 |
| LDAISDGS | ASSLRDSHNSPLY | 0.65794253 |
| LGGFAAML | ASSLRDSHNSPLY | 0.57331574 |
| LGGKNPMI | ASSLRDSHNSPLY | 0.5299298 |
| LGPKGVAAIL | ASSLRDSHNSPLY | 0.63886195 |
| LMVGVSI | ASSLRDSHNSPLY | 0.64420474 |
| LVGAAFVL | ASSLRDSHNSPLY | 0.57810116 |
| LVLAVAPL | ASSLRDSHNSPLY | 0.60105985 |
| MAAIAVIDAA | ASSLRDSHNSPLY | 0.59694064 |
| NNGINAELQ | ASSLRDSHNSPLY | 0.58505327 |
| RGEPLVLVHG | ASSLRDSHNSPLY | 0.6332794 |
| RGGAGGAAGLI | ASSLRDSHNSPLY | 0.54963154 |
| RSYRMHPAV | ASSLRDSHNSPLY | 0.6563344 |
| SAAHLLAGQLA | ASSLRDSHNSPLY | 0.53911495 |
| SAAVGSY | ASSLRDSHNSPLY | 0.64169216 |
| SPHAVLHRI | ASSLRDSHNSPLY | 0.6344711 |
| SQIAKVL | ASSLRDSHNSPLY | 0.64540815 |
| TALGHLVP | ASSLRDSHNSPLY | 0.5905307 |
| TAPTTSGAA | ASSLRDSHNSPLY | 0.5912014 |
| TDGDDGNGGSPGT | ASSLRDSHNSPLY | 0.5635993 |
| TDGLGEN | ASSLRDSHNSPLY | 0.41777495 |
| TGGPPLPGQA | ASSLRDSHNSPLY | 0.6560989 |
| TIPAITI | ASSLRDSHNSPLY | 0.58356345 |
| TIPGFTI | ASSLRDSHNSPLY | 0.65662134 |
| TSIASGALA | ASSLRDSHNSPLY | 0.6446963 |
| TVTLPGAAI | ASSLRDSHNSPLY | 0.5778137 |
| VASGLSAALP | ASSLRDSHNSPLY | 0.6379162 |
| VGAGVIGI | ASSLRDSHNSPLY | 0.580969 |
| VGPKRFGV | ASSLRDSHNSPLY | 0.574261 |
| VNVAFPD | ASSLRDSHNSPLY | 0.63578475 |
| VSMTNSGVS | ASSLRDSHNSPLY | 0.649667 |
| VTDVLGDLSV | ASSLRDSHNSPLY | 0.5546042 |
| VVVVGAGFAGL | ASSLRDSHNSPLY | 0.63486254 |
| VYTVGTVV | ASSLRDSHNSPLY | 0.63073015 |
| WTTFSGP | ASSLRDSHNSPLY | 0.5716287 |
| YDELVEAF | ASSLRDSHNSPLY | 0.6311743 |
| YGTGGAGGNGG | ASSLRDSHNSPLY | 0.6547413 |
| YTNVMAV | ASSLRDSHNSPLY | 0.6313522 |

| | | |
|---|---|---|
| FLEQVHQL | ASSLRDSHNSPLY | 0.56795806 |
| GLSLLAVAV | ASSLRDSHNSPLY | 0.6448232 |
| LEQVHQL | ASSLRDSHNSPLY | 0.5571078 |
| LNAAGAI | ASSLRDSHNSPLY | 0.6374408 |
| PAVHASL | ASSLRDSHNSPLY | 0.62272465 |
| TELEDDA | ASSLRDSHNSPLY | 0.5931213 |
| TPPGPPN | ASSLRDSHNSPLY | 0.651924 |
| TPPPPPPP | ASSLRDSHNSPLY | 0.57437754 |
| TQSPALI | ASGDAKTGGAAETLY | 0.55298233 |
| EFAAAFPQ | ASGDAKTGGAAETLY | 0.60170627 |
| LLSGNLDVL | ASGDAKTGGAAETLY | 0.636395 |
| GPANLGNYNI | ASGDAKTGGAAETLY | 0.65102565 |
| TLGPTLP | ASGDAKTGGAAETLY | 0.5748398 |
| VYAYSSCCK | ASGDAKTGGAAETLY | 0.6275861 |
| APAPAPAPNI | ASGDAKTGGAAETLY | 0.6016356 |
| PGGPVPIP | ASGDAKTGGAAETLY | 0.5645561 |
| FLTVALEP | ASGDAKTGGAAETLY | 0.61407137 |
| SQYFGANL | ASGDAKTGGAAETLY | 0.62015265 |
| GGVKPAA | ASGDAKTGGAAETLY | 0.56037915 |
| GGTGGAGGAGGW | ASGDAKTGGAAETLY | 0.64640117 |
| GVGGVAGLGGAGG | ASGDAKTGGAAETLY | 0.61453176 |
| VLLVLATTI | ASGDAKTGGAAETLY | 0.64788 |
| IGIFAAML | ASGDAKTGGAAETLY | 0.65926766 |
| AVGDLID | ASGDAKTGGAAETLY | 0.5925456 |
| NISPLPL | ASGDAKTGGAAETLY | 0.63957083 |
| VTAVFVAI | ASGDAKTGGAAETLY | 0.64090365 |
| SGDDNDAAL | ASGDAKTGGAAETLY | 0.6389556 |
| AVAALAHS | ASGDAKTGGAAETLY | 0.6288091 |
| GGVTVMATP | ASGDAKTGGAAETLY | 0.5970671 |
| AGLAANVRD | ASGDAKTGGAAETLY | 0.6612543 |
| TNRPDLIDPALLRPG | ASGDAKTGGAAETLY | 0.65116113 |
| LANLGVTAL | ASGDAKTGGAAETLY | 0.6430389 |
| TGERIEL | ASGDAKTGGAAETLY | 0.6454754 |
| NGEELAA | ASGDAKTGGAAETLY | 0.6469783 |
| VLSLVWG | ASGDAKTGGAAETLY | 0.64312685 |
| IGYALWL | ASGDAKTGGAAETLY | 0.5885284 |
| LSLLLAPL | ASGDAKTGGAAETLY | 0.5243266 |
| TLDEALK | ASGDAKTGGAAETLY | 0.62944585 |
| TDGAGTAEGG | ASGDAKTGGAAETLY | 0.6174829 |
| NAPGGAGGNGG | ASGDAKTGGAAETLY | 0.611225 |
| VSVFMGMGGNF | ASGDAKTGGAAETLY | 0.46954772 |
| LASYLAAL | ASGDAKTGGAAETLY | 0.657048 |
| LVGGTPAL | ASGDAKTGGAAETLY | 0.66954136 |
| VRNADVL | ASGDAKTGGAAETLY | 0.60209537 |
| TCSSPFLVSAM | ASGDAKTGGAAETLY | 0.6655904 |
| VHITDAE | ASGDAKTGGAAETLY | 0.53153765 |
| EYAAPSF | ASGDAKTGGAAETLY | 0.6504095 |
| VVDAFHGGL | ASGDAKTGGAAETLY | 0.65466106 |
| ASVVDPQ | ASGDAKTGGAAETLY | 0.55781144 |
| SYNEFGGLS | ASGDAKTGGAAETLY | 0.6599947 |
| DPDGSVY | ASGDAKTGGAAETLY | 0.5862113 |

| | | |
|---|---|---|
| MVSTIHATE | ASGDAKTGGAAETLY | 0.6243191 |
| LAAGVADPV | ASGDAKTGGAAETLY | 0.6575151 |
| TKEQIAA | ASGDAKTGGAAETLY | 0.6290394 |
| SLASPAAL | ASGDAKTGGAAETLY | 0.6401442 |
| IGLVFYGV | ASGDAKTGGAAETLY | 0.63518894 |
| FLLDASNL | ASGDAKTGGAAETLY | 0.6019813 |
| YQVGNFP | ASGDAKTGGAAETLY | 0.6399042 |
| ESLLSQST | ASGDAKTGGAAETLY | 0.6537943 |
| LFTVGFAAVG | ASGDAKTGGAAETLY | 0.55995333 |
| ATPPTPT | ASGDAKTGGAAETLY | 0.66822374 |
| TDLATLG | ASGDAKTGGAAETLY | 0.5706452 |
| SAGGTGR | ASGDAKTGGAAETLY | 0.67013 |
| GFSNSGS | ASGDAKTGGAAETLY | 0.6438255 |
| GGNGGSGGTSV | ASGDAKTGGAAETLY | 0.61979127 |
| GTSVATGGAGN | ASGDAKTGGAAETLY | 0.60736036 |
| NGGNGGSGGTS | ASGDAKTGGAAETLY | 0.654772 |
| AVVKFPH | ASGDAKTGGAAETLY | 0.61053646 |
| TPPPPPP | ASGDAKTGGAAETLY | 0.64138144 |
| AAPAAAAAPA | ASGDAKTGGAAETLY | 0.664534 |
| IALNEDL | ASGDAKTGGAAETLY | 0.5054345 |
| IVNVSSM | ASGDAKTGGAAETLY | 0.7640467 |
| TGIPPAPR | ASGDAKTGGAAETLY | 0.66445017 |
| FDLGGGTF | ASGDAKTGGAAETLY | 0.73942554 |
| FDLGGGTFDVS | ASGDAKTGGAAETLY | 0.49314803 |
| NEPTAAA | ASGDAKTGGAAETLY | 0.29637095 |
| NEPTAAALA | ASGDAKTGGAAETLY | 0.6416378 |
| GSIINISSI | ASGDAKTGGAAETLY | 0.53454417 |
| VAPGNAGTA | ASGDAKTGGAAETLY | 0.7731879 |
| TGGGASLE | ASGDAKTGGAAETLY | 0.75398386 |
| VAVYNHYKRI | ASGDAKTGGAAETLY | 0.515153 |
| TDPAPATT | ASGDAKTGGAAETLY | 0.78261364 |
| TNRPDILDPALLRPG | ASGDAKTGGAAETLY | 0.7714799 |
| LTTTTTS | ASGDAKTGGAAETLY | 0.40824217 |
| PTGSGKTL | ASGDAKTGGAAETLY | 0.7124403 |
| TGSGKTL | ASGDAKTGGAAETLY | 0.35796535 |
| TIGHVDHGK | ASGDAKTGGAAETLY | 0.65376234 |
| TIGHVDHGKTT | ASGDAKTGGAAETLY | 0.25724518 |
| GHVDHGKTT | ASGDAKTGGAAETLY | 0.71948683 |
| AQFEHTLL | ASGDAKTGGAAETLY | 0.72194684 |
| IVNQDEL | ASGDAKTGGAAETLY | 0.36610106 |
| TNGSQFF | ASGDAKTGGAAETLY | 0.7400814 |
| TGKGIVE | ASGDAKTGGAAETLY | 0.70612836 |
| LISNASDA | ASGDAKTGGAAETLY | 0.73535645 |
| AELRTLGELTKIAKAHGAQ | ASGDAKTGGAAETLY | 0.76907706 |
| APNAVAAAEPSW | ASGDAKTGGAAETLY | 0.50022537 |
| APQVAGAATAVL | ASGDAKTGGAAETLY | 0.65907526 |
| ARRDPAAAAAILPTDA | ASGDAKTGGAAETLY | 0.7529707 |
| DAAARDVLAEAGLAEYF | ASGDAKTGGAAETLY | 0.78201485 |
| DDQIVLSAKANLAKVEQ | ASGDAKTGGAAETLY | 0.6051942 |
| DELLPDTGEDSDEDGH | ASGDAKTGGAAETLY | 0.7248863 |
| ELPEAPANRATVPQD | ASGDAKTGGAAETLY | 0.7787124 |

| | | |
|---|---|---|
| FGQSKIERVTGIG | ASGDAKTGGAAETLY | 0.6262844 |
| GAGGTGSTAGGAGGAGGAGGLY | ASGDAKTGGAAETLY | 0.75181144 |
| GHGGTGAAGGNGGTGGAGGSGI | ASGDAKTGGAAETLY | 0.7624513 |
| GNGGNGGAGGEGGPGGAGGAGA | ASGDAKTGGAAETLY | 0.44937575 |
| GSTAGNGGNGGNATNGTIAGQP | ASGDAKTGGAAETLY | 0.7776189 |
| GTGGDGGNGGNGGTGG | ASGDAKTGGAAETLY | 0.77783656 |
| GVRRPAGIPADAPV | ASGDAKTGGAAETLY | 0.6465361 |
| GYRQVTDAHLLTLARR | ASGDAKTGGAAETLY | 0.6913583 |
| IAKMGQDGHDRG | ASGDAKTGGAAETLY | 0.4359281 |
| KAEPSIEIQYT | ASGDAKTGGAAETLY | 0.65899503 |
| KPVRIRMAADEDTVAVT | ASGDAKTGGAAETLY | 0.28776956 |
| MPPRIAGMRLLVIKPEP | ASGDAKTGGAAETLY | 0.7383757 |
| MKFYAVALFRSRGVPPTR | ASGDAKTGGAAETLY | 0.7541065 |
| NAGGFGILGAVAHSP | ASGDAKTGGAAETLY | 0.70469916 |
| NEGREITLAMAEM | ASGDAKTGGAAETLY | 0.42209634 |
| NELVEWSHPLGHRWRWE | ASGDAKTGGAAETLY | 0.39306748 |
| NLGAGGTGGNGGNPG | ASGDAKTGGAAETLY | 0.6819562 |
| NTEWPKWETVMAADPVD | ASGDAKTGGAAETLY | 0.56541675 |
| PASAAPAETQP | ASGDAKTGGAAETLY | 0.43249738 |
| QQSSDGKNGVRAAEITGE | ASGDAKTGGAAETLY | 0.34196514 |
| RAPDAVTTSWWKE | ASGDAKTGGAAETLY | 0.7652414 |
| SGFGNVGAHTSGWFNQ | ASGDAKTGGAAETLY | 0.76579666 |
| SGVAGAVGQAASVGGLKV | ASGDAKTGGAAETLY | 0.66232073 |
| SNGGAGTGSAGGNGGT | ASGDAKTGGAAETLY | 0.7671622 |
| TGGAGGEGGAAG | ASGDAKTGGAAETLY | 0.77020574 |
| TGGDGGAAGAGGAPGHG | ASGDAKTGGAAETLY | 0.580518 |
| TMPPLLAAIANAGRHGV | ASGDAKTGGAAETLY | 0.63435006 |
| TSSTAGNSGGA | ASGDAKTGGAAETLY | 0.41278774 |
| VPATVPVGILC | ASGDAKTGGAAETLY | 0.78537 |
| VQRPDSGGVVLTGRIS | ASGDAKTGGAAETLY | 0.649132 |
| WTPNPVPATKGVPAGLVQAT | ASGDAKTGGAAETLY | 0.7374241 |
| YALKRALAKFSDRFDVVII | ASGDAKTGGAAETLY | 0.7443996 |
| YLIGSVIV | ASGDAKTGGAAETLY | 0.7042684 |
| YGQLVAVID | ASGDAKTGGAAETLY | 0.60536766 |
| VTVGGMTLPA | ASGDAKTGGAAETLY | 0.7302526 |
| VLALPAV | ASGDAKTGGAAETLY | 0.69584954 |
| VIPPTLNYET | ASGDAKTGGAAETLY | 0.55479896 |
| VGMGGMHPSSG | ASGDAKTGGAAETLY | 0.2927465 |
| VGLAGGAATA | ASGDAKTGGAAETLY | 0.7288744 |
| VGGAGGAGGSGP | ASGDAKTGGAAETLY | 0.7227512 |
| VFAAFMV | ASGDAKTGGAAETLY | 0.781134 |
| TLYDKLR | ASGDAKTGGAAETLY | 0.61717886 |
| TLAVFTVL | ASGDAKTGGAAETLY | 0.55566895 |
| TGSVQPARV | ASGDAKTGGAAETLY | 0.77407855 |
| TGGGAGSGGT | ASGDAKTGGAAETLY | 0.68224037 |
| SHLETSL | ASGDAKTGGAAETLY | 0.6651286 |
| SFTTGMT | ASGDAKTGGAAETLY | 0.71206 |
| QVEVAKRMGVRQPTVS | ASGDAKTGGAAETLY | 0.66699433 |
| QGSGGPALAM | ASGDAKTGGAAETLY | 0.6774068 |
| MLAPSNP | ASGDAKTGGAAETLY | 0.7292583 |
| LVNNAAI | ASGDAKTGGAAETLY | 0.6859218 |

| | | |
|---|---|---|
| LTGEINTLA | ASGDAKTGGAAETLY | 0.74563414 |
| LSIFGPV | ASGDAKTGGAAETLY | 0.6528541 |
| LLALLANL | ASGDAKTGGAAETLY | 0.64611197 |
| LISNASDALDKLR | ASGDAKTGGAAETLY | 0.38521928 |
| LINEAVP | ASGDAKTGGAAETLY | 0.5038394 |
| LGYAFSGL | ASGDAKTGGAAETLY | 0.7438544 |
| LDFVAVAL | ASGDAKTGGAAETLY | 0.7301731 |
| LAQPGAE | ASGDAKTGGAAETLY | 0.7673439 |
| ISNLGAVTGF | ASGDAKTGGAAETLY | 0.7056517 |
| HQEDPGRI | ASGDAKTGGAAETLY | 0.59676486 |
| GTPGEPTP | ASGDAKTGGAAETLY | 0.75454605 |
| GFFNSGS | ASGDAKTGGAAETLY | 0.77594745 |
| GAGGGGLTAGAN | ASGDAKTGGAAETLY | 0.34835842 |
| FVDALVNM | ASGDAKTGGAAETLY | 0.654211 |
| FLTVAPEL | ASGDAKTGGAAETLY | 0.78276646 |
| FETLTAL | ASGDAKTGGAAETLY | 0.7653711 |
| DVINPIEL | ASGDAKTGGAAETLY | 0.64651406 |
| APVSGAPV | ASGDAKTGGAAETLY | 0.6884208 |
| ALVGGLAW | ASGDAKTGGAAETLY | 0.79027426 |
| ALAAAAATAI | ASGDAKTGGAAETLY | 0.7425519 |
| AFGLGLLL | ASGDAKTGGAAETLY | 0.78648674 |
| AFFAAASL | ASGDAKTGGAAETLY | 0.3925221 |
| AAAALPAFSPP | ASGDAKTGGAAETLY | 0.52913636 |
| VVLAPAL | ASGDAKTGGAAETLY | 0.31490567 |
| TIGHVDHGKTTL | ASGDAKTGGAAETLY | 0.24528143 |
| LISNASDALDK | ASGDAKTGGAAETLY | 0.74352777 |
| LISNASDAL | ASGDAKTGGAAETLY | 0.6222224 |
| ISNASDALDK | ASGDAKTGGAAETLY | 0.771321 |
| NQAETLVYQTE | ASGDAKTGGAAETLY | 0.61333627 |
| TGGTGGTAGAG | ASGDAKTGGAAETLY | 0.75955504 |
| TGSTAGGAGGA | ASGDAKTGGAAETLY | 0.61110806 |
| DPLAVPVAAGPV | ASGDAKTGGAAETLY | 0.6879543 |
| GGTGGTPGNGGN | ASGDAKTGGAAETLY | 0.4484174 |
| PAGLAGVDPATL | ASGDAKTGGAAETLY | 0.74077404 |
| GNNLVVEAPHAGASV | ASGDAKTGGAAETLY | 0.7746627 |
| KVTRIMSASTLEQPA | ASGDAKTGGAAETLY | 0.64793956 |
| LGFALPIALPRRTVV | ASGDAKTGGAAETLY | 0.4714572 |
| TRLGNLTARKLVVLV | ASGDAKTGGAAETLY | 0.750003 |
| WSQQRRLIYFPSAGP | ASGDAKTGGAAETLY | 0.6693421 |
| GPNGNPDPMAAAVDIR | ASGDAKTGGAAETLY | 0.276383 |
| TQRCVSVADLRAAASR | ASGDAKTGGAAETLY | 0.4503423 |
| QFVLDVAEHPVEVLTETV | ASGDAKTGGAAETLY | 0.611861 |
| GGAGGTGGAGGNGGTQAGDG | ASGDAKTGGAAETLY | 0.5790156 |
| PLLATLRAIAVDVDAKPAQV | ASGDAKTGGAAETLY | 0.77231103 |
| APKFRTGGYMIDFWGVGYQVAK | ASGDAKTGGAAETLY | 0.7655874 |
| GAGGAGGAGGAAGGPGGTGGTG | ASGDAKTGGAAETLY | 0.6458184 |
| IGALGLLATDVAPAGSVAFGVA | ASGDAKTGGAAETLY | 0.74057925 |
| AGGAGGIGGNGGAGGAGGVLMG | ASGDAKTGGAAETLY | 0.7371837 |
| ADELIGLL | ASGDAKTGGAAETLY | 0.19351524 |
| ADRALKA | ASGDAKTGGAAETLY | 0.44767228 |
| AGAVGQVGGAGGA | ASGDAKTGGAAETLY | 0.44549626 |

| | | |
|---|---|---|
| AGGGGGLGGAP | ASGDAKTGGAAETLY | 0.763754 |
| AGGLLYGNGGAG | ASGDAKTGGAAETLY | 0.5427455 |
| AGLESIGL | ASGDAKTGGAAETLY | 0.68486035 |
| AGLKTGII | ASGDAKTGGAAETLY | 0.71067953 |
| ASLAALLL | ASGDAKTGGAAETLY | 0.76503456 |
| ASVINISSI | ASGDAKTGGAAETLY | 0.68463945 |
| ATPPGPA | ASGDAKTGGAAETLY | 0.38883847 |
| ATVASALV | ASGDAKTGGAAETLY | 0.2515574 |
| ATVASLIL | ASGDAKTGGAAETLY | 0.7414426 |
| AVNGPVLIRH | ASGDAKTGGAAETLY | 0.41451722 |
| DGAAGAVGI | ASGDAKTGGAAETLY | 0.68370724 |
| DPLLHGI | ASGDAKTGGAAETLY | 0.50849986 |
| ELGAALGAL | ASGDAKTGGAAETLY | 0.6929132 |
| ELGRTQAV | ASGDAKTGGAAETLY | 0.31699902 |
| FVAGGYLS | ASGDAKTGGAAETLY | 0.722883 |
| GASGGAGGAGGW | ASGDAKTGGAAETLY | 0.47847748 |
| GGAGAGGDGGAGGI | ASGDAKTGGAAETLY | 0.7149246 |
| GTGHNGSQPGMGGNG | ASGDAKTGGAAETLY | 0.7685554 |
| GYAAASASAS | ASGDAKTGGAAETLY | 0.76299226 |
| HQVDVAAVQQG | ASGDAKTGGAAETLY | 0.60772336 |
| IGAAIAHT | ASGDAKTGGAAETLY | 0.67271376 |
| IGMIEVAQL | ASGDAKTGGAAETLY | 0.4177331 |
| IRLAPEHTSAMD | ASGDAKTGGAAETLY | 0.61187875 |
| IYNVSVL | ASGDAKTGGAAETLY | 0.6442456 |
| KQGFLGM | ASGDAKTGGAAETLY | 0.7834989 |
| LESVITQL | ASGDAKTGGAAETLY | 0.53757656 |
| LGLGALVVV | ASGDAKTGGAAETLY | 0.39787158 |
| LIGNGGNGGSGGI | ASGDAKTGGAAETLY | 0.52712464 |
| LNAINAPTL | ASGDAKTGGAAETLY | 0.7702545 |
| LQLLNTGV | ASGDAKTGGAAETLY | 0.7483593 |
| LQTSIEF | ASGDAKTGGAAETLY | 0.59542894 |
| MSFAVSEL | ASGDAKTGGAAETLY | 0.44948357 |
| NTGALGL | ASGDAKTGGAAETLY | 0.65832436 |
| PPGAGGAGGAGTT | ASGDAKTGGAAETLY | 0.7110161 |
| QLVGLGI | ASGDAKTGGAAETLY | 0.48046196 |
| RFLEQVHQL | ASGDAKTGGAAETLY | 0.5017468 |
| RGVGYVLR | ASGDAKTGGAAETLY | 0.68199974 |
| SATGVFAGVI | ASGDAKTGGAAETLY | 0.791518 |
| SLPAITLP | ASGDAKTGGAAETLY | 0.6347625 |
| STALAFTV | ASGDAKTGGAAETLY | 0.23245122 |
| SVLTVGLGK | ASGDAKTGGAAETLY | 0.72876567 |
| TALAGAGSGI | ASGDAKTGGAAETLY | 0.53718746 |
| TGLPFTGI | ASGDAKTGGAAETLY | 0.6134528 |
| VAEHVTV | ASGDAKTGGAAETLY | 0.6551572 |
| VDIVLNSL | ASGDAKTGGAAETLY | 0.40890402 |
| VLTGGGAVL | ASGDAKTGGAAETLY | 0.69605213 |
| VNVGLAYGI | ASGDAKTGGAAETLY | 0.7440418 |
| VSSAAVGAV | ASGDAKTGGAAETLY | 0.7330945 |
| VTSVYEAGV | ASGDAKTGGAAETLY | 0.7060429 |
| WVALDRGA | ASGDAKTGGAAETLY | 0.7744483 |
| YLVGSVVI | ASGDAKTGGAAETLY | 0.7514976 |

| | | |
|---|---|---|
| GPPGTGKTLL | ASGDAKTGGAAETLY | 0.7787343 |
| GVGMTKF | ASGDAKTGGAAETLY | 0.77719027 |
| IGIGGSDL | ASGDAKTGGAAETLY | 0.61643595 |
| IGTLLAEL | ASGDAKTGGAAETLY | 0.60314906 |
| IYTSGTT | ASGDAKTGGAAETLY | 0.74915206 |
| LSFPAGV | ASGDAKTGGAAETLY | 0.73093534 |
| LYTSGTT | ASGDAKTGGAAETLY | 0.6819989 |
| VGAFPFGF | ASGDAKTGGAAETLY | 0.5274786 |
| VGAGTVEF | ASGDAKTGGAAETLY | 0.7787719 |
| VGKSTLLL | ASGDAKTGGAAETLY | 0.59250987 |
| AALAHAGGGIV | ASGDAKTGGAAETLY | 0.7827518 |
| AALVAVSLL | ASGDAKTGGAAETLY | 0.3076647 |
| AAVSRAVAAL | ASGDAKTGGAAETLY | 0.583316 |
| AGGSSGIGGF | ASGDAKTGGAAETLY | 0.75980747 |
| AGLTVANANG | ASGDAKTGGAAETLY | 0.20598735 |
| AGLVGAVL | ASGDAKTGGAAETLY | 0.5231721 |
| ALSQAFTV | ASGDAKTGGAAETLY | 0.66018414 |
| ALTFSVSL | ASGDAKTGGAAETLY | 0.7247108 |
| ALTGAALVAL | ASGDAKTGGAAETLY | 0.20841713 |
| ASFSIGAGVL | ASGDAKTGGAAETLY | 0.6984408 |
| DGSVYKI | ASGDAKTGGAAETLY | 0.36953345 |
| EQLAEIV | ASGDAKTGGAAETLY | 0.702381 |
| FAVSELTVM | ASGDAKTGGAAETLY | 0.69247925 |
| GAAPLAAGL | ASGDAKTGGAAETLY | 0.2682895 |
| GADGGDGGSGGAGG | ASGDAKTGGAAETLY | 0.75415576 |
| GFGLDTGGAGG | ASGDAKTGGAAETLY | 0.26566207 |
| GLASGGLL | ASGDAKTGGAAETLY | 0.61107063 |
| GRIAGAAVLL | ASGDAKTGGAAETLY | 0.746858 |
| GSAGSLNTSF | ASGDAKTGGAAETLY | 0.7869878 |
| GTGQIGI | ASGDAKTGGAAETLY | 0.50289595 |
| GTLIALIL | ASGDAKTGGAAETLY | 0.7632153 |
| IAATMGFVVG | ASGDAKTGGAAETLY | 0.46954417 |
| IFAQALL | ASGDAKTGGAAETLY | 0.7701093 |
| IGAAVALGF | ASGDAKTGGAAETLY | 0.5924574 |
| IGLTPLAL | ASGDAKTGGAAETLY | 0.2977818 |
| ILGSNTSTL | ASGDAKTGGAAETLY | 0.76934993 |
| IVFSLTVL | ASGDAKTGGAAETLY | 0.6389149 |
| LASKATGF | ASGDAKTGGAAETLY | 0.67943436 |
| LASVALIL | ASGDAKTGGAAETLY | 0.7725598 |
| LFAVGLTL | ASGDAKTGGAAETLY | 0.47886592 |
| LFNGGIGTYI | ASGDAKTGGAAETLY | 0.70215034 |
| LTLGAGII | ASGDAKTGGAAETLY | 0.77900904 |
| LVAAALGDI | ASGDAKTGGAAETLY | 0.7638141 |
| NTAFVQLGI | ASGDAKTGGAAETLY | 0.45526522 |
| QLFSGAEL | ASGDAKTGGAAETLY | 0.7186961 |
| SAATFITL | ASGDAKTGGAAETLY | 0.73969597 |
| SNYGLANLGV | ASGDAKTGGAAETLY | 0.5133935 |
| SYGGAYAVM | ASGDAKTGGAAETLY | 0.7411784 |
| TTGVGGLTLG | ASGDAKTGGAAETLY | 0.53488344 |
| TTQTIYI | ASGDAKTGGAAETLY | 0.7444211 |
| VAAGAGLGAVI | ASGDAKTGGAAETLY | 0.688961 |

| | | |
|---|---|---|
| VAAVVTLL | ASGDAKTGGAAETLY | 0.78230107 |
| VGFLGGLI | ASGDAKTGGAAETLY | 0.31080696 |
| VGVIPILL | ASGDAKTGGAAETLY | 0.56568664 |
| VIFASSLI | ASGDAKTGGAAETLY | 0.6901345 |
| VKGGDGGAAAG | ASGDAKTGGAAETLY | 0.7532606 |
| VTKTFTNI | ASGDAKTGGAAETLY | 0.44357383 |
| VVAAAILL | ASGDAKTGGAAETLY | 0.46558 |
| YAALDVEL | ASGDAKTGGAAETLY | 0.67740804 |
| LASVALLL | ASGDAKTGGAAETLY | 0.77356124 |
| LDAVHRV | ASGDAKTGGAAETLY | 0.59266293 |
| QGLSSSIF | ASGDAKTGGAAETLY | 0.7196805 |
| RGTFANIRL | ASGDAKTGGAAETLY | 0.78516805 |
| RQLWWGHRI | ASGDAKTGGAAETLY | 0.51040137 |
| YGPPGTGKTLL | ASGDAKTGGAAETLY | 0.7065078 |
| AALVGAPANVA | ASGDAKTGGAAETLY | 0.75009763 |
| AATFRRTE | ASGDAKTGGAAETLY | 0.77835953 |
| AGALGGGI | ASGDAKTGGAAETLY | 0.56150234 |
| AGGAGGAGGIGAG | ASGDAKTGGAAETLY | 0.76676625 |
| AGSGALGL | ASGDAKTGGAAETLY | 0.5881668 |
| AQIADQI | ASGDAKTGGAAETLY | 0.7726054 |
| ASAVAVTYL | ASGDAKTGGAAETLY | 0.4075692 |
| ASTVLGGTGG | ASGDAKTGGAAETLY | 0.7168546 |
| AVGGEPAK | ASGDAKTGGAAETLY | 0.5915238 |
| CEQGTFGGTC | ASGDAKTGGAAETLY | 0.75980353 |
| DEGKAPAG | ASGDAKTGGAAETLY | 0.7490175 |
| DTVVFPAL | ASGDAKTGGAAETLY | 0.7742591 |
| FEGITMAG | ASGDAKTGGAAETLY | 0.45300466 |
| FPLVVQD | ASGDAKTGGAAETLY | 0.7603688 |
| GAFTTPQ | ASGDAKTGGAAETLY | 0.7753066 |
| GFGGFGTAGGVGGA | ASGDAKTGGAAETLY | 0.79261553 |
| GFLAQFL | ASGDAKTGGAAETLY | 0.5576934 |
| GGAGGAGGNAGVA | ASGDAKTGGAAETLY | 0.7597373 |
| GGVGTAGAGGAGGAGG | ASGDAKTGGAAETLY | 0.703099 |
| GNGGGGDAGNAGSGGN | ASGDAKTGGAAETLY | 0.57471675 |
| GTGGAGGAGNDAGS | ASGDAKTGGAAETLY | 0.46728003 |
| GTTNTGWFNS | ASGDAKTGGAAETLY | 0.4587682 |
| IFMGPLFTL | ASGDAKTGGAAETLY | 0.68871987 |
| ILAHAIEAI | ASGDAKTGGAAETLY | 0.6439309 |
| IRSAIREAL | ASGDAKTGGAAETLY | 0.7796835 |
| ITFAGGG | ASGDAKTGGAAETLY | 0.771869 |
| ITGMSGHQEV | ASGDAKTGGAAETLY | 0.75362533 |
| IVAGLACTL | ASGDAKTGGAAETLY | 0.7791776 |
| IVGGGLAGA | ASGDAKTGGAAETLY | 0.76082313 |
| IVTGVLNI | ASGDAKTGGAAETLY | 0.70509374 |
| LAQQGCI | ASGDAKTGGAAETLY | 0.5994798 |
| LCLHVEN | ASGDAKTGGAAETLY | 0.66520715 |
| LDAISDGS | ASGDAKTGGAAETLY | 0.43742228 |
| LGGFAAML | ASGDAKTGGAAETLY | 0.6771321 |
| LGGKNPMI | ASGDAKTGGAAETLY | 0.5923606 |
| LGPKGVAAIL | ASGDAKTGGAAETLY | 0.73690784 |
| LMVGVSI | ASGDAKTGGAAETLY | 0.20607042 |

| | | |
|---|---|---|
| LVGAAFVL | ASGDAKTGGAAETLY | 0.7598354 |
| LVLAVAPL | ASGDAKTGGAAETLY | 0.7853725 |
| MAAIAVIDAA | ASGDAKTGGAAETLY | 0.58630604 |
| NNGINAELQ | ASGDAKTGGAAETLY | 0.733083 |
| RGEPLVLVHG | ASGDAKTGGAAETLY | 0.72905827 |
| RGGAGGAAGLI | ASGDAKTGGAAETLY | 0.47510618 |
| RSYRMHPAV | ASGDAKTGGAAETLY | 0.76486313 |
| SAAHLLAGQLA | ASGDAKTGGAAETLY | 0.5266199 |
| SAAVGSY | ASGDAKTGGAAETLY | 0.75283015 |
| SPHAVLHRI | ASGDAKTGGAAETLY | 0.742282 |
| SQIAKVL | ASGDAKTGGAAETLY | 0.7100555 |
| TALGHLVP | ASGDAKTGGAAETLY | 0.6598288 |
| TAPTTSGAA | ASGDAKTGGAAETLY | 0.7083421 |
| TDGDDGNGGSPGT | ASGDAKTGGAAETLY | 0.29199496 |
| TDGLGEN | ASGDAKTGGAAETLY | 0.70751226 |
| TGGPPLPGQA | ASGDAKTGGAAETLY | 0.44666627 |
| TIPAITI | ASGDAKTGGAAETLY | 0.68186486 |
| TIPGFTI | ASGDAKTGGAAETLY | 0.36613518 |
| TSIASGALA | ASGDAKTGGAAETLY | 0.7725184 |
| TVTLPGAAI | ASGDAKTGGAAETLY | 0.74032027 |
| VASGLSAALP | ASGDAKTGGAAETLY | 0.7083372 |
| VGAGVIGI | ASGDAKTGGAAETLY | 0.48911947 |
| VGPKRFGV | ASGDAKTGGAAETLY | 0.75712824 |
| VNVAFPD | ASGDAKTGGAAETLY | 0.73816043 |
| VSMTNSGVS | ASGDAKTGGAAETLY | 0.5255191 |
| VTDVLGDLSV | ASGDAKTGGAAETLY | 0.63624215 |
| VVVVGAGFAGL | ASGDAKTGGAAETLY | 0.5427346 |
| VYTVGTVV | ASGDAKTGGAAETLY | 0.5070834 |
| WTTFSGP | ASGDAKTGGAAETLY | 0.7623793 |
| YDELVEAF | ASGDAKTGGAAETLY | 0.73257226 |
| YGTGGAGGNGG | ASGDAKTGGAAETLY | 0.25019076 |
| YTNVMAV | ASGDAKTGGAAETLY | 0.6570502 |
| FLEQVHQL | ASGDAKTGGAAETLY | 0.72158253 |
| GLSLLAVAV | ASGDAKTGGAAETLY | 0.50246173 |
| LEQVHQL | ASGDAKTGGAAETLY | 0.22566259 |
| LNAAGAI | ASGDAKTGGAAETLY | 0.69011545 |
| PAVHASL | ASGDAKTGGAAETLY | 0.5084478 |
| TELEDDA | ASGDAKTGGAAETLY | 0.38769537 |
| TPPGPPN | ASGDAKTGGAAETLY | 0.6084529 |
| TPPPPPPP | ASGDAKTGGAAETLY | 0.7778323 |
| TQSPALI | ASSIRTGEYEQY | 0.5376437 |
| EFAAAFPQ | ASSIRTGEYEQY | 0.74089575 |
| LLSGNLDVL | ASSIRTGEYEQY | 0.7558229 |
| GPANLGNYNI | ASSIRTGEYEQY | 0.7467479 |
| TLGPTLP | ASSIRTGEYEQY | 0.7790471 |
| VYAYSSCCK | ASSIRTGEYEQY | 0.73598355 |
| APAPAPAPNI | ASSIRTGEYEQY | 0.4218425 |
| PGGPVPIP | ASSIRTGEYEQY | 0.72397625 |
| FLTVALEP | ASSIRTGEYEQY | 0.54245865 |
| SQYFGANL | ASSIRTGEYEQY | 0.6993519 |
| GGVKPAA | ASSIRTGEYEQY | 0.55040336 |

| | | |
|---|---|---|
| GGTGGAGGAGGW | ASSIRTGEYEQY | 0.47052073 |
| GVGGVAGLGGAGG | ASSIRTGEYEQY | 0.6202259 |
| VLLVLATTI | ASSIRTGEYEQY | 0.713624 |
| IGIFAAML | ASSIRTGEYEQY | 0.75151324 |
| AVGDLID | ASSIRTGEYEQY | 0.55599093 |
| NISPLPL | ASSIRTGEYEQY | 0.36162388 |
| VTAVFVAI | ASSIRTGEYEQY | 0.67860126 |
| SGDDNDAAL | ASSIRTGEYEQY | 0.3363515 |
| AVAALAHS | ASSIRTGEYEQY | 0.76148224 |
| GGVTVMATP | ASSIRTGEYEQY | 0.48907268 |
| AGLAANVRD | ASSIRTGEYEQY | 0.7919209 |
| TNRPDLIDPALLRPG | ASSIRTGEYEQY | 0.7705623 |
| LANLGVTAL | ASSIRTGEYEQY | 0.6818656 |
| TGERIEL | ASSIRTGEYEQY | 0.7657429 |
| NGEELAA | ASSIRTGEYEQY | 0.45892957 |
| VLSLVWG | ASSIRTGEYEQY | 0.5923718 |
| IGYALWL | ASSIRTGEYEQY | 0.60267437 |
| LSLLLAPL | ASSIRTGEYEQY | 0.65388536 |
| TLDEALK | ASSIRTGEYEQY | 0.70696497 |
| TDGAGTAEGG | ASSIRTGEYEQY | 0.64950985 |
| NAPGGAGGNGG | ASSIRTGEYEQY | 0.25246656 |
| VSVFMGMGGNF | ASSIRTGEYEQY | 0.774817 |
| LASYLAAL | ASSIRTGEYEQY | 0.5777168 |
| LVGGTPAL | ASSIRTGEYEQY | 0.41603988 |
| VRNADVL | ASSIRTGEYEQY | 0.59871894 |
| TCSSPFLVSAM | ASSIRTGEYEQY | 0.7812574 |
| VHITDAE | ASSIRTGEYEQY | 0.7816032 |
| EYAAPSF | ASSIRTGEYEQY | 0.6378139 |
| VVDAFHGGL | ASSIRTGEYEQY | 0.49449283 |
| ASVVDPQ | ASSIRTGEYEQY | 0.76836187 |
| SYNEFGGLS | ASSIRTGEYEQY | 0.5888085 |
| DPDGSVY | ASSIRTGEYEQY | 0.6340276 |
| MVSTIHATE | ASSIRTGEYEQY | 0.7062805 |
| LAAGVADPV | ASSIRTGEYEQY | 0.6582368 |
| TKEQIAA | ASSIRTGEYEQY | 0.52794874 |
| SLASPAAL | ASSIRTGEYEQY | 0.73977447 |
| IGLVFYGV | ASSIRTGEYEQY | 0.62683916 |
| FLLDASNL | ASSIRTGEYEQY | 0.62686324 |
| YQVGNFP | ASSIRTGEYEQY | 0.21254534 |
| ESLLSQST | ASSIRTGEYEQY | 0.6652564 |
| LFTVGFAAVG | ASSIRTGEYEQY | 0.66012335 |
| ATPPTPT | ASSIRTGEYEQY | 0.37135005 |
| TDLATLG | ASSIRTGEYEQY | 0.24123031 |
| SAGGTGR | ASSIRTGEYEQY | 0.6287502 |
| GFSNSGS | ASSIRTGEYEQY | 0.6969681 |
| GGNGGSGGTSV | ASSIRTGEYEQY | 0.77247834 |
| GTSVATGGAGN | ASSIRTGEYEQY | 0.6709672 |
| NGGNGGSGGTS | ASSIRTGEYEQY | 0.76349986 |
| AVVKFPH | ASSIRTGEYEQY | 0.65458167 |
| TPPPPPP | ASSIRTGEYEQY | 0.67075706 |
| AAPAAAAPA | ASSIRTGEYEQY | 0.30666977 |

| | | |
|---|---|---|
| IALNEDL | ASSIRTGEYEQY | 0.7260872 |
| IVNVSSM | ASSIRTGEYEQY | 0.73632956 |
| TGIPPAPR | ASSIRTGEYEQY | 0.7185197 |
| FDLGGGTF | ASSIRTGEYEQY | 0.6933756 |
| FDLGGGTFDVS | ASSIRTGEYEQY | 0.49944562 |
| NEPTAAA | ASSIRTGEYEQY | 0.74283075 |
| NEPTAAALA | ASSIRTGEYEQY | 0.72515756 |
| GSIINISSI | ASSIRTGEYEQY | 0.44962794 |
| VAPGNAGTA | ASSIRTGEYEQY | 0.44772673 |
| TGGGASLE | ASSIRTGEYEQY | 0.52609146 |
| VAVYNHYKRI | ASSIRTGEYEQY | 0.29141808 |
| TDPAPATT | ASSIRTGEYEQY | 0.7441237 |
| TNRPDILDPALLRPG | ASSIRTGEYEQY | 0.6686964 |
| LTTTTTS | ASSIRTGEYEQY | 0.7142932 |
| PTGSGKTL | ASSIRTGEYEQY | 0.6714821 |
| TGSGKTL | ASSIRTGEYEQY | 0.6720309 |
| TIGHVDHGK | ASSIRTGEYEQY | 0.6867842 |
| TIGHVDHGKTT | ASSIRTGEYEQY | 0.72373414 |
| GHVDHGKTT | ASSIRTGEYEQY | 0.6078489 |
| AQFEHTLL | ASSIRTGEYEQY | 0.73667634 |
| IVNQDEL | ASSIRTGEYEQY | 0.48108757 |
| TNGSQFF | ASSIRTGEYEQY | 0.59795594 |
| TGKGIVE | ASSIRTGEYEQY | 0.78368986 |
| LISNASDA | ASSIRTGEYEQY | 0.3973068 |
| AELRTLGELTKIAKAHGAQ | ASSIRTGEYEQY | 0.6125995 |
| APNAVAAAEPSW | ASSIRTGEYEQY | 0.73432803 |
| APQVAGAATAVL | ASSIRTGEYEQY | 0.5685493 |
| ARRDPAAAAAILPTDA | ASSIRTGEYEQY | 0.7612264 |
| DAAARDVLAEAGLAEYF | ASSIRTGEYEQY | 0.7492031 |
| DDQIVLSAKANLAKVEQ | ASSIRTGEYEQY | 0.7414596 |
| DELLPDTGEDSDEDGH | ASSIRTGEYEQY | 0.74285007 |
| ELPEAPANRATVPQD | ASSIRTGEYEQY | 0.7828585 |
| FGQSKIERVTGIG | ASSIRTGEYEQY | 0.76627207 |
| GAGGTGSTAGGAGGAGGAGGLY | ASSIRTGEYEQY | 0.5734139 |
| GHGGTGAAGGNGGTGGAGGSGI | ASSIRTGEYEQY | 0.7821039 |
| GNGGNGGAGGEGGPGGAGGAGA | ASSIRTGEYEQY | 0.41797268 |
| GSTAGNGGNGGNATNGTIAGQP | ASSIRTGEYEQY | 0.7579533 |
| GTGGDGGNGGNGGTGG | ASSIRTGEYEQY | 0.76364636 |
| GVRRPAGIPADAPV | ASSIRTGEYEQY | 0.4290743 |
| GYRQVTDAHLLTLARR | ASSIRTGEYEQY | 0.70956135 |
| IAKMGQDGHDRG | ASSIRTGEYEQY | 0.74463725 |
| KAEPSIEIQYT | ASSIRTGEYEQY | 0.78048825 |
| KPVRIRMAADEDTVAVT | ASSIRTGEYEQY | 0.4849481 |
| MPPRIAGMRLLVIKPEP | ASSIRTGEYEQY | 0.6500241 |
| MKFYAVALFRSRGVPPTR | ASSIRTGEYEQY | 0.5695008 |
| NAGGFGILGAVAHSP | ASSIRTGEYEQY | 0.3721221 |
| NEGREITLAMAEM | ASSIRTGEYEQY | 0.377272 |
| NELVEWSHPLGHRWRWE | ASSIRTGEYEQY | 0.7320392 |
| NLGAGGTGGNGGNPG | ASSIRTGEYEQY | 0.67112994 |
| NTEWPKWETVMAADPVD | ASSIRTGEYEQY | 0.7756064 |
| PASAAPAETQP | ASSIRTGEYEQY | 0.6935822 |

| | | |
|---|---|---|
| QQSSDGKNGVRAAEITGE | ASSIRTGEYEQY | 0.7625402 |
| RAPDAVTTSWWKE | ASSIRTGEYEQY | 0.5570556 |
| SGFGNVGAHTSGWFNQ | ASSIRTGEYEQY | 0.7317164 |
| SGVAGAVGQAASVGGLKV | ASSIRTGEYEQY | 0.2912179 |
| SNGGAGTGSAGGNGGT | ASSIRTGEYEQY | 0.37303567 |
| TGGAGGEGGAAG | ASSIRTGEYEQY | 0.3676588 |
| TGGDGGAAGAGGAPGHG | ASSIRTGEYEQY | 0.7291038 |
| TMPPLLAAIANAGRHGV | ASSIRTGEYEQY | 0.23818146 |
| TSSTAGNSGGA | ASSIRTGEYEQY | 0.7555095 |
| VPATVPVGILC | ASSIRTGEYEQY | 0.71785176 |
| VQRPDSGGVVLTGRIS | ASSIRTGEYEQY | 0.7324449 |
| WTPNPVPATKGVPAGLVQAT | ASSIRTGEYEQY | 0.61479557 |
| YALKRALAKFSDRFDVVII | ASSIRTGEYEQY | 0.57962966 |
| YLIGSVIV | ASSIRTGEYEQY | 0.5897525 |
| YGQLVAVID | ASSIRTGEYEQY | 0.7800617 |
| VTVGGMTLPA | ASSIRTGEYEQY | 0.42786413 |
| VLALPAV | ASSIRTGEYEQY | 0.25386146 |
| VIPPTLNYET | ASSIRTGEYEQY | 0.45291987 |
| VGMGGMHPSSG | ASSIRTGEYEQY | 0.35692483 |
| VGLAGGAATA | ASSIRTGEYEQY | 0.768872 |
| VGGAGGAGGSGP | ASSIRTGEYEQY | 0.75067306 |
| VFAAFMV | ASSIRTGEYEQY | 0.7468133 |
| TLYDKLR | ASSIRTGEYEQY | 0.78695166 |
| TLAVFTVL | ASSIRTGEYEQY | 0.2134217 |
| TGSVQPARV | ASSIRTGEYEQY | 0.6326293 |
| TGGGAGSGGT | ASSIRTGEYEQY | 0.7848977 |
| SHLETSL | ASSIRTGEYEQY | 0.65723836 |
| SFTTGMT | ASSIRTGEYEQY | 0.6885714 |
| QVEVAKRMGVRQPTVS | ASSIRTGEYEQY | 0.7692061 |
| QGSGGPALAM | ASSIRTGEYEQY | 0.68338287 |
| MLAPSNP | ASSIRTGEYEQY | 0.5702523 |
| LVNNAAI | ASSIRTGEYEQY | 0.7336992 |
| LTGEINTLA | ASSIRTGEYEQY | 0.6886121 |
| LSIFGPV | ASSIRTGEYEQY | 0.4435845 |
| LLALLANL | ASSIRTGEYEQY | 0.35898215 |
| LISNASDALDKLR | ASSIRTGEYEQY | 0.74506545 |
| LINEAVP | ASSIRTGEYEQY | 0.7881311 |
| LGYAFSGL | ASSIRTGEYEQY | 0.7369946 |
| LDFVAVAL | ASSIRTGEYEQY | 0.5826026 |
| LAQPGAE | ASSIRTGEYEQY | 0.7297207 |
| ISNLGAVTGF | ASSIRTGEYEQY | 0.7251481 |
| HQEDPGRI | ASSIRTGEYEQY | 0.556926 |
| GTPGEPTP | ASSIRTGEYEQY | 0.5369376 |
| GFFNSGS | ASSIRTGEYEQY | 0.763401 |
| GAGGGGLTAGAN | ASSIRTGEYEQY | 0.51275325 |
| FVDALVNM | ASSIRTGEYEQY | 0.71102226 |
| FLTVAPEL | ASSIRTGEYEQY | 0.51135087 |
| FETLTAL | ASSIRTGEYEQY | 0.78719443 |
| DVINPIEL | ASSIRTGEYEQY | 0.7374212 |
| APVSGAPV | ASSIRTGEYEQY | 0.73638123 |
| ALVGGLAW | ASSIRTGEYEQY | 0.6621076 |

| | | |
|---|---|---|
| ALAAAAATAI | ASSIRTGEYEQY | 0.59032464 |
| AFGLGLLL | ASSIRTGEYEQY | 0.7061237 |
| AFFAAASL | ASSIRTGEYEQY | 0.24693349 |
| AAAALPAFSPP | ASSIRTGEYEQY | 0.77383876 |
| VVLAPAL | ASSIRTGEYEQY | 0.5797822 |
| TIGHVDHGKTTL | ASSIRTGEYEQY | 0.58032864 |
| LISNASDALDK | ASSIRTGEYEQY | 0.7545434 |
| LISNASDAL | ASSIRTGEYEQY | 0.7852357 |
| ISNASDALDK | ASSIRTGEYEQY | 0.5605279 |
| NQAETLVYQTE | ASSIRTGEYEQY | 0.75383186 |
| TGGTGGTAGAG | ASSIRTGEYEQY | 0.65300727 |
| TGSTAGGAGGA | ASSIRTGEYEQY | 0.7296483 |
| DPLAVPVAAGPV | ASSIRTGEYEQY | 0.78163266 |
| GGTGGTPGNGGN | ASSIRTGEYEQY | 0.72370577 |
| PAGLAGVDPATL | ASSIRTGEYEQY | 0.7890856 |
| GNNLVVEAPHAGASV | ASSIRTGEYEQY | 0.6011734 |
| KVTRIMSASTLEQPA | ASSIRTGEYEQY | 0.3176283 |
| LGFALPIALPRRTVV | ASSIRTGEYEQY | 0.6260228 |
| TRLGNLTARKLVVLV | ASSIRTGEYEQY | 0.70247877 |
| WSQQRRLIYFPSAGP | ASSIRTGEYEQY | 0.7492488 |
| GPNGNPDPMAAAVDIR | ASSIRTGEYEQY | 0.7388526 |
| TQRCVSVADLRAAASR | ASSIRTGEYEQY | 0.7413309 |
| QFVLDVAEHPVEVLTETV | ASSIRTGEYEQY | 0.47373974 |
| GGAGGTGGAGGNGGTQAGDG | ASSIRTGEYEQY | 0.7687408 |
| PLLATLRAIAVDVDAKPAQV | ASSIRTGEYEQY | 0.7487142 |
| APKFRTGGYMIDFWGVGYQVAK | ASSIRTGEYEQY | 0.75520504 |
| GAGGAGGAGGAAGGPGGTGGTG | ASSIRTGEYEQY | 0.50654733 |
| IGALGLLATDVAPAGSVAFGVA | ASSIRTGEYEQY | 0.49222597 |
| AGGAGGIGGNGGAGGAGGVLMG | ASSIRTGEYEQY | 0.6740009 |
| ADELIGLL | ASSIRTGEYEQY | 0.7282134 |
| ADRALKA | ASSIRTGEYEQY | 0.42000222 |
| AGAVGQVGGAGGA | ASSIRTGEYEQY | 0.7191855 |
| AGGGGGLGGAP | ASSIRTGEYEQY | 0.743459 |
| AGGLLYGNGGAG | ASSIRTGEYEQY | 0.33210376 |
| AGLESIGL | ASSIRTGEYEQY | 0.50911224 |
| AGLKTGII | ASSIRTGEYEQY | 0.7709322 |
| ASLAALLL | ASSIRTGEYEQY | 0.6273181 |
| ASVINISSI | ASSIRTGEYEQY | 0.57653373 |
| ATPPGPA | ASSIRTGEYEQY | 0.7753431 |
| ATVASALV | ASSIRTGEYEQY | 0.784408 |
| ATVASLIL | ASSIRTGEYEQY | 0.7645416 |
| AVNGPVLIRH | ASSIRTGEYEQY | 0.5004136 |
| DGAAGAVGI | ASSIRTGEYEQY | 0.21796513 |
| DPLLHGI | ASSIRTGEYEQY | 0.75036466 |
| ELGAALGAL | ASSIRTGEYEQY | 0.7797061 |
| ELGRTQAV | ASSIRTGEYEQY | 0.71036756 |
| FVAGGYLS | ASSIRTGEYEQY | 0.68073386 |
| GASGGAGGAGGW | ASSIRTGEYEQY | 0.632126 |
| GGAGAGGDGGAGGI | ASSIRTGEYEQY | 0.7517766 |
| GTGHNGSQPGMGGNG | ASSIRTGEYEQY | 0.52990246 |
| GYAAASASAS | ASSIRTGEYEQY | 0.27197775 |

| | | |
|---|---|---|
| HQVDVAAVQQG | ASSIRTGEYEQY | 0.6515179 |
| IGAAIAHT | ASSIRTGEYEQY | 0.7670661 |
| IGMIEVAQL | ASSIRTGEYEQY | 0.62777543 |
| IRLAPEHTSAMD | ASSIRTGEYEQY | 0.6670946 |
| IYNVSVL | ASSIRTGEYEQY | 0.7505454 |
| KQGFLGM | ASSIRTGEYEQY | 0.53400326 |
| LESVITQL | ASSIRTGEYEQY | 0.7046392 |
| LGLGALVVV | ASSIRTGEYEQY | 0.5291115 |
| LIGNGGNGGSGGI | ASSIRTGEYEQY | 0.76126766 |
| LNAINAPTL | ASSIRTGEYEQY | 0.4637193 |
| LQLLNTGV | ASSIRTGEYEQY | 0.6796434 |
| LQTSIEF | ASSIRTGEYEQY | 0.7604656 |
| MSFAVSEL | ASSIRTGEYEQY | 0.7017475 |
| NTGALGL | ASSIRTGEYEQY | 0.63981473 |
| PPGAGGAGGAGTT | ASSIRTGEYEQY | 0.5780753 |
| QLVGLGI | ASSIRTGEYEQY | 0.35547042 |
| RFLEQVHQL | ASSIRTGEYEQY | 0.5638734 |
| RGVGYVLR | ASSIRTGEYEQY | 0.77660275 |
| SATGVFAGVI | ASSIRTGEYEQY | 0.48878723 |
| SLPAITLP | ASSIRTGEYEQY | 0.7757617 |
| STALAFTV | ASSIRTGEYEQY | 0.70304155 |
| SVLTVGLGK | ASSIRTGEYEQY | 0.57502556 |
| TALAGAGSGI | ASSIRTGEYEQY | 0.6431474 |
| TGLPFTGI | ASSIRTGEYEQY | 0.51505256 |
| VAEHVTV | ASSIRTGEYEQY | 0.72996473 |
| VDIVLNSL | ASSIRTGEYEQY | 0.58454645 |
| VLTGGGAVL | ASSIRTGEYEQY | 0.7424747 |
| VNVGLAYGI | ASSIRTGEYEQY | 0.5492619 |
| VSSAAVGAV | ASSIRTGEYEQY | 0.64211214 |
| VTSVYEAGV | ASSIRTGEYEQY | 0.71106446 |
| WVALDRGA | ASSIRTGEYEQY | 0.46470684 |
| YLVGSVVI | ASSIRTGEYEQY | 0.688125 |
| GPPGTGKTLL | ASSIRTGEYEQY | 0.7756829 |
| GVGMTKF | ASSIRTGEYEQY | 0.7030021 |
| IGIGGSDL | ASSIRTGEYEQY | 0.41334555 |
| IGTLLAEL | ASSIRTGEYEQY | 0.7249254 |
| IYTSGTT | ASSIRTGEYEQY | 0.56828547 |
| LSFPAGV | ASSIRTGEYEQY | 0.7337224 |
| LYTSGTT | ASSIRTGEYEQY | 0.6680589 |
| VGAFPFGF | ASSIRTGEYEQY | 0.65510136 |
| VGAGTVEF | ASSIRTGEYEQY | 0.67159355 |
| VGKSTLLL | ASSIRTGEYEQY | 0.2602686 |
| AALAHAGGGIV | ASSIRTGEYEQY | 0.4846056 |
| AALVAVSLL | ASSIRTGEYEQY | 0.5712209 |
| AAVSRAVAAL | ASSIRTGEYEQY | 0.7608406 |
| AGGSSGIGGF | ASSIRTGEYEQY | 0.7568506 |
| AGLTVANANG | ASSIRTGEYEQY | 0.51244867 |
| AGLVGAVL | ASSIRTGEYEQY | 0.70461124 |
| ALSQAFTV | ASSIRTGEYEQY | 0.5491767 |
| ALTFSVSL | ASSIRTGEYEQY | 0.45997667 |
| ALTGAALVAL | ASSIRTGEYEQY | 0.5980605 |

| | | |
|---|---|---|
| ASFSIGAGVL | ASSIRTGEYEQY | 0.4455725 |
| DGSVYKI | ASSIRTGEYEQY | 0.41902563 |
| EQLAEIV | ASSIRTGEYEQY | 0.7250015 |
| FAVSELTVM | ASSIRTGEYEQY | 0.52021235 |
| GAAPLAAGL | ASSIRTGEYEQY | 0.7081555 |
| GADGGDGGSGGAGG | ASSIRTGEYEQY | 0.7682659 |
| GFGLDTGGAGG | ASSIRTGEYEQY | 0.4958334 |
| GLASGGLL | ASSIRTGEYEQY | 0.6340935 |
| GRIAGAAVLL | ASSIRTGEYEQY | 0.6787121 |
| GSAGSLNTSF | ASSIRTGEYEQY | 0.7495365 |
| GTGQIGI | ASSIRTGEYEQY | 0.7309284 |
| GTLIALIL | ASSIRTGEYEQY | 0.49240467 |
| IAATMGFVVG | ASSIRTGEYEQY | 0.7371499 |
| IFAQALL | ASSIRTGEYEQY | 0.75209653 |
| IGAAVALGF | ASSIRTGEYEQY | 0.7491527 |
| IGLTPLAL | ASSIRTGEYEQY | 0.70692927 |
| ILGSNTSTL | ASSIRTGEYEQY | 0.7777252 |
| IVFSLTVL | ASSIRTGEYEQY | 0.7552198 |
| LASKATGF | ASSIRTGEYEQY | 0.6660615 |
| LASVALIL | ASSIRTGEYEQY | 0.27738088 |
| LFAVGLTL | ASSIRTGEYEQY | 0.7245872 |
| LFNGGIGTYI | ASSIRTGEYEQY | 0.7286838 |
| LTLGAGII | ASSIRTGEYEQY | 0.51502216 |
| LVAAALGDI | ASSIRTGEYEQY | 0.277476 |
| NTAFVQLGI | ASSIRTGEYEQY | 0.7760383 |
| QLFSGAEL | ASSIRTGEYEQY | 0.7856725 |
| SAATFITL | ASSIRTGEYEQY | 0.7082106 |
| SNYGLANLGV | ASSIRTGEYEQY | 0.76660705 |
| SYGGAYAVM | ASSIRTGEYEQY | 0.31736827 |
| TTGVGGLTLG | ASSIRTGEYEQY | 0.7456155 |
| TTQTIYI | ASSIRTGEYEQY | 0.7803876 |
| VAAGAGLGAVI | ASSIRTGEYEQY | 0.58249885 |
| VAAVVTLL | ASSIRTGEYEQY | 0.74869764 |
| VGFLGGLI | ASSIRTGEYEQY | 0.32638687 |
| VGVIPILL | ASSIRTGEYEQY | 0.4962604 |
| VIFASSLI | ASSIRTGEYEQY | 0.78211975 |
| VKGGDGGAAAG | ASSIRTGEYEQY | 0.7345849 |
| VTKTFTNI | ASSIRTGEYEQY | 0.7037877 |
| VVAAAILL | ASSIRTGEYEQY | 0.7345412 |
| YAALDVEL | ASSIRTGEYEQY | 0.4794978 |
| LASVALLL | ASSIRTGEYEQY | 0.67567104 |
| LDAVHRV | ASSIRTGEYEQY | 0.7740861 |
| QGLSSSIF | ASSIRTGEYEQY | 0.6928649 |
| RGTFANIRL | ASSIRTGEYEQY | 0.73696387 |
| RQLWWGHRI | ASSIRTGEYEQY | 0.5649244 |
| YGPPGTGKTLL | ASSIRTGEYEQY | 0.7787288 |
| AALVGAPANVA | ASSIRTGEYEQY | 0.7869221 |
| AATFRRTE | ASSIRTGEYEQY | 0.7295674 |
| AGALGGGI | ASSIRTGEYEQY | 0.4830662 |
| AGGAGGAGGIGAG | ASSIRTGEYEQY | 0.7777003 |
| AGSGALGL | ASSIRTGEYEQY | 0.7321882 |

| | | |
|---|---|---|
| AQIADQI | ASSIRTGEYEQY | 0.72625625 |
| ASAVAVTYL | ASSIRTGEYEQY | 0.7303279 |
| ASTVLGGTGG | ASSIRTGEYEQY | 0.13913845 |
| AVGGEPAK | ASSIRTGEYEQY | 0.49492925 |
| CEQGTFGGTC | ASSIRTGEYEQY | 0.2386939 |
| DEGKAPAG | ASSIRTGEYEQY | 0.20602915 |
| DTVVFPAL | ASSIRTGEYEQY | 0.15899086 |
| FEGITMAG | ASSIRTGEYEQY | 0.12290337 |
| FPLVVQD | ASSIRTGEYEQY | 0.16625163 |
| GAFTTPQ | ASSIRTGEYEQY | 0.1455316 |
| GFGGFGTAGGVGGA | ASSIRTGEYEQY | 0.65893686 |
| GFLAQFL | ASSIRTGEYEQY | 0.21849902 |
| GGAGGAGGNAGVA | ASSIRTGEYEQY | 0.16366263 |
| GGVGTAGAGGAGGAGG | ASSIRTGEYEQY | 0.35399222 |
| GNGGGGDAGNAGSGGN | ASSIRTGEYEQY | 0.35022303 |
| GTGGAGGAGNDAGS | ASSIRTGEYEQY | 0.12223943 |
| GTTNTGWFNS | ASSIRTGEYEQY | 0.18509568 |
| IFMGPLFTL | ASSIRTGEYEQY | 0.13935767 |
| ILAHAIEAI | ASSIRTGEYEQY | 0.16981892 |
| IRSAIREAL | ASSIRTGEYEQY | 0.10363155 |
| ITFAGGG | ASSIRTGEYEQY | 0.21073979 |
| ITGMSGHQEV | ASSIRTGEYEQY | 0.14583796 |
| IVAGLACTL | ASSIRTGEYEQY | 0.09604841 |
| IVGGGLAGA | ASSIRTGEYEQY | 0.20433858 |
| IVTGVLNI | ASSIRTGEYEQY | 0.16769864 |
| LAQQGCI | ASSIRTGEYEQY | 0.19506411 |
| LCLHVEN | ASSIRTGEYEQY | 0.34932497 |
| LDAISDGS | ASSIRTGEYEQY | 0.12564166 |
| LGGFAAML | ASSIRTGEYEQY | 0.15567607 |
| LGGKNPMI | ASSIRTGEYEQY | 0.26942056 |
| LGPKGVAAIL | ASSIRTGEYEQY | 0.42992476 |
| LMVGVSI | ASSIRTGEYEQY | 0.14613608 |
| LVGAAFVL | ASSIRTGEYEQY | 0.20937894 |
| LVLAVAPL | ASSIRTGEYEQY | 0.45761675 |
| MAAIAVIDAA | ASSIRTGEYEQY | 0.21178375 |
| NNGINAELQ | ASSIRTGEYEQY | 0.23886791 |
| RGEPLVLVHG | ASSIRTGEYEQY | 0.28431576 |
| RGGAGGAAGLI | ASSIRTGEYEQY | 0.12875636 |
| RSYRMHPAV | ASSIRTGEYEQY | 0.33788866 |
| SAAHLLAGQLA | ASSIRTGEYEQY | 0.4585968 |
| SAAVGSY | ASSIRTGEYEQY | 0.19187464 |
| SPHAVLHRI | ASSIRTGEYEQY | 0.17392288 |
| SQIAKVL | ASSIRTGEYEQY | 0.13744335 |
| TALGHLVP | ASSIRTGEYEQY | 0.15772463 |
| TAPTTSGAA | ASSIRTGEYEQY | 0.110847734 |
| TDGDDGNGGSPGT | ASSIRTGEYEQY | 0.252073 |
| TDGLGEN | ASSIRTGEYEQY | 0.3089425 |
| TGGPPLPGQA | ASSIRTGEYEQY | 0.16237672 |
| TIPAITI | ASSIRTGEYEQY | 0.11434635 |
| TIPGFTI | ASSIRTGEYEQY | 0.120635316 |
| TSIASGALA | ASSIRTGEYEQY | 0.18345553 |

| | | |
|---|---|---|
| TVTLPGAAI | ASSIRTGEYEQY | 0.14967243 |
| VASGLSAALP | ASSIRTGEYEQY | 0.11278745 |
| VGAGVIGI | ASSIRTGEYEQY | 0.14013293 |
| VGPKRFGV | ASSIRTGEYEQY | 0.29301944 |
| VNVAFPD | ASSIRTGEYEQY | 0.3031816 |
| VSMTNSGVS | ASSIRTGEYEQY | 0.12980907 |
| VTDVLGDLSV | ASSIRTGEYEQY | 0.45337215 |
| VVVVGAGFAGL | ASSIRTGEYEQY | 0.38708222 |
| VYTVGTVV | ASSIRTGEYEQY | 0.37765032 |
| WTTFSGP | ASSIRTGEYEQY | 0.17591502 |
| YDELVEAF | ASSIRTGEYEQY | 0.14734152 |
| YGTGGAGGNGG | ASSIRTGEYEQY | 0.70716476 |
| YTNVMAV | ASSIRTGEYEQY | 0.17949249 |
| FLEQVHQL | ASSIRTGEYEQY | 0.16413787 |
| GLSLLAVAV | ASSIRTGEYEQY | 0.2656049 |
| LEQVHQL | ASSIRTGEYEQY | 0.21342134 |
| LNAAGAI | ASSIRTGEYEQY | 0.14405043 |
| PAVHASL | ASSIRTGEYEQY | 0.325537 |
| TELEDDA | ASSIRTGEYEQY | 0.23502523 |
| TPPGPPN | ASSIRTGEYEQY | 0.11649102 |
| TPPPPPPP | ASSIRTGEYEQY | 0.15542658 |
| TQSPALI | TCSADRGQAPL | 0.12852207 |
| EFAAAFPQ | TCSADRGQAPL | 0.19175601 |
| LLSGNLDVL | TCSADRGQAPL | 0.36982572 |
| GPANLGNYNI | TCSADRGQAPL | 0.20340262 |
| TLGPTLP | TCSADRGQAPL | 0.17291099 |
| VYAYSSCCK | TCSADRGQAPL | 0.28561798 |
| APAPAPAPNI | TCSADRGQAPL | 0.2439119 |
| PGGPVPIP | TCSADRGQAPL | 0.18949707 |
| FLTVALEP | TCSADRGQAPL | 0.23932132 |
| SQYFGANL | TCSADRGQAPL | 0.15315253 |
| GGVKPAA | TCSADRGQAPL | 0.24220821 |
| GGTGGAGGAGGW | TCSADRGQAPL | 0.22349335 |
| GVGGVAGLGGAGG | TCSADRGQAPL | 0.2178676 |
| VLLVLATTI | TCSADRGQAPL | 0.22943923 |
| IGIFAAML | TCSADRGQAPL | 0.31896934 |
| AVGDLID | TCSADRGQAPL | 0.1862025 |
| NISPLPL | TCSADRGQAPL | 0.1124488 |
| VTAVFVAI | TCSADRGQAPL | 0.13682811 |
| SGDDNDAAL | TCSADRGQAPL | 0.2457254 |
| AVAALAHS | TCSADRGQAPL | 0.22577542 |
| GGVTVMATP | TCSADRGQAPL | 0.23266286 |
| AGLAANVRD | TCSADRGQAPL | 0.21859625 |
| TNRPDLIDPALLRPG | TCSADRGQAPL | 0.13437116 |
| LANLGVTAL | TCSADRGQAPL | 0.24485555 |
| TGERIEL | TCSADRGQAPL | 0.34720486 |
| NGEELAA | TCSADRGQAPL | 0.11433891 |
| VLSLVWG | TCSADRGQAPL | 0.166064 |
| IGYALWL | TCSADRGQAPL | 0.6170111 |
| LSLLLAPL | TCSADRGQAPL | 0.35332412 |
| TLDEALK | TCSADRGQAPL | 0.16769835 |

| | | |
|---|---|---|
| TDGAGTAEGG | TCSADRGQAPL | 0.16202614 |
| NAPGGAGGNGG | TCSADRGQAPL | 0.7665 |
| VSVFMGMGGNF | TCSADRGQAPL | 0.13862185 |
| LASYLAAL | TCSADRGQAPL | 0.43011513 |
| LVGGTPAL | TCSADRGQAPL | 0.10625977 |
| VRNADVL | TCSADRGQAPL | 0.16871282 |
| TCSSPFLVSAM | TCSADRGQAPL | 0.13645458 |
| VHITDAE | TCSADRGQAPL | 0.10959183 |
| EYAAPSF | TCSADRGQAPL | 0.23018718 |
| VVDAFHGGL | TCSADRGQAPL | 0.16030826 |
| ASVVDPQ | TCSADRGQAPL | 0.30015653 |
| SYNEFGGLS | TCSADRGQAPL | 0.14896041 |
| DPDGSVY | TCSADRGQAPL | 0.27297407 |
| MVSTIHATE | TCSADRGQAPL | 0.24792996 |
| LAAGVADPV | TCSADRGQAPL | 0.20052737 |
| TKEQIAA | TCSADRGQAPL | 0.15512711 |
| SLASPAAL | TCSADRGQAPL | 0.21122642 |
| IGLVFYGV | TCSADRGQAPL | 0.26682073 |
| FLLDASNL | TCSADRGQAPL | 0.22875309 |
| YQVGNFP | TCSADRGQAPL | 0.12543713 |
| ESLLSQST | TCSADRGQAPL | 0.15147704 |
| LFTVGFAAVG | TCSADRGQAPL | 0.17565277 |
| ATPPTPT | TCSADRGQAPL | 0.1126274 |
| TDLATLG | TCSADRGQAPL | 0.101872444 |
| SAGGTGR | TCSADRGQAPL | 0.15087758 |
| GFSNSGS | TCSADRGQAPL | 0.13917439 |
| GGNGGSGGTSV | TCSADRGQAPL | 0.29715568 |
| GTSVATGGAGN | TCSADRGQAPL | 0.35108742 |
| NGGNGGSGGTS | TCSADRGQAPL | 0.16677836 |
| AVVKFPH | TCSADRGQAPL | 0.34449485 |
| TPPPPPP | TCSADRGQAPL | 0.28492537 |
| AAPAAAAAPA | TCSADRGQAPL | 0.09398835 |
| IALNEDL | TCSADRGQAPL | 0.14622247 |
| IVNVSSM | TCSADRGQAPL | 0.121165276 |
| TGIPPAPR | TCSADRGQAPL | 0.23438352 |
| FDLGGGTF | TCSADRGQAPL | 0.13400695 |
| FDLGGGTFDVS | TCSADRGQAPL | 0.13027082 |
| NEPTAAA | TCSADRGQAPL | 0.1961025 |
| NEPTAAALA | TCSADRGQAPL | 0.5348643 |
| GSIINISSI | TCSADRGQAPL | 0.17365739 |
| VAPGNAGTA | TCSADRGQAPL | 0.13107517 |
| TGGGASLE | TCSADRGQAPL | 0.10106909 |
| VAVYNHYKRI | TCSADRGQAPL | 0.24388374 |
| TDPAPATT | TCSADRGQAPL | 0.14429215 |
| TNRPDILDPALLRPG | TCSADRGQAPL | 0.15528613 |
| LTTTTTS | TCSADRGQAPL | 0.1075478 |
| PTGSGKTL | TCSADRGQAPL | 0.16952693 |
| TGSGKTL | TCSADRGQAPL | 0.12017463 |
| TIGHVDHGK | TCSADRGQAPL | 0.25077322 |
| TIGHVDHGKTT | TCSADRGQAPL | 0.15795581 |
| GHVDHGKTT | TCSADRGQAPL | 0.15659189 |

| | | |
|---|---|---|
| AQFEHTLL | TCSADRGQAPL | 0.42983234 |
| IVNQDEL | TCSADRGQAPL | 0.6254602 |
| TNGSQFF | TCSADRGQAPL | 0.16540585 |
| TGKGIVE | TCSADRGQAPL | 0.19166976 |
| LISNASDA | TCSADRGQAPL | 0.133312 |
| AELRTLGELTKIAKAHGAQ | TCSADRGQAPL | 0.15348434 |
| APNAVAAAEPSW | TCSADRGQAPL | 0.15434068 |
| APQVAGAATAVL | TCSADRGQAPL | 0.41020972 |
| ARRDPAAAAAILPTDA | TCSADRGQAPL | 0.16125077 |
| DAAARDVLAEAGLAEYF | TCSADRGQAPL | 0.12690696 |
| DDQIVLSAKANLAKVEQ | TCSADRGQAPL | 0.204147 |
| DELLPDTGEDSDEDGH | TCSADRGQAPL | 0.31927252 |
| ELPEAPANRATVPQD | TCSADRGQAPL | 0.6073692 |
| FGQSKIERVTGIG | TCSADRGQAPL | 0.14250737 |
| GAGGTGSTAGGAGGAGGAGGLY | TCSADRGQAPL | 0.16275238 |
| GHGGTGAAGGNGGTGGAGGSGI | TCSADRGQAPL | 0.17531365 |
| GNGGNGGAGGEGGPGGAGGAGA | TCSADRGQAPL | 0.17695072 |
| GSTAGNGGNGGNATNGTIAGQP | TCSADRGQAPL | 0.14949614 |
| GTGGDGGNGGNGGTGG | TCSADRGQAPL | 0.1497636 |
| GVRRPAGIPADAPV | TCSADRGQAPL | 0.22854605 |
| GYRQVTDAHLLTLARR | TCSADRGQAPL | 0.7592139 |
| IAKMGQDGHDRG | TCSADRGQAPL | 0.16218847 |
| KAEPSIEIQYT | TCSADRGQAPL | 0.10646241 |
| KPVRIRMAADEDTVAVT | TCSADRGQAPL | 0.18709862 |
| MPPRIAGMRLLVIKPEP | TCSADRGQAPL | 0.12845132 |
| MKFYAVALFRSRGVPPTR | TCSADRGQAPL | 0.14963967 |
| NAGGFGILGAVAHSP | TCSADRGQAPL | 0.22993676 |
| NEGREITLAMAEM | TCSADRGQAPL | 0.11246487 |
| NELVEWSHPLGHRWRWE | TCSADRGQAPL | 0.15453786 |
| NLGAGGTGGNGGNPG | TCSADRGQAPL | 0.6710986 |
| NTEWPKWETVMAADPVD | TCSADRGQAPL | 0.19401896 |
| PASAAPAETQP | TCSADRGQAPL | 0.27605206 |
| QQSSDGKNGVRAAEITGE | TCSADRGQAPL | 0.42542422 |
| RAPDAVTTSWWKE | TCSADRGQAPL | 0.22880006 |
| SGFGNVGAHTSGWFNQ | TCSADRGQAPL | 0.47688904 |
| SGVAGAVGQAASVGGLKV | TCSADRGQAPL | 0.42691883 |
| SNGGAGTGSAGGNGGT | TCSADRGQAPL | 0.19114217 |
| TGGAGGEGGAAG | TCSADRGQAPL | 0.17411152 |
| TGGDGGAAGAGGAPGHG | TCSADRGQAPL | 0.2197161 |
| TMPPLLAAIANAGRHGV | TCSADRGQAPL | 0.18984294 |
| TSSTAGNSGGA | TCSADRGQAPL | 0.19744131 |
| VPATVPVGILC | TCSADRGQAPL | 0.13926391 |
| VQRPDSGGVVLTGRIS | TCSADRGQAPL | 0.32602155 |
| WTPNPVPATKGVPAGLVQAT | TCSADRGQAPL | 0.18245651 |
| YALKRALAKFSDRFDVVII | TCSADRGQAPL | 0.40309873 |
| YLIGSVIV | TCSADRGQAPL | 0.12426193 |
| YGQLVAVID | TCSADRGQAPL | 0.14740357 |
| VTVGGMTLPA | TCSADRGQAPL | 0.3438129 |
| VLALPAV | TCSADRGQAPL | 0.10111553 |
| VIPPTLNYET | TCSADRGQAPL | 0.106193125 |
| VGMGGMHPSSG | TCSADRGQAPL | 0.19092473 |

| | | |
|---|---|---|
| VGLAGGAATA | TCSADRGQAPL | 0.43941343 |
| VGGAGGAGGSGP | TCSADRGQAPL | 0.101123326 |
| VFAAFMV | TCSADRGQAPL | 0.15162504 |
| TLYDKLR | TCSADRGQAPL | 0.1264149 |
| TLAVFTVL | TCSADRGQAPL | 0.23908678 |
| TGSVQPARV | TCSADRGQAPL | 0.20919536 |
| TGGGAGSGGT | TCSADRGQAPL | 0.115024276 |
| SHLETSL | TCSADRGQAPL | 0.23238175 |
| SFTTGMT | TCSADRGQAPL | 0.10688576 |
| QVEVAKRMGVRQPTVS | TCSADRGQAPL | 0.1517247 |
| QGSGGPALAM | TCSADRGQAPL | 0.24889809 |
| MLAPSNP | TCSADRGQAPL | 0.64046717 |
| LVNNAAI | TCSADRGQAPL | 0.1282169 |
| LTGEINTLA | TCSADRGQAPL | 0.28710756 |
| LSIFGPV | TCSADRGQAPL | 0.12330676 |
| LLALLANL | TCSADRGQAPL | 0.7359942 |
| LISNASDALDKLR | TCSADRGQAPL | 0.16733095 |
| LINEAVP | TCSADRGQAPL | 0.11591975 |
| LGYAFSGL | TCSADRGQAPL | 0.32179618 |
| LDFVAVAL | TCSADRGQAPL | 0.24886292 |
| LAQPGAE | TCSADRGQAPL | 0.21006128 |
| ISNLGAVTGF | TCSADRGQAPL | 0.49256617 |
| HQEDPGRI | TCSADRGQAPL | 0.16667764 |
| GTPGEPTP | TCSADRGQAPL | 0.17612849 |
| GFFNSGS | TCSADRGQAPL | 0.3783599 |
| GAGGGGLTAGAN | TCSADRGQAPL | 0.3241291 |
| FVDALVNM | TCSADRGQAPL | 0.15611628 |
| FLTVAPEL | TCSADRGQAPL | 0.2587124 |
| FETLTAL | TCSADRGQAPL | 0.2427354 |
| DVINPIEL | TCSADRGQAPL | 0.15318066 |
| APVSGAPV | TCSADRGQAPL | 0.35696787 |
| ALVGGLAW | TCSADRGQAPL | 0.18423837 |
| ALAAAAATAI | TCSADRGQAPL | 0.29142416 |
| AFGLGLLL | TCSADRGQAPL | 0.1965592 |
| AFFAAASL | TCSADRGQAPL | 0.7669087 |
| AAAALPAFSPP | TCSADRGQAPL | 0.12811874 |
| VVLAPAL | TCSADRGQAPL | 0.1933122 |
| TIGHVDHGKTTL | TCSADRGQAPL | 0.22443151 |
| LISNASDALDK | TCSADRGQAPL | 0.25112614 |
| LISNASDAL | TCSADRGQAPL | 0.15595773 |
| ISNASDALDK | TCSADRGQAPL | 0.14417984 |
| NQAETLVYQTE | TCSADRGQAPL | 0.19538538 |
| TGGTGGTAGAG | TCSADRGQAPL | 0.42926997 |
| TGSTAGGAGGA | TCSADRGQAPL | 0.17320853 |
| DPLAVPVAAGPV | TCSADRGQAPL | 0.2004372 |
| GGTGGTPGNGGN | TCSADRGQAPL | 0.5875245 |
| PAGLAGVDPATL | TCSADRGQAPL | 0.15272151 |
| GNNLVVEAPHAGASV | TCSADRGQAPL | 0.20802741 |
| KVTRIMSASTLEQPA | TCSADRGQAPL | 0.26062524 |
| LGFALPIALPRRTVV | TCSADRGQAPL | 0.31585416 |
| TRLGNLTARKLVVLV | TCSADRGQAPL | 0.16895483 |

| | | |
|---|---|---|
| WSQQRRLIYFPSAGP | TCSADRGQAPL | 0.38075173 |
| GPNGNPDPMAAAVDIR | TCSADRGQAPL | 0.13337234 |
| TQRCVSVADLRAAASR | TCSADRGQAPL | 0.37679726 |
| QFVLDVAEHPVEVLTETV | TCSADRGQAPL | 0.12522732 |
| GGAGGTGGAGGNGGTQAGDG | TCSADRGQAPL | 0.1899753 |
| PLLATLRAIAVDVDAKPAQV | TCSADRGQAPL | 0.13932014 |
| APKFRTGGYMIDFWGVGYQVAK | TCSADRGQAPL | 0.29604357 |
| GAGGAGGAGGAAGGPGGTGGTG | TCSADRGQAPL | 0.38870168 |
| IGALGLLATDVAPAGSVAFGVA | TCSADRGQAPL | 0.31030357 |
| AGGAGGIGGNGGAGGAGGVLMG | TCSADRGQAPL | 0.10305769 |
| ADELIGLL | TCSADRGQAPL | 0.36221153 |
| ADRALKA | TCSADRGQAPL | 0.36101654 |
| AGAVGQVGGAGGA | TCSADRGQAPL | 0.78537416 |
| AGGGGGLGGAP | TCSADRGQAPL | 0.18740965 |
| AGGLLYGNGGAG | TCSADRGQAPL | 0.24868615 |
| AGLESIGL | TCSADRGQAPL | 0.20616993 |
| AGLKTGII | TCSADRGQAPL | 0.17131454 |
| ASLAALLL | TCSADRGQAPL | 0.11919972 |
| ASVINISSI | TCSADRGQAPL | 0.17891096 |
| ATPPGPA | TCSADRGQAPL | 0.15812245 |
| ATVASALV | TCSADRGQAPL | 0.14301035 |
| ATVASLIL | TCSADRGQAPL | 0.6689371 |
| AVNGPVLIRH | TCSADRGQAPL | 0.35361332 |
| DGAAGAVGI | TCSADRGQAPL | 0.4269015 |
| DPLLHGI | TCSADRGQAPL | 0.4517738 |
| ELGAALGAL | TCSADRGQAPL | 0.5195819 |
| ELGRTQAV | TCSADRGQAPL | 0.23320462 |
| FVAGGYLS | TCSADRGQAPL | 0.15140574 |
| GASGGAGGAGGW | TCSADRGQAPL | 0.15564185 |
| GGAGAGGDGGAGGI | TCSADRGQAPL | 0.11894145 |
| GTGHNGSQPGMGGNG | TCSADRGQAPL | 0.14216553 |
| GYAAASASAS | TCSADRGQAPL | 0.15787841 |
| HQVDVAAVQQG | TCSADRGQAPL | 0.19282483 |
| IGAAIAHT | TCSADRGQAPL | 0.101418406 |
| IGMIEVAQL | TCSADRGQAPL | 0.33230016 |
| IRLAPEHTSAMD | TCSADRGQAPL | 0.44142497 |
| IYNVSVL | TCSADRGQAPL | 0.25381505 |
| KQGFLGM | TCSADRGQAPL | 0.23555389 |
| LESVITQL | TCSADRGQAPL | 0.19806191 |
| LGLGALVVV | TCSADRGQAPL | 0.13195424 |
| LIGNGGNGGSGGI | TCSADRGQAPL | 0.33333418 |
| LNAINAPTL | TCSADRGQAPL | 0.13239688 |
| LQLLNTGV | TCSADRGQAPL | 0.23435819 |
| LQTSIEF | TCSADRGQAPL | 0.31982896 |
| MSFAVSEL | TCSADRGQAPL | 0.20540176 |
| NTGALGL | TCSADRGQAPL | 0.18080449 |
| PPGAGGAGGAGTT | TCSADRGQAPL | 0.18418439 |
| QLVGLGI | TCSADRGQAPL | 0.12088199 |
| RFLEQVHQL | TCSADRGQAPL | 0.1948638 |
| RGVGYVLR | TCSADRGQAPL | 0.12323188 |
| SATGVFAGVI | TCSADRGQAPL | 0.18275476 |

| | | |
|---|---|---|
| SLPAITLP | TCSADRGQAPL | 0.13167837 |
| STALAFTV | TCSADRGQAPL | 0.23494285 |
| SVLTVGLGK | TCSADRGQAPL | 0.23484197 |
| TALAGAGSGI | TCSADRGQAPL | 0.2273249 |
| TGLPFTGI | TCSADRGQAPL | 0.13173768 |
| VAEHVTV | TCSADRGQAPL | 0.17820309 |
| VDIVLNSL | TCSADRGQAPL | 0.16991563 |
| VLTGGGAVL | TCSADRGQAPL | 0.12990026 |
| VNVGLAYGI | TCSADRGQAPL | 0.2514651 |
| VSSAAVGAV | TCSADRGQAPL | 0.16279137 |
| VTSVYEAGV | TCSADRGQAPL | 0.16133796 |
| WVALDRGA | TCSADRGQAPL | 0.27085513 |
| YLVGSVVI | TCSADRGQAPL | 0.22670837 |
| GPPGTGKTLL | TCSADRGQAPL | 0.116972856 |
| GVGMTKF | TCSADRGQAPL | 0.20949383 |
| IGIGGSDL | TCSADRGQAPL | 0.18061453 |
| IGTLLAEL | TCSADRGQAPL | 0.16984668 |
| IYTSGTT | TCSADRGQAPL | 0.09768333 |
| LSFPAGV | TCSADRGQAPL | 0.2131607 |
| LYTSGTT | TCSADRGQAPL | 0.15500766 |
| VGAFPFGF | TCSADRGQAPL | 0.13041161 |
| VGAGTVEF | TCSADRGQAPL | 0.15780137 |
| VGKSTLLL | TCSADRGQAPL | 0.39826506 |
| AALAHAGGGIV | TCSADRGQAPL | 0.31452206 |
| AALVAVSLL | TCSADRGQAPL | 0.18322587 |
| AAVSRAVAAL | TCSADRGQAPL | 0.28098625 |
| AGGSSGIGGF | TCSADRGQAPL | 0.26033399 |
| AGLTVANANG | TCSADRGQAPL | 0.49973238 |
| AGLVGAVL | TCSADRGQAPL | 0.30668008 |
| ALSQAFTV | TCSADRGQAPL | 0.15858649 |
| ALTFSVSL | TCSADRGQAPL | 0.20837618 |
| ALTGAALVAL | TCSADRGQAPL | 0.16427007 |
| ASFSIGAGVL | TCSADRGQAPL | 0.10777575 |
| DGSVYKI | TCSADRGQAPL | 0.108578995 |
| EQLAEIV | TCSADRGQAPL | 0.16522776 |
| FAVSELTVM | TCSADRGQAPL | 0.10792173 |
| GAAPLAAGL | TCSADRGQAPL | 0.20026518 |
| GADGGDGGSGGAGG | TCSADRGQAPL | 0.25842342 |
| GFGLDTGGAGG | TCSADRGQAPL | 0.17392564 |
| GLASGGLL | TCSADRGQAPL | 0.121865205 |
| GRIAGAAVLL | TCSADRGQAPL | 0.14906691 |
| GSAGSLNTSF | TCSADRGQAPL | 0.1452196 |
| GTGQIGI | TCSADRGQAPL | 0.3079293 |
| GTLIALIL | TCSADRGQAPL | 0.18919761 |
| IAATMGFVVG | TCSADRGQAPL | 0.798125 |
| IFAQALL | TCSADRGQAPL | 0.40980643 |
| IGAAVALGF | TCSADRGQAPL | 0.1720705 |
| IGLTPLAL | TCSADRGQAPL | 0.31441328 |
| ILGSNTSTL | TCSADRGQAPL | 0.14271359 |
| IVFSLTVL | TCSADRGQAPL | 0.1479093 |
| LASKATGF | TCSADRGQAPL | 0.16609178 |

| | | |
|---|---|---|
| LASVALIL | TCSADRGQAPL | 0.2334145 |
| LFAVGLTL | TCSADRGQAPL | 0.22517473 |
| LFNGGIGTYI | TCSADRGQAPL | 0.16136356 |
| LTLGAGII | TCSADRGQAPL | 0.12073574 |
| LVAAALGDI | TCSADRGQAPL | 0.27989945 |
| NTAFVQLGI | TCSADRGQAPL | 0.22404441 |
| QLFSGAEL | TCSADRGQAPL | 0.11393965 |
| SAATFITL | TCSADRGQAPL | 0.14155325 |
| SNYGLANLGV | TCSADRGQAPL | 0.64466894 |
| SYGGAYAVM | TCSADRGQAPL | 0.54851234 |
| TTGVGGLTLG | TCSADRGQAPL | 0.13509107 |
| TTQTIYI | TCSADRGQAPL | 0.14306907 |
| VAAGAGLGAVI | TCSADRGQAPL | 0.5369495 |
| VAAVVTLL | TCSADRGQAPL | 0.1264162 |
| VGFLGGLI | TCSADRGQAPL | 0.1961393 |
| VGVIPILL | TCSADRGQAPL | 0.21436048 |
| VIFASSLI | TCSADRGQAPL | 0.30114943 |
| VKGGDGGAAAG | TCSADRGQAPL | 0.15306585 |
| VTKTFTNI | TCSADRGQAPL | 0.20483167 |
| VVAAAILL | TCSADRGQAPL | 0.22912362 |
| YAALDVEL | TCSADRGQAPL | 0.22122233 |
| LASVALLL | TCSADRGQAPL | 0.101080395 |
| LDAVHRV | TCSADRGQAPL | 0.2074933 |
| QGLSSSIF | TCSADRGQAPL | 0.12995836 |
| RGTFANIRL | TCSADRGQAPL | 0.14904061 |
| RQLWWGHRI | TCSADRGQAPL | 0.11261976 |
| YGPPGTGKTLL | TCSADRGQAPL | 0.16048142 |
| AALVGAPANVA | TCSADRGQAPL | 0.49117887 |
| AATFRRTE | TCSADRGQAPL | 0.44213337 |
| AGALGGGI | TCSADRGQAPL | 0.121891424 |
| AGGAGGAGGIGAG | TCSADRGQAPL | 0.33115786 |
| AGSGALGL | TCSADRGQAPL | 0.5889815 |
| AQIADQI | TCSADRGQAPL | 0.13867 |
| ASAVAVTYL | TCSADRGQAPL | 0.119165435 |
| ASTVLGGTGG | TCSADRGQAPL | 0.25569955 |
| AVGGEPAK | TCSADRGQAPL | 0.17686711 |
| CEQGTFGGTC | TCSADRGQAPL | 0.2391748 |
| DEGKAPAG | TCSADRGQAPL | 0.57710814 |
| DTVVFPAL | TCSADRGQAPL | 0.117973186 |
| FEGITMAG | TCSADRGQAPL | 0.23680894 |
| FPLVVQD | TCSADRGQAPL | 0.34027603 |
| GAFTTPQ | TCSADRGQAPL | 0.1342622 |
| GFGGFGTAGGVGGA | TCSADRGQAPL | 0.12967463 |
| GFLAQFL | TCSADRGQAPL | 0.14363737 |
| GGAGGAGGNAGVA | TCSADRGQAPL | 0.10781918 |
| GGVGTAGAGGAGGAGG | TCSADRGQAPL | 0.2657107 |
| GNGGGDAGNAGSGGN | TCSADRGQAPL | 0.14023593 |
| GTGGAGGAGNDAGS | TCSADRGQAPL | 0.17873438 |
| GTTNTGWFNS | TCSADRGQAPL | 0.17486948 |
| IFMGPLFTL | TCSADRGQAPL | 0.17236952 |
| ILAHAIEAI | TCSADRGQAPL | 0.1781217 |

| | | |
|---|---|---|
| IRSAIREAL | TCSADRGQAPL | 0.18770742 |
| ITFAGGG | TCSADRGQAPL | 0.19965655 |
| ITGMSGHQEV | TCSADRGQAPL | 0.24941412 |
| IVAGLACTL | TCSADRGQAPL | 0.15651302 |
| IVGGGLAGA | TCSADRGQAPL | 0.17864078 |
| IVTGVLNI | TCSADRGQAPL | 0.4850818 |
| LAQQGCI | TCSADRGQAPL | 0.14374642 |
| LCLHVEN | TCSADRGQAPL | 0.1462822 |
| LDAISDGS | TCSADRGQAPL | 0.32874975 |
| LGGFAAML | TCSADRGQAPL | 0.12472392 |
| LGGKNPMI | TCSADRGQAPL | 0.31031662 |
| LGPKGVAAIL | TCSADRGQAPL | 0.25601512 |
| LMVGVSI | TCSADRGQAPL | 0.23308474 |
| LVGAAFVL | TCSADRGQAPL | 0.19457197 |
| LVLAVAPL | TCSADRGQAPL | 0.61648446 |
| MAAIAVIDAA | TCSADRGQAPL | 0.2554971 |
| NNGINAELQ | TCSADRGQAPL | 0.15258664 |
| RGEPLVLVHG | TCSADRGQAPL | 0.52034783 |
| RGGAGGAAGLI | TCSADRGQAPL | 0.14366324 |
| RSYRMHPAV | TCSADRGQAPL | 0.26990384 |
| SAAHLLAGQLA | TCSADRGQAPL | 0.24548951 |
| SAAVGSY | TCSADRGQAPL | 0.12798063 |
| SPHAVLHRI | TCSADRGQAPL | 0.17806545 |
| SQIAKVL | TCSADRGQAPL | 0.30579317 |
| TALGHLVP | TCSADRGQAPL | 0.45583594 |
| TAPTTSGAA | TCSADRGQAPL | 0.11636865 |
| TDGDDGNGGSPGT | TCSADRGQAPL | 0.15277165 |
| TDGLGEN | TCSADRGQAPL | 0.12679662 |
| TGGPPLPGQA | TCSADRGQAPL | 0.17137091 |
| TIPAITI | TCSADRGQAPL | 0.14712109 |
| TIPGFTI | TCSADRGQAPL | 0.27607507 |
| TSIASGALA | TCSADRGQAPL | 0.16685341 |
| TVTLPGAAI | TCSADRGQAPL | 0.5492957 |
| VASGLSAALP | TCSADRGQAPL | 0.20488048 |
| VGAGVIGI | TCSADRGQAPL | 0.28364477 |
| VGPKRFGV | TCSADRGQAPL | 0.1494347 |
| VNVAFPD | TCSADRGQAPL | 0.19916666 |
| VSMTNSGVS | TCSADRGQAPL | 0.1241486 |
| VTDVLGDLSV | TCSADRGQAPL | 0.14483292 |
| VVVVGAGFAGL | TCSADRGQAPL | 0.13350657 |
| VYTVGTVV | TCSADRGQAPL | 0.19196138 |
| WTTFSGP | TCSADRGQAPL | 0.11012702 |
| YDELVEAF | TCSADRGQAPL | 0.40148705 |
| YGTGGAGGNGG | TCSADRGQAPL | 0.18013582 |
| YTNVMAV | TCSADRGQAPL | 0.26039255 |
| FLEQVHQL | TCSADRGQAPL | 0.1622424 |
| GLSLLAVAV | TCSADRGQAPL | 0.15654239 |
| LEQVHQL | TCSADRGQAPL | 0.16470307 |
| LNAAGAI | TCSADRGQAPL | 0.35120142 |
| PAVHASL | TCSADRGQAPL | 0.12970877 |
| TELEDDA | TCSADRGQAPL | 0.109275855 |

| | | |
|---|---|---|
| TPPGPPN | TCSADRGQAPL | 0.14674783 |
| TPPPPPPP | TCSADRGQAPL | 0.1083154 |
| TQSPALI | ASGDGTGRNTEVF | 0.44262898 |
| EFAAAFPQ | ASGDGTGRNTEVF | 0.31316018 |
| LLSGNLDVL | ASGDGTGRNTEVF | 0.19777909 |
| GPANLGNYNI | ASGDGTGRNTEVF | 0.4048652 |
| TLGPTLP | ASGDGTGRNTEVF | 0.10379749 |
| VYAYSSCCK | ASGDGTGRNTEVF | 0.17895465 |
| APAPAPAPNI | ASGDGTGRNTEVF | 0.5189533 |
| PGGPVPIP | ASGDGTGRNTEVF | 0.1964509 |
| FLTVALEP | ASGDGTGRNTEVF | 0.1829434 |
| SQYFGANL | ASGDGTGRNTEVF | 0.37020364 |
| GGVKPAA | ASGDGTGRNTEVF | 0.16625114 |
| GGTGGAGGAGGW | ASGDGTGRNTEVF | 0.17172599 |
| GVGGVAGLGGAGG | ASGDGTGRNTEVF | 0.22456387 |
| VLLVLATTI | ASGDGTGRNTEVF | 0.20273896 |
| IGIFAAML | ASGDGTGRNTEVF | 0.1829366 |
| AVGDLID | ASGDGTGRNTEVF | 0.13283122 |
| NISPLPL | ASGDGTGRNTEVF | 0.22095913 |
| VTAVFVAI | ASGDGTGRNTEVF | 0.76024234 |
| SGDDNDAAL | ASGDGTGRNTEVF | 0.17963131 |
| AVAALAHS | ASGDGTGRNTEVF | 0.37163523 |
| GGVTVMATP | ASGDGTGRNTEVF | 0.22839834 |
| AGLAANVRD | ASGDGTGRNTEVF | 0.37943798 |
| TNRPDLIDPALLRPG | ASGDGTGRNTEVF | 0.14527057 |
| LANLGVTAL | ASGDGTGRNTEVF | 0.13308842 |
| TGERIEL | ASGDGTGRNTEVF | 0.16238429 |
| NGEELAA | ASGDGTGRNTEVF | 0.12134977 |
| VLSLVWG | ASGDGTGRNTEVF | 0.18276584 |
| IGYALWL | ASGDGTGRNTEVF | 0.15661696 |
| LSLLLAPL | ASGDGTGRNTEVF | 0.81868494 |
| TLDEALK | ASGDGTGRNTEVF | 0.25174466 |
| TDGAGTAEGG | ASGDGTGRNTEVF | 0.31958762 |
| NAPGGAGGNGG | ASGDGTGRNTEVF | 0.13992153 |
| VSVFMGMGGNF | ASGDGTGRNTEVF | 0.17482977 |
| LASYLAAL | ASGDGTGRNTEVF | 0.21043941 |
| LVGGTPAL | ASGDGTGRNTEVF | 0.12418296 |
| VRNADVL | ASGDGTGRNTEVF | 0.4364708 |
| TCSSPFLVSAM | ASGDGTGRNTEVF | 0.14148846 |
| VHITDAE | ASGDGTGRNTEVF | 0.1786201 |
| EYAAPSF | ASGDGTGRNTEVF | 0.28784218 |
| VVDAFHGGL | ASGDGTGRNTEVF | 0.71340185 |
| ASVVDPQ | ASGDGTGRNTEVF | 0.1595258 |
| SYNEFGGLS | ASGDGTGRNTEVF | 0.18070471 |
| DPDGSVY | ASGDGTGRNTEVF | 0.1505419 |
| MVSTIHATE | ASGDGTGRNTEVF | 0.28188166 |
| LAAGVADPV | ASGDGTGRNTEVF | 0.4697124 |
| TKEQIAA | ASGDGTGRNTEVF | 0.29760897 |
| SLASPAAL | ASGDGTGRNTEVF | 0.73324835 |
| IGLVFYGV | ASGDGTGRNTEVF | 0.18602726 |
| FLLDASNL | ASGDGTGRNTEVF | 0.138778 |

| | | |
|---|---|---|
| YQVGNFP | ASGDGTGRNTEVF | 0.18859829 |
| ESLLSQST | ASGDGTGRNTEVF | 0.2717932 |
| LFTVGFAAVG | ASGDGTGRNTEVF | 0.474826 |
| ATPPTPT | ASGDGTGRNTEVF | 0.32507756 |
| TDLATLG | ASGDGTGRNTEVF | 0.21543176 |
| SAGGTGR | ASGDGTGRNTEVF | 0.14288686 |
| GFSNSGS | ASGDGTGRNTEVF | 0.35668415 |
| GGNGGSGGTSV | ASGDGTGRNTEVF | 0.4785151 |
| GTSVATGGAGN | ASGDGTGRNTEVF | 0.32845336 |
| NGGNGGSGGTS | ASGDGTGRNTEVF | 0.16073222 |
| AVVKFPH | ASGDGTGRNTEVF | 0.16211033 |
| TPPPPPP | ASGDGTGRNTEVF | 0.18016142 |
| AAPAAAAAPA | ASGDGTGRNTEVF | 0.4600305 |
| IALNEDL | ASGDGTGRNTEVF | 0.14049059 |
| IVNVSSM | ASGDGTGRNTEVF | 0.19740297 |
| TGIPPAPR | ASGDGTGRNTEVF | 0.20880216 |
| FDLGGGTF | ASGDGTGRNTEVF | 0.120227836 |
| FDLGGGTFDVS | ASGDGTGRNTEVF | 0.2174942 |
| NEPTAAA | ASGDGTGRNTEVF | 0.4807054 |
| NEPTAAALA | ASGDGTGRNTEVF | 0.16849649 |
| GSIINISSI | ASGDGTGRNTEVF | 0.13534266 |
| VAPGNAGTA | ASGDGTGRNTEVF | 0.48256737 |
| TGGGASLE | ASGDGTGRNTEVF | 0.7763941 |
| VAVYNHYKRI | ASGDGTGRNTEVF | 0.22292879 |
| TDPAPATT | ASGDGTGRNTEVF | 0.123733826 |
| TNRPDILDPALLRPG | ASGDGTGRNTEVF | 0.10131602 |
| LTTTTTS | ASGDGTGRNTEVF | 0.20909235 |
| PTGSGKTL | ASGDGTGRNTEVF | 0.57752836 |
| TGSGKTL | ASGDGTGRNTEVF | 0.1809233 |
| TIGHVDHGK | ASGDGTGRNTEVF | 0.1535444 |
| TIGHVDHGKTT | ASGDGTGRNTEVF | 0.19393146 |
| GHVDHGKTT | ASGDGTGRNTEVF | 0.31059092 |
| AQFEHTLL | ASGDGTGRNTEVF | 0.14622132 |
| IVNQDEL | ASGDGTGRNTEVF | 0.13511236 |
| TNGSQFF | ASGDGTGRNTEVF | 0.1726814 |
| TGKGIVE | ASGDGTGRNTEVF | 0.29556912 |
| LISNASDA | ASGDGTGRNTEVF | 0.13216107 |
| AELRTLGELTKIAKAHGAQ | ASGDGTGRNTEVF | 0.1659657 |
| APNAVAAAEPSW | ASGDGTGRNTEVF | 0.18671511 |
| APQVAGAATAVL | ASGDGTGRNTEVF | 0.15733908 |
| ARRDPAAAAAILPTDA | ASGDGTGRNTEVF | 0.22548452 |
| DAAARDVLAEAGLAEYF | ASGDGTGRNTEVF | 0.13779214 |
| DDQIVLSAKANLAKVEQ | ASGDGTGRNTEVF | 0.5525094 |
| DELLPDTGEDSDEDGH | ASGDGTGRNTEVF | 0.12297017 |
| ELPEAPANRATVPQD | ASGDGTGRNTEVF | 0.25970647 |
| FGQSKIERVTGIG | ASGDGTGRNTEVF | 0.24274656 |
| GAGGTGSTAGGAGGAGGAGGLY | ASGDGTGRNTEVF | 0.2144439 |
| GHGGTGAAGGNGGTGGAGGSGI | ASGDGTGRNTEVF | 0.16515318 |
| GNGGNGGAGGEGGPGGAGGAGA | ASGDGTGRNTEVF | 0.1311122 |
| GSTAGNGGNGGNATNGTIAGQP | ASGDGTGRNTEVF | 0.12443401 |
| GTGGDGGNGGNGGTGG | ASGDGTGRNTEVF | 0.11256728 |

| | | |
|---|---|---|
| GVRRPAGIPADAPV | ASGDGTGRNTEVF | 0.33178955 |
| GYRQVTDAHLLTLARR | ASGDGTGRNTEVF | 0.16087681 |
| IAKMGQDGHDRG | ASGDGTGRNTEVF | 0.3705243 |
| KAEPSIEIQYT | ASGDGTGRNTEVF | 0.19348228 |
| KPVRIRMAADEDTVAVT | ASGDGTGRNTEVF | 0.14829959 |
| MPPRIAGMRLLVIKPEP | ASGDGTGRNTEVF | 0.21020661 |
| MKFYAVALFRSRGVPPTR | ASGDGTGRNTEVF | 0.14280124 |
| NAGGFGILGAVAHSP | ASGDGTGRNTEVF | 0.12668073 |
| NEGREITLAMAEM | ASGDGTGRNTEVF | 0.32419187 |
| NELVEWSHPLGHRWRWE | ASGDGTGRNTEVF | 0.2308203 |
| NLGAGGTGGNGGNPG | ASGDGTGRNTEVF | 0.1421587 |
| NTEWPKWETVMAADPVD | ASGDGTGRNTEVF | 0.19548965 |
| PASAAPAETQP | ASGDGTGRNTEVF | 0.20422187 |
| QQSSDGKNGVRAAEITGE | ASGDGTGRNTEVF | 0.1399583 |
| RAPDAVTTSWWKE | ASGDGTGRNTEVF | 0.17287458 |
| SGFGNVGAHTSGWFNQ | ASGDGTGRNTEVF | 0.3876146 |
| SGVAGAVGQAASVGGLKV | ASGDGTGRNTEVF | 0.19236034 |
| SNGGAGTGSAGGNGGT | ASGDGTGRNTEVF | 0.12656479 |
| TGGAGGEGGAAG | ASGDGTGRNTEVF | 0.28571135 |
| TGGDGGAAGAGGAPGHG | ASGDGTGRNTEVF | 0.13068573 |
| TMPPLLAAIANAGRHGV | ASGDGTGRNTEVF | 0.28169647 |
| TSSTAGNSGGA | ASGDGTGRNTEVF | 0.21305215 |
| VPATVPVGILC | ASGDGTGRNTEVF | 0.18240069 |
| VQRPDSGGVVLTGRIS | ASGDGTGRNTEVF | 0.29217023 |
| WTPNPVPATKGVPAGLVQAT | ASGDGTGRNTEVF | 0.123103276 |
| YALKRALAKFSDRFDVVII | ASGDGTGRNTEVF | 0.13812506 |
| YLIGSVIV | ASGDGTGRNTEVF | 0.16182522 |
| YGQLVAVID | ASGDGTGRNTEVF | 0.24115083 |
| VTVGGMTLPA | ASGDGTGRNTEVF | 0.29602432 |
| VLALPAV | ASGDGTGRNTEVF | 0.1248275 |
| VIPPTLNYET | ASGDGTGRNTEVF | 0.20891856 |
| VGMGGMHPSSG | ASGDGTGRNTEVF | 0.15410689 |
| VGLAGGAATA | ASGDGTGRNTEVF | 0.1394506 |
| VGGAGGAGGSGP | ASGDGTGRNTEVF | 0.17688216 |
| VFAAFMV | ASGDGTGRNTEVF | 0.14502825 |
| TLYDKLR | ASGDGTGRNTEVF | 0.14569387 |
| TLAVFTVL | ASGDGTGRNTEVF | 0.21276115 |
| TGSVQPARV | ASGDGTGRNTEVF | 0.1631355 |
| TGGGAGSGGT | ASGDGTGRNTEVF | 0.22783361 |
| SHLETSL | ASGDGTGRNTEVF | 0.26659644 |
| SFTTGMT | ASGDGTGRNTEVF | 0.1420524 |
| QVEVAKRMGVRQPTVS | ASGDGTGRNTEVF | 0.2228121 |
| QGSGGPALAM | ASGDGTGRNTEVF | 0.2398526 |
| MLAPSNP | ASGDGTGRNTEVF | 0.26758918 |
| LVNNAAI | ASGDGTGRNTEVF | 0.18203779 |
| LTGEINTLA | ASGDGTGRNTEVF | 0.13171884 |
| LSIFGPV | ASGDGTGRNTEVF | 0.28850812 |
| LLALLANL | ASGDGTGRNTEVF | 0.32298398 |
| LISNASDALDKLR | ASGDGTGRNTEVF | 0.2741903 |
| LINEAVP | ASGDGTGRNTEVF | 0.23554286 |
| LGYAFSGL | ASGDGTGRNTEVF | 0.27535656 |

| | | |
|---|---|---|
| LDFVAVAL | ASGDGTGRNTEVF | 0.2820242 |
| LAQPGAE | ASGDGTGRNTEVF | 0.1566561 |
| ISNLGAVTGF | ASGDGTGRNTEVF | 0.11957479 |
| HQEDPGRI | ASGDGTGRNTEVF | 0.19812825 |
| GTPGEPTP | ASGDGTGRNTEVF | 0.19397327 |
| GFFNSGS | ASGDGTGRNTEVF | 0.1484062 |
| GAGGGGLTAGAN | ASGDGTGRNTEVF | 0.12797387 |
| FVDALVNM | ASGDGTGRNTEVF | 0.39160156 |
| FLTVAPEL | ASGDGTGRNTEVF | 0.7732313 |
| FETLTAL | ASGDGTGRNTEVF | 0.16893627 |
| DVINPIEL | ASGDGTGRNTEVF | 0.3513067 |
| APVSGAPV | ASGDGTGRNTEVF | 0.11268792 |
| ALVGGLAW | ASGDGTGRNTEVF | 0.3112682 |
| ALAAAAATAI | ASGDGTGRNTEVF | 0.40101853 |
| AFGLGLLL | ASGDGTGRNTEVF | 0.13525933 |
| AFFAAASL | ASGDGTGRNTEVF | 0.24774835 |
| AAAALPAFSPP | ASGDGTGRNTEVF | 0.13059103 |
| VVLAPAL | ASGDGTGRNTEVF | 0.15248406 |
| TIGHVDHGKTTL | ASGDGTGRNTEVF | 0.4631044 |
| LISNASDALDK | ASGDGTGRNTEVF | 0.19490975 |
| LISNASDAL | ASGDGTGRNTEVF | 0.18734574 |
| ISNASDALDK | ASGDGTGRNTEVF | 0.19109416 |
| NQAETLVYQTE | ASGDGTGRNTEVF | 0.15197031 |
| TGGTGGTAGAG | ASGDGTGRNTEVF | 0.20054138 |
| TGSTAGGAGGA | ASGDGTGRNTEVF | 0.32740355 |
| DPLAVPVAAGPV | ASGDGTGRNTEVF | 0.13504785 |
| GGTGGTPGNGGN | ASGDGTGRNTEVF | 0.42181575 |
| PAGLAGVDPATL | ASGDGTGRNTEVF | 0.13924941 |
| GNNLVVEAPHAGASV | ASGDGTGRNTEVF | 0.6569176 |
| KVTRIMSASTLEQPA | ASGDGTGRNTEVF | 0.6770066 |
| LGFALPIALPRRTVV | ASGDGTGRNTEVF | 0.22895849 |
| TRLGNLTARKLVVLV | ASGDGTGRNTEVF | 0.16406327 |
| WSQQRRLIYFPSAGP | ASGDGTGRNTEVF | 0.38645256 |
| GPNGNPDPMAAAVDIR | ASGDGTGRNTEVF | 0.18372728 |
| TQRCVSVADLRAAASR | ASGDGTGRNTEVF | 0.22530717 |
| QFVLDVAEHPVEVLTETV | ASGDGTGRNTEVF | 0.23749387 |
| GGAGGTGGAGGNGGTQAGDG | ASGDGTGRNTEVF | 0.17683734 |
| PLLATLRAIAVDVDAKPAQV | ASGDGTGRNTEVF | 0.8417765 |
| APKFRTGGYMIDFWGVGYQVAK | ASGDGTGRNTEVF | 0.7241819 |
| GAGGAGGAGGAAGGPGGTGGTG | ASGDGTGRNTEVF | 0.53241646 |
| IGALGLLATDVAPAGSVAFGVA | ASGDGTGRNTEVF | 0.3566875 |
| AGGAGGIGGNGGAGGAGGVLMG | ASGDGTGRNTEVF | 0.19323516 |
| ADELIGLL | ASGDGTGRNTEVF | 0.38531885 |
| ADRALKA | ASGDGTGRNTEVF | 0.2527936 |
| AGAVGQVGGAGGA | ASGDGTGRNTEVF | 0.8176667 |
| AGGGGGLGGAP | ASGDGTGRNTEVF | 0.6747441 |
| AGGLLYGNGGAG | ASGDGTGRNTEVF | 0.34721035 |
| AGLESIGL | ASGDGTGRNTEVF | 0.7300249 |
| AGLKTGII | ASGDGTGRNTEVF | 0.7699925 |
| ASLAALLL | ASGDGTGRNTEVF | 0.17870574 |
| ASVINISSI | ASGDGTGRNTEVF | 0.49252582 |

| | | |
|---|---|---|
| ATPPGPA | ASGDGTGRNTEVF | 0.42116657 |
| ATVASALV | ASGDGTGRNTEVF | 0.37291312 |
| ATVASLIL | ASGDGTGRNTEVF | 0.13500139 |
| AVNGPVLIRH | ASGDGTGRNTEVF | 0.72481894 |
| DGAAGAVGI | ASGDGTGRNTEVF | 0.45999637 |
| DPLLHGI | ASGDGTGRNTEVF | 0.10058658 |
| ELGAALGAL | ASGDGTGRNTEVF | 0.53468245 |
| ELGRTQAV | ASGDGTGRNTEVF | 0.4044664 |
| FVAGGYLS | ASGDGTGRNTEVF | 0.3666287 |
| GASGGAGGAGGW | ASGDGTGRNTEVF | 0.6933963 |
| GGAGAGGDGGAGGI | ASGDGTGRNTEVF | 0.5287338 |
| GTGHNGSQPGMGGNG | ASGDGTGRNTEVF | 0.32567763 |
| GYAAASASAS | ASGDGTGRNTEVF | 0.74150705 |
| HQVDVAAVQQG | ASGDGTGRNTEVF | 0.7589953 |
| IGAAIAHT | ASGDGTGRNTEVF | 0.26509526 |
| IGMIEVAQL | ASGDGTGRNTEVF | 0.59368587 |
| IRLAPEHTSAMD | ASGDGTGRNTEVF | 0.8290826 |
| IYNVSVL | ASGDGTGRNTEVF | 0.5193586 |
| KQGFLGM | ASGDGTGRNTEVF | 0.7875386 |
| LESVITQL | ASGDGTGRNTEVF | 0.80640435 |
| LGLGALVVV | ASGDGTGRNTEVF | 0.4127959 |
| LIGNGGNGGSGGI | ASGDGTGRNTEVF | 0.7665164 |
| LNAINAPTL | ASGDGTGRNTEVF | 0.8536361 |
| LQLLNTGV | ASGDGTGRNTEVF | 0.5119834 |
| LQTSIEF | ASGDGTGRNTEVF | 0.44604415 |
| MSFAVSEL | ASGDGTGRNTEVF | 0.18527524 |
| NTGALGL | ASGDGTGRNTEVF | 0.5245676 |
| PPGAGGAGGAGTT | ASGDGTGRNTEVF | 0.27576822 |
| QLVGLGI | ASGDGTGRNTEVF | 0.69803625 |
| RFLEQVHQL | ASGDGTGRNTEVF | 0.8254445 |
| RGVGYVLR | ASGDGTGRNTEVF | 0.45232046 |
| SATGVFAGVI | ASGDGTGRNTEVF | 0.1829257 |
| SLPAITLP | ASGDGTGRNTEVF | 0.21056522 |
| STALAFTV | ASGDGTGRNTEVF | 0.68138784 |
| SVLTVGLGK | ASGDGTGRNTEVF | 0.23670846 |
| TALAGAGSGI | ASGDGTGRNTEVF | 0.17091164 |
| TGLPFTGI | ASGDGTGRNTEVF | 0.42542157 |
| VAEHVTV | ASGDGTGRNTEVF | 0.75540113 |
| VDIVLNSL | ASGDGTGRNTEVF | 0.7630544 |
| VLTGGGAVL | ASGDGTGRNTEVF | 0.18959337 |
| VNVGLAYGI | ASGDGTGRNTEVF | 0.8294692 |
| VSSAAVGAV | ASGDGTGRNTEVF | 0.80002123 |
| VTSVYEAGV | ASGDGTGRNTEVF | 0.8571371 |
| WVALDRGA | ASGDGTGRNTEVF | 0.5682771 |
| YLVGSVVI | ASGDGTGRNTEVF | 0.18910263 |
| GPPGTGKTLL | ASGDGTGRNTEVF | 0.8259746 |
| GVGMTKF | ASGDGTGRNTEVF | 0.41681033 |
| IGIGGSDL | ASGDGTGRNTEVF | 0.49570653 |
| IGTLLAEL | ASGDGTGRNTEVF | 0.6613104 |
| IYTSGTT | ASGDGTGRNTEVF | 0.51209617 |
| LSFPAGV | ASGDGTGRNTEVF | 0.30268753 |

| | | |
|---|---|---|
| LYTSGTT | ASGDGTGRNTEVF | 0.7522526 |
| VGAFPFGF | ASGDGTGRNTEVF | 0.7728691 |
| VGAGTVEF | ASGDGTGRNTEVF | 0.28620017 |
| VGKSTLLL | ASGDGTGRNTEVF | 0.2945894 |
| AALAHAGGGIV | ASGDGTGRNTEVF | 0.39767653 |
| AALVAVSLL | ASGDGTGRNTEVF | 0.5114106 |
| AAVSRAVAAL | ASGDGTGRNTEVF | 0.7426579 |
| AGGSSGIGGF | ASGDGTGRNTEVF | 0.6026108 |
| AGLTVANANG | ASGDGTGRNTEVF | 0.602679 |
| AGLVGAVL | ASGDGTGRNTEVF | 0.68395835 |
| ALSQAFTV | ASGDGTGRNTEVF | 0.73249334 |
| ALTFSVSL | ASGDGTGRNTEVF | 0.5168276 |
| ALTGAALVAL | ASGDGTGRNTEVF | 0.6921462 |
| ASFSIGAGVL | ASGDGTGRNTEVF | 0.35189766 |
| DGSVYKI | ASGDGTGRNTEVF | 0.7551652 |
| EQLAEIV | ASGDGTGRNTEVF | 0.70600563 |
| FAVSELTVM | ASGDGTGRNTEVF | 0.705903 |
| GAAPLAAGL | ASGDGTGRNTEVF | 0.78304565 |
| GADGGDGGSGGAGG | ASGDGTGRNTEVF | 0.7467315 |
| GFGLDTGGAGG | ASGDGTGRNTEVF | 0.63216686 |
| GLASGGLL | ASGDGTGRNTEVF | 0.15797073 |
| GRIAGAAVLL | ASGDGTGRNTEVF | 0.4645623 |
| GSAGSLNTSF | ASGDGTGRNTEVF | 0.54458183 |
| GTGQIGI | ASGDGTGRNTEVF | 0.66787326 |
| GTLIALIL | ASGDGTGRNTEVF | 0.66681373 |
| IAATMGFVVG | ASGDGTGRNTEVF | 0.7359858 |
| IFAQALL | ASGDGTGRNTEVF | 0.37832862 |
| IGAAVALGF | ASGDGTGRNTEVF | 0.7563528 |
| IGLTPLAL | ASGDGTGRNTEVF | 0.8101258 |
| ILGSNTSTL | ASGDGTGRNTEVF | 0.20967403 |
| IVFSLTVL | ASGDGTGRNTEVF | 0.32507563 |
| LASKATGF | ASGDGTGRNTEVF | 0.8376736 |
| LASVALIL | ASGDGTGRNTEVF | 0.68450403 |
| LFAVGLTL | ASGDGTGRNTEVF | 0.40773058 |
| LFNGGIGTYI | ASGDGTGRNTEVF | 0.40158075 |
| LTLGAGII | ASGDGTGRNTEVF | 0.8557318 |
| LVAAALGDI | ASGDGTGRNTEVF | 0.43492308 |
| NTAFVQLGI | ASGDGTGRNTEVF | 0.8232413 |
| QLFSGAEL | ASGDGTGRNTEVF | 0.13501726 |
| SAATFITL | ASGDGTGRNTEVF | 0.47256297 |
| SNYGLANLGV | ASGDGTGRNTEVF | 0.40815192 |
| SYGGAYAVM | ASGDGTGRNTEVF | 0.10431445 |
| TTGVGGLTLG | ASGDGTGRNTEVF | 0.7298207 |
| TTQTIYI | ASGDGTGRNTEVF | 0.2549741 |
| VAAGAGLGAVI | ASGDGTGRNTEVF | 0.6578216 |
| VAAVVTLL | ASGDGTGRNTEVF | 0.49911433 |
| VGFLGGLI | ASGDGTGRNTEVF | 0.71597725 |
| VGVIPILL | ASGDGTGRNTEVF | 0.666025 |
| VIFASSLI | ASGDGTGRNTEVF | 0.45269853 |
| VKGGDGGAAAG | ASGDGTGRNTEVF | 0.2509445 |
| VTKTFTNI | ASGDGTGRNTEVF | 0.6396152 |

| | | |
|---|---|---|
| VVAAAILL | ASGDGTGRNTEVF | 0.7395188 |
| YAALDVEL | ASGDGTGRNTEVF | 0.7120523 |
| LASVALLL | ASGDGTGRNTEVF | 0.17941132 |
| LDAVHRV | ASGDGTGRNTEVF | 0.5420145 |
| QGLSSSIF | ASGDGTGRNTEVF | 0.4851887 |
| RGTFANIRL | ASGDGTGRNTEVF | 0.15191713 |
| RQLWWGHRI | ASGDGTGRNTEVF | 0.102984026 |
| YGPPGTGKTLL | ASGDGTGRNTEVF | 0.5052622 |
| AALVGAPANVA | ASGDGTGRNTEVF | 0.5553381 |
| AATFRRTE | ASGDGTGRNTEVF | 0.68730253 |
| AGALGGGI | ASGDGTGRNTEVF | 0.7123418 |
| AGGAGGAGGIGAG | ASGDGTGRNTEVF | 0.32297105 |
| AGSGALGL | ASGDGTGRNTEVF | 0.76605123 |
| AQIADQI | ASGDGTGRNTEVF | 0.5322763 |
| ASAVAVTYL | ASGDGTGRNTEVF | 0.0863214 |
| ASTVLGGTGG | ASGDGTGRNTEVF | 0.48517177 |
| AVGGEPAK | ASGDGTGRNTEVF | 0.16726348 |
| CEQGTFGGTC | ASGDGTGRNTEVF | 0.79870576 |
| DEGKAPAG | ASGDGTGRNTEVF | 0.19311064 |
| DTVVFPAL | ASGDGTGRNTEVF | 0.3791171 |
| FEGITMAG | ASGDGTGRNTEVF | 0.41082513 |
| FPLVVQD | ASGDGTGRNTEVF | 0.8071627 |
| GAFTTPQ | ASGDGTGRNTEVF | 0.6509267 |
| GFGGFGTAGGVGGA | ASGDGTGRNTEVF | 0.362005 |
| GFLAQFL | ASGDGTGRNTEVF | 0.1364326 |
| GGAGGAGGNAGVA | ASGDGTGRNTEVF | 0.6186406 |
| GGVGTAGAGGAGGAGG | ASGDGTGRNTEVF | 0.56647176 |
| GNGGGGDAGNAGSGGN | ASGDGTGRNTEVF | 0.25398406 |
| GTGGAGGAGNDAGS | ASGDGTGRNTEVF | 0.15211242 |
| GTTNTGWFNS | ASGDGTGRNTEVF | 0.19434205 |
| IFMGPLFTL | ASGDGTGRNTEVF | 0.22109018 |
| ILAHAIEAI | ASGDGTGRNTEVF | 0.75586796 |
| IRSAIREAL | ASGDGTGRNTEVF | 0.28460062 |
| ITFAGGG | ASGDGTGRNTEVF | 0.37744793 |
| ITGMSGHQEV | ASGDGTGRNTEVF | 0.8513216 |
| IVAGLACTL | ASGDGTGRNTEVF | 0.77122825 |
| IVGGGLAGA | ASGDGTGRNTEVF | 0.43041295 |
| IVTGVLNI | ASGDGTGRNTEVF | 0.39061314 |
| LAQQGCI | ASGDGTGRNTEVF | 0.24890386 |
| LCLHVEN | ASGDGTGRNTEVF | 0.29053771 |
| LDAISDGS | ASGDGTGRNTEVF | 0.3424846 |
| LGGFAAML | ASGDGTGRNTEVF | 0.7358904 |
| LGGKNPMI | ASGDGTGRNTEVF | 0.43999898 |
| LGPKGVAAIL | ASGDGTGRNTEVF | 0.20161866 |
| LMVGVSI | ASGDGTGRNTEVF | 0.7942964 |
| LVGAAFVL | ASGDGTGRNTEVF | 0.8230532 |
| LVLAVAPL | ASGDGTGRNTEVF | 0.85772926 |
| MAAIAVIDAA | ASGDGTGRNTEVF | 0.22220817 |
| NNGINAELQ | ASGDGTGRNTEVF | 0.5447703 |
| RGEPLVLVHG | ASGDGTGRNTEVF | 0.3719257 |
| RGGAGGAAGLI | ASGDGTGRNTEVF | 0.4596092 |

| | | |
|---|---|---|
| RSYRMHPAV | ASGDGTGRNTEVF | 0.2994141 |
| SAAHLLAGQLA | ASGDGTGRNTEVF | 0.24507925 |
| SAAVGSY | ASGDGTGRNTEVF | 0.7568002 |
| SPHAVLHRI | ASGDGTGRNTEVF | 0.8677933 |
| SQIAKVL | ASGDGTGRNTEVF | 0.43212166 |
| TALGHLVP | ASGDGTGRNTEVF | 0.2164132 |
| TAPTTSGAA | ASGDGTGRNTEVF | 0.4137333 |
| TDGDDGNGGSPGT | ASGDGTGRNTEVF | 0.3757446 |
| TDGLGEN | ASGDGTGRNTEVF | 0.26524633 |
| TGGPPLPGQA | ASGDGTGRNTEVF | 0.51846486 |
| TIPAITI | ASGDGTGRNTEVF | 0.15574265 |
| TIPGFTI | ASGDGTGRNTEVF | 0.47579825 |
| TSIASGALA | ASGDGTGRNTEVF | 0.8616314 |
| TVTLPGAAI | ASGDGTGRNTEVF | 0.6489661 |
| VASGLSAALP | ASGDGTGRNTEVF | 0.63530123 |
| VGAGVIGI | ASGDGTGRNTEVF | 0.8040825 |
| VGPKRFGV | ASGDGTGRNTEVF | 0.70066637 |
| VNVAFPD | ASGDGTGRNTEVF | 0.7786882 |
| VSMTNSGVS | ASGDGTGRNTEVF | 0.8068248 |
| VTDVLGDLSV | ASGDGTGRNTEVF | 0.43055955 |
| VVVVGAGFAGL | ASGDGTGRNTEVF | 0.65132535 |
| VYTVGTVV | ASGDGTGRNTEVF | 0.71672547 |
| WTTFSGP | ASGDGTGRNTEVF | 0.39905465 |
| YDELVEAF | ASGDGTGRNTEVF | 0.5029367 |
| YGTGGAGGNGG | ASGDGTGRNTEVF | 0.51246595 |
| YTNVMAV | ASGDGTGRNTEVF | 0.750044 |
| FLEQVHQL | ASGDGTGRNTEVF | 0.6539552 |
| GLSLLAVAV | ASGDGTGRNTEVF | 0.8260072 |
| LEQVHQL | ASGDGTGRNTEVF | 0.13204867 |
| LNAAGAI | ASGDGTGRNTEVF | 0.26127172 |
| PAVHASL | ASGDGTGRNTEVF | 0.78469205 |
| TELEDDA | ASGDGTGRNTEVF | 0.112256005 |
| TPPGPPN | ASGDGTGRNTEVF | 0.1401813 |
| TPPPPPPP | ASGDGTGRNTEVF | 0.5717841 |
| TQSPALI | ASSSGEDTQY | 0.8462547 |
| EFAAAFPQ | ASSSGEDTQY | 0.11227563 |
| LLSGNLDVL | ASSSGEDTQY | 0.3455115 |
| GPANLGNYNI | ASSSGEDTQY | 0.17915799 |
| TLGPTLP | ASSSGEDTQY | 0.8118042 |
| VYAYSSCCK | ASSSGEDTQY | 0.53439385 |
| APAPAPAPNI | ASSSGEDTQY | 0.16850609 |
| PGGPVPIP | ASSSGEDTQY | 0.66368484 |
| FLTVALEP | ASSSGEDTQY | 0.17704551 |
| SQYFGANL | ASSSGEDTQY | 0.44929636 |
| GGVKPAA | ASSSGEDTQY | 0.7239456 |
| GGTGGAGGAGGW | ASSSGEDTQY | 0.83162844 |
| GVGGVAGLGGAGG | ASSSGEDTQY | 0.15276037 |
| VLLVLATTI | ASSSGEDTQY | 0.7185546 |
| IGIFAAML | ASSSGEDTQY | 0.45760053 |
| AVGDLID | ASSSGEDTQY | 0.8722654 |
| NISPLPL | ASSSGEDTQY | 0.41335085 |

| | | |
|---|---|---|
| VTAVFVAI | ASSSGEDTQY | 0.17292286 |
| SGDDNDAAL | ASSSGEDTQY | 0.74218106 |
| AVAALAHS | ASSSGEDTQY | 0.6895252 |
| GGVTVMATP | ASSSGEDTQY | 0.4688028 |
| AGLAANVRD | ASSSGEDTQY | 0.7858678 |
| TNRPDLIDPALLRPG | ASSSGEDTQY | 0.60130364 |
| LANLGVTAL | ASSSGEDTQY | 0.48861516 |
| TGERIEL | ASSSGEDTQY | 0.73903465 |
| NGEELAA | ASSSGEDTQY | 0.66383404 |
| VLSLVWG | ASSSGEDTQY | 0.28714702 |
| IGYALWL | ASSSGEDTQY | 0.7168156 |
| LSLLLAPL | ASSSGEDTQY | 0.4869539 |
| TLDEALK | ASSSGEDTQY | 0.20063657 |
| TDGAGTAEGG | ASSSGEDTQY | 0.85964835 |
| NAPGGAGGNGG | ASSSGEDTQY | 0.54807466 |
| VSVFMGMGGNF | ASSSGEDTQY | 0.7575558 |
| LASYLAAL | ASSSGEDTQY | 0.69197726 |
| LVGGTPAL | ASSSGEDTQY | 0.852374 |
| VRNADVL | ASSSGEDTQY | 0.23160458 |
| TCSSPFLVSAM | ASSSGEDTQY | 0.5986762 |
| VHITDAE | ASSSGEDTQY | 0.58474547 |
| EYAAPSF | ASSSGEDTQY | 0.7564399 |
| VVDAFHGGL | ASSSGEDTQY | 0.59236693 |
| ASVVDPQ | ASSSGEDTQY | 0.64230555 |
| SYNEFGGLS | ASSSGEDTQY | 0.67254996 |
| DPDGSVY | ASSSGEDTQY | 0.793948 |
| MVSTIHATE | ASSSGEDTQY | 0.5106047 |
| LAAGVADPV | ASSSGEDTQY | 0.6558852 |
| TKEQIAA | ASSSGEDTQY | 0.81435204 |
| SLASPAAL | ASSSGEDTQY | 0.18739423 |
| IGLVFYGV | ASSSGEDTQY | 0.48080173 |
| FLLDASNL | ASSSGEDTQY | 0.74860615 |
| YQVGNFP | ASSSGEDTQY | 0.7340337 |
| ESLLSQST | ASSSGEDTQY | 0.47977376 |
| LFTVGFAAVG | ASSSGEDTQY | 0.76753676 |
| ATPPTPT | ASSSGEDTQY | 0.40164676 |
| TDLATLG | ASSSGEDTQY | 0.6598572 |
| SAGGTGR | ASSSGEDTQY | 0.5551499 |
| GFSNSGS | ASSSGEDTQY | 0.5396173 |
| GGNGGSGGTSV | ASSSGEDTQY | 0.25575036 |
| GTSVATGGAGN | ASSSGEDTQY | 0.6485064 |
| NGGNGGSGGTS | ASSSGEDTQY | 0.7744117 |
| AVVKFPH | ASSSGEDTQY | 0.70948005 |
| TPPPPPP | ASSSGEDTQY | 0.11410508 |
| AAPAAAAAPA | ASSSGEDTQY | 0.6789638 |
| IALNEDL | ASSSGEDTQY | 0.7146641 |
| IVNVSSM | ASSSGEDTQY | 0.84381795 |
| TGIPPAPR | ASSSGEDTQY | 0.63616526 |
| FDLGGGTF | ASSSGEDTQY | 0.660059 |
| FDLGGGTFDVS | ASSSGEDTQY | 0.37549472 |
| NEPTAAA | ASSSGEDTQY | 0.59062684 |

| | | |
|---|---|---|
| NEPTAAALA | ASSSGEDTQY | 0.23546265 |
| GSIINISSI | ASSSGEDTQY | 0.2734211 |
| VAPGNAGTA | ASSSGEDTQY | 0.43732136 |
| TGGGASLE | ASSSGEDTQY | 0.26761088 |
| VAVYNHYKRI | ASSSGEDTQY | 0.818782 |
| TDPAPATT | ASSSGEDTQY | 0.7473563 |
| TNRPDILDPALLRPG | ASSSGEDTQY | 0.8068243 |
| LTTTTTS | ASSSGEDTQY | 0.81788385 |
| PTGSGKTL | ASSSGEDTQY | 0.8274236 |
| TGSGKTL | ASSSGEDTQY | 0.6583703 |
| TIGHVDHGK | ASSSGEDTQY | 0.3566453 |
| TIGHVDHGKTT | ASSSGEDTQY | 0.5283865 |
| GHVDHGKTT | ASSSGEDTQY | 0.14841333 |
| AQFEHTLL | ASSSGEDTQY | 0.44448256 |
| IVNQDEL | ASSSGEDTQY | 0.46262124 |
| TNGSQFF | ASSSGEDTQY | 0.49245274 |
| TGKGIVE | ASSSGEDTQY | 0.10315816 |
| LISNASDA | ASSSGEDTQY | 0.76857185 |
| AELRTLGELTKIAKAHGAQ | ASSSGEDTQY | 0.74880695 |
| APNAVAAAEPSW | ASSSGEDTQY | 0.5048307 |
| APQVAGAATAVL | ASSSGEDTQY | 0.7192601 |
| ARRDPAAAAAILPTDA | ASSSGEDTQY | 0.7337384 |
| DAAARDVLAEAGLAEYF | ASSSGEDTQY | 0.3801825 |
| DDQIVLSAKANLAKVEQ | ASSSGEDTQY | 0.75664115 |
| DELLPDTGEDSDEDGH | ASSSGEDTQY | 0.3781666 |
| ELPEAPANRATVPQD | ASSSGEDTQY | 0.7055069 |
| FGQSKIERVTGIG | ASSSGEDTQY | 0.844293 |
| GAGGTGSTAGGAGGAGGAGGLY | ASSSGEDTQY | 0.6559608 |
| GHGGTGAAGGNGGTGGAGGSGI | ASSSGEDTQY | 0.52163744 |
| GNGGNGGAGGEGGPGGAGGAGA | ASSSGEDTQY | 0.50670576 |
| GSTAGNGGNGGNATNGTIAGQP | ASSSGEDTQY | 0.2025164 |
| GTGGDGGNGGNGGTGG | ASSSGEDTQY | 0.5455735 |
| GVRRPAGIPADAPV | ASSSGEDTQY | 0.10162664 |
| GYRQVTDAHLLTLARR | ASSSGEDTQY | 0.66525686 |
| IAKMGQDGHDRG | ASSSGEDTQY | 0.3045851 |
| KAEPSIEIQYT | ASSSGEDTQY | 0.5474882 |
| KPVRIRMAADEDTVAVT | ASSSGEDTQY | 0.44710666 |
| MPPRIAGMRLLVIKPEP | ASSSGEDTQY | 0.6434432 |
| MKFYAVALFRSRGVPPTR | ASSSGEDTQY | 0.28831232 |
| NAGGFGILGAVAHSP | ASSSGEDTQY | 0.6182984 |
| NEGREITLAMAEM | ASSSGEDTQY | 0.6406098 |
| NELVEWSHPLGHRWRWE | ASSSGEDTQY | 0.2291594 |
| NLGAGGTGGNGGNPG | ASSSGEDTQY | 0.7215621 |
| NTEWPKWETVMAADPVD | ASSSGEDTQY | 0.49387532 |
| PASAAPAETQP | ASSSGEDTQY | 0.59781194 |
| QQSSDGKNGVRAAEITGE | ASSSGEDTQY | 0.8040498 |
| RAPDAVTTSWWKE | ASSSGEDTQY | 0.57612383 |
| SGFGNVGAHTSGWFNQ | ASSSGEDTQY | 0.195894 |
| SGVAGAVGQAASVGGLKV | ASSSGEDTQY | 0.56087345 |
| SNGGAGTGSAGGNGGT | ASSSGEDTQY | 0.5584965 |
| TGGAGGEGGAAG | ASSSGEDTQY | 0.45813206 |

| | | |
|---|---|---|
| TGGDGGAAGAGGAPGHG | ASSSGEDTQY | 0.102231465 |
| TMPPLLAAIANAGRHGV | ASSSGEDTQY | 0.7304165 |
| TSSTAGNSGGA | ASSSGEDTQY | 0.20573372 |
| VPATVPVGILC | ASSSGEDTQY | 0.42395175 |
| VQRPDSGGVVLTGRIS | ASSSGEDTQY | 0.30342078 |
| WTPNPVPATKGVPAGLVQAT | ASSSGEDTQY | 0.7290028 |
| YALKRALAKFSDRFDVVII | ASSSGEDTQY | 0.68619657 |
| YLIGSVIV | ASSSGEDTQY | 0.5668779 |
| YGQLVAVID | ASSSGEDTQY | 0.66062915 |
| VTVGGMTLPA | ASSSGEDTQY | 0.59210074 |
| VLALPAV | ASSSGEDTQY | 0.76390624 |
| VIPPTLNYET | ASSSGEDTQY | 0.66332877 |
| VGMGGMHPSSG | ASSSGEDTQY | 0.61380327 |
| VGLAGGAATA | ASSSGEDTQY | 0.61965746 |
| VGGAGGAGGSGP | ASSSGEDTQY | 0.49080417 |
| VFAAFMV | ASSSGEDTQY | 0.15107232 |
| TLYDKLR | ASSSGEDTQY | 0.26884958 |
| TLAVFTVL | ASSSGEDTQY | 0.52268744 |
| TGSVQPARV | ASSSGEDTQY | 0.1569736 |
| TGGGAGSGGT | ASSSGEDTQY | 0.6489751 |
| SHLETSL | ASSSGEDTQY | 0.79069585 |
| SFTTGMT | ASSSGEDTQY | 0.7411709 |
| QVEVAKRMGVRQPTVS | ASSSGEDTQY | 0.15549095 |
| QGSGGPALAM | ASSSGEDTQY | 0.40866512 |
| MLAPSNP | ASSSGEDTQY | 0.45174158 |
| LVNNAAI | ASSSGEDTQY | 0.6411152 |
| LTGEINTLA | ASSSGEDTQY | 0.808702 |
| LSIFGPV | ASSSGEDTQY | 0.8495395 |
| LLALLANL | ASSSGEDTQY | 0.7525765 |
| LISNASDALDKLR | ASSSGEDTQY | 0.33855543 |
| LINEAVP | ASSSGEDTQY | 0.66544944 |
| LGYAFSGL | ASSSGEDTQY | 0.4671176 |
| LDFVAVAL | ASSSGEDTQY | 0.47164387 |
| LAQPGAE | ASSSGEDTQY | 0.5028408 |
| ISNLGAVTGF | ASSSGEDTQY | 0.71148473 |
| HQEDPGRI | ASSSGEDTQY | 0.47114652 |
| GTPGEPTP | ASSSGEDTQY | 0.31655195 |
| GFFNSGS | ASSSGEDTQY | 0.21772671 |
| GAGGGGLTAGAN | ASSSGEDTQY | 0.6867738 |
| FVDALVNM | ASSSGEDTQY | 0.51423067 |
| FLTVAPEL | ASSSGEDTQY | 0.26157838 |
| FETLTAL | ASSSGEDTQY | 0.4532621 |
| DVINPIEL | ASSSGEDTQY | 0.8395669 |
| APVSGAPV | ASSSGEDTQY | 0.8383851 |
| ALVGGLAW | ASSSGEDTQY | 0.3762268 |
| ALAAAAATAI | ASSSGEDTQY | 0.34056693 |
| AFGLGLLL | ASSSGEDTQY | 0.7950207 |
| AFFAAASL | ASSSGEDTQY | 0.23496005 |
| AAAALPAFSPP | ASSSGEDTQY | 0.6329846 |
| VVLAPAL | ASSSGEDTQY | 0.6422401 |
| TIGHVDHGKTTL | ASSSGEDTQY | 0.49539888 |

| | | |
|---|---|---|
| LISNASDALDK | ASSSGEDTQY | 0.29905945 |
| LISNASDAL | ASSSGEDTQY | 0.7001387 |
| ISNASDALDK | ASSSGEDTQY | 0.43403617 |
| NQAETLVYQTE | ASSSGEDTQY | 0.6970446 |
| TGGTGGTAGAG | ASSSGEDTQY | 0.1360888 |
| TGSTAGGAGGA | ASSSGEDTQY | 0.7221309 |
| DPLAVPVAAGPV | ASSSGEDTQY | 0.2248469 |
| GGTGGTPGNGGN | ASSSGEDTQY | 0.46514356 |
| PAGLAGVDPATL | ASSSGEDTQY | 0.13502586 |
| GNNLVVEAPHAGASV | ASSSGEDTQY | 0.34674066 |
| KVTRIMSASTLEQPA | ASSSGEDTQY | 0.7945815 |
| LGFALPIALPRRTVV | ASSSGEDTQY | 0.8049727 |
| TRLGNLTARKLVVLV | ASSSGEDTQY | 0.3570029 |
| WSQQRRLIYFPSAGP | ASSSGEDTQY | 0.6633229 |
| GPNGNPDPMAAAVDIR | ASSSGEDTQY | 0.8568676 |
| TQRCVSVADLRAAASR | ASSSGEDTQY | 0.37957272 |
| QFVLDVAEHPVEVLTETV | ASSSGEDTQY | 0.13344067 |
| GGAGGTGGAGGNGGTQAGDG | ASSSGEDTQY | 0.6112397 |
| PLLATLRAIAVDVDAKPAQV | ASSSGEDTQY | 0.6961491 |
| APKFRTGGYMIDFWGVGYQVAK | ASSSGEDTQY | 0.7994345 |
| GAGGAGGAGGAAGGPGGTGGTG | ASSSGEDTQY | 0.811463 |
| IGALGLLATDVAPAGSVAFGVA | ASSSGEDTQY | 0.16176379 |
| AGGAGGIGGNGGAGGAGGVLMG | ASSSGEDTQY | 0.75146544 |
| ADELIGLL | ASSSGEDTQY | 0.78656393 |
| ADRALKA | ASSSGEDTQY | 0.2608088 |
| AGAVGQVGGAGGA | ASSSGEDTQY | 0.2425479 |
| AGGGGGLGGAP | ASSSGEDTQY | 0.26466405 |
| AGGLLYGNGGAG | ASSSGEDTQY | 0.15816018 |
| AGLESIGL | ASSSGEDTQY | 0.6664821 |
| AGLKTGII | ASSSGEDTQY | 0.44556907 |
| ASLAALLL | ASSSGEDTQY | 0.4995587 |
| ASVINISSI | ASSSGEDTQY | 0.42550635 |
| ATPPGPA | ASSSGEDTQY | 0.38820955 |
| ATVASALV | ASSSGEDTQY | 0.3359427 |
| ATVASLIL | ASSSGEDTQY | 0.3693169 |
| AVNGPVLIRH | ASSSGEDTQY | 0.7112786 |
| DGAAGAVGI | ASSSGEDTQY | 0.7438067 |
| DPLLHGI | ASSSGEDTQY | 0.70234984 |
| ELGAALGAL | ASSSGEDTQY | 0.56243587 |
| ELGRTQAV | ASSSGEDTQY | 0.80298644 |
| FVAGGYLS | ASSSGEDTQY | 0.2591468 |
| GASGGAGGAGGW | ASSSGEDTQY | 0.26469153 |
| GGAGAGGDGGAGGI | ASSSGEDTQY | 0.7219678 |
| GTGHNGSQPGMGGNG | ASSSGEDTQY | 0.34948903 |
| GYAAASASAS | ASSSGEDTQY | 0.7814322 |
| HQVDVAAVQQG | ASSSGEDTQY | 0.5330905 |
| IGAAIAHT | ASSSGEDTQY | 0.58660275 |
| IGMIEVAQL | ASSSGEDTQY | 0.7539568 |
| IRLAPEHTSAMD | ASSSGEDTQY | 0.8302936 |
| IYNVSVL | ASSSGEDTQY | 0.6368058 |
| KQGFLGM | ASSSGEDTQY | 0.35133874 |

| | | |
|---|---|---|
| LESVITQL | ASSSGEDTQY | 0.75786006 |
| LGLGALVVV | ASSSGEDTQY | 0.20783207 |
| LIGNGGNGGSGGI | ASSSGEDTQY | 0.6755917 |
| LNAINAPTL | ASSSGEDTQY | 0.7178927 |
| LQLLNTGV | ASSSGEDTQY | 0.24959612 |
| LQTSIEF | ASSSGEDTQY | 0.49962223 |
| MSFAVSEL | ASSSGEDTQY | 0.8034471 |
| NTGALGL | ASSSGEDTQY | 0.70209324 |
| PPGAGGAGGAGTT | ASSSGEDTQY | 0.23085108 |
| QLVGLGI | ASSSGEDTQY | 0.28854504 |
| RFLEQVHQL | ASSSGEDTQY | 0.26185387 |
| RGVGYVLR | ASSSGEDTQY | 0.5855489 |
| SATGVFAGVI | ASSSGEDTQY | 0.1634161 |
| SLPAITLP | ASSSGEDTQY | 0.7425744 |
| STALAFTV | ASSSGEDTQY | 0.397188 |
| SVLTVGLGK | ASSSGEDTQY | 0.8147384 |
| TALAGAGSGI | ASSSGEDTQY | 0.6151845 |
| TGLPFTGI | ASSSGEDTQY | 0.7565096 |
| VAEHVTV | ASSSGEDTQY | 0.40297198 |
| VDIVLNSL | ASSSGEDTQY | 0.3948455 |
| VLTGGGAVL | ASSSGEDTQY | 0.17079745 |
| VNVGLAYGI | ASSSGEDTQY | 0.35315314 |
| VSSAAVGAV | ASSSGEDTQY | 0.28567624 |
| VTSVYEAGV | ASSSGEDTQY | 0.5138498 |
| WVALDRGA | ASSSGEDTQY | 0.19244993 |
| YLVGSVVI | ASSSGEDTQY | 0.8496674 |
| GPPGTGKTLL | ASSSGEDTQY | 0.3998943 |
| GVGMTKF | ASSSGEDTQY | 0.6568179 |
| IGIGGSDL | ASSSGEDTQY | 0.4913932 |
| IGTLLAEL | ASSSGEDTQY | 0.20507899 |
| IYTSGTT | ASSSGEDTQY | 0.3412149 |
| LSFPAGV | ASSSGEDTQY | 0.732791 |
| LYTSGTT | ASSSGEDTQY | 0.18353811 |
| VGAFPFGF | ASSSGEDTQY | 0.14671521 |
| VGAGTVEF | ASSSGEDTQY | 0.35992104 |
| VGKSTLLL | ASSSGEDTQY | 0.2741753 |
| AALAHAGGGIV | ASSSGEDTQY | 0.83978045 |
| AALVAVSLL | ASSSGEDTQY | 0.7964735 |
| AAVSRAVAAL | ASSSGEDTQY | 0.61502373 |
| AGGSSGIGGF | ASSSGEDTQY | 0.8377081 |
| AGLTVANANG | ASSSGEDTQY | 0.10533451 |
| AGLVGAVL | ASSSGEDTQY | 0.48606277 |
| ALSQAFTV | ASSSGEDTQY | 0.8046975 |
| ALTFSVSL | ASSSGEDTQY | 0.72613937 |
| ALTGAALVAL | ASSSGEDTQY | 0.44984758 |
| ASFSIGAGVL | ASSSGEDTQY | 0.74380755 |
| DGSVYKI | ASSSGEDTQY | 0.30853456 |
| EQLAEIV | ASSSGEDTQY | 0.46294466 |
| FAVSELTVM | ASSSGEDTQY | 0.4684786 |
| GAAPLAAGL | ASSSGEDTQY | 0.5890238 |
| GADGGDGGSGGAGG | ASSSGEDTQY | 0.61619914 |

| | | |
|---|---|---|
| GFGLDTGGAGG | ASSSGEDTQY | 0.17358012 |
| GLASGGLL | ASSSGEDTQY | 0.6331199 |
| GRIAGAAVLL | ASSSGEDTQY | 0.8599069 |
| GSAGSLNTSF | ASSSGEDTQY | 0.38676512 |
| GTGQIGI | ASSSGEDTQY | 0.78971684 |
| GTLIALIL | ASSSGEDTQY | 0.58696604 |
| IAATMGFVVG | ASSSGEDTQY | 0.8259011 |
| IFAQALL | ASSSGEDTQY | 0.3281446 |
| IGAAVALGF | ASSSGEDTQY | 0.19896078 |
| IGLTPLAL | ASSSGEDTQY | 0.62499297 |
| ILGSNTSTL | ASSSGEDTQY | 0.212571 |
| IVFSLTVL | ASSSGEDTQY | 0.5669322 |
| LASKATGF | ASSSGEDTQY | 0.48912472 |
| LASVALIL | ASSSGEDTQY | 0.8745211 |
| LFAVGLTL | ASSSGEDTQY | 0.7386733 |
| LFNGGIGTYI | ASSSGEDTQY | 0.5872606 |
| LTLGAGII | ASSSGEDTQY | 0.22982278 |
| LVAAALGDI | ASSSGEDTQY | 0.5908905 |
| NTAFVQLGI | ASSSGEDTQY | 0.7401719 |
| QLFSGAEL | ASSSGEDTQY | 0.21780632 |
| SAATFITL | ASSSGEDTQY | 0.79749715 |
| SNYGLANLGV | ASSSGEDTQY | 0.26105866 |
| SYGGAYAVM | ASSSGEDTQY | 0.71837735 |
| TTGVGGLTLG | ASSSGEDTQY | 0.6951045 |
| TTQTIYI | ASSSGEDTQY | 0.8383373 |
| VAAGAGLGAVI | ASSSGEDTQY | 0.43784916 |
| VAAVVTLL | ASSSGEDTQY | 0.60654306 |
| VGFLGGLI | ASSSGEDTQY | 0.20051679 |
| VGVIPILL | ASSSGEDTQY | 0.7898151 |
| VIFASSLI | ASSSGEDTQY | 0.77227676 |
| VKGGDGGAAAG | ASSSGEDTQY | 0.7605623 |
| VTKTFTNI | ASSSGEDTQY | 0.84511685 |
| VVAAAILL | ASSSGEDTQY | 0.767433 |
| YAALDVEL | ASSSGEDTQY | 0.15549369 |
| LASVALLL | ASSSGEDTQY | 0.42029527 |
| LDAVHRV | ASSSGEDTQY | 0.7565832 |
| QGLSSSIF | ASSSGEDTQY | 0.81569695 |
| RGTFANIRL | ASSSGEDTQY | 0.8324041 |
| RQLWWGHRI | ASSSGEDTQY | 0.5767076 |
| YGPPGTGKTLL | ASSSGEDTQY | 0.47161838 |
| AALVGAPANVA | ASSSGEDTQY | 0.8220315 |
| AATFRRTE | ASSSGEDTQY | 0.8811412 |
| AGALGGGI | ASSSGEDTQY | 0.6984312 |
| AGGAGGAGGIGAG | ASSSGEDTQY | 0.42901465 |
| AGSGALGL | ASSSGEDTQY | 0.4635654 |
| AQIADQI | ASSSGEDTQY | 0.6506547 |
| ASAVAVTYL | ASSSGEDTQY | 0.76624525 |
| ASTVLGGTGG | ASSSGEDTQY | 0.22397593 |
| AVGGEPAK | ASSSGEDTQY | 0.5591427 |
| CEQGTFGGTC | ASSSGEDTQY | 0.5618664 |
| DEGKAPAG | ASSSGEDTQY | 0.12389858 |

| | | |
|---|---|---|
| DTVVFPAL | ASSSGEDTQY | 0.78045726 |
| FEGITMAG | ASSSGEDTQY | 0.8016249 |
| FPLVVQD | ASSSGEDTQY | 0.5141261 |
| GAFTTPQ | ASSSGEDTQY | 0.23062274 |
| GFGGFGTAGGVGGA | ASSSGEDTQY | 0.80580974 |
| GFLAQFL | ASSSGEDTQY | 0.8489691 |
| GGAGGAGGNAGVA | ASSSGEDTQY | 0.7767173 |
| GGVGTAGAGGAGGAGG | ASSSGEDTQY | 0.19139577 |
| GNGGGGDAGNAGSGGN | ASSSGEDTQY | 0.11425925 |
| GTGGAGGAGNDAGS | ASSSGEDTQY | 0.32366982 |
| GTTNTGWFNS | ASSSGEDTQY | 0.8337538 |
| IFMGPLFTL | ASSSGEDTQY | 0.49372983 |
| ILAHAIEAI | ASSSGEDTQY | 0.3572176 |
| IRSAIREAL | ASSSGEDTQY | 0.57514626 |
| ITFAGGG | ASSSGEDTQY | 0.7831859 |
| ITGMSGHQEV | ASSSGEDTQY | 0.18086472 |
| IVAGLACTL | ASSSGEDTQY | 0.2851048 |
| IVGGGLAGA | ASSSGEDTQY | 0.6304133 |
| IVTGVLNI | ASSSGEDTQY | 0.7656925 |
| LAQQGCI | ASSSGEDTQY | 0.2377109 |
| LCLHVEN | ASSSGEDTQY | 0.4154398 |
| LDAISDGS | ASSSGEDTQY | 0.49305224 |
| LGGFAAML | ASSSGEDTQY | 0.71204084 |
| LGGKNPMI | ASSSGEDTQY | 0.6618413 |
| LGPKGVAAIL | ASSSGEDTQY | 0.22423571 |
| LMVGVSI | ASSSGEDTQY | 0.8038317 |
| LVGAAFVL | ASSSGEDTQY | 0.18841696 |
| LVLAVAPL | ASSSGEDTQY | 0.74524546 |
| MAAIAVIDAA | ASSSGEDTQY | 0.50905895 |
| NNGINAELQ | ASSSGEDTQY | 0.6169355 |
| RGEPLVLVHG | ASSSGEDTQY | 0.40167966 |
| RGGAGGAAGLI | ASSSGEDTQY | 0.32159847 |
| RSYRMHPAV | ASSSGEDTQY | 0.23335426 |
| SAAHLLAGQLA | ASSSGEDTQY | 0.25414476 |
| SAAVGSY | ASSSGEDTQY | 0.74552405 |
| SPHAVLHRI | ASSSGEDTQY | 0.18085986 |
| SQIAKVL | ASSSGEDTQY | 0.6823145 |
| TALGHLVP | ASSSGEDTQY | 0.59031904 |
| TAPTTSGAA | ASSSGEDTQY | 0.21198253 |
| TDGDDGNGGSPGT | ASSSGEDTQY | 0.66458124 |
| TDGLGEN | ASSSGEDTQY | 0.21098363 |
| TGGPPLPGQA | ASSSGEDTQY | 0.41559273 |
| TIPAITI | ASSSGEDTQY | 0.78486943 |
| TIPGFTI | ASSSGEDTQY | 0.6029199 |
| TSIASGALA | ASSSGEDTQY | 0.3088946 |
| TVTLPGAAI | ASSSGEDTQY | 0.6081959 |
| VASGLSAALP | ASSSGEDTQY | 0.69114065 |
| VGAGVIGI | ASSSGEDTQY | 0.33345017 |
| VGPKRFGV | ASSSGEDTQY | 0.46580172 |
| VNVAFPD | ASSSGEDTQY | 0.79332167 |
| VSMTNSGVS | ASSSGEDTQY | 0.5105908 |

| | | |
|---|---|---|
| VTDVLGDLSV | ASSSGEDTQY | 0.16867393 |
| VVVVGAGFAGL | ASSSGEDTQY | 0.7333687 |
| VYTVGTVV | ASSSGEDTQY | 0.3317504 |
| WTTFSGP | ASSSGEDTQY | 0.6290035 |
| YDELVEAF | ASSSGEDTQY | 0.45712966 |
| YGTGGAGGNGG | ASSSGEDTQY | 0.4981112 |
| YTNVMAV | ASSSGEDTQY | 0.75399256 |
| FLEQVHQL | ASSSGEDTQY | 0.3861869 |
| GLSLLAVAV | ASSSGEDTQY | 0.39026502 |
| LEQVHQL | ASSSGEDTQY | 0.42265105 |
| LNAAGAI | ASSSGEDTQY | 0.4753095 |
| PAVHASL | ASSSGEDTQY | 0.70030427 |
| TELEDDA | ASSSGEDTQY | 0.22811443 |
| TPPGPPN | ASSSGEDTQY | 0.66833436 |
| TPPPPPPP | ASSSGEDTQY | 0.48358697 |
| TQSPALI | ASSSLGDEQY | 0.28167787 |
| EFAAAFPQ | ASSSLGDEQY | 0.48902038 |
| LLSGNLDVL | ASSSLGDEQY | 0.4336323 |
| GPANLGNYNI | ASSSLGDEQY | 0.18976039 |
| TLGPTLP | ASSSLGDEQY | 0.6753831 |
| VYAYSSCCK | ASSSLGDEQY | 0.29758525 |
| APAPAPAPNI | ASSSLGDEQY | 0.6904253 |
| PGGPVPIP | ASSSLGDEQY | 0.8343289 |
| FLTVALEP | ASSSLGDEQY | 0.20674062 |
| SQYFGANL | ASSSLGDEQY | 0.707813 |
| GGVKPAA | ASSSLGDEQY | 0.64782494 |
| GGTGGAGGAGGW | ASSSLGDEQY | 0.59825003 |
| GVGGVAGLGGAGG | ASSSLGDEQY | 0.43634403 |
| VLLVLATTI | ASSSLGDEQY | 0.3199947 |
| IGIFAAML | ASSSLGDEQY | 0.7981825 |
| AVGDLID | ASSSLGDEQY | 0.6535367 |
| NISPLPL | ASSSLGDEQY | 0.6635089 |
| VTAVFVAI | ASSSLGDEQY | 0.7166305 |
| SGDDNDAAL | ASSSLGDEQY | 0.70953333 |
| AVAALAHS | ASSSLGDEQY | 0.6719576 |
| GGVTVMATP | ASSSLGDEQY | 0.34392694 |
| AGLAANVRD | ASSSLGDEQY | 0.18147555 |
| TNRPDLIDPALLRPG | ASSSLGDEQY | 0.42917913 |
| LANLGVTAL | ASSSLGDEQY | 0.48769096 |
| TGERIEL | ASSSLGDEQY | 0.2628001 |
| NGEELAA | ASSSLGDEQY | 0.22252749 |
| VLSLVWG | ASSSLGDEQY | 0.7565503 |
| IGYALWL | ASSSLGDEQY | 0.8612062 |
| LSLLLAPL | ASSSLGDEQY | 0.4274284 |
| TLDEALK | ASSSLGDEQY | 0.85544306 |
| TDGAGTAEGG | ASSSLGDEQY | 0.20631772 |
| NAPGGAGGNGG | ASSSLGDEQY | 0.7568615 |
| VSVFMGMGGNF | ASSSLGDEQY | 0.78122705 |
| LASYLAAL | ASSSLGDEQY | 0.2608389 |
| LVGGTPAL | ASSSLGDEQY | 0.65642965 |
| VRNADVL | ASSSLGDEQY | 0.34056246 |

| | | |
|---|---|---|
| TCSSPFLVSAM | ASSSLGDEQY | 0.2548489 |
| VHITDAE | ASSSLGDEQY | 0.71705484 |
| EYAAPSF | ASSSLGDEQY | 0.47667676 |
| VVDAFHGGL | ASSSLGDEQY | 0.6260599 |
| ASVVDPQ | ASSSLGDEQY | 0.42213094 |
| SYNEFGGLS | ASSSLGDEQY | 0.25804752 |
| DPDGSVY | ASSSLGDEQY | 0.53372353 |
| MVSTIHATE | ASSSLGDEQY | 0.6983546 |
| LAAGVADPV | ASSSLGDEQY | 0.54888666 |
| TKEQIAA | ASSSLGDEQY | 0.8769076 |
| SLASPAAL | ASSSLGDEQY | 0.2570985 |
| IGLVFYGV | ASSSLGDEQY | 0.85766685 |
| FLLDASNL | ASSSLGDEQY | 0.82267654 |
| YQVGNFP | ASSSLGDEQY | 0.6098839 |
| ESLLSQST | ASSSLGDEQY | 0.4652419 |
| LFTVGFAAVG | ASSSLGDEQY | 0.80100185 |
| ATPPTPT | ASSSLGDEQY | 0.43008435 |
| TDLATLG | ASSSLGDEQY | 0.6700363 |
| SAGGTGR | ASSSLGDEQY | 0.77038693 |
| GFSNSGS | ASSSLGDEQY | 0.28611356 |
| GGNGGSGGTSV | ASSSLGDEQY | 0.7057431 |
| GTSVATGGAGN | ASSSLGDEQY | 0.5841788 |
| NGGNGGSGGTS | ASSSLGDEQY | 0.57463515 |
| AVVKFPH | ASSSLGDEQY | 0.41171917 |
| TPPPPPP | ASSSLGDEQY | 0.27989352 |
| AAPAAAAAPA | ASSSLGDEQY | 0.41950187 |
| IALNEDL | ASSSLGDEQY | 0.35762456 |
| IVNVSSM | ASSSLGDEQY | 0.71148306 |
| TGIPPAPR | ASSSLGDEQY | 0.5569922 |
| FDLGGGTF | ASSSLGDEQY | 0.40305543 |
| FDLGGGTFDVS | ASSSLGDEQY | 0.6686009 |
| NEPTAAA | ASSSLGDEQY | 0.69543105 |
| NEPTAAALA | ASSSLGDEQY | 0.29360017 |
| GSIINISSI | ASSSLGDEQY | 0.4287332 |
| VAPGNAGTA | ASSSLGDEQY | 0.36797255 |
| TGGGASLE | ASSSLGDEQY | 0.4839417 |
| VAVYNHYKRI | ASSSLGDEQY | 0.28544325 |
| TDPAPATT | ASSSLGDEQY | 0.5406889 |
| TNRPDILDPALLRPG | ASSSLGDEQY | 0.33219627 |
| LTTTTTS | ASSSLGDEQY | 0.22341116 |
| PTGSGKTL | ASSSLGDEQY | 0.59040534 |
| TGSGKTL | ASSSLGDEQY | 0.40745318 |
| TIGHVDHGK | ASSSLGDEQY | 0.40478706 |
| TIGHVDHGKTT | ASSSLGDEQY | 0.57499295 |
| GHVDHGKTT | ASSSLGDEQY | 0.38929066 |
| AQFEHTLL | ASSSLGDEQY | 0.36240396 |
| IVNQDEL | ASSSLGDEQY | 0.62393713 |
| TNGSQFF | ASSSLGDEQY | 0.60353935 |
| TGKGIVE | ASSSLGDEQY | 0.32755798 |
| LISNASDA | ASSSLGDEQY | 0.4767803 |
| AELRTLGELTKIAKAHGAQ | ASSSLGDEQY | 0.71369946 |

| | | |
|---|---|---|
| APNAVAAAEPSW | ASSSLGDEQY | 0.5252167 |
| APQVAGAATAVL | ASSSLGDEQY | 0.61863524 |
| ARRDPAAAAAILPTDA | ASSSLGDEQY | 0.6421211 |
| DAAARDVLAEAGLAEYF | ASSSLGDEQY | 0.43186373 |
| DDQIVLSAKANLAKVEQ | ASSSLGDEQY | 0.6218029 |
| DELLPDTGEDSDEDGH | ASSSLGDEQY | 0.6799561 |
| ELPEAPANRATVPQD | ASSSLGDEQY | 0.46781665 |
| FGQSKIERVTGIG | ASSSLGDEQY | 0.5024651 |
| GAGGTGSTAGGAGGAGGAGGLY | ASSSLGDEQY | 0.27668476 |
| GHGGTGAAGGNGGTGGAGGSGI | ASSSLGDEQY | 0.54717475 |
| GNGGNGGAGGEGGPGGAGGAGA | ASSSLGDEQY | 0.27501413 |
| GSTAGNGGNGGNATNGTIAGQP | ASSSLGDEQY | 0.63736755 |
| GTGGDGGNGGNGGTGG | ASSSLGDEQY | 0.6613546 |
| GVRRPAGIPADAPV | ASSSLGDEQY | 0.4964798 |
| GYRQVTDAHLLTLARR | ASSSLGDEQY | 0.27537245 |
| IAKMGQDGHDRG | ASSSLGDEQY | 0.25868922 |
| KAEPSIEIQYT | ASSSLGDEQY | 0.5789768 |
| KPVRIRMAADEDTVAVT | ASSSLGDEQY | 0.27373955 |
| MPPRIAGMRLLVIKPEP | ASSSLGDEQY | 0.24446154 |
| MKFYAVALFRSRGVPPTR | ASSSLGDEQY | 0.50699395 |
| NAGGFGILGAVAHSP | ASSSLGDEQY | 0.6813919 |
| NEGREITLAMAEM | ASSSLGDEQY | 0.6872927 |
| NELVEWSHPLGHRWRWE | ASSSLGDEQY | 0.4333009 |
| NLGAGGTGGNGGNPG | ASSSLGDEQY | 0.67266464 |
| NTEWPKWETVMAADPVD | ASSSLGDEQY | 0.6607293 |
| PASAAPAETQP | ASSSLGDEQY | 0.6454191 |
| QQSSDGKNGVRAAEITGE | ASSSLGDEQY | 0.42299455 |
| RAPDAVTTSWWKE | ASSSLGDEQY | 0.2966231 |
| SGFGNVGAHTSGWFNQ | ASSSLGDEQY | 0.69621253 |
| SGVAGAVGQAASVGGLKV | ASSSLGDEQY | 0.52754736 |
| SNGGAGTGSAGGNGGT | ASSSLGDEQY | 0.5924445 |
| TGGAGGEGGAAG | ASSSLGDEQY | 0.6588656 |
| TGGDGGAAGAGGAPGHG | ASSSLGDEQY | 0.4500221 |
| TMPPLLAAIANAGRHGV | ASSSLGDEQY | 0.34779978 |
| TSSTAGNSGGA | ASSSLGDEQY | 0.628973 |
| VPATVPVGILC | ASSSLGDEQY | 0.58621436 |
| VQRPDSGGVVLTGRIS | ASSSLGDEQY | 0.38172436 |
| WTPNPVPATKGVPAGLVQAT | ASSSLGDEQY | 0.31490132 |
| YALKRALAKFSDRFDVVII | ASSSLGDEQY | 0.47204226 |
| YLIGSVIV | ASSSLGDEQY | 0.5504239 |
| YGQLVAVID | ASSSLGDEQY | 0.6227132 |
| VTVGGMTLPA | ASSSLGDEQY | 0.51377296 |
| VLALPAV | ASSSLGDEQY | 0.49622345 |
| VIPPTLNYET | ASSSLGDEQY | 0.5202548 |
| VGMGGMHPSSG | ASSSLGDEQY | 0.59365946 |
| VGLAGGAATA | ASSSLGDEQY | 0.60439515 |
| VGGAGGAGGSGP | ASSSLGDEQY | 0.6548836 |
| VFAAFMV | ASSSLGDEQY | 0.46933493 |
| TLYDKLR | ASSSLGDEQY | 0.5879075 |
| TLAVFTVL | ASSSLGDEQY | 0.63886154 |
| TGSVQPARV | ASSSLGDEQY | 0.5078411 |

| | | |
|---|---|---|
| TGGGAGSGGT | ASSSLGDEQY | 0.61128294 |
| SHLETSL | ASSSLGDEQY | 0.5426141 |
| SFTTGMT | ASSSLGDEQY | 0.5000811 |
| QVEVAKRMGVRQPTVS | ASSSLGDEQY | 0.31486648 |
| QGSGGPALAM | ASSSLGDEQY | 0.47208935 |
| MLAPSNP | ASSSLGDEQY | 0.5327685 |
| LVNNAAI | ASSSLGDEQY | 0.5482968 |
| LTGEINTLA | ASSSLGDEQY | 0.64195466 |
| LSIFGPV | ASSSLGDEQY | 0.49039388 |
| LLALLANL | ASSSLGDEQY | 0.4068299 |
| LISNASDALDKLR | ASSSLGDEQY | 0.6944437 |
| LINEAVP | ASSSLGDEQY | 0.68118584 |
| LGYAFSGL | ASSSLGDEQY | 0.3057804 |
| LDFVAVAL | ASSSLGDEQY | 0.5164419 |
| LAQPGAE | ASSSLGDEQY | 0.7232998 |
| ISNLGAVTGF | ASSSLGDEQY | 0.5916586 |
| HQEDPGRI | ASSSLGDEQY | 0.46106106 |
| GTPGEPTP | ASSSLGDEQY | 0.46070004 |
| GFFNSGS | ASSSLGDEQY | 0.7151724 |
| GAGGGGLTAGAN | ASSSLGDEQY | 0.39863092 |
| FVDALVNM | ASSSLGDEQY | 0.6894198 |
| FLTVAPEL | ASSSLGDEQY | 0.26852244 |
| FETLTAL | ASSSLGDEQY | 0.32601225 |
| DVINPIEL | ASSSLGDEQY | 0.3593243 |
| APVSGAPV | ASSSLGDEQY | 0.22662713 |
| ALVGGLAW | ASSSLGDEQY | 0.6746267 |
| ALAAAAATAI | ASSSLGDEQY | 0.34309983 |
| AFGLGLLL | ASSSLGDEQY | 0.6159117 |
| AFFAAASL | ASSSLGDEQY | 0.38847667 |
| AAAALPAFSPP | ASSSLGDEQY | 0.6461126 |
| VVLAPAL | ASSSLGDEQY | 0.5988851 |
| TIGHVDHGKTTL | ASSSLGDEQY | 0.57180864 |
| LISNASDALDK | ASSSLGDEQY | 0.33755758 |
| LISNASDAL | ASSSLGDEQY | 0.6020283 |
| ISNASDALDK | ASSSLGDEQY | 0.6891483 |
| NQAETLVYQTE | ASSSLGDEQY | 0.5183023 |
| TGGTGGTAGAG | ASSSLGDEQY | 0.27460653 |
| TGSTAGGAGGA | ASSSLGDEQY | 0.56387436 |
| DPLAVPVAAGPV | ASSSLGDEQY | 0.51105154 |
| GGTGGTPGNGGN | ASSSLGDEQY | 0.31027898 |
| PAGLAGVDPATL | ASSSLGDEQY | 0.23082547 |
| GNNLVVEAPHAGASV | ASSSLGDEQY | 0.37824687 |
| KVTRIMSASTLEQPA | ASSSLGDEQY | 0.5474591 |
| LGFALPIALPRRTVV | ASSSLGDEQY | 0.6448985 |
| TRLGNLTARKLVVLV | ASSSLGDEQY | 0.66575146 |
| WSQQRRLIYFPSAGP | ASSSLGDEQY | 0.40782756 |
| GPNGNPDPMAAAVDIR | ASSSLGDEQY | 0.6565585 |
| TQRCVSVADLRAAASR | ASSSLGDEQY | 0.6285651 |
| QFVLDVAEHPVEVLTETV | ASSSLGDEQY | 0.20929839 |
| GGAGGTGGAGGNGGTQAGDG | ASSSLGDEQY | 0.44621307 |
| PLLATLRAIAVDVDAKPAQV | ASSSLGDEQY | 0.28293797 |

| | | |
|---|---|---|
| APKFRTGGYMIDFWGVGYQVAK | ASSSLGDEQY | 0.63082254 |
| GAGGAGGAGGAAGGPGGTGGTG | ASSSLGDEQY | 0.35440314 |
| IGALGLLATDVAPAGSVAFGVA | ASSSLGDEQY | 0.5194758 |
| AGGAGGIGGNGGAGGAGGVLMG | ASSSLGDEQY | 0.5111437 |
| ADELIGLL | ASSSLGDEQY | 0.66216844 |
| ADRALKA | ASSSLGDEQY | 0.5848789 |
| AGAVGQVGGAGGA | ASSSLGDEQY | 0.3267644 |
| AGGGGGLGGAP | ASSSLGDEQY | 0.29425523 |
| AGGLLYGNGGAG | ASSSLGDEQY | 0.62759745 |
| AGLESIGL | ASSSLGDEQY | 0.46984917 |
| AGLKTGII | ASSSLGDEQY | 0.3269999 |
| ASLAALLL | ASSSLGDEQY | 0.40849376 |
| ASVINISSI | ASSSLGDEQY | 0.6287153 |
| ATPPGPA | ASSSLGDEQY | 0.28606907 |
| ATVASALV | ASSSLGDEQY | 0.56533265 |
| ATVASLIL | ASSSLGDEQY | 0.2651319 |
| AVNGPVLIRH | ASSSLGDEQY | 0.50859034 |
| DGAAGAVGI | ASSSLGDEQY | 0.7120583 |
| DPLLHGI | ASSSLGDEQY | 0.67211246 |
| ELGAALGAL | ASSSLGDEQY | 0.53626835 |
| ELGRTQAV | ASSSLGDEQY | 0.43075168 |
| FVAGGYLS | ASSSLGDEQY | 0.3498309 |
| GASGGAGGAGGW | ASSSLGDEQY | 0.3987857 |
| GGAGAGGDGGAGGI | ASSSLGDEQY | 0.3775449 |
| GTGHNGSQPGMGGNG | ASSSLGDEQY | 0.6828407 |
| GYAAASASAS | ASSSLGDEQY | 0.43948632 |
| HQVDVAAVQQG | ASSSLGDEQY | 0.32758614 |
| IGAAIAHT | ASSSLGDEQY | 0.53888726 |
| IGMIEVAQL | ASSSLGDEQY | 0.69447744 |
| IRLAPEHTSAMD | ASSSLGDEQY | 0.71142185 |
| IYNVSVL | ASSSLGDEQY | 0.3603744 |
| KQGFLGM | ASSSLGDEQY | 0.48395085 |
| LESVITQL | ASSSLGDEQY | 0.44086698 |
| LGLGALVVV | ASSSLGDEQY | 0.5645007 |
| LIGNGGNGGSGGI | ASSSLGDEQY | 0.340778 |
| LNAINAPTL | ASSSLGDEQY | 0.34379125 |
| LQLLNTGV | ASSSLGDEQY | 0.5456065 |
| LQTSIEF | ASSSLGDEQY | 0.73580503 |
| MSFAVSEL | ASSSLGDEQY | 0.5037168 |
| NTGALGL | ASSSLGDEQY | 0.28057376 |
| PPGAGGAGGAGTT | ASSSLGDEQY | 0.50999266 |
| QLVGLGI | ASSSLGDEQY | 0.31803215 |
| RFLEQVHQL | ASSSLGDEQY | 0.39920503 |
| RGVGYVLR | ASSSLGDEQY | 0.42837822 |
| SATGVFAGVI | ASSSLGDEQY | 0.26372197 |
| SLPAITLP | ASSSLGDEQY | 0.41484356 |
| STALAFTV | ASSSLGDEQY | 0.71083546 |
| SVLTVGLGK | ASSSLGDEQY | 0.5612663 |
| TALAGAGSGI | ASSSLGDEQY | 0.44195586 |
| TGLPFTGI | ASSSLGDEQY | 0.6794097 |
| VAEHVTV | ASSSLGDEQY | 0.55612785 |

| | | |
|---|---|---|
| VDIVLNSL | ASSSLGDEQY | 0.6609267 |
| VLTGGGAVL | ASSSLGDEQY | 0.66777515 |
| VNVGLAYGI | ASSSLGDEQY | 0.5511065 |
| VSSAAVGAV | ASSSLGDEQY | 0.50430393 |
| VTSVYEAGV | ASSSLGDEQY | 0.6358783 |
| WVALDRGA | ASSSLGDEQY | 0.50791776 |
| YLVGSVVI | ASSSLGDEQY | 0.5456365 |
| GPPGTGKTLL | ASSSLGDEQY | 0.46858823 |
| GVGMTKF | ASSSLGDEQY | 0.6765269 |
| IGIGGSDL | ASSSLGDEQY | 0.48031074 |
| IGTLLAEL | ASSSLGDEQY | 0.70305836 |
| IYTSGTT | ASSSLGDEQY | 0.23416819 |
| LSFPAGV | ASSSLGDEQY | 0.3401489 |
| LYTSGTT | ASSSLGDEQY | 0.67009807 |
| VGAFPFGF | ASSSLGDEQY | 0.22484152 |
| VGAGTVEF | ASSSLGDEQY | 0.23656255 |
| VGKSTLLL | ASSSLGDEQY | 0.36649486 |
| AALAHAGGGIV | ASSSLGDEQY | 0.7008439 |
| AALVAVSLL | ASSSLGDEQY | 0.22596523 |
| AAVSRAVAAL | ASSSLGDEQY | 0.42319405 |
| AGGSSGIGGF | ASSSLGDEQY | 0.24601276 |
| AGLTVANANG | ASSSLGDEQY | 0.63975066 |
| AGLVGAVL | ASSSLGDEQY | 0.50822496 |
| ALSQAFTV | ASSSLGDEQY | 0.2652228 |
| ALTFSVSL | ASSSLGDEQY | 0.57800364 |
| ALTGAALVAL | ASSSLGDEQY | 0.31577826 |
| ASFSIGAGVL | ASSSLGDEQY | 0.5972754 |
| DGSVYKI | ASSSLGDEQY | 0.6294291 |
| EQLAEIV | ASSSLGDEQY | 0.7131508 |
| FAVSELTVM | ASSSLGDEQY | 0.2770361 |
| GAAPLAAGL | ASSSLGDEQY | 0.51934123 |
| GADGGDGGSGGAGG | ASSSLGDEQY | 0.3463936 |
| GFGLDTGGAGG | ASSSLGDEQY | 0.7340305 |
| GLASGGLL | ASSSLGDEQY | 0.46422422 |
| GRIAGAAVLL | ASSSLGDEQY | 0.24488279 |
| GSAGSLNTSF | ASSSLGDEQY | 0.5975217 |
| GTGQIGI | ASSSLGDEQY | 0.5322141 |
| GTLIALIL | ASSSLGDEQY | 0.43636143 |
| IAATMGFVVG | ASSSLGDEQY | 0.65556675 |
| IFAQALL | ASSSLGDEQY | 0.42830434 |
| IGAAVALGF | ASSSLGDEQY | 0.43824583 |
| IGLTPLAL | ASSSLGDEQY | 0.6677289 |
| ILGSNTSTL | ASSSLGDEQY | 0.64980507 |
| IVFSLTVL | ASSSLGDEQY | 0.37545282 |
| LASKATGF | ASSSLGDEQY | 0.63181984 |
| LASVALIL | ASSSLGDEQY | 0.59418666 |
| LFAVGLTL | ASSSLGDEQY | 0.31751034 |
| LFNGGIGTYI | ASSSLGDEQY | 0.7299012 |
| LTLGAGII | ASSSLGDEQY | 0.37397042 |
| LVAAALGDI | ASSSLGDEQY | 0.64642894 |
| NTAFVQLGI | ASSSLGDEQY | 0.5775525 |

| | | |
|---|---|---|
| QLFSGAEL | ASSSLGDEQY | 0.7200124 |
| SAATFITL | ASSSLGDEQY | 0.3218936 |
| SNYGLANLGV | ASSSLGDEQY | 0.6357552 |
| SYGGAYAVM | ASSSLGDEQY | 0.60510206 |
| TTGVGGLTLG | ASSSLGDEQY | 0.577821 |
| TTQTIYI | ASSSLGDEQY | 0.37378567 |
| VAAGAGLGAVI | ASSSLGDEQY | 0.60107034 |
| VAAVVTLL | ASSSLGDEQY | 0.5786582 |
| VGFLGGLI | ASSSLGDEQY | 0.68143195 |
| VGVIPILL | ASSSLGDEQY | 0.4644891 |
| VIFASSLI | ASSSLGDEQY | 0.62197113 |
| VKGGDGGAAAG | ASSSLGDEQY | 0.701297 |
| VTKTFTNI | ASSSLGDEQY | 0.26849034 |
| VVAAAILL | ASSSLGDEQY | 0.54363775 |
| YAALDVEL | ASSSLGDEQY | 0.6012629 |
| LASVALLL | ASSSLGDEQY | 0.67859054 |
| LDAVHRV | ASSSLGDEQY | 0.42284685 |
| QGLSSSIF | ASSSLGDEQY | 0.6824939 |
| RGTFANIRL | ASSSLGDEQY | 0.42963362 |
| RQLWWGHRI | ASSSLGDEQY | 0.64141536 |
| YGPPGTGKTLL | ASSSLGDEQY | 0.40847382 |
| AALVGAPANVA | ASSSLGDEQY | 0.47883427 |
| AATFRRTE | ASSSLGDEQY | 0.3453906 |
| AGALGGGI | ASSSLGDEQY | 0.6163805 |
| AGGAGGAGGIGAG | ASSSLGDEQY | 0.65829515 |
| AGSGALGL | ASSSLGDEQY | 0.64071804 |
| AQIADQI | ASSSLGDEQY | 0.25915486 |
| ASAVAVTYL | ASSSLGDEQY | 0.59335834 |
| ASTVLGGTGG | ASSSLGDEQY | 0.6685883 |
| AVGGEPAK | ASSSLGDEQY | 0.71247935 |
| CEQGTFGGTC | ASSSLGDEQY | 0.4938785 |
| DEGKAPAG | ASSSLGDEQY | 0.61107445 |
| DTVVFPAL | ASSSLGDEQY | 0.50322074 |
| FEGITMAG | ASSSLGDEQY | 0.46821004 |
| FPLVVQD | ASSSLGDEQY | 0.3297275 |
| GAFTTPQ | ASSSLGDEQY | 0.3056811 |
| GFGGFGTAGGVGGA | ASSSLGDEQY | 0.48097962 |
| GFLAQFL | ASSSLGDEQY | 0.40275943 |
| GGAGGAGGNAGVA | ASSSLGDEQY | 0.6969784 |
| GGVGTAGAGGAGGAGG | ASSSLGDEQY | 0.680075 |
| GNGGGGDAGNAGSGGN | ASSSLGDEQY | 0.65368783 |
| GTGGAGGAGNDAGS | ASSSLGDEQY | 0.6663376 |
| GTTNTGWFNS | ASSSLGDEQY | 0.6904144 |
| IFMGPLFTL | ASSSLGDEQY | 0.5093239 |
| ILAHAIEAI | ASSSLGDEQY | 0.4582237 |
| IRSAIREAL | ASSSLGDEQY | 0.5111795 |
| ITFAGGG | ASSSLGDEQY | 0.28306255 |
| ITGMSGHQEV | ASSSLGDEQY | 0.35582715 |
| IVAGLACTL | ASSSLGDEQY | 0.43788642 |
| IVGGGLAGA | ASSSLGDEQY | 0.49047378 |
| IVTGVLNI | ASSSLGDEQY | 0.21495488 |

| | | |
|---|---|---|
| LAQQGCI | ASSSLGDEQY | 0.64930713 |
| LCLHVEN | ASSSLGDEQY | 0.68475527 |
| LDAISDGS | ASSSLGDEQY | 0.57462984 |
| LGGFAAML | ASSSLGDEQY | 0.65520835 |
| LGGKNPMI | ASSSLGDEQY | 0.5349257 |
| LGPKGVAAIL | ASSSLGDEQY | 0.37520665 |
| LMVGVSI | ASSSLGDEQY | 0.6207019 |
| LVGAAFVL | ASSSLGDEQY | 0.3331677 |
| LVLAVAPL | ASSSLGDEQY | 0.574717 |
| MAAIAVIDAA | ASSSLGDEQY | 0.6880863 |
| NNGINAELQ | ASSSLGDEQY | 0.5810077 |
| RGEPLVLVHG | ASSSLGDEQY | 0.51453125 |
| RGGAGGAAGLI | ASSSLGDEQY | 0.4939785 |
| RSYRMHPAV | ASSSLGDEQY | 0.29766914 |
| SAAHLLAGQLA | ASSSLGDEQY | 0.53097767 |
| SAAVGSY | ASSSLGDEQY | 0.32987958 |
| SPHAVLHRI | ASSSLGDEQY | 0.52909195 |
| SQIAKVL | ASSSLGDEQY | 0.31247592 |
| TALGHLVP | ASSSLGDEQY | 0.566978 |
| TAPTTSGAA | ASSSLGDEQY | 0.5960449 |
| TDGDDGNGGSPGT | ASSSLGDEQY | 0.54371107 |
| TDGLGEN | ASSSLGDEQY | 0.38611573 |
| TGGPPLPGQA | ASSSLGDEQY | 0.5184878 |
| TIPAITI | ASSSLGDEQY | 0.61463535 |
| TIPGFTI | ASSSLGDEQY | 0.30105698 |
| TSIASGALA | ASSSLGDEQY | 0.57377815 |
| TVTLPGAAI | ASSSLGDEQY | 0.36116767 |
| VASGLSAALP | ASSSLGDEQY | 0.5798241 |
| VGAGVIGI | ASSSLGDEQY | 0.67476904 |
| VGPKRFGV | ASSSLGDEQY | 0.43986624 |
| VNVAFPD | ASSSLGDEQY | 0.29927364 |
| VSMTNSGVS | ASSSLGDEQY | 0.47658727 |
| VTDVLGDLSV | ASSSLGDEQY | 0.4673376 |
| VVVVGAGFAGL | ASSSLGDEQY | 0.46043104 |
| VYTVGTVV | ASSSLGDEQY | 0.23002829 |
| WTTFSGP | ASSSLGDEQY | 0.6543613 |
| YDELVEAF | ASSSLGDEQY | 0.32303828 |
| YGTGGAGGNGG | ASSSLGDEQY | 0.43606532 |
| YTNVMAV | ASSSLGDEQY | 0.4589669 |
| FLEQVHQL | ASSSLGDEQY | 0.5929402 |
| GLSLLAVAV | ASSSLGDEQY | 0.54351056 |
| LEQVHQL | ASSSLGDEQY | 0.42768866 |
| LNAAGAI | ASSSLGDEQY | 0.5826756 |
| PAVHASL | ASSSLGDEQY | 0.5852757 |
| TELEDDA | ASSSLGDEQY | 0.675451 |
| TPPGPPN | ASSSLGDEQY | 0.6214392 |
| TPPPPPPP | ASSSLGDEQY | 0.48087847 |
| TQSPALI | SSSQPGTNSDYT | 0.6643994 |
| EFAAAFPQ | SSSQPGTNSDYT | 0.46396878 |
| LLSGNLDVL | SSSQPGTNSDYT | 0.27309278 |
| GPANLGNYNI | SSSQPGTNSDYT | 0.29140633 |

| | | |
|---|---|---|
| TLGPTLP | SSSQPGTNSDYT | 0.3257855 |
| VYAYSSCCK | SSSQPGTNSDYT | 0.28588933 |
| APAPAPAPNI | SSSQPGTNSDYT | 0.6067555 |
| PGGPVPIP | SSSQPGTNSDYT | 0.6041268 |
| FLTVALEP | SSSQPGTNSDYT | 0.4951207 |
| SQYFGANL | SSSQPGTNSDYT | 0.25858173 |
| GGVKPAA | SSSQPGTNSDYT | 0.39196646 |
| GGTGGAGGAGGW | SSSQPGTNSDYT | 0.4274006 |
| GVGGVAGLGGAGG | SSSQPGTNSDYT | 0.5722972 |
| VLLVLATTI | SSSQPGTNSDYT | 0.62808776 |
| IGIFAAML | SSSQPGTNSDYT | 0.72346103 |
| AVGDLID | SSSQPGTNSDYT | 0.68311346 |
| NISPLPL | SSSQPGTNSDYT | 0.51320624 |
| VTAVFVAI | SSSQPGTNSDYT | 0.61846155 |
| SGDDNDAAL | SSSQPGTNSDYT | 0.41337568 |
| AVAALAHS | SSSQPGTNSDYT | 0.48013198 |
| GGVTVMATP | SSSQPGTNSDYT | 0.47708106 |
| AGLAANVRD | SSSQPGTNSDYT | 0.57758224 |
| TNRPDLIDPALLRPG | SSSQPGTNSDYT | 0.43653536 |
| LANLGVTAL | SSSQPGTNSDYT | 0.4149037 |
| TGERIEL | SSSQPGTNSDYT | 0.30716035 |
| NGEELAA | SSSQPGTNSDYT | 0.512123 |
| VLSLVWG | SSSQPGTNSDYT | 0.36373478 |
| IGYALWL | SSSQPGTNSDYT | 0.25539368 |
| LSLLLAPL | SSSQPGTNSDYT | 0.50313735 |
| TLDEALK | SSSQPGTNSDYT | 0.68579876 |
| TDGAGTAEGG | SSSQPGTNSDYT | 0.69034183 |
| NAPGGAGGNGG | SSSQPGTNSDYT | 0.3813359 |
| VSVFMGMGGNF | SSSQPGTNSDYT | 0.35808086 |
| LASYLAAL | SSSQPGTNSDYT | 0.6711636 |
| LVGGTPAL | SSSQPGTNSDYT | 0.39733562 |
| VRNADVL | SSSQPGTNSDYT | 0.5356661 |
| TCSSPFLVSAM | SSSQPGTNSDYT | 0.5316573 |
| VHITDAE | SSSQPGTNSDYT | 0.5999853 |
| EYAAPSF | SSSQPGTNSDYT | 0.36614662 |
| VVDAFHGGL | SSSQPGTNSDYT | 0.5736232 |
| ASVVDPQ | SSSQPGTNSDYT | 0.5586828 |
| SYNEFGGLS | SSSQPGTNSDYT | 0.48403805 |
| DPDGSVY | SSSQPGTNSDYT | 0.24611713 |
| MVSTIHATE | SSSQPGTNSDYT | 0.6381438 |
| LAAGVADPV | SSSQPGTNSDYT | 0.38883033 |
| TKEQIAA | SSSQPGTNSDYT | 0.31653327 |
| SLASPAAL | SSSQPGTNSDYT | 0.23323517 |
| IGLVFYGV | SSSQPGTNSDYT | 0.5440695 |
| FLLDASNL | SSSQPGTNSDYT | 0.60694635 |
| YQVGNFP | SSSQPGTNSDYT | 0.67959416 |
| ESLLSQST | SSSQPGTNSDYT | 0.46503025 |
| LFTVGFAAVG | SSSQPGTNSDYT | 0.6376458 |
| ATPPTPT | SSSQPGTNSDYT | 0.6674018 |
| TDLATLG | SSSQPGTNSDYT | 0.44437486 |
| SAGGTGR | SSSQPGTNSDYT | 0.26239154 |

| | | |
|---|---|---|
| GFSNSGS | SSSQPGTNSDYT | 0.52090514 |
| GGNGGSGGTSV | SSSQPGTNSDYT | 0.6030544 |
| GTSVATGGAGN | SSSQPGTNSDYT | 0.65791774 |
| NGGNGGSGGTS | SSSQPGTNSDYT | 0.6950829 |
| AVVKFPH | SSSQPGTNSDYT | 0.30212253 |
| TPPPPPP | SSSQPGTNSDYT | 0.5795755 |
| AAPAAAAAPA | SSSQPGTNSDYT | 0.54329395 |
| IALNEDL | SSSQPGTNSDYT | 0.3067968 |
| IVNVSSM | SSSQPGTNSDYT | 0.40427113 |
| TGIPPAPR | SSSQPGTNSDYT | 0.47051954 |
| FDLGGGTF | SSSQPGTNSDYT | 0.28468037 |
| FDLGGGTFDVS | SSSQPGTNSDYT | 0.6022873 |
| NEPTAAA | SSSQPGTNSDYT | 0.4814685 |
| NEPTAAALA | SSSQPGTNSDYT | 0.4147365 |
| GSIINISSI | SSSQPGTNSDYT | 0.44269586 |
| VAPGNAGTA | SSSQPGTNSDYT | 0.3655565 |
| TGGGASLE | SSSQPGTNSDYT | 0.4015504 |
| VAVYNHYKRI | SSSQPGTNSDYT | 0.4232723 |
| TDPAPATT | SSSQPGTNSDYT | 0.6631045 |
| TNRPDILDPALLRPG | SSSQPGTNSDYT | 0.5905255 |
| LTTTTTS | SSSQPGTNSDYT | 0.5490963 |
| PTGSGKTL | SSSQPGTNSDYT | 0.50966644 |
| TGSGKTL | SSSQPGTNSDYT | 0.68714887 |
| TIGHVDHGK | SSSQPGTNSDYT | 0.28213108 |
| TIGHVDHGKTT | SSSQPGTNSDYT | 0.41393358 |
| GHVDHGKTT | SSSQPGTNSDYT | 0.5470011 |
| AQFEHTLL | SSSQPGTNSDYT | 0.37872046 |
| IVNQDEL | SSSQPGTNSDYT | 0.6273995 |
| TNGSQFF | SSSQPGTNSDYT | 0.6418762 |
| TGKGIVE | SSSQPGTNSDYT | 0.5569882 |
| LISNASDA | SSSQPGTNSDYT | 0.48189148 |
| AELRTLGELTKIAKAHGAQ | SSSQPGTNSDYT | 0.69893825 |
| APNAVAAAEPSW | SSSQPGTNSDYT | 0.5121734 |
| APQVAGAATAVL | SSSQPGTNSDYT | 0.44209665 |
| ARRDPAAAAAILPTDA | SSSQPGTNSDYT | 0.62316763 |
| DAAARDVLAEAGLAEYF | SSSQPGTNSDYT | 0.36439893 |
| DDQIVLSAKANLAKVEQ | SSSQPGTNSDYT | 0.63969445 |
| DELLPDTGEDSDEDGH | SSSQPGTNSDYT | 0.56465983 |
| ELPEAPANRATVPQD | SSSQPGTNSDYT | 0.3344307 |
| FGQSKIERVTGIG | SSSQPGTNSDYT | 0.45605892 |
| GAGGTGSTAGGAGGAGGAGGLY | SSSQPGTNSDYT | 0.66678417 |
| GHGGTGAAGGNGGTGGAGGSGI | SSSQPGTNSDYT | 0.64100415 |
| GNGGNGGAGGEGGPGGAGGAGA | SSSQPGTNSDYT | 0.31549028 |
| GSTAGNGGNGGNATNGTIAGQP | SSSQPGTNSDYT | 0.40314257 |
| GTGGDGGNGGNGGTGG | SSSQPGTNSDYT | 0.3567198 |
| GVRRPAGIPADAPV | SSSQPGTNSDYT | 0.3849798 |
| GYRQVTDAHLLTLARR | SSSQPGTNSDYT | 0.28415897 |
| IAKMGQDGHDRG | SSSQPGTNSDYT | 0.6003417 |
| KAEPSIEIQYT | SSSQPGTNSDYT | 0.44277567 |
| KPVRIRMAADEDTVAVT | SSSQPGTNSDYT | 0.69198596 |
| MPPRIAGMRLLVIKPEP | SSSQPGTNSDYT | 0.5837421 |

| | | |
|---|---|---|
| MKFYAVALFRSRGVPPTR | SSSQPGTNSDYT | 0.6780977 |
| NAGGFGILGAVAHSP | SSSQPGTNSDYT | 0.36070266 |
| NEGREITLAMAEM | SSSQPGTNSDYT | 0.56183887 |
| NELVEWSHPLGHRWRWE | SSSQPGTNSDYT | 0.24891631 |
| NLGAGGTGGNGGNPG | SSSQPGTNSDYT | 0.44687888 |
| NTEWPKWETVMAADPVD | SSSQPGTNSDYT | 0.28336608 |
| PASAAPAETQP | SSSQPGTNSDYT | 0.531811 |
| QQSSDGKNGVRAAEITGE | SSSQPGTNSDYT | 0.26491502 |
| RAPDAVTTSWWKE | SSSQPGTNSDYT | 0.72897846 |
| SGFGNVGAHTSGWFNQ | SSSQPGTNSDYT | 0.35001397 |
| SGVAGAVGQAASVGGLKV | SSSQPGTNSDYT | 0.6274879 |
| SNGGAGTGSAGGNGGT | SSSQPGTNSDYT | 0.43576366 |
| TGGAGGEGGAAG | SSSQPGTNSDYT | 0.29929405 |
| TGGDGGAAGAGGAPGHG | SSSQPGTNSDYT | 0.44477078 |
| TMPPLLAAIANAGRHGV | SSSQPGTNSDYT | 0.5628718 |
| TSSTAGNSGGA | SSSQPGTNSDYT | 0.33569542 |
| VPATVPVGILC | SSSQPGTNSDYT | 0.25190738 |
| VQRPDSGGVVLTGRIS | SSSQPGTNSDYT | 0.3910971 |
| WTPNPVPATKGVPAGLVQAT | SSSQPGTNSDYT | 0.40452856 |
| YALKRALAKFSDRFDVVII | SSSQPGTNSDYT | 0.7040621 |
| YLIGSVIV | SSSQPGTNSDYT | 0.66921914 |
| YGQLVAVID | SSSQPGTNSDYT | 0.4620446 |
| VTVGGMTLPA | SSSQPGTNSDYT | 0.71674526 |
| VLALPAV | SSSQPGTNSDYT | 0.21795498 |
| VIPPTLNYET | SSSQPGTNSDYT | 0.47793823 |
| VGMGGMHPSSG | SSSQPGTNSDYT | 0.716459 |
| VGLAGGAATA | SSSQPGTNSDYT | 0.54250836 |
| VGGAGGAGGSGP | SSSQPGTNSDYT | 0.47837204 |
| VFAAFMV | SSSQPGTNSDYT | 0.6222899 |
| TLYDKLR | SSSQPGTNSDYT | 0.4649719 |
| TLAVFTVL | SSSQPGTNSDYT | 0.47197968 |
| TGSVQPARV | SSSQPGTNSDYT | 0.45506176 |
| TGGGAGSGGT | SSSQPGTNSDYT | 0.577938 |
| SHLETSL | SSSQPGTNSDYT | 0.42360485 |
| SFTTGMT | SSSQPGTNSDYT | 0.2897077 |
| QVEVAKRMGVRQPTVS | SSSQPGTNSDYT | 0.5112329 |
| QGSGGPALAM | SSSQPGTNSDYT | 0.72062194 |
| MLAPSNP | SSSQPGTNSDYT | 0.41623053 |
| LVNNAAI | SSSQPGTNSDYT | 0.63375425 |
| LTGEINTLA | SSSQPGTNSDYT | 0.48890793 |
| LSIFGPV | SSSQPGTNSDYT | 0.48362166 |
| LLALLANL | SSSQPGTNSDYT | 0.38713664 |
| LISNASDALDKLR | SSSQPGTNSDYT | 0.31679368 |
| LINEAVP | SSSQPGTNSDYT | 0.5614195 |
| LGYAFSGL | SSSQPGTNSDYT | 0.3097294 |
| LDFVAVAL | SSSQPGTNSDYT | 0.42295945 |
| LAQPGAE | SSSQPGTNSDYT | 0.50055707 |
| ISNLGAVTGF | SSSQPGTNSDYT | 0.7302637 |
| HQEDPGRI | SSSQPGTNSDYT | 0.6439688 |
| GTPGEPTP | SSSQPGTNSDYT | 0.517537 |
| GFFNSGS | SSSQPGTNSDYT | 0.41616136 |

| | | |
|---|---|---|
| GAGGGGLTAGAN | SSSQPGTNSDYT | 0.46009138 |
| FVDALVNM | SSSQPGTNSDYT | 0.5909742 |
| FLTVAPEL | SSSQPGTNSDYT | 0.3018411 |
| FETLTAL | SSSQPGTNSDYT | 0.6978413 |
| DVINPIEL | SSSQPGTNSDYT | 0.3124601 |
| APVSGAPV | SSSQPGTNSDYT | 0.613201 |
| ALVGGLAW | SSSQPGTNSDYT | 0.61092854 |
| ALAAAAATAI | SSSQPGTNSDYT | 0.70730615 |
| AFGLGLLL | SSSQPGTNSDYT | 0.49392527 |
| AFFAAASL | SSSQPGTNSDYT | 0.57432276 |
| AAAALPAFSPP | SSSQPGTNSDYT | 0.36851886 |
| VVLAPAL | SSSQPGTNSDYT | 0.61485714 |
| TIGHVDHGKTTL | SSSQPGTNSDYT | 0.6746373 |
| LISNASDALDK | SSSQPGTNSDYT | 0.59256065 |
| LISNASDAL | SSSQPGTNSDYT | 0.7239915 |
| ISNASDALDK | SSSQPGTNSDYT | 0.561133 |
| NQAETLVYQTE | SSSQPGTNSDYT | 0.25871676 |
| TGGTGGTAGAG | SSSQPGTNSDYT | 0.41107512 |
| TGSTAGGAGGA | SSSQPGTNSDYT | 0.646286 |
| DPLAVPVAAGPV | SSSQPGTNSDYT | 0.6840652 |
| GGTGGTPGNGGN | SSSQPGTNSDYT | 0.6636219 |
| PAGLAGVDPATL | SSSQPGTNSDYT | 0.57050526 |
| GNNLVVEAPHAGASV | SSSQPGTNSDYT | 0.45662332 |
| KVTRIMSASTLEQPA | SSSQPGTNSDYT | 0.68649566 |
| LGFALPIALPRRTVV | SSSQPGTNSDYT | 0.7301849 |
| TRLGNLTARKLVVLV | SSSQPGTNSDYT | 0.55950284 |
| WSQQRRLIYFPSAGP | SSSQPGTNSDYT | 0.3751471 |
| GPNGNPDPMAAAVDIR | SSSQPGTNSDYT | 0.3624411 |
| TQRCVSVADLRAAASR | SSSQPGTNSDYT | 0.5992364 |
| QFVLDVAEHPVEVLTETV | SSSQPGTNSDYT | 0.6870432 |
| GGAGGTGGAGGNGGTQAGDG | SSSQPGTNSDYT | 0.26007256 |
| PLLATLRAIAVDVDAKPAQV | SSSQPGTNSDYT | 0.4730988 |
| APKFRTGGYMIDFWGVGYQVAK | SSSQPGTNSDYT | 0.58242416 |
| GAGGAGGAGGAAGGPGGTGGTG | SSSQPGTNSDYT | 0.24267632 |
| IGALGLLATDVAPAGSVAFGVA | SSSQPGTNSDYT | 0.5628029 |
| AGGAGGIGGNGGAGGAGGVLMG | SSSQPGTNSDYT | 0.68199414 |
| ADELIGLL | SSSQPGTNSDYT | 0.49910614 |
| ADRALKA | SSSQPGTNSDYT | 0.26151904 |
| AGAVGQVGGAGGA | SSSQPGTNSDYT | 0.6959641 |
| AGGGGGLGGAP | SSSQPGTNSDYT | 0.6791324 |
| AGGLLYGNGGAG | SSSQPGTNSDYT | 0.62747896 |
| AGLESIGL | SSSQPGTNSDYT | 0.29018816 |
| AGLKTGII | SSSQPGTNSDYT | 0.22483142 |
| ASLAALLL | SSSQPGTNSDYT | 0.5297688 |
| ASVINISSI | SSSQPGTNSDYT | 0.656821 |
| ATPPGPA | SSSQPGTNSDYT | 0.4766789 |
| ATVASALV | SSSQPGTNSDYT | 0.32202283 |
| ATVASLIL | SSSQPGTNSDYT | 0.41044736 |
| AVNGPVLIRH | SSSQPGTNSDYT | 0.6769162 |
| DGAAGAVGI | SSSQPGTNSDYT | 0.27372986 |
| DPLLHGI | SSSQPGTNSDYT | 0.29233953 |

| | | |
|---|---|---|
| ELGAALGAL | SSSQPGTNSDYT | 0.5733751 |
| ELGRTQAV | SSSQPGTNSDYT | 0.5800339 |
| FVAGGYLS | SSSQPGTNSDYT | 0.35419515 |
| GASGGAGGAGGW | SSSQPGTNSDYT | 0.4134007 |
| GGAGAGGDGGAGGI | SSSQPGTNSDYT | 0.6173531 |
| GTGHNGSQPGMGGNG | SSSQPGTNSDYT | 0.44001824 |
| GYAAASASAS | SSSQPGTNSDYT | 0.53328884 |
| HQVDVAAVQQG | SSSQPGTNSDYT | 0.28231317 |
| IGAAIAHT | SSSQPGTNSDYT | 0.696159 |
| IGMIEVAQL | SSSQPGTNSDYT | 0.2915126 |
| IRLAPEHTSAMD | SSSQPGTNSDYT | 0.6141134 |
| IYNVSVL | SSSQPGTNSDYT | 0.5457468 |
| KQGFLGM | SSSQPGTNSDYT | 0.56081516 |
| LESVITQL | SSSQPGTNSDYT | 0.46278447 |
| LGLGALVVV | SSSQPGTNSDYT | 0.2963242 |
| LIGNGGNGGSGGI | SSSQPGTNSDYT | 0.3185703 |
| LNAINAPTL | SSSQPGTNSDYT | 0.28300208 |
| LQLLNTGV | SSSQPGTNSDYT | 0.6903462 |
| LQTSIEF | SSSQPGTNSDYT | 0.5847012 |
| MSFAVSEL | SSSQPGTNSDYT | 0.6019498 |
| NTGALGL | SSSQPGTNSDYT | 0.50914985 |
| PPGAGGAGGAGTT | SSSQPGTNSDYT | 0.35666338 |
| QLVGLGI | SSSQPGTNSDYT | 0.56134593 |
| RFLEQVHQL | SSSQPGTNSDYT | 0.27914262 |
| RGVGYVLR | SSSQPGTNSDYT | 0.45181853 |
| SATGVFAGVI | SSSQPGTNSDYT | 0.45310563 |
| SLPAITLP | SSSQPGTNSDYT | 0.55748785 |
| STALAFTV | SSSQPGTNSDYT | 0.36346927 |
| SVLTVGLGK | SSSQPGTNSDYT | 0.4940943 |
| TALAGAGSGI | SSSQPGTNSDYT | 0.5803633 |
| TGLPFTGI | SSSQPGTNSDYT | 0.34946632 |
| VAEHVTV | SSSQPGTNSDYT | 0.5034144 |
| VDIVLNSL | SSSQPGTNSDYT | 0.688986 |
| VLTGGGAVL | SSSQPGTNSDYT | 0.533563 |
| VNVGLAYGI | SSSQPGTNSDYT | 0.26027787 |
| VSSAAVGAV | SSSQPGTNSDYT | 0.567905 |
| VTSVYEAGV | SSSQPGTNSDYT | 0.41756505 |
| WVALDRGA | SSSQPGTNSDYT | 0.59808 |
| YLVGSVVI | SSSQPGTNSDYT | 0.563601 |
| GPPGTGKTLL | SSSQPGTNSDYT | 0.51909417 |
| GVGMTKF | SSSQPGTNSDYT | 0.5394835 |
| IGIGGSDL | SSSQPGTNSDYT | 0.39215484 |
| IGTLLAEL | SSSQPGTNSDYT | 0.38400584 |
| IYTSGTT | SSSQPGTNSDYT | 0.44200498 |
| LSFPAGV | SSSQPGTNSDYT | 0.56597596 |
| LYTSGTT | SSSQPGTNSDYT | 0.57864046 |
| VGAFPFGF | SSSQPGTNSDYT | 0.35806572 |
| VGAGTVEF | SSSQPGTNSDYT | 0.5253191 |
| VGKSTLLL | SSSQPGTNSDYT | 0.43424675 |
| AALAHAGGGIV | SSSQPGTNSDYT | 0.3729267 |
| AALVAVSLL | SSSQPGTNSDYT | 0.4148951 |

| | | |
|---|---|---|
| AAVSRAVAAL | SSSQPGTNSDYT | 0.41345546 |
| AGGSSGIGGF | SSSQPGTNSDYT | 0.2510944 |
| AGLTVANANG | SSSQPGTNSDYT | 0.63383317 |
| AGLVGAVL | SSSQPGTNSDYT | 0.3297884 |
| ALSQAFTV | SSSQPGTNSDYT | 0.49892354 |
| ALTFSVSL | SSSQPGTNSDYT | 0.6681992 |
| ALTGAALVAL | SSSQPGTNSDYT | 0.28434148 |
| ASFSIGAGVL | SSSQPGTNSDYT | 0.55182254 |
| DGSVYKI | SSSQPGTNSDYT | 0.48579228 |
| EQLAEIV | SSSQPGTNSDYT | 0.54757375 |
| FAVSELTVM | SSSQPGTNSDYT | 0.48199758 |
| GAAPLAAGL | SSSQPGTNSDYT | 0.3790574 |
| GADGGDGGSGGAGG | SSSQPGTNSDYT | 0.6595409 |
| GFGLDTGGAGG | SSSQPGTNSDYT | 0.56063175 |
| GLASGGLL | SSSQPGTNSDYT | 0.62104356 |
| GRIAGAAVLL | SSSQPGTNSDYT | 0.6363152 |
| GSAGSLNTSF | SSSQPGTNSDYT | 0.6629033 |
| GTGQIGI | SSSQPGTNSDYT | 0.60909826 |
| GTLIALIL | SSSQPGTNSDYT | 0.41065764 |
| IAATMGFVVG | SSSQPGTNSDYT | 0.2465587 |
| IFAQALL | SSSQPGTNSDYT | 0.5558971 |
| IGAAVALGF | SSSQPGTNSDYT | 0.5497153 |
| IGLTPLAL | SSSQPGTNSDYT | 0.32211116 |
| ILGSNTSTL | SSSQPGTNSDYT | 0.258477 |
| IVFSLTVL | SSSQPGTNSDYT | 0.6897727 |
| LASKATGF | SSSQPGTNSDYT | 0.72403 |
| LASVALIL | SSSQPGTNSDYT | 0.38080177 |
| LFAVGLTL | SSSQPGTNSDYT | 0.6986908 |
| LFNGGIGTYI | SSSQPGTNSDYT | 0.27375674 |
| LTLGAGII | SSSQPGTNSDYT | 0.5878662 |
| LVAAALGDI | SSSQPGTNSDYT | 0.6903056 |
| NTAFVQLGI | SSSQPGTNSDYT | 0.29698014 |
| QLFSGAEL | SSSQPGTNSDYT | 0.6379789 |
| SAATFITL | SSSQPGTNSDYT | 0.36445522 |
| SNYGLANLGV | SSSQPGTNSDYT | 0.34311265 |
| SYGGAYAVM | SSSQPGTNSDYT | 0.666219 |
| TTGVGGLTLG | SSSQPGTNSDYT | 0.47509092 |
| TTQTIYI | SSSQPGTNSDYT | 0.59215844 |
| VAAGAGLGAVI | SSSQPGTNSDYT | 0.39436802 |
| VAAVVTLL | SSSQPGTNSDYT | 0.38073993 |
| VGFLGGLI | SSSQPGTNSDYT | 0.5717735 |
| VGVIPILL | SSSQPGTNSDYT | 0.6745663 |
| VIFASSLI | SSSQPGTNSDYT | 0.4335065 |
| VKGGDGGAAAG | SSSQPGTNSDYT | 0.71554077 |
| VTKTFTNI | SSSQPGTNSDYT | 0.46176547 |
| VVAAAILL | SSSQPGTNSDYT | 0.7277409 |
| YAALDVEL | SSSQPGTNSDYT | 0.69615453 |
| LASVALLL | SSSQPGTNSDYT | 0.5710846 |
| LDAVHRV | SSSQPGTNSDYT | 0.5007465 |
| QGLSSSIF | SSSQPGTNSDYT | 0.6797502 |
| RGTFANIRL | SSSQPGTNSDYT | 0.39687127 |

| | | |
|---|---|---|
| RQLWWGHRI | SSSQPGTNSDYT | 0.6165198 |
| YGPPGTGKTLL | SSSQPGTNSDYT | 0.5872426 |
| AALVGAPANVA | SSSQPGTNSDYT | 0.58200586 |
| AATFRRTE | SSSQPGTNSDYT | 0.84762716 |
| AGALGGGI | SSSQPGTNSDYT | 0.8139663 |
| AGGAGGAGGIGAG | SSSQPGTNSDYT | 0.7496256 |
| AGSGALGL | SSSQPGTNSDYT | 0.68262964 |
| AQIADQI | SSSQPGTNSDYT | 0.5008933 |
| ASAVAVTYL | SSSQPGTNSDYT | 0.74979854 |
| ASTVLGGTGG | SSSQPGTNSDYT | 0.5960033 |
| AVGGEPAK | SSSQPGTNSDYT | 0.82780254 |
| CEQGTFGGTC | SSSQPGTNSDYT | 0.8385457 |
| DEGKAPAG | SSSQPGTNSDYT | 0.67670393 |
| DTVVFPAL | SSSQPGTNSDYT | 0.84676874 |
| FEGITMAG | SSSQPGTNSDYT | 0.82456446 |
| FPLVVQD | SSSQPGTNSDYT | 0.56708187 |
| GAFTTPQ | SSSQPGTNSDYT | 0.7688287 |
| GFGGFGTAGGVGGA | SSSQPGTNSDYT | 0.7439241 |
| GFLAQFL | SSSQPGTNSDYT | 0.71349055 |
| GGAGGAGGNAGVA | SSSQPGTNSDYT | 0.5603462 |
| GGVGTAGAGGAGGAGG | SSSQPGTNSDYT | 0.8151802 |
| GNGGGGDAGNAGSGGN | SSSQPGTNSDYT | 0.7258784 |
| GTGGAGGAGNDAGS | SSSQPGTNSDYT | 0.27358824 |
| GTTNTGWFNS | SSSQPGTNSDYT | 0.8128382 |
| IFMGPLFTL | SSSQPGTNSDYT | 0.7723413 |
| ILAHAIEAI | SSSQPGTNSDYT | 0.74832875 |
| IRSAIREAL | SSSQPGTNSDYT | 0.83548945 |
| ITFAGGG | SSSQPGTNSDYT | 0.64804655 |
| ITGMSGHQEV | SSSQPGTNSDYT | 0.73265815 |
| IVAGLACTL | SSSQPGTNSDYT | 0.84010327 |
| IVGGGLAGA | SSSQPGTNSDYT | 0.8454336 |
| IVTGVLNI | SSSQPGTNSDYT | 0.5715031 |
| LAQQGCI | SSSQPGTNSDYT | 0.8025726 |
| LCLHVEN | SSSQPGTNSDYT | 0.83849823 |
| LDAISDGS | SSSQPGTNSDYT | 0.8004058 |
| LGGFAAML | SSSQPGTNSDYT | 0.8408556 |
| LGGKNPMI | SSSQPGTNSDYT | 0.844486 |
| LGPKGVAAIL | SSSQPGTNSDYT | 0.7522558 |
| LMVGVSI | SSSQPGTNSDYT | 0.84645593 |
| LVGAAFVL | SSSQPGTNSDYT | 0.85368335 |
| LVLAVAPL | SSSQPGTNSDYT | 0.76041126 |
| MAAIAVIDAA | SSSQPGTNSDYT | 0.7507132 |
| NNGINAELQ | SSSQPGTNSDYT | 0.5892921 |
| RGEPLVLVHG | SSSQPGTNSDYT | 0.79318607 |
| RGGAGGAAGLI | SSSQPGTNSDYT | 0.5426447 |
| RSYRMHPAV | SSSQPGTNSDYT | 0.81347835 |
| SAAHLLAGQLA | SSSQPGTNSDYT | 0.8378298 |
| SAAVGSY | SSSQPGTNSDYT | 0.74924403 |
| SPHAVLHRI | SSSQPGTNSDYT | 0.59580666 |
| SQIAKVL | SSSQPGTNSDYT | 0.5708907 |
| TALGHLVP | SSSQPGTNSDYT | 0.7930448 |

| | | |
|---|---|---|
| TAPTTSGAA | SSSQPGTNSDYT | 0.5941602 |
| TDGDDGNGGSPGT | SSSQPGTNSDYT | 0.3942866 |
| TDGLGEN | SSSQPGTNSDYT | 0.75308245 |
| TGGPPLPGQA | SSSQPGTNSDYT | 0.8350197 |
| TIPAITI | SSSQPGTNSDYT | 0.8266357 |
| TIPGFTI | SSSQPGTNSDYT | 0.5630005 |
| TSIASGALA | SSSQPGTNSDYT | 0.85063696 |
| TVTLPGAAI | SSSQPGTNSDYT | 0.82774144 |
| VASGLSAALP | SSSQPGTNSDYT | 0.8330418 |
| VGAGVIGI | SSSQPGTNSDYT | 0.7601071 |
| VGPKRFGV | SSSQPGTNSDYT | 0.5654118 |
| VNVAFPD | SSSQPGTNSDYT | 0.83660877 |
| VSMTNSGVS | SSSQPGTNSDYT | 0.7697263 |
| VTDVLGDLSV | SSSQPGTNSDYT | 0.7519511 |
| VVVVGAGFAGL | SSSQPGTNSDYT | 0.82448673 |
| VYTVGTVV | SSSQPGTNSDYT | 0.77945316 |
| WTTFSGP | SSSQPGTNSDYT | 0.6970476 |
| YDELVEAF | SSSQPGTNSDYT | 0.76587045 |
| YGTGGAGGNGG | SSSQPGTNSDYT | 0.8273722 |
| YTNVMAV | SSSQPGTNSDYT | 0.7167965 |
| FLEQVHQL | SSSQPGTNSDYT | 0.47626498 |
| GLSLLAVAV | SSSQPGTNSDYT | 0.7081001 |
| LEQVHQL | SSSQPGTNSDYT | 0.77752244 |
| LNAAGAI | SSSQPGTNSDYT | 0.8441225 |
| PAVHASL | SSSQPGTNSDYT | 0.8039214 |
| TELEDDA | SSSQPGTNSDYT | 0.72126055 |
| TPPGPPN | SSSQPGTNSDYT | 0.8233087 |
| TPPPPPPP | SSSQPGTNSDYT | 0.7941059 |
| TQSPALI | ASGIGQTYEQY | 0.76448274 |
| EFAAAFPQ | ASGIGQTYEQY | 0.8228779 |
| LLSGNLDVL | ASGIGQTYEQY | 0.7027813 |
| GPANLGNYNI | ASGIGQTYEQY | 0.8296803 |
| TLGPTLP | ASGIGQTYEQY | 0.8084005 |
| VYAYSSCCK | ASGIGQTYEQY | 0.8189325 |
| APAPAPAPNI | ASGIGQTYEQY | 0.84205747 |
| PGGPVPIP | ASGIGQTYEQY | 0.81905234 |
| FLTVALEP | ASGIGQTYEQY | 0.76650363 |
| SQYFGANL | ASGIGQTYEQY | 0.44043568 |
| GGVKPAA | ASGIGQTYEQY | 0.7835882 |
| GGTGGAGGAGGW | ASGIGQTYEQY | 0.813742 |
| GVGGVAGLGGAGG | ASGIGQTYEQY | 0.8093884 |
| VLLVLATTI | ASGIGQTYEQY | 0.8360765 |
| IGIFAAML | ASGIGQTYEQY | 0.801166 |
| AVGDLID | ASGIGQTYEQY | 0.73129284 |
| NISPLPL | ASGIGQTYEQY | 0.8407134 |
| VTAVFVAI | ASGIGQTYEQY | 0.8341023 |
| SGDDNDAAL | ASGIGQTYEQY | 0.48772064 |
| AVAALAHS | ASGIGQTYEQY | 0.77565193 |
| GGVTVMATP | ASGIGQTYEQY | 0.85419106 |
| AGLAANVRD | ASGIGQTYEQY | 0.80453765 |
| TNRPDLIDPALLRPG | ASGIGQTYEQY | 0.7680614 |

| | | |
|---|---|---|
| LANLGVTAL | ASGIGQTYEQY | 0.65246135 |
| TGERIEL | ASGIGQTYEQY | 0.8402498 |
| NGEELAA | ASGIGQTYEQY | 0.72779626 |
| VLSLVWG | ASGIGQTYEQY | 0.85213566 |
| IGYALWL | ASGIGQTYEQY | 0.28081077 |
| LSLLLAPL | ASGIGQTYEQY | 0.6552237 |
| TLDEALK | ASGIGQTYEQY | 0.5488732 |
| TDGAGTAEGG | ASGIGQTYEQY | 0.25706467 |
| NAPGGAGGNGG | ASGIGQTYEQY | 0.8375103 |
| VSVFMGMGGNF | ASGIGQTYEQY | 0.662265 |
| LASYLAAL | ASGIGQTYEQY | 0.8186397 |
| LVGGTPAL | ASGIGQTYEQY | 0.7910235 |
| VRNADVL | ASGIGQTYEQY | 0.8408201 |
| TCSSPFLVSAM | ASGIGQTYEQY | 0.7929019 |
| VHITDAE | ASGIGQTYEQY | 0.781278 |
| EYAAPSF | ASGIGQTYEQY | 0.57340324 |
| VVDAFHGGL | ASGIGQTYEQY | 0.831043 |
| ASVVDPQ | ASGIGQTYEQY | 0.8500515 |
| SYNEFGGLS | ASGIGQTYEQY | 0.7905623 |
| DPDGSVY | ASGIGQTYEQY | 0.50867546 |
| MVSTIHATE | ASGIGQTYEQY | 0.7924459 |
| LAAGVADPV | ASGIGQTYEQY | 0.76752824 |
| TKEQIAA | ASGIGQTYEQY | 0.43013906 |
| SLASPAAL | ASGIGQTYEQY | 0.2742536 |
| IGLVFYGV | ASGIGQTYEQY | 0.7238172 |
| FLLDASNL | ASGIGQTYEQY | 0.76586866 |
| YQVGNFP | ASGIGQTYEQY | 0.8412156 |
| ESLLSQST | ASGIGQTYEQY | 0.8415729 |
| LFTVGFAAVG | ASGIGQTYEQY | 0.7028804 |
| ATPPTPT | ASGIGQTYEQY | 0.8288541 |
| TDLATLG | ASGIGQTYEQY | 0.81161183 |
| SAGGTGR | ASGIGQTYEQY | 0.14080858 |
| GFSNSGS | ASGIGQTYEQY | 0.7106844 |
| GGNGGSGGTSV | ASGIGQTYEQY | 0.5861037 |
| GTSVATGGAGN | ASGIGQTYEQY | 0.84832484 |
| NGGNGGSGGTS | ASGIGQTYEQY | 0.7314705 |
| AVVKFPH | ASGIGQTYEQY | 0.7665423 |
| TPPPPPP | ASGIGQTYEQY | 0.75046515 |
| AAPAAAAAPA | ASGIGQTYEQY | 0.8072815 |
| IALNEDL | ASGIGQTYEQY | 0.7779241 |
| IVNVSSM | ASGIGQTYEQY | 0.59363437 |
| TGIPPAPR | ASGIGQTYEQY | 0.39231753 |
| FDLGGGTF | ASGIGQTYEQY | 0.8214833 |
| FDLGGGTFDVS | ASGIGQTYEQY | 0.71173525 |
| NEPTAAA | ASGIGQTYEQY | 0.67158586 |
| NEPTAAALA | ASGIGQTYEQY | 0.50871986 |
| GSIINISSI | ASGIGQTYEQY | 0.80370295 |
| VAPGNAGTA | ASGIGQTYEQY | 0.45585102 |
| TGGGASLE | ASGIGQTYEQY | 0.82701945 |
| VAVYNHYKRI | ASGIGQTYEQY | 0.73673075 |
| TDPAPATT | ASGIGQTYEQY | 0.7892246 |

| | | |
|---|---|---|
| TNRPDILDPALLRPG | ASGIGQTYEQY | 0.8551222 |
| LTTTTTS | ASGIGQTYEQY | 0.7986202 |
| PTGSGKTL | ASGIGQTYEQY | 0.7411122 |
| TGSGKTL | ASGIGQTYEQY | 0.6782607 |
| TIGHVDHGK | ASGIGQTYEQY | 0.60062265 |
| TIGHVDHGKTT | ASGIGQTYEQY | 0.6677724 |
| GHVDHGKTT | ASGIGQTYEQY | 0.7012421 |
| AQFEHTLL | ASGIGQTYEQY | 0.85024863 |
| IVNQDEL | ASGIGQTYEQY | 0.71415615 |
| TNGSQFF | ASGIGQTYEQY | 0.54397535 |
| TGKGIVE | ASGIGQTYEQY | 0.7396599 |
| LISNASDA | ASGIGQTYEQY | 0.8472537 |
| AELRTLGELTKIAKAHGAQ | ASGIGQTYEQY | 0.8529013 |
| APNAVAAAEPSW | ASGIGQTYEQY | 0.7211734 |
| APQVAGAATAVL | ASGIGQTYEQY | 0.68223834 |
| ARRDPAAAAAILPTDA | ASGIGQTYEQY | 0.6770888 |
| DAAARDVLAEAGLAEYF | ASGIGQTYEQY | 0.78497803 |
| DDQIVLSAKANLAKVEQ | ASGIGQTYEQY | 0.6188471 |
| DELLPDTGEDSDEDGH | ASGIGQTYEQY | 0.5304481 |
| ELPEAPANRATVPQD | ASGIGQTYEQY | 0.8074156 |
| FGQSKIERVTGIG | ASGIGQTYEQY | 0.84880656 |
| GAGGTGSTAGGAGGAGGAGGLY | ASGIGQTYEQY | 0.7370887 |
| GHGGTGAAGGNGGTGGAGGSGI | ASGIGQTYEQY | 0.6209518 |
| GNGGNGGAGGEGGPGGAGGAGA | ASGIGQTYEQY | 0.78086066 |
| GSTAGNGGNGGNATNGTIAGQP | ASGIGQTYEQY | 0.6958034 |
| GTGGDGGNGGNGGTGG | ASGIGQTYEQY | 0.6539638 |
| GVRRPAGIPADAPV | ASGIGQTYEQY | 0.77917564 |
| GYRQVTDAHLLTLARR | ASGIGQTYEQY | 0.44862914 |
| IAKMGQDGHDRG | ASGIGQTYEQY | 0.76438826 |
| KAEPSIEIQYT | ASGIGQTYEQY | 0.8493102 |
| KPVRIRMAADEDTVAVT | ASGIGQTYEQY | 0.7774193 |
| MPPRIAGMRLLVIKPEP | ASGIGQTYEQY | 0.79993737 |
| MKFYAVALFRSRGVPPTR | ASGIGQTYEQY | 0.844968 |
| NAGGFGILGAVAHSP | ASGIGQTYEQY | 0.8209068 |
| NEGREITLAMAEM | ASGIGQTYEQY | 0.8431833 |
| NELVEWSHPLGHRWRWE | ASGIGQTYEQY | 0.8373315 |
| NLGAGGTGGNGGNPG | ASGIGQTYEQY | 0.7384743 |
| NTEWPKWETVMAADPVD | ASGIGQTYEQY | 0.7576816 |
| PASAAPAETQP | ASGIGQTYEQY | 0.8397413 |
| QQSSDGKNGVRAAEITGE | ASGIGQTYEQY | 0.78623337 |
| RAPDAVTTSWWKE | ASGIGQTYEQY | 0.76362634 |
| SGFGNVGAHTSGWFNQ | ASGIGQTYEQY | 0.69914556 |
| SGVAGAVGQAASVGGLKV | ASGIGQTYEQY | 0.8392737 |
| SNGGAGTGSAGGNGGT | ASGIGQTYEQY | 0.7136128 |
| TGGAGGEGGAAG | ASGIGQTYEQY | 0.8329979 |
| TGGDGGAAGAGGAPGHG | ASGIGQTYEQY | 0.28216687 |
| TMPPLLAAIANAGRHGV | ASGIGQTYEQY | 0.54767406 |
| TSSTAGNSGGA | ASGIGQTYEQY | 0.8340287 |
| VPATVPVGILC | ASGIGQTYEQY | 0.1509471 |
| VQRPDSGGVVLTGRIS | ASGIGQTYEQY | 0.2662163 |
| WTPNPVPATKGVPAGLVQAT | ASGIGQTYEQY | 0.7647638 |

| | | |
|---|---|---|
| YALKRALAKFSDRFDVVII | ASGIGQTYEQY | 0.83716166 |
| YLIGSVIV | ASGIGQTYEQY | 0.21131125 |
| YGQLVAVID | ASGIGQTYEQY | 0.6948793 |
| VTVGGMTLPA | ASGIGQTYEQY | 0.4180591 |
| VLALPAV | ASGIGQTYEQY | 0.7987664 |
| VIPPTLNYET | ASGIGQTYEQY | 0.7777163 |
| VGMGGMHPSSG | ASGIGQTYEQY | 0.41754958 |
| VGLAGGAATA | ASGIGQTYEQY | 0.8171538 |
| VGGAGGAGGSGP | ASGIGQTYEQY | 0.45394963 |
| VFAAFMV | ASGIGQTYEQY | 0.731147 |
| TLYDKLR | ASGIGQTYEQY | 0.8433811 |
| TLAVFTVL | ASGIGQTYEQY | 0.8472067 |
| TGSVQPARV | ASGIGQTYEQY | 0.658091 |
| TGGGAGSGGT | ASGIGQTYEQY | 0.82940114 |
| SHLETSL | ASGIGQTYEQY | 0.7039702 |
| SFTTGMT | ASGIGQTYEQY | 0.856254 |
| QVEVAKRMGVRQPTVS | ASGIGQTYEQY | 0.75000864 |
| QGSGGPALAM | ASGIGQTYEQY | 0.4115914 |
| MLAPSNP | ASGIGQTYEQY | 0.83217335 |
| LVNNAAI | ASGIGQTYEQY | 0.8047499 |
| LTGEINTLA | ASGIGQTYEQY | 0.7806072 |
| LSIFGPV | ASGIGQTYEQY | 0.8242456 |
| LLALLANL | ASGIGQTYEQY | 0.6166092 |
| LISNASDALDKLR | ASGIGQTYEQY | 0.7446533 |
| LINEAVP | ASGIGQTYEQY | 0.8408706 |
| LGYAFSGL | ASGIGQTYEQY | 0.8385531 |
| LDFVAVAL | ASGIGQTYEQY | 0.57670903 |
| LAQPGAE | ASGIGQTYEQY | 0.83014745 |
| ISNLGAVTGF | ASGIGQTYEQY | 0.77408767 |
| HQEDPGRI | ASGIGQTYEQY | 0.64013726 |
| GTPGEPTP | ASGIGQTYEQY | 0.852514 |
| GFFNSGS | ASGIGQTYEQY | 0.73467135 |
| GAGGGGLTAGAN | ASGIGQTYEQY | 0.824216 |
| FVDALVNM | ASGIGQTYEQY | 0.8106609 |
| FLTVAPEL | ASGIGQTYEQY | 0.82574296 |
| FETLTAL | ASGIGQTYEQY | 0.6294072 |
| DVINPIEL | ASGIGQTYEQY | 0.819909 |
| APVSGAPV | ASGIGQTYEQY | 0.793233 |
| ALVGGLAW | ASGIGQTYEQY | 0.8321692 |
| ALAAAAATAI | ASGIGQTYEQY | 0.77341926 |
| AFGLGLLL | ASGIGQTYEQY | 0.8029947 |
| AFFAAASL | ASGIGQTYEQY | 0.8073643 |
| AAAALPAFSPP | ASGIGQTYEQY | 0.8437371 |
| VVLAPAL | ASGIGQTYEQY | 0.7739967 |
| TIGHVDHGKTTL | ASGIGQTYEQY | 0.8216291 |
| LISNASDALDK | ASGIGQTYEQY | 0.84853494 |
| LISNASDAL | ASGIGQTYEQY | 0.62688184 |
| ISNASDALDK | ASGIGQTYEQY | 0.74565804 |
| NQAETLVYQTE | ASGIGQTYEQY | 0.82931256 |
| TGGTGGTAGAG | ASGIGQTYEQY | 0.8439977 |
| TGSTAGGAGGA | ASGIGQTYEQY | 0.7081666 |

| | | |
|---|---|---|
| DPLAVPVAAGPV | ASGIGQTYEQY | 0.84017813 |
| GGTGGTPGNGGN | ASGIGQTYEQY | 0.77056074 |
| PAGLAGVDPATL | ASGIGQTYEQY | 0.83005714 |
| GNNLVVEAPHAGASV | ASGIGQTYEQY | 0.78044593 |
| KVTRIMSASTLEQPA | ASGIGQTYEQY | 0.7983564 |
| LGFALPIALPRRTVV | ASGIGQTYEQY | 0.7039622 |
| TRLGNLTARKLVVLV | ASGIGQTYEQY | 0.83603567 |
| WSQQRRLIYFPSAGP | ASGIGQTYEQY | 0.8351399 |
| GPNGNPDPMAAAVDIR | ASGIGQTYEQY | 0.83026576 |
| TQRCVSVADLRAAASR | ASGIGQTYEQY | 0.26161778 |
| QFVLDVAEHPVEVLTETV | ASGIGQTYEQY | 0.8291446 |
| GGAGGTGGAGGNGGTQAGDG | ASGIGQTYEQY | 0.81718826 |
| PLLATLRAIAVDVDAKPAQV | ASGIGQTYEQY | 0.8499966 |
| APKFRTGGYMIDFWGVGYQVAK | ASGIGQTYEQY | 0.72106755 |
| GAGGAGGAGGAAGGPGGTGGTG | ASGIGQTYEQY | 0.7950053 |
| IGALGLLATDVAPAGSVAFGVA | ASGIGQTYEQY | 0.7780897 |
| AGGAGGIGGNGGAGGAGGVLMG | ASGIGQTYEQY | 0.73936415 |
| ADELIGLL | ASGIGQTYEQY | 0.5071977 |
| ADRALKA | ASGIGQTYEQY | 0.7467587 |
| AGAVGQVGGAGGA | ASGIGQTYEQY | 0.7743945 |
| AGGGGGLGGAP | ASGIGQTYEQY | 0.6827954 |
| AGGLLYGNGGAG | ASGIGQTYEQY | 0.8289848 |
| AGLESIGL | ASGIGQTYEQY | 0.8412776 |
| AGLKTGII | ASGIGQTYEQY | 0.8429022 |
| ASLAALLL | ASGIGQTYEQY | 0.8443916 |
| ASVINISSI | ASGIGQTYEQY | 0.84508574 |
| ATPPGPA | ASGIGQTYEQY | 0.8242979 |
| ATVASALV | ASGIGQTYEQY | 0.70943725 |
| ATVASLIL | ASGIGQTYEQY | 0.80971587 |
| AVNGPVLIRH | ASGIGQTYEQY | 0.5416987 |
| DGAAGAVGI | ASGIGQTYEQY | 0.7784674 |
| DPLLHGI | ASGIGQTYEQY | 0.71870327 |
| ELGAALGAL | ASGIGQTYEQY | 0.7962344 |
| ELGRTQAV | ASGIGQTYEQY | 0.15156472 |
| FVAGGYLS | ASGIGQTYEQY | 0.8074093 |
| GASGGAGGAGGW | ASGIGQTYEQY | 0.85152215 |
| GGAGAGGDGGAGGI | ASGIGQTYEQY | 0.76820344 |
| GTGHNGSQPGMGGNG | ASGIGQTYEQY | 0.80010235 |
| GYAAASASAS | ASGIGQTYEQY | 0.82093865 |
| HQVDVAAVQQG | ASGIGQTYEQY | 0.70548874 |
| IGAAIAHT | ASGIGQTYEQY | 0.82864404 |
| IGMIEVAQL | ASGIGQTYEQY | 0.6655715 |
| IRLAPEHTSAMD | ASGIGQTYEQY | 0.81274605 |
| IYNVSVL | ASGIGQTYEQY | 0.8470049 |
| KQGFLGM | ASGIGQTYEQY | 0.7998965 |
| LESVITQL | ASGIGQTYEQY | 0.8052395 |
| LGLGALVVV | ASGIGQTYEQY | 0.72643626 |
| LIGNGGNGGSGGI | ASGIGQTYEQY | 0.50140953 |
| LNAINAPTL | ASGIGQTYEQY | 0.8201112 |
| LQLLNTGV | ASGIGQTYEQY | 0.6116238 |
| LQTSIEF | ASGIGQTYEQY | 0.78700364 |

| | | |
|---|---|---|
| MSFAVSEL | ASGIGQTYEQY | 0.61085 |
| NTGALGL | ASGIGQTYEQY | 0.817832 |
| PPGAGGAGGAGTT | ASGIGQTYEQY | 0.8054583 |
| QLVGLGI | ASGIGQTYEQY | 0.80605775 |
| RFLEQVHQL | ASGIGQTYEQY | 0.7001064 |
| RGVGYVLR | ASGIGQTYEQY | 0.801031 |
| SATGVFAGVI | ASGIGQTYEQY | 0.7674309 |
| SLPAITLP | ASGIGQTYEQY | 0.6031051 |
| STALAFTV | ASGIGQTYEQY | 0.8328476 |
| SVLTVGLGK | ASGIGQTYEQY | 0.71357155 |
| TALAGAGSGI | ASGIGQTYEQY | 0.7696664 |
| TGLPFTGI | ASGIGQTYEQY | 0.8384056 |
| VAEHVTV | ASGIGQTYEQY | 0.8014436 |
| VDIVLNSL | ASGIGQTYEQY | 0.46929294 |
| VLTGGGAVL | ASGIGQTYEQY | 0.7752118 |
| VNVGLAYGI | ASGIGQTYEQY | 0.79519475 |
| VSSAAVGAV | ASGIGQTYEQY | 0.7438624 |
| VTSVYEAGV | ASGIGQTYEQY | 0.23664415 |
| WVALDRGA | ASGIGQTYEQY | 0.8240231 |
| YLVGSVVI | ASGIGQTYEQY | 0.6378228 |
| GPPGTGKTLL | ASGIGQTYEQY | 0.7157667 |
| GVGMTKF | ASGIGQTYEQY | 0.72222805 |
| IGIGGSDL | ASGIGQTYEQY | 0.8322675 |
| IGTLLAEL | ASGIGQTYEQY | 0.75647247 |
| IYTSGTT | ASGIGQTYEQY | 0.67593575 |
| LSFPAGV | ASGIGQTYEQY | 0.8262975 |
| LYTSGTT | ASGIGQTYEQY | 0.82442015 |
| VGAFPFGF | ASGIGQTYEQY | 0.84180486 |
| VGAGTVEF | ASGIGQTYEQY | 0.8134402 |
| VGKSTLLL | ASGIGQTYEQY | 0.7289963 |
| AALAHAGGGIV | ASGIGQTYEQY | 0.80811846 |
| AALVAVSLL | ASGIGQTYEQY | 0.7116272 |
| AAVSRAVAAL | ASGIGQTYEQY | 0.6001177 |
| AGGSSGIGGF | ASGIGQTYEQY | 0.42819345 |
| AGLTVANANG | ASGIGQTYEQY | 0.74970543 |
| AGLVGAVL | ASGIGQTYEQY | 0.44699806 |
| ALSQAFTV | ASGIGQTYEQY | 0.81115055 |
| ALTFSVSL | ASGIGQTYEQY | 0.83755153 |
| ALTGAALVAL | ASGIGQTYEQY | 0.7636681 |
| ASFSIGAGVL | ASGIGQTYEQY | 0.47964683 |
| DGSVYKI | ASGIGQTYEQY | 0.7764336 |
| EQLAEIV | ASGIGQTYEQY | 0.7638645 |
| FAVSELTVM | ASGIGQTYEQY | 0.82694685 |
| GAAPLAAGL | ASGIGQTYEQY | 0.8088745 |
| GADGGDGGSGGAGG | ASGIGQTYEQY | 0.8487219 |
| GFGLDTGGAGG | ASGIGQTYEQY | 0.8381228 |
| GLASGGLL | ASGIGQTYEQY | 0.7314556 |
| GRIAGAAVLL | ASGIGQTYEQY | 0.8207507 |
| GSAGSLNTSF | ASGIGQTYEQY | 0.69004345 |
| GTGQIGI | ASGIGQTYEQY | 0.80227304 |
| GTLIALIL | ASGIGQTYEQY | 0.7789048 |

| | | |
|---|---|---|
| IAATMGFVVG | ASGIGQTYEQY | 0.7888218 |
| IFAQALL | ASGIGQTYEQY | 0.7615314 |
| IGAAVALGF | ASGIGQTYEQY | 0.7516027 |
| IGLTPLAL | ASGIGQTYEQY | 0.3938765 |
| ILGSNTSTL | ASGIGQTYEQY | 0.8068756 |
| IVFSLTVL | ASGIGQTYEQY | 0.647364 |
| LASKATGF | ASGIGQTYEQY | 0.39706874 |
| LASVALIL | ASGIGQTYEQY | 0.7388165 |
| LFAVGLTL | ASGIGQTYEQY | 0.8548281 |
| LFNGGIGTYI | ASGIGQTYEQY | 0.84730655 |
| LTLGAGII | ASGIGQTYEQY | 0.7180699 |
| LVAAALGDI | ASGIGQTYEQY | 0.65250176 |
| NTAFVQLGI | ASGIGQTYEQY | 0.8172468 |
| QLFSGAEL | ASGIGQTYEQY | 0.47220856 |
| SAATFITL | ASGIGQTYEQY | 0.79305816 |
| SNYGLANLGV | ASGIGQTYEQY | 0.79366183 |
| SYGGAYAVM | ASGIGQTYEQY | 0.78225565 |
| TTGVGGLTLG | ASGIGQTYEQY | 0.5651994 |
| TTQTIYI | ASGIGQTYEQY | 0.81588125 |
| VAAGAGLGAVI | ASGIGQTYEQY | 0.7824547 |
| VAAVVTLL | ASGIGQTYEQY | 0.7833315 |
| VGFLGGLI | ASGIGQTYEQY | 0.52086943 |
| VGVIPILL | ASGIGQTYEQY | 0.8394467 |
| VIFASSLI | ASGIGQTYEQY | 0.716696 |
| VKGGDGGAAAG | ASGIGQTYEQY | 0.5014364 |
| VTKTFTNI | ASGIGQTYEQY | 0.2686035 |
| VVAAAILL | ASGIGQTYEQY | 0.7595478 |
| YAALDVEL | ASGIGQTYEQY | 0.83525825 |
| LASVALLL | ASGIGQTYEQY | 0.8411073 |
| LDAVHRV | ASGIGQTYEQY | 0.7162595 |
| QGLSSSIF | ASGIGQTYEQY | 0.84319496 |
| RGTFANIRL | ASGIGQTYEQY | 0.84738225 |
| RQLWWGHRI | ASGIGQTYEQY | 0.7663071 |
| YGPPGTGKTLL | ASGIGQTYEQY | 0.39933705 |
| AALVGAPANVA | ASGIGQTYEQY | 0.78869665 |
| AATFRRTE | ASGIGQTYEQY | 0.79749024 |
| AGALGGGI | ASGIGQTYEQY | 0.8392197 |
| AGGAGGAGGIGAG | ASGIGQTYEQY | 0.82685655 |
| AGSGALGL | ASGIGQTYEQY | 0.57282984 |
| AQIADQI | ASGIGQTYEQY | 0.81408083 |
| ASAVAVTYL | ASGIGQTYEQY | 0.82983685 |
| ASTVLGGTGG | ASGIGQTYEQY | 0.5840008 |
| AVGGEPAK | ASGIGQTYEQY | 0.7158266 |
| CEQGTFGGTC | ASGIGQTYEQY | 0.6750961 |
| DEGKAPAG | ASGIGQTYEQY | 0.53061736 |
| DTVVFPAL | ASGIGQTYEQY | 0.82874835 |
| FEGITMAG | ASGIGQTYEQY | 0.7580586 |
| FPLVVQD | ASGIGQTYEQY | 0.72954786 |
| GAFTTPQ | ASGIGQTYEQY | 0.73827565 |
| GFGGFGTAGGVGGA | ASGIGQTYEQY | 0.7069789 |
| GFLAQFL | ASGIGQTYEQY | 0.65252215 |

| | | |
|---|---|---|
| GGAGGAGGNAGVA | ASGIGQTYEQY | 0.73630905 |
| GGVGTAGAGGAGGAGG | ASGIGQTYEQY | 0.8283987 |
| GNGGGGDAGNAGSGGN | ASGIGQTYEQY | 0.8230171 |
| GTGGAGGAGNDAGS | ASGIGQTYEQY | 0.78983974 |
| GTTNTGWFNS | ASGIGQTYEQY | 0.745991 |
| IFMGPLFTL | ASGIGQTYEQY | 0.84854245 |
| ILAHAIEAI | ASGIGQTYEQY | 0.5745399 |
| IRSAIREAL | ASGIGQTYEQY | 0.654462 |
| ITFAGGG | ASGIGQTYEQY | 0.81120306 |
| ITGMSGHQEV | ASGIGQTYEQY | 0.5965661 |
| IVAGLACTL | ASGIGQTYEQY | 0.831047 |
| IVGGGLAGA | ASGIGQTYEQY | 0.8179283 |
| IVTGVLNI | ASGIGQTYEQY | 0.80669016 |
| LAQQGCI | ASGIGQTYEQY | 0.78193897 |
| LCLHVEN | ASGIGQTYEQY | 0.8563997 |
| LDAISDGS | ASGIGQTYEQY | 0.82356095 |
| LGGFAAML | ASGIGQTYEQY | 0.7399831 |
| LGGKNPMI | ASGIGQTYEQY | 0.82772386 |
| LGPKGVAAIL | ASGIGQTYEQY | 0.5170634 |
| LMVGVSI | ASGIGQTYEQY | 0.8419876 |
| LVGAAFVL | ASGIGQTYEQY | 0.8130819 |
| LVLAVAPL | ASGIGQTYEQY | 0.6241178 |
| MAAIAVIDAA | ASGIGQTYEQY | 0.7660879 |
| NNGINAELQ | ASGIGQTYEQY | 0.8398616 |
| RGEPLVLVHG | ASGIGQTYEQY | 0.8408843 |
| RGGAGGAAGLI | ASGIGQTYEQY | 0.47538507 |
| RSYRMHPAV | ASGIGQTYEQY | 0.7010468 |
| SAAHLLAGQLA | ASGIGQTYEQY | 0.4422206 |
| SAAVGSY | ASGIGQTYEQY | 0.5353866 |
| SPHAVLHRI | ASGIGQTYEQY | 0.42319208 |
| SQIAKVL | ASGIGQTYEQY | 0.8104798 |
| TALGHLVP | ASGIGQTYEQY | 0.74592656 |
| TAPTTSGAA | ASGIGQTYEQY | 0.8137492 |
| TDGDDGNGGSPGT | ASGIGQTYEQY | 0.799324 |
| TDGLGEN | ASGIGQTYEQY | 0.8408523 |
| TGGPPLPGQA | ASGIGQTYEQY | 0.6981567 |
| TIPAITI | ASGIGQTYEQY | 0.7935534 |
| TIPGFTI | ASGIGQTYEQY | 0.37409437 |
| TSIASGALA | ASGIGQTYEQY | 0.6111623 |
| TVTLPGAAI | ASGIGQTYEQY | 0.6041179 |
| VASGLSAALP | ASGIGQTYEQY | 0.7672329 |
| VGAGVIGI | ASGIGQTYEQY | 0.53470576 |
| VGPKRFGV | ASGIGQTYEQY | 0.83911395 |
| VNVAFPD | ASGIGQTYEQY | 0.7703221 |
| VSMTNSGVS | ASGIGQTYEQY | 0.8116927 |
| VTDVLGDLSV | ASGIGQTYEQY | 0.7584034 |
| VVVVGAGFAGL | ASGIGQTYEQY | 0.67077243 |
| VYTVGTVV | ASGIGQTYEQY | 0.694944 |
| WTTFSGP | ASGIGQTYEQY | 0.82786393 |
| YDELVEAF | ASGIGQTYEQY | 0.68571043 |
| YGTGGAGGNGG | ASGIGQTYEQY | 0.34886438 |

| | | |
|---|---|---|
| YTNVMAV | ASGIGQTYEQY | 0.63943607 |
| FLEQVHQL | ASGIGQTYEQY | 0.699718 |
| GLSLLAVAV | ASGIGQTYEQY | 0.8464329 |
| LEQVHQL | ASGIGQTYEQY | 0.83856165 |
| LNAAGAI | ASGIGQTYEQY | 0.79666215 |
| PAVHASL | ASGIGQTYEQY | 0.841865 |
| TELEDDA | ASGIGQTYEQY | 0.1750556 |
| TPPGPPN | ASGIGQTYEQY | 0.78081274 |
| TPPPPPPP | ASGIGQTYEQY | 0.84570247 |
| TQSPALI | AWSLRDFNYAEQF | 0.8093722 |
| EFAAAFPQ | AWSLRDFNYAEQF | 0.77332693 |
| LLSGNLDVL | AWSLRDFNYAEQF | 0.84053767 |
| GPANLGNYNI | AWSLRDFNYAEQF | 0.70491934 |
| TLGPTLP | AWSLRDFNYAEQF | 0.7425017 |
| VYAYSSCCK | AWSLRDFNYAEQF | 0.7955289 |
| APAPAPAPNI | AWSLRDFNYAEQF | 0.8179677 |
| PGGPVPIP | AWSLRDFNYAEQF | 0.78398275 |
| FLTVALEP | AWSLRDFNYAEQF | 0.52842176 |
| SQYFGANL | AWSLRDFNYAEQF | 0.7707216 |
| GGVKPAA | AWSLRDFNYAEQF | 0.85899436 |
| GGTGGAGGAGGW | AWSLRDFNYAEQF | 0.77135885 |
| GVGGVAGLGGAGG | AWSLRDFNYAEQF | 0.8383547 |
| VLLVLATTI | AWSLRDFNYAEQF | 0.7937813 |
| IGIFAAML | AWSLRDFNYAEQF | 0.7884308 |
| AVGDLID | AWSLRDFNYAEQF | 0.6382115 |
| NISPLPL | AWSLRDFNYAEQF | 0.660409 |
| VTAVFVAI | AWSLRDFNYAEQF | 0.8074219 |
| SGDDNDAAL | AWSLRDFNYAEQF | 0.67035735 |
| AVAALAHS | AWSLRDFNYAEQF | 0.76002055 |
| GGVTVMATP | AWSLRDFNYAEQF | 0.7151337 |
| AGLAANVRD | AWSLRDFNYAEQF | 0.8567894 |
| TNRPDLIDPALLRPG | AWSLRDFNYAEQF | 0.81001115 |
| LANLGVTAL | AWSLRDFNYAEQF | 0.8107798 |
| TGERIEL | AWSLRDFNYAEQF | 0.6438896 |
| NGEELAA | AWSLRDFNYAEQF | 0.7848445 |
| VLSLVWG | AWSLRDFNYAEQF | 0.81946576 |
| IGYALWL | AWSLRDFNYAEQF | 0.45155734 |
| LSLLLAPL | AWSLRDFNYAEQF | 0.8448329 |
| TLDEALK | AWSLRDFNYAEQF | 0.62222224 |
| TDGAGTAEGG | AWSLRDFNYAEQF | 0.77325416 |
| NAPGGAGGNGG | AWSLRDFNYAEQF | 0.8085733 |
| VSVFMGMGGNF | AWSLRDFNYAEQF | 0.8397397 |
| LASYLAAL | AWSLRDFNYAEQF | 0.7099489 |
| LVGGTPAL | AWSLRDFNYAEQF | 0.7700286 |
| VRNADVL | AWSLRDFNYAEQF | 0.60038066 |
| TCSSPFLVSAM | AWSLRDFNYAEQF | 0.8094801 |
| VHITDAE | AWSLRDFNYAEQF | 0.8422905 |
| EYAAPSF | AWSLRDFNYAEQF | 0.77007174 |
| VVDAFHGGL | AWSLRDFNYAEQF | 0.8509513 |
| ASVVDPQ | AWSLRDFNYAEQF | 0.79725087 |
| SYNEFGGLS | AWSLRDFNYAEQF | 0.31618485 |

| | | |
|---|---|---|
| DPDGSVY | AWSLRDFNYAEQF | 0.7467339 |
| MVSTIHATE | AWSLRDFNYAEQF | 0.84639 |
| LAAGVADPV | AWSLRDFNYAEQF | 0.84400374 |
| TKEQIAA | AWSLRDFNYAEQF | 0.8516948 |
| SLASPAAL | AWSLRDFNYAEQF | 0.8356559 |
| IGLVFYGV | AWSLRDFNYAEQF | 0.7606642 |
| FLLDASNL | AWSLRDFNYAEQF | 0.84378076 |
| YQVGNFP | AWSLRDFNYAEQF | 0.8545536 |
| ESLLSQST | AWSLRDFNYAEQF | 0.8199036 |
| LFTVGFAAVG | AWSLRDFNYAEQF | 0.5906398 |
| ATPPTPT | AWSLRDFNYAEQF | 0.5325556 |
| TDLATLG | AWSLRDFNYAEQF | 0.8011632 |
| SAGGTGR | AWSLRDFNYAEQF | 0.8333133 |
| GFSNSGS | AWSLRDFNYAEQF | 0.48668662 |
| GGNGGSGGTSV | AWSLRDFNYAEQF | 0.79292727 |
| GTSVATGGAGN | AWSLRDFNYAEQF | 0.8011093 |
| NGGNGGSGGTS | AWSLRDFNYAEQF | 0.43691415 |
| AVVKFPH | AWSLRDFNYAEQF | 0.82712317 |
| TPPPPPP | AWSLRDFNYAEQF | 0.85146934 |
| AAPAAAAAPA | AWSLRDFNYAEQF | 0.75940156 |
| IALNEDL | AWSLRDFNYAEQF | 0.6212952 |
| IVNVSSM | AWSLRDFNYAEQF | 0.8437016 |
| TGIPPAPR | AWSLRDFNYAEQF | 0.85644376 |
| FDLGGGTF | AWSLRDFNYAEQF | 0.8422173 |
| FDLGGGTFDVS | AWSLRDFNYAEQF | 0.45681086 |
| NEPTAAA | AWSLRDFNYAEQF | 0.19922784 |
| NEPTAAALA | AWSLRDFNYAEQF | 0.7753123 |
| GSIINISSI | AWSLRDFNYAEQF | 0.8199574 |
| VAPGNAGTA | AWSLRDFNYAEQF | 0.75395805 |
| TGGGASLE | AWSLRDFNYAEQF | 0.69252884 |
| VAVYNHYKRI | AWSLRDFNYAEQF | 0.8198968 |
| TDPAPATT | AWSLRDFNYAEQF | 0.8325779 |
| TNRPDILDPALLRPG | AWSLRDFNYAEQF | 0.60546124 |
| LTTTTTS | AWSLRDFNYAEQF | 0.4467048 |
| PTGSGKTL | AWSLRDFNYAEQF | 0.83423483 |
| TGSGKTL | AWSLRDFNYAEQF | 0.81905055 |
| TIGHVDHGK | AWSLRDFNYAEQF | 0.6735335 |
| TIGHVDHGKTT | AWSLRDFNYAEQF | 0.75415134 |
| GHVDHGKTT | AWSLRDFNYAEQF | 0.79060316 |
| AQFEHTLL | AWSLRDFNYAEQF | 0.7406601 |
| IVNQDEL | AWSLRDFNYAEQF | 0.8103976 |
| TNGSQFF | AWSLRDFNYAEQF | 0.5357152 |
| TGKGIVE | AWSLRDFNYAEQF | 0.85082877 |
| LISNASDA | AWSLRDFNYAEQF | 0.51253664 |
| AELRTLGELTKIAKAHGAQ | AWSLRDFNYAEQF | 0.8223948 |
| APNAVAAAEPSW | AWSLRDFNYAEQF | 0.81169784 |
| APQVAGAATAVL | AWSLRDFNYAEQF | 0.8075339 |
| ARRDPAAAAAILPTDA | AWSLRDFNYAEQF | 0.7509609 |
| DAAARDVLAEAGLAEYF | AWSLRDFNYAEQF | 0.6916615 |
| DDQIVLSAKANLAKVEQ | AWSLRDFNYAEQF | 0.57898366 |
| DELLPDTGEDSDEDGH | AWSLRDFNYAEQF | 0.43881255 |

| | | |
|---|---|---|
| ELPEAPANRATVPQD | AWSLRDFNYAEQF | 0.84608245 |
| FGQSKIERVTGIG | AWSLRDFNYAEQF | 0.79859203 |
| GAGGTGSTAGGAGGAGGAGGLY | AWSLRDFNYAEQF | 0.83699805 |
| GHGGTGAAGGNGGTGGAGGSGI | AWSLRDFNYAEQF | 0.808533 |
| GNGGNGGAGGEGGPGGAGGAGA | AWSLRDFNYAEQF | 0.6647779 |
| GSTAGNGGNGGNATNGTIAGQP | AWSLRDFNYAEQF | 0.7964829 |
| GTGGDGGNGGNGGTGG | AWSLRDFNYAEQF | 0.61653334 |
| GVRRPAGIPADAPV | AWSLRDFNYAEQF | 0.71283627 |
| GYRQVTDAHLLTLARR | AWSLRDFNYAEQF | 0.7846683 |
| IAKMGQDGHDRG | AWSLRDFNYAEQF | 0.8061423 |
| KAEPSIEIQYT | AWSLRDFNYAEQF | 0.710694 |
| KPVRIRMAADEDTVAVT | AWSLRDFNYAEQF | 0.7838938 |
| MPPRIAGMRLLVIKPEP | AWSLRDFNYAEQF | 0.8052447 |
| MKFYAVALFRSRGVPPTR | AWSLRDFNYAEQF | 0.59498847 |
| NAGGFGILGAVAHSP | AWSLRDFNYAEQF | 0.75999415 |
| NEGREITLAMAEM | AWSLRDFNYAEQF | 0.84198284 |
| NELVEWSHPLGHRWRWE | AWSLRDFNYAEQF | 0.7660452 |
| NLGAGGTGGNGGNPG | AWSLRDFNYAEQF | 0.5981998 |
| NTEWPKWETVMAADPVD | AWSLRDFNYAEQF | 0.8274608 |
| PASAAPAETQP | AWSLRDFNYAEQF | 0.64688456 |
| QQSSDGKNGVRAAEITGE | AWSLRDFNYAEQF | 0.809828 |
| RAPDAVTTSWWKE | AWSLRDFNYAEQF | 0.78687024 |
| SGFGNVGAHTSGWFNQ | AWSLRDFNYAEQF | 0.69651115 |
| SGVAGAVGQAASVGGLKV | AWSLRDFNYAEQF | 0.82724893 |
| SNGGAGTGSAGGNGGT | AWSLRDFNYAEQF | 0.573761 |
| TGGAGGEGGAAG | AWSLRDFNYAEQF | 0.68973285 |
| TGGDGGAAGAGGAPGHG | AWSLRDFNYAEQF | 0.7115022 |
| TMPPLLAAIANAGRHGV | AWSLRDFNYAEQF | 0.78249395 |
| TSSTAGNSGGA | AWSLRDFNYAEQF | 0.81913614 |
| VPATVPVGILC | AWSLRDFNYAEQF | 0.5992135 |
| VQRPDSGGVVLTGRIS | AWSLRDFNYAEQF | 0.7880034 |
| WTPNPVPATKGVPAGLVQAT | AWSLRDFNYAEQF | 0.73052186 |
| YALKRALAKFSDRFDVVII | AWSLRDFNYAEQF | 0.62604195 |
| YLIGSVIV | AWSLRDFNYAEQF | 0.6904851 |
| YGQLVAVID | AWSLRDFNYAEQF | 0.7188742 |
| VTVGGMTLPA | AWSLRDFNYAEQF | 0.5539714 |
| VLALPAV | AWSLRDFNYAEQF | 0.82281125 |
| VIPPTLNYET | AWSLRDFNYAEQF | 0.6696431 |
| VGMGGMHPSSG | AWSLRDFNYAEQF | 0.8198782 |
| VGLAGGAATA | AWSLRDFNYAEQF | 0.8468059 |
| VGGAGGAGGSGP | AWSLRDFNYAEQF | 0.53139484 |
| VFAAFMV | AWSLRDFNYAEQF | 0.81077033 |
| TLYDKLR | AWSLRDFNYAEQF | 0.81797963 |
| TLAVFTVL | AWSLRDFNYAEQF | 0.81952566 |
| TGSVQPARV | AWSLRDFNYAEQF | 0.7648201 |
| TGGGAGSGGT | AWSLRDFNYAEQF | 0.7151855 |
| SHLETSL | AWSLRDFNYAEQF | 0.8485065 |
| SFTTGMT | AWSLRDFNYAEQF | 0.82145256 |
| QVEVAKRMGVRQPTVS | AWSLRDFNYAEQF | 0.81524014 |
| QGSGGPALAM | AWSLRDFNYAEQF | 0.82347804 |
| MLAPSNP | AWSLRDFNYAEQF | 0.8249117 |

| | | |
|---|---|---|
| LVNNAAI | AWSLRDFNYAEQF | 0.8209782 |
| LTGEINTLA | AWSLRDFNYAEQF | 0.7175907 |
| LSIFGPV | AWSLRDFNYAEQF | 0.37552112 |
| LLALLANL | AWSLRDFNYAEQF | 0.75985026 |
| LISNASDALDKLR | AWSLRDFNYAEQF | 0.74120027 |
| LINEAVP | AWSLRDFNYAEQF | 0.6594161 |
| LGYAFSGL | AWSLRDFNYAEQF | 0.3094622 |
| LDFVAVAL | AWSLRDFNYAEQF | 0.83592594 |
| LAQPGAE | AWSLRDFNYAEQF | 0.8575909 |
| ISNLGAVTGF | AWSLRDFNYAEQF | 0.74624294 |
| HQEDPGRI | AWSLRDFNYAEQF | 0.85321367 |
| GTPGEPTP | AWSLRDFNYAEQF | 0.48567438 |
| GFFNSGS | AWSLRDFNYAEQF | 0.8261085 |
| GAGGGGLTAGAN | AWSLRDFNYAEQF | 0.83631206 |
| FVDALVNM | AWSLRDFNYAEQF | 0.6243029 |
| FLTVAPEL | AWSLRDFNYAEQF | 0.8220093 |
| FETLTAL | AWSLRDFNYAEQF | 0.7328182 |
| DVINPIEL | AWSLRDFNYAEQF | 0.73298776 |
| APVSGAPV | AWSLRDFNYAEQF | 0.8448959 |
| ALVGGLAW | AWSLRDFNYAEQF | 0.7530935 |
| ALAAAAATAI | AWSLRDFNYAEQF | 0.80652744 |
| AFGLGLLL | AWSLRDFNYAEQF | 0.7462261 |
| AFFAAASL | AWSLRDFNYAEQF | 0.5934465 |
| AAAALPAFSPP | AWSLRDFNYAEQF | 0.8015624 |
| VVLAPAL | AWSLRDFNYAEQF | 0.83463067 |
| TIGHVDHGKTTL | AWSLRDFNYAEQF | 0.71608984 |
| LISNASDALDK | AWSLRDFNYAEQF | 0.850222 |
| LISNASDAL | AWSLRDFNYAEQF | 0.7574473 |
| ISNASDALDK | AWSLRDFNYAEQF | 0.8586227 |
| NQAETLVYQTE | AWSLRDFNYAEQF | 0.8405066 |
| TGGTGGTAGAG | AWSLRDFNYAEQF | 0.7985254 |
| TGSTAGGAGGA | AWSLRDFNYAEQF | 0.69648194 |
| DPLAVPVAAGPV | AWSLRDFNYAEQF | 0.84374887 |
| GGTGGTPGNGGN | AWSLRDFNYAEQF | 0.7389209 |
| PAGLAGVDPATL | AWSLRDFNYAEQF | 0.82942194 |
| GNNLVVEAPHAGASV | AWSLRDFNYAEQF | 0.8438159 |
| KVTRIMSASTLEQPA | AWSLRDFNYAEQF | 0.3898164 |
| LGFALPIALPRRTVV | AWSLRDFNYAEQF | 0.8596921 |
| TRLGNLTARKLVVLV | AWSLRDFNYAEQF | 0.79486555 |
| WSQQRRLIYFPSAGP | AWSLRDFNYAEQF | 0.74035734 |
| GPNGNPDPMAAAVDIR | AWSLRDFNYAEQF | 0.5783501 |
| TQRCVSVADLRAAASR | AWSLRDFNYAEQF | 0.31580627 |
| QFVLDVAEHPVEVLTETV | AWSLRDFNYAEQF | 0.6706576 |
| GGAGGTGGAGGNGGTQAGDG | AWSLRDFNYAEQF | 0.45129243 |
| PLLATLRAIAVDVDAKPAQV | AWSLRDFNYAEQF | 0.8602333 |
| APKFRTGGYMIDFWGVGYQVAK | AWSLRDFNYAEQF | 0.82831687 |
| GAGGAGGAGGAAGGPGGTGGTG | AWSLRDFNYAEQF | 0.5431666 |
| IGALGLLATDVAPAGSVAFGVA | AWSLRDFNYAEQF | 0.83893436 |
| AGGAGGIGGNGGAGGAGGVLMG | AWSLRDFNYAEQF | 0.8109783 |
| ADELIGLL | AWSLRDFNYAEQF | 0.35155815 |
| ADRALKA | AWSLRDFNYAEQF | 0.71101344 |

| | | |
|---|---|---|
| AGAVGQVGGAGGA | AWSLRDFNYAEQF | 0.708728 |
| AGGGGGLGGAP | AWSLRDFNYAEQF | 0.68013126 |
| AGGLLYGNGGAG | AWSLRDFNYAEQF | 0.35862547 |
| AGLESIGL | AWSLRDFNYAEQF | 0.8091399 |
| AGLKTGII | AWSLRDFNYAEQF | 0.6838987 |
| ASLAALLL | AWSLRDFNYAEQF | 0.1470154 |
| ASVINISSI | AWSLRDFNYAEQF | 0.7799068 |
| ATPPGPA | AWSLRDFNYAEQF | 0.67562217 |
| ATVASALV | AWSLRDFNYAEQF | 0.58814836 |
| ATVASLIL | AWSLRDFNYAEQF | 0.8383138 |
| AVNGPVLIRH | AWSLRDFNYAEQF | 0.6696926 |
| DGAAGAVGI | AWSLRDFNYAEQF | 0.5122816 |
| DPLLHGI | AWSLRDFNYAEQF | 0.8190694 |
| ELGAALGAL | AWSLRDFNYAEQF | 0.8369179 |
| ELGRTQAV | AWSLRDFNYAEQF | 0.34609997 |
| FVAGGYLS | AWSLRDFNYAEQF | 0.7881705 |
| GASGGAGGAGGW | AWSLRDFNYAEQF | 0.85212946 |
| GGAGAGGDGGAGGI | AWSLRDFNYAEQF | 0.7055706 |
| GTGHNGSQPGMGGNG | AWSLRDFNYAEQF | 0.86433035 |
| GYAAASASAS | AWSLRDFNYAEQF | 0.8670389 |
| HQVDVAAVQQG | AWSLRDFNYAEQF | 0.6123533 |
| IGAAIAHT | AWSLRDFNYAEQF | 0.8509308 |
| IGMIEVAQL | AWSLRDFNYAEQF | 0.864902 |
| IRLAPEHTSAMD | AWSLRDFNYAEQF | 0.64172184 |
| IYNVSVL | AWSLRDFNYAEQF | 0.72018623 |
| KQGFLGM | AWSLRDFNYAEQF | 0.35930496 |
| LESVITQL | AWSLRDFNYAEQF | 0.68929136 |
| LGLGALVVV | AWSLRDFNYAEQF | 0.37266326 |
| LIGNGGNGGSGGI | AWSLRDFNYAEQF | 0.8082662 |
| LNAINAPTL | AWSLRDFNYAEQF | 0.84518886 |
| LQLLNTGV | AWSLRDFNYAEQF | 0.6720613 |
| LQTSIEF | AWSLRDFNYAEQF | 0.38837928 |
| MSFAVSEL | AWSLRDFNYAEQF | 0.41603917 |
| NTGALGL | AWSLRDFNYAEQF | 0.78361076 |
| PPGAGGAGGAGTT | AWSLRDFNYAEQF | 0.44318026 |
| QLVGLGI | AWSLRDFNYAEQF | 0.3221954 |
| RFLEQVHQL | AWSLRDFNYAEQF | 0.6935075 |
| RGVGYVLR | AWSLRDFNYAEQF | 0.81117386 |
| SATGVFAGVI | AWSLRDFNYAEQF | 0.81819093 |
| SLPAITLP | AWSLRDFNYAEQF | 0.49983284 |
| STALAFTV | AWSLRDFNYAEQF | 0.8734678 |
| SVLTVGLGK | AWSLRDFNYAEQF | 0.85474986 |
| TALAGAGSGI | AWSLRDFNYAEQF | 0.8665715 |
| TGLPFTGI | AWSLRDFNYAEQF | 0.69795924 |
| VAEHVTV | AWSLRDFNYAEQF | 0.4093905 |
| VDIVLNSL | AWSLRDFNYAEQF | 0.8623097 |
| VLTGGGAVL | AWSLRDFNYAEQF | 0.65419924 |
| VNVGLAYGI | AWSLRDFNYAEQF | 0.73334503 |
| VSSAAVGAV | AWSLRDFNYAEQF | 0.8260562 |
| VTSVYEAGV | AWSLRDFNYAEQF | 0.62761503 |
| WVALDRGA | AWSLRDFNYAEQF | 0.5913913 |

| | | |
|---|---|---|
| YLVGSVVI | AWSLRDFNYAEQF | 0.77816117 |
| GPPGTGKTLL | AWSLRDFNYAEQF | 0.857739 |
| GVGMTKF | AWSLRDFNYAEQF | 0.55018896 |
| IGIGGSDL | AWSLRDFNYAEQF | 0.39434087 |
| IGTLLAEL | AWSLRDFNYAEQF | 0.6286292 |
| IYTSGTT | AWSLRDFNYAEQF | 0.67659533 |
| LSFPAGV | AWSLRDFNYAEQF | 0.839249 |
| LYTSGTT | AWSLRDFNYAEQF | 0.76558197 |
| VGAFPFGF | AWSLRDFNYAEQF | 0.74065864 |
| VGAGTVEF | AWSLRDFNYAEQF | 0.79579604 |
| VGKSTLLL | AWSLRDFNYAEQF | 0.8049615 |
| AALAHAGGGIV | AWSLRDFNYAEQF | 0.7440933 |
| AALVAVSLL | AWSLRDFNYAEQF | 0.7412456 |
| AAVSRAVAAL | AWSLRDFNYAEQF | 0.62699735 |
| AGGSSGIGGF | AWSLRDFNYAEQF | 0.8616644 |
| AGLTVANANG | AWSLRDFNYAEQF | 0.8282634 |
| AGLVGAVL | AWSLRDFNYAEQF | 0.83394134 |
| ALSQAFTV | AWSLRDFNYAEQF | 0.8646724 |
| ALTFSVSL | AWSLRDFNYAEQF | 0.75830024 |
| ALTGAALVAL | AWSLRDFNYAEQF | 0.73796606 |
| ASFSIGAGVL | AWSLRDFNYAEQF | 0.31253895 |
| DGSVYKI | AWSLRDFNYAEQF | 0.6178774 |
| EQLAEIV | AWSLRDFNYAEQF | 0.75646937 |
| FAVSELTVM | AWSLRDFNYAEQF | 0.8038404 |
| GAAPLAAGL | AWSLRDFNYAEQF | 0.82979214 |
| GADGGDGGSGGAGG | AWSLRDFNYAEQF | 0.7728169 |
| GFGLDTGGAGG | AWSLRDFNYAEQF | 0.60771275 |
| GLASGGLL | AWSLRDFNYAEQF | 0.8202245 |
| GRIAGAAVLL | AWSLRDFNYAEQF | 0.8491691 |
| GSAGSLNTSF | AWSLRDFNYAEQF | 0.412457 |
| GTGQIGI | AWSLRDFNYAEQF | 0.6666064 |
| GTLIALIL | AWSLRDFNYAEQF | 0.87766075 |
| IAATMGFVVG | AWSLRDFNYAEQF | 0.82393765 |
| IFAQALL | AWSLRDFNYAEQF | 0.63938403 |
| IGAAVALGF | AWSLRDFNYAEQF | 0.6397637 |
| IGLTPLAL | AWSLRDFNYAEQF | 0.8696735 |
| ILGSNTSTL | AWSLRDFNYAEQF | 0.72335637 |
| IVFSLTVL | AWSLRDFNYAEQF | 0.8627535 |
| LASKATGF | AWSLRDFNYAEQF | 0.25014746 |
| LASVALIL | AWSLRDFNYAEQF | 0.6553129 |
| LFAVGLTL | AWSLRDFNYAEQF | 0.5508759 |
| LFNGGIGTYI | AWSLRDFNYAEQF | 0.17474389 |
| LTLGAGII | AWSLRDFNYAEQF | 0.8235956 |
| LVAAALGDI | AWSLRDFNYAEQF | 0.50007415 |
| NTAFVQLGI | AWSLRDFNYAEQF | 0.80925846 |
| QLFSGAEL | AWSLRDFNYAEQF | 0.67289686 |
| SAATFITL | AWSLRDFNYAEQF | 0.8499756 |
| SNYGLANLGV | AWSLRDFNYAEQF | 0.81231916 |
| SYGGAYAVM | AWSLRDFNYAEQF | 0.68796927 |
| TTGVGGLTLG | AWSLRDFNYAEQF | 0.5088376 |
| TTQTIYI | AWSLRDFNYAEQF | 0.7846776 |

| | | |
|---|---|---|
| VAAGAGLGAVI | AWSLRDFNYAEQF | 0.8414022 |
| VAAVVTLL | AWSLRDFNYAEQF | 0.8449528 |
| VGFLGGLI | AWSLRDFNYAEQF | 0.24780464 |
| VGVIPILL | AWSLRDFNYAEQF | 0.80195016 |
| VIFASSLI | AWSLRDFNYAEQF | 0.6276151 |
| VKGGDGGAAAG | AWSLRDFNYAEQF | 0.2628284 |
| VTKTFTNI | AWSLRDFNYAEQF | 0.20316657 |
| VVAAAILL | AWSLRDFNYAEQF | 0.65464723 |
| YAALDVEL | AWSLRDFNYAEQF | 0.7291025 |
| LASVALLL | AWSLRDFNYAEQF | 0.8113222 |
| LDAVHRV | AWSLRDFNYAEQF | 0.8270604 |
| QGLSSSIF | AWSLRDFNYAEQF | 0.6227808 |
| RGTFANIRL | AWSLRDFNYAEQF | 0.8242226 |
| RQLWWGHRI | AWSLRDFNYAEQF | 0.68104273 |
| YGPPGTGKTLL | AWSLRDFNYAEQF | 0.10202045 |
| AALVGAPANVA | AWSLRDFNYAEQF | 0.6041749 |
| AATFRRTE | AWSLRDFNYAEQF | 0.34338528 |
| AGALGGGI | AWSLRDFNYAEQF | 0.8684721 |
| AGGAGGAGGIGAG | AWSLRDFNYAEQF | 0.5277184 |
| AGSGALGL | AWSLRDFNYAEQF | 0.6992898 |
| AQIADQI | AWSLRDFNYAEQF | 0.6898892 |
| ASAVAVTYL | AWSLRDFNYAEQF | 0.8279675 |
| ASTVLGGTGG | AWSLRDFNYAEQF | 0.7523906 |
| AVGGEPAK | AWSLRDFNYAEQF | 0.53685176 |
| CEQGTFGGTC | AWSLRDFNYAEQF | 0.23217557 |
| DEGKAPAG | AWSLRDFNYAEQF | 0.7760509 |
| DTVVFPAL | AWSLRDFNYAEQF | 0.5554056 |
| FEGITMAG | AWSLRDFNYAEQF | 0.39812756 |
| FPLVVQD | AWSLRDFNYAEQF | 0.48106074 |
| GAFTTPQ | AWSLRDFNYAEQF | 0.4886895 |
| GFGGFGTAGGVGGA | AWSLRDFNYAEQF | 0.2962307 |
| GFLAQFL | AWSLRDFNYAEQF | 0.8425946 |
| GGAGGAGGNAGVA | AWSLRDFNYAEQF | 0.5687929 |
| GGVGTAGAGGAGGAGG | AWSLRDFNYAEQF | 0.6882302 |
| GNGGGGDAGNAGSGGN | AWSLRDFNYAEQF | 0.87581635 |
| GTGGAGGAGNDAGS | AWSLRDFNYAEQF | 0.80675226 |
| GTTNTGWFNS | AWSLRDFNYAEQF | 0.70577645 |
| IFMGPLFTL | AWSLRDFNYAEQF | 0.52107394 |
| ILAHAIEAI | AWSLRDFNYAEQF | 0.5021411 |
| IRSAIREAL | AWSLRDFNYAEQF | 0.45789567 |
| ITFAGGG | AWSLRDFNYAEQF | 0.49132168 |
| ITGMSGHQEV | AWSLRDFNYAEQF | 0.8582525 |
| IVAGLACTL | AWSLRDFNYAEQF | 0.7329432 |
| IVGGGLAGA | AWSLRDFNYAEQF | 0.40033525 |
| IVTGVLNI | AWSLRDFNYAEQF | 0.8078151 |
| LAQQGCI | AWSLRDFNYAEQF | 0.8656677 |
| LCLHVEN | AWSLRDFNYAEQF | 0.878918 |
| LDAISDGS | AWSLRDFNYAEQF | 0.51053953 |
| LGGFAAML | AWSLRDFNYAEQF | 0.6156212 |
| LGGKNPMI | AWSLRDFNYAEQF | 0.6097606 |
| LGPKGVAAIL | AWSLRDFNYAEQF | 0.7219276 |

| | | |
|---|---|---|
| LMVGVSI | AWSLRDFNYAEQF | 0.5214062 |
| LVGAAFVL | AWSLRDFNYAEQF | 0.42419064 |
| LVLAVAPL | AWSLRDFNYAEQF | 0.8411021 |
| MAAIAVIDAA | AWSLRDFNYAEQF | 0.88317883 |
| NNGINAELQ | AWSLRDFNYAEQF | 0.66003525 |
| RGEPLVLVHG | AWSLRDFNYAEQF | 0.37463355 |
| RGGAGGAAGLI | AWSLRDFNYAEQF | 0.6740861 |
| RSYRMHPAV | AWSLRDFNYAEQF | 0.56956816 |
| SAAHLLAGQLA | AWSLRDFNYAEQF | 0.53010046 |
| SAAVGSY | AWSLRDFNYAEQF | 0.6337641 |
| SPHAVLHRI | AWSLRDFNYAEQF | 0.32506692 |
| SQIAKVL | AWSLRDFNYAEQF | 0.6823888 |
| TALGHLVP | AWSLRDFNYAEQF | 0.86243486 |
| TAPTTSGAA | AWSLRDFNYAEQF | 0.7499802 |
| TDGDDGNGGSPGT | AWSLRDFNYAEQF | 0.7916849 |
| TDGLGEN | AWSLRDFNYAEQF | 0.86028063 |
| TGGPPLPGQA | AWSLRDFNYAEQF | 0.8210937 |
| TIPAITI | AWSLRDFNYAEQF | 0.8583361 |
| TIPGFTI | AWSLRDFNYAEQF | 0.863551 |
| TSIASGALA | AWSLRDFNYAEQF | 0.72112334 |
| TVTLPGAAI | AWSLRDFNYAEQF | 0.73697734 |
| VASGLSAALP | AWSLRDFNYAEQF | 0.84586024 |
| VGAGVIGI | AWSLRDFNYAEQF | 0.6540058 |
| VGPKRFGV | AWSLRDFNYAEQF | 0.7205956 |
| VNVAFPD | AWSLRDFNYAEQF | 0.70883226 |
| VSMTNSGVS | AWSLRDFNYAEQF | 0.8128143 |
| VTDVLGDLSV | AWSLRDFNYAEQF | 0.68330336 |
| VVVVGAGFAGL | AWSLRDFNYAEQF | 0.86944187 |
| VYTVGTVV | AWSLRDFNYAEQF | 0.22075227 |
| WTTFSGP | AWSLRDFNYAEQF | 0.40688765 |
| YDELVEAF | AWSLRDFNYAEQF | 0.85905397 |
| YGTGGAGGNGG | AWSLRDFNYAEQF | 0.13086581 |
| YTNVMAV | AWSLRDFNYAEQF | 0.24047205 |
| FLEQVHQL | AWSLRDFNYAEQF | 0.7530927 |
| GLSLLAVAV | AWSLRDFNYAEQF | 0.8539946 |
| LEQVHQL | AWSLRDFNYAEQF | 0.1437064 |
| LNAAGAI | AWSLRDFNYAEQF | 0.56573594 |
| PAVHASL | AWSLRDFNYAEQF | 0.3443539 |
| TELEDDA | AWSLRDFNYAEQF | 0.82508427 |
| TPPGPPN | AWSLRDFNYAEQF | 0.68472016 |
| TPPPPPPP | AWSLRDFNYAEQF | 0.3249661 |
| TQSPALI | ASRGQGAQAPL | 0.82322496 |
| EFAAAFPQ | ASRGQGAQAPL | 0.4696784 |
| LLSGNLDVL | ASRGQGAQAPL | 0.74589217 |
| GPANLGNYNI | ASRGQGAQAPL | 0.8482164 |
| TLGPTLP | ASRGQGAQAPL | 0.8776337 |
| VYAYSSCCK | ASRGQGAQAPL | 0.39493084 |
| APAPAPAPNI | ASRGQGAQAPL | 0.81556535 |
| PGGPVPIP | ASRGQGAQAPL | 0.65648377 |
| FLTVALEP | ASRGQGAQAPL | 0.8848072 |
| SQYFGANL | ASRGQGAQAPL | 0.6340517 |

| | | |
|---|---|---|
| GGVKPAA | ASRGQGAQAPL | 0.3209446 |
| GGTGGAGGAGGW | ASRGQGAQAPL | 0.8321619 |
| GVGGVAGLGGAGG | ASRGQGAQAPL | 0.8422905 |
| VLLVLATTI | ASRGQGAQAPL | 0.5925382 |
| IGIFAAML | ASRGQGAQAPL | 0.85574293 |
| AVGDLID | ASRGQGAQAPL | 0.6700386 |
| NISPLPL | ASRGQGAQAPL | 0.6750727 |
| VTAVFVAI | ASRGQGAQAPL | 0.8445973 |
| SGDDNDAAL | ASRGQGAQAPL | 0.83055043 |
| AVAALAHS | ASRGQGAQAPL | 0.45213753 |
| GGVTVMATP | ASRGQGAQAPL | 0.8388026 |
| AGLAANVRD | ASRGQGAQAPL | 0.71420276 |
| TNRPDLIDPALLRPG | ASRGQGAQAPL | 0.40709424 |
| LANLGVTAL | ASRGQGAQAPL | 0.87260306 |
| TGERIEL | ASRGQGAQAPL | 0.72536194 |
| NGEELAA | ASRGQGAQAPL | 0.82554394 |
| VLSLVWG | ASRGQGAQAPL | 0.7686431 |
| IGYALWL | ASRGQGAQAPL | 0.8685495 |
| LSLLLAPL | ASRGQGAQAPL | 0.42922997 |
| TLDEALK | ASRGQGAQAPL | 0.7414225 |
| TDGAGTAEGG | ASRGQGAQAPL | 0.70314497 |
| NAPGGAGGNGG | ASRGQGAQAPL | 0.8405938 |
| VSVFMGMGGNF | ASRGQGAQAPL | 0.7551543 |
| LASYLAAL | ASRGQGAQAPL | 0.72298527 |
| LVGGTPAL | ASRGQGAQAPL | 0.8066225 |
| VRNADVL | ASRGQGAQAPL | 0.84642684 |
| TCSSPFLVSAM | ASRGQGAQAPL | 0.75599307 |
| VHITDAE | ASRGQGAQAPL | 0.7154983 |
| EYAAPSF | ASRGQGAQAPL | 0.86906403 |
| VVDAFHGGL | ASRGQGAQAPL | 0.403906 |
| ASVVDPQ | ASRGQGAQAPL | 0.6897191 |
| SYNEFGGLS | ASRGQGAQAPL | 0.8242213 |
| DPDGSVY | ASRGQGAQAPL | 0.8363347 |
| MVSTIHATE | ASRGQGAQAPL | 0.7311988 |
| LAAGVADPV | ASRGQGAQAPL | 0.860121 |
| TKEQIAA | ASRGQGAQAPL | 0.6628459 |
| SLASPAAL | ASRGQGAQAPL | 0.82093745 |
| IGLVFYGV | ASRGQGAQAPL | 0.7543394 |
| FLLDASNL | ASRGQGAQAPL | 0.69504035 |
| YQVGNFP | ASRGQGAQAPL | 0.45259786 |
| ESLLSQST | ASRGQGAQAPL | 0.80068314 |
| LFTVGFAAVG | ASRGQGAQAPL | 0.8399266 |
| ATPPTPT | ASRGQGAQAPL | 0.8187372 |
| TDLATLG | ASRGQGAQAPL | 0.21101673 |
| SAGGTGR | ASRGQGAQAPL | 0.81797683 |
| GFSNSGS | ASRGQGAQAPL | 0.8283433 |
| GGNGGSGGTSV | ASRGQGAQAPL | 0.8830308 |
| GTSVATGGAGN | ASRGQGAQAPL | 0.6944511 |
| NGGNGGSGGTS | ASRGQGAQAPL | 0.78290963 |
| AVVKFPH | ASRGQGAQAPL | 0.61677194 |
| TPPPPPP | ASRGQGAQAPL | 0.7587991 |

| | | |
|---|---|---|
| AAPAAAAAPA | ASRGQGAQAPL | 0.35397106 |
| IALNEDL | ASRGQGAQAPL | 0.4661479 |
| IVNVSSM | ASRGQGAQAPL | 0.6953318 |
| TGIPPAPR | ASRGQGAQAPL | 0.5526569 |
| FDLGGGTF | ASRGQGAQAPL | 0.84197056 |
| FDLGGGTFDVS | ASRGQGAQAPL | 0.84010863 |
| NEPTAAA | ASRGQGAQAPL | 0.83144504 |
| NEPTAAALA | ASRGQGAQAPL | 0.851086 |
| GSIINISSI | ASRGQGAQAPL | 0.8692566 |
| VAPGNAGTA | ASRGQGAQAPL | 0.8476083 |
| TGGGASLE | ASRGQGAQAPL | 0.6078366 |
| VAVYNHYKRI | ASRGQGAQAPL | 0.7257533 |
| TDPAPATT | ASRGQGAQAPL | 0.34234113 |
| TNRPDILDPALLRPG | ASRGQGAQAPL | 0.6854353 |
| LTTTTTS | ASRGQGAQAPL | 0.6175418 |
| PTGSGKTL | ASRGQGAQAPL | 0.7443968 |
| TGSGKTL | ASRGQGAQAPL | 0.12396903 |
| TIGHVDHGK | ASRGQGAQAPL | 0.8201431 |
| TIGHVDHGKTT | ASRGQGAQAPL | 0.86377454 |
| GHVDHGKTT | ASRGQGAQAPL | 0.674796 |
| AQFEHTLL | ASRGQGAQAPL | 0.7727753 |
| IVNQDEL | ASRGQGAQAPL | 0.82923687 |
| TNGSQFF | ASRGQGAQAPL | 0.658981 |
| TGKGIVE | ASRGQGAQAPL | 0.7989028 |
| LISNASDA | ASRGQGAQAPL | 0.594886 |
| AELRTLGELTKIAKAHGAQ | ASRGQGAQAPL | 0.8309766 |
| APNAVAAAEPSW | ASRGQGAQAPL | 0.8537831 |
| APQVAGAATAVL | ASRGQGAQAPL | 0.7740797 |
| ARRDPAAAAAILPTDA | ASRGQGAQAPL | 0.6244744 |
| DAAARDVLAEAGLAEYF | ASRGQGAQAPL | 0.7064959 |
| DDQIVLSAKANLAKVEQ | ASRGQGAQAPL | 0.39712942 |
| DELLPDTGEDSDEDGH | ASRGQGAQAPL | 0.698459 |
| ELPEAPANRATVPQD | ASRGQGAQAPL | 0.1685898 |
| FGQSKIERVTGIG | ASRGQGAQAPL | 0.77421206 |
| GAGGTGSTAGGAGGAGGAGGLY | ASRGQGAQAPL | 0.4823055 |
| GHGGTGAAGGNGGTGGAGGSGI | ASRGQGAQAPL | 0.75156075 |
| GNGGNGGAGGEGGPGGAGGAGA | ASRGQGAQAPL | 0.7280371 |
| GSTAGNGGNGGNATNGTIAGQP | ASRGQGAQAPL | 0.7876117 |
| GTGGDGGNGGNGGTGG | ASRGQGAQAPL | 0.51516867 |
| GVRRPAGIPADAPV | ASRGQGAQAPL | 0.81474257 |
| GYRQVTDAHLLTLARR | ASRGQGAQAPL | 0.7670864 |
| IAKMGQDGHDRG | ASRGQGAQAPL | 0.3229637 |
| KAEPSIEIQYT | ASRGQGAQAPL | 0.85181415 |
| KPVRIRMAADEDTVAVT | ASRGQGAQAPL | 0.6735747 |
| MPPRIAGMRLLVIKPEP | ASRGQGAQAPL | 0.7643084 |
| MKFYAVALFRSRGVPPTR | ASRGQGAQAPL | 0.830115 |
| NAGGFGILGAVAHSP | ASRGQGAQAPL | 0.7140057 |
| NEGREITLAMAEM | ASRGQGAQAPL | 0.4792058 |
| NELVEWSHPLGHRWRWE | ASRGQGAQAPL | 0.69877565 |
| NLGAGGTGGNGGNPG | ASRGQGAQAPL | 0.7360732 |
| NTEWPKWETVMAADPVD | ASRGQGAQAPL | 0.6382512 |

| | | |
|---|---|---|
| PASAAPAETQP | ASRGQGAQAPL | 0.17815435 |
| QQSSDGKNGVRAAEITGE | ASRGQGAQAPL | 0.83183014 |
| RAPDAVTTSWWKE | ASRGQGAQAPL | 0.40443754 |
| SGFGNVGAHTSGWFNQ | ASRGQGAQAPL | 0.6519699 |
| SGVAGAVGQAASVGGLKV | ASRGQGAQAPL | 0.6532595 |
| SNGGAGTGSAGGNGGT | ASRGQGAQAPL | 0.83096814 |
| TGGAGGEGGAAG | ASRGQGAQAPL | 0.7576292 |
| TGGDGGAAGAGGAPGHG | ASRGQGAQAPL | 0.521026 |
| TMPPLLAAIANAGRHGV | ASRGQGAQAPL | 0.8062199 |
| TSSTAGNSGGA | ASRGQGAQAPL | 0.7007861 |
| VPATVPVGILC | ASRGQGAQAPL | 0.8335918 |
| VQRPDSGGVVLTGRIS | ASRGQGAQAPL | 0.70703983 |
| WTPNPVPATKGVPAGLVQAT | ASRGQGAQAPL | 0.72897124 |
| YALKRALAKFSDRFDVVII | ASRGQGAQAPL | 0.74114966 |
| YLIGSVIV | ASRGQGAQAPL | 0.55635697 |
| YGQLVAVID | ASRGQGAQAPL | 0.452671 |
| VTVGGMTLPA | ASRGQGAQAPL | 0.4135275 |
| VLALPAV | ASRGQGAQAPL | 0.6921071 |
| VIPPTLNYET | ASRGQGAQAPL | 0.350195 |
| VGMGGMHPSSG | ASRGQGAQAPL | 0.76406276 |
| VGLAGGAATA | ASRGQGAQAPL | 0.86478907 |
| VGGAGGAGGSGP | ASRGQGAQAPL | 0.7641635 |
| VFAAFMV | ASRGQGAQAPL | 0.30379125 |
| TLYDKLR | ASRGQGAQAPL | 0.6546326 |
| TLAVFTVL | ASRGQGAQAPL | 0.74962723 |
| TGSVQPARV | ASRGQGAQAPL | 0.8169991 |
| TGGGAGSGGT | ASRGQGAQAPL | 0.83875173 |
| SHLETSL | ASRGQGAQAPL | 0.8818429 |
| SFTTGMT | ASRGQGAQAPL | 0.82681626 |
| QVEVAKRMGVRQPTVS | ASRGQGAQAPL | 0.61146474 |
| QGSGGPALAM | ASRGQGAQAPL | 0.8251231 |
| MLAPSNP | ASRGQGAQAPL | 0.7068144 |
| LVNNAAI | ASRGQGAQAPL | 0.7760538 |
| LTGEINTLA | ASRGQGAQAPL | 0.75756514 |
| LSIFGPV | ASRGQGAQAPL | 0.82126456 |
| LLALLANL | ASRGQGAQAPL | 0.5791836 |
| LISNASDALDKLR | ASRGQGAQAPL | 0.6645422 |
| LINEAVP | ASRGQGAQAPL | 0.39091104 |
| LGYAFSGL | ASRGQGAQAPL | 0.83973575 |
| LDFVAVAL | ASRGQGAQAPL | 0.6040852 |
| LAQPGAE | ASRGQGAQAPL | 0.49149436 |
| ISNLGAVTGF | ASRGQGAQAPL | 0.6715038 |
| HQEDPGRI | ASRGQGAQAPL | 0.8656883 |
| GTPGEPTP | ASRGQGAQAPL | 0.8550464 |
| GFFNSGS | ASRGQGAQAPL | 0.59358346 |
| GAGGGGLTAGAN | ASRGQGAQAPL | 0.5417367 |
| FVDALVNM | ASRGQGAQAPL | 0.8271375 |
| FLTVAPEL | ASRGQGAQAPL | 0.4819949 |
| FETLTAL | ASRGQGAQAPL | 0.8242846 |
| DVINPIEL | ASRGQGAQAPL | 0.76731384 |
| APVSGAPV | ASRGQGAQAPL | 0.74919647 |

| | | |
|---|---|---|
| ALVGGLAW | ASRGQGAQAPL | 0.50877845 |
| ALAAAAATAI | ASRGQGAQAPL | 0.8198452 |
| AFGLGLLL | ASRGQGAQAPL | 0.6919973 |
| AFFAAASL | ASRGQGAQAPL | 0.80954874 |
| AAAALPAFSPP | ASRGQGAQAPL | 0.1990307 |
| VVLAPAL | ASRGQGAQAPL | 0.81972575 |
| TIGHVDHGKTTL | ASRGQGAQAPL | 0.5211638 |
| LISNASDALDK | ASRGQGAQAPL | 0.48965046 |
| LISNASDAL | ASRGQGAQAPL | 0.23903522 |
| ISNASDALDK | ASRGQGAQAPL | 0.69283223 |
| NQAETLVYQTE | ASRGQGAQAPL | 0.83529687 |
| TGGTGGTAGAG | ASRGQGAQAPL | 0.86043376 |
| TGSTAGGAGGA | ASRGQGAQAPL | 0.7496691 |
| DPLAVPVAAGPV | ASRGQGAQAPL | 0.82182163 |
| GGTGGTPGNGGN | ASRGQGAQAPL | 0.86699545 |
| PAGLAGVDPATL | ASRGQGAQAPL | 0.66421616 |
| GNNLVVEAPHAGASV | ASRGQGAQAPL | 0.22536205 |
| KVTRIMSASTLEQPA | ASRGQGAQAPL | 0.7283814 |
| LGFALPIALPRRTVV | ASRGQGAQAPL | 0.7949294 |
| TRLGNLTARKLVVLV | ASRGQGAQAPL | 0.85643363 |
| WSQQRRLIYFPSAGP | ASRGQGAQAPL | 0.80835223 |
| GPNGNPDPMAAAVDIR | ASRGQGAQAPL | 0.40472603 |
| TQRCVSVADLRAAASR | ASRGQGAQAPL | 0.7444103 |
| QFVLDVAEHPVEVLTETV | ASRGQGAQAPL | 0.8258781 |
| GGAGGTGGAGGNGGTQAGDG | ASRGQGAQAPL | 0.45682216 |
| PLLATLRAIAVDVDAKPAQV | ASRGQGAQAPL | 0.5349225 |
| APKFRTGGYMIDFWGVGYQVAK | ASRGQGAQAPL | 0.5606781 |
| GAGGAGGAGGAAGGPGGTGGTG | ASRGQGAQAPL | 0.37145072 |
| IGALGLLATDVAPAGSVAFGVA | ASRGQGAQAPL | 0.754305 |
| AGGAGGIGGNGGAGGAGGVLMG | ASRGQGAQAPL | 0.6835581 |
| ADELIGLL | ASRGQGAQAPL | 0.56228566 |
| ADRALKA | ASRGQGAQAPL | 0.71505547 |
| AGAVGQVGGAGGA | ASRGQGAQAPL | 0.47394076 |
| AGGGGGLGGAP | ASRGQGAQAPL | 0.522125 |
| AGGLLYGNGGAG | ASRGQGAQAPL | 0.5941203 |
| AGLESIGL | ASRGQGAQAPL | 0.7876355 |
| AGLKTGII | ASRGQGAQAPL | 0.8390822 |
| ASLAALLL | ASRGQGAQAPL | 0.7422744 |
| ASVINISSI | ASRGQGAQAPL | 0.7559388 |
| ATPPGPA | ASRGQGAQAPL | 0.8565241 |
| ATVASALV | ASRGQGAQAPL | 0.34869918 |
| ATVASLIL | ASRGQGAQAPL | 0.5218643 |
| AVNGPVLIRH | ASRGQGAQAPL | 0.808076 |
| DGAAGAVGI | ASRGQGAQAPL | 0.5558461 |
| DPLLHGI | ASRGQGAQAPL | 0.85470897 |
| ELGAALGAL | ASRGQGAQAPL | 0.754745 |
| ELGRTQAV | ASRGQGAQAPL | 0.78518045 |
| FVAGGYLS | ASRGQGAQAPL | 0.79830015 |
| GASGGAGGAGGW | ASRGQGAQAPL | 0.87940705 |
| GGAGAGGDGGAGGI | ASRGQGAQAPL | 0.7923805 |
| GTGHNGSQPGMGGNG | ASRGQGAQAPL | 0.6842826 |

| | | |
|---|---|---|
| GYAAASASAS | ASRGQGAQAPL | 0.8477434 |
| HQVDVAAVQQG | ASRGQGAQAPL | 0.3628923 |
| IGAAIAHT | ASRGQGAQAPL | 0.83894145 |
| IGMIEVAQL | ASRGQGAQAPL | 0.8163115 |
| IRLAPEHTSAMD | ASRGQGAQAPL | 0.5023613 |
| IYNVSVL | ASRGQGAQAPL | 0.6907044 |
| KQGFLGM | ASRGQGAQAPL | 0.8651389 |
| LESVITQL | ASRGQGAQAPL | 0.85916436 |
| LGLGALVVV | ASRGQGAQAPL | 0.46509778 |
| LIGNGGNGGSGGI | ASRGQGAQAPL | 0.598384 |
| LNAINAPTL | ASRGQGAQAPL | 0.39116147 |
| LQLLNTGV | ASRGQGAQAPL | 0.54502 |
| LQTSIEF | ASRGQGAQAPL | 0.28975633 |
| MSFAVSEL | ASRGQGAQAPL | 0.8017998 |
| NTGALGL | ASRGQGAQAPL | 0.6172739 |
| PPGAGGAGGAGTT | ASRGQGAQAPL | 0.84222615 |
| QLVGLGI | ASRGQGAQAPL | 0.7616843 |
| RFLEQVHQL | ASRGQGAQAPL | 0.8314233 |
| RGVGYVLR | ASRGQGAQAPL | 0.57688415 |
| SATGVFAGVI | ASRGQGAQAPL | 0.6213652 |
| SLPAITLP | ASRGQGAQAPL | 0.29324615 |
| STALAFTV | ASRGQGAQAPL | 0.34851444 |
| SVLTVGLGK | ASRGQGAQAPL | 0.58675647 |
| TALAGAGSGI | ASRGQGAQAPL | 0.6173625 |
| TGLPFTGI | ASRGQGAQAPL | 0.23731437 |
| VAEHVTV | ASRGQGAQAPL | 0.87041074 |
| VDIVLNSL | ASRGQGAQAPL | 0.6696332 |
| VLTGGGAVL | ASRGQGAQAPL | 0.79003584 |
| VNVGLAYGI | ASRGQGAQAPL | 0.57775235 |
| VSSAAVGAV | ASRGQGAQAPL | 0.42184746 |
| VTSVYEAGV | ASRGQGAQAPL | 0.57890016 |
| WVALDRGA | ASRGQGAQAPL | 0.79331934 |
| YLVGSVVI | ASRGQGAQAPL | 0.4772363 |
| GPPGTGKTLL | ASRGQGAQAPL | 0.33748114 |
| GVGMTKF | ASRGQGAQAPL | 0.47252333 |
| IGIGGSDL | ASRGQGAQAPL | 0.7341622 |
| IGTLLAEL | ASRGQGAQAPL | 0.87719125 |
| IYTSGTT | ASRGQGAQAPL | 0.8492242 |
| LSFPAGV | ASRGQGAQAPL | 0.7597964 |
| LYTSGTT | ASRGQGAQAPL | 0.8726007 |
| VGAFPFGF | ASRGQGAQAPL | 0.15797757 |
| VGAGTVEF | ASRGQGAQAPL | 0.63891375 |
| VGKSTLLL | ASRGQGAQAPL | 0.8577813 |
| AALAHAGGGIV | ASRGQGAQAPL | 0.80047226 |
| AALVAVSLL | ASRGQGAQAPL | 0.6711657 |
| AAVSRAVAAL | ASRGQGAQAPL | 0.8496659 |
| AGGSSGIGGF | ASRGQGAQAPL | 0.5115848 |
| AGLTVANANG | ASRGQGAQAPL | 0.6269612 |
| AGLVGAVL | ASRGQGAQAPL | 0.67959136 |
| ALSQAFTV | ASRGQGAQAPL | 0.8253321 |
| ALTFSVSL | ASRGQGAQAPL | 0.8140905 |

| | | |
|---|---|---|
| ALTGAALVAL | ASRGQGAQAPL | 0.3788436 |
| ASFSIGAGVL | ASRGQGAQAPL | 0.765144 |
| DGSVYKI | ASRGQGAQAPL | 0.8850507 |
| EQLAEIV | ASRGQGAQAPL | 0.6210871 |
| FAVSELTVM | ASRGQGAQAPL | 0.8533656 |
| GAAPLAAGL | ASRGQGAQAPL | 0.7491152 |
| GADGGDGGSGGAGG | ASRGQGAQAPL | 0.7391765 |
| GFGLDTGGAGG | ASRGQGAQAPL | 0.5314119 |
| GLASGGLL | ASRGQGAQAPL | 0.49716553 |
| GRIAGAAVLL | ASRGQGAQAPL | 0.8120024 |
| GSAGSLNTSF | ASRGQGAQAPL | 0.5263064 |
| GTGQIGI | ASRGQGAQAPL | 0.70394087 |
| GTLIALIL | ASRGQGAQAPL | 0.720127 |
| IAATMGFVVG | ASRGQGAQAPL | 0.8840165 |
| IFAQALL | ASRGQGAQAPL | 0.79986453 |
| IGAAVALGF | ASRGQGAQAPL | 0.8012284 |
| IGLTPLAL | ASRGQGAQAPL | 0.48981568 |
| ILGSNTSTL | ASRGQGAQAPL | 0.79830885 |
| IVFSLTVL | ASRGQGAQAPL | 0.8243638 |
| LASKATGF | ASRGQGAQAPL | 0.4676519 |
| LASVALIL | ASRGQGAQAPL | 0.84849095 |
| LFAVGLTL | ASRGQGAQAPL | 0.47111475 |
| LFNGGIGTYI | ASRGQGAQAPL | 0.77351314 |
| LTLGAGII | ASRGQGAQAPL | 0.78625876 |
| LVAAALGDI | ASRGQGAQAPL | 0.8604462 |
| NTAFVQLGI | ASRGQGAQAPL | 0.6870736 |
| QLFSGAEL | ASRGQGAQAPL | 0.8058113 |
| SAATFITL | ASRGQGAQAPL | 0.4294268 |
| SNYGLANLGV | ASRGQGAQAPL | 0.83267283 |
| SYGGAYAVM | ASRGQGAQAPL | 0.8466021 |
| TTGVGGLTLG | ASRGQGAQAPL | 0.80604696 |
| TTQTIYI | ASRGQGAQAPL | 0.8841047 |
| VAAGAGLGAVI | ASRGQGAQAPL | 0.78495234 |
| VAAVVTLL | ASRGQGAQAPL | 0.24056703 |
| VGFLGGLI | ASRGQGAQAPL | 0.6798423 |
| VGVIPILL | ASRGQGAQAPL | 0.85236496 |
| VIFASSLI | ASRGQGAQAPL | 0.8685497 |
| VKGGDGGAAAG | ASRGQGAQAPL | 0.8462132 |
| VTKTFTNI | ASRGQGAQAPL | 0.8024365 |
| VVAAAILL | ASRGQGAQAPL | 0.67866063 |
| YAALDVEL | ASRGQGAQAPL | 0.8651769 |
| LASVALLL | ASRGQGAQAPL | 0.88762826 |
| LDAVHRV | ASRGQGAQAPL | 0.7956275 |
| QGLSSSIF | ASRGQGAQAPL | 0.5152236 |
| RGTFANIRL | ASRGQGAQAPL | 0.4709319 |
| RQLWWGHRI | ASRGQGAQAPL | 0.78813267 |
| YGPPGTGKTLL | ASRGQGAQAPL | 0.8410989 |
| AALVGAPANVA | ASRGQGAQAPL | 0.39105207 |
| AATFRRTE | ASRGQGAQAPL | 0.7186909 |
| AGALGGGI | ASRGQGAQAPL | 0.7334674 |
| AGGAGGAGGIGAG | ASRGQGAQAPL | 0.2130866 |

| | | |
|---|---|---|
| AGSGALGL | ASRGQGAQAPL | 0.8414354 |
| AQIADQI | ASRGQGAQAPL | 0.8691299 |
| ASAVAVTYL | ASRGQGAQAPL | 0.7757266 |
| ASTVLGGTGG | ASRGQGAQAPL | 0.4477731 |
| AVGGEPAK | ASRGQGAQAPL | 0.85053414 |
| CEQGTFGGTC | ASRGQGAQAPL | 0.86919075 |
| DEGKAPAG | ASRGQGAQAPL | 0.8682564 |
| DTVVFPAL | ASRGQGAQAPL | 0.29099053 |
| FEGITMAG | ASRGQGAQAPL | 0.16805945 |
| FPLVVQD | ASRGQGAQAPL | 0.7065395 |
| GAFTTPQ | ASRGQGAQAPL | 0.8294995 |
| GFGGFGTAGGVGGA | ASRGQGAQAPL | 0.57218194 |
| GFLAQFL | ASRGQGAQAPL | 0.4501826 |
| GGAGGAGGNAGVA | ASRGQGAQAPL | 0.7871599 |
| GGVGTAGAGGAGGAGG | ASRGQGAQAPL | 0.85018003 |
| GNGGGGDAGNAGSGGN | ASRGQGAQAPL | 0.36691573 |
| GTGGAGGAGNDAGS | ASRGQGAQAPL | 0.38692117 |
| GTTNTGWFNS | ASRGQGAQAPL | 0.838828 |
| IFMGPLFTL | ASRGQGAQAPL | 0.82882965 |
| ILAHAIEAI | ASRGQGAQAPL | 0.5536024 |
| IRSAIREAL | ASRGQGAQAPL | 0.67194116 |
| ITFAGGG | ASRGQGAQAPL | 0.7551048 |
| ITGMSGHQEV | ASRGQGAQAPL | 0.81168085 |
| IVAGLACTL | ASRGQGAQAPL | 0.7967627 |
| IVGGGLAGA | ASRGQGAQAPL | 0.35431 |
| IVTGVLNI | ASRGQGAQAPL | 0.85735446 |
| LAQQGCI | ASRGQGAQAPL | 0.39546233 |
| LCLHVEN | ASRGQGAQAPL | 0.83964694 |
| LDAISDGS | ASRGQGAQAPL | 0.74544835 |
| LGGFAAML | ASRGQGAQAPL | 0.7846974 |
| LGGKNPMI | ASRGQGAQAPL | 0.62645084 |
| LGPKGVAAIL | ASRGQGAQAPL | 0.53557515 |
| LMVGVSI | ASRGQGAQAPL | 0.4611048 |
| LVGAAFVL | ASRGQGAQAPL | 0.35605407 |
| LVLAVAPL | ASRGQGAQAPL | 0.8484253 |
| MAAIAVIDAA | ASRGQGAQAPL | 0.30012816 |
| NNGINAELQ | ASRGQGAQAPL | 0.84199136 |
| RGEPLVLVHG | ASRGQGAQAPL | 0.72248566 |
| RGGAGGAAGLI | ASRGQGAQAPL | 0.5275508 |
| RSYRMHPAV | ASRGQGAQAPL | 0.8178098 |
| SAAHLLAGQLA | ASRGQGAQAPL | 0.31602666 |
| SAAVGSY | ASRGQGAQAPL | 0.7489659 |
| SPHAVLHRI | ASRGQGAQAPL | 0.76153517 |
| SQIAKVL | ASRGQGAQAPL | 0.8085463 |
| TALGHLVP | ASRGQGAQAPL | 0.57190883 |
| TAPTTSGAA | ASRGQGAQAPL | 0.809459 |
| TDGDDGNGGSPGT | ASRGQGAQAPL | 0.79411995 |
| TDGLGEN | ASRGQGAQAPL | 0.4560562 |
| TGGPPLPGQA | ASRGQGAQAPL | 0.7515862 |
| TIPAITI | ASRGQGAQAPL | 0.8452746 |
| TIPGFTI | ASRGQGAQAPL | 0.7669305 |

| | | |
|---|---|---|
| TSIASGALA | ASRGQGAQAPL | 0.3456784 |
| TVTLPGAAI | ASRGQGAQAPL | 0.76893187 |
| VASGLSAALP | ASRGQGAQAPL | 0.57296956 |
| VGAGVIGI | ASRGQGAQAPL | 0.79467654 |
| VGPKRFGV | ASRGQGAQAPL | 0.7725191 |
| VNVAFPD | ASRGQGAQAPL | 0.5417035 |
| VSMTNSGVS | ASRGQGAQAPL | 0.7975614 |
| VTDVLGDLSV | ASRGQGAQAPL | 0.36736155 |
| VVVVGAGFAGL | ASRGQGAQAPL | 0.5653919 |
| VYTVGTVV | ASRGQGAQAPL | 0.62630945 |
| WTTFSGP | ASRGQGAQAPL | 0.70019436 |
| YDELVEAF | ASRGQGAQAPL | 0.8151053 |
| YGTGGAGGNGG | ASRGQGAQAPL | 0.45615876 |
| YTNVMAV | ASRGQGAQAPL | 0.7679813 |
| FLEQVHQL | ASRGQGAQAPL | 0.6499259 |
| GLSLLAVAV | ASRGQGAQAPL | 0.48492348 |
| LEQVHQL | ASRGQGAQAPL | 0.7397561 |
| LNAAGAI | ASRGQGAQAPL | 0.6334716 |
| PAVHASL | ASRGQGAQAPL | 0.3795175 |
| TELEDDA | ASRGQGAQAPL | 0.82327515 |
| TPPGPPN | ASRGQGAQAPL | 0.41258103 |
| TPPPPPPP | ASRGQGAQAPL | 0.7267138 |
| TQSPALI | ASSPSRYQAPL | 0.8824171 |
| EFAAAFPQ | ASSPSRYQAPL | 0.329418 |
| LLSGNLDVL | ASSPSRYQAPL | 0.8279609 |
| GPANLGNYNI | ASSPSRYQAPL | 0.7704325 |
| TLGPTLP | ASSPSRYQAPL | 0.7832986 |
| VYAYSSCCK | ASSPSRYQAPL | 0.64858997 |
| APAPAPAPNI | ASSPSRYQAPL | 0.5658635 |
| PGGPVPIP | ASSPSRYQAPL | 0.8581687 |
| FLTVALEP | ASSPSRYQAPL | 0.7726308 |
| SQYFGANL | ASSPSRYQAPL | 0.72773665 |
| GGVKPAA | ASSPSRYQAPL | 0.80280674 |
| GGTGGAGGAGGW | ASSPSRYQAPL | 0.8466664 |
| GVGGVAGLGGAGG | ASSPSRYQAPL | 0.82839644 |
| VLLVLATTI | ASSPSRYQAPL | 0.59174734 |
| IGIFAAML | ASSPSRYQAPL | 0.2880834 |
| AVGDLID | ASSPSRYQAPL | 0.6254786 |
| NISPLPL | ASSPSRYQAPL | 0.6650959 |
| VTAVFVAI | ASSPSRYQAPL | 0.5881789 |
| SGDDNDAAL | ASSPSRYQAPL | 0.30336434 |
| AVAALAHS | ASSPSRYQAPL | 0.84099746 |
| GGVTVMATP | ASSPSRYQAPL | 0.88289595 |
| AGLAANVRD | ASSPSRYQAPL | 0.5542803 |
| TNRPDLIDPALLRPG | ASSPSRYQAPL | 0.88688016 |
| LANLGVTAL | ASSPSRYQAPL | 0.40070122 |
| TGERIEL | ASSPSRYQAPL | 0.7698583 |
| NGEELAA | ASSPSRYQAPL | 0.8410104 |
| VLSLVWG | ASSPSRYQAPL | 0.4874493 |
| IGYALWL | ASSPSRYQAPL | 0.754241 |
| LSLLLAPL | ASSPSRYQAPL | 0.56641585 |

| | | |
|---|---|---|
| TLDEALK | ASSPSRYQAPL | 0.5600109 |
| TDGAGTAEGG | ASSPSRYQAPL | 0.8420553 |
| NAPGGAGGNGG | ASSPSRYQAPL | 0.51677 |
| VSVFMGMGGNF | ASSPSRYQAPL | 0.78139997 |
| LASYLAAL | ASSPSRYQAPL | 0.57733965 |
| LVGGTPAL | ASSPSRYQAPL | 0.47581223 |
| VRNADVL | ASSPSRYQAPL | 0.79644245 |
| TCSSPFLVSAM | ASSPSRYQAPL | 0.82229733 |
| VHITDAE | ASSPSRYQAPL | 0.75318086 |
| EYAAPSF | ASSPSRYQAPL | 0.88644665 |
| VVDAFHGGL | ASSPSRYQAPL | 0.5366603 |
| ASVVDPQ | ASSPSRYQAPL | 0.8822341 |
| SYNEFGGLS | ASSPSRYQAPL | 0.8656677 |
| DPDGSVY | ASSPSRYQAPL | 0.7682221 |
| MVSTIHATE | ASSPSRYQAPL | 0.6557893 |
| LAAGVADPV | ASSPSRYQAPL | 0.8304445 |
| TKEQIAA | ASSPSRYQAPL | 0.59739923 |
| SLASPAAL | ASSPSRYQAPL | 0.7368115 |
| IGLVFYGV | ASSPSRYQAPL | 0.86813456 |
| FLLDASNL | ASSPSRYQAPL | 0.8139443 |
| YQVGNFP | ASSPSRYQAPL | 0.8864354 |
| ESLLSQST | ASSPSRYQAPL | 0.87042475 |
| LFTVGFAAVG | ASSPSRYQAPL | 0.87774605 |
| ATPPTPT | ASSPSRYQAPL | 0.832167 |
| TDLATLG | ASSPSRYQAPL | 0.68654895 |
| SAGGTGR | ASSPSRYQAPL | 0.8209225 |
| GFSNSGS | ASSPSRYQAPL | 0.7642471 |
| GGNGGSGGTSV | ASSPSRYQAPL | 0.87026703 |
| GTSVATGGAGN | ASSPSRYQAPL | 0.90321267 |
| NGGNGGSGGTS | ASSPSRYQAPL | 0.8275513 |
| AVVKFPH | ASSPSRYQAPL | 0.89113736 |
| TPPPPPP | ASSPSRYQAPL | 0.8931957 |
| AAPAAAAAPA | ASSPSRYQAPL | 0.68093085 |
| IALNEDL | ASSPSRYQAPL | 0.8267664 |
| IVNVSSM | ASSPSRYQAPL | 0.7920283 |
| TGIPPAPR | ASSPSRYQAPL | 0.8682183 |
| FDLGGGTF | ASSPSRYQAPL | 0.43881232 |
| FDLGGGTFDVS | ASSPSRYQAPL | 0.8955327 |
| NEPTAAA | ASSPSRYQAPL | 0.75739706 |
| NEPTAAALA | ASSPSRYQAPL | 0.29068825 |
| GSIINISSI | ASSPSRYQAPL | 0.8658758 |
| VAPGNAGTA | ASSPSRYQAPL | 0.841272 |
| TGGGASLE | ASSPSRYQAPL | 0.86209905 |
| VAVYNHYKRI | ASSPSRYQAPL | 0.89676094 |
| TDPAPATT | ASSPSRYQAPL | 0.70284355 |
| TNRPDILDPALLRPG | ASSPSRYQAPL | 0.7651601 |
| LTTTTTS | ASSPSRYQAPL | 0.8672454 |
| PTGSGKTL | ASSPSRYQAPL | 0.90635973 |
| TGSGKTL | ASSPSRYQAPL | 0.74296874 |
| TIGHVDHGK | ASSPSRYQAPL | 0.87432766 |
| TIGHVDHGKTT | ASSPSRYQAPL | 0.8917348 |

| | | |
|---|---|---|
| GHVDHGKTT | ASSPSRYQAPL | 0.86585957 |
| AQFEHTLL | ASSPSRYQAPL | 0.9077088 |
| IVNQDEL | ASSPSRYQAPL | 0.9124435 |
| TNGSQFF | ASSPSRYQAPL | 0.776491 |
| TGKGIVE | ASSPSRYQAPL | 0.9049748 |
| LISNASDA | ASSPSRYQAPL | 0.8969587 |
| AELRTLGELTKIAKAHGAQ | ASSPSRYQAPL | 0.84459794 |
| APNAVAAAEPSW | ASSPSRYQAPL | 0.8446974 |
| APQVAGAATAVL | ASSPSRYQAPL | 0.79298735 |
| ARRDPAAAAAILPTDA | ASSPSRYQAPL | 0.83730435 |
| DAAARDVLAEAGLAEYF | ASSPSRYQAPL | 0.6913599 |
| DDQIVLSAKANLAKVEQ | ASSPSRYQAPL | 0.84521663 |
| DELLPDTGEDSDEDGH | ASSPSRYQAPL | 0.9008492 |
| ELPEAPANRATVPQD | ASSPSRYQAPL | 0.81450176 |
| FGQSKIERVTGIG | ASSPSRYQAPL | 0.73942953 |
| GAGGTGSTAGGAGGAGGAGGLY | ASSPSRYQAPL | 0.73810047 |
| GHGGTGAAGGNGGTGGAGGSGI | ASSPSRYQAPL | 0.8439728 |
| GNGGNGGAGGEGGPGGAGGAGA | ASSPSRYQAPL | 0.84841764 |
| GSTAGNGGNGGNATNGTIAGQP | ASSPSRYQAPL | 0.62192625 |
| GTGGDGGNGGNGGTGG | ASSPSRYQAPL | 0.82478946 |
| GVRRPAGIPADAPV | ASSPSRYQAPL | 0.886919 |
| GYRQVTDAHLLTLARR | ASSPSRYQAPL | 0.8890562 |
| IAKMGQDGHDRG | ASSPSRYQAPL | 0.64055586 |
| KAEPSIEIQYT | ASSPSRYQAPL | 0.9082797 |
| KPVRIRMAADEDTVAVT | ASSPSRYQAPL | 0.83847886 |
| MPPRIAGMRLLVIKPEP | ASSPSRYQAPL | 0.8952491 |
| MKFYAVALFRSRGVPPTR | ASSPSRYQAPL | 0.8534785 |
| NAGGFGILGAVAHSP | ASSPSRYQAPL | 0.77155066 |
| NEGREITLAMAEM | ASSPSRYQAPL | 0.87394315 |
| NELVEWSHPLGHRWRWE | ASSPSRYQAPL | 0.807952 |
| NLGAGGTGGNGGNPG | ASSPSRYQAPL | 0.8074504 |
| NTEWPKWETVMAADPVD | ASSPSRYQAPL | 0.8929932 |
| PASAAPAETQP | ASSPSRYQAPL | 0.8288411 |
| QQSSDGKNGVRAAEITGE | ASSPSRYQAPL | 0.7603806 |
| RAPDAVTTSWWKE | ASSPSRYQAPL | 0.71865857 |
| SGFGNVGAHTSGWFNQ | ASSPSRYQAPL | 0.9018054 |
| SGVAGAVGQAASVGGLKV | ASSPSRYQAPL | 0.68286955 |
| SNGGAGTGSAGGNGGT | ASSPSRYQAPL | 0.84086025 |
| TGGAGGEGGAAG | ASSPSRYQAPL | 0.74924606 |
| TGGDGGAAGAGGAPGHG | ASSPSRYQAPL | 0.8323495 |
| TMPPLLAAIANAGRHGV | ASSPSRYQAPL | 0.9005605 |
| TSSTAGNSGGA | ASSPSRYQAPL | 0.887177 |
| VPATVPVGILC | ASSPSRYQAPL | 0.7869228 |
| VQRPDSGGVVLTGRIS | ASSPSRYQAPL | 0.8822663 |
| WTPNPVPATKGVPAGLVQAT | ASSPSRYQAPL | 0.88035 |
| YALKRALAKFSDRFDVVII | ASSPSRYQAPL | 0.8640841 |
| YLIGSVIV | ASSPSRYQAPL | 0.8636425 |
| YGQLVAVID | ASSPSRYQAPL | 0.7383211 |
| VTVGGMTLPA | ASSPSRYQAPL | 0.89638925 |
| VLALPAV | ASSPSRYQAPL | 0.88096964 |
| VIPPTLNYET | ASSPSRYQAPL | 0.907202 |

| | | |
|---|---|---|
| VGMGGMHPSSG | ASSPSRYQAPL | 0.9031083 |
| VGLAGGAATA | ASSPSRYQAPL | 0.8503568 |
| VGGAGGAGGSGP | ASSPSRYQAPL | 0.8600449 |
| VFAAFMV | ASSPSRYQAPL | 0.634627 |
| TLYDKLR | ASSPSRYQAPL | 0.73608047 |
| TLAVFTVL | ASSPSRYQAPL | 0.8733604 |
| TGSVQPARV | ASSPSRYQAPL | 0.85335624 |
| TGGGAGSGGT | ASSPSRYQAPL | 0.89065766 |
| SHLETSL | ASSPSRYQAPL | 0.8917135 |
| SFTTGMT | ASSPSRYQAPL | 0.7958176 |
| QVEVAKRMGVRQPTVS | ASSPSRYQAPL | 0.88913727 |
| QGSGGPALAM | ASSPSRYQAPL | 0.8994633 |
| MLAPSNP | ASSPSRYQAPL | 0.6543002 |
| LVNNAAI | ASSPSRYQAPL | 0.82822776 |
| LTGEINTLA | ASSPSRYQAPL | 0.91213816 |
| LSIFGPV | ASSPSRYQAPL | 0.8397559 |
| LLALLANL | ASSPSRYQAPL | 0.8372427 |
| LISNASDALDKLR | ASSPSRYQAPL | 0.8651006 |
| LINEAVP | ASSPSRYQAPL | 0.8913944 |
| LGYAFSGL | ASSPSRYQAPL | 0.82663953 |
| LDFVAVAL | ASSPSRYQAPL | 0.90036714 |
| LAQPGAE | ASSPSRYQAPL | 0.50661707 |
| ISNLGAVTGF | ASSPSRYQAPL | 0.8732103 |
| HQEDPGRI | ASSPSRYQAPL | 0.7843548 |
| GTPGEPTP | ASSPSRYQAPL | 0.7156105 |
| GFFNSGS | ASSPSRYQAPL | 0.89472985 |
| GAGGGGLTAGAN | ASSPSRYQAPL | 0.8535404 |
| FVDALVNM | ASSPSRYQAPL | 0.8885987 |
| FLTVAPEL | ASSPSRYQAPL | 0.8362797 |
| FETLTAL | ASSPSRYQAPL | 0.90033567 |
| DVINPIEL | ASSPSRYQAPL | 0.8281853 |
| APVSGAPV | ASSPSRYQAPL | 0.85827494 |
| ALVGGLAW | ASSPSRYQAPL | 0.8351983 |
| ALAAAAATAI | ASSPSRYQAPL | 0.8864091 |
| AFGLGLLL | ASSPSRYQAPL | 0.9009727 |
| AFFAAASL | ASSPSRYQAPL | 0.8940227 |
| AAAALPAFSPP | ASSPSRYQAPL | 0.60817647 |
| VVLAPAL | ASSPSRYQAPL | 0.82071155 |
| TIGHVDHGKTTL | ASSPSRYQAPL | 0.8564187 |
| LISNASDALDK | ASSPSRYQAPL | 0.41099766 |
| LISNASDAL | ASSPSRYQAPL | 0.55181354 |
| ISNASDALDK | ASSPSRYQAPL | 0.78706026 |
| NQAETLVYQTE | ASSPSRYQAPL | 0.7940608 |
| TGGTGGTAGAG | ASSPSRYQAPL | 0.88846254 |
| TGSTAGGAGGA | ASSPSRYQAPL | 0.85712487 |
| DPLAVPVAAGPV | ASSPSRYQAPL | 0.8388163 |
| GGTGGTPGNGGN | ASSPSRYQAPL | 0.87237716 |
| PAGLAGVDPATL | ASSPSRYQAPL | 0.83345234 |
| GNNLVVEAPHAGASV | ASSPSRYQAPL | 0.18454784 |
| KVTRIMSASTLEQPA | ASSPSRYQAPL | 0.7966809 |
| LGFALPIALPRRTVV | ASSPSRYQAPL | 0.70226586 |

| | | |
|---|---|---|
| TRLGNLTARKLVVLV | ASSPSRYQAPL | 0.90763354 |
| WSQQRRLIYFPSAGP | ASSPSRYQAPL | 0.8244924 |
| GPNGNPDPMAAAVDIR | ASSPSRYQAPL | 0.7266758 |
| TQRCVSVADLRAAASR | ASSPSRYQAPL | 0.8422903 |
| QFVLDVAEHPVEVLTETV | ASSPSRYQAPL | 0.82076997 |
| GGAGGTGGAGGNGGTQAGDG | ASSPSRYQAPL | 0.818694 |
| PLLATLRAIAVDVDAKPAQV | ASSPSRYQAPL | 0.7168241 |
| APKFRTGGYMIDFWGVGYQVAK | ASSPSRYQAPL | 0.45492142 |
| GAGGAGGAGGAAGGPGGTGGTG | ASSPSRYQAPL | 0.8828412 |
| IGALGLLATDVAPAGSVAFGVA | ASSPSRYQAPL | 0.6849102 |
| AGGAGGIGGNGGAGGAGGVLMG | ASSPSRYQAPL | 0.79555434 |
| ADELIGLL | ASSPSRYQAPL | 0.5765494 |
| ADRALKA | ASSPSRYQAPL | 0.7253952 |
| AGAVGQVGGAGGA | ASSPSRYQAPL | 0.66131693 |
| AGGGGGLGGAP | ASSPSRYQAPL | 0.90540755 |
| AGGLLYGNGGAG | ASSPSRYQAPL | 0.8219279 |
| AGLESIGL | ASSPSRYQAPL | 0.788594 |
| AGLKTGII | ASSPSRYQAPL | 0.9019218 |
| ASLAALLL | ASSPSRYQAPL | 0.79436916 |
| ASVINISSI | ASSPSRYQAPL | 0.8194462 |
| ATPPGPA | ASSPSRYQAPL | 0.81462073 |
| ATVASALV | ASSPSRYQAPL | 0.8153715 |
| ATVASLIL | ASSPSRYQAPL | 0.79467773 |
| AVNGPVLIRH | ASSPSRYQAPL | 0.81190157 |
| DGAAGAVGI | ASSPSRYQAPL | 0.90330327 |
| DPLLHGI | ASSPSRYQAPL | 0.8802056 |
| ELGAALGAL | ASSPSRYQAPL | 0.70681083 |
| ELGRTQAV | ASSPSRYQAPL | 0.8912879 |
| FVAGGYLS | ASSPSRYQAPL | 0.9043318 |
| GASGGAGGAGGW | ASSPSRYQAPL | 0.90964246 |
| GGAGAGGDGGAGGI | ASSPSRYQAPL | 0.7774954 |
| GTGHNGSQPGMGGNG | ASSPSRYQAPL | 0.7904167 |
| GYAAASASAS | ASSPSRYQAPL | 0.840577 |
| HQVDVAAVQQG | ASSPSRYQAPL | 0.80922896 |
| IGAAIAHT | ASSPSRYQAPL | 0.8270029 |
| IGMIEVAQL | ASSPSRYQAPL | 0.77127665 |
| IRLAPEHTSAMD | ASSPSRYQAPL | 0.8862786 |
| IYNVSVL | ASSPSRYQAPL | 0.9068092 |
| KQGFLGM | ASSPSRYQAPL | 0.8349843 |
| LESVITQL | ASSPSRYQAPL | 0.5978324 |
| LGLGALVVV | ASSPSRYQAPL | 0.8236571 |
| LIGNGGNGGSGGI | ASSPSRYQAPL | 0.7405566 |
| LNAINAPTL | ASSPSRYQAPL | 0.78851146 |
| LQLLNTGV | ASSPSRYQAPL | 0.86038816 |
| LQTSIEF | ASSPSRYQAPL | 0.6731779 |
| MSFAVSEL | ASSPSRYQAPL | 0.83559895 |
| NTGALGL | ASSPSRYQAPL | 0.9078417 |
| PPGAGGAGGAGTT | ASSPSRYQAPL | 0.79266083 |
| QLVGLGI | ASSPSRYQAPL | 0.87340415 |
| RFLEQVHQL | ASSPSRYQAPL | 0.89608604 |
| RGVGYVLR | ASSPSRYQAPL | 0.884787 |

| | | |
|---|---|---|
| SATGVFAGVI | ASSPSRYQAPL | 0.9092236 |
| SLPAITLP | ASSPSRYQAPL | 0.8771819 |
| STALAFTV | ASSPSRYQAPL | 0.87929046 |
| SVLTVGLGK | ASSPSRYQAPL | 0.84403324 |
| TALAGAGSGI | ASSPSRYQAPL | 0.8623018 |
| TGLPFTGI | ASSPSRYQAPL | 0.83779526 |
| VAEHVTV | ASSPSRYQAPL | 0.8436845 |
| VDIVLNSL | ASSPSRYQAPL | 0.8338957 |
| VLTGGGAVL | ASSPSRYQAPL | 0.8955463 |
| VNVGLAYGI | ASSPSRYQAPL | 0.8400248 |
| VSSAAVGAV | ASSPSRYQAPL | 0.8938302 |
| VTSVYEAGV | ASSPSRYQAPL | 0.77167696 |
| WVALDRGA | ASSPSRYQAPL | 0.778648 |
| YLVGSVVI | ASSPSRYQAPL | 0.89741343 |
| GPPGTGKTLL | ASSPSRYQAPL | 0.17595534 |
| GVGMTKF | ASSPSRYQAPL | 0.629505 |
| IGIGGSDL | ASSPSRYQAPL | 0.84865403 |
| IGTLLAEL | ASSPSRYQAPL | 0.8970228 |
| IYTSGTT | ASSPSRYQAPL | 0.23317672 |
| LSFPAGV | ASSPSRYQAPL | 0.726411 |
| LYTSGTT | ASSPSRYQAPL | 0.76720643 |
| VGAFPFGF | ASSPSRYQAPL | 0.8940258 |
| VGAGTVEF | ASSPSRYQAPL | 0.8346008 |
| VGKSTLLL | ASSPSRYQAPL | 0.6244549 |
| AALAHAGGGIV | ASSPSRYQAPL | 0.9032107 |
| AALVAVSLL | ASSPSRYQAPL | 0.59840035 |
| AAVSRAVAAL | ASSPSRYQAPL | 0.75072354 |
| AGGSSGIGGF | ASSPSRYQAPL | 0.9000132 |
| AGLTVANANG | ASSPSRYQAPL | 0.89898443 |
| AGLVGAVL | ASSPSRYQAPL | 0.80259466 |
| ALSQAFTV | ASSPSRYQAPL | 0.895694 |
| ALTFSVSL | ASSPSRYQAPL | 0.7798403 |
| ALTGAALVAL | ASSPSRYQAPL | 0.9159387 |
| ASFSIGAGVL | ASSPSRYQAPL | 0.8473451 |
| DGSVYKI | ASSPSRYQAPL | 0.76871514 |
| EQLAEIV | ASSPSRYQAPL | 0.8961339 |
| FAVSELTVM | ASSPSRYQAPL | 0.893085 |
| GAAPLAAGL | ASSPSRYQAPL | 0.8736106 |
| GADGGDGGSGGAGG | ASSPSRYQAPL | 0.86839634 |
| GFGLDTGGAGG | ASSPSRYQAPL | 0.8571885 |
| GLASGGLL | ASSPSRYQAPL | 0.82797796 |
| GRIAGAAVLL | ASSPSRYQAPL | 0.8986007 |
| GSAGSLNTSF | ASSPSRYQAPL | 0.8967512 |
| GTGQIGI | ASSPSRYQAPL | 0.7654056 |
| GTLIALIL | ASSPSRYQAPL | 0.9019694 |
| IAATMGFVVG | ASSPSRYQAPL | 0.8070433 |
| IFAQALL | ASSPSRYQAPL | 0.7876042 |
| IGAAVALGF | ASSPSRYQAPL | 0.90881443 |
| IGLTPLAL | ASSPSRYQAPL | 0.876806 |
| ILGSNTSTL | ASSPSRYQAPL | 0.8905741 |
| IVFSLTVL | ASSPSRYQAPL | 0.8968215 |

| | | |
|---|---|---|
| LASKATGF | ASSPSRYQAPL | 0.8768852 |
| LASVALIL | ASSPSRYQAPL | 0.7357603 |
| LFAVGLTL | ASSPSRYQAPL | 0.8348031 |
| LFNGGIGTYI | ASSPSRYQAPL | 0.8329134 |
| LTLGAGII | ASSPSRYQAPL | 0.8957554 |
| LVAAALGDI | ASSPSRYQAPL | 0.8703717 |
| NTAFVQLGI | ASSPSRYQAPL | 0.7494937 |
| QLFSGAEL | ASSPSRYQAPL | 0.89321065 |
| SAATFITL | ASSPSRYQAPL | 0.8980659 |
| SNYGLANLGV | ASSPSRYQAPL | 0.85831493 |
| SYGGAYAVM | ASSPSRYQAPL | 0.8523526 |
| TTGVGGLTLG | ASSPSRYQAPL | 0.91037714 |
| TTQTIYI | ASSPSRYQAPL | 0.83096004 |
| VAAGAGLGAVI | ASSPSRYQAPL | 0.8288492 |
| VAAVVTLL | ASSPSRYQAPL | 0.8924445 |
| VGFLGGLI | ASSPSRYQAPL | 0.89715195 |
| VGVIPILL | ASSPSRYQAPL | 0.8805927 |
| VIFASSLI | ASSPSRYQAPL | 0.8957279 |
| VKGGDGGAAAG | ASSPSRYQAPL | 0.72559184 |
| VTKTFTNI | ASSPSRYQAPL | 0.8815146 |
| VVAAAILL | ASSPSRYQAPL | 0.7305622 |
| YAALDVEL | ASSPSRYQAPL | 0.8554299 |
| LASVALLL | ASSPSRYQAPL | 0.7334864 |
| LDAVHRV | ASSPSRYQAPL | 0.86224985 |
| QGLSSSIF | ASSPSRYQAPL | 0.87874126 |
| RGTFANIRL | ASSPSRYQAPL | 0.87710404 |
| RQLWWGHRI | ASSPSRYQAPL | 0.52481365 |
| YGPPGTGKTLL | ASSPSRYQAPL | 0.87942255 |
| AALVGAPANVA | ASSPSRYQAPL | 0.8732206 |
| AATFRRTE | ASSPSRYQAPL | 0.91229534 |
| AGALGGGI | ASSPSRYQAPL | 0.8430751 |
| AGGAGGAGGIGAG | ASSPSRYQAPL | 0.88491213 |
| AGSGALGL | ASSPSRYQAPL | 0.7962021 |
| AQIADQI | ASSPSRYQAPL | 0.86572015 |
| ASAVAVTYL | ASSPSRYQAPL | 0.6998774 |
| ASTVLGGTGG | ASSPSRYQAPL | 0.78853416 |
| AVGGEPAK | ASSPSRYQAPL | 0.75619453 |
| CEQGTFGGTC | ASSPSRYQAPL | 0.77896667 |
| DEGKAPAG | ASSPSRYQAPL | 0.8680725 |
| DTVVFPAL | ASSPSRYQAPL | 0.8890481 |
| FEGITMAG | ASSPSRYQAPL | 0.8801535 |
| FPLVVQD | ASSPSRYQAPL | 0.88786805 |
| GAFTTPQ | ASSPSRYQAPL | 0.8999026 |
| GFGGFGTAGGVGGA | ASSPSRYQAPL | 0.91179335 |
| GFLAQFL | ASSPSRYQAPL | 0.79859716 |
| GGAGGAGGNAGVA | ASSPSRYQAPL | 0.8162248 |
| GGVGTAGAGGAGGAGG | ASSPSRYQAPL | 0.72028625 |
| GNGGGGDAGNAGSGGN | ASSPSRYQAPL | 0.8232066 |
| GTGGAGGAGNDAGS | ASSPSRYQAPL | 0.8082895 |
| GTTNTGWFNS | ASSPSRYQAPL | 0.8625684 |
| IFMGPLFTL | ASSPSRYQAPL | 0.18087122 |

| | | |
|---|---|---|
| ILAHAIEAI | ASSPSRYQAPL | 0.7931154 |
| IRSAIREAL | ASSPSRYQAPL | 0.90807486 |
| ITFAGGG | ASSPSRYQAPL | 0.7438438 |
| ITGMSGHQEV | ASSPSRYQAPL | 0.86305207 |
| IVAGLACTL | ASSPSRYQAPL | 0.8459015 |
| IVGGGLAGA | ASSPSRYQAPL | 0.7911495 |
| IVTGVLNI | ASSPSRYQAPL | 0.87051535 |
| LAQQGCI | ASSPSRYQAPL | 0.8012136 |
| LCLHVEN | ASSPSRYQAPL | 0.90094143 |
| LDAISDGS | ASSPSRYQAPL | 0.9095957 |
| LGGFAAML | ASSPSRYQAPL | 0.8596823 |
| LGGKNPMI | ASSPSRYQAPL | 0.84384084 |
| LGPKGVAAIL | ASSPSRYQAPL | 0.87539935 |
| LMVGVSI | ASSPSRYQAPL | 0.6855017 |
| LVGAAFVL | ASSPSRYQAPL | 0.8458094 |
| LVLAVAPL | ASSPSRYQAPL | 0.5138916 |
| MAAIAVIDAA | ASSPSRYQAPL | 0.83403194 |
| NNGINAELQ | ASSPSRYQAPL | 0.83575237 |
| RGEPLVLVHG | ASSPSRYQAPL | 0.8681165 |
| RGGAGGAAGLI | ASSPSRYQAPL | 0.83055294 |
| RSYRMHPAV | ASSPSRYQAPL | 0.8467771 |
| SAAHLLAGQLA | ASSPSRYQAPL | 0.7390753 |
| SAAVGSY | ASSPSRYQAPL | 0.84714055 |
| SPHAVLHRI | ASSPSRYQAPL | 0.78522766 |
| SQIAKVL | ASSPSRYQAPL | 0.72302336 |
| TALGHLVP | ASSPSRYQAPL | 0.8701234 |
| TAPTTSGAA | ASSPSRYQAPL | 0.86390585 |
| TDGDDGNGGSPGT | ASSPSRYQAPL | 0.7877772 |
| TDGLGEN | ASSPSRYQAPL | 0.89288616 |
| TGGPPLPGQA | ASSPSRYQAPL | 0.8878561 |
| TIPAITI | ASSPSRYQAPL | 0.6169918 |
| TIPGFTI | ASSPSRYQAPL | 0.7924495 |
| TSIASGALA | ASSPSRYQAPL | 0.85015 |
| TVTLPGAAI | ASSPSRYQAPL | 0.860612 |
| VASGLSAALP | ASSPSRYQAPL | 0.2453756 |
| VGAGVIGI | ASSPSRYQAPL | 0.8785611 |
| VGPKRFGV | ASSPSRYQAPL | 0.7877839 |
| VNVAFPD | ASSPSRYQAPL | 0.6735729 |
| VSMTNSGVS | ASSPSRYQAPL | 0.8133574 |
| VTDVLGDLSV | ASSPSRYQAPL | 0.8935125 |
| VVVVGAGFAGL | ASSPSRYQAPL | 0.7820586 |
| VYTVGTVV | ASSPSRYQAPL | 0.8680444 |
| WTTFSGP | ASSPSRYQAPL | 0.87991166 |
| YDELVEAF | ASSPSRYQAPL | 0.86601144 |
| YGTGGAGGNGG | ASSPSRYQAPL | 0.89956594 |
| YTNVMAV | ASSPSRYQAPL | 0.87714887 |
| FLEQVHQL | ASSPSRYQAPL | 0.84776545 |
| GLSLLAVAV | ASSPSRYQAPL | 0.8777695 |
| LEQVHQL | ASSPSRYQAPL | 0.83082944 |
| LNAAGAI | ASSPSRYQAPL | 0.71351606 |
| PAVHASL | ASSPSRYQAPL | 0.639149 |

| | | |
|---|---|---|
| TELEDDA | ASSPSRYQAPL | 0.8420888 |
| TPPGPPN | ASSPSRYQAPL | 0.6730105 |
| TPPPPPPP | ASSPSRYQAPL | 0.86891973 |
| TQSPALI | ASGDSGQGSERLF | 0.9123507 |
| EFAAAFPQ | ASGDSGQGSERLF | 0.845948 |
| LLSGNLDVL | ASGDSGQGSERLF | 0.73737127 |
| GPANLGNYNI | ASGDSGQGSERLF | 0.8171387 |
| TLGPTLP | ASGDSGQGSERLF | 0.8040006 |
| VYAYSSCCK | ASGDSGQGSERLF | 0.90047854 |
| APAPAPAPNI | ASGDSGQGSERLF | 0.8977562 |
| PGGPVPIP | ASGDSGQGSERLF | 0.9033476 |
| FLTVALEP | ASGDSGQGSERLF | 0.85665905 |
| SQYFGANL | ASGDSGQGSERLF | 0.7724909 |
| GGVKPAA | ASGDSGQGSERLF | 0.8974551 |
| GGTGGAGGAGGW | ASGDSGQGSERLF | 0.832947 |
| GVGGVAGLGGAGG | ASGDSGQGSERLF | 0.81038713 |
| VLLVLATTI | ASGDSGQGSERLF | 0.8682418 |
| IGIFAAML | ASGDSGQGSERLF | 0.854638 |
| AVGDLID | ASGDSGQGSERLF | 0.85618556 |
| NISPLPL | ASGDSGQGSERLF | 0.84894663 |
| VTAVFVAI | ASGDSGQGSERLF | 0.7142769 |
| SGDDNDAAL | ASGDSGQGSERLF | 0.8634358 |
| AVAALAHS | ASGDSGQGSERLF | 0.86085707 |
| GGVTVMATP | ASGDSGQGSERLF | 0.67370474 |
| AGLAANVRD | ASGDSGQGSERLF | 0.81673384 |
| TNRPDLIDPALLRPG | ASGDSGQGSERLF | 0.91030943 |
| LANLGVTAL | ASGDSGQGSERLF | 0.8912802 |
| TGERIEL | ASGDSGQGSERLF | 0.7292776 |
| NGEELAA | ASGDSGQGSERLF | 0.79265445 |
| VLSLVWG | ASGDSGQGSERLF | 0.8053364 |
| IGYALWL | ASGDSGQGSERLF | 0.66562474 |
| LSLLLAPL | ASGDSGQGSERLF | 0.89951515 |
| TLDEALK | ASGDSGQGSERLF | 0.81006193 |
| TDGAGTAEGG | ASGDSGQGSERLF | 0.80099416 |
| NAPGGAGGNGG | ASGDSGQGSERLF | 0.830907 |
| VSVFMGMGGNF | ASGDSGQGSERLF | 0.8559879 |
| LASYLAAL | ASGDSGQGSERLF | 0.76864094 |
| LVGGTPAL | ASGDSGQGSERLF | 0.8953452 |
| VRNADVL | ASGDSGQGSERLF | 0.22760682 |
| TCSSPFLVSAM | ASGDSGQGSERLF | 0.88207316 |
| VHITDAE | ASGDSGQGSERLF | 0.7404469 |
| EYAAPSF | ASGDSGQGSERLF | 0.8003178 |
| VVDAFHGGL | ASGDSGQGSERLF | 0.6154978 |
| ASVVDPQ | ASGDSGQGSERLF | 0.8270282 |
| SYNEFGGLS | ASGDSGQGSERLF | 0.8812293 |
| DPDGSVY | ASGDSGQGSERLF | 0.8928553 |
| MVSTIHATE | ASGDSGQGSERLF | 0.7675736 |
| LAAGVADPV | ASGDSGQGSERLF | 0.8852642 |
| TKEQIAA | ASGDSGQGSERLF | 0.90112245 |
| SLASPAAL | ASGDSGQGSERLF | 0.79814637 |
| IGLVFYGV | ASGDSGQGSERLF | 0.6055007 |

| | | |
|---|---|---|
| FLLDASNL | ASGDSGQGSERLF | 0.8826435 |
| YQVGNFP | ASGDSGQGSERLF | 0.7526298 |
| ESLLSQST | ASGDSGQGSERLF | 0.8913648 |
| LFTVGFAAVG | ASGDSGQGSERLF | 0.8694575 |
| ATPPTPT | ASGDSGQGSERLF | 0.7032734 |
| TDLATLG | ASGDSGQGSERLF | 0.89615905 |
| SAGGTGR | ASGDSGQGSERLF | 0.89167297 |
| GFSNSGS | ASGDSGQGSERLF | 0.7481147 |
| GGNGGSGGTSV | ASGDSGQGSERLF | 0.7864497 |
| GTSVATGGAGN | ASGDSGQGSERLF | 0.7911732 |
| NGGNGGSGGTS | ASGDSGQGSERLF | 0.6716269 |
| AVVKFPH | ASGDSGQGSERLF | 0.8661436 |
| TPPPPPP | ASGDSGQGSERLF | 0.7618326 |
| AAPAAAAAPA | ASGDSGQGSERLF | 0.7833779 |
| IALNEDL | ASGDSGQGSERLF | 0.85399604 |
| IVNVSSM | ASGDSGQGSERLF | 0.80847883 |
| TGIPPAPR | ASGDSGQGSERLF | 0.8244201 |
| FDLGGGTF | ASGDSGQGSERLF | 0.82861066 |
| FDLGGGTFDVS | ASGDSGQGSERLF | 0.84598374 |
| NEPTAAA | ASGDSGQGSERLF | 0.8646922 |
| NEPTAAALA | ASGDSGQGSERLF | 0.80134964 |
| GSIINISSI | ASGDSGQGSERLF | 0.82158756 |
| VAPGNAGTA | ASGDSGQGSERLF | 0.8982035 |
| TGGGASLE | ASGDSGQGSERLF | 0.6820301 |
| VAVYNHYKRI | ASGDSGQGSERLF | 0.74819267 |
| TDPAPATT | ASGDSGQGSERLF | 0.882117 |
| TNRPDILDPALLRPG | ASGDSGQGSERLF | 0.72467285 |
| LTTTTTS | ASGDSGQGSERLF | 0.8905416 |
| PTGSGKTL | ASGDSGQGSERLF | 0.83065295 |
| TGSGKTL | ASGDSGQGSERLF | 0.8646154 |
| TIGHVDHGK | ASGDSGQGSERLF | 0.7969988 |
| TIGHVDHGKTT | ASGDSGQGSERLF | 0.9129595 |
| GHVDHGKTT | ASGDSGQGSERLF | 0.8994993 |
| AQFEHTLL | ASGDSGQGSERLF | 0.84834373 |
| IVNQDEL | ASGDSGQGSERLF | 0.8616996 |
| TNGSQFF | ASGDSGQGSERLF | 0.6995011 |
| TGKGIVE | ASGDSGQGSERLF | 0.8942746 |
| LISNASDA | ASGDSGQGSERLF | 0.89642644 |
| AELRTLGELTKIAKAHGAQ | ASGDSGQGSERLF | 0.70836544 |
| APNAVAAAEPSW | ASGDSGQGSERLF | 0.830474 |
| APQVAGAATAVL | ASGDSGQGSERLF | 0.9000093 |
| ARRDPAAAAAILPTDA | ASGDSGQGSERLF | 0.90287817 |
| DAAARDVLAEAGLAEYF | ASGDSGQGSERLF | 0.725585 |
| DDQIVLSAKANLAKVEQ | ASGDSGQGSERLF | 0.7901755 |
| DELLPDTGEDSDEDGH | ASGDSGQGSERLF | 0.6107989 |
| ELPEAPANRATVPQD | ASGDSGQGSERLF | 0.77064335 |
| FGQSKIERVTGIG | ASGDSGQGSERLF | 0.729234 |
| GAGGTGSTAGGAGGAGGAGGLY | ASGDSGQGSERLF | 0.861861 |
| GHGGTGAAGGNGGTGGAGGSGI | ASGDSGQGSERLF | 0.8257978 |
| GNGGNGGAGGEGGPGGAGGAGA | ASGDSGQGSERLF | 0.86220765 |
| GSTAGNGGNGGNATNGTIAGQP | ASGDSGQGSERLF | 0.8678754 |

| | | |
|---|---|---|
| GTGGDGGNGGNGGTGG | ASGDSGQGSERLF | 0.9019199 |
| GVRRPAGIPADAPV | ASGDSGQGSERLF | 0.8202301 |
| GYRQVTDAHLLTLARR | ASGDSGQGSERLF | 0.83579636 |
| IAKMGQDGHDRG | ASGDSGQGSERLF | 0.75690585 |
| KAEPSIEIQYT | ASGDSGQGSERLF | 0.70972633 |
| KPVRIRMAADEDTVAVT | ASGDSGQGSERLF | 0.8050451 |
| MPPRIAGMRLLVIKPEP | ASGDSGQGSERLF | 0.79276407 |
| MKFYAVALFRSRGVPPTR | ASGDSGQGSERLF | 0.4579426 |
| NAGGFGILGAVAHSP | ASGDSGQGSERLF | 0.8990402 |
| NEGREITLAMAEM | ASGDSGQGSERLF | 0.8747674 |
| NELVEWSHPLGHRWRWE | ASGDSGQGSERLF | 0.8788401 |
| NLGAGGTGGNGGNPG | ASGDSGQGSERLF | 0.78100836 |
| NTEWPKWETVMAADPVD | ASGDSGQGSERLF | 0.8482828 |
| PASAAPAETQP | ASGDSGQGSERLF | 0.7845539 |
| QQSSDGKNGVRAAEITGE | ASGDSGQGSERLF | 0.8860886 |
| RAPDAVTTSWWKE | ASGDSGQGSERLF | 0.8040224 |
| SGFGNVGAHTSGWFNQ | ASGDSGQGSERLF | 0.5668688 |
| SGVAGAVGQAASVGGLKV | ASGDSGQGSERLF | 0.754945 |
| SNGGAGTGSAGGNGGT | ASGDSGQGSERLF | 0.6114727 |
| TGGAGGEGGAAG | ASGDSGQGSERLF | 0.9015801 |
| TGGDGGAAGAGGAPGHG | ASGDSGQGSERLF | 0.8918564 |
| TMPPLLAAIANAGRHGV | ASGDSGQGSERLF | 0.8719976 |
| TSSTAGNSGGA | ASGDSGQGSERLF | 0.90705943 |
| VPATVPVGILC | ASGDSGQGSERLF | 0.39702934 |
| VQRPDSGGVVLTGRIS | ASGDSGQGSERLF | 0.73737884 |
| WTPNPVPATKGVPAGLVQAT | ASGDSGQGSERLF | 0.8985716 |
| YALKRALAKFSDRFDVVII | ASGDSGQGSERLF | 0.86934316 |
| YLIGSVIV | ASGDSGQGSERLF | 0.83773506 |
| YGQLVAVID | ASGDSGQGSERLF | 0.91222286 |
| VTVGGMTLPA | ASGDSGQGSERLF | 0.7565094 |
| VLALPAV | ASGDSGQGSERLF | 0.8568237 |
| VIPPTLNYET | ASGDSGQGSERLF | 0.8675481 |
| VGMGGMHPSSG | ASGDSGQGSERLF | 0.89254427 |
| VGLAGGAATA | ASGDSGQGSERLF | 0.87237096 |
| VGGAGGAGGSGP | ASGDSGQGSERLF | 0.7807481 |
| VFAAFMV | ASGDSGQGSERLF | 0.88454527 |
| TLYDKLR | ASGDSGQGSERLF | 0.9043814 |
| TLAVFTVL | ASGDSGQGSERLF | 0.82240534 |
| TGSVQPARV | ASGDSGQGSERLF | 0.8827416 |
| TGGGAGSGGT | ASGDSGQGSERLF | 0.8851698 |
| SHLETSL | ASGDSGQGSERLF | 0.8320205 |
| SFTTGMT | ASGDSGQGSERLF | 0.790622 |
| QVEVAKRMGVRQPTVS | ASGDSGQGSERLF | 0.77222955 |
| QGSGGPALAM | ASGDSGQGSERLF | 0.84629506 |
| MLAPSNP | ASGDSGQGSERLF | 0.7019522 |
| LVNNAAI | ASGDSGQGSERLF | 0.8688209 |
| LTGEINTLA | ASGDSGQGSERLF | 0.8365427 |
| LSIFGPV | ASGDSGQGSERLF | 0.9178111 |
| LLALLANL | ASGDSGQGSERLF | 0.87294 |
| LISNASDALDKLR | ASGDSGQGSERLF | 0.88748896 |
| LINEAVP | ASGDSGQGSERLF | 0.7169954 |

| | | |
|---|---|---|
| LGYAFSGL | ASGDSGQGSERLF | 0.8602521 |
| LDFVAVAL | ASGDSGQGSERLF | 0.8601859 |
| LAQPGAE | ASGDSGQGSERLF | 0.746986 |
| ISNLGAVTGF | ASGDSGQGSERLF | 0.89573437 |
| HQEDPGRI | ASGDSGQGSERLF | 0.7839079 |
| GTPGEPTP | ASGDSGQGSERLF | 0.80302 |
| GFFNSGS | ASGDSGQGSERLF | 0.8786122 |
| GAGGGGLTAGAN | ASGDSGQGSERLF | 0.8805508 |
| FVDALVNM | ASGDSGQGSERLF | 0.85921 |
| FLTVAPEL | ASGDSGQGSERLF | 0.8215075 |
| FETLTAL | ASGDSGQGSERLF | 0.7685281 |
| DVINPIEL | ASGDSGQGSERLF | 0.8889043 |
| APVSGAPV | ASGDSGQGSERLF | 0.9002681 |
| ALVGGLAW | ASGDSGQGSERLF | 0.8030945 |
| ALAAAAATAI | ASGDSGQGSERLF | 0.908057 |
| AFGLGLLL | ASGDSGQGSERLF | 0.89437664 |
| AFFAAASL | ASGDSGQGSERLF | 0.65307623 |
| AAAALPAFSPP | ASGDSGQGSERLF | 0.8680692 |
| VVLAPAL | ASGDSGQGSERLF | 0.89852977 |
| TIGHVDHGKTTL | ASGDSGQGSERLF | 0.882447 |
| LISNASDALDK | ASGDSGQGSERLF | 0.90740705 |
| LISNASDAL | ASGDSGQGSERLF | 0.8865966 |
| ISNASDALDK | ASGDSGQGSERLF | 0.7955362 |
| NQAETLVYQTE | ASGDSGQGSERLF | 0.8976664 |
| TGGTGGTAGAG | ASGDSGQGSERLF | 0.9116173 |
| TGSTAGGAGGA | ASGDSGQGSERLF | 0.8974496 |
| DPLAVPVAAGPV | ASGDSGQGSERLF | 0.80544055 |
| GGTGGTPGNGGN | ASGDSGQGSERLF | 0.7837305 |
| PAGLAGVDPATL | ASGDSGQGSERLF | 0.8695816 |
| GNNLVVEAPHAGASV | ASGDSGQGSERLF | 0.89523876 |
| KVTRIMSASTLEQPA | ASGDSGQGSERLF | 0.8273104 |
| LGFALPIALPRRTVV | ASGDSGQGSERLF | 0.86653817 |
| TRLGNLTARKLVVLV | ASGDSGQGSERLF | 0.8492735 |
| WSQQRRLIYFPSAGP | ASGDSGQGSERLF | 0.65012693 |
| GPNGNPDPMAAAVDIR | ASGDSGQGSERLF | 0.8676479 |
| TQRCVSVADLRAAASR | ASGDSGQGSERLF | 0.8942048 |
| QFVLDVAEHPVEVLTETV | ASGDSGQGSERLF | 0.88327646 |
| GGAGGTGGAGGNGGTQAGDG | ASGDSGQGSERLF | 0.81695956 |
| PLLATLRAIAVDVDAKPAQV | ASGDSGQGSERLF | 0.8891643 |
| APKFRTGGYMIDFWGVGYQVAK | ASGDSGQGSERLF | 0.91313934 |
| GAGGAGGAGGAAGGPGGTGGTG | ASGDSGQGSERLF | 0.91522074 |
| IGALGLLATDVAPAGSVAFGVA | ASGDSGQGSERLF | 0.6417097 |
| AGGAGGIGGNGGAGGAGGVLMG | ASGDSGQGSERLF | 0.45068982 |
| ADELIGLL | ASGDSGQGSERLF | 0.8229703 |
| ADRALKA | ASGDSGQGSERLF | 0.85903233 |
| AGAVGQVGGAGGA | ASGDSGQGSERLF | 0.83193517 |
| AGGGGGLGGAP | ASGDSGQGSERLF | 0.8339005 |
| AGGLLYGNGGAG | ASGDSGQGSERLF | 0.8608899 |
| AGLESIGL | ASGDSGQGSERLF | 0.89421415 |
| AGLKTGII | ASGDSGQGSERLF | 0.8178586 |
| ASLAALLL | ASGDSGQGSERLF | 0.716466 |

| | | |
|---|---|---|
| ASVINISSI | ASGDSGQGSERLF | 0.87249625 |
| ATPPGPA | ASGDSGQGSERLF | 0.89755356 |
| ATVASALV | ASGDSGQGSERLF | 0.722081 |
| ATVASLIL | ASGDSGQGSERLF | 0.8327588 |
| AVNGPVLIRH | ASGDSGQGSERLF | 0.8147348 |
| DGAAGAVGI | ASGDSGQGSERLF | 0.77776366 |
| DPLLHGI | ASGDSGQGSERLF | 0.8272269 |
| ELGAALGAL | ASGDSGQGSERLF | 0.73868144 |
| ELGRTQAV | ASGDSGQGSERLF | 0.8883704 |
| FVAGGYLS | ASGDSGQGSERLF | 0.74594915 |
| GASGGAGGAGGW | ASGDSGQGSERLF | 0.8798897 |
| GGAGAGGDGGAGGI | ASGDSGQGSERLF | 0.8711128 |
| GTGHNGSQPGMGGNG | ASGDSGQGSERLF | 0.87298983 |
| GYAAASASAS | ASGDSGQGSERLF | 0.8357489 |
| HQVDVAAVQQG | ASGDSGQGSERLF | 0.7820033 |
| IGAAIAHT | ASGDSGQGSERLF | 0.7570506 |
| IGMIEVAQL | ASGDSGQGSERLF | 0.5850639 |
| IRLAPEHTSAMD | ASGDSGQGSERLF | 0.9001312 |
| IYNVSVL | ASGDSGQGSERLF | 0.7601156 |
| KQGFLGM | ASGDSGQGSERLF | 0.90818083 |
| LESVITQL | ASGDSGQGSERLF | 0.89042246 |
| LGLGALVVV | ASGDSGQGSERLF | 0.7830081 |
| LIGNGGNGGSGGI | ASGDSGQGSERLF | 0.87971437 |
| LNAINAPTL | ASGDSGQGSERLF | 0.76004946 |
| LQLLNTGV | ASGDSGQGSERLF | 0.8102256 |
| LQTSIEF | ASGDSGQGSERLF | 0.86434186 |
| MSFAVSEL | ASGDSGQGSERLF | 0.89540184 |
| NTGALGL | ASGDSGQGSERLF | 0.791127 |
| PPGAGGAGGAGTT | ASGDSGQGSERLF | 0.8958281 |
| QLVGLGI | ASGDSGQGSERLF | 0.8542808 |
| RFLEQVHQL | ASGDSGQGSERLF | 0.7533722 |
| RGVGYVLR | ASGDSGQGSERLF | 0.8824355 |
| SATGVFAGVI | ASGDSGQGSERLF | 0.89450216 |
| SLPAITLP | ASGDSGQGSERLF | 0.89525783 |
| STALAFTV | ASGDSGQGSERLF | 0.8169104 |
| SVLTVGLGK | ASGDSGQGSERLF | 0.8640568 |
| TALAGAGSGI | ASGDSGQGSERLF | 0.75344765 |
| TGLPFTGI | ASGDSGQGSERLF | 0.85028267 |
| VAEHVTV | ASGDSGQGSERLF | 0.8636831 |
| VDIVLNSL | ASGDSGQGSERLF | 0.8242382 |
| VLTGGGAVL | ASGDSGQGSERLF | 0.8957188 |
| VNVGLAYGI | ASGDSGQGSERLF | 0.63839734 |
| VSSAAVGAV | ASGDSGQGSERLF | 0.74754953 |
| VTSVYEAGV | ASGDSGQGSERLF | 0.8254521 |
| WVALDRGA | ASGDSGQGSERLF | 0.86251235 |
| YLVGSVVI | ASGDSGQGSERLF | 0.8950435 |
| GPPGTGKTLL | ASGDSGQGSERLF | 0.7334025 |
| GVGMTKF | ASGDSGQGSERLF | 0.8407147 |
| IGIGGSDL | ASGDSGQGSERLF | 0.8435881 |
| IGTLLAEL | ASGDSGQGSERLF | 0.80022264 |
| IYTSGTT | ASGDSGQGSERLF | 0.8960495 |

| | | |
|---|---|---|
| LSFPAGV | ASGDSGQGSERLF | 0.84214103 |
| LYTSGTT | ASGDSGQGSERLF | 0.7965019 |
| VGAFPFGF | ASGDSGQGSERLF | 0.881801 |
| VGAGTVEF | ASGDSGQGSERLF | 0.8399221 |
| VGKSTLLL | ASGDSGQGSERLF | 0.8696877 |
| AALAHAGGGIV | ASGDSGQGSERLF | 0.8942939 |
| AALVAVSLL | ASGDSGQGSERLF | 0.8001982 |
| AAVSRAVAAL | ASGDSGQGSERLF | 0.88357645 |
| AGGSSGIGGF | ASGDSGQGSERLF | 0.8704381 |
| AGLTVANANG | ASGDSGQGSERLF | 0.88365054 |
| AGLVGAVL | ASGDSGQGSERLF | 0.8295508 |
| ALSQAFTV | ASGDSGQGSERLF | 0.78777266 |
| ALTFSVSL | ASGDSGQGSERLF | 0.90468323 |
| ALTGAALVAL | ASGDSGQGSERLF | 0.8760036 |
| ASFSIGAGVL | ASGDSGQGSERLF | 0.87089396 |
| DGSVYKI | ASGDSGQGSERLF | 0.8910859 |
| EQLAEIV | ASGDSGQGSERLF | 0.86564296 |
| FAVSELTVM | ASGDSGQGSERLF | 0.88890374 |
| GAAPLAAGL | ASGDSGQGSERLF | 0.75764 |
| GADGGDGGSGGAGG | ASGDSGQGSERLF | 0.74992204 |
| GFGLDTGGAGG | ASGDSGQGSERLF | 0.8255785 |
| GLASGGLL | ASGDSGQGSERLF | 0.81327534 |
| GRIAGAAVLL | ASGDSGQGSERLF | 0.84384084 |
| GSAGSLNTSF | ASGDSGQGSERLF | 0.63872606 |
| GTGQIGI | ASGDSGQGSERLF | 0.88951147 |
| GTLIALIL | ASGDSGQGSERLF | 0.9177654 |
| IAATMGFVVG | ASGDSGQGSERLF | 0.7596123 |
| IFAQALL | ASGDSGQGSERLF | 0.9159893 |
| IGAAVALGF | ASGDSGQGSERLF | 0.74416703 |
| IGLTPLAL | ASGDSGQGSERLF | 0.8778337 |
| ILGSNTSTL | ASGDSGQGSERLF | 0.89557827 |
| IVFSLTVL | ASGDSGQGSERLF | 0.730397 |
| LASKATGF | ASGDSGQGSERLF | 0.8610184 |
| LASVALIL | ASGDSGQGSERLF | 0.81351733 |
| LFAVGLTL | ASGDSGQGSERLF | 0.8486978 |
| LFNGGIGTYI | ASGDSGQGSERLF | 0.9067868 |
| LTLGAGII | ASGDSGQGSERLF | 0.8098041 |
| LVAAALGDI | ASGDSGQGSERLF | 0.8706627 |
| NTAFVQLGI | ASGDSGQGSERLF | 0.86782426 |
| QLFSGAEL | ASGDSGQGSERLF | 0.81032753 |
| SAATFITL | ASGDSGQGSERLF | 0.88992643 |
| SNYGLANLGV | ASGDSGQGSERLF | 0.89573646 |
| SYGGAYAVM | ASGDSGQGSERLF | 0.78850234 |
| TTGVGGLTLG | ASGDSGQGSERLF | 0.90723026 |
| TTQTIYI | ASGDSGQGSERLF | 0.77698684 |
| VAAGAGLGAVI | ASGDSGQGSERLF | 0.90746903 |
| VAAVVTLL | ASGDSGQGSERLF | 0.8849827 |
| VGFLGGLI | ASGDSGQGSERLF | 0.8296727 |
| VGVIPILL | ASGDSGQGSERLF | 0.83543813 |
| VIFASSLI | ASGDSGQGSERLF | 0.89033675 |
| VKGGDGGAAAG | ASGDSGQGSERLF | 0.84724665 |

| | | |
|---|---|---|
| VTKTFTNI | ASGDSGQGSERLF | 0.8607898 |
| VVAAAILL | ASGDSGQGSERLF | 0.9128115 |
| YAALDVEL | ASGDSGQGSERLF | 0.41208103 |
| LASVALLL | ASGDSGQGSERLF | 0.6334521 |
| LDAVHRV | ASGDSGQGSERLF | 0.5863331 |
| QGLSSSIF | ASGDSGQGSERLF | 0.5094861 |
| RGTFANIRL | ASGDSGQGSERLF | 0.4353265 |
| RQLWWGHRI | ASGDSGQGSERLF | 0.36266258 |
| YGPPGTGKTLL | ASGDSGQGSERLF | 0.5164179 |
| AALVGAPANVA | ASGDSGQGSERLF | 0.4216334 |
| AATFRRTE | ASGDSGQGSERLF | 0.6208551 |
| AGALGGGI | ASGDSGQGSERLF | 0.5951294 |
| AGGAGGAGGIGAG | ASGDSGQGSERLF | 0.42132387 |
| AGSGALGL | ASGDSGQGSERLF | 0.64650077 |
| AQIADQI | ASGDSGQGSERLF | 0.6130842 |
| ASAVAVTYL | ASGDSGQGSERLF | 0.38870776 |
| ASTVLGGTGG | ASGDSGQGSERLF | 0.4907815 |
| AVGGEPAK | ASGDSGQGSERLF | 0.42618132 |
| CEQGTFGGTC | ASGDSGQGSERLF | 0.47794706 |
| DEGKAPAG | ASGDSGQGSERLF | 0.35357666 |
| DTVVFPAL | ASGDSGQGSERLF | 0.5588028 |
| FEGITMAG | ASGDSGQGSERLF | 0.4797225 |
| FPLVVQD | ASGDSGQGSERLF | 0.3293018 |
| GAFTTPQ | ASGDSGQGSERLF | 0.60475343 |
| GFGGFGTAGGVGGA | ASGDSGQGSERLF | 0.44998562 |
| GFLAQFL | ASGDSGQGSERLF | 0.49434984 |
| GGAGGAGGNAGVA | ASGDSGQGSERLF | 0.61973447 |
| GGVGTAGAGGAGGAGG | ASGDSGQGSERLF | 0.5042715 |
| GNGGGGDAGNAGSGGN | ASGDSGQGSERLF | 0.45184666 |
| GTGGAGGAGNDAGS | ASGDSGQGSERLF | 0.6182564 |
| GTTNTGWFNS | ASGDSGQGSERLF | 0.6404686 |
| IFMGPLFTL | ASGDSGQGSERLF | 0.42043084 |
| ILAHAIEAI | ASGDSGQGSERLF | 0.53958476 |
| IRSAIREAL | ASGDSGQGSERLF | 0.6426009 |
| ITFAGGG | ASGDSGQGSERLF | 0.6085454 |
| ITGMSGHQEV | ASGDSGQGSERLF | 0.62812203 |
| IVAGLACTL | ASGDSGQGSERLF | 0.63831145 |
| IVGGGLAGA | ASGDSGQGSERLF | 0.42893767 |
| IVTGVLNI | ASGDSGQGSERLF | 0.62524945 |
| LAQQGCI | ASGDSGQGSERLF | 0.6436765 |
| LCLHVEN | ASGDSGQGSERLF | 0.5335776 |
| LDAISDGS | ASGDSGQGSERLF | 0.54254794 |
| LGGFAAML | ASGDSGQGSERLF | 0.44517958 |
| LGGKNPMI | ASGDSGQGSERLF | 0.5601697 |
| LGPKGVAAIL | ASGDSGQGSERLF | 0.36211568 |
| LMVGVSI | ASGDSGQGSERLF | 0.58630925 |
| LVGAAFVL | ASGDSGQGSERLF | 0.63517576 |
| LVLAVAPL | ASGDSGQGSERLF | 0.53220177 |
| MAAIAVIDAA | ASGDSGQGSERLF | 0.3993112 |
| NNGINAELQ | ASGDSGQGSERLF | 0.3590065 |
| RGEPLVLVHG | ASGDSGQGSERLF | 0.5558214 |

| | | |
|---|---|---|
| RGGAGGAAGLI | ASGDSGQGSERLF | 0.40204185 |
| RSYRMHPAV | ASGDSGQGSERLF | 0.3744945 |
| SAAHLLAGQLA | ASGDSGQGSERLF | 0.50219595 |
| SAAVGSY | ASGDSGQGSERLF | 0.6387737 |
| SPHAVLHRI | ASGDSGQGSERLF | 0.62621117 |
| SQIAKVL | ASGDSGQGSERLF | 0.4859468 |
| TALGHLVP | ASGDSGQGSERLF | 0.63842887 |
| TAPTTSGAA | ASGDSGQGSERLF | 0.61091244 |
| TDGDDGNGGSPGT | ASGDSGQGSERLF | 0.59792536 |
| TDGLGEN | ASGDSGQGSERLF | 0.4503407 |
| TGGPPLPGQA | ASGDSGQGSERLF | 0.43808508 |
| TIPAITI | ASGDSGQGSERLF | 0.6193226 |
| TIPGFTI | ASGDSGQGSERLF | 0.56385374 |
| TSIASGALA | ASGDSGQGSERLF | 0.5920763 |
| TVTLPGAAI | ASGDSGQGSERLF | 0.619554 |
| VASGLSAALP | ASGDSGQGSERLF | 0.565493 |
| VGAGVIGI | ASGDSGQGSERLF | 0.508112 |
| VGPKRFGV | ASGDSGQGSERLF | 0.5601369 |
| VNVAFPD | ASGDSGQGSERLF | 0.6131796 |
| VSMTNSGVS | ASGDSGQGSERLF | 0.47227636 |
| VTDVLGDLSV | ASGDSGQGSERLF | 0.39666846 |
| VVVVGAGFAGL | ASGDSGQGSERLF | 0.47545612 |
| VYTVGTVV | ASGDSGQGSERLF | 0.54849654 |
| WTTFSGP | ASGDSGQGSERLF | 0.6356553 |
| YDELVEAF | ASGDSGQGSERLF | 0.54859924 |
| YGTGGAGGNGG | ASGDSGQGSERLF | 0.53797925 |
| YTNVMAV | ASGDSGQGSERLF | 0.5795584 |
| FLEQVHQL | ASGDSGQGSERLF | 0.58240557 |
| GLSLLAVAV | ASGDSGQGSERLF | 0.5396174 |
| LEQVHQL | ASGDSGQGSERLF | 0.63123643 |
| LNAAGAI | ASGDSGQGSERLF | 0.5147785 |
| PAVHASL | ASGDSGQGSERLF | 0.610514 |
| TELEDDA | ASGDSGQGSERLF | 0.58694094 |
| TPPGPPN | ASGDSGQGSERLF | 0.590534 |
| TPPPPPPP | ASGDSGQGSERLF | 0.62593156 |
| TQSPALI | AWSPTGTNTGQLY | 0.5976795 |
| EFAAAFPQ | AWSPTGTNTGQLY | 0.48906654 |
| LLSGNLDVL | AWSPTGTNTGQLY | 0.40314358 |
| GPANLGNYNI | AWSPTGTNTGQLY | 0.43596935 |
| TLGPTLP | AWSPTGTNTGQLY | 0.6052843 |
| VYAYSSCCK | AWSPTGTNTGQLY | 0.57373154 |
| APAPAPAPNI | AWSPTGTNTGQLY | 0.6246178 |
| PGGPVPIP | AWSPTGTNTGQLY | 0.5870921 |
| FLTVALEP | AWSPTGTNTGQLY | 0.5002392 |
| SQYFGANL | AWSPTGTNTGQLY | 0.63773537 |
| GGVKPAA | AWSPTGTNTGQLY | 0.6278471 |
| GGTGGAGGAGGW | AWSPTGTNTGQLY | 0.36172435 |
| GVGGVAGLGGAGG | AWSPTGTNTGQLY | 0.47411796 |
| VLLVLATTI | AWSPTGTNTGQLY | 0.6436801 |
| IGIFAAML | AWSPTGTNTGQLY | 0.59929425 |
| AVGDLID | AWSPTGTNTGQLY | 0.5054538 |

| | | |
|---|---|---|
| NISPLPL | AWSPTGTNTGQLY | 0.4612168 |
| VTAVFVAI | AWSPTGTNTGQLY | 0.6395305 |
| SGDDNDAAL | AWSPTGTNTGQLY | 0.52214926 |
| AVAALAHS | AWSPTGTNTGQLY | 0.64356375 |
| GGVTVMATP | AWSPTGTNTGQLY | 0.34008536 |
| AGLAANVRD | AWSPTGTNTGQLY | 0.37921387 |
| TNRPDLIDPALLRPG | AWSPTGTNTGQLY | 0.4268195 |
| LANLGVTAL | AWSPTGTNTGQLY | 0.32774535 |
| TGERIEL | AWSPTGTNTGQLY | 0.61971295 |
| NGEELAA | AWSPTGTNTGQLY | 0.44515517 |
| VLSLVWG | AWSPTGTNTGQLY | 0.6060015 |
| IGYALWL | AWSPTGTNTGQLY | 0.5111207 |
| LSLLLAPL | AWSPTGTNTGQLY | 0.61969376 |
| TLDEALK | AWSPTGTNTGQLY | 0.5605135 |
| TDGAGTAEGG | AWSPTGTNTGQLY | 0.5669535 |
| NAPGGAGGNGG | AWSPTGTNTGQLY | 0.44502342 |
| VSVFMGMGGNF | AWSPTGTNTGQLY | 0.6086898 |
| LASYLAAL | AWSPTGTNTGQLY | 0.6427901 |
| LVGGTPAL | AWSPTGTNTGQLY | 0.57827663 |
| VRNADVL | AWSPTGTNTGQLY | 0.37738508 |
| TCSSPFLVSAM | AWSPTGTNTGQLY | 0.59796035 |
| VHITDAE | AWSPTGTNTGQLY | 0.5500008 |
| EYAAPSF | AWSPTGTNTGQLY | 0.37309596 |
| VVDAFHGGL | AWSPTGTNTGQLY | 0.35998282 |
| ASVVDPQ | AWSPTGTNTGQLY | 0.524078 |
| SYNEFGGLS | AWSPTGTNTGQLY | 0.5380197 |
| DPDGSVY | AWSPTGTNTGQLY | 0.61547816 |
| MVSTIHATE | AWSPTGTNTGQLY | 0.6279851 |
| LAAGVADPV | AWSPTGTNTGQLY | 0.44850925 |
| TKEQIAA | AWSPTGTNTGQLY | 0.61584264 |
| SLASPAAL | AWSPTGTNTGQLY | 0.6149443 |
| IGLVFYGV | AWSPTGTNTGQLY | 0.3004843 |
| FLLDASNL | AWSPTGTNTGQLY | 0.4510953 |
| YQVGNFP | AWSPTGTNTGQLY | 0.39459822 |
| ESLLSQST | AWSPTGTNTGQLY | 0.62141705 |
| LFTVGFAAVG | AWSPTGTNTGQLY | 0.38608152 |
| ATPPTPT | AWSPTGTNTGQLY | 0.52060807 |
| TDLATLG | AWSPTGTNTGQLY | 0.52017874 |
| SAGGTGR | AWSPTGTNTGQLY | 0.59123456 |
| GFSNSGS | AWSPTGTNTGQLY | 0.5399914 |
| GGNGGSGGTSV | AWSPTGTNTGQLY | 0.43343294 |
| GTSVATGGAGN | AWSPTGTNTGQLY | 0.363288 |
| NGGNGGSGGTS | AWSPTGTNTGQLY | 0.6312384 |
| AVVKFPH | AWSPTGTNTGQLY | 0.5003759 |
| TPPPPPP | AWSPTGTNTGQLY | 0.43204856 |
| AAPAAAAAPA | AWSPTGTNTGQLY | 0.4522168 |
| IALNEDL | AWSPTGTNTGQLY | 0.61756355 |
| IVNVSSM | AWSPTGTNTGQLY | 0.3356869 |
| TGIPPAPR | AWSPTGTNTGQLY | 0.6205675 |
| FDLGGGTF | AWSPTGTNTGQLY | 0.48087293 |
| FDLGGGTFDVS | AWSPTGTNTGQLY | 0.56828964 |

| | | |
|---|---|---|
| NEPTAAA | AWSPTGTNTGQLY | 0.6501694 |
| NEPTAAALA | AWSPTGTNTGQLY | 0.59205955 |
| GSIINISSI | AWSPTGTNTGQLY | 0.53881025 |
| VAPGNAGTA | AWSPTGTNTGQLY | 0.5114982 |
| TGGGASLE | AWSPTGTNTGQLY | 0.37314898 |
| VAVYNHYKRI | AWSPTGTNTGQLY | 0.47313792 |
| TDPAPATT | AWSPTGTNTGQLY | 0.46670395 |
| TNRPDILDPALLRPG | AWSPTGTNTGQLY | 0.6470237 |
| LTTTTTS | AWSPTGTNTGQLY | 0.43157846 |
| PTGSGKTL | AWSPTGTNTGQLY | 0.39190394 |
| TGSGKTL | AWSPTGTNTGQLY | 0.43405008 |
| TIGHVDHGK | AWSPTGTNTGQLY | 0.6345506 |
| TIGHVDHGKTT | AWSPTGTNTGQLY | 0.6473561 |
| GHVDHGKTT | AWSPTGTNTGQLY | 0.5114606 |
| AQFEHTLL | AWSPTGTNTGQLY | 0.47300825 |
| IVNQDEL | AWSPTGTNTGQLY | 0.43821824 |
| TNGSQFF | AWSPTGTNTGQLY | 0.56364435 |
| TGKGIVE | AWSPTGTNTGQLY | 0.48450404 |
| LISNASDA | AWSPTGTNTGQLY | 0.4340948 |
| AELRTLGELTKIAKAHGAQ | AWSPTGTNTGQLY | 0.5909859 |
| APNAVAAAEPSW | AWSPTGTNTGQLY | 0.65243673 |
| APQVAGAATAVL | AWSPTGTNTGQLY | 0.4770592 |
| ARRDPAAAAAILPTDA | AWSPTGTNTGQLY | 0.34171492 |
| DAAARDVLAEAGLAEYF | AWSPTGTNTGQLY | 0.5000851 |
| DDQIVLSAKANLAKVEQ | AWSPTGTNTGQLY | 0.44800943 |
| DELLPDTGEDSDEDGH | AWSPTGTNTGQLY | 0.4363693 |
| ELPEAPANRATVPQD | AWSPTGTNTGQLY | 0.5668931 |
| FGQSKIERVTGIG | AWSPTGTNTGQLY | 0.34584045 |
| GAGGTGSTAGGAGGAGGAGGLY | AWSPTGTNTGQLY | 0.47641763 |
| GHGGTGAAGGNGGTGGAGGSGI | AWSPTGTNTGQLY | 0.6504084 |
| GNGGNGGAGGEGGPGGAGGAGA | AWSPTGTNTGQLY | 0.501176 |
| GSTAGNGGNGGNATNGTIAGQP | AWSPTGTNTGQLY | 0.5336128 |
| GTGGDGGNGGNGGTGG | AWSPTGTNTGQLY | 0.63178384 |
| GVRRPAGIPADAPV | AWSPTGTNTGQLY | 0.58030164 |
| GYRQVTDAHLLTLARR | AWSPTGTNTGQLY | 0.6268118 |
| IAKMGQDGHDRG | AWSPTGTNTGQLY | 0.6266285 |
| KAEPSIEIQYT | AWSPTGTNTGQLY | 0.54294187 |
| KPVRIRMAADEDTVAVT | AWSPTGTNTGQLY | 0.55780137 |
| MPPRIAGMRLLVIKPEP | AWSPTGTNTGQLY | 0.60701925 |
| MKFYAVALFRSRGVPPTR | AWSPTGTNTGQLY | 0.56555796 |
| NAGGFGILGAVAHSP | AWSPTGTNTGQLY | 0.5135143 |
| NEGREITLAMAEM | AWSPTGTNTGQLY | 0.46510428 |
| NELVEWSHPLGHRWRWE | AWSPTGTNTGQLY | 0.60163194 |
| NLGAGGTGGNGGNPG | AWSPTGTNTGQLY | 0.47314388 |
| NTEWPKWETVMAADPVD | AWSPTGTNTGQLY | 0.60992026 |
| PASAAPAETQP | AWSPTGTNTGQLY | 0.33857813 |
| QQSSDGKNGVRAAEITGE | AWSPTGTNTGQLY | 0.42062137 |
| RAPDAVTTSWWKE | AWSPTGTNTGQLY | 0.6110408 |
| SGFGNVGAHTSGWFNQ | AWSPTGTNTGQLY | 0.32809576 |
| SGVAGAVGQAASVGGLKV | AWSPTGTNTGQLY | 0.32902467 |
| SNGGAGTGSAGGNGGT | AWSPTGTNTGQLY | 0.5337223 |

| | | |
|---|---|---|
| TGGAGGEGGAAG | AWSPTGTNTGQLY | 0.6245862 |
| TGGDGGAAGAGGAPGHG | AWSPTGTNTGQLY | 0.34417874 |
| TMPPLLAAIANAGRHGV | AWSPTGTNTGQLY | 0.44439667 |
| TSSTAGNSGGA | AWSPTGTNTGQLY | 0.3792546 |
| VPATVPVGILC | AWSPTGTNTGQLY | 0.59548527 |
| VQRPDSGGVVLTGRIS | AWSPTGTNTGQLY | 0.5793054 |
| WTPNPVPATKGVPAGLVQAT | AWSPTGTNTGQLY | 0.32465595 |
| YALKRALAKFSDRFDVVII | AWSPTGTNTGQLY | 0.5625365 |
| YLIGSVIV | AWSPTGTNTGQLY | 0.35180104 |
| YGQLVAVID | AWSPTGTNTGQLY | 0.58786756 |
| VTVGGMTLPA | AWSPTGTNTGQLY | 0.6163461 |
| VLALPAV | AWSPTGTNTGQLY | 0.6456229 |
| VIPPTLNYET | AWSPTGTNTGQLY | 0.3644942 |
| VGMGGMHPSSG | AWSPTGTNTGQLY | 0.60711515 |
| VGLAGGAATA | AWSPTGTNTGQLY | 0.45247963 |
| VGGAGGAGGSGP | AWSPTGTNTGQLY | 0.65316105 |
| VFAAFMV | AWSPTGTNTGQLY | 0.4909994 |
| TLYDKLR | AWSPTGTNTGQLY | 0.3407835 |
| TLAVFTVL | AWSPTGTNTGQLY | 0.5948383 |
| TGSVQPARV | AWSPTGTNTGQLY | 0.5571385 |
| TGGGAGSGGT | AWSPTGTNTGQLY | 0.56961936 |
| SHLETSL | AWSPTGTNTGQLY | 0.6031627 |
| SFTTGMT | AWSPTGTNTGQLY | 0.40071473 |
| QVEVAKRMGVRQPTVS | AWSPTGTNTGQLY | 0.4477773 |
| QGSGGPALAM | AWSPTGTNTGQLY | 0.622236 |
| MLAPSNP | AWSPTGTNTGQLY | 0.6108618 |
| LVNNAAI | AWSPTGTNTGQLY | 0.4767225 |
| LTGEINTLA | AWSPTGTNTGQLY | 0.6353708 |
| LSIFGPV | AWSPTGTNTGQLY | 0.5803765 |
| LLALLANL | AWSPTGTNTGQLY | 0.471967 |
| LISNASDALDKLR | AWSPTGTNTGQLY | 0.64166063 |
| LINEAVP | AWSPTGTNTGQLY | 0.5362764 |
| LGYAFSGL | AWSPTGTNTGQLY | 0.6135228 |
| LDFVAVAL | AWSPTGTNTGQLY | 0.55887514 |
| LAQPGAE | AWSPTGTNTGQLY | 0.6400603 |
| ISNLGAVTGF | AWSPTGTNTGQLY | 0.38637012 |
| HQEDPGRI | AWSPTGTNTGQLY | 0.5994835 |
| GTPGEPTP | AWSPTGTNTGQLY | 0.5504443 |
| GFFNSGS | AWSPTGTNTGQLY | 0.6159538 |
| GAGGGGLTAGAN | AWSPTGTNTGQLY | 0.5131484 |
| FVDALVNM | AWSPTGTNTGQLY | 0.6000137 |
| FLTVAPEL | AWSPTGTNTGQLY | 0.5409328 |
| FETLTAL | AWSPTGTNTGQLY | 0.6424203 |
| DVINPIEL | AWSPTGTNTGQLY | 0.54721475 |
| APVSGAPV | AWSPTGTNTGQLY | 0.6029787 |
| ALVGGLAW | AWSPTGTNTGQLY | 0.6377865 |
| ALAAAAATAI | AWSPTGTNTGQLY | 0.4219861 |
| AFGLGLLL | AWSPTGTNTGQLY | 0.57637954 |
| AFFAAASL | AWSPTGTNTGQLY | 0.6124746 |
| AAAALPAFSPP | AWSPTGTNTGQLY | 0.64693683 |
| VVLAPAL | AWSPTGTNTGQLY | 0.42736188 |

| | | |
|---|---|---|
| TIGHVDHGKTTL | AWSPTGTNTGQLY | 0.638704 |
| LISNASDALDK | AWSPTGTNTGQLY | 0.4749558 |
| LISNASDAL | AWSPTGTNTGQLY | 0.6234962 |
| ISNASDALDK | AWSPTGTNTGQLY | 0.5319433 |
| NQAETLVYQTE | AWSPTGTNTGQLY | 0.5668658 |
| TGGTGGTAGAG | AWSPTGTNTGQLY | 0.5118375 |
| TGSTAGGAGGA | AWSPTGTNTGQLY | 0.61758345 |
| DPLAVPVAAGPV | AWSPTGTNTGQLY | 0.62196636 |
| GGTGGTPGNGGN | AWSPTGTNTGQLY | 0.62472 |
| PAGLAGVDPATL | AWSPTGTNTGQLY | 0.364059 |
| GNNLVVEAPHAGASV | AWSPTGTNTGQLY | 0.61270154 |
| KVTRIMSASTLEQPA | AWSPTGTNTGQLY | 0.62968135 |
| LGFALPIALPRRTVV | AWSPTGTNTGQLY | 0.64473736 |
| TRLGNLTARKLVVLV | AWSPTGTNTGQLY | 0.5456015 |
| WSQQRRLIYFPSAGP | AWSPTGTNTGQLY | 0.5533499 |
| GPNGNPDPMAAAVDIR | AWSPTGTNTGQLY | 0.5806453 |
| TQRCVSVADLRAAASR | AWSPTGTNTGQLY | 0.4694381 |
| QFVLDVAEHPVEVLTETV | AWSPTGTNTGQLY | 0.3973359 |
| GGAGGTGGAGGNGGTQAGDG | AWSPTGTNTGQLY | 0.5645995 |
| PLLATLRAIAVDVDAKPAQV | AWSPTGTNTGQLY | 0.5450528 |
| APKFRTGGYMIDFWGVGYQVAK | AWSPTGTNTGQLY | 0.483164 |
| GAGGAGGAGGAAGGPGGTGGTG | AWSPTGTNTGQLY | 0.6150476 |
| IGALGLLATDVAPAGSVAFGVA | AWSPTGTNTGQLY | 0.6486642 |
| AGGAGGIGGNGGAGGAGGVLMG | AWSPTGTNTGQLY | 0.6403743 |
| ADELIGLL | AWSPTGTNTGQLY | 0.61150026 |
| ADRALKA | AWSPTGTNTGQLY | 0.635966 |
| AGAVGQVGGAGGA | AWSPTGTNTGQLY | 0.5872341 |
| AGGGGGLGGAP | AWSPTGTNTGQLY | 0.48279622 |
| AGGLLYGNGGAG | AWSPTGTNTGQLY | 0.5020823 |
| AGLESIGL | AWSPTGTNTGQLY | 0.40966126 |
| AGLKTGII | AWSPTGTNTGQLY | 0.43112487 |
| ASLAALLL | AWSPTGTNTGQLY | 0.4726219 |
| ASVINISSI | AWSPTGTNTGQLY | 0.53362274 |
| ATPPGPA | AWSPTGTNTGQLY | 0.31249532 |
| ATVASALV | AWSPTGTNTGQLY | 0.5834805 |
| ATVASLIL | AWSPTGTNTGQLY | 0.6494447 |
| AVNGPVLIRH | AWSPTGTNTGQLY | 0.58712566 |
| DGAAGAVGI | AWSPTGTNTGQLY | 0.5548984 |
| DPLLHGI | AWSPTGTNTGQLY | 0.57798815 |
| ELGAALGAL | AWSPTGTNTGQLY | 0.47839868 |
| ELGRTQAV | AWSPTGTNTGQLY | 0.5892904 |
| FVAGGYLS | AWSPTGTNTGQLY | 0.4568843 |
| GASGGAGGAGGW | AWSPTGTNTGQLY | 0.5842688 |
| GGAGAGGDGGAGGI | AWSPTGTNTGQLY | 0.6341346 |
| GTGHNGSQPGMGGNG | AWSPTGTNTGQLY | 0.54441977 |
| GYAAASASAS | AWSPTGTNTGQLY | 0.5885213 |
| HQVDVAAVQQG | AWSPTGTNTGQLY | 0.48049325 |
| IGAAIAHT | AWSPTGTNTGQLY | 0.36421847 |
| IGMIEVAQL | AWSPTGTNTGQLY | 0.6052741 |
| IRLAPEHTSAMD | AWSPTGTNTGQLY | 0.522735 |
| IYNVSVL | AWSPTGTNTGQLY | 0.54076666 |

| | | |
|---|---|---|
| KQGFLGM | AWSPTGTNTGQLY | 0.38407084 |
| LESVITQL | AWSPTGTNTGQLY | 0.5750231 |
| LGLGALVVV | AWSPTGTNTGQLY | 0.6013013 |
| LIGNGGNGGSGGI | AWSPTGTNTGQLY | 0.5884521 |
| LNAINAPTL | AWSPTGTNTGQLY | 0.41055158 |
| LQLLNTGV | AWSPTGTNTGQLY | 0.5792645 |
| LQTSIEF | AWSPTGTNTGQLY | 0.5891702 |
| MSFAVSEL | AWSPTGTNTGQLY | 0.39541635 |
| NTGALGL | AWSPTGTNTGQLY | 0.62397647 |
| PPGAGGAGGAGTT | AWSPTGTNTGQLY | 0.4882366 |
| QLVGLGI | AWSPTGTNTGQLY | 0.5548662 |
| RFLEQVHQL | AWSPTGTNTGQLY | 0.632774 |
| RGVGYVLR | AWSPTGTNTGQLY | 0.5714381 |
| SATGVFAGVI | AWSPTGTNTGQLY | 0.33904108 |
| SLPAITLP | AWSPTGTNTGQLY | 0.5225651 |
| STALAFTV | AWSPTGTNTGQLY | 0.5088953 |
| SVLTVGLGK | AWSPTGTNTGQLY | 0.4950085 |
| TALAGAGSGI | AWSPTGTNTGQLY | 0.32432625 |
| TGLPFTGI | AWSPTGTNTGQLY | 0.60271263 |
| VAEHVTV | AWSPTGTNTGQLY | 0.4763737 |
| VDIVLNSL | AWSPTGTNTGQLY | 0.45432895 |
| VLTGGGAVL | AWSPTGTNTGQLY | 0.44242573 |
| VNVGLAYGI | AWSPTGTNTGQLY | 0.608833 |
| VSSAAVGAV | AWSPTGTNTGQLY | 0.61719966 |
| VTSVYEAGV | AWSPTGTNTGQLY | 0.5224614 |
| WVALDRGA | AWSPTGTNTGQLY | 0.62937874 |
| YLVGSVVI | AWSPTGTNTGQLY | 0.63757926 |
| GPPGTGKTLL | AWSPTGTNTGQLY | 0.642908 |
| GVGMTKF | AWSPTGTNTGQLY | 0.6135436 |
| IGIGGSDL | AWSPTGTNTGQLY | 0.5546001 |
| IGTLLAEL | AWSPTGTNTGQLY | 0.57985663 |
| IYTSGTT | AWSPTGTNTGQLY | 0.5279604 |
| LSFPAGV | AWSPTGTNTGQLY | 0.39257553 |
| LYTSGTT | AWSPTGTNTGQLY | 0.38460138 |
| VGAFPFGF | AWSPTGTNTGQLY | 0.45350718 |
| VGAGTVEF | AWSPTGTNTGQLY | 0.38744414 |
| VGKSTLLL | AWSPTGTNTGQLY | 0.5408628 |
| AALAHAGGGIV | AWSPTGTNTGQLY | 0.6194826 |
| AALVAVSLL | AWSPTGTNTGQLY | 0.3975277 |
| AAVSRAVAAL | AWSPTGTNTGQLY | 0.3914612 |
| AGGSSGIGGF | AWSPTGTNTGQLY | 0.4967905 |
| AGLTVANANG | AWSPTGTNTGQLY | 0.4351506 |
| AGLVGAVL | AWSPTGTNTGQLY | 0.6060765 |
| ALSQAFTV | AWSPTGTNTGQLY | 0.58296657 |
| ALTFSVSL | AWSPTGTNTGQLY | 0.64987975 |
| ALTGAALVAL | AWSPTGTNTGQLY | 0.6380756 |
| ASFSIGAGVL | AWSPTGTNTGQLY | 0.5386557 |
| DGSVYKI | AWSPTGTNTGQLY | 0.5866461 |
| EQLAEIV | AWSPTGTNTGQLY | 0.44101244 |
| FAVSELTVM | AWSPTGTNTGQLY | 0.57077914 |
| GAAPLAAGL | AWSPTGTNTGQLY | 0.5459324 |

| | | |
|---|---|---|
| GADGGDGGSGGAGG | AWSPTGTNTGQLY | 0.5889293 |
| GFGLDTGGAGG | AWSPTGTNTGQLY | 0.55804396 |
| GLASGGLL | AWSPTGTNTGQLY | 0.5369127 |
| GRIAGAAVLL | AWSPTGTNTGQLY | 0.3384695 |
| GSAGSLNTSF | AWSPTGTNTGQLY | 0.614939 |
| GTGQIGI | AWSPTGTNTGQLY | 0.5299283 |
| GTLIALIL | AWSPTGTNTGQLY | 0.36327413 |
| IAATMGFVVG | AWSPTGTNTGQLY | 0.53422976 |
| IFAQALL | AWSPTGTNTGQLY | 0.6479709 |
| IGAAVALGF | AWSPTGTNTGQLY | 0.64505976 |
| IGLTPLAL | AWSPTGTNTGQLY | 0.51076376 |
| ILGSNTSTL | AWSPTGTNTGQLY | 0.43834502 |
| IVFSLTVL | AWSPTGTNTGQLY | 0.59742093 |
| LASKATGF | AWSPTGTNTGQLY | 0.4170174 |
| LASVALIL | AWSPTGTNTGQLY | 0.48426318 |
| LFAVGLTL | AWSPTGTNTGQLY | 0.57206434 |
| LFNGGIGTYI | AWSPTGTNTGQLY | 0.56520736 |
| LTLGAGII | AWSPTGTNTGQLY | 0.38242865 |
| LVAAALGDI | AWSPTGTNTGQLY | 0.57632107 |
| NTAFVQLGI | AWSPTGTNTGQLY | 0.5705029 |
| QLFSGAEL | AWSPTGTNTGQLY | 0.59638053 |
| SAATFITL | AWSPTGTNTGQLY | 0.34955937 |
| SNYGLANLGV | AWSPTGTNTGQLY | 0.5931937 |
| SYGGAYAVM | AWSPTGTNTGQLY | 0.5047878 |
| TTGVGGLTLG | AWSPTGTNTGQLY | 0.42347622 |
| TTQTIYI | AWSPTGTNTGQLY | 0.34320244 |
| VAAGAGLGAVI | AWSPTGTNTGQLY | 0.54875 |
| VAAVVTLL | AWSPTGTNTGQLY | 0.608081 |
| VGFLGGLI | AWSPTGTNTGQLY | 0.62791026 |
| VGVIPILL | AWSPTGTNTGQLY | 0.55866504 |
| VIFASSLI | AWSPTGTNTGQLY | 0.62454134 |
| VKGGDGGAAAG | AWSPTGTNTGQLY | 0.64187664 |
| VTKTFTNI | AWSPTGTNTGQLY | 0.5489971 |
| VVAAAILL | AWSPTGTNTGQLY | 0.37743258 |
| YAALDVEL | AWSPTGTNTGQLY | 0.5861315 |
| LASVALLL | AWSPTGTNTGQLY | 0.5701794 |
| LDAVHRV | AWSPTGTNTGQLY | 0.61988056 |
| QGLSSSIF | AWSPTGTNTGQLY | 0.6377398 |
| RGTFANIRL | AWSPTGTNTGQLY | 0.4143536 |
| RQLWWGHRI | AWSPTGTNTGQLY | 0.56190974 |
| YGPPGTGKTLL | AWSPTGTNTGQLY | 0.6185886 |
| AALVGAPANVA | AWSPTGTNTGQLY | 0.40489012 |
| AATFRRTE | AWSPTGTNTGQLY | 0.47566724 |
| AGALGGGI | AWSPTGTNTGQLY | 0.4788308 |
| AGGAGGAGGIGAG | AWSPTGTNTGQLY | 0.43375665 |
| AGSGALGL | AWSPTGTNTGQLY | 0.62776095 |
| AQIADQI | AWSPTGTNTGQLY | 0.5277754 |
| ASAVAVTYL | AWSPTGTNTGQLY | 0.49390724 |
| ASTVLGGTGG | AWSPTGTNTGQLY | 0.49110195 |
| AVGGEPAK | AWSPTGTNTGQLY | 0.4380469 |
| CEQGTFGGTC | AWSPTGTNTGQLY | 0.49886328 |

| | | |
|---|---|---|
| DEGKAPAG | AWSPTGTNTGQLY | 0.5224621 |
| DTVVFPAL | AWSPTGTNTGQLY | 0.51940954 |
| FEGITMAG | AWSPTGTNTGQLY | 0.5909624 |
| FPLVVQD | AWSPTGTNTGQLY | 0.47386727 |
| GAFTTPQ | AWSPTGTNTGQLY | 0.523373 |
| GFGGFGTAGGVGGA | AWSPTGTNTGQLY | 0.6434008 |
| GFLAQFL | AWSPTGTNTGQLY | 0.41024327 |
| GGAGGAGGNAGVA | AWSPTGTNTGQLY | 0.43721047 |
| GGVGTAGAGGAGGAGG | AWSPTGTNTGQLY | 0.55451447 |
| GNGGGGDAGNAGSGGN | AWSPTGTNTGQLY | 0.43456116 |
| GTGGAGGAGNDAGS | AWSPTGTNTGQLY | 0.59978116 |
| GTTNTGWFNS | AWSPTGTNTGQLY | 0.59843576 |
| IFMGPLFTL | AWSPTGTNTGQLY | 0.5565544 |
| ILAHAIEAI | AWSPTGTNTGQLY | 0.5289458 |
| IRSAIREAL | AWSPTGTNTGQLY | 0.65469086 |
| ITFAGGG | AWSPTGTNTGQLY | 0.5962935 |
| ITGMSGHQEV | AWSPTGTNTGQLY | 0.48077837 |
| IVAGLACTL | AWSPTGTNTGQLY | 0.6082071 |
| IVGGGLAGA | AWSPTGTNTGQLY | 0.41043615 |
| IVTGVLNI | AWSPTGTNTGQLY | 0.62269247 |
| LAQQGCI | AWSPTGTNTGQLY | 0.5702203 |
| LCLHVEN | AWSPTGTNTGQLY | 0.44118083 |
| LDAISDGS | AWSPTGTNTGQLY | 0.46105915 |
| LGGFAAML | AWSPTGTNTGQLY | 0.6335216 |
| LGGKNPMI | AWSPTGTNTGQLY | 0.62568355 |
| LGPKGVAAIL | AWSPTGTNTGQLY | 0.38333672 |
| LMVGVSI | AWSPTGTNTGQLY | 0.49158314 |
| LVGAAFVL | AWSPTGTNTGQLY | 0.47727627 |
| LVLAVAPL | AWSPTGTNTGQLY | 0.45141324 |
| MAAIAVIDAA | AWSPTGTNTGQLY | 0.3994214 |
| NNGINAELQ | AWSPTGTNTGQLY | 0.59145033 |
| RGEPLVLVHG | AWSPTGTNTGQLY | 0.4890171 |
| RGGAGGAAGLI | AWSPTGTNTGQLY | 0.61487174 |
| RSYRMHPAV | AWSPTGTNTGQLY | 0.54785806 |
| SAAHLLAGQLA | AWSPTGTNTGQLY | 0.61793065 |
| SAAVGSY | AWSPTGTNTGQLY | 0.40484804 |
| SPHAVLHRI | AWSPTGTNTGQLY | 0.5855767 |
| SQIAKVL | AWSPTGTNTGQLY | 0.36407894 |
| TALGHLVP | AWSPTGTNTGQLY | 0.6013303 |
| TAPTTSGAA | AWSPTGTNTGQLY | 0.4357001 |
| TDGDDGNGGSPGT | AWSPTGTNTGQLY | 0.50374675 |
| TDGLGEN | AWSPTGTNTGQLY | 0.44266135 |
| TGGPPLPGQA | AWSPTGTNTGQLY | 0.634724 |
| TIPAITI | AWSPTGTNTGQLY | 0.52860874 |
| TIPGFTI | AWSPTGTNTGQLY | 0.6146411 |
| TSIASGALA | AWSPTGTNTGQLY | 0.51621306 |
| TVTLPGAAI | AWSPTGTNTGQLY | 0.4960101 |
| VASGLSAALP | AWSPTGTNTGQLY | 0.4795788 |
| VGAGVIGI | AWSPTGTNTGQLY | 0.5847238 |
| VGPKRFGV | AWSPTGTNTGQLY | 0.37271103 |
| VNVAFPD | AWSPTGTNTGQLY | 0.32485765 |

| | | |
|---|---|---|
| VSMTNSGVS | AWSPTGTNTGQLY | 0.46401608 |
| VTDVLGDLSV | AWSPTGTNTGQLY | 0.39945412 |
| VVVVGAGFAGL | AWSPTGTNTGQLY | 0.6255778 |
| VYTVGTVV | AWSPTGTNTGQLY | 0.63020295 |
| WTTFSGP | AWSPTGTNTGQLY | 0.5070472 |
| YDELVEAF | AWSPTGTNTGQLY | 0.60747886 |
| YGTGGAGGNGG | AWSPTGTNTGQLY | 0.3194659 |
| YTNVMAV | AWSPTGTNTGQLY | 0.5592198 |
| FLEQVHQL | AWSPTGTNTGQLY | 0.6449238 |
| GLSLLAVAV | AWSPTGTNTGQLY | 0.59747165 |
| LEQVHQL | AWSPTGTNTGQLY | 0.51595247 |
| LNAAGAI | AWSPTGTNTGQLY | 0.62663 |
| PAVHASL | AWSPTGTNTGQLY | 0.5331739 |
| TELEDDA | AWSPTGTNTGQLY | 0.49043393 |
| TPPGPPN | AWSPTGTNTGQLY | 0.5722463 |
| TPPPPPPP | AWSPTGTNTGQLY | 0.5976781 |
| TQSPALI | ASSRDTGNSNTGQLY | 0.5717794 |
| EFAAAFPQ | ASSRDTGNSNTGQLY | 0.42585683 |
| LLSGNLDVL | ASSRDTGNSNTGQLY | 0.51465625 |
| GPANLGNYNI | ASSRDTGNSNTGQLY | 0.65633595 |
| TLGPTLP | ASSRDTGNSNTGQLY | 0.4854324 |
| VYAYSSCCK | ASSRDTGNSNTGQLY | 0.62080777 |
| APAPAPAPNI | ASSRDTGNSNTGQLY | 0.57591236 |
| PGGPVPIP | ASSRDTGNSNTGQLY | 0.6164224 |
| FLTVALEP | ASSRDTGNSNTGQLY | 0.41395098 |
| SQYFGANL | ASSRDTGNSNTGQLY | 0.46541512 |
| GGVKPAA | ASSRDTGNSNTGQLY | 0.5917101 |
| GGTGGAGGAGGW | ASSRDTGNSNTGQLY | 0.4504554 |
| GVGGVAGLGGAGG | ASSRDTGNSNTGQLY | 0.45256907 |
| VLLVLATTI | ASSRDTGNSNTGQLY | 0.44944942 |
| IGIFAAML | ASSRDTGNSNTGQLY | 0.65829337 |
| AVGDLID | ASSRDTGNSNTGQLY | 0.600158 |
| NISPLPL | ASSRDTGNSNTGQLY | 0.5996363 |
| VTAVFVAI | ASSRDTGNSNTGQLY | 0.5022188 |
| SGDDNDAAL | ASSRDTGNSNTGQLY | 0.54805636 |
| AVAALAHS | ASSRDTGNSNTGQLY | 0.58920425 |
| GGVTVMATP | ASSRDTGNSNTGQLY | 0.34876457 |
| AGLAANVRD | ASSRDTGNSNTGQLY | 0.6395182 |
| TNRPDLIDPALLRPG | ASSRDTGNSNTGQLY | 0.46766886 |
| LANLGVTAL | ASSRDTGNSNTGQLY | 0.5350438 |
| TGERIEL | ASSRDTGNSNTGQLY | 0.56922954 |
| NGEELAA | ASSRDTGNSNTGQLY | 0.6394398 |
| VLSLVWG | ASSRDTGNSNTGQLY | 0.49331158 |
| IGYALWL | ASSRDTGNSNTGQLY | 0.5889658 |
| LSLLLAPL | ASSRDTGNSNTGQLY | 0.46652797 |
| TLDEALK | ASSRDTGNSNTGQLY | 0.5751053 |
| TDGAGTAEGG | ASSRDTGNSNTGQLY | 0.62118685 |
| NAPGGAGGNGG | ASSRDTGNSNTGQLY | 0.5802882 |
| VSVFMGMGGNF | ASSRDTGNSNTGQLY | 0.6459483 |
| LASYLAAL | ASSRDTGNSNTGQLY | 0.54453176 |
| LVGGTPAL | ASSRDTGNSNTGQLY | 0.36522168 |

| | | |
|---|---|---|
| VRNADVL | ASSRDTGNSNTGQLY | 0.5143273 |
| TCSSPFLVSAM | ASSRDTGNSNTGQLY | 0.61813676 |
| VHITDAE | ASSRDTGNSNTGQLY | 0.6300527 |
| EYAAPSF | ASSRDTGNSNTGQLY | 0.59899366 |
| VVDAFHGGL | ASSRDTGNSNTGQLY | 0.61442935 |
| ASVVDPQ | ASSRDTGNSNTGQLY | 0.52842456 |
| SYNEFGGLS | ASSRDTGNSNTGQLY | 0.6341682 |
| DPDGSVY | ASSRDTGNSNTGQLY | 0.65266174 |
| MVSTIHATE | ASSRDTGNSNTGQLY | 0.60068744 |
| LAAGVADPV | ASSRDTGNSNTGQLY | 0.43552306 |
| TKEQIAA | ASSRDTGNSNTGQLY | 0.39040524 |
| SLASPAAL | ASSRDTGNSNTGQLY | 0.54720885 |
| IGLVFYGV | ASSRDTGNSNTGQLY | 0.6177397 |
| FLLDASNL | ASSRDTGNSNTGQLY | 0.36280274 |
| YQVGNFP | ASSRDTGNSNTGQLY | 0.5393734 |
| ESLLSQST | ASSRDTGNSNTGQLY | 0.5651655 |
| LFTVGFAAVG | ASSRDTGNSNTGQLY | 0.3753378 |
| ATPPTPT | ASSRDTGNSNTGQLY | 0.579897 |
| TDLATLG | ASSRDTGNSNTGQLY | 0.6468044 |
| SAGGTGR | ASSRDTGNSNTGQLY | 0.4876776 |
| GFSNSGS | ASSRDTGNSNTGQLY | 0.37728217 |
| GGNGGSGGTSV | ASSRDTGNSNTGQLY | 0.6402257 |
| GTSVATGGAGN | ASSRDTGNSNTGQLY | 0.65418476 |
| NGGNGGSGGTS | ASSRDTGNSNTGQLY | 0.60548574 |
| AVVKFPH | ASSRDTGNSNTGQLY | 0.3641577 |
| TPPPPPP | ASSRDTGNSNTGQLY | 0.3145427 |
| AAPAAAAAPA | ASSRDTGNSNTGQLY | 0.60222954 |
| IALNEDL | ASSRDTGNSNTGQLY | 0.61723113 |
| IVNVSSM | ASSRDTGNSNTGQLY | 0.5261669 |
| TGIPPAPR | ASSRDTGNSNTGQLY | 0.45494437 |
| FDLGGGTF | ASSRDTGNSNTGQLY | 0.5768991 |
| FDLGGGTFDVS | ASSRDTGNSNTGQLY | 0.6272377 |
| NEPTAAA | ASSRDTGNSNTGQLY | 0.41343856 |
| NEPTAAALA | ASSRDTGNSNTGQLY | 0.40988538 |
| GSIINISSI | ASSRDTGNSNTGQLY | 0.59957904 |
| VAPGNAGTA | ASSRDTGNSNTGQLY | 0.5495998 |
| TGGGASLE | ASSRDTGNSNTGQLY | 0.46681792 |
| VAVYNHYKRI | ASSRDTGNSNTGQLY | 0.49260312 |
| TDPAPATT | ASSRDTGNSNTGQLY | 0.6213571 |
| TNRPDILDPALLRPG | ASSRDTGNSNTGQLY | 0.57797587 |
| LTTTTTS | ASSRDTGNSNTGQLY | 0.58070517 |
| PTGSGKTL | ASSRDTGNSNTGQLY | 0.34671345 |
| TGSGKTL | ASSRDTGNSNTGQLY | 0.6408636 |
| TIGHVDHGK | ASSRDTGNSNTGQLY | 0.3582102 |
| TIGHVDHGKTT | ASSRDTGNSNTGQLY | 0.6182435 |
| GHVDHGKTT | ASSRDTGNSNTGQLY | 0.5983123 |
| AQFEHTLL | ASSRDTGNSNTGQLY | 0.5821622 |
| IVNQDEL | ASSRDTGNSNTGQLY | 0.4942546 |
| TNGSQFF | ASSRDTGNSNTGQLY | 0.39946994 |
| TGKGIVE | ASSRDTGNSNTGQLY | 0.36533287 |
| LISNASDA | ASSRDTGNSNTGQLY | 0.3827535 |

| | | |
|---|---|---|
| AELRTLGELTKIAKAHGAQ | ASSRDTGNSNTGQLY | 0.6443769 |
| APNAVAAAEPSW | ASSRDTGNSNTGQLY | 0.5907866 |
| APQVAGAATAVL | ASSRDTGNSNTGQLY | 0.6243293 |
| ARRDPAAAAAILPTDA | ASSRDTGNSNTGQLY | 0.58492446 |
| DAAARDVLAEAGLAEYF | ASSRDTGNSNTGQLY | 0.49179173 |
| DDQIVLSAKANLAKVEQ | ASSRDTGNSNTGQLY | 0.5823029 |
| DELLPDTGEDSDEDGH | ASSRDTGNSNTGQLY | 0.4529234 |
| ELPEAPANRATVPQD | ASSRDTGNSNTGQLY | 0.5215222 |
| FGQSKIERVTGIG | ASSRDTGNSNTGQLY | 0.5583474 |
| GAGGTGSTAGGAGGAGGAGGLY | ASSRDTGNSNTGQLY | 0.54236335 |
| GHGGTGAAGGNGGTGGAGGSGI | ASSRDTGNSNTGQLY | 0.4075754 |
| GNGGNGGAGGEGGPGGAGGAGA | ASSRDTGNSNTGQLY | 0.5223602 |
| GSTAGNGGNGGNATNGTIAGQP | ASSRDTGNSNTGQLY | 0.56550235 |
| GTGGDGGNGGNGGTGG | ASSRDTGNSNTGQLY | 0.37788203 |
| GVRRPAGIPADAPV | ASSRDTGNSNTGQLY | 0.4560693 |
| GYRQVTDAHLLTLARR | ASSRDTGNSNTGQLY | 0.63493437 |
| IAKMGQDGHDRG | ASSRDTGNSNTGQLY | 0.51923734 |
| KAEPSIEIQYT | ASSRDTGNSNTGQLY | 0.36007032 |
| KPVRIRMAADEDTVAVT | ASSRDTGNSNTGQLY | 0.619529 |
| MPPRIAGMRLLVIKPEP | ASSRDTGNSNTGQLY | 0.5114486 |
| MKFYAVALFRSRGVPPTR | ASSRDTGNSNTGQLY | 0.56994027 |
| NAGGFGILGAVAHSP | ASSRDTGNSNTGQLY | 0.5886466 |
| NEGREITLAMAEM | ASSRDTGNSNTGQLY | 0.45362097 |
| NELVEWSHPLGHRWRWE | ASSRDTGNSNTGQLY | 0.61899626 |
| NLGAGGTGGNGGNPG | ASSRDTGNSNTGQLY | 0.40889823 |
| NTEWPKWETVMAADPVD | ASSRDTGNSNTGQLY | 0.49137545 |
| PASAAPAETQP | ASSRDTGNSNTGQLY | 0.54807866 |
| QQSSDGKNGVRAAEITGE | ASSRDTGNSNTGQLY | 0.60386693 |
| RAPDAVTTSWWKE | ASSRDTGNSNTGQLY | 0.603012 |
| SGFGNVGAHTSGWFNQ | ASSRDTGNSNTGQLY | 0.46624523 |
| SGVAGAVGQAASVGGLKV | ASSRDTGNSNTGQLY | 0.5396719 |
| SNGGAGTGSAGGNGGT | ASSRDTGNSNTGQLY | 0.4631718 |
| TGGAGGEGGAAG | ASSRDTGNSNTGQLY | 0.39895186 |
| TGGDGGAAGAGGAPGHG | ASSRDTGNSNTGQLY | 0.40995523 |
| TMPPLLAAIANAGRHGV | ASSRDTGNSNTGQLY | 0.50548977 |
| TSSTAGNSGGA | ASSRDTGNSNTGQLY | 0.38779384 |
| VPATVPVGILC | ASSRDTGNSNTGQLY | 0.6199248 |
| VQRPDSGGVVLTGRIS | ASSRDTGNSNTGQLY | 0.48248285 |
| WTPNPVPATKGVPAGLVQAT | ASSRDTGNSNTGQLY | 0.58899456 |
| YALKRALAKFSDRFDVVII | ASSRDTGNSNTGQLY | 0.6338396 |
| YLIGSVIV | ASSRDTGNSNTGQLY | 0.49332827 |
| YGQLVAVID | ASSRDTGNSNTGQLY | 0.6010984 |
| VTVGGMTLPA | ASSRDTGNSNTGQLY | 0.5852489 |
| VLALPAV | ASSRDTGNSNTGQLY | 0.6105289 |
| VIPPTLNYET | ASSRDTGNSNTGQLY | 0.5162804 |
| VGMGGMHPSSG | ASSRDTGNSNTGQLY | 0.3813664 |
| VGLAGGAATA | ASSRDTGNSNTGQLY | 0.6049656 |
| VGGAGGAGGSGP | ASSRDTGNSNTGQLY | 0.6207769 |
| VFAAFMV | ASSRDTGNSNTGQLY | 0.6040484 |
| TLYDKLR | ASSRDTGNSNTGQLY | 0.61971635 |
| TLAVFTVL | ASSRDTGNSNTGQLY | 0.6134863 |

| | | |
|---|---|---|
| TGSVQPARV | ASSRDTGNSNTGQLY | 0.6012712 |
| TGGGAGSGGT | ASSRDTGNSNTGQLY | 0.5432755 |
| SHLETSL | ASSRDTGNSNTGQLY | 0.36949348 |
| SFTTGMT | ASSRDTGNSNTGQLY | 0.5712006 |
| QVEVAKRMGVRQPTVS | ASSRDTGNSNTGQLY | 0.5827747 |
| QGSGGPALAM | ASSRDTGNSNTGQLY | 0.44218087 |
| MLAPSNP | ASSRDTGNSNTGQLY | 0.33924866 |
| LVNNAAI | ASSRDTGNSNTGQLY | 0.6250815 |
| LTGEINTLA | ASSRDTGNSNTGQLY | 0.6560781 |
| LSIFGPV | ASSRDTGNSNTGQLY | 0.48024005 |
| LLALLANL | ASSRDTGNSNTGQLY | 0.63018525 |
| LISNASDALDKLR | ASSRDTGNSNTGQLY | 0.36137494 |
| LINEAVP | ASSRDTGNSNTGQLY | 0.6203167 |
| LGYAFSGL | ASSRDTGNSNTGQLY | 0.6290209 |
| LDFVAVAL | ASSRDTGNSNTGQLY | 0.45301616 |
| LAQPGAE | ASSRDTGNSNTGQLY | 0.61065197 |
| ISNLGAVTGF | ASSRDTGNSNTGQLY | 0.40319097 |
| HQEDPGRI | ASSRDTGNSNTGQLY | 0.43350047 |
| GTPGEPTP | ASSRDTGNSNTGQLY | 0.6388019 |
| GFFNSGS | ASSRDTGNSNTGQLY | 0.48624265 |
| GAGGGGLTAGAN | ASSRDTGNSNTGQLY | 0.5868289 |
| FVDALVNM | ASSRDTGNSNTGQLY | 0.48956627 |
| FLTVAPEL | ASSRDTGNSNTGQLY | 0.43635157 |
| FETLTAL | ASSRDTGNSNTGQLY | 0.56762165 |
| DVINPIEL | ASSRDTGNSNTGQLY | 0.6275763 |
| APVSGAPV | ASSRDTGNSNTGQLY | 0.50013614 |
| ALVGGLAW | ASSRDTGNSNTGQLY | 0.64651966 |
| ALAAAAATAI | ASSRDTGNSNTGQLY | 0.53822637 |
| AFGLGLLL | ASSRDTGNSNTGQLY | 0.6360447 |
| AFFAAASL | ASSRDTGNSNTGQLY | 0.6175011 |
| AAAALPAFSPP | ASSRDTGNSNTGQLY | 0.5609891 |
| VVLAPAL | ASSRDTGNSNTGQLY | 0.5035311 |
| TIGHVDHGKTTL | ASSRDTGNSNTGQLY | 0.6219436 |
| LISNASDALDK | ASSRDTGNSNTGQLY | 0.45214704 |
| LISNASDAL | ASSRDTGNSNTGQLY | 0.63084626 |
| ISNASDALDK | ASSRDTGNSNTGQLY | 0.614309 |
| NQAETLVYQTE | ASSRDTGNSNTGQLY | 0.6525052 |
| TGGTGGTAGAG | ASSRDTGNSNTGQLY | 0.7491423 |
| TGSTAGGAGGA | ASSRDTGNSNTGQLY | 0.7105189 |
| DPLAVPVAAGPV | ASSRDTGNSNTGQLY | 0.7300908 |
| GGTGGTPGNGGN | ASSRDTGNSNTGQLY | 0.66793084 |
| PAGLAGVDPATL | ASSRDTGNSNTGQLY | 0.61410874 |
| GNNLVVEAPHAGASV | ASSRDTGNSNTGQLY | 0.6948189 |
| KVTRIMSASTLEQPA | ASSRDTGNSNTGQLY | 0.60788596 |
| LGFALPIALPRRTVV | ASSRDTGNSNTGQLY | 0.75353134 |
| TRLGNLTARKLVVLV | ASSRDTGNSNTGQLY | 0.7576113 |
| WSQQRRLIYFPSAGP | ASSRDTGNSNTGQLY | 0.6541023 |
| GPNGNPDPMAAAVDIR | ASSRDTGNSNTGQLY | 0.7611939 |
| TQRCVSVADLRAAASR | ASSRDTGNSNTGQLY | 0.74126816 |
| QFVLDVAEHPVEVLTETV | ASSRDTGNSNTGQLY | 0.587186 |
| GGAGGTGGAGGNGGTQAGDG | ASSRDTGNSNTGQLY | 0.69028217 |

| | | |
|---|---|---|
| PLLATLRAIAVDVDAKPAQV | ASSRDTGNSNTGQLY | 0.70572126 |
| APKFRTGGYMIDFWGVGYQVAK | ASSRDTGNSNTGQLY | 0.7235817 |
| GAGGAGGAGGAAGGPGGTGGTG | ASSRDTGNSNTGQLY | 0.51227874 |
| IGALGLLATDVAPAGSVAFGVA | ASSRDTGNSNTGQLY | 0.7524226 |
| AGGAGGIGGNGGAGGAGGVLMG | ASSRDTGNSNTGQLY | 0.6219974 |
| ADELIGLL | ASSRDTGNSNTGQLY | 0.34273887 |
| ADRALKA | ASSRDTGNSNTGQLY | 0.74292505 |
| AGAVGQVGGAGGA | ASSRDTGNSNTGQLY | 0.68224275 |
| AGGGGGLGGAP | ASSRDTGNSNTGQLY | 0.7072915 |
| AGGLLYGNGGAG | ASSRDTGNSNTGQLY | 0.7533926 |
| AGLESIGL | ASSRDTGNSNTGQLY | 0.57846105 |
| AGLKTGII | ASSRDTGNSNTGQLY | 0.6369908 |
| ASLAALLL | ASSRDTGNSNTGQLY | 0.7182559 |
| ASVINISSI | ASSRDTGNSNTGQLY | 0.7500105 |
| ATPPGPA | ASSRDTGNSNTGQLY | 0.6295471 |
| ATVASALV | ASSRDTGNSNTGQLY | 0.7093911 |
| ATVASLIL | ASSRDTGNSNTGQLY | 0.7526165 |
| AVNGPVLIRH | ASSRDTGNSNTGQLY | 0.73078895 |
| DGAAGAVGI | ASSRDTGNSNTGQLY | 0.76634324 |
| DPLLHGI | ASSRDTGNSNTGQLY | 0.7698055 |
| ELGAALGAL | ASSRDTGNSNTGQLY | 0.65896374 |
| ELGRTQAV | ASSRDTGNSNTGQLY | 0.7545827 |
| FVAGGYLS | ASSRDTGNSNTGQLY | 0.7722902 |
| GASGGAGGAGGW | ASSRDTGNSNTGQLY | 0.6858758 |
| GGAGAGGDGGAGGI | ASSRDTGNSNTGQLY | 0.7017789 |
| GTGHNGSQPGMGGNG | ASSRDTGNSNTGQLY | 0.6256428 |
| GYAAASASAS | ASSRDTGNSNTGQLY | 0.6947624 |
| HQVDVAAVQQG | ASSRDTGNSNTGQLY | 0.5894157 |
| IGAAIAHT | ASSRDTGNSNTGQLY | 0.69824576 |
| IGMIEVAQL | ASSRDTGNSNTGQLY | 0.7685927 |
| IRLAPEHTSAMD | ASSRDTGNSNTGQLY | 0.6732545 |
| IYNVSVL | ASSRDTGNSNTGQLY | 0.57879627 |
| KQGFLGM | ASSRDTGNSNTGQLY | 0.55020297 |
| LESVITQL | ASSRDTGNSNTGQLY | 0.7155029 |
| LGLGALVVV | ASSRDTGNSNTGQLY | 0.67843926 |
| LIGNGGNGGSGGI | ASSRDTGNSNTGQLY | 0.563529 |
| LNAINAPTL | ASSRDTGNSNTGQLY | 0.6702707 |
| LQLLNTGV | ASSRDTGNSNTGQLY | 0.7516158 |
| LQTSIEF | ASSRDTGNSNTGQLY | 0.7502105 |
| MSFAVSEL | ASSRDTGNSNTGQLY | 0.5691624 |
| NTGALGL | ASSRDTGNSNTGQLY | 0.76619023 |
| PPGAGGAGGAGTT | ASSRDTGNSNTGQLY | 0.7167903 |
| QLVGLGI | ASSRDTGNSNTGQLY | 0.74895024 |
| RFLEQVHQL | ASSRDTGNSNTGQLY | 0.69704115 |
| RGVGYVLR | ASSRDTGNSNTGQLY | 0.64787775 |
| SATGVFAGVI | ASSRDTGNSNTGQLY | 0.7429132 |
| SLPAITLP | ASSRDTGNSNTGQLY | 0.6106008 |
| STALAFTV | ASSRDTGNSNTGQLY | 0.68289775 |
| SVLTVGLGK | ASSRDTGNSNTGQLY | 0.7579533 |
| TALAGAGSGI | ASSRDTGNSNTGQLY | 0.6925702 |
| TGLPFTGI | ASSRDTGNSNTGQLY | 0.61680114 |

| | | |
|---|---|---|
| VAEHVTV | ASSRDTGNSNTGQLY | 0.61181515 |
| VDIVLNSL | ASSRDTGNSNTGQLY | 0.76454383 |
| VLTGGGAVL | ASSRDTGNSNTGQLY | 0.55154604 |
| VNVGLAYGI | ASSRDTGNSNTGQLY | 0.64983714 |
| VSSAAVGAV | ASSRDTGNSNTGQLY | 0.6237592 |
| VTSVYEAGV | ASSRDTGNSNTGQLY | 0.6640432 |
| WVALDRGA | ASSRDTGNSNTGQLY | 0.75596833 |
| YLVGSVVI | ASSRDTGNSNTGQLY | 0.745528 |
| GPPGTGKTLL | ASSRDTGNSNTGQLY | 0.6463244 |
| GVGMTKF | ASSRDTGNSNTGQLY | 0.7306235 |
| IGIGGSDL | ASSRDTGNSNTGQLY | 0.7384745 |
| IGTLLAEL | ASSRDTGNSNTGQLY | 0.71181977 |
| IYTSGTT | ASSRDTGNSNTGQLY | 0.6951262 |
| LSFPAGV | ASSRDTGNSNTGQLY | 0.57770914 |
| LYTSGTT | ASSRDTGNSNTGQLY | 0.75520146 |
| VGAFPFGF | ASSRDTGNSNTGQLY | 0.7471473 |
| VGAGTVEF | ASSRDTGNSNTGQLY | 0.7614131 |
| VGKSTLLL | ASSRDTGNSNTGQLY | 0.7572226 |
| AALAHAGGGIV | ASSRDTGNSNTGQLY | 0.70473707 |
| AALVAVSLL | ASSRDTGNSNTGQLY | 0.7082124 |
| AAVSRAVAAL | ASSRDTGNSNTGQLY | 0.5430968 |
| AGGSSGIGGF | ASSRDTGNSNTGQLY | 0.6378207 |
| AGLTVANANG | ASSRDTGNSNTGQLY | 0.69113886 |
| AGLVGAVL | ASSRDTGNSNTGQLY | 0.7123918 |
| ALSQAFTV | ASSRDTGNSNTGQLY | 0.7433656 |
| ALTFSVSL | ASSRDTGNSNTGQLY | 0.75044954 |
| ALTGAALVAL | ASSRDTGNSNTGQLY | 0.60660774 |
| ASFSIGAGVL | ASSRDTGNSNTGQLY | 0.7464214 |
| DGSVYKI | ASSRDTGNSNTGQLY | 0.752905 |
| EQLAEIV | ASSRDTGNSNTGQLY | 0.56374645 |
| FAVSELTVM | ASSRDTGNSNTGQLY | 0.66569173 |
| GAAPLAAGL | ASSRDTGNSNTGQLY | 0.77855825 |
| GADGGDGGSGGAGG | ASSRDTGNSNTGQLY | 0.71126384 |
| GFGLDTGGAGG | ASSRDTGNSNTGQLY | 0.67205983 |
| GLASGGLL | ASSRDTGNSNTGQLY | 0.7063653 |
| GRIAGAAVLL | ASSRDTGNSNTGQLY | 0.7651999 |
| GSAGSLNTSF | ASSRDTGNSNTGQLY | 0.6831702 |
| GTGQIGI | ASSRDTGNSNTGQLY | 0.7742089 |
| GTLIALIL | ASSRDTGNSNTGQLY | 0.49373776 |
| IAATMGFVVG | ASSRDTGNSNTGQLY | 0.71394247 |
| IFAQALL | ASSRDTGNSNTGQLY | 0.6637655 |
| IGAAVALGF | ASSRDTGNSNTGQLY | 0.46658736 |
| IGLTPLAL | ASSRDTGNSNTGQLY | 0.7495025 |
| ILGSNTSTL | ASSRDTGNSNTGQLY | 0.69213057 |
| IVFSLTVL | ASSRDTGNSNTGQLY | 0.7444525 |
| LASKATGF | ASSRDTGNSNTGQLY | 0.6768714 |
| LASVALIL | ASSRDTGNSNTGQLY | 0.7588379 |
| LFAVGLTL | ASSRDTGNSNTGQLY | 0.68681633 |
| LFNGGIGTYI | ASSRDTGNSNTGQLY | 0.6891722 |
| LTLGAGII | ASSRDTGNSNTGQLY | 0.71170235 |
| LVAAALGDI | ASSRDTGNSNTGQLY | 0.7424161 |

| | | |
|---|---|---|
| NTAFVQLGI | ASSRDTGNSNTGQLY | 0.75906265 |
| QLFSGAEL | ASSRDTGNSNTGQLY | 0.76555544 |
| SAATFITL | ASSRDTGNSNTGQLY | 0.5550138 |
| SNYGLANLGV | ASSRDTGNSNTGQLY | 0.69854754 |
| SYGGAYAVM | ASSRDTGNSNTGQLY | 0.6960351 |
| TTGVGGLTLG | ASSRDTGNSNTGQLY | 0.459138 |
| TTQTIYI | ASSRDTGNSNTGQLY | 0.5052221 |
| VAAGAGLGAVI | ASSRDTGNSNTGQLY | 0.6692794 |
| VAAVVTLL | ASSRDTGNSNTGQLY | 0.66119313 |
| VGFLGGLI | ASSRDTGNSNTGQLY | 0.7354637 |
| VGVIPILL | ASSRDTGNSNTGQLY | 0.7225628 |
| VIFASSLI | ASSRDTGNSNTGQLY | 0.6817187 |
| VKGGDGGAAAG | ASSRDTGNSNTGQLY | 0.7009125 |
| VTKTFTNI | ASSRDTGNSNTGQLY | 0.6840335 |
| VVAAAILL | ASSRDTGNSNTGQLY | 0.21557026 |
| YAALDVEL | ASSRDTGNSNTGQLY | 0.626504 |
| LASVALLL | ASSRDTGNSNTGQLY | 0.5990438 |
| LDAVHRV | ASSRDTGNSNTGQLY | 0.77186024 |
| QGLSSSIF | ASSRDTGNSNTGQLY | 0.652395 |
| RGTFANIRL | ASSRDTGNSNTGQLY | 0.6170145 |
| RQLWWGHRI | ASSRDTGNSNTGQLY | 0.69608295 |
| YGPPGTGKTLL | ASSRDTGNSNTGQLY | 0.6908713 |
| AALVGAPANVA | ASSRDTGNSNTGQLY | 0.66431457 |
| AATFRRTE | ASSRDTGNSNTGQLY | 0.59653085 |
| AGALGGGI | ASSRDTGNSNTGQLY | 0.49525097 |
| AGGAGGAGGIGAG | ASSRDTGNSNTGQLY | 0.75033665 |
| AGSGALGL | ASSRDTGNSNTGQLY | 0.6293662 |
| AQIADQI | ASSRDTGNSNTGQLY | 0.63460344 |
| ASAVAVTYL | ASSRDTGNSNTGQLY | 0.5560633 |
| ASTVLGGTGG | ASSRDTGNSNTGQLY | 0.6302319 |
| AVGGEPAK | ASSRDTGNSNTGQLY | 0.5681341 |
| CEQGTFGGTC | ASSRDTGNSNTGQLY | 0.76711607 |
| DEGKAPAG | ASSRDTGNSNTGQLY | 0.64467496 |
| DTVVFPAL | ASSRDTGNSNTGQLY | 0.6360264 |
| FEGITMAG | ASSRDTGNSNTGQLY | 0.7670326 |
| FPLVVQD | ASSRDTGNSNTGQLY | 0.6615974 |
| GAFTTPQ | ASSRDTGNSNTGQLY | 0.6629536 |
| GFGGFGTAGGVGGA | ASSRDTGNSNTGQLY | 0.6884446 |
| GFLAQFL | ASSRDTGNSNTGQLY | 0.6441719 |
| GGAGGAGGNAGVA | ASSRDTGNSNTGQLY | 0.6465592 |
| GGVGTAGAGGAGGAGG | ASSRDTGNSNTGQLY | 0.66567075 |
| GNGGGGDAGNAGSGGN | ASSRDTGNSNTGQLY | 0.7673843 |
| GTGGAGGAGNDAGS | ASSRDTGNSNTGQLY | 0.7326211 |
| GTTNTGWFNS | ASSRDTGNSNTGQLY | 0.6504477 |
| IFMGPLFTL | ASSRDTGNSNTGQLY | 0.7480094 |
| ILAHAIEAI | ASSRDTGNSNTGQLY | 0.7699379 |
| IRSAIREAL | ASSRDTGNSNTGQLY | 0.76243716 |
| ITFAGGG | ASSRDTGNSNTGQLY | 0.60958934 |
| ITGMSGHQEV | ASSRDTGNSNTGQLY | 0.66215944 |
| IVAGLACTL | ASSRDTGNSNTGQLY | 0.6705849 |
| IVGGGLAGA | ASSRDTGNSNTGQLY | 0.6418244 |

| | | |
|---|---|---|
| IVTGVLNI | ASSRDTGNSNTGQLY | 0.6661537 |
| LAQQGCI | ASSRDTGNSNTGQLY | 0.65377736 |
| LCLHVEN | ASSRDTGNSNTGQLY | 0.7451278 |
| LDAISDGS | ASSRDTGNSNTGQLY | 0.77040386 |
| LGGFAAML | ASSRDTGNSNTGQLY | 0.68489385 |
| LGGKNPMI | ASSRDTGNSNTGQLY | 0.572626 |
| LGPKGVAAIL | ASSRDTGNSNTGQLY | 0.64944375 |
| LMVGVSI | ASSRDTGNSNTGQLY | 0.58597744 |
| LVGAAFVL | ASSRDTGNSNTGQLY | 0.61028475 |
| LVLAVAPL | ASSRDTGNSNTGQLY | 0.6660557 |
| MAAIAVIDAA | ASSRDTGNSNTGQLY | 0.54805505 |
| NNGINAELQ | ASSRDTGNSNTGQLY | 0.7010658 |
| RGEPLVLVHG | ASSRDTGNSNTGQLY | 0.75965214 |
| RGGAGGAAGLI | ASSRDTGNSNTGQLY | 0.6258858 |
| RSYRMHPAV | ASSRDTGNSNTGQLY | 0.73879397 |
| SAAHLLAGQLA | ASSRDTGNSNTGQLY | 0.7665546 |
| SAAVGSY | ASSRDTGNSNTGQLY | 0.74335206 |
| SPHAVLHRI | ASSRDTGNSNTGQLY | 0.7613573 |
| SQIAKVL | ASSRDTGNSNTGQLY | 0.72918344 |
| TALGHLVP | ASSRDTGNSNTGQLY | 0.75158685 |
| TAPTTSGAA | ASSRDTGNSNTGQLY | 0.69236904 |
| TDGDDGNGGSPGT | ASSRDTGNSNTGQLY | 0.73549855 |
| TDGLGEN | ASSRDTGNSNTGQLY | 0.66573757 |
| TGGPPLPGQA | ASSRDTGNSNTGQLY | 0.6940968 |
| TIPAITI | ASSRDTGNSNTGQLY | 0.6711951 |
| TIPGFTI | ASSRDTGNSNTGQLY | 0.7312767 |
| TSIASGALA | ASSRDTGNSNTGQLY | 0.65532494 |
| TVTLPGAAI | ASSRDTGNSNTGQLY | 0.76481664 |
| VASGLSAALP | ASSRDTGNSNTGQLY | 0.5185699 |
| VGAGVIGI | ASSRDTGNSNTGQLY | 0.6331912 |
| VGPKRFGV | ASSRDTGNSNTGQLY | 0.751637 |
| VNVAFPD | ASSRDTGNSNTGQLY | 0.2633182 |
| VSMTNSGVS | ASSRDTGNSNTGQLY | 0.51793647 |
| VTDVLGDLSV | ASSRDTGNSNTGQLY | 0.704867 |
| VVVVGAGFAGL | ASSRDTGNSNTGQLY | 0.7437059 |
| VYTVGTVV | ASSRDTGNSNTGQLY | 0.29713407 |
| WTTFSGP | ASSRDTGNSNTGQLY | 0.6682801 |
| YDELVEAF | ASSRDTGNSNTGQLY | 0.59993386 |
| YGTGGAGGNGG | ASSRDTGNSNTGQLY | 0.7627127 |
| YTNVMAV | ASSRDTGNSNTGQLY | 0.69627655 |
| FLEQVHQL | ASSRDTGNSNTGQLY | 0.5324696 |
| GLSLLAVAV | ASSRDTGNSNTGQLY | 0.7607878 |
| LEQVHQL | ASSRDTGNSNTGQLY | 0.57322824 |
| LNAAGAI | ASSRDTGNSNTGQLY | 0.639492 |
| PAVHASL | ASSRDTGNSNTGQLY | 0.75946903 |
| TELEDDA | ASSRDTGNSNTGQLY | 0.7634005 |
| TPPGPPN | ASSRDTGNSNTGQLY | 0.62927353 |
| TPPPPPPP | ASSRDTGNSNTGQLY | 0.7367251 |
| TQSPALI | TCSGTGQDTQY | 0.62585914 |
| EFAAAFPQ | TCSGTGQDTQY | 0.769716 |
| LLSGNLDVL | TCSGTGQDTQY | 0.6824087 |

| | | |
|---|---|---|
| GPANLGNYNI | TCSGTGQDTQY | 0.61602867 |
| TLGPTLP | TCSGTGQDTQY | 0.75521725 |
| VYAYSSCCK | TCSGTGQDTQY | 0.7503368 |
| APAPAPAPNI | TCSGTGQDTQY | 0.72336435 |
| PGGPVPIP | TCSGTGQDTQY | 0.72441715 |
| FLTVALEP | TCSGTGQDTQY | 0.69211006 |
| SQYFGANL | TCSGTGQDTQY | 0.65233594 |
| GGVKPAA | TCSGTGQDTQY | 0.7510792 |
| GGTGGAGGAGGW | TCSGTGQDTQY | 0.73989373 |
| GVGGVAGLGGAGG | TCSGTGQDTQY | 0.66695035 |
| VLLVLATTI | TCSGTGQDTQY | 0.7665267 |
| IGIFAAML | TCSGTGQDTQY | 0.69106126 |
| AVGDLID | TCSGTGQDTQY | 0.6691878 |
| NISPLPL | TCSGTGQDTQY | 0.7689489 |
| VTAVFVAI | TCSGTGQDTQY | 0.7153527 |
| SGDDNDAAL | TCSGTGQDTQY | 0.7415477 |
| AVAALAHS | TCSGTGQDTQY | 0.75291955 |
| GGVTVMATP | TCSGTGQDTQY | 0.7352855 |
| AGLAANVRD | TCSGTGQDTQY | 0.6492197 |
| TNRPDLIDPALLRPG | TCSGTGQDTQY | 0.68362504 |
| LANLGVTAL | TCSGTGQDTQY | 0.67400205 |
| TGERIEL | TCSGTGQDTQY | 0.74243 |
| NGEELAA | TCSGTGQDTQY | 0.71965533 |
| VLSLVWG | TCSGTGQDTQY | 0.65521586 |
| IGYALWL | TCSGTGQDTQY | 0.7535572 |
| LSLLLAPL | TCSGTGQDTQY | 0.7662236 |
| TLDEALK | TCSGTGQDTQY | 0.72476673 |
| TDGAGTAEGG | TCSGTGQDTQY | 0.7081626 |
| NAPGGAGGNGG | TCSGTGQDTQY | 0.7660196 |
| VSVFMGMGGNF | TCSGTGQDTQY | 0.6867365 |
| LASYLAAL | TCSGTGQDTQY | 0.70824575 |
| LVGGTPAL | TCSGTGQDTQY | 0.74631286 |
| VRNADVL | TCSGTGQDTQY | 0.75457126 |
| TCSSPFLVSAM | TCSGTGQDTQY | 0.7175035 |
| VHITDAE | TCSGTGQDTQY | 0.7427403 |
| EYAAPSF | TCSGTGQDTQY | 0.6158098 |
| VVDAFHGGL | TCSGTGQDTQY | 0.72313493 |
| ASVVDPQ | TCSGTGQDTQY | 0.5737958 |
| SYNEFGGLS | TCSGTGQDTQY | 0.71305835 |
| DPDGSVY | TCSGTGQDTQY | 0.5798015 |
| MVSTIHATE | TCSGTGQDTQY | 0.733179 |
| LAAGVADPV | TCSGTGQDTQY | 0.72750247 |
| TKEQIAA | TCSGTGQDTQY | 0.74650055 |
| SLASPAAL | TCSGTGQDTQY | 0.47824702 |
| IGLVFYGV | TCSGTGQDTQY | 0.71760136 |
| FLLDASNL | TCSGTGQDTQY | 0.74212426 |
| YQVGNFP | TCSGTGQDTQY | 0.77485394 |
| ESLLSQST | TCSGTGQDTQY | 0.6645187 |
| LFTVGFAAVG | TCSGTGQDTQY | 0.7248369 |
| ATPPTPT | TCSGTGQDTQY | 0.63944644 |
| TDLATLG | TCSGTGQDTQY | 0.7131398 |

| | | |
|---|---|---|
| SAGGTGR | TCSGTGQDTQY | 0.52159727 |
| GFSNSGS | TCSGTGQDTQY | 0.64496404 |
| GGNGGSGGTSV | TCSGTGQDTQY | 0.6150385 |
| GTSVATGGAGN | TCSGTGQDTQY | 0.5857885 |
| NGGNGGSGGTS | TCSGTGQDTQY | 0.7360971 |
| AVVKFPH | TCSGTGQDTQY | 0.7500945 |
| TPPPPPP | TCSGTGQDTQY | 0.7417054 |
| AAPAAAAAPA | TCSGTGQDTQY | 0.7413258 |
| IALNEDL | TCSGTGQDTQY | 0.7551422 |
| IVNVSSM | TCSGTGQDTQY | 0.7699029 |
| TGIPPAPR | TCSGTGQDTQY | 0.6487546 |
| FDLGGGTF | TCSGTGQDTQY | 0.69049644 |
| FDLGGGTFDVS | TCSGTGQDTQY | 0.57926583 |
| NEPTAAA | TCSGTGQDTQY | 0.66574776 |
| NEPTAAALA | TCSGTGQDTQY | 0.6667865 |
| GSIINISSI | TCSGTGQDTQY | 0.704304 |
| VAPGNAGTA | TCSGTGQDTQY | 0.28983137 |
| TGGGASLE | TCSGTGQDTQY | 0.6756455 |
| VAVYNHYKRI | TCSGTGQDTQY | 0.7722906 |
| TDPAPATT | TCSGTGQDTQY | 0.5730492 |
| TNRPDILDPALLRPG | TCSGTGQDTQY | 0.7197099 |
| LTTTTTS | TCSGTGQDTQY | 0.7231637 |
| PTGSGKTL | TCSGTGQDTQY | 0.656728 |
| TGSGKTL | TCSGTGQDTQY | 0.7155662 |
| TIGHVDHGK | TCSGTGQDTQY | 0.6645807 |
| TIGHVDHGKTT | TCSGTGQDTQY | 0.7582298 |
| GHVDHGKTT | TCSGTGQDTQY | 0.7656964 |
| AQFEHTLL | TCSGTGQDTQY | 0.7168011 |
| IVNQDEL | TCSGTGQDTQY | 0.66499025 |
| TNGSQFF | TCSGTGQDTQY | 0.7186018 |
| TGKGIVE | TCSGTGQDTQY | 0.62436986 |
| LISNASDA | TCSGTGQDTQY | 0.6896925 |
| AELRTLGELTKIAKAHGAQ | TCSGTGQDTQY | 0.5348041 |
| APNAVAAAEPSW | TCSGTGQDTQY | 0.65882474 |
| APQVAGAATAVL | TCSGTGQDTQY | 0.6772535 |
| ARRDPAAAAAILPTDA | TCSGTGQDTQY | 0.71590924 |
| DAAARDVLAEAGLAEYF | TCSGTGQDTQY | 0.69563025 |
| DDQIVLSAKANLAKVEQ | TCSGTGQDTQY | 0.71250206 |
| DELLPDTGEDSDEDGH | TCSGTGQDTQY | 0.58846295 |
| ELPEAPANRATVPQD | TCSGTGQDTQY | 0.7151511 |
| FGQSKIERVTGIG | TCSGTGQDTQY | 0.6533536 |
| GAGGTGSTAGGAGGAGGAGGLY | TCSGTGQDTQY | 0.58532953 |
| GHGGTGAAGGNGGTGGAGGSGI | TCSGTGQDTQY | 0.7472097 |
| GNGGNGGAGGEGGPGGAGGAGA | TCSGTGQDTQY | 0.7224626 |
| GSTAGNGGNGGNATNGTIAGQP | TCSGTGQDTQY | 0.6351304 |
| GTGGDGGNGGNGGTGG | TCSGTGQDTQY | 0.75618666 |
| GVRRPAGIPADAPV | TCSGTGQDTQY | 0.7332361 |
| GYRQVTDAHLLTLARR | TCSGTGQDTQY | 0.6260002 |
| IAKMGQDGHDRG | TCSGTGQDTQY | 0.5989544 |
| KAEPSIEIQYT | TCSGTGQDTQY | 0.69734734 |
| KPVRIRMAADEDTVAVT | TCSGTGQDTQY | 0.6884492 |

| | | |
|---|---|---|
| MPPRIAGMRLLVIKPEP | TCSGTGQDTQY | 0.35111463 |
| MKFYAVALFRSRGVPPTR | TCSGTGQDTQY | 0.7367266 |
| NAGGFGILGAVAHSP | TCSGTGQDTQY | 0.6682271 |
| NEGREITLAMAEM | TCSGTGQDTQY | 0.5732728 |
| NELVEWSHPLGHRWRWE | TCSGTGQDTQY | 0.68494666 |
| NLGAGGTGGNGGNPG | TCSGTGQDTQY | 0.7478982 |
| NTEWPKWETVMAADPVD | TCSGTGQDTQY | 0.61513543 |
| PASAAPAETQP | TCSGTGQDTQY | 0.7112304 |
| QQSSDGKNGVRAAEITGE | TCSGTGQDTQY | 0.7099911 |
| RAPDAVTTSWWKE | TCSGTGQDTQY | 0.72555995 |
| SGFGNVGAHTSGWFNQ | TCSGTGQDTQY | 0.75729984 |
| SGVAGAVGQAASVGGLKV | TCSGTGQDTQY | 0.7257042 |
| SNGGAGTGSAGGNGGT | TCSGTGQDTQY | 0.710039 |
| TGGAGGEGGAAG | TCSGTGQDTQY | 0.7336366 |
| TGGDGGAAGAGGAPGHG | TCSGTGQDTQY | 0.7022625 |
| TMPPLLAAIANAGRHGV | TCSGTGQDTQY | 0.6177472 |
| TSSTAGNSGGA | TCSGTGQDTQY | 0.53889686 |
| VPATVPVGILC | TCSGTGQDTQY | 0.66856956 |
| VQRPDSGGVVLTGRIS | TCSGTGQDTQY | 0.61210126 |
| WTPNPVPATKGVPAGLVQAT | TCSGTGQDTQY | 0.71424985 |
| YALKRALAKFSDRFDVVII | TCSGTGQDTQY | 0.7739577 |
| YLIGSVIV | TCSGTGQDTQY | 0.68514323 |
| YGQLVAVID | TCSGTGQDTQY | 0.55981356 |
| VTVGGMTLPA | TCSGTGQDTQY | 0.6650998 |
| VLALPAV | TCSGTGQDTQY | 0.68420696 |
| VIPPTLNYET | TCSGTGQDTQY | 0.73928094 |
| VGMGGMHPSSG | TCSGTGQDTQY | 0.7545224 |
| VGLAGGAATA | TCSGTGQDTQY | 0.7728161 |
| VGGAGGAGGSGP | TCSGTGQDTQY | 0.7331334 |
| VFAAFMV | TCSGTGQDTQY | 0.6338043 |
| TLYDKLR | TCSGTGQDTQY | 0.74637234 |
| TLAVFTVL | TCSGTGQDTQY | 0.67735577 |
| TGSVQPARV | TCSGTGQDTQY | 0.70742893 |
| TGGGAGSGGT | TCSGTGQDTQY | 0.73341906 |
| SHLETSL | TCSGTGQDTQY | 0.72596234 |
| SFTTGMT | TCSGTGQDTQY | 0.70188236 |
| QVEVAKRMGVRQPTVS | TCSGTGQDTQY | 0.7128153 |
| QGSGGPALAM | TCSGTGQDTQY | 0.58282113 |
| MLAPSNP | TCSGTGQDTQY | 0.7509043 |
| LVNNAAI | TCSGTGQDTQY | 0.6799879 |
| LTGEINTLA | TCSGTGQDTQY | 0.58728385 |
| LSIFGPV | TCSGTGQDTQY | 0.6642573 |
| LLALLANL | TCSGTGQDTQY | 0.7586885 |
| LISNASDALDKLR | TCSGTGQDTQY | 0.7437554 |
| LINEAVP | TCSGTGQDTQY | 0.59268373 |
| LGYAFSGL | TCSGTGQDTQY | 0.6402057 |
| LDFVAVAL | TCSGTGQDTQY | 0.69105285 |
| LAQPGAE | TCSGTGQDTQY | 0.584981 |
| ISNLGAVTGF | TCSGTGQDTQY | 0.7520428 |
| HQEDPGRI | TCSGTGQDTQY | 0.6431741 |
| GTPGEPTP | TCSGTGQDTQY | 0.66217935 |

| | | |
|---|---|---|
| GFFNSGS | TCSGTGQDTQY | 0.65662146 |
| GAGGGGLTAGAN | TCSGTGQDTQY | 0.7309899 |
| FVDALVNM | TCSGTGQDTQY | 0.6382409 |
| FLTVAPEL | TCSGTGQDTQY | 0.7462818 |
| FETLTAL | TCSGTGQDTQY | 0.343757 |
| DVINPIEL | TCSGTGQDTQY | 0.7552209 |
| APVSGAPV | TCSGTGQDTQY | 0.5851379 |
| ALVGGLAW | TCSGTGQDTQY | 0.6281578 |
| ALAAAAATAI | TCSGTGQDTQY | 0.4501575 |
| AFGLGLLL | TCSGTGQDTQY | 0.7184339 |
| AFFAAASL | TCSGTGQDTQY | 0.71973056 |
| AAAALPAFSPP | TCSGTGQDTQY | 0.7548699 |
| VVLAPAL | TCSGTGQDTQY | 0.6744214 |
| TIGHVDHGKTTL | TCSGTGQDTQY | 0.7395125 |
| LISNASDALDK | TCSGTGQDTQY | 0.7437848 |
| LISNASDAL | TCSGTGQDTQY | 0.64337075 |
| ISNASDALDK | TCSGTGQDTQY | 0.49899143 |
| NQAETLVYQTE | TCSGTGQDTQY | 0.7244743 |
| TGGTGGTAGAG | TCSGTGQDTQY | 0.6552428 |
| TGSTAGGAGGA | TCSGTGQDTQY | 0.7522931 |
| DPLAVPVAAGPV | TCSGTGQDTQY | 0.6880014 |
| GGTGGTPGNGGN | TCSGTGQDTQY | 0.52911043 |
| PAGLAGVDPATL | TCSGTGQDTQY | 0.73934925 |
| GNNLVVEAPHAGASV | TCSGTGQDTQY | 0.73184586 |
| KVTRIMSASTLEQPA | TCSGTGQDTQY | 0.5711888 |
| LGFALPIALPRRTVV | TCSGTGQDTQY | 0.65520424 |
| TRLGNLTARKLVVLV | TCSGTGQDTQY | 0.69075847 |
| WSQQRRLIYFPSAGP | TCSGTGQDTQY | 0.55112636 |
| GPNGNPDPMAAAVDIR | TCSGTGQDTQY | 0.73481315 |
| TQRCVSVADLRAAASR | TCSGTGQDTQY | 0.6215297 |
| QFVLDVAEHPVEVLTETV | TCSGTGQDTQY | 0.65456235 |
| GGAGGTGGAGGNGGTQAGDG | TCSGTGQDTQY | 0.7199969 |
| PLLATLRAIAVDVDAKPAQV | TCSGTGQDTQY | 0.6835471 |
| APKFRTGGYMIDFWGVGYQVAK | TCSGTGQDTQY | 0.7032025 |
| GAGGAGGAGGAAGGPGGTGGTG | TCSGTGQDTQY | 0.696648 |
| IGALGLLATDVAPAGSVAFGVA | TCSGTGQDTQY | 0.74759585 |
| AGGAGGIGGNGGAGGAGGVLMG | TCSGTGQDTQY | 0.73688376 |
| ADELIGLL | TCSGTGQDTQY | 0.6838399 |
| ADRALKA | TCSGTGQDTQY | 0.7080188 |
| AGAVGQVGGAGGA | TCSGTGQDTQY | 0.75971717 |
| AGGGGGLGGAP | TCSGTGQDTQY | 0.58481884 |
| AGGLLYGNGGAG | TCSGTGQDTQY | 0.65197706 |
| AGLESIGL | TCSGTGQDTQY | 0.7307533 |
| AGLKTGII | TCSGTGQDTQY | 0.6223965 |
| ASLAALLL | TCSGTGQDTQY | 0.74945927 |
| ASVINISSI | TCSGTGQDTQY | 0.6962495 |
| ATPPGPA | TCSGTGQDTQY | 0.7173743 |
| ATVASALV | TCSGTGQDTQY | 0.6644344 |
| ATVASLIL | TCSGTGQDTQY | 0.7683149 |
| AVNGPVLIRH | TCSGTGQDTQY | 0.7449912 |
| DGAAGAVGI | TCSGTGQDTQY | 0.723047 |

| | | |
|---|---|---|
| DPLLHGI | TCSGTGQDTQY | 0.75268614 |
| ELGAALGAL | TCSGTGQDTQY | 0.59309053 |
| ELGRTQAV | TCSGTGQDTQY | 0.74978316 |
| FVAGGYLS | TCSGTGQDTQY | 0.740061 |
| GASGGAGGAGGW | TCSGTGQDTQY | 0.55605567 |
| GGAGAGGDGGAGGI | TCSGTGQDTQY | 0.6515655 |
| GTGHNGSQPGMGGNG | TCSGTGQDTQY | 0.76030016 |
| GYAAASASAS | TCSGTGQDTQY | 0.762761 |
| HQVDVAAVQQG | TCSGTGQDTQY | 0.61682606 |
| IGAAIAHT | TCSGTGQDTQY | 0.6558148 |
| IGMIEVAQL | TCSGTGQDTQY | 0.545542 |
| IRLAPEHTSAMD | TCSGTGQDTQY | 0.6067635 |
| IYNVSVL | TCSGTGQDTQY | 0.632467 |
| KQGFLGM | TCSGTGQDTQY | 0.72750914 |
| LESVITQL | TCSGTGQDTQY | 0.6667255 |
| LGLGALVVV | TCSGTGQDTQY | 0.7234888 |
| LIGNGGNGGSGGI | TCSGTGQDTQY | 0.71480626 |
| LNAINAPTL | TCSGTGQDTQY | 0.76235056 |
| LQLLNTGV | TCSGTGQDTQY | 0.6413659 |
| LQTSIEF | TCSGTGQDTQY | 0.6879689 |
| MSFAVSEL | TCSGTGQDTQY | 0.5946677 |
| NTGALGL | TCSGTGQDTQY | 0.62526995 |
| PPGAGGAGGAGTT | TCSGTGQDTQY | 0.6697201 |
| QLVGLGI | TCSGTGQDTQY | 0.6694594 |
| RFLEQVHQL | TCSGTGQDTQY | 0.45462894 |
| RGVGYVLR | TCSGTGQDTQY | 0.7699473 |
| SATGVFAGVI | TCSGTGQDTQY | 0.6987344 |
| SLPAITLP | TCSGTGQDTQY | 0.7486087 |
| STALAFTV | TCSGTGQDTQY | 0.6778867 |
| SVLTVGLGK | TCSGTGQDTQY | 0.6919056 |
| TALAGAGSGI | TCSGTGQDTQY | 0.6445965 |
| TGLPFTGI | TCSGTGQDTQY | 0.7220744 |
| VAEHVTV | TCSGTGQDTQY | 0.6197637 |
| VDIVLNSL | TCSGTGQDTQY | 0.57704604 |
| VLTGGGAVL | TCSGTGQDTQY | 0.66368973 |
| VNVGLAYGI | TCSGTGQDTQY | 0.62757623 |
| VSSAAVGAV | TCSGTGQDTQY | 0.76463634 |
| VTSVYEAGV | TCSGTGQDTQY | 0.7551749 |
| WVALDRGA | TCSGTGQDTQY | 0.70787823 |
| YLVGSVVI | TCSGTGQDTQY | 0.7697669 |
| GPPGTGKTLL | TCSGTGQDTQY | 0.33209267 |
| GVGMTKF | TCSGTGQDTQY | 0.67142713 |
| IGIGGSDL | TCSGTGQDTQY | 0.75873345 |
| IGTLLAEL | TCSGTGQDTQY | 0.7230046 |
| IYTSGTT | TCSGTGQDTQY | 0.6976079 |
| LSFPAGV | TCSGTGQDTQY | 0.76360947 |
| LYTSGTT | TCSGTGQDTQY | 0.6185584 |
| VGAFPFGF | TCSGTGQDTQY | 0.68579614 |
| VGAGTVEF | TCSGTGQDTQY | 0.7169977 |
| VGKSTLLL | TCSGTGQDTQY | 0.7624887 |
| AALAHAGGGIV | TCSGTGQDTQY | 0.73168445 |

| | | |
|---|---|---|
| AALVAVSLL | TCSGTGQDTQY | 0.6089871 |
| AAVSRAVAAL | TCSGTGQDTQY | 0.7317912 |
| AGGSSGIGGF | TCSGTGQDTQY | 0.77636397 |
| AGLTVANANG | TCSGTGQDTQY | 0.64383024 |
| AGLVGAVL | TCSGTGQDTQY | 0.73742485 |
| ALSQAFTV | TCSGTGQDTQY | 0.7255334 |
| ALTFSVSL | TCSGTGQDTQY | 0.6088986 |
| ALTGAALVAL | TCSGTGQDTQY | 0.60807896 |
| ASFSIGAGVL | TCSGTGQDTQY | 0.6240985 |
| DGSVYKI | TCSGTGQDTQY | 0.70913285 |
| EQLAEIV | TCSGTGQDTQY | 0.56490517 |
| FAVSELTVM | TCSGTGQDTQY | 0.7132903 |
| GAAPLAAGL | TCSGTGQDTQY | 0.7020581 |
| GADGGDGGSGGAGG | TCSGTGQDTQY | 0.7743822 |
| GFGLDTGGAGG | TCSGTGQDTQY | 0.722984 |
| GLASGGLL | TCSGTGQDTQY | 0.73789126 |
| GRIAGAAVLL | TCSGTGQDTQY | 0.56828517 |
| GSAGSLNTSF | TCSGTGQDTQY | 0.72047067 |
| GTGQIGI | TCSGTGQDTQY | 0.7180501 |
| GTLIALIL | TCSGTGQDTQY | 0.63743055 |
| IAATMGFVVG | TCSGTGQDTQY | 0.74419844 |
| IFAQALL | TCSGTGQDTQY | 0.6462845 |
| IGAAVALGF | TCSGTGQDTQY | 0.66206115 |
| IGLTPLAL | TCSGTGQDTQY | 0.7494335 |
| ILGSNTSTL | TCSGTGQDTQY | 0.76278436 |
| IVFSLTVL | TCSGTGQDTQY | 0.7071574 |
| LASKATGF | TCSGTGQDTQY | 0.71751195 |
| LASVALIL | TCSGTGQDTQY | 0.6313283 |
| LFAVGLTL | TCSGTGQDTQY | 0.746305 |
| LFNGGIGTYI | TCSGTGQDTQY | 0.75033295 |
| LTLGAGII | TCSGTGQDTQY | 0.6846366 |
| LVAAALGDI | TCSGTGQDTQY | 0.7758005 |
| NTAFVQLGI | TCSGTGQDTQY | 0.74955 |
| QLFSGAEL | TCSGTGQDTQY | 0.5399015 |
| SAATFITL | TCSGTGQDTQY | 0.7098082 |
| SNYGLANLGV | TCSGTGQDTQY | 0.7567069 |
| SYGGAYAVM | TCSGTGQDTQY | 0.74871016 |
| TTGVGGLTLG | TCSGTGQDTQY | 0.7737967 |
| TTQTIYI | TCSGTGQDTQY | 0.72134 |
| VAAGAGLGAVI | TCSGTGQDTQY | 0.6081194 |
| VAAVVTLL | TCSGTGQDTQY | 0.7595215 |
| VGFLGGLI | TCSGTGQDTQY | 0.7744436 |
| VGVIPILL | TCSGTGQDTQY | 0.74706763 |
| VIFASSLI | TCSGTGQDTQY | 0.64665073 |
| VKGGDGGAAAG | TCSGTGQDTQY | 0.6442122 |
| VTKTFTNI | TCSGTGQDTQY | 0.7408669 |
| VVAAAILL | TCSGTGQDTQY | 0.7471702 |
| YAALDVEL | TCSGTGQDTQY | 0.6528423 |
| LASVALLL | TCSGTGQDTQY | 0.7005539 |
| LDAVHRV | TCSGTGQDTQY | 0.69608945 |
| QGLSSSIF | TCSGTGQDTQY | 0.53880984 |

| | | |
|---|---|---|
| RGTFANIRL | TCSGTGQDTQY | 0.7285713 |
| RQLWWGHRI | TCSGTGQDTQY | 0.74903905 |
| YGPPGTGKTLL | TCSGTGQDTQY | 0.75540346 |
| AALVGAPANVA | TCSGTGQDTQY | 0.62021375 |
| AATFRRTE | TCSGTGQDTQY | 0.73875797 |
| AGALGGGI | TCSGTGQDTQY | 0.760841 |
| AGGAGGAGGIGAG | TCSGTGQDTQY | 0.775696 |
| AGSGALGL | TCSGTGQDTQY | 0.5835073 |
| AQIADQI | TCSGTGQDTQY | 0.37294507 |
| ASAVAVTYL | TCSGTGQDTQY | 0.6953628 |
| ASTVLGGTGG | TCSGTGQDTQY | 0.71820515 |
| AVGGEPAK | TCSGTGQDTQY | 0.6885983 |
| CEQGTFGGTC | TCSGTGQDTQY | 0.6539458 |
| DEGKAPAG | TCSGTGQDTQY | 0.7094784 |
| DTVVFPAL | TCSGTGQDTQY | 0.7500297 |
| FEGITMAG | TCSGTGQDTQY | 0.67065877 |
| FPLVVQD | TCSGTGQDTQY | 0.58021426 |
| GAFTTPQ | TCSGTGQDTQY | 0.7476975 |
| GFGGFGTAGGVGGA | TCSGTGQDTQY | 0.74744225 |
| GFLAQFL | TCSGTGQDTQY | 0.6210413 |
| GGAGGAGGNAGVA | TCSGTGQDTQY | 0.65203696 |
| GGVGTAGAGGAGGAGG | TCSGTGQDTQY | 0.70625544 |
| GNGGGGDAGNAGSGGN | TCSGTGQDTQY | 0.6618341 |
| GTGGAGGAGNDAGS | TCSGTGQDTQY | 0.6935565 |
| GTTNTGWFNS | TCSGTGQDTQY | 0.5354472 |
| IFMGPLFTL | TCSGTGQDTQY | 0.7604043 |
| ILAHAIEAI | TCSGTGQDTQY | 0.59678346 |
| IRSAIREAL | TCSGTGQDTQY | 0.73188776 |
| ITFAGGG | TCSGTGQDTQY | 0.71589595 |
| ITGMSGHQEV | TCSGTGQDTQY | 0.7354416 |
| IVAGLACTL | TCSGTGQDTQY | 0.6691927 |
| IVGGGLAGA | TCSGTGQDTQY | 0.5993356 |
| IVTGVLNI | TCSGTGQDTQY | 0.6118194 |
| LAQQGCI | TCSGTGQDTQY | 0.49242508 |
| LCLHVEN | TCSGTGQDTQY | 0.7532069 |
| LDAISDGS | TCSGTGQDTQY | 0.6461102 |
| LGGFAAML | TCSGTGQDTQY | 0.7670315 |
| LGGKNPMI | TCSGTGQDTQY | 0.7401732 |
| LGPKGVAAIL | TCSGTGQDTQY | 0.686721 |
| LMVGVSI | TCSGTGQDTQY | 0.7299669 |
| LVGAAFVL | TCSGTGQDTQY | 0.58214736 |
| LVLAVAPL | TCSGTGQDTQY | 0.6993508 |
| MAAIAVIDAA | TCSGTGQDTQY | 0.67702204 |
| NNGINAELQ | TCSGTGQDTQY | 0.75699604 |
| RGEPLVLVHG | TCSGTGQDTQY | 0.64864653 |
| RGGAGGAAGLI | TCSGTGQDTQY | 0.7536356 |
| RSYRMHPAV | TCSGTGQDTQY | 0.7082387 |
| SAAHLLAGQLA | TCSGTGQDTQY | 0.59907997 |
| SAAVGSY | TCSGTGQDTQY | 0.741664 |
| SPHAVLHRI | TCSGTGQDTQY | 0.7454649 |
| SQIAKVL | TCSGTGQDTQY | 0.7522014 |

| | | |
|---|---|---|
| TALGHLVP | TCSGTGQDTQY | 0.5885265 |
| TAPTTSGAA | TCSGTGQDTQY | 0.7211422 |
| TDGDDGNGGSPGT | TCSGTGQDTQY | 0.60198176 |
| TDGLGEN | TCSGTGQDTQY | 0.7062365 |
| TGGPPLPGQA | TCSGTGQDTQY | 0.7171265 |
| TIPAITI | TCSGTGQDTQY | 0.67359674 |
| TIPGFTI | TCSGTGQDTQY | 0.75326335 |
| TSIASGALA | TCSGTGQDTQY | 0.5810526 |
| TVTLPGAAI | TCSGTGQDTQY | 0.61414266 |
| VASGLSAALP | TCSGTGQDTQY | 0.6886033 |
| VGAGVIGI | TCSGTGQDTQY | 0.7177628 |
| VGPKRFGV | TCSGTGQDTQY | 0.7411083 |
| VNVAFPD | TCSGTGQDTQY | 0.5674111 |
| VSMTNSGVS | TCSGTGQDTQY | 0.6783077 |
| VTDVLGDLSV | TCSGTGQDTQY | 0.7028197 |
| VVVVGAGFAGL | TCSGTGQDTQY | 0.62516713 |
| VYTVGTVV | TCSGTGQDTQY | 0.74453014 |
| WTTFSGP | TCSGTGQDTQY | 0.7053967 |
| YDELVEAF | TCSGTGQDTQY | 0.5922414 |
| YGTGGAGGNGG | TCSGTGQDTQY | 0.75316125 |
| YTNVMAV | TCSGTGQDTQY | 0.6849132 |
| FLEQVHQL | TCSGTGQDTQY | 0.72953385 |
| GLSLLAVAV | TCSGTGQDTQY | 0.7586224 |
| LEQVHQL | TCSGTGQDTQY | 0.52301913 |
| LNAAGAI | TCSGTGQDTQY | 0.7333565 |
| PAVHASL | TCSGTGQDTQY | 0.70627844 |
| TELEDDA | TCSGTGQDTQY | 0.70255655 |
| TPPGPPN | TCSGTGQDTQY | 0.69168246 |
| TPPPPPPP | TCSGTGQDTQY | 0.6558932 |
| TQSPALI | ASSSPGTTNTEVF | 0.76241744 |
| EFAAAFPQ | ASSSPGTTNTEVF | 0.73724055 |
| LLSGNLDVL | ASSSPGTTNTEVF | 0.7232695 |
| GPANLGNYNI | ASSSPGTTNTEVF | 0.735018 |
| TLGPTLP | ASSSPGTTNTEVF | 0.7378074 |
| VYAYSSCCK | ASSSPGTTNTEVF | 0.7529771 |
| APAPAPAPNI | ASSSPGTTNTEVF | 0.58077896 |
| PGGPVPIP | ASSSPGTTNTEVF | 0.5594974 |
| FLTVALEP | ASSSPGTTNTEVF | 0.6700969 |
| SQYFGANL | ASSSPGTTNTEVF | 0.65953875 |
| GGVKPAA | ASSSPGTTNTEVF | 0.7000507 |
| GGTGGAGGAGGW | ASSSPGTTNTEVF | 0.53103244 |
| GVGGVAGLGGAGG | ASSSPGTTNTEVF | 0.7480494 |
| VLLVLATTI | ASSSPGTTNTEVF | 0.7734589 |
| IGIFAAML | ASSSPGTTNTEVF | 0.63417935 |
| AVGDLID | ASSSPGTTNTEVF | 0.7790665 |
| NISPLPL | ASSSPGTTNTEVF | 0.6019721 |
| VTAVFVAI | ASSSPGTTNTEVF | 0.7334981 |
| SGDDNDAAL | ASSSPGTTNTEVF | 0.75703686 |
| AVAALAHS | ASSSPGTTNTEVF | 0.5782171 |
| GGVTVMATP | ASSSPGTTNTEVF | 0.72678185 |
| AGLAANVRD | ASSSPGTTNTEVF | 0.67699087 |

| | | |
|---|---|---|
| TNRPDLIDPALLRPG | ASSSPGTTNTEVF | 0.686169 |
| LANLGVTAL | ASSSPGTTNTEVF | 0.7539979 |
| TGERIEL | ASSSPGTTNTEVF | 0.6779755 |
| NGEELAA | ASSSPGTTNTEVF | 0.75027233 |
| VLSLVWG | ASSSPGTTNTEVF | 0.7186089 |
| IGYALWL | ASSSPGTTNTEVF | 0.6557152 |
| LSLLLAPL | ASSSPGTTNTEVF | 0.755104 |
| TLDEALK | ASSSPGTTNTEVF | 0.7523873 |
| TDGAGTAEGG | ASSSPGTTNTEVF | 0.6708363 |
| NAPGGAGGNGG | ASSSPGTTNTEVF | 0.7719237 |
| VSVFMGMGGNF | ASSSPGTTNTEVF | 0.6364577 |
| LASYLAAL | ASSSPGTTNTEVF | 0.77673924 |
| LVGGTPAL | ASSSPGTTNTEVF | 0.7485279 |
| VRNADVL | ASSSPGTTNTEVF | 0.6757397 |
| TCSSPFLVSAM | ASSSPGTTNTEVF | 0.69629705 |
| VHITDAE | ASSSPGTTNTEVF | 0.72977203 |
| EYAAPSF | ASSSPGTTNTEVF | 0.677104 |
| VVDAFHGGL | ASSSPGTTNTEVF | 0.7188053 |
| ASVVDPQ | ASSSPGTTNTEVF | 0.7719108 |
| SYNEFGGLS | ASSSPGTTNTEVF | 0.7391553 |
| DPDGSVY | ASSSPGTTNTEVF | 0.8605773 |
| MVSTIHATE | ASSSPGTTNTEVF | 0.8178403 |
| LAAGVADPV | ASSSPGTTNTEVF | 0.8270827 |
| TKEQIAA | ASSSPGTTNTEVF | 0.7582283 |
| SLASPAAL | ASSSPGTTNTEVF | 0.5733978 |
| IGLVFYGV | ASSSPGTTNTEVF | 0.8112609 |
| FLLDASNL | ASSSPGTTNTEVF | 0.7151075 |
| YQVGNFP | ASSSPGTTNTEVF | 0.77861416 |
| ESLLSQST | ASSSPGTTNTEVF | 0.85255593 |
| LFTVGFAAVG | ASSSPGTTNTEVF | 0.7578393 |
| ATPPTPT | ASSSPGTTNTEVF | 0.86295366 |
| TDLATLG | ASSSPGTTNTEVF | 0.86637896 |
| SAGGTGR | ASSSPGTTNTEVF | 0.56352663 |
| GFSNSGS | ASSSPGTTNTEVF | 0.75475895 |
| GGNGGSGGTSV | ASSSPGTTNTEVF | 0.7236712 |
| GTSVATGGAGN | ASSSPGTTNTEVF | 0.7938472 |
| NGGNGGSGGTS | ASSSPGTTNTEVF | 0.3756197 |
| AVVKFPH | ASSSPGTTNTEVF | 0.8530065 |
| TPPPPPP | ASSSPGTTNTEVF | 0.69243336 |
| AAPAAAAAPA | ASSSPGTTNTEVF | 0.2629177 |
| IALNEDL | ASSSPGTTNTEVF | 0.8206624 |
| IVNVSSM | ASSSPGTTNTEVF | 0.7755226 |
| TGIPPAPR | ASSSPGTTNTEVF | 0.7851759 |
| FDLGGGTF | ASSSPGTTNTEVF | 0.84169173 |
| FDLGGGTFDVS | ASSSPGTTNTEVF | 0.6006649 |
| NEPTAAA | ASSSPGTTNTEVF | 0.7353625 |
| NEPTAAALA | ASSSPGTTNTEVF | 0.8323642 |
| GSIINISSI | ASSSPGTTNTEVF | 0.8713965 |
| VAPGNAGTA | ASSSPGTTNTEVF | 0.6975208 |
| TGGGASLE | ASSSPGTTNTEVF | 0.8109054 |
| VAVYNHYKRI | ASSSPGTTNTEVF | 0.868269 |

| | | |
|---|---|---|
| TDPAPATT | ASSSPGTTNTEVF | 0.8405344 |
| TNRPDILDPALLRPG | ASSSPGTTNTEVF | 0.86146456 |
| LTTTTTS | ASSSPGTTNTEVF | 0.8740936 |
| PTGSGKTL | ASSSPGTTNTEVF | 0.7307432 |
| TGSGKTL | ASSSPGTTNTEVF | 0.8537958 |
| TIGHVDHGK | ASSSPGTTNTEVF | 0.8685796 |
| TIGHVDHGKTT | ASSSPGTTNTEVF | 0.7933426 |
| GHVDHGKTT | ASSSPGTTNTEVF | 0.81506985 |
| AQFEHTLL | ASSSPGTTNTEVF | 0.7385827 |
| IVNQDEL | ASSSPGTTNTEVF | 0.76617074 |
| TNGSQFF | ASSSPGTTNTEVF | 0.42114168 |
| TGKGIVE | ASSSPGTTNTEVF | 0.8042766 |
| LISNASDA | ASSSPGTTNTEVF | 0.85835147 |
| AELRTLGELTKIAKAHGAQ | ASSSPGTTNTEVF | 0.75154006 |
| APNAVAAAEPSW | ASSSPGTTNTEVF | 0.6185637 |
| APQVAGAATAVL | ASSSPGTTNTEVF | 0.6714242 |
| ARRDPAAAAAILPTDA | ASSSPGTTNTEVF | 0.7632787 |
| DAAARDVLAEAGLAEYF | ASSSPGTTNTEVF | 0.7947466 |
| DDQIVLSAKANLAKVEQ | ASSSPGTTNTEVF | 0.5778366 |
| DELLPDTGEDSDEDGH | ASSSPGTTNTEVF | 0.65884644 |
| ELPEAPANRATVPQD | ASSSPGTTNTEVF | 0.8603428 |
| FGQSKIERVTGIG | ASSSPGTTNTEVF | 0.8665444 |
| GAGGTGSTAGGAGGAGGAGGLY | ASSSPGTTNTEVF | 0.49829313 |
| GHGGTGAAGGNGGTGGAGGSGI | ASSSPGTTNTEVF | 0.8541169 |
| GNGGNGGAGGEGGPGGAGGAGA | ASSSPGTTNTEVF | 0.8352421 |
| GSTAGNGGNGGNATNGTIAGQP | ASSSPGTTNTEVF | 0.8686769 |
| GTGGDGGNGGNGGTGG | ASSSPGTTNTEVF | 0.82601035 |
| GVRRPAGIPADAPV | ASSSPGTTNTEVF | 0.70475644 |
| GYRQVTDAHLLTLARR | ASSSPGTTNTEVF | 0.7984412 |
| IAKMGQDGHDRG | ASSSPGTTNTEVF | 0.72403795 |
| KAEPSIEIQYT | ASSSPGTTNTEVF | 0.7613528 |
| KPVRIRMAADEDTVAVT | ASSSPGTTNTEVF | 0.85359275 |
| MPPRIAGMRLLVIKPEP | ASSSPGTTNTEVF | 0.77885985 |
| MKFYAVALFRSRGVPPTR | ASSSPGTTNTEVF | 0.7180651 |
| NAGGFGILGAVAHSP | ASSSPGTTNTEVF | 0.6549667 |
| NEGREITLAMAEM | ASSSPGTTNTEVF | 0.87078524 |
| NELVEWSHPLGHRWRWE | ASSSPGTTNTEVF | 0.56215715 |
| NLGAGGTGGNGGNPG | ASSSPGTTNTEVF | 0.713068 |
| NTEWPKWETVMAADPVD | ASSSPGTTNTEVF | 0.711325 |
| PASAAPAETQP | ASSSPGTTNTEVF | 0.80021405 |
| QQSSDGKNGVRAAEITGE | ASSSPGTTNTEVF | 0.858432 |
| RAPDAVTTSWWKE | ASSSPGTTNTEVF | 0.8420844 |
| SGFGNVGAHTSGWFNQ | ASSSPGTTNTEVF | 0.7419828 |
| SGVAGAVGQAASVGGLKV | ASSSPGTTNTEVF | 0.83184206 |
| SNGGAGTGSAGGNGGT | ASSSPGTTNTEVF | 0.8485134 |
| TGGAGGEGGAAG | ASSSPGTTNTEVF | 0.82667273 |
| TGGDGGAAGAGGAPGHG | ASSSPGTTNTEVF | 0.8448864 |
| TMPPLLAAIANAGRHGV | ASSSPGTTNTEVF | 0.6690106 |
| TSSTAGNSGGA | ASSSPGTTNTEVF | 0.8682484 |
| VPATVPVGILC | ASSSPGTTNTEVF | 0.83587736 |
| VQRPDSGGVVLTGRIS | ASSSPGTTNTEVF | 0.86003643 |

| | | |
|---|---|---|
| WTPNPVPATKGVPAGLVQAT | ASSSPGTTNTEVF | 0.85083205 |
| YALKRALAKFSDRFDVVII | ASSSPGTTNTEVF | 0.820934 |
| YLIGSVIV | ASSSPGTTNTEVF | 0.81762713 |
| YGQLVAVID | ASSSPGTTNTEVF | 0.49445382 |
| VTVGGMTLPA | ASSSPGTTNTEVF | 0.75297487 |
| VLALPAV | ASSSPGTTNTEVF | 0.8165716 |
| VIPPTLNYET | ASSSPGTTNTEVF | 0.79487056 |
| VGMGGMHPSSG | ASSSPGTTNTEVF | 0.83972377 |
| VGLAGGAATA | ASSSPGTTNTEVF | 0.8554624 |
| VGGAGGAGGSGP | ASSSPGTTNTEVF | 0.7217118 |
| VFAAFMV | ASSSPGTTNTEVF | 0.86829615 |
| TLYDKLR | ASSSPGTTNTEVF | 0.8433001 |
| TLAVFTVL | ASSSPGTTNTEVF | 0.55427027 |
| TGSVQPARV | ASSSPGTTNTEVF | 0.7726838 |
| TGGGAGSGGT | ASSSPGTTNTEVF | 0.8665266 |
| SHLETSL | ASSSPGTTNTEVF | 0.8001523 |
| SFTTGMT | ASSSPGTTNTEVF | 0.7777294 |
| QVEVAKRMGVRQPTVS | ASSSPGTTNTEVF | 0.7932284 |
| QGSGGPALAM | ASSSPGTTNTEVF | 0.8610178 |
| MLAPSNP | ASSSPGTTNTEVF | 0.76717967 |
| LVNNAAI | ASSSPGTTNTEVF | 0.8724865 |
| LTGEINTLA | ASSSPGTTNTEVF | 0.2925494 |
| LSIFGPV | ASSSPGTTNTEVF | 0.7780043 |
| LLALLANL | ASSSPGTTNTEVF | 0.7408925 |
| LISNASDALDKLR | ASSSPGTTNTEVF | 0.5778534 |
| LINEAVP | ASSSPGTTNTEVF | 0.86249405 |
| LGYAFSGL | ASSSPGTTNTEVF | 0.7919808 |
| LDFVAVAL | ASSSPGTTNTEVF | 0.84839463 |
| LAQPGAE | ASSSPGTTNTEVF | 0.773993 |
| ISNLGAVTGF | ASSSPGTTNTEVF | 0.85828644 |
| HQEDPGRI | ASSSPGTTNTEVF | 0.78794396 |
| GTPGEPTP | ASSSPGTTNTEVF | 0.8098565 |
| GFFNSGS | ASSSPGTTNTEVF | 0.7546624 |
| GAGGGGLTAGAN | ASSSPGTTNTEVF | 0.8525667 |
| FVDALVNM | ASSSPGTTNTEVF | 0.87400436 |
| FLTVAPEL | ASSSPGTTNTEVF | 0.85484934 |
| FETLTAL | ASSSPGTTNTEVF | 0.65483654 |
| DVINPIEL | ASSSPGTTNTEVF | 0.77028733 |
| APVSGAPV | ASSSPGTTNTEVF | 0.8359048 |
| ALVGGLAW | ASSSPGTTNTEVF | 0.36673543 |
| ALAAAAATAI | ASSSPGTTNTEVF | 0.4300424 |
| AFGLGLLL | ASSSPGTTNTEVF | 0.7085067 |
| AFFAAASL | ASSSPGTTNTEVF | 0.6213156 |
| AAAALPAFSPP | ASSSPGTTNTEVF | 0.8584251 |
| VVLAPAL | ASSSPGTTNTEVF | 0.8375498 |
| TIGHVDHGKTTL | ASSSPGTTNTEVF | 0.7766714 |
| LISNASDALDK | ASSSPGTTNTEVF | 0.8378761 |
| LISNASDAL | ASSSPGTTNTEVF | 0.817249 |
| ISNASDALDK | ASSSPGTTNTEVF | 0.1367305 |
| NQAETLVYQTE | ASSSPGTTNTEVF | 0.7408973 |
| TGGTGGTAGAG | ASSSPGTTNTEVF | 0.5885885 |

| | | |
|---|---|---|
| TGSTAGGAGGA | ASSSPGTTNTEVF | 0.87237394 |
| DPLAVPVAAGPV | ASSSPGTTNTEVF | 0.71856666 |
| GGTGGTPGNGGN | ASSSPGTTNTEVF | 0.6130979 |
| PAGLAGVDPATL | ASSSPGTTNTEVF | 0.78085876 |
| GNNLVVEAPHAGASV | ASSSPGTTNTEVF | 0.7390339 |
| KVTRIMSASTLEQPA | ASSSPGTTNTEVF | 0.77656245 |
| LGFALPIALPRRTVV | ASSSPGTTNTEVF | 0.68712384 |
| TRLGNLTARKLVVLV | ASSSPGTTNTEVF | 0.24097867 |
| WSQQRRLIYFPSAGP | ASSSPGTTNTEVF | 0.8550284 |
| GPNGNPDPMAAAVDIR | ASSSPGTTNTEVF | 0.5299963 |
| TQRCVSVADLRAAASR | ASSSPGTTNTEVF | 0.7436546 |
| QFVLDVAEHPVEVLTETV | ASSSPGTTNTEVF | 0.42926592 |
| GGAGGTGGAGGNGGTQAGDG | ASSSPGTTNTEVF | 0.44621173 |
| PLLATLRAIAVDVDAKPAQV | ASSSPGTTNTEVF | 0.6079625 |
| APKFRTGGYMIDFWGVGYQVAK | ASSSPGTTNTEVF | 0.86459655 |
| GAGGAGGAGGAAGGPGGTGGTG | ASSSPGTTNTEVF | 0.7103821 |
| IGALGLLATDVAPAGSVAFGVA | ASSSPGTTNTEVF | 0.7002984 |
| AGGAGGIGGNGGAGGAGGVLMG | ASSSPGTTNTEVF | 0.85848546 |
| ADELIGLL | ASSSPGTTNTEVF | 0.7420393 |
| ADRALKA | ASSSPGTTNTEVF | 0.7188778 |
| AGAVGQVGGAGGA | ASSSPGTTNTEVF | 0.7587085 |
| AGGGGGLGGAP | ASSSPGTTNTEVF | 0.6877892 |
| AGGLLYGNGGAG | ASSSPGTTNTEVF | 0.73851466 |
| AGLESIGL | ASSSPGTTNTEVF | 0.73382854 |
| AGLKTGII | ASSSPGTTNTEVF | 0.8754516 |
| ASLAALLL | ASSSPGTTNTEVF | 0.7798267 |
| ASVINISSI | ASSSPGTTNTEVF | 0.63987386 |
| ATPPGPA | ASSSPGTTNTEVF | 0.8047357 |
| ATVASALV | ASSSPGTTNTEVF | 0.8634807 |
| ATVASLIL | ASSSPGTTNTEVF | 0.8783265 |
| AVNGPVLIRH | ASSSPGTTNTEVF | 0.67741185 |
| DGAAGAVGI | ASSSPGTTNTEVF | 0.78039813 |
| DPLLHGI | ASSSPGTTNTEVF | 0.7970336 |
| ELGAALGAL | ASSSPGTTNTEVF | 0.7915618 |
| ELGRTQAV | ASSSPGTTNTEVF | 0.7779702 |
| FVAGGYLS | ASSSPGTTNTEVF | 0.6650048 |
| GASGGAGGAGGW | ASSSPGTTNTEVF | 0.8563873 |
| GGAGAGGDGGAGGI | ASSSPGTTNTEVF | 0.87393093 |
| GTGHNGSQPGMGGNG | ASSSPGTTNTEVF | 0.8003573 |
| GYAAASASAS | ASSSPGTTNTEVF | 0.58284056 |
| HQVDVAAVQQG | ASSSPGTTNTEVF | 0.7805084 |
| IGAAIAHT | ASSSPGTTNTEVF | 0.65246534 |
| IGMIEVAQL | ASSSPGTTNTEVF | 0.7443503 |
| IRLAPEHTSAMD | ASSSPGTTNTEVF | 0.8327167 |
| IYNVSVL | ASSSPGTTNTEVF | 0.5032975 |
| KQGFLGM | ASSSPGTTNTEVF | 0.7234744 |
| LESVITQL | ASSSPGTTNTEVF | 0.8775256 |
| LGLGALVVV | ASSSPGTTNTEVF | 0.69369036 |
| LIGNGGNGGSGGI | ASSSPGTTNTEVF | 0.7970512 |
| LNAINAPTL | ASSSPGTTNTEVF | 0.84810853 |
| LQLLNTGV | ASSSPGTTNTEVF | 0.8187022 |

| | | |
|---|---|---|
| LQTSIEF | ASSSPGTTNTEVF | 0.8618902 |
| MSFAVSEL | ASSSPGTTNTEVF | 0.839466 |
| NTGALGL | ASSSPGTTNTEVF | 0.82197404 |
| PPGAGGAGGAGTT | ASSSPGTTNTEVF | 0.8086368 |
| QLVGLGI | ASSSPGTTNTEVF | 0.76002073 |
| RFLEQVHQL | ASSSPGTTNTEVF | 0.8029464 |
| RGVGYVLR | ASSSPGTTNTEVF | 0.77205086 |
| SATGVFAGVI | ASSSPGTTNTEVF | 0.7280144 |
| SLPAITLP | ASSSPGTTNTEVF | 0.8528083 |
| STALAFTV | ASSSPGTTNTEVF | 0.8109892 |
| SVLTVGLGK | ASSSPGTTNTEVF | 0.84374154 |
| TALAGAGSGI | ASSSPGTTNTEVF | 0.7001852 |
| TGLPFTGI | ASSSPGTTNTEVF | 0.7034342 |
| VAEHVTV | ASSSPGTTNTEVF | 0.8426944 |
| VDIVLNSL | ASSSPGTTNTEVF | 0.2547129 |
| VLTGGGAVL | ASSSPGTTNTEVF | 0.45627755 |
| VNVGLAYGI | ASSSPGTTNTEVF | 0.8139012 |
| VSSAAVGAV | ASSSPGTTNTEVF | 0.85944724 |
| VTSVYEAGV | ASSSPGTTNTEVF | 0.26110715 |
| WVALDRGA | ASSSPGTTNTEVF | 0.71663916 |
| YLVGSVVI | ASSSPGTTNTEVF | 0.5712535 |
| GPPGTGKTLL | ASSSPGTTNTEVF | 0.8430287 |
| GVGMTKF | ASSSPGTTNTEVF | 0.80286306 |
| IGIGGSDL | ASSSPGTTNTEVF | 0.55638576 |
| IGTLLAEL | ASSSPGTTNTEVF | 0.8462852 |
| IYTSGTT | ASSSPGTTNTEVF | 0.4723251 |
| LSFPAGV | ASSSPGTTNTEVF | 0.52332896 |
| LYTSGTT | ASSSPGTTNTEVF | 0.8564292 |
| VGAFPFGF | ASSSPGTTNTEVF | 0.85471696 |
| VGAGTVEF | ASSSPGTTNTEVF | 0.68195426 |
| VGKSTLLL | ASSSPGTTNTEVF | 0.85525954 |
| AALAHAGGGIV | ASSSPGTTNTEVF | 0.70936245 |
| AALVAVSLL | ASSSPGTTNTEVF | 0.8760286 |
| AAVSRAVAAL | ASSSPGTTNTEVF | 0.7696192 |
| AGGSSGIGGF | ASSSPGTTNTEVF | 0.58441037 |
| AGLTVANANG | ASSSPGTTNTEVF | 0.8410422 |
| AGLVGAVL | ASSSPGTTNTEVF | 0.80396444 |
| ALSQAFTV | ASSSPGTTNTEVF | 0.85087943 |
| ALTFSVSL | ASSSPGTTNTEVF | 0.8353288 |
| ALTGAALVAL | ASSSPGTTNTEVF | 0.7448523 |
| ASFSIGAGVL | ASSSPGTTNTEVF | 0.73536265 |
| DGSVYKI | ASSSPGTTNTEVF | 0.85151494 |
| EQLAEIV | ASSSPGTTNTEVF | 0.84754574 |
| FAVSELTVM | ASSSPGTTNTEVF | 0.66232395 |
| GAAPLAAGL | ASSSPGTTNTEVF | 0.8583662 |
| GADGGDGGSGGAGG | ASSSPGTTNTEVF | 0.77764356 |
| GFGLDTGGAGG | ASSSPGTTNTEVF | 0.72313696 |
| GLASGGLL | ASSSPGTTNTEVF | 0.87464374 |
| GRIAGAAVLL | ASSSPGTTNTEVF | 0.82692873 |
| GSAGSLNTSF | ASSSPGTTNTEVF | 0.84255886 |
| GTGQIGI | ASSSPGTTNTEVF | 0.85996383 |

| | | |
|---|---|---|
| GTLIALIL | ASSSPGTTNTEVF | 0.8600386 |
| IAATMGFVVG | ASSSPGTTNTEVF | 0.6429153 |
| IFAQALL | ASSSPGTTNTEVF | 0.82965696 |
| IGAAVALGF | ASSSPGTTNTEVF | 0.81224114 |
| IGLTPLAL | ASSSPGTTNTEVF | 0.8453301 |
| ILGSNTSTL | ASSSPGTTNTEVF | 0.79660076 |
| IVFSLTVL | ASSSPGTTNTEVF | 0.64672923 |
| LASKATGF | ASSSPGTTNTEVF | 0.8265717 |
| LASVALIL | ASSSPGTTNTEVF | 0.86122465 |
| LFAVGLTL | ASSSPGTTNTEVF | 0.82297313 |
| LFNGGIGTYI | ASSSPGTTNTEVF | 0.8288541 |
| LTLGAGII | ASSSPGTTNTEVF | 0.8673065 |
| LVAAALGDI | ASSSPGTTNTEVF | 0.7583379 |
| NTAFVQLGI | ASSSPGTTNTEVF | 0.7892233 |
| QLFSGAEL | ASSSPGTTNTEVF | 0.8582659 |
| SAATFITL | ASSSPGTTNTEVF | 0.86148995 |
| SNYGLANLGV | ASSSPGTTNTEVF | 0.80050004 |
| SYGGAYAVM | ASSSPGTTNTEVF | 0.8691434 |
| TTGVGGLTLG | ASSSPGTTNTEVF | 0.7179549 |
| TTQTIYI | ASSSPGTTNTEVF | 0.8350459 |
| VAAGAGLGAVI | ASSSPGTTNTEVF | 0.6620226 |
| VAAVVTLL | ASSSPGTTNTEVF | 0.8222891 |
| VGFLGGLI | ASSSPGTTNTEVF | 0.62239707 |
| VGVIPILL | ASSSPGTTNTEVF | 0.8376269 |
| VIFASSLI | ASSSPGTTNTEVF | 0.8419604 |
| VKGGDGGAAAG | ASSSPGTTNTEVF | 0.8375774 |
| VTKTFTNI | ASSSPGTTNTEVF | 0.42129973 |
| VVAAAILL | ASSSPGTTNTEVF | 0.84670043 |
| YAALDVEL | ASSSPGTTNTEVF | 0.842113 |
| LASVALLL | ASSSPGTTNTEVF | 0.8734791 |
| LDAVHRV | ASSSPGTTNTEVF | 0.81576556 |
| QGLSSSIF | ASSSPGTTNTEVF | 0.85247946 |
| RGTFANIRL | ASSSPGTTNTEVF | 0.7723317 |
| RQLWWGHRI | ASSSPGTTNTEVF | 0.8030966 |
| YGPPGTGKTLL | ASSSPGTTNTEVF | 0.63517535 |
| AALVGAPANVA | ASSSPGTTNTEVF | 0.7690989 |
| AATFRRTE | ASSSPGTTNTEVF | 0.72234905 |
| AGALGGGI | ASSSPGTTNTEVF | 0.66403186 |
| AGGAGGAGGIGAG | ASSSPGTTNTEVF | 0.83626425 |
| AGSGALGL | ASSSPGTTNTEVF | 0.8596902 |
| AQIADQI | ASSSPGTTNTEVF | 0.8475282 |
| ASAVAVTYL | ASSSPGTTNTEVF | 0.8563173 |
| ASTVLGGTGG | ASSSPGTTNTEVF | 0.85572684 |
| AVGGEPAK | ASSSPGTTNTEVF | 0.8681119 |
| CEQGTFGGTC | ASSSPGTTNTEVF | 0.75041395 |
| DEGKAPAG | ASSSPGTTNTEVF | 0.7484191 |
| DTVVFPAL | ASSSPGTTNTEVF | 0.6307479 |
| FEGITMAG | ASSSPGTTNTEVF | 0.7370288 |
| FPLVVQD | ASSSPGTTNTEVF | 0.7492237 |
| GAFTTPQ | ASSSPGTTNTEVF | 0.80759025 |
| GFGGFGTAGGVGGA | ASSSPGTTNTEVF | 0.20491111 |

| | | |
|---|---|---|
| GFLAQFL | ASSSPGTTNTEVF | 0.7370805 |
| GGAGGAGGNAGVA | ASSSPGTTNTEVF | 0.869 |
| GGVGTAGAGGAGGAGG | ASSSPGTTNTEVF | 0.6397041 |
| GNGGGGDAGNAGSGGN | ASSSPGTTNTEVF | 0.8390969 |
| GTGGAGGAGNDAGS | ASSSPGTTNTEVF | 0.7171587 |
| GTTNTGWFNS | ASSSPGTTNTEVF | 0.71671945 |
| IFMGPLFTL | ASSSPGTTNTEVF | 0.84670746 |
| ILAHAIEAI | ASSSPGTTNTEVF | 0.7135701 |
| IRSAIREAL | ASSSPGTTNTEVF | 0.84538 |
| ITFAGGG | ASSSPGTTNTEVF | 0.86908895 |
| ITGMSGHQEV | ASSSPGTTNTEVF | 0.8097501 |
| IVAGLACTL | ASSSPGTTNTEVF | 0.8206531 |
| IVGGGLAGA | ASSSPGTTNTEVF | 0.82259357 |
| IVTGVLNI | ASSSPGTTNTEVF | 0.58638495 |
| LAQQGCI | ASSSPGTTNTEVF | 0.8354914 |
| LCLHVEN | ASSSPGTTNTEVF | 0.18913388 |
| LDAISDGS | ASSSPGTTNTEVF | 0.75383514 |
| LGGFAAML | ASSSPGTTNTEVF | 0.7071018 |
| LGGKNPMI | ASSSPGTTNTEVF | 0.82331276 |
| LGPKGVAAIL | ASSSPGTTNTEVF | 0.7919522 |
| LMVGVSI | ASSSPGTTNTEVF | 0.84350556 |
| LVGAAFVL | ASSSPGTTNTEVF | 0.65489024 |
| LVLAVAPL | ASSSPGTTNTEVF | 0.77858794 |
| MAAIAVIDAA | ASSSPGTTNTEVF | 0.728678 |
| NNGINAELQ | ASSSPGTTNTEVF | 0.6565228 |
| RGEPLVLVHG | ASSSPGTTNTEVF | 0.8347346 |
| RGGAGGAAGLI | ASSSPGTTNTEVF | 0.8205682 |
| RSYRMHPAV | ASSSPGTTNTEVF | 0.69857866 |
| SAAHLLAGQLA | ASSSPGTTNTEVF | 0.861838 |
| SAAVGSY | ASSSPGTTNTEVF | 0.8203231 |
| SPHAVLHRI | ASSSPGTTNTEVF | 0.61995727 |
| SQIAKVL | ASSSPGTTNTEVF | 0.7355548 |
| TALGHLVP | ASSSPGTTNTEVF | 0.8011055 |
| TAPTTSGAA | ASSSPGTTNTEVF | 0.79492515 |
| TDGDDGNGGSPGT | ASSSPGTTNTEVF | 0.1820731 |
| TDGLGEN | ASSSPGTTNTEVF | 0.8112967 |
| TGGPPLPGQA | ASSSPGTTNTEVF | 0.7269578 |
| TIPAITI | ASSSPGTTNTEVF | 0.61603206 |
| TIPGFTI | ASSSPGTTNTEVF | 0.7440063 |
| TSIASGALA | ASSSPGTTNTEVF | 0.8319204 |
| TVTLPGAAI | ASSSPGTTNTEVF | 0.7037459 |
| VASGLSAALP | ASSSPGTTNTEVF | 0.82202005 |
| VGAGVIGI | ASSSPGTTNTEVF | 0.8411936 |
| VGPKRFGV | ASSSPGTTNTEVF | 0.8619919 |
| VNVAFPD | ASSSPGTTNTEVF | 0.8697995 |
| VSMTNSGVS | ASSSPGTTNTEVF | 0.82968235 |
| VTDVLGDLSV | ASSSPGTTNTEVF | 0.7952118 |
| VVVVGAGFAGL | ASSSPGTTNTEVF | 0.8406168 |
| VYTVGTVV | ASSSPGTTNTEVF | 0.8259971 |
| WTTFSGP | ASSSPGTTNTEVF | 0.5569442 |
| YDELVEAF | ASSSPGTTNTEVF | 0.4762959 |

| | | |
|---|---|---|
| YGTGGAGGNGG | ASSSPGTTNTEVF | 0.81024635 |
| YTNVMAV | ASSSPGTTNTEVF | 0.51962256 |
| FLEQVHQL | ASSSPGTTNTEVF | 0.81581604 |
| GLSLLAVAV | ASSSPGTTNTEVF | 0.8713944 |
| LEQVHQL | ASSSPGTTNTEVF | 0.7956496 |
| LNAAGAI | ASSSPGTTNTEVF | 0.6680949 |
| PAVHASL | ASSSPGTTNTEVF | 0.7582499 |
| TELEDDA | ASSSPGTTNTEVF | 0.7175802 |
| TPPGPPN | ASSSPGTTNTEVF | 0.83901834 |
| TPPPPPPP | ASSSPGTTNTEVF | 0.8547906 |
| TQSPALI | ASGPTGGQDTQY | 0.86376387 |
| EFAAAFPQ | ASGPTGGQDTQY | 0.85281336 |
| LLSGNLDVL | ASGPTGGQDTQY | 0.741228 |
| GPANLGNYNI | ASGPTGGQDTQY | 0.82751393 |
| TLGPTLP | ASGPTGGQDTQY | 0.74070895 |
| VYAYSSCCK | ASGPTGGQDTQY | 0.74272573 |
| APAPAPAPNI | ASGPTGGQDTQY | 0.8047629 |
| PGGPVPIP | ASGPTGGQDTQY | 0.82323706 |
| FLTVALEP | ASGPTGGQDTQY | 0.8028736 |
| SQYFGANL | ASGPTGGQDTQY | 0.76697344 |
| GGVKPAA | ASGPTGGQDTQY | 0.6151988 |
| GGTGGAGGAGGW | ASGPTGGQDTQY | 0.84538716 |
| GVGGVAGLGGAGG | ASGPTGGQDTQY | 0.80339575 |
| VLLVLATTI | ASGPTGGQDTQY | 0.6442653 |
| IGIFAAML | ASGPTGGQDTQY | 0.73163736 |
| AVGDLID | ASGPTGGQDTQY | 0.8762796 |
| NISPLPL | ASGPTGGQDTQY | 0.8708515 |
| VTAVFVAI | ASGPTGGQDTQY | 0.6850076 |
| SGDDNDAAL | ASGPTGGQDTQY | 0.7402234 |
| AVAALAHS | ASGPTGGQDTQY | 0.75308824 |
| GGVTVMATP | ASGPTGGQDTQY | 0.4996946 |
| AGLAANVRD | ASGPTGGQDTQY | 0.8332209 |
| TNRPDLIDPALLRPG | ASGPTGGQDTQY | 0.76609516 |
| LANLGVTAL | ASGPTGGQDTQY | 0.7461866 |
| TGERIEL | ASGPTGGQDTQY | 0.76985025 |
| NGEELAA | ASGPTGGQDTQY | 0.785702 |
| VLSLVWG | ASGPTGGQDTQY | 0.7258729 |
| IGYALWL | ASGPTGGQDTQY | 0.86216605 |
| LSLLLAPL | ASGPTGGQDTQY | 0.21942651 |
| TLDEALK | ASGPTGGQDTQY | 0.8419435 |
| TDGAGTAEGG | ASGPTGGQDTQY | 0.6177653 |
| NAPGGAGGNGG | ASGPTGGQDTQY | 0.7451522 |
| VSVFMGMGGNF | ASGPTGGQDTQY | 0.604056 |
| LASYLAAL | ASGPTGGQDTQY | 0.7902775 |
| LVGGTPAL | ASGPTGGQDTQY | 0.84803814 |
| VRNADVL | ASGPTGGQDTQY | 0.8476176 |
| TCSSPFLVSAM | ASGPTGGQDTQY | 0.64900756 |
| VHITDAE | ASGPTGGQDTQY | 0.8516545 |
| EYAAPSF | ASGPTGGQDTQY | 0.88091636 |
| VVDAFHGGL | ASGPTGGQDTQY | 0.69562054 |
| ASVVDPQ | ASGPTGGQDTQY | 0.5698144 |

| | | |
|---|---|---|
| SYNEFGGLS | ASGPTGGQDTQY | 0.84019077 |
| DPDGSVY | ASGPTGGQDTQY | 0.7572965 |
| MVSTIHATE | ASGPTGGQDTQY | 0.8380185 |
| LAAGVADPV | ASGPTGGQDTQY | 0.8280241 |
| TKEQIAA | ASGPTGGQDTQY | 0.5486233 |
| SLASPAAL | ASGPTGGQDTQY | 0.8700947 |
| IGLVFYGV | ASGPTGGQDTQY | 0.8511934 |
| FLLDASNL | ASGPTGGQDTQY | 0.70103467 |
| YQVGNFP | ASGPTGGQDTQY | 0.7482921 |
| ESLLSQST | ASGPTGGQDTQY | 0.7402551 |
| LFTVGFAAVG | ASGPTGGQDTQY | 0.44381893 |
| ATPPTPT | ASGPTGGQDTQY | 0.85926974 |
| TDLATLG | ASGPTGGQDTQY | 0.63274145 |
| SAGGTGR | ASGPTGGQDTQY | 0.76896256 |
| GFSNSGS | ASGPTGGQDTQY | 0.8038415 |
| GGNGGSGGTSV | ASGPTGGQDTQY | 0.78395736 |
| GTSVATGGAGN | ASGPTGGQDTQY | 0.7726099 |
| NGGNGGSGGTS | ASGPTGGQDTQY | 0.7686664 |
| AVVKFPH | ASGPTGGQDTQY | 0.81814855 |
| TPPPPPP | ASGPTGGQDTQY | 0.8006791 |
| AAPAAAAAPA | ASGPTGGQDTQY | 0.7823013 |
| IALNEDL | ASGPTGGQDTQY | 0.7958202 |
| IVNVSSM | ASGPTGGQDTQY | 0.8585471 |
| TGIPPAPR | ASGPTGGQDTQY | 0.61578983 |
| FDLGGGTF | ASGPTGGQDTQY | 0.73232514 |
| FDLGGGTFDVS | ASGPTGGQDTQY | 0.82857734 |
| NEPTAAA | ASGPTGGQDTQY | 0.59700614 |
| NEPTAAALA | ASGPTGGQDTQY | 0.8355205 |
| GSIINISSI | ASGPTGGQDTQY | 0.8024148 |
| VAPGNAGTA | ASGPTGGQDTQY | 0.79689926 |
| TGGGASLE | ASGPTGGQDTQY | 0.7746759 |
| VAVYNHYKRI | ASGPTGGQDTQY | 0.8769705 |
| TDPAPATT | ASGPTGGQDTQY | 0.84533584 |
| TNRPDILDPALLRPG | ASGPTGGQDTQY | 0.8024509 |
| LTTTTTS | ASGPTGGQDTQY | 0.7898693 |
| PTGSGKTL | ASGPTGGQDTQY | 0.7235267 |
| TGSGKTL | ASGPTGGQDTQY | 0.8562343 |
| TIGHVDHGK | ASGPTGGQDTQY | 0.84281814 |
| TIGHVDHGKTT | ASGPTGGQDTQY | 0.5931742 |
| GHVDHGKTT | ASGPTGGQDTQY | 0.74174714 |
| AQFEHTLL | ASGPTGGQDTQY | 0.87359166 |
| IVNQDEL | ASGPTGGQDTQY | 0.85459405 |
| TNGSQFF | ASGPTGGQDTQY | 0.5949667 |
| TGKGIVE | ASGPTGGQDTQY | 0.74142224 |
| LISNASDA | ASGPTGGQDTQY | 0.54063195 |
| AELRTLGELTKIAKAHGAQ | ASGPTGGQDTQY | 0.7797107 |
| APNAVAAAEPSW | ASGPTGGQDTQY | 0.68015873 |
| APQVAGAATAVL | ASGPTGGQDTQY | 0.8307202 |
| ARRDPAAAAAILPTDA | ASGPTGGQDTQY | 0.7641589 |
| DAAARDVLAEAGLAEYF | ASGPTGGQDTQY | 0.8113207 |
| DDQIVLSAKANLAKVEQ | ASGPTGGQDTQY | 0.8145716 |

| | | |
|---|---|---|
| DELLPDTGEDSDEDGH | ASGPTGGQDTQY | 0.87165695 |
| ELPEAPANRATVPQD | ASGPTGGQDTQY | 0.7345653 |
| FGQSKIERVTGIG | ASGPTGGQDTQY | 0.77888584 |
| GAGGTGSTAGGAGGAGGAGGLY | ASGPTGGQDTQY | 0.6033337 |
| GHGGTGAAGGNGGTGGAGGSGI | ASGPTGGQDTQY | 0.6587244 |
| GNGGNGGAGGEGGPGGAGGAGA | ASGPTGGQDTQY | 0.74229896 |
| GSTAGNGGNGGNATNGTIAGQP | ASGPTGGQDTQY | 0.78597736 |
| GTGGDGGNGGNGGTGG | ASGPTGGQDTQY | 0.32437027 |
| GVRRPAGIPADAPV | ASGPTGGQDTQY | 0.85916615 |
| GYRQVTDAHLLTLARR | ASGPTGGQDTQY | 0.81924 |
| IAKMGQDGHDRG | ASGPTGGQDTQY | 0.85158026 |
| KAEPSIEIQYT | ASGPTGGQDTQY | 0.7891202 |
| KPVRIRMAADEDTVAVT | ASGPTGGQDTQY | 0.79343766 |
| MPPRIAGMRLLVIKPEP | ASGPTGGQDTQY | 0.7338015 |
| MKFYAVALFRSRGVPPTR | ASGPTGGQDTQY | 0.84190416 |
| NAGGFGILGAVAHSP | ASGPTGGQDTQY | 0.70118254 |
| NEGREITLAMAEM | ASGPTGGQDTQY | 0.5612511 |
| NELVEWSHPLGHRWRWE | ASGPTGGQDTQY | 0.77133536 |
| NLGAGGTGGNGGNPG | ASGPTGGQDTQY | 0.59295434 |
| NTEWPKWETVMAADPVD | ASGPTGGQDTQY | 0.86055446 |
| PASAAPAETQP | ASGPTGGQDTQY | 0.8536353 |
| QQSSDGKNGVRAAEITGE | ASGPTGGQDTQY | 0.8153004 |
| RAPDAVTTSWWKE | ASGPTGGQDTQY | 0.8528042 |
| SGFGNVGAHTSGWFNQ | ASGPTGGQDTQY | 0.41133878 |
| SGVAGAVGQAASVGGLKV | ASGPTGGQDTQY | 0.75583607 |
| SNGGAGTGSAGGNGGT | ASGPTGGQDTQY | 0.87228894 |
| TGGAGGEGGAAG | ASGPTGGQDTQY | 0.8276242 |
| TGGDGGAAGAGGAPGHG | ASGPTGGQDTQY | 0.7701382 |
| TMPPLLAAIANAGRHGV | ASGPTGGQDTQY | 0.86382645 |
| TSSTAGNSGGA | ASGPTGGQDTQY | 0.72275907 |
| VPATVPVGILC | ASGPTGGQDTQY | 0.7996167 |
| VQRPDSGGVVLTGRIS | ASGPTGGQDTQY | 0.7983502 |
| WTPNPVPATKGVPAGLVQAT | ASGPTGGQDTQY | 0.8460964 |
| YALKRALAKFSDRFDVVII | ASGPTGGQDTQY | 0.8337016 |
| YLIGSVIV | ASGPTGGQDTQY | 0.7027267 |
| YGQLVAVID | ASGPTGGQDTQY | 0.8446021 |
| VTVGGMTLPA | ASGPTGGQDTQY | 0.8745917 |
| VLALPAV | ASGPTGGQDTQY | 0.747638 |
| VIPPTLNYET | ASGPTGGQDTQY | 0.84076285 |
| VGMGGMHPSSG | ASGPTGGQDTQY | 0.826641 |
| VGLAGGAATA | ASGPTGGQDTQY | 0.8057519 |
| VGGAGGAGGSGP | ASGPTGGQDTQY | 0.6961181 |
| VFAAFMV | ASGPTGGQDTQY | 0.7295242 |
| TLYDKLR | ASGPTGGQDTQY | 0.8031925 |
| TLAVFTVL | ASGPTGGQDTQY | 0.63263285 |
| TGSVQPARV | ASGPTGGQDTQY | 0.80236566 |
| TGGGAGSGGT | ASGPTGGQDTQY | 0.76839894 |
| SHLETSL | ASGPTGGQDTQY | 0.8830323 |
| SFTTGMT | ASGPTGGQDTQY | 0.8273653 |
| QVEVAKRMGVRQPTVS | ASGPTGGQDTQY | 0.8247549 |
| QGSGGPALAM | ASGPTGGQDTQY | 0.7059172 |

| | | |
|---|---|---|
| MLAPSNP | ASGPTGGQDTQY | 0.8364874 |
| LVNNAAI | ASGPTGGQDTQY | 0.8263874 |
| LTGEINTLA | ASGPTGGQDTQY | 0.6315274 |
| LSIFGPV | ASGPTGGQDTQY | 0.8664365 |
| LLALLANL | ASGPTGGQDTQY | 0.7067095 |
| LISNASDALDKLR | ASGPTGGQDTQY | 0.73423284 |
| LINEAVP | ASGPTGGQDTQY | 0.8494477 |
| LGYAFSGL | ASGPTGGQDTQY | 0.8515069 |
| LDFVAVAL | ASGPTGGQDTQY | 0.7770089 |
| LAQPGAE | ASGPTGGQDTQY | 0.7737808 |
| ISNLGAVTGF | ASGPTGGQDTQY | 0.70210546 |
| HQEDPGRI | ASGPTGGQDTQY | 0.8569104 |
| GTPGEPTP | ASGPTGGQDTQY | 0.85393625 |
| GFFNSGS | ASGPTGGQDTQY | 0.7881926 |
| GAGGGGLTAGAN | ASGPTGGQDTQY | 0.8627972 |
| FVDALVNM | ASGPTGGQDTQY | 0.84024614 |
| FLTVAPEL | ASGPTGGQDTQY | 0.5820198 |
| FETLTAL | ASGPTGGQDTQY | 0.8118335 |
| DVINPIEL | ASGPTGGQDTQY | 0.8473946 |
| APVSGAPV | ASGPTGGQDTQY | 0.83876127 |
| ALVGGLAW | ASGPTGGQDTQY | 0.8807155 |
| ALAAAAATAI | ASGPTGGQDTQY | 0.8312908 |
| AFGLGLLL | ASGPTGGQDTQY | 0.75035995 |
| AFFAAASL | ASGPTGGQDTQY | 0.8628328 |
| AAAALPAFSPP | ASGPTGGQDTQY | 0.87932825 |
| VVLAPAL | ASGPTGGQDTQY | 0.8522171 |
| TIGHVDHGKTTL | ASGPTGGQDTQY | 0.7834936 |
| LISNASDALDK | ASGPTGGQDTQY | 0.76081276 |
| LISNASDAL | ASGPTGGQDTQY | 0.8247515 |
| ISNASDALDK | ASGPTGGQDTQY | 0.849095 |
| NQAETLVYQTE | ASGPTGGQDTQY | 0.7226249 |
| TGGTGGTAGAG | ASGPTGGQDTQY | 0.77347255 |
| TGSTAGGAGGA | ASGPTGGQDTQY | 0.8131975 |
| DPLAVPVAAGPV | ASGPTGGQDTQY | 0.5836124 |
| GGTGGTPGNGGN | ASGPTGGQDTQY | 0.8302039 |
| PAGLAGVDPATL | ASGPTGGQDTQY | 0.86018515 |
| GNNLVVEAPHAGASV | ASGPTGGQDTQY | 0.7964053 |
| KVTRIMSASTLEQPA | ASGPTGGQDTQY | 0.7354809 |
| LGFALPIALPRRTVV | ASGPTGGQDTQY | 0.85353893 |
| TRLGNLTARKLVVLV | ASGPTGGQDTQY | 0.88044065 |
| WSQQRRLIYFPSAGP | ASGPTGGQDTQY | 0.8637709 |
| GPNGNPDPMAAAVDIR | ASGPTGGQDTQY | 0.62108463 |
| TQRCVSVADLRAAASR | ASGPTGGQDTQY | 0.3946367 |
| QFVLDVAEHPVEVLTETV | ASGPTGGQDTQY | 0.7374537 |
| GGAGGTGGAGGNGGTQAGDG | ASGPTGGQDTQY | 0.8244329 |
| PLLATLRAIAVDVDAKPAQV | ASGPTGGQDTQY | 0.77252316 |
| APKFRTGGYMIDFWGVGYQVAK | ASGPTGGQDTQY | 0.74606395 |
| GAGGAGGAGGAAGGPGGTGGTG | ASGPTGGQDTQY | 0.7839228 |
| IGALGLLATDVAPAGSVAFGVA | ASGPTGGQDTQY | 0.8564441 |
| AGGAGGIGGNGGAGGAGGVLMG | ASGPTGGQDTQY | 0.74687093 |
| ADELIGLL | ASGPTGGQDTQY | 0.6548993 |

| | | |
|---|---|---|
| ADRALKA | ASGPTGGQDTQY | 0.80343676 |
| AGAVGQVGGAGGA | ASGPTGGQDTQY | 0.8451116 |
| AGGGGGLGGAP | ASGPTGGQDTQY | 0.6637583 |
| AGGLLYGNGGAG | ASGPTGGQDTQY | 0.7548896 |
| AGLESIGL | ASGPTGGQDTQY | 0.7086161 |
| AGLKTGII | ASGPTGGQDTQY | 0.677267 |
| ASLAALLL | ASGPTGGQDTQY | 0.8018373 |
| ASVINISSI | ASGPTGGQDTQY | 0.70671237 |
| ATPPGPA | ASGPTGGQDTQY | 0.85125136 |
| ATVASALV | ASGPTGGQDTQY | 0.6746109 |
| ATVASLIL | ASGPTGGQDTQY | 0.8540056 |
| AVNGPVLIRH | ASGPTGGQDTQY | 0.82662034 |
| DGAAGAVGI | ASGPTGGQDTQY | 0.8418909 |
| DPLLHGI | ASGPTGGQDTQY | 0.75030893 |
| ELGAALGAL | ASGPTGGQDTQY | 0.66696244 |
| ELGRTQAV | ASGPTGGQDTQY | 0.63276255 |
| FVAGGYLS | ASGPTGGQDTQY | 0.48060596 |
| GASGGAGGAGGW | ASGPTGGQDTQY | 0.861607 |
| GGAGAGGDGGAGGI | ASGPTGGQDTQY | 0.36996767 |
| GTGHNGSQPGMGGNG | ASGPTGGQDTQY | 0.8591028 |
| GYAAASASAS | ASGPTGGQDTQY | 0.8512443 |
| HQVDVAAVQQG | ASGPTGGQDTQY | 0.685062 |
| IGAAIAHT | ASGPTGGQDTQY | 0.83481896 |
| IGMIEVAQL | ASGPTGGQDTQY | 0.64728713 |
| IRLAPEHTSAMD | ASGPTGGQDTQY | 0.75378925 |
| IYNVSVL | ASGPTGGQDTQY | 0.8537884 |
| KQGFLGM | ASGPTGGQDTQY | 0.84045815 |
| LESVITQL | ASGPTGGQDTQY | 0.71283257 |
| LGLGALVVV | ASGPTGGQDTQY | 0.8248962 |
| LIGNGGNGGSGGI | ASGPTGGQDTQY | 0.81117797 |
| LNAINAPTL | ASGPTGGQDTQY | 0.7409543 |
| LQLLNTGV | ASGPTGGQDTQY | 0.81212735 |
| LQTSIEF | ASGPTGGQDTQY | 0.84682864 |
| MSFAVSEL | ASGPTGGQDTQY | 0.8415782 |
| NTGALGL | ASGPTGGQDTQY | 0.7195794 |
| PPGAGGAGGAGTT | ASGPTGGQDTQY | 0.83800274 |
| QLVGLGI | ASGPTGGQDTQY | 0.6049414 |
| RFLEQVHQL | ASGPTGGQDTQY | 0.8153168 |
| RGVGYVLR | ASGPTGGQDTQY | 0.79576486 |
| SATGVFAGVI | ASGPTGGQDTQY | 0.8051759 |
| SLPAITLP | ASGPTGGQDTQY | 0.8304171 |
| STALAFTV | ASGPTGGQDTQY | 0.6647689 |
| SVLTVGLGK | ASGPTGGQDTQY | 0.7094859 |
| TALAGAGSGI | ASGPTGGQDTQY | 0.785706 |
| TGLPFTGI | ASGPTGGQDTQY | 0.8290197 |
| VAEHVTV | ASGPTGGQDTQY | 0.8438949 |
| VDIVLNSL | ASGPTGGQDTQY | 0.68469626 |
| VLTGGGAVL | ASGPTGGQDTQY | 0.8219292 |
| VNVGLAYGI | ASGPTGGQDTQY | 0.7751343 |
| VSSAAVGAV | ASGPTGGQDTQY | 0.6915815 |
| VTSVYEAGV | ASGPTGGQDTQY | 0.81386614 |

| | | |
|---|---|---|
| WVALDRGA | ASGPTGGQDTQY | 0.7941474 |
| YLVGSVVI | ASGPTGGQDTQY | 0.74352336 |
| GPPGTGKTLL | ASGPTGGQDTQY | 0.8355047 |
| GVGMTKF | ASGPTGGQDTQY | 0.70658296 |
| IGIGGSDL | ASGPTGGQDTQY | 0.8468168 |
| IGTLLAEL | ASGPTGGQDTQY | 0.878186 |
| IYTSGTT | ASGPTGGQDTQY | 0.77398086 |
| LSFPAGV | ASGPTGGQDTQY | 0.8530523 |
| LYTSGTT | ASGPTGGQDTQY | 0.82483864 |
| VGAFPFGF | ASGPTGGQDTQY | 0.8112956 |
| VGAGTVEF | ASGPTGGQDTQY | 0.78440595 |
| VGKSTLLL | ASGPTGGQDTQY | 0.70226836 |
| AALAHAGGGIV | ASGPTGGQDTQY | 0.8534935 |
| AALVAVSLL | ASGPTGGQDTQY | 0.84394026 |
| AAVSRAVAAL | ASGPTGGQDTQY | 0.85978657 |
| AGGSSGIGGF | ASGPTGGQDTQY | 0.8148706 |
| AGLTVANANG | ASGPTGGQDTQY | 0.8277262 |
| AGLVGAVL | ASGPTGGQDTQY | 0.8381675 |
| ALSQAFTV | ASGPTGGQDTQY | 0.7061088 |
| ALTFSVSL | ASGPTGGQDTQY | 0.5713589 |
| ALTGAALVAL | ASGPTGGQDTQY | 0.76302356 |
| ASFSIGAGVL | ASGPTGGQDTQY | 0.7039708 |
| DGSVYKI | ASGPTGGQDTQY | 0.7803432 |
| EQLAEIV | ASGPTGGQDTQY | 0.6151319 |
| FAVSELTVM | ASGPTGGQDTQY | 0.84764624 |
| GAAPLAAGL | ASGPTGGQDTQY | 0.8824697 |
| GADGGDGGSGGAGG | ASGPTGGQDTQY | 0.7767004 |
| GFGLDTGGAGG | ASGPTGGQDTQY | 0.87045985 |
| GLASGGLL | ASGPTGGQDTQY | 0.62619716 |
| GRIAGAAVLL | ASGPTGGQDTQY | 0.8543019 |
| GSAGSLNTSF | ASGPTGGQDTQY | 0.86157876 |
| GTGQIGI | ASGPTGGQDTQY | 0.66464347 |
| GTLIALIL | ASGPTGGQDTQY | 0.84644186 |
| IAATMGFVVG | ASGPTGGQDTQY | 0.73047304 |
| IFAQALL | ASGPTGGQDTQY | 0.77716714 |
| IGAAVALGF | ASGPTGGQDTQY | 0.8685759 |
| IGLTPLAL | ASGPTGGQDTQY | 0.7884706 |
| ILGSNTSTL | ASGPTGGQDTQY | 0.8237524 |
| IVFSLTVL | ASGPTGGQDTQY | 0.81148756 |
| LASKATGF | ASGPTGGQDTQY | 0.7390124 |
| LASVALIL | ASGPTGGQDTQY | 0.82155037 |
| LFAVGLTL | ASGPTGGQDTQY | 0.865193 |
| LFNGGIGTYI | ASGPTGGQDTQY | 0.7563634 |
| LTLGAGII | ASGPTGGQDTQY | 0.8774855 |
| LVAAALGDI | ASGPTGGQDTQY | 0.73667246 |
| NTAFVQLGI | ASGPTGGQDTQY | 0.869745 |
| QLFSGAEL | ASGPTGGQDTQY | 0.8295082 |
| SAATFITL | ASGPTGGQDTQY | 0.7818134 |
| SNYGLANLGV | ASGPTGGQDTQY | 0.76789725 |
| SYGGAYAVM | ASGPTGGQDTQY | 0.85770285 |
| TTGVGGLTLG | ASGPTGGQDTQY | 0.7779523 |

| | | |
|---|---|---|
| TTQTIYI | ASGPTGGQDTQY | 0.84884024 |
| VAAGAGLGAVI | ASGPTGGQDTQY | 0.8690495 |
| VAAVVTLL | ASGPTGGQDTQY | 0.3110767 |
| VGFLGGLI | ASGPTGGQDTQY | 0.6569822 |
| VGVIPILL | ASGPTGGQDTQY | 0.49579394 |
| VIFASSLI | ASGPTGGQDTQY | 0.44668898 |
| VKGGDGGAAAG | ASGPTGGQDTQY | 0.38979173 |
| VTKTFTNI | ASGPTGGQDTQY | 0.27638823 |
| VVAAAILL | ASGPTGGQDTQY | 0.36493686 |
| YAALDVEL | ASGPTGGQDTQY | 0.36672217 |
| LASVALLL | ASGPTGGQDTQY | 0.73715293 |
| LDAVHRV | ASGPTGGQDTQY | 0.45847523 |
| QGLSSSIF | ASGPTGGQDTQY | 0.38862497 |
| RGTFANIRL | ASGPTGGQDTQY | 0.6881939 |
| RQLWWGHRI | ASGPTGGQDTQY | 0.6355734 |
| YGPPGTGKTLL | ASGPTGGQDTQY | 0.2705648 |
| AALVGAPANVA | ASGPTGGQDTQY | 0.47722197 |
| AATFRRTE | ASGPTGGQDTQY | 0.29627946 |
| AGALGGGI | ASGPTGGQDTQY | 0.3702172 |
| AGGAGGAGGIGAG | ASGPTGGQDTQY | 0.22774164 |
| AGSGALGL | ASGPTGGQDTQY | 0.48591822 |
| AQIADQI | ASGPTGGQDTQY | 0.5739784 |
| ASAVAVTYL | ASGPTGGQDTQY | 0.24437658 |
| ASTVLGGTGG | ASGPTGGQDTQY | 0.42937297 |
| AVGGEPAK | ASGPTGGQDTQY | 0.44385144 |
| CEQGTFGGTC | ASGPTGGQDTQY | 0.4655001 |
| DEGKAPAG | ASGPTGGQDTQY | 0.64993465 |
| DTVVFPAL | ASGPTGGQDTQY | 0.30097002 |
| FEGITMAG | ASGPTGGQDTQY | 0.38965288 |
| FPLVVQD | ASGPTGGQDTQY | 0.6152815 |
| GAFTTPQ | ASGPTGGQDTQY | 0.7009541 |
| GFGGFGTAGGVGGA | ASGPTGGQDTQY | 0.31823096 |
| GFLAQFL | ASGPTGGQDTQY | 0.53589493 |
| GGAGGAGGNAGVA | ASGPTGGQDTQY | 0.70469224 |
| GGVGTAGAGGAGGAGG | ASGPTGGQDTQY | 0.48465592 |
| GNGGGGDAGNAGSGGN | ASGPTGGQDTQY | 0.49509382 |
| GTGGAGGAGNDAGS | ASGPTGGQDTQY | 0.56257457 |
| GTTNTGWFNS | ASGPTGGQDTQY | 0.30852664 |
| IFMGPLFTL | ASGPTGGQDTQY | 0.63611114 |
| ILAHAIEAI | ASGPTGGQDTQY | 0.6626082 |
| IRSAIREAL | ASGPTGGQDTQY | 0.41165763 |
| ITFAGGG | ASGPTGGQDTQY | 0.41753 |
| ITGMSGHQEV | ASGPTGGQDTQY | 0.30335414 |
| IVAGLACTL | ASGPTGGQDTQY | 0.36612418 |
| IVGGGLAGA | ASGPTGGQDTQY | 0.26392907 |
| IVTGVLNI | ASGPTGGQDTQY | 0.62369907 |
| LAQQGCI | ASGPTGGQDTQY | 0.6077744 |
| LCLHVEN | ASGPTGGQDTQY | 0.48812562 |
| LDAISDGS | ASGPTGGQDTQY | 0.2646003 |
| LGGFAAML | ASGPTGGQDTQY | 0.32631388 |
| LGGKNPMI | ASGPTGGQDTQY | 0.4341898 |

| | | |
|---|---|---|
| LGPKGVAAIL | ASGPTGGQDTQY | 0.32754815 |
| LMVGVSI | ASGPTGGQDTQY | 0.28375596 |
| LVGAAFVL | ASGPTGGQDTQY | 0.28199378 |
| LVLAVAPL | ASGPTGGQDTQY | 0.6603769 |
| MAAIAVIDAA | ASGPTGGQDTQY | 0.6895157 |
| NNGINAELQ | ASGPTGGQDTQY | 0.38565814 |
| RGEPLVLVHG | ASGPTGGQDTQY | 0.6005367 |
| RGGAGGAAGLI | ASGPTGGQDTQY | 0.7260531 |
| RSYRMHPAV | ASGPTGGQDTQY | 0.57268167 |
| SAAHLLAGQLA | ASGPTGGQDTQY | 0.42219755 |
| SAAVGSY | ASGPTGGQDTQY | 0.29012907 |
| SPHAVLHRI | ASGPTGGQDTQY | 0.73737276 |
| SQIAKVL | ASGPTGGQDTQY | 0.5399734 |
| TALGHLVP | ASGPTGGQDTQY | 0.5309454 |
| TAPTTSGAA | ASGPTGGQDTQY | 0.52615315 |
| TDGDDGNGGSPGT | ASGPTGGQDTQY | 0.49433517 |
| TDGLGEN | ASGPTGGQDTQY | 0.365968 |
| TGGPPLPGQA | ASGPTGGQDTQY | 0.6655601 |
| TIPAITI | ASGPTGGQDTQY | 0.4807478 |
| TIPGFTI | ASGPTGGQDTQY | 0.32960442 |
| TSIASGALA | ASGPTGGQDTQY | 0.30177632 |
| TVTLPGAAI | ASGPTGGQDTQY | 0.5725812 |
| VASGLSAALP | ASGPTGGQDTQY | 0.59142965 |
| VGAGVIGI | ASGPTGGQDTQY | 0.6644908 |
| VGPKRFGV | ASGPTGGQDTQY | 0.40451574 |
| VNVAFPD | ASGPTGGQDTQY | 0.32548597 |
| VSMTNSGVS | ASGPTGGQDTQY | 0.63454115 |
| VTDVLGDLSV | ASGPTGGQDTQY | 0.50523484 |
| VVVVGAGFAGL | ASGPTGGQDTQY | 0.44627076 |
| VYTVGTVV | ASGPTGGQDTQY | 0.5664393 |
| WTTFSGP | ASGPTGGQDTQY | 0.4411837 |
| YDELVEAF | ASGPTGGQDTQY | 0.45770228 |
| YGTGGAGGNGG | ASGPTGGQDTQY | 0.52460665 |
| YTNVMAV | ASGPTGGQDTQY | 0.40907294 |
| FLEQVHQL | ASGPTGGQDTQY | 0.47538382 |
| GLSLLAVAV | ASGPTGGQDTQY | 0.47450924 |
| LEQVHQL | ASGPTGGQDTQY | 0.41159344 |
| LNAAGAI | ASGPTGGQDTQY | 0.31593156 |
| PAVHASL | ASGPTGGQDTQY | 0.34770507 |
| TELEDDA | ASGPTGGQDTQY | 0.55194217 |
| TPPGPPN | ASGPTGGQDTQY | 0.5472311 |
| TPPPPPPP | ASGPTGGQDTQY | 0.5747727 |
| TQSPALI | ASSQDQTGSQNTLY | 0.40400794 |
| EFAAAFPQ | ASSQDQTGSQNTLY | 0.38850418 |
| LLSGNLDVL | ASSQDQTGSQNTLY | 0.60019433 |
| GPANLGNYNI | ASSQDQTGSQNTLY | 0.651285 |
| TLGPTLP | ASSQDQTGSQNTLY | 0.26780388 |
| VYAYSSCCK | ASSQDQTGSQNTLY | 0.4403925 |
| APAPAPAPNI | ASSQDQTGSQNTLY | 0.6789688 |
| PGGPVPIP | ASSQDQTGSQNTLY | 0.711254 |
| FLTVALEP | ASSQDQTGSQNTLY | 0.46600592 |

| | | |
|---|---|---|
| SQYFGANL | ASSQDQTGSQNTLY | 0.3479826 |
| GGVKPAA | ASSQDQTGSQNTLY | 0.75414705 |
| GGTGGAGGAGGW | ASSQDQTGSQNTLY | 0.49472108 |
| GVGGVAGLGGAGG | ASSQDQTGSQNTLY | 0.68655324 |
| VLLVLATTI | ASSQDQTGSQNTLY | 0.25816107 |
| IGIFAAML | ASSQDQTGSQNTLY | 0.34536466 |
| AVGDLID | ASSQDQTGSQNTLY | 0.31777382 |
| NISPLPL | ASSQDQTGSQNTLY | 0.24098384 |
| VTAVFVAI | ASSQDQTGSQNTLY | 0.619434 |
| SGDDNDAAL | ASSQDQTGSQNTLY | 0.34210798 |
| AVAALAHS | ASSQDQTGSQNTLY | 0.5748233 |
| GGVTVMATP | ASSQDQTGSQNTLY | 0.38450068 |
| AGLAANVRD | ASSQDQTGSQNTLY | 0.5204202 |
| TNRPDLIDPALLRPG | ASSQDQTGSQNTLY | 0.39145872 |
| LANLGVTAL | ASSQDQTGSQNTLY | 0.41321522 |
| TGERIEL | ASSQDQTGSQNTLY | 0.30066395 |
| NGEELAA | ASSQDQTGSQNTLY | 0.6348529 |
| VLSLVWG | ASSQDQTGSQNTLY | 0.6031804 |
| IGYALWL | ASSQDQTGSQNTLY | 0.4062578 |
| LSLLLAPL | ASSQDQTGSQNTLY | 0.28516424 |
| TLDEALK | ASSQDQTGSQNTLY | 0.5879285 |
| TDGAGTAEGG | ASSQDQTGSQNTLY | 0.49914813 |
| NAPGGAGGNGG | ASSQDQTGSQNTLY | 0.2648424 |
| VSVFMGMGGNF | ASSQDQTGSQNTLY | 0.27077347 |
| LASYLAAL | ASSQDQTGSQNTLY | 0.39241135 |
| LVGGTPAL | ASSQDQTGSQNTLY | 0.35458007 |
| VRNADVL | ASSQDQTGSQNTLY | 0.6713186 |
| TCSSPFLVSAM | ASSQDQTGSQNTLY | 0.6527504 |
| VHITDAE | ASSQDQTGSQNTLY | 0.43673015 |
| EYAAPSF | ASSQDQTGSQNTLY | 0.6303574 |
| VVDAFHGGL | ASSQDQTGSQNTLY | 0.6099509 |
| ASVVDPQ | ASSQDQTGSQNTLY | 0.2195565 |
| SYNEFGGLS | ASSQDQTGSQNTLY | 0.36129713 |
| DPDGSVY | ASSQDQTGSQNTLY | 0.2723068 |
| MVSTIHATE | ASSQDQTGSQNTLY | 0.4874845 |
| LAAGVADPV | ASSQDQTGSQNTLY | 0.32158628 |
| TKEQIAA | ASSQDQTGSQNTLY | 0.41223708 |
| SLASPAAL | ASSQDQTGSQNTLY | 0.5112226 |
| IGLVFYGV | ASSQDQTGSQNTLY | 0.7081566 |
| FLLDASNL | ASSQDQTGSQNTLY | 0.53138226 |
| YQVGNFP | ASSQDQTGSQNTLY | 0.27924138 |
| ESLLSQST | ASSQDQTGSQNTLY | 0.24043572 |
| LFTVGFAAVG | ASSQDQTGSQNTLY | 0.601969 |
| ATPPTPT | ASSQDQTGSQNTLY | 0.29587567 |
| TDLATLG | ASSQDQTGSQNTLY | 0.34270394 |
| SAGGTGR | ASSQDQTGSQNTLY | 0.2567986 |
| GFSNSGS | ASSQDQTGSQNTLY | 0.3548231 |
| GGNGGSGGTSV | ASSQDQTGSQNTLY | 0.26580402 |
| GTSVATGGAGN | ASSQDQTGSQNTLY | 0.42323053 |
| NGGNGGSGGTS | ASSQDQTGSQNTLY | 0.3528692 |
| AVVKFPH | ASSQDQTGSQNTLY | 0.48682532 |

| | | |
|---|---|---|
| TPPPPPP | ASSQDQTGSQNTLY | 0.67500633 |
| AAPAAAAAPA | ASSQDQTGSQNTLY | 0.71375763 |
| IALNEDL | ASSQDQTGSQNTLY | 0.41215497 |
| IVNVSSM | ASSQDQTGSQNTLY | 0.4097999 |
| TGIPPAPR | ASSQDQTGSQNTLY | 0.30103418 |
| FDLGGGTF | ASSQDQTGSQNTLY | 0.3775576 |
| FDLGGGTFDVS | ASSQDQTGSQNTLY | 0.35636923 |
| NEPTAAA | ASSQDQTGSQNTLY | 0.6643982 |
| NEPTAAALA | ASSQDQTGSQNTLY | 0.32630482 |
| GSIINISSI | ASSQDQTGSQNTLY | 0.27034587 |
| VAPGNAGTA | ASSQDQTGSQNTLY | 0.3626327 |
| TGGGASLE | ASSQDQTGSQNTLY | 0.6457895 |
| VAVYNHYKRI | ASSQDQTGSQNTLY | 0.7034568 |
| TDPAPATT | ASSQDQTGSQNTLY | 0.3247872 |
| TNRPDILDPALLRPG | ASSQDQTGSQNTLY | 0.40956032 |
| LTTTTTS | ASSQDQTGSQNTLY | 0.37431598 |
| PTGSGKTL | ASSQDQTGSQNTLY | 0.55822474 |
| TGSGKTL | ASSQDQTGSQNTLY | 0.30623794 |
| TIGHVDHGK | ASSQDQTGSQNTLY | 0.3108424 |
| TIGHVDHGKTT | ASSQDQTGSQNTLY | 0.46243924 |
| GHVDHGKTT | ASSQDQTGSQNTLY | 0.75948936 |
| AQFEHTLL | ASSQDQTGSQNTLY | 0.3827576 |
| IVNQDEL | ASSQDQTGSQNTLY | 0.23935798 |
| TNGSQFF | ASSQDQTGSQNTLY | 0.5707042 |
| TGKGIVE | ASSQDQTGSQNTLY | 0.35246468 |
| LISNASDA | ASSQDQTGSQNTLY | 0.42150158 |
| AELRTLGELTKIAKAHGAQ | ASSQDQTGSQNTLY | 0.5009623 |
| APNAVAAAEPSW | ASSQDQTGSQNTLY | 0.2521865 |
| APQVAGAATAVL | ASSQDQTGSQNTLY | 0.40824962 |
| ARRDPAAAAAILPTDA | ASSQDQTGSQNTLY | 0.648597 |
| DAAARDVLAEAGLAEYF | ASSQDQTGSQNTLY | 0.6721982 |
| DDQIVLSAKANLAKVEQ | ASSQDQTGSQNTLY | 0.41991737 |
| DELLPDTGEDSDEDGH | ASSQDQTGSQNTLY | 0.6506582 |
| ELPEAPANRATVPQD | ASSQDQTGSQNTLY | 0.5675265 |
| FGQSKIERVTGIG | ASSQDQTGSQNTLY | 0.70040655 |
| GAGGTGSTAGGAGGAGGAGGLY | ASSQDQTGSQNTLY | 0.72894776 |
| GHGGTGAAGGNGGTGGAGGSGI | ASSQDQTGSQNTLY | 0.34785706 |
| GNGGNGGAGGEGGPGGAGGAGA | ASSQDQTGSQNTLY | 0.4043565 |
| GSTAGNGGNGGNATNGTIAGQP | ASSQDQTGSQNTLY | 0.5907934 |
| GTGGDGGNGGNGGTGG | ASSQDQTGSQNTLY | 0.5604254 |
| GVRRPAGIPADAPV | ASSQDQTGSQNTLY | 0.5208324 |
| GYRQVTDAHLLTLARR | ASSQDQTGSQNTLY | 0.32966414 |
| IAKMGQDGHDRG | ASSQDQTGSQNTLY | 0.70336246 |
| KAEPSIEIQYT | ASSQDQTGSQNTLY | 0.45685136 |
| KPVRIRMAADEDTVAVT | ASSQDQTGSQNTLY | 0.7277095 |
| MPPRIAGMRLLVIKPEP | ASSQDQTGSQNTLY | 0.27077165 |
| MKFYAVALFRSRGVPPTR | ASSQDQTGSQNTLY | 0.32966563 |
| NAGGFGILGAVAHSP | ASSQDQTGSQNTLY | 0.6155667 |
| NEGREITLAMAEM | ASSQDQTGSQNTLY | 0.23581983 |
| NELVEWSHPLGHRWRWE | ASSQDQTGSQNTLY | 0.29066685 |
| NLGAGGTGGNGGNPG | ASSQDQTGSQNTLY | 0.40858203 |

| | | |
|---|---|---|
| NTEWPKWETVMAADPVD | ASSQDQTGSQNTLY | 0.6344328 |
| PASAAPAETQP | ASSQDQTGSQNTLY | 0.23157512 |
| QQSSDGKNGVRAAEITGE | ASSQDQTGSQNTLY | 0.4087309 |
| RAPDAVTTSWWKE | ASSQDQTGSQNTLY | 0.27318552 |
| SGFGNVGAHTSGWFNQ | ASSQDQTGSQNTLY | 0.4209525 |
| SGVAGAVGQAASVGGLKV | ASSQDQTGSQNTLY | 0.47697204 |
| SNGGAGTGSAGGNGGT | ASSQDQTGSQNTLY | 0.26173913 |
| TGGAGGEGGAAG | ASSQDQTGSQNTLY | 0.49688014 |
| TGGDGGAAGAGGAPGHG | ASSQDQTGSQNTLY | 0.23919478 |
| TMPPLLAAIANAGRHGV | ASSQDQTGSQNTLY | 0.36922416 |
| TSSTAGNSGGA | ASSQDQTGSQNTLY | 0.5026529 |
| VPATVPVGILC | ASSQDQTGSQNTLY | 0.7416694 |
| VQRPDSGGVVLTGRIS | ASSQDQTGSQNTLY | 0.30685428 |
| WTPNPVPATKGVPAGLVQAT | ASSQDQTGSQNTLY | 0.5845437 |
| YALKRALAKFSDRFDVVII | ASSQDQTGSQNTLY | 0.30485383 |
| YLIGSVIV | ASSQDQTGSQNTLY | 0.72717243 |
| YGQLVAVID | ASSQDQTGSQNTLY | 0.4385273 |
| VTVGGMTLPA | ASSQDQTGSQNTLY | 0.26202822 |
| VLALPAV | ASSQDQTGSQNTLY | 0.62650925 |
| VIPPTLNYET | ASSQDQTGSQNTLY | 0.38028544 |
| VGMGGMHPSSG | ASSQDQTGSQNTLY | 0.40838897 |
| VGLAGGAATA | ASSQDQTGSQNTLY | 0.7246163 |
| VGGAGGAGGSGP | ASSQDQTGSQNTLY | 0.33204427 |
| VFAAFMV | ASSQDQTGSQNTLY | 0.500689 |
| TLYDKLR | ASSQDQTGSQNTLY | 0.681394 |
| TLAVFTVL | ASSQDQTGSQNTLY | 0.5720511 |
| TGSVQPARV | ASSQDQTGSQNTLY | 0.32632133 |
| TGGGAGSGGT | ASSQDQTGSQNTLY | 0.4249437 |
| SHLETSL | ASSQDQTGSQNTLY | 0.5951184 |
| SFTTGMT | ASSQDQTGSQNTLY | 0.34157228 |
| QVEVAKRMGVRQPTVS | ASSQDQTGSQNTLY | 0.65606785 |
| QGSGGPALAM | ASSQDQTGSQNTLY | 0.33598244 |
| MLAPSNP | ASSQDQTGSQNTLY | 0.58554673 |
| LVNNAAI | ASSQDQTGSQNTLY | 0.51317537 |
| LTGEINTLA | ASSQDQTGSQNTLY | 0.75607526 |
| LSIFGPV | ASSQDQTGSQNTLY | 0.2966663 |
| LLALLANL | ASSQDQTGSQNTLY | 0.3893175 |
| LISNASDALDKLR | ASSQDQTGSQNTLY | 0.5364636 |
| LINEAVP | ASSQDQTGSQNTLY | 0.51704955 |
| LGYAFSGL | ASSQDQTGSQNTLY | 0.31566775 |
| LDFVAVAL | ASSQDQTGSQNTLY | 0.33624914 |
| LAQPGAE | ASSQDQTGSQNTLY | 0.4202967 |
| ISNLGAVTGF | ASSQDQTGSQNTLY | 0.6688266 |
| HQEDPGRI | ASSQDQTGSQNTLY | 0.35563505 |
| GTPGEPTP | ASSQDQTGSQNTLY | 0.47457144 |
| GFFNSGS | ASSQDQTGSQNTLY | 0.72553015 |
| GAGGGGLTAGAN | ASSQDQTGSQNTLY | 0.32964835 |
| FVDALVNM | ASSQDQTGSQNTLY | 0.47215796 |
| FLTVAPEL | ASSQDQTGSQNTLY | 0.5126028 |
| FETLTAL | ASSQDQTGSQNTLY | 0.63546324 |
| DVINPIEL | ASSQDQTGSQNTLY | 0.34117192 |

| | | |
|---|---|---|
| APVSGAPV | ASSQDQTGSQNTLY | 0.59806395 |
| ALVGGLAW | ASSQDQTGSQNTLY | 0.34425694 |
| ALAAAAATAI | ASSQDQTGSQNTLY | 0.5882814 |
| AFGLGLLL | ASSQDQTGSQNTLY | 0.30755523 |
| AFFAAASL | ASSQDQTGSQNTLY | 0.49510354 |
| AAAALPAFSPP | ASSQDQTGSQNTLY | 0.38736382 |
| VVLAPAL | ASSQDQTGSQNTLY | 0.64825964 |
| TIGHVDHGKTTL | ASSQDQTGSQNTLY | 0.63736105 |
| LISNASDALDK | ASSQDQTGSQNTLY | 0.63434464 |
| LISNASDAL | ASSQDQTGSQNTLY | 0.25718975 |
| ISNASDALDK | ASSQDQTGSQNTLY | 0.6507062 |
| NQAETLVYQTE | ASSQDQTGSQNTLY | 0.6434274 |
| TGGTGGTAGAG | ASSQDQTGSQNTLY | 0.7544899 |
| TGSTAGGAGGA | ASSQDQTGSQNTLY | 0.43608898 |
| DPLAVPVAAGPV | ASSQDQTGSQNTLY | 0.5280895 |
| GGTGGTPGNGGN | ASSQDQTGSQNTLY | 0.4990918 |
| PAGLAGVDPATL | ASSQDQTGSQNTLY | 0.37231284 |
| GNNLVVEAPHAGASV | ASSQDQTGSQNTLY | 0.32831582 |
| KVTRIMSASTLEQPA | ASSQDQTGSQNTLY | 0.37391588 |
| LGFALPIALPRRTVV | ASSQDQTGSQNTLY | 0.47815606 |
| TRLGNLTARKLVVLV | ASSQDQTGSQNTLY | 0.38941583 |
| WSQQRRLIYFPSAGP | ASSQDQTGSQNTLY | 0.73511314 |
| GPNGNPDPMAAAVDIR | ASSQDQTGSQNTLY | 0.63046324 |
| TQRCVSVADLRAAASR | ASSQDQTGSQNTLY | 0.5806657 |
| QFVLDVAEHPVEVLTETV | ASSQDQTGSQNTLY | 0.70975614 |
| GGAGGTGGAGGNGGTQAGDG | ASSQDQTGSQNTLY | 0.65969193 |
| PLLATLRAIAVDVDAKPAQV | ASSQDQTGSQNTLY | 0.4154198 |
| APKFRTGGYMIDFWGVGYQVAK | ASSQDQTGSQNTLY | 0.4193089 |
| GAGGAGGAGGAAGGPGGTGGTG | ASSQDQTGSQNTLY | 0.43622136 |
| IGALGLLATDVAPAGSVAFGVA | ASSQDQTGSQNTLY | 0.29053667 |
| AGGAGGIGGNGGAGGAGGVLMG | ASSQDQTGSQNTLY | 0.37867633 |
| ADELIGLL | ASSQDQTGSQNTLY | 0.38460317 |
| ADRALKA | ASSQDQTGSQNTLY | 0.531631 |
| AGAVGQVGGAGGA | ASSQDQTGSQNTLY | 0.22730497 |
| AGGGGGLGGAP | ASSQDQTGSQNTLY | 0.6972064 |
| AGGLLYGNGGAG | ASSQDQTGSQNTLY | 0.6625458 |
| AGLESIGL | ASSQDQTGSQNTLY | 0.49315077 |
| AGLKTGII | ASSQDQTGSQNTLY | 0.59023625 |
| ASLAALLL | ASSQDQTGSQNTLY | 0.48314196 |
| ASVINISSI | ASSQDQTGSQNTLY | 0.29900286 |
| ATPPGPA | ASSQDQTGSQNTLY | 0.6628678 |
| ATVASALV | ASSQDQTGSQNTLY | 0.309858 |
| ATVASLIL | ASSQDQTGSQNTLY | 0.45480365 |
| AVNGPVLIRH | ASSQDQTGSQNTLY | 0.6513604 |
| DGAAGAVGI | ASSQDQTGSQNTLY | 0.46732056 |
| DPLLHGI | ASSQDQTGSQNTLY | 0.46353987 |
| ELGAALGAL | ASSQDQTGSQNTLY | 0.3789939 |
| ELGRTQAV | ASSQDQTGSQNTLY | 0.27975476 |
| FVAGGYLS | ASSQDQTGSQNTLY | 0.4905141 |
| GASGGAGGAGGW | ASSQDQTGSQNTLY | 0.266817 |
| GGAGAGGDGGAGGI | ASSQDQTGSQNTLY | 0.50699663 |

| | | |
|---|---|---|
| GTGHNGSQPGMGGNG | ASSQDQTGSQNTLY | 0.29972795 |
| GYAAASASAS | ASSQDQTGSQNTLY | 0.612267 |
| HQVDVAAVQQG | ASSQDQTGSQNTLY | 0.57729673 |
| IGAAIAHT | ASSQDQTGSQNTLY | 0.46722728 |
| IGMIEVAQL | ASSQDQTGSQNTLY | 0.36286455 |
| IRLAPEHTSAMD | ASSQDQTGSQNTLY | 0.6074569 |
| IYNVSVL | ASSQDQTGSQNTLY | 0.6166061 |
| KQGFLGM | ASSQDQTGSQNTLY | 0.31069925 |
| LESVITQL | ASSQDQTGSQNTLY | 0.39414686 |
| LGLGALVVV | ASSQDQTGSQNTLY | 0.34352142 |
| LIGNGGNGGSGGI | ASSQDQTGSQNTLY | 0.3597172 |
| LNAINAPTL | ASSQDQTGSQNTLY | 0.6441256 |
| LQLLNTGV | ASSQDQTGSQNTLY | 0.4346197 |
| LQTSIEF | ASSQDQTGSQNTLY | 0.24795826 |
| MSFAVSEL | ASSQDQTGSQNTLY | 0.5637264 |
| NTGALGL | ASSQDQTGSQNTLY | 0.5152837 |
| PPGAGGAGGAGTT | ASSQDQTGSQNTLY | 0.44831136 |
| QLVGLGI | ASSQDQTGSQNTLY | 0.22972749 |
| RFLEQVHQL | ASSQDQTGSQNTLY | 0.4222062 |
| RGVGYVLR | ASSQDQTGSQNTLY | 0.33659977 |
| SATGVFAGVI | ASSQDQTGSQNTLY | 0.31442648 |
| SLPAITLP | ASSQDQTGSQNTLY | 0.32983485 |
| STALAFTV | ASSQDQTGSQNTLY | 0.6753243 |
| SVLTVGLGK | ASSQDQTGSQNTLY | 0.45153713 |
| TALAGAGSGI | ASSQDQTGSQNTLY | 0.51158774 |
| TGLPFTGI | ASSQDQTGSQNTLY | 0.5839322 |
| VAEHVTV | ASSQDQTGSQNTLY | 0.54925615 |
| VDIVLNSL | ASSQDQTGSQNTLY | 0.69881654 |
| VLTGGGAVL | ASSQDQTGSQNTLY | 0.53013897 |
| VNVGLAYGI | ASSQDQTGSQNTLY | 0.36372653 |
| VSSAAVGAV | ASSQDQTGSQNTLY | 0.4563486 |
| VTSVYEAGV | ASSQDQTGSQNTLY | 0.3796076 |
| WVALDRGA | ASSQDQTGSQNTLY | 0.3778838 |
| YLVGSVVI | ASSQDQTGSQNTLY | 0.3061662 |
| GPPGTGKTLL | ASSQDQTGSQNTLY | 0.43156648 |
| GVGMTKF | ASSQDQTGSQNTLY | 0.35126653 |
| IGIGGSDL | ASSQDQTGSQNTLY | 0.56649494 |
| IGTLLAEL | ASSQDQTGSQNTLY | 0.4416127 |
| IYTSGTT | ASSQDQTGSQNTLY | 0.4767474 |
| LSFPAGV | ASSQDQTGSQNTLY | 0.28288338 |
| LYTSGTT | ASSQDQTGSQNTLY | 0.40226287 |
| VGAFPFGF | ASSQDQTGSQNTLY | 0.3179512 |
| VGAGTVEF | ASSQDQTGSQNTLY | 0.63234234 |
| VGKSTLLL | ASSQDQTGSQNTLY | 0.49362776 |
| AALAHAGGGIV | ASSQDQTGSQNTLY | 0.75241387 |
| AALVAVSLL | ASSQDQTGSQNTLY | 0.6914791 |
| AAVSRAVAAL | ASSQDQTGSQNTLY | 0.48522487 |
| AGGSSGIGGF | ASSQDQTGSQNTLY | 0.6370528 |
| AGLTVANANG | ASSQDQTGSQNTLY | 0.31648445 |
| AGLVGAVL | ASSQDQTGSQNTLY | 0.3257506 |
| ALSQAFTV | ASSQDQTGSQNTLY | 0.3293642 |

| | | |
|---|---|---|
| ALTFSVSL | ASSQDQTGSQNTLY | 0.45054582 |
| ALTGAALVAL | ASSQDQTGSQNTLY | 0.4360814 |
| ASFSIGAGVL | ASSQDQTGSQNTLY | 0.31766003 |
| DGSVYKI | ASSQDQTGSQNTLY | 0.26624426 |
| EQLAEIV | ASSQDQTGSQNTLY | 0.6576522 |
| FAVSELTVM | ASSQDQTGSQNTLY | 0.5552739 |
| GAAPLAAGL | ASSQDQTGSQNTLY | 0.30388582 |
| GADGGDGGSGGAGG | ASSQDQTGSQNTLY | 0.32662615 |
| GFGLDTGGAGG | ASSQDQTGSQNTLY | 0.70413715 |
| GLASGGLL | ASSQDQTGSQNTLY | 0.72830355 |
| GRIAGAAVLL | ASSQDQTGSQNTLY | 0.40128762 |
| GSAGSLNTSF | ASSQDQTGSQNTLY | 0.3447251 |
| GTGQIGI | ASSQDQTGSQNTLY | 0.7063011 |
| GTLIALIL | ASSQDQTGSQNTLY | 0.3538099 |
| IAATMGFVVG | ASSQDQTGSQNTLY | 0.3644762 |
| IFAQALL | ASSQDQTGSQNTLY | 0.56507 |
| IGAAVALGF | ASSQDQTGSQNTLY | 0.41131973 |
| IGLTPLAL | ASSQDQTGSQNTLY | 0.32653564 |
| ILGSNTSTL | ASSQDQTGSQNTLY | 0.5200906 |
| IVFSLTVL | ASSQDQTGSQNTLY | 0.44813758 |
| LASKATGF | ASSQDQTGSQNTLY | 0.3810552 |
| LASVALIL | ASSQDQTGSQNTLY | 0.23069713 |
| LFAVGLTL | ASSQDQTGSQNTLY | 0.39134124 |
| LFNGGIGTYI | ASSQDQTGSQNTLY | 0.38835394 |
| LTLGAGII | ASSQDQTGSQNTLY | 0.3111262 |
| LVAAALGDI | ASSQDQTGSQNTLY | 0.25076693 |
| NTAFVQLGI | ASSQDQTGSQNTLY | 0.36023474 |
| QLFSGAEL | ASSQDQTGSQNTLY | 0.64725214 |
| SAATFITL | ASSQDQTGSQNTLY | 0.67941016 |
| SNYGLANLGV | ASSQDQTGSQNTLY | 0.4138071 |
| SYGGAYAVM | ASSQDQTGSQNTLY | 0.59766763 |
| TTGVGGLTLG | ASSQDQTGSQNTLY | 0.60946083 |
| TTQTIYI | ASSQDQTGSQNTLY | 0.3949807 |
| VAAGAGLGAVI | ASSQDQTGSQNTLY | 0.26663354 |
| VAAVVTLL | ASSQDQTGSQNTLY | 0.53825986 |
| VGFLGGLI | ASSQDQTGSQNTLY | 0.5776682 |
| VGVIPILL | ASSQDQTGSQNTLY | 0.48118055 |
| VIFASSLI | ASSQDQTGSQNTLY | 0.6782533 |
| VKGGDGGAAAG | ASSQDQTGSQNTLY | 0.30591694 |
| VTKTFTNI | ASSQDQTGSQNTLY | 0.51991546 |
| VVAAAILL | ASSQDQTGSQNTLY | 0.47020912 |
| YAALDVEL | ASSQDQTGSQNTLY | 0.30203512 |
| LASVALLL | ASSQDQTGSQNTLY | 0.28721014 |
| LDAVHRV | ASSQDQTGSQNTLY | 0.29913417 |
| QGLSSSIF | ASSQDQTGSQNTLY | 0.24928263 |
| RGTFANIRL | ASSQDQTGSQNTLY | 0.49381483 |
| RQLWWGHRI | ASSQDQTGSQNTLY | 0.42760703 |
| YGPPGTGKTLL | ASSQDQTGSQNTLY | 0.42856908 |
| AALVGAPANVA | ASSQDQTGSQNTLY | 0.392753 |
| AATFRRTE | ASSQDQTGSQNTLY | 0.37241536 |
| AGALGGGI | ASSQDQTGSQNTLY | 0.40750387 |

| | | |
|---|---|---|
| AGGAGGAGGIGAG | ASSQDQTGSQNTLY | 0.46168822 |
| AGSGALGL | ASSQDQTGSQNTLY | 0.42260543 |
| AQIADQI | ASSQDQTGSQNTLY | 0.53042 |
| ASAVAVTYL | ASSQDQTGSQNTLY | 0.34095424 |
| ASTVLGGTGG | ASSQDQTGSQNTLY | 0.34290558 |
| AVGGEPAK | ASSQDQTGSQNTLY | 0.71239114 |
| CEQGTFGGTC | ASSQDQTGSQNTLY | 0.30317527 |
| DEGKAPAG | ASSQDQTGSQNTLY | 0.30636805 |
| DTVVFPAL | ASSQDQTGSQNTLY | 0.5371877 |
| FEGITMAG | ASSQDQTGSQNTLY | 0.37351102 |
| FPLVVQD | ASSQDQTGSQNTLY | 0.58793235 |
| GAFTTPQ | ASSQDQTGSQNTLY | 0.59936523 |
| GFGGFGTAGGVGGA | ASSQDQTGSQNTLY | 0.5784013 |
| GFLAQFL | ASSQDQTGSQNTLY | 0.6664871 |
| GGAGGAGGNAGVA | ASSQDQTGSQNTLY | 0.67744064 |
| GGVGTAGAGGAGGAGG | ASSQDQTGSQNTLY | 0.640007 |
| GNGGGGDAGNAGSGGN | ASSQDQTGSQNTLY | 0.31633648 |
| GTGGAGGAGNDAGS | ASSQDQTGSQNTLY | 0.73427486 |
| GTTNTGWFNS | ASSQDQTGSQNTLY | 0.2971797 |
| IFMGPLFTL | ASSQDQTGSQNTLY | 0.52475697 |
| ILAHAIEAI | ASSQDQTGSQNTLY | 0.5476617 |
| IRSAIREAL | ASSQDQTGSQNTLY | 0.32870537 |
| ITFAGGG | ASSQDQTGSQNTLY | 0.5149963 |
| ITGMSGHQEV | ASSQDQTGSQNTLY | 0.56310046 |
| IVAGLACTL | ASSQDQTGSQNTLY | 0.6129922 |
| IVGGGLAGA | ASSQDQTGSQNTLY | 0.34938675 |
| IVTGVLNI | ASSQDQTGSQNTLY | 0.35654154 |
| LAQQGCI | ASSQDQTGSQNTLY | 0.35924247 |
| LCLHVEN | ASSQDQTGSQNTLY | 0.34121522 |
| LDAISDGS | ASSQDQTGSQNTLY | 0.29991502 |
| LGGFAAML | ASSQDQTGSQNTLY | 0.5448035 |
| LGGKNPMI | ASSQDQTGSQNTLY | 0.45673606 |
| LGPKGVAAIL | ASSQDQTGSQNTLY | 0.73319614 |
| LMVGVSI | ASSQDQTGSQNTLY | 0.56740814 |
| LVGAAFVL | ASSQDQTGSQNTLY | 0.6873354 |
| LVLAVAPL | ASSQDQTGSQNTLY | 0.3978874 |
| MAAIAVIDAA | ASSQDQTGSQNTLY | 0.51985157 |
| NNGINAELQ | ASSQDQTGSQNTLY | 0.26203465 |
| RGEPLVLVHG | ASSQDQTGSQNTLY | 0.3401311 |
| RGGAGGAAGLI | ASSQDQTGSQNTLY | 0.29438174 |
| RSYRMHPAV | ASSQDQTGSQNTLY | 0.4598124 |
| SAAHLLAGQLA | ASSQDQTGSQNTLY | 0.24764507 |
| SAAVGSY | ASSQDQTGSQNTLY | 0.66126335 |
| SPHAVLHRI | ASSQDQTGSQNTLY | 0.5323105 |
| SQIAKVL | ASSQDQTGSQNTLY | 0.6181748 |
| TALGHLVP | ASSQDQTGSQNTLY | 0.3450235 |
| TAPTTSGAA | ASSQDQTGSQNTLY | 0.31351048 |
| TDGDDGNGGSPGT | ASSQDQTGSQNTLY | 0.41023678 |
| TDGLGEN | ASSQDQTGSQNTLY | 0.68873614 |
| TGGPPLPGQA | ASSQDQTGSQNTLY | 0.32088473 |
| TIPAITI | ASSQDQTGSQNTLY | 0.24404481 |

| | | |
|---|---|---|
| TIPGFTI | ASSQDQTGSQNTLY | 0.32642877 |
| TSIASGALA | ASSQDQTGSQNTLY | 0.25324383 |
| TVTLPGAAI | ASSQDQTGSQNTLY | 0.6443643 |
| VASGLSAALP | ASSQDQTGSQNTLY | 0.58015805 |
| VGAGVIGI | ASSQDQTGSQNTLY | 0.549386 |
| VGPKRFGV | ASSQDQTGSQNTLY | 0.729301 |
| VNVAFPD | ASSQDQTGSQNTLY | 0.23498136 |
| VSMTNSGVS | ASSQDQTGSQNTLY | 0.34958583 |
| VTDVLGDLSV | ASSQDQTGSQNTLY | 0.71792704 |
| VVVVGAGFAGL | ASSQDQTGSQNTLY | 0.5117315 |
| VYTVGTVV | ASSQDQTGSQNTLY | 0.4815498 |
| WTTFSGP | ASSQDQTGSQNTLY | 0.6400567 |
| YDELVEAF | ASSQDQTGSQNTLY | 0.4203781 |
| YGTGGAGGNGG | ASSQDQTGSQNTLY | 0.46992317 |
| YTNVMAV | ASSQDQTGSQNTLY | 0.50170094 |
| FLEQVHQL | ASSQDQTGSQNTLY | 0.3806527 |
| GLSLLAVAV | ASSQDQTGSQNTLY | 0.33638245 |
| LEQVHQL | ASSQDQTGSQNTLY | 0.28246704 |
| LNAAGAI | ASSQDQTGSQNTLY | 0.4491148 |
| PAVHASL | ASSQDQTGSQNTLY | 0.7311871 |
| TELEDDA | ASSQDQTGSQNTLY | 0.54239964 |
| TPPGPPN | ASSQDQTGSQNTLY | 0.46236855 |
| TPPPPPPP | ASSQDQTGSQNTLY | 0.4848474 |
| TQSPALI | AWSLGVGGSTSAETLY | 0.72241545 |
| EFAAAFPQ | AWSLGVGGSTSAETLY | 0.37864512 |
| LLSGNLDVL | AWSLGVGGSTSAETLY | 0.3303418 |
| GPANLGNYNI | AWSLGVGGSTSAETLY | 0.64577657 |
| TLGPTLP | AWSLGVGGSTSAETLY | 0.30754393 |
| VYAYSSCCK | AWSLGVGGSTSAETLY | 0.40348127 |
| APAPAPAPNI | AWSLGVGGSTSAETLY | 0.32069984 |
| PGGPVPIP | AWSLGVGGSTSAETLY | 0.74263275 |
| FLTVALEP | AWSLGVGGSTSAETLY | 0.57378864 |
| SQYFGANL | AWSLGVGGSTSAETLY | 0.4575726 |
| GGVKPAA | AWSLGVGGSTSAETLY | 0.43924016 |
| GGTGGAGGAGGW | AWSLGVGGSTSAETLY | 0.3760274 |
| GVGGVAGLGGAGG | AWSLGVGGSTSAETLY | 0.5249767 |
| VLLVLATTI | AWSLGVGGSTSAETLY | 0.26247656 |
| IGIFAAML | AWSLGVGGSTSAETLY | 0.6951649 |
| AVGDLID | AWSLGVGGSTSAETLY | 0.34882176 |
| NISPLPL | AWSLGVGGSTSAETLY | 0.40220058 |
| VTAVFVAI | AWSLGVGGSTSAETLY | 0.5427282 |
| SGDDNDAAL | AWSLGVGGSTSAETLY | 0.74914896 |
| AVAALAHS | AWSLGVGGSTSAETLY | 0.34562868 |
| GGVTVMATP | AWSLGVGGSTSAETLY | 0.6267158 |
| AGLAANVRD | AWSLGVGGSTSAETLY | 0.35871312 |
| TNRPDLIDPALLRPG | AWSLGVGGSTSAETLY | 0.54540837 |
| LANLGVTAL | AWSLGVGGSTSAETLY | 0.71281004 |
| TGERIEL | AWSLGVGGSTSAETLY | 0.70358706 |
| NGEELAA | AWSLGVGGSTSAETLY | 0.746981 |
| VLSLVWG | AWSLGVGGSTSAETLY | 0.36544958 |
| IGYALWL | AWSLGVGGSTSAETLY | 0.28676832 |

| | | |
|---|---|---|
| LSLLLAPL | AWSLGVGGSTSAETLY | 0.3848847 |
| TLDEALK | AWSLGVGGSTSAETLY | 0.42546707 |
| TDGAGTAEGG | AWSLGVGGSTSAETLY | 0.61241835 |
| NAPGGAGGNGG | AWSLGVGGSTSAETLY | 0.6294913 |
| VSVFMGMGGNF | AWSLGVGGSTSAETLY | 0.44094282 |
| LASYLAAL | AWSLGVGGSTSAETLY | 0.37414122 |
| LVGGTPAL | AWSLGVGGSTSAETLY | 0.6645426 |
| VRNADVL | AWSLGVGGSTSAETLY | 0.65712106 |
| TCSSPFLVSAM | AWSLGVGGSTSAETLY | 0.6079297 |
| VHITDAE | AWSLGVGGSTSAETLY | 0.35346633 |
| EYAAPSF | AWSLGVGGSTSAETLY | 0.33644688 |
| VVDAFHGGL | AWSLGVGGSTSAETLY | 0.4145368 |
| ASVVDPQ | AWSLGVGGSTSAETLY | 0.62466025 |
| SYNEFGGLS | AWSLGVGGSTSAETLY | 0.30389512 |
| DPDGSVY | AWSLGVGGSTSAETLY | 0.482772 |
| MVSTIHATE | AWSLGVGGSTSAETLY | 0.63716996 |
| LAAGVADPV | AWSLGVGGSTSAETLY | 0.2763236 |
| TKEQIAA | AWSLGVGGSTSAETLY | 0.36095202 |
| SLASPAAL | AWSLGVGGSTSAETLY | 0.617501 |
| IGLVFYGV | AWSLGVGGSTSAETLY | 0.340229 |
| FLLDASNL | AWSLGVGGSTSAETLY | 0.34352913 |
| YQVGNFP | AWSLGVGGSTSAETLY | 0.6932646 |
| ESLLSQST | AWSLGVGGSTSAETLY | 0.72547215 |
| LFTVGFAAVG | AWSLGVGGSTSAETLY | 0.5379265 |
| ATPPTPT | AWSLGVGGSTSAETLY | 0.27033848 |
| TDLATLG | AWSLGVGGSTSAETLY | 0.22764085 |
| SAGGTGR | AWSLGVGGSTSAETLY | 0.53710985 |
| GFSNSGS | AWSLGVGGSTSAETLY | 0.73066175 |
| GGNGGSGGTSV | AWSLGVGGSTSAETLY | 0.4138555 |
| GTSVATGGAGN | AWSLGVGGSTSAETLY | 0.41484648 |
| NGGNGGSGGTS | AWSLGVGGSTSAETLY | 0.40155235 |
| AVVKFPH | AWSLGVGGSTSAETLY | 0.6036953 |
| TPPPPPP | AWSLGVGGSTSAETLY | 0.336738 |
| AAPAAAAAPA | AWSLGVGGSTSAETLY | 0.285861 |
| IALNEDL | AWSLGVGGSTSAETLY | 0.3507732 |
| IVNVSSM | AWSLGVGGSTSAETLY | 0.59748524 |
| TGIPPAPR | AWSLGVGGSTSAETLY | 0.30005318 |
| FDLGGGTF | AWSLGVGGSTSAETLY | 0.48389727 |
| FDLGGGTFDVS | AWSLGVGGSTSAETLY | 0.58858514 |
| NEPTAAA | AWSLGVGGSTSAETLY | 0.330588 |
| NEPTAAALA | AWSLGVGGSTSAETLY | 0.5160521 |
| GSIINISSI | AWSLGVGGSTSAETLY | 0.3464755 |
| VAPGNAGTA | AWSLGVGGSTSAETLY | 0.64963496 |
| TGGGASLE | AWSLGVGGSTSAETLY | 0.2975161 |
| VAVYNHYKRI | AWSLGVGGSTSAETLY | 0.4715185 |
| TDPAPATT | AWSLGVGGSTSAETLY | 0.5339343 |
| TNRPDILDPALLRPG | AWSLGVGGSTSAETLY | 0.40821427 |
| LTTTTTS | AWSLGVGGSTSAETLY | 0.4691326 |
| PTGSGKTL | AWSLGVGGSTSAETLY | 0.35558963 |
| TGSGKTL | AWSLGVGGSTSAETLY | 0.31298214 |
| TIGHVDHGK | AWSLGVGGSTSAETLY | 0.32107064 |

| | | |
|---|---|---|
| TIGHVDHGKTT | AWSLGVGGSTSAETLY | 0.62522876 |
| GHVDHGKTT | AWSLGVGGSTSAETLY | 0.35904557 |
| AQFEHTLL | AWSLGVGGSTSAETLY | 0.5603807 |
| IVNQDEL | AWSLGVGGSTSAETLY | 0.56643176 |
| TNGSQFF | AWSLGVGGSTSAETLY | 0.30815816 |
| TGKGIVE | AWSLGVGGSTSAETLY | 0.4447843 |
| LISNASDA | AWSLGVGGSTSAETLY | 0.36513078 |
| AELRTLGELTKIAKAHGAQ | AWSLGVGGSTSAETLY | 0.5350198 |
| APNAVAAAEPSW | AWSLGVGGSTSAETLY | 0.52358603 |
| APQVAGAATAVL | AWSLGVGGSTSAETLY | 0.4352423 |
| ARRDPAAAAAILPTDA | AWSLGVGGSTSAETLY | 0.3687926 |
| DAAARDVLAEAGLAEYF | AWSLGVGGSTSAETLY | 0.38533106 |
| DDQIVLSAKANLAKVEQ | AWSLGVGGSTSAETLY | 0.5018404 |
| DELLPDTGEDSDEDGH | AWSLGVGGSTSAETLY | 0.3566834 |
| ELPEAPANRATVPQD | AWSLGVGGSTSAETLY | 0.35973766 |
| FGQSKIERVTGIG | AWSLGVGGSTSAETLY | 0.6777996 |
| GAGGTGSTAGGAGGAGGAGGLY | AWSLGVGGSTSAETLY | 0.38717985 |
| GHGGTGAAGGNGGTGGAGGSGI | AWSLGVGGSTSAETLY | 0.32470286 |
| GNGGNGGAGGEGGPGGAGGAGA | AWSLGVGGSTSAETLY | 0.5167841 |
| GSTAGNGGNGGNATNGTIAGQP | AWSLGVGGSTSAETLY | 0.30157313 |
| GTGGDGGNGGNGGTGG | AWSLGVGGSTSAETLY | 0.58995914 |
| GVRRPAGIPADAPV | AWSLGVGGSTSAETLY | 0.35372916 |
| GYRQVTDAHLLTLARR | AWSLGVGGSTSAETLY | 0.42912972 |
| IAKMGQDGHDRG | AWSLGVGGSTSAETLY | 0.39681283 |
| KAEPSIEIQYT | AWSLGVGGSTSAETLY | 0.26968852 |
| KPVRIRMAADEDTVAVT | AWSLGVGGSTSAETLY | 0.34008408 |
| MPPRIAGMRLLVIKPEP | AWSLGVGGSTSAETLY | 0.34170374 |
| MKFYAVALFRSRGVPPTR | AWSLGVGGSTSAETLY | 0.5780467 |
| NAGGFGILGAVAHSP | AWSLGVGGSTSAETLY | 0.5889044 |
| NEGREITLAMAEM | AWSLGVGGSTSAETLY | 0.33894515 |
| NELVEWSHPLGHRWRWE | AWSLGVGGSTSAETLY | 0.48879355 |
| NLGAGGTGGNGGNPG | AWSLGVGGSTSAETLY | 0.37736315 |
| NTEWPKWETVMAADPVD | AWSLGVGGSTSAETLY | 0.3443181 |
| PASAAPAETQP | AWSLGVGGSTSAETLY | 0.3509587 |
| QQSSDGKNGVRAAEITGE | AWSLGVGGSTSAETLY | 0.3274227 |
| RAPDAVTTSWWKE | AWSLGVGGSTSAETLY | 0.3155783 |
| SGFGNVGAHTSGWFNQ | AWSLGVGGSTSAETLY | 0.39608836 |
| SGVAGAVGQAASVGGLKV | AWSLGVGGSTSAETLY | 0.32681805 |
| SNGGAGTGSAGGNGGT | AWSLGVGGSTSAETLY | 0.45471665 |
| TGGAGGEGGAAG | AWSLGVGGSTSAETLY | 0.5833063 |
| TGGDGGAAGAGGAPGHG | AWSLGVGGSTSAETLY | 0.34652478 |
| TMPPLLAAIANAGRHGV | AWSLGVGGSTSAETLY | 0.43882203 |
| TSSTAGNSGGA | AWSLGVGGSTSAETLY | 0.47286287 |
| VPATVPVGILC | AWSLGVGGSTSAETLY | 0.53994274 |
| VQRPDSGGVVLTGRIS | AWSLGVGGSTSAETLY | 0.49458045 |
| WTPNPVPATKGVPAGLVQAT | AWSLGVGGSTSAETLY | 0.3273012 |
| YALKRALAKFSDRFDVVII | AWSLGVGGSTSAETLY | 0.4935981 |
| YLIGSVIV | AWSLGVGGSTSAETLY | 0.5941454 |
| YGQLVAVID | AWSLGVGGSTSAETLY | 0.5355244 |
| VTVGGMTLPA | AWSLGVGGSTSAETLY | 0.48809844 |
| VLALPAV | AWSLGVGGSTSAETLY | 0.7085733 |

| | | |
|---|---|---|
| VIPPTLNYET | AWSLGVGGSTSAETLY | 0.5670361 |
| VGMGGMHPSSG | AWSLGVGGSTSAETLY | 0.4479433 |
| VGLAGGAATA | AWSLGVGGSTSAETLY | 0.250673 |
| VGGAGGAGGSGP | AWSLGVGGSTSAETLY | 0.51705647 |
| VFAAFMV | AWSLGVGGSTSAETLY | 0.5521053 |
| TLYDKLR | AWSLGVGGSTSAETLY | 0.30598935 |
| TLAVFTVL | AWSLGVGGSTSAETLY | 0.26836514 |
| TGSVQPARV | AWSLGVGGSTSAETLY | 0.6806264 |
| TGGGAGSGGT | AWSLGVGGSTSAETLY | 0.7251867 |
| SHLETSL | AWSLGVGGSTSAETLY | 0.38818192 |
| SFTTGMT | AWSLGVGGSTSAETLY | 0.6627817 |
| QVEVAKRMGVRQPTVS | AWSLGVGGSTSAETLY | 0.26078445 |
| QGSGGPALAM | AWSLGVGGSTSAETLY | 0.580183 |
| MLAPSNP | AWSLGVGGSTSAETLY | 0.6630978 |
| LVNNAAI | AWSLGVGGSTSAETLY | 0.3302302 |
| LTGEINTLA | AWSLGVGGSTSAETLY | 0.57242715 |
| LSIFGPV | AWSLGVGGSTSAETLY | 0.31360972 |
| LLALLANL | AWSLGVGGSTSAETLY | 0.33494565 |
| LISNASDALDKLR | AWSLGVGGSTSAETLY | 0.6729809 |
| LINEAVP | AWSLGVGGSTSAETLY | 0.47400683 |
| LGYAFSGL | AWSLGVGGSTSAETLY | 0.37678534 |
| LDFVAVAL | AWSLGVGGSTSAETLY | 0.44432133 |
| LAQPGAE | AWSLGVGGSTSAETLY | 0.340434 |
| ISNLGAVTGF | AWSLGVGGSTSAETLY | 0.35601133 |
| HQEDPGRI | AWSLGVGGSTSAETLY | 0.5653854 |
| GTPGEPTP | AWSLGVGGSTSAETLY | 0.5475081 |
| GFFNSGS | AWSLGVGGSTSAETLY | 0.63413453 |
| GAGGGGLTAGAN | AWSLGVGGSTSAETLY | 0.3403817 |
| FVDALVNM | AWSLGVGGSTSAETLY | 0.691939 |
| FLTVAPEL | AWSLGVGGSTSAETLY | 0.73298407 |
| FETLTAL | AWSLGVGGSTSAETLY | 0.5930679 |
| DVINPIEL | AWSLGVGGSTSAETLY | 0.3242146 |
| APVSGAPV | AWSLGVGGSTSAETLY | 0.70397246 |
| ALVGGLAW | AWSLGVGGSTSAETLY | 0.4887976 |
| ALAAAAATAI | AWSLGVGGSTSAETLY | 0.5396044 |
| AFGLGLLL | AWSLGVGGSTSAETLY | 0.43722713 |
| AFFAAASL | AWSLGVGGSTSAETLY | 0.5069058 |
| AAAALPAFSPP | AWSLGVGGSTSAETLY | 0.6512629 |
| VVLAPAL | AWSLGVGGSTSAETLY | 0.61972857 |
| TIGHVDHGKTTL | AWSLGVGGSTSAETLY | 0.62559265 |
| LISNASDALDK | AWSLGVGGSTSAETLY | 0.57129735 |
| LISNASDAL | AWSLGVGGSTSAETLY | 0.50537735 |
| ISNASDALDK | AWSLGVGGSTSAETLY | 0.5825113 |
| NQAETLVYQTE | AWSLGVGGSTSAETLY | 0.539992 |
| TGGTGGTAGAG | AWSLGVGGSTSAETLY | 0.65801364 |
| TGSTAGGAGGA | AWSLGVGGSTSAETLY | 0.65295994 |
| DPLAVPVAAGPV | AWSLGVGGSTSAETLY | 0.5584536 |
| GGTGGTPGNGGN | AWSLGVGGSTSAETLY | 0.6533821 |
| PAGLAGVDPATL | AWSLGVGGSTSAETLY | 0.63937366 |
| GNNLVVEAPHAGASV | AWSLGVGGSTSAETLY | 0.46058756 |
| KVTRIMSASTLEQPA | AWSLGVGGSTSAETLY | 0.58929724 |

| | | |
|---|---|---|
| LGFALPIALPRRTVV | AWSLGVGGSTSAETLY | 0.56787014 |
| TRLGNLTARKLVVLV | AWSLGVGGSTSAETLY | 0.60643023 |
| WSQQRRLIYFPSAGP | AWSLGVGGSTSAETLY | 0.44635016 |
| GPNGNPDPMAAAVDIR | AWSLGVGGSTSAETLY | 0.64303005 |
| TQRCVSVADLRAAASR | AWSLGVGGSTSAETLY | 0.5554092 |
| QFVLDVAEHPVEVLTETV | AWSLGVGGSTSAETLY | 0.36812174 |
| GGAGGTGGAGGNGGTQAGDG | AWSLGVGGSTSAETLY | 0.62580234 |
| PLLATLRAIAVDVDAKPAQV | AWSLGVGGSTSAETLY | 0.6093787 |
| APKFRTGGYMIDFWGVGYQVAK | AWSLGVGGSTSAETLY | 0.6060639 |
| GAGGAGGAGGAAGGPGGTGGTG | AWSLGVGGSTSAETLY | 0.644753 |
| IGALGLLATDVAPAGSVAFGVA | AWSLGVGGSTSAETLY | 0.5573666 |
| AGGAGGIGGNGGAGGAGGVLMG | AWSLGVGGSTSAETLY | 0.54873073 |
| ADELIGLL | AWSLGVGGSTSAETLY | 0.6290704 |
| ADRALKA | AWSLGVGGSTSAETLY | 0.6507344 |
| AGAVGQVGGAGGA | AWSLGVGGSTSAETLY | 0.54606664 |
| AGGGGGLGGAP | AWSLGVGGSTSAETLY | 0.6149049 |
| AGGLLYGNGGAG | AWSLGVGGSTSAETLY | 0.6545004 |
| AGLESIGL | AWSLGVGGSTSAETLY | 0.6175805 |
| AGLKTGII | AWSLGVGGSTSAETLY | 0.6565362 |
| ASLAALLL | AWSLGVGGSTSAETLY | 0.66117543 |
| ASVINISSI | AWSLGVGGSTSAETLY | 0.56597334 |
| ATPPGPA | AWSLGVGGSTSAETLY | 0.6520416 |
| ATVASALV | AWSLGVGGSTSAETLY | 0.65319717 |
| ATVASLIL | AWSLGVGGSTSAETLY | 0.58311534 |
| AVNGPVLIRH | AWSLGVGGSTSAETLY | 0.60937136 |
| DGAAGAVGI | AWSLGVGGSTSAETLY | 0.4804573 |
| DPLLHGI | AWSLGVGGSTSAETLY | 0.5973032 |
| ELGAALGAL | AWSLGVGGSTSAETLY | 0.5226288 |
| ELGRTQAV | AWSLGVGGSTSAETLY | 0.624413 |
| FVAGGYLS | AWSLGVGGSTSAETLY | 0.65851915 |
| GASGGAGGAGGW | AWSLGVGGSTSAETLY | 0.5930147 |
| GGAGAGGDGGAGGI | AWSLGVGGSTSAETLY | 0.5114558 |
| GTGHNGSQPGMGGNG | AWSLGVGGSTSAETLY | 0.48376912 |
| GYAAASASAS | AWSLGVGGSTSAETLY | 0.62024295 |
| HQVDVAAVQQG | AWSLGVGGSTSAETLY | 0.5235422 |
| IGAAIAHT | AWSLGVGGSTSAETLY | 0.43521038 |
| IGMIEVAQL | AWSLGVGGSTSAETLY | 0.5976763 |
| IRLAPEHTSAMD | AWSLGVGGSTSAETLY | 0.65037906 |
| IYNVSVL | AWSLGVGGSTSAETLY | 0.65013206 |
| KQGFLGM | AWSLGVGGSTSAETLY | 0.52130955 |
| LESVITQL | AWSLGVGGSTSAETLY | 0.6589687 |
| LGLGALVVV | AWSLGVGGSTSAETLY | 0.6402106 |
| LIGNGGNGGSGGI | AWSLGVGGSTSAETLY | 0.6443502 |
| LNAINAPTL | AWSLGVGGSTSAETLY | 0.6055503 |
| LQLLNTGV | AWSLGVGGSTSAETLY | 0.504492 |
| LQTSIEF | AWSLGVGGSTSAETLY | 0.65361917 |
| MSFAVSEL | AWSLGVGGSTSAETLY | 0.5799726 |
| NTGALGL | AWSLGVGGSTSAETLY | 0.6172151 |
| PPGAGGAGGAGTT | AWSLGVGGSTSAETLY | 0.64750683 |
| QLVGLGI | AWSLGVGGSTSAETLY | 0.60551554 |
| RFLEQVHQL | AWSLGVGGSTSAETLY | 0.5729596 |

| | | |
|---|---|---|
| RGVGYVLR | AWSLGVGGSTSAETLY | 0.59753144 |
| SATGVFAGVI | AWSLGVGGSTSAETLY | 0.6520561 |
| SLPAITLP | AWSLGVGGSTSAETLY | 0.5443908 |
| STALAFTV | AWSLGVGGSTSAETLY | 0.53951174 |
| SVLTVGLGK | AWSLGVGGSTSAETLY | 0.59260005 |
| TALAGAGSGI | AWSLGVGGSTSAETLY | 0.60792327 |
| TGLPFTGI | AWSLGVGGSTSAETLY | 0.6524708 |
| VAEHVTV | AWSLGVGGSTSAETLY | 0.6283511 |
| VDIVLNSL | AWSLGVGGSTSAETLY | 0.5802562 |
| VLTGGGAVL | AWSLGVGGSTSAETLY | 0.6296916 |
| VNVGLAYGI | AWSLGVGGSTSAETLY | 0.6378642 |
| VSSAAVGAV | AWSLGVGGSTSAETLY | 0.6223813 |
| VTSVYEAGV | AWSLGVGGSTSAETLY | 0.6149638 |
| WVALDRGA | AWSLGVGGSTSAETLY | 0.5633593 |
| YLVGSVVI | AWSLGVGGSTSAETLY | 0.645311 |
| GPPGTGKTLL | AWSLGVGGSTSAETLY | 0.6405242 |
| GVGMTKF | AWSLGVGGSTSAETLY | 0.6456271 |
| IGIGGSDL | AWSLGVGGSTSAETLY | 0.65384024 |
| IGTLLAEL | AWSLGVGGSTSAETLY | 0.63028175 |
| IYTSGTT | AWSLGVGGSTSAETLY | 0.607577 |
| LSFPAGV | AWSLGVGGSTSAETLY | 0.43574756 |
| LYTSGTT | AWSLGVGGSTSAETLY | 0.60097784 |
| VGAFPFGF | AWSLGVGGSTSAETLY | 0.6093828 |
| VGAGTVEF | AWSLGVGGSTSAETLY | 0.62186414 |
| VGKSTLLL | AWSLGVGGSTSAETLY | 0.6421666 |
| AALAHAGGGIV | AWSLGVGGSTSAETLY | 0.6359503 |
| AALVAVSLL | AWSLGVGGSTSAETLY | 0.5761208 |
| AAVSRAVAAL | AWSLGVGGSTSAETLY | 0.6497258 |
| AGGSSGIGGF | AWSLGVGGSTSAETLY | 0.65111375 |
| AGLTVANANG | AWSLGVGGSTSAETLY | 0.49290547 |
| AGLVGAVL | AWSLGVGGSTSAETLY | 0.59958494 |
| ALSQAFTV | AWSLGVGGSTSAETLY | 0.6651781 |
| ALTFSVSL | AWSLGVGGSTSAETLY | 0.62396044 |
| ALTGAALVAL | AWSLGVGGSTSAETLY | 0.5863304 |
| ASFSIGAGVL | AWSLGVGGSTSAETLY | 0.5953603 |
| DGSVYKI | AWSLGVGGSTSAETLY | 0.6638303 |
| EQLAEIV | AWSLGVGGSTSAETLY | 0.60024875 |
| FAVSELTVM | AWSLGVGGSTSAETLY | 0.65621644 |
| GAAPLAAGL | AWSLGVGGSTSAETLY | 0.4017975 |
| GADGGDGGSGGAGG | AWSLGVGGSTSAETLY | 0.57358664 |
| GFGLDTGGAGG | AWSLGVGGSTSAETLY | 0.50957733 |
| GLASGGLL | AWSLGVGGSTSAETLY | 0.37871182 |
| GRIAGAAVLL | AWSLGVGGSTSAETLY | 0.6471746 |
| GSAGSLNTSF | AWSLGVGGSTSAETLY | 0.54641324 |
| GTGQIGI | AWSLGVGGSTSAETLY | 0.63209695 |
| GTLIALIL | AWSLGVGGSTSAETLY | 0.59263474 |
| IAATMGFVVG | AWSLGVGGSTSAETLY | 0.65451837 |
| IFAQALL | AWSLGVGGSTSAETLY | 0.59792733 |
| IGAAVALGF | AWSLGVGGSTSAETLY | 0.6113137 |
| IGLTPLAL | AWSLGVGGSTSAETLY | 0.5599943 |
| ILGSNTSTL | AWSLGVGGSTSAETLY | 0.6362308 |

| | | |
|---|---|---|
| IVFSLTVL | AWSLGVGGSTSAETLY | 0.65587455 |
| LASKATGF | AWSLGVGGSTSAETLY | 0.6462537 |
| LASVALIL | AWSLGVGGSTSAETLY | 0.4462576 |
| LFAVGLTL | AWSLGVGGSTSAETLY | 0.61478776 |
| LFNGGIGTYI | AWSLGVGGSTSAETLY | 0.60223854 |
| LTLGAGII | AWSLGVGGSTSAETLY | 0.48427236 |
| LVAAALGDI | AWSLGVGGSTSAETLY | 0.41643527 |
| NTAFVQLGI | AWSLGVGGSTSAETLY | 0.5989248 |
| QLFSGAEL | AWSLGVGGSTSAETLY | 0.5918378 |
| SAATFITL | AWSLGVGGSTSAETLY | 0.6346699 |
| SNYGLANLGV | AWSLGVGGSTSAETLY | 0.6325887 |
| SYGGAYAVM | AWSLGVGGSTSAETLY | 0.58663046 |
| TTGVGGLTLG | AWSLGVGGSTSAETLY | 0.62563753 |
| TTQTIYI | AWSLGVGGSTSAETLY | 0.6100271 |
| VAAGAGLGAVI | AWSLGVGGSTSAETLY | 0.30597612 |
| VAAVVTLL | AWSLGVGGSTSAETLY | 0.55575794 |
| VGFLGGLI | AWSLGVGGSTSAETLY | 0.48156434 |
| VGVIPILL | AWSLGVGGSTSAETLY | 0.6624415 |
| VIFASSLI | AWSLGVGGSTSAETLY | 0.57825255 |
| VKGGDGGAAAG | AWSLGVGGSTSAETLY | 0.5791716 |
| VTKTFTNI | AWSLGVGGSTSAETLY | 0.61057734 |
| VVAAAILL | AWSLGVGGSTSAETLY | 0.62107736 |
| YAALDVEL | AWSLGVGGSTSAETLY | 0.6056944 |
| LASVALLL | AWSLGVGGSTSAETLY | 0.53137505 |
| LDAVHRV | AWSLGVGGSTSAETLY | 0.44349095 |
| QGLSSSIF | AWSLGVGGSTSAETLY | 0.6404705 |
| RGTFANIRL | AWSLGVGGSTSAETLY | 0.56299376 |
| RQLWWGHRI | AWSLGVGGSTSAETLY | 0.53901714 |
| YGPPGTGKTLL | AWSLGVGGSTSAETLY | 0.49675182 |
| AALVGAPANVA | AWSLGVGGSTSAETLY | 0.6174902 |
| AATFRRTE | AWSLGVGGSTSAETLY | 0.45860726 |
| AGALGGGI | AWSLGVGGSTSAETLY | 0.64682007 |
| AGGAGGAGGIGAG | AWSLGVGGSTSAETLY | 0.541975 |
| AGSGALGL | AWSLGVGGSTSAETLY | 0.59709847 |
| AQIADQI | AWSLGVGGSTSAETLY | 0.66332865 |
| ASAVAVTYL | AWSLGVGGSTSAETLY | 0.62464935 |
| ASTVLGGTGG | AWSLGVGGSTSAETLY | 0.5956396 |
| AVGGEPAK | AWSLGVGGSTSAETLY | 0.567834 |
| CEQGTFGGTC | AWSLGVGGSTSAETLY | 0.5469364 |
| DEGKAPAG | AWSLGVGGSTSAETLY | 0.5540039 |
| DTVVFPAL | AWSLGVGGSTSAETLY | 0.58233356 |
| FEGITMAG | AWSLGVGGSTSAETLY | 0.6568488 |
| FPLVVQD | AWSLGVGGSTSAETLY | 0.60860455 |
| GAFTTPQ | AWSLGVGGSTSAETLY | 0.49628997 |
| GFGGFGTAGGVGGA | AWSLGVGGSTSAETLY | 0.623295 |
| GFLAQFL | AWSLGVGGSTSAETLY | 0.6614151 |
| GGAGGAGGNAGVA | AWSLGVGGSTSAETLY | 0.65250814 |
| GGVGTAGAGGAGGAGG | AWSLGVGGSTSAETLY | 0.54320985 |
| GNGGGGDAGNAGSGGN | AWSLGVGGSTSAETLY | 0.5549622 |
| GTGGAGGAGNDAGS | AWSLGVGGSTSAETLY | 0.5610893 |
| GTTNTGWFNS | AWSLGVGGSTSAETLY | 0.595484 |

| | | |
|---|---|---|
| IFMGPLFTL | AWSLGVGGSTSAETLY | 0.53034806 |
| ILAHAIEAI | AWSLGVGGSTSAETLY | 0.55202156 |
| IRSAIREAL | AWSLGVGGSTSAETLY | 0.63616216 |
| ITFAGGG | AWSLGVGGSTSAETLY | 0.6725365 |
| ITGMSGHQEV | AWSLGVGGSTSAETLY | 0.599562 |
| IVAGLACTL | AWSLGVGGSTSAETLY | 0.44575223 |
| IVGGGLAGA | AWSLGVGGSTSAETLY | 0.5953417 |
| IVTGVLNI | AWSLGVGGSTSAETLY | 0.5482583 |
| LAQQGCI | AWSLGVGGSTSAETLY | 0.5615268 |
| LCLHVEN | AWSLGVGGSTSAETLY | 0.57888114 |
| LDAISDGS | AWSLGVGGSTSAETLY | 0.46339905 |
| LGGFAAML | AWSLGVGGSTSAETLY | 0.61607236 |
| LGGKNPMI | AWSLGVGGSTSAETLY | 0.65706146 |
| LGPKGVAAIL | AWSLGVGGSTSAETLY | 0.57592136 |
| LMVGVSI | AWSLGVGGSTSAETLY | 0.61490506 |
| LVGAAFVL | AWSLGVGGSTSAETLY | 0.6526681 |
| LVLAVAPL | AWSLGVGGSTSAETLY | 0.63294417 |
| MAAIAVIDAA | AWSLGVGGSTSAETLY | 0.6508503 |
| NNGINAELQ | AWSLGVGGSTSAETLY | 0.6513911 |
| RGEPLVLVHG | AWSLGVGGSTSAETLY | 0.6350205 |
| RGGAGGAAGLI | AWSLGVGGSTSAETLY | 0.5976203 |
| RSYRMHPAV | AWSLGVGGSTSAETLY | 0.6434142 |
| SAAHLLAGQLA | AWSLGVGGSTSAETLY | 0.5979873 |
| SAAVGSY | AWSLGVGGSTSAETLY | 0.6043032 |
| SPHAVLHRI | AWSLGVGGSTSAETLY | 0.56731504 |
| SQIAKVL | AWSLGVGGSTSAETLY | 0.64552 |
| TALGHLVP | AWSLGVGGSTSAETLY | 0.57797694 |
| TAPTTSGAA | AWSLGVGGSTSAETLY | 0.662272 |
| TDGDDGNGGSPGT | AWSLGVGGSTSAETLY | 0.39790356 |
| TDGLGEN | AWSLGVGGSTSAETLY | 0.5382637 |
| TGGPPLPGQA | AWSLGVGGSTSAETLY | 0.6461396 |
| TIPAITI | AWSLGVGGSTSAETLY | 0.31781498 |
| TIPGFTI | AWSLGVGGSTSAETLY | 0.4003673 |
| TSIASGALA | AWSLGVGGSTSAETLY | 0.57672834 |
| TVTLPGAAI | AWSLGVGGSTSAETLY | 0.64753675 |
| VASGLSAALP | AWSLGVGGSTSAETLY | 0.3340592 |
| VGAGVIGI | AWSLGVGGSTSAETLY | 0.5662432 |
| VGPKRFGV | AWSLGVGGSTSAETLY | 0.47802073 |
| VNVAFPD | AWSLGVGGSTSAETLY | 0.6495429 |
| VSMTNSGVS | AWSLGVGGSTSAETLY | 0.6216475 |
| VTDVLGDLSV | AWSLGVGGSTSAETLY | 0.49609053 |
| VVVVGAGFAGL | AWSLGVGGSTSAETLY | 0.6439864 |
| VYTVGTVV | AWSLGVGGSTSAETLY | 0.45584175 |
| WTTFSGP | AWSLGVGGSTSAETLY | 0.62530017 |
| YDELVEAF | AWSLGVGGSTSAETLY | 0.6475623 |
| YGTGGAGGNGG | AWSLGVGGSTSAETLY | 0.6642346 |
| YTNVMAV | AWSLGVGGSTSAETLY | 0.53322387 |
| FLEQVHQL | AWSLGVGGSTSAETLY | 0.63565195 |
| GLSLLAVAV | AWSLGVGGSTSAETLY | 0.57395554 |
| LEQVHQL | AWSLGVGGSTSAETLY | 0.66384244 |
| LNAAGAI | AWSLGVGGSTSAETLY | 0.58889896 |

| | | |
|---|---|---|
| PAVHASL | AWSLGVGGSTSAETLY | 0.48634413 |
| TELEDDA | AWSLGVGGSTSAETLY | 0.64323103 |
| TPPGPPN | AWSLGVGGSTSAETLY | 0.63188076 |
| TPPPPPPP | AWSLGVGGSTSAETLY | 0.5912708 |
| TQSPALI | ASSLLLGGYAEQF | 0.64046466 |
| EFAAAFPQ | ASSLLLGGYAEQF | 0.5915412 |
| LLSGNLDVL | ASSLLLGGYAEQF | 0.59100866 |
| GPANLGNYNI | ASSLLLGGYAEQF | 0.64524424 |
| TLGPTLP | ASSLLLGGYAEQF | 0.64204824 |
| VYAYSSCCK | ASSLLLGGYAEQF | 0.5723098 |
| APAPAPAPNI | ASSLLLGGYAEQF | 0.6508411 |
| PGGPVPIP | ASSLLLGGYAEQF | 0.6053961 |
| FLTVALEP | ASSLLLGGYAEQF | 0.5236548 |
| SQYFGANL | ASSLLLGGYAEQF | 0.6613486 |
| GGVKPAA | ASSLLLGGYAEQF | 0.5911958 |
| GGTGGAGGAGGW | ASSLLLGGYAEQF | 0.6365017 |
| GVGGVAGLGGAGG | ASSLLLGGYAEQF | 0.64506567 |
| VLLVLATTI | ASSLLLGGYAEQF | 0.6600903 |
| IGIFAAML | ASSLLLGGYAEQF | 0.5536951 |
| AVGDLID | ASSLLLGGYAEQF | 0.6038648 |
| NISPLPL | ASSLLLGGYAEQF | 0.59871316 |
| VTAVFVAI | ASSLLLGGYAEQF | 0.64438033 |
| SGDDNDAAL | ASSLLLGGYAEQF | 0.5985445 |
| AVAALAHS | ASSLLLGGYAEQF | 0.59325075 |
| GGVTVMATP | ASSLLLGGYAEQF | 0.6379721 |
| AGLAANVRD | ASSLLLGGYAEQF | 0.6549641 |
| TNRPDLIDPALLRPG | ASSLLLGGYAEQF | 0.6097396 |
| LANLGVTAL | ASSLLLGGYAEQF | 0.61466324 |
| TGERIEL | ASSLLLGGYAEQF | 0.65665865 |
| NGEELAA | ASSLLLGGYAEQF | 0.5152972 |
| VLSLVWG | ASSLLLGGYAEQF | 0.6068932 |
| IGYALWL | ASSLLLGGYAEQF | 0.6299182 |
| LSLLLAPL | ASSLLLGGYAEQF | 0.648921 |
| TLDEALK | ASSLLLGGYAEQF | 0.60403746 |
| TDGAGTAEGG | ASSLLLGGYAEQF | 0.6496402 |
| NAPGGAGGNGG | ASSLLLGGYAEQF | 0.58084327 |
| VSVFMGMGGNF | ASSLLLGGYAEQF | 0.6385197 |
| LASYLAAL | ASSLLLGGYAEQF | 0.56765056 |
| LVGGTPAL | ASSLLLGGYAEQF | 0.5999255 |
| VRNADVL | ASSLLLGGYAEQF | 0.55364436 |
| TCSSPFLVSAM | ASSLLLGGYAEQF | 0.6219488 |
| VHITDAE | ASSLLLGGYAEQF | 0.6357451 |
| EYAAPSF | ASSLLLGGYAEQF | 0.641471 |
| VVDAFHGGL | ASSLLLGGYAEQF | 0.40126637 |
| ASVVDPQ | ASSLLLGGYAEQF | 0.6358538 |
| SYNEFGGLS | ASSLLLGGYAEQF | 0.64513785 |
| DPDGSVY | ASSLLLGGYAEQF | 0.66363347 |
| MVSTIHATE | ASSLLLGGYAEQF | 0.5792622 |
| LAAGVADPV | ASSLLLGGYAEQF | 0.6235104 |
| TKEQIAA | ASSLLLGGYAEQF | 0.5592886 |
| SLASPAAL | ASSLLLGGYAEQF | 0.6131352 |

| | | |
|---|---|---|
| IGLVFYGV | ASSLLLGGYAEQF | 0.49922147 |
| FLLDASNL | ASSLLLGGYAEQF | 0.5215953 |
| YQVGNFP | ASSLLLGGYAEQF | 0.5860255 |
| ESLLSQST | ASSLLLGGYAEQF | 0.5531726 |
| LFTVGFAAVG | ASSLLLGGYAEQF | 0.6437963 |
| ATPPTPT | ASSLLLGGYAEQF | 0.6472503 |
| TDLATLG | ASSLLLGGYAEQF | 0.648708 |
| SAGGTGR | ASSLLLGGYAEQF | 0.6463618 |
| GFSNSGS | ASSLLLGGYAEQF | 0.6494845 |
| GGNGGSGGTSV | ASSLLLGGYAEQF | 0.65420043 |
| GTSVATGGAGN | ASSLLLGGYAEQF | 0.58148545 |
| NGGNGGSGGTS | ASSLLLGGYAEQF | 0.62035626 |
| AVVKFPH | ASSLLLGGYAEQF | 0.50007015 |
| TPPPPPP | ASSLLLGGYAEQF | 0.59961164 |
| AAPAAAAAPA | ASSLLLGGYAEQF | 0.59265625 |
| IALNEDL | ASSLLLGGYAEQF | 0.624516 |
| IVNVSSM | ASSLLLGGYAEQF | 0.32321924 |
| TGIPPAPR | ASSLLLGGYAEQF | 0.6293591 |
| FDLGGGTF | ASSLLLGGYAEQF | 0.660088 |
| FDLGGGTFDVS | ASSLLLGGYAEQF | 0.5657778 |
| NEPTAAA | ASSLLLGGYAEQF | 0.63121146 |
| NEPTAAALA | ASSLLLGGYAEQF | 0.6314065 |
| GSIINISSI | ASSLLLGGYAEQF | 0.5686946 |
| VAPGNAGTA | ASSLLLGGYAEQF | 0.62833977 |
| TGGGASLE | ASSLLLGGYAEQF | 0.5778281 |
| VAVYNHYKRI | ASSLLLGGYAEQF | 0.63961315 |
| TDPAPATT | ASSLLLGGYAEQF | 0.65795255 |
| TNRPDILDPALLRPG | ASSLLLGGYAEQF | 0.62273073 |
| LTTTTTS | ASSLLLGGYAEQF | 0.5605184 |
| PTGSGKTL | ASSLLLGGYAEQF | 0.61685187 |
| TGSGKTL | ASSLLLGGYAEQF | 0.5025766 |
| TIGHVDHGK | ASSLLLGGYAEQF | 0.5988486 |
| TIGHVDHGKTT | ASSLLLGGYAEQF | 0.5448735 |
| GHVDHGKTT | ASSLLLGGYAEQF | 0.59880376 |
| AQFEHTLL | ASSLLLGGYAEQF | 0.56648093 |
| IVNQDEL | ASSLLLGGYAEQF | 0.62794113 |
| TNGSQFF | ASSLLLGGYAEQF | 0.61660516 |
| TGKGIVE | ASSLLLGGYAEQF | 0.6111729 |
| LISNASDA | ASSLLLGGYAEQF | 0.5131993 |
| AELRTLGELTKIAKAHGAQ | ASSLLLGGYAEQF | 0.6351286 |
| APNAVAAAEPSW | ASSLLLGGYAEQF | 0.6044993 |
| APQVAGAATAVL | ASSLLLGGYAEQF | 0.49915224 |
| ARRDPAAAAAILPTDA | ASSLLLGGYAEQF | 0.6394902 |
| DAAARDVLAEAGLAEYF | ASSLLLGGYAEQF | 0.5768295 |
| DDQIVLSAKANLAKVEQ | ASSLLLGGYAEQF | 0.6010066 |
| DELLPDTGEDSDEDGH | ASSLLLGGYAEQF | 0.6524628 |
| ELPEAPANRATVPQD | ASSLLLGGYAEQF | 0.6194563 |
| FGQSKIERVTGIG | ASSLLLGGYAEQF | 0.45377705 |
| GAGGTGSTAGGAGGAGGAGGLY | ASSLLLGGYAEQF | 0.57350296 |
| GHGGTGAAGGNGGTGGAGGSGI | ASSLLLGGYAEQF | 0.60231096 |
| GNGGNGGAGGEGGPGGAGGAGA | ASSLLLGGYAEQF | 0.59433335 |

| | | |
|---|---|---|
| GSTAGNGGNGGNATNGTIAGQP | ASSLLLGGYAEQF | 0.36183932 |
| GTGGDGGNGGNGGTGG | ASSLLLGGYAEQF | 0.6373929 |
| GVRRPAGIPADAPV | ASSLLLGGYAEQF | 0.5201001 |
| GYRQVTDAHLLTLARR | ASSLLLGGYAEQF | 0.54715157 |
| IAKMGQDGHDRG | ASSLLLGGYAEQF | 0.59077895 |
| KAEPSIEIQYT | ASSLLLGGYAEQF | 0.64135015 |
| KPVRIRMAADEDTVAVT | ASSLLLGGYAEQF | 0.57598007 |
| MPPRIAGMRLLVIKPEP | ASSLLLGGYAEQF | 0.61290294 |
| MKFYAVALFRSRGVPPTR | ASSLLLGGYAEQF | 0.61721313 |
| NAGGFGILGAVAHSP | ASSLLLGGYAEQF | 0.6305617 |
| NEGREITLAMAEM | ASSLLLGGYAEQF | 0.6524453 |
| NELVEWSHPLGHRWRWE | ASSLLLGGYAEQF | 0.6272944 |
| NLGAGGTGGNGGNPG | ASSLLLGGYAEQF | 0.5925556 |
| NTEWPKWETVMAADPVD | ASSLLLGGYAEQF | 0.6216409 |
| PASAAPAETQP | ASSLLLGGYAEQF | 0.5728896 |
| QQSSDGKNGVRAAEITGE | ASSLLLGGYAEQF | 0.5452228 |
| RAPDAVTTSWWKE | ASSLLLGGYAEQF | 0.48911 |
| SGFGNVGAHTSGWFNQ | ASSLLLGGYAEQF | 0.53520966 |
| SGVAGAVGQAASVGGLKV | ASSLLLGGYAEQF | 0.5090049 |
| SNGGAGTGSAGGNGGT | ASSLLLGGYAEQF | 0.61905015 |
| TGGAGGEGGAAG | ASSLLLGGYAEQF | 0.6570181 |
| TGGDGGAAGAGGAPGHG | ASSLLLGGYAEQF | 0.60447955 |
| TMPPLLAAIANAGRHGV | ASSLLLGGYAEQF | 0.45043588 |
| TSSTAGNSGGA | ASSLLLGGYAEQF | 0.57465357 |
| VPATVPVGILC | ASSLLLGGYAEQF | 0.60827327 |
| VQRPDSGGVVLTGRIS | ASSLLLGGYAEQF | 0.63564885 |
| WTPNPVPATKGVPAGLVQAT | ASSLLLGGYAEQF | 0.645478 |
| YALKRALAKFSDRFDVVII | ASSLLLGGYAEQF | 0.67164177 |
| YLIGSVIV | ASSLLLGGYAEQF | 0.645594 |
| YGQLVAVID | ASSLLLGGYAEQF | 0.569293 |
| VTVGGMTLPA | ASSLLLGGYAEQF | 0.6389674 |
| VLALPAV | ASSLLLGGYAEQF | 0.5912849 |
| VIPPTLNYET | ASSLLLGGYAEQF | 0.60722476 |
| VGMGGMHPSSG | ASSLLLGGYAEQF | 0.61575115 |
| VGLAGGAATA | ASSLLLGGYAEQF | 0.61228275 |
| VGGAGGAGGSGP | ASSLLLGGYAEQF | 0.58604276 |
| VFAAFMV | ASSLLLGGYAEQF | 0.59864515 |
| TLYDKLR | ASSLLLGGYAEQF | 0.4435961 |
| TLAVFTVL | ASSLLLGGYAEQF | 0.63570946 |
| TGSVQPARV | ASSLLLGGYAEQF | 0.47588804 |
| TGGGAGSGGT | ASSLLLGGYAEQF | 0.467067 |
| SHLETSL | ASSLLLGGYAEQF | 0.59136075 |
| SFTTGMT | ASSLLLGGYAEQF | 0.65849704 |
| QVEVAKRMGVRQPTVS | ASSLLLGGYAEQF | 0.64939874 |
| QGSGGPALAM | ASSLLLGGYAEQF | 0.5596862 |
| MLAPSNP | ASSLLLGGYAEQF | 0.5596777 |
| LVNNAAI | ASSLLLGGYAEQF | 0.624678 |
| LTGEINTLA | ASSLLLGGYAEQF | 0.5112529 |
| LSIFGPV | ASSLLLGGYAEQF | 0.63691646 |
| LLALLANL | ASSLLLGGYAEQF | 0.60380936 |
| LISNASDALDKLR | ASSLLLGGYAEQF | 0.6137192 |

| | | |
|---|---|---|
| LINEAVP | ASSLLLGGYAEQF | 0.5235747 |
| LGYAFSGL | ASSLLLGGYAEQF | 0.6313288 |
| LDFVAVAL | ASSLLLGGYAEQF | 0.58319193 |
| LAQPGAE | ASSLLLGGYAEQF | 0.6326653 |
| ISNLGAVTGF | ASSLLLGGYAEQF | 0.37952456 |
| HQEDPGRI | ASSLLLGGYAEQF | 0.65286773 |
| GTPGEPTP | ASSLLLGGYAEQF | 0.5526428 |
| GFFNSGS | ASSLLLGGYAEQF | 0.53409195 |
| GAGGGGLTAGAN | ASSLLLGGYAEQF | 0.37385923 |
| FVDALVNM | ASSLLLGGYAEQF | 0.6100099 |
| FLTVAPEL | ASSLLLGGYAEQF | 0.6161363 |
| FETLTAL | ASSLLLGGYAEQF | 0.64570016 |
| DVINPIEL | ASSLLLGGYAEQF | 0.5946155 |
| APVSGAPV | ASSLLLGGYAEQF | 0.63993114 |
| ALVGGLAW | ASSLLLGGYAEQF | 0.63507926 |
| ALAAAAATAI | ASSLLLGGYAEQF | 0.58691776 |
| AFGLGLLL | ASSLLLGGYAEQF | 0.42395926 |
| AFFAAASL | ASSLLLGGYAEQF | 0.6241936 |
| AAAALPAFSPP | ASSLLLGGYAEQF | 0.5982625 |
| VVLAPAL | ASSLLLGGYAEQF | 0.65014845 |
| TIGHVDHGKTTL | ASSLLLGGYAEQF | 0.6138552 |
| LISNASDALDK | ASSLLLGGYAEQF | 0.5048014 |
| LISNASDAL | ASSLLLGGYAEQF | 0.6282581 |
| ISNASDALDK | ASSLLLGGYAEQF | 0.6404474 |
| NQAETLVYQTE | ASSLLLGGYAEQF | 0.5030215 |
| TGGTGGTAGAG | ASSLLLGGYAEQF | 0.5289345 |
| TGSTAGGAGGA | ASSLLLGGYAEQF | 0.56486833 |
| DPLAVPVAAGPV | ASSLLLGGYAEQF | 0.48684657 |
| GGTGGTPGNGGN | ASSLLLGGYAEQF | 0.63838863 |
| PAGLAGVDPATL | ASSLLLGGYAEQF | 0.5841359 |
| GNNLVVEAPHAGASV | ASSLLLGGYAEQF | 0.603755 |
| KVTRIMSASTLEQPA | ASSLLLGGYAEQF | 0.6087389 |
| LGFALPIALPRRTVV | ASSLLLGGYAEQF | 0.5904692 |
| TRLGNLTARKLVVLV | ASSLLLGGYAEQF | 0.5856346 |
| WSQQRRLIYFPSAGP | ASSLLLGGYAEQF | 0.59751135 |
| GPNGNPDPMAAAVDIR | ASSLLLGGYAEQF | 0.6412325 |
| TQRCVSVADLRAAASR | ASSLLLGGYAEQF | 0.63654494 |
| QFVLDVAEHPVEVLTETV | ASSLLLGGYAEQF | 0.60431004 |
| GGAGGTGGAGGNGGTQAGDG | ASSLLLGGYAEQF | 0.6019669 |
| PLLATLRAIAVDVDAKPAQV | ASSLLLGGYAEQF | 0.6540902 |
| APKFRTGGYMIDFWGVGYQVAK | ASSLLLGGYAEQF | 0.4918183 |
| GAGGAGGAGGAAGGPGGTGGTG | ASSLLLGGYAEQF | 0.58500767 |
| IGALGLLATDVAPAGSVAFGVA | ASSLLLGGYAEQF | 0.6306007 |
| AGGAGGIGGNGGAGGAGGVLMG | ASSLLLGGYAEQF | 0.55292034 |
| ADELIGLL | ASSLLLGGYAEQF | 0.64504075 |
| ADRALKA | ASSLLLGGYAEQF | 0.6178658 |
| AGAVGQVGGAGGA | ASSLLLGGYAEQF | 0.62993133 |
| AGGGGGLGGAP | ASSLLLGGYAEQF | 0.60701096 |
| AGGLLYGNGGAG | ASSLLLGGYAEQF | 0.65743005 |
| AGLESIGL | ASSLLLGGYAEQF | 0.6344423 |
| AGLKTGII | ASSLLLGGYAEQF | 0.60045314 |

| | | |
|---|---|---|
| ASLAALLL | ASSLLLGGYAEQF | 0.65031546 |
| ASVINISSI | ASSLLLGGYAEQF | 0.47757328 |
| ATPPGPA | ASSLLLGGYAEQF | 0.6451661 |
| ATVASALV | ASSLLLGGYAEQF | 0.6271872 |
| ATVASLIL | ASSLLLGGYAEQF | 0.52607155 |
| AVNGPVLIRH | ASSLLLGGYAEQF | 0.5935656 |
| DGAAGAVGI | ASSLLLGGYAEQF | 0.6510977 |
| DPLLHGI | ASSLLLGGYAEQF | 0.6451692 |
| ELGAALGAL | ASSLLLGGYAEQF | 0.54081136 |
| ELGRTQAV | ASSLLLGGYAEQF | 0.57288635 |
| FVAGGYLS | ASSLLLGGYAEQF | 0.47486654 |
| GASGGAGGAGGW | ASSLLLGGYAEQF | 0.45387608 |
| GGAGAGGDGGAGGI | ASSLLLGGYAEQF | 0.46103853 |
| GTGHNGSQPGMGGNG | ASSLLLGGYAEQF | 0.6278286 |
| GYAAASASAS | ASSLLLGGYAEQF | 0.57574224 |
| HQVDVAAVQQG | ASSLLLGGYAEQF | 0.6479374 |
| IGAAIAHT | ASSLLLGGYAEQF | 0.6197589 |
| IGMIEVAQL | ASSLLLGGYAEQF | 0.6538515 |
| IRLAPEHTSAMD | ASSLLLGGYAEQF | 0.5531404 |
| IYNVSVL | ASSLLLGGYAEQF | 0.6013702 |
| KQGFLGM | ASSLLLGGYAEQF | 0.41679767 |
| LESVITQL | ASSLLLGGYAEQF | 0.56058204 |
| LGLGALVVV | ASSLLLGGYAEQF | 0.53307307 |
| LIGNGGNGGSGGI | ASSLLLGGYAEQF | 0.61223084 |
| LNAINAPTL | ASSLLLGGYAEQF | 0.4532231 |
| LQLLNTGV | ASSLLLGGYAEQF | 0.66209155 |
| LQTSIEF | ASSLLLGGYAEQF | 0.5890285 |
| MSFAVSEL | ASSLLLGGYAEQF | 0.6368104 |
| NTGALGL | ASSLLLGGYAEQF | 0.57634544 |
| PPGAGGAGGAGTT | ASSLLLGGYAEQF | 0.5386895 |
| QLVGLGI | ASSLLLGGYAEQF | 0.5828866 |
| RFLEQVHQL | ASSLLLGGYAEQF | 0.6301079 |
| RGVGYVLR | ASSLLLGGYAEQF | 0.5649766 |
| SATGVFAGVI | ASSLLLGGYAEQF | 0.38086098 |
| SLPAITLP | ASSLLLGGYAEQF | 0.54039806 |
| STALAFTV | ASSLLLGGYAEQF | 0.53301716 |
| SVLTVGLGK | ASSLLLGGYAEQF | 0.6552353 |
| TALAGAGSGI | ASSLLLGGYAEQF | 0.6487983 |
| TGLPFTGI | ASSLLLGGYAEQF | 0.61405927 |
| VAEHVTV | ASSLLLGGYAEQF | 0.65684104 |
| VDIVLNSL | ASSLLLGGYAEQF | 0.32973924 |
| VLTGGGAVL | ASSLLLGGYAEQF | 0.58951116 |
| VNVGLAYGI | ASSLLLGGYAEQF | 0.6565838 |
| VSSAAVGAV | ASSLLLGGYAEQF | 0.62314105 |
| VTSVYEAGV | ASSLLLGGYAEQF | 0.5968063 |
| WVALDRGA | ASSLLLGGYAEQF | 0.65173817 |
| YLVGSVVI | ASSLLLGGYAEQF | 0.5625093 |
| GPPGTGKTLL | ASSLLLGGYAEQF | 0.5956634 |
| GVGMTKF | ASSLLLGGYAEQF | 0.6180321 |
| IGIGGSDL | ASSLLLGGYAEQF | 0.64267147 |
| IGTLLAEL | ASSLLLGGYAEQF | 0.6102564 |

| | | |
|---|---|---|
| IYTSGTT | ASSLLLGGYAEQF | 0.48966143 |
| LSFPAGV | ASSLLLGGYAEQF | 0.6265127 |
| LYTSGTT | ASSLLLGGYAEQF | 0.6669398 |
| VGAFPFGF | ASSLLLGGYAEQF | 0.5757527 |
| VGAGTVEF | ASSLLLGGYAEQF | 0.63430214 |
| VGKSTLLL | ASSLLLGGYAEQF | 0.6177571 |
| AALAHAGGGIV | ASSLLLGGYAEQF | 0.5689329 |
| AALVAVSLL | ASSLLLGGYAEQF | 0.54334 |
| AAVSRAVAAL | ASSLLLGGYAEQF | 0.5360638 |
| AGGSSGIGGF | ASSLLLGGYAEQF | 0.63715667 |
| AGLTVANANG | ASSLLLGGYAEQF | 0.5417914 |
| AGLVGAVL | ASSLLLGGYAEQF | 0.5941772 |
| ALSQAFTV | ASSLLLGGYAEQF | 0.60448575 |
| ALTFSVSL | ASSLLLGGYAEQF | 0.6651322 |
| ALTGAALVAL | ASSLLLGGYAEQF | 0.6257494 |
| ASFSIGAGVL | ASSLLLGGYAEQF | 0.63495207 |
| DGSVYKI | ASSLLLGGYAEQF | 0.5534381 |
| EQLAEIV | ASSLLLGGYAEQF | 0.6109081 |
| FAVSELTVM | ASSLLLGGYAEQF | 0.63280344 |
| GAAPLAAGL | ASSLLLGGYAEQF | 0.46812236 |
| GADGGDGGSGGAGG | ASSLLLGGYAEQF | 0.65067303 |
| GFGLDTGGAGG | ASSLLLGGYAEQF | 0.5419817 |
| GLASGGLL | ASSLLLGGYAEQF | 0.5971466 |
| GRIAGAAVLL | ASSLLLGGYAEQF | 0.63995045 |
| GSAGSLNTSF | ASSLLLGGYAEQF | 0.6641799 |
| GTGQIGI | ASSLLLGGYAEQF | 0.60497963 |
| GTLIALIL | ASSLLLGGYAEQF | 0.6300899 |
| IAATMGFVVG | ASSLLLGGYAEQF | 0.5464238 |
| IFAQALL | ASSLLLGGYAEQF | 0.6409273 |
| IGAAVALGF | ASSLLLGGYAEQF | 0.6461005 |
| IGLTPLAL | ASSLLLGGYAEQF | 0.6154169 |
| ILGSNTSTL | ASSLLLGGYAEQF | 0.6663101 |
| IVFSLTVL | ASSLLLGGYAEQF | 0.64387393 |
| LASKATGF | ASSLLLGGYAEQF | 0.40860063 |
| LASVALIL | ASSLLLGGYAEQF | 0.6083724 |
| LFAVGLTL | ASSLLLGGYAEQF | 0.65416485 |
| LFNGGIGTYI | ASSLLLGGYAEQF | 0.64987767 |
| LTLGAGII | ASSLLLGGYAEQF | 0.6558473 |
| LVAAALGDI | ASSLLLGGYAEQF | 0.6228356 |
| NTAFVQLGI | ASSLLLGGYAEQF | 0.5626502 |
| QLFSGAEL | ASSLLLGGYAEQF | 0.6555559 |
| SAATFITL | ASSLLLGGYAEQF | 0.66543466 |
| SNYGLANLGV | ASSLLLGGYAEQF | 0.63332105 |
| SYGGAYAVM | ASSLLLGGYAEQF | 0.5344489 |
| TTGVGGLTLG | ASSLLLGGYAEQF | 0.4947723 |
| TTQTIYI | ASSLLLGGYAEQF | 0.6341022 |
| VAAGAGLGAVI | ASSLLLGGYAEQF | 0.6445228 |
| VAAVVTLL | ASSLLLGGYAEQF | 0.46604547 |
| VGFLGGLI | ASSLLLGGYAEQF | 0.61974454 |
| VGVIPILL | ASSLLLGGYAEQF | 0.6186446 |
| VIFASSLI | ASSLLLGGYAEQF | 0.40517834 |

| | | |
|---|---|---|
| VKGGDGGAAAG | ASSLLLGGYAEQF | 0.61869353 |
| VTKTFTNI | ASSLLLGGYAEQF | 0.6457222 |
| VVAAAILL | ASSLLLGGYAEQF | 0.63003623 |
| YAALDVEL | ASSLLLGGYAEQF | 0.5338433 |
| LASVALLL | ASSLLLGGYAEQF | 0.6472886 |
| LDAVHRV | ASSLLLGGYAEQF | 0.653867 |
| QGLSSSIF | ASSLLLGGYAEQF | 0.66106343 |
| RGTFANIRL | ASSLLLGGYAEQF | 0.4685675 |
| RQLWWGHRI | ASSLLLGGYAEQF | 0.35492918 |
| YGPPGTGKTLL | ASSLLLGGYAEQF | 0.6076056 |
| AALVGAPANVA | ASSLLLGGYAEQF | 0.64102376 |
| AATFRRTE | ASSLLLGGYAEQF | 0.60646117 |
| AGALGGGI | ASSLLLGGYAEQF | 0.5557332 |
| AGGAGGAGGIGAG | ASSLLLGGYAEQF | 0.6185423 |
| AGSGALGL | ASSLLLGGYAEQF | 0.6502202 |
| AQIADQI | ASSLLLGGYAEQF | 0.5062707 |
| ASAVAVTYL | ASSLLLGGYAEQF | 0.46475843 |
| ASTVLGGTGG | ASSLLLGGYAEQF | 0.63621414 |
| AVGGEPAK | ASSLLLGGYAEQF | 0.6360468 |
| CEQGTFGGTC | ASSLLLGGYAEQF | 0.5552375 |
| DEGKAPAG | ASSLLLGGYAEQF | 0.5804157 |
| DTVVFPAL | ASSLLLGGYAEQF | 0.6201606 |
| FEGITMAG | ASSLLLGGYAEQF | 0.6090106 |
| FPLVVQD | ASSLLLGGYAEQF | 0.6194921 |
| GAFTTPQ | ASSLLLGGYAEQF | 0.48606288 |
| GFGGFGTAGGVGGA | ASSLLLGGYAEQF | 0.6444123 |
| GFLAQFL | ASSLLLGGYAEQF | 0.5046515 |
| GGAGGAGGNAGVA | ASSLLLGGYAEQF | 0.62793255 |
| GGVGTAGAGGAGGAGG | ASSLLLGGYAEQF | 0.61046594 |
| GNGGGGDAGNAGSGGN | ASSLLLGGYAEQF | 0.6189661 |
| GTGGAGGAGNDAGS | ASSLLLGGYAEQF | 0.58849615 |
| GTTNTGWFNS | ASSLLLGGYAEQF | 0.54523116 |
| IFMGPLFTL | ASSLLLGGYAEQF | 0.5297966 |
| ILAHAIEAI | ASSLLLGGYAEQF | 0.46998006 |
| IRSAIREAL | ASSLLLGGYAEQF | 0.65064365 |
| ITFAGGG | ASSLLLGGYAEQF | 0.61488086 |
| ITGMSGHQEV | ASSLLLGGYAEQF | 0.64507663 |
| IVAGLACTL | ASSLLLGGYAEQF | 0.62779933 |
| IVGGGLAGA | ASSLLLGGYAEQF | 0.5649579 |
| IVTGVLNI | ASSLLLGGYAEQF | 0.6285341 |
| LAQQGCI | ASSLLLGGYAEQF | 0.5066067 |
| LCLHVEN | ASSLLLGGYAEQF | 0.61217576 |
| LDAISDGS | ASSLLLGGYAEQF | 0.56214464 |
| LGGFAAML | ASSLLLGGYAEQF | 0.638137 |
| LGGKNPMI | ASSLLLGGYAEQF | 0.5328855 |
| LGPKGVAAIL | ASSLLLGGYAEQF | 0.63264126 |
| LMVGVSI | ASSLLLGGYAEQF | 0.61895573 |
| LVGAAFVL | ASSLLLGGYAEQF | 0.52317375 |
| LVLAVAPL | ASSLLLGGYAEQF | 0.63069254 |
| MAAIAVIDAA | ASSLLLGGYAEQF | 0.6431204 |
| NNGINAELQ | ASSLLLGGYAEQF | 0.6251832 |

| | | |
|---|---|---|
| RGEPLVLVHG | ASSLLLGGYAEQF | 0.5045556 |
| RGGAGGAAGLI | ASSLLLGGYAEQF | 0.6282958 |
| RSYRMHPAV | ASSLLLGGYAEQF | 0.553694 |
| SAAHLLAGQLA | ASSLLLGGYAEQF | 0.61667746 |
| SAAVGSY | ASSLLLGGYAEQF | 0.6179276 |
| SPHAVLHRI | ASSLLLGGYAEQF | 0.5497607 |
| SQIAKVL | ASSLLLGGYAEQF | 0.6398866 |
| TALGHLVP | ASSLLLGGYAEQF | 0.4849145 |
| TAPTTSGAA | ASSLLLGGYAEQF | 0.53947264 |
| TDGDDGNGGSPGT | ASSLLLGGYAEQF | 0.57474077 |
| TDGLGEN | ASSLLLGGYAEQF | 0.60819906 |
| TGGPPLPGQA | ASSLLLGGYAEQF | 0.63807523 |
| TIPAITI | ASSLLLGGYAEQF | 0.5306746 |
| TIPGFTI | ASSLLLGGYAEQF | 0.6072981 |
| TSIASGALA | ASSLLLGGYAEQF | 0.602309 |
| TVTLPGAAI | ASSLLLGGYAEQF | 0.55152404 |
| VASGLSAALP | ASSLLLGGYAEQF | 0.6172448 |
| VGAGVIGI | ASSLLLGGYAEQF | 0.5918618 |
| VGPKRFGV | ASSLLLGGYAEQF | 0.45815533 |
| VNVAFPD | ASSLLLGGYAEQF | 0.6381839 |
| VSMTNSGVS | ASSLLLGGYAEQF | 0.5526542 |
| VTDVLGDLSV | ASSLLLGGYAEQF | 0.6238687 |
| VVVVGAGFAGL | ASSLLLGGYAEQF | 0.6483071 |
| VYTVGTVV | ASSLLLGGYAEQF | 0.4646275 |
| WTTFSGP | ASSLLLGGYAEQF | 0.6241745 |
| YDELVEAF | ASSLLLGGYAEQF | 0.61745095 |
| YGTGGAGGNGG | ASSLLLGGYAEQF | 0.6187964 |
| YTNVMAV | ASSLLLGGYAEQF | 0.6125066 |
| FLEQVHQL | ASSLLLGGYAEQF | 0.57476264 |
| GLSLLAVAV | ASSLLLGGYAEQF | 0.6582243 |
| LEQVHQL | ASSLLLGGYAEQF | 0.6414968 |
| LNAAGAI | ASSLLLGGYAEQF | 0.6069393 |
| PAVHASL | ASSLLLGGYAEQF | 0.6337689 |
| TELEDDA | ASSLLLGGYAEQF | 0.6430661 |
| TPPGPPN | ASSLLLGGYAEQF | 0.6442224 |
| TPPPPPPP | ASSLLLGGYAEQF | 0.56777406 |
| TQSPALI | ASSLRDTNSGNTLY | 0.4496755 |
| EFAAAFPQ | ASSLRDTNSGNTLY | 0.5959858 |
| LLSGNLDVL | ASSLRDTNSGNTLY | 0.58952284 |
| GPANLGNYNI | ASSLRDTNSGNTLY | 0.57808137 |
| TLGPTLP | ASSLRDTNSGNTLY | 0.43680543 |
| VYAYSSCCK | ASSLRDTNSGNTLY | 0.6486375 |
| APAPAPAPNI | ASSLRDTNSGNTLY | 0.66455644 |
| PGGPVPIP | ASSLRDTNSGNTLY | 0.5689426 |
| FLTVALEP | ASSLRDTNSGNTLY | 0.6667527 |
| SQYFGANL | ASSLRDTNSGNTLY | 0.50479716 |
| GGVKPAA | ASSLRDTNSGNTLY | 0.64141786 |
| GGTGGAGGAGGW | ASSLRDTNSGNTLY | 0.65267 |
| GVGGVAGLGGAGG | ASSLRDTNSGNTLY | 0.52616405 |
| VLLVLATTI | ASSLRDTNSGNTLY | 0.6448737 |
| IGIFAAML | ASSLRDTNSGNTLY | 0.5758291 |

| | | |
|---|---|---|
| AVGDLID | ASSLRDTNSGNTLY | 0.5842116 |
| NISPLPL | ASSLRDTNSGNTLY | 0.6437531 |
| VTAVFVAI | ASSLRDTNSGNTLY | 0.5772589 |
| SGDDNDAAL | ASSLRDTNSGNTLY | 0.6362466 |
| AVAALAHS | ASSLRDTNSGNTLY | 0.6028142 |
| GGVTVMATP | ASSLRDTNSGNTLY | 0.5643095 |
| AGLAANVRD | ASSLRDTNSGNTLY | 0.63490963 |
| TNRPDLIDPALLRPG | ASSLRDTNSGNTLY | 0.6354312 |
| LANLGVTAL | ASSLRDTNSGNTLY | 0.5931357 |
| TGERIEL | ASSLRDTNSGNTLY | 0.6643861 |
| NGEELAA | ASSLRDTNSGNTLY | 0.5480524 |
| VLSLVWG | ASSLRDTNSGNTLY | 0.66018337 |
| IGYALWL | ASSLRDTNSGNTLY | 0.6457234 |
| LSLLLAPL | ASSLRDTNSGNTLY | 0.6098335 |
| TLDEALK | ASSLRDTNSGNTLY | 0.5861569 |
| TDGAGTAEGG | ASSLRDTNSGNTLY | 0.6445559 |
| NAPGGAGGNGG | ASSLRDTNSGNTLY | 0.58342457 |
| VSVFMGMGGNF | ASSLRDTNSGNTLY | 0.6186315 |
| LASYLAAL | ASSLRDTNSGNTLY | 0.66055006 |
| LVGGTPAL | ASSLRDTNSGNTLY | 0.39819664 |
| VRNADVL | ASSLRDTNSGNTLY | 0.81169784 |
| TCSSPFLVSAM | ASSLRDTNSGNTLY | 0.681783 |
| VHITDAE | ASSLRDTNSGNTLY | 0.7501864 |
| EYAAPSF | ASSLRDTNSGNTLY | 0.46707103 |
| VVDAFHGGL | ASSLRDTNSGNTLY | 0.2832215 |
| ASVVDPQ | ASSLRDTNSGNTLY | 0.59888965 |
| SYNEFGGLS | ASSLRDTNSGNTLY | 0.4782716 |
| DPDGSVY | ASSLRDTNSGNTLY | 0.72083616 |
| MVSTIHATE | ASSLRDTNSGNTLY | 0.7650664 |
| LAAGVADPV | ASSLRDTNSGNTLY | 0.5164675 |
| TKEQIAA | ASSLRDTNSGNTLY | 0.7579421 |
| SLASPAAL | ASSLRDTNSGNTLY | 0.8109189 |
| IGLVFYGV | ASSLRDTNSGNTLY | 0.28257024 |
| FLLDASNL | ASSLRDTNSGNTLY | 0.6168647 |
| YQVGNFP | ASSLRDTNSGNTLY | 0.3800435 |
| ESLLSQST | ASSLRDTNSGNTLY | 0.5844287 |
| LFTVGFAAVG | ASSLRDTNSGNTLY | 0.17505226 |
| ATPPTPT | ASSLRDTNSGNTLY | 0.75779724 |
| TDLATLG | ASSLRDTNSGNTLY | 0.4121427 |
| SAGGTGR | ASSLRDTNSGNTLY | 0.20248409 |
| GFSNSGS | ASSLRDTNSGNTLY | 0.70230746 |
| GGNGGSGGTSV | ASSLRDTNSGNTLY | 0.60508746 |
| GTSVATGGAGN | ASSLRDTNSGNTLY | 0.6511869 |
| NGGNGGSGGTS | ASSLRDTNSGNTLY | 0.77701175 |
| AVVKFPH | ASSLRDTNSGNTLY | 0.59538996 |
| TPPPPPP | ASSLRDTNSGNTLY | 0.51748073 |
| AAPAAAAAPA | ASSLRDTNSGNTLY | 0.6694124 |
| IALNEDL | ASSLRDTNSGNTLY | 0.80150545 |
| IVNVSSM | ASSLRDTNSGNTLY | 0.5519479 |
| TGIPPAPR | ASSLRDTNSGNTLY | 0.6539046 |
| FDLGGGTF | ASSLRDTNSGNTLY | 0.844151 |

| | | |
|---|---|---|
| FDLGGGTFDVS | ASSLRDTNSGNTLY | 0.6569529 |
| NEPTAAA | ASSLRDTNSGNTLY | 0.81414545 |
| NEPTAAALA | ASSLRDTNSGNTLY | 0.8317613 |
| GSIINISSI | ASSLRDTNSGNTLY | 0.4045324 |
| VAPGNAGTA | ASSLRDTNSGNTLY | 0.7562517 |
| TGGGASLE | ASSLRDTNSGNTLY | 0.7606799 |
| VAVYNHYKRI | ASSLRDTNSGNTLY | 0.6401441 |
| TDPAPATT | ASSLRDTNSGNTLY | 0.66021895 |
| TNRPDILDPALLRPG | ASSLRDTNSGNTLY | 0.37656453 |
| LTTTTTS | ASSLRDTNSGNTLY | 0.6052668 |
| PTGSGKTL | ASSLRDTNSGNTLY | 0.27239737 |
| TGSGKTL | ASSLRDTNSGNTLY | 0.66172236 |
| TIGHVDHGK | ASSLRDTNSGNTLY | 0.8251067 |
| TIGHVDHGKTT | ASSLRDTNSGNTLY | 0.6243076 |
| GHVDHGKTT | ASSLRDTNSGNTLY | 0.34486753 |
| AQFEHTLL | ASSLRDTNSGNTLY | 0.26254153 |
| IVNQDEL | ASSLRDTNSGNTLY | 0.7150543 |
| TNGSQFF | ASSLRDTNSGNTLY | 0.46430513 |
| TGKGIVE | ASSLRDTNSGNTLY | 0.3703813 |
| LISNASDA | ASSLRDTNSGNTLY | 0.36825305 |
| AELRTLGELTKIAKAHGAQ | ASSLRDTNSGNTLY | 0.8017442 |
| APNAVAAAEPSW | ASSLRDTNSGNTLY | 0.8092917 |
| APQVAGAATAVL | ASSLRDTNSGNTLY | 0.5226568 |
| ARRDPAAAAAILPTDA | ASSLRDTNSGNTLY | 0.79873735 |
| DAAARDVLAEAGLAEYF | ASSLRDTNSGNTLY | 0.72610795 |
| DDQIVLSAKANLAKVEQ | ASSLRDTNSGNTLY | 0.7670698 |
| DELLPDTGEDSDEDGH | ASSLRDTNSGNTLY | 0.678151 |
| ELPEAPANRATVPQD | ASSLRDTNSGNTLY | 0.3772207 |
| FGQSKIERVTGIG | ASSLRDTNSGNTLY | 0.7745046 |
| GAGGTGSTAGGAGGAGGAGGLY | ASSLRDTNSGNTLY | 0.5302297 |
| GHGGTGAAGGNGGTGGAGGSGI | ASSLRDTNSGNTLY | 0.7651732 |
| GNGGNGGAGGEGGPGGAGGAGA | ASSLRDTNSGNTLY | 0.7608305 |
| GSTAGNGGNGGNATNGTIAGQP | ASSLRDTNSGNTLY | 0.6782961 |
| GTGGDGGNGGNGGTGG | ASSLRDTNSGNTLY | 0.4968398 |
| GVRRPAGIPADAPV | ASSLRDTNSGNTLY | 0.6200238 |
| GYRQVTDAHLLTLARR | ASSLRDTNSGNTLY | 0.8179595 |
| IAKMGQDGHDRG | ASSLRDTNSGNTLY | 0.36274636 |
| KAEPSIEIQYT | ASSLRDTNSGNTLY | 0.41295147 |
| KPVRIRMAADEDTVAVT | ASSLRDTNSGNTLY | 0.65835524 |
| MPPRIAGMRLLVIKPEP | ASSLRDTNSGNTLY | 0.6917485 |
| MKFYAVALFRSRGVPPTR | ASSLRDTNSGNTLY | 0.8095571 |
| NAGGFGILGAVAHSP | ASSLRDTNSGNTLY | 0.7218181 |
| NEGREITLAMAEM | ASSLRDTNSGNTLY | 0.6012088 |
| NELVEWSHPLGHRWRWE | ASSLRDTNSGNTLY | 0.7374709 |
| NLGAGGTGGNGGNPG | ASSLRDTNSGNTLY | 0.78763175 |
| NTEWPKWETVMAADPVD | ASSLRDTNSGNTLY | 0.6955509 |
| PASAAPAETQP | ASSLRDTNSGNTLY | 0.74883264 |
| QQSSDGKNGVRAAEITGE | ASSLRDTNSGNTLY | 0.5016159 |
| RAPDAVTTSWWKE | ASSLRDTNSGNTLY | 0.7910454 |
| SGFGNVGAHTSGWFNQ | ASSLRDTNSGNTLY | 0.7806696 |
| SGVAGAVGQAASVGGLKV | ASSLRDTNSGNTLY | 0.7649492 |

| | | |
|---|---|---|
| SNGGAGTGSAGGNGGT | ASSLRDTNSGNTLY | 0.79498327 |
| TGGAGGEGGAAG | ASSLRDTNSGNTLY | 0.75660825 |
| TGGDGGAAGAGGAPGHG | ASSLRDTNSGNTLY | 0.7003646 |
| TMPPLLAAIANAGRHGV | ASSLRDTNSGNTLY | 0.27994654 |
| TSSTAGNSGGA | ASSLRDTNSGNTLY | 0.4211212 |
| VPATVPVGILC | ASSLRDTNSGNTLY | 0.66629803 |
| VQRPDSGGVVLTGRIS | ASSLRDTNSGNTLY | 0.6680816 |
| WTPNPVPATKGVPAGLVQAT | ASSLRDTNSGNTLY | 0.7178928 |
| YALKRALAKFSDRFDVVII | ASSLRDTNSGNTLY | 0.7958019 |
| YLIGSVIV | ASSLRDTNSGNTLY | 0.49769258 |
| YGQLVAVID | ASSLRDTNSGNTLY | 0.8226661 |
| VTVGGMTLPA | ASSLRDTNSGNTLY | 0.80178595 |
| VLALPAV | ASSLRDTNSGNTLY | 0.29438496 |
| VIPPTLNYET | ASSLRDTNSGNTLY | 0.5276637 |
| VGMGGMHPSSG | ASSLRDTNSGNTLY | 0.7911153 |
| VGLAGGAATA | ASSLRDTNSGNTLY | 0.6719755 |
| VGGAGGAGGSGP | ASSLRDTNSGNTLY | 0.571997 |
| VFAAFMV | ASSLRDTNSGNTLY | 0.6572093 |
| TLYDKLR | ASSLRDTNSGNTLY | 0.7941286 |
| TLAVFTVL | ASSLRDTNSGNTLY | 0.674183 |
| TGSVQPARV | ASSLRDTNSGNTLY | 0.807887 |
| TGGGAGSGGT | ASSLRDTNSGNTLY | 0.21437587 |
| SHLETSL | ASSLRDTNSGNTLY | 0.6350695 |
| SFTTGMT | ASSLRDTNSGNTLY | 0.54568887 |
| QVEVAKRMGVRQPTVS | ASSLRDTNSGNTLY | 0.17631204 |
| QGSGGPALAM | ASSLRDTNSGNTLY | 0.7669345 |
| MLAPSNP | ASSLRDTNSGNTLY | 0.5816066 |
| LVNNAAI | ASSLRDTNSGNTLY | 0.73931885 |
| LTGEINTLA | ASSLRDTNSGNTLY | 0.48703638 |
| LSIFGPV | ASSLRDTNSGNTLY | 0.77075195 |
| LLALLANL | ASSLRDTNSGNTLY | 0.63017905 |
| LISNASDALDKLR | ASSLRDTNSGNTLY | 0.5944024 |
| LINEAVP | ASSLRDTNSGNTLY | 0.4375875 |
| LGYAFSGL | ASSLRDTNSGNTLY | 0.72595984 |
| LDFVAVAL | ASSLRDTNSGNTLY | 0.81156254 |
| LAQPGAE | ASSLRDTNSGNTLY | 0.80814016 |
| ISNLGAVTGF | ASSLRDTNSGNTLY | 0.30724007 |
| HQEDPGRI | ASSLRDTNSGNTLY | 0.7182768 |
| GTPGEPTP | ASSLRDTNSGNTLY | 0.6286973 |
| GFFNSGS | ASSLRDTNSGNTLY | 0.38155395 |
| GAGGGGLTAGAN | ASSLRDTNSGNTLY | 0.2543517 |
| FVDALVNM | ASSLRDTNSGNTLY | 0.53839713 |
| FLTVAPEL | ASSLRDTNSGNTLY | 0.59085953 |
| FETLTAL | ASSLRDTNSGNTLY | 0.7320199 |
| DVINPIEL | ASSLRDTNSGNTLY | 0.7297426 |
| APVSGAPV | ASSLRDTNSGNTLY | 0.57229257 |
| ALVGGLAW | ASSLRDTNSGNTLY | 0.7416501 |
| ALAAAAATAI | ASSLRDTNSGNTLY | 0.73169947 |
| AFGLGLLL | ASSLRDTNSGNTLY | 0.08714202 |
| AFFAAASL | ASSLRDTNSGNTLY | 0.47891706 |
| AAAALPAFSPP | ASSLRDTNSGNTLY | 0.2848399 |

| | | |
|---|---|---|
| VVLAPAL | ASSLRDTNSGNTLY | 0.8232744 |
| TIGHVDHGKTTL | ASSLRDTNSGNTLY | 0.37749252 |
| LISNASDALDK | ASSLRDTNSGNTLY | 0.52677405 |
| LISNASDAL | ASSLRDTNSGNTLY | 0.6470547 |
| ISNASDALDK | ASSLRDTNSGNTLY | 0.6633565 |
| NQAETLVYQTE | ASSLRDTNSGNTLY | 0.6687213 |
| TGGTGGTAGAG | ASSLRDTNSGNTLY | 0.5057126 |
| TGSTAGGAGGA | ASSLRDTNSGNTLY | 0.2565852 |
| DPLAVPVAAGPV | ASSLRDTNSGNTLY | 0.75473666 |
| GGTGGTPGNGGN | ASSLRDTNSGNTLY | 0.5275694 |
| PAGLAGVDPATL | ASSLRDTNSGNTLY | 0.5423163 |
| GNNLVVEAPHAGASV | ASSLRDTNSGNTLY | 0.41405624 |
| KVTRIMSASTLEQPA | ASSLRDTNSGNTLY | 0.70812666 |
| LGFALPIALPRRTVV | ASSLRDTNSGNTLY | 0.309264 |
| TRLGNLTARKLVVLV | ASSLRDTNSGNTLY | 0.80899477 |
| WSQQRRLIYFPSAGP | ASSLRDTNSGNTLY | 0.3521614 |
| GPNGNPDPMAAAVDIR | ASSLRDTNSGNTLY | 0.59347874 |
| TQRCVSVADLRAAASR | ASSLRDTNSGNTLY | 0.8080966 |
| QFVLDVAEHPVEVLTETV | ASSLRDTNSGNTLY | 0.6839607 |
| GGAGGTGGAGGNGGTQAGDG | ASSLRDTNSGNTLY | 0.52277195 |
| PLLATLRAIAVDVDAKPAQV | ASSLRDTNSGNTLY | 0.62370074 |
| APKFRTGGYMIDFWGVGYQVAK | ASSLRDTNSGNTLY | 0.35068244 |
| GAGGAGGAGGAAGGPGGTGGTG | ASSLRDTNSGNTLY | 0.5046484 |
| IGALGLLATDVAPAGSVAFGVA | ASSLRDTNSGNTLY | 0.6003851 |
| AGGAGGIGGNGGAGGAGGVLMG | ASSLRDTNSGNTLY | 0.8152741 |
| ADELIGLL | ASSLRDTNSGNTLY | 0.6428113 |
| ADRALKA | ASSLRDTNSGNTLY | 0.3600347 |
| AGAVGQVGGAGGA | ASSLRDTNSGNTLY | 0.7838831 |
| AGGGGGLGGAP | ASSLRDTNSGNTLY | 0.82271266 |
| AGGLLYGNGGAG | ASSLRDTNSGNTLY | 0.8038089 |
| AGLESIGL | ASSLRDTNSGNTLY | 0.39549404 |
| AGLKTGII | ASSLRDTNSGNTLY | 0.6001554 |
| ASLAALLL | ASSLRDTNSGNTLY | 0.6166013 |
| ASVINISSI | ASSLRDTNSGNTLY | 0.61029375 |
| ATPPGPA | ASSLRDTNSGNTLY | 0.5577325 |
| ATVASALV | ASSLRDTNSGNTLY | 0.5230474 |
| ATVASLIL | ASSLRDTNSGNTLY | 0.7832085 |
| AVNGPVLIRH | ASSLRDTNSGNTLY | 0.85584235 |
| DGAAGAVGI | ASSLRDTNSGNTLY | 0.64671284 |
| DPLLHGI | ASSLRDTNSGNTLY | 0.17825876 |
| ELGAALGAL | ASSLRDTNSGNTLY | 0.61604106 |
| ELGRTQAV | ASSLRDTNSGNTLY | 0.37885028 |
| FVAGGYLS | ASSLRDTNSGNTLY | 0.5013556 |
| GASGGAGGAGGW | ASSLRDTNSGNTLY | 0.64141953 |
| GGAGAGGDGGAGGI | ASSLRDTNSGNTLY | 0.28138632 |
| GTGHNGSQPGMGGNG | ASSLRDTNSGNTLY | 0.65864235 |
| GYAAASASAS | ASSLRDTNSGNTLY | 0.8444754 |
| HQVDVAAVQQG | ASSLRDTNSGNTLY | 0.3910991 |
| IGAAIAHT | ASSLRDTNSGNTLY | 0.68287814 |
| IGMIEVAQL | ASSLRDTNSGNTLY | 0.77770865 |
| IRLAPEHTSAMD | ASSLRDTNSGNTLY | 0.69798136 |

| | | |
|---|---|---|
| IYNVSVL | ASSLRDTNSGNTLY | 0.8038471 |
| KQGFLGM | ASSLRDTNSGNTLY | 0.73806477 |
| LESVITQL | ASSLRDTNSGNTLY | 0.67841125 |
| LGLGALVVV | ASSLRDTNSGNTLY | 0.67778784 |
| LIGNGGNGGSGGI | ASSLRDTNSGNTLY | 0.7425996 |
| LNAINAPTL | ASSLRDTNSGNTLY | 0.61874425 |
| LQLLNTGV | ASSLRDTNSGNTLY | 0.6325092 |
| LQTSIEF | ASSLRDTNSGNTLY | 0.51247525 |
| MSFAVSEL | ASSLRDTNSGNTLY | 0.7476501 |
| NTGALGL | ASSLRDTNSGNTLY | 0.6851396 |
| PPGAGGAGGAGTT | ASSLRDTNSGNTLY | 0.7897525 |
| QLVGLGI | ASSLRDTNSGNTLY | 0.2596812 |
| RFLEQVHQL | ASSLRDTNSGNTLY | 0.5273836 |
| RGVGYVLR | ASSLRDTNSGNTLY | 0.7788011 |
| SATGVFAGVI | ASSLRDTNSGNTLY | 0.1533985 |
| SLPAITLP | ASSLRDTNSGNTLY | 0.36099923 |
| STALAFTV | ASSLRDTNSGNTLY | 0.6551514 |
| SVLTVGLGK | ASSLRDTNSGNTLY | 0.79852307 |
| TALAGAGSGI | ASSLRDTNSGNTLY | 0.17306945 |
| TGLPFTGI | ASSLRDTNSGNTLY | 0.5974853 |
| VAEHVTV | ASSLRDTNSGNTLY | 0.30793187 |
| VDIVLNSL | ASSLRDTNSGNTLY | 0.82087374 |
| VLTGGGAVL | ASSLRDTNSGNTLY | 0.6811344 |
| VNVGLAYGI | ASSLRDTNSGNTLY | 0.20200427 |
| VSSAAVGAV | ASSLRDTNSGNTLY | 0.7832634 |
| VTSVYEAGV | ASSLRDTNSGNTLY | 0.24380419 |
| WVALDRGA | ASSLRDTNSGNTLY | 0.65050244 |
| YLVGSVVI | ASSLRDTNSGNTLY | 0.7353543 |
| GPPGTGKTLL | ASSLRDTNSGNTLY | 0.8141854 |
| GVGMTKF | ASSLRDTNSGNTLY | 0.33730292 |
| IGIGGSDL | ASSLRDTNSGNTLY | 0.7407035 |
| IGTLLAEL | ASSLRDTNSGNTLY | 0.53865 |
| IYTSGTT | ASSLRDTNSGNTLY | 0.83463776 |
| LSFPAGV | ASSLRDTNSGNTLY | 0.5141992 |
| LYTSGTT | ASSLRDTNSGNTLY | 0.31469822 |
| VGAFPFGF | ASSLRDTNSGNTLY | 0.7855116 |
| VGAGTVEF | ASSLRDTNSGNTLY | 0.68326473 |
| VGKSTLLL | ASSLRDTNSGNTLY | 0.75193024 |
| AALAHAGGGIV | ASSLRDTNSGNTLY | 0.7108913 |
| AALVAVSLL | ASSLRDTNSGNTLY | 0.64416134 |
| AAVSRAVAAL | ASSLRDTNSGNTLY | 0.5730265 |
| AGGSSGIGGF | ASSLRDTNSGNTLY | 0.72931564 |
| AGLTVANANG | ASSLRDTNSGNTLY | 0.73807836 |
| AGLVGAVL | ASSLRDTNSGNTLY | 0.55676067 |
| ALSQAFTV | ASSLRDTNSGNTLY | 0.79926264 |
| ALTFSVSL | ASSLRDTNSGNTLY | 0.71220094 |
| ALTGAALVAL | ASSLRDTNSGNTLY | 0.4903356 |
| ASFSIGAGVL | ASSLRDTNSGNTLY | 0.828429 |
| DGSVYKI | ASSLRDTNSGNTLY | 0.66237384 |
| EQLAEIV | ASSLRDTNSGNTLY | 0.760025 |
| FAVSELTVM | ASSLRDTNSGNTLY | 0.74483955 |

| | | |
|---|---|---|
| GAAPLAAGL | ASSLRDTNSGNTLY | 0.80805737 |
| GADGGDGGSGGAGG | ASSLRDTNSGNTLY | 0.28687465 |
| GFGLDTGGAGG | ASSLRDTNSGNTLY | 0.69077754 |
| GLASGGLL | ASSLRDTNSGNTLY | 0.67271286 |
| GRIAGAAVLL | ASSLRDTNSGNTLY | 0.7544384 |
| GSAGSLNTSF | ASSLRDTNSGNTLY | 0.6135191 |
| GTGQIGI | ASSLRDTNSGNTLY | 0.58684736 |
| GTLIALIL | ASSLRDTNSGNTLY | 0.7251432 |
| IAATMGFVVG | ASSLRDTNSGNTLY | 0.8291301 |
| IFAQALL | ASSLRDTNSGNTLY | 0.6541425 |
| IGAAVALGF | ASSLRDTNSGNTLY | 0.727996 |
| IGLTPLAL | ASSLRDTNSGNTLY | 0.83324254 |
| ILGSNTSTL | ASSLRDTNSGNTLY | 0.40569162 |
| IVFSLTVL | ASSLRDTNSGNTLY | 0.7221378 |
| LASKATGF | ASSLRDTNSGNTLY | 0.7812189 |
| LASVALIL | ASSLRDTNSGNTLY | 0.80778867 |
| LFAVGLTL | ASSLRDTNSGNTLY | 0.6308319 |
| LFNGGIGTYI | ASSLRDTNSGNTLY | 0.7982868 |
| LTLGAGII | ASSLRDTNSGNTLY | 0.42483526 |
| LVAAALGDI | ASSLRDTNSGNTLY | 0.71409416 |
| NTAFVQLGI | ASSLRDTNSGNTLY | 0.49503735 |
| QLFSGAEL | ASSLRDTNSGNTLY | 0.5593226 |
| SAATFITL | ASSLRDTNSGNTLY | 0.4354294 |
| SNYGLANLGV | ASSLRDTNSGNTLY | 0.6111719 |
| SYGGAYAVM | ASSLRDTNSGNTLY | 0.7413972 |
| TTGVGGLTLG | ASSLRDTNSGNTLY | 0.70836186 |
| TTQTIYI | ASSLRDTNSGNTLY | 0.28933045 |
| VAAGAGLGAVI | ASSLRDTNSGNTLY | 0.7358993 |
| VAAVVTLL | ASSLRDTNSGNTLY | 0.7313943 |
| VGFLGGLI | ASSLRDTNSGNTLY | 0.8330747 |
| VGVIPILL | ASSLRDTNSGNTLY | 0.6196909 |
| VIFASSLI | ASSLRDTNSGNTLY | 0.77445567 |
| VKGGDGGAAAG | ASSLRDTNSGNTLY | 0.5825192 |
| VTKTFTNI | ASSLRDTNSGNTLY | 0.6709803 |
| VVAAAILL | ASSLRDTNSGNTLY | 0.38384438 |
| YAALDVEL | ASSLRDTNSGNTLY | 0.47541714 |
| LASVALLL | ASSLRDTNSGNTLY | 0.5763725 |
| LDAVHRV | ASSLRDTNSGNTLY | 0.4528797 |
| QGLSSSIF | ASSLRDTNSGNTLY | 0.76425934 |
| RGTFANIRL | ASSLRDTNSGNTLY | 0.78720844 |
| RQLWWGHRI | ASSLRDTNSGNTLY | 0.7857181 |
| YGPPGTGKTLL | ASSLRDTNSGNTLY | 0.7978915 |
| AALVGAPANVA | ASSLRDTNSGNTLY | 0.79529846 |
| AATFRRTE | ASSLRDTNSGNTLY | 0.7908597 |
| AGALGGGI | ASSLRDTNSGNTLY | 0.5562802 |
| AGGAGGAGGIGAG | ASSLRDTNSGNTLY | 0.46591675 |
| AGSGALGL | ASSLRDTNSGNTLY | 0.31852394 |
| AQIADQI | ASSLRDTNSGNTLY | 0.5934616 |
| ASAVAVTYL | ASSLRDTNSGNTLY | 0.5779767 |
| ASTVLGGTGG | ASSLRDTNSGNTLY | 0.6908051 |
| AVGGEPAK | ASSLRDTNSGNTLY | 0.13012516 |

| | | |
|---|---|---|
| CEQGTFGGTC | ASSLRDTNSGNTLY | 0.6568518 |
| DEGKAPAG | ASSLRDTNSGNTLY | 0.81871295 |
| DTVVFPAL | ASSLRDTNSGNTLY | 0.50759685 |
| FEGITMAG | ASSLRDTNSGNTLY | 0.76642215 |
| FPLVVQD | ASSLRDTNSGNTLY | 0.7030001 |
| GAFTTPQ | ASSLRDTNSGNTLY | 0.46056083 |
| GFGGFGTAGGVGGA | ASSLRDTNSGNTLY | 0.7627081 |
| GFLAQFL | ASSLRDTNSGNTLY | 0.52857774 |
| GGAGGAGGNAGVA | ASSLRDTNSGNTLY | 0.7816228 |
| GGVGTAGAGGAGGAGG | ASSLRDTNSGNTLY | 0.8021194 |
| GNGGGGDAGNAGSGGN | ASSLRDTNSGNTLY | 0.7089244 |
| GTGGAGGAGNDAGS | ASSLRDTNSGNTLY | 0.6412028 |
| GTTNTGWFNS | ASSLRDTNSGNTLY | 0.7219255 |
| IFMGPLFTL | ASSLRDTNSGNTLY | 0.24852043 |
| ILAHAIEAI | ASSLRDTNSGNTLY | 0.65504754 |
| IRSAIREAL | ASSLRDTNSGNTLY | 0.31341198 |
| ITFAGGG | ASSLRDTNSGNTLY | 0.6210481 |
| ITGMSGHQEV | ASSLRDTNSGNTLY | 0.39213783 |
| IVAGLACTL | ASSLRDTNSGNTLY | 0.7246244 |
| IVGGGLAGA | ASSLRDTNSGNTLY | 0.7110944 |
| IVTGVLNI | ASSLRDTNSGNTLY | 0.7164912 |
| LAQQGCI | ASSLRDTNSGNTLY | 0.30595356 |
| LCLHVEN | ASSLRDTNSGNTLY | 0.73987204 |
| LDAISDGS | ASSLRDTNSGNTLY | 0.70820695 |
| LGGFAAML | ASSLRDTNSGNTLY | 0.3802297 |
| LGGKNPMI | ASSLRDTNSGNTLY | 0.70747256 |
| LGPKGVAAIL | ASSLRDTNSGNTLY | 0.6760274 |
| LMVGVSI | ASSLRDTNSGNTLY | 0.5471571 |
| LVGAAFVL | ASSLRDTNSGNTLY | 0.7907037 |
| LVLAVAPL | ASSLRDTNSGNTLY | 0.7199857 |
| MAAIAVIDAA | ASSLRDTNSGNTLY | 0.21013889 |
| NNGINAELQ | ASSLRDTNSGNTLY | 0.5254656 |
| RGEPLVLVHG | ASSLRDTNSGNTLY | 0.64501613 |
| RGGAGGAAGLI | ASSLRDTNSGNTLY | 0.5092554 |
| RSYRMHPAV | ASSLRDTNSGNTLY | 0.17324896 |
| SAAHLLAGQLA | ASSLRDTNSGNTLY | 0.68232185 |
| SAAVGSY | ASSLRDTNSGNTLY | 0.504442 |
| SPHAVLHRI | ASSLRDTNSGNTLY | 0.31504384 |
| SQIAKVL | ASSLRDTNSGNTLY | 0.56052834 |
| TALGHLVP | ASSLRDTNSGNTLY | 0.7851192 |
| TAPTTSGAA | ASSLRDTNSGNTLY | 0.63139486 |
| TDGDDGNGGSPGT | ASSLRDTNSGNTLY | 0.74529886 |
| TDGLGEN | ASSLRDTNSGNTLY | 0.7197749 |
| TGGPPLPGQA | ASSLRDTNSGNTLY | 0.77939206 |
| TIPAITI | ASSLRDTNSGNTLY | 0.7981969 |
| TIPGFTI | ASSLRDTNSGNTLY | 0.75303584 |
| TSIASGALA | ASSLRDTNSGNTLY | 0.67603153 |
| TVTLPGAAI | ASSLRDTNSGNTLY | 0.74135286 |
| VASGLSAALP | ASSLRDTNSGNTLY | 0.6109377 |
| VGAGVIGI | ASSLRDTNSGNTLY | 0.48970813 |
| VGPKRFGV | ASSLRDTNSGNTLY | 0.44982713 |

| | | |
|---|---|---|
| VNVAFPD | ASSLRDTNSGNTLY | 0.40546477 |
| VSMTNSGVS | ASSLRDTNSGNTLY | 0.43787622 |
| VTDVLGDLSV | ASSLRDTNSGNTLY | 0.6811923 |
| VVVVGAGFAGL | ASSLRDTNSGNTLY | 0.8249092 |
| VYTVGTVV | ASSLRDTNSGNTLY | 0.5800755 |
| WTTFSGP | ASSLRDTNSGNTLY | 0.2985854 |
| YDELVEAF | ASSLRDTNSGNTLY | 0.5772209 |
| YGTGGAGGNGG | ASSLRDTNSGNTLY | 0.26220688 |
| YTNVMAV | ASSLRDTNSGNTLY | 0.7338016 |
| FLEQVHQL | ASSLRDTNSGNTLY | 0.7373935 |
| GLSLLAVAV | ASSLRDTNSGNTLY | 0.82611626 |
| LEQVHQL | ASSLRDTNSGNTLY | 0.75970006 |
| LNAAGAI | ASSLRDTNSGNTLY | 0.610344 |
| PAVHASL | ASSLRDTNSGNTLY | 0.7613104 |
| TELEDDA | ASSLRDTNSGNTLY | 0.4979114 |
| TPPGPPN | ASSLRDTNSGNTLY | 0.62882596 |
| TPPPPPPP | ASSLRDTNSGNTLY | 0.61783993 |
| TQSPALI | ASGDEGRGQNTLY | 0.6666974 |
| EFAAAFPQ | ASGDEGRGQNTLY | 0.67744255 |
| LLSGNLDVL | ASGDEGRGQNTLY | 0.56115204 |
| GPANLGNYNI | ASGDEGRGQNTLY | 0.22174254 |
| TLGPTLP | ASGDEGRGQNTLY | 0.72310865 |
| VYAYSSCCK | ASGDEGRGQNTLY | 0.32314545 |
| APAPAPAPNI | ASGDEGRGQNTLY | 0.4861285 |
| PGGPVPIP | ASGDEGRGQNTLY | 0.5676503 |
| FLTVALEP | ASGDEGRGQNTLY | 0.8097297 |
| SQYFGANL | ASGDEGRGQNTLY | 0.83958155 |
| GGVKPAA | ASGDEGRGQNTLY | 0.54533905 |
| GGTGGAGGAGGW | ASGDEGRGQNTLY | 0.46287376 |
| GVGGVAGLGGAGG | ASGDEGRGQNTLY | 0.7036247 |
| VLLVLATTI | ASGDEGRGQNTLY | 0.45630226 |
| IGIFAAML | ASGDEGRGQNTLY | 0.7349144 |
| AVGDLID | ASGDEGRGQNTLY | 0.61206245 |
| NISPLPL | ASGDEGRGQNTLY | 0.63313526 |
| VTAVFVAI | ASGDEGRGQNTLY | 0.53076434 |
| SGDDNDAAL | ASGDEGRGQNTLY | 0.7110585 |
| AVAALAHS | ASGDEGRGQNTLY | 0.5335333 |
| GGVTVMATP | ASGDEGRGQNTLY | 0.7726184 |
| AGLAANVRD | ASGDEGRGQNTLY | 0.17165545 |
| TNRPDLIDPALLRPG | ASGDEGRGQNTLY | 0.74452734 |
| LANLGVTAL | ASGDEGRGQNTLY | 0.47440726 |
| TGERIEL | ASGDEGRGQNTLY | 0.5037137 |
| NGEELAA | ASGDEGRGQNTLY | 0.22960271 |
| VLSLVWG | ASGDEGRGQNTLY | 0.6387135 |
| IGYALWL | ASGDEGRGQNTLY | 0.6541112 |
| LSLLLAPL | ASGDEGRGQNTLY | 0.78172666 |
| TLDEALK | ASGDEGRGQNTLY | 0.58125365 |
| TDGAGTAEGG | ASGDEGRGQNTLY | 0.7058979 |
| NAPGGAGGNGG | ASGDEGRGQNTLY | 0.72185826 |
| VSVFMGMGGNF | ASGDEGRGQNTLY | 0.55152774 |
| LASYLAAL | ASGDEGRGQNTLY | 0.23679797 |

| | | |
|---|---|---|
| LVGGTPAL | ASGDEGRGQNTLY | 0.74975735 |
| VRNADVL | ASGDEGRGQNTLY | 0.51105064 |
| TCSSPFLVSAM | ASGDEGRGQNTLY | 0.78778136 |
| VHITDAE | ASGDEGRGQNTLY | 0.72608495 |
| EYAAPSF | ASGDEGRGQNTLY | 0.29344034 |
| VVDAFHGGL | ASGDEGRGQNTLY | 0.8098079 |
| ASVVDPQ | ASGDEGRGQNTLY | 0.72600305 |
| SYNEFGGLS | ASGDEGRGQNTLY | 0.37024528 |
| DPDGSVY | ASGDEGRGQNTLY | 0.4552985 |
| MVSTIHATE | ASGDEGRGQNTLY | 0.52175516 |
| LAAGVADPV | ASGDEGRGQNTLY | 0.3198812 |
| TKEQIAA | ASGDEGRGQNTLY | 0.7856358 |
| SLASPAAL | ASGDEGRGQNTLY | 0.54461086 |
| IGLVFYGV | ASGDEGRGQNTLY | 0.68943846 |
| FLLDASNL | ASGDEGRGQNTLY | 0.64606816 |
| YQVGNFP | ASGDEGRGQNTLY | 0.6550646 |
| ESLLSQST | ASGDEGRGQNTLY | 0.63205075 |
| LFTVGFAAVG | ASGDEGRGQNTLY | 0.6570628 |
| ATPPTPT | ASGDEGRGQNTLY | 0.6593017 |
| TDLATLG | ASGDEGRGQNTLY | 0.72433174 |
| SAGGTGR | ASGDEGRGQNTLY | 0.5804124 |
| GFSNSGS | ASGDEGRGQNTLY | 0.6577173 |
| GGNGGSGGTSV | ASGDEGRGQNTLY | 0.7996755 |
| GTSVATGGAGN | ASGDEGRGQNTLY | 0.38130787 |
| NGGNGGSGGTS | ASGDEGRGQNTLY | 0.5818796 |
| AVVKFPH | ASGDEGRGQNTLY | 0.7753233 |
| TPPPPPP | ASGDEGRGQNTLY | 0.5458094 |
| AAPAAAAAPA | ASGDEGRGQNTLY | 0.7856572 |
| IALNEDL | ASGDEGRGQNTLY | 0.7431564 |
| IVNVSSM | ASGDEGRGQNTLY | 0.7126013 |
| TGIPPAPR | ASGDEGRGQNTLY | 0.5677073 |
| FDLGGGTF | ASGDEGRGQNTLY | 0.8013413 |
| FDLGGGTFDVS | ASGDEGRGQNTLY | 0.7559709 |
| NEPTAAA | ASGDEGRGQNTLY | 0.5952582 |
| NEPTAAALA | ASGDEGRGQNTLY | 0.70872426 |
| GSIINISSI | ASGDEGRGQNTLY | 0.44427866 |
| VAPGNAGTA | ASGDEGRGQNTLY | 0.71382916 |
| TGGGASLE | ASGDEGRGQNTLY | 0.7668899 |
| VAVYNHYKRI | ASGDEGRGQNTLY | 0.32462546 |
| TDPAPATT | ASGDEGRGQNTLY | 0.58065593 |
| TNRPDILDPALLRPG | ASGDEGRGQNTLY | 0.8214948 |
| LTTTTTS | ASGDEGRGQNTLY | 0.7433483 |
| PTGSGKTL | ASGDEGRGQNTLY | 0.45691502 |
| TGSGKTL | ASGDEGRGQNTLY | 0.52101827 |
| TIGHVDHGK | ASGDEGRGQNTLY | 0.5528529 |
| TIGHVDHGKTT | ASGDEGRGQNTLY | 0.6067877 |
| GHVDHGKTT | ASGDEGRGQNTLY | 0.3740567 |
| AQFEHTLL | ASGDEGRGQNTLY | 0.7392833 |
| IVNQDEL | ASGDEGRGQNTLY | 0.55767035 |
| TNGSQFF | ASGDEGRGQNTLY | 0.7740072 |
| TGKGIVE | ASGDEGRGQNTLY | 0.6739003 |

| | | |
|---|---|---|
| LISNASDA | ASGDEGRGQNTLY | 0.80312276 |
| AELRTLGELTKIAKAHGAQ | ASGDEGRGQNTLY | 0.5335969 |
| APNAVAAAEPSW | ASGDEGRGQNTLY | 0.67570674 |
| APQVAGAATAVL | ASGDEGRGQNTLY | 0.27304715 |
| ARRDPAAAAAILPTDA | ASGDEGRGQNTLY | 0.4578208 |
| DAAARDVLAEAGLAEYF | ASGDEGRGQNTLY | 0.44371426 |
| DDQIVLSAKANLAKVEQ | ASGDEGRGQNTLY | 0.68490875 |
| DELLPDTGEDSDEDGH | ASGDEGRGQNTLY | 0.23981968 |
| ELPEAPANRATVPQD | ASGDEGRGQNTLY | 0.82586676 |
| FGQSKIERVTGIG | ASGDEGRGQNTLY | 0.602407 |
| GAGGTGSTAGGAGGAGGAGGLY | ASGDEGRGQNTLY | 0.74949455 |
| GHGGTGAAGGNGGTGGAGGSGI | ASGDEGRGQNTLY | 0.6254151 |
| GNGGNGGAGGEGGPGGAGGAGA | ASGDEGRGQNTLY | 0.50191855 |
| GSTAGNGGNGGNATNGTIAGQP | ASGDEGRGQNTLY | 0.5596047 |
| GTGGDGGNGGNGGTGG | ASGDEGRGQNTLY | 0.77393293 |
| GVRRPAGIPADAPV | ASGDEGRGQNTLY | 0.2901603 |
| GYRQVTDAHLLTLARR | ASGDEGRGQNTLY | 0.16980655 |
| IAKMGQDGHDRG | ASGDEGRGQNTLY | 0.47031498 |
| KAEPSIEIQYT | ASGDEGRGQNTLY | 0.2769689 |
| KPVRIRMAADEDTVAVT | ASGDEGRGQNTLY | 0.81454027 |
| MPPRIAGMRLLVIKPEP | ASGDEGRGQNTLY | 0.7821171 |
| MKFYAVALFRSRGVPPTR | ASGDEGRGQNTLY | 0.6492737 |
| NAGGFGILGAVAHSP | ASGDEGRGQNTLY | 0.80659 |
| NEGREITLAMAEM | ASGDEGRGQNTLY | 0.14065994 |
| NELVEWSHPLGHRWRWE | ASGDEGRGQNTLY | 0.6008089 |
| NLGAGGTGGNGGNPG | ASGDEGRGQNTLY | 0.8244699 |
| NTEWPKWETVMAADPVD | ASGDEGRGQNTLY | 0.67302585 |
| PASAAPAETQP | ASGDEGRGQNTLY | 0.5970622 |
| QQSSDGKNGVRAAEITGE | ASGDEGRGQNTLY | 0.7881112 |
| RAPDAVTTSWWKE | ASGDEGRGQNTLY | 0.6133096 |
| SGFGNVGAHTSGWFNQ | ASGDEGRGQNTLY | 0.5427115 |
| SGVAGAVGQAASVGGLKV | ASGDEGRGQNTLY | 0.7164794 |
| SNGGAGTGSAGGNGGT | ASGDEGRGQNTLY | 0.7207042 |
| TGGAGGEGGAAG | ASGDEGRGQNTLY | 0.6999688 |
| TGGDGGAAGAGGAPGHG | ASGDEGRGQNTLY | 0.27723527 |
| TMPPLLAAIANAGRHGV | ASGDEGRGQNTLY | 0.76433384 |
| TSSTAGNSGGA | ASGDEGRGQNTLY | 0.81965554 |
| VPATVPVGILC | ASGDEGRGQNTLY | 0.55742747 |
| VQRPDSGGVVLTGRIS | ASGDEGRGQNTLY | 0.70102286 |
| WTPNPVPATKGVPAGLVQAT | ASGDEGRGQNTLY | 0.7002224 |
| YALKRALAKFSDRFDVVII | ASGDEGRGQNTLY | 0.45428538 |
| YLIGSVIV | ASGDEGRGQNTLY | 0.42686024 |
| YGQLVAVID | ASGDEGRGQNTLY | 0.388659 |
| VTVGGMTLPA | ASGDEGRGQNTLY | 0.75525194 |
| VLALPAV | ASGDEGRGQNTLY | 0.35243148 |
| VIPPTLNYET | ASGDEGRGQNTLY | 0.6894262 |
| VGMGGMHPSSG | ASGDEGRGQNTLY | 0.63776934 |
| VGLAGGAATA | ASGDEGRGQNTLY | 0.8523842 |
| VGGAGGAGGSGP | ASGDEGRGQNTLY | 0.72950983 |
| VFAAFMV | ASGDEGRGQNTLY | 0.7267576 |
| TLYDKLR | ASGDEGRGQNTLY | 0.51510483 |

| | | |
|---|---|---|
| TLAVFTVL | ASGDEGRGQNTLY | 0.67032933 |
| TGSVQPARV | ASGDEGRGQNTLY | 0.6999607 |
| TGGGAGSGGT | ASGDEGRGQNTLY | 0.22001323 |
| SHLETSL | ASGDEGRGQNTLY | 0.8141079 |
| SFTTGMT | ASGDEGRGQNTLY | 0.44173443 |
| QVEVAKRMGVRQPTVS | ASGDEGRGQNTLY | 0.6258644 |
| QGSGGPALAM | ASGDEGRGQNTLY | 0.75061715 |
| MLAPSNP | ASGDEGRGQNTLY | 0.77702385 |
| LVNNAAI | ASGDEGRGQNTLY | 0.62805873 |
| LTGEINTLA | ASGDEGRGQNTLY | 0.7628757 |
| LSIFGPV | ASGDEGRGQNTLY | 0.5158063 |
| LLALLANL | ASGDEGRGQNTLY | 0.77009505 |
| LISNASDALDKLR | ASGDEGRGQNTLY | 0.7849846 |
| LINEAVP | ASGDEGRGQNTLY | 0.4802528 |
| LGYAFSGL | ASGDEGRGQNTLY | 0.79805505 |
| LDFVAVAL | ASGDEGRGQNTLY | 0.78633934 |
| LAQPGAE | ASGDEGRGQNTLY | 0.32728192 |
| ISNLGAVTGF | ASGDEGRGQNTLY | 0.6162653 |
| HQEDPGRI | ASGDEGRGQNTLY | 0.752568 |
| GTPGEPTP | ASGDEGRGQNTLY | 0.7684773 |
| GFFNSGS | ASGDEGRGQNTLY | 0.7728522 |
| GAGGGGLTAGAN | ASGDEGRGQNTLY | 0.7326678 |
| FVDALVNM | ASGDEGRGQNTLY | 0.41684717 |
| FLTVAPEL | ASGDEGRGQNTLY | 0.8063108 |
| FETLTAL | ASGDEGRGQNTLY | 0.86514354 |
| DVINPIEL | ASGDEGRGQNTLY | 0.7378927 |
| APVSGAPV | ASGDEGRGQNTLY | 0.5483926 |
| ALVGGLAW | ASGDEGRGQNTLY | 0.5895724 |
| ALAAAAATAI | ASGDEGRGQNTLY | 0.7547063 |
| AFGLGLLL | ASGDEGRGQNTLY | 0.76378834 |
| AFFAAASL | ASGDEGRGQNTLY | 0.41107512 |
| AAAALPAFSPP | ASGDEGRGQNTLY | 0.6685566 |
| VVLAPAL | ASGDEGRGQNTLY | 0.7192811 |
| TIGHVDHGKTTL | ASGDEGRGQNTLY | 0.2402932 |
| LISNASDALDK | ASGDEGRGQNTLY | 0.7004738 |
| LISNASDAL | ASGDEGRGQNTLY | 0.77163625 |
| ISNASDALDK | ASGDEGRGQNTLY | 0.7097243 |
| NQAETLVYQTE | ASGDEGRGQNTLY | 0.43352315 |
| TGGTGGTAGAG | ASGDEGRGQNTLY | 0.81238997 |
| TGSTAGGAGGA | ASGDEGRGQNTLY | 0.7895908 |
| DPLAVPVAAGPV | ASGDEGRGQNTLY | 0.8262466 |
| GGTGGTPGNGGN | ASGDEGRGQNTLY | 0.36075562 |
| PAGLAGVDPATL | ASGDEGRGQNTLY | 0.18119627 |
| GNNLVVEAPHAGASV | ASGDEGRGQNTLY | 0.6909466 |
| KVTRIMSASTLEQPA | ASGDEGRGQNTLY | 0.7440367 |
| LGFALPIALPRRTVV | ASGDEGRGQNTLY | 0.6744989 |
| TRLGNLTARKLVVLV | ASGDEGRGQNTLY | 0.51369107 |
| WSQQRRLIYFPSAGP | ASGDEGRGQNTLY | 0.6568034 |
| GPNGNPDPMAAAVDIR | ASGDEGRGQNTLY | 0.789999 |
| TQRCVSVADLRAAASR | ASGDEGRGQNTLY | 0.4142195 |
| QFVLDVAEHPVEVLTETV | ASGDEGRGQNTLY | 0.37095857 |

| | | |
|---|---|---|
| GGAGGTGGAGGNGGTQAGDG | ASGDEGRGQNTLY | 0.71830964 |
| PLLATLRAIAVDVDAKPAQV | ASGDEGRGQNTLY | 0.7856878 |
| APKFRTGGYMIDFWGVGYQVAK | ASGDEGRGQNTLY | 0.38393736 |
| GAGGAGGAGGAAGGPGGTGGTG | ASGDEGRGQNTLY | 0.5139149 |
| IGALGLLATDVAPAGSVAFGVA | ASGDEGRGQNTLY | 0.71069056 |
| AGGAGGIGGNGGAGGAGGVLMG | ASGDEGRGQNTLY | 0.72899294 |
| ADELIGLL | ASGDEGRGQNTLY | 0.64705265 |
| ADRALKA | ASGDEGRGQNTLY | 0.41732064 |
| AGAVGQVGGAGGA | ASGDEGRGQNTLY | 0.68798774 |
| AGGGGGLGGAP | ASGDEGRGQNTLY | 0.3349049 |
| AGGLLYGNGGAG | ASGDEGRGQNTLY | 0.7686629 |
| AGLESIGL | ASGDEGRGQNTLY | 0.6976488 |
| AGLKTGII | ASGDEGRGQNTLY | 0.69229543 |
| ASLAALLL | ASGDEGRGQNTLY | 0.5825539 |
| ASVINISSI | ASGDEGRGQNTLY | 0.3679583 |
| ATPPGPA | ASGDEGRGQNTLY | 0.277739 |
| ATVASALV | ASGDEGRGQNTLY | 0.4436262 |
| ATVASLIL | ASGDEGRGQNTLY | 0.7871275 |
| AVNGPVLIRH | ASGDEGRGQNTLY | 0.5634401 |
| DGAAGAVGI | ASGDEGRGQNTLY | 0.759704 |
| DPLLHGI | ASGDEGRGQNTLY | 0.6874552 |
| ELGAALGAL | ASGDEGRGQNTLY | 0.40253752 |
| ELGRTQAV | ASGDEGRGQNTLY | 0.7208569 |
| FVAGGYLS | ASGDEGRGQNTLY | 0.42498827 |
| GASGGAGGAGGW | ASGDEGRGQNTLY | 0.61673546 |
| GGAGAGGDGGAGGI | ASGDEGRGQNTLY | 0.61118114 |
| GTGHNGSQPGMGGNG | ASGDEGRGQNTLY | 0.7674035 |
| GYAAASASAS | ASGDEGRGQNTLY | 0.44236976 |
| HQVDVAAVQQG | ASGDEGRGQNTLY | 0.74043393 |
| IGAAIAHT | ASGDEGRGQNTLY | 0.70405835 |
| IGMIEVAQL | ASGDEGRGQNTLY | 0.40947902 |
| IRLAPEHTSAMD | ASGDEGRGQNTLY | 0.712912 |
| IYNVSVL | ASGDEGRGQNTLY | 0.772956 |
| KQGFLGM | ASGDEGRGQNTLY | 0.73614895 |
| LESVITQL | ASGDEGRGQNTLY | 0.24574348 |
| LGLGALVVV | ASGDEGRGQNTLY | 0.73609877 |
| LIGNGGNGGSGGI | ASGDEGRGQNTLY | 0.47752088 |
| LNAINAPTL | ASGDEGRGQNTLY | 0.6962098 |
| LQLLNTGV | ASGDEGRGQNTLY | 0.6459435 |
| LQTSIEF | ASGDEGRGQNTLY | 0.6864147 |
| MSFAVSEL | ASGDEGRGQNTLY | 0.77381825 |
| NTGALGL | ASGDEGRGQNTLY | 0.35152927 |
| PPGAGGAGGAGTT | ASGDEGRGQNTLY | 0.46320286 |
| QLVGLGI | ASGDEGRGQNTLY | 0.5645444 |
| RFLEQVHQL | ASGDEGRGQNTLY | 0.73412216 |
| RGVGYVLR | ASGDEGRGQNTLY | 0.7517239 |
| SATGVFAGVI | ASGDEGRGQNTLY | 0.42016053 |
| SLPAITLP | ASGDEGRGQNTLY | 0.6657536 |
| STALAFTV | ASGDEGRGQNTLY | 0.54976225 |
| SVLTVGLGK | ASGDEGRGQNTLY | 0.40580556 |
| TALAGAGSGI | ASGDEGRGQNTLY | 0.68248415 |

| | | |
|---|---|---|
| TGLPFTGI | ASGDEGRGQNTLY | 0.55916905 |
| VAEHVTV | ASGDEGRGQNTLY | 0.36254972 |
| VDIVLNSL | ASGDEGRGQNTLY | 0.72682667 |
| VLTGGGAVL | ASGDEGRGQNTLY | 0.57001185 |
| VNVGLAYGI | ASGDEGRGQNTLY | 0.77727807 |
| VSSAAVGAV | ASGDEGRGQNTLY | 0.7712078 |
| VTSVYEAGV | ASGDEGRGQNTLY | 0.46585578 |
| WVALDRGA | ASGDEGRGQNTLY | 0.74613154 |
| YLVGSVVI | ASGDEGRGQNTLY | 0.6099256 |
| GPPGTGKTLL | ASGDEGRGQNTLY | 0.68357646 |
| GVGMTKF | ASGDEGRGQNTLY | 0.68330973 |
| IGIGGSDL | ASGDEGRGQNTLY | 0.37062958 |
| IGTLLAEL | ASGDEGRGQNTLY | 0.77064717 |
| IYTSGTT | ASGDEGRGQNTLY | 0.79038644 |
| LSFPAGV | ASGDEGRGQNTLY | 0.74963593 |
| LYTSGTT | ASGDEGRGQNTLY | 0.7119616 |
| VGAFPFGF | ASGDEGRGQNTLY | 0.71440476 |
| VGAGTVEF | ASGDEGRGQNTLY | 0.7874509 |
| VGKSTLLL | ASGDEGRGQNTLY | 0.52775776 |
| AALAHAGGGIV | ASGDEGRGQNTLY | 0.26480234 |
| AALVAVSLL | ASGDEGRGQNTLY | 0.6765584 |
| AAVSRAVAAL | ASGDEGRGQNTLY | 0.6075847 |
| AGGSSGIGGF | ASGDEGRGQNTLY | 0.4812373 |
| AGLTVANANG | ASGDEGRGQNTLY | 0.4060037 |
| AGLVGAVL | ASGDEGRGQNTLY | 0.75491154 |
| ALSQAFTV | ASGDEGRGQNTLY | 0.84225 |
| ALTFSVSL | ASGDEGRGQNTLY | 0.60792494 |
| ALTGAALVAL | ASGDEGRGQNTLY | 0.8302976 |
| ASFSIGAGVL | ASGDEGRGQNTLY | 0.33379304 |
| DGSVYKI | ASGDEGRGQNTLY | 0.8017576 |
| EQLAEIV | ASGDEGRGQNTLY | 0.7994108 |
| FAVSELTVM | ASGDEGRGQNTLY | 0.39739618 |
| GAAPLAAGL | ASGDEGRGQNTLY | 0.74724317 |
| GADGGDGGSGGAGG | ASGDEGRGQNTLY | 0.29780722 |
| GFGLDTGGAGG | ASGDEGRGQNTLY | 0.42138433 |
| GLASGGLL | ASGDEGRGQNTLY | 0.7847316 |
| GRIAGAAVLL | ASGDEGRGQNTLY | 0.6953515 |
| GSAGSLNTSF | ASGDEGRGQNTLY | 0.7238665 |
| GTGQIGI | ASGDEGRGQNTLY | 0.66798943 |
| GTLIALIL | ASGDEGRGQNTLY | 0.49816674 |
| IAATMGFVVG | ASGDEGRGQNTLY | 0.66497695 |
| IFAQALL | ASGDEGRGQNTLY | 0.7976109 |
| IGAAVALGF | ASGDEGRGQNTLY | 0.66944945 |
| IGLTPLAL | ASGDEGRGQNTLY | 0.86722004 |
| ILGSNTSTL | ASGDEGRGQNTLY | 0.41859856 |
| IVFSLTVL | ASGDEGRGQNTLY | 0.78196704 |
| LASKATGF | ASGDEGRGQNTLY | 0.79374313 |
| LASVALIL | ASGDEGRGQNTLY | 0.6599781 |
| LFAVGLTL | ASGDEGRGQNTLY | 0.59843016 |
| LFNGGIGTYI | ASGDEGRGQNTLY | 0.7720014 |
| LTLGAGII | ASGDEGRGQNTLY | 0.6365329 |

| | | |
|---|---|---|
| LVAAALGDI | ASGDEGRGQNTLY | 0.75861 |
| NTAFVQLGI | ASGDEGRGQNTLY | 0.8054453 |
| QLFSGAEL | ASGDEGRGQNTLY | 0.35623452 |
| SAATFITL | ASGDEGRGQNTLY | 0.56116575 |
| SNYGLANLGV | ASGDEGRGQNTLY | 0.488512 |
| SYGGAYAVM | ASGDEGRGQNTLY | 0.5260128 |
| TTGVGGLTLG | ASGDEGRGQNTLY | 0.4305586 |
| TTQTIYI | ASGDEGRGQNTLY | 0.39775005 |
| VAAGAGLGAVI | ASGDEGRGQNTLY | 0.4631253 |
| VAAVVTLL | ASGDEGRGQNTLY | 0.4428224 |
| VGFLGGLI | ASGDEGRGQNTLY | 0.64551145 |
| VGVIPILL | ASGDEGRGQNTLY | 0.5424845 |
| VIFASSLI | ASGDEGRGQNTLY | 0.44001383 |
| VKGGDGGAAAG | ASGDEGRGQNTLY | 0.59578407 |
| VTKTFTNI | ASGDEGRGQNTLY | 0.59430826 |
| VVAAAILL | ASGDEGRGQNTLY | 0.36616346 |
| YAALDVEL | ASGDEGRGQNTLY | 0.447672 |
| LASVALLL | ASGDEGRGQNTLY | 0.47248703 |
| LDAVHRV | ASGDEGRGQNTLY | 0.41053882 |
| QGLSSSIF | ASGDEGRGQNTLY | 0.31957498 |
| RGTFANIRL | ASGDEGRGQNTLY | 0.4961549 |
| RQLWWGHRI | ASGDEGRGQNTLY | 0.51677155 |
| YGPPGTGKTLL | ASGDEGRGQNTLY | 0.37843126 |
| AALVGAPANVA | ASGDEGRGQNTLY | 0.5304959 |
| AATFRRTE | ASGDEGRGQNTLY | 0.44651687 |
| AGALGGGI | ASGDEGRGQNTLY | 0.49501592 |
| AGGAGGAGGIGAG | ASGDEGRGQNTLY | 0.5666438 |
| AGSGALGL | ASGDEGRGQNTLY | 0.46766296 |
| AQIADQI | ASGDEGRGQNTLY | 0.50178486 |
| ASAVAVTYL | ASGDEGRGQNTLY | 0.5487107 |
| ASTVLGGTGG | ASGDEGRGQNTLY | 0.6199821 |
| AVGGEPAK | ASGDEGRGQNTLY | 0.45413017 |
| CEQGTFGGTC | ASGDEGRGQNTLY | 0.4436882 |
| DEGKAPAG | ASGDEGRGQNTLY | 0.62700385 |
| DTVVFPAL | ASGDEGRGQNTLY | 0.5110922 |
| FEGITMAG | ASGDEGRGQNTLY | 0.5977816 |
| FPLVVQD | ASGDEGRGQNTLY | 0.6020236 |
| GAFTTPQ | ASGDEGRGQNTLY | 0.41450334 |
| GFGGFGTAGGVGGA | ASGDEGRGQNTLY | 0.59050727 |
| GFLAQFL | ASGDEGRGQNTLY | 0.5939703 |
| GGAGGAGGNAGVA | ASGDEGRGQNTLY | 0.48543346 |
| GGVGTAGAGGAGGAGG | ASGDEGRGQNTLY | 0.5001423 |
| GNGGGGDAGNAGSGGN | ASGDEGRGQNTLY | 0.36629912 |
| GTGGAGGAGNDAGS | ASGDEGRGQNTLY | 0.4641287 |
| GTTNTGWFNS | ASGDEGRGQNTLY | 0.39627972 |
| IFMGPLFTL | ASGDEGRGQNTLY | 0.52968395 |
| ILAHAIEAI | ASGDEGRGQNTLY | 0.5977491 |
| IRSAIREAL | ASGDEGRGQNTLY | 0.5135564 |
| ITFAGGG | ASGDEGRGQNTLY | 0.3303608 |
| ITGMSGHQEV | ASGDEGRGQNTLY | 0.3846577 |
| IVAGLACTL | ASGDEGRGQNTLY | 0.42133296 |

| | | |
|---|---|---|
| IVGGGLAGA | ASGDEGRGQNTLY | 0.36351714 |
| IVTGVLNI | ASGDEGRGQNTLY | 0.45091522 |
| LAQQGCI | ASGDEGRGQNTLY | 0.3679114 |
| LCLHVEN | ASGDEGRGQNTLY | 0.61869454 |
| LDAISDGS | ASGDEGRGQNTLY | 0.6107521 |
| LGGFAAML | ASGDEGRGQNTLY | 0.4893732 |
| LGGKNPMI | ASGDEGRGQNTLY | 0.5391167 |
| LGPKGVAAIL | ASGDEGRGQNTLY | 0.6280541 |
| LMVGVSI | ASGDEGRGQNTLY | 0.5799514 |
| LVGAAFVL | ASGDEGRGQNTLY | 0.47643745 |
| LVLAVAPL | ASGDEGRGQNTLY | 0.34928545 |
| MAAIAVIDAA | ASGDEGRGQNTLY | 0.6323608 |
| NNGINAELQ | ASGDEGRGQNTLY | 0.456202 |
| RGEPLVLVHG | ASGDEGRGQNTLY | 0.6325674 |
| RGGAGGAAGLI | ASGDEGRGQNTLY | 0.58100235 |
| RSYRMHPAV | ASGDEGRGQNTLY | 0.5309615 |
| SAAHLLAGQLA | ASGDEGRGQNTLY | 0.41128814 |
| SAAVGSY | ASGDEGRGQNTLY | 0.5875056 |
| SPHAVLHRI | ASGDEGRGQNTLY | 0.5350574 |
| SQIAKVL | ASGDEGRGQNTLY | 0.3794878 |
| TALGHLVP | ASGDEGRGQNTLY | 0.37169036 |
| TAPTTSGAA | ASGDEGRGQNTLY | 0.5875911 |
| TDGDDGNGGSPGT | ASGDEGRGQNTLY | 0.4942152 |
| TDGLGEN | ASGDEGRGQNTLY | 0.6254605 |
| TGGPPLPGQA | ASGDEGRGQNTLY | 0.46121943 |
| TIPAITI | ASGDEGRGQNTLY | 0.3969795 |
| TIPGFTI | ASGDEGRGQNTLY | 0.57149124 |
| TSIASGALA | ASGDEGRGQNTLY | 0.50619483 |
| TVTLPGAAI | ASGDEGRGQNTLY | 0.4970174 |
| VASGLSAALP | ASGDEGRGQNTLY | 0.57660747 |
| VGAGVIGI | ASGDEGRGQNTLY | 0.4426982 |
| VGPKRFGV | ASGDEGRGQNTLY | 0.5061193 |
| VNVAFPD | ASGDEGRGQNTLY | 0.57678276 |
| VSMTNSGVS | ASGDEGRGQNTLY | 0.4903496 |
| VTDVLGDLSV | ASGDEGRGQNTLY | 0.5866775 |
| VVVVGAGFAGL | ASGDEGRGQNTLY | 0.4931464 |
| VYTVGTVV | ASGDEGRGQNTLY | 0.47376007 |
| WTTFSGP | ASGDEGRGQNTLY | 0.45314652 |
| YDELVEAF | ASGDEGRGQNTLY | 0.48906758 |
| YGTGGAGGNGG | ASGDEGRGQNTLY | 0.5374963 |
| YTNVMAV | ASGDEGRGQNTLY | 0.49336246 |
| FLEQVHQL | ASGDEGRGQNTLY | 0.5955738 |
| GLSLLAVAV | ASGDEGRGQNTLY | 0.54162246 |
| LEQVHQL | ASGDEGRGQNTLY | 0.452711 |
| LNAAGAI | ASGDEGRGQNTLY | 0.6190918 |
| PAVHASL | ASGDEGRGQNTLY | 0.585588 |
| TELEDDA | ASGDEGRGQNTLY | 0.3884379 |
| TPPGPPN | ASGDEGRGQNTLY | 0.49529195 |
| TPPPPPPP | ASGDEGRGQNTLY | 0.56359184 |
| TQSPALI | TCSADPDNSPLY | 0.61504 |
| EFAAAFPQ | TCSADPDNSPLY | 0.48753148 |

| | | |
|---|---|---|
| LLSGNLDVL | TCSADPDNSPLY | 0.48889786 |
| GPANLGNYNI | TCSADPDNSPLY | 0.6504352 |
| TLGPTLP | TCSADPDNSPLY | 0.6084657 |
| VYAYSSCCK | TCSADPDNSPLY | 0.60813075 |
| APAPAPAPNI | TCSADPDNSPLY | 0.3895245 |
| PGGPVPIP | TCSADPDNSPLY | 0.42158362 |
| FLTVALEP | TCSADPDNSPLY | 0.39803854 |
| SQYFGANL | TCSADPDNSPLY | 0.31319323 |
| GGVKPAA | TCSADPDNSPLY | 0.60366327 |
| GGTGGAGGAGGW | TCSADPDNSPLY | 0.42589527 |
| GVGGVAGLGGAGG | TCSADPDNSPLY | 0.53246665 |
| VLLVLATTI | TCSADPDNSPLY | 0.47218478 |
| IGIFAAML | TCSADPDNSPLY | 0.5672519 |
| AVGDLID | TCSADPDNSPLY | 0.50472736 |
| NISPLPL | TCSADPDNSPLY | 0.43114242 |
| VTAVFVAI | TCSADPDNSPLY | 0.37811953 |
| SGDDNDAAL | TCSADPDNSPLY | 0.580065 |
| AVAALAHS | TCSADPDNSPLY | 0.6261089 |
| GGVTVMATP | TCSADPDNSPLY | 0.52733815 |
| AGLAANVRD | TCSADPDNSPLY | 0.38359532 |
| TNRPDLIDPALLRPG | TCSADPDNSPLY | 0.6160699 |
| LANLGVTAL | TCSADPDNSPLY | 0.5424266 |
| TGERIEL | TCSADPDNSPLY | 0.39653933 |
| NGEELAA | TCSADPDNSPLY | 0.41836008 |
| VLSLVWG | TCSADPDNSPLY | 0.39874122 |
| IGYALWL | TCSADPDNSPLY | 0.38412288 |
| LSLLLAPL | TCSADPDNSPLY | 0.570022 |
| TLDEALK | TCSADPDNSPLY | 0.59247714 |
| TDGAGTAEGG | TCSADPDNSPLY | 0.43268347 |
| NAPGGAGGNGG | TCSADPDNSPLY | 0.54114497 |
| VSVFMGMGGNF | TCSADPDNSPLY | 0.53124607 |
| LASYLAAL | TCSADPDNSPLY | 0.30564764 |
| LVGGTPAL | TCSADPDNSPLY | 0.42289257 |
| VRNADVL | TCSADPDNSPLY | 0.3819545 |
| TCSSPFLVSAM | TCSADPDNSPLY | 0.58020645 |
| VHITDAE | TCSADPDNSPLY | 0.38589942 |
| EYAAPSF | TCSADPDNSPLY | 0.5035641 |
| VVDAFHGGL | TCSADPDNSPLY | 0.511091 |
| ASVVDPQ | TCSADPDNSPLY | 0.6130151 |
| SYNEFGGLS | TCSADPDNSPLY | 0.5386544 |
| DPDGSVY | TCSADPDNSPLY | 0.3627894 |
| MVSTIHATE | TCSADPDNSPLY | 0.3629432 |
| LAAGVADPV | TCSADPDNSPLY | 0.57847285 |
| TKEQIAA | TCSADPDNSPLY | 0.53368706 |
| SLASPAAL | TCSADPDNSPLY | 0.47377363 |
| IGLVFYGV | TCSADPDNSPLY | 0.47218513 |
| FLLDASNL | TCSADPDNSPLY | 0.5961432 |
| YQVGNFP | TCSADPDNSPLY | 0.3577561 |
| ESLLSQST | TCSADPDNSPLY | 0.55981046 |
| LFTVGFAAVG | TCSADPDNSPLY | 0.37454933 |
| ATPPTPT | TCSADPDNSPLY | 0.48954082 |

| | | |
|---|---|---|
| TDLATLG | TCSADPDNSPLY | 0.601455 |
| SAGGTGR | TCSADPDNSPLY | 0.6080213 |
| GFSNSGS | TCSADPDNSPLY | 0.45416412 |
| GGNGGSGGTSV | TCSADPDNSPLY | 0.46873882 |
| GTSVATGGAGN | TCSADPDNSPLY | 0.3889157 |
| NGGNGGSGGTS | TCSADPDNSPLY | 0.4721165 |
| AVVKFPH | TCSADPDNSPLY | 0.4799155 |
| TPPPPPP | TCSADPDNSPLY | 0.60962343 |
| AAPAAAAAPA | TCSADPDNSPLY | 0.44428304 |
| IALNEDL | TCSADPDNSPLY | 0.36539552 |
| IVNVSSM | TCSADPDNSPLY | 0.45624465 |
| TGIPPAPR | TCSADPDNSPLY | 0.58951825 |
| FDLGGGTF | TCSADPDNSPLY | 0.6077336 |
| FDLGGGTFDVS | TCSADPDNSPLY | 0.40052766 |
| NEPTAAA | TCSADPDNSPLY | 0.44532347 |
| NEPTAAALA | TCSADPDNSPLY | 0.45023173 |
| GSIINISSI | TCSADPDNSPLY | 0.50799626 |
| VAPGNAGTA | TCSADPDNSPLY | 0.37750968 |
| TGGGASLE | TCSADPDNSPLY | 0.43686393 |
| VAVYNHYKRI | TCSADPDNSPLY | 0.50663114 |
| TDPAPATT | TCSADPDNSPLY | 0.6612219 |
| TNRPDILDPALLRPG | TCSADPDNSPLY | 0.4561935 |
| LTTTTTS | TCSADPDNSPLY | 0.34299055 |
| PTGSGKTL | TCSADPDNSPLY | 0.53505963 |
| TGSGKTL | TCSADPDNSPLY | 0.39396033 |
| TIGHVDHGK | TCSADPDNSPLY | 0.42756194 |
| TIGHVDHGKTT | TCSADPDNSPLY | 0.51396775 |
| GHVDHGKTT | TCSADPDNSPLY | 0.341072 |
| AQFEHTLL | TCSADPDNSPLY | 0.47970396 |
| IVNQDEL | TCSADPDNSPLY | 0.60378104 |
| TNGSQFF | TCSADPDNSPLY | 0.5710632 |
| TGKGIVE | TCSADPDNSPLY | 0.44131437 |
| LISNASDA | TCSADPDNSPLY | 0.5796369 |
| AELRTLGELTKIAKAHGAQ | TCSADPDNSPLY | 0.49964198 |
| APNAVAAAEPSW | TCSADPDNSPLY | 0.5912634 |
| APQVAGAATAVL | TCSADPDNSPLY | 0.6418725 |
| ARRDPAAAAAILPTDA | TCSADPDNSPLY | 0.45203063 |
| DAAARDVLAEAGLAEYF | TCSADPDNSPLY | 0.43560427 |
| DDQIVLSAKANLAKVEQ | TCSADPDNSPLY | 0.5330079 |
| DELLPDTGEDSDEDGH | TCSADPDNSPLY | 0.5250309 |
| ELPEAPANRATVPQD | TCSADPDNSPLY | 0.5234975 |
| FGQSKIERVTGIG | TCSADPDNSPLY | 0.44466355 |
| GAGGTGSTAGGAGGAGGAGGLY | TCSADPDNSPLY | 0.6033615 |
| GHGGTGAAGGNGGTGGAGGSGI | TCSADPDNSPLY | 0.4425099 |
| GNGGNGGAGGEGGPGGAGGAGA | TCSADPDNSPLY | 0.6431979 |
| GSTAGNGGNGGNATNGTIAGQP | TCSADPDNSPLY | 0.33169517 |
| GTGGDGGNGGNGGTGG | TCSADPDNSPLY | 0.43899703 |
| GVRRPAGIPADAPV | TCSADPDNSPLY | 0.5637676 |
| GYRQVTDAHLLTLARR | TCSADPDNSPLY | 0.3345207 |
| IAKMGQDGHDRG | TCSADPDNSPLY | 0.43597525 |
| KAEPSIEIQYT | TCSADPDNSPLY | 0.44268888 |

| | | |
|---|---|---|
| KPVRIRMAADEDTVAVT | TCSADPDNSPLY | 0.51968443 |
| MPPRIAGMRLLVIKPEP | TCSADPDNSPLY | 0.3255139 |
| MKFYAVALFRSRGVPPTR | TCSADPDNSPLY | 0.54957694 |
| NAGGFGILGAVAHSP | TCSADPDNSPLY | 0.35950568 |
| NEGREITLAMAEM | TCSADPDNSPLY | 0.46402496 |
| NELVEWSHPLGHRWRWE | TCSADPDNSPLY | 0.5074584 |
| NLGAGGTGGNGGNPG | TCSADPDNSPLY | 0.38428885 |
| NTEWPKWETVMAADPVD | TCSADPDNSPLY | 0.52054393 |
| PASAAPAETQP | TCSADPDNSPLY | 0.35871437 |
| QQSSDGKNGVRAAEITGE | TCSADPDNSPLY | 0.55635303 |
| RAPDAVTTSWWKE | TCSADPDNSPLY | 0.5747021 |
| SGFGNVGAHTSGWFNQ | TCSADPDNSPLY | 0.6517495 |
| SGVAGAVGQAASVGGLKV | TCSADPDNSPLY | 0.37545097 |
| SNGGAGTGSAGGNGGT | TCSADPDNSPLY | 0.5657194 |
| TGGAGGEGGAAG | TCSADPDNSPLY | 0.38487616 |
| TGGDGGAAGAGGAPGHG | TCSADPDNSPLY | 0.6384996 |
| TMPPLLAAIANAGRHGV | TCSADPDNSPLY | 0.48092481 |
| TSSTAGNSGGA | TCSADPDNSPLY | 0.34160218 |
| VPATVPVGILC | TCSADPDNSPLY | 0.56075406 |
| VQRPDSGGVVLTGRIS | TCSADPDNSPLY | 0.4226964 |
| WTPNPVPATKGVPAGLVQAT | TCSADPDNSPLY | 0.4967476 |
| YALKRALAKFSDRFDVVII | TCSADPDNSPLY | 0.62398076 |
| YLIGSVIV | TCSADPDNSPLY | 0.38562685 |
| YGQLVAVID | TCSADPDNSPLY | 0.48233336 |
| VTVGGMTLPA | TCSADPDNSPLY | 0.5525488 |
| VLALPAV | TCSADPDNSPLY | 0.55301267 |
| VIPPTLNYET | TCSADPDNSPLY | 0.44777638 |
| VGMGGMHPSSG | TCSADPDNSPLY | 0.569406 |
| VGLAGGAATA | TCSADPDNSPLY | 0.55538857 |
| VGGAGGAGGSGP | TCSADPDNSPLY | 0.39968356 |
| VFAAFMV | TCSADPDNSPLY | 0.6094819 |
| TLYDKLR | TCSADPDNSPLY | 0.43453616 |
| TLAVFTVL | TCSADPDNSPLY | 0.52743185 |
| TGSVQPARV | TCSADPDNSPLY | 0.5361677 |
| TGGGAGSGGT | TCSADPDNSPLY | 0.6604497 |
| SHLETSL | TCSADPDNSPLY | 0.48273838 |
| SFTTGMT | TCSADPDNSPLY | 0.4761707 |
| QVEVAKRMGVRQPTVS | TCSADPDNSPLY | 0.557765 |
| QGSGGPALAM | TCSADPDNSPLY | 0.55337024 |
| MLAPSNP | TCSADPDNSPLY | 0.3941218 |
| LVNNAAI | TCSADPDNSPLY | 0.58588254 |
| LTGEINTLA | TCSADPDNSPLY | 0.4902686 |
| LSIFGPV | TCSADPDNSPLY | 0.59897065 |
| LLALLANL | TCSADPDNSPLY | 0.42346007 |
| LISNASDALDKLR | TCSADPDNSPLY | 0.5552814 |
| LINEAVP | TCSADPDNSPLY | 0.620039 |
| LGYAFSGL | TCSADPDNSPLY | 0.37919956 |
| LDFVAVAL | TCSADPDNSPLY | 0.5445801 |
| LAQPGAE | TCSADPDNSPLY | 0.5402066 |
| ISNLGAVTGF | TCSADPDNSPLY | 0.6233729 |
| HQEDPGRI | TCSADPDNSPLY | 0.40529123 |

| | | |
|---|---|---|
| GTPGEPTP | TCSADPDNSPLY | 0.55963916 |
| GFFNSGS | TCSADPDNSPLY | 0.5101038 |
| GAGGGGLTAGAN | TCSADPDNSPLY | 0.55428225 |
| FVDALVNM | TCSADPDNSPLY | 0.3813128 |
| FLTVAPEL | TCSADPDNSPLY | 0.45898658 |
| FETLTAL | TCSADPDNSPLY | 0.4204257 |
| DVINPIEL | TCSADPDNSPLY | 0.5241016 |
| APVSGAPV | TCSADPDNSPLY | 0.5596885 |
| ALVGGLAW | TCSADPDNSPLY | 0.5555304 |
| ALAAAAATAI | TCSADPDNSPLY | 0.43160427 |
| AFGLGLLL | TCSADPDNSPLY | 0.57047856 |
| AFFAAASL | TCSADPDNSPLY | 0.5742528 |
| AAAALPAFSPP | TCSADPDNSPLY | 0.6489883 |
| VVLAPAL | TCSADPDNSPLY | 0.44440943 |
| TIGHVDHGKTTL | TCSADPDNSPLY | 0.5910842 |
| LISNASDALDK | TCSADPDNSPLY | 0.5454182 |
| LISNASDAL | TCSADPDNSPLY | 0.44488543 |
| ISNASDALDK | TCSADPDNSPLY | 0.35394886 |
| NQAETLVYQTE | TCSADPDNSPLY | 0.39887825 |
| TGGTGGTAGAG | TCSADPDNSPLY | 0.5174695 |
| TGSTAGGAGGA | TCSADPDNSPLY | 0.4026918 |
| DPLAVPVAAGPV | TCSADPDNSPLY | 0.62561893 |
| GGTGGTPGNGGN | TCSADPDNSPLY | 0.6130089 |
| PAGLAGVDPATL | TCSADPDNSPLY | 0.5981151 |
| GNNLVVEAPHAGASV | TCSADPDNSPLY | 0.59891486 |
| KVTRIMSASTLEQPA | TCSADPDNSPLY | 0.5445061 |
| LGFALPIALPRRTVV | TCSADPDNSPLY | 0.5149879 |
| TRLGNLTARKLVVLV | TCSADPDNSPLY | 0.46612138 |
| WSQQRRLIYFPSAGP | TCSADPDNSPLY | 0.51324195 |
| GPNGNPDPMAAAVDIR | TCSADPDNSPLY | 0.3398227 |
| TQRCVSVADLRAAASR | TCSADPDNSPLY | 0.42229488 |
| QFVLDVAEHPVEVLTETV | TCSADPDNSPLY | 0.44106007 |
| GGAGGTGGAGGNGGTQAGDG | TCSADPDNSPLY | 0.5201492 |
| PLLATLRAIAVDVDAKPAQV | TCSADPDNSPLY | 0.3251934 |
| APKFRTGGYMIDFWGVGYQVAK | TCSADPDNSPLY | 0.60380554 |
| GAGGAGGAGGAAGGPGGTGGTG | TCSADPDNSPLY | 0.6000459 |
| IGALGLLATDVAPAGSVAFGVA | TCSADPDNSPLY | 0.55767393 |
| AGGAGGIGGNGGAGGAGGVLMG | TCSADPDNSPLY | 0.57170993 |
| ADELIGLL | TCSADPDNSPLY | 0.49987137 |
| ADRALKA | TCSADPDNSPLY | 0.36002544 |
| AGAVGQVGGAGGA | TCSADPDNSPLY | 0.5615477 |
| AGGGGGLGGAP | TCSADPDNSPLY | 0.416314 |
| AGGLLYGNGGAG | TCSADPDNSPLY | 0.55683947 |
| AGLESIGL | TCSADPDNSPLY | 0.5830866 |
| AGLKTGII | TCSADPDNSPLY | 0.54710734 |
| ASLAALLL | TCSADPDNSPLY | 0.5325285 |
| ASVINISSI | TCSADPDNSPLY | 0.511575 |
| ATPPGPA | TCSADPDNSPLY | 0.4867826 |
| ATVASALV | TCSADPDNSPLY | 0.54369885 |
| ATVASLIL | TCSADPDNSPLY | 0.43223193 |
| AVNGPVLIRH | TCSADPDNSPLY | 0.51267016 |

| | | |
|---|---|---|
| DGAAGAVGI | TCSADPDNSPLY | 0.39063546 |
| DPLLHGI | TCSADPDNSPLY | 0.5880078 |
| ELGAALGAL | TCSADPDNSPLY | 0.5467701 |
| ELGRTQAV | TCSADPDNSPLY | 0.48070866 |
| FVAGGYLS | TCSADPDNSPLY | 0.45517328 |
| GASGGAGGAGGW | TCSADPDNSPLY | 0.62356204 |
| GGAGAGGDGGAGGI | TCSADPDNSPLY | 0.61818117 |
| GTGHNGSQPGMGGNG | TCSADPDNSPLY | 0.4314489 |
| GYAAASASAS | TCSADPDNSPLY | 0.44229388 |
| HQVDVAAVQQG | TCSADPDNSPLY | 0.4155192 |
| IGAAIAHT | TCSADPDNSPLY | 0.44491443 |
| IGMIEVAQL | TCSADPDNSPLY | 0.60585505 |
| IRLAPEHTSAMD | TCSADPDNSPLY | 0.51847875 |
| IYNVSVL | TCSADPDNSPLY | 0.40328744 |
| KQGFLGM | TCSADPDNSPLY | 0.44708905 |
| LESVITQL | TCSADPDNSPLY | 0.52631676 |
| LGLGALVVV | TCSADPDNSPLY | 0.4698971 |
| LIGNGGNGGSGGI | TCSADPDNSPLY | 0.34368467 |
| LNAINAPTL | TCSADPDNSPLY | 0.4527214 |
| LQLLNTGV | TCSADPDNSPLY | 0.4006951 |
| LQTSIEF | TCSADPDNSPLY | 0.42220867 |
| MSFAVSEL | TCSADPDNSPLY | 0.42125574 |
| NTGALGL | TCSADPDNSPLY | 0.5919304 |
| PPGAGGAGGAGTT | TCSADPDNSPLY | 0.58277965 |
| QLVGLGI | TCSADPDNSPLY | 0.50888014 |
| RFLEQVHQL | TCSADPDNSPLY | 0.5716555 |
| RGVGYVLR | TCSADPDNSPLY | 0.58455265 |
| SATGVFAGVI | TCSADPDNSPLY | 0.5790353 |
| SLPAITLP | TCSADPDNSPLY | 0.5748077 |
| STALAFTV | TCSADPDNSPLY | 0.43819314 |
| SVLTVGLGK | TCSADPDNSPLY | 0.52584445 |
| TALAGAGSGI | TCSADPDNSPLY | 0.42294964 |
| TGLPFTGI | TCSADPDNSPLY | 0.5294207 |
| VAEHVTV | TCSADPDNSPLY | 0.5267943 |
| VDIVLNSL | TCSADPDNSPLY | 0.44604695 |
| VLTGGGAVL | TCSADPDNSPLY | 0.55320334 |
| VNVGLAYGI | TCSADPDNSPLY | 0.5663953 |
| VSSAAVGAV | TCSADPDNSPLY | 0.5768005 |
| VTSVYEAGV | TCSADPDNSPLY | 0.39789882 |
| WVALDRGA | TCSADPDNSPLY | 0.34815258 |
| YLVGSVVI | TCSADPDNSPLY | 0.42113158 |
| GPPGTGKTLL | TCSADPDNSPLY | 0.4762398 |
| GVGMTKF | TCSADPDNSPLY | 0.5705036 |
| IGIGGSDL | TCSADPDNSPLY | 0.5008347 |
| IGTLLAEL | TCSADPDNSPLY | 0.6529633 |
| IYTSGTT | TCSADPDNSPLY | 0.60096014 |
| LSFPAGV | TCSADPDNSPLY | 0.4885112 |
| LYTSGTT | TCSADPDNSPLY | 0.5641027 |
| VGAFPFGF | TCSADPDNSPLY | 0.39307758 |
| VGAGTVEF | TCSADPDNSPLY | 0.41232002 |
| VGKSTLLL | TCSADPDNSPLY | 0.4302687 |

| | | |
|---|---|---|
| AALAHAGGGIV | TCSADPDNSPLY | 0.42025816 |
| AALVAVSLL | TCSADPDNSPLY | 0.46279028 |
| AAVSRAVAAL | TCSADPDNSPLY | 0.41208524 |
| AGGSSGIGGF | TCSADPDNSPLY | 0.36605301 |
| AGLTVANANG | TCSADPDNSPLY | 0.61496425 |
| AGLVGAVL | TCSADPDNSPLY | 0.50439626 |
| ALSQAFTV | TCSADPDNSPLY | 0.46147686 |
| ALTFSVSL | TCSADPDNSPLY | 0.4481948 |
| ALTGAALVAL | TCSADPDNSPLY | 0.62163126 |
| ASFSIGAGVL | TCSADPDNSPLY | 0.62836164 |
| DGSVYKI | TCSADPDNSPLY | 0.44106895 |
| EQLAEIV | TCSADPDNSPLY | 0.4513597 |
| FAVSELTVM | TCSADPDNSPLY | 0.6062328 |
| GAAPLAAGL | TCSADPDNSPLY | 0.50120443 |
| GADGGDGGSGGAGG | TCSADPDNSPLY | 0.45782876 |
| GFGLDTGGAGG | TCSADPDNSPLY | 0.5232562 |
| GLASGGLL | TCSADPDNSPLY | 0.47188932 |
| GRIAGAAVLL | TCSADPDNSPLY | 0.41644007 |
| GSAGSLNTSF | TCSADPDNSPLY | 0.54037404 |
| GTGQIGI | TCSADPDNSPLY | 0.48617023 |
| GTLIALIL | TCSADPDNSPLY | 0.4733385 |
| IAATMGFVVG | TCSADPDNSPLY | 0.38175687 |
| IFAQALL | TCSADPDNSPLY | 0.551837 |
| IGAAVALGF | TCSADPDNSPLY | 0.40536064 |
| IGLTPLAL | TCSADPDNSPLY | 0.37119612 |
| ILGSNTSTL | TCSADPDNSPLY | 0.3235492 |
| IVFSLTVL | TCSADPDNSPLY | 0.45488048 |
| LASKATGF | TCSADPDNSPLY | 0.534734 |
| LASVALIL | TCSADPDNSPLY | 0.59055483 |
| LFAVGLTL | TCSADPDNSPLY | 0.59007066 |
| LFNGGIGTYI | TCSADPDNSPLY | 0.5469592 |
| LTLGAGII | TCSADPDNSPLY | 0.56412625 |
| LVAAALGDI | TCSADPDNSPLY | 0.44858074 |
| NTAFVQLGI | TCSADPDNSPLY | 0.380788 |
| QLFSGAEL | TCSADPDNSPLY | 0.5290178 |
| SAATFITL | TCSADPDNSPLY | 0.58507544 |
| SNYGLANLGV | TCSADPDNSPLY | 0.53053147 |
| SYGGAYAVM | TCSADPDNSPLY | 0.56896245 |
| TTGVGGLTLG | TCSADPDNSPLY | 0.35777515 |
| TTQTIYI | TCSADPDNSPLY | 0.582907 |
| VAAGAGLGAVI | TCSADPDNSPLY | 0.5551438 |
| VAAVVTLL | TCSADPDNSPLY | 0.35063672 |
| VGFLGGLI | TCSADPDNSPLY | 0.4065526 |
| VGVIPILL | TCSADPDNSPLY | 0.3906109 |
| VIFASSLI | TCSADPDNSPLY | 0.36820796 |
| VKGGDGGAAAG | TCSADPDNSPLY | 0.5667363 |
| VTKTFTNI | TCSADPDNSPLY | 0.43143898 |
| VVAAAILL | TCSADPDNSPLY | 0.5672204 |
| YAALDVEL | TCSADPDNSPLY | 0.42105666 |
| LASVALLL | TCSADPDNSPLY | 0.5018415 |
| LDAVHRV | TCSADPDNSPLY | 0.5022519 |

| | | |
|---|---|---|
| QGLSSSIF | TCSADPDNSPLY | 0.5337373 |
| RGTFANIRL | TCSADPDNSPLY | 0.5563717 |
| RQLWWGHRI | TCSADPDNSPLY | 0.5356133 |
| YGPPGTGKTLL | TCSADPDNSPLY | 0.41293272 |
| AALVGAPANVA | TCSADPDNSPLY | 0.44560698 |
| AATFRRTE | TCSADPDNSPLY | 0.6129848 |
| AGALGGGI | TCSADPDNSPLY | 0.39792213 |
| AGGAGGAGGIGAG | TCSADPDNSPLY | 0.43407413 |
| AGSGALGL | TCSADPDNSPLY | 0.48170918 |
| AQIADQI | TCSADPDNSPLY | 0.5338669 |
| ASAVAVTYL | TCSADPDNSPLY | 0.5752105 |
| ASTVLGGTGG | TCSADPDNSPLY | 0.5643211 |
| AVGGEPAK | TCSADPDNSPLY | 0.52962893 |
| CEQGTFGGTC | TCSADPDNSPLY | 0.59788704 |
| DEGKAPAG | TCSADPDNSPLY | 0.59604275 |
| DTVVFPAL | TCSADPDNSPLY | 0.554413 |
| FEGITMAG | TCSADPDNSPLY | 0.40824735 |
| FPLVVQD | TCSADPDNSPLY | 0.6226506 |
| GAFTTPQ | TCSADPDNSPLY | 0.36818662 |
| GFGGFGTAGGVGGA | TCSADPDNSPLY | 0.52512985 |
| GFLAQFL | TCSADPDNSPLY | 0.53469104 |
| GGAGGAGGNAGVA | TCSADPDNSPLY | 0.39881766 |
| GGVGTAGAGGAGGAGG | TCSADPDNSPLY | 0.49260867 |
| GNGGGGDAGNAGSGGN | TCSADPDNSPLY | 0.5895101 |
| GTGGAGGAGNDAGS | TCSADPDNSPLY | 0.5561105 |
| GTTNTGWFNS | TCSADPDNSPLY | 0.53330266 |
| IFMGPLFTL | TCSADPDNSPLY | 0.4058471 |
| ILAHAIEAI | TCSADPDNSPLY | 0.47737622 |
| IRSAIREAL | TCSADPDNSPLY | 0.38449353 |
| ITFAGGG | TCSADPDNSPLY | 0.36161503 |
| ITGMSGHQEV | TCSADPDNSPLY | 0.5290208 |
| IVAGLACTL | TCSADPDNSPLY | 0.5045561 |
| IVGGGLAGA | TCSADPDNSPLY | 0.63827264 |
| IVTGVLNI | TCSADPDNSPLY | 0.55494845 |
| LAQQGCI | TCSADPDNSPLY | 0.6135954 |
| LCLHVEN | TCSADPDNSPLY | 0.42459047 |
| LDAISDGS | TCSADPDNSPLY | 0.52613676 |
| LGGFAAML | TCSADPDNSPLY | 0.34254265 |
| LGGKNPMI | TCSADPDNSPLY | 0.537479 |
| LGPKGVAAIL | TCSADPDNSPLY | 0.38771427 |
| LMVGVSI | TCSADPDNSPLY | 0.5501552 |
| LVGAAFVL | TCSADPDNSPLY | 0.37158367 |
| LVLAVAPL | TCSADPDNSPLY | 0.5551522 |
| MAAIAVIDAA | TCSADPDNSPLY | 0.44632548 |
| NNGINAELQ | TCSADPDNSPLY | 0.5795733 |
| RGEPLVLVHG | TCSADPDNSPLY | 0.48708695 |
| RGGAGGAAGLI | TCSADPDNSPLY | 0.39179456 |
| RSYRMHPAV | TCSADPDNSPLY | 0.45721766 |
| SAAHLLAGQLA | TCSADPDNSPLY | 0.5953499 |
| SAAVGSY | TCSADPDNSPLY | 0.38207185 |
| SPHAVLHRI | TCSADPDNSPLY | 0.33682412 |

| | | |
|---|---|---|
| SQIAKVL | TCSADPDNSPLY | 0.40199056 |
| TALGHLVP | TCSADPDNSPLY | 0.49282566 |
| TAPTTSGAA | TCSADPDNSPLY | 0.5906681 |
| TDGDDGNGGSPGT | TCSADPDNSPLY | 0.57618845 |
| TDGLGEN | TCSADPDNSPLY | 0.48856992 |
| TGGPPLPGQA | TCSADPDNSPLY | 0.64225364 |
| TIPAITI | TCSADPDNSPLY | 0.31330457 |
| TIPGFTI | TCSADPDNSPLY | 0.48992482 |
| TSIASGALA | TCSADPDNSPLY | 0.6352629 |
| TVTLPGAAI | TCSADPDNSPLY | 0.49845052 |
| VASGLSAALP | TCSADPDNSPLY | 0.514855 |
| VGAGVIGI | TCSADPDNSPLY | 0.58276546 |
| VGPKRFGV | TCSADPDNSPLY | 0.47554415 |
| VNVAFPD | TCSADPDNSPLY | 0.49642903 |
| VSMTNSGVS | TCSADPDNSPLY | 0.5408412 |
| VTDVLGDLSV | TCSADPDNSPLY | 0.47771648 |
| VVVVGAGFAGL | TCSADPDNSPLY | 0.3973205 |
| VYTVGTVV | TCSADPDNSPLY | 0.35852385 |
| WTTFSGP | TCSADPDNSPLY | 0.54466397 |
| YDELVEAF | TCSADPDNSPLY | 0.6207609 |
| YGTGGAGGNGG | TCSADPDNSPLY | 0.48518938 |
| YTNVMAV | TCSADPDNSPLY | 0.58013654 |
| FLEQVHQL | TCSADPDNSPLY | 0.5068831 |
| GLSLLAVAV | TCSADPDNSPLY | 0.5944731 |
| LEQVHQL | TCSADPDNSPLY | 0.43096521 |
| LNAAGAI | TCSADPDNSPLY | 0.36434415 |
| PAVHASL | TCSADPDNSPLY | 0.63094974 |
| TELEDDA | TCSADPDNSPLY | 0.37915972 |
| TPPGPPN | TCSADPDNSPLY | 0.46568564 |
| TPPPPPPP | TCSADPDNSPLY | 0.45313805 |
| TQSPALI | ASSRGQNNSPLY | 0.63267756 |
| EFAAAFPQ | ASSRGQNNSPLY | 0.5162412 |
| LLSGNLDVL | ASSRGQNNSPLY | 0.5206769 |
| GPANLGNYNI | ASSRGQNNSPLY | 0.48661682 |
| TLGPTLP | ASSRGQNNSPLY | 0.46515846 |
| VYAYSSCCK | ASSRGQNNSPLY | 0.5436124 |
| APAPAPAPNI | ASSRGQNNSPLY | 0.36528245 |
| PGGPVPIP | ASSRGQNNSPLY | 0.59547746 |
| FLTVALEP | ASSRGQNNSPLY | 0.3938376 |
| SQYFGANL | ASSRGQNNSPLY | 0.41622016 |
| GGVKPAA | ASSRGQNNSPLY | 0.48939383 |
| GGTGGAGGAGGW | ASSRGQNNSPLY | 0.6452037 |
| GVGGVAGLGGAGG | ASSRGQNNSPLY | 0.44046047 |
| VLLVLATTI | ASSRGQNNSPLY | 0.62921095 |
| IGIFAAML | ASSRGQNNSPLY | 0.43614662 |
| AVGDLID | ASSRGQNNSPLY | 0.5096445 |
| NISPLPL | ASSRGQNNSPLY | 0.61352026 |
| VTAVFVAI | ASSRGQNNSPLY | 0.604106 |
| SGDDNDAAL | ASSRGQNNSPLY | 0.64437544 |
| AVAALAHS | ASSRGQNNSPLY | 0.48344535 |
| GGVTVMATP | ASSRGQNNSPLY | 0.34277096 |

| | | |
|---|---|---|
| AGLAANVRD | ASSRGQNNSPLY | 0.4583399 |
| TNRPDLIDPALLRPG | ASSRGQNNSPLY | 0.5504333 |
| LANLGVTAL | ASSRGQNNSPLY | 0.549374 |
| TGERIEL | ASSRGQNNSPLY | 0.5900551 |
| NGEELAA | ASSRGQNNSPLY | 0.5455121 |
| VLSLVWG | ASSRGQNNSPLY | 0.44830126 |
| IGYALWL | ASSRGQNNSPLY | 0.5902349 |
| LSLLLAPL | ASSRGQNNSPLY | 0.6326645 |
| TLDEALK | ASSRGQNNSPLY | 0.56605494 |
| TDGAGTAEGG | ASSRGQNNSPLY | 0.3808296 |
| NAPGGAGGNGG | ASSRGQNNSPLY | 0.37525627 |
| VSVFMGMGGNF | ASSRGQNNSPLY | 0.5325932 |
| LASYLAAL | ASSRGQNNSPLY | 0.5270799 |
| LVGGTPAL | ASSRGQNNSPLY | 0.37215447 |
| VRNADVL | ASSRGQNNSPLY | 0.46975303 |
| TCSSPFLVSAM | ASSRGQNNSPLY | 0.5288386 |
| VHITDAE | ASSRGQNNSPLY | 0.33893445 |
| EYAAPSF | ASSRGQNNSPLY | 0.46456206 |
| VVDAFHGGL | ASSRGQNNSPLY | 0.5785781 |
| ASVVDPQ | ASSRGQNNSPLY | 0.44761428 |
| SYNEFGGLS | ASSRGQNNSPLY | 0.36874363 |
| DPDGSVY | ASSRGQNNSPLY | 0.6010175 |
| MVSTIHATE | ASSRGQNNSPLY | 0.6175111 |
| LAAGVADPV | ASSRGQNNSPLY | 0.57698345 |
| TKEQIAA | ASSRGQNNSPLY | 0.39142913 |
| SLASPAAL | ASSRGQNNSPLY | 0.31126693 |
| IGLVFYGV | ASSRGQNNSPLY | 0.58480096 |
| FLLDASNL | ASSRGQNNSPLY | 0.62981135 |
| YQVGNFP | ASSRGQNNSPLY | 0.5197927 |
| ESLLSQST | ASSRGQNNSPLY | 0.44315302 |
| LFTVGFAAVG | ASSRGQNNSPLY | 0.40660784 |
| ATPPTPT | ASSRGQNNSPLY | 0.544612 |
| TDLATLG | ASSRGQNNSPLY | 0.38380772 |
| SAGGTGR | ASSRGQNNSPLY | 0.35324666 |
| GFSNSGS | ASSRGQNNSPLY | 0.4127354 |
| GGNGGSGGTSV | ASSRGQNNSPLY | 0.551561 |
| GTSVATGGAGN | ASSRGQNNSPLY | 0.37846982 |
| NGGNGGSGGTS | ASSRGQNNSPLY | 0.4136477 |
| AVVKFPH | ASSRGQNNSPLY | 0.6099242 |
| TPPPPPP | ASSRGQNNSPLY | 0.5230425 |
| AAPAAAAAPA | ASSRGQNNSPLY | 0.51943725 |
| IALNEDL | ASSRGQNNSPLY | 0.4059369 |
| IVNVSSM | ASSRGQNNSPLY | 0.5364881 |
| TGIPPAPR | ASSRGQNNSPLY | 0.38877654 |
| FDLGGGTF | ASSRGQNNSPLY | 0.53108454 |
| FDLGGGTFDVS | ASSRGQNNSPLY | 0.5355566 |
| NEPTAAA | ASSRGQNNSPLY | 0.5230891 |
| NEPTAAALA | ASSRGQNNSPLY | 0.46421063 |
| GSIINISSI | ASSRGQNNSPLY | 0.4013994 |
| VAPGNAGTA | ASSRGQNNSPLY | 0.39079657 |
| TGGGASLE | ASSRGQNNSPLY | 0.46996993 |

| | | |
|---|---|---|
| VAVYNHYKRI | ASSRGQNNSPLY | 0.61346745 |
| TDPAPATT | ASSRGQNNSPLY | 0.56225157 |
| TNRPDILDPALLRPG | ASSRGQNNSPLY | 0.56210923 |
| LTTTTTS | ASSRGQNNSPLY | 0.5651648 |
| PTGSGKTL | ASSRGQNNSPLY | 0.3893299 |
| TGSGKTL | ASSRGQNNSPLY | 0.4687815 |
| TIGHVDHGK | ASSRGQNNSPLY | 0.37369418 |
| TIGHVDHGKTT | ASSRGQNNSPLY | 0.62108326 |
| GHVDHGKTT | ASSRGQNNSPLY | 0.3925341 |
| AQFEHTLL | ASSRGQNNSPLY | 0.49421394 |
| IVNQDEL | ASSRGQNNSPLY | 0.47795042 |
| TNGSQFF | ASSRGQNNSPLY | 0.45524174 |
| TGKGIVE | ASSRGQNNSPLY | 0.55520314 |
| LISNASDA | ASSRGQNNSPLY | 0.4356779 |
| AELRTLGELTKIAKAHGAQ | ASSRGQNNSPLY | 0.48922664 |
| APNAVAAAEPSW | ASSRGQNNSPLY | 0.5750275 |
| APQVAGAATAVL | ASSRGQNNSPLY | 0.49492565 |
| ARRDPAAAAAILPTDA | ASSRGQNNSPLY | 0.36110097 |
| DAAARDVLAEAGLAEYF | ASSRGQNNSPLY | 0.5647179 |
| DDQIVLSAKANLAKVEQ | ASSRGQNNSPLY | 0.39073014 |
| DELLPDTGEDSDEDGH | ASSRGQNNSPLY | 0.5342705 |
| ELPEAPANRATVPQD | ASSRGQNNSPLY | 0.46766013 |
| FGQSKIERVTGIG | ASSRGQNNSPLY | 0.46452945 |
| GAGGTGSTAGGAGGAGGAGGLY | ASSRGQNNSPLY | 0.49193794 |
| GHGGTGAAGGNGGTGGAGGSGI | ASSRGQNNSPLY | 0.36327428 |
| GNGGNGGAGGEGGPGGAGGAGA | ASSRGQNNSPLY | 0.4356977 |
| GSTAGNGGNGGNATNGTIAGQP | ASSRGQNNSPLY | 0.42934075 |
| GTGGDGGNGGNGGTGG | ASSRGQNNSPLY | 0.58146495 |
| GVRRPAGIPADAPV | ASSRGQNNSPLY | 0.54366803 |
| GYRQVTDAHLLTLARR | ASSRGQNNSPLY | 0.37906283 |
| IAKMGQDGHDRG | ASSRGQNNSPLY | 0.54682314 |
| KAEPSIEIQYT | ASSRGQNNSPLY | 0.44939917 |
| KPVRIRMAADEDTVAVT | ASSRGQNNSPLY | 0.4421257 |
| MPPRIAGMRLLVIKPEP | ASSRGQNNSPLY | 0.4342314 |
| MKFYAVALFRSRGVPPTR | ASSRGQNNSPLY | 0.40300226 |
| NAGGFGILGAVAHSP | ASSRGQNNSPLY | 0.36180404 |
| NEGREITLAMAEM | ASSRGQNNSPLY | 0.49531627 |
| NELVEWSHPLGHRWRWE | ASSRGQNNSPLY | 0.42937785 |
| NLGAGGTGGNGGNPG | ASSRGQNNSPLY | 0.53061587 |
| NTEWPKWETVMAADPVD | ASSRGQNNSPLY | 0.5338373 |
| PASAAPAETQP | ASSRGQNNSPLY | 0.39629218 |
| QQSSDGKNGVRAAEITGE | ASSRGQNNSPLY | 0.4983356 |
| RAPDAVTTSWWKE | ASSRGQNNSPLY | 0.46446127 |
| SGFGNVGAHTSGWFNQ | ASSRGQNNSPLY | 0.5322208 |
| SGVAGAVGQAASVGGLKV | ASSRGQNNSPLY | 0.5526364 |
| SNGGAGTGSAGGNGGT | ASSRGQNNSPLY | 0.40982947 |
| TGGAGGEGGAAG | ASSRGQNNSPLY | 0.52720547 |
| TGGDGGAAGAGGAPGHG | ASSRGQNNSPLY | 0.597903 |
| TMPPLLAAIANAGRHGV | ASSRGQNNSPLY | 0.55369884 |
| TSSTAGNSGGA | ASSRGQNNSPLY | 0.47222692 |
| VPATVPVGILC | ASSRGQNNSPLY | 0.6301217 |

| | | |
|---|---|---|
| VQRPDSGGVVLTGRIS | ASSRGQNNSPLY | 0.5707221 |
| WTPNPVPATKGVPAGLVQAT | ASSRGQNNSPLY | 0.4532865 |
| YALKRALAKFSDRFDVVII | ASSRGQNNSPLY | 0.326928 |
| YLIGSVIV | ASSRGQNNSPLY | 0.5282347 |
| YGQLVAVID | ASSRGQNNSPLY | 0.53023016 |
| VTVGGMTLPA | ASSRGQNNSPLY | 0.39150488 |
| VLALPAV | ASSRGQNNSPLY | 0.3476455 |
| VIPPTLNYET | ASSRGQNNSPLY | 0.569405 |
| VGMGGMHPSSG | ASSRGQNNSPLY | 0.627559 |
| VGLAGGAATA | ASSRGQNNSPLY | 0.52861446 |
| VGGAGGAGGSGP | ASSRGQNNSPLY | 0.57723296 |
| VFAAFMV | ASSRGQNNSPLY | 0.34613568 |
| TLYDKLR | ASSRGQNNSPLY | 0.5890399 |
| TLAVFTVL | ASSRGQNNSPLY | 0.5746785 |
| TGSVQPARV | ASSRGQNNSPLY | 0.39284843 |
| TGGGAGSGGT | ASSRGQNNSPLY | 0.55492467 |
| SHLETSL | ASSRGQNNSPLY | 0.45166036 |
| SFTTGMT | ASSRGQNNSPLY | 0.4107944 |
| QVEVAKRMGVRQPTVS | ASSRGQNNSPLY | 0.58450174 |
| QGSGGPALAM | ASSRGQNNSPLY | 0.55053985 |
| MLAPSNP | ASSRGQNNSPLY | 0.50190425 |
| LVNNAAI | ASSRGQNNSPLY | 0.5021111 |
| LTGEINTLA | ASSRGQNNSPLY | 0.42297062 |
| LSIFGPV | ASSRGQNNSPLY | 0.45048264 |
| LLALLANL | ASSRGQNNSPLY | 0.55785966 |
| LISNASDALDKLR | ASSRGQNNSPLY | 0.6038264 |
| LINEAVP | ASSRGQNNSPLY | 0.6175773 |
| LGYAFSGL | ASSRGQNNSPLY | 0.43182456 |
| LDFVAVAL | ASSRGQNNSPLY | 0.62098265 |
| LAQPGAE | ASSRGQNNSPLY | 0.6325903 |
| ISNLGAVTGF | ASSRGQNNSPLY | 0.51992476 |
| HQEDPGRI | ASSRGQNNSPLY | 0.3961531 |
| GTPGEPTP | ASSRGQNNSPLY | 0.61785966 |
| GFFNSGS | ASSRGQNNSPLY | 0.4985343 |
| GAGGGGLTAGAN | ASSRGQNNSPLY | 0.5855328 |
| FVDALVNM | ASSRGQNNSPLY | 0.5623739 |
| FLTVAPEL | ASSRGQNNSPLY | 0.24434164 |
| FETLTAL | ASSRGQNNSPLY | 0.7943847 |
| DVINPIEL | ASSRGQNNSPLY | 0.6780838 |
| APVSGAPV | ASSRGQNNSPLY | 0.43069994 |
| ALVGGLAW | ASSRGQNNSPLY | 0.23480451 |
| ALAAAAATAI | ASSRGQNNSPLY | 0.1598566 |
| AFGLGLLL | ASSRGQNNSPLY | 0.38072756 |
| AFFAAASL | ASSRGQNNSPLY | 0.24480718 |
| AAAALPAFSPP | ASSRGQNNSPLY | 0.773955 |
| VVLAPAL | ASSRGQNNSPLY | 0.67564607 |
| TIGHVDHGKTTL | ASSRGQNNSPLY | 0.256663 |
| LISNASDALDK | ASSRGQNNSPLY | 0.7185437 |
| LISNASDAL | ASSRGQNNSPLY | 0.5168367 |
| ISNASDALDK | ASSRGQNNSPLY | 0.20081383 |
| NQAETLVYQTE | ASSRGQNNSPLY | 0.52873516 |

| | | |
|---|---|---|
| TGGTGGTAGAG | ASSRGQNNSPLY | 0.2261921 |
| TGSTAGGAGGA | ASSRGQNNSPLY | 0.39213127 |
| DPLAVPVAAGPV | ASSRGQNNSPLY | 0.23544133 |
| GGTGGTPGNGGN | ASSRGQNNSPLY | 0.6004848 |
| PAGLAGVDPATL | ASSRGQNNSPLY | 0.39862865 |
| GNNLVVEAPHAGASV | ASSRGQNNSPLY | 0.16487673 |
| KVTRIMSASTLEQPA | ASSRGQNNSPLY | 0.6391562 |
| LGFALPIALPRRTVV | ASSRGQNNSPLY | 0.3912058 |
| TRLGNLTARKLVVLV | ASSRGQNNSPLY | 0.4031852 |
| WSQQRRLIYFPSAGP | ASSRGQNNSPLY | 0.6741808 |
| GPNGNPDPMAAAVDIR | ASSRGQNNSPLY | 0.33865902 |
| TQRCVSVADLRAAASR | ASSRGQNNSPLY | 0.36432624 |
| QFVLDVAEHPVEVLTETV | ASSRGQNNSPLY | 0.7462945 |
| GGAGGTGGAGGNGGTQAGDG | ASSRGQNNSPLY | 0.68087643 |
| PLLATLRAIAVDVDAKPAQV | ASSRGQNNSPLY | 0.24327418 |
| APKFRTGGYMIDFWGVGYQVAK | ASSRGQNNSPLY | 0.51095855 |
| GAGGAGGAGGAAGGPGGTGGTG | ASSRGQNNSPLY | 0.67130226 |
| IGALGLLATDVAPAGSVAFGVA | ASSRGQNNSPLY | 0.4137196 |
| AGGAGGIGGNGGAGGAGGVLMG | ASSRGQNNSPLY | 0.6555939 |
| ADELIGLL | ASSRGQNNSPLY | 0.66070807 |
| ADRALKA | ASSRGQNNSPLY | 0.29444098 |
| AGAVGQVGGAGGA | ASSRGQNNSPLY | 0.72135156 |
| AGGGGGLGGAP | ASSRGQNNSPLY | 0.81885237 |
| AGGLLYGNGGAG | ASSRGQNNSPLY | 0.3896176 |
| AGLESIGL | ASSRGQNNSPLY | 0.40740803 |
| AGLKTGII | ASSRGQNNSPLY | 0.21670663 |
| ASLAALLL | ASSRGQNNSPLY | 0.5206549 |
| ASVINISSI | ASSRGQNNSPLY | 0.18371952 |
| ATPPGPA | ASSRGQNNSPLY | 0.62472117 |
| ATVASALV | ASSRGQNNSPLY | 0.6520116 |
| ATVASLIL | ASSRGQNNSPLY | 0.43193275 |
| AVNGPVLIRH | ASSRGQNNSPLY | 0.25779712 |
| DGAAGAVGI | ASSRGQNNSPLY | 0.1966008 |
| DPLLHGI | ASSRGQNNSPLY | 0.5953966 |
| ELGAALGAL | ASSRGQNNSPLY | 0.20745896 |
| ELGRTQAV | ASSRGQNNSPLY | 0.19737734 |
| FVAGGYLS | ASSRGQNNSPLY | 0.34362227 |
| GASGGAGGAGGW | ASSRGQNNSPLY | 0.52685106 |
| GGAGAGGDGGAGGI | ASSRGQNNSPLY | 0.52135456 |
| GTGHNGSQPGMGGNG | ASSRGQNNSPLY | 0.327817 |
| GYAAASASAS | ASSRGQNNSPLY | 0.7797447 |
| HQVDVAAVQQG | ASSRGQNNSPLY | 0.7953714 |
| IGAAIAHT | ASSRGQNNSPLY | 0.62277025 |
| IGMIEVAQL | ASSRGQNNSPLY | 0.33291638 |
| IRLAPEHTSAMD | ASSRGQNNSPLY | 0.2644734 |
| IYNVSVL | ASSRGQNNSPLY | 0.810885 |
| KQGFLGM | ASSRGQNNSPLY | 0.31216556 |
| LESVITQL | ASSRGQNNSPLY | 0.57634264 |
| LGLGALVVV | ASSRGQNNSPLY | 0.5698345 |
| LIGNGGNGGSGGI | ASSRGQNNSPLY | 0.5057342 |
| LNAINAPTL | ASSRGQNNSPLY | 0.29601032 |

| | | |
|---|---|---|
| LQLLNTGV | ASSRGQNNSPLY | 0.6766817 |
| LQTSIEF | ASSRGQNNSPLY | 0.72989243 |
| MSFAVSEL | ASSRGQNNSPLY | 0.29293567 |
| NTGALGL | ASSRGQNNSPLY | 0.20287624 |
| PPGAGGAGGAGTT | ASSRGQNNSPLY | 0.34929532 |
| QLVGLGI | ASSRGQNNSPLY | 0.42463005 |
| RFLEQVHQL | ASSRGQNNSPLY | 0.65536976 |
| RGVGYVLR | ASSRGQNNSPLY | 0.51889336 |
| SATGVFAGVI | ASSRGQNNSPLY | 0.4035815 |
| SLPAITLP | ASSRGQNNSPLY | 0.638544 |
| STALAFTV | ASSRGQNNSPLY | 0.44159687 |
| SVLTVGLGK | ASSRGQNNSPLY | 0.4365833 |
| TALAGAGSGI | ASSRGQNNSPLY | 0.6589227 |
| TGLPFTGI | ASSRGQNNSPLY | 0.37700635 |
| VAEHVTV | ASSRGQNNSPLY | 0.68347085 |
| VDIVLNSL | ASSRGQNNSPLY | 0.6002752 |
| VLTGGGAVL | ASSRGQNNSPLY | 0.4765094 |
| VNVGLAYGI | ASSRGQNNSPLY | 0.6836904 |
| VSSAAVGAV | ASSRGQNNSPLY | 0.46802932 |
| VTSVYEAGV | ASSRGQNNSPLY | 0.44019014 |
| WVALDRGA | ASSRGQNNSPLY | 0.16444537 |
| YLVGSVVI | ASSRGQNNSPLY | 0.22327499 |
| GPPGTGKTLL | ASSRGQNNSPLY | 0.65307534 |
| GVGMTKF | ASSRGQNNSPLY | 0.6161388 |
| IGIGGSDL | ASSRGQNNSPLY | 0.70145154 |
| IGTLLAEL | ASSRGQNNSPLY | 0.47483575 |
| IYTSGTT | ASSRGQNNSPLY | 0.38841134 |
| LSFPAGV | ASSRGQNNSPLY | 0.73889303 |
| LYTSGTT | ASSRGQNNSPLY | 0.6736326 |
| VGAFPFGF | ASSRGQNNSPLY | 0.17400372 |
| VGAGTVEF | ASSRGQNNSPLY | 0.42449257 |
| VGKSTLLL | ASSRGQNNSPLY | 0.81455934 |
| AALAHAGGGIV | ASSRGQNNSPLY | 0.6803387 |
| AALVAVSLL | ASSRGQNNSPLY | 0.3391133 |
| AAVSRAVAAL | ASSRGQNNSPLY | 0.34356418 |
| AGGSSGIGGF | ASSRGQNNSPLY | 0.8237885 |
| AGLTVANANG | ASSRGQNNSPLY | 0.35322696 |
| AGLVGAVL | ASSRGQNNSPLY | 0.82698697 |
| ALSQAFTV | ASSRGQNNSPLY | 0.15922467 |
| ALTFSVSL | ASSRGQNNSPLY | 0.3435288 |
| ALTGAALVAL | ASSRGQNNSPLY | 0.2898681 |
| ASFSIGAGVL | ASSRGQNNSPLY | 0.15222248 |
| DGSVYKI | ASSRGQNNSPLY | 0.64721966 |
| EQLAEIV | ASSRGQNNSPLY | 0.33069223 |
| FAVSELTVM | ASSRGQNNSPLY | 0.47907197 |
| GAAPLAAGL | ASSRGQNNSPLY | 0.4270386 |
| GADGGDGGSGGAGG | ASSRGQNNSPLY | 0.6897452 |
| GFGLDTGGAGG | ASSRGQNNSPLY | 0.6256701 |
| GLASGGLL | ASSRGQNNSPLY | 0.40418577 |
| GRIAGAAVLL | ASSRGQNNSPLY | 0.24838176 |
| GSAGSLNTSF | ASSRGQNNSPLY | 0.51125133 |

| | | |
|---|---|---|
| GTGQIGI | ASSRGQNNSPLY | 0.7841765 |
| GTLIALIL | ASSRGQNNSPLY | 0.5696046 |
| IAATMGFVVG | ASSRGQNNSPLY | 0.18148486 |
| IFAQALL | ASSRGQNNSPLY | 0.7042418 |
| IGAAVALGF | ASSRGQNNSPLY | 0.32841888 |
| IGLTPLAL | ASSRGQNNSPLY | 0.2765472 |
| ILGSNTSTL | ASSRGQNNSPLY | 0.14731471 |
| IVFSLTVL | ASSRGQNNSPLY | 0.39368582 |
| LASKATGF | ASSRGQNNSPLY | 0.360711 |
| LASVALIL | ASSRGQNNSPLY | 0.72748095 |
| LFAVGLTL | ASSRGQNNSPLY | 0.76054454 |
| LFNGGIGTYI | ASSRGQNNSPLY | 0.320435 |
| LTLGAGII | ASSRGQNNSPLY | 0.69711745 |
| LVAAALGDI | ASSRGQNNSPLY | 0.6409355 |
| NTAFVQLGI | ASSRGQNNSPLY | 0.11663349 |
| QLFSGAEL | ASSRGQNNSPLY | 0.2548597 |
| SAATFITL | ASSRGQNNSPLY | 0.19281806 |
| SNYGLANLGV | ASSRGQNNSPLY | 0.7149244 |
| SYGGAYAVM | ASSRGQNNSPLY | 0.3245328 |
| TTGVGGLTLG | ASSRGQNNSPLY | 0.47605112 |
| TTQTIYI | ASSRGQNNSPLY | 0.3695048 |
| VAAGAGLGAVI | ASSRGQNNSPLY | 0.7809853 |
| VAAVVTLL | ASSRGQNNSPLY | 0.44524372 |
| VGFLGGLI | ASSRGQNNSPLY | 0.23181732 |
| VGVIPILL | ASSRGQNNSPLY | 0.18583897 |
| VIFASSLI | ASSRGQNNSPLY | 0.4638148 |
| VKGGDGGAAAG | ASSRGQNNSPLY | 0.39355212 |
| VTKTFTNI | ASSRGQNNSPLY | 0.27447924 |
| VVAAAILL | ASSRGQNNSPLY | 0.32658863 |
| YAALDVEL | ASSRGQNNSPLY | 0.7863923 |
| LASVALLL | ASSRGQNNSPLY | 0.18579343 |
| LDAVHRV | ASSRGQNNSPLY | 0.58730507 |
| QGLSSSIF | ASSRGQNNSPLY | 0.22188497 |
| RGTFANIRL | ASSRGQNNSPLY | 0.47072637 |
| RQLWWGHRI | ASSRGQNNSPLY | 0.78740144 |
| YGPPGTGKTLL | ASSRGQNNSPLY | 0.7612581 |
| AALVGAPANVA | ASSRGQNNSPLY | 0.30470163 |
| AATFRRTE | ASSRGQNNSPLY | 0.29494292 |
| AGALGGGI | ASSRGQNNSPLY | 0.24218519 |
| AGGAGGAGGIGAG | ASSRGQNNSPLY | 0.3787905 |
| AGSGALGL | ASSRGQNNSPLY | 0.36940515 |
| AQIADQI | ASSRGQNNSPLY | 0.7083236 |
| ASAVAVTYL | ASSRGQNNSPLY | 0.4354303 |
| ASTVLGGTGG | ASSRGQNNSPLY | 0.26776707 |
| AVGGEPAK | ASSRGQNNSPLY | 0.51440555 |
| CEQGTFGGTC | ASSRGQNNSPLY | 0.7803236 |
| DEGKAPAG | ASSRGQNNSPLY | 0.8123022 |
| DTVVFPAL | ASSRGQNNSPLY | 0.28851426 |
| FEGITMAG | ASSRGQNNSPLY | 0.29755747 |
| FPLVVQD | ASSRGQNNSPLY | 0.30170813 |
| GAFTTPQ | ASSRGQNNSPLY | 0.5231389 |

| | | |
|---|---|---|
| GFGGFGTAGGVGGA | ASSRGQNNSPLY | 0.2908943 |
| GFLAQFL | ASSRGQNNSPLY | 0.2683004 |
| GGAGGAGGNAGVA | ASSRGQNNSPLY | 0.6434165 |
| GGVGTAGAGGAGGAGG | ASSRGQNNSPLY | 0.83961886 |
| GNGGGGDAGNAGSGGN | ASSRGQNNSPLY | 0.37143433 |
| GTGGAGGAGNDAGS | ASSRGQNNSPLY | 0.19399892 |
| GTTNTGWFNS | ASSRGQNNSPLY | 0.4201557 |
| IFMGPLFTL | ASSRGQNNSPLY | 0.27165574 |
| ILAHAIEAI | ASSRGQNNSPLY | 0.27667946 |
| IRSAIREAL | ASSRGQNNSPLY | 0.35115993 |
| ITFAGGG | ASSRGQNNSPLY | 0.18567935 |
| ITGMSGHQEV | ASSRGQNNSPLY | 0.46163586 |
| IVAGLACTL | ASSRGQNNSPLY | 0.7310142 |
| IVGGGLAGA | ASSRGQNNSPLY | 0.36968362 |
| IVTGVLNI | ASSRGQNNSPLY | 0.50953674 |
| LAQQGCI | ASSRGQNNSPLY | 0.80164623 |
| LCLHVEN | ASSRGQNNSPLY | 0.59643465 |
| LDAISDGS | ASSRGQNNSPLY | 0.75542164 |
| LGGFAAML | ASSRGQNNSPLY | 0.7936153 |
| LGGKNPMI | ASSRGQNNSPLY | 0.39544326 |
| LGPKGVAAIL | ASSRGQNNSPLY | 0.43310738 |
| LMVGVSI | ASSRGQNNSPLY | 0.77972806 |
| LVGAAFVL | ASSRGQNNSPLY | 0.398362 |
| LVLAVAPL | ASSRGQNNSPLY | 0.39968088 |
| MAAIAVIDAA | ASSRGQNNSPLY | 0.34808815 |
| NNGINAELQ | ASSRGQNNSPLY | 0.7764498 |
| RGEPLVLVHG | ASSRGQNNSPLY | 0.30808008 |
| RGGAGGAAGLI | ASSRGQNNSPLY | 0.78064173 |
| RSYRMHPAV | ASSRGQNNSPLY | 0.16285643 |
| SAAHLLAGQLA | ASSRGQNNSPLY | 0.25479874 |
| SAAVGSY | ASSRGQNNSPLY | 0.7550155 |
| SPHAVLHRI | ASSRGQNNSPLY | 0.1279975 |
| SQIAKVL | ASSRGQNNSPLY | 0.16674985 |
| TALGHLVP | ASSRGQNNSPLY | 0.5099455 |
| TAPTTSGAA | ASSRGQNNSPLY | 0.73811096 |
| TDGDDGNGGSPGT | ASSRGQNNSPLY | 0.1473075 |
| TDGLGEN | ASSRGQNNSPLY | 0.30241454 |
| TGGPPLPGQA | ASSRGQNNSPLY | 0.1745962 |
| TIPAITI | ASSRGQNNSPLY | 0.45258862 |
| TIPGFTI | ASSRGQNNSPLY | 0.4222517 |
| TSIASGALA | ASSRGQNNSPLY | 0.14161283 |
| TVTLPGAAI | ASSRGQNNSPLY | 0.5685029 |
| VASGLSAALP | ASSRGQNNSPLY | 0.18885365 |
| VGAGVIGI | ASSRGQNNSPLY | 0.40426907 |
| VGPKRFGV | ASSRGQNNSPLY | 0.68167496 |
| VNVAFPD | ASSRGQNNSPLY | 0.82687235 |
| VSMTNSGVS | ASSRGQNNSPLY | 0.25345224 |
| VTDVLGDLSV | ASSRGQNNSPLY | 0.540691 |
| VVVVGAGFAGL | ASSRGQNNSPLY | 0.38772798 |
| VYTVGTVV | ASSRGQNNSPLY | 0.8460566 |
| WTTFSGP | ASSRGQNNSPLY | 0.25911194 |

| | | |
|---|---|---|
| YDELVEAF | ASSRGQNNSPLY | 0.1898686 |
| YGTGGAGGNGG | ASSRGQNNSPLY | 0.7335547 |
| YTNVMAV | ASSRGQNNSPLY | 0.4594946 |
| FLEQVHQL | ASSRGQNNSPLY | 0.36592716 |
| GLSLLAVAV | ASSRGQNNSPLY | 0.7827759 |
| LEQVHQL | ASSRGQNNSPLY | 0.31381488 |
| LNAAGAI | ASSRGQNNSPLY | 0.34002918 |
| PAVHASL | ASSRGQNNSPLY | 0.69798476 |
| TELEDDA | ASSRGQNNSPLY | 0.7372568 |
| TPPGPPN | ASSRGQNNSPLY | 0.30321842 |
| TPPPPPPP | ASSRGQNNSPLY | 0.6585793 |
| TQSPALI | ASSSGSSYEQY | 0.4838881 |
| EFAAAFPQ | ASSSGSSYEQY | 0.23451516 |
| LLSGNLDVL | ASSSGSSYEQY | 0.7835325 |
| GPANLGNYNI | ASSSGSSYEQY | 0.32098606 |
| TLGPTLP | ASSSGSSYEQY | 0.65907896 |
| VYAYSSCCK | ASSSGSSYEQY | 0.4895661 |
| APAPAPAPNI | ASSSGSSYEQY | 0.7742934 |
| PGGPVPIP | ASSSGSSYEQY | 0.18336973 |
| FLTVALEP | ASSSGSSYEQY | 0.6856932 |
| SQYFGANL | ASSSGSSYEQY | 0.36875716 |
| GGVKPAA | ASSSGSSYEQY | 0.56982386 |
| GGTGGAGGAGGW | ASSSGSSYEQY | 0.5132126 |
| GVGGVAGLGGAGG | ASSSGSSYEQY | 0.57147086 |
| VLLVLATTI | ASSSGSSYEQY | 0.58038896 |
| IGIFAAML | ASSSGSSYEQY | 0.7358676 |
| AVGDLID | ASSSGSSYEQY | 0.44728434 |
| NISPLPL | ASSSGSSYEQY | 0.6041838 |
| VTAVFVAI | ASSSGSSYEQY | 0.79506624 |
| SGDDNDAAL | ASSSGSSYEQY | 0.24462742 |
| AVAALAHS | ASSSGSSYEQY | 0.4269097 |
| GGVTVMATP | ASSSGSSYEQY | 0.6897987 |
| AGLAANVRD | ASSSGSSYEQY | 0.730611 |
| TNRPDLIDPALLRPG | ASSSGSSYEQY | 0.38423252 |
| LANLGVTAL | ASSSGSSYEQY | 0.80166465 |
| TGERIEL | ASSSGSSYEQY | 0.29631615 |
| NGEELAA | ASSSGSSYEQY | 0.74866223 |
| VLSLVWG | ASSSGSSYEQY | 0.5387608 |
| IGYALWL | ASSSGSSYEQY | 0.32226878 |
| LSLLLAPL | ASSSGSSYEQY | 0.33279523 |
| TLDEALK | ASSSGSSYEQY | 0.7281097 |
| TDGAGTAEGG | ASSSGSSYEQY | 0.7112162 |
| NAPGGAGGNGG | ASSSGSSYEQY | 0.6655756 |
| VSVFMGMGGNF | ASSSGSSYEQY | 0.1837281 |
| LASYLAAL | ASSSGSSYEQY | 0.59885865 |
| LVGGTPAL | ASSSGSSYEQY | 0.6727201 |
| VRNADVL | ASSSGSSYEQY | 0.8439805 |
| TCSSPFLVSAM | ASSSGSSYEQY | 0.30845034 |
| VHITDAE | ASSSGSSYEQY | 0.48421994 |
| EYAAPSF | ASSSGSSYEQY | 0.5264845 |
| VVDAFHGGL | ASSSGSSYEQY | 0.36112148 |

| | | |
|---|---|---|
| ASVVDPQ | ASSSGSSYEQY | 0.18616311 |
| SYNEFGGLS | ASSSGSSYEQY | 0.29838046 |
| DPDGSVY | ASSSGSSYEQY | 0.3827423 |
| MVSTIHATE | ASSSGSSYEQY | 0.2959208 |
| LAAGVADPV | ASSSGSSYEQY | 0.80811656 |
| TKEQIAA | ASSSGSSYEQY | 0.7810402 |
| SLASPAAL | ASSSGSSYEQY | 0.6811641 |
| IGLVFYGV | ASSSGSSYEQY | 0.74311256 |
| FLLDASNL | ASSSGSSYEQY | 0.75963885 |
| YQVGNFP | ASSSGSSYEQY | 0.48782945 |
| ESLLSQST | ASSSGSSYEQY | 0.3508327 |
| LFTVGFAAVG | ASSSGSSYEQY | 0.26950774 |
| ATPPTPT | ASSSGSSYEQY | 0.2534805 |
| TDLATLG | ASSSGSSYEQY | 0.39581805 |
| SAGGTGR | ASSSGSSYEQY | 0.32333624 |
| GFSNSGS | ASSSGSSYEQY | 0.42493588 |
| GGNGGSGGTSV | ASSSGSSYEQY | 0.12269393 |
| GTSVATGGAGN | ASSSGSSYEQY | 0.758482 |
| NGGNGGSGGTS | ASSSGSSYEQY | 0.7488959 |
| AVVKFPH | ASSSGSSYEQY | 0.4811669 |
| TPPPPPP | ASSSGSSYEQY | 0.5167229 |
| AAPAAAAAPA | ASSSGSSYEQY | 0.62066424 |
| IALNEDL | ASSSGSSYEQY | 0.32282305 |
| IVNVSSM | ASSSGSSYEQY | 0.59683895 |
| TGIPPAPR | ASSSGSSYEQY | 0.30849752 |
| FDLGGGTF | ASSSGSSYEQY | 0.5963749 |
| FDLGGGTFDVS | ASSSGSSYEQY | 0.648641 |
| NEPTAAA | ASSSGSSYEQY | 0.57171404 |
| NEPTAAALA | ASSSGSSYEQY | 0.4811015 |
| GSIINISSI | ASSSGSSYEQY | 0.3836432 |
| VAPGNAGTA | ASSSGSSYEQY | 0.16271809 |
| TGGGASLE | ASSSGSSYEQY | 0.5431249 |
| VAVYNHYKRI | ASSSGSSYEQY | 0.21998456 |
| TDPAPATT | ASSSGSSYEQY | 0.46140403 |
| TNRPDILDPALLRPG | ASSSGSSYEQY | 0.22712211 |
| LTTTTTS | ASSSGSSYEQY | 0.5625781 |
| PTGSGKTL | ASSSGSSYEQY | 0.562057 |
| TGSGKTL | ASSSGSSYEQY | 0.56474173 |
| TIGHVDHGK | ASSSGSSYEQY | 0.23899949 |
| TIGHVDHGKTT | ASSSGSSYEQY | 0.56252503 |
| GHVDHGKTT | ASSSGSSYEQY | 0.6857582 |
| AQFEHTLL | ASSSGSSYEQY | 0.23345226 |
| IVNQDEL | ASSSGSSYEQY | 0.68766326 |
| TNGSQFF | ASSSGSSYEQY | 0.43909845 |
| TGKGIVE | ASSSGSSYEQY | 0.33320302 |
| LISNASDA | ASSSGSSYEQY | 0.63424325 |
| AELRTLGELTKIAKAHGAQ | ASSSGSSYEQY | 0.41552848 |
| APNAVAAAEPSW | ASSSGSSYEQY | 0.15872017 |
| APQVAGAATAVL | ASSSGSSYEQY | 0.36109853 |
| ARRDPAAAAAILPTDA | ASSSGSSYEQY | 0.5484987 |
| DAAARDVLAEAGLAEYF | ASSSGSSYEQY | 0.24314077 |

| | | |
|---|---|---|
| DDQIVLSAKANLAKVEQ | ASSSGSSYEQY | 0.16035235 |
| DELLPDTGEDSDEDGH | ASSSGSSYEQY | 0.6719355 |
| ELPEAPANRATVPQD | ASSSGSSYEQY | 0.23410924 |
| FGQSKIERVTGIG | ASSSGSSYEQY | 0.21717662 |
| GAGGTGSTAGGAGGAGGAGGLY | ASSSGSSYEQY | 0.417552 |
| GHGGTGAAGGNGGTGGAGGSGI | ASSSGSSYEQY | 0.6575277 |
| GNGGNGGAGGEGGPGGAGGAGA | ASSSGSSYEQY | 0.26494476 |
| GSTAGNGGNGGNATNGTIAGQP | ASSSGSSYEQY | 0.30196834 |
| GTGGDGGNGGNGGTGG | ASSSGSSYEQY | 0.6740942 |
| GVRRPAGIPADAPV | ASSSGSSYEQY | 0.5947336 |
| GYRQVTDAHLLTLARR | ASSSGSSYEQY | 0.57282627 |
| IAKMGQDGHDRG | ASSSGSSYEQY | 0.582935 |
| KAEPSIEIQYT | ASSSGSSYEQY | 0.3946434 |
| KPVRIRMAADEDTVAVT | ASSSGSSYEQY | 0.57718575 |
| MPPRIAGMRLLVIKPEP | ASSSGSSYEQY | 0.3608666 |
| MKFYAVALFRSRGVPPTR | ASSSGSSYEQY | 0.29654014 |
| NAGGFGILGAVAHSP | ASSSGSSYEQY | 0.23088679 |
| NEGREITLAMAEM | ASSSGSSYEQY | 0.3368595 |
| NELVEWSHPLGHRWRWE | ASSSGSSYEQY | 0.2620281 |
| NLGAGGTGGNGGNPG | ASSSGSSYEQY | 0.37132883 |
| NTEWPKWETVMAADPVD | ASSSGSSYEQY | 0.6440967 |
| PASAAPAETQP | ASSSGSSYEQY | 0.33861235 |
| QQSSDGKNGVRAAEITGE | ASSSGSSYEQY | 0.20143333 |
| RAPDAVTTSWWKE | ASSSGSSYEQY | 0.52840626 |
| SGFGNVGAHTSGWFNQ | ASSSGSSYEQY | 0.19509247 |
| SGVAGAVGQAASVGGLKV | ASSSGSSYEQY | 0.596859 |
| SNGGAGTGSAGGNGGT | ASSSGSSYEQY | 0.5117866 |
| TGGAGGEGGAAG | ASSSGSSYEQY | 0.8449863 |
| TGGDGGAAGAGGAPGHG | ASSSGSSYEQY | 0.7770644 |
| TMPPLLAAIANAGRHGV | ASSSGSSYEQY | 0.46412358 |
| TSSTAGNSGGA | ASSSGSSYEQY | 0.6918899 |
| VPATVPVGILC | ASSSGSSYEQY | 0.24636303 |
| VQRPDSGGVVLTGRIS | ASSSGSSYEQY | 0.59154564 |
| WTPNPVPATKGVPAGLVQAT | ASSSGSSYEQY | 0.48897833 |
| YALKRALAKFSDRFDVVII | ASSSGSSYEQY | 0.5937115 |
| YLIGSVIV | ASSSGSSYEQY | 0.3808024 |
| YGQLVAVID | ASSSGSSYEQY | 0.38184315 |
| VTVGGMTLPA | ASSSGSSYEQY | 0.15618923 |
| VLALPAV | ASSSGSSYEQY | 0.7366422 |
| VIPPTLNYET | ASSSGSSYEQY | 0.21631998 |
| VGMGGMHPSSG | ASSSGSSYEQY | 0.17849544 |
| VGLAGGAATA | ASSSGSSYEQY | 0.32232392 |
| VGGAGGAGGSGP | ASSSGSSYEQY | 0.8181861 |
| VFAAFMV | ASSSGSSYEQY | 0.7250799 |
| TLYDKLR | ASSSGSSYEQY | 0.3460091 |
| TLAVFTVL | ASSSGSSYEQY | 0.27420515 |
| TGSVQPARV | ASSSGSSYEQY | 0.793674 |
| TGGGAGSGGT | ASSSGSSYEQY | 0.25101936 |
| SHLETSL | ASSSGSSYEQY | 0.5670324 |
| SFTTGMT | ASSSGSSYEQY | 0.44776887 |
| QVEVAKRMGVRQPTVS | ASSSGSSYEQY | 0.441997 |

| | | |
|---|---|---|
| QGSGGPALAM | ASSSGSSYEQY | 0.24729477 |
| MLAPSNP | ASSSGSSYEQY | 0.68978757 |
| LVNNAAI | ASSSGSSYEQY | 0.45201385 |
| LTGEINTLA | ASSSGSSYEQY | 0.45345485 |
| LSIFGPV | ASSSGSSYEQY | 0.23113425 |
| LLALLANL | ASSSGSSYEQY | 0.6853702 |
| LISNASDALDKLR | ASSSGSSYEQY | 0.32472298 |
| LINEAVP | ASSSGSSYEQY | 0.22047919 |
| LGYAFSGL | ASSSGSSYEQY | 0.18141785 |
| LDFVAVAL | ASSSGSSYEQY | 0.42578632 |
| LAQPGAE | ASSSGSSYEQY | 0.72057474 |
| ISNLGAVTGF | ASSSGSSYEQY | 0.7751088 |
| HQEDPGRI | ASSSGSSYEQY | 0.4427871 |
| GTPGEPTP | ASSSGSSYEQY | 0.66709757 |
| GFFNSGS | ASSSGSSYEQY | 0.7198975 |
| GAGGGGLTAGAN | ASSSGSSYEQY | 0.41781777 |
| FVDALVNM | ASSSGSSYEQY | 0.17640042 |
| FLTVAPEL | ASSSGSSYEQY | 0.43144038 |
| FETLTAL | ASSSGSSYEQY | 0.72420895 |
| DVINPIEL | ASSSGSSYEQY | 0.6843258 |
| APVSGAPV | ASSSGSSYEQY | 0.7396213 |
| ALVGGLAW | ASSSGSSYEQY | 0.19457251 |
| ALAAAAATAI | ASSSGSSYEQY | 0.4809 |
| AFGLGLLL | ASSSGSSYEQY | 0.59386855 |
| AFFAAASL | ASSSGSSYEQY | 0.2061567 |
| AAAALPAFSPP | ASSSGSSYEQY | 0.4036255 |
| VVLAPAL | ASSSGSSYEQY | 0.3379054 |
| TIGHVDHGKTTL | ASSSGSSYEQY | 0.30846286 |
| LISNASDALDK | ASSSGSSYEQY | 0.64731395 |
| LISNASDAL | ASSSGSSYEQY | 0.43531826 |
| ISNASDALDK | ASSSGSSYEQY | 0.3628434 |
| NQAETLVYQTE | ASSSGSSYEQY | 0.36616048 |
| TGGTGGTAGAG | ASSSGSSYEQY | 0.37632382 |
| TGSTAGGAGGA | ASSSGSSYEQY | 0.29424405 |
| DPLAVPVAAGPV | ASSSGSSYEQY | 0.38676423 |
| GGTGGTPGNGGN | ASSSGSSYEQY | 0.48860145 |
| PAGLAGVDPATL | ASSSGSSYEQY | 0.7195367 |
| GNNLVVEAPHAGASV | ASSSGSSYEQY | 0.52371377 |
| KVTRIMSASTLEQPA | ASSSGSSYEQY | 0.38199568 |
| LGFALPIALPRRTVV | ASSSGSSYEQY | 0.7959022 |
| TRLGNLTARKLVVLV | ASSSGSSYEQY | 0.20897985 |
| WSQQRRLIYFPSAGP | ASSSGSSYEQY | 0.25513256 |
| GPNGNPDPMAAAVDIR | ASSSGSSYEQY | 0.56267816 |
| TQRCVSVADLRAAASR | ASSSGSSYEQY | 0.31295526 |
| QFVLDVAEHPVEVLTETV | ASSSGSSYEQY | 0.67348784 |
| GGAGGTGGAGGNGGTQAGDG | ASSSGSSYEQY | 0.6655307 |
| PLLATLRAIAVDVDAKPAQV | ASSSGSSYEQY | 0.5109288 |
| APKFRTGGYMIDFWGVGYQVAK | ASSSGSSYEQY | 0.5696651 |
| GAGGAGGAGGAAGGPGGTGGTG | ASSSGSSYEQY | 0.8194407 |
| IGALGLLATDVAPAGSVAFGVA | ASSSGSSYEQY | 0.6254569 |
| AGGAGGIGGNGGAGGAGGVLMG | ASSSGSSYEQY | 0.41964984 |

| | | |
|---|---|---|
| ADELIGLL | ASSSGSSYEQY | 0.8099445 |
| ADRALKA | ASSSGSSYEQY | 0.24533321 |
| AGAVGQVGGAGGA | ASSSGSSYEQY | 0.6959627 |
| AGGGGGLGGAP | ASSSGSSYEQY | 0.6584288 |
| AGGLLYGNGGAG | ASSSGSSYEQY | 0.18004213 |
| AGLESIGL | ASSSGSSYEQY | 0.36828196 |
| AGLKTGII | ASSSGSSYEQY | 0.73446727 |
| ASLAALLL | ASSSGSSYEQY | 0.7403681 |
| ASVINISSI | ASSSGSSYEQY | 0.21242268 |
| ATPPGPA | ASSSGSSYEQY | 0.33776852 |
| ATVASALV | ASSSGSSYEQY | 0.22431844 |
| ATVASLIL | ASSSGSSYEQY | 0.28192484 |
| AVNGPVLIRH | ASSSGSSYEQY | 0.20006482 |
| DGAAGAVGI | ASSSGSSYEQY | 0.46136165 |
| DPLLHGI | ASSSGSSYEQY | 0.3457866 |
| ELGAALGAL | ASSSGSSYEQY | 0.7701153 |
| ELGRTQAV | ASSSGSSYEQY | 0.37300742 |
| FVAGGYLS | ASSSGSSYEQY | 0.63146424 |
| GASGGAGGAGGW | ASSSGSSYEQY | 0.34721375 |
| GGAGAGGDGGAGGI | ASSSGSSYEQY | 0.5317245 |
| GTGHNGSQPGMGGNG | ASSSGSSYEQY | 0.17793044 |
| GYAAASASAS | ASSSGSSYEQY | 0.2009594 |
| HQVDVAAVQQG | ASSSGSSYEQY | 0.28311765 |
| IGAAIAHT | ASSSGSSYEQY | 0.46863887 |
| IGMIEVAQL | ASSSGSSYEQY | 0.17884381 |
| IRLAPEHTSAMD | ASSSGSSYEQY | 0.70642555 |
| IYNVSVL | ASSSGSSYEQY | 0.3427032 |
| KQGFLGM | ASSSGSSYEQY | 0.46816722 |
| LESVITQL | ASSSGSSYEQY | 0.38002154 |
| LGLGALVVV | ASSSGSSYEQY | 0.3060131 |
| LIGNGGNGGSGGI | ASSSGSSYEQY | 0.32876584 |
| LNAINAPTL | ASSSGSSYEQY | 0.67171454 |
| LQLLNTGV | ASSSGSSYEQY | 0.24212733 |
| LQTSIEF | ASSSGSSYEQY | 0.16889575 |
| MSFAVSEL | ASSSGSSYEQY | 0.3016076 |
| NTGALGL | ASSSGSSYEQY | 0.24747968 |
| PPGAGGAGGAGTT | ASSSGSSYEQY | 0.8029995 |
| QLVGLGI | ASSSGSSYEQY | 0.72313726 |
| RFLEQVHQL | ASSSGSSYEQY | 0.4570373 |
| RGVGYVLR | ASSSGSSYEQY | 0.7774037 |
| SATGVFAGVI | ASSSGSSYEQY | 0.13568217 |
| SLPAITLP | ASSSGSSYEQY | 0.4371704 |
| STALAFTV | ASSSGSSYEQY | 0.6606862 |
| SVLTVGLGK | ASSSGSSYEQY | 0.42540497 |
| TALAGAGSGI | ASSSGSSYEQY | 0.35607103 |
| TGLPFTGI | ASSSGSSYEQY | 0.71580774 |
| VAEHVTV | ASSSGSSYEQY | 0.35321057 |
| VDIVLNSL | ASSSGSSYEQY | 0.23857872 |
| VLTGGGAVL | ASSSGSSYEQY | 0.5738467 |
| VNVGLAYGI | ASSSGSSYEQY | 0.64561415 |
| VSSAAVGAV | ASSSGSSYEQY | 0.6322563 |

| | | |
|---|---|---|
| VTSVYEAGV | ASSSGSSYEQY | 0.23313646 |
| WVALDRGA | ASSSGSSYEQY | 0.42117903 |
| YLVGSVVI | ASSSGSSYEQY | 0.8587502 |
| GPPGTGKTLL | ASSSGSSYEQY | 0.38577762 |
| GVGMTKF | ASSSGSSYEQY | 0.5880735 |
| IGIGGSDL | ASSSGSSYEQY | 0.39080727 |
| IGTLLAEL | ASSSGSSYEQY | 0.40325797 |
| IYTSGTT | ASSSGSSYEQY | 0.24092203 |
| LSFPAGV | ASSSGSSYEQY | 0.23089786 |
| LYTSGTT | ASSSGSSYEQY | 0.63353395 |
| VGAFPFGF | ASSSGSSYEQY | 0.19506502 |
| VGAGTVEF | ASSSGSSYEQY | 0.45547694 |
| VGKSTLLL | ASSSGSSYEQY | 0.34994316 |
| AALAHAGGGIV | ASSSGSSYEQY | 0.85483307 |
| AALVAVSLL | ASSSGSSYEQY | 0.65486693 |
| AAVSRAVAAL | ASSSGSSYEQY | 0.49965233 |
| AGGSSGIGGF | ASSSGSSYEQY | 0.3512951 |
| AGLTVANANG | ASSSGSSYEQY | 0.4879767 |
| AGLVGAVL | ASSSGSSYEQY | 0.6407891 |
| ALSQAFTV | ASSSGSSYEQY | 0.16686659 |
| ALTFSVSL | ASSSGSSYEQY | 0.71048635 |
| ALTGAALVAL | ASSSGSSYEQY | 0.23134044 |
| ASFSIGAGVL | ASSSGSSYEQY | 0.47867993 |
| DGSVYKI | ASSSGSSYEQY | 0.38382617 |
| EQLAEIV | ASSSGSSYEQY | 0.83200586 |
| FAVSELTVM | ASSSGSSYEQY | 0.4282437 |
| GAAPLAAGL | ASSSGSSYEQY | 0.5515138 |
| GADGGDGGSGGAGG | ASSSGSSYEQY | 0.2843231 |
| GFGLDTGGAGG | ASSSGSSYEQY | 0.4623646 |
| GLASGGLL | ASSSGSSYEQY | 0.7460688 |
| GRIAGAAVLL | ASSSGSSYEQY | 0.37840146 |
| GSAGSLNTSF | ASSSGSSYEQY | 0.838766 |
| GTGQIGI | ASSSGSSYEQY | 0.46017003 |
| GTLIALIL | ASSSGSSYEQY | 0.16903754 |
| IAATMGFVVG | ASSSGSSYEQY | 0.2935167 |
| IFAQALL | ASSSGSSYEQY | 0.7059614 |
| IGAAVALGF | ASSSGSSYEQY | 0.79278016 |
| IGLTPLAL | ASSSGSSYEQY | 0.69579804 |
| ILGSNTSTL | ASSSGSSYEQY | 0.73753804 |
| IVFSLTVL | ASSSGSSYEQY | 0.34131336 |
| LASKATGF | ASSSGSSYEQY | 0.7474108 |
| LASVALIL | ASSSGSSYEQY | 0.8294362 |
| LFAVGLTL | ASSSGSSYEQY | 0.43117648 |
| LFNGGIGTYI | ASSSGSSYEQY | 0.20303711 |
| LTLGAGII | ASSSGSSYEQY | 0.19498456 |
| LVAAALGDI | ASSSGSSYEQY | 0.62121636 |
| NTAFVQLGI | ASSSGSSYEQY | 0.7409793 |
| QLFSGAEL | ASSSGSSYEQY | 0.21209434 |
| SAATFITL | ASSSGSSYEQY | 0.46354133 |
| SNYGLANLGV | ASSSGSSYEQY | 0.49536592 |
| SYGGAYAVM | ASSSGSSYEQY | 0.16823834 |

| | | |
|---|---|---|
| TTGVGGLTLG | ASSSGSSYEQY | 0.68185395 |
| TTQTIYI | ASSSGSSYEQY | 0.822322 |
| VAAGAGLGAVI | ASSSGSSYEQY | 0.46070552 |
| VAAVVTLL | ASSSGSSYEQY | 0.2011055 |
| VGFLGGLI | ASSSGSSYEQY | 0.7519403 |
| VGVIPILL | ASSSGSSYEQY | 0.84183663 |
| VIFASSLI | ASSSGSSYEQY | 0.7254476 |
| VKGGDGGAAAG | ASSSGSSYEQY | 0.18589528 |
| VTKTFTNI | ASSSGSSYEQY | 0.13448396 |
| VVAAAILL | ASSSGSSYEQY | 0.55548126 |
| YAALDVEL | ASSSGSSYEQY | 0.7476827 |
| LASVALLL | ASSSGSSYEQY | 0.4596492 |
| LDAVHRV | ASSSGSSYEQY | 0.29725394 |
| QGLSSSIF | ASSSGSSYEQY | 0.5956675 |
| RGTFANIRL | ASSSGSSYEQY | 0.66630495 |
| RQLWWGHRI | ASSSGSSYEQY | 0.23192315 |
| YGPPGTGKTLL | ASSSGSSYEQY | 0.19651066 |
| AALVGAPANVA | ASSSGSSYEQY | 0.72754467 |
| AATFRRTE | ASSSGSSYEQY | 0.63522995 |
| AGALGGGI | ASSSGSSYEQY | 0.23489311 |
| AGGAGGAGGIGAG | ASSSGSSYEQY | 0.40270865 |
| AGSGALGL | ASSSGSSYEQY | 0.5254471 |
| AQIADQI | ASSSGSSYEQY | 0.45757762 |
| ASAVAVTYL | ASSSGSSYEQY | 0.5173942 |
| ASTVLGGTGG | ASSSGSSYEQY | 0.17756994 |
| AVGGEPAK | ASSSGSSYEQY | 0.7817701 |
| CEQGTFGGTC | ASSSGSSYEQY | 0.19843152 |
| DEGKAPAG | ASSSGSSYEQY | 0.6435814 |
| DTVVFPAL | ASSSGSSYEQY | 0.5949619 |
| FEGITMAG | ASSSGSSYEQY | 0.55555284 |
| FPLVVQD | ASSSGSSYEQY | 0.38756996 |
| GAFTTPQ | ASSSGSSYEQY | 0.20662028 |
| GFGGFGTAGGVGGA | ASSSGSSYEQY | 0.20034112 |
| GFLAQFL | ASSSGSSYEQY | 0.1949006 |
| GGAGGAGGNAGVA | ASSSGSSYEQY | 0.7693995 |
| GGVGTAGAGGAGGAGG | ASSSGSSYEQY | 0.5202874 |
| GNGGGGDAGNAGSGGN | ASSSGSSYEQY | 0.685352 |
| GTGGAGGAGNDAGS | ASSSGSSYEQY | 0.41287285 |
| GTTNTGWFNS | ASSSGSSYEQY | 0.24756625 |
| IFMGPLFTL | ASSSGSSYEQY | 0.5509116 |
| ILAHAIEAI | ASSSGSSYEQY | 0.23743705 |
| IRSAIREAL | ASSSGSSYEQY | 0.44143015 |
| ITFAGGG | ASSSGSSYEQY | 0.32313466 |
| ITGMSGHQEV | ASSSGSSYEQY | 0.45042002 |
| IVAGLACTL | ASSSGSSYEQY | 0.27207524 |
| IVGGGLAGA | ASSSGSSYEQY | 0.4983546 |
| IVTGVLNI | ASSSGSSYEQY | 0.55071485 |
| LAQQGCI | ASSSGSSYEQY | 0.2345407 |
| LCLHVEN | ASSSGSSYEQY | 0.4620698 |
| LDAISDGS | ASSSGSSYEQY | 0.7923195 |
| LGGFAAML | ASSSGSSYEQY | 0.5039744 |

| | | |
|---|---|---|
| LGGKNPMI | ASSSGSSYEQY | 0.17654896 |
| LGPKGVAAIL | ASSSGSSYEQY | 0.6153137 |
| LMVGVSI | ASSSGSSYEQY | 0.26290435 |
| LVGAAFVL | ASSSGSSYEQY | 0.575137 |
| LVLAVAPL | ASSSGSSYEQY | 0.53751147 |
| MAAIAVIDAA | ASSSGSSYEQY | 0.33003947 |
| NNGINAELQ | ASSSGSSYEQY | 0.5453 |
| RGEPLVLVHG | ASSSGSSYEQY | 0.29790556 |
| RGGAGGAAGLI | ASSSGSSYEQY | 0.37256807 |
| RSYRMHPAV | ASSSGSSYEQY | 0.3179083 |
| SAAHLLAGQLA | ASSSGSSYEQY | 0.5470001 |
| SAAVGSY | ASSSGSSYEQY | 0.5131567 |
| SPHAVLHRI | ASSSGSSYEQY | 0.22585788 |
| SQIAKVL | ASSSGSSYEQY | 0.46811724 |
| TALGHLVP | ASSSGSSYEQY | 0.271744 |
| TAPTTSGAA | ASSSGSSYEQY | 0.20052807 |
| TDGDDGNGGSPGT | ASSSGSSYEQY | 0.42845201 |
| TDGLGEN | ASSSGSSYEQY | 0.41231337 |
| TGGPPLPGQA | ASSSGSSYEQY | 0.19507869 |
| TIPAITI | ASSSGSSYEQY | 0.62132835 |
| TIPGFTI | ASSSGSSYEQY | 0.32413596 |
| TSIASGALA | ASSSGSSYEQY | 0.56000614 |
| TVTLPGAAI | ASSSGSSYEQY | 0.80000705 |
| VASGLSAALP | ASSSGSSYEQY | 0.21600497 |
| VGAGVIGI | ASSSGSSYEQY | 0.5485631 |
| VGPKRFGV | ASSSGSSYEQY | 0.41979244 |
| VNVAFPD | ASSSGSSYEQY | 0.63960767 |
| VSMTNSGVS | ASSSGSSYEQY | 0.4729096 |
| VTDVLGDLSV | ASSSGSSYEQY | 0.23966645 |
| VVVVGAGFAGL | ASSSGSSYEQY | 0.74978495 |
| VYTVGTVV | ASSSGSSYEQY | 0.510163 |
| WTTFSGP | ASSSGSSYEQY | 0.39584303 |
| YDELVEAF | ASSSGSSYEQY | 0.57581496 |
| YGTGGAGGNGG | ASSSGSSYEQY | 0.7920289 |
| YTNVMAV | ASSSGSSYEQY | 0.58926666 |
| FLEQVHQL | ASSSGSSYEQY | 0.3197893 |
| GLSLLAVAV | ASSSGSSYEQY | 0.16826646 |
| LEQVHQL | ASSSGSSYEQY | 0.4921369 |
| LNAAGAI | ASSSGSSYEQY | 0.52803683 |
| PAVHASL | ASSSGSSYEQY | 0.28479114 |
| TELEDDA | ASSSGSSYEQY | 0.1573566 |
| TPPGPPN | ASSSGSSYEQY | 0.62430453 |
| TPPPPPPP | ASSSGSSYEQY | 0.83288145 |
| TQSPALI | ASSQEGNQAPL | 0.3385098 |
| EFAAAFPQ | ASSQEGNQAPL | 0.81546676 |
| LLSGNLDVL | ASSQEGNQAPL | 0.2008335 |
| GPANLGNYNI | ASSQEGNQAPL | 0.53331506 |
| TLGPTLP | ASSQEGNQAPL | 0.690555 |
| VYAYSSCCK | ASSQEGNQAPL | 0.19411287 |
| APAPAPAPNI | ASSQEGNQAPL | 0.40158406 |
| PGGPVPIP | ASSQEGNQAPL | 0.19963916 |

| | | |
|---|---|---|
| FLTVALEP | ASSQEGNQAPL | 0.24871832 |
| SQYFGANL | ASSQEGNQAPL | 0.75288284 |
| GGVKPAA | ASSQEGNQAPL | 0.35896206 |
| GGTGGAGGAGGW | ASSQEGNQAPL | 0.569481 |
| GVGGVAGLGGAGG | ASSQEGNQAPL | 0.35524926 |
| VLLVLATTI | ASSQEGNQAPL | 0.26458558 |
| IGIFAAML | ASSQEGNQAPL | 0.51887536 |
| AVGDLID | ASSQEGNQAPL | 0.66471934 |
| NISPLPL | ASSQEGNQAPL | 0.28212395 |
| VTAVFVAI | ASSQEGNQAPL | 0.79925746 |
| SGDDNDAAL | ASSQEGNQAPL | 0.32509333 |
| AVAALAHS | ASSQEGNQAPL | 0.8391994 |
| GGVTVMATP | ASSQEGNQAPL | 0.7888435 |
| AGLAANVRD | ASSQEGNQAPL | 0.5967924 |
| TNRPDLIDPALLRPG | ASSQEGNQAPL | 0.32093948 |
| LANLGVTAL | ASSQEGNQAPL | 0.77505064 |
| TGERIEL | ASSQEGNQAPL | 0.32886812 |
| NGEELAA | ASSQEGNQAPL | 0.6510976 |
| VLSLVWG | ASSQEGNQAPL | 0.6589296 |
| IGYALWL | ASSQEGNQAPL | 0.69349605 |
| LSLLLAPL | ASSQEGNQAPL | 0.73410773 |
| TLDEALK | ASSQEGNQAPL | 0.7189327 |
| TDGAGTAEGG | ASSQEGNQAPL | 0.7137269 |
| NAPGGAGGNGG | ASSQEGNQAPL | 0.69948167 |
| VSVFMGMGGNF | ASSQEGNQAPL | 0.5749349 |
| LASYLAAL | ASSQEGNQAPL | 0.7171676 |
| LVGGTPAL | ASSQEGNQAPL | 0.67681634 |
| VRNADVL | ASSQEGNQAPL | 0.7273082 |
| TCSSPFLVSAM | ASSQEGNQAPL | 0.72916126 |
| VHITDAE | ASSQEGNQAPL | 0.6834857 |
| EYAAPSF | ASSQEGNQAPL | 0.73494 |
| VVDAFHGGL | ASSQEGNQAPL | 0.7322741 |
| ASVVDPQ | ASSQEGNQAPL | 0.63275015 |
| SYNEFGGLS | ASSQEGNQAPL | 0.70667917 |
| DPDGSVY | ASSQEGNQAPL | 0.68849754 |
| MVSTIHATE | ASSQEGNQAPL | 0.7112948 |
| LAAGVADPV | ASSQEGNQAPL | 0.5311184 |
| TKEQIAA | ASSQEGNQAPL | 0.7302616 |
| SLASPAAL | ASSQEGNQAPL | 0.6741976 |
| IGLVFYGV | ASSQEGNQAPL | 0.29308352 |
| FLLDASNL | ASSQEGNQAPL | 0.7135736 |
| YQVGNFP | ASSQEGNQAPL | 0.71073693 |
| ESLLSQST | ASSQEGNQAPL | 0.71330106 |
| LFTVGFAAVG | ASSQEGNQAPL | 0.7290142 |
| ATPPTPT | ASSQEGNQAPL | 0.52349854 |
| TDLATLG | ASSQEGNQAPL | 0.68141115 |
| SAGGTGR | ASSQEGNQAPL | 0.7295537 |
| GFSNSGS | ASSQEGNQAPL | 0.7331059 |
| GGNGGSGGTSV | ASSQEGNQAPL | 0.6478429 |
| GTSVATGGAGN | ASSQEGNQAPL | 0.71902406 |
| NGGNGGSGGTS | ASSQEGNQAPL | 0.73172367 |

| | | |
|---|---|---|
| AVVKFPH | ASSQEGNQAPL | 0.72222745 |
| TPPPPPP | ASSQEGNQAPL | 0.7339815 |
| AAPAAAAAPA | ASSQEGNQAPL | 0.7363374 |
| IALNEDL | ASSQEGNQAPL | 0.6456825 |
| IVNVSSM | ASSQEGNQAPL | 0.73480463 |
| TGIPPAPR | ASSQEGNQAPL | 0.73660606 |
| FDLGGGTF | ASSQEGNQAPL | 0.7103814 |
| FDLGGGTFDVS | ASSQEGNQAPL | 0.71219563 |
| NEPTAAA | ASSQEGNQAPL | 0.67298853 |
| NEPTAAALA | ASSQEGNQAPL | 0.697433 |
| GSIINISSI | ASSQEGNQAPL | 0.55697113 |
| VAPGNAGTA | ASSQEGNQAPL | 0.71774614 |
| TGGGASLE | ASSQEGNQAPL | 0.7342731 |
| VAVYNHYKRI | ASSQEGNQAPL | 0.69707143 |
| TDPAPATT | ASSQEGNQAPL | 0.6513432 |
| TNRPDILDPALLRPG | ASSQEGNQAPL | 0.6798407 |
| LTTTTTS | ASSQEGNQAPL | 0.6813282 |
| PTGSGKTL | ASSQEGNQAPL | 0.7023946 |
| TGSGKTL | ASSQEGNQAPL | 0.5178211 |
| TIGHVDHGK | ASSQEGNQAPL | 0.65332973 |
| TIGHVDHGKTT | ASSQEGNQAPL | 0.73140216 |
| GHVDHGKTT | ASSQEGNQAPL | 0.73226345 |
| AQFEHTLL | ASSQEGNQAPL | 0.57238984 |
| IVNQDEL | ASSQEGNQAPL | 0.73442245 |
| TNGSQFF | ASSQEGNQAPL | 0.7261833 |
| TGKGIVE | ASSQEGNQAPL | 0.7367884 |
| LISNASDA | ASSQEGNQAPL | 0.71853393 |
| AELRTLGELTKIAKAHGAQ | ASSQEGNQAPL | 0.67237663 |
| APNAVAAAEPSW | ASSQEGNQAPL | 0.7229635 |
| APQVAGAATAVL | ASSQEGNQAPL | 0.6823958 |
| ARRDPAAAAAILPTDA | ASSQEGNQAPL | 0.62650645 |
| DAAARDVLAEAGLAEYF | ASSQEGNQAPL | 0.72890836 |
| DDQIVLSAKANLAKVEQ | ASSQEGNQAPL | 0.70333993 |
| DELLPDTGEDSDEDGH | ASSQEGNQAPL | 0.6691052 |
| ELPEAPANRATVPQD | ASSQEGNQAPL | 0.675308 |
| FGQSKIERVTGIG | ASSQEGNQAPL | 0.73239326 |
| GAGGTGSTAGGAGGAGGAGGLY | ASSQEGNQAPL | 0.6484585 |
| GHGGTGAAGGNGGTGGAGGSGI | ASSQEGNQAPL | 0.61438525 |
| GNGGNGGAGGEGGPGGAGGAGA | ASSQEGNQAPL | 0.65066916 |
| GSTAGNGGNGGNATNGTIAGQP | ASSQEGNQAPL | 0.70801675 |
| GTGGDGGNGGNGGTGG | ASSQEGNQAPL | 0.7309538 |
| GVRRPAGIPADAPV | ASSQEGNQAPL | 0.7296544 |
| GYRQVTDAHLLTLARR | ASSQEGNQAPL | 0.6674088 |
| IAKMGQDGHDRG | ASSQEGNQAPL | 0.72259814 |
| KAEPSIEIQYT | ASSQEGNQAPL | 0.72618294 |
| KPVRIRMAADEDTVAVT | ASSQEGNQAPL | 0.71231735 |
| MPPRIAGMRLLVIKPEP | ASSQEGNQAPL | 0.7085026 |
| MKFYAVALFRSRGVPPTR | ASSQEGNQAPL | 0.66888595 |
| NAGGFGILGAVAHSP | ASSQEGNQAPL | 0.73328805 |
| NEGREITLAMAEM | ASSQEGNQAPL | 0.7217128 |
| NELVEWSHPLGHRWRWE | ASSQEGNQAPL | 0.7337597 |

| | | |
|---|---|---|
| NLGAGGTGGNGGNPG | ASSQEGNQAPL | 0.73371494 |
| NTEWPKWETVMAADPVD | ASSQEGNQAPL | 0.70560104 |
| PASAAPAETQP | ASSQEGNQAPL | 0.71556795 |
| QQSSDGKNGVRAAEITGE | ASSQEGNQAPL | 0.5248525 |
| RAPDAVTTSWWKE | ASSQEGNQAPL | 0.65301704 |
| SGFGNVGAHTSGWFNQ | ASSQEGNQAPL | 0.7143613 |
| SGVAGAVGQAASVGGLKV | ASSQEGNQAPL | 0.71803045 |
| SNGGAGTGSAGGNGGT | ASSQEGNQAPL | 0.7303062 |
| TGGAGGEGGAAG | ASSQEGNQAPL | 0.72825223 |
| TGGDGGAAGAGGAPGHG | ASSQEGNQAPL | 0.6684823 |
| TMPPLLAAIANAGRHGV | ASSQEGNQAPL | 0.7290079 |
| TSSTAGNSGGA | ASSQEGNQAPL | 0.73260796 |
| VPATVPVGILC | ASSQEGNQAPL | 0.60760605 |
| VQRPDSGGVVLTGRIS | ASSQEGNQAPL | 0.68155456 |
| WTPNPVPATKGVPAGLVQAT | ASSQEGNQAPL | 0.7386 |
| YALKRALAKFSDRFDVVII | ASSQEGNQAPL | 0.71169424 |
| YLIGSVIV | ASSQEGNQAPL | 0.70165443 |
| YGQLVAVID | ASSQEGNQAPL | 0.6969656 |
| VTVGGMTLPA | ASSQEGNQAPL | 0.73783016 |
| VLALPAV | ASSQEGNQAPL | 0.67399824 |
| VIPPTLNYET | ASSQEGNQAPL | 0.73672134 |
| VGMGGMHPSSG | ASSQEGNQAPL | 0.42860252 |
| VGLAGGAATA | ASSQEGNQAPL | 0.7080333 |
| VGGAGGAGGSGP | ASSQEGNQAPL | 0.6740833 |
| VFAAFMV | ASSQEGNQAPL | 0.49299937 |
| TLYDKLR | ASSQEGNQAPL | 0.73220223 |
| TLAVFTVL | ASSQEGNQAPL | 0.7152462 |
| TGSVQPARV | ASSQEGNQAPL | 0.7284192 |
| TGGGAGSGGT | ASSQEGNQAPL | 0.7124847 |
| SHLETSL | ASSQEGNQAPL | 0.7344366 |
| SFTTGMT | ASSQEGNQAPL | 0.7129227 |
| QVEVAKRMGVRQPTVS | ASSQEGNQAPL | 0.709064 |
| QGSGGPALAM | ASSQEGNQAPL | 0.64947045 |
| MLAPSNP | ASSQEGNQAPL | 0.7297224 |
| LVNNAAI | ASSQEGNQAPL | 0.7323493 |
| LTGEINTLA | ASSQEGNQAPL | 0.7299837 |
| LSIFGPV | ASSQEGNQAPL | 0.6003463 |
| LLALLANL | ASSQEGNQAPL | 0.68544513 |
| LISNASDALDKLR | ASSQEGNQAPL | 0.72257566 |
| LINEAVP | ASSQEGNQAPL | 0.32791147 |
| LGYAFSGL | ASSQEGNQAPL | 0.38210016 |
| LDFVAVAL | ASSQEGNQAPL | 0.6811627 |
| LAQPGAE | ASSQEGNQAPL | 0.6450577 |
| ISNLGAVTGF | ASSQEGNQAPL | 0.7329721 |
| HQEDPGRI | ASSQEGNQAPL | 0.72397155 |
| GTPGEPTP | ASSQEGNQAPL | 0.7060446 |
| GFFNSGS | ASSQEGNQAPL | 0.7144134 |
| GAGGGGLTAGAN | ASSQEGNQAPL | 0.7012737 |
| FVDALVNM | ASSQEGNQAPL | 0.25387412 |
| FLTVAPEL | ASSQEGNQAPL | 0.6725739 |
| FETLTAL | ASSQEGNQAPL | 0.64270747 |

| | | |
|---|---|---|
| DVINPIEL | ASSQEGNQAPL | 0.7382317 |
| APVSGAPV | ASSQEGNQAPL | 0.6652154 |
| ALVGGLAW | ASSQEGNQAPL | 0.6554796 |
| ALAAAAATAI | ASSQEGNQAPL | 0.70873344 |
| AFGLGLLL | ASSQEGNQAPL | 0.7119658 |
| AFFAAASL | ASSQEGNQAPL | 0.7040541 |
| AAAALPAFSPP | ASSQEGNQAPL | 0.66716206 |
| VVLAPAL | ASSQEGNQAPL | 0.3441927 |
| TIGHVDHGKTTL | ASSQEGNQAPL | 0.7298405 |
| LISNASDALDK | ASSQEGNQAPL | 0.57163894 |
| LISNASDAL | ASSQEGNQAPL | 0.6711414 |
| ISNASDALDK | ASSQEGNQAPL | 0.42078537 |
| NQAETLVYQTE | ASSQEGNQAPL | 0.57111704 |
| TGGTGGTAGAG | ASSQEGNQAPL | 0.6045747 |
| TGSTAGGAGGA | ASSQEGNQAPL | 0.73109394 |
| DPLAVPVAAGPV | ASSQEGNQAPL | 0.701494 |
| GGTGGTPGNGGN | ASSQEGNQAPL | 0.6878667 |
| PAGLAGVDPATL | ASSQEGNQAPL | 0.7381269 |
| GNNLVVEAPHAGASV | ASSQEGNQAPL | 0.7069113 |
| KVTRIMSASTLEQPA | ASSQEGNQAPL | 0.68501073 |
| LGFALPIALPRRTVV | ASSQEGNQAPL | 0.6940361 |
| TRLGNLTARKLVVLV | ASSQEGNQAPL | 0.6684021 |
| WSQQRRLIYFPSAGP | ASSQEGNQAPL | 0.662345 |
| GPNGNPDPMAAAVDIR | ASSQEGNQAPL | 0.68441457 |
| TQRCVSVADLRAAASR | ASSQEGNQAPL | 0.7362396 |
| QFVLDVAEHPVEVLTETV | ASSQEGNQAPL | 0.70757693 |
| GGAGGTGGAGGNGGTQAGDG | ASSQEGNQAPL | 0.6698068 |
| PLLATLRAIAVDVDAKPAQV | ASSQEGNQAPL | 0.7191561 |
| APKFRTGGYMIDFWGVGYQVAK | ASSQEGNQAPL | 0.73659706 |
| GAGGAGGAGGAAGGPGGTGGTG | ASSQEGNQAPL | 0.73775756 |
| IGALGLLATDVAPAGSVAFGVA | ASSQEGNQAPL | 0.65721756 |
| AGGAGGIGGNGGAGGAGGVLMG | ASSQEGNQAPL | 0.7078346 |
| ADELIGLL | ASSQEGNQAPL | 0.695022 |
| ADRALKA | ASSQEGNQAPL | 0.7036693 |
| AGAVGQVGGAGGA | ASSQEGNQAPL | 0.68657315 |
| AGGGGGLGGAP | ASSQEGNQAPL | 0.6176418 |
| AGGLLYGNGGAG | ASSQEGNQAPL | 0.7276586 |
| AGLESIGL | ASSQEGNQAPL | 0.7391254 |
| AGLKTGII | ASSQEGNQAPL | 0.67632484 |
| ASLAALLL | ASSQEGNQAPL | 0.6523683 |
| ASVINISSI | ASSQEGNQAPL | 0.7014761 |
| ATPPGPA | ASSQEGNQAPL | 0.6715451 |
| ATVASALV | ASSQEGNQAPL | 0.6849703 |
| ATVASLIL | ASSQEGNQAPL | 0.70464265 |
| AVNGPVLIRH | ASSQEGNQAPL | 0.6188146 |
| DGAAGAVGI | ASSQEGNQAPL | 0.70723057 |
| DPLLHGI | ASSQEGNQAPL | 0.73771685 |
| ELGAALGAL | ASSQEGNQAPL | 0.69982743 |
| ELGRTQAV | ASSQEGNQAPL | 0.7215054 |
| FVAGGYLS | ASSQEGNQAPL | 0.7345437 |
| GASGGAGGAGGW | ASSQEGNQAPL | 0.7250265 |

| | | |
|---|---|---|
| GGAGAGGDGGAGGI | ASSQEGNQAPL | 0.7332263 |
| GTGHNGSQPGMGGNG | ASSQEGNQAPL | 0.7286502 |
| GYAAASASAS | ASSQEGNQAPL | 0.7086252 |
| HQVDVAAVQQG | ASSQEGNQAPL | 0.7110058 |
| IGAAIAHT | ASSQEGNQAPL | 0.7216923 |
| IGMIEVAQL | ASSQEGNQAPL | 0.7079407 |
| IRLAPEHTSAMD | ASSQEGNQAPL | 0.69916034 |
| IYNVSVL | ASSQEGNQAPL | 0.68007374 |
| KQGFLGM | ASSQEGNQAPL | 0.7340536 |
| LESVITQL | ASSQEGNQAPL | 0.69798386 |
| LGLGALVVV | ASSQEGNQAPL | 0.7319622 |
| LIGNGGNGGSGGI | ASSQEGNQAPL | 0.59400034 |
| LNAINAPTL | ASSQEGNQAPL | 0.629686 |
| LQLLNTGV | ASSQEGNQAPL | 0.7286806 |
| LQTSIEF | ASSQEGNQAPL | 0.25964427 |
| MSFAVSEL | ASSQEGNQAPL | 0.4182184 |
| NTGALGL | ASSQEGNQAPL | 0.71969086 |
| PPGAGGAGGAGTT | ASSQEGNQAPL | 0.731606 |
| QLVGLGI | ASSQEGNQAPL | 0.26804352 |
| RFLEQVHQL | ASSQEGNQAPL | 0.6450058 |
| RGVGYVLR | ASSQEGNQAPL | 0.6340107 |
| SATGVFAGVI | ASSQEGNQAPL | 0.7249377 |
| SLPAITLP | ASSQEGNQAPL | 0.71541774 |
| STALAFTV | ASSQEGNQAPL | 0.54300576 |
| SVLTVGLGK | ASSQEGNQAPL | 0.7303686 |
| TALAGAGSGI | ASSQEGNQAPL | 0.5643779 |
| TGLPFTGI | ASSQEGNQAPL | 0.5588694 |
| VAEHVTV | ASSQEGNQAPL | 0.73217475 |
| VDIVLNSL | ASSQEGNQAPL | 0.733669 |
| VLTGGGAVL | ASSQEGNQAPL | 0.6285577 |
| VNVGLAYGI | ASSQEGNQAPL | 0.7290139 |
| VSSAAVGAV | ASSQEGNQAPL | 0.6806683 |
| VTSVYEAGV | ASSQEGNQAPL | 0.7383206 |
| WVALDRGA | ASSQEGNQAPL | 0.70069826 |
| YLVGSVVI | ASSQEGNQAPL | 0.6111027 |
| GPPGTGKTLL | ASSQEGNQAPL | 0.7323226 |
| GVGMTKF | ASSQEGNQAPL | 0.7239225 |
| IGIGGSDL | ASSQEGNQAPL | 0.71149987 |
| IGTLLAEL | ASSQEGNQAPL | 0.7283721 |
| IYTSGTT | ASSQEGNQAPL | 0.6579446 |
| LSFPAGV | ASSQEGNQAPL | 0.7099017 |
| LYTSGTT | ASSQEGNQAPL | 0.73514587 |
| VGAFPFGF | ASSQEGNQAPL | 0.7254425 |
| VGAGTVEF | ASSQEGNQAPL | 0.6123567 |
| VGKSTLLL | ASSQEGNQAPL | 0.7307828 |
| AALAHAGGGIV | ASSQEGNQAPL | 0.69370234 |
| AALVAVSLL | ASSQEGNQAPL | 0.6860037 |
| AAVSRAVAAL | ASSQEGNQAPL | 0.7372551 |
| AGGSSGIGGF | ASSQEGNQAPL | 0.7123014 |
| AGLTVANANG | ASSQEGNQAPL | 0.72764254 |
| AGLVGAVL | ASSQEGNQAPL | 0.7341898 |

| | | |
|---|---|---|
| ALSQAFTV | ASSQEGNQAPL | 0.73540807 |
| ALTFSVSL | ASSQEGNQAPL | 0.5516711 |
| ALTGAALVAL | ASSQEGNQAPL | 0.7086187 |
| ASFSIGAGVL | ASSQEGNQAPL | 0.70137894 |
| DGSVYKI | ASSQEGNQAPL | 0.73196304 |
| EQLAEIV | ASSQEGNQAPL | 0.7182636 |
| FAVSELTVM | ASSQEGNQAPL | 0.6502665 |
| GAAPLAAGL | ASSQEGNQAPL | 0.7246684 |
| GADGGDGGSGGAGG | ASSQEGNQAPL | 0.73347837 |
| GFGLDTGGAGG | ASSQEGNQAPL | 0.712922 |
| GLASGGLL | ASSQEGNQAPL | 0.71222186 |
| GRIAGAAVLL | ASSQEGNQAPL | 0.73505265 |
| GSAGSLNTSF | ASSQEGNQAPL | 0.70510197 |
| GTGQIGI | ASSQEGNQAPL | 0.6977984 |
| GTLIALIL | ASSQEGNQAPL | 0.73223144 |
| IAATMGFVVG | ASSQEGNQAPL | 0.73000145 |
| IFAQALL | ASSQEGNQAPL | 0.7109951 |
| IGAAVALGF | ASSQEGNQAPL | 0.72973686 |
| IGLTPLAL | ASSQEGNQAPL | 0.6187144 |
| ILGSNTSTL | ASSQEGNQAPL | 0.7251059 |
| IVFSLTVL | ASSQEGNQAPL | 0.6653589 |
| LASKATGF | ASSQEGNQAPL | 0.72354543 |
| LASVALIL | ASSQEGNQAPL | 0.6469391 |
| LFAVGLTL | ASSQEGNQAPL | 0.7285496 |
| LFNGGIGTYI | ASSQEGNQAPL | 0.7268641 |
| LTLGAGII | ASSQEGNQAPL | 0.7304957 |
| LVAAALGDI | ASSQEGNQAPL | 0.3972793 |
| NTAFVQLGI | ASSQEGNQAPL | 0.7288422 |
| QLFSGAEL | ASSQEGNQAPL | 0.73075294 |
| SAATFITL | ASSQEGNQAPL | 0.73815244 |
| SNYGLANLGV | ASSQEGNQAPL | 0.70365095 |
| SYGGAYAVM | ASSQEGNQAPL | 0.7215959 |
| TTGVGGLTLG | ASSQEGNQAPL | 0.68057853 |
| TTQTIYI | ASSQEGNQAPL | 0.7035835 |
| VAAGAGLGAVI | ASSQEGNQAPL | 0.63993645 |
| VAAVVTLL | ASSQEGNQAPL | 0.69745195 |
| VGFLGGLI | ASSQEGNQAPL | 0.6698141 |
| VGVIPILL | ASSQEGNQAPL | 0.6766999 |
| VIFASSLI | ASSQEGNQAPL | 0.72233915 |
| VKGGDGGAAAG | ASSQEGNQAPL | 0.7284645 |
| VTKTFTNI | ASSQEGNQAPL | 0.7288909 |
| VVAAAILL | ASSQEGNQAPL | 0.73509073 |
| YAALDVEL | ASSQEGNQAPL | 0.7321239 |
| LASVALLL | ASSQEGNQAPL | 0.73483837 |
| LDAVHRV | ASSQEGNQAPL | 0.6870431 |
| QGLSSSIF | ASSQEGNQAPL | 0.6930417 |
| RGTFANIRL | ASSQEGNQAPL | 0.65379274 |
| RQLWWGHRI | ASSQEGNQAPL | 0.7047212 |
| YGPPGTGKTLL | ASSQEGNQAPL | 0.69167626 |
| AALVGAPANVA | ASSQEGNQAPL | 0.716998 |
| AATFRRTE | ASSQEGNQAPL | 0.26691654 |

| | | |
|---|---|---|
| AGALGGGI | ASSQEGNQAPL | 0.7020813 |
| AGGAGGAGGIGAG | ASSQEGNQAPL | 0.73605967 |
| AGSGALGL | ASSQEGNQAPL | 0.63787115 |
| AQIADQI | ASSQEGNQAPL | 0.7222208 |
| ASAVAVTYL | ASSQEGNQAPL | 0.70992327 |
| ASTVLGGTGG | ASSQEGNQAPL | 0.6819206 |
| AVGGEPAK | ASSQEGNQAPL | 0.727972 |
| CEQGTFGGTC | ASSQEGNQAPL | 0.6745463 |
| DEGKAPAG | ASSQEGNQAPL | 0.7260963 |
| DTVVFPAL | ASSQEGNQAPL | 0.7370968 |
| FEGITMAG | ASSQEGNQAPL | 0.72233653 |
| FPLVVQD | ASSQEGNQAPL | 0.6988778 |
| GAFTTPQ | ASSQEGNQAPL | 0.7119255 |
| GFGGFGTAGGVGGA | ASSQEGNQAPL | 0.5285774 |
| GFLAQFL | ASSQEGNQAPL | 0.7155514 |
| GGAGGAGGNAGVA | ASSQEGNQAPL | 0.293553 |
| GGVGTAGAGGAGGAGG | ASSQEGNQAPL | 0.71056116 |
| GNGGGGDAGNAGSGGN | ASSQEGNQAPL | 0.66594195 |
| GTGGAGGAGNDAGS | ASSQEGNQAPL | 0.71740586 |
| GTTNTGWFNS | ASSQEGNQAPL | 0.7047434 |
| IFMGPLFTL | ASSQEGNQAPL | 0.7192027 |
| ILAHAIEAI | ASSQEGNQAPL | 0.67737055 |
| IRSAIREAL | ASSQEGNQAPL | 0.7056693 |
| ITFAGGG | ASSQEGNQAPL | 0.6644714 |
| ITGMSGHQEV | ASSQEGNQAPL | 0.6431291 |
| IVAGLACTL | ASSQEGNQAPL | 0.72659934 |
| IVGGGLAGA | ASSQEGNQAPL | 0.713551 |
| IVTGVLNI | ASSQEGNQAPL | 0.68108726 |
| LAQQGCI | ASSQEGNQAPL | 0.7349602 |
| LCLHVEN | ASSQEGNQAPL | 0.7219287 |
| LDAISDGS | ASSQEGNQAPL | 0.5383457 |
| LGGFAAML | ASSQEGNQAPL | 0.69412065 |
| LGGKNPMI | ASSQEGNQAPL | 0.7135496 |
| LGPKGVAAIL | ASSQEGNQAPL | 0.7067233 |
| LMVGVSI | ASSQEGNQAPL | 0.27932894 |
| LVGAAFVL | ASSQEGNQAPL | 0.71353966 |
| LVLAVAPL | ASSQEGNQAPL | 0.6861252 |
| MAAIAVIDAA | ASSQEGNQAPL | 0.6596355 |
| NNGINAELQ | ASSQEGNQAPL | 0.6838602 |
| RGEPLVLVHG | ASSQEGNQAPL | 0.72843325 |
| RGGAGGAAGLI | ASSQEGNQAPL | 0.590706 |
| RSYRMHPAV | ASSQEGNQAPL | 0.71807206 |
| SAAHLLAGQLA | ASSQEGNQAPL | 0.72237796 |
| SAAVGSY | ASSQEGNQAPL | 0.72369474 |
| SPHAVLHRI | ASSQEGNQAPL | 0.73424995 |
| SQIAKVL | ASSQEGNQAPL | 0.71452796 |
| TALGHLVP | ASSQEGNQAPL | 0.70282376 |
| TAPTTSGAA | ASSQEGNQAPL | 0.71871114 |
| TDGDDGNGGSPGT | ASSQEGNQAPL | 0.70160234 |
| TDGLGEN | ASSQEGNQAPL | 0.5451811 |
| TGGPPLPGQA | ASSQEGNQAPL | 0.470533 |

| | | |
|---|---|---|
| TIPAITI | ASSQEGNQAPL | 0.72010624 |
| TIPGFTI | ASSQEGNQAPL | 0.48880792 |
| TSIASGALA | ASSQEGNQAPL | 0.71742105 |
| TVTLPGAAI | ASSQEGNQAPL | 0.73674035 |
| VASGLSAALP | ASSQEGNQAPL | 0.683715 |
| VGAGVIGI | ASSQEGNQAPL | 0.6173488 |
| VGPKRFGV | ASSQEGNQAPL | 0.6985266 |
| VNVAFPD | ASSQEGNQAPL | 0.68333626 |
| VSMTNSGVS | ASSQEGNQAPL | 0.7275306 |
| VTDVLGDLSV | ASSQEGNQAPL | 0.7298523 |
| VVVVGAGFAGL | ASSQEGNQAPL | 0.73642004 |
| VYTVGTVV | ASSQEGNQAPL | 0.72940755 |
| WTTFSGP | ASSQEGNQAPL | 0.6849899 |
| YDELVEAF | ASSQEGNQAPL | 0.72191364 |
| YGTGGAGGNGG | ASSQEGNQAPL | 0.70203525 |
| YTNVMAV | ASSQEGNQAPL | 0.7046038 |
| FLEQVHQL | ASSQEGNQAPL | 0.71273804 |
| GLSLLAVAV | ASSQEGNQAPL | 0.7152512 |
| LEQVHQL | ASSQEGNQAPL | 0.70496565 |
| LNAAGAI | ASSQEGNQAPL | 0.7028384 |
| PAVHASL | ASSQEGNQAPL | 0.6100403 |
| TELEDDA | ASSQEGNQAPL | 0.7262204 |
| TPPGPPN | ASSQEGNQAPL | 0.71750736 |
| TPPPPPPP | ASSQEGNQAPL | 0.52383494 |
| TQSPALI | ASSQSDRGLANTEVF | 0.69127 |
| EFAAAFPQ | ASSQSDRGLANTEVF | 0.73616993 |
| LLSGNLDVL | ASSQSDRGLANTEVF | 0.73310673 |
| GPANLGNYNI | ASSQSDRGLANTEVF | 0.67247283 |
| TLGPTLP | ASSQSDRGLANTEVF | 0.68568444 |
| VYAYSSCCK | ASSQSDRGLANTEVF | 0.7000525 |
| APAPAPAPNI | ASSQSDRGLANTEVF | 0.5230243 |
| PGGPVPIP | ASSQSDRGLANTEVF | 0.7212024 |
| FLTVALEP | ASSQSDRGLANTEVF | 0.7048656 |
| SQYFGANL | ASSQSDRGLANTEVF | 0.69477916 |
| GGVKPAA | ASSQSDRGLANTEVF | 0.67795813 |
| GGTGGAGGAGGW | ASSQSDRGLANTEVF | 0.7163844 |
| GVGGVAGLGGAGG | ASSQSDRGLANTEVF | 0.6793784 |
| VLLVLATTI | ASSQSDRGLANTEVF | 0.7253971 |
| IGIFAAML | ASSQSDRGLANTEVF | 0.27058142 |
| AVGDLID | ASSQSDRGLANTEVF | 0.7243495 |
| NISPLPL | ASSQSDRGLANTEVF | 0.64719045 |
| VTAVFVAI | ASSQSDRGLANTEVF | 0.6446948 |
| SGDDNDAAL | ASSQSDRGLANTEVF | 0.47188473 |
| AVAALAHS | ASSQSDRGLANTEVF | 0.7106432 |
| GGVTVMATP | ASSQSDRGLANTEVF | 0.730212 |
| AGLAANVRD | ASSQSDRGLANTEVF | 0.7284978 |
| TNRPDLIDPALLRPG | ASSQSDRGLANTEVF | 0.6148596 |
| LANLGVTAL | ASSQSDRGLANTEVF | 0.73064005 |
| TGERIEL | ASSQSDRGLANTEVF | 0.7336179 |
| NGEELAA | ASSQSDRGLANTEVF | 0.6871549 |
| VLSLVWG | ASSQSDRGLANTEVF | 0.42073107 |

| | | |
|---|---|---|
| IGYALWL | ASSQSDRGLANTEVF | 0.7261368 |
| LSLLLAPL | ASSQSDRGLANTEVF | 0.69863206 |
| TLDEALK | ASSQSDRGLANTEVF | 0.7290902 |
| TDGAGTAEGG | ASSQSDRGLANTEVF | 0.7149224 |
| NAPGGAGGNGG | ASSQSDRGLANTEVF | 0.6398849 |
| VSVFMGMGGNF | ASSQSDRGLANTEVF | 0.72963953 |
| LASYLAAL | ASSQSDRGLANTEVF | 0.7342005 |
| LVGGTPAL | ASSQSDRGLANTEVF | 0.65490496 |
| VRNADVL | ASSQSDRGLANTEVF | 0.66229033 |
| TCSSPFLVSAM | ASSQSDRGLANTEVF | 0.6749923 |
| VHITDAE | ASSQSDRGLANTEVF | 0.5405718 |
| EYAAPSF | ASSQSDRGLANTEVF | 0.72803897 |
| VVDAFHGGL | ASSQSDRGLANTEVF | 0.67910445 |
| ASVVDPQ | ASSQSDRGLANTEVF | 0.68929183 |
| SYNEFGGLS | ASSQSDRGLANTEVF | 0.7193058 |
| DPDGSVY | ASSQSDRGLANTEVF | 0.6917095 |
| MVSTIHATE | ASSQSDRGLANTEVF | 0.6969703 |
| LAAGVADPV | ASSQSDRGLANTEVF | 0.70091236 |
| TKEQIAA | ASSQSDRGLANTEVF | 0.7226018 |
| SLASPAAL | ASSQSDRGLANTEVF | 0.71822625 |
| IGLVFYGV | ASSQSDRGLANTEVF | 0.7086909 |
| FLLDASNL | ASSQSDRGLANTEVF | 0.7182332 |
| YQVGNFP | ASSQSDRGLANTEVF | 0.73439074 |
| ESLLSQST | ASSQSDRGLANTEVF | 0.60562503 |
| LFTVGFAAVG | ASSQSDRGLANTEVF | 0.6687368 |
| ATPPTPT | ASSQSDRGLANTEVF | 0.71833754 |
| TDLATLG | ASSQSDRGLANTEVF | 0.61188936 |
| SAGGTGR | ASSQSDRGLANTEVF | 0.7243716 |
| GFSNSGS | ASSQSDRGLANTEVF | 0.7020739 |
| GGNGGSGGTSV | ASSQSDRGLANTEVF | 0.71564734 |
| GTSVATGGAGN | ASSQSDRGLANTEVF | 0.6978338 |
| NGGNGGSGGTS | ASSQSDRGLANTEVF | 0.7378514 |
| AVVKFPH | ASSQSDRGLANTEVF | 0.73224115 |
| TPPPPPP | ASSQSDRGLANTEVF | 0.71910995 |
| AAPAAAAAPA | ASSQSDRGLANTEVF | 0.72164404 |
| IALNEDL | ASSQSDRGLANTEVF | 0.6185943 |
| IVNVSSM | ASSQSDRGLANTEVF | 0.7314859 |
| TGIPPAPR | ASSQSDRGLANTEVF | 0.7261537 |
| FDLGGGTF | ASSQSDRGLANTEVF | 0.65543604 |
| FDLGGGTFDVS | ASSQSDRGLANTEVF | 0.71200156 |
| NEPTAAA | ASSQSDRGLANTEVF | 0.73459667 |
| NEPTAAALA | ASSQSDRGLANTEVF | 0.7338675 |
| GSIINISSI | ASSQSDRGLANTEVF | 0.5758679 |
| VAPGNAGTA | ASSQSDRGLANTEVF | 0.6974453 |
| TGGGASLE | ASSQSDRGLANTEVF | 0.4533411 |
| VAVYNHYKRI | ASSQSDRGLANTEVF | 0.6475394 |
| TDPAPATT | ASSQSDRGLANTEVF | 0.6400159 |
| TNRPDILDPALLRPG | ASSQSDRGLANTEVF | 0.720275 |
| LTTTTTS | ASSQSDRGLANTEVF | 0.6968069 |
| PTGSGKTL | ASSQSDRGLANTEVF | 0.7189661 |
| TGSGKTL | ASSQSDRGLANTEVF | 0.7149309 |

| | | |
|---|---|---|
| TIGHVDHGK | ASSQSDRGLANTEVF | 0.7353076 |
| TIGHVDHGKTT | ASSQSDRGLANTEVF | 0.6938555 |
| GHVDHGKTT | ASSQSDRGLANTEVF | 0.70446765 |
| AQFEHTLL | ASSQSDRGLANTEVF | 0.5854099 |
| IVNQDEL | ASSQSDRGLANTEVF | 0.58463645 |
| TNGSQFF | ASSQSDRGLANTEVF | 0.6777313 |
| TGKGIVE | ASSQSDRGLANTEVF | 0.7030587 |
| LISNASDA | ASSQSDRGLANTEVF | 0.3435921 |
| AELRTLGELTKIAKAHGAQ | ASSQSDRGLANTEVF | 0.7338976 |
| APNAVAAAEPSW | ASSQSDRGLANTEVF | 0.7182248 |
| APQVAGAATAVL | ASSQSDRGLANTEVF | 0.72802544 |
| ARRDPAAAAAILPTDA | ASSQSDRGLANTEVF | 0.681993 |
| DAAARDVLAEAGLAEYF | ASSQSDRGLANTEVF | 0.7052372 |
| DDQIVLSAKANLAKVEQ | ASSQSDRGLANTEVF | 0.6802393 |
| DELLPDTGEDSDEDGH | ASSQSDRGLANTEVF | 0.7241524 |
| ELPEAPANRATVPQD | ASSQSDRGLANTEVF | 0.64415693 |
| FGQSKIERVTGIG | ASSQSDRGLANTEVF | 0.6240444 |
| GAGGTGSTAGGAGGAGGAGGLY | ASSQSDRGLANTEVF | 0.6964736 |
| GHGGTGAAGGNGGTGGAGGSGI | ASSQSDRGLANTEVF | 0.6256505 |
| GNGGNGGAGGEGGPGGAGGAGA | ASSQSDRGLANTEVF | 0.7344417 |
| GSTAGNGGNGGNATNGTIAGQP | ASSQSDRGLANTEVF | 0.7340567 |
| GTGGDGGNGGNGGTGG | ASSQSDRGLANTEVF | 0.7231704 |
| GVRRPAGIPADAPV | ASSQSDRGLANTEVF | 0.73223424 |
| GYRQVTDAHLLTLARR | ASSQSDRGLANTEVF | 0.31859046 |
| IAKMGQDGHDRG | ASSQSDRGLANTEVF | 0.67779803 |
| KAEPSIEIQYT | ASSQSDRGLANTEVF | 0.73475647 |
| KPVRIRMAADEDTVAVT | ASSQSDRGLANTEVF | 0.7218751 |
| MPPRIAGMRLLVIKPEP | ASSQSDRGLANTEVF | 0.7028141 |
| MKFYAVALFRSRGVPPTR | ASSQSDRGLANTEVF | 0.7334526 |
| NAGGFGILGAVAHSP | ASSQSDRGLANTEVF | 0.67617035 |
| NEGREITLAMAEM | ASSQSDRGLANTEVF | 0.70688903 |
| NELVEWSHPLGHRWRWE | ASSQSDRGLANTEVF | 0.71189344 |
| NLGAGGTGGNGGNPG | ASSQSDRGLANTEVF | 0.7262409 |
| NTEWPKWETVMAADPVD | ASSQSDRGLANTEVF | 0.7224913 |
| PASAAPAETQP | ASSQSDRGLANTEVF | 0.6555215 |
| QQSSDGKNGVRAAEITGE | ASSQSDRGLANTEVF | 0.7235136 |
| RAPDAVTTSWWKE | ASSQSDRGLANTEVF | 0.73920256 |
| SGFGNVGAHTSGWFNQ | ASSQSDRGLANTEVF | 0.7024636 |
| SGVAGAVGQAASVGGLKV | ASSQSDRGLANTEVF | 0.7274029 |
| SNGGAGTGSAGGNGGT | ASSQSDRGLANTEVF | 0.72470963 |
| TGGAGGEGGAAG | ASSQSDRGLANTEVF | 0.72164845 |
| TGGDGGAAGAGGAPGHG | ASSQSDRGLANTEVF | 0.6806272 |
| TMPPLLAAIANAGRHGV | ASSQSDRGLANTEVF | 0.67627823 |
| TSSTAGNSGGA | ASSQSDRGLANTEVF | 0.7063846 |
| VPATVPVGILC | ASSQSDRGLANTEVF | 0.6308091 |
| VQRPDSGGVVLTGRIS | ASSQSDRGLANTEVF | 0.7137197 |
| WTPNPVPATKGVPAGLVQAT | ASSQSDRGLANTEVF | 0.7007458 |
| YALKRALAKFSDRFDVVII | ASSQSDRGLANTEVF | 0.7406492 |
| YLIGSVIV | ASSQSDRGLANTEVF | 0.7230855 |
| YGQLVAVID | ASSQSDRGLANTEVF | 0.7206806 |
| VTVGGMTLPA | ASSQSDRGLANTEVF | 0.6563654 |

| | | |
|---|---|---|
| VLALPAV | ASSQSDRGLANTEVF | 0.7231325 |
| VIPPTLNYET | ASSQSDRGLANTEVF | 0.7217737 |
| VGMGGMHPSSG | ASSQSDRGLANTEVF | 0.6334356 |
| VGLAGGAATA | ASSQSDRGLANTEVF | 0.731395 |
| VGGAGGAGGSGP | ASSQSDRGLANTEVF | 0.6716006 |
| VFAAFMV | ASSQSDRGLANTEVF | 0.6932136 |
| TLYDKLR | ASSQSDRGLANTEVF | 0.7292472 |
| TLAVFTVL | ASSQSDRGLANTEVF | 0.7346051 |
| TGSVQPARV | ASSQSDRGLANTEVF | 0.6945719 |
| TGGGAGSGGT | ASSQSDRGLANTEVF | 0.6760757 |
| SHLETSL | ASSQSDRGLANTEVF | 0.6707487 |
| SFTTGMT | ASSQSDRGLANTEVF | 0.7284887 |
| QVEVAKRMGVRQPTVS | ASSQSDRGLANTEVF | 0.7317426 |
| QGSGGPALAM | ASSQSDRGLANTEVF | 0.70945936 |
| MLAPSNP | ASSQSDRGLANTEVF | 0.7363174 |
| LVNNAAI | ASSQSDRGLANTEVF | 0.72557074 |
| LTGEINTLA | ASSQSDRGLANTEVF | 0.5621363 |
| LSIFGPV | ASSQSDRGLANTEVF | 0.7095664 |
| LLALLANL | ASSQSDRGLANTEVF | 0.73135173 |
| LISNASDALDKLR | ASSQSDRGLANTEVF | 0.7284605 |
| LINEAVP | ASSQSDRGLANTEVF | 0.7391555 |
| LGYAFSGL | ASSQSDRGLANTEVF | 0.72331154 |
| LDFVAVAL | ASSQSDRGLANTEVF | 0.68992186 |
| LAQPGAE | ASSQSDRGLANTEVF | 0.7358624 |
| ISNLGAVTGF | ASSQSDRGLANTEVF | 0.7396106 |
| HQEDPGRI | ASSQSDRGLANTEVF | 0.72911656 |
| GTPGEPTP | ASSQSDRGLANTEVF | 0.6888918 |
| GFFNSGS | ASSQSDRGLANTEVF | 0.6628181 |
| GAGGGGLTAGAN | ASSQSDRGLANTEVF | 0.7212454 |
| FVDALVNM | ASSQSDRGLANTEVF | 0.7267842 |
| FLTVAPEL | ASSQSDRGLANTEVF | 0.6684549 |
| FETLTAL | ASSQSDRGLANTEVF | 0.7115439 |
| DVINPIEL | ASSQSDRGLANTEVF | 0.71582854 |
| APVSGAPV | ASSQSDRGLANTEVF | 0.6139486 |
| ALVGGLAW | ASSQSDRGLANTEVF | 0.7288723 |
| ALAAAAATAI | ASSQSDRGLANTEVF | 0.730979 |
| AFGLGLLL | ASSQSDRGLANTEVF | 0.71452117 |
| AFFAAASL | ASSQSDRGLANTEVF | 0.69249743 |
| AAAALPAFSPP | ASSQSDRGLANTEVF | 0.7288085 |
| VVLAPAL | ASSQSDRGLANTEVF | 0.74035513 |
| TIGHVDHGKTTL | ASSQSDRGLANTEVF | 0.7365526 |
| LISNASDALDK | ASSQSDRGLANTEVF | 0.58546025 |
| LISNASDAL | ASSQSDRGLANTEVF | 0.44662845 |
| ISNASDALDK | ASSQSDRGLANTEVF | 0.6789848 |
| NQAETLVYQTE | ASSQSDRGLANTEVF | 0.7265659 |
| TGGTGGTAGAG | ASSQSDRGLANTEVF | 0.7053888 |
| TGSTAGGAGGA | ASSQSDRGLANTEVF | 0.69757295 |
| DPLAVPVAAGPV | ASSQSDRGLANTEVF | 0.71706223 |
| GGTGGTPGNGGN | ASSQSDRGLANTEVF | 0.7339004 |
| PAGLAGVDPATL | ASSQSDRGLANTEVF | 0.7006973 |
| GNNLVVEAPHAGASV | ASSQSDRGLANTEVF | 0.5807906 |

| | | |
|---|---|---|
| KVTRIMSASTLEQPA | ASSQSDRGLANTEVF | 0.72407115 |
| LGFALPIALPRRTVV | ASSQSDRGLANTEVF | 0.7281709 |
| TRLGNLTARKLVVLV | ASSQSDRGLANTEVF | 0.6788625 |
| WSQQRRLIYFPSAGP | ASSQSDRGLANTEVF | 0.70332575 |
| GPNGNPDPMAAAVDIR | ASSQSDRGLANTEVF | 0.6704545 |
| TQRCVSVADLRAAASR | ASSQSDRGLANTEVF | 0.6448605 |
| QFVLDVAEHPVEVLTETV | ASSQSDRGLANTEVF | 0.7123903 |
| GGAGGTGGAGGNGGTQAGDG | ASSQSDRGLANTEVF | 0.6633681 |
| PLLATLRAIAVDVDAKPAQV | ASSQSDRGLANTEVF | 0.73140675 |
| APKFRTGGYMIDFWGVGYQVAK | ASSQSDRGLANTEVF | 0.67002416 |
| GAGGAGGAGGAAGGPGGTGGTG | ASSQSDRGLANTEVF | 0.7153356 |
| IGALGLLATDVAPAGSVAFGVA | ASSQSDRGLANTEVF | 0.7203107 |
| AGGAGGIGGNGGAGGAGGVLMG | ASSQSDRGLANTEVF | 0.7260436 |
| ADELIGLL | ASSQSDRGLANTEVF | 0.69531107 |
| ADRALKA | ASSQSDRGLANTEVF | 0.68825233 |
| AGAVGQVGGAGGA | ASSQSDRGLANTEVF | 0.63253826 |
| AGGGGGLGGAP | ASSQSDRGLANTEVF | 0.48185873 |
| AGGLLYGNGGAG | ASSQSDRGLANTEVF | 0.7315665 |
| AGLESIGL | ASSQSDRGLANTEVF | 0.52695715 |
| AGLKTGII | ASSQSDRGLANTEVF | 0.73414296 |
| ASLAALLL | ASSQSDRGLANTEVF | 0.73037946 |
| ASVINISSI | ASSQSDRGLANTEVF | 0.66353905 |
| ATPPGPA | ASSQSDRGLANTEVF | 0.7234528 |
| ATVASALV | ASSQSDRGLANTEVF | 0.63266975 |
| ATVASLIL | ASSQSDRGLANTEVF | 0.65717095 |
| AVNGPVLIRH | ASSQSDRGLANTEVF | 0.7258275 |
| DGAAGAVGI | ASSQSDRGLANTEVF | 0.72836137 |
| DPLLHGI | ASSQSDRGLANTEVF | 0.66754645 |
| ELGAALGAL | ASSQSDRGLANTEVF | 0.71894586 |
| ELGRTQAV | ASSQSDRGLANTEVF | 0.7190063 |
| FVAGGYLS | ASSQSDRGLANTEVF | 0.66163814 |
| GASGGAGGAGGW | ASSQSDRGLANTEVF | 0.7186165 |
| GGAGAGGDGGAGGI | ASSQSDRGLANTEVF | 0.73300743 |
| GTGHNGSQPGMGGNG | ASSQSDRGLANTEVF | 0.7200039 |
| GYAAASASAS | ASSQSDRGLANTEVF | 0.6753589 |
| HQVDVAAVQQG | ASSQSDRGLANTEVF | 0.72654384 |
| IGAAIAHT | ASSQSDRGLANTEVF | 0.65186214 |
| IGMIEVAQL | ASSQSDRGLANTEVF | 0.720337 |
| IRLAPEHTSAMD | ASSQSDRGLANTEVF | 0.70004416 |
| IYNVSVL | ASSQSDRGLANTEVF | 0.69849336 |
| KQGFLGM | ASSQSDRGLANTEVF | 0.7280036 |
| LESVITQL | ASSQSDRGLANTEVF | 0.5838253 |
| LGLGALVVV | ASSQSDRGLANTEVF | 0.66542083 |
| LIGNGGNGGSGGI | ASSQSDRGLANTEVF | 0.69088584 |
| LNAINAPTL | ASSQSDRGLANTEVF | 0.71667933 |
| LQLLNTGV | ASSQSDRGLANTEVF | 0.73050934 |
| LQTSIEF | ASSQSDRGLANTEVF | 0.6706058 |
| MSFAVSEL | ASSQSDRGLANTEVF | 0.71391 |
| NTGALGL | ASSQSDRGLANTEVF | 0.70457965 |
| PPGAGGAGGAGTT | ASSQSDRGLANTEVF | 0.6750804 |
| QLVGLGI | ASSQSDRGLANTEVF | 0.71830046 |

| | | |
|---|---|---|
| RFLEQVHQL | ASSQSDRGLANTEVF | 0.71037567 |
| RGVGYVLR | ASSQSDRGLANTEVF | 0.6452333 |
| SATGVFAGVI | ASSQSDRGLANTEVF | 0.7246276 |
| SLPAITLP | ASSQSDRGLANTEVF | 0.6666341 |
| STALAFTV | ASSQSDRGLANTEVF | 0.71912485 |
| SVLTVGLGK | ASSQSDRGLANTEVF | 0.7373555 |
| TALAGAGSGI | ASSQSDRGLANTEVF | 0.66206557 |
| TGLPFTGI | ASSQSDRGLANTEVF | 0.721985 |
| VAEHVTV | ASSQSDRGLANTEVF | 0.7211679 |
| VDIVLNSL | ASSQSDRGLANTEVF | 0.71919405 |
| VLTGGGAVL | ASSQSDRGLANTEVF | 0.70569646 |
| VNVGLAYGI | ASSQSDRGLANTEVF | 0.68154377 |
| VSSAAVGAV | ASSQSDRGLANTEVF | 0.73558605 |
| VTSVYEAGV | ASSQSDRGLANTEVF | 0.7243278 |
| WVALDRGA | ASSQSDRGLANTEVF | 0.7214002 |
| YLVGSVVI | ASSQSDRGLANTEVF | 0.722456 |
| GPPGTGKTLL | ASSQSDRGLANTEVF | 0.7234953 |
| GVGMTKF | ASSQSDRGLANTEVF | 0.72539616 |
| IGIGGSDL | ASSQSDRGLANTEVF | 0.6733031 |
| IGTLLAEL | ASSQSDRGLANTEVF | 0.5851612 |
| IYTSGTT | ASSQSDRGLANTEVF | 0.7018247 |
| LSFPAGV | ASSQSDRGLANTEVF | 0.70230734 |
| LYTSGTT | ASSQSDRGLANTEVF | 0.70919806 |
| VGAFPFGF | ASSQSDRGLANTEVF | 0.5020407 |
| VGAGTVEF | ASSQSDRGLANTEVF | 0.7322842 |
| VGKSTLLL | ASSQSDRGLANTEVF | 0.73897177 |
| AALAHAGGGIV | ASSQSDRGLANTEVF | 0.6725662 |
| AALVAVSLL | ASSQSDRGLANTEVF | 0.738253 |
| AAVSRAVAAL | ASSQSDRGLANTEVF | 0.6467142 |
| AGGSSGIGGF | ASSQSDRGLANTEVF | 0.72505766 |
| AGLTVANANG | ASSQSDRGLANTEVF | 0.73289955 |
| AGLVGAVL | ASSQSDRGLANTEVF | 0.6517279 |
| ALSQAFTV | ASSQSDRGLANTEVF | 0.72883993 |
| ALTFSVSL | ASSQSDRGLANTEVF | 0.66495687 |
| ALTGAALVAL | ASSQSDRGLANTEVF | 0.6942638 |
| ASFSIGAGVL | ASSQSDRGLANTEVF | 0.73403245 |
| DGSVYKI | ASSQSDRGLANTEVF | 0.69410396 |
| EQLAEIV | ASSQSDRGLANTEVF | 0.71988386 |
| FAVSELTVM | ASSQSDRGLANTEVF | 0.7170359 |
| GAAPLAAGL | ASSQSDRGLANTEVF | 0.6629008 |
| GADGGDGGSGGAGG | ASSQSDRGLANTEVF | 0.72165835 |
| GFGLDTGGAGG | ASSQSDRGLANTEVF | 0.7308719 |
| GLASGGLL | ASSQSDRGLANTEVF | 0.66007423 |
| GRIAGAAVLL | ASSQSDRGLANTEVF | 0.7395032 |
| GSAGSLNTSF | ASSQSDRGLANTEVF | 0.66635334 |
| GTGQIGI | ASSQSDRGLANTEVF | 0.74034727 |
| GTLIALIL | ASSQSDRGLANTEVF | 0.7208244 |
| IAATMGFVVG | ASSQSDRGLANTEVF | 0.70887 |
| IFAQALL | ASSQSDRGLANTEVF | 0.68901074 |
| IGAAVALGF | ASSQSDRGLANTEVF | 0.7320938 |
| IGLTPLAL | ASSQSDRGLANTEVF | 0.70641106 |

| | | |
|---|---|---|
| ILGSNTSTL | ASSQSDRGLANTEVF | 0.7150663 |
| IVFSLTVL | ASSQSDRGLANTEVF | 0.73773617 |
| LASKATGF | ASSQSDRGLANTEVF | 0.8320295 |
| LASVALIL | ASSQSDRGLANTEVF | 0.8937038 |
| LFAVGLTL | ASSQSDRGLANTEVF | 0.86171514 |
| LFNGGIGTYI | ASSQSDRGLANTEVF | 0.8775606 |
| LTLGAGII | ASSQSDRGLANTEVF | 0.80606186 |
| LVAAALGDI | ASSQSDRGLANTEVF | 0.4694734 |
| NTAFVQLGI | ASSQSDRGLANTEVF | 0.87261117 |
| QLFSGAEL | ASSQSDRGLANTEVF | 0.7870798 |
| SAATFITL | ASSQSDRGLANTEVF | 0.87026674 |
| SNYGLANLGV | ASSQSDRGLANTEVF | 0.89401805 |
| SYGGAYAVM | ASSQSDRGLANTEVF | 0.8200355 |
| TTGVGGLTLG | ASSQSDRGLANTEVF | 0.8950085 |
| TTQTIYI | ASSQSDRGLANTEVF | 0.8897728 |
| VAAGAGLGAVI | ASSQSDRGLANTEVF | 0.7647722 |
| VAAVVTLL | ASSQSDRGLANTEVF | 0.83645195 |
| VGFLGGLI | ASSQSDRGLANTEVF | 0.7831051 |
| VGVIPILL | ASSQSDRGLANTEVF | 0.865553 |
| VIFASSLI | ASSQSDRGLANTEVF | 0.7238017 |
| VKGGDGGAAAG | ASSQSDRGLANTEVF | 0.8801425 |
| VTKTFTNI | ASSQSDRGLANTEVF | 0.7850384 |
| VVAAAILL | ASSQSDRGLANTEVF | 0.49026078 |
| YAALDVEL | ASSQSDRGLANTEVF | 0.88018227 |
| LASVALLL | ASSQSDRGLANTEVF | 0.86170787 |
| LDAVHRV | ASSQSDRGLANTEVF | 0.8506502 |
| QGLSSSIF | ASSQSDRGLANTEVF | 0.89074504 |
| RGTFANIRL | ASSQSDRGLANTEVF | 0.73221666 |
| RQLWWGHRI | ASSQSDRGLANTEVF | 0.8187018 |
| YGPPGTGKTLL | ASSQSDRGLANTEVF | 0.8822939 |
| AALVGAPANVA | ASSQSDRGLANTEVF | 0.89834666 |
| AATFRRTE | ASSQSDRGLANTEVF | 0.7806004 |
| AGALGGGI | ASSQSDRGLANTEVF | 0.86529475 |
| AGGAGGAGGIGAG | ASSQSDRGLANTEVF | 0.8941062 |
| AGSGALGL | ASSQSDRGLANTEVF | 0.8719058 |
| AQIADQI | ASSQSDRGLANTEVF | 0.8971348 |
| ASAVAVTYL | ASSQSDRGLANTEVF | 0.89768636 |
| ASTVLGGTGG | ASSQSDRGLANTEVF | 0.81794846 |
| AVGGEPAK | ASSQSDRGLANTEVF | 0.89477116 |
| CEQGTFGGTC | ASSQSDRGLANTEVF | 0.9032593 |
| DEGKAPAG | ASSQSDRGLANTEVF | 0.85451806 |
| DTVVFPAL | ASSQSDRGLANTEVF | 0.86605483 |
| FEGITMAG | ASSQSDRGLANTEVF | 0.7871219 |
| FPLVVQD | ASSQSDRGLANTEVF | 0.84174716 |
| GAFTTPQ | ASSQSDRGLANTEVF | 0.4698904 |
| GFGGFGTAGGVGGA | ASSQSDRGLANTEVF | 0.86071205 |
| GFLAQFL | ASSQSDRGLANTEVF | 0.89384687 |
| GGAGGAGGNAGVA | ASSQSDRGLANTEVF | 0.8437362 |
| GGVGTAGAGGAGGAGG | ASSQSDRGLANTEVF | 0.79508305 |
| GNGGGGDAGNAGSGGN | ASSQSDRGLANTEVF | 0.7636702 |
| GTGGAGGAGNDAGS | ASSQSDRGLANTEVF | 0.8577629 |

| | | |
|---|---|---|
| GTTNTGWFNS | ASSQSDRGLANTEVF | 0.793347 |
| IFMGPLFTL | ASSQSDRGLANTEVF | 0.63819957 |
| ILAHAIEAI | ASSQSDRGLANTEVF | 0.8163407 |
| IRSAIREAL | ASSQSDRGLANTEVF | 0.8900758 |
| ITFAGGG | ASSQSDRGLANTEVF | 0.88853306 |
| ITGMSGHQEV | ASSQSDRGLANTEVF | 0.76088583 |
| IVAGLACTL | ASSQSDRGLANTEVF | 0.9014475 |
| IVGGGLAGA | ASSQSDRGLANTEVF | 0.8543854 |
| IVTGVLNI | ASSQSDRGLANTEVF | 0.88644236 |
| LAQQGCI | ASSQSDRGLANTEVF | 0.85518265 |
| LCLHVEN | ASSQSDRGLANTEVF | 0.81709373 |
| LDAISDGS | ASSQSDRGLANTEVF | 0.8901339 |
| LGGFAAML | ASSQSDRGLANTEVF | 0.8311453 |
| LGGKNPMI | ASSQSDRGLANTEVF | 0.842288 |
| LGPKGVAAIL | ASSQSDRGLANTEVF | 0.8819722 |
| LMVGVSI | ASSQSDRGLANTEVF | 0.8533567 |
| LVGAAFVL | ASSQSDRGLANTEVF | 0.8084769 |
| LVLAVAPL | ASSQSDRGLANTEVF | 0.7944251 |
| MAAIAVIDAA | ASSQSDRGLANTEVF | 0.8905812 |
| NNGINAELQ | ASSQSDRGLANTEVF | 0.77153194 |
| RGEPLVLVHG | ASSQSDRGLANTEVF | 0.68736386 |
| RGGAGGAAGLI | ASSQSDRGLANTEVF | 0.7732223 |
| RSYRMHPAV | ASSQSDRGLANTEVF | 0.8481467 |
| SAAHLLAGQLA | ASSQSDRGLANTEVF | 0.8978981 |
| SAAVGSY | ASSQSDRGLANTEVF | 0.8794546 |
| SPHAVLHRI | ASSQSDRGLANTEVF | 0.83987284 |
| SQIAKVL | ASSQSDRGLANTEVF | 0.88197213 |
| TALGHLVP | ASSQSDRGLANTEVF | 0.8730071 |
| TAPTTSGAA | ASSQSDRGLANTEVF | 0.8616464 |
| TDGDDGNGGSPGT | ASSQSDRGLANTEVF | 0.883325 |
| TDGLGEN | ASSQSDRGLANTEVF | 0.8087395 |
| TGGPPLPGQA | ASSQSDRGLANTEVF | 0.88810855 |
| TIPAITI | ASSQSDRGLANTEVF | 0.8823287 |
| TIPGFTI | ASSQSDRGLANTEVF | 0.8887234 |
| TSIASGALA | ASSQSDRGLANTEVF | 0.8940493 |
| TVTLPGAAI | ASSQSDRGLANTEVF | 0.8678534 |
| VASGLSAALP | ASSQSDRGLANTEVF | 0.85004824 |
| VGAGVIGI | ASSQSDRGLANTEVF | 0.5692463 |
| VGPKRFGV | ASSQSDRGLANTEVF | 0.7886777 |
| VNVAFPD | ASSQSDRGLANTEVF | 0.8790735 |
| VSMTNSGVS | ASSQSDRGLANTEVF | 0.85162807 |
| VTDVLGDLSV | ASSQSDRGLANTEVF | 0.8861921 |
| VVVVGAGFAGL | ASSQSDRGLANTEVF | 0.8831447 |
| VYTVGTVV | ASSQSDRGLANTEVF | 0.8188698 |
| WTTFSGP | ASSQSDRGLANTEVF | 0.89349663 |
| YDELVEAF | ASSQSDRGLANTEVF | 0.89178747 |
| YGTGGAGGNGG | ASSQSDRGLANTEVF | 0.70697105 |
| YTNVMAV | ASSQSDRGLANTEVF | 0.83596283 |
| FLEQVHQL | ASSQSDRGLANTEVF | 0.9035099 |
| GLSLLAVAV | ASSQSDRGLANTEVF | 0.85464746 |
| LEQVHQL | ASSQSDRGLANTEVF | 0.83993834 |

| | | |
|---|---|---|
| LNAAGAI | ASSQSDRGLANTEVF | 0.8525395 |
| PAVHASL | ASSQSDRGLANTEVF | 0.89266956 |
| TELEDDA | ASSQSDRGLANTEVF | 0.8227798 |
| TPPGPPN | ASSQSDRGLANTEVF | 0.90216494 |
| TPPPPPPP | ASSQSDRGLANTEVF | 0.6101886 |
| TQSPALI | ASSRTGLSYEQY | 0.8507688 |
| EFAAAFPQ | ASSRTGLSYEQY | 0.7791639 |
| LLSGNLDVL | ASSRTGLSYEQY | 0.50775176 |
| GPANLGNYNI | ASSRTGLSYEQY | 0.8849615 |
| TLGPTLP | ASSRTGLSYEQY | 0.86990714 |
| VYAYSSCCK | ASSRTGLSYEQY | 0.88487595 |
| APAPAPAPNI | ASSRTGLSYEQY | 0.8470423 |
| PGGPVPIP | ASSRTGLSYEQY | 0.89101267 |
| FLTVALEP | ASSRTGLSYEQY | 0.86769557 |
| SQYFGANL | ASSRTGLSYEQY | 0.85688937 |
| GGVKPAA | ASSRTGLSYEQY | 0.827861 |
| GGTGGAGGAGGW | ASSRTGLSYEQY | 0.87504864 |
| GVGGVAGLGGAGG | ASSRTGLSYEQY | 0.89888144 |
| VLLVLATTI | ASSRTGLSYEQY | 0.88884735 |
| IGIFAAML | ASSRTGLSYEQY | 0.60366535 |
| AVGDLID | ASSRTGLSYEQY | 0.845418 |
| NISPLPL | ASSRTGLSYEQY | 0.84897244 |
| VTAVFVAI | ASSRTGLSYEQY | 0.46573865 |
| SGDDNDAAL | ASSRTGLSYEQY | 0.44020432 |
| AVAALAHS | ASSRTGLSYEQY | 0.7843794 |
| GGVTVMATP | ASSRTGLSYEQY | 0.81801385 |
| AGLAANVRD | ASSRTGLSYEQY | 0.8867193 |
| TNRPDLIDPALLRPG | ASSRTGLSYEQY | 0.8864932 |
| LANLGVTAL | ASSRTGLSYEQY | 0.8500212 |
| TGERIEL | ASSRTGLSYEQY | 0.88536996 |
| NGEELAA | ASSRTGLSYEQY | 0.8703003 |
| VLSLVWG | ASSRTGLSYEQY | 0.11123209 |
| IGYALWL | ASSRTGLSYEQY | 0.8301458 |
| LSLLLAPL | ASSRTGLSYEQY | 0.7813916 |
| TLDEALK | ASSRTGLSYEQY | 0.89477825 |
| TDGAGTAEGG | ASSRTGLSYEQY | 0.8217437 |
| NAPGGAGGNGG | ASSRTGLSYEQY | 0.83004874 |
| VSVFMGMGGNF | ASSRTGLSYEQY | 0.860607 |
| LASYLAAL | ASSRTGLSYEQY | 0.85178614 |
| LVGGTPAL | ASSRTGLSYEQY | 0.8397839 |
| VRNADVL | ASSRTGLSYEQY | 0.7590792 |
| TCSSPFLVSAM | ASSRTGLSYEQY | 0.4748683 |
| VHITDAE | ASSRTGLSYEQY | 0.8822209 |
| EYAAPSF | ASSRTGLSYEQY | 0.7517903 |
| VVDAFHGGL | ASSRTGLSYEQY | 0.80334926 |
| ASVVDPQ | ASSRTGLSYEQY | 0.7763122 |
| SYNEFGGLS | ASSRTGLSYEQY | 0.8284323 |
| DPDGSVY | ASSRTGLSYEQY | 0.6829329 |
| MVSTIHATE | ASSRTGLSYEQY | 0.8958991 |
| LAAGVADPV | ASSRTGLSYEQY | 0.8052782 |
| TKEQIAA | ASSRTGLSYEQY | 0.8444113 |

| | | |
|---|---|---|
| SLASPAAL | ASSRTGLSYEQY | 0.9019626 |
| IGLVFYGV | ASSRTGLSYEQY | 0.8430138 |
| FLLDASNL | ASSRTGLSYEQY | 0.8427695 |
| YQVGNFP | ASSRTGLSYEQY | 0.8318481 |
| ESLLSQST | ASSRTGLSYEQY | 0.80474627 |
| LFTVGFAAVG | ASSRTGLSYEQY | 0.8212863 |
| ATPPTPT | ASSRTGLSYEQY | 0.8255179 |
| TDLATLG | ASSRTGLSYEQY | 0.8985854 |
| SAGGTGR | ASSRTGLSYEQY | 0.8605613 |
| GFSNSGS | ASSRTGLSYEQY | 0.82956386 |
| GGNGGSGGTSV | ASSRTGLSYEQY | 0.8751687 |
| GTSVATGGAGN | ASSRTGLSYEQY | 0.8958539 |
| NGGNGGSGGTS | ASSRTGLSYEQY | 0.9003047 |
| AVVKFPH | ASSRTGLSYEQY | 0.8122208 |
| TPPPPPP | ASSRTGLSYEQY | 0.8337514 |
| AAPAAAAAPA | ASSRTGLSYEQY | 0.8543695 |
| IALNEDL | ASSRTGLSYEQY | 0.8424438 |
| IVNVSSM | ASSRTGLSYEQY | 0.84374905 |
| TGIPPAPR | ASSRTGLSYEQY | 0.7953244 |
| FDLGGGTF | ASSRTGLSYEQY | 0.881645 |
| FDLGGGTFDVS | ASSRTGLSYEQY | 0.8977699 |
| NEPTAAA | ASSRTGLSYEQY | 0.84655464 |
| NEPTAAALA | ASSRTGLSYEQY | 0.68035686 |
| GSIINISSI | ASSRTGLSYEQY | 0.8448937 |
| VAPGNAGTA | ASSRTGLSYEQY | 0.78429866 |
| TGGGASLE | ASSRTGLSYEQY | 0.82038367 |
| VAVYNHYKRI | ASSRTGLSYEQY | 0.86794996 |
| TDPAPATT | ASSRTGLSYEQY | 0.74176866 |
| TNRPDILDPALLRPG | ASSRTGLSYEQY | 0.85332686 |
| LTTTTTS | ASSRTGLSYEQY | 0.90041375 |
| PTGSGKTL | ASSRTGLSYEQY | 0.8226616 |
| TGSGKTL | ASSRTGLSYEQY | 0.8848908 |
| TIGHVDHGK | ASSRTGLSYEQY | 0.89709127 |
| TIGHVDHGKTT | ASSRTGLSYEQY | 0.8765732 |
| GHVDHGKTT | ASSRTGLSYEQY | 0.89724255 |
| AQFEHTLL | ASSRTGLSYEQY | 0.87956035 |
| IVNQDEL | ASSRTGLSYEQY | 0.8583056 |
| TNGSQFF | ASSRTGLSYEQY | 0.8500266 |
| TGKGIVE | ASSRTGLSYEQY | 0.8838599 |
| LISNASDA | ASSRTGLSYEQY | 0.8554115 |
| AELRTLGELTKIAKAHGAQ | ASSRTGLSYEQY | 0.8550023 |
| APNAVAAAEPSW | ASSRTGLSYEQY | 0.8397097 |
| APQVAGAATAVL | ASSRTGLSYEQY | 0.891611 |
| ARRDPAAAAAILPTDA | ASSRTGLSYEQY | 0.8374601 |
| DAAARDVLAEAGLAEYF | ASSRTGLSYEQY | 0.87771475 |
| DDQIVLSAKANLAKVEQ | ASSRTGLSYEQY | 0.56285703 |
| DELLPDTGEDSDEDGH | ASSRTGLSYEQY | 0.78459775 |
| ELPEAPANRATVPQD | ASSRTGLSYEQY | 0.8825772 |
| FGQSKIERVTGIG | ASSRTGLSYEQY | 0.1502606 |
| GAGGTGSTAGGAGGAGGAGGLY | ASSRTGLSYEQY | 0.53399616 |
| GHGGTGAAGGNGGTGGAGGSGI | ASSRTGLSYEQY | 0.86480254 |

| | | |
|---|---|---|
| GNGGNGGAGGEGGPGGAGGAGA | ASSRTGLSYEQY | 0.8876682 |
| GSTAGNGGNGGNATNGTIAGQP | ASSRTGLSYEQY | 0.18474111 |
| GTGGDGGNGGNGGTGG | ASSRTGLSYEQY | 0.81529117 |
| GVRRPAGIPADAPV | ASSRTGLSYEQY | 0.6784211 |
| GYRQVTDAHLLTLARR | ASSRTGLSYEQY | 0.8751128 |
| IAKMGQDGHDRG | ASSRTGLSYEQY | 0.8584754 |
| KAEPSIEIQYT | ASSRTGLSYEQY | 0.4543695 |
| KPVRIRMAADEDTVAVT | ASSRTGLSYEQY | 0.8869246 |
| MPPRIAGMRLLVIKPEP | ASSRTGLSYEQY | 0.742006 |
| MKFYAVALFRSRGVPPTR | ASSRTGLSYEQY | 0.7943065 |
| NAGGFGILGAVAHSP | ASSRTGLSYEQY | 0.8946474 |
| NEGREITLAMAEM | ASSRTGLSYEQY | 0.8933351 |
| NELVEWSHPLGHRWRWE | ASSRTGLSYEQY | 0.8166825 |
| NLGAGGTGGNGGNPG | ASSRTGLSYEQY | 0.8854325 |
| NTEWPKWETVMAADPVD | ASSRTGLSYEQY | 0.81855065 |
| PASAAPAETQP | ASSRTGLSYEQY | 0.9036868 |
| QQSSDGKNGVRAAEITGE | ASSRTGLSYEQY | 0.82396436 |
| RAPDAVTTSWWKE | ASSRTGLSYEQY | 0.7559261 |
| SGFGNVGAHTSGWFNQ | ASSRTGLSYEQY | 0.88684094 |
| SGVAGAVGQAASVGGLKV | ASSRTGLSYEQY | 0.8795911 |
| SNGGAGTGSAGGNGGT | ASSRTGLSYEQY | 0.8644904 |
| TGGAGGEGGAAG | ASSRTGLSYEQY | 0.8726245 |
| TGGDGGAAGAGGAPGHG | ASSRTGLSYEQY | 0.8152176 |
| TMPPLLAAIANAGRHGV | ASSRTGLSYEQY | 0.84350896 |
| TSSTAGNSGGA | ASSRTGLSYEQY | 0.8950842 |
| VPATVPVGILC | ASSRTGLSYEQY | 0.8898884 |
| VQRPDSGGVVLTGRIS | ASSRTGLSYEQY | 0.76662934 |
| WTPNPVPATKGVPAGLVQAT | ASSRTGLSYEQY | 0.89771074 |
| YALKRALAKFSDRFDVVII | ASSRTGLSYEQY | 0.8553171 |
| YLIGSVIV | ASSRTGLSYEQY | 0.80357945 |
| YGQLVAVID | ASSRTGLSYEQY | 0.8977276 |
| VTVGGMTLPA | ASSRTGLSYEQY | 0.8662425 |
| VLALPAV | ASSRTGLSYEQY | 0.8872576 |
| VIPPTLNYET | ASSRTGLSYEQY | 0.8749901 |
| VGMGGMHPSSG | ASSRTGLSYEQY | 0.871198 |
| VGLAGGAATA | ASSRTGLSYEQY | 0.51902294 |
| VGGAGGAGGSGP | ASSRTGLSYEQY | 0.88349736 |
| VFAAFMV | ASSRTGLSYEQY | 0.8492354 |
| TLYDKLR | ASSRTGLSYEQY | 0.8891657 |
| TLAVFTVL | ASSRTGLSYEQY | 0.8722764 |
| TGSVQPARV | ASSRTGLSYEQY | 0.8209982 |
| TGGGAGSGGT | ASSRTGLSYEQY | 0.8730582 |
| SHLETSL | ASSRTGLSYEQY | 0.8977885 |
| SFTTGMT | ASSRTGLSYEQY | 0.8605597 |
| QVEVAKRMGVRQPTVS | ASSRTGLSYEQY | 0.8839981 |
| QGSGGPALAM | ASSRTGLSYEQY | 0.9000236 |
| MLAPSNP | ASSRTGLSYEQY | 0.81869906 |
| LVNNAAI | ASSRTGLSYEQY | 0.8626659 |
| LTGEINTLA | ASSRTGLSYEQY | 0.88686955 |
| LSIFGPV | ASSRTGLSYEQY | 0.8974537 |
| LLALLANL | ASSRTGLSYEQY | 0.8625933 |

| | | |
|---|---|---|
| LISNASDALDKLR | ASSRTGLSYEQY | 0.89894044 |
| LINEAVP | ASSRTGLSYEQY | 0.7501529 |
| LGYAFSGL | ASSRTGLSYEQY | 0.8909819 |
| LDFVAVAL | ASSRTGLSYEQY | 0.8034563 |
| LAQPGAE | ASSRTGLSYEQY | 0.8649505 |
| ISNLGAVTGF | ASSRTGLSYEQY | 0.7667258 |
| HQEDPGRI | ASSRTGLSYEQY | 0.8918754 |
| GTPGEPTP | ASSRTGLSYEQY | 0.88517785 |
| GFFNSGS | ASSRTGLSYEQY | 0.88731307 |
| GAGGGGLTAGAN | ASSRTGLSYEQY | 0.6121073 |
| FVDALVNM | ASSRTGLSYEQY | 0.8869138 |
| FLTVAPEL | ASSRTGLSYEQY | 0.8898703 |
| FETLTAL | ASSRTGLSYEQY | 0.90130204 |
| DVINPIEL | ASSRTGLSYEQY | 0.83208764 |
| APVSGAPV | ASSRTGLSYEQY | 0.88463897 |
| ALVGGLAW | ASSRTGLSYEQY | 0.85295796 |
| ALAAAAATAI | ASSRTGLSYEQY | 0.8510726 |
| AFGLGLLL | ASSRTGLSYEQY | 0.7442424 |
| AFFAAASL | ASSRTGLSYEQY | 0.8337839 |
| AAAALPAFSPP | ASSRTGLSYEQY | 0.8356248 |
| VVLAPAL | ASSRTGLSYEQY | 0.81070185 |
| TIGHVDHGKTTL | ASSRTGLSYEQY | 0.88202345 |
| LISNASDALDK | ASSRTGLSYEQY | 0.8951775 |
| LISNASDAL | ASSRTGLSYEQY | 0.89264965 |
| ISNASDALDK | ASSRTGLSYEQY | 0.88596666 |
| NQAETLVYQTE | ASSRTGLSYEQY | 0.89595926 |
| TGGTGGTAGAG | ASSRTGLSYEQY | 0.89096427 |
| TGSTAGGAGGA | ASSRTGLSYEQY | 0.8280902 |
| DPLAVPVAAGPV | ASSRTGLSYEQY | 0.83507735 |
| GGTGGTPGNGGN | ASSRTGLSYEQY | 0.78474975 |
| PAGLAGVDPATL | ASSRTGLSYEQY | 0.85956395 |
| GNNLVVEAPHAGASV | ASSRTGLSYEQY | 0.822321 |
| KVTRIMSASTLEQPA | ASSRTGLSYEQY | 0.87849027 |
| LGFALPIALPRRTVV | ASSRTGLSYEQY | 0.11195622 |
| TRLGNLTARKLVVLV | ASSRTGLSYEQY | 0.85160464 |
| WSQQRRLIYFPSAGP | ASSRTGLSYEQY | 0.90054566 |
| GPNGNPDPMAAAVDIR | ASSRTGLSYEQY | 0.76513463 |
| TQRCVSVADLRAAASR | ASSRTGLSYEQY | 0.86614066 |
| QFVLDVAEHPVEVLTETV | ASSRTGLSYEQY | 0.8813536 |
| GGAGGTGGAGGNGGTQAGDG | ASSRTGLSYEQY | 0.79950464 |
| PLLATLRAIAVDVDAKPAQV | ASSRTGLSYEQY | 0.87273204 |
| APKFRTGGYMIDFWGVGYQVAK | ASSRTGLSYEQY | 0.8153791 |
| GAGGAGGAGGAAGGPGGTGGTG | ASSRTGLSYEQY | 0.8868668 |
| IGALGLLATDVAPAGSVAFGVA | ASSRTGLSYEQY | 0.8970911 |
| AGGAGGIGGNGGAGGAGGVLMG | ASSRTGLSYEQY | 0.86815643 |
| ADELIGLL | ASSRTGLSYEQY | 0.86269015 |
| ADRALKA | ASSRTGLSYEQY | 0.8681171 |
| AGAVGQVGGAGGA | ASSRTGLSYEQY | 0.38816094 |
| AGGGGGLGGAP | ASSRTGLSYEQY | 0.87106234 |
| AGGLLYGNGGAG | ASSRTGLSYEQY | 0.5688836 |
| AGLESIGL | ASSRTGLSYEQY | 0.8433769 |

| | | |
|---|---|---|
| AGLKTGII | ASSRTGLSYEQY | 0.77010393 |
| ASLAALLL | ASSRTGLSYEQY | 0.87397134 |
| ASVINISSI | ASSRTGLSYEQY | 0.86491454 |
| ATPPGPA | ASSRTGLSYEQY | 0.87105393 |
| ATVASALV | ASSRTGLSYEQY | 0.7844016 |
| ATVASLIL | ASSRTGLSYEQY | 0.85620004 |
| AVNGPVLIRH | ASSRTGLSYEQY | 0.83693343 |
| DGAAGAVGI | ASSRTGLSYEQY | 0.75391895 |
| DPLLHGI | ASSRTGLSYEQY | 0.8917004 |
| ELGAALGAL | ASSRTGLSYEQY | 0.8606626 |
| ELGRTQAV | ASSRTGLSYEQY | 0.8362887 |
| FVAGGYLS | ASSRTGLSYEQY | 0.8889013 |
| GASGGAGGAGGW | ASSRTGLSYEQY | 0.87137324 |
| GGAGAGGDGGAGGI | ASSRTGLSYEQY | 0.58055085 |
| GTGHNGSQPGMGGNG | ASSRTGLSYEQY | 0.82335424 |
| GYAAASASAS | ASSRTGLSYEQY | 0.85819125 |
| HQVDVAAVQQG | ASSRTGLSYEQY | 0.8046315 |
| IGAAIAHT | ASSRTGLSYEQY | 0.35593513 |
| IGMIEVAQL | ASSRTGLSYEQY | 0.87902063 |
| IRLAPEHTSAMD | ASSRTGLSYEQY | 0.80474144 |
| IYNVSVL | ASSRTGLSYEQY | 0.7773867 |
| KQGFLGM | ASSRTGLSYEQY | 0.837075 |
| LESVITQL | ASSRTGLSYEQY | 0.88827556 |
| LGLGALVVV | ASSRTGLSYEQY | 0.63926405 |
| LIGNGGNGGSGGI | ASSRTGLSYEQY | 0.8508165 |
| LNAINAPTL | ASSRTGLSYEQY | 0.8848472 |
| LQLLNTGV | ASSRTGLSYEQY | 0.88049746 |
| LQTSIEF | ASSRTGLSYEQY | 0.89484614 |
| MSFAVSEL | ASSRTGLSYEQY | 0.88146627 |
| NTGALGL | ASSRTGLSYEQY | 0.8435145 |
| PPGAGGAGGAGTT | ASSRTGLSYEQY | 0.8771421 |
| QLVGLGI | ASSRTGLSYEQY | 0.8588315 |
| RFLEQVHQL | ASSRTGLSYEQY | 0.7714206 |
| RGVGYVLR | ASSRTGLSYEQY | 0.71097875 |
| SATGVFAGVI | ASSRTGLSYEQY | 0.8398546 |
| SLPAITLP | ASSRTGLSYEQY | 0.6541076 |
| STALAFTV | ASSRTGLSYEQY | 0.8644002 |
| SVLTVGLGK | ASSRTGLSYEQY | 0.89872575 |
| TALAGAGSGI | ASSRTGLSYEQY | 0.79415846 |
| TGLPFTGI | ASSRTGLSYEQY | 0.759204 |
| VAEHVTV | ASSRTGLSYEQY | 0.8377175 |
| VDIVLNSL | ASSRTGLSYEQY | 0.7282605 |
| VLTGGGAVL | ASSRTGLSYEQY | 0.87913436 |
| VNVGLAYGI | ASSRTGLSYEQY | 0.87006634 |
| VSSAAVGAV | ASSRTGLSYEQY | 0.8976533 |
| VTSVYEAGV | ASSRTGLSYEQY | 0.8888997 |
| WVALDRGA | ASSRTGLSYEQY | 0.83585703 |
| YLVGSVVI | ASSRTGLSYEQY | 0.89048195 |
| GPPGTGKTLL | ASSRTGLSYEQY | 0.8248013 |
| GVGMTKF | ASSRTGLSYEQY | 0.88231015 |
| IGIGGSDL | ASSRTGLSYEQY | 0.8617381 |

| | | |
|---|---|---|
| IGTLLAEL | ASSRTGLSYEQY | 0.864658 |
| IYTSGTT | ASSRTGLSYEQY | 0.85557806 |
| LSFPAGV | ASSRTGLSYEQY | 0.85027343 |
| LYTSGTT | ASSRTGLSYEQY | 0.5878961 |
| VGAFPFGF | ASSRTGLSYEQY | 0.8714398 |
| VGAGTVEF | ASSRTGLSYEQY | 0.7275382 |
| VGKSTLLL | ASSRTGLSYEQY | 0.6120102 |
| AALAHAGGGIV | ASSRTGLSYEQY | 0.82170475 |
| AALVAVSLL | ASSRTGLSYEQY | 0.9029015 |
| AAVSRAVAAL | ASSRTGLSYEQY | 0.8934852 |
| AGGSSGIGGF | ASSRTGLSYEQY | 0.7938582 |
| AGLTVANANG | ASSRTGLSYEQY | 0.7936371 |
| AGLVGAVL | ASSRTGLSYEQY | 0.86071205 |
| ALSQAFTV | ASSRTGLSYEQY | 0.7372966 |
| ALTFSVSL | ASSRTGLSYEQY | 0.88006616 |
| ALTGAALVAL | ASSRTGLSYEQY | 0.8479173 |
| ASFSIGAGVL | ASSRTGLSYEQY | 0.84784377 |
| DGSVYKI | ASSRTGLSYEQY | 0.8268292 |
| EQLAEIV | ASSRTGLSYEQY | 0.86038995 |
| FAVSELTVM | ASSRTGLSYEQY | 0.840098 |
| GAAPLAAGL | ASSRTGLSYEQY | 0.8822674 |
| GADGGDGGSGGAGG | ASSRTGLSYEQY | 0.2569965 |
| GFGLDTGGAGG | ASSRTGLSYEQY | 0.8844001 |
| GLASGGLL | ASSRTGLSYEQY | 0.80672956 |
| GRIAGAAVLL | ASSRTGLSYEQY | 0.75966954 |
| GSAGSLNTSF | ASSRTGLSYEQY | 0.70193386 |
| GTGQIGI | ASSRTGLSYEQY | 0.8746773 |
| GTLIALIL | ASSRTGLSYEQY | 0.8795928 |
| IAATMGFVVG | ASSRTGLSYEQY | 0.88403136 |
| IFAQALL | ASSRTGLSYEQY | 0.7918565 |
| IGAAVALGF | ASSRTGLSYEQY | 0.8898553 |
| IGLTPLAL | ASSRTGLSYEQY | 0.88494265 |
| ILGSNTSTL | ASSRTGLSYEQY | 0.8453401 |
| IVFSLTVL | ASSRTGLSYEQY | 0.46892452 |
| LASKATGF | ASSRTGLSYEQY | 0.8761161 |
| LASVALIL | ASSRTGLSYEQY | 0.85079664 |
| LFAVGLTL | ASSRTGLSYEQY | 0.8854941 |
| LFNGGIGTYI | ASSRTGLSYEQY | 0.87720263 |
| LTLGAGII | ASSRTGLSYEQY | 0.7626883 |
| LVAAALGDI | ASSRTGLSYEQY | 0.8854461 |
| NTAFVQLGI | ASSRTGLSYEQY | 0.89518595 |
| QLFSGAEL | ASSRTGLSYEQY | 0.7570778 |
| SAATFITL | ASSRTGLSYEQY | 0.8581748 |
| SNYGLANLGV | ASSRTGLSYEQY | 0.8531659 |
| SYGGAYAVM | ASSRTGLSYEQY | 0.77207476 |
| TTGVGGLTLG | ASSRTGLSYEQY | 0.8817805 |
| TTQTIYI | ASSRTGLSYEQY | 0.8168541 |
| VAAGAGLGAVI | ASSRTGLSYEQY | 0.83112806 |
| VAAVVTLL | ASSRTGLSYEQY | 0.8562083 |
| VGFLGGLI | ASSRTGLSYEQY | 0.83535767 |
| VGVIPILL | ASSRTGLSYEQY | 0.8354375 |

| | | |
|---|---|---|
| VIFASSLI | ASSRTGLSYEQY | 0.84900504 |
| VKGGDGGAAAG | ASSRTGLSYEQY | 0.8610009 |
| VTKTFTNI | ASSRTGLSYEQY | 0.8671542 |
| VVAAAILL | ASSRTGLSYEQY | 0.84952533 |
| YAALDVEL | ASSRTGLSYEQY | 0.8712063 |
| LASVALLL | ASSRTGLSYEQY | 0.9008655 |
| LDAVHRV | ASSRTGLSYEQY | 0.69728243 |
| QGLSSSIF | ASSRTGLSYEQY | 0.79916954 |
| RGTFANIRL | ASSRTGLSYEQY | 0.8708141 |
| RQLWWGHRI | ASSRTGLSYEQY | 0.75116545 |
| YGPPGTGKTLL | ASSRTGLSYEQY | 0.8912927 |
| AALVGAPANVA | ASSRTGLSYEQY | 0.8700131 |
| AATFRRTE | ASSRTGLSYEQY | 0.8801613 |
| AGALGGGI | ASSRTGLSYEQY | 0.81210494 |
| AGGAGGAGGIGAG | ASSRTGLSYEQY | 0.9034317 |
| AGSGALGL | ASSRTGLSYEQY | 0.8858391 |
| AQIADQI | ASSRTGLSYEQY | 0.8734514 |
| ASAVAVTYL | ASSRTGLSYEQY | 0.88543177 |
| ASTVLGGTGG | ASSRTGLSYEQY | 0.8084803 |
| AVGGEPAK | ASSRTGLSYEQY | 0.88630056 |
| CEQGTFGGTC | ASSRTGLSYEQY | 0.88085586 |
| DEGKAPAG | ASSRTGLSYEQY | 0.7445643 |
| DTVVFPAL | ASSRTGLSYEQY | 0.8437072 |
| FEGITMAG | ASSRTGLSYEQY | 0.88986844 |
| FPLVVQD | ASSRTGLSYEQY | 0.8973812 |
| GAFTTPQ | ASSRTGLSYEQY | 0.6205857 |
| GFGGFGTAGGVGGA | ASSRTGLSYEQY | 0.83617455 |
| GFLAQFL | ASSRTGLSYEQY | 0.6516694 |
| GGAGGAGGNAGVA | ASSRTGLSYEQY | 0.7945256 |
| GGVGTAGAGGAGGAGG | ASSRTGLSYEQY | 0.68435895 |
| GNGGGGDAGNAGSGGN | ASSRTGLSYEQY | 0.8655703 |
| GTGGAGGAGNDAGS | ASSRTGLSYEQY | 0.84137356 |
| GTTNTGWFNS | ASSRTGLSYEQY | 0.86707675 |
| IFMGPLFTL | ASSRTGLSYEQY | 0.8650887 |
| ILAHAIEAI | ASSRTGLSYEQY | 0.89170784 |
| IRSAIREAL | ASSRTGLSYEQY | 0.81923044 |
| ITFAGGG | ASSRTGLSYEQY | 0.85999846 |
| ITGMSGHQEV | ASSRTGLSYEQY | 0.7552154 |
| IVAGLACTL | ASSRTGLSYEQY | 0.526989 |
| IVGGGLAGA | ASSRTGLSYEQY | 0.8002515 |
| IVTGVLNI | ASSRTGLSYEQY | 0.82501554 |
| LAQQGCI | ASSRTGLSYEQY | 0.29526854 |
| LCLHVEN | ASSRTGLSYEQY | 0.8950157 |
| LDAISDGS | ASSRTGLSYEQY | 0.8656845 |
| LGGFAAML | ASSRTGLSYEQY | 0.88074064 |
| LGGKNPMI | ASSRTGLSYEQY | 0.8549508 |
| LGPKGVAAIL | ASSRTGLSYEQY | 0.86464024 |
| LMVGVSI | ASSRTGLSYEQY | 0.8194639 |
| LVGAAFVL | ASSRTGLSYEQY | 0.88233477 |
| LVLAVAPL | ASSRTGLSYEQY | 0.7632213 |
| MAAIAVIDAA | ASSRTGLSYEQY | 0.70107496 |

| | | |
|---|---|---|
| NNGINAELQ | ASSRTGLSYEQY | 0.84906274 |
| RGEPLVLVHG | ASSRTGLSYEQY | 0.7741649 |
| RGGAGGAAGLI | ASSRTGLSYEQY | 0.89136153 |
| RSYRMHPAV | ASSRTGLSYEQY | 0.89033854 |
| SAAHLLAGQLA | ASSRTGLSYEQY | 0.8670387 |
| SAAVGSY | ASSRTGLSYEQY | 0.8901895 |
| SPHAVLHRI | ASSRTGLSYEQY | 0.2524083 |
| SQIAKVL | ASSRTGLSYEQY | 0.84828526 |
| TALGHLVP | ASSRTGLSYEQY | 0.89701647 |
| TAPTTSGAA | ASSRTGLSYEQY | 0.86808544 |
| TDGDDGNGGSPGT | ASSRTGLSYEQY | 0.8515253 |
| TDGLGEN | ASSRTGLSYEQY | 0.8998717 |
| TGGPPLPGQA | ASSRTGLSYEQY | 0.8168385 |
| TIPAITI | ASSRTGLSYEQY | 0.8089042 |
| TIPGFTI | ASSRTGLSYEQY | 0.86734533 |
| TSIASGALA | ASSRTGLSYEQY | 0.8817277 |
| TVTLPGAAI | ASSRTGLSYEQY | 0.87765324 |
| VASGLSAALP | ASSRTGLSYEQY | 0.7560214 |
| VGAGVIGI | ASSRTGLSYEQY | 0.88244367 |
| VGPKRFGV | ASSRTGLSYEQY | 0.9051078 |
| VNVAFPD | ASSRTGLSYEQY | 0.84084785 |
| VSMTNSGVS | ASSRTGLSYEQY | 0.8814911 |
| VTDVLGDLSV | ASSRTGLSYEQY | 0.87928486 |
| VVVVGAGFAGL | ASSRTGLSYEQY | 0.80452085 |
| VYTVGTVV | ASSRTGLSYEQY | 0.8085071 |
| WTTFSGP | ASSRTGLSYEQY | 0.7773613 |
| YDELVEAF | ASSRTGLSYEQY | 0.8543177 |
| YGTGGAGGNGG | ASSRTGLSYEQY | 0.78132343 |
| YTNVMAV | ASSRTGLSYEQY | 0.8655055 |
| FLEQVHQL | ASSRTGLSYEQY | 0.85695463 |
| GLSLLAVAV | ASSRTGLSYEQY | 0.90645546 |
| LEQVHQL | ASSRTGLSYEQY | 0.8771938 |
| LNAAGAI | ASSRTGLSYEQY | 0.88433427 |
| PAVHASL | ASSRTGLSYEQY | 0.78876865 |
| TELEDDA | ASSRTGLSYEQY | 0.8604021 |
| TPPGPPN | ASSRTGLSYEQY | 0.8683895 |
| TPPPPPPP | ASSRTGLSYEQY | 0.58395773 |
| TQSPALI | ASSDAGNTLY | 0.89514506 |
| EFAAAFPQ | ASSDAGNTLY | 0.80046207 |
| LLSGNLDVL | ASSDAGNTLY | 0.8355628 |
| GPANLGNYNI | ASSDAGNTLY | 0.87408257 |
| TLGPTLP | ASSDAGNTLY | 0.88643146 |
| VYAYSSCCK | ASSDAGNTLY | 0.86399186 |
| APAPAPAPNI | ASSDAGNTLY | 0.8523944 |
| PGGPVPIP | ASSDAGNTLY | 0.7977562 |
| FLTVALEP | ASSDAGNTLY | 0.86895865 |
| SQYFGANL | ASSDAGNTLY | 0.89294994 |
| GGVKPAA | ASSDAGNTLY | 0.80276847 |
| GGTGGAGGAGGW | ASSDAGNTLY | 0.89333236 |
| GVGGVAGLGGAGG | ASSDAGNTLY | 0.87334335 |
| VLLVLATTI | ASSDAGNTLY | 0.5434146 |

| | | |
|---|---|---|
| IGIFAAML | ASSDAGNTLY | 0.8385416 |
| AVGDLID | ASSDAGNTLY | 0.89386207 |
| NISPLPL | ASSDAGNTLY | 0.8797134 |
| VTAVFVAI | ASSDAGNTLY | 0.8977796 |
| SGDDNDAAL | ASSDAGNTLY | 0.89151204 |
| AVAALAHS | ASSDAGNTLY | 0.808202 |
| GGVTVMATP | ASSDAGNTLY | 0.89373153 |
| AGLAANVRD | ASSDAGNTLY | 0.9015895 |
| TNRPDLIDPALLRPG | ASSDAGNTLY | 0.8894743 |
| LANLGVTAL | ASSDAGNTLY | 0.8022013 |
| TGERIEL | ASSDAGNTLY | 0.7096158 |
| NGEELAA | ASSDAGNTLY | 0.87448287 |
| VLSLVWG | ASSDAGNTLY | 0.886914 |
| IGYALWL | ASSDAGNTLY | 0.79079485 |
| LSLLLAPL | ASSDAGNTLY | 0.8726832 |
| TLDEALK | ASSDAGNTLY | 0.8575318 |
| TDGAGTAEGG | ASSDAGNTLY | 0.63927734 |
| NAPGGAGGNGG | ASSDAGNTLY | 0.88731194 |
| VSVFMGMGGNF | ASSDAGNTLY | 0.89951503 |
| LASYLAAL | ASSDAGNTLY | 0.87649703 |
| LVGGTPAL | ASSDAGNTLY | 0.7692114 |
| VRNADVL | ASSDAGNTLY | 0.89727294 |
| TCSSPFLVSAM | ASSDAGNTLY | 0.90382224 |
| VHITDAE | ASSDAGNTLY | 0.89697057 |
| EYAAPSF | ASSDAGNTLY | 0.7130905 |
| VVDAFHGGL | ASSDAGNTLY | 0.48180756 |
| ASVVDPQ | ASSDAGNTLY | 0.8430614 |
| SYNEFGGLS | ASSDAGNTLY | 0.87580454 |
| DPDGSVY | ASSDAGNTLY | 0.8431121 |
| MVSTIHATE | ASSDAGNTLY | 0.820791 |
| LAAGVADPV | ASSDAGNTLY | 0.8706994 |
| TKEQIAA | ASSDAGNTLY | 0.8910068 |
| SLASPAAL | ASSDAGNTLY | 0.80972767 |
| IGLVFYGV | ASSDAGNTLY | 0.6368059 |
| FLLDASNL | ASSDAGNTLY | 0.8908652 |
| YQVGNFP | ASSDAGNTLY | 0.88751394 |
| ESLLSQST | ASSDAGNTLY | 0.80801713 |
| LFTVGFAAVG | ASSDAGNTLY | 0.83162034 |
| ATPPTPT | ASSDAGNTLY | 0.8477141 |
| TDLATLG | ASSDAGNTLY | 0.8048283 |
| SAGGTGR | ASSDAGNTLY | 0.84937066 |
| GFSNSGS | ASSDAGNTLY | 0.74810505 |
| GGNGGSGGTSV | ASSDAGNTLY | 0.89861196 |
| GTSVATGGAGN | ASSDAGNTLY | 0.7797039 |
| NGGNGGSGGTS | ASSDAGNTLY | 0.87969595 |
| AVVKFPH | ASSDAGNTLY | 0.87725365 |
| TPPPPPP | ASSDAGNTLY | 0.8746236 |
| AAPAAAAAPA | ASSDAGNTLY | 0.84812534 |
| IALNEDL | ASSDAGNTLY | 0.8142056 |
| IVNVSSM | ASSDAGNTLY | 0.7063497 |
| TGIPPAPR | ASSDAGNTLY | 0.64924777 |

| | | |
|---|---|---|
| FDLGGGTF | ASSDAGNTLY | 0.89666307 |
| FDLGGGTFDVS | ASSDAGNTLY | 0.7712613 |
| NEPTAAA | ASSDAGNTLY | 0.8957801 |
| NEPTAAALA | ASSDAGNTLY | 0.87585783 |
| GSIINISSI | ASSDAGNTLY | 0.8110416 |
| VAPGNAGTA | ASSDAGNTLY | 0.87532926 |
| TGGGASLE | ASSDAGNTLY | 0.64497095 |
| VAVYNHYKRI | ASSDAGNTLY | 0.80367017 |
| TDPAPATT | ASSDAGNTLY | 0.8507581 |
| TNRPDILDPALLRPG | ASSDAGNTLY | 0.88460296 |
| LTTTTTS | ASSDAGNTLY | 0.8231033 |
| PTGSGKTL | ASSDAGNTLY | 0.8830006 |
| TGSGKTL | ASSDAGNTLY | 0.86360264 |
| TIGHVDHGK | ASSDAGNTLY | 0.77245545 |
| TIGHVDHGKTT | ASSDAGNTLY | 0.87830657 |
| GHVDHGKTT | ASSDAGNTLY | 0.8933295 |
| AQFEHTLL | ASSDAGNTLY | 0.88181436 |
| IVNQDEL | ASSDAGNTLY | 0.74308974 |
| TNGSQFF | ASSDAGNTLY | 0.88210934 |
| TGKGIVE | ASSDAGNTLY | 0.7946878 |
| LISNASDA | ASSDAGNTLY | 0.8670688 |
| AELRTLGELTKIAKAHGAQ | ASSDAGNTLY | 0.86897004 |
| APNAVAAAEPSW | ASSDAGNTLY | 0.83578014 |
| APQVAGAATAVL | ASSDAGNTLY | 0.8881054 |
| ARRDPAAAAAILPTDA | ASSDAGNTLY | 0.76505494 |
| DAAARDVLAEAGLAEYF | ASSDAGNTLY | 0.8025416 |
| DDQIVLSAKANLAKVEQ | ASSDAGNTLY | 0.8499211 |
| DELLPDTGEDSDEDGH | ASSDAGNTLY | 0.8760696 |
| ELPEAPANRATVPQD | ASSDAGNTLY | 0.88941425 |
| FGQSKIERVTGIG | ASSDAGNTLY | 0.76342416 |
| GAGGTGSTAGGAGGAGGAGGLY | ASSDAGNTLY | 0.84859127 |
| GHGGTGAAGGNGGTGGAGGSGI | ASSDAGNTLY | 0.8011664 |
| GNGGNGGAGGEGGPGGAGGAGA | ASSDAGNTLY | 0.7492788 |
| GSTAGNGGNGGNATNGTIAGQP | ASSDAGNTLY | 0.87529933 |
| GTGGDGGNGGNGGTGG | ASSDAGNTLY | 0.866606 |
| GVRRPAGIPADAPV | ASSDAGNTLY | 0.75143075 |
| GYRQVTDAHLLTLARR | ASSDAGNTLY | 0.8761309 |
| IAKMGQDGHDRG | ASSDAGNTLY | 0.81191653 |
| KAEPSIEIQYT | ASSDAGNTLY | 0.87930065 |
| KPVRIRMAADEDTVAVT | ASSDAGNTLY | 0.8818437 |
| MPPRIAGMRLLVIKPEP | ASSDAGNTLY | 0.78902096 |
| MKFYAVALFRSRGVPPTR | ASSDAGNTLY | 0.879236 |
| NAGGFGILGAVAHSP | ASSDAGNTLY | 0.8747556 |
| NEGREITLAMAEM | ASSDAGNTLY | 0.8822926 |
| NELVEWSHPLGHRWRWE | ASSDAGNTLY | 0.8421116 |
| NLGAGGTGGNGGNPG | ASSDAGNTLY | 0.7648934 |
| NTEWPKWETVMAADPVD | ASSDAGNTLY | 0.8986161 |
| PASAAPAETQP | ASSDAGNTLY | 0.8879578 |
| QQSSDGKNGVRAAEITGE | ASSDAGNTLY | 0.88021666 |
| RAPDAVTTSWWKE | ASSDAGNTLY | 0.8846529 |
| SGFGNVGAHTSGWFNQ | ASSDAGNTLY | 0.88584954 |

| | | |
|---|---|---|
| SGVAGAVGQAASVGGLKV | ASSDAGNTLY | 0.8897346 |
| SNGGAGTGSAGGNGGT | ASSDAGNTLY | 0.81841564 |
| TGGAGGEGGAAG | ASSDAGNTLY | 0.6891989 |
| TGGDGGAAGAGGAPGHG | ASSDAGNTLY | 0.8533756 |
| TMPPLLAAIANAGRHGV | ASSDAGNTLY | 0.8414339 |
| TSSTAGNSGGA | ASSDAGNTLY | 0.8634131 |
| VPATVPVGILC | ASSDAGNTLY | 0.49800104 |
| VQRPDSGGVVLTGRIS | ASSDAGNTLY | 0.89166534 |
| WTPNPVPATKGVPAGLVQAT | ASSDAGNTLY | 0.9072496 |
| YALKRALAKFSDRFDVVII | ASSDAGNTLY | 0.80124104 |
| YLIGSVIV | ASSDAGNTLY | 0.89865196 |
| YGQLVAVID | ASSDAGNTLY | 0.7881227 |
| VTVGGMTLPA | ASSDAGNTLY | 0.8835266 |
| VLALPAV | ASSDAGNTLY | 0.89650106 |
| VIPPTLNYET | ASSDAGNTLY | 0.75758827 |
| VGMGGMHPSSG | ASSDAGNTLY | 0.8697808 |
| VGLAGGAATA | ASSDAGNTLY | 0.7533037 |
| VGGAGGAGGSGP | ASSDAGNTLY | 0.81952596 |
| VFAAFMV | ASSDAGNTLY | 0.8988672 |
| TLYDKLR | ASSDAGNTLY | 0.83673483 |
| TLAVFTVL | ASSDAGNTLY | 0.86937976 |
| TGSVQPARV | ASSDAGNTLY | 0.8537396 |
| TGGGAGSGGT | ASSDAGNTLY | 0.79244214 |
| SHLETSL | ASSDAGNTLY | 0.87810075 |
| SFTTGMT | ASSDAGNTLY | 0.8928658 |
| QVEVAKRMGVRQPTVS | ASSDAGNTLY | 0.76006955 |
| QGSGGPALAM | ASSDAGNTLY | 0.9000441 |
| MLAPSNP | ASSDAGNTLY | 0.8300481 |
| LVNNAAI | ASSDAGNTLY | 0.9050846 |
| LTGEINTLA | ASSDAGNTLY | 0.891003 |
| LSIFGPV | ASSDAGNTLY | 0.84604144 |
| LLALLANL | ASSDAGNTLY | 0.8423737 |
| LISNASDALDKLR | ASSDAGNTLY | 0.8904997 |
| LINEAVP | ASSDAGNTLY | 0.84603775 |
| LGYAFSGL | ASSDAGNTLY | 0.8898976 |
| LDFVAVAL | ASSDAGNTLY | 0.8964709 |
| LAQPGAE | ASSDAGNTLY | 0.5442074 |
| ISNLGAVTGF | ASSDAGNTLY | 0.6605315 |
| HQEDPGRI | ASSDAGNTLY | 0.6316922 |
| GTPGEPTP | ASSDAGNTLY | 0.59688854 |
| GFFNSGS | ASSDAGNTLY | 0.5572628 |
| GAGGGGLTAGAN | ASSDAGNTLY | 0.44688946 |
| FVDALVNM | ASSDAGNTLY | 0.6176683 |
| FLTVAPEL | ASSDAGNTLY | 0.5286752 |
| FETLTAL | ASSDAGNTLY | 0.65039194 |
| DVINPIEL | ASSDAGNTLY | 0.65429556 |
| APVSGAPV | ASSDAGNTLY | 0.5445454 |
| ALVGGLAW | ASSDAGNTLY | 0.66399693 |
| ALAAAAATAI | ASSDAGNTLY | 0.6398882 |
| AFGLGLLL | ASSDAGNTLY | 0.47621828 |
| AFFAAASL | ASSDAGNTLY | 0.58669347 |

| | | |
|---|---|---|
| AAAALPAFSPP | ASSDAGNTLY | 0.5925661 |
| VVLAPAL | ASSDAGNTLY | 0.60406804 |
| TIGHVDHGKTTL | ASSDAGNTLY | 0.48780435 |
| LISNASDALDK | ASSDAGNTLY | 0.64962196 |
| LISNASDAL | ASSDAGNTLY | 0.59015775 |
| ISNASDALDK | ASSDAGNTLY | 0.33833277 |
| NQAETLVYQTE | ASSDAGNTLY | 0.6247911 |
| TGGTGGTAGAG | ASSDAGNTLY | 0.6201596 |
| TGSTAGGAGGA | ASSDAGNTLY | 0.6080572 |
| DPLAVPVAAGPV | ASSDAGNTLY | 0.6574264 |
| GGTGGTPGNGGN | ASSDAGNTLY | 0.56874835 |
| PAGLAGVDPATL | ASSDAGNTLY | 0.5603885 |
| GNNLVVEAPHAGASV | ASSDAGNTLY | 0.6532823 |
| KVTRIMSASTLEQPA | ASSDAGNTLY | 0.6630398 |
| LGFALPIALPRRTVV | ASSDAGNTLY | 0.50155413 |
| TRLGNLTARKLVVLV | ASSDAGNTLY | 0.6379276 |
| WSQQRRLIYFPSAGP | ASSDAGNTLY | 0.6509206 |
| GPNGNPDPMAAAVDIR | ASSDAGNTLY | 0.63860667 |
| TQRCVSVADLRAAASR | ASSDAGNTLY | 0.6630691 |
| QFVLDVAEHPVEVLTETV | ASSDAGNTLY | 0.66468865 |
| GGAGGTGGAGGNGGTQAGDG | ASSDAGNTLY | 0.54591215 |
| PLLATLRAIAVDVDAKPAQV | ASSDAGNTLY | 0.6615811 |
| APKFRTGGYMIDFWGVGYQVAK | ASSDAGNTLY | 0.6659664 |
| GAGGAGGAGGAAGGPGGTGGTG | ASSDAGNTLY | 0.5785986 |
| IGALGLLATDVAPAGSVAFGVA | ASSDAGNTLY | 0.62694204 |
| AGGAGGIGGNGGAGGAGGVLMG | ASSDAGNTLY | 0.5232002 |
| ADELIGLL | ASSDAGNTLY | 0.60826075 |
| ADRALKA | ASSDAGNTLY | 0.41394895 |
| AGAVGQVGGAGGA | ASSDAGNTLY | 0.6436746 |
| AGGGGGLGGAP | ASSDAGNTLY | 0.66217184 |
| AGGLLYGNGGAG | ASSDAGNTLY | 0.609395 |
| AGLESIGL | ASSDAGNTLY | 0.5570205 |
| AGLKTGII | ASSDAGNTLY | 0.54197705 |
| ASLAALLL | ASSDAGNTLY | 0.606891 |
| ASVINISSI | ASSDAGNTLY | 0.5745409 |
| ATPPGPA | ASSDAGNTLY | 0.4178094 |
| ATVASALV | ASSDAGNTLY | 0.5939505 |
| ATVASLIL | ASSDAGNTLY | 0.6548758 |
| AVNGPVLIRH | ASSDAGNTLY | 0.6541789 |
| DGAAGAVGI | ASSDAGNTLY | 0.43693286 |
| DPLLHGI | ASSDAGNTLY | 0.6633047 |
| ELGAALGAL | ASSDAGNTLY | 0.64303297 |
| ELGRTQAV | ASSDAGNTLY | 0.65844005 |
| FVAGGYLS | ASSDAGNTLY | 0.6111374 |
| GASGGAGGAGGW | ASSDAGNTLY | 0.54790217 |
| GGAGAGGDGGAGGI | ASSDAGNTLY | 0.65275794 |
| GTGHNGSQPGMGGNG | ASSDAGNTLY | 0.6033533 |
| GYAAASASAS | ASSDAGNTLY | 0.592935 |
| HQVDVAAVQQG | ASSDAGNTLY | 0.65282184 |
| IGAAIAHT | ASSDAGNTLY | 0.620388 |
| IGMIEVAQL | ASSDAGNTLY | 0.57728845 |

| | | |
|---|---|---|
| IRLAPEHTSAMD | ASSDAGNTLY | 0.5965758 |
| IYNVSVL | ASSDAGNTLY | 0.65563476 |
| KQGFLGM | ASSDAGNTLY | 0.5420212 |
| LESVITQL | ASSDAGNTLY | 0.5283435 |
| LGLGALVVV | ASSDAGNTLY | 0.550683 |
| LIGNGGNGGSGGI | ASSDAGNTLY | 0.6214043 |
| LNAINAPTL | ASSDAGNTLY | 0.6600557 |
| LQLLNTGV | ASSDAGNTLY | 0.6444597 |
| LQTSIEF | ASSDAGNTLY | 0.5966698 |
| MSFAVSEL | ASSDAGNTLY | 0.6446716 |
| NTGALGL | ASSDAGNTLY | 0.64275014 |
| PPGAGGAGGAGTT | ASSDAGNTLY | 0.6278686 |
| QLVGLGI | ASSDAGNTLY | 0.62979746 |
| RFLEQVHQL | ASSDAGNTLY | 0.5868754 |
| RGVGYVLR | ASSDAGNTLY | 0.655055 |
| SATGVFAGVI | ASSDAGNTLY | 0.6421503 |
| SLPAITLP | ASSDAGNTLY | 0.6547935 |
| STALAFTV | ASSDAGNTLY | 0.6593747 |
| SVLTVGLGK | ASSDAGNTLY | 0.6366243 |
| TALAGAGSGI | ASSDAGNTLY | 0.62170905 |
| TGLPFTGI | ASSDAGNTLY | 0.38092184 |
| VAEHVTV | ASSDAGNTLY | 0.55813885 |
| VDIVLNSL | ASSDAGNTLY | 0.6348386 |
| VLTGGGAVL | ASSDAGNTLY | 0.6333357 |
| VNVGLAYGI | ASSDAGNTLY | 0.65034246 |
| VSSAAVGAV | ASSDAGNTLY | 0.63529646 |
| VTSVYEAGV | ASSDAGNTLY | 0.57103765 |
| WVALDRGA | ASSDAGNTLY | 0.6475106 |
| YLVGSVVI | ASSDAGNTLY | 0.6548611 |
| GPPGTGKTLL | ASSDAGNTLY | 0.46043083 |
| GVGMTKF | ASSDAGNTLY | 0.59098923 |
| IGIGGSDL | ASSDAGNTLY | 0.669564 |
| IGTLLAEL | ASSDAGNTLY | 0.63013923 |
| IYTSGTT | ASSDAGNTLY | 0.60045826 |
| LSFPAGV | ASSDAGNTLY | 0.5723658 |
| LYTSGTT | ASSDAGNTLY | 0.6640717 |
| VGAFPFGF | ASSDAGNTLY | 0.6063627 |
| VGAGTVEF | ASSDAGNTLY | 0.66158783 |
| VGKSTLLL | ASSDAGNTLY | 0.34216365 |
| AALAHAGGGIV | ASSDAGNTLY | 0.5615076 |
| AALVAVSLL | ASSDAGNTLY | 0.47161338 |
| AAVSRAVAAL | ASSDAGNTLY | 0.40358067 |
| AGGSSGIGGF | ASSDAGNTLY | 0.65703386 |
| AGLTVANANG | ASSDAGNTLY | 0.54286885 |
| AGLVGAVL | ASSDAGNTLY | 0.6388265 |
| ALSQAFTV | ASSDAGNTLY | 0.6366802 |
| ALTFSVSL | ASSDAGNTLY | 0.6586557 |
| ALTGAALVAL | ASSDAGNTLY | 0.6325412 |
| ASFSIGAGVL | ASSDAGNTLY | 0.6193677 |
| DGSVYKI | ASSDAGNTLY | 0.5685921 |
| EQLAEIV | ASSDAGNTLY | 0.64806646 |

| | | |
|---|---|---|
| FAVSELTVM | ASSDAGNTLY | 0.6548368 |
| GAAPLAAGL | ASSDAGNTLY | 0.64607406 |
| GADGGDGGSGGAGG | ASSDAGNTLY | 0.4409893 |
| GFGLDTGGAGG | ASSDAGNTLY | 0.62162364 |
| GLASGGLL | ASSDAGNTLY | 0.6198095 |
| GRIAGAAVLL | ASSDAGNTLY | 0.45583776 |
| GSAGSLNTSF | ASSDAGNTLY | 0.3841578 |
| GTGQIGI | ASSDAGNTLY | 0.5809789 |
| GTLIALIL | ASSDAGNTLY | 0.5903149 |
| IAATMGFVVG | ASSDAGNTLY | 0.6546402 |
| IFAQALL | ASSDAGNTLY | 0.65408117 |
| IGAAVALGF | ASSDAGNTLY | 0.6076971 |
| IGLTPLAL | ASSDAGNTLY | 0.64589953 |
| ILGSNTSTL | ASSDAGNTLY | 0.62081766 |
| IVFSLTVL | ASSDAGNTLY | 0.30979043 |
| LASKATGF | ASSDAGNTLY | 0.5637901 |
| LASVALIL | ASSDAGNTLY | 0.49399722 |
| LFAVGLTL | ASSDAGNTLY | 0.6633017 |
| LFNGGIGTYI | ASSDAGNTLY | 0.56612366 |
| LTLGAGII | ASSDAGNTLY | 0.5716377 |
| LVAAALGDI | ASSDAGNTLY | 0.62142044 |
| NTAFVQLGI | ASSDAGNTLY | 0.6294141 |
| QLFSGAEL | ASSDAGNTLY | 0.62225336 |
| SAATFITL | ASSDAGNTLY | 0.54247063 |
| SNYGLANLGV | ASSDAGNTLY | 0.37469608 |
| SYGGAYAVM | ASSDAGNTLY | 0.6494952 |
| TTGVGGLTLG | ASSDAGNTLY | 0.5334048 |
| TTQTIYI | ASSDAGNTLY | 0.5203331 |
| VAAGAGLGAVI | ASSDAGNTLY | 0.4125913 |
| VAAVVTLL | ASSDAGNTLY | 0.6307457 |
| VGFLGGLI | ASSDAGNTLY | 0.42454255 |
| VGVIPILL | ASSDAGNTLY | 0.65776396 |
| VIFASSLI | ASSDAGNTLY | 0.604475 |
| VKGGDGGAAAG | ASSDAGNTLY | 0.60727805 |
| VTKTFTNI | ASSDAGNTLY | 0.6682368 |
| VVAAAILL | ASSDAGNTLY | 0.6267377 |
| YAALDVEL | ASSDAGNTLY | 0.6048661 |
| LASVALLL | ASSDAGNTLY | 0.54387707 |
| LDAVHRV | ASSDAGNTLY | 0.52414346 |
| QGLSSSIF | ASSDAGNTLY | 0.5571854 |
| RGTFANIRL | ASSDAGNTLY | 0.5623806 |
| RQLWWGHRI | ASSDAGNTLY | 0.66513026 |
| YGPPGTGKTLL | ASSDAGNTLY | 0.5904193 |
| AALVGAPANVA | ASSDAGNTLY | 0.50148505 |
| AATFRRTE | ASSDAGNTLY | 0.58095205 |
| AGALGGGI | ASSDAGNTLY | 0.6641975 |
| AGGAGGAGGIGAG | ASSDAGNTLY | 0.6688626 |
| AGSGALGL | ASSDAGNTLY | 0.57089764 |
| AQIADQI | ASSDAGNTLY | 0.5614839 |
| ASAVAVTYL | ASSDAGNTLY | 0.5583211 |
| ASTVLGGTGG | ASSDAGNTLY | 0.6212537 |

| | | |
|---|---|---|
| AVGGEPAK | ASSDAGNTLY | 0.5667231 |
| CEQGTFGGTC | ASSDAGNTLY | 0.48234162 |
| DEGKAPAG | ASSDAGNTLY | 0.6470793 |
| DTVVFPAL | ASSDAGNTLY | 0.66816974 |
| FEGITMAG | ASSDAGNTLY | 0.5824716 |
| FPLVVQD | ASSDAGNTLY | 0.5410906 |
| GAFTTPQ | ASSDAGNTLY | 0.6136433 |
| GFGGFGTAGGVGGA | ASSDAGNTLY | 0.56804043 |
| GFLAQFL | ASSDAGNTLY | 0.5938115 |
| GGAGGAGGNAGVA | ASSDAGNTLY | 0.6158799 |
| GGVGTAGAGGAGGAGG | ASSDAGNTLY | 0.44694614 |
| GNGGGGDAGNAGSGGN | ASSDAGNTLY | 0.60931814 |
| GTGGAGGAGNDAGS | ASSDAGNTLY | 0.6685592 |
| GTTNTGWFNS | ASSDAGNTLY | 0.5984534 |
| IFMGPLFTL | ASSDAGNTLY | 0.630402 |
| ILAHAIEAI | ASSDAGNTLY | 0.66422534 |
| IRSAIREAL | ASSDAGNTLY | 0.64844173 |
| ITFAGGG | ASSDAGNTLY | 0.66209745 |
| ITGMSGHQEV | ASSDAGNTLY | 0.65294564 |
| IVAGLACTL | ASSDAGNTLY | 0.6139668 |
| IVGGGLAGA | ASSDAGNTLY | 0.6164697 |
| IVTGVLNI | ASSDAGNTLY | 0.64955086 |
| LAQQGCI | ASSDAGNTLY | 0.62246025 |
| LCLHVEN | ASSDAGNTLY | 0.6072283 |
| LDAISDGS | ASSDAGNTLY | 0.5581512 |
| LGGFAAML | ASSDAGNTLY | 0.6540044 |
| LGGKNPMI | ASSDAGNTLY | 0.6013191 |
| LGPKGVAAIL | ASSDAGNTLY | 0.6567477 |
| LMVGVSI | ASSDAGNTLY | 0.4288826 |
| LVGAAFVL | ASSDAGNTLY | 0.5186957 |
| LVLAVAPL | ASSDAGNTLY | 0.6531158 |
| MAAIAVIDAA | ASSDAGNTLY | 0.3106118 |
| NNGINAELQ | ASSDAGNTLY | 0.3618086 |
| RGEPLVLVHG | ASSDAGNTLY | 0.61271966 |
| RGGAGGAAGLI | ASSDAGNTLY | 0.65808624 |
| RSYRMHPAV | ASSDAGNTLY | 0.32648107 |
| SAAHLLAGQLA | ASSDAGNTLY | 0.53419614 |
| SAAVGSY | ASSDAGNTLY | 0.4913015 |
| SPHAVLHRI | ASSDAGNTLY | 0.64231443 |
| SQIAKVL | ASSDAGNTLY | 0.6247506 |
| TALGHLVP | ASSDAGNTLY | 0.42159933 |
| TAPTTSGAA | ASSDAGNTLY | 0.6481388 |
| TDGDDGNGGSPGT | ASSDAGNTLY | 0.48451257 |
| TDGLGEN | ASSDAGNTLY | 0.59860843 |
| TGGPPLPGQA | ASSDAGNTLY | 0.65747374 |
| TIPAITI | ASSDAGNTLY | 0.66293097 |
| TIPGFTI | ASSDAGNTLY | 0.51438 |
| TSIASGALA | ASSDAGNTLY | 0.6520031 |
| TVTLPGAAI | ASSDAGNTLY | 0.58883953 |
| VASGLSAALP | ASSDAGNTLY | 0.67085385 |
| VGAGVIGI | ASSDAGNTLY | 0.5913328 |

| | | |
|---|---|---|
| VGPKRFGV | ASSDAGNTLY | 0.4872367 |
| VNVAFPD | ASSDAGNTLY | 0.6537767 |
| VSMTNSGVS | ASSDAGNTLY | 0.63460875 |
| VTDVLGDLSV | ASSDAGNTLY | 0.577018 |
| VVVVGAGFAGL | ASSDAGNTLY | 0.6482143 |
| VYTVGTVV | ASSDAGNTLY | 0.55661553 |
| WTTFSGP | ASSDAGNTLY | 0.61407936 |
| YDELVEAF | ASSDAGNTLY | 0.66267735 |
| YGTGGAGGNGG | ASSDAGNTLY | 0.6520357 |
| YTNVMAV | ASSDAGNTLY | 0.50739586 |
| FLEQVHQL | ASSDAGNTLY | 0.65616876 |
| GLSLLAVAV | ASSDAGNTLY | 0.60965776 |
| LEQVHQL | ASSDAGNTLY | 0.53172445 |
| LNAAGAI | ASSDAGNTLY | 0.66477376 |
| PAVHASL | ASSDAGNTLY | 0.6237296 |
| TELEDDA | ASSDAGNTLY | 0.65244484 |
| TPPGPPN | ASSDAGNTLY | 0.6471275 |
| TPPPPPPP | ASSDAGNTLY | 0.65153724 |
| TQSPALI | ASSQDPLGNTLY | 0.45730382 |
| EFAAAFPQ | ASSQDPLGNTLY | 0.610707 |
| LLSGNLDVL | ASSQDPLGNTLY | 0.6087251 |
| GPANLGNYNI | ASSQDPLGNTLY | 0.6558326 |
| TLGPTLP | ASSQDPLGNTLY | 0.6171281 |
| VYAYSSCCK | ASSQDPLGNTLY | 0.61716324 |
| APAPAPAPNI | ASSQDPLGNTLY | 0.64399815 |
| PGGPVPIP | ASSQDPLGNTLY | 0.66081774 |
| FLTVALEP | ASSQDPLGNTLY | 0.6145649 |
| SQYFGANL | ASSQDPLGNTLY | 0.62289447 |
| GGVKPAA | ASSQDPLGNTLY | 0.6646424 |
| GGTGGAGGAGGW | ASSQDPLGNTLY | 0.53892946 |
| GVGGVAGLGGAGG | ASSQDPLGNTLY | 0.5976286 |
| VLLVLATTI | ASSQDPLGNTLY | 0.63962734 |
| IGIFAAML | ASSQDPLGNTLY | 0.65394187 |
| AVGDLID | ASSQDPLGNTLY | 0.60496706 |
| NISPLPL | ASSQDPLGNTLY | 0.6610359 |
| VTAVFVAI | ASSQDPLGNTLY | 0.5554465 |
| SGDDNDAAL | ASSQDPLGNTLY | 0.6525049 |
| AVAALAHS | ASSQDPLGNTLY | 0.59286505 |
| GGVTVMATP | ASSQDPLGNTLY | 0.6388642 |
| AGLAANVRD | ASSQDPLGNTLY | 0.5663301 |
| TNRPDLIDPALLRPG | ASSQDPLGNTLY | 0.6455574 |
| LANLGVTAL | ASSQDPLGNTLY | 0.65666634 |
| TGERIEL | ASSQDPLGNTLY | 0.6526228 |
| NGEELAA | ASSQDPLGNTLY | 0.35965797 |
| VLSLVWG | ASSQDPLGNTLY | 0.6577342 |
| IGYALWL | ASSQDPLGNTLY | 0.65373135 |
| LSLLLAPL | ASSQDPLGNTLY | 0.66905487 |
| TLDEALK | ASSQDPLGNTLY | 0.60581946 |
| TDGAGTAEGG | ASSQDPLGNTLY | 0.62119913 |
| NAPGGAGGNGG | ASSQDPLGNTLY | 0.5952696 |
| VSVFMGMGGNF | ASSQDPLGNTLY | 0.61641514 |

| | | |
|---|---|---|
| LASYLAAL | ASSQDPLGNTLY | 0.51156604 |
| LVGGTPAL | ASSQDPLGNTLY | 0.58018726 |
| VRNADVL | ASSQDPLGNTLY | 0.6093128 |
| TCSSPFLVSAM | ASSQDPLGNTLY | 0.5820738 |
| VHITDAE | ASSQDPLGNTLY | 0.65067416 |
| EYAAPSF | ASSQDPLGNTLY | 0.6594404 |
| VVDAFHGGL | ASSQDPLGNTLY | 0.65524405 |
| ASVVDPQ | ASSQDPLGNTLY | 0.6589935 |
| SYNEFGGLS | ASSQDPLGNTLY | 0.66125464 |
| DPDGSVY | ASSQDPLGNTLY | 0.65738326 |
| MVSTIHATE | ASSQDPLGNTLY | 0.5760101 |
| LAAGVADPV | ASSQDPLGNTLY | 0.61740494 |
| TKEQIAA | ASSQDPLGNTLY | 0.52254516 |
| SLASPAAL | ASSQDPLGNTLY | 0.6116395 |
| IGLVFYGV | ASSQDPLGNTLY | 0.5919122 |
| FLLDASNL | ASSQDPLGNTLY | 0.63981336 |
| YQVGNFP | ASSQDPLGNTLY | 0.3143229 |
| ESLLSQST | ASSQDPLGNTLY | 0.6321814 |
| LFTVGFAAVG | ASSQDPLGNTLY | 0.6651245 |
| ATPPTPT | ASSQDPLGNTLY | 0.5803604 |
| TDLATLG | ASSQDPLGNTLY | 0.6308418 |
| SAGGTGR | ASSQDPLGNTLY | 0.64014614 |
| GFSNSGS | ASSQDPLGNTLY | 0.58206165 |
| GGNGGSGGTSV | ASSQDPLGNTLY | 0.646903 |
| GTSVATGGAGN | ASSQDPLGNTLY | 0.5582877 |
| NGGNGGSGGTS | ASSQDPLGNTLY | 0.64827347 |
| AVVKFPH | ASSQDPLGNTLY | 0.66424835 |
| TPPPPPP | ASSQDPLGNTLY | 0.63311267 |
| AAPAAAAAPA | ASSQDPLGNTLY | 0.58555675 |
| IALNEDL | ASSQDPLGNTLY | 0.61137676 |
| IVNVSSM | ASSQDPLGNTLY | 0.43199325 |
| TGIPPAPR | ASSQDPLGNTLY | 0.6237143 |
| FDLGGGTF | ASSQDPLGNTLY | 0.5027752 |
| FDLGGGTFDVS | ASSQDPLGNTLY | 0.6193034 |
| NEPTAAA | ASSQDPLGNTLY | 0.54598767 |
| NEPTAAALA | ASSQDPLGNTLY | 0.63597214 |
| GSIINISSI | ASSQDPLGNTLY | 0.626064 |
| VAPGNAGTA | ASSQDPLGNTLY | 0.62472355 |
| TGGGASLE | ASSQDPLGNTLY | 0.53336716 |
| VAVYNHYKRI | ASSQDPLGNTLY | 0.64068425 |
| TDPAPATT | ASSQDPLGNTLY | 0.6111069 |
| TNRPDILDPALLRPG | ASSQDPLGNTLY | 0.49352568 |
| LTTTTTS | ASSQDPLGNTLY | 0.65201974 |
| PTGSGKTL | ASSQDPLGNTLY | 0.5801845 |
| TGSGKTL | ASSQDPLGNTLY | 0.614318 |
| TIGHVDHGK | ASSQDPLGNTLY | 0.65801984 |
| TIGHVDHGKTT | ASSQDPLGNTLY | 0.6225306 |
| GHVDHGKTT | ASSQDPLGNTLY | 0.44090524 |
| AQFEHTLL | ASSQDPLGNTLY | 0.6132528 |
| IVNQDEL | ASSQDPLGNTLY | 0.6081985 |
| TNGSQFF | ASSQDPLGNTLY | 0.6032533 |

| | | |
|---|---|---|
| TGKGIVE | ASSQDPLGNTLY | 0.35330608 |
| LISNASDA | ASSQDPLGNTLY | 0.6437003 |
| AELRTLGELTKIAKAHGAQ | ASSQDPLGNTLY | 0.5297909 |
| APNAVAAAEPSW | ASSQDPLGNTLY | 0.56485844 |
| APQVAGAATAVL | ASSQDPLGNTLY | 0.5755776 |
| ARRDPAAAAAILPTDA | ASSQDPLGNTLY | 0.65363765 |
| DAAARDVLAEAGLAEYF | ASSQDPLGNTLY | 0.613151 |
| DDQIVLSAKANLAKVEQ | ASSQDPLGNTLY | 0.6174252 |
| DELLPDTGEDSDEDGH | ASSQDPLGNTLY | 0.646896 |
| ELPEAPANRATVPQD | ASSQDPLGNTLY | 0.6423415 |
| FGQSKIERVTGIG | ASSQDPLGNTLY | 0.6609775 |
| GAGGTGSTAGGAGGAGGAGGLY | ASSQDPLGNTLY | 0.62779343 |
| GHGGTGAAGGNGGTGGAGGSGI | ASSQDPLGNTLY | 0.5774934 |
| GNGGNGGAGGEGGPGGAGGAGA | ASSQDPLGNTLY | 0.61887294 |
| GSTAGNGGNGGNATNGTIAGQP | ASSQDPLGNTLY | 0.6031579 |
| GTGGDGGNGGNGGTGG | ASSQDPLGNTLY | 0.53459907 |
| GVRRPAGIPADAPV | ASSQDPLGNTLY | 0.4683764 |
| GYRQVTDAHLLTLARR | ASSQDPLGNTLY | 0.6264496 |
| IAKMGQDGHDRG | ASSQDPLGNTLY | 0.43311536 |
| KAEPSIEIQYT | ASSQDPLGNTLY | 0.61686933 |
| KPVRIRMAADEDTVAVT | ASSQDPLGNTLY | 0.6597913 |
| MPPRIAGMRLLVIKPEP | ASSQDPLGNTLY | 0.5950928 |
| MKFYAVALFRSRGVPPTR | ASSQDPLGNTLY | 0.46947512 |
| NAGGFGILGAVAHSP | ASSQDPLGNTLY | 0.5862178 |
| NEGREITLAMAEM | ASSQDPLGNTLY | 0.59664357 |
| NELVEWSHPLGHRWRWE | ASSQDPLGNTLY | 0.6553198 |
| NLGAGGTGGNGGNPG | ASSQDPLGNTLY | 0.63944423 |
| NTEWPKWETVMAADPVD | ASSQDPLGNTLY | 0.67011476 |
| PASAAPAETQP | ASSQDPLGNTLY | 0.656812 |
| QQSSDGKNGVRAAEITGE | ASSQDPLGNTLY | 0.579835 |
| RAPDAVTTSWWKE | ASSQDPLGNTLY | 0.6456498 |
| SGFGNVGAHTSGWFNQ | ASSQDPLGNTLY | 0.599859 |
| SGVAGAVGQAASVGGLKV | ASSQDPLGNTLY | 0.62764204 |
| SNGGAGTGSAGGNGGT | ASSQDPLGNTLY | 0.62545264 |
| TGGAGGEGGAAG | ASSQDPLGNTLY | 0.6239195 |
| TGGDGGAAGAGGAPGHG | ASSQDPLGNTLY | 0.59747237 |
| TMPPLLAAIANAGRHGV | ASSQDPLGNTLY | 0.59232193 |
| TSSTAGNSGGA | ASSQDPLGNTLY | 0.4337267 |
| VPATVPVGILC | ASSQDPLGNTLY | 0.6356921 |
| VQRPDSGGVVLTGRIS | ASSQDPLGNTLY | 0.598298 |
| WTPNPVPATKGVPAGLVQAT | ASSQDPLGNTLY | 0.47393152 |
| YALKRALAKFSDRFDVVII | ASSQDPLGNTLY | 0.5850219 |
| YLIGSVIV | ASSQDPLGNTLY | 0.66558635 |
| YGQLVAVID | ASSQDPLGNTLY | 0.6613834 |
| VTVGGMTLPA | ASSQDPLGNTLY | 0.5842615 |
| VLALPAV | ASSQDPLGNTLY | 0.55200386 |
| VIPPTLNYET | ASSQDPLGNTLY | 0.6285743 |
| VGMGGMHPSSG | ASSQDPLGNTLY | 0.405875 |
| VGLAGGAATA | ASSQDPLGNTLY | 0.6325715 |
| VGGAGGAGGSGP | ASSQDPLGNTLY | 0.62816495 |
| VFAAFMV | ASSQDPLGNTLY | 0.6254099 |

| | | |
|---|---|---|
| TLYDKLR | ASSQDPLGNTLY | 0.5087813 |
| TLAVFTVL | ASSQDPLGNTLY | 0.6325099 |
| TGSVQPARV | ASSQDPLGNTLY | 0.60177326 |
| TGGGAGSGGT | ASSQDPLGNTLY | 0.64559937 |
| SHLETSL | ASSQDPLGNTLY | 0.37388384 |
| SFTTGMT | ASSQDPLGNTLY | 0.64527154 |
| QVEVAKRMGVRQPTVS | ASSQDPLGNTLY | 0.55080354 |
| QGSGGPALAM | ASSQDPLGNTLY | 0.5524615 |
| MLAPSNP | ASSQDPLGNTLY | 0.38967958 |
| LVNNAAI | ASSQDPLGNTLY | 0.6107692 |
| LTGEINTLA | ASSQDPLGNTLY | 0.6543719 |
| LSIFGPV | ASSQDPLGNTLY | 0.6535622 |
| LLALLANL | ASSQDPLGNTLY | 0.583517 |
| LISNASDALDKLR | ASSQDPLGNTLY | 0.6531334 |
| LINEAVP | ASSQDPLGNTLY | 0.6657322 |
| LGYAFSGL | ASSQDPLGNTLY | 0.6138382 |
| LDFVAVAL | ASSQDPLGNTLY | 0.40661085 |
| LAQPGAE | ASSQDPLGNTLY | 0.6374991 |
| ISNLGAVTGF | ASSQDPLGNTLY | 0.6109724 |
| HQEDPGRI | ASSQDPLGNTLY | 0.6545652 |
| GTPGEPTP | ASSQDPLGNTLY | 0.63200086 |
| GFFNSGS | ASSQDPLGNTLY | 0.5244637 |
| GAGGGGLTAGAN | ASSQDPLGNTLY | 0.6312421 |
| FVDALVNM | ASSQDPLGNTLY | 0.656906 |
| FLTVAPEL | ASSQDPLGNTLY | 0.5384195 |
| FETLTAL | ASSQDPLGNTLY | 0.5525789 |
| DVINPIEL | ASSQDPLGNTLY | 0.5592488 |
| APVSGAPV | ASSQDPLGNTLY | 0.4838884 |
| ALVGGLAW | ASSQDPLGNTLY | 0.6409136 |
| ALAAAAATAI | ASSQDPLGNTLY | 0.58709407 |
| AFGLGLLL | ASSQDPLGNTLY | 0.58532476 |
| AFFAAASL | ASSQDPLGNTLY | 0.62200284 |
| AAAALPAFSPP | ASSQDPLGNTLY | 0.56284213 |
| VVLAPAL | ASSQDPLGNTLY | 0.5711751 |
| TIGHVDHGKTTL | ASSQDPLGNTLY | 0.59060395 |
| LISNASDALDK | ASSQDPLGNTLY | 0.63508546 |
| LISNASDAL | ASSQDPLGNTLY | 0.6400953 |
| ISNASDALDK | ASSQDPLGNTLY | 0.61709905 |
| NQAETLVYQTE | ASSQDPLGNTLY | 0.621444 |
| TGGTGGTAGAG | ASSQDPLGNTLY | 0.6659894 |
| TGSTAGGAGGA | ASSQDPLGNTLY | 0.50820374 |
| DPLAVPVAAGPV | ASSQDPLGNTLY | 0.5452424 |
| GGTGGTPGNGGN | ASSQDPLGNTLY | 0.63872015 |
| PAGLAGVDPATL | ASSQDPLGNTLY | 0.49666303 |
| GNNLVVEAPHAGASV | ASSQDPLGNTLY | 0.6531493 |
| KVTRIMSASTLEQPA | ASSQDPLGNTLY | 0.6184119 |
| LGFALPIALPRRTVV | ASSQDPLGNTLY | 0.6391027 |
| TRLGNLTARKLVVLV | ASSQDPLGNTLY | 0.6254288 |
| WSQQRRLIYFPSAGP | ASSQDPLGNTLY | 0.66639185 |
| GPNGNPDPMAAAVDIR | ASSQDPLGNTLY | 0.65473515 |
| TQRCVSVADLRAAASR | ASSQDPLGNTLY | 0.6104293 |

| | | |
|---|---|---|
| QFVLDVAEHPVEVLTETV | ASSQDPLGNTLY | 0.6517913 |
| GGAGGTGGAGGNGGTQAGDG | ASSQDPLGNTLY | 0.4671679 |
| PLLATLRAIAVDVDAKPAQV | ASSQDPLGNTLY | 0.6548931 |
| APKFRTGGYMIDFWGVGYQVAK | ASSQDPLGNTLY | 0.6340316 |
| GAGGAGGAGGAAGGPGGTGGTG | ASSQDPLGNTLY | 0.55048954 |
| IGALGLLATDVAPAGSVAFGVA | ASSQDPLGNTLY | 0.6169856 |
| AGGAGGIGGNGGAGGAGGVLMG | ASSQDPLGNTLY | 0.6606608 |
| ADELIGLL | ASSQDPLGNTLY | 0.65736824 |
| ADRALKA | ASSQDPLGNTLY | 0.46778873 |
| AGAVGQVGGAGGA | ASSQDPLGNTLY | 0.60219014 |
| AGGGGGLGGAP | ASSQDPLGNTLY | 0.39893007 |
| AGGLLYGNGGAG | ASSQDPLGNTLY | 0.4734426 |
| AGLESIGL | ASSQDPLGNTLY | 0.44607267 |
| AGLKTGII | ASSQDPLGNTLY | 0.64570796 |
| ASLAALLL | ASSQDPLGNTLY | 0.58800656 |
| ASVINISSI | ASSQDPLGNTLY | 0.6382491 |
| ATPPGPA | ASSQDPLGNTLY | 0.6229999 |
| ATVASALV | ASSQDPLGNTLY | 0.660334 |
| ATVASLIL | ASSQDPLGNTLY | 0.54179 |
| AVNGPVLIRH | ASSQDPLGNTLY | 0.6056018 |
| DGAAGAVGI | ASSQDPLGNTLY | 0.42796317 |
| DPLLHGI | ASSQDPLGNTLY | 0.54869986 |
| ELGAALGAL | ASSQDPLGNTLY | 0.57009274 |
| ELGRTQAV | ASSQDPLGNTLY | 0.5997411 |
| FVAGGYLS | ASSQDPLGNTLY | 0.4446628 |
| GASGGAGGAGGW | ASSQDPLGNTLY | 0.66126275 |
| GGAGAGGDGGAGGI | ASSQDPLGNTLY | 0.6311108 |
| GTGHNGSQPGMGGNG | ASSQDPLGNTLY | 0.6458603 |
| GYAAASASAS | ASSQDPLGNTLY | 0.5684375 |
| HQVDVAAVQQG | ASSQDPLGNTLY | 0.6001979 |
| IGAAIAHT | ASSQDPLGNTLY | 0.5802389 |
| IGMIEVAQL | ASSQDPLGNTLY | 0.6394574 |
| IRLAPEHTSAMD | ASSQDPLGNTLY | 0.544156 |
| IYNVSVL | ASSQDPLGNTLY | 0.4345502 |
| KQGFLGM | ASSQDPLGNTLY | 0.5796124 |
| LESVITQL | ASSQDPLGNTLY | 0.55340636 |
| LGLGALVVV | ASSQDPLGNTLY | 0.6607542 |
| LIGNGGNGGSGGI | ASSQDPLGNTLY | 0.6595302 |
| LNAINAPTL | ASSQDPLGNTLY | 0.6375969 |
| LQLLNTGV | ASSQDPLGNTLY | 0.65213877 |
| LQTSIEF | ASSQDPLGNTLY | 0.34098417 |
| MSFAVSEL | ASSQDPLGNTLY | 0.587097 |
| NTGALGL | ASSQDPLGNTLY | 0.66150177 |
| PPGAGGAGGAGTT | ASSQDPLGNTLY | 0.6436533 |
| QLVGLGI | ASSQDPLGNTLY | 0.6032221 |
| RFLEQVHQL | ASSQDPLGNTLY | 0.6632537 |
| RGVGYVLR | ASSQDPLGNTLY | 0.5458875 |
| SATGVFAGVI | ASSQDPLGNTLY | 0.5932951 |
| SLPAITLP | ASSQDPLGNTLY | 0.6363468 |
| STALAFTV | ASSQDPLGNTLY | 0.645973 |
| SVLTVGLGK | ASSQDPLGNTLY | 0.6402601 |

| | | |
|---|---|---|
| TALAGAGSGI | ASSQDPLGNTLY | 0.52499765 |
| TGLPFTGI | ASSQDPLGNTLY | 0.6260723 |
| VAEHVTV | ASSQDPLGNTLY | 0.67244315 |
| VDIVLNSL | ASSQDPLGNTLY | 0.6055062 |
| VLTGGGAVL | ASSQDPLGNTLY | 0.65777785 |
| VNVGLAYGI | ASSQDPLGNTLY | 0.6369938 |
| VSSAAVGAV | ASSQDPLGNTLY | 0.63347477 |
| VTSVYEAGV | ASSQDPLGNTLY | 0.5439763 |
| WVALDRGA | ASSQDPLGNTLY | 0.57805794 |
| YLVGSVVI | ASSQDPLGNTLY | 0.6384424 |
| GPPGTGKTLL | ASSQDPLGNTLY | 0.57220984 |
| GVGMTKF | ASSQDPLGNTLY | 0.5942763 |
| IGIGGSDL | ASSQDPLGNTLY | 0.5852482 |
| IGTLLAEL | ASSQDPLGNTLY | 0.6730329 |
| IYTSGTT | ASSQDPLGNTLY | 0.6371341 |
| LSFPAGV | ASSQDPLGNTLY | 0.65180343 |
| LYTSGTT | ASSQDPLGNTLY | 0.5496579 |
| VGAFPFGF | ASSQDPLGNTLY | 0.6293169 |
| VGAGTVEF | ASSQDPLGNTLY | 0.6468445 |
| VGKSTLLL | ASSQDPLGNTLY | 0.4619736 |
| AALAHAGGGIV | ASSQDPLGNTLY | 0.66037786 |
| AALVAVSLL | ASSQDPLGNTLY | 0.56172466 |
| AAVSRAVAAL | ASSQDPLGNTLY | 0.60505474 |
| AGGSSGIGGF | ASSQDPLGNTLY | 0.6452671 |
| AGLTVANANG | ASSQDPLGNTLY | 0.6616089 |
| AGLVGAVL | ASSQDPLGNTLY | 0.59950244 |
| ALSQAFTV | ASSQDPLGNTLY | 0.6307731 |
| ALTFSVSL | ASSQDPLGNTLY | 0.53984326 |
| ALTGAALVAL | ASSQDPLGNTLY | 0.6487229 |
| ASFSIGAGVL | ASSQDPLGNTLY | 0.65857995 |
| DGSVYKI | ASSQDPLGNTLY | 0.62567353 |
| EQLAEIV | ASSQDPLGNTLY | 0.66736203 |
| FAVSELTVM | ASSQDPLGNTLY | 0.6363932 |
| GAAPLAAGL | ASSQDPLGNTLY | 0.39158309 |
| GADGGDGGSGGAGG | ASSQDPLGNTLY | 0.616284 |
| GFGLDTGGAGG | ASSQDPLGNTLY | 0.65488577 |
| GLASGGLL | ASSQDPLGNTLY | 0.65791917 |
| GRIAGAAVLL | ASSQDPLGNTLY | 0.6639342 |
| GSAGSLNTSF | ASSQDPLGNTLY | 0.6302805 |
| GTGQIGI | ASSQDPLGNTLY | 0.5977653 |
| GTLIALIL | ASSQDPLGNTLY | 0.66416323 |
| IAATMGFVVG | ASSQDPLGNTLY | 0.66712916 |
| IFAQALL | ASSQDPLGNTLY | 0.64649457 |
| IGAAVALGF | ASSQDPLGNTLY | 0.56270057 |
| IGLTPLAL | ASSQDPLGNTLY | 0.5045258 |
| ILGSNTSTL | ASSQDPLGNTLY | 0.63094467 |
| IVFSLTVL | ASSQDPLGNTLY | 0.6543305 |
| LASKATGF | ASSQDPLGNTLY | 0.4833972 |
| LASVALIL | ASSQDPLGNTLY | 0.6332562 |
| LFAVGLTL | ASSQDPLGNTLY | 0.6343484 |
| LFNGGIGTYI | ASSQDPLGNTLY | 0.41735336 |

| | | |
|---|---|---|
| LTLGAGII | ASSQDPLGNTLY | 0.6468024 |
| LVAAALGDI | ASSQDPLGNTLY | 0.66072875 |
| NTAFVQLGI | ASSQDPLGNTLY | 0.62317324 |
| QLFSGAEL | ASSQDPLGNTLY | 0.57968485 |
| SAATFITL | ASSQDPLGNTLY | 0.65790033 |
| SNYGLANLGV | ASSQDPLGNTLY | 0.6715567 |
| SYGGAYAVM | ASSQDPLGNTLY | 0.66363263 |
| TTGVGGLTLG | ASSQDPLGNTLY | 0.43332532 |
| TTQTIYI | ASSQDPLGNTLY | 0.37308782 |
| VAAGAGLGAVI | ASSQDPLGNTLY | 0.59131587 |
| VAAVVTLL | ASSQDPLGNTLY | 0.64004683 |
| VGFLGGLI | ASSQDPLGNTLY | 0.59902614 |
| VGVIPILL | ASSQDPLGNTLY | 0.57844234 |
| VIFASSLI | ASSQDPLGNTLY | 0.6399181 |
| VKGGDGGAAAG | ASSQDPLGNTLY | 0.6589382 |
| VTKTFTNI | ASSQDPLGNTLY | 0.51691157 |
| VVAAAILL | ASSQDPLGNTLY | 0.4539009 |
| YAALDVEL | ASSQDPLGNTLY | 0.6469259 |
| LASVALLL | ASSQDPLGNTLY | 0.6387795 |
| LDAVHRV | ASSQDPLGNTLY | 0.5699617 |
| QGLSSSIF | ASSQDPLGNTLY | 0.6111492 |
| RGTFANIRL | ASSQDPLGNTLY | 0.5883668 |
| RQLWWGHRI | ASSQDPLGNTLY | 0.6085274 |
| YGPPGTGKTLL | ASSQDPLGNTLY | 0.63619274 |
| AALVGAPANVA | ASSQDPLGNTLY | 0.49990672 |
| AATFRRTE | ASSQDPLGNTLY | 0.6611054 |
| AGALGGGI | ASSQDPLGNTLY | 0.52755535 |
| AGGAGGAGGIGAG | ASSQDPLGNTLY | 0.6445811 |
| AGSGALGL | ASSQDPLGNTLY | 0.6303181 |
| AQIADQI | ASSQDPLGNTLY | 0.6361085 |
| ASAVAVTYL | ASSQDPLGNTLY | 0.5917218 |
| ASTVLGGTGG | ASSQDPLGNTLY | 0.5877905 |
| AVGGEPAK | ASSQDPLGNTLY | 0.48198557 |
| CEQGTFGGTC | ASSQDPLGNTLY | 0.44099683 |
| DEGKAPAG | ASSQDPLGNTLY | 0.6586246 |
| DTVVFPAL | ASSQDPLGNTLY | 0.6189245 |
| FEGITMAG | ASSQDPLGNTLY | 0.65799445 |
| FPLVVQD | ASSQDPLGNTLY | 0.64785963 |
| GAFTTPQ | ASSQDPLGNTLY | 0.5618583 |
| GFGGFGTAGGVGGA | ASSQDPLGNTLY | 0.6395952 |
| GFLAQFL | ASSQDPLGNTLY | 0.5339217 |
| GGAGGAGGNAGVA | ASSQDPLGNTLY | 0.5952157 |
| GGVGTAGAGGAGGAGG | ASSQDPLGNTLY | 0.6318631 |
| GNGGGGDAGNAGSGGN | ASSQDPLGNTLY | 0.6433278 |
| GTGGAGGAGNDAGS | ASSQDPLGNTLY | 0.5444437 |
| GTTNTGWFNS | ASSQDPLGNTLY | 0.63672507 |
| IFMGPLFTL | ASSQDPLGNTLY | 0.6398103 |
| ILAHAIEAI | ASSQDPLGNTLY | 0.55027777 |
| IRSAIREAL | ASSQDPLGNTLY | 0.6160474 |
| ITFAGGG | ASSQDPLGNTLY | 0.6602692 |
| ITGMSGHQEV | ASSQDPLGNTLY | 0.62986004 |

| | | |
|---|---|---|
| IVAGLACTL | ASSQDPLGNTLY | 0.5495901 |
| IVGGGLAGA | ASSQDPLGNTLY | 0.6508149 |
| IVTGVLNI | ASSQDPLGNTLY | 0.55943227 |
| LAQQGCI | ASSQDPLGNTLY | 0.63803196 |
| LCLHVEN | ASSQDPLGNTLY | 0.62606657 |
| LDAISDGS | ASSQDPLGNTLY | 0.5617188 |
| LGGFAAML | ASSQDPLGNTLY | 0.6522862 |
| LGGKNPMI | ASSQDPLGNTLY | 0.50053465 |
| LGPKGVAAIL | ASSQDPLGNTLY | 0.55436105 |
| LMVGVSI | ASSQDPLGNTLY | 0.59501106 |
| LVGAAFVL | ASSQDPLGNTLY | 0.6224146 |
| LVLAVAPL | ASSQDPLGNTLY | 0.6543508 |
| MAAIAVIDAA | ASSQDPLGNTLY | 0.55777836 |
| NNGINAELQ | ASSQDPLGNTLY | 0.63247716 |
| RGEPLVLVHG | ASSQDPLGNTLY | 0.5777223 |
| RGGAGGAAGLI | ASSQDPLGNTLY | 0.52942467 |
| RSYRMHPAV | ASSQDPLGNTLY | 0.61153275 |
| SAAHLLAGQLA | ASSQDPLGNTLY | 0.6168031 |
| SAAVGSY | ASSQDPLGNTLY | 0.55519533 |
| SPHAVLHRI | ASSQDPLGNTLY | 0.6418042 |
| SQIAKVL | ASSQDPLGNTLY | 0.5542523 |
| TALGHLVP | ASSQDPLGNTLY | 0.62400764 |
| TAPTTSGAA | ASSQDPLGNTLY | 0.6670267 |
| TDGDDGNGGSPGT | ASSQDPLGNTLY | 0.5468342 |
| TDGLGEN | ASSQDPLGNTLY | 0.6452677 |
| TGGPPLPGQA | ASSQDPLGNTLY | 0.6418216 |
| TIPAITI | ASSQDPLGNTLY | 0.6446819 |
| TIPGFTI | ASSQDPLGNTLY | 0.5925387 |
| TSIASGALA | ASSQDPLGNTLY | 0.5702708 |
| TVTLPGAAI | ASSQDPLGNTLY | 0.65897757 |
| VASGLSAALP | ASSQDPLGNTLY | 0.6505995 |
| VGAGVIGI | ASSQDPLGNTLY | 0.60921204 |
| VGPKRFGV | ASSQDPLGNTLY | 0.64449626 |
| VNVAFPD | ASSQDPLGNTLY | 0.6492655 |
| VSMTNSGVS | ASSQDPLGNTLY | 0.6481651 |
| VTDVLGDLSV | ASSQDPLGNTLY | 0.59535027 |
| VVVVGAGFAGL | ASSQDPLGNTLY | 0.46892285 |
| VYTVGTVV | ASSQDPLGNTLY | 0.60777056 |
| WTTFSGP | ASSQDPLGNTLY | 0.597595 |
| YDELVEAF | ASSQDPLGNTLY | 0.6058785 |
| YGTGGAGGNGG | ASSQDPLGNTLY | 0.4208971 |
| YTNVMAV | ASSQDPLGNTLY | 0.65527594 |
| FLEQVHQL | ASSQDPLGNTLY | 0.67033994 |
| GLSLLAVAV | ASSQDPLGNTLY | 0.5681464 |
| LEQVHQL | ASSQDPLGNTLY | 0.6669972 |
| LNAAGAI | ASSQDPLGNTLY | 0.51198035 |
| PAVHASL | ASSQDPLGNTLY | 0.64625424 |
| TELEDDA | ASSQDPLGNTLY | 0.6569801 |
| TPPGPPN | ASSQDPLGNTLY | 0.5408223 |
| TPPPPPPP | ASSQDPLGNTLY | 0.64961684 |
| TQSPALI | ASSQEDRNTEVF | 0.5639279 |

| | | |
|---|---|---|
| EFAAAFPQ | ASSQEDRNTEVF | 0.5982709 |
| LLSGNLDVL | ASSQEDRNTEVF | 0.6604897 |
| GPANLGNYNI | ASSQEDRNTEVF | 0.5621599 |
| TLGPTLP | ASSQEDRNTEVF | 0.6356099 |
| VYAYSSCCK | ASSQEDRNTEVF | 0.60212797 |
| APAPAPAPNI | ASSQEDRNTEVF | 0.52858615 |
| PGGPVPIP | ASSQEDRNTEVF | 0.63530475 |
| FLTVALEP | ASSQEDRNTEVF | 0.6439208 |
| SQYFGANL | ASSQEDRNTEVF | 0.5846873 |
| GGVKPAA | ASSQEDRNTEVF | 0.66693014 |
| GGTGGAGGAGGW | ASSQEDRNTEVF | 0.5507734 |
| GVGGVAGLGGAGG | ASSQEDRNTEVF | 0.6667803 |
| VLLVLATTI | ASSQEDRNTEVF | 0.64498127 |
| IGIFAAML | ASSQEDRNTEVF | 0.6264658 |
| AVGDLID | ASSQEDRNTEVF | 0.58872736 |
| NISPLPL | ASSQEDRNTEVF | 0.65868866 |
| VTAVFVAI | ASSQEDRNTEVF | 0.57868886 |
| SGDDNDAAL | ASSQEDRNTEVF | 0.6345325 |
| AVAALAHS | ASSQEDRNTEVF | 0.66423124 |
| GGVTVMATP | ASSQEDRNTEVF | 0.5506636 |
| AGLAANVRD | ASSQEDRNTEVF | 0.83011246 |
| TNRPDLIDPALLRPG | ASSQEDRNTEVF | 0.76142144 |
| LANLGVTAL | ASSQEDRNTEVF | 0.8066998 |
| TGERIEL | ASSQEDRNTEVF | 0.6483495 |
| NGEELAA | ASSQEDRNTEVF | 0.39605492 |
| VLSLVWG | ASSQEDRNTEVF | 0.7573323 |
| IGYALWL | ASSQEDRNTEVF | 0.5624463 |
| LSLLLAPL | ASSQEDRNTEVF | 0.7864687 |
| TLDEALK | ASSQEDRNTEVF | 0.81429607 |
| TDGAGTAEGG | ASSQEDRNTEVF | 0.62189007 |
| NAPGGAGGNGG | ASSQEDRNTEVF | 0.8210131 |
| VSVFMGMGGNF | ASSQEDRNTEVF | 0.83938754 |
| LASYLAAL | ASSQEDRNTEVF | 0.39753103 |
| LVGGTPAL | ASSQEDRNTEVF | 0.69665676 |
| VRNADVL | ASSQEDRNTEVF | 0.68169665 |
| TCSSPFLVSAM | ASSQEDRNTEVF | 0.70973974 |
| VHITDAE | ASSQEDRNTEVF | 0.24557021 |
| EYAAPSF | ASSQEDRNTEVF | 0.8079717 |
| VVDAFHGGL | ASSQEDRNTEVF | 0.645011 |
| ASVVDPQ | ASSQEDRNTEVF | 0.15721196 |
| SYNEFGGLS | ASSQEDRNTEVF | 0.7742171 |
| DPDGSVY | ASSQEDRNTEVF | 0.6782467 |
| MVSTIHATE | ASSQEDRNTEVF | 0.6586618 |
| LAAGVADPV | ASSQEDRNTEVF | 0.821664 |
| TKEQIAA | ASSQEDRNTEVF | 0.5022526 |
| SLASPAAL | ASSQEDRNTEVF | 0.5337132 |
| IGLVFYGV | ASSQEDRNTEVF | 0.77076936 |
| FLLDASNL | ASSQEDRNTEVF | 0.8399018 |
| YQVGNFP | ASSQEDRNTEVF | 0.50103414 |
| ESLLSQST | ASSQEDRNTEVF | 0.7594102 |
| LFTVGFAAVG | ASSQEDRNTEVF | 0.8597007 |

| | | |
|---|---|---|
| ATPPTPT | ASSQEDRNTEVF | 0.77118206 |
| TDLATLG | ASSQEDRNTEVF | 0.8393569 |
| SAGGTGR | ASSQEDRNTEVF | 0.85667735 |
| GFSNSGS | ASSQEDRNTEVF | 0.6343948 |
| GGNGGSGGTSV | ASSQEDRNTEVF | 0.8174968 |
| GTSVATGGAGN | ASSQEDRNTEVF | 0.8392739 |
| NGGNGGSGGTS | ASSQEDRNTEVF | 0.667187 |
| AVVKFPH | ASSQEDRNTEVF | 0.7439306 |
| TPPPPPP | ASSQEDRNTEVF | 0.5979419 |
| AAPAAAAAPA | ASSQEDRNTEVF | 0.70181036 |
| IALNEDL | ASSQEDRNTEVF | 0.2852136 |
| IVNVSSM | ASSQEDRNTEVF | 0.7486771 |
| TGIPPAPR | ASSQEDRNTEVF | 0.8328972 |
| FDLGGGTF | ASSQEDRNTEVF | 0.7053234 |
| FDLGGGTFDVS | ASSQEDRNTEVF | 0.47704542 |
| NEPTAAA | ASSQEDRNTEVF | 0.42476293 |
| NEPTAAALA | ASSQEDRNTEVF | 0.7169149 |
| GSIINISSI | ASSQEDRNTEVF | 0.6320495 |
| VAPGNAGTA | ASSQEDRNTEVF | 0.3387974 |
| TGGGASLE | ASSQEDRNTEVF | 0.62990373 |
| VAVYNHYKRI | ASSQEDRNTEVF | 0.8348532 |
| TDPAPATT | ASSQEDRNTEVF | 0.84422696 |
| TNRPDILDPALLRPG | ASSQEDRNTEVF | 0.48900807 |
| LTTTTTS | ASSQEDRNTEVF | 0.84277624 |
| PTGSGKTL | ASSQEDRNTEVF | 0.80581206 |
| TGSGKTL | ASSQEDRNTEVF | 0.855036 |
| TIGHVDHGK | ASSQEDRNTEVF | 0.7691641 |
| TIGHVDHGKTT | ASSQEDRNTEVF | 0.5319755 |
| GHVDHGKTT | ASSQEDRNTEVF | 0.8358589 |
| AQFEHTLL | ASSQEDRNTEVF | 0.66456026 |
| IVNQDEL | ASSQEDRNTEVF | 0.6621239 |
| TNGSQFF | ASSQEDRNTEVF | 0.82530004 |
| TGKGIVE | ASSQEDRNTEVF | 0.712024 |
| LISNASDA | ASSQEDRNTEVF | 0.6002211 |
| AELRTLGELTKIAKAHGAQ | ASSQEDRNTEVF | 0.69156766 |
| APNAVAAAEPSW | ASSQEDRNTEVF | 0.84965444 |
| APQVAGAATAVL | ASSQEDRNTEVF | 0.5228846 |
| ARRDPAAAAAILPTDA | ASSQEDRNTEVF | 0.46409753 |
| DAAARDVLAEAGLAEYF | ASSQEDRNTEVF | 0.6219306 |
| DDQIVLSAKANLAKVEQ | ASSQEDRNTEVF | 0.73881733 |
| DELLPDTGEDSDEDGH | ASSQEDRNTEVF | 0.83443916 |
| ELPEAPANRATVPQD | ASSQEDRNTEVF | 0.7942511 |
| FGQSKIERVTGIG | ASSQEDRNTEVF | 0.6576721 |
| GAGGTGSTAGGAGGAGGAGGLY | ASSQEDRNTEVF | 0.7751899 |
| GHGGTGAAGGNGGTGGAGGSGI | ASSQEDRNTEVF | 0.8082572 |
| GNGGNGGAGGEGGPGGAGGAGA | ASSQEDRNTEVF | 0.7692968 |
| GSTAGNGGNGGNATNGTIAGQP | ASSQEDRNTEVF | 0.81679755 |
| GTGGDGGNGGNGGTGG | ASSQEDRNTEVF | 0.57330805 |
| GVRRPAGIPADAPV | ASSQEDRNTEVF | 0.8386973 |
| GYRQVTDAHLLTLARR | ASSQEDRNTEVF | 0.82478976 |
| IAKMGQDGHDRG | ASSQEDRNTEVF | 0.8164203 |

| | | |
|---|---|---|
| KAEPSIEIQYT | ASSQEDRNTEVF | 0.8218031 |
| KPVRIRMAADEDTVAVT | ASSQEDRNTEVF | 0.7381826 |
| MPPRIAGMRLLVIKPEP | ASSQEDRNTEVF | 0.77188826 |
| MKFYAVALFRSRGVPPTR | ASSQEDRNTEVF | 0.39768034 |
| NAGGFGILGAVAHSP | ASSQEDRNTEVF | 0.72256553 |
| NEGREITLAMAEM | ASSQEDRNTEVF | 0.74943984 |
| NELVEWSHPLGHRWRWE | ASSQEDRNTEVF | 0.7559838 |
| NLGAGGTGGNGGNPG | ASSQEDRNTEVF | 0.78204036 |
| NTEWPKWETVMAADPVD | ASSQEDRNTEVF | 0.8232837 |
| PASAAPAETQP | ASSQEDRNTEVF | 0.61931896 |
| QQSSDGKNGVRAAEITGE | ASSQEDRNTEVF | 0.84599304 |
| RAPDAVTTSWWKE | ASSQEDRNTEVF | 0.8286978 |
| SGFGNVGAHTSGWFNQ | ASSQEDRNTEVF | 0.4618822 |
| SGVAGAVGQAASVGGLKV | ASSQEDRNTEVF | 0.7083658 |
| SNGGAGTGSAGGNGGT | ASSQEDRNTEVF | 0.8369238 |
| TGGAGGEGGAAG | ASSQEDRNTEVF | 0.78625196 |
| TGGDGGAAGAGGAPGHG | ASSQEDRNTEVF | 0.6749638 |
| TMPPLLAAIANAGRHGV | ASSQEDRNTEVF | 0.74230826 |
| TSSTAGNSGGA | ASSQEDRNTEVF | 0.85821784 |
| VPATVPVGILC | ASSQEDRNTEVF | 0.6657259 |
| VQRPDSGGVVLTGRIS | ASSQEDRNTEVF | 0.8527531 |
| WTPNPVPATKGVPAGLVQAT | ASSQEDRNTEVF | 0.24748254 |
| YALKRALAKFSDRFDVVII | ASSQEDRNTEVF | 0.7159053 |
| YLIGSVIV | ASSQEDRNTEVF | 0.6908228 |
| YGQLVAVID | ASSQEDRNTEVF | 0.23427238 |
| VTVGGMTLPA | ASSQEDRNTEVF | 0.8213476 |
| VLALPAV | ASSQEDRNTEVF | 0.70852864 |
| VIPPTLNYET | ASSQEDRNTEVF | 0.81366134 |
| VGMGGMHPSSG | ASSQEDRNTEVF | 0.7143668 |
| VGLAGGAATA | ASSQEDRNTEVF | 0.83290267 |
| VGGAGGAGGSGP | ASSQEDRNTEVF | 0.72993994 |
| VFAAFMV | ASSQEDRNTEVF | 0.7358282 |
| TLYDKLR | ASSQEDRNTEVF | 0.50896 |
| TLAVFTVL | ASSQEDRNTEVF | 0.81005543 |
| TGSVQPARV | ASSQEDRNTEVF | 0.8425447 |
| TGGGAGSGGT | ASSQEDRNTEVF | 0.838902 |
| SHLETSL | ASSQEDRNTEVF | 0.3322119 |
| SFTTGMT | ASSQEDRNTEVF | 0.70637316 |
| QVEVAKRMGVRQPTVS | ASSQEDRNTEVF | 0.7526031 |
| QGSGGPALAM | ASSQEDRNTEVF | 0.2584089 |
| MLAPSNP | ASSQEDRNTEVF | 0.22523086 |
| LVNNAAI | ASSQEDRNTEVF | 0.6219939 |
| LTGEINTLA | ASSQEDRNTEVF | 0.6474914 |
| LSIFGPV | ASSQEDRNTEVF | 0.80891776 |
| LLALLANL | ASSQEDRNTEVF | 0.7898933 |
| LISNASDALDKLR | ASSQEDRNTEVF | 0.6938914 |
| LINEAVP | ASSQEDRNTEVF | 0.79981613 |
| LGYAFSGL | ASSQEDRNTEVF | 0.74033296 |
| LDFVAVAL | ASSQEDRNTEVF | 0.09336738 |
| LAQPGAE | ASSQEDRNTEVF | 0.6524993 |
| ISNLGAVTGF | ASSQEDRNTEVF | 0.4160694 |

| | | |
|---|---|---|
| HQEDPGRI | ASSQEDRNTEVF | 0.848783 |
| GTPGEPTP | ASSQEDRNTEVF | 0.5833355 |
| GFFNSGS | ASSQEDRNTEVF | 0.62844193 |
| GAGGGGLTAGAN | ASSQEDRNTEVF | 0.73760873 |
| FVDALVNM | ASSQEDRNTEVF | 0.74814487 |
| FLTVAPEL | ASSQEDRNTEVF | 0.7440871 |
| FETLTAL | ASSQEDRNTEVF | 0.5598016 |
| DVINPIEL | ASSQEDRNTEVF | 0.19034095 |
| APVSGAPV | ASSQEDRNTEVF | 0.822438 |
| ALVGGLAW | ASSQEDRNTEVF | 0.40244383 |
| ALAAAAATAI | ASSQEDRNTEVF | 0.5245746 |
| AFGLGLLL | ASSQEDRNTEVF | 0.38704497 |
| AFFAAASL | ASSQEDRNTEVF | 0.21992931 |
| AAAALPAFSPP | ASSQEDRNTEVF | 0.37623674 |
| VVLAPAL | ASSQEDRNTEVF | 0.8316679 |
| TIGHVDHGKTTL | ASSQEDRNTEVF | 0.6106426 |
| LISNASDALDK | ASSQEDRNTEVF | 0.6833257 |
| LISNASDAL | ASSQEDRNTEVF | 0.8406927 |
| ISNASDALDK | ASSQEDRNTEVF | 0.73142785 |
| NQAETLVYQTE | ASSQEDRNTEVF | 0.6882875 |
| TGGTGGTAGAG | ASSQEDRNTEVF | 0.6112115 |
| TGSTAGGAGGA | ASSQEDRNTEVF | 0.53254974 |
| DPLAVPVAAGPV | ASSQEDRNTEVF | 0.58483934 |
| GGTGGTPGNGGN | ASSQEDRNTEVF | 0.6032392 |
| PAGLAGVDPATL | ASSQEDRNTEVF | 0.84955823 |
| GNNLVVEAPHAGASV | ASSQEDRNTEVF | 0.72029305 |
| KVTRIMSASTLEQPA | ASSQEDRNTEVF | 0.5011977 |
| LGFALPIALPRRTVV | ASSQEDRNTEVF | 0.78792465 |
| TRLGNLTARKLVVLV | ASSQEDRNTEVF | 0.8432926 |
| WSQQRRLIYFPSAGP | ASSQEDRNTEVF | 0.8651104 |
| GPNGNPDPMAAAVDIR | ASSQEDRNTEVF | 0.54048234 |
| TQRCVSVADLRAAASR | ASSQEDRNTEVF | 0.73799753 |
| QFVLDVAEHPVEVLTETV | ASSQEDRNTEVF | 0.72999114 |
| GGAGGTGGAGGNGGTQAGDG | ASSQEDRNTEVF | 0.7123518 |
| PLLATLRAIAVDVDAKPAQV | ASSQEDRNTEVF | 0.6702707 |
| APKFRTGGYMIDFWGVGYQVAK | ASSQEDRNTEVF | 0.5169635 |
| GAGGAGGAGGAAGGPGGTGGTG | ASSQEDRNTEVF | 0.8258636 |
| IGALGLLATDVAPAGSVAFGVA | ASSQEDRNTEVF | 0.8746213 |
| AGGAGGIGGNGGAGGAGGVLMG | ASSQEDRNTEVF | 0.7158357 |
| ADELIGLL | ASSQEDRNTEVF | 0.46406543 |
| ADRALKA | ASSQEDRNTEVF | 0.69857115 |
| AGAVGQVGGAGGA | ASSQEDRNTEVF | 0.534819 |
| AGGGGGLGGAP | ASSQEDRNTEVF | 0.6300471 |
| AGGLLYGNGGAG | ASSQEDRNTEVF | 0.7270696 |
| AGLESIGL | ASSQEDRNTEVF | 0.41083747 |
| AGLKTGII | ASSQEDRNTEVF | 0.72463524 |
| ASLAALLL | ASSQEDRNTEVF | 0.865412 |
| ASVINISSI | ASSQEDRNTEVF | 0.66261566 |
| ATPPGPA | ASSQEDRNTEVF | 0.786415 |
| ATVASALV | ASSQEDRNTEVF | 0.82461095 |
| ATVASLIL | ASSQEDRNTEVF | 0.7875073 |

| | | |
|---|---|---|
| AVNGPVLIRH | ASSQEDRNTEVF | 0.8437926 |
| DGAAGAVGI | ASSQEDRNTEVF | 0.81056035 |
| DPLLHGI | ASSQEDRNTEVF | 0.7373618 |
| ELGAALGAL | ASSQEDRNTEVF | 0.7550791 |
| ELGRTQAV | ASSQEDRNTEVF | 0.7663864 |
| FVAGGYLS | ASSQEDRNTEVF | 0.7171383 |
| GASGGAGGAGGW | ASSQEDRNTEVF | 0.7301786 |
| GGAGAGGDGGAGGI | ASSQEDRNTEVF | 0.6725769 |
| GTGHNGSQPGMGGNG | ASSQEDRNTEVF | 0.8132882 |
| GYAAASASAS | ASSQEDRNTEVF | 0.7554362 |
| HQVDVAAVQQG | ASSQEDRNTEVF | 0.8455411 |
| IGAAIAHT | ASSQEDRNTEVF | 0.32702547 |
| IGMIEVAQL | ASSQEDRNTEVF | 0.505914 |
| IRLAPEHTSAMD | ASSQEDRNTEVF | 0.8150029 |
| IYNVSVL | ASSQEDRNTEVF | 0.15699948 |
| KQGFLGM | ASSQEDRNTEVF | 0.25977966 |
| LESVITQL | ASSQEDRNTEVF | 0.74779844 |
| LGLGALVVV | ASSQEDRNTEVF | 0.84463125 |
| LIGNGGNGGSGGI | ASSQEDRNTEVF | 0.14488263 |
| LNAINAPTL | ASSQEDRNTEVF | 0.58527434 |
| LQLLNTGV | ASSQEDRNTEVF | 0.39666122 |
| LQTSIEF | ASSQEDRNTEVF | 0.8163285 |
| MSFAVSEL | ASSQEDRNTEVF | 0.72975636 |
| NTGALGL | ASSQEDRNTEVF | 0.32582402 |
| PPGAGGAGGAGTT | ASSQEDRNTEVF | 0.8140705 |
| QLVGLGI | ASSQEDRNTEVF | 0.3971871 |
| RFLEQVHQL | ASSQEDRNTEVF | 0.55027187 |
| RGVGYVLR | ASSQEDRNTEVF | 0.82495093 |
| SATGVFAGVI | ASSQEDRNTEVF | 0.84610164 |
| SLPAITLP | ASSQEDRNTEVF | 0.5101825 |
| STALAFTV | ASSQEDRNTEVF | 0.7966596 |
| SVLTVGLGK | ASSQEDRNTEVF | 0.65368795 |
| TALAGAGSGI | ASSQEDRNTEVF | 0.87132573 |
| TGLPFTGI | ASSQEDRNTEVF | 0.69692373 |
| VAEHVTV | ASSQEDRNTEVF | 0.3794202 |
| VDIVLNSL | ASSQEDRNTEVF | 0.8192109 |
| VLTGGGAVL | ASSQEDRNTEVF | 0.80030525 |
| VNVGLAYGI | ASSQEDRNTEVF | 0.77347016 |
| VSSAAVGAV | ASSQEDRNTEVF | 0.8131452 |
| VTSVYEAGV | ASSQEDRNTEVF | 0.6687806 |
| WVALDRGA | ASSQEDRNTEVF | 0.6798079 |
| YLVGSVVI | ASSQEDRNTEVF | 0.8122219 |
| GPPGTGKTLL | ASSQEDRNTEVF | 0.8002299 |
| GVGMTKF | ASSQEDRNTEVF | 0.4952532 |
| IGIGGSDL | ASSQEDRNTEVF | 0.8427522 |
| IGTLLAEL | ASSQEDRNTEVF | 0.7390616 |
| IYTSGTT | ASSQEDRNTEVF | 0.6357726 |
| LSFPAGV | ASSQEDRNTEVF | 0.8638011 |
| LYTSGTT | ASSQEDRNTEVF | 0.74691546 |
| VGAFPFGF | ASSQEDRNTEVF | 0.8179019 |
| VGAGTVEF | ASSQEDRNTEVF | 0.82257116 |

| | | |
|---|---|---|
| VGKSTLLL | ASSQEDRNTEVF | 0.85437506 |
| AALAHAGGGIV | ASSQEDRNTEVF | 0.4515552 |
| AALVAVSLL | ASSQEDRNTEVF | 0.8092882 |
| AAVSRAVAAL | ASSQEDRNTEVF | 0.75440097 |
| AGGSSGIGGF | ASSQEDRNTEVF | 0.80821127 |
| AGLTVANANG | ASSQEDRNTEVF | 0.73238707 |
| AGLVGAVL | ASSQEDRNTEVF | 0.6078273 |
| ALSQAFTV | ASSQEDRNTEVF | 0.7925793 |
| ALTFSVSL | ASSQEDRNTEVF | 0.84515953 |
| ALTGAALVAL | ASSQEDRNTEVF | 0.7595607 |
| ASFSIGAGVL | ASSQEDRNTEVF | 0.77956855 |
| DGSVYKI | ASSQEDRNTEVF | 0.854815 |
| EQLAEIV | ASSQEDRNTEVF | 0.5676993 |
| FAVSELTVM | ASSQEDRNTEVF | 0.7437215 |
| GAAPLAAGL | ASSQEDRNTEVF | 0.8245485 |
| GADGGDGGSGGAGG | ASSQEDRNTEVF | 0.83445084 |
| GFGLDTGGAGG | ASSQEDRNTEVF | 0.72686785 |
| GLASGGLL | ASSQEDRNTEVF | 0.8427802 |
| GRIAGAAVLL | ASSQEDRNTEVF | 0.6295273 |
| GSAGSLNTSF | ASSQEDRNTEVF | 0.77981365 |
| GTGQIGI | ASSQEDRNTEVF | 0.63241214 |
| GTLIALIL | ASSQEDRNTEVF | 0.7383722 |
| IAATMGFVVG | ASSQEDRNTEVF | 0.4509216 |
| IFAQALL | ASSQEDRNTEVF | 0.76273966 |
| IGAAVALGF | ASSQEDRNTEVF | 0.81014854 |
| IGLTPLAL | ASSQEDRNTEVF | 0.78720814 |
| ILGSNTSTL | ASSQEDRNTEVF | 0.24882104 |
| IVFSLTVL | ASSQEDRNTEVF | 0.79061115 |
| LASKATGF | ASSQEDRNTEVF | 0.8014515 |
| LASVALIL | ASSQEDRNTEVF | 0.8722656 |
| LFAVGLTL | ASSQEDRNTEVF | 0.74183446 |
| LFNGGIGTYI | ASSQEDRNTEVF | 0.828205 |
| LTLGAGII | ASSQEDRNTEVF | 0.6316798 |
| LVAAALGDI | ASSQEDRNTEVF | 0.735159 |
| NTAFVQLGI | ASSQEDRNTEVF | 0.5000668 |
| QLFSGAEL | ASSQEDRNTEVF | 0.6912328 |
| SAATFITL | ASSQEDRNTEVF | 0.6592574 |
| SNYGLANLGV | ASSQEDRNTEVF | 0.5582444 |
| SYGGAYAVM | ASSQEDRNTEVF | 0.8107405 |
| TTGVGGLTLG | ASSQEDRNTEVF | 0.8298379 |
| TTQTIYI | ASSQEDRNTEVF | 0.8257544 |
| VAAGAGLGAVI | ASSQEDRNTEVF | 0.8303774 |
| VAAVVTLL | ASSQEDRNTEVF | 0.8476058 |
| VGFLGGLI | ASSQEDRNTEVF | 0.8351761 |
| VGVIPILL | ASSQEDRNTEVF | 0.64118534 |
| VIFASSLI | ASSQEDRNTEVF | 0.71042156 |
| VKGGDGGAAAG | ASSQEDRNTEVF | 0.47589836 |
| VTKTFTNI | ASSQEDRNTEVF | 0.6793373 |
| VVAAAILL | ASSQEDRNTEVF | 0.65159684 |
| YAALDVEL | ASSQEDRNTEVF | 0.76564693 |
| LASVALLL | ASSQEDRNTEVF | 0.12658547 |

| | | |
|---|---|---|
| LDAVHRV | ASSQEDRNTEVF | 0.75774205 |
| QGLSSSIF | ASSQEDRNTEVF | 0.8462868 |
| RGTFANIRL | ASSQEDRNTEVF | 0.59253085 |
| RQLWWGHRI | ASSQEDRNTEVF | 0.80420923 |
| YGPPGTGKTLL | ASSQEDRNTEVF | 0.7755729 |
| AALVGAPANVA | ASSQEDRNTEVF | 0.6302674 |
| AATFRRTE | ASSQEDRNTEVF | 0.80026925 |
| AGALGGGI | ASSQEDRNTEVF | 0.59904265 |
| AGGAGGAGGIGAG | ASSQEDRNTEVF | 0.8242444 |
| AGSGALGL | ASSQEDRNTEVF | 0.8585892 |
| AQIADQI | ASSQEDRNTEVF | 0.7666689 |
| ASAVAVTYL | ASSQEDRNTEVF | 0.7414466 |
| ASTVLGGTGG | ASSQEDRNTEVF | 0.78141606 |
| AVGGEPAK | ASSQEDRNTEVF | 0.37654358 |
| CEQGTFGGTC | ASSQEDRNTEVF | 0.75199753 |
| DEGKAPAG | ASSQEDRNTEVF | 0.112922505 |
| DTVVFPAL | ASSQEDRNTEVF | 0.7131463 |
| FEGITMAG | ASSQEDRNTEVF | 0.52540624 |
| FPLVVQD | ASSQEDRNTEVF | 0.7656168 |
| GAFTTPQ | ASSQEDRNTEVF | 0.73785853 |
| GFGGFGTAGGVGGA | ASSQEDRNTEVF | 0.79287165 |
| GFLAQFL | ASSQEDRNTEVF | 0.54519415 |
| GGAGGAGGNAGVA | ASSQEDRNTEVF | 0.70612615 |
| GGVGTAGAGGAGGAGG | ASSQEDRNTEVF | 0.66645724 |
| GNGGGGDAGNAGSGGN | ASSQEDRNTEVF | 0.45410237 |
| GTGGAGGAGNDAGS | ASSQEDRNTEVF | 0.8266927 |
| GTTNTGWFNS | ASSQEDRNTEVF | 0.72701776 |
| IFMGPLFTL | ASSQEDRNTEVF | 0.69115835 |
| ILAHAIEAI | ASSQEDRNTEVF | 0.82776177 |
| IRSAIREAL | ASSQEDRNTEVF | 0.7589508 |
| ITFAGGG | ASSQEDRNTEVF | 0.39798203 |
| ITGMSGHQEV | ASSQEDRNTEVF | 0.69309515 |
| IVAGLACTL | ASSQEDRNTEVF | 0.7367039 |
| IVGGGLAGA | ASSQEDRNTEVF | 0.68704176 |
| IVTGVLNI | ASSQEDRNTEVF | 0.1198588 |
| LAQQGCI | ASSQEDRNTEVF | 0.7754793 |
| LCLHVEN | ASSQEDRNTEVF | 0.6346691 |
| LDAISDGS | ASSQEDRNTEVF | 0.55101883 |
| LGGFAAML | ASSQEDRNTEVF | 0.6755611 |
| LGGKNPMI | ASSQEDRNTEVF | 0.81874025 |
| LGPKGVAAIL | ASSQEDRNTEVF | 0.6030555 |
| LMVGVSI | ASSQEDRNTEVF | 0.725104 |
| LVGAAFVL | ASSQEDRNTEVF | 0.8097837 |
| LVLAVAPL | ASSQEDRNTEVF | 0.79223067 |
| MAAIAVIDAA | ASSQEDRNTEVF | 0.8438417 |
| NNGINAELQ | ASSQEDRNTEVF | 0.77846944 |
| RGEPLVLVHG | ASSQEDRNTEVF | 0.76086605 |
| RGGAGGAAGLI | ASSQEDRNTEVF | 0.7938272 |
| RSYRMHPAV | ASSQEDRNTEVF | 0.7565115 |
| SAAHLLAGQLA | ASSQEDRNTEVF | 0.38388568 |
| SAAVGSY | ASSQEDRNTEVF | 0.3421119 |

| | | |
|---|---|---|
| SPHAVLHRI | ASSQEDRNTEVF | 0.67358696 |
| SQIAKVL | ASSQEDRNTEVF | 0.33229053 |
| TALGHLVP | ASSQEDRNTEVF | 0.7728249 |
| TAPTTSGAA | ASSQEDRNTEVF | 0.8502811 |
| TDGDDGNGGSPGT | ASSQEDRNTEVF | 0.7472427 |
| TDGLGEN | ASSQEDRNTEVF | 0.4370102 |
| TGGPPLPGQA | ASSQEDRNTEVF | 0.697 |
| TIPAITI | ASSQEDRNTEVF | 0.6026215 |
| TIPGFTI | ASSQEDRNTEVF | 0.8022903 |
| TSIASGALA | ASSQEDRNTEVF | 0.82209814 |
| TVTLPGAAI | ASSQEDRNTEVF | 0.85548776 |
| VASGLSAALP | ASSQEDRNTEVF | 0.8159983 |
| VGAGVIGI | ASSQEDRNTEVF | 0.64511496 |
| VGPKRFGV | ASSQEDRNTEVF | 0.8127824 |
| VNVAFPD | ASSQEDRNTEVF | 0.6862631 |
| VSMTNSGVS | ASSQEDRNTEVF | 0.7647554 |
| VTDVLGDLSV | ASSQEDRNTEVF | 0.7296677 |
| VVVVGAGFAGL | ASSQEDRNTEVF | 0.7627096 |
| VYTVGTVV | ASSQEDRNTEVF | 0.6698755 |
| WTTFSGP | ASSQEDRNTEVF | 0.6729928 |
| YDELVEAF | ASSQEDRNTEVF | 0.3693633 |
| YGTGGAGGNGG | ASSQEDRNTEVF | 0.79980016 |
| YTNVMAV | ASSQEDRNTEVF | 0.67783606 |
| FLEQVHQL | ASSQEDRNTEVF | 0.40818986 |
| GLSLLAVAV | ASSQEDRNTEVF | 0.6604407 |
| LEQVHQL | ASSQEDRNTEVF | 0.8518645 |
| LNAAGAI | ASSQEDRNTEVF | 0.8483828 |
| PAVHASL | ASSQEDRNTEVF | 0.620775 |
| TELEDDA | ASSQEDRNTEVF | 0.5708874 |
| TPPGPPN | ASSQEDRNTEVF | 0.7653184 |
| TPPPPPPP | ASSQEDRNTEVF | 0.462513 |
| TQSPALI | ASSQSPGTGDTGQLY | 0.79340255 |
| EFAAAFPQ | ASSQSPGTGDTGQLY | 0.6971538 |
| LLSGNLDVL | ASSQSPGTGDTGQLY | 0.665925 |
| GPANLGNYNI | ASSQSPGTGDTGQLY | 0.67420256 |
| TLGPTLP | ASSQSPGTGDTGQLY | 0.7600261 |
| VYAYSSCCK | ASSQSPGTGDTGQLY | 0.64141333 |
| APAPAPAPNI | ASSQSPGTGDTGQLY | 0.8260041 |
| PGGPVPIP | ASSQSPGTGDTGQLY | 0.15350716 |
| FLTVALEP | ASSQSPGTGDTGQLY | 0.80202013 |
| SQYFGANL | ASSQSPGTGDTGQLY | 0.53572804 |
| GGVKPAA | ASSQSPGTGDTGQLY | 0.6026723 |
| GGTGGAGGAGGW | ASSQSPGTGDTGQLY | 0.29313973 |
| GVGGVAGLGGAGG | ASSQSPGTGDTGQLY | 0.7763555 |
| VLLVLATTI | ASSQSPGTGDTGQLY | 0.80725634 |
| IGIFAAML | ASSQSPGTGDTGQLY | 0.82380116 |
| AVGDLID | ASSQSPGTGDTGQLY | 0.5857444 |
| NISPLPL | ASSQSPGTGDTGQLY | 0.7986362 |
| VTAVFVAI | ASSQSPGTGDTGQLY | 0.84488803 |
| SGDDNDAAL | ASSQSPGTGDTGQLY | 0.65759176 |
| AVAALAHS | ASSQSPGTGDTGQLY | 0.30621088 |

| | | |
|---|---|---|
| GGVTVMATP | ASSQSPGTGDTGQLY | 0.7812741 |
| AGLAANVRD | ASSQSPGTGDTGQLY | 0.72497714 |
| TNRPDLIDPALLRPG | ASSQSPGTGDTGQLY | 0.8029522 |
| LANLGVTAL | ASSQSPGTGDTGQLY | 0.7990363 |
| TGERIEL | ASSQSPGTGDTGQLY | 0.4465382 |
| NGEELAA | ASSQSPGTGDTGQLY | 0.82912326 |
| VLSLVWG | ASSQSPGTGDTGQLY | 0.8061218 |
| IGYALWL | ASSQSPGTGDTGQLY | 0.5321486 |
| LSLLLAPL | ASSQSPGTGDTGQLY | 0.68778455 |
| TLDEALK | ASSQSPGTGDTGQLY | 0.646086 |
| TDGAGTAEGG | ASSQSPGTGDTGQLY | 0.35730863 |
| NAPGGAGGNGG | ASSQSPGTGDTGQLY | 0.8014795 |
| VSVFMGMGGNF | ASSQSPGTGDTGQLY | 0.6259431 |
| LASYLAAL | ASSQSPGTGDTGQLY | 0.65176845 |
| LVGGTPAL | ASSQSPGTGDTGQLY | 0.7290288 |
| VRNADVL | ASSQSPGTGDTGQLY | 0.63974285 |
| TCSSPFLVSAM | ASSQSPGTGDTGQLY | 0.6449077 |
| VHITDAE | ASSQSPGTGDTGQLY | 0.67447925 |
| EYAAPSF | ASSQSPGTGDTGQLY | 0.7385751 |
| VVDAFHGGL | ASSQSPGTGDTGQLY | 0.77531636 |
| ASVVDPQ | ASSQSPGTGDTGQLY | 0.7077343 |
| SYNEFGGLS | ASSQSPGTGDTGQLY | 0.7805391 |
| DPDGSVY | ASSQSPGTGDTGQLY | 0.83539397 |
| MVSTIHATE | ASSQSPGTGDTGQLY | 0.40958732 |
| LAAGVADPV | ASSQSPGTGDTGQLY | 0.57949823 |
| TKEQIAA | ASSQSPGTGDTGQLY | 0.79547644 |
| SLASPAAL | ASSQSPGTGDTGQLY | 0.51019907 |
| IGLVFYGV | ASSQSPGTGDTGQLY | 0.8092418 |
| FLLDASNL | ASSQSPGTGDTGQLY | 0.74162245 |
| YQVGNFP | ASSQSPGTGDTGQLY | 0.78254795 |
| ESLLSQST | ASSQSPGTGDTGQLY | 0.7364061 |
| LFTVGFAAVG | ASSQSPGTGDTGQLY | 0.8579022 |
| ATPPTPT | ASSQSPGTGDTGQLY | 0.8068335 |
| TDLATLG | ASSQSPGTGDTGQLY | 0.7577509 |
| SAGGTGR | ASSQSPGTGDTGQLY | 0.76684535 |
| GFSNSGS | ASSQSPGTGDTGQLY | 0.56327075 |
| GGNGGSGGTSV | ASSQSPGTGDTGQLY | 0.80014247 |
| GTSVATGGAGN | ASSQSPGTGDTGQLY | 0.81078565 |
| NGGNGGSGGTS | ASSQSPGTGDTGQLY | 0.5104555 |
| AVVKFPH | ASSQSPGTGDTGQLY | 0.6838417 |
| TPPPPPP | ASSQSPGTGDTGQLY | 0.8530484 |
| AAPAAAAAPA | ASSQSPGTGDTGQLY | 0.8287281 |
| IALNEDL | ASSQSPGTGDTGQLY | 0.38910347 |
| IVNVSSM | ASSQSPGTGDTGQLY | 0.6468147 |
| TGIPPAPR | ASSQSPGTGDTGQLY | 0.45878047 |
| FDLGGGTF | ASSQSPGTGDTGQLY | 0.71170324 |
| FDLGGGTFDVS | ASSQSPGTGDTGQLY | 0.4342267 |
| NEPTAAA | ASSQSPGTGDTGQLY | 0.7909647 |
| NEPTAAALA | ASSQSPGTGDTGQLY | 0.65352917 |
| GSIINISSI | ASSQSPGTGDTGQLY | 0.81776965 |
| VAPGNAGTA | ASSQSPGTGDTGQLY | 0.7669181 |

| | | |
|---|---|---|
| TGGGASLE | ASSQSPGTGDTGQLY | 0.84595656 |
| VAVYNHYKRI | ASSQSPGTGDTGQLY | 0.640177 |
| TDPAPATT | ASSQSPGTGDTGQLY | 0.6661275 |
| TNRPDILDPALLRPG | ASSQSPGTGDTGQLY | 0.3490085 |
| LTTTTTS | ASSQSPGTGDTGQLY | 0.40719962 |
| PTGSGKTL | ASSQSPGTGDTGQLY | 0.58964527 |
| TGSGKTL | ASSQSPGTGDTGQLY | 0.67313725 |
| TIGHVDHGK | ASSQSPGTGDTGQLY | 0.18845338 |
| TIGHVDHGKTT | ASSQSPGTGDTGQLY | 0.8399701 |
| GHVDHGKTT | ASSQSPGTGDTGQLY | 0.72247076 |
| AQFEHTLL | ASSQSPGTGDTGQLY | 0.8185134 |
| IVNQDEL | ASSQSPGTGDTGQLY | 0.636413 |
| TNGSQFF | ASSQSPGTGDTGQLY | 0.64763427 |
| TGKGIVE | ASSQSPGTGDTGQLY | 0.6368946 |
| LISNASDA | ASSQSPGTGDTGQLY | 0.8073985 |
| AELRTLGELTKIAKAHGAQ | ASSQSPGTGDTGQLY | 0.5262562 |
| APNAVAAAEPSW | ASSQSPGTGDTGQLY | 0.29997498 |
| APQVAGAATAVL | ASSQSPGTGDTGQLY | 0.6758683 |
| ARRDPAAAAAILPTDA | ASSQSPGTGDTGQLY | 0.52479386 |
| DAAARDVLAEAGLAEYF | ASSQSPGTGDTGQLY | 0.8610908 |
| DDQIVLSAKANLAKVEQ | ASSQSPGTGDTGQLY | 0.8212307 |
| DELLPDTGEDSDEDGH | ASSQSPGTGDTGQLY | 0.74522626 |
| ELPEAPANRATVPQD | ASSQSPGTGDTGQLY | 0.8499788 |
| FGQSKIERVTGIG | ASSQSPGTGDTGQLY | 0.18739088 |
| GAGGTGSTAGGAGGAGGAGGLY | ASSQSPGTGDTGQLY | 0.66685545 |
| GHGGTGAAGGNGGTGGAGGSGI | ASSQSPGTGDTGQLY | 0.8583407 |
| GNGGNGGAGGEGGPGGAGGAGA | ASSQSPGTGDTGQLY | 0.7708236 |
| GSTAGNGGNGGNATNGTIAGQP | ASSQSPGTGDTGQLY | 0.7029438 |
| GTGGDGGNGGNGGTGG | ASSQSPGTGDTGQLY | 0.8276341 |
| GVRRPAGIPADAPV | ASSQSPGTGDTGQLY | 0.5913174 |
| GYRQVTDAHLLTLARR | ASSQSPGTGDTGQLY | 0.7024767 |
| IAKMGQDGHDRG | ASSQSPGTGDTGQLY | 0.7246724 |
| KAEPSIEIQYT | ASSQSPGTGDTGQLY | 0.8007715 |
| KPVRIRMAADEDTVAVT | ASSQSPGTGDTGQLY | 0.7829757 |
| MPPRIAGMRLLVIKPEP | ASSQSPGTGDTGQLY | 0.47480083 |
| MKFYAVALFRSRGVPPTR | ASSQSPGTGDTGQLY | 0.8120338 |
| NAGGFGILGAVAHSP | ASSQSPGTGDTGQLY | 0.86416954 |
| NEGREITLAMAEM | ASSQSPGTGDTGQLY | 0.6586678 |
| NELVEWSHPLGHRWRWE | ASSQSPGTGDTGQLY | 0.79581964 |
| NLGAGGTGGNGGNPG | ASSQSPGTGDTGQLY | 0.7753465 |
| NTEWPKWETVMAADPVD | ASSQSPGTGDTGQLY | 0.7459407 |
| PASAAPAETQP | ASSQSPGTGDTGQLY | 0.58114386 |
| QQSSDGKNGVRAAEITGE | ASSQSPGTGDTGQLY | 0.5719334 |
| RAPDAVTTSWWKE | ASSQSPGTGDTGQLY | 0.7291615 |
| SGFGNVGAHTSGWFNQ | ASSQSPGTGDTGQLY | 0.50367415 |
| SGVAGAVGQAASVGGLKV | ASSQSPGTGDTGQLY | 0.74738634 |
| SNGGAGTGSAGGNGGT | ASSQSPGTGDTGQLY | 0.71731496 |
| TGGAGGEGGAAG | ASSQSPGTGDTGQLY | 0.87856275 |
| TGGDGGAAGAGGAPGHG | ASSQSPGTGDTGQLY | 0.7990734 |
| TMPPLLAAIANAGRHGV | ASSQSPGTGDTGQLY | 0.7734816 |
| TSSTAGNSGGA | ASSQSPGTGDTGQLY | 0.54698145 |

| | | |
|---|---|---|
| VPATVPVGILC | ASSQSPGTGDTGQLY | 0.76757395 |
| VQRPDSGGVVLTGRIS | ASSQSPGTGDTGQLY | 0.7661655 |
| WTPNPVPATKGVPAGLVQAT | ASSQSPGTGDTGQLY | 0.3888185 |
| YALKRALAKFSDRFDVVII | ASSQSPGTGDTGQLY | 0.84557164 |
| YLIGSVIV | ASSQSPGTGDTGQLY | 0.55325365 |
| YGQLVAVID | ASSQSPGTGDTGQLY | 0.6961937 |
| VTVGGMTLPA | ASSQSPGTGDTGQLY | 0.80463314 |
| VLALPAV | ASSQSPGTGDTGQLY | 0.83293784 |
| VIPPTLNYET | ASSQSPGTGDTGQLY | 0.709951 |
| VGMGGMHPSSG | ASSQSPGTGDTGQLY | 0.7435788 |
| VGLAGGAATA | ASSQSPGTGDTGQLY | 0.52891505 |
| VGGAGGAGGSGP | ASSQSPGTGDTGQLY | 0.81770706 |
| VFAAFMV | ASSQSPGTGDTGQLY | 0.8221921 |
| TLYDKLR | ASSQSPGTGDTGQLY | 0.7305906 |
| TLAVFTVL | ASSQSPGTGDTGQLY | 0.8551953 |
| TGSVQPARV | ASSQSPGTGDTGQLY | 0.77483845 |
| TGGGAGSGGT | ASSQSPGTGDTGQLY | 0.3673851 |
| SHLETSL | ASSQSPGTGDTGQLY | 0.7316706 |
| SFTTGMT | ASSQSPGTGDTGQLY | 0.8338524 |
| QVEVAKRMGVRQPTVS | ASSQSPGTGDTGQLY | 0.82564044 |
| QGSGGPALAM | ASSQSPGTGDTGQLY | 0.83777416 |
| MLAPSNP | ASSQSPGTGDTGQLY | 0.7958629 |
| LVNNAAI | ASSQSPGTGDTGQLY | 0.6406673 |
| LTGEINTLA | ASSQSPGTGDTGQLY | 0.83941364 |
| LSIFGPV | ASSQSPGTGDTGQLY | 0.8794354 |
| LLALLANL | ASSQSPGTGDTGQLY | 0.82313365 |
| LISNASDALDKLR | ASSQSPGTGDTGQLY | 0.6421212 |
| LINEAVP | ASSQSPGTGDTGQLY | 0.6143714 |
| LGYAFSGL | ASSQSPGTGDTGQLY | 0.79527545 |
| LDFVAVAL | ASSQSPGTGDTGQLY | 0.8102545 |
| LAQPGAE | ASSQSPGTGDTGQLY | 0.54675394 |
| ISNLGAVTGF | ASSQSPGTGDTGQLY | 0.74606025 |
| HQEDPGRI | ASSQSPGTGDTGQLY | 0.7672955 |
| GTPGEPTP | ASSQSPGTGDTGQLY | 0.33244225 |
| GFFNSGS | ASSQSPGTGDTGQLY | 0.79078114 |
| GAGGGGLTAGAN | ASSQSPGTGDTGQLY | 0.8387724 |
| FVDALVNM | ASSQSPGTGDTGQLY | 0.7775248 |
| FLTVAPEL | ASSQSPGTGDTGQLY | 0.5683222 |
| FETLTAL | ASSQSPGTGDTGQLY | 0.83249265 |
| DVINPIEL | ASSQSPGTGDTGQLY | 0.8627305 |
| APVSGAPV | ASSQSPGTGDTGQLY | 0.84842575 |
| ALVGGLAW | ASSQSPGTGDTGQLY | 0.32300457 |
| ALAAAAATAI | ASSQSPGTGDTGQLY | 0.22544748 |
| AFGLGLLL | ASSQSPGTGDTGQLY | 0.6941983 |
| AFFAAASL | ASSQSPGTGDTGQLY | 0.8229612 |
| AAAALPAFSPP | ASSQSPGTGDTGQLY | 0.66828126 |
| VVLAPAL | ASSQSPGTGDTGQLY | 0.5811626 |
| TIGHVDHGKTTL | ASSQSPGTGDTGQLY | 0.76221335 |
| LISNASDALDK | ASSQSPGTGDTGQLY | 0.8287296 |
| LISNASDAL | ASSQSPGTGDTGQLY | 0.57095045 |
| ISNASDALDK | ASSQSPGTGDTGQLY | 0.42213 |

| | | |
|---|---|---|
| NQAETLVYQTE | ASSQSPGTGDTGQLY | 0.81662935 |
| TGGTGGTAGAG | ASSQSPGTGDTGQLY | 0.8199549 |
| TGSTAGGAGGA | ASSQSPGTGDTGQLY | 0.52079177 |
| DPLAVPVAAGPV | ASSQSPGTGDTGQLY | 0.6765946 |
| GGTGGTPGNGGN | ASSQSPGTGDTGQLY | 0.72283816 |
| PAGLAGVDPATL | ASSQSPGTGDTGQLY | 0.65368444 |
| GNNLVVEAPHAGASV | ASSQSPGTGDTGQLY | 0.7482952 |
| KVTRIMSASTLEQPA | ASSQSPGTGDTGQLY | 0.48881844 |
| LGFALPIALPRRTVV | ASSQSPGTGDTGQLY | 0.79817 |
| TRLGNLTARKLVVLV | ASSQSPGTGDTGQLY | 0.48476005 |
| WSQQRRLIYFPSAGP | ASSQSPGTGDTGQLY | 0.83059514 |
| GPNGNPDPMAAAVDIR | ASSQSPGTGDTGQLY | 0.75273263 |
| TQRCVSVADLRAAASR | ASSQSPGTGDTGQLY | 0.77958095 |
| QFVLDVAEHPVEVLTETV | ASSQSPGTGDTGQLY | 0.6744679 |
| GGAGGTGGAGGNGGTQAGDG | ASSQSPGTGDTGQLY | 0.56308955 |
| PLLATLRAIAVDVDAKPAQV | ASSQSPGTGDTGQLY | 0.5317108 |
| APKFRTGGYMIDFWGVGYQVAK | ASSQSPGTGDTGQLY | 0.35800838 |
| GAGGAGGAGGAAGGPGGTGGTG | ASSQSPGTGDTGQLY | 0.828475 |
| IGALGLLATDVAPAGSVAFGVA | ASSQSPGTGDTGQLY | 0.1808248 |
| AGGAGGIGGNGGAGGAGGVLMG | ASSQSPGTGDTGQLY | 0.8217542 |
| ADELIGLL | ASSQSPGTGDTGQLY | 0.7890186 |
| ADRALKA | ASSQSPGTGDTGQLY | 0.5136787 |
| AGAVGQVGGAGGA | ASSQSPGTGDTGQLY | 0.7900604 |
| AGGGGGLGGAP | ASSQSPGTGDTGQLY | 0.41732287 |
| AGGLLYGNGGAG | ASSQSPGTGDTGQLY | 0.6682996 |
| AGLESIGL | ASSQSPGTGDTGQLY | 0.7976258 |
| AGLKTGII | ASSQSPGTGDTGQLY | 0.8191344 |
| ASLAALLL | ASSQSPGTGDTGQLY | 0.5990345 |
| ASVINISSI | ASSQSPGTGDTGQLY | 0.7952807 |
| ATPPGPA | ASSQSPGTGDTGQLY | 0.77416974 |
| ATVASALV | ASSQSPGTGDTGQLY | 0.62885433 |
| ATVASLIL | ASSQSPGTGDTGQLY | 0.77256054 |
| AVNGPVLIRH | ASSQSPGTGDTGQLY | 0.8123957 |
| DGAAGAVGI | ASSQSPGTGDTGQLY | 0.79661644 |
| DPLLHGI | ASSQSPGTGDTGQLY | 0.50474215 |
| ELGAALGAL | ASSQSPGTGDTGQLY | 0.78707314 |
| ELGRTQAV | ASSQSPGTGDTGQLY | 0.5855752 |
| FVAGGYLS | ASSQSPGTGDTGQLY | 0.7631503 |
| GASGGAGGAGGW | ASSQSPGTGDTGQLY | 0.7180414 |
| GGAGAGGDGGAGGI | ASSQSPGTGDTGQLY | 0.7166163 |
| GTGHNGSQPGMGGNG | ASSQSPGTGDTGQLY | 0.8071066 |
| GYAAASASAS | ASSQSPGTGDTGQLY | 0.5686587 |
| HQVDVAAVQQG | ASSQSPGTGDTGQLY | 0.6100707 |
| IGAAIAHT | ASSQSPGTGDTGQLY | 0.67519313 |
| IGMIEVAQL | ASSQSPGTGDTGQLY | 0.7466268 |
| IRLAPEHTSAMD | ASSQSPGTGDTGQLY | 0.8069443 |
| IYNVSVL | ASSQSPGTGDTGQLY | 0.5299848 |
| KQGFLGM | ASSQSPGTGDTGQLY | 0.74068105 |
| LESVITQL | ASSQSPGTGDTGQLY | 0.68679535 |
| LGLGALVVV | ASSQSPGTGDTGQLY | 0.56161445 |
| LIGNGGNGGSGGI | ASSQSPGTGDTGQLY | 0.7703957 |

| | | |
|---|---|---|
| LNAINAPTL | ASSQSPGTGDTGQLY | 0.70997727 |
| LQLLNTGV | ASSQSPGTGDTGQLY | 0.48451236 |
| LQTSIEF | ASSQSPGTGDTGQLY | 0.7904736 |
| MSFAVSEL | ASSQSPGTGDTGQLY | 0.5853931 |
| NTGALGL | ASSQSPGTGDTGQLY | 0.7958969 |
| PPGAGGAGGAGTT | ASSQSPGTGDTGQLY | 0.8441209 |
| QLVGLGI | ASSQSPGTGDTGQLY | 0.6282445 |
| RFLEQVHQL | ASSQSPGTGDTGQLY | 0.8200111 |
| RGVGYVLR | ASSQSPGTGDTGQLY | 0.75423396 |
| SATGVFAGVI | ASSQSPGTGDTGQLY | 0.7610857 |
| SLPAITLP | ASSQSPGTGDTGQLY | 0.6970926 |
| STALAFTV | ASSQSPGTGDTGQLY | 0.6001904 |
| SVLTVGLGK | ASSQSPGTGDTGQLY | 0.8338834 |
| TALAGAGSGI | ASSQSPGTGDTGQLY | 0.8114664 |
| TGLPFTGI | ASSQSPGTGDTGQLY | 0.817798 |
| VAEHVTV | ASSQSPGTGDTGQLY | 0.7693435 |
| VDIVLNSL | ASSQSPGTGDTGQLY | 0.8203618 |
| VLTGGGAVL | ASSQSPGTGDTGQLY | 0.8185197 |
| VNVGLAYGI | ASSQSPGTGDTGQLY | 0.58722186 |
| VSSAAVGAV | ASSQSPGTGDTGQLY | 0.36977398 |
| VTSVYEAGV | ASSQSPGTGDTGQLY | 0.6668819 |
| WVALDRGA | ASSQSPGTGDTGQLY | 0.6516662 |
| YLVGSVVI | ASSQSPGTGDTGQLY | 0.6817398 |
| GPPGTGKTLL | ASSQSPGTGDTGQLY | 0.3548812 |
| GVGMTKF | ASSQSPGTGDTGQLY | 0.8131499 |
| IGIGGSDL | ASSQSPGTGDTGQLY | 0.86622137 |
| IGTLLAEL | ASSQSPGTGDTGQLY | 0.58324754 |
| IYTSGTT | ASSQSPGTGDTGQLY | 0.8593519 |
| LSFPAGV | ASSQSPGTGDTGQLY | 0.49393553 |
| LYTSGTT | ASSQSPGTGDTGQLY | 0.8158592 |
| VGAFPFGF | ASSQSPGTGDTGQLY | 0.83160293 |
| VGAGTVEF | ASSQSPGTGDTGQLY | 0.51729876 |
| VGKSTLLL | ASSQSPGTGDTGQLY | 0.7853198 |
| AALAHAGGGIV | ASSQSPGTGDTGQLY | 0.64646584 |
| AALVAVSLL | ASSQSPGTGDTGQLY | 0.6809764 |
| AAVSRAVAAL | ASSQSPGTGDTGQLY | 0.8307072 |
| AGGSSGIGGF | ASSQSPGTGDTGQLY | 0.66723514 |
| AGLTVANANG | ASSQSPGTGDTGQLY | 0.77758616 |
| AGLVGAVL | ASSQSPGTGDTGQLY | 0.68746215 |
| ALSQAFTV | ASSQSPGTGDTGQLY | 0.5574615 |
| ALTFSVSL | ASSQSPGTGDTGQLY | 0.758039 |
| ALTGAALVAL | ASSQSPGTGDTGQLY | 0.8240031 |
| ASFSIGAGVL | ASSQSPGTGDTGQLY | 0.66156983 |
| DGSVYKI | ASSQSPGTGDTGQLY | 0.8772346 |
| EQLAEIV | ASSQSPGTGDTGQLY | 0.568747 |
| FAVSELTVM | ASSQSPGTGDTGQLY | 0.84537363 |
| GAAPLAAGL | ASSQSPGTGDTGQLY | 0.82813585 |
| GADGGDGGSGGAGG | ASSQSPGTGDTGQLY | 0.7232977 |
| GFGLDTGGAGG | ASSQSPGTGDTGQLY | 0.68225664 |
| GLASGGLL | ASSQSPGTGDTGQLY | 0.81385237 |
| GRIAGAAVLL | ASSQSPGTGDTGQLY | 0.7007178 |

| | | |
|---|---|---|
| GSAGSLNTSF | ASSQSPGTGDTGQLY | 0.81297946 |
| GTGQIGI | ASSQSPGTGDTGQLY | 0.8409663 |
| GTLIALIL | ASSQSPGTGDTGQLY | 0.5203572 |
| IAATMGFVVG | ASSQSPGTGDTGQLY | 0.6444331 |
| IFAQALL | ASSQSPGTGDTGQLY | 0.6120718 |
| IGAAVALGF | ASSQSPGTGDTGQLY | 0.6294495 |
| IGLTPLAL | ASSQSPGTGDTGQLY | 0.5585152 |
| ILGSNTSTL | ASSQSPGTGDTGQLY | 0.501569 |
| IVFSLTVL | ASSQSPGTGDTGQLY | 0.6150258 |
| LASKATGF | ASSQSPGTGDTGQLY | 0.51652795 |
| LASVALIL | ASSQSPGTGDTGQLY | 0.6460513 |
| LFAVGLTL | ASSQSPGTGDTGQLY | 0.65038264 |
| LFNGGIGTYI | ASSQSPGTGDTGQLY | 0.54094106 |
| LTLGAGII | ASSQSPGTGDTGQLY | 0.6672325 |
| LVAAALGDI | ASSQSPGTGDTGQLY | 0.65583163 |
| NTAFVQLGI | ASSQSPGTGDTGQLY | 0.4621492 |
| QLFSGAEL | ASSQSPGTGDTGQLY | 0.56593996 |
| SAATFITL | ASSQSPGTGDTGQLY | 0.5996395 |
| SNYGLANLGV | ASSQSPGTGDTGQLY | 0.59397715 |
| SYGGAYAVM | ASSQSPGTGDTGQLY | 0.4015508 |
| TTGVGGLTLG | ASSQSPGTGDTGQLY | 0.6456419 |
| TTQTIYI | ASSQSPGTGDTGQLY | 0.5827441 |
| VAAGAGLGAVI | ASSQSPGTGDTGQLY | 0.3598983 |
| VAAVVTLL | ASSQSPGTGDTGQLY | 0.6365589 |
| VGFLGGLI | ASSQSPGTGDTGQLY | 0.60161793 |
| VGVIPILL | ASSQSPGTGDTGQLY | 0.575638 |
| VIFASSLI | ASSQSPGTGDTGQLY | 0.65899545 |
| VKGGDGGAAAG | ASSQSPGTGDTGQLY | 0.448677 |
| VTKTFTNI | ASSQSPGTGDTGQLY | 0.53551495 |
| VVAAAILL | ASSQSPGTGDTGQLY | 0.6474987 |
| YAALDVEL | ASSQSPGTGDTGQLY | 0.66796637 |
| LASVALLL | ASSQSPGTGDTGQLY | 0.5098696 |
| LDAVHRV | ASSQSPGTGDTGQLY | 0.628559 |
| QGLSSSIF | ASSQSPGTGDTGQLY | 0.6612939 |
| RGTFANIRL | ASSQSPGTGDTGQLY | 0.64438397 |
| RQLWWGHRI | ASSQSPGTGDTGQLY | 0.665067 |
| YGPPGTGKTLL | ASSQSPGTGDTGQLY | 0.6694434 |
| AALVGAPANVA | ASSQSPGTGDTGQLY | 0.58264 |
| AATFRRTE | ASSQSPGTGDTGQLY | 0.6597655 |
| AGALGGGI | ASSQSPGTGDTGQLY | 0.67060876 |
| AGGAGGAGGIGAG | ASSQSPGTGDTGQLY | 0.55315185 |
| AGSGALGL | ASSQSPGTGDTGQLY | 0.63256466 |
| AQIADQI | ASSQSPGTGDTGQLY | 0.5479438 |
| ASAVAVTYL | ASSQSPGTGDTGQLY | 0.61395746 |
| ASTVLGGTGG | ASSQSPGTGDTGQLY | 0.39955145 |
| AVGGEPAK | ASSQSPGTGDTGQLY | 0.63224256 |
| CEQGTFGGTC | ASSQSPGTGDTGQLY | 0.6593724 |
| DEGKAPAG | ASSQSPGTGDTGQLY | 0.6046237 |
| DTVVFPAL | ASSQSPGTGDTGQLY | 0.51363754 |
| FEGITMAG | ASSQSPGTGDTGQLY | 0.5099652 |
| FPLVVQD | ASSQSPGTGDTGQLY | 0.57272756 |

| | | |
|---|---|---|
| GAFTTPQ | ASSQSPGTGDTGQLY | 0.57604146 |
| GFGGFGTAGGVGGA | ASSQSPGTGDTGQLY | 0.45679584 |
| GFLAQFL | ASSQSPGTGDTGQLY | 0.57239366 |
| GGAGGAGGNAGVA | ASSQSPGTGDTGQLY | 0.65981185 |
| GGVGTAGAGGAGGAGG | ASSQSPGTGDTGQLY | 0.6572541 |
| GNGGGGDAGNAGSGGN | ASSQSPGTGDTGQLY | 0.46967173 |
| GTGGAGGAGNDAGS | ASSQSPGTGDTGQLY | 0.66214454 |
| GTTNTGWFNS | ASSQSPGTGDTGQLY | 0.632563 |
| IFMGPLFTL | ASSQSPGTGDTGQLY | 0.66379696 |
| ILAHAIEAI | ASSQSPGTGDTGQLY | 0.6160987 |
| IRSAIREAL | ASSQSPGTGDTGQLY | 0.4915806 |
| ITFAGGG | ASSQSPGTGDTGQLY | 0.654117 |
| ITGMSGHQEV | ASSQSPGTGDTGQLY | 0.5981398 |
| IVAGLACTL | ASSQSPGTGDTGQLY | 0.6057408 |
| IVGGGLAGA | ASSQSPGTGDTGQLY | 0.65982616 |
| IVTGVLNI | ASSQSPGTGDTGQLY | 0.60120624 |
| LAQQGCI | ASSQSPGTGDTGQLY | 0.5455966 |
| LCLHVEN | ASSQSPGTGDTGQLY | 0.56462514 |
| LDAISDGS | ASSQSPGTGDTGQLY | 0.6449752 |
| LGGFAAML | ASSQSPGTGDTGQLY | 0.5076232 |
| LGGKNPMI | ASSQSPGTGDTGQLY | 0.4966477 |
| LGPKGVAAIL | ASSQSPGTGDTGQLY | 0.559969 |
| LMVGVSI | ASSQSPGTGDTGQLY | 0.6187013 |
| LVGAAFVL | ASSQSPGTGDTGQLY | 0.6618421 |
| LVLAVAPL | ASSQSPGTGDTGQLY | 0.62235457 |
| MAAIAVIDAA | ASSQSPGTGDTGQLY | 0.56635284 |
| NNGINAELQ | ASSQSPGTGDTGQLY | 0.64395237 |
| RGEPLVLVHG | ASSQSPGTGDTGQLY | 0.62454325 |
| RGGAGGAAGLI | ASSQSPGTGDTGQLY | 0.6101401 |
| RSYRMHPAV | ASSQSPGTGDTGQLY | 0.6453241 |
| SAAHLLAGQLA | ASSQSPGTGDTGQLY | 0.5458522 |
| SAAVGSY | ASSQSPGTGDTGQLY | 0.64032286 |
| SPHAVLHRI | ASSQSPGTGDTGQLY | 0.6449177 |
| SQIAKVL | ASSQSPGTGDTGQLY | 0.6445931 |
| TALGHLVP | ASSQSPGTGDTGQLY | 0.6562894 |
| TAPTTSGAA | ASSQSPGTGDTGQLY | 0.59945774 |
| TDGDDGNGGSPGT | ASSQSPGTGDTGQLY | 0.6076331 |
| TDGLGEN | ASSQSPGTGDTGQLY | 0.41946274 |
| TGGPPLPGQA | ASSQSPGTGDTGQLY | 0.63391465 |
| TIPAITI | ASSQSPGTGDTGQLY | 0.64232785 |
| TIPGFTI | ASSQSPGTGDTGQLY | 0.6188494 |
| TSIASGALA | ASSQSPGTGDTGQLY | 0.6545589 |
| TVTLPGAAI | ASSQSPGTGDTGQLY | 0.63499355 |
| VASGLSAALP | ASSQSPGTGDTGQLY | 0.5589402 |
| VGAGVIGI | ASSQSPGTGDTGQLY | 0.66177356 |
| VGPKRFGV | ASSQSPGTGDTGQLY | 0.6586213 |
| VNVAFPD | ASSQSPGTGDTGQLY | 0.4647002 |
| VSMTNSGVS | ASSQSPGTGDTGQLY | 0.59037286 |
| VTDVLGDLSV | ASSQSPGTGDTGQLY | 0.6597398 |
| VVVVGAGFAGL | ASSQSPGTGDTGQLY | 0.623997 |
| VYTVGTVV | ASSQSPGTGDTGQLY | 0.6059099 |

| | | |
|---|---|---|
| WTTFSGP | ASSQSPGTGDTGQLY | 0.60750353 |
| YDELVEAF | ASSQSPGTGDTGQLY | 0.6650057 |
| YGTGGAGGNGG | ASSQSPGTGDTGQLY | 0.60605365 |
| YTNVMAV | ASSQSPGTGDTGQLY | 0.6681767 |
| FLEQVHQL | ASSQSPGTGDTGQLY | 0.36829096 |
| GLSLLAVAV | ASSQSPGTGDTGQLY | 0.54319876 |
| LEQVHQL | ASSQSPGTGDTGQLY | 0.4799778 |
| LNAAGAI | ASSQSPGTGDTGQLY | 0.3803273 |
| PAVHASL | ASSQSPGTGDTGQLY | 0.6557946 |
| TELEDDA | ASSQSPGTGDTGQLY | 0.5979893 |
| TPPGPPN | ASSQSPGTGDTGQLY | 0.64880365 |
| TPPPPPPP | ASSQSPGTGDTGQLY | 0.6498469 |
| TQSPALI | ASSLDQGNYAEQF | 0.65171593 |
| EFAAAFPQ | ASSLDQGNYAEQF | 0.6144348 |
| LLSGNLDVL | ASSLDQGNYAEQF | 0.6238143 |
| GPANLGNYNI | ASSLDQGNYAEQF | 0.52727467 |
| TLGPTLP | ASSLDQGNYAEQF | 0.64914215 |
| VYAYSSCCK | ASSLDQGNYAEQF | 0.667891 |
| APAPAPAPNI | ASSLDQGNYAEQF | 0.6278973 |
| PGGPVPIP | ASSLDQGNYAEQF | 0.45632684 |
| FLTVALEP | ASSLDQGNYAEQF | 0.5972614 |
| SQYFGANL | ASSLDQGNYAEQF | 0.6107156 |
| GGVKPAA | ASSLDQGNYAEQF | 0.3977871 |
| GGTGGAGGAGGW | ASSLDQGNYAEQF | 0.41503155 |
| GVGGVAGLGGAGG | ASSLDQGNYAEQF | 0.54353195 |
| VLLVLATTI | ASSLDQGNYAEQF | 0.5272825 |
| IGIFAAML | ASSLDQGNYAEQF | 0.6556325 |
| AVGDLID | ASSLDQGNYAEQF | 0.6481543 |
| NISPLPL | ASSLDQGNYAEQF | 0.6003659 |
| VTAVFVAI | ASSLDQGNYAEQF | 0.65245986 |
| SGDDNDAAL | ASSLDQGNYAEQF | 0.61656857 |
| AVAALAHS | ASSLDQGNYAEQF | 0.30435553 |
| GGVTVMATP | ASSLDQGNYAEQF | 0.57520914 |
| AGLAANVRD | ASSLDQGNYAEQF | 0.4971229 |
| TNRPDLIDPALLRPG | ASSLDQGNYAEQF | 0.6603186 |
| LANLGVTAL | ASSLDQGNYAEQF | 0.55127203 |
| TGERIEL | ASSLDQGNYAEQF | 0.5645563 |
| NGEELAA | ASSLDQGNYAEQF | 0.61337435 |
| VLSLVWG | ASSLDQGNYAEQF | 0.61294746 |
| IGYALWL | ASSLDQGNYAEQF | 0.612162 |
| LSLLLAPL | ASSLDQGNYAEQF | 0.47376764 |
| TLDEALK | ASSLDQGNYAEQF | 0.3483921 |
| TDGAGTAEGG | ASSLDQGNYAEQF | 0.65207803 |
| NAPGGAGGNGG | ASSLDQGNYAEQF | 0.46874422 |
| VSVFMGMGGNF | ASSLDQGNYAEQF | 0.5116643 |
| LASYLAAL | ASSLDQGNYAEQF | 0.4316298 |
| LVGGTPAL | ASSLDQGNYAEQF | 0.56193835 |
| VRNADVL | ASSLDQGNYAEQF | 0.4238401 |
| TCSSPFLVSAM | ASSLDQGNYAEQF | 0.66007686 |
| VHITDAE | ASSLDQGNYAEQF | 0.57662845 |
| EYAAPSF | ASSLDQGNYAEQF | 0.5908616 |

| | | |
|---|---|---|
| VVDAFHGGL | ASSLDQGNYAEQF | 0.66649294 |
| ASVVDPQ | ASSLDQGNYAEQF | 0.6111496 |
| SYNEFGGLS | ASSLDQGNYAEQF | 0.59225535 |
| DPDGSVY | ASSLDQGNYAEQF | 0.53877234 |
| MVSTIHATE | ASSLDQGNYAEQF | 0.5169816 |
| LAAGVADPV | ASSLDQGNYAEQF | 0.54346305 |
| TKEQIAA | ASSLDQGNYAEQF | 0.54870385 |
| SLASPAAL | ASSLDQGNYAEQF | 0.6697359 |
| IGLVFYGV | ASSLDQGNYAEQF | 0.5590757 |
| FLLDASNL | ASSLDQGNYAEQF | 0.47665173 |
| YQVGNFP | ASSLDQGNYAEQF | 0.5194887 |
| ESLLSQST | ASSLDQGNYAEQF | 0.66797185 |
| LFTVGFAAVG | ASSLDQGNYAEQF | 0.6742073 |
| ATPPTPT | ASSLDQGNYAEQF | 0.5534551 |
| TDLATLG | ASSLDQGNYAEQF | 0.55649287 |
| SAGGTGR | ASSLDQGNYAEQF | 0.5847426 |
| GFSNSGS | ASSLDQGNYAEQF | 0.6055135 |
| GGNGGSGGTSV | ASSLDQGNYAEQF | 0.55501175 |
| GTSVATGGAGN | ASSLDQGNYAEQF | 0.48737508 |
| NGGNGGSGGTS | ASSLDQGNYAEQF | 0.6320335 |
| AVVKFPH | ASSLDQGNYAEQF | 0.67266595 |
| TPPPPPP | ASSLDQGNYAEQF | 0.57051027 |
| AAPAAAAAPA | ASSLDQGNYAEQF | 0.4704596 |
| IALNEDL | ASSLDQGNYAEQF | 0.60223854 |
| IVNVSSM | ASSLDQGNYAEQF | 0.5311049 |
| TGIPPAPR | ASSLDQGNYAEQF | 0.5714367 |
| FDLGGGTF | ASSLDQGNYAEQF | 0.6247374 |
| FDLGGGTFDVS | ASSLDQGNYAEQF | 0.45215544 |
| NEPTAAA | ASSLDQGNYAEQF | 0.60059315 |
| NEPTAAALA | ASSLDQGNYAEQF | 0.6684135 |
| GSIINISSI | ASSLDQGNYAEQF | 0.5851852 |
| VAPGNAGTA | ASSLDQGNYAEQF | 0.62956196 |
| TGGGASLE | ASSLDQGNYAEQF | 0.66501176 |
| VAVYNHYKRI | ASSLDQGNYAEQF | 0.6446302 |
| TDPAPATT | ASSLDQGNYAEQF | 0.6652036 |
| TNRPDILDPALLRPG | ASSLDQGNYAEQF | 0.64562476 |
| LTTTTTS | ASSLDQGNYAEQF | 0.6074878 |
| PTGSGKTL | ASSLDQGNYAEQF | 0.60113925 |
| TGSGKTL | ASSLDQGNYAEQF | 0.6453723 |
| TIGHVDHGK | ASSLDQGNYAEQF | 0.61811715 |
| TIGHVDHGKTT | ASSLDQGNYAEQF | 0.61260056 |
| GHVDHGKTT | ASSLDQGNYAEQF | 0.55416375 |
| AQFEHTLL | ASSLDQGNYAEQF | 0.65783805 |
| IVNQDEL | ASSLDQGNYAEQF | 0.57734495 |
| TNGSQFF | ASSLDQGNYAEQF | 0.65464395 |
| TGKGIVE | ASSLDQGNYAEQF | 0.47152716 |
| LISNASDA | ASSLDQGNYAEQF | 0.5173723 |
| AELRTLGELTKIAKAHGAQ | ASSLDQGNYAEQF | 0.64283544 |
| APNAVAAAEPSW | ASSLDQGNYAEQF | 0.31810653 |
| APQVAGAATAVL | ASSLDQGNYAEQF | 0.38430527 |
| ARRDPAAAAAILPTDA | ASSLDQGNYAEQF | 0.60618365 |

| | | |
|---|---|---|
| DAAARDVLAEAGLAEYF | ASSLDQGNYAEQF | 0.65875274 |
| DDQIVLSAKANLAKVEQ | ASSLDQGNYAEQF | 0.32100832 |
| DELLPDTGEDSDEDGH | ASSLDQGNYAEQF | 0.5359249 |
| ELPEAPANRATVPQD | ASSLDQGNYAEQF | 0.49579063 |
| FGQSKIERVTGIG | ASSLDQGNYAEQF | 0.60530865 |
| GAGGTGSTAGGAGGAGGAGGLY | ASSLDQGNYAEQF | 0.6139096 |
| GHGGTGAAGGNGGTGGAGGSGI | ASSLDQGNYAEQF | 0.46107987 |
| GNGGNGGAGGEGGPGGAGGAGA | ASSLDQGNYAEQF | 0.64333963 |
| GSTAGNGGNGGNATNGTIAGQP | ASSLDQGNYAEQF | 0.4346261 |
| GTGGDGGNGGNGGTGG | ASSLDQGNYAEQF | 0.50302017 |
| GVRRPAGIPADAPV | ASSLDQGNYAEQF | 0.6524514 |
| GYRQVTDAHLLTLARR | ASSLDQGNYAEQF | 0.6588763 |
| IAKMGQDGHDRG | ASSLDQGNYAEQF | 0.56323063 |
| KAEPSIEIQYT | ASSLDQGNYAEQF | 0.65222806 |
| KPVRIRMAADEDTVAVT | ASSLDQGNYAEQF | 0.5220965 |
| MPPRIAGMRLLVIKPEP | ASSLDQGNYAEQF | 0.6783447 |
| MKFYAVALFRSRGVPPTR | ASSLDQGNYAEQF | 0.59667027 |
| NAGGFGILGAVAHSP | ASSLDQGNYAEQF | 0.4586413 |
| NEGREITLAMAEM | ASSLDQGNYAEQF | 0.6437908 |
| NELVEWSHPLGHRWRWE | ASSLDQGNYAEQF | 0.6262171 |
| NLGAGGTGGNGGNPG | ASSLDQGNYAEQF | 0.6092474 |
| NTEWPKWETVMAADPVD | ASSLDQGNYAEQF | 0.6414779 |
| PASAAPAETQP | ASSLDQGNYAEQF | 0.48201275 |
| QQSSDGKNGVRAAEITGE | ASSLDQGNYAEQF | 0.59837264 |
| RAPDAVTTSWWKE | ASSLDQGNYAEQF | 0.65970975 |
| SGFGNVGAHTSGWFNQ | ASSLDQGNYAEQF | 0.649127 |
| SGVAGAVGQAASVGGLKV | ASSLDQGNYAEQF | 0.5174577 |
| SNGGAGTGSAGGNGGT | ASSLDQGNYAEQF | 0.6578898 |
| TGGAGGEGGAAG | ASSLDQGNYAEQF | 0.5993235 |
| TGGDGGAAGAGGAPGHG | ASSLDQGNYAEQF | 0.5339424 |
| TMPPLLAAIANAGRHGV | ASSLDQGNYAEQF | 0.66442823 |
| TSSTAGNSGGA | ASSLDQGNYAEQF | 0.6096901 |
| VPATVPVGILC | ASSLDQGNYAEQF | 0.6558392 |
| VQRPDSGGVVLTGRIS | ASSLDQGNYAEQF | 0.6470776 |
| WTPNPVPATKGVPAGLVQAT | ASSLDQGNYAEQF | 0.6596652 |
| YALKRALAKFSDRFDVVII | ASSLDQGNYAEQF | 0.5602953 |
| YLIGSVIV | ASSLDQGNYAEQF | 0.63708323 |
| YGQLVAVID | ASSLDQGNYAEQF | 0.6268162 |
| VTVGGMTLPA | ASSLDQGNYAEQF | 0.6483479 |
| VLALPAV | ASSLDQGNYAEQF | 0.58181864 |
| VIPPTLNYET | ASSLDQGNYAEQF | 0.55765 |
| VGMGGMHPSSG | ASSLDQGNYAEQF | 0.63923883 |
| VGLAGGAATA | ASSLDQGNYAEQF | 0.6625787 |
| VGGAGGAGGSGP | ASSLDQGNYAEQF | 0.59049344 |
| VFAAFMV | ASSLDQGNYAEQF | 0.62758195 |
| TLYDKLR | ASSLDQGNYAEQF | 0.667075 |
| TLAVFTVL | ASSLDQGNYAEQF | 0.5483273 |
| TGSVQPARV | ASSLDQGNYAEQF | 0.60236037 |
| TGGGAGSGGT | ASSLDQGNYAEQF | 0.62812674 |
| SHLETSL | ASSLDQGNYAEQF | 0.66556036 |
| SFTTGMT | ASSLDQGNYAEQF | 0.5714097 |

| | | |
|---|---|---|
| QVEVAKRMGVRQPTVS | ASSLDQGNYAEQF | 0.66542757 |
| QGSGGPALAM | ASSLDQGNYAEQF | 0.58240306 |
| MLAPSNP | ASSLDQGNYAEQF | 0.6486194 |
| LVNNAAI | ASSLDQGNYAEQF | 0.5866721 |
| LTGEINTLA | ASSLDQGNYAEQF | 0.6367274 |
| LSIFGPV | ASSLDQGNYAEQF | 0.5299515 |
| LLALLANL | ASSLDQGNYAEQF | 0.6440009 |
| LISNASDALDKLR | ASSLDQGNYAEQF | 0.6554729 |
| LINEAVP | ASSLDQGNYAEQF | 0.65414095 |
| LGYAFSGL | ASSLDQGNYAEQF | 0.39789525 |
| LDFVAVAL | ASSLDQGNYAEQF | 0.6479062 |
| LAQPGAE | ASSLDQGNYAEQF | 0.64427286 |
| ISNLGAVTGF | ASSLDQGNYAEQF | 0.673432 |
| HQEDPGRI | ASSLDQGNYAEQF | 0.6018039 |
| GTPGEPTP | ASSLDQGNYAEQF | 0.6430342 |
| GFFNSGS | ASSLDQGNYAEQF | 0.5841014 |
| GAGGGGLTAGAN | ASSLDQGNYAEQF | 0.58169633 |
| FVDALVNM | ASSLDQGNYAEQF | 0.5006155 |
| FLTVAPEL | ASSLDQGNYAEQF | 0.5799693 |
| FETLTAL | ASSLDQGNYAEQF | 0.6084173 |
| DVINPIEL | ASSLDQGNYAEQF | 0.5461331 |
| APVSGAPV | ASSLDQGNYAEQF | 0.6500088 |
| ALVGGLAW | ASSLDQGNYAEQF | 0.6672503 |
| ALAAAAATAI | ASSLDQGNYAEQF | 0.64975965 |
| AFGLGLLL | ASSLDQGNYAEQF | 0.6580822 |
| AFFAAASL | ASSLDQGNYAEQF | 0.6626284 |
| AAAALPAFSPP | ASSLDQGNYAEQF | 0.6499753 |
| VVLAPAL | ASSLDQGNYAEQF | 0.55582595 |
| TIGHVDHGKTTL | ASSLDQGNYAEQF | 0.63893175 |
| LISNASDALDK | ASSLDQGNYAEQF | 0.46659577 |
| LISNASDAL | ASSLDQGNYAEQF | 0.5968721 |
| ISNASDALDK | ASSLDQGNYAEQF | 0.5682124 |
| NQAETLVYQTE | ASSLDQGNYAEQF | 0.6398221 |
| TGGTGGTAGAG | ASSLDQGNYAEQF | 0.3108922 |
| TGSTAGGAGGA | ASSLDQGNYAEQF | 0.6205251 |
| DPLAVPVAAGPV | ASSLDQGNYAEQF | 0.66990584 |
| GGTGGTPGNGGN | ASSLDQGNYAEQF | 0.56721306 |
| PAGLAGVDPATL | ASSLDQGNYAEQF | 0.6402404 |
| GNNLVVEAPHAGASV | ASSLDQGNYAEQF | 0.63543844 |
| KVTRIMSASTLEQPA | ASSLDQGNYAEQF | 0.53127503 |
| LGFALPIALPRRTVV | ASSLDQGNYAEQF | 0.64762753 |
| TRLGNLTARKLVVLV | ASSLDQGNYAEQF | 0.55021 |
| WSQQRRLIYFPSAGP | ASSLDQGNYAEQF | 0.6501447 |
| GPNGNPDPMAAAVDIR | ASSLDQGNYAEQF | 0.66427433 |
| TQRCVSVADLRAAASR | ASSLDQGNYAEQF | 0.6314959 |
| QFVLDVAEHPVEVLTETV | ASSLDQGNYAEQF | 0.63169616 |
| GGAGGTGGAGGNGGTQAGDG | ASSLDQGNYAEQF | 0.6295869 |
| PLLATLRAIAVDVDAKPAQV | ASSLDQGNYAEQF | 0.50870574 |
| APKFRTGGYMIDFWGVGYQVAK | ASSLDQGNYAEQF | 0.6361636 |
| GAGGAGGAGGAAGGPGGTGGTG | ASSLDQGNYAEQF | 0.396524 |
| IGALGLLATDVAPAGSVAFGVA | ASSLDQGNYAEQF | 0.6027524 |

| | | |
|---|---|---|
| AGGAGGIGGNGGAGGAGGVLMG | ASSLDQGNYAEQF | 0.5334049 |
| ADELIGLL | ASSLDQGNYAEQF | 0.6361327 |
| ADRALKA | ASSLDQGNYAEQF | 0.61283696 |
| AGAVGQVGGAGGA | ASSLDQGNYAEQF | 0.61782694 |
| AGGGGGLGGAP | ASSLDQGNYAEQF | 0.5320203 |
| AGGLLYGNGGAG | ASSLDQGNYAEQF | 0.6244265 |
| AGLESIGL | ASSLDQGNYAEQF | 0.6005862 |
| AGLKTGII | ASSLDQGNYAEQF | 0.4788543 |
| ASLAALLL | ASSLDQGNYAEQF | 0.6538439 |
| ASVINISSI | ASSLDQGNYAEQF | 0.56462204 |
| ATPPGPA | ASSLDQGNYAEQF | 0.5679614 |
| ATVASALV | ASSLDQGNYAEQF | 0.66049963 |
| ATVASLIL | ASSLDQGNYAEQF | 0.6240904 |
| AVNGPVLIRH | ASSLDQGNYAEQF | 0.4427572 |
| DGAAGAVGI | ASSLDQGNYAEQF | 0.60269034 |
| DPLLHGI | ASSLDQGNYAEQF | 0.6163632 |
| ELGAALGAL | ASSLDQGNYAEQF | 0.6129808 |
| ELGRTQAV | ASSLDQGNYAEQF | 0.3292596 |
| FVAGGYLS | ASSLDQGNYAEQF | 0.6122589 |
| GASGGAGGAGGW | ASSLDQGNYAEQF | 0.535845 |
| GGAGAGGDGGAGGI | ASSLDQGNYAEQF | 0.51923406 |
| GTGHNGSQPGMGGNG | ASSLDQGNYAEQF | 0.5478083 |
| GYAAASASAS | ASSLDQGNYAEQF | 0.6539123 |
| HQVDVAAVQQG | ASSLDQGNYAEQF | 0.58236873 |
| IGAAIAHT | ASSLDQGNYAEQF | 0.6118605 |
| IGMIEVAQL | ASSLDQGNYAEQF | 0.6514993 |
| IRLAPEHTSAMD | ASSLDQGNYAEQF | 0.6468811 |
| IYNVSVL | ASSLDQGNYAEQF | 0.6698539 |
| KQGFLGM | ASSLDQGNYAEQF | 0.61305034 |
| LESVITQL | ASSLDQGNYAEQF | 0.5478136 |
| LGLGALVVV | ASSLDQGNYAEQF | 0.63566947 |
| LIGNGGNGGSGGI | ASSLDQGNYAEQF | 0.60742855 |
| LNAINAPTL | ASSLDQGNYAEQF | 0.52476007 |
| LQLLNTGV | ASSLDQGNYAEQF | 0.45163485 |
| LQTSIEF | ASSLDQGNYAEQF | 0.61650074 |
| MSFAVSEL | ASSLDQGNYAEQF | 0.47933757 |
| NTGALGL | ASSLDQGNYAEQF | 0.6238281 |
| PPGAGGAGGAGTT | ASSLDQGNYAEQF | 0.6547808 |
| QLVGLGI | ASSLDQGNYAEQF | 0.58732307 |
| RFLEQVHQL | ASSLDQGNYAEQF | 0.48933995 |
| RGVGYVLR | ASSLDQGNYAEQF | 0.5659865 |
| SATGVFAGVI | ASSLDQGNYAEQF | 0.61901504 |
| SLPAITLP | ASSLDQGNYAEQF | 0.6603652 |
| STALAFTV | ASSLDQGNYAEQF | 0.6253103 |
| SVLTVGLGK | ASSLDQGNYAEQF | 0.66794527 |
| TALAGAGSGI | ASSLDQGNYAEQF | 0.64533323 |
| TGLPFTGI | ASSLDQGNYAEQF | 0.5649012 |
| VAEHVTV | ASSLDQGNYAEQF | 0.6471263 |
| VDIVLNSL | ASSLDQGNYAEQF | 0.5775279 |
| VLTGGGAVL | ASSLDQGNYAEQF | 0.6167374 |
| VNVGLAYGI | ASSLDQGNYAEQF | 0.6248764 |

| | | |
|---|---|---|
| VSSAAVGAV | ASSLDQGNYAEQF | 0.5634513 |
| VTSVYEAGV | ASSLDQGNYAEQF | 0.58305264 |
| WVALDRGA | ASSLDQGNYAEQF | 0.60348326 |
| YLVGSVVI | ASSLDQGNYAEQF | 0.42237517 |
| GPPGTGKTLL | ASSLDQGNYAEQF | 0.62721807 |
| GVGMTKF | ASSLDQGNYAEQF | 0.6335301 |
| IGIGGSDL | ASSLDQGNYAEQF | 0.46673113 |
| IGTLLAEL | ASSLDQGNYAEQF | 0.5650531 |
| IYTSGTT | ASSLDQGNYAEQF | 0.67191166 |
| LSFPAGV | ASSLDQGNYAEQF | 0.6670779 |
| LYTSGTT | ASSLDQGNYAEQF | 0.54936373 |
| VGAFPFGF | ASSLDQGNYAEQF | 0.5352871 |
| VGAGTVEF | ASSLDQGNYAEQF | 0.6116221 |
| VGKSTLLL | ASSLDQGNYAEQF | 0.45593885 |
| AALAHAGGGIV | ASSLDQGNYAEQF | 0.60409904 |
| AALVAVSLL | ASSLDQGNYAEQF | 0.5973597 |
| AAVSRAVAAL | ASSLDQGNYAEQF | 0.5877056 |
| AGGSSGIGGF | ASSLDQGNYAEQF | 0.5609086 |
| AGLTVANANG | ASSLDQGNYAEQF | 0.6200533 |
| AGLVGAVL | ASSLDQGNYAEQF | 0.544547 |
| ALSQAFTV | ASSLDQGNYAEQF | 0.63246846 |
| ALTFSVSL | ASSLDQGNYAEQF | 0.3304223 |
| ALTGAALVAL | ASSLDQGNYAEQF | 0.6189177 |
| ASFSIGAGVL | ASSLDQGNYAEQF | 0.54085636 |
| DGSVYKI | ASSLDQGNYAEQF | 0.51874954 |
| EQLAEIV | ASSLDQGNYAEQF | 0.42175698 |
| FAVSELTVM | ASSLDQGNYAEQF | 0.61139184 |
| GAAPLAAGL | ASSLDQGNYAEQF | 0.6526822 |
| GADGGDGGSGGAGG | ASSLDQGNYAEQF | 0.64766264 |
| GFGLDTGGAGG | ASSLDQGNYAEQF | 0.5730003 |
| GLASGGLL | ASSLDQGNYAEQF | 0.6530847 |
| GRIAGAAVLL | ASSLDQGNYAEQF | 0.66975796 |
| GSAGSLNTSF | ASSLDQGNYAEQF | 0.6063057 |
| GTGQIGI | ASSLDQGNYAEQF | 0.4778127 |
| GTLIALIL | ASSLDQGNYAEQF | 0.63576925 |
| IAATMGFVVG | ASSLDQGNYAEQF | 0.5832164 |
| IFAQALL | ASSLDQGNYAEQF | 0.62676775 |
| IGAAVALGF | ASSLDQGNYAEQF | 0.6429439 |
| IGLTPLAL | ASSLDQGNYAEQF | 0.48762107 |
| ILGSNTSTL | ASSLDQGNYAEQF | 0.6577674 |
| IVFSLTVL | ASSLDQGNYAEQF | 0.6508332 |
| LASKATGF | ASSLDQGNYAEQF | 0.53006893 |
| LASVALIL | ASSLDQGNYAEQF | 0.5617283 |
| LFAVGLTL | ASSLDQGNYAEQF | 0.5422826 |
| LFNGGIGTYI | ASSLDQGNYAEQF | 0.3951991 |
| LTLGAGII | ASSLDQGNYAEQF | 0.643596 |
| LVAAALGDI | ASSLDQGNYAEQF | 0.56506073 |
| NTAFVQLGI | ASSLDQGNYAEQF | 0.5855427 |
| QLFSGAEL | ASSLDQGNYAEQF | 0.6056368 |
| SAATFITL | ASSLDQGNYAEQF | 0.5696303 |
| SNYGLANLGV | ASSLDQGNYAEQF | 0.58139175 |

| | | |
|---|---|---|
| SYGGAYAVM | ASSLDQGNYAEQF | 0.59585077 |
| TTGVGGLTLG | ASSLDQGNYAEQF | 0.60292333 |
| TTQTIYI | ASSLDQGNYAEQF | 0.622653 |
| VAAGAGLGAVI | ASSLDQGNYAEQF | 0.5805886 |
| VAAVVTLL | ASSLDQGNYAEQF | 0.61826366 |
| VGFLGGLI | ASSLDQGNYAEQF | 0.67055696 |
| VGVIPILL | ASSLDQGNYAEQF | 0.48473817 |
| VIFASSLI | ASSLDQGNYAEQF | 0.53689826 |
| VKGGDGGAAAG | ASSLDQGNYAEQF | 0.61326015 |
| VTKTFTNI | ASSLDQGNYAEQF | 0.5029712 |
| VVAAAILL | ASSLDQGNYAEQF | 0.65073466 |
| YAALDVEL | ASSLDQGNYAEQF | 0.6086241 |
| LASVALLL | ASSLDQGNYAEQF | 0.6378619 |
| LDAVHRV | ASSLDQGNYAEQF | 0.633092 |
| QGLSSSIF | ASSLDQGNYAEQF | 0.67259854 |
| RGTFANIRL | ASSLDQGNYAEQF | 0.65204173 |
| RQLWWGHRI | ASSLDQGNYAEQF | 0.6031129 |
| YGPPGTGKTLL | ASSLDQGNYAEQF | 0.65242016 |
| AALVGAPANVA | ASSLDQGNYAEQF | 0.4754726 |
| AATFRRTE | ASSLDQGNYAEQF | 0.6563017 |
| AGALGGGI | ASSLDQGNYAEQF | 0.636291 |
| AGGAGGAGGIGAG | ASSLDQGNYAEQF | 0.52081156 |
| AGSGALGL | ASSLDQGNYAEQF | 0.59895164 |
| AQIADQI | ASSLDQGNYAEQF | 0.6651033 |
| ASAVAVTYL | ASSLDQGNYAEQF | 0.6603263 |
| ASTVLGGTGG | ASSLDQGNYAEQF | 0.48598063 |
| AVGGEPAK | ASSLDQGNYAEQF | 0.57553136 |
| CEQGTFGGTC | ASSLDQGNYAEQF | 0.43925774 |
| DEGKAPAG | ASSLDQGNYAEQF | 0.4808104 |
| DTVVFPAL | ASSLDQGNYAEQF | 0.4626629 |
| FEGITMAG | ASSLDQGNYAEQF | 0.6434841 |
| FPLVVQD | ASSLDQGNYAEQF | 0.60177255 |
| GAFTTPQ | ASSLDQGNYAEQF | 0.6398177 |
| GFGGFGTAGGVGGA | ASSLDQGNYAEQF | 0.62454295 |
| GFLAQFL | ASSLDQGNYAEQF | 0.6615776 |
| GGAGGAGGNAGVA | ASSLDQGNYAEQF | 0.54723203 |
| GGVGTAGAGGAGGAGG | ASSLDQGNYAEQF | 0.5902969 |
| GNGGGGDAGNAGSGGN | ASSLDQGNYAEQF | 0.43325058 |
| GTGGAGGAGNDAGS | ASSLDQGNYAEQF | 0.54586416 |
| GTTNTGWFNS | ASSLDQGNYAEQF | 0.5647685 |
| IFMGPLFTL | ASSLDQGNYAEQF | 0.5804086 |
| ILAHAIEAI | ASSLDQGNYAEQF | 0.3838662 |
| IRSAIREAL | ASSLDQGNYAEQF | 0.6487908 |
| ITFAGGG | ASSLDQGNYAEQF | 0.62601 |
| ITGMSGHQEV | ASSLDQGNYAEQF | 0.64985657 |
| IVAGLACTL | ASSLDQGNYAEQF | 0.5689666 |
| IVGGGLAGA | ASSLDQGNYAEQF | 0.6032894 |
| IVTGVLNI | ASSLDQGNYAEQF | 0.55219805 |
| LAQQGCI | ASSLDQGNYAEQF | 0.64496076 |
| LCLHVEN | ASSLDQGNYAEQF | 0.54439694 |
| LDAISDGS | ASSLDQGNYAEQF | 0.39331102 |

| | | |
|---|---|---|
| LGGFAAML | ASSLDQGNYAEQF | 0.57874525 |
| LGGKNPMI | ASSLDQGNYAEQF | 0.4923133 |
| LGPKGVAAIL | ASSLDQGNYAEQF | 0.6620211 |
| LMVGVSI | ASSLDQGNYAEQF | 0.65940654 |
| LVGAAFVL | ASSLDQGNYAEQF | 0.6297764 |
| LVLAVAPL | ASSLDQGNYAEQF | 0.65303946 |
| MAAIAVIDAA | ASSLDQGNYAEQF | 0.3652955 |
| NNGINAELQ | ASSLDQGNYAEQF | 0.5746187 |
| RGEPLVLVHG | ASSLDQGNYAEQF | 0.66654325 |
| RGGAGGAAGLI | ASSLDQGNYAEQF | 0.6328425 |
| RSYRMHPAV | ASSLDQGNYAEQF | 0.6122399 |
| SAAHLLAGQLA | ASSLDQGNYAEQF | 0.66512996 |
| SAAVGSY | ASSLDQGNYAEQF | 0.54217625 |
| SPHAVLHRI | ASSLDQGNYAEQF | 0.6148367 |
| SQIAKVL | ASSLDQGNYAEQF | 0.6188551 |
| TALGHLVP | ASSLDQGNYAEQF | 0.64687943 |
| TAPTTSGAA | ASSLDQGNYAEQF | 0.6166538 |
| TDGDDGNGGSPGT | ASSLDQGNYAEQF | 0.5373121 |
| TDGLGEN | ASSLDQGNYAEQF | 0.6425278 |
| TGGPPLPGQA | ASSLDQGNYAEQF | 0.6744526 |
| TIPAITI | ASSLDQGNYAEQF | 0.5919702 |
| TIPGFTI | ASSLDQGNYAEQF | 0.6554089 |
| TSIASGALA | ASSLDQGNYAEQF | 0.64216363 |
| TVTLPGAAI | ASSLDQGNYAEQF | 0.62678564 |
| VASGLSAALP | ASSLDQGNYAEQF | 0.54229957 |
| VGAGVIGI | ASSLDQGNYAEQF | 0.55593383 |
| VGPKRFGV | ASSLDQGNYAEQF | 0.62689406 |
| VNVAFPD | ASSLDQGNYAEQF | 0.55956286 |
| VSMTNSGVS | ASSLDQGNYAEQF | 0.6047946 |
| VTDVLGDLSV | ASSLDQGNYAEQF | 0.5777113 |
| VVVVGAGFAGL | ASSLDQGNYAEQF | 0.6784331 |
| VYTVGTVV | ASSLDQGNYAEQF | 0.6450434 |
| WTTFSGP | ASSLDQGNYAEQF | 0.638766 |
| YDELVEAF | ASSLDQGNYAEQF | 0.51776 |
| YGTGGAGGNGG | ASSLDQGNYAEQF | 0.612608 |
| YTNVMAV | ASSLDQGNYAEQF | 0.63143563 |
| FLEQVHQL | ASSLDQGNYAEQF | 0.45363444 |
| GLSLLAVAV | ASSLDQGNYAEQF | 0.6684097 |
| LEQVHQL | ASSLDQGNYAEQF | 0.560663 |
| LNAAGAI | ASSLDQGNYAEQF | 0.5334581 |
| PAVHASL | ASSLDQGNYAEQF | 0.6264994 |
| TELEDDA | ASSLDQGNYAEQF | 0.65820426 |
| TPPGPPN | ASSLDQGNYAEQF | 0.61169285 |
| TPPPPPPP | ASSLDQGNYAEQF | 0.63292664 |
| TQSPALI | ASSSTGGAYTQY | 0.5363876 |
| EFAAAFPQ | ASSSTGGAYTQY | 0.6434177 |
| LLSGNLDVL | ASSSTGGAYTQY | 0.6594121 |
| GPANLGNYNI | ASSSTGGAYTQY | 0.6149616 |
| TLGPTLP | ASSSTGGAYTQY | 0.66784847 |
| VYAYSSCCK | ASSSTGGAYTQY | 0.5811769 |
| APAPAPAPNI | ASSSTGGAYTQY | 0.40796983 |

| | | |
|---|---|---|
| PGGPVPIP | ASSSTGGAYTQY | 0.6144444 |
| FLTVALEP | ASSSTGGAYTQY | 0.6525136 |
| SQYFGANL | ASSSTGGAYTQY | 0.64970654 |
| GGVKPAA | ASSSTGGAYTQY | 0.66419494 |
| GGTGGAGGAGGW | ASSSTGGAYTQY | 0.64405996 |
| GVGGVAGLGGAGG | ASSSTGGAYTQY | 0.5631014 |
| VLLVLATTI | ASSSTGGAYTQY | 0.6657373 |
| IGIFAAML | ASSSTGGAYTQY | 0.6710286 |
| AVGDLID | ASSSTGGAYTQY | 0.6516833 |
| NISPLPL | ASSSTGGAYTQY | 0.53025633 |
| VTAVFVAI | ASSSTGGAYTQY | 0.4663246 |
| SGDDNDAAL | ASSSTGGAYTQY | 0.63253087 |
| AVAALAHS | ASSSTGGAYTQY | 0.63228035 |
| GGVTVMATP | ASSSTGGAYTQY | 0.50142604 |
| AGLAANVRD | ASSSTGGAYTQY | 0.6408116 |
| TNRPDLIDPALLRPG | ASSSTGGAYTQY | 0.6304255 |
| LANLGVTAL | ASSSTGGAYTQY | 0.43325478 |
| TGERIEL | ASSSTGGAYTQY | 0.6188976 |
| NGEELAA | ASSSTGGAYTQY | 0.66465306 |
| VLSLVWG | ASSSTGGAYTQY | 0.6143515 |
| IGYALWL | ASSSTGGAYTQY | 0.5870781 |
| LSLLLAPL | ASSSTGGAYTQY | 0.6643823 |
| TLDEALK | ASSSTGGAYTQY | 0.67387885 |
| TDGAGTAEGG | ASSSTGGAYTQY | 0.66751254 |
| NAPGGAGGNGG | ASSSTGGAYTQY | 0.44708678 |
| VSVFMGMGGNF | ASSSTGGAYTQY | 0.35459659 |
| LASYLAAL | ASSSTGGAYTQY | 0.5968654 |
| LVGGTPAL | ASSSTGGAYTQY | 0.6347766 |
| VRNADVL | ASSSTGGAYTQY | 0.5840602 |
| TCSSPFLVSAM | ASSSTGGAYTQY | 0.5326508 |
| VHITDAE | ASSSTGGAYTQY | 0.6357479 |
| EYAAPSF | ASSSTGGAYTQY | 0.6562178 |
| VVDAFHGGL | ASSSTGGAYTQY | 0.5380892 |
| ASVVDPQ | ASSSTGGAYTQY | 0.42434645 |
| SYNEFGGLS | ASSSTGGAYTQY | 0.6434532 |
| DPDGSVY | ASSSTGGAYTQY | 0.63972414 |
| MVSTIHATE | ASSSTGGAYTQY | 0.54597425 |
| LAAGVADPV | ASSSTGGAYTQY | 0.58871824 |
| TKEQIAA | ASSSTGGAYTQY | 0.6018685 |
| SLASPAAL | ASSSTGGAYTQY | 0.4819809 |
| IGLVFYGV | ASSSTGGAYTQY | 0.6146726 |
| FLLDASNL | ASSSTGGAYTQY | 0.4626835 |
| YQVGNFP | ASSSTGGAYTQY | 0.6433031 |
| ESLLSQST | ASSSTGGAYTQY | 0.5145531 |
| LFTVGFAAVG | ASSSTGGAYTQY | 0.64917874 |
| ATPPTPT | ASSSTGGAYTQY | 0.6307421 |
| TDLATLG | ASSSTGGAYTQY | 0.6348804 |
| SAGGTGR | ASSSTGGAYTQY | 0.6034383 |
| GFSNSGS | ASSSTGGAYTQY | 0.5687453 |
| GGNGGSGGTSV | ASSSTGGAYTQY | 0.5227751 |
| GTSVATGGAGN | ASSSTGGAYTQY | 0.4166465 |

| | | |
|---|---|---|
| NGGNGGSGGTS | ASSSTGGAYTQY | 0.6687643 |
| AVVKFPH | ASSSTGGAYTQY | 0.5436162 |
| TPPPPPP | ASSSTGGAYTQY | 0.6576808 |
| AAPAAAAAPA | ASSSTGGAYTQY | 0.6519669 |
| IALNEDL | ASSSTGGAYTQY | 0.5319757 |
| IVNVSSM | ASSSTGGAYTQY | 0.62227994 |
| TGIPPAPR | ASSSTGGAYTQY | 0.4921593 |
| FDLGGGTF | ASSSTGGAYTQY | 0.6088525 |
| FDLGGGTFDVS | ASSSTGGAYTQY | 0.6281843 |
| NEPTAAA | ASSSTGGAYTQY | 0.6356685 |
| NEPTAAALA | ASSSTGGAYTQY | 0.56314087 |
| GSIINISSI | ASSSTGGAYTQY | 0.62964416 |
| VAPGNAGTA | ASSSTGGAYTQY | 0.6303588 |
| TGGGASLE | ASSSTGGAYTQY | 0.59290427 |
| VAVYNHYKRI | ASSSTGGAYTQY | 0.6167682 |
| TDPAPATT | ASSSTGGAYTQY | 0.65519565 |
| TNRPDILDPALLRPG | ASSSTGGAYTQY | 0.6387427 |
| LTTTTTS | ASSSTGGAYTQY | 0.5746204 |
| PTGSGKTL | ASSSTGGAYTQY | 0.6442747 |
| TGSGKTL | ASSSTGGAYTQY | 0.54874843 |
| TIGHVDHGK | ASSSTGGAYTQY | 0.62718827 |
| TIGHVDHGKTT | ASSSTGGAYTQY | 0.6017495 |
| GHVDHGKTT | ASSSTGGAYTQY | 0.5778918 |
| AQFEHTLL | ASSSTGGAYTQY | 0.6420871 |
| IVNQDEL | ASSSTGGAYTQY | 0.449486 |
| TNGSQFF | ASSSTGGAYTQY | 0.5596789 |
| TGKGIVE | ASSSTGGAYTQY | 0.59914076 |
| LISNASDA | ASSSTGGAYTQY | 0.6250952 |
| AELRTLGELTKIAKAHGAQ | ASSSTGGAYTQY | 0.65289015 |
| APNAVAAAEPSW | ASSSTGGAYTQY | 0.51512015 |
| APQVAGAATAVL | ASSSTGGAYTQY | 0.62106 |
| ARRDPAAAAAILPTDA | ASSSTGGAYTQY | 0.5983137 |
| DAAARDVLAEAGLAEYF | ASSSTGGAYTQY | 0.5412152 |
| DDQIVLSAKANLAKVEQ | ASSSTGGAYTQY | 0.58274364 |
| DELLPDTGEDSDEDGH | ASSSTGGAYTQY | 0.60660017 |
| ELPEAPANRATVPQD | ASSSTGGAYTQY | 0.5434923 |
| FGQSKIERVTGIG | ASSSTGGAYTQY | 0.6436549 |
| GAGGTGSTAGGAGGAGGAGGLY | ASSSTGGAYTQY | 0.5338494 |
| GHGGTGAAGGNGGTGGAGGSGI | ASSSTGGAYTQY | 0.6264482 |
| GNGGNGGAGGEGGPGGAGGAGA | ASSSTGGAYTQY | 0.65602064 |
| GSTAGNGGNGGNATNGTIAGQP | ASSSTGGAYTQY | 0.58342606 |
| GTGGDGGNGGNGGTGG | ASSSTGGAYTQY | 0.6424096 |
| GVRRPAGIPADAPV | ASSSTGGAYTQY | 0.6115915 |
| GYRQVTDAHLLTLARR | ASSSTGGAYTQY | 0.6498523 |
| IAKMGQDGHDRG | ASSSTGGAYTQY | 0.6010999 |
| KAEPSIEIQYT | ASSSTGGAYTQY | 0.58458763 |
| KPVRIRMAADEDTVAVT | ASSSTGGAYTQY | 0.64867514 |
| MPPRIAGMRLLVIKPEP | ASSSTGGAYTQY | 0.65039086 |
| MKFYAVALFRSRGVPPTR | ASSSTGGAYTQY | 0.66181546 |
| NAGGFGILGAVAHSP | ASSSTGGAYTQY | 0.61692995 |
| NEGREITLAMAEM | ASSSTGGAYTQY | 0.6488372 |

| | | |
|---|---|---|
| NELVEWSHPLGHRWRWE | ASSSTGGAYTQY | 0.64871526 |
| NLGAGGTGGNGGNPG | ASSSTGGAYTQY | 0.56777614 |
| NTEWPKWETVMAADPVD | ASSSTGGAYTQY | 0.43493903 |
| PASAAPAETQP | ASSSTGGAYTQY | 0.58869857 |
| QQSSDGKNGVRAAEITGE | ASSSTGGAYTQY | 0.6025416 |
| RAPDAVTTSWWKE | ASSSTGGAYTQY | 0.5963052 |
| SGFGNVGAHTSGWFNQ | ASSSTGGAYTQY | 0.3985211 |
| SGVAGAVGQAASVGGLKV | ASSSTGGAYTQY | 0.6548018 |
| SNGGAGTGSAGGNGGT | ASSSTGGAYTQY | 0.6761281 |
| TGGAGGEGGAAG | ASSSTGGAYTQY | 0.5594772 |
| TGGDGGAAGAGGAPGHG | ASSSTGGAYTQY | 0.66753 |
| TMPPLLAAIANAGRHGV | ASSSTGGAYTQY | 0.48064536 |
| TSSTAGNSGGA | ASSSTGGAYTQY | 0.6506822 |
| VPATVPVGILC | ASSSTGGAYTQY | 0.66518116 |
| VQRPDSGGVVLTGRIS | ASSSTGGAYTQY | 0.52107215 |
| WTPNPVPATKGVPAGLVQAT | ASSSTGGAYTQY | 0.63751817 |
| YALKRALAKFSDRFDVVII | ASSSTGGAYTQY | 0.63182086 |
| YLIGSVIV | ASSSTGGAYTQY | 0.6140481 |
| YGQLVAVID | ASSSTGGAYTQY | 0.66871744 |
| VTVGGMTLPA | ASSSTGGAYTQY | 0.5861956 |
| VLALPAV | ASSSTGGAYTQY | 0.64540815 |
| VIPPTLNYET | ASSSTGGAYTQY | 0.58704436 |
| VGMGGMHPSSG | ASSSTGGAYTQY | 0.5209664 |
| VGLAGGAATA | ASSSTGGAYTQY | 0.6319912 |
| VGGAGGAGGSGP | ASSSTGGAYTQY | 0.6551476 |
| VFAAFMV | ASSSTGGAYTQY | 0.5796777 |
| TLYDKLR | ASSSTGGAYTQY | 0.6695307 |
| TLAVFTVL | ASSSTGGAYTQY | 0.55846876 |
| TGSVQPARV | ASSSTGGAYTQY | 0.66794074 |
| TGGGAGSGGT | ASSSTGGAYTQY | 0.64670026 |
| SHLETSL | ASSSTGGAYTQY | 0.6023006 |
| SFTTGMT | ASSSTGGAYTQY | 0.5748459 |
| QVEVAKRMGVRQPTVS | ASSSTGGAYTQY | 0.66382134 |
| QGSGGPALAM | ASSSTGGAYTQY | 0.58329034 |
| MLAPSNP | ASSSTGGAYTQY | 0.65344405 |
| LVNNAAI | ASSSTGGAYTQY | 0.6587437 |
| LTGEINTLA | ASSSTGGAYTQY | 0.35678995 |
| LSIFGPV | ASSSTGGAYTQY | 0.66266704 |
| LLALLANL | ASSSTGGAYTQY | 0.62318784 |
| LISNASDALDKLR | ASSSTGGAYTQY | 0.6146518 |
| LINEAVP | ASSSTGGAYTQY | 0.49000442 |
| LGYAFSGL | ASSSTGGAYTQY | 0.35749832 |
| LDFVAVAL | ASSSTGGAYTQY | 0.5337032 |
| LAQPGAE | ASSSTGGAYTQY | 0.44422972 |
| ISNLGAVTGF | ASSSTGGAYTQY | 0.69662297 |
| HQEDPGRI | ASSSTGGAYTQY | 0.6401095 |
| GTPGEPTP | ASSSTGGAYTQY | 0.4755005 |
| GFFNSGS | ASSSTGGAYTQY | 0.6367912 |
| GAGGGGLTAGAN | ASSSTGGAYTQY | 0.66595715 |
| FVDALVNM | ASSSTGGAYTQY | 0.35105476 |
| FLTVAPEL | ASSSTGGAYTQY | 0.5257236 |

| | | |
|---|---|---|
| FETLTAL | ASSSTGGAYTQY | 0.44989687 |
| DVINPIEL | ASSSTGGAYTQY | 0.50369227 |
| APVSGAPV | ASSSTGGAYTQY | 0.29906195 |
| ALVGGLAW | ASSSTGGAYTQY | 0.63307375 |
| ALAAAAATAI | ASSSTGGAYTQY | 0.49764293 |
| AFGLGLLL | ASSSTGGAYTQY | 0.28568104 |
| AFFAAASL | ASSSTGGAYTQY | 0.6072209 |
| AAAALPAFSPP | ASSSTGGAYTQY | 0.4938279 |
| VVLAPAL | ASSSTGGAYTQY | 0.5078471 |
| TIGHVDHGKTTL | ASSSTGGAYTQY | 0.63963556 |
| LISNASDALDK | ASSSTGGAYTQY | 0.55254406 |
| LISNASDAL | ASSSTGGAYTQY | 0.4566986 |
| ISNASDALDK | ASSSTGGAYTQY | 0.6201479 |
| NQAETLVYQTE | ASSSTGGAYTQY | 0.6597754 |
| TGGTGGTAGAG | ASSSTGGAYTQY | 0.40612045 |
| TGSTAGGAGGA | ASSSTGGAYTQY | 0.5961633 |
| DPLAVPVAAGPV | ASSSTGGAYTQY | 0.68527484 |
| GGTGGTPGNGGN | ASSSTGGAYTQY | 0.57093143 |
| PAGLAGVDPATL | ASSSTGGAYTQY | 0.6728559 |
| GNNLVVEAPHAGASV | ASSSTGGAYTQY | 0.6839712 |
| KVTRIMSASTLEQPA | ASSSTGGAYTQY | 0.51235396 |
| LGFALPIALPRRTVV | ASSSTGGAYTQY | 0.6535106 |
| TRLGNLTARKLVVLV | ASSSTGGAYTQY | 0.6871362 |
| WSQQRRLIYFPSAGP | ASSSTGGAYTQY | 0.4868089 |
| GPNGNPDPMAAAVDIR | ASSSTGGAYTQY | 0.5415945 |
| TQRCVSVADLRAAASR | ASSSTGGAYTQY | 0.39464423 |
| QFVLDVAEHPVEVLTETV | ASSSTGGAYTQY | 0.56270766 |
| GGAGGTGGAGGNGGTQAGDG | ASSSTGGAYTQY | 0.44389212 |
| PLLATLRAIAVDVDAKPAQV | ASSSTGGAYTQY | 0.6121206 |
| APKFRTGGYMIDFWGVGYQVAK | ASSSTGGAYTQY | 0.6766502 |
| GAGGAGGAGGAAGGPGGTGGTG | ASSSTGGAYTQY | 0.5414385 |
| IGALGLLATDVAPAGSVAFGVA | ASSSTGGAYTQY | 0.35040158 |
| AGGAGGIGGNGGAGGAGGVLMG | ASSSTGGAYTQY | 0.36626998 |
| ADELIGLL | ASSSTGGAYTQY | 0.5924065 |
| ADRALKA | ASSSTGGAYTQY | 0.39844745 |
| AGAVGQVGGAGGA | ASSSTGGAYTQY | 0.36918104 |
| AGGGGGLGGAP | ASSSTGGAYTQY | 0.42236102 |
| AGGLLYGNGGAG | ASSSTGGAYTQY | 0.6517683 |
| AGLESIGL | ASSSTGGAYTQY | 0.65269005 |
| AGLKTGII | ASSSTGGAYTQY | 0.47174627 |
| ASLAALLL | ASSSTGGAYTQY | 0.67773557 |
| ASVINISSI | ASSSTGGAYTQY | 0.67859185 |
| ATPPGPA | ASSSTGGAYTQY | 0.69464433 |
| ATVASALV | ASSSTGGAYTQY | 0.55024517 |
| ATVASLIL | ASSSTGGAYTQY | 0.3624845 |
| AVNGPVLIRH | ASSSTGGAYTQY | 0.681318 |
| DGAAGAVGI | ASSSTGGAYTQY | 0.5151207 |
| DPLLHGI | ASSSTGGAYTQY | 0.60601115 |
| ELGAALGAL | ASSSTGGAYTQY | 0.63267684 |
| ELGRTQAV | ASSSTGGAYTQY | 0.5413156 |
| FVAGGYLS | ASSSTGGAYTQY | 0.45270717 |

| | | |
|---|---|---|
| GASGGAGGAGGW | ASSSTGGAYTQY | 0.6310178 |
| GGAGAGGDGGAGGI | ASSSTGGAYTQY | 0.6833955 |
| GTGHNGSQPGMGGNG | ASSSTGGAYTQY | 0.40908015 |
| GYAAASASAS | ASSSTGGAYTQY | 0.40286395 |
| HQVDVAAVQQG | ASSSTGGAYTQY | 0.54021376 |
| IGAAIAHT | ASSSTGGAYTQY | 0.54230183 |
| IGMIEVAQL | ASSSTGGAYTQY | 0.6604749 |
| IRLAPEHTSAMD | ASSSTGGAYTQY | 0.5845613 |
| IYNVSVL | ASSSTGGAYTQY | 0.529631 |
| KQGFLGM | ASSSTGGAYTQY | 0.62312156 |
| LESVITQL | ASSSTGGAYTQY | 0.6302901 |
| LGLGALVVV | ASSSTGGAYTQY | 0.59161204 |
| LIGNGGNGGSGGI | ASSSTGGAYTQY | 0.64980924 |
| LNAINAPTL | ASSSTGGAYTQY | 0.44679075 |
| LQLLNTGV | ASSSTGGAYTQY | 0.6693902 |
| LQTSIEF | ASSSTGGAYTQY | 0.64582443 |
| MSFAVSEL | ASSSTGGAYTQY | 0.6363517 |
| NTGALGL | ASSSTGGAYTQY | 0.6757252 |
| PPGAGGAGGAGTT | ASSSTGGAYTQY | 0.59479636 |
| QLVGLGI | ASSSTGGAYTQY | 0.5967161 |
| RFLEQVHQL | ASSSTGGAYTQY | 0.39701045 |
| RGVGYVLR | ASSSTGGAYTQY | 0.5216818 |
| SATGVFAGVI | ASSSTGGAYTQY | 0.598044 |
| SLPAITLP | ASSSTGGAYTQY | 0.5950356 |
| STALAFTV | ASSSTGGAYTQY | 0.66313416 |
| SVLTVGLGK | ASSSTGGAYTQY | 0.6374514 |
| TALAGAGSGI | ASSSTGGAYTQY | 0.49340746 |
| TGLPFTGI | ASSSTGGAYTQY | 0.6882071 |
| VAEHVTV | ASSSTGGAYTQY | 0.6685724 |
| VDIVLNSL | ASSSTGGAYTQY | 0.4119351 |
| VLTGGGAVL | ASSSTGGAYTQY | 0.4934552 |
| VNVGLAYGI | ASSSTGGAYTQY | 0.6799668 |
| VSSAAVGAV | ASSSTGGAYTQY | 0.65592057 |
| VTSVYEAGV | ASSSTGGAYTQY | 0.5019623 |
| WVALDRGA | ASSSTGGAYTQY | 0.5791129 |
| YLVGSVVI | ASSSTGGAYTQY | 0.70357406 |
| GPPGTGKTLL | ASSSTGGAYTQY | 0.57093686 |
| GVGMTKF | ASSSTGGAYTQY | 0.6787096 |
| IGIGGSDL | ASSSTGGAYTQY | 0.324121 |
| IGTLLAEL | ASSSTGGAYTQY | 0.5506039 |
| IYTSGTT | ASSSTGGAYTQY | 0.4920006 |
| LSFPAGV | ASSSTGGAYTQY | 0.2846264 |
| LYTSGTT | ASSSTGGAYTQY | 0.6504045 |
| VGAFPFGF | ASSSTGGAYTQY | 0.5092144 |
| VGAGTVEF | ASSSTGGAYTQY | 0.6392286 |
| VGKSTLLL | ASSSTGGAYTQY | 0.47807887 |
| AALAHAGGGIV | ASSSTGGAYTQY | 0.65612483 |
| AALVAVSLL | ASSSTGGAYTQY | 0.6078994 |
| AAVSRAVAAL | ASSSTGGAYTQY | 0.5257291 |
| AGGSSGIGGF | ASSSTGGAYTQY | 0.40498525 |
| AGLTVANANG | ASSSTGGAYTQY | 0.60826445 |

| | | |
|---|---|---|
| AGLVGAVL | ASSSTGGAYTQY | 0.6902224 |
| ALSQAFTV | ASSSTGGAYTQY | 0.67526793 |
| ALTFSVSL | ASSSTGGAYTQY | 0.33520707 |
| ALTGAALVAL | ASSSTGGAYTQY | 0.6125573 |
| ASFSIGAGVL | ASSSTGGAYTQY | 0.5463058 |
| DGSVYKI | ASSSTGGAYTQY | 0.38899362 |
| EQLAEIV | ASSSTGGAYTQY | 0.30859584 |
| FAVSELTVM | ASSSTGGAYTQY | 0.4773806 |
| GAAPLAAGL | ASSSTGGAYTQY | 0.5606154 |
| GADGGDGGSGGAGG | ASSSTGGAYTQY | 0.61364746 |
| GFGLDTGGAGG | ASSSTGGAYTQY | 0.6299528 |
| GLASGGLL | ASSSTGGAYTQY | 0.47663748 |
| GRIAGAAVLL | ASSSTGGAYTQY | 0.6324129 |
| GSAGSLNTSF | ASSSTGGAYTQY | 0.56954247 |
| GTGQIGI | ASSSTGGAYTQY | 0.25581148 |
| GTLIALIL | ASSSTGGAYTQY | 0.45494232 |
| IAATMGFVVG | ASSSTGGAYTQY | 0.34902453 |
| IFAQALL | ASSSTGGAYTQY | 0.6852288 |
| IGAAVALGF | ASSSTGGAYTQY | 0.3918789 |
| IGLTPLAL | ASSSTGGAYTQY | 0.5119871 |
| ILGSNTSTL | ASSSTGGAYTQY | 0.55247307 |
| IVFSLTVL | ASSSTGGAYTQY | 0.6594267 |
| LASKATGF | ASSSTGGAYTQY | 0.5838699 |
| LASVALIL | ASSSTGGAYTQY | 0.45220345 |
| LFAVGLTL | ASSSTGGAYTQY | 0.37984326 |
| LFNGGIGTYI | ASSSTGGAYTQY | 0.61037105 |
| LTLGAGII | ASSSTGGAYTQY | 0.5469936 |
| LVAAALGDI | ASSSTGGAYTQY | 0.42189234 |
| NTAFVQLGI | ASSSTGGAYTQY | 0.41959864 |
| QLFSGAEL | ASSSTGGAYTQY | 0.61453503 |
| SAATFITL | ASSSTGGAYTQY | 0.34447452 |
| SNYGLANLGV | ASSSTGGAYTQY | 0.6632576 |
| SYGGAYAVM | ASSSTGGAYTQY | 0.3977474 |
| TTGVGGLTLG | ASSSTGGAYTQY | 0.4865541 |
| TTQTIYI | ASSSTGGAYTQY | 0.6849321 |
| VAAGAGLGAVI | ASSSTGGAYTQY | 0.6465583 |
| VAAVVTLL | ASSSTGGAYTQY | 0.5051949 |
| VGFLGGLI | ASSSTGGAYTQY | 0.4354297 |
| VGVIPILL | ASSSTGGAYTQY | 0.4023912 |
| VIFASSLI | ASSSTGGAYTQY | 0.4525885 |
| VKGGDGGAAAG | ASSSTGGAYTQY | 0.4507041 |
| VTKTFTNI | ASSSTGGAYTQY | 0.6662508 |
| VVAAAILL | ASSSTGGAYTQY | 0.5993015 |
| YAALDVEL | ASSSTGGAYTQY | 0.38476214 |
| LASVALLL | ASSSTGGAYTQY | 0.6489199 |
| LDAVHRV | ASSSTGGAYTQY | 0.6775122 |
| QGLSSSIF | ASSSTGGAYTQY | 0.69761133 |
| RGTFANIRL | ASSSTGGAYTQY | 0.39438838 |
| RQLWWGHRI | ASSSTGGAYTQY | 0.5190031 |
| YGPPGTGKTLL | ASSSTGGAYTQY | 0.5564439 |
| AALVGAPANVA | ASSSTGGAYTQY | 0.5480652 |

| | | |
|---|---|---|
| AATFRRTE | ASSSTGGAYTQY | 0.4425676 |
| AGALGGGI | ASSSTGGAYTQY | 0.43443945 |
| AGGAGGAGGIGAG | ASSSTGGAYTQY | 0.6599233 |
| AGSGALGL | ASSSTGGAYTQY | 0.71190083 |
| AQIADQI | ASSSTGGAYTQY | 0.53598166 |
| ASAVAVTYL | ASSSTGGAYTQY | 0.33781624 |
| ASTVLGGTGG | ASSSTGGAYTQY | 0.5357952 |
| AVGGEPAK | ASSSTGGAYTQY | 0.4350673 |
| CEQGTFGGTC | ASSSTGGAYTQY | 0.45645136 |
| DEGKAPAG | ASSSTGGAYTQY | 0.5426655 |
| DTVVFPAL | ASSSTGGAYTQY | 0.34596452 |
| FEGITMAG | ASSSTGGAYTQY | 0.5205531 |
| FPLVVQD | ASSSTGGAYTQY | 0.68594474 |
| GAFTTPQ | ASSSTGGAYTQY | 0.5638108 |
| GFGGFGTAGGVGGA | ASSSTGGAYTQY | 0.5999166 |
| GFLAQFL | ASSSTGGAYTQY | 0.66424584 |
| GGAGGAGGNAGVA | ASSSTGGAYTQY | 0.62020046 |
| GGVGTAGAGGAGGAGG | ASSSTGGAYTQY | 0.67345595 |
| GNGGGDAGNAGSGGN | ASSSTGGAYTQY | 0.6862329 |
| GTGGAGGAGNDAGS | ASSSTGGAYTQY | 0.5457541 |
| GTTNTGWFNS | ASSSTGGAYTQY | 0.5745442 |
| IFMGPLFTL | ASSSTGGAYTQY | 0.666479 |
| ILAHAIEAI | ASSSTGGAYTQY | 0.532778 |
| IRSAIREAL | ASSSTGGAYTQY | 0.56975454 |
| ITFAGGG | ASSSTGGAYTQY | 0.53156304 |
| ITGMSGHQEV | ASSSTGGAYTQY | 0.6326312 |
| IVAGLACTL | ASSSTGGAYTQY | 0.5314793 |
| IVGGGLAGA | ASSSTGGAYTQY | 0.7005526 |
| IVTGVLNI | ASSSTGGAYTQY | 0.30463344 |
| LAQQGCI | ASSSTGGAYTQY | 0.42250383 |
| LCLHVEN | ASSSTGGAYTQY | 0.66732824 |
| LDAISDGS | ASSSTGGAYTQY | 0.2857527 |
| LGGFAAML | ASSSTGGAYTQY | 0.33256775 |
| LGGKNPMI | ASSSTGGAYTQY | 0.58930546 |
| LGPKGVAAIL | ASSSTGGAYTQY | 0.67703676 |
| LMVGVSI | ASSSTGGAYTQY | 0.28864905 |
| LVGAAFVL | ASSSTGGAYTQY | 0.5092789 |
| LVLAVAPL | ASSSTGGAYTQY | 0.3535235 |
| MAAIAVIDAA | ASSSTGGAYTQY | 0.6573345 |
| NNGINAELQ | ASSSTGGAYTQY | 0.54580724 |
| RGEPLVLVHG | ASSSTGGAYTQY | 0.3425615 |
| RGGAGGAAGLI | ASSSTGGAYTQY | 0.647449 |
| RSYRMHPAV | ASSSTGGAYTQY | 0.3586556 |
| SAAHLLAGQLA | ASSSTGGAYTQY | 0.61768544 |
| SAAVGSY | ASSSTGGAYTQY | 0.651085 |
| SPHAVLHRI | ASSSTGGAYTQY | 0.70348907 |
| SQIAKVL | ASSSTGGAYTQY | 0.35888115 |
| TALGHLVP | ASSSTGGAYTQY | 0.6247289 |
| TAPTTSGAA | ASSSTGGAYTQY | 0.49089533 |
| TDGDDGNGGSPGT | ASSSTGGAYTQY | 0.7048157 |
| TDGLGEN | ASSSTGGAYTQY | 0.4880317 |

| | | |
|---|---|---|
| TGGPPLPGQA | ASSSTGGAYTQY | 0.3479371 |
| TIPAITI | ASSSTGGAYTQY | 0.6521196 |
| TIPGFTI | ASSSTGGAYTQY | 0.6533958 |
| TSIASGALA | ASSSTGGAYTQY | 0.57106286 |
| TVTLPGAAI | ASSSTGGAYTQY | 0.674044 |
| VASGLSAALP | ASSSTGGAYTQY | 0.5563064 |
| VGAGVIGI | ASSSTGGAYTQY | 0.52136886 |
| VGPKRFGV | ASSSTGGAYTQY | 0.6293196 |
| VNVAFPD | ASSSTGGAYTQY | 0.63343596 |
| VSMTNSGVS | ASSSTGGAYTQY | 0.46662533 |
| VTDVLGDLSV | ASSSTGGAYTQY | 0.679423 |
| VVVVGAGFAGL | ASSSTGGAYTQY | 0.58403146 |
| VYTVGTVV | ASSSTGGAYTQY | 0.4125718 |
| WTTFSGP | ASSSTGGAYTQY | 0.69519114 |
| YDELVEAF | ASSSTGGAYTQY | 0.5630237 |
| YGTGGAGGNGG | ASSSTGGAYTQY | 0.63991034 |
| YTNVMAV | ASSSTGGAYTQY | 0.62379485 |
| FLEQVHQL | ASSSTGGAYTQY | 0.7117153 |
| GLSLLAVAV | ASSSTGGAYTQY | 0.40948948 |
| LEQVHQL | ASSSTGGAYTQY | 0.64430964 |
| LNAAGAI | ASSSTGGAYTQY | 0.57844424 |
| PAVHASL | ASSSTGGAYTQY | 0.6547716 |
| TELEDDA | ASSSTGGAYTQY | 0.6206815 |
| TPPGPPN | ASSSTGGAYTQY | 0.5886684 |
| TPPPPPPP | ASSSTGGAYTQY | 0.6298931 |
| TQSPALI | ASSFWGNYAEQF | 0.66277045 |
| EFAAAFPQ | ASSFWGNYAEQF | 0.5503399 |
| LLSGNLDVL | ASSFWGNYAEQF | 0.63222957 |
| GPANLGNYNI | ASSFWGNYAEQF | 0.68640935 |
| TLGPTLP | ASSFWGNYAEQF | 0.41839167 |
| VYAYSSCCK | ASSFWGNYAEQF | 0.57789475 |
| APAPAPAPNI | ASSFWGNYAEQF | 0.6559479 |
| PGGPVPIP | ASSFWGNYAEQF | 0.6812161 |
| FLTVALEP | ASSFWGNYAEQF | 0.5420511 |
| SQYFGANL | ASSFWGNYAEQF | 0.6723621 |
| GGVKPAA | ASSFWGNYAEQF | 0.51760733 |
| GGTGGAGGAGGW | ASSFWGNYAEQF | 0.6102662 |
| GVGGVAGLGGAGG | ASSFWGNYAEQF | 0.5703958 |
| VLLVLATTI | ASSFWGNYAEQF | 0.4953087 |
| IGIFAAML | ASSFWGNYAEQF | 0.40633637 |
| AVGDLID | ASSFWGNYAEQF | 0.5873842 |
| NISPLPL | ASSFWGNYAEQF | 0.6453856 |
| VTAVFVAI | ASSFWGNYAEQF | 0.60828704 |
| SGDDNDAAL | ASSFWGNYAEQF | 0.31315312 |
| AVAALAHS | ASSFWGNYAEQF | 0.6199671 |
| GGVTVMATP | ASSFWGNYAEQF | 0.6219592 |
| AGLAANVRD | ASSFWGNYAEQF | 0.701818 |
| TNRPDLIDPALLRPG | ASSFWGNYAEQF | 0.5425563 |
| LANLGVTAL | ASSFWGNYAEQF | 0.66248834 |
| TGERIEL | ASSFWGNYAEQF | 0.51908493 |
| NGEELAA | ASSFWGNYAEQF | 0.5975499 |

| | | |
|---|---|---|
| VLSLVWG | ASSFWGNYAEQF | 0.35996047 |
| IGYALWL | ASSFWGNYAEQF | 0.4916417 |
| LSLLLAPL | ASSFWGNYAEQF | 0.53512394 |
| TLDEALK | ASSFWGNYAEQF | 0.43882456 |
| TDGAGTAEGG | ASSFWGNYAEQF | 0.6688118 |
| NAPGGAGGNGG | ASSFWGNYAEQF | 0.67504835 |
| VSVFMGMGGNF | ASSFWGNYAEQF | 0.6500364 |
| LASYLAAL | ASSFWGNYAEQF | 0.6654495 |
| LVGGTPAL | ASSFWGNYAEQF | 0.68213844 |
| VRNADVL | ASSFWGNYAEQF | 0.6688509 |
| TCSSPFLVSAM | ASSFWGNYAEQF | 0.52588403 |
| VHITDAE | ASSFWGNYAEQF | 0.5335636 |
| EYAAPSF | ASSFWGNYAEQF | 0.3468099 |
| VVDAFHGGL | ASSFWGNYAEQF | 0.48822805 |
| ASVVDPQ | ASSFWGNYAEQF | 0.5018644 |
| SYNEFGGLS | ASSFWGNYAEQF | 0.54937446 |
| DPDGSVY | ASSFWGNYAEQF | 0.27742535 |
| MVSTIHATE | ASSFWGNYAEQF | 0.6468277 |
| LAAGVADPV | ASSFWGNYAEQF | 0.66757977 |
| TKEQIAA | ASSFWGNYAEQF | 0.55019253 |
| SLASPAAL | ASSFWGNYAEQF | 0.633156 |
| IGLVFYGV | ASSFWGNYAEQF | 0.6327983 |
| FLLDASNL | ASSFWGNYAEQF | 0.46663934 |
| YQVGNFP | ASSFWGNYAEQF | 0.62773526 |
| ESLLSQST | ASSFWGNYAEQF | 0.4812284 |
| LFTVGFAAVG | ASSFWGNYAEQF | 0.6480205 |
| ATPPTPT | ASSFWGNYAEQF | 0.6727227 |
| TDLATLG | ASSFWGNYAEQF | 0.6068518 |
| SAGGTGR | ASSFWGNYAEQF | 0.6158563 |
| GFSNSGS | ASSFWGNYAEQF | 0.5989504 |
| GGNGGSGGTSV | ASSFWGNYAEQF | 0.37715718 |
| GTSVATGGAGN | ASSFWGNYAEQF | 0.625509 |
| NGGNGGSGGTS | ASSFWGNYAEQF | 0.47794273 |
| AVVKFPH | ASSFWGNYAEQF | 0.5930437 |
| TPPPPPP | ASSFWGNYAEQF | 0.44460222 |
| AAPAAAAAPA | ASSFWGNYAEQF | 0.6104408 |
| IALNEDL | ASSFWGNYAEQF | 0.5637784 |
| IVNVSSM | ASSFWGNYAEQF | 0.60691273 |
| TGIPPAPR | ASSFWGNYAEQF | 0.42813432 |
| FDLGGGTF | ASSFWGNYAEQF | 0.63283634 |
| FDLGGGTFDVS | ASSFWGNYAEQF | 0.6024728 |
| NEPTAAA | ASSFWGNYAEQF | 0.36913458 |
| NEPTAAALA | ASSFWGNYAEQF | 0.6660011 |
| GSIINISSI | ASSFWGNYAEQF | 0.5474221 |
| VAPGNAGTA | ASSFWGNYAEQF | 0.5710717 |
| TGGGASLE | ASSFWGNYAEQF | 0.65741116 |
| VAVYNHYKRI | ASSFWGNYAEQF | 0.57040423 |
| TDPAPATT | ASSFWGNYAEQF | 0.36727586 |
| TNRPDILDPALLRPG | ASSFWGNYAEQF | 0.51570076 |
| LTTTTTS | ASSFWGNYAEQF | 0.6015531 |
| PTGSGKTL | ASSFWGNYAEQF | 0.51873803 |

| | | |
|---|---|---|
| TGSGKTL | ASSFWGNYAEQF | 0.30823362 |
| TIGHVDHGK | ASSFWGNYAEQF | 0.60509324 |
| TIGHVDHGKTT | ASSFWGNYAEQF | 0.411114 |
| GHVDHGKTT | ASSFWGNYAEQF | 0.47564113 |
| AQFEHTLL | ASSFWGNYAEQF | 0.5141062 |
| IVNQDEL | ASSFWGNYAEQF | 0.6502564 |
| TNGSQFF | ASSFWGNYAEQF | 0.6336707 |
| TGKGIVE | ASSFWGNYAEQF | 0.55946016 |
| LISNASDA | ASSFWGNYAEQF | 0.6434642 |
| AELRTLGELTKIAKAHGAQ | ASSFWGNYAEQF | 0.58697695 |
| APNAVAAAEPSW | ASSFWGNYAEQF | 0.6531849 |
| APQVAGAATAVL | ASSFWGNYAEQF | 0.6171179 |
| ARRDPAAAAAILPTDA | ASSFWGNYAEQF | 0.59248453 |
| DAAARDVLAEAGLAEYF | ASSFWGNYAEQF | 0.62267125 |
| DDQIVLSAKANLAKVEQ | ASSFWGNYAEQF | 0.47929248 |
| DELLPDTGEDSDEDGH | ASSFWGNYAEQF | 0.4005737 |
| ELPEAPANRATVPQD | ASSFWGNYAEQF | 0.4005046 |
| FGQSKIERVTGIG | ASSFWGNYAEQF | 0.44978982 |
| GAGGTGSTAGGAGGAGGAGGLY | ASSFWGNYAEQF | 0.39066643 |
| GHGGTGAAGGNGGTGGAGGSGI | ASSFWGNYAEQF | 0.61269134 |
| GNGGNGGAGGEGGPGGAGGAGA | ASSFWGNYAEQF | 0.6928433 |
| GSTAGNGGNGGNATNGTIAGQP | ASSFWGNYAEQF | 0.57758844 |
| GTGGDGGNGGNGGTGG | ASSFWGNYAEQF | 0.34591755 |
| GVRRPAGIPADAPV | ASSFWGNYAEQF | 0.48504722 |
| GYRQVTDAHLLTLARR | ASSFWGNYAEQF | 0.50018376 |
| IAKMGQDGHDRG | ASSFWGNYAEQF | 0.6356232 |
| KAEPSIEIQYT | ASSFWGNYAEQF | 0.67297554 |
| KPVRIRMAADEDTVAVT | ASSFWGNYAEQF | 0.7042171 |
| MPPRIAGMRLLVIKPEP | ASSFWGNYAEQF | 0.6393266 |
| MKFYAVALFRSRGVPPTR | ASSFWGNYAEQF | 0.5059997 |
| NAGGFGILGAVAHSP | ASSFWGNYAEQF | 0.6453651 |
| NEGREITLAMAEM | ASSFWGNYAEQF | 0.5086255 |
| NELVEWSHPLGHRWRWE | ASSFWGNYAEQF | 0.59957397 |
| NLGAGGTGGNGGNPG | ASSFWGNYAEQF | 0.5507133 |
| NTEWPKWETVMAADPVD | ASSFWGNYAEQF | 0.6414515 |
| PASAAPAETQP | ASSFWGNYAEQF | 0.5290806 |
| QQSSDGKNGVRAAEITGE | ASSFWGNYAEQF | 0.4959364 |
| RAPDAVTTSWWKE | ASSFWGNYAEQF | 0.36075762 |
| SGFGNVGAHTSGWFNQ | ASSFWGNYAEQF | 0.672957 |
| SGVAGAVGQAASVGGLKV | ASSFWGNYAEQF | 0.43388683 |
| SNGGAGTGSAGGNGGT | ASSFWGNYAEQF | 0.41220307 |
| TGGAGGEGGAAG | ASSFWGNYAEQF | 0.51506245 |
| TGGDGGAAGAGGAPGHG | ASSFWGNYAEQF | 0.6885834 |
| TMPPLLAAIANAGRHGV | ASSFWGNYAEQF | 0.6822183 |
| TSSTAGNSGGA | ASSFWGNYAEQF | 0.48078448 |
| VPATVPVGILC | ASSFWGNYAEQF | 0.46452704 |
| VQRPDSGGVVLTGRIS | ASSFWGNYAEQF | 0.651305 |
| WTPNPVPATKGVPAGLVQAT | ASSFWGNYAEQF | 0.49574232 |
| YALKRALAKFSDRFDVVII | ASSFWGNYAEQF | 0.64280653 |
| YLIGSVIV | ASSFWGNYAEQF | 0.5808365 |
| YGQLVAVID | ASSFWGNYAEQF | 0.5374974 |

| | | |
|---|---|---|
| VTVGGMTLPA | ASSFWGNYAEQF | 0.49987692 |
| VLALPAV | ASSFWGNYAEQF | 0.62487733 |
| VIPPTLNYET | ASSFWGNYAEQF | 0.5175554 |
| VGMGGMHPSSG | ASSFWGNYAEQF | 0.6435977 |
| VGLAGGAATA | ASSFWGNYAEQF | 0.3510066 |
| VGGAGGAGGSGP | ASSFWGNYAEQF | 0.6517598 |
| VFAAFMV | ASSFWGNYAEQF | 0.4103833 |
| TLYDKLR | ASSFWGNYAEQF | 0.44086546 |
| TLAVFTVL | ASSFWGNYAEQF | 0.3047221 |
| TGSVQPARV | ASSFWGNYAEQF | 0.5390256 |
| TGGGAGSGGT | ASSFWGNYAEQF | 0.64827627 |
| SHLETSL | ASSFWGNYAEQF | 0.6709498 |
| SFTTGMT | ASSFWGNYAEQF | 0.55571574 |
| QVEVAKRMGVRQPTVS | ASSFWGNYAEQF | 0.63460827 |
| QGSGGPALAM | ASSFWGNYAEQF | 0.6763759 |
| MLAPSNP | ASSFWGNYAEQF | 0.44710243 |
| LVNNAAI | ASSFWGNYAEQF | 0.31495035 |
| LTGEINTLA | ASSFWGNYAEQF | 0.59537125 |
| LSIFGPV | ASSFWGNYAEQF | 0.6397749 |
| LLALLANL | ASSFWGNYAEQF | 0.67500496 |
| LISNASDALDKLR | ASSFWGNYAEQF | 0.66390747 |
| LINEAVP | ASSFWGNYAEQF | 0.350618 |
| LGYAFSGL | ASSFWGNYAEQF | 0.6692556 |
| LDFVAVAL | ASSFWGNYAEQF | 0.5908878 |
| LAQPGAE | ASSFWGNYAEQF | 0.37201074 |
| ISNLGAVTGF | ASSFWGNYAEQF | 0.46161988 |
| HQEDPGRI | ASSFWGNYAEQF | 0.42738876 |
| GTPGEPTP | ASSFWGNYAEQF | 0.40232316 |
| GFFNSGS | ASSFWGNYAEQF | 0.6257489 |
| GAGGGGLTAGAN | ASSFWGNYAEQF | 0.48335147 |
| FVDALVNM | ASSFWGNYAEQF | 0.5403023 |
| FLTVAPEL | ASSFWGNYAEQF | 0.53904486 |
| FETLTAL | ASSFWGNYAEQF | 0.4729585 |
| DVINPIEL | ASSFWGNYAEQF | 0.49092397 |
| APVSGAPV | ASSFWGNYAEQF | 0.5014887 |
| ALVGGLAW | ASSFWGNYAEQF | 0.6197095 |
| ALAAAAATAI | ASSFWGNYAEQF | 0.6485507 |
| AFGLGLLL | ASSFWGNYAEQF | 0.5671389 |
| AFFAAASL | ASSFWGNYAEQF | 0.63520426 |
| AAAALPAFSPP | ASSFWGNYAEQF | 0.65220773 |
| VVLAPAL | ASSFWGNYAEQF | 0.3878886 |
| TIGHVDHGKTTL | ASSFWGNYAEQF | 0.43741193 |
| LISNASDALDK | ASSFWGNYAEQF | 0.63423276 |
| LISNASDAL | ASSFWGNYAEQF | 0.50175226 |
| ISNASDALDK | ASSFWGNYAEQF | 0.6681845 |
| NQAETLVYQTE | ASSFWGNYAEQF | 0.5880489 |
| TGGTGGTAGAG | ASSFWGNYAEQF | 0.5899974 |
| TGSTAGGAGGA | ASSFWGNYAEQF | 0.6096293 |
| DPLAVPVAAGPV | ASSFWGNYAEQF | 0.6936381 |
| GGTGGTPGNGGN | ASSFWGNYAEQF | 0.6054642 |
| PAGLAGVDPATL | ASSFWGNYAEQF | 0.5226157 |

| | | |
|---|---|---|
| GNNLVVEAPHAGASV | ASSFWGNYAEQF | 0.6902659 |
| KVTRIMSASTLEQPA | ASSFWGNYAEQF | 0.38576347 |
| LGFALPIALPRRTVV | ASSFWGNYAEQF | 0.64136636 |
| TRLGNLTARKLVVLV | ASSFWGNYAEQF | 0.64737356 |
| WSQQRRLIYFPSAGP | ASSFWGNYAEQF | 0.415529 |
| GPNGNPDPMAAAVDIR | ASSFWGNYAEQF | 0.52337694 |
| TQRCVSVADLRAAASR | ASSFWGNYAEQF | 0.68275195 |
| QFVLDVAEHPVEVLTETV | ASSFWGNYAEQF | 0.66702497 |
| GGAGGTGGAGGNGGTQAGDG | ASSFWGNYAEQF | 0.46751797 |
| PLLATLRAIAVDVDAKPAQV | ASSFWGNYAEQF | 0.4455134 |
| APKFRTGGYMIDFWGVGYQVAK | ASSFWGNYAEQF | 0.43242598 |
| GAGGAGGAGGAAGGPGGTGGTG | ASSFWGNYAEQF | 0.47834718 |
| IGALGLLATDVAPAGSVAFGVA | ASSFWGNYAEQF | 0.35118496 |
| AGGAGGIGGNGGAGGAGGVLMG | ASSFWGNYAEQF | 0.62595963 |
| ADELIGLL | ASSFWGNYAEQF | 0.4857849 |
| ADRALKA | ASSFWGNYAEQF | 0.6893709 |
| AGAVGQVGGAGGA | ASSFWGNYAEQF | 0.60918236 |
| AGGGGGLGGAP | ASSFWGNYAEQF | 0.6604396 |
| AGGLLYGNGGAG | ASSFWGNYAEQF | 0.51346064 |
| AGLESIGL | ASSFWGNYAEQF | 0.51782155 |
| AGLKTGII | ASSFWGNYAEQF | 0.33446625 |
| ASLAALLL | ASSFWGNYAEQF | 0.4421252 |
| ASVINISSI | ASSFWGNYAEQF | 0.43652514 |
| ATPPGPA | ASSFWGNYAEQF | 0.5520612 |
| ATVASALV | ASSFWGNYAEQF | 0.35355902 |
| ATVASLIL | ASSFWGNYAEQF | 0.6856878 |
| AVNGPVLIRH | ASSFWGNYAEQF | 0.517159 |
| DGAAGAVGI | ASSFWGNYAEQF | 0.62820464 |
| DPLLHGI | ASSFWGNYAEQF | 0.5074104 |
| ELGAALGAL | ASSFWGNYAEQF | 0.43960473 |
| ELGRTQAV | ASSFWGNYAEQF | 0.5150199 |
| FVAGGYLS | ASSFWGNYAEQF | 0.6448618 |
| GASGGAGGAGGW | ASSFWGNYAEQF | 0.36030775 |
| GGAGAGGDGGAGGI | ASSFWGNYAEQF | 0.32264328 |
| GTGHNGSQPGMGGNG | ASSFWGNYAEQF | 0.44394106 |
| GYAAASASAS | ASSFWGNYAEQF | 0.43292743 |
| HQVDVAAVQQG | ASSFWGNYAEQF | 0.695477 |
| IGAAIAHT | ASSFWGNYAEQF | 0.66519487 |
| IGMIEVAQL | ASSFWGNYAEQF | 0.56327355 |
| IRLAPEHTSAMD | ASSFWGNYAEQF | 0.69899064 |
| IYNVSVL | ASSFWGNYAEQF | 0.27849713 |
| KQGFLGM | ASSFWGNYAEQF | 0.52253574 |
| LESVITQL | ASSFWGNYAEQF | 0.6817539 |
| LGLGALVVV | ASSFWGNYAEQF | 0.6308733 |
| LIGNGGNGGSGGI | ASSFWGNYAEQF | 0.51999974 |
| LNAINAPTL | ASSFWGNYAEQF | 0.6565417 |
| LQLLNTGV | ASSFWGNYAEQF | 0.45740086 |
| LQTSIEF | ASSFWGNYAEQF | 0.52963173 |
| MSFAVSEL | ASSFWGNYAEQF | 0.554325 |
| NTGALGL | ASSFWGNYAEQF | 0.6239526 |
| PPGAGGAGGAGTT | ASSFWGNYAEQF | 0.6468879 |

| | | |
|---|---|---|
| QLVGLGI | ASSFWGNYAEQF | 0.3459929 |
| RFLEQVHQL | ASSFWGNYAEQF | 0.6309669 |
| RGVGYVLR | ASSFWGNYAEQF | 0.6937187 |
| SATGVFAGVI | ASSFWGNYAEQF | 0.5085985 |
| SLPAITLP | ASSFWGNYAEQF | 0.6437408 |
| STALAFTV | ASSFWGNYAEQF | 0.5901483 |
| SVLTVGLGK | ASSFWGNYAEQF | 0.60846776 |
| TALAGAGSGI | ASSFWGNYAEQF | 0.44899336 |
| TGLPFTGI | ASSFWGNYAEQF | 0.38615853 |
| VAEHVTV | ASSFWGNYAEQF | 0.6310034 |
| VDIVLNSL | ASSFWGNYAEQF | 0.37498927 |
| VLTGGGAVL | ASSFWGNYAEQF | 0.5576919 |
| VNVGLAYGI | ASSFWGNYAEQF | 0.5716219 |
| VSSAAVGAV | ASSFWGNYAEQF | 0.70280176 |
| VTSVYEAGV | ASSFWGNYAEQF | 0.62478673 |
| WVALDRGA | ASSFWGNYAEQF | 0.60674417 |
| YLVGSVVI | ASSFWGNYAEQF | 0.47836357 |
| GPPGTGKTLL | ASSFWGNYAEQF | 0.6102444 |
| GVGMTKF | ASSFWGNYAEQF | 0.6323067 |
| IGIGGSDL | ASSFWGNYAEQF | 0.37151214 |
| IGTLLAEL | ASSFWGNYAEQF | 0.6557021 |
| IYTSGTT | ASSFWGNYAEQF | 0.3898805 |
| LSFPAGV | ASSFWGNYAEQF | 0.5812923 |
| LYTSGTT | ASSFWGNYAEQF | 0.5768045 |
| VGAFPFGF | ASSFWGNYAEQF | 0.69474685 |
| VGAGTVEF | ASSFWGNYAEQF | 0.5379803 |
| VGKSTLLL | ASSFWGNYAEQF | 0.62489617 |
| AALAHAGGGIV | ASSFWGNYAEQF | 0.42609015 |
| AALVAVSLL | ASSFWGNYAEQF | 0.6551133 |
| AAVSRAVAAL | ASSFWGNYAEQF | 0.66921055 |
| AGGSSGIGGF | ASSFWGNYAEQF | 0.678941 |
| AGLTVANANG | ASSFWGNYAEQF | 0.70147204 |
| AGLVGAVL | ASSFWGNYAEQF | 0.64493567 |
| ALSQAFTV | ASSFWGNYAEQF | 0.32447398 |
| ALTFSVSL | ASSFWGNYAEQF | 0.5387547 |
| ALTGAALVAL | ASSFWGNYAEQF | 0.6676675 |
| ASFSIGAGVL | ASSFWGNYAEQF | 0.67679685 |
| DGSVYKI | ASSFWGNYAEQF | 0.6758994 |
| EQLAEIV | ASSFWGNYAEQF | 0.6829032 |
| FAVSELTVM | ASSFWGNYAEQF | 0.53001904 |
| GAAPLAAGL | ASSFWGNYAEQF | 0.6721517 |
| GADGGDGGSGGAGG | ASSFWGNYAEQF | 0.71240455 |
| GFGLDTGGAGG | ASSFWGNYAEQF | 0.6395776 |
| GLASGGLL | ASSFWGNYAEQF | 0.43219057 |
| GRIAGAAVLL | ASSFWGNYAEQF | 0.4189193 |
| GSAGSLNTSF | ASSFWGNYAEQF | 0.6302997 |
| GTGQIGI | ASSFWGNYAEQF | 0.65922815 |
| GTLIALIL | ASSFWGNYAEQF | 0.38362107 |
| IAATMGFVVG | ASSFWGNYAEQF | 0.5317092 |
| IFAQALL | ASSFWGNYAEQF | 0.5695317 |
| IGAAVALGF | ASSFWGNYAEQF | 0.3120245 |

| | | |
|---|---|---|
| IGLTPLAL | ASSFWGNYAEQF | 0.64229167 |
| ILGSNTSTL | ASSFWGNYAEQF | 0.6805757 |
| IVFSLTVL | ASSFWGNYAEQF | 0.5942687 |
| LASKATGF | ASSFWGNYAEQF | 0.36989206 |
| LASVALIL | ASSFWGNYAEQF | 0.6634345 |
| LFAVGLTL | ASSFWGNYAEQF | 0.6729698 |
| LFNGGIGTYI | ASSFWGNYAEQF | 0.68296105 |
| LTLGAGII | ASSFWGNYAEQF | 0.3445191 |
| LVAAALGDI | ASSFWGNYAEQF | 0.28383985 |
| NTAFVQLGI | ASSFWGNYAEQF | 0.58905137 |
| QLFSGAEL | ASSFWGNYAEQF | 0.68063694 |
| SAATFITL | ASSFWGNYAEQF | 0.5057266 |
| SNYGLANLGV | ASSFWGNYAEQF | 0.45327845 |
| SYGGAYAVM | ASSFWGNYAEQF | 0.60177135 |
| TTGVGGLTLG | ASSFWGNYAEQF | 0.6571876 |
| TTQTIYI | ASSFWGNYAEQF | 0.40719688 |
| VAAGAGLGAVI | ASSFWGNYAEQF | 0.40457135 |
| VAAVVTLL | ASSFWGNYAEQF | 0.64741844 |
| VGFLGGLI | ASSFWGNYAEQF | 0.662875 |
| VGVIPILL | ASSFWGNYAEQF | 0.40515792 |
| VIFASSLI | ASSFWGNYAEQF | 0.5192152 |
| VKGGDGGAAAG | ASSFWGNYAEQF | 0.6029721 |
| VTKTFTNI | ASSFWGNYAEQF | 0.6172559 |
| VVAAAILL | ASSFWGNYAEQF | 0.59221727 |
| YAALDVEL | ASSFWGNYAEQF | 0.39015383 |
| LASVALLL | ASSFWGNYAEQF | 0.64108634 |
| LDAVHRV | ASSFWGNYAEQF | 0.36094674 |
| QGLSSSIF | ASSFWGNYAEQF | 0.6527852 |
| RGTFANIRL | ASSFWGNYAEQF | 0.60534865 |
| RQLWWGHRI | ASSFWGNYAEQF | 0.6131739 |
| YGPPGTGKTLL | ASSFWGNYAEQF | 0.5012636 |
| AALVGAPANVA | ASSFWGNYAEQF | 0.4121796 |
| AATFRRTE | ASSFWGNYAEQF | 0.40000737 |
| AGALGGGI | ASSFWGNYAEQF | 0.40027875 |
| AGGAGGAGGIGAG | ASSFWGNYAEQF | 0.6744964 |
| AGSGALGL | ASSFWGNYAEQF | 0.6168283 |
| AQIADQI | ASSFWGNYAEQF | 0.6622871 |
| ASAVAVTYL | ASSFWGNYAEQF | 0.57140964 |
| ASTVLGGTGG | ASSFWGNYAEQF | 0.39471114 |
| AVGGEPAK | ASSFWGNYAEQF | 0.63294756 |
| CEQGTFGGTC | ASSFWGNYAEQF | 0.3885526 |
| DEGKAPAG | ASSFWGNYAEQF | 0.58560216 |
| DTVVFPAL | ASSFWGNYAEQF | 0.6105371 |
| FEGITMAG | ASSFWGNYAEQF | 0.64547896 |
| FPLVVQD | ASSFWGNYAEQF | 0.468868 |
| GAFTTPQ | ASSFWGNYAEQF | 0.6153702 |
| GFGGFGTAGGVGGA | ASSFWGNYAEQF | 0.6160915 |
| GFLAQFL | ASSFWGNYAEQF | 0.39884177 |
| GGAGGAGGNAGVA | ASSFWGNYAEQF | 0.6076344 |
| GGVGTAGAGGAGGAGG | ASSFWGNYAEQF | 0.63741666 |
| GNGGGGDAGNAGSGGN | ASSFWGNYAEQF | 0.6173042 |

| | | |
|---|---|---|
| GTGGAGGAGNDAGS | ASSFWGNYAEQF | 0.32686356 |
| GTTNTGWFNS | ASSFWGNYAEQF | 0.5861281 |
| IFMGPLFTL | ASSFWGNYAEQF | 0.45169294 |
| ILAHAIEAI | ASSFWGNYAEQF | 0.6060521 |
| IRSAIREAL | ASSFWGNYAEQF | 0.546767 |
| ITFAGGG | ASSFWGNYAEQF | 0.4703613 |
| ITGMSGHQEV | ASSFWGNYAEQF | 0.61812717 |
| IVAGLACTL | ASSFWGNYAEQF | 0.40542805 |
| IVGGGLAGA | ASSFWGNYAEQF | 0.496606 |
| IVTGVLNI | ASSFWGNYAEQF | 0.52394843 |
| LAQQGCI | ASSFWGNYAEQF | 0.587652 |
| LCLHVEN | ASSFWGNYAEQF | 0.62492454 |
| LDAISDGS | ASSFWGNYAEQF | 0.39583597 |
| LGGFAAML | ASSFWGNYAEQF | 0.5999239 |
| LGGKNPMI | ASSFWGNYAEQF | 0.51567054 |
| LGPKGVAAIL | ASSFWGNYAEQF | 0.41849497 |
| LMVGVSI | ASSFWGNYAEQF | 0.56539214 |
| LVGAAFVL | ASSFWGNYAEQF | 0.50063694 |
| LVLAVAPL | ASSFWGNYAEQF | 0.35968304 |
| MAAIAVIDAA | ASSFWGNYAEQF | 0.6269663 |
| NNGINAELQ | ASSFWGNYAEQF | 0.4191982 |
| RGEPLVLVHG | ASSFWGNYAEQF | 0.6092216 |
| RGGAGGAAGLI | ASSFWGNYAEQF | 0.615036 |
| RSYRMHPAV | ASSFWGNYAEQF | 0.40496594 |
| SAAHLLAGQLA | ASSFWGNYAEQF | 0.6416389 |
| SAAVGSY | ASSFWGNYAEQF | 0.57336277 |
| SPHAVLHRI | ASSFWGNYAEQF | 0.6040926 |
| SQIAKVL | ASSFWGNYAEQF | 0.5414213 |
| TALGHLVP | ASSFWGNYAEQF | 0.42196378 |
| TAPTTSGAA | ASSFWGNYAEQF | 0.6619801 |
| TDGDDGNGGSPGT | ASSFWGNYAEQF | 0.6169522 |
| TDGLGEN | ASSFWGNYAEQF | 0.64878553 |
| TGGPPLPGQA | ASSFWGNYAEQF | 0.63988006 |
| TIPAITI | ASSFWGNYAEQF | 0.66675997 |
| TIPGFTI | ASSFWGNYAEQF | 0.6516448 |
| TSIASGALA | ASSFWGNYAEQF | 0.45805937 |
| TVTLPGAAI | ASSFWGNYAEQF | 0.33217043 |
| VASGLSAALP | ASSFWGNYAEQF | 0.5138247 |
| VGAGVIGI | ASSFWGNYAEQF | 0.5195743 |
| VGPKRFGV | ASSFWGNYAEQF | 0.45278564 |
| VNVAFPD | ASSFWGNYAEQF | 0.35047606 |
| VSMTNSGVS | ASSFWGNYAEQF | 0.63912976 |
| VTDVLGDLSV | ASSFWGNYAEQF | 0.70382094 |
| VVVVGAGFAGL | ASSFWGNYAEQF | 0.4944608 |
| VYTVGTVV | ASSFWGNYAEQF | 0.70483863 |
| WTTFSGP | ASSFWGNYAEQF | 0.3797825 |
| YDELVEAF | ASSFWGNYAEQF | 0.636733 |
| YGTGGAGGNGG | ASSFWGNYAEQF | 0.6492094 |
| YTNVMAV | ASSFWGNYAEQF | 0.4062092 |
| FLEQVHQL | ASSFWGNYAEQF | 0.6042718 |
| GLSLLAVAV | ASSFWGNYAEQF | 0.40865907 |

| | | |
|---|---|---|
| LEQVHQL | ASSFWGNYAEQF | 0.438437 |
| LNAAGAI | ASSFWGNYAEQF | 0.65535355 |
| PAVHASL | ASSFWGNYAEQF | 0.5277763 |
| TELEDDA | ASSFWGNYAEQF | 0.6084824 |
| TPPGPPN | ASSFWGNYAEQF | 0.536041 |
| TPPPPPPP | ASSFWGNYAEQF | 0.4497489 |
| TQSPALI | ASSLGQKYAEQF | 0.5881582 |
| EFAAAFPQ | ASSLGQKYAEQF | 0.6543198 |
| LLSGNLDVL | ASSLGQKYAEQF | 0.5804318 |
| GPANLGNYNI | ASSLGQKYAEQF | 0.7103736 |
| TLGPTLP | ASSLGQKYAEQF | 0.43420702 |
| VYAYSSCCK | ASSLGQKYAEQF | 0.6835035 |
| APAPAPAPNI | ASSLGQKYAEQF | 0.68615496 |
| PGGPVPIP | ASSLGQKYAEQF | 0.5949639 |
| FLTVALEP | ASSLGQKYAEQF | 0.5010235 |
| SQYFGANL | ASSLGQKYAEQF | 0.6513028 |
| GGVKPAA | ASSLGQKYAEQF | 0.56456435 |
| GGTGGAGGAGGW | ASSLGQKYAEQF | 0.65161 |
| GVGGVAGLGGAGG | ASSLGQKYAEQF | 0.6786145 |
| VLLVLATTI | ASSLGQKYAEQF | 0.47590533 |
| IGIFAAML | ASSLGQKYAEQF | 0.64685065 |
| AVGDLID | ASSLGQKYAEQF | 0.61756504 |
| NISPLPL | ASSLGQKYAEQF | 0.6144065 |
| VTAVFVAI | ASSLGQKYAEQF | 0.53526 |
| SGDDNDAAL | ASSLGQKYAEQF | 0.4870446 |
| AVAALAHS | ASSLGQKYAEQF | 0.5738922 |
| GGVTVMATP | ASSLGQKYAEQF | 0.51671124 |
| AGLAANVRD | ASSLGQKYAEQF | 0.6250703 |
| TNRPDLIDPALLRPG | ASSLGQKYAEQF | 0.6348062 |
| LANLGVTAL | ASSLGQKYAEQF | 0.5403092 |
| TGERIEL | ASSLGQKYAEQF | 0.64330757 |
| NGEELAA | ASSLGQKYAEQF | 0.6360301 |
| VLSLVWG | ASSLGQKYAEQF | 0.43304566 |
| IGYALWL | ASSLGQKYAEQF | 0.5841999 |
| LSLLLAPL | ASSLGQKYAEQF | 0.55304617 |
| TLDEALK | ASSLGQKYAEQF | 0.566599 |
| TDGAGTAEGG | ASSLGQKYAEQF | 0.42545992 |
| NAPGGAGGNGG | ASSLGQKYAEQF | 0.61622405 |
| VSVFMGMGGNF | ASSLGQKYAEQF | 0.52826345 |
| LASYLAAL | ASSLGQKYAEQF | 0.34272835 |
| LVGGTPAL | ASSLGQKYAEQF | 0.61695784 |
| VRNADVL | ASSLGQKYAEQF | 0.5815555 |
| TCSSPFLVSAM | ASSLGQKYAEQF | 0.57701755 |
| VHITDAE | ASSLGQKYAEQF | 0.62487614 |
| EYAAPSF | ASSLGQKYAEQF | 0.5564314 |
| VVDAFHGGL | ASSLGQKYAEQF | 0.5682575 |
| ASVVDPQ | ASSLGQKYAEQF | 0.62199104 |
| SYNEFGGLS | ASSLGQKYAEQF | 0.6415181 |
| DPDGSVY | ASSLGQKYAEQF | 0.5600334 |
| MVSTIHATE | ASSLGQKYAEQF | 0.58119416 |
| LAAGVADPV | ASSLGQKYAEQF | 0.6445792 |

| | | |
|---|---|---|
| TKEQIAA | ASSLGQKYAEQF | 0.60326767 |
| SLASPAAL | ASSLGQKYAEQF | 0.6420994 |
| IGLVFYGV | ASSLGQKYAEQF | 0.64817 |
| FLLDASNL | ASSLGQKYAEQF | 0.5386195 |
| YQVGNFP | ASSLGQKYAEQF | 0.6337315 |
| ESLLSQST | ASSLGQKYAEQF | 0.6504449 |
| LFTVGFAAVG | ASSLGQKYAEQF | 0.59590614 |
| ATPPTPT | ASSLGQKYAEQF | 0.59325176 |
| TDLATLG | ASSLGQKYAEQF | 0.44880348 |
| SAGGTGR | ASSLGQKYAEQF | 0.59116393 |
| GFSNSGS | ASSLGQKYAEQF | 0.43470156 |
| GGNGGSGGTSV | ASSLGQKYAEQF | 0.59675056 |
| GTSVATGGAGN | ASSLGQKYAEQF | 0.64414954 |
| NGGNGGSGGTS | ASSLGQKYAEQF | 0.5590675 |
| AVVKFPH | ASSLGQKYAEQF | 0.4847448 |
| TPPPPPP | ASSLGQKYAEQF | 0.46118274 |
| AAPAAAAAPA | ASSLGQKYAEQF | 0.60027766 |
| IALNEDL | ASSLGQKYAEQF | 0.5268761 |
| IVNVSSM | ASSLGQKYAEQF | 0.40843987 |
| TGIPPAPR | ASSLGQKYAEQF | 0.59698224 |
| FDLGGGTF | ASSLGQKYAEQF | 0.6383983 |
| FDLGGGTFDVS | ASSLGQKYAEQF | 0.6414777 |
| NEPTAAA | ASSLGQKYAEQF | 0.47943786 |
| NEPTAAALA | ASSLGQKYAEQF | 0.64646256 |
| GSIINISSI | ASSLGQKYAEQF | 0.6073078 |
| VAPGNAGTA | ASSLGQKYAEQF | 0.64888823 |
| TGGGASLE | ASSLGQKYAEQF | 0.6012437 |
| VAVYNHYKRI | ASSLGQKYAEQF | 0.51699924 |
| TDPAPATT | ASSLGQKYAEQF | 0.61742777 |
| TNRPDILDPALLRPG | ASSLGQKYAEQF | 0.5709631 |
| LTTTTTS | ASSLGQKYAEQF | 0.6138783 |
| PTGSGKTL | ASSLGQKYAEQF | 0.63223124 |
| TGSGKTL | ASSLGQKYAEQF | 0.61041147 |
| TIGHVDHGK | ASSLGQKYAEQF | 0.54154897 |
| TIGHVDHGKTT | ASSLGQKYAEQF | 0.5635268 |
| GHVDHGKTT | ASSLGQKYAEQF | 0.63690066 |
| AQFEHTLL | ASSLGQKYAEQF | 0.51247364 |
| IVNQDEL | ASSLGQKYAEQF | 0.52544445 |
| TNGSQFF | ASSLGQKYAEQF | 0.55213565 |
| TGKGIVE | ASSLGQKYAEQF | 0.59402287 |
| LISNASDA | ASSLGQKYAEQF | 0.6382465 |
| AELRTLGELTKIAKAHGAQ | ASSLGQKYAEQF | 0.62315965 |
| APNAVAAAEPSW | ASSLGQKYAEQF | 0.5742839 |
| APQVAGAATAVL | ASSLGQKYAEQF | 0.6165541 |
| ARRDPAAAAAILPTDA | ASSLGQKYAEQF | 0.6268453 |
| DAAARDVLAEAGLAEYF | ASSLGQKYAEQF | 0.60306346 |
| DDQIVLSAKANLAKVEQ | ASSLGQKYAEQF | 0.6050427 |
| DELLPDTGEDSDEDGH | ASSLGQKYAEQF | 0.5374767 |
| ELPEAPANRATVPQD | ASSLGQKYAEQF | 0.62908584 |
| FGQSKIERVTGIG | ASSLGQKYAEQF | 0.62064725 |
| GAGGTGSTAGGAGGAGGAGGLY | ASSLGQKYAEQF | 0.628915 |

| | | |
|---|---|---|
| GHGGTGAAGGNGGTGGAGGSGI | ASSLGQKYAEQF | 0.6352717 |
| GNGGNGGAGGEGGPGGAGGAGA | ASSLGQKYAEQF | 0.6372782 |
| GSTAGNGGNGGNATNGTIAGQP | ASSLGQKYAEQF | 0.59637743 |
| GTGGDGGNGGNGGTGG | ASSLGQKYAEQF | 0.41313735 |
| GVRRPAGIPADAPV | ASSLGQKYAEQF | 0.56727946 |
| GYRQVTDAHLLTLARR | ASSLGQKYAEQF | 0.6066675 |
| IAKMGQDGHDRG | ASSLGQKYAEQF | 0.6039605 |
| KAEPSIEIQYT | ASSLGQKYAEQF | 0.6270987 |
| KPVRIRMAADEDTVAVT | ASSLGQKYAEQF | 0.63495505 |
| MPPRIAGMRLLVIKPEP | ASSLGQKYAEQF | 0.5530492 |
| MKFYAVALFRSRGVPPTR | ASSLGQKYAEQF | 0.63906235 |
| NAGGFGILGAVAHSP | ASSLGQKYAEQF | 0.637431 |
| NEGREITLAMAEM | ASSLGQKYAEQF | 0.41133565 |
| NELVEWSHPLGHRWRWE | ASSLGQKYAEQF | 0.5714265 |
| NLGAGGTGGNGGNPG | ASSLGQKYAEQF | 0.6538274 |
| NTEWPKWETVMAADPVD | ASSLGQKYAEQF | 0.59681547 |
| PASAAPAETQP | ASSLGQKYAEQF | 0.5672853 |
| QQSSDGKNGVRAAEITGE | ASSLGQKYAEQF | 0.5774107 |
| RAPDAVTTSWWKE | ASSLGQKYAEQF | 0.645285 |
| SGFGNVGAHTSGWFNQ | ASSLGQKYAEQF | 0.57330024 |
| SGVAGAVGQAASVGGLKV | ASSLGQKYAEQF | 0.6497055 |
| SNGGAGTGSAGGNGGT | ASSLGQKYAEQF | 0.37246442 |
| TGGAGGEGGAAG | ASSLGQKYAEQF | 0.5372069 |
| TGGDGGAAGAGGAPGHG | ASSLGQKYAEQF | 0.5120147 |
| TMPPLLAAIANAGRHGV | ASSLGQKYAEQF | 0.36694402 |
| TSSTAGNSGGA | ASSLGQKYAEQF | 0.6296307 |
| VPATVPVGILC | ASSLGQKYAEQF | 0.55957866 |
| VQRPDSGGVVLTGRIS | ASSLGQKYAEQF | 0.6262948 |
| WTPNPVPATKGVPAGLVQAT | ASSLGQKYAEQF | 0.587503 |
| YALKRALAKFSDRFDVVII | ASSLGQKYAEQF | 0.6353118 |
| YLIGSVIV | ASSLGQKYAEQF | 0.5866828 |
| YGQLVAVID | ASSLGQKYAEQF | 0.59985346 |
| VTVGGMTLPA | ASSLGQKYAEQF | 0.53870946 |
| VLALPAV | ASSLGQKYAEQF | 0.6274729 |
| VIPPTLNYET | ASSLGQKYAEQF | 0.6454445 |
| VGMGGMHPSSG | ASSLGQKYAEQF | 0.6323596 |
| VGLAGGAATA | ASSLGQKYAEQF | 0.50670224 |
| VGGAGGAGGSGP | ASSLGQKYAEQF | 0.60356987 |
| VFAAFMV | ASSLGQKYAEQF | 0.5929434 |
| TLYDKLR | ASSLGQKYAEQF | 0.422666 |
| TLAVFTVL | ASSLGQKYAEQF | 0.37014306 |
| TGSVQPARV | ASSLGQKYAEQF | 0.5893112 |
| TGGGAGSGGT | ASSLGQKYAEQF | 0.5597295 |
| SHLETSL | ASSLGQKYAEQF | 0.62542695 |
| SFTTGMT | ASSLGQKYAEQF | 0.62859446 |
| QVEVAKRMGVRQPTVS | ASSLGQKYAEQF | 0.55995584 |
| QGSGGPALAM | ASSLGQKYAEQF | 0.62455106 |
| MLAPSNP | ASSLGQKYAEQF | 0.61388254 |
| LVNNAAI | ASSLGQKYAEQF | 0.31042764 |
| LTGEINTLA | ASSLGQKYAEQF | 0.54181886 |
| LSIFGPV | ASSLGQKYAEQF | 0.4557803 |

| | | |
|---|---|---|
| LLALLANL | ASSLGQKYAEQF | 0.6449386 |
| LISNASDALDKLR | ASSLGQKYAEQF | 0.52202594 |
| LINEAVP | ASSLGQKYAEQF | 0.52856374 |
| LGYAFSGL | ASSLGQKYAEQF | 0.58627284 |
| LDFVAVAL | ASSLGQKYAEQF | 0.58952343 |
| LAQPGAE | ASSLGQKYAEQF | 0.5882143 |
| ISNLGAVTGF | ASSLGQKYAEQF | 0.5199114 |
| HQEDPGRI | ASSLGQKYAEQF | 0.4244038 |
| GTPGEPTP | ASSLGQKYAEQF | 0.6330584 |
| GFFNSGS | ASSLGQKYAEQF | 0.54578876 |
| GAGGGGLTAGAN | ASSLGQKYAEQF | 0.5665604 |
| FVDALVNM | ASSLGQKYAEQF | 0.43644074 |
| FLTVAPEL | ASSLGQKYAEQF | 0.5837527 |
| FETLTAL | ASSLGQKYAEQF | 0.46326292 |
| DVINPIEL | ASSLGQKYAEQF | 0.641542 |
| APVSGAPV | ASSLGQKYAEQF | 0.553403 |
| ALVGGLAW | ASSLGQKYAEQF | 0.57555425 |
| ALAAAAATAI | ASSLGQKYAEQF | 0.650739 |
| AFGLGLLL | ASSLGQKYAEQF | 0.5905791 |
| AFFAAASL | ASSLGQKYAEQF | 0.57671154 |
| AAAALPAFSPP | ASSLGQKYAEQF | 0.5902978 |
| VVLAPAL | ASSLGQKYAEQF | 0.46874368 |
| TIGHVDHGKTTL | ASSLGQKYAEQF | 0.5552486 |
| LISNASDALDK | ASSLGQKYAEQF | 0.58371145 |
| LISNASDAL | ASSLGQKYAEQF | 0.6491003 |
| ISNASDALDK | ASSLGQKYAEQF | 0.5702359 |
| NQAETLVYQTE | ASSLGQKYAEQF | 0.47218883 |
| TGGTGGTAGAG | ASSLGQKYAEQF | 0.6061538 |
| TGSTAGGAGGA | ASSLGQKYAEQF | 0.6435826 |
| DPLAVPVAAGPV | ASSLGQKYAEQF | 0.65062475 |
| GGTGGTPGNGGN | ASSLGQKYAEQF | 0.5208676 |
| PAGLAGVDPATL | ASSLGQKYAEQF | 0.5886581 |
| GNNLVVEAPHAGASV | ASSLGQKYAEQF | 0.57688475 |
| KVTRIMSASTLEQPA | ASSLGQKYAEQF | 0.57477355 |
| LGFALPIALPRRTVV | ASSLGQKYAEQF | 0.55921423 |
| TRLGNLTARKLVVLV | ASSLGQKYAEQF | 0.51033944 |
| WSQQRRLIYFPSAGP | ASSLGQKYAEQF | 0.6197007 |
| GPNGNPDPMAAAVDIR | ASSLGQKYAEQF | 0.64787245 |
| TQRCVSVADLRAAASR | ASSLGQKYAEQF | 0.5815904 |
| QFVLDVAEHPVEVLTETV | ASSLGQKYAEQF | 0.4335819 |
| GGAGGTGGAGGNGGTQAGDG | ASSLGQKYAEQF | 0.5830913 |
| PLLATLRAIAVDVDAKPAQV | ASSLGQKYAEQF | 0.5249284 |
| APKFRTGGYMIDFWGVGYQVAK | ASSLGQKYAEQF | 0.5280305 |
| GAGGAGGAGGAAGGPGGTGGTG | ASSLGQKYAEQF | 0.5993526 |
| IGALGLLATDVAPAGSVAFGVA | ASSLGQKYAEQF | 0.40631944 |
| AGGAGGIGGNGGAGGAGGVLMG | ASSLGQKYAEQF | 0.59288347 |
| ADELIGLL | ASSLGQKYAEQF | 0.6537482 |
| ADRALKA | ASSLGQKYAEQF | 0.5519395 |
| AGAVGQVGGAGGA | ASSLGQKYAEQF | 0.62064993 |
| AGGGGGLGGAP | ASSLGQKYAEQF | 0.6360954 |
| AGGLLYGNGGAG | ASSLGQKYAEQF | 0.6080522 |

| | | |
|---|---|---|
| AGLESIGL | ASSLGQKYAEQF | 0.62592983 |
| AGLKTGII | ASSLGQKYAEQF | 0.6157986 |
| ASLAALLL | ASSLGQKYAEQF | 0.6178133 |
| ASVINISSI | ASSLGQKYAEQF | 0.6009872 |
| ATPPGPA | ASSLGQKYAEQF | 0.6148488 |
| ATVASALV | ASSLGQKYAEQF | 0.5875726 |
| ATVASLIL | ASSLGQKYAEQF | 0.5827712 |
| AVNGPVLIRH | ASSLGQKYAEQF | 0.53363323 |
| DGAAGAVGI | ASSLGQKYAEQF | 0.62527454 |
| DPLLHGI | ASSLGQKYAEQF | 0.57837516 |
| ELGAALGAL | ASSLGQKYAEQF | 0.6287209 |
| ELGRTQAV | ASSLGQKYAEQF | 0.42121446 |
| FVAGGYLS | ASSLGQKYAEQF | 0.5409602 |
| GASGGAGGAGGW | ASSLGQKYAEQF | 0.6237687 |
| GGAGAGGDGGAGGI | ASSLGQKYAEQF | 0.33888268 |
| GTGHNGSQPGMGGNG | ASSLGQKYAEQF | 0.3836091 |
| GYAAASASAS | ASSLGQKYAEQF | 0.57313746 |
| HQVDVAAVQQG | ASSLGQKYAEQF | 0.6381619 |
| IGAAIAHT | ASSLGQKYAEQF | 0.34526694 |
| IGMIEVAQL | ASSLGQKYAEQF | 0.5464395 |
| IRLAPEHTSAMD | ASSLGQKYAEQF | 0.47244632 |
| IYNVSVL | ASSLGQKYAEQF | 0.6449115 |
| KQGFLGM | ASSLGQKYAEQF | 0.60535026 |
| LESVITQL | ASSLGQKYAEQF | 0.422627 |
| LGLGALVVV | ASSLGQKYAEQF | 0.6166986 |
| LIGNGGNGGSGGI | ASSLGQKYAEQF | 0.4212453 |
| LNAINAPTL | ASSLGQKYAEQF | 0.5674526 |
| LQLLNTGV | ASSLGQKYAEQF | 0.63343483 |
| LQTSIEF | ASSLGQKYAEQF | 0.63681024 |
| MSFAVSEL | ASSLGQKYAEQF | 0.4676085 |
| NTGALGL | ASSLGQKYAEQF | 0.62585706 |
| PPGAGGAGGAGTT | ASSLGQKYAEQF | 0.5319288 |
| QLVGLGI | ASSLGQKYAEQF | 0.65536153 |
| RFLEQVHQL | ASSLGQKYAEQF | 0.5755217 |
| RGVGYVLR | ASSLGQKYAEQF | 0.41288346 |
| SATGVFAGVI | ASSLGQKYAEQF | 0.6287063 |
| SLPAITLP | ASSLGQKYAEQF | 0.60746026 |
| STALAFTV | ASSLGQKYAEQF | 0.6139359 |
| SVLTVGLGK | ASSLGQKYAEQF | 0.60290515 |
| TALAGAGSGI | ASSLGQKYAEQF | 0.5682608 |
| TGLPFTGI | ASSLGQKYAEQF | 0.55309683 |
| VAEHVTV | ASSLGQKYAEQF | 0.629581 |
| VDIVLNSL | ASSLGQKYAEQF | 0.6244966 |
| VLTGGGAVL | ASSLGQKYAEQF | 0.54785883 |
| VNVGLAYGI | ASSLGQKYAEQF | 0.6460306 |
| VSSAAVGAV | ASSLGQKYAEQF | 0.5978805 |
| VTSVYEAGV | ASSLGQKYAEQF | 0.50729483 |
| WVALDRGA | ASSLGQKYAEQF | 0.6489122 |
| YLVGSVVI | ASSLGQKYAEQF | 0.56372136 |
| GPPGTGKTLL | ASSLGQKYAEQF | 0.623918 |
| GVGMTKF | ASSLGQKYAEQF | 0.6229111 |

| | | |
|---|---|---|
| IGIGGSDL | ASSLGQKYAEQF | 0.63939595 |
| IGTLLAEL | ASSLGQKYAEQF | 0.5021341 |
| IYTSGTT | ASSLGQKYAEQF | 0.6024234 |
| LSFPAGV | ASSLGQKYAEQF | 0.5957501 |
| LYTSGTT | ASSLGQKYAEQF | 0.62746775 |
| VGAFPFGF | ASSLGQKYAEQF | 0.5985314 |
| VGAGTVEF | ASSLGQKYAEQF | 0.61024165 |
| VGKSTLLL | ASSLGQKYAEQF | 0.6183648 |
| AALAHAGGGIV | ASSLGQKYAEQF | 0.64583504 |
| AALVAVSLL | ASSLGQKYAEQF | 0.5920732 |
| AAVSRAVAAL | ASSLGQKYAEQF | 0.60486215 |
| AGGSSGIGGF | ASSLGQKYAEQF | 0.6328909 |
| AGLTVANANG | ASSLGQKYAEQF | 0.48025227 |
| AGLVGAVL | ASSLGQKYAEQF | 0.6032779 |
| ALSQAFTV | ASSLGQKYAEQF | 0.62843144 |
| ALTFSVSL | ASSLGQKYAEQF | 0.64049 |
| ALTGAALVAL | ASSLGQKYAEQF | 0.5719469 |
| ASFSIGAGVL | ASSLGQKYAEQF | 0.636221 |
| DGSVYKI | ASSLGQKYAEQF | 0.5558453 |
| EQLAEIV | ASSLGQKYAEQF | 0.6234388 |
| FAVSELTVM | ASSLGQKYAEQF | 0.558154 |
| GAAPLAAGL | ASSLGQKYAEQF | 0.60371614 |
| GADGGDGGSGGAGG | ASSLGQKYAEQF | 0.56439894 |
| GFGLDTGGAGG | ASSLGQKYAEQF | 0.62083435 |
| GLASGGLL | ASSLGQKYAEQF | 0.62366784 |
| GRIAGAAVLL | ASSLGQKYAEQF | 0.63048685 |
| GSAGSLNTSF | ASSLGQKYAEQF | 0.40121782 |
| GTGQIGI | ASSLGQKYAEQF | 0.61427015 |
| GTLIALIL | ASSLGQKYAEQF | 0.6353929 |
| IAATMGFVVG | ASSLGQKYAEQF | 0.63956386 |
| IFAQALL | ASSLGQKYAEQF | 0.5936009 |
| IGAAVALGF | ASSLGQKYAEQF | 0.62511873 |
| IGLTPLAL | ASSLGQKYAEQF | 0.58515084 |
| ILGSNTSTL | ASSLGQKYAEQF | 0.57232386 |
| IVFSLTVL | ASSLGQKYAEQF | 0.48649216 |
| LASKATGF | ASSLGQKYAEQF | 0.5309645 |
| LASVALIL | ASSLGQKYAEQF | 0.5620471 |
| LFAVGLTL | ASSLGQKYAEQF | 0.5033302 |
| LFNGGIGTYI | ASSLGQKYAEQF | 0.6242041 |
| LTLGAGII | ASSLGQKYAEQF | 0.6371122 |
| LVAAALGDI | ASSLGQKYAEQF | 0.6438352 |
| NTAFVQLGI | ASSLGQKYAEQF | 0.6330416 |
| QLFSGAEL | ASSLGQKYAEQF | 0.636391 |
| SAATFITL | ASSLGQKYAEQF | 0.6336999 |
| SNYGLANLGV | ASSLGQKYAEQF | 0.56140804 |
| SYGGAYAVM | ASSLGQKYAEQF | 0.5950101 |
| TTGVGGLTLG | ASSLGQKYAEQF | 0.4669095 |
| TTQTIYI | ASSLGQKYAEQF | 0.5531186 |
| VAAGAGLGAVI | ASSLGQKYAEQF | 0.56416774 |
| VAAVVTLL | ASSLGQKYAEQF | 0.5967579 |
| VGFLGGLI | ASSLGQKYAEQF | 0.34007305 |

| | | |
|---|---|---|
| VGVIPILL | ASSLGQKYAEQF | 0.5780145 |
| VIFASSLI | ASSLGQKYAEQF | 0.6499606 |
| VKGGDGGAAAG | ASSLGQKYAEQF | 0.58251923 |
| VTKTFTNI | ASSLGQKYAEQF | 0.6200642 |
| VVAAAILL | ASSLGQKYAEQF | 0.6018387 |
| YAALDVEL | ASSLGQKYAEQF | 0.549773 |
| LASVALLL | ASSLGQKYAEQF | 0.62118053 |
| LDAVHRV | ASSLGQKYAEQF | 0.5519289 |
| QGLSSSIF | ASSLGQKYAEQF | 0.6244356 |
| RGTFANIRL | ASSLGQKYAEQF | 0.64788693 |
| RQLWWGHRI | ASSLGQKYAEQF | 0.6086529 |
| YGPPGTGKTLL | ASSLGQKYAEQF | 0.57952195 |
| AALVGAPANVA | ASSLGQKYAEQF | 0.60228735 |
| AATFRRTE | ASSLGQKYAEQF | 0.45626116 |
| AGALGGGI | ASSLGQKYAEQF | 0.6033753 |
| AGGAGGAGGIGAG | ASSLGQKYAEQF | 0.5118196 |
| AGSGALGL | ASSLGQKYAEQF | 0.57256526 |
| AQIADQI | ASSLGQKYAEQF | 0.5037086 |
| ASAVAVTYL | ASSLGQKYAEQF | 0.6165251 |
| ASTVLGGTGG | ASSLGQKYAEQF | 0.60932577 |
| AVGGEPAK | ASSLGQKYAEQF | 0.6088561 |
| CEQGTFGGTC | ASSLGQKYAEQF | 0.4979052 |
| DEGKAPAG | ASSLGQKYAEQF | 0.6063164 |
| DTVVFPAL | ASSLGQKYAEQF | 0.60283405 |
| FEGITMAG | ASSLGQKYAEQF | 0.5163738 |
| FPLVVQD | ASSLGQKYAEQF | 0.6294243 |
| GAFTTPQ | ASSLGQKYAEQF | 0.5774709 |
| GFGGFGTAGGVGGA | ASSLGQKYAEQF | 0.5725953 |
| GFLAQFL | ASSLGQKYAEQF | 0.63953626 |
| GGAGGAGGNAGVA | ASSLGQKYAEQF | 0.6103109 |
| GGVGTAGAGGAGGAGG | ASSLGQKYAEQF | 0.42267573 |
| GNGGGGDAGNAGSGGN | ASSLGQKYAEQF | 0.5665123 |
| GTGGAGGAGNDAGS | ASSLGQKYAEQF | 0.58345157 |
| GTTNTGWFNS | ASSLGQKYAEQF | 0.5782696 |
| IFMGPLFTL | ASSLGQKYAEQF | 0.3508569 |
| ILAHAIEAI | ASSLGQKYAEQF | 0.6289188 |
| IRSAIREAL | ASSLGQKYAEQF | 0.5124363 |
| ITFAGGG | ASSLGQKYAEQF | 0.51981604 |
| ITGMSGHQEV | ASSLGQKYAEQF | 0.5400971 |
| IVAGLACTL | ASSLGQKYAEQF | 0.61926794 |
| IVGGGLAGA | ASSLGQKYAEQF | 0.6095997 |
| IVTGVLNI | ASSLGQKYAEQF | 0.6130353 |
| LAQQGCI | ASSLGQKYAEQF | 0.60522544 |
| LCLHVEN | ASSLGQKYAEQF | 0.63817096 |
| LDAISDGS | ASSLGQKYAEQF | 0.6431856 |
| LGGFAAML | ASSLGQKYAEQF | 0.63341236 |
| LGGKNPMI | ASSLGQKYAEQF | 0.5890902 |
| LGPKGVAAIL | ASSLGQKYAEQF | 0.6204303 |
| LMVGVSI | ASSLGQKYAEQF | 0.5931602 |
| LVGAAFVL | ASSLGQKYAEQF | 0.45061803 |
| LVLAVAPL | ASSLGQKYAEQF | 0.4452613 |

| | | |
|---|---|---|
| MAAIAVIDAA | ASSLGQKYAEQF | 0.5782565 |
| NNGINAELQ | ASSLGQKYAEQF | 0.4564669 |
| RGEPLVLVHG | ASSLGQKYAEQF | 0.6035573 |
| RGGAGGAAGLI | ASSLGQKYAEQF | 0.64344585 |
| RSYRMHPAV | ASSLGQKYAEQF | 0.5931522 |
| SAAHLLAGQLA | ASSLGQKYAEQF | 0.42007014 |
| SAAVGSY | ASSLGQKYAEQF | 0.56553113 |
| SPHAVLHRI | ASSLGQKYAEQF | 0.5349045 |
| SQIAKVL | ASSLGQKYAEQF | 0.61762166 |
| TALGHLVP | ASSLGQKYAEQF | 0.6373923 |
| TAPTTSGAA | ASSLGQKYAEQF | 0.6476939 |
| TDGDDGNGGSPGT | ASSLGQKYAEQF | 0.6369997 |
| TDGLGEN | ASSLGQKYAEQF | 0.57139015 |
| TGGPPLPGQA | ASSLGQKYAEQF | 0.6135317 |
| TIPAITI | ASSLGQKYAEQF | 0.5278616 |
| TIPGFTI | ASSLGQKYAEQF | 0.601578 |
| TSIASGALA | ASSLGQKYAEQF | 0.59869623 |
| TVTLPGAAI | ASSLGQKYAEQF | 0.60457194 |
| VASGLSAALP | ASSLGQKYAEQF | 0.60928476 |
| VGAGVIGI | ASSLGQKYAEQF | 0.58747816 |
| VGPKRFGV | ASSLGQKYAEQF | 0.4515478 |
| VNVAFPD | ASSLGQKYAEQF | 0.6207997 |
| VSMTNSGVS | ASSLGQKYAEQF | 0.580793 |
| VTDVLGDLSV | ASSLGQKYAEQF | 0.42022663 |
| VVVVGAGFAGL | ASSLGQKYAEQF | 0.5715401 |
| VYTVGTVV | ASSLGQKYAEQF | 0.64703745 |
| WTTFSGP | ASSLGQKYAEQF | 0.64319634 |
| YDELVEAF | ASSLGQKYAEQF | 0.52051866 |
| YGTGGAGGNGG | ASSLGQKYAEQF | 0.5320146 |
| YTNVMAV | ASSLGQKYAEQF | 0.588277 |
| FLEQVHQL | ASSLGQKYAEQF | 0.44965255 |
| GLSLLAVAV | ASSLGQKYAEQF | 0.6056136 |
| LEQVHQL | ASSLGQKYAEQF | 0.5774381 |
| LNAAGAI | ASSLGQKYAEQF | 0.5906633 |
| PAVHASL | ASSLGQKYAEQF | 0.54732394 |
| TELEDDA | ASSLGQKYAEQF | 0.61258745 |
| TPPGPPN | ASSLGQKYAEQF | 0.591969 |
| TPPPPPPP | ASSLGQKYAEQF | 0.6195304 |
| TQSPALI | ASGDAQKDTQY | 0.34807062 |
| EFAAAFPQ | ASGDAQKDTQY | 0.634974 |
| LLSGNLDVL | ASGDAQKDTQY | 0.5221251 |
| GPANLGNYNI | ASGDAQKDTQY | 0.5277856 |
| TLGPTLP | ASGDAQKDTQY | 0.39005703 |
| VYAYSSCCK | ASGDAQKDTQY | 0.58925915 |
| APAPAPAPNI | ASGDAQKDTQY | 0.6261889 |
| PGGPVPIP | ASGDAQKDTQY | 0.63171697 |
| FLTVALEP | ASGDAQKDTQY | 0.55486465 |
| SQYFGANL | ASGDAQKDTQY | 0.62878966 |
| GGVKPAA | ASGDAQKDTQY | 0.6491283 |
| GGTGGAGGAGGW | ASGDAQKDTQY | 0.5595757 |
| GVGGVAGLGGAGG | ASGDAQKDTQY | 0.44638556 |

| | | |
|---|---|---|
| VLLVLATTI | ASGDAQKDTQY | 0.6194823 |
| IGIFAAML | ASGDAQKDTQY | 0.5656184 |
| AVGDLID | ASGDAQKDTQY | 0.6340227 |
| NISPLPL | ASGDAQKDTQY | 0.62774426 |
| VTAVFVAI | ASGDAQKDTQY | 0.4452513 |
| SGDDNDAAL | ASGDAQKDTQY | 0.630716 |
| AVAALAHS | ASGDAQKDTQY | 0.6384535 |
| GGVTVMATP | ASGDAQKDTQY | 0.5214187 |
| AGLAANVRD | ASGDAQKDTQY | 0.4855176 |
| TNRPDLIDPALLRPG | ASGDAQKDTQY | 0.54108304 |
| LANLGVTAL | ASGDAQKDTQY | 0.4771225 |
| TGERIEL | ASGDAQKDTQY | 0.63808864 |
| NGEELAA | ASGDAQKDTQY | 0.5529344 |
| VLSLVWG | ASGDAQKDTQY | 0.5985929 |
| IGYALWL | ASGDAQKDTQY | 0.57481325 |
| LSLLLAPL | ASGDAQKDTQY | 0.5976429 |
| TLDEALK | ASGDAQKDTQY | 0.57886344 |
| TDGAGTAEGG | ASGDAQKDTQY | 0.59704316 |
| NAPGGAGGNGG | ASGDAQKDTQY | 0.62862694 |
| VSVFMGMGGNF | ASGDAQKDTQY | 0.61455786 |
| LASYLAAL | ASGDAQKDTQY | 0.5790819 |
| LVGGTPAL | ASGDAQKDTQY | 0.58886206 |
| VRNADVL | ASGDAQKDTQY | 0.6415657 |
| TCSSPFLVSAM | ASGDAQKDTQY | 0.5434513 |
| VHITDAE | ASGDAQKDTQY | 0.5557556 |
| EYAAPSF | ASGDAQKDTQY | 0.6148497 |
| VVDAFHGGL | ASGDAQKDTQY | 0.512624 |
| ASVVDPQ | ASGDAQKDTQY | 0.6190626 |
| SYNEFGGLS | ASGDAQKDTQY | 0.6169013 |
| DPDGSVY | ASGDAQKDTQY | 0.60095054 |
| MVSTIHATE | ASGDAQKDTQY | 0.5847169 |
| LAAGVADPV | ASGDAQKDTQY | 0.64797527 |
| TKEQIAA | ASGDAQKDTQY | 0.6174056 |
| SLASPAAL | ASGDAQKDTQY | 0.57267445 |
| IGLVFYGV | ASGDAQKDTQY | 0.61906105 |
| FLLDASNL | ASGDAQKDTQY | 0.4824322 |
| YQVGNFP | ASGDAQKDTQY | 0.635425 |
| ESLLSQST | ASGDAQKDTQY | 0.62175554 |
| LFTVGFAAVG | ASGDAQKDTQY | 0.4841985 |
| ATPPTPT | ASGDAQKDTQY | 0.55902267 |
| TDLATLG | ASGDAQKDTQY | 0.643558 |
| SAGGTGR | ASGDAQKDTQY | 0.63210255 |
| GFSNSGS | ASGDAQKDTQY | 0.4907817 |
| GGNGGSGGTSV | ASGDAQKDTQY | 0.5432053 |
| GTSVATGGAGN | ASGDAQKDTQY | 0.4747222 |
| NGGNGGSGGTS | ASGDAQKDTQY | 0.5130643 |
| AVVKFPH | ASGDAQKDTQY | 0.460661 |
| TPPPPPP | ASGDAQKDTQY | 0.61285543 |
| AAPAAAAAPA | ASGDAQKDTQY | 0.5643527 |
| IALNEDL | ASGDAQKDTQY | 0.6164505 |
| IVNVSSM | ASGDAQKDTQY | 0.59954536 |

| | | |
|---|---|---|
| TGIPPAPR | ASGDAQKDTQY | 0.6407093 |
| FDLGGGTF | ASGDAQKDTQY | 0.52383 |
| FDLGGGTFDVS | ASGDAQKDTQY | 0.6060453 |
| NEPTAAA | ASGDAQKDTQY | 0.440974 |
| NEPTAAALA | ASGDAQKDTQY | 0.5948886 |
| GSIINISSI | ASGDAQKDTQY | 0.5282586 |
| VAPGNAGTA | ASGDAQKDTQY | 0.6123281 |
| TGGGASLE | ASGDAQKDTQY | 0.41356125 |
| VAVYNHYKRI | ASGDAQKDTQY | 0.64733124 |
| TDPAPATT | ASGDAQKDTQY | 0.5676578 |
| TNRPDILDPALLRPG | ASGDAQKDTQY | 0.6249448 |
| LTTTTTS | ASGDAQKDTQY | 0.5862542 |
| PTGSGKTL | ASGDAQKDTQY | 0.55885017 |
| TGSGKTL | ASGDAQKDTQY | 0.5529976 |
| TIGHVDHGK | ASGDAQKDTQY | 0.6207804 |
| TIGHVDHGKTT | ASGDAQKDTQY | 0.5050471 |
| GHVDHGKTT | ASGDAQKDTQY | 0.37381834 |
| AQFEHTLL | ASGDAQKDTQY | 0.56426203 |
| IVNQDEL | ASGDAQKDTQY | 0.47383764 |
| TNGSQFF | ASGDAQKDTQY | 0.6395952 |
| TGKGIVE | ASGDAQKDTQY | 0.63736135 |
| LISNASDA | ASGDAQKDTQY | 0.58373994 |
| AELRTLGELTKIAKAHGAQ | ASGDAQKDTQY | 0.62720054 |
| APNAVAAAEPSW | ASGDAQKDTQY | 0.33317226 |
| APQVAGAATAVL | ASGDAQKDTQY | 0.58950603 |
| ARRDPAAAAAILPTDA | ASGDAQKDTQY | 0.6450947 |
| DAAARDVLAEAGLAEYF | ASGDAQKDTQY | 0.6080869 |
| DDQIVLSAKANLAKVEQ | ASGDAQKDTQY | 0.5859911 |
| DELLPDTGEDSDEDGH | ASGDAQKDTQY | 0.6370998 |
| ELPEAPANRATVPQD | ASGDAQKDTQY | 0.5737881 |
| FGQSKIERVTGIG | ASGDAQKDTQY | 0.5885997 |
| GAGGTGSTAGGAGGAGGAGGLY | ASGDAQKDTQY | 0.6026641 |
| GHGGTGAAGGNGGTGGAGGSGI | ASGDAQKDTQY | 0.63121593 |
| GNGGNGGAGGEGGPGGAGGAGA | ASGDAQKDTQY | 0.6156668 |
| GSTAGNGGNGGNATNGTIAGQP | ASGDAQKDTQY | 0.5152835 |
| GTGGDGGNGGNGGTGG | ASGDAQKDTQY | 0.6165752 |
| GVRRPAGIPADAPV | ASGDAQKDTQY | 0.6538263 |
| GYRQVTDAHLLTLARR | ASGDAQKDTQY | 0.5542373 |
| IAKMGQDGHDRG | ASGDAQKDTQY | 0.63271296 |
| KAEPSIEIQYT | ASGDAQKDTQY | 0.6034845 |
| KPVRIRMAADEDTVAVT | ASGDAQKDTQY | 0.6157354 |
| MPPRIAGMRLLVIKPEP | ASGDAQKDTQY | 0.5102239 |
| MKFYAVALFRSRGVPPTR | ASGDAQKDTQY | 0.5193928 |
| NAGGFGILGAVAHSP | ASGDAQKDTQY | 0.6148418 |
| NEGREITLAMAEM | ASGDAQKDTQY | 0.51574314 |
| NELVEWSHPLGHRWRWE | ASGDAQKDTQY | 0.58623105 |
| NLGAGGTGGNGGNPG | ASGDAQKDTQY | 0.55600524 |
| NTEWPKWETVMAADPVD | ASGDAQKDTQY | 0.6588121 |
| PASAAPAETQP | ASGDAQKDTQY | 0.61627865 |
| QQSSDGKNGVRAAEITGE | ASGDAQKDTQY | 0.6147014 |
| RAPDAVTTSWWKE | ASGDAQKDTQY | 0.52854306 |

| | | |
|---|---|---|
| SGFGNVGAHTSGWFNQ | ASGDAQKDTQY | 0.60153747 |
| SGVAGAVGQAASVGGLKV | ASGDAQKDTQY | 0.6144313 |
| SNGGAGTGSAGGNGGT | ASGDAQKDTQY | 0.47342426 |
| TGGAGGEGGAAG | ASGDAQKDTQY | 0.64058465 |
| TGGDGGAAGAGGAPGHG | ASGDAQKDTQY | 0.53224015 |
| TMPPLLAAIANAGRHGV | ASGDAQKDTQY | 0.5904761 |
| TSSTAGNSGGA | ASGDAQKDTQY | 0.6332102 |
| VPATVPVGILC | ASGDAQKDTQY | 0.63560486 |
| VQRPDSGGVVLTGRIS | ASGDAQKDTQY | 0.5693207 |
| WTPNPVPATKGVPAGLVQAT | ASGDAQKDTQY | 0.60587084 |
| YALKRALAKFSDRFDVVII | ASGDAQKDTQY | 0.5211803 |
| YLIGSVIV | ASGDAQKDTQY | 0.61913997 |
| YGQLVAVID | ASGDAQKDTQY | 0.6260001 |
| VTVGGMTLPA | ASGDAQKDTQY | 0.59958696 |
| VLALPAV | ASGDAQKDTQY | 0.64617854 |
| VIPPTLNYET | ASGDAQKDTQY | 0.62856245 |
| VGMGGMHPSSG | ASGDAQKDTQY | 0.4065723 |
| VGLAGGAATA | ASGDAQKDTQY | 0.5910759 |
| VGGAGGAGGSGP | ASGDAQKDTQY | 0.6368463 |
| VFAAFMV | ASGDAQKDTQY | 0.6333416 |
| TLYDKLR | ASGDAQKDTQY | 0.65108573 |
| TLAVFTVL | ASGDAQKDTQY | 0.6243236 |
| TGSVQPARV | ASGDAQKDTQY | 0.5615722 |
| TGGGAGSGGT | ASGDAQKDTQY | 0.63898784 |
| SHLETSL | ASGDAQKDTQY | 0.6559168 |
| SFTTGMT | ASGDAQKDTQY | 0.62107635 |
| QVEVAKRMGVRQPTVS | ASGDAQKDTQY | 0.54966265 |
| QGSGGPALAM | ASGDAQKDTQY | 0.5318238 |
| MLAPSNP | ASGDAQKDTQY | 0.6218545 |
| LVNNAAI | ASGDAQKDTQY | 0.62332773 |
| LTGEINTLA | ASGDAQKDTQY | 0.47325754 |
| LSIFGPV | ASGDAQKDTQY | 0.59856427 |
| LLALLANL | ASGDAQKDTQY | 0.60615945 |
| LISNASDALDKLR | ASGDAQKDTQY | 0.40520105 |
| LINEAVP | ASGDAQKDTQY | 0.626659 |
| LGYAFSGL | ASGDAQKDTQY | 0.6330211 |
| LDFVAVAL | ASGDAQKDTQY | 0.59286773 |
| LAQPGAE | ASGDAQKDTQY | 0.5156413 |
| ISNLGAVTGF | ASGDAQKDTQY | 0.63699055 |
| HQEDPGRI | ASGDAQKDTQY | 0.6488027 |
| GTPGEPTP | ASGDAQKDTQY | 0.635043 |
| GFFNSGS | ASGDAQKDTQY | 0.47439617 |
| GAGGGGLTAGAN | ASGDAQKDTQY | 0.3438042 |
| FVDALVNM | ASGDAQKDTQY | 0.59548265 |
| FLTVAPEL | ASGDAQKDTQY | 0.6266748 |
| FETLTAL | ASGDAQKDTQY | 0.6035863 |
| DVINPIEL | ASGDAQKDTQY | 0.5422994 |
| APVSGAPV | ASGDAQKDTQY | 0.59893477 |
| ALVGGLAW | ASGDAQKDTQY | 0.6297741 |
| ALAAAAATAI | ASGDAQKDTQY | 0.47736266 |
| AFGLGLLL | ASGDAQKDTQY | 0.47340256 |

| | | |
|---|---|---|
| AFFAAASL | ASGDAQKDTQY | 0.6231321 |
| AAAALPAFSPP | ASGDAQKDTQY | 0.6172757 |
| VVLAPAL | ASGDAQKDTQY | 0.5279367 |
| TIGHVDHGKTTL | ASGDAQKDTQY | 0.5415709 |
| LISNASDALDK | ASGDAQKDTQY | 0.60294306 |
| LISNASDAL | ASGDAQKDTQY | 0.5815706 |
| ISNASDALDK | ASGDAQKDTQY | 0.60378003 |
| NQAETLVYQTE | ASGDAQKDTQY | 0.47419325 |
| TGGTGGTAGAG | ASGDAQKDTQY | 0.6410717 |
| TGSTAGGAGGA | ASGDAQKDTQY | 0.4337331 |
| DPLAVPVAAGPV | ASGDAQKDTQY | 0.6204275 |
| GGTGGTPGNGGN | ASGDAQKDTQY | 0.6141871 |
| PAGLAGVDPATL | ASGDAQKDTQY | 0.6184749 |
| GNNLVVEAPHAGASV | ASGDAQKDTQY | 0.5584457 |
| KVTRIMSASTLEQPA | ASGDAQKDTQY | 0.51924443 |
| LGFALPIALPRRTVV | ASGDAQKDTQY | 0.47456056 |
| TRLGNLTARKLVVLV | ASGDAQKDTQY | 0.45505136 |
| WSQQRRLIYFPSAGP | ASGDAQKDTQY | 0.6415715 |
| GPNGNPDPMAAAVDIR | ASGDAQKDTQY | 0.58225507 |
| TQRCVSVADLRAAASR | ASGDAQKDTQY | 0.6348965 |
| QFVLDVAEHPVEVLTETV | ASGDAQKDTQY | 0.61078376 |
| GGAGGTGGAGGNGGTQAGDG | ASGDAQKDTQY | 0.55434835 |
| PLLATLRAIAVDVDAKPAQV | ASGDAQKDTQY | 0.6123098 |
| APKFRTGGYMIDFWGVGYQVAK | ASGDAQKDTQY | 0.51571494 |
| GAGGAGGAGGAAGGPGGTGGTG | ASGDAQKDTQY | 0.5499823 |
| IGALGLLATDVAPAGSVAFGVA | ASGDAQKDTQY | 0.61120015 |
| AGGAGGIGGNGGAGGAGGVLMG | ASGDAQKDTQY | 0.6168643 |
| ADELIGLL | ASGDAQKDTQY | 0.5060903 |
| ADRALKA | ASGDAQKDTQY | 0.61057454 |
| AGAVGQVGGAGGA | ASGDAQKDTQY | 0.6070585 |
| AGGGGGLGGAP | ASGDAQKDTQY | 0.54042304 |
| AGGLLYGNGGAG | ASGDAQKDTQY | 0.6090163 |
| AGLESIGL | ASGDAQKDTQY | 0.63822013 |
| AGLKTGII | ASGDAQKDTQY | 0.6136495 |
| ASLAALLL | ASGDAQKDTQY | 0.47530568 |
| ASVINISSI | ASGDAQKDTQY | 0.62859404 |
| ATPPGPA | ASGDAQKDTQY | 0.51220846 |
| ATVASALV | ASGDAQKDTQY | 0.5940404 |
| ATVASLIL | ASGDAQKDTQY | 0.6064381 |
| AVNGPVLIRH | ASGDAQKDTQY | 0.57907665 |
| DGAAGAVGI | ASGDAQKDTQY | 0.6387201 |
| DPLLHGI | ASGDAQKDTQY | 0.5536483 |
| ELGAALGAL | ASGDAQKDTQY | 0.53076595 |
| ELGRTQAV | ASGDAQKDTQY | 0.5716648 |
| FVAGGYLS | ASGDAQKDTQY | 0.6205809 |
| GASGGAGGAGGW | ASGDAQKDTQY | 0.61828154 |
| GGAGAGGDGGAGGI | ASGDAQKDTQY | 0.5005779 |
| GTGHNGSQPGMGGNG | ASGDAQKDTQY | 0.59970105 |
| GYAAASASAS | ASGDAQKDTQY | 0.5768512 |
| HQVDVAAVQQG | ASGDAQKDTQY | 0.52933526 |
| IGAAIAHT | ASGDAQKDTQY | 0.5760957 |

| | | |
|---|---|---|
| IGMIEVAQL | ASGDAQKDTQY | 0.59129906 |
| IRLAPEHTSAMD | ASGDAQKDTQY | 0.46966508 |
| IYNVSVL | ASGDAQKDTQY | 0.62710243 |
| KQGFLGM | ASGDAQKDTQY | 0.57758987 |
| LESVITQL | ASGDAQKDTQY | 0.63144743 |
| LGLGALVVV | ASGDAQKDTQY | 0.6396894 |
| LIGNGGNGGSGGI | ASGDAQKDTQY | 0.50465494 |
| LNAINAPTL | ASGDAQKDTQY | 0.61283565 |
| LQLLNTGV | ASGDAQKDTQY | 0.6082151 |
| LQTSIEF | ASGDAQKDTQY | 0.6127233 |
| MSFAVSEL | ASGDAQKDTQY | 0.6041224 |
| NTGALGL | ASGDAQKDTQY | 0.54094374 |
| PPGAGGAGGAGTT | ASGDAQKDTQY | 0.634203 |
| QLVGLGI | ASGDAQKDTQY | 0.63561845 |
| RFLEQVHQL | ASGDAQKDTQY | 0.61199665 |
| RGVGYVLR | ASGDAQKDTQY | 0.6294838 |
| SATGVFAGVI | ASGDAQKDTQY | 0.6180552 |
| SLPAITLP | ASGDAQKDTQY | 0.61818254 |
| STALAFTV | ASGDAQKDTQY | 0.5556957 |
| SVLTVGLGK | ASGDAQKDTQY | 0.47070912 |
| TALAGAGSGI | ASGDAQKDTQY | 0.60144854 |
| TGLPFTGI | ASGDAQKDTQY | 0.5813985 |
| VAEHVTV | ASGDAQKDTQY | 0.5449687 |
| VDIVLNSL | ASGDAQKDTQY | 0.43159637 |
| VLTGGGAVL | ASGDAQKDTQY | 0.6332192 |
| VNVGLAYGI | ASGDAQKDTQY | 0.65409493 |
| VSSAAVGAV | ASGDAQKDTQY | 0.5816288 |
| VTSVYEAGV | ASGDAQKDTQY | 0.6465788 |
| WVALDRGA | ASGDAQKDTQY | 0.4464758 |
| YLVGSVVI | ASGDAQKDTQY | 0.6378176 |
| GPPGTGKTLL | ASGDAQKDTQY | 0.6402254 |
| GVGMTKF | ASGDAQKDTQY | 0.511938 |
| IGIGGSDL | ASGDAQKDTQY | 0.6358202 |
| IGTLLAEL | ASGDAQKDTQY | 0.5465643 |
| IYTSGTT | ASGDAQKDTQY | 0.5377355 |
| LSFPAGV | ASGDAQKDTQY | 0.64027774 |
| LYTSGTT | ASGDAQKDTQY | 0.6004511 |
| VGAFPFGF | ASGDAQKDTQY | 0.62341684 |
| VGAGTVEF | ASGDAQKDTQY | 0.57575035 |
| VGKSTLLL | ASGDAQKDTQY | 0.5312902 |
| AALAHAGGGIV | ASGDAQKDTQY | 0.6173387 |
| AALVAVSLL | ASGDAQKDTQY | 0.63410383 |
| AAVSRAVAAL | ASGDAQKDTQY | 0.55057895 |
| AGGSSGIGGF | ASGDAQKDTQY | 0.655406 |
| AGLTVANANG | ASGDAQKDTQY | 0.5392739 |
| AGLVGAVL | ASGDAQKDTQY | 0.6462339 |
| ALSQAFTV | ASGDAQKDTQY | 0.61459893 |
| ALTFSVSL | ASGDAQKDTQY | 0.57526064 |
| ALTGAALVAL | ASGDAQKDTQY | 0.5722551 |
| ASFSIGAGVL | ASGDAQKDTQY | 0.6305416 |
| DGSVYKI | ASGDAQKDTQY | 0.5755293 |

| | | |
|---|---|---|
| EQLAEIV | ASGDAQKDTQY | 0.614212 |
| FAVSELTVM | ASGDAQKDTQY | 0.6437089 |
| GAAPLAAGL | ASGDAQKDTQY | 0.25599378 |
| GADGGDGGSGGAGG | ASGDAQKDTQY | 0.7938523 |
| GFGLDTGGAGG | ASGDAQKDTQY | 0.47454178 |
| GLASGGLL | ASGDAQKDTQY | 0.42981434 |
| GRIAGAAVLL | ASGDAQKDTQY | 0.27641723 |
| GSAGSLNTSF | ASGDAQKDTQY | 0.1748531 |
| GTGQIGI | ASGDAQKDTQY | 0.37811756 |
| GTLIALIL | ASGDAQKDTQY | 0.21335554 |
| IAATMGFVVG | ASGDAQKDTQY | 0.6412358 |
| IFAQALL | ASGDAQKDTQY | 0.6297714 |
| IGAAVALGF | ASGDAQKDTQY | 0.2590718 |
| IGLTPLAL | ASGDAQKDTQY | 0.7655283 |
| ILGSNTSTL | ASGDAQKDTQY | 0.65741473 |
| IVFSLTVL | ASGDAQKDTQY | 0.18437384 |
| LASKATGF | ASGDAQKDTQY | 0.3468936 |
| LASVALIL | ASGDAQKDTQY | 0.3074773 |
| LFAVGLTL | ASGDAQKDTQY | 0.39293855 |
| LFNGGIGTYI | ASGDAQKDTQY | 0.16269402 |
| LTLGAGII | ASGDAQKDTQY | 0.53433055 |
| LVAAALGDI | ASGDAQKDTQY | 0.25928277 |
| NTAFVQLGI | ASGDAQKDTQY | 0.07953507 |
| QLFSGAEL | ASGDAQKDTQY | 0.5207229 |
| SAATFITL | ASGDAQKDTQY | 0.34812516 |
| SNYGLANLGV | ASGDAQKDTQY | 0.3176153 |
| SYGGAYAVM | ASGDAQKDTQY | 0.7037777 |
| TTGVGGLTLG | ASGDAQKDTQY | 0.12119122 |
| TTQTIYI | ASGDAQKDTQY | 0.24048519 |
| VAAGAGLGAVI | ASGDAQKDTQY | 0.69612384 |
| VAAVVTLL | ASGDAQKDTQY | 0.6979787 |
| VGFLGGLI | ASGDAQKDTQY | 0.20315969 |
| VGVIPILL | ASGDAQKDTQY | 0.5147689 |
| VIFASSLI | ASGDAQKDTQY | 0.7443608 |
| VKGGDGGAAAG | ASGDAQKDTQY | 0.4793321 |
| VTKTFTNI | ASGDAQKDTQY | 0.613633 |
| VVAAAILL | ASGDAQKDTQY | 0.65740025 |
| YAALDVEL | ASGDAQKDTQY | 0.23766631 |
| LASVALLL | ASGDAQKDTQY | 0.7097349 |
| LDAVHRV | ASGDAQKDTQY | 0.8272754 |
| QGLSSSIF | ASGDAQKDTQY | 0.35401416 |
| RGTFANIRL | ASGDAQKDTQY | 0.4175279 |
| RQLWWGHRI | ASGDAQKDTQY | 0.26156205 |
| YGPPGTGKTLL | ASGDAQKDTQY | 0.39444548 |
| AALVGAPANVA | ASGDAQKDTQY | 0.10493859 |
| AATFRRTE | ASGDAQKDTQY | 0.59385467 |
| AGALGGGI | ASGDAQKDTQY | 0.5999732 |
| AGGAGGAGGIGAG | ASGDAQKDTQY | 0.34342033 |
| AGSGALGL | ASGDAQKDTQY | 0.20013861 |
| AQIADQI | ASGDAQKDTQY | 0.15945666 |
| ASAVAVTYL | ASGDAQKDTQY | 0.24139458 |

| | | |
|---|---|---|
| ASTVLGGTGG | ASGDAQKDTQY | 0.26879463 |
| AVGGEPAK | ASGDAQKDTQY | 0.10682537 |
| CEQGTFGGTC | ASGDAQKDTQY | 0.3941365 |
| DEGKAPAG | ASGDAQKDTQY | 0.60484064 |
| DTVVFPAL | ASGDAQKDTQY | 0.6161191 |
| FEGITMAG | ASGDAQKDTQY | 0.15803926 |
| FPLVVQD | ASGDAQKDTQY | 0.8273065 |
| GAFTTPQ | ASGDAQKDTQY | 0.5450908 |
| GFGGFGTAGGVGGA | ASGDAQKDTQY | 0.6240745 |
| GFLAQFL | ASGDAQKDTQY | 0.3430005 |
| GGAGGAGGNAGVA | ASGDAQKDTQY | 0.26579136 |
| GGVGTAGAGGAGGAGG | ASGDAQKDTQY | 0.76224613 |
| GNGGGGDAGNAGSGGN | ASGDAQKDTQY | 0.38178337 |
| GTGGAGGAGNDAGS | ASGDAQKDTQY | 0.35382032 |
| GTTNTGWFNS | ASGDAQKDTQY | 0.6844009 |
| IFMGPLFTL | ASGDAQKDTQY | 0.33911535 |
| ILAHAIEAI | ASGDAQKDTQY | 0.25465554 |
| IRSAIREAL | ASGDAQKDTQY | 0.4793733 |
| ITFAGGG | ASGDAQKDTQY | 0.5131278 |
| ITGMSGHQEV | ASGDAQKDTQY | 0.2187892 |
| IVAGLACTL | ASGDAQKDTQY | 0.22241463 |
| IVGGGLAGA | ASGDAQKDTQY | 0.17583989 |
| IVTGVLNI | ASGDAQKDTQY | 0.38015372 |
| LAQQGCI | ASGDAQKDTQY | 0.6847736 |
| LCLHVEN | ASGDAQKDTQY | 0.45503503 |
| LDAISDGS | ASGDAQKDTQY | 0.27014297 |
| LGGFAAML | ASGDAQKDTQY | 0.5859575 |
| LGGKNPMI | ASGDAQKDTQY | 0.48521104 |
| LGPKGVAAIL | ASGDAQKDTQY | 0.4477484 |
| LMVGVSI | ASGDAQKDTQY | 0.62372625 |
| LVGAAFVL | ASGDAQKDTQY | 0.28815496 |
| LVLAVAPL | ASGDAQKDTQY | 0.57258713 |
| MAAIAVIDAA | ASGDAQKDTQY | 0.5865002 |
| NNGINAELQ | ASGDAQKDTQY | 0.50139445 |
| RGEPLVLVHG | ASGDAQKDTQY | 0.58572656 |
| RGGAGGAAGLI | ASGDAQKDTQY | 0.42752397 |
| RSYRMHPAV | ASGDAQKDTQY | 0.39890718 |
| SAAHLLAGQLA | ASGDAQKDTQY | 0.12979122 |
| SAAVGSY | ASGDAQKDTQY | 0.32194164 |
| SPHAVLHRI | ASGDAQKDTQY | 0.54326445 |
| SQIAKVL | ASGDAQKDTQY | 0.5199931 |
| TALGHLVP | ASGDAQKDTQY | 0.6418164 |
| TAPTTSGAA | ASGDAQKDTQY | 0.4235737 |
| TDGDDGNGGSPGT | ASGDAQKDTQY | 0.25491256 |
| TDGLGEN | ASGDAQKDTQY | 0.62215436 |
| TGGPPLPGQA | ASGDAQKDTQY | 0.7264066 |
| TIPAITI | ASGDAQKDTQY | 0.1712377 |
| TIPGFTI | ASGDAQKDTQY | 0.36698747 |
| TSIASGALA | ASGDAQKDTQY | 0.8492385 |
| TVTLPGAAI | ASGDAQKDTQY | 0.6350657 |
| VASGLSAALP | ASGDAQKDTQY | 0.355786 |

| | | |
|---|---|---|
| VGAGVIGI | ASGDAQKDTQY | 0.31591424 |
| VGPKRFGV | ASGDAQKDTQY | 0.76012754 |
| VNVAFPD | ASGDAQKDTQY | 0.2109383 |
| VSMTNSGVS | ASGDAQKDTQY | 0.8103126 |
| VTDVLGDLSV | ASGDAQKDTQY | 0.10485838 |
| VVVVGAGFAGL | ASGDAQKDTQY | 0.24398746 |
| VYTVGTVV | ASGDAQKDTQY | 0.20434318 |
| WTTFSGP | ASGDAQKDTQY | 0.12973338 |
| YDELVEAF | ASGDAQKDTQY | 0.6856248 |
| YGTGGAGGNGG | ASGDAQKDTQY | 0.2698809 |
| YTNVMAV | ASGDAQKDTQY | 0.6377715 |
| FLEQVHQL | ASGDAQKDTQY | 0.3729335 |
| GLSLLAVAV | ASGDAQKDTQY | 0.68158054 |
| LEQVHQL | ASGDAQKDTQY | 0.59292775 |
| LNAAGAI | ASGDAQKDTQY | 0.36888087 |
| PAVHASL | ASGDAQKDTQY | 0.26323226 |
| TELEDDA | ASGDAQKDTQY | 0.5947956 |
| TPPGPPN | ASGDAQKDTQY | 0.6651027 |
| TPPPPPPP | ASGDAQKDTQY | 0.41460726 |
| TQSPALI | ASSLANSGNTLY | 0.15870917 |
| EFAAAFPQ | ASSLANSGNTLY | 0.36271116 |
| LLSGNLDVL | ASSLANSGNTLY | 0.3867724 |
| GPANLGNYNI | ASSLANSGNTLY | 0.11772183 |
| TLGPTLP | ASSLANSGNTLY | 0.092185125 |
| VYAYSSCCK | ASSLANSGNTLY | 0.19925788 |
| APAPAPAPNI | ASSLANSGNTLY | 0.28865194 |
| PGGPVPIP | ASSLANSGNTLY | 0.7177249 |
| FLTVALEP | ASSLANSGNTLY | 0.70550597 |
| SQYFGANL | ASSLANSGNTLY | 0.3487776 |
| GGVKPAA | ASSLANSGNTLY | 0.6023486 |
| GGTGGAGGAGGW | ASSLANSGNTLY | 0.47120202 |
| GVGGVAGLGGAGG | ASSLANSGNTLY | 0.06901138 |
| VLLVLATTI | ASSLANSGNTLY | 0.27944022 |
| IGIFAAML | ASSLANSGNTLY | 0.18225622 |
| AVGDLID | ASSLANSGNTLY | 0.58801603 |
| NISPLPL | ASSLANSGNTLY | 0.2729652 |
| VTAVFVAI | ASSLANSGNTLY | 0.3745434 |
| SGDDNDAAL | ASSLANSGNTLY | 0.39684725 |
| AVAALAHS | ASSLANSGNTLY | 0.63179344 |
| GGVTVMATP | ASSLANSGNTLY | 0.46506706 |
| AGLAANVRD | ASSLANSGNTLY | 0.18722066 |
| TNRPDLIDPALLRPG | ASSLANSGNTLY | 0.08408444 |
| LANLGVTAL | ASSLANSGNTLY | 0.5794051 |
| TGERIEL | ASSLANSGNTLY | 0.15642184 |
| NGEELAA | ASSLANSGNTLY | 0.22139354 |
| VLSLVWG | ASSLANSGNTLY | 0.14205462 |
| IGYALWL | ASSLANSGNTLY | 0.09289713 |
| LSLLLAPL | ASSLANSGNTLY | 0.15310779 |
| TLDEALK | ASSLANSGNTLY | 0.5415111 |
| TDGAGTAEGG | ASSLANSGNTLY | 0.33728403 |
| NAPGGAGGNGG | ASSLANSGNTLY | 0.42393047 |

| | | |
|---|---|---|
| VSVFMGMGGNF | ASSLANSGNTLY | 0.8301111 |
| LASYLAAL | ASSLANSGNTLY | 0.63221055 |
| LVGGTPAL | ASSLANSGNTLY | 0.40628284 |
| VRNADVL | ASSLANSGNTLY | 0.27978507 |
| TCSSPFLVSAM | ASSLANSGNTLY | 0.23614357 |
| VHITDAE | ASSLANSGNTLY | 0.2434871 |
| EYAAPSF | ASSLANSGNTLY | 0.22501999 |
| VVDAFHGGL | ASSLANSGNTLY | 0.78010774 |
| ASVVDPQ | ASSLANSGNTLY | 0.27363735 |
| SYNEFGGLS | ASSLANSGNTLY | 0.24370217 |
| DPDGSVY | ASSLANSGNTLY | 0.27409878 |
| MVSTIHATE | ASSLANSGNTLY | 0.7472962 |
| LAAGVADPV | ASSLANSGNTLY | 0.80977356 |
| TKEQIAA | ASSLANSGNTLY | 0.24614373 |
| SLASPAAL | ASSLANSGNTLY | 0.32890287 |
| IGLVFYGV | ASSLANSGNTLY | 0.34045112 |
| FLLDASNL | ASSLANSGNTLY | 0.47021574 |
| YQVGNFP | ASSLANSGNTLY | 0.30378118 |
| ESLLSQST | ASSLANSGNTLY | 0.20863456 |
| LFTVGFAAVG | ASSLANSGNTLY | 0.49181664 |
| ATPPTPT | ASSLANSGNTLY | 0.76839805 |
| TDLATLG | ASSLANSGNTLY | 0.32666972 |
| SAGGTGR | ASSLANSGNTLY | 0.18534769 |
| GFSNSGS | ASSLANSGNTLY | 0.380928 |
| GGNGGSGGTSV | ASSLANSGNTLY | 0.2012994 |
| GTSVATGGAGN | ASSLANSGNTLY | 0.27278483 |
| NGGNGGSGGTS | ASSLANSGNTLY | 0.32970762 |
| AVVKFPH | ASSLANSGNTLY | 0.17986056 |
| TPPPPPP | ASSLANSGNTLY | 0.35588995 |
| AAPAAAAAPA | ASSLANSGNTLY | 0.796315 |
| IALNEDL | ASSLANSGNTLY | 0.36526504 |
| IVNVSSM | ASSLANSGNTLY | 0.6318868 |
| TGIPPAPR | ASSLANSGNTLY | 0.78002197 |
| FDLGGGTF | ASSLANSGNTLY | 0.6205009 |
| FDLGGGTFDVS | ASSLANSGNTLY | 0.7700332 |
| NEPTAAA | ASSLANSGNTLY | 0.6102288 |
| NEPTAAALA | ASSLANSGNTLY | 0.43939188 |
| GSIINISSI | ASSLANSGNTLY | 0.42053744 |
| VAPGNAGTA | ASSLANSGNTLY | 0.5133159 |
| TGGGASLE | ASSLANSGNTLY | 0.38090205 |
| VAVYNHYKRI | ASSLANSGNTLY | 0.40139687 |
| TDPAPATT | ASSLANSGNTLY | 0.33010578 |
| TNRPDILDPALLRPG | ASSLANSGNTLY | 0.7229882 |
| LTTTTTS | ASSLANSGNTLY | 0.40219253 |
| PTGSGKTL | ASSLANSGNTLY | 0.6452984 |
| TGSGKTL | ASSLANSGNTLY | 0.17203045 |
| TIGHVDHGK | ASSLANSGNTLY | 0.20141216 |
| TIGHVDHGKTT | ASSLANSGNTLY | 0.70078063 |
| GHVDHGKTT | ASSLANSGNTLY | 0.08475955 |
| AQFEHTLL | ASSLANSGNTLY | 0.09360616 |
| IVNQDEL | ASSLANSGNTLY | 0.3503667 |

| | | |
|---|---|---|
| TNGSQFF | ASSLANSGNTLY | 0.74482083 |
| TGKGIVE | ASSLANSGNTLY | 0.080727905 |
| LISNASDA | ASSLANSGNTLY | 0.23795915 |
| AELRTLGELTKIAKAHGAQ | ASSLANSGNTLY | 0.1694542 |
| APNAVAAAEPSW | ASSLANSGNTLY | 0.39834142 |
| APQVAGAATAVL | ASSLANSGNTLY | 0.4443403 |
| ARRDPAAAAAILPTDA | ASSLANSGNTLY | 0.15708515 |
| DAAARDVLAEAGLAEYF | ASSLANSGNTLY | 0.5714507 |
| DDQIVLSAKANLAKVEQ | ASSLANSGNTLY | 0.15313047 |
| DELLPDTGEDSDEDGH | ASSLANSGNTLY | 0.14628953 |
| ELPEAPANRATVPQD | ASSLANSGNTLY | 0.6785385 |
| FGQSKIERVTGIG | ASSLANSGNTLY | 0.7413517 |
| GAGGTGSTAGGAGGAGGAGGLY | ASSLANSGNTLY | 0.23533046 |
| GHGGTGAAGGNGGTGGAGGSGI | ASSLANSGNTLY | 0.62022597 |
| GNGGNGGAGGEGGPGGAGGAGA | ASSLANSGNTLY | 0.19712201 |
| GSTAGNGGNGGNATNGTIAGQP | ASSLANSGNTLY | 0.8309224 |
| GTGGDGGNGGNGGTGG | ASSLANSGNTLY | 0.31938425 |
| GVRRPAGIPADAPV | ASSLANSGNTLY | 0.15610602 |
| GYRQVTDAHLLTLARR | ASSLANSGNTLY | 0.6440449 |
| IAKMGQDGHDRG | ASSLANSGNTLY | 0.70878464 |
| KAEPSIEIQYT | ASSLANSGNTLY | 0.41914633 |
| KPVRIRMAADEDTVAVT | ASSLANSGNTLY | 0.585104 |
| MPPRIAGMRLLVIKPEP | ASSLANSGNTLY | 0.21609531 |
| MKFYAVALFRSRGVPPTR | ASSLANSGNTLY | 0.33435848 |
| NAGGFGILGAVAHSP | ASSLANSGNTLY | 0.73978555 |
| NEGREITLAMAEM | ASSLANSGNTLY | 0.7084093 |
| NELVEWSHPLGHRWRWE | ASSLANSGNTLY | 0.21375935 |
| NLGAGGTGGNGGNPG | ASSLANSGNTLY | 0.6300659 |
| NTEWPKWETVMAADPVD | ASSLANSGNTLY | 0.40117586 |
| PASAAPAETQP | ASSLANSGNTLY | 0.2850797 |
| QQSSDGKNGVRAAEITGE | ASSLANSGNTLY | 0.7881733 |
| RAPDAVTTSWWKE | ASSLANSGNTLY | 0.34077084 |
| SGFGNVGAHTSGWFNQ | ASSLANSGNTLY | 0.6687025 |
| SGVAGAVGQAASVGGLKV | ASSLANSGNTLY | 0.5096683 |
| SNGGAGTGSAGGNGGT | ASSLANSGNTLY | 0.536085 |
| TGGAGGEGGAAG | ASSLANSGNTLY | 0.2226156 |
| TGGDGGAAGAGGAPGHG | ASSLANSGNTLY | 0.5671016 |
| TMPPLLAAIANAGRHGV | ASSLANSGNTLY | 0.47585443 |
| TSSTAGNSGGA | ASSLANSGNTLY | 0.5682126 |
| VPATVPVGILC | ASSLANSGNTLY | 0.27767375 |
| VQRPDSGGVVLTGRIS | ASSLANSGNTLY | 0.1978673 |
| WTPNPVPATKGVPAGLVQAT | ASSLANSGNTLY | 0.51729894 |
| YALKRALAKFSDRFDVVII | ASSLANSGNTLY | 0.73738647 |
| YLIGSVIV | ASSLANSGNTLY | 0.41899812 |
| YGQLVAVID | ASSLANSGNTLY | 0.5044379 |
| VTVGGMTLPA | ASSLANSGNTLY | 0.79948735 |
| VLALPAV | ASSLANSGNTLY | 0.2440167 |
| VIPPTLNYET | ASSLANSGNTLY | 0.36564848 |
| VGMGGMHPSSG | ASSLANSGNTLY | 0.5907238 |
| VGLAGGAATA | ASSLANSGNTLY | 0.7102694 |
| VGGAGGAGGSGP | ASSLANSGNTLY | 0.3212738 |

| | | |
|---|---|---|
| VFAAFMV | ASSLANSGNTLY | 0.7441226 |
| TLYDKLR | ASSLANSGNTLY | 0.25067082 |
| TLAVFTVL | ASSLANSGNTLY | 0.6453992 |
| TGSVQPARV | ASSLANSGNTLY | 0.20580456 |
| TGGGAGSGGT | ASSLANSGNTLY | 0.48767552 |
| SHLETSL | ASSLANSGNTLY | 0.20288603 |
| SFTTGMT | ASSLANSGNTLY | 0.6992738 |
| QVEVAKRMGVRQPTVS | ASSLANSGNTLY | 0.68107116 |
| QGSGGPALAM | ASSLANSGNTLY | 0.7122023 |
| MLAPSNP | ASSLANSGNTLY | 0.11065022 |
| LVNNAAI | ASSLANSGNTLY | 0.57315236 |
| LTGEINTLA | ASSLANSGNTLY | 0.6288763 |
| LSIFGPV | ASSLANSGNTLY | 0.828619 |
| LLALLANL | ASSLANSGNTLY | 0.37095594 |
| LISNASDALDKLR | ASSLANSGNTLY | 0.5561135 |
| LINEAVP | ASSLANSGNTLY | 0.29418588 |
| LGYAFSGL | ASSLANSGNTLY | 0.3006841 |
| LDFVAVAL | ASSLANSGNTLY | 0.16460228 |
| LAQPGAE | ASSLANSGNTLY | 0.34849328 |
| ISNLGAVTGF | ASSLANSGNTLY | 0.33447236 |
| HQEDPGRI | ASSLANSGNTLY | 0.24870291 |
| GTPGEPTP | ASSLANSGNTLY | 0.7153263 |
| GFFNSGS | ASSLANSGNTLY | 0.75455356 |
| GAGGGGLTAGAN | ASSLANSGNTLY | 0.75620687 |
| FVDALVNM | ASSLANSGNTLY | 0.7178416 |
| FLTVAPEL | ASSLANSGNTLY | 0.78218234 |
| FETLTAL | ASSLANSGNTLY | 0.5023724 |
| DVINPIEL | ASSLANSGNTLY | 0.27011615 |
| APVSGAPV | ASSLANSGNTLY | 0.3696652 |
| ALVGGLAW | ASSLANSGNTLY | 0.18836455 |
| ALAAAAATAI | ASSLANSGNTLY | 0.35075277 |
| AFGLGLLL | ASSLANSGNTLY | 0.26675665 |
| AFFAAASL | ASSLANSGNTLY | 0.44734806 |
| AAAALPAFSPP | ASSLANSGNTLY | 0.07644109 |
| VVLAPAL | ASSLANSGNTLY | 0.58634615 |
| TIGHVDHGKTTL | ASSLANSGNTLY | 0.80488205 |
| LISNASDALDK | ASSLANSGNTLY | 0.22451979 |
| LISNASDAL | ASSLANSGNTLY | 0.51325685 |
| ISNASDALDK | ASSLANSGNTLY | 0.6890476 |
| NQAETLVYQTE | ASSLANSGNTLY | 0.26454443 |
| TGGTGGTAGAG | ASSLANSGNTLY | 0.6141834 |
| TGSTAGGAGGA | ASSLANSGNTLY | 0.1974255 |
| DPLAVPVAAGPV | ASSLANSGNTLY | 0.5727226 |
| GGTGGTPGNGGN | ASSLANSGNTLY | 0.7888448 |
| PAGLAGVDPATL | ASSLANSGNTLY | 0.509128 |
| GNNLVVEAPHAGASV | ASSLANSGNTLY | 0.38986427 |
| KVTRIMSASTLEQPA | ASSLANSGNTLY | 0.39496082 |
| LGFALPIALPRRTVV | ASSLANSGNTLY | 0.19699484 |
| TRLGNLTARKLVVLV | ASSLANSGNTLY | 0.46679717 |
| WSQQRRLIYFPSAGP | ASSLANSGNTLY | 0.07502967 |
| GPNGNPDPMAAAVDIR | ASSLANSGNTLY | 0.48301616 |

| | | |
|---|---|---|
| TQRCVSVADLRAAASR | ASSLANSGNTLY | 0.20904987 |
| QFVLDVAEHPVEVLTETV | ASSLANSGNTLY | 0.46348494 |
| GGAGGTGGAGGNGGTQAGDG | ASSLANSGNTLY | 0.47399238 |
| PLLATLRAIAVDVDAKPAQV | ASSLANSGNTLY | 0.5174612 |
| APKFRTGGYMIDFWGVGYQVAK | ASSLANSGNTLY | 0.219578 |
| GAGGAGGAGGAAGGPGGTGGTG | ASSLANSGNTLY | 0.3641883 |
| IGALGLLATDVAPAGSVAFGVA | ASSLANSGNTLY | 0.25104284 |
| AGGAGGIGGNGGAGGAGGVLMG | ASSLANSGNTLY | 0.17679565 |
| ADELIGLL | ASSLANSGNTLY | 0.7364696 |
| ADRALKA | ASSLANSGNTLY | 0.29101345 |
| AGAVGQVGGAGGA | ASSLANSGNTLY | 0.42238787 |
| AGGGGGLGGAP | ASSLANSGNTLY | 0.63921213 |
| AGGLLYGNGGAG | ASSLANSGNTLY | 0.46417952 |
| AGLESIGL | ASSLANSGNTLY | 0.18659717 |
| AGLKTGII | ASSLANSGNTLY | 0.45390645 |
| ASLAALLL | ASSLANSGNTLY | 0.35787007 |
| ASVINISSI | ASSLANSGNTLY | 0.2945045 |
| ATPPGPA | ASSLANSGNTLY | 0.0768209 |
| ATVASALV | ASSLANSGNTLY | 0.5481931 |
| ATVASLIL | ASSLANSGNTLY | 0.29021814 |
| AVNGPVLIRH | ASSLANSGNTLY | 0.30892676 |
| DGAAGAVGI | ASSLANSGNTLY | 0.37838352 |
| DPLLHGI | ASSLANSGNTLY | 0.6733085 |
| ELGAALGAL | ASSLANSGNTLY | 0.18352586 |
| ELGRTQAV | ASSLANSGNTLY | 0.2754725 |
| FVAGGYLS | ASSLANSGNTLY | 0.68527985 |
| GASGGAGGAGGW | ASSLANSGNTLY | 0.5176183 |
| GGAGAGGDGGAGGI | ASSLANSGNTLY | 0.7430872 |
| GTGHNGSQPGMGGNG | ASSLANSGNTLY | 0.42963186 |
| GYAAASASAS | ASSLANSGNTLY | 0.30049214 |
| HQVDVAAVQQG | ASSLANSGNTLY | 0.4890146 |
| IGAAIAHT | ASSLANSGNTLY | 0.35256037 |
| IGMIEVAQL | ASSLANSGNTLY | 0.11879081 |
| IRLAPEHTSAMD | ASSLANSGNTLY | 0.105431154 |
| IYNVSVL | ASSLANSGNTLY | 0.29564348 |
| KQGFLGM | ASSLANSGNTLY | 0.11485514 |
| LESVITQL | ASSLANSGNTLY | 0.4555151 |
| LGLGALVVV | ASSLANSGNTLY | 0.5782702 |
| LIGNGGNGGSGGI | ASSLANSGNTLY | 0.28843126 |
| LNAINAPTL | ASSLANSGNTLY | 0.19517352 |
| LQLLNTGV | ASSLANSGNTLY | 0.31070325 |
| LQTSIEF | ASSLANSGNTLY | 0.33202145 |
| MSFAVSEL | ASSLANSGNTLY | 0.6030294 |
| NTGALGL | ASSLANSGNTLY | 0.30239105 |
| PPGAGGAGGAGTT | ASSLANSGNTLY | 0.7886066 |
| QLVGLGI | ASSLANSGNTLY | 0.70265555 |
| RFLEQVHQL | ASSLANSGNTLY | 0.3100003 |
| RGVGYVLR | ASSLANSGNTLY | 0.6704359 |
| SATGVFAGVI | ASSLANSGNTLY | 0.33532855 |
| SLPAITLP | ASSLANSGNTLY | 0.5117997 |
| STALAFTV | ASSLANSGNTLY | 0.44469833 |

| | | |
|---|---|---|
| SVLTVGLGK | ASSLANSGNTLY | 0.34997934 |
| TALAGAGSGI | ASSLANSGNTLY | 0.3261406 |
| TGLPFTGI | ASSLANSGNTLY | 0.41371867 |
| VAEHVTV | ASSLANSGNTLY | 0.16716188 |
| VDIVLNSL | ASSLANSGNTLY | 0.34154868 |
| VLTGGGAVL | ASSLANSGNTLY | 0.27773273 |
| VNVGLAYGI | ASSLANSGNTLY | 0.105731994 |
| VSSAAVGAV | ASSLANSGNTLY | 0.33296666 |
| VTSVYEAGV | ASSLANSGNTLY | 0.82890654 |
| WVALDRGA | ASSLANSGNTLY | 0.73179126 |
| YLVGSVVI | ASSLANSGNTLY | 0.21618679 |
| GPPGTGKTLL | ASSLANSGNTLY | 0.21585882 |
| GVGMTKF | ASSLANSGNTLY | 0.6748459 |
| IGIGGSDL | ASSLANSGNTLY | 0.14624718 |
| IGTLLAEL | ASSLANSGNTLY | 0.418523 |
| IYTSGTT | ASSLANSGNTLY | 0.4682177 |
| LSFPAGV | ASSLANSGNTLY | 0.4621951 |
| LYTSGTT | ASSLANSGNTLY | 0.254435 |
| VGAFPFGF | ASSLANSGNTLY | 0.5495257 |
| VGAGTVEF | ASSLANSGNTLY | 0.35583997 |
| VGKSTLLL | ASSLANSGNTLY | 0.43861902 |
| AALAHAGGGIV | ASSLANSGNTLY | 0.081212595 |
| AALVAVSLL | ASSLANSGNTLY | 0.5569294 |
| AAVSRAVAAL | ASSLANSGNTLY | 0.24854928 |
| AGGSSGIGGF | ASSLANSGNTLY | 0.22708072 |
| AGLTVANANG | ASSLANSGNTLY | 0.13065054 |
| AGLVGAVL | ASSLANSGNTLY | 0.4296043 |
| ALSQAFTV | ASSLANSGNTLY | 0.6409863 |
| ALTFSVSL | ASSLANSGNTLY | 0.7410017 |
| ALTGAALVAL | ASSLANSGNTLY | 0.18337235 |
| ASFSIGAGVL | ASSLANSGNTLY | 0.6905619 |
| DGSVYKI | ASSLANSGNTLY | 0.7641218 |
| EQLAEIV | ASSLANSGNTLY | 0.41169754 |
| FAVSELTVM | ASSLANSGNTLY | 0.18139535 |
| GAAPLAAGL | ASSLANSGNTLY | 0.43532217 |
| GADGGDGGSGGAGG | ASSLANSGNTLY | 0.36042202 |
| GFGLDTGGAGG | ASSLANSGNTLY | 0.5200832 |
| GLASGGLL | ASSLANSGNTLY | 0.63657784 |
| GRIAGAAVLL | ASSLANSGNTLY | 0.17932802 |
| GSAGSLNTSF | ASSLANSGNTLY | 0.49494624 |
| GTGQIGI | ASSLANSGNTLY | 0.6888002 |
| GTLIALIL | ASSLANSGNTLY | 0.21102306 |
| IAATMGFVVG | ASSLANSGNTLY | 0.32839024 |
| IFAQALL | ASSLANSGNTLY | 0.32387716 |
| IGAAVALGF | ASSLANSGNTLY | 0.10905413 |
| IGLTPLAL | ASSLANSGNTLY | 0.54385775 |
| ILGSNTSTL | ASSLANSGNTLY | 0.3458978 |
| IVFSLTVL | ASSLANSGNTLY | 0.256881 |
| LASKATGF | ASSLANSGNTLY | 0.38150436 |
| LASVALIL | ASSLANSGNTLY | 0.25854397 |
| LFAVGLTL | ASSLANSGNTLY | 0.29180318 |

| | | |
|---|---|---|
| LFNGGIGTYI | ASSLANSGNTLY | 0.31262144 |
| LTLGAGII | ASSLANSGNTLY | 0.44186515 |
| LVAAALGDI | ASSLANSGNTLY | 0.5848728 |
| NTAFVQLGI | ASSLANSGNTLY | 0.4565922 |
| QLFSGAEL | ASSLANSGNTLY | 0.5009558 |
| SAATFITL | ASSLANSGNTLY | 0.81396174 |
| SNYGLANLGV | ASSLANSGNTLY | 0.16688429 |
| SYGGAYAVM | ASSLANSGNTLY | 0.23064579 |
| TTGVGGLTLG | ASSLANSGNTLY | 0.5475963 |
| TTQTIYI | ASSLANSGNTLY | 0.15405652 |
| VAAGAGLGAVI | ASSLANSGNTLY | 0.5940739 |
| VAAVVTLL | ASSLANSGNTLY | 0.49920702 |
| VGFLGGLI | ASSLANSGNTLY | 0.514645 |
| VGVIPILL | ASSLANSGNTLY | 0.3732711 |
| VIFASSLI | ASSLANSGNTLY | 0.84908515 |
| VKGGDGGAAAG | ASSLANSGNTLY | 0.59145796 |
| VTKTFTNI | ASSLANSGNTLY | 0.39210123 |
| VVAAAILL | ASSLANSGNTLY | 0.572884 |
| YAALDVEL | ASSLANSGNTLY | 0.23793092 |
| LASVALLL | ASSLANSGNTLY | 0.67935175 |
| LDAVHRV | ASSLANSGNTLY | 0.521111 |
| QGLSSSIF | ASSLANSGNTLY | 0.20319197 |
| RGTFANIRL | ASSLANSGNTLY | 0.3551554 |
| RQLWWGHRI | ASSLANSGNTLY | 0.7011412 |
| YGPPGTGKTLL | ASSLANSGNTLY | 0.77631533 |
| AALVGAPANVA | ASSLANSGNTLY | 0.12879369 |
| AATFRRTE | ASSLANSGNTLY | 0.30196485 |
| AGALGGGI | ASSLANSGNTLY | 0.17015946 |
| AGGAGGAGGIGAG | ASSLANSGNTLY | 0.28073704 |
| AGSGALGL | ASSLANSGNTLY | 0.25313926 |
| AQIADQI | ASSLANSGNTLY | 0.5874463 |
| ASAVAVTYL | ASSLANSGNTLY | 0.3361186 |
| ASTVLGGTGG | ASSLANSGNTLY | 0.65538764 |
| AVGGEPAK | ASSLANSGNTLY | 0.44551116 |
| CEQGTFGGTC | ASSLANSGNTLY | 0.7028015 |
| DEGKAPAG | ASSLANSGNTLY | 0.2533813 |
| DTVVFPAL | ASSLANSGNTLY | 0.36238396 |
| FEGITMAG | ASSLANSGNTLY | 0.15343575 |
| FPLVVQD | ASSLANSGNTLY | 0.15526743 |
| GAFTTPQ | ASSLANSGNTLY | 0.19500253 |
| GFGGFGTAGGVGGA | ASSLANSGNTLY | 0.32748562 |
| GFLAQFL | ASSLANSGNTLY | 0.09301203 |
| GGAGGAGGNAGVA | ASSLANSGNTLY | 0.71054363 |
| GGVGTAGAGGAGGAGG | ASSLANSGNTLY | 0.4035529 |
| GNGGGGDAGNAGSGGN | ASSLANSGNTLY | 0.5719816 |
| GTGGAGGAGNDAGS | ASSLANSGNTLY | 0.32436788 |
| GTTNTGWFNS | ASSLANSGNTLY | 0.32323575 |
| IFMGPLFTL | ASSLANSGNTLY | 0.2736208 |
| ILAHAIEAI | ASSLANSGNTLY | 0.57688636 |
| IRSAIREAL | ASSLANSGNTLY | 0.23585233 |
| ITFAGGG | ASSLANSGNTLY | 0.15196109 |

| | | |
|---|---|---|
| ITGMSGHQEV | ASSLANSGNTLY | 0.28985164 |
| IVAGLACTL | ASSLANSGNTLY | 0.23435321 |
| IVGGGLAGA | ASSLANSGNTLY | 0.74366844 |
| IVTGVLNI | ASSLANSGNTLY | 0.65283275 |
| LAQQGCI | ASSLANSGNTLY | 0.479829 |
| LCLHVEN | ASSLANSGNTLY | 0.67051756 |
| LDAISDGS | ASSLANSGNTLY | 0.10959753 |
| LGGFAAML | ASSLANSGNTLY | 0.35105482 |
| LGGKNPMI | ASSLANSGNTLY | 0.7601404 |
| LGPKGVAAIL | ASSLANSGNTLY | 0.39375842 |
| LMVGVSI | ASSLANSGNTLY | 0.39506334 |
| LVGAAFVL | ASSLANSGNTLY | 0.6877628 |
| LVLAVAPL | ASSLANSGNTLY | 0.26999727 |
| MAAIAVIDAA | ASSLANSGNTLY | 0.28565824 |
| NNGINAELQ | ASSLANSGNTLY | 0.38720083 |
| RGEPLVLVHG | ASSLANSGNTLY | 0.58256364 |
| RGGAGGAAGLI | ASSLANSGNTLY | 0.28022152 |
| RSYRMHPAV | ASSLANSGNTLY | 0.23450282 |
| SAAHLLAGQLA | ASSLANSGNTLY | 0.4712022 |
| SAAVGSY | ASSLANSGNTLY | 0.8719866 |
| SPHAVLHRI | ASSLANSGNTLY | 0.33955905 |
| SQIAKVL | ASSLANSGNTLY | 0.6271486 |
| TALGHLVP | ASSLANSGNTLY | 0.5039346 |
| TAPTTSGAA | ASSLANSGNTLY | 0.31181568 |
| TDGDDGNGGSPGT | ASSLANSGNTLY | 0.2283677 |
| TDGLGEN | ASSLANSGNTLY | 0.25195685 |
| TGGPPLPGQA | ASSLANSGNTLY | 0.37462735 |
| TIPAITI | ASSLANSGNTLY | 0.18732 |
| TIPGFTI | ASSLANSGNTLY | 0.34635952 |
| TSIASGALA | ASSLANSGNTLY | 0.34935528 |
| TVTLPGAAI | ASSLANSGNTLY | 0.8643515 |
| VASGLSAALP | ASSLANSGNTLY | 0.619395 |
| VGAGVIGI | ASSLANSGNTLY | 0.50316447 |
| VGPKRFGV | ASSLANSGNTLY | 0.1982547 |
| VNVAFPD | ASSLANSGNTLY | 0.37853572 |
| VSMTNSGVS | ASSLANSGNTLY | 0.50755364 |
| VTDVLGDLSV | ASSLANSGNTLY | 0.19744645 |
| VVVVGAGFAGL | ASSLANSGNTLY | 0.74835396 |
| VYTVGTVV | ASSLANSGNTLY | 0.24288988 |
| WTTFSGP | ASSLANSGNTLY | 0.37631914 |
| YDELVEAF | ASSLANSGNTLY | 0.5610074 |
| YGTGGAGGNGG | ASSLANSGNTLY | 0.7399403 |
| YTNVMAV | ASSLANSGNTLY | 0.3790325 |
| FLEQVHQL | ASSLANSGNTLY | 0.40532172 |
| GLSLLAVAV | ASSLANSGNTLY | 0.22534914 |
| LEQVHQL | ASSLANSGNTLY | 0.5899598 |
| LNAAGAI | ASSLANSGNTLY | 0.729056 |
| PAVHASL | ASSLANSGNTLY | 0.2178701 |
| TELEDDA | ASSLANSGNTLY | 0.7940757 |
| TPPGPPN | ASSLANSGNTLY | 0.33976918 |
| TPPPPPPP | ASSLANSGNTLY | 0.19273789 |

| | | |
|---|---|---|
| TQSPALI | SSSQRQYEQY | 0.36107105 |
| EFAAAFPQ | SSSQRQYEQY | 0.7547358 |
| LLSGNLDVL | SSSQRQYEQY | 0.74962133 |
| GPANLGNYNI | SSSQRQYEQY | 0.6427707 |
| TLGPTLP | SSSQRQYEQY | 0.54492223 |
| VYAYSSCCK | SSSQRQYEQY | 0.32551017 |
| APAPAPAPNI | SSSQRQYEQY | 0.7515453 |
| PGGPVPIP | SSSQRQYEQY | 0.6985668 |
| FLTVALEP | SSSQRQYEQY | 0.53534913 |
| SQYFGANL | SSSQRQYEQY | 0.27433863 |
| GGVKPAA | SSSQRQYEQY | 0.23165527 |
| GGTGGAGGAGGW | SSSQRQYEQY | 0.4488706 |
| GVGGVAGLGGAGG | SSSQRQYEQY | 0.6806925 |
| VLLVLATTI | SSSQRQYEQY | 0.18315578 |
| IGIFAAML | SSSQRQYEQY | 0.4459055 |
| AVGDLID | SSSQRQYEQY | 0.461605 |
| NISPLPL | SSSQRQYEQY | 0.17861514 |
| VTAVFVAI | SSSQRQYEQY | 0.51609933 |
| SGDDNDAAL | SSSQRQYEQY | 0.8292528 |
| AVAALAHS | SSSQRQYEQY | 0.43868315 |
| GGVTVMATP | SSSQRQYEQY | 0.21333009 |
| AGLAANVRD | SSSQRQYEQY | 0.7639615 |
| TNRPDLIDPALLRPG | SSSQRQYEQY | 0.8473518 |
| LANLGVTAL | SSSQRQYEQY | 0.6123351 |
| TGERIEL | SSSQRQYEQY | 0.17936154 |
| NGEELAA | SSSQRQYEQY | 0.101824574 |
| VLSLVWG | SSSQRQYEQY | 0.37968898 |
| IGYALWL | SSSQRQYEQY | 0.5987593 |
| LSLLLAPL | SSSQRQYEQY | 0.27769518 |
| TLDEALK | SSSQRQYEQY | 0.22479004 |
| TDGAGTAEGG | SSSQRQYEQY | 0.5986628 |
| NAPGGAGGNGG | SSSQRQYEQY | 0.6973208 |
| VSVFMGMGGNF | SSSQRQYEQY | 0.22803102 |
| LASYLAAL | SSSQRQYEQY | 0.17424802 |
| LVGGTPAL | SSSQRQYEQY | 0.6589179 |
| VRNADVL | SSSQRQYEQY | 0.5628474 |
| TCSSPFLVSAM | SSSQRQYEQY | 0.25772476 |
| VHITDAE | SSSQRQYEQY | 0.3427674 |
| EYAAPSF | SSSQRQYEQY | 0.425332 |
| VVDAFHGGL | SSSQRQYEQY | 0.13959268 |
| ASVVDPQ | SSSQRQYEQY | 0.5255659 |
| SYNEFGGLS | SSSQRQYEQY | 0.17254317 |
| DPDGSVY | SSSQRQYEQY | 0.8210606 |
| MVSTIHATE | SSSQRQYEQY | 0.20096485 |
| LAAGVADPV | SSSQRQYEQY | 0.5865692 |
| TKEQIAA | SSSQRQYEQY | 0.49840495 |
| SLASPAAL | SSSQRQYEQY | 0.52466834 |
| IGLVFYGV | SSSQRQYEQY | 0.35811943 |
| FLLDASNL | SSSQRQYEQY | 0.22789654 |
| YQVGNFP | SSSQRQYEQY | 0.20233326 |
| ESLLSQST | SSSQRQYEQY | 0.102024704 |

| | | |
|---|---|---|
| LFTVGFAAVG | SSSQRQYEQY | 0.75958234 |
| ATPPTPT | SSSQRQYEQY | 0.08399094 |
| TDLATLG | SSSQRQYEQY | 0.7398358 |
| SAGGTGR | SSSQRQYEQY | 0.52246284 |
| GFSNSGS | SSSQRQYEQY | 0.2690679 |
| GGNGGSGGTSV | SSSQRQYEQY | 0.47319007 |
| GTSVATGGAGN | SSSQRQYEQY | 0.16310008 |
| NGGNGGSGGTS | SSSQRQYEQY | 0.17752108 |
| AVVKFPH | SSSQRQYEQY | 0.5792798 |
| TPPPPPP | SSSQRQYEQY | 0.5728772 |
| AAPAAAAAPA | SSSQRQYEQY | 0.27164876 |
| IALNEDL | SSSQRQYEQY | 0.44448626 |
| IVNVSSM | SSSQRQYEQY | 0.5463647 |
| TGIPPAPR | SSSQRQYEQY | 0.22132647 |
| FDLGGGTF | SSSQRQYEQY | 0.39388472 |
| FDLGGGTFDVS | SSSQRQYEQY | 0.74871165 |
| NEPTAAA | SSSQRQYEQY | 0.49341837 |
| NEPTAAALA | SSSQRQYEQY | 0.18999091 |
| GSIINISSI | SSSQRQYEQY | 0.60900676 |
| VAPGNAGTA | SSSQRQYEQY | 0.23119394 |
| TGGGASLE | SSSQRQYEQY | 0.5379334 |
| VAVYNHYKRI | SSSQRQYEQY | 0.4676128 |
| TDPAPATT | SSSQRQYEQY | 0.32219127 |
| TNRPDILDPALLRPG | SSSQRQYEQY | 0.6425004 |
| LTTTTTS | SSSQRQYEQY | 0.21632752 |
| PTGSGKTL | SSSQRQYEQY | 0.24196096 |
| TGSGKTL | SSSQRQYEQY | 0.30175632 |
| TIGHVDHGK | SSSQRQYEQY | 0.41918045 |
| TIGHVDHGKTT | SSSQRQYEQY | 0.5699016 |
| GHVDHGKTT | SSSQRQYEQY | 0.22146702 |
| AQFEHTLL | SSSQRQYEQY | 0.45087326 |
| IVNQDEL | SSSQRQYEQY | 0.26061475 |
| TNGSQFF | SSSQRQYEQY | 0.20216522 |
| TGKGIVE | SSSQRQYEQY | 0.3374833 |
| LISNASDA | SSSQRQYEQY | 0.31459722 |
| AELRTLGELTKIAKAHGAQ | SSSQRQYEQY | 0.19729117 |
| APNAVAAAEPSW | SSSQRQYEQY | 0.6150038 |
| APQVAGAATAVL | SSSQRQYEQY | 0.25631917 |
| ARRDPAAAAAILPTDA | SSSQRQYEQY | 0.47441518 |
| DAAARDVLAEAGLAEYF | SSSQRQYEQY | 0.73656094 |
| DDQIVLSAKANLAKVEQ | SSSQRQYEQY | 0.23997529 |
| DELLPDTGEDSDEDGH | SSSQRQYEQY | 0.53154707 |
| ELPEAPANRATVPQD | SSSQRQYEQY | 0.47217515 |
| FGQSKIERVTGIG | SSSQRQYEQY | 0.59932303 |
| GAGGTGSTAGGAGGAGGAGGLY | SSSQRQYEQY | 0.35042256 |
| GHGGTGAAGGNGGTGGAGGSGI | SSSQRQYEQY | 0.2546838 |
| GNGGNGGAGGEGGPGGAGGAGA | SSSQRQYEQY | 0.772554 |
| GSTAGNGGNGGNATNGTIAGQP | SSSQRQYEQY | 0.54238474 |
| GTGGDGGNGGNGGTGG | SSSQRQYEQY | 0.5845536 |
| GVRRPAGIPADAPV | SSSQRQYEQY | 0.5423665 |
| GYRQVTDAHLLTLARR | SSSQRQYEQY | 0.68767124 |

| | | |
|---|---|---|
| IAKMGQDGHDRG | SSSQRQYEQY | 0.61272544 |
| KAEPSIEIQYT | SSSQRQYEQY | 0.23826408 |
| KPVRIRMAADEDTVAVT | SSSQRQYEQY | 0.15013085 |
| MPPRIAGMRLLVIKPEP | SSSQRQYEQY | 0.34685093 |
| MKFYAVALFRSRGVPPTR | SSSQRQYEQY | 0.35419124 |
| NAGGFGILGAVAHSP | SSSQRQYEQY | 0.30856606 |
| NEGREITLAMAEM | SSSQRQYEQY | 0.13966282 |
| NELVEWSHPLGHRWRWE | SSSQRQYEQY | 0.7314629 |
| NLGAGGTGGNGGNPG | SSSQRQYEQY | 0.85550374 |
| NTEWPKWETVMAADPVD | SSSQRQYEQY | 0.25511312 |
| PASAAPAETQP | SSSQRQYEQY | 0.58931065 |
| QQSSDGKNGVRAAEITGE | SSSQRQYEQY | 0.1642533 |
| RAPDAVTTSWWKE | SSSQRQYEQY | 0.55581677 |
| SGFGNVGAHTSGWFNQ | SSSQRQYEQY | 0.73281896 |
| SGVAGAVGQAASVGGLKV | SSSQRQYEQY | 0.20558189 |
| SNGGAGTGSAGGNGGT | SSSQRQYEQY | 0.50730693 |
| TGGAGGEGGAAG | SSSQRQYEQY | 0.28195268 |
| TGGDGGAAGAGGAPGHG | SSSQRQYEQY | 0.32070005 |
| TMPPLLAAIANAGRHGV | SSSQRQYEQY | 0.7379035 |
| TSSTAGNSGGA | SSSQRQYEQY | 0.28032303 |
| VPATVPVGILC | SSSQRQYEQY | 0.57957494 |
| VQRPDSGGVVLTGRIS | SSSQRQYEQY | 0.31749895 |
| WTPNPVPATKGVPAGLVQAT | SSSQRQYEQY | 0.22870894 |
| YALKRALAKFSDRFDVVII | SSSQRQYEQY | 0.53754795 |
| YLIGSVIV | SSSQRQYEQY | 0.6832204 |
| YGQLVAVID | SSSQRQYEQY | 0.19758525 |
| VTVGGMTLPA | SSSQRQYEQY | 0.66261995 |
| VLALPAV | SSSQRQYEQY | 0.3211468 |
| VIPPTLNYET | SSSQRQYEQY | 0.86329794 |
| VGMGGMHPSSG | SSSQRQYEQY | 0.7656684 |
| VGLAGGAATA | SSSQRQYEQY | 0.48157224 |
| VGGAGGAGGSGP | SSSQRQYEQY | 0.31842732 |
| VFAAFMV | SSSQRQYEQY | 0.7348968 |
| TLYDKLR | SSSQRQYEQY | 0.27112347 |
| TLAVFTVL | SSSQRQYEQY | 0.58527607 |
| TGSVQPARV | SSSQRQYEQY | 0.5783888 |
| TGGGAGSGGT | SSSQRQYEQY | 0.37690097 |
| SHLETSL | SSSQRQYEQY | 0.8466761 |
| SFTTGMT | SSSQRQYEQY | 0.75912035 |
| QVEVAKRMGVRQPTVS | SSSQRQYEQY | 0.79636854 |
| QGSGGPALAM | SSSQRQYEQY | 0.6399284 |
| MLAPSNP | SSSQRQYEQY | 0.3597885 |
| LVNNAAI | SSSQRQYEQY | 0.6709639 |
| LTGEINTLA | SSSQRQYEQY | 0.50904906 |
| LSIFGPV | SSSQRQYEQY | 0.8346495 |
| LLALLANL | SSSQRQYEQY | 0.83272654 |
| LISNASDALDKLR | SSSQRQYEQY | 0.6002436 |
| LINEAVP | SSSQRQYEQY | 0.81192565 |
| LGYAFSGL | SSSQRQYEQY | 0.80272865 |
| LDFVAVAL | SSSQRQYEQY | 0.38197947 |
| LAQPGAE | SSSQRQYEQY | 0.73819363 |

| | | |
|---|---|---|
| ISNLGAVTGF | SSSQRQYEQY | 0.66958547 |
| HQEDPGRI | SSSQRQYEQY | 0.6960787 |
| GTPGEPTP | SSSQRQYEQY | 0.29305947 |
| GFFNSGS | SSSQRQYEQY | 0.80693537 |
| GAGGGGLTAGAN | SSSQRQYEQY | 0.64076114 |
| FVDALVNM | SSSQRQYEQY | 0.15510353 |
| FLTVAPEL | SSSQRQYEQY | 0.7834145 |
| FETLTAL | SSSQRQYEQY | 0.6795519 |
| DVINPIEL | SSSQRQYEQY | 0.5859581 |
| APVSGAPV | SSSQRQYEQY | 0.81487167 |
| ALVGGLAW | SSSQRQYEQY | 0.56741124 |
| ALAAAAATAI | SSSQRQYEQY | 0.59720546 |
| AFGLGLLL | SSSQRQYEQY | 0.8004488 |
| AFFAAASL | SSSQRQYEQY | 0.83445436 |
| AAAALPAFSPP | SSSQRQYEQY | 0.47907057 |
| VVLAPAL | SSSQRQYEQY | 0.7531836 |
| TIGHVDHGKTTL | SSSQRQYEQY | 0.84978354 |
| LISNASDALDK | SSSQRQYEQY | 0.71521264 |
| LISNASDAL | SSSQRQYEQY | 0.8564829 |
| ISNASDALDK | SSSQRQYEQY | 0.8625667 |
| NQAETLVYQTE | SSSQRQYEQY | 0.6610886 |
| TGGTGGTAGAG | SSSQRQYEQY | 0.83756995 |
| TGSTAGGAGGA | SSSQRQYEQY | 0.836178 |
| DPLAVPVAAGPV | SSSQRQYEQY | 0.6990525 |
| GGTGGTPGNGGN | SSSQRQYEQY | 0.7267886 |
| PAGLAGVDPATL | SSSQRQYEQY | 0.3633053 |
| GNNLVVEAPHAGASV | SSSQRQYEQY | 0.7310693 |
| KVTRIMSASTLEQPA | SSSQRQYEQY | 0.42594355 |
| LGFALPIALPRRTVV | SSSQRQYEQY | 0.7697513 |
| TRLGNLTARKLVVLV | SSSQRQYEQY | 0.8544453 |
| WSQQRRLIYFPSAGP | SSSQRQYEQY | 0.71492493 |
| GPNGNPDPMAAAVDIR | SSSQRQYEQY | 0.36084723 |
| TQRCVSVADLRAAASR | SSSQRQYEQY | 0.46938357 |
| QFVLDVAEHPVEVLTETV | SSSQRQYEQY | 0.7906073 |
| GGAGGTGGAGGNGGTQAGDG | SSSQRQYEQY | 0.44587857 |
| PLLATLRAIAVDVDAKPAQV | SSSQRQYEQY | 0.43535292 |
| APKFRTGGYMIDFWGVGYQVAK | SSSQRQYEQY | 0.61664265 |
| GAGGAGGAGGAAGGPGGTGGTG | SSSQRQYEQY | 0.81883633 |
| IGALGLLATDVAPAGSVAFGVA | SSSQRQYEQY | 0.83249605 |
| AGGAGGIGGNGGAGGAGGVLMG | SSSQRQYEQY | 0.53655887 |
| ADELIGLL | SSSQRQYEQY | 0.85393405 |
| ADRALKA | SSSQRQYEQY | 0.8250863 |
| AGAVGQVGGAGGA | SSSQRQYEQY | 0.86214864 |
| AGGGGGLGGAP | SSSQRQYEQY | 0.7590768 |
| AGGLLYGNGGAG | SSSQRQYEQY | 0.4272342 |
| AGLESIGL | SSSQRQYEQY | 0.8326643 |
| AGLKTGII | SSSQRQYEQY | 0.6306879 |
| ASLAALLL | SSSQRQYEQY | 0.7747899 |
| ASVINISSI | SSSQRQYEQY | 0.8162348 |
| ATPPGPA | SSSQRQYEQY | 0.67638093 |
| ATVASALV | SSSQRQYEQY | 0.60469854 |

| | | |
|---|---|---|
| ATVASLIL | SSSQRQYEQY | 0.7442379 |
| AVNGPVLIRH | SSSQRQYEQY | 0.8591187 |
| DGAAGAVGI | SSSQRQYEQY | 0.55859816 |
| DPLLHGI | SSSQRQYEQY | 0.4487911 |
| ELGAALGAL | SSSQRQYEQY | 0.68407786 |
| ELGRTQAV | SSSQRQYEQY | 0.68270206 |
| FVAGGYLS | SSSQRQYEQY | 0.8427621 |
| GASGGAGGAGGW | SSSQRQYEQY | 0.80278635 |
| GGAGAGGDGGAGGI | SSSQRQYEQY | 0.73940235 |
| GTGHNGSQPGMGGNG | SSSQRQYEQY | 0.77861947 |
| GYAAASASAS | SSSQRQYEQY | 0.817909 |
| HQVDVAAVQQG | SSSQRQYEQY | 0.7568926 |
| IGAAIAHT | SSSQRQYEQY | 0.7505202 |
| IGMIEVAQL | SSSQRQYEQY | 0.5933485 |
| IRLAPEHTSAMD | SSSQRQYEQY | 0.8530793 |
| IYNVSVL | SSSQRQYEQY | 0.8232933 |
| KQGFLGM | SSSQRQYEQY | 0.8314476 |
| LESVITQL | SSSQRQYEQY | 0.8513439 |
| LGLGALVVV | SSSQRQYEQY | 0.77657557 |
| LIGNGGNGGSGGI | SSSQRQYEQY | 0.77403235 |
| LNAINAPTL | SSSQRQYEQY | 0.41076392 |
| LQLLNTGV | SSSQRQYEQY | 0.56179446 |
| LQTSIEF | SSSQRQYEQY | 0.7356511 |
| MSFAVSEL | SSSQRQYEQY | 0.78092 |
| NTGALGL | SSSQRQYEQY | 0.8117925 |
| PPGAGGAGGAGTT | SSSQRQYEQY | 0.82324433 |
| QLVGLGI | SSSQRQYEQY | 0.6461105 |
| RFLEQVHQL | SSSQRQYEQY | 0.84140515 |
| RGVGYVLR | SSSQRQYEQY | 0.83996993 |
| SATGVFAGVI | SSSQRQYEQY | 0.4908781 |
| SLPAITLP | SSSQRQYEQY | 0.6357488 |
| STALAFTV | SSSQRQYEQY | 0.85291 |
| SVLTVGLGK | SSSQRQYEQY | 0.8042493 |
| TALAGAGSGI | SSSQRQYEQY | 0.6878904 |
| TGLPFTGI | SSSQRQYEQY | 0.71864986 |
| VAEHVTV | SSSQRQYEQY | 0.8581527 |
| VDIVLNSL | SSSQRQYEQY | 0.72354156 |
| VLTGGGAVL | SSSQRQYEQY | 0.8490844 |
| VNVGLAYGI | SSSQRQYEQY | 0.27692637 |
| VSSAAVGAV | SSSQRQYEQY | 0.7260205 |
| VTSVYEAGV | SSSQRQYEQY | 0.6698087 |
| WVALDRGA | SSSQRQYEQY | 0.16854061 |
| YLVGSVVI | SSSQRQYEQY | 0.8287418 |
| GPPGTGKTLL | SSSQRQYEQY | 0.5616611 |
| GVGMTKF | SSSQRQYEQY | 0.7882501 |
| IGIGGSDL | SSSQRQYEQY | 0.63744074 |
| IGTLLAEL | SSSQRQYEQY | 0.84134 |
| IYTSGTT | SSSQRQYEQY | 0.78135586 |
| LSFPAGV | SSSQRQYEQY | 0.6568053 |
| LYTSGTT | SSSQRQYEQY | 0.56477714 |
| VGAFPFGF | SSSQRQYEQY | 0.78090984 |

| | | |
|---|---|---|
| VGAGTVEF | SSSQRQYEQY | 0.850587 |
| VGKSTLLL | SSSQRQYEQY | 0.85703504 |
| AALAHAGGGIV | SSSQRQYEQY | 0.32774454 |
| AALVAVSLL | SSSQRQYEQY | 0.7826981 |
| AAVSRAVAAL | SSSQRQYEQY | 0.70502484 |
| AGGSSGIGGF | SSSQRQYEQY | 0.2520126 |
| AGLTVANANG | SSSQRQYEQY | 0.22407681 |
| AGLVGAVL | SSSQRQYEQY | 0.68217677 |
| ALSQAFTV | SSSQRQYEQY | 0.7040091 |
| ALTFSVSL | SSSQRQYEQY | 0.7922584 |
| ALTGAALVAL | SSSQRQYEQY | 0.8238175 |
| ASFSIGAGVL | SSSQRQYEQY | 0.65117776 |
| DGSVYKI | SSSQRQYEQY | 0.81077117 |
| EQLAEIV | SSSQRQYEQY | 0.69556195 |
| FAVSELTVM | SSSQRQYEQY | 0.0984662 |
| GAAPLAAGL | SSSQRQYEQY | 0.6049225 |
| GADGGDGGSGGAGG | SSSQRQYEQY | 0.37079912 |
| GFGLDTGGAGG | SSSQRQYEQY | 0.8584156 |
| GLASGGLL | SSSQRQYEQY | 0.42360604 |
| GRIAGAAVLL | SSSQRQYEQY | 0.6807344 |
| GSAGSLNTSF | SSSQRQYEQY | 0.72597563 |
| GTGQIGI | SSSQRQYEQY | 0.80103457 |
| GTLIALIL | SSSQRQYEQY | 0.755545 |
| IAATMGFVVG | SSSQRQYEQY | 0.6083869 |
| IFAQALL | SSSQRQYEQY | 0.23720805 |
| IGAAVALGF | SSSQRQYEQY | 0.7997099 |
| IGLTPLAL | SSSQRQYEQY | 0.673564 |
| ILGSNTSTL | SSSQRQYEQY | 0.50111854 |
| IVFSLTVL | SSSQRQYEQY | 0.46490467 |
| LASKATGF | SSSQRQYEQY | 0.6466116 |
| LASVALIL | SSSQRQYEQY | 0.2964413 |
| LFAVGLTL | SSSQRQYEQY | 0.84349316 |
| LFNGGIGTYI | SSSQRQYEQY | 0.56436896 |
| LTLGAGII | SSSQRQYEQY | 0.6806622 |
| LVAAALGDI | SSSQRQYEQY | 0.8602319 |
| NTAFVQLGI | SSSQRQYEQY | 0.7656393 |
| QLFSGAEL | SSSQRQYEQY | 0.71031594 |
| SAATFITL | SSSQRQYEQY | 0.579352 |
| SNYGLANLGV | SSSQRQYEQY | 0.5312947 |
| SYGGAYAVM | SSSQRQYEQY | 0.5486015 |
| TTGVGGLTLG | SSSQRQYEQY | 0.58718526 |
| TTQTIYI | SSSQRQYEQY | 0.84519744 |
| VAAGAGLGAVI | SSSQRQYEQY | 0.774735 |
| VAAVVTLL | SSSQRQYEQY | 0.40034193 |
| VGFLGGLI | SSSQRQYEQY | 0.8344849 |
| VGVIPILL | SSSQRQYEQY | 0.8495964 |
| VIFASSLI | SSSQRQYEQY | 0.8611531 |
| VKGGDGGAAAG | SSSQRQYEQY | 0.51319766 |
| VTKTFTNI | SSSQRQYEQY | 0.6655395 |
| VVAAAILL | SSSQRQYEQY | 0.69163096 |
| YAALDVEL | SSSQRQYEQY | 0.69573194 |

| | | |
|---|---|---|
| LASVALLL | SSSQRQYEQY | 0.48196763 |
| LDAVHRV | SSSQRQYEQY | 0.49713677 |
| QGLSSSIF | SSSQRQYEQY | 0.8393393 |
| RGTFANIRL | SSSQRQYEQY | 0.8686662 |
| RQLWWGHRI | SSSQRQYEQY | 0.7131692 |
| YGPPGTGKTLL | SSSQRQYEQY | 0.28888854 |
| AALVGAPANVA | SSSQRQYEQY | 0.696475 |
| AATFRRTE | SSSQRQYEQY | 0.614043 |
| AGALGGGI | SSSQRQYEQY | 0.5956999 |
| AGGAGGAGGIGAG | SSSQRQYEQY | 0.6625999 |
| AGSGALGL | SSSQRQYEQY | 0.3579721 |
| AQIADQI | SSSQRQYEQY | 0.7311059 |
| ASAVAVTYL | SSSQRQYEQY | 0.86016405 |
| ASTVLGGTGG | SSSQRQYEQY | 0.7193318 |
| AVGGEPAK | SSSQRQYEQY | 0.784052 |
| CEQGTFGGTC | SSSQRQYEQY | 0.8435265 |
| DEGKAPAG | SSSQRQYEQY | 0.79721606 |
| DTVVFPAL | SSSQRQYEQY | 0.8328657 |
| FEGITMAG | SSSQRQYEQY | 0.83600795 |
| FPLVVQD | SSSQRQYEQY | 0.7566553 |
| GAFTTPQ | SSSQRQYEQY | 0.7618383 |
| GFGGFGTAGGVGGA | SSSQRQYEQY | 0.8078152 |
| GFLAQFL | SSSQRQYEQY | 0.67456234 |
| GGAGGAGGNAGVA | SSSQRQYEQY | 0.751213 |
| GGVGTAGAGGAGGAGG | SSSQRQYEQY | 0.73543566 |
| GNGGGGDAGNAGSGGN | SSSQRQYEQY | 0.79751444 |
| GTGGAGGAGNDAGS | SSSQRQYEQY | 0.7175094 |
| GTTNTGWFNS | SSSQRQYEQY | 0.8622216 |
| IFMGPLFTL | SSSQRQYEQY | 0.21705043 |
| ILAHAIEAI | SSSQRQYEQY | 0.4835077 |
| IRSAIREAL | SSSQRQYEQY | 0.8329644 |
| ITFAGGG | SSSQRQYEQY | 0.15256155 |
| ITGMSGHQEV | SSSQRQYEQY | 0.30327666 |
| IVAGLACTL | SSSQRQYEQY | 0.77328616 |
| IVGGGLAGA | SSSQRQYEQY | 0.8421574 |
| IVTGVLNI | SSSQRQYEQY | 0.1624926 |
| LAQQGCI | SSSQRQYEQY | 0.68167436 |
| LCLHVEN | SSSQRQYEQY | 0.4248485 |
| LDAISDGS | SSSQRQYEQY | 0.8432269 |
| LGGFAAML | SSSQRQYEQY | 0.70330083 |
| LGGKNPMI | SSSQRQYEQY | 0.35562396 |
| LGPKGVAAIL | SSSQRQYEQY | 0.81639034 |
| LMVGVSI | SSSQRQYEQY | 0.45885277 |
| LVGAAFVL | SSSQRQYEQY | 0.743994 |
| LVLAVAPL | SSSQRQYEQY | 0.84102774 |
| MAAIAVIDAA | SSSQRQYEQY | 0.8611075 |
| NNGINAELQ | SSSQRQYEQY | 0.4098646 |
| RGEPLVLVHG | SSSQRQYEQY | 0.817537 |
| RGGAGGAAGLI | SSSQRQYEQY | 0.6692531 |
| RSYRMHPAV | SSSQRQYEQY | 0.8677388 |
| SAAHLLAGQLA | SSSQRQYEQY | 0.66983116 |

| | | |
|---|---|---|
| SAAVGSY | SSSQRQYEQY | 0.31865823 |
| SPHAVLHRI | SSSQRQYEQY | 0.8281256 |
| SQIAKVL | SSSQRQYEQY | 0.81198454 |
| TALGHLVP | SSSQRQYEQY | 0.68833363 |
| TAPTTSGAA | SSSQRQYEQY | 0.8214838 |
| TDGDDGNGGSPGT | SSSQRQYEQY | 0.72891754 |
| TDGLGEN | SSSQRQYEQY | 0.6888626 |
| TGGPPLPGQA | SSSQRQYEQY | 0.8156338 |
| TIPAITI | SSSQRQYEQY | 0.8038577 |
| TIPGFTI | SSSQRQYEQY | 0.5583969 |
| TSIASGALA | SSSQRQYEQY | 0.8573264 |
| TVTLPGAAI | SSSQRQYEQY | 0.72526467 |
| VASGLSAALP | SSSQRQYEQY | 0.4639448 |
| VGAGVIGI | SSSQRQYEQY | 0.8668341 |
| VGPKRFGV | SSSQRQYEQY | 0.7446847 |
| VNVAFPD | SSSQRQYEQY | 0.8137947 |
| VSMTNSGVS | SSSQRQYEQY | 0.8061171 |
| VTDVLGDLSV | SSSQRQYEQY | 0.86188793 |
| VVVVGAGFAGL | SSSQRQYEQY | 0.45605493 |
| VYTVGTVV | SSSQRQYEQY | 0.69673276 |
| WTTFSGP | SSSQRQYEQY | 0.7076322 |
| YDELVEAF | SSSQRQYEQY | 0.83248067 |
| YGTGGAGGNGG | SSSQRQYEQY | 0.7284099 |
| YTNVMAV | SSSQRQYEQY | 0.661911 |
| FLEQVHQL | SSSQRQYEQY | 0.81495523 |
| GLSLLAVAV | SSSQRQYEQY | 0.84717107 |
| LEQVHQL | SSSQRQYEQY | 0.76082826 |
| LNAAGAI | SSSQRQYEQY | 0.759511 |
| PAVHASL | SSSQRQYEQY | 0.85190177 |
| TELEDDA | SSSQRQYEQY | 0.47261062 |
| TPPGPPN | SSSQRQYEQY | 0.73846275 |
| TPPPPPPP | SSSQRQYEQY | 0.8309256 |
| TQSPALI | ASSRRDERLF | 0.83731544 |
| EFAAAFPQ | ASSRRDERLF | 0.7417289 |
| LLSGNLDVL | ASSRRDERLF | 0.84464926 |
| GPANLGNYNI | ASSRRDERLF | 0.68594724 |
| TLGPTLP | ASSRRDERLF | 0.7794518 |
| VYAYSSCCK | ASSRRDERLF | 0.70598817 |
| APAPAPAPNI | ASSRRDERLF | 0.6708857 |
| PGGPVPIP | ASSRRDERLF | 0.4884324 |
| FLTVALEP | ASSRRDERLF | 0.7537682 |
| SQYFGANL | ASSRRDERLF | 0.8213444 |
| GGVKPAA | ASSRRDERLF | 0.7766145 |
| GGTGGAGGAGGW | ASSRRDERLF | 0.22604176 |
| GVGGVAGLGGAGG | ASSRRDERLF | 0.7875385 |
| VLLVLATTI | ASSRRDERLF | 0.79006886 |
| IGIFAAML | ASSRRDERLF | 0.8619365 |
| AVGDLID | ASSRRDERLF | 0.7200744 |
| NISPLPL | ASSRRDERLF | 0.8205812 |
| VTAVFVAI | ASSRRDERLF | 0.6550004 |
| SGDDNDAAL | ASSRRDERLF | 0.77742684 |

| | | |
|---|---|---|
| AVAALAHS | ASSRRDERLF | 0.40154177 |
| GGVTVMATP | ASSRRDERLF | 0.4722185 |
| AGLAANVRD | ASSRRDERLF | 0.6854732 |
| TNRPDLIDPALLRPG | ASSRRDERLF | 0.5533564 |
| LANLGVTAL | ASSRRDERLF | 0.8165436 |
| TGERIEL | ASSRRDERLF | 0.8276626 |
| NGEELAA | ASSRRDERLF | 0.8347173 |
| VLSLVWG | ASSRRDERLF | 0.83630145 |
| IGYALWL | ASSRRDERLF | 0.8467287 |
| LSLLLAPL | ASSRRDERLF | 0.8488685 |
| TLDEALK | ASSRRDERLF | 0.6381575 |
| TDGAGTAEGG | ASSRRDERLF | 0.6683332 |
| NAPGGAGGNGG | ASSRRDERLF | 0.30002558 |
| VSVFMGMGGNF | ASSRRDERLF | 0.66970813 |
| LASYLAAL | ASSRRDERLF | 0.64650905 |
| LVGGTPAL | ASSRRDERLF | 0.7445799 |
| VRNADVL | ASSRRDERLF | 0.14400525 |
| TCSSPFLVSAM | ASSRRDERLF | 0.76967967 |
| VHITDAE | ASSRRDERLF | 0.847875 |
| EYAAPSF | ASSRRDERLF | 0.65949583 |
| VVDAFHGGL | ASSRRDERLF | 0.7923709 |
| ASVVDPQ | ASSRRDERLF | 0.8002277 |
| SYNEFGGLS | ASSRRDERLF | 0.64683735 |
| DPDGSVY | ASSRRDERLF | 0.80387634 |
| MVSTIHATE | ASSRRDERLF | 0.6773674 |
| LAAGVADPV | ASSRRDERLF | 0.8275283 |
| TKEQIAA | ASSRRDERLF | 0.85213137 |
| SLASPAAL | ASSRRDERLF | 0.7897358 |
| IGLVFYGV | ASSRRDERLF | 0.7215926 |
| FLLDASNL | ASSRRDERLF | 0.7787137 |
| YQVGNFP | ASSRRDERLF | 0.5101696 |
| ESLLSQST | ASSRRDERLF | 0.7510092 |
| LFTVGFAAVG | ASSRRDERLF | 0.27941516 |
| ATPPTPT | ASSRRDERLF | 0.7551845 |
| TDLATLG | ASSRRDERLF | 0.5959571 |
| SAGGTGR | ASSRRDERLF | 0.77702 |
| GFSNSGS | ASSRRDERLF | 0.7300651 |
| GGNGGSGGTSV | ASSRRDERLF | 0.8020499 |
| GTSVATGGAGN | ASSRRDERLF | 0.5608376 |
| NGGNGGSGGTS | ASSRRDERLF | 0.79608786 |
| AVVKFPH | ASSRRDERLF | 0.7638078 |
| TPPPPPP | ASSRRDERLF | 0.39637995 |
| AAPAAAAAPA | ASSRRDERLF | 0.8252523 |
| IALNEDL | ASSRRDERLF | 0.7547611 |
| IVNVSSM | ASSRRDERLF | 0.71367395 |
| TGIPPAPR | ASSRRDERLF | 0.82668996 |
| FDLGGGTF | ASSRRDERLF | 0.7523296 |
| FDLGGGTFDVS | ASSRRDERLF | 0.4532394 |
| NEPTAAA | ASSRRDERLF | 0.66078717 |
| NEPTAAALA | ASSRRDERLF | 0.7630607 |
| GSIINISSI | ASSRRDERLF | 0.6844635 |

| | | |
|---|---|---|
| VAPGNAGTA | ASSRRDERLF | 0.16898082 |
| TGGGASLE | ASSRRDERLF | 0.8131614 |
| VAVYNHYKRI | ASSRRDERLF | 0.4683088 |
| TDPAPATT | ASSRRDERLF | 0.63463354 |
| TNRPDILDPALLRPG | ASSRRDERLF | 0.62652457 |
| LTTTTTS | ASSRRDERLF | 0.8193227 |
| PTGSGKTL | ASSRRDERLF | 0.7486557 |
| TGSGKTL | ASSRRDERLF | 0.5924207 |
| TIGHVDHGK | ASSRRDERLF | 0.77798927 |
| TIGHVDHGKTT | ASSRRDERLF | 0.7773826 |
| GHVDHGKTT | ASSRRDERLF | 0.8170434 |
| AQFEHTLL | ASSRRDERLF | 0.7272902 |
| IVNQDEL | ASSRRDERLF | 0.77139425 |
| TNGSQFF | ASSRRDERLF | 0.77850443 |
| TGKGIVE | ASSRRDERLF | 0.5967253 |
| LISNASDA | ASSRRDERLF | 0.5210972 |
| AELRTLGELTKIAKAHGAQ | ASSRRDERLF | 0.546535 |
| APNAVAAAEPSW | ASSRRDERLF | 0.62209535 |
| APQVAGAATAVL | ASSRRDERLF | 0.49619734 |
| ARRDPAAAAAILPTDA | ASSRRDERLF | 0.7845237 |
| DAAARDVLAEAGLAEYF | ASSRRDERLF | 0.8639698 |
| DDQIVLSAKANLAKVEQ | ASSRRDERLF | 0.7673068 |
| DELLPDTGEDSDEDGH | ASSRRDERLF | 0.33652484 |
| ELPEAPANRATVPQD | ASSRRDERLF | 0.705914 |
| FGQSKIERVTGIG | ASSRRDERLF | 0.6999835 |
| GAGGTGSTAGGAGGAGGAGGLY | ASSRRDERLF | 0.79629374 |
| GHGGTGAAGGNGGTGGAGGSGI | ASSRRDERLF | 0.8373569 |
| GNGGNGGAGGEGGPGGAGGAGA | ASSRRDERLF | 0.8598448 |
| GSTAGNGGNGGNATNGTIAGQP | ASSRRDERLF | 0.8146707 |
| GTGGDGGNGGNGGTGG | ASSRRDERLF | 0.6317899 |
| GVRRPAGIPADAPV | ASSRRDERLF | 0.81798816 |
| GYRQVTDAHLLTLARR | ASSRRDERLF | 0.701814 |
| IAKMGQDGHDRG | ASSRRDERLF | 0.7224468 |
| KAEPSIEIQYT | ASSRRDERLF | 0.7760155 |
| KPVRIRMAADEDTVAVT | ASSRRDERLF | 0.8055422 |
| MPPRIAGMRLLVIKPEP | ASSRRDERLF | 0.6597184 |
| MKFYAVALFRSRGVPPTR | ASSRRDERLF | 0.69532466 |
| NAGGFGILGAVAHSP | ASSRRDERLF | 0.41973144 |
| NEGREITLAMAEM | ASSRRDERLF | 0.83764195 |
| NELVEWSHPLGHRWRWE | ASSRRDERLF | 0.4841881 |
| NLGAGGTGGNGGNPG | ASSRRDERLF | 0.5755964 |
| NTEWPKWETVMAADPVD | ASSRRDERLF | 0.7053267 |
| PASAAPAETQP | ASSRRDERLF | 0.8461852 |
| QQSSDGKNGVRAAEITGE | ASSRRDERLF | 0.8531649 |
| RAPDAVTTSWWKE | ASSRRDERLF | 0.6464815 |
| SGFGNVGAHTSGWFNQ | ASSRRDERLF | 0.6030681 |
| SGVAGAVGQAASVGGLKV | ASSRRDERLF | 0.7968284 |
| SNGGAGTGSAGGNGGT | ASSRRDERLF | 0.5914905 |
| TGGAGGEGGAAG | ASSRRDERLF | 0.82759583 |
| TGGDGGAAGAGGAPGHG | ASSRRDERLF | 0.73100626 |
| TMPPLLAAIANAGRHGV | ASSRRDERLF | 0.69915634 |

| | | |
|---|---|---|
| TSSTAGNSGGA | ASSRRDERLF | 0.6391057 |
| VPATVPVGILC | ASSRRDERLF | 0.8092203 |
| VQRPDSGGVVLTGRIS | ASSRRDERLF | 0.67801994 |
| WTPNPVPATKGVPAGLVQAT | ASSRRDERLF | 0.81901205 |
| YALKRALAKFSDRFDVVII | ASSRRDERLF | 0.2299913 |
| YLIGSVIV | ASSRRDERLF | 0.82211447 |
| YGQLVAVID | ASSRRDERLF | 0.51581746 |
| VTVGGMTLPA | ASSRRDERLF | 0.57773876 |
| VLALPAV | ASSRRDERLF | 0.21939448 |
| VIPPTLNYET | ASSRRDERLF | 0.69597113 |
| VGMGGMHPSSG | ASSRRDERLF | 0.7887068 |
| VGLAGGAATA | ASSRRDERLF | 0.8448099 |
| VGGAGGAGGSGP | ASSRRDERLF | 0.74914813 |
| VFAAFMV | ASSRRDERLF | 0.79567796 |
| TLYDKLR | ASSRRDERLF | 0.8463 |
| TLAVFTVL | ASSRRDERLF | 0.64728546 |
| TGSVQPARV | ASSRRDERLF | 0.24586117 |
| TGGGAGSGGT | ASSRRDERLF | 0.7469783 |
| SHLETSL | ASSRRDERLF | 0.76059455 |
| SFTTGMT | ASSRRDERLF | 0.8431757 |
| QVEVAKRMGVRQPTVS | ASSRRDERLF | 0.78259814 |
| QGSGGPALAM | ASSRRDERLF | 0.37742513 |
| MLAPSNP | ASSRRDERLF | 0.79864746 |
| LVNNAAI | ASSRRDERLF | 0.8143586 |
| LTGEINTLA | ASSRRDERLF | 0.48517364 |
| LSIFGPV | ASSRRDERLF | 0.52784055 |
| LLALLANL | ASSRRDERLF | 0.5366735 |
| LISNASDALDKLR | ASSRRDERLF | 0.31075442 |
| LINEAVP | ASSRRDERLF | 0.80001605 |
| LGYAFSGL | ASSRRDERLF | 0.6804487 |
| LDFVAVAL | ASSRRDERLF | 0.68967515 |
| LAQPGAE | ASSRRDERLF | 0.7175742 |
| ISNLGAVTGF | ASSRRDERLF | 0.6110264 |
| HQEDPGRI | ASSRRDERLF | 0.64203465 |
| GTPGEPTP | ASSRRDERLF | 0.6590526 |
| GFFNSGS | ASSRRDERLF | 0.7733274 |
| GAGGGGLTAGAN | ASSRRDERLF | 0.8247911 |
| FVDALVNM | ASSRRDERLF | 0.6935668 |
| FLTVAPEL | ASSRRDERLF | 0.75465333 |
| FETLTAL | ASSRRDERLF | 0.83979577 |
| DVINPIEL | ASSRRDERLF | 0.42522907 |
| APVSGAPV | ASSRRDERLF | 0.58298606 |
| ALVGGLAW | ASSRRDERLF | 0.80365163 |
| ALAAAAATAI | ASSRRDERLF | 0.6402063 |
| AFGLGLLL | ASSRRDERLF | 0.8369962 |
| AFFAAASL | ASSRRDERLF | 0.7592914 |
| AAAALPAFSPP | ASSRRDERLF | 0.7742679 |
| VVLAPAL | ASSRRDERLF | 0.7528919 |
| TIGHVDHGKTTL | ASSRRDERLF | 0.8626704 |
| LISNASDALDK | ASSRRDERLF | 0.789258 |
| LISNASDAL | ASSRRDERLF | 0.6685735 |

| | | |
|---|---|---|
| ISNASDALDK | ASSRRDERLF | 0.76974624 |
| NQAETLVYQTE | ASSRRDERLF | 0.40366873 |
| TGGTGGTAGAG | ASSRRDERLF | 0.8180388 |
| TGSTAGGAGGA | ASSRRDERLF | 0.8177688 |
| DPLAVPVAAGPV | ASSRRDERLF | 0.539817 |
| GGTGGTPGNGGN | ASSRRDERLF | 0.69637835 |
| PAGLAGVDPATL | ASSRRDERLF | 0.86043066 |
| GNNLVVEAPHAGASV | ASSRRDERLF | 0.83188176 |
| KVTRIMSASTLEQPA | ASSRRDERLF | 0.6066896 |
| LGFALPIALPRRTVV | ASSRRDERLF | 0.59306777 |
| TRLGNLTARKLVVLV | ASSRRDERLF | 0.55101216 |
| WSQQRRLIYFPSAGP | ASSRRDERLF | 0.6561787 |
| GPNGNPDPMAAAVDIR | ASSRRDERLF | 0.32145298 |
| TQRCVSVADLRAAASR | ASSRRDERLF | 0.80202913 |
| QFVLDVAEHPVEVLTETV | ASSRRDERLF | 0.6815071 |
| GGAGGTGGAGGNGGTQAGDG | ASSRRDERLF | 0.8249618 |
| PLLATLRAIAVDVDAKPAQV | ASSRRDERLF | 0.78450036 |
| APKFRTGGYMIDFWGVGYQVAK | ASSRRDERLF | 0.83942056 |
| GAGGAGGAGGAAGGPGGTGGTG | ASSRRDERLF | 0.6736481 |
| IGALGLLATDVAPAGSVAFGVA | ASSRRDERLF | 0.66123605 |
| AGGAGGIGGNGGAGGAGGVLMG | ASSRRDERLF | 0.34092677 |
| ADELIGLL | ASSRRDERLF | 0.39640486 |
| ADRALKA | ASSRRDERLF | 0.62807226 |
| AGAVGQVGGAGGA | ASSRRDERLF | 0.6994672 |
| AGGGGGLGGAP | ASSRRDERLF | 0.30255687 |
| AGGLLYGNGGAG | ASSRRDERLF | 0.85807824 |
| AGLESIGL | ASSRRDERLF | 0.6460359 |
| AGLKTGII | ASSRRDERLF | 0.8076192 |
| ASLAALLL | ASSRRDERLF | 0.6842489 |
| ASVINISSI | ASSRRDERLF | 0.39897192 |
| ATPPGPA | ASSRRDERLF | 0.6113889 |
| ATVASALV | ASSRRDERLF | 0.7978148 |
| ATVASLIL | ASSRRDERLF | 0.37150773 |
| AVNGPVLIRH | ASSRRDERLF | 0.33961505 |
| DGAAGAVGI | ASSRRDERLF | 0.5230676 |
| DPLLHGI | ASSRRDERLF | 0.7000911 |
| ELGAALGAL | ASSRRDERLF | 0.8603312 |
| ELGRTQAV | ASSRRDERLF | 0.84230435 |
| FVAGGYLS | ASSRRDERLF | 0.75090945 |
| GASGGAGGAGGW | ASSRRDERLF | 0.8685376 |
| GGAGAGGDGGAGGI | ASSRRDERLF | 0.15622398 |
| GTGHNGSQPGMGGNG | ASSRRDERLF | 0.69402033 |
| GYAAASASAS | ASSRRDERLF | 0.84447 |
| HQVDVAAVQQG | ASSRRDERLF | 0.7924817 |
| IGAAIAHT | ASSRRDERLF | 0.7113886 |
| IGMIEVAQL | ASSRRDERLF | 0.83261377 |
| IRLAPEHTSAMD | ASSRRDERLF | 0.5706368 |
| IYNVSVL | ASSRRDERLF | 0.68044937 |
| KQGFLGM | ASSRRDERLF | 0.6491498 |
| LESVITQL | ASSRRDERLF | 0.8312942 |
| LGLGALVVV | ASSRRDERLF | 0.78821635 |

| | | |
|---|---|---|
| LIGNGGNGGSGGI | ASSRRDERLF | 0.36270657 |
| LNAINAPTL | ASSRRDERLF | 0.81424904 |
| LQLLNTGV | ASSRRDERLF | 0.86692774 |
| LQTSIEF | ASSRRDERLF | 0.63094026 |
| MSFAVSEL | ASSRRDERLF | 0.82767975 |
| NTGALGL | ASSRRDERLF | 0.74714434 |
| PPGAGGAGGAGTT | ASSRRDERLF | 0.6563705 |
| QLVGLGI | ASSRRDERLF | 0.6007379 |
| RFLEQVHQL | ASSRRDERLF | 0.4924739 |
| RGVGYVLR | ASSRRDERLF | 0.79739 |
| SATGVFAGVI | ASSRRDERLF | 0.4985515 |
| SLPAITLP | ASSRRDERLF | 0.7329047 |
| STALAFTV | ASSRRDERLF | 0.7298013 |
| SVLTVGLGK | ASSRRDERLF | 0.87298787 |
| TALAGAGSGI | ASSRRDERLF | 0.78800476 |
| TGLPFTGI | ASSRRDERLF | 0.78704536 |
| VAEHVTV | ASSRRDERLF | 0.5670061 |
| VDIVLNSL | ASSRRDERLF | 0.8002007 |
| VLTGGGAVL | ASSRRDERLF | 0.80034864 |
| VNVGLAYGI | ASSRRDERLF | 0.4608667 |
| VSSAAVGAV | ASSRRDERLF | 0.8369151 |
| VTSVYEAGV | ASSRRDERLF | 0.5275483 |
| WVALDRGA | ASSRRDERLF | 0.7720132 |
| YLVGSVVI | ASSRRDERLF | 0.78813857 |
| GPPGTGKTLL | ASSRRDERLF | 0.83883363 |
| GVGMTKF | ASSRRDERLF | 0.70991004 |
| IGIGGSDL | ASSRRDERLF | 0.80174863 |
| IGTLLAEL | ASSRRDERLF | 0.48953304 |
| IYTSGTT | ASSRRDERLF | 0.82520366 |
| LSFPAGV | ASSRRDERLF | 0.8321027 |
| LYTSGTT | ASSRRDERLF | 0.69592214 |
| VGAFPFGF | ASSRRDERLF | 0.8687583 |
| VGAGTVEF | ASSRRDERLF | 0.8024403 |
| VGKSTLLL | ASSRRDERLF | 0.2740633 |
| AALAHAGGGIV | ASSRRDERLF | 0.734231 |
| AALVAVSLL | ASSRRDERLF | 0.84111917 |
| AAVSRAVAAL | ASSRRDERLF | 0.84581 |
| AGGSSGIGGF | ASSRRDERLF | 0.82972705 |
| AGLTVANANG | ASSRRDERLF | 0.78414434 |
| AGLVGAVL | ASSRRDERLF | 0.7051465 |
| ALSQAFTV | ASSRRDERLF | 0.8484733 |
| ALTFSVSL | ASSRRDERLF | 0.88407737 |
| ALTGAALVAL | ASSRRDERLF | 0.7998599 |
| ASFSIGAGVL | ASSRRDERLF | 0.5841611 |
| DGSVYKI | ASSRRDERLF | 0.5489907 |
| EQLAEIV | ASSRRDERLF | 0.8119975 |
| FAVSELTVM | ASSRRDERLF | 0.8083999 |
| GAAPLAAGL | ASSRRDERLF | 0.46629688 |
| GADGGDGGSGGAGG | ASSRRDERLF | 0.70484066 |
| GFGLDTGGAGG | ASSRRDERLF | 0.7518828 |
| GLASGGLL | ASSRRDERLF | 0.22514221 |

| | | |
|---|---|---|
| GRIAGAAVLL | ASSRRDERLF | 0.8021009 |
| GSAGSLNTSF | ASSRRDERLF | 0.8446673 |
| GTGQIGI | ASSRRDERLF | 0.80095834 |
| GTLIALIL | ASSRRDERLF | 0.42128444 |
| IAATMGFVVG | ASSRRDERLF | 0.8359413 |
| IFAQALL | ASSRRDERLF | 0.85141546 |
| IGAAVALGF | ASSRRDERLF | 0.855629 |
| IGLTPLAL | ASSRRDERLF | 0.39034328 |
| ILGSNTSTL | ASSRRDERLF | 0.16698985 |
| IVFSLTVL | ASSRRDERLF | 0.7319963 |
| LASKATGF | ASSRRDERLF | 0.8081227 |
| LASVALIL | ASSRRDERLF | 0.6533978 |
| LFAVGLTL | ASSRRDERLF | 0.48034477 |
| LFNGGIGTYI | ASSRRDERLF | 0.70231223 |
| LTLGAGII | ASSRRDERLF | 0.83376074 |
| LVAAALGDI | ASSRRDERLF | 0.4094418 |
| NTAFVQLGI | ASSRRDERLF | 0.49569497 |
| QLFSGAEL | ASSRRDERLF | 0.77820647 |
| SAATFITL | ASSRRDERLF | 0.83275634 |
| SNYGLANLGV | ASSRRDERLF | 0.58011186 |
| SYGGAYAVM | ASSRRDERLF | 0.65634334 |
| TTGVGGLTLG | ASSRRDERLF | 0.7762687 |
| TTQTIYI | ASSRRDERLF | 0.80767095 |
| VAAGAGLGAVI | ASSRRDERLF | 0.77473706 |
| VAAVVTLL | ASSRRDERLF | 0.44213808 |
| VGFLGGLI | ASSRRDERLF | 0.8323494 |
| VGVIPILL | ASSRRDERLF | 0.46590245 |
| VIFASSLI | ASSRRDERLF | 0.8211776 |
| VKGGDGGAAAG | ASSRRDERLF | 0.73999614 |
| VTKTFTNI | ASSRRDERLF | 0.7955748 |
| VVAAAILL | ASSRRDERLF | 0.63742375 |
| YAALDVEL | ASSRRDERLF | 0.529519 |
| LASVALLL | ASSRRDERLF | 0.44868058 |
| LDAVHRV | ASSRRDERLF | 0.49185342 |
| QGLSSSIF | ASSRRDERLF | 0.83122885 |
| RGTFANIRL | ASSRRDERLF | 0.31445763 |
| RQLWWGHRI | ASSRRDERLF | 0.8213867 |
| YGPPGTGKTLL | ASSRRDERLF | 0.7511605 |
| AALVGAPANVA | ASSRRDERLF | 0.56245244 |
| AATFRRTE | ASSRRDERLF | 0.810179 |
| AGALGGGI | ASSRRDERLF | 0.33116806 |
| AGGAGGAGGIGAG | ASSRRDERLF | 0.7733655 |
| AGSGALGL | ASSRRDERLF | 0.69372857 |
| AQIADQI | ASSRRDERLF | 0.79815865 |
| ASAVAVTYL | ASSRRDERLF | 0.6363997 |
| ASTVLGGTGG | ASSRRDERLF | 0.8148525 |
| AVGGEPAK | ASSRRDERLF | 0.7904217 |
| CEQGTFGGTC | ASSRRDERLF | 0.4825579 |
| DEGKAPAG | ASSRRDERLF | 0.7931626 |
| DTVVFPAL | ASSRRDERLF | 0.8275796 |
| FEGITMAG | ASSRRDERLF | 0.8049785 |

| | | |
|---|---|---|
| FPLVVQD | ASSRRDERLF | 0.29401675 |
| GAFTTPQ | ASSRRDERLF | 0.7880007 |
| GFGGFGTAGGVGGA | ASSRRDERLF | 0.5695652 |
| GFLAQFL | ASSRRDERLF | 0.77076256 |
| GGAGGAGGNAGVA | ASSRRDERLF | 0.75015974 |
| GGVGTAGAGGAGGAGG | ASSRRDERLF | 0.6359093 |
| GNGGGGDAGNAGSGGN | ASSRRDERLF | 0.81575644 |
| GTGGAGGAGNDAGS | ASSRRDERLF | 0.38114658 |
| GTTNTGWFNS | ASSRRDERLF | 0.65283084 |
| IFMGPLFTL | ASSRRDERLF | 0.6546439 |
| ILAHAIEAI | ASSRRDERLF | 0.75357467 |
| IRSAIREAL | ASSRRDERLF | 0.7999109 |
| ITFAGGG | ASSRRDERLF | 0.47001833 |
| ITGMSGHQEV | ASSRRDERLF | 0.7781757 |
| IVAGLACTL | ASSRRDERLF | 0.7034288 |
| IVGGGLAGA | ASSRRDERLF | 0.5492188 |
| IVTGVLNI | ASSRRDERLF | 0.78264725 |
| LAQQGCI | ASSRRDERLF | 0.63294876 |
| LCLHVEN | ASSRRDERLF | 0.3677982 |
| LDAISDGS | ASSRRDERLF | 0.8219725 |
| LGGFAAML | ASSRRDERLF | 0.46455395 |
| LGGKNPMI | ASSRRDERLF | 0.78823626 |
| LGPKGVAAIL | ASSRRDERLF | 0.852795 |
| LMVGVSI | ASSRRDERLF | 0.31963545 |
| LVGAAFVL | ASSRRDERLF | 0.8180429 |
| LVLAVAPL | ASSRRDERLF | 0.7384664 |
| MAAIAVIDAA | ASSRRDERLF | 0.7521089 |
| NNGINAELQ | ASSRRDERLF | 0.69869685 |
| RGEPLVLVHG | ASSRRDERLF | 0.5713601 |
| RGGAGGAAGLI | ASSRRDERLF | 0.8384006 |
| RSYRMHPAV | ASSRRDERLF | 0.8072159 |
| SAAHLLAGQLA | ASSRRDERLF | 0.7167569 |
| SAAVGSY | ASSRRDERLF | 0.78015673 |
| SPHAVLHRI | ASSRRDERLF | 0.8055334 |
| SQIAKVL | ASSRRDERLF | 0.8244091 |
| TALGHLVP | ASSRRDERLF | 0.6247061 |
| TAPTTSGAA | ASSRRDERLF | 0.3690472 |
| TDGDDGNGGSPGT | ASSRRDERLF | 0.6837354 |
| TDGLGEN | ASSRRDERLF | 0.6585789 |
| TGGPPLPGQA | ASSRRDERLF | 0.56878096 |
| TIPAITI | ASSRRDERLF | 0.37416428 |
| TIPGFTI | ASSRRDERLF | 0.81128764 |
| TSIASGALA | ASSRRDERLF | 0.8715118 |
| TVTLPGAAI | ASSRRDERLF | 0.66434914 |
| VASGLSAALP | ASSRRDERLF | 0.8709974 |
| VGAGVIGI | ASSRRDERLF | 0.3894801 |
| VGPKRFGV | ASSRRDERLF | 0.81019926 |
| VNVAFPD | ASSRRDERLF | 0.82440174 |
| VSMTNSGVS | ASSRRDERLF | 0.53933036 |
| VTDVLGDLSV | ASSRRDERLF | 0.7601515 |
| VVVVGAGFAGL | ASSRRDERLF | 0.5185343 |

| | | |
|---|---|---|
| VYTVGTVV | ASSRRDERLF | 0.5252211 |
| WTTFSGP | ASSRRDERLF | 0.8241178 |
| YDELVEAF | ASSRRDERLF | 0.6134312 |
| YGTGGAGGNGG | ASSRRDERLF | 0.8025271 |
| YTNVMAV | ASSRRDERLF | 0.5935534 |
| FLEQVHQL | ASSRRDERLF | 0.5341307 |
| GLSLLAVAV | ASSRRDERLF | 0.79645455 |
| LEQVHQL | ASSRRDERLF | 0.8128731 |
| LNAAGAI | ASSRRDERLF | 0.75059134 |
| PAVHASL | ASSRRDERLF | 0.88186294 |
| TELEDDA | ASSRRDERLF | 0.61940396 |
| TPPGPPN | ASSRRDERLF | 0.85933375 |
| TPPPPPPP | ASSRRDERLF | 0.8516077 |
| TQSPALI | ASSMTGRNTEVF | 0.7528491 |
| EFAAAFPQ | ASSMTGRNTEVF | 0.70402586 |
| LLSGNLDVL | ASSMTGRNTEVF | 0.821129 |
| GPANLGNYNI | ASSMTGRNTEVF | 0.6492529 |
| TLGPTLP | ASSMTGRNTEVF | 0.75755584 |
| VYAYSSCCK | ASSMTGRNTEVF | 0.85931075 |
| APAPAPAPNI | ASSMTGRNTEVF | 0.5353502 |
| PGGPVPIP | ASSMTGRNTEVF | 0.6966665 |
| FLTVALEP | ASSMTGRNTEVF | 0.6415827 |
| SQYFGANL | ASSMTGRNTEVF | 0.6393672 |
| GGVKPAA | ASSMTGRNTEVF | 0.5881962 |
| GGTGGAGGAGGW | ASSMTGRNTEVF | 0.44104457 |
| GVGGVAGLGGAGG | ASSMTGRNTEVF | 0.61743015 |
| VLLVLATTI | ASSMTGRNTEVF | 0.54692006 |
| IGIFAAML | ASSMTGRNTEVF | 0.678971 |
| AVGDLID | ASSMTGRNTEVF | 0.6750531 |
| NISPLPL | ASSMTGRNTEVF | 0.6016438 |
| VTAVFVAI | ASSMTGRNTEVF | 0.6951914 |
| SGDDNDAAL | ASSMTGRNTEVF | 0.6998052 |
| AVAALAHS | ASSMTGRNTEVF | 0.44834495 |
| GGVTVMATP | ASSMTGRNTEVF | 0.6298865 |
| AGLAANVRD | ASSMTGRNTEVF | 0.50608474 |
| TNRPDLIDPALLRPG | ASSMTGRNTEVF | 0.5993844 |
| LANLGVTAL | ASSMTGRNTEVF | 0.32201347 |
| TGERIEL | ASSMTGRNTEVF | 0.67088807 |
| NGEELAA | ASSMTGRNTEVF | 0.5479965 |
| VLSLVWG | ASSMTGRNTEVF | 0.27605218 |
| IGYALWL | ASSMTGRNTEVF | 0.63552356 |
| LSLLLAPL | ASSMTGRNTEVF | 0.62047595 |
| TLDEALK | ASSMTGRNTEVF | 0.61365217 |
| TDGAGTAEGG | ASSMTGRNTEVF | 0.677334 |
| NAPGGAGGNGG | ASSMTGRNTEVF | 0.4880613 |
| VSVFMGMGGNF | ASSMTGRNTEVF | 0.5781954 |
| LASYLAAL | ASSMTGRNTEVF | 0.66523314 |
| LVGGTPAL | ASSMTGRNTEVF | 0.7063722 |
| VRNADVL | ASSMTGRNTEVF | 0.54141927 |
| TCSSPFLVSAM | ASSMTGRNTEVF | 0.63970125 |
| VHITDAE | ASSMTGRNTEVF | 0.7040365 |

| | | |
|---|---|---|
| EYAAPSF | ASSMTGRNTEVF | 0.6412808 |
| VVDAFHGGL | ASSMTGRNTEVF | 0.6833615 |
| ASVVDPQ | ASSMTGRNTEVF | 0.7039875 |
| SYNEFGGLS | ASSMTGRNTEVF | 0.5269479 |
| DPDGSVY | ASSMTGRNTEVF | 0.69539726 |
| MVSTIHATE | ASSMTGRNTEVF | 0.70939624 |
| LAAGVADPV | ASSMTGRNTEVF | 0.6308528 |
| TKEQIAA | ASSMTGRNTEVF | 0.6088782 |
| SLASPAAL | ASSMTGRNTEVF | 0.526757 |
| IGLVFYGV | ASSMTGRNTEVF | 0.6127633 |
| FLLDASNL | ASSMTGRNTEVF | 0.38439468 |
| YQVGNFP | ASSMTGRNTEVF | 0.6283175 |
| ESLLSQST | ASSMTGRNTEVF | 0.686949 |
| LFTVGFAAVG | ASSMTGRNTEVF | 0.5828341 |
| ATPPTPT | ASSMTGRNTEVF | 0.43767276 |
| TDLATLG | ASSMTGRNTEVF | 0.5281055 |
| SAGGTGR | ASSMTGRNTEVF | 0.62038356 |
| GFSNSGS | ASSMTGRNTEVF | 0.5783054 |
| GGNGGSGGTSV | ASSMTGRNTEVF | 0.38178775 |
| GTSVATGGAGN | ASSMTGRNTEVF | 0.53101104 |
| NGGNGGSGGTS | ASSMTGRNTEVF | 0.68751043 |
| AVVKFPH | ASSMTGRNTEVF | 0.6931142 |
| TPPPPPP | ASSMTGRNTEVF | 0.45324862 |
| AAPAAAAAPA | ASSMTGRNTEVF | 0.69978946 |
| IALNEDL | ASSMTGRNTEVF | 0.6713984 |
| IVNVSSM | ASSMTGRNTEVF | 0.7134486 |
| TGIPPAPR | ASSMTGRNTEVF | 0.6533875 |
| FDLGGGTF | ASSMTGRNTEVF | 0.53720605 |
| FDLGGGTFDVS | ASSMTGRNTEVF | 0.676896 |
| NEPTAAA | ASSMTGRNTEVF | 0.57632077 |
| NEPTAAALA | ASSMTGRNTEVF | 0.52889156 |
| GSIINISSI | ASSMTGRNTEVF | 0.67657053 |
| VAPGNAGTA | ASSMTGRNTEVF | 0.6366898 |
| TGGGASLE | ASSMTGRNTEVF | 0.53595525 |
| VAVYNHYKRI | ASSMTGRNTEVF | 0.55577487 |
| TDPAPATT | ASSMTGRNTEVF | 0.6975452 |
| TNRPDILDPALLRPG | ASSMTGRNTEVF | 0.4619428 |
| LTTTTTS | ASSMTGRNTEVF | 0.563267 |
| PTGSGKTL | ASSMTGRNTEVF | 0.5391389 |
| TGSGKTL | ASSMTGRNTEVF | 0.6344491 |
| TIGHVDHGK | ASSMTGRNTEVF | 0.6985097 |
| TIGHVDHGKTT | ASSMTGRNTEVF | 0.6688233 |
| GHVDHGKTT | ASSMTGRNTEVF | 0.5626836 |
| AQFEHTLL | ASSMTGRNTEVF | 0.6648636 |
| IVNQDEL | ASSMTGRNTEVF | 0.6636573 |
| TNGSQFF | ASSMTGRNTEVF | 0.63578635 |
| TGKGIVE | ASSMTGRNTEVF | 0.6737321 |
| LISNASDA | ASSMTGRNTEVF | 0.520539 |
| AELRTLGELTKIAKAHGAQ | ASSMTGRNTEVF | 0.68871385 |
| APNAVAAAEPSW | ASSMTGRNTEVF | 0.65352106 |
| APQVAGAATAVL | ASSMTGRNTEVF | 0.68106717 |

| | | |
|---|---|---|
| ARRDPAAAAAILPTDA | ASSMTGRNTEVF | 0.680776 |
| DAAARDVLAEAGLAEYF | ASSMTGRNTEVF | 0.67303103 |
| DDQIVLSAKANLAKVEQ | ASSMTGRNTEVF | 0.64529 |
| DELLPDTGEDSDEDGH | ASSMTGRNTEVF | 0.39568052 |
| ELPEAPANRATVPQD | ASSMTGRNTEVF | 0.55042535 |
| FGQSKIERVTGIG | ASSMTGRNTEVF | 0.6527699 |
| GAGGTGSTAGGAGGAGGAGGLY | ASSMTGRNTEVF | 0.6367881 |
| GHGGTGAAGGNGGTGGAGGSGI | ASSMTGRNTEVF | 0.6705415 |
| GNGGNGGAGGEGGPGGAGGAGA | ASSMTGRNTEVF | 0.671944 |
| GSTAGNGGNGGNATNGTIAGQP | ASSMTGRNTEVF | 0.55221677 |
| GTGGDGGNGGNGGTGG | ASSMTGRNTEVF | 0.7006635 |
| GVRRPAGIPADAPV | ASSMTGRNTEVF | 0.6988827 |
| GYRQVTDAHLLTLARR | ASSMTGRNTEVF | 0.40441412 |
| IAKMGQDGHDRG | ASSMTGRNTEVF | 0.57421035 |
| KAEPSIEIQYT | ASSMTGRNTEVF | 0.71094006 |
| KPVRIRMAADEDTVAVT | ASSMTGRNTEVF | 0.6334988 |
| MPPRIAGMRLLVIKPEP | ASSMTGRNTEVF | 0.6174476 |
| MKFYAVALFRSRGVPPTR | ASSMTGRNTEVF | 0.57799745 |
| NAGGFGILGAVAHSP | ASSMTGRNTEVF | 0.7134648 |
| NEGREITLAMAEM | ASSMTGRNTEVF | 0.5279928 |
| NELVEWSHPLGHRWRWE | ASSMTGRNTEVF | 0.71234816 |
| NLGAGGTGGNGGNPG | ASSMTGRNTEVF | 0.3265334 |
| NTEWPKWETVMAADPVD | ASSMTGRNTEVF | 0.59527284 |
| PASAAPAETQP | ASSMTGRNTEVF | 0.5678182 |
| QQSSDGKNGVRAAEITGE | ASSMTGRNTEVF | 0.36657405 |
| RAPDAVTTSWWKE | ASSMTGRNTEVF | 0.6735183 |
| SGFGNVGAHTSGWFNQ | ASSMTGRNTEVF | 0.6142507 |
| SGVAGAVGQAASVGGLKV | ASSMTGRNTEVF | 0.67169106 |
| SNGGAGTGSAGGNGGT | ASSMTGRNTEVF | 0.6021428 |
| TGGAGGEGGAAG | ASSMTGRNTEVF | 0.67925406 |
| TGGDGGAAGAGGAPGHG | ASSMTGRNTEVF | 0.6187108 |
| TMPPLLAAIANAGRHGV | ASSMTGRNTEVF | 0.6448123 |
| TSSTAGNSGGA | ASSMTGRNTEVF | 0.53804004 |
| VPATVPVGILC | ASSMTGRNTEVF | 0.6689432 |
| VQRPDSGGVVLTGRIS | ASSMTGRNTEVF | 0.7032641 |
| WTPNPVPATKGVPAGLVQAT | ASSMTGRNTEVF | 0.6748591 |
| YALKRALAKFSDRFDVVII | ASSMTGRNTEVF | 0.47769395 |
| YLIGSVIV | ASSMTGRNTEVF | 0.54300046 |
| YGQLVAVID | ASSMTGRNTEVF | 0.64049053 |
| VTVGGMTLPA | ASSMTGRNTEVF | 0.36369446 |
| VLALPAV | ASSMTGRNTEVF | 0.32756823 |
| VIPPTLNYET | ASSMTGRNTEVF | 0.6003773 |
| VGMGGMHPSSG | ASSMTGRNTEVF | 0.53434557 |
| VGLAGGAATA | ASSMTGRNTEVF | 0.6913613 |
| VGGAGGAGGSGP | ASSMTGRNTEVF | 0.66759473 |
| VFAAFMV | ASSMTGRNTEVF | 0.5974949 |
| TLYDKLR | ASSMTGRNTEVF | 0.6782538 |
| TLAVFTVL | ASSMTGRNTEVF | 0.6306812 |
| TGSVQPARV | ASSMTGRNTEVF | 0.25314173 |
| TGGGAGSGGT | ASSMTGRNTEVF | 0.56920844 |
| SHLETSL | ASSMTGRNTEVF | 0.45778358 |

| | | |
|---|---|---|
| SFTTGMT | ASSMTGRNTEVF | 0.70384526 |
| QVEVAKRMGVRQPTVS | ASSMTGRNTEVF | 0.51180637 |
| QGSGGPALAM | ASSMTGRNTEVF | 0.50598145 |
| MLAPSNP | ASSMTGRNTEVF | 0.62333494 |
| LVNNAAI | ASSMTGRNTEVF | 0.6202061 |
| LTGEINTLA | ASSMTGRNTEVF | 0.60356283 |
| LSIFGPV | ASSMTGRNTEVF | 0.51509327 |
| LLALLANL | ASSMTGRNTEVF | 0.3162055 |
| LISNASDALDKLR | ASSMTGRNTEVF | 0.6759352 |
| LINEAVP | ASSMTGRNTEVF | 0.46631092 |
| LGYAFSGL | ASSMTGRNTEVF | 0.56180096 |
| LDFVAVAL | ASSMTGRNTEVF | 0.33601096 |
| LAQPGAE | ASSMTGRNTEVF | 0.4905262 |
| ISNLGAVTGF | ASSMTGRNTEVF | 0.41789386 |
| HQEDPGRI | ASSMTGRNTEVF | 0.67172754 |
| GTPGEPTP | ASSMTGRNTEVF | 0.6072726 |
| GFFNSGS | ASSMTGRNTEVF | 0.5733526 |
| GAGGGGLTAGAN | ASSMTGRNTEVF | 0.69901556 |
| FVDALVNM | ASSMTGRNTEVF | 0.6319613 |
| FLTVAPEL | ASSMTGRNTEVF | 0.5710211 |
| FETLTAL | ASSMTGRNTEVF | 0.6154976 |
| DVINPIEL | ASSMTGRNTEVF | 0.5033318 |
| APVSGAPV | ASSMTGRNTEVF | 0.56961256 |
| ALVGGLAW | ASSMTGRNTEVF | 0.564116 |
| ALAAAAATAI | ASSMTGRNTEVF | 0.7095751 |
| AFGLGLLL | ASSMTGRNTEVF | 0.5724708 |
| AFFAAASL | ASSMTGRNTEVF | 0.44607112 |
| AAAALPAFSPP | ASSMTGRNTEVF | 0.6422156 |
| VVLAPAL | ASSMTGRNTEVF | 0.7040758 |
| TIGHVDHGKTTL | ASSMTGRNTEVF | 0.7214443 |
| LISNASDALDK | ASSMTGRNTEVF | 0.5313715 |
| LISNASDAL | ASSMTGRNTEVF | 0.6306016 |
| ISNASDALDK | ASSMTGRNTEVF | 0.6184788 |
| NQAETLVYQTE | ASSMTGRNTEVF | 0.59725696 |
| TGGTGGTAGAG | ASSMTGRNTEVF | 0.54497564 |
| TGSTAGGAGGA | ASSMTGRNTEVF | 0.5062977 |
| DPLAVPVAAGPV | ASSMTGRNTEVF | 0.6796686 |
| GGTGGTPGNGGN | ASSMTGRNTEVF | 0.71965575 |
| PAGLAGVDPATL | ASSMTGRNTEVF | 0.5975817 |
| GNNLVVEAPHAGASV | ASSMTGRNTEVF | 0.3473354 |
| KVTRIMSASTLEQPA | ASSMTGRNTEVF | 0.62854224 |
| LGFALPIALPRRTVV | ASSMTGRNTEVF | 0.51897144 |
| TRLGNLTARKLVVLV | ASSMTGRNTEVF | 0.57393175 |
| WSQQRRLIYFPSAGP | ASSMTGRNTEVF | 0.64102584 |
| GPNGNPDPMAAAVDIR | ASSMTGRNTEVF | 0.3765483 |
| TQRCVSVADLRAAASR | ASSMTGRNTEVF | 0.5876547 |
| QFVLDVAEHPVEVLTETV | ASSMTGRNTEVF | 0.7126218 |
| GGAGGTGGAGGNGGTQAGDG | ASSMTGRNTEVF | 0.59430516 |
| PLLATLRAIAVDVDAKPAQV | ASSMTGRNTEVF | 0.670104 |
| APKFRTGGYMIDFWGVGYQVAK | ASSMTGRNTEVF | 0.6850356 |
| GAGGAGGAGGAAGGPGGTGGTG | ASSMTGRNTEVF | 0.6534723 |

| | | |
|---|---|---|
| IGALGLLATDVAPAGSVAFGVA | ASSMTGRNTEVF | 0.6853692 |
| AGGAGGIGGNGGAGGAGGVLMG | ASSMTGRNTEVF | 0.6802912 |
| ADELIGLL | ASSMTGRNTEVF | 0.6275854 |
| ADRALKA | ASSMTGRNTEVF | 0.62458616 |
| AGAVGQVGGAGGA | ASSMTGRNTEVF | 0.67487633 |
| AGGGGGLGGAP | ASSMTGRNTEVF | 0.6359167 |
| AGGLLYGNGGAG | ASSMTGRNTEVF | 0.6248361 |
| AGLESIGL | ASSMTGRNTEVF | 0.52618635 |
| AGLKTGII | ASSMTGRNTEVF | 0.69474787 |
| ASLAALLL | ASSMTGRNTEVF | 0.6233705 |
| ASVINISSI | ASSMTGRNTEVF | 0.6959628 |
| ATPPGPA | ASSMTGRNTEVF | 0.46084672 |
| ATVASALV | ASSMTGRNTEVF | 0.5363822 |
| ATVASLIL | ASSMTGRNTEVF | 0.6679814 |
| AVNGPVLIRH | ASSMTGRNTEVF | 0.30887818 |
| DGAAGAVGI | ASSMTGRNTEVF | 0.3440402 |
| DPLLHGI | ASSMTGRNTEVF | 0.6522515 |
| ELGAALGAL | ASSMTGRNTEVF | 0.7030305 |
| ELGRTQAV | ASSMTGRNTEVF | 0.29130188 |
| FVAGGYLS | ASSMTGRNTEVF | 0.56541616 |
| GASGGAGGAGGW | ASSMTGRNTEVF | 0.4866417 |
| GGAGAGGDGGAGGI | ASSMTGRNTEVF | 0.6779511 |
| GTGHNGSQPGMGGNG | ASSMTGRNTEVF | 0.62973845 |
| GYAAASASAS | ASSMTGRNTEVF | 0.4231773 |
| HQVDVAAVQQG | ASSMTGRNTEVF | 0.6660085 |
| IGAAIAHT | ASSMTGRNTEVF | 0.33441588 |
| IGMIEVAQL | ASSMTGRNTEVF | 0.51702195 |
| IRLAPEHTSAMD | ASSMTGRNTEVF | 0.67229474 |
| IYNVSVL | ASSMTGRNTEVF | 0.69802046 |
| KQGFLGM | ASSMTGRNTEVF | 0.4785999 |
| LESVITQL | ASSMTGRNTEVF | 0.6794789 |
| LGLGALVVV | ASSMTGRNTEVF | 0.5312552 |
| LIGNGGNGGSGGI | ASSMTGRNTEVF | 0.7228553 |
| LNAINAPTL | ASSMTGRNTEVF | 0.6165724 |
| LQLLNTGV | ASSMTGRNTEVF | 0.41030416 |
| LQTSIEF | ASSMTGRNTEVF | 0.6710234 |
| MSFAVSEL | ASSMTGRNTEVF | 0.6653338 |
| NTGALGL | ASSMTGRNTEVF | 0.66738766 |
| PPGAGGAGGAGTT | ASSMTGRNTEVF | 0.6717645 |
| QLVGLGI | ASSMTGRNTEVF | 0.5836325 |
| RFLEQVHQL | ASSMTGRNTEVF | 0.6096397 |
| RGVGYVLR | ASSMTGRNTEVF | 0.68831277 |
| SATGVFAGVI | ASSMTGRNTEVF | 0.6709582 |
| SLPAITLP | ASSMTGRNTEVF | 0.535447 |
| STALAFTV | ASSMTGRNTEVF | 0.6820333 |
| SVLTVGLGK | ASSMTGRNTEVF | 0.61205775 |
| TALAGAGSGI | ASSMTGRNTEVF | 0.58056325 |
| TGLPFTGI | ASSMTGRNTEVF | 0.69939405 |
| VAEHVTV | ASSMTGRNTEVF | 0.6203556 |
| VDIVLNSL | ASSMTGRNTEVF | 0.67500687 |
| VLTGGGAVL | ASSMTGRNTEVF | 0.6789979 |

| | | |
|---|---|---|
| VNVGLAYGI | ASSMTGRNTEVF | 0.71967965 |
| VSSAAVGAV | ASSMTGRNTEVF | 0.46560454 |
| VTSVYEAGV | ASSMTGRNTEVF | 0.67360485 |
| WVALDRGA | ASSMTGRNTEVF | 0.6419246 |
| YLVGSVVI | ASSMTGRNTEVF | 0.6859255 |
| GPPGTGKTLL | ASSMTGRNTEVF | 0.6132577 |
| GVGMTKF | ASSMTGRNTEVF | 0.5263845 |
| IGIGGSDL | ASSMTGRNTEVF | 0.6539902 |
| IGTLLAEL | ASSMTGRNTEVF | 0.6919248 |
| IYTSGTT | ASSMTGRNTEVF | 0.6242156 |
| LSFPAGV | ASSMTGRNTEVF | 0.6439757 |
| LYTSGTT | ASSMTGRNTEVF | 0.7005304 |
| VGAFPFGF | ASSMTGRNTEVF | 0.59276634 |
| VGAGTVEF | ASSMTGRNTEVF | 0.60970354 |
| VGKSTLLL | ASSMTGRNTEVF | 0.67356104 |
| AALAHAGGGIV | ASSMTGRNTEVF | 0.69351065 |
| AALVAVSLL | ASSMTGRNTEVF | 0.58878917 |
| AAVSRAVAAL | ASSMTGRNTEVF | 0.70364785 |
| AGGSSGIGGF | ASSMTGRNTEVF | 0.5266902 |
| AGLTVANANG | ASSMTGRNTEVF | 0.6748978 |
| AGLVGAVL | ASSMTGRNTEVF | 0.55718696 |
| ALSQAFTV | ASSMTGRNTEVF | 0.6451972 |
| ALTFSVSL | ASSMTGRNTEVF | 0.5411394 |
| ALTGAALVAL | ASSMTGRNTEVF | 0.66770244 |
| ASFSIGAGVL | ASSMTGRNTEVF | 0.6760385 |
| DGSVYKI | ASSMTGRNTEVF | 0.6799572 |
| EQLAEIV | ASSMTGRNTEVF | 0.31314155 |
| FAVSELTVM | ASSMTGRNTEVF | 0.6672027 |
| GAAPLAAGL | ASSMTGRNTEVF | 0.6877351 |
| GADGGDGGSGGAGG | ASSMTGRNTEVF | 0.71075594 |
| GFGLDTGGAGG | ASSMTGRNTEVF | 0.6138418 |
| GLASGGLL | ASSMTGRNTEVF | 0.6858718 |
| GRIAGAAVLL | ASSMTGRNTEVF | 0.6097679 |
| GSAGSLNTSF | ASSMTGRNTEVF | 0.5922984 |
| GTGQIGI | ASSMTGRNTEVF | 0.4818383 |
| GTLIALIL | ASSMTGRNTEVF | 0.64857125 |
| IAATMGFVVG | ASSMTGRNTEVF | 0.55199534 |
| IFAQALL | ASSMTGRNTEVF | 0.5343292 |
| IGAAVALGF | ASSMTGRNTEVF | 0.67521715 |
| IGLTPLAL | ASSMTGRNTEVF | 0.68396 |
| ILGSNTSTL | ASSMTGRNTEVF | 0.6839932 |
| IVFSLTVL | ASSMTGRNTEVF | 0.70577717 |
| LASKATGF | ASSMTGRNTEVF | 0.71047914 |
| LASVALIL | ASSMTGRNTEVF | 0.68926686 |
| LFAVGLTL | ASSMTGRNTEVF | 0.5779687 |
| LFNGGIGTYI | ASSMTGRNTEVF | 0.609694 |
| LTLGAGII | ASSMTGRNTEVF | 0.4316401 |
| LVAAALGDI | ASSMTGRNTEVF | 0.5791538 |
| NTAFVQLGI | ASSMTGRNTEVF | 0.56895375 |
| QLFSGAEL | ASSMTGRNTEVF | 0.6390314 |
| SAATFITL | ASSMTGRNTEVF | 0.29216614 |

| | | |
|---|---|---|
| SNYGLANLGV | ASSMTGRNTEVF | 0.6078367 |
| SYGGAYAVM | ASSMTGRNTEVF | 0.7062341 |
| TTGVGGLTLG | ASSMTGRNTEVF | 0.5376549 |
| TTQTIYI | ASSMTGRNTEVF | 0.66491807 |
| VAAGAGLGAVI | ASSMTGRNTEVF | 0.6467482 |
| VAAVVTLL | ASSMTGRNTEVF | 0.5220064 |
| VGFLGGLI | ASSMTGRNTEVF | 0.6916894 |
| VGVIPILL | ASSMTGRNTEVF | 0.5163995 |
| VIFASSLI | ASSMTGRNTEVF | 0.65242016 |
| VKGGDGGAAAG | ASSMTGRNTEVF | 0.70210457 |
| VTKTFTNI | ASSMTGRNTEVF | 0.64696306 |
| VVAAAILL | ASSMTGRNTEVF | 0.6366854 |
| YAALDVEL | ASSMTGRNTEVF | 0.62613773 |
| LASVALLL | ASSMTGRNTEVF | 0.43385023 |
| LDAVHRV | ASSMTGRNTEVF | 0.6626619 |
| QGLSSSIF | ASSMTGRNTEVF | 0.4253086 |
| RGTFANIRL | ASSMTGRNTEVF | 0.61640674 |
| RQLWWGHRI | ASSMTGRNTEVF | 0.5170044 |
| YGPPGTGKTLL | ASSMTGRNTEVF | 0.66569984 |
| AALVGAPANVA | ASSMTGRNTEVF | 0.6262194 |
| AATFRRTE | ASSMTGRNTEVF | 0.6636764 |
| AGALGGGI | ASSMTGRNTEVF | 0.53628707 |
| AGGAGGAGGIGAG | ASSMTGRNTEVF | 0.5824094 |
| AGSGALGL | ASSMTGRNTEVF | 0.54002726 |
| AQIADQI | ASSMTGRNTEVF | 0.5106418 |
| ASAVAVTYL | ASSMTGRNTEVF | 0.67651033 |
| ASTVLGGTGG | ASSMTGRNTEVF | 0.61235607 |
| AVGGEPAK | ASSMTGRNTEVF | 0.5631677 |
| CEQGTFGGTC | ASSMTGRNTEVF | 0.6893651 |
| DEGKAPAG | ASSMTGRNTEVF | 0.6478078 |
| DTVVFPAL | ASSMTGRNTEVF | 0.38741922 |
| FEGITMAG | ASSMTGRNTEVF | 0.6196228 |
| FPLVVQD | ASSMTGRNTEVF | 0.64330065 |
| GAFTTPQ | ASSMTGRNTEVF | 0.624004 |
| GFGGFGTAGGVGGA | ASSMTGRNTEVF | 0.2860708 |
| GFLAQFL | ASSMTGRNTEVF | 0.6580943 |
| GGAGGAGGNAGVA | ASSMTGRNTEVF | 0.57680106 |
| GGVGTAGAGGAGGAGG | ASSMTGRNTEVF | 0.52599514 |
| GNGGGGDAGNAGSGGN | ASSMTGRNTEVF | 0.54572654 |
| GTGGAGGAGNDAGS | ASSMTGRNTEVF | 0.68203485 |
| GTTNTGWFNS | ASSMTGRNTEVF | 0.58592725 |
| IFMGPLFTL | ASSMTGRNTEVF | 0.65011024 |
| ILAHAIEAI | ASSMTGRNTEVF | 0.667218 |
| IRSAIREAL | ASSMTGRNTEVF | 0.6799704 |
| ITFAGGG | ASSMTGRNTEVF | 0.698752 |
| ITGMSGHQEV | ASSMTGRNTEVF | 0.6855942 |
| IVAGLACTL | ASSMTGRNTEVF | 0.62533796 |
| IVGGGLAGA | ASSMTGRNTEVF | 0.6719687 |
| IVTGVLNI | ASSMTGRNTEVF | 0.6037362 |
| LAQQGCI | ASSMTGRNTEVF | 0.3773491 |
| LCLHVEN | ASSMTGRNTEVF | 0.3574991 |

| | | |
|---|---|---|
| LDAISDGS | ASSMTGRNTEVF | 0.6421641 |
| LGGFAAML | ASSMTGRNTEVF | 0.3787853 |
| LGGKNPMI | ASSMTGRNTEVF | 0.65814567 |
| LGPKGVAAIL | ASSMTGRNTEVF | 0.69690925 |
| LMVGVSI | ASSMTGRNTEVF | 0.6368991 |
| LVGAAFVL | ASSMTGRNTEVF | 0.47548574 |
| LVLAVAPL | ASSMTGRNTEVF | 0.60526276 |
| MAAIAVIDAA | ASSMTGRNTEVF | 0.5480764 |
| NNGINAELQ | ASSMTGRNTEVF | 0.6723851 |
| RGEPLVLVHG | ASSMTGRNTEVF | 0.6883312 |
| RGGAGGAAGLI | ASSMTGRNTEVF | 0.7093954 |
| RSYRMHPAV | ASSMTGRNTEVF | 0.68842655 |
| SAAHLLAGQLA | ASSMTGRNTEVF | 0.56886816 |
| SAAVGSY | ASSMTGRNTEVF | 0.667528 |
| SPHAVLHRI | ASSMTGRNTEVF | 0.5411696 |
| SQIAKVL | ASSMTGRNTEVF | 0.6200386 |
| TALGHLVP | ASSMTGRNTEVF | 0.59650904 |
| TAPTTSGAA | ASSMTGRNTEVF | 0.64889896 |
| TDGDDGNGGSPGT | ASSMTGRNTEVF | 0.640102 |
| TDGLGEN | ASSMTGRNTEVF | 0.56755424 |
| TGGPPLPGQA | ASSMTGRNTEVF | 0.41010118 |
| TIPAITI | ASSMTGRNTEVF | 0.68480843 |
| TIPGFTI | ASSMTGRNTEVF | 0.676838 |
| TSIASGALA | ASSMTGRNTEVF | 0.38140926 |
| TVTLPGAAI | ASSMTGRNTEVF | 0.5577764 |
| VASGLSAALP | ASSMTGRNTEVF | 0.7171299 |
| VGAGVIGI | ASSMTGRNTEVF | 0.7144002 |
| VGPKRFGV | ASSMTGRNTEVF | 0.5366365 |
| VNVAFPD | ASSMTGRNTEVF | 0.55779403 |
| VSMTNSGVS | ASSMTGRNTEVF | 0.61928695 |
| VTDVLGDLSV | ASSMTGRNTEVF | 0.44332477 |
| VVVVGAGFAGL | ASSMTGRNTEVF | 0.63945436 |
| VYTVGTVV | ASSMTGRNTEVF | 0.5804566 |
| WTTFSGP | ASSMTGRNTEVF | 0.58986974 |
| YDELVEAF | ASSMTGRNTEVF | 0.575175 |
| YGTGGAGGNGG | ASSMTGRNTEVF | 0.6284921 |
| YTNVMAV | ASSMTGRNTEVF | 0.58496994 |
| FLEQVHQL | ASSMTGRNTEVF | 0.6771772 |
| GLSLLAVAV | ASSMTGRNTEVF | 0.28743425 |
| LEQVHQL | ASSMTGRNTEVF | 0.64956677 |
| LNAAGAI | ASSMTGRNTEVF | 0.5161005 |
| PAVHASL | ASSMTGRNTEVF | 0.5650928 |
| TELEDDA | ASSMTGRNTEVF | 0.38953528 |
| TPPGPPN | ASSMTGRNTEVF | 0.60243964 |
| TPPPPPPP | ASSMTGRNTEVF | 0.6948821 |
| TQSPALI | ASSHGTGGSAETLY | 0.6793753 |
| EFAAAFPQ | ASSHGTGGSAETLY | 0.45423174 |
| LLSGNLDVL | ASSHGTGGSAETLY | 0.6821563 |
| GPANLGNYNI | ASSHGTGGSAETLY | 0.7177728 |
| TLGPTLP | ASSHGTGGSAETLY | 0.5390147 |
| VYAYSSCCK | ASSHGTGGSAETLY | 0.4108037 |

| | | |
|---|---|---|
| APAPAPAPNI | ASSHGTGGSAETLY | 0.658548 |
| PGGPVPIP | ASSHGTGGSAETLY | 0.6129142 |
| FLTVALEP | ASSHGTGGSAETLY | 0.66826683 |
| SQYFGANL | ASSHGTGGSAETLY | 0.69860756 |
| GGVKPAA | ASSHGTGGSAETLY | 0.45338833 |
| GGTGGAGGAGGW | ASSHGTGGSAETLY | 0.6985767 |
| GVGGVAGLGGAGG | ASSHGTGGSAETLY | 0.6991664 |
| VLLVLATTI | ASSHGTGGSAETLY | 0.5236925 |
| IGIFAAML | ASSHGTGGSAETLY | 0.49384585 |
| AVGDLID | ASSHGTGGSAETLY | 0.52679294 |
| NISPLPL | ASSHGTGGSAETLY | 0.34885094 |
| VTAVFVAI | ASSHGTGGSAETLY | 0.6824254 |
| SGDDNDAAL | ASSHGTGGSAETLY | 0.53723705 |
| AVAALAHS | ASSHGTGGSAETLY | 0.59526306 |
| GGVTVMATP | ASSHGTGGSAETLY | 0.619923 |
| AGLAANVRD | ASSHGTGGSAETLY | 0.6068824 |
| TNRPDLIDPALLRPG | ASSHGTGGSAETLY | 0.5925686 |
| LANLGVTAL | ASSHGTGGSAETLY | 0.61094886 |
| TGERIEL | ASSHGTGGSAETLY | 0.6384233 |
| NGEELAA | ASSHGTGGSAETLY | 0.6426048 |
| VLSLVWG | ASSHGTGGSAETLY | 0.5837057 |
| IGYALWL | ASSHGTGGSAETLY | 0.63069046 |
| LSLLLAPL | ASSHGTGGSAETLY | 0.694152 |
| TLDEALK | ASSHGTGGSAETLY | 0.5171654 |
| TDGAGTAEGG | ASSHGTGGSAETLY | 0.54743075 |
| NAPGGAGGNGG | ASSHGTGGSAETLY | 0.6630325 |
| VSVFMGMGGNF | ASSHGTGGSAETLY | 0.44821298 |
| LASYLAAL | ASSHGTGGSAETLY | 0.6641861 |
| LVGGTPAL | ASSHGTGGSAETLY | 0.62871015 |
| VRNADVL | ASSHGTGGSAETLY | 0.6429979 |
| TCSSPFLVSAM | ASSHGTGGSAETLY | 0.6356956 |
| VHITDAE | ASSHGTGGSAETLY | 0.7137369 |
| EYAAPSF | ASSHGTGGSAETLY | 0.66528594 |
| VVDAFHGGL | ASSHGTGGSAETLY | 0.579888 |
| ASVVDPQ | ASSHGTGGSAETLY | 0.67753065 |
| SYNEFGGLS | ASSHGTGGSAETLY | 0.54064566 |
| DPDGSVY | ASSHGTGGSAETLY | 0.68468654 |
| MVSTIHATE | ASSHGTGGSAETLY | 0.68646085 |
| LAAGVADPV | ASSHGTGGSAETLY | 0.49667913 |
| TKEQIAA | ASSHGTGGSAETLY | 0.61889404 |
| SLASPAAL | ASSHGTGGSAETLY | 0.6985271 |
| IGLVFYGV | ASSHGTGGSAETLY | 0.6863054 |
| FLLDASNL | ASSHGTGGSAETLY | 0.41025305 |
| YQVGNFP | ASSHGTGGSAETLY | 0.5540203 |
| ESLLSQST | ASSHGTGGSAETLY | 0.45573658 |
| LFTVGFAAVG | ASSHGTGGSAETLY | 0.61564624 |
| ATPPTPT | ASSHGTGGSAETLY | 0.5213956 |
| TDLATLG | ASSHGTGGSAETLY | 0.6621182 |
| SAGGTGR | ASSHGTGGSAETLY | 0.6075591 |
| GFSNSGS | ASSHGTGGSAETLY | 0.6650362 |
| GGNGGSGGTSV | ASSHGTGGSAETLY | 0.6483919 |

| | | |
|---|---|---|
| GTSVATGGAGN | ASSHGTGGSAETLY | 0.69518304 |
| NGGNGGSGGTS | ASSHGTGGSAETLY | 0.5900078 |
| AVVKFPH | ASSHGTGGSAETLY | 0.62167746 |
| TPPPPPP | ASSHGTGGSAETLY | 0.4232153 |
| AAPAAAAAPA | ASSHGTGGSAETLY | 0.5357593 |
| IALNEDL | ASSHGTGGSAETLY | 0.5267801 |
| IVNVSSM | ASSHGTGGSAETLY | 0.6118189 |
| TGIPPAPR | ASSHGTGGSAETLY | 0.33610418 |
| FDLGGGTF | ASSHGTGGSAETLY | 0.7014031 |
| FDLGGGTFDVS | ASSHGTGGSAETLY | 0.62246716 |
| NEPTAAA | ASSHGTGGSAETLY | 0.6698549 |
| NEPTAAALA | ASSHGTGGSAETLY | 0.5951087 |
| GSIINISSI | ASSHGTGGSAETLY | 0.57298374 |
| VAPGNAGTA | ASSHGTGGSAETLY | 0.5656919 |
| TGGGASLE | ASSHGTGGSAETLY | 0.6867137 |
| VAVYNHYKRI | ASSHGTGGSAETLY | 0.50098825 |
| TDPAPATT | ASSHGTGGSAETLY | 0.3675525 |
| TNRPDILDPALLRPG | ASSHGTGGSAETLY | 0.5548815 |
| LTTTTTS | ASSHGTGGSAETLY | 0.36823872 |
| PTGSGKTL | ASSHGTGGSAETLY | 0.70999914 |
| TGSGKTL | ASSHGTGGSAETLY | 0.6846624 |
| TIGHVDHGK | ASSHGTGGSAETLY | 0.6471176 |
| TIGHVDHGKTT | ASSHGTGGSAETLY | 0.70053667 |
| GHVDHGKTT | ASSHGTGGSAETLY | 0.33766884 |
| AQFEHTLL | ASSHGTGGSAETLY | 0.60220575 |
| IVNQDEL | ASSHGTGGSAETLY | 0.7066822 |
| TNGSQFF | ASSHGTGGSAETLY | 0.6431621 |
| TGKGIVE | ASSHGTGGSAETLY | 0.6262451 |
| LISNASDA | ASSHGTGGSAETLY | 0.68666863 |
| AELRTLGELTKIAKAHGAQ | ASSHGTGGSAETLY | 0.5561486 |
| APNAVAAAEPSW | ASSHGTGGSAETLY | 0.6385643 |
| APQVAGAATAVL | ASSHGTGGSAETLY | 0.6188264 |
| ARRDPAAAAAILPTDA | ASSHGTGGSAETLY | 0.65675056 |
| DAAARDVLAEAGLAEYF | ASSHGTGGSAETLY | 0.6401843 |
| DDQIVLSAKANLAKVEQ | ASSHGTGGSAETLY | 0.49367064 |
| DELLPDTGEDSDEDGH | ASSHGTGGSAETLY | 0.6702026 |
| ELPEAPANRATVPQD | ASSHGTGGSAETLY | 0.71776915 |
| FGQSKIERVTGIG | ASSHGTGGSAETLY | 0.60447764 |
| GAGGTGSTAGGAGGAGGAGGLY | ASSHGTGGSAETLY | 0.6793712 |
| GHGGTGAAGGNGGTGGAGGSGI | ASSHGTGGSAETLY | 0.6516845 |
| GNGGNGGAGGEGGPGGAGGAGA | ASSHGTGGSAETLY | 0.6635207 |
| GSTAGNGGNGGNATNGTIAGQP | ASSHGTGGSAETLY | 0.5544173 |
| GTGGDGGNGGNGGTGG | ASSHGTGGSAETLY | 0.5344825 |
| GVRRPAGIPADAPV | ASSHGTGGSAETLY | 0.577808 |
| GYRQVTDAHLLTLARR | ASSHGTGGSAETLY | 0.46391585 |
| IAKMGQDGHDRG | ASSHGTGGSAETLY | 0.62694985 |
| KAEPSIEIQYT | ASSHGTGGSAETLY | 0.5492823 |
| KPVRIRMAADEDTVAVT | ASSHGTGGSAETLY | 0.72895676 |
| MPPRIAGMRLLVIKPEP | ASSHGTGGSAETLY | 0.661812 |
| MKFYAVALFRSRGVPPTR | ASSHGTGGSAETLY | 0.6523556 |
| NAGGFGILGAVAHSP | ASSHGTGGSAETLY | 0.53769845 |

| | | |
|---|---|---|
| NEGREITLAMAEM | ASSHGTGGSAETLY | 0.6416397 |
| NELVEWSHPLGHRWRWE | ASSHGTGGSAETLY | 0.6287589 |
| NLGAGGTGGNGGNPG | ASSHGTGGSAETLY | 0.41081688 |
| NTEWPKWETVMAADPVD | ASSHGTGGSAETLY | 0.69921815 |
| PASAAPAETQP | ASSHGTGGSAETLY | 0.54239696 |
| QQSSDGKNGVRAAEITGE | ASSHGTGGSAETLY | 0.58704776 |
| RAPDAVTTSWWKE | ASSHGTGGSAETLY | 0.6749408 |
| SGFGNVGAHTSGWFNQ | ASSHGTGGSAETLY | 0.69247824 |
| SGVAGAVGQAASVGGLKV | ASSHGTGGSAETLY | 0.575269 |
| SNGGAGTGSAGGNGGT | ASSHGTGGSAETLY | 0.5677599 |
| TGGAGGEGGAAG | ASSHGTGGSAETLY | 0.547619 |
| TGGDGGAAGAGGAPGHG | ASSHGTGGSAETLY | 0.68028444 |
| TMPPLLAAIANAGRHGV | ASSHGTGGSAETLY | 0.6903777 |
| TSSTAGNSGGA | ASSHGTGGSAETLY | 0.66002643 |
| VPATVPVGILC | ASSHGTGGSAETLY | 0.7111324 |
| VQRPDSGGVVLTGRIS | ASSHGTGGSAETLY | 0.6467851 |
| WTPNPVPATKGVPAGLVQAT | ASSHGTGGSAETLY | 0.49033386 |
| YALKRALAKFSDRFDVVII | ASSHGTGGSAETLY | 0.6521427 |
| YLIGSVIV | ASSHGTGGSAETLY | 0.675616 |
| YGQLVAVID | ASSHGTGGSAETLY | 0.6883818 |
| VTVGGMTLPA | ASSHGTGGSAETLY | 0.71394664 |
| VLALPAV | ASSHGTGGSAETLY | 0.67659426 |
| VIPPTLNYET | ASSHGTGGSAETLY | 0.5828042 |
| VGMGGMHPSSG | ASSHGTGGSAETLY | 0.69850343 |
| VGLAGGAATA | ASSHGTGGSAETLY | 0.72389805 |
| VGGAGGAGGSGP | ASSHGTGGSAETLY | 0.67412347 |
| VFAAFMV | ASSHGTGGSAETLY | 0.58875144 |
| TLYDKLR | ASSHGTGGSAETLY | 0.5828557 |
| TLAVFTVL | ASSHGTGGSAETLY | 0.6370591 |
| TGSVQPARV | ASSHGTGGSAETLY | 0.67057717 |
| TGGGAGSGGT | ASSHGTGGSAETLY | 0.5231435 |
| SHLETSL | ASSHGTGGSAETLY | 0.62678695 |
| SFTTGMT | ASSHGTGGSAETLY | 0.6548325 |
| QVEVAKRMGVRQPTVS | ASSHGTGGSAETLY | 0.45176488 |
| QGSGGPALAM | ASSHGTGGSAETLY | 0.6670183 |
| MLAPSNP | ASSHGTGGSAETLY | 0.7028154 |
| LVNNAAI | ASSHGTGGSAETLY | 0.589612 |
| LTGEINTLA | ASSHGTGGSAETLY | 0.5571743 |
| LSIFGPV | ASSHGTGGSAETLY | 0.6937442 |
| LLALLANL | ASSHGTGGSAETLY | 0.7258752 |
| LISNASDALDKLR | ASSHGTGGSAETLY | 0.6904755 |
| LINEAVP | ASSHGTGGSAETLY | 0.44685623 |
| LGYAFSGL | ASSHGTGGSAETLY | 0.31219557 |
| LDFVAVAL | ASSHGTGGSAETLY | 0.5560701 |
| LAQPGAE | ASSHGTGGSAETLY | 0.68244696 |
| ISNLGAVTGF | ASSHGTGGSAETLY | 0.6136544 |
| HQEDPGRI | ASSHGTGGSAETLY | 0.5784097 |
| GTPGEPTP | ASSHGTGGSAETLY | 0.6339444 |
| GFFNSGS | ASSHGTGGSAETLY | 0.6846994 |
| GAGGGGLTAGAN | ASSHGTGGSAETLY | 0.59625065 |
| FVDALVNM | ASSHGTGGSAETLY | 0.525836 |

| | | |
|---|---|---|
| FLTVAPEL | ASSHGTGGSAETLY | 0.64157045 |
| FETLTAL | ASSHGTGGSAETLY | 0.69004786 |
| DVINPIEL | ASSHGTGGSAETLY | 0.50151795 |
| APVSGAPV | ASSHGTGGSAETLY | 0.5899972 |
| ALVGGLAW | ASSHGTGGSAETLY | 0.58429205 |
| ALAAAAATAI | ASSHGTGGSAETLY | 0.5546643 |
| AFGLGLLL | ASSHGTGGSAETLY | 0.6191903 |
| AFFAAASL | ASSHGTGGSAETLY | 0.5490548 |
| AAAALPAFSPP | ASSHGTGGSAETLY | 0.69123864 |
| VVLAPAL | ASSHGTGGSAETLY | 0.4829887 |
| TIGHVDHGKTTL | ASSHGTGGSAETLY | 0.66644275 |
| LISNASDALDK | ASSHGTGGSAETLY | 0.653229 |
| LISNASDAL | ASSHGTGGSAETLY | 0.65966475 |
| ISNASDALDK | ASSHGTGGSAETLY | 0.6013152 |
| NQAETLVYQTE | ASSHGTGGSAETLY | 0.5449316 |
| TGGTGGTAGAG | ASSHGTGGSAETLY | 0.4730128 |
| TGSTAGGAGGA | ASSHGTGGSAETLY | 0.39850312 |
| DPLAVPVAAGPV | ASSHGTGGSAETLY | 0.68395066 |
| GGTGGTPGNGGN | ASSHGTGGSAETLY | 0.51225126 |
| PAGLAGVDPATL | ASSHGTGGSAETLY | 0.6893662 |
| GNNLVVEAPHAGASV | ASSHGTGGSAETLY | 0.65806353 |
| KVTRIMSASTLEQPA | ASSHGTGGSAETLY | 0.53491956 |
| LGFALPIALPRRTVV | ASSHGTGGSAETLY | 0.64501405 |
| TRLGNLTARKLVVLV | ASSHGTGGSAETLY | 0.5407906 |
| WSQQRRLIYFPSAGP | ASSHGTGGSAETLY | 0.5069362 |
| GPNGNPDPMAAAVDIR | ASSHGTGGSAETLY | 0.69130236 |
| TQRCVSVADLRAAASR | ASSHGTGGSAETLY | 0.6506864 |
| QFVLDVAEHPVEVLTETV | ASSHGTGGSAETLY | 0.557385 |
| GGAGGTGGAGGNGGTQAGDG | ASSHGTGGSAETLY | 0.6441843 |
| PLLATLRAIAVDVDAKPAQV | ASSHGTGGSAETLY | 0.6473932 |
| APKFRTGGYMIDFWGVGYQVAK | ASSHGTGGSAETLY | 0.57012904 |
| GAGGAGGAGGAAGGPGGTGGTG | ASSHGTGGSAETLY | 0.6201973 |
| IGALGLLATDVAPAGSVAFGVA | ASSHGTGGSAETLY | 0.6940618 |
| AGGAGGIGGNGGAGGAGGVLMG | ASSHGTGGSAETLY | 0.6388008 |
| ADELIGLL | ASSHGTGGSAETLY | 0.49946293 |
| ADRALKA | ASSHGTGGSAETLY | 0.65956986 |
| AGAVGQVGGAGGA | ASSHGTGGSAETLY | 0.5070766 |
| AGGGGGLGGAP | ASSHGTGGSAETLY | 0.640728 |
| AGGLLYGNGGAG | ASSHGTGGSAETLY | 0.6091797 |
| AGLESIGL | ASSHGTGGSAETLY | 0.6256135 |
| AGLKTGII | ASSHGTGGSAETLY | 0.67440146 |
| ASLAALLL | ASSHGTGGSAETLY | 0.46381584 |
| ASVINISSI | ASSHGTGGSAETLY | 0.5346662 |
| ATPPGPA | ASSHGTGGSAETLY | 0.573158 |
| ATVASALV | ASSHGTGGSAETLY | 0.65593857 |
| ATVASLIL | ASSHGTGGSAETLY | 0.6717938 |
| AVNGPVLIRH | ASSHGTGGSAETLY | 0.5003423 |
| DGAAGAVGI | ASSHGTGGSAETLY | 0.65139246 |
| DPLLHGI | ASSHGTGGSAETLY | 0.609726 |
| ELGAALGAL | ASSHGTGGSAETLY | 0.55859166 |
| ELGRTQAV | ASSHGTGGSAETLY | 0.6102442 |

| | | |
|---|---|---|
| FVAGGYLS | ASSHGTGGSAETLY | 0.5617547 |
| GASGGAGGAGGW | ASSHGTGGSAETLY | 0.5221298 |
| GGAGAGGDGGAGGI | ASSHGTGGSAETLY | 0.6576358 |
| GTGHNGSQPGMGGNG | ASSHGTGGSAETLY | 0.57859224 |
| GYAAASASAS | ASSHGTGGSAETLY | 0.6683058 |
| HQVDVAAVQQG | ASSHGTGGSAETLY | 0.659212 |
| IGAAIAHT | ASSHGTGGSAETLY | 0.557872 |
| IGMIEVAQL | ASSHGTGGSAETLY | 0.65846324 |
| IRLAPEHTSAMD | ASSHGTGGSAETLY | 0.6438123 |
| IYNVSVL | ASSHGTGGSAETLY | 0.65525126 |
| KQGFLGM | ASSHGTGGSAETLY | 0.61640453 |
| LESVITQL | ASSHGTGGSAETLY | 0.5284764 |
| LGLGALVVV | ASSHGTGGSAETLY | 0.67918825 |
| LIGNGGNGGSGGI | ASSHGTGGSAETLY | 0.6692779 |
| LNAINAPTL | ASSHGTGGSAETLY | 0.68158656 |
| LQLLNTGV | ASSHGTGGSAETLY | 0.6589656 |
| LQTSIEF | ASSHGTGGSAETLY | 0.6517339 |
| MSFAVSEL | ASSHGTGGSAETLY | 0.6663469 |
| NTGALGL | ASSHGTGGSAETLY | 0.5642192 |
| PPGAGGAGGAGTT | ASSHGTGGSAETLY | 0.43270674 |
| QLVGLGI | ASSHGTGGSAETLY | 0.6136339 |
| RFLEQVHQL | ASSHGTGGSAETLY | 0.61833286 |
| RGVGYVLR | ASSHGTGGSAETLY | 0.5472413 |
| SATGVFAGVI | ASSHGTGGSAETLY | 0.44065523 |
| SLPAITLP | ASSHGTGGSAETLY | 0.6849341 |
| STALAFTV | ASSHGTGGSAETLY | 0.7262484 |
| SVLTVGLGK | ASSHGTGGSAETLY | 0.5855101 |
| TALAGAGSGI | ASSHGTGGSAETLY | 0.71727437 |
| TGLPFTGI | ASSHGTGGSAETLY | 0.40306544 |
| VAEHVTV | ASSHGTGGSAETLY | 0.67231923 |
| VDIVLNSL | ASSHGTGGSAETLY | 0.69351983 |
| VLTGGGAVL | ASSHGTGGSAETLY | 0.4977754 |
| VNVGLAYGI | ASSHGTGGSAETLY | 0.68116045 |
| VSSAAVGAV | ASSHGTGGSAETLY | 0.50107723 |
| VTSVYEAGV | ASSHGTGGSAETLY | 0.5608766 |
| WVALDRGA | ASSHGTGGSAETLY | 0.70001084 |
| YLVGSVVI | ASSHGTGGSAETLY | 0.6220684 |
| GPPGTGKTLL | ASSHGTGGSAETLY | 0.65006363 |
| GVGMTKF | ASSHGTGGSAETLY | 0.61724085 |
| IGIGGSDL | ASSHGTGGSAETLY | 0.5516918 |
| IGTLLAEL | ASSHGTGGSAETLY | 0.6310923 |
| IYTSGTT | ASSHGTGGSAETLY | 0.701446 |
| LSFPAGV | ASSHGTGGSAETLY | 0.52131706 |
| LYTSGTT | ASSHGTGGSAETLY | 0.7230177 |
| VGAFPFGF | ASSHGTGGSAETLY | 0.5420584 |
| VGAGTVEF | ASSHGTGGSAETLY | 0.7134999 |
| VGKSTLLL | ASSHGTGGSAETLY | 0.6764062 |
| AALAHAGGGIV | ASSHGTGGSAETLY | 0.61063147 |
| AALVAVSLL | ASSHGTGGSAETLY | 0.5720482 |
| AAVSRAVAAL | ASSHGTGGSAETLY | 0.68710595 |
| AGGSSGIGGF | ASSHGTGGSAETLY | 0.63091284 |

| | | |
|---|---|---|
| AGLTVANANG | ASSHGTGGSAETLY | 0.65584993 |
| AGLVGAVL | ASSHGTGGSAETLY | 0.6961074 |
| ALSQAFTV | ASSHGTGGSAETLY | 0.6988951 |
| ALTFSVSL | ASSHGTGGSAETLY | 0.8233944 |
| ALTGAALVAL | ASSHGTGGSAETLY | 0.79076934 |
| ASFSIGAGVL | ASSHGTGGSAETLY | 0.8098112 |
| DGSVYKI | ASSHGTGGSAETLY | 0.6789291 |
| EQLAEIV | ASSHGTGGSAETLY | 0.48505646 |
| FAVSELTVM | ASSHGTGGSAETLY | 0.77798986 |
| GAAPLAAGL | ASSHGTGGSAETLY | 0.64322746 |
| GADGGDGGSGGAGG | ASSHGTGGSAETLY | 0.81836736 |
| GFGLDTGGAGG | ASSHGTGGSAETLY | 0.8222164 |
| GLASGGLL | ASSHGTGGSAETLY | 0.6940656 |
| GRIAGAAVLL | ASSHGTGGSAETLY | 0.80415106 |
| GSAGSLNTSF | ASSHGTGGSAETLY | 0.8205606 |
| GTGQIGI | ASSHGTGGSAETLY | 0.53273433 |
| GTLIALIL | ASSHGTGGSAETLY | 0.7847893 |
| IAATMGFVVG | ASSHGTGGSAETLY | 0.6921834 |
| IFAQALL | ASSHGTGGSAETLY | 0.7789629 |
| IGAAVALGF | ASSHGTGGSAETLY | 0.48449314 |
| IGLTPLAL | ASSHGTGGSAETLY | 0.8131536 |
| ILGSNTSTL | ASSHGTGGSAETLY | 0.75152373 |
| IVFSLTVL | ASSHGTGGSAETLY | 0.3034476 |
| LASKATGF | ASSHGTGGSAETLY | 0.7879677 |
| LASVALIL | ASSHGTGGSAETLY | 0.74701834 |
| LFAVGLTL | ASSHGTGGSAETLY | 0.7831593 |
| LFNGGIGTYI | ASSHGTGGSAETLY | 0.818972 |
| LTLGAGII | ASSHGTGGSAETLY | 0.7401964 |
| LVAAALGDI | ASSHGTGGSAETLY | 0.68056226 |
| NTAFVQLGI | ASSHGTGGSAETLY | 0.7985027 |
| QLFSGAEL | ASSHGTGGSAETLY | 0.81685466 |
| SAATFITL | ASSHGTGGSAETLY | 0.61821264 |
| SNYGLANLGV | ASSHGTGGSAETLY | 0.79803973 |
| SYGGAYAVM | ASSHGTGGSAETLY | 0.8232145 |
| TTGVGGLTLG | ASSHGTGGSAETLY | 0.7336755 |
| TTQTIYI | ASSHGTGGSAETLY | 0.8300342 |
| VAAGAGLGAVI | ASSHGTGGSAETLY | 0.832591 |
| VAAVVTLL | ASSHGTGGSAETLY | 0.7463168 |
| VGFLGGLI | ASSHGTGGSAETLY | 0.81883717 |
| VGVIPILL | ASSHGTGGSAETLY | 0.8163826 |
| VIFASSLI | ASSHGTGGSAETLY | 0.7731237 |
| VKGGDGGAAAG | ASSHGTGGSAETLY | 0.7648814 |
| VTKTFTNI | ASSHGTGGSAETLY | 0.6489316 |
| VVAAAILL | ASSHGTGGSAETLY | 0.7893005 |
| YAALDVEL | ASSHGTGGSAETLY | 0.55450225 |
| LASVALLL | ASSHGTGGSAETLY | 0.7869446 |
| LDAVHRV | ASSHGTGGSAETLY | 0.82501495 |
| QGLSSSIF | ASSHGTGGSAETLY | 0.7412969 |
| RGTFANIRL | ASSHGTGGSAETLY | 0.63196373 |
| RQLWWGHRI | ASSHGTGGSAETLY | 0.6214553 |
| YGPPGTGKTLL | ASSHGTGGSAETLY | 0.8059152 |

| | | |
|---|---|---|
| AALVGAPANVA | ASSHGTGGSAETLY | 0.7472904 |
| AATFRRTE | ASSHGTGGSAETLY | 0.64582825 |
| AGALGGGI | ASSHGTGGSAETLY | 0.75607276 |
| AGGAGGAGGIGAG | ASSHGTGGSAETLY | 0.8029478 |
| AGSGALGL | ASSHGTGGSAETLY | 0.8142298 |
| AQIADQI | ASSHGTGGSAETLY | 0.57281786 |
| ASAVAVTYL | ASSHGTGGSAETLY | 0.8254264 |
| ASTVLGGTGG | ASSHGTGGSAETLY | 0.81413436 |
| AVGGEPAK | ASSHGTGGSAETLY | 0.82426214 |
| CEQGTFGGTC | ASSHGTGGSAETLY | 0.7952935 |
| DEGKAPAG | ASSHGTGGSAETLY | 0.69444627 |
| DTVVFPAL | ASSHGTGGSAETLY | 0.8133943 |
| FEGITMAG | ASSHGTGGSAETLY | 0.7027284 |
| FPLVVQD | ASSHGTGGSAETLY | 0.7901141 |
| GAFTTPQ | ASSHGTGGSAETLY | 0.81073904 |
| GFGGFGTAGGVGGA | ASSHGTGGSAETLY | 0.7266079 |
| GFLAQFL | ASSHGTGGSAETLY | 0.65915674 |
| GGAGGAGGNAGVA | ASSHGTGGSAETLY | 0.77764857 |
| GGVGTAGAGGAGGAGG | ASSHGTGGSAETLY | 0.82946527 |
| GNGGGGDAGNAGSGGN | ASSHGTGGSAETLY | 0.66049165 |
| GTGGAGGAGNDAGS | ASSHGTGGSAETLY | 0.6196312 |
| GTTNTGWFNS | ASSHGTGGSAETLY | 0.77584577 |
| IFMGPLFTL | ASSHGTGGSAETLY | 0.73610675 |
| ILAHAIEAI | ASSHGTGGSAETLY | 0.81919557 |
| IRSAIREAL | ASSHGTGGSAETLY | 0.8155809 |
| ITFAGGG | ASSHGTGGSAETLY | 0.7793499 |
| ITGMSGHQEV | ASSHGTGGSAETLY | 0.8078735 |
| IVAGLACTL | ASSHGTGGSAETLY | 0.81726897 |
| IVGGGLAGA | ASSHGTGGSAETLY | 0.7862458 |
| IVTGVLNI | ASSHGTGGSAETLY | 0.81121874 |
| LAQQGCI | ASSHGTGGSAETLY | 0.71430695 |
| LCLHVEN | ASSHGTGGSAETLY | 0.8261163 |
| LDAISDGS | ASSHGTGGSAETLY | 0.8070234 |
| LGGFAAML | ASSHGTGGSAETLY | 0.827322 |
| LGGKNPMI | ASSHGTGGSAETLY | 0.82741034 |
| LGPKGVAAIL | ASSHGTGGSAETLY | 0.80742997 |
| LMVGVSI | ASSHGTGGSAETLY | 0.80299985 |
| LVGAAFVL | ASSHGTGGSAETLY | 0.41392437 |
| LVLAVAPL | ASSHGTGGSAETLY | 0.71975654 |
| MAAIAVIDAA | ASSHGTGGSAETLY | 0.7974515 |
| NNGINAELQ | ASSHGTGGSAETLY | 0.79801553 |
| RGEPLVLVHG | ASSHGTGGSAETLY | 0.81349695 |
| RGGAGGAAGLI | ASSHGTGGSAETLY | 0.82526565 |
| RSYRMHPAV | ASSHGTGGSAETLY | 0.717949 |
| SAAHLLAGQLA | ASSHGTGGSAETLY | 0.8262032 |
| SAAVGSY | ASSHGTGGSAETLY | 0.8215451 |
| SPHAVLHRI | ASSHGTGGSAETLY | 0.561738 |
| SQIAKVL | ASSHGTGGSAETLY | 0.75800854 |
| TALGHLVP | ASSHGTGGSAETLY | 0.82853526 |
| TAPTTSGAA | ASSHGTGGSAETLY | 0.804078 |
| TDGDDGNGGSPGT | ASSHGTGGSAETLY | 0.7060152 |

| | | |
|---|---|---|
| TDGLGEN | ASSHGTGGSAETLY | 0.7989192 |
| TGGPPLPGQA | ASSHGTGGSAETLY | 0.8250766 |
| TIPAITI | ASSHGTGGSAETLY | 0.8085191 |
| TIPGFTI | ASSHGTGGSAETLY | 0.8237666 |
| TSIASGALA | ASSHGTGGSAETLY | 0.3880734 |
| TVTLPGAAI | ASSHGTGGSAETLY | 0.8060398 |
| VASGLSAALP | ASSHGTGGSAETLY | 0.7797642 |
| VGAGVIGI | ASSHGTGGSAETLY | 0.32026175 |
| VGPKRFGV | ASSHGTGGSAETLY | 0.8053416 |
| VNVAFPD | ASSHGTGGSAETLY | 0.7644409 |
| VSMTNSGVS | ASSHGTGGSAETLY | 0.8228194 |
| VTDVLGDLSV | ASSHGTGGSAETLY | 0.75318134 |
| VVVVGAGFAGL | ASSHGTGGSAETLY | 0.81889856 |
| VYTVGTVV | ASSHGTGGSAETLY | 0.7779249 |
| WTTFSGP | ASSHGTGGSAETLY | 0.7871572 |
| YDELVEAF | ASSHGTGGSAETLY | 0.74659157 |
| YGTGGAGGNGG | ASSHGTGGSAETLY | 0.77925956 |
| YTNVMAV | ASSHGTGGSAETLY | 0.8262891 |
| FLEQVHQL | ASSHGTGGSAETLY | 0.8297328 |
| GLSLLAVAV | ASSHGTGGSAETLY | 0.40781102 |
| LEQVHQL | ASSHGTGGSAETLY | 0.81425726 |
| LNAAGAI | ASSHGTGGSAETLY | 0.74019086 |
| PAVHASL | ASSHGTGGSAETLY | 0.32363495 |
| TELEDDA | ASSHGTGGSAETLY | 0.55355376 |
| TPPGPPN | ASSHGTGGSAETLY | 0.7660079 |
| TPPPPPPP | ASSHGTGGSAETLY | 0.76578635 |
| TQSPALI | ASSLNGENTEVF | 0.7907071 |
| EFAAAFPQ | ASSLNGENTEVF | 0.80052567 |
| LLSGNLDVL | ASSLNGENTEVF | 0.7112242 |
| GPANLGNYNI | ASSLNGENTEVF | 0.7997207 |
| TLGPTLP | ASSLNGENTEVF | 0.7591675 |
| VYAYSSCCK | ASSLNGENTEVF | 0.16581692 |
| APAPAPAPNI | ASSLNGENTEVF | 0.74671596 |
| PGGPVPIP | ASSLNGENTEVF | 0.54112804 |
| FLTVALEP | ASSLNGENTEVF | 0.8312886 |
| SQYFGANL | ASSLNGENTEVF | 0.7215267 |
| GGVKPAA | ASSLNGENTEVF | 0.6944872 |
| GGTGGAGGAGGW | ASSLNGENTEVF | 0.7483525 |
| GVGGVAGLGGAGG | ASSLNGENTEVF | 0.79832965 |
| VLLVLATTI | ASSLNGENTEVF | 0.77667105 |
| IGIFAAML | ASSLNGENTEVF | 0.7511976 |
| AVGDLID | ASSLNGENTEVF | 0.4013049 |
| NISPLPL | ASSLNGENTEVF | 0.79035413 |
| VTAVFVAI | ASSLNGENTEVF | 0.73069584 |
| SGDDNDAAL | ASSLNGENTEVF | 0.7049267 |
| AVAALAHS | ASSLNGENTEVF | 0.687999 |
| GGVTVMATP | ASSLNGENTEVF | 0.78415155 |
| AGLAANVRD | ASSLNGENTEVF | 0.5312567 |
| TNRPDLIDPALLRPG | ASSLNGENTEVF | 0.82751155 |
| LANLGVTAL | ASSLNGENTEVF | 0.75062704 |
| TGERIEL | ASSLNGENTEVF | 0.70777965 |

| | | |
|---|---|---|
| NGEELAA | ASSLNGENTEVF | 0.8211172 |
| VLSLVWG | ASSLNGENTEVF | 0.78313756 |
| IGYALWL | ASSLNGENTEVF | 0.7500154 |
| LSLLLAPL | ASSLNGENTEVF | 0.73805416 |
| TLDEALK | ASSLNGENTEVF | 0.6619526 |
| TDGAGTAEGG | ASSLNGENTEVF | 0.6675547 |
| NAPGGAGGNGG | ASSLNGENTEVF | 0.70987225 |
| VSVFMGMGGNF | ASSLNGENTEVF | 0.82661116 |
| LASYLAAL | ASSLNGENTEVF | 0.81224763 |
| LVGGTPAL | ASSLNGENTEVF | 0.65061045 |
| VRNADVL | ASSLNGENTEVF | 0.82601583 |
| TCSSPFLVSAM | ASSLNGENTEVF | 0.8236738 |
| VHITDAE | ASSLNGENTEVF | 0.82379544 |
| EYAAPSF | ASSLNGENTEVF | 0.67832214 |
| VVDAFHGGL | ASSLNGENTEVF | 0.77784085 |
| ASVVDPQ | ASSLNGENTEVF | 0.78764725 |
| SYNEFGGLS | ASSLNGENTEVF | 0.74167454 |
| DPDGSVY | ASSLNGENTEVF | 0.7673329 |
| MVSTIHATE | ASSLNGENTEVF | 0.6487608 |
| LAAGVADPV | ASSLNGENTEVF | 0.8244166 |
| TKEQIAA | ASSLNGENTEVF | 0.833401 |
| SLASPAAL | ASSLNGENTEVF | 0.77562976 |
| IGLVFYGV | ASSLNGENTEVF | 0.5596429 |
| FLLDASNL | ASSLNGENTEVF | 0.7285056 |
| YQVGNFP | ASSLNGENTEVF | 0.69377637 |
| ESLLSQST | ASSLNGENTEVF | 0.6752534 |
| LFTVGFAAVG | ASSLNGENTEVF | 0.7615371 |
| ATPPTPT | ASSLNGENTEVF | 0.55700266 |
| TDLATLG | ASSLNGENTEVF | 0.77449876 |
| SAGGTGR | ASSLNGENTEVF | 0.8231328 |
| GFSNSGS | ASSLNGENTEVF | 0.74329424 |
| GGNGGSGGTSV | ASSLNGENTEVF | 0.8131231 |
| GTSVATGGAGN | ASSLNGENTEVF | 0.82012063 |
| NGGNGGSGGTS | ASSLNGENTEVF | 0.8073499 |
| AVVKFPH | ASSLNGENTEVF | 0.82326037 |
| TPPPPPP | ASSLNGENTEVF | 0.8216331 |
| AAPAAAAAPA | ASSLNGENTEVF | 0.79670095 |
| IALNEDL | ASSLNGENTEVF | 0.79792625 |
| IVNVSSM | ASSLNGENTEVF | 0.80580664 |
| TGIPPAPR | ASSLNGENTEVF | 0.73741007 |
| FDLGGGTF | ASSLNGENTEVF | 0.7475946 |
| FDLGGGTFDVS | ASSLNGENTEVF | 0.77733576 |
| NEPTAAA | ASSLNGENTEVF | 0.79810405 |
| NEPTAAALA | ASSLNGENTEVF | 0.78250265 |
| GSIINISSI | ASSLNGENTEVF | 0.82816726 |
| VAPGNAGTA | ASSLNGENTEVF | 0.4673318 |
| TGGGASLE | ASSLNGENTEVF | 0.64753795 |
| VAVYNHYKRI | ASSLNGENTEVF | 0.8207596 |
| TDPAPATT | ASSLNGENTEVF | 0.22401465 |
| TNRPDILDPALLRPG | ASSLNGENTEVF | 0.67001516 |
| LTTTTTS | ASSLNGENTEVF | 0.79953885 |

| | | |
|---|---|---|
| PTGSGKTL | ASSLNGENTEVF | 0.8147678 |
| TGSGKTL | ASSLNGENTEVF | 0.25049734 |
| TIGHVDHGK | ASSLNGENTEVF | 0.71141195 |
| TIGHVDHGKTT | ASSLNGENTEVF | 0.6473701 |
| GHVDHGKTT | ASSLNGENTEVF | 0.8316761 |
| AQFEHTLL | ASSLNGENTEVF | 0.7486571 |
| IVNQDEL | ASSLNGENTEVF | 0.47809982 |
| TNGSQFF | ASSLNGENTEVF | 0.82181805 |
| TGKGIVE | ASSLNGENTEVF | 0.57358783 |
| LISNASDA | ASSLNGENTEVF | 0.78293633 |
| AELRTLGELTKIAKAHGAQ | ASSLNGENTEVF | 0.81714547 |
| APNAVAAAEPSW | ASSLNGENTEVF | 0.8313868 |
| APQVAGAATAVL | ASSLNGENTEVF | 0.6417558 |
| ARRDPAAAAAILPTDA | ASSLNGENTEVF | 0.8144736 |
| DAAARDVLAEAGLAEYF | ASSLNGENTEVF | 0.77128905 |
| DDQIVLSAKANLAKVEQ | ASSLNGENTEVF | 0.82652485 |
| DELLPDTGEDSDEDGH | ASSLNGENTEVF | 0.70225143 |
| ELPEAPANRATVPQD | ASSLNGENTEVF | 0.59240615 |
| FGQSKIERVTGIG | ASSLNGENTEVF | 0.8209199 |
| GAGGTGSTAGGAGGAGGAGGLY | ASSLNGENTEVF | 0.8176626 |
| GHGGTGAAGGNGGTGGAGGSGI | ASSLNGENTEVF | 0.79428756 |
| GNGGNGGAGGEGGPGGAGGAGA | ASSLNGENTEVF | 0.8143322 |
| GSTAGNGGNGGNATNGTIAGQP | ASSLNGENTEVF | 0.8095813 |
| GTGGDGGNGGNGGTGG | ASSLNGENTEVF | 0.7547617 |
| GVRRPAGIPADAPV | ASSLNGENTEVF | 0.81099665 |
| GYRQVTDAHLLTLARR | ASSLNGENTEVF | 0.80914915 |
| IAKMGQDGHDRG | ASSLNGENTEVF | 0.7072306 |
| KAEPSIEIQYT | ASSLNGENTEVF | 0.82465035 |
| KPVRIRMAADEDTVAVT | ASSLNGENTEVF | 0.7563423 |
| MPPRIAGMRLLVIKPEP | ASSLNGENTEVF | 0.7024515 |
| MKFYAVALFRSRGVPPTR | ASSLNGENTEVF | 0.8229968 |
| NAGGFGILGAVAHSP | ASSLNGENTEVF | 0.80795574 |
| NEGREITLAMAEM | ASSLNGENTEVF | 0.80833703 |
| NELVEWSHPLGHRWRWE | ASSLNGENTEVF | 0.8094997 |
| NLGAGGTGGNGGNPG | ASSLNGENTEVF | 0.828968 |
| NTEWPKWETVMAADPVD | ASSLNGENTEVF | 0.5422389 |
| PASAAPAETQP | ASSLNGENTEVF | 0.79146445 |
| QQSSDGKNGVRAAEITGE | ASSLNGENTEVF | 0.76219827 |
| RAPDAVTTSWWKE | ASSLNGENTEVF | 0.8218081 |
| SGFGNVGAHTSGWFNQ | ASSLNGENTEVF | 0.8010094 |
| SGVAGAVGQAASVGGLKV | ASSLNGENTEVF | 0.73743355 |
| SNGGAGTGSAGGNGGT | ASSLNGENTEVF | 0.815729 |
| TGGAGGEGGAAG | ASSLNGENTEVF | 0.81804293 |
| TGGDGGAAGAGGAPGHG | ASSLNGENTEVF | 0.80312395 |
| TMPPLLAAIANAGRHGV | ASSLNGENTEVF | 0.8056506 |
| TSSTAGNSGGA | ASSLNGENTEVF | 0.8272369 |
| VPATVPVGILC | ASSLNGENTEVF | 0.70101655 |
| VQRPDSGGVVLTGRIS | ASSLNGENTEVF | 0.77274334 |
| WTPNPVPATKGVPAGLVQAT | ASSLNGENTEVF | 0.816931 |
| YALKRALAKFSDRFDVVII | ASSLNGENTEVF | 0.82531166 |
| YLIGSVIV | ASSLNGENTEVF | 0.8062739 |

| | | |
|---|---|---|
| YGQLVAVID | ASSLNGENTEVF | 0.8256071 |
| VTVGGMTLPA | ASSLNGENTEVF | 0.75497687 |
| VLALPAV | ASSLNGENTEVF | 0.79549366 |
| VIPPTLNYET | ASSLNGENTEVF | 0.7527056 |
| VGMGGMHPSSG | ASSLNGENTEVF | 0.7787061 |
| VGLAGGAATA | ASSLNGENTEVF | 0.68497425 |
| VGGAGGAGGSGP | ASSLNGENTEVF | 0.7749912 |
| VFAAFMV | ASSLNGENTEVF | 0.805223 |
| TLYDKLR | ASSLNGENTEVF | 0.7956989 |
| TLAVFTVL | ASSLNGENTEVF | 0.5074802 |
| TGSVQPARV | ASSLNGENTEVF | 0.7960336 |
| TGGGAGSGGT | ASSLNGENTEVF | 0.8064292 |
| SHLETSL | ASSLNGENTEVF | 0.832034 |
| SFTTGMT | ASSLNGENTEVF | 0.77689195 |
| QVEVAKRMGVRQPTVS | ASSLNGENTEVF | 0.8168279 |
| QGSGGPALAM | ASSLNGENTEVF | 0.7010951 |
| MLAPSNP | ASSLNGENTEVF | 0.80524576 |
| LVNNAAI | ASSLNGENTEVF | 0.63801825 |
| LTGEINTLA | ASSLNGENTEVF | 0.7281744 |
| LSIFGPV | ASSLNGENTEVF | 0.7172367 |
| LLALLANL | ASSLNGENTEVF | 0.6745255 |
| LISNASDALDKLR | ASSLNGENTEVF | 0.8070372 |
| LINEAVP | ASSLNGENTEVF | 0.8194152 |
| LGYAFSGL | ASSLNGENTEVF | 0.8050731 |
| LDFVAVAL | ASSLNGENTEVF | 0.8180223 |
| LAQPGAE | ASSLNGENTEVF | 0.8184483 |
| ISNLGAVTGF | ASSLNGENTEVF | 0.83234376 |
| HQEDPGRI | ASSLNGENTEVF | 0.74156463 |
| GTPGEPTP | ASSLNGENTEVF | 0.7505816 |
| GFFNSGS | ASSLNGENTEVF | 0.45820612 |
| GAGGGGLTAGAN | ASSLNGENTEVF | 0.74936265 |
| FVDALVNM | ASSLNGENTEVF | 0.74430335 |
| FLTVAPEL | ASSLNGENTEVF | 0.76876485 |
| FETLTAL | ASSLNGENTEVF | 0.22128597 |
| DVINPIEL | ASSLNGENTEVF | 0.79247314 |
| APVSGAPV | ASSLNGENTEVF | 0.82748854 |
| ALVGGLAW | ASSLNGENTEVF | 0.7134134 |
| ALAAAAATAI | ASSLNGENTEVF | 0.80441815 |
| AFGLGLLL | ASSLNGENTEVF | 0.79296094 |
| AFFAAASL | ASSLNGENTEVF | 0.76056993 |
| AAAALPAFSPP | ASSLNGENTEVF | 0.8004515 |
| VVLAPAL | ASSLNGENTEVF | 0.7512506 |
| TIGHVDHGKTTL | ASSLNGENTEVF | 0.8234234 |
| LISNASDALDK | ASSLNGENTEVF | 0.82492614 |
| LISNASDAL | ASSLNGENTEVF | 0.79813033 |
| ISNASDALDK | ASSLNGENTEVF | 0.7864917 |
| NQAETLVYQTE | ASSLNGENTEVF | 0.80825025 |
| TGGTGGTAGAG | ASSLNGENTEVF | 0.53768 |
| TGSTAGGAGGA | ASSLNGENTEVF | 0.79454714 |
| DPLAVPVAAGPV | ASSLNGENTEVF | 0.6544373 |
| GGTGGTPGNGGN | ASSLNGENTEVF | 0.778746 |

| | | |
|---|---|---|
| PAGLAGVDPATL | ASSLNGENTEVF | 0.7181071 |
| GNNLVVEAPHAGASV | ASSLNGENTEVF | 0.79075426 |
| KVTRIMSASTLEQPA | ASSLNGENTEVF | 0.7572439 |
| LGFALPIALPRRTVV | ASSLNGENTEVF | 0.8081304 |
| TRLGNLTARKLVVLV | ASSLNGENTEVF | 0.62574947 |
| WSQQRRLIYFPSAGP | ASSLNGENTEVF | 0.8199946 |
| GPNGNPDPMAAAVDIR | ASSLNGENTEVF | 0.7978641 |
| TQRCVSVADLRAAASR | ASSLNGENTEVF | 0.589241 |
| QFVLDVAEHPVEVLTETV | ASSLNGENTEVF | 0.8157541 |
| GGAGGTGGAGGNGGTQAGDG | ASSLNGENTEVF | 0.8025893 |
| PLLATLRAIAVDVDAKPAQV | ASSLNGENTEVF | 0.7639767 |
| APKFRTGGYMIDFWGVGYQVAK | ASSLNGENTEVF | 0.8125838 |
| GAGGAGGAGGAAGGPGGTGGTG | ASSLNGENTEVF | 0.79736066 |
| IGALGLLATDVAPAGSVAFGVA | ASSLNGENTEVF | 0.56158274 |
| AGGAGGIGGNGGAGGAGGVLMG | ASSLNGENTEVF | 0.7144811 |
| ADELIGLL | ASSLNGENTEVF | 0.7824049 |
| ADRALKA | ASSLNGENTEVF | 0.7101733 |
| AGAVGQVGGAGGA | ASSLNGENTEVF | 0.45594913 |
| AGGGGGLGGAP | ASSLNGENTEVF | 0.81369114 |
| AGGLLYGNGGAG | ASSLNGENTEVF | 0.7060635 |
| AGLESIGL | ASSLNGENTEVF | 0.7155465 |
| AGLKTGII | ASSLNGENTEVF | 0.75546026 |
| ASLAALLL | ASSLNGENTEVF | 0.8177726 |
| ASVINISSI | ASSLNGENTEVF | 0.77124524 |
| ATPPGPA | ASSLNGENTEVF | 0.8138032 |
| ATVASALV | ASSLNGENTEVF | 0.80602956 |
| ATVASLIL | ASSLNGENTEVF | 0.7618005 |
| AVNGPVLIRH | ASSLNGENTEVF | 0.8129203 |
| DGAAGAVGI | ASSLNGENTEVF | 0.80396086 |
| DPLLHGI | ASSLNGENTEVF | 0.7989435 |
| ELGAALGAL | ASSLNGENTEVF | 0.80799854 |
| ELGRTQAV | ASSLNGENTEVF | 0.757468 |
| FVAGGYLS | ASSLNGENTEVF | 0.7515739 |
| GASGGAGGAGGW | ASSLNGENTEVF | 0.7484592 |
| GGAGAGGDGGAGGI | ASSLNGENTEVF | 0.716804 |
| GTGHNGSQPGMGGNG | ASSLNGENTEVF | 0.6751971 |
| GYAAASASAS | ASSLNGENTEVF | 0.78030765 |
| HQVDVAAVQQG | ASSLNGENTEVF | 0.833437 |
| IGAAIAHT | ASSLNGENTEVF | 0.80108726 |
| IGMIEVAQL | ASSLNGENTEVF | 0.6187321 |
| IRLAPEHTSAMD | ASSLNGENTEVF | 0.7728267 |
| IYNVSVL | ASSLNGENTEVF | 0.70406187 |
| KQGFLGM | ASSLNGENTEVF | 0.81557834 |
| LESVITQL | ASSLNGENTEVF | 0.8199202 |
| LGLGALVVV | ASSLNGENTEVF | 0.8289186 |
| LIGNGGNGGSGGI | ASSLNGENTEVF | 0.80653584 |
| LNAINAPTL | ASSLNGENTEVF | 0.72430813 |
| LQLLNTGV | ASSLNGENTEVF | 0.8172108 |
| LQTSIEF | ASSLNGENTEVF | 0.7693567 |
| MSFAVSEL | ASSLNGENTEVF | 0.79832554 |
| NTGALGL | ASSLNGENTEVF | 0.796761 |

| | | |
|---|---|---|
| PPGAGGAGGAGTT | ASSLNGENTEVF | 0.80438244 |
| QLVGLGI | ASSLNGENTEVF | 0.79676294 |
| RFLEQVHQL | ASSLNGENTEVF | 0.79552454 |
| RGVGYVLR | ASSLNGENTEVF | 0.5694198 |
| SATGVFAGVI | ASSLNGENTEVF | 0.8203553 |
| SLPAITLP | ASSLNGENTEVF | 0.5730481 |
| STALAFTV | ASSLNGENTEVF | 0.75892186 |
| SVLTVGLGK | ASSLNGENTEVF | 0.76000106 |
| TALAGAGSGI | ASSLNGENTEVF | 0.81831 |
| TGLPFTGI | ASSLNGENTEVF | 0.8212037 |
| VAEHVTV | ASSLNGENTEVF | 0.7193072 |
| VDIVLNSL | ASSLNGENTEVF | 0.7138456 |
| VLTGGGAVL | ASSLNGENTEVF | 0.8004308 |
| VNVGLAYGI | ASSLNGENTEVF | 0.6174873 |
| VSSAAVGAV | ASSLNGENTEVF | 0.8228382 |
| VTSVYEAGV | ASSLNGENTEVF | 0.7587944 |
| WVALDRGA | ASSLNGENTEVF | 0.76049036 |
| YLVGSVVI | ASSLNGENTEVF | 0.7745689 |
| GPPGTGKTLL | ASSLNGENTEVF | 0.8107809 |
| GVGMTKF | ASSLNGENTEVF | 0.7637749 |
| IGIGGSDL | ASSLNGENTEVF | 0.8252655 |
| IGTLLAEL | ASSLNGENTEVF | 0.269731 |
| IYTSGTT | ASSLNGENTEVF | 0.81630284 |
| LSFPAGV | ASSLNGENTEVF | 0.5870793 |
| LYTSGTT | ASSLNGENTEVF | 0.75014305 |
| VGAFPFGF | ASSLNGENTEVF | 0.43752792 |
| VGAGTVEF | ASSLNGENTEVF | 0.7894666 |
| VGKSTLLL | ASSLNGENTEVF | 0.7954789 |
| AALAHAGGGIV | ASSLNGENTEVF | 0.82181776 |
| AALVAVSLL | ASSLNGENTEVF | 0.7941946 |
| AAVSRAVAAL | ASSLNGENTEVF | 0.81104666 |
| AGGSSGIGGF | ASSLNGENTEVF | 0.81795937 |
| AGLTVANANG | ASSLNGENTEVF | 0.71679175 |
| AGLVGAVL | ASSLNGENTEVF | 0.35481164 |
| ALSQAFTV | ASSLNGENTEVF | 0.79886234 |
| ALTFSVSL | ASSLNGENTEVF | 0.7839986 |
| ALTGAALVAL | ASSLNGENTEVF | 0.82506114 |
| ASFSIGAGVL | ASSLNGENTEVF | 0.7880069 |
| DGSVYKI | ASSLNGENTEVF | 0.56924725 |
| EQLAEIV | ASSLNGENTEVF | 0.81958747 |
| FAVSELTVM | ASSLNGENTEVF | 0.8125658 |
| GAAPLAAGL | ASSLNGENTEVF | 0.6502198 |
| GADGGDGGSGGAGG | ASSLNGENTEVF | 0.73194575 |
| GFGLDTGGAGG | ASSLNGENTEVF | 0.75238204 |
| GLASGGLL | ASSLNGENTEVF | 0.58740133 |
| GRIAGAAVLL | ASSLNGENTEVF | 0.7930072 |
| GSAGSLNTSF | ASSLNGENTEVF | 0.72721326 |
| GTGQIGI | ASSLNGENTEVF | 0.754153 |
| GTLIALIL | ASSLNGENTEVF | 0.77363753 |
| IAATMGFVVG | ASSLNGENTEVF | 0.71828485 |
| IFAQALL | ASSLNGENTEVF | 0.70851284 |

| | | |
|---|---|---|
| IGAAVALGF | ASSLNGENTEVF | 0.7154839 |
| IGLTPLAL | ASSLNGENTEVF | 0.7948475 |
| ILGSNTSTL | ASSLNGENTEVF | 0.81337947 |
| IVFSLTVL | ASSLNGENTEVF | 0.76672953 |
| LASKATGF | ASSLNGENTEVF | 0.80239534 |
| LASVALIL | ASSLNGENTEVF | 0.8148297 |
| LFAVGLTL | ASSLNGENTEVF | 0.60937905 |
| LFNGGIGTYI | ASSLNGENTEVF | 0.71312654 |
| LTLGAGII | ASSLNGENTEVF | 0.8179318 |
| LVAAALGDI | ASSLNGENTEVF | 0.7545018 |
| NTAFVQLGI | ASSLNGENTEVF | 0.82079613 |
| QLFSGAEL | ASSLNGENTEVF | 0.77822965 |
| SAATFITL | ASSLNGENTEVF | 0.77890444 |
| SNYGLANLGV | ASSLNGENTEVF | 0.7950926 |
| SYGGAYAVM | ASSLNGENTEVF | 0.82879245 |
| TTGVGGLTLG | ASSLNGENTEVF | 0.81411934 |
| TTQTIYI | ASSLNGENTEVF | 0.78650093 |
| VAAGAGLGAVI | ASSLNGENTEVF | 0.8114255 |
| VAAVVTLL | ASSLNGENTEVF | 0.69943464 |
| VGFLGGLI | ASSLNGENTEVF | 0.8168621 |
| VGVIPILL | ASSLNGENTEVF | 0.82083476 |
| VIFASSLI | ASSLNGENTEVF | 0.6571963 |
| VKGGDGGAAAG | ASSLNGENTEVF | 0.75558376 |
| VTKTFTNI | ASSLNGENTEVF | 0.82767564 |
| VVAAAILL | ASSLNGENTEVF | 0.82587576 |
| YAALDVEL | ASSLNGENTEVF | 0.7481422 |
| LASVALLL | ASSLNGENTEVF | 0.7228565 |
| LDAVHRV | ASSLNGENTEVF | 0.63120055 |
| QGLSSSIF | ASSLNGENTEVF | 0.78600615 |
| RGTFANIRL | ASSLNGENTEVF | 0.5614612 |
| RQLWWGHRI | ASSLNGENTEVF | 0.80328286 |
| YGPPGTGKTLL | ASSLNGENTEVF | 0.69909227 |
| AALVGAPANVA | ASSLNGENTEVF | 0.8171403 |
| AATFRRTE | ASSLNGENTEVF | 0.77176434 |
| AGALGGGI | ASSLNGENTEVF | 0.81131685 |
| AGGAGGAGGIGAG | ASSLNGENTEVF | 0.7453909 |
| AGSGALGL | ASSLNGENTEVF | 0.74401116 |
| AQIADQI | ASSLNGENTEVF | 0.65190375 |
| ASAVAVTYL | ASSLNGENTEVF | 0.5432179 |
| ASTVLGGTGG | ASSLNGENTEVF | 0.75803447 |
| AVGGEPAK | ASSLNGENTEVF | 0.7621128 |
| CEQGTFGGTC | ASSLNGENTEVF | 0.40313688 |
| DEGKAPAG | ASSLNGENTEVF | 0.8268534 |
| DTVVFPAL | ASSLNGENTEVF | 0.7854787 |
| FEGITMAG | ASSLNGENTEVF | 0.7899288 |
| FPLVVQD | ASSLNGENTEVF | 0.7135894 |
| GAFTTPQ | ASSLNGENTEVF | 0.7221801 |
| GFGGFGTAGGVGGA | ASSLNGENTEVF | 0.7173152 |
| GFLAQFL | ASSLNGENTEVF | 0.81393385 |
| GGAGGAGGNAGVA | ASSLNGENTEVF | 0.6603204 |
| GGVGTAGAGGAGGAGG | ASSLNGENTEVF | 0.4744434 |

| | | |
|---|---|---|
| GNGGGGDAGNAGSGGN | ASSLNGENTEVF | 0.72146225 |
| GTGGAGGAGNDAGS | ASSLNGENTEVF | 0.6927719 |
| GTTNTGWFNS | ASSLNGENTEVF | 0.82564527 |
| IFMGPLFTL | ASSLNGENTEVF | 0.81719655 |
| ILAHAIEAI | ASSLNGENTEVF | 0.7966608 |
| IRSAIREAL | ASSLNGENTEVF | 0.82994616 |
| ITFAGGG | ASSLNGENTEVF | 0.22931042 |
| ITGMSGHQEV | ASSLNGENTEVF | 0.72233486 |
| IVAGLACTL | ASSLNGENTEVF | 0.82169384 |
| IVGGGLAGA | ASSLNGENTEVF | 0.798997 |
| IVTGVLNI | ASSLNGENTEVF | 0.7273846 |
| LAQQGCI | ASSLNGENTEVF | 0.81888044 |
| LCLHVEN | ASSLNGENTEVF | 0.70094603 |
| LDAISDGS | ASSLNGENTEVF | 0.72467256 |
| LGGFAAML | ASSLNGENTEVF | 0.79010105 |
| LGGKNPMI | ASSLNGENTEVF | 0.8164686 |
| LGPKGVAAIL | ASSLNGENTEVF | 0.8070067 |
| LMVGVSI | ASSLNGENTEVF | 0.5387112 |
| LVGAAFVL | ASSLNGENTEVF | 0.81563544 |
| LVLAVAPL | ASSLNGENTEVF | 0.8259266 |
| MAAIAVIDAA | ASSLNGENTEVF | 0.7234396 |
| NNGINAELQ | ASSLNGENTEVF | 0.794055 |
| RGEPLVLVHG | ASSLNGENTEVF | 0.79758155 |
| RGGAGGAAGLI | ASSLNGENTEVF | 0.7549218 |
| RSYRMHPAV | ASSLNGENTEVF | 0.6706722 |
| SAAHLLAGQLA | ASSLNGENTEVF | 0.6417163 |
| SAAVGSY | ASSLNGENTEVF | 0.8196855 |
| SPHAVLHRI | ASSLNGENTEVF | 0.5975261 |
| SQIAKVL | ASSLNGENTEVF | 0.80216914 |
| TALGHLVP | ASSLNGENTEVF | 0.76825285 |
| TAPTTSGAA | ASSLNGENTEVF | 0.82997376 |
| TDGDDGNGGSPGT | ASSLNGENTEVF | 0.8002598 |
| TDGLGEN | ASSLNGENTEVF | 0.80595934 |
| TGGPPLPGQA | ASSLNGENTEVF | 0.64787805 |
| TIPAITI | ASSLNGENTEVF | 0.8091351 |
| TIPGFTI | ASSLNGENTEVF | 0.797256 |
| TSIASGALA | ASSLNGENTEVF | 0.5559196 |
| TVTLPGAAI | ASSLNGENTEVF | 0.8144781 |
| VASGLSAALP | ASSLNGENTEVF | 0.61797136 |
| VGAGVIGI | ASSLNGENTEVF | 0.7912748 |
| VGPKRFGV | ASSLNGENTEVF | 0.8164675 |
| VNVAFPD | ASSLNGENTEVF | 0.8220797 |
| VSMTNSGVS | ASSLNGENTEVF | 0.74040174 |
| VTDVLGDLSV | ASSLNGENTEVF | 0.80801934 |
| VVVVGAGFAGL | ASSLNGENTEVF | 0.6370619 |
| VYTVGTVV | ASSLNGENTEVF | 0.81599015 |
| WTTFSGP | ASSLNGENTEVF | 0.81482255 |
| YDELVEAF | ASSLNGENTEVF | 0.79108924 |
| YGTGGAGGNGG | ASSLNGENTEVF | 0.8318011 |
| YTNVMAV | ASSLNGENTEVF | 0.8248318 |
| FLEQVHQL | ASSLNGENTEVF | 0.50755703 |

| | | |
|---|---|---|
| GLSLLAVAV | ASSLNGENTEVF | 0.7992566 |
| LEQVHQL | ASSLNGENTEVF | 0.81683475 |
| LNAAGAI | ASSLNGENTEVF | 0.82017773 |
| PAVHASL | ASSLNGENTEVF | 0.8175559 |
| TELEDDA | ASSLNGENTEVF | 0.8199835 |
| TPPGPPN | ASSLNGENTEVF | 0.75330174 |
| TPPPPPPP | ASSLNGENTEVF | 0.8229437 |
| TQSPALI | ASSLSGTDAEQF | 0.8354715 |
| EFAAAFPQ | ASSLSGTDAEQF | 0.80766565 |
| LLSGNLDVL | ASSLSGTDAEQF | 0.7479613 |
| GPANLGNYNI | ASSLSGTDAEQF | 0.76356137 |
| TLGPTLP | ASSLSGTDAEQF | 0.8093774 |
| VYAYSSCCK | ASSLSGTDAEQF | 0.8152518 |
| APAPAPAPNI | ASSLSGTDAEQF | 0.72918653 |
| PGGPVPIP | ASSLSGTDAEQF | 0.7557463 |
| FLTVALEP | ASSLSGTDAEQF | 0.7580879 |
| SQYFGANL | ASSLSGTDAEQF | 0.4496978 |
| GGVKPAA | ASSLSGTDAEQF | 0.8040806 |
| GGTGGAGGAGGW | ASSLSGTDAEQF | 0.82106954 |
| GVGGVAGLGGAGG | ASSLSGTDAEQF | 0.8136897 |
| VLLVLATTI | ASSLSGTDAEQF | 0.7212901 |
| IGIFAAML | ASSLSGTDAEQF | 0.81261116 |
| AVGDLID | ASSLSGTDAEQF | 0.81788504 |
| NISPLPL | ASSLSGTDAEQF | 0.8313131 |
| VTAVFVAI | ASSLSGTDAEQF | 0.52569747 |
| SGDDNDAAL | ASSLSGTDAEQF | 0.28590065 |
| AVAALAHS | ASSLSGTDAEQF | 0.77888817 |
| GGVTVMATP | ASSLSGTDAEQF | 0.8048576 |
| AGLAANVRD | ASSLSGTDAEQF | 0.74602723 |
| TNRPDLIDPALLRPG | ASSLSGTDAEQF | 0.7485845 |
| LANLGVTAL | ASSLSGTDAEQF | 0.7911259 |
| TGERIEL | ASSLSGTDAEQF | 0.81413573 |
| NGEELAA | ASSLSGTDAEQF | 0.6866878 |
| VLSLVWG | ASSLSGTDAEQF | 0.69863266 |
| IGYALWL | ASSLSGTDAEQF | 0.8071079 |
| LSLLLAPL | ASSLSGTDAEQF | 0.82250917 |
| TLDEALK | ASSLSGTDAEQF | 0.7005672 |
| TDGAGTAEGG | ASSLSGTDAEQF | 0.7468786 |
| NAPGGAGGNGG | ASSLSGTDAEQF | 0.7400776 |
| VSVFMGMGGNF | ASSLSGTDAEQF | 0.80674815 |
| LASYLAAL | ASSLSGTDAEQF | 0.7817108 |
| LVGGTPAL | ASSLSGTDAEQF | 0.62015754 |
| VRNADVL | ASSLSGTDAEQF | 0.81208926 |
| TCSSPFLVSAM | ASSLSGTDAEQF | 0.534477 |
| VHITDAE | ASSLSGTDAEQF | 0.8138644 |
| EYAAPSF | ASSLSGTDAEQF | 0.8011317 |
| VVDAFHGGL | ASSLSGTDAEQF | 0.80456537 |
| ASVVDPQ | ASSLSGTDAEQF | 0.67615616 |
| SYNEFGGLS | ASSLSGTDAEQF | 0.67557347 |
| DPDGSVY | ASSLSGTDAEQF | 0.5802972 |
| MVSTIHATE | ASSLSGTDAEQF | 0.7177267 |

| | | |
|---|---|---|
| LAAGVADPV | ASSLSGTDAEQF | 0.819839 |
| TKEQIAA | ASSLSGTDAEQF | 0.7712976 |
| SLASPAAL | ASSLSGTDAEQF | 0.82220733 |
| IGLVFYGV | ASSLSGTDAEQF | 0.7782662 |
| FLLDASNL | ASSLSGTDAEQF | 0.68803036 |
| YQVGNFP | ASSLSGTDAEQF | 0.81338507 |
| ESLLSQST | ASSLSGTDAEQF | 0.66853833 |
| LFTVGFAAVG | ASSLSGTDAEQF | 0.794399 |
| ATPPTPT | ASSLSGTDAEQF | 0.7775042 |
| TDLATLG | ASSLSGTDAEQF | 0.8195837 |
| SAGGTGR | ASSLSGTDAEQF | 0.6682103 |
| GFSNSGS | ASSLSGTDAEQF | 0.8209338 |
| GGNGGSGGTSV | ASSLSGTDAEQF | 0.799999 |
| GTSVATGGAGN | ASSLSGTDAEQF | 0.72785497 |
| NGGNGGSGGTS | ASSLSGTDAEQF | 0.81305057 |
| AVVKFPH | ASSLSGTDAEQF | 0.8146312 |
| TPPPPPP | ASSLSGTDAEQF | 0.8142854 |
| AAPAAAAAPA | ASSLSGTDAEQF | 0.5905894 |
| IALNEDL | ASSLSGTDAEQF | 0.78860754 |
| IVNVSSM | ASSLSGTDAEQF | 0.72359115 |
| TGIPPAPR | ASSLSGTDAEQF | 0.7868413 |
| FDLGGGTF | ASSLSGTDAEQF | 0.787246 |
| FDLGGGTFDVS | ASSLSGTDAEQF | 0.76660043 |
| NEPTAAA | ASSLSGTDAEQF | 0.81784713 |
| NEPTAAALA | ASSLSGTDAEQF | 0.66943085 |
| GSIINISSI | ASSLSGTDAEQF | 0.7331218 |
| VAPGNAGTA | ASSLSGTDAEQF | 0.7395861 |
| TGGGASLE | ASSLSGTDAEQF | 0.7612586 |
| VAVYNHYKRI | ASSLSGTDAEQF | 0.8085058 |
| TDPAPATT | ASSLSGTDAEQF | 0.6029608 |
| TNRPDILDPALLRPG | ASSLSGTDAEQF | 0.7953163 |
| LTTTTTS | ASSLSGTDAEQF | 0.77040815 |
| PTGSGKTL | ASSLSGTDAEQF | 0.65645707 |
| TGSGKTL | ASSLSGTDAEQF | 0.8162425 |
| TIGHVDHGK | ASSLSGTDAEQF | 0.77260447 |
| TIGHVDHGKTT | ASSLSGTDAEQF | 0.6630262 |
| GHVDHGKTT | ASSLSGTDAEQF | 0.812387 |
| AQFEHTLL | ASSLSGTDAEQF | 0.73475313 |
| IVNQDEL | ASSLSGTDAEQF | 0.8050388 |
| TNGSQFF | ASSLSGTDAEQF | 0.8289126 |
| TGKGIVE | ASSLSGTDAEQF | 0.6627476 |
| LISNASDA | ASSLSGTDAEQF | 0.81254685 |
| AELRTLGELTKIAKAHGAQ | ASSLSGTDAEQF | 0.78742313 |
| APNAVAAAEPSW | ASSLSGTDAEQF | 0.8033946 |
| APQVAGAATAVL | ASSLSGTDAEQF | 0.7291688 |
| ARRDPAAAAAILPTDA | ASSLSGTDAEQF | 0.71999973 |
| DAAARDVLAEAGLAEYF | ASSLSGTDAEQF | 0.81689656 |
| DDQIVLSAKANLAKVEQ | ASSLSGTDAEQF | 0.7912178 |
| DELLPDTGEDSDEDGH | ASSLSGTDAEQF | 0.81349355 |
| ELPEAPANRATVPQD | ASSLSGTDAEQF | 0.8162237 |
| FGQSKIERVTGIG | ASSLSGTDAEQF | 0.8107035 |

| | | |
|---|---|---|
| GAGGTGSTAGGAGGAGGAGGLY | ASSLSGTDAEQF | 0.8193896 |
| GHGGTGAAGGNGGTGGAGGSGI | ASSLSGTDAEQF | 0.70222354 |
| GNGGNGGAGGEGGPGGAGGAGA | ASSLSGTDAEQF | 0.64255434 |
| GSTAGNGGNGGNATNGTIAGQP | ASSLSGTDAEQF | 0.75959367 |
| GTGGDGGNGGNGGTGG | ASSLSGTDAEQF | 0.7397051 |
| GVRRPAGIPADAPV | ASSLSGTDAEQF | 0.7462287 |
| GYRQVTDAHLLTLARR | ASSLSGTDAEQF | 0.680735 |
| IAKMGQDGHDRG | ASSLSGTDAEQF | 0.810783 |
| KAEPSIEIQYT | ASSLSGTDAEQF | 0.8290453 |
| KPVRIRMAADEDTVAVT | ASSLSGTDAEQF | 0.7245566 |
| MPPRIAGMRLLVIKPEP | ASSLSGTDAEQF | 0.8341534 |
| MKFYAVALFRSRGVPPTR | ASSLSGTDAEQF | 0.65238416 |
| NAGGFGILGAVAHSP | ASSLSGTDAEQF | 0.8003587 |
| NEGREITLAMAEM | ASSLSGTDAEQF | 0.8179477 |
| NELVEWSHPLGHRWRWE | ASSLSGTDAEQF | 0.62695444 |
| NLGAGGTGGNGGNPG | ASSLSGTDAEQF | 0.7996566 |
| NTEWPKWETVMAADPVD | ASSLSGTDAEQF | 0.66706187 |
| PASAAPAETQP | ASSLSGTDAEQF | 0.6910844 |
| QQSSDGKNGVRAAEITGE | ASSLSGTDAEQF | 0.818758 |
| RAPDAVTTSWWKE | ASSLSGTDAEQF | 0.71284974 |
| SGFGNVGAHTSGWFNQ | ASSLSGTDAEQF | 0.81179154 |
| SGVAGAVGQAASVGGLKV | ASSLSGTDAEQF | 0.79001606 |
| SNGGAGTGSAGGNGGT | ASSLSGTDAEQF | 0.6785923 |
| TGGAGGEGGAAG | ASSLSGTDAEQF | 0.80683666 |
| TGGDGGAAGAGGAPGHG | ASSLSGTDAEQF | 0.823506 |
| TMPPLLAAIANAGRHGV | ASSLSGTDAEQF | 0.79961294 |
| TSSTAGNSGGA | ASSLSGTDAEQF | 0.8360412 |
| VPATVPVGILC | ASSLSGTDAEQF | 0.6668781 |
| VQRPDSGGVVLTGRIS | ASSLSGTDAEQF | 0.81335074 |
| WTPNPVPATKGVPAGLVQAT | ASSLSGTDAEQF | 0.82222426 |
| YALKRALAKFSDRFDVVII | ASSLSGTDAEQF | 0.77210844 |
| YLIGSVIV | ASSLSGTDAEQF | 0.77319884 |
| YGQLVAVID | ASSLSGTDAEQF | 0.8060174 |
| VTVGGMTLPA | ASSLSGTDAEQF | 0.7890823 |
| VLALPAV | ASSLSGTDAEQF | 0.8094873 |
| VIPPTLNYET | ASSLSGTDAEQF | 0.8322362 |
| VGMGGMHPSSG | ASSLSGTDAEQF | 0.35660818 |
| VGLAGGAATA | ASSLSGTDAEQF | 0.7862531 |
| VGGAGGAGGSGP | ASSLSGTDAEQF | 0.74438787 |
| VFAAFMV | ASSLSGTDAEQF | 0.76413786 |
| TLYDKLR | ASSLSGTDAEQF | 0.56683266 |
| TLAVFTVL | ASSLSGTDAEQF | 0.41550413 |
| TGSVQPARV | ASSLSGTDAEQF | 0.6505277 |
| TGGGAGSGGT | ASSLSGTDAEQF | 0.5610932 |
| SHLETSL | ASSLSGTDAEQF | 0.8044002 |
| SFTTGMT | ASSLSGTDAEQF | 0.7575315 |
| QVEVAKRMGVRQPTVS | ASSLSGTDAEQF | 0.5627231 |
| QGSGGPALAM | ASSLSGTDAEQF | 0.7747944 |
| MLAPSNP | ASSLSGTDAEQF | 0.7900232 |
| LVNNAAI | ASSLSGTDAEQF | 0.35488468 |
| LTGEINTLA | ASSLSGTDAEQF | 0.6814177 |

| | | |
|---|---|---|
| LSIFGPV | ASSLSGTDAEQF | 0.67971516 |
| LLALLANL | ASSLSGTDAEQF | 0.6064609 |
| LISNASDALDKLR | ASSLSGTDAEQF | 0.28224683 |
| LINEAVP | ASSLSGTDAEQF | 0.7296746 |
| LGYAFSGL | ASSLSGTDAEQF | 0.6286799 |
| LDFVAVAL | ASSLSGTDAEQF | 0.41238922 |
| LAQPGAE | ASSLSGTDAEQF | 0.7485203 |
| ISNLGAVTGF | ASSLSGTDAEQF | 0.57737523 |
| HQEDPGRI | ASSLSGTDAEQF | 0.7016224 |
| GTPGEPTP | ASSLSGTDAEQF | 0.78222084 |
| GFFNSGS | ASSLSGTDAEQF | 0.77088976 |
| GAGGGGLTAGAN | ASSLSGTDAEQF | 0.67335117 |
| FVDALVNM | ASSLSGTDAEQF | 0.7389601 |
| FLTVAPEL | ASSLSGTDAEQF | 0.7954341 |
| FETLTAL | ASSLSGTDAEQF | 0.6297186 |
| DVINPIEL | ASSLSGTDAEQF | 0.6997268 |
| APVSGAPV | ASSLSGTDAEQF | 0.8080811 |
| ALVGGLAW | ASSLSGTDAEQF | 0.6920091 |
| ALAAAAATAI | ASSLSGTDAEQF | 0.809766 |
| AFGLGLLL | ASSLSGTDAEQF | 0.81074923 |
| AFFAAASL | ASSLSGTDAEQF | 0.57132053 |
| AAAALPAFSPP | ASSLSGTDAEQF | 0.7871686 |
| VVLAPAL | ASSLSGTDAEQF | 0.7875528 |
| TIGHVDHGKTTL | ASSLSGTDAEQF | 0.6848329 |
| LISNASDALDK | ASSLSGTDAEQF | 0.69961905 |
| LISNASDAL | ASSLSGTDAEQF | 0.37639952 |
| ISNASDALDK | ASSLSGTDAEQF | 0.6911694 |
| NQAETLVYQTE | ASSLSGTDAEQF | 0.5944161 |
| TGGTGGTAGAG | ASSLSGTDAEQF | 0.7546295 |
| TGSTAGGAGGA | ASSLSGTDAEQF | 0.7997843 |
| DPLAVPVAAGPV | ASSLSGTDAEQF | 0.7071992 |
| GGTGGTPGNGGN | ASSLSGTDAEQF | 0.37218338 |
| PAGLAGVDPATL | ASSLSGTDAEQF | 0.37055528 |
| GNNLVVEAPHAGASV | ASSLSGTDAEQF | 0.7468399 |
| KVTRIMSASTLEQPA | ASSLSGTDAEQF | 0.40237227 |
| LGFALPIALPRRTVV | ASSLSGTDAEQF | 0.52458763 |
| TRLGNLTARKLVVLV | ASSLSGTDAEQF | 0.5693877 |
| WSQQRRLIYFPSAGP | ASSLSGTDAEQF | 0.7941151 |
| GPNGNPDPMAAAVDIR | ASSLSGTDAEQF | 0.7990571 |
| TQRCVSVADLRAAASR | ASSLSGTDAEQF | 0.6183444 |
| QFVLDVAEHPVEVLTETV | ASSLSGTDAEQF | 0.7874652 |
| GGAGGTGGAGGNGGTQAGDG | ASSLSGTDAEQF | 0.7929162 |
| PLLATLRAIAVDVDAKPAQV | ASSLSGTDAEQF | 0.792367 |
| APKFRTGGYMIDFWGVGYQVAK | ASSLSGTDAEQF | 0.70639586 |
| GAGGAGGAGGAAGGPGGTGGTG | ASSLSGTDAEQF | 0.3757615 |
| IGALGLLATDVAPAGSVAFGVA | ASSLSGTDAEQF | 0.7978412 |
| AGGAGGIGGNGGAGGAGGVLMG | ASSLSGTDAEQF | 0.6538479 |
| ADELIGLL | ASSLSGTDAEQF | 0.8100413 |
| ADRALKA | ASSLSGTDAEQF | 0.7857305 |
| AGAVGQVGGAGGA | ASSLSGTDAEQF | 0.7333975 |
| AGGGGGLGGAP | ASSLSGTDAEQF | 0.55353224 |

| | | |
|---|---|---|
| AGGLLYGNGGAG | ASSLSGTDAEQF | 0.75743115 |
| AGLESIGL | ASSLSGTDAEQF | 0.79385173 |
| AGLKTGII | ASSLSGTDAEQF | 0.47186983 |
| ASLAALLL | ASSLSGTDAEQF | 0.38480306 |
| ASVINISSI | ASSLSGTDAEQF | 0.76547337 |
| ATPPGPA | ASSLSGTDAEQF | 0.7152616 |
| ATVASALV | ASSLSGTDAEQF | 0.80226386 |
| ATVASLIL | ASSLSGTDAEQF | 0.7168589 |
| AVNGPVLIRH | ASSLSGTDAEQF | 0.68047833 |
| DGAAGAVGI | ASSLSGTDAEQF | 0.7717948 |
| DPLLHGI | ASSLSGTDAEQF | 0.7710358 |
| ELGAALGAL | ASSLSGTDAEQF | 0.7019158 |
| ELGRTQAV | ASSLSGTDAEQF | 0.7636789 |
| FVAGGYLS | ASSLSGTDAEQF | 0.60783076 |
| GASGGAGGAGGW | ASSLSGTDAEQF | 0.7840096 |
| GGAGAGGDGGAGGI | ASSLSGTDAEQF | 0.7870927 |
| GTGHNGSQPGMGGNG | ASSLSGTDAEQF | 0.77092266 |
| GYAAASASAS | ASSLSGTDAEQF | 0.7966239 |
| HQVDVAAVQQG | ASSLSGTDAEQF | 0.739605 |
| IGAAIAHT | ASSLSGTDAEQF | 0.71400243 |
| IGMIEVAQL | ASSLSGTDAEQF | 0.504467 |
| IRLAPEHTSAMD | ASSLSGTDAEQF | 0.65730095 |
| IYNVSVL | ASSLSGTDAEQF | 0.7112383 |
| KQGFLGM | ASSLSGTDAEQF | 0.74459594 |
| LESVITQL | ASSLSGTDAEQF | 0.771462 |
| LGLGALVVV | ASSLSGTDAEQF | 0.7814102 |
| LIGNGGNGGSGGI | ASSLSGTDAEQF | 0.61203086 |
| LNAINAPTL | ASSLSGTDAEQF | 0.80526674 |
| LQLLNTGV | ASSLSGTDAEQF | 0.7962217 |
| LQTSIEF | ASSLSGTDAEQF | 0.46147758 |
| MSFAVSEL | ASSLSGTDAEQF | 0.6829345 |
| NTGALGL | ASSLSGTDAEQF | 0.78993344 |
| PPGAGGAGGAGTT | ASSLSGTDAEQF | 0.7869186 |
| QLVGLGI | ASSLSGTDAEQF | 0.6689974 |
| RFLEQVHQL | ASSLSGTDAEQF | 0.7268958 |
| RGVGYVLR | ASSLSGTDAEQF | 0.809433 |
| SATGVFAGVI | ASSLSGTDAEQF | 0.788961 |
| SLPAITLP | ASSLSGTDAEQF | 0.8007897 |
| STALAFTV | ASSLSGTDAEQF | 0.38047725 |
| SVLTVGLGK | ASSLSGTDAEQF | 0.6593268 |
| TALAGAGSGI | ASSLSGTDAEQF | 0.56583774 |
| TGLPFTGI | ASSLSGTDAEQF | 0.20596594 |
| VAEHVTV | ASSLSGTDAEQF | 0.7936703 |
| VDIVLNSL | ASSLSGTDAEQF | 0.5308137 |
| VLTGGGAVL | ASSLSGTDAEQF | 0.7694562 |
| VNVGLAYGI | ASSLSGTDAEQF | 0.57321715 |
| VSSAAVGAV | ASSLSGTDAEQF | 0.78876853 |
| VTSVYEAGV | ASSLSGTDAEQF | 0.72453076 |
| WVALDRGA | ASSLSGTDAEQF | 0.68762 |
| YLVGSVVI | ASSLSGTDAEQF | 0.5014877 |
| GPPGTGKTLL | ASSLSGTDAEQF | 0.76210463 |

| | | |
|---|---|---|
| GVGMTKF | ASSLSGTDAEQF | 0.8019947 |
| IGIGGSDL | ASSLSGTDAEQF | 0.79911876 |
| IGTLLAEL | ASSLSGTDAEQF | 0.41483438 |
| IYTSGTT | ASSLSGTDAEQF | 0.8039454 |
| LSFPAGV | ASSLSGTDAEQF | 0.6947528 |
| LYTSGTT | ASSLSGTDAEQF | 0.6179424 |
| VGAFPFGF | ASSLSGTDAEQF | 0.5331199 |
| VGAGTVEF | ASSLSGTDAEQF | 0.63744175 |
| VGKSTLLL | ASSLSGTDAEQF | 0.6951122 |
| AALAHAGGGIV | ASSLSGTDAEQF | 0.7579452 |
| AALVAVSLL | ASSLSGTDAEQF | 0.78709865 |
| AAVSRAVAAL | ASSLSGTDAEQF | 0.56768167 |
| AGGSSGIGGF | ASSLSGTDAEQF | 0.7648921 |
| AGLTVANANG | ASSLSGTDAEQF | 0.69494665 |
| AGLVGAVL | ASSLSGTDAEQF | 0.16451131 |
| ALSQAFTV | ASSLSGTDAEQF | 0.5751529 |
| ALTFSVSL | ASSLSGTDAEQF | 0.3582005 |
| ALTGAALVAL | ASSLSGTDAEQF | 0.800929 |
| ASFSIGAGVL | ASSLSGTDAEQF | 0.48242587 |
| DGSVYKI | ASSLSGTDAEQF | 0.67336816 |
| EQLAEIV | ASSLSGTDAEQF | 0.668589 |
| FAVSELTVM | ASSLSGTDAEQF | 0.7742634 |
| GAAPLAAGL | ASSLSGTDAEQF | 0.7492094 |
| GADGGDGGSGGAGG | ASSLSGTDAEQF | 0.54472727 |
| GFGLDTGGAGG | ASSLSGTDAEQF | 0.61099166 |
| GLASGGLL | ASSLSGTDAEQF | 0.7563196 |
| GRIAGAAVLL | ASSLSGTDAEQF | 0.7816292 |
| GSAGSLNTSF | ASSLSGTDAEQF | 0.6435187 |
| GTGQIGI | ASSLSGTDAEQF | 0.6726296 |
| GTLIALIL | ASSLSGTDAEQF | 0.80612266 |
| IAATMGFVVG | ASSLSGTDAEQF | 0.34058353 |
| IFAQALL | ASSLSGTDAEQF | 0.7966161 |
| IGAAVALGF | ASSLSGTDAEQF | 0.5005675 |
| IGLTPLAL | ASSLSGTDAEQF | 0.64411646 |
| ILGSNTSTL | ASSLSGTDAEQF | 0.7946085 |
| IVFSLTVL | ASSLSGTDAEQF | 0.7649467 |
| LASKATGF | ASSLSGTDAEQF | 0.6728538 |
| LASVALIL | ASSLSGTDAEQF | 0.66391444 |
| LFAVGLTL | ASSLSGTDAEQF | 0.47661674 |
| LFNGGIGTYI | ASSLSGTDAEQF | 0.6185428 |
| LTLGAGII | ASSLSGTDAEQF | 0.64566034 |
| LVAAALGDI | ASSLSGTDAEQF | 0.80315 |
| NTAFVQLGI | ASSLSGTDAEQF | 0.71259797 |
| QLFSGAEL | ASSLSGTDAEQF | 0.37866896 |
| SAATFITL | ASSLSGTDAEQF | 0.78020334 |
| SNYGLANLGV | ASSLSGTDAEQF | 0.7989917 |
| SYGGAYAVM | ASSLSGTDAEQF | 0.8018417 |
| TTGVGGLTLG | ASSLSGTDAEQF | 0.46379006 |
| TTQTIYI | ASSLSGTDAEQF | 0.64411044 |
| VAAGAGLGAVI | ASSLSGTDAEQF | 0.683746 |
| VAAVVTLL | ASSLSGTDAEQF | 0.6796924 |

| | | |
|---|---|---|
| VGFLGGLI | ASSLSGTDAEQF | 0.5717845 |
| VGVIPILL | ASSLSGTDAEQF | 0.5848593 |
| VIFASSLI | ASSLSGTDAEQF | 0.7859589 |
| VKGGDGGAAAG | ASSLSGTDAEQF | 0.82163733 |
| VTKTFTNI | ASSLSGTDAEQF | 0.6375195 |
| VVAAAILL | ASSLSGTDAEQF | 0.30367792 |
| YAALDVEL | ASSLSGTDAEQF | 0.68686223 |
| LASVALLL | ASSLSGTDAEQF | 0.5316285 |
| LDAVHRV | ASSLSGTDAEQF | 0.54313314 |
| QGLSSSIF | ASSLSGTDAEQF | 0.6824447 |
| RGTFANIRL | ASSLSGTDAEQF | 0.3262555 |
| RQLWWGHRI | ASSLSGTDAEQF | 0.70087457 |
| YGPPGTGKTLL | ASSLSGTDAEQF | 0.8069055 |
| AALVGAPANVA | ASSLSGTDAEQF | 0.6755409 |
| AATFRRTE | ASSLSGTDAEQF | 0.70264137 |
| AGALGGGI | ASSLSGTDAEQF | 0.7870698 |
| AGGAGGAGGIGAG | ASSLSGTDAEQF | 0.7556107 |
| AGSGALGL | ASSLSGTDAEQF | 0.795611 |
| AQIADQI | ASSLSGTDAEQF | 0.79977024 |
| ASAVAVTYL | ASSLSGTDAEQF | 0.70933235 |
| ASTVLGGTGG | ASSLSGTDAEQF | 0.72055745 |
| AVGGEPAK | ASSLSGTDAEQF | 0.78516793 |
| CEQGTFGGTC | ASSLSGTDAEQF | 0.6729339 |
| DEGKAPAG | ASSLSGTDAEQF | 0.7191925 |
| DTVVFPAL | ASSLSGTDAEQF | 0.6893059 |
| FEGITMAG | ASSLSGTDAEQF | 0.75124127 |
| FPLVVQD | ASSLSGTDAEQF | 0.7319697 |
| GAFTTPQ | ASSLSGTDAEQF | 0.8066232 |
| GFGGFGTAGGVGGA | ASSLSGTDAEQF | 0.23604429 |
| GFLAQFL | ASSLSGTDAEQF | 0.6191566 |
| GGAGGAGGNAGVA | ASSLSGTDAEQF | 0.7962955 |
| GGVGTAGAGGAGGAGG | ASSLSGTDAEQF | 0.2572008 |
| GNGGGGDAGNAGSGGN | ASSLSGTDAEQF | 0.6025296 |
| GTGGAGGAGNDAGS | ASSLSGTDAEQF | 0.68652475 |
| GTTNTGWFNS | ASSLSGTDAEQF | 0.79455864 |
| IFMGPLFTL | ASSLSGTDAEQF | 0.34229994 |
| ILAHAIEAI | ASSLSGTDAEQF | 0.73307025 |
| IRSAIREAL | ASSLSGTDAEQF | 0.33254248 |
| ITFAGGG | ASSLSGTDAEQF | 0.79915595 |
| ITGMSGHQEV | ASSLSGTDAEQF | 0.68967617 |
| IVAGLACTL | ASSLSGTDAEQF | 0.38212937 |
| IVGGGLAGA | ASSLSGTDAEQF | 0.7836211 |
| IVTGVLNI | ASSLSGTDAEQF | 0.47939733 |
| LAQQGCI | ASSLSGTDAEQF | 0.7883843 |
| LCLHVEN | ASSLSGTDAEQF | 0.7827598 |
| LDAISDGS | ASSLSGTDAEQF | 0.8180093 |
| LGGFAAML | ASSLSGTDAEQF | 0.40907714 |
| LGGKNPMI | ASSLSGTDAEQF | 0.7815571 |
| LGPKGVAAIL | ASSLSGTDAEQF | 0.64342177 |
| LMVGVSI | ASSLSGTDAEQF | 0.81415606 |
| LVGAAFVL | ASSLSGTDAEQF | 0.6325232 |

| | | |
|---|---|---|
| LVLAVAPL | ASSLSGTDAEQF | 0.31100562 |
| MAAIAVIDAA | ASSLSGTDAEQF | 0.7849241 |
| NNGINAELQ | ASSLSGTDAEQF | 0.7220495 |
| RGEPLVLVHG | ASSLSGTDAEQF | 0.68799466 |
| RGGAGGAAGLI | ASSLSGTDAEQF | 0.78339076 |
| RSYRMHPAV | ASSLSGTDAEQF | 0.67414105 |
| SAAHLLAGQLA | ASSLSGTDAEQF | 0.6403892 |
| SAAVGSY | ASSLSGTDAEQF | 0.7683116 |
| SPHAVLHRI | ASSLSGTDAEQF | 0.7539873 |
| SQIAKVL | ASSLSGTDAEQF | 0.6726219 |
| TALGHLVP | ASSLSGTDAEQF | 0.7928641 |
| TAPTTSGAA | ASSLSGTDAEQF | 0.73001754 |
| TDGDDGNGGSPGT | ASSLSGTDAEQF | 0.5091358 |
| TDGLGEN | ASSLSGTDAEQF | 0.80599016 |
| TGGPPLPGQA | ASSLSGTDAEQF | 0.6682781 |
| TIPAITI | ASSLSGTDAEQF | 0.7729709 |
| TIPGFTI | ASSLSGTDAEQF | 0.7402692 |
| TSIASGALA | ASSLSGTDAEQF | 0.81656355 |
| TVTLPGAAI | ASSLSGTDAEQF | 0.66531646 |
| VASGLSAALP | ASSLSGTDAEQF | 0.7540417 |
| VGAGVIGI | ASSLSGTDAEQF | 0.7291728 |
| VGPKRFGV | ASSLSGTDAEQF | 0.7862878 |
| VNVAFPD | ASSLSGTDAEQF | 0.690716 |
| VSMTNSGVS | ASSLSGTDAEQF | 0.79003114 |
| VTDVLGDLSV | ASSLSGTDAEQF | 0.74263966 |
| VVVVGAGFAGL | ASSLSGTDAEQF | 0.7989763 |
| VYTVGTVV | ASSLSGTDAEQF | 0.6422105 |
| WTTFSGP | ASSLSGTDAEQF | 0.7341837 |
| YDELVEAF | ASSLSGTDAEQF | 0.8099414 |
| YGTGGAGGNGG | ASSLSGTDAEQF | 0.4241153 |
| YTNVMAV | ASSLSGTDAEQF | 0.724918 |
| FLEQVHQL | ASSLSGTDAEQF | 0.780339 |
| GLSLLAVAV | ASSLSGTDAEQF | 0.7988141 |
| LEQVHQL | ASSLSGTDAEQF | 0.6618651 |
| LNAAGAI | ASSLSGTDAEQF | 0.78763306 |
| PAVHASL | ASSLSGTDAEQF | 0.6937182 |
| TELEDDA | ASSLSGTDAEQF | 0.747496 |
| TPPGPPN | ASSLSGTDAEQF | 0.6751783 |
| TPPPPPPP | ASSLSGTDAEQF | 0.6613009 |
| TQSPALI | ASSLVDRDGTGQLY | 0.62620056 |
| EFAAAFPQ | ASSLVDRDGTGQLY | 0.6948708 |
| LLSGNLDVL | ASSLVDRDGTGQLY | 0.77348083 |
| GPANLGNYNI | ASSLVDRDGTGQLY | 0.7508265 |
| TLGPTLP | ASSLVDRDGTGQLY | 0.55399525 |
| VYAYSSCCK | ASSLVDRDGTGQLY | 0.7559805 |
| APAPAPAPNI | ASSLVDRDGTGQLY | 0.7765535 |
| PGGPVPIP | ASSLVDRDGTGQLY | 0.81610477 |
| FLTVALEP | ASSLVDRDGTGQLY | 0.7255821 |
| SQYFGANL | ASSLVDRDGTGQLY | 0.78347886 |
| GGVKPAA | ASSLVDRDGTGQLY | 0.7174945 |
| GGTGGAGGAGGW | ASSLVDRDGTGQLY | 0.7350592 |

| | | |
|---|---|---|
| GVGGVAGLGGAGG | ASSLVDRDGTGQLY | 0.4320169 |
| VLLVLATTI | ASSLVDRDGTGQLY | 0.5137651 |
| IGIFAAML | ASSLVDRDGTGQLY | 0.71316016 |
| AVGDLID | ASSLVDRDGTGQLY | 0.55022454 |
| NISPLPL | ASSLVDRDGTGQLY | 0.78338075 |
| VTAVFVAI | ASSLVDRDGTGQLY | 0.79492044 |
| SGDDNDAAL | ASSLVDRDGTGQLY | 0.7881799 |
| AVAALAHS | ASSLVDRDGTGQLY | 0.79849565 |
| GGVTVMATP | ASSLVDRDGTGQLY | 0.7906449 |
| AGLAANVRD | ASSLVDRDGTGQLY | 0.7827128 |
| TNRPDLIDPALLRPG | ASSLVDRDGTGQLY | 0.63412213 |
| LANLGVTAL | ASSLVDRDGTGQLY | 0.6911348 |
| TGERIEL | ASSLVDRDGTGQLY | 0.3422196 |
| NGEELAA | ASSLVDRDGTGQLY | 0.5675981 |
| VLSLVWG | ASSLVDRDGTGQLY | 0.61326325 |
| IGYALWL | ASSLVDRDGTGQLY | 0.7131667 |
| LSLLLAPL | ASSLVDRDGTGQLY | 0.25201106 |
| TLDEALK | ASSLVDRDGTGQLY | 0.76351047 |
| TDGAGTAEGG | ASSLVDRDGTGQLY | 0.80145246 |
| NAPGGAGGNGG | ASSLVDRDGTGQLY | 0.7477355 |
| VSVFMGMGGNF | ASSLVDRDGTGQLY | 0.7766166 |
| LASYLAAL | ASSLVDRDGTGQLY | 0.7450103 |
| LVGGTPAL | ASSLVDRDGTGQLY | 0.5734736 |
| VRNADVL | ASSLVDRDGTGQLY | 0.7781092 |
| TCSSPFLVSAM | ASSLVDRDGTGQLY | 0.658598 |
| VHITDAE | ASSLVDRDGTGQLY | 0.79737794 |
| EYAAPSF | ASSLVDRDGTGQLY | 0.7942126 |
| VVDAFHGGL | ASSLVDRDGTGQLY | 0.74088347 |
| ASVVDPQ | ASSLVDRDGTGQLY | 0.6917917 |
| SYNEFGGLS | ASSLVDRDGTGQLY | 0.7652372 |
| DPDGSVY | ASSLVDRDGTGQLY | 0.68197525 |
| MVSTIHATE | ASSLVDRDGTGQLY | 0.71590877 |
| LAAGVADPV | ASSLVDRDGTGQLY | 0.72406936 |
| TKEQIAA | ASSLVDRDGTGQLY | 0.7261918 |
| SLASPAAL | ASSLVDRDGTGQLY | 0.47621942 |
| IGLVFYGV | ASSLVDRDGTGQLY | 0.7675955 |
| FLLDASNL | ASSLVDRDGTGQLY | 0.7159099 |
| YQVGNFP | ASSLVDRDGTGQLY | 0.7317072 |
| ESLLSQST | ASSLVDRDGTGQLY | 0.5368992 |
| LFTVGFAAVG | ASSLVDRDGTGQLY | 0.8069364 |
| ATPPTPT | ASSLVDRDGTGQLY | 0.7910748 |
| TDLATLG | ASSLVDRDGTGQLY | 0.5081165 |
| SAGGTGR | ASSLVDRDGTGQLY | 0.7437071 |
| GFSNSGS | ASSLVDRDGTGQLY | 0.6440336 |
| GGNGGSGGTSV | ASSLVDRDGTGQLY | 0.67961025 |
| GTSVATGGAGN | ASSLVDRDGTGQLY | 0.78928506 |
| NGGNGGSGGTS | ASSLVDRDGTGQLY | 0.7326143 |
| AVVKFPH | ASSLVDRDGTGQLY | 0.52608347 |
| TPPPPPP | ASSLVDRDGTGQLY | 0.62507546 |
| AAPAAAAAPA | ASSLVDRDGTGQLY | 0.72806835 |
| IALNEDL | ASSLVDRDGTGQLY | 0.6472528 |

| | | |
|---|---|---|
| IVNVSSM | ASSLVDRDGTGQLY | 0.32726315 |
| TGIPPAPR | ASSLVDRDGTGQLY | 0.72795844 |
| FDLGGGTF | ASSLVDRDGTGQLY | 0.5134176 |
| FDLGGGTFDVS | ASSLVDRDGTGQLY | 0.5661957 |
| NEPTAAA | ASSLVDRDGTGQLY | 0.59011376 |
| NEPTAAALA | ASSLVDRDGTGQLY | 0.7888736 |
| GSIINISSI | ASSLVDRDGTGQLY | 0.79458404 |
| VAPGNAGTA | ASSLVDRDGTGQLY | 0.7154288 |
| TGGGASLE | ASSLVDRDGTGQLY | 0.75353247 |
| VAVYNHYKRI | ASSLVDRDGTGQLY | 0.75338924 |
| TDPAPATT | ASSLVDRDGTGQLY | 0.78225195 |
| TNRPDILDPALLRPG | ASSLVDRDGTGQLY | 0.76519865 |
| LTTTTTS | ASSLVDRDGTGQLY | 0.7173246 |
| PTGSGKTL | ASSLVDRDGTGQLY | 0.76706207 |
| TGSGKTL | ASSLVDRDGTGQLY | 0.6692609 |
| TIGHVDHGK | ASSLVDRDGTGQLY | 0.7320671 |
| TIGHVDHGKTT | ASSLVDRDGTGQLY | 0.72908926 |
| GHVDHGKTT | ASSLVDRDGTGQLY | 0.52571666 |
| AQFEHTLL | ASSLVDRDGTGQLY | 0.697131 |
| IVNQDEL | ASSLVDRDGTGQLY | 0.7460302 |
| TNGSQFF | ASSLVDRDGTGQLY | 0.80394435 |
| TGKGIVE | ASSLVDRDGTGQLY | 0.7120955 |
| LISNASDA | ASSLVDRDGTGQLY | 0.3396756 |
| AELRTLGELTKIAKAHGAQ | ASSLVDRDGTGQLY | 0.6164237 |
| APNAVAAAEPSW | ASSLVDRDGTGQLY | 0.6992768 |
| APQVAGAATAVL | ASSLVDRDGTGQLY | 0.78600776 |
| ARRDPAAAAAILPTDA | ASSLVDRDGTGQLY | 0.78363717 |
| DAAARDVLAEAGLAEYF | ASSLVDRDGTGQLY | 0.8116685 |
| DDQIVLSAKANLAKVEQ | ASSLVDRDGTGQLY | 0.77804375 |
| DELLPDTGEDSDEDGH | ASSLVDRDGTGQLY | 0.6344701 |
| ELPEAPANRATVPQD | ASSLVDRDGTGQLY | 0.7800888 |
| FGQSKIERVTGIG | ASSLVDRDGTGQLY | 0.61028135 |
| GAGGTGSTAGGAGGAGGAGGLY | ASSLVDRDGTGQLY | 0.7199898 |
| GHGGTGAAGGNGGTGGAGGSGI | ASSLVDRDGTGQLY | 0.6884662 |
| GNGGNGGAGGEGGPGGAGGAGA | ASSLVDRDGTGQLY | 0.72607154 |
| GSTAGNGGNGGNATNGTIAGQP | ASSLVDRDGTGQLY | 0.65481746 |
| GTGGDGGNGGNGGTGG | ASSLVDRDGTGQLY | 0.6675167 |
| GVRRPAGIPADAPV | ASSLVDRDGTGQLY | 0.38532913 |
| GYRQVTDAHLLTLARR | ASSLVDRDGTGQLY | 0.7637991 |
| IAKMGQDGHDRG | ASSLVDRDGTGQLY | 0.69396436 |
| KAEPSIEIQYT | ASSLVDRDGTGQLY | 0.7158576 |
| KPVRIRMAADEDTVAVT | ASSLVDRDGTGQLY | 0.6734559 |
| MPPRIAGMRLLVIKPEP | ASSLVDRDGTGQLY | 0.8004814 |
| MKFYAVALFRSRGVPPTR | ASSLVDRDGTGQLY | 0.8083994 |
| NAGGFGILGAVAHSP | ASSLVDRDGTGQLY | 0.6448777 |
| NEGREITLAMAEM | ASSLVDRDGTGQLY | 0.58664536 |
| NELVEWSHPLGHRWRWE | ASSLVDRDGTGQLY | 0.7810584 |
| NLGAGGTGGNGGNPG | ASSLVDRDGTGQLY | 0.665382 |
| NTEWPKWETVMAADPVD | ASSLVDRDGTGQLY | 0.75152075 |
| PASAAPAETQP | ASSLVDRDGTGQLY | 0.72492695 |
| QQSSDGKNGVRAAEITGE | ASSLVDRDGTGQLY | 0.66088253 |

| | | |
|---|---|---|
| RAPDAVTTSWWKE | ASSLVDRDGTGQLY | 0.58211935 |
| SGFGNVGAHTSGWFNQ | ASSLVDRDGTGQLY | 0.77277064 |
| SGVAGAVGQAASVGGLKV | ASSLVDRDGTGQLY | 0.63784146 |
| SNGGAGTGSAGGNGGT | ASSLVDRDGTGQLY | 0.75426435 |
| TGGAGGEGGAAG | ASSLVDRDGTGQLY | 0.62219965 |
| TGGDGGAAGAGGAPGHG | ASSLVDRDGTGQLY | 0.78452444 |
| TMPPLLAAIANAGRHGV | ASSLVDRDGTGQLY | 0.49361187 |
| TSSTAGNSGGA | ASSLVDRDGTGQLY | 0.5794029 |
| VPATVPVGILC | ASSLVDRDGTGQLY | 0.23847426 |
| VQRPDSGGVVLTGRIS | ASSLVDRDGTGQLY | 0.67199945 |
| WTPNPVPATKGVPAGLVQAT | ASSLVDRDGTGQLY | 0.75530833 |
| YALKRALAKFSDRFDVVII | ASSLVDRDGTGQLY | 0.80214703 |
| YLIGSVIV | ASSLVDRDGTGQLY | 0.7843693 |
| YGQLVAVID | ASSLVDRDGTGQLY | 0.749267 |
| VTVGGMTLPA | ASSLVDRDGTGQLY | 0.80129975 |
| VLALPAV | ASSLVDRDGTGQLY | 0.62513995 |
| VIPPTLNYET | ASSLVDRDGTGQLY | 0.36464682 |
| VGMGGMHPSSG | ASSLVDRDGTGQLY | 0.74500483 |
| VGLAGGAATA | ASSLVDRDGTGQLY | 0.7133223 |
| VGGAGGAGGSGP | ASSLVDRDGTGQLY | 0.8016851 |
| VFAAFMV | ASSLVDRDGTGQLY | 0.77637935 |
| TLYDKLR | ASSLVDRDGTGQLY | 0.36351925 |
| TLAVFTVL | ASSLVDRDGTGQLY | 0.7527498 |
| TGSVQPARV | ASSLVDRDGTGQLY | 0.74302745 |
| TGGGAGSGGT | ASSLVDRDGTGQLY | 0.43418902 |
| SHLETSL | ASSLVDRDGTGQLY | 0.5530561 |
| SFTTGMT | ASSLVDRDGTGQLY | 0.56344247 |
| QVEVAKRMGVRQPTVS | ASSLVDRDGTGQLY | 0.5973109 |
| QGSGGPALAM | ASSLVDRDGTGQLY | 0.7254135 |
| MLAPSNP | ASSLVDRDGTGQLY | 0.61625594 |
| LVNNAAI | ASSLVDRDGTGQLY | 0.7575058 |
| LTGEINTLA | ASSLVDRDGTGQLY | 0.6118925 |
| LSIFGPV | ASSLVDRDGTGQLY | 0.65651464 |
| LLALLANL | ASSLVDRDGTGQLY | 0.696205 |
| LISNASDALDKLR | ASSLVDRDGTGQLY | 0.71652424 |
| LINEAVP | ASSLVDRDGTGQLY | 0.76146424 |
| LGYAFSGL | ASSLVDRDGTGQLY | 0.7796221 |
| LDFVAVAL | ASSLVDRDGTGQLY | 0.6348588 |
| LAQPGAE | ASSLVDRDGTGQLY | 0.7160903 |
| ISNLGAVTGF | ASSLVDRDGTGQLY | 0.7915792 |
| HQEDPGRI | ASSLVDRDGTGQLY | 0.5492004 |
| GTPGEPTP | ASSLVDRDGTGQLY | 0.5787332 |
| GFFNSGS | ASSLVDRDGTGQLY | 0.76670337 |
| GAGGGGLTAGAN | ASSLVDRDGTGQLY | 0.7395742 |
| FVDALVNM | ASSLVDRDGTGQLY | 0.7900031 |
| FLTVAPEL | ASSLVDRDGTGQLY | 0.7341265 |
| FETLTAL | ASSLVDRDGTGQLY | 0.7359451 |
| DVINPIEL | ASSLVDRDGTGQLY | 0.75002956 |
| APVSGAPV | ASSLVDRDGTGQLY | 0.79937315 |
| ALVGGLAW | ASSLVDRDGTGQLY | 0.7702044 |
| ALAAAAATAI | ASSLVDRDGTGQLY | 0.55834377 |

| | | |
|---|---|---|
| AFGLGLLL | ASSLVDRDGTGQLY | 0.76852477 |
| AFFAAASL | ASSLVDRDGTGQLY | 0.40610304 |
| AAAALPAFSPP | ASSLVDRDGTGQLY | 0.7616993 |
| VVLAPAL | ASSLVDRDGTGQLY | 0.7808682 |
| TIGHVDHGKTTL | ASSLVDRDGTGQLY | 0.53375906 |
| LISNASDALDK | ASSLVDRDGTGQLY | 0.65338254 |
| LISNASDAL | ASSLVDRDGTGQLY | 0.80381155 |
| ISNASDALDK | ASSLVDRDGTGQLY | 0.7780243 |
| NQAETLVYQTE | ASSLVDRDGTGQLY | 0.72186434 |
| TGGTGGTAGAG | ASSLVDRDGTGQLY | 0.5227917 |
| TGSTAGGAGGA | ASSLVDRDGTGQLY | 0.68720925 |
| DPLAVPVAAGPV | ASSLVDRDGTGQLY | 0.6376289 |
| GGTGGTPGNGGN | ASSLVDRDGTGQLY | 0.3192604 |
| PAGLAGVDPATL | ASSLVDRDGTGQLY | 0.7699125 |
| GNNLVVEAPHAGASV | ASSLVDRDGTGQLY | 0.67493516 |
| KVTRIMSASTLEQPA | ASSLVDRDGTGQLY | 0.79984736 |
| LGFALPIALPRRTVV | ASSLVDRDGTGQLY | 0.7433465 |
| TRLGNLTARKLVVLV | ASSLVDRDGTGQLY | 0.7961191 |
| WSQQRRLIYFPSAGP | ASSLVDRDGTGQLY | 0.62618273 |
| GPNGNPDPMAAAVDIR | ASSLVDRDGTGQLY | 0.709625 |
| TQRCVSVADLRAAASR | ASSLVDRDGTGQLY | 0.2752593 |
| QFVLDVAEHPVEVLTETV | ASSLVDRDGTGQLY | 0.74099517 |
| GGAGGTGGAGGNGGTQAGDG | ASSLVDRDGTGQLY | 0.48921686 |
| PLLATLRAIAVDVDAKPAQV | ASSLVDRDGTGQLY | 0.7475009 |
| APKFRTGGYMIDFWGVGYQVAK | ASSLVDRDGTGQLY | 0.63073194 |
| GAGGAGGAGGAAGGPGGTGGTG | ASSLVDRDGTGQLY | 0.79884696 |
| IGALGLLATDVAPAGSVAFGVA | ASSLVDRDGTGQLY | 0.646582 |
| AGGAGGIGGNGGAGGAGGVLMG | ASSLVDRDGTGQLY | 0.76753426 |
| ADELIGLL | ASSLVDRDGTGQLY | 0.6267791 |
| ADRALKA | ASSLVDRDGTGQLY | 0.45264915 |
| AGAVGQVGGAGGA | ASSLVDRDGTGQLY | 0.63613296 |
| AGGGGGLGGAP | ASSLVDRDGTGQLY | 0.7888582 |
| AGGLLYGNGGAG | ASSLVDRDGTGQLY | 0.4378876 |
| AGLESIGL | ASSLVDRDGTGQLY | 0.27283636 |
| AGLKTGII | ASSLVDRDGTGQLY | 0.5611774 |
| ASLAALLL | ASSLVDRDGTGQLY | 0.72695184 |
| ASVINISSI | ASSLVDRDGTGQLY | 0.8017759 |
| ATPPGPA | ASSLVDRDGTGQLY | 0.7877164 |
| ATVASALV | ASSLVDRDGTGQLY | 0.7221271 |
| ATVASLIL | ASSLVDRDGTGQLY | 0.8086046 |
| AVNGPVLIRH | ASSLVDRDGTGQLY | 0.18349005 |
| DGAAGAVGI | ASSLVDRDGTGQLY | 0.65565693 |
| DPLLHGI | ASSLVDRDGTGQLY | 0.8046988 |
| ELGAALGAL | ASSLVDRDGTGQLY | 0.73379385 |
| ELGRTQAV | ASSLVDRDGTGQLY | 0.6848116 |
| FVAGGYLS | ASSLVDRDGTGQLY | 0.7915896 |
| GASGGAGGAGGW | ASSLVDRDGTGQLY | 0.62266684 |
| GGAGAGGDGGAGGI | ASSLVDRDGTGQLY | 0.6587563 |
| GTGHNGSQPGMGGNG | ASSLVDRDGTGQLY | 0.7544248 |
| GYAAASASAS | ASSLVDRDGTGQLY | 0.7545015 |
| HQVDVAAVQQG | ASSLVDRDGTGQLY | 0.75482583 |

| | | |
|---|---|---|
| IGAAIAHT | ASSLVDRDGTGQLY | 0.31548172 |
| IGMIEVAQL | ASSLVDRDGTGQLY | 0.7755008 |
| IRLAPEHTSAMD | ASSLVDRDGTGQLY | 0.8046955 |
| IYNVSVL | ASSLVDRDGTGQLY | 0.6522584 |
| KQGFLGM | ASSLVDRDGTGQLY | 0.79304963 |
| LESVITQL | ASSLVDRDGTGQLY | 0.7231345 |
| LGLGALVVV | ASSLVDRDGTGQLY | 0.7666391 |
| LIGNGGNGGSGGI | ASSLVDRDGTGQLY | 0.5729482 |
| LNAINAPTL | ASSLVDRDGTGQLY | 0.47793677 |
| LQLLNTGV | ASSLVDRDGTGQLY | 0.8093186 |
| LQTSIEF | ASSLVDRDGTGQLY | 0.5289413 |
| MSFAVSEL | ASSLVDRDGTGQLY | 0.7304405 |
| NTGALGL | ASSLVDRDGTGQLY | 0.6727158 |
| PPGAGGAGGAGTT | ASSLVDRDGTGQLY | 0.8155401 |
| QLVGLGI | ASSLVDRDGTGQLY | 0.75410473 |
| RFLEQVHQL | ASSLVDRDGTGQLY | 0.7601338 |
| RGVGYVLR | ASSLVDRDGTGQLY | 0.63593495 |
| SATGVFAGVI | ASSLVDRDGTGQLY | 0.7058026 |
| SLPAITLP | ASSLVDRDGTGQLY | 0.76965725 |
| STALAFTV | ASSLVDRDGTGQLY | 0.38739026 |
| SVLTVGLGK | ASSLVDRDGTGQLY | 0.7949522 |
| TALAGAGSGI | ASSLVDRDGTGQLY | 0.4983313 |
| TGLPFTGI | ASSLVDRDGTGQLY | 0.72019243 |
| VAEHVTV | ASSLVDRDGTGQLY | 0.7456385 |
| VDIVLNSL | ASSLVDRDGTGQLY | 0.8022224 |
| VLTGGGAVL | ASSLVDRDGTGQLY | 0.679578 |
| VNVGLAYGI | ASSLVDRDGTGQLY | 0.80795527 |
| VSSAAVGAV | ASSLVDRDGTGQLY | 0.5858841 |
| VTSVYEAGV | ASSLVDRDGTGQLY | 0.7815573 |
| WVALDRGA | ASSLVDRDGTGQLY | 0.7974895 |
| YLVGSVVI | ASSLVDRDGTGQLY | 0.7590576 |
| GPPGTGKTLL | ASSLVDRDGTGQLY | 0.8113657 |
| GVGMTKF | ASSLVDRDGTGQLY | 0.7816863 |
| IGIGGSDL | ASSLVDRDGTGQLY | 0.29252502 |
| IGTLLAEL | ASSLVDRDGTGQLY | 0.70113957 |
| IYTSGTT | ASSLVDRDGTGQLY | 0.7862028 |
| LSFPAGV | ASSLVDRDGTGQLY | 0.7964396 |
| LYTSGTT | ASSLVDRDGTGQLY | 0.7877694 |
| VGAFPFGF | ASSLVDRDGTGQLY | 0.7831081 |
| VGAGTVEF | ASSLVDRDGTGQLY | 0.66980875 |
| VGKSTLLL | ASSLVDRDGTGQLY | 0.7972001 |
| AALAHAGGGIV | ASSLVDRDGTGQLY | 0.82463765 |
| AALVAVSLL | ASSLVDRDGTGQLY | 0.7715552 |
| AAVSRAVAAL | ASSLVDRDGTGQLY | 0.5970627 |
| AGGSSGIGGF | ASSLVDRDGTGQLY | 0.60410595 |
| AGLTVANANG | ASSLVDRDGTGQLY | 0.76146203 |
| AGLVGAVL | ASSLVDRDGTGQLY | 0.77426946 |
| ALSQAFTV | ASSLVDRDGTGQLY | 0.43777373 |
| ALTFSVSL | ASSLVDRDGTGQLY | 0.6797911 |
| ALTGAALVAL | ASSLVDRDGTGQLY | 0.73866963 |
| ASFSIGAGVL | ASSLVDRDGTGQLY | 0.2749884 |

| | | |
|---|---|---|
| DGSVYKI | ASSLVDRDGTGQLY | 0.74058044 |
| EQLAEIV | ASSLVDRDGTGQLY | 0.78629136 |
| FAVSELTVM | ASSLVDRDGTGQLY | 0.73703885 |
| GAAPLAAGL | ASSLVDRDGTGQLY | 0.43484676 |
| GADGGDGGSGGAGG | ASSLVDRDGTGQLY | 0.7919949 |
| GFGLDTGGAGG | ASSLVDRDGTGQLY | 0.80097806 |
| GLASGGLL | ASSLVDRDGTGQLY | 0.8057147 |
| GRIAGAAVLL | ASSLVDRDGTGQLY | 0.476152 |
| GSAGSLNTSF | ASSLVDRDGTGQLY | 0.20509091 |
| GTGQIGI | ASSLVDRDGTGQLY | 0.7750628 |
| GTLIALIL | ASSLVDRDGTGQLY | 0.7841536 |
| IAATMGFVVG | ASSLVDRDGTGQLY | 0.686515 |
| IFAQALL | ASSLVDRDGTGQLY | 0.5565233 |
| IGAAVALGF | ASSLVDRDGTGQLY | 0.72688 |
| IGLTPLAL | ASSLVDRDGTGQLY | 0.77362895 |
| ILGSNTSTL | ASSLVDRDGTGQLY | 0.43182984 |
| IVFSLTVL | ASSLVDRDGTGQLY | 0.4455708 |
| LASKATGF | ASSLVDRDGTGQLY | 0.73659235 |
| LASVALIL | ASSLVDRDGTGQLY | 0.79318625 |
| LFAVGLTL | ASSLVDRDGTGQLY | 0.48419148 |
| LFNGGIGTYI | ASSLVDRDGTGQLY | 0.6014272 |
| LTLGAGII | ASSLVDRDGTGQLY | 0.77752316 |
| LVAAALGDI | ASSLVDRDGTGQLY | 0.8039911 |
| NTAFVQLGI | ASSLVDRDGTGQLY | 0.7343257 |
| QLFSGAEL | ASSLVDRDGTGQLY | 0.45695853 |
| SAATFITL | ASSLVDRDGTGQLY | 0.7477248 |
| SNYGLANLGV | ASSLVDRDGTGQLY | 0.43938223 |
| SYGGAYAVM | ASSLVDRDGTGQLY | 0.77524954 |
| TTGVGGLTLG | ASSLVDRDGTGQLY | 0.73801625 |
| TTQTIYI | ASSLVDRDGTGQLY | 0.71566474 |
| VAAGAGLGAVI | ASSLVDRDGTGQLY | 0.6697554 |
| VAAVVTLL | ASSLVDRDGTGQLY | 0.47567764 |
| VGFLGGLI | ASSLVDRDGTGQLY | 0.42922246 |
| VGVIPILL | ASSLVDRDGTGQLY | 0.69876164 |
| VIFASSLI | ASSLVDRDGTGQLY | 0.7946752 |
| VKGGDGGAAAG | ASSLVDRDGTGQLY | 0.79077846 |
| VTKTFTNI | ASSLVDRDGTGQLY | 0.7672187 |
| VVAAAILL | ASSLVDRDGTGQLY | 0.7483808 |
| YAALDVEL | ASSLVDRDGTGQLY | 0.51737964 |
| LASVALLL | ASSLVDRDGTGQLY | 0.7520846 |
| LDAVHRV | ASSLVDRDGTGQLY | 0.4932645 |
| QGLSSSIF | ASSLVDRDGTGQLY | 0.78636837 |
| RGTFANIRL | ASSLVDRDGTGQLY | 0.67178476 |
| RQLWWGHRI | ASSLVDRDGTGQLY | 0.7646873 |
| YGPPGTGKTLL | ASSLVDRDGTGQLY | 0.63319856 |
| AALVGAPANVA | ASSLVDRDGTGQLY | 0.7507234 |
| AATFRRTE | ASSLVDRDGTGQLY | 0.75040144 |
| AGALGGGI | ASSLVDRDGTGQLY | 0.5156648 |
| AGGAGGAGGIGAG | ASSLVDRDGTGQLY | 0.7169169 |
| AGSGALGL | ASSLVDRDGTGQLY | 0.78720963 |
| AQIADQI | ASSLVDRDGTGQLY | 0.77356565 |

| | | |
|---|---|---|
| ASAVAVTYL | ASSLVDRDGTGQLY | 0.35449326 |
| ASTVLGGTGG | ASSLVDRDGTGQLY | 0.7584715 |
| AVGGEPAK | ASSLVDRDGTGQLY | 0.5699216 |
| CEQGTFGGTC | ASSLVDRDGTGQLY | 0.7336661 |
| DEGKAPAG | ASSLVDRDGTGQLY | 0.7208371 |
| DTVVFPAL | ASSLVDRDGTGQLY | 0.7225505 |
| FEGITMAG | ASSLVDRDGTGQLY | 0.74465656 |
| FPLVVQD | ASSLVDRDGTGQLY | 0.4649191 |
| GAFTTPQ | ASSLVDRDGTGQLY | 0.60615826 |
| GFGGFGTAGGVGGA | ASSLVDRDGTGQLY | 0.63961375 |
| GFLAQFL | ASSLVDRDGTGQLY | 0.7624551 |
| GGAGGAGGNAGVA | ASSLVDRDGTGQLY | 0.76583815 |
| GGVGTAGAGGAGGAGG | ASSLVDRDGTGQLY | 0.47777906 |
| GNGGGGDAGNAGSGGN | ASSLVDRDGTGQLY | 0.7388146 |
| GTGGAGGAGNDAGS | ASSLVDRDGTGQLY | 0.66556394 |
| GTTNTGWFNS | ASSLVDRDGTGQLY | 0.5379542 |
| IFMGPLFTL | ASSLVDRDGTGQLY | 0.69345534 |
| ILAHAIEAI | ASSLVDRDGTGQLY | 0.6158135 |
| IRSAIREAL | ASSLVDRDGTGQLY | 0.37232938 |
| ITFAGGG | ASSLVDRDGTGQLY | 0.7608727 |
| ITGMSGHQEV | ASSLVDRDGTGQLY | 0.6073453 |
| IVAGLACTL | ASSLVDRDGTGQLY | 0.75222445 |
| IVGGGLAGA | ASSLVDRDGTGQLY | 0.79449344 |
| IVTGVLNI | ASSLVDRDGTGQLY | 0.39945138 |
| LAQQGCI | ASSLVDRDGTGQLY | 0.7617072 |
| LCLHVEN | ASSLVDRDGTGQLY | 0.68013376 |
| LDAISDGS | ASSLVDRDGTGQLY | 0.73170674 |
| LGGFAAML | ASSLVDRDGTGQLY | 0.7407156 |
| LGGKNPMI | ASSLVDRDGTGQLY | 0.5293443 |
| LGPKGVAAIL | ASSLVDRDGTGQLY | 0.78030145 |
| LMVGVSI | ASSLVDRDGTGQLY | 0.78181183 |
| LVGAAFVL | ASSLVDRDGTGQLY | 0.75865734 |
| LVLAVAPL | ASSLVDRDGTGQLY | 0.7229309 |
| MAAIAVIDAA | ASSLVDRDGTGQLY | 0.78592163 |
| NNGINAELQ | ASSLVDRDGTGQLY | 0.78900015 |
| RGEPLVLVHG | ASSLVDRDGTGQLY | 0.6433078 |
| RGGAGGAAGLI | ASSLVDRDGTGQLY | 0.3071853 |
| RSYRMHPAV | ASSLVDRDGTGQLY | 0.72986 |
| SAAHLLAGQLA | ASSLVDRDGTGQLY | 0.72903746 |
| SAAVGSY | ASSLVDRDGTGQLY | 0.50840664 |
| SPHAVLHRI | ASSLVDRDGTGQLY | 0.41207555 |
| SQIAKVL | ASSLVDRDGTGQLY | 0.77618533 |
| TALGHLVP | ASSLVDRDGTGQLY | 0.8108469 |
| TAPTTSGAA | ASSLVDRDGTGQLY | 0.7370439 |
| TDGDDGNGGSPGT | ASSLVDRDGTGQLY | 0.81816435 |
| TDGLGEN | ASSLVDRDGTGQLY | 0.38160706 |
| TGGPPLPGQA | ASSLVDRDGTGQLY | 0.78595495 |
| TIPAITI | ASSLVDRDGTGQLY | 0.7965652 |
| TIPGFTI | ASSLVDRDGTGQLY | 0.53714436 |
| TSIASGALA | ASSLVDRDGTGQLY | 0.7533729 |
| TVTLPGAAI | ASSLVDRDGTGQLY | 0.64407086 |

| | | |
|---|---|---|
| VASGLSAALP | ASSLVDRDGTGQLY | 0.5337321 |
| VGAGVIGI | ASSLVDRDGTGQLY | 0.7887284 |
| VGPKRFGV | ASSLVDRDGTGQLY | 0.7353511 |
| VNVAFPD | ASSLVDRDGTGQLY | 0.76475644 |
| VSMTNSGVS | ASSLVDRDGTGQLY | 0.6942229 |
| VTDVLGDLSV | ASSLVDRDGTGQLY | 0.5388423 |
| VVVVGAGFAGL | ASSLVDRDGTGQLY | 0.7347651 |
| VYTVGTVV | ASSLVDRDGTGQLY | 0.7842911 |
| WTTFSGP | ASSLVDRDGTGQLY | 0.7918232 |
| YDELVEAF | ASSLVDRDGTGQLY | 0.82323766 |
| YGTGGAGGNGG | ASSLVDRDGTGQLY | 0.61352205 |
| YTNVMAV | ASSLVDRDGTGQLY | 0.7959821 |
| FLEQVHQL | ASSLVDRDGTGQLY | 0.803292 |
| GLSLLAVAV | ASSLVDRDGTGQLY | 0.7273835 |
| LEQVHQL | ASSLVDRDGTGQLY | 0.6876646 |
| LNAAGAI | ASSLVDRDGTGQLY | 0.7929148 |
| PAVHASL | ASSLVDRDGTGQLY | 0.7206048 |
| TELEDDA | ASSLVDRDGTGQLY | 0.7594507 |
| TPPGPPN | ASSLVDRDGTGQLY | 0.7968612 |
| TPPPPPPP | ASSLVDRDGTGQLY | 0.2089971 |
| TQSPALI | ASSLLQANTEVF | 0.56678635 |
| EFAAAFPQ | ASSLLQANTEVF | 0.38000715 |
| LLSGNLDVL | ASSLLQANTEVF | 0.488382 |
| GPANLGNYNI | ASSLLQANTEVF | 0.2641065 |
| TLGPTLP | ASSLLQANTEVF | 0.23457012 |
| VYAYSSCCK | ASSLLQANTEVF | 0.29546544 |
| APAPAPAPNI | ASSLLQANTEVF | 0.22669949 |
| PGGPVPIP | ASSLLQANTEVF | 0.63155025 |
| FLTVALEP | ASSLLQANTEVF | 0.4576428 |
| SQYFGANL | ASSLLQANTEVF | 0.2675404 |
| GGVKPAA | ASSLLQANTEVF | 0.44374055 |
| GGTGGAGGAGGW | ASSLLQANTEVF | 0.5133547 |
| GVGGVAGLGGAGG | ASSLLQANTEVF | 0.17806181 |
| VLLVLATTI | ASSLLQANTEVF | 0.3273065 |
| IGIFAAML | ASSLLQANTEVF | 0.25682858 |
| AVGDLID | ASSLLQANTEVF | 0.30174977 |
| NISPLPL | ASSLLQANTEVF | 0.12557417 |
| VTAVFVAI | ASSLLQANTEVF | 0.44119707 |
| SGDDNDAAL | ASSLLQANTEVF | 0.21643877 |
| AVAALAHS | ASSLLQANTEVF | 0.14197294 |
| GGVTVMATP | ASSLLQANTEVF | 0.34886613 |
| AGLAANVRD | ASSLLQANTEVF | 0.26046222 |
| TNRPDLIDPALLRPG | ASSLLQANTEVF | 0.27787632 |
| LANLGVTAL | ASSLLQANTEVF | 0.46662274 |
| TGERIEL | ASSLLQANTEVF | 0.37178215 |
| NGEELAA | ASSLLQANTEVF | 0.21455503 |
| VLSLVWG | ASSLLQANTEVF | 0.34419933 |
| IGYALWL | ASSLLQANTEVF | 0.51229924 |
| LSLLLAPL | ASSLLQANTEVF | 0.24728113 |
| TLDEALK | ASSLLQANTEVF | 0.3040625 |
| TDGAGTAEGG | ASSLLQANTEVF | 0.6539473 |

| | | |
|---|---|---|
| NAPGGAGGNGG | ASSLLQANTEVF | 0.35298386 |
| VSVFMGMGGNF | ASSLLQANTEVF | 0.60261464 |
| LASYLAAL | ASSLLQANTEVF | 0.651151 |
| LVGGTPAL | ASSLLQANTEVF | 0.2813734 |
| VRNADVL | ASSLLQANTEVF | 0.48811713 |
| TCSSPFLVSAM | ASSLLQANTEVF | 0.45991668 |
| VHITDAE | ASSLLQANTEVF | 0.39503124 |
| EYAAPSF | ASSLLQANTEVF | 0.30882403 |
| VVDAFHGGL | ASSLLQANTEVF | 0.20738727 |
| ASVVDPQ | ASSLLQANTEVF | 0.31446922 |
| SYNEFGGLS | ASSLLQANTEVF | 0.2521558 |
| DPDGSVY | ASSLLQANTEVF | 0.36670297 |
| MVSTIHATE | ASSLLQANTEVF | 0.5955938 |
| LAAGVADPV | ASSLLQANTEVF | 0.28153312 |
| TKEQIAA | ASSLLQANTEVF | 0.15616275 |
| SLASPAAL | ASSLLQANTEVF | 0.15616407 |
| IGLVFYGV | ASSLLQANTEVF | 0.42983824 |
| FLLDASNL | ASSLLQANTEVF | 0.21878164 |
| YQVGNFP | ASSLLQANTEVF | 0.28958932 |
| ESLLSQST | ASSLLQANTEVF | 0.22372547 |
| LFTVGFAAVG | ASSLLQANTEVF | 0.50524664 |
| ATPPTPT | ASSLLQANTEVF | 0.5430274 |
| TDLATLG | ASSLLQANTEVF | 0.215506 |
| SAGGTGR | ASSLLQANTEVF | 0.5362097 |
| GFSNSGS | ASSLLQANTEVF | 0.55830956 |
| GGNGGSGGTSV | ASSLLQANTEVF | 0.6120623 |
| GTSVATGGAGN | ASSLLQANTEVF | 0.38886473 |
| NGGNGGSGGTS | ASSLLQANTEVF | 0.19307494 |
| AVVKFPH | ASSLLQANTEVF | 0.57914275 |
| TPPPPPP | ASSLLQANTEVF | 0.2440402 |
| AAPAAAAAPA | ASSLLQANTEVF | 0.53855217 |
| IALNEDL | ASSLLQANTEVF | 0.46413994 |
| IVNVSSM | ASSLLQANTEVF | 0.35659397 |
| TGIPPAPR | ASSLLQANTEVF | 0.21385509 |
| FDLGGGTF | ASSLLQANTEVF | 0.43489587 |
| FDLGGGTFDVS | ASSLLQANTEVF | 0.57512003 |
| NEPTAAA | ASSLLQANTEVF | 0.1517084 |
| NEPTAAALA | ASSLLQANTEVF | 0.2368897 |
| GSIINISSI | ASSLLQANTEVF | 0.4055684 |
| VAPGNAGTA | ASSLLQANTEVF | 0.3036472 |
| TGGGASLE | ASSLLQANTEVF | 0.53050274 |
| VAVYNHYKRI | ASSLLQANTEVF | 0.47175813 |
| TDPAPATT | ASSLLQANTEVF | 0.32698208 |
| TNRPDILDPALLRPG | ASSLLQANTEVF | 0.44182375 |
| LTTTTTS | ASSLLQANTEVF | 0.51766247 |
| PTGSGKTL | ASSLLQANTEVF | 0.36935994 |
| TGSGKTL | ASSLLQANTEVF | 0.3728704 |
| TIGHVDHGK | ASSLLQANTEVF | 0.1973296 |
| TIGHVDHGKTT | ASSLLQANTEVF | 0.55330956 |
| GHVDHGKTT | ASSLLQANTEVF | 0.5298345 |
| AQFEHTLL | ASSLLQANTEVF | 0.48530492 |

| | | |
|---|---|---|
| IVNQDEL | ASSLLQANTEVF | 0.5649281 |
| TNGSQFF | ASSLLQANTEVF | 0.40603992 |
| TGKGIVE | ASSLLQANTEVF | 0.42422482 |
| LISNASDA | ASSLLQANTEVF | 0.1634308 |
| AELRTLGELTKIAKAHGAQ | ASSLLQANTEVF | 0.31092295 |
| APNAVAAAEPSW | ASSLLQANTEVF | 0.2561582 |
| APQVAGAATAVL | ASSLLQANTEVF | 0.38693437 |
| ARRDPAAAAAILPTDA | ASSLLQANTEVF | 0.3860517 |
| DAAARDVLAEAGLAEYF | ASSLLQANTEVF | 0.5778011 |
| DDQIVLSAKANLAKVEQ | ASSLLQANTEVF | 0.23445807 |
| DELLPDTGEDSDEDGH | ASSLLQANTEVF | 0.5473934 |
| ELPEAPANRATVPQD | ASSLLQANTEVF | 0.62224627 |
| FGQSKIERVTGIG | ASSLLQANTEVF | 0.18216676 |
| GAGGTGSTAGGAGGAGGAGGLY | ASSLLQANTEVF | 0.27738166 |
| GHGGTGAAGGNGGTGGAGGSGI | ASSLLQANTEVF | 0.5368433 |
| GNGGNGGAGGEGGPGGAGGAGA | ASSLLQANTEVF | 0.46766886 |
| GSTAGNGGNGGNATNGTIAGQP | ASSLLQANTEVF | 0.2783973 |
| GTGGDGGNGGNGGTGG | ASSLLQANTEVF | 0.38878077 |
| GVRRPAGIPADAPV | ASSLLQANTEVF | 0.63395256 |
| GYRQVTDAHLLTLARR | ASSLLQANTEVF | 0.44722942 |
| IAKMGQDGHDRG | ASSLLQANTEVF | 0.60833484 |
| KAEPSIEIQYT | ASSLLQANTEVF | 0.15989597 |
| KPVRIRMAADEDTVAVT | ASSLLQANTEVF | 0.41593856 |
| MPPRIAGMRLLVIKPEP | ASSLLQANTEVF | 0.34376913 |
| MKFYAVALFRSRGVPPTR | ASSLLQANTEVF | 0.116999365 |
| NAGGFGILGAVAHSP | ASSLLQANTEVF | 0.4763742 |
| NEGREITLAMAEM | ASSLLQANTEVF | 0.2830722 |
| NELVEWSHPLGHRWRWE | ASSLLQANTEVF | 0.42974603 |
| NLGAGGTGGNGGNPG | ASSLLQANTEVF | 0.27045903 |
| NTEWPKWETVMAADPVD | ASSLLQANTEVF | 0.48375905 |
| PASAAPAETQP | ASSLLQANTEVF | 0.29224312 |
| QQSSDGKNGVRAAEITGE | ASSLLQANTEVF | 0.29317918 |
| RAPDAVTTSWWKE | ASSLLQANTEVF | 0.27451405 |
| SGFGNVGAHTSGWFNQ | ASSLLQANTEVF | 0.38534963 |
| SGVAGAVGQAASVGGLKV | ASSLLQANTEVF | 0.5182019 |
| SNGGAGTGSAGGNGGT | ASSLLQANTEVF | 0.6280101 |
| TGGAGGEGGAAG | ASSLLQANTEVF | 0.1581087 |
| TGGDGGAAGAGGAPGHG | ASSLLQANTEVF | 0.5118791 |
| TMPPLLAAIANAGRHGV | ASSLLQANTEVF | 0.32263637 |
| TSSTAGNSGGA | ASSLLQANTEVF | 0.21111178 |
| VPATVPVGILC | ASSLLQANTEVF | 0.18721983 |
| VQRPDSGGVVLTGRIS | ASSLLQANTEVF | 0.23560706 |
| WTPNPVPATKGVPAGLVQAT | ASSLLQANTEVF | 0.263486 |
| YALKRALAKFSDRFDVVII | ASSLLQANTEVF | 0.33454743 |
| YLIGSVIV | ASSLLQANTEVF | 0.36180246 |
| YGQLVAVID | ASSLLQANTEVF | 0.25445473 |
| VTVGGMTLPA | ASSLLQANTEVF | 0.38993746 |
| VLALPAV | ASSLLQANTEVF | 0.31719536 |
| VIPPTLNYET | ASSLLQANTEVF | 0.0942855 |
| VGMGGMHPSSG | ASSLLQANTEVF | 0.2751492 |
| VGLAGGAATA | ASSLLQANTEVF | 0.17199335 |

| | | |
|---|---|---|
| VGGAGGAGGSGP | ASSLLQANTEVF | 0.6150225 |
| VFAAFMV | ASSLLQANTEVF | 0.18519905 |
| TLYDKLR | ASSLLQANTEVF | 0.21517205 |
| TLAVFTVL | ASSLLQANTEVF | 0.28274652 |
| TGSVQPARV | ASSLLQANTEVF | 0.51152945 |
| TGGGAGSGGT | ASSLLQANTEVF | 0.35067475 |
| SHLETSL | ASSLLQANTEVF | 0.25012428 |
| SFTTGMT | ASSLLQANTEVF | 0.18079104 |
| QVEVAKRMGVRQPTVS | ASSLLQANTEVF | 0.42429292 |
| QGSGGPALAM | ASSLLQANTEVF | 0.3798189 |
| MLAPSNP | ASSLLQANTEVF | 0.27315006 |
| LVNNAAI | ASSLLQANTEVF | 0.22600818 |
| LTGEINTLA | ASSLLQANTEVF | 0.6421065 |
| LSIFGPV | ASSLLQANTEVF | 0.19760375 |
| LLALLANL | ASSLLQANTEVF | 0.5932215 |
| LISNASDALDKLR | ASSLLQANTEVF | 0.19727731 |
| LINEAVP | ASSLLQANTEVF | 0.22775039 |
| LGYAFSGL | ASSLLQANTEVF | 0.62502474 |
| LDFVAVAL | ASSLLQANTEVF | 0.46860301 |
| LAQPGAE | ASSLLQANTEVF | 0.27202746 |
| ISNLGAVTGF | ASSLLQANTEVF | 0.33072528 |
| HQEDPGRI | ASSLLQANTEVF | 0.21613613 |
| GTPGEPTP | ASSLLQANTEVF | 0.2210087 |
| GFFNSGS | ASSLLQANTEVF | 0.29493472 |
| GAGGGGLTAGAN | ASSLLQANTEVF | 0.5046668 |
| FVDALVNM | ASSLLQANTEVF | 0.40845987 |
| FLTVAPEL | ASSLLQANTEVF | 0.19502537 |
| FETLTAL | ASSLLQANTEVF | 0.6282979 |
| DVINPIEL | ASSLLQANTEVF | 0.63004565 |
| APVSGAPV | ASSLLQANTEVF | 0.5938316 |
| ALVGGLAW | ASSLLQANTEVF | 0.19711226 |
| ALAAAAATAI | ASSLLQANTEVF | 0.36267078 |
| AFGLGLLL | ASSLLQANTEVF | 0.32334208 |
| AFFAAASL | ASSLLQANTEVF | 0.24826202 |
| AAAALPAFSPP | ASSLLQANTEVF | 0.28009135 |
| VVLAPAL | ASSLLQANTEVF | 0.27232528 |
| TIGHVDHGKTTL | ASSLLQANTEVF | 0.5595532 |
| LISNASDALDK | ASSLLQANTEVF | 0.7451193 |
| LISNASDAL | ASSLLQANTEVF | 0.31388518 |
| ISNASDALDK | ASSLLQANTEVF | 0.12925607 |
| NQAETLVYQTE | ASSLLQANTEVF | 0.25698078 |
| TGGTGGTAGAG | ASSLLQANTEVF | 0.17865005 |
| TGSTAGGAGGA | ASSLLQANTEVF | 0.1919892 |
| DPLAVPVAAGPV | ASSLLQANTEVF | 0.29309678 |
| GGTGGTPGNGGN | ASSLLQANTEVF | 0.15152301 |
| PAGLAGVDPATL | ASSLLQANTEVF | 0.32480344 |
| GNNLVVEAPHAGASV | ASSLLQANTEVF | 0.65793097 |
| KVTRIMSASTLEQPA | ASSLLQANTEVF | 0.23528232 |
| LGFALPIALPRRTVV | ASSLLQANTEVF | 0.44453362 |
| TRLGNLTARKLVVLV | ASSLLQANTEVF | 0.57163334 |
| WSQQRRLIYFPSAGP | ASSLLQANTEVF | 0.39304274 |

| | | |
|---|---|---|
| GPNGNPDPMAAAVDIR | ASSLLQANTEVF | 0.5567973 |
| TQRCVSVADLRAAASR | ASSLLQANTEVF | 0.56885874 |
| QFVLDVAEHPVEVLTETV | ASSLLQANTEVF | 0.3842696 |
| GGAGGTGGAGGNGGTQAGDG | ASSLLQANTEVF | 0.39013585 |
| PLLATLRAIAVDVDAKPAQV | ASSLLQANTEVF | 0.5682418 |
| APKFRTGGYMIDFWGVGYQVAK | ASSLLQANTEVF | 0.24783942 |
| GAGGAGGAGGAAGGPGGTGGTG | ASSLLQANTEVF | 0.3370753 |
| IGALGLLATDVAPAGSVAFGVA | ASSLLQANTEVF | 0.35050538 |
| AGGAGGIGGNGGAGGAGGVLMG | ASSLLQANTEVF | 0.4655952 |
| ADELIGLL | ASSLLQANTEVF | 0.36996764 |
| ADRALKA | ASSLLQANTEVF | 0.66999084 |
| AGAVGQVGGAGGA | ASSLLQANTEVF | 0.15786812 |
| AGGGGGLGGAP | ASSLLQANTEVF | 0.23637378 |
| AGGLLYGNGGAG | ASSLLQANTEVF | 0.5518951 |
| AGLESIGL | ASSLLQANTEVF | 0.13105395 |
| AGLKTGII | ASSLLQANTEVF | 0.2646939 |
| ASLAALLL | ASSLLQANTEVF | 0.3612535 |
| ASVINISSI | ASSLLQANTEVF | 0.5735723 |
| ATPPGPA | ASSLLQANTEVF | 0.13223961 |
| ATVASALV | ASSLLQANTEVF | 0.33536673 |
| ATVASLIL | ASSLLQANTEVF | 0.19270647 |
| AVNGPVLIRH | ASSLLQANTEVF | 0.64357144 |
| DGAAGAVGI | ASSLLQANTEVF | 0.32270205 |
| DPLLHGI | ASSLLQANTEVF | 0.18177147 |
| ELGAALGAL | ASSLLQANTEVF | 0.49060664 |
| ELGRTQAV | ASSLLQANTEVF | 0.14731514 |
| FVAGGYLS | ASSLLQANTEVF | 0.5196239 |
| GASGGAGGAGGW | ASSLLQANTEVF | 0.43000543 |
| GGAGAGGDGGAGGI | ASSLLQANTEVF | 0.6727679 |
| GTGHNGSQPGMGGNG | ASSLLQANTEVF | 0.18744962 |
| GYAAASASAS | ASSLLQANTEVF | 0.40946603 |
| HQVDVAAVQQG | ASSLLQANTEVF | 0.26749146 |
| IGAAIAHT | ASSLLQANTEVF | 0.6888416 |
| IGMIEVAQL | ASSLLQANTEVF | 0.2862345 |
| IRLAPEHTSAMD | ASSLLQANTEVF | 0.2081224 |
| IYNVSVL | ASSLLQANTEVF | 0.5161334 |
| KQGFLGM | ASSLLQANTEVF | 0.43600538 |
| LESVITQL | ASSLLQANTEVF | 0.40466142 |
| LGLGALVVV | ASSLLQANTEVF | 0.5506593 |
| LIGNGGNGGSGGI | ASSLLQANTEVF | 0.41995567 |
| LNAINAPTL | ASSLLQANTEVF | 0.25951084 |
| LQLLNTGV | ASSLLQANTEVF | 0.44397295 |
| LQTSIEF | ASSLLQANTEVF | 0.34294662 |
| MSFAVSEL | ASSLLQANTEVF | 0.28487316 |
| NTGALGL | ASSLLQANTEVF | 0.51031625 |
| PPGAGGAGGAGTT | ASSLLQANTEVF | 0.31048253 |
| QLVGLGI | ASSLLQANTEVF | 0.23180383 |
| RFLEQVHQL | ASSLLQANTEVF | 0.6339433 |
| RGVGYVLR | ASSLLQANTEVF | 0.3618118 |
| SATGVFAGVI | ASSLLQANTEVF | 0.4595727 |
| SLPAITLP | ASSLLQANTEVF | 0.45608068 |

| | | |
|---|---|---|
| STALAFTV | ASSLLQANTEVF | 0.6765872 |
| SVLTVGLGK | ASSLLQANTEVF | 0.25701848 |
| TALAGAGSGI | ASSLLQANTEVF | 0.36125648 |
| TGLPFTGI | ASSLLQANTEVF | 0.3281946 |
| VAEHVTV | ASSLLQANTEVF | 0.5036398 |
| VDIVLNSL | ASSLLQANTEVF | 0.3046176 |
| VLTGGGAVL | ASSLLQANTEVF | 0.43565863 |
| VNVGLAYGI | ASSLLQANTEVF | 0.47487885 |
| VSSAAVGAV | ASSLLQANTEVF | 0.58797175 |
| VTSVYEAGV | ASSLLQANTEVF | 0.3487102 |
| WVALDRGA | ASSLLQANTEVF | 0.33888808 |
| YLVGSVVI | ASSLLQANTEVF | 0.6279883 |
| GPPGTGKTLL | ASSLLQANTEVF | 0.23733127 |
| GVGMTKF | ASSLLQANTEVF | 0.34391388 |
| IGIGGSDL | ASSLLQANTEVF | 0.5239167 |
| IGTLLAEL | ASSLLQANTEVF | 0.47917417 |
| IYTSGTT | ASSLLQANTEVF | 0.4001265 |
| LSFPAGV | ASSLLQANTEVF | 0.511203 |
| LYTSGTT | ASSLLQANTEVF | 0.26447698 |
| VGAFPFGF | ASSLLQANTEVF | 0.3437825 |
| VGAGTVEF | ASSLLQANTEVF | 0.3226822 |
| VGKSTLLL | ASSLLQANTEVF | 0.23285031 |
| AALAHAGGGIV | ASSLLQANTEVF | 0.1892012 |
| AALVAVSLL | ASSLLQANTEVF | 0.25075603 |
| AAVSRAVAAL | ASSLLQANTEVF | 0.4360016 |
| AGGSSGIGGF | ASSLLQANTEVF | 0.36081207 |
| AGLTVANANG | ASSLLQANTEVF | 0.18735749 |
| AGLVGAVL | ASSLLQANTEVF | 0.3954367 |
| ALSQAFTV | ASSLLQANTEVF | 0.41930443 |
| ALTFSVSL | ASSLLQANTEVF | 0.71269417 |
| ALTGAALVAL | ASSLLQANTEVF | 0.37246382 |
| ASFSIGAGVL | ASSLLQANTEVF | 0.5279864 |
| DGSVYKI | ASSLLQANTEVF | 0.23409785 |
| EQLAEIV | ASSLLQANTEVF | 0.38048014 |
| FAVSELTVM | ASSLLQANTEVF | 0.15309188 |
| GAAPLAAGL | ASSLLQANTEVF | 0.25100896 |
| GADGGDGGSGGAGG | ASSLLQANTEVF | 0.2714835 |
| GFGLDTGGAGG | ASSLLQANTEVF | 0.19587389 |
| GLASGGLL | ASSLLQANTEVF | 0.5802504 |
| GRIAGAAVLL | ASSLLQANTEVF | 0.49281967 |
| GSAGSLNTSF | ASSLLQANTEVF | 0.52274793 |
| GTGQIGI | ASSLLQANTEVF | 0.6065649 |
| GTLIALIL | ASSLLQANTEVF | 0.59048074 |
| IAATMGFVVG | ASSLLQANTEVF | 0.50462204 |
| IFAQALL | ASSLLQANTEVF | 0.2402998 |
| IGAAVALGF | ASSLLQANTEVF | 0.29296303 |
| IGLTPLAL | ASSLLQANTEVF | 0.14717302 |
| ILGSNTSTL | ASSLLQANTEVF | 0.28045076 |
| IVFSLTVL | ASSLLQANTEVF | 0.26536134 |
| LASKATGF | ASSLLQANTEVF | 0.3388864 |
| LASVALIL | ASSLLQANTEVF | 0.12811342 |

| | | |
|---|---|---|
| LFAVGLTL | ASSLLQANTEVF | 0.47840452 |
| LFNGGIGTYI | ASSLLQANTEVF | 0.51603776 |
| LTLGAGII | ASSLLQANTEVF | 0.2583138 |
| LVAAALGDI | ASSLLQANTEVF | 0.47104734 |
| NTAFVQLGI | ASSLLQANTEVF | 0.43258232 |
| QLFSGAEL | ASSLLQANTEVF | 0.20889184 |
| SAATFITL | ASSLLQANTEVF | 0.48457363 |
| SNYGLANLGV | ASSLLQANTEVF | 0.26907024 |
| SYGGAYAVM | ASSLLQANTEVF | 0.5864285 |
| TTGVGGLTLG | ASSLLQANTEVF | 0.5760943 |
| TTQTIYI | ASSLLQANTEVF | 0.42644915 |
| VAAGAGLGAVI | ASSLLQANTEVF | 0.24646752 |
| VAAVVTLL | ASSLLQANTEVF | 0.46026853 |
| VGFLGGLI | ASSLLQANTEVF | 0.23977627 |
| VGVIPILL | ASSLLQANTEVF | 0.26126173 |
| VIFASSLI | ASSLLQANTEVF | 0.27061507 |
| VKGGDGGAAAG | ASSLLQANTEVF | 0.3405552 |
| VTKTFTNI | ASSLLQANTEVF | 0.25296757 |
| VVAAAILL | ASSLLQANTEVF | 0.3705701 |
| YAALDVEL | ASSLLQANTEVF | 0.30571097 |
| LASVALLL | ASSLLQANTEVF | 0.382379 |
| LDAVHRV | ASSLLQANTEVF | 0.17584187 |
| QGLSSSIF | ASSLLQANTEVF | 0.5574038 |
| RGTFANIRL | ASSLLQANTEVF | 0.5613169 |
| RQLWWGHRI | ASSLLQANTEVF | 0.19828181 |
| YGPPGTGKTLL | ASSLLQANTEVF | 0.41361308 |
| AALVGAPANVA | ASSLLQANTEVF | 0.3852029 |
| AATFRRTE | ASSLLQANTEVF | 0.25095412 |
| AGALGGGI | ASSLLQANTEVF | 0.5414057 |
| AGGAGGAGGIGAG | ASSLLQANTEVF | 0.35464856 |
| AGSGALGL | ASSLLQANTEVF | 0.15853739 |
| AQIADQI | ASSLLQANTEVF | 0.19936469 |
| ASAVAVTYL | ASSLLQANTEVF | 0.40082535 |
| ASTVLGGTGG | ASSLLQANTEVF | 0.29081205 |
| AVGGEPAK | ASSLLQANTEVF | 0.14433867 |
| CEQGTFGGTC | ASSLLQANTEVF | 0.38169485 |
| DEGKAPAG | ASSLLQANTEVF | 0.23803456 |
| DTVVFPAL | ASSLLQANTEVF | 0.1739573 |
| FEGITMAG | ASSLLQANTEVF | 0.247121 |
| FPLVVQD | ASSLLQANTEVF | 0.5098728 |
| GAFTTPQ | ASSLLQANTEVF | 0.47205907 |
| GFGGFGTAGGVGGA | ASSLLQANTEVF | 0.41371688 |
| GFLAQFL | ASSLLQANTEVF | 0.33951548 |
| GGAGGAGGNAGVA | ASSLLQANTEVF | 0.4281625 |
| GGVGTAGAGGAGGAGG | ASSLLQANTEVF | 0.5047119 |
| GNGGGGDAGNAGSGGN | ASSLLQANTEVF | 0.41231787 |
| GTGGAGGAGNDAGS | ASSLLQANTEVF | 0.40461993 |
| GTTNTGWFNS | ASSLLQANTEVF | 0.43117374 |
| IFMGPLFTL | ASSLLQANTEVF | 0.3364227 |
| ILAHAIEAI | ASSLLQANTEVF | 0.29485407 |
| IRSAIREAL | ASSLLQANTEVF | 0.30109835 |

| | | |
|---|---|---|
| ITFAGGG | ASSLLQANTEVF | 0.19675368 |
| ITGMSGHQEV | ASSLLQANTEVF | 0.29563951 |
| IVAGLACTL | ASSLLQANTEVF | 0.40546653 |
| IVGGGLAGA | ASSLLQANTEVF | 0.6286343 |
| IVTGVLNI | ASSLLQANTEVF | 0.40887052 |
| LAQQGCI | ASSLLQANTEVF | 0.1895383 |
| LCLHVEN | ASSLLQANTEVF | 0.29244882 |
| LDAISDGS | ASSLLQANTEVF | 0.31551093 |
| LGGFAAML | ASSLLQANTEVF | 0.46677786 |
| LGGKNPMI | ASSLLQANTEVF | 0.55657613 |
| LGPKGVAAIL | ASSLLQANTEVF | 0.6975948 |
| LMVGVSI | ASSLLQANTEVF | 0.46642187 |
| LVGAAFVL | ASSLLQANTEVF | 0.22537987 |
| LVLAVAPL | ASSLLQANTEVF | 0.4520795 |
| MAAIAVIDAA | ASSLLQANTEVF | 0.288801 |
| NNGINAELQ | ASSLLQANTEVF | 0.32864225 |
| RGEPLVLVHG | ASSLLQANTEVF | 0.3386892 |
| RGGAGGAAGLI | ASSLLQANTEVF | 0.3656145 |
| RSYRMHPAV | ASSLLQANTEVF | 0.28425384 |
| SAAHLLAGQLA | ASSLLQANTEVF | 0.3082095 |
| SAAVGSY | ASSLLQANTEVF | 0.18915261 |
| SPHAVLHRI | ASSLLQANTEVF | 0.44984674 |
| SQIAKVL | ASSLLQANTEVF | 0.19880669 |
| TALGHLVP | ASSLLQANTEVF | 0.35297406 |
| TAPTTSGAA | ASSLLQANTEVF | 0.270742 |
| TDGDDGNGGSPGT | ASSLLQANTEVF | 0.60346544 |
| TDGLGEN | ASSLLQANTEVF | 0.6533015 |
| TGGPPLPGQA | ASSLLQANTEVF | 0.23890324 |
| TIPAITI | ASSLLQANTEVF | 0.25696045 |
| TIPGFTI | ASSLLQANTEVF | 0.49523526 |
| TSIASGALA | ASSLLQANTEVF | 0.2288845 |
| TVTLPGAAI | ASSLLQANTEVF | 0.4791755 |
| VASGLSAALP | ASSLLQANTEVF | 0.30535603 |
| VGAGVIGI | ASSLLQANTEVF | 0.25552192 |
| VGPKRFGV | ASSLLQANTEVF | 0.28418115 |
| VNVAFPD | ASSLLQANTEVF | 0.45722073 |
| VSMTNSGVS | ASSLLQANTEVF | 0.18628785 |
| VTDVLGDLSV | ASSLLQANTEVF | 0.48687983 |
| VVVVGAGFAGL | ASSLLQANTEVF | 0.13801895 |
| VYTVGTVV | ASSLLQANTEVF | 0.45646697 |
| WTTFSGP | ASSLLQANTEVF | 0.16716892 |
| YDELVEAF | ASSLLQANTEVF | 0.2907228 |
| YGTGGAGGNGG | ASSLLQANTEVF | 0.14373975 |
| YTNVMAV | ASSLLQANTEVF | 0.29420334 |
| FLEQVHQL | ASSLLQANTEVF | 0.36682647 |
| GLSLLAVAV | ASSLLQANTEVF | 0.5823625 |
| LEQVHQL | ASSLLQANTEVF | 0.45044303 |
| LNAAGAI | ASSLLQANTEVF | 0.34010413 |
| PAVHASL | ASSLLQANTEVF | 0.45235127 |
| TELEDDA | ASSLLQANTEVF | 0.2285024 |
| TPPGPPN | ASSLLQANTEVF | 0.16896957 |

| | | |
|---|---|---|
| TPPPPPPP | ASSLLQANTEVF | 0.39340368 |
| TQSPALI | ASSLGQSETLY | 0.2737188 |
| EFAAAFPQ | ASSLGQSETLY | 0.5534014 |
| LLSGNLDVL | ASSLGQSETLY | 0.42470154 |
| GPANLGNYNI | ASSLGQSETLY | 0.13884555 |
| TLGPTLP | ASSLGQSETLY | 0.5247934 |
| VYAYSSCCK | ASSLGQSETLY | 0.48874784 |
| APAPAPAPNI | ASSLGQSETLY | 0.19528978 |
| PGGPVPIP | ASSLGQSETLY | 0.22466117 |
| FLTVALEP | ASSLGQSETLY | 0.26221532 |
| SQYFGANL | ASSLGQSETLY | 0.18770672 |
| GGVKPAA | ASSLGQSETLY | 0.41534576 |
| GGTGGAGGAGGW | ASSLGQSETLY | 0.22169353 |
| GVGGVAGLGGAGG | ASSLGQSETLY | 0.36512664 |
| VLLVLATTI | ASSLGQSETLY | 0.29484427 |
| IGIFAAML | ASSLGQSETLY | 0.3222557 |
| AVGDLID | ASSLGQSETLY | 0.33612406 |
| NISPLPL | ASSLGQSETLY | 0.32771546 |
| VTAVFVAI | ASSLGQSETLY | 0.3736673 |
| SGDDNDAAL | ASSLGQSETLY | 0.47664332 |
| AVAALAHS | ASSLGQSETLY | 0.22412246 |
| GGVTVMATP | ASSLGQSETLY | 0.34882224 |
| AGLAANVRD | ASSLGQSETLY | 0.56787556 |
| TNRPDLIDPALLRPG | ASSLGQSETLY | 0.1835382 |
| LANLGVTAL | ASSLGQSETLY | 0.23997992 |
| TGERIEL | ASSLGQSETLY | 0.44791597 |
| NGEELAA | ASSLGQSETLY | 0.3534333 |
| VLSLVWG | ASSLGQSETLY | 0.57037354 |
| IGYALWL | ASSLGQSETLY | 0.34802812 |
| LSLLLAPL | ASSLGQSETLY | 0.36430624 |
| TLDEALK | ASSLGQSETLY | 0.43247646 |
| TDGAGTAEGG | ASSLGQSETLY | 0.5691029 |
| NAPGGAGGNGG | ASSLGQSETLY | 0.43929148 |
| VSVFMGMGGNF | ASSLGQSETLY | 0.31427664 |
| LASYLAAL | ASSLGQSETLY | 0.4317559 |
| LVGGTPAL | ASSLGQSETLY | 0.24039234 |
| VRNADVL | ASSLGQSETLY | 0.3556219 |
| TCSSPFLVSAM | ASSLGQSETLY | 0.5344089 |
| VHITDAE | ASSLGQSETLY | 0.17209396 |
| EYAAPSF | ASSLGQSETLY | 0.2645044 |
| VVDAFHGGL | ASSLGQSETLY | 0.628203 |
| ASVVDPQ | ASSLGQSETLY | 0.43228054 |
| SYNEFGGLS | ASSLGQSETLY | 0.32994235 |
| DPDGSVY | ASSLGQSETLY | 0.22743177 |
| MVSTIHATE | ASSLGQSETLY | 0.31678393 |
| LAAGVADPV | ASSLGQSETLY | 0.39416087 |
| TKEQIAA | ASSLGQSETLY | 0.21035516 |
| SLASPAAL | ASSLGQSETLY | 0.46006322 |
| IGLVFYGV | ASSLGQSETLY | 0.28112733 |
| FLLDASNL | ASSLGQSETLY | 0.6209854 |
| YQVGNFP | ASSLGQSETLY | 0.38481706 |

| | | |
|---|---|---|
| ESLLSQST | ASSLGQSETLY | 0.5188278 |
| LFTVGFAAVG | ASSLGQSETLY | 0.29354364 |
| ATPPTPT | ASSLGQSETLY | 0.31480277 |
| TDLATLG | ASSLGQSETLY | 0.20221226 |
| SAGGTGR | ASSLGQSETLY | 0.251634 |
| GFSNSGS | ASSLGQSETLY | 0.25219128 |
| GGNGGSGGTSV | ASSLGQSETLY | 0.3445341 |
| GTSVATGGAGN | ASSLGQSETLY | 0.17038415 |
| NGGNGGSGGTS | ASSLGQSETLY | 0.61946607 |
| AVVKFPH | ASSLGQSETLY | 0.26994175 |
| TPPPPPP | ASSLGQSETLY | 0.4387287 |
| AAPAAAAAPA | ASSLGQSETLY | 0.32659212 |
| IALNEDL | ASSLGQSETLY | 0.2245842 |
| IVNVSSM | ASSLGQSETLY | 0.23027016 |
| TGIPPAPR | ASSLGQSETLY | 0.5293069 |
| FDLGGGTF | ASSLGQSETLY | 0.1691283 |
| FDLGGGTFDVS | ASSLGQSETLY | 0.121988215 |
| NEPTAAA | ASSLGQSETLY | 0.24556303 |
| NEPTAAALA | ASSLGQSETLY | 0.16742782 |
| GSIINISSI | ASSLGQSETLY | 0.6405238 |
| VAPGNAGTA | ASSLGQSETLY | 0.55201536 |
| TGGGASLE | ASSLGQSETLY | 0.34285828 |
| VAVYNHYKRI | ASSLGQSETLY | 0.68340915 |
| TDPAPATT | ASSLGQSETLY | 0.11655307 |
| TNRPDILDPALLRPG | ASSLGQSETLY | 0.32186258 |
| LTTTTTS | ASSLGQSETLY | 0.58544976 |
| PTGSGKTL | ASSLGQSETLY | 0.37332898 |
| TGSGKTL | ASSLGQSETLY | 0.31060404 |
| TIGHVDHGK | ASSLGQSETLY | 0.4662276 |
| TIGHVDHGKTT | ASSLGQSETLY | 0.23988365 |
| GHVDHGKTT | ASSLGQSETLY | 0.30129054 |
| AQFEHTLL | ASSLGQSETLY | 0.32861722 |
| IVNQDEL | ASSLGQSETLY | 0.44519073 |
| TNGSQFF | ASSLGQSETLY | 0.48435947 |
| TGKGIVE | ASSLGQSETLY | 0.16295677 |
| LISNASDA | ASSLGQSETLY | 0.5207293 |
| AELRTLGELTKIAKAHGAQ | ASSLGQSETLY | 0.65981376 |
| APNAVAAAEPSW | ASSLGQSETLY | 0.1994395 |
| APQVAGAATAVL | ASSLGQSETLY | 0.34923947 |
| ARRDPAAAAAILPTDA | ASSLGQSETLY | 0.34127834 |
| DAAARDVLAEAGLAEYF | ASSLGQSETLY | 0.21540385 |
| DDQIVLSAKANLAKVEQ | ASSLGQSETLY | 0.22489844 |
| DELLPDTGEDSDEDGH | ASSLGQSETLY | 0.16102558 |
| ELPEAPANRATVPQD | ASSLGQSETLY | 0.57149035 |
| FGQSKIERVTGIG | ASSLGQSETLY | 0.17964497 |
| GAGGTGSTAGGAGGAGGAGGLY | ASSLGQSETLY | 0.4377063 |
| GHGGTGAAGGNGGTGGAGGSGI | ASSLGQSETLY | 0.3596947 |
| GNGGNGGAGGEGGPGGAGGAGA | ASSLGQSETLY | 0.71528155 |
| GSTAGNGGNGGNATNGTIAGQP | ASSLGQSETLY | 0.4166032 |
| GTGGDGGNGGNGGTGG | ASSLGQSETLY | 0.37498415 |
| GVRRPAGIPADAPV | ASSLGQSETLY | 0.19437242 |

| | | |
|---|---|---|
| GYRQVTDAHLLTLARR | ASSLGQSETLY | 0.36357993 |
| IAKMGQDGHDRG | ASSLGQSETLY | 0.46723333 |
| KAEPSIEIQYT | ASSLGQSETLY | 0.19162352 |
| KPVRIRMAADEDTVAVT | ASSLGQSETLY | 0.5708237 |
| MPPRIAGMRLLVIKPEP | ASSLGQSETLY | 0.21665195 |
| MKFYAVALFRSRGVPPTR | ASSLGQSETLY | 0.40428573 |
| NAGGFGILGAVAHSP | ASSLGQSETLY | 0.44801182 |
| NEGREITLAMAEM | ASSLGQSETLY | 0.6434084 |
| NELVEWSHPLGHRWRWE | ASSLGQSETLY | 0.3316469 |
| NLGAGGTGGNGGNPG | ASSLGQSETLY | 0.58310765 |
| NTEWPKWETVMAADPVD | ASSLGQSETLY | 0.20742695 |
| PASAAPAETQP | ASSLGQSETLY | 0.52758753 |
| QQSSDGKNGVRAAEITGE | ASSLGQSETLY | 0.5150745 |
| RAPDAVTTSWWKE | ASSLGQSETLY | 0.44894528 |
| SGFGNVGAHTSGWFNQ | ASSLGQSETLY | 0.6775 |
| SGVAGAVGQAASVGGLKV | ASSLGQSETLY | 0.5054144 |
| SNGGAGTGSAGGNGGT | ASSLGQSETLY | 0.20675556 |
| TGGAGGEGGAAG | ASSLGQSETLY | 0.35088614 |
| TGGDGGAAGAGGAPGHG | ASSLGQSETLY | 0.46873498 |
| TMPPLLAAIANAGRHGV | ASSLGQSETLY | 0.59543586 |
| TSSTAGNSGGA | ASSLGQSETLY | 0.4355818 |
| VPATVPVGILC | ASSLGQSETLY | 0.41117537 |
| VQRPDSGGVVLTGRIS | ASSLGQSETLY | 0.23646824 |
| WTPNPVPATKGVPAGLVQAT | ASSLGQSETLY | 0.61644447 |
| YALKRALAKFSDRFDVVII | ASSLGQSETLY | 0.79809994 |
| YLIGSVIV | ASSLGQSETLY | 0.41320536 |
| YGQLVAVID | ASSLGQSETLY | 0.2477653 |
| VTVGGMTLPA | ASSLGQSETLY | 0.36031032 |
| VLALPAV | ASSLGQSETLY | 0.48088884 |
| VIPPTLNYET | ASSLGQSETLY | 0.43421865 |
| VGMGGMHPSSG | ASSLGQSETLY | 0.25657606 |
| VGLAGGAATA | ASSLGQSETLY | 0.32032654 |
| VGGAGGAGGSGP | ASSLGQSETLY | 0.35862395 |
| VFAAFMV | ASSLGQSETLY | 0.1410966 |
| TLYDKLR | ASSLGQSETLY | 0.44547623 |
| TLAVFTVL | ASSLGQSETLY | 0.5231088 |
| TGSVQPARV | ASSLGQSETLY | 0.45344454 |
| TGGGAGSGGT | ASSLGQSETLY | 0.21383609 |
| SHLETSL | ASSLGQSETLY | 0.56522375 |
| SFTTGMT | ASSLGQSETLY | 0.60955966 |
| QVEVAKRMGVRQPTVS | ASSLGQSETLY | 0.66247636 |
| QGSGGPALAM | ASSLGQSETLY | 0.1980513 |
| MLAPSNP | ASSLGQSETLY | 0.12344625 |
| LVNNAAI | ASSLGQSETLY | 0.35749584 |
| LTGEINTLA | ASSLGQSETLY | 0.55836 |
| LSIFGPV | ASSLGQSETLY | 0.33739853 |
| LLALLANL | ASSLGQSETLY | 0.21927999 |
| LISNASDALDKLR | ASSLGQSETLY | 0.3114657 |
| LINEAVP | ASSLGQSETLY | 0.52919376 |
| LGYAFSGL | ASSLGQSETLY | 0.23510028 |
| LDFVAVAL | ASSLGQSETLY | 0.21182622 |

| | | |
|---|---|---|
| LAQPGAE | ASSLGQSETLY | 0.3461837 |
| ISNLGAVTGF | ASSLGQSETLY | 0.5825132 |
| HQEDPGRI | ASSLGQSETLY | 0.16116133 |
| GTPGEPTP | ASSLGQSETLY | 0.19617188 |
| GFFNSGS | ASSLGQSETLY | 0.40525052 |
| GAGGGGLTAGAN | ASSLGQSETLY | 0.5986903 |
| FVDALVNM | ASSLGQSETLY | 0.35152477 |
| FLTVAPEL | ASSLGQSETLY | 0.22273041 |
| FETLTAL | ASSLGQSETLY | 0.35063988 |
| DVINPIEL | ASSLGQSETLY | 0.17141713 |
| APVSGAPV | ASSLGQSETLY | 0.4908918 |
| ALVGGLAW | ASSLGQSETLY | 0.27479506 |
| ALAAAAATAI | ASSLGQSETLY | 0.39923325 |
| AFGLGLLL | ASSLGQSETLY | 0.2677135 |
| AFFAAASL | ASSLGQSETLY | 0.17576335 |
| AAAALPAFSPP | ASSLGQSETLY | 0.22115144 |
| VVLAPAL | ASSLGQSETLY | 0.22406967 |
| TIGHVDHGKTTL | ASSLGQSETLY | 0.4725474 |
| LISNASDALDK | ASSLGQSETLY | 0.46812117 |
| LISNASDAL | ASSLGQSETLY | 0.3992504 |
| ISNASDALDK | ASSLGQSETLY | 0.35091117 |
| NQAETLVYQTE | ASSLGQSETLY | 0.22254664 |
| TGGTGGTAGAG | ASSLGQSETLY | 0.42997977 |
| TGSTAGGAGGA | ASSLGQSETLY | 0.16749288 |
| DPLAVPVAAGPV | ASSLGQSETLY | 0.5492979 |
| GGTGGTPGNGGN | ASSLGQSETLY | 0.35035658 |
| PAGLAGVDPATL | ASSLGQSETLY | 0.48113197 |
| GNNLVVEAPHAGASV | ASSLGQSETLY | 0.24719478 |
| KVTRIMSASTLEQPA | ASSLGQSETLY | 0.49870095 |
| LGFALPIALPRRTVV | ASSLGQSETLY | 0.44411984 |
| TRLGNLTARKLVVLV | ASSLGQSETLY | 0.26231146 |
| WSQQRRLIYFPSAGP | ASSLGQSETLY | 0.44164628 |
| GPNGNPDPMAAAVDIR | ASSLGQSETLY | 0.5505885 |
| TQRCVSVADLRAAASR | ASSLGQSETLY | 0.4680547 |
| QFVLDVAEHPVEVLTETV | ASSLGQSETLY | 0.15876219 |
| GGAGGTGGAGGNGGTQAGDG | ASSLGQSETLY | 0.39896825 |
| PLLATLRAIAVDVDAKPAQV | ASSLGQSETLY | 0.20022643 |
| APKFRTGGYMIDFWGVGYQVAK | ASSLGQSETLY | 0.36721504 |
| GAGGAGGAGGAAGGPGGTGGTG | ASSLGQSETLY | 0.30675477 |
| IGALGLLATDVAPAGSVAFGVA | ASSLGQSETLY | 0.37238553 |
| AGGAGGIGGNGGAGGAGGVLMG | ASSLGQSETLY | 0.47973904 |
| ADELIGLL | ASSLGQSETLY | 0.2271235 |
| ADRALKA | ASSLGQSETLY | 0.21777152 |
| AGAVGQVGGAGGA | ASSLGQSETLY | 0.30942184 |
| AGGGGGLGGAP | ASSLGQSETLY | 0.4169895 |
| AGGLLYGNGGAG | ASSLGQSETLY | 0.4507188 |
| AGLESIGL | ASSLGQSETLY | 0.20142813 |
| AGLKTGII | ASSLGQSETLY | 0.39354402 |
| ASLAALLL | ASSLGQSETLY | 0.33451325 |
| ASVINISSI | ASSLGQSETLY | 0.2343056 |
| ATPPGPA | ASSLGQSETLY | 0.46319285 |

| | | |
|---|---|---|
| ATVASALV | ASSLGQSETLY | 0.29305544 |
| ATVASLIL | ASSLGQSETLY | 0.1856912 |
| AVNGPVLIRH | ASSLGQSETLY | 0.46250385 |
| DGAAGAVGI | ASSLGQSETLY | 0.300785 |
| DPLLHGI | ASSLGQSETLY | 0.4837338 |
| ELGAALGAL | ASSLGQSETLY | 0.55750936 |
| ELGRTQAV | ASSLGQSETLY | 0.17483632 |
| FVAGGYLS | ASSLGQSETLY | 0.4685309 |
| GASGGAGGAGGW | ASSLGQSETLY | 0.29647645 |
| GGAGAGGDGGAGGI | ASSLGQSETLY | 0.28472883 |
| GTGHNGSQPGMGGNG | ASSLGQSETLY | 0.33667442 |
| GYAAASASAS | ASSLGQSETLY | 0.2535153 |
| HQVDVAAVQQG | ASSLGQSETLY | 0.5229727 |
| IGAAIAHT | ASSLGQSETLY | 0.46693248 |
| IGMIEVAQL | ASSLGQSETLY | 0.39151388 |
| IRLAPEHTSAMD | ASSLGQSETLY | 0.3307035 |
| IYNVSVL | ASSLGQSETLY | 0.45403352 |
| KQGFLGM | ASSLGQSETLY | 0.49497047 |
| LESVITQL | ASSLGQSETLY | 0.19233093 |
| LGLGALVVV | ASSLGQSETLY | 0.16017087 |
| LIGNGGNGGSGGI | ASSLGQSETLY | 0.3458714 |
| LNAINAPTL | ASSLGQSETLY | 0.34370115 |
| LQLLNTGV | ASSLGQSETLY | 0.24019292 |
| LQTSIEF | ASSLGQSETLY | 0.25188056 |
| MSFAVSEL | ASSLGQSETLY | 0.4633162 |
| NTGALGL | ASSLGQSETLY | 0.65681285 |
| PPGAGGAGGAGTT | ASSLGQSETLY | 0.25114796 |
| QLVGLGI | ASSLGQSETLY | 0.7172477 |
| RFLEQVHQL | ASSLGQSETLY | 0.1939809 |
| RGVGYVLR | ASSLGQSETLY | 0.4878807 |
| SATGVFAGVI | ASSLGQSETLY | 0.5479366 |
| SLPAITLP | ASSLGQSETLY | 0.17051437 |
| STALAFTV | ASSLGQSETLY | 0.39460942 |
| SVLTVGLGK | ASSLGQSETLY | 0.25089613 |
| TALAGAGSGI | ASSLGQSETLY | 0.24184941 |
| TGLPFTGI | ASSLGQSETLY | 0.47275987 |
| VAEHVTV | ASSLGQSETLY | 0.3498829 |
| VDIVLNSL | ASSLGQSETLY | 0.4552156 |
| VLTGGGAVL | ASSLGQSETLY | 0.30544588 |
| VNVGLAYGI | ASSLGQSETLY | 0.22620179 |
| VSSAAVGAV | ASSLGQSETLY | 0.3837838 |
| VTSVYEAGV | ASSLGQSETLY | 0.44506162 |
| WVALDRGA | ASSLGQSETLY | 0.49724233 |
| YLVGSVVI | ASSLGQSETLY | 0.79234314 |
| GPPGTGKTLL | ASSLGQSETLY | 0.20270823 |
| GVGMTKF | ASSLGQSETLY | 0.5088269 |
| IGIGGSDL | ASSLGQSETLY | 0.6286298 |
| IGTLLAEL | ASSLGQSETLY | 0.29822505 |
| IYTSGTT | ASSLGQSETLY | 0.30292177 |
| LSFPAGV | ASSLGQSETLY | 0.5328001 |
| LYTSGTT | ASSLGQSETLY | 0.28183624 |

| | | |
|---|---|---|
| VGAFPFGF | ASSLGQSETLY | 0.387978 |
| VGAGTVEF | ASSLGQSETLY | 0.59486747 |
| VGKSTLLL | ASSLGQSETLY | 0.1788244 |
| AALAHAGGGIV | ASSLGQSETLY | 0.73267174 |
| AALVAVSLL | ASSLGQSETLY | 0.41125092 |
| AAVSRAVAAL | ASSLGQSETLY | 0.4099794 |
| AGGSSGIGGF | ASSLGQSETLY | 0.22864766 |
| AGLTVANANG | ASSLGQSETLY | 0.14801364 |
| AGLVGAVL | ASSLGQSETLY | 0.29245284 |
| ALSQAFTV | ASSLGQSETLY | 0.19787905 |
| ALTFSVSL | ASSLGQSETLY | 0.75994486 |
| ALTGAALVAL | ASSLGQSETLY | 0.5085344 |
| ASFSIGAGVL | ASSLGQSETLY | 0.241963 |
| DGSVYKI | ASSLGQSETLY | 0.70082575 |
| EQLAEIV | ASSLGQSETLY | 0.71153927 |
| FAVSELTVM | ASSLGQSETLY | 0.14460477 |
| GAAPLAAGL | ASSLGQSETLY | 0.2585282 |
| GADGGDGGSGGAGG | ASSLGQSETLY | 0.2293532 |
| GFGLDTGGAGG | ASSLGQSETLY | 0.267776 |
| GLASGGLL | ASSLGQSETLY | 0.10495825 |
| GRIAGAAVLL | ASSLGQSETLY | 0.4845263 |
| GSAGSLNTSF | ASSLGQSETLY | 0.24493288 |
| GTGQIGI | ASSLGQSETLY | 0.089350715 |
| GTLIALIL | ASSLGQSETLY | 0.31613395 |
| IAATMGFVVG | ASSLGQSETLY | 0.28832835 |
| IFAQALL | ASSLGQSETLY | 0.30292016 |
| IGAAVALGF | ASSLGQSETLY | 0.69606245 |
| IGLTPLAL | ASSLGQSETLY | 0.18998215 |
| ILGSNTSTL | ASSLGQSETLY | 0.19436714 |
| IVFSLTVL | ASSLGQSETLY | 0.5157443 |
| LASKATGF | ASSLGQSETLY | 0.75787306 |
| LASVALIL | ASSLGQSETLY | 0.1934837 |
| LFAVGLTL | ASSLGQSETLY | 0.47337022 |
| LFNGGIGTYI | ASSLGQSETLY | 0.80381274 |
| LTLGAGII | ASSLGQSETLY | 0.45998216 |
| LVAAALGDI | ASSLGQSETLY | 0.6419022 |
| NTAFVQLGI | ASSLGQSETLY | 0.6979683 |
| QLFSGAEL | ASSLGQSETLY | 0.18196028 |
| SAATFITL | ASSLGQSETLY | 0.6631598 |
| SNYGLANLGV | ASSLGQSETLY | 0.7869003 |
| SYGGAYAVM | ASSLGQSETLY | 0.26526332 |
| TTGVGGLTLG | ASSLGQSETLY | 0.32282436 |
| TTQTIYI | ASSLGQSETLY | 0.20505103 |
| VAAGAGLGAVI | ASSLGQSETLY | 0.28721106 |
| VAAVVTLL | ASSLGQSETLY | 0.11734409 |
| VGFLGGLI | ASSLGQSETLY | 0.56390667 |
| VGVIPILL | ASSLGQSETLY | 0.6435133 |
| VIFASSLI | ASSLGQSETLY | 0.32131198 |
| VKGGDGGAAAG | ASSLGQSETLY | 0.14142218 |
| VTKTFTNI | ASSLGQSETLY | 0.13108343 |
| VVAAAILL | ASSLGQSETLY | 0.31557208 |

| | | |
|---|---|---|
| YAALDVEL | ASSLGQSETLY | 0.23465234 |
| LASVALLL | ASSLGQSETLY | 0.12541884 |
| LDAVHRV | ASSLGQSETLY | 0.21761173 |
| QGLSSSIF | ASSLGQSETLY | 0.64579964 |
| RGTFANIRL | ASSLGQSETLY | 0.6705178 |
| RQLWWGHRI | ASSLGQSETLY | 0.15725785 |
| YGPPGTGKTLL | ASSLGQSETLY | 0.79601824 |
| AALVGAPANVA | ASSLGQSETLY | 0.6479123 |
| AATFRRTE | ASSLGQSETLY | 0.53956324 |
| AGALGGGI | ASSLGQSETLY | 0.30166778 |
| AGGAGGAGGIGAG | ASSLGQSETLY | 0.18775915 |
| AGSGALGL | ASSLGQSETLY | 0.8000697 |
| AQIADQI | ASSLGQSETLY | 0.3723194 |
| ASAVAVTYL | ASSLGQSETLY | 0.27563664 |
| ASTVLGGTGG | ASSLGQSETLY | 0.5098115 |
| AVGGEPAK | ASSLGQSETLY | 0.36154488 |
| CEQGTFGGTC | ASSLGQSETLY | 0.23516756 |
| DEGKAPAG | ASSLGQSETLY | 0.5956095 |
| DTVVFPAL | ASSLGQSETLY | 0.4273023 |
| FEGITMAG | ASSLGQSETLY | 0.15588933 |
| FPLVVQD | ASSLGQSETLY | 0.1929335 |
| GAFTTPQ | ASSLGQSETLY | 0.19402853 |
| GFGGFGTAGGVGGA | ASSLGQSETLY | 0.41866153 |
| GFLAQFL | ASSLGQSETLY | 0.7080523 |
| GGAGGAGGNAGVA | ASSLGQSETLY | 0.35482997 |
| GGVGTAGAGGAGGAGG | ASSLGQSETLY | 0.26368204 |
| GNGGGGDAGNAGSGGN | ASSLGQSETLY | 0.63007617 |
| GTGGAGGAGNDAGS | ASSLGQSETLY | 0.51409024 |
| GTTNTGWFNS | ASSLGQSETLY | 0.33590612 |
| IFMGPLFTL | ASSLGQSETLY | 0.57615083 |
| ILAHAIEAI | ASSLGQSETLY | 0.29003596 |
| IRSAIREAL | ASSLGQSETLY | 0.42298442 |
| ITFAGGG | ASSLGQSETLY | 0.46951687 |
| ITGMSGHQEV | ASSLGQSETLY | 0.49218807 |
| IVAGLACTL | ASSLGQSETLY | 0.5814455 |
| IVGGGLAGA | ASSLGQSETLY | 0.45594466 |
| IVTGVLNI | ASSLGQSETLY | 0.29038638 |
| LAQQGCI | ASSLGQSETLY | 0.12303993 |
| LCLHVEN | ASSLGQSETLY | 0.26022345 |
| LDAISDGS | ASSLGQSETLY | 0.4639828 |
| LGGFAAML | ASSLGQSETLY | 0.4351029 |
| LGGKNPMI | ASSLGQSETLY | 0.55074227 |
| LGPKGVAAIL | ASSLGQSETLY | 0.44235787 |
| LMVGVSI | ASSLGQSETLY | 0.20334552 |
| LVGAAFVL | ASSLGQSETLY | 0.6283455 |
| LVLAVAPL | ASSLGQSETLY | 0.7162938 |
| MAAIAVIDAA | ASSLGQSETLY | 0.14949594 |
| NNGINAELQ | ASSLGQSETLY | 0.30901593 |
| RGEPLVLVHG | ASSLGQSETLY | 0.79751366 |
| RGGAGGAAGLI | ASSLGQSETLY | 0.6933915 |
| RSYRMHPAV | ASSLGQSETLY | 0.28371698 |

| | | |
|---|---|---|
| SAAHLLAGQLA | ASSLGQSETLY | 0.33377093 |
| SAAVGSY | ASSLGQSETLY | 0.8195894 |
| SPHAVLHRI | ASSLGQSETLY | 0.2591626 |
| SQIAKVL | ASSLGQSETLY | 0.726562 |
| TALGHLVP | ASSLGQSETLY | 0.10856593 |
| TAPTTSGAA | ASSLGQSETLY | 0.2413931 |
| TDGDDGNGGSPGT | ASSLGQSETLY | 0.19375718 |
| TDGLGEN | ASSLGQSETLY | 0.1080762 |
| TGGPPLPGQA | ASSLGQSETLY | 0.6081747 |
| TIPAITI | ASSLGQSETLY | 0.22230527 |
| TIPGFTI | ASSLGQSETLY | 0.6363299 |
| TSIASGALA | ASSLGQSETLY | 0.29892156 |
| TVTLPGAAI | ASSLGQSETLY | 0.59888107 |
| VASGLSAALP | ASSLGQSETLY | 0.38542566 |
| VGAGVIGI | ASSLGQSETLY | 0.32510853 |
| VGPKRFGV | ASSLGQSETLY | 0.19187109 |
| VNVAFPD | ASSLGQSETLY | 0.53156054 |
| VSMTNSGVS | ASSLGQSETLY | 0.6249546 |
| VTDVLGDLSV | ASSLGQSETLY | 0.37047344 |
| VVVVGAGFAGL | ASSLGQSETLY | 0.14426866 |
| VYTVGTVV | ASSLGQSETLY | 0.29476446 |
| WTTFSGP | ASSLGQSETLY | 0.34351647 |
| YDELVEAF | ASSLGQSETLY | 0.11811826 |
| YGTGGAGGNGG | ASSLGQSETLY | 0.1003779 |
| YTNVMAV | ASSLGQSETLY | 0.17261152 |
| FLEQVHQL | ASSLGQSETLY | 0.32437783 |
| GLSLLAVAV | ASSLGQSETLY | 0.60119927 |
| LEQVHQL | ASSLGQSETLY | 0.6234728 |
| LNAAGAI | ASSLGQSETLY | 0.26487988 |
| PAVHASL | ASSLGQSETLY | 0.6068306 |
| TELEDDA | ASSLGQSETLY | 0.5196303 |
| TPPGPPN | ASSLGQSETLY | 0.08223943 |
| TPPPPPPP | ASSLGQSETLY | 0.25310296 |
| TQSPALI | ASSQDRGASNSDYT | 0.14598006 |
| EFAAAFPQ | ASSQDRGASNSDYT | 0.55555284 |
| LLSGNLDVL | ASSQDRGASNSDYT | 0.18856248 |
| GPANLGNYNI | ASSQDRGASNSDYT | 0.2631692 |
| TLGPTLP | ASSQDRGASNSDYT | 0.3450758 |
| VYAYSSCCK | ASSQDRGASNSDYT | 0.7179683 |
| APAPAPAPNI | ASSQDRGASNSDYT | 0.44833457 |
| PGGPVPIP | ASSQDRGASNSDYT | 0.16770795 |
| FLTVALEP | ASSQDRGASNSDYT | 0.101477414 |
| SQYFGANL | ASSQDRGASNSDYT | 0.53666145 |
| GGVKPAA | ASSQDRGASNSDYT | 0.1944634 |
| GGTGGAGGAGGW | ASSQDRGASNSDYT | 0.19377601 |
| GVGGVAGLGGAGG | ASSQDRGASNSDYT | 0.12309734 |
| VLLVLATTI | ASSQDRGASNSDYT | 0.4172421 |
| IGIFAAML | ASSQDRGASNSDYT | 0.14766693 |
| AVGDLID | ASSQDRGASNSDYT | 0.55984116 |
| NISPLPL | ASSQDRGASNSDYT | 0.22512728 |
| VTAVFVAI | ASSQDRGASNSDYT | 0.32116288 |

| | | |
|---|---|---|
| SGDDNDAAL | ASSQDRGASNSDYT | 0.7838526 |
| AVAALAHS | ASSQDRGASNSDYT | 0.738313 |
| GGVTVMATP | ASSQDRGASNSDYT | 0.29360732 |
| AGLAANVRD | ASSQDRGASNSDYT | 0.23790297 |
| TNRPDLIDPALLRPG | ASSQDRGASNSDYT | 0.17063469 |
| LANLGVTAL | ASSQDRGASNSDYT | 0.19862671 |
| TGERIEL | ASSQDRGASNSDYT | 0.20760077 |
| NGEELAA | ASSQDRGASNSDYT | 0.7365185 |
| VLSLVWG | ASSQDRGASNSDYT | 0.25684264 |
| IGYALWL | ASSQDRGASNSDYT | 0.16353184 |
| LSLLLAPL | ASSQDRGASNSDYT | 0.3674627 |
| TLDEALK | ASSQDRGASNSDYT | 0.7335317 |
| TDGAGTAEGG | ASSQDRGASNSDYT | 0.8109381 |
| NAPGGAGGNGG | ASSQDRGASNSDYT | 0.17060229 |
| VSVFMGMGGNF | ASSQDRGASNSDYT | 0.26771414 |
| LASYLAAL | ASSQDRGASNSDYT | 0.26850334 |
| LVGGTPAL | ASSQDRGASNSDYT | 0.4033469 |
| VRNADVL | ASSQDRGASNSDYT | 0.24737653 |
| TCSSPFLVSAM | ASSQDRGASNSDYT | 0.19155994 |
| VHITDAE | ASSQDRGASNSDYT | 0.41457096 |
| EYAAPSF | ASSQDRGASNSDYT | 0.85332847 |
| VVDAFHGGL | ASSQDRGASNSDYT | 0.2592906 |
| ASVVDPQ | ASSQDRGASNSDYT | 0.13473031 |
| SYNEFGGLS | ASSQDRGASNSDYT | 0.30364245 |
| DPDGSVY | ASSQDRGASNSDYT | 0.17809969 |
| MVSTIHATE | ASSQDRGASNSDYT | 0.22440904 |
| LAAGVADPV | ASSQDRGASNSDYT | 0.42137665 |
| TKEQIAA | ASSQDRGASNSDYT | 0.13947147 |
| SLASPAAL | ASSQDRGASNSDYT | 0.29528874 |
| IGLVFYGV | ASSQDRGASNSDYT | 0.8035907 |
| FLLDASNL | ASSQDRGASNSDYT | 0.36291012 |
| YQVGNFP | ASSQDRGASNSDYT | 0.5274117 |
| ESLLSQST | ASSQDRGASNSDYT | 0.7203798 |
| LFTVGFAAVG | ASSQDRGASNSDYT | 0.5031878 |
| ATPPTPT | ASSQDRGASNSDYT | 0.78075486 |
| TDLATLG | ASSQDRGASNSDYT | 0.7304721 |
| SAGGTGR | ASSQDRGASNSDYT | 0.2880552 |
| GFSNSGS | ASSQDRGASNSDYT | 0.32398713 |
| GGNGGSGGTSV | ASSQDRGASNSDYT | 0.68365645 |
| GTSVATGGAGN | ASSQDRGASNSDYT | 0.33628064 |
| NGGNGGSGGTS | ASSQDRGASNSDYT | 0.33284712 |
| AVVKFPH | ASSQDRGASNSDYT | 0.24220227 |
| TPPPPPP | ASSQDRGASNSDYT | 0.69070953 |
| AAPAAAAAPA | ASSQDRGASNSDYT | 0.3650036 |
| IALNEDL | ASSQDRGASNSDYT | 0.687538 |
| IVNVSSM | ASSQDRGASNSDYT | 0.13829857 |
| TGIPPAPR | ASSQDRGASNSDYT | 0.18593465 |
| FDLGGGTF | ASSQDRGASNSDYT | 0.7146242 |
| FDLGGGTFDVS | ASSQDRGASNSDYT | 0.09452691 |
| NEPTAAA | ASSQDRGASNSDYT | 0.1140306 |
| NEPTAAALA | ASSQDRGASNSDYT | 0.26792735 |

| | | |
|---|---|---|
| GSIINISSI | ASSQDRGASNSDYT | 0.77277005 |
| VAPGNAGTA | ASSQDRGASNSDYT | 0.090371095 |
| TGGGASLE | ASSQDRGASNSDYT | 0.21705167 |
| VAVYNHYKRI | ASSQDRGASNSDYT | 0.14398165 |
| TDPAPATT | ASSQDRGASNSDYT | 0.44948792 |
| TNRPDILDPALLRPG | ASSQDRGASNSDYT | 0.4172601 |
| LTTTTTS | ASSQDRGASNSDYT | 0.13443081 |
| PTGSGKTL | ASSQDRGASNSDYT | 0.53206027 |
| TGSGKTL | ASSQDRGASNSDYT | 0.1255044 |
| TIGHVDHGK | ASSQDRGASNSDYT | 0.24873807 |
| TIGHVDHGKTT | ASSQDRGASNSDYT | 0.5552345 |
| GHVDHGKTT | ASSQDRGASNSDYT | 0.8057921 |
| AQFEHTLL | ASSQDRGASNSDYT | 0.16511285 |
| IVNQDEL | ASSQDRGASNSDYT | 0.62259257 |
| TNGSQFF | ASSQDRGASNSDYT | 0.21596731 |
| TGKGIVE | ASSQDRGASNSDYT | 0.8338249 |
| LISNASDA | ASSQDRGASNSDYT | 0.25227386 |
| AELRTLGELTKIAKAHGAQ | ASSQDRGASNSDYT | 0.13213664 |
| APNAVAAAEPSW | ASSQDRGASNSDYT | 0.6172902 |
| APQVAGAATAVL | ASSQDRGASNSDYT | 0.5035732 |
| ARRDPAAAAAILPTDA | ASSQDRGASNSDYT | 0.3481419 |
| DAAARDVLAEAGLAEYF | ASSQDRGASNSDYT | 0.67946213 |
| DDQIVLSAKANLAKVEQ | ASSQDRGASNSDYT | 0.26207542 |
| DELLPDTGEDSDEDGH | ASSQDRGASNSDYT | 0.26831 |
| ELPEAPANRATVPQD | ASSQDRGASNSDYT | 0.69519776 |
| FGQSKIERVTGIG | ASSQDRGASNSDYT | 0.53514636 |
| GAGGTGSTAGGAGGAGGAGGLY | ASSQDRGASNSDYT | 0.20413299 |
| GHGGTGAAGGNGGTGGAGGSGI | ASSQDRGASNSDYT | 0.5956439 |
| GNGGNGGAGGEGGPGGAGGAGA | ASSQDRGASNSDYT | 0.45261356 |
| GSTAGNGGNGGNATNGTIAGQP | ASSQDRGASNSDYT | 0.21151173 |
| GTGGDGGNGGNGGTGG | ASSQDRGASNSDYT | 0.79879785 |
| GVRRPAGIPADAPV | ASSQDRGASNSDYT | 0.34203985 |
| GYRQVTDAHLLTLARR | ASSQDRGASNSDYT | 0.61776423 |
| IAKMGQDGHDRG | ASSQDRGASNSDYT | 0.45113248 |
| KAEPSIEIQYT | ASSQDRGASNSDYT | 0.82382286 |
| KPVRIRMAADEDTVAVT | ASSQDRGASNSDYT | 0.16181108 |
| MPPRIAGMRLLVIKPEP | ASSQDRGASNSDYT | 0.5621773 |
| MKFYAVALFRSRGVPPTR | ASSQDRGASNSDYT | 0.37409204 |
| NAGGFGILGAVAHSP | ASSQDRGASNSDYT | 0.56781256 |
| NEGREITLAMAEM | ASSQDRGASNSDYT | 0.2970431 |
| NELVEWSHPLGHRWRWE | ASSQDRGASNSDYT | 0.28827578 |
| NLGAGGTGGNGGNPG | ASSQDRGASNSDYT | 0.38447693 |
| NTEWPKWETVMAADPVD | ASSQDRGASNSDYT | 0.7078278 |
| PASAAPAETQP | ASSQDRGASNSDYT | 0.2680678 |
| QQSSDGKNGVRAAEITGE | ASSQDRGASNSDYT | 0.48884976 |
| RAPDAVTTSWWKE | ASSQDRGASNSDYT | 0.81958604 |
| SGFGNVGAHTSGWFNQ | ASSQDRGASNSDYT | 0.18196037 |
| SGVAGAVGQAASVGGLKV | ASSQDRGASNSDYT | 0.35900438 |
| SNGGAGTGSAGGNGGT | ASSQDRGASNSDYT | 0.44215402 |
| TGGAGGEGGAAG | ASSQDRGASNSDYT | 0.66099155 |
| TGGDGGAAGAGGAPGHG | ASSQDRGASNSDYT | 0.29954445 |

| | | |
|---|---|---|
| TMPPLLAAIANAGRHGV | ASSQDRGASNSDYT | 0.6761982 |
| TSSTAGNSGGA | ASSQDRGASNSDYT | 0.2504232 |
| VPATVPVGILC | ASSQDRGASNSDYT | 0.5431775 |
| VQRPDSGGVVLTGRIS | ASSQDRGASNSDYT | 0.2269291 |
| WTPNPVPATKGVPAGLVQAT | ASSQDRGASNSDYT | 0.43735605 |
| YALKRALAKFSDRFDVVII | ASSQDRGASNSDYT | 0.1927913 |
| YLIGSVIV | ASSQDRGASNSDYT | 0.53363305 |
| YGQLVAVID | ASSQDRGASNSDYT | 0.6686182 |
| VTVGGMTLPA | ASSQDRGASNSDYT | 0.63818455 |
| VLALPAV | ASSQDRGASNSDYT | 0.10283125 |
| VIPPTLNYET | ASSQDRGASNSDYT | 0.6461483 |
| VGMGGMHPSSG | ASSQDRGASNSDYT | 0.602654 |
| VGLAGGAATA | ASSQDRGASNSDYT | 0.8546255 |
| VGGAGGAGGSGP | ASSQDRGASNSDYT | 0.34474868 |
| VFAAFMV | ASSQDRGASNSDYT | 0.52215815 |
| TLYDKLR | ASSQDRGASNSDYT | 0.27968043 |
| TLAVFTVL | ASSQDRGASNSDYT | 0.317576 |
| TGSVQPARV | ASSQDRGASNSDYT | 0.15364674 |
| TGGGAGSGGT | ASSQDRGASNSDYT | 0.3330394 |
| SHLETSL | ASSQDRGASNSDYT | 0.3762707 |
| SFTTGMT | ASSQDRGASNSDYT | 0.20633483 |
| QVEVAKRMGVRQPTVS | ASSQDRGASNSDYT | 0.7814528 |
| QGSGGPALAM | ASSQDRGASNSDYT | 0.7016139 |
| MLAPSNP | ASSQDRGASNSDYT | 0.6805218 |
| LVNNAAI | ASSQDRGASNSDYT | 0.7113075 |
| LTGEINTLA | ASSQDRGASNSDYT | 0.77569956 |
| LSIFGPV | ASSQDRGASNSDYT | 0.5163857 |
| LLALLANL | ASSQDRGASNSDYT | 0.2581573 |
| LISNASDALDKLR | ASSQDRGASNSDYT | 0.30850035 |
| LINEAVP | ASSQDRGASNSDYT | 0.15219438 |
| LGYAFSGL | ASSQDRGASNSDYT | 0.26398858 |
| LDFVAVAL | ASSQDRGASNSDYT | 0.262905 |
| LAQPGAE | ASSQDRGASNSDYT | 0.4259836 |
| ISNLGAVTGF | ASSQDRGASNSDYT | 0.09231438 |
| HQEDPGRI | ASSQDRGASNSDYT | 0.6640066 |
| GTPGEPTP | ASSQDRGASNSDYT | 0.75906783 |
| GFFNSGS | ASSQDRGASNSDYT | 0.3260107 |
| GAGGGGLTAGAN | ASSQDRGASNSDYT | 0.5206474 |
| FVDALVNM | ASSQDRGASNSDYT | 0.5563456 |
| FLTVAPEL | ASSQDRGASNSDYT | 0.20519006 |
| FETLTAL | ASSQDRGASNSDYT | 0.63800764 |
| DVINPIEL | ASSQDRGASNSDYT | 0.19375741 |
| APVSGAPV | ASSQDRGASNSDYT | 0.59381294 |
| ALVGGLAW | ASSQDRGASNSDYT | 0.757375 |
| ALAAAAATAI | ASSQDRGASNSDYT | 0.35236114 |
| AFGLGLLL | ASSQDRGASNSDYT | 0.3555062 |
| AFFAAASL | ASSQDRGASNSDYT | 0.3864771 |
| AAAALPAFSPP | ASSQDRGASNSDYT | 0.14988878 |
| VVLAPAL | ASSQDRGASNSDYT | 0.37285334 |
| TIGHVDHGKTTL | ASSQDRGASNSDYT | 0.1257959 |
| LISNASDALDK | ASSQDRGASNSDYT | 0.37232742 |

| | | |
|---|---|---|
| LISNASDAL | ASSQDRGASNSDYT | 0.16527112 |
| ISNASDALDK | ASSQDRGASNSDYT | 0.42401296 |
| NQAETLVYQTE | ASSQDRGASNSDYT | 0.43576393 |
| TGGTGGTAGAG | ASSQDRGASNSDYT | 0.3358624 |
| TGSTAGGAGGA | ASSQDRGASNSDYT | 0.17553133 |
| DPLAVPVAAGPV | ASSQDRGASNSDYT | 0.39031082 |
| GGTGGTPGNGGN | ASSQDRGASNSDYT | 0.24378508 |
| PAGLAGVDPATL | ASSQDRGASNSDYT | 0.15268077 |
| GNNLVVEAPHAGASV | ASSQDRGASNSDYT | 0.6048757 |
| KVTRIMSASTLEQPA | ASSQDRGASNSDYT | 0.23445962 |
| LGFALPIALPRRTVV | ASSQDRGASNSDYT | 0.377024 |
| TRLGNLTARKLVVLV | ASSQDRGASNSDYT | 0.640299 |
| WSQQRRLIYFPSAGP | ASSQDRGASNSDYT | 0.4137954 |
| GPNGNPDPMAAAVDIR | ASSQDRGASNSDYT | 0.13769248 |
| TQRCVSVADLRAAASR | ASSQDRGASNSDYT | 0.3808427 |
| QFVLDVAEHPVEVLTETV | ASSQDRGASNSDYT | 0.30146253 |
| GGAGGTGGAGGNGGTQAGDG | ASSQDRGASNSDYT | 0.26487315 |
| PLLATLRAIAVDVDAKPAQV | ASSQDRGASNSDYT | 0.09131033 |
| APKFRTGGYMIDFWGVGYQVAK | ASSQDRGASNSDYT | 0.41175342 |
| GAGGAGGAGGAAGGPGGTGGTG | ASSQDRGASNSDYT | 0.21477005 |
| IGALGLLATDVAPAGSVAFGVA | ASSQDRGASNSDYT | 0.19598165 |
| AGGAGGIGGNGGAGGAGGVLMG | ASSQDRGASNSDYT | 0.27928823 |
| ADELIGLL | ASSQDRGASNSDYT | 0.6830243 |
| ADRALKA | ASSQDRGASNSDYT | 0.38424304 |
| AGAVGQVGGAGGA | ASSQDRGASNSDYT | 0.4231978 |
| AGGGGGLGGAP | ASSQDRGASNSDYT | 0.6195158 |
| AGGLLYGNGGAG | ASSQDRGASNSDYT | 0.49585193 |
| AGLESIGL | ASSQDRGASNSDYT | 0.7766005 |
| AGLKTGII | ASSQDRGASNSDYT | 0.47386557 |
| ASLAALLL | ASSQDRGASNSDYT | 0.27753597 |
| ASVINISSI | ASSQDRGASNSDYT | 0.37765253 |
| ATPPGPA | ASSQDRGASNSDYT | 0.33066624 |
| ATVASALV | ASSQDRGASNSDYT | 0.11476113 |
| ATVASLIL | ASSQDRGASNSDYT | 0.11553218 |
| AVNGPVLIRH | ASSQDRGASNSDYT | 0.22834036 |
| DGAAGAVGI | ASSQDRGASNSDYT | 0.11259364 |
| DPLLHGI | ASSQDRGASNSDYT | 0.40041786 |
| ELGAALGAL | ASSQDRGASNSDYT | 0.5565983 |
| ELGRTQAV | ASSQDRGASNSDYT | 0.3031717 |
| FVAGGYLS | ASSQDRGASNSDYT | 0.14989746 |
| GASGGAGGAGGW | ASSQDRGASNSDYT | 0.22224376 |
| GGAGAGGDGGAGGI | ASSQDRGASNSDYT | 0.23530984 |
| GTGHNGSQPGMGGNG | ASSQDRGASNSDYT | 0.6658548 |
| GYAAASASAS | ASSQDRGASNSDYT | 0.32094288 |
| HQVDVAAVQQG | ASSQDRGASNSDYT | 0.83642715 |
| IGAAIAHT | ASSQDRGASNSDYT | 0.7031636 |
| IGMIEVAQL | ASSQDRGASNSDYT | 0.29459506 |
| IRLAPEHTSAMD | ASSQDRGASNSDYT | 0.60791457 |
| IYNVSVL | ASSQDRGASNSDYT | 0.24164715 |
| KQGFLGM | ASSQDRGASNSDYT | 0.35725597 |
| LESVITQL | ASSQDRGASNSDYT | 0.2659523 |

| | | |
|---|---|---|
| LGLGALVVV | ASSQDRGASNSDYT | 0.3351487 |
| LIGNGGNGGSGGI | ASSQDRGASNSDYT | 0.27969405 |
| LNAINAPTL | ASSQDRGASNSDYT | 0.22469673 |
| LQLLNTGV | ASSQDRGASNSDYT | 0.13903163 |
| LQTSIEF | ASSQDRGASNSDYT | 0.31845555 |
| MSFAVSEL | ASSQDRGASNSDYT | 0.21658672 |
| NTGALGL | ASSQDRGASNSDYT | 0.12781963 |
| PPGAGGAGGAGTT | ASSQDRGASNSDYT | 0.21540965 |
| QLVGLGI | ASSQDRGASNSDYT | 0.8319255 |
| RFLEQVHQL | ASSQDRGASNSDYT | 0.80544806 |
| RGVGYVLR | ASSQDRGASNSDYT | 0.20288622 |
| SATGVFAGVI | ASSQDRGASNSDYT | 0.19423029 |
| SLPAITLP | ASSQDRGASNSDYT | 0.68995005 |
| STALAFTV | ASSQDRGASNSDYT | 0.14707309 |
| SVLTVGLGK | ASSQDRGASNSDYT | 0.3583389 |
| TALAGAGSGI | ASSQDRGASNSDYT | 0.5243999 |
| TGLPFTGI | ASSQDRGASNSDYT | 0.4076706 |
| VAEHVTV | ASSQDRGASNSDYT | 0.20143093 |
| VDIVLNSL | ASSQDRGASNSDYT | 0.42480397 |
| VLTGGGAVL | ASSQDRGASNSDYT | 0.3310219 |
| VNVGLAYGI | ASSQDRGASNSDYT | 0.45477492 |
| VSSAAVGAV | ASSQDRGASNSDYT | 0.103565685 |
| VTSVYEAGV | ASSQDRGASNSDYT | 0.5255225 |
| WVALDRGA | ASSQDRGASNSDYT | 0.192159 |
| YLVGSVVI | ASSQDRGASNSDYT | 0.2200632 |
| GPPGTGKTLL | ASSQDRGASNSDYT | 0.11208082 |
| GVGMTKF | ASSQDRGASNSDYT | 0.28822488 |
| IGIGGSDL | ASSQDRGASNSDYT | 0.62950015 |
| IGTLLAEL | ASSQDRGASNSDYT | 0.69829285 |
| IYTSGTT | ASSQDRGASNSDYT | 0.16298336 |
| LSFPAGV | ASSQDRGASNSDYT | 0.65290797 |
| LYTSGTT | ASSQDRGASNSDYT | 0.73515964 |
| VGAFPFGF | ASSQDRGASNSDYT | 0.30981138 |
| VGAGTVEF | ASSQDRGASNSDYT | 0.13128495 |
| VGKSTLLL | ASSQDRGASNSDYT | 0.56638265 |
| AALAHAGGGIV | ASSQDRGASNSDYT | 0.39166135 |
| AALVAVSLL | ASSQDRGASNSDYT | 0.6065468 |
| AAVSRAVAAL | ASSQDRGASNSDYT | 0.6791866 |
| AGGSSGIGGF | ASSQDRGASNSDYT | 0.13934678 |
| AGLTVANANG | ASSQDRGASNSDYT | 0.61313903 |
| AGLVGAVL | ASSQDRGASNSDYT | 0.46097177 |
| ALSQAFTV | ASSQDRGASNSDYT | 0.15495892 |
| ALTFSVSL | ASSQDRGASNSDYT | 0.19721252 |
| ALTGAALVAL | ASSQDRGASNSDYT | 0.22086 |
| ASFSIGAGVL | ASSQDRGASNSDYT | 0.117571585 |
| DGSVYKI | ASSQDRGASNSDYT | 0.43413728 |
| EQLAEIV | ASSQDRGASNSDYT | 0.24863873 |
| FAVSELTVM | ASSQDRGASNSDYT | 0.29698187 |
| GAAPLAAGL | ASSQDRGASNSDYT | 0.2888464 |
| GADGGDGGSGGAGG | ASSQDRGASNSDYT | 0.22852397 |
| GFGLDTGGAGG | ASSQDRGASNSDYT | 0.26721048 |

| | | |
|---|---|---|
| GLASGGLL | ASSQDRGASNSDYT | 0.278607 |
| GRIAGAAVLL | ASSQDRGASNSDYT | 0.39345646 |
| GSAGSLNTSF | ASSQDRGASNSDYT | 0.48105875 |
| GTGQIGI | ASSQDRGASNSDYT | 0.39961046 |
| GTLIALIL | ASSQDRGASNSDYT | 0.47225145 |
| IAATMGFVVG | ASSQDRGASNSDYT | 0.7754458 |
| IFAQALL | ASSQDRGASNSDYT | 0.16534527 |
| IGAAVALGF | ASSQDRGASNSDYT | 0.20094928 |
| IGLTPLAL | ASSQDRGASNSDYT | 0.5669702 |
| ILGSNTSTL | ASSQDRGASNSDYT | 0.15425166 |
| IVFSLTVL | ASSQDRGASNSDYT | 0.67960596 |
| LASKATGF | ASSQDRGASNSDYT | 0.44203904 |
| LASVALIL | ASSQDRGASNSDYT | 0.4946813 |
| LFAVGLTL | ASSQDRGASNSDYT | 0.4320646 |
| LFNGGIGTYI | ASSQDRGASNSDYT | 0.8281519 |
| LTLGAGII | ASSQDRGASNSDYT | 0.5705579 |
| LVAAALGDI | ASSQDRGASNSDYT | 0.24967909 |
| NTAFVQLGI | ASSQDRGASNSDYT | 0.67920095 |
| QLFSGAEL | ASSQDRGASNSDYT | 0.18150268 |
| SAATFITL | ASSQDRGASNSDYT | 0.59223896 |
| SNYGLANLGV | ASSQDRGASNSDYT | 0.53610986 |
| SYGGAYAVM | ASSQDRGASNSDYT | 0.18185417 |
| TTGVGGLTLG | ASSQDRGASNSDYT | 0.27481928 |
| TTQTIYI | ASSQDRGASNSDYT | 0.59096074 |
| VAAGAGLGAVI | ASSQDRGASNSDYT | 0.7119075 |
| VAAVVTLL | ASSQDRGASNSDYT | 0.12577517 |
| VGFLGGLI | ASSQDRGASNSDYT | 0.23482075 |
| VGVIPILL | ASSQDRGASNSDYT | 0.15077579 |
| VIFASSLI | ASSQDRGASNSDYT | 0.22920573 |
| VKGGDGGAAAG | ASSQDRGASNSDYT | 0.17668752 |
| VTKTFTNI | ASSQDRGASNSDYT | 0.522354 |
| VVAAAILL | ASSQDRGASNSDYT | 0.2669161 |
| YAALDVEL | ASSQDRGASNSDYT | 0.77365506 |
| LASVALLL | ASSQDRGASNSDYT | 0.41932443 |
| LDAVHRV | ASSQDRGASNSDYT | 0.6739569 |
| QGLSSSIF | ASSQDRGASNSDYT | 0.18849087 |
| RGTFANIRL | ASSQDRGASNSDYT | 0.31894186 |
| RQLWWGHRI | ASSQDRGASNSDYT | 0.13475575 |
| YGPPGTGKTLL | ASSQDRGASNSDYT | 0.16701558 |
| AALVGAPANVA | ASSQDRGASNSDYT | 0.16546716 |
| AATFRRTE | ASSQDRGASNSDYT | 0.32358494 |
| AGALGGGI | ASSQDRGASNSDYT | 0.10999685 |
| AGGAGGAGGIGAG | ASSQDRGASNSDYT | 0.7636601 |
| AGSGALGL | ASSQDRGASNSDYT | 0.3902922 |
| AQIADQI | ASSQDRGASNSDYT | 0.5573312 |
| ASAVAVTYL | ASSQDRGASNSDYT | 0.26719627 |
| ASTVLGGTGG | ASSQDRGASNSDYT | 0.2141041 |
| AVGGEPAK | ASSQDRGASNSDYT | 0.25772876 |
| CEQGTFGGTC | ASSQDRGASNSDYT | 0.66976964 |
| DEGKAPAG | ASSQDRGASNSDYT | 0.15698296 |
| DTVVFPAL | ASSQDRGASNSDYT | 0.11203458 |

| | | |
|---|---|---|
| FEGITMAG | ASSQDRGASNSDYT | 0.24371433 |
| FPLVVQD | ASSQDRGASNSDYT | 0.1555409 |
| GAFTTPQ | ASSQDRGASNSDYT | 0.7571607 |
| GFGGFGTAGGVGGA | ASSQDRGASNSDYT | 0.6394676 |
| GFLAQFL | ASSQDRGASNSDYT | 0.3627232 |
| GGAGGAGGNAGVA | ASSQDRGASNSDYT | 0.64035 |
| GGVGTAGAGGAGGAGG | ASSQDRGASNSDYT | 0.09954953 |
| GNGGGGDAGNAGSGGN | ASSQDRGASNSDYT | 0.29701778 |
| GTGGAGGAGNDAGS | ASSQDRGASNSDYT | 0.80208784 |
| GTTNTGWFNS | ASSQDRGASNSDYT | 0.41452897 |
| IFMGPLFTL | ASSQDRGASNSDYT | 0.30232012 |
| ILAHAIEAI | ASSQDRGASNSDYT | 0.7248682 |
| IRSAIREAL | ASSQDRGASNSDYT | 0.24020791 |
| ITFAGGG | ASSQDRGASNSDYT | 0.26545924 |
| ITGMSGHQEV | ASSQDRGASNSDYT | 0.41858882 |
| IVAGLACTL | ASSQDRGASNSDYT | 0.33059877 |
| IVGGGLAGA | ASSQDRGASNSDYT | 0.31653798 |
| IVTGVLNI | ASSQDRGASNSDYT | 0.1485945 |
| LAQQGCI | ASSQDRGASNSDYT | 0.4719444 |
| LCLHVEN | ASSQDRGASNSDYT | 0.85367084 |
| LDAISDGS | ASSQDRGASNSDYT | 0.29278588 |
| LGGFAAML | ASSQDRGASNSDYT | 0.63232714 |
| LGGKNPMI | ASSQDRGASNSDYT | 0.5547575 |
| LGPKGVAAIL | ASSQDRGASNSDYT | 0.6477971 |
| LMVGVSI | ASSQDRGASNSDYT | 0.19433203 |
| LVGAAFVL | ASSQDRGASNSDYT | 0.17960884 |
| LVLAVAPL | ASSQDRGASNSDYT | 0.37690642 |
| MAAIAVIDAA | ASSQDRGASNSDYT | 0.18629552 |
| NNGINAELQ | ASSQDRGASNSDYT | 0.34905678 |
| RGEPLVLVHG | ASSQDRGASNSDYT | 0.26659462 |
| RGGAGGAAGLI | ASSQDRGASNSDYT | 0.8652596 |
| RSYRMHPAV | ASSQDRGASNSDYT | 0.61189914 |
| SAAHLLAGQLA | ASSQDRGASNSDYT | 0.58345616 |
| SAAVGSY | ASSQDRGASNSDYT | 0.18870842 |
| SPHAVLHRI | ASSQDRGASNSDYT | 0.25897336 |
| SQIAKVL | ASSQDRGASNSDYT | 0.5554295 |
| TALGHLVP | ASSQDRGASNSDYT | 0.14662082 |
| TAPTTSGAA | ASSQDRGASNSDYT | 0.73629665 |
| TDGDDGNGGSPGT | ASSQDRGASNSDYT | 0.21455336 |
| TDGLGEN | ASSQDRGASNSDYT | 0.33513868 |
| TGGPPLPGQA | ASSQDRGASNSDYT | 0.5482895 |
| TIPAITI | ASSQDRGASNSDYT | 0.80796057 |
| TIPGFTI | ASSQDRGASNSDYT | 0.2497712 |
| TSIASGALA | ASSQDRGASNSDYT | 0.34950095 |
| TVTLPGAAI | ASSQDRGASNSDYT | 0.19121192 |
| VASGLSAALP | ASSQDRGASNSDYT | 0.519541 |
| VGAGVIGI | ASSQDRGASNSDYT | 0.7323477 |
| VGPKRFGV | ASSQDRGASNSDYT | 0.4049116 |
| VNVAFPD | ASSQDRGASNSDYT | 0.80421305 |
| VSMTNSGVS | ASSQDRGASNSDYT | 0.446411 |
| VTDVLGDLSV | ASSQDRGASNSDYT | 0.15339157 |

| | | |
|---|---|---|
| VVVVGAGFAGL | ASSQDRGASNSDYT | 0.29472327 |
| VYTVGTVV | ASSQDRGASNSDYT | 0.6350039 |
| WTTFSGP | ASSQDRGASNSDYT | 0.749076 |
| YDELVEAF | ASSQDRGASNSDYT | 0.6590854 |
| YGTGGAGGNGG | ASSQDRGASNSDYT | 0.5434229 |
| YTNVMAV | ASSQDRGASNSDYT | 0.20655625 |
| FLEQVHQL | ASSQDRGASNSDYT | 0.72249186 |
| GLSLLAVAV | ASSQDRGASNSDYT | 0.7667807 |
| LEQVHQL | ASSQDRGASNSDYT | 0.6563815 |
| LNAAGAI | ASSQDRGASNSDYT | 0.22695048 |
| PAVHASL | ASSQDRGASNSDYT | 0.22753179 |
| TELEDDA | ASSQDRGASNSDYT | 0.3645985 |
| TPPGPPN | ASSQDRGASNSDYT | 0.7467382 |
| TPPPPPPP | ASSQDRGASNSDYT | 0.17626666 |
| TQSPALI | ASSSDPDREDTQY | 0.45338646 |
| EFAAAFPQ | ASSSDPDREDTQY | 0.4631354 |
| LLSGNLDVL | ASSSDPDREDTQY | 0.13317604 |
| GPANLGNYNI | ASSSDPDREDTQY | 0.40743047 |
| TLGPTLP | ASSSDPDREDTQY | 0.79734135 |
| VYAYSSCCK | ASSSDPDREDTQY | 0.32717648 |
| APAPAPAPNI | ASSSDPDREDTQY | 0.20794496 |
| PGGPVPIP | ASSSDPDREDTQY | 0.74595594 |
| FLTVALEP | ASSSDPDREDTQY | 0.8470247 |
| SQYFGANL | ASSSDPDREDTQY | 0.6313433 |
| GGVKPAA | ASSSDPDREDTQY | 0.14898516 |
| GGTGGAGGAGGW | ASSSDPDREDTQY | 0.101092115 |
| GVGGVAGLGGAGG | ASSSDPDREDTQY | 0.313711 |
| VLLVLATTI | ASSSDPDREDTQY | 0.7217443 |
| IGIFAAML | ASSSDPDREDTQY | 0.28258085 |
| AVGDLID | ASSSDPDREDTQY | 0.22771099 |
| NISPLPL | ASSSDPDREDTQY | 0.584916 |
| VTAVFVAI | ASSSDPDREDTQY | 0.6693646 |
| SGDDNDAAL | ASSSDPDREDTQY | 0.17832795 |
| AVAALAHS | ASSSDPDREDTQY | 0.15908496 |
| GGVTVMATP | ASSSDPDREDTQY | 0.5669223 |
| AGLAANVRD | ASSSDPDREDTQY | 0.6054288 |
| TNRPDLIDPALLRPG | ASSSDPDREDTQY | 0.1640837 |
| LANLGVTAL | ASSSDPDREDTQY | 0.25475022 |
| TGERIEL | ASSSDPDREDTQY | 0.32784647 |
| NGEELAA | ASSSDPDREDTQY | 0.29253352 |
| VLSLVWG | ASSSDPDREDTQY | 0.46517032 |
| IGYALWL | ASSSDPDREDTQY | 0.17022774 |
| LSLLLAPL | ASSSDPDREDTQY | 0.72773314 |
| TLDEALK | ASSSDPDREDTQY | 0.15675405 |
| TDGAGTAEGG | ASSSDPDREDTQY | 0.5101018 |
| NAPGGAGGNGG | ASSSDPDREDTQY | 0.3712187 |
| VSVFMGMGGNF | ASSSDPDREDTQY | 0.32493943 |
| LASYLAAL | ASSSDPDREDTQY | 0.2904207 |
| LVGGTPAL | ASSSDPDREDTQY | 0.20948133 |
| VRNADVL | ASSSDPDREDTQY | 0.15851846 |
| TCSSPFLVSAM | ASSSDPDREDTQY | 0.11279524 |

| | | |
|---|---|---|
| VHITDAE | ASSSDPDREDTQY | 0.69889534 |
| EYAAPSF | ASSSDPDREDTQY | 0.27658778 |
| VVDAFHGGL | ASSSDPDREDTQY | 0.5784892 |
| ASVVDPQ | ASSSDPDREDTQY | 0.4748851 |
| SYNEFGGLS | ASSSDPDREDTQY | 0.17988554 |
| DPDGSVY | ASSSDPDREDTQY | 0.38299116 |
| MVSTIHATE | ASSSDPDREDTQY | 0.19918633 |
| LAAGVADPV | ASSSDPDREDTQY | 0.18267037 |
| TKEQIAA | ASSSDPDREDTQY | 0.5023674 |
| SLASPAAL | ASSSDPDREDTQY | 0.5710413 |
| IGLVFYGV | ASSSDPDREDTQY | 0.19926761 |
| FLLDASNL | ASSSDPDREDTQY | 0.42199087 |
| YQVGNFP | ASSSDPDREDTQY | 0.38496152 |
| ESLLSQST | ASSSDPDREDTQY | 0.18421055 |
| LFTVGFAAVG | ASSSDPDREDTQY | 0.31314537 |
| ATPPTPT | ASSSDPDREDTQY | 0.6859498 |
| TDLATLG | ASSSDPDREDTQY | 0.40587354 |
| SAGGTGR | ASSSDPDREDTQY | 0.16001222 |
| GFSNSGS | ASSSDPDREDTQY | 0.55649674 |
| GGNGGSGGTSV | ASSSDPDREDTQY | 0.21067294 |
| GTSVATGGAGN | ASSSDPDREDTQY | 0.46085486 |
| NGGNGGSGGTS | ASSSDPDREDTQY | 0.32452762 |
| AVVKFPH | ASSSDPDREDTQY | 0.27716622 |
| TPPPPPP | ASSSDPDREDTQY | 0.55478036 |
| AAPAAAAAPA | ASSSDPDREDTQY | 0.17204927 |
| IALNEDL | ASSSDPDREDTQY | 0.19809788 |
| IVNVSSM | ASSSDPDREDTQY | 0.25850707 |
| TGIPPAPR | ASSSDPDREDTQY | 0.37778863 |
| FDLGGGTF | ASSSDPDREDTQY | 0.65139675 |
| FDLGGGTFDVS | ASSSDPDREDTQY | 0.18440631 |
| NEPTAAA | ASSSDPDREDTQY | 0.32888556 |
| NEPTAAALA | ASSSDPDREDTQY | 0.22768424 |
| GSIINISSI | ASSSDPDREDTQY | 0.18103613 |
| VAPGNAGTA | ASSSDPDREDTQY | 0.30366105 |
| TGGGASLE | ASSSDPDREDTQY | 0.25251037 |
| VAVYNHYKRI | ASSSDPDREDTQY | 0.18088336 |
| TDPAPATT | ASSSDPDREDTQY | 0.38598248 |
| TNRPDILDPALLRPG | ASSSDPDREDTQY | 0.23337427 |
| LTTTTTS | ASSSDPDREDTQY | 0.37793946 |
| PTGSGKTL | ASSSDPDREDTQY | 0.6610408 |
| TGSGKTL | ASSSDPDREDTQY | 0.21134518 |
| TIGHVDHGK | ASSSDPDREDTQY | 0.4284147 |
| TIGHVDHGKTT | ASSSDPDREDTQY | 0.54787326 |
| GHVDHGKTT | ASSSDPDREDTQY | 0.50136846 |
| AQFEHTLL | ASSSDPDREDTQY | 0.2948749 |
| IVNQDEL | ASSSDPDREDTQY | 0.19087803 |
| TNGSQFF | ASSSDPDREDTQY | 0.72988975 |
| TGKGIVE | ASSSDPDREDTQY | 0.578298 |
| LISNASDA | ASSSDPDREDTQY | 0.64976776 |
| AELRTLGELTKIAKAHGAQ | ASSSDPDREDTQY | 0.51806945 |
| APNAVAAAEPSW | ASSSDPDREDTQY | 0.68594706 |

| | | |
|---|---|---|
| APQVAGAATAVL | ASSSDPDREDTQY | 0.5706717 |
| ARRDPAAAAAILPTDA | ASSSDPDREDTQY | 0.2730651 |
| DAAARDVLAEAGLAEYF | ASSSDPDREDTQY | 0.13768159 |
| DDQIVLSAKANLAKVEQ | ASSSDPDREDTQY | 0.31766856 |
| DELLPDTGEDSDEDGH | ASSSDPDREDTQY | 0.2519801 |
| ELPEAPANRATVPQD | ASSSDPDREDTQY | 0.21514028 |
| FGQSKIERVTGIG | ASSSDPDREDTQY | 0.14701182 |
| GAGGTGSTAGGAGGAGGAGGLY | ASSSDPDREDTQY | 0.7247351 |
| GHGGTGAAGGNGGTGGAGGSGI | ASSSDPDREDTQY | 0.855809 |
| GNGGNGGAGGEGGPGGAGGAGA | ASSSDPDREDTQY | 0.23268795 |
| GSTAGNGGNGGNATNGTIAGQP | ASSSDPDREDTQY | 0.7715908 |
| GTGGDGGNGGNGGTGG | ASSSDPDREDTQY | 0.14321344 |
| GVRRPAGIPADAPV | ASSSDPDREDTQY | 0.6187334 |
| GYRQVTDAHLLTLARR | ASSSDPDREDTQY | 0.73540366 |
| IAKMGQDGHDRG | ASSSDPDREDTQY | 0.19031484 |
| KAEPSIEIQYT | ASSSDPDREDTQY | 0.51177 |
| KPVRIRMAADEDTVAVT | ASSSDPDREDTQY | 0.18850768 |
| MPPRIAGMRLLVIKPEP | ASSSDPDREDTQY | 0.25094625 |
| MKFYAVALFRSRGVPPTR | ASSSDPDREDTQY | 0.717674 |
| NAGGFGILGAVAHSP | ASSSDPDREDTQY | 0.30077758 |
| NEGREITLAMAEM | ASSSDPDREDTQY | 0.364183 |
| NELVEWSHPLGHRWRWE | ASSSDPDREDTQY | 0.34069866 |
| NLGAGGTGGNGGNPG | ASSSDPDREDTQY | 0.2023097 |
| NTEWPKWETVMAADPVD | ASSSDPDREDTQY | 0.31899825 |
| PASAAPAETQP | ASSSDPDREDTQY | 0.6284172 |
| QQSSDGKNGVRAAEITGE | ASSSDPDREDTQY | 0.19747338 |
| RAPDAVTTSWWKE | ASSSDPDREDTQY | 0.7096525 |
| SGFGNVGAHTSGWFNQ | ASSSDPDREDTQY | 0.17313841 |
| SGVAGAVGQAASVGGLKV | ASSSDPDREDTQY | 0.8147435 |
| SNGGAGTGSAGGNGGT | ASSSDPDREDTQY | 0.79884994 |
| TGGAGGEGGAAG | ASSSDPDREDTQY | 0.42311466 |
| TGGDGGAAGAGGAPGHG | ASSSDPDREDTQY | 0.25258008 |
| TMPPLLAAIANAGRHGV | ASSSDPDREDTQY | 0.75936687 |
| TSSTAGNSGGA | ASSSDPDREDTQY | 0.313689 |
| VPATVPVGILC | ASSSDPDREDTQY | 0.5635519 |
| VQRPDSGGVVLTGRIS | ASSSDPDREDTQY | 0.5326804 |
| WTPNPVPATKGVPAGLVQAT | ASSSDPDREDTQY | 0.16172704 |
| YALKRALAKFSDRFDVVII | ASSSDPDREDTQY | 0.63856286 |
| YLIGSVIV | ASSSDPDREDTQY | 0.47747123 |
| YGQLVAVID | ASSSDPDREDTQY | 0.50346565 |
| VTVGGMTLPA | ASSSDPDREDTQY | 0.20988654 |
| VLALPAV | ASSSDPDREDTQY | 0.1769877 |
| VIPPTLNYET | ASSSDPDREDTQY | 0.25400293 |
| VGMGGMHPSSG | ASSSDPDREDTQY | 0.18337518 |
| VGLAGGAATA | ASSSDPDREDTQY | 0.7757149 |
| VGGAGGAGGSGP | ASSSDPDREDTQY | 0.45675385 |
| VFAAFMV | ASSSDPDREDTQY | 0.20743325 |
| TLYDKLR | ASSSDPDREDTQY | 0.454449 |
| TLAVFTVL | ASSSDPDREDTQY | 0.6008374 |
| TGSVQPARV | ASSSDPDREDTQY | 0.15061381 |
| TGGGAGSGGT | ASSSDPDREDTQY | 0.32600248 |

| | | |
|---|---|---|
| SHLETSL | ASSSDPDREDTQY | 0.2555392 |
| SFTTGMT | ASSSDPDREDTQY | 0.26320857 |
| QVEVAKRMGVRQPTVS | ASSSDPDREDTQY | 0.12733585 |
| QGSGGPALAM | ASSSDPDREDTQY | 0.45961088 |
| MLAPSNP | ASSSDPDREDTQY | 0.21419546 |
| LVNNAAI | ASSSDPDREDTQY | 0.13128355 |
| LTGEINTLA | ASSSDPDREDTQY | 0.36966032 |
| LSIFGPV | ASSSDPDREDTQY | 0.22571388 |
| LLALLANL | ASSSDPDREDTQY | 0.24569306 |
| LISNASDALDKLR | ASSSDPDREDTQY | 0.53405046 |
| LINEAVP | ASSSDPDREDTQY | 0.46435288 |
| LGYAFSGL | ASSSDPDREDTQY | 0.23555571 |
| LDFVAVAL | ASSSDPDREDTQY | 0.45087016 |
| LAQPGAE | ASSSDPDREDTQY | 0.5986255 |
| ISNLGAVTGF | ASSSDPDREDTQY | 0.21433172 |
| HQEDPGRI | ASSSDPDREDTQY | 0.3794024 |
| GTPGEPTP | ASSSDPDREDTQY | 0.7376337 |
| GFFNSGS | ASSSDPDREDTQY | 0.27788043 |
| GAGGGGLTAGAN | ASSSDPDREDTQY | 0.7045393 |
| FVDALVNM | ASSSDPDREDTQY | 0.72749376 |
| FLTVAPEL | ASSSDPDREDTQY | 0.24910483 |
| FETLTAL | ASSSDPDREDTQY | 0.58985054 |
| DVINPIEL | ASSSDPDREDTQY | 0.5804532 |
| APVSGAPV | ASSSDPDREDTQY | 0.33818316 |
| ALVGGLAW | ASSSDPDREDTQY | 0.31919795 |
| ALAAAAATAI | ASSSDPDREDTQY | 0.15635292 |
| AFGLGLLL | ASSSDPDREDTQY | 0.33553118 |
| AFFAAASL | ASSSDPDREDTQY | 0.19757277 |
| AAAALPAFSPP | ASSSDPDREDTQY | 0.5158835 |
| VVLAPAL | ASSSDPDREDTQY | 0.6773672 |
| TIGHVDHGKTTL | ASSSDPDREDTQY | 0.2929502 |
| LISNASDALDK | ASSSDPDREDTQY | 0.14265022 |
| LISNASDAL | ASSSDPDREDTQY | 0.15299094 |
| ISNASDALDK | ASSSDPDREDTQY | 0.4299889 |
| NQAETLVYQTE | ASSSDPDREDTQY | 0.17399205 |
| TGGTGGTAGAG | ASSSDPDREDTQY | 0.21089366 |
| TGSTAGGAGGA | ASSSDPDREDTQY | 0.18891153 |
| DPLAVPVAAGPV | ASSSDPDREDTQY | 0.5233766 |
| GGTGGTPGNGGN | ASSSDPDREDTQY | 0.56810963 |
| PAGLAGVDPATL | ASSSDPDREDTQY | 0.19322845 |
| GNNLVVEAPHAGASV | ASSSDPDREDTQY | 0.6085616 |
| KVTRIMSASTLEQPA | ASSSDPDREDTQY | 0.7222384 |
| LGFALPIALPRRTVV | ASSSDPDREDTQY | 0.7634069 |
| TRLGNLTARKLVVLV | ASSSDPDREDTQY | 0.36780465 |
| WSQQRRLIYFPSAGP | ASSSDPDREDTQY | 0.1515625 |
| GPNGNPDPMAAAVDIR | ASSSDPDREDTQY | 0.69084495 |
| TQRCVSVADLRAAASR | ASSSDPDREDTQY | 0.21371633 |
| QFVLDVAEHPVEVLTETV | ASSSDPDREDTQY | 0.6647246 |
| GGAGGTGGAGGNGGTQAGDG | ASSSDPDREDTQY | 0.51614505 |
| PLLATLRAIAVDVDAKPAQV | ASSSDPDREDTQY | 0.30755305 |
| APKFRTGGYMIDFWGVGYQVAK | ASSSDPDREDTQY | 0.19131452 |

| | | |
|---|---|---|
| GAGGAGGAGGAAGGPGGTGGTG | ASSSDPDREDTQY | 0.64402246 |
| IGALGLLATDVAPAGSVAFGVA | ASSSDPDREDTQY | 0.62250376 |
| AGGAGGIGGNGGAGGAGGVLMG | ASSSDPDREDTQY | 0.16241279 |
| ADELIGLL | ASSSDPDREDTQY | 0.18070917 |
| ADRALKA | ASSSDPDREDTQY | 0.4711334 |
| AGAVGQVGGAGGA | ASSSDPDREDTQY | 0.28564474 |
| AGGGGGLGGAP | ASSSDPDREDTQY | 0.6147618 |
| AGGLLYGNGGAG | ASSSDPDREDTQY | 0.40812838 |
| AGLESIGL | ASSSDPDREDTQY | 0.41478282 |
| AGLKTGII | ASSSDPDREDTQY | 0.49180105 |
| ASLAALLL | ASSSDPDREDTQY | 0.54966563 |
| ASVINISSI | ASSSDPDREDTQY | 0.37169448 |
| ATPPGPA | ASSSDPDREDTQY | 0.45850757 |
| ATVASALV | ASSSDPDREDTQY | 0.20046005 |
| ATVASLIL | ASSSDPDREDTQY | 0.60882187 |
| AVNGPVLIRH | ASSSDPDREDTQY | 0.58615834 |
| DGAAGAVGI | ASSSDPDREDTQY | 0.50892603 |
| DPLLHGI | ASSSDPDREDTQY | 0.68855596 |
| ELGAALGAL | ASSSDPDREDTQY | 0.39578292 |
| ELGRTQAV | ASSSDPDREDTQY | 0.44775826 |
| FVAGGYLS | ASSSDPDREDTQY | 0.16408585 |
| GASGGAGGAGGW | ASSSDPDREDTQY | 0.2577185 |
| GGAGAGGDGGAGGI | ASSSDPDREDTQY | 0.27615008 |
| GTGHNGSQPGMGGNG | ASSSDPDREDTQY | 0.45069373 |
| GYAAASASAS | ASSSDPDREDTQY | 0.5096799 |
| HQVDVAAVQQG | ASSSDPDREDTQY | 0.5499551 |
| IGAAIAHT | ASSSDPDREDTQY | 0.227413 |
| IGMIEVAQL | ASSSDPDREDTQY | 0.64778984 |
| IRLAPEHTSAMD | ASSSDPDREDTQY | 0.71478975 |
| IYNVSVL | ASSSDPDREDTQY | 0.16493955 |
| KQGFLGM | ASSSDPDREDTQY | 0.24484138 |
| LESVITQL | ASSSDPDREDTQY | 0.55522966 |
| LGLGALVVV | ASSSDPDREDTQY | 0.56986797 |
| LIGNGGNGGSGGI | ASSSDPDREDTQY | 0.23872977 |
| LNAINAPTL | ASSSDPDREDTQY | 0.36645475 |
| LQLLNTGV | ASSSDPDREDTQY | 0.7517467 |
| LQTSIEF | ASSSDPDREDTQY | 0.57279336 |
| MSFAVSEL | ASSSDPDREDTQY | 0.6807407 |
| NTGALGL | ASSSDPDREDTQY | 0.14497183 |
| PPGAGGAGGAGTT | ASSSDPDREDTQY | 0.33623454 |
| QLVGLGI | ASSSDPDREDTQY | 0.27349404 |
| RFLEQVHQL | ASSSDPDREDTQY | 0.11874677 |
| RGVGYVLR | ASSSDPDREDTQY | 0.5027713 |
| SATGVFAGVI | ASSSDPDREDTQY | 0.21566096 |
| SLPAITLP | ASSSDPDREDTQY | 0.4732411 |
| STALAFTV | ASSSDPDREDTQY | 0.22701326 |
| SVLTVGLGK | ASSSDPDREDTQY | 0.5643395 |
| TALAGAGSGI | ASSSDPDREDTQY | 0.45811713 |
| TGLPFTGI | ASSSDPDREDTQY | 0.28481436 |
| VAEHVTV | ASSSDPDREDTQY | 0.19977653 |
| VDIVLNSL | ASSSDPDREDTQY | 0.33380714 |

| | | |
|---|---|---|
| VLTGGGAVL | ASSSDPDREDTQY | 0.62321246 |
| VNVGLAYGI | ASSSDPDREDTQY | 0.634357 |
| VSSAAVGAV | ASSSDPDREDTQY | 0.15598993 |
| VTSVYEAGV | ASSSDPDREDTQY | 0.6838914 |
| WVALDRGA | ASSSDPDREDTQY | 0.2655904 |
| YLVGSVVI | ASSSDPDREDTQY | 0.16174673 |
| GPPGTGKTLL | ASSSDPDREDTQY | 0.1509298 |
| GVGMTKF | ASSSDPDREDTQY | 0.22953865 |
| IGIGGSDL | ASSSDPDREDTQY | 0.25520322 |
| IGTLLAEL | ASSSDPDREDTQY | 0.35357565 |
| IYTSGTT | ASSSDPDREDTQY | 0.522928 |
| LSFPAGV | ASSSDPDREDTQY | 0.20558101 |
| LYTSGTT | ASSSDPDREDTQY | 0.5418755 |
| VGAFPFGF | ASSSDPDREDTQY | 0.2772372 |
| VGAGTVEF | ASSSDPDREDTQY | 0.10545204 |
| VGKSTLLL | ASSSDPDREDTQY | 0.24816495 |
| AALAHAGGGIV | ASSSDPDREDTQY | 0.1423946 |
| AALVAVSLL | ASSSDPDREDTQY | 0.6652392 |
| AAVSRAVAAL | ASSSDPDREDTQY | 0.16109334 |
| AGGSSGIGGF | ASSSDPDREDTQY | 0.24058351 |
| AGLTVANANG | ASSSDPDREDTQY | 0.24999061 |
| AGLVGAVL | ASSSDPDREDTQY | 0.70901585 |
| ALSQAFTV | ASSSDPDREDTQY | 0.42961812 |
| ALTFSVSL | ASSSDPDREDTQY | 0.22892827 |
| ALTGAALVAL | ASSSDPDREDTQY | 0.21714355 |
| ASFSIGAGVL | ASSSDPDREDTQY | 0.36768347 |
| DGSVYKI | ASSSDPDREDTQY | 0.5012696 |
| EQLAEIV | ASSSDPDREDTQY | 0.22702102 |
| FAVSELTVM | ASSSDPDREDTQY | 0.21732207 |
| GAAPLAAGL | ASSSDPDREDTQY | 0.7718425 |
| GADGGDGGSGGAGG | ASSSDPDREDTQY | 0.16633222 |
| GFGLDTGGAGG | ASSSDPDREDTQY | 0.6532231 |
| GLASGGLL | ASSSDPDREDTQY | 0.19112173 |
| GRIAGAAVLL | ASSSDPDREDTQY | 0.21467188 |
| GSAGSLNTSF | ASSSDPDREDTQY | 0.7154763 |
| GTGQIGI | ASSSDPDREDTQY | 0.6427231 |
| GTLIALIL | ASSSDPDREDTQY | 0.2800217 |
| IAATMGFVVG | ASSSDPDREDTQY | 0.28230911 |
| IFAQALL | ASSSDPDREDTQY | 0.17880547 |
| IGAAVALGF | ASSSDPDREDTQY | 0.1998809 |
| IGLTPLAL | ASSSDPDREDTQY | 0.24760929 |
| ILGSNTSTL | ASSSDPDREDTQY | 0.5807042 |
| IVFSLTVL | ASSSDPDREDTQY | 0.3874638 |
| LASKATGF | ASSSDPDREDTQY | 0.16743672 |
| LASVALIL | ASSSDPDREDTQY | 0.6601573 |
| LFAVGLTL | ASSSDPDREDTQY | 0.7018831 |
| LFNGGIGTYI | ASSSDPDREDTQY | 0.7341357 |
| LTLGAGII | ASSSDPDREDTQY | 0.17095417 |
| LVAAALGDI | ASSSDPDREDTQY | 0.3183195 |
| NTAFVQLGI | ASSSDPDREDTQY | 0.3392335 |
| QLFSGAEL | ASSSDPDREDTQY | 0.27580085 |

| | | |
|---|---|---|
| SAATFITL | ASSSDPDREDTQY | 0.22895142 |
| SNYGLANLGV | ASSSDPDREDTQY | 0.21071264 |
| SYGGAYAVM | ASSSDPDREDTQY | 0.61714154 |
| TTGVGGLTLG | ASSSDPDREDTQY | 0.8314102 |
| TTQTIYI | ASSSDPDREDTQY | 0.28684336 |
| VAAGAGLGAVI | ASSSDPDREDTQY | 0.13007522 |
| VAAVVTLL | ASSSDPDREDTQY | 0.2444661 |
| VGFLGGLI | ASSSDPDREDTQY | 0.20342551 |
| VGVIPILL | ASSSDPDREDTQY | 0.1741351 |
| VIFASSLI | ASSSDPDREDTQY | 0.27222922 |
| VKGGDGGAAAG | ASSSDPDREDTQY | 0.13859563 |
| VTKTFTNI | ASSSDPDREDTQY | 0.3025785 |
| VVAAAILL | ASSSDPDREDTQY | 0.7208869 |
| YAALDVEL | ASSSDPDREDTQY | 0.27869368 |
| LASVALLL | ASSSDPDREDTQY | 0.4692696 |
| LDAVHRV | ASSSDPDREDTQY | 0.6765921 |
| QGLSSSIF | ASSSDPDREDTQY | 0.45525151 |
| RGTFANIRL | ASSSDPDREDTQY | 0.66089493 |
| RQLWWGHRI | ASSSDPDREDTQY | 0.7509576 |
| YGPPGTGKTLL | ASSSDPDREDTQY | 0.29997253 |
| AALVGAPANVA | ASSSDPDREDTQY | 0.48182803 |
| AATFRRTE | ASSSDPDREDTQY | 0.61025405 |
| AGALGGGI | ASSSDPDREDTQY | 0.24941029 |
| AGGAGGAGGIGAG | ASSSDPDREDTQY | 0.32711095 |
| AGSGALGL | ASSSDPDREDTQY | 0.3541423 |
| AQIADQI | ASSSDPDREDTQY | 0.43770748 |
| ASAVAVTYL | ASSSDPDREDTQY | 0.43741116 |
| ASTVLGGTGG | ASSSDPDREDTQY | 0.7557184 |
| AVGGEPAK | ASSSDPDREDTQY | 0.13359761 |
| CEQGTFGGTC | ASSSDPDREDTQY | 0.20782933 |
| DEGKAPAG | ASSSDPDREDTQY | 0.7139398 |
| DTVVFPAL | ASSSDPDREDTQY | 0.12694922 |
| FEGITMAG | ASSSDPDREDTQY | 0.1959391 |
| FPLVVQD | ASSSDPDREDTQY | 0.3523413 |
| GAFTTPQ | ASSSDPDREDTQY | 0.7300296 |
| GFGGFGTAGGVGGA | ASSSDPDREDTQY | 0.12706603 |
| GFLAQFL | ASSSDPDREDTQY | 0.35592118 |
| GGAGGAGGNAGVA | ASSSDPDREDTQY | 0.15991746 |
| GGVGTAGAGGAGGAGG | ASSSDPDREDTQY | 0.7002994 |
| GNGGGGDAGNAGSGGN | ASSSDPDREDTQY | 0.2666721 |
| GTGGAGGAGNDAGS | ASSSDPDREDTQY | 0.15481338 |
| GTTNTGWFNS | ASSSDPDREDTQY | 0.5309678 |
| IFMGPLFTL | ASSSDPDREDTQY | 0.16058987 |
| ILAHAIEAI | ASSSDPDREDTQY | 0.5731277 |
| IRSAIREAL | ASSSDPDREDTQY | 0.557521 |
| ITFAGGG | ASSSDPDREDTQY | 0.7976338 |
| ITGMSGHQEV | ASSSDPDREDTQY | 0.15347633 |
| IVAGLACTL | ASSSDPDREDTQY | 0.5066985 |
| IVGGGLAGA | ASSSDPDREDTQY | 0.28735888 |
| IVTGVLNI | ASSSDPDREDTQY | 0.8209698 |
| LAQQGCI | ASSSDPDREDTQY | 0.2393716 |

| | | |
|---|---|---|
| LCLHVEN | ASSSDPDREDTQY | 0.15898265 |
| LDAISDGS | ASSSDPDREDTQY | 0.5839075 |
| LGGFAAML | ASSSDPDREDTQY | 0.50357527 |
| LGGKNPMI | ASSSDPDREDTQY | 0.298794 |
| LGPKGVAAIL | ASSSDPDREDTQY | 0.6789435 |
| LMVGVSI | ASSSDPDREDTQY | 0.36321738 |
| LVGAAFVL | ASSSDPDREDTQY | 0.27140144 |
| LVLAVAPL | ASSSDPDREDTQY | 0.4932282 |
| MAAIAVIDAA | ASSSDPDREDTQY | 0.4031375 |
| NNGINAELQ | ASSSDPDREDTQY | 0.2333312 |
| RGEPLVLVHG | ASSSDPDREDTQY | 0.5700304 |
| RGGAGGAAGLI | ASSSDPDREDTQY | 0.30262285 |
| RSYRMHPAV | ASSSDPDREDTQY | 0.17434001 |
| SAAHLLAGQLA | ASSSDPDREDTQY | 0.74532866 |
| SAAVGSY | ASSSDPDREDTQY | 0.364023 |
| SPHAVLHRI | ASSSDPDREDTQY | 0.5233764 |
| SQIAKVL | ASSSDPDREDTQY | 0.40895352 |
| TALGHLVP | ASSSDPDREDTQY | 0.795997 |
| TAPTTSGAA | ASSSDPDREDTQY | 0.21036169 |
| TDGDDGNGGSPGT | ASSSDPDREDTQY | 0.38186428 |
| TDGLGEN | ASSSDPDREDTQY | 0.33963686 |
| TGGPPLPGQA | ASSSDPDREDTQY | 0.5901282 |
| TIPAITI | ASSSDPDREDTQY | 0.36296526 |
| TIPGFTI | ASSSDPDREDTQY | 0.5904747 |
| TSIASGALA | ASSSDPDREDTQY | 0.4793555 |
| TVTLPGAAI | ASSSDPDREDTQY | 0.5928067 |
| VASGLSAALP | ASSSDPDREDTQY | 0.28399363 |
| VGAGVIGI | ASSSDPDREDTQY | 0.41099083 |
| VGPKRFGV | ASSSDPDREDTQY | 0.73533016 |
| VNVAFPD | ASSSDPDREDTQY | 0.17111814 |
| VSMTNSGVS | ASSSDPDREDTQY | 0.3217474 |
| VTDVLGDLSV | ASSSDPDREDTQY | 0.5468401 |
| VVVVGAGFAGL | ASSSDPDREDTQY | 0.57185364 |
| VYTVGTVV | ASSSDPDREDTQY | 0.34592813 |
| WTTFSGP | ASSSDPDREDTQY | 0.6370262 |
| YDELVEAF | ASSSDPDREDTQY | 0.26484406 |
| YGTGGAGGNGG | ASSSDPDREDTQY | 0.4005487 |
| YTNVMAV | ASSSDPDREDTQY | 0.37416017 |
| FLEQVHQL | ASSSDPDREDTQY | 0.2456756 |
| GLSLLAVAV | ASSSDPDREDTQY | 0.18015438 |
| LEQVHQL | ASSSDPDREDTQY | 0.26252833 |
| LNAAGAI | ASSSDPDREDTQY | 0.58150876 |
| PAVHASL | ASSSDPDREDTQY | 0.4031638 |
| TELEDDA | ASSSDPDREDTQY | 0.16981831 |
| TPPGPPN | ASSSDPDREDTQY | 0.39375663 |
| TPPPPPPP | ASSSDPDREDTQY | 0.48768133 |
| TQSPALI | ASGDEGEVF | 0.7996594 |
| EFAAAFPQ | ASGDEGEVF | 0.45339793 |
| LLSGNLDVL | ASGDEGEVF | 0.5993682 |
| GPANLGNYNI | ASGDEGEVF | 0.26069576 |
| TLGPTLP | ASGDEGEVF | 0.35987103 |

| | | |
|---|---|---|
| VYAYSSCCK | ASGDEGEVF | 0.15523377 |
| APAPAPAPNI | ASGDEGEVF | 0.19825606 |
| PGGPVPIP | ASGDEGEVF | 0.32820964 |
| FLTVALEP | ASGDEGEVF | 0.1805429 |
| SQYFGANL | ASGDEGEVF | 0.69210166 |
| GGVKPAA | ASGDEGEVF | 0.6141521 |
| GGTGGAGGAGGW | ASGDEGEVF | 0.5080893 |
| GVGGVAGLGGAGG | ASGDEGEVF | 0.72478163 |
| VLLVLATTI | ASGDEGEVF | 0.70997393 |
| IGIFAAML | ASGDEGEVF | 0.52277845 |
| AVGDLID | ASGDEGEVF | 0.20983797 |
| NISPLPL | ASGDEGEVF | 0.26305667 |
| VTAVFVAI | ASGDEGEVF | 0.13390522 |
| SGDDNDAAL | ASGDEGEVF | 0.25008765 |
| AVAALAHS | ASGDEGEVF | 0.24547963 |
| GGVTVMATP | ASGDEGEVF | 0.30920482 |
| AGLAANVRD | ASGDEGEVF | 0.124042936 |
| TNRPDLIDPALLRPG | ASGDEGEVF | 0.6673178 |
| LANLGVTAL | ASGDEGEVF | 0.5781549 |
| TGERIEL | ASGDEGEVF | 0.39724028 |
| NGEELAA | ASGDEGEVF | 0.53008246 |
| VLSLVWG | ASGDEGEVF | 0.49800682 |
| IGYALWL | ASGDEGEVF | 0.21395524 |
| LSLLLAPL | ASGDEGEVF | 0.55966955 |
| TLDEALK | ASGDEGEVF | 0.29069233 |
| TDGAGTAEGG | ASGDEGEVF | 0.6701648 |
| NAPGGAGGNGG | ASGDEGEVF | 0.55706125 |
| VSVFMGMGGNF | ASGDEGEVF | 0.4853248 |
| LASYLAAL | ASGDEGEVF | 0.30965763 |
| LVGGTPAL | ASGDEGEVF | 0.49911922 |
| VRNADVL | ASGDEGEVF | 0.21906373 |
| TCSSPFLVSAM | ASGDEGEVF | 0.33100966 |
| VHITDAE | ASGDEGEVF | 0.31614864 |
| EYAAPSF | ASGDEGEVF | 0.44201392 |
| VVDAFHGGL | ASGDEGEVF | 0.194504 |
| ASVVDPQ | ASGDEGEVF | 0.40903622 |
| SYNEFGGLS | ASGDEGEVF | 0.2693285 |
| DPDGSVY | ASGDEGEVF | 0.41606852 |
| MVSTIHATE | ASGDEGEVF | 0.1670122 |
| LAAGVADPV | ASGDEGEVF | 0.68499416 |
| TKEQIAA | ASGDEGEVF | 0.7271485 |
| SLASPAAL | ASGDEGEVF | 0.18152162 |
| IGLVFYGV | ASGDEGEVF | 0.5340535 |
| FLLDASNL | ASGDEGEVF | 0.30038473 |
| YQVGNFP | ASGDEGEVF | 0.26143062 |
| ESLLSQST | ASGDEGEVF | 0.592898 |
| LFTVGFAAVG | ASGDEGEVF | 0.31364852 |
| ATPPTPT | ASGDEGEVF | 0.16977842 |
| TDLATLG | ASGDEGEVF | 0.21769398 |
| SAGGTGR | ASGDEGEVF | 0.4098704 |
| GFSNSGS | ASGDEGEVF | 0.26004633 |

| | | |
|---|---|---|
| GGNGGSGGTSV | ASGDEGEVF | 0.15014641 |
| GTSVATGGAGN | ASGDEGEVF | 0.37439227 |
| NGGNGGSGGTS | ASGDEGEVF | 0.17739986 |
| AVVKFPH | ASGDEGEVF | 0.19479539 |
| TPPPPPP | ASGDEGEVF | 0.19839947 |
| AAPAAAAAPA | ASGDEGEVF | 0.60088325 |
| IALNEDL | ASGDEGEVF | 0.72997606 |
| IVNVSSM | ASGDEGEVF | 0.31424248 |
| TGIPPAPR | ASGDEGEVF | 0.3982535 |
| FDLGGGTF | ASGDEGEVF | 0.3516709 |
| FDLGGGTFDVS | ASGDEGEVF | 0.5052671 |
| NEPTAAA | ASGDEGEVF | 0.38319832 |
| NEPTAAALA | ASGDEGEVF | 0.4094068 |
| GSIINISSI | ASGDEGEVF | 0.49565274 |
| VAPGNAGTA | ASGDEGEVF | 0.29299295 |
| TGGGASLE | ASGDEGEVF | 0.27302676 |
| VAVYNHYKRI | ASGDEGEVF | 0.37061328 |
| TDPAPATT | ASGDEGEVF | 0.20100938 |
| TNRPDILDPALLRPG | ASGDEGEVF | 0.22475913 |
| LTTTTTS | ASGDEGEVF | 0.40847665 |
| PTGSGKTL | ASGDEGEVF | 0.7024971 |
| TGSGKTL | ASGDEGEVF | 0.45963523 |
| TIGHVDHGK | ASGDEGEVF | 0.14746834 |
| TIGHVDHGKTT | ASGDEGEVF | 0.26324773 |
| GHVDHGKTT | ASGDEGEVF | 0.27649638 |
| AQFEHTLL | ASGDEGEVF | 0.501973 |
| IVNQDEL | ASGDEGEVF | 0.6756723 |
| TNGSQFF | ASGDEGEVF | 0.8122076 |
| TGKGIVE | ASGDEGEVF | 0.50775707 |
| LISNASDA | ASGDEGEVF | 0.20789517 |
| AELRTLGELTKIAKAHGAQ | ASGDEGEVF | 0.53296196 |
| APNAVAAAEPSW | ASGDEGEVF | 0.29011565 |
| APQVAGAATAVL | ASGDEGEVF | 0.3358625 |
| ARRDPAAAAAILPTDA | ASGDEGEVF | 0.33636692 |
| DAAARDVLAEAGLAEYF | ASGDEGEVF | 0.4129253 |
| DDQIVLSAKANLAKVEQ | ASGDEGEVF | 0.259822 |
| DELLPDTGEDSDEDGH | ASGDEGEVF | 0.25345954 |
| ELPEAPANRATVPQD | ASGDEGEVF | 0.1681776 |
| FGQSKIERVTGIG | ASGDEGEVF | 0.42824972 |
| GAGGTGSTAGGAGGAGGAGGLY | ASGDEGEVF | 0.24503821 |
| GHGGTGAAGGNGGTGGAGGSGI | ASGDEGEVF | 0.3214541 |
| GNGGNGGAGGEGGPGGAGGAGA | ASGDEGEVF | 0.3042253 |
| GSTAGNGGNGGNATNGTIAGQP | ASGDEGEVF | 0.713995 |
| GTGGDGGNGGNGGTGG | ASGDEGEVF | 0.7549428 |
| GVRRPAGIPADAPV | ASGDEGEVF | 0.26415575 |
| GYRQVTDAHLLTLARR | ASGDEGEVF | 0.22222503 |
| IAKMGQDGHDRG | ASGDEGEVF | 0.6675517 |
| KAEPSIEIQYT | ASGDEGEVF | 0.22226757 |
| KPVRIRMAADEDTVAVT | ASGDEGEVF | 0.49837542 |
| MPPRIAGMRLLVIKPEP | ASGDEGEVF | 0.3624972 |
| MKFYAVALFRSRGVPPTR | ASGDEGEVF | 0.26746142 |

| | | |
|---|---|---|
| NAGGFGILGAVAHSP | ASGDEGEVF | 0.2647122 |
| NEGREITLAMAEM | ASGDEGEVF | 0.5394969 |
| NELVEWSHPLGHRWRWE | ASGDEGEVF | 0.20011954 |
| NLGAGGTGGNGGNPG | ASGDEGEVF | 0.5300501 |
| NTEWPKWETVMAADPVD | ASGDEGEVF | 0.13764428 |
| PASAAPAETQP | ASGDEGEVF | 0.51057994 |
| QQSSDGKNGVRAAEITGE | ASGDEGEVF | 0.15753886 |
| RAPDAVTTSWWKE | ASGDEGEVF | 0.27566063 |
| SGFGNVGAHTSGWFNQ | ASGDEGEVF | 0.12869485 |
| SGVAGAVGQAASVGGLKV | ASGDEGEVF | 0.25257605 |
| SNGGAGTGSAGGNGGT | ASGDEGEVF | 0.5365708 |
| TGGAGGEGGAAG | ASGDEGEVF | 0.70321465 |
| TGGDGGAAGAGGAPGHG | ASGDEGEVF | 0.5646801 |
| TMPPLLAAIANAGRHGV | ASGDEGEVF | 0.3676718 |
| TSSTAGNSGGA | ASGDEGEVF | 0.749264 |
| VPATVPVGILC | ASGDEGEVF | 0.20186163 |
| VQRPDSGGVVLTGRIS | ASGDEGEVF | 0.14084326 |
| WTPNPVPATKGVPAGLVQAT | ASGDEGEVF | 0.35197 |
| YALKRALAKFSDRFDVVII | ASGDEGEVF | 0.4076172 |
| YLIGSVIV | ASGDEGEVF | 0.65238786 |
| YGQLVAVID | ASGDEGEVF | 0.5803752 |
| VTVGGMTLPA | ASGDEGEVF | 0.13425265 |
| VLALPAV | ASGDEGEVF | 0.53241634 |
| VIPPTLNYET | ASGDEGEVF | 0.55111074 |
| VGMGGMHPSSG | ASGDEGEVF | 0.18423603 |
| VGLAGGAATA | ASGDEGEVF | 0.18857157 |
| VGGAGGAGGSGP | ASGDEGEVF | 0.2036886 |
| VFAAFMV | ASGDEGEVF | 0.21220623 |
| TLYDKLR | ASGDEGEVF | 0.38888425 |
| TLAVFTVL | ASGDEGEVF | 0.23225448 |
| TGSVQPARV | ASGDEGEVF | 0.40463173 |
| TGGGAGSGGT | ASGDEGEVF | 0.25942683 |
| SHLETSL | ASGDEGEVF | 0.26803026 |
| SFTTGMT | ASGDEGEVF | 0.2896449 |
| QVEVAKRMGVRQPTVS | ASGDEGEVF | 0.2940062 |
| QGSGGPALAM | ASGDEGEVF | 0.40485024 |
| MLAPSNP | ASGDEGEVF | 0.5831958 |
| LVNNAAI | ASGDEGEVF | 0.26386124 |
| LTGEINTLA | ASGDEGEVF | 0.36523482 |
| LSIFGPV | ASGDEGEVF | 0.58817816 |
| LLALLANL | ASGDEGEVF | 0.18298794 |
| LISNASDALDKLR | ASGDEGEVF | 0.21313998 |
| LINEAVP | ASGDEGEVF | 0.50013316 |
| LGYAFSGL | ASGDEGEVF | 0.42034772 |
| LDFVAVAL | ASGDEGEVF | 0.7264936 |
| LAQPGAE | ASGDEGEVF | 0.33057064 |
| ISNLGAVTGF | ASGDEGEVF | 0.3448704 |
| HQEDPGRI | ASGDEGEVF | 0.57763016 |
| GTPGEPTP | ASGDEGEVF | 0.6746057 |
| GFFNSGS | ASGDEGEVF | 0.40705478 |
| GAGGGGLTAGAN | ASGDEGEVF | 0.24407607 |

| | | |
|---|---|---|
| FVDALVNM | ASGDEGEVF | 0.6012914 |
| FLTVAPEL | ASGDEGEVF | 0.17834409 |
| FETLTAL | ASGDEGEVF | 0.46539798 |
| DVINPIEL | ASGDEGEVF | 0.54938734 |
| APVSGAPV | ASGDEGEVF | 0.16289285 |
| ALVGGLAW | ASGDEGEVF | 0.2794607 |
| ALAAAAATAI | ASGDEGEVF | 0.7003836 |
| AFGLGLLL | ASGDEGEVF | 0.4910776 |
| AFFAAASL | ASGDEGEVF | 0.35616916 |
| AAAALPAFSPP | ASGDEGEVF | 0.19873291 |
| VVLAPAL | ASGDEGEVF | 0.3048359 |
| TIGHVDHGKTTL | ASGDEGEVF | 0.41267782 |
| LISNASDALDK | ASGDEGEVF | 0.14947598 |
| LISNASDAL | ASGDEGEVF | 0.5487944 |
| ISNASDALDK | ASGDEGEVF | 0.24014832 |
| NQAETLVYQTE | ASGDEGEVF | 0.75817704 |
| TGGTGGTAGAG | ASGDEGEVF | 0.38753673 |
| TGSTAGGAGGA | ASGDEGEVF | 0.5076237 |
| DPLAVPVAAGPV | ASGDEGEVF | 0.26669502 |
| GGTGGTPGNGGN | ASGDEGEVF | 0.28466567 |
| PAGLAGVDPATL | ASGDEGEVF | 0.15704814 |
| GNNLVVEAPHAGASV | ASGDEGEVF | 0.28409386 |
| KVTRIMSASTLEQPA | ASGDEGEVF | 0.20600694 |
| LGFALPIALPRRTVV | ASGDEGEVF | 0.4046237 |
| TRLGNLTARKLVVLV | ASGDEGEVF | 0.1656917 |
| WSQQRRLIYFPSAGP | ASGDEGEVF | 0.70988226 |
| GPNGNPDPMAAAVDIR | ASGDEGEVF | 0.2424199 |
| TQRCVSVADLRAAASR | ASGDEGEVF | 0.3948244 |
| QFVLDVAEHPVEVLTETV | ASGDEGEVF | 0.28675783 |
| GGAGGTGGAGGNGGTQAGDG | ASGDEGEVF | 0.16855453 |
| PLLATLRAIAVDVDAKPAQV | ASGDEGEVF | 0.20437968 |
| APKFRTGGYMIDFWGVGYQVAK | ASGDEGEVF | 0.6118723 |
| GAGGAGGAGGAAGGPGGTGGTG | ASGDEGEVF | 0.15293159 |
| IGALGLLATDVAPAGSVAFGVA | ASGDEGEVF | 0.13710365 |
| AGGAGGIGGNGGAGGAGGVLMG | ASGDEGEVF | 0.21487251 |
| ADELIGLL | ASGDEGEVF | 0.29136848 |
| ADRALKA | ASGDEGEVF | 0.7652427 |
| AGAVGQVGGAGGA | ASGDEGEVF | 0.6780876 |
| AGGGGGLGGAP | ASGDEGEVF | 0.362976 |
| AGGLLYGNGGAG | ASGDEGEVF | 0.76650655 |
| AGLESIGL | ASGDEGEVF | 0.1186915 |
| AGLKTGII | ASGDEGEVF | 0.28070414 |
| ASLAALLL | ASGDEGEVF | 0.68831646 |
| ASVINISSI | ASGDEGEVF | 0.44677722 |
| ATPPGPA | ASGDEGEVF | 0.26022035 |
| ATVASALV | ASGDEGEVF | 0.56649256 |
| ATVASLIL | ASGDEGEVF | 0.21436104 |
| AVNGPVLIRH | ASGDEGEVF | 0.26332206 |
| DGAAGAVGI | ASGDEGEVF | 0.33012486 |
| DPLLHGI | ASGDEGEVF | 0.4413793 |
| ELGAALGAL | ASGDEGEVF | 0.45429322 |

| | | |
|---|---|---|
| ELGRTQAV | ASGDEGEVF | 0.14125414 |
| FVAGGYLS | ASGDEGEVF | 0.5801015 |
| GASGGAGGAGGW | ASGDEGEVF | 0.7730863 |
| GGAGAGGDGGAGGI | ASGDEGEVF | 0.20765436 |
| GTGHNGSQPGMGGNG | ASGDEGEVF | 0.6143904 |
| GYAAASASAS | ASGDEGEVF | 0.32139546 |
| HQVDVAAVQQG | ASGDEGEVF | 0.42411274 |
| IGAAIAHT | ASGDEGEVF | 0.2033267 |
| IGMIEVAQL | ASGDEGEVF | 0.14981684 |
| IRLAPEHTSAMD | ASGDEGEVF | 0.7083849 |
| IYNVSVL | ASGDEGEVF | 0.17643733 |
| KQGFLGM | ASGDEGEVF | 0.40712208 |
| LESVITQL | ASGDEGEVF | 0.35341775 |
| LGLGALVVV | ASGDEGEVF | 0.8221581 |
| LIGNGGNGGSGGI | ASGDEGEVF | 0.4810214 |
| LNAINAPTL | ASGDEGEVF | 0.35911202 |
| LQLLNTGV | ASGDEGEVF | 0.20004366 |
| LQTSIEF | ASGDEGEVF | 0.38124454 |
| MSFAVSEL | ASGDEGEVF | 0.5886973 |
| NTGALGL | ASGDEGEVF | 0.17143351 |
| PPGAGGAGGAGTT | ASGDEGEVF | 0.65717727 |
| QLVGLGI | ASGDEGEVF | 0.17858662 |
| RFLEQVHQL | ASGDEGEVF | 0.4861896 |
| RGVGYVLR | ASGDEGEVF | 0.4441071 |
| SATGVFAGVI | ASGDEGEVF | 0.78412896 |
| SLPAITLP | ASGDEGEVF | 0.29764137 |
| STALAFTV | ASGDEGEVF | 0.74237216 |
| SVLTVGLGK | ASGDEGEVF | 0.18251601 |
| TALAGAGSGI | ASGDEGEVF | 0.651765 |
| TGLPFTGI | ASGDEGEVF | 0.61939245 |
| VAEHVTV | ASGDEGEVF | 0.7129849 |
| VDIVLNSL | ASGDEGEVF | 0.78377247 |
| VLTGGGAVL | ASGDEGEVF | 0.56140006 |
| VNVGLAYGI | ASGDEGEVF | 0.15398976 |
| VSSAAVGAV | ASGDEGEVF | 0.32495257 |
| VTSVYEAGV | ASGDEGEVF | 0.606261 |
| WVALDRGA | ASGDEGEVF | 0.7481898 |
| YLVGSVVI | ASGDEGEVF | 0.4855204 |
| GPPGTGKTLL | ASGDEGEVF | 0.42434818 |
| GVGMTKF | ASGDEGEVF | 0.3420807 |
| IGIGGSDL | ASGDEGEVF | 0.72121495 |
| IGTLLAEL | ASGDEGEVF | 0.8386909 |
| IYTSGTT | ASGDEGEVF | 0.46330127 |
| LSFPAGV | ASGDEGEVF | 0.2577092 |
| LYTSGTT | ASGDEGEVF | 0.31335825 |
| VGAFPFGF | ASGDEGEVF | 0.52337515 |
| VGAGTVEF | ASGDEGEVF | 0.45309758 |
| VGKSTLLL | ASGDEGEVF | 0.19701895 |
| AALAHAGGGIV | ASGDEGEVF | 0.26609817 |
| AALVAVSLL | ASGDEGEVF | 0.3303819 |
| AAVSRAVAAL | ASGDEGEVF | 0.12635887 |

| | | |
|---|---|---|
| AGGSSGIGGF | ASGDEGEVF | 0.6318938 |
| AGLTVANANG | ASGDEGEVF | 0.6436065 |
| AGLVGAVL | ASGDEGEVF | 0.45098627 |
| ALSQAFTV | ASGDEGEVF | 0.17948437 |
| ALTFSVSL | ASGDEGEVF | 0.6601633 |
| ALTGAALVAL | ASGDEGEVF | 0.68298006 |
| ASFSIGAGVL | ASGDEGEVF | 0.6951519 |
| DGSVYKI | ASGDEGEVF | 0.17466234 |
| EQLAEIV | ASGDEGEVF | 0.11970959 |
| FAVSELTVM | ASGDEGEVF | 0.39251605 |
| GAAPLAAGL | ASGDEGEVF | 0.6458566 |
| GADGGDGGSGGAGG | ASGDEGEVF | 0.31525478 |
| GFGLDTGGAGG | ASGDEGEVF | 0.19405314 |
| GLASGGLL | ASGDEGEVF | 0.3623755 |
| GRIAGAAVLL | ASGDEGEVF | 0.60280716 |
| GSAGSLNTSF | ASGDEGEVF | 0.16951951 |
| GTGQIGI | ASGDEGEVF | 0.18334357 |
| GTLIALIL | ASGDEGEVF | 0.35157397 |
| IAATMGFVVG | ASGDEGEVF | 0.64801097 |
| IFAQALL | ASGDEGEVF | 0.16799821 |
| IGAAVALGF | ASGDEGEVF | 0.21423863 |
| IGLTPLAL | ASGDEGEVF | 0.41810638 |
| ILGSNTSTL | ASGDEGEVF | 0.715514 |
| IVFSLTVL | ASGDEGEVF | 0.44918203 |
| LASKATGF | ASGDEGEVF | 0.17819718 |
| LASVALIL | ASGDEGEVF | 0.39330974 |
| LFAVGLTL | ASGDEGEVF | 0.15593696 |
| LFNGGIGTYI | ASGDEGEVF | 0.6005259 |
| LTLGAGII | ASGDEGEVF | 0.2974846 |
| LVAAALGDI | ASGDEGEVF | 0.3974964 |
| NTAFVQLGI | ASGDEGEVF | 0.22786224 |
| QLFSGAEL | ASGDEGEVF | 0.16823608 |
| SAATFITL | ASGDEGEVF | 0.17369506 |
| SNYGLANLGV | ASGDEGEVF | 0.28807962 |
| SYGGAYAVM | ASGDEGEVF | 0.591103 |
| TTGVGGLTLG | ASGDEGEVF | 0.6607486 |
| TTQTIYI | ASGDEGEVF | 0.4314296 |
| VAAGAGLGAVI | ASGDEGEVF | 0.2850902 |
| VAAVVTLL | ASGDEGEVF | 0.17616348 |
| VGFLGGLI | ASGDEGEVF | 0.43053418 |
| VGVIPILL | ASGDEGEVF | 0.15377766 |
| VIFASSLI | ASGDEGEVF | 0.54328895 |
| VKGGDGGAAAG | ASGDEGEVF | 0.43031088 |
| VTKTFTNI | ASGDEGEVF | 0.49867308 |
| VVAAAILL | ASGDEGEVF | 0.23834497 |
| YAALDVEL | ASGDEGEVF | 0.47874808 |
| LASVALLL | ASGDEGEVF | 0.5181517 |
| LDAVHRV | ASGDEGEVF | 0.22387236 |
| QGLSSSIF | ASGDEGEVF | 0.41103196 |
| RGTFANIRL | ASGDEGEVF | 0.64871395 |
| RQLWWGHRI | ASGDEGEVF | 0.46514422 |

| | | |
|---|---|---|
| YGPPGTGKTLL | ASGDEGEVF | 0.1367129 |
| AALVGAPANVA | ASGDEGEVF | 0.34350824 |
| AATFRRTE | ASGDEGEVF | 0.21008097 |
| AGALGGGI | ASGDEGEVF | 0.43910542 |
| AGGAGGAGGIGAG | ASGDEGEVF | 0.28813362 |
| AGSGALGL | ASGDEGEVF | 0.36386287 |
| AQIADQI | ASGDEGEVF | 0.42695767 |
| ASAVAVTYL | ASGDEGEVF | 0.19439425 |
| ASTVLGGTGG | ASGDEGEVF | 0.23818284 |
| AVGGEPAK | ASGDEGEVF | 0.2951105 |
| CEQGTFGGTC | ASGDEGEVF | 0.38861793 |
| DEGKAPAG | ASGDEGEVF | 0.46849263 |
| DTVVFPAL | ASGDEGEVF | 0.16717899 |
| FEGITMAG | ASGDEGEVF | 0.49368352 |
| FPLVVQD | ASGDEGEVF | 0.3033244 |
| GAFTTPQ | ASGDEGEVF | 0.18802518 |
| GFGGFGTAGGVGGA | ASGDEGEVF | 0.38908055 |
| GFLAQFL | ASGDEGEVF | 0.26868135 |
| GGAGGAGGNAGVA | ASGDEGEVF | 0.16264585 |
| GGVGTAGAGGAGGAGG | ASGDEGEVF | 0.5036328 |
| GNGGGGDAGNAGSGGN | ASGDEGEVF | 0.21908492 |
| GTGGAGGAGNDAGS | ASGDEGEVF | 0.49127346 |
| GTTNTGWFNS | ASGDEGEVF | 0.7986928 |
| IFMGPLFTL | ASGDEGEVF | 0.1507913 |
| ILAHAIEAI | ASGDEGEVF | 0.5359256 |
| IRSAIREAL | ASGDEGEVF | 0.2600105 |
| ITFAGGG | ASGDEGEVF | 0.33754623 |
| ITGMSGHQEV | ASGDEGEVF | 0.3123356 |
| IVAGLACTL | ASGDEGEVF | 0.2065622 |
| IVGGGLAGA | ASGDEGEVF | 0.5576463 |
| IVTGVLNI | ASGDEGEVF | 0.43893087 |
| LAQQGCI | ASGDEGEVF | 0.38731664 |
| LCLHVEN | ASGDEGEVF | 0.38581964 |
| LDAISDGS | ASGDEGEVF | 0.5182019 |
| LGGFAAML | ASGDEGEVF | 0.5916395 |
| LGGKNPMI | ASGDEGEVF | 0.21355404 |
| LGPKGVAAIL | ASGDEGEVF | 0.14999591 |
| LMVGVSI | ASGDEGEVF | 0.32320738 |
| LVGAAFVL | ASGDEGEVF | 0.31974342 |
| LVLAVAPL | ASGDEGEVF | 0.19639558 |
| MAAIAVIDAA | ASGDEGEVF | 0.19243248 |
| NNGINAELQ | ASGDEGEVF | 0.54273224 |
| RGEPLVLVHG | ASGDEGEVF | 0.76167154 |
| RGGAGGAAGLI | ASGDEGEVF | 0.32028747 |
| RSYRMHPAV | ASGDEGEVF | 0.7970439 |
| SAAHLLAGQLA | ASGDEGEVF | 0.16238815 |
| SAAVGSY | ASGDEGEVF | 0.49674606 |
| SPHAVLHRI | ASGDEGEVF | 0.607875 |
| SQIAKVL | ASGDEGEVF | 0.1682162 |
| TALGHLVP | ASGDEGEVF | 0.39447376 |
| TAPTTSGAA | ASGDEGEVF | 0.19183001 |

| | | |
|---|---|---|
| TDGDDGNGGSPGT | ASGDEGEVF | 0.1831066 |
| TDGLGEN | ASGDEGEVF | 0.53234774 |
| TGGPPLPGQA | ASGDEGEVF | 0.35769257 |
| TIPAITI | ASGDEGEVF | 0.4689212 |
| TIPGFTI | ASGDEGEVF | 0.25384626 |
| TSIASGALA | ASGDEGEVF | 0.18206468 |
| TVTLPGAAI | ASGDEGEVF | 0.36467987 |
| VASGLSAALP | ASGDEGEVF | 0.5261049 |
| VGAGVIGI | ASGDEGEVF | 0.6242222 |
| VGPKRFGV | ASGDEGEVF | 0.83567464 |
| VNVAFPD | ASGDEGEVF | 0.18303093 |
| VSMTNSGVS | ASGDEGEVF | 0.73104656 |
| VTDVLGDLSV | ASGDEGEVF | 0.771863 |
| VVVVGAGFAGL | ASGDEGEVF | 0.4019454 |
| VYTVGTVV | ASGDEGEVF | 0.32784116 |
| WTTFSGP | ASGDEGEVF | 0.6289253 |
| YDELVEAF | ASGDEGEVF | 0.28109056 |
| YGTGGAGGNGG | ASGDEGEVF | 0.44784153 |
| YTNVMAV | ASGDEGEVF | 0.6492286 |
| FLEQVHQL | ASGDEGEVF | 0.7480869 |
| GLSLLAVAV | ASGDEGEVF | 0.8271295 |
| LEQVHQL | ASGDEGEVF | 0.80321467 |
| LNAAGAI | ASGDEGEVF | 0.8095781 |
| PAVHASL | ASGDEGEVF | 0.75848436 |
| TELEDDA | ASGDEGEVF | 0.5341756 |
| TPPGPPN | ASGDEGEVF | 0.8014666 |
| TPPPPPPP | ASGDEGEVF | 0.6797234 |
| TQSPALI | AWSPSGTGENYAEQF | 0.83834785 |
| EFAAAFPQ | AWSPSGTGENYAEQF | 0.83227 |
| LLSGNLDVL | AWSPSGTGENYAEQF | 0.7397027 |
| GPANLGNYNI | AWSPSGTGENYAEQF | 0.82665884 |
| TLGPTLP | AWSPSGTGENYAEQF | 0.81015617 |
| VYAYSSCCK | AWSPSGTGENYAEQF | 0.6812146 |
| APAPAPAPNI | AWSPSGTGENYAEQF | 0.78534436 |
| PGGPVPIP | AWSPSGTGENYAEQF | 0.7791711 |
| FLTVALEP | AWSPSGTGENYAEQF | 0.8116127 |
| SQYFGANL | AWSPSGTGENYAEQF | 0.6580627 |
| GGVKPAA | AWSPSGTGENYAEQF | 0.8289002 |
| GGTGGAGGAGGW | AWSPSGTGENYAEQF | 0.76728606 |
| GVGGVAGLGGAGG | AWSPSGTGENYAEQF | 0.37451738 |
| VLLVLATTI | AWSPSGTGENYAEQF | 0.82100403 |
| IGIFAAML | AWSPSGTGENYAEQF | 0.7969752 |
| AVGDLID | AWSPSGTGENYAEQF | 0.7945311 |
| NISPLPL | AWSPSGTGENYAEQF | 0.82958746 |
| VTAVFVAI | AWSPSGTGENYAEQF | 0.7515815 |
| SGDDNDAAL | AWSPSGTGENYAEQF | 0.6768593 |
| AVAALAHS | AWSPSGTGENYAEQF | 0.8130851 |
| GGVTVMATP | AWSPSGTGENYAEQF | 0.82964796 |
| AGLAANVRD | AWSPSGTGENYAEQF | 0.60665786 |
| TNRPDLIDPALLRPG | AWSPSGTGENYAEQF | 0.81375116 |
| LANLGVTAL | AWSPSGTGENYAEQF | 0.83054733 |

| | | |
|---|---|---|
| TGERIEL | AWSPSGTGENYAEQF | 0.80062664 |
| NGEELAA | AWSPSGTGENYAEQF | 0.83806574 |
| VLSLVWG | AWSPSGTGENYAEQF | 0.84045124 |
| IGYALWL | AWSPSGTGENYAEQF | 0.75358784 |
| LSLLLAPL | AWSPSGTGENYAEQF | 0.8311542 |
| TLDEALK | AWSPSGTGENYAEQF | 0.83438337 |
| TDGAGTAEGG | AWSPSGTGENYAEQF | 0.75896424 |
| NAPGGAGGNGG | AWSPSGTGENYAEQF | 0.79825 |
| VSVFMGMGGNF | AWSPSGTGENYAEQF | 0.7256942 |
| LASYLAAL | AWSPSGTGENYAEQF | 0.7970666 |
| LVGGTPAL | AWSPSGTGENYAEQF | 0.5907519 |
| VRNADVL | AWSPSGTGENYAEQF | 0.8079622 |
| TCSSPFLVSAM | AWSPSGTGENYAEQF | 0.834069 |
| VHITDAE | AWSPSGTGENYAEQF | 0.7819844 |
| EYAAPSF | AWSPSGTGENYAEQF | 0.7138276 |
| VVDAFHGGL | AWSPSGTGENYAEQF | 0.68273574 |
| ASVVDPQ | AWSPSGTGENYAEQF | 0.80041337 |
| SYNEFGGLS | AWSPSGTGENYAEQF | 0.75956714 |
| DPDGSVY | AWSPSGTGENYAEQF | 0.5841493 |
| MVSTIHATE | AWSPSGTGENYAEQF | 0.79027224 |
| LAAGVADPV | AWSPSGTGENYAEQF | 0.8237874 |
| TKEQIAA | AWSPSGTGENYAEQF | 0.8287387 |
| SLASPAAL | AWSPSGTGENYAEQF | 0.6571194 |
| IGLVFYGV | AWSPSGTGENYAEQF | 0.83616596 |
| FLLDASNL | AWSPSGTGENYAEQF | 0.8364521 |
| YQVGNFP | AWSPSGTGENYAEQF | 0.8288406 |
| ESLLSQST | AWSPSGTGENYAEQF | 0.78576773 |
| LFTVGFAAVG | AWSPSGTGENYAEQF | 0.72190464 |
| ATPPTPT | AWSPSGTGENYAEQF | 0.83632106 |
| TDLATLG | AWSPSGTGENYAEQF | 0.7571149 |
| SAGGTGR | AWSPSGTGENYAEQF | 0.800315 |
| GFSNSGS | AWSPSGTGENYAEQF | 0.8312839 |
| GGNGGSGGTSV | AWSPSGTGENYAEQF | 0.7432952 |
| GTSVATGGAGN | AWSPSGTGENYAEQF | 0.73414326 |
| NGGNGGSGGTS | AWSPSGTGENYAEQF | 0.8015239 |
| AVVKFPH | AWSPSGTGENYAEQF | 0.8399818 |
| TPPPPPP | AWSPSGTGENYAEQF | 0.70383847 |
| AAPAAAAAPA | AWSPSGTGENYAEQF | 0.68499243 |
| IALNEDL | AWSPSGTGENYAEQF | 0.7657201 |
| IVNVSSM | AWSPSGTGENYAEQF | 0.786191 |
| TGIPPAPR | AWSPSGTGENYAEQF | 0.82901573 |
| FDLGGGTF | AWSPSGTGENYAEQF | 0.82368815 |
| FDLGGGTFDVS | AWSPSGTGENYAEQF | 0.7742579 |
| NEPTAAA | AWSPSGTGENYAEQF | 0.8178302 |
| NEPTAAALA | AWSPSGTGENYAEQF | 0.8188139 |
| GSIINISSI | AWSPSGTGENYAEQF | 0.80950916 |
| VAPGNAGTA | AWSPSGTGENYAEQF | 0.8228875 |
| TGGGASLE | AWSPSGTGENYAEQF | 0.72991335 |
| VAVYNHYKRI | AWSPSGTGENYAEQF | 0.83749783 |
| TDPAPATT | AWSPSGTGENYAEQF | 0.82587373 |
| TNRPDILDPALLRPG | AWSPSGTGENYAEQF | 0.83346176 |

| | | |
|---|---|---|
| LTTTTTS | AWSPSGTGENYAEQF | 0.83636886 |
| PTGSGKTL | AWSPSGTGENYAEQF | 0.76736057 |
| TGSGKTL | AWSPSGTGENYAEQF | 0.8087486 |
| TIGHVDHGK | AWSPSGTGENYAEQF | 0.5668148 |
| TIGHVDHGKTT | AWSPSGTGENYAEQF | 0.72025836 |
| GHVDHGKTT | AWSPSGTGENYAEQF | 0.8084736 |
| AQFEHTLL | AWSPSGTGENYAEQF | 0.8071815 |
| IVNQDEL | AWSPSGTGENYAEQF | 0.82682544 |
| TNGSQFF | AWSPSGTGENYAEQF | 0.81968856 |
| TGKGIVE | AWSPSGTGENYAEQF | 0.7621927 |
| LISNASDA | AWSPSGTGENYAEQF | 0.8311382 |
| AELRTLGELTKIAKAHGAQ | AWSPSGTGENYAEQF | 0.83117306 |
| APNAVAAAEPSW | AWSPSGTGENYAEQF | 0.6799525 |
| APQVAGAATAVL | AWSPSGTGENYAEQF | 0.7770686 |
| ARRDPAAAAAILPTDA | AWSPSGTGENYAEQF | 0.84142584 |
| DAAARDVLAEAGLAEYF | AWSPSGTGENYAEQF | 0.8225543 |
| DDQIVLSAKANLAKVEQ | AWSPSGTGENYAEQF | 0.76876414 |
| DELLPDTGEDSDEDGH | AWSPSGTGENYAEQF | 0.7973249 |
| ELPEAPANRATVPQD | AWSPSGTGENYAEQF | 0.8432044 |
| FGQSKIERVTGIG | AWSPSGTGENYAEQF | 0.8116053 |
| GAGGTGSTAGGAGGAGGAGGLY | AWSPSGTGENYAEQF | 0.8382145 |
| GHGGTGAAGGNGGTGGAGGSGI | AWSPSGTGENYAEQF | 0.4856304 |
| GNGGNGGAGGEGGPGGAGGAGA | AWSPSGTGENYAEQF | 0.79376173 |
| GSTAGNGGNGGNATNGTIAGQP | AWSPSGTGENYAEQF | 0.70048374 |
| GTGGDGGNGGNGGTGG | AWSPSGTGENYAEQF | 0.44653016 |
| GVRRPAGIPADAPV | AWSPSGTGENYAEQF | 0.8272958 |
| GYRQVTDAHLLTLARR | AWSPSGTGENYAEQF | 0.7696501 |
| IAKMGQDGHDRG | AWSPSGTGENYAEQF | 0.81762314 |
| KAEPSIEIQYT | AWSPSGTGENYAEQF | 0.78672385 |
| KPVRIRMAADEDTVAVT | AWSPSGTGENYAEQF | 0.83370185 |
| MPPRIAGMRLLVIKPEP | AWSPSGTGENYAEQF | 0.7996588 |
| MKFYAVALFRSRGVPPTR | AWSPSGTGENYAEQF | 0.8080194 |
| NAGGFGILGAVAHSP | AWSPSGTGENYAEQF | 0.7825628 |
| NEGREITLAMAEM | AWSPSGTGENYAEQF | 0.8141608 |
| NELVEWSHPLGHRWRWE | AWSPSGTGENYAEQF | 0.8347094 |
| NLGAGGTGGNGGNPG | AWSPSGTGENYAEQF | 0.8363085 |
| NTEWPKWETVMAADPVD | AWSPSGTGENYAEQF | 0.45442617 |
| PASAAPAETQP | AWSPSGTGENYAEQF | 0.8200817 |
| QQSSDGKNGVRAAEITGE | AWSPSGTGENYAEQF | 0.7841554 |
| RAPDAVTTSWWKE | AWSPSGTGENYAEQF | 0.44355968 |
| SGFGNVGAHTSGWFNQ | AWSPSGTGENYAEQF | 0.6242213 |
| SGVAGAVGQAASVGGLKV | AWSPSGTGENYAEQF | 0.7564953 |
| SNGGAGTGSAGGNGGT | AWSPSGTGENYAEQF | 0.777442 |
| TGGAGGEGGAAG | AWSPSGTGENYAEQF | 0.81824756 |
| TGGDGGAAGAGGAPGHG | AWSPSGTGENYAEQF | 0.8156407 |
| TMPPLLAAIANAGRHGV | AWSPSGTGENYAEQF | 0.7784169 |
| TSSTAGNSGGA | AWSPSGTGENYAEQF | 0.81308794 |
| VPATVPVGILC | AWSPSGTGENYAEQF | 0.7721628 |
| VQRPDSGGVVLTGRIS | AWSPSGTGENYAEQF | 0.16318037 |
| WTPNPVPATKGVPAGLVQAT | AWSPSGTGENYAEQF | 0.7287589 |
| YALKRALAKFSDRFDVVII | AWSPSGTGENYAEQF | 0.70145386 |

| | | |
|---|---|---|
| YLIGSVIV | AWSPSGTGENYAEQF | 0.84030646 |
| YGQLVAVID | AWSPSGTGENYAEQF | 0.77095944 |
| VTVGGMTLPA | AWSPSGTGENYAEQF | 0.7618817 |
| VLALPAV | AWSPSGTGENYAEQF | 0.7960675 |
| VIPPTLNYET | AWSPSGTGENYAEQF | 0.8200896 |
| VGMGGMHPSSG | AWSPSGTGENYAEQF | 0.7929853 |
| VGLAGGAATA | AWSPSGTGENYAEQF | 0.72656643 |
| VGGAGGAGGSGP | AWSPSGTGENYAEQF | 0.42264807 |
| VFAAFMV | AWSPSGTGENYAEQF | 0.81592345 |
| TLYDKLR | AWSPSGTGENYAEQF | 0.6613315 |
| TLAVFTVL | AWSPSGTGENYAEQF | 0.66502786 |
| TGSVQPARV | AWSPSGTGENYAEQF | 0.72304076 |
| TGGGAGSGGT | AWSPSGTGENYAEQF | 0.6042496 |
| SHLETSL | AWSPSGTGENYAEQF | 0.5093751 |
| SFTTGMT | AWSPSGTGENYAEQF | 0.8345673 |
| QVEVAKRMGVRQPTVS | AWSPSGTGENYAEQF | 0.746956 |
| QGSGGPALAM | AWSPSGTGENYAEQF | 0.7686589 |
| MLAPSNP | AWSPSGTGENYAEQF | 0.83717406 |
| LVNNAAI | AWSPSGTGENYAEQF | 0.8105606 |
| LTGEINTLA | AWSPSGTGENYAEQF | 0.77489495 |
| LSIFGPV | AWSPSGTGENYAEQF | 0.71091545 |
| LLALLANL | AWSPSGTGENYAEQF | 0.7627501 |
| LISNASDALDKLR | AWSPSGTGENYAEQF | 0.72378135 |
| LINEAVP | AWSPSGTGENYAEQF | 0.71731126 |
| LGYAFSGL | AWSPSGTGENYAEQF | 0.8321747 |
| LDFVAVAL | AWSPSGTGENYAEQF | 0.81398237 |
| LAQPGAE | AWSPSGTGENYAEQF | 0.7073592 |
| ISNLGAVTGF | AWSPSGTGENYAEQF | 0.8206867 |
| HQEDPGRI | AWSPSGTGENYAEQF | 0.83694124 |
| GTPGEPTP | AWSPSGTGENYAEQF | 0.8380404 |
| GFFNSGS | AWSPSGTGENYAEQF | 0.7688411 |
| GAGGGGLTAGAN | AWSPSGTGENYAEQF | 0.7547574 |
| FVDALVNM | AWSPSGTGENYAEQF | 0.76538736 |
| FLTVAPEL | AWSPSGTGENYAEQF | 0.78183746 |
| FETLTAL | AWSPSGTGENYAEQF | 0.75922513 |
| DVINPIEL | AWSPSGTGENYAEQF | 0.69166625 |
| APVSGAPV | AWSPSGTGENYAEQF | 0.8337213 |
| ALVGGLAW | AWSPSGTGENYAEQF | 0.8474604 |
| ALAAAAATAI | AWSPSGTGENYAEQF | 0.78653204 |
| AFGLGLLL | AWSPSGTGENYAEQF | 0.711174 |
| AFFAAASL | AWSPSGTGENYAEQF | 0.7816576 |
| AAAALPAFSPP | AWSPSGTGENYAEQF | 0.70871735 |
| VVLAPAL | AWSPSGTGENYAEQF | 0.75881267 |
| TIGHVDHGKTTL | AWSPSGTGENYAEQF | 0.74835813 |
| LISNASDALDK | AWSPSGTGENYAEQF | 0.6523519 |
| LISNASDAL | AWSPSGTGENYAEQF | 0.78649426 |
| ISNASDALDK | AWSPSGTGENYAEQF | 0.83575904 |
| NQAETLVYQTE | AWSPSGTGENYAEQF | 0.7848327 |
| TGGTGGTAGAG | AWSPSGTGENYAEQF | 0.80916846 |
| TGSTAGGAGGA | AWSPSGTGENYAEQF | 0.83398664 |
| DPLAVPVAAGPV | AWSPSGTGENYAEQF | 0.8239895 |

| | | |
|---|---|---|
| GGTGGTPGNGGN | AWSPSGTGENYAEQF | 0.8375833 |
| PAGLAGVDPATL | AWSPSGTGENYAEQF | 0.83906925 |
| GNNLVVEAPHAGASV | AWSPSGTGENYAEQF | 0.8201256 |
| KVTRIMSASTLEQPA | AWSPSGTGENYAEQF | 0.7905522 |
| LGFALPIALPRRTVV | AWSPSGTGENYAEQF | 0.8218997 |
| TRLGNLTARKLVVLV | AWSPSGTGENYAEQF | 0.79254895 |
| WSQQRRLIYFPSAGP | AWSPSGTGENYAEQF | 0.7906663 |
| GPNGNPDPMAAAVDIR | AWSPSGTGENYAEQF | 0.7816681 |
| TQRCVSVADLRAAASR | AWSPSGTGENYAEQF | 0.8255208 |
| QFVLDVAEHPVEVLTETV | AWSPSGTGENYAEQF | 0.77858365 |
| GGAGGTGGAGGNGGTQAGDG | AWSPSGTGENYAEQF | 0.84246486 |
| PLLATLRAIAVDVDAKPAQV | AWSPSGTGENYAEQF | 0.48618913 |
| APKFRTGGYMIDFWGVGYQVAK | AWSPSGTGENYAEQF | 0.6791161 |
| GAGGAGGAGGAAGGPGGTGGTG | AWSPSGTGENYAEQF | 0.8284637 |
| IGALGLLATDVAPAGSVAFGVA | AWSPSGTGENYAEQF | 0.20213947 |
| AGGAGGIGGNGGAGGAGGVLMG | AWSPSGTGENYAEQF | 0.64982915 |
| ADELIGLL | AWSPSGTGENYAEQF | 0.8085439 |
| ADRALKA | AWSPSGTGENYAEQF | 0.8261967 |
| AGAVGQVGGAGGA | AWSPSGTGENYAEQF | 0.23578346 |
| AGGGGGLGGAP | AWSPSGTGENYAEQF | 0.70432216 |
| AGGLLYGNGGAG | AWSPSGTGENYAEQF | 0.6298063 |
| AGLESIGL | AWSPSGTGENYAEQF | 0.82690686 |
| AGLKTGII | AWSPSGTGENYAEQF | 0.79044473 |
| ASLAALLL | AWSPSGTGENYAEQF | 0.56481004 |
| ASVINISSI | AWSPSGTGENYAEQF | 0.8332331 |
| ATPPGPA | AWSPSGTGENYAEQF | 0.6708317 |
| ATVASALV | AWSPSGTGENYAEQF | 0.78318346 |
| ATVASLIL | AWSPSGTGENYAEQF | 0.83214945 |
| AVNGPVLIRH | AWSPSGTGENYAEQF | 0.8432101 |
| DGAAGAVGI | AWSPSGTGENYAEQF | 0.7389703 |
| DPLLHGI | AWSPSGTGENYAEQF | 0.81793505 |
| ELGAALGAL | AWSPSGTGENYAEQF | 0.7665932 |
| ELGRTQAV | AWSPSGTGENYAEQF | 0.8401917 |
| FVAGGYLS | AWSPSGTGENYAEQF | 0.7672833 |
| GASGGAGGAGGW | AWSPSGTGENYAEQF | 0.6979235 |
| GGAGAGGDGGAGGI | AWSPSGTGENYAEQF | 0.8304894 |
| GTGHNGSQPGMGGNG | AWSPSGTGENYAEQF | 0.8256587 |
| GYAAASASAS | AWSPSGTGENYAEQF | 0.77548885 |
| HQVDVAAVQQG | AWSPSGTGENYAEQF | 0.8361939 |
| IGAAIAHT | AWSPSGTGENYAEQF | 0.7833752 |
| IGMIEVAQL | AWSPSGTGENYAEQF | 0.7823209 |
| IRLAPEHTSAMD | AWSPSGTGENYAEQF | 0.8276542 |
| IYNVSVL | AWSPSGTGENYAEQF | 0.8295163 |
| KQGFLGM | AWSPSGTGENYAEQF | 0.66316825 |
| LESVITQL | AWSPSGTGENYAEQF | 0.8344426 |
| LGLGALVVV | AWSPSGTGENYAEQF | 0.7776465 |
| LIGNGGNGGSGGI | AWSPSGTGENYAEQF | 0.72097707 |
| LNAINAPTL | AWSPSGTGENYAEQF | 0.83835185 |
| LQLLNTGV | AWSPSGTGENYAEQF | 0.80938125 |
| LQTSIEF | AWSPSGTGENYAEQF | 0.82302254 |
| MSFAVSEL | AWSPSGTGENYAEQF | 0.82441676 |

| | | |
|---|---|---|
| NTGALGL | AWSPSGTGENYAEQF | 0.84134716 |
| PPGAGGAGGAGTT | AWSPSGTGENYAEQF | 0.66780776 |
| QLVGLGI | AWSPSGTGENYAEQF | 0.8064602 |
| RFLEQVHQL | AWSPSGTGENYAEQF | 0.7890464 |
| RGVGYVLR | AWSPSGTGENYAEQF | 0.83096886 |
| SATGVFAGVI | AWSPSGTGENYAEQF | 0.8104981 |
| SLPAITLP | AWSPSGTGENYAEQF | 0.75529146 |
| STALAFTV | AWSPSGTGENYAEQF | 0.8290658 |
| SVLTVGLGK | AWSPSGTGENYAEQF | 0.8315291 |
| TALAGAGSGI | AWSPSGTGENYAEQF | 0.81524867 |
| TGLPFTGI | AWSPSGTGENYAEQF | 0.8123771 |
| VAEHVTV | AWSPSGTGENYAEQF | 0.84005284 |
| VDIVLNSL | AWSPSGTGENYAEQF | 0.7535496 |
| VLTGGGAVL | AWSPSGTGENYAEQF | 0.78400683 |
| VNVGLAYGI | AWSPSGTGENYAEQF | 0.8294977 |
| VSSAAVGAV | AWSPSGTGENYAEQF | 0.8301287 |
| VTSVYEAGV | AWSPSGTGENYAEQF | 0.80840087 |
| WVALDRGA | AWSPSGTGENYAEQF | 0.8377358 |
| YLVGSVVI | AWSPSGTGENYAEQF | 0.72828335 |
| GPPGTGKTLL | AWSPSGTGENYAEQF | 0.8148366 |
| GVGMTKF | AWSPSGTGENYAEQF | 0.7207471 |
| IGIGGSDL | AWSPSGTGENYAEQF | 0.78474295 |
| IGTLLAEL | AWSPSGTGENYAEQF | 0.68133384 |
| IYTSGTT | AWSPSGTGENYAEQF | 0.8155417 |
| LSFPAGV | AWSPSGTGENYAEQF | 0.8212986 |
| LYTSGTT | AWSPSGTGENYAEQF | 0.81877494 |
| VGAFPFGF | AWSPSGTGENYAEQF | 0.48636788 |
| VGAGTVEF | AWSPSGTGENYAEQF | 0.8193841 |
| VGKSTLLL | AWSPSGTGENYAEQF | 0.82231176 |
| AALAHAGGGIV | AWSPSGTGENYAEQF | 0.8455509 |
| AALVAVSLL | AWSPSGTGENYAEQF | 0.783934 |
| AAVSRAVAAL | AWSPSGTGENYAEQF | 0.8128052 |
| AGGSSGIGGF | AWSPSGTGENYAEQF | 0.73489785 |
| AGLTVANANG | AWSPSGTGENYAEQF | 0.81633663 |
| AGLVGAVL | AWSPSGTGENYAEQF | 0.68677986 |
| ALSQAFTV | AWSPSGTGENYAEQF | 0.72053975 |
| ALTFSVSL | AWSPSGTGENYAEQF | 0.77155626 |
| ALTGAALVAL | AWSPSGTGENYAEQF | 0.7488746 |
| ASFSIGAGVL | AWSPSGTGENYAEQF | 0.826934 |
| DGSVYKI | AWSPSGTGENYAEQF | 0.8326796 |
| EQLAEIV | AWSPSGTGENYAEQF | 0.81988573 |
| FAVSELTVM | AWSPSGTGENYAEQF | 0.8244784 |
| GAAPLAAGL | AWSPSGTGENYAEQF | 0.8328441 |
| GADGGDGGSGGAGG | AWSPSGTGENYAEQF | 0.8380739 |
| GFGLDTGGAGG | AWSPSGTGENYAEQF | 0.75751925 |
| GLASGGLL | AWSPSGTGENYAEQF | 0.78581107 |
| GRIAGAAVLL | AWSPSGTGENYAEQF | 0.69466114 |
| GSAGSLNTSF | AWSPSGTGENYAEQF | 0.7841939 |
| GTGQIGI | AWSPSGTGENYAEQF | 0.74963737 |
| GTLIALIL | AWSPSGTGENYAEQF | 0.80902743 |
| IAATMGFVVG | AWSPSGTGENYAEQF | 0.18796492 |

| | | |
|---|---|---|
| IFAQALL | AWSPSGTGENYAEQF | 0.8179021 |
| IGAAVALGF | AWSPSGTGENYAEQF | 0.8328694 |
| IGLTPLAL | AWSPSGTGENYAEQF | 0.71149206 |
| ILGSNTSTL | AWSPSGTGENYAEQF | 0.8148762 |
| IVFSLTVL | AWSPSGTGENYAEQF | 0.810529 |
| LASKATGF | AWSPSGTGENYAEQF | 0.767933 |
| LASVALIL | AWSPSGTGENYAEQF | 0.80856025 |
| LFAVGLTL | AWSPSGTGENYAEQF | 0.7342066 |
| LFNGGIGTYI | AWSPSGTGENYAEQF | 0.82980514 |
| LTLGAGII | AWSPSGTGENYAEQF | 0.83634573 |
| LVAAALGDI | AWSPSGTGENYAEQF | 0.8075568 |
| NTAFVQLGI | AWSPSGTGENYAEQF | 0.77831864 |
| QLFSGAEL | AWSPSGTGENYAEQF | 0.80940616 |
| SAATFITL | AWSPSGTGENYAEQF | 0.57649183 |
| SNYGLANLGV | AWSPSGTGENYAEQF | 0.8016936 |
| SYGGAYAVM | AWSPSGTGENYAEQF | 0.2858328 |
| TTGVGGLTLG | AWSPSGTGENYAEQF | 0.7944383 |
| TTQTIYI | AWSPSGTGENYAEQF | 0.73338807 |
| VAAGAGLGAVI | AWSPSGTGENYAEQF | 0.80913985 |
| VAAVVTLL | AWSPSGTGENYAEQF | 0.7792798 |
| VGFLGGLI | AWSPSGTGENYAEQF | 0.81871283 |
| VGVIPILL | AWSPSGTGENYAEQF | 0.72805345 |
| VIFASSLI | AWSPSGTGENYAEQF | 0.82276064 |
| VKGGDGGAAAG | AWSPSGTGENYAEQF | 0.7962769 |
| VTKTFTNI | AWSPSGTGENYAEQF | 0.58100903 |
| VVAAAILL | AWSPSGTGENYAEQF | 0.8296247 |
| YAALDVEL | AWSPSGTGENYAEQF | 0.80619544 |
| LASVALLL | AWSPSGTGENYAEQF | 0.7774076 |
| LDAVHRV | AWSPSGTGENYAEQF | 0.82757056 |
| QGLSSSIF | AWSPSGTGENYAEQF | 0.80942535 |
| RGTFANIRL | AWSPSGTGENYAEQF | 0.6627367 |
| RQLWWGHRI | AWSPSGTGENYAEQF | 0.76464725 |
| YGPPGTGKTLL | AWSPSGTGENYAEQF | 0.7903526 |
| AALVGAPANVA | AWSPSGTGENYAEQF | 0.77312565 |
| AATFRRTE | AWSPSGTGENYAEQF | 0.26021323 |
| AGALGGGI | AWSPSGTGENYAEQF | 0.8286525 |
| AGGAGGAGGIGAG | AWSPSGTGENYAEQF | 0.71357775 |
| AGSGALGL | AWSPSGTGENYAEQF | 0.72781813 |
| AQIADQI | AWSPSGTGENYAEQF | 0.77876884 |
| ASAVAVTYL | AWSPSGTGENYAEQF | 0.82820666 |
| ASTVLGGTGG | AWSPSGTGENYAEQF | 0.71480465 |
| AVGGEPAK | AWSPSGTGENYAEQF | 0.790949 |
| CEQGTFGGTC | AWSPSGTGENYAEQF | 0.82218474 |
| DEGKAPAG | AWSPSGTGENYAEQF | 0.7939135 |
| DTVVFPAL | AWSPSGTGENYAEQF | 0.8259728 |
| FEGITMAG | AWSPSGTGENYAEQF | 0.80032456 |
| FPLVVQD | AWSPSGTGENYAEQF | 0.7874844 |
| GAFTTPQ | AWSPSGTGENYAEQF | 0.8109338 |
| GFGGFGTAGGVGGA | AWSPSGTGENYAEQF | 0.76028866 |
| GFLAQFL | AWSPSGTGENYAEQF | 0.78633785 |
| GGAGGAGGNAGVA | AWSPSGTGENYAEQF | 0.7323108 |

| | | |
|---|---|---|
| GGVGTAGAGGAGGAGG | AWSPSGTGENYAEQF | 0.7668929 |
| GNGGGGDAGNAGSGGN | AWSPSGTGENYAEQF | 0.7366345 |
| GTGGAGGAGNDAGS | AWSPSGTGENYAEQF | 0.8032111 |
| GTTNTGWFNS | AWSPSGTGENYAEQF | 0.84035313 |
| IFMGPLFTL | AWSPSGTGENYAEQF | 0.7953687 |
| ILAHAIEAI | AWSPSGTGENYAEQF | 0.65217495 |
| IRSAIREAL | AWSPSGTGENYAEQF | 0.7755959 |
| ITFAGGG | AWSPSGTGENYAEQF | 0.7812427 |
| ITGMSGHQEV | AWSPSGTGENYAEQF | 0.8242357 |
| IVAGLACTL | AWSPSGTGENYAEQF | 0.82163024 |
| IVGGGLAGA | AWSPSGTGENYAEQF | 0.8459202 |
| IVTGVLNI | AWSPSGTGENYAEQF | 0.82351106 |
| LAQQGCI | AWSPSGTGENYAEQF | 0.7385819 |
| LCLHVEN | AWSPSGTGENYAEQF | 0.8281623 |
| LDAISDGS | AWSPSGTGENYAEQF | 0.78804064 |
| LGGFAAML | AWSPSGTGENYAEQF | 0.79574156 |
| LGGKNPMI | AWSPSGTGENYAEQF | 0.81910765 |
| LGPKGVAAIL | AWSPSGTGENYAEQF | 0.81954455 |
| LMVGVSI | AWSPSGTGENYAEQF | 0.77266496 |
| LVGAAFVL | AWSPSGTGENYAEQF | 0.80117404 |
| LVLAVAPL | AWSPSGTGENYAEQF | 0.62175 |
| MAAIAVIDAA | AWSPSGTGENYAEQF | 0.83603406 |
| NNGINAELQ | AWSPSGTGENYAEQF | 0.63859266 |
| RGEPLVLVHG | AWSPSGTGENYAEQF | 0.7568533 |
| RGGAGGAAGLI | AWSPSGTGENYAEQF | 0.77628833 |
| RSYRMHPAV | AWSPSGTGENYAEQF | 0.8369062 |
| SAAHLLAGQLA | AWSPSGTGENYAEQF | 0.827075 |
| SAAVGSY | AWSPSGTGENYAEQF | 0.7395625 |
| SPHAVLHRI | AWSPSGTGENYAEQF | 0.7106457 |
| SQIAKVL | AWSPSGTGENYAEQF | 0.8139564 |
| TALGHLVP | AWSPSGTGENYAEQF | 0.6655599 |
| TAPTTSGAA | AWSPSGTGENYAEQF | 0.82915103 |
| TDGDDGNGGSPGT | AWSPSGTGENYAEQF | 0.7829772 |
| TDGLGEN | AWSPSGTGENYAEQF | 0.7699765 |
| TGGPPLPGQA | AWSPSGTGENYAEQF | 0.7443394 |
| TIPAITI | AWSPSGTGENYAEQF | 0.8170701 |
| TIPGFTI | AWSPSGTGENYAEQF | 0.7439975 |
| TSIASGALA | AWSPSGTGENYAEQF | 0.8302275 |
| TVTLPGAAI | AWSPSGTGENYAEQF | 0.320719 |
| VASGLSAALP | AWSPSGTGENYAEQF | 0.82661104 |
| VGAGVIGI | AWSPSGTGENYAEQF | 0.7161881 |
| VGPKRFGV | AWSPSGTGENYAEQF | 0.72832835 |
| VNVAFPD | AWSPSGTGENYAEQF | 0.53597903 |
| VSMTNSGVS | AWSPSGTGENYAEQF | 0.8070364 |
| VTDVLGDLSV | AWSPSGTGENYAEQF | 0.8114471 |
| VVVVGAGFAGL | AWSPSGTGENYAEQF | 0.8319796 |
| VYTVGTVV | AWSPSGTGENYAEQF | 0.810749 |
| WTTFSGP | AWSPSGTGENYAEQF | 0.8203338 |
| YDELVEAF | AWSPSGTGENYAEQF | 0.82492185 |
| YGTGGAGGNGG | AWSPSGTGENYAEQF | 0.76794034 |
| YTNVMAV | AWSPSGTGENYAEQF | 0.5035028 |

| | | |
|---|---|---|
| FLEQVHQL | AWSPSGTGENYAEQF | 0.8006122 |
| GLSLLAVAV | AWSPSGTGENYAEQF | 0.81406784 |
| LEQVHQL | AWSPSGTGENYAEQF | 0.8331077 |
| LNAAGAI | AWSPSGTGENYAEQF | 0.80305326 |
| PAVHASL | AWSPSGTGENYAEQF | 0.7060188 |
| TELEDDA | AWSPSGTGENYAEQF | 0.8130338 |
| TPPGPPN | AWSPSGTGENYAEQF | 0.8214549 |
| TPPPPPPP | AWSPSGTGENYAEQF | 0.6781004 |
| TQSPALI | ASSPPTNSGNTLY | 0.7709291 |
| EFAAAFPQ | ASSPPTNSGNTLY | 0.7712718 |
| LLSGNLDVL | ASSPPTNSGNTLY | 0.6129185 |
| GPANLGNYNI | ASSPPTNSGNTLY | 0.8087801 |
| TLGPTLP | ASSPPTNSGNTLY | 0.752383 |
| VYAYSSCCK | ASSPPTNSGNTLY | 0.7318927 |
| APAPAPAPNI | ASSPPTNSGNTLY | 0.8052083 |
| PGGPVPIP | ASSPPTNSGNTLY | 0.73177564 |
| FLTVALEP | ASSPPTNSGNTLY | 0.74288225 |
| SQYFGANL | ASSPPTNSGNTLY | 0.73450696 |
| GGVKPAA | ASSPPTNSGNTLY | 0.8079583 |
| GGTGGAGGAGGW | ASSPPTNSGNTLY | 0.8225422 |
| GVGGVAGLGGAGG | ASSPPTNSGNTLY | 0.7954186 |
| VLLVLATTI | ASSPPTNSGNTLY | 0.8139587 |
| IGIFAAML | ASSPPTNSGNTLY | 0.8306712 |
| AVGDLID | ASSPPTNSGNTLY | 0.5488385 |
| NISPLPL | ASSPPTNSGNTLY | 0.7180485 |
| VTAVFVAI | ASSPPTNSGNTLY | 0.8217279 |
| SGDDNDAAL | ASSPPTNSGNTLY | 0.70675594 |
| AVAALAHS | ASSPPTNSGNTLY | 0.83191645 |
| GGVTVMATP | ASSPPTNSGNTLY | 0.77537334 |
| AGLAANVRD | ASSPPTNSGNTLY | 0.8135071 |
| TNRPDLIDPALLRPG | ASSPPTNSGNTLY | 0.8036274 |
| LANLGVTAL | ASSPPTNSGNTLY | 0.83590585 |
| TGERIEL | ASSPPTNSGNTLY | 0.8255412 |
| NGEELAA | ASSPPTNSGNTLY | 0.803892 |
| VLSLVWG | ASSPPTNSGNTLY | 0.83080924 |
| IGYALWL | ASSPPTNSGNTLY | 0.65595675 |
| LSLLLAPL | ASSPPTNSGNTLY | 0.8236779 |
| TLDEALK | ASSPPTNSGNTLY | 0.82311773 |
| TDGAGTAEGG | ASSPPTNSGNTLY | 0.6897754 |
| NAPGGAGGNGG | ASSPPTNSGNTLY | 0.78471357 |
| VSVFMGMGGNF | ASSPPTNSGNTLY | 0.83792746 |
| LASYLAAL | ASSPPTNSGNTLY | 0.8346895 |
| LVGGTPAL | ASSPPTNSGNTLY | 0.7286923 |
| VRNADVL | ASSPPTNSGNTLY | 0.7856755 |
| TCSSPFLVSAM | ASSPPTNSGNTLY | 0.5712253 |
| VHITDAE | ASSPPTNSGNTLY | 0.6894009 |
| EYAAPSF | ASSPPTNSGNTLY | 0.5313632 |
| VVDAFHGGL | ASSPPTNSGNTLY | 0.820794 |
| ASVVDPQ | ASSPPTNSGNTLY | 0.75929296 |
| SYNEFGGLS | ASSPPTNSGNTLY | 0.8353859 |
| DPDGSVY | ASSPPTNSGNTLY | 0.80812335 |

| | | |
|---|---|---|
| MVSTIHATE | ASSPPTNSGNTLY | 0.83421916 |
| LAAGVADPV | ASSPPTNSGNTLY | 0.7536813 |
| TKEQIAA | ASSPPTNSGNTLY | 0.74871325 |
| SLASPAAL | ASSPPTNSGNTLY | 0.6455816 |
| IGLVFYGV | ASSPPTNSGNTLY | 0.5491133 |
| FLLDASNL | ASSPPTNSGNTLY | 0.7684185 |
| YQVGNFP | ASSPPTNSGNTLY | 0.7487179 |
| ESLLSQST | ASSPPTNSGNTLY | 0.34637776 |
| LFTVGFAAVG | ASSPPTNSGNTLY | 0.8358375 |
| ATPPTPT | ASSPPTNSGNTLY | 0.8047352 |
| TDLATLG | ASSPPTNSGNTLY | 0.8183359 |
| SAGGTGR | ASSPPTNSGNTLY | 0.726936 |
| GFSNSGS | ASSPPTNSGNTLY | 0.7587359 |
| GGNGGSGGTSV | ASSPPTNSGNTLY | 0.7440382 |
| GTSVATGGAGN | ASSPPTNSGNTLY | 0.8168255 |
| NGGNGGSGGTS | ASSPPTNSGNTLY | 0.7501699 |
| AVVKFPH | ASSPPTNSGNTLY | 0.6485331 |
| TPPPPPP | ASSPPTNSGNTLY | 0.7381364 |
| AAPAAAAAPA | ASSPPTNSGNTLY | 0.76102 |
| IALNEDL | ASSPPTNSGNTLY | 0.83814013 |
| IVNVSSM | ASSPPTNSGNTLY | 0.8270369 |
| TGIPPAPR | ASSPPTNSGNTLY | 0.81285584 |
| FDLGGGTF | ASSPPTNSGNTLY | 0.8412825 |
| FDLGGGTFDVS | ASSPPTNSGNTLY | 0.2669491 |
| NEPTAAA | ASSPPTNSGNTLY | 0.74354947 |
| NEPTAAALA | ASSPPTNSGNTLY | 0.8291501 |
| GSIINISSI | ASSPPTNSGNTLY | 0.8155863 |
| VAPGNAGTA | ASSPPTNSGNTLY | 0.7738559 |
| TGGGASLE | ASSPPTNSGNTLY | 0.8342618 |
| VAVYNHYKRI | ASSPPTNSGNTLY | 0.68979526 |
| TDPAPATT | ASSPPTNSGNTLY | 0.7705275 |
| TNRPDILDPALLRPG | ASSPPTNSGNTLY | 0.80687356 |
| LTTTTTS | ASSPPTNSGNTLY | 0.8344668 |
| PTGSGKTL | ASSPPTNSGNTLY | 0.81317246 |
| TGSGKTL | ASSPPTNSGNTLY | 0.72833014 |
| TIGHVDHGK | ASSPPTNSGNTLY | 0.81734943 |
| TIGHVDHGKTT | ASSPPTNSGNTLY | 0.84090006 |
| GHVDHGKTT | ASSPPTNSGNTLY | 0.7749891 |
| AQFEHTLL | ASSPPTNSGNTLY | 0.82234746 |
| IVNQDEL | ASSPPTNSGNTLY | 0.8089447 |
| TNGSQFF | ASSPPTNSGNTLY | 0.7577489 |
| TGKGIVE | ASSPPTNSGNTLY | 0.73531806 |
| LISNASDA | ASSPPTNSGNTLY | 0.74377406 |
| AELRTLGELTKIAKAHGAQ | ASSPPTNSGNTLY | 0.8251976 |
| APNAVAAAEPSW | ASSPPTNSGNTLY | 0.7117276 |
| APQVAGAATAVL | ASSPPTNSGNTLY | 0.79413414 |
| ARRDPAAAAAILPTDA | ASSPPTNSGNTLY | 0.7985079 |
| DAAARDVLAEAGLAEYF | ASSPPTNSGNTLY | 0.84147847 |
| DDQIVLSAKANLAKVEQ | ASSPPTNSGNTLY | 0.816074 |
| DELLPDTGEDSDEDGH | ASSPPTNSGNTLY | 0.816234 |
| ELPEAPANRATVPQD | ASSPPTNSGNTLY | 0.7191639 |

| | | |
|---|---|---|
| FGQSKIERVTGIG | ASSPPTNSGNTLY | 0.8194781 |
| GAGGTGSTAGGAGGAGGAGGLY | ASSPPTNSGNTLY | 0.81266546 |
| GHGGTGAAGGNGGTGGAGGSGI | ASSPPTNSGNTLY | 0.6628201 |
| GNGGNGGAGGEGGPGGAGGAGA | ASSPPTNSGNTLY | 0.826788 |
| GSTAGNGGNGGNATNGTIAGQP | ASSPPTNSGNTLY | 0.6895951 |
| GTGGDGGNGGNGGTGG | ASSPPTNSGNTLY | 0.78380734 |
| GVRRPAGIPADAPV | ASSPPTNSGNTLY | 0.81358445 |
| GYRQVTDAHLLTLARR | ASSPPTNSGNTLY | 0.8435295 |
| IAKMGQDGHDRG | ASSPPTNSGNTLY | 0.79587245 |
| KAEPSIEIQYT | ASSPPTNSGNTLY | 0.82468116 |
| KPVRIRMAADEDTVAVT | ASSPPTNSGNTLY | 0.71458286 |
| MPPRIAGMRLLVIKPEP | ASSPPTNSGNTLY | 0.8293015 |
| MKFYAVALFRSRGVPPTR | ASSPPTNSGNTLY | 0.83356965 |
| NAGGFGILGAVAHSP | ASSPPTNSGNTLY | 0.8097459 |
| NEGREITLAMAEM | ASSPPTNSGNTLY | 0.84499836 |
| NELVEWSHPLGHRWRWE | ASSPPTNSGNTLY | 0.8216826 |
| NLGAGGTGGNGGNPG | ASSPPTNSGNTLY | 0.4524451 |
| NTEWPKWETVMAADPVD | ASSPPTNSGNTLY | 0.8069408 |
| PASAAPAETQP | ASSPPTNSGNTLY | 0.83238965 |
| QQSSDGKNGVRAAEITGE | ASSPPTNSGNTLY | 0.8286776 |
| RAPDAVTTSWWKE | ASSPPTNSGNTLY | 0.83391577 |
| SGFGNVGAHTSGWFNQ | ASSPPTNSGNTLY | 0.81875235 |
| SGVAGAVGQAASVGGLKV | ASSPPTNSGNTLY | 0.7697448 |
| SNGGAGTGSAGGNGGT | ASSPPTNSGNTLY | 0.83393264 |
| TGGAGGEGGAAG | ASSPPTNSGNTLY | 0.840236 |
| TGGDGGAAGAGGAPGHG | ASSPPTNSGNTLY | 0.8226632 |
| TMPPLLAAIANAGRHGV | ASSPPTNSGNTLY | 0.7620446 |
| TSSTAGNSGGA | ASSPPTNSGNTLY | 0.6856209 |
| VPATVPVGILC | ASSPPTNSGNTLY | 0.81581986 |
| VQRPDSGGVVLTGRIS | ASSPPTNSGNTLY | 0.8261137 |
| WTPNPVPATKGVPAGLVQAT | ASSPPTNSGNTLY | 0.70384467 |
| YALKRALAKFSDRFDVVII | ASSPPTNSGNTLY | 0.79864335 |
| YLIGSVIV | ASSPPTNSGNTLY | 0.7970549 |
| YGQLVAVID | ASSPPTNSGNTLY | 0.4874639 |
| VTVGGMTLPA | ASSPPTNSGNTLY | 0.8117403 |
| VLALPAV | ASSPPTNSGNTLY | 0.8311655 |
| VIPPTLNYET | ASSPPTNSGNTLY | 0.822016 |
| VGMGGMHPSSG | ASSPPTNSGNTLY | 0.7252158 |
| VGLAGGAATA | ASSPPTNSGNTLY | 0.8261844 |
| VGGAGGAGGSGP | ASSPPTNSGNTLY | 0.8371054 |
| VFAAFMV | ASSPPTNSGNTLY | 0.8412717 |
| TLYDKLR | ASSPPTNSGNTLY | 0.5633944 |
| TLAVFTVL | ASSPPTNSGNTLY | 0.4041454 |
| TGSVQPARV | ASSPPTNSGNTLY | 0.7852787 |
| TGGGAGSGGT | ASSPPTNSGNTLY | 0.82797104 |
| SHLETSL | ASSPPTNSGNTLY | 0.74807906 |
| SFTTGMT | ASSPPTNSGNTLY | 0.75640357 |
| QVEVAKRMGVRQPTVS | ASSPPTNSGNTLY | 0.80786514 |
| QGSGGPALAM | ASSPPTNSGNTLY | 0.82916546 |
| MLAPSNP | ASSPPTNSGNTLY | 0.7274867 |
| LVNNAAI | ASSPPTNSGNTLY | 0.628813 |

| | | |
|---|---|---|
| LTGEINTLA | ASSPPTNSGNTLY | 0.8214379 |
| LSIFGPV | ASSPPTNSGNTLY | 0.8296082 |
| LLALLANL | ASSPPTNSGNTLY | 0.7369244 |
| LISNASDALDKLR | ASSPPTNSGNTLY | 0.7895856 |
| LINEAVP | ASSPPTNSGNTLY | 0.77579933 |
| LGYAFSGL | ASSPPTNSGNTLY | 0.790678 |
| LDFVAVAL | ASSPPTNSGNTLY | 0.7973535 |
| LAQPGAE | ASSPPTNSGNTLY | 0.6600912 |
| ISNLGAVTGF | ASSPPTNSGNTLY | 0.8296281 |
| HQEDPGRI | ASSPPTNSGNTLY | 0.7011837 |
| GTPGEPTP | ASSPPTNSGNTLY | 0.8284329 |
| GFFNSGS | ASSPPTNSGNTLY | 0.80517685 |
| GAGGGGLTAGAN | ASSPPTNSGNTLY | 0.82272196 |
| FVDALVNM | ASSPPTNSGNTLY | 0.7696357 |
| FLTVAPEL | ASSPPTNSGNTLY | 0.752518 |
| FETLTAL | ASSPPTNSGNTLY | 0.6860924 |
| DVINPIEL | ASSPPTNSGNTLY | 0.66176796 |
| APVSGAPV | ASSPPTNSGNTLY | 0.8306407 |
| ALVGGLAW | ASSPPTNSGNTLY | 0.4314995 |
| ALAAAAATAI | ASSPPTNSGNTLY | 0.8340726 |
| AFGLGLLL | ASSPPTNSGNTLY | 0.8127023 |
| AFFAAASL | ASSPPTNSGNTLY | 0.76455736 |
| AAAALPAFSPP | ASSPPTNSGNTLY | 0.8262686 |
| VVLAPAL | ASSPPTNSGNTLY | 0.63164973 |
| TIGHVDHGKTTL | ASSPPTNSGNTLY | 0.8222954 |
| LISNASDALDK | ASSPPTNSGNTLY | 0.78507173 |
| LISNASDAL | ASSPPTNSGNTLY | 0.8295835 |
| ISNASDALDK | ASSPPTNSGNTLY | 0.74457747 |
| NQAETLVYQTE | ASSPPTNSGNTLY | 0.8244659 |
| TGGTGGTAGAG | ASSPPTNSGNTLY | 0.8138839 |
| TGSTAGGAGGA | ASSPPTNSGNTLY | 0.67236954 |
| DPLAVPVAAGPV | ASSPPTNSGNTLY | 0.82030654 |
| GGTGGTPGNGGN | ASSPPTNSGNTLY | 0.8275101 |
| PAGLAGVDPATL | ASSPPTNSGNTLY | 0.82443106 |
| GNNLVVEAPHAGASV | ASSPPTNSGNTLY | 0.7111083 |
| KVTRIMSASTLEQPA | ASSPPTNSGNTLY | 0.8030592 |
| LGFALPIALPRRTVV | ASSPPTNSGNTLY | 0.75597084 |
| TRLGNLTARKLVVLV | ASSPPTNSGNTLY | 0.8078953 |
| WSQQRRLIYFPSAGP | ASSPPTNSGNTLY | 0.8163067 |
| GPNGNPDPMAAAVDIR | ASSPPTNSGNTLY | 0.7405275 |
| TQRCVSVADLRAAASR | ASSPPTNSGNTLY | 0.822764 |
| QFVLDVAEHPVEVLTETV | ASSPPTNSGNTLY | 0.55826104 |
| GGAGGTGGAGGNGGTQAGDG | ASSPPTNSGNTLY | 0.72569454 |
| PLLATLRAIAVDVDAKPAQV | ASSPPTNSGNTLY | 0.78137696 |
| APKFRTGGYMIDFWGVGYQVAK | ASSPPTNSGNTLY | 0.7798425 |
| GAGGAGGAGGAAGGPGGTGGTG | ASSPPTNSGNTLY | 0.8235196 |
| IGALGLLATDVAPAGSVAFGVA | ASSPPTNSGNTLY | 0.7093223 |
| AGGAGGIGGNGGAGGAGGVLMG | ASSPPTNSGNTLY | 0.8016397 |
| ADELIGLL | ASSPPTNSGNTLY | 0.76023936 |
| ADRALKA | ASSPPTNSGNTLY | 0.7026475 |
| AGAVGQVGGAGGA | ASSPPTNSGNTLY | 0.8192607 |

| | | |
|---|---|---|
| AGGGGGLGGAP | ASSPPTNSGNTLY | 0.7830645 |
| AGGLLYGNGGAG | ASSPPTNSGNTLY | 0.6906005 |
| AGLESIGL | ASSPPTNSGNTLY | 0.8299017 |
| AGLKTGII | ASSPPTNSGNTLY | 0.7146828 |
| ASLAALLL | ASSPPTNSGNTLY | 0.79520977 |
| ASVINISSI | ASSPPTNSGNTLY | 0.83540773 |
| ATPPGPA | ASSPPTNSGNTLY | 0.67858624 |
| ATVASALV | ASSPPTNSGNTLY | 0.82827675 |
| ATVASLIL | ASSPPTNSGNTLY | 0.8051151 |
| AVNGPVLIRH | ASSPPTNSGNTLY | 0.8145213 |
| DGAAGAVGI | ASSPPTNSGNTLY | 0.7536267 |
| DPLLHGI | ASSPPTNSGNTLY | 0.74685645 |
| ELGAALGAL | ASSPPTNSGNTLY | 0.83579206 |
| ELGRTQAV | ASSPPTNSGNTLY | 0.81156886 |
| FVAGGYLS | ASSPPTNSGNTLY | 0.7917355 |
| GASGGAGGAGGW | ASSPPTNSGNTLY | 0.8187041 |
| GGAGAGGDGGAGGI | ASSPPTNSGNTLY | 0.8350809 |
| GTGHNGSQPGMGGNG | ASSPPTNSGNTLY | 0.8308337 |
| GYAAASASAS | ASSPPTNSGNTLY | 0.74629647 |
| HQVDVAAVQQG | ASSPPTNSGNTLY | 0.60328174 |
| IGAAIAHT | ASSPPTNSGNTLY | 0.76334333 |
| IGMIEVAQL | ASSPPTNSGNTLY | 0.7352214 |
| IRLAPEHTSAMD | ASSPPTNSGNTLY | 0.7943424 |
| IYNVSVL | ASSPPTNSGNTLY | 0.5630729 |
| KQGFLGM | ASSPPTNSGNTLY | 0.8232024 |
| LESVITQL | ASSPPTNSGNTLY | 0.8423362 |
| LGLGALVVV | ASSPPTNSGNTLY | 0.67879665 |
| LIGNGGNGGSGGI | ASSPPTNSGNTLY | 0.84244573 |
| LNAINAPTL | ASSPPTNSGNTLY | 0.67556894 |
| LQLLNTGV | ASSPPTNSGNTLY | 0.81212723 |
| LQTSIEF | ASSPPTNSGNTLY | 0.8261657 |
| MSFAVSEL | ASSPPTNSGNTLY | 0.67095107 |
| NTGALGL | ASSPPTNSGNTLY | 0.7929846 |
| PPGAGGAGGAGTT | ASSPPTNSGNTLY | 0.76145464 |
| QLVGLGI | ASSPPTNSGNTLY | 0.76686805 |
| RFLEQVHQL | ASSPPTNSGNTLY | 0.8273902 |
| RGVGYVLR | ASSPPTNSGNTLY | 0.67936647 |
| SATGVFAGVI | ASSPPTNSGNTLY | 0.82271063 |
| SLPAITLP | ASSPPTNSGNTLY | 0.8003178 |
| STALAFTV | ASSPPTNSGNTLY | 0.73476505 |
| SVLTVGLGK | ASSPPTNSGNTLY | 0.8284619 |
| TALAGAGSGI | ASSPPTNSGNTLY | 0.8202629 |
| TGLPFTGI | ASSPPTNSGNTLY | 0.8145142 |
| VAEHVTV | ASSPPTNSGNTLY | 0.8394475 |
| VDIVLNSL | ASSPPTNSGNTLY | 0.7256311 |
| VLTGGGAVL | ASSPPTNSGNTLY | 0.8394052 |
| VNVGLAYGI | ASSPPTNSGNTLY | 0.8329624 |
| VSSAAVGAV | ASSPPTNSGNTLY | 0.7982726 |
| VTSVYEAGV | ASSPPTNSGNTLY | 0.79077184 |
| WVALDRGA | ASSPPTNSGNTLY | 0.82471263 |
| YLVGSVVI | ASSPPTNSGNTLY | 0.7851811 |

| | | |
|---|---|---|
| GPPGTGKTLL | ASSPPTNSGNTLY | 0.8190413 |
| GVGMTKF | ASSPPTNSGNTLY | 0.8406098 |
| IGIGGSDL | ASSPPTNSGNTLY | 0.33471486 |
| IGTLLAEL | ASSPPTNSGNTLY | 0.6439038 |
| IYTSGTT | ASSPPTNSGNTLY | 0.58502233 |
| LSFPAGV | ASSPPTNSGNTLY | 0.60273623 |
| LYTSGTT | ASSPPTNSGNTLY | 0.45166525 |
| VGAFPFGF | ASSPPTNSGNTLY | 0.35134217 |
| VGAGTVEF | ASSPPTNSGNTLY | 0.5235687 |
| VGKSTLLL | ASSPPTNSGNTLY | 0.4051603 |
| AALAHAGGGIV | ASSPPTNSGNTLY | 0.67328453 |
| AALVAVSLL | ASSPPTNSGNTLY | 0.6277056 |
| AAVSRAVAAL | ASSPPTNSGNTLY | 0.43369988 |
| AGGSSGIGGF | ASSPPTNSGNTLY | 0.5987551 |
| AGLTVANANG | ASSPPTNSGNTLY | 0.62053096 |
| AGLVGAVL | ASSPPTNSGNTLY | 0.34587356 |
| ALSQAFTV | ASSPPTNSGNTLY | 0.5200438 |
| ALTFSVSL | ASSPPTNSGNTLY | 0.5063053 |
| ALTGAALVAL | ASSPPTNSGNTLY | 0.49136952 |
| ASFSIGAGVL | ASSPPTNSGNTLY | 0.3139367 |
| DGSVYKI | ASSPPTNSGNTLY | 0.61110884 |
| EQLAEIV | ASSPPTNSGNTLY | 0.45455176 |
| FAVSELTVM | ASSPPTNSGNTLY | 0.2967536 |
| GAAPLAAGL | ASSPPTNSGNTLY | 0.58137625 |
| GADGGDGGSGGAGG | ASSPPTNSGNTLY | 0.46292126 |
| GFGLDTGGAGG | ASSPPTNSGNTLY | 0.4644925 |
| GLASGGLL | ASSPPTNSGNTLY | 0.6108825 |
| GRIAGAAVLL | ASSPPTNSGNTLY | 0.6071639 |
| GSAGSLNTSF | ASSPPTNSGNTLY | 0.47232133 |
| GTGQIGI | ASSPPTNSGNTLY | 0.59104633 |
| GTLIALIL | ASSPPTNSGNTLY | 0.6520985 |
| IAATMGFVVG | ASSPPTNSGNTLY | 0.40736854 |
| IFAQALL | ASSPPTNSGNTLY | 0.5474322 |
| IGAAVALGF | ASSPPTNSGNTLY | 0.6742667 |
| IGLTPLAL | ASSPPTNSGNTLY | 0.56164974 |
| ILGSNTSTL | ASSPPTNSGNTLY | 0.66743696 |
| IVFSLTVL | ASSPPTNSGNTLY | 0.676267 |
| LASKATGF | ASSPPTNSGNTLY | 0.4892512 |
| LASVALIL | ASSPPTNSGNTLY | 0.6395881 |
| LFAVGLTL | ASSPPTNSGNTLY | 0.6357385 |
| LFNGGIGTYI | ASSPPTNSGNTLY | 0.5151009 |
| LTLGAGII | ASSPPTNSGNTLY | 0.5360417 |
| LVAAALGDI | ASSPPTNSGNTLY | 0.32960704 |
| NTAFVQLGI | ASSPPTNSGNTLY | 0.516719 |
| QLFSGAEL | ASSPPTNSGNTLY | 0.40545285 |
| SAATFITL | ASSPPTNSGNTLY | 0.59204173 |
| SNYGLANLGV | ASSPPTNSGNTLY | 0.6591573 |
| SYGGAYAVM | ASSPPTNSGNTLY | 0.54835254 |
| TTGVGGLTLG | ASSPPTNSGNTLY | 0.31681883 |
| TTQTIYI | ASSPPTNSGNTLY | 0.36312306 |
| VAAGAGLGAVI | ASSPPTNSGNTLY | 0.5574677 |

| | | |
|---|---|---|
| VAAVVTLL | ASSPPTNSGNTLY | 0.3624266 |
| VGFLGGLI | ASSPPTNSGNTLY | 0.38083944 |
| VGVIPILL | ASSPPTNSGNTLY | 0.4185708 |
| VIFASSLI | ASSPPTNSGNTLY | 0.64608896 |
| VKGGDGGAAAG | ASSPPTNSGNTLY | 0.65501887 |
| VTKTFTNI | ASSPPTNSGNTLY | 0.4678738 |
| VVAAAILL | ASSPPTNSGNTLY | 0.64926213 |
| YAALDVEL | ASSPPTNSGNTLY | 0.6662625 |
| LASVALLL | ASSPPTNSGNTLY | 0.6733421 |
| LDAVHRV | ASSPPTNSGNTLY | 0.5653556 |
| QGLSSSIF | ASSPPTNSGNTLY | 0.3347172 |
| RGTFANIRL | ASSPPTNSGNTLY | 0.65881383 |
| RQLWWGHRI | ASSPPTNSGNTLY | 0.46865398 |
| YGPPGTGKTLL | ASSPPTNSGNTLY | 0.6483396 |
| AALVGAPANVA | ASSPPTNSGNTLY | 0.6218592 |
| AATFRRTE | ASSPPTNSGNTLY | 0.49517554 |
| AGALGGGI | ASSPPTNSGNTLY | 0.4445109 |
| AGGAGGAGGIGAG | ASSPPTNSGNTLY | 0.63177127 |
| AGSGALGL | ASSPPTNSGNTLY | 0.66184837 |
| AQIADQI | ASSPPTNSGNTLY | 0.3971581 |
| ASAVAVTYL | ASSPPTNSGNTLY | 0.3831092 |
| ASTVLGGTGG | ASSPPTNSGNTLY | 0.56288874 |
| AVGGEPAK | ASSPPTNSGNTLY | 0.54278284 |
| CEQGTFGGTC | ASSPPTNSGNTLY | 0.65809923 |
| DEGKAPAG | ASSPPTNSGNTLY | 0.5995604 |
| DTVVFPAL | ASSPPTNSGNTLY | 0.55426013 |
| FEGITMAG | ASSPPTNSGNTLY | 0.59168553 |
| FPLVVQD | ASSPPTNSGNTLY | 0.62807846 |
| GAFTTPQ | ASSPPTNSGNTLY | 0.57044894 |
| GFGGFGTAGGVGGA | ASSPPTNSGNTLY | 0.55756116 |
| GFLAQFL | ASSPPTNSGNTLY | 0.4363826 |
| GGAGGAGGNAGVA | ASSPPTNSGNTLY | 0.6448176 |
| GGVGTAGAGGAGGAGG | ASSPPTNSGNTLY | 0.64523077 |
| GNGGGGDAGNAGSGGN | ASSPPTNSGNTLY | 0.63752544 |
| GTGGAGGAGNDAGS | ASSPPTNSGNTLY | 0.6638936 |
| GTTNTGWFNS | ASSPPTNSGNTLY | 0.6084986 |
| IFMGPLFTL | ASSPPTNSGNTLY | 0.5732907 |
| ILAHAIEAI | ASSPPTNSGNTLY | 0.41953963 |
| IRSAIREAL | ASSPPTNSGNTLY | 0.50630707 |
| ITFAGGG | ASSPPTNSGNTLY | 0.5085623 |
| ITGMSGHQEV | ASSPPTNSGNTLY | 0.5767149 |
| IVAGLACTL | ASSPPTNSGNTLY | 0.6272035 |
| IVGGGLAGA | ASSPPTNSGNTLY | 0.6235515 |
| IVTGVLNI | ASSPPTNSGNTLY | 0.4601807 |
| LAQQGCI | ASSPPTNSGNTLY | 0.6549554 |
| LCLHVEN | ASSPPTNSGNTLY | 0.65585446 |
| LDAISDGS | ASSPPTNSGNTLY | 0.4192375 |
| LGGFAAML | ASSPPTNSGNTLY | 0.49072427 |
| LGGKNPMI | ASSPPTNSGNTLY | 0.64582705 |
| LGPKGVAAIL | ASSPPTNSGNTLY | 0.63489586 |
| LMVGVSI | ASSPPTNSGNTLY | 0.48422176 |

| | | |
|---|---|---|
| LVGAAFVL | ASSPPTNSGNTLY | 0.53837436 |
| LVLAVAPL | ASSPPTNSGNTLY | 0.6813089 |
| MAAIAVIDAA | ASSPPTNSGNTLY | 0.60910964 |
| NNGINAELQ | ASSPPTNSGNTLY | 0.6491029 |
| RGEPLVLVHG | ASSPPTNSGNTLY | 0.35467958 |
| RGGAGGAAGLI | ASSPPTNSGNTLY | 0.5159115 |
| RSYRMHPAV | ASSPPTNSGNTLY | 0.45028386 |
| SAAHLLAGQLA | ASSPPTNSGNTLY | 0.28097028 |
| SAAVGSY | ASSPPTNSGNTLY | 0.6395687 |
| SPHAVLHRI | ASSPPTNSGNTLY | 0.44903168 |
| SQIAKVL | ASSPPTNSGNTLY | 0.59010804 |
| TALGHLVP | ASSPPTNSGNTLY | 0.43979886 |
| TAPTTSGAA | ASSPPTNSGNTLY | 0.6540462 |
| TDGDDGNGGSPGT | ASSPPTNSGNTLY | 0.60344434 |
| TDGLGEN | ASSPPTNSGNTLY | 0.52483046 |
| TGGPPLPGQA | ASSPPTNSGNTLY | 0.4233792 |
| TIPAITI | ASSPPTNSGNTLY | 0.5892696 |
| TIPGFTI | ASSPPTNSGNTLY | 0.6543193 |
| TSIASGALA | ASSPPTNSGNTLY | 0.6654143 |
| TVTLPGAAI | ASSPPTNSGNTLY | 0.35696405 |
| VASGLSAALP | ASSPPTNSGNTLY | 0.62681246 |
| VGAGVIGI | ASSPPTNSGNTLY | 0.5548463 |
| VGPKRFGV | ASSPPTNSGNTLY | 0.3593002 |
| VNVAFPD | ASSPPTNSGNTLY | 0.30549118 |
| VSMTNSGVS | ASSPPTNSGNTLY | 0.4153849 |
| VTDVLGDLSV | ASSPPTNSGNTLY | 0.48462477 |
| VVVVGAGFAGL | ASSPPTNSGNTLY | 0.57576203 |
| VYTVGTVV | ASSPPTNSGNTLY | 0.6396053 |
| WTTFSGP | ASSPPTNSGNTLY | 0.47327825 |
| YDELVEAF | ASSPPTNSGNTLY | 0.6131655 |
| YGTGGAGGNGG | ASSPPTNSGNTLY | 0.5342651 |
| YTNVMAV | ASSPPTNSGNTLY | 0.26502234 |
| FLEQVHQL | ASSPPTNSGNTLY | 0.4605924 |
| GLSLLAVAV | ASSPPTNSGNTLY | 0.3351673 |
| LEQVHQL | ASSPPTNSGNTLY | 0.6726461 |
| LNAAGAI | ASSPPTNSGNTLY | 0.37110835 |
| PAVHASL | ASSPPTNSGNTLY | 0.49968076 |
| TELEDDA | ASSPPTNSGNTLY | 0.54789186 |
| TPPGPPN | ASSPPTNSGNTLY | 0.650743 |
| TPPPPPPP | ASSPPTNSGNTLY | 0.57375234 |
| TQSPALI | ASSGTGEAETLY | 0.45473665 |
| EFAAAFPQ | ASSGTGEAETLY | 0.34019017 |
| LLSGNLDVL | ASSGTGEAETLY | 0.6164502 |
| GPANLGNYNI | ASSGTGEAETLY | 0.6230493 |
| TLGPTLP | ASSGTGEAETLY | 0.41852775 |
| VYAYSSCCK | ASSGTGEAETLY | 0.44136792 |
| APAPAPAPNI | ASSGTGEAETLY | 0.51453876 |
| PGGPVPIP | ASSGTGEAETLY | 0.319921 |
| FLTVALEP | ASSGTGEAETLY | 0.65745443 |
| SQYFGANL | ASSGTGEAETLY | 0.371703 |
| GGVKPAA | ASSGTGEAETLY | 0.48126638 |

| | | |
|---|---|---|
| GGTGGAGGAGGW | ASSGTGEAETLY | 0.66078824 |
| GVGGVAGLGGAGG | ASSGTGEAETLY | 0.6393108 |
| VLLVLATTI | ASSGTGEAETLY | 0.50976205 |
| IGIFAAML | ASSGTGEAETLY | 0.45666122 |
| AVGDLID | ASSGTGEAETLY | 0.3901478 |
| NISPLPL | ASSGTGEAETLY | 0.42633075 |
| VTAVFVAI | ASSGTGEAETLY | 0.43559998 |
| SGDDNDAAL | ASSGTGEAETLY | 0.6364846 |
| AVAALAHS | ASSGTGEAETLY | 0.56191975 |
| GGVTVMATP | ASSGTGEAETLY | 0.37254673 |
| AGLAANVRD | ASSGTGEAETLY | 0.63516885 |
| TNRPDLIDPALLRPG | ASSGTGEAETLY | 0.6496942 |
| LANLGVTAL | ASSGTGEAETLY | 0.66788447 |
| TGERIEL | ASSGTGEAETLY | 0.4073357 |
| NGEELAA | ASSGTGEAETLY | 0.5268557 |
| VLSLVWG | ASSGTGEAETLY | 0.513888 |
| IGYALWL | ASSGTGEAETLY | 0.521316 |
| LSLLLAPL | ASSGTGEAETLY | 0.43872887 |
| TLDEALK | ASSGTGEAETLY | 0.42341462 |
| TDGAGTAEGG | ASSGTGEAETLY | 0.6423098 |
| NAPGGAGGNGG | ASSGTGEAETLY | 0.69973445 |
| VSVFMGMGGNF | ASSGTGEAETLY | 0.529799 |
| LASYLAAL | ASSGTGEAETLY | 0.30940422 |
| LVGGTPAL | ASSGTGEAETLY | 0.5271635 |
| VRNADVL | ASSGTGEAETLY | 0.44089586 |
| TCSSPFLVSAM | ASSGTGEAETLY | 0.43916884 |
| VHITDAE | ASSGTGEAETLY | 0.53118217 |
| EYAAPSF | ASSGTGEAETLY | 0.3368222 |
| VVDAFHGGL | ASSGTGEAETLY | 0.5225648 |
| ASVVDPQ | ASSGTGEAETLY | 0.67229044 |
| SYNEFGGLS | ASSGTGEAETLY | 0.5414306 |
| DPDGSVY | ASSGTGEAETLY | 0.5239345 |
| MVSTIHATE | ASSGTGEAETLY | 0.6378591 |
| LAAGVADPV | ASSGTGEAETLY | 0.5862137 |
| TKEQIAA | ASSGTGEAETLY | 0.64614445 |
| SLASPAAL | ASSGTGEAETLY | 0.67120576 |
| IGLVFYGV | ASSGTGEAETLY | 0.5500023 |
| FLLDASNL | ASSGTGEAETLY | 0.5617472 |
| YQVGNFP | ASSGTGEAETLY | 0.61188585 |
| ESLLSQST | ASSGTGEAETLY | 0.51258206 |
| LFTVGFAAVG | ASSGTGEAETLY | 0.5673348 |
| ATPPTPT | ASSGTGEAETLY | 0.5265867 |
| TDLATLG | ASSGTGEAETLY | 0.58836883 |
| SAGGTGR | ASSGTGEAETLY | 0.5314743 |
| GFSNSGS | ASSGTGEAETLY | 0.6832372 |
| GGNGGSGGTSV | ASSGTGEAETLY | 0.29031774 |
| GTSVATGGAGN | ASSGTGEAETLY | 0.41430563 |
| NGGNGGSGGTS | ASSGTGEAETLY | 0.6461613 |
| AVVKFPH | ASSGTGEAETLY | 0.2923175 |
| TPPPPPP | ASSGTGEAETLY | 0.35035682 |
| AAPAAAAAPA | ASSGTGEAETLY | 0.544201 |

| | | |
|---|---|---|
| IALNEDL | ASSGTGEAETLY | 0.6599392 |
| IVNVSSM | ASSGTGEAETLY | 0.29514092 |
| TGIPPAPR | ASSGTGEAETLY | 0.55468446 |
| FDLGGGTF | ASSGTGEAETLY | 0.34561723 |
| FDLGGGTFDVS | ASSGTGEAETLY | 0.65766907 |
| NEPTAAA | ASSGTGEAETLY | 0.5401302 |
| NEPTAAALA | ASSGTGEAETLY | 0.37080064 |
| GSIINISSI | ASSGTGEAETLY | 0.6298506 |
| VAPGNAGTA | ASSGTGEAETLY | 0.37623072 |
| TGGGASLE | ASSGTGEAETLY | 0.6196568 |
| VAVYNHYKRI | ASSGTGEAETLY | 0.6412623 |
| TDPAPATT | ASSGTGEAETLY | 0.688643 |
| TNRPDILDPALLRPG | ASSGTGEAETLY | 0.34323093 |
| LTTTTTS | ASSGTGEAETLY | 0.6332355 |
| PTGSGKTL | ASSGTGEAETLY | 0.479298 |
| TGSGKTL | ASSGTGEAETLY | 0.6800839 |
| TIGHVDHGK | ASSGTGEAETLY | 0.47713816 |
| TIGHVDHGKTT | ASSGTGEAETLY | 0.32868946 |
| GHVDHGKTT | ASSGTGEAETLY | 0.62610674 |
| AQFEHTLL | ASSGTGEAETLY | 0.58651304 |
| IVNQDEL | ASSGTGEAETLY | 0.50974786 |
| TNGSQFF | ASSGTGEAETLY | 0.6508473 |
| TGKGIVE | ASSGTGEAETLY | 0.5497291 |
| LISNASDA | ASSGTGEAETLY | 0.5158466 |
| AELRTLGELTKIAKAHGAQ | ASSGTGEAETLY | 0.6203902 |
| APNAVAAAEPSW | ASSGTGEAETLY | 0.6043098 |
| APQVAGAATAVL | ASSGTGEAETLY | 0.4481709 |
| ARRDPAAAAAILPTDA | ASSGTGEAETLY | 0.6557851 |
| DAAARDVLAEAGLAEYF | ASSGTGEAETLY | 0.5591965 |
| DDQIVLSAKANLAKVEQ | ASSGTGEAETLY | 0.37989646 |
| DELLPDTGEDSDEDGH | ASSGTGEAETLY | 0.67278576 |
| ELPEAPANRATVPQD | ASSGTGEAETLY | 0.5672213 |
| FGQSKIERVTGIG | ASSGTGEAETLY | 0.61437005 |
| GAGGTGSTAGGAGGAGGAGGLY | ASSGTGEAETLY | 0.6104138 |
| GHGGTGAAGGNGGTGGAGGSGI | ASSGTGEAETLY | 0.69962895 |
| GNGGNGGAGGEGGPGGAGGAGA | ASSGTGEAETLY | 0.4434814 |
| GSTAGNGGNGGNATNGTIAGQP | ASSGTGEAETLY | 0.5532462 |
| GTGGDGGNGGNGGTGG | ASSGTGEAETLY | 0.5539918 |
| GVRRPAGIPADAPV | ASSGTGEAETLY | 0.6496285 |
| GYRQVTDAHLLTLARR | ASSGTGEAETLY | 0.5703672 |
| IAKMGQDGHDRG | ASSGTGEAETLY | 0.5986662 |
| KAEPSIEIQYT | ASSGTGEAETLY | 0.5964746 |
| KPVRIRMAADEDTVAVT | ASSGTGEAETLY | 0.64245206 |
| MPPRIAGMRLLVIKPEP | ASSGTGEAETLY | 0.5276959 |
| MKFYAVALFRSRGVPPTR | ASSGTGEAETLY | 0.54641366 |
| NAGGFGILGAVAHSP | ASSGTGEAETLY | 0.6674541 |
| NEGREITLAMAEM | ASSGTGEAETLY | 0.3585802 |
| NELVEWSHPLGHRWRWE | ASSGTGEAETLY | 0.571404 |
| NLGAGGTGGNGGNPG | ASSGTGEAETLY | 0.63676894 |
| NTEWPKWETVMAADPVD | ASSGTGEAETLY | 0.63777876 |
| PASAAPAETQP | ASSGTGEAETLY | 0.5177753 |

| | | |
|---|---|---|
| QQSSDGKNGVRAAEITGE | ASSGTGEAETLY | 0.64746225 |
| RAPDAVTTSWWKE | ASSGTGEAETLY | 0.4977888 |
| SGFGNVGAHTSGWFNQ | ASSGTGEAETLY | 0.58185905 |
| SGVAGAVGQAASVGGLKV | ASSGTGEAETLY | 0.5415036 |
| SNGGAGTGSAGGNGGT | ASSGTGEAETLY | 0.47658187 |
| TGGAGGEGGAAG | ASSGTGEAETLY | 0.3670215 |
| TGGDGGAAGAGGAPGHG | ASSGTGEAETLY | 0.49952447 |
| TMPPLLAAIANAGRHGV | ASSGTGEAETLY | 0.6185492 |
| TSSTAGNSGGA | ASSGTGEAETLY | 0.5689265 |
| VPATVPVGILC | ASSGTGEAETLY | 0.33437124 |
| VQRPDSGGVVLTGRIS | ASSGTGEAETLY | 0.58893645 |
| WTPNPVPATKGVPAGLVQAT | ASSGTGEAETLY | 0.60362655 |
| YALKRALAKFSDRFDVVII | ASSGTGEAETLY | 0.6794728 |
| YLIGSVIV | ASSGTGEAETLY | 0.54930544 |
| YGQLVAVID | ASSGTGEAETLY | 0.632507 |
| VTVGGMTLPA | ASSGTGEAETLY | 0.53026444 |
| VLALPAV | ASSGTGEAETLY | 0.58164334 |
| VIPPTLNYET | ASSGTGEAETLY | 0.34686685 |
| VGMGGMHPSSG | ASSGTGEAETLY | 0.36759618 |
| VGLAGGAATA | ASSGTGEAETLY | 0.55585873 |
| VGGAGGAGGSGP | ASSGTGEAETLY | 0.39724514 |
| VFAAFMV | ASSGTGEAETLY | 0.6430887 |
| TLYDKLR | ASSGTGEAETLY | 0.6305752 |
| TLAVFTVL | ASSGTGEAETLY | 0.6343947 |
| TGSVQPARV | ASSGTGEAETLY | 0.64869654 |
| TGGGAGSGGT | ASSGTGEAETLY | 0.64725053 |
| SHLETSL | ASSGTGEAETLY | 0.65472263 |
| SFTTGMT | ASSGTGEAETLY | 0.44170028 |
| QVEVAKRMGVRQPTVS | ASSGTGEAETLY | 0.5058059 |
| QGSGGPALAM | ASSGTGEAETLY | 0.3183853 |
| MLAPSNP | ASSGTGEAETLY | 0.4527617 |
| LVNNAAI | ASSGTGEAETLY | 0.44089618 |
| LTGEINTLA | ASSGTGEAETLY | 0.54605836 |
| LSIFGPV | ASSGTGEAETLY | 0.28909314 |
| LLALLANL | ASSGTGEAETLY | 0.6269157 |
| LISNASDALDKLR | ASSGTGEAETLY | 0.63190776 |
| LINEAVP | ASSGTGEAETLY | 0.56296015 |
| LGYAFSGL | ASSGTGEAETLY | 0.61723506 |
| LDFVAVAL | ASSGTGEAETLY | 0.5785093 |
| LAQPGAE | ASSGTGEAETLY | 0.4317458 |
| ISNLGAVTGF | ASSGTGEAETLY | 0.6121222 |
| HQEDPGRI | ASSGTGEAETLY | 0.5102007 |
| GTPGEPTP | ASSGTGEAETLY | 0.64563906 |
| GFFNSGS | ASSGTGEAETLY | 0.6468239 |
| GAGGGGLTAGAN | ASSGTGEAETLY | 0.59187114 |
| FVDALVNM | ASSGTGEAETLY | 0.50065815 |
| FLTVAPEL | ASSGTGEAETLY | 0.60253996 |
| FETLTAL | ASSGTGEAETLY | 0.46364594 |
| DVINPIEL | ASSGTGEAETLY | 0.5322169 |
| APVSGAPV | ASSGTGEAETLY | 0.34015614 |
| ALVGGLAW | ASSGTGEAETLY | 0.57821435 |

| | | |
|---|---|---|
| ALAAAAATAI | ASSGTGEAETLY | 0.40869576 |
| AFGLGLLL | ASSGTGEAETLY | 0.5936425 |
| AFFAAASL | ASSGTGEAETLY | 0.5309416 |
| AAAALPAFSPP | ASSGTGEAETLY | 0.60318774 |
| VVLAPAL | ASSGTGEAETLY | 0.41806293 |
| TIGHVDHGKTTL | ASSGTGEAETLY | 0.64972734 |
| LISNASDALDK | ASSGTGEAETLY | 0.63216674 |
| LISNASDAL | ASSGTGEAETLY | 0.36414796 |
| ISNASDALDK | ASSGTGEAETLY | 0.6167762 |
| NQAETLVYQTE | ASSGTGEAETLY | 0.51013106 |
| TGGTGGTAGAG | ASSGTGEAETLY | 0.5097199 |
| TGSTAGGAGGA | ASSGTGEAETLY | 0.6418154 |
| DPLAVPVAAGPV | ASSGTGEAETLY | 0.552792 |
| GGTGGTPGNGGN | ASSGTGEAETLY | 0.38706473 |
| PAGLAGVDPATL | ASSGTGEAETLY | 0.4887469 |
| GNNLVVEAPHAGASV | ASSGTGEAETLY | 0.5714896 |
| KVTRIMSASTLEQPA | ASSGTGEAETLY | 0.50543565 |
| LGFALPIALPRRTVV | ASSGTGEAETLY | 0.32409245 |
| TRLGNLTARKLVVLV | ASSGTGEAETLY | 0.59253645 |
| WSQQRRLIYFPSAGP | ASSGTGEAETLY | 0.38085228 |
| GPNGNPDPMAAAVDIR | ASSGTGEAETLY | 0.45500642 |
| TQRCVSVADLRAAASR | ASSGTGEAETLY | 0.44923598 |
| QFVLDVAEHPVEVLTETV | ASSGTGEAETLY | 0.6335986 |
| GGAGGTGGAGGNGGTQAGDG | ASSGTGEAETLY | 0.6685506 |
| PLLATLRAIAVDVDAKPAQV | ASSGTGEAETLY | 0.4936191 |
| APKFRTGGYMIDFWGVGYQVAK | ASSGTGEAETLY | 0.56400967 |
| GAGGAGGAGGAAGGPGGTGGTG | ASSGTGEAETLY | 0.5913886 |
| IGALGLLATDVAPAGSVAFGVA | ASSGTGEAETLY | 0.629004 |
| AGGAGGIGGNGGAGGAGGVLMG | ASSGTGEAETLY | 0.5496175 |
| ADELIGLL | ASSGTGEAETLY | 0.54406184 |
| ADRALKA | ASSGTGEAETLY | 0.5989474 |
| AGAVGQVGGAGGA | ASSGTGEAETLY | 0.5295849 |
| AGGGGGLGGAP | ASSGTGEAETLY | 0.41226465 |
| AGGLLYGNGGAG | ASSGTGEAETLY | 0.49150646 |
| AGLESIGL | ASSGTGEAETLY | 0.44946957 |
| AGLKTGII | ASSGTGEAETLY | 0.44004667 |
| ASLAALLL | ASSGTGEAETLY | 0.5911847 |
| ASVINISSI | ASSGTGEAETLY | 0.6808211 |
| ATPPGPA | ASSGTGEAETLY | 0.5922751 |
| ATVASALV | ASSGTGEAETLY | 0.31505537 |
| ATVASLIL | ASSGTGEAETLY | 0.45977443 |
| AVNGPVLIRH | ASSGTGEAETLY | 0.55953765 |
| DGAAGAVGI | ASSGTGEAETLY | 0.60028696 |
| DPLLHGI | ASSGTGEAETLY | 0.6483651 |
| ELGAALGAL | ASSGTGEAETLY | 0.68530923 |
| ELGRTQAV | ASSGTGEAETLY | 0.62209475 |
| FVAGGYLS | ASSGTGEAETLY | 0.46328926 |
| GASGGAGGAGGW | ASSGTGEAETLY | 0.6248754 |
| GGAGAGGDGGAGGI | ASSGTGEAETLY | 0.49974316 |
| GTGHNGSQPGMGGNG | ASSGTGEAETLY | 0.48226416 |
| GYAAASASAS | ASSGTGEAETLY | 0.54719186 |

| | | |
|---|---|---|
| HQVDVAAVQQG | ASSGTGEAETLY | 0.6206498 |
| IGAAIAHT | ASSGTGEAETLY | 0.46735123 |
| IGMIEVAQL | ASSGTGEAETLY | 0.48286182 |
| IRLAPEHTSAMD | ASSGTGEAETLY | 0.3675577 |
| IYNVSVL | ASSGTGEAETLY | 0.6453335 |
| KQGFLGM | ASSGTGEAETLY | 0.46895695 |
| LESVITQL | ASSGTGEAETLY | 0.47857663 |
| LGLGALVVV | ASSGTGEAETLY | 0.5230503 |
| LIGNGGNGGSGGI | ASSGTGEAETLY | 0.6607424 |
| LNAINAPTL | ASSGTGEAETLY | 0.6681224 |
| LQLLNTGV | ASSGTGEAETLY | 0.49323773 |
| LQTSIEF | ASSGTGEAETLY | 0.42814612 |
| MSFAVSEL | ASSGTGEAETLY | 0.6272203 |
| NTGALGL | ASSGTGEAETLY | 0.49263608 |
| PPGAGGAGGAGTT | ASSGTGEAETLY | 0.61789274 |
| QLVGLGI | ASSGTGEAETLY | 0.58430815 |
| RFLEQVHQL | ASSGTGEAETLY | 0.5302768 |
| RGVGYVLR | ASSGTGEAETLY | 0.47153413 |
| SATGVFAGVI | ASSGTGEAETLY | 0.6135204 |
| SLPAITLP | ASSGTGEAETLY | 0.49664944 |
| STALAFTV | ASSGTGEAETLY | 0.6326905 |
| SVLTVGLGK | ASSGTGEAETLY | 0.38700393 |
| TALAGAGSGI | ASSGTGEAETLY | 0.6470835 |
| TGLPFTGI | ASSGTGEAETLY | 0.38913202 |
| VAEHVTV | ASSGTGEAETLY | 0.4786618 |
| VDIVLNSL | ASSGTGEAETLY | 0.29856735 |
| VLTGGGAVL | ASSGTGEAETLY | 0.52761215 |
| VNVGLAYGI | ASSGTGEAETLY | 0.6013802 |
| VSSAAVGAV | ASSGTGEAETLY | 0.6511417 |
| VTSVYEAGV | ASSGTGEAETLY | 0.58792526 |
| WVALDRGA | ASSGTGEAETLY | 0.5864553 |
| YLVGSVVI | ASSGTGEAETLY | 0.6635777 |
| GPPGTGKTLL | ASSGTGEAETLY | 0.42222667 |
| GVGMTKF | ASSGTGEAETLY | 0.31361237 |
| IGIGGSDL | ASSGTGEAETLY | 0.5776753 |
| IGTLLAEL | ASSGTGEAETLY | 0.6169213 |
| IYTSGTT | ASSGTGEAETLY | 0.6586034 |
| LSFPAGV | ASSGTGEAETLY | 0.62089574 |
| LYTSGTT | ASSGTGEAETLY | 0.3238742 |
| VGAFPFGF | ASSGTGEAETLY | 0.58812994 |
| VGAGTVEF | ASSGTGEAETLY | 0.60059726 |
| VGKSTLLL | ASSGTGEAETLY | 0.36820397 |
| AALAHAGGGIV | ASSGTGEAETLY | 0.39675602 |
| AALVAVSLL | ASSGTGEAETLY | 0.4192623 |
| AAVSRAVAAL | ASSGTGEAETLY | 0.37348682 |
| AGGSSGIGGF | ASSGTGEAETLY | 0.60911727 |
| AGLTVANANG | ASSGTGEAETLY | 0.4641687 |
| AGLVGAVL | ASSGTGEAETLY | 0.5688889 |
| ALSQAFTV | ASSGTGEAETLY | 0.4920662 |
| ALTFSVSL | ASSGTGEAETLY | 0.448358 |
| ALTGAALVAL | ASSGTGEAETLY | 0.5371525 |

| | | |
|---|---|---|
| ASFSIGAGVL | ASSGTGEAETLY | 0.49827987 |
| DGSVYKI | ASSGTGEAETLY | 0.6034274 |
| EQLAEIV | ASSGTGEAETLY | 0.62368906 |
| FAVSELTVM | ASSGTGEAETLY | 0.55031157 |
| GAAPLAAGL | ASSGTGEAETLY | 0.59717566 |
| GADGGDGGSGGAGG | ASSGTGEAETLY | 0.6307726 |
| GFGLDTGGAGG | ASSGTGEAETLY | 0.3804345 |
| GLASGGLL | ASSGTGEAETLY | 0.45010468 |
| GRIAGAAVLL | ASSGTGEAETLY | 0.61560315 |
| GSAGSLNTSF | ASSGTGEAETLY | 0.5140437 |
| GTGQIGI | ASSGTGEAETLY | 0.65193 |
| GTLIALIL | ASSGTGEAETLY | 0.56671536 |
| IAATMGFVVG | ASSGTGEAETLY | 0.57996976 |
| IFAQALL | ASSGTGEAETLY | 0.58919716 |
| IGAAVALGF | ASSGTGEAETLY | 0.6535006 |
| IGLTPLAL | ASSGTGEAETLY | 0.57766753 |
| ILGSNTSTL | ASSGTGEAETLY | 0.47783697 |
| IVFSLTVL | ASSGTGEAETLY | 0.63209033 |
| LASKATGF | ASSGTGEAETLY | 0.36298573 |
| LASVALIL | ASSGTGEAETLY | 0.6133514 |
| LFAVGLTL | ASSGTGEAETLY | 0.6236049 |
| LFNGGIGTYI | ASSGTGEAETLY | 0.39708346 |
| LTLGAGII | ASSGTGEAETLY | 0.5173584 |
| LVAAALGDI | ASSGTGEAETLY | 0.66153884 |
| NTAFVQLGI | ASSGTGEAETLY | 0.621661 |
| QLFSGAEL | ASSGTGEAETLY | 0.49465775 |
| SAATFITL | ASSGTGEAETLY | 0.4527594 |
| SNYGLANLGV | ASSGTGEAETLY | 0.46180147 |
| SYGGAYAVM | ASSGTGEAETLY | 0.43666303 |
| TTGVGGLTLG | ASSGTGEAETLY | 0.3285666 |
| TTQTIYI | ASSGTGEAETLY | 0.61050665 |
| VAAGAGLGAVI | ASSGTGEAETLY | 0.49048162 |
| VAAVVTLL | ASSGTGEAETLY | 0.67224944 |
| VGFLGGLI | ASSGTGEAETLY | 0.58736396 |
| VGVIPILL | ASSGTGEAETLY | 0.65001744 |
| VIFASSLI | ASSGTGEAETLY | 0.4756377 |
| VKGGDGGAAAG | ASSGTGEAETLY | 0.51064456 |
| VTKTFTNI | ASSGTGEAETLY | 0.33040538 |
| VVAAAILL | ASSGTGEAETLY | 0.4502688 |
| YAALDVEL | ASSGTGEAETLY | 0.4163006 |
| LASVALLL | ASSGTGEAETLY | 0.5487392 |
| LDAVHRV | ASSGTGEAETLY | 0.34713456 |
| QGLSSSIF | ASSGTGEAETLY | 0.66732126 |
| RGTFANIRL | ASSGTGEAETLY | 0.46602505 |
| RQLWWGHRI | ASSGTGEAETLY | 0.62567174 |
| YGPPGTGKTLL | ASSGTGEAETLY | 0.51824933 |
| AALVGAPANVA | ASSGTGEAETLY | 0.36094755 |
| AATFRRTE | ASSGTGEAETLY | 0.45526356 |
| AGALGGGI | ASSGTGEAETLY | 0.61890805 |
| AGGAGGAGGIGAG | ASSGTGEAETLY | 0.3500841 |
| AGSGALGL | ASSGTGEAETLY | 0.32170767 |

| | | |
|---|---|---|
| AQIADQI | ASSGTGEAETLY | 0.4708557 |
| ASAVAVTYL | ASSGTGEAETLY | 0.5524608 |
| ASTVLGGTGG | ASSGTGEAETLY | 0.6689118 |
| AVGGEPAK | ASSGTGEAETLY | 0.64182323 |
| CEQGTFGGTC | ASSGTGEAETLY | 0.5388315 |
| DEGKAPAG | ASSGTGEAETLY | 0.6820115 |
| DTVVFPAL | ASSGTGEAETLY | 0.27675864 |
| FEGITMAG | ASSGTGEAETLY | 0.5573823 |
| FPLVVQD | ASSGTGEAETLY | 0.66740686 |
| GAFTTPQ | ASSGTGEAETLY | 0.589487 |
| GFGGFGTAGGVGGA | ASSGTGEAETLY | 0.5027744 |
| GFLAQFL | ASSGTGEAETLY | 0.6436918 |
| GGAGGAGGNAGVA | ASSGTGEAETLY | 0.4482953 |
| GGVGTAGAGGAGGAGG | ASSGTGEAETLY | 0.51643103 |
| GNGGGGDAGNAGSGGN | ASSGTGEAETLY | 0.5351751 |
| GTGGAGGAGNDAGS | ASSGTGEAETLY | 0.60893583 |
| GTTNTGWFNS | ASSGTGEAETLY | 0.5925462 |
| IFMGPLFTL | ASSGTGEAETLY | 0.33649078 |
| ILAHAIEAI | ASSGTGEAETLY | 0.6318903 |
| IRSAIREAL | ASSGTGEAETLY | 0.6564282 |
| ITFAGGG | ASSGTGEAETLY | 0.46320683 |
| ITGMSGHQEV | ASSGTGEAETLY | 0.6490108 |
| IVAGLACTL | ASSGTGEAETLY | 0.56449676 |
| IVGGGLAGA | ASSGTGEAETLY | 0.6015092 |
| IVTGVLNI | ASSGTGEAETLY | 0.44375035 |
| LAQQGCI | ASSGTGEAETLY | 0.35631692 |
| LCLHVEN | ASSGTGEAETLY | 0.6520586 |
| LDAISDGS | ASSGTGEAETLY | 0.43352973 |
| LGGFAAML | ASSGTGEAETLY | 0.5467969 |
| LGGKNPMI | ASSGTGEAETLY | 0.56839573 |
| LGPKGVAAIL | ASSGTGEAETLY | 0.67508346 |
| LMVGVSI | ASSGTGEAETLY | 0.59149444 |
| LVGAAFVL | ASSGTGEAETLY | 0.5877816 |
| LVLAVAPL | ASSGTGEAETLY | 0.46590298 |
| MAAIAVIDAA | ASSGTGEAETLY | 0.5767532 |
| NNGINAELQ | ASSGTGEAETLY | 0.62263846 |
| RGEPLVLVHG | ASSGTGEAETLY | 0.38310418 |
| RGGAGGAAGLI | ASSGTGEAETLY | 0.6437636 |
| RSYRMHPAV | ASSGTGEAETLY | 0.4020361 |
| SAAHLLAGQLA | ASSGTGEAETLY | 0.5823697 |
| SAAVGSY | ASSGTGEAETLY | 0.5960212 |
| SPHAVLHRI | ASSGTGEAETLY | 0.68077755 |
| SQIAKVL | ASSGTGEAETLY | 0.530012 |
| TALGHLVP | ASSGTGEAETLY | 0.6599258 |
| TAPTTSGAA | ASSGTGEAETLY | 0.39771077 |
| TDGDDGNGGSPGT | ASSGTGEAETLY | 0.6380634 |
| TDGLGEN | ASSGTGEAETLY | 0.64378333 |
| TGGPPLPGQA | ASSGTGEAETLY | 0.64904183 |
| TIPAITI | ASSGTGEAETLY | 0.6818524 |
| TIPGFTI | ASSGTGEAETLY | 0.63892996 |
| TSIASGALA | ASSGTGEAETLY | 0.31446388 |

| | | |
|---|---|---|
| TVTLPGAAI | ASSGTGEAETLY | 0.52937514 |
| VASGLSAALP | ASSGTGEAETLY | 0.65533924 |
| VGAGVIGI | ASSGTGEAETLY | 0.6492249 |
| VGPKRFGV | ASSGTGEAETLY | 0.64042956 |
| VNVAFPD | ASSGTGEAETLY | 0.5924286 |
| VSMTNSGVS | ASSGTGEAETLY | 0.50860214 |
| VTDVLGDLSV | ASSGTGEAETLY | 0.65592146 |
| VVVVGAGFAGL | ASSGTGEAETLY | 0.6974808 |
| VYTVGTVV | ASSGTGEAETLY | 0.6231588 |
| WTTFSGP | ASSGTGEAETLY | 0.47187382 |
| YDELVEAF | ASSGTGEAETLY | 0.42259216 |
| YGTGGAGGNGG | ASSGTGEAETLY | 0.60189795 |
| YTNVMAV | ASSGTGEAETLY | 0.618484 |
| FLEQVHQL | ASSGTGEAETLY | 0.3679682 |
| GLSLLAVAV | ASSGTGEAETLY | 0.52577853 |
| LEQVHQL | ASSGTGEAETLY | 0.56281793 |
| LNAAGAI | ASSGTGEAETLY | 0.289334 |
| PAVHASL | ASSGTGEAETLY | 0.6189326 |
| TELEDDA | ASSGTGEAETLY | 0.6301801 |
| TPPGPPN | ASSGTGEAETLY | 0.5817576 |
| TPPPPPPP | ASSGTGEAETLY | 0.3581083 |
| TQSPALI | AWIQGGAEQF | 0.6412959 |
| EFAAAFPQ | AWIQGGAEQF | 0.62144685 |
| LLSGNLDVL | AWIQGGAEQF | 0.6646129 |
| GPANLGNYNI | AWIQGGAEQF | 0.3790507 |
| TLGPTLP | AWIQGGAEQF | 0.28260207 |
| VYAYSSCCK | AWIQGGAEQF | 0.5971783 |
| APAPAPAPNI | AWIQGGAEQF | 0.6561188 |
| PGGPVPIP | AWIQGGAEQF | 0.49356455 |
| FLTVALEP | AWIQGGAEQF | 0.40231007 |
| SQYFGANL | AWIQGGAEQF | 0.49927974 |
| GGVKPAA | AWIQGGAEQF | 0.6401911 |
| GGTGGAGGAGGW | AWIQGGAEQF | 0.34294853 |
| GVGGVAGLGGAGG | AWIQGGAEQF | 0.386964 |
| VLLVLATTI | AWIQGGAEQF | 0.54382664 |
| IGIFAAML | AWIQGGAEQF | 0.6467592 |
| AVGDLID | AWIQGGAEQF | 0.42141253 |
| NISPLPL | AWIQGGAEQF | 0.4520519 |
| VTAVFVAI | AWIQGGAEQF | 0.6226349 |
| SGDDNDAAL | AWIQGGAEQF | 0.6586832 |
| AVAALAHS | AWIQGGAEQF | 0.5814991 |
| GGVTVMATP | AWIQGGAEQF | 0.38471228 |
| AGLAANVRD | AWIQGGAEQF | 0.5923171 |
| TNRPDLIDPALLRPG | AWIQGGAEQF | 0.3812076 |
| LANLGVTAL | AWIQGGAEQF | 0.6306277 |
| TGERIEL | AWIQGGAEQF | 0.5315484 |
| NGEELAA | AWIQGGAEQF | 0.5725019 |
| VLSLVWG | AWIQGGAEQF | 0.47814375 |
| IGYALWL | AWIQGGAEQF | 0.3965934 |
| LSLLLAPL | AWIQGGAEQF | 0.37374952 |
| TLDEALK | AWIQGGAEQF | 0.43810084 |

| | | |
|---|---|---|
| TDGAGTAEGG | AWIQGGAEQF | 0.6336829 |
| NAPGGAGGNGG | AWIQGGAEQF | 0.48783213 |
| VSVFMGMGGNF | AWIQGGAEQF | 0.6002076 |
| LASYLAAL | AWIQGGAEQF | 0.5658666 |
| LVGGTPAL | AWIQGGAEQF | 0.39006573 |
| VRNADVL | AWIQGGAEQF | 0.605842 |
| TCSSPFLVSAM | AWIQGGAEQF | 0.34294972 |
| VHITDAE | AWIQGGAEQF | 0.6098033 |
| EYAAPSF | AWIQGGAEQF | 0.5362892 |
| VVDAFHGGL | AWIQGGAEQF | 0.6180877 |
| ASVVDPQ | AWIQGGAEQF | 0.4538151 |
| SYNEFGGLS | AWIQGGAEQF | 0.59716713 |
| DPDGSVY | AWIQGGAEQF | 0.59997314 |
| MVSTIHATE | AWIQGGAEQF | 0.40878737 |
| LAAGVADPV | AWIQGGAEQF | 0.5693209 |
| TKEQIAA | AWIQGGAEQF | 0.64047235 |
| SLASPAAL | AWIQGGAEQF | 0.59042525 |
| IGLVFYGV | AWIQGGAEQF | 0.3132357 |
| FLLDASNL | AWIQGGAEQF | 0.5778028 |
| YQVGNFP | AWIQGGAEQF | 0.4340982 |
| ESLLSQST | AWIQGGAEQF | 0.5848341 |
| LFTVGFAAVG | AWIQGGAEQF | 0.52915263 |
| ATPPTPT | AWIQGGAEQF | 0.46270126 |
| TDLATLG | AWIQGGAEQF | 0.5981419 |
| SAGGTGR | AWIQGGAEQF | 0.41396356 |
| GFSNSGS | AWIQGGAEQF | 0.4557922 |
| GGNGGSGGTSV | AWIQGGAEQF | 0.5160982 |
| GTSVATGGAGN | AWIQGGAEQF | 0.57478565 |
| NGGNGGSGGTS | AWIQGGAEQF | 0.6125111 |
| AVVKFPH | AWIQGGAEQF | 0.3817382 |
| TPPPPPP | AWIQGGAEQF | 0.60110736 |
| AAPAAAAAPA | AWIQGGAEQF | 0.50620973 |
| IALNEDL | AWIQGGAEQF | 0.4183014 |
| IVNVSSM | AWIQGGAEQF | 0.5537758 |
| TGIPPAPR | AWIQGGAEQF | 0.5033585 |
| FDLGGGTF | AWIQGGAEQF | 0.34448755 |
| FDLGGGTFDVS | AWIQGGAEQF | 0.61870277 |
| NEPTAAA | AWIQGGAEQF | 0.37052375 |
| NEPTAAALA | AWIQGGAEQF | 0.6038461 |
| GSIINISSI | AWIQGGAEQF | 0.669423 |
| VAPGNAGTA | AWIQGGAEQF | 0.33748448 |
| TGGGASLE | AWIQGGAEQF | 0.6199254 |
| VAVYNHYKRI | AWIQGGAEQF | 0.53425133 |
| TDPAPATT | AWIQGGAEQF | 0.53847116 |
| TNRPDILDPALLRPG | AWIQGGAEQF | 0.5279073 |
| LTTTTTS | AWIQGGAEQF | 0.4425223 |
| PTGSGKTL | AWIQGGAEQF | 0.64937735 |
| TGSGKTL | AWIQGGAEQF | 0.6215528 |
| TIGHVDHGK | AWIQGGAEQF | 0.555982 |
| TIGHVDHGKTT | AWIQGGAEQF | 0.5519273 |
| GHVDHGKTT | AWIQGGAEQF | 0.64164835 |

| | | |
|---|---|---|
| AQFEHTLL | AWIQGGAEQF | 0.6411492 |
| IVNQDEL | AWIQGGAEQF | 0.4815383 |
| TNGSQFF | AWIQGGAEQF | 0.33868366 |
| TGKGIVE | AWIQGGAEQF | 0.5243067 |
| LISNASDA | AWIQGGAEQF | 0.5219493 |
| AELRTLGELTKIAKAHGAQ | AWIQGGAEQF | 0.44037366 |
| APNAVAAAEPSW | AWIQGGAEQF | 0.36102954 |
| APQVAGAATAVL | AWIQGGAEQF | 0.62623155 |
| ARRDPAAAAAILPTDA | AWIQGGAEQF | 0.6799915 |
| DAAARDVLAEAGLAEYF | AWIQGGAEQF | 0.5389756 |
| DDQIVLSAKANLAKVEQ | AWIQGGAEQF | 0.685228 |
| DELLPDTGEDSDEDGH | AWIQGGAEQF | 0.35706663 |
| ELPEAPANRATVPQD | AWIQGGAEQF | 0.6356776 |
| FGQSKIERVTGIG | AWIQGGAEQF | 0.6352215 |
| GAGGTGSTAGGAGGAGGAGGLY | AWIQGGAEQF | 0.41080135 |
| GHGGTGAAGGNGGTGGAGGSGI | AWIQGGAEQF | 0.6020067 |
| GNGGNGGAGGEGGPGGAGGAGA | AWIQGGAEQF | 0.41014284 |
| GSTAGNGGNGGNATNGTIAGQP | AWIQGGAEQF | 0.4364201 |
| GTGGDGGNGGNGGTGG | AWIQGGAEQF | 0.6220735 |
| GVRRPAGIPADAPV | AWIQGGAEQF | 0.53306293 |
| GYRQVTDAHLLTLARR | AWIQGGAEQF | 0.5898624 |
| IAKMGQDGHDRG | AWIQGGAEQF | 0.49080455 |
| KAEPSIEIQYT | AWIQGGAEQF | 0.4090457 |
| KPVRIRMAADEDTVAVT | AWIQGGAEQF | 0.5878484 |
| MPPRIAGMRLLVIKPEP | AWIQGGAEQF | 0.58929676 |
| MKFYAVALFRSRGVPPTR | AWIQGGAEQF | 0.6142825 |
| NAGGFGILGAVAHSP | AWIQGGAEQF | 0.69405556 |
| NEGREITLAMAEM | AWIQGGAEQF | 0.39831862 |
| NELVEWSHPLGHRWRWE | AWIQGGAEQF | 0.65739566 |
| NLGAGGTGGNGGNPG | AWIQGGAEQF | 0.6649996 |
| NTEWPKWETVMAADPVD | AWIQGGAEQF | 0.5807003 |
| PASAAPAETQP | AWIQGGAEQF | 0.5185752 |
| QQSSDGKNGVRAAEITGE | AWIQGGAEQF | 0.6390853 |
| RAPDAVTTSWWKE | AWIQGGAEQF | 0.5190701 |
| SGFGNVGAHTSGWFNQ | AWIQGGAEQF | 0.55000454 |
| SGVAGAVGQAASVGGLKV | AWIQGGAEQF | 0.67056733 |
| SNGGAGTGSAGGNGGT | AWIQGGAEQF | 0.5846888 |
| TGGAGGEGGAAG | AWIQGGAEQF | 0.86647135 |
| TGGDGGAAGAGGAPGHG | AWIQGGAEQF | 0.8092172 |
| TMPPLLAAIANAGRHGV | AWIQGGAEQF | 0.71680236 |
| TSSTAGNSGGA | AWIQGGAEQF | 0.66872174 |
| VPATVPVGILC | AWIQGGAEQF | 0.31204545 |
| VQRPDSGGVVLTGRIS | AWIQGGAEQF | 0.67121047 |
| WTPNPVPATKGVPAGLVQAT | AWIQGGAEQF | 0.5704179 |
| YALKRALAKFSDRFDVVII | AWIQGGAEQF | 0.7651391 |
| YLIGSVIV | AWIQGGAEQF | 0.8066299 |
| YGQLVAVID | AWIQGGAEQF | 0.62975174 |
| VTVGGMTLPA | AWIQGGAEQF | 0.8799814 |
| VLALPAV | AWIQGGAEQF | 0.858043 |
| VIPPTLNYET | AWIQGGAEQF | 0.4310147 |
| VGMGGMHPSSG | AWIQGGAEQF | 0.7263099 |

| | | |
|---|---|---|
| VGLAGGAATA | AWIQGGAEQF | 0.56257254 |
| VGGAGGAGGSGP | AWIQGGAEQF | 0.7189064 |
| VFAAFMV | AWIQGGAEQF | 0.29467463 |
| TLYDKLR | AWIQGGAEQF | 0.84623235 |
| TLAVFTVL | AWIQGGAEQF | 0.59536 |
| TGSVQPARV | AWIQGGAEQF | 0.115974076 |
| TGGGAGSGGT | AWIQGGAEQF | 0.7484125 |
| SHLETSL | AWIQGGAEQF | 0.73309726 |
| SFTTGMT | AWIQGGAEQF | 0.7238684 |
| QVEVAKRMGVRQPTVS | AWIQGGAEQF | 0.8587202 |
| QGSGGPALAM | AWIQGGAEQF | 0.19490513 |
| MLAPSNP | AWIQGGAEQF | 0.5258802 |
| LVNNAAI | AWIQGGAEQF | 0.8524431 |
| LTGEINTLA | AWIQGGAEQF | 0.8747053 |
| LSIFGPV | AWIQGGAEQF | 0.42931512 |
| LLALLANL | AWIQGGAEQF | 0.82421404 |
| LISNASDALDKLR | AWIQGGAEQF | 0.84317756 |
| LINEAVP | AWIQGGAEQF | 0.7983637 |
| LGYAFSGL | AWIQGGAEQF | 0.8383307 |
| LDFVAVAL | AWIQGGAEQF | 0.8627199 |
| LAQPGAE | AWIQGGAEQF | 0.50423014 |
| ISNLGAVTGF | AWIQGGAEQF | 0.86904955 |
| HQEDPGRI | AWIQGGAEQF | 0.875554 |
| GTPGEPTP | AWIQGGAEQF | 0.6509905 |
| GFFNSGS | AWIQGGAEQF | 0.80495393 |
| GAGGGGLTAGAN | AWIQGGAEQF | 0.51988196 |
| FVDALVNM | AWIQGGAEQF | 0.57224846 |
| FLTVAPEL | AWIQGGAEQF | 0.2239104 |
| FETLTAL | AWIQGGAEQF | 0.8248993 |
| DVINPIEL | AWIQGGAEQF | 0.8555051 |
| APVSGAPV | AWIQGGAEQF | 0.7369211 |
| ALVGGLAW | AWIQGGAEQF | 0.5358182 |
| ALAAAAATAI | AWIQGGAEQF | 0.5318125 |
| AFGLGLLL | AWIQGGAEQF | 0.6507044 |
| AFFAAASL | AWIQGGAEQF | 0.7673606 |
| AAAALPAFSPP | AWIQGGAEQF | 0.20964947 |
| VVLAPAL | AWIQGGAEQF | 0.6024613 |
| TIGHVDHGKTTL | AWIQGGAEQF | 0.8641306 |
| LISNASDALDK | AWIQGGAEQF | 0.8427174 |
| LISNASDAL | AWIQGGAEQF | 0.3207708 |
| ISNASDALDK | AWIQGGAEQF | 0.8740685 |
| NQAETLVYQTE | AWIQGGAEQF | 0.80414766 |
| TGGTGGTAGAG | AWIQGGAEQF | 0.8421157 |
| TGSTAGGAGGA | AWIQGGAEQF | 0.5965135 |
| DPLAVPVAAGPV | AWIQGGAEQF | 0.56660664 |
| GGTGGTPGNGGN | AWIQGGAEQF | 0.81112707 |
| PAGLAGVDPATL | AWIQGGAEQF | 0.74150896 |
| GNNLVVEAPHAGASV | AWIQGGAEQF | 0.37648803 |
| KVTRIMSASTLEQPA | AWIQGGAEQF | 0.85007215 |
| LGFALPIALPRRTVV | AWIQGGAEQF | 0.7020665 |
| TRLGNLTARKLVVLV | AWIQGGAEQF | 0.71241724 |

| | | |
|---|---|---|
| WSQQRRLIYFPSAGP | AWIQGGAEQF | 0.62723726 |
| GPNGNPDPMAAAVDIR | AWIQGGAEQF | 0.79293823 |
| TQRCVSVADLRAAASR | AWIQGGAEQF | 0.55851454 |
| QFVLDVAEHPVEVLTETV | AWIQGGAEQF | 0.5546893 |
| GGAGGTGGAGGNGGTQAGDG | AWIQGGAEQF | 0.36833 |
| PLLATLRAIAVDVDAKPAQV | AWIQGGAEQF | 0.76606286 |
| APKFRTGGYMIDFWGVGYQVAK | AWIQGGAEQF | 0.8666728 |
| GAGGAGGAGGAAGGPGGTGGTG | AWIQGGAEQF | 0.7301732 |
| IGALGLLATDVAPAGSVAFGVA | AWIQGGAEQF | 0.6542575 |
| AGGAGGIGGNGGAGGAGGVLMG | AWIQGGAEQF | 0.8376175 |
| ADELIGLL | AWIQGGAEQF | 0.76856065 |
| ADRALKA | AWIQGGAEQF | 0.7326008 |
| AGAVGQVGGAGGA | AWIQGGAEQF | 0.8003262 |
| AGGGGGLGGAP | AWIQGGAEQF | 0.66397965 |
| AGGLLYGNGGAG | AWIQGGAEQF | 0.80944324 |
| AGLESIGL | AWIQGGAEQF | 0.7926862 |
| AGLKTGII | AWIQGGAEQF | 0.8316642 |
| ASLAALLL | AWIQGGAEQF | 0.8354805 |
| ASVINISSI | AWIQGGAEQF | 0.781655 |
| ATPPGPA | AWIQGGAEQF | 0.6873007 |
| ATVASALV | AWIQGGAEQF | 0.19699307 |
| ATVASLIL | AWIQGGAEQF | 0.5145936 |
| AVNGPVLIRH | AWIQGGAEQF | 0.8313568 |
| DGAAGAVGI | AWIQGGAEQF | 0.80728245 |
| DPLLHGI | AWIQGGAEQF | 0.83663386 |
| ELGAALGAL | AWIQGGAEQF | 0.73577404 |
| ELGRTQAV | AWIQGGAEQF | 0.58540666 |
| FVAGGYLS | AWIQGGAEQF | 0.804426 |
| GASGGAGGAGGW | AWIQGGAEQF | 0.829498 |
| GGAGAGGDGGAGGI | AWIQGGAEQF | 0.40410015 |
| GTGHNGSQPGMGGNG | AWIQGGAEQF | 0.6487594 |
| GYAAASASAS | AWIQGGAEQF | 0.8862518 |
| HQVDVAAVQQG | AWIQGGAEQF | 0.7687277 |
| IGAAIAHT | AWIQGGAEQF | 0.7528316 |
| IGMIEVAQL | AWIQGGAEQF | 0.56962883 |
| IRLAPEHTSAMD | AWIQGGAEQF | 0.862248 |
| IYNVSVL | AWIQGGAEQF | 0.5447052 |
| KQGFLGM | AWIQGGAEQF | 0.870041 |
| LESVITQL | AWIQGGAEQF | 0.16967122 |
| LGLGALVVV | AWIQGGAEQF | 0.5286933 |
| LIGNGGNGGSGGI | AWIQGGAEQF | 0.373194 |
| LNAINAPTL | AWIQGGAEQF | 0.37024516 |
| LQLLNTGV | AWIQGGAEQF | 0.84931344 |
| LQTSIEF | AWIQGGAEQF | 0.58027685 |
| MSFAVSEL | AWIQGGAEQF | 0.8495701 |
| NTGALGL | AWIQGGAEQF | 0.7643206 |
| PPGAGGAGGAGTT | AWIQGGAEQF | 0.84582984 |
| QLVGLGI | AWIQGGAEQF | 0.7927761 |
| RFLEQVHQL | AWIQGGAEQF | 0.7934277 |
| RGVGYVLR | AWIQGGAEQF | 0.5122701 |
| SATGVFAGVI | AWIQGGAEQF | 0.84389657 |

| | | |
|---|---|---|
| SLPAITLP | AWIQGGAEQF | 0.8413287 |
| STALAFTV | AWIQGGAEQF | 0.69392025 |
| SVLTVGLGK | AWIQGGAEQF | 0.41939953 |
| TALAGAGSGI | AWIQGGAEQF | 0.58466685 |
| TGLPFTGI | AWIQGGAEQF | 0.7808808 |
| VAEHVTV | AWIQGGAEQF | 0.17796287 |
| VDIVLNSL | AWIQGGAEQF | 0.16439848 |
| VLTGGGAVL | AWIQGGAEQF | 0.6691901 |
| VNVGLAYGI | AWIQGGAEQF | 0.49989635 |
| VSSAAVGAV | AWIQGGAEQF | 0.87290525 |
| VTSVYEAGV | AWIQGGAEQF | 0.8615968 |
| WVALDRGA | AWIQGGAEQF | 0.7732675 |
| YLVGSVVI | AWIQGGAEQF | 0.80557436 |
| GPPGTGKTLL | AWIQGGAEQF | 0.79646647 |
| GVGMTKF | AWIQGGAEQF | 0.10596511 |
| IGIGGSDL | AWIQGGAEQF | 0.5433425 |
| IGTLLAEL | AWIQGGAEQF | 0.50143546 |
| IYTSGTT | AWIQGGAEQF | 0.86823475 |
| LSFPAGV | AWIQGGAEQF | 0.56518424 |
| LYTSGTT | AWIQGGAEQF | 0.6189441 |
| VGAFPFGF | AWIQGGAEQF | 0.8092587 |
| VGAGTVEF | AWIQGGAEQF | 0.7689069 |
| VGKSTLLL | AWIQGGAEQF | 0.71064126 |
| AALAHAGGGIV | AWIQGGAEQF | 0.49697423 |
| AALVAVSLL | AWIQGGAEQF | 0.115604155 |
| AAVSRAVAAL | AWIQGGAEQF | 0.8573794 |
| AGGSSGIGGF | AWIQGGAEQF | 0.2506933 |
| AGLTVANANG | AWIQGGAEQF | 0.502815 |
| AGLVGAVL | AWIQGGAEQF | 0.17451037 |
| ALSQAFTV | AWIQGGAEQF | 0.13235755 |
| ALTFSVSL | AWIQGGAEQF | 0.3360109 |
| ALTGAALVAL | AWIQGGAEQF | 0.8332219 |
| ASFSIGAGVL | AWIQGGAEQF | 0.73734736 |
| DGSVYKI | AWIQGGAEQF | 0.7055206 |
| EQLAEIV | AWIQGGAEQF | 0.8758851 |
| FAVSELTVM | AWIQGGAEQF | 0.78031874 |
| GAAPLAAGL | AWIQGGAEQF | 0.72584915 |
| GADGGDGGSGGAGG | AWIQGGAEQF | 0.6164541 |
| GFGLDTGGAGG | AWIQGGAEQF | 0.55456936 |
| GLASGGLL | AWIQGGAEQF | 0.5175079 |
| GRIAGAAVLL | AWIQGGAEQF | 0.5132612 |
| GSAGSLNTSF | AWIQGGAEQF | 0.88376594 |
| GTGQIGI | AWIQGGAEQF | 0.55159295 |
| GTLIALIL | AWIQGGAEQF | 0.3922425 |
| IAATMGFVVG | AWIQGGAEQF | 0.5729459 |
| IFAQALL | AWIQGGAEQF | 0.87022793 |
| IGAAVALGF | AWIQGGAEQF | 0.8834721 |
| IGLTPLAL | AWIQGGAEQF | 0.639869 |
| ILGSNTSTL | AWIQGGAEQF | 0.4726295 |
| IVFSLTVL | AWIQGGAEQF | 0.5620478 |
| LASKATGF | AWIQGGAEQF | 0.80021214 |

| | | |
|---|---|---|
| LASVALIL | AWIQGGAEQF | 0.52663106 |
| LFAVGLTL | AWIQGGAEQF | 0.42729867 |
| LFNGGIGTYI | AWIQGGAEQF | 0.77330434 |
| LTLGAGII | AWIQGGAEQF | 0.8747692 |
| LVAAALGDI | AWIQGGAEQF | 0.59404814 |
| NTAFVQLGI | AWIQGGAEQF | 0.34331304 |
| QLFSGAEL | AWIQGGAEQF | 0.7816768 |
| SAATFITL | AWIQGGAEQF | 0.6257279 |
| SNYGLANLGV | AWIQGGAEQF | 0.7391974 |
| SYGGAYAVM | AWIQGGAEQF | 0.7564452 |
| TTGVGGLTLG | AWIQGGAEQF | 0.3962918 |
| TTQTIYI | AWIQGGAEQF | 0.5788648 |
| VAAGAGLGAVI | AWIQGGAEQF | 0.87312627 |
| VAAVVTLL | AWIQGGAEQF | 0.68504506 |
| VGFLGGLI | AWIQGGAEQF | 0.7668694 |
| VGVIPILL | AWIQGGAEQF | 0.8630805 |
| VIFASSLI | AWIQGGAEQF | 0.8333746 |
| VKGGDGGAAAG | AWIQGGAEQF | 0.8739666 |
| VTKTFTNI | AWIQGGAEQF | 0.83043945 |
| VVAAAILL | AWIQGGAEQF | 0.67464685 |
| YAALDVEL | AWIQGGAEQF | 0.65020245 |
| LASVALLL | AWIQGGAEQF | 0.7615386 |
| LDAVHRV | AWIQGGAEQF | 0.8214407 |
| QGLSSSIF | AWIQGGAEQF | 0.7512815 |
| RGTFANIRL | AWIQGGAEQF | 0.5079984 |
| RQLWWGHRI | AWIQGGAEQF | 0.85954595 |
| YGPPGTGKTLL | AWIQGGAEQF | 0.57918334 |
| AALVGAPANVA | AWIQGGAEQF | 0.81277 |
| AATFRRTE | AWIQGGAEQF | 0.5298436 |
| AGALGGGI | AWIQGGAEQF | 0.4763277 |
| AGGAGGAGGIGAG | AWIQGGAEQF | 0.8536389 |
| AGSGALGL | AWIQGGAEQF | 0.10622227 |
| AQIADQI | AWIQGGAEQF | 0.17003347 |
| ASAVAVTYL | AWIQGGAEQF | 0.6946622 |
| ASTVLGGTGG | AWIQGGAEQF | 0.85942966 |
| AVGGEPAK | AWIQGGAEQF | 0.12308218 |
| CEQGTFGGTC | AWIQGGAEQF | 0.4526105 |
| DEGKAPAG | AWIQGGAEQF | 0.4092668 |
| DTVVFPAL | AWIQGGAEQF | 0.75231445 |
| FEGITMAG | AWIQGGAEQF | 0.76766634 |
| FPLVVQD | AWIQGGAEQF | 0.35631946 |
| GAFTTPQ | AWIQGGAEQF | 0.83125865 |
| GFGGFGTAGGVGGA | AWIQGGAEQF | 0.34712207 |
| GFLAQFL | AWIQGGAEQF | 0.29075348 |
| GGAGGAGGNAGVA | AWIQGGAEQF | 0.83513904 |
| GGVGTAGAGGAGGAGG | AWIQGGAEQF | 0.8497013 |
| GNGGGGDAGNAGSGGN | AWIQGGAEQF | 0.45682588 |
| GTGGAGGAGNDAGS | AWIQGGAEQF | 0.8491456 |
| GTTNTGWFNS | AWIQGGAEQF | 0.52692276 |
| IFMGPLFTL | AWIQGGAEQF | 0.87641287 |
| ILAHAIEAI | AWIQGGAEQF | 0.68084764 |

| | | |
|---|---|---|
| IRSAIREAL | AWIQGGAEQF | 0.3392425 |
| ITFAGGG | AWIQGGAEQF | 0.8311477 |
| ITGMSGHQEV | AWIQGGAEQF | 0.7917545 |
| IVAGLACTL | AWIQGGAEQF | 0.694141 |
| IVGGGLAGA | AWIQGGAEQF | 0.82589126 |
| IVTGVLNI | AWIQGGAEQF | 0.4737302 |
| LAQQGCI | AWIQGGAEQF | 0.76528335 |
| LCLHVEN | AWIQGGAEQF | 0.8742073 |
| LDAISDGS | AWIQGGAEQF | 0.86202425 |
| LGGFAAML | AWIQGGAEQF | 0.49087575 |
| LGGKNPMI | AWIQGGAEQF | 0.8071529 |
| LGPKGVAAIL | AWIQGGAEQF | 0.7859492 |
| LMVGVSI | AWIQGGAEQF | 0.595184 |
| LVGAAFVL | AWIQGGAEQF | 0.86290675 |
| LVLAVAPL | AWIQGGAEQF | 0.70785916 |
| MAAIAVIDAA | AWIQGGAEQF | 0.83659196 |
| NNGINAELQ | AWIQGGAEQF | 0.79507804 |
| RGEPLVLVHG | AWIQGGAEQF | 0.82220376 |
| RGGAGGAAGLI | AWIQGGAEQF | 0.36581543 |
| RSYRMHPAV | AWIQGGAEQF | 0.7304964 |
| SAAHLLAGQLA | AWIQGGAEQF | 0.72087765 |
| SAAVGSY | AWIQGGAEQF | 0.8357183 |
| SPHAVLHRI | AWIQGGAEQF | 0.698001 |
| SQIAKVL | AWIQGGAEQF | 0.27726933 |
| TALGHLVP | AWIQGGAEQF | 0.7610967 |
| TAPTTSGAA | AWIQGGAEQF | 0.85798866 |
| TDGDDGNGGSPGT | AWIQGGAEQF | 0.7214126 |
| TDGLGEN | AWIQGGAEQF | 0.7321394 |
| TGGPPLPGQA | AWIQGGAEQF | 0.88111055 |
| TIPAITI | AWIQGGAEQF | 0.57796353 |
| TIPGFTI | AWIQGGAEQF | 0.7580416 |
| TSIASGALA | AWIQGGAEQF | 0.7665176 |
| TVTLPGAAI | AWIQGGAEQF | 0.84246784 |
| VASGLSAALP | AWIQGGAEQF | 0.64939535 |
| VGAGVIGI | AWIQGGAEQF | 0.8579396 |
| VGPKRFGV | AWIQGGAEQF | 0.42891476 |
| VNVAFPD | AWIQGGAEQF | 0.861346 |
| VSMTNSGVS | AWIQGGAEQF | 0.560961 |
| VTDVLGDLSV | AWIQGGAEQF | 0.83340305 |
| VVVVGAGFAGL | AWIQGGAEQF | 0.6713705 |
| VYTVGTVV | AWIQGGAEQF | 0.8582701 |
| WTTFSGP | AWIQGGAEQF | 0.85117906 |
| YDELVEAF | AWIQGGAEQF | 0.8646352 |
| YGTGGAGGNGG | AWIQGGAEQF | 0.14958002 |
| YTNVMAV | AWIQGGAEQF | 0.8601806 |
| FLEQVHQL | AWIQGGAEQF | 0.8646797 |
| GLSLLAVAV | AWIQGGAEQF | 0.8835862 |
| LEQVHQL | AWIQGGAEQF | 0.63635856 |
| LNAAGAI | AWIQGGAEQF | 0.75902283 |
| PAVHASL | AWIQGGAEQF | 0.7256425 |
| TELEDDA | AWIQGGAEQF | 0.641503 |

| | | |
|---|---|---|
| TPPGPPN | AWIQGGAEQF | 0.61275274 |
| TPPPPPPP | AWIQGGAEQF | 0.6328961 |
| TQSPALI | TCSAEGENEQY | 0.745965 |
| EFAAAFPQ | TCSAEGENEQY | 0.68026376 |
| LLSGNLDVL | TCSAEGENEQY | 0.83664834 |
| GPANLGNYNI | TCSAEGENEQY | 0.8460587 |
| TLGPTLP | TCSAEGENEQY | 0.8442801 |
| VYAYSSCCK | TCSAEGENEQY | 0.86124194 |
| APAPAPAPNI | TCSAEGENEQY | 0.85268956 |
| PGGPVPIP | TCSAEGENEQY | 0.83049214 |
| FLTVALEP | TCSAEGENEQY | 0.5954615 |
| SQYFGANL | TCSAEGENEQY | 0.68215144 |
| GGVKPAA | TCSAEGENEQY | 0.5653728 |
| GGTGGAGGAGGW | TCSAEGENEQY | 0.6495423 |
| GVGGVAGLGGAGG | TCSAEGENEQY | 0.6730549 |
| VLLVLATTI | TCSAEGENEQY | 0.80155176 |
| IGIFAAML | TCSAEGENEQY | 0.107862 |
| AVGDLID | TCSAEGENEQY | 0.72366726 |
| NISPLPL | TCSAEGENEQY | 0.8778652 |
| VTAVFVAI | TCSAEGENEQY | 0.5874905 |
| SGDDNDAAL | TCSAEGENEQY | 0.74932426 |
| AVAALAHS | TCSAEGENEQY | 0.68892753 |
| GGVTVMATP | TCSAEGENEQY | 0.64454204 |
| AGLAANVRD | TCSAEGENEQY | 0.84327865 |
| TNRPDLIDPALLRPG | TCSAEGENEQY | 0.5395929 |
| LANLGVTAL | TCSAEGENEQY | 0.8029436 |
| TGERIEL | TCSAEGENEQY | 0.86463225 |
| NGEELAA | TCSAEGENEQY | 0.74517894 |
| VLSLVWG | TCSAEGENEQY | 0.6945219 |
| IGYALWL | TCSAEGENEQY | 0.65751755 |
| LSLLLAPL | TCSAEGENEQY | 0.33973044 |
| TLDEALK | TCSAEGENEQY | 0.7851933 |
| TDGAGTAEGG | TCSAEGENEQY | 0.1022252 |
| NAPGGAGGNGG | TCSAEGENEQY | 0.76107335 |
| VSVFMGMGGNF | TCSAEGENEQY | 0.50270855 |
| LASYLAAL | TCSAEGENEQY | 0.8278047 |
| LVGGTPAL | TCSAEGENEQY | 0.80697787 |
| VRNADVL | TCSAEGENEQY | 0.7731943 |
| TCSSPFLVSAM | TCSAEGENEQY | 0.6342164 |
| VHITDAE | TCSAEGENEQY | 0.6974669 |
| EYAAPSF | TCSAEGENEQY | 0.44584376 |
| VVDAFHGGL | TCSAEGENEQY | 0.39311084 |
| ASVVDPQ | TCSAEGENEQY | 0.82148093 |
| SYNEFGGLS | TCSAEGENEQY | 0.5816059 |
| DPDGSVY | TCSAEGENEQY | 0.654863 |
| MVSTIHATE | TCSAEGENEQY | 0.85735893 |
| LAAGVADPV | TCSAEGENEQY | 0.761942 |
| TKEQIAA | TCSAEGENEQY | 0.3500423 |
| SLASPAAL | TCSAEGENEQY | 0.75716907 |
| IGLVFYGV | TCSAEGENEQY | 0.758888 |
| FLLDASNL | TCSAEGENEQY | 0.71850014 |

| | | |
|---|---|---|
| YQVGNFP | TCSAEGENEQY | 0.10028469 |
| ESLLSQST | TCSAEGENEQY | 0.7733494 |
| LFTVGFAAVG | TCSAEGENEQY | 0.62222403 |
| ATPPTPT | TCSAEGENEQY | 0.55685496 |
| TDLATLG | TCSAEGENEQY | 0.58210164 |
| SAGGTGR | TCSAEGENEQY | 0.8540635 |
| GFSNSGS | TCSAEGENEQY | 0.25858656 |
| GGNGGSGGTSV | TCSAEGENEQY | 0.7776737 |
| GTSVATGGAGN | TCSAEGENEQY | 0.8351441 |
| NGGNGGSGGTS | TCSAEGENEQY | 0.83799136 |
| AVVKFPH | TCSAEGENEQY | 0.87807393 |
| TPPPPPP | TCSAEGENEQY | 0.80066526 |
| AAPAAAAAPA | TCSAEGENEQY | 0.461208 |
| IALNEDL | TCSAEGENEQY | 0.7419187 |
| IVNVSSM | TCSAEGENEQY | 0.5792974 |
| TGIPPAPR | TCSAEGENEQY | 0.20859043 |
| FDLGGGTF | TCSAEGENEQY | 0.17355391 |
| FDLGGGTFDVS | TCSAEGENEQY | 0.80944043 |
| NEPTAAA | TCSAEGENEQY | 0.1982317 |
| NEPTAAALA | TCSAEGENEQY | 0.76002955 |
| GSIINISSI | TCSAEGENEQY | 0.8353454 |
| VAPGNAGTA | TCSAEGENEQY | 0.5501446 |
| TGGGASLE | TCSAEGENEQY | 0.3899914 |
| VAVYNHYKRI | TCSAEGENEQY | 0.6744666 |
| TDPAPATT | TCSAEGENEQY | 0.7087022 |
| TNRPDILDPALLRPG | TCSAEGENEQY | 0.84548336 |
| LTTTTTS | TCSAEGENEQY | 0.72782016 |
| PTGSGKTL | TCSAEGENEQY | 0.8714324 |
| TGSGKTL | TCSAEGENEQY | 0.8535887 |
| TIGHVDHGK | TCSAEGENEQY | 0.7202707 |
| TIGHVDHGKTT | TCSAEGENEQY | 0.83114815 |
| GHVDHGKTT | TCSAEGENEQY | 0.65320706 |
| AQFEHTLL | TCSAEGENEQY | 0.6212015 |
| IVNQDEL | TCSAEGENEQY | 0.6716832 |
| TNGSQFF | TCSAEGENEQY | 0.7161618 |
| TGKGIVE | TCSAEGENEQY | 0.7504328 |
| LISNASDA | TCSAEGENEQY | 0.63712025 |
| AELRTLGELTKIAKAHGAQ | TCSAEGENEQY | 0.32462347 |
| APNAVAAAEPSW | TCSAEGENEQY | 0.7747917 |
| APQVAGAATAVL | TCSAEGENEQY | 0.7615508 |
| ARRDPAAAAAILPTDA | TCSAEGENEQY | 0.23301895 |
| DAAARDVLAEAGLAEYF | TCSAEGENEQY | 0.62935174 |
| DDQIVLSAKANLAKVEQ | TCSAEGENEQY | 0.8870307 |
| DELLPDTGEDSDEDGH | TCSAEGENEQY | 0.85987735 |
| ELPEAPANRATVPQD | TCSAEGENEQY | 0.61401165 |
| FGQSKIERVTGIG | TCSAEGENEQY | 0.5871426 |
| GAGGTGSTAGGAGGAGGAGGLY | TCSAEGENEQY | 0.7262057 |
| GHGGTGAAGGNGGTGGAGGSGI | TCSAEGENEQY | 0.23862626 |
| GNGGNGGAGGEGGPGGAGGAGA | TCSAEGENEQY | 0.74532646 |
| GSTAGNGGNGGNATNGTIAGQP | TCSAEGENEQY | 0.788857 |
| GTGGDGGNGGNGGTGG | TCSAEGENEQY | 0.78778934 |

| | | |
|---|---|---|
| GVRRPAGIPADAPV | TCSAEGENEQY | 0.46029636 |
| GYRQVTDAHLLTLARR | TCSAEGENEQY | 0.76392233 |
| IAKMGQDGHDRG | TCSAEGENEQY | 0.68748116 |
| KAEPSIEIQYT | TCSAEGENEQY | 0.7706722 |
| KPVRIRMAADEDTVAVT | TCSAEGENEQY | 0.13015977 |
| MPPRIAGMRLLVIKPEP | TCSAEGENEQY | 0.7567212 |
| MKFYAVALFRSRGVPPTR | TCSAEGENEQY | 0.6443687 |
| NAGGFGILGAVAHSP | TCSAEGENEQY | 0.44749293 |
| NEGREITLAMAEM | TCSAEGENEQY | 0.33460322 |
| NELVEWSHPLGHRWRWE | TCSAEGENEQY | 0.6786992 |
| NLGAGGTGGNGGNPG | TCSAEGENEQY | 0.8396672 |
| NTEWPKWETVMAADPVD | TCSAEGENEQY | 0.8468931 |
| PASAAPAETQP | TCSAEGENEQY | 0.31573045 |
| QQSSDGKNGVRAAEITGE | TCSAEGENEQY | 0.8623796 |
| RAPDAVTTSWWKE | TCSAEGENEQY | 0.87455475 |
| SGFGNVGAHTSGWFNQ | TCSAEGENEQY | 0.6697045 |
| SGVAGAVGQAASVGGLKV | TCSAEGENEQY | 0.34479254 |
| SNGGAGTGSAGGNGGT | TCSAEGENEQY | 0.8238692 |
| TGGAGGEGGAAG | TCSAEGENEQY | 0.584349 |
| TGGDGGAAGAGGAPGHG | TCSAEGENEQY | 0.814888 |
| TMPPLLAAIANAGRHGV | TCSAEGENEQY | 0.78029203 |
| TSSTAGNSGGA | TCSAEGENEQY | 0.5612485 |
| VPATVPVGILC | TCSAEGENEQY | 0.80863345 |
| VQRPDSGGVVLTGRIS | TCSAEGENEQY | 0.84377503 |
| WTPNPVPATKGVPAGLVQAT | TCSAEGENEQY | 0.6124166 |
| YALKRALAKFSDRFDVVII | TCSAEGENEQY | 0.46808973 |
| YLIGSVIV | TCSAEGENEQY | 0.52467024 |
| YGQLVAVID | TCSAEGENEQY | 0.19193101 |
| VTVGGMTLPA | TCSAEGENEQY | 0.81952643 |
| VLALPAV | TCSAEGENEQY | 0.6600629 |
| VIPPTLNYET | TCSAEGENEQY | 0.57432365 |
| VGMGGMHPSSG | TCSAEGENEQY | 0.8350477 |
| VGLAGGAATA | TCSAEGENEQY | 0.5517844 |
| VGGAGGAGGSGP | TCSAEGENEQY | 0.6784916 |
| VFAAFMV | TCSAEGENEQY | 0.6223203 |
| TLYDKLR | TCSAEGENEQY | 0.7535864 |
| TLAVFTVL | TCSAEGENEQY | 0.79076415 |
| TGSVQPARV | TCSAEGENEQY | 0.76167023 |
| TGGGAGSGGT | TCSAEGENEQY | 0.6670318 |
| SHLETSL | TCSAEGENEQY | 0.8754338 |
| SFTTGMT | TCSAEGENEQY | 0.4443671 |
| QVEVAKRMGVRQPTVS | TCSAEGENEQY | 0.6335303 |
| QGSGGPALAM | TCSAEGENEQY | 0.7995889 |
| MLAPSNP | TCSAEGENEQY | 0.2997398 |
| LVNNAAI | TCSAEGENEQY | 0.8340173 |
| LTGEINTLA | TCSAEGENEQY | 0.79159987 |
| LSIFGPV | TCSAEGENEQY | 0.8046708 |
| LLALLANL | TCSAEGENEQY | 0.51564145 |
| LISNASDALDKLR | TCSAEGENEQY | 0.8851887 |
| LINEAVP | TCSAEGENEQY | 0.85430205 |
| LGYAFSGL | TCSAEGENEQY | 0.68255186 |

| | | |
|---|---|---|
| LDFVAVAL | TCSAEGENEQY | 0.75865114 |
| LAQPGAE | TCSAEGENEQY | 0.43138802 |
| ISNLGAVTGF | TCSAEGENEQY | 0.8541231 |
| HQEDPGRI | TCSAEGENEQY | 0.7889118 |
| GTPGEPTP | TCSAEGENEQY | 0.55461764 |
| GFFNSGS | TCSAEGENEQY | 0.7696283 |
| GAGGGGLTAGAN | TCSAEGENEQY | 0.84039116 |
| FVDALVNM | TCSAEGENEQY | 0.863705 |
| FLTVAPEL | TCSAEGENEQY | 0.21577603 |
| FETLTAL | TCSAEGENEQY | 0.7624285 |
| DVINPIEL | TCSAEGENEQY | 0.19759673 |
| APVSGAPV | TCSAEGENEQY | 0.3897546 |
| ALVGGLAW | TCSAEGENEQY | 0.5157722 |
| ALAAAAATAI | TCSAEGENEQY | 0.8011699 |
| AFGLGLLL | TCSAEGENEQY | 0.72414804 |
| AFFAAASL | TCSAEGENEQY | 0.7751391 |
| AAAALPAFSPP | TCSAEGENEQY | 0.7914896 |
| VVLAPAL | TCSAEGENEQY | 0.86850595 |
| TIGHVDHGKTTL | TCSAEGENEQY | 0.55402565 |
| LISNASDALDK | TCSAEGENEQY | 0.7652627 |
| LISNASDAL | TCSAEGENEQY | 0.37085888 |
| ISNASDALDK | TCSAEGENEQY | 0.27769583 |
| NQAETLVYQTE | TCSAEGENEQY | 0.49191692 |
| TGGTGGTAGAG | TCSAEGENEQY | 0.7156265 |
| TGSTAGGAGGA | TCSAEGENEQY | 0.18806782 |
| DPLAVPVAAGPV | TCSAEGENEQY | 0.8470541 |
| GGTGGTPGNGGN | TCSAEGENEQY | 0.8137915 |
| PAGLAGVDPATL | TCSAEGENEQY | 0.84551334 |
| GNNLVVEAPHAGASV | TCSAEGENEQY | 0.5762357 |
| KVTRIMSASTLEQPA | TCSAEGENEQY | 0.7055199 |
| LGFALPIALPRRTVV | TCSAEGENEQY | 0.6451268 |
| TRLGNLTARKLVVLV | TCSAEGENEQY | 0.82910144 |
| WSQQRRLIYFPSAGP | TCSAEGENEQY | 0.5690191 |
| GPNGNPDPMAAAVDIR | TCSAEGENEQY | 0.3276317 |
| TQRCVSVADLRAAASR | TCSAEGENEQY | 0.49602866 |
| QFVLDVAEHPVEVLTETV | TCSAEGENEQY | 0.4828343 |
| GGAGGTGGAGGNGGTQAGDG | TCSAEGENEQY | 0.8516391 |
| PLLATLRAIAVDVDAKPAQV | TCSAEGENEQY | 0.8463713 |
| APKFRTGGYMIDFWGVGYQVAK | TCSAEGENEQY | 0.824203 |
| GAGGAGGAGGAAGGPGGTGGTG | TCSAEGENEQY | 0.8167554 |
| IGALGLLATDVAPAGSVAFGVA | TCSAEGENEQY | 0.22999252 |
| AGGAGGIGGNGGAGGAGGVLMG | TCSAEGENEQY | 0.61371386 |
| ADELIGLL | TCSAEGENEQY | 0.8821486 |
| ADRALKA | TCSAEGENEQY | 0.757775 |
| AGAVGQVGGAGGA | TCSAEGENEQY | 0.74841547 |
| AGGGGGLGGAP | TCSAEGENEQY | 0.8720634 |
| AGGLLYGNGGAG | TCSAEGENEQY | 0.66266966 |
| AGLESIGL | TCSAEGENEQY | 0.7126384 |
| AGLKTGII | TCSAEGENEQY | 0.73533535 |
| ASLAALLL | TCSAEGENEQY | 0.76733094 |
| ASVINISSI | TCSAEGENEQY | 0.7028114 |

| | | |
|---|---|---|
| ATPPGPA | TCSAEGENEQY | 0.5543978 |
| ATVASALV | TCSAEGENEQY | 0.74346775 |
| ATVASLIL | TCSAEGENEQY | 0.88634074 |
| AVNGPVLIRH | TCSAEGENEQY | 0.7702147 |
| DGAAGAVGI | TCSAEGENEQY | 0.8583388 |
| DPLLHGI | TCSAEGENEQY | 0.81622773 |
| ELGAALGAL | TCSAEGENEQY | 0.8044992 |
| ELGRTQAV | TCSAEGENEQY | 0.63133466 |
| FVAGGYLS | TCSAEGENEQY | 0.73747635 |
| GASGGAGGAGGW | TCSAEGENEQY | 0.68763757 |
| GGAGAGGDGGAGGI | TCSAEGENEQY | 0.6724031 |
| GTGHNGSQPGMGGNG | TCSAEGENEQY | 0.6772473 |
| GYAAASASAS | TCSAEGENEQY | 0.6003899 |
| HQVDVAAVQQG | TCSAEGENEQY | 0.88952535 |
| IGAAIAHT | TCSAEGENEQY | 0.80403703 |
| IGMIEVAQL | TCSAEGENEQY | 0.81750906 |
| IRLAPEHTSAMD | TCSAEGENEQY | 0.59129024 |
| IYNVSVL | TCSAEGENEQY | 0.715739 |
| KQGFLGM | TCSAEGENEQY | 0.82462746 |
| LESVITQL | TCSAEGENEQY | 0.39469594 |
| LGLGALVVV | TCSAEGENEQY | 0.8704497 |
| LIGNGGNGGSGGI | TCSAEGENEQY | 0.69235647 |
| LNAINAPTL | TCSAEGENEQY | 0.5422434 |
| LQLLNTGV | TCSAEGENEQY | 0.84796584 |
| LQTSIEF | TCSAEGENEQY | 0.86182976 |
| MSFAVSEL | TCSAEGENEQY | 0.6066816 |
| NTGALGL | TCSAEGENEQY | 0.60833305 |
| PPGAGGAGGAGTT | TCSAEGENEQY | 0.6770178 |
| QLVGLGI | TCSAEGENEQY | 0.8029077 |
| RFLEQVHQL | TCSAEGENEQY | 0.8684442 |
| RGVGYVLR | TCSAEGENEQY | 0.51978004 |
| SATGVFAGVI | TCSAEGENEQY | 0.8683448 |
| SLPAITLP | TCSAEGENEQY | 0.6765371 |
| STALAFTV | TCSAEGENEQY | 0.36518547 |
| SVLTVGLGK | TCSAEGENEQY | 0.72139114 |
| TALAGAGSGI | TCSAEGENEQY | 0.844162 |
| TGLPFTGI | TCSAEGENEQY | 0.83918107 |
| VAEHVTV | TCSAEGENEQY | 0.8873135 |
| VDIVLNSL | TCSAEGENEQY | 0.78115267 |
| VLTGGGAVL | TCSAEGENEQY | 0.5663376 |
| VNVGLAYGI | TCSAEGENEQY | 0.8704383 |
| VSSAAVGAV | TCSAEGENEQY | 0.85850847 |
| VTSVYEAGV | TCSAEGENEQY | 0.84830105 |
| WVALDRGA | TCSAEGENEQY | 0.6138793 |
| YLVGSVVI | TCSAEGENEQY | 0.46223423 |
| GPPGTGKTLL | TCSAEGENEQY | 0.6772291 |
| GVGMTKF | TCSAEGENEQY | 0.8480153 |
| IGIGGSDL | TCSAEGENEQY | 0.48339257 |
| IGTLLAEL | TCSAEGENEQY | 0.77280426 |
| IYTSGTT | TCSAEGENEQY | 0.80720073 |
| LSFPAGV | TCSAEGENEQY | 0.4023653 |

| | | |
|---|---|---|
| LYTSGTT | TCSAEGENEQY | 0.7703949 |
| VGAFPFGF | TCSAEGENEQY | 0.86368144 |
| VGAGTVEF | TCSAEGENEQY | 0.6258625 |
| VGKSTLLL | TCSAEGENEQY | 0.67764735 |
| AALAHAGGGIV | TCSAEGENEQY | 0.84991574 |
| AALVAVSLL | TCSAEGENEQY | 0.893156 |
| AAVSRAVAAL | TCSAEGENEQY | 0.8534697 |
| AGGSSGIGGF | TCSAEGENEQY | 0.3999543 |
| AGLTVANANG | TCSAEGENEQY | 0.22548191 |
| AGLVGAVL | TCSAEGENEQY | 0.5963962 |
| ALSQAFTV | TCSAEGENEQY | 0.8115435 |
| ALTFSVSL | TCSAEGENEQY | 0.61899596 |
| ALTGAALVAL | TCSAEGENEQY | 0.6284958 |
| ASFSIGAGVL | TCSAEGENEQY | 0.823449 |
| DGSVYKI | TCSAEGENEQY | 0.86308694 |
| EQLAEIV | TCSAEGENEQY | 0.5402963 |
| FAVSELTVM | TCSAEGENEQY | 0.34917632 |
| GAAPLAAGL | TCSAEGENEQY | 0.7580212 |
| GADGGDGGSGGAGG | TCSAEGENEQY | 0.8074479 |
| GFGLDTGGAGG | TCSAEGENEQY | 0.70587057 |
| GLASGGLL | TCSAEGENEQY | 0.7583256 |
| GRIAGAAVLL | TCSAEGENEQY | 0.4924077 |
| GSAGSLNTSF | TCSAEGENEQY | 0.2435572 |
| GTGQIGI | TCSAEGENEQY | 0.828966 |
| GTLIALIL | TCSAEGENEQY | 0.52988887 |
| IAATMGFVVG | TCSAEGENEQY | 0.87170863 |
| IFAQALL | TCSAEGENEQY | 0.58248734 |
| IGAAVALGF | TCSAEGENEQY | 0.80085063 |
| IGLTPLAL | TCSAEGENEQY | 0.8340082 |
| ILGSNTSTL | TCSAEGENEQY | 0.79723066 |
| IVFSLTVL | TCSAEGENEQY | 0.7121316 |
| LASKATGF | TCSAEGENEQY | 0.60678875 |
| LASVALIL | TCSAEGENEQY | 0.4180268 |
| LFAVGLTL | TCSAEGENEQY | 0.20506462 |
| LFNGGIGTYI | TCSAEGENEQY | 0.8575313 |
| LTLGAGII | TCSAEGENEQY | 0.13002433 |
| LVAAALGDI | TCSAEGENEQY | 0.8701154 |
| NTAFVQLGI | TCSAEGENEQY | 0.83111334 |
| QLFSGAEL | TCSAEGENEQY | 0.6129974 |
| SAATFITL | TCSAEGENEQY | 0.78173167 |
| SNYGLANLGV | TCSAEGENEQY | 0.6070518 |
| SYGGAYAVM | TCSAEGENEQY | 0.3695953 |
| TTGVGGLTLG | TCSAEGENEQY | 0.8324548 |
| TTQTIYI | TCSAEGENEQY | 0.8277136 |
| VAAGAGLGAVI | TCSAEGENEQY | 0.61835235 |
| VAAVVTLL | TCSAEGENEQY | 0.75939333 |
| VGFLGGLI | TCSAEGENEQY | 0.77520025 |
| VGVIPILL | TCSAEGENEQY | 0.46589488 |
| VIFASSLI | TCSAEGENEQY | 0.6396357 |
| VKGGDGGAAAG | TCSAEGENEQY | 0.8528969 |
| VTKTFTNI | TCSAEGENEQY | 0.7084262 |

| | | |
|---|---|---|
| VVAAAILL | TCSAEGENEQY | 0.61892414 |
| YAALDVEL | TCSAEGENEQY | 0.80824566 |
| LASVALLL | TCSAEGENEQY | 0.52786297 |
| LDAVHRV | TCSAEGENEQY | 0.84700036 |
| QGLSSSIF | TCSAEGENEQY | 0.7366682 |
| RGTFANIRL | TCSAEGENEQY | 0.67933655 |
| RQLWWGHRI | TCSAEGENEQY | 0.8089645 |
| YGPPGTGKTLL | TCSAEGENEQY | 0.3753515 |
| AALVGAPANVA | TCSAEGENEQY | 0.42484054 |
| AATFRRTE | TCSAEGENEQY | 0.6776798 |
| AGALGGGI | TCSAEGENEQY | 0.7921721 |
| AGGAGGAGGIGAG | TCSAEGENEQY | 0.85076165 |
| AGSGALGL | TCSAEGENEQY | 0.64364123 |
| AQIADQI | TCSAEGENEQY | 0.7638921 |
| ASAVAVTYL | TCSAEGENEQY | 0.64713496 |
| ASTVLGGTGG | TCSAEGENEQY | 0.5439331 |
| AVGGEPAK | TCSAEGENEQY | 0.614438 |
| CEQGTFGGTC | TCSAEGENEQY | 0.5492302 |
| DEGKAPAG | TCSAEGENEQY | 0.6132863 |
| DTVVFPAL | TCSAEGENEQY | 0.8039272 |
| FEGITMAG | TCSAEGENEQY | 0.5615803 |
| FPLVVQD | TCSAEGENEQY | 0.7020045 |
| GAFTTPQ | TCSAEGENEQY | 0.84513086 |
| GFGGFGTAGGVGGA | TCSAEGENEQY | 0.6976763 |
| GFLAQFL | TCSAEGENEQY | 0.7700772 |
| GGAGGAGGNAGVA | TCSAEGENEQY | 0.8182821 |
| GGVGTAGAGGAGGAGG | TCSAEGENEQY | 0.82907766 |
| GNGGGGDAGNAGSGGN | TCSAEGENEQY | 0.65995944 |
| GTGGAGGAGNDAGS | TCSAEGENEQY | 0.6891681 |
| GTTNTGWFNS | TCSAEGENEQY | 0.8709112 |
| IFMGPLFTL | TCSAEGENEQY | 0.8358854 |
| ILAHAIEAI | TCSAEGENEQY | 0.78021526 |
| IRSAIREAL | TCSAEGENEQY | 0.82833016 |
| ITFAGGG | TCSAEGENEQY | 0.79014945 |
| ITGMSGHQEV | TCSAEGENEQY | 0.82595503 |
| IVAGLACTL | TCSAEGENEQY | 0.7467749 |
| IVGGGLAGA | TCSAEGENEQY | 0.41687405 |
| IVTGVLNI | TCSAEGENEQY | 0.64876205 |
| LAQQGCI | TCSAEGENEQY | 0.7877732 |
| LCLHVEN | TCSAEGENEQY | 0.6111441 |
| LDAISDGS | TCSAEGENEQY | 0.34444162 |
| LGGFAAML | TCSAEGENEQY | 0.86993706 |
| LGGKNPMI | TCSAEGENEQY | 0.8904512 |
| LGPKGVAAIL | TCSAEGENEQY | 0.5254145 |
| LMVGVSI | TCSAEGENEQY | 0.8733096 |
| LVGAAFVL | TCSAEGENEQY | 0.37333348 |
| LVLAVAPL | TCSAEGENEQY | 0.80765045 |
| MAAIAVIDAA | TCSAEGENEQY | 0.8656837 |
| NNGINAELQ | TCSAEGENEQY | 0.6318927 |
| RGEPLVLVHG | TCSAEGENEQY | 0.82737243 |
| RGGAGGAAGLI | TCSAEGENEQY | 0.5489819 |

| | | |
|---|---|---|
| RSYRMHPAV | TCSAEGENEQY | 0.67069465 |
| SAAHLLAGQLA | TCSAEGENEQY | 0.87197125 |
| SAAVGSY | TCSAEGENEQY | 0.6559384 |
| SPHAVLHRI | TCSAEGENEQY | 0.74236166 |
| SQIAKVL | TCSAEGENEQY | 0.7271502 |
| TALGHLVP | TCSAEGENEQY | 0.57583743 |
| TAPTTSGAA | TCSAEGENEQY | 0.7632269 |
| TDGDDGNGGSPGT | TCSAEGENEQY | 0.8542887 |
| TDGLGEN | TCSAEGENEQY | 0.37627694 |
| TGGPPLPGQA | TCSAEGENEQY | 0.85356504 |
| TIPAITI | TCSAEGENEQY | 0.59592086 |
| TIPGFTI | TCSAEGENEQY | 0.88909954 |
| TSIASGALA | TCSAEGENEQY | 0.8211427 |
| TVTLPGAAI | TCSAEGENEQY | 0.8073102 |
| VASGLSAALP | TCSAEGENEQY | 0.6404671 |
| VGAGVIGI | TCSAEGENEQY | 0.87086356 |
| VGPKRFGV | TCSAEGENEQY | 0.7597414 |
| VNVAFPD | TCSAEGENEQY | 0.80073774 |
| VSMTNSGVS | TCSAEGENEQY | 0.8591809 |
| VTDVLGDLSV | TCSAEGENEQY | 0.7233654 |
| VVVVGAGFAGL | TCSAEGENEQY | 0.8058078 |
| VYTVGTVV | TCSAEGENEQY | 0.7709409 |
| WTTFSGP | TCSAEGENEQY | 0.7644662 |
| YDELVEAF | TCSAEGENEQY | 0.7314888 |
| YGTGGAGGNGG | TCSAEGENEQY | 0.62566733 |
| YTNVMAV | TCSAEGENEQY | 0.7668745 |
| FLEQVHQL | TCSAEGENEQY | 0.67671454 |
| GLSLLAVAV | TCSAEGENEQY | 0.7494845 |
| LEQVHQL | TCSAEGENEQY | 0.8010653 |
| LNAAGAI | TCSAEGENEQY | 0.7230843 |
| PAVHASL | TCSAEGENEQY | 0.8011731 |
| TELEDDA | TCSAEGENEQY | 0.78527594 |
| TPPGPPN | TCSAEGENEQY | 0.647001 |
| TPPPPPPP | TCSAEGENEQY | 0.7457304 |
| TQSPALI | ASSLAEVF | 0.74266374 |
| EFAAAFPQ | ASSLAEVF | 0.76058996 |
| LLSGNLDVL | ASSLAEVF | 0.61265826 |
| GPANLGNYNI | ASSLAEVF | 0.79365295 |
| TLGPTLP | ASSLAEVF | 0.6913877 |
| VYAYSSCCK | ASSLAEVF | 0.32640845 |
| APAPAPAPNI | ASSLAEVF | 0.7871747 |
| PGGPVPIP | ASSLAEVF | 0.7545777 |
| FLTVALEP | ASSLAEVF | 0.742551 |
| SQYFGANL | ASSLAEVF | 0.79942083 |
| GGVKPAA | ASSLAEVF | 0.62014735 |
| GGTGGAGGAGGW | ASSLAEVF | 0.681276 |
| GVGGVAGLGGAGG | ASSLAEVF | 0.7745397 |
| VLLVLATTI | ASSLAEVF | 0.8070128 |
| IGIFAAML | ASSLAEVF | 0.66024846 |
| AVGDLID | ASSLAEVF | 0.7662709 |
| NISPLPL | ASSLAEVF | 0.79334515 |

| | | |
|---|---|---|
| VTAVFVAI | ASSLAEVF | 0.7946231 |
| SGDDNDAAL | ASSLAEVF | 0.8118641 |
| AVAALAHS | ASSLAEVF | 0.81597143 |
| GGVTVMATP | ASSLAEVF | 0.7031468 |
| AGLAANVRD | ASSLAEVF | 0.8012317 |
| TNRPDLIDPALLRPG | ASSLAEVF | 0.8036042 |
| LANLGVTAL | ASSLAEVF | 0.76026773 |
| TGERIEL | ASSLAEVF | 0.75772077 |
| NGEELAA | ASSLAEVF | 0.7336498 |
| VLSLVWG | ASSLAEVF | 0.72172886 |
| IGYALWL | ASSLAEVF | 0.51152295 |
| LSLLLAPL | ASSLAEVF | 0.76027614 |
| TLDEALK | ASSLAEVF | 0.8085459 |
| TDGAGTAEGG | ASSLAEVF | 0.72621006 |
| NAPGGAGGNGG | ASSLAEVF | 0.6648098 |
| VSVFMGMGGNF | ASSLAEVF | 0.6365507 |
| LASYLAAL | ASSLAEVF | 0.76436615 |
| LVGGTPAL | ASSLAEVF | 0.7318243 |
| VRNADVL | ASSLAEVF | 0.5738876 |
| TCSSPFLVSAM | ASSLAEVF | 0.74956936 |
| VHITDAE | ASSLAEVF | 0.7932924 |
| EYAAPSF | ASSLAEVF | 0.797789 |
| VVDAFHGGL | ASSLAEVF | 0.52773994 |
| ASVVDPQ | ASSLAEVF | 0.80814505 |
| SYNEFGGLS | ASSLAEVF | 0.7624738 |
| DPDGSVY | ASSLAEVF | 0.7970709 |
| MVSTIHATE | ASSLAEVF | 0.7832804 |
| LAAGVADPV | ASSLAEVF | 0.7331002 |
| TKEQIAA | ASSLAEVF | 0.78344804 |
| SLASPAAL | ASSLAEVF | 0.70762116 |
| IGLVFYGV | ASSLAEVF | 0.6693497 |
| FLLDASNL | ASSLAEVF | 0.79931426 |
| YQVGNFP | ASSLAEVF | 0.73452663 |
| ESLLSQST | ASSLAEVF | 0.6942695 |
| LFTVGFAAVG | ASSLAEVF | 0.64069706 |
| ATPPTPT | ASSLAEVF | 0.8145632 |
| TDLATLG | ASSLAEVF | 0.6115593 |
| SAGGTGR | ASSLAEVF | 0.7216204 |
| GFSNSGS | ASSLAEVF | 0.6162014 |
| GGNGGSGGTSV | ASSLAEVF | 0.7438182 |
| GTSVATGGAGN | ASSLAEVF | 0.8012091 |
| NGGNGGSGGTS | ASSLAEVF | 0.7987629 |
| AVVKFPH | ASSLAEVF | 0.7269477 |
| TPPPPPP | ASSLAEVF | 0.7761202 |
| AAPAAAAAPA | ASSLAEVF | 0.7932798 |
| IALNEDL | ASSLAEVF | 0.7699868 |
| IVNVSSM | ASSLAEVF | 0.79146934 |
| TGIPPAPR | ASSLAEVF | 0.66037816 |
| FDLGGGTF | ASSLAEVF | 0.80931276 |
| FDLGGGTFDVS | ASSLAEVF | 0.7872645 |
| NEPTAAA | ASSLAEVF | 0.8047482 |

| | | |
|---|---|---|
| NEPTAAALA | ASSLAEVF | 0.8071701 |
| GSIINISSI | ASSLAEVF | 0.77187806 |
| VAPGNAGTA | ASSLAEVF | 0.7743791 |
| TGGGASLE | ASSLAEVF | 0.50731844 |
| VAVYNHYKRI | ASSLAEVF | 0.7385521 |
| TDPAPATT | ASSLAEVF | 0.77068526 |
| TNRPDILDPALLRPG | ASSLAEVF | 0.7697842 |
| LTTTTTS | ASSLAEVF | 0.7852255 |
| PTGSGKTL | ASSLAEVF | 0.79000926 |
| TGSGKTL | ASSLAEVF | 0.70865566 |
| TIGHVDHGK | ASSLAEVF | 0.78904706 |
| TIGHVDHGKTT | ASSLAEVF | 0.7929368 |
| GHVDHGKTT | ASSLAEVF | 0.62257695 |
| AQFEHTLL | ASSLAEVF | 0.74205315 |
| IVNQDEL | ASSLAEVF | 0.8175776 |
| TNGSQFF | ASSLAEVF | 0.73579496 |
| TGKGIVE | ASSLAEVF | 0.7371479 |
| LISNASDA | ASSLAEVF | 0.73814404 |
| AELRTLGELTKIAKAHGAQ | ASSLAEVF | 0.79994154 |
| APNAVAAAEPSW | ASSLAEVF | 0.6783717 |
| APQVAGAATAVL | ASSLAEVF | 0.81180346 |
| ARRDPAAAAAILPTDA | ASSLAEVF | 0.36117667 |
| DAAARDVLAEAGLAEYF | ASSLAEVF | 0.7645072 |
| DDQIVLSAKANLAKVEQ | ASSLAEVF | 0.7368721 |
| DELLPDTGEDSDEDGH | ASSLAEVF | 0.6166166 |
| ELPEAPANRATVPQD | ASSLAEVF | 0.80247384 |
| FGQSKIERVTGIG | ASSLAEVF | 0.7480775 |
| GAGGTGSTAGGAGGAGGAGGLY | ASSLAEVF | 0.77872676 |
| GHGGTGAAGGNGGTGGAGGSGI | ASSLAEVF | 0.77820003 |
| GNGGNGGAGGEGGPGGAGGAGA | ASSLAEVF | 0.8076837 |
| GSTAGNGGNGGNATNGTIAGQP | ASSLAEVF | 0.73560065 |
| GTGGDGGNGGNGGTGG | ASSLAEVF | 0.7549759 |
| GVRRPAGIPADAPV | ASSLAEVF | 0.76513594 |
| GYRQVTDAHLLTLARR | ASSLAEVF | 0.79333204 |
| IAKMGQDGHDRG | ASSLAEVF | 0.799425 |
| KAEPSIEIQYT | ASSLAEVF | 0.8076172 |
| KPVRIRMAADEDTVAVT | ASSLAEVF | 0.55632544 |
| MPPRIAGMRLLVIKPEP | ASSLAEVF | 0.72422874 |
| MKFYAVALFRSRGVPPTR | ASSLAEVF | 0.7683012 |
| NAGGFGILGAVAHSP | ASSLAEVF | 0.4787985 |
| NEGREITLAMAEM | ASSLAEVF | 0.4422625 |
| NELVEWSHPLGHRWRWE | ASSLAEVF | 0.715084 |
| NLGAGGTGGNGGNPG | ASSLAEVF | 0.71703243 |
| NTEWPKWETVMAADPVD | ASSLAEVF | 0.79826796 |
| PASAAPAETQP | ASSLAEVF | 0.77786136 |
| QQSSDGKNGVRAAEITGE | ASSLAEVF | 0.7383992 |
| RAPDAVTTSWWKE | ASSLAEVF | 0.7826717 |
| SGFGNVGAHTSGWFNQ | ASSLAEVF | 0.730245 |
| SGVAGAVGQAASVGGLKV | ASSLAEVF | 0.21370445 |
| SNGGAGTGSAGGNGGT | ASSLAEVF | 0.7269792 |
| TGGAGGEGGAAG | ASSLAEVF | 0.6460717 |

| | | |
|---|---|---|
| TGGDGGAAGAGGAPGHG | ASSLAEVF | 0.8093213 |
| TMPPLLAAIANAGRHGV | ASSLAEVF | 0.74234706 |
| TSSTAGNSGGA | ASSLAEVF | 0.68817323 |
| VPATVPVGILC | ASSLAEVF | 0.75145644 |
| VQRPDSGGVVLTGRIS | ASSLAEVF | 0.731131 |
| WTPNPVPATKGVPAGLVQAT | ASSLAEVF | 0.74296755 |
| YALKRALAKFSDRFDVVII | ASSLAEVF | 0.67528343 |
| YLIGSVIV | ASSLAEVF | 0.35936606 |
| YGQLVAVID | ASSLAEVF | 0.79394907 |
| VTVGGMTLPA | ASSLAEVF | 0.5949961 |
| VLALPAV | ASSLAEVF | 0.67368317 |
| VIPPTLNYET | ASSLAEVF | 0.5546849 |
| VGMGGMHPSSG | ASSLAEVF | 0.6153226 |
| VGLAGGAATA | ASSLAEVF | 0.56456476 |
| VGGAGGAGGSGP | ASSLAEVF | 0.81005126 |
| VFAAFMV | ASSLAEVF | 0.7078272 |
| TLYDKLR | ASSLAEVF | 0.7171394 |
| TLAVFTVL | ASSLAEVF | 0.8099273 |
| TGSVQPARV | ASSLAEVF | 0.6730521 |
| TGGGAGSGGT | ASSLAEVF | 0.73058486 |
| SHLETSL | ASSLAEVF | 0.7254261 |
| SFTTGMT | ASSLAEVF | 0.7017846 |
| QVEVAKRMGVRQPTVS | ASSLAEVF | 0.7125598 |
| QGSGGPALAM | ASSLAEVF | 0.6969282 |
| MLAPSNP | ASSLAEVF | 0.8083816 |
| LVNNAAI | ASSLAEVF | 0.7658078 |
| LTGEINTLA | ASSLAEVF | 0.7362298 |
| LSIFGPV | ASSLAEVF | 0.7821812 |
| LLALLANL | ASSLAEVF | 0.8065221 |
| LISNASDALDKLR | ASSLAEVF | 0.81151944 |
| LINEAVP | ASSLAEVF | 0.7161682 |
| LGYAFSGL | ASSLAEVF | 0.72457266 |
| LDFVAVAL | ASSLAEVF | 0.7267592 |
| LAQPGAE | ASSLAEVF | 0.724639 |
| ISNLGAVTGF | ASSLAEVF | 0.75923485 |
| HQEDPGRI | ASSLAEVF | 0.68066335 |
| GTPGEPTP | ASSLAEVF | 0.803189 |
| GFFNSGS | ASSLAEVF | 0.80190444 |
| GAGGGGLTAGAN | ASSLAEVF | 0.763516 |
| FVDALVNM | ASSLAEVF | 0.63803184 |
| FLTVAPEL | ASSLAEVF | 0.7307951 |
| FETLTAL | ASSLAEVF | 0.6387426 |
| DVINPIEL | ASSLAEVF | 0.7058517 |
| APVSGAPV | ASSLAEVF | 0.7571609 |
| ALVGGLAW | ASSLAEVF | 0.61818 |
| ALAAAAATAI | ASSLAEVF | 0.75268763 |
| AFGLGLLL | ASSLAEVF | 0.81090844 |
| AFFAAASL | ASSLAEVF | 0.6575171 |
| AAAALPAFSPP | ASSLAEVF | 0.76885426 |
| VVLAPAL | ASSLAEVF | 0.80797195 |
| TIGHVDHGKTTL | ASSLAEVF | 0.7904339 |

| | | |
|---|---|---|
| LISNASDALDK | ASSLAEVF | 0.8082427 |
| LISNASDAL | ASSLAEVF | 0.7794118 |
| ISNASDALDK | ASSLAEVF | 0.7876904 |
| NQAETLVYQTE | ASSLAEVF | 0.7563826 |
| TGGTGGTAGAG | ASSLAEVF | 0.76779085 |
| TGSTAGGAGGA | ASSLAEVF | 0.74716204 |
| DPLAVPVAAGPV | ASSLAEVF | 0.7443023 |
| GGTGGTPGNGGN | ASSLAEVF | 0.71445006 |
| PAGLAGVDPATL | ASSLAEVF | 0.79358053 |
| GNNLVVEAPHAGASV | ASSLAEVF | 0.76948166 |
| KVTRIMSASTLEQPA | ASSLAEVF | 0.79154074 |
| LGFALPIALPRRTVV | ASSLAEVF | 0.67654073 |
| TRLGNLTARKLVVLV | ASSLAEVF | 0.68564457 |
| WSQQRRLIYFPSAGP | ASSLAEVF | 0.7944293 |
| GPNGNPDPMAAAVDIR | ASSLAEVF | 0.234527 |
| TQRCVSVADLRAAASR | ASSLAEVF | 0.47329402 |
| QFVLDVAEHPVEVLTETV | ASSLAEVF | 0.7775458 |
| GGAGGTGGAGGNGGTQAGDG | ASSLAEVF | 0.7947441 |
| PLLATLRAIAVDVDAKPAQV | ASSLAEVF | 0.31157705 |
| APKFRTGGYMIDFWGVGYQVAK | ASSLAEVF | 0.6658293 |
| GAGGAGGAGGAAGGPGGTGGTG | ASSLAEVF | 0.6311147 |
| IGALGLLATDVAPAGSVAFGVA | ASSLAEVF | 0.800367 |
| AGGAGGIGGNGGAGGAGGVLMG | ASSLAEVF | 0.74640393 |
| ADELIGLL | ASSLAEVF | 0.6214529 |
| ADRALKA | ASSLAEVF | 0.79475456 |
| AGAVGQVGGAGGA | ASSLAEVF | 0.5919616 |
| AGGGGGLGGAP | ASSLAEVF | 0.6461774 |
| AGGLLYGNGGAG | ASSLAEVF | 0.8078377 |
| AGLESIGL | ASSLAEVF | 0.80128765 |
| AGLKTGII | ASSLAEVF | 0.73289037 |
| ASLAALLL | ASSLAEVF | 0.7925545 |
| ASVINISSI | ASSLAEVF | 0.70148075 |
| ATPPGPA | ASSLAEVF | 0.8136852 |
| ATVASALV | ASSLAEVF | 0.73754823 |
| ATVASLIL | ASSLAEVF | 0.6874371 |
| AVNGPVLIRH | ASSLAEVF | 0.79315645 |
| DGAAGAVGI | ASSLAEVF | 0.780878 |
| DPLLHGI | ASSLAEVF | 0.7649061 |
| ELGAALGAL | ASSLAEVF | 0.77881163 |
| ELGRTQAV | ASSLAEVF | 0.7509029 |
| FVAGGYLS | ASSLAEVF | 0.7265315 |
| GASGGAGGAGGW | ASSLAEVF | 0.7949733 |
| GGAGAGGDGGAGGI | ASSLAEVF | 0.79982436 |
| GTGHNGSQPGMGGNG | ASSLAEVF | 0.68405485 |
| GYAAASASAS | ASSLAEVF | 0.8103516 |
| HQVDVAAVQQG | ASSLAEVF | 0.7263303 |
| IGAAIAHT | ASSLAEVF | 0.70643294 |
| IGMIEVAQL | ASSLAEVF | 0.816514 |
| IRLAPEHTSAMD | ASSLAEVF | 0.7873895 |
| IYNVSVL | ASSLAEVF | 0.79329693 |
| KQGFLGM | ASSLAEVF | 0.8017945 |

| | | |
|---|---|---|
| LESVITQL | ASSLAEVF | 0.7731269 |
| LGLGALVVV | ASSLAEVF | 0.70246506 |
| LIGNGGNGGSGGI | ASSLAEVF | 0.77089393 |
| LNAINAPTL | ASSLAEVF | 0.7488196 |
| LQLLNTGV | ASSLAEVF | 0.79934955 |
| LQTSIEF | ASSLAEVF | 0.77492476 |
| MSFAVSEL | ASSLAEVF | 0.6427493 |
| NTGALGL | ASSLAEVF | 0.7873463 |
| PPGAGGAGGAGTT | ASSLAEVF | 0.80662024 |
| QLVGLGI | ASSLAEVF | 0.7839754 |
| RFLEQVHQL | ASSLAEVF | 0.77603084 |
| RGVGYVLR | ASSLAEVF | 0.80949724 |
| SATGVFAGVI | ASSLAEVF | 0.75416034 |
| SLPAITLP | ASSLAEVF | 0.7449759 |
| STALAFTV | ASSLAEVF | 0.7972467 |
| SVLTVGLGK | ASSLAEVF | 0.7931438 |
| TALAGAGSGI | ASSLAEVF | 0.76883626 |
| TGLPFTGI | ASSLAEVF | 0.8137231 |
| VAEHVTV | ASSLAEVF | 0.69554967 |
| VDIVLNSL | ASSLAEVF | 0.77665937 |
| VLTGGGAVL | ASSLAEVF | 0.6740259 |
| VNVGLAYGI | ASSLAEVF | 0.7610318 |
| VSSAAVGAV | ASSLAEVF | 0.6402233 |
| VTSVYEAGV | ASSLAEVF | 0.7700975 |
| WVALDRGA | ASSLAEVF | 0.7898372 |
| YLVGSVVI | ASSLAEVF | 0.7837516 |
| GPPGTGKTLL | ASSLAEVF | 0.43904856 |
| GVGMTKF | ASSLAEVF | 0.7933876 |
| IGIGGSDL | ASSLAEVF | 0.76216245 |
| IGTLLAEL | ASSLAEVF | 0.8144904 |
| IYTSGTT | ASSLAEVF | 0.75357825 |
| LSFPAGV | ASSLAEVF | 0.77045095 |
| LYTSGTT | ASSLAEVF | 0.7092837 |
| VGAFPFGF | ASSLAEVF | 0.7678851 |
| VGAGTVEF | ASSLAEVF | 0.61693096 |
| VGKSTLLL | ASSLAEVF | 0.70360196 |
| AALAHAGGGIV | ASSLAEVF | 0.7103697 |
| AALVAVSLL | ASSLAEVF | 0.6581867 |
| AAVSRAVAAL | ASSLAEVF | 0.7751491 |
| AGGSSGIGGF | ASSLAEVF | 0.80321395 |
| AGLTVANANG | ASSLAEVF | 0.78559375 |
| AGLVGAVL | ASSLAEVF | 0.7935226 |
| ALSQAFTV | ASSLAEVF | 0.8030275 |
| ALTFSVSL | ASSLAEVF | 0.81230986 |
| ALTGAALVAL | ASSLAEVF | 0.71229327 |
| ASFSIGAGVL | ASSLAEVF | 0.7503277 |
| DGSVYKI | ASSLAEVF | 0.6482705 |
| EQLAEIV | ASSLAEVF | 0.7467856 |
| FAVSELTVM | ASSLAEVF | 0.7222557 |
| GAAPLAAGL | ASSLAEVF | 0.7722477 |
| GADGGDGGSGGAGG | ASSLAEVF | 0.23660828 |

| | | |
|---|---|---|
| GFGLDTGGAGG | ASSLAEVF | 0.724738 |
| GLASGGLL | ASSLAEVF | 0.80980444 |
| GRIAGAAVLL | ASSLAEVF | 0.5828605 |
| GSAGSLNTSF | ASSLAEVF | 0.7742848 |
| GTGQIGI | ASSLAEVF | 0.7578634 |
| GTLIALIL | ASSLAEVF | 0.7126127 |
| IAATMGFVVG | ASSLAEVF | 0.7763894 |
| IFAQALL | ASSLAEVF | 0.71232474 |
| IGAAVALGF | ASSLAEVF | 0.7931586 |
| IGLTPLAL | ASSLAEVF | 0.8113693 |
| ILGSNTSTL | ASSLAEVF | 0.7699922 |
| IVFSLTVL | ASSLAEVF | 0.732289 |
| LASKATGF | ASSLAEVF | 0.7657002 |
| LASVALIL | ASSLAEVF | 0.6277637 |
| LFAVGLTL | ASSLAEVF | 0.75506 |
| LFNGGIGTYI | ASSLAEVF | 0.31844872 |
| LTLGAGII | ASSLAEVF | 0.73985267 |
| LVAAALGDI | ASSLAEVF | 0.73456097 |
| NTAFVQLGI | ASSLAEVF | 0.7642035 |
| QLFSGAEL | ASSLAEVF | 0.73987055 |
| SAATFITL | ASSLAEVF | 0.77903557 |
| SNYGLANLGV | ASSLAEVF | 0.6296929 |
| SYGGAYAVM | ASSLAEVF | 0.73563594 |
| TTGVGGLTLG | ASSLAEVF | 0.6566723 |
| TTQTIYI | ASSLAEVF | 0.6293829 |
| VAAGAGLGAVI | ASSLAEVF | 0.8012047 |
| VAAVVTLL | ASSLAEVF | 0.7806281 |
| VGFLGGLI | ASSLAEVF | 0.730176 |
| VGVIPILL | ASSLAEVF | 0.7974472 |
| VIFASSLI | ASSLAEVF | 0.7716731 |
| VKGGDGGAAAG | ASSLAEVF | 0.6374635 |
| VTKTFTNI | ASSLAEVF | 0.69585556 |
| VVAAAILL | ASSLAEVF | 0.751365 |
| YAALDVEL | ASSLAEVF | 0.75164783 |
| LASVALLL | ASSLAEVF | 0.33717474 |
| LDAVHRV | ASSLAEVF | 0.7934119 |
| QGLSSSIF | ASSLAEVF | 0.70851827 |
| RGTFANIRL | ASSLAEVF | 0.63803935 |
| RQLWWGHRI | ASSLAEVF | 0.7395226 |
| YGPPGTGKTLL | ASSLAEVF | 0.7909379 |
| AALVGAPANVA | ASSLAEVF | 0.6433934 |
| AATFRRTE | ASSLAEVF | 0.7360566 |
| AGALGGGI | ASSLAEVF | 0.7903143 |
| AGGAGGAGGIGAG | ASSLAEVF | 0.7711427 |
| AGSGALGL | ASSLAEVF | 0.8042819 |
| AQIADQI | ASSLAEVF | 0.78009945 |
| ASAVAVTYL | ASSLAEVF | 0.7678535 |
| ASTVLGGTGG | ASSLAEVF | 0.77144915 |
| AVGGEPAK | ASSLAEVF | 0.7755221 |
| CEQGTFGGTC | ASSLAEVF | 0.6074748 |
| DEGKAPAG | ASSLAEVF | 0.5208375 |

| | | |
|---|---|---|
| DTVVFPAL | ASSLAEVF | 0.7279626 |
| FEGITMAG | ASSLAEVF | 0.5560639 |
| FPLVVQD | ASSLAEVF | 0.7668949 |
| GAFTTPQ | ASSLAEVF | 0.8133365 |
| GFGGFGTAGGVGGA | ASSLAEVF | 0.7698714 |
| GFLAQFL | ASSLAEVF | 0.67620194 |
| GGAGGAGGNAGVA | ASSLAEVF | 0.7371943 |
| GGVGTAGAGGAGGAGG | ASSLAEVF | 0.7484401 |
| GNGGGGDAGNAGSGGN | ASSLAEVF | 0.79368454 |
| GTGGAGGAGNDAGS | ASSLAEVF | 0.8000858 |
| GTTNTGWFNS | ASSLAEVF | 0.8016495 |
| IFMGPLFTL | ASSLAEVF | 0.78869015 |
| ILAHAIEAI | ASSLAEVF | 0.6787223 |
| IRSAIREAL | ASSLAEVF | 0.7927563 |
| ITFAGGG | ASSLAEVF | 0.73884237 |
| ITGMSGHQEV | ASSLAEVF | 0.7411237 |
| IVAGLACTL | ASSLAEVF | 0.7774793 |
| IVGGGLAGA | ASSLAEVF | 0.77253413 |
| IVTGVLNI | ASSLAEVF | 0.7701928 |
| LAQQGCI | ASSLAEVF | 0.74314076 |
| LCLHVEN | ASSLAEVF | 0.56173563 |
| LDAISDGS | ASSLAEVF | 0.777002 |
| LGGFAAML | ASSLAEVF | 0.7179444 |
| LGGKNPMI | ASSLAEVF | 0.5970975 |
| LGPKGVAAIL | ASSLAEVF | 0.72257483 |
| LMVGVSI | ASSLAEVF | 0.8160307 |
| LVGAAFVL | ASSLAEVF | 0.7996886 |
| LVLAVAPL | ASSLAEVF | 0.6275949 |
| MAAIAVIDAA | ASSLAEVF | 0.6942116 |
| NNGINAELQ | ASSLAEVF | 0.73582804 |
| RGEPLVLVHG | ASSLAEVF | 0.5547557 |
| RGGAGGAAGLI | ASSLAEVF | 0.7959599 |
| RSYRMHPAV | ASSLAEVF | 0.72149056 |
| SAAHLLAGQLA | ASSLAEVF | 0.7202539 |
| SAAVGSY | ASSLAEVF | 0.70236856 |
| SPHAVLHRI | ASSLAEVF | 0.7712556 |
| SQIAKVL | ASSLAEVF | 0.67702866 |
| TALGHLVP | ASSLAEVF | 0.8041335 |
| TAPTTSGAA | ASSLAEVF | 0.28874248 |
| TDGDDGNGGSPGT | ASSLAEVF | 0.80288416 |
| TDGLGEN | ASSLAEVF | 0.66002434 |
| TGGPPLPGQA | ASSLAEVF | 0.74120456 |
| TIPAITI | ASSLAEVF | 0.6459129 |
| TIPGFTI | ASSLAEVF | 0.7677287 |
| TSIASGALA | ASSLAEVF | 0.77316767 |
| TVTLPGAAI | ASSLAEVF | 0.7979034 |
| VASGLSAALP | ASSLAEVF | 0.66710603 |
| VGAGVIGI | ASSLAEVF | 0.7903951 |
| VGPKRFGV | ASSLAEVF | 0.798488 |
| VNVAFPD | ASSLAEVF | 0.7235762 |
| VSMTNSGVS | ASSLAEVF | 0.6763475 |

| | | |
|---|---|---|
| VTDVLGDLSV | ASSLAEVF | 0.77828497 |
| VVVVGAGFAGL | ASSLAEVF | 0.7023234 |
| VYTVGTVV | ASSLAEVF | 0.7973347 |
| WTTFSGP | ASSLAEVF | 0.74599445 |
| YDELVEAF | ASSLAEVF | 0.60618067 |
| YGTGGAGGNGG | ASSLAEVF | 0.77242774 |
| YTNVMAV | ASSLAEVF | 0.79682976 |
| FLEQVHQL | ASSLAEVF | 0.6689833 |
| GLSLLAVAV | ASSLAEVF | 0.7507489 |
| LEQVHQL | ASSLAEVF | 0.7483089 |
| LNAAGAI | ASSLAEVF | 0.59836245 |
| PAVHASL | ASSLAEVF | 0.77175176 |
| TELEDDA | ASSLAEVF | 0.68395144 |
| TPPGPPN | ASSLAEVF | 0.7020142 |
| TPPPPPPP | ASSLAEVF | 0.76054823 |
| TQSPALI | SSSHRTGEDTQY | 0.7256999 |
| EFAAAFPQ | SSSHRTGEDTQY | 0.73319674 |
| LLSGNLDVL | SSSHRTGEDTQY | 0.7241827 |
| GPANLGNYNI | SSSHRTGEDTQY | 0.777244 |
| TLGPTLP | SSSHRTGEDTQY | 0.78019434 |
| VYAYSSCCK | SSSHRTGEDTQY | 0.7449077 |
| APAPAPAPNI | SSSHRTGEDTQY | 0.75079674 |
| PGGPVPIP | SSSHRTGEDTQY | 0.80789745 |
| FLTVALEP | SSSHRTGEDTQY | 0.57577324 |
| SQYFGANL | SSSHRTGEDTQY | 0.68094647 |
| GGVKPAA | SSSHRTGEDTQY | 0.78558 |
| GGTGGAGGAGGW | SSSHRTGEDTQY | 0.59642947 |
| GVGGVAGLGGAGG | SSSHRTGEDTQY | 0.79519296 |
| VLLVLATTI | SSSHRTGEDTQY | 0.730496 |
| IGIFAAML | SSSHRTGEDTQY | 0.7714338 |
| AVGDLID | SSSHRTGEDTQY | 0.73626053 |
| NISPLPL | SSSHRTGEDTQY | 0.8133418 |
| VTAVFVAI | SSSHRTGEDTQY | 0.79566765 |
| SGDDNDAAL | SSSHRTGEDTQY | 0.7729544 |
| AVAALAHS | SSSHRTGEDTQY | 0.78449583 |
| GGVTVMATP | SSSHRTGEDTQY | 0.71005845 |
| AGLAANVRD | SSSHRTGEDTQY | 0.79451704 |
| TNRPDLIDPALLRPG | SSSHRTGEDTQY | 0.78888124 |
| LANLGVTAL | SSSHRTGEDTQY | 0.63107735 |
| TGERIEL | SSSHRTGEDTQY | 0.72602123 |
| NGEELAA | SSSHRTGEDTQY | 0.81230843 |
| VLSLVWG | SSSHRTGEDTQY | 0.80882007 |
| IGYALWL | SSSHRTGEDTQY | 0.56622356 |
| LSLLLAPL | SSSHRTGEDTQY | 0.7275211 |
| TLDEALK | SSSHRTGEDTQY | 0.5576202 |
| TDGAGTAEGG | SSSHRTGEDTQY | 0.7316749 |
| NAPGGAGGNGG | SSSHRTGEDTQY | 0.68371236 |
| VSVFMGMGGNF | SSSHRTGEDTQY | 0.783846 |
| LASYLAAL | SSSHRTGEDTQY | 0.72219896 |
| LVGGTPAL | SSSHRTGEDTQY | 0.74300015 |
| VRNADVL | SSSHRTGEDTQY | 0.7716061 |

| | | |
|---|---|---|
| TCSSPFLVSAM | SSSHRTGEDTQY | 0.80889904 |
| VHITDAE | SSSHRTGEDTQY | 0.7133811 |
| EYAAPSF | SSSHRTGEDTQY | 0.727044 |
| VVDAFHGGL | SSSHRTGEDTQY | 0.64527404 |
| ASVVDPQ | SSSHRTGEDTQY | 0.5355316 |
| SYNEFGGLS | SSSHRTGEDTQY | 0.7509968 |
| DPDGSVY | SSSHRTGEDTQY | 0.71943927 |
| MVSTIHATE | SSSHRTGEDTQY | 0.39382136 |
| LAAGVADPV | SSSHRTGEDTQY | 0.80926883 |
| TKEQIAA | SSSHRTGEDTQY | 0.759424 |
| SLASPAAL | SSSHRTGEDTQY | 0.7947188 |
| IGLVFYGV | SSSHRTGEDTQY | 0.7135102 |
| FLLDASNL | SSSHRTGEDTQY | 0.7738526 |
| YQVGNFP | SSSHRTGEDTQY | 0.7043716 |
| ESLLSQST | SSSHRTGEDTQY | 0.78061277 |
| LFTVGFAAVG | SSSHRTGEDTQY | 0.71770275 |
| ATPPTPT | SSSHRTGEDTQY | 0.57751715 |
| TDLATLG | SSSHRTGEDTQY | 0.75264 |
| SAGGTGR | SSSHRTGEDTQY | 0.6591681 |
| GFSNSGS | SSSHRTGEDTQY | 0.8051133 |
| GGNGGSGGTSV | SSSHRTGEDTQY | 0.79852283 |
| GTSVATGGAGN | SSSHRTGEDTQY | 0.76261103 |
| NGGNGGSGGTS | SSSHRTGEDTQY | 0.8042801 |
| AVVKFPH | SSSHRTGEDTQY | 0.42042536 |
| TPPPPPP | SSSHRTGEDTQY | 0.7135012 |
| AAPAAAAAPA | SSSHRTGEDTQY | 0.79920065 |
| IALNEDL | SSSHRTGEDTQY | 0.7834151 |
| IVNVSSM | SSSHRTGEDTQY | 0.74431443 |
| TGIPPAPR | SSSHRTGEDTQY | 0.8105823 |
| FDLGGGTF | SSSHRTGEDTQY | 0.63763666 |
| FDLGGGTFDVS | SSSHRTGEDTQY | 0.7504721 |
| NEPTAAA | SSSHRTGEDTQY | 0.77040625 |
| NEPTAAALA | SSSHRTGEDTQY | 0.80109525 |
| GSIINISSI | SSSHRTGEDTQY | 0.7933394 |
| VAPGNAGTA | SSSHRTGEDTQY | 0.7228793 |
| TGGGASLE | SSSHRTGEDTQY | 0.7773048 |
| VAVYNHYKRI | SSSHRTGEDTQY | 0.81861806 |
| TDPAPATT | SSSHRTGEDTQY | 0.70220757 |
| TNRPDILDPALLRPG | SSSHRTGEDTQY | 0.79017526 |
| LTTTTTS | SSSHRTGEDTQY | 0.77702785 |
| PTGSGKTL | SSSHRTGEDTQY | 0.618716 |
| TGSGKTL | SSSHRTGEDTQY | 0.66927975 |
| TIGHVDHGK | SSSHRTGEDTQY | 0.6921065 |
| TIGHVDHGKTT | SSSHRTGEDTQY | 0.7253844 |
| GHVDHGKTT | SSSHRTGEDTQY | 0.66331196 |
| AQFEHTLL | SSSHRTGEDTQY | 0.76627403 |
| IVNQDEL | SSSHRTGEDTQY | 0.75797606 |
| TNGSQFF | SSSHRTGEDTQY | 0.82001185 |
| TGKGIVE | SSSHRTGEDTQY | 0.77428406 |
| LISNASDA | SSSHRTGEDTQY | 0.7943654 |
| AELRTLGELTKIAKAHGAQ | SSSHRTGEDTQY | 0.6484296 |

| | | |
|---|---|---|
| APNAVAAAEPSW | SSSHRTGEDTQY | 0.7818535 |
| APQVAGAATAVL | SSSHRTGEDTQY | 0.7751279 |
| ARRDPAAAAAILPTDA | SSSHRTGEDTQY | 0.6065272 |
| DAAARDVLAEAGLAEYF | SSSHRTGEDTQY | 0.8042605 |
| DDQIVLSAKANLAKVEQ | SSSHRTGEDTQY | 0.70572466 |
| DELLPDTGEDSDEDGH | SSSHRTGEDTQY | 0.7163478 |
| ELPEAPANRATVPQD | SSSHRTGEDTQY | 0.78977853 |
| FGQSKIERVTGIG | SSSHRTGEDTQY | 0.777334 |
| GAGGTGSTAGGAGGAGGAGGLY | SSSHRTGEDTQY | 0.76326835 |
| GHGGTGAAGGNGGTGGAGGSGI | SSSHRTGEDTQY | 0.7161524 |
| GNGGNGGAGGEGGPGGAGGAGA | SSSHRTGEDTQY | 0.67697966 |
| GSTAGNGGNGGNATNGTIAGQP | SSSHRTGEDTQY | 0.79674053 |
| GTGGDGGNGGNGGTGG | SSSHRTGEDTQY | 0.7978057 |
| GVRRPAGIPADAPV | SSSHRTGEDTQY | 0.716578 |
| GYRQVTDAHLLTLARR | SSSHRTGEDTQY | 0.8098264 |
| IAKMGQDGHDRG | SSSHRTGEDTQY | 0.80155116 |
| KAEPSIEIQYT | SSSHRTGEDTQY | 0.62443197 |
| KPVRIRMAADEDTVAVT | SSSHRTGEDTQY | 0.7639004 |
| MPPRIAGMRLLVIKPEP | SSSHRTGEDTQY | 0.80685973 |
| MKFYAVALFRSRGVPPTR | SSSHRTGEDTQY | 0.79978955 |
| NAGGFGILGAVAHSP | SSSHRTGEDTQY | 0.8111605 |
| NEGREITLAMAEM | SSSHRTGEDTQY | 0.78620565 |
| NELVEWSHPLGHRWRWE | SSSHRTGEDTQY | 0.6933086 |
| NLGAGGTGGNGGNPG | SSSHRTGEDTQY | 0.80526006 |
| NTEWPKWETVMAADPVD | SSSHRTGEDTQY | 0.81674117 |
| PASAAPAETQP | SSSHRTGEDTQY | 0.7923204 |
| QQSSDGKNGVRAAEITGE | SSSHRTGEDTQY | 0.7252113 |
| RAPDAVTTSWWKE | SSSHRTGEDTQY | 0.7056875 |
| SGFGNVGAHTSGWFNQ | SSSHRTGEDTQY | 0.7814447 |
| SGVAGAVGQAASVGGLKV | SSSHRTGEDTQY | 0.7978535 |
| SNGGAGTGSAGGNGGT | SSSHRTGEDTQY | 0.7128658 |
| TGGAGGEGGAAG | SSSHRTGEDTQY | 0.7612928 |
| TGGDGGAAGAGGAPGHG | SSSHRTGEDTQY | 0.75166464 |
| TMPPLLAAIANAGRHGV | SSSHRTGEDTQY | 0.59486246 |
| TSSTAGNSGGA | SSSHRTGEDTQY | 0.7802095 |
| VPATVPVGILC | SSSHRTGEDTQY | 0.80370355 |
| VQRPDSGGVVLTGRIS | SSSHRTGEDTQY | 0.7794748 |
| WTPNPVPATKGVPAGLVQAT | SSSHRTGEDTQY | 0.71476424 |
| YALKRALAKFSDRFDVVII | SSSHRTGEDTQY | 0.8003235 |
| YLIGSVIV | SSSHRTGEDTQY | 0.8171188 |
| YGQLVAVID | SSSHRTGEDTQY | 0.8116116 |
| VTVGGMTLPA | SSSHRTGEDTQY | 0.6135612 |
| VLALPAV | SSSHRTGEDTQY | 0.46571362 |
| VIPPTLNYET | SSSHRTGEDTQY | 0.7167287 |
| VGMGGMHPSSG | SSSHRTGEDTQY | 0.75435036 |
| VGLAGGAATA | SSSHRTGEDTQY | 0.7166149 |
| VGGAGGAGGSGP | SSSHRTGEDTQY | 0.69673973 |
| VFAAFMV | SSSHRTGEDTQY | 0.77576786 |
| TLYDKLR | SSSHRTGEDTQY | 0.80026984 |
| TLAVFTVL | SSSHRTGEDTQY | 0.71925193 |
| TGSVQPARV | SSSHRTGEDTQY | 0.6788919 |

| | | |
|---|---|---|
| TGGGAGSGGT | SSSHRTGEDTQY | 0.7896888 |
| SHLETSL | SSSHRTGEDTQY | 0.7926893 |
| SFTTGMT | SSSHRTGEDTQY | 0.6663772 |
| QVEVAKRMGVRQPTVS | SSSHRTGEDTQY | 0.72133726 |
| QGSGGPALAM | SSSHRTGEDTQY | 0.69426286 |
| MLAPSNP | SSSHRTGEDTQY | 0.6706548 |
| LVNNAAI | SSSHRTGEDTQY | 0.7574035 |
| LTGEINTLA | SSSHRTGEDTQY | 0.6602006 |
| LSIFGPV | SSSHRTGEDTQY | 0.80396897 |
| LLALLANL | SSSHRTGEDTQY | 0.68372464 |
| LISNASDALDKLR | SSSHRTGEDTQY | 0.8001944 |
| LINEAVP | SSSHRTGEDTQY | 0.7687479 |
| LGYAFSGL | SSSHRTGEDTQY | 0.7899699 |
| LDFVAVAL | SSSHRTGEDTQY | 0.7321664 |
| LAQPGAE | SSSHRTGEDTQY | 0.6868676 |
| ISNLGAVTGF | SSSHRTGEDTQY | 0.68267226 |
| HQEDPGRI | SSSHRTGEDTQY | 0.5061332 |
| GTPGEPTP | SSSHRTGEDTQY | 0.80083466 |
| GFFNSGS | SSSHRTGEDTQY | 0.5308484 |
| GAGGGGLTAGAN | SSSHRTGEDTQY | 0.8064299 |
| FVDALVNM | SSSHRTGEDTQY | 0.7959047 |
| FLTVAPEL | SSSHRTGEDTQY | 0.7197447 |
| FETLTAL | SSSHRTGEDTQY | 0.79104245 |
| DVINPIEL | SSSHRTGEDTQY | 0.65239733 |
| APVSGAPV | SSSHRTGEDTQY | 0.6892309 |
| ALVGGLAW | SSSHRTGEDTQY | 0.7494511 |
| ALAAAAATAI | SSSHRTGEDTQY | 0.79271984 |
| AFGLGLLL | SSSHRTGEDTQY | 0.71078604 |
| AFFAAASL | SSSHRTGEDTQY | 0.7938435 |
| AAAALPAFSPP | SSSHRTGEDTQY | 0.77371496 |
| VVLAPAL | SSSHRTGEDTQY | 0.70610154 |
| TIGHVDHGKTTL | SSSHRTGEDTQY | 0.7739085 |
| LISNASDALDK | SSSHRTGEDTQY | 0.788882 |
| LISNASDAL | SSSHRTGEDTQY | 0.7913601 |
| ISNASDALDK | SSSHRTGEDTQY | 0.7053187 |
| NQAETLVYQTE | SSSHRTGEDTQY | 0.77814806 |
| TGGTGGTAGAG | SSSHRTGEDTQY | 0.66116714 |
| TGSTAGGAGGA | SSSHRTGEDTQY | 0.74982905 |
| DPLAVPVAAGPV | SSSHRTGEDTQY | 0.77840495 |
| GGTGGTPGNGGN | SSSHRTGEDTQY | 0.73504794 |
| PAGLAGVDPATL | SSSHRTGEDTQY | 0.79291195 |
| GNNLVVEAPHAGASV | SSSHRTGEDTQY | 0.71089196 |
| KVTRIMSASTLEQPA | SSSHRTGEDTQY | 0.6795551 |
| LGFALPIALPRRTVV | SSSHRTGEDTQY | 0.7311004 |
| TRLGNLTARKLVVLV | SSSHRTGEDTQY | 0.76155794 |
| WSQQRRLIYFPSAGP | SSSHRTGEDTQY | 0.7963287 |
| GPNGNPDPMAAAVDIR | SSSHRTGEDTQY | 0.6249094 |
| TQRCVSVADLRAAASR | SSSHRTGEDTQY | 0.75968635 |
| QFVLDVAEHPVEVLTETV | SSSHRTGEDTQY | 0.7461747 |
| GGAGGTGGAGGNGGTQAGDG | SSSHRTGEDTQY | 0.6819896 |
| PLLATLRAIAVDVDAKPAQV | SSSHRTGEDTQY | 0.78498304 |

| | | |
|---|---|---|
| APKFRTGGYMIDFWGVGYQVAK | SSSHRTGEDTQY | 0.7762031 |
| GAGGAGGAGGAAGGPGGTGGTG | SSSHRTGEDTQY | 0.6838686 |
| IGALGLLATDVAPAGSVAFGVA | SSSHRTGEDTQY | 0.78886044 |
| AGGAGGIGGNGGAGGAGGVLMG | SSSHRTGEDTQY | 0.71395844 |
| ADELIGLL | SSSHRTGEDTQY | 0.77668566 |
| ADRALKA | SSSHRTGEDTQY | 0.8147351 |
| AGAVGQVGGAGGA | SSSHRTGEDTQY | 0.7378239 |
| AGGGGGLGGAP | SSSHRTGEDTQY | 0.7999898 |
| AGGLLYGNGGAG | SSSHRTGEDTQY | 0.7693174 |
| AGLESIGL | SSSHRTGEDTQY | 0.7819566 |
| AGLKTGII | SSSHRTGEDTQY | 0.73838955 |
| ASLAALLL | SSSHRTGEDTQY | 0.7149848 |
| ASVINISSI | SSSHRTGEDTQY | 0.8107119 |
| ATPPGPA | SSSHRTGEDTQY | 0.7850679 |
| ATVASALV | SSSHRTGEDTQY | 0.78375185 |
| ATVASLIL | SSSHRTGEDTQY | 0.7793619 |
| AVNGPVLIRH | SSSHRTGEDTQY | 0.7758081 |
| DGAAGAVGI | SSSHRTGEDTQY | 0.7915739 |
| DPLLHGI | SSSHRTGEDTQY | 0.6688183 |
| ELGAALGAL | SSSHRTGEDTQY | 0.6244663 |
| ELGRTQAV | SSSHRTGEDTQY | 0.70762926 |
| FVAGGYLS | SSSHRTGEDTQY | 0.7153866 |
| GASGGAGGAGGW | SSSHRTGEDTQY | 0.7749567 |
| GGAGAGGDGGAGGI | SSSHRTGEDTQY | 0.6286594 |
| GTGHNGSQPGMGGNG | SSSHRTGEDTQY | 0.7886803 |
| GYAAASASAS | SSSHRTGEDTQY | 0.81951475 |
| HQVDVAAVQQG | SSSHRTGEDTQY | 0.6928395 |
| IGAAIAHT | SSSHRTGEDTQY | 0.81665146 |
| IGMIEVAQL | SSSHRTGEDTQY | 0.6757343 |
| IRLAPEHTSAMD | SSSHRTGEDTQY | 0.786991 |
| IYNVSVL | SSSHRTGEDTQY | 0.80158937 |
| KQGFLGM | SSSHRTGEDTQY | 0.6515889 |
| LESVITQL | SSSHRTGEDTQY | 0.7768522 |
| LGLGALVVV | SSSHRTGEDTQY | 0.74565154 |
| LIGNGGNGGSGGI | SSSHRTGEDTQY | 0.7662605 |
| LNAINAPTL | SSSHRTGEDTQY | 0.8028889 |
| LQLLNTGV | SSSHRTGEDTQY | 0.70088536 |
| LQTSIEF | SSSHRTGEDTQY | 0.7751794 |
| MSFAVSEL | SSSHRTGEDTQY | 0.75675833 |
| NTGALGL | SSSHRTGEDTQY | 0.7081096 |
| PPGAGGAGGAGTT | SSSHRTGEDTQY | 0.7942153 |
| QLVGLGI | SSSHRTGEDTQY | 0.79335344 |
| RFLEQVHQL | SSSHRTGEDTQY | 0.6949399 |
| RGVGYVLR | SSSHRTGEDTQY | 0.8142284 |
| SATGVFAGVI | SSSHRTGEDTQY | 0.677914 |
| SLPAITLP | SSSHRTGEDTQY | 0.8170011 |
| STALAFTV | SSSHRTGEDTQY | 0.7913719 |
| SVLTVGLGK | SSSHRTGEDTQY | 0.7358013 |
| TALAGAGSGI | SSSHRTGEDTQY | 0.75058925 |
| TGLPFTGI | SSSHRTGEDTQY | 0.79322934 |
| VAEHVTV | SSSHRTGEDTQY | 0.72173834 |

| | | |
|---|---|---|
| VDIVLNSL | SSSHRTGEDTQY | 0.7907307 |
| VLTGGGAVL | SSSHRTGEDTQY | 0.81292695 |
| VNVGLAYGI | SSSHRTGEDTQY | 0.4797666 |
| VSSAAVGAV | SSSHRTGEDTQY | 0.7514699 |
| VTSVYEAGV | SSSHRTGEDTQY | 0.6294435 |
| WVALDRGA | SSSHRTGEDTQY | 0.6922292 |
| YLVGSVVI | SSSHRTGEDTQY | 0.52865374 |
| GPPGTGKTLL | SSSHRTGEDTQY | 0.43851718 |
| GVGMTKF | SSSHRTGEDTQY | 0.6059575 |
| IGIGGSDL | SSSHRTGEDTQY | 0.4693238 |
| IGTLLAEL | SSSHRTGEDTQY | 0.6964623 |
| IYTSGTT | SSSHRTGEDTQY | 0.7440226 |
| LSFPAGV | SSSHRTGEDTQY | 0.5192669 |
| LYTSGTT | SSSHRTGEDTQY | 0.7384667 |
| VGAFPFGF | SSSHRTGEDTQY | 0.7416212 |
| VGAGTVEF | SSSHRTGEDTQY | 0.37906852 |
| VGKSTLLL | SSSHRTGEDTQY | 0.5883838 |
| AALAHAGGGIV | SSSHRTGEDTQY | 0.51664597 |
| AALVAVSLL | SSSHRTGEDTQY | 0.60657084 |
| AAVSRAVAAL | SSSHRTGEDTQY | 0.35158262 |
| AGGSSGIGGF | SSSHRTGEDTQY | 0.70940703 |
| AGLTVANANG | SSSHRTGEDTQY | 0.44506437 |
| AGLVGAVL | SSSHRTGEDTQY | 0.27844542 |
| ALSQAFTV | SSSHRTGEDTQY | 0.7006855 |
| ALTFSVSL | SSSHRTGEDTQY | 0.58841556 |
| ALTGAALVAL | SSSHRTGEDTQY | 0.5483894 |
| ASFSIGAGVL | SSSHRTGEDTQY | 0.7089443 |
| DGSVYKI | SSSHRTGEDTQY | 0.5605875 |
| EQLAEIV | SSSHRTGEDTQY | 0.4798492 |
| FAVSELTVM | SSSHRTGEDTQY | 0.683258 |
| GAAPLAAGL | SSSHRTGEDTQY | 0.7556317 |
| GADGGDGGSGGAGG | SSSHRTGEDTQY | 0.47343293 |
| GFGLDTGGAGG | SSSHRTGEDTQY | 0.6150893 |
| GLASGGLL | SSSHRTGEDTQY | 0.7645459 |
| GRIAGAAVLL | SSSHRTGEDTQY | 0.66154325 |
| GSAGSLNTSF | SSSHRTGEDTQY | 0.76884496 |
| GTGQIGI | SSSHRTGEDTQY | 0.7843671 |
| GTLIALIL | SSSHRTGEDTQY | 0.5982512 |
| IAATMGFVVG | SSSHRTGEDTQY | 0.73378354 |
| IFAQALL | SSSHRTGEDTQY | 0.7415539 |
| IGAAVALGF | SSSHRTGEDTQY | 0.62110543 |
| IGLTPLAL | SSSHRTGEDTQY | 0.6373766 |
| ILGSNTSTL | SSSHRTGEDTQY | 0.4585949 |
| IVFSLTVL | SSSHRTGEDTQY | 0.6349936 |
| LASKATGF | SSSHRTGEDTQY | 0.42353952 |
| LASVALIL | SSSHRTGEDTQY | 0.6307514 |
| LFAVGLTL | SSSHRTGEDTQY | 0.75981414 |
| LFNGGIGTYI | SSSHRTGEDTQY | 0.5669685 |
| LTLGAGII | SSSHRTGEDTQY | 0.41910246 |
| LVAAALGDI | SSSHRTGEDTQY | 0.38476932 |
| NTAFVQLGI | SSSHRTGEDTQY | 0.6746876 |

| | | |
|---|---|---|
| QLFSGAEL | SSSHRTGEDTQY | 0.46717292 |
| SAATFITL | SSSHRTGEDTQY | 0.4577545 |
| SNYGLANLGV | SSSHRTGEDTQY | 0.58038366 |
| SYGGAYAVM | SSSHRTGEDTQY | 0.73535186 |
| TTGVGGLTLG | SSSHRTGEDTQY | 0.7291012 |
| TTQTIYI | SSSHRTGEDTQY | 0.481608 |
| VAAGAGLGAVI | SSSHRTGEDTQY | 0.7480599 |
| VAAVVTLL | SSSHRTGEDTQY | 0.6608424 |
| VGFLGGLI | SSSHRTGEDTQY | 0.7458533 |
| VGVIPILL | SSSHRTGEDTQY | 0.6474599 |
| VIFASSLI | SSSHRTGEDTQY | 0.48230997 |
| VKGGDGGAAAG | SSSHRTGEDTQY | 0.69406307 |
| VTKTFTNI | SSSHRTGEDTQY | 0.54850286 |
| VVAAAILL | SSSHRTGEDTQY | 0.72493935 |
| YAALDVEL | SSSHRTGEDTQY | 0.7352142 |
| LASVALLL | SSSHRTGEDTQY | 0.5780252 |
| LDAVHRV | SSSHRTGEDTQY | 0.5128017 |
| QGLSSSIF | SSSHRTGEDTQY | 0.51344496 |
| RGTFANIRL | SSSHRTGEDTQY | 0.77027375 |
| RQLWWGHRI | SSSHRTGEDTQY | 0.44112873 |
| YGPPGTGKTLL | SSSHRTGEDTQY | 0.4786703 |
| AALVGAPANVA | SSSHRTGEDTQY | 0.57875705 |
| AATFRRTE | SSSHRTGEDTQY | 0.6073833 |
| AGALGGGI | SSSHRTGEDTQY | 0.7499688 |
| AGGAGGAGGIGAG | SSSHRTGEDTQY | 0.71189755 |
| AGSGALGL | SSSHRTGEDTQY | 0.60186225 |
| AQIADQI | SSSHRTGEDTQY | 0.6707471 |
| ASAVAVTYL | SSSHRTGEDTQY | 0.716164 |
| ASTVLGGTGG | SSSHRTGEDTQY | 0.6639105 |
| AVGGEPAK | SSSHRTGEDTQY | 0.7240672 |
| CEQGTFGGTC | SSSHRTGEDTQY | 0.49316645 |
| DEGKAPAG | SSSHRTGEDTQY | 0.7516978 |
| DTVVFPAL | SSSHRTGEDTQY | 0.71296453 |
| FEGITMAG | SSSHRTGEDTQY | 0.7171445 |
| FPLVVQD | SSSHRTGEDTQY | 0.7514081 |
| GAFTTPQ | SSSHRTGEDTQY | 0.6864648 |
| GFGGFGTAGGVGGA | SSSHRTGEDTQY | 0.66305274 |
| GFLAQFL | SSSHRTGEDTQY | 0.38913292 |
| GGAGGAGGNAGVA | SSSHRTGEDTQY | 0.6374209 |
| GGVGTAGAGGAGGAGG | SSSHRTGEDTQY | 0.62616926 |
| GNGGGGDAGNAGSGGN | SSSHRTGEDTQY | 0.63641083 |
| GTGGAGGAGNDAGS | SSSHRTGEDTQY | 0.69836056 |
| GTTNTGWFNS | SSSHRTGEDTQY | 0.7180452 |
| IFMGPLFTL | SSSHRTGEDTQY | 0.5167084 |
| ILAHAIEAI | SSSHRTGEDTQY | 0.76543796 |
| IRSAIREAL | SSSHRTGEDTQY | 0.7462244 |
| ITFAGGG | SSSHRTGEDTQY | 0.38542342 |
| ITGMSGHQEV | SSSHRTGEDTQY | 0.586393 |
| IVAGLACTL | SSSHRTGEDTQY | 0.7584879 |
| IVGGGLAGA | SSSHRTGEDTQY | 0.6269479 |
| IVTGVLNI | SSSHRTGEDTQY | 0.5847679 |

| | | |
|---|---|---|
| LAQQGCI | SSSHRTGEDTQY | 0.6065548 |
| LCLHVEN | SSSHRTGEDTQY | 0.72446126 |
| LDAISDGS | SSSHRTGEDTQY | 0.6509586 |
| LGGFAAML | SSSHRTGEDTQY | 0.7815746 |
| LGGKNPMI | SSSHRTGEDTQY | 0.29208383 |
| LGPKGVAAIL | SSSHRTGEDTQY | 0.5866307 |
| LMVGVSI | SSSHRTGEDTQY | 0.5824388 |
| LVGAAFVL | SSSHRTGEDTQY | 0.29901645 |
| LVLAVAPL | SSSHRTGEDTQY | 0.73247504 |
| MAAIAVIDAA | SSSHRTGEDTQY | 0.5802544 |
| NNGINAELQ | SSSHRTGEDTQY | 0.7098098 |
| RGEPLVLVHG | SSSHRTGEDTQY | 0.5731523 |
| RGGAGGAAGLI | SSSHRTGEDTQY | 0.7542822 |
| RSYRMHPAV | SSSHRTGEDTQY | 0.59131896 |
| SAAHLLAGQLA | SSSHRTGEDTQY | 0.61965895 |
| SAAVGSY | SSSHRTGEDTQY | 0.5844055 |
| SPHAVLHRI | SSSHRTGEDTQY | 0.6883396 |
| SQIAKVL | SSSHRTGEDTQY | 0.77461565 |
| TALGHLVP | SSSHRTGEDTQY | 0.75995874 |
| TAPTTSGAA | SSSHRTGEDTQY | 0.38473186 |
| TDGDDGNGGSPGT | SSSHRTGEDTQY | 0.67368764 |
| TDGLGEN | SSSHRTGEDTQY | 0.6313458 |
| TGGPPLPGQA | SSSHRTGEDTQY | 0.4039747 |
| TIPAITI | SSSHRTGEDTQY | 0.37429574 |
| TIPGFTI | SSSHRTGEDTQY | 0.54198605 |
| TSIASGALA | SSSHRTGEDTQY | 0.535557 |
| TVTLPGAAI | SSSHRTGEDTQY | 0.6816277 |
| VASGLSAALP | SSSHRTGEDTQY | 0.6881462 |
| VGAGVIGI | SSSHRTGEDTQY | 0.5404169 |
| VGPKRFGV | SSSHRTGEDTQY | 0.69748205 |
| VNVAFPD | SSSHRTGEDTQY | 0.66841114 |
| VSMTNSGVS | SSSHRTGEDTQY | 0.16997102 |
| VTDVLGDLSV | SSSHRTGEDTQY | 0.5577949 |
| VVVVGAGFAGL | SSSHRTGEDTQY | 0.3794128 |
| VYTVGTVV | SSSHRTGEDTQY | 0.7673061 |
| WTTFSGP | SSSHRTGEDTQY | 0.4992506 |
| YDELVEAF | SSSHRTGEDTQY | 0.5097488 |
| YGTGGAGGNGG | SSSHRTGEDTQY | 0.6020541 |
| YTNVMAV | SSSHRTGEDTQY | 0.5873312 |
| FLEQVHQL | SSSHRTGEDTQY | 0.5956384 |
| GLSLLAVAV | SSSHRTGEDTQY | 0.50018275 |
| LEQVHQL | SSSHRTGEDTQY | 0.38295087 |
| LNAAGAI | SSSHRTGEDTQY | 0.7206088 |
| PAVHASL | SSSHRTGEDTQY | 0.54903364 |
| TELEDDA | SSSHRTGEDTQY | 0.5220156 |
| TPPGPPN | SSSHRTGEDTQY | 0.5173615 |
| TPPPPPPP | SSSHRTGEDTQY | 0.679178 |
| TQSPALI | ASSLAGGSAETLY | 0.34721124 |
| EFAAAFPQ | ASSLAGGSAETLY | 0.75096506 |
| LLSGNLDVL | ASSLAGGSAETLY | 0.44424355 |
| GPANLGNYNI | ASSLAGGSAETLY | 0.5832294 |

| | | |
|---|---|---|
| TLGPTLP | ASSLAGGSAETLY | 0.76898044 |
| VYAYSSCCK | ASSLAGGSAETLY | 0.57192105 |
| APAPAPAPNI | ASSLAGGSAETLY | 0.591187 |
| PGGPVPIP | ASSLAGGSAETLY | 0.57970554 |
| FLTVALEP | ASSLAGGSAETLY | 0.45732346 |
| SQYFGANL | ASSLAGGSAETLY | 0.50287443 |
| GGVKPAA | ASSLAGGSAETLY | 0.5150299 |
| GGTGGAGGAGGW | ASSLAGGSAETLY | 0.76855797 |
| GVGGVAGLGGAGG | ASSLAGGSAETLY | 0.6252967 |
| VLLVLATTI | ASSLAGGSAETLY | 0.48271975 |
| IGIFAAML | ASSLAGGSAETLY | 0.67294997 |
| AVGDLID | ASSLAGGSAETLY | 0.7669443 |
| NISPLPL | ASSLAGGSAETLY | 0.7606211 |
| VTAVFVAI | ASSLAGGSAETLY | 0.5043604 |
| SGDDNDAAL | ASSLAGGSAETLY | 0.6030015 |
| AVAALAHS | ASSLAGGSAETLY | 0.6090257 |
| GGVTVMATP | ASSLAGGSAETLY | 0.5725196 |
| AGLAANVRD | ASSLAGGSAETLY | 0.55255854 |
| TNRPDLIDPALLRPG | ASSLAGGSAETLY | 0.48870406 |
| LANLGVTAL | ASSLAGGSAETLY | 0.7307765 |
| TGERIEL | ASSLAGGSAETLY | 0.762825 |
| NGEELAA | ASSLAGGSAETLY | 0.63210595 |
| VLSLVWG | ASSLAGGSAETLY | 0.31498176 |
| IGYALWL | ASSLAGGSAETLY | 0.5918766 |
| LSLLLAPL | ASSLAGGSAETLY | 0.42753598 |
| TLDEALK | ASSLAGGSAETLY | 0.48888683 |
| TDGAGTAEGG | ASSLAGGSAETLY | 0.5918554 |
| NAPGGAGGNGG | ASSLAGGSAETLY | 0.33919248 |
| VSVFMGMGGNF | ASSLAGGSAETLY | 0.61404395 |
| LASYLAAL | ASSLAGGSAETLY | 0.76993626 |
| LVGGTPAL | ASSLAGGSAETLY | 0.46827844 |
| VRNADVL | ASSLAGGSAETLY | 0.64582425 |
| TCSSPFLVSAM | ASSLAGGSAETLY | 0.74460196 |
| VHITDAE | ASSLAGGSAETLY | 0.6745647 |
| EYAAPSF | ASSLAGGSAETLY | 0.72861314 |
| VVDAFHGGL | ASSLAGGSAETLY | 0.6837464 |
| ASVVDPQ | ASSLAGGSAETLY | 0.71359503 |
| SYNEFGGLS | ASSLAGGSAETLY | 0.66059875 |
| DPDGSVY | ASSLAGGSAETLY | 0.6932239 |
| MVSTIHATE | ASSLAGGSAETLY | 0.597911 |
| LAAGVADPV | ASSLAGGSAETLY | 0.6167116 |
| TKEQIAA | ASSLAGGSAETLY | 0.6071499 |
| SLASPAAL | ASSLAGGSAETLY | 0.6937984 |
| IGLVFYGV | ASSLAGGSAETLY | 0.6042619 |
| FLLDASNL | ASSLAGGSAETLY | 0.73419774 |
| YQVGNFP | ASSLAGGSAETLY | 0.3383994 |
| ESLLSQST | ASSLAGGSAETLY | 0.460738 |
| LFTVGFAAVG | ASSLAGGSAETLY | 0.7061261 |
| ATPPTPT | ASSLAGGSAETLY | 0.24548633 |
| TDLATLG | ASSLAGGSAETLY | 0.41091534 |
| SAGGTGR | ASSLAGGSAETLY | 0.6110227 |

| | | |
|---|---|---|
| GFSNSGS | ASSLAGGSAETLY | 0.73443526 |
| GGNGGSGGTSV | ASSLAGGSAETLY | 0.2509374 |
| GTSVATGGAGN | ASSLAGGSAETLY | 0.5752971 |
| NGGNGGSGGTS | ASSLAGGSAETLY | 0.3909305 |
| AVVKFPH | ASSLAGGSAETLY | 0.7534519 |
| TPPPPPP | ASSLAGGSAETLY | 0.60481876 |
| AAPAAAAAPA | ASSLAGGSAETLY | 0.40729704 |
| IALNEDL | ASSLAGGSAETLY | 0.71649516 |
| IVNVSSM | ASSLAGGSAETLY | 0.34274596 |
| TGIPPAPR | ASSLAGGSAETLY | 0.6369115 |
| FDLGGGTF | ASSLAGGSAETLY | 0.73811716 |
| FDLGGGTFDVS | ASSLAGGSAETLY | 0.7401488 |
| NEPTAAA | ASSLAGGSAETLY | 0.46368158 |
| NEPTAAALA | ASSLAGGSAETLY | 0.68674153 |
| GSIINISSI | ASSLAGGSAETLY | 0.52545667 |
| VAPGNAGTA | ASSLAGGSAETLY | 0.7856369 |
| TGGGASLE | ASSLAGGSAETLY | 0.5881017 |
| VAVYNHYKRI | ASSLAGGSAETLY | 0.40911454 |
| TDPAPATT | ASSLAGGSAETLY | 0.70747364 |
| TNRPDILDPALLRPG | ASSLAGGSAETLY | 0.66671264 |
| LTTTTTS | ASSLAGGSAETLY | 0.67864007 |
| PTGSGKTL | ASSLAGGSAETLY | 0.65553176 |
| TGSGKTL | ASSLAGGSAETLY | 0.5832923 |
| TIGHVDHGK | ASSLAGGSAETLY | 0.5446582 |
| TIGHVDHGKTT | ASSLAGGSAETLY | 0.6989689 |
| GHVDHGKTT | ASSLAGGSAETLY | 0.70698595 |
| AQFEHTLL | ASSLAGGSAETLY | 0.5128132 |
| IVNQDEL | ASSLAGGSAETLY | 0.76507145 |
| TNGSQFF | ASSLAGGSAETLY | 0.6116158 |
| TGKGIVE | ASSLAGGSAETLY | 0.5360371 |
| LISNASDA | ASSLAGGSAETLY | 0.77235407 |
| AELRTLGELTKIAKAHGAQ | ASSLAGGSAETLY | 0.62087005 |
| APNAVAAAEPSW | ASSLAGGSAETLY | 0.7083598 |
| APQVAGAATAVL | ASSLAGGSAETLY | 0.71101606 |
| ARRDPAAAAAILPTDA | ASSLAGGSAETLY | 0.7197254 |
| DAAARDVLAEAGLAEYF | ASSLAGGSAETLY | 0.51495105 |
| DDQIVLSAKANLAKVEQ | ASSLAGGSAETLY | 0.6919867 |
| DELLPDTGEDSDEDGH | ASSLAGGSAETLY | 0.63546795 |
| ELPEAPANRATVPQD | ASSLAGGSAETLY | 0.72246736 |
| FGQSKIERVTGIG | ASSLAGGSAETLY | 0.60454005 |
| GAGGTGSTAGGAGGAGGAGGLY | ASSLAGGSAETLY | 0.5835955 |
| GHGGTGAAGGNGGTGGAGGSGI | ASSLAGGSAETLY | 0.71358025 |
| GNGGNGGAGGEGGPGGAGGAGA | ASSLAGGSAETLY | 0.7582327 |
| GSTAGNGGNGGNATNGTIAGQP | ASSLAGGSAETLY | 0.6852349 |
| GTGGDGGNGGNGGTGG | ASSLAGGSAETLY | 0.7028089 |
| GVRRPAGIPADAPV | ASSLAGGSAETLY | 0.7447597 |
| GYRQVTDAHLLTLARR | ASSLAGGSAETLY | 0.49160734 |
| IAKMGQDGHDRG | ASSLAGGSAETLY | 0.6378353 |
| KAEPSIEIQYT | ASSLAGGSAETLY | 0.72769237 |
| KPVRIRMAADEDTVAVT | ASSLAGGSAETLY | 0.7581497 |
| MPPRIAGMRLLVIKPEP | ASSLAGGSAETLY | 0.61917716 |

| | | |
|---|---|---|
| MKFYAVALFRSRGVPPTR | ASSLAGGSAETLY | 0.75098395 |
| NAGGFGILGAVAHSP | ASSLAGGSAETLY | 0.59728175 |
| NEGREITLAMAEM | ASSLAGGSAETLY | 0.6541458 |
| NELVEWSHPLGHRWRWE | ASSLAGGSAETLY | 0.50740266 |
| NLGAGGTGGNGGNPG | ASSLAGGSAETLY | 0.5886099 |
| NTEWPKWETVMAADPVD | ASSLAGGSAETLY | 0.43614313 |
| PASAAPAETQP | ASSLAGGSAETLY | 0.66441023 |
| QQSSDGKNGVRAAEITGE | ASSLAGGSAETLY | 0.6863723 |
| RAPDAVTTSWWKE | ASSLAGGSAETLY | 0.6846186 |
| SGFGNVGAHTSGWFNQ | ASSLAGGSAETLY | 0.38603762 |
| SGVAGAVGQAASVGGLKV | ASSLAGGSAETLY | 0.67708254 |
| SNGGAGTGSAGGNGGT | ASSLAGGSAETLY | 0.6906582 |
| TGGAGGEGGAAG | ASSLAGGSAETLY | 0.7467778 |
| TGGDGGAAGAGGAPGHG | ASSLAGGSAETLY | 0.63398725 |
| TMPPLLAAIANAGRHGV | ASSLAGGSAETLY | 0.71428597 |
| TSSTAGNSGGA | ASSLAGGSAETLY | 0.5739852 |
| VPATVPVGILC | ASSLAGGSAETLY | 0.6508712 |
| VQRPDSGGVVLTGRIS | ASSLAGGSAETLY | 0.38379684 |
| WTPNPVPATKGVPAGLVQAT | ASSLAGGSAETLY | 0.5443755 |
| YALKRALAKFSDRFDVVII | ASSLAGGSAETLY | 0.56431305 |
| YLIGSVIV | ASSLAGGSAETLY | 0.45394412 |
| YGQLVAVID | ASSLAGGSAETLY | 0.6920735 |
| VTVGGMTLPA | ASSLAGGSAETLY | 0.7555453 |
| VLALPAV | ASSLAGGSAETLY | 0.73390657 |
| VIPPTLNYET | ASSLAGGSAETLY | 0.7234929 |
| VGMGGMHPSSG | ASSLAGGSAETLY | 0.73365515 |
| VGLAGGAATA | ASSLAGGSAETLY | 0.74556005 |
| VGGAGGAGGSGP | ASSLAGGSAETLY | 0.5202117 |
| VFAAFMV | ASSLAGGSAETLY | 0.61330825 |
| TLYDKLR | ASSLAGGSAETLY | 0.36208817 |
| TLAVFTVL | ASSLAGGSAETLY | 0.57786745 |
| TGSVQPARV | ASSLAGGSAETLY | 0.5305105 |
| TGGGAGSGGT | ASSLAGGSAETLY | 0.65191257 |
| SHLETSL | ASSLAGGSAETLY | 0.21554965 |
| SFTTGMT | ASSLAGGSAETLY | 0.57657933 |
| QVEVAKRMGVRQPTVS | ASSLAGGSAETLY | 0.77014655 |
| QGSGGPALAM | ASSLAGGSAETLY | 0.53807676 |
| MLAPSNP | ASSLAGGSAETLY | 0.66393685 |
| LVNNAAI | ASSLAGGSAETLY | 0.67322564 |
| LTGEINTLA | ASSLAGGSAETLY | 0.5261545 |
| LSIFGPV | ASSLAGGSAETLY | 0.6857953 |
| LLALLANL | ASSLAGGSAETLY | 0.5344391 |
| LISNASDALDKLR | ASSLAGGSAETLY | 0.7306823 |
| LINEAVP | ASSLAGGSAETLY | 0.7557618 |
| LGYAFSGL | ASSLAGGSAETLY | 0.6616467 |
| LDFVAVAL | ASSLAGGSAETLY | 0.6201921 |
| LAQPGAE | ASSLAGGSAETLY | 0.68546367 |
| ISNLGAVTGF | ASSLAGGSAETLY | 0.45623866 |
| HQEDPGRI | ASSLAGGSAETLY | 0.6613276 |
| GTPGEPTP | ASSLAGGSAETLY | 0.4950254 |
| GFFNSGS | ASSLAGGSAETLY | 0.56121105 |

| | | |
|---|---|---|
| GAGGGGLTAGAN | ASSLAGGSAETLY | 0.519697 |
| FVDALVNM | ASSLAGGSAETLY | 0.684086 |
| FLTVAPEL | ASSLAGGSAETLY | 0.66586185 |
| FETLTAL | ASSLAGGSAETLY | 0.69302624 |
| DVINPIEL | ASSLAGGSAETLY | 0.43007267 |
| APVSGAPV | ASSLAGGSAETLY | 0.7058399 |
| ALVGGLAW | ASSLAGGSAETLY | 0.67330366 |
| ALAAAAATAI | ASSLAGGSAETLY | 0.4308086 |
| AFGLGLLL | ASSLAGGSAETLY | 0.7320465 |
| AFFAAASL | ASSLAGGSAETLY | 0.6269734 |
| AAAALPAFSPP | ASSLAGGSAETLY | 0.5306811 |
| VVLAPAL | ASSLAGGSAETLY | 0.7354639 |
| TIGHVDHGKTTL | ASSLAGGSAETLY | 0.6508616 |
| LISNASDALDK | ASSLAGGSAETLY | 0.379034 |
| LISNASDAL | ASSLAGGSAETLY | 0.47562534 |
| ISNASDALDK | ASSLAGGSAETLY | 0.6226974 |
| NQAETLVYQTE | ASSLAGGSAETLY | 0.5849749 |
| TGGTGGTAGAG | ASSLAGGSAETLY | 0.28453907 |
| TGSTAGGAGGA | ASSLAGGSAETLY | 0.7224236 |
| DPLAVPVAAGPV | ASSLAGGSAETLY | 0.54627967 |
| GGTGGTPGNGGN | ASSLAGGSAETLY | 0.45656455 |
| PAGLAGVDPATL | ASSLAGGSAETLY | 0.54755044 |
| GNNLVVEAPHAGASV | ASSLAGGSAETLY | 0.70273924 |
| KVTRIMSASTLEQPA | ASSLAGGSAETLY | 0.6109391 |
| LGFALPIALPRRTVV | ASSLAGGSAETLY | 0.5495976 |
| TRLGNLTARKLVVLV | ASSLAGGSAETLY | 0.6563076 |
| WSQQRRLIYFPSAGP | ASSLAGGSAETLY | 0.7264613 |
| GPNGNPDPMAAAVDIR | ASSLAGGSAETLY | 0.7586297 |
| TQRCVSVADLRAAASR | ASSLAGGSAETLY | 0.6747218 |
| QFVLDVAEHPVEVLTETV | ASSLAGGSAETLY | 0.6645982 |
| GGAGGTGGAGGNGGTQAGDG | ASSLAGGSAETLY | 0.71796435 |
| PLLATLRAIAVDVDAKPAQV | ASSLAGGSAETLY | 0.61564296 |
| APKFRTGGYMIDFWGVGYQVAK | ASSLAGGSAETLY | 0.5282676 |
| GAGGAGGAGGAAGGPGGTGGTG | ASSLAGGSAETLY | 0.4792191 |
| IGALGLLATDVAPAGSVAFGVA | ASSLAGGSAETLY | 0.4625547 |
| AGGAGGIGGNGGAGGAGGVLMG | ASSLAGGSAETLY | 0.51337993 |
| ADELIGLL | ASSLAGGSAETLY | 0.651512 |
| ADRALKA | ASSLAGGSAETLY | 0.7532617 |
| AGAVGQVGGAGGA | ASSLAGGSAETLY | 0.60853297 |
| AGGGGGLGGAP | ASSLAGGSAETLY | 0.42848662 |
| AGGLLYGNGGAG | ASSLAGGSAETLY | 0.57674617 |
| AGLESIGL | ASSLAGGSAETLY | 0.5829614 |
| AGLKTGII | ASSLAGGSAETLY | 0.6883592 |
| ASLAALLL | ASSLAGGSAETLY | 0.7227833 |
| ASVINISSI | ASSLAGGSAETLY | 0.7435458 |
| ATPPGPA | ASSLAGGSAETLY | 0.7285829 |
| ATVASALV | ASSLAGGSAETLY | 0.54393345 |
| ATVASLIL | ASSLAGGSAETLY | 0.6995105 |
| AVNGPVLIRH | ASSLAGGSAETLY | 0.5611064 |
| DGAAGAVGI | ASSLAGGSAETLY | 0.6621773 |
| DPLLHGI | ASSLAGGSAETLY | 0.67293864 |

| | | |
|---|---|---|
| ELGAALGAL | ASSLAGGSAETLY | 0.64312613 |
| ELGRTQAV | ASSLAGGSAETLY | 0.6202643 |
| FVAGGYLS | ASSLAGGSAETLY | 0.62400764 |
| GASGGAGGAGGW | ASSLAGGSAETLY | 0.3675779 |
| GGAGAGGDGGAGGI | ASSLAGGSAETLY | 0.68421584 |
| GTGHNGSQPGMGGNG | ASSLAGGSAETLY | 0.45050368 |
| GYAAASASAS | ASSLAGGSAETLY | 0.5061864 |
| HQVDVAAVQQG | ASSLAGGSAETLY | 0.5755014 |
| IGAAIAHT | ASSLAGGSAETLY | 0.76290584 |
| IGMIEVAQL | ASSLAGGSAETLY | 0.74713063 |
| IRLAPEHTSAMD | ASSLAGGSAETLY | 0.49840102 |
| IYNVSVL | ASSLAGGSAETLY | 0.5070809 |
| KQGFLGM | ASSLAGGSAETLY | 0.6109251 |
| LESVITQL | ASSLAGGSAETLY | 0.473908 |
| LGLGALVVV | ASSLAGGSAETLY | 0.7041664 |
| LIGNGGNGGSGGI | ASSLAGGSAETLY | 0.5707549 |
| LNAINAPTL | ASSLAGGSAETLY | 0.59714144 |
| LQLLNTGV | ASSLAGGSAETLY | 0.5791762 |
| LQTSIEF | ASSLAGGSAETLY | 0.68423 |
| MSFAVSEL | ASSLAGGSAETLY | 0.555836 |
| NTGALGL | ASSLAGGSAETLY | 0.7182302 |
| PPGAGGAGGAGTT | ASSLAGGSAETLY | 0.22925037 |
| QLVGLGI | ASSLAGGSAETLY | 0.74891967 |
| RFLEQVHQL | ASSLAGGSAETLY | 0.4540798 |
| RGVGYVLR | ASSLAGGSAETLY | 0.5361957 |
| SATGVFAGVI | ASSLAGGSAETLY | 0.3206209 |
| SLPAITLP | ASSLAGGSAETLY | 0.66948414 |
| STALAFTV | ASSLAGGSAETLY | 0.6365149 |
| SVLTVGLGK | ASSLAGGSAETLY | 0.7221556 |
| TALAGAGSGI | ASSLAGGSAETLY | 0.6462103 |
| TGLPFTGI | ASSLAGGSAETLY | 0.69277215 |
| VAEHVTV | ASSLAGGSAETLY | 0.7154542 |
| VDIVLNSL | ASSLAGGSAETLY | 0.5492512 |
| VLTGGGAVL | ASSLAGGSAETLY | 0.40887433 |
| VNVGLAYGI | ASSLAGGSAETLY | 0.6538519 |
| VSSAAVGAV | ASSLAGGSAETLY | 0.5576829 |
| VTSVYEAGV | ASSLAGGSAETLY | 0.7285872 |
| WVALDRGA | ASSLAGGSAETLY | 0.6833283 |
| YLVGSVVI | ASSLAGGSAETLY | 0.35946986 |
| GPPGTGKTLL | ASSLAGGSAETLY | 0.7164308 |
| GVGMTKF | ASSLAGGSAETLY | 0.700401 |
| IGIGGSDL | ASSLAGGSAETLY | 0.42952302 |
| IGTLLAEL | ASSLAGGSAETLY | 0.58396125 |
| IYTSGTT | ASSLAGGSAETLY | 0.6153742 |
| LSFPAGV | ASSLAGGSAETLY | 0.45897943 |
| LYTSGTT | ASSLAGGSAETLY | 0.71833944 |
| VGAFPFGF | ASSLAGGSAETLY | 0.51603717 |
| VGAGTVEF | ASSLAGGSAETLY | 0.56637484 |
| VGKSTLLL | ASSLAGGSAETLY | 0.6045486 |
| AALAHAGGGIV | ASSLAGGSAETLY | 0.5573054 |
| AALVAVSLL | ASSLAGGSAETLY | 0.5717845 |

| | | |
|---|---|---|
| AAVSRAVAAL | ASSLAGGSAETLY | 0.5863526 |
| AGGSSGIGGF | ASSLAGGSAETLY | 0.6749516 |
| AGLTVANANG | ASSLAGGSAETLY | 0.6883607 |
| AGLVGAVL | ASSLAGGSAETLY | 0.5793231 |
| ALSQAFTV | ASSLAGGSAETLY | 0.6262964 |
| ALTFSVSL | ASSLAGGSAETLY | 0.75318444 |
| ALTGAALVAL | ASSLAGGSAETLY | 0.40138197 |
| ASFSIGAGVL | ASSLAGGSAETLY | 0.579309 |
| DGSVYKI | ASSLAGGSAETLY | 0.6657229 |
| EQLAEIV | ASSLAGGSAETLY | 0.5611228 |
| FAVSELTVM | ASSLAGGSAETLY | 0.72539586 |
| GAAPLAAGL | ASSLAGGSAETLY | 0.68090457 |
| GADGGDGGSGGAGG | ASSLAGGSAETLY | 0.65812886 |
| GFGLDTGGAGG | ASSLAGGSAETLY | 0.57170695 |
| GLASGGLL | ASSLAGGSAETLY | 0.7677045 |
| GRIAGAAVLL | ASSLAGGSAETLY | 0.67906713 |
| GSAGSLNTSF | ASSLAGGSAETLY | 0.6349089 |
| GTGQIGI | ASSLAGGSAETLY | 0.66420925 |
| GTLIALIL | ASSLAGGSAETLY | 0.5287941 |
| IAATMGFVVG | ASSLAGGSAETLY | 0.7270477 |
| IFAQALL | ASSLAGGSAETLY | 0.66999817 |
| IGAAVALGF | ASSLAGGSAETLY | 0.41731024 |
| IGLTPLAL | ASSLAGGSAETLY | 0.5552881 |
| ILGSNTSTL | ASSLAGGSAETLY | 0.7808927 |
| IVFSLTVL | ASSLAGGSAETLY | 0.71406627 |
| LASKATGF | ASSLAGGSAETLY | 0.58105993 |
| LASVALIL | ASSLAGGSAETLY | 0.5254851 |
| LFAVGLTL | ASSLAGGSAETLY | 0.5556761 |
| LFNGGIGTYI | ASSLAGGSAETLY | 0.591163 |
| LTLGAGII | ASSLAGGSAETLY | 0.5070306 |
| LVAAALGDI | ASSLAGGSAETLY | 0.6795069 |
| NTAFVQLGI | ASSLAGGSAETLY | 0.578222 |
| QLFSGAEL | ASSLAGGSAETLY | 0.66422915 |
| SAATFITL | ASSLAGGSAETLY | 0.6500673 |
| SNYGLANLGV | ASSLAGGSAETLY | 0.7434157 |
| SYGGAYAVM | ASSLAGGSAETLY | 0.5275903 |
| TTGVGGLTLG | ASSLAGGSAETLY | 0.62904596 |
| TTQTIYI | ASSLAGGSAETLY | 0.36306754 |
| VAAGAGLGAVI | ASSLAGGSAETLY | 0.554742 |
| VAAVVTLL | ASSLAGGSAETLY | 0.55538756 |
| VGFLGGLI | ASSLAGGSAETLY | 0.5978565 |
| VGVIPILL | ASSLAGGSAETLY | 0.32344088 |
| VIFASSLI | ASSLAGGSAETLY | 0.75832164 |
| VKGGDGGAAAG | ASSLAGGSAETLY | 0.5597008 |
| VTKTFTNI | ASSLAGGSAETLY | 0.70054406 |
| VVAAAILL | ASSLAGGSAETLY | 0.5925716 |
| YAALDVEL | ASSLAGGSAETLY | 0.5387934 |
| LASVALLL | ASSLAGGSAETLY | 0.5089297 |
| LDAVHRV | ASSLAGGSAETLY | 0.69884884 |
| QGLSSSIF | ASSLAGGSAETLY | 0.46392506 |
| RGTFANIRL | ASSLAGGSAETLY | 0.28836042 |

| | | |
|---|---|---|
| RQLWWGHRI | ASSLAGGSAETLY | 0.5389784 |
| YGPPGTGKTLL | ASSLAGGSAETLY | 0.4356066 |
| AALVGAPANVA | ASSLAGGSAETLY | 0.7579915 |
| AATFRRTE | ASSLAGGSAETLY | 0.73237836 |
| AGALGGGI | ASSLAGGSAETLY | 0.6192444 |
| AGGAGGAGGIGAG | ASSLAGGSAETLY | 0.75514245 |
| AGSGALGL | ASSLAGGSAETLY | 0.2496612 |
| AQIADQI | ASSLAGGSAETLY | 0.60258174 |
| ASAVAVTYL | ASSLAGGSAETLY | 0.7573932 |
| ASTVLGGTGG | ASSLAGGSAETLY | 0.677389 |
| AVGGEPAK | ASSLAGGSAETLY | 0.6179453 |
| CEQGTFGGTC | ASSLAGGSAETLY | 0.7346025 |
| DEGKAPAG | ASSLAGGSAETLY | 0.5089805 |
| DTVVFPAL | ASSLAGGSAETLY | 0.5933089 |
| FEGITMAG | ASSLAGGSAETLY | 0.6133287 |
| FPLVVQD | ASSLAGGSAETLY | 0.743222 |
| GAFTTPQ | ASSLAGGSAETLY | 0.6838427 |
| GFGGFGTAGGVGGA | ASSLAGGSAETLY | 0.4593813 |
| GFLAQFL | ASSLAGGSAETLY | 0.6959648 |
| GGAGGAGGNAGVA | ASSLAGGSAETLY | 0.7853285 |
| GGVGTAGAGGAGGAGG | ASSLAGGSAETLY | 0.4954226 |
| GNGGGGDAGNAGSGGN | ASSLAGGSAETLY | 0.6654955 |
| GTGGAGGAGNDAGS | ASSLAGGSAETLY | 0.62828285 |
| GTTNTGWFNS | ASSLAGGSAETLY | 0.53650117 |
| IFMGPLFTL | ASSLAGGSAETLY | 0.46773008 |
| ILAHAIEAI | ASSLAGGSAETLY | 0.43441465 |
| IRSAIREAL | ASSLAGGSAETLY | 0.7200565 |
| ITFAGGG | ASSLAGGSAETLY | 0.466561 |
| ITGMSGHQEV | ASSLAGGSAETLY | 0.62758905 |
| IVAGLACTL | ASSLAGGSAETLY | 0.6024805 |
| IVGGGLAGA | ASSLAGGSAETLY | 0.7945088 |
| IVTGVLNI | ASSLAGGSAETLY | 0.670174 |
| LAQQGCI | ASSLAGGSAETLY | 0.6857444 |
| LCLHVEN | ASSLAGGSAETLY | 0.48029393 |
| LDAISDGS | ASSLAGGSAETLY | 0.6908076 |
| LGGFAAML | ASSLAGGSAETLY | 0.67442507 |
| LGGKNPMI | ASSLAGGSAETLY | 0.39436662 |
| LGPKGVAAIL | ASSLAGGSAETLY | 0.7563011 |
| LMVGVSI | ASSLAGGSAETLY | 0.47292125 |
| LVGAAFVL | ASSLAGGSAETLY | 0.6086133 |
| LVLAVAPL | ASSLAGGSAETLY | 0.7119258 |
| MAAIAVIDAA | ASSLAGGSAETLY | 0.71537215 |
| NNGINAELQ | ASSLAGGSAETLY | 0.6379 |
| RGEPLVLVHG | ASSLAGGSAETLY | 0.7291281 |
| RGGAGGAAGLI | ASSLAGGSAETLY | 0.47221762 |
| RSYRMHPAV | ASSLAGGSAETLY | 0.70018023 |
| SAAHLLAGQLA | ASSLAGGSAETLY | 0.71014947 |
| SAAVGSY | ASSLAGGSAETLY | 0.5810085 |
| SPHAVLHRI | ASSLAGGSAETLY | 0.75966346 |
| SQIAKVL | ASSLAGGSAETLY | 0.70357805 |
| TALGHLVP | ASSLAGGSAETLY | 0.42179114 |

| | | |
|---|---|---|
| TAPTTSGAA | ASSLAGGSAETLY | 0.6424827 |
| TDGDDGNGGSPGT | ASSLAGGSAETLY | 0.74875355 |
| TDGLGEN | ASSLAGGSAETLY | 0.72631735 |
| TGGPPLPGQA | ASSLAGGSAETLY | 0.7294561 |
| TIPAITI | ASSLAGGSAETLY | 0.6758112 |
| TIPGFTI | ASSLAGGSAETLY | 0.5501497 |
| TSIASGALA | ASSLAGGSAETLY | 0.7540939 |
| TVTLPGAAI | ASSLAGGSAETLY | 0.7999501 |
| VASGLSAALP | ASSLAGGSAETLY | 0.68228906 |
| VGAGVIGI | ASSLAGGSAETLY | 0.47478124 |
| VGPKRFGV | ASSLAGGSAETLY | 0.49869543 |
| VNVAFPD | ASSLAGGSAETLY | 0.7159753 |
| VSMTNSGVS | ASSLAGGSAETLY | 0.7099973 |
| VTDVLGDLSV | ASSLAGGSAETLY | 0.44094682 |
| VVVVGAGFAGL | ASSLAGGSAETLY | 0.641835 |
| VYTVGTVV | ASSLAGGSAETLY | 0.6390066 |
| WTTFSGP | ASSLAGGSAETLY | 0.34906837 |
| YDELVEAF | ASSLAGGSAETLY | 0.6855844 |
| YGTGGAGGNGG | ASSLAGGSAETLY | 0.73256344 |
| YTNVMAV | ASSLAGGSAETLY | 0.7028314 |
| FLEQVHQL | ASSLAGGSAETLY | 0.4538281 |
| GLSLLAVAV | ASSLAGGSAETLY | 0.7491048 |
| LEQVHQL | ASSLAGGSAETLY | 0.7250521 |
| LNAAGAI | ASSLAGGSAETLY | 0.77292466 |
| PAVHASL | ASSLAGGSAETLY | 0.40431345 |
| TELEDDA | ASSLAGGSAETLY | 0.25059593 |
| TPPGPPN | ASSLAGGSAETLY | 0.6678836 |
| TPPPPPPP | ASSLAGGSAETLY | 0.65580964 |
| TQSPALI | ASSARGDNQAPL | 0.563381 |
| EFAAAFPQ | ASSARGDNQAPL | 0.46273413 |
| LLSGNLDVL | ASSARGDNQAPL | 0.6194078 |
| GPANLGNYNI | ASSARGDNQAPL | 0.7347989 |
| TLGPTLP | ASSARGDNQAPL | 0.47418275 |
| VYAYSSCCK | ASSARGDNQAPL | 0.44759455 |
| APAPAPAPNI | ASSARGDNQAPL | 0.7070998 |
| PGGPVPIP | ASSARGDNQAPL | 0.69622225 |
| FLTVALEP | ASSARGDNQAPL | 0.46979252 |
| SQYFGANL | ASSARGDNQAPL | 0.5142692 |
| GGVKPAA | ASSARGDNQAPL | 0.6966813 |
| GGTGGAGGAGGW | ASSARGDNQAPL | 0.6586754 |
| GVGGVAGLGGAGG | ASSARGDNQAPL | 0.63380253 |
| VLLVLATTI | ASSARGDNQAPL | 0.40645024 |
| IGIFAAML | ASSARGDNQAPL | 0.7328898 |
| AVGDLID | ASSARGDNQAPL | 0.39162397 |
| NISPLPL | ASSARGDNQAPL | 0.7269684 |
| VTAVFVAI | ASSARGDNQAPL | 0.636994 |
| SGDDNDAAL | ASSARGDNQAPL | 0.68990934 |
| AVAALAHS | ASSARGDNQAPL | 0.5680075 |
| GGVTVMATP | ASSARGDNQAPL | 0.40962356 |
| AGLAANVRD | ASSARGDNQAPL | 0.45699736 |
| TNRPDLIDPALLRPG | ASSARGDNQAPL | 0.41288176 |

| | | |
|---|---|---|
| LANLGVTAL | ASSARGDNQAPL | 0.7549008 |
| TGERIEL | ASSARGDNQAPL | 0.6305171 |
| NGEELAA | ASSARGDNQAPL | 0.7128897 |
| VLSLVWG | ASSARGDNQAPL | 0.6542302 |
| IGYALWL | ASSARGDNQAPL | 0.5384055 |
| LSLLLAPL | ASSARGDNQAPL | 0.69347274 |
| TLDEALK | ASSARGDNQAPL | 0.37251484 |
| TDGAGTAEGG | ASSARGDNQAPL | 0.67613643 |
| NAPGGAGGNGG | ASSARGDNQAPL | 0.6075027 |
| VSVFMGMGGNF | ASSARGDNQAPL | 0.7073579 |
| LASYLAAL | ASSARGDNQAPL | 0.5247151 |
| LVGGTPAL | ASSARGDNQAPL | 0.7026667 |
| VRNADVL | ASSARGDNQAPL | 0.66816646 |
| TCSSPFLVSAM | ASSARGDNQAPL | 0.5065646 |
| VHITDAE | ASSARGDNQAPL | 0.69840777 |
| EYAAPSF | ASSARGDNQAPL | 0.72681975 |
| VVDAFHGGL | ASSARGDNQAPL | 0.71535826 |
| ASVVDPQ | ASSARGDNQAPL | 0.36530155 |
| SYNEFGGLS | ASSARGDNQAPL | 0.67574006 |
| DPDGSVY | ASSARGDNQAPL | 0.46996742 |
| MVSTIHATE | ASSARGDNQAPL | 0.63322824 |
| LAAGVADPV | ASSARGDNQAPL | 0.66234064 |
| TKEQIAA | ASSARGDNQAPL | 0.5920628 |
| SLASPAAL | ASSARGDNQAPL | 0.70928085 |
| IGLVFYGV | ASSARGDNQAPL | 0.46900952 |
| FLLDASNL | ASSARGDNQAPL | 0.50655967 |
| YQVGNFP | ASSARGDNQAPL | 0.5729522 |
| ESLLSQST | ASSARGDNQAPL | 0.6904905 |
| LFTVGFAAVG | ASSARGDNQAPL | 0.6944441 |
| ATPPTPT | ASSARGDNQAPL | 0.43747675 |
| TDLATLG | ASSARGDNQAPL | 0.6252144 |
| SAGGTGR | ASSARGDNQAPL | 0.6120955 |
| GFSNSGS | ASSARGDNQAPL | 0.47380576 |
| GGNGGSGGTSV | ASSARGDNQAPL | 0.6620591 |
| GTSVATGGAGN | ASSARGDNQAPL | 0.6195254 |
| NGGNGGSGGTS | ASSARGDNQAPL | 0.3775682 |
| AVVKFPH | ASSARGDNQAPL | 0.72974354 |
| TPPPPPP | ASSARGDNQAPL | 0.51069653 |
| AAPAAAAAPA | ASSARGDNQAPL | 0.7226228 |
| IALNEDL | ASSARGDNQAPL | 0.76655203 |
| IVNVSSM | ASSARGDNQAPL | 0.43499264 |
| TGIPPAPR | ASSARGDNQAPL | 0.7200852 |
| FDLGGGTF | ASSARGDNQAPL | 0.6232422 |
| FDLGGGTFDVS | ASSARGDNQAPL | 0.63396895 |
| NEPTAAA | ASSARGDNQAPL | 0.6088664 |
| NEPTAAALA | ASSARGDNQAPL | 0.52366996 |
| GSIINISSI | ASSARGDNQAPL | 0.7434895 |
| VAPGNAGTA | ASSARGDNQAPL | 0.72086823 |
| TGGGASLE | ASSARGDNQAPL | 0.7163909 |
| VAVYNHYKRI | ASSARGDNQAPL | 0.6903141 |
| TDPAPATT | ASSARGDNQAPL | 0.6865592 |

| | | |
|---|---|---|
| TNRPDILDPALLRPG | ASSARGDNQAPL | 0.71659106 |
| LTTTTTS | ASSARGDNQAPL | 0.48119003 |
| PTGSGKTL | ASSARGDNQAPL | 0.35020077 |
| TGSGKTL | ASSARGDNQAPL | 0.6114317 |
| TIGHVDHGK | ASSARGDNQAPL | 0.5871179 |
| TIGHVDHGKTT | ASSARGDNQAPL | 0.5504269 |
| GHVDHGKTT | ASSARGDNQAPL | 0.42638284 |
| AQFEHTLL | ASSARGDNQAPL | 0.7209973 |
| IVNQDEL | ASSARGDNQAPL | 0.7837045 |
| TNGSQFF | ASSARGDNQAPL | 0.5372124 |
| TGKGIVE | ASSARGDNQAPL | 0.7755204 |
| LISNASDA | ASSARGDNQAPL | 0.41075838 |
| AELRTLGELTKIAKAHGAQ | ASSARGDNQAPL | 0.7337839 |
| APNAVAAAEPSW | ASSARGDNQAPL | 0.7570567 |
| APQVAGAATAVL | ASSARGDNQAPL | 0.41766113 |
| ARRDPAAAAAILPTDA | ASSARGDNQAPL | 0.6978207 |
| DAAARDVLAEAGLAEYF | ASSARGDNQAPL | 0.5201227 |
| DDQIVLSAKANLAKVEQ | ASSARGDNQAPL | 0.55729187 |
| DELLPDTGEDSDEDGH | ASSARGDNQAPL | 0.7371847 |
| ELPEAPANRATVPQD | ASSARGDNQAPL | 0.5755132 |
| FGQSKIERVTGIG | ASSARGDNQAPL | 0.7386808 |
| GAGGTGSTAGGAGGAGGAGGLY | ASSARGDNQAPL | 0.55538887 |
| GHGGTGAAGGNGGTGGAGGSGI | ASSARGDNQAPL | 0.4970366 |
| GNGGNGGAGGEGGPGGAGGAGA | ASSARGDNQAPL | 0.71388984 |
| GSTAGNGGNGGNATNGTIAGQP | ASSARGDNQAPL | 0.7238568 |
| GTGGDGGNGGNGGTGG | ASSARGDNQAPL | 0.63298863 |
| GVRRPAGIPADAPV | ASSARGDNQAPL | 0.79515356 |
| GYRQVTDAHLLTLARR | ASSARGDNQAPL | 0.5281222 |
| IAKMGQDGHDRG | ASSARGDNQAPL | 0.7497747 |
| KAEPSIEIQYT | ASSARGDNQAPL | 0.70114386 |
| KPVRIRMAADEDTVAVT | ASSARGDNQAPL | 0.57642454 |
| MPPRIAGMRLLVIKPEP | ASSARGDNQAPL | 0.6029108 |
| MKFYAVALFRSRGVPPTR | ASSARGDNQAPL | 0.69555974 |
| NAGGFGILGAVAHSP | ASSARGDNQAPL | 0.53419685 |
| NEGREITLAMAEM | ASSARGDNQAPL | 0.7269639 |
| NELVEWSHPLGHRWRWE | ASSARGDNQAPL | 0.76598823 |
| NLGAGGTGGNGGNPG | ASSARGDNQAPL | 0.35475838 |
| NTEWPKWETVMAADPVD | ASSARGDNQAPL | 0.6322154 |
| PASAAPAETQP | ASSARGDNQAPL | 0.52295095 |
| QQSSDGKNGVRAAEITGE | ASSARGDNQAPL | 0.6292014 |
| RAPDAVTTSWWKE | ASSARGDNQAPL | 0.4222028 |
| SGFGNVGAHTSGWFNQ | ASSARGDNQAPL | 0.31315583 |
| SGVAGAVGQAASVGGLKV | ASSARGDNQAPL | 0.53355217 |
| SNGGAGTGSAGGNGGT | ASSARGDNQAPL | 0.38215718 |
| TGGAGGEGGAAG | ASSARGDNQAPL | 0.66282964 |
| TGGDGGAAGAGGAPGHG | ASSARGDNQAPL | 0.6320199 |
| TMPPLLAAIANAGRHGV | ASSARGDNQAPL | 0.41432512 |
| TSSTAGNSGGA | ASSARGDNQAPL | 0.6158776 |
| VPATVPVGILC | ASSARGDNQAPL | 0.65414757 |
| VQRPDSGGVVLTGRIS | ASSARGDNQAPL | 0.37167525 |
| WTPNPVPATKGVPAGLVQAT | ASSARGDNQAPL | 0.43991533 |

| | | |
|---|---|---|
| YALKRALAKFSDRFDVVII | ASSARGDNQAPL | 0.4308049 |
| YLIGSVIV | ASSARGDNQAPL | 0.5279887 |
| YGQLVAVID | ASSARGDNQAPL | 0.30371362 |
| VTVGGMTLPA | ASSARGDNQAPL | 0.56265646 |
| VLALPAV | ASSARGDNQAPL | 0.40400285 |
| VIPPTLNYET | ASSARGDNQAPL | 0.268574 |
| VGMGGMHPSSG | ASSARGDNQAPL | 0.59621155 |
| VGLAGGAATA | ASSARGDNQAPL | 0.4417831 |
| VGGAGGAGGSGP | ASSARGDNQAPL | 0.48837838 |
| VFAAFMV | ASSARGDNQAPL | 0.6306251 |
| TLYDKLR | ASSARGDNQAPL | 0.52693665 |
| TLAVFTVL | ASSARGDNQAPL | 0.4008659 |
| TGSVQPARV | ASSARGDNQAPL | 0.5477875 |
| TGGGAGSGGT | ASSARGDNQAPL | 0.67440397 |
| SHLETSL | ASSARGDNQAPL | 0.3709616 |
| SFTTGMT | ASSARGDNQAPL | 0.57502127 |
| QVEVAKRMGVRQPTVS | ASSARGDNQAPL | 0.66526926 |
| QGSGGPALAM | ASSARGDNQAPL | 0.54353017 |
| MLAPSNP | ASSARGDNQAPL | 0.67283833 |
| LVNNAAI | ASSARGDNQAPL | 0.675648 |
| LTGEINTLA | ASSARGDNQAPL | 0.42684582 |
| LSIFGPV | ASSARGDNQAPL | 0.652328 |
| LLALLANL | ASSARGDNQAPL | 0.6464394 |
| LISNASDALDKLR | ASSARGDNQAPL | 0.48147598 |
| LINEAVP | ASSARGDNQAPL | 0.5375544 |
| LGYAFSGL | ASSARGDNQAPL | 0.35645866 |
| LDFVAVAL | ASSARGDNQAPL | 0.5002639 |
| LAQPGAE | ASSARGDNQAPL | 0.32737955 |
| ISNLGAVTGF | ASSARGDNQAPL | 0.5475068 |
| HQEDPGRI | ASSARGDNQAPL | 0.63211566 |
| GTPGEPTP | ASSARGDNQAPL | 0.5261959 |
| GFFNSGS | ASSARGDNQAPL | 0.3159635 |
| GAGGGGLTAGAN | ASSARGDNQAPL | 0.31475854 |
| FVDALVNM | ASSARGDNQAPL | 0.48259464 |
| FLTVAPEL | ASSARGDNQAPL | 0.38161868 |
| FETLTAL | ASSARGDNQAPL | 0.3221594 |
| DVINPIEL | ASSARGDNQAPL | 0.4499038 |
| APVSGAPV | ASSARGDNQAPL | 0.6397358 |
| ALVGGLAW | ASSARGDNQAPL | 0.6518287 |
| ALAAAAATAI | ASSARGDNQAPL | 0.3993451 |
| AFGLGLLL | ASSARGDNQAPL | 0.67719245 |
| AFFAAASL | ASSARGDNQAPL | 0.6376759 |
| AAAALPAFSPP | ASSARGDNQAPL | 0.6621864 |
| VVLAPAL | ASSARGDNQAPL | 0.4787117 |
| TIGHVDHGKTTL | ASSARGDNQAPL | 0.35318 |
| LISNASDALDK | ASSARGDNQAPL | 0.66719514 |
| LISNASDAL | ASSARGDNQAPL | 0.49538225 |
| ISNASDALDK | ASSARGDNQAPL | 0.60316694 |
| NQAETLVYQTE | ASSARGDNQAPL | 0.645301 |
| TGGTGGTAGAG | ASSARGDNQAPL | 0.44086456 |
| TGSTAGGAGGA | ASSARGDNQAPL | 0.41304106 |

| | | |
|---|---|---|
| DPLAVPVAAGPV | ASSARGDNQAPL | 0.5996269 |
| GGTGGTPGNGGN | ASSARGDNQAPL | 0.61792326 |
| PAGLAGVDPATL | ASSARGDNQAPL | 0.34273967 |
| GNNLVVEAPHAGASV | ASSARGDNQAPL | 0.37348855 |
| KVTRIMSASTLEQPA | ASSARGDNQAPL | 0.4653942 |
| LGFALPIALPRRTVV | ASSARGDNQAPL | 0.48298255 |
| TRLGNLTARKLVVLV | ASSARGDNQAPL | 0.679693 |
| WSQQRRLIYFPSAGP | ASSARGDNQAPL | 0.5473318 |
| GPNGNPDPMAAAVDIR | ASSARGDNQAPL | 0.4812051 |
| TQRCVSVADLRAAASR | ASSARGDNQAPL | 0.6196761 |
| QFVLDVAEHPVEVLTETV | ASSARGDNQAPL | 0.58631444 |
| GGAGGTGGAGGNGGTQAGDG | ASSARGDNQAPL | 0.5276807 |
| PLLATLRAIAVDVDAKPAQV | ASSARGDNQAPL | 0.5571175 |
| APKFRTGGYMIDFWGVGYQVAK | ASSARGDNQAPL | 0.42853856 |
| GAGGAGGAGGAAGGPGGTGGTG | ASSARGDNQAPL | 0.60340154 |
| IGALGLLATDVAPAGSVAFGVA | ASSARGDNQAPL | 0.62259096 |
| AGGAGGIGGNGGAGGAGGVLMG | ASSARGDNQAPL | 0.63436204 |
| ADELIGLL | ASSARGDNQAPL | 0.6585951 |
| ADRALKA | ASSARGDNQAPL | 0.53513426 |
| AGAVGQVGGAGGA | ASSARGDNQAPL | 0.5043992 |
| AGGGGGLGGAP | ASSARGDNQAPL | 0.35291165 |
| AGGLLYGNGGAG | ASSARGDNQAPL | 0.4721634 |
| AGLESIGL | ASSARGDNQAPL | 0.52974033 |
| AGLKTGII | ASSARGDNQAPL | 0.5296021 |
| ASLAALLL | ASSARGDNQAPL | 0.61796135 |
| ASVINISSI | ASSARGDNQAPL | 0.6097786 |
| ATPPGPA | ASSARGDNQAPL | 0.4035207 |
| ATVASALV | ASSARGDNQAPL | 0.6006002 |
| ATVASLIL | ASSARGDNQAPL | 0.6469037 |
| AVNGPVLIRH | ASSARGDNQAPL | 0.37539178 |
| DGAAGAVGI | ASSARGDNQAPL | 0.54047024 |
| DPLLHGI | ASSARGDNQAPL | 0.6729644 |
| ELGAALGAL | ASSARGDNQAPL | 0.6492889 |
| ELGRTQAV | ASSARGDNQAPL | 0.43956408 |
| FVAGGYLS | ASSARGDNQAPL | 0.55232596 |
| GASGGAGGAGGW | ASSARGDNQAPL | 0.6903454 |
| GGAGAGGDGGAGGI | ASSARGDNQAPL | 0.551143 |
| GTGHNGSQPGMGGNG | ASSARGDNQAPL | 0.6245499 |
| GYAAASASAS | ASSARGDNQAPL | 0.33403087 |
| HQVDVAAVQQG | ASSARGDNQAPL | 0.50124574 |
| IGAAIAHT | ASSARGDNQAPL | 0.3633747 |
| IGMIEVAQL | ASSARGDNQAPL | 0.2811986 |
| IRLAPEHTSAMD | ASSARGDNQAPL | 0.64442545 |
| IYNVSVL | ASSARGDNQAPL | 0.44956386 |
| KQGFLGM | ASSARGDNQAPL | 0.63295656 |
| LESVITQL | ASSARGDNQAPL | 0.4055819 |
| LGLGALVVV | ASSARGDNQAPL | 0.6699945 |
| LIGNGGNGGSGGI | ASSARGDNQAPL | 0.5903177 |
| LNAINAPTL | ASSARGDNQAPL | 0.5584607 |
| LQLLNTGV | ASSARGDNQAPL | 0.44329262 |
| LQTSIEF | ASSARGDNQAPL | 0.57559824 |

| | | |
|---|---|---|
| MSFAVSEL | ASSARGDNQAPL | 0.6189473 |
| NTGALGL | ASSARGDNQAPL | 0.6110431 |
| PPGAGGAGGAGTT | ASSARGDNQAPL | 0.3162502 |
| QLVGLGI | ASSARGDNQAPL | 0.5740464 |
| RFLEQVHQL | ASSARGDNQAPL | 0.51439655 |
| RGVGYVLR | ASSARGDNQAPL | 0.3028378 |
| SATGVFAGVI | ASSARGDNQAPL | 0.28187275 |
| SLPAITLP | ASSARGDNQAPL | 0.34183624 |
| STALAFTV | ASSARGDNQAPL | 0.44773877 |
| SVLTVGLGK | ASSARGDNQAPL | 0.58058834 |
| TALAGAGSGI | ASSARGDNQAPL | 0.61303353 |
| TGLPFTGI | ASSARGDNQAPL | 0.47004026 |
| VAEHVTV | ASSARGDNQAPL | 0.57351255 |
| VDIVLNSL | ASSARGDNQAPL | 0.50233936 |
| VLTGGGAVL | ASSARGDNQAPL | 0.2541771 |
| VNVGLAYGI | ASSARGDNQAPL | 0.40335354 |
| VSSAAVGAV | ASSARGDNQAPL | 0.3742212 |
| VTSVYEAGV | ASSARGDNQAPL | 0.65995806 |
| WVALDRGA | ASSARGDNQAPL | 0.4036728 |
| YLVGSVVI | ASSARGDNQAPL | 0.48210245 |
| GPPGTGKTLL | ASSARGDNQAPL | 0.5142151 |
| GVGMTKF | ASSARGDNQAPL | 0.63758016 |
| IGIGGSDL | ASSARGDNQAPL | 0.5346163 |
| IGTLLAEL | ASSARGDNQAPL | 0.34821147 |
| IYTSGTT | ASSARGDNQAPL | 0.28539848 |
| LSFPAGV | ASSARGDNQAPL | 0.6122398 |
| LYTSGTT | ASSARGDNQAPL | 0.533352 |
| VGAFPFGF | ASSARGDNQAPL | 0.40819305 |
| VGAGTVEF | ASSARGDNQAPL | 0.4495558 |
| VGKSTLLL | ASSARGDNQAPL | 0.41036594 |
| AALAHAGGGIV | ASSARGDNQAPL | 0.30627552 |
| AALVAVSLL | ASSARGDNQAPL | 0.641945 |
| AAVSRAVAAL | ASSARGDNQAPL | 0.39294538 |
| AGGSSGIGGF | ASSARGDNQAPL | 0.4662138 |
| AGLTVANANG | ASSARGDNQAPL | 0.64539754 |
| AGLVGAVL | ASSARGDNQAPL | 0.6306692 |
| ALSQAFTV | ASSARGDNQAPL | 0.5096759 |
| ALTFSVSL | ASSARGDNQAPL | 0.38769838 |
| ALTGAALVAL | ASSARGDNQAPL | 0.41165045 |
| ASFSIGAGVL | ASSARGDNQAPL | 0.3687061 |
| DGSVYKI | ASSARGDNQAPL | 0.35711944 |
| EQLAEIV | ASSARGDNQAPL | 0.6564572 |
| FAVSELTVM | ASSARGDNQAPL | 0.5704455 |
| GAAPLAAGL | ASSARGDNQAPL | 0.39342317 |
| GADGGDGGSGGAGG | ASSARGDNQAPL | 0.60759 |
| GFGLDTGGAGG | ASSARGDNQAPL | 0.63714576 |
| GLASGGLL | ASSARGDNQAPL | 0.66066337 |
| GRIAGAAVLL | ASSARGDNQAPL | 0.44292992 |
| GSAGSLNTSF | ASSARGDNQAPL | 0.43604425 |
| GTGQIGI | ASSARGDNQAPL | 0.54939836 |
| GTLIALIL | ASSARGDNQAPL | 0.50895965 |

| | | |
|---|---|---|
| IAATMGFVVG | ASSARGDNQAPL | 0.43399394 |
| IFAQALL | ASSARGDNQAPL | 0.39366135 |
| IGAAVALGF | ASSARGDNQAPL | 0.59522796 |
| IGLTPLAL | ASSARGDNQAPL | 0.6968291 |
| ILGSNTSTL | ASSARGDNQAPL | 0.4743828 |
| IVFSLTVL | ASSARGDNQAPL | 0.30587575 |
| LASKATGF | ASSARGDNQAPL | 0.43815637 |
| LASVALIL | ASSARGDNQAPL | 0.36384147 |
| LFAVGLTL | ASSARGDNQAPL | 0.41100338 |
| LFNGGIGTYI | ASSARGDNQAPL | 0.5760616 |
| LTLGAGII | ASSARGDNQAPL | 0.36405945 |
| LVAAALGDI | ASSARGDNQAPL | 0.44050246 |
| NTAFVQLGI | ASSARGDNQAPL | 0.66784096 |
| QLFSGAEL | ASSARGDNQAPL | 0.5424189 |
| SAATFITL | ASSARGDNQAPL | 0.5842614 |
| SNYGLANLGV | ASSARGDNQAPL | 0.6377611 |
| SYGGAYAVM | ASSARGDNQAPL | 0.5892441 |
| TTGVGGLTLG | ASSARGDNQAPL | 0.66956526 |
| TTQTIYI | ASSARGDNQAPL | 0.6659543 |
| VAAGAGLGAVI | ASSARGDNQAPL | 0.54153395 |
| VAAVVTLL | ASSARGDNQAPL | 0.46475518 |
| VGFLGGLI | ASSARGDNQAPL | 0.5825787 |
| VGVIPILL | ASSARGDNQAPL | 0.47578588 |
| VIFASSLI | ASSARGDNQAPL | 0.52249265 |
| VKGGDGGAAAG | ASSARGDNQAPL | 0.47459385 |
| VTKTFTNI | ASSARGDNQAPL | 0.59938204 |
| VVAAAILL | ASSARGDNQAPL | 0.43505913 |
| YAALDVEL | ASSARGDNQAPL | 0.67839766 |
| LASVALLL | ASSARGDNQAPL | 0.29459295 |
| LDAVHRV | ASSARGDNQAPL | 0.3961176 |
| QGLSSSIF | ASSARGDNQAPL | 0.6585251 |
| RGTFANIRL | ASSARGDNQAPL | 0.26937494 |
| RQLWWGHRI | ASSARGDNQAPL | 0.3150907 |
| YGPPGTGKTLL | ASSARGDNQAPL | 0.46469605 |
| AALVGAPANVA | ASSARGDNQAPL | 0.6438674 |
| AATFRRTE | ASSARGDNQAPL | 0.268506 |
| AGALGGGI | ASSARGDNQAPL | 0.52757317 |
| AGGAGGAGGIGAG | ASSARGDNQAPL | 0.34560356 |
| AGSGALGL | ASSARGDNQAPL | 0.6288657 |
| AQIADQI | ASSARGDNQAPL | 0.510819 |
| ASAVAVTYL | ASSARGDNQAPL | 0.33777532 |
| ASTVLGGTGG | ASSARGDNQAPL | 0.6507715 |
| AVGGEPAK | ASSARGDNQAPL | 0.35183552 |
| CEQGTFGGTC | ASSARGDNQAPL | 0.53780663 |
| DEGKAPAG | ASSARGDNQAPL | 0.6633914 |
| DTVVFPAL | ASSARGDNQAPL | 0.6921378 |
| FEGITMAG | ASSARGDNQAPL | 0.38406968 |
| FPLVVQD | ASSARGDNQAPL | 0.64421535 |
| GAFTTPQ | ASSARGDNQAPL | 0.41235608 |
| GFGGFGTAGGVGGA | ASSARGDNQAPL | 0.6805103 |
| GFLAQFL | ASSARGDNQAPL | 0.4429203 |

| | | |
|---|---|---|
| GGAGGAGGNAGVA | ASSARGDNQAPL | 0.3157526 |
| GGVGTAGAGGAGGAGG | ASSARGDNQAPL | 0.630423 |
| GNGGGGDAGNAGSGGN | ASSARGDNQAPL | 0.62499774 |
| GTGGAGGAGNDAGS | ASSARGDNQAPL | 0.5289524 |
| GTTNTGWFNS | ASSARGDNQAPL | 0.6388006 |
| IFMGPLFTL | ASSARGDNQAPL | 0.53295976 |
| ILAHAIEAI | ASSARGDNQAPL | 0.44963303 |
| IRSAIREAL | ASSARGDNQAPL | 0.64717174 |
| ITFAGGG | ASSARGDNQAPL | 0.6002592 |
| ITGMSGHQEV | ASSARGDNQAPL | 0.40912306 |
| IVAGLACTL | ASSARGDNQAPL | 0.65714955 |
| IVGGGLAGA | ASSARGDNQAPL | 0.51602006 |
| IVTGVLNI | ASSARGDNQAPL | 0.38579583 |
| LAQQGCI | ASSARGDNQAPL | 0.6720916 |
| LCLHVEN | ASSARGDNQAPL | 0.55862427 |
| LDAISDGS | ASSARGDNQAPL | 0.5911081 |
| LGGFAAML | ASSARGDNQAPL | 0.5447451 |
| LGGKNPMI | ASSARGDNQAPL | 0.6875793 |
| LGPKGVAAIL | ASSARGDNQAPL | 0.3869299 |
| LMVGVSI | ASSARGDNQAPL | 0.54351354 |
| LVGAAFVL | ASSARGDNQAPL | 0.5047742 |
| LVLAVAPL | ASSARGDNQAPL | 0.6417637 |
| MAAIAVIDAA | ASSARGDNQAPL | 0.5310044 |
| NNGINAELQ | ASSARGDNQAPL | 0.5058025 |
| RGEPLVLVHG | ASSARGDNQAPL | 0.57732487 |
| RGGAGGAAGLI | ASSARGDNQAPL | 0.61585605 |
| RSYRMHPAV | ASSARGDNQAPL | 0.4509253 |
| SAAHLLAGQLA | ASSARGDNQAPL | 0.5170926 |
| SAAVGSY | ASSARGDNQAPL | 0.68770576 |
| SPHAVLHRI | ASSARGDNQAPL | 0.3592433 |
| SQIAKVL | ASSARGDNQAPL | 0.5426134 |
| TALGHLVP | ASSARGDNQAPL | 0.5712619 |
| TAPTTSGAA | ASSARGDNQAPL | 0.62176895 |
| TDGDDGNGGSPGT | ASSARGDNQAPL | 0.5495979 |
| TDGLGEN | ASSARGDNQAPL | 0.64578754 |
| TGGPPLPGQA | ASSARGDNQAPL | 0.46799132 |
| TIPAITI | ASSARGDNQAPL | 0.6023089 |
| TIPGFTI | ASSARGDNQAPL | 0.47812176 |
| TSIASGALA | ASSARGDNQAPL | 0.532445 |
| TVTLPGAAI | ASSARGDNQAPL | 0.35869512 |
| VASGLSAALP | ASSARGDNQAPL | 0.51309735 |
| VGAGVIGI | ASSARGDNQAPL | 0.5929351 |
| VGPKRFGV | ASSARGDNQAPL | 0.58702284 |
| VNVAFPD | ASSARGDNQAPL | 0.34163016 |
| VSMTNSGVS | ASSARGDNQAPL | 0.59238577 |
| VTDVLGDLSV | ASSARGDNQAPL | 0.5923388 |
| VVVVGAGFAGL | ASSARGDNQAPL | 0.6938169 |
| VYTVGTVV | ASSARGDNQAPL | 0.46259975 |
| WTTFSGP | ASSARGDNQAPL | 0.65903366 |
| YDELVEAF | ASSARGDNQAPL | 0.45689687 |
| YGTGGAGGNGG | ASSARGDNQAPL | 0.5464435 |

| | | |
|---|---|---|
| YTNVMAV | ASSARGDNQAPL | 0.308526 |
| FLEQVHQL | ASSARGDNQAPL | 0.3782181 |
| GLSLLAVAV | ASSARGDNQAPL | 0.5298818 |
| LEQVHQL | ASSARGDNQAPL | 0.43082392 |
| LNAAGAI | ASSARGDNQAPL | 0.65858984 |
| PAVHASL | ASSARGDNQAPL | 0.6355578 |
| TELEDDA | ASSARGDNQAPL | 0.6178595 |
| TPPGPPN | ASSARGDNQAPL | 0.62863314 |
| TPPPPPPP | ASSARGDNQAPL | 0.6507531 |
| TQSPALI | ASSERVGGRDEQY | 0.6612849 |
| EFAAAFPQ | ASSERVGGRDEQY | 0.4333305 |
| LLSGNLDVL | ASSERVGGRDEQY | 0.48825005 |
| GPANLGNYNI | ASSERVGGRDEQY | 0.32392722 |
| TLGPTLP | ASSERVGGRDEQY | 0.40888575 |
| VYAYSSCCK | ASSERVGGRDEQY | 0.41835433 |
| APAPAPAPNI | ASSERVGGRDEQY | 0.5384171 |
| PGGPVPIP | ASSERVGGRDEQY | 0.27221975 |
| FLTVALEP | ASSERVGGRDEQY | 0.6103039 |
| SQYFGANL | ASSERVGGRDEQY | 0.65941536 |
| GGVKPAA | ASSERVGGRDEQY | 0.4980212 |
| GGTGGAGGAGGW | ASSERVGGRDEQY | 0.58379894 |
| GVGGVAGLGGAGG | ASSERVGGRDEQY | 0.58932173 |
| VLLVLATTI | ASSERVGGRDEQY | 0.3678531 |
| IGIFAAML | ASSERVGGRDEQY | 0.56100386 |
| AVGDLID | ASSERVGGRDEQY | 0.3979051 |
| NISPLPL | ASSERVGGRDEQY | 0.6576315 |
| VTAVFVAI | ASSERVGGRDEQY | 0.6509391 |
| SGDDNDAAL | ASSERVGGRDEQY | 0.5346468 |
| AVAALAHS | ASSERVGGRDEQY | 0.42469585 |
| GGVTVMATP | ASSERVGGRDEQY | 0.6049346 |
| AGLAANVRD | ASSERVGGRDEQY | 0.39612323 |
| TNRPDLIDPALLRPG | ASSERVGGRDEQY | 0.45446953 |
| LANLGVTAL | ASSERVGGRDEQY | 0.2857316 |
| TGERIEL | ASSERVGGRDEQY | 0.5472554 |
| NGEELAA | ASSERVGGRDEQY | 0.36016136 |
| VLSLVWG | ASSERVGGRDEQY | 0.6080802 |
| IGYALWL | ASSERVGGRDEQY | 0.50465524 |
| LSLLLAPL | ASSERVGGRDEQY | 0.5094856 |
| TLDEALK | ASSERVGGRDEQY | 0.36609867 |
| TDGAGTAEGG | ASSERVGGRDEQY | 0.62322605 |
| NAPGGAGGNGG | ASSERVGGRDEQY | 0.57251024 |
| VSVFMGMGGNF | ASSERVGGRDEQY | 0.3132927 |
| LASYLAAL | ASSERVGGRDEQY | 0.63917637 |
| LVGGTPAL | ASSERVGGRDEQY | 0.44229177 |
| VRNADVL | ASSERVGGRDEQY | 0.5184355 |
| TCSSPFLVSAM | ASSERVGGRDEQY | 0.6010925 |
| VHITDAE | ASSERVGGRDEQY | 0.5691206 |
| EYAAPSF | ASSERVGGRDEQY | 0.35164642 |
| VVDAFHGGL | ASSERVGGRDEQY | 0.43363154 |
| ASVVDPQ | ASSERVGGRDEQY | 0.47189635 |
| SYNEFGGLS | ASSERVGGRDEQY | 0.45546186 |

| | | |
|---|---|---|
| DPDGSVY | ASSERVGGRDEQY | 0.29716066 |
| MVSTIHATE | ASSERVGGRDEQY | 0.5963917 |
| LAAGVADPV | ASSERVGGRDEQY | 0.38699514 |
| TKEQIAA | ASSERVGGRDEQY | 0.41453564 |
| SLASPAAL | ASSERVGGRDEQY | 0.5053079 |
| IGLVFYGV | ASSERVGGRDEQY | 0.6641016 |
| FLLDASNL | ASSERVGGRDEQY | 0.5918188 |
| YQVGNFP | ASSERVGGRDEQY | 0.47210076 |
| ESLLSQST | ASSERVGGRDEQY | 0.59971565 |
| LFTVGFAAVG | ASSERVGGRDEQY | 0.53138834 |
| ATPPTPT | ASSERVGGRDEQY | 0.6606342 |
| TDLATLG | ASSERVGGRDEQY | 0.5284075 |
| SAGGTGR | ASSERVGGRDEQY | 0.43095607 |
| GFSNSGS | ASSERVGGRDEQY | 0.62593603 |
| GGNGGSGGTSV | ASSERVGGRDEQY | 0.4573618 |
| GTSVATGGAGN | ASSERVGGRDEQY | 0.3627255 |
| NGGNGGSGGTS | ASSERVGGRDEQY | 0.39042044 |
| AVVKFPH | ASSERVGGRDEQY | 0.40064865 |
| TPPPPPP | ASSERVGGRDEQY | 0.32666203 |
| AAPAAAAAPA | ASSERVGGRDEQY | 0.5692497 |
| IALNEDL | ASSERVGGRDEQY | 0.67605954 |
| IVNVSSM | ASSERVGGRDEQY | 0.5322888 |
| TGIPPAPR | ASSERVGGRDEQY | 0.31733513 |
| FDLGGGTF | ASSERVGGRDEQY | 0.38570148 |
| FDLGGGTFDVS | ASSERVGGRDEQY | 0.4939675 |
| NEPTAAA | ASSERVGGRDEQY | 0.617767 |
| NEPTAAALA | ASSERVGGRDEQY | 0.55435485 |
| GSIINISSI | ASSERVGGRDEQY | 0.68804693 |
| VAPGNAGTA | ASSERVGGRDEQY | 0.6183804 |
| TGGGASLE | ASSERVGGRDEQY | 0.42487782 |
| VAVYNHYKRI | ASSERVGGRDEQY | 0.65630597 |
| TDPAPATT | ASSERVGGRDEQY | 0.46961334 |
| TNRPDILDPALLRPG | ASSERVGGRDEQY | 0.47749704 |
| LTTTTTS | ASSERVGGRDEQY | 0.518381 |
| PTGSGKTL | ASSERVGGRDEQY | 0.5426746 |
| TGSGKTL | ASSERVGGRDEQY | 0.44240516 |
| TIGHVDHGK | ASSERVGGRDEQY | 0.48422962 |
| TIGHVDHGKTT | ASSERVGGRDEQY | 0.33605832 |
| GHVDHGKTT | ASSERVGGRDEQY | 0.5753636 |
| AQFEHTLL | ASSERVGGRDEQY | 0.3835666 |
| IVNQDEL | ASSERVGGRDEQY | 0.3821931 |
| TNGSQFF | ASSERVGGRDEQY | 0.48139673 |
| TGKGIVE | ASSERVGGRDEQY | 0.6837138 |
| LISNASDA | ASSERVGGRDEQY | 0.65655965 |
| AELRTLGELTKIAKAHGAQ | ASSERVGGRDEQY | 0.42225057 |
| APNAVAAAEPSW | ASSERVGGRDEQY | 0.3872977 |
| APQVAGAATAVL | ASSERVGGRDEQY | 0.61740744 |
| ARRDPAAAAAILPTDA | ASSERVGGRDEQY | 0.445399 |
| DAAARDVLAEAGLAEYF | ASSERVGGRDEQY | 0.6318029 |
| DDQIVLSAKANLAKVEQ | ASSERVGGRDEQY | 0.55917585 |
| DELLPDTGEDSDEDGH | ASSERVGGRDEQY | 0.50484014 |

| | | |
|---|---|---|
| ELPEAPANRATVPQD | ASSERVGGRDEQY | 0.45158917 |
| FGQSKIERVTGIG | ASSERVGGRDEQY | 0.5673643 |
| GAGGTGSTAGGAGGAGGAGGLY | ASSERVGGRDEQY | 0.44223705 |
| GHGGTGAAGGNGGTGGAGGSGI | ASSERVGGRDEQY | 0.6198153 |
| GNGGNGGAGGEGGPGGAGGAGA | ASSERVGGRDEQY | 0.31268376 |
| GSTAGNGGNGGNATNGTIAGQP | ASSERVGGRDEQY | 0.65750545 |
| GTGGDGGNGGNGGTGG | ASSERVGGRDEQY | 0.38005888 |
| GVRRPAGIPADAPV | ASSERVGGRDEQY | 0.36854813 |
| GYRQVTDAHLLTLARR | ASSERVGGRDEQY | 0.27836326 |
| IAKMGQDGHDRG | ASSERVGGRDEQY | 0.5534681 |
| KAEPSIEIQYT | ASSERVGGRDEQY | 0.5575625 |
| KPVRIRMAADEDTVAVT | ASSERVGGRDEQY | 0.63862467 |
| MPPRIAGMRLLVIKPEP | ASSERVGGRDEQY | 0.5258387 |
| MKFYAVALFRSRGVPPTR | ASSERVGGRDEQY | 0.5881691 |
| NAGGFGILGAVAHSP | ASSERVGGRDEQY | 0.64818597 |
| NEGREITLAMAEM | ASSERVGGRDEQY | 0.43149504 |
| NELVEWSHPLGHRWRWE | ASSERVGGRDEQY | 0.30954498 |
| NLGAGGTGGNGGNPG | ASSERVGGRDEQY | 0.6256911 |
| NTEWPKWETVMAADPVD | ASSERVGGRDEQY | 0.60300857 |
| PASAAPAETQP | ASSERVGGRDEQY | 0.65537107 |
| QQSSDGKNGVRAAEITGE | ASSERVGGRDEQY | 0.598801 |
| RAPDAVTTSWWKE | ASSERVGGRDEQY | 0.3457985 |
| SGFGNVGAHTSGWFNQ | ASSERVGGRDEQY | 0.6298059 |
| SGVAGAVGQAASVGGLKV | ASSERVGGRDEQY | 0.62456197 |
| SNGGAGTGSAGGNGGT | ASSERVGGRDEQY | 0.32945246 |
| TGGAGGEGGAAG | ASSERVGGRDEQY | 0.42539808 |
| TGGDGGAAGAGGAPGHG | ASSERVGGRDEQY | 0.43876994 |
| TMPPLLAAIANAGRHGV | ASSERVGGRDEQY | 0.3386936 |
| TSSTAGNSGGA | ASSERVGGRDEQY | 0.4898548 |
| VPATVPVGILC | ASSERVGGRDEQY | 0.41251564 |
| VQRPDSGGVVLTGRIS | ASSERVGGRDEQY | 0.5156014 |
| WTPNPVPATKGVPAGLVQAT | ASSERVGGRDEQY | 0.47254366 |
| YALKRALAKFSDRFDVVII | ASSERVGGRDEQY | 0.36396256 |
| YLIGSVIV | ASSERVGGRDEQY | 0.52512753 |
| YGQLVAVID | ASSERVGGRDEQY | 0.48343313 |
| VTVGGMTLPA | ASSERVGGRDEQY | 0.576155 |
| VLALPAV | ASSERVGGRDEQY | 0.5849642 |
| VIPPTLNYET | ASSERVGGRDEQY | 0.48842704 |
| VGMGGMHPSSG | ASSERVGGRDEQY | 0.62345535 |
| VGLAGGAATA | ASSERVGGRDEQY | 0.6486629 |
| VGGAGGAGGSGP | ASSERVGGRDEQY | 0.33967513 |
| VFAAFMV | ASSERVGGRDEQY | 0.4082696 |
| TLYDKLR | ASSERVGGRDEQY | 0.6116283 |
| TLAVFTVL | ASSERVGGRDEQY | 0.43039814 |
| TGSVQPARV | ASSERVGGRDEQY | 0.6636469 |
| TGGGAGSGGT | ASSERVGGRDEQY | 0.48779237 |
| SHLETSL | ASSERVGGRDEQY | 0.55830735 |
| SFTTGMT | ASSERVGGRDEQY | 0.5386986 |
| QVEVAKRMGVRQPTVS | ASSERVGGRDEQY | 0.6720042 |
| QGSGGPALAM | ASSERVGGRDEQY | 0.6083587 |
| MLAPSNP | ASSERVGGRDEQY | 0.48944175 |

| | | |
|---|---|---|
| LVNNAAI | ASSERVGGRDEQY | 0.6323886 |
| LTGEINTLA | ASSERVGGRDEQY | 0.3522672 |
| LSIFGPV | ASSERVGGRDEQY | 0.5963747 |
| LLALLANL | ASSERVGGRDEQY | 0.6252887 |
| LISNASDALDKLR | ASSERVGGRDEQY | 0.37428048 |
| LINEAVP | ASSERVGGRDEQY | 0.44552118 |
| LGYAFSGL | ASSERVGGRDEQY | 0.63320005 |
| LDFVAVAL | ASSERVGGRDEQY | 0.65767884 |
| LAQPGAE | ASSERVGGRDEQY | 0.3688937 |
| ISNLGAVTGF | ASSERVGGRDEQY | 0.5061269 |
| HQEDPGRI | ASSERVGGRDEQY | 0.35455805 |
| GTPGEPTP | ASSERVGGRDEQY | 0.36767584 |
| GFFNSGS | ASSERVGGRDEQY | 0.32631207 |
| GAGGGGLTAGAN | ASSERVGGRDEQY | 0.5819951 |
| FVDALVNM | ASSERVGGRDEQY | 0.4383912 |
| FLTVAPEL | ASSERVGGRDEQY | 0.6627846 |
| FETLTAL | ASSERVGGRDEQY | 0.57724845 |
| DVINPIEL | ASSERVGGRDEQY | 0.65686965 |
| APVSGAPV | ASSERVGGRDEQY | 0.38176233 |
| ALVGGLAW | ASSERVGGRDEQY | 0.48492458 |
| ALAAAAATAI | ASSERVGGRDEQY | 0.32807827 |
| AFGLGLLL | ASSERVGGRDEQY | 0.3585644 |
| AFFAAASL | ASSERVGGRDEQY | 0.34218383 |
| AAAALPAFSPP | ASSERVGGRDEQY | 0.50315386 |
| VVLAPAL | ASSERVGGRDEQY | 0.2984367 |
| TIGHVDHGKTTL | ASSERVGGRDEQY | 0.6660193 |
| LISNASDALDK | ASSERVGGRDEQY | 0.51848364 |
| LISNASDAL | ASSERVGGRDEQY | 0.6125886 |
| ISNASDALDK | ASSERVGGRDEQY | 0.4635374 |
| NQAETLVYQTE | ASSERVGGRDEQY | 0.36295834 |
| TGGTGGTAGAG | ASSERVGGRDEQY | 0.44380122 |
| TGSTAGGAGGA | ASSERVGGRDEQY | 0.6404328 |
| DPLAVPVAAGPV | ASSERVGGRDEQY | 0.36445153 |
| GGTGGTPGNGGN | ASSERVGGRDEQY | 0.29920155 |
| PAGLAGVDPATL | ASSERVGGRDEQY | 0.39885288 |
| GNNLVVEAPHAGASV | ASSERVGGRDEQY | 0.42060935 |
| KVTRIMSASTLEQPA | ASSERVGGRDEQY | 0.66407967 |
| LGFALPIALPRRTVV | ASSERVGGRDEQY | 0.6239022 |
| TRLGNLTARKLVVLV | ASSERVGGRDEQY | 0.54664433 |
| WSQQRRLIYFPSAGP | ASSERVGGRDEQY | 0.67946446 |
| GPNGNPDPMAAAVDIR | ASSERVGGRDEQY | 0.27341574 |
| TQRCVSVADLRAAASR | ASSERVGGRDEQY | 0.50029355 |
| QFVLDVAEHPVEVLTETV | ASSERVGGRDEQY | 0.68511283 |
| GGAGGTGGAGGNGGTQAGDG | ASSERVGGRDEQY | 0.52783036 |
| PLLATLRAIAVDVDAKPAQV | ASSERVGGRDEQY | 0.45162165 |
| APKFRTGGYMIDFWGVGYQVAK | ASSERVGGRDEQY | 0.6663717 |
| GAGGAGGAGGAAGGPGGTGGTG | ASSERVGGRDEQY | 0.39736322 |
| IGALGLLATDVAPAGSVAFGVA | ASSERVGGRDEQY | 0.4584291 |
| AGGAGGIGGNGGAGGAGGVLMG | ASSERVGGRDEQY | 0.534971 |
| ADELIGLL | ASSERVGGRDEQY | 0.5918275 |
| ADRALKA | ASSERVGGRDEQY | 0.49639323 |

| | | |
|---|---|---|
| AGAVGQVGGAGGA | ASSERVGGRDEQY | 0.33796614 |
| AGGGGGLGGAP | ASSERVGGRDEQY | 0.649912 |
| AGGLLYGNGGAG | ASSERVGGRDEQY | 0.67353034 |
| AGLESIGL | ASSERVGGRDEQY | 0.5055587 |
| AGLKTGII | ASSERVGGRDEQY | 0.62011683 |
| ASLAALLL | ASSERVGGRDEQY | 0.60335773 |
| ASVINISSI | ASSERVGGRDEQY | 0.46210814 |
| ATPPGPA | ASSERVGGRDEQY | 0.397064 |
| ATVASALV | ASSERVGGRDEQY | 0.35543773 |
| ATVASLIL | ASSERVGGRDEQY | 0.6188185 |
| AVNGPVLIRH | ASSERVGGRDEQY | 0.40409976 |
| DGAAGAVGI | ASSERVGGRDEQY | 0.5463648 |
| DPLLHGI | ASSERVGGRDEQY | 0.54496205 |
| ELGAALGAL | ASSERVGGRDEQY | 0.6855415 |
| ELGRTQAV | ASSERVGGRDEQY | 0.5561085 |
| FVAGGYLS | ASSERVGGRDEQY | 0.62077725 |
| GASGGAGGAGGW | ASSERVGGRDEQY | 0.42872378 |
| GGAGAGGDGGAGGI | ASSERVGGRDEQY | 0.49780867 |
| GTGHNGSQPGMGGNG | ASSERVGGRDEQY | 0.5767892 |
| GYAAASASAS | ASSERVGGRDEQY | 0.35014004 |
| HQVDVAAVQQG | ASSERVGGRDEQY | 0.63016754 |
| IGAAIAHT | ASSERVGGRDEQY | 0.3878972 |
| IGMIEVAQL | ASSERVGGRDEQY | 0.539403 |
| IRLAPEHTSAMD | ASSERVGGRDEQY | 0.5803158 |
| IYNVSVL | ASSERVGGRDEQY | 0.67442435 |
| KQGFLGM | ASSERVGGRDEQY | 0.5197923 |
| LESVITQL | ASSERVGGRDEQY | 0.6066499 |
| LGLGALVVV | ASSERVGGRDEQY | 0.40107805 |
| LIGNGGNGGSGGI | ASSERVGGRDEQY | 0.6130415 |
| LNAINAPTL | ASSERVGGRDEQY | 0.6679381 |
| LQLLNTGV | ASSERVGGRDEQY | 0.58963287 |
| LQTSIEF | ASSERVGGRDEQY | 0.6887555 |
| MSFAVSEL | ASSERVGGRDEQY | 0.615764 |
| NTGALGL | ASSERVGGRDEQY | 0.31209382 |
| PPGAGGAGGAGTT | ASSERVGGRDEQY | 0.55286545 |
| QLVGLGI | ASSERVGGRDEQY | 0.6783303 |
| RFLEQVHQL | ASSERVGGRDEQY | 0.6399447 |
| RGVGYVLR | ASSERVGGRDEQY | 0.62274873 |
| SATGVFAGVI | ASSERVGGRDEQY | 0.5218591 |
| SLPAITLP | ASSERVGGRDEQY | 0.43787152 |
| STALAFTV | ASSERVGGRDEQY | 0.6509825 |
| SVLTVGLGK | ASSERVGGRDEQY | 0.6918787 |
| TALAGAGSGI | ASSERVGGRDEQY | 0.66251516 |
| TGLPFTGI | ASSERVGGRDEQY | 0.43069276 |
| VAEHVTV | ASSERVGGRDEQY | 0.38391092 |
| VDIVLNSL | ASSERVGGRDEQY | 0.53475124 |
| VLTGGGAVL | ASSERVGGRDEQY | 0.6458031 |
| VNVGLAYGI | ASSERVGGRDEQY | 0.36799246 |
| VSSAAVGAV | ASSERVGGRDEQY | 0.52314246 |
| VTSVYEAGV | ASSERVGGRDEQY | 0.47140127 |
| WVALDRGA | ASSERVGGRDEQY | 0.2900073 |

| | | |
|---|---|---|
| YLVGSVVI | ASSERVGGRDEQY | 0.58199394 |
| GPPGTGKTLL | ASSERVGGRDEQY | 0.6536313 |
| GVGMTKF | ASSERVGGRDEQY | 0.5847226 |
| IGIGGSDL | ASSERVGGRDEQY | 0.362801 |
| IGTLLAEL | ASSERVGGRDEQY | 0.62926173 |
| IYTSGTT | ASSERVGGRDEQY | 0.64982474 |
| LSFPAGV | ASSERVGGRDEQY | 0.6670257 |
| LYTSGTT | ASSERVGGRDEQY | 0.3543076 |
| VGAFPFGF | ASSERVGGRDEQY | 0.27039185 |
| VGAGTVEF | ASSERVGGRDEQY | 0.5849289 |
| VGKSTLLL | ASSERVGGRDEQY | 0.65545696 |
| AALAHAGGGIV | ASSERVGGRDEQY | 0.39291653 |
| AALVAVSLL | ASSERVGGRDEQY | 0.40417033 |
| AAVSRAVAAL | ASSERVGGRDEQY | 0.5383121 |
| AGGSSGIGGF | ASSERVGGRDEQY | 0.6309997 |
| AGLTVANANG | ASSERVGGRDEQY | 0.34717712 |
| AGLVGAVL | ASSERVGGRDEQY | 0.31551522 |
| ALSQAFTV | ASSERVGGRDEQY | 0.5617415 |
| ALTFSVSL | ASSERVGGRDEQY | 0.6628804 |
| ALTGAALVAL | ASSERVGGRDEQY | 0.37107965 |
| ASFSIGAGVL | ASSERVGGRDEQY | 0.4503321 |
| DGSVYKI | ASSERVGGRDEQY | 0.5810264 |
| EQLAEIV | ASSERVGGRDEQY | 0.6007501 |
| FAVSELTVM | ASSERVGGRDEQY | 0.52395797 |
| GAAPLAAGL | ASSERVGGRDEQY | 0.33574864 |
| GADGGDGGSGGAGG | ASSERVGGRDEQY | 0.606454 |
| GFGLDTGGAGG | ASSERVGGRDEQY | 0.34382877 |
| GLASGGLL | ASSERVGGRDEQY | 0.5942691 |
| GRIAGAAVLL | ASSERVGGRDEQY | 0.5356375 |
| GSAGSLNTSF | ASSERVGGRDEQY | 0.555835 |
| GTGQIGI | ASSERVGGRDEQY | 0.45755464 |
| GTLIALIL | ASSERVGGRDEQY | 0.38287276 |
| IAATMGFVVG | ASSERVGGRDEQY | 0.345253 |
| IFAQALL | ASSERVGGRDEQY | 0.3343323 |
| IGAAVALGF | ASSERVGGRDEQY | 0.63451684 |
| IGLTPLAL | ASSERVGGRDEQY | 0.34332386 |
| ILGSNTSTL | ASSERVGGRDEQY | 0.63077974 |
| IVFSLTVL | ASSERVGGRDEQY | 0.5721745 |
| LASKATGF | ASSERVGGRDEQY | 0.41891155 |
| LASVALIL | ASSERVGGRDEQY | 0.5428149 |
| LFAVGLTL | ASSERVGGRDEQY | 0.3381777 |
| LFNGGIGTYI | ASSERVGGRDEQY | 0.55619335 |
| LTLGAGII | ASSERVGGRDEQY | 0.5007241 |
| LVAAALGDI | ASSERVGGRDEQY | 0.64449096 |
| NTAFVQLGI | ASSERVGGRDEQY | 0.4220274 |
| QLFSGAEL | ASSERVGGRDEQY | 0.6127498 |
| SAATFITL | ASSERVGGRDEQY | 0.56293464 |
| SNYGLANLGV | ASSERVGGRDEQY | 0.34849572 |
| SYGGAYAVM | ASSERVGGRDEQY | 0.57413787 |
| TTGVGGLTLG | ASSERVGGRDEQY | 0.6383674 |
| TTQTIYI | ASSERVGGRDEQY | 0.6045593 |

| | | |
|---|---|---|
| VAAGAGLGAVI | ASSERVGGRDEQY | 0.33091247 |
| VAAVVTLL | ASSERVGGRDEQY | 0.544901 |
| VGFLGGLI | ASSERVGGRDEQY | 0.4590685 |
| VGVIPILL | ASSERVGGRDEQY | 0.54924995 |
| VIFASSLI | ASSERVGGRDEQY | 0.5551026 |
| VKGGDGGAAAG | ASSERVGGRDEQY | 0.41886163 |
| VTKTFTNI | ASSERVGGRDEQY | 0.60697806 |
| VVAAAILL | ASSERVGGRDEQY | 0.346133 |
| YAALDVEL | ASSERVGGRDEQY | 0.3847848 |
| LASVALLL | ASSERVGGRDEQY | 0.5228549 |
| LDAVHRV | ASSERVGGRDEQY | 0.5684585 |
| QGLSSSIF | ASSERVGGRDEQY | 0.65056676 |
| RGTFANIRL | ASSERVGGRDEQY | 0.37322313 |
| RQLWWGHRI | ASSERVGGRDEQY | 0.55022365 |
| YGPPGTGKTLL | ASSERVGGRDEQY | 0.4085811 |
| AALVGAPANVA | ASSERVGGRDEQY | 0.37857854 |
| AATFRRTE | ASSERVGGRDEQY | 0.5735236 |
| AGALGGGI | ASSERVGGRDEQY | 0.42438826 |
| AGGAGGAGGIGAG | ASSERVGGRDEQY | 0.35356066 |
| AGSGALGL | ASSERVGGRDEQY | 0.5987594 |
| AQIADQI | ASSERVGGRDEQY | 0.36956492 |
| ASAVAVTYL | ASSERVGGRDEQY | 0.49846953 |
| ASTVLGGTGG | ASSERVGGRDEQY | 0.68093956 |
| AVGGEPAK | ASSERVGGRDEQY | 0.3526963 |
| CEQGTFGGTC | ASSERVGGRDEQY | 0.5709416 |
| DEGKAPAG | ASSERVGGRDEQY | 0.5806907 |
| DTVVFPAL | ASSERVGGRDEQY | 0.5643014 |
| FEGITMAG | ASSERVGGRDEQY | 0.47197425 |
| FPLVVQD | ASSERVGGRDEQY | 0.38784042 |
| GAFTTPQ | ASSERVGGRDEQY | 0.67457855 |
| GFGGFGTAGGVGGA | ASSERVGGRDEQY | 0.6109205 |
| GFLAQFL | ASSERVGGRDEQY | 0.58360136 |
| GGAGGAGGNAGVA | ASSERVGGRDEQY | 0.5900047 |
| GGVGTAGAGGAGGAGG | ASSERVGGRDEQY | 0.6596147 |
| GNGGGGDAGNAGSGGN | ASSERVGGRDEQY | 0.6673117 |
| GTGGAGGAGNDAGS | ASSERVGGRDEQY | 0.4402449 |
| GTTNTGWFNS | ASSERVGGRDEQY | 0.33538675 |
| IFMGPLFTL | ASSERVGGRDEQY | 0.5108413 |
| ILAHAIEAI | ASSERVGGRDEQY | 0.503914 |
| IRSAIREAL | ASSERVGGRDEQY | 0.46476305 |
| ITFAGGG | ASSERVGGRDEQY | 0.32090026 |
| ITGMSGHQEV | ASSERVGGRDEQY | 0.64926124 |
| IVAGLACTL | ASSERVGGRDEQY | 0.6979393 |
| IVGGGLAGA | ASSERVGGRDEQY | 0.45955884 |
| IVTGVLNI | ASSERVGGRDEQY | 0.68272305 |
| LAQQGCI | ASSERVGGRDEQY | 0.32342368 |
| LCLHVEN | ASSERVGGRDEQY | 0.5913892 |
| LDAISDGS | ASSERVGGRDEQY | 0.66060257 |
| LGGFAAML | ASSERVGGRDEQY | 0.3730452 |
| LGGKNPMI | ASSERVGGRDEQY | 0.5788545 |
| LGPKGVAAIL | ASSERVGGRDEQY | 0.36922616 |

| | | |
|---|---|---|
| LMVGVSI | ASSERVGGRDEQY | 0.43647018 |
| LVGAAFVL | ASSERVGGRDEQY | 0.6516733 |
| LVLAVAPL | ASSERVGGRDEQY | 0.47396803 |
| MAAIAVIDAA | ASSERVGGRDEQY | 0.54804736 |
| NNGINAELQ | ASSERVGGRDEQY | 0.5236962 |
| RGEPLVLVHG | ASSERVGGRDEQY | 0.40444064 |
| RGGAGGAAGLI | ASSERVGGRDEQY | 0.6045835 |
| RSYRMHPAV | ASSERVGGRDEQY | 0.62061924 |
| SAAHLLAGQLA | ASSERVGGRDEQY | 0.5307034 |
| SAAVGSY | ASSERVGGRDEQY | 0.6898097 |
| SPHAVLHRI | ASSERVGGRDEQY | 0.35813272 |
| SQIAKVL | ASSERVGGRDEQY | 0.67291224 |
| TALGHLVP | ASSERVGGRDEQY | 0.6829794 |
| TAPTTSGAA | ASSERVGGRDEQY | 0.5385765 |
| TDGDDGNGGSPGT | ASSERVGGRDEQY | 0.4739299 |
| TDGLGEN | ASSERVGGRDEQY | 0.6472007 |
| TGGPPLPGQA | ASSERVGGRDEQY | 0.52625334 |
| TIPAITI | ASSERVGGRDEQY | 0.5305369 |
| TIPGFTI | ASSERVGGRDEQY | 0.6682874 |
| TSIASGALA | ASSERVGGRDEQY | 0.38003772 |
| TVTLPGAAI | ASSERVGGRDEQY | 0.8110107 |
| VASGLSAALP | ASSERVGGRDEQY | 0.78804827 |
| VGAGVIGI | ASSERVGGRDEQY | 0.68296504 |
| VGPKRFGV | ASSERVGGRDEQY | 0.6073526 |
| VNVAFPD | ASSERVGGRDEQY | 0.3560678 |
| VSMTNSGVS | ASSERVGGRDEQY | 0.6176511 |
| VTDVLGDLSV | ASSERVGGRDEQY | 0.5444246 |
| VVVVGAGFAGL | ASSERVGGRDEQY | 0.8146719 |
| VYTVGTVV | ASSERVGGRDEQY | 0.77747416 |
| WTTFSGP | ASSERVGGRDEQY | 0.56521404 |
| YDELVEAF | ASSERVGGRDEQY | 0.78779924 |
| YGTGGAGGNGG | ASSERVGGRDEQY | 0.7802955 |
| YTNVMAV | ASSERVGGRDEQY | 0.3708453 |
| FLEQVHQL | ASSERVGGRDEQY | 0.73303604 |
| GLSLLAVAV | ASSERVGGRDEQY | 0.5552627 |
| LEQVHQL | ASSERVGGRDEQY | 0.6125896 |
| LNAAGAI | ASSERVGGRDEQY | 0.28621674 |
| PAVHASL | ASSERVGGRDEQY | 0.7511054 |
| TELEDDA | ASSERVGGRDEQY | 0.7290262 |
| TPPGPPN | ASSERVGGRDEQY | 0.19979984 |
| TPPPPPPP | ASSERVGGRDEQY | 0.75005555 |
| TQSPALI | ASSGDNYEQY | 0.6447717 |
| EFAAAFPQ | ASSGDNYEQY | 0.66340196 |
| LLSGNLDVL | ASSGDNYEQY | 0.78618383 |
| GPANLGNYNI | ASSGDNYEQY | 0.6755605 |
| TLGPTLP | ASSGDNYEQY | 0.6414179 |
| VYAYSSCCK | ASSGDNYEQY | 0.78709114 |
| APAPAPAPNI | ASSGDNYEQY | 0.8008957 |
| PGGPVPIP | ASSGDNYEQY | 0.46406418 |
| FLTVALEP | ASSGDNYEQY | 0.75374657 |
| SQYFGANL | ASSGDNYEQY | 0.8166858 |

| | | |
|---|---|---|
| GGVKPAA | ASSGDNYEQY | 0.74187773 |
| GGTGGAGGAGGW | ASSGDNYEQY | 0.80237216 |
| GVGGVAGLGGAGG | ASSGDNYEQY | 0.80814743 |
| VLLVLATTI | ASSGDNYEQY | 0.5168883 |
| IGIFAAML | ASSGDNYEQY | 0.80064106 |
| AVGDLID | ASSGDNYEQY | 0.8122169 |
| NISPLPL | ASSGDNYEQY | 0.6630938 |
| VTAVFVAI | ASSGDNYEQY | 0.6474255 |
| SGDDNDAAL | ASSGDNYEQY | 0.46842223 |
| AVAALAHS | ASSGDNYEQY | 0.72383887 |
| GGVTVMATP | ASSGDNYEQY | 0.45556417 |
| AGLAANVRD | ASSGDNYEQY | 0.7689171 |
| TNRPDLIDPALLRPG | ASSGDNYEQY | 0.81240034 |
| LANLGVTAL | ASSGDNYEQY | 0.6873603 |
| TGERIEL | ASSGDNYEQY | 0.4076956 |
| NGEELAA | ASSGDNYEQY | 0.47082275 |
| VLSLVWG | ASSGDNYEQY | 0.75304055 |
| IGYALWL | ASSGDNYEQY | 0.42986676 |
| LSLLLAPL | ASSGDNYEQY | 0.36201978 |
| TLDEALK | ASSGDNYEQY | 0.52626157 |
| TDGAGTAEGG | ASSGDNYEQY | 0.7902195 |
| NAPGGAGGNGG | ASSGDNYEQY | 0.7928959 |
| VSVFMGMGGNF | ASSGDNYEQY | 0.5656051 |
| LASYLAAL | ASSGDNYEQY | 0.81045556 |
| LVGGTPAL | ASSGDNYEQY | 0.81026185 |
| VRNADVL | ASSGDNYEQY | 0.82052976 |
| TCSSPFLVSAM | ASSGDNYEQY | 0.72642195 |
| VHITDAE | ASSGDNYEQY | 0.4561576 |
| EYAAPSF | ASSGDNYEQY | 0.80938745 |
| VVDAFHGGL | ASSGDNYEQY | 0.6975035 |
| ASVVDPQ | ASSGDNYEQY | 0.7292032 |
| SYNEFGGLS | ASSGDNYEQY | 0.7749666 |
| DPDGSVY | ASSGDNYEQY | 0.72562265 |
| MVSTIHATE | ASSGDNYEQY | 0.58223665 |
| LAAGVADPV | ASSGDNYEQY | 0.7729657 |
| TKEQIAA | ASSGDNYEQY | 0.8162759 |
| SLASPAAL | ASSGDNYEQY | 0.52796936 |
| IGLVFYGV | ASSGDNYEQY | 0.4223138 |
| FLLDASNL | ASSGDNYEQY | 0.67991257 |
| YQVGNFP | ASSGDNYEQY | 0.7295995 |
| ESLLSQST | ASSGDNYEQY | 0.80055565 |
| LFTVGFAAVG | ASSGDNYEQY | 0.74134743 |
| ATPPTPT | ASSGDNYEQY | 0.71896684 |
| TDLATLG | ASSGDNYEQY | 0.77502656 |
| SAGGTGR | ASSGDNYEQY | 0.7788254 |
| GFSNSGS | ASSGDNYEQY | 0.72065085 |
| GGNGGSGGTSV | ASSGDNYEQY | 0.7852521 |
| GTSVATGGAGN | ASSGDNYEQY | 0.61170936 |
| NGGNGGSGGTS | ASSGDNYEQY | 0.814881 |
| AVVKFPH | ASSGDNYEQY | 0.7727579 |
| TPPPPPP | ASSGDNYEQY | 0.7550205 |

| | | |
|---|---|---|
| AAPAAAAAPA | ASSGDNYEQY | 0.80024475 |
| IALNEDL | ASSGDNYEQY | 0.7865151 |
| IVNVSSM | ASSGDNYEQY | 0.74864566 |
| TGIPPAPR | ASSGDNYEQY | 0.4928425 |
| FDLGGGTF | ASSGDNYEQY | 0.57580453 |
| FDLGGGTFDVS | ASSGDNYEQY | 0.741951 |
| NEPTAAA | ASSGDNYEQY | 0.77205235 |
| NEPTAAALA | ASSGDNYEQY | 0.7861329 |
| GSIINISSI | ASSGDNYEQY | 0.7586036 |
| VAPGNAGTA | ASSGDNYEQY | 0.599882 |
| TGGGASLE | ASSGDNYEQY | 0.8086864 |
| VAVYNHYKRI | ASSGDNYEQY | 0.8131423 |
| TDPAPATT | ASSGDNYEQY | 0.45085308 |
| TNRPDILDPALLRPG | ASSGDNYEQY | 0.6605783 |
| LTTTTTS | ASSGDNYEQY | 0.8134135 |
| PTGSGKTL | ASSGDNYEQY | 0.7901683 |
| TGSGKTL | ASSGDNYEQY | 0.6489283 |
| TIGHVDHGK | ASSGDNYEQY | 0.60421276 |
| TIGHVDHGKTT | ASSGDNYEQY | 0.8209369 |
| GHVDHGKTT | ASSGDNYEQY | 0.66816723 |
| AQFEHTLL | ASSGDNYEQY | 0.81725264 |
| IVNQDEL | ASSGDNYEQY | 0.28719127 |
| TNGSQFF | ASSGDNYEQY | 0.6692207 |
| TGKGIVE | ASSGDNYEQY | 0.662848 |
| LISNASDA | ASSGDNYEQY | 0.20956258 |
| AELRTLGELTKIAKAHGAQ | ASSGDNYEQY | 0.7771009 |
| APNAVAAAEPSW | ASSGDNYEQY | 0.46986902 |
| APQVAGAATAVL | ASSGDNYEQY | 0.77207756 |
| ARRDPAAAAAILPTDA | ASSGDNYEQY | 0.63981044 |
| DAAARDVLAEAGLAEYF | ASSGDNYEQY | 0.78914213 |
| DDQIVLSAKANLAKVEQ | ASSGDNYEQY | 0.77191496 |
| DELLPDTGEDSDEDGH | ASSGDNYEQY | 0.6832672 |
| ELPEAPANRATVPQD | ASSGDNYEQY | 0.46965736 |
| FGQSKIERVTGIG | ASSGDNYEQY | 0.75949776 |
| GAGGTGSTAGGAGGAGGAGGLY | ASSGDNYEQY | 0.8068504 |
| GHGGTGAAGGNGGTGGAGGSGI | ASSGDNYEQY | 0.8002236 |
| GNGGNGGAGGEGGPGGAGGAGA | ASSGDNYEQY | 0.33669272 |
| GSTAGNGGNGGNATNGTIAGQP | ASSGDNYEQY | 0.7736231 |
| GTGGDGGNGGNGGTGG | ASSGDNYEQY | 0.6740422 |
| GVRRPAGIPADAPV | ASSGDNYEQY | 0.43063438 |
| GYRQVTDAHLLTLARR | ASSGDNYEQY | 0.24843204 |
| IAKMGQDGHDRG | ASSGDNYEQY | 0.71002007 |
| KAEPSIEIQYT | ASSGDNYEQY | 0.70764065 |
| KPVRIRMAADEDTVAVT | ASSGDNYEQY | 0.7739898 |
| MPPRIAGMRLLVIKPEP | ASSGDNYEQY | 0.7974422 |
| MKFYAVALFRSRGVPPTR | ASSGDNYEQY | 0.6301833 |
| NAGGFGILGAVAHSP | ASSGDNYEQY | 0.798347 |
| NEGREITLAMAEM | ASSGDNYEQY | 0.74234164 |
| NELVEWSHPLGHRWRWE | ASSGDNYEQY | 0.15710524 |
| NLGAGGTGGNGGNPG | ASSGDNYEQY | 0.6227337 |
| NTEWPKWETVMAADPVD | ASSGDNYEQY | 0.35214284 |

| | | |
|---|---|---|
| PASAAPAETQP | ASSGDNYEQY | 0.79040533 |
| QQSSDGKNGVRAAEITGE | ASSGDNYEQY | 0.4926056 |
| RAPDAVTTSWWKE | ASSGDNYEQY | 0.61486065 |
| SGFGNVGAHTSGWFNQ | ASSGDNYEQY | 0.70586205 |
| SGVAGAVGQAASVGGLKV | ASSGDNYEQY | 0.79030365 |
| SNGGAGTGSAGGNGGT | ASSGDNYEQY | 0.7315496 |
| TGGAGGEGGAAG | ASSGDNYEQY | 0.6346688 |
| TGGDGGAAGAGGAPGHG | ASSGDNYEQY | 0.39887536 |
| TMPPLLAAIANAGRHGV | ASSGDNYEQY | 0.7598452 |
| TSSTAGNSGGA | ASSGDNYEQY | 0.6913652 |
| VPATVPVGILC | ASSGDNYEQY | 0.4896373 |
| VQRPDSGGVVLTGRIS | ASSGDNYEQY | 0.42741787 |
| WTPNPVPATKGVPAGLVQAT | ASSGDNYEQY | 0.7480041 |
| YALKRALAKFSDRFDVVII | ASSGDNYEQY | 0.37069708 |
| YLIGSVIV | ASSGDNYEQY | 0.7900709 |
| YGQLVAVID | ASSGDNYEQY | 0.55161583 |
| VTVGGMTLPA | ASSGDNYEQY | 0.6558807 |
| VLALPAV | ASSGDNYEQY | 0.8141566 |
| VIPPTLNYET | ASSGDNYEQY | 0.782428 |
| VGMGGMHPSSG | ASSGDNYEQY | 0.6667603 |
| VGLAGGAATA | ASSGDNYEQY | 0.62534696 |
| VGGAGGAGGSGP | ASSGDNYEQY | 0.45537612 |
| VFAAFMV | ASSGDNYEQY | 0.64021444 |
| TLYDKLR | ASSGDNYEQY | 0.5835191 |
| TLAVFTVL | ASSGDNYEQY | 0.80561006 |
| TGSVQPARV | ASSGDNYEQY | 0.63598776 |
| TGGGAGSGGT | ASSGDNYEQY | 0.37208402 |
| SHLETSL | ASSGDNYEQY | 0.72922754 |
| SFTTGMT | ASSGDNYEQY | 0.8144112 |
| QVEVAKRMGVRQPTVS | ASSGDNYEQY | 0.8262137 |
| QGSGGPALAM | ASSGDNYEQY | 0.50476545 |
| MLAPSNP | ASSGDNYEQY | 0.6515402 |
| LVNNAAI | ASSGDNYEQY | 0.5908344 |
| LTGEINTLA | ASSGDNYEQY | 0.69824684 |
| LSIFGPV | ASSGDNYEQY | 0.5770833 |
| LLALLANL | ASSGDNYEQY | 0.49640188 |
| LISNASDALDKLR | ASSGDNYEQY | 0.81078905 |
| LINEAVP | ASSGDNYEQY | 0.82528675 |
| LGYAFSGL | ASSGDNYEQY | 0.6500348 |
| LDFVAVAL | ASSGDNYEQY | 0.39067554 |
| LAQPGAE | ASSGDNYEQY | 0.70285726 |
| ISNLGAVTGF | ASSGDNYEQY | 0.62023056 |
| HQEDPGRI | ASSGDNYEQY | 0.6089331 |
| GTPGEPTP | ASSGDNYEQY | 0.7100438 |
| GFFNSGS | ASSGDNYEQY | 0.34615442 |
| GAGGGGLTAGAN | ASSGDNYEQY | 0.6918604 |
| FVDALVNM | ASSGDNYEQY | 0.8203516 |
| FLTVAPEL | ASSGDNYEQY | 0.7494718 |
| FETLTAL | ASSGDNYEQY | 0.7440078 |
| DVINPIEL | ASSGDNYEQY | 0.8077351 |
| APVSGAPV | ASSGDNYEQY | 0.7736055 |

| | | |
|---|---|---|
| ALVGGLAW | ASSGDNYEQY | 0.80314416 |
| ALAAAAATAI | ASSGDNYEQY | 0.8108558 |
| AFGLGLLL | ASSGDNYEQY | 0.6315894 |
| AFFAAASL | ASSGDNYEQY | 0.7434266 |
| AAAALPAFSPP | ASSGDNYEQY | 0.8023158 |
| VVLAPAL | ASSGDNYEQY | 0.6806561 |
| TIGHVDHGKTTL | ASSGDNYEQY | 0.730682 |
| LISNASDALDK | ASSGDNYEQY | 0.63777786 |
| LISNASDAL | ASSGDNYEQY | 0.78517467 |
| ISNASDALDK | ASSGDNYEQY | 0.7485167 |
| NQAETLVYQTE | ASSGDNYEQY | 0.8168867 |
| TGGTGGTAGAG | ASSGDNYEQY | 0.23194365 |
| TGSTAGGAGGA | ASSGDNYEQY | 0.50981027 |
| DPLAVPVAAGPV | ASSGDNYEQY | 0.79895926 |
| GGTGGTPGNGGN | ASSGDNYEQY | 0.29277375 |
| PAGLAGVDPATL | ASSGDNYEQY | 0.28538114 |
| GNNLVVEAPHAGASV | ASSGDNYEQY | 0.78757745 |
| KVTRIMSASTLEQPA | ASSGDNYEQY | 0.8123722 |
| LGFALPIALPRRTVV | ASSGDNYEQY | 0.23718417 |
| TRLGNLTARKLVVLV | ASSGDNYEQY | 0.6599783 |
| WSQQRRLIYFPSAGP | ASSGDNYEQY | 0.3481317 |
| GPNGNPDPMAAAVDIR | ASSGDNYEQY | 0.76765954 |
| TQRCVSVADLRAAASR | ASSGDNYEQY | 0.71235317 |
| QFVLDVAEHPVEVLTETV | ASSGDNYEQY | 0.30335936 |
| GGAGGTGGAGGNGGTQAGDG | ASSGDNYEQY | 0.75261855 |
| PLLATLRAIAVDVDAKPAQV | ASSGDNYEQY | 0.39172953 |
| APKFRTGGYMIDFWGVGYQVAK | ASSGDNYEQY | 0.73294026 |
| GAGGAGGAGGAAGGPGGTGGTG | ASSGDNYEQY | 0.7956765 |
| IGALGLLATDVAPAGSVAFGVA | ASSGDNYEQY | 0.82111645 |
| AGGAGGIGGNGGAGGAGGVLMG | ASSGDNYEQY | 0.38490105 |
| ADELIGLL | ASSGDNYEQY | 0.7822675 |
| ADRALKA | ASSGDNYEQY | 0.64722186 |
| AGAVGQVGGAGGA | ASSGDNYEQY | 0.8290279 |
| AGGGGGLGGAP | ASSGDNYEQY | 0.62995815 |
| AGGLLYGNGGAG | ASSGDNYEQY | 0.33117694 |
| AGLESIGL | ASSGDNYEQY | 0.79503953 |
| AGLKTGII | ASSGDNYEQY | 0.77157885 |
| ASLAALLL | ASSGDNYEQY | 0.6766287 |
| ASVINISSI | ASSGDNYEQY | 0.80688536 |
| ATPPGPA | ASSGDNYEQY | 0.62562037 |
| ATVASALV | ASSGDNYEQY | 0.6956521 |
| ATVASLIL | ASSGDNYEQY | 0.77764076 |
| AVNGPVLIRH | ASSGDNYEQY | 0.77682155 |
| DGAAGAVGI | ASSGDNYEQY | 0.53238004 |
| DPLLHGI | ASSGDNYEQY | 0.797969 |
| ELGAALGAL | ASSGDNYEQY | 0.72577477 |
| ELGRTQAV | ASSGDNYEQY | 0.48967028 |
| FVAGGYLS | ASSGDNYEQY | 0.8172585 |
| GASGGAGGAGGW | ASSGDNYEQY | 0.7219728 |
| GGAGAGGDGGAGGI | ASSGDNYEQY | 0.79447603 |
| GTGHNGSQPGMGGNG | ASSGDNYEQY | 0.7602171 |

| | | |
|---|---|---|
| GYAAASASAS | ASSGDNYEQY | 0.824155 |
| HQVDVAAVQQG | ASSGDNYEQY | 0.37804824 |
| IGAAIAHT | ASSGDNYEQY | 0.744235 |
| IGMIEVAQL | ASSGDNYEQY | 0.7629497 |
| IRLAPEHTSAMD | ASSGDNYEQY | 0.7896169 |
| IYNVSVL | ASSGDNYEQY | 0.65832865 |
| KQGFLGM | ASSGDNYEQY | 0.76158786 |
| LESVITQL | ASSGDNYEQY | 0.7450938 |
| LGLGALVVV | ASSGDNYEQY | 0.80980355 |
| LIGNGGNGGSGGI | ASSGDNYEQY | 0.754742 |
| LNAINAPTL | ASSGDNYEQY | 0.7462722 |
| LQLLNTGV | ASSGDNYEQY | 0.8135607 |
| LQTSIEF | ASSGDNYEQY | 0.52290833 |
| MSFAVSEL | ASSGDNYEQY | 0.7033336 |
| NTGALGL | ASSGDNYEQY | 0.8057381 |
| PPGAGGAGGAGTT | ASSGDNYEQY | 0.8019558 |
| QLVGLGI | ASSGDNYEQY | 0.66158414 |
| RFLEQVHQL | ASSGDNYEQY | 0.81102914 |
| RGVGYVLR | ASSGDNYEQY | 0.63356245 |
| SATGVFAGVI | ASSGDNYEQY | 0.77308905 |
| SLPAITLP | ASSGDNYEQY | 0.67974997 |
| STALAFTV | ASSGDNYEQY | 0.68985355 |
| SVLTVGLGK | ASSGDNYEQY | 0.5899507 |
| TALAGAGSGI | ASSGDNYEQY | 0.74944997 |
| TGLPFTGI | ASSGDNYEQY | 0.80266523 |
| VAEHVTV | ASSGDNYEQY | 0.77203375 |
| VDIVLNSL | ASSGDNYEQY | 0.25848708 |
| VLTGGGAVL | ASSGDNYEQY | 0.77376735 |
| VNVGLAYGI | ASSGDNYEQY | 0.7729343 |
| VSSAAVGAV | ASSGDNYEQY | 0.8215131 |
| VTSVYEAGV | ASSGDNYEQY | 0.7403055 |
| WVALDRGA | ASSGDNYEQY | 0.73646265 |
| YLVGSVVI | ASSGDNYEQY | 0.71092 |
| GPPGTGKTLL | ASSGDNYEQY | 0.72550523 |
| GVGMTKF | ASSGDNYEQY | 0.49478516 |
| IGIGGSDL | ASSGDNYEQY | 0.6543261 |
| IGTLLAEL | ASSGDNYEQY | 0.690359 |
| IYTSGTT | ASSGDNYEQY | 0.5771525 |
| LSFPAGV | ASSGDNYEQY | 0.8043845 |
| LYTSGTT | ASSGDNYEQY | 0.81079626 |
| VGAFPFGF | ASSGDNYEQY | 0.7989597 |
| VGAGTVEF | ASSGDNYEQY | 0.81717134 |
| VGKSTLLL | ASSGDNYEQY | 0.81680524 |
| AALAHAGGGIV | ASSGDNYEQY | 0.78353 |
| AALVAVSLL | ASSGDNYEQY | 0.6475618 |
| AAVSRAVAAL | ASSGDNYEQY | 0.67068994 |
| AGGSSGIGGF | ASSGDNYEQY | 0.38908073 |
| AGLTVANANG | ASSGDNYEQY | 0.64073217 |
| AGLVGAVL | ASSGDNYEQY | 0.6641314 |
| ALSQAFTV | ASSGDNYEQY | 0.70310354 |
| ALTFSVSL | ASSGDNYEQY | 0.18831624 |

| | | |
|---|---|---|
| ALTGAALVAL | ASSGDNYEQY | 0.78134376 |
| ASFSIGAGVL | ASSGDNYEQY | 0.80896735 |
| DGSVYKI | ASSGDNYEQY | 0.75496274 |
| EQLAEIV | ASSGDNYEQY | 0.77681196 |
| FAVSELTVM | ASSGDNYEQY | 0.7750489 |
| GAAPLAAGL | ASSGDNYEQY | 0.6501373 |
| GADGGDGGSGGAGG | ASSGDNYEQY | 0.8010271 |
| GFGLDTGGAGG | ASSGDNYEQY | 0.5613647 |
| GLASGGLL | ASSGDNYEQY | 0.7708856 |
| GRIAGAAVLL | ASSGDNYEQY | 0.8083453 |
| GSAGSLNTSF | ASSGDNYEQY | 0.7580668 |
| GTGQIGI | ASSGDNYEQY | 0.73226154 |
| GTLIALIL | ASSGDNYEQY | 0.65791166 |
| IAATMGFVVG | ASSGDNYEQY | 0.3331764 |
| IFAQALL | ASSGDNYEQY | 0.74529785 |
| IGAAVALGF | ASSGDNYEQY | 0.6632047 |
| IGLTPLAL | ASSGDNYEQY | 0.74522316 |
| ILGSNTSTL | ASSGDNYEQY | 0.48899296 |
| IVFSLTVL | ASSGDNYEQY | 0.75182116 |
| LASKATGF | ASSGDNYEQY | 0.7221513 |
| LASVALIL | ASSGDNYEQY | 0.79510653 |
| LFAVGLTL | ASSGDNYEQY | 0.53809905 |
| LFNGGIGTYI | ASSGDNYEQY | 0.76645106 |
| LTLGAGII | ASSGDNYEQY | 0.7409637 |
| LVAAALGDI | ASSGDNYEQY | 0.46824563 |
| NTAFVQLGI | ASSGDNYEQY | 0.79539526 |
| QLFSGAEL | ASSGDNYEQY | 0.7393069 |
| SAATFITL | ASSGDNYEQY | 0.6968379 |
| SNYGLANLGV | ASSGDNYEQY | 0.79346573 |
| SYGGAYAVM | ASSGDNYEQY | 0.7469562 |
| TTGVGGLTLG | ASSGDNYEQY | 0.4043694 |
| TTQTIYI | ASSGDNYEQY | 0.72392154 |
| VAAGAGLGAVI | ASSGDNYEQY | 0.74574685 |
| VAAVVTLL | ASSGDNYEQY | 0.6851308 |
| VGFLGGLI | ASSGDNYEQY | 0.23707725 |
| VGVIPILL | ASSGDNYEQY | 0.75133896 |
| VIFASSLI | ASSGDNYEQY | 0.47739547 |
| VKGGDGGAAAG | ASSGDNYEQY | 0.63327456 |
| VTKTFTNI | ASSGDNYEQY | 0.66665304 |
| VVAAAILL | ASSGDNYEQY | 0.7965834 |
| YAALDVEL | ASSGDNYEQY | 0.79241717 |
| LASVALLL | ASSGDNYEQY | 0.6280086 |
| LDAVHRV | ASSGDNYEQY | 0.79003656 |
| QGLSSSIF | ASSGDNYEQY | 0.74237746 |
| RGTFANIRL | ASSGDNYEQY | 0.79549456 |
| RQLWWGHRI | ASSGDNYEQY | 0.77099746 |
| YGPPGTGKTLL | ASSGDNYEQY | 0.75959706 |
| AALVGAPANVA | ASSGDNYEQY | 0.7268693 |
| AATFRRTE | ASSGDNYEQY | 0.6876121 |
| AGALGGGI | ASSGDNYEQY | 0.40593937 |
| AGGAGGAGGIGAG | ASSGDNYEQY | 0.44127375 |

| | | |
|---|---|---|
| AGSGALGL | ASSGDNYEQY | 0.69504404 |
| AQIADQI | ASSGDNYEQY | 0.4242974 |
| ASAVAVTYL | ASSGDNYEQY | 0.7432959 |
| ASTVLGGTGG | ASSGDNYEQY | 0.8018973 |
| AVGGEPAK | ASSGDNYEQY | 0.75632906 |
| CEQGTFGGTC | ASSGDNYEQY | 0.33050874 |
| DEGKAPAG | ASSGDNYEQY | 0.7061044 |
| DTVVFPAL | ASSGDNYEQY | 0.5387168 |
| FEGITMAG | ASSGDNYEQY | 0.7770245 |
| FPLVVQD | ASSGDNYEQY | 0.79136294 |
| GAFTTPQ | ASSGDNYEQY | 0.817855 |
| GFGGFGTAGGVGGA | ASSGDNYEQY | 0.7940616 |
| GFLAQFL | ASSGDNYEQY | 0.639162 |
| GGAGGAGGNAGVA | ASSGDNYEQY | 0.78641534 |
| GGVGTAGAGGAGGAGG | ASSGDNYEQY | 0.6014368 |
| GNGGGGDAGNAGSGGN | ASSGDNYEQY | 0.7009624 |
| GTGGAGGAGNDAGS | ASSGDNYEQY | 0.67167413 |
| GTTNTGWFNS | ASSGDNYEQY | 0.78704154 |
| IFMGPLFTL | ASSGDNYEQY | 0.7063489 |
| ILAHAIEAI | ASSGDNYEQY | 0.5817262 |
| IRSAIREAL | ASSGDNYEQY | 0.39261746 |
| ITFAGGG | ASSGDNYEQY | 0.80979776 |
| ITGMSGHQEV | ASSGDNYEQY | 0.73616266 |
| IVAGLACTL | ASSGDNYEQY | 0.45183626 |
| IVGGGLAGA | ASSGDNYEQY | 0.5560708 |
| IVTGVLNI | ASSGDNYEQY | 0.81908286 |
| LAQQGCI | ASSGDNYEQY | 0.81998694 |
| LCLHVEN | ASSGDNYEQY | 0.6404424 |
| LDAISDGS | ASSGDNYEQY | 0.5729599 |
| LGGFAAML | ASSGDNYEQY | 0.7950181 |
| LGGKNPMI | ASSGDNYEQY | 0.5849701 |
| LGPKGVAAIL | ASSGDNYEQY | 0.7484882 |
| LMVGVSI | ASSGDNYEQY | 0.72858 |
| LVGAAFVL | ASSGDNYEQY | 0.72472 |
| LVLAVAPL | ASSGDNYEQY | 0.5075722 |
| MAAIAVIDAA | ASSGDNYEQY | 0.79064035 |
| NNGINAELQ | ASSGDNYEQY | 0.7199643 |
| RGEPLVLVHG | ASSGDNYEQY | 0.78063977 |
| RGGAGGAAGLI | ASSGDNYEQY | 0.30792817 |
| RSYRMHPAV | ASSGDNYEQY | 0.769193 |
| SAAHLLAGQLA | ASSGDNYEQY | 0.47337785 |
| SAAVGSY | ASSGDNYEQY | 0.62022173 |
| SPHAVLHRI | ASSGDNYEQY | 0.27143294 |
| SQIAKVL | ASSGDNYEQY | 0.64975613 |
| TALGHLVP | ASSGDNYEQY | 0.81440866 |
| TAPTTSGAA | ASSGDNYEQY | 0.81313384 |
| TDGDDGNGGSPGT | ASSGDNYEQY | 0.67323554 |
| TDGLGEN | ASSGDNYEQY | 0.7909968 |
| TGGPPLPGQA | ASSGDNYEQY | 0.8267399 |
| TIPAITI | ASSGDNYEQY | 0.5820509 |
| TIPGFTI | ASSGDNYEQY | 0.25463286 |

| | | |
|---|---|---|
| TSIASGALA | ASSGDNYEQY | 0.72088367 |
| TVTLPGAAI | ASSGDNYEQY | 0.77681625 |
| VASGLSAALP | ASSGDNYEQY | 0.7940601 |
| VGAGVIGI | ASSGDNYEQY | 0.81453055 |
| VGPKRFGV | ASSGDNYEQY | 0.45912465 |
| VNVAFPD | ASSGDNYEQY | 0.7279203 |
| VSMTNSGVS | ASSGDNYEQY | 0.7734529 |
| VTDVLGDLSV | ASSGDNYEQY | 0.47865105 |
| VVVVGAGFAGL | ASSGDNYEQY | 0.5297884 |
| VYTVGTVV | ASSGDNYEQY | 0.49262542 |
| WTTFSGP | ASSGDNYEQY | 0.40624 |
| YDELVEAF | ASSGDNYEQY | 0.7684351 |
| YGTGGAGGNGG | ASSGDNYEQY | 0.6371012 |
| YTNVMAV | ASSGDNYEQY | 0.6599321 |
| FLEQVHQL | ASSGDNYEQY | 0.62894475 |
| GLSLLAVAV | ASSGDNYEQY | 0.59049886 |
| LEQVHQL | ASSGDNYEQY | 0.6078353 |
| LNAAGAI | ASSGDNYEQY | 0.6716193 |
| PAVHASL | ASSGDNYEQY | 0.76368546 |
| TELEDDA | ASSGDNYEQY | 0.79963857 |
| TPPGPPN | ASSGDNYEQY | 0.73150724 |
| TPPPPPPP | ASSGDNYEQY | 0.71553385 |
| TQSPALI | ASSLEGASERLF | 0.8084358 |
| EFAAAFPQ | ASSLEGASERLF | 0.5356297 |
| LLSGNLDVL | ASSLEGASERLF | 0.5481469 |
| GPANLGNYNI | ASSLEGASERLF | 0.79050255 |
| TLGPTLP | ASSLEGASERLF | 0.5913354 |
| VYAYSSCCK | ASSLEGASERLF | 0.79671866 |
| APAPAPAPNI | ASSLEGASERLF | 0.73959136 |
| PGGPVPIP | ASSLEGASERLF | 0.7511635 |
| FLTVALEP | ASSLEGASERLF | 0.7875028 |
| SQYFGANL | ASSLEGASERLF | 0.8218847 |
| GGVKPAA | ASSLEGASERLF | 0.75734985 |
| GGTGGAGGAGGW | ASSLEGASERLF | 0.5939765 |
| GVGGVAGLGGAGG | ASSLEGASERLF | 0.8054602 |
| VLLVLATTI | ASSLEGASERLF | 0.43241847 |
| IGIFAAML | ASSLEGASERLF | 0.79481 |
| AVGDLID | ASSLEGASERLF | 0.7899809 |
| NISPLPL | ASSLEGASERLF | 0.5473555 |
| VTAVFVAI | ASSLEGASERLF | 0.7034669 |
| SGDDNDAAL | ASSLEGASERLF | 0.8204862 |
| AVAALAHS | ASSLEGASERLF | 0.8014422 |
| GGVTVMATP | ASSLEGASERLF | 0.48633105 |
| AGLAANVRD | ASSLEGASERLF | 0.5650877 |
| TNRPDLIDPALLRPG | ASSLEGASERLF | 0.6164379 |
| LANLGVTAL | ASSLEGASERLF | 0.7310884 |
| TGERIEL | ASSLEGASERLF | 0.30498084 |
| NGEELAA | ASSLEGASERLF | 0.78759104 |
| VLSLVWG | ASSLEGASERLF | 0.6382727 |
| IGYALWL | ASSLEGASERLF | 0.80974483 |
| LSLLLAPL | ASSLEGASERLF | 0.75091577 |

| | | |
|---|---|---|
| TLDEALK | ASSLEGASERLF | 0.7951451 |
| TDGAGTAEGG | ASSLEGASERLF | 0.66445756 |
| NAPGGAGGNGG | ASSLEGASERLF | 0.65916336 |
| VSVFMGMGGNF | ASSLEGASERLF | 0.30074975 |
| LASYLAAL | ASSLEGASERLF | 0.635303 |
| LVGGTPAL | ASSLEGASERLF | 0.58004844 |
| VRNADVL | ASSLEGASERLF | 0.7108044 |
| TCSSPFLVSAM | ASSLEGASERLF | 0.41165623 |
| VHITDAE | ASSLEGASERLF | 0.8130369 |
| EYAAPSF | ASSLEGASERLF | 0.6385608 |
| VVDAFHGGL | ASSLEGASERLF | 0.7746817 |
| ASVVDPQ | ASSLEGASERLF | 0.6017286 |
| SYNEFGGLS | ASSLEGASERLF | 0.514505 |
| DPDGSVY | ASSLEGASERLF | 0.6500026 |
| MVSTIHATE | ASSLEGASERLF | 0.7922477 |
| LAAGVADPV | ASSLEGASERLF | 0.421542 |
| TKEQIAA | ASSLEGASERLF | 0.28117856 |
| SLASPAAL | ASSLEGASERLF | 0.673122 |
| IGLVFYGV | ASSLEGASERLF | 0.51551414 |
| FLLDASNL | ASSLEGASERLF | 0.82008743 |
| YQVGNFP | ASSLEGASERLF | 0.8051113 |
| ESLLSQST | ASSLEGASERLF | 0.72380775 |
| LFTVGFAAVG | ASSLEGASERLF | 0.8234138 |
| ATPPTPT | ASSLEGASERLF | 0.19377832 |
| TDLATLG | ASSLEGASERLF | 0.62555635 |
| SAGGTGR | ASSLEGASERLF | 0.8081633 |
| GFSNSGS | ASSLEGASERLF | 0.7802226 |
| GGNGGSGGTSV | ASSLEGASERLF | 0.68611467 |
| GTSVATGGAGN | ASSLEGASERLF | 0.79557943 |
| NGGNGGSGGTS | ASSLEGASERLF | 0.5723023 |
| AVVKFPH | ASSLEGASERLF | 0.6850946 |
| TPPPPPP | ASSLEGASERLF | 0.73640597 |
| AAPAAAAAPA | ASSLEGASERLF | 0.7422882 |
| IALNEDL | ASSLEGASERLF | 0.76179796 |
| IVNVSSM | ASSLEGASERLF | 0.37021536 |
| TGIPPAPR | ASSLEGASERLF | 0.69009006 |
| FDLGGGTF | ASSLEGASERLF | 0.8257541 |
| FDLGGGTFDVS | ASSLEGASERLF | 0.6794356 |
| NEPTAAA | ASSLEGASERLF | 0.80636144 |
| NEPTAAALA | ASSLEGASERLF | 0.72475445 |
| GSIINISSI | ASSLEGASERLF | 0.80156106 |
| VAPGNAGTA | ASSLEGASERLF | 0.5712955 |
| TGGGASLE | ASSLEGASERLF | 0.50712067 |
| VAVYNHYKRI | ASSLEGASERLF | 0.7578098 |
| TDPAPATT | ASSLEGASERLF | 0.4523248 |
| TNRPDILDPALLRPG | ASSLEGASERLF | 0.69791186 |
| LTTTTTS | ASSLEGASERLF | 0.63815737 |
| PTGSGKTL | ASSLEGASERLF | 0.82935154 |
| TGSGKTL | ASSLEGASERLF | 0.7834234 |
| TIGHVDHGK | ASSLEGASERLF | 0.76244974 |
| TIGHVDHGKTT | ASSLEGASERLF | 0.61990726 |

| | | |
|---|---|---|
| GHVDHGKTT | ASSLEGASERLF | 0.7791611 |
| AQFEHTLL | ASSLEGASERLF | 0.7753785 |
| IVNQDEL | ASSLEGASERLF | 0.37680125 |
| TNGSQFF | ASSLEGASERLF | 0.8062064 |
| TGKGIVE | ASSLEGASERLF | 0.4812281 |
| LISNASDA | ASSLEGASERLF | 0.73827815 |
| AELRTLGELTKIAKAHGAQ | ASSLEGASERLF | 0.7565483 |
| APNAVAAAEPSW | ASSLEGASERLF | 0.8131647 |
| APQVAGAATAVL | ASSLEGASERLF | 0.6563224 |
| ARRDPAAAAAILPTDA | ASSLEGASERLF | 0.75392514 |
| DAAARDVLAEAGLAEYF | ASSLEGASERLF | 0.5431038 |
| DDQIVLSAKANLAKVEQ | ASSLEGASERLF | 0.8012237 |
| DELLPDTGEDSDEDGH | ASSLEGASERLF | 0.8034146 |
| ELPEAPANRATVPQD | ASSLEGASERLF | 0.79907584 |
| FGQSKIERVTGIG | ASSLEGASERLF | 0.82369816 |
| GAGGTGSTAGGAGGAGGAGGLY | ASSLEGASERLF | 0.7274573 |
| GHGGTGAAGGNGGTGGAGGSGI | ASSLEGASERLF | 0.30449885 |
| GNGGNGGAGGEGGPGGAGGAGA | ASSLEGASERLF | 0.6736719 |
| GSTAGNGGNGGNATNGTIAGQP | ASSLEGASERLF | 0.79991126 |
| GTGGDGGNGGNGGTGG | ASSLEGASERLF | 0.81635964 |
| GVRRPAGIPADAPV | ASSLEGASERLF | 0.80267566 |
| GYRQVTDAHLLTLARR | ASSLEGASERLF | 0.7756524 |
| IAKMGQDGHDRG | ASSLEGASERLF | 0.7042905 |
| KAEPSIEIQYT | ASSLEGASERLF | 0.81219184 |
| KPVRIRMAADEDTVAVT | ASSLEGASERLF | 0.8290082 |
| MPPRIAGMRLLVIKPEP | ASSLEGASERLF | 0.77107555 |
| MKFYAVALFRSRGVPPTR | ASSLEGASERLF | 0.64760864 |
| NAGGFGILGAVAHSP | ASSLEGASERLF | 0.6275182 |
| NEGREITLAMAEM | ASSLEGASERLF | 0.77522665 |
| NELVEWSHPLGHRWRWE | ASSLEGASERLF | 0.7953455 |
| NLGAGGTGGNGGNPG | ASSLEGASERLF | 0.44885963 |
| NTEWPKWETVMAADPVD | ASSLEGASERLF | 0.7129494 |
| PASAAPAETQP | ASSLEGASERLF | 0.7440999 |
| QQSSDGKNGVRAAEITGE | ASSLEGASERLF | 0.29273403 |
| RAPDAVTTSWWKE | ASSLEGASERLF | 0.77729946 |
| SGFGNVGAHTSGWFNQ | ASSLEGASERLF | 0.8153054 |
| SGVAGAVGQAASVGGLKV | ASSLEGASERLF | 0.6793556 |
| SNGGAGTGSAGGNGGT | ASSLEGASERLF | 0.4410494 |
| TGGAGGEGGAAG | ASSLEGASERLF | 0.81028813 |
| TGGDGGAAGAGGAPGHG | ASSLEGASERLF | 0.8217652 |
| TMPPLLAAIANAGRHGV | ASSLEGASERLF | 0.78991884 |
| TSSTAGNSGGA | ASSLEGASERLF | 0.31418186 |
| VPATVPVGILC | ASSLEGASERLF | 0.21406816 |
| VQRPDSGGVVLTGRIS | ASSLEGASERLF | 0.7300105 |
| WTPNPVPATKGVPAGLVQAT | ASSLEGASERLF | 0.8072337 |
| YALKRALAKFSDRFDVVII | ASSLEGASERLF | 0.6697024 |
| YLIGSVIV | ASSLEGASERLF | 0.603014 |
| YGQLVAVID | ASSLEGASERLF | 0.738163 |
| VTVGGMTLPA | ASSLEGASERLF | 0.7883285 |
| VLALPAV | ASSLEGASERLF | 0.42382485 |
| VIPPTLNYET | ASSLEGASERLF | 0.61855876 |

| | | |
|---|---|---|
| VGMGGMHPSSG | ASSLEGASERLF | 0.74173415 |
| VGLAGGAATA | ASSLEGASERLF | 0.7982277 |
| VGGAGGAGGSGP | ASSLEGASERLF | 0.51342833 |
| VFAAFMV | ASSLEGASERLF | 0.6952226 |
| TLYDKLR | ASSLEGASERLF | 0.75316334 |
| TLAVFTVL | ASSLEGASERLF | 0.75401103 |
| TGSVQPARV | ASSLEGASERLF | 0.7507165 |
| TGGGAGSGGT | ASSLEGASERLF | 0.500242 |
| SHLETSL | ASSLEGASERLF | 0.80319977 |
| SFTTGMT | ASSLEGASERLF | 0.40923196 |
| QVEVAKRMGVRQPTVS | ASSLEGASERLF | 0.8005927 |
| QGSGGPALAM | ASSLEGASERLF | 0.737067 |
| MLAPSNP | ASSLEGASERLF | 0.7444688 |
| LVNNAAI | ASSLEGASERLF | 0.6396383 |
| LTGEINTLA | ASSLEGASERLF | 0.5540137 |
| LSIFGPV | ASSLEGASERLF | 0.45942876 |
| LLALLANL | ASSLEGASERLF | 0.5971329 |
| LISNASDALDKLR | ASSLEGASERLF | 0.80385435 |
| LINEAVP | ASSLEGASERLF | 0.7592652 |
| LGYAFSGL | ASSLEGASERLF | 0.79132646 |
| LDFVAVAL | ASSLEGASERLF | 0.73182535 |
| LAQPGAE | ASSLEGASERLF | 0.50677955 |
| ISNLGAVTGF | ASSLEGASERLF | 0.7881192 |
| HQEDPGRI | ASSLEGASERLF | 0.48931473 |
| GTPGEPTP | ASSLEGASERLF | 0.66790164 |
| GFFNSGS | ASSLEGASERLF | 0.78696775 |
| GAGGGGLTAGAN | ASSLEGASERLF | 0.7400924 |
| FVDALVNM | ASSLEGASERLF | 0.55075437 |
| FLTVAPEL | ASSLEGASERLF | 0.7518146 |
| FETLTAL | ASSLEGASERLF | 0.7687716 |
| DVINPIEL | ASSLEGASERLF | 0.5219604 |
| APVSGAPV | ASSLEGASERLF | 0.62913835 |
| ALVGGLAW | ASSLEGASERLF | 0.7930382 |
| ALAAAAATAI | ASSLEGASERLF | 0.69465786 |
| AFGLGLLL | ASSLEGASERLF | 0.38322794 |
| AFFAAASL | ASSLEGASERLF | 0.7632523 |
| AAAALPAFSPP | ASSLEGASERLF | 0.55462915 |
| VVLAPAL | ASSLEGASERLF | 0.76605606 |
| TIGHVDHGKTTL | ASSLEGASERLF | 0.705775 |
| LISNASDALDK | ASSLEGASERLF | 0.678769 |
| LISNASDAL | ASSLEGASERLF | 0.77803105 |
| ISNASDALDK | ASSLEGASERLF | 0.53696954 |
| NQAETLVYQTE | ASSLEGASERLF | 0.6698034 |
| TGGTGGTAGAG | ASSLEGASERLF | 0.57509315 |
| TGSTAGGAGGA | ASSLEGASERLF | 0.73682964 |
| DPLAVPVAAGPV | ASSLEGASERLF | 0.75907755 |
| GGTGGTPGNGGN | ASSLEGASERLF | 0.43355432 |
| PAGLAGVDPATL | ASSLEGASERLF | 0.7656026 |
| GNNLVVEAPHAGASV | ASSLEGASERLF | 0.64860725 |
| KVTRIMSASTLEQPA | ASSLEGASERLF | 0.4769282 |
| LGFALPIALPRRTVV | ASSLEGASERLF | 0.6451709 |

| | | |
|---|---|---|
| TRLGNLTARKLVVLV | ASSLEGASERLF | 0.6571649 |
| WSQQRRLIYFPSAGP | ASSLEGASERLF | 0.36868954 |
| GPNGNPDPMAAAVDIR | ASSLEGASERLF | 0.7670654 |
| TQRCVSVADLRAAASR | ASSLEGASERLF | 0.56421053 |
| QFVLDVAEHPVEVLTETV | ASSLEGASERLF | 0.7564217 |
| GGAGGTGGAGGNGGTQAGDG | ASSLEGASERLF | 0.7641454 |
| PLLATLRAIAVDVDAKPAQV | ASSLEGASERLF | 0.493325 |
| APKFRTGGYMIDFWGVGYQVAK | ASSLEGASERLF | 0.7997177 |
| GAGGAGGAGGAAGGPGGTGGTG | ASSLEGASERLF | 0.7539002 |
| IGALGLLATDVAPAGSVAFGVA | ASSLEGASERLF | 0.7474497 |
| AGGAGGIGGNGGAGGAGGVLMG | ASSLEGASERLF | 0.69710666 |
| ADELIGLL | ASSLEGASERLF | 0.49875137 |
| ADRALKA | ASSLEGASERLF | 0.7901554 |
| AGAVGQVGGAGGA | ASSLEGASERLF | 0.77868927 |
| AGGGGGLGGAP | ASSLEGASERLF | 0.77134717 |
| AGGLLYGNGGAG | ASSLEGASERLF | 0.734829 |
| AGLESIGL | ASSLEGASERLF | 0.7983453 |
| AGLKTGII | ASSLEGASERLF | 0.7829468 |
| ASLAALLL | ASSLEGASERLF | 0.629755 |
| ASVINISSI | ASSLEGASERLF | 0.32180417 |
| ATPPGPA | ASSLEGASERLF | 0.6795442 |
| ATVASALV | ASSLEGASERLF | 0.69690216 |
| ATVASLIL | ASSLEGASERLF | 0.507524 |
| AVNGPVLIRH | ASSLEGASERLF | 0.4141248 |
| DGAAGAVGI | ASSLEGASERLF | 0.7820529 |
| DPLLHGI | ASSLEGASERLF | 0.82677025 |
| ELGAALGAL | ASSLEGASERLF | 0.6848521 |
| ELGRTQAV | ASSLEGASERLF | 0.8222853 |
| FVAGGYLS | ASSLEGASERLF | 0.3433159 |
| GASGGAGGAGGW | ASSLEGASERLF | 0.790495 |
| GGAGAGGDGGAGGI | ASSLEGASERLF | 0.7938482 |
| GTGHNGSQPGMGGNG | ASSLEGASERLF | 0.5590732 |
| GYAAASASAS | ASSLEGASERLF | 0.7458097 |
| HQVDVAAVQQG | ASSLEGASERLF | 0.47786534 |
| IGAAIAHT | ASSLEGASERLF | 0.58100194 |
| IGMIEVAQL | ASSLEGASERLF | 0.796303 |
| IRLAPEHTSAMD | ASSLEGASERLF | 0.62128615 |
| IYNVSVL | ASSLEGASERLF | 0.73404706 |
| KQGFLGM | ASSLEGASERLF | 0.65165937 |
| LESVITQL | ASSLEGASERLF | 0.55250615 |
| LGLGALVVV | ASSLEGASERLF | 0.7078917 |
| LIGNGGNGGSGGI | ASSLEGASERLF | 0.7986764 |
| LNAINAPTL | ASSLEGASERLF | 0.73619664 |
| LQLLNTGV | ASSLEGASERLF | 0.828525 |
| LQTSIEF | ASSLEGASERLF | 0.5642917 |
| MSFAVSEL | ASSLEGASERLF | 0.82039124 |
| NTGALGL | ASSLEGASERLF | 0.81690645 |
| PPGAGGAGGAGTT | ASSLEGASERLF | 0.74568963 |
| QLVGLGI | ASSLEGASERLF | 0.6275469 |
| RFLEQVHQL | ASSLEGASERLF | 0.7962403 |
| RGVGYVLR | ASSLEGASERLF | 0.662459 |

| | | |
|---|---|---|
| SATGVFAGVI | ASSLEGASERLF | 0.7721914 |
| SLPAITLP | ASSLEGASERLF | 0.790235 |
| STALAFTV | ASSLEGASERLF | 0.18253252 |
| SVLTVGLGK | ASSLEGASERLF | 0.44521606 |
| TALAGAGSGI | ASSLEGASERLF | 0.32480597 |
| TGLPFTGI | ASSLEGASERLF | 0.27083182 |
| VAEHVTV | ASSLEGASERLF | 0.202667 |
| VDIVLNSL | ASSLEGASERLF | 0.167784 |
| VLTGGGAVL | ASSLEGASERLF | 0.25191998 |
| VNVGLAYGI | ASSLEGASERLF | 0.18440847 |
| VSSAAVGAV | ASSLEGASERLF | 0.734672 |
| VTSVYEAGV | ASSLEGASERLF | 0.3179824 |
| WVALDRGA | ASSLEGASERLF | 0.20007922 |
| YLVGSVVI | ASSLEGASERLF | 0.38596788 |
| GPPGTGKTLL | ASSLEGASERLF | 0.3464982 |
| GVGMTKF | ASSLEGASERLF | 0.14575446 |
| IGIGGSDL | ASSLEGASERLF | 0.24753162 |
| IGTLLAEL | ASSLEGASERLF | 0.2022527 |
| IYTSGTT | ASSLEGASERLF | 0.20672317 |
| LSFPAGV | ASSLEGASERLF | 0.11706488 |
| LYTSGTT | ASSLEGASERLF | 0.26285115 |
| VGAFPFGF | ASSLEGASERLF | 0.28173384 |
| VGAGTVEF | ASSLEGASERLF | 0.12455292 |
| VGKSTLLL | ASSLEGASERLF | 0.28676957 |
| AALAHAGGGIV | ASSLEGASERLF | 0.20704491 |
| AALVAVSLL | ASSLEGASERLF | 0.22594872 |
| AAVSRAVAAL | ASSLEGASERLF | 0.32392022 |
| AGGSSGIGGF | ASSLEGASERLF | 0.24348637 |
| AGLTVANANG | ASSLEGASERLF | 0.21329011 |
| AGLVGAVL | ASSLEGASERLF | 0.304213 |
| ALSQAFTV | ASSLEGASERLF | 0.41076896 |
| ALTFSVSL | ASSLEGASERLF | 0.18281478 |
| ALTGAALVAL | ASSLEGASERLF | 0.22665974 |
| ASFSIGAGVL | ASSLEGASERLF | 0.48655108 |
| DGSVYKI | ASSLEGASERLF | 0.25091165 |
| EQLAEIV | ASSLEGASERLF | 0.34877893 |
| FAVSELTVM | ASSLEGASERLF | 0.39506376 |
| GAAPLAAGL | ASSLEGASERLF | 0.18489265 |
| GADGGDGGSGGAGG | ASSLEGASERLF | 0.37998447 |
| GFGLDTGGAGG | ASSLEGASERLF | 0.36932233 |
| GLASGGLL | ASSLEGASERLF | 0.25735617 |
| GRIAGAAVLL | ASSLEGASERLF | 0.21479185 |
| GSAGSLNTSF | ASSLEGASERLF | 0.1847336 |
| GTGQIGI | ASSLEGASERLF | 0.24721256 |
| GTLIALIL | ASSLEGASERLF | 0.17100526 |
| IAATMGFVVG | ASSLEGASERLF | 0.27806002 |
| IFAQALL | ASSLEGASERLF | 0.4642777 |
| IGAAVALGF | ASSLEGASERLF | 0.20554386 |
| IGLTPLAL | ASSLEGASERLF | 0.14229478 |
| ILGSNTSTL | ASSLEGASERLF | 0.15235403 |
| IVFSLTVL | ASSLEGASERLF | 0.25193942 |

| | | |
|---|---|---|
| LASKATGF | ASSLEGASERLF | 0.16140813 |
| LASVALIL | ASSLEGASERLF | 0.1917629 |
| LFAVGLTL | ASSLEGASERLF | 0.18091993 |
| LFNGGIGTYI | ASSLEGASERLF | 0.42790383 |
| LTLGAGII | ASSLEGASERLF | 0.41496003 |
| LVAAALGDI | ASSLEGASERLF | 0.1819074 |
| NTAFVQLGI | ASSLEGASERLF | 0.37624428 |
| QLFSGAEL | ASSLEGASERLF | 0.6405953 |
| SAATFITL | ASSLEGASERLF | 0.41306233 |
| SNYGLANLGV | ASSLEGASERLF | 0.27871042 |
| SYGGAYAVM | ASSLEGASERLF | 0.18074155 |
| TTGVGGLTLG | ASSLEGASERLF | 0.6974134 |
| TTQTIYI | ASSLEGASERLF | 0.19802386 |
| VAAGAGLGAVI | ASSLEGASERLF | 0.3541942 |
| VAAVVTLL | ASSLEGASERLF | 0.32061058 |
| VGFLGGLI | ASSLEGASERLF | 0.31311694 |
| VGVIPILL | ASSLEGASERLF | 0.16609485 |
| VIFASSLI | ASSLEGASERLF | 0.43726745 |
| VKGGDGGAAAG | ASSLEGASERLF | 0.3990872 |
| VTKTFTNI | ASSLEGASERLF | 0.14106423 |
| VVAAAILL | ASSLEGASERLF | 0.1867137 |
| YAALDVEL | ASSLEGASERLF | 0.26352245 |
| LASVALLL | ASSLEGASERLF | 0.2366272 |
| LDAVHRV | ASSLEGASERLF | 0.4291093 |
| QGLSSSIF | ASSLEGASERLF | 0.29458284 |
| RGTFANIRL | ASSLEGASERLF | 0.18925476 |
| RQLWWGHRI | ASSLEGASERLF | 0.30557668 |
| YGPPGTGKTLL | ASSLEGASERLF | 0.29771203 |
| AALVGAPANVA | ASSLEGASERLF | 0.23289187 |
| AATFRRTE | ASSLEGASERLF | 0.33926278 |
| AGALGGGI | ASSLEGASERLF | 0.1738543 |
| AGGAGGAGGIGAG | ASSLEGASERLF | 0.39556837 |
| AGSGALGL | ASSLEGASERLF | 0.2803795 |
| AQIADQI | ASSLEGASERLF | 0.29454657 |
| ASAVAVTYL | ASSLEGASERLF | 0.34595412 |
| ASTVLGGTGG | ASSLEGASERLF | 0.33166987 |
| AVGGEPAK | ASSLEGASERLF | 0.2638085 |
| CEQGTFGGTC | ASSLEGASERLF | 0.16463783 |
| DEGKAPAG | ASSLEGASERLF | 0.20174992 |
| DTVVFPAL | ASSLEGASERLF | 0.2518202 |
| FEGITMAG | ASSLEGASERLF | 0.26465252 |
| FPLVVQD | ASSLEGASERLF | 0.32007697 |
| GAFTTPQ | ASSLEGASERLF | 0.31074852 |
| GFGGFGTAGGVGGA | ASSLEGASERLF | 0.18233332 |
| GFLAQFL | ASSLEGASERLF | 0.40086958 |
| GGAGGAGGNAGVA | ASSLEGASERLF | 0.4204229 |
| GGVGTAGAGGAGGAGG | ASSLEGASERLF | 0.13343619 |
| GNGGGGDAGNAGSGGN | ASSLEGASERLF | 0.22130656 |
| GTGGAGGAGNDAGS | ASSLEGASERLF | 0.46002954 |
| GTTNTGWFNS | ASSLEGASERLF | 0.5086577 |
| IFMGPLFTL | ASSLEGASERLF | 0.23881927 |

| | | |
|---|---|---|
| ILAHAIEAI | ASSLEGASERLF | 0.21963276 |
| IRSAIREAL | ASSLEGASERLF | 0.75176775 |
| ITFAGGG | ASSLEGASERLF | 0.2730111 |
| ITGMSGHQEV | ASSLEGASERLF | 0.46055126 |
| IVAGLACTL | ASSLEGASERLF | 0.13270414 |
| IVGGGLAGA | ASSLEGASERLF | 0.24980728 |
| IVTGVLNI | ASSLEGASERLF | 0.2163609 |
| LAQQGCI | ASSLEGASERLF | 0.11898447 |
| LCLHVEN | ASSLEGASERLF | 0.3287203 |
| LDAISDGS | ASSLEGASERLF | 0.23722193 |
| LGGFAAML | ASSLEGASERLF | 0.2897628 |
| LGGKNPMI | ASSLEGASERLF | 0.23211084 |
| LGPKGVAAIL | ASSLEGASERLF | 0.32839057 |
| LMVGVSI | ASSLEGASERLF | 0.21491165 |
| LVGAAFVL | ASSLEGASERLF | 0.23174097 |
| LVLAVAPL | ASSLEGASERLF | 0.20121694 |
| MAAIAVIDAA | ASSLEGASERLF | 0.31976897 |
| NNGINAELQ | ASSLEGASERLF | 0.39561224 |
| RGEPLVLVHG | ASSLEGASERLF | 0.350116 |
| RGGAGGAAGLI | ASSLEGASERLF | 0.151089 |
| RSYRMHPAV | ASSLEGASERLF | 0.38148266 |
| SAAHLLAGQLA | ASSLEGASERLF | 0.2350738 |
| SAAVGSY | ASSLEGASERLF | 0.18226255 |
| SPHAVLHRI | ASSLEGASERLF | 0.16128242 |
| SQIAKVL | ASSLEGASERLF | 0.2325879 |
| TALGHLVP | ASSLEGASERLF | 0.18721862 |
| TAPTTSGAA | ASSLEGASERLF | 0.31808728 |
| TDGDDGNGGSPGT | ASSLEGASERLF | 0.38294214 |
| TDGLGEN | ASSLEGASERLF | 0.20098642 |
| TGGPPLPGQA | ASSLEGASERLF | 0.35448232 |
| TIPAITI | ASSLEGASERLF | 0.2479939 |
| TIPGFTI | ASSLEGASERLF | 0.103505775 |
| TSIASGALA | ASSLEGASERLF | 0.18687005 |
| TVTLPGAAI | ASSLEGASERLF | 0.14494976 |
| VASGLSAALP | ASSLEGASERLF | 0.34110928 |
| VGAGVIGI | ASSLEGASERLF | 0.16805595 |
| VGPKRFGV | ASSLEGASERLF | 0.18366599 |
| VNVAFPD | ASSLEGASERLF | 0.23308581 |
| VSMTNSGVS | ASSLEGASERLF | 0.5785428 |
| VTDVLGDLSV | ASSLEGASERLF | 0.22274531 |
| VVVVGAGFAGL | ASSLEGASERLF | 0.17565769 |
| VYTVGTVV | ASSLEGASERLF | 0.14920373 |
| WTTFSGP | ASSLEGASERLF | 0.3251933 |
| YDELVEAF | ASSLEGASERLF | 0.2485915 |
| YGTGGAGGNGG | ASSLEGASERLF | 0.20699258 |
| YTNVMAV | ASSLEGASERLF | 0.17301688 |
| FLEQVHQL | ASSLEGASERLF | 0.36390078 |
| GLSLLAVAV | ASSLEGASERLF | 0.14691804 |
| LEQVHQL | ASSLEGASERLF | 0.32132483 |
| LNAAGAI | ASSLEGASERLF | 0.19100673 |
| PAVHASL | ASSLEGASERLF | 0.22135201 |

| | | |
|---|---|---|
| TELEDDA | ASSLEGASERLF | 0.49491218 |
| TPPGPPN | ASSLEGASERLF | 0.56118757 |
| TPPPPPPP | ASSLEGASERLF | 0.19529419 |
| TQSPALI | ASSETDTEVF | 0.24499157 |
| EFAAAFPQ | ASSETDTEVF | 0.17198734 |
| LLSGNLDVL | ASSETDTEVF | 0.20196137 |
| GPANLGNYNI | ASSETDTEVF | 0.20368332 |
| TLGPTLP | ASSETDTEVF | 0.41173515 |
| VYAYSSCCK | ASSETDTEVF | 0.22809523 |
| APAPAPAPNI | ASSETDTEVF | 0.16976257 |
| PGGPVPIP | ASSETDTEVF | 0.3063325 |
| FLTVALEP | ASSETDTEVF | 0.4374727 |
| SQYFGANL | ASSETDTEVF | 0.5509348 |
| GGVKPAA | ASSETDTEVF | 0.16845839 |
| GGTGGAGGAGGW | ASSETDTEVF | 0.23631093 |
| GVGGVAGLGGAGG | ASSETDTEVF | 0.23662302 |
| VLLVLATTI | ASSETDTEVF | 0.2141805 |
| IGIFAAML | ASSETDTEVF | 0.20867732 |
| AVGDLID | ASSETDTEVF | 0.18567173 |
| NISPLPL | ASSETDTEVF | 0.35478216 |
| VTAVFVAI | ASSETDTEVF | 0.76929986 |
| SGDDNDAAL | ASSETDTEVF | 0.21573247 |
| AVAALAHS | ASSETDTEVF | 0.1327019 |
| GGVTVMATP | ASSETDTEVF | 0.2509645 |
| AGLAANVRD | ASSETDTEVF | 0.15816295 |
| TNRPDLIDPALLRPG | ASSETDTEVF | 0.17685473 |
| LANLGVTAL | ASSETDTEVF | 0.24506924 |
| TGERIEL | ASSETDTEVF | 0.12708482 |
| NGEELAA | ASSETDTEVF | 0.23333506 |
| VLSLVWG | ASSETDTEVF | 0.5137292 |
| IGYALWL | ASSETDTEVF | 0.3684513 |
| LSLLLAPL | ASSETDTEVF | 0.30790284 |
| TLDEALK | ASSETDTEVF | 0.40919107 |
| TDGAGTAEGG | ASSETDTEVF | 0.27573538 |
| NAPGGAGGNGG | ASSETDTEVF | 0.38979083 |
| VSVFMGMGGNF | ASSETDTEVF | 0.63508415 |
| LASYLAAL | ASSETDTEVF | 0.24807912 |
| LVGGTPAL | ASSETDTEVF | 0.24216828 |
| VRNADVL | ASSETDTEVF | 0.4051281 |
| TCSSPFLVSAM | ASSETDTEVF | 0.23217727 |
| VHITDAE | ASSETDTEVF | 0.2723734 |
| EYAAPSF | ASSETDTEVF | 0.19514401 |
| VVDAFHGGL | ASSETDTEVF | 0.38084316 |
| ASVVDPQ | ASSETDTEVF | 0.25244772 |
| SYNEFGGLS | ASSETDTEVF | 0.6446899 |
| DPDGSVY | ASSETDTEVF | 0.13830593 |
| MVSTIHATE | ASSETDTEVF | 0.17794372 |
| LAAGVADPV | ASSETDTEVF | 0.33922255 |
| TKEQIAA | ASSETDTEVF | 0.13386565 |
| SLASPAAL | ASSETDTEVF | 0.16732018 |
| IGLVFYGV | ASSETDTEVF | 0.30060947 |

| | | |
|---|---|---|
| FLLDASNL | ASSETDTEVF | 0.3337998 |
| YQVGNFP | ASSETDTEVF | 0.13605504 |
| ESLLSQST | ASSETDTEVF | 0.246225 |
| LFTVGFAAVG | ASSETDTEVF | 0.16436574 |
| ATPPTPT | ASSETDTEVF | 0.31792745 |
| TDLATLG | ASSETDTEVF | 0.24670017 |
| SAGGTGR | ASSETDTEVF | 0.14376593 |
| GFSNSGS | ASSETDTEVF | 0.27742425 |
| GGNGGSGGTSV | ASSETDTEVF | 0.13101201 |
| GTSVATGGAGN | ASSETDTEVF | 0.3108719 |
| NGGNGGSGGTS | ASSETDTEVF | 0.3320665 |
| AVVKFPH | ASSETDTEVF | 0.72200155 |
| TPPPPPP | ASSETDTEVF | 0.1707025 |
| AAPAAAAAPA | ASSETDTEVF | 0.3008842 |
| IALNEDL | ASSETDTEVF | 0.16421202 |
| IVNVSSM | ASSETDTEVF | 0.66954315 |
| TGIPPAPR | ASSETDTEVF | 0.25424734 |
| FDLGGGTF | ASSETDTEVF | 0.15006779 |
| FDLGGGTFDVS | ASSETDTEVF | 0.37867147 |
| NEPTAAA | ASSETDTEVF | 0.2748859 |
| NEPTAAALA | ASSETDTEVF | 0.28124395 |
| GSIINISSI | ASSETDTEVF | 0.6487893 |
| VAPGNAGTA | ASSETDTEVF | 0.21436709 |
| TGGGASLE | ASSETDTEVF | 0.20865913 |
| VAVYNHYKRI | ASSETDTEVF | 0.33721656 |
| TDPAPATT | ASSETDTEVF | 0.30192524 |
| TNRPDILDPALLRPG | ASSETDTEVF | 0.20419489 |
| LTTTTTS | ASSETDTEVF | 0.3695837 |
| PTGSGKTL | ASSETDTEVF | 0.26112422 |
| TGSGKTL | ASSETDTEVF | 0.21084902 |
| TIGHVDHGK | ASSETDTEVF | 0.5034464 |
| TIGHVDHGKTT | ASSETDTEVF | 0.24848548 |
| GHVDHGKTT | ASSETDTEVF | 0.31777185 |
| AQFEHTLL | ASSETDTEVF | 0.29756224 |
| IVNQDEL | ASSETDTEVF | 0.78322875 |
| TNGSQFF | ASSETDTEVF | 0.18039776 |
| TGKGIVE | ASSETDTEVF | 0.28896385 |
| LISNASDA | ASSETDTEVF | 0.28854337 |
| AELRTLGELTKIAKAHGAQ | ASSETDTEVF | 0.33015934 |
| APNAVAAAEPSW | ASSETDTEVF | 0.20210366 |
| APQVAGAATAVL | ASSETDTEVF | 0.3294369 |
| ARRDPAAAAAILPTDA | ASSETDTEVF | 0.26286384 |
| DAAARDVLAEAGLAEYF | ASSETDTEVF | 0.49084187 |
| DDQIVLSAKANLAKVEQ | ASSETDTEVF | 0.2551344 |
| DELLPDTGEDSDEDGH | ASSETDTEVF | 0.26024207 |
| ELPEAPANRATVPQD | ASSETDTEVF | 0.47900268 |
| FGQSKIERVTGIG | ASSETDTEVF | 0.22330275 |
| GAGGTGSTAGGAGGAGGAGGLY | ASSETDTEVF | 0.2348952 |
| GHGGTGAAGGNGGTGGAGGSGI | ASSETDTEVF | 0.3968825 |
| GNGGNGGAGGEGGPGGAGGAGA | ASSETDTEVF | 0.39662388 |
| GSTAGNGGNGGNATNGTIAGQP | ASSETDTEVF | 0.21796381 |

| | | |
|---|---|---|
| GTGGDGGNGGNGGTGG | ASSETDTEVF | 0.42754436 |
| GVRRPAGIPADAPV | ASSETDTEVF | 0.19805023 |
| GYRQVTDAHLLTLARR | ASSETDTEVF | 0.29465353 |
| IAKMGQDGHDRG | ASSETDTEVF | 0.1895422 |
| KAEPSIEIQYT | ASSETDTEVF | 0.20746829 |
| KPVRIRMAADEDTVAVT | ASSETDTEVF | 0.18735416 |
| MPPRIAGMRLLVIKPEP | ASSETDTEVF | 0.25464702 |
| MKFYAVALFRSRGVPPTR | ASSETDTEVF | 0.39687073 |
| NAGGFGILGAVAHSP | ASSETDTEVF | 0.2935337 |
| NEGREITLAMAEM | ASSETDTEVF | 0.15322939 |
| NELVEWSHPLGHRWRWE | ASSETDTEVF | 0.3586514 |
| NLGAGGTGGNGGNPG | ASSETDTEVF | 0.3071652 |
| NTEWPKWETVMAADPVD | ASSETDTEVF | 0.7450359 |
| PASAAPAETQP | ASSETDTEVF | 0.23289298 |
| QQSSDGKNGVRAAEITGE | ASSETDTEVF | 0.32054538 |
| RAPDAVTTSWWKE | ASSETDTEVF | 0.26584733 |
| SGFGNVGAHTSGWFNQ | ASSETDTEVF | 0.22995955 |
| SGVAGAVGQAASVGGLKV | ASSETDTEVF | 0.13396086 |
| SNGGAGTGSAGGNGGT | ASSETDTEVF | 0.23530135 |
| TGGAGGEGGAAG | ASSETDTEVF | 0.19926718 |
| TGGDGGAAGAGGAPGHG | ASSETDTEVF | 0.16483945 |
| TMPPLLAAIANAGRHGV | ASSETDTEVF | 0.6440592 |
| TSSTAGNSGGA | ASSETDTEVF | 0.43290645 |
| VPATVPVGILC | ASSETDTEVF | 0.43750638 |
| VQRPDSGGVVLTGRIS | ASSETDTEVF | 0.5134047 |
| WTPNPVPATKGVPAGLVQAT | ASSETDTEVF | 0.40617868 |
| YALKRALAKFSDRFDVVII | ASSETDTEVF | 0.3276768 |
| YLIGSVIV | ASSETDTEVF | 0.192795 |
| YGQLVAVID | ASSETDTEVF | 0.22262399 |
| VTVGGMTLPA | ASSETDTEVF | 0.13855731 |
| VLALPAV | ASSETDTEVF | 0.20671707 |
| VIPPTLNYET | ASSETDTEVF | 0.19575104 |
| VGMGGMHPSSG | ASSETDTEVF | 0.25405946 |
| VGLAGGAATA | ASSETDTEVF | 0.12732509 |
| VGGAGGAGGSGP | ASSETDTEVF | 0.57509124 |
| VFAAFMV | ASSETDTEVF | 0.42816374 |
| TLYDKLR | ASSETDTEVF | 0.30633265 |
| TLAVFTVL | ASSETDTEVF | 0.31490186 |
| TGSVQPARV | ASSETDTEVF | 0.29372194 |
| TGGGAGSGGT | ASSETDTEVF | 0.1615968 |
| SHLETSL | ASSETDTEVF | 0.377899 |
| SFTTGMT | ASSETDTEVF | 0.16473283 |
| QVEVAKRMGVRQPTVS | ASSETDTEVF | 0.28491914 |
| QGSGGPALAM | ASSETDTEVF | 0.39314312 |
| MLAPSNP | ASSETDTEVF | 0.2825872 |
| LVNNAAI | ASSETDTEVF | 0.25452194 |
| LTGEINTLA | ASSETDTEVF | 0.24451824 |
| LSIFGPV | ASSETDTEVF | 0.168312 |
| LLALLANL | ASSETDTEVF | 0.27430817 |
| LISNASDALDKLR | ASSETDTEVF | 0.24850126 |
| LINEAVP | ASSETDTEVF | 0.24582437 |

| | | |
|---|---|---|
| LGYAFSGL | ASSETDTEVF | 0.17032333 |
| LDFVAVAL | ASSETDTEVF | 0.33081293 |
| LAQPGAE | ASSETDTEVF | 0.27720004 |
| ISNLGAVTGF | ASSETDTEVF | 0.30822727 |
| HQEDPGRI | ASSETDTEVF | 0.16917561 |
| GTPGEPTP | ASSETDTEVF | 0.36321765 |
| GFFNSGS | ASSETDTEVF | 0.39139694 |
| GAGGGGLTAGAN | ASSETDTEVF | 0.16592446 |
| FVDALVNM | ASSETDTEVF | 0.2887418 |
| FLTVAPEL | ASSETDTEVF | 0.26337135 |
| FETLTAL | ASSETDTEVF | 0.19124019 |
| DVINPIEL | ASSETDTEVF | 0.3777434 |
| APVSGAPV | ASSETDTEVF | 0.2702084 |
| ALVGGLAW | ASSETDTEVF | 0.14968412 |
| ALAAAAATAI | ASSETDTEVF | 0.22119728 |
| AFGLGLLL | ASSETDTEVF | 0.29203948 |
| AFFAAASL | ASSETDTEVF | 0.22396246 |
| AAAALPAFSPP | ASSETDTEVF | 0.1403213 |
| VVLAPAL | ASSETDTEVF | 0.2688536 |
| TIGHVDHGKTTL | ASSETDTEVF | 0.2124683 |
| LISNASDALDK | ASSETDTEVF | 0.16226974 |
| LISNASDAL | ASSETDTEVF | 0.19212005 |
| ISNASDALDK | ASSETDTEVF | 0.37638748 |
| NQAETLVYQTE | ASSETDTEVF | 0.4459291 |
| TGGTGGTAGAG | ASSETDTEVF | 0.23949279 |
| TGSTAGGAGGA | ASSETDTEVF | 0.34098333 |
| DPLAVPVAAGPV | ASSETDTEVF | 0.35192215 |
| GGTGGTPGNGGN | ASSETDTEVF | 0.37221518 |
| PAGLAGVDPATL | ASSETDTEVF | 0.39032385 |
| GNNLVVEAPHAGASV | ASSETDTEVF | 0.26215002 |
| KVTRIMSASTLEQPA | ASSETDTEVF | 0.28049517 |
| LGFALPIALPRRTVV | ASSETDTEVF | 0.2550198 |
| TRLGNLTARKLVVLV | ASSETDTEVF | 0.20379783 |
| WSQQRRLIYFPSAGP | ASSETDTEVF | 0.17718025 |
| GPNGNPDPMAAAVDIR | ASSETDTEVF | 0.1863269 |
| TQRCVSVADLRAAASR | ASSETDTEVF | 0.19120814 |
| QFVLDVAEHPVEVLTETV | ASSETDTEVF | 0.28614715 |
| GGAGGTGGAGGNGGTQAGDG | ASSETDTEVF | 0.3603549 |
| PLLATLRAIAVDVDAKPAQV | ASSETDTEVF | 0.26047224 |
| APKFRTGGYMIDFWGVGYQVAK | ASSETDTEVF | 0.16444364 |
| GAGGAGGAGGAAGGPGGTGGTG | ASSETDTEVF | 0.23818882 |
| IGALGLLATDVAPAGSVAFGVA | ASSETDTEVF | 0.19010656 |
| AGGAGGIGGNGGAGGAGGVLMG | ASSETDTEVF | 0.29506612 |
| ADELIGLL | ASSETDTEVF | 0.3295965 |
| ADRALKA | ASSETDTEVF | 0.7460025 |
| AGAVGQVGGAGGA | ASSETDTEVF | 0.462861 |
| AGGGGGLGGAP | ASSETDTEVF | 0.18384585 |
| AGGLLYGNGGAG | ASSETDTEVF | 0.31984222 |
| AGLESIGL | ASSETDTEVF | 0.1756087 |
| AGLKTGII | ASSETDTEVF | 0.24415775 |
| ASLAALLL | ASSETDTEVF | 0.22916763 |

| | | |
|---|---|---|
| ASVINISSI | ASSETDTEVF | 0.27577847 |
| ATPPGPA | ASSETDTEVF | 0.26714265 |
| ATVASALV | ASSETDTEVF | 0.23565343 |
| ATVASLIL | ASSETDTEVF | 0.14065419 |
| AVNGPVLIRH | ASSETDTEVF | 0.5179045 |
| DGAAGAVGI | ASSETDTEVF | 0.263792 |
| DPLLHGI | ASSETDTEVF | 0.1827929 |
| ELGAALGAL | ASSETDTEVF | 0.17026317 |
| ELGRTQAV | ASSETDTEVF | 0.53764045 |
| FVAGGYLS | ASSETDTEVF | 0.5861412 |
| GASGGAGGAGGW | ASSETDTEVF | 0.17859802 |
| GGAGAGGDGGAGGI | ASSETDTEVF | 0.18320721 |
| GTGHNGSQPGMGGNG | ASSETDTEVF | 0.6354319 |
| GYAAASASAS | ASSETDTEVF | 0.18621022 |
| HQVDVAAVQQG | ASSETDTEVF | 0.26360348 |
| IGAAIAHT | ASSETDTEVF | 0.22208652 |
| IGMIEVAQL | ASSETDTEVF | 0.2254472 |
| IRLAPEHTSAMD | ASSETDTEVF | 0.2134069 |
| IYNVSVL | ASSETDTEVF | 0.29887524 |
| KQGFLGM | ASSETDTEVF | 0.21820523 |
| LESVITQL | ASSETDTEVF | 0.31209397 |
| LGLGALVVV | ASSETDTEVF | 0.123811476 |
| LIGNGGNGGSGGI | ASSETDTEVF | 0.29542586 |
| LNAINAPTL | ASSETDTEVF | 0.15336804 |
| LQLLNTGV | ASSETDTEVF | 0.18406887 |
| LQTSIEF | ASSETDTEVF | 0.13792843 |
| MSFAVSEL | ASSETDTEVF | 0.23757288 |
| NTGALGL | ASSETDTEVF | 0.4079279 |
| PPGAGGAGGAGTT | ASSETDTEVF | 0.46499956 |
| QLVGLGI | ASSETDTEVF | 0.25817457 |
| RFLEQVHQL | ASSETDTEVF | 0.32787633 |
| RGVGYVLR | ASSETDTEVF | 0.45064294 |
| SATGVFAGVI | ASSETDTEVF | 0.1833153 |
| SLPAITLP | ASSETDTEVF | 0.15832898 |
| STALAFTV | ASSETDTEVF | 0.27062386 |
| SVLTVGLGK | ASSETDTEVF | 0.42341956 |
| TALAGAGSGI | ASSETDTEVF | 0.3007045 |
| TGLPFTGI | ASSETDTEVF | 0.45332563 |
| VAEHVTV | ASSETDTEVF | 0.1407249 |
| VDIVLNSL | ASSETDTEVF | 0.29294506 |
| VLTGGGAVL | ASSETDTEVF | 0.33704674 |
| VNVGLAYGI | ASSETDTEVF | 0.15145116 |
| VSSAAVGAV | ASSETDTEVF | 0.18991768 |
| VTSVYEAGV | ASSETDTEVF | 0.20588346 |
| WVALDRGA | ASSETDTEVF | 0.16411793 |
| YLVGSVVI | ASSETDTEVF | 0.33184543 |
| GPPGTGKTLL | ASSETDTEVF | 0.17670009 |
| GVGMTKF | ASSETDTEVF | 0.24532175 |
| IGIGGSDL | ASSETDTEVF | 0.19649118 |
| IGTLLAEL | ASSETDTEVF | 0.22420141 |
| IYTSGTT | ASSETDTEVF | 0.22744437 |

| | | |
|---|---|---|
| LSFPAGV | ASSETDTEVF | 0.28828287 |
| LYTSGTT | ASSETDTEVF | 0.28500155 |
| VGAFPFGF | ASSETDTEVF | 0.3350489 |
| VGAGTVEF | ASSETDTEVF | 0.18944225 |
| VGKSTLLL | ASSETDTEVF | 0.23272914 |
| AALAHAGGGIV | ASSETDTEVF | 0.48287988 |
| AALVAVSLL | ASSETDTEVF | 0.175077 |
| AAVSRAVAAL | ASSETDTEVF | 0.17235166 |
| AGGSSGIGGF | ASSETDTEVF | 0.30369854 |
| AGLTVANANG | ASSETDTEVF | 0.220602 |
| AGLVGAVL | ASSETDTEVF | 0.32230383 |
| ALSQAFTV | ASSETDTEVF | 0.27539203 |
| ALTFSVSL | ASSETDTEVF | 0.27720633 |
| ALTGAALVAL | ASSETDTEVF | 0.519986 |
| ASFSIGAGVL | ASSETDTEVF | 0.5170269 |
| DGSVYKI | ASSETDTEVF | 0.30468383 |
| EQLAEIV | ASSETDTEVF | 0.21651423 |
| FAVSELTVM | ASSETDTEVF | 0.6553037 |
| GAAPLAAGL | ASSETDTEVF | 0.20290135 |
| GADGGDGGSGGAGG | ASSETDTEVF | 0.3158285 |
| GFGLDTGGAGG | ASSETDTEVF | 0.343007 |
| GLASGGLL | ASSETDTEVF | 0.15392616 |
| GRIAGAAVLL | ASSETDTEVF | 0.21205583 |
| GSAGSLNTSF | ASSETDTEVF | 0.46169195 |
| GTGQIGI | ASSETDTEVF | 0.3668041 |
| GTLIALIL | ASSETDTEVF | 0.20107979 |
| IAATMGFVVG | ASSETDTEVF | 0.16491075 |
| IFAQALL | ASSETDTEVF | 0.23546523 |
| IGAAVALGF | ASSETDTEVF | 0.25211886 |
| IGLTPLAL | ASSETDTEVF | 0.18202543 |
| ILGSNTSTL | ASSETDTEVF | 0.31324896 |
| IVFSLTVL | ASSETDTEVF | 0.2383957 |
| LASKATGF | ASSETDTEVF | 0.665723 |
| LASVALIL | ASSETDTEVF | 0.30202398 |
| LFAVGLTL | ASSETDTEVF | 0.39075518 |
| LFNGGIGTYI | ASSETDTEVF | 0.22414537 |
| LTLGAGII | ASSETDTEVF | 0.24936755 |
| LVAAALGDI | ASSETDTEVF | 0.14481401 |
| NTAFVQLGI | ASSETDTEVF | 0.28601807 |
| QLFSGAEL | ASSETDTEVF | 0.2108928 |
| SAATFITL | ASSETDTEVF | 0.25922328 |
| SNYGLANLGV | ASSETDTEVF | 0.15923053 |
| SYGGAYAVM | ASSETDTEVF | 0.40255362 |
| TTGVGGLTLG | ASSETDTEVF | 0.19469231 |
| TTQTIYI | ASSETDTEVF | 0.35037294 |
| VAAGAGLGAVI | ASSETDTEVF | 0.2096351 |
| VAAVVTLL | ASSETDTEVF | 0.2161439 |
| VGFLGGLI | ASSETDTEVF | 0.20013528 |
| VGVIPILL | ASSETDTEVF | 0.3946728 |
| VIFASSLI | ASSETDTEVF | 0.15917474 |
| VKGGDGGAAAG | ASSETDTEVF | 0.1258412 |

| | | |
|---|---|---|
| VTKTFTNI | ASSETDTEVF | 0.210028 |
| VVAAAILL | ASSETDTEVF | 0.14349352 |
| YAALDVEL | ASSETDTEVF | 0.4971949 |
| LASVALLL | ASSETDTEVF | 0.38244992 |
| LDAVHRV | ASSETDTEVF | 0.24363461 |
| QGLSSSIF | ASSETDTEVF | 0.7016846 |
| RGTFANIRL | ASSETDTEVF | 0.11730812 |
| RQLWWGHRI | ASSETDTEVF | 0.22160113 |
| YGPPGTGKTLL | ASSETDTEVF | 0.4703682 |
| AALVGAPANVA | ASSETDTEVF | 0.29486796 |
| AATFRRTE | ASSETDTEVF | 0.26576146 |
| AGALGGGI | ASSETDTEVF | 0.33495522 |
| AGGAGGAGGIGAG | ASSETDTEVF | 0.18178132 |
| AGSGALGL | ASSETDTEVF | 0.23216432 |
| AQIADQI | ASSETDTEVF | 0.2649151 |
| ASAVAVTYL | ASSETDTEVF | 0.2951946 |
| ASTVLGGTGG | ASSETDTEVF | 0.2508356 |
| AVGGEPAK | ASSETDTEVF | 0.16756329 |
| CEQGTFGGTC | ASSETDTEVF | 0.29737833 |
| DEGKAPAG | ASSETDTEVF | 0.6594055 |
| DTVVFPAL | ASSETDTEVF | 0.22275838 |
| FEGITMAG | ASSETDTEVF | 0.29094243 |
| FPLVVQD | ASSETDTEVF | 0.2454571 |
| GAFTTPQ | ASSETDTEVF | 0.5652784 |
| GFGGFGTAGGVGGA | ASSETDTEVF | 0.18615994 |
| GFLAQFL | ASSETDTEVF | 0.15167782 |
| GGAGGAGGNAGVA | ASSETDTEVF | 0.35935962 |
| GGVGTAGAGGAGGAGG | ASSETDTEVF | 0.13209799 |
| GNGGGGDAGNAGSGGN | ASSETDTEVF | 0.27038616 |
| GTGGAGGAGNDAGS | ASSETDTEVF | 0.18705143 |
| GTTNTGWFNS | ASSETDTEVF | 0.678499 |
| IFMGPLFTL | ASSETDTEVF | 0.2880006 |
| ILAHAIEAI | ASSETDTEVF | 0.2787305 |
| IRSAIREAL | ASSETDTEVF | 0.18731993 |
| ITFAGGG | ASSETDTEVF | 0.26566467 |
| ITGMSGHQEV | ASSETDTEVF | 0.27230644 |
| IVAGLACTL | ASSETDTEVF | 0.14626399 |
| IVGGGLAGA | ASSETDTEVF | 0.41704577 |
| IVTGVLNI | ASSETDTEVF | 0.16977702 |
| LAQQGCI | ASSETDTEVF | 0.23076499 |
| LCLHVEN | ASSETDTEVF | 0.28322697 |
| LDAISDGS | ASSETDTEVF | 0.70176744 |
| LGGFAAML | ASSETDTEVF | 0.20282704 |
| LGGKNPMI | ASSETDTEVF | 0.35485375 |
| LGPKGVAAIL | ASSETDTEVF | 0.18012829 |
| LMVGVSI | ASSETDTEVF | 0.33256447 |
| LVGAAFVL | ASSETDTEVF | 0.4638809 |
| LVLAVAPL | ASSETDTEVF | 0.55824256 |
| MAAIAVIDAA | ASSETDTEVF | 0.7558445 |
| NNGINAELQ | ASSETDTEVF | 0.27201182 |
| RGEPLVLVHG | ASSETDTEVF | 0.17618178 |

| | | |
|---|---|---|
| RGGAGGAAGLI | ASSETDTEVF | 0.25190458 |
| RSYRMHPAV | ASSETDTEVF | 0.34588668 |
| SAAHLLAGQLA | ASSETDTEVF | 0.43002617 |
| SAAVGSY | ASSETDTEVF | 0.36448336 |
| SPHAVLHRI | ASSETDTEVF | 0.2987762 |
| SQIAKVL | ASSETDTEVF | 0.19454733 |
| TALGHLVP | ASSETDTEVF | 0.4013059 |
| TAPTTSGAA | ASSETDTEVF | 0.5828904 |
| TDGDDGNGGSPGT | ASSETDTEVF | 0.2972644 |
| TDGLGEN | ASSETDTEVF | 0.17743388 |
| TGGPPLPGQA | ASSETDTEVF | 0.21454594 |
| TIPAITI | ASSETDTEVF | 0.2871596 |
| TIPGFTI | ASSETDTEVF | 0.32935262 |
| TSIASGALA | ASSETDTEVF | 0.18370669 |
| TVTLPGAAI | ASSETDTEVF | 0.22038944 |
| VASGLSAALP | ASSETDTEVF | 0.2791452 |
| VGAGVIGI | ASSETDTEVF | 0.15727618 |
| VGPKRFGV | ASSETDTEVF | 0.28184733 |
| VNVAFPD | ASSETDTEVF | 0.47071594 |
| VSMTNSGVS | ASSETDTEVF | 0.24885815 |
| VTDVLGDLSV | ASSETDTEVF | 0.16112958 |
| VVVVGAGFAGL | ASSETDTEVF | 0.48467875 |
| VYTVGTVV | ASSETDTEVF | 0.5728315 |
| WTTFSGP | ASSETDTEVF | 0.33731642 |
| YDELVEAF | ASSETDTEVF | 0.14865415 |
| YGTGGAGGNGG | ASSETDTEVF | 0.11358886 |
| YTNVMAV | ASSETDTEVF | 0.31480876 |
| FLEQVHQL | ASSETDTEVF | 0.6808309 |
| GLSLLAVAV | ASSETDTEVF | 0.23450449 |
| LEQVHQL | ASSETDTEVF | 0.18750024 |
| LNAAGAI | ASSETDTEVF | 0.2265553 |
| PAVHASL | ASSETDTEVF | 0.32171643 |
| TELEDDA | ASSETDTEVF | 0.21496381 |
| TPPGPPN | ASSETDTEVF | 0.18546106 |
| TPPPPPPP | ASSETDTEVF | 0.26244593 |
| TQSPALI | ASSLGTGRVF | 0.356963 |
| EFAAAFPQ | ASSLGTGRVF | 0.14275727 |
| LLSGNLDVL | ASSLGTGRVF | 0.19083032 |
| GPANLGNYNI | ASSLGTGRVF | 0.4078692 |
| TLGPTLP | ASSLGTGRVF | 0.33026922 |
| VYAYSSCCK | ASSLGTGRVF | 0.23120038 |
| APAPAPAPNI | ASSLGTGRVF | 0.1670123 |
| PGGPVPIP | ASSLGTGRVF | 0.38642144 |
| FLTVALEP | ASSLGTGRVF | 0.1528155 |
| SQYFGANL | ASSLGTGRVF | 0.33535397 |
| GGVKPAA | ASSLGTGRVF | 0.25401142 |
| GGTGGAGGAGGW | ASSLGTGRVF | 0.2779694 |
| GVGGVAGLGGAGG | ASSLGTGRVF | 0.22007392 |
| VLLVLATTI | ASSLGTGRVF | 0.15561229 |
| IGIFAAML | ASSLGTGRVF | 0.16330194 |
| AVGDLID | ASSLGTGRVF | 0.1743602 |

| | | |
|---|---|---|
| NISPLPL | ASSLGTGRVF | 0.36733586 |
| VTAVFVAI | ASSLGTGRVF | 0.37335244 |
| SGDDNDAAL | ASSLGTGRVF | 0.33265743 |
| AVAALAHS | ASSLGTGRVF | 0.27318722 |
| GGVTVMATP | ASSLGTGRVF | 0.1792215 |
| AGLAANVRD | ASSLGTGRVF | 0.2882044 |
| TNRPDLIDPALLRPG | ASSLGTGRVF | 0.16545507 |
| LANLGVTAL | ASSLGTGRVF | 0.28608754 |
| TGERIEL | ASSLGTGRVF | 0.25262582 |
| NGEELAA | ASSLGTGRVF | 0.25647074 |
| VLSLVWG | ASSLGTGRVF | 0.20128684 |
| IGYALWL | ASSLGTGRVF | 0.2820176 |
| LSLLLAPL | ASSLGTGRVF | 0.28243282 |
| TLDEALK | ASSLGTGRVF | 0.21656168 |
| TDGAGTAEGG | ASSLGTGRVF | 0.24803992 |
| NAPGGAGGNGG | ASSLGTGRVF | 0.37973088 |
| VSVFMGMGGNF | ASSLGTGRVF | 0.2651509 |
| LASYLAAL | ASSLGTGRVF | 0.15754694 |
| LVGGTPAL | ASSLGTGRVF | 0.3189858 |
| VRNADVL | ASSLGTGRVF | 0.15909928 |
| TCSSPFLVSAM | ASSLGTGRVF | 0.2896133 |
| VHITDAE | ASSLGTGRVF | 0.23077118 |
| EYAAPSF | ASSLGTGRVF | 0.23929217 |
| VVDAFHGGL | ASSLGTGRVF | 0.3257234 |
| ASVVDPQ | ASSLGTGRVF | 0.17971367 |
| SYNEFGGLS | ASSLGTGRVF | 0.1863346 |
| DPDGSVY | ASSLGTGRVF | 0.18329285 |
| MVSTIHATE | ASSLGTGRVF | 0.37024915 |
| LAAGVADPV | ASSLGTGRVF | 0.28626394 |
| TKEQIAA | ASSLGTGRVF | 0.15169258 |
| SLASPAAL | ASSLGTGRVF | 0.2741881 |
| IGLVFYGV | ASSLGTGRVF | 0.22357428 |
| FLLDASNL | ASSLGTGRVF | 0.17499106 |
| YQVGNFP | ASSLGTGRVF | 0.24748369 |
| ESLLSQST | ASSLGTGRVF | 0.22147608 |
| LFTVGFAAVG | ASSLGTGRVF | 0.15852323 |
| ATPPTPT | ASSLGTGRVF | 0.28657424 |
| TDLATLG | ASSLGTGRVF | 0.2225352 |
| SAGGTGR | ASSLGTGRVF | 0.3132951 |
| GFSNSGS | ASSLGTGRVF | 0.3547112 |
| GGNGGSGGTSV | ASSLGTGRVF | 0.17407292 |
| GTSVATGGAGN | ASSLGTGRVF | 0.32724804 |
| NGGNGGSGGTS | ASSLGTGRVF | 0.24711768 |
| AVVKFPH | ASSLGTGRVF | 0.2595341 |
| TPPPPPP | ASSLGTGRVF | 0.28205723 |
| AAPAAAAAPA | ASSLGTGRVF | 0.1710179 |
| IALNEDL | ASSLGTGRVF | 0.35416964 |
| IVNVSSM | ASSLGTGRVF | 0.3866533 |
| TGIPPAPR | ASSLGTGRVF | 0.30981737 |
| FDLGGGTF | ASSLGTGRVF | 0.24680328 |
| FDLGGGTFDVS | ASSLGTGRVF | 0.5380669 |

| | | |
|---|---|---|
| NEPTAAA | ASSLGTGRVF | 0.29717588 |
| NEPTAAALA | ASSLGTGRVF | 0.17978688 |
| GSIINISSI | ASSLGTGRVF | 0.13180557 |
| VAPGNAGTA | ASSLGTGRVF | 0.26816973 |
| TGGGASLE | ASSLGTGRVF | 0.30343276 |
| VAVYNHYKRI | ASSLGTGRVF | 0.20818602 |
| TDPAPATT | ASSLGTGRVF | 0.15771644 |
| TNRPDILDPALLRPG | ASSLGTGRVF | 0.31483248 |
| LTTTTTS | ASSLGTGRVF | 0.6039704 |
| PTGSGKTL | ASSLGTGRVF | 0.2476953 |
| TGSGKTL | ASSLGTGRVF | 0.45062986 |
| TIGHVDHGK | ASSLGTGRVF | 0.14663444 |
| TIGHVDHGKTT | ASSLGTGRVF | 0.36862093 |
| GHVDHGKTT | ASSLGTGRVF | 0.355228 |
| AQFEHTLL | ASSLGTGRVF | 0.15428568 |
| IVNQDEL | ASSLGTGRVF | 0.2963074 |
| TNGSQFF | ASSLGTGRVF | 0.18807153 |
| TGKGIVE | ASSLGTGRVF | 0.19463369 |
| LISNASDA | ASSLGTGRVF | 0.3980459 |
| AELRTLGELTKIAKAHGAQ | ASSLGTGRVF | 0.26353022 |
| APNAVAAAEPSW | ASSLGTGRVF | 0.27725577 |
| APQVAGAATAVL | ASSLGTGRVF | 0.22442062 |
| ARRDPAAAAAILPTDA | ASSLGTGRVF | 0.18423185 |
| DAAARDVLAEAGLAEYF | ASSLGTGRVF | 0.25615618 |
| DDQIVLSAKANLAKVEQ | ASSLGTGRVF | 0.3514732 |
| DELLPDTGEDSDEDGH | ASSLGTGRVF | 0.2870064 |
| ELPEAPANRATVPQD | ASSLGTGRVF | 0.52515745 |
| FGQSKIERVTGIG | ASSLGTGRVF | 0.18932967 |
| GAGGTGSTAGGAGGAGGAGGLY | ASSLGTGRVF | 0.54061633 |
| GHGGTGAAGGNGGTGGAGGSGI | ASSLGTGRVF | 0.73196137 |
| GNGGNGGAGGEGGPGGAGGAGA | ASSLGTGRVF | 0.23263332 |
| GSTAGNGGNGGNATNGTIAGQP | ASSLGTGRVF | 0.2017945 |
| GTGGDGGNGGNGGTGG | ASSLGTGRVF | 0.43975928 |
| GVRRPAGIPADAPV | ASSLGTGRVF | 0.23768395 |
| GYRQVTDAHLLTLARR | ASSLGTGRVF | 0.30686495 |
| IAKMGQDGHDRG | ASSLGTGRVF | 0.33932757 |
| KAEPSIEIQYT | ASSLGTGRVF | 0.7744262 |
| KPVRIRMAADEDTVAVT | ASSLGTGRVF | 0.9051502 |
| MPPRIAGMRLLVIKPEP | ASSLGTGRVF | 0.87378293 |
| MKFYAVALFRSRGVPPTR | ASSLGTGRVF | 0.87995934 |
| NAGGFGILGAVAHSP | ASSLGTGRVF | 0.80717075 |
| NEGREITLAMAEM | ASSLGTGRVF | 0.5712261 |
| NELVEWSHPLGHRWRWE | ASSLGTGRVF | 0.85959446 |
| NLGAGGTGGNGGNPG | ASSLGTGRVF | 0.7604 |
| NTEWPKWETVMAADPVD | ASSLGTGRVF | 0.8855731 |
| PASAAPAETQP | ASSLGTGRVF | 0.90154004 |
| QQSSDGKNGVRAAEITGE | ASSLGTGRVF | 0.8370848 |
| RAPDAVTTSWWKE | ASSLGTGRVF | 0.9013152 |
| SGFGNVGAHTSGWFNQ | ASSLGTGRVF | 0.90362567 |
| SGVAGAVGQAASVGGLKV | ASSLGTGRVF | 0.6791752 |
| SNGGAGTGSAGGNGGT | ASSLGTGRVF | 0.8608996 |

| | | |
|---|---|---|
| TGGAGGEGGAAG | ASSLGTGRVF | 0.8200517 |
| TGGDGGAAGAGGAPGHG | ASSLGTGRVF | 0.8593297 |
| TMPPLLAAIANAGRHGV | ASSLGTGRVF | 0.5597979 |
| TSSTAGNSGGA | ASSLGTGRVF | 0.8958911 |
| VPATVPVGILC | ASSLGTGRVF | 0.8130011 |
| VQRPDSGGVVLTGRIS | ASSLGTGRVF | 0.36087972 |
| WTPNPVPATKGVPAGLVQAT | ASSLGTGRVF | 0.8712188 |
| YALKRALAKFSDRFDVVII | ASSLGTGRVF | 0.8715963 |
| YLIGSVIV | ASSLGTGRVF | 0.8496721 |
| YGQLVAVID | ASSLGTGRVF | 0.89748704 |
| VTVGGMTLPA | ASSLGTGRVF | 0.75622654 |
| VLALPAV | ASSLGTGRVF | 0.8144968 |
| VIPPTLNYET | ASSLGTGRVF | 0.89493495 |
| VGMGGMHPSSG | ASSLGTGRVF | 0.9020947 |
| VGLAGGAATA | ASSLGTGRVF | 0.75046825 |
| VGGAGGAGGSGP | ASSLGTGRVF | 0.87999606 |
| VFAAFMV | ASSLGTGRVF | 0.90151143 |
| TLYDKLR | ASSLGTGRVF | 0.8751442 |
| TLAVFTVL | ASSLGTGRVF | 0.89979076 |
| TGSVQPARV | ASSLGTGRVF | 0.90462816 |
| TGGGAGSGGT | ASSLGTGRVF | 0.84913504 |
| SHLETSL | ASSLGTGRVF | 0.90203536 |
| SFTTGMT | ASSLGTGRVF | 0.90731037 |
| QVEVAKRMGVRQPTVS | ASSLGTGRVF | 0.8683716 |
| QGSGGPALAM | ASSLGTGRVF | 0.8640005 |
| MLAPSNP | ASSLGTGRVF | 0.6942005 |
| LVNNAAI | ASSLGTGRVF | 0.8560459 |
| LTGEINTLA | ASSLGTGRVF | 0.64647996 |
| LSIFGPV | ASSLGTGRVF | 0.87378097 |
| LLALLANL | ASSLGTGRVF | 0.9031249 |
| LISNASDALDKLR | ASSLGTGRVF | 0.85111743 |
| LINEAVP | ASSLGTGRVF | 0.7661697 |
| LGYAFSGL | ASSLGTGRVF | 0.7920396 |
| LDFVAVAL | ASSLGTGRVF | 0.87411 |
| LAQPGAE | ASSLGTGRVF | 0.83727884 |
| ISNLGAVTGF | ASSLGTGRVF | 0.6002891 |
| HQEDPGRI | ASSLGTGRVF | 0.8480324 |
| GTPGEPTP | ASSLGTGRVF | 0.89797705 |
| GFFNSGS | ASSLGTGRVF | 0.8969191 |
| GAGGGGLTAGAN | ASSLGTGRVF | 0.7389124 |
| FVDALVNM | ASSLGTGRVF | 0.9077847 |
| FLTVAPEL | ASSLGTGRVF | 0.88908696 |
| FETLTAL | ASSLGTGRVF | 0.89862573 |
| DVINPIEL | ASSLGTGRVF | 0.8734357 |
| APVSGAPV | ASSLGTGRVF | 0.7275423 |
| ALVGGLAW | ASSLGTGRVF | 0.89510345 |
| ALAAAAATAI | ASSLGTGRVF | 0.8364799 |
| AFGLGLLL | ASSLGTGRVF | 0.8191432 |
| AFFAAASL | ASSLGTGRVF | 0.89219344 |
| AAAALPAFSPP | ASSLGTGRVF | 0.8508897 |
| VVLAPAL | ASSLGTGRVF | 0.8164737 |

| | | |
|---|---|---|
| TIGHVDHGKTTL | ASSLGTGRVF | 0.83125496 |
| LISNASDALDK | ASSLGTGRVF | 0.90113485 |
| LISNASDAL | ASSLGTGRVF | 0.7955979 |
| ISNASDALDK | ASSLGTGRVF | 0.7093014 |
| NQAETLVYQTE | ASSLGTGRVF | 0.82484865 |
| TGGTGGTAGAG | ASSLGTGRVF | 0.8613926 |
| TGSTAGGAGGA | ASSLGTGRVF | 0.901 |
| DPLAVPVAAGPV | ASSLGTGRVF | 0.88953143 |
| GGTGGTPGNGGN | ASSLGTGRVF | 0.8435925 |
| PAGLAGVDPATL | ASSLGTGRVF | 0.8807421 |
| GNNLVVEAPHAGASV | ASSLGTGRVF | 0.8885598 |
| KVTRIMSASTLEQPA | ASSLGTGRVF | 0.87844676 |
| LGFALPIALPRRTVV | ASSLGTGRVF | 0.88156176 |
| TRLGNLTARKLVVLV | ASSLGTGRVF | 0.80457383 |
| WSQQRRLIYFPSAGP | ASSLGTGRVF | 0.89933145 |
| GPNGNPDPMAAAVDIR | ASSLGTGRVF | 0.88821095 |
| TQRCVSVADLRAAASR | ASSLGTGRVF | 0.8986281 |
| QFVLDVAEHPVEVLTETV | ASSLGTGRVF | 0.9019469 |
| GGAGGTGGAGGNGGTQAGDG | ASSLGTGRVF | 0.88195825 |
| PLLATLRAIAVDVDAKPAQV | ASSLGTGRVF | 0.87754333 |
| APKFRTGGYMIDFWGVGYQVAK | ASSLGTGRVF | 0.5587858 |
| GAGGAGGAGGAAGGPGGTGGTG | ASSLGTGRVF | 0.85076326 |
| IGALGLLATDVAPAGSVAFGVA | ASSLGTGRVF | 0.87410545 |
| AGGAGGIGGNGGAGGAGGVLMG | ASSLGTGRVF | 0.87712985 |
| ADELIGLL | ASSLGTGRVF | 0.8912581 |
| ADRALKA | ASSLGTGRVF | 0.89136875 |
| AGAVGQVGGAGGA | ASSLGTGRVF | 0.8340851 |
| AGGGGGLGGAP | ASSLGTGRVF | 0.89757085 |
| AGGLLYGNGGAG | ASSLGTGRVF | 0.8999876 |
| AGLESIGL | ASSLGTGRVF | 0.6784463 |
| AGLKTGII | ASSLGTGRVF | 0.84778726 |
| ASLAALLL | ASSLGTGRVF | 0.9098058 |
| ASVINISSI | ASSLGTGRVF | 0.87428117 |
| ATPPGPA | ASSLGTGRVF | 0.85734326 |
| ATVASALV | ASSLGTGRVF | 0.84314626 |
| ATVASLIL | ASSLGTGRVF | 0.9050653 |
| AVNGPVLIRH | ASSLGTGRVF | 0.8395064 |
| DGAAGAVGI | ASSLGTGRVF | 0.9058236 |
| DPLLHGI | ASSLGTGRVF | 0.465765 |
| ELGAALGAL | ASSLGTGRVF | 0.85482144 |
| ELGRTQAV | ASSLGTGRVF | 0.72380173 |
| FVAGGYLS | ASSLGTGRVF | 0.39320317 |
| GASGGAGGAGGW | ASSLGTGRVF | 0.89603996 |
| GGAGAGGDGGAGGI | ASSLGTGRVF | 0.8600658 |
| GTGHNGSQPGMGGNG | ASSLGTGRVF | 0.89151895 |
| GYAAASASAS | ASSLGTGRVF | 0.8636397 |
| HQVDVAAVQQG | ASSLGTGRVF | 0.9000353 |
| IGAAIAHT | ASSLGTGRVF | 0.8689274 |
| IGMIEVAQL | ASSLGTGRVF | 0.8753324 |
| IRLAPEHTSAMD | ASSLGTGRVF | 0.755245 |
| IYNVSVL | ASSLGTGRVF | 0.892416 |

| | | |
|---|---|---|
| KQGFLGM | ASSLGTGRVF | 0.90329957 |
| LESVITQL | ASSLGTGRVF | 0.89109087 |
| LGLGALVVV | ASSLGTGRVF | 0.677869 |
| LIGNGGNGGSGGI | ASSLGTGRVF | 0.8476219 |
| LNAINAPTL | ASSLGTGRVF | 0.86914337 |
| LQLLNTGV | ASSLGTGRVF | 0.2836068 |
| LQTSIEF | ASSLGTGRVF | 0.29054803 |
| MSFAVSEL | ASSLGTGRVF | 0.8352897 |
| NTGALGL | ASSLGTGRVF | 0.83751714 |
| PPGAGGAGGAGTT | ASSLGTGRVF | 0.8979124 |
| QLVGLGI | ASSLGTGRVF | 0.8924172 |
| RFLEQVHQL | ASSLGTGRVF | 0.8585273 |
| RGVGYVLR | ASSLGTGRVF | 0.8903322 |
| SATGVFAGVI | ASSLGTGRVF | 0.8754777 |
| SLPAITLP | ASSLGTGRVF | 0.11267939 |
| STALAFTV | ASSLGTGRVF | 0.8403224 |
| SVLTVGLGK | ASSLGTGRVF | 0.6742925 |
| TALAGAGSGI | ASSLGTGRVF | 0.9087031 |
| TGLPFTGI | ASSLGTGRVF | 0.8137799 |
| VAEHVTV | ASSLGTGRVF | 0.8259212 |
| VDIVLNSL | ASSLGTGRVF | 0.8664293 |
| VLTGGGAVL | ASSLGTGRVF | 0.875387 |
| VNVGLAYGI | ASSLGTGRVF | 0.8662312 |
| VSSAAVGAV | ASSLGTGRVF | 0.7801785 |
| VTSVYEAGV | ASSLGTGRVF | 0.26053077 |
| WVALDRGA | ASSLGTGRVF | 0.89010143 |
| YLVGSVVI | ASSLGTGRVF | 0.72736746 |
| GPPGTGKTLL | ASSLGTGRVF | 0.7922783 |
| GVGMTKF | ASSLGTGRVF | 0.5825095 |
| IGIGGSDL | ASSLGTGRVF | 0.4164032 |
| IGTLLAEL | ASSLGTGRVF | 0.74268824 |
| IYTSGTT | ASSLGTGRVF | 0.89279795 |
| LSFPAGV | ASSLGTGRVF | 0.84060013 |
| LYTSGTT | ASSLGTGRVF | 0.8528919 |
| VGAFPFGF | ASSLGTGRVF | 0.90856254 |
| VGAGTVEF | ASSLGTGRVF | 0.8603102 |
| VGKSTLLL | ASSLGTGRVF | 0.84612274 |
| AALAHAGGGIV | ASSLGTGRVF | 0.8175565 |
| AALVAVSLL | ASSLGTGRVF | 0.7970663 |
| AAVSRAVAAL | ASSLGTGRVF | 0.78738916 |
| AGGSSGIGGF | ASSLGTGRVF | 0.8186141 |
| AGLTVANANG | ASSLGTGRVF | 0.90668654 |
| AGLVGAVL | ASSLGTGRVF | 0.8568225 |
| ALSQAFTV | ASSLGTGRVF | 0.7415726 |
| ALTFSVSL | ASSLGTGRVF | 0.8786136 |
| ALTGAALVAL | ASSLGTGRVF | 0.9049481 |
| ASFSIGAGVL | ASSLGTGRVF | 0.90676117 |
| DGSVYKI | ASSLGTGRVF | 0.78853995 |
| EQLAEIV | ASSLGTGRVF | 0.81135976 |
| FAVSELTVM | ASSLGTGRVF | 0.8487042 |
| GAAPLAAGL | ASSLGTGRVF | 0.85527563 |

| | | |
|---|---|---|
| GADGGDGGSGGAGG | ASSLGTGRVF | 0.7951388 |
| GFGLDTGGAGG | ASSLGTGRVF | 0.734925 |
| GLASGGLL | ASSLGTGRVF | 0.8944205 |
| GRIAGAAVLL | ASSLGTGRVF | 0.90987873 |
| GSAGSLNTSF | ASSLGTGRVF | 0.86329997 |
| GTGQIGI | ASSLGTGRVF | 0.6370287 |
| GTLIALIL | ASSLGTGRVF | 0.8597227 |
| IAATMGFVVG | ASSLGTGRVF | 0.8117449 |
| IFAQALL | ASSLGTGRVF | 0.82403326 |
| IGAAVALGF | ASSLGTGRVF | 0.86365056 |
| IGLTPLAL | ASSLGTGRVF | 0.7156603 |
| ILGSNTSTL | ASSLGTGRVF | 0.86894596 |
| IVFSLTVL | ASSLGTGRVF | 0.9060325 |
| LASKATGF | ASSLGTGRVF | 0.83424044 |
| LASVALIL | ASSLGTGRVF | 0.8888542 |
| LFAVGLTL | ASSLGTGRVF | 0.9016272 |
| LFNGGIGTYI | ASSLGTGRVF | 0.8891595 |
| LTLGAGII | ASSLGTGRVF | 0.90114963 |
| LVAAALGDI | ASSLGTGRVF | 0.89302194 |
| NTAFVQLGI | ASSLGTGRVF | 0.8674396 |
| QLFSGAEL | ASSLGTGRVF | 0.8625258 |
| SAATFITL | ASSLGTGRVF | 0.8840601 |
| SNYGLANLGV | ASSLGTGRVF | 0.8672145 |
| SYGGAYAVM | ASSLGTGRVF | 0.8596779 |
| TTGVGGLTLG | ASSLGTGRVF | 0.830676 |
| TTQTIYI | ASSLGTGRVF | 0.9003147 |
| VAAGAGLGAVI | ASSLGTGRVF | 0.8408549 |
| VAAVVTLL | ASSLGTGRVF | 0.9000431 |
| VGFLGGLI | ASSLGTGRVF | 0.59103584 |
| VGVIPILL | ASSLGTGRVF | 0.7284236 |
| VIFASSLI | ASSLGTGRVF | 0.89705503 |
| VKGGDGGAAAG | ASSLGTGRVF | 0.12831536 |
| VTKTFTNI | ASSLGTGRVF | 0.44270602 |
| VVAAAILL | ASSLGTGRVF | 0.87165934 |
| YAALDVEL | ASSLGTGRVF | 0.9012015 |
| LASVALLL | ASSLGTGRVF | 0.14788522 |
| LDAVHRV | ASSLGTGRVF | 0.80700374 |
| QGLSSSIF | ASSLGTGRVF | 0.70682955 |
| RGTFANIRL | ASSLGTGRVF | 0.8928464 |
| RQLWWGHRI | ASSLGTGRVF | 0.8664802 |
| YGPPGTGKTLL | ASSLGTGRVF | 0.6686741 |
| AALVGAPANVA | ASSLGTGRVF | 0.8948072 |
| AATFRRTE | ASSLGTGRVF | 0.5898745 |
| AGALGGGI | ASSLGTGRVF | 0.76207393 |
| AGGAGGAGGIGAG | ASSLGTGRVF | 0.89759827 |
| AGSGALGL | ASSLGTGRVF | 0.9006041 |
| AQIADQI | ASSLGTGRVF | 0.790466 |
| ASAVAVTYL | ASSLGTGRVF | 0.89293087 |
| ASTVLGGTGG | ASSLGTGRVF | 0.84034157 |
| AVGGEPAK | ASSLGTGRVF | 0.9069032 |
| CEQGTFGGTC | ASSLGTGRVF | 0.8343091 |

| | | |
|---|---|---|
| DEGKAPAG | ASSLGTGRVF | 0.7101799 |
| DTVVFPAL | ASSLGTGRVF | 0.8994334 |
| FEGITMAG | ASSLGTGRVF | 0.892691 |
| FPLVVQD | ASSLGTGRVF | 0.87004185 |
| GAFTTPQ | ASSLGTGRVF | 0.8871983 |
| GFGGFGTAGGVGGA | ASSLGTGRVF | 0.8273047 |
| GFLAQFL | ASSLGTGRVF | 0.8568668 |
| GGAGGAGGNAGVA | ASSLGTGRVF | 0.89921045 |
| GGVGTAGAGGAGGAGG | ASSLGTGRVF | 0.89580697 |
| GNGGGGDAGNAGSGGN | ASSLGTGRVF | 0.70779514 |
| GTGGAGGAGNDAGS | ASSLGTGRVF | 0.90065336 |
| GTTNTGWFNS | ASSLGTGRVF | 0.8602581 |
| IFMGPLFTL | ASSLGTGRVF | 0.74932885 |
| ILAHAIEAI | ASSLGTGRVF | 0.90738666 |
| IRSAIREAL | ASSLGTGRVF | 0.87065136 |
| ITFAGGG | ASSLGTGRVF | 0.8909479 |
| ITGMSGHQEV | ASSLGTGRVF | 0.89435357 |
| IVAGLACTL | ASSLGTGRVF | 0.90221053 |
| IVGGGLAGA | ASSLGTGRVF | 0.681417 |
| IVTGVLNI | ASSLGTGRVF | 0.8682759 |
| LAQQGCI | ASSLGTGRVF | 0.8648912 |
| LCLHVEN | ASSLGTGRVF | 0.8964588 |
| LDAISDGS | ASSLGTGRVF | 0.8707652 |
| LGGFAAML | ASSLGTGRVF | 0.7956815 |
| LGGKNPMI | ASSLGTGRVF | 0.8865653 |
| LGPKGVAAIL | ASSLGTGRVF | 0.90301245 |
| LMVGVSI | ASSLGTGRVF | 0.87138593 |
| LVGAAFVL | ASSLGTGRVF | 0.8792616 |
| LVLAVAPL | ASSLGTGRVF | 0.90428245 |
| MAAIAVIDAA | ASSLGTGRVF | 0.7622113 |
| NNGINAELQ | ASSLGTGRVF | 0.8622086 |
| RGEPLVLVHG | ASSLGTGRVF | 0.896286 |
| RGGAGGAAGLI | ASSLGTGRVF | 0.900195 |
| RSYRMHPAV | ASSLGTGRVF | 0.8629575 |
| SAAHLLAGQLA | ASSLGTGRVF | 0.90182817 |
| SAAVGSY | ASSLGTGRVF | 0.8169657 |
| SPHAVLHRI | ASSLGTGRVF | 0.894225 |
| SQIAKVL | ASSLGTGRVF | 0.8214355 |
| TALGHLVP | ASSLGTGRVF | 0.8764144 |
| TAPTTSGAA | ASSLGTGRVF | 0.7731855 |
| TDGDDGNGGSPGT | ASSLGTGRVF | 0.8887133 |
| TDGLGEN | ASSLGTGRVF | 0.89201427 |
| TGGPPLPGQA | ASSLGTGRVF | 0.8938732 |
| TIPAITI | ASSLGTGRVF | 0.3267847 |
| TIPGFTI | ASSLGTGRVF | 0.890443 |
| TSIASGALA | ASSLGTGRVF | 0.8948215 |
| TVTLPGAAI | ASSLGTGRVF | 0.90825754 |
| VASGLSAALP | ASSLGTGRVF | 0.8393458 |
| VGAGVIGI | ASSLGTGRVF | 0.8911842 |
| VGPKRFGV | ASSLGTGRVF | 0.82459694 |
| VNVAFPD | ASSLGTGRVF | 0.85561514 |

| | | |
|---|---|---|
| VSMTNSGVS | ASSLGTGRVF | 0.7424711 |
| VTDVLGDLSV | ASSLGTGRVF | 0.78965604 |
| VVVVGAGFAGL | ASSLGTGRVF | 0.8346955 |
| VYTVGTVV | ASSLGTGRVF | 0.80321944 |
| WTTFSGP | ASSLGTGRVF | 0.89026165 |
| YDELVEAF | ASSLGTGRVF | 0.8887159 |
| YGTGGAGGNGG | ASSLGTGRVF | 0.8990451 |
| YTNVMAV | ASSLGTGRVF | 0.90122867 |
| FLEQVHQL | ASSLGTGRVF | 0.90253437 |
| GLSLLAVAV | ASSLGTGRVF | 0.90061593 |
| LEQVHQL | ASSLGTGRVF | 0.8058052 |
| LNAAGAI | ASSLGTGRVF | 0.8703829 |
| PAVHASL | ASSLGTGRVF | 0.7164162 |
| TELEDDA | ASSLGTGRVF | 0.8686569 |
| TPPGPPN | ASSLGTGRVF | 0.827422 |
| TPPPPPPP | ASSLGTGRVF | 0.87677324 |
| TQSPALI | ASSQGDYAEQF | 0.12145966 |
| EFAAAFPQ | ASSQGDYAEQF | 0.86677873 |
| LLSGNLDVL | ASSQGDYAEQF | 0.90768343 |
| GPANLGNYNI | ASSQGDYAEQF | 0.7873776 |
| TLGPTLP | ASSQGDYAEQF | 0.8840022 |
| VYAYSSCCK | ASSQGDYAEQF | 0.8840459 |
| APAPAPAPNI | ASSQGDYAEQF | 0.82578385 |
| PGGPVPIP | ASSQGDYAEQF | 0.89034677 |
| FLTVALEP | ASSQGDYAEQF | 0.8241031 |
| SQYFGANL | ASSQGDYAEQF | 0.88879114 |
| GGVKPAA | ASSQGDYAEQF | 0.90455663 |
| GGTGGAGGAGGW | ASSQGDYAEQF | 0.88275105 |
| GVGGVAGLGGAGG | ASSQGDYAEQF | 0.8579364 |
| VLLVLATTI | ASSQGDYAEQF | 0.86860895 |
| IGIFAAML | ASSQGDYAEQF | 0.4382164 |
| AVGDLID | ASSQGDYAEQF | 0.87405956 |
| NISPLPL | ASSQGDYAEQF | 0.12956475 |
| VTAVFVAI | ASSQGDYAEQF | 0.8657901 |
| SGDDNDAAL | ASSQGDYAEQF | 0.7873868 |
| AVAALAHS | ASSQGDYAEQF | 0.87904525 |
| GGVTVMATP | ASSQGDYAEQF | 0.8717444 |
| AGLAANVRD | ASSQGDYAEQF | 0.884578 |
| TNRPDLIDPALLRPG | ASSQGDYAEQF | 0.80014837 |
| LANLGVTAL | ASSQGDYAEQF | 0.8681487 |
| TGERIEL | ASSQGDYAEQF | 0.83215195 |
| NGEELAA | ASSQGDYAEQF | 0.75188774 |
| VLSLVWG | ASSQGDYAEQF | 0.89155805 |
| IGYALWL | ASSQGDYAEQF | 0.8590721 |
| LSLLLAPL | ASSQGDYAEQF | 0.85375357 |
| TLDEALK | ASSQGDYAEQF | 0.9023028 |
| TDGAGTAEGG | ASSQGDYAEQF | 0.87557024 |
| NAPGGAGGNGG | ASSQGDYAEQF | 0.58401513 |
| VSVFMGMGGNF | ASSQGDYAEQF | 0.84271073 |
| LASYLAAL | ASSQGDYAEQF | 0.874952 |
| LVGGTPAL | ASSQGDYAEQF | 0.8346014 |

| | | |
|---|---|---|
| VRNADVL | ASSQGDYAEQF | 0.17132606 |
| TCSSPFLVSAM | ASSQGDYAEQF | 0.89093685 |
| VHITDAE | ASSQGDYAEQF | 0.75420696 |
| EYAAPSF | ASSQGDYAEQF | 0.79543525 |
| VVDAFHGGL | ASSQGDYAEQF | 0.8318563 |
| ASVVDPQ | ASSQGDYAEQF | 0.8937983 |
| SYNEFGGLS | ASSQGDYAEQF | 0.80344594 |
| DPDGSVY | ASSQGDYAEQF | 0.85328865 |
| MVSTIHATE | ASSQGDYAEQF | 0.8830687 |
| LAAGVADPV | ASSQGDYAEQF | 0.889694 |
| TKEQIAA | ASSQGDYAEQF | 0.90112287 |
| SLASPAAL | ASSQGDYAEQF | 0.88211656 |
| IGLVFYGV | ASSQGDYAEQF | 0.8344873 |
| FLLDASNL | ASSQGDYAEQF | 0.89118 |
| YQVGNFP | ASSQGDYAEQF | 0.83607626 |
| ESLLSQST | ASSQGDYAEQF | 0.73811847 |
| LFTVGFAAVG | ASSQGDYAEQF | 0.6270101 |
| ATPPTPT | ASSQGDYAEQF | 0.8551782 |
| TDLATLG | ASSQGDYAEQF | 0.66764176 |
| SAGGTGR | ASSQGDYAEQF | 0.877594 |
| GFSNSGS | ASSQGDYAEQF | 0.905033 |
| GGNGGSGGTSV | ASSQGDYAEQF | 0.8652029 |
| GTSVATGGAGN | ASSQGDYAEQF | 0.6390293 |
| NGGNGGSGGTS | ASSQGDYAEQF | 0.8547847 |
| AVVKFPH | ASSQGDYAEQF | 0.82862616 |
| TPPPPPP | ASSQGDYAEQF | 0.88416016 |
| AAPAAAAAPA | ASSQGDYAEQF | 0.8902103 |
| IALNEDL | ASSQGDYAEQF | 0.9089452 |
| IVNVSSM | ASSQGDYAEQF | 0.8980462 |
| TGIPPAPR | ASSQGDYAEQF | 0.8334176 |
| FDLGGGTF | ASSQGDYAEQF | 0.89046025 |
| FDLGGGTFDVS | ASSQGDYAEQF | 0.8455198 |
| NEPTAAA | ASSQGDYAEQF | 0.8646439 |
| NEPTAAALA | ASSQGDYAEQF | 0.8664222 |
| GSIINISSI | ASSQGDYAEQF | 0.87597775 |
| VAPGNAGTA | ASSQGDYAEQF | 0.8521271 |
| TGGGASLE | ASSQGDYAEQF | 0.8318854 |
| VAVYNHYKRI | ASSQGDYAEQF | 0.689595 |
| TDPAPATT | ASSQGDYAEQF | 0.89162827 |
| TNRPDILDPALLRPG | ASSQGDYAEQF | 0.8473923 |
| LTTTTTS | ASSQGDYAEQF | 0.63411504 |
| PTGSGKTL | ASSQGDYAEQF | 0.85308295 |
| TGSGKTL | ASSQGDYAEQF | 0.9070306 |
| TIGHVDHGK | ASSQGDYAEQF | 0.9025431 |
| TIGHVDHGKTT | ASSQGDYAEQF | 0.81755847 |
| GHVDHGKTT | ASSQGDYAEQF | 0.80172384 |
| AQFEHTLL | ASSQGDYAEQF | 0.87350845 |
| IVNQDEL | ASSQGDYAEQF | 0.6800024 |
| TNGSQFF | ASSQGDYAEQF | 0.8887441 |
| TGKGIVE | ASSQGDYAEQF | 0.8598436 |
| LISNASDA | ASSQGDYAEQF | 0.8600247 |

| | | |
|---|---|---|
| AELRTLGELTKIAKAHGAQ | ASSQGDYAEQF | 0.83813894 |
| APNAVAAAEPSW | ASSQGDYAEQF | 0.8788152 |
| APQVAGAATAVL | ASSQGDYAEQF | 0.84026384 |
| ARRDPAAAAAILPTDA | ASSQGDYAEQF | 0.8889721 |
| DAAARDVLAEAGLAEYF | ASSQGDYAEQF | 0.20285502 |
| DDQIVLSAKANLAKVEQ | ASSQGDYAEQF | 0.89353865 |
| DELLPDTGEDSDEDGH | ASSQGDYAEQF | 0.8129302 |
| ELPEAPANRATVPQD | ASSQGDYAEQF | 0.7209163 |
| FGQSKIERVTGIG | ASSQGDYAEQF | 0.42451262 |
| GAGGTGSTAGGAGGAGGAGGLY | ASSQGDYAEQF | 0.8606202 |
| GHGGTGAAGGNGGTGGAGGSGI | ASSQGDYAEQF | 0.8856012 |
| GNGGNGGAGGEGGPGGAGGAGA | ASSQGDYAEQF | 0.89543396 |
| GSTAGNGGNGGNATNGTIAGQP | ASSQGDYAEQF | 0.77757955 |
| GTGGDGGNGGNGGTGG | ASSQGDYAEQF | 0.89652115 |
| GVRRPAGIPADAPV | ASSQGDYAEQF | 0.89744455 |
| GYRQVTDAHLLTLARR | ASSQGDYAEQF | 0.8423853 |
| IAKMGQDGHDRG | ASSQGDYAEQF | 0.3993858 |
| KAEPSIEIQYT | ASSQGDYAEQF | 0.8841554 |
| KPVRIRMAADEDTVAVT | ASSQGDYAEQF | 0.8419972 |
| MPPRIAGMRLLVIKPEP | ASSQGDYAEQF | 0.8955307 |
| MKFYAVALFRSRGVPPTR | ASSQGDYAEQF | 0.8747444 |
| NAGGFGILGAVAHSP | ASSQGDYAEQF | 0.7468077 |
| NEGREITLAMAEM | ASSQGDYAEQF | 0.9029286 |
| NELVEWSHPLGHRWRWE | ASSQGDYAEQF | 0.90175885 |
| NLGAGGTGGNGGNPG | ASSQGDYAEQF | 0.79840314 |
| NTEWPKWETVMAADPVD | ASSQGDYAEQF | 0.7876606 |
| PASAAPAETQP | ASSQGDYAEQF | 0.8152114 |
| QQSSDGKNGVRAAEITGE | ASSQGDYAEQF | 0.5559121 |
| RAPDAVTTSWWKE | ASSQGDYAEQF | 0.89813083 |
| SGFGNVGAHTSGWFNQ | ASSQGDYAEQF | 0.83845055 |
| SGVAGAVGQAASVGGLKV | ASSQGDYAEQF | 0.8309114 |
| SNGGAGTGSAGGNGGT | ASSQGDYAEQF | 0.8733785 |
| TGGAGGEGGAAG | ASSQGDYAEQF | 0.8471801 |
| TGGDGGAAGAGGAPGHG | ASSQGDYAEQF | 0.8096617 |
| TMPPLLAAIANAGRHGV | ASSQGDYAEQF | 0.8400793 |
| TSSTAGNSGGA | ASSQGDYAEQF | 0.8748664 |
| VPATVPVGILC | ASSQGDYAEQF | 0.8851607 |
| VQRPDSGGVVLTGRIS | ASSQGDYAEQF | 0.8631373 |
| WTPNPVPATKGVPAGLVQAT | ASSQGDYAEQF | 0.87557435 |
| YALKRALAKFSDRFDVVII | ASSQGDYAEQF | 0.90339434 |
| YLIGSVIV | ASSQGDYAEQF | 0.68631446 |
| YGQLVAVID | ASSQGDYAEQF | 0.82168937 |
| VTVGGMTLPA | ASSQGDYAEQF | 0.8869035 |
| VLALPAV | ASSQGDYAEQF | 0.7337557 |
| VIPPTLNYET | ASSQGDYAEQF | 0.89734185 |
| VGMGGMHPSSG | ASSQGDYAEQF | 0.8772712 |
| VGLAGGAATA | ASSQGDYAEQF | 0.88032234 |
| VGGAGGAGGSGP | ASSQGDYAEQF | 0.845201 |
| VFAAFMV | ASSQGDYAEQF | 0.908983 |
| TLYDKLR | ASSQGDYAEQF | 0.8941848 |
| TLAVFTVL | ASSQGDYAEQF | 0.8748385 |

| | | |
|---|---|---|
| TGSVQPARV | ASSQGDYAEQF | 0.8850791 |
| TGGGAGSGGT | ASSQGDYAEQF | 0.7332604 |
| SHLETSL | ASSQGDYAEQF | 0.9003056 |
| SFTTGMT | ASSQGDYAEQF | 0.89059776 |
| QVEVAKRMGVRQPTVS | ASSQGDYAEQF | 0.74476194 |
| QGSGGPALAM | ASSQGDYAEQF | 0.86246043 |
| MLAPSNP | ASSQGDYAEQF | 0.90073025 |
| LVNNAAI | ASSQGDYAEQF | 0.9002167 |
| LTGEINTLA | ASSQGDYAEQF | 0.64841366 |
| LSIFGPV | ASSQGDYAEQF | 0.8318512 |
| LLALLANL | ASSQGDYAEQF | 0.61324 |
| LISNASDALDKLR | ASSQGDYAEQF | 0.7778046 |
| LINEAVP | ASSQGDYAEQF | 0.6095966 |
| LGYAFSGL | ASSQGDYAEQF | 0.88404757 |
| LDFVAVAL | ASSQGDYAEQF | 0.8479745 |
| LAQPGAE | ASSQGDYAEQF | 0.8814151 |
| ISNLGAVTGF | ASSQGDYAEQF | 0.87311363 |
| HQEDPGRI | ASSQGDYAEQF | 0.9010834 |
| GTPGEPTP | ASSQGDYAEQF | 0.8397171 |
| GFFNSGS | ASSQGDYAEQF | 0.8580215 |
| GAGGGGLTAGAN | ASSQGDYAEQF | 0.67607236 |
| FVDALVNM | ASSQGDYAEQF | 0.61346066 |
| FLTVAPEL | ASSQGDYAEQF | 0.80013674 |
| FETLTAL | ASSQGDYAEQF | 0.85824245 |
| DVINPIEL | ASSQGDYAEQF | 0.32160288 |
| APVSGAPV | ASSQGDYAEQF | 0.90530276 |
| ALVGGLAW | ASSQGDYAEQF | 0.8723117 |
| ALAAAAATAI | ASSQGDYAEQF | 0.88428986 |
| AFGLGLLL | ASSQGDYAEQF | 0.85991156 |
| AFFAAASL | ASSQGDYAEQF | 0.8143383 |
| AAAALPAFSPP | ASSQGDYAEQF | 0.8098445 |
| VVLAPAL | ASSQGDYAEQF | 0.88879645 |
| TIGHVDHGKTTL | ASSQGDYAEQF | 0.7693859 |
| LISNASDALDK | ASSQGDYAEQF | 0.56618714 |
| LISNASDAL | ASSQGDYAEQF | 0.7704339 |
| ISNASDALDK | ASSQGDYAEQF | 0.73389804 |
| NQAETLVYQTE | ASSQGDYAEQF | 0.90360737 |
| TGGTGGTAGAG | ASSQGDYAEQF | 0.8994388 |
| TGSTAGGAGGA | ASSQGDYAEQF | 0.87805307 |
| DPLAVPVAAGPV | ASSQGDYAEQF | 0.9032352 |
| GGTGGTPGNGGN | ASSQGDYAEQF | 0.22400784 |
| PAGLAGVDPATL | ASSQGDYAEQF | 0.8575913 |
| GNNLVVEAPHAGASV | ASSQGDYAEQF | 0.90487474 |
| KVTRIMSASTLEQPA | ASSQGDYAEQF | 0.87651014 |
| LGFALPIALPRRTVV | ASSQGDYAEQF | 0.8647085 |
| TRLGNLTARKLVVLV | ASSQGDYAEQF | 0.8991109 |
| WSQQRRLIYFPSAGP | ASSQGDYAEQF | 0.8187401 |
| GPNGNPDPMAAAVDIR | ASSQGDYAEQF | 0.8404971 |
| TQRCVSVADLRAAASR | ASSQGDYAEQF | 0.8574992 |
| QFVLDVAEHPVEVLTETV | ASSQGDYAEQF | 0.89125824 |
| GGAGGTGGAGGNGGTQAGDG | ASSQGDYAEQF | 0.859977 |

| | | |
|---|---|---|
| PLLATLRAIAVDVDAKPAQV | ASSQGDYAEQF | 0.74683684 |
| APKFRTGGYMIDFWGVGYQVAK | ASSQGDYAEQF | 0.8882121 |
| GAGGAGGAGGAAGGPGGTGGTG | ASSQGDYAEQF | 0.9091911 |
| IGALGLLATDVAPAGSVAFGVA | ASSQGDYAEQF | 0.8367774 |
| AGGAGGIGGNGGAGGAGGVLMG | ASSQGDYAEQF | 0.89105487 |
| ADELIGLL | ASSQGDYAEQF | 0.885211 |
| ADRALKA | ASSQGDYAEQF | 0.875721 |
| AGAVGQVGGAGGA | ASSQGDYAEQF | 0.8107181 |
| AGGGGGLGGAP | ASSQGDYAEQF | 0.8027019 |
| AGGLLYGNGGAG | ASSQGDYAEQF | 0.86822724 |
| AGLESIGL | ASSQGDYAEQF | 0.79231894 |
| AGLKTGII | ASSQGDYAEQF | 0.86333203 |
| ASLAALLL | ASSQGDYAEQF | 0.861637 |
| ASVINISSI | ASSQGDYAEQF | 0.91096836 |
| ATPPGPA | ASSQGDYAEQF | 0.8824167 |
| ATVASALV | ASSQGDYAEQF | 0.88424444 |
| ATVASLIL | ASSQGDYAEQF | 0.78814495 |
| AVNGPVLIRH | ASSQGDYAEQF | 0.878453 |
| DGAAGAVGI | ASSQGDYAEQF | 0.88656455 |
| DPLLHGI | ASSQGDYAEQF | 0.69402057 |
| ELGAALGAL | ASSQGDYAEQF | 0.9018425 |
| ELGRTQAV | ASSQGDYAEQF | 0.8045094 |
| FVAGGYLS | ASSQGDYAEQF | 0.8624983 |
| GASGGAGGAGGW | ASSQGDYAEQF | 0.8950153 |
| GGAGAGGDGGAGGI | ASSQGDYAEQF | 0.9013363 |
| GTGHNGSQPGMGGNG | ASSQGDYAEQF | 0.85306275 |
| GYAAASASAS | ASSQGDYAEQF | 0.8533367 |
| HQVDVAAVQQG | ASSQGDYAEQF | 0.76971257 |
| IGAAIAHT | ASSQGDYAEQF | 0.89367276 |
| IGMIEVAQL | ASSQGDYAEQF | 0.8973092 |
| IRLAPEHTSAMD | ASSQGDYAEQF | 0.83688307 |
| IYNVSVL | ASSQGDYAEQF | 0.90373486 |
| KQGFLGM | ASSQGDYAEQF | 0.88839436 |
| LESVITQL | ASSQGDYAEQF | 0.46205407 |
| LGLGALVVV | ASSQGDYAEQF | 0.86155635 |
| LIGNGGNGGSGGI | ASSQGDYAEQF | 0.90097713 |
| LNAINAPTL | ASSQGDYAEQF | 0.8962153 |
| LQLLNTGV | ASSQGDYAEQF | 0.90761393 |
| LQTSIEF | ASSQGDYAEQF | 0.8872767 |
| MSFAVSEL | ASSQGDYAEQF | 0.8409638 |
| NTGALGL | ASSQGDYAEQF | 0.9037912 |
| PPGAGGAGGAGTT | ASSQGDYAEQF | 0.9101759 |
| QLVGLGI | ASSQGDYAEQF | 0.8938647 |
| RFLEQVHQL | ASSQGDYAEQF | 0.81024325 |
| RGVGYVLR | ASSQGDYAEQF | 0.77810127 |
| SATGVFAGVI | ASSQGDYAEQF | 0.88281447 |
| SLPAITLP | ASSQGDYAEQF | 0.8909688 |
| STALAFTV | ASSQGDYAEQF | 0.7552508 |
| SVLTVGLGK | ASSQGDYAEQF | 0.8761345 |
| TALAGAGSGI | ASSQGDYAEQF | 0.87579656 |
| TGLPFTGI | ASSQGDYAEQF | 0.4670853 |

| | | |
|---|---|---|
| VAEHVTV | ASSQGDYAEQF | 0.8902216 |
| VDIVLNSL | ASSQGDYAEQF | 0.90293473 |
| VLTGGGAVL | ASSQGDYAEQF | 0.8775347 |
| VNVGLAYGI | ASSQGDYAEQF | 0.83014596 |
| VSSAAVGAV | ASSQGDYAEQF | 0.8992478 |
| VTSVYEAGV | ASSQGDYAEQF | 0.90829533 |
| WVALDRGA | ASSQGDYAEQF | 0.90422225 |
| YLVGSVVI | ASSQGDYAEQF | 0.5999667 |
| GPPGTGKTLL | ASSQGDYAEQF | 0.37578517 |
| GVGMTKF | ASSQGDYAEQF | 0.84219235 |
| IGIGGSDL | ASSQGDYAEQF | 0.88971364 |
| IGTLLAEL | ASSQGDYAEQF | 0.85999346 |
| IYTSGTT | ASSQGDYAEQF | 0.8344534 |
| LSFPAGV | ASSQGDYAEQF | 0.88021976 |
| LYTSGTT | ASSQGDYAEQF | 0.9001664 |
| VGAFPFGF | ASSQGDYAEQF | 0.75167 |
| VGAGTVEF | ASSQGDYAEQF | 0.57713705 |
| VGKSTLLL | ASSQGDYAEQF | 0.88762623 |
| AALAHAGGGIV | ASSQGDYAEQF | 0.8916485 |
| AALVAVSLL | ASSQGDYAEQF | 0.79941845 |
| AAVSRAVAAL | ASSQGDYAEQF | 0.84075433 |
| AGGSSGIGGF | ASSQGDYAEQF | 0.8496586 |
| AGLTVANANG | ASSQGDYAEQF | 0.84345967 |
| AGLVGAVL | ASSQGDYAEQF | 0.8766002 |
| ALSQAFTV | ASSQGDYAEQF | 0.7780248 |
| ALTFSVSL | ASSQGDYAEQF | 0.90253115 |
| ALTGAALVAL | ASSQGDYAEQF | 0.7786796 |
| ASFSIGAGVL | ASSQGDYAEQF | 0.88762355 |
| DGSVYKI | ASSQGDYAEQF | 0.8788661 |
| EQLAEIV | ASSQGDYAEQF | 0.8862885 |
| FAVSELTVM | ASSQGDYAEQF | 0.84718853 |
| GAAPLAAGL | ASSQGDYAEQF | 0.82893944 |
| GADGGDGGSGGAGG | ASSQGDYAEQF | 0.7623684 |
| GFGLDTGGAGG | ASSQGDYAEQF | 0.62320423 |
| GLASGGLL | ASSQGDYAEQF | 0.89635056 |
| GRIAGAAVLL | ASSQGDYAEQF | 0.42096233 |
| GSAGSLNTSF | ASSQGDYAEQF | 0.89980006 |
| GTGQIGI | ASSQGDYAEQF | 0.8814538 |
| GTLIALIL | ASSQGDYAEQF | 0.75099033 |
| IAATMGFVVG | ASSQGDYAEQF | 0.8803989 |
| IFAQALL | ASSQGDYAEQF | 0.6991453 |
| IGAAVALGF | ASSQGDYAEQF | 0.8248811 |
| IGLTPLAL | ASSQGDYAEQF | 0.87821877 |
| ILGSNTSTL | ASSQGDYAEQF | 0.8924992 |
| IVFSLTVL | ASSQGDYAEQF | 0.813365 |
| LASKATGF | ASSQGDYAEQF | 0.8858677 |
| LASVALIL | ASSQGDYAEQF | 0.88483435 |
| LFAVGLTL | ASSQGDYAEQF | 0.8035133 |
| LFNGGIGTYI | ASSQGDYAEQF | 0.88396215 |
| LTLGAGII | ASSQGDYAEQF | 0.8993285 |
| LVAAALGDI | ASSQGDYAEQF | 0.8812635 |

| | | |
|---|---|---|
| NTAFVQLGI | ASSQGDYAEQF | 0.74331796 |
| QLFSGAEL | ASSQGDYAEQF | 0.88925254 |
| SAATFITL | ASSQGDYAEQF | 0.80283463 |
| SNYGLANLGV | ASSQGDYAEQF | 0.88102174 |
| SYGGAYAVM | ASSQGDYAEQF | 0.8641404 |
| TTGVGGLTLG | ASSQGDYAEQF | 0.847686 |
| TTQTIYI | ASSQGDYAEQF | 0.8953737 |
| VAAGAGLGAVI | ASSQGDYAEQF | 0.6286415 |
| VAAVVTLL | ASSQGDYAEQF | 0.79825544 |
| VGFLGGLI | ASSQGDYAEQF | 0.84401274 |
| VGVIPILL | ASSQGDYAEQF | 0.86943465 |
| VIFASSLI | ASSQGDYAEQF | 0.8946279 |
| VKGGDGGAAAG | ASSQGDYAEQF | 0.7733998 |
| VTKTFTNI | ASSQGDYAEQF | 0.8782646 |
| VVAAAILL | ASSQGDYAEQF | 0.83387786 |
| YAALDVEL | ASSQGDYAEQF | 0.7358321 |
| LASVALLL | ASSQGDYAEQF | 0.87457424 |
| LDAVHRV | ASSQGDYAEQF | 0.84647596 |
| QGLSSSIF | ASSQGDYAEQF | 0.7722469 |
| RGTFANIRL | ASSQGDYAEQF | 0.88876987 |
| RQLWWGHRI | ASSQGDYAEQF | 0.7586769 |
| YGPPGTGKTLL | ASSQGDYAEQF | 0.89027673 |
| AALVGAPANVA | ASSQGDYAEQF | 0.9037516 |
| AATFRRTE | ASSQGDYAEQF | 0.76097554 |
| AGALGGGI | ASSQGDYAEQF | 0.8863642 |
| AGGAGGAGGIGAG | ASSQGDYAEQF | 0.8834479 |
| AGSGALGL | ASSQGDYAEQF | 0.88144374 |
| AQIADQI | ASSQGDYAEQF | 0.85505867 |
| ASAVAVTYL | ASSQGDYAEQF | 0.8417316 |
| ASTVLGGTGG | ASSQGDYAEQF | 0.9055933 |
| AVGGEPAK | ASSQGDYAEQF | 0.8892654 |
| CEQGTFGGTC | ASSQGDYAEQF | 0.8923284 |
| DEGKAPAG | ASSQGDYAEQF | 0.89281964 |
| DTVVFPAL | ASSQGDYAEQF | 0.88687044 |
| FEGITMAG | ASSQGDYAEQF | 0.89172155 |
| FPLVVQD | ASSQGDYAEQF | 0.8262757 |
| GAFTTPQ | ASSQGDYAEQF | 0.69093734 |
| GFGGFGTAGGVGGA | ASSQGDYAEQF | 0.8428623 |
| GFLAQFL | ASSQGDYAEQF | 0.8598562 |
| GGAGGAGGNAGVA | ASSQGDYAEQF | 0.8398356 |
| GGVGTAGAGGAGGAGG | ASSQGDYAEQF | 0.56000555 |
| GNGGGGDAGNAGSGGN | ASSQGDYAEQF | 0.8975528 |
| GTGGAGGAGNDAGS | ASSQGDYAEQF | 0.9102015 |
| GTTNTGWFNS | ASSQGDYAEQF | 0.8265838 |
| IFMGPLFTL | ASSQGDYAEQF | 0.90924156 |
| ILAHAIEAI | ASSQGDYAEQF | 0.76785153 |
| IRSAIREAL | ASSQGDYAEQF | 0.8899501 |
| ITFAGGG | ASSQGDYAEQF | 0.89778817 |
| ITGMSGHQEV | ASSQGDYAEQF | 0.77937746 |
| IVAGLACTL | ASSQGDYAEQF | 0.89498407 |
| IVGGGLAGA | ASSQGDYAEQF | 0.79798096 |

| | | |
|---|---|---|
| IVTGVLNI | ASSQGDYAEQF | 0.83049583 |
| LAQQGCI | ASSQGDYAEQF | 0.9006762 |
| LCLHVEN | ASSQGDYAEQF | 0.8552656 |
| LDAISDGS | ASSQGDYAEQF | 0.88320816 |
| LGGFAAML | ASSQGDYAEQF | 0.8565865 |
| LGGKNPMI | ASSQGDYAEQF | 0.75606036 |
| LGPKGVAAIL | ASSQGDYAEQF | 0.88047516 |
| LMVGVSI | ASSQGDYAEQF | 0.90201837 |
| LVGAAFVL | ASSQGDYAEQF | 0.81365734 |
| LVLAVAPL | ASSQGDYAEQF | 0.9092913 |
| MAAIAVIDAA | ASSQGDYAEQF | 0.80599 |
| NNGINAELQ | ASSQGDYAEQF | 0.91051376 |
| RGEPLVLVHG | ASSQGDYAEQF | 0.8981433 |
| RGGAGGAAGLI | ASSQGDYAEQF | 0.865549 |
| RSYRMHPAV | ASSQGDYAEQF | 0.83830094 |
| SAAHLLAGQLA | ASSQGDYAEQF | 0.8973214 |
| SAAVGSY | ASSQGDYAEQF | 0.8485583 |
| SPHAVLHRI | ASSQGDYAEQF | 0.87834793 |
| SQIAKVL | ASSQGDYAEQF | 0.9061514 |
| TALGHLVP | ASSQGDYAEQF | 0.7161974 |
| TAPTTSGAA | ASSQGDYAEQF | 0.9065481 |
| TDGDDGNGGSPGT | ASSQGDYAEQF | 0.84831834 |
| TDGLGEN | ASSQGDYAEQF | 0.8481614 |
| TGGPPLPGQA | ASSQGDYAEQF | 0.7359675 |
| TIPAITI | ASSQGDYAEQF | 0.35064808 |
| TIPGFTI | ASSQGDYAEQF | 0.84495413 |
| TSIASGALA | ASSQGDYAEQF | 0.6710602 |
| TVTLPGAAI | ASSQGDYAEQF | 0.8557885 |
| VASGLSAALP | ASSQGDYAEQF | 0.8925694 |
| VGAGVIGI | ASSQGDYAEQF | 0.79693586 |
| VGPKRFGV | ASSQGDYAEQF | 0.90751654 |
| VNVAFPD | ASSQGDYAEQF | 0.9054666 |
| VSMTNSGVS | ASSQGDYAEQF | 0.58010113 |
| VTDVLGDLSV | ASSQGDYAEQF | 0.83052707 |
| VVVVGAGFAGL | ASSQGDYAEQF | 0.8087796 |
| VYTVGTVV | ASSQGDYAEQF | 0.82152617 |
| WTTFSGP | ASSQGDYAEQF | 0.5016108 |
| YDELVEAF | ASSQGDYAEQF | 0.88997716 |
| YGTGGAGGNGG | ASSQGDYAEQF | 0.7491925 |
| YTNVMAV | ASSQGDYAEQF | 0.15444429 |
| FLEQVHQL | ASSQGDYAEQF | 0.8343576 |
| GLSLLAVAV | ASSQGDYAEQF | 0.8609762 |
| LEQVHQL | ASSQGDYAEQF | 0.7958989 |
| LNAAGAI | ASSQGDYAEQF | 0.8985741 |
| PAVHASL | ASSQGDYAEQF | 0.39326096 |
| TELEDDA | ASSQGDYAEQF | 0.76371413 |
| TPPGPPN | ASSQGDYAEQF | 0.89714706 |
| TPPPPPPP | ASSQGDYAEQF | 0.90562344 |
| TQSPALI | ASGPRQGANSDYT | 0.61393654 |
| EFAAAFPQ | ASGPRQGANSDYT | 0.87465525 |
| LLSGNLDVL | ASGPRQGANSDYT | 0.89943016 |

| | | |
|---|---|---|
| GPANLGNYNI | ASGPRQGANSDYT | 0.8617313 |
| TLGPTLP | ASGPRQGANSDYT | 0.8948493 |
| VYAYSSCCK | ASGPRQGANSDYT | 0.9024088 |
| APAPAPAPNI | ASGPRQGANSDYT | 0.7715241 |
| PGGPVPIP | ASGPRQGANSDYT | 0.90393627 |
| FLTVALEP | ASGPRQGANSDYT | 0.91091686 |
| SQYFGANL | ASGPRQGANSDYT | 0.8331766 |
| GGVKPAA | ASGPRQGANSDYT | 0.8566401 |
| GGTGGAGGAGGW | ASGPRQGANSDYT | 0.6697923 |
| GVGGVAGLGGAGG | ASGPRQGANSDYT | 0.80050874 |
| VLLVLATTI | ASGPRQGANSDYT | 0.30871806 |
| IGIFAAML | ASGPRQGANSDYT | 0.8725603 |
| AVGDLID | ASGPRQGANSDYT | 0.8946527 |
| NISPLPL | ASGPRQGANSDYT | 0.82446504 |
| VTAVFVAI | ASGPRQGANSDYT | 0.71385634 |
| SGDDNDAAL | ASGPRQGANSDYT | 0.75571483 |
| AVAALAHS | ASGPRQGANSDYT | 0.8070502 |
| GGVTVMATP | ASGPRQGANSDYT | 0.8125722 |
| AGLAANVRD | ASGPRQGANSDYT | 0.32898408 |
| TNRPDLIDPALLRPG | ASGPRQGANSDYT | 0.71121514 |
| LANLGVTAL | ASGPRQGANSDYT | 0.89974564 |
| TGERIEL | ASGPRQGANSDYT | 0.8947844 |
| NGEELAA | ASGPRQGANSDYT | 0.51883405 |
| VLSLVWG | ASGPRQGANSDYT | 0.90879184 |
| IGYALWL | ASGPRQGANSDYT | 0.87016046 |
| LSLLLAPL | ASGPRQGANSDYT | 0.89578825 |
| TLDEALK | ASGPRQGANSDYT | 0.84944296 |
| TDGAGTAEGG | ASGPRQGANSDYT | 0.6950928 |
| NAPGGAGGNGG | ASGPRQGANSDYT | 0.88508034 |
| VSVFMGMGGNF | ASGPRQGANSDYT | 0.81789684 |
| LASYLAAL | ASGPRQGANSDYT | 0.6234962 |
| LVGGTPAL | ASGPRQGANSDYT | 0.88600296 |
| VRNADVL | ASGPRQGANSDYT | 0.7730537 |
| TCSSPFLVSAM | ASGPRQGANSDYT | 0.7906846 |
| VHITDAE | ASGPRQGANSDYT | 0.78554595 |
| EYAAPSF | ASGPRQGANSDYT | 0.8847181 |
| VVDAFHGGL | ASGPRQGANSDYT | 0.74790263 |
| ASVVDPQ | ASGPRQGANSDYT | 0.502841 |
| SYNEFGGLS | ASGPRQGANSDYT | 0.67528605 |
| DPDGSVY | ASGPRQGANSDYT | 0.8493259 |
| MVSTIHATE | ASGPRQGANSDYT | 0.9027433 |
| LAAGVADPV | ASGPRQGANSDYT | 0.8591616 |
| TKEQIAA | ASGPRQGANSDYT | 0.77869344 |
| SLASPAAL | ASGPRQGANSDYT | 0.87651455 |
| IGLVFYGV | ASGPRQGANSDYT | 0.86508536 |
| FLLDASNL | ASGPRQGANSDYT | 0.86050665 |
| YQVGNFP | ASGPRQGANSDYT | 0.8656697 |
| ESLLSQST | ASGPRQGANSDYT | 0.7855989 |
| LFTVGFAAVG | ASGPRQGANSDYT | 0.88718295 |
| ATPPTPT | ASGPRQGANSDYT | 0.8769159 |
| TDLATLG | ASGPRQGANSDYT | 0.89170086 |

| | | |
|---|---|---|
| SAGGTGR | ASGPRQGANSDYT | 0.89688826 |
| GFSNSGS | ASGPRQGANSDYT | 0.84734404 |
| GGNGGSGGTSV | ASGPRQGANSDYT | 0.8441824 |
| GTSVATGGAGN | ASGPRQGANSDYT | 0.29520884 |
| NGGNGGSGGTS | ASGPRQGANSDYT | 0.79667765 |
| AVVKFPH | ASGPRQGANSDYT | 0.86314815 |
| TPPPPPP | ASGPRQGANSDYT | 0.8629819 |
| AAPAAAAAPA | ASGPRQGANSDYT | 0.88521826 |
| IALNEDL | ASGPRQGANSDYT | 0.8590517 |
| IVNVSSM | ASGPRQGANSDYT | 0.7863438 |
| TGIPPAPR | ASGPRQGANSDYT | 0.8816387 |
| FDLGGGTF | ASGPRQGANSDYT | 0.89535475 |
| FDLGGGTFDVS | ASGPRQGANSDYT | 0.56195647 |
| NEPTAAA | ASGPRQGANSDYT | 0.8150994 |
| NEPTAAALA | ASGPRQGANSDYT | 0.91320276 |
| GSIINISSI | ASGPRQGANSDYT | 0.86137867 |
| VAPGNAGTA | ASGPRQGANSDYT | 0.84351707 |
| TGGGASLE | ASGPRQGANSDYT | 0.7449418 |
| VAVYNHYKRI | ASGPRQGANSDYT | 0.9017216 |
| TDPAPATT | ASGPRQGANSDYT | 0.7329674 |
| TNRPDILDPALLRPG | ASGPRQGANSDYT | 0.90743244 |
| LTTTTTS | ASGPRQGANSDYT | 0.20877221 |
| PTGSGKTL | ASGPRQGANSDYT | 0.74884784 |
| TGSGKTL | ASGPRQGANSDYT | 0.46495402 |
| TIGHVDHGK | ASGPRQGANSDYT | 0.3072436 |
| TIGHVDHGKTT | ASGPRQGANSDYT | 0.8944943 |
| GHVDHGKTT | ASGPRQGANSDYT | 0.8020657 |
| AQFEHTLL | ASGPRQGANSDYT | 0.887656 |
| IVNQDEL | ASGPRQGANSDYT | 0.85602605 |
| TNGSQFF | ASGPRQGANSDYT | 0.8970586 |
| TGKGIVE | ASGPRQGANSDYT | 0.86431575 |
| LISNASDA | ASGPRQGANSDYT | 0.856481 |
| AELRTLGELTKIAKAHGAQ | ASGPRQGANSDYT | 0.59566903 |
| APNAVAAAEPSW | ASGPRQGANSDYT | 0.89062136 |
| APQVAGAATAVL | ASGPRQGANSDYT | 0.8975722 |
| ARRDPAAAAAILPTDA | ASGPRQGANSDYT | 0.8484447 |
| DAAARDVLAEAGLAEYF | ASGPRQGANSDYT | 0.5734085 |
| DDQIVLSAKANLAKVEQ | ASGPRQGANSDYT | 0.7674822 |
| DELLPDTGEDSDEDGH | ASGPRQGANSDYT | 0.86118066 |
| ELPEAPANRATVPQD | ASGPRQGANSDYT | 0.19381541 |
| FGQSKIERVTGIG | ASGPRQGANSDYT | 0.16843274 |
| GAGGTGSTAGGAGGAGGAGGLY | ASGPRQGANSDYT | 0.7569545 |
| GHGGTGAAGGNGGTGGAGGSGI | ASGPRQGANSDYT | 0.74451375 |
| GNGGNGGAGGEGGPGGAGGAGA | ASGPRQGANSDYT | 0.90307796 |
| GSTAGNGGNGGNATNGTIAGQP | ASGPRQGANSDYT | 0.89791244 |
| GTGGDGGNGGNGGTGG | ASGPRQGANSDYT | 0.8469678 |
| GVRRPAGIPADAPV | ASGPRQGANSDYT | 0.8819496 |
| GYRQVTDAHLLTLARR | ASGPRQGANSDYT | 0.8618958 |
| IAKMGQDGHDRG | ASGPRQGANSDYT | 0.08614885 |
| KAEPSIEIQYT | ASGPRQGANSDYT | 0.78664994 |
| KPVRIRMAADEDTVAVT | ASGPRQGANSDYT | 0.6151953 |

| | | |
|---|---|---|
| MPPRIAGMRLLVIKPEP | ASGPRQGANSDYT | 0.9088445 |
| MKFYAVALFRSRGVPPTR | ASGPRQGANSDYT | 0.7237853 |
| NAGGFGILGAVAHSP | ASGPRQGANSDYT | 0.7983881 |
| NEGREITLAMAEM | ASGPRQGANSDYT | 0.8642688 |
| NELVEWSHPLGHRWRWE | ASGPRQGANSDYT | 0.85472924 |
| NLGAGGTGGNGGNPG | ASGPRQGANSDYT | 0.84856844 |
| NTEWPKWETVMAADPVD | ASGPRQGANSDYT | 0.63755834 |
| PASAAPAETQP | ASGPRQGANSDYT | 0.12348131 |
| QQSSDGKNGVRAAEITGE | ASGPRQGANSDYT | 0.8884302 |
| RAPDAVTTSWWKE | ASGPRQGANSDYT | 0.45587817 |
| SGFGNVGAHTSGWFNQ | ASGPRQGANSDYT | 0.7028459 |
| SGVAGAVGQAASVGGLKV | ASGPRQGANSDYT | 0.28226215 |
| SNGGAGTGSAGGNGGT | ASGPRQGANSDYT | 0.1677627 |
| TGGAGGEGGAAG | ASGPRQGANSDYT | 0.5483285 |
| TGGDGGAAGAGGAPGHG | ASGPRQGANSDYT | 0.87528396 |
| TMPPLLAAIANAGRHGV | ASGPRQGANSDYT | 0.8276086 |
| TSSTAGNSGGA | ASGPRQGANSDYT | 0.8330544 |
| VPATVPVGILC | ASGPRQGANSDYT | 0.9115719 |
| VQRPDSGGVVLTGRIS | ASGPRQGANSDYT | 0.84764516 |
| WTPNPVPATKGVPAGLVQAT | ASGPRQGANSDYT | 0.8326306 |
| YALKRALAKFSDRFDVVII | ASGPRQGANSDYT | 0.72961676 |
| YLIGSVIV | ASGPRQGANSDYT | 0.75162804 |
| YGQLVAVID | ASGPRQGANSDYT | 0.68102837 |
| VTVGGMTLPA | ASGPRQGANSDYT | 0.6991315 |
| VLALPAV | ASGPRQGANSDYT | 0.91110635 |
| VIPPTLNYET | ASGPRQGANSDYT | 0.75401413 |
| VGMGGMHPSSG | ASGPRQGANSDYT | 0.6200199 |
| VGLAGGAATA | ASGPRQGANSDYT | 0.80960476 |
| VGGAGGAGGSGP | ASGPRQGANSDYT | 0.9049014 |
| VFAAFMV | ASGPRQGANSDYT | 0.91060835 |
| TLYDKLR | ASGPRQGANSDYT | 0.73471415 |
| TLAVFTVL | ASGPRQGANSDYT | 0.6959152 |
| TGSVQPARV | ASGPRQGANSDYT | 0.7966262 |
| TGGGAGSGGT | ASGPRQGANSDYT | 0.8561591 |
| SHLETSL | ASGPRQGANSDYT | 0.6753963 |
| SFTTGMT | ASGPRQGANSDYT | 0.5264318 |
| QVEVAKRMGVRQPTVS | ASGPRQGANSDYT | 0.8751864 |
| QGSGGPALAM | ASGPRQGANSDYT | 0.90968496 |
| MLAPSNP | ASGPRQGANSDYT | 0.79855305 |
| LVNNAAI | ASGPRQGANSDYT | 0.70367014 |
| LTGEINTLA | ASGPRQGANSDYT | 0.85293615 |
| LSIFGPV | ASGPRQGANSDYT | 0.77292985 |
| LLALLANL | ASGPRQGANSDYT | 0.81903243 |
| LISNASDALDKLR | ASGPRQGANSDYT | 0.85906374 |
| LINEAVP | ASGPRQGANSDYT | 0.6495779 |
| LGYAFSGL | ASGPRQGANSDYT | 0.8247112 |
| LDFVAVAL | ASGPRQGANSDYT | 0.9076294 |
| LAQPGAE | ASGPRQGANSDYT | 0.7997315 |
| ISNLGAVTGF | ASGPRQGANSDYT | 0.8723576 |
| HQEDPGRI | ASGPRQGANSDYT | 0.9018444 |
| GTPGEPTP | ASGPRQGANSDYT | 0.8839687 |

| | | |
|---|---|---|
| GFFNSGS | ASGPRQGANSDYT | 0.9028877 |
| GAGGGGLTAGAN | ASGPRQGANSDYT | 0.88215846 |
| FVDALVNM | ASGPRQGANSDYT | 0.7804818 |
| FLTVAPEL | ASGPRQGANSDYT | 0.80380815 |
| FETLTAL | ASGPRQGANSDYT | 0.86377406 |
| DVINPIEL | ASGPRQGANSDYT | 0.86688095 |
| APVSGAPV | ASGPRQGANSDYT | 0.8400764 |
| ALVGGLAW | ASGPRQGANSDYT | 0.7534098 |
| ALAAAAATAI | ASGPRQGANSDYT | 0.90365505 |
| AFGLGLLL | ASGPRQGANSDYT | 0.8029125 |
| AFFAAASL | ASGPRQGANSDYT | 0.886418 |
| AAAALPAFSPP | ASGPRQGANSDYT | 0.5237412 |
| VVLAPAL | ASGPRQGANSDYT | 0.5929141 |
| TIGHVDHGKTTL | ASGPRQGANSDYT | 0.8929266 |
| LISNASDALDK | ASGPRQGANSDYT | 0.106731534 |
| LISNASDAL | ASGPRQGANSDYT | 0.22021727 |
| ISNASDALDK | ASGPRQGANSDYT | 0.84299487 |
| NQAETLVYQTE | ASGPRQGANSDYT | 0.9015433 |
| TGGTGGTAGAG | ASGPRQGANSDYT | 0.112511516 |
| TGSTAGGAGGA | ASGPRQGANSDYT | 0.72424954 |
| DPLAVPVAAGPV | ASGPRQGANSDYT | 0.62945926 |
| GGTGGTPGNGGN | ASGPRQGANSDYT | 0.85481155 |
| PAGLAGVDPATL | ASGPRQGANSDYT | 0.8496726 |
| GNNLVVEAPHAGASV | ASGPRQGANSDYT | 0.5139077 |
| KVTRIMSASTLEQPA | ASGPRQGANSDYT | 0.8846705 |
| LGFALPIALPRRTVV | ASGPRQGANSDYT | 0.5317246 |
| TRLGNLTARKLVVLV | ASGPRQGANSDYT | 0.47085953 |
| WSQQRRLIYFPSAGP | ASGPRQGANSDYT | 0.89343244 |
| GPNGNPDPMAAAVDIR | ASGPRQGANSDYT | 0.89345723 |
| TQRCVSVADLRAAASR | ASGPRQGANSDYT | 0.72556025 |
| QFVLDVAEHPVEVLTETV | ASGPRQGANSDYT | 0.89476 |
| GGAGGTGGAGGNGGTQAGDG | ASGPRQGANSDYT | 0.78759235 |
| PLLATLRAIAVDVDAKPAQV | ASGPRQGANSDYT | 0.90738326 |
| APKFRTGGYMIDFWGVGYQVAK | ASGPRQGANSDYT | 0.77970755 |
| GAGGAGGAGGAAGGPGGTGGTG | ASGPRQGANSDYT | 0.4663306 |
| IGALGLLATDVAPAGSVAFGVA | ASGPRQGANSDYT | 0.8945726 |
| AGGAGGIGGNGGAGGAGGVLMG | ASGPRQGANSDYT | 0.8758399 |
| ADELIGLL | ASGPRQGANSDYT | 0.82630944 |
| ADRALKA | ASGPRQGANSDYT | 0.8738406 |
| AGAVGQVGGAGGA | ASGPRQGANSDYT | 0.66660476 |
| AGGGGGLGGAP | ASGPRQGANSDYT | 0.8494092 |
| AGGLLYGNGGAG | ASGPRQGANSDYT | 0.90399075 |
| AGLESIGL | ASGPRQGANSDYT | 0.89369345 |
| AGLKTGII | ASGPRQGANSDYT | 0.53067756 |
| ASLAALLL | ASGPRQGANSDYT | 0.8829525 |
| ASVINISSI | ASGPRQGANSDYT | 0.8444394 |
| ATPPGPA | ASGPRQGANSDYT | 0.6874173 |
| ATVASALV | ASGPRQGANSDYT | 0.9072551 |
| ATVASLIL | ASGPRQGANSDYT | 0.8327357 |
| AVNGPVLIRH | ASGPRQGANSDYT | 0.88683796 |
| DGAAGAVGI | ASGPRQGANSDYT | 0.8753771 |

| | | |
|---|---|---|
| DPLLHGI | ASGPRQGANSDYT | 0.88756037 |
| ELGAALGAL | ASGPRQGANSDYT | 0.39082044 |
| ELGRTQAV | ASGPRQGANSDYT | 0.82046086 |
| FVAGGYLS | ASGPRQGANSDYT | 0.84604174 |
| GASGGAGGAGGW | ASGPRQGANSDYT | 0.8929014 |
| GGAGAGGDGGAGGI | ASGPRQGANSDYT | 0.8378778 |
| GTGHNGSQPGMGGNG | ASGPRQGANSDYT | 0.59054565 |
| GYAAASASAS | ASGPRQGANSDYT | 0.8614061 |
| HQVDVAAVQQG | ASGPRQGANSDYT | 0.9012855 |
| IGAAIAHT | ASGPRQGANSDYT | 0.8171117 |
| IGMIEVAQL | ASGPRQGANSDYT | 0.85557795 |
| IRLAPEHTSAMD | ASGPRQGANSDYT | 0.9080495 |
| IYNVSVL | ASGPRQGANSDYT | 0.68208736 |
| KQGFLGM | ASGPRQGANSDYT | 0.83093274 |
| LESVITQL | ASGPRQGANSDYT | 0.8799819 |
| LGLGALVVV | ASGPRQGANSDYT | 0.8966038 |
| LIGNGGNGGSGGI | ASGPRQGANSDYT | 0.8011173 |
| LNAINAPTL | ASGPRQGANSDYT | 0.89990455 |
| LQLLNTGV | ASGPRQGANSDYT | 0.71984434 |
| LQTSIEF | ASGPRQGANSDYT | 0.8947413 |
| MSFAVSEL | ASGPRQGANSDYT | 0.7610986 |
| NTGALGL | ASGPRQGANSDYT | 0.8812783 |
| PPGAGGAGGAGTT | ASGPRQGANSDYT | 0.7047941 |
| QLVGLGI | ASGPRQGANSDYT | 0.8944884 |
| RFLEQVHQL | ASGPRQGANSDYT | 0.8927413 |
| RGVGYVLR | ASGPRQGANSDYT | 0.8985183 |
| SATGVFAGVI | ASGPRQGANSDYT | 0.17839205 |
| SLPAITLP | ASGPRQGANSDYT | 0.8958148 |
| STALAFTV | ASGPRQGANSDYT | 0.898207 |
| SVLTVGLGK | ASGPRQGANSDYT | 0.9111654 |
| TALAGAGSGI | ASGPRQGANSDYT | 0.77070916 |
| TGLPFTGI | ASGPRQGANSDYT | 0.87946516 |
| VAEHVTV | ASGPRQGANSDYT | 0.81383514 |
| VDIVLNSL | ASGPRQGANSDYT | 0.78267115 |
| VLTGGGAVL | ASGPRQGANSDYT | 0.7047254 |
| VNVGLAYGI | ASGPRQGANSDYT | 0.74934757 |
| VSSAAVGAV | ASGPRQGANSDYT | 0.8294305 |
| VTSVYEAGV | ASGPRQGANSDYT | 0.7770529 |
| WVALDRGA | ASGPRQGANSDYT | 0.88452077 |
| YLVGSVVI | ASGPRQGANSDYT | 0.8829408 |
| GPPGTGKTLL | ASGPRQGANSDYT | 0.89434266 |
| GVGMTKF | ASGPRQGANSDYT | 0.90326285 |
| IGIGGSDL | ASGPRQGANSDYT | 0.900501 |
| IGTLLAEL | ASGPRQGANSDYT | 0.8892781 |
| IYTSGTT | ASGPRQGANSDYT | 0.72375023 |
| LSFPAGV | ASGPRQGANSDYT | 0.851187 |
| LYTSGTT | ASGPRQGANSDYT | 0.6922013 |
| VGAFPFGF | ASGPRQGANSDYT | 0.84874004 |
| VGAGTVEF | ASGPRQGANSDYT | 0.74154615 |
| VGKSTLLL | ASGPRQGANSDYT | 0.87645113 |
| AALAHAGGGIV | ASGPRQGANSDYT | 0.09758294 |

| | | |
|---|---|---|
| AALVAVSLL | ASGPRQGANSDYT | 0.84736055 |
| AAVSRAVAAL | ASGPRQGANSDYT | 0.91030014 |
| AGGSSGIGGF | ASGPRQGANSDYT | 0.7015098 |
| AGLTVANANG | ASGPRQGANSDYT | 0.8676952 |
| AGLVGAVL | ASGPRQGANSDYT | 0.85261273 |
| ALSQAFTV | ASGPRQGANSDYT | 0.75544685 |
| ALTFSVSL | ASGPRQGANSDYT | 0.8909336 |
| ALTGAALVAL | ASGPRQGANSDYT | 0.7022948 |
| ASFSIGAGVL | ASGPRQGANSDYT | 0.86946756 |
| DGSVYKI | ASGPRQGANSDYT | 0.9054304 |
| EQLAEIV | ASGPRQGANSDYT | 0.8648262 |
| FAVSELTVM | ASGPRQGANSDYT | 0.815735 |
| GAAPLAAGL | ASGPRQGANSDYT | 0.8188375 |
| GADGGDGGSGGAGG | ASGPRQGANSDYT | 0.3244384 |
| GFGLDTGGAGG | ASGPRQGANSDYT | 0.8560312 |
| GLASGGLL | ASGPRQGANSDYT | 0.09081344 |
| GRIAGAAVLL | ASGPRQGANSDYT | 0.8513047 |
| GSAGSLNTSF | ASGPRQGANSDYT | 0.6069636 |
| GTGQIGI | ASGPRQGANSDYT | 0.8707099 |
| GTLIALIL | ASGPRQGANSDYT | 0.86304295 |
| IAATMGFVVG | ASGPRQGANSDYT | 0.86793506 |
| IFAQALL | ASGPRQGANSDYT | 0.7664927 |
| IGAAVALGF | ASGPRQGANSDYT | 0.8185284 |
| IGLTPLAL | ASGPRQGANSDYT | 0.7052394 |
| ILGSNTSTL | ASGPRQGANSDYT | 0.6149167 |
| IVFSLTVL | ASGPRQGANSDYT | 0.88206506 |
| LASKATGF | ASGPRQGANSDYT | 0.7508785 |
| LASVALIL | ASGPRQGANSDYT | 0.81702405 |
| LFAVGLTL | ASGPRQGANSDYT | 0.90144825 |
| LFNGGIGTYI | ASGPRQGANSDYT | 0.8491058 |
| LTLGAGII | ASGPRQGANSDYT | 0.5007625 |
| LVAAALGDI | ASGPRQGANSDYT | 0.8309666 |
| NTAFVQLGI | ASGPRQGANSDYT | 0.860744 |
| QLFSGAEL | ASGPRQGANSDYT | 0.7718495 |
| SAATFITL | ASGPRQGANSDYT | 0.09979302 |
| SNYGLANLGV | ASGPRQGANSDYT | 0.8637927 |
| SYGGAYAVM | ASGPRQGANSDYT | 0.6951007 |
| TTGVGGLTLG | ASGPRQGANSDYT | 0.7756164 |
| TTQTIYI | ASGPRQGANSDYT | 0.75659996 |
| VAAGAGLGAVI | ASGPRQGANSDYT | 0.8926134 |
| VAAVVTLL | ASGPRQGANSDYT | 0.46438497 |
| VGFLGGLI | ASGPRQGANSDYT | 0.8186128 |
| VGVIPILL | ASGPRQGANSDYT | 0.8792112 |
| VIFASSLI | ASGPRQGANSDYT | 0.8882236 |
| VKGGDGGAAAG | ASGPRQGANSDYT | 0.90509397 |
| VTKTFTNI | ASGPRQGANSDYT | 0.8470838 |
| VVAAAILL | ASGPRQGANSDYT | 0.6756244 |
| YAALDVEL | ASGPRQGANSDYT | 0.8733734 |
| LASVALLL | ASGPRQGANSDYT | 0.81714326 |
| LDAVHRV | ASGPRQGANSDYT | 0.44448128 |
| QGLSSSIF | ASGPRQGANSDYT | 0.3213821 |

| | | |
|---|---|---|
| RGTFANIRL | ASGPRQGANSDYT | 0.86312306 |
| RQLWWGHRI | ASGPRQGANSDYT | 0.35710043 |
| YGPPGTGKTLL | ASGPRQGANSDYT | 0.85437894 |
| AALVGAPANVA | ASGPRQGANSDYT | 0.89871466 |
| AATFRRTE | ASGPRQGANSDYT | 0.7978925 |
| AGALGGGI | ASGPRQGANSDYT | 0.50750166 |
| AGGAGGAGGIGAG | ASGPRQGANSDYT | 0.81879574 |
| AGSGALGL | ASGPRQGANSDYT | 0.7779341 |
| AQIADQI | ASGPRQGANSDYT | 0.8815854 |
| ASAVAVTYL | ASGPRQGANSDYT | 0.8296951 |
| ASTVLGGTGG | ASGPRQGANSDYT | 0.9075822 |
| AVGGEPAK | ASGPRQGANSDYT | 0.90112424 |
| CEQGTFGGTC | ASGPRQGANSDYT | 0.80791765 |
| DEGKAPAG | ASGPRQGANSDYT | 0.8861778 |
| DTVVFPAL | ASGPRQGANSDYT | 0.8027687 |
| FEGITMAG | ASGPRQGANSDYT | 0.8155835 |
| FPLVVQD | ASGPRQGANSDYT | 0.8140595 |
| GAFTTPQ | ASGPRQGANSDYT | 0.8338796 |
| GFGGFGTAGGVGGA | ASGPRQGANSDYT | 0.7800386 |
| GFLAQFL | ASGPRQGANSDYT | 0.7459854 |
| GGAGGAGGNAGVA | ASGPRQGANSDYT | 0.56601703 |
| GGVGTAGAGGAGGAGG | ASGPRQGANSDYT | 0.86665094 |
| GNGGGGDAGNAGSGGN | ASGPRQGANSDYT | 0.83951604 |
| GTGGAGGAGNDAGS | ASGPRQGANSDYT | 0.3395727 |
| GTTNTGWFNS | ASGPRQGANSDYT | 0.8099166 |
| IFMGPLFTL | ASGPRQGANSDYT | 0.91133976 |
| ILAHAIEAI | ASGPRQGANSDYT | 0.9046636 |
| IRSAIREAL | ASGPRQGANSDYT | 0.7597831 |
| ITFAGGG | ASGPRQGANSDYT | 0.6988679 |
| ITGMSGHQEV | ASGPRQGANSDYT | 0.84899414 |
| IVAGLACTL | ASGPRQGANSDYT | 0.36859864 |
| IVGGGLAGA | ASGPRQGANSDYT | 0.86003196 |
| IVTGVLNI | ASGPRQGANSDYT | 0.8558574 |
| LAQQGCI | ASGPRQGANSDYT | 0.8500345 |
| LCLHVEN | ASGPRQGANSDYT | 0.7170291 |
| LDAISDGS | ASGPRQGANSDYT | 0.8544697 |
| LGGFAAML | ASGPRQGANSDYT | 0.8052937 |
| LGGKNPMI | ASGPRQGANSDYT | 0.86442673 |
| LGPKGVAAIL | ASGPRQGANSDYT | 0.1349473 |
| LMVGVSI | ASGPRQGANSDYT | 0.86967385 |
| LVGAAFVL | ASGPRQGANSDYT | 0.7798546 |
| LVLAVAPL | ASGPRQGANSDYT | 0.5812306 |
| MAAIAVIDAA | ASGPRQGANSDYT | 0.26544887 |
| NNGINAELQ | ASGPRQGANSDYT | 0.84364814 |
| RGEPLVLVHG | ASGPRQGANSDYT | 0.8860314 |
| RGGAGGAAGLI | ASGPRQGANSDYT | 0.8907907 |
| RSYRMHPAV | ASGPRQGANSDYT | 0.51975703 |
| SAAHLLAGQLA | ASGPRQGANSDYT | 0.8985146 |
| SAAVGSY | ASGPRQGANSDYT | 0.9031993 |
| SPHAVLHRI | ASGPRQGANSDYT | 0.82370645 |
| SQIAKVL | ASGPRQGANSDYT | 0.28633592 |

| | | |
|---|---|---|
| TALGHLVP | ASGPRQGANSDYT | 0.8773964 |
| TAPTTSGAA | ASGPRQGANSDYT | 0.7903192 |
| TDGDDGNGGSPGT | ASGPRQGANSDYT | 0.8853186 |
| TDGLGEN | ASGPRQGANSDYT | 0.8646848 |
| TGGPPLPGQA | ASGPRQGANSDYT | 0.69947416 |
| TIPAITI | ASGPRQGANSDYT | 0.89779073 |
| TIPGFTI | ASGPRQGANSDYT | 0.9027761 |
| TSIASGALA | ASGPRQGANSDYT | 0.75271696 |
| TVTLPGAAI | ASGPRQGANSDYT | 0.72874427 |
| VASGLSAALP | ASGPRQGANSDYT | 0.7081419 |
| VGAGVIGI | ASGPRQGANSDYT | 0.27603662 |
| VGPKRFGV | ASGPRQGANSDYT | 0.88991594 |
| VNVAFPD | ASGPRQGANSDYT | 0.80938244 |
| VSMTNSGVS | ASGPRQGANSDYT | 0.7669319 |
| VTDVLGDLSV | ASGPRQGANSDYT | 0.86850226 |
| VVVVGAGFAGL | ASGPRQGANSDYT | 0.75510633 |
| VYTVGTVV | ASGPRQGANSDYT | 0.76678693 |
| WTTFSGP | ASGPRQGANSDYT | 0.79315066 |
| YDELVEAF | ASGPRQGANSDYT | 0.85170233 |
| YGTGGAGGNGG | ASGPRQGANSDYT | 0.8670969 |
| YTNVMAV | ASGPRQGANSDYT | 0.8509993 |
| FLEQVHQL | ASGPRQGANSDYT | 0.8591707 |
| GLSLLAVAV | ASGPRQGANSDYT | 0.9078114 |
| LEQVHQL | ASGPRQGANSDYT | 0.5241949 |
| LNAAGAI | ASGPRQGANSDYT | 0.7530458 |
| PAVHASL | ASGPRQGANSDYT | 0.87111187 |
| TELEDDA | ASGPRQGANSDYT | 0.43274134 |
| TPPGPPN | ASGPRQGANSDYT | 0.89036846 |
| TPPPPPPP | ASGPRQGANSDYT | 0.86456966 |
| TQSPALI | ASSQEDRENTEVF | 0.8774466 |
| EFAAAFPQ | ASSQEDRENTEVF | 0.7906983 |
| LLSGNLDVL | ASSQEDRENTEVF | 0.91303086 |
| GPANLGNYNI | ASSQEDRENTEVF | 0.8959062 |
| TLGPTLP | ASSQEDRENTEVF | 0.83903986 |
| VYAYSSCCK | ASSQEDRENTEVF | 0.85427594 |
| APAPAPAPNI | ASSQEDRENTEVF | 0.5686474 |
| PGGPVPIP | ASSQEDRENTEVF | 0.899559 |
| FLTVALEP | ASSQEDRENTEVF | 0.87952316 |
| SQYFGANL | ASSQEDRENTEVF | 0.6985634 |
| GGVKPAA | ASSQEDRENTEVF | 0.8572508 |
| GGTGGAGGAGGW | ASSQEDRENTEVF | 0.89230514 |
| GVGGVAGLGGAGG | ASSQEDRENTEVF | 0.89886373 |
| VLLVLATTI | ASSQEDRENTEVF | 0.3131997 |
| IGIFAAML | ASSQEDRENTEVF | 0.8226925 |
| AVGDLID | ASSQEDRENTEVF | 0.2816247 |
| NISPLPL | ASSQEDRENTEVF | 0.60930765 |
| VTAVFVAI | ASSQEDRENTEVF | 0.535961 |
| SGDDNDAAL | ASSQEDRENTEVF | 0.8698816 |
| AVAALAHS | ASSQEDRENTEVF | 0.831325 |
| GGVTVMATP | ASSQEDRENTEVF | 0.8594976 |
| AGLAANVRD | ASSQEDRENTEVF | 0.8525223 |

| | | |
|---|---|---|
| TNRPDLIDPALLRPG | ASSQEDRENTEVF | 0.90212303 |
| LANLGVTAL | ASSQEDRENTEVF | 0.79441655 |
| TGERIEL | ASSQEDRENTEVF | 0.8323734 |
| NGEELAA | ASSQEDRENTEVF | 0.4456039 |
| VLSLVWG | ASSQEDRENTEVF | 0.34170127 |
| IGYALWL | ASSQEDRENTEVF | 0.6687069 |
| LSLLLAPL | ASSQEDRENTEVF | 0.8278213 |
| TLDEALK | ASSQEDRENTEVF | 0.18553059 |
| TDGAGTAEGG | ASSQEDRENTEVF | 0.9012544 |
| NAPGGAGGNGG | ASSQEDRENTEVF | 0.8716449 |
| VSVFMGMGGNF | ASSQEDRENTEVF | 0.87887335 |
| LASYLAAL | ASSQEDRENTEVF | 0.8123528 |
| LVGGTPAL | ASSQEDRENTEVF | 0.7687605 |
| VRNADVL | ASSQEDRENTEVF | 0.73601186 |
| TCSSPFLVSAM | ASSQEDRENTEVF | 0.882537 |
| VHITDAE | ASSQEDRENTEVF | 0.6804286 |
| EYAAPSF | ASSQEDRENTEVF | 0.5940823 |
| VVDAFHGGL | ASSQEDRENTEVF | 0.72973454 |
| ASVVDPQ | ASSQEDRENTEVF | 0.73822886 |
| SYNEFGGLS | ASSQEDRENTEVF | 0.9001153 |
| DPDGSVY | ASSQEDRENTEVF | 0.8968272 |
| MVSTIHATE | ASSQEDRENTEVF | 0.87669736 |
| LAAGVADPV | ASSQEDRENTEVF | 0.89313674 |
| TKEQIAA | ASSQEDRENTEVF | 0.20952056 |
| SLASPAAL | ASSQEDRENTEVF | 0.8236616 |
| IGLVFYGV | ASSQEDRENTEVF | 0.90962803 |
| FLLDASNL | ASSQEDRENTEVF | 0.8435748 |
| YQVGNFP | ASSQEDRENTEVF | 0.8461494 |
| ESLLSQST | ASSQEDRENTEVF | 0.90025663 |
| LFTVGFAAVG | ASSQEDRENTEVF | 0.78079146 |
| ATPPTPT | ASSQEDRENTEVF | 0.78186095 |
| TDLATLG | ASSQEDRENTEVF | 0.82656586 |
| SAGGTGR | ASSQEDRENTEVF | 0.8667536 |
| GFSNSGS | ASSQEDRENTEVF | 0.79518956 |
| GGNGGSGGTSV | ASSQEDRENTEVF | 0.6650162 |
| GTSVATGGAGN | ASSQEDRENTEVF | 0.86983716 |
| NGGNGGSGGTS | ASSQEDRENTEVF | 0.91396224 |
| AVVKFPH | ASSQEDRENTEVF | 0.8267759 |
| TPPPPPP | ASSQEDRENTEVF | 0.8928417 |
| AAPAAAAAPA | ASSQEDRENTEVF | 0.87957567 |
| IALNEDL | ASSQEDRENTEVF | 0.8698785 |
| IVNVSSM | ASSQEDRENTEVF | 0.7722062 |
| TGIPPAPR | ASSQEDRENTEVF | 0.79048306 |
| FDLGGGTF | ASSQEDRENTEVF | 0.8215115 |
| FDLGGGTFDVS | ASSQEDRENTEVF | 0.7726701 |
| NEPTAAA | ASSQEDRENTEVF | 0.8189126 |
| NEPTAAALA | ASSQEDRENTEVF | 0.82114613 |
| GSIINISSI | ASSQEDRENTEVF | 0.9147632 |
| VAPGNAGTA | ASSQEDRENTEVF | 0.87187153 |
| TGGGASLE | ASSQEDRENTEVF | 0.8710186 |
| VAVYNHYKRI | ASSQEDRENTEVF | 0.7495891 |

| | | |
|---|---|---|
| TDPAPATT | ASSQEDRENTEVF | 0.8361523 |
| TNRPDILDPALLRPG | ASSQEDRENTEVF | 0.87554806 |
| LTTTTTS | ASSQEDRENTEVF | 0.5873921 |
| PTGSGKTL | ASSQEDRENTEVF | 0.9043424 |
| TGSGKTL | ASSQEDRENTEVF | 0.77271557 |
| TIGHVDHGK | ASSQEDRENTEVF | 0.80750597 |
| TIGHVDHGKTT | ASSQEDRENTEVF | 0.88906705 |
| GHVDHGKTT | ASSQEDRENTEVF | 0.8989022 |
| AQFEHTLL | ASSQEDRENTEVF | 0.8054918 |
| IVNQDEL | ASSQEDRENTEVF | 0.78349066 |
| TNGSQFF | ASSQEDRENTEVF | 0.7248816 |
| TGKGIVE | ASSQEDRENTEVF | 0.87891 |
| LISNASDA | ASSQEDRENTEVF | 0.89949155 |
| AELRTLGELTKIAKAHGAQ | ASSQEDRENTEVF | 0.73340017 |
| APNAVAAAEPSW | ASSQEDRENTEVF | 0.8998101 |
| APQVAGAATAVL | ASSQEDRENTEVF | 0.85465425 |
| ARRDPAAAAAILPTDA | ASSQEDRENTEVF | 0.36595857 |
| DAAARDVLAEAGLAEYF | ASSQEDRENTEVF | 0.8294678 |
| DDQIVLSAKANLAKVEQ | ASSQEDRENTEVF | 0.89507675 |
| DELLPDTGEDSDEDGH | ASSQEDRENTEVF | 0.8921173 |
| ELPEAPANRATVPQD | ASSQEDRENTEVF | 0.91096956 |
| FGQSKIERVTGIG | ASSQEDRENTEVF | 0.8597344 |
| GAGGTGSTAGGAGGAGGAGGLY | ASSQEDRENTEVF | 0.8064548 |
| GHGGTGAAGGNGGTGGAGGSGI | ASSQEDRENTEVF | 0.9062556 |
| GNGGNGGAGGEGGPGGAGGAGA | ASSQEDRENTEVF | 0.90981674 |
| GSTAGNGGNGGNATNGTIAGQP | ASSQEDRENTEVF | 0.8958858 |
| GTGGDGGNGGNGGTGG | ASSQEDRENTEVF | 0.7660159 |
| GVRRPAGIPADAPV | ASSQEDRENTEVF | 0.60130906 |
| GYRQVTDAHLLTLARR | ASSQEDRENTEVF | 0.8432499 |
| IAKMGQDGHDRG | ASSQEDRENTEVF | 0.8855721 |
| KAEPSIEIQYT | ASSQEDRENTEVF | 0.6183935 |
| KPVRIRMAADEDTVAVT | ASSQEDRENTEVF | 0.86186856 |
| MPPRIAGMRLLVIKPEP | ASSQEDRENTEVF | 0.8661917 |
| MKFYAVALFRSRGVPPTR | ASSQEDRENTEVF | 0.41509974 |
| NAGGFGILGAVAHSP | ASSQEDRENTEVF | 0.879159 |
| NEGREITLAMAEM | ASSQEDRENTEVF | 0.9022781 |
| NELVEWSHPLGHRWRWE | ASSQEDRENTEVF | 0.8128625 |
| NLGAGGTGGNGGNPG | ASSQEDRENTEVF | 0.8039144 |
| NTEWPKWETVMAADPVD | ASSQEDRENTEVF | 0.89750594 |
| PASAAPAETQP | ASSQEDRENTEVF | 0.9141762 |
| QQSSDGKNGVRAAEITGE | ASSQEDRENTEVF | 0.89978886 |
| RAPDAVTTSWWKE | ASSQEDRENTEVF | 0.46727857 |
| SGFGNVGAHTSGWFNQ | ASSQEDRENTEVF | 0.30173957 |
| SGVAGAVGQAASVGGLKV | ASSQEDRENTEVF | 0.75379133 |
| SNGGAGTGSAGGNGGT | ASSQEDRENTEVF | 0.87619555 |
| TGGAGGEGGAAG | ASSQEDRENTEVF | 0.8084904 |
| TGGDGGAAGAGGAPGHG | ASSQEDRENTEVF | 0.7596599 |
| TMPPLLAAIANAGRHGV | ASSQEDRENTEVF | 0.86962163 |
| TSSTAGNSGGA | ASSQEDRENTEVF | 0.9020261 |
| VPATVPVGILC | ASSQEDRENTEVF | 0.6562189 |
| VQRPDSGGVVLTGRIS | ASSQEDRENTEVF | 0.3649554 |

| | | |
|---|---|---|
| WTPNPVPATKGVPAGLVQAT | ASSQEDRENTEVF | 0.8717952 |
| YALKRALAKFSDRFDVVII | ASSQEDRENTEVF | 0.88535064 |
| YLIGSVIV | ASSQEDRENTEVF | 0.76406235 |
| YGQLVAVID | ASSQEDRENTEVF | 0.82497144 |
| VTVGGMTLPA | ASSQEDRENTEVF | 0.7241629 |
| VLALPAV | ASSQEDRENTEVF | 0.6485619 |
| VIPPTLNYET | ASSQEDRENTEVF | 0.8716316 |
| VGMGGMHPSSG | ASSQEDRENTEVF | 0.7112301 |
| VGLAGGAATA | ASSQEDRENTEVF | 0.9065946 |
| VGGAGGAGGSGP | ASSQEDRENTEVF | 0.7669742 |
| VFAAFMV | ASSQEDRENTEVF | 0.86970735 |
| TLYDKLR | ASSQEDRENTEVF | 0.87111974 |
| TLAVFTVL | ASSQEDRENTEVF | 0.87238735 |
| TGSVQPARV | ASSQEDRENTEVF | 0.8253508 |
| TGGGAGSGGT | ASSQEDRENTEVF | 0.8158431 |
| SHLETSL | ASSQEDRENTEVF | 0.615493 |
| SFTTGMT | ASSQEDRENTEVF | 0.3416616 |
| QVEVAKRMGVRQPTVS | ASSQEDRENTEVF | 0.8948544 |
| QGSGGPALAM | ASSQEDRENTEVF | 0.15616564 |
| MLAPSNP | ASSQEDRENTEVF | 0.898399 |
| LVNNAAI | ASSQEDRENTEVF | 0.8782537 |
| LTGEINTLA | ASSQEDRENTEVF | 0.67271674 |
| LSIFGPV | ASSQEDRENTEVF | 0.849725 |
| LLALLANL | ASSQEDRENTEVF | 0.6049676 |
| LISNASDALDKLR | ASSQEDRENTEVF | 0.61961484 |
| LINEAVP | ASSQEDRENTEVF | 0.8670466 |
| LGYAFSGL | ASSQEDRENTEVF | 0.8730561 |
| LDFVAVAL | ASSQEDRENTEVF | 0.7875196 |
| LAQPGAE | ASSQEDRENTEVF | 0.8525683 |
| ISNLGAVTGF | ASSQEDRENTEVF | 0.868113 |
| HQEDPGRI | ASSQEDRENTEVF | 0.7679031 |
| GTPGEPTP | ASSQEDRENTEVF | 0.8366641 |
| GFFNSGS | ASSQEDRENTEVF | 0.9020176 |
| GAGGGGLTAGAN | ASSQEDRENTEVF | 0.84627974 |
| FVDALVNM | ASSQEDRENTEVF | 0.7262658 |
| FLTVAPEL | ASSQEDRENTEVF | 0.88404953 |
| FETLTAL | ASSQEDRENTEVF | 0.7280651 |
| DVINPIEL | ASSQEDRENTEVF | 0.8813041 |
| APVSGAPV | ASSQEDRENTEVF | 0.821387 |
| ALVGGLAW | ASSQEDRENTEVF | 0.8091593 |
| ALAAAAATAI | ASSQEDRENTEVF | 0.87335014 |
| AFGLGLLL | ASSQEDRENTEVF | 0.494936 |
| AFFAAASL | ASSQEDRENTEVF | 0.7407104 |
| AAAALPAFSPP | ASSQEDRENTEVF | 0.79312205 |
| VVLAPAL | ASSQEDRENTEVF | 0.8570815 |
| TIGHVDHGKTTL | ASSQEDRENTEVF | 0.895627 |
| LISNASDALDK | ASSQEDRENTEVF | 0.71800417 |
| LISNASDAL | ASSQEDRENTEVF | 0.8637835 |
| ISNASDALDK | ASSQEDRENTEVF | 0.7326244 |
| NQAETLVYQTE | ASSQEDRENTEVF | 0.61156595 |
| TGGTGGTAGAG | ASSQEDRENTEVF | 0.8046739 |

| | | |
|---|---|---|
| TGSTAGGAGGA | ASSQEDRENTEVF | 0.75850075 |
| DPLAVPVAAGPV | ASSQEDRENTEVF | 0.755005 |
| GGTGGTPGNGGN | ASSQEDRENTEVF | 0.8737651 |
| PAGLAGVDPATL | ASSQEDRENTEVF | 0.55446506 |
| GNNLVVEAPHAGASV | ASSQEDRENTEVF | 0.85930216 |
| KVTRIMSASTLEQPA | ASSQEDRENTEVF | 0.899993 |
| LGFALPIALPRRTVV | ASSQEDRENTEVF | 0.7554153 |
| TRLGNLTARKLVVLV | ASSQEDRENTEVF | 0.8632251 |
| WSQQRRLIYFPSAGP | ASSQEDRENTEVF | 0.8796047 |
| GPNGNPDPMAAAVDIR | ASSQEDRENTEVF | 0.8712325 |
| TQRCVSVADLRAAASR | ASSQEDRENTEVF | 0.815279 |
| QFVLDVAEHPVEVLTETV | ASSQEDRENTEVF | 0.81667554 |
| GGAGGTGGAGGNGGTQAGDG | ASSQEDRENTEVF | 0.9064888 |
| PLLATLRAIAVDVDAKPAQV | ASSQEDRENTEVF | 0.88570136 |
| APKFRTGGYMIDFWGVGYQVAK | ASSQEDRENTEVF | 0.8887189 |
| GAGGAGGAGGAAGGPGGTGGTG | ASSQEDRENTEVF | 0.88270754 |
| IGALGLLATDVAPAGSVAFGVA | ASSQEDRENTEVF | 0.8758068 |
| AGGAGGIGGNGGAGGAGGVLMG | ASSQEDRENTEVF | 0.8850004 |
| ADELIGLL | ASSQEDRENTEVF | 0.8198655 |
| ADRALKA | ASSQEDRENTEVF | 0.5290362 |
| AGAVGQVGGAGGA | ASSQEDRENTEVF | 0.7937948 |
| AGGGGGLGGAP | ASSQEDRENTEVF | 0.84678614 |
| AGGLLYGNGGAG | ASSQEDRENTEVF | 0.80239767 |
| AGLESIGL | ASSQEDRENTEVF | 0.3696427 |
| AGLKTGII | ASSQEDRENTEVF | 0.90208954 |
| ASLAALLL | ASSQEDRENTEVF | 0.9149349 |
| ASVINISSI | ASSQEDRENTEVF | 0.76404995 |
| ATPPGPA | ASSQEDRENTEVF | 0.90854204 |
| ATVASALV | ASSQEDRENTEVF | 0.62343264 |
| ATVASLIL | ASSQEDRENTEVF | 0.8757829 |
| AVNGPVLIRH | ASSQEDRENTEVF | 0.89951175 |
| DGAAGAVGI | ASSQEDRENTEVF | 0.73467696 |
| DPLLHGI | ASSQEDRENTEVF | 0.8915434 |
| ELGAALGAL | ASSQEDRENTEVF | 0.7118553 |
| ELGRTQAV | ASSQEDRENTEVF | 0.7867885 |
| FVAGGYLS | ASSQEDRENTEVF | 0.9017776 |
| GASGGAGGAGGW | ASSQEDRENTEVF | 0.8170333 |
| GGAGAGGDGGAGGI | ASSQEDRENTEVF | 0.85093856 |
| GTGHNGSQPGMGGNG | ASSQEDRENTEVF | 0.79832685 |
| GYAAASASAS | ASSQEDRENTEVF | 0.6023707 |
| HQVDVAAVQQG | ASSQEDRENTEVF | 0.8437595 |
| IGAAIAHT | ASSQEDRENTEVF | 0.9025358 |
| IGMIEVAQL | ASSQEDRENTEVF | 0.6020608 |
| IRLAPEHTSAMD | ASSQEDRENTEVF | 0.90923977 |
| IYNVSVL | ASSQEDRENTEVF | 0.7761048 |
| KQGFLGM | ASSQEDRENTEVF | 0.91605043 |
| LESVITQL | ASSQEDRENTEVF | 0.88902736 |
| LGLGALVVV | ASSQEDRENTEVF | 0.85775954 |
| LIGNGGNGGSGGI | ASSQEDRENTEVF | 0.76889336 |
| LNAINAPTL | ASSQEDRENTEVF | 0.90324056 |
| LQLLNTGV | ASSQEDRENTEVF | 0.8208674 |

| | | |
|---|---|---|
| LQTSIEF | ASSQEDRENTEVF | 0.86294997 |
| MSFAVSEL | ASSQEDRENTEVF | 0.90418875 |
| NTGALGL | ASSQEDRENTEVF | 0.35764897 |
| PPGAGGAGGAGTT | ASSQEDRENTEVF | 0.59204566 |
| QLVGLGI | ASSQEDRENTEVF | 0.5098331 |
| RFLEQVHQL | ASSQEDRENTEVF | 0.48699573 |
| RGVGYVLR | ASSQEDRENTEVF | 0.45134664 |
| SATGVFAGVI | ASSQEDRENTEVF | 0.37270835 |
| SLPAITLP | ASSQEDRENTEVF | 0.4138138 |
| STALAFTV | ASSQEDRENTEVF | 0.43263614 |
| SVLTVGLGK | ASSQEDRENTEVF | 0.5892417 |
| TALAGAGSGI | ASSQEDRENTEVF | 0.5023388 |
| TGLPFTGI | ASSQEDRENTEVF | 0.43399328 |
| VAEHVTV | ASSQEDRENTEVF | 0.56846666 |
| VDIVLNSL | ASSQEDRENTEVF | 0.5662795 |
| VLTGGGAVL | ASSQEDRENTEVF | 0.37403893 |
| VNVGLAYGI | ASSQEDRENTEVF | 0.45558947 |
| VSSAAVGAV | ASSQEDRENTEVF | 0.39757967 |
| VTSVYEAGV | ASSQEDRENTEVF | 0.44618958 |
| WVALDRGA | ASSQEDRENTEVF | 0.3338578 |
| YLVGSVVI | ASSQEDRENTEVF | 0.5119087 |
| GPPGTGKTLL | ASSQEDRENTEVF | 0.34705913 |
| GVGMTKF | ASSQEDRENTEVF | 0.31987292 |
| IGIGGSDL | ASSQEDRENTEVF | 0.52142435 |
| IGTLLAEL | ASSQEDRENTEVF | 0.40067676 |
| IYTSGTT | ASSQEDRENTEVF | 0.44303864 |
| LSFPAGV | ASSQEDRENTEVF | 0.5023253 |
| LYTSGTT | ASSQEDRENTEVF | 0.42927653 |
| VGAFPFGF | ASSQEDRENTEVF | 0.41146567 |
| VGAGTVEF | ASSQEDRENTEVF | 0.53476775 |
| VGKSTLLL | ASSQEDRENTEVF | 0.52649087 |
| AALAHAGGGIV | ASSQEDRENTEVF | 0.40151036 |
| AALVAVSLL | ASSQEDRENTEVF | 0.42479303 |
| AAVSRAVAAL | ASSQEDRENTEVF | 0.57291 |
| AGGSSGIGGF | ASSQEDRENTEVF | 0.48819804 |
| AGLTVANANG | ASSQEDRENTEVF | 0.50411576 |
| AGLVGAVL | ASSQEDRENTEVF | 0.51398367 |
| ALSQAFTV | ASSQEDRENTEVF | 0.43384254 |
| ALTFSVSL | ASSQEDRENTEVF | 0.54099727 |
| ALTGAALVAL | ASSQEDRENTEVF | 0.5594127 |
| ASFSIGAGVL | ASSQEDRENTEVF | 0.46768185 |
| DGSVYKI | ASSQEDRENTEVF | 0.47454625 |
| EQLAEIV | ASSQEDRENTEVF | 0.34236792 |
| FAVSELTVM | ASSQEDRENTEVF | 0.47141415 |
| GAAPLAAGL | ASSQEDRENTEVF | 0.40982017 |
| GADGGDGGSGGAGG | ASSQEDRENTEVF | 0.5087837 |
| GFGLDTGGAGG | ASSQEDRENTEVF | 0.5733789 |
| GLASGGLL | ASSQEDRENTEVF | 0.48495817 |
| GRIAGAAVLL | ASSQEDRENTEVF | 0.3412884 |
| GSAGSLNTSF | ASSQEDRENTEVF | 0.3580588 |
| GTGQIGI | ASSQEDRENTEVF | 0.5116743 |

| | | |
|---|---|---|
| GTLIALIL | ASSQEDRENTEVF | 0.34087747 |
| IAATMGFVVG | ASSQEDRENTEVF | 0.3521122 |
| IFAQALL | ASSQEDRENTEVF | 0.4284183 |
| IGAAVALGF | ASSQEDRENTEVF | 0.59196997 |
| IGLTPLAL | ASSQEDRENTEVF | 0.573199 |
| ILGSNTSTL | ASSQEDRENTEVF | 0.47588134 |
| IVFSLTVL | ASSQEDRENTEVF | 0.5313206 |
| LASKATGF | ASSQEDRENTEVF | 0.5490328 |
| LASVALIL | ASSQEDRENTEVF | 0.53709316 |
| LFAVGLTL | ASSQEDRENTEVF | 0.41220024 |
| LFNGGIGTYI | ASSQEDRENTEVF | 0.37817678 |
| LTLGAGII | ASSQEDRENTEVF | 0.56408644 |
| LVAAALGDI | ASSQEDRENTEVF | 0.4533981 |
| NTAFVQLGI | ASSQEDRENTEVF | 0.58146346 |
| QLFSGAEL | ASSQEDRENTEVF | 0.5533219 |
| SAATFITL | ASSQEDRENTEVF | 0.45907694 |
| SNYGLANLGV | ASSQEDRENTEVF | 0.3953147 |
| SYGGAYAVM | ASSQEDRENTEVF | 0.52435005 |
| TTGVGGLTLG | ASSQEDRENTEVF | 0.54558694 |
| TTQTIYI | ASSQEDRENTEVF | 0.4032481 |
| VAAGAGLGAVI | ASSQEDRENTEVF | 0.38857412 |
| VAAVVTLL | ASSQEDRENTEVF | 0.48489064 |
| VGFLGGLI | ASSQEDRENTEVF | 0.47434533 |
| VGVIPILL | ASSQEDRENTEVF | 0.5640747 |
| VIFASSLI | ASSQEDRENTEVF | 0.4734512 |
| VKGGDGGAAAG | ASSQEDRENTEVF | 0.4143226 |
| VTKTFTNI | ASSQEDRENTEVF | 0.48023835 |
| VVAAAILL | ASSQEDRENTEVF | 0.52528083 |
| YAALDVEL | ASSQEDRENTEVF | 0.50203353 |
| LASVALLL | ASSQEDRENTEVF | 0.47302294 |
| LDAVHRV | ASSQEDRENTEVF | 0.44573998 |
| QGLSSSIF | ASSQEDRENTEVF | 0.52550364 |
| RGTFANIRL | ASSQEDRENTEVF | 0.5346468 |
| RQLWWGHRI | ASSQEDRENTEVF | 0.4411245 |
| YGPPGTGKTLL | ASSQEDRENTEVF | 0.49428815 |
| AALVGAPANVA | ASSQEDRENTEVF | 0.47618836 |
| AATFRRTE | ASSQEDRENTEVF | 0.4591714 |
| AGALGGGI | ASSQEDRENTEVF | 0.39884603 |
| AGGAGGAGGIGAG | ASSQEDRENTEVF | 0.39695966 |
| AGSGALGL | ASSQEDRENTEVF | 0.42400187 |
| AQIADQI | ASSQEDRENTEVF | 0.48860943 |
| ASAVAVTYL | ASSQEDRENTEVF | 0.5172883 |
| ASTVLGGTGG | ASSQEDRENTEVF | 0.4689536 |
| AVGGEPAK | ASSQEDRENTEVF | 0.43317765 |
| CEQGTFGGTC | ASSQEDRENTEVF | 0.5495355 |
| DEGKAPAG | ASSQEDRENTEVF | 0.53594494 |
| DTVVFPAL | ASSQEDRENTEVF | 0.38327166 |
| FEGITMAG | ASSQEDRENTEVF | 0.468093 |
| FPLVVQD | ASSQEDRENTEVF | 0.6081421 |
| GAFTTPQ | ASSQEDRENTEVF | 0.49628466 |
| GFGGFGTAGGVGGA | ASSQEDRENTEVF | 0.45027703 |

| | | |
|---|---|---|
| GFLAQFL | ASSQEDRENTEVF | 0.42187408 |
| GGAGGAGGNAGVA | ASSQEDRENTEVF | 0.6037228 |
| GGVGTAGAGGAGGAGG | ASSQEDRENTEVF | 0.47459307 |
| GNGGGGDAGNAGSGGN | ASSQEDRENTEVF | 0.5880522 |
| GTGGAGGAGNDAGS | ASSQEDRENTEVF | 0.35004997 |
| GTTNTGWFNS | ASSQEDRENTEVF | 0.38165447 |
| IFMGPLFTL | ASSQEDRENTEVF | 0.4101218 |
| ILAHAIEAI | ASSQEDRENTEVF | 0.31917658 |
| IRSAIREAL | ASSQEDRENTEVF | 0.58308494 |
| ITFAGGG | ASSQEDRENTEVF | 0.3882671 |
| ITGMSGHQEV | ASSQEDRENTEVF | 0.4955756 |
| IVAGLACTL | ASSQEDRENTEVF | 0.391943 |
| IVGGGLAGA | ASSQEDRENTEVF | 0.55914754 |
| IVTGVLNI | ASSQEDRENTEVF | 0.49955928 |
| LAQQGCI | ASSQEDRENTEVF | 0.48630154 |
| LCLHVEN | ASSQEDRENTEVF | 0.41294613 |
| LDAISDGS | ASSQEDRENTEVF | 0.54565555 |
| LGGFAAML | ASSQEDRENTEVF | 0.5475195 |
| LGGKNPMI | ASSQEDRENTEVF | 0.49919242 |
| LGPKGVAAIL | ASSQEDRENTEVF | 0.36553097 |
| LMVGVSI | ASSQEDRENTEVF | 0.52985436 |
| LVGAAFVL | ASSQEDRENTEVF | 0.5017406 |
| LVLAVAPL | ASSQEDRENTEVF | 0.39055383 |
| MAAIAVIDAA | ASSQEDRENTEVF | 0.33590433 |
| NNGINAELQ | ASSQEDRENTEVF | 0.43968436 |
| RGEPLVLVHG | ASSQEDRENTEVF | 0.42710644 |
| RGGAGGAAGLI | ASSQEDRENTEVF | 0.5370019 |
| RSYRMHPAV | ASSQEDRENTEVF | 0.56367195 |
| SAAHLLAGQLA | ASSQEDRENTEVF | 0.41263536 |
| SAAVGSY | ASSQEDRENTEVF | 0.4757538 |
| SPHAVLHRI | ASSQEDRENTEVF | 0.5269974 |
| SQIAKVL | ASSQEDRENTEVF | 0.30303144 |
| TALGHLVP | ASSQEDRENTEVF | 0.3849436 |
| TAPTTSGAA | ASSQEDRENTEVF | 0.36562783 |
| TDGDDGNGGSPGT | ASSQEDRENTEVF | 0.53498274 |
| TDGLGEN | ASSQEDRENTEVF | 0.39044544 |
| TGGPPLPGQA | ASSQEDRENTEVF | 0.475778 |
| TIPAITI | ASSQEDRENTEVF | 0.49010044 |
| TIPGFTI | ASSQEDRENTEVF | 0.52635884 |
| TSIASGALA | ASSQEDRENTEVF | 0.48257512 |
| TVTLPGAAI | ASSQEDRENTEVF | 0.37235695 |
| VASGLSAALP | ASSQEDRENTEVF | 0.40486106 |
| VGAGVIGI | ASSQEDRENTEVF | 0.5569267 |
| VGPKRFGV | ASSQEDRENTEVF | 0.525803 |
| VNVAFPD | ASSQEDRENTEVF | 0.40893435 |
| VSMTNSGVS | ASSQEDRENTEVF | 0.41664368 |
| VTDVLGDLSV | ASSQEDRENTEVF | 0.57969016 |
| VVVVGAGFAGL | ASSQEDRENTEVF | 0.34335226 |
| VYTVGTVV | ASSQEDRENTEVF | 0.4575807 |
| WTTFSGP | ASSQEDRENTEVF | 0.35652885 |
| YDELVEAF | ASSQEDRENTEVF | 0.47061446 |

| | | |
|---|---|---|
| YGTGGAGGNGG | ASSQEDRENTEVF | 0.5959555 |
| YTNVMAV | ASSQEDRENTEVF | 0.5644659 |
| FLEQVHQL | ASSQEDRENTEVF | 0.47288325 |
| GLSLLAVAV | ASSQEDRENTEVF | 0.45035103 |
| LEQVHQL | ASSQEDRENTEVF | 0.3819913 |
| LNAAGAI | ASSQEDRENTEVF | 0.42779052 |
| PAVHASL | ASSQEDRENTEVF | 0.42450598 |
| TELEDDA | ASSQEDRENTEVF | 0.5775915 |
| TPPGPPN | ASSQEDRENTEVF | 0.44342798 |
| TPPPPPPP | ASSQEDRENTEVF | 0.373117 |
| TQSPALI | ASSDEGQGLNERLF | 0.4984301 |
| EFAAAFPQ | ASSDEGQGLNERLF | 0.57378733 |
| LLSGNLDVL | ASSDEGQGLNERLF | 0.5695508 |
| GPANLGNYNI | ASSDEGQGLNERLF | 0.40840223 |
| TLGPTLP | ASSDEGQGLNERLF | 0.40963656 |
| VYAYSSCCK | ASSDEGQGLNERLF | 0.3978294 |
| APAPAPAPNI | ASSDEGQGLNERLF | 0.4904322 |
| PGGPVPIP | ASSDEGQGLNERLF | 0.35607418 |
| FLTVALEP | ASSDEGQGLNERLF | 0.4028029 |
| SQYFGANL | ASSDEGQGLNERLF | 0.51823384 |
| GGVKPAA | ASSDEGQGLNERLF | 0.62515557 |
| GGTGGAGGAGGW | ASSDEGQGLNERLF | 0.46095705 |
| GVGGVAGLGGAGG | ASSDEGQGLNERLF | 0.32671115 |
| VLLVLATTI | ASSDEGQGLNERLF | 0.4828499 |
| IGIFAAML | ASSDEGQGLNERLF | 0.3762746 |
| AVGDLID | ASSDEGQGLNERLF | 0.4049098 |
| NISPLPL | ASSDEGQGLNERLF | 0.4025111 |
| VTAVFVAI | ASSDEGQGLNERLF | 0.3434808 |
| SGDDNDAAL | ASSDEGQGLNERLF | 0.4415023 |
| AVAALAHS | ASSDEGQGLNERLF | 0.600083 |
| GGVTVMATP | ASSDEGQGLNERLF | 0.4820898 |
| AGLAANVRD | ASSDEGQGLNERLF | 0.4205517 |
| TNRPDLIDPALLRPG | ASSDEGQGLNERLF | 0.5762354 |
| LANLGVTAL | ASSDEGQGLNERLF | 0.5150839 |
| TGERIEL | ASSDEGQGLNERLF | 0.5423718 |
| NGEELAA | ASSDEGQGLNERLF | 0.52455026 |
| VLSLVWG | ASSDEGQGLNERLF | 0.52086115 |
| IGYALWL | ASSDEGQGLNERLF | 0.4477395 |
| LSLLLAPL | ASSDEGQGLNERLF | 0.56581783 |
| TLDEALK | ASSDEGQGLNERLF | 0.47006506 |
| TDGAGTAEGG | ASSDEGQGLNERLF | 0.49934345 |
| NAPGGAGGNGG | ASSDEGQGLNERLF | 0.40893018 |
| VSVFMGMGGNF | ASSDEGQGLNERLF | 0.56788063 |
| LASYLAAL | ASSDEGQGLNERLF | 0.40465063 |
| LVGGTPAL | ASSDEGQGLNERLF | 0.5790394 |
| VRNADVL | ASSDEGQGLNERLF | 0.3204047 |
| TCSSPFLVSAM | ASSDEGQGLNERLF | 0.3956609 |
| VHITDAE | ASSDEGQGLNERLF | 0.55726504 |
| EYAAPSF | ASSDEGQGLNERLF | 0.3197293 |
| VVDAFHGGL | ASSDEGQGLNERLF | 0.34200612 |
| ASVVDPQ | ASSDEGQGLNERLF | 0.4160758 |

| | | |
|---|---|---|
| SYNEFGGLS | ASSDEGQGLNERLF | 0.56027144 |
| DPDGSVY | ASSDEGQGLNERLF | 0.32188326 |
| MVSTIHATE | ASSDEGQGLNERLF | 0.45782933 |
| LAAGVADPV | ASSDEGQGLNERLF | 0.3457185 |
| TKEQIAA | ASSDEGQGLNERLF | 0.51933 |
| SLASPAAL | ASSDEGQGLNERLF | 0.46272832 |
| IGLVFYGV | ASSDEGQGLNERLF | 0.35640076 |
| FLLDASNL | ASSDEGQGLNERLF | 0.52383006 |
| YQVGNFP | ASSDEGQGLNERLF | 0.35892084 |
| ESLLSQST | ASSDEGQGLNERLF | 0.5430523 |
| LFTVGFAAVG | ASSDEGQGLNERLF | 0.5257377 |
| ATPPTPT | ASSDEGQGLNERLF | 0.5803835 |
| TDLATLG | ASSDEGQGLNERLF | 0.35245243 |
| SAGGTGR | ASSDEGQGLNERLF | 0.45809564 |
| GFSNSGS | ASSDEGQGLNERLF | 0.38372135 |
| GGNGGSGGTSV | ASSDEGQGLNERLF | 0.6137326 |
| GTSVATGGAGN | ASSDEGQGLNERLF | 0.45850226 |
| NGGNGGSGGTS | ASSDEGQGLNERLF | 0.3298769 |
| AVVKFPH | ASSDEGQGLNERLF | 0.53412294 |
| TPPPPPP | ASSDEGQGLNERLF | 0.46505845 |
| AAPAAAAAPA | ASSDEGQGLNERLF | 0.45495036 |
| IALNEDL | ASSDEGQGLNERLF | 0.5282586 |
| IVNVSSM | ASSDEGQGLNERLF | 0.41234365 |
| TGIPPAPR | ASSDEGQGLNERLF | 0.44733885 |
| FDLGGGTF | ASSDEGQGLNERLF | 0.543651 |
| FDLGGGTFDVS | ASSDEGQGLNERLF | 0.5080904 |
| NEPTAAA | ASSDEGQGLNERLF | 0.43336886 |
| NEPTAAALA | ASSDEGQGLNERLF | 0.5221576 |
| GSIINISSI | ASSDEGQGLNERLF | 0.53324836 |
| VAPGNAGTA | ASSDEGQGLNERLF | 0.38521448 |
| TGGGASLE | ASSDEGQGLNERLF | 0.6055876 |
| VAVYNHYKRI | ASSDEGQGLNERLF | 0.3734942 |
| TDPAPATT | ASSDEGQGLNERLF | 0.50880724 |
| TNRPDILDPALLRPG | ASSDEGQGLNERLF | 0.53514403 |
| LTTTTTS | ASSDEGQGLNERLF | 0.59941775 |
| PTGSGKTL | ASSDEGQGLNERLF | 0.39786285 |
| TGSGKTL | ASSDEGQGLNERLF | 0.46423882 |
| TIGHVDHGK | ASSDEGQGLNERLF | 0.4780863 |
| TIGHVDHGKTT | ASSDEGQGLNERLF | 0.48018777 |
| GHVDHGKTT | ASSDEGQGLNERLF | 0.37944293 |
| AQFEHTLL | ASSDEGQGLNERLF | 0.56529915 |
| IVNQDEL | ASSDEGQGLNERLF | 0.5214718 |
| TNGSQFF | ASSDEGQGLNERLF | 0.56574464 |
| TGKGIVE | ASSDEGQGLNERLF | 0.4374432 |
| LISNASDA | ASSDEGQGLNERLF | 0.45993897 |
| AELRTLGELTKIAKAHGAQ | ASSDEGQGLNERLF | 0.5911439 |
| APNAVAAAEPSW | ASSDEGQGLNERLF | 0.35654244 |
| APQVAGAATAVL | ASSDEGQGLNERLF | 0.5241799 |
| ARRDPAAAAAILPTDA | ASSDEGQGLNERLF | 0.518273 |
| DAAARDVLAEAGLAEYF | ASSDEGQGLNERLF | 0.53669155 |
| DDQIVLSAKANLAKVEQ | ASSDEGQGLNERLF | 0.41671383 |

| | | |
|---|---|---|
| DELLPDTGEDSDEDGH | ASSDEGQGLNERLF | 0.56486577 |
| ELPEAPANRATVPQD | ASSDEGQGLNERLF | 0.41215998 |
| FGQSKIERVTGIG | ASSDEGQGLNERLF | 0.54872024 |
| GAGGTGSTAGGAGGAGGAGGLY | ASSDEGQGLNERLF | 0.42145878 |
| GHGGTGAAGGNGGTGGAGGSGI | ASSDEGQGLNERLF | 0.39697096 |
| GNGGNGGAGGEGGPGGAGGAGA | ASSDEGQGLNERLF | 0.42747626 |
| GSTAGNGGNGGNATNGTIAGQP | ASSDEGQGLNERLF | 0.52407676 |
| GTGGDGGNGGNGGTGG | ASSDEGQGLNERLF | 0.51567984 |
| GVRRPAGIPADAPV | ASSDEGQGLNERLF | 0.5397037 |
| GYRQVTDAHLLTLARR | ASSDEGQGLNERLF | 0.35675183 |
| IAKMGQDGHDRG | ASSDEGQGLNERLF | 0.5308262 |
| KAEPSIEIQYT | ASSDEGQGLNERLF | 0.56973577 |
| KPVRIRMAADEDTVAVT | ASSDEGQGLNERLF | 0.59517086 |
| MPPRIAGMRLLVIKPEP | ASSDEGQGLNERLF | 0.42311436 |
| MKFYAVALFRSRGVPPTR | ASSDEGQGLNERLF | 0.49250218 |
| NAGGFGILGAVAHSP | ASSDEGQGLNERLF | 0.49185488 |
| NEGREITLAMAEM | ASSDEGQGLNERLF | 0.4688223 |
| NELVEWSHPLGHRWRWE | ASSDEGQGLNERLF | 0.34118265 |
| NLGAGGTGGNGGNPG | ASSDEGQGLNERLF | 0.3752117 |
| NTEWPKWETVMAADPVD | ASSDEGQGLNERLF | 0.44482562 |
| PASAAPAETQP | ASSDEGQGLNERLF | 0.42791462 |
| QQSSDGKNGVRAAEITGE | ASSDEGQGLNERLF | 0.5800428 |
| RAPDAVTTSWWKE | ASSDEGQGLNERLF | 0.52890646 |
| SGFGNVGAHTSGWFNQ | ASSDEGQGLNERLF | 0.53664756 |
| SGVAGAVGQAASVGGLKV | ASSDEGQGLNERLF | 0.5537071 |
| SNGGAGTGSAGGNGGT | ASSDEGQGLNERLF | 0.55231124 |
| TGGAGGEGGAAG | ASSDEGQGLNERLF | 0.44016847 |
| TGGDGGAAGAGGAPGHG | ASSDEGQGLNERLF | 0.47164595 |
| TMPPLLAAIANAGRHGV | ASSDEGQGLNERLF | 0.42324823 |
| TSSTAGNSGGA | ASSDEGQGLNERLF | 0.35114613 |
| VPATVPVGILC | ASSDEGQGLNERLF | 0.37833259 |
| VQRPDSGGVVLTGRIS | ASSDEGQGLNERLF | 0.46092612 |
| WTPNPVPATKGVPAGLVQAT | ASSDEGQGLNERLF | 0.44812214 |
| YALKRALAKFSDRFDVVII | ASSDEGQGLNERLF | 0.31422108 |
| YLIGSVIV | ASSDEGQGLNERLF | 0.5319318 |
| YGQLVAVID | ASSDEGQGLNERLF | 0.5712249 |
| VTVGGMTLPA | ASSDEGQGLNERLF | 0.5278927 |
| VLALPAV | ASSDEGQGLNERLF | 0.52639127 |
| VIPPTLNYET | ASSDEGQGLNERLF | 0.45883662 |
| VGMGGMHPSSG | ASSDEGQGLNERLF | 0.38275325 |
| VGLAGGAATA | ASSDEGQGLNERLF | 0.50091225 |
| VGGAGGAGGSGP | ASSDEGQGLNERLF | 0.40813404 |
| VFAAFMV | ASSDEGQGLNERLF | 0.49892747 |
| TLYDKLR | ASSDEGQGLNERLF | 0.5551547 |
| TLAVFTVL | ASSDEGQGLNERLF | 0.5166521 |
| TGSVQPARV | ASSDEGQGLNERLF | 0.3880262 |
| TGGGAGSGGT | ASSDEGQGLNERLF | 0.44396096 |
| SHLETSL | ASSDEGQGLNERLF | 0.40754572 |
| SFTTGMT | ASSDEGQGLNERLF | 0.41600773 |
| QVEVAKRMGVRQPTVS | ASSDEGQGLNERLF | 0.42993134 |
| QGSGGPALAM | ASSDEGQGLNERLF | 0.48389953 |

| | | |
|---|---|---|
| MLAPSNP | ASSDEGQGLNERLF | 0.41360432 |
| LVNNAAI | ASSDEGQGLNERLF | 0.55039257 |
| LTGEINTLA | ASSDEGQGLNERLF | 0.53186744 |
| LSIFGPV | ASSDEGQGLNERLF | 0.4781229 |
| LLALLANL | ASSDEGQGLNERLF | 0.40032807 |
| LISNASDALDKLR | ASSDEGQGLNERLF | 0.5516904 |
| LINEAVP | ASSDEGQGLNERLF | 0.57755697 |
| LGYAFSGL | ASSDEGQGLNERLF | 0.3806944 |
| LDFVAVAL | ASSDEGQGLNERLF | 0.45379424 |
| LAQPGAE | ASSDEGQGLNERLF | 0.41144064 |
| ISNLGAVTGF | ASSDEGQGLNERLF | 0.47965497 |
| HQEDPGRI | ASSDEGQGLNERLF | 0.58514154 |
| GTPGEPTP | ASSDEGQGLNERLF | 0.4471522 |
| GFFNSGS | ASSDEGQGLNERLF | 0.38649517 |
| GAGGGGLTAGAN | ASSDEGQGLNERLF | 0.40040326 |
| FVDALVNM | ASSDEGQGLNERLF | 0.48128858 |
| FLTVAPEL | ASSDEGQGLNERLF | 0.47056133 |
| FETLTAL | ASSDEGQGLNERLF | 0.32994545 |
| DVINPIEL | ASSDEGQGLNERLF | 0.54787076 |
| APVSGAPV | ASSDEGQGLNERLF | 0.38604683 |
| ALVGGLAW | ASSDEGQGLNERLF | 0.4382404 |
| ALAAAAATAI | ASSDEGQGLNERLF | 0.4453553 |
| AFGLGLLL | ASSDEGQGLNERLF | 0.53509 |
| AFFAAASL | ASSDEGQGLNERLF | 0.5131022 |
| AAAALPAFSPP | ASSDEGQGLNERLF | 0.43662995 |
| VVLAPAL | ASSDEGQGLNERLF | 0.44435266 |
| TIGHVDHGKTTL | ASSDEGQGLNERLF | 0.50391495 |
| LISNASDALDK | ASSDEGQGLNERLF | 0.5260321 |
| LISNASDAL | ASSDEGQGLNERLF | 0.5620119 |
| ISNASDALDK | ASSDEGQGLNERLF | 0.46938503 |
| NQAETLVYQTE | ASSDEGQGLNERLF | 0.54273456 |
| TGGTGGTAGAG | ASSDEGQGLNERLF | 0.38967845 |
| TGSTAGGAGGA | ASSDEGQGLNERLF | 0.4050846 |
| DPLAVPVAAGPV | ASSDEGQGLNERLF | 0.43728325 |
| GGTGGTPGNGGN | ASSDEGQGLNERLF | 0.3785643 |
| PAGLAGVDPATL | ASSDEGQGLNERLF | 0.40093178 |
| GNNLVVEAPHAGASV | ASSDEGQGLNERLF | 0.54177004 |
| KVTRIMSASTLEQPA | ASSDEGQGLNERLF | 0.4936242 |
| LGFALPIALPRRTVV | ASSDEGQGLNERLF | 0.408266 |
| TRLGNLTARKLVVLV | ASSDEGQGLNERLF | 0.33713064 |
| WSQQRRLIYFPSAGP | ASSDEGQGLNERLF | 0.4203345 |
| GPNGNPDPMAAAVDIR | ASSDEGQGLNERLF | 0.4285152 |
| TQRCVSVADLRAAASR | ASSDEGQGLNERLF | 0.5052391 |
| QFVLDVAEHPVEVLTETV | ASSDEGQGLNERLF | 0.5615094 |
| GGAGGTGGAGGNGGTQAGDG | ASSDEGQGLNERLF | 0.61229753 |
| PLLATLRAIAVDVDAKPAQV | ASSDEGQGLNERLF | 0.5941965 |
| APKFRTGGYMIDFWGVGYQVAK | ASSDEGQGLNERLF | 0.49070603 |
| GAGGAGGAGGAAGGPGGTGGTG | ASSDEGQGLNERLF | 0.5450536 |
| IGALGLLATDVAPAGSVAFGVA | ASSDEGQGLNERLF | 0.38554642 |
| AGGAGGIGGNGGAGGAGGVLMG | ASSDEGQGLNERLF | 0.43523175 |
| ADELIGLL | ASSDEGQGLNERLF | 0.46101174 |

| | | |
|---|---|---|
| ADRALKA | ASSDEGQGLNERLF | 0.48874983 |
| AGAVGQVGGAGGA | ASSDEGQGLNERLF | 0.4008761 |
| AGGGGGLGGAP | ASSDEGQGLNERLF | 0.43058568 |
| AGGLLYGNGGAG | ASSDEGQGLNERLF | 0.35105005 |
| AGLESIGL | ASSDEGQGLNERLF | 0.4845478 |
| AGLKTGII | ASSDEGQGLNERLF | 0.4110288 |
| ASLAALLL | ASSDEGQGLNERLF | 0.34492886 |
| ASVINISSI | ASSDEGQGLNERLF | 0.45733368 |
| ATPPGPA | ASSDEGQGLNERLF | 0.5480518 |
| ATVASALV | ASSDEGQGLNERLF | 0.55202144 |
| ATVASLIL | ASSDEGQGLNERLF | 0.44587985 |
| AVNGPVLIRH | ASSDEGQGLNERLF | 0.4445297 |
| DGAAGAVGI | ASSDEGQGLNERLF | 0.5439798 |
| DPLLHGI | ASSDEGQGLNERLF | 0.45970175 |
| ELGAALGAL | ASSDEGQGLNERLF | 0.47621164 |
| ELGRTQAV | ASSDEGQGLNERLF | 0.46485126 |
| FVAGGYLS | ASSDEGQGLNERLF | 0.51041853 |
| GASGGAGGAGGW | ASSDEGQGLNERLF | 0.3712936 |
| GGAGAGGDGGAGGI | ASSDEGQGLNERLF | 0.5234083 |
| GTGHNGSQPGMGGNG | ASSDEGQGLNERLF | 0.5149234 |
| GYAAASASAS | ASSDEGQGLNERLF | 0.41615444 |
| HQVDVAAVQQG | ASSDEGQGLNERLF | 0.34968406 |
| IGAAIAHT | ASSDEGQGLNERLF | 0.57147646 |
| IGMIEVAQL | ASSDEGQGLNERLF | 0.40844715 |
| IRLAPEHTSAMD | ASSDEGQGLNERLF | 0.371932 |
| IYNVSVL | ASSDEGQGLNERLF | 0.3147215 |
| KQGFLGM | ASSDEGQGLNERLF | 0.46754792 |
| LESVITQL | ASSDEGQGLNERLF | 0.46217883 |
| LGLGALVVV | ASSDEGQGLNERLF | 0.5685935 |
| LIGNGGNGGSGGI | ASSDEGQGLNERLF | 0.48906654 |
| LNAINAPTL | ASSDEGQGLNERLF | 0.5207272 |
| LQLLNTGV | ASSDEGQGLNERLF | 0.54030967 |
| LQTSIEF | ASSDEGQGLNERLF | 0.42109892 |
| MSFAVSEL | ASSDEGQGLNERLF | 0.35719082 |
| NTGALGL | ASSDEGQGLNERLF | 0.45970616 |
| PPGAGGAGGAGTT | ASSDEGQGLNERLF | 0.45505008 |
| QLVGLGI | ASSDEGQGLNERLF | 0.5526577 |
| RFLEQVHQL | ASSDEGQGLNERLF | 0.55950665 |
| RGVGYVLR | ASSDEGQGLNERLF | 0.35659117 |
| SATGVFAGVI | ASSDEGQGLNERLF | 0.48591 |
| SLPAITLP | ASSDEGQGLNERLF | 0.4998362 |
| STALAFTV | ASSDEGQGLNERLF | 0.34416094 |
| SVLTVGLGK | ASSDEGQGLNERLF | 0.38063517 |
| TALAGAGSGI | ASSDEGQGLNERLF | 0.42771345 |
| TGLPFTGI | ASSDEGQGLNERLF | 0.36236656 |
| VAEHVTV | ASSDEGQGLNERLF | 0.51480114 |
| VDIVLNSL | ASSDEGQGLNERLF | 0.454726 |
| VLTGGGAVL | ASSDEGQGLNERLF | 0.44603857 |
| VNVGLAYGI | ASSDEGQGLNERLF | 0.4783442 |
| VSSAAVGAV | ASSDEGQGLNERLF | 0.42120913 |
| VTSVYEAGV | ASSDEGQGLNERLF | 0.42496023 |

| | | |
|---|---|---|
| WVALDRGA | ASSDEGQGLNERLF | 0.4490235 |
| YLVGSVVI | ASSDEGQGLNERLF | 0.5572072 |
| GPPGTGKTLL | ASSDEGQGLNERLF | 0.5290122 |
| GVGMTKF | ASSDEGQGLNERLF | 0.43273538 |
| IGIGGSDL | ASSDEGQGLNERLF | 0.42408478 |
| IGTLLAEL | ASSDEGQGLNERLF | 0.57853055 |
| IYTSGTT | ASSDEGQGLNERLF | 0.38134456 |
| LSFPAGV | ASSDEGQGLNERLF | 0.43110704 |
| LYTSGTT | ASSDEGQGLNERLF | 0.476192 |
| VGAFPFGF | ASSDEGQGLNERLF | 0.5034051 |
| VGAGTVEF | ASSDEGQGLNERLF | 0.52402306 |
| VGKSTLLL | ASSDEGQGLNERLF | 0.5589172 |
| AALAHAGGGIV | ASSDEGQGLNERLF | 0.4777244 |
| AALVAVSLL | ASSDEGQGLNERLF | 0.40486717 |
| AAVSRAVAAL | ASSDEGQGLNERLF | 0.55977935 |
| AGGSSGIGGF | ASSDEGQGLNERLF | 0.4950267 |
| AGLTVANANG | ASSDEGQGLNERLF | 0.4163704 |
| AGLVGAVL | ASSDEGQGLNERLF | 0.47498852 |
| ALSQAFTV | ASSDEGQGLNERLF | 0.37438476 |
| ALTFSVSL | ASSDEGQGLNERLF | 0.53170073 |
| ALTGAALVAL | ASSDEGQGLNERLF | 0.49349123 |
| ASFSIGAGVL | ASSDEGQGLNERLF | 0.40015477 |
| DGSVYKI | ASSDEGQGLNERLF | 0.45947796 |
| EQLAEIV | ASSDEGQGLNERLF | 0.55696553 |
| FAVSELTVM | ASSDEGQGLNERLF | 0.5304911 |
| GAAPLAAGL | ASSDEGQGLNERLF | 0.4704936 |
| GADGGDGGSGGAGG | ASSDEGQGLNERLF | 0.43090564 |
| GFGLDTGGAGG | ASSDEGQGLNERLF | 0.40425044 |
| GLASGGLL | ASSDEGQGLNERLF | 0.39242214 |
| GRIAGAAVLL | ASSDEGQGLNERLF | 0.3726182 |
| GSAGSLNTSF | ASSDEGQGLNERLF | 0.45341623 |
| GTGQIGI | ASSDEGQGLNERLF | 0.43632844 |
| GTLIALIL | ASSDEGQGLNERLF | 0.56164074 |
| IAATMGFVVG | ASSDEGQGLNERLF | 0.5438599 |
| IFAQALL | ASSDEGQGLNERLF | 0.58815855 |
| IGAAVALGF | ASSDEGQGLNERLF | 0.4078142 |
| IGLTPLAL | ASSDEGQGLNERLF | 0.5304513 |
| ILGSNTSTL | ASSDEGQGLNERLF | 0.3317392 |
| IVFSLTVL | ASSDEGQGLNERLF | 0.5152025 |
| LASKATGF | ASSDEGQGLNERLF | 0.35543185 |
| LASVALIL | ASSDEGQGLNERLF | 0.50634634 |
| LFAVGLTL | ASSDEGQGLNERLF | 0.36045817 |
| LFNGGIGTYI | ASSDEGQGLNERLF | 0.603566 |
| LTLGAGII | ASSDEGQGLNERLF | 0.3719509 |
| LVAAALGDI | ASSDEGQGLNERLF | 0.5535095 |
| NTAFVQLGI | ASSDEGQGLNERLF | 0.43235824 |
| QLFSGAEL | ASSDEGQGLNERLF | 0.35058036 |
| SAATFITL | ASSDEGQGLNERLF | 0.46666214 |
| SNYGLANLGV | ASSDEGQGLNERLF | 0.50813127 |
| SYGGAYAVM | ASSDEGQGLNERLF | 0.3747876 |
| TTGVGGLTLG | ASSDEGQGLNERLF | 0.32844058 |

| | | |
|---|---|---|
| TTQTIYI | ASSDEGQGLNERLF | 0.3683716 |
| VAAGAGLGAVI | ASSDEGQGLNERLF | 0.40914464 |
| VAAVVTLL | ASSDEGQGLNERLF | 0.5935544 |
| VGFLGGLI | ASSDEGQGLNERLF | 0.5719956 |
| VGVIPILL | ASSDEGQGLNERLF | 0.4721985 |
| VIFASSLI | ASSDEGQGLNERLF | 0.5740813 |
| VKGGDGGAAAG | ASSDEGQGLNERLF | 0.30795968 |
| VTKTFTNI | ASSDEGQGLNERLF | 0.46298152 |
| VVAAAILL | ASSDEGQGLNERLF | 0.5912824 |
| YAALDVEL | ASSDEGQGLNERLF | 0.50914335 |
| LASVALLL | ASSDEGQGLNERLF | 0.45643 |
| LDAVHRV | ASSDEGQGLNERLF | 0.5210493 |
| QGLSSSIF | ASSDEGQGLNERLF | 0.4651521 |
| RGTFANIRL | ASSDEGQGLNERLF | 0.4772084 |
| RQLWWGHRI | ASSDEGQGLNERLF | 0.4631545 |
| YGPPGTGKTLL | ASSDEGQGLNERLF | 0.51662785 |
| AALVGAPANVA | ASSDEGQGLNERLF | 0.41459864 |
| AATFRRTE | ASSDEGQGLNERLF | 0.34816778 |
| AGALGGGI | ASSDEGQGLNERLF | 0.45525703 |
| AGGAGGAGGIGAG | ASSDEGQGLNERLF | 0.6049507 |
| AGSGALGL | ASSDEGQGLNERLF | 0.4345672 |
| AQIADQI | ASSDEGQGLNERLF | 0.52103543 |
| ASAVAVTYL | ASSDEGQGLNERLF | 0.42354053 |
| ASTVLGGTGG | ASSDEGQGLNERLF | 0.4952973 |
| AVGGEPAK | ASSDEGQGLNERLF | 0.41442877 |
| CEQGTFGGTC | ASSDEGQGLNERLF | 0.36179852 |
| DEGKAPAG | ASSDEGQGLNERLF | 0.564294 |
| DTVVFPAL | ASSDEGQGLNERLF | 0.35730803 |
| FEGITMAG | ASSDEGQGLNERLF | 0.42015648 |
| FPLVVQD | ASSDEGQGLNERLF | 0.43916023 |
| GAFTTPQ | ASSDEGQGLNERLF | 0.6211261 |
| GFGGFGTAGGVGGA | ASSDEGQGLNERLF | 0.49237347 |
| GFLAQFL | ASSDEGQGLNERLF | 0.46236038 |
| GGAGGAGGNAGVA | ASSDEGQGLNERLF | 0.4535584 |
| GGVGTAGAGGAGGAGG | ASSDEGQGLNERLF | 0.4434446 |
| GNGGGGDAGNAGSGGN | ASSDEGQGLNERLF | 0.5226346 |
| GTGGAGGAGNDAGS | ASSDEGQGLNERLF | 0.35421026 |
| GTTNTGWFNS | ASSDEGQGLNERLF | 0.55137503 |
| IFMGPLFTL | ASSDEGQGLNERLF | 0.37093753 |
| ILAHAIEAI | ASSDEGQGLNERLF | 0.50492054 |
| IRSAIREAL | ASSDEGQGLNERLF | 0.4871464 |
| ITFAGGG | ASSDEGQGLNERLF | 0.60057247 |
| ITGMSGHQEV | ASSDEGQGLNERLF | 0.44614953 |
| IVAGLACTL | ASSDEGQGLNERLF | 0.5604786 |
| IVGGGLAGA | ASSDEGQGLNERLF | 0.4314775 |
| IVTGVLNI | ASSDEGQGLNERLF | 0.4689952 |
| LAQQGCI | ASSDEGQGLNERLF | 0.5868201 |
| LCLHVEN | ASSDEGQGLNERLF | 0.4127994 |
| LDAISDGS | ASSDEGQGLNERLF | 0.6062151 |
| LGGFAAML | ASSDEGQGLNERLF | 0.46231404 |
| LGGKNPMI | ASSDEGQGLNERLF | 0.34671375 |

| | | |
|---|---|---|
| LGPKGVAAIL | ASSDEGQGLNERLF | 0.44625056 |
| LMVGVSI | ASSDEGQGLNERLF | 0.53860825 |
| LVGAAFVL | ASSDEGQGLNERLF | 0.5666952 |
| LVLAVAPL | ASSDEGQGLNERLF | 0.53961456 |
| MAAIAVIDAA | ASSDEGQGLNERLF | 0.4445958 |
| NNGINAELQ | ASSDEGQGLNERLF | 0.42760158 |
| RGEPLVLVHG | ASSDEGQGLNERLF | 0.5816808 |
| RGGAGGAAGLI | ASSDEGQGLNERLF | 0.6227378 |
| RSYRMHPAV | ASSDEGQGLNERLF | 0.46465915 |
| SAAHLLAGQLA | ASSDEGQGLNERLF | 0.350721 |
| SAAVGSY | ASSDEGQGLNERLF | 0.36629814 |
| SPHAVLHRI | ASSDEGQGLNERLF | 0.536511 |
| SQIAKVL | ASSDEGQGLNERLF | 0.5750072 |
| TALGHLVP | ASSDEGQGLNERLF | 0.3384342 |
| TAPTTSGAA | ASSDEGQGLNERLF | 0.4164737 |
| TDGDDGNGGSPGT | ASSDEGQGLNERLF | 0.5093002 |
| TDGLGEN | ASSDEGQGLNERLF | 0.33233878 |
| TGGPPLPGQA | ASSDEGQGLNERLF | 0.46949053 |
| TIPAITI | ASSDEGQGLNERLF | 0.545432 |
| TIPGFTI | ASSDEGQGLNERLF | 0.45210728 |
| TSIASGALA | ASSDEGQGLNERLF | 0.33663064 |
| TVTLPGAAI | ASSDEGQGLNERLF | 0.54630584 |
| VASGLSAALP | ASSDEGQGLNERLF | 0.56869525 |
| VGAGVIGI | ASSDEGQGLNERLF | 0.5546899 |
| VGPKRFGV | ASSDEGQGLNERLF | 0.36299977 |
| VNVAFPD | ASSDEGQGLNERLF | 0.31087482 |
| VSMTNSGVS | ASSDEGQGLNERLF | 0.5354646 |
| VTDVLGDLSV | ASSDEGQGLNERLF | 0.5123299 |
| VVVVGAGFAGL | ASSDEGQGLNERLF | 0.46165678 |
| VYTVGTVV | ASSDEGQGLNERLF | 0.3881381 |
| WTTFSGP | ASSDEGQGLNERLF | 0.38903576 |
| YDELVEAF | ASSDEGQGLNERLF | 0.57350117 |
| YGTGGAGGNGG | ASSDEGQGLNERLF | 0.36244935 |
| YTNVMAV | ASSDEGQGLNERLF | 0.36120418 |
| FLEQVHQL | ASSDEGQGLNERLF | 0.45543614 |
| GLSLLAVAV | ASSDEGQGLNERLF | 0.5241494 |
| LEQVHQL | ASSDEGQGLNERLF | 0.38351315 |
| LNAAGAI | ASSDEGQGLNERLF | 0.44002324 |
| PAVHASL | ASSDEGQGLNERLF | 0.5732224 |
| TELEDDA | ASSDEGQGLNERLF | 0.55611455 |
| TPPGPPN | ASSDEGQGLNERLF | 0.48719257 |
| TPPPPPPP | ASSDEGQGLNERLF | 0.36467335 |
| TQSPALI | ASSPTPYEQY | 0.5719779 |
| EFAAAFPQ | ASSPTPYEQY | 0.35930333 |
| LLSGNLDVL | ASSPTPYEQY | 0.47595337 |
| GPANLGNYNI | ASSPTPYEQY | 0.47399986 |
| TLGPTLP | ASSPTPYEQY | 0.50536656 |
| VYAYSSCCK | ASSPTPYEQY | 0.44427037 |
| APAPAPAPNI | ASSPTPYEQY | 0.3401345 |
| PGGPVPIP | ASSPTPYEQY | 0.3838891 |
| FLTVALEP | ASSPTPYEQY | 0.37677205 |

| | | |
|---|---|---|
| SQYFGANL | ASSPTPYEQY | 0.5457901 |
| GGVKPAA | ASSPTPYEQY | 0.54376864 |
| GGTGGAGGAGGW | ASSPTPYEQY | 0.47955394 |
| GVGGVAGLGGAGG | ASSPTPYEQY | 0.4965577 |
| VLLVLATTI | ASSPTPYEQY | 0.42975384 |
| IGIFAAML | ASSPTPYEQY | 0.4883998 |
| AVGDLID | ASSPTPYEQY | 0.37132323 |
| NISPLPL | ASSPTPYEQY | 0.48874366 |
| VTAVFVAI | ASSPTPYEQY | 0.3661845 |
| SGDDNDAAL | ASSPTPYEQY | 0.46182346 |
| AVAALAHS | ASSPTPYEQY | 0.40968555 |
| GGVTVMATP | ASSPTPYEQY | 0.4487505 |
| AGLAANVRD | ASSPTPYEQY | 0.5194095 |
| TNRPDLIDPALLRPG | ASSPTPYEQY | 0.3662677 |
| LANLGVTAL | ASSPTPYEQY | 0.46536863 |
| TGERIEL | ASSPTPYEQY | 0.58341587 |
| NGEELAA | ASSPTPYEQY | 0.46357638 |
| VLSLVWG | ASSPTPYEQY | 0.33189335 |
| IGYALWL | ASSPTPYEQY | 0.493823 |
| LSLLLAPL | ASSPTPYEQY | 0.402241 |
| TLDEALK | ASSPTPYEQY | 0.52858144 |
| TDGAGTAEGG | ASSPTPYEQY | 0.5082529 |
| NAPGGAGGNGG | ASSPTPYEQY | 0.42160758 |
| VSVFMGMGGNF | ASSPTPYEQY | 0.4960541 |
| LASYLAAL | ASSPTPYEQY | 0.36513048 |
| LVGGTPAL | ASSPTPYEQY | 0.38241127 |
| VRNADVL | ASSPTPYEQY | 0.41437936 |
| TCSSPFLVSAM | ASSPTPYEQY | 0.5464657 |
| VHITDAE | ASSPTPYEQY | 0.48286986 |
| EYAAPSF | ASSPTPYEQY | 0.39013714 |
| VVDAFHGGL | ASSPTPYEQY | 0.4816544 |
| ASVVDPQ | ASSPTPYEQY | 0.4780686 |
| SYNEFGGLS | ASSPTPYEQY | 0.4286056 |
| DPDGSVY | ASSPTPYEQY | 0.42018577 |
| MVSTIHATE | ASSPTPYEQY | 0.37443382 |
| LAAGVADPV | ASSPTPYEQY | 0.32989764 |
| TKEQIAA | ASSPTPYEQY | 0.53153646 |
| SLASPAAL | ASSPTPYEQY | 0.4087154 |
| IGLVFYGV | ASSPTPYEQY | 0.476022 |
| FLLDASNL | ASSPTPYEQY | 0.52133334 |
| YQVGNFP | ASSPTPYEQY | 0.32950538 |
| ESLLSQST | ASSPTPYEQY | 0.44735202 |
| LFTVGFAAVG | ASSPTPYEQY | 0.44538125 |
| ATPPTPT | ASSPTPYEQY | 0.43287438 |
| TDLATLG | ASSPTPYEQY | 0.48747733 |
| SAGGTGR | ASSPTPYEQY | 0.40523934 |
| GFSNSGS | ASSPTPYEQY | 0.5309628 |
| GGNGGSGGTSV | ASSPTPYEQY | 0.521287 |
| GTSVATGGAGN | ASSPTPYEQY | 0.48167828 |
| NGGNGGSGGTS | ASSPTPYEQY | 0.50486106 |
| AVVKFPH | ASSPTPYEQY | 0.51799566 |

| | | |
|---|---|---|
| TPPPPPP | ASSPTPYEQY | 0.53197974 |
| AAPAAAAAPA | ASSPTPYEQY | 0.4070026 |
| IALNEDL | ASSPTPYEQY | 0.33156073 |
| IVNVSSM | ASSPTPYEQY | 0.5317299 |
| TGIPPAPR | ASSPTPYEQY | 0.5300162 |
| FDLGGGTF | ASSPTPYEQY | 0.3712505 |
| FDLGGGTFDVS | ASSPTPYEQY | 0.33448786 |
| NEPTAAA | ASSPTPYEQY | 0.5526133 |
| NEPTAAALA | ASSPTPYEQY | 0.60028994 |
| GSIINISSI | ASSPTPYEQY | 0.41442657 |
| VAPGNAGTA | ASSPTPYEQY | 0.5966807 |
| TGGGASLE | ASSPTPYEQY | 0.33961588 |
| VAVYNHYKRI | ASSPTPYEQY | 0.5121825 |
| TDPAPATT | ASSPTPYEQY | 0.54841614 |
| TNRPDILDPALLRPG | ASSPTPYEQY | 0.3607797 |
| LTTTTTS | ASSPTPYEQY | 0.5313675 |
| PTGSGKTL | ASSPTPYEQY | 0.36953813 |
| TGSGKTL | ASSPTPYEQY | 0.37541094 |
| TIGHVDHGK | ASSPTPYEQY | 0.5398011 |
| TIGHVDHGKTT | ASSPTPYEQY | 0.46794716 |
| GHVDHGKTT | ASSPTPYEQY | 0.51536804 |
| AQFEHTLL | ASSPTPYEQY | 0.4243247 |
| IVNQDEL | ASSPTPYEQY | 0.43568614 |
| TNGSQFF | ASSPTPYEQY | 0.5049241 |
| TGKGIVE | ASSPTPYEQY | 0.5390134 |
| LISNASDA | ASSPTPYEQY | 0.43670583 |
| AELRTLGELTKIAKAHGAQ | ASSPTPYEQY | 0.6038855 |
| APNAVAAAEPSW | ASSPTPYEQY | 0.40755358 |
| APQVAGAATAVL | ASSPTPYEQY | 0.61146474 |
| ARRDPAAAAAILPTDA | ASSPTPYEQY | 0.5049676 |
| DAAARDVLAEAGLAEYF | ASSPTPYEQY | 0.5201559 |
| DDQIVLSAKANLAKVEQ | ASSPTPYEQY | 0.42882663 |
| DELLPDTGEDSDEDGH | ASSPTPYEQY | 0.5709201 |
| ELPEAPANRATVPQD | ASSPTPYEQY | 0.45596242 |
| FGQSKIERVTGIG | ASSPTPYEQY | 0.4383222 |
| GAGGTGSTAGGAGGAGGAGGLY | ASSPTPYEQY | 0.49428314 |
| GHGGTGAAGGNGGTGGAGGSGI | ASSPTPYEQY | 0.61498684 |
| GNGGNGGAGGEGGPGGAGGAGA | ASSPTPYEQY | 0.8850632 |
| GSTAGNGGNGGNATNGTIAGQP | ASSPTPYEQY | 0.8207834 |
| GTGGDGGNGGNGGTGG | ASSPTPYEQY | 0.8060498 |
| GVRRPAGIPADAPV | ASSPTPYEQY | 0.66887605 |
| GYRQVTDAHLLTLARR | ASSPTPYEQY | 0.3305088 |
| IAKMGQDGHDRG | ASSPTPYEQY | 0.8070094 |
| KAEPSIEIQYT | ASSPTPYEQY | 0.5902101 |
| KPVRIRMAADEDTVAVT | ASSPTPYEQY | 0.85582626 |
| MPPRIAGMRLLVIKPEP | ASSPTPYEQY | 0.8541207 |
| MKFYAVALFRSRGVPPTR | ASSPTPYEQY | 0.69878227 |
| NAGGFGILGAVAHSP | ASSPTPYEQY | 0.8727269 |
| NEGREITLAMAEM | ASSPTPYEQY | 0.8798069 |
| NELVEWSHPLGHRWRWE | ASSPTPYEQY | 0.42947716 |
| NLGAGGTGGNGGNPG | ASSPTPYEQY | 0.76311153 |

| | | |
|---|---|---|
| NTEWPKWETVMAADPVD | ASSPTPYEQY | 0.7294514 |
| PASAAPAETQP | ASSPTPYEQY | 0.74584365 |
| QQSSDGKNGVRAAEITGE | ASSPTPYEQY | 0.44081682 |
| RAPDAVTTSWWKE | ASSPTPYEQY | 0.8528625 |
| SGFGNVGAHTSGWFNQ | ASSPTPYEQY | 0.7225138 |
| SGVAGAVGQAASVGGLKV | ASSPTPYEQY | 0.13042875 |
| SNGGAGTGSAGGNGGT | ASSPTPYEQY | 0.77366483 |
| TGGAGGEGGAAG | ASSPTPYEQY | 0.7746744 |
| TGGDGGAAGAGGAPGHG | ASSPTPYEQY | 0.70522785 |
| TMPPLLAAIANAGRHGV | ASSPTPYEQY | 0.86756206 |
| TSSTAGNSGGA | ASSPTPYEQY | 0.44861847 |
| VPATVPVGILC | ASSPTPYEQY | 0.65660334 |
| VQRPDSGGVVLTGRIS | ASSPTPYEQY | 0.85685635 |
| WTPNPVPATKGVPAGLVQAT | ASSPTPYEQY | 0.8761173 |
| YALKRALAKFSDRFDVVII | ASSPTPYEQY | 0.5101627 |
| YLIGSVIV | ASSPTPYEQY | 0.82838124 |
| YGQLVAVID | ASSPTPYEQY | 0.88096607 |
| VTVGGMTLPA | ASSPTPYEQY | 0.8309375 |
| VLALPAV | ASSPTPYEQY | 0.8769678 |
| VIPPTLNYET | ASSPTPYEQY | 0.8836068 |
| VGMGGMHPSSG | ASSPTPYEQY | 0.6787436 |
| VGLAGGAATA | ASSPTPYEQY | 0.86814845 |
| VGGAGGAGGSGP | ASSPTPYEQY | 0.8853583 |
| VFAAFMV | ASSPTPYEQY | 0.7397396 |
| TLYDKLR | ASSPTPYEQY | 0.78690195 |
| TLAVFTVL | ASSPTPYEQY | 0.63453734 |
| TGSVQPARV | ASSPTPYEQY | 0.75061786 |
| TGGGAGSGGT | ASSPTPYEQY | 0.29085857 |
| SHLETSL | ASSPTPYEQY | 0.83368236 |
| SFTTGMT | ASSPTPYEQY | 0.87385666 |
| QVEVAKRMGVRQPTVS | ASSPTPYEQY | 0.75177467 |
| QGSGGPALAM | ASSPTPYEQY | 0.5748488 |
| MLAPSNP | ASSPTPYEQY | 0.5918956 |
| LVNNAAI | ASSPTPYEQY | 0.78968287 |
| LTGEINTLA | ASSPTPYEQY | 0.72687185 |
| LSIFGPV | ASSPTPYEQY | 0.30297822 |
| LLALLANL | ASSPTPYEQY | 0.71571565 |
| LISNASDALDKLR | ASSPTPYEQY | 0.86931944 |
| LINEAVP | ASSPTPYEQY | 0.87281406 |
| LGYAFSGL | ASSPTPYEQY | 0.38679916 |
| LDFVAVAL | ASSPTPYEQY | 0.88267565 |
| LAQPGAE | ASSPTPYEQY | 0.8451601 |
| ISNLGAVTGF | ASSPTPYEQY | 0.8871208 |
| HQEDPGRI | ASSPTPYEQY | 0.8223553 |
| GTPGEPTP | ASSPTPYEQY | 0.6395012 |
| GFFNSGS | ASSPTPYEQY | 0.8598318 |
| GAGGGGLTAGAN | ASSPTPYEQY | 0.749751 |
| FVDALVNM | ASSPTPYEQY | 0.5872262 |
| FLTVAPEL | ASSPTPYEQY | 0.8532008 |
| FETLTAL | ASSPTPYEQY | 0.72173196 |
| DVINPIEL | ASSPTPYEQY | 0.70058095 |

| | | |
|---|---|---|
| APVSGAPV | ASSPTPYEQY | 0.75574327 |
| ALVGGLAW | ASSPTPYEQY | 0.86809725 |
| ALAAAAATAI | ASSPTPYEQY | 0.6190723 |
| AFGLGLLL | ASSPTPYEQY | 0.51229 |
| AFFAAASL | ASSPTPYEQY | 0.5999786 |
| AAAALPAFSPP | ASSPTPYEQY | 0.79499567 |
| VVLAPAL | ASSPTPYEQY | 0.8705089 |
| TIGHVDHGKTTL | ASSPTPYEQY | 0.82639337 |
| LISNASDALDK | ASSPTPYEQY | 0.7617909 |
| LISNASDAL | ASSPTPYEQY | 0.8405282 |
| ISNASDALDK | ASSPTPYEQY | 0.8505202 |
| NQAETLVYQTE | ASSPTPYEQY | 0.8030489 |
| TGGTGGTAGAG | ASSPTPYEQY | 0.8516951 |
| TGSTAGGAGGA | ASSPTPYEQY | 0.70008236 |
| DPLAVPVAAGPV | ASSPTPYEQY | 0.86392987 |
| GGTGGTPGNGGN | ASSPTPYEQY | 0.83788157 |
| PAGLAGVDPATL | ASSPTPYEQY | 0.8667458 |
| GNNLVVEAPHAGASV | ASSPTPYEQY | 0.86858404 |
| KVTRIMSASTLEQPA | ASSPTPYEQY | 0.8339406 |
| LGFALPIALPRRTVV | ASSPTPYEQY | 0.8002414 |
| TRLGNLTARKLVVLV | ASSPTPYEQY | 0.25305238 |
| WSQQRRLIYFPSAGP | ASSPTPYEQY | 0.746619 |
| GPNGNPDPMAAAVDIR | ASSPTPYEQY | 0.8204484 |
| TQRCVSVADLRAAASR | ASSPTPYEQY | 0.8267292 |
| QFVLDVAEHPVEVLTETV | ASSPTPYEQY | 0.8420866 |
| GGAGGTGGAGGNGGTQAGDG | ASSPTPYEQY | 0.849085 |
| PLLATLRAIAVDVDAKPAQV | ASSPTPYEQY | 0.6736665 |
| APKFRTGGYMIDFWGVGYQVAK | ASSPTPYEQY | 0.8653454 |
| GAGGAGGAGGAAGGPGGTGGTG | ASSPTPYEQY | 0.86956215 |
| IGALGLLATDVAPAGSVAFGVA | ASSPTPYEQY | 0.44179106 |
| AGGAGGIGGNGGAGGAGGVLMG | ASSPTPYEQY | 0.73236084 |
| ADELIGLL | ASSPTPYEQY | 0.89255464 |
| ADRALKA | ASSPTPYEQY | 0.8265176 |
| AGAVGQVGGAGGA | ASSPTPYEQY | 0.75058687 |
| AGGGGGLGGAP | ASSPTPYEQY | 0.7024352 |
| AGGLLYGNGGAG | ASSPTPYEQY | 0.8860525 |
| AGLESIGL | ASSPTPYEQY | 0.6859995 |
| AGLKTGII | ASSPTPYEQY | 0.8810226 |
| ASLAALLL | ASSPTPYEQY | 0.17646891 |
| ASVINISSI | ASSPTPYEQY | 0.7079119 |
| ATPPGPA | ASSPTPYEQY | 0.57604194 |
| ATVASALV | ASSPTPYEQY | 0.24125707 |
| ATVASLIL | ASSPTPYEQY | 0.8621482 |
| AVNGPVLIRH | ASSPTPYEQY | 0.73276484 |
| DGAAGAVGI | ASSPTPYEQY | 0.857079 |
| DPLLHGI | ASSPTPYEQY | 0.8164522 |
| ELGAALGAL | ASSPTPYEQY | 0.8620157 |
| ELGRTQAV | ASSPTPYEQY | 0.83296376 |
| FVAGGYLS | ASSPTPYEQY | 0.8000324 |
| GASGGAGGAGGW | ASSPTPYEQY | 0.56373286 |
| GGAGAGGDGGAGGI | ASSPTPYEQY | 0.8465751 |

| | | |
|---|---|---|
| GTGHNGSQPGMGGNG | ASSPTPYEQY | 0.8669156 |
| GYAAASASAS | ASSPTPYEQY | 0.84107673 |
| HQVDVAAVQQG | ASSPTPYEQY | 0.39594245 |
| IGAAIAHT | ASSPTPYEQY | 0.7216994 |
| IGMIEVAQL | ASSPTPYEQY | 0.77839875 |
| IRLAPEHTSAMD | ASSPTPYEQY | 0.23988225 |
| IYNVSVL | ASSPTPYEQY | 0.16681565 |
| KQGFLGM | ASSPTPYEQY | 0.67539036 |
| LESVITQL | ASSPTPYEQY | 0.72581404 |
| LGLGALVVV | ASSPTPYEQY | 0.8663 |
| LIGNGGNGGSGGI | ASSPTPYEQY | 0.86236745 |
| LNAINAPTL | ASSPTPYEQY | 0.75572765 |
| LQLLNTGV | ASSPTPYEQY | 0.8606354 |
| LQTSIEF | ASSPTPYEQY | 0.8117707 |
| MSFAVSEL | ASSPTPYEQY | 0.09584936 |
| NTGALGL | ASSPTPYEQY | 0.7832817 |
| PPGAGGAGGAGTT | ASSPTPYEQY | 0.45158917 |
| QLVGLGI | ASSPTPYEQY | 0.88424855 |
| RFLEQVHQL | ASSPTPYEQY | 0.59542376 |
| RGVGYVLR | ASSPTPYEQY | 0.7060219 |
| SATGVFAGVI | ASSPTPYEQY | 0.79359204 |
| SLPAITLP | ASSPTPYEQY | 0.8212346 |
| STALAFTV | ASSPTPYEQY | 0.8133146 |
| SVLTVGLGK | ASSPTPYEQY | 0.59982944 |
| TALAGAGSGI | ASSPTPYEQY | 0.14470717 |
| TGLPFTGI | ASSPTPYEQY | 0.8534243 |
| VAEHVTV | ASSPTPYEQY | 0.56780636 |
| VDIVLNSL | ASSPTPYEQY | 0.49575615 |
| VLTGGGAVL | ASSPTPYEQY | 0.28081992 |
| VNVGLAYGI | ASSPTPYEQY | 0.22051805 |
| VSSAAVGAV | ASSPTPYEQY | 0.4217669 |
| VTSVYEAGV | ASSPTPYEQY | 0.8691032 |
| WVALDRGA | ASSPTPYEQY | 0.76518893 |
| YLVGSVVI | ASSPTPYEQY | 0.74838674 |
| GPPGTGKTLL | ASSPTPYEQY | 0.8853645 |
| GVGMTKF | ASSPTPYEQY | 0.8043116 |
| IGIGGSDL | ASSPTPYEQY | 0.74622536 |
| IGTLLAEL | ASSPTPYEQY | 0.65539455 |
| IYTSGTT | ASSPTPYEQY | 0.6035348 |
| LSFPAGV | ASSPTPYEQY | 0.6001364 |
| LYTSGTT | ASSPTPYEQY | 0.56960654 |
| VGAFPFGF | ASSPTPYEQY | 0.8841125 |
| VGAGTVEF | ASSPTPYEQY | 0.71681285 |
| VGKSTLLL | ASSPTPYEQY | 0.5516355 |
| AALAHAGGGIV | ASSPTPYEQY | 0.83979154 |
| AALVAVSLL | ASSPTPYEQY | 0.88180774 |
| AAVSRAVAAL | ASSPTPYEQY | 0.8959435 |
| AGGSSGIGGF | ASSPTPYEQY | 0.59541667 |
| AGLTVANANG | ASSPTPYEQY | 0.78828394 |
| AGLVGAVL | ASSPTPYEQY | 0.74421024 |
| ALSQAFTV | ASSPTPYEQY | 0.79011333 |

| | | |
|---|---|---|
| ALTFSVSL | ASSPTPYEQY | 0.7141175 |
| ALTGAALVAL | ASSPTPYEQY | 0.4194215 |
| ASFSIGAGVL | ASSPTPYEQY | 0.8532266 |
| DGSVYKI | ASSPTPYEQY | 0.89044833 |
| EQLAEIV | ASSPTPYEQY | 0.71907645 |
| FAVSELTVM | ASSPTPYEQY | 0.66694057 |
| GAAPLAAGL | ASSPTPYEQY | 0.765889 |
| GADGGDGGSGGAGG | ASSPTPYEQY | 0.6837563 |
| GFGLDTGGAGG | ASSPTPYEQY | 0.712498 |
| GLASGGLL | ASSPTPYEQY | 0.8268756 |
| GRIAGAAVLL | ASSPTPYEQY | 0.44692993 |
| GSAGSLNTSF | ASSPTPYEQY | 0.7460355 |
| GTGQIGI | ASSPTPYEQY | 0.89207286 |
| GTLIALIL | ASSPTPYEQY | 0.7608546 |
| IAATMGFVVG | ASSPTPYEQY | 0.8385224 |
| IFAQALL | ASSPTPYEQY | 0.87114745 |
| IGAAVALGF | ASSPTPYEQY | 0.83987635 |
| IGLTPLAL | ASSPTPYEQY | 0.8781695 |
| ILGSNTSTL | ASSPTPYEQY | 0.859148 |
| IVFSLTVL | ASSPTPYEQY | 0.7362431 |
| LASKATGF | ASSPTPYEQY | 0.7872478 |
| LASVALIL | ASSPTPYEQY | 0.82347137 |
| LFAVGLTL | ASSPTPYEQY | 0.7889567 |
| LFNGGIGTYI | ASSPTPYEQY | 0.77519786 |
| LTLGAGII | ASSPTPYEQY | 0.7091822 |
| LVAAALGDI | ASSPTPYEQY | 0.86823285 |
| NTAFVQLGI | ASSPTPYEQY | 0.8185979 |
| QLFSGAEL | ASSPTPYEQY | 0.8635414 |
| SAATFITL | ASSPTPYEQY | 0.39678478 |
| SNYGLANLGV | ASSPTPYEQY | 0.495781 |
| SYGGAYAVM | ASSPTPYEQY | 0.8644091 |
| TTGVGGLTLG | ASSPTPYEQY | 0.13252634 |
| TTQTIYI | ASSPTPYEQY | 0.2013384 |
| VAAGAGLGAVI | ASSPTPYEQY | 0.8091999 |
| VAAVVTLL | ASSPTPYEQY | 0.88328004 |
| VGFLGGLI | ASSPTPYEQY | 0.14808294 |
| VGVIPILL | ASSPTPYEQY | 0.58264256 |
| VIFASSLI | ASSPTPYEQY | 0.52623415 |
| VKGGDGGAAAG | ASSPTPYEQY | 0.86749935 |
| VTKTFTNI | ASSPTPYEQY | 0.7987318 |
| VVAAAILL | ASSPTPYEQY | 0.36364037 |
| YAALDVEL | ASSPTPYEQY | 0.84599125 |
| LASVALLL | ASSPTPYEQY | 0.47837937 |
| LDAVHRV | ASSPTPYEQY | 0.47907278 |
| QGLSSSIF | ASSPTPYEQY | 0.86223316 |
| RGTFANIRL | ASSPTPYEQY | 0.8718519 |
| RQLWWGHRI | ASSPTPYEQY | 0.52546334 |
| YGPPGTGKTLL | ASSPTPYEQY | 0.8647056 |
| AALVGAPANVA | ASSPTPYEQY | 0.7306515 |
| AATFRRTE | ASSPTPYEQY | 0.8950009 |
| AGALGGGI | ASSPTPYEQY | 0.706758 |

| | | |
|---|---|---|
| AGGAGGAGGIGAG | ASSPTPYEQY | 0.35916984 |
| AGSGALGL | ASSPTPYEQY | 0.8609288 |
| AQIADQI | ASSPTPYEQY | 0.8478081 |
| ASAVAVTYL | ASSPTPYEQY | 0.7763374 |
| ASTVLGGTGG | ASSPTPYEQY | 0.85245585 |
| AVGGEPAK | ASSPTPYEQY | 0.72061527 |
| CEQGTFGGTC | ASSPTPYEQY | 0.771924 |
| DEGKAPAG | ASSPTPYEQY | 0.87180096 |
| DTVVFPAL | ASSPTPYEQY | 0.8569799 |
| FEGITMAG | ASSPTPYEQY | 0.4418586 |
| FPLVVQD | ASSPTPYEQY | 0.8585326 |
| GAFTTPQ | ASSPTPYEQY | 0.78226584 |
| GFGGFGTAGGVGGA | ASSPTPYEQY | 0.63842666 |
| GFLAQFL | ASSPTPYEQY | 0.89109284 |
| GGAGGAGGNAGVA | ASSPTPYEQY | 0.8178045 |
| GGVGTAGAGGAGGAGG | ASSPTPYEQY | 0.8614334 |
| GNGGGGDAGNAGSGGN | ASSPTPYEQY | 0.8448268 |
| GTGGAGGAGNDAGS | ASSPTPYEQY | 0.8734646 |
| GTTNTGWFNS | ASSPTPYEQY | 0.36719212 |
| IFMGPLFTL | ASSPTPYEQY | 0.8313518 |
| ILAHAIEAI | ASSPTPYEQY | 0.79358983 |
| IRSAIREAL | ASSPTPYEQY | 0.86364007 |
| ITFAGGG | ASSPTPYEQY | 0.8197496 |
| ITGMSGHQEV | ASSPTPYEQY | 0.54143655 |
| IVAGLACTL | ASSPTPYEQY | 0.8156996 |
| IVGGGLAGA | ASSPTPYEQY | 0.8772838 |
| IVTGVLNI | ASSPTPYEQY | 0.76995826 |
| LAQQGCI | ASSPTPYEQY | 0.83314633 |
| LCLHVEN | ASSPTPYEQY | 0.8821826 |
| LDAISDGS | ASSPTPYEQY | 0.6153667 |
| LGGFAAML | ASSPTPYEQY | 0.77093625 |
| LGGKNPMI | ASSPTPYEQY | 0.8468503 |
| LGPKGVAAIL | ASSPTPYEQY | 0.86641073 |
| LMVGVSI | ASSPTPYEQY | 0.7547213 |
| LVGAAFVL | ASSPTPYEQY | 0.8797588 |
| LVLAVAPL | ASSPTPYEQY | 0.70282316 |
| MAAIAVIDAA | ASSPTPYEQY | 0.85174274 |
| NNGINAELQ | ASSPTPYEQY | 0.72192436 |
| RGEPLVLVHG | ASSPTPYEQY | 0.84083855 |
| RGGAGGAAGLI | ASSPTPYEQY | 0.55584466 |
| RSYRMHPAV | ASSPTPYEQY | 0.84443575 |
| SAAHLLAGQLA | ASSPTPYEQY | 0.8686483 |
| SAAVGSY | ASSPTPYEQY | 0.8602304 |
| SPHAVLHRI | ASSPTPYEQY | 0.19765128 |
| SQIAKVL | ASSPTPYEQY | 0.8611028 |
| TALGHLVP | ASSPTPYEQY | 0.85833764 |
| TAPTTSGAA | ASSPTPYEQY | 0.8872453 |
| TDGDDGNGGSPGT | ASSPTPYEQY | 0.7786428 |
| TDGLGEN | ASSPTPYEQY | 0.8432533 |
| TGGPPLPGQA | ASSPTPYEQY | 0.74083686 |
| TIPAITI | ASSPTPYEQY | 0.77499783 |

| | | |
|---|---|---|
| TIPGFTI | ASSPTPYEQY | 0.5963994 |
| TSIASGALA | ASSPTPYEQY | 0.7566327 |
| TVTLPGAAI | ASSPTPYEQY | 0.7539586 |
| VASGLSAALP | ASSPTPYEQY | 0.6459745 |
| VGAGVIGI | ASSPTPYEQY | 0.85743463 |
| VGPKRFGV | ASSPTPYEQY | 0.8630846 |
| VNVAFPD | ASSPTPYEQY | 0.8665496 |
| VSMTNSGVS | ASSPTPYEQY | 0.8749528 |
| VTDVLGDLSV | ASSPTPYEQY | 0.88007355 |
| VVVVGAGFAGL | ASSPTPYEQY | 0.8690908 |
| VYTVGTVV | ASSPTPYEQY | 0.698186 |
| WTTFSGP | ASSPTPYEQY | 0.794671 |
| YDELVEAF | ASSPTPYEQY | 0.5440996 |
| YGTGGAGGNGG | ASSPTPYEQY | 0.76695967 |
| YTNVMAV | ASSPTPYEQY | 0.70841706 |
| FLEQVHQL | ASSPTPYEQY | 0.8211793 |
| GLSLLAVAV | ASSPTPYEQY | 0.12648554 |
| LEQVHQL | ASSPTPYEQY | 0.8124835 |
| LNAAGAI | ASSPTPYEQY | 0.88461906 |
| PAVHASL | ASSPTPYEQY | 0.72075427 |
| TELEDDA | ASSPTPYEQY | 0.845052 |
| TPPGPPN | ASSPTPYEQY | 0.8177552 |
| TPPPPPPP | ASSPTPYEQY | 0.6880679 |
| TQSPALI | ASSLVQANTEVF | 0.85778415 |
| EFAAAFPQ | ASSLVQANTEVF | 0.61644614 |
| LLSGNLDVL | ASSLVQANTEVF | 0.84572583 |
| GPANLGNYNI | ASSLVQANTEVF | 0.8882108 |
| TLGPTLP | ASSLVQANTEVF | 0.81269944 |
| VYAYSSCCK | ASSLVQANTEVF | 0.79967976 |
| APAPAPAPNI | ASSLVQANTEVF | 0.78080636 |
| PGGPVPIP | ASSLVQANTEVF | 0.33515066 |
| FLTVALEP | ASSLVQANTEVF | 0.8140779 |
| SQYFGANL | ASSLVQANTEVF | 0.113530375 |
| GGVKPAA | ASSLVQANTEVF | 0.81288815 |
| GGTGGAGGAGGW | ASSLVQANTEVF | 0.5412239 |
| GVGGVAGLGGAGG | ASSLVQANTEVF | 0.80394125 |
| VLLVLATTI | ASSLVQANTEVF | 0.8023257 |
| IGIFAAML | ASSLVQANTEVF | 0.83498955 |
| AVGDLID | ASSLVQANTEVF | 0.6391751 |
| NISPLPL | ASSLVQANTEVF | 0.7823103 |
| VTAVFVAI | ASSLVQANTEVF | 0.67081136 |
| SGDDNDAAL | ASSLVQANTEVF | 0.46371296 |
| AVAALAHS | ASSLVQANTEVF | 0.86627585 |
| GGVTVMATP | ASSLVQANTEVF | 0.7039546 |
| AGLAANVRD | ASSLVQANTEVF | 0.7910308 |
| TNRPDLIDPALLRPG | ASSLVQANTEVF | 0.8745611 |
| LANLGVTAL | ASSLVQANTEVF | 0.809049 |
| TGERIEL | ASSLVQANTEVF | 0.46203995 |
| NGEELAA | ASSLVQANTEVF | 0.80364317 |
| VLSLVWG | ASSLVQANTEVF | 0.7919949 |
| IGYALWL | ASSLVQANTEVF | 0.7227223 |

| | | |
|---|---|---|
| LSLLLAPL | ASSLVQANTEVF | 0.12247847 |
| TLDEALK | ASSLVQANTEVF | 0.8370331 |
| TDGAGTAEGG | ASSLVQANTEVF | 0.638664 |
| NAPGGAGGNGG | ASSLVQANTEVF | 0.72372997 |
| VSVFMGMGGNF | ASSLVQANTEVF | 0.677034 |
| LASYLAAL | ASSLVQANTEVF | 0.8593582 |
| LVGGTPAL | ASSLVQANTEVF | 0.5231527 |
| VRNADVL | ASSLVQANTEVF | 0.75768876 |
| TCSSPFLVSAM | ASSLVQANTEVF | 0.8526896 |
| VHITDAE | ASSLVQANTEVF | 0.84105635 |
| EYAAPSF | ASSLVQANTEVF | 0.8783508 |
| VVDAFHGGL | ASSLVQANTEVF | 0.82466507 |
| ASVVDPQ | ASSLVQANTEVF | 0.7593597 |
| SYNEFGGLS | ASSLVQANTEVF | 0.81970686 |
| DPDGSVY | ASSLVQANTEVF | 0.83010733 |
| MVSTIHATE | ASSLVQANTEVF | 0.2555598 |
| LAAGVADPV | ASSLVQANTEVF | 0.278072 |
| TKEQIAA | ASSLVQANTEVF | 0.82964647 |
| SLASPAAL | ASSLVQANTEVF | 0.24137056 |
| IGLVFYGV | ASSLVQANTEVF | 0.81463885 |
| FLLDASNL | ASSLVQANTEVF | 0.8811727 |
| YQVGNFP | ASSLVQANTEVF | 0.82541394 |
| ESLLSQST | ASSLVQANTEVF | 0.4597577 |
| LFTVGFAAVG | ASSLVQANTEVF | 0.73770356 |
| ATPPTPT | ASSLVQANTEVF | 0.74562865 |
| TDLATLG | ASSLVQANTEVF | 0.85685205 |
| SAGGTGR | ASSLVQANTEVF | 0.8486102 |
| GFSNSGS | ASSLVQANTEVF | 0.87901866 |
| GGNGGSGGTSV | ASSLVQANTEVF | 0.86627203 |
| GTSVATGGAGN | ASSLVQANTEVF | 0.7258656 |
| NGGNGGSGGTS | ASSLVQANTEVF | 0.85061246 |
| AVVKFPH | ASSLVQANTEVF | 0.74558556 |
| TPPPPPP | ASSLVQANTEVF | 0.797175 |
| AAPAAAAAPA | ASSLVQANTEVF | 0.75384027 |
| IALNEDL | ASSLVQANTEVF | 0.83441013 |
| IVNVSSM | ASSLVQANTEVF | 0.75270736 |
| TGIPPAPR | ASSLVQANTEVF | 0.65275323 |
| FDLGGGTF | ASSLVQANTEVF | 0.44077653 |
| FDLGGGTFDVS | ASSLVQANTEVF | 0.8345169 |
| NEPTAAA | ASSLVQANTEVF | 0.8229281 |
| NEPTAAALA | ASSLVQANTEVF | 0.384628 |
| GSIINISSI | ASSLVQANTEVF | 0.6919838 |
| VAPGNAGTA | ASSLVQANTEVF | 0.8882515 |
| TGGGASLE | ASSLVQANTEVF | 0.8840992 |
| VAVYNHYKRI | ASSLVQANTEVF | 0.6348492 |
| TDPAPATT | ASSLVQANTEVF | 0.6234146 |
| TNRPDILDPALLRPG | ASSLVQANTEVF | 0.81998694 |
| LTTTTTS | ASSLVQANTEVF | 0.32916248 |
| PTGSGKTL | ASSLVQANTEVF | 0.82700086 |
| TGSGKTL | ASSLVQANTEVF | 0.7981738 |
| TIGHVDHGK | ASSLVQANTEVF | 0.79491055 |

| | | |
|---|---|---|
| TIGHVDHGKTT | ASSLVQANTEVF | 0.62432253 |
| GHVDHGKTT | ASSLVQANTEVF | 0.8180597 |
| AQFEHTLL | ASSLVQANTEVF | 0.7575051 |
| IVNQDEL | ASSLVQANTEVF | 0.856493 |
| TNGSQFF | ASSLVQANTEVF | 0.16106908 |
| TGKGIVE | ASSLVQANTEVF | 0.83660686 |
| LISNASDA | ASSLVQANTEVF | 0.6416912 |
| AELRTLGELTKIAKAHGAQ | ASSLVQANTEVF | 0.7104274 |
| APNAVAAAEPSW | ASSLVQANTEVF | 0.29316327 |
| APQVAGAATAVL | ASSLVQANTEVF | 0.8076446 |
| ARRDPAAAAAILPTDA | ASSLVQANTEVF | 0.871987 |
| DAAARDVLAEAGLAEYF | ASSLVQANTEVF | 0.86974156 |
| DDQIVLSAKANLAKVEQ | ASSLVQANTEVF | 0.49252015 |
| DELLPDTGEDSDEDGH | ASSLVQANTEVF | 0.8604175 |
| ELPEAPANRATVPQD | ASSLVQANTEVF | 0.891242 |
| FGQSKIERVTGIG | ASSLVQANTEVF | 0.7511194 |
| GAGGTGSTAGGAGGAGGAGGLY | ASSLVQANTEVF | 0.25104117 |
| GHGGTGAAGGNGGTGGAGGSGI | ASSLVQANTEVF | 0.8407117 |
| GNGGNGGAGGEGGPGGAGGAGA | ASSLVQANTEVF | 0.8048543 |
| GSTAGNGGNGGNATNGTIAGQP | ASSLVQANTEVF | 0.86676234 |
| GTGGDGGNGGNGGTGG | ASSLVQANTEVF | 0.85726506 |
| GVRRPAGIPADAPV | ASSLVQANTEVF | 0.522011 |
| GYRQVTDAHLLTLARR | ASSLVQANTEVF | 0.86820644 |
| IAKMGQDGHDRG | ASSLVQANTEVF | 0.8680665 |
| KAEPSIEIQYT | ASSLVQANTEVF | 0.5926628 |
| KPVRIRMAADEDTVAVT | ASSLVQANTEVF | 0.72021484 |
| MPPRIAGMRLLVIKPEP | ASSLVQANTEVF | 0.7087043 |
| MKFYAVALFRSRGVPPTR | ASSLVQANTEVF | 0.30168226 |
| NAGGFGILGAVAHSP | ASSLVQANTEVF | 0.84780705 |
| NEGREITLAMAEM | ASSLVQANTEVF | 0.7248339 |
| NELVEWSHPLGHRWRWE | ASSLVQANTEVF | 0.7101668 |
| NLGAGGTGGNGGNPG | ASSLVQANTEVF | 0.7957767 |
| NTEWPKWETVMAADPVD | ASSLVQANTEVF | 0.6319242 |
| PASAAPAETQP | ASSLVQANTEVF | 0.66620743 |
| QQSSDGKNGVRAAEITGE | ASSLVQANTEVF | 0.69072795 |
| RAPDAVTTSWWKE | ASSLVQANTEVF | 0.82033324 |
| SGFGNVGAHTSGWFNQ | ASSLVQANTEVF | 0.8365969 |
| SGVAGAVGQAASVGGLKV | ASSLVQANTEVF | 0.8293267 |
| SNGGAGTGSAGGNGGT | ASSLVQANTEVF | 0.84204865 |
| TGGAGGEGGAAG | ASSLVQANTEVF | 0.87900126 |
| TGGDGGAAGAGGAPGHG | ASSLVQANTEVF | 0.46503824 |
| TMPPLLAAIANAGRHGV | ASSLVQANTEVF | 0.6326959 |
| TSSTAGNSGGA | ASSLVQANTEVF | 0.85359204 |
| VPATVPVGILC | ASSLVQANTEVF | 0.42847294 |
| VQRPDSGGVVLTGRIS | ASSLVQANTEVF | 0.8607293 |
| WTPNPVPATKGVPAGLVQAT | ASSLVQANTEVF | 0.80228394 |
| YALKRALAKFSDRFDVVII | ASSLVQANTEVF | 0.8281498 |
| YLIGSVIV | ASSLVQANTEVF | 0.8181509 |
| YGQLVAVID | ASSLVQANTEVF | 0.89360416 |
| VTVGGMTLPA | ASSLVQANTEVF | 0.85783994 |
| VLALPAV | ASSLVQANTEVF | 0.7757804 |

| | | |
|---|---|---|
| VIPPTLNYET | ASSLVQANTEVF | 0.82691365 |
| VGMGGMHPSSG | ASSLVQANTEVF | 0.5954202 |
| VGLAGGAATA | ASSLVQANTEVF | 0.8603755 |
| VGGAGGAGGSGP | ASSLVQANTEVF | 0.85464585 |
| VFAAFMV | ASSLVQANTEVF | 0.62674147 |
| TLYDKLR | ASSLVQANTEVF | 0.7754594 |
| TLAVFTVL | ASSLVQANTEVF | 0.8756943 |
| TGSVQPARV | ASSLVQANTEVF | 0.86844325 |
| TGGGAGSGGT | ASSLVQANTEVF | 0.28878728 |
| SHLETSL | ASSLVQANTEVF | 0.7232429 |
| SFTTGMT | ASSLVQANTEVF | 0.31024155 |
| QVEVAKRMGVRQPTVS | ASSLVQANTEVF | 0.6916329 |
| QGSGGPALAM | ASSLVQANTEVF | 0.48297098 |
| MLAPSNP | ASSLVQANTEVF | 0.84564674 |
| LVNNAAI | ASSLVQANTEVF | 0.74154675 |
| LTGEINTLA | ASSLVQANTEVF | 0.8371663 |
| LSIFGPV | ASSLVQANTEVF | 0.8090802 |
| LLALLANL | ASSLVQANTEVF | 0.8719614 |
| LISNASDALDKLR | ASSLVQANTEVF | 0.67712986 |
| LINEAVP | ASSLVQANTEVF | 0.7367275 |
| LGYAFSGL | ASSLVQANTEVF | 0.34653232 |
| LDFVAVAL | ASSLVQANTEVF | 0.41772997 |
| LAQPGAE | ASSLVQANTEVF | 0.62344927 |
| ISNLGAVTGF | ASSLVQANTEVF | 0.7388922 |
| HQEDPGRI | ASSLVQANTEVF | 0.28728026 |
| GTPGEPTP | ASSLVQANTEVF | 0.8871459 |
| GFFNSGS | ASSLVQANTEVF | 0.81538826 |
| GAGGGGLTAGAN | ASSLVQANTEVF | 0.8480738 |
| FVDALVNM | ASSLVQANTEVF | 0.72021556 |
| FLTVAPEL | ASSLVQANTEVF | 0.71748704 |
| FETLTAL | ASSLVQANTEVF | 0.7017483 |
| DVINPIEL | ASSLVQANTEVF | 0.8461045 |
| APVSGAPV | ASSLVQANTEVF | 0.55817163 |
| ALVGGLAW | ASSLVQANTEVF | 0.4403945 |
| ALAAAAATAI | ASSLVQANTEVF | 0.7884817 |
| AFGLGLLL | ASSLVQANTEVF | 0.653424 |
| AFFAAASL | ASSLVQANTEVF | 0.8847191 |
| AAAALPAFSPP | ASSLVQANTEVF | 0.86772877 |
| VVLAPAL | ASSLVQANTEVF | 0.8228288 |
| TIGHVDHGKTTL | ASSLVQANTEVF | 0.86904407 |
| LISNASDALDK | ASSLVQANTEVF | 0.20483412 |
| LISNASDAL | ASSLVQANTEVF | 0.7655463 |
| ISNASDALDK | ASSLVQANTEVF | 0.8831667 |
| NQAETLVYQTE | ASSLVQANTEVF | 0.8436408 |
| TGGTGGTAGAG | ASSLVQANTEVF | 0.7698331 |
| TGSTAGGAGGA | ASSLVQANTEVF | 0.87650293 |
| DPLAVPVAAGPV | ASSLVQANTEVF | 0.6534555 |
| GGTGGTPGNGGN | ASSLVQANTEVF | 0.7361529 |
| PAGLAGVDPATL | ASSLVQANTEVF | 0.780926 |
| GNNLVVEAPHAGASV | ASSLVQANTEVF | 0.80809486 |
| KVTRIMSASTLEQPA | ASSLVQANTEVF | 0.8224125 |

| | | |
|---|---|---|
| LGFALPIALPRRTVV | ASSLVQANTEVF | 0.55414957 |
| TRLGNLTARKLVVLV | ASSLVQANTEVF | 0.83144784 |
| WSQQRRLIYFPSAGP | ASSLVQANTEVF | 0.8944124 |
| GPNGNPDPMAAAVDIR | ASSLVQANTEVF | 0.75333536 |
| TQRCVSVADLRAAASR | ASSLVQANTEVF | 0.8720393 |
| QFVLDVAEHPVEVLTETV | ASSLVQANTEVF | 0.842779 |
| GGAGGTGGAGGNGGTQAGDG | ASSLVQANTEVF | 0.8387771 |
| PLLATLRAIAVDVDAKPAQV | ASSLVQANTEVF | 0.6534543 |
| APKFRTGGYMIDFWGVGYQVAK | ASSLVQANTEVF | 0.6746716 |
| GAGGAGGAGGAAGGPGGTGGTG | ASSLVQANTEVF | 0.7757194 |
| IGALGLLATDVAPAGSVAFGVA | ASSLVQANTEVF | 0.6503304 |
| AGGAGGIGGNGGAGGAGGVLMG | ASSLVQANTEVF | 0.80115235 |
| ADELIGLL | ASSLVQANTEVF | 0.73639 |
| ADRALKA | ASSLVQANTEVF | 0.89835435 |
| AGAVGQVGGAGGA | ASSLVQANTEVF | 0.8432473 |
| AGGGGGLGGAP | ASSLVQANTEVF | 0.84128135 |
| AGGLLYGNGGAG | ASSLVQANTEVF | 0.58484703 |
| AGLESIGL | ASSLVQANTEVF | 0.804087 |
| AGLKTGII | ASSLVQANTEVF | 0.8353749 |
| ASLAALLL | ASSLVQANTEVF | 0.47229403 |
| ASVINISSI | ASSLVQANTEVF | 0.8771135 |
| ATPPGPA | ASSLVQANTEVF | 0.6566653 |
| ATVASALV | ASSLVQANTEVF | 0.79140157 |
| ATVASLIL | ASSLVQANTEVF | 0.8590394 |
| AVNGPVLIRH | ASSLVQANTEVF | 0.86321837 |
| DGAAGAVGI | ASSLVQANTEVF | 0.7333553 |
| DPLLHGI | ASSLVQANTEVF | 0.7396529 |
| ELGAALGAL | ASSLVQANTEVF | 0.57889867 |
| ELGRTQAV | ASSLVQANTEVF | 0.86349744 |
| FVAGGYLS | ASSLVQANTEVF | 0.86999285 |
| GASGGAGGAGGW | ASSLVQANTEVF | 0.7809701 |
| GGAGAGGDGGAGGI | ASSLVQANTEVF | 0.8793175 |
| GTGHNGSQPGMGGNG | ASSLVQANTEVF | 0.8592675 |
| GYAAASASAS | ASSLVQANTEVF | 0.36838308 |
| HQVDVAAVQQG | ASSLVQANTEVF | 0.76248306 |
| IGAAIAHT | ASSLVQANTEVF | 0.8649553 |
| IGMIEVAQL | ASSLVQANTEVF | 0.8723284 |
| IRLAPEHTSAMD | ASSLVQANTEVF | 0.88602495 |
| IYNVSVL | ASSLVQANTEVF | 0.8407876 |
| KQGFLGM | ASSLVQANTEVF | 0.7335663 |
| LESVITQL | ASSLVQANTEVF | 0.8807341 |
| LGLGALVVV | ASSLVQANTEVF | 0.899408 |
| LIGNGGNGGSGGI | ASSLVQANTEVF | 0.86523765 |
| LNAINAPTL | ASSLVQANTEVF | 0.71694577 |
| LQLLNTGV | ASSLVQANTEVF | 0.65576804 |
| LQTSIEF | ASSLVQANTEVF | 0.7949145 |
| MSFAVSEL | ASSLVQANTEVF | 0.86257845 |
| NTGALGL | ASSLVQANTEVF | 0.5713105 |
| PPGAGGAGGAGTT | ASSLVQANTEVF | 0.8110542 |
| QLVGLGI | ASSLVQANTEVF | 0.8193091 |
| RFLEQVHQL | ASSLVQANTEVF | 0.3814181 |

| | | |
|---|---|---|
| RGVGYVLR | ASSLVQANTEVF | 0.8542708 |
| SATGVFAGVI | ASSLVQANTEVF | 0.8801094 |
| SLPAITLP | ASSLVQANTEVF | 0.7545591 |
| STALAFTV | ASSLVQANTEVF | 0.711663 |
| SVLTVGLGK | ASSLVQANTEVF | 0.8725976 |
| TALAGAGSGI | ASSLVQANTEVF | 0.89568514 |
| TGLPFTGI | ASSLVQANTEVF | 0.8731205 |
| VAEHVTV | ASSLVQANTEVF | 0.37872946 |
| VDIVLNSL | ASSLVQANTEVF | 0.24562505 |
| VLTGGGAVL | ASSLVQANTEVF | 0.63651645 |
| VNVGLAYGI | ASSLVQANTEVF | 0.853055 |
| VSSAAVGAV | ASSLVQANTEVF | 0.71804374 |
| VTSVYEAGV | ASSLVQANTEVF | 0.6458153 |
| WVALDRGA | ASSLVQANTEVF | 0.83775854 |
| YLVGSVVI | ASSLVQANTEVF | 0.8746793 |
| GPPGTGKTLL | ASSLVQANTEVF | 0.5751748 |
| GVGMTKF | ASSLVQANTEVF | 0.5226494 |
| IGIGGSDL | ASSLVQANTEVF | 0.86290276 |
| IGTLLAEL | ASSLVQANTEVF | 0.86398447 |
| IYTSGTT | ASSLVQANTEVF | 0.6345055 |
| LSFPAGV | ASSLVQANTEVF | 0.7495545 |
| LYTSGTT | ASSLVQANTEVF | 0.65732527 |
| VGAFPFGF | ASSLVQANTEVF | 0.6780632 |
| VGAGTVEF | ASSLVQANTEVF | 0.8243266 |
| VGKSTLLL | ASSLVQANTEVF | 0.5596063 |
| AALAHAGGGIV | ASSLVQANTEVF | 0.88307554 |
| AALVAVSLL | ASSLVQANTEVF | 0.571038 |
| AAVSRAVAAL | ASSLVQANTEVF | 0.8553344 |
| AGGSSGIGGF | ASSLVQANTEVF | 0.8125415 |
| AGLTVANANG | ASSLVQANTEVF | 0.8124155 |
| AGLVGAVL | ASSLVQANTEVF | 0.7365546 |
| ALSQAFTV | ASSLVQANTEVF | 0.722435 |
| ALTFSVSL | ASSLVQANTEVF | 0.5111326 |
| ALTGAALVAL | ASSLVQANTEVF | 0.31661683 |
| ASFSIGAGVL | ASSLVQANTEVF | 0.86721563 |
| DGSVYKI | ASSLVQANTEVF | 0.192131 |
| EQLAEIV | ASSLVQANTEVF | 0.8637276 |
| FAVSELTVM | ASSLVQANTEVF | 0.8409532 |
| GAAPLAAGL | ASSLVQANTEVF | 0.57904404 |
| GADGGDGGSGGAGG | ASSLVQANTEVF | 0.8283131 |
| GFGLDTGGAGG | ASSLVQANTEVF | 0.51997256 |
| GLASGGLL | ASSLVQANTEVF | 0.52695864 |
| GRIAGAAVLL | ASSLVQANTEVF | 0.85750103 |
| GSAGSLNTSF | ASSLVQANTEVF | 0.8403586 |
| GTGQIGI | ASSLVQANTEVF | 0.6605441 |
| GTLIALIL | ASSLVQANTEVF | 0.8227709 |
| IAATMGFVVG | ASSLVQANTEVF | 0.8243332 |
| IFAQALL | ASSLVQANTEVF | 0.69164324 |
| IGAAVALGF | ASSLVQANTEVF | 0.7420107 |
| IGLTPLAL | ASSLVQANTEVF | 0.87189335 |
| ILGSNTSTL | ASSLVQANTEVF | 0.80029804 |

| | | |
|---|---|---|
| IVFSLTVL | ASSLVQANTEVF | 0.5743228 |
| LASKATGF | ASSLVQANTEVF | 0.856009 |
| LASVALIL | ASSLVQANTEVF | 0.6202872 |
| LFAVGLTL | ASSLVQANTEVF | 0.83140993 |
| LFNGGIGTYI | ASSLVQANTEVF | 0.77650964 |
| LTLGAGII | ASSLVQANTEVF | 0.75096 |
| LVAAALGDI | ASSLVQANTEVF | 0.85365725 |
| NTAFVQLGI | ASSLVQANTEVF | 0.686936 |
| QLFSGAEL | ASSLVQANTEVF | 0.680095 |
| SAATFITL | ASSLVQANTEVF | 0.7030908 |
| SNYGLANLGV | ASSLVQANTEVF | 0.77746946 |
| SYGGAYAVM | ASSLVQANTEVF | 0.8653331 |
| TTGVGGLTLG | ASSLVQANTEVF | 0.59494364 |
| TTQTIYI | ASSLVQANTEVF | 0.8183564 |
| VAAGAGLGAVI | ASSLVQANTEVF | 0.71764004 |
| VAAVVTLL | ASSLVQANTEVF | 0.55443156 |
| VGFLGGLI | ASSLVQANTEVF | 0.7634796 |
| VGVIPILL | ASSLVQANTEVF | 0.7651423 |
| VIFASSLI | ASSLVQANTEVF | 0.6813179 |
| VKGGDGGAAAG | ASSLVQANTEVF | 0.82682955 |
| VTKTFTNI | ASSLVQANTEVF | 0.504675 |
| VVAAAILL | ASSLVQANTEVF | 0.8197279 |
| YAALDVEL | ASSLVQANTEVF | 0.88810384 |
| LASVALLL | ASSLVQANTEVF | 0.7036979 |
| LDAVHRV | ASSLVQANTEVF | 0.84877825 |
| QGLSSSIF | ASSLVQANTEVF | 0.84191304 |
| RGTFANIRL | ASSLVQANTEVF | 0.8312821 |
| RQLWWGHRI | ASSLVQANTEVF | 0.71274275 |
| YGPPGTGKTLL | ASSLVQANTEVF | 0.69939363 |
| AALVGAPANVA | ASSLVQANTEVF | 0.8814893 |
| AATFRRTE | ASSLVQANTEVF | 0.85254633 |
| AGALGGGI | ASSLVQANTEVF | 0.86289984 |
| AGGAGGAGGIGAG | ASSLVQANTEVF | 0.8546396 |
| AGSGALGL | ASSLVQANTEVF | 0.8487102 |
| AQIADQI | ASSLVQANTEVF | 0.8515223 |
| ASAVAVTYL | ASSLVQANTEVF | 0.72165155 |
| ASTVLGGTGG | ASSLVQANTEVF | 0.42358032 |
| AVGGEPAK | ASSLVQANTEVF | 0.7202661 |
| CEQGTFGGTC | ASSLVQANTEVF | 0.72629005 |
| DEGKAPAG | ASSLVQANTEVF | 0.74619174 |
| DTVVFPAL | ASSLVQANTEVF | 0.38751692 |
| FEGITMAG | ASSLVQANTEVF | 0.8708058 |
| FPLVVQD | ASSLVQANTEVF | 0.895572 |
| GAFTTPQ | ASSLVQANTEVF | 0.68812525 |
| GFGGFGTAGGVGGA | ASSLVQANTEVF | 0.8897731 |
| GFLAQFL | ASSLVQANTEVF | 0.4908864 |
| GGAGGAGGNAGVA | ASSLVQANTEVF | 0.8471452 |
| GGVGTAGAGGAGGAGG | ASSLVQANTEVF | 0.87568516 |
| GNGGGGDAGNAGSGGN | ASSLVQANTEVF | 0.6491397 |
| GTGGAGGAGNDAGS | ASSLVQANTEVF | 0.8524616 |
| GTTNTGWFNS | ASSLVQANTEVF | 0.6270145 |

| | | |
|---|---|---|
| IFMGPLFTL | ASSLVQANTEVF | 0.742851 |
| ILAHAIEAI | ASSLVQANTEVF | 0.8735398 |
| IRSAIREAL | ASSLVQANTEVF | 0.71049607 |
| ITFAGGG | ASSLVQANTEVF | 0.80775934 |
| ITGMSGHQEV | ASSLVQANTEVF | 0.71782285 |
| IVAGLACTL | ASSLVQANTEVF | 0.56325006 |
| IVGGGLAGA | ASSLVQANTEVF | 0.78509164 |
| IVTGVLNI | ASSLVQANTEVF | 0.8723359 |
| LAQQGCI | ASSLVQANTEVF | 0.63185036 |
| LCLHVEN | ASSLVQANTEVF | 0.89816266 |
| LDAISDGS | ASSLVQANTEVF | 0.6223893 |
| LGGFAAML | ASSLVQANTEVF | 0.8928777 |
| LGGKNPMI | ASSLVQANTEVF | 0.86160374 |
| LGPKGVAAIL | ASSLVQANTEVF | 0.8051865 |
| LMVGVSI | ASSLVQANTEVF | 0.72129834 |
| LVGAAFVL | ASSLVQANTEVF | 0.8704393 |
| LVLAVAPL | ASSLVQANTEVF | 0.730966 |
| MAAIAVIDAA | ASSLVQANTEVF | 0.8464757 |
| NNGINAELQ | ASSLVQANTEVF | 0.87862444 |
| RGEPLVLVHG | ASSLVQANTEVF | 0.50110006 |
| RGGAGGAAGLI | ASSLVQANTEVF | 0.853552 |
| RSYRMHPAV | ASSLVQANTEVF | 0.75937605 |
| SAAHLLAGQLA | ASSLVQANTEVF | 0.7993492 |
| SAAVGSY | ASSLVQANTEVF | 0.5526073 |
| SPHAVLHRI | ASSLVQANTEVF | 0.4519853 |
| SQIAKVL | ASSLVQANTEVF | 0.651136 |
| TALGHLVP | ASSLVQANTEVF | 0.5136269 |
| TAPTTSGAA | ASSLVQANTEVF | 0.79701644 |
| TDGDDGNGGSPGT | ASSLVQANTEVF | 0.83423984 |
| TDGLGEN | ASSLVQANTEVF | 0.5947263 |
| TGGPPLPGQA | ASSLVQANTEVF | 0.8245647 |
| TIPAITI | ASSLVQANTEVF | 0.84375536 |
| TIPGFTI | ASSLVQANTEVF | 0.40628192 |
| TSIASGALA | ASSLVQANTEVF | 0.6770218 |
| TVTLPGAAI | ASSLVQANTEVF | 0.60945934 |
| VASGLSAALP | ASSLVQANTEVF | 0.69957435 |
| VGAGVIGI | ASSLVQANTEVF | 0.25524205 |
| VGPKRFGV | ASSLVQANTEVF | 0.8105332 |
| VNVAFPD | ASSLVQANTEVF | 0.5686604 |
| VSMTNSGVS | ASSLVQANTEVF | 0.21830556 |
| VTDVLGDLSV | ASSLVQANTEVF | 0.7731172 |
| VVVVGAGFAGL | ASSLVQANTEVF | 0.6571194 |
| VYTVGTVV | ASSLVQANTEVF | 0.66881216 |
| WTTFSGP | ASSLVQANTEVF | 0.83076733 |
| YDELVEAF | ASSLVQANTEVF | 0.660864 |
| YGTGGAGGNGG | ASSLVQANTEVF | 0.5253148 |
| YTNVMAV | ASSLVQANTEVF | 0.76581407 |
| FLEQVHQL | ASSLVQANTEVF | 0.84164965 |
| GLSLLAVAV | ASSLVQANTEVF | 0.5363953 |
| LEQVHQL | ASSLVQANTEVF | 0.7435616 |
| LNAAGAI | ASSLVQANTEVF | 0.8538908 |

| | | |
|---|---|---|
| PAVHASL | ASSLVQANTEVF | 0.76542985 |
| TELEDDA | ASSLVQANTEVF | 0.8582453 |
| TPPGPPN | ASSLVQANTEVF | 0.8676615 |
| TPPPPPPP | ASSLVQANTEVF | 0.59965026 |
| TQSPALI | ASSLSGGALAEQF | 0.8294679 |
| EFAAAFPQ | ASSLSGGALAEQF | 0.8423816 |
| LLSGNLDVL | ASSLSGGALAEQF | 0.7052754 |
| GPANLGNYNI | ASSLSGGALAEQF | 0.7397566 |
| TLGPTLP | ASSLSGGALAEQF | 0.42538708 |
| VYAYSSCCK | ASSLSGGALAEQF | 0.6796832 |
| APAPAPAPNI | ASSLSGGALAEQF | 0.4320479 |
| PGGPVPIP | ASSLSGGALAEQF | 0.7355623 |
| FLTVALEP | ASSLSGGALAEQF | 0.8613378 |
| SQYFGANL | ASSLSGGALAEQF | 0.6718031 |
| GGVKPAA | ASSLSGGALAEQF | 0.44180667 |
| GGTGGAGGAGGW | ASSLSGGALAEQF | 0.37997374 |
| GVGGVAGLGGAGG | ASSLSGGALAEQF | 0.7625059 |
| VLLVLATTI | ASSLSGGALAEQF | 0.55842686 |
| IGIFAAML | ASSLSGGALAEQF | 0.5350484 |
| AVGDLID | ASSLSGGALAEQF | 0.63189954 |
| NISPLPL | ASSLSGGALAEQF | 0.8195243 |
| VTAVFVAI | ASSLSGGALAEQF | 0.8396524 |
| SGDDNDAAL | ASSLSGGALAEQF | 0.49579608 |
| AVAALAHS | ASSLSGGALAEQF | 0.85597265 |
| GGVTVMATP | ASSLSGGALAEQF | 0.78604037 |
| AGLAANVRD | ASSLSGGALAEQF | 0.8632493 |
| TNRPDLIDPALLRPG | ASSLSGGALAEQF | 0.74772894 |
| LANLGVTAL | ASSLSGGALAEQF | 0.5275263 |
| TGERIEL | ASSLSGGALAEQF | 0.8119651 |
| NGEELAA | ASSLSGGALAEQF | 0.6268309 |
| VLSLVWG | ASSLSGGALAEQF | 0.7769784 |
| IGYALWL | ASSLSGGALAEQF | 0.8198621 |
| LSLLLAPL | ASSLSGGALAEQF | 0.68894774 |
| TLDEALK | ASSLSGGALAEQF | 0.59121764 |
| TDGAGTAEGG | ASSLSGGALAEQF | 0.62960887 |
| NAPGGAGGNGG | ASSLSGGALAEQF | 0.86840135 |
| VSVFMGMGGNF | ASSLSGGALAEQF | 0.4489297 |
| LASYLAAL | ASSLSGGALAEQF | 0.5113056 |
| LVGGTPAL | ASSLSGGALAEQF | 0.67477876 |
| VRNADVL | ASSLSGGALAEQF | 0.6958152 |
| TCSSPFLVSAM | ASSLSGGALAEQF | 0.8411634 |
| VHITDAE | ASSLSGGALAEQF | 0.81303376 |
| EYAAPSF | ASSLSGGALAEQF | 0.7226725 |
| VVDAFHGGL | ASSLSGGALAEQF | 0.7596986 |
| ASVVDPQ | ASSLSGGALAEQF | 0.83283556 |
| SYNEFGGLS | ASSLSGGALAEQF | 0.7571103 |
| DPDGSVY | ASSLSGGALAEQF | 0.704549 |
| MVSTIHATE | ASSLSGGALAEQF | 0.5657599 |
| LAAGVADPV | ASSLSGGALAEQF | 0.85753787 |
| TKEQIAA | ASSLSGGALAEQF | 0.8182839 |
| SLASPAAL | ASSLSGGALAEQF | 0.8365553 |

| | | |
|---|---|---|
| IGLVFYGV | ASSLSGGALAEQF | 0.83846915 |
| FLLDASNL | ASSLSGGALAEQF | 0.80744606 |
| YQVGNFP | ASSLSGGALAEQF | 0.77172804 |
| ESLLSQST | ASSLSGGALAEQF | 0.33461162 |
| LFTVGFAAVG | ASSLSGGALAEQF | 0.6092339 |
| ATPPTPT | ASSLSGGALAEQF | 0.6495844 |
| TDLATLG | ASSLSGGALAEQF | 0.7603127 |
| SAGGTGR | ASSLSGGALAEQF | 0.7490884 |
| GFSNSGS | ASSLSGGALAEQF | 0.84935206 |
| GGNGGSGGTSV | ASSLSGGALAEQF | 0.56328136 |
| GTSVATGGAGN | ASSLSGGALAEQF | 0.79791385 |
| NGGNGGSGGTS | ASSLSGGALAEQF | 0.8494954 |
| AVVKFPH | ASSLSGGALAEQF | 0.39243484 |
| TPPPPPP | ASSLSGGALAEQF | 0.6678752 |
| AAPAAAAAPA | ASSLSGGALAEQF | 0.8752483 |
| IALNEDL | ASSLSGGALAEQF | 0.7481721 |
| IVNVSSM | ASSLSGGALAEQF | 0.6261438 |
| TGIPPAPR | ASSLSGGALAEQF | 0.71355176 |
| FDLGGGTF | ASSLSGGALAEQF | 0.84852755 |
| FDLGGGTFDVS | ASSLSGGALAEQF | 0.7557609 |
| NEPTAAA | ASSLSGGALAEQF | 0.85190535 |
| NEPTAAALA | ASSLSGGALAEQF | 0.27664605 |
| GSIINISSI | ASSLSGGALAEQF | 0.73691046 |
| VAPGNAGTA | ASSLSGGALAEQF | 0.6029843 |
| TGGGASLE | ASSLSGGALAEQF | 0.26190716 |
| VAVYNHYKRI | ASSLSGGALAEQF | 0.8177624 |
| TDPAPATT | ASSLSGGALAEQF | 0.6849942 |
| TNRPDILDPALLRPG | ASSLSGGALAEQF | 0.80571836 |
| LTTTTTS | ASSLSGGALAEQF | 0.6476725 |
| PTGSGKTL | ASSLSGGALAEQF | 0.8335836 |
| TGSGKTL | ASSLSGGALAEQF | 0.71859527 |
| TIGHVDHGK | ASSLSGGALAEQF | 0.697977 |
| TIGHVDHGKTT | ASSLSGGALAEQF | 0.6572678 |
| GHVDHGKTT | ASSLSGGALAEQF | 0.7707403 |
| AQFEHTLL | ASSLSGGALAEQF | 0.8125725 |
| IVNQDEL | ASSLSGGALAEQF | 0.87574935 |
| TNGSQFF | ASSLSGGALAEQF | 0.37559247 |
| TGKGIVE | ASSLSGGALAEQF | 0.76430583 |
| LISNASDA | ASSLSGGALAEQF | 0.68212295 |
| AELRTLGELTKIAKAHGAQ | ASSLSGGALAEQF | 0.35393384 |
| APNAVAAAEPSW | ASSLSGGALAEQF | 0.29500806 |
| APQVAGAATAVL | ASSLSGGALAEQF | 0.6320229 |
| ARRDPAAAAAILPTDA | ASSLSGGALAEQF | 0.65754735 |
| DAAARDVLAEAGLAEYF | ASSLSGGALAEQF | 0.7743911 |
| DDQIVLSAKANLAKVEQ | ASSLSGGALAEQF | 0.77242017 |
| DELLPDTGEDSDEDGH | ASSLSGGALAEQF | 0.6559675 |
| ELPEAPANRATVPQD | ASSLSGGALAEQF | 0.7760484 |
| FGQSKIERVTGIG | ASSLSGGALAEQF | 0.7406025 |
| GAGGTGSTAGGAGGAGGAGGLY | ASSLSGGALAEQF | 0.10417815 |
| GHGGTGAAGGNGGTGGAGGSGI | ASSLSGGALAEQF | 0.6343691 |
| GNGGNGGAGGEGGPGGAGGAGA | ASSLSGGALAEQF | 0.3939247 |

| | | |
|---|---|---|
| GSTAGNGGNGGNATNGTIAGQP | ASSLSGGALAEQF | 0.8627913 |
| GTGGDGGNGGNGGTGG | ASSLSGGALAEQF | 0.532048 |
| GVRRPAGIPADAPV | ASSLSGGALAEQF | 0.5519742 |
| GYRQVTDAHLLTLARR | ASSLSGGALAEQF | 0.6887936 |
| IAKMGQDGHDRG | ASSLSGGALAEQF | 0.73458755 |
| KAEPSIEIQYT | ASSLSGGALAEQF | 0.7214854 |
| KPVRIRMAADEDTVAVT | ASSLSGGALAEQF | 0.56380165 |
| MPPRIAGMRLLVIKPEP | ASSLSGGALAEQF | 0.2777782 |
| MKFYAVALFRSRGVPPTR | ASSLSGGALAEQF | 0.79446125 |
| NAGGFGILGAVAHSP | ASSLSGGALAEQF | 0.6042603 |
| NEGREITLAMAEM | ASSLSGGALAEQF | 0.5631987 |
| NELVEWSHPLGHRWRWE | ASSLSGGALAEQF | 0.37810117 |
| NLGAGGTGGNGGNPG | ASSLSGGALAEQF | 0.7081413 |
| NTEWPKWETVMAADPVD | ASSLSGGALAEQF | 0.45667058 |
| PASAAPAETQP | ASSLSGGALAEQF | 0.85862184 |
| QQSSDGKNGVRAAEITGE | ASSLSGGALAEQF | 0.57476753 |
| RAPDAVTTSWWKE | ASSLSGGALAEQF | 0.628188 |
| SGFGNVGAHTSGWFNQ | ASSLSGGALAEQF | 0.8633995 |
| SGVAGAVGQAASVGGLKV | ASSLSGGALAEQF | 0.68720084 |
| SNGGAGTGSAGGNGGT | ASSLSGGALAEQF | 0.660934 |
| TGGAGGEGGAAG | ASSLSGGALAEQF | 0.6490729 |
| TGGDGGAAGAGGAPGHG | ASSLSGGALAEQF | 0.4811867 |
| TMPPLLAAIANAGRHGV | ASSLSGGALAEQF | 0.53884643 |
| TSSTAGNSGGA | ASSLSGGALAEQF | 0.60280657 |
| VPATVPVGILC | ASSLSGGALAEQF | 0.8507771 |
| VQRPDSGGVVLTGRIS | ASSLSGGALAEQF | 0.7566985 |
| WTPNPVPATKGVPAGLVQAT | ASSLSGGALAEQF | 0.4706622 |
| YALKRALAKFSDRFDVVII | ASSLSGGALAEQF | 0.8399565 |
| YLIGSVIV | ASSLSGGALAEQF | 0.8534754 |
| YGQLVAVID | ASSLSGGALAEQF | 0.86459863 |
| VTVGGMTLPA | ASSLSGGALAEQF | 0.511522 |
| VLALPAV | ASSLSGGALAEQF | 0.7290466 |
| VIPPTLNYET | ASSLSGGALAEQF | 0.702736 |
| VGMGGMHPSSG | ASSLSGGALAEQF | 0.65839684 |
| VGLAGGAATA | ASSLSGGALAEQF | 0.67255193 |
| VGGAGGAGGSGP | ASSLSGGALAEQF | 0.5576354 |
| VFAAFMV | ASSLSGGALAEQF | 0.8458974 |
| TLYDKLR | ASSLSGGALAEQF | 0.87682307 |
| TLAVFTVL | ASSLSGGALAEQF | 0.70074886 |
| TGSVQPARV | ASSLSGGALAEQF | 0.29624623 |
| TGGGAGSGGT | ASSLSGGALAEQF | 0.63398385 |
| SHLETSL | ASSLSGGALAEQF | 0.5028038 |
| SFTTGMT | ASSLSGGALAEQF | 0.5543497 |
| QVEVAKRMGVRQPTVS | ASSLSGGALAEQF | 0.72429425 |
| QGSGGPALAM | ASSLSGGALAEQF | 0.3888958 |
| MLAPSNP | ASSLSGGALAEQF | 0.70588017 |
| LVNNAAI | ASSLSGGALAEQF | 0.86831003 |
| LTGEINTLA | ASSLSGGALAEQF | 0.5390618 |
| LSIFGPV | ASSLSGGALAEQF | 0.8024379 |
| LLALLANL | ASSLSGGALAEQF | 0.85211504 |
| LISNASDALDKLR | ASSLSGGALAEQF | 0.7923232 |

| | | |
|---|---|---|
| LINEAVP | ASSLSGGALAEQF | 0.85121167 |
| LGYAFSGL | ASSLSGGALAEQF | 0.7934933 |
| LDFVAVAL | ASSLSGGALAEQF | 0.7614022 |
| LAQPGAE | ASSLSGGALAEQF | 0.7322536 |
| ISNLGAVTGF | ASSLSGGALAEQF | 0.7794 |
| HQEDPGRI | ASSLSGGALAEQF | 0.6400586 |
| GTPGEPTP | ASSLSGGALAEQF | 0.7023682 |
| GFFNSGS | ASSLSGGALAEQF | 0.66174865 |
| GAGGGGLTAGAN | ASSLSGGALAEQF | 0.78966117 |
| FVDALVNM | ASSLSGGALAEQF | 0.71025884 |
| FLTVAPEL | ASSLSGGALAEQF | 0.8472414 |
| FETLTAL | ASSLSGGALAEQF | 0.3329709 |
| DVINPIEL | ASSLSGGALAEQF | 0.527212 |
| APVSGAPV | ASSLSGGALAEQF | 0.8421979 |
| ALVGGLAW | ASSLSGGALAEQF | 0.1714361 |
| ALAAAAATAI | ASSLSGGALAEQF | 0.42584097 |
| AFGLGLLL | ASSLSGGALAEQF | 0.7544035 |
| AFFAAASL | ASSLSGGALAEQF | 0.8324889 |
| AAAALPAFSPP | ASSLSGGALAEQF | 0.20109235 |
| VVLAPAL | ASSLSGGALAEQF | 0.6222795 |
| TIGHVDHGKTTL | ASSLSGGALAEQF | 0.4216161 |
| LISNASDALDK | ASSLSGGALAEQF | 0.86914575 |
| LISNASDAL | ASSLSGGALAEQF | 0.73438114 |
| ISNASDALDK | ASSLSGGALAEQF | 0.4066139 |
| NQAETLVYQTE | ASSLSGGALAEQF | 0.82765275 |
| TGGTGGTAGAG | ASSLSGGALAEQF | 0.33744368 |
| TGSTAGGAGGA | ASSLSGGALAEQF | 0.6791152 |
| DPLAVPVAAGPV | ASSLSGGALAEQF | 0.81244844 |
| GGTGGTPGNGGN | ASSLSGGALAEQF | 0.84730184 |
| PAGLAGVDPATL | ASSLSGGALAEQF | 0.4527476 |
| GNNLVVEAPHAGASV | ASSLSGGALAEQF | 0.810398 |
| KVTRIMSASTLEQPA | ASSLSGGALAEQF | 0.63258135 |
| LGFALPIALPRRTVV | ASSLSGGALAEQF | 0.8799256 |
| TRLGNLTARKLVVLV | ASSLSGGALAEQF | 0.60065246 |
| WSQQRRLIYFPSAGP | ASSLSGGALAEQF | 0.4523664 |
| GPNGNPDPMAAAVDIR | ASSLSGGALAEQF | 0.83381814 |
| TQRCVSVADLRAAASR | ASSLSGGALAEQF | 0.8091344 |
| QFVLDVAEHPVEVLTETV | ASSLSGGALAEQF | 0.7630398 |
| GGAGGTGGAGGNGGTQAGDG | ASSLSGGALAEQF | 0.80537045 |
| PLLATLRAIAVDVDAKPAQV | ASSLSGGALAEQF | 0.7517996 |
| APKFRTGGYMIDFWGVGYQVAK | ASSLSGGALAEQF | 0.6521592 |
| GAGGAGGAGGAAGGPGGTGGTG | ASSLSGGALAEQF | 0.82536125 |
| IGALGLLATDVAPAGSVAFGVA | ASSLSGGALAEQF | 0.7629619 |
| AGGAGGIGGNGGAGGAGGVLMG | ASSLSGGALAEQF | 0.6051578 |
| ADELIGLL | ASSLSGGALAEQF | 0.86405843 |
| ADRALKA | ASSLSGGALAEQF | 0.7250453 |
| AGAVGQVGGAGGA | ASSLSGGALAEQF | 0.50527024 |
| AGGGGGLGGAP | ASSLSGGALAEQF | 0.8712367 |
| AGGLLYGNGGAG | ASSLSGGALAEQF | 0.78794205 |
| AGLESIGL | ASSLSGGALAEQF | 0.8029654 |
| AGLKTGII | ASSLSGGALAEQF | 0.8077982 |

| | | |
|---|---|---|
| ASLAALLL | ASSLSGGALAEQF | 0.8517197 |
| ASVINISSI | ASSLSGGALAEQF | 0.50986385 |
| ATPPGPA | ASSLSGGALAEQF | 0.6905292 |
| ATVASALV | ASSLSGGALAEQF | 0.71685386 |
| ATVASLIL | ASSLSGGALAEQF | 0.8135456 |
| AVNGPVLIRH | ASSLSGGALAEQF | 0.76690376 |
| DGAAGAVGI | ASSLSGGALAEQF | 0.6107421 |
| DPLLHGI | ASSLSGGALAEQF | 0.7962115 |
| ELGAALGAL | ASSLSGGALAEQF | 0.85592425 |
| ELGRTQAV | ASSLSGGALAEQF | 0.7310755 |
| FVAGGYLS | ASSLSGGALAEQF | 0.69935846 |
| GASGGAGGAGGW | ASSLSGGALAEQF | 0.8605463 |
| GGAGAGGDGGAGGI | ASSLSGGALAEQF | 0.50281596 |
| GTGHNGSQPGMGGNG | ASSLSGGALAEQF | 0.75263333 |
| GYAAASASAS | ASSLSGGALAEQF | 0.82522666 |
| HQVDVAAVQQG | ASSLSGGALAEQF | 0.8023717 |
| IGAAIAHT | ASSLSGGALAEQF | 0.75084054 |
| IGMIEVAQL | ASSLSGGALAEQF | 0.835659 |
| IRLAPEHTSAMD | ASSLSGGALAEQF | 0.568895 |
| IYNVSVL | ASSLSGGALAEQF | 0.79668725 |
| KQGFLGM | ASSLSGGALAEQF | 0.6087339 |
| LESVITQL | ASSLSGGALAEQF | 0.71206284 |
| LGLGALVVV | ASSLSGGALAEQF | 0.484756 |
| LIGNGGNGGSGGI | ASSLSGGALAEQF | 0.7190466 |
| LNAINAPTL | ASSLSGGALAEQF | 0.80660176 |
| LQLLNTGV | ASSLSGGALAEQF | 0.7848098 |
| LQTSIEF | ASSLSGGALAEQF | 0.30465955 |
| MSFAVSEL | ASSLSGGALAEQF | 0.7980015 |
| NTGALGL | ASSLSGGALAEQF | 0.80073434 |
| PPGAGGAGGAGTT | ASSLSGGALAEQF | 0.8760179 |
| QLVGLGI | ASSLSGGALAEQF | 0.6988541 |
| RFLEQVHQL | ASSLSGGALAEQF | 0.82105 |
| RGVGYVLR | ASSLSGGALAEQF | 0.5921402 |
| SATGVFAGVI | ASSLSGGALAEQF | 0.7760673 |
| SLPAITLP | ASSLSGGALAEQF | 0.40431285 |
| STALAFTV | ASSLSGGALAEQF | 0.5295359 |
| SVLTVGLGK | ASSLSGGALAEQF | 0.621416 |
| TALAGAGSGI | ASSLSGGALAEQF | 0.51824 |
| TGLPFTGI | ASSLSGGALAEQF | 0.79811597 |
| VAEHVTV | ASSLSGGALAEQF | 0.79514825 |
| VDIVLNSL | ASSLSGGALAEQF | 0.83079624 |
| VLTGGGAVL | ASSLSGGALAEQF | 0.8384204 |
| VNVGLAYGI | ASSLSGGALAEQF | 0.8540051 |
| VSSAAVGAV | ASSLSGGALAEQF | 0.85846215 |
| VTSVYEAGV | ASSLSGGALAEQF | 0.5651638 |
| WVALDRGA | ASSLSGGALAEQF | 0.67719257 |
| YLVGSVVI | ASSLSGGALAEQF | 0.39515838 |
| GPPGTGKTLL | ASSLSGGALAEQF | 0.65352243 |
| GVGMTKF | ASSLSGGALAEQF | 0.64949214 |
| IGIGGSDL | ASSLSGGALAEQF | 0.7402576 |
| IGTLLAEL | ASSLSGGALAEQF | 0.18184794 |

| | | |
|---|---|---|
| IYTSGTT | ASSLSGGALAEQF | 0.7200222 |
| LSFPAGV | ASSLSGGALAEQF | 0.8543124 |
| LYTSGTT | ASSLSGGALAEQF | 0.61091435 |
| VGAFPFGF | ASSLSGGALAEQF | 0.7856835 |
| VGAGTVEF | ASSLSGGALAEQF | 0.7561813 |
| VGKSTLLL | ASSLSGGALAEQF | 0.6165575 |
| AALAHAGGGIV | ASSLSGGALAEQF | 0.7882824 |
| AALVAVSLL | ASSLSGGALAEQF | 0.6298635 |
| AAVSRAVAAL | ASSLSGGALAEQF | 0.8407917 |
| AGGSSGIGGF | ASSLSGGALAEQF | 0.84712625 |
| AGLTVANANG | ASSLSGGALAEQF | 0.7690764 |
| AGLVGAVL | ASSLSGGALAEQF | 0.6304847 |
| ALSQAFTV | ASSLSGGALAEQF | 0.77407634 |
| ALTFSVSL | ASSLSGGALAEQF | 0.4103654 |
| ALTGAALVAL | ASSLSGGALAEQF | 0.64371675 |
| ASFSIGAGVL | ASSLSGGALAEQF | 0.4205889 |
| DGSVYKI | ASSLSGGALAEQF | 0.70646274 |
| EQLAEIV | ASSLSGGALAEQF | 0.5072933 |
| FAVSELTVM | ASSLSGGALAEQF | 0.7493987 |
| GAAPLAAGL | ASSLSGGALAEQF | 0.7285575 |
| GADGGDGGSGGAGG | ASSLSGGALAEQF | 0.7569947 |
| GFGLDTGGAGG | ASSLSGGALAEQF | 0.42736626 |
| GLASGGLL | ASSLSGGALAEQF | 0.82137364 |
| GRIAGAAVLL | ASSLSGGALAEQF | 0.75860333 |
| GSAGSLNTSF | ASSLSGGALAEQF | 0.44376314 |
| GTGQIGI | ASSLSGGALAEQF | 0.8101444 |
| GTLIALIL | ASSLSGGALAEQF | 0.74456453 |
| IAATMGFVVG | ASSLSGGALAEQF | 0.6567229 |
| IFAQALL | ASSLSGGALAEQF | 0.8326527 |
| IGAAVALGF | ASSLSGGALAEQF | 0.7750414 |
| IGLTPLAL | ASSLSGGALAEQF | 0.38794684 |
| ILGSNTSTL | ASSLSGGALAEQF | 0.5872495 |
| IVFSLTVL | ASSLSGGALAEQF | 0.72285247 |
| LASKATGF | ASSLSGGALAEQF | 0.61838526 |
| LASVALIL | ASSLSGGALAEQF | 0.21058966 |
| LFAVGLTL | ASSLSGGALAEQF | 0.78253186 |
| LFNGGIGTYI | ASSLSGGALAEQF | 0.5145594 |
| LTLGAGII | ASSLSGGALAEQF | 0.4607353 |
| LVAAALGDI | ASSLSGGALAEQF | 0.6459028 |
| NTAFVQLGI | ASSLSGGALAEQF | 0.8216264 |
| QLFSGAEL | ASSLSGGALAEQF | 0.753569 |
| SAATFITL | ASSLSGGALAEQF | 0.69956917 |
| SNYGLANLGV | ASSLSGGALAEQF | 0.7507167 |
| SYGGAYAVM | ASSLSGGALAEQF | 0.7883117 |
| TTGVGGLTLG | ASSLSGGALAEQF | 0.84972817 |
| TTQTIYI | ASSLSGGALAEQF | 0.75464386 |
| VAAGAGLGAVI | ASSLSGGALAEQF | 0.70419234 |
| VAAVVTLL | ASSLSGGALAEQF | 0.80064094 |
| VGFLGGLI | ASSLSGGALAEQF | 0.7463356 |
| VGVIPILL | ASSLSGGALAEQF | 0.5948384 |
| VIFASSLI | ASSLSGGALAEQF | 0.5137284 |

| | | |
|---|---|---|
| VKGGDGGAAAG | ASSLSGGALAEQF | 0.5708169 |
| VTKTFTNI | ASSLSGGALAEQF | 0.5311946 |
| VVAAAILL | ASSLSGGALAEQF | 0.72336435 |
| YAALDVEL | ASSLSGGALAEQF | 0.8809815 |
| LASVALLL | ASSLSGGALAEQF | 0.74202484 |
| LDAVHRV | ASSLSGGALAEQF | 0.3441223 |
| QGLSSSIF | ASSLSGGALAEQF | 0.63322484 |
| RGTFANIRL | ASSLSGGALAEQF | 0.661358 |
| RQLWWGHRI | ASSLSGGALAEQF | 0.7906608 |
| YGPPGTGKTLL | ASSLSGGALAEQF | 0.81784105 |
| AALVGAPANVA | ASSLSGGALAEQF | 0.860146 |
| AATFRRTE | ASSLSGGALAEQF | 0.7967546 |
| AGALGGGI | ASSLSGGALAEQF | 0.61530364 |
| AGGAGGAGGIGAG | ASSLSGGALAEQF | 0.79957086 |
| AGSGALGL | ASSLSGGALAEQF | 0.6819801 |
| AQIADQI | ASSLSGGALAEQF | 0.67809296 |
| ASAVAVTYL | ASSLSGGALAEQF | 0.7366521 |
| ASTVLGGTGG | ASSLSGGALAEQF | 0.78402567 |
| AVGGEPAK | ASSLSGGALAEQF | 0.68559706 |
| CEQGTFGGTC | ASSLSGGALAEQF | 0.62908113 |
| DEGKAPAG | ASSLSGGALAEQF | 0.40244395 |
| DTVVFPAL | ASSLSGGALAEQF | 0.8159585 |
| FEGITMAG | ASSLSGGALAEQF | 0.63995767 |
| FPLVVQD | ASSLSGGALAEQF | 0.64100343 |
| GAFTTPQ | ASSLSGGALAEQF | 0.61500734 |
| GFGGFGTAGGVGGA | ASSLSGGALAEQF | 0.8568795 |
| GFLAQFL | ASSLSGGALAEQF | 0.85152626 |
| GGAGGAGGNAGVA | ASSLSGGALAEQF | 0.59622025 |
| GGVGTAGAGGAGGAGG | ASSLSGGALAEQF | 0.5740911 |
| GNGGGGDAGNAGSGGN | ASSLSGGALAEQF | 0.73086405 |
| GTGGAGGAGNDAGS | ASSLSGGALAEQF | 0.48451936 |
| GTTNTGWFNS | ASSLSGGALAEQF | 0.8291577 |
| IFMGPLFTL | ASSLSGGALAEQF | 0.6920941 |
| ILAHAIEAI | ASSLSGGALAEQF | 0.70078397 |
| IRSAIREAL | ASSLSGGALAEQF | 0.6599994 |
| ITFAGGG | ASSLSGGALAEQF | 0.78255713 |
| ITGMSGHQEV | ASSLSGGALAEQF | 0.6276978 |
| IVAGLACTL | ASSLSGGALAEQF | 0.8370849 |
| IVGGGLAGA | ASSLSGGALAEQF | 0.19197655 |
| IVTGVLNI | ASSLSGGALAEQF | 0.8256373 |
| LAQQGCI | ASSLSGGALAEQF | 0.47780842 |
| LCLHVEN | ASSLSGGALAEQF | 0.62638694 |
| LDAISDGS | ASSLSGGALAEQF | 0.305259 |
| LGGFAAML | ASSLSGGALAEQF | 0.6693039 |
| LGGKNPMI | ASSLSGGALAEQF | 0.78579843 |
| LGPKGVAAIL | ASSLSGGALAEQF | 0.84002215 |
| LMVGVSI | ASSLSGGALAEQF | 0.67348325 |
| LVGAAFVL | ASSLSGGALAEQF | 0.76116985 |
| LVLAVAPL | ASSLSGGALAEQF | 0.84669316 |
| MAAIAVIDAA | ASSLSGGALAEQF | 0.62825763 |
| NNGINAELQ | ASSLSGGALAEQF | 0.31616253 |

| | | |
|---|---|---|
| RGEPLVLVHG | ASSLSGGALAEQF | 0.79862654 |
| RGGAGGAAGLI | ASSLSGGALAEQF | 0.60317194 |
| RSYRMHPAV | ASSLSGGALAEQF | 0.8391497 |
| SAAHLLAGQLA | ASSLSGGALAEQF | 0.7612819 |
| SAAVGSY | ASSLSGGALAEQF | 0.37756318 |
| SPHAVLHRI | ASSLSGGALAEQF | 0.8383705 |
| SQIAKVL | ASSLSGGALAEQF | 0.8156183 |
| TALGHLVP | ASSLSGGALAEQF | 0.48560464 |
| TAPTTSGAA | ASSLSGGALAEQF | 0.46633118 |
| TDGDDGNGGSPGT | ASSLSGGALAEQF | 0.58482057 |
| TDGLGEN | ASSLSGGALAEQF | 0.4362092 |
| TGGPPLPGQA | ASSLSGGALAEQF | 0.8096194 |
| TIPAITI | ASSLSGGALAEQF | 0.600781 |
| TIPGFTI | ASSLSGGALAEQF | 0.68535006 |
| TSIASGALA | ASSLSGGALAEQF | 0.7280261 |
| TVTLPGAAI | ASSLSGGALAEQF | 0.6511142 |
| VASGLSAALP | ASSLSGGALAEQF | 0.6624952 |
| VGAGVIGI | ASSLSGGALAEQF | 0.69252324 |
| VGPKRFGV | ASSLSGGALAEQF | 0.76141524 |
| VNVAFPD | ASSLSGGALAEQF | 0.7990557 |
| VSMTNSGVS | ASSLSGGALAEQF | 0.6804333 |
| VTDVLGDLSV | ASSLSGGALAEQF | 0.7565993 |
| VVVVGAGFAGL | ASSLSGGALAEQF | 0.846632 |
| VYTVGTVV | ASSLSGGALAEQF | 0.5013294 |
| WTTFSGP | ASSLSGGALAEQF | 0.6121047 |
| YDELVEAF | ASSLSGGALAEQF | 0.8183755 |
| YGTGGAGGNGG | ASSLSGGALAEQF | 0.62925106 |
| YTNVMAV | ASSLSGGALAEQF | 0.83884203 |
| FLEQVHQL | ASSLSGGALAEQF | 0.7397475 |
| GLSLLAVAV | ASSLSGGALAEQF | 0.756063 |
| LEQVHQL | ASSLSGGALAEQF | 0.69032025 |
| LNAAGAI | ASSLSGGALAEQF | 0.86075366 |
| PAVHASL | ASSLSGGALAEQF | 0.8049809 |
| TELEDDA | ASSLSGGALAEQF | 0.6966271 |
| TPPGPPN | ASSLSGGALAEQF | 0.788854 |
| TPPPPPPP | ASSLSGGALAEQF | 0.4585135 |
| TQSPALI | ASSFRDIQDTQY | 0.78980565 |
| EFAAAFPQ | ASSFRDIQDTQY | 0.80094385 |
| LLSGNLDVL | ASSFRDIQDTQY | 0.47330695 |
| GPANLGNYNI | ASSFRDIQDTQY | 0.6526047 |
| TLGPTLP | ASSFRDIQDTQY | 0.8572503 |
| VYAYSSCCK | ASSFRDIQDTQY | 0.83701545 |
| APAPAPAPNI | ASSFRDIQDTQY | 0.54937166 |
| PGGPVPIP | ASSFRDIQDTQY | 0.63195795 |
| FLTVALEP | ASSFRDIQDTQY | 0.49866685 |
| SQYFGANL | ASSFRDIQDTQY | 0.648474 |
| GGVKPAA | ASSFRDIQDTQY | 0.4327132 |
| GGTGGAGGAGGW | ASSFRDIQDTQY | 0.7676624 |
| GVGGVAGLGGAGG | ASSFRDIQDTQY | 0.6197833 |
| VLLVLATTI | ASSFRDIQDTQY | 0.7917167 |
| IGIFAAML | ASSFRDIQDTQY | 0.7274597 |

| | | |
|---|---|---|
| AVGDLID | ASSFRDIQDTQY | 0.8325007 |
| NISPLPL | ASSFRDIQDTQY | 0.5911524 |
| VTAVFVAI | ASSFRDIQDTQY | 0.6570298 |
| SGDDNDAAL | ASSFRDIQDTQY | 0.43253186 |
| AVAALAHS | ASSFRDIQDTQY | 0.5324257 |
| GGVTVMATP | ASSFRDIQDTQY | 0.5995749 |
| AGLAANVRD | ASSFRDIQDTQY | 0.67637104 |
| TNRPDLIDPALLRPG | ASSFRDIQDTQY | 0.32738483 |
| LANLGVTAL | ASSFRDIQDTQY | 0.8689809 |
| TGERIEL | ASSFRDIQDTQY | 0.6977526 |
| NGEELAA | ASSFRDIQDTQY | 0.8000605 |
| VLSLVWG | ASSFRDIQDTQY | 0.66870046 |
| IGYALWL | ASSFRDIQDTQY | 0.5573592 |
| LSLLLAPL | ASSFRDIQDTQY | 0.6043293 |
| TLDEALK | ASSFRDIQDTQY | 0.7741329 |
| TDGAGTAEGG | ASSFRDIQDTQY | 0.47362643 |
| NAPGGAGGNGG | ASSFRDIQDTQY | 0.2584965 |
| VSVFMGMGGNF | ASSFRDIQDTQY | 0.61464274 |
| LASYLAAL | ASSFRDIQDTQY | 0.39867103 |
| LVGGTPAL | ASSFRDIQDTQY | 0.86364883 |
| VRNADVL | ASSFRDIQDTQY | 0.836957 |
| TCSSPFLVSAM | ASSFRDIQDTQY | 0.7551552 |
| VHITDAE | ASSFRDIQDTQY | 0.8616463 |
| EYAAPSF | ASSFRDIQDTQY | 0.20617236 |
| VVDAFHGGL | ASSFRDIQDTQY | 0.70296955 |
| ASVVDPQ | ASSFRDIQDTQY | 0.85707694 |
| SYNEFGGLS | ASSFRDIQDTQY | 0.78670794 |
| DPDGSVY | ASSFRDIQDTQY | 0.6708322 |
| MVSTIHATE | ASSFRDIQDTQY | 0.83897626 |
| LAAGVADPV | ASSFRDIQDTQY | 0.5931051 |
| TKEQIAA | ASSFRDIQDTQY | 0.627211 |
| SLASPAAL | ASSFRDIQDTQY | 0.7377922 |
| IGLVFYGV | ASSFRDIQDTQY | 0.8110647 |
| FLLDASNL | ASSFRDIQDTQY | 0.81540966 |
| YQVGNFP | ASSFRDIQDTQY | 0.41863957 |
| ESLLSQST | ASSFRDIQDTQY | 0.8161373 |
| LFTVGFAAVG | ASSFRDIQDTQY | 0.8627388 |
| ATPPTPT | ASSFRDIQDTQY | 0.6033311 |
| TDLATLG | ASSFRDIQDTQY | 0.79296994 |
| SAGGTGR | ASSFRDIQDTQY | 0.7695728 |
| GFSNSGS | ASSFRDIQDTQY | 0.69519526 |
| GGNGGSGGTSV | ASSFRDIQDTQY | 0.52602077 |
| GTSVATGGAGN | ASSFRDIQDTQY | 0.48627377 |
| NGGNGGSGGTS | ASSFRDIQDTQY | 0.8154491 |
| AVVKFPH | ASSFRDIQDTQY | 0.49715626 |
| TPPPPPP | ASSFRDIQDTQY | 0.73620605 |
| AAPAAAAAPA | ASSFRDIQDTQY | 0.7116116 |
| IALNEDL | ASSFRDIQDTQY | 0.8860987 |
| IVNVSSM | ASSFRDIQDTQY | 0.75417596 |
| TGIPPAPR | ASSFRDIQDTQY | 0.8304531 |
| FDLGGGTF | ASSFRDIQDTQY | 0.51399326 |

| | | |
|---|---|---|
| FDLGGGTFDVS | ASSFRDIQDTQY | 0.77699393 |
| NEPTAAA | ASSFRDIQDTQY | 0.7871207 |
| NEPTAAALA | ASSFRDIQDTQY | 0.44897115 |
| GSIINISSI | ASSFRDIQDTQY | 0.84398925 |
| VAPGNAGTA | ASSFRDIQDTQY | 0.54824615 |
| TGGGASLE | ASSFRDIQDTQY | 0.7382194 |
| VAVYNHYKRI | ASSFRDIQDTQY | 0.8321084 |
| TDPAPATT | ASSFRDIQDTQY | 0.8172232 |
| TNRPDILDPALLRPG | ASSFRDIQDTQY | 0.70196605 |
| LTTTTTS | ASSFRDIQDTQY | 0.80361485 |
| PTGSGKTL | ASSFRDIQDTQY | 0.5352729 |
| TGSGKTL | ASSFRDIQDTQY | 0.8216065 |
| TIGHVDHGK | ASSFRDIQDTQY | 0.8263999 |
| TIGHVDHGKTT | ASSFRDIQDTQY | 0.7358136 |
| GHVDHGKTT | ASSFRDIQDTQY | 0.85250074 |
| AQFEHTLL | ASSFRDIQDTQY | 0.8407147 |
| IVNQDEL | ASSFRDIQDTQY | 0.37378103 |
| TNGSQFF | ASSFRDIQDTQY | 0.6928089 |
| TGKGIVE | ASSFRDIQDTQY | 0.83652765 |
| LISNASDA | ASSFRDIQDTQY | 0.8360759 |
| AELRTLGELTKIAKAHGAQ | ASSFRDIQDTQY | 0.8202501 |
| APNAVAAAEPSW | ASSFRDIQDTQY | 0.74392104 |
| APQVAGAATAVL | ASSFRDIQDTQY | 0.59819525 |
| ARRDPAAAAAILPTDA | ASSFRDIQDTQY | 0.85605156 |
| DAAARDVLAEAGLAEYF | ASSFRDIQDTQY | 0.8949597 |
| DDQIVLSAKANLAKVEQ | ASSFRDIQDTQY | 0.826851 |
| DELLPDTGEDSDEDGH | ASSFRDIQDTQY | 0.60421675 |
| ELPEAPANRATVPQD | ASSFRDIQDTQY | 0.6370744 |
| FGQSKIERVTGIG | ASSFRDIQDTQY | 0.8032943 |
| GAGGTGSTAGGAGGAGGAGGLY | ASSFRDIQDTQY | 0.8269533 |
| GHGGTGAAGGNGGTGGAGGSGI | ASSFRDIQDTQY | 0.5182692 |
| GNGGNGGAGGEGGPGGAGGAGA | ASSFRDIQDTQY | 0.73873323 |
| GSTAGNGGNGGNATNGTIAGQP | ASSFRDIQDTQY | 0.731365 |
| GTGGDGGNGGNGGTGG | ASSFRDIQDTQY | 0.26132867 |
| GVRRPAGIPADAPV | ASSFRDIQDTQY | 0.78449917 |
| GYRQVTDAHLLTLARR | ASSFRDIQDTQY | 0.8259169 |
| IAKMGQDGHDRG | ASSFRDIQDTQY | 0.8044201 |
| KAEPSIEIQYT | ASSFRDIQDTQY | 0.56716704 |
| KPVRIRMAADEDTVAVT | ASSFRDIQDTQY | 0.83061254 |
| MPPRIAGMRLLVIKPEP | ASSFRDIQDTQY | 0.8404467 |
| MKFYAVALFRSRGVPPTR | ASSFRDIQDTQY | 0.87629515 |
| NAGGFGILGAVAHSP | ASSFRDIQDTQY | 0.37962157 |
| NEGREITLAMAEM | ASSFRDIQDTQY | 0.21070915 |
| NELVEWSHPLGHRWRWE | ASSFRDIQDTQY | 0.700389 |
| NLGAGGTGGNGGNPG | ASSFRDIQDTQY | 0.7846767 |
| NTEWPKWETVMAADPVD | ASSFRDIQDTQY | 0.67624533 |
| PASAAPAETQP | ASSFRDIQDTQY | 0.58696336 |
| QQSSDGKNGVRAAEITGE | ASSFRDIQDTQY | 0.759521 |
| RAPDAVTTSWWKE | ASSFRDIQDTQY | 0.8402017 |
| SGFGNVGAHTSGWFNQ | ASSFRDIQDTQY | 0.48018098 |
| SGVAGAVGQAASVGGLKV | ASSFRDIQDTQY | 0.47234213 |

| | | |
|---|---|---|
| SNGGAGTGSAGGNGGT | ASSFRDIQDTQY | 0.7872918 |
| TGGAGGEGGAAG | ASSFRDIQDTQY | 0.8280325 |
| TGGDGGAAGAGGAPGHG | ASSFRDIQDTQY | 0.5303056 |
| TMPPLLAAIANAGRHGV | ASSFRDIQDTQY | 0.6167065 |
| TSSTAGNSGGA | ASSFRDIQDTQY | 0.73268205 |
| VPATVPVGILC | ASSFRDIQDTQY | 0.7788633 |
| VQRPDSGGVVLTGRIS | ASSFRDIQDTQY | 0.7372987 |
| WTPNPVPATKGVPAGLVQAT | ASSFRDIQDTQY | 0.4610785 |
| YALKRALAKFSDRFDVVII | ASSFRDIQDTQY | 0.8206023 |
| YLIGSVIV | ASSFRDIQDTQY | 0.4311026 |
| YGQLVAVID | ASSFRDIQDTQY | 0.8219922 |
| VTVGGMTLPA | ASSFRDIQDTQY | 0.6910914 |
| VLALPAV | ASSFRDIQDTQY | 0.77698934 |
| VIPPTLNYET | ASSFRDIQDTQY | 0.625499 |
| VGMGGMHPSSG | ASSFRDIQDTQY | 0.47459495 |
| VGLAGGAATA | ASSFRDIQDTQY | 0.50247604 |
| VGGAGGAGGSGP | ASSFRDIQDTQY | 0.44390902 |
| VFAAFMV | ASSFRDIQDTQY | 0.79285705 |
| TLYDKLR | ASSFRDIQDTQY | 0.703494 |
| TLAVFTVL | ASSFRDIQDTQY | 0.8168274 |
| TGSVQPARV | ASSFRDIQDTQY | 0.7518396 |
| TGGGAGSGGT | ASSFRDIQDTQY | 0.5785212 |
| SHLETSL | ASSFRDIQDTQY | 0.7942338 |
| SFTTGMT | ASSFRDIQDTQY | 0.4667838 |
| QVEVAKRMGVRQPTVS | ASSFRDIQDTQY | 0.73964345 |
| QGSGGPALAM | ASSFRDIQDTQY | 0.75517774 |
| MLAPSNP | ASSFRDIQDTQY | 0.82512236 |
| LVNNAAI | ASSFRDIQDTQY | 0.5089681 |
| LTGEINTLA | ASSFRDIQDTQY | 0.8158281 |
| LSIFGPV | ASSFRDIQDTQY | 0.77643037 |
| LLALLANL | ASSFRDIQDTQY | 0.54099417 |
| LISNASDALDKLR | ASSFRDIQDTQY | 0.79685736 |
| LINEAVP | ASSFRDIQDTQY | 0.8364723 |
| LGYAFSGL | ASSFRDIQDTQY | 0.79310226 |
| LDFVAVAL | ASSFRDIQDTQY | 0.396494 |
| LAQPGAE | ASSFRDIQDTQY | 0.78695 |
| ISNLGAVTGF | ASSFRDIQDTQY | 0.530436 |
| HQEDPGRI | ASSFRDIQDTQY | 0.7544899 |
| GTPGEPTP | ASSFRDIQDTQY | 0.74705553 |
| GFFNSGS | ASSFRDIQDTQY | 0.6837565 |
| GAGGGGLTAGAN | ASSFRDIQDTQY | 0.8374312 |
| FVDALVNM | ASSFRDIQDTQY | 0.56157947 |
| FLTVAPEL | ASSFRDIQDTQY | 0.5851448 |
| FETLTAL | ASSFRDIQDTQY | 0.70049626 |
| DVINPIEL | ASSFRDIQDTQY | 0.75752383 |
| APVSGAPV | ASSFRDIQDTQY | 0.8347153 |
| ALVGGLAW | ASSFRDIQDTQY | 0.4897404 |
| ALAAAAATAI | ASSFRDIQDTQY | 0.756137 |
| AFGLGLLL | ASSFRDIQDTQY | 0.63946855 |
| AFFAAASL | ASSFRDIQDTQY | 0.45456636 |
| AAAALPAFSPP | ASSFRDIQDTQY | 0.79345423 |

| | | |
|---|---|---|
| VVLAPAL | ASSFRDIQDTQY | 0.7098541 |
| TIGHVDHGKTTL | ASSFRDIQDTQY | 0.5255926 |
| LISNASDALDK | ASSFRDIQDTQY | 0.7931644 |
| LISNASDAL | ASSFRDIQDTQY | 0.5763119 |
| ISNASDALDK | ASSFRDIQDTQY | 0.8129798 |
| NQAETLVYQTE | ASSFRDIQDTQY | 0.869377 |
| TGGTGGTAGAG | ASSFRDIQDTQY | 0.43216997 |
| TGSTAGGAGGA | ASSFRDIQDTQY | 0.8195751 |
| DPLAVPVAAGPV | ASSFRDIQDTQY | 0.75397444 |
| GGTGGTPGNGGN | ASSFRDIQDTQY | 0.699033 |
| PAGLAGVDPATL | ASSFRDIQDTQY | 0.6862725 |
| GNNLVVEAPHAGASV | ASSFRDIQDTQY | 0.575506 |
| KVTRIMSASTLEQPA | ASSFRDIQDTQY | 0.85275376 |
| LGFALPIALPRRTVV | ASSFRDIQDTQY | 0.82488376 |
| TRLGNLTARKLVVLV | ASSFRDIQDTQY | 0.7861967 |
| WSQQRRLIYFPSAGP | ASSFRDIQDTQY | 0.78226197 |
| GPNGNPDPMAAAVDIR | ASSFRDIQDTQY | 0.76418054 |
| TQRCVSVADLRAAASR | ASSFRDIQDTQY | 0.82796884 |
| QFVLDVAEHPVEVLTETV | ASSFRDIQDTQY | 0.55301756 |
| GGAGGTGGAGGNGGTQAGDG | ASSFRDIQDTQY | 0.37072814 |
| PLLATLRAIAVDVDAKPAQV | ASSFRDIQDTQY | 0.68156517 |
| APKFRTGGYMIDFWGVGYQVAK | ASSFRDIQDTQY | 0.65848446 |
| GAGGAGGAGGAAGGPGGTGGTG | ASSFRDIQDTQY | 0.5794798 |
| IGALGLLATDVAPAGSVAFGVA | ASSFRDIQDTQY | 0.48929513 |
| AGGAGGIGGNGGAGGAGGVLMG | ASSFRDIQDTQY | 0.8260033 |
| ADELIGLL | ASSFRDIQDTQY | 0.87148786 |
| ADRALKA | ASSFRDIQDTQY | 0.6194765 |
| AGAVGQVGGAGGA | ASSFRDIQDTQY | 0.8854798 |
| AGGGGGLGGAP | ASSFRDIQDTQY | 0.4878006 |
| AGGLLYGNGGAG | ASSFRDIQDTQY | 0.8207885 |
| AGLESIGL | ASSFRDIQDTQY | 0.8411129 |
| AGLKTGII | ASSFRDIQDTQY | 0.5094036 |
| ASLAALLL | ASSFRDIQDTQY | 0.80008197 |
| ASVINISSI | ASSFRDIQDTQY | 0.51344436 |
| ATPPGPA | ASSFRDIQDTQY | 0.6492123 |
| ATVASALV | ASSFRDIQDTQY | 0.8359891 |
| ATVASLIL | ASSFRDIQDTQY | 0.6901455 |
| AVNGPVLIRH | ASSFRDIQDTQY | 0.7897242 |
| DGAAGAVGI | ASSFRDIQDTQY | 0.6823107 |
| DPLLHGI | ASSFRDIQDTQY | 0.5176766 |
| ELGAALGAL | ASSFRDIQDTQY | 0.7656847 |
| ELGRTQAV | ASSFRDIQDTQY | 0.8136995 |
| FVAGGYLS | ASSFRDIQDTQY | 0.7393464 |
| GASGGAGGAGGW | ASSFRDIQDTQY | 0.8932885 |
| GGAGAGGDGGAGGI | ASSFRDIQDTQY | 0.48452434 |
| GTGHNGSQPGMGGNG | ASSFRDIQDTQY | 0.84864736 |
| GYAAASASAS | ASSFRDIQDTQY | 0.83955556 |
| HQVDVAAVQQG | ASSFRDIQDTQY | 0.7123432 |
| IGAAIAHT | ASSFRDIQDTQY | 0.6916704 |
| IGMIEVAQL | ASSFRDIQDTQY | 0.8078637 |
| IRLAPEHTSAMD | ASSFRDIQDTQY | 0.6351737 |

| | | |
|---|---|---|
| IYNVSVL | ASSFRDIQDTQY | 0.716273 |
| KQGFLGM | ASSFRDIQDTQY | 0.8636353 |
| LESVITQL | ASSFRDIQDTQY | 0.3551225 |
| LGLGALVVV | ASSFRDIQDTQY | 0.6735156 |
| LIGNGGNGGSGGI | ASSFRDIQDTQY | 0.54747915 |
| LNAINAPTL | ASSFRDIQDTQY | 0.6119456 |
| LQLLNTGV | ASSFRDIQDTQY | 0.4849993 |
| LQTSIEF | ASSFRDIQDTQY | 0.2896439 |
| MSFAVSEL | ASSFRDIQDTQY | 0.5914762 |
| NTGALGL | ASSFRDIQDTQY | 0.41388178 |
| PPGAGGAGGAGTT | ASSFRDIQDTQY | 0.6943855 |
| QLVGLGI | ASSFRDIQDTQY | 0.67329216 |
| RFLEQVHQL | ASSFRDIQDTQY | 0.4709506 |
| RGVGYVLR | ASSFRDIQDTQY | 0.6803676 |
| SATGVFAGVI | ASSFRDIQDTQY | 0.70569026 |
| SLPAITLP | ASSFRDIQDTQY | 0.3171439 |
| STALAFTV | ASSFRDIQDTQY | 0.41519976 |
| SVLTVGLGK | ASSFRDIQDTQY | 0.42205322 |
| TALAGAGSGI | ASSFRDIQDTQY | 0.53276724 |
| TGLPFTGI | ASSFRDIQDTQY | 0.30431533 |
| VAEHVTV | ASSFRDIQDTQY | 0.61908984 |
| VDIVLNSL | ASSFRDIQDTQY | 0.39964473 |
| VLTGGGAVL | ASSFRDIQDTQY | 0.24170086 |
| VNVGLAYGI | ASSFRDIQDTQY | 0.60680604 |
| VSSAAVGAV | ASSFRDIQDTQY | 0.5373674 |
| VTSVYEAGV | ASSFRDIQDTQY | 0.47399515 |
| WVALDRGA | ASSFRDIQDTQY | 0.6627972 |
| YLVGSVVI | ASSFRDIQDTQY | 0.30228853 |
| GPPGTGKTLL | ASSFRDIQDTQY | 0.39974374 |
| GVGMTKF | ASSFRDIQDTQY | 0.6148473 |
| IGIGGSDL | ASSFRDIQDTQY | 0.68921405 |
| IGTLLAEL | ASSFRDIQDTQY | 0.35473865 |
| IYTSGTT | ASSFRDIQDTQY | 0.6247577 |
| LSFPAGV | ASSFRDIQDTQY | 0.69304806 |
| LYTSGTT | ASSFRDIQDTQY | 0.5354062 |
| VGAFPFGF | ASSFRDIQDTQY | 0.68688804 |
| VGAGTVEF | ASSFRDIQDTQY | 0.698175 |
| VGKSTLLL | ASSFRDIQDTQY | 0.47209477 |
| AALAHAGGGIV | ASSFRDIQDTQY | 0.6821967 |
| AALVAVSLL | ASSFRDIQDTQY | 0.698912 |
| AAVSRAVAAL | ASSFRDIQDTQY | 0.44049925 |
| AGGSSGIGGF | ASSFRDIQDTQY | 0.60104233 |
| AGLTVANANG | ASSFRDIQDTQY | 0.3941862 |
| AGLVGAVL | ASSFRDIQDTQY | 0.581136 |
| ALSQAFTV | ASSFRDIQDTQY | 0.28088605 |
| ALTFSVSL | ASSFRDIQDTQY | 0.6111162 |
| ALTGAALVAL | ASSFRDIQDTQY | 0.6445434 |
| ASFSIGAGVL | ASSFRDIQDTQY | 0.5549033 |
| DGSVYKI | ASSFRDIQDTQY | 0.32047513 |
| EQLAEIV | ASSFRDIQDTQY | 0.30478725 |
| FAVSELTVM | ASSFRDIQDTQY | 0.44453904 |

| | | |
|---|---|---|
| GAAPLAAGL | ASSFRDIQDTQY | 0.3988288 |
| GADGGDGGSGGAGG | ASSFRDIQDTQY | 0.28865802 |
| GFGLDTGGAGG | ASSFRDIQDTQY | 0.4404932 |
| GLASGGLL | ASSFRDIQDTQY | 0.65517586 |
| GRIAGAAVLL | ASSFRDIQDTQY | 0.67328274 |
| GSAGSLNTSF | ASSFRDIQDTQY | 0.30411166 |
| GTGQIGI | ASSFRDIQDTQY | 0.7177585 |
| GTLIALIL | ASSFRDIQDTQY | 0.66620326 |
| IAATMGFVVG | ASSFRDIQDTQY | 0.69823587 |
| IFAQALL | ASSFRDIQDTQY | 0.52850753 |
| IGAAVALGF | ASSFRDIQDTQY | 0.34231865 |
| IGLTPLAL | ASSFRDIQDTQY | 0.708066 |
| ILGSNTSTL | ASSFRDIQDTQY | 0.5048606 |
| IVFSLTVL | ASSFRDIQDTQY | 0.5220971 |
| LASKATGF | ASSFRDIQDTQY | 0.66135335 |
| LASVALIL | ASSFRDIQDTQY | 0.42344838 |
| LFAVGLTL | ASSFRDIQDTQY | 0.43914503 |
| LFNGGIGTYI | ASSFRDIQDTQY | 0.6312277 |
| LTLGAGII | ASSFRDIQDTQY | 0.65205926 |
| LVAAALGDI | ASSFRDIQDTQY | 0.35226813 |
| NTAFVQLGI | ASSFRDIQDTQY | 0.35096323 |
| QLFSGAEL | ASSFRDIQDTQY | 0.36117908 |
| SAATFITL | ASSFRDIQDTQY | 0.506315 |
| SNYGLANLGV | ASSFRDIQDTQY | 0.6884607 |
| SYGGAYAVM | ASSFRDIQDTQY | 0.5853956 |
| TTGVGGLTLG | ASSFRDIQDTQY | 0.4306389 |
| TTQTIYI | ASSFRDIQDTQY | 0.650895 |
| VAAGAGLGAVI | ASSFRDIQDTQY | 0.5799761 |
| VAAVVTLL | ASSFRDIQDTQY | 0.5831132 |
| VGFLGGLI | ASSFRDIQDTQY | 0.6979516 |
| VGVIPILL | ASSFRDIQDTQY | 0.46696252 |
| VIFASSLI | ASSFRDIQDTQY | 0.63955665 |
| VKGGDGGAAAG | ASSFRDIQDTQY | 0.62464285 |
| VTKTFTNI | ASSFRDIQDTQY | 0.65785575 |
| VVAAAILL | ASSFRDIQDTQY | 0.68372977 |
| YAALDVEL | ASSFRDIQDTQY | 0.5280056 |
| LASVALLL | ASSFRDIQDTQY | 0.56470776 |
| LDAVHRV | ASSFRDIQDTQY | 0.28305984 |
| QGLSSSIF | ASSFRDIQDTQY | 0.49537987 |
| RGTFANIRL | ASSFRDIQDTQY | 0.66557586 |
| RQLWWGHRI | ASSFRDIQDTQY | 0.5396203 |
| YGPPGTGKTLL | ASSFRDIQDTQY | 0.6932048 |
| AALVGAPANVA | ASSFRDIQDTQY | 0.64196694 |
| AATFRRTE | ASSFRDIQDTQY | 0.44854584 |
| AGALGGGI | ASSFRDIQDTQY | 0.70869267 |
| AGGAGGAGGIGAG | ASSFRDIQDTQY | 0.67631364 |
| AGSGALGL | ASSFRDIQDTQY | 0.37628534 |
| AQIADQI | ASSFRDIQDTQY | 0.5300392 |
| ASAVAVTYL | ASSFRDIQDTQY | 0.7150835 |
| ASTVLGGTGG | ASSFRDIQDTQY | 0.685933 |
| AVGGEPAK | ASSFRDIQDTQY | 0.49586272 |

| | | |
|---|---|---|
| CEQGTFGGTC | ASSFRDIQDTQY | 0.57979476 |
| DEGKAPAG | ASSFRDIQDTQY | 0.7232194 |
| DTVVFPAL | ASSFRDIQDTQY | 0.5107594 |
| FEGITMAG | ASSFRDIQDTQY | 0.6897745 |
| FPLVVQD | ASSFRDIQDTQY | 0.26695463 |
| GAFTTPQ | ASSFRDIQDTQY | 0.5124093 |
| GFGGFGTAGGVGGA | ASSFRDIQDTQY | 0.34339768 |
| GFLAQFL | ASSFRDIQDTQY | 0.26112738 |
| GGAGGAGGNAGVA | ASSFRDIQDTQY | 0.66999745 |
| GGVGTAGAGGAGGAGG | ASSFRDIQDTQY | 0.52426296 |
| GNGGGGDAGNAGSGGN | ASSFRDIQDTQY | 0.6442802 |
| GTGGAGGAGNDAGS | ASSFRDIQDTQY | 0.49442473 |
| GTTNTGWFNS | ASSFRDIQDTQY | 0.6895009 |
| IFMGPLFTL | ASSFRDIQDTQY | 0.65659064 |
| ILAHAIEAI | ASSFRDIQDTQY | 0.5643261 |
| IRSAIREAL | ASSFRDIQDTQY | 0.3803891 |
| ITFAGGG | ASSFRDIQDTQY | 0.5563043 |
| ITGMSGHQEV | ASSFRDIQDTQY | 0.7139696 |
| IVAGLACTL | ASSFRDIQDTQY | 0.5855241 |
| IVGGGLAGA | ASSFRDIQDTQY | 0.2854812 |
| IVTGVLNI | ASSFRDIQDTQY | 0.58129144 |
| LAQQGCI | ASSFRDIQDTQY | 0.52030873 |
| LCLHVEN | ASSFRDIQDTQY | 0.2659488 |
| LDAISDGS | ASSFRDIQDTQY | 0.2562928 |
| LGGFAAML | ASSFRDIQDTQY | 0.31447014 |
| LGGKNPMI | ASSFRDIQDTQY | 0.45798397 |
| LGPKGVAAIL | ASSFRDIQDTQY | 0.66385484 |
| LMVGVSI | ASSFRDIQDTQY | 0.65464103 |
| LVGAAFVL | ASSFRDIQDTQY | 0.4840625 |
| LVLAVAPL | ASSFRDIQDTQY | 0.63558394 |
| MAAIAVIDAA | ASSFRDIQDTQY | 0.5229467 |
| NNGINAELQ | ASSFRDIQDTQY | 0.2337846 |
| RGEPLVLVHG | ASSFRDIQDTQY | 0.38254222 |
| RGGAGGAAGLI | ASSFRDIQDTQY | 0.3552036 |
| RSYRMHPAV | ASSFRDIQDTQY | 0.6752355 |
| SAAHLLAGQLA | ASSFRDIQDTQY | 0.4495616 |
| SAAVGSY | ASSFRDIQDTQY | 0.49914345 |
| SPHAVLHRI | ASSFRDIQDTQY | 0.580375 |
| SQIAKVL | ASSFRDIQDTQY | 0.6768751 |
| TALGHLVP | ASSFRDIQDTQY | 0.600561 |
| TAPTTSGAA | ASSFRDIQDTQY | 0.31592044 |
| TDGDDGNGGSPGT | ASSFRDIQDTQY | 0.24661279 |
| TDGLGEN | ASSFRDIQDTQY | 0.60326827 |
| TGGPPLPGQA | ASSFRDIQDTQY | 0.39732906 |
| TIPAITI | ASSFRDIQDTQY | 0.33632493 |
| TIPGFTI | ASSFRDIQDTQY | 0.35012257 |
| TSIASGALA | ASSFRDIQDTQY | 0.2835948 |
| TVTLPGAAI | ASSFRDIQDTQY | 0.2972732 |
| VASGLSAALP | ASSFRDIQDTQY | 0.6709463 |
| VGAGVIGI | ASSFRDIQDTQY | 0.40534717 |
| VGPKRFGV | ASSFRDIQDTQY | 0.49694502 |

| | | |
|---|---|---|
| VNVAFPD | ASSFRDIQDTQY | 0.7037866 |
| VSMTNSGVS | ASSFRDIQDTQY | 0.66961575 |
| VTDVLGDLSV | ASSFRDIQDTQY | 0.5566913 |
| VVVVGAGFAGL | ASSFRDIQDTQY | 0.35859194 |
| VYTVGTVV | ASSFRDIQDTQY | 0.41492492 |
| WTTFSGP | ASSFRDIQDTQY | 0.395176 |
| YDELVEAF | ASSFRDIQDTQY | 0.36543834 |
| YGTGGAGGNGG | ASSFRDIQDTQY | 0.7027786 |
| YTNVMAV | ASSFRDIQDTQY | 0.5401056 |
| FLEQVHQL | ASSFRDIQDTQY | 0.40161324 |
| GLSLLAVAV | ASSFRDIQDTQY | 0.60660994 |
| LEQVHQL | ASSFRDIQDTQY | 0.68621457 |
| LNAAGAI | ASSFRDIQDTQY | 0.7066996 |
| PAVHASL | ASSFRDIQDTQY | 0.38653544 |
| TELEDDA | ASSFRDIQDTQY | 0.4773586 |
| TPPGPPN | ASSFRDIQDTQY | 0.5585247 |
| TPPPPPPP | ASSFRDIQDTQY | 0.5693798 |
| TQSPALI | TCSAAQGTEVF | 0.38110757 |
| EFAAAFPQ | TCSAAQGTEVF | 0.3944417 |
| LLSGNLDVL | TCSAAQGTEVF | 0.6288984 |
| GPANLGNYNI | TCSAAQGTEVF | 0.7243962 |
| TLGPTLP | TCSAAQGTEVF | 0.50855213 |
| VYAYSSCCK | TCSAAQGTEVF | 0.35153133 |
| APAPAPAPNI | TCSAAQGTEVF | 0.4751001 |
| PGGPVPIP | TCSAAQGTEVF | 0.3699525 |
| FLTVALEP | TCSAAQGTEVF | 0.49411708 |
| SQYFGANL | TCSAAQGTEVF | 0.57509524 |
| GGVKPAA | TCSAAQGTEVF | 0.34592122 |
| GGTGGAGGAGGW | TCSAAQGTEVF | 0.5272415 |
| GVGGVAGLGGAGG | TCSAAQGTEVF | 0.71653974 |
| VLLVLATTI | TCSAAQGTEVF | 0.6185457 |
| IGIFAAML | TCSAAQGTEVF | 0.64986587 |
| AVGDLID | TCSAAQGTEVF | 0.69052637 |
| NISPLPL | TCSAAQGTEVF | 0.6272235 |
| VTAVFVAI | TCSAAQGTEVF | 0.70030224 |
| SGDDNDAAL | TCSAAQGTEVF | 0.6963005 |
| AVAALAHS | TCSAAQGTEVF | 0.5176718 |
| GGVTVMATP | TCSAAQGTEVF | 0.5053476 |
| AGLAANVRD | TCSAAQGTEVF | 0.66016054 |
| TNRPDLIDPALLRPG | TCSAAQGTEVF | 0.58398664 |
| LANLGVTAL | TCSAAQGTEVF | 0.53941435 |
| TGERIEL | TCSAAQGTEVF | 0.55181897 |
| NGEELAA | TCSAAQGTEVF | 0.6837673 |
| VLSLVWG | TCSAAQGTEVF | 0.43222874 |
| IGYALWL | TCSAAQGTEVF | 0.7033303 |
| LSLLLAPL | TCSAAQGTEVF | 0.2779326 |
| TLDEALK | TCSAAQGTEVF | 0.35926586 |
| TDGAGTAEGG | TCSAAQGTEVF | 0.6738541 |
| NAPGGAGGNGG | TCSAAQGTEVF | 0.24003598 |
| VSVFMGMGGNF | TCSAAQGTEVF | 0.27099705 |
| LASYLAAL | TCSAAQGTEVF | 0.5257392 |

| | | |
|---|---|---|
| LVGGTPAL | TCSAAQGTEVF | 0.6912875 |
| VRNADVL | TCSAAQGTEVF | 0.24277353 |
| TCSSPFLVSAM | TCSAAQGTEVF | 0.4183463 |
| VHITDAE | TCSAAQGTEVF | 0.29794702 |
| EYAAPSF | TCSAAQGTEVF | 0.6229508 |
| VVDAFHGGL | TCSAAQGTEVF | 0.5860355 |
| ASVVDPQ | TCSAAQGTEVF | 0.30705544 |
| SYNEFGGLS | TCSAAQGTEVF | 0.656289 |
| DPDGSVY | TCSAAQGTEVF | 0.39045942 |
| MVSTIHATE | TCSAAQGTEVF | 0.43398005 |
| LAAGVADPV | TCSAAQGTEVF | 0.6867771 |
| TKEQIAA | TCSAAQGTEVF | 0.7157156 |
| SLASPAAL | TCSAAQGTEVF | 0.40002203 |
| IGLVFYGV | TCSAAQGTEVF | 0.667295 |
| FLLDASNL | TCSAAQGTEVF | 0.44400173 |
| YQVGNFP | TCSAAQGTEVF | 0.7120279 |
| ESLLSQST | TCSAAQGTEVF | 0.5086567 |
| LFTVGFAAVG | TCSAAQGTEVF | 0.2940729 |
| ATPPTPT | TCSAAQGTEVF | 0.6716929 |
| TDLATLG | TCSAAQGTEVF | 0.6713066 |
| SAGGTGR | TCSAAQGTEVF | 0.5470822 |
| GFSNSGS | TCSAAQGTEVF | 0.67771816 |
| GGNGGSGGTSV | TCSAAQGTEVF | 0.46364665 |
| GTSVATGGAGN | TCSAAQGTEVF | 0.51594967 |
| NGGNGGSGGTS | TCSAAQGTEVF | 0.6891482 |
| AVVKFPH | TCSAAQGTEVF | 0.68199944 |
| TPPPPPP | TCSAAQGTEVF | 0.36687547 |
| AAPAAAAAPA | TCSAAQGTEVF | 0.7046289 |
| IALNEDL | TCSAAQGTEVF | 0.50106746 |
| IVNVSSM | TCSAAQGTEVF | 0.37933514 |
| TGIPPAPR | TCSAAQGTEVF | 0.7071904 |
| FDLGGGTF | TCSAAQGTEVF | 0.5755629 |
| FDLGGGTFDVS | TCSAAQGTEVF | 0.6563189 |
| NEPTAAA | TCSAAQGTEVF | 0.5943128 |
| NEPTAAALA | TCSAAQGTEVF | 0.7089059 |
| GSIINISSI | TCSAAQGTEVF | 0.3278392 |
| VAPGNAGTA | TCSAAQGTEVF | 0.6280234 |
| TGGGASLE | TCSAAQGTEVF | 0.5690243 |
| VAVYNHYKRI | TCSAAQGTEVF | 0.67217904 |
| TDPAPATT | TCSAAQGTEVF | 0.53325313 |
| TNRPDILDPALLRPG | TCSAAQGTEVF | 0.34177345 |
| LTTTTTS | TCSAAQGTEVF | 0.63587236 |
| PTGSGKTL | TCSAAQGTEVF | 0.67637986 |
| TGSGKTL | TCSAAQGTEVF | 0.4451205 |
| TIGHVDHGK | TCSAAQGTEVF | 0.67571354 |
| TIGHVDHGKTT | TCSAAQGTEVF | 0.71275485 |
| GHVDHGKTT | TCSAAQGTEVF | 0.38529313 |
| AQFEHTLL | TCSAAQGTEVF | 0.55708104 |
| IVNQDEL | TCSAAQGTEVF | 0.57295835 |
| TNGSQFF | TCSAAQGTEVF | 0.7108535 |
| TGKGIVE | TCSAAQGTEVF | 0.5405335 |

| | | |
|---|---|---|
| LISNASDA | TCSAAQGTEVF | 0.7018201 |
| AELRTLGELTKIAKAHGAQ | TCSAAQGTEVF | 0.510775 |
| APNAVAAAEPSW | TCSAAQGTEVF | 0.6496099 |
| APQVAGAATAVL | TCSAAQGTEVF | 0.46292943 |
| ARRDPAAAAAILPTDA | TCSAAQGTEVF | 0.592405 |
| DAAARDVLAEAGLAEYF | TCSAAQGTEVF | 0.30633962 |
| DDQIVLSAKANLAKVEQ | TCSAAQGTEVF | 0.63521755 |
| DELLPDTGEDSDEDGH | TCSAAQGTEVF | 0.6553473 |
| ELPEAPANRATVPQD | TCSAAQGTEVF | 0.6612013 |
| FGQSKIERVTGIG | TCSAAQGTEVF | 0.27385807 |
| GAGGTGSTAGGAGGAGGAGGLY | TCSAAQGTEVF | 0.6608577 |
| GHGGTGAAGGNGGTGGAGGSGI | TCSAAQGTEVF | 0.65556324 |
| GNGGNGGAGGEGGPGGAGGAGA | TCSAAQGTEVF | 0.72293997 |
| GSTAGNGGNGGNATNGTIAGQP | TCSAAQGTEVF | 0.46056998 |
| GTGGDGGNGGNGGTGG | TCSAAQGTEVF | 0.67772865 |
| GVRRPAGIPADAPV | TCSAAQGTEVF | 0.42088956 |
| GYRQVTDAHLLTLARR | TCSAAQGTEVF | 0.5512137 |
| IAKMGQDGHDRG | TCSAAQGTEVF | 0.29537594 |
| KAEPSIEIQYT | TCSAAQGTEVF | 0.44144517 |
| KPVRIRMAADEDTVAVT | TCSAAQGTEVF | 0.5494885 |
| MPPRIAGMRLLVIKPEP | TCSAAQGTEVF | 0.43374574 |
| MKFYAVALFRSRGVPPTR | TCSAAQGTEVF | 0.69869256 |
| NAGGFGILGAVAHSP | TCSAAQGTEVF | 0.7049837 |
| NEGREITLAMAEM | TCSAAQGTEVF | 0.6837424 |
| NELVEWSHPLGHRWRWE | TCSAAQGTEVF | 0.65320724 |
| NLGAGGTGGNGGNPG | TCSAAQGTEVF | 0.7041 |
| NTEWPKWETVMAADPVD | TCSAAQGTEVF | 0.6747657 |
| PASAAPAETQP | TCSAAQGTEVF | 0.5110253 |
| QQSSDGKNGVRAAEITGE | TCSAAQGTEVF | 0.5377002 |
| RAPDAVTTSWWKE | TCSAAQGTEVF | 0.3135395 |
| SGFGNVGAHTSGWFNQ | TCSAAQGTEVF | 0.5091244 |
| SGVAGAVGQAASVGGLKV | TCSAAQGTEVF | 0.43817523 |
| SNGGAGTGSAGGNGGT | TCSAAQGTEVF | 0.62228245 |
| TGGAGGEGGAAG | TCSAAQGTEVF | 0.24153528 |
| TGGDGGAAGAGGAPGHG | TCSAAQGTEVF | 0.66063535 |
| TMPPLLAAIANAGRHGV | TCSAAQGTEVF | 0.7054182 |
| TSSTAGNSGGA | TCSAAQGTEVF | 0.39011988 |
| VPATVPVGILC | TCSAAQGTEVF | 0.6351802 |
| VQRPDSGGVVLTGRIS | TCSAAQGTEVF | 0.66939586 |
| WTPNPVPATKGVPAGLVQAT | TCSAAQGTEVF | 0.34503126 |
| YALKRALAKFSDRFDVVII | TCSAAQGTEVF | 0.58496845 |
| YLIGSVIV | TCSAAQGTEVF | 0.38425502 |
| YGQLVAVID | TCSAAQGTEVF | 0.66965044 |
| VTVGGMTLPA | TCSAAQGTEVF | 0.6971762 |
| VLALPAV | TCSAAQGTEVF | 0.6100068 |
| VIPPTLNYET | TCSAAQGTEVF | 0.65550137 |
| VGMGGMHPSSG | TCSAAQGTEVF | 0.6314412 |
| VGLAGGAATA | TCSAAQGTEVF | 0.30757046 |
| VGGAGGAGGSGP | TCSAAQGTEVF | 0.66217494 |
| VFAAFMV | TCSAAQGTEVF | 0.24464238 |
| TLYDKLR | TCSAAQGTEVF | 0.60521257 |

| | | |
|---|---|---|
| TLAVFTVL | TCSAAQGTEVF | 0.3837701 |
| TGSVQPARV | TCSAAQGTEVF | 0.5677213 |
| TGGGAGSGGT | TCSAAQGTEVF | 0.5291989 |
| SHLETSL | TCSAAQGTEVF | 0.5181016 |
| SFTTGMT | TCSAAQGTEVF | 0.43182996 |
| QVEVAKRMGVRQPTVS | TCSAAQGTEVF | 0.5971196 |
| QGSGGPALAM | TCSAAQGTEVF | 0.4947987 |
| MLAPSNP | TCSAAQGTEVF | 0.30124167 |
| LVNNAAI | TCSAAQGTEVF | 0.6947967 |
| LTGEINTLA | TCSAAQGTEVF | 0.4209489 |
| LSIFGPV | TCSAAQGTEVF | 0.60588706 |
| LLALLANL | TCSAAQGTEVF | 0.65396583 |
| LISNASDALDKLR | TCSAAQGTEVF | 0.5782963 |
| LINEAVP | TCSAAQGTEVF | 0.33389544 |
| LGYAFSGL | TCSAAQGTEVF | 0.5162951 |
| LDFVAVAL | TCSAAQGTEVF | 0.52522033 |
| LAQPGAE | TCSAAQGTEVF | 0.49708092 |
| ISNLGAVTGF | TCSAAQGTEVF | 0.24960974 |
| HQEDPGRI | TCSAAQGTEVF | 0.6279941 |
| GTPGEPTP | TCSAAQGTEVF | 0.3668564 |
| GFFNSGS | TCSAAQGTEVF | 0.4873035 |
| GAGGGGLTAGAN | TCSAAQGTEVF | 0.5371207 |
| FVDALVNM | TCSAAQGTEVF | 0.6845242 |
| FLTVAPEL | TCSAAQGTEVF | 0.32763508 |
| FETLTAL | TCSAAQGTEVF | 0.47029752 |
| DVINPIEL | TCSAAQGTEVF | 0.6898561 |
| APVSGAPV | TCSAAQGTEVF | 0.5885612 |
| ALVGGLAW | TCSAAQGTEVF | 0.69376993 |
| ALAAAAATAI | TCSAAQGTEVF | 0.5951043 |
| AFGLGLLL | TCSAAQGTEVF | 0.44799477 |
| AFFAAASL | TCSAAQGTEVF | 0.63658214 |
| AAAALPAFSPP | TCSAAQGTEVF | 0.4164831 |
| VVLAPAL | TCSAAQGTEVF | 0.30818036 |
| TIGHVDHGKTTL | TCSAAQGTEVF | 0.30913258 |
| LISNASDALDK | TCSAAQGTEVF | 0.47930893 |
| LISNASDAL | TCSAAQGTEVF | 0.27465013 |
| ISNASDALDK | TCSAAQGTEVF | 0.6301087 |
| NQAETLVYQTE | TCSAAQGTEVF | 0.69716364 |
| TGGTGGTAGAG | TCSAAQGTEVF | 0.5016198 |
| TGSTAGGAGGA | TCSAAQGTEVF | 0.32097572 |
| DPLAVPVAAGPV | TCSAAQGTEVF | 0.40426272 |
| GGTGGTPGNGGN | TCSAAQGTEVF | 0.45179307 |
| PAGLAGVDPATL | TCSAAQGTEVF | 0.65082616 |
| GNNLVVEAPHAGASV | TCSAAQGTEVF | 0.5460365 |
| KVTRIMSASTLEQPA | TCSAAQGTEVF | 0.7178271 |
| LGFALPIALPRRTVV | TCSAAQGTEVF | 0.66693455 |
| TRLGNLTARKLVVLV | TCSAAQGTEVF | 0.45797187 |
| WSQQRRLIYFPSAGP | TCSAAQGTEVF | 0.6719311 |
| GPNGNPDPMAAAVDIR | TCSAAQGTEVF | 0.51812446 |
| TQRCVSVADLRAAASR | TCSAAQGTEVF | 0.5531995 |
| QFVLDVAEHPVEVLTETV | TCSAAQGTEVF | 0.5750241 |

| | | |
|---|---|---|
| GGAGGTGGAGGNGGTQAGDG | TCSAAQGTEVF | 0.51504004 |
| PLLATLRAIAVDVDAKPAQV | TCSAAQGTEVF | 0.5590193 |
| APKFRTGGYMIDFWGVGYQVAK | TCSAAQGTEVF | 0.5052423 |
| GAGGAGGAGGAAGGPGGTGGTG | TCSAAQGTEVF | 0.32387683 |
| IGALGLLATDVAPAGSVAFGVA | TCSAAQGTEVF | 0.5748127 |
| AGGAGGIGGNGGAGGAGGVLMG | TCSAAQGTEVF | 0.3628453 |
| ADELIGLL | TCSAAQGTEVF | 0.30509365 |
| ADRALKA | TCSAAQGTEVF | 0.5133678 |
| AGAVGQVGGAGGA | TCSAAQGTEVF | 0.7127167 |
| AGGGGGLGGAP | TCSAAQGTEVF | 0.68377084 |
| AGGLLYGNGGAG | TCSAAQGTEVF | 0.40933323 |
| AGLESIGL | TCSAAQGTEVF | 0.37134212 |
| AGLKTGII | TCSAAQGTEVF | 0.66910005 |
| ASLAALLL | TCSAAQGTEVF | 0.34474617 |
| ASVINISSI | TCSAAQGTEVF | 0.63056505 |
| ATPPGPA | TCSAAQGTEVF | 0.60767883 |
| ATVASALV | TCSAAQGTEVF | 0.5689608 |
| ATVASLIL | TCSAAQGTEVF | 0.4729606 |
| AVNGPVLIRH | TCSAAQGTEVF | 0.5589475 |
| DGAAGAVGI | TCSAAQGTEVF | 0.48333862 |
| DPLLHGI | TCSAAQGTEVF | 0.6585232 |
| ELGAALGAL | TCSAAQGTEVF | 0.25432956 |
| ELGRTQAV | TCSAAQGTEVF | 0.6457951 |
| FVAGGYLS | TCSAAQGTEVF | 0.3677345 |
| GASGGAGGAGGW | TCSAAQGTEVF | 0.34895682 |
| GGAGAGGDGGAGGI | TCSAAQGTEVF | 0.25697967 |
| GTGHNGSQPGMGGNG | TCSAAQGTEVF | 0.60397196 |
| GYAAASASAS | TCSAAQGTEVF | 0.62943316 |
| HQVDVAAVQQG | TCSAAQGTEVF | 0.65152705 |
| IGAAIAHT | TCSAAQGTEVF | 0.47425663 |
| IGMIEVAQL | TCSAAQGTEVF | 0.69330937 |
| IRLAPEHTSAMD | TCSAAQGTEVF | 0.6898565 |
| IYNVSVL | TCSAAQGTEVF | 0.5195686 |
| KQGFLGM | TCSAAQGTEVF | 0.27725518 |
| LESVITQL | TCSAAQGTEVF | 0.6368951 |
| LGLGALVVV | TCSAAQGTEVF | 0.6351403 |
| LIGNGGNGGSGGI | TCSAAQGTEVF | 0.66333354 |
| LNAINAPTL | TCSAAQGTEVF | 0.64932084 |
| LQLLNTGV | TCSAAQGTEVF | 0.3511927 |
| LQTSIEF | TCSAAQGTEVF | 0.68443704 |
| MSFAVSEL | TCSAAQGTEVF | 0.64378 |
| NTGALGL | TCSAAQGTEVF | 0.33706445 |
| PPGAGGAGGAGTT | TCSAAQGTEVF | 0.52796537 |
| QLVGLGI | TCSAAQGTEVF | 0.47463268 |
| RFLEQVHQL | TCSAAQGTEVF | 0.2802884 |
| RGVGYVLR | TCSAAQGTEVF | 0.59500253 |
| SATGVFAGVI | TCSAAQGTEVF | 0.44182187 |
| SLPAITLP | TCSAAQGTEVF | 0.48793066 |
| STALAFTV | TCSAAQGTEVF | 0.48780406 |
| SVLTVGLGK | TCSAAQGTEVF | 0.40259838 |
| TALAGAGSGI | TCSAAQGTEVF | 0.4378399 |

| | | |
|---|---|---|
| TGLPFTGI | TCSAAQGTEVF | 0.4463429 |
| VAEHVTV | TCSAAQGTEVF | 0.5842298 |
| VDIVLNSL | TCSAAQGTEVF | 0.60166717 |
| VLTGGGAVL | TCSAAQGTEVF | 0.5817319 |
| VNVGLAYGI | TCSAAQGTEVF | 0.67294157 |
| VSSAAVGAV | TCSAAQGTEVF | 0.69864595 |
| VTSVYEAGV | TCSAAQGTEVF | 0.31664333 |
| WVALDRGA | TCSAAQGTEVF | 0.40236947 |
| YLVGSVVI | TCSAAQGTEVF | 0.6350532 |
| GPPGTGKTLL | TCSAAQGTEVF | 0.3466992 |
| GVGMTKF | TCSAAQGTEVF | 0.68098855 |
| IGIGGSDL | TCSAAQGTEVF | 0.49627212 |
| IGTLLAEL | TCSAAQGTEVF | 0.614835 |
| IYTSGTT | TCSAAQGTEVF | 0.5423513 |
| LSFPAGV | TCSAAQGTEVF | 0.7238444 |
| LYTSGTT | TCSAAQGTEVF | 0.64917874 |
| VGAFPFGF | TCSAAQGTEVF | 0.57459795 |
| VGAGTVEF | TCSAAQGTEVF | 0.624634 |
| VGKSTLLL | TCSAAQGTEVF | 0.38235447 |
| AALAHAGGGIV | TCSAAQGTEVF | 0.6803196 |
| AALVAVSLL | TCSAAQGTEVF | 0.65317416 |
| AAVSRAVAAL | TCSAAQGTEVF | 0.40851468 |
| AGGSSGIGGF | TCSAAQGTEVF | 0.5118921 |
| AGLTVANANG | TCSAAQGTEVF | 0.6666981 |
| AGLVGAVL | TCSAAQGTEVF | 0.6945044 |
| ALSQAFTV | TCSAAQGTEVF | 0.3077283 |
| ALTFSVSL | TCSAAQGTEVF | 0.46742404 |
| ALTGAALVAL | TCSAAQGTEVF | 0.27502775 |
| ASFSIGAGVL | TCSAAQGTEVF | 0.35111004 |
| DGSVYKI | TCSAAQGTEVF | 0.29824072 |
| EQLAEIV | TCSAAQGTEVF | 0.6151913 |
| FAVSELTVM | TCSAAQGTEVF | 0.51847756 |
| GAAPLAAGL | TCSAAQGTEVF | 0.6915742 |
| GADGGDGGSGGAGG | TCSAAQGTEVF | 0.623728 |
| GFGLDTGGAGG | TCSAAQGTEVF | 0.68791777 |
| GLASGGLL | TCSAAQGTEVF | 0.4337566 |
| GRIAGAAVLL | TCSAAQGTEVF | 0.46436864 |
| GSAGSLNTSF | TCSAAQGTEVF | 0.30751574 |
| GTGQIGI | TCSAAQGTEVF | 0.2899633 |
| GTLIALIL | TCSAAQGTEVF | 0.3189482 |
| IAATMGFVVG | TCSAAQGTEVF | 0.46347576 |
| IFAQALL | TCSAAQGTEVF | 0.2586434 |
| IGAAVALGF | TCSAAQGTEVF | 0.69388974 |
| IGLTPLAL | TCSAAQGTEVF | 0.5688246 |
| ILGSNTSTL | TCSAAQGTEVF | 0.602008 |
| IVFSLTVL | TCSAAQGTEVF | 0.50348103 |
| LASKATGF | TCSAAQGTEVF | 0.44711602 |
| LASVALIL | TCSAAQGTEVF | 0.49683523 |
| LFAVGLTL | TCSAAQGTEVF | 0.66304207 |
| LFNGGIGTYI | TCSAAQGTEVF | 0.35811982 |
| LTLGAGII | TCSAAQGTEVF | 0.31333366 |

| | | |
|---|---|---|
| LVAAALGDI | TCSAAQGTEVF | 0.38223767 |
| NTAFVQLGI | TCSAAQGTEVF | 0.4025551 |
| QLFSGAEL | TCSAAQGTEVF | 0.70984125 |
| SAATFITL | TCSAAQGTEVF | 0.66276485 |
| SNYGLANLGV | TCSAAQGTEVF | 0.58521545 |
| SYGGAYAVM | TCSAAQGTEVF | 0.7008903 |
| TTGVGGLTLG | TCSAAQGTEVF | 0.25136614 |
| TTQTIYI | TCSAAQGTEVF | 0.53885484 |
| VAAGAGLGAVI | TCSAAQGTEVF | 0.71506405 |
| VAAVVTLL | TCSAAQGTEVF | 0.58594835 |
| VGFLGGLI | TCSAAQGTEVF | 0.526614 |
| VGVIPILL | TCSAAQGTEVF | 0.6970612 |
| VIFASSLI | TCSAAQGTEVF | 0.407418 |
| VKGGDGGAAAG | TCSAAQGTEVF | 0.50401276 |
| VTKTFTNI | TCSAAQGTEVF | 0.54131156 |
| VVAAAILL | TCSAAQGTEVF | 0.6440038 |
| YAALDVEL | TCSAAQGTEVF | 0.4592449 |
| LASVALLL | TCSAAQGTEVF | 0.3498295 |
| LDAVHRV | TCSAAQGTEVF | 0.6615728 |
| QGLSSSIF | TCSAAQGTEVF | 0.7172353 |
| RGTFANIRL | TCSAAQGTEVF | 0.48976237 |
| RQLWWGHRI | TCSAAQGTEVF | 0.6643696 |
| YGPPGTGKTLL | TCSAAQGTEVF | 0.6399689 |
| AALVGAPANVA | TCSAAQGTEVF | 0.50815594 |
| AATFRRTE | TCSAAQGTEVF | 0.47913912 |
| AGALGGGI | TCSAAQGTEVF | 0.35643703 |
| AGGAGGAGGIGAG | TCSAAQGTEVF | 0.5744902 |
| AGSGALGL | TCSAAQGTEVF | 0.36913183 |
| AQIADQI | TCSAAQGTEVF | 0.56211174 |
| ASAVAVTYL | TCSAAQGTEVF | 0.57336575 |
| ASTVLGGTGG | TCSAAQGTEVF | 0.72272706 |
| AVGGEPAK | TCSAAQGTEVF | 0.6351238 |
| CEQGTFGGTC | TCSAAQGTEVF | 0.64608586 |
| DEGKAPAG | TCSAAQGTEVF | 0.39451182 |
| DTVVFPAL | TCSAAQGTEVF | 0.51097816 |
| FEGITMAG | TCSAAQGTEVF | 0.6143804 |
| FPLVVQD | TCSAAQGTEVF | 0.34663948 |
| GAFTTPQ | TCSAAQGTEVF | 0.66286373 |
| GFGGFGTAGGVGGA | TCSAAQGTEVF | 0.3911821 |
| GFLAQFL | TCSAAQGTEVF | 0.59406984 |
| GGAGGAGGNAGVA | TCSAAQGTEVF | 0.5732233 |
| GGVGTAGAGGAGGAGG | TCSAAQGTEVF | 0.7090789 |
| GNGGGGDAGNAGSGGN | TCSAAQGTEVF | 0.58880717 |
| GTGGAGGAGNDAGS | TCSAAQGTEVF | 0.54908025 |
| GTTNTGWFNS | TCSAAQGTEVF | 0.34304118 |
| IFMGPLFTL | TCSAAQGTEVF | 0.63335526 |
| ILAHAIEAI | TCSAAQGTEVF | 0.6877048 |
| IRSAIREAL | TCSAAQGTEVF | 0.5329745 |
| ITFAGGG | TCSAAQGTEVF | 0.71778053 |
| ITGMSGHQEV | TCSAAQGTEVF | 0.5932584 |
| IVAGLACTL | TCSAAQGTEVF | 0.28316724 |

| | | |
|---|---|---|
| IVGGGLAGA | TCSAAQGTEVF | 0.5020788 |
| IVTGVLNI | TCSAAQGTEVF | 0.7006475 |
| LAQQGCI | TCSAAQGTEVF | 0.66879153 |
| LCLHVEN | TCSAAQGTEVF | 0.6895765 |
| LDAISDGS | TCSAAQGTEVF | 0.67124486 |
| LGGFAAML | TCSAAQGTEVF | 0.50765836 |
| LGGKNPMI | TCSAAQGTEVF | 0.68068904 |
| LGPKGVAAIL | TCSAAQGTEVF | 0.71953213 |
| LMVGVSI | TCSAAQGTEVF | 0.6857718 |
| LVGAAFVL | TCSAAQGTEVF | 0.39253986 |
| LVLAVAPL | TCSAAQGTEVF | 0.32755312 |
| MAAIAVIDAA | TCSAAQGTEVF | 0.62309664 |
| NNGINAELQ | TCSAAQGTEVF | 0.6862174 |
| RGEPLVLVHG | TCSAAQGTEVF | 0.37754557 |
| RGGAGGAAGLI | TCSAAQGTEVF | 0.5917329 |
| RSYRMHPAV | TCSAAQGTEVF | 0.46833175 |
| SAAHLLAGQLA | TCSAAQGTEVF | 0.2786817 |
| SAAVGSY | TCSAAQGTEVF | 0.5664344 |
| SPHAVLHRI | TCSAAQGTEVF | 0.71298945 |
| SQIAKVL | TCSAAQGTEVF | 0.6163964 |
| TALGHLVP | TCSAAQGTEVF | 0.33652496 |
| TAPTTSGAA | TCSAAQGTEVF | 0.6826165 |
| TDGDDGNGGSPGT | TCSAAQGTEVF | 0.7191069 |
| TDGLGEN | TCSAAQGTEVF | 0.67469305 |
| TGGPPLPGQA | TCSAAQGTEVF | 0.31909695 |
| TIPAITI | TCSAAQGTEVF | 0.25406417 |
| TIPGFTI | TCSAAQGTEVF | 0.5181797 |
| TSIASGALA | TCSAAQGTEVF | 0.69833076 |
| TVTLPGAAI | TCSAAQGTEVF | 0.43035805 |
| VASGLSAALP | TCSAAQGTEVF | 0.3977922 |
| VGAGVIGI | TCSAAQGTEVF | 0.5711033 |
| VGPKRFGV | TCSAAQGTEVF | 0.6711372 |
| VNVAFPD | TCSAAQGTEVF | 0.37375808 |
| VSMTNSGVS | TCSAAQGTEVF | 0.28940225 |
| VTDVLGDLSV | TCSAAQGTEVF | 0.58432937 |
| VVVVGAGFAGL | TCSAAQGTEVF | 0.67314386 |
| VYTVGTVV | TCSAAQGTEVF | 0.36114573 |
| WTTFSGP | TCSAAQGTEVF | 0.49791253 |
| YDELVEAF | TCSAAQGTEVF | 0.43795893 |
| YGTGGAGGNGG | TCSAAQGTEVF | 0.4155562 |
| YTNVMAV | TCSAAQGTEVF | 0.5497682 |
| FLEQVHQL | TCSAAQGTEVF | 0.32538694 |
| GLSLLAVAV | TCSAAQGTEVF | 0.6987723 |
| LEQVHQL | TCSAAQGTEVF | 0.36669666 |
| LNAAGAI | TCSAAQGTEVF | 0.6295301 |
| PAVHASL | TCSAAQGTEVF | 0.6227779 |
| TELEDDA | TCSAAQGTEVF | 0.63073474 |
| TPPGPPN | TCSAAQGTEVF | 0.502961 |
| TPPPPPPP | TCSAAQGTEVF | 0.47635072 |
| TQSPALI | ASSQSSYEQY | 0.33325112 |
| EFAAAFPQ | ASSQSSYEQY | 0.27380043 |

| | | |
|---|---|---|
| LLSGNLDVL | ASSQSSYEQY | 0.7088715 |
| GPANLGNYNI | ASSQSSYEQY | 0.25299746 |
| TLGPTLP | ASSQSSYEQY | 0.69916415 |
| VYAYSSCCK | ASSQSSYEQY | 0.6019705 |
| APAPAPAPNI | ASSQSSYEQY | 0.3564038 |
| PGGPVPIP | ASSQSSYEQY | 0.5677327 |
| FLTVALEP | ASSQSSYEQY | 0.29877412 |
| SQYFGANL | ASSQSSYEQY | 0.46281004 |
| GGVKPAA | ASSQSSYEQY | 0.56777394 |
| GGTGGAGGAGGW | ASSQSSYEQY | 0.66464627 |
| GVGGVAGLGGAGG | ASSQSSYEQY | 0.5200311 |
| VLLVLATTI | ASSQSSYEQY | 0.60559356 |
| IGIFAAML | ASSQSSYEQY | 0.62337524 |
| AVGDLID | ASSQSSYEQY | 0.3432239 |
| NISPLPL | ASSQSSYEQY | 0.61668247 |
| VTAVFVAI | ASSQSSYEQY | 0.6847849 |
| SGDDNDAAL | ASSQSSYEQY | 0.64710295 |
| AVAALAHS | ASSQSSYEQY | 0.33847484 |
| GGVTVMATP | ASSQSSYEQY | 0.65411747 |
| AGLAANVRD | ASSQSSYEQY | 0.42718154 |
| TNRPDLIDPALLRPG | ASSQSSYEQY | 0.59173214 |
| LANLGVTAL | ASSQSSYEQY | 0.56011623 |
| TGERIEL | ASSQSSYEQY | 0.5001456 |
| NGEELAA | ASSQSSYEQY | 0.65040076 |
| VLSLVWG | ASSQSSYEQY | 0.30680355 |
| IGYALWL | ASSQSSYEQY | 0.45894617 |
| LSLLLAPL | ASSQSSYEQY | 0.5468274 |
| TLDEALK | ASSQSSYEQY | 0.5240119 |
| TDGAGTAEGG | ASSQSSYEQY | 0.6755743 |
| NAPGGAGGNGG | ASSQSSYEQY | 0.38774827 |
| VSVFMGMGGNF | ASSQSSYEQY | 0.5990556 |
| LASYLAAL | ASSQSSYEQY | 0.41424277 |
| LVGGTPAL | ASSQSSYEQY | 0.41158837 |
| VRNADVL | ASSQSSYEQY | 0.601068 |
| TCSSPFLVSAM | ASSQSSYEQY | 0.42895025 |
| VHITDAE | ASSQSSYEQY | 0.34616905 |
| EYAAPSF | ASSQSSYEQY | 0.62605345 |
| VVDAFHGGL | ASSQSSYEQY | 0.35188624 |
| ASVVDPQ | ASSQSSYEQY | 0.56619245 |
| SYNEFGGLS | ASSQSSYEQY | 0.69741714 |
| DPDGSVY | ASSQSSYEQY | 0.43511826 |
| MVSTIHATE | ASSQSSYEQY | 0.5977378 |
| LAAGVADPV | ASSQSSYEQY | 0.627698 |
| TKEQIAA | ASSQSSYEQY | 0.66776013 |
| SLASPAAL | ASSQSSYEQY | 0.3987503 |
| IGLVFYGV | ASSQSSYEQY | 0.3772904 |
| FLLDASNL | ASSQSSYEQY | 0.70219475 |
| YQVGNFP | ASSQSSYEQY | 0.6529857 |
| ESLLSQST | ASSQSSYEQY | 0.69207394 |
| LFTVGFAAVG | ASSQSSYEQY | 0.6586464 |
| ATPPTPT | ASSQSSYEQY | 0.6957319 |

| | | |
|---|---|---|
| TDLATLG | ASSQSSYEQY | 0.6742935 |
| SAGGTGR | ASSQSSYEQY | 0.47455814 |
| GFSNSGS | ASSQSSYEQY | 0.30293587 |
| GGNGGSGGTSV | ASSQSSYEQY | 0.5099491 |
| GTSVATGGAGN | ASSQSSYEQY | 0.39061323 |
| NGGNGGSGGTS | ASSQSSYEQY | 0.52612567 |
| AVVKFPH | ASSQSSYEQY | 0.2818964 |
| TPPPPPP | ASSQSSYEQY | 0.689131 |
| AAPAAAAAPA | ASSQSSYEQY | 0.7274426 |
| IALNEDL | ASSQSSYEQY | 0.43147933 |
| IVNVSSM | ASSQSSYEQY | 0.7077179 |
| TGIPPAPR | ASSQSSYEQY | 0.31280452 |
| FDLGGGTF | ASSQSSYEQY | 0.6545005 |
| FDLGGGTFDVS | ASSQSSYEQY | 0.68993866 |
| NEPTAAA | ASSQSSYEQY | 0.3881018 |
| NEPTAAALA | ASSQSSYEQY | 0.63418305 |
| GSIINISSI | ASSQSSYEQY | 0.39012334 |
| VAPGNAGTA | ASSQSSYEQY | 0.45563608 |
| TGGGASLE | ASSQSSYEQY | 0.70291054 |
| VAVYNHYKRI | ASSQSSYEQY | 0.41750282 |
| TDPAPATT | ASSQSSYEQY | 0.6023896 |
| TNRPDILDPALLRPG | ASSQSSYEQY | 0.49140096 |
| LTTTTTS | ASSQSSYEQY | 0.3699169 |
| PTGSGKTL | ASSQSSYEQY | 0.61957645 |
| TGSGKTL | ASSQSSYEQY | 0.6925039 |
| TIGHVDHGK | ASSQSSYEQY | 0.40081263 |
| TIGHVDHGKTT | ASSQSSYEQY | 0.71848464 |
| GHVDHGKTT | ASSQSSYEQY | 0.43224552 |
| AQFEHTLL | ASSQSSYEQY | 0.71891016 |
| IVNQDEL | ASSQSSYEQY | 0.7107214 |
| TNGSQFF | ASSQSSYEQY | 0.5514418 |
| TGKGIVE | ASSQSSYEQY | 0.46151114 |
| LISNASDA | ASSQSSYEQY | 0.69255877 |
| AELRTLGELTKIAKAHGAQ | ASSQSSYEQY | 0.51656556 |
| APNAVAAAEPSW | ASSQSSYEQY | 0.69035447 |
| APQVAGAATAVL | ASSQSSYEQY | 0.68778944 |
| ARRDPAAAAAILPTDA | ASSQSSYEQY | 0.3450382 |
| DAAARDVLAEAGLAEYF | ASSQSSYEQY | 0.6741303 |
| DDQIVLSAKANLAKVEQ | ASSQSSYEQY | 0.63708144 |
| DELLPDTGEDSDEDGH | ASSQSSYEQY | 0.5232361 |
| ELPEAPANRATVPQD | ASSQSSYEQY | 0.4673811 |
| FGQSKIERVTGIG | ASSQSSYEQY | 0.3307121 |
| GAGGTGSTAGGAGGAGGAGGLY | ASSQSSYEQY | 0.41042754 |
| GHGGTGAAGGNGGTGGAGGSGI | ASSQSSYEQY | 0.43091834 |
| GNGGNGGAGGEGGPGGAGGAGA | ASSQSSYEQY | 0.6655173 |
| GSTAGNGGNGGNATNGTIAGQP | ASSQSSYEQY | 0.6027982 |
| GTGGDGGNGGNGGTGG | ASSQSSYEQY | 0.4489839 |
| GVRRPAGIPADAPV | ASSQSSYEQY | 0.65261686 |
| GYRQVTDAHLLTLARR | ASSQSSYEQY | 0.62792134 |
| IAKMGQDGHDRG | ASSQSSYEQY | 0.3241651 |
| KAEPSIEIQYT | ASSQSSYEQY | 0.54790705 |

| | | |
|---|---|---|
| KPVRIRMAADEDTVAVT | ASSQSSYEQY | 0.3598333 |
| MPPRIAGMRLLVIKPEP | ASSQSSYEQY | 0.45199537 |
| MKFYAVALFRSRGVPPTR | ASSQSSYEQY | 0.28556585 |
| NAGGFGILGAVAHSP | ASSQSSYEQY | 0.6007849 |
| NEGREITLAMAEM | ASSQSSYEQY | 0.54453695 |
| NELVEWSHPLGHRWRWE | ASSQSSYEQY | 0.27703825 |
| NLGAGGTGGNGGNPG | ASSQSSYEQY | 0.5337978 |
| NTEWPKWETVMAADPVD | ASSQSSYEQY | 0.50541127 |
| PASAAPAETQP | ASSQSSYEQY | 0.5284684 |
| QQSSDGKNGVRAAEITGE | ASSQSSYEQY | 0.6442939 |
| RAPDAVTTSWWKE | ASSQSSYEQY | 0.40028727 |
| SGFGNVGAHTSGWFNQ | ASSQSSYEQY | 0.44428158 |
| SGVAGAVGQAASVGGLKV | ASSQSSYEQY | 0.63199306 |
| SNGGAGTGSAGGNGGT | ASSQSSYEQY | 0.6794003 |
| TGGAGGEGGAAG | ASSQSSYEQY | 0.3966689 |
| TGGDGGAAGAGGAPGHG | ASSQSSYEQY | 0.5924134 |
| TMPPLLAAIANAGRHGV | ASSQSSYEQY | 0.6690309 |
| TSSTAGNSGGA | ASSQSSYEQY | 0.59390146 |
| VPATVPVGILC | ASSQSSYEQY | 0.6138678 |
| VQRPDSGGVVLTGRIS | ASSQSSYEQY | 0.65230596 |
| WTPNPVPATKGVPAGLVQAT | ASSQSSYEQY | 0.3681714 |
| YALKRALAKFSDRFDVVII | ASSQSSYEQY | 0.6677426 |
| YLIGSVIV | ASSQSSYEQY | 0.67071587 |
| YGQLVAVID | ASSQSSYEQY | 0.4918518 |
| VTVGGMTLPA | ASSQSSYEQY | 0.48604876 |
| VLALPAV | ASSQSSYEQY | 0.34526435 |
| VIPPTLNYET | ASSQSSYEQY | 0.47780535 |
| VGMGGMHPSSG | ASSQSSYEQY | 0.3529382 |
| VGLAGGAATA | ASSQSSYEQY | 0.61302674 |
| VGGAGGAGGSGP | ASSQSSYEQY | 0.6663477 |
| VFAAFMV | ASSQSSYEQY | 0.52923805 |
| TLYDKLR | ASSQSSYEQY | 0.33852738 |
| TLAVFTVL | ASSQSSYEQY | 0.3612485 |
| TGSVQPARV | ASSQSSYEQY | 0.5444819 |
| TGGGAGSGGT | ASSQSSYEQY | 0.42333156 |
| SHLETSL | ASSQSSYEQY | 0.31820837 |
| SFTTGMT | ASSQSSYEQY | 0.39797089 |
| QVEVAKRMGVRQPTVS | ASSQSSYEQY | 0.6508303 |
| QGSGGPALAM | ASSQSSYEQY | 0.6501154 |
| MLAPSNP | ASSQSSYEQY | 0.4141755 |
| LVNNAAI | ASSQSSYEQY | 0.678666 |
| LTGEINTLA | ASSQSSYEQY | 0.6615557 |
| LSIFGPV | ASSQSSYEQY | 0.6621001 |
| LLALLANL | ASSQSSYEQY | 0.47822422 |
| LISNASDALDKLR | ASSQSSYEQY | 0.36924762 |
| LINEAVP | ASSQSSYEQY | 0.66941714 |
| LGYAFSGL | ASSQSSYEQY | 0.5678721 |
| LDFVAVAL | ASSQSSYEQY | 0.53564584 |
| LAQPGAE | ASSQSSYEQY | 0.608303 |
| ISNLGAVTGF | ASSQSSYEQY | 0.5603094 |
| HQEDPGRI | ASSQSSYEQY | 0.46346548 |

| | | |
|---|---|---|
| GTPGEPTP | ASSQSSYEQY | 0.6129184 |
| GFFNSGS | ASSQSSYEQY | 0.6189728 |
| GAGGGGLTAGAN | ASSQSSYEQY | 0.37089735 |
| FVDALVNM | ASSQSSYEQY | 0.42367572 |
| FLTVAPEL | ASSQSSYEQY | 0.4774112 |
| FETLTAL | ASSQSSYEQY | 0.5761119 |
| DVINPIEL | ASSQSSYEQY | 0.6702253 |
| APVSGAPV | ASSQSSYEQY | 0.5156555 |
| ALVGGLAW | ASSQSSYEQY | 0.4980421 |
| ALAAAAATAI | ASSQSSYEQY | 0.63021874 |
| AFGLGLLL | ASSQSSYEQY | 0.59603906 |
| AFFAAASL | ASSQSSYEQY | 0.52112705 |
| AAAALPAFSPP | ASSQSSYEQY | 0.57891756 |
| VVLAPAL | ASSQSSYEQY | 0.48131564 |
| TIGHVDHGKTTL | ASSQSSYEQY | 0.6198066 |
| LISNASDALDK | ASSQSSYEQY | 0.57726717 |
| LISNASDAL | ASSQSSYEQY | 0.56515265 |
| ISNASDALDK | ASSQSSYEQY | 0.6114719 |
| NQAETLVYQTE | ASSQSSYEQY | 0.63750505 |
| TGGTGGTAGAG | ASSQSSYEQY | 0.5202683 |
| TGSTAGGAGGA | ASSQSSYEQY | 0.34148443 |
| DPLAVPVAAGPV | ASSQSSYEQY | 0.39298177 |
| GGTGGTPGNGGN | ASSQSSYEQY | 0.62137914 |
| PAGLAGVDPATL | ASSQSSYEQY | 0.60929674 |
| GNNLVVEAPHAGASV | ASSQSSYEQY | 0.64010835 |
| KVTRIMSASTLEQPA | ASSQSSYEQY | 0.54486346 |
| LGFALPIALPRRTVV | ASSQSSYEQY | 0.45437446 |
| TRLGNLTARKLVVLV | ASSQSSYEQY | 0.634852 |
| WSQQRRLIYFPSAGP | ASSQSSYEQY | 0.6604741 |
| GPNGNPDPMAAAVDIR | ASSQSSYEQY | 0.32665175 |
| TQRCVSVADLRAAASR | ASSQSSYEQY | 0.46478155 |
| QFVLDVAEHPVEVLTETV | ASSQSSYEQY | 0.68190956 |
| GGAGGTGGAGGNGGTQAGDG | ASSQSSYEQY | 0.64151263 |
| PLLATLRAIAVDVDAKPAQV | ASSQSSYEQY | 0.50339985 |
| APKFRTGGYMIDFWGVGYQVAK | ASSQSSYEQY | 0.4265374 |
| GAGGAGGAGGAAGGPGGTGGTG | ASSQSSYEQY | 0.6878196 |
| IGALGLLATDVAPAGSVAFGVA | ASSQSSYEQY | 0.47943944 |
| AGGAGGIGGNGGAGGAGGVLMG | ASSQSSYEQY | 0.6772356 |
| ADELIGLL | ASSQSSYEQY | 0.29888457 |
| ADRALKA | ASSQSSYEQY | 0.41437432 |
| AGAVGQVGGAGGA | ASSQSSYEQY | 0.39207375 |
| AGGGGGLGGAP | ASSQSSYEQY | 0.30311757 |
| AGGLLYGNGGAG | ASSQSSYEQY | 0.626807 |
| AGLESIGL | ASSQSSYEQY | 0.38642123 |
| AGLKTGII | ASSQSSYEQY | 0.6115162 |
| ASLAALLL | ASSQSSYEQY | 0.43876565 |
| ASVINISSI | ASSQSSYEQY | 0.6379274 |
| ATPPGPA | ASSQSSYEQY | 0.5828847 |
| ATVASALV | ASSQSSYEQY | 0.5184547 |
| ATVASLIL | ASSQSSYEQY | 0.38039684 |
| AVNGPVLIRH | ASSQSSYEQY | 0.6182984 |

| | | |
|---|---|---|
| DGAAGAVGI | ASSQSSYEQY | 0.6598259 |
| DPLLHGI | ASSQSSYEQY | 0.55162776 |
| ELGAALGAL | ASSQSSYEQY | 0.34916738 |
| ELGRTQAV | ASSQSSYEQY | 0.5500007 |
| FVAGGYLS | ASSQSSYEQY | 0.53879666 |
| GASGGAGGAGGW | ASSQSSYEQY | 0.3240287 |
| GGAGAGGDGGAGGI | ASSQSSYEQY | 0.29148936 |
| GTGHNGSQPGMGGNG | ASSQSSYEQY | 0.5144904 |
| GYAAASASAS | ASSQSSYEQY | 0.45179424 |
| HQVDVAAVQQG | ASSQSSYEQY | 0.6506521 |
| IGAAIAHT | ASSQSSYEQY | 0.6443029 |
| IGMIEVAQL | ASSQSSYEQY | 0.51290023 |
| IRLAPEHTSAMD | ASSQSSYEQY | 0.6511001 |
| IYNVSVL | ASSQSSYEQY | 0.61766124 |
| KQGFLGM | ASSQSSYEQY | 0.2675625 |
| LESVITQL | ASSQSSYEQY | 0.39110452 |
| LGLGALVVV | ASSQSSYEQY | 0.3280647 |
| LIGNGGNGGSGGI | ASSQSSYEQY | 0.60003376 |
| LNAINAPTL | ASSQSSYEQY | 0.3736282 |
| LQLLNTGV | ASSQSSYEQY | 0.41395783 |
| LQTSIEF | ASSQSSYEQY | 0.5808484 |
| MSFAVSEL | ASSQSSYEQY | 0.6478111 |
| NTGALGL | ASSQSSYEQY | 0.5371287 |
| PPGAGGAGGAGTT | ASSQSSYEQY | 0.38866585 |
| QLVGLGI | ASSQSSYEQY | 0.29688516 |
| RFLEQVHQL | ASSQSSYEQY | 0.62296426 |
| RGVGYVLR | ASSQSSYEQY | 0.4116559 |
| SATGVFAGVI | ASSQSSYEQY | 0.40703508 |
| SLPAITLP | ASSQSSYEQY | 0.30656624 |
| STALAFTV | ASSQSSYEQY | 0.5038513 |
| SVLTVGLGK | ASSQSSYEQY | 0.3120873 |
| TALAGAGSGI | ASSQSSYEQY | 0.59495056 |
| TGLPFTGI | ASSQSSYEQY | 0.48867658 |
| VAEHVTV | ASSQSSYEQY | 0.50583243 |
| VDIVLNSL | ASSQSSYEQY | 0.6831301 |
| VLTGGGAVL | ASSQSSYEQY | 0.6441103 |
| VNVGLAYGI | ASSQSSYEQY | 0.5135752 |
| VSSAAVGAV | ASSQSSYEQY | 0.48592418 |
| VTSVYEAGV | ASSQSSYEQY | 0.36557737 |
| WVALDRGA | ASSQSSYEQY | 0.454512 |
| YLVGSVVI | ASSQSSYEQY | 0.43929315 |
| GPPGTGKTLL | ASSQSSYEQY | 0.66640216 |
| GVGMTKF | ASSQSSYEQY | 0.40511534 |
| IGIGGSDL | ASSQSSYEQY | 0.31951445 |
| IGTLLAEL | ASSQSSYEQY | 0.44826648 |
| IYTSGTT | ASSQSSYEQY | 0.66591835 |
| LSFPAGV | ASSQSSYEQY | 0.6949618 |
| LYTSGTT | ASSQSSYEQY | 0.41378924 |
| VGAFPFGF | ASSQSSYEQY | 0.43168753 |
| VGAGTVEF | ASSQSSYEQY | 0.4461095 |
| VGKSTLLL | ASSQSSYEQY | 0.54485035 |

| | | |
|---|---|---|
| AALAHAGGGIV | ASSQSSYEQY | 0.40005976 |
| AALVAVSLL | ASSQSSYEQY | 0.37239096 |
| AAVSRAVAAL | ASSQSSYEQY | 0.6013523 |
| AGGSSGIGGF | ASSQSSYEQY | 0.6937307 |
| AGLTVANANG | ASSQSSYEQY | 0.48500335 |
| AGLVGAVL | ASSQSSYEQY | 0.290877 |
| ALSQAFTV | ASSQSSYEQY | 0.5765374 |
| ALTFSVSL | ASSQSSYEQY | 0.4512338 |
| ALTGAALVAL | ASSQSSYEQY | 0.4909773 |
| ASFSIGAGVL | ASSQSSYEQY | 0.58326197 |
| DGSVYKI | ASSQSSYEQY | 0.3085366 |
| EQLAEIV | ASSQSSYEQY | 0.49507013 |
| FAVSELTVM | ASSQSSYEQY | 0.69052374 |
| GAAPLAAGL | ASSQSSYEQY | 0.60900676 |
| GADGGDGGSGGAGG | ASSQSSYEQY | 0.6116471 |
| GFGLDTGGAGG | ASSQSSYEQY | 0.66582596 |
| GLASGGLL | ASSQSSYEQY | 0.6222912 |
| GRIAGAAVLL | ASSQSSYEQY | 0.66693485 |
| GSAGSLNTSF | ASSQSSYEQY | 0.66591835 |
| GTGQIGI | ASSQSSYEQY | 0.47315136 |
| GTLIALIL | ASSQSSYEQY | 0.51879287 |
| IAATMGFVVG | ASSQSSYEQY | 0.6346935 |
| IFAQALL | ASSQSSYEQY | 0.5723246 |
| IGAAVALGF | ASSQSSYEQY | 0.56343126 |
| IGLTPLAL | ASSQSSYEQY | 0.40010452 |
| ILGSNTSTL | ASSQSSYEQY | 0.6470432 |
| IVFSLTVL | ASSQSSYEQY | 0.5088311 |
| LASKATGF | ASSQSSYEQY | 0.6763874 |
| LASVALIL | ASSQSSYEQY | 0.32085055 |
| LFAVGLTL | ASSQSSYEQY | 0.40643466 |
| LFNGGIGTYI | ASSQSSYEQY | 0.6493946 |
| LTLGAGII | ASSQSSYEQY | 0.28803793 |
| LVAAALGDI | ASSQSSYEQY | 0.2993192 |
| NTAFVQLGI | ASSQSSYEQY | 0.55244714 |
| QLFSGAEL | ASSQSSYEQY | 0.67426986 |
| SAATFITL | ASSQSSYEQY | 0.2852843 |
| SNYGLANLGV | ASSQSSYEQY | 0.4176664 |
| SYGGAYAVM | ASSQSSYEQY | 0.34516877 |
| TTGVGGLTLG | ASSQSSYEQY | 0.5451082 |
| TTQTIYI | ASSQSSYEQY | 0.56356966 |
| VAAGAGLGAVI | ASSQSSYEQY | 0.32100272 |
| VAAVVTLL | ASSQSSYEQY | 0.5898945 |
| VGFLGGLI | ASSQSSYEQY | 0.29458234 |
| VGVIPILL | ASSQSSYEQY | 0.46683884 |
| VIFASSLI | ASSQSSYEQY | 0.6276579 |
| VKGGDGGAAAG | ASSQSSYEQY | 0.68411136 |
| VTKTFTNI | ASSQSSYEQY | 0.33946642 |
| VVAAAILL | ASSQSSYEQY | 0.64516544 |
| YAALDVEL | ASSQSSYEQY | 0.3858904 |
| LASVALLL | ASSQSSYEQY | 0.69926345 |
| LDAVHRV | ASSQSSYEQY | 0.5002473 |

| | | |
|---|---|---|
| QGLSSSIF | ASSQSSYEQY | 0.31506637 |
| RGTFANIRL | ASSQSSYEQY | 0.6463789 |
| RQLWWGHRI | ASSQSSYEQY | 0.57960397 |
| YGPPGTGKTLL | ASSQSSYEQY | 0.52803546 |
| AALVGAPANVA | ASSQSSYEQY | 0.66651344 |
| AATFRRTE | ASSQSSYEQY | 0.40702283 |
| AGALGGGI | ASSQSSYEQY | 0.5548732 |
| AGGAGGAGGIGAG | ASSQSSYEQY | 0.64693475 |
| AGSGALGL | ASSQSSYEQY | 0.63939476 |
| AQIADQI | ASSQSSYEQY | 0.37595922 |
| ASAVAVTYL | ASSQSSYEQY | 0.6263286 |
| ASTVLGGTGG | ASSQSSYEQY | 0.58568287 |
| AVGGEPAK | ASSQSSYEQY | 0.38276494 |
| CEQGTFGGTC | ASSQSSYEQY | 0.6748625 |
| DEGKAPAG | ASSQSSYEQY | 0.46325058 |
| DTVVFPAL | ASSQSSYEQY | 0.64209914 |
| FEGITMAG | ASSQSSYEQY | 0.5971206 |
| FPLVVQD | ASSQSSYEQY | 0.6918314 |
| GAFTTPQ | ASSQSSYEQY | 0.34654176 |
| GFGGFGTAGGVGGA | ASSQSSYEQY | 0.54345876 |
| GFLAQFL | ASSQSSYEQY | 0.57188845 |
| GGAGGAGGNAGVA | ASSQSSYEQY | 0.61732423 |
| GGVGTAGAGGAGGAGG | ASSQSSYEQY | 0.4464709 |
| GNGGGGDAGNAGSGGN | ASSQSSYEQY | 0.49863607 |
| GTGGAGGAGNDAGS | ASSQSSYEQY | 0.5561731 |
| GTTNTGWFNS | ASSQSSYEQY | 0.66871095 |
| IFMGPLFTL | ASSQSSYEQY | 0.49394363 |
| ILAHAIEAI | ASSQSSYEQY | 0.5363705 |
| IRSAIREAL | ASSQSSYEQY | 0.6792213 |
| ITFAGGG | ASSQSSYEQY | 0.3940468 |
| ITGMSGHQEV | ASSQSSYEQY | 0.54771286 |
| IVAGLACTL | ASSQSSYEQY | 0.64112973 |
| IVGGGLAGA | ASSQSSYEQY | 0.6606952 |
| IVTGVLNI | ASSQSSYEQY | 0.43135154 |
| LAQQGCI | ASSQSSYEQY | 0.6591325 |
| LCLHVEN | ASSQSSYEQY | 0.43780595 |
| LDAISDGS | ASSQSSYEQY | 0.63973194 |
| LGGFAAML | ASSQSSYEQY | 0.45971712 |
| LGGKNPMI | ASSQSSYEQY | 0.52016026 |
| LGPKGVAAIL | ASSQSSYEQY | 0.45505583 |
| LMVGVSI | ASSQSSYEQY | 0.61867046 |
| LVGAAFVL | ASSQSSYEQY | 0.6566791 |
| LVLAVAPL | ASSQSSYEQY | 0.63201904 |
| MAAIAVIDAA | ASSQSSYEQY | 0.28843316 |
| NNGINAELQ | ASSQSSYEQY | 0.6444761 |
| RGEPLVLVHG | ASSQSSYEQY | 0.62631726 |
| RGGAGGAAGLI | ASSQSSYEQY | 0.689304 |
| RSYRMHPAV | ASSQSSYEQY | 0.53106457 |
| SAAHLLAGQLA | ASSQSSYEQY | 0.5829764 |
| SAAVGSY | ASSQSSYEQY | 0.54153764 |
| SPHAVLHRI | ASSQSSYEQY | 0.49192542 |

| | | |
|---|---|---|
| SQIAKVL | ASSQSSYEQY | 0.37685737 |
| TALGHLVP | ASSQSSYEQY | 0.4893576 |
| TAPTTSGAA | ASSQSSYEQY | 0.5248729 |
| TDGDDGNGGSPGT | ASSQSSYEQY | 0.4579268 |
| TDGLGEN | ASSQSSYEQY | 0.669215 |
| TGGPPLPGQA | ASSQSSYEQY | 0.65612745 |
| TIPAITI | ASSQSSYEQY | 0.66579443 |
| TIPGFTI | ASSQSSYEQY | 0.6760011 |
| TSIASGALA | ASSQSSYEQY | 0.685428 |
| TVTLPGAAI | ASSQSSYEQY | 0.59211516 |
| VASGLSAALP | ASSQSSYEQY | 0.4935209 |
| VGAGVIGI | ASSQSSYEQY | 0.4570571 |
| VGPKRFGV | ASSQSSYEQY | 0.334538 |
| VNVAFPD | ASSQSSYEQY | 0.42133138 |
| VSMTNSGVS | ASSQSSYEQY | 0.49565798 |
| VTDVLGDLSV | ASSQSSYEQY | 0.5507491 |
| VVVVGAGFAGL | ASSQSSYEQY | 0.28287068 |
| VYTVGTVV | ASSQSSYEQY | 0.6308452 |
| WTTFSGP | ASSQSSYEQY | 0.67459476 |
| YDELVEAF | ASSQSSYEQY | 0.6037341 |
| YGTGGAGGNGG | ASSQSSYEQY | 0.5988663 |
| YTNVMAV | ASSQSSYEQY | 0.58441573 |
| FLEQVHQL | ASSQSSYEQY | 0.42520106 |
| GLSLLAVAV | ASSQSSYEQY | 0.6523817 |
| LEQVHQL | ASSQSSYEQY | 0.42807636 |
| LNAAGAI | ASSQSSYEQY | 0.5803175 |
| PAVHASL | ASSQSSYEQY | 0.67197263 |
| TELEDDA | ASSQSSYEQY | 0.57467675 |
| TPPGPPN | ASSQSSYEQY | 0.55949074 |
| TPPPPPPP | ASSQSSYEQY | 0.50204325 |
| TQSPALI | GGRDQGSGNTLY | 0.3232037 |
| EFAAAFPQ | GGRDQGSGNTLY | 0.5828923 |
| LLSGNLDVL | GGRDQGSGNTLY | 0.4048373 |
| GPANLGNYNI | GGRDQGSGNTLY | 0.5368389 |
| TLGPTLP | GGRDQGSGNTLY | 0.39772755 |
| VYAYSSCCK | GGRDQGSGNTLY | 0.6101892 |
| APAPAPAPNI | GGRDQGSGNTLY | 0.5930776 |
| PGGPVPIP | GGRDQGSGNTLY | 0.60295725 |
| FLTVALEP | GGRDQGSGNTLY | 0.41294968 |
| SQYFGANL | GGRDQGSGNTLY | 0.600662 |
| GGVKPAA | GGRDQGSGNTLY | 0.5604521 |
| GGTGGAGGAGGW | GGRDQGSGNTLY | 0.3620018 |
| GVGGVAGLGGAGG | GGRDQGSGNTLY | 0.6090805 |
| VLLVLATTI | GGRDQGSGNTLY | 0.47425523 |
| IGIFAAML | GGRDQGSGNTLY | 0.52201205 |
| AVGDLID | GGRDQGSGNTLY | 0.6523454 |
| NISPLPL | GGRDQGSGNTLY | 0.59943736 |
| VTAVFVAI | GGRDQGSGNTLY | 0.30869928 |
| SGDDNDAAL | GGRDQGSGNTLY | 0.59586346 |
| AVAALAHS | GGRDQGSGNTLY | 0.56162626 |
| GGVTVMATP | GGRDQGSGNTLY | 0.5593981 |

| | | |
|---|---|---|
| AGLAANVRD | GGRDQGSGNTLY | 0.27534518 |
| TNRPDLIDPALLRPG | GGRDQGSGNTLY | 0.57003903 |
| LANLGVTAL | GGRDQGSGNTLY | 0.38504934 |
| TGERIEL | GGRDQGSGNTLY | 0.4230308 |
| NGEELAA | GGRDQGSGNTLY | 0.42722207 |
| VLSLVWG | GGRDQGSGNTLY | 0.65673554 |
| IGYALWL | GGRDQGSGNTLY | 0.6170904 |
| LSLLLAPL | GGRDQGSGNTLY | 0.48871458 |
| TLDEALK | GGRDQGSGNTLY | 0.6123581 |
| TDGAGTAEGG | GGRDQGSGNTLY | 0.6174927 |
| NAPGGAGGNGG | GGRDQGSGNTLY | 0.6711694 |
| VSVFMGMGGNF | GGRDQGSGNTLY | 0.62231374 |
| LASYLAAL | GGRDQGSGNTLY | 0.49150786 |
| LVGGTPAL | GGRDQGSGNTLY | 0.5094489 |
| VRNADVL | GGRDQGSGNTLY | 0.42464176 |
| TCSSPFLVSAM | GGRDQGSGNTLY | 0.33066788 |
| VHITDAE | GGRDQGSGNTLY | 0.33151934 |
| EYAAPSF | GGRDQGSGNTLY | 0.5404511 |
| VVDAFHGGL | GGRDQGSGNTLY | 0.33193487 |
| ASVVDPQ | GGRDQGSGNTLY | 0.5921606 |
| SYNEFGGLS | GGRDQGSGNTLY | 0.59936416 |
| DPDGSVY | GGRDQGSGNTLY | 0.5103478 |
| MVSTIHATE | GGRDQGSGNTLY | 0.3257035 |
| LAAGVADPV | GGRDQGSGNTLY | 0.48101008 |
| TKEQIAA | GGRDQGSGNTLY | 0.40216047 |
| SLASPAAL | GGRDQGSGNTLY | 0.6361358 |
| IGLVFYGV | GGRDQGSGNTLY | 0.5956844 |
| FLLDASNL | GGRDQGSGNTLY | 0.68706274 |
| YQVGNFP | GGRDQGSGNTLY | 0.652141 |
| ESLLSQST | GGRDQGSGNTLY | 0.51920617 |
| LFTVGFAAVG | GGRDQGSGNTLY | 0.6405822 |
| ATPPTPT | GGRDQGSGNTLY | 0.4073679 |
| TDLATLG | GGRDQGSGNTLY | 0.41337213 |
| SAGGTGR | GGRDQGSGNTLY | 0.43912426 |
| GFSNSGS | GGRDQGSGNTLY | 0.59234035 |
| GGNGGSGGTSV | GGRDQGSGNTLY | 0.5763395 |
| GTSVATGGAGN | GGRDQGSGNTLY | 0.41547656 |
| NGGNGGSGGTS | GGRDQGSGNTLY | 0.31594828 |
| AVVKFPH | GGRDQGSGNTLY | 0.65382874 |
| TPPPPPP | GGRDQGSGNTLY | 0.60083663 |
| AAPAAAAAPA | GGRDQGSGNTLY | 0.33156046 |
| IALNEDL | GGRDQGSGNTLY | 0.4189561 |
| IVNVSSM | GGRDQGSGNTLY | 0.6947439 |
| TGIPPAPR | GGRDQGSGNTLY | 0.6857826 |
| FDLGGGTF | GGRDQGSGNTLY | 0.43682098 |
| FDLGGGTFDVS | GGRDQGSGNTLY | 0.42975715 |
| NEPTAAA | GGRDQGSGNTLY | 0.64689493 |
| NEPTAAALA | GGRDQGSGNTLY | 0.4078547 |
| GSIINISSI | GGRDQGSGNTLY | 0.50165457 |
| VAPGNAGTA | GGRDQGSGNTLY | 0.5908401 |
| TGGGASLE | GGRDQGSGNTLY | 0.5969666 |

| | | |
|---|---|---|
| VAVYNHYKRI | GGRDQGSGNTLY | 0.3944515 |
| TDPAPATT | GGRDQGSGNTLY | 0.60417616 |
| TNRPDILDPALLRPG | GGRDQGSGNTLY | 0.565602 |
| LTTTTTS | GGRDQGSGNTLY | 0.5492734 |
| PTGSGKTL | GGRDQGSGNTLY | 0.28886634 |
| TGSGKTL | GGRDQGSGNTLY | 0.57323194 |
| TIGHVDHGK | GGRDQGSGNTLY | 0.42417008 |
| TIGHVDHGKTT | GGRDQGSGNTLY | 0.40663442 |
| GHVDHGKTT | GGRDQGSGNTLY | 0.3114951 |
| AQFEHTLL | GGRDQGSGNTLY | 0.41070354 |
| IVNQDEL | GGRDQGSGNTLY | 0.66881466 |
| TNGSQFF | GGRDQGSGNTLY | 0.66975224 |
| TGKGIVE | GGRDQGSGNTLY | 0.42255503 |
| LISNASDA | GGRDQGSGNTLY | 0.64739525 |
| AELRTLGELTKIAKAHGAQ | GGRDQGSGNTLY | 0.6921687 |
| APNAVAAAEPSW | GGRDQGSGNTLY | 0.4284274 |
| APQVAGAATAVL | GGRDQGSGNTLY | 0.30971724 |
| ARRDPAAAAAILPTDA | GGRDQGSGNTLY | 0.5971003 |
| DAAARDVLAEAGLAEYF | GGRDQGSGNTLY | 0.5515331 |
| DDQIVLSAKANLAKVEQ | GGRDQGSGNTLY | 0.6058782 |
| DELLPDTGEDSDEDGH | GGRDQGSGNTLY | 0.67805475 |
| ELPEAPANRATVPQD | GGRDQGSGNTLY | 0.363828 |
| FGQSKIERVTGIG | GGRDQGSGNTLY | 0.60149306 |
| GAGGTGSTAGGAGGAGGAGGLY | GGRDQGSGNTLY | 0.64957595 |
| GHGGTGAAGGNGGTGGAGGSGI | GGRDQGSGNTLY | 0.40154243 |
| GNGGNGGAGGEGGPGGAGGAGA | GGRDQGSGNTLY | 0.33524013 |
| GSTAGNGGNGGNATNGTIAGQP | GGRDQGSGNTLY | 0.35889894 |
| GTGGDGGNGGNGGTGG | GGRDQGSGNTLY | 0.31435224 |
| GVRRPAGIPADAPV | GGRDQGSGNTLY | 0.6290841 |
| GYRQVTDAHLLTLARR | GGRDQGSGNTLY | 0.4687131 |
| IAKMGQDGHDRG | GGRDQGSGNTLY | 0.49204248 |
| KAEPSIEIQYT | GGRDQGSGNTLY | 0.5088465 |
| KPVRIRMAADEDTVAVT | GGRDQGSGNTLY | 0.4705251 |
| MPPRIAGMRLLVIKPEP | GGRDQGSGNTLY | 0.4652177 |
| MKFYAVALFRSRGVPPTR | GGRDQGSGNTLY | 0.49674988 |
| NAGGFGILGAVAHSP | GGRDQGSGNTLY | 0.57519746 |
| NEGREITLAMAEM | GGRDQGSGNTLY | 0.63412833 |
| NELVEWSHPLGHRWRWE | GGRDQGSGNTLY | 0.4720111 |
| NLGAGGTGGNGGNPG | GGRDQGSGNTLY | 0.45505625 |
| NTEWPKWETVMAADPVD | GGRDQGSGNTLY | 0.67427635 |
| PASAAPAETQP | GGRDQGSGNTLY | 0.38128638 |
| QQSSDGKNGVRAAEITGE | GGRDQGSGNTLY | 0.40883207 |
| RAPDAVTTSWWKE | GGRDQGSGNTLY | 0.6309952 |
| SGFGNVGAHTSGWFNQ | GGRDQGSGNTLY | 0.40326756 |
| SGVAGAVGQAASVGGLKV | GGRDQGSGNTLY | 0.6376279 |
| SNGGAGTGSAGGNGGT | GGRDQGSGNTLY | 0.60864973 |
| TGGAGGEGGAAG | GGRDQGSGNTLY | 0.60266984 |
| TGGDGGAAGAGGAPGHG | GGRDQGSGNTLY | 0.5806182 |
| TMPPLLAAIANAGRHGV | GGRDQGSGNTLY | 0.6878519 |
| TSSTAGNSGGA | GGRDQGSGNTLY | 0.6186116 |
| VPATVPVGILC | GGRDQGSGNTLY | 0.3870178 |

| | | |
|---|---|---|
| VQRPDSGGVVLTGRIS | GGRDQGSGNTLY | 0.6581311 |
| WTPNPVPATKGVPAGLVQAT | GGRDQGSGNTLY | 0.35319015 |
| YALKRALAKFSDRFDVVII | GGRDQGSGNTLY | 0.63532746 |
| YLIGSVIV | GGRDQGSGNTLY | 0.6117386 |
| YGQLVAVID | GGRDQGSGNTLY | 0.41594377 |
| VTVGGMTLPA | GGRDQGSGNTLY | 0.5593286 |
| VLALPAV | GGRDQGSGNTLY | 0.6657726 |
| VIPPTLNYET | GGRDQGSGNTLY | 0.6648887 |
| VGMGGMHPSSG | GGRDQGSGNTLY | 0.35725892 |
| VGLAGGAATA | GGRDQGSGNTLY | 0.4344374 |
| VGGAGGAGGSGP | GGRDQGSGNTLY | 0.37962738 |
| VFAAFMV | GGRDQGSGNTLY | 0.42969143 |
| TLYDKLR | GGRDQGSGNTLY | 0.34223077 |
| TLAVFTVL | GGRDQGSGNTLY | 0.63385093 |
| TGSVQPARV | GGRDQGSGNTLY | 0.50034773 |
| TGGGAGSGGT | GGRDQGSGNTLY | 0.6641462 |
| SHLETSL | GGRDQGSGNTLY | 0.59950316 |
| SFTTGMT | GGRDQGSGNTLY | 0.6609342 |
| QVEVAKRMGVRQPTVS | GGRDQGSGNTLY | 0.4460592 |
| QGSGGPALAM | GGRDQGSGNTLY | 0.53814745 |
| MLAPSNP | GGRDQGSGNTLY | 0.3226469 |
| LVNNAAI | GGRDQGSGNTLY | 0.43998298 |
| LTGEINTLA | GGRDQGSGNTLY | 0.37325686 |
| LSIFGPV | GGRDQGSGNTLY | 0.53009266 |
| LLALLANL | GGRDQGSGNTLY | 0.30764166 |
| LISNASDALDKLR | GGRDQGSGNTLY | 0.66447556 |
| LINEAVP | GGRDQGSGNTLY | 0.5278911 |
| LGYAFSGL | GGRDQGSGNTLY | 0.6310107 |
| LDFVAVAL | GGRDQGSGNTLY | 0.45481437 |
| LAQPGAE | GGRDQGSGNTLY | 0.4110546 |
| ISNLGAVTGF | GGRDQGSGNTLY | 0.5030194 |
| HQEDPGRI | GGRDQGSGNTLY | 0.65402067 |
| GTPGEPTP | GGRDQGSGNTLY | 0.3718002 |
| GFFNSGS | GGRDQGSGNTLY | 0.2943702 |
| GAGGGGLTAGAN | GGRDQGSGNTLY | 0.3933821 |
| FVDALVNM | GGRDQGSGNTLY | 0.3015869 |
| FLTVAPEL | GGRDQGSGNTLY | 0.6824806 |
| FETLTAL | GGRDQGSGNTLY | 0.6564127 |
| DVINPIEL | GGRDQGSGNTLY | 0.57747966 |
| APVSGAPV | GGRDQGSGNTLY | 0.6833006 |
| ALVGGLAW | GGRDQGSGNTLY | 0.28432864 |
| ALAAAAATAI | GGRDQGSGNTLY | 0.48333845 |
| AFGLGLLL | GGRDQGSGNTLY | 0.6722932 |
| AFFAAASL | GGRDQGSGNTLY | 0.60262203 |
| AAAALPAFSPP | GGRDQGSGNTLY | 0.5350902 |
| VVLAPAL | GGRDQGSGNTLY | 0.6639615 |
| TIGHVDHGKTTL | GGRDQGSGNTLY | 0.4837779 |
| LISNASDALDK | GGRDQGSGNTLY | 0.55652946 |
| LISNASDAL | GGRDQGSGNTLY | 0.569401 |
| ISNASDALDK | GGRDQGSGNTLY | 0.52413034 |
| NQAETLVYQTE | GGRDQGSGNTLY | 0.4989078 |

| | | |
|---|---|---|
| TGGTGGTAGAG | GGRDQGSGNTLY | 0.3432224 |
| TGSTAGGAGGA | GGRDQGSGNTLY | 0.5407406 |
| DPLAVPVAAGPV | GGRDQGSGNTLY | 0.6945821 |
| GGTGGTPGNGGN | GGRDQGSGNTLY | 0.551792 |
| PAGLAGVDPATL | GGRDQGSGNTLY | 0.6579927 |
| GNNLVVEAPHAGASV | GGRDQGSGNTLY | 0.5942101 |
| KVTRIMSASTLEQPA | GGRDQGSGNTLY | 0.6422705 |
| LGFALPIALPRRTVV | GGRDQGSGNTLY | 0.4302582 |
| TRLGNLTARKLVVLV | GGRDQGSGNTLY | 0.4157955 |
| WSQQRRLIYFPSAGP | GGRDQGSGNTLY | 0.5866008 |
| GPNGNPDPMAAAVDIR | GGRDQGSGNTLY | 0.4090569 |
| TQRCVSVADLRAAASR | GGRDQGSGNTLY | 0.48080927 |
| QFVLDVAEHPVEVLTETV | GGRDQGSGNTLY | 0.38910663 |
| GGAGGTGGAGGNGGTQAGDG | GGRDQGSGNTLY | 0.7015312 |
| PLLATLRAIAVDVDAKPAQV | GGRDQGSGNTLY | 0.6192093 |
| APKFRTGGYMIDFWGVGYQVAK | GGRDQGSGNTLY | 0.62779343 |
| GAGGAGGAGGAAGGPGGTGGTG | GGRDQGSGNTLY | 0.46563104 |
| IGALGLLATDVAPAGSVAFGVA | GGRDQGSGNTLY | 0.53313434 |
| AGGAGGIGGNGGAGGAGGVLMG | GGRDQGSGNTLY | 0.6235018 |
| ADELIGLL | GGRDQGSGNTLY | 0.318131 |
| ADRALKA | GGRDQGSGNTLY | 0.6669868 |
| AGAVGQVGGAGGA | GGRDQGSGNTLY | 0.4317312 |
| AGGGGGLGGAP | GGRDQGSGNTLY | 0.5181731 |
| AGGLLYGNGGAG | GGRDQGSGNTLY | 0.6054225 |
| AGLESIGL | GGRDQGSGNTLY | 0.67485964 |
| AGLKTGII | GGRDQGSGNTLY | 0.47599667 |
| ASLAALLL | GGRDQGSGNTLY | 0.5798706 |
| ASVINISSI | GGRDQGSGNTLY | 0.433387 |
| ATPPGPA | GGRDQGSGNTLY | 0.63654995 |
| ATVASALV | GGRDQGSGNTLY | 0.6717205 |
| ATVASLIL | GGRDQGSGNTLY | 0.62882054 |
| AVNGPVLIRH | GGRDQGSGNTLY | 0.6920967 |
| DGAAGAVGI | GGRDQGSGNTLY | 0.46757096 |
| DPLLHGI | GGRDQGSGNTLY | 0.32701346 |
| ELGAALGAL | GGRDQGSGNTLY | 0.51024246 |
| ELGRTQAV | GGRDQGSGNTLY | 0.6410396 |
| FVAGGYLS | GGRDQGSGNTLY | 0.6768035 |
| GASGGAGGAGGW | GGRDQGSGNTLY | 0.6727021 |
| GGAGAGGDGGAGGI | GGRDQGSGNTLY | 0.5936415 |
| GTGHNGSQPGMGGNG | GGRDQGSGNTLY | 0.48052835 |
| GYAAASASAS | GGRDQGSGNTLY | 0.66543937 |
| HQVDVAAVQQG | GGRDQGSGNTLY | 0.68619275 |
| IGAAIAHT | GGRDQGSGNTLY | 0.6383158 |
| IGMIEVAQL | GGRDQGSGNTLY | 0.45502687 |
| IRLAPEHTSAMD | GGRDQGSGNTLY | 0.43663344 |
| IYNVSVL | GGRDQGSGNTLY | 0.56789184 |
| KQGFLGM | GGRDQGSGNTLY | 0.6588179 |
| LESVITQL | GGRDQGSGNTLY | 0.36417037 |
| LGLGALVVV | GGRDQGSGNTLY | 0.5767297 |
| LIGNGGNGGSGGI | GGRDQGSGNTLY | 0.59932995 |
| LNAINAPTL | GGRDQGSGNTLY | 0.31543753 |

| | | |
|---|---|---|
| LQLLNTGV | GGRDQGSGNTLY | 0.53827226 |
| LQTSIEF | GGRDQGSGNTLY | 0.66928923 |
| MSFAVSEL | GGRDQGSGNTLY | 0.44661856 |
| NTGALGL | GGRDQGSGNTLY | 0.37644225 |
| PPGAGGAGGAGTT | GGRDQGSGNTLY | 0.67329097 |
| QLVGLGI | GGRDQGSGNTLY | 0.69723463 |
| RFLEQVHQL | GGRDQGSGNTLY | 0.6121983 |
| RGVGYVLR | GGRDQGSGNTLY | 0.32702458 |
| SATGVFAGVI | GGRDQGSGNTLY | 0.28105056 |
| SLPAITLP | GGRDQGSGNTLY | 0.58256054 |
| STALAFTV | GGRDQGSGNTLY | 0.6676892 |
| SVLTVGLGK | GGRDQGSGNTLY | 0.5109507 |
| TALAGAGSGI | GGRDQGSGNTLY | 0.47683668 |
| TGLPFTGI | GGRDQGSGNTLY | 0.5787986 |
| VAEHVTV | GGRDQGSGNTLY | 0.63686705 |
| VDIVLNSL | GGRDQGSGNTLY | 0.3652333 |
| VLTGGGAVL | GGRDQGSGNTLY | 0.38099948 |
| VNVGLAYGI | GGRDQGSGNTLY | 0.46834573 |
| VSSAAVGAV | GGRDQGSGNTLY | 0.61840326 |
| VTSVYEAGV | GGRDQGSGNTLY | 0.41288137 |
| WVALDRGA | GGRDQGSGNTLY | 0.5321312 |
| YLVGSVVI | GGRDQGSGNTLY | 0.5884736 |
| GPPGTGKTLL | GGRDQGSGNTLY | 0.52191657 |
| GVGMTKF | GGRDQGSGNTLY | 0.6064447 |
| IGIGGSDL | GGRDQGSGNTLY | 0.41091248 |
| IGTLLAEL | GGRDQGSGNTLY | 0.66926086 |
| IYTSGTT | GGRDQGSGNTLY | 0.3490821 |
| LSFPAGV | GGRDQGSGNTLY | 0.62410784 |
| LYTSGTT | GGRDQGSGNTLY | 0.6122304 |
| VGAFPFGF | GGRDQGSGNTLY | 0.5638819 |
| VGAGTVEF | GGRDQGSGNTLY | 0.52333903 |
| VGKSTLLL | GGRDQGSGNTLY | 0.4017157 |
| AALAHAGGGIV | GGRDQGSGNTLY | 0.35128903 |
| AALVAVSLL | GGRDQGSGNTLY | 0.37796122 |
| AAVSRAVAAL | GGRDQGSGNTLY | 0.6612923 |
| AGGSSGIGGF | GGRDQGSGNTLY | 0.5143935 |
| AGLTVANANG | GGRDQGSGNTLY | 0.6541387 |
| AGLVGAVL | GGRDQGSGNTLY | 0.5854239 |
| ALSQAFTV | GGRDQGSGNTLY | 0.41589642 |
| ALTFSVSL | GGRDQGSGNTLY | 0.5882003 |
| ALTGAALVAL | GGRDQGSGNTLY | 0.4012958 |
| ASFSIGAGVL | GGRDQGSGNTLY | 0.4345535 |
| DGSVYKI | GGRDQGSGNTLY | 0.6555972 |
| EQLAEIV | GGRDQGSGNTLY | 0.5555742 |
| FAVSELTVM | GGRDQGSGNTLY | 0.3973933 |
| GAAPLAAGL | GGRDQGSGNTLY | 0.5264106 |
| GADGGDGGSGGAGG | GGRDQGSGNTLY | 0.58418834 |
| GFGLDTGGAGG | GGRDQGSGNTLY | 0.36472216 |
| GLASGGLL | GGRDQGSGNTLY | 0.4484229 |
| GRIAGAAVLL | GGRDQGSGNTLY | 0.6498222 |
| GSAGSLNTSF | GGRDQGSGNTLY | 0.513687 |

| | | |
|---|---|---|
| GTGQIGI | GGRDQGSGNTLY | 0.34808442 |
| GTLIALIL | GGRDQGSGNTLY | 0.63023376 |
| IAATMGFVVG | GGRDQGSGNTLY | 0.39951736 |
| IFAQALL | GGRDQGSGNTLY | 0.61476475 |
| IGAAVALGF | GGRDQGSGNTLY | 0.53926694 |
| IGLTPLAL | GGRDQGSGNTLY | 0.4843487 |
| ILGSNTSTL | GGRDQGSGNTLY | 0.6426754 |
| IVFSLTVL | GGRDQGSGNTLY | 0.33763325 |
| LASKATGF | GGRDQGSGNTLY | 0.42157903 |
| LASVALIL | GGRDQGSGNTLY | 0.4722882 |
| LFAVGLTL | GGRDQGSGNTLY | 0.57765627 |
| LFNGGIGTYI | GGRDQGSGNTLY | 0.6275685 |
| LTLGAGII | GGRDQGSGNTLY | 0.39171493 |
| LVAAALGDI | GGRDQGSGNTLY | 0.58076483 |
| NTAFVQLGI | GGRDQGSGNTLY | 0.49246216 |
| QLFSGAEL | GGRDQGSGNTLY | 0.41537708 |
| SAATFITL | GGRDQGSGNTLY | 0.43487263 |
| SNYGLANLGV | GGRDQGSGNTLY | 0.40843043 |
| SYGGAYAVM | GGRDQGSGNTLY | 0.3686369 |
| TTGVGGLTLG | GGRDQGSGNTLY | 0.5641245 |
| TTQTIYI | GGRDQGSGNTLY | 0.41794008 |
| VAAGAGLGAVI | GGRDQGSGNTLY | 0.5953862 |
| VAAVVTLL | GGRDQGSGNTLY | 0.59033644 |
| VGFLGGLI | GGRDQGSGNTLY | 0.3807238 |
| VGVIPILL | GGRDQGSGNTLY | 0.6018667 |
| VIFASSLI | GGRDQGSGNTLY | 0.6133703 |
| VKGGDGGAAAG | GGRDQGSGNTLY | 0.63235307 |
| VTKTFTNI | GGRDQGSGNTLY | 0.5201434 |
| VVAAAILL | GGRDQGSGNTLY | 0.3742648 |
| YAALDVEL | GGRDQGSGNTLY | 0.64636976 |
| LASVALLL | GGRDQGSGNTLY | 0.6430146 |
| LDAVHRV | GGRDQGSGNTLY | 0.6127793 |
| QGLSSSIF | GGRDQGSGNTLY | 0.5914773 |
| RGTFANIRL | GGRDQGSGNTLY | 0.6360709 |
| RQLWWGHRI | GGRDQGSGNTLY | 0.6130248 |
| YGPPGTGKTLL | GGRDQGSGNTLY | 0.5358528 |
| AALVGAPANVA | GGRDQGSGNTLY | 0.33738154 |
| AATFRRTE | GGRDQGSGNTLY | 0.5411718 |
| AGALGGGI | GGRDQGSGNTLY | 0.5387118 |
| AGGAGGAGGIGAG | GGRDQGSGNTLY | 0.37482283 |
| AGSGALGL | GGRDQGSGNTLY | 0.33095032 |
| AQIADQI | GGRDQGSGNTLY | 0.64684033 |
| ASAVAVTYL | GGRDQGSGNTLY | 0.70082164 |
| ASTVLGGTGG | GGRDQGSGNTLY | 0.48032087 |
| AVGGEPAK | GGRDQGSGNTLY | 0.68220425 |
| CEQGTFGGTC | GGRDQGSGNTLY | 0.3165347 |
| DEGKAPAG | GGRDQGSGNTLY | 0.65420496 |
| DTVVFPAL | GGRDQGSGNTLY | 0.6594816 |
| FEGITMAG | GGRDQGSGNTLY | 0.43936336 |
| FPLVVQD | GGRDQGSGNTLY | 0.6063466 |
| GAFTTPQ | GGRDQGSGNTLY | 0.3579026 |

| | | |
|---|---|---|
| GFGGFGTAGGVGGA | GGRDQGSGNTLY | 0.42113578 |
| GFLAQFL | GGRDQGSGNTLY | 0.6753725 |
| GGAGGAGGNAGVA | GGRDQGSGNTLY | 0.5109056 |
| GGVGTAGAGGAGGAGG | GGRDQGSGNTLY | 0.4945508 |
| GNGGGGDAGNAGSGGN | GGRDQGSGNTLY | 0.5176226 |
| GTGGAGGAGNDAGS | GGRDQGSGNTLY | 0.4394401 |
| GTTNTGWFNS | GGRDQGSGNTLY | 0.513685 |
| IFMGPLFTL | GGRDQGSGNTLY | 0.64569235 |
| ILAHAIEAI | GGRDQGSGNTLY | 0.49367034 |
| IRSAIREAL | GGRDQGSGNTLY | 0.682099 |
| ITFAGGG | GGRDQGSGNTLY | 0.42857593 |
| ITGMSGHQEV | GGRDQGSGNTLY | 0.68229324 |
| IVAGLACTL | GGRDQGSGNTLY | 0.67952853 |
| IVGGGLAGA | GGRDQGSGNTLY | 0.5991843 |
| IVTGVLNI | GGRDQGSGNTLY | 0.43935537 |
| LAQQGCI | GGRDQGSGNTLY | 0.65895534 |
| LCLHVEN | GGRDQGSGNTLY | 0.519833 |
| LDAISDGS | GGRDQGSGNTLY | 0.5905286 |
| LGGFAAML | GGRDQGSGNTLY | 0.6065358 |
| LGGKNPMI | GGRDQGSGNTLY | 0.122831434 |
| LGPKGVAAIL | GGRDQGSGNTLY | 0.12625976 |
| LMVGVSI | GGRDQGSGNTLY | 0.12379559 |
| LVGAAFVL | GGRDQGSGNTLY | 0.12914605 |
| LVLAVAPL | GGRDQGSGNTLY | 0.12667999 |
| MAAIAVIDAA | GGRDQGSGNTLY | 0.12309323 |
| NNGINAELQ | GGRDQGSGNTLY | 0.12451492 |
| RGEPLVLVHG | GGRDQGSGNTLY | 0.124384075 |
| RGGAGGAAGLI | GGRDQGSGNTLY | 0.12436919 |
| RSYRMHPAV | GGRDQGSGNTLY | 0.12630984 |
| SAAHLLAGQLA | GGRDQGSGNTLY | 0.12655993 |
| SAAVGSY | GGRDQGSGNTLY | 0.13330822 |
| SPHAVLHRI | GGRDQGSGNTLY | 0.12970702 |
| SQIAKVL | GGRDQGSGNTLY | 0.12300731 |
| TALGHLVP | GGRDQGSGNTLY | 0.12373601 |
| TAPTTSGAA | GGRDQGSGNTLY | 0.12457414 |
| TDGDDGNGGSPGT | GGRDQGSGNTLY | 0.1255403 |
| TDGLGEN | GGRDQGSGNTLY | 0.12201501 |
| TGGPPLPGQA | GGRDQGSGNTLY | 0.12630406 |
| TIPAITI | GGRDQGSGNTLY | 0.12222225 |
| TIPGFTI | GGRDQGSGNTLY | 0.12189822 |
| TSIASGALA | GGRDQGSGNTLY | 0.12628081 |
| TVTLPGAAI | GGRDQGSGNTLY | 0.12555444 |
| VASGLSAALP | GGRDQGSGNTLY | 0.12427721 |
| VGAGVIGI | GGRDQGSGNTLY | 0.12770182 |
| VGPKRFGV | GGRDQGSGNTLY | 0.123139225 |
| VNVAFPD | GGRDQGSGNTLY | 0.12424287 |
| VSMTNSGVS | GGRDQGSGNTLY | 0.12656671 |
| VTDVLGDLSV | GGRDQGSGNTLY | 0.13369496 |
| VVVVGAGFAGL | GGRDQGSGNTLY | 0.12455298 |
| VYTVGTVV | GGRDQGSGNTLY | 0.123909466 |
| WTTFSGP | GGRDQGSGNTLY | 0.13399023 |

| | | |
|---|---|---|
| YDELVEAF | GGRDQGSGNTLY | 0.12768912 |
| YGTGGAGGNGG | GGRDQGSGNTLY | 0.12758943 |
| YTNVMAV | GGRDQGSGNTLY | 0.12864147 |
| FLEQVHQL | GGRDQGSGNTLY | 0.122800075 |
| GLSLLAVAV | GGRDQGSGNTLY | 0.13037351 |
| LEQVHQL | GGRDQGSGNTLY | 0.12727761 |
| LNAAGAI | GGRDQGSGNTLY | 0.12577412 |
| PAVHASL | GGRDQGSGNTLY | 0.12833658 |
| TELEDDA | GGRDQGSGNTLY | 0.12321879 |
| TPPGPPN | GGRDQGSGNTLY | 0.12375167 |
| TPPPPPPP | GGRDQGSGNTLY | 0.122099616 |
| TQSPALI | TCSAEGQTNERLF | 0.124703795 |
| EFAAAFPQ | TCSAEGQTNERLF | 0.12870649 |
| LLSGNLDVL | TCSAEGQTNERLF | 0.12766849 |
| GPANLGNYNI | TCSAEGQTNERLF | 0.122545615 |
| TLGPTLP | TCSAEGQTNERLF | 0.123450436 |
| VYAYSSCCK | TCSAEGQTNERLF | 0.122193135 |
| APAPAPAPNI | TCSAEGQTNERLF | 0.12364329 |
| PGGPVPIP | TCSAEGQTNERLF | 0.12232484 |
| FLTVALEP | TCSAEGQTNERLF | 0.1229011 |
| SQYFGANL | TCSAEGQTNERLF | 0.13547687 |
| GGVKPAA | TCSAEGQTNERLF | 0.13210608 |
| GGTGGAGGAGGW | TCSAEGQTNERLF | 0.12617904 |
| GVGGVAGLGGAGG | TCSAEGQTNERLF | 0.12832516 |
| VLLVLATTI | TCSAEGQTNERLF | 0.12285865 |
| IGIFAAML | TCSAEGQTNERLF | 0.12710515 |
| AVGDLID | TCSAEGQTNERLF | 0.12583931 |
| NISPLPL | TCSAEGQTNERLF | 0.12267913 |
| VTAVFVAI | TCSAEGQTNERLF | 0.1273685 |
| SGDDNDAAL | TCSAEGQTNERLF | 0.12571198 |
| AVAALAHS | TCSAEGQTNERLF | 0.124476045 |
| GGVTVMATP | TCSAEGQTNERLF | 0.13087517 |
| AGLAANVRD | TCSAEGQTNERLF | 0.12510799 |
| TNRPDLIDPALLRPG | TCSAEGQTNERLF | 0.124638975 |
| LANLGVTAL | TCSAEGQTNERLF | 0.12275968 |
| TGERIEL | TCSAEGQTNERLF | 0.124923736 |
| NGEELAA | TCSAEGQTNERLF | 0.122935005 |
| VLSLVWG | TCSAEGQTNERLF | 0.124384075 |
| IGYALWL | TCSAEGQTNERLF | 0.12286437 |
| LSLLLAPL | TCSAEGQTNERLF | 0.12615299 |
| TLDEALK | TCSAEGQTNERLF | 0.13627954 |
| TDGAGTAEGG | TCSAEGQTNERLF | 0.12748188 |
| NAPGGAGGNGG | TCSAEGQTNERLF | 0.122880615 |
| VSVFMGMGGNF | TCSAEGQTNERLF | 0.12661669 |
| LASYLAAL | TCSAEGQTNERLF | 0.12736635 |
| LVGGTPAL | TCSAEGQTNERLF | 0.12595387 |
| VRNADVL | TCSAEGQTNERLF | 0.1238222 |
| TCSSPFLVSAM | TCSAEGQTNERLF | 0.12338711 |
| VHITDAE | TCSAEGQTNERLF | 0.12492376 |
| EYAAPSF | TCSAEGQTNERLF | 0.13230409 |
| VVDAFHGGL | TCSAEGQTNERLF | 0.12781924 |

| | | |
|---|---|---|
| ASVVDPQ | TCSAEGQTNERLF | 0.12587921 |
| SYNEFGGLS | TCSAEGQTNERLF | 0.12514853 |
| DPDGSVY | TCSAEGQTNERLF | 0.12428026 |
| MVSTIHATE | TCSAEGQTNERLF | 0.124803565 |
| LAAGVADPV | TCSAEGQTNERLF | 0.123865776 |
| TKEQIAA | TCSAEGQTNERLF | 0.12432478 |
| SLASPAAL | TCSAEGQTNERLF | 0.12435902 |
| IGLVFYGV | TCSAEGQTNERLF | 0.12499609 |
| FLLDASNL | TCSAEGQTNERLF | 0.12607946 |
| YQVGNFP | TCSAEGQTNERLF | 0.12321997 |
| ESLLSQST | TCSAEGQTNERLF | 0.12463931 |
| LFTVGFAAVG | TCSAEGQTNERLF | 0.12843156 |
| ATPPTPT | TCSAEGQTNERLF | 0.12373721 |
| TDLATLG | TCSAEGQTNERLF | 0.12453192 |
| SAGGTGR | TCSAEGQTNERLF | 0.12885873 |
| GFSNSGS | TCSAEGQTNERLF | 0.12442996 |
| GGNGGSGGTSV | TCSAEGQTNERLF | 0.12684818 |
| GTSVATGGAGN | TCSAEGQTNERLF | 0.12605494 |
| NGGNGGSGGTS | TCSAEGQTNERLF | 0.12673603 |
| AVVKFPH | TCSAEGQTNERLF | 0.12267168 |
| TPPPPPP | TCSAEGQTNERLF | 0.12719716 |
| AAPAAAAAPA | TCSAEGQTNERLF | 0.123933546 |
| IALNEDL | TCSAEGQTNERLF | 0.123836756 |
| IVNVSSM | TCSAEGQTNERLF | 0.122514375 |
| TGIPPAPR | TCSAEGQTNERLF | 0.12211429 |
| FDLGGGTF | TCSAEGQTNERLF | 0.1317027 |
| FDLGGGTFDVS | TCSAEGQTNERLF | 0.12342842 |
| NEPTAAA | TCSAEGQTNERLF | 0.12678684 |
| NEPTAAALA | TCSAEGQTNERLF | 0.124442816 |
| GSIINISSI | TCSAEGQTNERLF | 0.13040784 |
| VAPGNAGTA | TCSAEGQTNERLF | 0.12573248 |
| TGGGASLE | TCSAEGQTNERLF | 0.12484503 |
| VAVYNHYKRI | TCSAEGQTNERLF | 0.12364714 |
| TDPAPATT | TCSAEGQTNERLF | 0.12958214 |
| TNRPDILDPALLRPG | TCSAEGQTNERLF | 0.12486957 |
| LTTTTTS | TCSAEGQTNERLF | 0.1260369 |
| PTGSGKTL | TCSAEGQTNERLF | 0.123404354 |
| TGSGKTL | TCSAEGQTNERLF | 0.12252956 |
| TIGHVDHGK | TCSAEGQTNERLF | 0.12762639 |
| TIGHVDHGKTT | TCSAEGQTNERLF | 0.12240857 |
| GHVDHGKTT | TCSAEGQTNERLF | 0.12187365 |
| AQFEHTLL | TCSAEGQTNERLF | 0.121976405 |
| IVNQDEL | TCSAEGQTNERLF | 0.12258584 |
| TNGSQFF | TCSAEGQTNERLF | 0.13029002 |
| TGKGIVE | TCSAEGQTNERLF | 0.12983502 |
| LISNASDA | TCSAEGQTNERLF | 0.12725595 |
| AELRTLGELTKIAKAHGAQ | TCSAEGQTNERLF | 0.1267645 |
| APNAVAAAEPSW | TCSAEGQTNERLF | 0.1265968 |
| APQVAGAATAVL | TCSAEGQTNERLF | 0.12173924 |
| ARRDPAAAAAILPTDA | TCSAEGQTNERLF | 0.124005534 |
| DAAARDVLAEAGLAEYF | TCSAEGQTNERLF | 0.12319205 |

| | | |
|---|---|---|
| DDQIVLSAKANLAKVEQ | TCSAEGQTNERLF | 0.12805302 |
| DELLPDTGEDSDEDGH | TCSAEGQTNERLF | 0.122710675 |
| ELPEAPANRATVPQD | TCSAEGQTNERLF | 0.12373916 |
| FGQSKIERVTGIG | TCSAEGQTNERLF | 0.13026787 |
| GAGGTGSTAGGAGGAGGAGGLY | TCSAEGQTNERLF | 0.12396176 |
| GHGGTGAAGGNGGTGGAGGSGI | TCSAEGQTNERLF | 0.12509438 |
| GNGGNGGAGGEGGPGGAGGAGA | TCSAEGQTNERLF | 0.122407936 |
| GSTAGNGGNGGNATNGTIAGQP | TCSAEGQTNERLF | 0.121892065 |
| GTGGDGGNGGNGGTGG | TCSAEGQTNERLF | 0.13354391 |
| GVRRPAGIPADAPV | TCSAEGQTNERLF | 0.1228161 |
| GYRQVTDAHLLTLARR | TCSAEGQTNERLF | 0.12357689 |
| IAKMGQDGHDRG | TCSAEGQTNERLF | 0.12223529 |
| KAEPSIEIQYT | TCSAEGQTNERLF | 0.123029545 |
| KPVRIRMAADEDTVAVT | TCSAEGQTNERLF | 0.12266411 |
| MPPRIAGMRLLVIKPEP | TCSAEGQTNERLF | 0.12715384 |
| MKFYAVALFRSRGVPPTR | TCSAEGQTNERLF | 0.12329927 |
| NAGGFGILGAVAHSP | TCSAEGQTNERLF | 0.12597206 |
| NEGREITLAMAEM | TCSAEGQTNERLF | 0.12834796 |
| NELVEWSHPLGHRWRWE | TCSAEGQTNERLF | 0.12599784 |
| NLGAGGTGGNGGNPG | TCSAEGQTNERLF | 0.12660919 |
| NTEWPKWETVMAADPVD | TCSAEGQTNERLF | 0.124930106 |
| PASAAPAETQP | TCSAEGQTNERLF | 0.12434533 |
| QQSSDGKNGVRAAEITGE | TCSAEGQTNERLF | 0.12383408 |
| RAPDAVTTSWWKE | TCSAEGQTNERLF | 0.12468593 |
| SGFGNVGAHTSGWFNQ | TCSAEGQTNERLF | 0.13243334 |
| SGVAGAVGQAASVGGLKV | TCSAEGQTNERLF | 0.124488436 |
| SNGGAGTGSAGGNGGT | TCSAEGQTNERLF | 0.122807875 |
| TGGAGGEGGAAG | TCSAEGQTNERLF | 0.124358624 |
| TGGDGGAAGAGGAPGHG | TCSAEGQTNERLF | 0.12943177 |
| TMPPLLAAIANAGRHGV | TCSAEGQTNERLF | 0.13086522 |
| TSSTAGNSGGA | TCSAEGQTNERLF | 0.123196386 |
| VPATVPVGILC | TCSAEGQTNERLF | 0.12464273 |
| VQRPDSGGVVLTGRIS | TCSAEGQTNERLF | 0.124745205 |
| WTPNPVPATKGVPAGLVQAT | TCSAEGQTNERLF | 0.12531877 |
| YALKRALAKFSDRFDVVII | TCSAEGQTNERLF | 0.12329724 |
| YLIGSVIV | TCSAEGQTNERLF | 0.12498699 |
| YGQLVAVID | TCSAEGQTNERLF | 0.12454113 |
| VTVGGMTLPA | TCSAEGQTNERLF | 0.13097823 |
| VLALPAV | TCSAEGQTNERLF | 0.12396194 |
| VIPPTLNYET | TCSAEGQTNERLF | 0.12258177 |
| VGMGGMHPSSG | TCSAEGQTNERLF | 0.12698582 |
| VGLAGGAATA | TCSAEGQTNERLF | 0.12306174 |
| VGGAGGAGGSGP | TCSAEGQTNERLF | 0.12561025 |
| VFAAFMV | TCSAEGQTNERLF | 0.1258959 |
| TLYDKLR | TCSAEGQTNERLF | 0.12280734 |
| TLAVFTVL | TCSAEGQTNERLF | 0.124822326 |
| TGSVQPARV | TCSAEGQTNERLF | 0.1317919 |
| TGGGAGSGGT | TCSAEGQTNERLF | 0.123890996 |
| SHLETSL | TCSAEGQTNERLF | 0.12478884 |
| SFTTGMT | TCSAEGQTNERLF | 0.13018589 |
| QVEVAKRMGVRQPTVS | TCSAEGQTNERLF | 0.12701528 |

| | | |
|---|---|---|
| QGSGGPALAM | TCSAEGQTNERLF | 0.12895764 |
| MLAPSNP | TCSAEGQTNERLF | 0.1233037 |
| LVNNAAI | TCSAEGQTNERLF | 0.12645052 |
| LTGEINTLA | TCSAEGQTNERLF | 0.12365571 |
| LSIFGPV | TCSAEGQTNERLF | 0.124106444 |
| LLALLANL | TCSAEGQTNERLF | 0.12833415 |
| LISNASDALDKLR | TCSAEGQTNERLF | 0.12837209 |
| LINEAVP | TCSAEGQTNERLF | 0.12470367 |
| LGYAFSGL | TCSAEGQTNERLF | 0.13147902 |
| LDFVAVAL | TCSAEGQTNERLF | 0.12388653 |
| LAQPGAE | TCSAEGQTNERLF | 0.12600875 |
| ISNLGAVTGF | TCSAEGQTNERLF | 0.12268116 |
| HQEDPGRI | TCSAEGQTNERLF | 0.12459269 |
| GTPGEPTP | TCSAEGQTNERLF | 0.12541431 |
| GFFNSGS | TCSAEGQTNERLF | 0.1223208 |
| GAGGGGLTAGAN | TCSAEGQTNERLF | 0.12194786 |
| FVDALVNM | TCSAEGQTNERLF | 0.123436786 |
| FLTVAPEL | TCSAEGQTNERLF | 0.12658781 |
| FETLTAL | TCSAEGQTNERLF | 0.12174542 |
| DVINPIEL | TCSAEGQTNERLF | 0.12459617 |
| APVSGAPV | TCSAEGQTNERLF | 0.122961484 |
| ALVGGLAW | TCSAEGQTNERLF | 0.12361869 |
| ALAAAAATAI | TCSAEGQTNERLF | 0.12802379 |
| AFGLGLLL | TCSAEGQTNERLF | 0.12345728 |
| AFFAAASL | TCSAEGQTNERLF | 0.12738347 |
| AAAALPAFSPP | TCSAEGQTNERLF | 0.12265114 |
| VVLAPAL | TCSAEGQTNERLF | 0.12364382 |
| TIGHVDHGKTTL | TCSAEGQTNERLF | 0.12836835 |
| LISNASDALDK | TCSAEGQTNERLF | 0.12569585 |
| LISNASDAL | TCSAEGQTNERLF | 0.12337563 |
| ISNASDALDK | TCSAEGQTNERLF | 0.12962662 |
| NQAETLVYQTE | TCSAEGQTNERLF | 0.12237748 |
| TGGTGGTAGAG | TCSAEGQTNERLF | 0.13102145 |
| TGSTAGGAGGA | TCSAEGQTNERLF | 0.12773216 |
| DPLAVPVAAGPV | TCSAEGQTNERLF | 0.12255075 |
| GGTGGTPGNGGN | TCSAEGQTNERLF | 0.12582648 |
| PAGLAGVDPATL | TCSAEGQTNERLF | 0.12524202 |
| GNNLVVEAPHAGASV | TCSAEGQTNERLF | 0.12592046 |
| KVTRIMSASTLEQPA | TCSAEGQTNERLF | 0.123668775 |
| LGFALPIALPRRTVV | TCSAEGQTNERLF | 0.12275424 |
| TRLGNLTARKLVVLV | TCSAEGQTNERLF | 0.12638052 |
| WSQQRRLIYFPSAGP | TCSAEGQTNERLF | 0.13164 |
| GPNGNPDPMAAAVDIR | TCSAEGQTNERLF | 0.12498475 |
| TQRCVSVADLRAAASR | TCSAEGQTNERLF | 0.124167494 |
| QFVLDVAEHPVEVLTETV | TCSAEGQTNERLF | 0.12985982 |
| GGAGGTGGAGGNGGTQAGDG | TCSAEGQTNERLF | 0.12645619 |
| PLLATLRAIAVDVDAKPAQV | TCSAEGQTNERLF | 0.12368101 |
| APKFRTGGYMIDFWGVGYQVAK | TCSAEGQTNERLF | 0.12857224 |
| GAGGAGGAGGAAGGPGGTGGTG | TCSAEGQTNERLF | 0.124570176 |
| IGALGLLATDVAPAGSVAFGVA | TCSAEGQTNERLF | 0.12998277 |
| AGGAGGIGGNGGAGGAGGVLMG | TCSAEGQTNERLF | 0.12820701 |

| | | |
|---|---|---|
| ADELIGLL | TCSAEGQTNERLF | 0.12481747 |
| ADRALKA | TCSAEGQTNERLF | 0.12337684 |
| AGAVGQVGGAGGA | TCSAEGQTNERLF | 0.12317917 |
| AGGGGGLGGAP | TCSAEGQTNERLF | 0.12522502 |
| AGGLLYGNGGAG | TCSAEGQTNERLF | 0.12701114 |
| AGLESIGL | TCSAEGQTNERLF | 0.122890905 |
| AGLKTGII | TCSAEGQTNERLF | 0.12334637 |
| ASLAALLL | TCSAEGQTNERLF | 0.12633231 |
| ASVINISSI | TCSAEGQTNERLF | 0.13181673 |
| ATPPGPA | TCSAEGQTNERLF | 0.12341047 |
| ATVASALV | TCSAEGQTNERLF | 0.1236537 |
| ATVASLIL | TCSAEGQTNERLF | 0.1334309 |
| AVNGPVLIRH | TCSAEGQTNERLF | 0.12316975 |
| DGAAGAVGI | TCSAEGQTNERLF | 0.12824048 |
| DPLLHGI | TCSAEGQTNERLF | 0.12584959 |
| ELGAALGAL | TCSAEGQTNERLF | 0.12660904 |
| ELGRTQAV | TCSAEGQTNERLF | 0.12478582 |
| FVAGGYLS | TCSAEGQTNERLF | 0.12652327 |
| GASGGAGGAGGW | TCSAEGQTNERLF | 0.12251085 |
| GGAGAGGDGGAGGI | TCSAEGQTNERLF | 0.12944554 |
| GTGHNGSQPGMGGNG | TCSAEGQTNERLF | 0.12195894 |
| GYAAASASAS | TCSAEGQTNERLF | 0.1273146 |
| HQVDVAAVQQG | TCSAEGQTNERLF | 0.12282834 |
| IGAAIAHT | TCSAEGQTNERLF | 0.12691237 |
| IGMIEVAQL | TCSAEGQTNERLF | 0.12740515 |
| IRLAPEHTSAMD | TCSAEGQTNERLF | 0.13060097 |
| IYNVSVL | TCSAEGQTNERLF | 0.12202865 |
| KQGFLGM | TCSAEGQTNERLF | 0.12979412 |
| LESVITQL | TCSAEGQTNERLF | 0.1282272 |
| LGLGALVVV | TCSAEGQTNERLF | 0.12910968 |
| LIGNGGNGGSGGI | TCSAEGQTNERLF | 0.124021985 |
| LNAINAPTL | TCSAEGQTNERLF | 0.12840715 |
| LQLLNTGV | TCSAEGQTNERLF | 0.1249833 |
| LQTSIEF | TCSAEGQTNERLF | 0.12507826 |
| MSFAVSEL | TCSAEGQTNERLF | 0.12278707 |
| NTGALGL | TCSAEGQTNERLF | 0.12284829 |
| PPGAGGAGGAGTT | TCSAEGQTNERLF | 0.12467645 |
| QLVGLGI | TCSAEGQTNERLF | 0.12476483 |
| RFLEQVHQL | TCSAEGQTNERLF | 0.12706974 |
| RGVGYVLR | TCSAEGQTNERLF | 0.12553184 |
| SATGVFAGVI | TCSAEGQTNERLF | 0.12868658 |
| SLPAITLP | TCSAEGQTNERLF | 0.12855883 |
| STALAFTV | TCSAEGQTNERLF | 0.12769152 |
| SVLTVGLGK | TCSAEGQTNERLF | 0.12606892 |
| TALAGAGSGI | TCSAEGQTNERLF | 0.12472316 |
| TGLPFTGI | TCSAEGQTNERLF | 0.1255168 |
| VAEHVTV | TCSAEGQTNERLF | 0.12328607 |
| VDIVLNSL | TCSAEGQTNERLF | 0.12424297 |
| VLTGGGAVL | TCSAEGQTNERLF | 0.12412568 |
| VNVGLAYGI | TCSAEGQTNERLF | 0.12968023 |
| VSSAAVGAV | TCSAEGQTNERLF | 0.121638246 |

| | | |
|---|---|---|
| VTSVYEAGV | TCSAEGQTNERLF | 0.122886755 |
| WVALDRGA | TCSAEGQTNERLF | 0.1329742 |
| YLVGSVVI | TCSAEGQTNERLF | 0.12292916 |
| GPPGTGKTLL | TCSAEGQTNERLF | 0.1267909 |
| GVGMTKF | TCSAEGQTNERLF | 0.12513693 |
| IGIGGSDL | TCSAEGQTNERLF | 0.12246927 |
| IGTLLAEL | TCSAEGQTNERLF | 0.12858306 |
| IYTSGTT | TCSAEGQTNERLF | 0.123458676 |
| LSFPAGV | TCSAEGQTNERLF | 0.12802787 |
| LYTSGTT | TCSAEGQTNERLF | 0.13053095 |
| VGAFPFGF | TCSAEGQTNERLF | 0.12581405 |
| VGAGTVEF | TCSAEGQTNERLF | 0.12302705 |
| VGKSTLLL | TCSAEGQTNERLF | 0.12665956 |
| AALAHAGGGIV | TCSAEGQTNERLF | 0.123715624 |
| AALVAVSLL | TCSAEGQTNERLF | 0.12334892 |
| AAVSRAVAAL | TCSAEGQTNERLF | 0.12241303 |
| AGGSSGIGGF | TCSAEGQTNERLF | 0.12519078 |
| AGLTVANANG | TCSAEGQTNERLF | 0.12330187 |
| AGLVGAVL | TCSAEGQTNERLF | 0.1297518 |
| ALSQAFTV | TCSAEGQTNERLF | 0.1271405 |
| ALTFSVSL | TCSAEGQTNERLF | 0.12436842 |
| ALTGAALVAL | TCSAEGQTNERLF | 0.124128945 |
| ASFSIGAGVL | TCSAEGQTNERLF | 0.12354313 |
| DGSVYKI | TCSAEGQTNERLF | 0.12292062 |
| EQLAEIV | TCSAEGQTNERLF | 0.123247035 |
| FAVSELTVM | TCSAEGQTNERLF | 0.12358964 |
| GAAPLAAGL | TCSAEGQTNERLF | 0.1236074 |
| GADGGDGGSGGAGG | TCSAEGQTNERLF | 0.123405084 |
| GFGLDTGGAGG | TCSAEGQTNERLF | 0.13039587 |
| GLASGGLL | TCSAEGQTNERLF | 0.12673776 |
| GRIAGAAVLL | TCSAEGQTNERLF | 0.12293316 |
| GSAGSLNTSF | TCSAEGQTNERLF | 0.12383742 |
| GTGQIGI | TCSAEGQTNERLF | 0.12488514 |
| GTLIALIL | TCSAEGQTNERLF | 0.12702668 |
| IAATMGFVVG | TCSAEGQTNERLF | 0.12172521 |
| IFAQALL | TCSAEGQTNERLF | 0.12370094 |
| IGAAVALGF | TCSAEGQTNERLF | 0.12378177 |
| IGLTPLAL | TCSAEGQTNERLF | 0.12295572 |
| ILGSNTSTL | TCSAEGQTNERLF | 0.12396878 |
| IVFSLTVL | TCSAEGQTNERLF | 0.13079527 |
| LASKATGF | TCSAEGQTNERLF | 0.12477273 |
| LASVALIL | TCSAEGQTNERLF | 0.124133356 |
| LFAVGLTL | TCSAEGQTNERLF | 0.12484781 |
| LFNGGIGTYI | TCSAEGQTNERLF | 0.1306626 |
| LTLGAGII | TCSAEGQTNERLF | 0.13376817 |
| LVAAALGDI | TCSAEGQTNERLF | 0.12650363 |
| NTAFVQLGI | TCSAEGQTNERLF | 0.12336864 |
| QLFSGAEL | TCSAEGQTNERLF | 0.12648384 |
| SAATFITL | TCSAEGQTNERLF | 0.12536818 |
| SNYGLANLGV | TCSAEGQTNERLF | 0.12233552 |
| SYGGAYAVM | TCSAEGQTNERLF | 0.12193943 |

| | | |
|---|---|---|
| TTGVGGLTLG | TCSAEGQTNERLF | 0.12632564 |
| TTQTIYI | TCSAEGQTNERLF | 0.12243031 |
| VAAGAGLGAVI | TCSAEGQTNERLF | 0.13010259 |
| VAAVVTLL | TCSAEGQTNERLF | 0.12869442 |
| VGFLGGLI | TCSAEGQTNERLF | 0.12273904 |
| VGVIPILL | TCSAEGQTNERLF | 0.12246979 |
| VIFASSLI | TCSAEGQTNERLF | 0.123122185 |
| VKGGDGGAAAG | TCSAEGQTNERLF | 0.12527905 |
| VTKTFTNI | TCSAEGQTNERLF | 0.12762073 |
| VVAAAILL | TCSAEGQTNERLF | 0.1230831 |
| YAALDVEL | TCSAEGQTNERLF | 0.12946552 |
| LASVALLL | TCSAEGQTNERLF | 0.12909243 |
| LDAVHRV | TCSAEGQTNERLF | 0.12479743 |
| QGLSSSIF | TCSAEGQTNERLF | 0.12854852 |
| RGTFANIRL | TCSAEGQTNERLF | 0.124588825 |
| RQLWWGHRI | TCSAEGQTNERLF | 0.1225961 |
| YGPPGTGKTLL | TCSAEGQTNERLF | 0.124906376 |
| AALVGAPANVA | TCSAEGQTNERLF | 0.1228592 |
| AATFRRTE | TCSAEGQTNERLF | 0.12313208 |
| AGALGGGI | TCSAEGQTNERLF | 0.12390236 |
| AGGAGGAGGIGAG | TCSAEGQTNERLF | 0.12339506 |
| AGSGALGL | TCSAEGQTNERLF | 0.12280359 |
| AQIADQI | TCSAEGQTNERLF | 0.12511708 |
| ASAVAVTYL | TCSAEGQTNERLF | 0.12219614 |
| ASTVLGGTGG | TCSAEGQTNERLF | 0.124895595 |
| AVGGEPAK | TCSAEGQTNERLF | 0.13166887 |
| CEQGTFGGTC | TCSAEGQTNERLF | 0.13327342 |
| DEGKAPAG | TCSAEGQTNERLF | 0.12364904 |
| DTVVFPAL | TCSAEGQTNERLF | 0.1238676 |
| FEGITMAG | TCSAEGQTNERLF | 0.12303667 |
| FPLVVQD | TCSAEGQTNERLF | 0.12644339 |
| GAFTTPQ | TCSAEGQTNERLF | 0.12504952 |
| GFGGFGTAGGVGGA | TCSAEGQTNERLF | 0.12516601 |
| GFLAQFL | TCSAEGQTNERLF | 0.124934174 |
| GGAGGAGGNAGVA | TCSAEGQTNERLF | 0.123575315 |
| GGVGTAGAGGAGGAGG | TCSAEGQTNERLF | 0.12389896 |
| GNGGGGDAGNAGSGGN | TCSAEGQTNERLF | 0.12358117 |
| GTGGAGGAGNDAGS | TCSAEGQTNERLF | 0.1257455 |
| GTTNTGWFNS | TCSAEGQTNERLF | 0.121848434 |
| IFMGPLFTL | TCSAEGQTNERLF | 0.124981605 |
| ILAHAIEAI | TCSAEGQTNERLF | 0.12407826 |
| IRSAIREAL | TCSAEGQTNERLF | 0.12313624 |
| ITFAGGG | TCSAEGQTNERLF | 0.12217381 |
| ITGMSGHQEV | TCSAEGQTNERLF | 0.124921195 |
| IVAGLACTL | TCSAEGQTNERLF | 0.124977514 |
| IVGGGLAGA | TCSAEGQTNERLF | 0.12716104 |
| IVTGVLNI | TCSAEGQTNERLF | 0.12254467 |
| LAQQGCI | TCSAEGQTNERLF | 0.1269609 |
| LCLHVEN | TCSAEGQTNERLF | 0.12781179 |
| LDAISDGS | TCSAEGQTNERLF | 0.12507522 |
| LGGFAAML | TCSAEGQTNERLF | 0.122778304 |

| | | |
|---|---|---|
| LGGKNPMI | TCSAEGQTNERLF | 0.12979314 |
| LGPKGVAAIL | TCSAEGQTNERLF | 0.12213527 |
| LMVGVSI | TCSAEGQTNERLF | 0.12426199 |
| LVGAAFVL | TCSAEGQTNERLF | 0.12656361 |
| LVLAVAPL | TCSAEGQTNERLF | 0.12319371 |
| MAAIAVIDAA | TCSAEGQTNERLF | 0.1285189 |
| NNGINAELQ | TCSAEGQTNERLF | 0.12966864 |
| RGEPLVLVHG | TCSAEGQTNERLF | 0.12313762 |
| RGGAGGAAGLI | TCSAEGQTNERLF | 0.12305729 |
| RSYRMHPAV | TCSAEGQTNERLF | 0.12345424 |
| SAAHLLAGQLA | TCSAEGQTNERLF | 0.12216042 |
| SAAVGSY | TCSAEGQTNERLF | 0.12741795 |
| SPHAVLHRI | TCSAEGQTNERLF | 0.123932205 |
| SQIAKVL | TCSAEGQTNERLF | 0.1256555 |
| TALGHLVP | TCSAEGQTNERLF | 0.12671173 |
| TAPTTSGAA | TCSAEGQTNERLF | 0.12474942 |
| TDGDDGNGGSPGT | TCSAEGQTNERLF | 0.12833765 |
| TDGLGEN | TCSAEGQTNERLF | 0.12712069 |
| TGGPPLPGQA | TCSAEGQTNERLF | 0.124416776 |
| TIPAITI | TCSAEGQTNERLF | 0.1238785 |
| TIPGFTI | TCSAEGQTNERLF | 0.1231597 |
| TSIASGALA | TCSAEGQTNERLF | 0.12396644 |
| TVTLPGAAI | TCSAEGQTNERLF | 0.13782187 |
| VASGLSAALP | TCSAEGQTNERLF | 0.12324871 |
| VGAGVIGI | TCSAEGQTNERLF | 0.12496561 |
| VGPKRFGV | TCSAEGQTNERLF | 0.124107614 |
| VNVAFPD | TCSAEGQTNERLF | 0.12268361 |
| VSMTNSGVS | TCSAEGQTNERLF | 0.12519392 |
| VTDVLGDLSV | TCSAEGQTNERLF | 0.12639736 |
| VVVVGAGFAGL | TCSAEGQTNERLF | 0.1296286 |
| VYTVGTVV | TCSAEGQTNERLF | 0.12252537 |
| WTTFSGP | TCSAEGQTNERLF | 0.12790525 |
| YDELVEAF | TCSAEGQTNERLF | 0.12813964 |
| YGTGGAGGNGG | TCSAEGQTNERLF | 0.123754404 |
| YTNVMAV | TCSAEGQTNERLF | 0.12260974 |
| FLEQVHQL | TCSAEGQTNERLF | 0.1227504 |
| GLSLLAVAV | TCSAEGQTNERLF | 0.12555484 |
| LEQVHQL | TCSAEGQTNERLF | 0.12663826 |
| LNAAGAI | TCSAEGQTNERLF | 0.12414275 |
| PAVHASL | TCSAEGQTNERLF | 0.12616931 |
| TELEDDA | TCSAEGQTNERLF | 0.12572344 |
| TPPGPPN | TCSAEGQTNERLF | 0.12776858 |
| TPPPPPPP | TCSAEGQTNERLF | 0.12244156 |
| TQSPALI | ASGLRLGGDAETLY | 0.1245614 |
| EFAAAFPQ | ASGLRLGGDAETLY | 0.12401077 |
| LLSGNLDVL | ASGLRLGGDAETLY | 0.1227911 |
| GPANLGNYNI | ASGLRLGGDAETLY | 0.12310863 |
| TLGPTLP | ASGLRLGGDAETLY | 0.1254373 |
| VYAYSSCCK | ASGLRLGGDAETLY | 0.12682688 |
| APAPAPAPNI | ASGLRLGGDAETLY | 0.12582816 |
| PGGPVPIP | ASGLRLGGDAETLY | 0.13096905 |

| | | |
|---|---|---|
| FLTVALEP | ASGLRLGGDAETLY | 0.13571717 |
| SQYFGANL | ASGLRLGGDAETLY | 0.123818696 |
| GGVKPAA | ASGLRLGGDAETLY | 0.12707679 |
| GGTGGAGGAGGW | ASGLRLGGDAETLY | 0.1225561 |
| GVGGVAGLGGAGG | ASGLRLGGDAETLY | 0.12362067 |
| VLLVLATTI | ASGLRLGGDAETLY | 0.12250175 |
| IGIFAAML | ASGLRLGGDAETLY | 0.12536782 |
| AVGDLID | ASGLRLGGDAETLY | 0.12205411 |
| NISPLPL | ASGLRLGGDAETLY | 0.12773001 |
| VTAVFVAI | ASGLRLGGDAETLY | 0.12486569 |
| SGDDNDAAL | ASGLRLGGDAETLY | 0.13146648 |
| AVAALAHS | ASGLRLGGDAETLY | 0.12551226 |
| GGVTVMATP | ASGLRLGGDAETLY | 0.12341607 |
| AGLAANVRD | ASGLRLGGDAETLY | 0.12476517 |
| TNRPDLIDPALLRPG | ASGLRLGGDAETLY | 0.12633725 |
| LANLGVTAL | ASGLRLGGDAETLY | 0.12288946 |
| TGERIEL | ASGLRLGGDAETLY | 0.12271441 |
| NGEELAA | ASGLRLGGDAETLY | 0.12392305 |
| VLSLVWG | ASGLRLGGDAETLY | 0.122794986 |
| IGYALWL | ASGLRLGGDAETLY | 0.12629367 |
| LSLLLAPL | ASGLRLGGDAETLY | 0.12894717 |
| TLDEALK | ASGLRLGGDAETLY | 0.12628964 |
| TDGAGTAEGG | ASGLRLGGDAETLY | 0.12678248 |
| NAPGGAGGNGG | ASGLRLGGDAETLY | 0.12196113 |
| VSVFMGMGGNF | ASGLRLGGDAETLY | 0.1253161 |
| LASYLAAL | ASGLRLGGDAETLY | 0.13775785 |
| LVGGTPAL | ASGLRLGGDAETLY | 0.12467857 |
| VRNADVL | ASGLRLGGDAETLY | 0.12762679 |
| TCSSPFLVSAM | ASGLRLGGDAETLY | 0.13220677 |
| VHITDAE | ASGLRLGGDAETLY | 0.12445788 |
| EYAAPSF | ASGLRLGGDAETLY | 0.1272102 |
| VVDAFHGGL | ASGLRLGGDAETLY | 0.124260396 |
| ASVVDPQ | ASGLRLGGDAETLY | 0.12596843 |
| SYNEFGGLS | ASGLRLGGDAETLY | 0.12294549 |
| DPDGSVY | ASGLRLGGDAETLY | 0.12322474 |
| MVSTIHATE | ASGLRLGGDAETLY | 0.12946522 |
| LAAGVADPV | ASGLRLGGDAETLY | 0.12958188 |
| TKEQIAA | ASGLRLGGDAETLY | 0.1256869 |
| SLASPAAL | ASGLRLGGDAETLY | 0.12943499 |
| IGLVFYGV | ASGLRLGGDAETLY | 0.12835523 |
| FLLDASNL | ASGLRLGGDAETLY | 0.124245726 |
| YQVGNFP | ASGLRLGGDAETLY | 0.1251567 |
| ESLLSQST | ASGLRLGGDAETLY | 0.12351475 |
| LFTVGFAAVG | ASGLRLGGDAETLY | 0.123336285 |
| ATPPTPT | ASGLRLGGDAETLY | 0.123424985 |
| TDLATLG | ASGLRLGGDAETLY | 0.12532863 |
| SAGGTGR | ASGLRLGGDAETLY | 0.12613782 |
| GFSNSGS | ASGLRLGGDAETLY | 0.13278973 |
| GGNGGSGGTSV | ASGLRLGGDAETLY | 0.1278398 |
| GTSVATGGAGN | ASGLRLGGDAETLY | 0.12856476 |
| NGGNGGSGGTS | ASGLRLGGDAETLY | 0.12374983 |

| | | |
|---|---|---|
| AVVKFPH | ASGLRLGGDAETLY | 0.12338999 |
| TPPPPPP | ASGLRLGGDAETLY | 0.12388599 |
| AAPAAAAAPA | ASGLRLGGDAETLY | 0.12379878 |
| IALNEDL | ASGLRLGGDAETLY | 0.13277404 |
| IVNVSSM | ASGLRLGGDAETLY | 0.12560372 |
| TGIPPAPR | ASGLRLGGDAETLY | 0.12298827 |
| FDLGGGTF | ASGLRLGGDAETLY | 0.12671313 |
| FDLGGGTFDVS | ASGLRLGGDAETLY | 0.12902828 |
| NEPTAAA | ASGLRLGGDAETLY | 0.1266648 |
| NEPTAAALA | ASGLRLGGDAETLY | 0.12384317 |
| GSIINISSI | ASGLRLGGDAETLY | 0.12480936 |
| VAPGNAGTA | ASGLRLGGDAETLY | 0.12577711 |
| TGGGASLE | ASGLRLGGDAETLY | 0.1354275 |
| VAVYNHYKRI | ASGLRLGGDAETLY | 0.12223391 |
| TDPAPATT | ASGLRLGGDAETLY | 0.12855035 |
| TNRPDILDPALLRPG | ASGLRLGGDAETLY | 0.12388784 |
| LTTTTTS | ASGLRLGGDAETLY | 0.122536875 |
| PTGSGKTL | ASGLRLGGDAETLY | 0.12534471 |
| TGSGKTL | ASGLRLGGDAETLY | 0.12736924 |
| TIGHVDHGK | ASGLRLGGDAETLY | 0.124869704 |
| TIGHVDHGKTT | ASGLRLGGDAETLY | 0.12789403 |
| GHVDHGKTT | ASGLRLGGDAETLY | 0.12232719 |
| AQFEHTLL | ASGLRLGGDAETLY | 0.12331994 |
| IVNQDEL | ASGLRLGGDAETLY | 0.13129784 |
| TNGSQFF | ASGLRLGGDAETLY | 0.12678453 |
| TGKGIVE | ASGLRLGGDAETLY | 0.13170406 |
| LISNASDA | ASGLRLGGDAETLY | 0.12383536 |
| AELRTLGELTKIAKAHGAQ | ASGLRLGGDAETLY | 0.12299147 |
| APNAVAAAEPSW | ASGLRLGGDAETLY | 0.1243525 |
| APQVAGAATAVL | ASGLRLGGDAETLY | 0.12603368 |
| ARRDPAAAAAILPTDA | ASGLRLGGDAETLY | 0.123397835 |
| DAAARDVLAEAGLAEYF | ASGLRLGGDAETLY | 0.1280642 |
| DDQIVLSAKANLAKVEQ | ASGLRLGGDAETLY | 0.12719938 |
| DELLPDTGEDSDEDGH | ASGLRLGGDAETLY | 0.12252026 |
| ELPEAPANRATVPQD | ASGLRLGGDAETLY | 0.124223314 |
| FGQSKIERVTGIG | ASGLRLGGDAETLY | 0.12609307 |
| GAGGTGSTAGGAGGAGGAGGLY | ASGLRLGGDAETLY | 0.12553817 |
| GHGGTGAAGGNGGTGGAGGSGI | ASGLRLGGDAETLY | 0.12283039 |
| GNGGNGGAGGEGGPGGAGGAGA | ASGLRLGGDAETLY | 0.13014218 |
| GSTAGNGGNGGNATNGTIAGQP | ASGLRLGGDAETLY | 0.13235235 |
| GTGGDGGNGGNGGTGG | ASGLRLGGDAETLY | 0.12839119 |
| GVRRPAGIPADAPV | ASGLRLGGDAETLY | 0.12342978 |
| GYRQVTDAHLLTLARR | ASGLRLGGDAETLY | 0.12203449 |
| IAKMGQDGHDRG | ASGLRLGGDAETLY | 0.12649152 |
| KAEPSIEIQYT | ASGLRLGGDAETLY | 0.12607387 |
| KPVRIRMAADEDTVAVT | ASGLRLGGDAETLY | 0.12551668 |
| MPPRIAGMRLLVIKPEP | ASGLRLGGDAETLY | 0.12329082 |
| MKFYAVALFRSRGVPPTR | ASGLRLGGDAETLY | 0.12315434 |
| NAGGFGILGAVAHSP | ASGLRLGGDAETLY | 0.13374501 |
| NEGREITLAMAEM | ASGLRLGGDAETLY | 0.12296602 |
| NELVEWSHPLGHRWRWE | ASGLRLGGDAETLY | 0.12246068 |

| | | |
|---|---|---|
| NLGAGGTGGNGGNPG | ASGLRLGGDAETLY | 0.12314266 |
| NTEWPKWETVMAADPVD | ASGLRLGGDAETLY | 0.12955856 |
| PASAAPAETQP | ASGLRLGGDAETLY | 0.12324012 |
| QQSSDGKNGVRAAEITGE | ASGLRLGGDAETLY | 0.12449161 |
| RAPDAVTTSWWKE | ASGLRLGGDAETLY | 0.12958808 |
| SGFGNVGAHTSGWFNQ | ASGLRLGGDAETLY | 0.12478304 |
| SGVAGAVGQAASVGGLKV | ASGLRLGGDAETLY | 0.12410976 |
| SNGGAGTGSAGGNGGT | ASGLRLGGDAETLY | 0.12375644 |
| TGGAGGEGGAAG | ASGLRLGGDAETLY | 0.12534897 |
| TGGDGGAAGAGGAPGHG | ASGLRLGGDAETLY | 0.12450019 |
| TMPPLLAAIANAGRHGV | ASGLRLGGDAETLY | 0.1244076 |
| TSSTAGNSGGA | ASGLRLGGDAETLY | 0.12542044 |
| VPATVPVGILC | ASGLRLGGDAETLY | 0.12590313 |
| VQRPDSGGVVLTGRIS | ASGLRLGGDAETLY | 0.12617898 |
| WTPNPVPATKGVPAGLVQAT | ASGLRLGGDAETLY | 0.12467712 |
| YALKRALAKFSDRFDVVII | ASGLRLGGDAETLY | 0.12323648 |
| YLIGSVIV | ASGLRLGGDAETLY | 0.12178974 |
| YGQLVAVID | ASGLRLGGDAETLY | 0.12639616 |
| VTVGGMTLPA | ASGLRLGGDAETLY | 0.12407386 |
| VLALPAV | ASGLRLGGDAETLY | 0.12759888 |
| VIPPTLNYET | ASGLRLGGDAETLY | 0.13044605 |
| VGMGGMHPSSG | ASGLRLGGDAETLY | 0.12400847 |
| VGLAGGAATA | ASGLRLGGDAETLY | 0.12491164 |
| VGGAGGAGGSGP | ASGLRLGGDAETLY | 0.12238723 |
| VFAAFMV | ASGLRLGGDAETLY | 0.12277474 |
| TLYDKLR | ASGLRLGGDAETLY | 0.12393908 |
| TLAVFTVL | ASGLRLGGDAETLY | 0.12610516 |
| TGSVQPARV | ASGLRLGGDAETLY | 0.12457759 |
| TGGGAGSGGT | ASGLRLGGDAETLY | 0.12649404 |
| SHLETSL | ASGLRLGGDAETLY | 0.12520568 |
| SFTTGMT | ASGLRLGGDAETLY | 0.12305353 |
| QVEVAKRMGVRQPTVS | ASGLRLGGDAETLY | 0.12551625 |
| QGSGGPALAM | ASGLRLGGDAETLY | 0.13089024 |
| MLAPSNP | ASGLRLGGDAETLY | 0.12747364 |
| LVNNAAI | ASGLRLGGDAETLY | 0.123525135 |
| LTGEINTLA | ASGLRLGGDAETLY | 0.12662007 |
| LSIFGPV | ASGLRLGGDAETLY | 0.12360645 |
| LLALLANL | ASGLRLGGDAETLY | 0.12674654 |
| LISNASDALDKLR | ASGLRLGGDAETLY | 0.12549025 |
| LINEAVP | ASGLRLGGDAETLY | 0.12570445 |
| LGYAFSGL | ASGLRLGGDAETLY | 0.12587318 |
| LDFVAVAL | ASGLRLGGDAETLY | 0.12259425 |
| LAQPGAE | ASGLRLGGDAETLY | 0.122896425 |
| ISNLGAVTGF | ASGLRLGGDAETLY | 0.12496638 |
| HQEDPGRI | ASGLRLGGDAETLY | 0.1295905 |
| GTPGEPTP | ASGLRLGGDAETLY | 0.1304144 |
| GFFNSGS | ASGLRLGGDAETLY | 0.12397156 |
| GAGGGGLTAGAN | ASGLRLGGDAETLY | 0.12478456 |
| FVDALVNM | ASGLRLGGDAETLY | 0.12468645 |
| FLTVAPEL | ASGLRLGGDAETLY | 0.12532589 |
| FETLTAL | ASGLRLGGDAETLY | 0.12550285 |

| | | |
|---|---|---|
| DVINPIEL | ASGLRLGGDAETLY | 0.12394576 |
| APVSGAPV | ASGLRLGGDAETLY | 0.12306789 |
| ALVGGLAW | ASGLRLGGDAETLY | 0.12622389 |
| ALAAAAATAI | ASGLRLGGDAETLY | 0.12349568 |
| AFGLGLLL | ASGLRLGGDAETLY | 0.12587774 |
| AFFAAASL | ASGLRLGGDAETLY | 0.12284438 |
| AAAALPAFSPP | ASGLRLGGDAETLY | 0.12313114 |
| VVLAPAL | ASGLRLGGDAETLY | 0.12458923 |
| TIGHVDHGKTTL | ASGLRLGGDAETLY | 0.1246871 |
| LISNASDALDK | ASGLRLGGDAETLY | 0.12490904 |
| LISNASDAL | ASGLRLGGDAETLY | 0.12670368 |
| ISNASDALDK | ASGLRLGGDAETLY | 0.12396225 |
| NQAETLVYQTE | ASGLRLGGDAETLY | 0.12769425 |
| TGGTGGTAGAG | ASGLRLGGDAETLY | 0.13452739 |
| TGSTAGGAGGA | ASGLRLGGDAETLY | 0.12660441 |
| DPLAVPVAAGPV | ASGLRLGGDAETLY | 0.12373128 |
| GGTGGTPGNGGN | ASGLRLGGDAETLY | 0.123453334 |
| PAGLAGVDPATL | ASGLRLGGDAETLY | 0.12867886 |
| GNNLVVEAPHAGASV | ASGLRLGGDAETLY | 0.1244752 |
| KVTRIMSASTLEQPA | ASGLRLGGDAETLY | 0.12231191 |
| LGFALPIALPRRTVV | ASGLRLGGDAETLY | 0.1265045 |
| TRLGNLTARKLVVLV | ASGLRLGGDAETLY | 0.12404897 |
| WSQQRRLIYFPSAGP | ASGLRLGGDAETLY | 0.12463862 |
| GPNGNPDPMAAAVDIR | ASGLRLGGDAETLY | 0.12245524 |
| TQRCVSVADLRAAASR | ASGLRLGGDAETLY | 0.13147883 |
| QFVLDVAEHPVEVLTETV | ASGLRLGGDAETLY | 0.13479893 |
| GGAGGTGGAGGNGGTQAGDG | ASGLRLGGDAETLY | 0.12429514 |
| PLLATLRAIAVDVDAKPAQV | ASGLRLGGDAETLY | 0.12537481 |
| APKFRTGGYMIDFWGVGYQVAK | ASGLRLGGDAETLY | 0.12249229 |
| GAGGAGGAGGAAGGPGGTGGTG | ASGLRLGGDAETLY | 0.12947908 |
| IGALGLLATDVAPAGSVAFGVA | ASGLRLGGDAETLY | 0.13148016 |
| AGGAGGIGGNGGAGGAGGVLMG | ASGLRLGGDAETLY | 0.12387742 |
| ADELIGLL | ASGLRLGGDAETLY | 0.12809792 |
| ADRALKA | ASGLRLGGDAETLY | 0.1239769 |
| AGAVGQVGGAGGA | ASGLRLGGDAETLY | 0.12469491 |
| AGGGGGLGGAP | ASGLRLGGDAETLY | 0.13065729 |
| AGGLLYGNGGAG | ASGLRLGGDAETLY | 0.12653522 |
| AGLESIGL | ASGLRLGGDAETLY | 0.12570776 |
| AGLKTGII | ASGLRLGGDAETLY | 0.12487632 |
| ASLAALLL | ASGLRLGGDAETLY | 0.124693915 |
| ASVINISSI | ASGLRLGGDAETLY | 0.12654649 |
| ATPPGPA | ASGLRLGGDAETLY | 0.12849699 |
| ATVASALV | ASGLRLGGDAETLY | 0.122675054 |
| ATVASLIL | ASGLRLGGDAETLY | 0.12682499 |
| AVNGPVLIRH | ASGLRLGGDAETLY | 0.123781085 |
| DGAAGAVGI | ASGLRLGGDAETLY | 0.12887074 |
| DPLLHGI | ASGLRLGGDAETLY | 0.12540136 |
| ELGAALGAL | ASGLRLGGDAETLY | 0.12595807 |
| ELGRTQAV | ASGLRLGGDAETLY | 0.12461211 |
| FVAGGYLS | ASGLRLGGDAETLY | 0.13373192 |
| GASGGAGGAGGW | ASGLRLGGDAETLY | 0.12467527 |

| | | |
|---|---|---|
| GGAGAGGDGGAGGI | ASGLRLGGDAETLY | 0.124155246 |
| GTGHNGSQPGMGGNG | ASGLRLGGDAETLY | 0.12755415 |
| GYAAASASAS | ASGLRLGGDAETLY | 0.45792812 |
| HQVDVAAVQQG | ASGLRLGGDAETLY | 0.86792773 |
| IGAAIAHT | ASGLRLGGDAETLY | 0.7975354 |
| IGMIEVAQL | ASGLRLGGDAETLY | 0.8083187 |
| IRLAPEHTSAMD | ASGLRLGGDAETLY | 0.6589556 |
| IYNVSVL | ASGLRLGGDAETLY | 0.38304266 |
| KQGFLGM | ASGLRLGGDAETLY | 0.6963175 |
| LESVITQL | ASGLRLGGDAETLY | 0.5612151 |
| LGLGALVVV | ASGLRLGGDAETLY | 0.8410969 |
| LIGNGGNGGSGGI | ASGLRLGGDAETLY | 0.859777 |
| LNAINAPTL | ASGLRLGGDAETLY | 0.61935943 |
| LQLLNTGV | ASGLRLGGDAETLY | 0.8280792 |
| LQTSIEF | ASGLRLGGDAETLY | 0.79326224 |
| MSFAVSEL | ASGLRLGGDAETLY | 0.4250176 |
| NTGALGL | ASGLRLGGDAETLY | 0.7859976 |
| PPGAGGAGGAGTT | ASGLRLGGDAETLY | 0.6714277 |
| QLVGLGI | ASGLRLGGDAETLY | 0.7508891 |
| RFLEQVHQL | ASGLRLGGDAETLY | 0.37208533 |
| RGVGYVLR | ASGLRLGGDAETLY | 0.8337842 |
| SATGVFAGVI | ASGLRLGGDAETLY | 0.6761916 |
| SLPAITLP | ASGLRLGGDAETLY | 0.18365112 |
| STALAFTV | ASGLRLGGDAETLY | 0.8161873 |
| SVLTVGLGK | ASGLRLGGDAETLY | 0.6861721 |
| TALAGAGSGI | ASGLRLGGDAETLY | 0.70490956 |
| TGLPFTGI | ASGLRLGGDAETLY | 0.84963405 |
| VAEHVTV | ASGLRLGGDAETLY | 0.4661548 |
| VDIVLNSL | ASGLRLGGDAETLY | 0.60229164 |
| VLTGGGAVL | ASGLRLGGDAETLY | 0.8328249 |
| VNVGLAYGI | ASGLRLGGDAETLY | 0.8537797 |
| VSSAAVGAV | ASGLRLGGDAETLY | 0.5140612 |
| VTSVYEAGV | ASGLRLGGDAETLY | 0.807006 |
| WVALDRGA | ASGLRLGGDAETLY | 0.86175835 |
| YLVGSVVI | ASGLRLGGDAETLY | 0.7138152 |
| GPPGTGKTLL | ASGLRLGGDAETLY | 0.8743479 |
| GVGMTKF | ASGLRLGGDAETLY | 0.8782624 |
| IGIGGSDL | ASGLRLGGDAETLY | 0.68255043 |
| IGTLLAEL | ASGLRLGGDAETLY | 0.8621248 |
| IYTSGTT | ASGLRLGGDAETLY | 0.83827776 |
| LSFPAGV | ASGLRLGGDAETLY | 0.6856783 |
| LYTSGTT | ASGLRLGGDAETLY | 0.769351 |
| VGAFPFGF | ASGLRLGGDAETLY | 0.37384513 |
| VGAGTVEF | ASGLRLGGDAETLY | 0.7523485 |
| VGKSTLLL | ASGLRLGGDAETLY | 0.4111484 |
| AALAHAGGGIV | ASGLRLGGDAETLY | 0.8089208 |
| AALVAVSLL | ASGLRLGGDAETLY | 0.8644935 |
| AAVSRAVAAL | ASGLRLGGDAETLY | 0.74493897 |
| AGGSSGIGGF | ASGLRLGGDAETLY | 0.47359365 |
| AGLTVANANG | ASGLRLGGDAETLY | 0.5155702 |
| AGLVGAVL | ASGLRLGGDAETLY | 0.8163707 |

| | | |
|---|---|---|
| ALSQAFTV | ASGLRLGGDAETLY | 0.49446726 |
| ALTFSVSL | ASGLRLGGDAETLY | 0.42911336 |
| ALTGAALVAL | ASGLRLGGDAETLY | 0.62993556 |
| ASFSIGAGVL | ASGLRLGGDAETLY | 0.82242 |
| DGSVYKI | ASGLRLGGDAETLY | 0.8328462 |
| EQLAEIV | ASGLRLGGDAETLY | 0.61238253 |
| FAVSELTVM | ASGLRLGGDAETLY | 0.8703884 |
| GAAPLAAGL | ASGLRLGGDAETLY | 0.8334888 |
| GADGGDGGSGGAGG | ASGLRLGGDAETLY | 0.8687885 |
| GFGLDTGGAGG | ASGLRLGGDAETLY | 0.74076974 |
| GLASGGLL | ASGLRLGGDAETLY | 0.5188162 |
| GRIAGAAVLL | ASGLRLGGDAETLY | 0.83967924 |
| GSAGSLNTSF | ASGLRLGGDAETLY | 0.6609729 |
| GTGQIGI | ASGLRLGGDAETLY | 0.8394537 |
| GTLIALIL | ASGLRLGGDAETLY | 0.84137577 |
| IAATMGFVVG | ASGLRLGGDAETLY | 0.6810211 |
| IFAQALL | ASGLRLGGDAETLY | 0.6259778 |
| IGAAVALGF | ASGLRLGGDAETLY | 0.7609831 |
| IGLTPLAL | ASGLRLGGDAETLY | 0.88050014 |
| ILGSNTSTL | ASGLRLGGDAETLY | 0.61600405 |
| IVFSLTVL | ASGLRLGGDAETLY | 0.39170396 |
| LASKATGF | ASGLRLGGDAETLY | 0.7192519 |
| LASVALIL | ASGLRLGGDAETLY | 0.6879512 |
| LFAVGLTL | ASGLRLGGDAETLY | 0.86730814 |
| LFNGGIGTYI | ASGLRLGGDAETLY | 0.8150658 |
| LTLGAGII | ASGLRLGGDAETLY | 0.7725992 |
| LVAAALGDI | ASGLRLGGDAETLY | 0.8158334 |
| NTAFVQLGI | ASGLRLGGDAETLY | 0.8165312 |
| QLFSGAEL | ASGLRLGGDAETLY | 0.7832569 |
| SAATFITL | ASGLRLGGDAETLY | 0.76380014 |
| SNYGLANLGV | ASGLRLGGDAETLY | 0.6669166 |
| SYGGAYAVM | ASGLRLGGDAETLY | 0.8757721 |
| TTGVGGLTLG | ASGLRLGGDAETLY | 0.8482101 |
| TTQTIYI | ASGLRLGGDAETLY | 0.8544996 |
| VAAGAGLGAVI | ASGLRLGGDAETLY | 0.87005377 |
| VAAVVTLL | ASGLRLGGDAETLY | 0.7813755 |
| VGFLGGLI | ASGLRLGGDAETLY | 0.79563904 |
| VGVIPILL | ASGLRLGGDAETLY | 0.3942158 |
| VIFASSLI | ASGLRLGGDAETLY | 0.48324853 |
| VKGGDGGAAAG | ASGLRLGGDAETLY | 0.770015 |
| VTKTFTNI | ASGLRLGGDAETLY | 0.8230958 |
| VVAAAILL | ASGLRLGGDAETLY | 0.8336457 |
| YAALDVEL | ASGLRLGGDAETLY | 0.8390252 |
| LASVALLL | ASGLRLGGDAETLY | 0.65858954 |
| LDAVHRV | ASGLRLGGDAETLY | 0.84497476 |
| QGLSSSIF | ASGLRLGGDAETLY | 0.8562829 |
| RGTFANIRL | ASGLRLGGDAETLY | 0.51773345 |
| RQLWWGHRI | ASGLRLGGDAETLY | 0.7221684 |
| YGPPGTGKTLL | ASGLRLGGDAETLY | 0.87355006 |
| AALVGAPANVA | ASGLRLGGDAETLY | 0.8152141 |
| AATFRRTE | ASGLRLGGDAETLY | 0.7131663 |

| | | |
|---|---|---|
| AGALGGGI | ASGLRLGGDAETLY | 0.73578453 |
| AGGAGGAGGIGAG | ASGLRLGGDAETLY | 0.86033446 |
| AGSGALGL | ASGLRLGGDAETLY | 0.79627335 |
| AQIADQI | ASGLRLGGDAETLY | 0.87422633 |
| ASAVAVTYL | ASGLRLGGDAETLY | 0.27078918 |
| ASTVLGGTGG | ASGLRLGGDAETLY | 0.7510569 |
| AVGGEPAK | ASGLRLGGDAETLY | 0.7407366 |
| CEQGTFGGTC | ASGLRLGGDAETLY | 0.18086953 |
| DEGKAPAG | ASGLRLGGDAETLY | 0.84854746 |
| DTVVFPAL | ASGLRLGGDAETLY | 0.58529365 |
| FEGITMAG | ASGLRLGGDAETLY | 0.7952931 |
| FPLVVQD | ASGLRLGGDAETLY | 0.6680589 |
| GAFTTPQ | ASGLRLGGDAETLY | 0.8682805 |
| GFGGFGTAGGVGGA | ASGLRLGGDAETLY | 0.81265503 |
| GFLAQFL | ASGLRLGGDAETLY | 0.6514394 |
| GGAGGAGGNAGVA | ASGLRLGGDAETLY | 0.5821099 |
| GGVGTAGAGGAGGAGG | ASGLRLGGDAETLY | 0.78487957 |
| GNGGGGDAGNAGSGGN | ASGLRLGGDAETLY | 0.85919994 |
| GTGGAGGAGNDAGS | ASGLRLGGDAETLY | 0.8745082 |
| GTTNTGWFNS | ASGLRLGGDAETLY | 0.34388787 |
| IFMGPLFTL | ASGLRLGGDAETLY | 0.82741797 |
| ILAHAIEAI | ASGLRLGGDAETLY | 0.71752334 |
| IRSAIREAL | ASGLRLGGDAETLY | 0.22107151 |
| ITFAGGG | ASGLRLGGDAETLY | 0.33838534 |
| ITGMSGHQEV | ASGLRLGGDAETLY | 0.7572477 |
| IVAGLACTL | ASGLRLGGDAETLY | 0.7210798 |
| IVGGGLAGA | ASGLRLGGDAETLY | 0.8109547 |
| IVTGVLNI | ASGLRLGGDAETLY | 0.85074246 |
| LAQQGCI | ASGLRLGGDAETLY | 0.6747187 |
| LCLHVEN | ASGLRLGGDAETLY | 0.83013225 |
| LDAISDGS | ASGLRLGGDAETLY | 0.7308214 |
| LGGFAAML | ASGLRLGGDAETLY | 0.0885135 |
| LGGKNPMI | ASGLRLGGDAETLY | 0.6056508 |
| LGPKGVAAIL | ASGLRLGGDAETLY | 0.40491658 |
| LMVGVSI | ASGLRLGGDAETLY | 0.8749234 |
| LVGAAFVL | ASGLRLGGDAETLY | 0.561153 |
| LVLAVAPL | ASGLRLGGDAETLY | 0.7245478 |
| MAAIAVIDAA | ASGLRLGGDAETLY | 0.75846624 |
| NNGINAELQ | ASGLRLGGDAETLY | 0.8085027 |
| RGEPLVLVHG | ASGLRLGGDAETLY | 0.7597245 |
| RGGAGGAAGLI | ASGLRLGGDAETLY | 0.6482744 |
| RSYRMHPAV | ASGLRLGGDAETLY | 0.31071383 |
| SAAHLLAGQLA | ASGLRLGGDAETLY | 0.8065355 |
| SAAVGSY | ASGLRLGGDAETLY | 0.6613903 |
| SPHAVLHRI | ASGLRLGGDAETLY | 0.59630203 |
| SQIAKVL | ASGLRLGGDAETLY | 0.56851375 |
| TALGHLVP | ASGLRLGGDAETLY | 0.8057035 |
| TAPTTSGAA | ASGLRLGGDAETLY | 0.2841081 |
| TDGDDGNGGSPGT | ASGLRLGGDAETLY | 0.85759956 |
| TDGLGEN | ASGLRLGGDAETLY | 0.6072663 |
| TGGPPLPGQA | ASGLRLGGDAETLY | 0.72548515 |

| | | |
|---|---|---|
| TIPAITI | ASGLRLGGDAETLY | 0.8753207 |
| TIPGFTI | ASGLRLGGDAETLY | 0.7845516 |
| TSIASGALA | ASGLRLGGDAETLY | 0.7499959 |
| TVTLPGAAI | ASGLRLGGDAETLY | 0.59426415 |
| VASGLSAALP | ASGLRLGGDAETLY | 0.5533694 |
| VGAGVIGI | ASGLRLGGDAETLY | 0.60052603 |
| VGPKRFGV | ASGLRLGGDAETLY | 0.6259777 |
| VNVAFPD | ASGLRLGGDAETLY | 0.86331296 |
| VSMTNSGVS | ASGLRLGGDAETLY | 0.8181807 |
| VTDVLGDLSV | ASGLRLGGDAETLY | 0.42641652 |
| VVVVGAGFAGL | ASGLRLGGDAETLY | 0.8534553 |
| VYTVGTVV | ASGLRLGGDAETLY | 0.8733921 |
| WTTFSGP | ASGLRLGGDAETLY | 0.8697605 |
| YDELVEAF | ASGLRLGGDAETLY | 0.5614178 |
| YGTGGAGGNGG | ASGLRLGGDAETLY | 0.6312653 |
| YTNVMAV | ASGLRLGGDAETLY | 0.70698524 |
| FLEQVHQL | ASGLRLGGDAETLY | 0.73499966 |
| GLSLLAVAV | ASGLRLGGDAETLY | 0.51523834 |
| LEQVHQL | ASGLRLGGDAETLY | 0.5390868 |
| LNAAGAI | ASGLRLGGDAETLY | 0.8662926 |
| PAVHASL | ASGLRLGGDAETLY | 0.8674351 |
| TELEDDA | ASGLRLGGDAETLY | 0.7257662 |
| TPPGPPN | ASGLRLGGDAETLY | 0.30594492 |
| TPPPPPPP | ASGLRLGGDAETLY | 0.7355348 |
| TQSPALI | ASGDNWGGGGAETLY | 0.60948503 |
| EFAAAFPQ | ASGDNWGGGGAETLY | 0.6475729 |
| LLSGNLDVL | ASGDNWGGGGAETLY | 0.60361373 |
| GPANLGNYNI | ASGDNWGGGGAETLY | 0.405235 |
| TLGPTLP | ASGDNWGGGGAETLY | 0.7572049 |
| VYAYSSCCK | ASGDNWGGGGAETLY | 0.8715472 |
| APAPAPAPNI | ASGDNWGGGGAETLY | 0.7191454 |
| PGGPVPIP | ASGDNWGGGGAETLY | 0.7855032 |
| FLTVALEP | ASGDNWGGGGAETLY | 0.8689017 |
| SQYFGANL | ASGDNWGGGGAETLY | 0.8357606 |
| GGVKPAA | ASGDNWGGGGAETLY | 0.8599599 |
| GGTGGAGGAGGW | ASGDNWGGGGAETLY | 0.8426988 |
| GVGGVAGLGGAGG | ASGDNWGGGGAETLY | 0.7625838 |
| VLLVLATTI | ASGDNWGGGGAETLY | 0.7874183 |
| IGIFAAML | ASGDNWGGGGAETLY | 0.84107137 |
| AVGDLID | ASGDNWGGGGAETLY | 0.7147368 |
| NISPLPL | ASGDNWGGGGAETLY | 0.77706754 |
| VTAVFVAI | ASGDNWGGGGAETLY | 0.7607385 |
| SGDDNDAAL | ASGDNWGGGGAETLY | 0.8137324 |
| AVAALAHS | ASGDNWGGGGAETLY | 0.7326786 |
| GGVTVMATP | ASGDNWGGGGAETLY | 0.8610343 |
| AGLAANVRD | ASGDNWGGGGAETLY | 0.20589323 |
| TNRPDLIDPALLRPG | ASGDNWGGGGAETLY | 0.52069414 |
| LANLGVTAL | ASGDNWGGGGAETLY | 0.8616466 |
| TGERIEL | ASGDNWGGGGAETLY | 0.13615076 |
| NGEELAA | ASGDNWGGGGAETLY | 0.39277983 |
| VLSLVWG | ASGDNWGGGGAETLY | 0.81198233 |

| | | |
|---|---|---|
| IGYALWL | ASGDNWGGGGAETLY | 0.8498082 |
| LSLLLAPL | ASGDNWGGGGAETLY | 0.14655694 |
| TLDEALK | ASGDNWGGGGAETLY | 0.7086064 |
| TDGAGTAEGG | ASGDNWGGGGAETLY | 0.39118809 |
| NAPGGAGGNGG | ASGDNWGGGGAETLY | 0.86123383 |
| VSVFMGMGGNF | ASGDNWGGGGAETLY | 0.7250929 |
| LASYLAAL | ASGDNWGGGGAETLY | 0.34692857 |
| LVGGTPAL | ASGDNWGGGGAETLY | 0.8500453 |
| VRNADVL | ASGDNWGGGGAETLY | 0.5211828 |
| TCSSPFLVSAM | ASGDNWGGGGAETLY | 0.779098 |
| VHITDAE | ASGDNWGGGGAETLY | 0.8628212 |
| EYAAPSF | ASGDNWGGGGAETLY | 0.8745215 |
| VVDAFHGGL | ASGDNWGGGGAETLY | 0.49364918 |
| ASVVDPQ | ASGDNWGGGGAETLY | 0.83355737 |
| SYNEFGGLS | ASGDNWGGGGAETLY | 0.73041046 |
| DPDGSVY | ASGDNWGGGGAETLY | 0.8807361 |
| MVSTIHATE | ASGDNWGGGGAETLY | 0.69668436 |
| LAAGVADPV | ASGDNWGGGGAETLY | 0.35227793 |
| TKEQIAA | ASGDNWGGGGAETLY | 0.8546246 |
| SLASPAAL | ASGDNWGGGGAETLY | 0.83540857 |
| IGLVFYGV | ASGDNWGGGGAETLY | 0.7047899 |
| FLLDASNL | ASGDNWGGGGAETLY | 0.83904123 |
| YQVGNFP | ASGDNWGGGGAETLY | 0.7558744 |
| ESLLSQST | ASGDNWGGGGAETLY | 0.73301876 |
| LFTVGFAAVG | ASGDNWGGGGAETLY | 0.8364192 |
| ATPPTPT | ASGDNWGGGGAETLY | 0.8407247 |
| TDLATLG | ASGDNWGGGGAETLY | 0.5436952 |
| SAGGTGR | ASGDNWGGGGAETLY | 0.8683733 |
| GFSNSGS | ASGDNWGGGGAETLY | 0.75638115 |
| GGNGGSGGTSV | ASGDNWGGGGAETLY | 0.51908946 |
| GTSVATGGAGN | ASGDNWGGGGAETLY | 0.87955403 |
| NGGNGGSGGTS | ASGDNWGGGGAETLY | 0.76504123 |
| AVVKFPH | ASGDNWGGGGAETLY | 0.8338873 |
| TPPPPPP | ASGDNWGGGGAETLY | 0.8178626 |
| AAPAAAAAPA | ASGDNWGGGGAETLY | 0.8386198 |
| IALNEDL | ASGDNWGGGGAETLY | 0.46642238 |
| IVNVSSM | ASGDNWGGGGAETLY | 0.642861 |
| TGIPPAPR | ASGDNWGGGGAETLY | 0.7200929 |
| FDLGGGTF | ASGDNWGGGGAETLY | 0.8537643 |
| FDLGGGTFDVS | ASGDNWGGGGAETLY | 0.7284012 |
| NEPTAAA | ASGDNWGGGGAETLY | 0.72954065 |
| NEPTAAALA | ASGDNWGGGGAETLY | 0.8435084 |
| GSIINISSI | ASGDNWGGGGAETLY | 0.8626262 |
| VAPGNAGTA | ASGDNWGGGGAETLY | 0.817788 |
| TGGGASLE | ASGDNWGGGGAETLY | 0.78566074 |
| VAVYNHYKRI | ASGDNWGGGGAETLY | 0.8768787 |
| TDPAPATT | ASGDNWGGGGAETLY | 0.53929967 |
| TNRPDILDPALLRPG | ASGDNWGGGGAETLY | 0.7697108 |
| LTTTTTS | ASGDNWGGGGAETLY | 0.854571 |
| PTGSGKTL | ASGDNWGGGGAETLY | 0.8515624 |
| TGSGKTL | ASGDNWGGGGAETLY | 0.7891047 |

| | | |
|---|---|---|
| TIGHVDHGK | ASGDNWGGGGAETLY | 0.8658039 |
| TIGHVDHGKTT | ASGDNWGGGGAETLY | 0.67983305 |
| GHVDHGKTT | ASGDNWGGGGAETLY | 0.7995563 |
| AQFEHTLL | ASGDNWGGGGAETLY | 0.7230988 |
| IVNQDEL | ASGDNWGGGGAETLY | 0.6606489 |
| TNGSQFF | ASGDNWGGGGAETLY | 0.5704386 |
| TGKGIVE | ASGDNWGGGGAETLY | 0.78963333 |
| LISNASDA | ASGDNWGGGGAETLY | 0.8387847 |
| AELRTLGELTKIAKAHGAQ | ASGDNWGGGGAETLY | 0.79861224 |
| APNAVAAAEPSW | ASGDNWGGGGAETLY | 0.29579872 |
| APQVAGAATAVL | ASGDNWGGGGAETLY | 0.8202829 |
| ARRDPAAAAAILPTDA | ASGDNWGGGGAETLY | 0.79004097 |
| DAAARDVLAEAGLAEYF | ASGDNWGGGGAETLY | 0.8768692 |
| DDQIVLSAKANLAKVEQ | ASGDNWGGGGAETLY | 0.73637605 |
| DELLPDTGEDSDEDGH | ASGDNWGGGGAETLY | 0.82767355 |
| ELPEAPANRATVPQD | ASGDNWGGGGAETLY | 0.6965428 |
| FGQSKIERVTGIG | ASGDNWGGGGAETLY | 0.79322714 |
| GAGGTGSTAGGAGGAGGAGGLY | ASGDNWGGGGAETLY | 0.42781305 |
| GHGGTGAAGGNGGTGGAGGSGI | ASGDNWGGGGAETLY | 0.52113414 |
| GNGGNGGAGGEGGPGGAGGAGA | ASGDNWGGGGAETLY | 0.7278782 |
| GSTAGNGGNGGNATNGTIAGQP | ASGDNWGGGGAETLY | 0.6258911 |
| GTGGDGGNGGNGGTGG | ASGDNWGGGGAETLY | 0.8379934 |
| GVRRPAGIPADAPV | ASGDNWGGGGAETLY | 0.8566753 |
| GYRQVTDAHLLTLARR | ASGDNWGGGGAETLY | 0.83602923 |
| IAKMGQDGHDRG | ASGDNWGGGGAETLY | 0.8571923 |
| KAEPSIEIQYT | ASGDNWGGGGAETLY | 0.8632245 |
| KPVRIRMAADEDTVAVT | ASGDNWGGGGAETLY | 0.8652548 |
| MPPRIAGMRLLVIKPEP | ASGDNWGGGGAETLY | 0.6648699 |
| MKFYAVALFRSRGVPPTR | ASGDNWGGGGAETLY | 0.6408702 |
| NAGGFGILGAVAHSP | ASGDNWGGGGAETLY | 0.35825124 |
| NEGREITLAMAEM | ASGDNWGGGGAETLY | 0.6966126 |
| NELVEWSHPLGHRWRWE | ASGDNWGGGGAETLY | 0.703582 |
| NLGAGGTGGNGGNPG | ASGDNWGGGGAETLY | 0.77521455 |
| NTEWPKWETVMAADPVD | ASGDNWGGGGAETLY | 0.12725891 |
| PASAAPAETQP | ASGDNWGGGGAETLY | 0.79071385 |
| QQSSDGKNGVRAAEITGE | ASGDNWGGGGAETLY | 0.8643521 |
| RAPDAVTTSWWKE | ASGDNWGGGGAETLY | 0.6531799 |
| SGFGNVGAHTSGWFNQ | ASGDNWGGGGAETLY | 0.7941765 |
| SGVAGAVGQAASVGGLKV | ASGDNWGGGGAETLY | 0.8163177 |
| SNGGAGTGSAGGNGGT | ASGDNWGGGGAETLY | 0.72622204 |
| TGGAGGEGGAAG | ASGDNWGGGGAETLY | 0.8169143 |
| TGGDGGAAGAGGAPGHG | ASGDNWGGGGAETLY | 0.6989746 |
| TMPPLLAAIANAGRHGV | ASGDNWGGGGAETLY | 0.8559953 |
| TSSTAGNSGGA | ASGDNWGGGGAETLY | 0.8560189 |
| VPATVPVGILC | ASGDNWGGGGAETLY | 0.8140645 |
| VQRPDSGGVVLTGRIS | ASGDNWGGGGAETLY | 0.7157563 |
| WTPNPVPATKGVPAGLVQAT | ASGDNWGGGGAETLY | 0.7935734 |
| YALKRALAKFSDRFDVVII | ASGDNWGGGGAETLY | 0.5241981 |
| YLIGSVIV | ASGDNWGGGGAETLY | 0.7712123 |
| YGQLVAVID | ASGDNWGGGGAETLY | 0.5452765 |
| VTVGGMTLPA | ASGDNWGGGGAETLY | 0.7751076 |

| | | |
|---|---|---|
| VLALPAV | ASGDNWGGGGAETLY | 0.62170434 |
| VIPPTLNYET | ASGDNWGGGGAETLY | 0.80906206 |
| VGMGGMHPSSG | ASGDNWGGGGAETLY | 0.7698945 |
| VGLAGGAATA | ASGDNWGGGGAETLY | 0.83479214 |
| VGGAGGAGGSGP | ASGDNWGGGGAETLY | 0.5780119 |
| VFAAFMV | ASGDNWGGGGAETLY | 0.8494852 |
| TLYDKLR | ASGDNWGGGGAETLY | 0.824926 |
| TLAVFTVL | ASGDNWGGGGAETLY | 0.45163321 |
| TGSVQPARV | ASGDNWGGGGAETLY | 0.84678173 |
| TGGGAGSGGT | ASGDNWGGGGAETLY | 0.79989505 |
| SHLETSL | ASGDNWGGGGAETLY | 0.7129965 |
| SFTTGMT | ASGDNWGGGGAETLY | 0.8390412 |
| QVEVAKRMGVRQPTVS | ASGDNWGGGGAETLY | 0.7797194 |
| QGSGGPALAM | ASGDNWGGGGAETLY | 0.47628585 |
| MLAPSNP | ASGDNWGGGGAETLY | 0.6515581 |
| LVNNAAI | ASGDNWGGGGAETLY | 0.7888832 |
| LTGEINTLA | ASGDNWGGGGAETLY | 0.6990775 |
| LSIFGPV | ASGDNWGGGGAETLY | 0.16762784 |
| LLALLANL | ASGDNWGGGGAETLY | 0.8395847 |
| LISNASDALDKLR | ASGDNWGGGGAETLY | 0.51769114 |
| LINEAVP | ASGDNWGGGGAETLY | 0.698562 |
| LGYAFSGL | ASGDNWGGGGAETLY | 0.74429506 |
| LDFVAVAL | ASGDNWGGGGAETLY | 0.8538604 |
| LAQPGAE | ASGDNWGGGGAETLY | 0.6597281 |
| ISNLGAVTGF | ASGDNWGGGGAETLY | 0.560586 |
| HQEDPGRI | ASGDNWGGGGAETLY | 0.8076661 |
| GTPGEPTP | ASGDNWGGGGAETLY | 0.7888355 |
| GFFNSGS | ASGDNWGGGGAETLY | 0.80241024 |
| GAGGGGLTAGAN | ASGDNWGGGGAETLY | 0.72978973 |
| FVDALVNM | ASGDNWGGGGAETLY | 0.8034948 |
| FLTVAPEL | ASGDNWGGGGAETLY | 0.77547485 |
| FETLTAL | ASGDNWGGGGAETLY | 0.5386872 |
| DVINPIEL | ASGDNWGGGGAETLY | 0.71575546 |
| APVSGAPV | ASGDNWGGGGAETLY | 0.68848664 |
| ALVGGLAW | ASGDNWGGGGAETLY | 0.6387931 |
| ALAAAAATAI | ASGDNWGGGGAETLY | 0.6137538 |
| AFGLGLLL | ASGDNWGGGGAETLY | 0.7978173 |
| AFFAAASL | ASGDNWGGGGAETLY | 0.8775324 |
| AAAALPAFSPP | ASGDNWGGGGAETLY | 0.76180756 |
| VVLAPAL | ASGDNWGGGGAETLY | 0.3917656 |
| TIGHVDHGKTTL | ASGDNWGGGGAETLY | 0.76162195 |
| LISNASDALDK | ASGDNWGGGGAETLY | 0.714434 |
| LISNASDAL | ASGDNWGGGGAETLY | 0.8312919 |
| ISNASDALDK | ASGDNWGGGGAETLY | 0.8522911 |
| NQAETLVYQTE | ASGDNWGGGGAETLY | 0.86280006 |
| TGGTGGTAGAG | ASGDNWGGGGAETLY | 0.83944094 |
| TGSTAGGAGGA | ASGDNWGGGGAETLY | 0.68230325 |
| DPLAVPVAAGPV | ASGDNWGGGGAETLY | 0.85193706 |
| GGTGGTPGNGGN | ASGDNWGGGGAETLY | 0.71700346 |
| PAGLAGVDPATL | ASGDNWGGGGAETLY | 0.7528026 |
| GNNLVVEAPHAGASV | ASGDNWGGGGAETLY | 0.81193584 |

| | | |
|---|---|---|
| KVTRIMSASTLEQPA | ASGDNWGGGGAETLY | 0.8296349 |
| LGFALPIALPRRTVV | ASGDNWGGGGAETLY | 0.6630951 |
| TRLGNLTARKLVVLV | ASGDNWGGGGAETLY | 0.742345 |
| WSQQRRLIYFPSAGP | ASGDNWGGGGAETLY | 0.4111398 |
| GPNGNPDPMAAAVDIR | ASGDNWGGGGAETLY | 0.84708536 |
| TQRCVSVADLRAAASR | ASGDNWGGGGAETLY | 0.437395 |
| QFVLDVAEHPVEVLTETV | ASGDNWGGGGAETLY | 0.570159 |
| GGAGGTGGAGGNGGTQAGDG | ASGDNWGGGGAETLY | 0.70688695 |
| PLLATLRAIAVDVDAKPAQV | ASGDNWGGGGAETLY | 0.8596408 |
| APKFRTGGYMIDFWGVGYQVAK | ASGDNWGGGGAETLY | 0.85948634 |
| GAGGAGGAGGAAGGPGGTGGTG | ASGDNWGGGGAETLY | 0.69065166 |
| IGALGLLATDVAPAGSVAFGVA | ASGDNWGGGGAETLY | 0.6408335 |
| AGGAGGIGGNGGAGGAGGVLMG | ASGDNWGGGGAETLY | 0.81207335 |
| ADELIGLL | ASGDNWGGGGAETLY | 0.59981334 |
| ADRALKA | ASGDNWGGGGAETLY | 0.8505218 |
| AGAVGQVGGAGGA | ASGDNWGGGGAETLY | 0.7536621 |
| AGGGGGLGGAP | ASGDNWGGGGAETLY | 0.7721378 |
| AGGLLYGNGGAG | ASGDNWGGGGAETLY | 0.6489247 |
| AGLESIGL | ASGDNWGGGGAETLY | 0.84250575 |
| AGLKTGII | ASGDNWGGGGAETLY | 0.73619395 |
| ASLAALLL | ASGDNWGGGGAETLY | 0.8313529 |
| ASVINISSI | ASGDNWGGGGAETLY | 0.17708552 |
| ATPPGPA | ASGDNWGGGGAETLY | 0.8540244 |
| ATVASALV | ASGDNWGGGGAETLY | 0.57058394 |
| ATVASLIL | ASGDNWGGGGAETLY | 0.61402345 |
| AVNGPVLIRH | ASGDNWGGGGAETLY | 0.22103241 |
| DGAAGAVGI | ASGDNWGGGGAETLY | 0.74149275 |
| DPLLHGI | ASGDNWGGGGAETLY | 0.79162925 |
| ELGAALGAL | ASGDNWGGGGAETLY | 0.8694615 |
| ELGRTQAV | ASGDNWGGGGAETLY | 0.803887 |
| FVAGGYLS | ASGDNWGGGGAETLY | 0.8011482 |
| GASGGAGGAGGW | ASGDNWGGGGAETLY | 0.84905577 |
| GGAGAGGDGGAGGI | ASGDNWGGGGAETLY | 0.70083296 |
| GTGHNGSQPGMGGNG | ASGDNWGGGGAETLY | 0.26610827 |
| GYAAASASAS | ASGDNWGGGGAETLY | 0.7657386 |
| HQVDVAAVQQG | ASGDNWGGGGAETLY | 0.77336633 |
| IGAAIAHT | ASGDNWGGGGAETLY | 0.86898005 |
| IGMIEVAQL | ASGDNWGGGGAETLY | 0.77852345 |
| IRLAPEHTSAMD | ASGDNWGGGGAETLY | 0.464221 |
| IYNVSVL | ASGDNWGGGGAETLY | 0.8104882 |
| KQGFLGM | ASGDNWGGGGAETLY | 0.80748355 |
| LESVITQL | ASGDNWGGGGAETLY | 0.49263242 |
| LGLGALVVV | ASGDNWGGGGAETLY | 0.5968652 |
| LIGNGGNGGSGGI | ASGDNWGGGGAETLY | 0.57440823 |
| LNAINAPTL | ASGDNWGGGGAETLY | 0.3331107 |
| LQLLNTGV | ASGDNWGGGGAETLY | 0.81231314 |
| LQTSIEF | ASGDNWGGGGAETLY | 0.7217002 |
| MSFAVSEL | ASGDNWGGGGAETLY | 0.69168174 |
| NTGALGL | ASGDNWGGGGAETLY | 0.77447844 |
| PPGAGGAGGAGTT | ASGDNWGGGGAETLY | 0.6318842 |
| QLVGLGI | ASGDNWGGGGAETLY | 0.67160946 |

| | | |
|---|---|---|
| RFLEQVHQL | ASGDNWGGGGAETLY | 0.68587685 |
| RGVGYVLR | ASGDNWGGGGAETLY | 0.7628094 |
| SATGVFAGVI | ASGDNWGGGGAETLY | 0.85392326 |
| SLPAITLP | ASGDNWGGGGAETLY | 0.70758754 |
| STALAFTV | ASGDNWGGGGAETLY | 0.75410366 |
| SVLTVGLGK | ASGDNWGGGGAETLY | 0.86360735 |
| TALAGAGSGI | ASGDNWGGGGAETLY | 0.4153769 |
| TGLPFTGI | ASGDNWGGGGAETLY | 0.6398377 |
| VAEHVTV | ASGDNWGGGGAETLY | 0.83168614 |
| VDIVLNSL | ASGDNWGGGGAETLY | 0.7445736 |
| VLTGGGAVL | ASGDNWGGGGAETLY | 0.861369 |
| VNVGLAYGI | ASGDNWGGGGAETLY | 0.7839957 |
| VSSAAVGAV | ASGDNWGGGGAETLY | 0.79592466 |
| VTSVYEAGV | ASGDNWGGGGAETLY | 0.7620791 |
| WVALDRGA | ASGDNWGGGGAETLY | 0.8700875 |
| YLVGSVVI | ASGDNWGGGGAETLY | 0.82409465 |
| GPPGTGKTLL | ASGDNWGGGGAETLY | 0.7214998 |
| GVGMTKF | ASGDNWGGGGAETLY | 0.81543624 |
| IGIGGSDL | ASGDNWGGGGAETLY | 0.3893962 |
| IGTLLAEL | ASGDNWGGGGAETLY | 0.83833236 |
| IYTSGTT | ASGDNWGGGGAETLY | 0.8365196 |
| LSFPAGV | ASGDNWGGGGAETLY | 0.58654845 |
| LYTSGTT | ASGDNWGGGGAETLY | 0.7412494 |
| VGAFPFGF | ASGDNWGGGGAETLY | 0.87988997 |
| VGAGTVEF | ASGDNWGGGGAETLY | 0.8581351 |
| VGKSTLLL | ASGDNWGGGGAETLY | 0.74044776 |
| AALAHAGGGIV | ASGDNWGGGGAETLY | 0.6659036 |
| AALVAVSLL | ASGDNWGGGGAETLY | 0.55061865 |
| AAVSRAVAAL | ASGDNWGGGGAETLY | 0.7072861 |
| AGGSSGIGGF | ASGDNWGGGGAETLY | 0.3389428 |
| AGLTVANANG | ASGDNWGGGGAETLY | 0.812762 |
| AGLVGAVL | ASGDNWGGGGAETLY | 0.6907514 |
| ALSQAFTV | ASGDNWGGGGAETLY | 0.818177 |
| ALTFSVSL | ASGDNWGGGGAETLY | 0.81185913 |
| ALTGAALVAL | ASGDNWGGGGAETLY | 0.8564127 |
| ASFSIGAGVL | ASGDNWGGGGAETLY | 0.6900104 |
| DGSVYKI | ASGDNWGGGGAETLY | 0.7049991 |
| EQLAEIV | ASGDNWGGGGAETLY | 0.38441914 |
| FAVSELTVM | ASGDNWGGGGAETLY | 0.35895917 |
| GAAPLAAGL | ASGDNWGGGGAETLY | 0.70007277 |
| GADGGDGGSGGAGG | ASGDNWGGGGAETLY | 0.7354709 |
| GFGLDTGGAGG | ASGDNWGGGGAETLY | 0.2183986 |
| GLASGGLL | ASGDNWGGGGAETLY | 0.87012905 |
| GRIAGAAVLL | ASGDNWGGGGAETLY | 0.6920042 |
| GSAGSLNTSF | ASGDNWGGGGAETLY | 0.8109277 |
| GTGQIGI | ASGDNWGGGGAETLY | 0.7028184 |
| GTLIALIL | ASGDNWGGGGAETLY | 0.40959394 |
| IAATMGFVVG | ASGDNWGGGGAETLY | 0.65946394 |
| IFAQALL | ASGDNWGGGGAETLY | 0.8224506 |
| IGAAVALGF | ASGDNWGGGGAETLY | 0.4667501 |
| IGLTPLAL | ASGDNWGGGGAETLY | 0.34338045 |

| | | |
|---|---|---|
| ILGSNTSTL | ASGDNWGGGGAETLY | 0.5033206 |
| IVFSLTVL | ASGDNWGGGGAETLY | 0.7493522 |
| LASKATGF | ASGDNWGGGGAETLY | 0.87578386 |
| LASVALIL | ASGDNWGGGGAETLY | 0.8600856 |
| LFAVGLTL | ASGDNWGGGGAETLY | 0.7961489 |
| LFNGGIGTYI | ASGDNWGGGGAETLY | 0.8743178 |
| LTLGAGII | ASGDNWGGGGAETLY | 0.14733681 |
| LVAAALGDI | ASGDNWGGGGAETLY | 0.7123569 |
| NTAFVQLGI | ASGDNWGGGGAETLY | 0.851259 |
| QLFSGAEL | ASGDNWGGGGAETLY | 0.8250003 |
| SAATFITL | ASGDNWGGGGAETLY | 0.7229819 |
| SNYGLANLGV | ASGDNWGGGGAETLY | 0.85689783 |
| SYGGAYAVM | ASGDNWGGGGAETLY | 0.6082396 |
| TTGVGGLTLG | ASGDNWGGGGAETLY | 0.6835214 |
| TTQTIYI | ASGDNWGGGGAETLY | 0.73085713 |
| VAAGAGLGAVI | ASGDNWGGGGAETLY | 0.8561721 |
| VAAVVTLL | ASGDNWGGGGAETLY | 0.8087511 |
| VGFLGGLI | ASGDNWGGGGAETLY | 0.3689788 |
| VGVIPILL | ASGDNWGGGGAETLY | 0.82169497 |
| VIFASSLI | ASGDNWGGGGAETLY | 0.88511384 |
| VKGGDGGAAAG | ASGDNWGGGGAETLY | 0.6760564 |
| VTKTFTNI | ASGDNWGGGGAETLY | 0.84617233 |
| VVAAAILL | ASGDNWGGGGAETLY | 0.7578792 |
| YAALDVEL | ASGDNWGGGGAETLY | 0.6098667 |
| LASVALLL | ASGDNWGGGGAETLY | 0.6297972 |
| LDAVHRV | ASGDNWGGGGAETLY | 0.5823446 |
| QGLSSSIF | ASGDNWGGGGAETLY | 0.85083145 |
| RGTFANIRL | ASGDNWGGGGAETLY | 0.5165488 |
| RQLWWGHRI | ASGDNWGGGGAETLY | 0.7619983 |
| YGPPGTGKTLL | ASGDNWGGGGAETLY | 0.75501335 |
| AALVGAPANVA | ASGDNWGGGGAETLY | 0.88764995 |
| AATFRRTE | ASGDNWGGGGAETLY | 0.8063129 |
| AGALGGGI | ASGDNWGGGGAETLY | 0.8229926 |
| AGGAGGAGGIGAG | ASGDNWGGGGAETLY | 0.6443128 |
| AGSGALGL | ASGDNWGGGGAETLY | 0.83611834 |
| AQIADQI | ASGDNWGGGGAETLY | 0.82786274 |
| ASAVAVTYL | ASGDNWGGGGAETLY | 0.46072337 |
| ASTVLGGTGG | ASGDNWGGGGAETLY | 0.8561617 |
| AVGGEPAK | ASGDNWGGGGAETLY | 0.5174444 |
| CEQGTFGGTC | ASGDNWGGGGAETLY | 0.79556084 |
| DEGKAPAG | ASGDNWGGGGAETLY | 0.7491118 |
| DTVVFPAL | ASGDNWGGGGAETLY | 0.8448561 |
| FEGITMAG | ASGDNWGGGGAETLY | 0.7415155 |
| FPLVVQD | ASGDNWGGGGAETLY | 0.85029954 |
| GAFTTPQ | ASGDNWGGGGAETLY | 0.58194077 |
| GFGGFGTAGGVGGA | ASGDNWGGGGAETLY | 0.8403846 |
| GFLAQFL | ASGDNWGGGGAETLY | 0.8542488 |
| GGAGGAGGNAGVA | ASGDNWGGGGAETLY | 0.7173785 |
| GGVGTAGAGGAGGAGG | ASGDNWGGGGAETLY | 0.87918496 |
| GNGGGGDAGNAGSGGN | ASGDNWGGGGAETLY | 0.81512266 |
| GTGGAGGAGNDAGS | ASGDNWGGGGAETLY | 0.27678135 |

| | | |
|---|---|---|
| GTTNTGWFNS | ASGDNWGGGGAETLY | 0.7520076 |
| IFMGPLFTL | ASGDNWGGGGAETLY | 0.8609761 |
| ILAHAIEAI | ASGDNWGGGGAETLY | 0.8687259 |
| IRSAIREAL | ASGDNWGGGGAETLY | 0.82046 |
| ITFAGGG | ASGDNWGGGGAETLY | 0.78149897 |
| ITGMSGHQEV | ASGDNWGGGGAETLY | 0.7524973 |
| IVAGLACTL | ASGDNWGGGGAETLY | 0.87143534 |
| IVGGGLAGA | ASGDNWGGGGAETLY | 0.8925253 |
| IVTGVLNI | ASGDNWGGGGAETLY | 0.8050538 |
| LAQQGCI | ASGDNWGGGGAETLY | 0.58049273 |
| LCLHVEN | ASGDNWGGGGAETLY | 0.5239421 |
| LDAISDGS | ASGDNWGGGGAETLY | 0.8362339 |
| LGGFAAML | ASGDNWGGGGAETLY | 0.8473114 |
| LGGKNPMI | ASGDNWGGGGAETLY | 0.51009715 |
| LGPKGVAAIL | ASGDNWGGGGAETLY | 0.7116207 |
| LMVGVSI | ASGDNWGGGGAETLY | 0.75466293 |
| LVGAAFVL | ASGDNWGGGGAETLY | 0.23350556 |
| LVLAVAPL | ASGDNWGGGGAETLY | 0.8191103 |
| MAAIAVIDAA | ASGDNWGGGGAETLY | 0.8568982 |
| NNGINAELQ | ASGDNWGGGGAETLY | 0.8416357 |
| RGEPLVLVHG | ASGDNWGGGGAETLY | 0.5157466 |
| RGGAGGAAGLI | ASGDNWGGGGAETLY | 0.8550043 |
| RSYRMHPAV | ASGDNWGGGGAETLY | 0.85947037 |
| SAAHLLAGQLA | ASGDNWGGGGAETLY | 0.8788342 |
| SAAVGSY | ASGDNWGGGGAETLY | 0.39711925 |
| SPHAVLHRI | ASGDNWGGGGAETLY | 0.16776106 |
| SQIAKVL | ASGDNWGGGGAETLY | 0.7827618 |
| TALGHLVP | ASGDNWGGGGAETLY | 0.8163365 |
| TAPTTSGAA | ASGDNWGGGGAETLY | 0.6861255 |
| TDGDDGNGGSPGT | ASGDNWGGGGAETLY | 0.5900927 |
| TDGLGEN | ASGDNWGGGGAETLY | 0.728786 |
| TGGPPLPGQA | ASGDNWGGGGAETLY | 0.8540653 |
| TIPAITI | ASGDNWGGGGAETLY | 0.47476768 |
| TIPGFTI | ASGDNWGGGGAETLY | 0.512763 |
| TSIASGALA | ASGDNWGGGGAETLY | 0.8008232 |
| TVTLPGAAI | ASGDNWGGGGAETLY | 0.8578331 |
| VASGLSAALP | ASGDNWGGGGAETLY | 0.6343615 |
| VGAGVIGI | ASGDNWGGGGAETLY | 0.71459466 |
| VGPKRFGV | ASGDNWGGGGAETLY | 0.805582 |
| VNVAFPD | ASGDNWGGGGAETLY | 0.792523 |
| VSMTNSGVS | ASGDNWGGGGAETLY | 0.8026403 |
| VTDVLGDLSV | ASGDNWGGGGAETLY | 0.4362843 |
| VVVVGAGFAGL | ASGDNWGGGGAETLY | 0.8612741 |
| VYTVGTVV | ASGDNWGGGGAETLY | 0.49435043 |
| WTTFSGP | ASGDNWGGGGAETLY | 0.84686226 |
| YDELVEAF | ASGDNWGGGGAETLY | 0.7692777 |
| YGTGGAGGNGG | ASGDNWGGGGAETLY | 0.82314444 |
| YTNVMAV | ASGDNWGGGGAETLY | 0.67555857 |
| FLEQVHQL | ASGDNWGGGGAETLY | 0.46897447 |
| GLSLLAVAV | ASGDNWGGGGAETLY | 0.48166782 |
| LEQVHQL | ASGDNWGGGGAETLY | 0.6109377 |

| | | |
|---|---|---|
| LNAAGAI | ASGDNWGGGGAETLY | 0.8535888 |
| PAVHASL | ASGDNWGGGGAETLY | 0.7188717 |
| TELEDDA | ASGDNWGGGGAETLY | 0.84925497 |
| TPPGPPN | ASGDNWGGGGAETLY | 0.7551223 |
| TPPPPPPP | ASGDNWGGGGAETLY | 0.6072737 |
| TQSPALI | ASSFYEQY | 0.8432861 |
| EFAAAFPQ | ASSFYEQY | 0.38951123 |
| LLSGNLDVL | ASSFYEQY | 0.79882276 |
| GPANLGNYNI | ASSFYEQY | 0.6446394 |
| TLGPTLP | ASSFYEQY | 0.83244026 |
| VYAYSSCCK | ASSFYEQY | 0.64070344 |
| APAPAPAPNI | ASSFYEQY | 0.8433839 |
| PGGPVPIP | ASSFYEQY | 0.81971955 |
| FLTVALEP | ASSFYEQY | 0.47810495 |
| SQYFGANL | ASSFYEQY | 0.8251815 |
| GGVKPAA | ASSFYEQY | 0.84997165 |
| GGTGGAGGAGGW | ASSFYEQY | 0.8332116 |
| GVGGVAGLGGAGG | ASSFYEQY | 0.31397218 |
| VLLVLATTI | ASSFYEQY | 0.7809926 |
| IGIFAAML | ASSFYEQY | 0.6098758 |
| AVGDLID | ASSFYEQY | 0.80811787 |
| NISPLPL | ASSFYEQY | 0.78771746 |
| VTAVFVAI | ASSFYEQY | 0.62697387 |
| SGDDNDAAL | ASSFYEQY | 0.8351512 |
| AVAALAHS | ASSFYEQY | 0.30523807 |
| GGVTVMATP | ASSFYEQY | 0.6996104 |
| AGLAANVRD | ASSFYEQY | 0.66929436 |
| TNRPDLIDPALLRPG | ASSFYEQY | 0.792076 |
| LANLGVTAL | ASSFYEQY | 0.8208666 |
| TGERIEL | ASSFYEQY | 0.48750535 |
| NGEELAA | ASSFYEQY | 0.7995751 |
| VLSLVWG | ASSFYEQY | 0.7328335 |
| IGYALWL | ASSFYEQY | 0.56232107 |
| LSLLLAPL | ASSFYEQY | 0.8178302 |
| TLDEALK | ASSFYEQY | 0.64648956 |
| TDGAGTAEGG | ASSFYEQY | 0.365291 |
| NAPGGAGGNGG | ASSFYEQY | 0.84907323 |
| VSVFMGMGGNF | ASSFYEQY | 0.5430281 |
| LASYLAAL | ASSFYEQY | 0.798307 |
| LVGGTPAL | ASSFYEQY | 0.8720453 |
| VRNADVL | ASSFYEQY | 0.30208927 |
| TCSSPFLVSAM | ASSFYEQY | 0.84645534 |
| VHITDAE | ASSFYEQY | 0.7869365 |
| EYAAPSF | ASSFYEQY | 0.7870757 |
| VVDAFHGGL | ASSFYEQY | 0.73270917 |
| ASVVDPQ | ASSFYEQY | 0.6046438 |
| SYNEFGGLS | ASSFYEQY | 0.8610542 |
| DPDGSVY | ASSFYEQY | 0.820075 |
| MVSTIHATE | ASSFYEQY | 0.6652057 |
| LAAGVADPV | ASSFYEQY | 0.8157231 |
| TKEQIAA | ASSFYEQY | 0.8263023 |

| | | |
|---|---|---|
| SLASPAAL | ASSFYEQY | 0.8554485 |
| IGLVFYGV | ASSFYEQY | 0.65198743 |
| FLLDASNL | ASSFYEQY | 0.38932794 |
| YQVGNFP | ASSFYEQY | 0.7350632 |
| ESLLSQST | ASSFYEQY | 0.6878864 |
| LFTVGFAAVG | ASSFYEQY | 0.60254264 |
| ATPPTPT | ASSFYEQY | 0.37886333 |
| TDLATLG | ASSFYEQY | 0.8222104 |
| SAGGTGR | ASSFYEQY | 0.8860563 |
| GFSNSGS | ASSFYEQY | 0.6705787 |
| GGNGGSGGTSV | ASSFYEQY | 0.88293016 |
| GTSVATGGAGN | ASSFYEQY | 0.4282814 |
| NGGNGGSGGTS | ASSFYEQY | 0.8153276 |
| AVVKFPH | ASSFYEQY | 0.8320582 |
| TPPPPPP | ASSFYEQY | 0.5305711 |
| AAPAAAAAPA | ASSFYEQY | 0.7502413 |
| IALNEDL | ASSFYEQY | 0.5226966 |
| IVNVSSM | ASSFYEQY | 0.56952465 |
| TGIPPAPR | ASSFYEQY | 0.8442255 |
| FDLGGGTF | ASSFYEQY | 0.6148175 |
| FDLGGGTFDVS | ASSFYEQY | 0.8245542 |
| NEPTAAA | ASSFYEQY | 0.67837226 |
| NEPTAAALA | ASSFYEQY | 0.5551913 |
| GSIINISSI | ASSFYEQY | 0.8279662 |
| VAPGNAGTA | ASSFYEQY | 0.8127619 |
| TGGGASLE | ASSFYEQY | 0.7871276 |
| VAVYNHYKRI | ASSFYEQY | 0.89116657 |
| TDPAPATT | ASSFYEQY | 0.65399444 |
| TNRPDILDPALLRPG | ASSFYEQY | 0.8793025 |
| LTTTTTS | ASSFYEQY | 0.8597308 |
| PTGSGKTL | ASSFYEQY | 0.7929039 |
| TGSGKTL | ASSFYEQY | 0.71720934 |
| TIGHVDHGK | ASSFYEQY | 0.8336338 |
| TIGHVDHGKTT | ASSFYEQY | 0.6338128 |
| GHVDHGKTT | ASSFYEQY | 0.78202057 |
| AQFEHTLL | ASSFYEQY | 0.87790215 |
| IVNQDEL | ASSFYEQY | 0.3512761 |
| TNGSQFF | ASSFYEQY | 0.7835808 |
| TGKGIVE | ASSFYEQY | 0.69910264 |
| LISNASDA | ASSFYEQY | 0.7117498 |
| AELRTLGELTKIAKAHGAQ | ASSFYEQY | 0.49974102 |
| APNAVAAAEPSW | ASSFYEQY | 0.39244276 |
| APQVAGAATAVL | ASSFYEQY | 0.6145089 |
| ARRDPAAAAAILPTDA | ASSFYEQY | 0.42197168 |
| DAAARDVLAEAGLAEYF | ASSFYEQY | 0.77115774 |
| DDQIVLSAKANLAKVEQ | ASSFYEQY | 0.7465683 |
| DELLPDTGEDSDEDGH | ASSFYEQY | 0.51948416 |
| ELPEAPANRATVPQD | ASSFYEQY | 0.708442 |
| FGQSKIERVTGIG | ASSFYEQY | 0.74538434 |
| GAGGTGSTAGGAGGAGGAGGLY | ASSFYEQY | 0.3341304 |
| GHGGTGAAGGNGGTGGAGGSGI | ASSFYEQY | 0.61687934 |

| | | |
|---|---|---|
| GNGGNGGAGGEGGPGGAGGAGA | ASSFYEQY | 0.6697151 |
| GSTAGNGGNGGNATNGTIAGQP | ASSFYEQY | 0.5915907 |
| GTGGDGGNGGNGGTGG | ASSFYEQY | 0.28715226 |
| GVRRPAGIPADAPV | ASSFYEQY | 0.72210467 |
| GYRQVTDAHLLTLARR | ASSFYEQY | 0.5634787 |
| IAKMGQDGHDRG | ASSFYEQY | 0.22190356 |
| KAEPSIEIQYT | ASSFYEQY | 0.6620406 |
| KPVRIRMAADEDTVAVT | ASSFYEQY | 0.5511744 |
| MPPRIAGMRLLVIKPEP | ASSFYEQY | 0.48549372 |
| MKFYAVALFRSRGVPPTR | ASSFYEQY | 0.7379465 |
| NAGGFGILGAVAHSP | ASSFYEQY | 0.6388495 |
| NEGREITLAMAEM | ASSFYEQY | 0.5195258 |
| NELVEWSHPLGHRWRWE | ASSFYEQY | 0.70096487 |
| NLGAGGTGGNGGNPG | ASSFYEQY | 0.73999834 |
| NTEWPKWETVMAADPVD | ASSFYEQY | 0.43332928 |
| PASAAPAETQP | ASSFYEQY | 0.63335156 |
| QQSSDGKNGVRAAEITGE | ASSFYEQY | 0.78796136 |
| RAPDAVTTSWWKE | ASSFYEQY | 0.6495433 |
| SGFGNVGAHTSGWFNQ | ASSFYEQY | 0.79785573 |
| SGVAGAVGQAASVGGLKV | ASSFYEQY | 0.8092935 |
| SNGGAGTGSAGGNGGT | ASSFYEQY | 0.5440952 |
| TGGAGGEGGAAG | ASSFYEQY | 0.7545158 |
| TGGDGGAAGAGGAPGHG | ASSFYEQY | 0.7554648 |
| TMPPLLAAIANAGRHGV | ASSFYEQY | 0.6405086 |
| TSSTAGNSGGA | ASSFYEQY | 0.626219 |
| VPATVPVGILC | ASSFYEQY | 0.37678555 |
| VQRPDSGGVVLTGRIS | ASSFYEQY | 0.63514173 |
| WTPNPVPATKGVPAGLVQAT | ASSFYEQY | 0.4158975 |
| YALKRALAKFSDRFDVVII | ASSFYEQY | 0.67648655 |
| YLIGSVIV | ASSFYEQY | 0.7867938 |
| YGQLVAVID | ASSFYEQY | 0.57505745 |
| VTVGGMTLPA | ASSFYEQY | 0.32400793 |
| VLALPAV | ASSFYEQY | 0.3370625 |
| VIPPTLNYET | ASSFYEQY | 0.72665787 |
| VGMGGMHPSSG | ASSFYEQY | 0.38142195 |
| VGLAGGAATA | ASSFYEQY | 0.45359746 |
| VGGAGGAGGSGP | ASSFYEQY | 0.59978646 |
| VFAAFMV | ASSFYEQY | 0.7295798 |
| TLYDKLR | ASSFYEQY | 0.756431 |
| TLAVFTVL | ASSFYEQY | 0.43344718 |
| TGSVQPARV | ASSFYEQY | 0.7701452 |
| TGGGAGSGGT | ASSFYEQY | 0.7448052 |
| SHLETSL | ASSFYEQY | 0.79098296 |
| SFTTGMT | ASSFYEQY | 0.6861194 |
| QVEVAKRMGVRQPTVS | ASSFYEQY | 0.35765284 |
| QGSGGPALAM | ASSFYEQY | 0.7632657 |
| MLAPSNP | ASSFYEQY | 0.53717893 |
| LVNNAAI | ASSFYEQY | 0.6990179 |
| LTGEINTLA | ASSFYEQY | 0.7294949 |
| LSIFGPV | ASSFYEQY | 0.6402285 |
| LLALLANL | ASSFYEQY | 0.475065 |

| | | |
|---|---|---|
| LISNASDALDKLR | ASSFYEQY | 0.6443865 |
| LINEAVP | ASSFYEQY | 0.80249923 |
| LGYAFSGL | ASSFYEQY | 0.3957314 |
| LDFVAVAL | ASSFYEQY | 0.43267393 |
| LAQPGAE | ASSFYEQY | 0.6024046 |
| ISNLGAVTGF | ASSFYEQY | 0.6018311 |
| HQEDPGRI | ASSFYEQY | 0.7539383 |
| GTPGEPTP | ASSFYEQY | 0.7321575 |
| GFFNSGS | ASSFYEQY | 0.645069 |
| GAGGGGLTAGAN | ASSFYEQY | 0.6798172 |
| FVDALVNM | ASSFYEQY | 0.7637106 |
| FLTVAPEL | ASSFYEQY | 0.68175423 |
| FETLTAL | ASSFYEQY | 0.6746919 |
| DVINPIEL | ASSFYEQY | 0.47560978 |
| APVSGAPV | ASSFYEQY | 0.7904929 |
| ALVGGLAW | ASSFYEQY | 0.7478466 |
| ALAAAAATAI | ASSFYEQY | 0.76286864 |
| AFGLGLLL | ASSFYEQY | 0.78677815 |
| AFFAAASL | ASSFYEQY | 0.71974003 |
| AAAALPAFSPP | ASSFYEQY | 0.6969173 |
| VVLAPAL | ASSFYEQY | 0.32468113 |
| TIGHVDHGKTTL | ASSFYEQY | 0.6280763 |
| LISNASDALDK | ASSFYEQY | 0.61148053 |
| LISNASDAL | ASSFYEQY | 0.68052137 |
| ISNASDALDK | ASSFYEQY | 0.72178257 |
| NQAETLVYQTE | ASSFYEQY | 0.7549098 |
| TGGTGGTAGAG | ASSFYEQY | 0.507856 |
| TGSTAGGAGGA | ASSFYEQY | 0.77308494 |
| DPLAVPVAAGPV | ASSFYEQY | 0.7753897 |
| GGTGGTPGNGGN | ASSFYEQY | 0.40744227 |
| PAGLAGVDPATL | ASSFYEQY | 0.529552 |
| GNNLVVEAPHAGASV | ASSFYEQY | 0.76876855 |
| KVTRIMSASTLEQPA | ASSFYEQY | 0.7041247 |
| LGFALPIALPRRTVV | ASSFYEQY | 0.56131226 |
| TRLGNLTARKLVVLV | ASSFYEQY | 0.6358841 |
| WSQQRRLIYFPSAGP | ASSFYEQY | 0.7758243 |
| GPNGNPDPMAAAVDIR | ASSFYEQY | 0.64209414 |
| TQRCVSVADLRAAASR | ASSFYEQY | 0.7833462 |
| QFVLDVAEHPVEVLTETV | ASSFYEQY | 0.26102674 |
| GGAGGTGGAGGNGGTQAGDG | ASSFYEQY | 0.6489909 |
| PLLATLRAIAVDVDAKPAQV | ASSFYEQY | 0.57370096 |
| APKFRTGGYMIDFWGVGYQVAK | ASSFYEQY | 0.20799947 |
| GAGGAGGAGGAAGGPGGTGGTG | ASSFYEQY | 0.72635716 |
| IGALGLLATDVAPAGSVAFGVA | ASSFYEQY | 0.50911486 |
| AGGAGGIGGNGGAGGAGGVLMG | ASSFYEQY | 0.7057117 |
| ADELIGLL | ASSFYEQY | 0.6120174 |
| ADRALKA | ASSFYEQY | 0.7467369 |
| AGAVGQVGGAGGA | ASSFYEQY | 0.68547404 |
| AGGGGGLGGAP | ASSFYEQY | 0.6026192 |
| AGGLLYGNGGAG | ASSFYEQY | 0.5048811 |
| AGLESIGL | ASSFYEQY | 0.6705452 |

| | | |
|---|---|---|
| AGLKTGII | ASSFYEQY | 0.77689314 |
| ASLAALLL | ASSFYEQY | 0.80520254 |
| ASVINISSI | ASSFYEQY | 0.31742963 |
| ATPPGPA | ASSFYEQY | 0.71164954 |
| ATVASALV | ASSFYEQY | 0.56177455 |
| ATVASLIL | ASSFYEQY | 0.5148683 |
| AVNGPVLIRH | ASSFYEQY | 0.27414164 |
| DGAAGAVGI | ASSFYEQY | 0.5784186 |
| DPLLHGI | ASSFYEQY | 0.6355326 |
| ELGAALGAL | ASSFYEQY | 0.6681714 |
| ELGRTQAV | ASSFYEQY | 0.7450386 |
| FVAGGYLS | ASSFYEQY | 0.5066787 |
| GASGGAGGAGGW | ASSFYEQY | 0.7344017 |
| GGAGAGGDGGAGGI | ASSFYEQY | 0.6110321 |
| GTGHNGSQPGMGGNG | ASSFYEQY | 0.14267254 |
| GYAAASASAS | ASSFYEQY | 0.6068145 |
| HQVDVAAVQQG | ASSFYEQY | 0.3132605 |
| IGAAIAHT | ASSFYEQY | 0.7946431 |
| IGMIEVAQL | ASSFYEQY | 0.3740121 |
| IRLAPEHTSAMD | ASSFYEQY | 0.52940327 |
| IYNVSVL | ASSFYEQY | 0.6009585 |
| KQGFLGM | ASSFYEQY | 0.7134861 |
| LESVITQL | ASSFYEQY | 0.6569891 |
| LGLGALVVV | ASSFYEQY | 0.51838565 |
| LIGNGGNGGSGGI | ASSFYEQY | 0.43251672 |
| LNAINAPTL | ASSFYEQY | 0.6970117 |
| LQLLNTGV | ASSFYEQY | 0.69015324 |
| LQTSIEF | ASSFYEQY | 0.42602128 |
| MSFAVSEL | ASSFYEQY | 0.48269463 |
| NTGALGL | ASSFYEQY | 0.76863885 |
| PPGAGGAGGAGTT | ASSFYEQY | 0.32850567 |
| QLVGLGI | ASSFYEQY | 0.79047257 |
| RFLEQVHQL | ASSFYEQY | 0.500765 |
| RGVGYVLR | ASSFYEQY | 0.5397552 |
| SATGVFAGVI | ASSFYEQY | 0.7925161 |
| SLPAITLP | ASSFYEQY | 0.6667385 |
| STALAFTV | ASSFYEQY | 0.60150284 |
| SVLTVGLGK | ASSFYEQY | 0.5421528 |
| TALAGAGSGI | ASSFYEQY | 0.44230458 |
| TGLPFTGI | ASSFYEQY | 0.47835636 |
| VAEHVTV | ASSFYEQY | 0.4965632 |
| VDIVLNSL | ASSFYEQY | 0.7727181 |
| VLTGGGAVL | ASSFYEQY | 0.6779896 |
| VNVGLAYGI | ASSFYEQY | 0.40510568 |
| VSSAAVGAV | ASSFYEQY | 0.7732528 |
| VTSVYEAGV | ASSFYEQY | 0.7858168 |
| WVALDRGA | ASSFYEQY | 0.78831345 |
| YLVGSVVI | ASSFYEQY | 0.40622565 |
| GPPGTGKTLL | ASSFYEQY | 0.6068408 |
| GVGMTKF | ASSFYEQY | 0.6005963 |
| IGIGGSDL | ASSFYEQY | 0.54233783 |

| | | |
|---|---|---|
| IGTLLAEL | ASSFYEQY | 0.5374781 |
| IYTSGTT | ASSFYEQY | 0.45075354 |
| LSFPAGV | ASSFYEQY | 0.7791765 |
| LYTSGTT | ASSFYEQY | 0.8167434 |
| VGAFPFGF | ASSFYEQY | 0.6345714 |
| VGAGTVEF | ASSFYEQY | 0.32332915 |
| VGKSTLLL | ASSFYEQY | 0.54368794 |
| AALAHAGGGIV | ASSFYEQY | 0.4593082 |
| AALVAVSLL | ASSFYEQY | 0.44306952 |
| AAVSRAVAAL | ASSFYEQY | 0.58489454 |
| AGGSSGIGGF | ASSFYEQY | 0.28723192 |
| AGLTVANANG | ASSFYEQY | 0.62043256 |
| AGLVGAVL | ASSFYEQY | 0.80013597 |
| ALSQAFTV | ASSFYEQY | 0.5708391 |
| ALTFSVSL | ASSFYEQY | 0.7307477 |
| ALTGAALVAL | ASSFYEQY | 0.7815043 |
| ASFSIGAGVL | ASSFYEQY | 0.7075218 |
| DGSVYKI | ASSFYEQY | 0.75533414 |
| EQLAEIV | ASSFYEQY | 0.7505433 |
| FAVSELTVM | ASSFYEQY | 0.67012656 |
| GAAPLAAGL | ASSFYEQY | 0.68316835 |
| GADGGDGGSGGAGG | ASSFYEQY | 0.74503195 |
| GFGLDTGGAGG | ASSFYEQY | 0.5480498 |
| GLASGGLL | ASSFYEQY | 0.63432133 |
| GRIAGAAVLL | ASSFYEQY | 0.65045273 |
| GSAGSLNTSF | ASSFYEQY | 0.6851838 |
| GTGQIGI | ASSFYEQY | 0.6943232 |
| GTLIALIL | ASSFYEQY | 0.8009391 |
| IAATMGFVVG | ASSFYEQY | 0.2627329 |
| IFAQALL | ASSFYEQY | 0.4502979 |
| IGAAVALGF | ASSFYEQY | 0.76095414 |
| IGLTPLAL | ASSFYEQY | 0.2155977 |
| ILGSNTSTL | ASSFYEQY | 0.3575306 |
| IVFSLTVL | ASSFYEQY | 0.69835675 |
| LASKATGF | ASSFYEQY | 0.77539 |
| LASVALIL | ASSFYEQY | 0.26989463 |
| LFAVGLTL | ASSFYEQY | 0.59849954 |
| LFNGGIGTYI | ASSFYEQY | 0.3776453 |
| LTLGAGII | ASSFYEQY | 0.7893847 |
| LVAAALGDI | ASSFYEQY | 0.5795331 |
| NTAFVQLGI | ASSFYEQY | 0.37907812 |
| QLFSGAEL | ASSFYEQY | 0.72576505 |
| SAATFITL | ASSFYEQY | 0.43108737 |
| SNYGLANLGV | ASSFYEQY | 0.7061281 |
| SYGGAYAVM | ASSFYEQY | 0.7475074 |
| TTGVGGLTLG | ASSFYEQY | 0.7989559 |
| TTQTIYI | ASSFYEQY | 0.35607842 |
| VAAGAGLGAVI | ASSFYEQY | 0.7172231 |
| VAAVVTLL | ASSFYEQY | 0.53741014 |
| VGFLGGLI | ASSFYEQY | 0.8138422 |
| VGVIPILL | ASSFYEQY | 0.562915 |

| | | |
|---|---|---|
| VIFASSLI | ASSFYEQY | 0.35084078 |
| VKGGDGGAAAG | ASSFYEQY | 0.7409452 |
| VTKTFTNI | ASSFYEQY | 0.7254555 |
| VVAAAILL | ASSFYEQY | 0.5900221 |
| YAALDVEL | ASSFYEQY | 0.74023235 |
| LASVALLL | ASSFYEQY | 0.6897484 |
| LDAVHRV | ASSFYEQY | 0.55823743 |
| QGLSSSIF | ASSFYEQY | 0.72734004 |
| RGTFANIRL | ASSFYEQY | 0.6969789 |
| RQLWWGHRI | ASSFYEQY | 0.55141366 |
| YGPPGTGKTLL | ASSFYEQY | 0.7823652 |
| AALVGAPANVA | ASSFYEQY | 0.5984528 |
| AATFRRTE | ASSFYEQY | 0.39653638 |
| AGALGGGI | ASSFYEQY | 0.79244876 |
| AGGAGGAGGIGAG | ASSFYEQY | 0.68398947 |
| AGSGALGL | ASSFYEQY | 0.74920344 |
| AQIADQI | ASSFYEQY | 0.71809375 |
| ASAVAVTYL | ASSFYEQY | 0.78833365 |
| ASTVLGGTGG | ASSFYEQY | 0.53227246 |
| AVGGEPAK | ASSFYEQY | 0.6656092 |
| CEQGTFGGTC | ASSFYEQY | 0.64960754 |
| DEGKAPAG | ASSFYEQY | 0.74513674 |
| DTVVFPAL | ASSFYEQY | 0.6569064 |
| FEGITMAG | ASSFYEQY | 0.7250142 |
| FPLVVQD | ASSFYEQY | 0.73048985 |
| GAFTTPQ | ASSFYEQY | 0.7835821 |
| GFGGFGTAGGVGGA | ASSFYEQY | 0.6697127 |
| GFLAQFL | ASSFYEQY | 0.66300476 |
| GGAGGAGGNAGVA | ASSFYEQY | 0.7728023 |
| GGVGTAGAGGAGGAGG | ASSFYEQY | 0.42705113 |
| GNGGGGDAGNAGSGGN | ASSFYEQY | 0.6363769 |
| GTGGAGGAGNDAGS | ASSFYEQY | 0.7734332 |
| GTTNTGWFNS | ASSFYEQY | 0.76291883 |
| IFMGPLFTL | ASSFYEQY | 0.6523443 |
| ILAHAIEAI | ASSFYEQY | 0.7771135 |
| IRSAIREAL | ASSFYEQY | 0.62157845 |
| ITFAGGG | ASSFYEQY | 0.69100785 |
| ITGMSGHQEV | ASSFYEQY | 0.5802416 |
| IVAGLACTL | ASSFYEQY | 0.5899171 |
| IVGGGLAGA | ASSFYEQY | 0.43202403 |
| IVTGVLNI | ASSFYEQY | 0.58585906 |
| LAQQGCI | ASSFYEQY | 0.74611163 |
| LCLHVEN | ASSFYEQY | 0.6530651 |
| LDAISDGS | ASSFYEQY | 0.31213528 |
| LGGFAAML | ASSFYEQY | 0.7060318 |
| LGGKNPMI | ASSFYEQY | 0.6961756 |
| LGPKGVAAIL | ASSFYEQY | 0.7997034 |
| LMVGVSI | ASSFYEQY | 0.67991126 |
| LVGAAFVL | ASSFYEQY | 0.7440381 |
| LVLAVAPL | ASSFYEQY | 0.6062366 |
| MAAIAVIDAA | ASSFYEQY | 0.6931227 |

| | | |
|---|---|---|
| NNGINAELQ | ASSFYEQY | 0.3345726 |
| RGEPLVLVHG | ASSFYEQY | 0.5001497 |
| RGGAGGAAGLI | ASSFYEQY | 0.60491836 |
| RSYRMHPAV | ASSFYEQY | 0.41130364 |
| SAAHLLAGQLA | ASSFYEQY | 0.72826874 |
| SAAVGSY | ASSFYEQY | 0.7433834 |
| SPHAVLHRI | ASSFYEQY | 0.75888366 |
| SQIAKVL | ASSFYEQY | 0.7683526 |
| TALGHLVP | ASSFYEQY | 0.7836384 |
| TAPTTSGAA | ASSFYEQY | 0.78217363 |
| TDGDDGNGGSPGT | ASSFYEQY | 0.49266887 |
| TDGLGEN | ASSFYEQY | 0.6377001 |
| TGGPPLPGQA | ASSFYEQY | 0.28673327 |
| TIPAITI | ASSFYEQY | 0.562085 |
| TIPGFTI | ASSFYEQY | 0.5375321 |
| TSIASGALA | ASSFYEQY | 0.6573286 |
| TVTLPGAAI | ASSFYEQY | 0.19935012 |
| VASGLSAALP | ASSFYEQY | 0.6876465 |
| VGAGVIGI | ASSFYEQY | 0.7827191 |
| VGPKRFGV | ASSFYEQY | 0.6076223 |
| VNVAFPD | ASSFYEQY | 0.7143439 |
| VSMTNSGVS | ASSFYEQY | 0.7325595 |
| VTDVLGDLSV | ASSFYEQY | 0.52698576 |
| VVVVGAGFAGL | ASSFYEQY | 0.72607493 |
| VYTVGTVV | ASSFYEQY | 0.58476156 |
| WTTFSGP | ASSFYEQY | 0.76250803 |
| YDELVEAF | ASSFYEQY | 0.7853623 |
| YGTGGAGGNGG | ASSFYEQY | 0.6916591 |
| YTNVMAV | ASSFYEQY | 0.6036217 |
| FLEQVHQL | ASSFYEQY | 0.7125747 |
| GLSLLAVAV | ASSFYEQY | 0.5019007 |
| LEQVHQL | ASSFYEQY | 0.61207116 |
| LNAAGAI | ASSFYEQY | 0.5928664 |
| PAVHASL | ASSFYEQY | 0.65582514 |
| TELEDDA | ASSFYEQY | 0.47400582 |
| TPPGPPN | ASSFYEQY | 0.64546764 |
| TPPPPPPP | ASSFYEQY | 0.61712056 |
| TQSPALI | ASSSKDTQY | 0.7238307 |
| EFAAAFPQ | ASSSKDTQY | 0.4128453 |
| LLSGNLDVL | ASSSKDTQY | 0.7338337 |
| GPANLGNYNI | ASSSKDTQY | 0.69218546 |
| TLGPTLP | ASSSKDTQY | 0.3658083 |
| VYAYSSCCK | ASSSKDTQY | 0.74650985 |
| APAPAPAPNI | ASSSKDTQY | 0.6751845 |
| PGGPVPIP | ASSSKDTQY | 0.6364498 |
| FLTVALEP | ASSSKDTQY | 0.7557245 |
| SQYFGANL | ASSSKDTQY | 0.6234007 |
| GGVKPAA | ASSSKDTQY | 0.47800094 |
| GGTGGAGGAGGW | ASSSKDTQY | 0.56682247 |
| GVGGVAGLGGAGG | ASSSKDTQY | 0.65868497 |
| VLLVLATTI | ASSSKDTQY | 0.58670115 |

| | | |
|---|---|---|
| IGIFAAML | ASSSKDTQY | 0.3066383 |
| AVGDLID | ASSSKDTQY | 0.7320075 |
| NISPLPL | ASSSKDTQY | 0.40394685 |
| VTAVFVAI | ASSSKDTQY | 0.53062683 |
| SGDDNDAAL | ASSSKDTQY | 0.4762467 |
| AVAALAHS | ASSSKDTQY | 0.7292138 |
| GGVTVMATP | ASSSKDTQY | 0.7103765 |
| AGLAANVRD | ASSSKDTQY | 0.60832846 |
| TNRPDLIDPALLRPG | ASSSKDTQY | 0.68766665 |
| LANLGVTAL | ASSSKDTQY | 0.6781556 |
| TGERIEL | ASSSKDTQY | 0.7498169 |
| NGEELAA | ASSSKDTQY | 0.6554499 |
| VLSLVWG | ASSSKDTQY | 0.68318295 |
| IGYALWL | ASSSKDTQY | 0.69154274 |
| LSLLLAPL | ASSSKDTQY | 0.6585357 |
| TLDEALK | ASSSKDTQY | 0.44564128 |
| TDGAGTAEGG | ASSSKDTQY | 0.50936145 |
| NAPGGAGGNGG | ASSSKDTQY | 0.56168973 |
| VSVFMGMGGNF | ASSSKDTQY | 0.4417532 |
| LASYLAAL | ASSSKDTQY | 0.6705867 |
| LVGGTPAL | ASSSKDTQY | 0.8060752 |
| VRNADVL | ASSSKDTQY | 0.7010029 |
| TCSSPFLVSAM | ASSSKDTQY | 0.3351599 |
| VHITDAE | ASSSKDTQY | 0.58048356 |
| EYAAPSF | ASSSKDTQY | 0.66269004 |
| VVDAFHGGL | ASSSKDTQY | 0.7282212 |
| ASVVDPQ | ASSSKDTQY | 0.7542729 |
| SYNEFGGLS | ASSSKDTQY | 0.8040916 |
| DPDGSVY | ASSSKDTQY | 0.7165202 |
| MVSTIHATE | ASSSKDTQY | 0.5086548 |
| LAAGVADPV | ASSSKDTQY | 0.72263026 |
| TKEQIAA | ASSSKDTQY | 0.6033957 |
| SLASPAAL | ASSSKDTQY | 0.6997228 |
| IGLVFYGV | ASSSKDTQY | 0.67959523 |
| FLLDASNL | ASSSKDTQY | 0.7274865 |
| YQVGNFP | ASSSKDTQY | 0.5779753 |
| ESLLSQST | ASSSKDTQY | 0.5830327 |
| LFTVGFAAVG | ASSSKDTQY | 0.39532536 |
| ATPPTPT | ASSSKDTQY | 0.7631502 |
| TDLATLG | ASSSKDTQY | 0.5633724 |
| SAGGTGR | ASSSKDTQY | 0.5697584 |
| GFSNSGS | ASSSKDTQY | 0.57389677 |
| GGNGGSGGTSV | ASSSKDTQY | 0.75030017 |
| GTSVATGGAGN | ASSSKDTQY | 0.7961157 |
| NGGNGGSGGTS | ASSSKDTQY | 0.49716228 |
| AVVKFPH | ASSSKDTQY | 0.48266953 |
| TPPPPPP | ASSSKDTQY | 0.7224066 |
| AAPAAAAAPA | ASSSKDTQY | 0.43866765 |
| IALNEDL | ASSSKDTQY | 0.7410581 |
| IVNVSSM | ASSSKDTQY | 0.6344143 |
| TGIPPAPR | ASSSKDTQY | 0.6132612 |

| | | |
|---|---|---|
| FDLGGGTF | ASSSKDTQY | 0.5421736 |
| FDLGGGTFDVS | ASSSKDTQY | 0.73114073 |
| NEPTAAA | ASSSKDTQY | 0.56641406 |
| NEPTAAALA | ASSSKDTQY | 0.75212383 |
| GSIINISSI | ASSSKDTQY | 0.40315124 |
| VAPGNAGTA | ASSSKDTQY | 0.72968054 |
| TGGGASLE | ASSSKDTQY | 0.3913874 |
| VAVYNHYKRI | ASSSKDTQY | 0.5812658 |
| TDPAPATT | ASSSKDTQY | 0.2947804 |
| TNRPDILDPALLRPG | ASSSKDTQY | 0.61224204 |
| LTTTTTS | ASSSKDTQY | 0.7282813 |
| PTGSGKTL | ASSSKDTQY | 0.77974397 |
| TGSGKTL | ASSSKDTQY | 0.6799243 |
| TIGHVDHGK | ASSSKDTQY | 0.677812 |
| TIGHVDHGKTT | ASSSKDTQY | 0.79173386 |
| GHVDHGKTT | ASSSKDTQY | 0.5226181 |
| AQFEHTLL | ASSSKDTQY | 0.3102459 |
| IVNQDEL | ASSSKDTQY | 0.6421064 |
| TNGSQFF | ASSSKDTQY | 0.6048585 |
| TGKGIVE | ASSSKDTQY | 0.7735437 |
| LISNASDA | ASSSKDTQY | 0.7116899 |
| AELRTLGELTKIAKAHGAQ | ASSSKDTQY | 0.2946325 |
| APNAVAAAEPSW | ASSSKDTQY | 0.690265 |
| APQVAGAATAVL | ASSSKDTQY | 0.755198 |
| ARRDPAAAAAILPTDA | ASSSKDTQY | 0.42495275 |
| DAAARDVLAEAGLAEYF | ASSSKDTQY | 0.50779986 |
| DDQIVLSAKANLAKVEQ | ASSSKDTQY | 0.50266707 |
| DELLPDTGEDSDEDGH | ASSSKDTQY | 0.4809633 |
| ELPEAPANRATVPQD | ASSSKDTQY | 0.7034973 |
| FGQSKIERVTGIG | ASSSKDTQY | 0.52765596 |
| GAGGTGSTAGGAGGAGGAGGLY | ASSSKDTQY | 0.610297 |
| GHGGTGAAGGNGGTGGAGGSGI | ASSSKDTQY | 0.5705115 |
| GNGGNGGAGGEGGPGGAGGAGA | ASSSKDTQY | 0.5159125 |
| GSTAGNGGNGGNATNGTIAGQP | ASSSKDTQY | 0.5639059 |
| GTGGDGGNGGNGGTGG | ASSSKDTQY | 0.59224725 |
| GVRRPAGIPADAPV | ASSSKDTQY | 0.70912635 |
| GYRQVTDAHLLTLARR | ASSSKDTQY | 0.74551487 |
| IAKMGQDGHDRG | ASSSKDTQY | 0.59787786 |
| KAEPSIEIQYT | ASSSKDTQY | 0.6944064 |
| KPVRIRMAADEDTVAVT | ASSSKDTQY | 0.77457106 |
| MPPRIAGMRLLVIKPEP | ASSSKDTQY | 0.4387288 |
| MKFYAVALFRSRGVPPTR | ASSSKDTQY | 0.49413913 |
| NAGGFGILGAVAHSP | ASSSKDTQY | 0.7293784 |
| NEGREITLAMAEM | ASSSKDTQY | 0.5531682 |
| NELVEWSHPLGHRWRWE | ASSSKDTQY | 0.76789665 |
| NLGAGGTGGNGGNPG | ASSSKDTQY | 0.6462518 |
| NTEWPKWETVMAADPVD | ASSSKDTQY | 0.6809419 |
| PASAAPAETQP | ASSSKDTQY | 0.68899107 |
| QQSSDGKNGVRAAEITGE | ASSSKDTQY | 0.7985616 |
| RAPDAVTTSWWKE | ASSSKDTQY | 0.69366366 |
| SGFGNVGAHTSGWFNQ | ASSSKDTQY | 0.5966292 |

| | | |
|---|---|---|
| SGVAGAVGQAASVGGLKV | ASSSKDTQY | 0.6993252 |
| SNGGAGTGSAGGNGGT | ASSSKDTQY | 0.40931815 |
| TGGAGGEGGAAG | ASSSKDTQY | 0.72889125 |
| TGGDGGAAGAGGAPGHG | ASSSKDTQY | 0.7415591 |
| TMPPLLAAIANAGRHGV | ASSSKDTQY | 0.43574724 |
| TSSTAGNSGGA | ASSSKDTQY | 0.5575462 |
| VPATVPVGILC | ASSSKDTQY | 0.804915 |
| VQRPDSGGVVLTGRIS | ASSSKDTQY | 0.7386953 |
| WTPNPVPATKGVPAGLVQAT | ASSSKDTQY | 0.52154684 |
| YALKRALAKFSDRFDVVII | ASSSKDTQY | 0.47178727 |
| YLIGSVIV | ASSSKDTQY | 0.54694706 |
| YGQLVAVID | ASSSKDTQY | 0.61533165 |
| VTVGGMTLPA | ASSSKDTQY | 0.34101343 |
| VLALPAV | ASSSKDTQY | 0.73206663 |
| VIPPTLNYET | ASSSKDTQY | 0.5601377 |
| VGMGGMHPSSG | ASSSKDTQY | 0.7519194 |
| VGLAGGAATA | ASSSKDTQY | 0.6683298 |
| VGGAGGAGGSGP | ASSSKDTQY | 0.74702936 |
| VFAAFMV | ASSSKDTQY | 0.54793286 |
| TLYDKLR | ASSSKDTQY | 0.57113945 |
| TLAVFTVL | ASSSKDTQY | 0.32419115 |
| TGSVQPARV | ASSSKDTQY | 0.5411731 |
| TGGGAGSGGT | ASSSKDTQY | 0.5709008 |
| SHLETSL | ASSSKDTQY | 0.5923793 |
| SFTTGMT | ASSSKDTQY | 0.45639044 |
| QVEVAKRMGVRQPTVS | ASSSKDTQY | 0.7918693 |
| QGSGGPALAM | ASSSKDTQY | 0.5634829 |
| MLAPSNP | ASSSKDTQY | 0.70065916 |
| LVNNAAI | ASSSKDTQY | 0.5958933 |
| LTGEINTLA | ASSSKDTQY | 0.46316803 |
| LSIFGPV | ASSSKDTQY | 0.50124586 |
| LLALLANL | ASSSKDTQY | 0.7158574 |
| LISNASDALDKLR | ASSSKDTQY | 0.3523067 |
| LINEAVP | ASSSKDTQY | 0.32430512 |
| LGYAFSGL | ASSSKDTQY | 0.5654366 |
| LDFVAVAL | ASSSKDTQY | 0.63465446 |
| LAQPGAE | ASSSKDTQY | 0.8014672 |
| ISNLGAVTGF | ASSSKDTQY | 0.7705228 |
| HQEDPGRI | ASSSKDTQY | 0.63511693 |
| GTPGEPTP | ASSSKDTQY | 0.81054604 |
| GFFNSGS | ASSSKDTQY | 0.20764633 |
| GAGGGGLTAGAN | ASSSKDTQY | 0.6073977 |
| FVDALVNM | ASSSKDTQY | 0.77024084 |
| FLTVAPEL | ASSSKDTQY | 0.694798 |
| FETLTAL | ASSSKDTQY | 0.6009684 |
| DVINPIEL | ASSSKDTQY | 0.7459445 |
| APVSGAPV | ASSSKDTQY | 0.4551975 |
| ALVGGLAW | ASSSKDTQY | 0.5889604 |
| ALAAAAATAI | ASSSKDTQY | 0.6041101 |
| AFGLGLLL | ASSSKDTQY | 0.7447215 |
| AFFAAASL | ASSSKDTQY | 0.762874 |

| | | |
|---|---|---|
| AAAALPAFSPP | ASSSKDTQY | 0.3344424 |
| VVLAPAL | ASSSKDTQY | 0.749112 |
| TIGHVDHGKTTL | ASSSKDTQY | 0.8044025 |
| LISNASDALDK | ASSSKDTQY | 0.4637591 |
| LISNASDAL | ASSSKDTQY | 0.7624206 |
| ISNASDALDK | ASSSKDTQY | 0.6336576 |
| NQAETLVYQTE | ASSSKDTQY | 0.6591687 |
| TGGTGGTAGAG | ASSSKDTQY | 0.4850065 |
| TGSTAGGAGGA | ASSSKDTQY | 0.37454882 |
| DPLAVPVAAGPV | ASSSKDTQY | 0.7223319 |
| GGTGGTPGNGGN | ASSSKDTQY | 0.47660568 |
| PAGLAGVDPATL | ASSSKDTQY | 0.67054707 |
| GNNLVVEAPHAGASV | ASSSKDTQY | 0.66387105 |
| KVTRIMSASTLEQPA | ASSSKDTQY | 0.8171924 |
| LGFALPIALPRRTVV | ASSSKDTQY | 0.7022968 |
| TRLGNLTARKLVVLV | ASSSKDTQY | 0.68635213 |
| WSQQRRLIYFPSAGP | ASSSKDTQY | 0.4056795 |
| GPNGNPDPMAAAVDIR | ASSSKDTQY | 0.71615183 |
| TQRCVSVADLRAAASR | ASSSKDTQY | 0.7167091 |
| QFVLDVAEHPVEVLTETV | ASSSKDTQY | 0.44886106 |
| GGAGGTGGAGGNGGTQAGDG | ASSSKDTQY | 0.7738802 |
| PLLATLRAIAVDVDAKPAQV | ASSSKDTQY | 0.4343906 |
| APKFRTGGYMIDFWGVGYQVAK | ASSSKDTQY | 0.69142365 |
| GAGGAGGAGGAAGGPGGTGGTG | ASSSKDTQY | 0.7344197 |
| IGALGLLATDVAPAGSVAFGVA | ASSSKDTQY | 0.7691325 |
| AGGAGGIGGNGGAGGAGGVLMG | ASSSKDTQY | 0.63484573 |
| ADELIGLL | ASSSKDTQY | 0.7574355 |
| ADRALKA | ASSSKDTQY | 0.37859228 |
| AGAVGQVGGAGGA | ASSSKDTQY | 0.76062226 |
| AGGGGGLGGAP | ASSSKDTQY | 0.7373145 |
| AGGLLYGNGGAG | ASSSKDTQY | 0.71330893 |
| AGLESIGL | ASSSKDTQY | 0.80660594 |
| AGLKTGII | ASSSKDTQY | 0.74660736 |
| ASLAALLL | ASSSKDTQY | 0.32559556 |
| ASVINISSI | ASSSKDTQY | 0.6483036 |
| ATPPGPA | ASSSKDTQY | 0.7419398 |
| ATVASALV | ASSSKDTQY | 0.7815872 |
| ATVASLIL | ASSSKDTQY | 0.7679616 |
| AVNGPVLIRH | ASSSKDTQY | 0.7281542 |
| DGAAGAVGI | ASSSKDTQY | 0.574391 |
| DPLLHGI | ASSSKDTQY | 0.7894132 |
| ELGAALGAL | ASSSKDTQY | 0.83559024 |
| ELGRTQAV | ASSSKDTQY | 0.7023256 |
| FVAGGYLS | ASSSKDTQY | 0.5225791 |
| GASGGAGGAGGW | ASSSKDTQY | 0.57791275 |
| GGAGAGGDGGAGGI | ASSSKDTQY | 0.7213372 |
| GTGHNGSQPGMGGNG | ASSSKDTQY | 0.7087232 |
| GYAAASASAS | ASSSKDTQY | 0.44266975 |
| HQVDVAAVQQG | ASSSKDTQY | 0.63653344 |
| IGAAIAHT | ASSSKDTQY | 0.6530972 |
| IGMIEVAQL | ASSSKDTQY | 0.23992419 |

| | | |
|---|---|---|
| IRLAPEHTSAMD | ASSSKDTQY | 0.73030066 |
| IYNVSVL | ASSSKDTQY | 0.77207506 |
| KQGFLGM | ASSSKDTQY | 0.72570765 |
| LESVITQL | ASSSKDTQY | 0.412053 |
| LGLGALVVV | ASSSKDTQY | 0.7730589 |
| LIGNGGNGGSGGI | ASSSKDTQY | 0.7806359 |
| LNAINAPTL | ASSSKDTQY | 0.7952117 |
| LQLLNTGV | ASSSKDTQY | 0.35775554 |
| LQTSIEF | ASSSKDTQY | 0.21507208 |
| MSFAVSEL | ASSSKDTQY | 0.66760397 |
| NTGALGL | ASSSKDTQY | 0.71266234 |
| PPGAGGAGGAGTT | ASSSKDTQY | 0.55607486 |
| QLVGLGI | ASSSKDTQY | 0.42939362 |
| RFLEQVHQL | ASSSKDTQY | 0.64991814 |
| RGVGYVLR | ASSSKDTQY | 0.76591134 |
| SATGVFAGVI | ASSSKDTQY | 0.38590458 |
| SLPAITLP | ASSSKDTQY | 0.4370194 |
| STALAFTV | ASSSKDTQY | 0.71918744 |
| SVLTVGLGK | ASSSKDTQY | 0.7610842 |
| TALAGAGSGI | ASSSKDTQY | 0.4252774 |
| TGLPFTGI | ASSSKDTQY | 0.4867291 |
| VAEHVTV | ASSSKDTQY | 0.6782645 |
| VDIVLNSL | ASSSKDTQY | 0.7424687 |
| VLTGGGAVL | ASSSKDTQY | 0.6735575 |
| VNVGLAYGI | ASSSKDTQY | 0.39464784 |
| VSSAAVGAV | ASSSKDTQY | 0.72566265 |
| VTSVYEAGV | ASSSKDTQY | 0.4048688 |
| WVALDRGA | ASSSKDTQY | 0.7675642 |
| YLVGSVVI | ASSSKDTQY | 0.61572254 |
| GPPGTGKTLL | ASSSKDTQY | 0.6989641 |
| GVGMTKF | ASSSKDTQY | 0.5571055 |
| IGIGGSDL | ASSSKDTQY | 0.43607965 |
| IGTLLAEL | ASSSKDTQY | 0.43136096 |
| IYTSGTT | ASSSKDTQY | 0.432775 |
| LSFPAGV | ASSSKDTQY | 0.7513397 |
| LYTSGTT | ASSSKDTQY | 0.7194083 |
| VGAFPFGF | ASSSKDTQY | 0.7347293 |
| VGAGTVEF | ASSSKDTQY | 0.6558909 |
| VGKSTLLL | ASSSKDTQY | 0.43424606 |
| AALAHAGGGIV | ASSSKDTQY | 0.74398255 |
| AALVAVSLL | ASSSKDTQY | 0.34972504 |
| AAVSRAVAAL | ASSSKDTQY | 0.71867144 |
| AGGSSGIGGF | ASSSKDTQY | 0.6479022 |
| AGLTVANANG | ASSSKDTQY | 0.7007927 |
| AGLVGAVL | ASSSKDTQY | 0.53997165 |
| ALSQAFTV | ASSSKDTQY | 0.75313586 |
| ALTFSVSL | ASSSKDTQY | 0.70654476 |
| ALTGAALVAL | ASSSKDTQY | 0.51084054 |
| ASFSIGAGVL | ASSSKDTQY | 0.6888604 |
| DGSVYKI | ASSSKDTQY | 0.7559943 |
| EQLAEIV | ASSSKDTQY | 0.7263863 |

| | | |
|---|---|---|
| FAVSELTVM | ASSSKDTQY | 0.31110835 |
| GAAPLAAGL | ASSSKDTQY | 0.69932914 |
| GADGGDGGSGGAGG | ASSSKDTQY | 0.4251042 |
| GFGLDTGGAGG | ASSSKDTQY | 0.6671972 |
| GLASGGLL | ASSSKDTQY | 0.65329075 |
| GRIAGAAVLL | ASSSKDTQY | 0.5973139 |
| GSAGSLNTSF | ASSSKDTQY | 0.7386651 |
| GTGQIGI | ASSSKDTQY | 0.47024056 |
| GTLIALIL | ASSSKDTQY | 0.53449893 |
| IAATMGFVVG | ASSSKDTQY | 0.57660395 |
| IFAQALL | ASSSKDTQY | 0.6631191 |
| IGAAVALGF | ASSSKDTQY | 0.7025969 |
| IGLTPLAL | ASSSKDTQY | 0.3727565 |
| ILGSNTSTL | ASSSKDTQY | 0.69631875 |
| IVFSLTVL | ASSSKDTQY | 0.5997767 |
| LASKATGF | ASSSKDTQY | 0.42344573 |
| LASVALIL | ASSSKDTQY | 0.69305557 |
| LFAVGLTL | ASSSKDTQY | 0.63725245 |
| LFNGGIGTYI | ASSSKDTQY | 0.3836542 |
| LTLGAGII | ASSSKDTQY | 0.73329604 |
| LVAAALGDI | ASSSKDTQY | 0.50752175 |
| NTAFVQLGI | ASSSKDTQY | 0.71699584 |
| QLFSGAEL | ASSSKDTQY | 0.7831818 |
| SAATFITL | ASSSKDTQY | 0.36151162 |
| SNYGLANLGV | ASSSKDTQY | 0.765075 |
| SYGGAYAVM | ASSSKDTQY | 0.60955364 |
| TTGVGGLTLG | ASSSKDTQY | 0.64698094 |
| TTQTIYI | ASSSKDTQY | 0.6037903 |
| VAAGAGLGAVI | ASSSKDTQY | 0.54318887 |
| VAAVVTLL | ASSSKDTQY | 0.7469735 |
| VGFLGGLI | ASSSKDTQY | 0.7285297 |
| VGVIPILL | ASSSKDTQY | 0.6970915 |
| VIFASSLI | ASSSKDTQY | 0.6916827 |
| VKGGDGGAAAG | ASSSKDTQY | 0.73682666 |
| VTKTFTNI | ASSSKDTQY | 0.7350112 |
| VVAAAILL | ASSSKDTQY | 0.49451 |
| YAALDVEL | ASSSKDTQY | 0.3379063 |
| LASVALLL | ASSSKDTQY | 0.5974984 |
| LDAVHRV | ASSSKDTQY | 0.6192163 |
| QGLSSSIF | ASSSKDTQY | 0.53910035 |
| RGTFANIRL | ASSSKDTQY | 0.42441577 |
| RQLWWGHRI | ASSSKDTQY | 0.71958935 |
| YGPPGTGKTLL | ASSSKDTQY | 0.79210067 |
| AALVGAPANVA | ASSSKDTQY | 0.5764526 |
| AATFRRTE | ASSSKDTQY | 0.8189504 |
| AGALGGGI | ASSSKDTQY | 0.39999935 |
| AGGAGGAGGIGAG | ASSSKDTQY | 0.72895014 |
| AGSGALGL | ASSSKDTQY | 0.7604912 |
| AQIADQI | ASSSKDTQY | 0.43413323 |
| ASAVAVTYL | ASSSKDTQY | 0.672765 |
| ASTVLGGTGG | ASSSKDTQY | 0.6317682 |

| | | |
|---|---|---|
| AVGGEPAK | ASSSKDTQY | 0.48015678 |
| CEQGTFGGTC | ASSSKDTQY | 0.74647504 |
| DEGKAPAG | ASSSKDTQY | 0.5540155 |
| DTVVFPAL | ASSSKDTQY | 0.7219036 |
| FEGITMAG | ASSSKDTQY | 0.5097325 |
| FPLVVQD | ASSSKDTQY | 0.42861253 |
| GAFTTPQ | ASSSKDTQY | 0.70093334 |
| GFGGFGTAGGVGGA | ASSSKDTQY | 0.7292055 |
| GFLAQFL | ASSSKDTQY | 0.69005185 |
| GGAGGAGGNAGVA | ASSSKDTQY | 0.8355577 |
| GGVGTAGAGGAGGAGG | ASSSKDTQY | 0.48799628 |
| GNGGGGDAGNAGSGGN | ASSSKDTQY | 0.789408 |
| GTGGAGGAGNDAGS | ASSSKDTQY | 0.780342 |
| GTTNTGWFNS | ASSSKDTQY | 0.62139773 |
| IFMGPLFTL | ASSSKDTQY | 0.6117609 |
| ILAHAIEAI | ASSSKDTQY | 0.72912586 |
| IRSAIREAL | ASSSKDTQY | 0.5336714 |
| ITFAGGG | ASSSKDTQY | 0.69841903 |
| ITGMSGHQEV | ASSSKDTQY | 0.80115604 |
| IVAGLACTL | ASSSKDTQY | 0.30937237 |
| IVGGGLAGA | ASSSKDTQY | 0.85047853 |
| IVTGVLNI | ASSSKDTQY | 0.7958908 |
| LAQQGCI | ASSSKDTQY | 0.7768862 |
| LCLHVEN | ASSSKDTQY | 0.5619637 |
| LDAISDGS | ASSSKDTQY | 0.2888509 |
| LGGFAAML | ASSSKDTQY | 0.6720923 |
| LGGKNPMI | ASSSKDTQY | 0.4963711 |
| LGPKGVAAIL | ASSSKDTQY | 0.85071266 |
| LMVGVSI | ASSSKDTQY | 0.8054174 |
| LVGAAFVL | ASSSKDTQY | 0.5484032 |
| LVLAVAPL | ASSSKDTQY | 0.8157809 |
| MAAIAVIDAA | ASSSKDTQY | 0.8154618 |
| NNGINAELQ | ASSSKDTQY | 0.38538712 |
| RGEPLVLVHG | ASSSKDTQY | 0.7216116 |
| RGGAGGAAGLI | ASSSKDTQY | 0.69235504 |
| RSYRMHPAV | ASSSKDTQY | 0.63962364 |
| SAAHLLAGQLA | ASSSKDTQY | 0.29236513 |
| SAAVGSY | ASSSKDTQY | 0.76550007 |
| SPHAVLHRI | ASSSKDTQY | 0.67815727 |
| SQIAKVL | ASSSKDTQY | 0.18631712 |
| TALGHLVP | ASSSKDTQY | 0.79721415 |
| TAPTTSGAA | ASSSKDTQY | 0.5705823 |
| TDGDDGNGGSPGT | ASSSKDTQY | 0.6216821 |
| TDGLGEN | ASSSKDTQY | 0.8177065 |
| TGGPPLPGQA | ASSSKDTQY | 0.72816336 |
| TIPAITI | ASSSKDTQY | 0.6271468 |
| TIPGFTI | ASSSKDTQY | 0.79386836 |
| TSIASGALA | ASSSKDTQY | 0.8505512 |
| TVTLPGAAI | ASSSKDTQY | 0.4846337 |
| VASGLSAALP | ASSSKDTQY | 0.73333824 |
| VGAGVIGI | ASSSKDTQY | 0.85515463 |

| | | |
|---|---|---|
| VGPKRFGV | ASSSKDTQY | 0.7629409 |
| VNVAFPD | ASSSKDTQY | 0.8571697 |
| VSMTNSGVS | ASSSKDTQY | 0.8547386 |
| VTDVLGDLSV | ASSSKDTQY | 0.543603 |
| VVVVGAGFAGL | ASSSKDTQY | 0.84118086 |
| VYTVGTVV | ASSSKDTQY | 0.8483857 |
| WTTFSGP | ASSSKDTQY | 0.67980623 |
| YDELVEAF | ASSSKDTQY | 0.7118915 |
| YGTGGAGGNGG | ASSSKDTQY | 0.33192247 |
| YTNVMAV | ASSSKDTQY | 0.71691185 |
| FLEQVHQL | ASSSKDTQY | 0.5103475 |
| GLSLLAVAV | ASSSKDTQY | 0.79330075 |
| LEQVHQL | ASSSKDTQY | 0.8433906 |
| LNAAGAI | ASSSKDTQY | 0.7202735 |
| PAVHASL | ASSSKDTQY | 0.3200546 |
| TELEDDA | ASSSKDTQY | 0.28906864 |
| TPPGPPN | ASSSKDTQY | 0.8298183 |
| TPPPPPPP | ASSSKDTQY | 0.3167721 |
| TQSPALI | ASSPTGGENSPLY | 0.39294684 |
| EFAAAFPQ | ASSPTGGENSPLY | 0.625119 |
| LLSGNLDVL | ASSPTGGENSPLY | 0.8332018 |
| GPANLGNYNI | ASSPTGGENSPLY | 0.8375766 |
| TLGPTLP | ASSPTGGENSPLY | 0.6183053 |
| VYAYSSCCK | ASSPTGGENSPLY | 0.84858286 |
| APAPAPAPNI | ASSPTGGENSPLY | 0.84284246 |
| PGGPVPIP | ASSPTGGENSPLY | 0.8469092 |
| FLTVALEP | ASSPTGGENSPLY | 0.7010728 |
| SQYFGANL | ASSPTGGENSPLY | 0.35938233 |
| GGVKPAA | ASSPTGGENSPLY | 0.8448193 |
| GGTGGAGGAGGW | ASSPTGGENSPLY | 0.6869079 |
| GVGGVAGLGGAGG | ASSPTGGENSPLY | 0.81591547 |
| VLLVLATTI | ASSPTGGENSPLY | 0.8138578 |
| IGIFAAML | ASSPTGGENSPLY | 0.72231066 |
| AVGDLID | ASSPTGGENSPLY | 0.5535275 |
| NISPLPL | ASSPTGGENSPLY | 0.7865193 |
| VTAVFVAI | ASSPTGGENSPLY | 0.8483356 |
| SGDDNDAAL | ASSPTGGENSPLY | 0.4545862 |
| AVAALAHS | ASSPTGGENSPLY | 0.35692424 |
| GGVTVMATP | ASSPTGGENSPLY | 0.71889377 |
| AGLAANVRD | ASSPTGGENSPLY | 0.6995543 |
| TNRPDLIDPALLRPG | ASSPTGGENSPLY | 0.8542595 |
| LANLGVTAL | ASSPTGGENSPLY | 0.74405444 |
| TGERIEL | ASSPTGGENSPLY | 0.740645 |
| NGEELAA | ASSPTGGENSPLY | 0.80565816 |
| VLSLVWG | ASSPTGGENSPLY | 0.80264103 |
| IGYALWL | ASSPTGGENSPLY | 0.7245564 |
| LSLLLAPL | ASSPTGGENSPLY | 0.7061072 |
| TLDEALK | ASSPTGGENSPLY | 0.63143957 |
| TDGAGTAEGG | ASSPTGGENSPLY | 0.8389137 |
| NAPGGAGGNGG | ASSPTGGENSPLY | 0.8223651 |
| VSVFMGMGGNF | ASSPTGGENSPLY | 0.7941103 |

| | | |
|---|---|---|
| LASYLAAL | ASSPTGGENSPLY | 0.8389468 |
| LVGGTPAL | ASSPTGGENSPLY | 0.77297604 |
| VRNADVL | ASSPTGGENSPLY | 0.73354954 |
| TCSSPFLVSAM | ASSPTGGENSPLY | 0.5095371 |
| VHITDAE | ASSPTGGENSPLY | 0.66592705 |
| EYAAPSF | ASSPTGGENSPLY | 0.7045256 |
| VVDAFHGGL | ASSPTGGENSPLY | 0.79446304 |
| ASVVDPQ | ASSPTGGENSPLY | 0.81192625 |
| SYNEFGGLS | ASSPTGGENSPLY | 0.7899823 |
| DPDGSVY | ASSPTGGENSPLY | 0.57909894 |
| MVSTIHATE | ASSPTGGENSPLY | 0.7955605 |
| LAAGVADPV | ASSPTGGENSPLY | 0.84341455 |
| TKEQIAA | ASSPTGGENSPLY | 0.460751 |
| SLASPAAL | ASSPTGGENSPLY | 0.72751486 |
| IGLVFYGV | ASSPTGGENSPLY | 0.8588567 |
| FLLDASNL | ASSPTGGENSPLY | 0.8092571 |
| YQVGNFP | ASSPTGGENSPLY | 0.67213005 |
| ESLLSQST | ASSPTGGENSPLY | 0.7005012 |
| LFTVGFAAVG | ASSPTGGENSPLY | 0.85268974 |
| ATPPTPT | ASSPTGGENSPLY | 0.7369729 |
| TDLATLG | ASSPTGGENSPLY | 0.848353 |
| SAGGTGR | ASSPTGGENSPLY | 0.28505886 |
| GFSNSGS | ASSPTGGENSPLY | 0.65752655 |
| GGNGGSGGTSV | ASSPTGGENSPLY | 0.64547527 |
| GTSVATGGAGN | ASSPTGGENSPLY | 0.15676945 |
| NGGNGGSGGTS | ASSPTGGENSPLY | 0.8398573 |
| AVVKFPH | ASSPTGGENSPLY | 0.52633977 |
| TPPPPPP | ASSPTGGENSPLY | 0.79608774 |
| AAPAAAAAPA | ASSPTGGENSPLY | 0.5880736 |
| IALNEDL | ASSPTGGENSPLY | 0.84448963 |
| IVNVSSM | ASSPTGGENSPLY | 0.78690255 |
| TGIPPAPR | ASSPTGGENSPLY | 0.66807127 |
| FDLGGGTF | ASSPTGGENSPLY | 0.51153517 |
| FDLGGGTFDVS | ASSPTGGENSPLY | 0.7841927 |
| NEPTAAA | ASSPTGGENSPLY | 0.81662095 |
| NEPTAAALA | ASSPTGGENSPLY | 0.8492376 |
| GSIINISSI | ASSPTGGENSPLY | 0.30117375 |
| VAPGNAGTA | ASSPTGGENSPLY | 0.814214 |
| TGGGASLE | ASSPTGGENSPLY | 0.7024852 |
| VAVYNHYKRI | ASSPTGGENSPLY | 0.49266335 |
| TDPAPATT | ASSPTGGENSPLY | 0.2445471 |
| TNRPDILDPALLRPG | ASSPTGGENSPLY | 0.6878168 |
| LTTTTTS | ASSPTGGENSPLY | 0.7512529 |
| PTGSGKTL | ASSPTGGENSPLY | 0.7948253 |
| TGSGKTL | ASSPTGGENSPLY | 0.84490716 |
| TIGHVDHGK | ASSPTGGENSPLY | 0.5313872 |
| TIGHVDHGKTT | ASSPTGGENSPLY | 0.8165076 |
| GHVDHGKTT | ASSPTGGENSPLY | 0.7217865 |
| AQFEHTLL | ASSPTGGENSPLY | 0.106985815 |
| IVNQDEL | ASSPTGGENSPLY | 0.62273186 |
| TNGSQFF | ASSPTGGENSPLY | 0.35160893 |

| | | |
|---|---|---|
| TGKGIVE | ASSPTGGENSPLY | 0.8280134 |
| LISNASDA | ASSPTGGENSPLY | 0.48052764 |
| AELRTLGELTKIAKAHGAQ | ASSPTGGENSPLY | 0.7238888 |
| APNAVAAAEPSW | ASSPTGGENSPLY | 0.67206347 |
| APQVAGAATAVL | ASSPTGGENSPLY | 0.81864166 |
| ARRDPAAAAAILPTDA | ASSPTGGENSPLY | 0.7678716 |
| DAAARDVLAEAGLAEYF | ASSPTGGENSPLY | 0.5327338 |
| DDQIVLSAKANLAKVEQ | ASSPTGGENSPLY | 0.47434962 |
| DELLPDTGEDSDEDGH | ASSPTGGENSPLY | 0.7933696 |
| ELPEAPANRATVPQD | ASSPTGGENSPLY | 0.7856997 |
| FGQSKIERVTGIG | ASSPTGGENSPLY | 0.54464024 |
| GAGGTGSTAGGAGGAGGAGGLY | ASSPTGGENSPLY | 0.6263459 |
| GHGGTGAAGGNGGTGGAGGSGI | ASSPTGGENSPLY | 0.8464793 |
| GNGGNGGAGGEGGPGGAGGAGA | ASSPTGGENSPLY | 0.25584084 |
| GSTAGNGGNGGNATNGTIAGQP | ASSPTGGENSPLY | 0.83423746 |
| GTGGDGGNGGNGGTGG | ASSPTGGENSPLY | 0.4424978 |
| GVRRPAGIPADAPV | ASSPTGGENSPLY | 0.6709264 |
| GYRQVTDAHLLTLARR | ASSPTGGENSPLY | 0.85164464 |
| IAKMGQDGHDRG | ASSPTGGENSPLY | 0.80486894 |
| KAEPSIEIQYT | ASSPTGGENSPLY | 0.70808375 |
| KPVRIRMAADEDTVAVT | ASSPTGGENSPLY | 0.6034887 |
| MPPRIAGMRLLVIKPEP | ASSPTGGENSPLY | 0.48348743 |
| MKFYAVALFRSRGVPPTR | ASSPTGGENSPLY | 0.57789415 |
| NAGGFGILGAVAHSP | ASSPTGGENSPLY | 0.5241778 |
| NEGREITLAMAEM | ASSPTGGENSPLY | 0.8370011 |
| NELVEWSHPLGHRWRWE | ASSPTGGENSPLY | 0.7310517 |
| NLGAGGTGGNGGNPG | ASSPTGGENSPLY | 0.3883659 |
| NTEWPKWETVMAADPVD | ASSPTGGENSPLY | 0.8409351 |
| PASAAPAETQP | ASSPTGGENSPLY | 0.8456219 |
| QQSSDGKNGVRAAEITGE | ASSPTGGENSPLY | 0.86177045 |
| RAPDAVTTSWWKE | ASSPTGGENSPLY | 0.5044249 |
| SGFGNVGAHTSGWFNQ | ASSPTGGENSPLY | 0.55545044 |
| SGVAGAVGQAASVGGLKV | ASSPTGGENSPLY | 0.67938864 |
| SNGGAGTGSAGGNGGT | ASSPTGGENSPLY | 0.6931244 |
| TGGAGGEGGAAG | ASSPTGGENSPLY | 0.4793773 |
| TGGDGGAAGAGGAPGHG | ASSPTGGENSPLY | 0.49753818 |
| TMPPLLAAIANAGRHGV | ASSPTGGENSPLY | 0.8378905 |
| TSSTAGNSGGA | ASSPTGGENSPLY | 0.87053496 |
| VPATVPVGILC | ASSPTGGENSPLY | 0.65617764 |
| VQRPDSGGVVLTGRIS | ASSPTGGENSPLY | 0.27282685 |
| WTPNPVPATKGVPAGLVQAT | ASSPTGGENSPLY | 0.6573946 |
| YALKRALAKFSDRFDVVII | ASSPTGGENSPLY | 0.52344334 |
| YLIGSVIV | ASSPTGGENSPLY | 0.5187329 |
| YGQLVAVID | ASSPTGGENSPLY | 0.7342703 |
| VTVGGMTLPA | ASSPTGGENSPLY | 0.31435114 |
| VLALPAV | ASSPTGGENSPLY | 0.66235876 |
| VIPPTLNYET | ASSPTGGENSPLY | 0.8579937 |
| VGMGGMHPSSG | ASSPTGGENSPLY | 0.7414385 |
| VGLAGGAATA | ASSPTGGENSPLY | 0.7431221 |
| VGGAGGAGGSGP | ASSPTGGENSPLY | 0.8442525 |
| VFAAFMV | ASSPTGGENSPLY | 0.78113234 |

| | | |
|---|---|---|
| TLYDKLR | ASSPTGGENSPLY | 0.84921086 |
| TLAVFTVL | ASSPTGGENSPLY | 0.8465115 |
| TGSVQPARV | ASSPTGGENSPLY | 0.7171665 |
| TGGGAGSGGT | ASSPTGGENSPLY | 0.7409634 |
| SHLETSL | ASSPTGGENSPLY | 0.8508656 |
| SFTTGMT | ASSPTGGENSPLY | 0.63739717 |
| QVEVAKRMGVRQPTVS | ASSPTGGENSPLY | 0.7509072 |
| QGSGGPALAM | ASSPTGGENSPLY | 0.7253175 |
| MLAPSNP | ASSPTGGENSPLY | 0.79546374 |
| LVNNAAI | ASSPTGGENSPLY | 0.7379434 |
| LTGEINTLA | ASSPTGGENSPLY | 0.85660744 |
| LSIFGPV | ASSPTGGENSPLY | 0.16726224 |
| LLALLANL | ASSPTGGENSPLY | 0.5163468 |
| LISNASDALDKLR | ASSPTGGENSPLY | 0.8497716 |
| LINEAVP | ASSPTGGENSPLY | 0.16669378 |
| LGYAFSGL | ASSPTGGENSPLY | 0.33644873 |
| LDFVAVAL | ASSPTGGENSPLY | 0.7653817 |
| LAQPGAE | ASSPTGGENSPLY | 0.8453446 |
| ISNLGAVTGF | ASSPTGGENSPLY | 0.24687316 |
| HQEDPGRI | ASSPTGGENSPLY | 0.7133608 |
| GTPGEPTP | ASSPTGGENSPLY | 0.26858854 |
| GFFNSGS | ASSPTGGENSPLY | 0.83821505 |
| GAGGGGLTAGAN | ASSPTGGENSPLY | 0.7064823 |
| FVDALVNM | ASSPTGGENSPLY | 0.30133593 |
| FLTVAPEL | ASSPTGGENSPLY | 0.8149545 |
| FETLTAL | ASSPTGGENSPLY | 0.462124 |
| DVINPIEL | ASSPTGGENSPLY | 0.81729245 |
| APVSGAPV | ASSPTGGENSPLY | 0.8483686 |
| ALVGGLAW | ASSPTGGENSPLY | 0.86982733 |
| ALAAAAATAI | ASSPTGGENSPLY | 0.36794984 |
| AFGLGLLL | ASSPTGGENSPLY | 0.8220249 |
| AFFAAASL | ASSPTGGENSPLY | 0.6037458 |
| AAAALPAFSPP | ASSPTGGENSPLY | 0.87339246 |
| VVLAPAL | ASSPTGGENSPLY | 0.6465306 |
| TIGHVDHGKTTL | ASSPTGGENSPLY | 0.24506855 |
| LISNASDALDK | ASSPTGGENSPLY | 0.8161324 |
| LISNASDAL | ASSPTGGENSPLY | 0.77087516 |
| ISNASDALDK | ASSPTGGENSPLY | 0.6461515 |
| NQAETLVYQTE | ASSPTGGENSPLY | 0.8403925 |
| TGGTGGTAGAG | ASSPTGGENSPLY | 0.70796883 |
| TGSTAGGAGGA | ASSPTGGENSPLY | 0.6066562 |
| DPLAVPVAAGPV | ASSPTGGENSPLY | 0.82649297 |
| GGTGGTPGNGGN | ASSPTGGENSPLY | 0.77362645 |
| PAGLAGVDPATL | ASSPTGGENSPLY | 0.62830454 |
| GNNLVVEAPHAGASV | ASSPTGGENSPLY | 0.8411669 |
| KVTRIMSASTLEQPA | ASSPTGGENSPLY | 0.7577678 |
| LGFALPIALPRRTVV | ASSPTGGENSPLY | 0.39871824 |
| TRLGNLTARKLVVLV | ASSPTGGENSPLY | 0.8681263 |
| WSQQRRLIYFPSAGP | ASSPTGGENSPLY | 0.6845578 |
| GPNGNPDPMAAAVDIR | ASSPTGGENSPLY | 0.815685 |
| TQRCVSVADLRAAASR | ASSPTGGENSPLY | 0.73734415 |

| | | |
|---|---|---|
| QFVLDVAEHPVEVLTETV | ASSPTGGENSPLY | 0.8594698 |
| GGAGGTGGAGGNGGTQAGDG | ASSPTGGENSPLY | 0.599668 |
| PLLATLRAIAVDVDAKPAQV | ASSPTGGENSPLY | 0.74780405 |
| APKFRTGGYMIDFWGVGYQVAK | ASSPTGGENSPLY | 0.7604404 |
| GAGGAGGAGGAAGGPGGTGGTG | ASSPTGGENSPLY | 0.81861913 |
| IGALGLLATDVAPAGSVAFGVA | ASSPTGGENSPLY | 0.71895856 |
| AGGAGGIGGNGGAGGAGGVLMG | ASSPTGGENSPLY | 0.8165215 |
| ADELIGLL | ASSPTGGENSPLY | 0.7820401 |
| ADRALKA | ASSPTGGENSPLY | 0.8382758 |
| AGAVGQVGGAGGA | ASSPTGGENSPLY | 0.71266824 |
| AGGGGGLGGAP | ASSPTGGENSPLY | 0.674962 |
| AGGLLYGNGGAG | ASSPTGGENSPLY | 0.8645506 |
| AGLESIGL | ASSPTGGENSPLY | 0.37278733 |
| AGLKTGII | ASSPTGGENSPLY | 0.73983115 |
| ASLAALLL | ASSPTGGENSPLY | 0.82878613 |
| ASVINISSI | ASSPTGGENSPLY | 0.82105863 |
| ATPPGPA | ASSPTGGENSPLY | 0.68012905 |
| ATVASALV | ASSPTGGENSPLY | 0.8276941 |
| ATVASLIL | ASSPTGGENSPLY | 0.7393271 |
| AVNGPVLIRH | ASSPTGGENSPLY | 0.8103869 |
| DGAAGAVGI | ASSPTGGENSPLY | 0.7499022 |
| DPLLHGI | ASSPTGGENSPLY | 0.673718 |
| ELGAALGAL | ASSPTGGENSPLY | 0.56303144 |
| ELGRTQAV | ASSPTGGENSPLY | 0.7295611 |
| FVAGGYLS | ASSPTGGENSPLY | 0.82715255 |
| GASGGAGGAGGW | ASSPTGGENSPLY | 0.78484213 |
| GGAGAGGDGGAGGI | ASSPTGGENSPLY | 0.33295503 |
| GTGHNGSQPGMGGNG | ASSPTGGENSPLY | 0.81251705 |
| GYAAASASAS | ASSPTGGENSPLY | 0.8111002 |
| HQVDVAAVQQG | ASSPTGGENSPLY | 0.8665943 |
| IGAAIAHT | ASSPTGGENSPLY | 0.73542833 |
| IGMIEVAQL | ASSPTGGENSPLY | 0.8193503 |
| IRLAPEHTSAMD | ASSPTGGENSPLY | 0.7592262 |
| IYNVSVL | ASSPTGGENSPLY | 0.7643503 |
| KQGFLGM | ASSPTGGENSPLY | 0.35006988 |
| LESVITQL | ASSPTGGENSPLY | 0.47141933 |
| LGLGALVVV | ASSPTGGENSPLY | 0.7404843 |
| LIGNGGNGGSGGI | ASSPTGGENSPLY | 0.5614519 |
| LNAINAPTL | ASSPTGGENSPLY | 0.835053 |
| LQLLNTGV | ASSPTGGENSPLY | 0.8241773 |
| LQTSIEF | ASSPTGGENSPLY | 0.8285382 |
| MSFAVSEL | ASSPTGGENSPLY | 0.84834546 |
| NTGALGL | ASSPTGGENSPLY | 0.8495651 |
| PPGAGGAGGAGTT | ASSPTGGENSPLY | 0.81208473 |
| QLVGLGI | ASSPTGGENSPLY | 0.6322601 |
| RFLEQVHQL | ASSPTGGENSPLY | 0.710814 |
| RGVGYVLR | ASSPTGGENSPLY | 0.32159153 |
| SATGVFAGVI | ASSPTGGENSPLY | 0.5164234 |
| SLPAITLP | ASSPTGGENSPLY | 0.6185647 |
| STALAFTV | ASSPTGGENSPLY | 0.72281235 |
| SVLTVGLGK | ASSPTGGENSPLY | 0.14449267 |

| | | |
|---|---|---|
| TALAGAGSGI | ASSPTGGENSPLY | 0.8166294 |
| TGLPFTGI | ASSPTGGENSPLY | 0.8419061 |
| VAEHVTV | ASSPTGGENSPLY | 0.7751219 |
| VDIVLNSL | ASSPTGGENSPLY | 0.79958045 |
| VLTGGGAVL | ASSPTGGENSPLY | 0.78158903 |
| VNVGLAYGI | ASSPTGGENSPLY | 0.6226752 |
| VSSAAVGAV | ASSPTGGENSPLY | 0.806632 |
| VTSVYEAGV | ASSPTGGENSPLY | 0.65459394 |
| WVALDRGA | ASSPTGGENSPLY | 0.8356091 |
| YLVGSVVI | ASSPTGGENSPLY | 0.8413056 |
| GPPGTGKTLL | ASSPTGGENSPLY | 0.76726854 |
| GVGMTKF | ASSPTGGENSPLY | 0.56338274 |
| IGIGGSDL | ASSPTGGENSPLY | 0.7662037 |
| IGTLLAEL | ASSPTGGENSPLY | 0.62227595 |
| IYTSGTT | ASSPTGGENSPLY | 0.5967071 |
| LSFPAGV | ASSPTGGENSPLY | 0.6622699 |
| LYTSGTT | ASSPTGGENSPLY | 0.7627555 |
| VGAFPFGF | ASSPTGGENSPLY | 0.4401365 |
| VGAGTVEF | ASSPTGGENSPLY | 0.79059935 |
| VGKSTLLL | ASSPTGGENSPLY | 0.73718274 |
| AALAHAGGGIV | ASSPTGGENSPLY | 0.7885239 |
| AALVAVSLL | ASSPTGGENSPLY | 0.5106828 |
| AAVSRAVAAL | ASSPTGGENSPLY | 0.8209651 |
| AGGSSGIGGF | ASSPTGGENSPLY | 0.7924366 |
| AGLTVANANG | ASSPTGGENSPLY | 0.36629117 |
| AGLVGAVL | ASSPTGGENSPLY | 0.79308736 |
| ALSQAFTV | ASSPTGGENSPLY | 0.68194497 |
| ALTFSVSL | ASSPTGGENSPLY | 0.7451345 |
| ALTGAALVAL | ASSPTGGENSPLY | 0.8249969 |
| ASFSIGAGVL | ASSPTGGENSPLY | 0.74145013 |
| DGSVYKI | ASSPTGGENSPLY | 0.5368308 |
| EQLAEIV | ASSPTGGENSPLY | 0.6277683 |
| FAVSELTVM | ASSPTGGENSPLY | 0.7428582 |
| GAAPLAAGL | ASSPTGGENSPLY | 0.67695606 |
| GADGGDGGSGGAGG | ASSPTGGENSPLY | 0.19218022 |
| GFGLDTGGAGG | ASSPTGGENSPLY | 0.79961014 |
| GLASGGLL | ASSPTGGENSPLY | 0.4031706 |
| GRIAGAAVLL | ASSPTGGENSPLY | 0.5989901 |
| GSAGSLNTSF | ASSPTGGENSPLY | 0.6542257 |
| GTGQIGI | ASSPTGGENSPLY | 0.8334297 |
| GTLIALIL | ASSPTGGENSPLY | 0.8129543 |
| IAATMGFVVG | ASSPTGGENSPLY | 0.6417111 |
| IFAQALL | ASSPTGGENSPLY | 0.7769929 |
| IGAAVALGF | ASSPTGGENSPLY | 0.7456509 |
| IGLTPLAL | ASSPTGGENSPLY | 0.82633865 |
| ILGSNTSTL | ASSPTGGENSPLY | 0.77601975 |
| IVFSLTVL | ASSPTGGENSPLY | 0.75661695 |
| LASKATGF | ASSPTGGENSPLY | 0.7872356 |
| LASVALIL | ASSPTGGENSPLY | 0.6166096 |
| LFAVGLTL | ASSPTGGENSPLY | 0.5442273 |
| LFNGGIGTYI | ASSPTGGENSPLY | 0.55258095 |

| | | |
|---|---|---|
| LTLGAGII | ASSPTGGENSPLY | 0.51816213 |
| LVAAALGDI | ASSPTGGENSPLY | 0.47406092 |
| NTAFVQLGI | ASSPTGGENSPLY | 0.7802019 |
| QLFSGAEL | ASSPTGGENSPLY | 0.8440577 |
| SAATFITL | ASSPTGGENSPLY | 0.72005475 |
| SNYGLANLGV | ASSPTGGENSPLY | 0.25694233 |
| SYGGAYAVM | ASSPTGGENSPLY | 0.6607153 |
| TTGVGGLTLG | ASSPTGGENSPLY | 0.7514249 |
| TTQTIYI | ASSPTGGENSPLY | 0.8243424 |
| VAAGAGLGAVI | ASSPTGGENSPLY | 0.84470737 |
| VAAVVTLL | ASSPTGGENSPLY | 0.8630084 |
| VGFLGGLI | ASSPTGGENSPLY | 0.82504106 |
| VGVIPILL | ASSPTGGENSPLY | 0.6448267 |
| VIFASSLI | ASSPTGGENSPLY | 0.8194858 |
| VKGGDGGAAAG | ASSPTGGENSPLY | 0.5838842 |
| VTKTFTNI | ASSPTGGENSPLY | 0.70373255 |
| VVAAAILL | ASSPTGGENSPLY | 0.7279892 |
| YAALDVEL | ASSPTGGENSPLY | 0.8141267 |
| LASVALLL | ASSPTGGENSPLY | 0.5856289 |
| LDAVHRV | ASSPTGGENSPLY | 0.6678681 |
| QGLSSSIF | ASSPTGGENSPLY | 0.35159123 |
| RGTFANIRL | ASSPTGGENSPLY | 0.82817036 |
| RQLWWGHRI | ASSPTGGENSPLY | 0.6453992 |
| YGPPGTGKTLL | ASSPTGGENSPLY | 0.49354982 |
| AALVGAPANVA | ASSPTGGENSPLY | 0.6666451 |
| AATFRRTE | ASSPTGGENSPLY | 0.8642567 |
| AGALGGGI | ASSPTGGENSPLY | 0.8594569 |
| AGGAGGAGGIGAG | ASSPTGGENSPLY | 0.6014222 |
| AGSGALGL | ASSPTGGENSPLY | 0.5402653 |
| AQIADQI | ASSPTGGENSPLY | 0.82863975 |
| ASAVAVTYL | ASSPTGGENSPLY | 0.6587451 |
| ASTVLGGTGG | ASSPTGGENSPLY | 0.7873425 |
| AVGGEPAK | ASSPTGGENSPLY | 0.7587149 |
| CEQGTFGGTC | ASSPTGGENSPLY | 0.7185689 |
| DEGKAPAG | ASSPTGGENSPLY | 0.5110634 |
| DTVVFPAL | ASSPTGGENSPLY | 0.8285486 |
| FEGITMAG | ASSPTGGENSPLY | 0.70537066 |
| FPLVVQD | ASSPTGGENSPLY | 0.7466668 |
| GAFTTPQ | ASSPTGGENSPLY | 0.61480856 |
| GFGGFGTAGGVGGA | ASSPTGGENSPLY | 0.8481529 |
| GFLAQFL | ASSPTGGENSPLY | 0.5100083 |
| GGAGGAGGNAGVA | ASSPTGGENSPLY | 0.53382343 |
| GGVGTAGAGGAGGAGG | ASSPTGGENSPLY | 0.2091673 |
| GNGGGGDAGNAGSGGN | ASSPTGGENSPLY | 0.6470202 |
| GTGGAGGAGNDAGS | ASSPTGGENSPLY | 0.82126665 |
| GTTNTGWFNS | ASSPTGGENSPLY | 0.860788 |
| IFMGPLFTL | ASSPTGGENSPLY | 0.7917568 |
| ILAHAIEAI | ASSPTGGENSPLY | 0.77108335 |
| IRSAIREAL | ASSPTGGENSPLY | 0.85394734 |
| ITFAGGG | ASSPTGGENSPLY | 0.6433071 |
| ITGMSGHQEV | ASSPTGGENSPLY | 0.2678393 |

| | | |
|---|---|---|
| IVAGLACTL | ASSPTGGENSPLY | 0.78268677 |
| IVGGGLAGA | ASSPTGGENSPLY | 0.7577263 |
| IVTGVLNI | ASSPTGGENSPLY | 0.85965776 |
| LAQQGCI | ASSPTGGENSPLY | 0.825277 |
| LCLHVEN | ASSPTGGENSPLY | 0.36030978 |
| LDAISDGS | ASSPTGGENSPLY | 0.73695856 |
| LGGFAAML | ASSPTGGENSPLY | 0.7733155 |
| LGGKNPMI | ASSPTGGENSPLY | 0.36450428 |
| LGPKGVAAIL | ASSPTGGENSPLY | 0.5488044 |
| LMVGVSI | ASSPTGGENSPLY | 0.5433836 |
| LVGAAFVL | ASSPTGGENSPLY | 0.5011046 |
| LVLAVAPL | ASSPTGGENSPLY | 0.72257185 |
| MAAIAVIDAA | ASSPTGGENSPLY | 0.62458384 |
| NNGINAELQ | ASSPTGGENSPLY | 0.72357714 |
| RGEPLVLVHG | ASSPTGGENSPLY | 0.5907094 |
| RGGAGGAAGLI | ASSPTGGENSPLY | 0.5065509 |
| RSYRMHPAV | ASSPTGGENSPLY | 0.62055314 |
| SAAHLLAGQLA | ASSPTGGENSPLY | 0.69102263 |
| SAAVGSY | ASSPTGGENSPLY | 0.81893146 |
| SPHAVLHRI | ASSPTGGENSPLY | 0.8374502 |
| SQIAKVL | ASSPTGGENSPLY | 0.68754864 |
| TALGHLVP | ASSPTGGENSPLY | 0.73489344 |
| TAPTTSGAA | ASSPTGGENSPLY | 0.8456507 |
| TDGDDGNGGSPGT | ASSPTGGENSPLY | 0.43365687 |
| TDGLGEN | ASSPTGGENSPLY | 0.49669692 |
| TGGPPLPGQA | ASSPTGGENSPLY | 0.8030697 |
| TIPAITI | ASSPTGGENSPLY | 0.6570436 |
| TIPGFTI | ASSPTGGENSPLY | 0.84362316 |
| TSIASGALA | ASSPTGGENSPLY | 0.76547176 |
| TVTLPGAAI | ASSPTGGENSPLY | 0.75914544 |
| VASGLSAALP | ASSPTGGENSPLY | 0.7868764 |
| VGAGVIGI | ASSPTGGENSPLY | 0.86261654 |
| VGPKRFGV | ASSPTGGENSPLY | 0.78964347 |
| VNVAFPD | ASSPTGGENSPLY | 0.5760958 |
| VSMTNSGVS | ASSPTGGENSPLY | 0.8074361 |
| VTDVLGDLSV | ASSPTGGENSPLY | 0.32997927 |
| VVVVGAGFAGL | ASSPTGGENSPLY | 0.79125404 |
| VYTVGTVV | ASSPTGGENSPLY | 0.8161932 |
| WTTFSGP | ASSPTGGENSPLY | 0.5277356 |
| YDELVEAF | ASSPTGGENSPLY | 0.60190856 |
| YGTGGAGGNGG | ASSPTGGENSPLY | 0.8604793 |
| YTNVMAV | ASSPTGGENSPLY | 0.8372737 |
| FLEQVHQL | ASSPTGGENSPLY | 0.53081477 |
| GLSLLAVAV | ASSPTGGENSPLY | 0.5562633 |
| LEQVHQL | ASSPTGGENSPLY | 0.63206357 |
| LNAAGAI | ASSPTGGENSPLY | 0.63676906 |
| PAVHASL | ASSPTGGENSPLY | 0.25302348 |
| TELEDDA | ASSPTGGENSPLY | 0.7990142 |
| TPPGPPN | ASSPTGGENSPLY | 0.6763436 |
| TPPPPPPP | ASSPTGGENSPLY | 0.8373718 |
| TQSPALI | ASSPKLRNTLY | 0.77898467 |

| | | |
|---|---|---|
| EFAAAFPQ | ASSPKLRNTLY | 0.8358341 |
| LLSGNLDVL | ASSPKLRNTLY | 0.57233846 |
| GPANLGNYNI | ASSPKLRNTLY | 0.7209131 |
| TLGPTLP | ASSPKLRNTLY | 0.25538653 |
| VYAYSSCCK | ASSPKLRNTLY | 0.64992386 |
| APAPAPAPNI | ASSPKLRNTLY | 0.5032532 |
| PGGPVPIP | ASSPKLRNTLY | 0.71548617 |
| FLTVALEP | ASSPKLRNTLY | 0.55602336 |
| SQYFGANL | ASSPKLRNTLY | 0.8594069 |
| GGVKPAA | ASSPKLRNTLY | 0.6230061 |
| GGTGGAGGAGGW | ASSPKLRNTLY | 0.8059927 |
| GVGGVAGLGGAGG | ASSPKLRNTLY | 0.6516324 |
| VLLVLATTI | ASSPKLRNTLY | 0.36287338 |
| IGIFAAML | ASSPKLRNTLY | 0.63168216 |
| AVGDLID | ASSPKLRNTLY | 0.82498777 |
| NISPLPL | ASSPKLRNTLY | 0.43108177 |
| VTAVFVAI | ASSPKLRNTLY | 0.26029724 |
| SGDDNDAAL | ASSPKLRNTLY | 0.5274313 |
| AVAALAHS | ASSPKLRNTLY | 0.7606704 |
| GGVTVMATP | ASSPKLRNTLY | 0.86533415 |
| AGLAANVRD | ASSPKLRNTLY | 0.83733857 |
| TNRPDLIDPALLRPG | ASSPKLRNTLY | 0.715287 |
| LANLGVTAL | ASSPKLRNTLY | 0.8678181 |
| TGERIEL | ASSPKLRNTLY | 0.132782 |
| NGEELAA | ASSPKLRNTLY | 0.72063136 |
| VLSLVWG | ASSPKLRNTLY | 0.85372156 |
| IGYALWL | ASSPKLRNTLY | 0.7852552 |
| LSLLLAPL | ASSPKLRNTLY | 0.6954733 |
| TLDEALK | ASSPKLRNTLY | 0.8520262 |
| TDGAGTAEGG | ASSPKLRNTLY | 0.5698089 |
| NAPGGAGGNGG | ASSPKLRNTLY | 0.6907828 |
| VSVFMGMGGNF | ASSPKLRNTLY | 0.7644144 |
| LASYLAAL | ASSPKLRNTLY | 0.7915907 |
| LVGGTPAL | ASSPKLRNTLY | 0.8194004 |
| VRNADVL | ASSPKLRNTLY | 0.27778596 |
| TCSSPFLVSAM | ASSPKLRNTLY | 0.7976904 |
| VHITDAE | ASSPKLRNTLY | 0.86620784 |
| EYAAPSF | ASSPKLRNTLY | 0.674023 |
| VVDAFHGGL | ASSPKLRNTLY | 0.85361785 |
| ASVVDPQ | ASSPKLRNTLY | 0.7769279 |
| SYNEFGGLS | ASSPKLRNTLY | 0.7746929 |
| DPDGSVY | ASSPKLRNTLY | 0.5653185 |
| MVSTIHATE | ASSPKLRNTLY | 0.459369 |
| LAAGVADPV | ASSPKLRNTLY | 0.8166716 |
| TKEQIAA | ASSPKLRNTLY | 0.5258174 |
| SLASPAAL | ASSPKLRNTLY | 0.7389506 |
| IGLVFYGV | ASSPKLRNTLY | 0.7208414 |
| FLLDASNL | ASSPKLRNTLY | 0.874617 |
| YQVGNFP | ASSPKLRNTLY | 0.78666735 |
| ESLLSQST | ASSPKLRNTLY | 0.823433 |
| LFTVGFAAVG | ASSPKLRNTLY | 0.5655329 |

| | | |
|---|---|---|
| ATPPTPT | ASSPKLRNTLY | 0.7774616 |
| TDLATLG | ASSPKLRNTLY | 0.8147434 |
| SAGGTGR | ASSPKLRNTLY | 0.37866583 |
| GFSNSGS | ASSPKLRNTLY | 0.8423492 |
| GGNGGSGGTSV | ASSPKLRNTLY | 0.47709405 |
| GTSVATGGAGN | ASSPKLRNTLY | 0.7787436 |
| NGGNGGSGGTS | ASSPKLRNTLY | 0.764402 |
| AVVKFPH | ASSPKLRNTLY | 0.8434221 |
| TPPPPPP | ASSPKLRNTLY | 0.70576906 |
| AAPAAAAAPA | ASSPKLRNTLY | 0.818917 |
| IALNEDL | ASSPKLRNTLY | 0.5215249 |
| IVNVSSM | ASSPKLRNTLY | 0.8116169 |
| TGIPPAPR | ASSPKLRNTLY | 0.8483013 |
| FDLGGGTF | ASSPKLRNTLY | 0.7586124 |
| FDLGGGTFDVS | ASSPKLRNTLY | 0.86879784 |
| NEPTAAA | ASSPKLRNTLY | 0.8206163 |
| NEPTAAALA | ASSPKLRNTLY | 0.22021063 |
| GSIINISSI | ASSPKLRNTLY | 0.69839334 |
| VAPGNAGTA | ASSPKLRNTLY | 0.8514509 |
| TGGGASLE | ASSPKLRNTLY | 0.85904443 |
| VAVYNHYKRI | ASSPKLRNTLY | 0.82262564 |
| TDPAPATT | ASSPKLRNTLY | 0.7966512 |
| TNRPDILDPALLRPG | ASSPKLRNTLY | 0.6801021 |
| LTTTTTS | ASSPKLRNTLY | 0.8501444 |
| PTGSGKTL | ASSPKLRNTLY | 0.88379925 |
| TGSGKTL | ASSPKLRNTLY | 0.82694674 |
| TIGHVDHGK | ASSPKLRNTLY | 0.5082239 |
| TIGHVDHGKTT | ASSPKLRNTLY | 0.5162108 |
| GHVDHGKTT | ASSPKLRNTLY | 0.8050814 |
| AQFEHTLL | ASSPKLRNTLY | 0.8390486 |
| IVNQDEL | ASSPKLRNTLY | 0.38950568 |
| TNGSQFF | ASSPKLRNTLY | 0.69334555 |
| TGKGIVE | ASSPKLRNTLY | 0.7392783 |
| LISNASDA | ASSPKLRNTLY | 0.1976384 |
| AELRTLGELTKIAKAHGAQ | ASSPKLRNTLY | 0.81793994 |
| APNAVAAAEPSW | ASSPKLRNTLY | 0.848997 |
| APQVAGAATAVL | ASSPKLRNTLY | 0.7435495 |
| ARRDPAAAAAILPTDA | ASSPKLRNTLY | 0.3409677 |
| DAAARDVLAEAGLAEYF | ASSPKLRNTLY | 0.8444419 |
| DDQIVLSAKANLAKVEQ | ASSPKLRNTLY | 0.85795045 |
| DELLPDTGEDSDEDGH | ASSPKLRNTLY | 0.8518587 |
| ELPEAPANRATVPQD | ASSPKLRNTLY | 0.360376 |
| FGQSKIERVTGIG | ASSPKLRNTLY | 0.14251772 |
| GAGGTGSTAGGAGGAGGAGGLY | ASSPKLRNTLY | 0.79582536 |
| GHGGTGAAGGNGGTGGAGGSGI | ASSPKLRNTLY | 0.8186966 |
| GNGGNGGAGGEGGPGGAGGAGA | ASSPKLRNTLY | 0.6021962 |
| GSTAGNGGNGGNATNGTIAGQP | ASSPKLRNTLY | 0.46316266 |
| GTGGDGGNGGNGGTGG | ASSPKLRNTLY | 0.75690186 |
| GVRRPAGIPADAPV | ASSPKLRNTLY | 0.82611585 |
| GYRQVTDAHLLTLARR | ASSPKLRNTLY | 0.34752268 |
| IAKMGQDGHDRG | ASSPKLRNTLY | 0.44475758 |

| | | |
|---|---|---|
| KAEPSIEIQYT | ASSPKLRNTLY | 0.7817043 |
| KPVRIRMAADEDTVAVT | ASSPKLRNTLY | 0.8399333 |
| MPPRIAGMRLLVIKPEP | ASSPKLRNTLY | 0.49167877 |
| MKFYAVALFRSRGVPPTR | ASSPKLRNTLY | 0.6152856 |
| NAGGFGILGAVAHSP | ASSPKLRNTLY | 0.81536365 |
| NEGREITLAMAEM | ASSPKLRNTLY | 0.83136773 |
| NELVEWSHPLGHRWRWE | ASSPKLRNTLY | 0.7625324 |
| NLGAGGTGGNGGNPG | ASSPKLRNTLY | 0.35714173 |
| NTEWPKWETVMAADPVD | ASSPKLRNTLY | 0.82976437 |
| PASAAPAETQP | ASSPKLRNTLY | 0.39371806 |
| QQSSDGKNGVRAAEITGE | ASSPKLRNTLY | 0.83501506 |
| RAPDAVTTSWWKE | ASSPKLRNTLY | 0.7395292 |
| SGFGNVGAHTSGWFNQ | ASSPKLRNTLY | 0.75371337 |
| SGVAGAVGQAASVGGLKV | ASSPKLRNTLY | 0.6732693 |
| SNGGAGTGSAGGNGGT | ASSPKLRNTLY | 0.4325578 |
| TGGAGGEGGAAG | ASSPKLRNTLY | 0.3945297 |
| TGGDGGAAGAGGAPGHG | ASSPKLRNTLY | 0.5274222 |
| TMPPLLAAIANAGRHGV | ASSPKLRNTLY | 0.8320483 |
| TSSTAGNSGGA | ASSPKLRNTLY | 0.8232126 |
| VPATVPVGILC | ASSPKLRNTLY | 0.7999399 |
| VQRPDSGGVVLTGRIS | ASSPKLRNTLY | 0.779469 |
| WTPNPVPATKGVPAGLVQAT | ASSPKLRNTLY | 0.5275531 |
| YALKRALAKFSDRFDVVII | ASSPKLRNTLY | 0.8051456 |
| YLIGSVIV | ASSPKLRNTLY | 0.39247072 |
| YGQLVAVID | ASSPKLRNTLY | 0.79920363 |
| VTVGGMTLPA | ASSPKLRNTLY | 0.62974995 |
| VLALPAV | ASSPKLRNTLY | 0.79075384 |
| VIPPTLNYET | ASSPKLRNTLY | 0.6289561 |
| VGMGGMHPSSG | ASSPKLRNTLY | 0.77931476 |
| VGLAGGAATA | ASSPKLRNTLY | 0.7761327 |
| VGGAGGAGGSGP | ASSPKLRNTLY | 0.43102348 |
| VFAAFMV | ASSPKLRNTLY | 0.7137461 |
| TLYDKLR | ASSPKLRNTLY | 0.8367709 |
| TLAVFTVL | ASSPKLRNTLY | 0.783713 |
| TGSVQPARV | ASSPKLRNTLY | 0.31870157 |
| TGGGAGSGGT | ASSPKLRNTLY | 0.7939795 |
| SHLETSL | ASSPKLRNTLY | 0.57843804 |
| SFTTGMT | ASSPKLRNTLY | 0.7642668 |
| QVEVAKRMGVRQPTVS | ASSPKLRNTLY | 0.76748216 |
| QGSGGPALAM | ASSPKLRNTLY | 0.62616134 |
| MLAPSNP | ASSPKLRNTLY | 0.79710466 |
| LVNNAAI | ASSPKLRNTLY | 0.40918198 |
| LTGEINTLA | ASSPKLRNTLY | 0.62821543 |
| LSIFGPV | ASSPKLRNTLY | 0.65978056 |
| LLALLANL | ASSPKLRNTLY | 0.7871755 |
| LISNASDALDKLR | ASSPKLRNTLY | 0.8268891 |
| LINEAVP | ASSPKLRNTLY | 0.4111339 |
| LGYAFSGL | ASSPKLRNTLY | 0.7645174 |
| LDFVAVAL | ASSPKLRNTLY | 0.6586926 |
| LAQPGAE | ASSPKLRNTLY | 0.5225018 |
| ISNLGAVTGF | ASSPKLRNTLY | 0.6883191 |

| | | |
|---|---|---|
| HQEDPGRI | ASSPKLRNTLY | 0.60009205 |
| GTPGEPTP | ASSPKLRNTLY | 0.32899925 |
| GFFNSGS | ASSPKLRNTLY | 0.8011116 |
| GAGGGGLTAGAN | ASSPKLRNTLY | 0.547294 |
| FVDALVNM | ASSPKLRNTLY | 0.75765526 |
| FLTVAPEL | ASSPKLRNTLY | 0.836596 |
| FETLTAL | ASSPKLRNTLY | 0.28134346 |
| DVINPIEL | ASSPKLRNTLY | 0.8234384 |
| APVSGAPV | ASSPKLRNTLY | 0.77149343 |
| ALVGGLAW | ASSPKLRNTLY | 0.7473476 |
| ALAAAAATAI | ASSPKLRNTLY | 0.7185399 |
| AFGLGLLL | ASSPKLRNTLY | 0.48378024 |
| AFFAAASL | ASSPKLRNTLY | 0.8524815 |
| AAAALPAFSPP | ASSPKLRNTLY | 0.8129105 |
| VVLAPAL | ASSPKLRNTLY | 0.76375705 |
| TIGHVDHGKTTL | ASSPKLRNTLY | 0.7831438 |
| LISNASDALDK | ASSPKLRNTLY | 0.8294625 |
| LISNASDAL | ASSPKLRNTLY | 0.82957023 |
| ISNASDALDK | ASSPKLRNTLY | 0.611745 |
| NQAETLVYQTE | ASSPKLRNTLY | 0.29365003 |
| TGGTGGTAGAG | ASSPKLRNTLY | 0.7353641 |
| TGSTAGGAGGA | ASSPKLRNTLY | 0.72890794 |
| DPLAVPVAAGPV | ASSPKLRNTLY | 0.4877811 |
| GGTGGTPGNGGN | ASSPKLRNTLY | 0.32079935 |
| PAGLAGVDPATL | ASSPKLRNTLY | 0.8251568 |
| GNNLVVEAPHAGASV | ASSPKLRNTLY | 0.8724388 |
| KVTRIMSASTLEQPA | ASSPKLRNTLY | 0.70622176 |
| LGFALPIALPRRTVV | ASSPKLRNTLY | 0.8765903 |
| TRLGNLTARKLVVLV | ASSPKLRNTLY | 0.32907867 |
| WSQQRRLIYFPSAGP | ASSPKLRNTLY | 0.8130758 |
| GPNGNPDPMAAAVDIR | ASSPKLRNTLY | 0.8424543 |
| TQRCVSVADLRAAASR | ASSPKLRNTLY | 0.52673614 |
| QFVLDVAEHPVEVLTETV | ASSPKLRNTLY | 0.7618814 |
| GGAGGTGGAGGNGGTQAGDG | ASSPKLRNTLY | 0.65507424 |
| PLLATLRAIAVDVDAKPAQV | ASSPKLRNTLY | 0.5905281 |
| APKFRTGGYMIDFWGVGYQVAK | ASSPKLRNTLY | 0.8343257 |
| GAGGAGGAGGAAGGPGGTGGTG | ASSPKLRNTLY | 0.666414 |
| IGALGLLATDVAPAGSVAFGVA | ASSPKLRNTLY | 0.7637808 |
| AGGAGGIGGNGGAGGAGGVLMG | ASSPKLRNTLY | 0.6046766 |
| ADELIGLL | ASSPKLRNTLY | 0.5139505 |
| ADRALKA | ASSPKLRNTLY | 0.7899721 |
| AGAVGQVGGAGGA | ASSPKLRNTLY | 0.7968462 |
| AGGGGGLGGAP | ASSPKLRNTLY | 0.7777225 |
| AGGLLYGNGGAG | ASSPKLRNTLY | 0.88244724 |
| AGLESIGL | ASSPKLRNTLY | 0.5882883 |
| AGLKTGII | ASSPKLRNTLY | 0.86900425 |
| ASLAALLL | ASSPKLRNTLY | 0.86305785 |
| ASVINISSI | ASSPKLRNTLY | 0.75666845 |
| ATPPGPA | ASSPKLRNTLY | 0.6759431 |
| ATVASALV | ASSPKLRNTLY | 0.841757 |
| ATVASLIL | ASSPKLRNTLY | 0.682016 |

| | | |
|---|---|---|
| AVNGPVLIRH | ASSPKLRNTLY | 0.7240325 |
| DGAAGAVGI | ASSPKLRNTLY | 0.83530146 |
| DPLLHGI | ASSPKLRNTLY | 0.16004936 |
| ELGAALGAL | ASSPKLRNTLY | 0.6119733 |
| ELGRTQAV | ASSPKLRNTLY | 0.3643618 |
| FVAGGYLS | ASSPKLRNTLY | 0.51563776 |
| GASGGAGGAGGW | ASSPKLRNTLY | 0.26572758 |
| GGAGAGGDGGAGGI | ASSPKLRNTLY | 0.17740455 |
| GTGHNGSQPGMGGNG | ASSPKLRNTLY | 0.31631345 |
| GYAAASASAS | ASSPKLRNTLY | 0.21941066 |
| HQVDVAAVQQG | ASSPKLRNTLY | 0.84450364 |
| IGAAIAHT | ASSPKLRNTLY | 0.493669 |
| IGMIEVAQL | ASSPKLRNTLY | 0.2769656 |
| IRLAPEHTSAMD | ASSPKLRNTLY | 0.6249466 |
| IYNVSVL | ASSPKLRNTLY | 0.63626957 |
| KQGFLGM | ASSPKLRNTLY | 0.14009832 |
| LESVITQL | ASSPKLRNTLY | 0.25087136 |
| LGLGALVVV | ASSPKLRNTLY | 0.23690704 |
| LIGNGGNGGSGGI | ASSPKLRNTLY | 0.35905102 |
| LNAINAPTL | ASSPKLRNTLY | 0.13686907 |
| LQLLNTGV | ASSPKLRNTLY | 0.41805398 |
| LQTSIEF | ASSPKLRNTLY | 0.19348374 |
| MSFAVSEL | ASSPKLRNTLY | 0.11127017 |
| NTGALGL | ASSPKLRNTLY | 0.44841838 |
| PPGAGGAGGAGTT | ASSPKLRNTLY | 0.29966244 |
| QLVGLGI | ASSPKLRNTLY | 0.2547741 |
| RFLEQVHQL | ASSPKLRNTLY | 0.5071676 |
| RGVGYVLR | ASSPKLRNTLY | 0.30093136 |
| SATGVFAGVI | ASSPKLRNTLY | 0.19464296 |
| SLPAITLP | ASSPKLRNTLY | 0.3547793 |
| STALAFTV | ASSPKLRNTLY | 0.6017784 |
| SVLTVGLGK | ASSPKLRNTLY | 0.20900795 |
| TALAGAGSGI | ASSPKLRNTLY | 0.40452588 |
| TGLPFTGI | ASSPKLRNTLY | 0.7616278 |
| VAEHVTV | ASSPKLRNTLY | 0.43317 |
| VDIVLNSL | ASSPKLRNTLY | 0.63838804 |
| VLTGGGAVL | ASSPKLRNTLY | 0.6743486 |
| VNVGLAYGI | ASSPKLRNTLY | 0.29380906 |
| VSSAAVGAV | ASSPKLRNTLY | 0.62102747 |
| VTSVYEAGV | ASSPKLRNTLY | 0.6730399 |
| WVALDRGA | ASSPKLRNTLY | 0.30978388 |
| YLVGSVVI | ASSPKLRNTLY | 0.46547613 |
| GPPGTGKTLL | ASSPKLRNTLY | 0.16621357 |
| GVGMTKF | ASSPKLRNTLY | 0.37464136 |
| IGIGGSDL | ASSPKLRNTLY | 0.17125842 |
| IGTLLAEL | ASSPKLRNTLY | 0.49785906 |
| IYTSGTT | ASSPKLRNTLY | 0.61349547 |
| LSFPAGV | ASSPKLRNTLY | 0.3323532 |
| LYTSGTT | ASSPKLRNTLY | 0.1273731 |
| VGAFPFGF | ASSPKLRNTLY | 0.139134 |
| VGAGTVEF | ASSPKLRNTLY | 0.45260108 |

| | | |
|---|---|---|
| VGKSTLLL | ASSPKLRNTLY | 0.18481363 |
| AALAHAGGGIV | ASSPKLRNTLY | 0.18403131 |
| AALVAVSLL | ASSPKLRNTLY | 0.25803608 |
| AAVSRAVAAL | ASSPKLRNTLY | 0.54433346 |
| AGGSSGIGGF | ASSPKLRNTLY | 0.66434914 |
| AGLTVANANG | ASSPKLRNTLY | 0.2175563 |
| AGLVGAVL | ASSPKLRNTLY | 0.73558235 |
| ALSQAFTV | ASSPKLRNTLY | 0.7542238 |
| ALTFSVSL | ASSPKLRNTLY | 0.7499054 |
| ALTGAALVAL | ASSPKLRNTLY | 0.33181196 |
| ASFSIGAGVL | ASSPKLRNTLY | 0.16161196 |
| DGSVYKI | ASSPKLRNTLY | 0.7997997 |
| EQLAEIV | ASSPKLRNTLY | 0.34259132 |
| FAVSELTVM | ASSPKLRNTLY | 0.57921386 |
| GAAPLAAGL | ASSPKLRNTLY | 0.52638686 |
| GADGGDGGSGGAGG | ASSPKLRNTLY | 0.2615195 |
| GFGLDTGGAGG | ASSPKLRNTLY | 0.21229051 |
| GLASGGLL | ASSPKLRNTLY | 0.6524971 |
| GRIAGAAVLL | ASSPKLRNTLY | 0.6193613 |
| GSAGSLNTSF | ASSPKLRNTLY | 0.14731072 |
| GTGQIGI | ASSPKLRNTLY | 0.19580568 |
| GTLIALIL | ASSPKLRNTLY | 0.29367673 |
| IAATMGFVVG | ASSPKLRNTLY | 0.31256726 |
| IFAQALL | ASSPKLRNTLY | 0.610053 |
| IGAAVALGF | ASSPKLRNTLY | 0.3872847 |
| IGLTPLAL | ASSPKLRNTLY | 0.3267836 |
| ILGSNTSTL | ASSPKLRNTLY | 0.5021262 |
| IVFSLTVL | ASSPKLRNTLY | 0.50187117 |
| LASKATGF | ASSPKLRNTLY | 0.44023645 |
| LASVALIL | ASSPKLRNTLY | 0.42097127 |
| LFAVGLTL | ASSPKLRNTLY | 0.26963076 |
| LFNGGIGTYI | ASSPKLRNTLY | 0.5598822 |
| LTLGAGII | ASSPKLRNTLY | 0.6263567 |
| LVAAALGDI | ASSPKLRNTLY | 0.49437445 |
| NTAFVQLGI | ASSPKLRNTLY | 0.5879351 |
| QLFSGAEL | ASSPKLRNTLY | 0.3630214 |
| SAATFITL | ASSPKLRNTLY | 0.4074837 |
| SNYGLANLGV | ASSPKLRNTLY | 0.16599706 |
| SYGGAYAVM | ASSPKLRNTLY | 0.39037693 |
| TTGVGGLTLG | ASSPKLRNTLY | 0.3699894 |
| TTQTIYI | ASSPKLRNTLY | 0.33460677 |
| VAAGAGLGAVI | ASSPKLRNTLY | 0.649918 |
| VAAVVTLL | ASSPKLRNTLY | 0.42229164 |
| VGFLGGLI | ASSPKLRNTLY | 0.2343122 |
| VGVIPILL | ASSPKLRNTLY | 0.57244056 |
| VIFASSLI | ASSPKLRNTLY | 0.7513143 |
| VKGGDGGAAAG | ASSPKLRNTLY | 0.20445076 |
| VTKTFTNI | ASSPKLRNTLY | 0.31181556 |
| VVAAAILL | ASSPKLRNTLY | 0.75263226 |
| YAALDVEL | ASSPKLRNTLY | 0.6921432 |
| LASVALLL | ASSPKLRNTLY | 0.2917466 |

| | | |
|---|---|---|
| LDAVHRV | ASSPKLRNTLY | 0.46357292 |
| QGLSSSIF | ASSPKLRNTLY | 0.8542677 |
| RGTFANIRL | ASSPKLRNTLY | 0.5032935 |
| RQLWWGHRI | ASSPKLRNTLY | 0.69322675 |
| YGPPGTGKTLL | ASSPKLRNTLY | 0.16170819 |
| AALVGAPANVA | ASSPKLRNTLY | 0.32004893 |
| AATFRRTE | ASSPKLRNTLY | 0.20888753 |
| AGALGGGI | ASSPKLRNTLY | 0.10971294 |
| AGGAGGAGGIGAG | ASSPKLRNTLY | 0.6847109 |
| AGSGALGL | ASSPKLRNTLY | 0.29224733 |
| AQIADQI | ASSPKLRNTLY | 0.540941 |
| ASAVAVTYL | ASSPKLRNTLY | 0.26868957 |
| ASTVLGGTGG | ASSPKLRNTLY | 0.6610526 |
| AVGGEPAK | ASSPKLRNTLY | 0.57621306 |
| CEQGTFGGTC | ASSPKLRNTLY | 0.3802792 |
| DEGKAPAG | ASSPKLRNTLY | 0.23482499 |
| DTVVFPAL | ASSPKLRNTLY | 0.48040283 |
| FEGITMAG | ASSPKLRNTLY | 0.6224284 |
| FPLVVQD | ASSPKLRNTLY | 0.5700729 |
| GAFTTPQ | ASSPKLRNTLY | 0.14940643 |
| GFGGFGTAGGVGGA | ASSPKLRNTLY | 0.65620273 |
| GFLAQFL | ASSPKLRNTLY | 0.37270653 |
| GGAGGAGGNAGVA | ASSPKLRNTLY | 0.16247241 |
| GGVGTAGAGGAGGAGG | ASSPKLRNTLY | 0.143922 |
| GNGGGGDAGNAGSGGN | ASSPKLRNTLY | 0.16133761 |
| GTGGAGGAGNDAGS | ASSPKLRNTLY | 0.3317623 |
| GTTNTGWFNS | ASSPKLRNTLY | 0.48838416 |
| IFMGPLFTL | ASSPKLRNTLY | 0.4365893 |
| ILAHAIEAI | ASSPKLRNTLY | 0.30429545 |
| IRSAIREAL | ASSPKLRNTLY | 0.5443978 |
| ITFAGGG | ASSPKLRNTLY | 0.4315737 |
| ITGMSGHQEV | ASSPKLRNTLY | 0.08913901 |
| IVAGLACTL | ASSPKLRNTLY | 0.26425704 |
| IVGGGLAGA | ASSPKLRNTLY | 0.14683679 |
| IVTGVLNI | ASSPKLRNTLY | 0.62128955 |
| LAQQGCI | ASSPKLRNTLY | 0.19731477 |
| LCLHVEN | ASSPKLRNTLY | 0.31128877 |
| LDAISDGS | ASSPKLRNTLY | 0.3617379 |
| LGGFAAML | ASSPKLRNTLY | 0.7535751 |
| LGGKNPMI | ASSPKLRNTLY | 0.48922023 |
| LGPKGVAAIL | ASSPKLRNTLY | 0.20128141 |
| LMVGVSI | ASSPKLRNTLY | 0.13065125 |
| LVGAAFVL | ASSPKLRNTLY | 0.47412193 |
| LVLAVAPL | ASSPKLRNTLY | 0.33967915 |
| MAAIAVIDAA | ASSPKLRNTLY | 0.18803675 |
| NNGINAELQ | ASSPKLRNTLY | 0.2296002 |
| RGEPLVLVHG | ASSPKLRNTLY | 0.43799466 |
| RGGAGGAAGLI | ASSPKLRNTLY | 0.1539327 |
| RSYRMHPAV | ASSPKLRNTLY | 0.5429405 |
| SAAHLLAGQLA | ASSPKLRNTLY | 0.1819548 |
| SAAVGSY | ASSPKLRNTLY | 0.24793579 |

| | | |
|---|---|---|
| SPHAVLHRI | ASSPKLRNTLY | 0.6352906 |
| SQIAKVL | ASSPKLRNTLY | 0.72928345 |
| TALGHLVP | ASSPKLRNTLY | 0.34753358 |
| TAPTTSGAA | ASSPKLRNTLY | 0.2397671 |
| TDGDDGNGGSPGT | ASSPKLRNTLY | 0.19377318 |
| TDGLGEN | ASSPKLRNTLY | 0.1962752 |
| TGGPPLPGQA | ASSPKLRNTLY | 0.21371605 |
| TIPAITI | ASSPKLRNTLY | 0.63671774 |
| TIPGFTI | ASSPKLRNTLY | 0.4378205 |
| TSIASGALA | ASSPKLRNTLY | 0.19919111 |
| TVTLPGAAI | ASSPKLRNTLY | 0.5675447 |
| VASGLSAALP | ASSPKLRNTLY | 0.74028623 |
| VGAGVIGI | ASSPKLRNTLY | 0.7679883 |
| VGPKRFGV | ASSPKLRNTLY | 0.17680281 |
| VNVAFPD | ASSPKLRNTLY | 0.3245886 |
| VSMTNSGVS | ASSPKLRNTLY | 0.37836343 |
| VTDVLGDLSV | ASSPKLRNTLY | 0.29052907 |
| VVVVGAGFAGL | ASSPKLRNTLY | 0.23757024 |
| VYTVGTVV | ASSPKLRNTLY | 0.24566938 |
| WTTFSGP | ASSPKLRNTLY | 0.5429163 |
| YDELVEAF | ASSPKLRNTLY | 0.8717146 |
| YGTGGAGGNGG | ASSPKLRNTLY | 0.3419594 |
| YTNVMAV | ASSPKLRNTLY | 0.12183072 |
| FLEQVHQL | ASSPKLRNTLY | 0.29309076 |
| GLSLLAVAV | ASSPKLRNTLY | 0.16212493 |
| LEQVHQL | ASSPKLRNTLY | 0.22911301 |
| LNAAGAI | ASSPKLRNTLY | 0.29880583 |
| PAVHASL | ASSPKLRNTLY | 0.14212541 |
| TELEDDA | ASSPKLRNTLY | 0.32735455 |
| TPPGPPN | ASSPKLRNTLY | 0.6985071 |
| TPPPPPPP | ASSPKLRNTLY | 0.37443614 |
| TQSPALI | ASSRDIANSDYT | 0.5067505 |
| EFAAAFPQ | ASSRDIANSDYT | 0.70059675 |
| LLSGNLDVL | ASSRDIANSDYT | 0.4327355 |
| GPANLGNYNI | ASSRDIANSDYT | 0.7072067 |
| TLGPTLP | ASSRDIANSDYT | 0.7350018 |
| VYAYSSCCK | ASSRDIANSDYT | 0.38057023 |
| APAPAPAPNI | ASSRDIANSDYT | 0.3144974 |
| PGGPVPIP | ASSRDIANSDYT | 0.64103127 |
| FLTVALEP | ASSRDIANSDYT | 0.4081818 |
| SQYFGANL | ASSRDIANSDYT | 0.37299705 |
| GGVKPAA | ASSRDIANSDYT | 0.5398841 |
| GGTGGAGGAGGW | ASSRDIANSDYT | 0.6320591 |
| GVGGVAGLGGAGG | ASSRDIANSDYT | 0.2772546 |
| VLLVLATTI | ASSRDIANSDYT | 0.8384371 |
| IGIFAAML | ASSRDIANSDYT | 0.12960573 |
| AVGDLID | ASSRDIANSDYT | 0.21440005 |
| NISPLPL | ASSRDIANSDYT | 0.6663588 |
| VTAVFVAI | ASSRDIANSDYT | 0.10347615 |
| SGDDNDAAL | ASSRDIANSDYT | 0.19609536 |
| AVAALAHS | ASSRDIANSDYT | 0.31877846 |

| | | |
|---|---|---|
| GGVTVMATP | ASSRDIANSDYT | 0.7236185 |
| AGLAANVRD | ASSRDIANSDYT | 0.109462075 |
| TNRPDLIDPALLRPG | ASSRDIANSDYT | 0.30109248 |
| LANLGVTAL | ASSRDIANSDYT | 0.15548617 |
| TGERIEL | ASSRDIANSDYT | 0.5079527 |
| NGEELAA | ASSRDIANSDYT | 0.38317123 |
| VLSLVWG | ASSRDIANSDYT | 0.1801263 |
| IGYALWL | ASSRDIANSDYT | 0.5616892 |
| LSLLLAPL | ASSRDIANSDYT | 0.15031236 |
| TLDEALK | ASSRDIANSDYT | 0.5648929 |
| TDGAGTAEGG | ASSRDIANSDYT | 0.60837275 |
| NAPGGAGGNGG | ASSRDIANSDYT | 0.8280127 |
| VSVFMGMGGNF | ASSRDIANSDYT | 0.1690937 |
| LASYLAAL | ASSRDIANSDYT | 0.48498762 |
| LVGGTPAL | ASSRDIANSDYT | 0.22609007 |
| VRNADVL | ASSRDIANSDYT | 0.8327627 |
| TCSSPFLVSAM | ASSRDIANSDYT | 0.26630616 |
| VHITDAE | ASSRDIANSDYT | 0.14865571 |
| EYAAPSF | ASSRDIANSDYT | 0.5139617 |
| VVDAFHGGL | ASSRDIANSDYT | 0.61167246 |
| ASVVDPQ | ASSRDIANSDYT | 0.2887843 |
| SYNEFGGLS | ASSRDIANSDYT | 0.74731004 |
| DPDGSVY | ASSRDIANSDYT | 0.3222962 |
| MVSTIHATE | ASSRDIANSDYT | 0.3195971 |
| LAAGVADPV | ASSRDIANSDYT | 0.716151 |
| TKEQIAA | ASSRDIANSDYT | 0.46363991 |
| SLASPAAL | ASSRDIANSDYT | 0.26119965 |
| IGLVFYGV | ASSRDIANSDYT | 0.6797373 |
| FLLDASNL | ASSRDIANSDYT | 0.38970572 |
| YQVGNFP | ASSRDIANSDYT | 0.17722704 |
| ESLLSQST | ASSRDIANSDYT | 0.8046939 |
| LFTVGFAAVG | ASSRDIANSDYT | 0.30989808 |
| ATPPTPT | ASSRDIANSDYT | 0.6257714 |
| TDLATLG | ASSRDIANSDYT | 0.48980242 |
| SAGGTGR | ASSRDIANSDYT | 0.86453146 |
| GFSNSGS | ASSRDIANSDYT | 0.21756607 |
| GGNGGSGGTSV | ASSRDIANSDYT | 0.47182038 |
| GTSVATGGAGN | ASSRDIANSDYT | 0.37485138 |
| NGGNGGSGGTS | ASSRDIANSDYT | 0.5152385 |
| AVVKFPH | ASSRDIANSDYT | 0.39674044 |
| TPPPPPP | ASSRDIANSDYT | 0.31423873 |
| AAPAAAAAPA | ASSRDIANSDYT | 0.5472989 |
| IALNEDL | ASSRDIANSDYT | 0.6218897 |
| IVNVSSM | ASSRDIANSDYT | 0.28404927 |
| TGIPPAPR | ASSRDIANSDYT | 0.31381848 |
| FDLGGGTF | ASSRDIANSDYT | 0.7581893 |
| FDLGGGTFDVS | ASSRDIANSDYT | 0.19014958 |
| NEPTAAA | ASSRDIANSDYT | 0.453105 |
| NEPTAAALA | ASSRDIANSDYT | 0.45221546 |
| GSIINISSI | ASSRDIANSDYT | 0.573226 |
| VAPGNAGTA | ASSRDIANSDYT | 0.32098407 |

| | | |
|---|---|---|
| TGGGASLE | ASSRDIANSDYT | 0.7109374 |
| VAVYNHYKRI | ASSRDIANSDYT | 0.3197173 |
| TDPAPATT | ASSRDIANSDYT | 0.5721883 |
| TNRPDILDPALLRPG | ASSRDIANSDYT | 0.40662846 |
| LTTTTTS | ASSRDIANSDYT | 0.3906548 |
| PTGSGKTL | ASSRDIANSDYT | 0.1442089 |
| TGSGKTL | ASSRDIANSDYT | 0.3527555 |
| TIGHVDHGK | ASSRDIANSDYT | 0.55760497 |
| TIGHVDHGKTT | ASSRDIANSDYT | 0.56550145 |
| GHVDHGKTT | ASSRDIANSDYT | 0.17029849 |
| AQFEHTLL | ASSRDIANSDYT | 0.53755903 |
| IVNQDEL | ASSRDIANSDYT | 0.6111913 |
| TNGSQFF | ASSRDIANSDYT | 0.8397251 |
| TGKGIVE | ASSRDIANSDYT | 0.30749565 |
| LISNASDA | ASSRDIANSDYT | 0.644048 |
| AELRTLGELTKIAKAHGAQ | ASSRDIANSDYT | 0.2280916 |
| APNAVAAAEPSW | ASSRDIANSDYT | 0.34704974 |
| APQVAGAATAVL | ASSRDIANSDYT | 0.14095236 |
| ARRDPAAAAAILPTDA | ASSRDIANSDYT | 0.18650168 |
| DAAARDVLAEAGLAEYF | ASSRDIANSDYT | 0.41554493 |
| DDQIVLSAKANLAKVEQ | ASSRDIANSDYT | 0.17896411 |
| DELLPDTGEDSDEDGH | ASSRDIANSDYT | 0.8022888 |
| ELPEAPANRATVPQD | ASSRDIANSDYT | 0.60524595 |
| FGQSKIERVTGIG | ASSRDIANSDYT | 0.49172044 |
| GAGGTGSTAGGAGGAGGAGGLY | ASSRDIANSDYT | 0.6433192 |
| GHGGTGAAGGNGGTGGAGGSGI | ASSRDIANSDYT | 0.75016046 |
| GNGGNGGAGGEGGPGGAGGAGA | ASSRDIANSDYT | 0.5854959 |
| GSTAGNGGNGGNATNGTIAGQP | ASSRDIANSDYT | 0.28760427 |
| GTGGDGGNGGNGGTGG | ASSRDIANSDYT | 0.41322345 |
| GVRRPAGIPADAPV | ASSRDIANSDYT | 0.12671879 |
| GYRQVTDAHLLTLARR | ASSRDIANSDYT | 0.26169586 |
| IAKMGQDGHDRG | ASSRDIANSDYT | 0.23630461 |
| KAEPSIEIQYT | ASSRDIANSDYT | 0.4114812 |
| KPVRIRMAADEDTVAVT | ASSRDIANSDYT | 0.10077288 |
| MPPRIAGMRLLVIKPEP | ASSRDIANSDYT | 0.6905693 |
| MKFYAVALFRSRGVPPTR | ASSRDIANSDYT | 0.6745037 |
| NAGGFGILGAVAHSP | ASSRDIANSDYT | 0.40309477 |
| NEGREITLAMAEM | ASSRDIANSDYT | 0.59923065 |
| NELVEWSHPLGHRWRWE | ASSRDIANSDYT | 0.58009136 |
| NLGAGGTGGNGGNPG | ASSRDIANSDYT | 0.17677462 |
| NTEWPKWETVMAADPVD | ASSRDIANSDYT | 0.49658126 |
| PASAAPAETQP | ASSRDIANSDYT | 0.23067051 |
| QQSSDGKNGVRAAEITGE | ASSRDIANSDYT | 0.6334915 |
| RAPDAVTTSWWKE | ASSRDIANSDYT | 0.68074 |
| SGFGNVGAHTSGWFNQ | ASSRDIANSDYT | 0.4761413 |
| SGVAGAVGQAASVGGLKV | ASSRDIANSDYT | 0.2618105 |
| SNGGAGTGSAGGNGGT | ASSRDIANSDYT | 0.5024925 |
| TGGAGGEGGAAG | ASSRDIANSDYT | 0.22174656 |
| TGGDGGAAGAGGAPGHG | ASSRDIANSDYT | 0.28441614 |
| TMPPLLAAIANAGRHGV | ASSRDIANSDYT | 0.177161 |
| TSSTAGNSGGA | ASSRDIANSDYT | 0.46122706 |

| | | |
|---|---|---|
| VPATVPVGILC | ASSRDIANSDYT | 0.20539047 |
| VQRPDSGGVVLTGRIS | ASSRDIANSDYT | 0.36631233 |
| WTPNPVPATKGVPAGLVQAT | ASSRDIANSDYT | 0.3530683 |
| YALKRALAKFSDRFDVVII | ASSRDIANSDYT | 0.3414096 |
| YLIGSVIV | ASSRDIANSDYT | 0.20524845 |
| YGQLVAVID | ASSRDIANSDYT | 0.6217829 |
| VTVGGMTLPA | ASSRDIANSDYT | 0.53579915 |
| VLALPAV | ASSRDIANSDYT | 0.15019459 |
| VIPPTLNYET | ASSRDIANSDYT | 0.59131944 |
| VGMGGMHPSSG | ASSRDIANSDYT | 0.2818755 |
| VGLAGGAATA | ASSRDIANSDYT | 0.40925407 |
| VGGAGGAGGSGP | ASSRDIANSDYT | 0.5768752 |
| VFAAFMV | ASSRDIANSDYT | 0.32357973 |
| TLYDKLR | ASSRDIANSDYT | 0.1926853 |
| TLAVFTVL | ASSRDIANSDYT | 0.23089795 |
| TGSVQPARV | ASSRDIANSDYT | 0.34441105 |
| TGGGAGSGGT | ASSRDIANSDYT | 0.28845567 |
| SHLETSL | ASSRDIANSDYT | 0.13359168 |
| SFTTGMT | ASSRDIANSDYT | 0.5458076 |
| QVEVAKRMGVRQPTVS | ASSRDIANSDYT | 0.1770485 |
| QGSGGPALAM | ASSRDIANSDYT | 0.2775757 |
| MLAPSNP | ASSRDIANSDYT | 0.2920981 |
| LVNNAAI | ASSRDIANSDYT | 0.5956816 |
| LTGEINTLA | ASSRDIANSDYT | 0.48635295 |
| LSIFGPV | ASSRDIANSDYT | 0.30623615 |
| LLALLANL | ASSRDIANSDYT | 0.46234137 |
| LISNASDALDKLR | ASSRDIANSDYT | 0.40660346 |
| LINEAVP | ASSRDIANSDYT | 0.71973056 |
| LGYAFSGL | ASSRDIANSDYT | 0.34531587 |
| LDFVAVAL | ASSRDIANSDYT | 0.27373663 |
| LAQPGAE | ASSRDIANSDYT | 0.53732955 |
| ISNLGAVTGF | ASSRDIANSDYT | 0.34052324 |
| HQEDPGRI | ASSRDIANSDYT | 0.24324128 |
| GTPGEPTP | ASSRDIANSDYT | 0.24789786 |
| GFFNSGS | ASSRDIANSDYT | 0.19487555 |
| GAGGGGLTAGAN | ASSRDIANSDYT | 0.24102828 |
| FVDALVNM | ASSRDIANSDYT | 0.49488616 |
| FLTVAPEL | ASSRDIANSDYT | 0.6627195 |
| FETLTAL | ASSRDIANSDYT | 0.4457156 |
| DVINPIEL | ASSRDIANSDYT | 0.1441283 |
| APVSGAPV | ASSRDIANSDYT | 0.19206196 |
| ALVGGLAW | ASSRDIANSDYT | 0.34120888 |
| ALAAAAATAI | ASSRDIANSDYT | 0.53988606 |
| AFGLGLLL | ASSRDIANSDYT | 0.47459096 |
| AFFAAASL | ASSRDIANSDYT | 0.8466102 |
| AAAALPAFSPP | ASSRDIANSDYT | 0.5782501 |
| VVLAPAL | ASSRDIANSDYT | 0.2669071 |
| TIGHVDHGKTTL | ASSRDIANSDYT | 0.5088726 |
| LISNASDALDK | ASSRDIANSDYT | 0.3421071 |
| LISNASDAL | ASSRDIANSDYT | 0.31668 |
| ISNASDALDK | ASSRDIANSDYT | 0.37059915 |

| | | |
|---|---|---|
| NQAETLVYQTE | ASSRDIANSDYT | 0.49409625 |
| TGGTGGTAGAG | ASSRDIANSDYT | 0.25224593 |
| TGSTAGGAGGA | ASSRDIANSDYT | 0.32102615 |
| DPLAVPVAAGPV | ASSRDIANSDYT | 0.16146466 |
| GGTGGTPGNGGN | ASSRDIANSDYT | 0.5344019 |
| PAGLAGVDPATL | ASSRDIANSDYT | 0.16411747 |
| GNNLVVEAPHAGASV | ASSRDIANSDYT | 0.23518609 |
| KVTRIMSASTLEQPA | ASSRDIANSDYT | 0.39516693 |
| LGFALPIALPRRTVV | ASSRDIANSDYT | 0.7660459 |
| TRLGNLTARKLVVLV | ASSRDIANSDYT | 0.6972678 |
| WSQQRRLIYFPSAGP | ASSRDIANSDYT | 0.23908615 |
| GPNGNPDPMAAAVDIR | ASSRDIANSDYT | 0.2190121 |
| TQRCVSVADLRAAASR | ASSRDIANSDYT | 0.6948019 |
| QFVLDVAEHPVEVLTETV | ASSRDIANSDYT | 0.23727646 |
| GGAGGTGGAGGNGGTQAGDG | ASSRDIANSDYT | 0.5165845 |
| PLLATLRAIAVDVDAKPAQV | ASSRDIANSDYT | 0.44899 |
| APKFRTGGYMIDFWGVGYQVAK | ASSRDIANSDYT | 0.41077948 |
| GAGGAGGAGGAAGGPGGTGGTG | ASSRDIANSDYT | 0.3087242 |
| IGALGLLATDVAPAGSVAFGVA | ASSRDIANSDYT | 0.4142787 |
| AGGAGGIGGNGGAGGAGGVLMG | ASSRDIANSDYT | 0.2926922 |
| ADELIGLL | ASSRDIANSDYT | 0.44249216 |
| ADRALKA | ASSRDIANSDYT | 0.14663814 |
| AGAVGQVGGAGGA | ASSRDIANSDYT | 0.61519086 |
| AGGGGGLGGAP | ASSRDIANSDYT | 0.17783634 |
| AGGLLYGNGGAG | ASSRDIANSDYT | 0.21823369 |
| AGLESIGL | ASSRDIANSDYT | 0.11811423 |
| AGLKTGII | ASSRDIANSDYT | 0.3179471 |
| ASLAALLL | ASSRDIANSDYT | 0.48136926 |
| ASVINISSI | ASSRDIANSDYT | 0.60306716 |
| ATPPGPA | ASSRDIANSDYT | 0.5578377 |
| ATVASALV | ASSRDIANSDYT | 0.5308014 |
| ATVASLIL | ASSRDIANSDYT | 0.6576002 |
| AVNGPVLIRH | ASSRDIANSDYT | 0.2805015 |
| DGAAGAVGI | ASSRDIANSDYT | 0.13264985 |
| DPLLHGI | ASSRDIANSDYT | 0.40905356 |
| ELGAALGAL | ASSRDIANSDYT | 0.57241625 |
| ELGRTQAV | ASSRDIANSDYT | 0.636371 |
| FVAGGYLS | ASSRDIANSDYT | 0.5986858 |
| GASGGAGGAGGW | ASSRDIANSDYT | 0.12826556 |
| GGAGAGGDGGAGGI | ASSRDIANSDYT | 0.52800244 |
| GTGHNGSQPGMGGNG | ASSRDIANSDYT | 0.4424231 |
| GYAAASASAS | ASSRDIANSDYT | 0.15245467 |
| HQVDVAAVQQG | ASSRDIANSDYT | 0.23385619 |
| IGAAIAHT | ASSRDIANSDYT | 0.27245227 |
| IGMIEVAQL | ASSRDIANSDYT | 0.1633635 |
| IRLAPEHTSAMD | ASSRDIANSDYT | 0.4101432 |
| IYNVSVL | ASSRDIANSDYT | 0.21886626 |
| KQGFLGM | ASSRDIANSDYT | 0.32845563 |
| LESVITQL | ASSRDIANSDYT | 0.2585959 |
| LGLGALVVV | ASSRDIANSDYT | 0.23138456 |
| LIGNGGNGGSGGI | ASSRDIANSDYT | 0.35419083 |

| | | |
|---|---|---|
| LNAINAPTL | ASSRDIANSDYT | 0.28949267 |
| LQLLNTGV | ASSRDIANSDYT | 0.6144916 |
| LQTSIEF | ASSRDIANSDYT | 0.44512516 |
| MSFAVSEL | ASSRDIANSDYT | 0.36557287 |
| NTGALGL | ASSRDIANSDYT | 0.47844237 |
| PPGAGGAGGAGTT | ASSRDIANSDYT | 0.6870213 |
| QLVGLGI | ASSRDIANSDYT | 0.16460845 |
| RFLEQVHQL | ASSRDIANSDYT | 0.30029124 |
| RGVGYVLR | ASSRDIANSDYT | 0.44620323 |
| SATGVFAGVI | ASSRDIANSDYT | 0.25001168 |
| SLPAITLP | ASSRDIANSDYT | 0.66729265 |
| STALAFTV | ASSRDIANSDYT | 0.3683619 |
| SVLTVGLGK | ASSRDIANSDYT | 0.44236124 |
| TALAGAGSGI | ASSRDIANSDYT | 0.3579437 |
| TGLPFTGI | ASSRDIANSDYT | 0.75145066 |
| VAEHVTV | ASSRDIANSDYT | 0.44931364 |
| VDIVLNSL | ASSRDIANSDYT | 0.3099187 |
| VLTGGGAVL | ASSRDIANSDYT | 0.65228295 |
| VNVGLAYGI | ASSRDIANSDYT | 0.21139707 |
| VSSAAVGAV | ASSRDIANSDYT | 0.5643863 |
| VTSVYEAGV | ASSRDIANSDYT | 0.5728652 |
| WVALDRGA | ASSRDIANSDYT | 0.19361112 |
| YLVGSVVI | ASSRDIANSDYT | 0.3100242 |
| GPPGTGKTLL | ASSRDIANSDYT | 0.6652845 |
| GVGMTKF | ASSRDIANSDYT | 0.6575796 |
| IGIGGSDL | ASSRDIANSDYT | 0.23374978 |
| IGTLLAEL | ASSRDIANSDYT | 0.19365926 |
| IYTSGTT | ASSRDIANSDYT | 0.18234879 |
| LSFPAGV | ASSRDIANSDYT | 0.21917313 |
| LYTSGTT | ASSRDIANSDYT | 0.16355352 |
| VGAFPFGF | ASSRDIANSDYT | 0.49804533 |
| VGAGTVEF | ASSRDIANSDYT | 0.312243 |
| VGKSTLLL | ASSRDIANSDYT | 0.8236597 |
| AALAHAGGGIV | ASSRDIANSDYT | 0.4592796 |
| AALVAVSLL | ASSRDIANSDYT | 0.66671526 |
| AAVSRAVAAL | ASSRDIANSDYT | 0.23814736 |
| AGGSSGIGGF | ASSRDIANSDYT | 0.29542577 |
| AGLTVANANG | ASSRDIANSDYT | 0.15181567 |
| AGLVGAVL | ASSRDIANSDYT | 0.1779821 |
| ALSQAFTV | ASSRDIANSDYT | 0.18492003 |
| ALTFSVSL | ASSRDIANSDYT | 0.32151926 |
| ALTGAALVAL | ASSRDIANSDYT | 0.14187151 |
| ASFSIGAGVL | ASSRDIANSDYT | 0.80578786 |
| DGSVYKI | ASSRDIANSDYT | 0.28579295 |
| EQLAEIV | ASSRDIANSDYT | 0.5109017 |
| FAVSELTVM | ASSRDIANSDYT | 0.2929524 |
| GAAPLAAGL | ASSRDIANSDYT | 0.18827468 |
| GADGGDGGSGGAGG | ASSRDIANSDYT | 0.27658033 |
| GFGLDTGGAGG | ASSRDIANSDYT | 0.53688455 |
| GLASGGLL | ASSRDIANSDYT | 0.16563572 |
| GRIAGAAVLL | ASSRDIANSDYT | 0.13094728 |

| | | |
|---|---|---|
| GSAGSLNTSF | ASSRDIANSDYT | 0.26008373 |
| GTGQIGI | ASSRDIANSDYT | 0.17775925 |
| GTLIALIL | ASSRDIANSDYT | 0.8147693 |
| IAATMGFVVG | ASSRDIANSDYT | 0.61211693 |
| IFAQALL | ASSRDIANSDYT | 0.34223306 |
| IGAAVALGF | ASSRDIANSDYT | 0.8338641 |
| IGLTPLAL | ASSRDIANSDYT | 0.10065098 |
| ILGSNTSTL | ASSRDIANSDYT | 0.33811527 |
| IVFSLTVL | ASSRDIANSDYT | 0.7746266 |
| LASKATGF | ASSRDIANSDYT | 0.37982357 |
| LASVALIL | ASSRDIANSDYT | 0.3181477 |
| LFAVGLTL | ASSRDIANSDYT | 0.6262847 |
| LFNGGIGTYI | ASSRDIANSDYT | 0.23134579 |
| LTLGAGII | ASSRDIANSDYT | 0.3106012 |
| LVAAALGDI | ASSRDIANSDYT | 0.291324 |
| NTAFVQLGI | ASSRDIANSDYT | 0.53208697 |
| QLFSGAEL | ASSRDIANSDYT | 0.4670136 |
| SAATFITL | ASSRDIANSDYT | 0.15492395 |
| SNYGLANLGV | ASSRDIANSDYT | 0.58633757 |
| SYGGAYAVM | ASSRDIANSDYT | 0.80181897 |
| TTGVGGLTLG | ASSRDIANSDYT | 0.22214817 |
| TTQTIYI | ASSRDIANSDYT | 0.4601243 |
| VAAGAGLGAVI | ASSRDIANSDYT | 0.423439 |
| VAAVVTLL | ASSRDIANSDYT | 0.191143 |
| VGFLGGLI | ASSRDIANSDYT | 0.24448329 |
| VGVIPILL | ASSRDIANSDYT | 0.14819376 |
| VIFASSLI | ASSRDIANSDYT | 0.66165245 |
| VKGGDGGAAAG | ASSRDIANSDYT | 0.23111562 |
| VTKTFTNI | ASSRDIANSDYT | 0.33167762 |
| VVAAAILL | ASSRDIANSDYT | 0.47780788 |
| YAALDVEL | ASSRDIANSDYT | 0.82277536 |
| LASVALLL | ASSRDIANSDYT | 0.6023333 |
| LDAVHRV | ASSRDIANSDYT | 0.437194 |
| QGLSSSIF | ASSRDIANSDYT | 0.19231457 |
| RGTFANIRL | ASSRDIANSDYT | 0.37736267 |
| RQLWWGHRI | ASSRDIANSDYT | 0.5123793 |
| YGPPGTGKTLL | ASSRDIANSDYT | 0.1775618 |
| AALVGAPANVA | ASSRDIANSDYT | 0.64657897 |
| AATFRRTE | ASSRDIANSDYT | 0.21156476 |
| AGALGGGI | ASSRDIANSDYT | 0.5183509 |
| AGGAGGAGGIGAG | ASSRDIANSDYT | 0.54795825 |
| AGSGALGL | ASSRDIANSDYT | 0.83595145 |
| AQIADQI | ASSRDIANSDYT | 0.4700052 |
| ASAVAVTYL | ASSRDIANSDYT | 0.62911105 |
| ASTVLGGTGG | ASSRDIANSDYT | 0.16827199 |
| AVGGEPAK | ASSRDIANSDYT | 0.56185275 |
| CEQGTFGGTC | ASSRDIANSDYT | 0.646279 |
| DEGKAPAG | ASSRDIANSDYT | 0.64009 |
| DTVVFPAL | ASSRDIANSDYT | 0.84175175 |
| FEGITMAG | ASSRDIANSDYT | 0.51878846 |
| FPLVVQD | ASSRDIANSDYT | 0.14655292 |

| | | |
|---|---|---|
| GAFTTPQ | ASSRDIANSDYT | 0.31145704 |
| GFGGFGTAGGVGGA | ASSRDIANSDYT | 0.6907263 |
| GFLAQFL | ASSRDIANSDYT | 0.6554992 |
| GGAGGAGGNAGVA | ASSRDIANSDYT | 0.62497914 |
| GGVGTAGAGGAGGAGG | ASSRDIANSDYT | 0.52161384 |
| GNGGGGDAGNAGSGGN | ASSRDIANSDYT | 0.2977398 |
| GTGGAGGAGNDAGS | ASSRDIANSDYT | 0.69382656 |
| GTTNTGWFNS | ASSRDIANSDYT | 0.8482324 |
| IFMGPLFTL | ASSRDIANSDYT | 0.5560911 |
| ILAHAIEAI | ASSRDIANSDYT | 0.23617887 |
| IRSAIREAL | ASSRDIANSDYT | 0.22966683 |
| ITFAGGG | ASSRDIANSDYT | 0.522616 |
| ITGMSGHQEV | ASSRDIANSDYT | 0.6590524 |
| IVAGLACTL | ASSRDIANSDYT | 0.18671885 |
| IVGGGLAGA | ASSRDIANSDYT | 0.40553415 |
| IVTGVLNI | ASSRDIANSDYT | 0.31347594 |
| LAQQGCI | ASSRDIANSDYT | 0.117340475 |
| LCLHVEN | ASSRDIANSDYT | 0.62515223 |
| LDAISDGS | ASSRDIANSDYT | 0.73754597 |
| LGGFAAML | ASSRDIANSDYT | 0.5219339 |
| LGGKNPMI | ASSRDIANSDYT | 0.16163424 |
| LGPKGVAAIL | ASSRDIANSDYT | 0.6852797 |
| LMVGVSI | ASSRDIANSDYT | 0.7308128 |
| LVGAAFVL | ASSRDIANSDYT | 0.70179856 |
| LVLAVAPL | ASSRDIANSDYT | 0.20329753 |
| MAAIAVIDAA | ASSRDIANSDYT | 0.103230685 |
| NNGINAELQ | ASSRDIANSDYT | 0.41229847 |
| RGEPLVLVHG | ASSRDIANSDYT | 0.7244836 |
| RGGAGGAAGLI | ASSRDIANSDYT | 0.24677767 |
| RSYRMHPAV | ASSRDIANSDYT | 0.182189 |
| SAAHLLAGQLA | ASSRDIANSDYT | 0.3416139 |
| SAAVGSY | ASSRDIANSDYT | 0.6398895 |
| SPHAVLHRI | ASSRDIANSDYT | 0.18217048 |
| SQIAKVL | ASSRDIANSDYT | 0.16931553 |
| TALGHLVP | ASSRDIANSDYT | 0.3630935 |
| TAPTTSGAA | ASSRDIANSDYT | 0.59139836 |
| TDGDDGNGGSPGT | ASSRDIANSDYT | 0.19644257 |
| TDGLGEN | ASSRDIANSDYT | 0.20400374 |
| TGGPPLPGQA | ASSRDIANSDYT | 0.37361485 |
| TIPAITI | ASSRDIANSDYT | 0.49285078 |
| TIPGFTI | ASSRDIANSDYT | 0.38580742 |
| TSIASGALA | ASSRDIANSDYT | 0.17918417 |
| TVTLPGAAI | ASSRDIANSDYT | 0.5848598 |
| VASGLSAALP | ASSRDIANSDYT | 0.17800593 |
| VGAGVIGI | ASSRDIANSDYT | 0.57800925 |
| VGPKRFGV | ASSRDIANSDYT | 0.33120787 |
| VNVAFPD | ASSRDIANSDYT | 0.40075988 |
| VSMTNSGVS | ASSRDIANSDYT | 0.32861874 |
| VTDVLGDLSV | ASSRDIANSDYT | 0.21864192 |
| VVVVGAGFAGL | ASSRDIANSDYT | 0.19461411 |
| VYTVGTVV | ASSRDIANSDYT | 0.1699581 |

| | | |
|---|---|---|
| WTTFSGP | ASSRDIANSDYT | 0.6024085 |
| YDELVEAF | ASSRDIANSDYT | 0.32414648 |
| YGTGGAGGNGG | ASSRDIANSDYT | 0.5647763 |
| YTNVMAV | ASSRDIANSDYT | 0.5352969 |
| FLEQVHQL | ASSRDIANSDYT | 0.18537739 |
| GLSLLAVAV | ASSRDIANSDYT | 0.43779206 |
| LEQVHQL | ASSRDIANSDYT | 0.14203832 |
| LNAAGAI | ASSRDIANSDYT | 0.6050283 |
| PAVHASL | ASSRDIANSDYT | 0.31561682 |
| TELEDDA | ASSRDIANSDYT | 0.5146465 |
| TPPGPPN | ASSRDIANSDYT | 0.3022022 |
| TPPPPPPP | ASSRDIANSDYT | 0.43087938 |
| TQSPALI | GARHRDRRGEQY | 0.5072423 |
| EFAAAFPQ | GARHRDRRGEQY | 0.19639803 |
| LLSGNLDVL | GARHRDRRGEQY | 0.51173997 |
| GPANLGNYNI | GARHRDRRGEQY | 0.6679404 |
| TLGPTLP | GARHRDRRGEQY | 0.5848578 |
| VYAYSSCCK | GARHRDRRGEQY | 0.13802868 |
| APAPAPAPNI | GARHRDRRGEQY | 0.4247962 |
| PGGPVPIP | GARHRDRRGEQY | 0.22903772 |
| FLTVALEP | GARHRDRRGEQY | 0.45032567 |
| SQYFGANL | GARHRDRRGEQY | 0.34848434 |
| GGVKPAA | GARHRDRRGEQY | 0.33113754 |
| GGTGGAGGAGGW | GARHRDRRGEQY | 0.39109385 |
| GVGGVAGLGGAGG | GARHRDRRGEQY | 0.27433506 |
| VLLVLATTI | GARHRDRRGEQY | 0.21066788 |
| IGIFAAML | GARHRDRRGEQY | 0.35386115 |
| AVGDLID | GARHRDRRGEQY | 0.40092662 |
| NISPLPL | GARHRDRRGEQY | 0.5204048 |
| VTAVFVAI | GARHRDRRGEQY | 0.19013803 |
| SGDDNDAAL | GARHRDRRGEQY | 0.423634 |
| AVAALAHS | GARHRDRRGEQY | 0.25592387 |
| GGVTVMATP | GARHRDRRGEQY | 0.23572206 |
| AGLAANVRD | GARHRDRRGEQY | 0.4223997 |
| TNRPDLIDPALLRPG | GARHRDRRGEQY | 0.34170765 |
| LANLGVTAL | GARHRDRRGEQY | 0.15047202 |
| TGERIEL | GARHRDRRGEQY | 0.52828723 |
| NGEELAA | GARHRDRRGEQY | 0.18620959 |
| VLSLVWG | GARHRDRRGEQY | 0.33748123 |
| IGYALWL | GARHRDRRGEQY | 0.7085426 |
| LSLLLAPL | GARHRDRRGEQY | 0.14614576 |
| TLDEALK | GARHRDRRGEQY | 0.4614901 |
| TDGAGTAEGG | GARHRDRRGEQY | 0.3605625 |
| NAPGGAGGNGG | GARHRDRRGEQY | 0.4327976 |
| VSVFMGMGGNF | GARHRDRRGEQY | 0.3022608 |
| LASYLAAL | GARHRDRRGEQY | 0.23973995 |
| LVGGTPAL | GARHRDRRGEQY | 0.6762452 |
| VRNADVL | GARHRDRRGEQY | 0.48324263 |
| TCSSPFLVSAM | GARHRDRRGEQY | 0.50446147 |
| VHITDAE | GARHRDRRGEQY | 0.44102287 |
| EYAAPSF | GARHRDRRGEQY | 0.7191126 |

| | | |
|---|---|---|
| VVDAFHGGL | GARHRDRRGEQY | 0.5789015 |
| ASVVDPQ | GARHRDRRGEQY | 0.24970418 |
| SYNEFGGLS | GARHRDRRGEQY | 0.13285585 |
| DPDGSVY | GARHRDRRGEQY | 0.3291107 |
| MVSTIHATE | GARHRDRRGEQY | 0.2925262 |
| LAAGVADPV | GARHRDRRGEQY | 0.26133645 |
| TKEQIAA | GARHRDRRGEQY | 0.17874777 |
| SLASPAAL | GARHRDRRGEQY | 0.71989787 |
| IGLVFYGV | GARHRDRRGEQY | 0.8155224 |
| FLLDASNL | GARHRDRRGEQY | 0.23360178 |
| YQVGNFP | GARHRDRRGEQY | 0.8282671 |
| ESLLSQST | GARHRDRRGEQY | 0.17462382 |
| LFTVGFAAVG | GARHRDRRGEQY | 0.5082972 |
| ATPPTPT | GARHRDRRGEQY | 0.75548023 |
| TDLATLG | GARHRDRRGEQY | 0.17328836 |
| SAGGTGR | GARHRDRRGEQY | 0.5184457 |
| GFSNSGS | GARHRDRRGEQY | 0.22710161 |
| GGNGGSGGTSV | GARHRDRRGEQY | 0.22301033 |
| GTSVATGGAGN | GARHRDRRGEQY | 0.67897695 |
| NGGNGGSGGTS | GARHRDRRGEQY | 0.28645223 |
| AVVKFPH | GARHRDRRGEQY | 0.4812047 |
| TPPPPPP | GARHRDRRGEQY | 0.3035646 |
| AAPAAAAAPA | GARHRDRRGEQY | 0.18459083 |
| IALNEDL | GARHRDRRGEQY | 0.4844771 |
| IVNVSSM | GARHRDRRGEQY | 0.64144 |
| TGIPPAPR | GARHRDRRGEQY | 0.47782102 |
| FDLGGGTF | GARHRDRRGEQY | 0.8286809 |
| FDLGGGTFDVS | GARHRDRRGEQY | 0.25067675 |
| NEPTAAA | GARHRDRRGEQY | 0.7843647 |
| NEPTAAALA | GARHRDRRGEQY | 0.806803 |
| GSIINISSI | GARHRDRRGEQY | 0.37947577 |
| VAPGNAGTA | GARHRDRRGEQY | 0.28457615 |
| TGGGASLE | GARHRDRRGEQY | 0.7146885 |
| VAVYNHYKRI | GARHRDRRGEQY | 0.3086565 |
| TDPAPATT | GARHRDRRGEQY | 0.3610144 |
| TNRPDILDPALLRPG | GARHRDRRGEQY | 0.6266633 |
| LTTTTTS | GARHRDRRGEQY | 0.12273507 |
| PTGSGKTL | GARHRDRRGEQY | 0.47300568 |
| TGSGKTL | GARHRDRRGEQY | 0.22349963 |
| TIGHVDHGK | GARHRDRRGEQY | 0.29889834 |
| TIGHVDHGKTT | GARHRDRRGEQY | 0.19938144 |
| GHVDHGKTT | GARHRDRRGEQY | 0.13655691 |
| AQFEHTLL | GARHRDRRGEQY | 0.20282854 |
| IVNQDEL | GARHRDRRGEQY | 0.15932883 |
| TNGSQFF | GARHRDRRGEQY | 0.82417595 |
| TGKGIVE | GARHRDRRGEQY | 0.315947 |
| LISNASDA | GARHRDRRGEQY | 0.20523977 |
| AELRTLGELTKIAKAHGAQ | GARHRDRRGEQY | 0.47158316 |
| APNAVAAAEPSW | GARHRDRRGEQY | 0.43954462 |
| APQVAGAATAVL | GARHRDRRGEQY | 0.11113636 |
| ARRDPAAAAAILPTDA | GARHRDRRGEQY | 0.1507831 |

| | | |
|---|---|---|
| DAAARDVLAEAGLAEYF | GARHRDRRGEQY | 0.17745581 |
| DDQIVLSAKANLAKVEQ | GARHRDRRGEQY | 0.23594075 |
| DELLPDTGEDSDEDGH | GARHRDRRGEQY | 0.10711243 |
| ELPEAPANRATVPQD | GARHRDRRGEQY | 0.27160487 |
| FGQSKIERVTGIG | GARHRDRRGEQY | 0.12744975 |
| GAGGTGSTAGGAGGAGGAGGLY | GARHRDRRGEQY | 0.09452681 |
| GHGGTGAAGGNGGTGGAGGSGI | GARHRDRRGEQY | 0.27171353 |
| GNGGNGGAGGEGGPGGAGGAGA | GARHRDRRGEQY | 0.2112497 |
| GSTAGNGGNGGNATNGTIAGQP | GARHRDRRGEQY | 0.17271022 |
| GTGGDGGNGGNGGTGG | GARHRDRRGEQY | 0.34892052 |
| GVRRPAGIPADAPV | GARHRDRRGEQY | 0.17013289 |
| GYRQVTDAHLLTLARR | GARHRDRRGEQY | 0.14239195 |
| IAKMGQDGHDRG | GARHRDRRGEQY | 0.263066 |
| KAEPSIEIQYT | GARHRDRRGEQY | 0.35909426 |
| KPVRIRMAADEDTVAVT | GARHRDRRGEQY | 0.15068997 |
| MPPRIAGMRLLVIKPEP | GARHRDRRGEQY | 0.24222225 |
| MKFYAVALFRSRGVPPTR | GARHRDRRGEQY | 0.56926703 |
| NAGGFGILGAVAHSP | GARHRDRRGEQY | 0.26657718 |
| NEGREITLAMAEM | GARHRDRRGEQY | 0.38609105 |
| NELVEWSHPLGHRWRWE | GARHRDRRGEQY | 0.4280516 |
| NLGAGGTGGNGGNPG | GARHRDRRGEQY | 0.1806153 |
| NTEWPKWETVMAADPVD | GARHRDRRGEQY | 0.43628833 |
| PASAAPAETQP | GARHRDRRGEQY | 0.5775424 |
| QQSSDGKNGVRAAEITGE | GARHRDRRGEQY | 0.21075131 |
| RAPDAVTTSWWKE | GARHRDRRGEQY | 0.30942246 |
| SGFGNVGAHTSGWFNQ | GARHRDRRGEQY | 0.13258506 |
| SGVAGAVGQAASVGGLKV | GARHRDRRGEQY | 0.22198838 |
| SNGGAGTGSAGGNGGT | GARHRDRRGEQY | 0.1304347 |
| TGGAGGEGGAAG | GARHRDRRGEQY | 0.31831232 |
| TGGDGGAAGAGGAPGHG | GARHRDRRGEQY | 0.37483895 |
| TMPPLLAAIANAGRHGV | GARHRDRRGEQY | 0.19776401 |
| TSSTAGNSGGA | GARHRDRRGEQY | 0.104378425 |
| VPATVPVGILC | GARHRDRRGEQY | 0.11206212 |
| VQRPDSGGVVLTGRIS | GARHRDRRGEQY | 0.27004853 |
| WTPNPVPATKGVPAGLVQAT | GARHRDRRGEQY | 0.13952073 |
| YALKRALAKFSDRFDVVII | GARHRDRRGEQY | 0.13053763 |
| YLIGSVIV | GARHRDRRGEQY | 0.17308362 |
| YGQLVAVID | GARHRDRRGEQY | 0.35962683 |
| VTVGGMTLPA | GARHRDRRGEQY | 0.44489434 |
| VLALPAV | GARHRDRRGEQY | 0.16002297 |
| VIPPTLNYET | GARHRDRRGEQY | 0.6136361 |
| VGMGGMHPSSG | GARHRDRRGEQY | 0.6589277 |
| VGLAGGAATA | GARHRDRRGEQY | 0.5954155 |
| VGGAGGAGGSGP | GARHRDRRGEQY | 0.20390433 |
| VFAAFMV | GARHRDRRGEQY | 0.12793249 |
| TLYDKLR | GARHRDRRGEQY | 0.77842164 |
| TLAVFTVL | GARHRDRRGEQY | 0.2374669 |
| TGSVQPARV | GARHRDRRGEQY | 0.4247641 |
| TGGGAGSGGT | GARHRDRRGEQY | 0.36967385 |
| SHLETSL | GARHRDRRGEQY | 0.19057913 |
| SFTTGMT | GARHRDRRGEQY | 0.15496321 |

| | | |
|---|---|---|
| QVEVAKRMGVRQPTVS | GARHRDRRGEQY | 0.5428869 |
| QGSGGPALAM | GARHRDRRGEQY | 0.3760922 |
| MLAPSNP | GARHRDRRGEQY | 0.119569525 |
| LVNNAAI | GARHRDRRGEQY | 0.15359266 |
| LTGEINTLA | GARHRDRRGEQY | 0.17420498 |
| LSIFGPV | GARHRDRRGEQY | 0.21100807 |
| LLALLANL | GARHRDRRGEQY | 0.37690455 |
| LISNASDALDKLR | GARHRDRRGEQY | 0.22384854 |
| LINEAVP | GARHRDRRGEQY | 0.18721107 |
| LGYAFSGL | GARHRDRRGEQY | 0.3043674 |
| LDFVAVAL | GARHRDRRGEQY | 0.29995713 |
| LAQPGAE | GARHRDRRGEQY | 0.2958391 |
| ISNLGAVTGF | GARHRDRRGEQY | 0.25164312 |
| HQEDPGRI | GARHRDRRGEQY | 0.19475217 |
| GTPGEPTP | GARHRDRRGEQY | 0.3432566 |
| GFFNSGS | GARHRDRRGEQY | 0.4425594 |
| GAGGGGLTAGAN | GARHRDRRGEQY | 0.28966185 |
| FVDALVNM | GARHRDRRGEQY | 0.33208764 |
| FLTVAPEL | GARHRDRRGEQY | 0.23210533 |
| FETLTAL | GARHRDRRGEQY | 0.2434521 |
| DVINPIEL | GARHRDRRGEQY | 0.12280747 |
| APVSGAPV | GARHRDRRGEQY | 0.22424608 |
| ALVGGLAW | GARHRDRRGEQY | 0.23918453 |
| ALAAAAATAI | GARHRDRRGEQY | 0.18672149 |
| AFGLGLLL | GARHRDRRGEQY | 0.44808766 |
| AFFAAASL | GARHRDRRGEQY | 0.24490908 |
| AAAALPAFSPP | GARHRDRRGEQY | 0.16129245 |
| VVLAPAL | GARHRDRRGEQY | 0.32375437 |
| TIGHVDHGKTTL | GARHRDRRGEQY | 0.5683787 |
| LISNASDALDK | GARHRDRRGEQY | 0.1475546 |
| LISNASDAL | GARHRDRRGEQY | 0.19243735 |
| ISNASDALDK | GARHRDRRGEQY | 0.6455009 |
| NQAETLVYQTE | GARHRDRRGEQY | 0.54051936 |
| TGGTGGTAGAG | GARHRDRRGEQY | 0.22590736 |
| TGSTAGGAGGA | GARHRDRRGEQY | 0.27663177 |
| DPLAVPVAAGPV | GARHRDRRGEQY | 0.8332248 |
| GGTGGTPGNGGN | GARHRDRRGEQY | 0.2820161 |
| PAGLAGVDPATL | GARHRDRRGEQY | 0.5108622 |
| GNNLVVEAPHAGASV | GARHRDRRGEQY | 0.11555833 |
| KVTRIMSASTLEQPA | GARHRDRRGEQY | 0.17986217 |
| LGFALPIALPRRTVV | GARHRDRRGEQY | 0.14544399 |
| TRLGNLTARKLVVLV | GARHRDRRGEQY | 0.09511728 |
| WSQQRRLIYFPSAGP | GARHRDRRGEQY | 0.5225986 |
| GPNGNPDPMAAAVDIR | GARHRDRRGEQY | 0.18398526 |
| TQRCVSVADLRAAASR | GARHRDRRGEQY | 0.36699802 |
| QFVLDVAEHPVEVLTETV | GARHRDRRGEQY | 0.18924399 |
| GGAGGTGGAGGNGGTQAGDG | GARHRDRRGEQY | 0.42154884 |
| PLLATLRAIAVDVDAKPAQV | GARHRDRRGEQY | 0.45653018 |
| APKFRTGGYMIDFWGVGYQVAK | GARHRDRRGEQY | 0.23015217 |
| GAGGAGGAGGAAGGPGGTGGTG | GARHRDRRGEQY | 0.16478947 |
| IGALGLLATDVAPAGSVAFGVA | GARHRDRRGEQY | 0.31848896 |

| | | |
|---|---|---|
| AGGAGGIGGNGGAGGAGGVLMG | GARHRDRRGEQY | 0.38884985 |
| ADELIGLL | GARHRDRRGEQY | 0.32411432 |
| ADRALKA | GARHRDRRGEQY | 0.119435325 |
| AGAVGQVGGAGGA | GARHRDRRGEQY | 0.43697459 |
| AGGGGGLGGAP | GARHRDRRGEQY | 0.23723876 |
| AGGLLYGNGGAG | GARHRDRRGEQY | 0.13160926 |
| AGLESIGL | GARHRDRRGEQY | 0.108595766 |
| AGLKTGII | GARHRDRRGEQY | 0.120343216 |
| ASLAALLL | GARHRDRRGEQY | 0.20719096 |
| ASVINISSI | GARHRDRRGEQY | 0.3382371 |
| ATPPGPA | GARHRDRRGEQY | 0.3470844 |
| ATVASALV | GARHRDRRGEQY | 0.2083034 |
| ATVASLIL | GARHRDRRGEQY | 0.36365646 |
| AVNGPVLIRH | GARHRDRRGEQY | 0.29509264 |
| DGAAGAVGI | GARHRDRRGEQY | 0.081170015 |
| DPLLHGI | GARHRDRRGEQY | 0.17644583 |
| ELGAALGAL | GARHRDRRGEQY | 0.116351835 |
| ELGRTQAV | GARHRDRRGEQY | 0.39964882 |
| FVAGGYLS | GARHRDRRGEQY | 0.13693008 |
| GASGGAGGAGGW | GARHRDRRGEQY | 0.2143124 |
| GGAGAGGDGGAGGI | GARHRDRRGEQY | 0.25629008 |
| GTGHNGSQPGMGGNG | GARHRDRRGEQY | 0.6994672 |
| GYAAASASAS | GARHRDRRGEQY | 0.31802112 |
| HQVDVAAVQQG | GARHRDRRGEQY | 0.13275516 |
| IGAAIAHT | GARHRDRRGEQY | 0.10702828 |
| IGMIEVAQL | GARHRDRRGEQY | 0.30870277 |
| IRLAPEHTSAMD | GARHRDRRGEQY | 0.20747808 |
| IYNVSVL | GARHRDRRGEQY | 0.1384127 |
| KQGFLGM | GARHRDRRGEQY | 0.14989929 |
| LESVITQL | GARHRDRRGEQY | 0.30812734 |
| LGLGALVVV | GARHRDRRGEQY | 0.12028907 |
| LIGNGGNGGSGGI | GARHRDRRGEQY | 0.31749102 |
| LNAINAPTL | GARHRDRRGEQY | 0.14595026 |
| LQLLNTGV | GARHRDRRGEQY | 0.19534011 |
| LQTSIEF | GARHRDRRGEQY | 0.53747773 |
| MSFAVSEL | GARHRDRRGEQY | 0.69356877 |
| NTGALGL | GARHRDRRGEQY | 0.2626147 |
| PPGAGGAGGAGTT | GARHRDRRGEQY | 0.17154434 |
| QLVGLGI | GARHRDRRGEQY | 0.14659756 |
| RFLEQVHQL | GARHRDRRGEQY | 0.14780287 |
| RGVGYVLR | GARHRDRRGEQY | 0.15860233 |
| SATGVFAGVI | GARHRDRRGEQY | 0.46501002 |
| SLPAITLP | GARHRDRRGEQY | 0.28058088 |
| STALAFTV | GARHRDRRGEQY | 0.14427157 |
| SVLTVGLGK | GARHRDRRGEQY | 0.3548643 |
| TALAGAGSGI | GARHRDRRGEQY | 0.6023723 |
| TGLPFTGI | GARHRDRRGEQY | 0.6312787 |
| VAEHVTV | GARHRDRRGEQY | 0.1289891 |
| VDIVLNSL | GARHRDRRGEQY | 0.1869076 |
| VLTGGGAVL | GARHRDRRGEQY | 0.21303575 |
| VNVGLAYGI | GARHRDRRGEQY | 0.19525556 |

| | | |
|---|---|---|
| VSSAAVGAV | GARHRDRRGEQY | 0.1602259 |
| VTSVYEAGV | GARHRDRRGEQY | 0.1717949 |
| WVALDRGA | GARHRDRRGEQY | 0.3057143 |
| YLVGSVVI | GARHRDRRGEQY | 0.86221176 |
| GPPGTGKTLL | GARHRDRRGEQY | 0.2187579 |
| GVGMTKF | GARHRDRRGEQY | 0.10239173 |
| IGIGGSDL | GARHRDRRGEQY | 0.21206436 |
| IGTLLAEL | GARHRDRRGEQY | 0.12055778 |
| IYTSGTT | GARHRDRRGEQY | 0.17375526 |
| LSFPAGV | GARHRDRRGEQY | 0.18236682 |
| LYTSGTT | GARHRDRRGEQY | 0.11118962 |
| VGAFPFGF | GARHRDRRGEQY | 0.21658465 |
| VGAGTVEF | GARHRDRRGEQY | 0.5424239 |
| VGKSTLLL | GARHRDRRGEQY | 0.28286147 |
| AALAHAGGGIV | GARHRDRRGEQY | 0.32195973 |
| AALVAVSLL | GARHRDRRGEQY | 0.58578455 |
| AAVSRAVAAL | GARHRDRRGEQY | 0.28190085 |
| AGGSSGIGGF | GARHRDRRGEQY | 0.5552588 |
| AGLTVANANG | GARHRDRRGEQY | 0.5482064 |
| AGLVGAVL | GARHRDRRGEQY | 0.24136522 |
| ALSQAFTV | GARHRDRRGEQY | 0.19459179 |
| ALTFSVSL | GARHRDRRGEQY | 0.5108969 |
| ALTGAALVAL | GARHRDRRGEQY | 0.27010325 |
| ASFSIGAGVL | GARHRDRRGEQY | 0.2756054 |
| DGSVYKI | GARHRDRRGEQY | 0.33588833 |
| EQLAEIV | GARHRDRRGEQY | 0.49640045 |
| FAVSELTVM | GARHRDRRGEQY | 0.18425296 |
| GAAPLAAGL | GARHRDRRGEQY | 0.798627 |
| GADGGDGGSGGAGG | GARHRDRRGEQY | 0.106902 |
| GFGLDTGGAGG | GARHRDRRGEQY | 0.15636322 |
| GLASGGLL | GARHRDRRGEQY | 0.45001587 |
| GRIAGAAVLL | GARHRDRRGEQY | 0.08905019 |
| GSAGSLNTSF | GARHRDRRGEQY | 0.130863 |
| GTGQIGI | GARHRDRRGEQY | 0.18580668 |
| GTLIALIL | GARHRDRRGEQY | 0.5947048 |
| IAATMGFVVG | GARHRDRRGEQY | 0.0914233 |
| IFAQALL | GARHRDRRGEQY | 0.18848053 |
| IGAAVALGF | GARHRDRRGEQY | 0.12314678 |
| IGLTPLAL | GARHRDRRGEQY | 0.2864727 |
| ILGSNTSTL | GARHRDRRGEQY | 0.24720329 |
| IVFSLTVL | GARHRDRRGEQY | 0.13589036 |
| LASKATGF | GARHRDRRGEQY | 0.33531225 |
| LASVALIL | GARHRDRRGEQY | 0.121060275 |
| LFAVGLTL | GARHRDRRGEQY | 0.35621452 |
| LFNGGIGTYI | GARHRDRRGEQY | 0.36584777 |
| LTLGAGII | GARHRDRRGEQY | 0.75695986 |
| LVAAALGDI | GARHRDRRGEQY | 0.12836482 |
| NTAFVQLGI | GARHRDRRGEQY | 0.28542736 |
| QLFSGAEL | GARHRDRRGEQY | 0.14288962 |
| SAATFITL | GARHRDRRGEQY | 0.7755642 |
| SNYGLANLGV | GARHRDRRGEQY | 0.20722121 |

| | | |
|---|---|---|
| SYGGAYAVM | GARHRDRRGEQY | 0.11200948 |
| TTGVGGLTLG | GARHRDRRGEQY | 0.31206554 |
| TTQTIYI | GARHRDRRGEQY | 0.47308457 |
| VAAGAGLGAVI | GARHRDRRGEQY | 0.20107248 |
| VAAVVTLL | GARHRDRRGEQY | 0.6236633 |
| VGFLGGLI | GARHRDRRGEQY | 0.18728326 |
| VGVIPILL | GARHRDRRGEQY | 0.2236228 |
| VIFASSLI | GARHRDRRGEQY | 0.58483297 |
| VKGGDGGAAAG | GARHRDRRGEQY | 0.31667197 |
| VTKTFTNI | GARHRDRRGEQY | 0.181719 |
| VVAAAILL | GARHRDRRGEQY | 0.45149708 |
| YAALDVEL | GARHRDRRGEQY | 0.25304455 |
| LASVALLL | GARHRDRRGEQY | 0.1358592 |
| LDAVHRV | GARHRDRRGEQY | 0.7171456 |
| QGLSSSIF | GARHRDRRGEQY | 0.18639988 |
| RGTFANIRL | GARHRDRRGEQY | 0.4427407 |
| RQLWWGHRI | GARHRDRRGEQY | 0.3159226 |
| YGPPGTGKTLL | GARHRDRRGEQY | 0.84224856 |
| AALVGAPANVA | GARHRDRRGEQY | 0.1560782 |
| AATFRRTE | GARHRDRRGEQY | 0.25887412 |
| AGALGGGI | GARHRDRRGEQY | 0.2396863 |
| AGGAGGAGGIGAG | GARHRDRRGEQY | 0.2778855 |
| AGSGALGL | GARHRDRRGEQY | 0.23319295 |
| AQIADQI | GARHRDRRGEQY | 0.20864514 |
| ASAVAVTYL | GARHRDRRGEQY | 0.37047994 |
| ASTVLGGTGG | GARHRDRRGEQY | 0.43907905 |
| AVGGEPAK | GARHRDRRGEQY | 0.18300267 |
| CEQGTFGGTC | GARHRDRRGEQY | 0.193792 |
| DEGKAPAG | GARHRDRRGEQY | 0.6527369 |
| DTVVFPAL | GARHRDRRGEQY | 0.13660865 |
| FEGITMAG | GARHRDRRGEQY | 0.29286367 |
| FPLVVQD | GARHRDRRGEQY | 0.27883047 |
| GAFTTPQ | GARHRDRRGEQY | 0.36914432 |
| GFGGFGTAGGVGGA | GARHRDRRGEQY | 0.1991528 |
| GFLAQFL | GARHRDRRGEQY | 0.5456332 |
| GGAGGAGGNAGVA | GARHRDRRGEQY | 0.19440143 |
| GGVGTAGAGGAGGAGG | GARHRDRRGEQY | 0.41967744 |
| GNGGGGDAGNAGSGGN | GARHRDRRGEQY | 0.2780542 |
| GTGGAGGAGNDAGS | GARHRDRRGEQY | 0.2495182 |
| GTTNTGWFNS | GARHRDRRGEQY | 0.1154529 |
| IFMGPLFTL | GARHRDRRGEQY | 0.24273582 |
| ILAHAIEAI | GARHRDRRGEQY | 0.38185865 |
| IRSAIREAL | GARHRDRRGEQY | 0.42374998 |
| ITFAGGG | GARHRDRRGEQY | 0.12201198 |
| ITGMSGHQEV | GARHRDRRGEQY | 0.3866424 |
| IVAGLACTL | GARHRDRRGEQY | 0.4314286 |
| IVGGGLAGA | GARHRDRRGEQY | 0.8245673 |
| IVTGVLNI | GARHRDRRGEQY | 0.19402185 |
| LAQQGCI | GARHRDRRGEQY | 0.38246247 |
| LCLHVEN | GARHRDRRGEQY | 0.17157954 |
| LDAISDGS | GARHRDRRGEQY | 0.20416951 |

| | | |
|---|---|---|
| LGGFAAML | GARHRDRRGEQY | 0.10715626 |
| LGGKNPMI | GARHRDRRGEQY | 0.14199013 |
| LGPKGVAAIL | GARHRDRRGEQY | 0.2708353 |
| LMVGVSI | GARHRDRRGEQY | 0.13397537 |
| LVGAAFVL | GARHRDRRGEQY | 0.7748133 |
| LVLAVAPL | GARHRDRRGEQY | 0.4028268 |
| MAAIAVIDAA | GARHRDRRGEQY | 0.36736786 |
| NNGINAELQ | GARHRDRRGEQY | 0.4294842 |
| RGEPLVLVHG | GARHRDRRGEQY | 0.64653015 |
| RGGAGGAAGLI | GARHRDRRGEQY | 0.32471657 |
| RSYRMHPAV | GARHRDRRGEQY | 0.20150916 |
| SAAHLLAGQLA | GARHRDRRGEQY | 0.26198894 |
| SAAVGSY | GARHRDRRGEQY | 0.10528661 |
| SPHAVLHRI | GARHRDRRGEQY | 0.18791018 |
| SQIAKVL | GARHRDRRGEQY | 0.1774933 |
| TALGHLVP | GARHRDRRGEQY | 0.29362386 |
| TAPTTSGAA | GARHRDRRGEQY | 0.08719598 |
| TDGDDGNGGSPGT | GARHRDRRGEQY | 0.6207293 |
| TDGLGEN | GARHRDRRGEQY | 0.5222907 |
| TGGPPLPGQA | GARHRDRRGEQY | 0.24847244 |
| TIPAITI | GARHRDRRGEQY | 0.4046632 |
| TIPGFTI | GARHRDRRGEQY | 0.46847337 |
| TSIASGALA | GARHRDRRGEQY | 0.12158576 |
| TVTLPGAAI | GARHRDRRGEQY | 0.3006323 |
| VASGLSAALP | GARHRDRRGEQY | 0.15074211 |
| VGAGVIGI | GARHRDRRGEQY | 0.39460358 |
| VGPKRFGV | GARHRDRRGEQY | 0.51065934 |
| VNVAFPD | GARHRDRRGEQY | 0.32588446 |
| VSMTNSGVS | GARHRDRRGEQY | 0.15808955 |
| VTDVLGDLSV | GARHRDRRGEQY | 0.28996593 |
| VVVVGAGFAGL | GARHRDRRGEQY | 0.16254343 |
| VYTVGTVV | GARHRDRRGEQY | 0.17112124 |
| WTTFSGP | GARHRDRRGEQY | 0.13317555 |
| YDELVEAF | GARHRDRRGEQY | 0.31887352 |
| YGTGGAGGNGG | GARHRDRRGEQY | 0.15522435 |
| YTNVMAV | GARHRDRRGEQY | 0.22137049 |
| FLEQVHQL | GARHRDRRGEQY | 0.26461598 |
| GLSLLAVAV | GARHRDRRGEQY | 0.20285359 |
| LEQVHQL | GARHRDRRGEQY | 0.15593381 |
| LNAAGAI | GARHRDRRGEQY | 0.37877652 |
| PAVHASL | GARHRDRRGEQY | 0.3759894 |
| TELEDDA | GARHRDRRGEQY | 0.1204666 |
| TPPGPPN | GARHRDRRGEQY | 0.39426515 |
| TPPPPPPP | GARHRDRRGEQY | 0.18044776 |
| TQSPALI | ASSQDRKNTLY | 0.2981069 |
| EFAAAFPQ | ASSQDRKNTLY | 0.4127605 |
| LLSGNLDVL | ASSQDRKNTLY | 0.22165416 |
| GPANLGNYNI | ASSQDRKNTLY | 0.14265804 |
| TLGPTLP | ASSQDRKNTLY | 0.17245328 |
| VYAYSSCCK | ASSQDRKNTLY | 0.22154263 |
| APAPAPAPNI | ASSQDRKNTLY | 0.20446447 |

| | | |
|---|---|---|
| PGGPVPIP | ASSQDRKNTLY | 0.104212455 |
| FLTVALEP | ASSQDRKNTLY | 0.36547613 |
| SQYFGANL | ASSQDRKNTLY | 0.13664249 |
| GGVKPAA | ASSQDRKNTLY | 0.20498449 |
| GGTGGAGGAGGW | ASSQDRKNTLY | 0.19004552 |
| GVGGVAGLGGAGG | ASSQDRKNTLY | 0.41257155 |
| VLLVLATTI | ASSQDRKNTLY | 0.2851746 |
| IGIFAAML | ASSQDRKNTLY | 0.206408 |
| AVGDLID | ASSQDRKNTLY | 0.3038054 |
| NISPLPL | ASSQDRKNTLY | 0.28635746 |
| VTAVFVAI | ASSQDRKNTLY | 0.55925155 |
| SGDDNDAAL | ASSQDRKNTLY | 0.22395581 |
| AVAALAHS | ASSQDRKNTLY | 0.18116431 |
| GGVTVMATP | ASSQDRKNTLY | 0.32289857 |
| AGLAANVRD | ASSQDRKNTLY | 0.2109701 |
| TNRPDLIDPALLRPG | ASSQDRKNTLY | 0.1558546 |
| LANLGVTAL | ASSQDRKNTLY | 0.1429319 |
| TGERIEL | ASSQDRKNTLY | 0.15477948 |
| NGEELAA | ASSQDRKNTLY | 0.1527659 |
| VLSLVWG | ASSQDRKNTLY | 0.3612113 |
| IGYALWL | ASSQDRKNTLY | 0.42946655 |
| LSLLLAPL | ASSQDRKNTLY | 0.222265 |
| TLDEALK | ASSQDRKNTLY | 0.11452854 |
| TDGAGTAEGG | ASSQDRKNTLY | 0.12929648 |
| NAPGGAGGNGG | ASSQDRKNTLY | 0.22107114 |
| VSVFMGMGGNF | ASSQDRKNTLY | 0.3708703 |
| LASYLAAL | ASSQDRKNTLY | 0.2930491 |
| LVGGTPAL | ASSQDRKNTLY | 0.84148896 |
| VRNADVL | ASSQDRKNTLY | 0.41994566 |
| TCSSPFLVSAM | ASSQDRKNTLY | 0.19354239 |
| VHITDAE | ASSQDRKNTLY | 0.350624 |
| EYAAPSF | ASSQDRKNTLY | 0.22157332 |
| VVDAFHGGL | ASSQDRKNTLY | 0.21740893 |
| ASVVDPQ | ASSQDRKNTLY | 0.22923225 |
| SYNEFGGLS | ASSQDRKNTLY | 0.28642452 |
| DPDGSVY | ASSQDRKNTLY | 0.15761551 |
| MVSTIHATE | ASSQDRKNTLY | 0.21007341 |
| LAAGVADPV | ASSQDRKNTLY | 0.13027255 |
| TKEQIAA | ASSQDRKNTLY | 0.32514095 |
| SLASPAAL | ASSQDRKNTLY | 0.13410684 |
| IGLVFYGV | ASSQDRKNTLY | 0.14437588 |
| FLLDASNL | ASSQDRKNTLY | 0.24023995 |
| YQVGNFP | ASSQDRKNTLY | 0.5680211 |
| ESLLSQST | ASSQDRKNTLY | 0.4592031 |
| LFTVGFAAVG | ASSQDRKNTLY | 0.15227416 |
| ATPPTPT | ASSQDRKNTLY | 0.15927954 |
| TDLATLG | ASSQDRKNTLY | 0.62512183 |
| SAGGTGR | ASSQDRKNTLY | 0.15426004 |
| GFSNSGS | ASSQDRKNTLY | 0.3103628 |
| GGNGGSGGTSV | ASSQDRKNTLY | 0.29912695 |
| GTSVATGGAGN | ASSQDRKNTLY | 0.3097444 |

| | | |
|---|---|---|
| NGGNGGSGGTS | ASSQDRKNTLY | 0.18621188 |
| AVVKFPH | ASSQDRKNTLY | 0.22207552 |
| TPPPPPP | ASSQDRKNTLY | 0.20970196 |
| AAPAAAAAPA | ASSQDRKNTLY | 0.23942731 |
| IALNEDL | ASSQDRKNTLY | 0.10821781 |
| IVNVSSM | ASSQDRKNTLY | 0.36129186 |
| TGIPPAPR | ASSQDRKNTLY | 0.14489116 |
| FDLGGGTF | ASSQDRKNTLY | 0.14748752 |
| FDLGGGTFDVS | ASSQDRKNTLY | 0.097589195 |
| NEPTAAA | ASSQDRKNTLY | 0.21043664 |
| NEPTAAALA | ASSQDRKNTLY | 0.31292385 |
| GSIINISSI | ASSQDRKNTLY | 0.37725988 |
| VAPGNAGTA | ASSQDRKNTLY | 0.33994025 |
| TGGGASLE | ASSQDRKNTLY | 0.33619323 |
| VAVYNHYKRI | ASSQDRKNTLY | 0.47810364 |
| TDPAPATT | ASSQDRKNTLY | 0.22545257 |
| TNRPDILDPALLRPG | ASSQDRKNTLY | 0.110098734 |
| LTTTTTS | ASSQDRKNTLY | 0.23812169 |
| PTGSGKTL | ASSQDRKNTLY | 0.42881322 |
| TGSGKTL | ASSQDRKNTLY | 0.37225336 |
| TIGHVDHGK | ASSQDRKNTLY | 0.42602956 |
| TIGHVDHGKTT | ASSQDRKNTLY | 0.10637577 |
| GHVDHGKTT | ASSQDRKNTLY | 0.3105631 |
| AQFEHTLL | ASSQDRKNTLY | 0.32079414 |
| IVNQDEL | ASSQDRKNTLY | 0.120432615 |
| TNGSQFF | ASSQDRKNTLY | 0.1647871 |
| TGKGIVE | ASSQDRKNTLY | 0.18889393 |
| LISNASDA | ASSQDRKNTLY | 0.11474549 |
| AELRTLGELTKIAKAHGAQ | ASSQDRKNTLY | 0.2805084 |
| APNAVAAAEPSW | ASSQDRKNTLY | 0.1656265 |
| APQVAGAATAVL | ASSQDRKNTLY | 0.20076433 |
| ARRDPAAAAAILPTDA | ASSQDRKNTLY | 0.18409762 |
| DAAARDVLAEAGLAEYF | ASSQDRKNTLY | 0.16722092 |
| DDQIVLSAKANLAKVEQ | ASSQDRKNTLY | 0.21548292 |
| DELLPDTGEDSDEDGH | ASSQDRKNTLY | 0.19928582 |
| ELPEAPANRATVPQD | ASSQDRKNTLY | 0.3746751 |
| FGQSKIERVTGIG | ASSQDRKNTLY | 0.24040955 |
| GAGGTGSTAGGAGGAGGAGGLY | ASSQDRKNTLY | 0.24851966 |
| GHGGTGAAGGNGGTGGAGGSGI | ASSQDRKNTLY | 0.27969772 |
| GNGGNGGAGGEGGPGGAGGAGA | ASSQDRKNTLY | 0.6161977 |
| GSTAGNGGNGGNATNGTIAGQP | ASSQDRKNTLY | 0.12804891 |
| GTGGDGGNGGNGGTGG | ASSQDRKNTLY | 0.18725066 |
| GVRRPAGIPADAPV | ASSQDRKNTLY | 0.24720669 |
| GYRQVTDAHLLTLARR | ASSQDRKNTLY | 0.17151974 |
| IAKMGQDGHDRG | ASSQDRKNTLY | 0.40305683 |
| KAEPSIEIQYT | ASSQDRKNTLY | 0.26094824 |
| KPVRIRMAADEDTVAVT | ASSQDRKNTLY | 0.315395 |
| MPPRIAGMRLLVIKPEP | ASSQDRKNTLY | 0.20996267 |
| MKFYAVALFRSRGVPPTR | ASSQDRKNTLY | 0.65534675 |
| NAGGFGILGAVAHSP | ASSQDRKNTLY | 0.2992891 |
| NEGREITLAMAEM | ASSQDRKNTLY | 0.21068087 |

| | | |
|---|---|---|
| NELVEWSHPLGHRWRWE | ASSQDRKNTLY | 0.49651647 |
| NLGAGGTGGNGGNPG | ASSQDRKNTLY | 0.15079625 |
| NTEWPKWETVMAADPVD | ASSQDRKNTLY | 0.38045144 |
| PASAAPAETQP | ASSQDRKNTLY | 0.33475912 |
| QQSSDGKNGVRAAEITGE | ASSQDRKNTLY | 0.14797482 |
| RAPDAVTTSWWKE | ASSQDRKNTLY | 0.22080153 |
| SGFGNVGAHTSGWFNQ | ASSQDRKNTLY | 0.44819817 |
| SGVAGAVGQAASVGGLKV | ASSQDRKNTLY | 0.53285944 |
| SNGGAGTGSAGGNGGT | ASSQDRKNTLY | 0.16089728 |
| TGGAGGEGGAAG | ASSQDRKNTLY | 0.14107437 |
| TGGDGGAAGAGGAPGHG | ASSQDRKNTLY | 0.13405213 |
| TMPPLLAAIANAGRHGV | ASSQDRKNTLY | 0.14704752 |
| TSSTAGNSGGA | ASSQDRKNTLY | 0.13079081 |
| VPATVPVGILC | ASSQDRKNTLY | 0.30106348 |
| VQRPDSGGVVLTGRIS | ASSQDRKNTLY | 0.23629618 |
| WTPNPVPATKGVPAGLVQAT | ASSQDRKNTLY | 0.77486223 |
| YALKRALAKFSDRFDVVII | ASSQDRKNTLY | 0.34167603 |
| YLIGSVIV | ASSQDRKNTLY | 0.48296335 |
| YGQLVAVID | ASSQDRKNTLY | 0.15916272 |
| VTVGGMTLPA | ASSQDRKNTLY | 0.21079084 |
| VLALPAV | ASSQDRKNTLY | 0.115457304 |
| VIPPTLNYET | ASSQDRKNTLY | 0.13614573 |
| VGMGGMHPSSG | ASSQDRKNTLY | 0.13319996 |
| VGLAGGAATA | ASSQDRKNTLY | 0.21156362 |
| VGGAGGAGGSGP | ASSQDRKNTLY | 0.11385639 |
| VFAAFMV | ASSQDRKNTLY | 0.7049202 |
| TLYDKLR | ASSQDRKNTLY | 0.19117574 |
| TLAVFTVL | ASSQDRKNTLY | 0.32986367 |
| TGSVQPARV | ASSQDRKNTLY | 0.20291245 |
| TGGGAGSGGT | ASSQDRKNTLY | 0.14015064 |
| SHLETSL | ASSQDRKNTLY | 0.19367239 |
| SFTTGMT | ASSQDRKNTLY | 0.30356157 |
| QVEVAKRMGVRQPTVS | ASSQDRKNTLY | 0.1225799 |
| QGSGGPALAM | ASSQDRKNTLY | 0.11450022 |
| MLAPSNP | ASSQDRKNTLY | 0.16802068 |
| LVNNAAI | ASSQDRKNTLY | 0.13929787 |
| LTGEINTLA | ASSQDRKNTLY | 0.7857604 |
| LSIFGPV | ASSQDRKNTLY | 0.43369484 |
| LLALLANL | ASSQDRKNTLY | 0.21774912 |
| LISNASDALDKLR | ASSQDRKNTLY | 0.782392 |
| LINEAVP | ASSQDRKNTLY | 0.08866017 |
| LGYAFSGL | ASSQDRKNTLY | 0.21190467 |
| LDFVAVAL | ASSQDRKNTLY | 0.6476989 |
| LAQPGAE | ASSQDRKNTLY | 0.24351719 |
| ISNLGAVTGF | ASSQDRKNTLY | 0.24395284 |
| HQEDPGRI | ASSQDRKNTLY | 0.41152835 |
| GTPGEPTP | ASSQDRKNTLY | 0.1647145 |
| GFFNSGS | ASSQDRKNTLY | 0.21299309 |
| GAGGGGLTAGAN | ASSQDRKNTLY | 0.1889513 |
| FVDALVNM | ASSQDRKNTLY | 0.335604 |
| FLTVAPEL | ASSQDRKNTLY | 0.26627174 |

| | | |
|---|---|---|
| FETLTAL | ASSQDRKNTLY | 0.1224184 |
| DVINPIEL | ASSQDRKNTLY | 0.3552941 |
| APVSGAPV | ASSQDRKNTLY | 0.7942214 |
| ALVGGLAW | ASSQDRKNTLY | 0.15200256 |
| ALAAAAATAI | ASSQDRKNTLY | 0.2870216 |
| AFGLGLLL | ASSQDRKNTLY | 0.2540995 |
| AFFAAASL | ASSQDRKNTLY | 0.13560857 |
| AAAALPAFSPP | ASSQDRKNTLY | 0.18791598 |
| VVLAPAL | ASSQDRKNTLY | 0.11828045 |
| TIGHVDHGKTTL | ASSQDRKNTLY | 0.44390303 |
| LISNASDALDK | ASSQDRKNTLY | 0.15278909 |
| LISNASDAL | ASSQDRKNTLY | 0.20322503 |
| ISNASDALDK | ASSQDRKNTLY | 0.3092175 |
| NQAETLVYQTE | ASSQDRKNTLY | 0.7922201 |
| TGGTGGTAGAG | ASSQDRKNTLY | 0.4364462 |
| TGSTAGGAGGA | ASSQDRKNTLY | 0.27131015 |
| DPLAVPVAAGPV | ASSQDRKNTLY | 0.13091461 |
| GGTGGTPGNGGN | ASSQDRKNTLY | 0.2149306 |
| PAGLAGVDPATL | ASSQDRKNTLY | 0.31439507 |
| GNNLVVEAPHAGASV | ASSQDRKNTLY | 0.14261031 |
| KVTRIMSASTLEQPA | ASSQDRKNTLY | 0.4713404 |
| LGFALPIALPRRTVV | ASSQDRKNTLY | 0.14981878 |
| TRLGNLTARKLVVLV | ASSQDRKNTLY | 0.3458886 |
| WSQQRRLIYFPSAGP | ASSQDRKNTLY | 0.33976018 |
| GPNGNPDPMAAAVDIR | ASSQDRKNTLY | 0.82864875 |
| TQRCVSVADLRAAASR | ASSQDRKNTLY | 0.32852682 |
| QFVLDVAEHPVEVLTETV | ASSQDRKNTLY | 0.49057245 |
| GGAGGTGGAGGNGGTQAGDG | ASSQDRKNTLY | 0.12674813 |
| PLLATLRAIAVDVDAKPAQV | ASSQDRKNTLY | 0.3402639 |
| APKFRTGGYMIDFWGVGYQVAK | ASSQDRKNTLY | 0.47868663 |
| GAGGAGGAGGAAGGPGGTGGTG | ASSQDRKNTLY | 0.42556316 |
| IGALGLLATDVAPAGSVAFGVA | ASSQDRKNTLY | 0.8262822 |
| AGGAGGIGGNGGAGGAGGVLMG | ASSQDRKNTLY | 0.3033871 |
| ADELIGLL | ASSQDRKNTLY | 0.11488146 |
| ADRALKA | ASSQDRKNTLY | 0.20207797 |
| AGAVGQVGGAGGA | ASSQDRKNTLY | 0.45240238 |
| AGGGGGLGGAP | ASSQDRKNTLY | 0.46821213 |
| AGGLLYGNGGAG | ASSQDRKNTLY | 0.40952373 |
| AGLESIGL | ASSQDRKNTLY | 0.3203513 |
| AGLKTGII | ASSQDRKNTLY | 0.18610787 |
| ASLAALLL | ASSQDRKNTLY | 0.5333586 |
| ASVINISSI | ASSQDRKNTLY | 0.780713 |
| ATPPGPA | ASSQDRKNTLY | 0.31520134 |
| ATVASALV | ASSQDRKNTLY | 0.15023261 |
| ATVASLIL | ASSQDRKNTLY | 0.15097566 |
| AVNGPVLIRH | ASSQDRKNTLY | 0.33782732 |
| DGAAGAVGI | ASSQDRKNTLY | 0.47458154 |
| DPLLHGI | ASSQDRKNTLY | 0.1301255 |
| ELGAALGAL | ASSQDRKNTLY | 0.28061455 |
| ELGRTQAV | ASSQDRKNTLY | 0.22002399 |
| FVAGGYLS | ASSQDRKNTLY | 0.101141036 |

| | | |
|---|---|---|
| GASGGAGGAGGW | ASSQDRKNTLY | 0.43172312 |
| GGAGAGGDGGAGGI | ASSQDRKNTLY | 0.63981926 |
| GTGHNGSQPGMGGNG | ASSQDRKNTLY | 0.33157828 |
| GYAAASASAS | ASSQDRKNTLY | 0.1209263 |
| HQVDVAAVQQG | ASSQDRKNTLY | 0.51851845 |
| IGAAIAHT | ASSQDRKNTLY | 0.6631759 |
| IGMIEVAQL | ASSQDRKNTLY | 0.46744275 |
| IRLAPEHTSAMD | ASSQDRKNTLY | 0.14239448 |
| IYNVSVL | ASSQDRKNTLY | 0.090752386 |
| KQGFLGM | ASSQDRKNTLY | 0.24527672 |
| LESVITQL | ASSQDRKNTLY | 0.6078646 |
| LGLGALVVV | ASSQDRKNTLY | 0.18039384 |
| LIGNGGNGGSGGI | ASSQDRKNTLY | 0.1308621 |
| LNAINAPTL | ASSQDRKNTLY | 0.20033862 |
| LQLLNTGV | ASSQDRKNTLY | 0.5027909 |
| LQTSIEF | ASSQDRKNTLY | 0.13160643 |
| MSFAVSEL | ASSQDRKNTLY | 0.12555231 |
| NTGALGL | ASSQDRKNTLY | 0.24699932 |
| PPGAGGAGGAGTT | ASSQDRKNTLY | 0.35211945 |
| QLVGLGI | ASSQDRKNTLY | 0.13825095 |
| RFLEQVHQL | ASSQDRKNTLY | 0.1438251 |
| RGVGYVLR | ASSQDRKNTLY | 0.2927756 |
| SATGVFAGVI | ASSQDRKNTLY | 0.2896917 |
| SLPAITLP | ASSQDRKNTLY | 0.24989979 |
| STALAFTV | ASSQDRKNTLY | 0.13205847 |
| SVLTVGLGK | ASSQDRKNTLY | 0.4335733 |
| TALAGAGSGI | ASSQDRKNTLY | 0.14478998 |
| TGLPFTGI | ASSQDRKNTLY | 0.3597355 |
| VAEHVTV | ASSQDRKNTLY | 0.21279047 |
| VDIVLNSL | ASSQDRKNTLY | 0.2570364 |
| VLTGGGAVL | ASSQDRKNTLY | 0.24446577 |
| VNVGLAYGI | ASSQDRKNTLY | 0.15426251 |
| VSSAAVGAV | ASSQDRKNTLY | 0.1496515 |
| VTSVYEAGV | ASSQDRKNTLY | 0.11063433 |
| WVALDRGA | ASSQDRKNTLY | 0.42568195 |
| YLVGSVVI | ASSQDRKNTLY | 0.2063486 |
| GPPGTGKTLL | ASSQDRKNTLY | 0.37980342 |
| GVGMTKF | ASSQDRKNTLY | 0.34444758 |
| IGIGGSDL | ASSQDRKNTLY | 0.14033946 |
| IGTLLAEL | ASSQDRKNTLY | 0.2577299 |
| IYTSGTT | ASSQDRKNTLY | 0.1128829 |
| LSFPAGV | ASSQDRKNTLY | 0.38617092 |
| LYTSGTT | ASSQDRKNTLY | 0.19442149 |
| VGAFPFGF | ASSQDRKNTLY | 0.29977205 |
| VGAGTVEF | ASSQDRKNTLY | 0.20565173 |
| VGKSTLLL | ASSQDRKNTLY | 0.26074558 |
| AALAHAGGGIV | ASSQDRKNTLY | 0.3458219 |
| AALVAVSLL | ASSQDRKNTLY | 0.13893309 |
| AAVSRAVAAL | ASSQDRKNTLY | 0.3423897 |
| AGGSSGIGGF | ASSQDRKNTLY | 0.52202535 |
| AGLTVANANG | ASSQDRKNTLY | 0.37722108 |

| | | |
|---|---|---|
| AGLVGAVL | ASSQDRKNTLY | 0.11103193 |
| ALSQAFTV | ASSQDRKNTLY | 0.31427938 |
| ALTFSVSL | ASSQDRKNTLY | 0.1510364 |
| ALTGAALVAL | ASSQDRKNTLY | 0.31223023 |
| ASFSIGAGVL | ASSQDRKNTLY | 0.23053253 |
| DGSVYKI | ASSQDRKNTLY | 0.21060531 |
| EQLAEIV | ASSQDRKNTLY | 0.27943653 |
| FAVSELTVM | ASSQDRKNTLY | 0.16934404 |
| GAAPLAAGL | ASSQDRKNTLY | 0.1438102 |
| GADGGDGGSGGAGG | ASSQDRKNTLY | 0.20986104 |
| GFGLDTGGAGG | ASSQDRKNTLY | 0.28279257 |
| GLASGGLL | ASSQDRKNTLY | 0.31747484 |
| GRIAGAAVLL | ASSQDRKNTLY | 0.14409351 |
| GSAGSLNTSF | ASSQDRKNTLY | 0.2678529 |
| GTGQIGI | ASSQDRKNTLY | 0.17598775 |
| GTLIALIL | ASSQDRKNTLY | 0.17116855 |
| IAATMGFVVG | ASSQDRKNTLY | 0.25098512 |
| IFAQALL | ASSQDRKNTLY | 0.20771623 |
| IGAAVALGF | ASSQDRKNTLY | 0.11742102 |
| IGLTPLAL | ASSQDRKNTLY | 0.3282149 |
| ILGSNTSTL | ASSQDRKNTLY | 0.1495184 |
| IVFSLTVL | ASSQDRKNTLY | 0.21987805 |
| LASKATGF | ASSQDRKNTLY | 0.5520431 |
| LASVALIL | ASSQDRKNTLY | 0.114689246 |
| LFAVGLTL | ASSQDRKNTLY | 0.26887423 |
| LFNGGIGTYI | ASSQDRKNTLY | 0.22456828 |
| LTLGAGII | ASSQDRKNTLY | 0.27990267 |
| LVAAALGDI | ASSQDRKNTLY | 0.2068342 |
| NTAFVQLGI | ASSQDRKNTLY | 0.17391898 |
| QLFSGAEL | ASSQDRKNTLY | 0.42990625 |
| SAATFITL | ASSQDRKNTLY | 0.3180599 |
| SNYGLANLGV | ASSQDRKNTLY | 0.31091392 |
| SYGGAYAVM | ASSQDRKNTLY | 0.27820128 |
| TTGVGGLTLG | ASSQDRKNTLY | 0.5865017 |
| TTQTIYI | ASSQDRKNTLY | 0.3572585 |
| VAAGAGLGAVI | ASSQDRKNTLY | 0.16745038 |
| VAAVVTLL | ASSQDRKNTLY | 0.10724783 |
| VGFLGGLI | ASSQDRKNTLY | 0.22077885 |
| VGVIPILL | ASSQDRKNTLY | 0.22208206 |
| VIFASSLI | ASSQDRKNTLY | 0.17890455 |
| VKGGDGGAAAG | ASSQDRKNTLY | 0.123242274 |
| VTKTFTNI | ASSQDRKNTLY | 0.58759624 |
| VVAAAILL | ASSQDRKNTLY | 0.7241165 |
| YAALDVEL | ASSQDRKNTLY | 0.15967175 |
| LASVALLL | ASSQDRKNTLY | 0.75350904 |
| LDAVHRV | ASSQDRKNTLY | 0.13207935 |
| QGLSSSIF | ASSQDRKNTLY | 0.31144 |
| RGTFANIRL | ASSQDRKNTLY | 0.61860824 |
| RQLWWGHRI | ASSQDRKNTLY | 0.1314361 |
| YGPPGTGKTLL | ASSQDRKNTLY | 0.30619073 |
| AALVGAPANVA | ASSQDRKNTLY | 0.1655687 |

| | | |
|---|---|---|
| AATFRRTE | ASSQDRKNTLY | 0.15450402 |
| AGALGGGI | ASSQDRKNTLY | 0.5035481 |
| AGGAGGAGGIGAG | ASSQDRKNTLY | 0.18636926 |
| AGSGALGL | ASSQDRKNTLY | 0.29001796 |
| AQIADQI | ASSQDRKNTLY | 0.19253579 |
| ASAVAVTYL | ASSQDRKNTLY | 0.1407133 |
| ASTVLGGTGG | ASSQDRKNTLY | 0.29853052 |
| AVGGEPAK | ASSQDRKNTLY | 0.45329165 |
| CEQGTFGGTC | ASSQDRKNTLY | 0.24559155 |
| DEGKAPAG | ASSQDRKNTLY | 0.7064258 |
| DTVVFPAL | ASSQDRKNTLY | 0.18006682 |
| FEGITMAG | ASSQDRKNTLY | 0.75297433 |
| FPLVVQD | ASSQDRKNTLY | 0.7231567 |
| GAFTTPQ | ASSQDRKNTLY | 0.23930907 |
| GFGGFGTAGGVGGA | ASSQDRKNTLY | 0.17165704 |
| GFLAQFL | ASSQDRKNTLY | 0.5971787 |
| GGAGGAGGNAGVA | ASSQDRKNTLY | 0.18079886 |
| GGVGTAGAGGAGGAGG | ASSQDRKNTLY | 0.2246797 |
| GNGGGGDAGNAGSGGN | ASSQDRKNTLY | 0.3807332 |
| GTGGAGGAGNDAGS | ASSQDRKNTLY | 0.5804344 |
| GTTNTGWFNS | ASSQDRKNTLY | 0.7074087 |
| IFMGPLFTL | ASSQDRKNTLY | 0.6953849 |
| ILAHAIEAI | ASSQDRKNTLY | 0.71541625 |
| IRSAIREAL | ASSQDRKNTLY | 0.5893879 |
| ITFAGGG | ASSQDRKNTLY | 0.48686135 |
| ITGMSGHQEV | ASSQDRKNTLY | 0.68554795 |
| IVAGLACTL | ASSQDRKNTLY | 0.55750763 |
| IVGGGLAGA | ASSQDRKNTLY | 0.72529376 |
| IVTGVLNI | ASSQDRKNTLY | 0.71850795 |
| LAQQGCI | ASSQDRKNTLY | 0.58454835 |
| LCLHVEN | ASSQDRKNTLY | 0.7123196 |
| LDAISDGS | ASSQDRKNTLY | 0.7186228 |
| LGGFAAML | ASSQDRKNTLY | 0.5422386 |
| LGGKNPMI | ASSQDRKNTLY | 0.6880863 |
| LGPKGVAAIL | ASSQDRKNTLY | 0.62064487 |
| LMVGVSI | ASSQDRKNTLY | 0.6861416 |
| LVGAAFVL | ASSQDRKNTLY | 0.41582114 |
| LVLAVAPL | ASSQDRKNTLY | 0.70887256 |
| MAAIAVIDAA | ASSQDRKNTLY | 0.67949164 |
| NNGINAELQ | ASSQDRKNTLY | 0.3666689 |
| RGEPLVLVHG | ASSQDRKNTLY | 0.70464396 |
| RGGAGGAAGLI | ASSQDRKNTLY | 0.6208584 |
| RSYRMHPAV | ASSQDRKNTLY | 0.67747676 |
| SAAHLLAGQLA | ASSQDRKNTLY | 0.71542835 |
| SAAVGSY | ASSQDRKNTLY | 0.6092155 |
| SPHAVLHRI | ASSQDRKNTLY | 0.60831773 |
| SQIAKVL | ASSQDRKNTLY | 0.6862014 |
| TALGHLVP | ASSQDRKNTLY | 0.7232633 |
| TAPTTSGAA | ASSQDRKNTLY | 0.5977471 |
| TDGDDGNGGSPGT | ASSQDRKNTLY | 0.6897942 |
| TDGLGEN | ASSQDRKNTLY | 0.7263199 |

| | | |
|---|---|---|
| TGGPPLPGQA | ASSQDRKNTLY | 0.6764306 |
| TIPAITI | ASSQDRKNTLY | 0.72919756 |
| TIPGFTI | ASSQDRKNTLY | 0.73203087 |
| TSIASGALA | ASSQDRKNTLY | 0.6603645 |
| TVTLPGAAI | ASSQDRKNTLY | 0.7180644 |
| VASGLSAALP | ASSQDRKNTLY | 0.7264575 |
| VGAGVIGI | ASSQDRKNTLY | 0.64924943 |
| VGPKRFGV | ASSQDRKNTLY | 0.6618644 |
| VNVAFPD | ASSQDRKNTLY | 0.5743779 |
| VSMTNSGVS | ASSQDRKNTLY | 0.6934018 |
| VTDVLGDLSV | ASSQDRKNTLY | 0.47225598 |
| VVVVGAGFAGL | ASSQDRKNTLY | 0.6829763 |
| VYTVGTVV | ASSQDRKNTLY | 0.7191174 |
| WTTFSGP | ASSQDRKNTLY | 0.6609552 |
| YDELVEAF | ASSQDRKNTLY | 0.51789284 |
| YGTGGAGGNGG | ASSQDRKNTLY | 0.45951593 |
| YTNVMAV | ASSQDRKNTLY | 0.67671704 |
| FLEQVHQL | ASSQDRKNTLY | 0.6524813 |
| GLSLLAVAV | ASSQDRKNTLY | 0.50830793 |
| LEQVHQL | ASSQDRKNTLY | 0.63998127 |
| LNAAGAI | ASSQDRKNTLY | 0.7113181 |
| PAVHASL | ASSQDRKNTLY | 0.71585995 |
| TELEDDA | ASSQDRKNTLY | 0.5713415 |
| TPPGPPN | ASSQDRKNTLY | 0.7209184 |
| TPPPPPPP | ASSQDRKNTLY | 0.717015 |
| TQSPALI | ASSHDRGISNERLF | 0.7218927 |
| EFAAAFPQ | ASSHDRGISNERLF | 0.67695034 |
| LLSGNLDVL | ASSHDRGISNERLF | 0.5130647 |
| GPANLGNYNI | ASSHDRGISNERLF | 0.72084105 |
| TLGPTLP | ASSHDRGISNERLF | 0.62139535 |
| VYAYSSCCK | ASSHDRGISNERLF | 0.69311976 |
| APAPAPAPNI | ASSHDRGISNERLF | 0.71916294 |
| PGGPVPIP | ASSHDRGISNERLF | 0.6540448 |
| FLTVALEP | ASSHDRGISNERLF | 0.5271513 |
| SQYFGANL | ASSHDRGISNERLF | 0.68135273 |
| GGVKPAA | ASSHDRGISNERLF | 0.72709286 |
| GGTGGAGGAGGW | ASSHDRGISNERLF | 0.50185025 |
| GVGGVAGLGGAGG | ASSHDRGISNERLF | 0.502759 |
| VLLVLATTI | ASSHDRGISNERLF | 0.66802573 |
| IGIFAAML | ASSHDRGISNERLF | 0.6616776 |
| AVGDLID | ASSHDRGISNERLF | 0.7227959 |
| NISPLPL | ASSHDRGISNERLF | 0.70500827 |
| VTAVFVAI | ASSHDRGISNERLF | 0.6652018 |
| SGDDNDAAL | ASSHDRGISNERLF | 0.7125711 |
| AVAALAHS | ASSHDRGISNERLF | 0.7023848 |
| GGVTVMATP | ASSHDRGISNERLF | 0.66273105 |
| AGLAANVRD | ASSHDRGISNERLF | 0.7143828 |
| TNRPDLIDPALLRPG | ASSHDRGISNERLF | 0.5859823 |
| LANLGVTAL | ASSHDRGISNERLF | 0.7203326 |
| TGERIEL | ASSHDRGISNERLF | 0.71275413 |
| NGEELAA | ASSHDRGISNERLF | 0.72301877 |

| | | |
|---|---|---|
| VLSLVWG | ASSHDRGISNERLF | 0.7237301 |
| IGYALWL | ASSHDRGISNERLF | 0.6691047 |
| LSLLLAPL | ASSHDRGISNERLF | 0.6932095 |
| TLDEALK | ASSHDRGISNERLF | 0.54790545 |
| TDGAGTAEGG | ASSHDRGISNERLF | 0.6704762 |
| NAPGGAGGNGG | ASSHDRGISNERLF | 0.70189154 |
| VSVFMGMGGNF | ASSHDRGISNERLF | 0.69320977 |
| LASYLAAL | ASSHDRGISNERLF | 0.7171966 |
| LVGGTPAL | ASSHDRGISNERLF | 0.71863115 |
| VRNADVL | ASSHDRGISNERLF | 0.5896533 |
| TCSSPFLVSAM | ASSHDRGISNERLF | 0.7293117 |
| VHITDAE | ASSHDRGISNERLF | 0.7228319 |
| EYAAPSF | ASSHDRGISNERLF | 0.5133332 |
| VVDAFHGGL | ASSHDRGISNERLF | 0.66424006 |
| ASVVDPQ | ASSHDRGISNERLF | 0.7211138 |
| SYNEFGGLS | ASSHDRGISNERLF | 0.7070464 |
| DPDGSVY | ASSHDRGISNERLF | 0.647914 |
| MVSTIHATE | ASSHDRGISNERLF | 0.7083961 |
| LAAGVADPV | ASSHDRGISNERLF | 0.72667754 |
| TKEQIAA | ASSHDRGISNERLF | 0.7001425 |
| SLASPAAL | ASSHDRGISNERLF | 0.7280855 |
| IGLVFYGV | ASSHDRGISNERLF | 0.505404 |
| FLLDASNL | ASSHDRGISNERLF | 0.69777477 |
| YQVGNFP | ASSHDRGISNERLF | 0.6343101 |
| ESLLSQST | ASSHDRGISNERLF | 0.34044206 |
| LFTVGFAAVG | ASSHDRGISNERLF | 0.70375323 |
| ATPPTPT | ASSHDRGISNERLF | 0.6757846 |
| TDLATLG | ASSHDRGISNERLF | 0.720997 |
| SAGGTGR | ASSHDRGISNERLF | 0.6788917 |
| GFSNSGS | ASSHDRGISNERLF | 0.7100832 |
| GGNGGSGGTSV | ASSHDRGISNERLF | 0.6608661 |
| GTSVATGGAGN | ASSHDRGISNERLF | 0.69988984 |
| NGGNGGSGGTS | ASSHDRGISNERLF | 0.62677574 |
| AVVKFPH | ASSHDRGISNERLF | 0.6959427 |
| TPPPPPP | ASSHDRGISNERLF | 0.72849524 |
| AAPAAAAAPA | ASSHDRGISNERLF | 0.7256006 |
| IALNEDL | ASSHDRGISNERLF | 0.4733264 |
| IVNVSSM | ASSHDRGISNERLF | 0.7039744 |
| TGIPPAPR | ASSHDRGISNERLF | 0.6599983 |
| FDLGGGTF | ASSHDRGISNERLF | 0.3565011 |
| FDLGGGTFDVS | ASSHDRGISNERLF | 0.5245218 |
| NEPTAAA | ASSHDRGISNERLF | 0.62156594 |
| NEPTAAALA | ASSHDRGISNERLF | 0.6302536 |
| GSIINISSI | ASSHDRGISNERLF | 0.69979393 |
| VAPGNAGTA | ASSHDRGISNERLF | 0.69898146 |
| TGGGASLE | ASSHDRGISNERLF | 0.64742744 |
| VAVYNHYKRI | ASSHDRGISNERLF | 0.7034408 |
| TDPAPATT | ASSHDRGISNERLF | 0.6534878 |
| TNRPDILDPALLRPG | ASSHDRGISNERLF | 0.24300419 |
| LTTTTTS | ASSHDRGISNERLF | 0.63671786 |
| PTGSGKTL | ASSHDRGISNERLF | 0.5630677 |

| | | |
|---|---|---|
| TGSGKTL | ASSHDRGISNERLF | 0.73018837 |
| TIGHVDHGK | ASSHDRGISNERLF | 0.6257113 |
| TIGHVDHGKTT | ASSHDRGISNERLF | 0.62475044 |
| GHVDHGKTT | ASSHDRGISNERLF | 0.65858316 |
| AQFEHTLL | ASSHDRGISNERLF | 0.69929165 |
| IVNQDEL | ASSHDRGISNERLF | 0.6794609 |
| TNGSQFF | ASSHDRGISNERLF | 0.5838677 |
| TGKGIVE | ASSHDRGISNERLF | 0.4210473 |
| LISNASDA | ASSHDRGISNERLF | 0.7014982 |
| AELRTLGELTKIAKAHGAQ | ASSHDRGISNERLF | 0.61458564 |
| APNAVAAAEPSW | ASSHDRGISNERLF | 0.6435061 |
| APQVAGAATAVL | ASSHDRGISNERLF | 0.64103323 |
| ARRDPAAAAAILPTDA | ASSHDRGISNERLF | 0.6603318 |
| DAAARDVLAEAGLAEYF | ASSHDRGISNERLF | 0.4929319 |
| DDQIVLSAKANLAKVEQ | ASSHDRGISNERLF | 0.728949 |
| DELLPDTGEDSDEDGH | ASSHDRGISNERLF | 0.6503537 |
| ELPEAPANRATVPQD | ASSHDRGISNERLF | 0.60315466 |
| FGQSKIERVTGIG | ASSHDRGISNERLF | 0.71427774 |
| GAGGTGSTAGGAGGAGGAGGLY | ASSHDRGISNERLF | 0.69150794 |
| GHGGTGAAGGNGGTGGAGGSGI | ASSHDRGISNERLF | 0.6466198 |
| GNGGNGGAGGEGGPGGAGGAGA | ASSHDRGISNERLF | 0.62630916 |
| GSTAGNGGNGGNATNGTIAGQP | ASSHDRGISNERLF | 0.6037545 |
| GTGGDGGNGGNGGTGG | ASSHDRGISNERLF | 0.5890627 |
| GVRRPAGIPADAPV | ASSHDRGISNERLF | 0.60971415 |
| GYRQVTDAHLLTLARR | ASSHDRGISNERLF | 0.72504914 |
| IAKMGQDGHDRG | ASSHDRGISNERLF | 0.7054466 |
| KAEPSIEIQYT | ASSHDRGISNERLF | 0.5445174 |
| KPVRIRMAADEDTVAVT | ASSHDRGISNERLF | 0.7146674 |
| MPPRIAGMRLLVIKPEP | ASSHDRGISNERLF | 0.7248575 |
| MKFYAVALFRSRGVPPTR | ASSHDRGISNERLF | 0.72187567 |
| NAGGFGILGAVAHSP | ASSHDRGISNERLF | 0.62470037 |
| NEGREITLAMAEM | ASSHDRGISNERLF | 0.65658414 |
| NELVEWSHPLGHRWRWE | ASSHDRGISNERLF | 0.68606937 |
| NLGAGGTGGNGGNPG | ASSHDRGISNERLF | 0.6348696 |
| NTEWPKWETVMAADPVD | ASSHDRGISNERLF | 0.67082655 |
| PASAAPAETQP | ASSHDRGISNERLF | 0.56423175 |
| QQSSDGKNGVRAAEITGE | ASSHDRGISNERLF | 0.72227854 |
| RAPDAVTTSWWKE | ASSHDRGISNERLF | 0.73566574 |
| SGFGNVGAHTSGWFNQ | ASSHDRGISNERLF | 0.63728863 |
| SGVAGAVGQAASVGGLKV | ASSHDRGISNERLF | 0.46999717 |
| SNGGAGTGSAGGNGGT | ASSHDRGISNERLF | 0.6464712 |
| TGGAGGEGGAAG | ASSHDRGISNERLF | 0.5634639 |
| TGGDGGAAGAGGAPGHG | ASSHDRGISNERLF | 0.57035327 |
| TMPPLLAAIANAGRHGV | ASSHDRGISNERLF | 0.6789552 |
| TSSTAGNSGGA | ASSHDRGISNERLF | 0.51943195 |
| VPATVPVGILC | ASSHDRGISNERLF | 0.66356564 |
| VQRPDSGGVVLTGRIS | ASSHDRGISNERLF | 0.7172134 |
| WTPNPVPATKGVPAGLVQAT | ASSHDRGISNERLF | 0.6639077 |
| YALKRALAKFSDRFDVVII | ASSHDRGISNERLF | 0.7106362 |
| YLIGSVIV | ASSHDRGISNERLF | 0.71697867 |
| YGQLVAVID | ASSHDRGISNERLF | 0.7037921 |

| | | |
|---|---|---|
| VTVGGMTLPA | ASSHDRGISNERLF | 0.71945834 |
| VLALPAV | ASSHDRGISNERLF | 0.7217339 |
| VIPPTLNYET | ASSHDRGISNERLF | 0.6984693 |
| VGMGGMHPSSG | ASSHDRGISNERLF | 0.6839571 |
| VGLAGGAATA | ASSHDRGISNERLF | 0.70833564 |
| VGGAGGAGGSGP | ASSHDRGISNERLF | 0.6457366 |
| VFAAFMV | ASSHDRGISNERLF | 0.67940295 |
| TLYDKLR | ASSHDRGISNERLF | 0.67078555 |
| TLAVFTVL | ASSHDRGISNERLF | 0.70286727 |
| TGSVQPARV | ASSHDRGISNERLF | 0.67789173 |
| TGGGAGSGGT | ASSHDRGISNERLF | 0.731014 |
| SHLETSL | ASSHDRGISNERLF | 0.4789859 |
| SFTTGMT | ASSHDRGISNERLF | 0.5974205 |
| QVEVAKRMGVRQPTVS | ASSHDRGISNERLF | 0.7130039 |
| QGSGGPALAM | ASSHDRGISNERLF | 0.3084037 |
| MLAPSNP | ASSHDRGISNERLF | 0.6207674 |
| LVNNAAI | ASSHDRGISNERLF | 0.69709194 |
| LTGEINTLA | ASSHDRGISNERLF | 0.7160789 |
| LSIFGPV | ASSHDRGISNERLF | 0.2903183 |
| LLALLANL | ASSHDRGISNERLF | 0.65847886 |
| LISNASDALDKLR | ASSHDRGISNERLF | 0.566972 |
| LINEAVP | ASSHDRGISNERLF | 0.7155197 |
| LGYAFSGL | ASSHDRGISNERLF | 0.65772206 |
| LDFVAVAL | ASSHDRGISNERLF | 0.4064596 |
| LAQPGAE | ASSHDRGISNERLF | 0.71824217 |
| ISNLGAVTGF | ASSHDRGISNERLF | 0.4842381 |
| HQEDPGRI | ASSHDRGISNERLF | 0.6430104 |
| GTPGEPTP | ASSHDRGISNERLF | 0.7136791 |
| GFFNSGS | ASSHDRGISNERLF | 0.7296617 |
| GAGGGGLTAGAN | ASSHDRGISNERLF | 0.60486287 |
| FVDALVNM | ASSHDRGISNERLF | 0.7125404 |
| FLTVAPEL | ASSHDRGISNERLF | 0.64318824 |
| FETLTAL | ASSHDRGISNERLF | 0.72963095 |
| DVINPIEL | ASSHDRGISNERLF | 0.63173485 |
| APVSGAPV | ASSHDRGISNERLF | 0.5340823 |
| ALVGGLAW | ASSHDRGISNERLF | 0.71058357 |
| ALAAAAATAI | ASSHDRGISNERLF | 0.7104838 |
| AFGLGLLL | ASSHDRGISNERLF | 0.7016456 |
| AFFAAASL | ASSHDRGISNERLF | 0.71343815 |
| AAAALPAFSPP | ASSHDRGISNERLF | 0.66549647 |
| VVLAPAL | ASSHDRGISNERLF | 0.646171 |
| TIGHVDHGKTTL | ASSHDRGISNERLF | 0.7077048 |
| LISNASDALDK | ASSHDRGISNERLF | 0.7064624 |
| LISNASDAL | ASSHDRGISNERLF | 0.5745606 |
| ISNASDALDK | ASSHDRGISNERLF | 0.7195812 |
| NQAETLVYQTE | ASSHDRGISNERLF | 0.67021346 |
| TGGTGGTAGAG | ASSHDRGISNERLF | 0.63573414 |
| TGSTAGGAGGA | ASSHDRGISNERLF | 0.7161762 |
| DPLAVPVAAGPV | ASSHDRGISNERLF | 0.6964859 |
| GGTGGTPGNGGN | ASSHDRGISNERLF | 0.70701325 |
| PAGLAGVDPATL | ASSHDRGISNERLF | 0.70202535 |

| | | |
|---|---|---|
| GNNLVVEAPHAGASV | ASSHDRGISNERLF | 0.7330482 |
| KVTRIMSASTLEQPA | ASSHDRGISNERLF | 0.53633976 |
| LGFALPIALPRRTVV | ASSHDRGISNERLF | 0.7064221 |
| TRLGNLTARKLVVLV | ASSHDRGISNERLF | 0.68467915 |
| WSQQRRLIYFPSAGP | ASSHDRGISNERLF | 0.7195683 |
| GPNGNPDPMAAAVDIR | ASSHDRGISNERLF | 0.69564795 |
| TQRCVSVADLRAAASR | ASSHDRGISNERLF | 0.6708183 |
| QFVLDVAEHPVEVLTETV | ASSHDRGISNERLF | 0.7136659 |
| GGAGGTGGAGGNGGTQAGDG | ASSHDRGISNERLF | 0.7140523 |
| PLLATLRAIAVDVDAKPAQV | ASSHDRGISNERLF | 0.6986439 |
| APKFRTGGYMIDFWGVGYQVAK | ASSHDRGISNERLF | 0.7046091 |
| GAGGAGGAGGAAGGPGGTGGTG | ASSHDRGISNERLF | 0.72337425 |
| IGALGLLATDVAPAGSVAFGVA | ASSHDRGISNERLF | 0.61294466 |
| AGGAGGIGGNGGAGGAGGVLMG | ASSHDRGISNERLF | 0.66899 |
| ADELIGLL | ASSHDRGISNERLF | 0.7174312 |
| ADRALKA | ASSHDRGISNERLF | 0.72557676 |
| AGAVGQVGGAGGA | ASSHDRGISNERLF | 0.6947533 |
| AGGGGGLGGAP | ASSHDRGISNERLF | 0.7243841 |
| AGGLLYGNGGAG | ASSHDRGISNERLF | 0.6283067 |
| AGLESIGL | ASSHDRGISNERLF | 0.6970202 |
| AGLKTGII | ASSHDRGISNERLF | 0.65107036 |
| ASLAALLL | ASSHDRGISNERLF | 0.6862229 |
| ASVINISSI | ASSHDRGISNERLF | 0.57245505 |
| ATPPGPA | ASSHDRGISNERLF | 0.6880597 |
| ATVASALV | ASSHDRGISNERLF | 0.6984028 |
| ATVASLIL | ASSHDRGISNERLF | 0.70407844 |
| AVNGPVLIRH | ASSHDRGISNERLF | 0.55702055 |
| DGAAGAVGI | ASSHDRGISNERLF | 0.68173254 |
| DPLLHGI | ASSHDRGISNERLF | 0.71029735 |
| ELGAALGAL | ASSHDRGISNERLF | 0.73217416 |
| ELGRTQAV | ASSHDRGISNERLF | 0.67969 |
| FVAGGYLS | ASSHDRGISNERLF | 0.71730566 |
| GASGGAGGAGGW | ASSHDRGISNERLF | 0.641937 |
| GGAGAGGDGGAGGI | ASSHDRGISNERLF | 0.6893717 |
| GTGHNGSQPGMGGNG | ASSHDRGISNERLF | 0.57192177 |
| GYAAASASAS | ASSHDRGISNERLF | 0.6583078 |
| HQVDVAAVQQG | ASSHDRGISNERLF | 0.6476375 |
| IGAAIAHT | ASSHDRGISNERLF | 0.6003669 |
| IGMIEVAQL | ASSHDRGISNERLF | 0.7163887 |
| IRLAPEHTSAMD | ASSHDRGISNERLF | 0.7227021 |
| IYNVSVL | ASSHDRGISNERLF | 0.69617903 |
| KQGFLGM | ASSHDRGISNERLF | 0.7171257 |
| LESVITQL | ASSHDRGISNERLF | 0.71630824 |
| LGLGALVVV | ASSHDRGISNERLF | 0.7286743 |
| LIGNGGNGGSGGI | ASSHDRGISNERLF | 0.60928154 |
| LNAINAPTL | ASSHDRGISNERLF | 0.68696755 |
| LQLLNTGV | ASSHDRGISNERLF | 0.46263507 |
| LQTSIEF | ASSHDRGISNERLF | 0.6281189 |
| MSFAVSEL | ASSHDRGISNERLF | 0.6232296 |
| NTGALGL | ASSHDRGISNERLF | 0.6689426 |
| PPGAGGAGGAGTT | ASSHDRGISNERLF | 0.2661877 |

| | | |
|---|---|---|
| QLVGLGI | ASSHDRGISNERLF | 0.70071125 |
| RFLEQVHQL | ASSHDRGISNERLF | 0.7269727 |
| RGVGYVLR | ASSHDRGISNERLF | 0.6046543 |
| SATGVFAGVI | ASSHDRGISNERLF | 0.70936286 |
| SLPAITLP | ASSHDRGISNERLF | 0.689208 |
| STALAFTV | ASSHDRGISNERLF | 0.6386887 |
| SVLTVGLGK | ASSHDRGISNERLF | 0.7032897 |
| TALAGAGSGI | ASSHDRGISNERLF | 0.6347177 |
| TGLPFTGI | ASSHDRGISNERLF | 0.72259223 |
| VAEHVTV | ASSHDRGISNERLF | 0.7195206 |
| VDIVLNSL | ASSHDRGISNERLF | 0.6927556 |
| VLTGGGAVL | ASSHDRGISNERLF | 0.6898637 |
| VNVGLAYGI | ASSHDRGISNERLF | 0.7098717 |
| VSSAAVGAV | ASSHDRGISNERLF | 0.5508524 |
| VTSVYEAGV | ASSHDRGISNERLF | 0.69256526 |
| WVALDRGA | ASSHDRGISNERLF | 0.5378436 |
| YLVGSVVI | ASSHDRGISNERLF | 0.6685872 |
| GPPGTGKTLL | ASSHDRGISNERLF | 0.5658071 |
| GVGMTKF | ASSHDRGISNERLF | 0.7003132 |
| IGIGGSDL | ASSHDRGISNERLF | 0.6471753 |
| IGTLLAEL | ASSHDRGISNERLF | 0.7092637 |
| IYTSGTT | ASSHDRGISNERLF | 0.5749377 |
| LSFPAGV | ASSHDRGISNERLF | 0.7095364 |
| LYTSGTT | ASSHDRGISNERLF | 0.69020873 |
| VGAFPFGF | ASSHDRGISNERLF | 0.50514966 |
| VGAGTVEF | ASSHDRGISNERLF | 0.71526617 |
| VGKSTLLL | ASSHDRGISNERLF | 0.68847334 |
| AALAHAGGGIV | ASSHDRGISNERLF | 0.5889646 |
| AALVAVSLL | ASSHDRGISNERLF | 0.7126167 |
| AAVSRAVAAL | ASSHDRGISNERLF | 0.69297844 |
| AGGSSGIGGF | ASSHDRGISNERLF | 0.49674323 |
| AGLTVANANG | ASSHDRGISNERLF | 0.6221787 |
| AGLVGAVL | ASSHDRGISNERLF | 0.6888895 |
| ALSQAFTV | ASSHDRGISNERLF | 0.6456953 |
| ALTFSVSL | ASSHDRGISNERLF | 0.29092264 |
| ALTGAALVAL | ASSHDRGISNERLF | 0.70057386 |
| ASFSIGAGVL | ASSHDRGISNERLF | 0.6356494 |
| DGSVYKI | ASSHDRGISNERLF | 0.53085005 |
| EQLAEIV | ASSHDRGISNERLF | 0.65585864 |
| FAVSELTVM | ASSHDRGISNERLF | 0.71647644 |
| GAAPLAAGL | ASSHDRGISNERLF | 0.67502415 |
| GADGGDGGSGGAGG | ASSHDRGISNERLF | 0.7030263 |
| GFGLDTGGAGG | ASSHDRGISNERLF | 0.70861906 |
| GLASGGLL | ASSHDRGISNERLF | 0.67324144 |
| GRIAGAAVLL | ASSHDRGISNERLF | 0.7211124 |
| GSAGSLNTSF | ASSHDRGISNERLF | 0.69908357 |
| GTGQIGI | ASSHDRGISNERLF | 0.68545306 |
| GTLIALIL | ASSHDRGISNERLF | 0.7130002 |
| IAATMGFVVG | ASSHDRGISNERLF | 0.68537056 |
| IFAQALL | ASSHDRGISNERLF | 0.6723151 |
| IGAAVALGF | ASSHDRGISNERLF | 0.64203584 |

| | | |
|---|---|---|
| IGLTPLAL | ASSHDRGISNERLF | 0.62084234 |
| ILGSNTSTL | ASSHDRGISNERLF | 0.66076505 |
| IVFSLTVL | ASSHDRGISNERLF | 0.69053143 |
| LASKATGF | ASSHDRGISNERLF | 0.73015344 |
| LASVALIL | ASSHDRGISNERLF | 0.69330263 |
| LFAVGLTL | ASSHDRGISNERLF | 0.50879794 |
| LFNGGIGTYI | ASSHDRGISNERLF | 0.67330027 |
| LTLGAGII | ASSHDRGISNERLF | 0.60765666 |
| LVAAALGDI | ASSHDRGISNERLF | 0.7159209 |
| NTAFVQLGI | ASSHDRGISNERLF | 0.7020016 |
| QLFSGAEL | ASSHDRGISNERLF | 0.7291265 |
| SAATFITL | ASSHDRGISNERLF | 0.6988497 |
| SNYGLANLGV | ASSHDRGISNERLF | 0.59607816 |
| SYGGAYAVM | ASSHDRGISNERLF | 0.7165839 |
| TTGVGGLTLG | ASSHDRGISNERLF | 0.6228915 |
| TTQTIYI | ASSHDRGISNERLF | 0.6926862 |
| VAAGAGLGAVI | ASSHDRGISNERLF | 0.69898105 |
| VAAVVTLL | ASSHDRGISNERLF | 0.68511415 |
| VGFLGGLI | ASSHDRGISNERLF | 0.68384045 |
| VGVIPILL | ASSHDRGISNERLF | 0.6967207 |
| VIFASSLI | ASSHDRGISNERLF | 0.4777542 |
| VKGGDGGAAAG | ASSHDRGISNERLF | 0.7254791 |
| VTKTFTNI | ASSHDRGISNERLF | 0.60324657 |
| VVAAAILL | ASSHDRGISNERLF | 0.62699986 |
| YAALDVEL | ASSHDRGISNERLF | 0.64256847 |
| LASVALLL | ASSHDRGISNERLF | 0.7235695 |
| LDAVHRV | ASSHDRGISNERLF | 0.72212094 |
| QGLSSSIF | ASSHDRGISNERLF | 0.5963464 |
| RGTFANIRL | ASSHDRGISNERLF | 0.5614455 |
| RQLWWGHRI | ASSHDRGISNERLF | 0.7058661 |
| YGPPGTGKTLL | ASSHDRGISNERLF | 0.6152885 |
| AALVGAPANVA | ASSHDRGISNERLF | 0.7145612 |
| AATFRRTE | ASSHDRGISNERLF | 0.6504717 |
| AGALGGGI | ASSHDRGISNERLF | 0.59903514 |
| AGGAGGAGGIGAG | ASSHDRGISNERLF | 0.6631048 |
| AGSGALGL | ASSHDRGISNERLF | 0.70921516 |
| AQIADQI | ASSHDRGISNERLF | 0.5822757 |
| ASAVAVTYL | ASSHDRGISNERLF | 0.71837026 |
| ASTVLGGTGG | ASSHDRGISNERLF | 0.3142579 |
| AVGGEPAK | ASSHDRGISNERLF | 0.6995944 |
| CEQGTFGGTC | ASSHDRGISNERLF | 0.5373487 |
| DEGKAPAG | ASSHDRGISNERLF | 0.6302748 |
| DTVVFPAL | ASSHDRGISNERLF | 0.44689563 |
| FEGITMAG | ASSHDRGISNERLF | 0.6860973 |
| FPLVVQD | ASSHDRGISNERLF | 0.69968295 |
| GAFTTPQ | ASSHDRGISNERLF | 0.7180395 |
| GFGGFGTAGGVGGA | ASSHDRGISNERLF | 0.69049346 |
| GFLAQFL | ASSHDRGISNERLF | 0.7108247 |
| GGAGGAGGNAGVA | ASSHDRGISNERLF | 0.7220385 |
| GGVGTAGAGGAGGAGG | ASSHDRGISNERLF | 0.60256064 |
| GNGGGGDAGNAGSGGN | ASSHDRGISNERLF | 0.478199 |

| | | |
|---|---|---|
| GTGGAGGAGNDAGS | ASSHDRGISNERLF | 0.69722307 |
| GTTNTGWFNS | ASSHDRGISNERLF | 0.6927334 |
| IFMGPLFTL | ASSHDRGISNERLF | 0.7141921 |
| ILAHAIEAI | ASSHDRGISNERLF | 0.7023126 |
| IRSAIREAL | ASSHDRGISNERLF | 0.5221574 |
| ITFAGGG | ASSHDRGISNERLF | 0.72067803 |
| ITGMSGHQEV | ASSHDRGISNERLF | 0.70217633 |
| IVAGLACTL | ASSHDRGISNERLF | 0.52219725 |
| IVGGGLAGA | ASSHDRGISNERLF | 0.6439866 |
| IVTGVLNI | ASSHDRGISNERLF | 0.6267824 |
| LAQQGCI | ASSHDRGISNERLF | 0.48455036 |
| LCLHVEN | ASSHDRGISNERLF | 0.67505825 |
| LDAISDGS | ASSHDRGISNERLF | 0.60796213 |
| LGGFAAML | ASSHDRGISNERLF | 0.66917586 |
| LGGKNPMI | ASSHDRGISNERLF | 0.6603993 |
| LGPKGVAAIL | ASSHDRGISNERLF | 0.64211875 |
| LMVGVSI | ASSHDRGISNERLF | 0.6754473 |
| LVGAAFVL | ASSHDRGISNERLF | 0.6607039 |
| LVLAVAPL | ASSHDRGISNERLF | 0.68899286 |
| MAAIAVIDAA | ASSHDRGISNERLF | 0.70756817 |
| NNGINAELQ | ASSHDRGISNERLF | 0.63745445 |
| RGEPLVLVHG | ASSHDRGISNERLF | 0.7067828 |
| RGGAGGAAGLI | ASSHDRGISNERLF | 0.7183436 |
| RSYRMHPAV | ASSHDRGISNERLF | 0.52356666 |
| SAAHLLAGQLA | ASSHDRGISNERLF | 0.5878602 |
| SAAVGSY | ASSHDRGISNERLF | 0.70613 |
| SPHAVLHRI | ASSHDRGISNERLF | 0.6464683 |
| SQIAKVL | ASSHDRGISNERLF | 0.71827734 |
| TALGHLVP | ASSHDRGISNERLF | 0.6661713 |
| TAPTTSGAA | ASSHDRGISNERLF | 0.68637335 |
| TDGDDGNGGSPGT | ASSHDRGISNERLF | 0.70899147 |
| TDGLGEN | ASSHDRGISNERLF | 0.7282978 |
| TGGPPLPGQA | ASSHDRGISNERLF | 0.7065891 |
| TIPAITI | ASSHDRGISNERLF | 0.65578985 |
| TIPGFTI | ASSHDRGISNERLF | 0.7151147 |
| TSIASGALA | ASSHDRGISNERLF | 0.5573528 |
| TVTLPGAAI | ASSHDRGISNERLF | 0.7116344 |
| VASGLSAALP | ASSHDRGISNERLF | 0.72155386 |
| VGAGVIGI | ASSHDRGISNERLF | 0.5105314 |
| VGPKRFGV | ASSHDRGISNERLF | 0.6486863 |
| VNVAFPD | ASSHDRGISNERLF | 0.72804934 |
| VSMTNSGVS | ASSHDRGISNERLF | 0.72056377 |
| VTDVLGDLSV | ASSHDRGISNERLF | 0.6315713 |
| VVVVGAGFAGL | ASSHDRGISNERLF | 0.6301062 |
| VYTVGTVV | ASSHDRGISNERLF | 0.6129668 |
| WTTFSGP | ASSHDRGISNERLF | 0.65788114 |
| YDELVEAF | ASSHDRGISNERLF | 0.5095414 |
| YGTGGAGGNGG | ASSHDRGISNERLF | 0.70667267 |
| YTNVMAV | ASSHDRGISNERLF | 0.6538421 |
| FLEQVHQL | ASSHDRGISNERLF | 0.72338164 |
| GLSLLAVAV | ASSHDRGISNERLF | 0.686785 |

| | | |
|---|---|---|
| LEQVHQL | ASSHDRGISNERLF | 0.7140228 |
| LNAAGAI | ASSHDRGISNERLF | 0.6445326 |
| PAVHASL | ASSHDRGISNERLF | 0.6517319 |
| TELEDDA | ASSHDRGISNERLF | 0.5462559 |
| TPPGPPN | ASSHDRGISNERLF | 0.54382 |
| TPPPPPPP | ASSHDRGISNERLF | 0.6537976 |
| TQSPALI | ASSLWDIYAEQF | 0.6637385 |
| EFAAAFPQ | ASSLWDIYAEQF | 0.36119455 |
| LLSGNLDVL | ASSLWDIYAEQF | 0.71661127 |
| GPANLGNYNI | ASSLWDIYAEQF | 0.6856324 |
| TLGPTLP | ASSLWDIYAEQF | 0.7056172 |
| VYAYSSCCK | ASSLWDIYAEQF | 0.6218749 |
| APAPAPAPNI | ASSLWDIYAEQF | 0.6592437 |
| PGGPVPIP | ASSLWDIYAEQF | 0.6121564 |
| FLTVALEP | ASSLWDIYAEQF | 0.7168156 |
| SQYFGANL | ASSLWDIYAEQF | 0.6054008 |
| GGVKPAA | ASSLWDIYAEQF | 0.38209942 |
| GGTGGAGGAGGW | ASSLWDIYAEQF | 0.65396535 |
| GVGGVAGLGGAGG | ASSLWDIYAEQF | 0.57263285 |
| VLLVLATTI | ASSLWDIYAEQF | 0.7252308 |
| IGIFAAML | ASSLWDIYAEQF | 0.7111731 |
| AVGDLID | ASSLWDIYAEQF | 0.6951721 |
| NISPLPL | ASSLWDIYAEQF | 0.7325361 |
| VTAVFVAI | ASSLWDIYAEQF | 0.2839401 |
| SGDDNDAAL | ASSLWDIYAEQF | 0.6490477 |
| AVAALAHS | ASSLWDIYAEQF | 0.72284675 |
| GGVTVMATP | ASSLWDIYAEQF | 0.6785785 |
| AGLAANVRD | ASSLWDIYAEQF | 0.6693971 |
| TNRPDLIDPALLRPG | ASSLWDIYAEQF | 0.72201073 |
| LANLGVTAL | ASSLWDIYAEQF | 0.5862596 |
| TGERIEL | ASSLWDIYAEQF | 0.659444 |
| NGEELAA | ASSLWDIYAEQF | 0.6865772 |
| VLSLVWG | ASSLWDIYAEQF | 0.71580344 |
| IGYALWL | ASSLWDIYAEQF | 0.70700425 |
| LSLLLAPL | ASSLWDIYAEQF | 0.5814556 |
| TLDEALK | ASSLWDIYAEQF | 0.71898717 |
| TDGAGTAEGG | ASSLWDIYAEQF | 0.72646797 |
| NAPGGAGGNGG | ASSLWDIYAEQF | 0.664263 |
| VSVFMGMGGNF | ASSLWDIYAEQF | 0.6914108 |
| LASYLAAL | ASSLWDIYAEQF | 0.6993669 |
| LVGGTPAL | ASSLWDIYAEQF | 0.6741004 |
| VRNADVL | ASSLWDIYAEQF | 0.5795989 |
| TCSSPFLVSAM | ASSLWDIYAEQF | 0.5670395 |
| VHITDAE | ASSLWDIYAEQF | 0.7079977 |
| EYAAPSF | ASSLWDIYAEQF | 0.5323441 |
| VVDAFHGGL | ASSLWDIYAEQF | 0.70865524 |
| ASVVDPQ | ASSLWDIYAEQF | 0.6515034 |
| SYNEFGGLS | ASSLWDIYAEQF | 0.7323203 |
| DPDGSVY | ASSLWDIYAEQF | 0.70631695 |
| MVSTIHATE | ASSLWDIYAEQF | 0.6956649 |
| LAAGVADPV | ASSLWDIYAEQF | 0.5760273 |

| | | |
|---|---|---|
| TKEQIAA | ASSLWDIYAEQF | 0.6996483 |
| SLASPAAL | ASSLWDIYAEQF | 0.68886995 |
| IGLVFYGV | ASSLWDIYAEQF | 0.44755697 |
| FLLDASNL | ASSLWDIYAEQF | 0.7162452 |
| YQVGNFP | ASSLWDIYAEQF | 0.5815164 |
| ESLLSQST | ASSLWDIYAEQF | 0.65628934 |
| LFTVGFAAVG | ASSLWDIYAEQF | 0.7078376 |
| ATPPTPT | ASSLWDIYAEQF | 0.7268192 |
| TDLATLG | ASSLWDIYAEQF | 0.67144966 |
| SAGGTGR | ASSLWDIYAEQF | 0.70432407 |
| GFSNSGS | ASSLWDIYAEQF | 0.5901473 |
| GGNGGSGGTSV | ASSLWDIYAEQF | 0.7143341 |
| GTSVATGGAGN | ASSLWDIYAEQF | 0.71439016 |
| NGGNGGSGGTS | ASSLWDIYAEQF | 0.7086525 |
| AVVKFPH | ASSLWDIYAEQF | 0.73246306 |
| TPPPPPP | ASSLWDIYAEQF | 0.703723 |
| AAPAAAAAPA | ASSLWDIYAEQF | 0.4454009 |
| IALNEDL | ASSLWDIYAEQF | 0.69617486 |
| IVNVSSM | ASSLWDIYAEQF | 0.7132776 |
| TGIPPAPR | ASSLWDIYAEQF | 0.7168277 |
| FDLGGGTF | ASSLWDIYAEQF | 0.71891797 |
| FDLGGGTFDVS | ASSLWDIYAEQF | 0.72162586 |
| NEPTAAA | ASSLWDIYAEQF | 0.6139958 |
| NEPTAAALA | ASSLWDIYAEQF | 0.7219469 |
| GSIINISSI | ASSLWDIYAEQF | 0.73288834 |
| VAPGNAGTA | ASSLWDIYAEQF | 0.7102245 |
| TGGGASLE | ASSLWDIYAEQF | 0.66923726 |
| VAVYNHYKRI | ASSLWDIYAEQF | 0.647367 |
| TDPAPATT | ASSLWDIYAEQF | 0.7056844 |
| TNRPDILDPALLRPG | ASSLWDIYAEQF | 0.69902694 |
| LTTTTTS | ASSLWDIYAEQF | 0.64572227 |
| PTGSGKTL | ASSLWDIYAEQF | 0.6702969 |
| TGSGKTL | ASSLWDIYAEQF | 0.6752582 |
| TIGHVDHGK | ASSLWDIYAEQF | 0.47897106 |
| TIGHVDHGKTT | ASSLWDIYAEQF | 0.69591683 |
| GHVDHGKTT | ASSLWDIYAEQF | 0.7194165 |
| AQFEHTLL | ASSLWDIYAEQF | 0.7054614 |
| IVNQDEL | ASSLWDIYAEQF | 0.6383209 |
| TNGSQFF | ASSLWDIYAEQF | 0.7161819 |
| TGKGIVE | ASSLWDIYAEQF | 0.7222025 |
| LISNASDA | ASSLWDIYAEQF | 0.72920215 |
| AELRTLGELTKIAKAHGAQ | ASSLWDIYAEQF | 0.57851803 |
| APNAVAAAEPSW | ASSLWDIYAEQF | 0.28981236 |
| APQVAGAATAVL | ASSLWDIYAEQF | 0.6976626 |
| ARRDPAAAAAILPTDA | ASSLWDIYAEQF | 0.71144634 |
| DAAARDVLAEAGLAEYF | ASSLWDIYAEQF | 0.64618886 |
| DDQIVLSAKANLAKVEQ | ASSLWDIYAEQF | 0.64072406 |
| DELLPDTGEDSDEDGH | ASSLWDIYAEQF | 0.68810713 |
| ELPEAPANRATVPQD | ASSLWDIYAEQF | 0.70693576 |
| FGQSKIERVTGIG | ASSLWDIYAEQF | 0.6131482 |
| GAGGTGSTAGGAGGAGGAGGLY | ASSLWDIYAEQF | 0.55431306 |

| | | |
|---|---|---|
| GHGGTGAAGGNGGTGGAGGSGI | ASSLWDIYAEQF | 0.7108958 |
| GNGGNGGAGGEGGPGGAGGAGA | ASSLWDIYAEQF | 0.7244985 |
| GSTAGNGGNGGNATNGTIAGQP | ASSLWDIYAEQF | 0.61355627 |
| GTGGDGGNGGNGGTGG | ASSLWDIYAEQF | 0.64689434 |
| GVRRPAGIPADAPV | ASSLWDIYAEQF | 0.69133914 |
| GYRQVTDAHLLTLARR | ASSLWDIYAEQF | 0.6606742 |
| IAKMGQDGHDRG | ASSLWDIYAEQF | 0.6591257 |
| KAEPSIEIQYT | ASSLWDIYAEQF | 0.496691 |
| KPVRIRMAADEDTVAVT | ASSLWDIYAEQF | 0.69779336 |
| MPPRIAGMRLLVIKPEP | ASSLWDIYAEQF | 0.50027466 |
| MKFYAVALFRSRGVPPTR | ASSLWDIYAEQF | 0.7174512 |
| NAGGFGILGAVAHSP | ASSLWDIYAEQF | 0.6993009 |
| NEGREITLAMAEM | ASSLWDIYAEQF | 0.69876254 |
| NELVEWSHPLGHRWRWE | ASSLWDIYAEQF | 0.6534684 |
| NLGAGGTGGNGGNPG | ASSLWDIYAEQF | 0.586373 |
| NTEWPKWETVMAADPVD | ASSLWDIYAEQF | 0.50917345 |
| PASAAPAETQP | ASSLWDIYAEQF | 0.62184846 |
| QQSSDGKNGVRAAEITGE | ASSLWDIYAEQF | 0.722998 |
| RAPDAVTTSWWKE | ASSLWDIYAEQF | 0.6183717 |
| SGFGNVGAHTSGWFNQ | ASSLWDIYAEQF | 0.71796525 |
| SGVAGAVGQAASVGGLKV | ASSLWDIYAEQF | 0.69332325 |
| SNGGAGTGSAGGNGGT | ASSLWDIYAEQF | 0.6079086 |
| TGGAGGEGGAAG | ASSLWDIYAEQF | 0.70750856 |
| TGGDGGAAGAGGAPGHG | ASSLWDIYAEQF | 0.5410466 |
| TMPPLLAAIANAGRHGV | ASSLWDIYAEQF | 0.70836747 |
| TSSTAGNSGGA | ASSLWDIYAEQF | 0.69638765 |
| VPATVPVGILC | ASSLWDIYAEQF | 0.7134524 |
| VQRPDSGGVVLTGRIS | ASSLWDIYAEQF | 0.62716615 |
| WTPNPVPATKGVPAGLVQAT | ASSLWDIYAEQF | 0.71660924 |
| YALKRALAKFSDRFDVVII | ASSLWDIYAEQF | 0.6903658 |
| YLIGSVIV | ASSLWDIYAEQF | 0.6616205 |
| YGQLVAVID | ASSLWDIYAEQF | 0.7057222 |
| VTVGGMTLPA | ASSLWDIYAEQF | 0.7097684 |
| VLALPAV | ASSLWDIYAEQF | 0.7142825 |
| VIPPTLNYET | ASSLWDIYAEQF | 0.6210209 |
| VGMGGMHPSSG | ASSLWDIYAEQF | 0.66895545 |
| VGLAGGAATA | ASSLWDIYAEQF | 0.64304286 |
| VGGAGGAGGSGP | ASSLWDIYAEQF | 0.67715275 |
| VFAAFMV | ASSLWDIYAEQF | 0.67693686 |
| TLYDKLR | ASSLWDIYAEQF | 0.68200785 |
| TLAVFTVL | ASSLWDIYAEQF | 0.70546365 |
| TGSVQPARV | ASSLWDIYAEQF | 0.50663406 |
| TGGGAGSGGT | ASSLWDIYAEQF | 0.61399174 |
| SHLETSL | ASSLWDIYAEQF | 0.67052436 |
| SFTTGMT | ASSLWDIYAEQF | 0.68510616 |
| QVEVAKRMGVRQPTVS | ASSLWDIYAEQF | 0.7058085 |
| QGSGGPALAM | ASSLWDIYAEQF | 0.48285127 |
| MLAPSNP | ASSLWDIYAEQF | 0.68884456 |
| LVNNAAI | ASSLWDIYAEQF | 0.6700095 |
| LTGEINTLA | ASSLWDIYAEQF | 0.58979154 |
| LSIFGPV | ASSLWDIYAEQF | 0.71299857 |

| | | |
|---|---|---|
| LLALLANL | ASSLWDIYAEQF | 0.66513973 |
| LISNASDALDKLR | ASSLWDIYAEQF | 0.6005477 |
| LINEAVP | ASSLWDIYAEQF | 0.7155048 |
| LGYAFSGL | ASSLWDIYAEQF | 0.63976824 |
| LDFVAVAL | ASSLWDIYAEQF | 0.68991065 |
| LAQPGAE | ASSLWDIYAEQF | 0.7061293 |
| ISNLGAVTGF | ASSLWDIYAEQF | 0.62489176 |
| HQEDPGRI | ASSLWDIYAEQF | 0.71271676 |
| GTPGEPTP | ASSLWDIYAEQF | 0.66659296 |
| GFFNSGS | ASSLWDIYAEQF | 0.7065357 |
| GAGGGGLTAGAN | ASSLWDIYAEQF | 0.67144585 |
| FVDALVNM | ASSLWDIYAEQF | 0.61112714 |
| FLTVAPEL | ASSLWDIYAEQF | 0.7095765 |
| FETLTAL | ASSLWDIYAEQF | 0.6967453 |
| DVINPIEL | ASSLWDIYAEQF | 0.7207497 |
| APVSGAPV | ASSLWDIYAEQF | 0.69930416 |
| ALVGGLAW | ASSLWDIYAEQF | 0.7218102 |
| ALAAAAATAI | ASSLWDIYAEQF | 0.7171926 |
| AFGLGLLL | ASSLWDIYAEQF | 0.59439194 |
| AFFAAASL | ASSLWDIYAEQF | 0.51024926 |
| AAAALPAFSPP | ASSLWDIYAEQF | 0.6577746 |
| VVLAPAL | ASSLWDIYAEQF | 0.6535631 |
| TIGHVDHGKTTL | ASSLWDIYAEQF | 0.6442476 |
| LISNASDALDK | ASSLWDIYAEQF | 0.5287013 |
| LISNASDAL | ASSLWDIYAEQF | 0.70954126 |
| ISNASDALDK | ASSLWDIYAEQF | 0.73003256 |
| NQAETLVYQTE | ASSLWDIYAEQF | 0.6494058 |
| TGGTGGTAGAG | ASSLWDIYAEQF | 0.7320248 |
| TGSTAGGAGGA | ASSLWDIYAEQF | 0.5165729 |
| DPLAVPVAAGPV | ASSLWDIYAEQF | 0.69769233 |
| GGTGGTPGNGGN | ASSLWDIYAEQF | 0.72340894 |
| PAGLAGVDPATL | ASSLWDIYAEQF | 0.5313457 |
| GNNLVVEAPHAGASV | ASSLWDIYAEQF | 0.6924565 |
| KVTRIMSASTLEQPA | ASSLWDIYAEQF | 0.6634102 |
| LGFALPIALPRRTVV | ASSLWDIYAEQF | 0.6320946 |
| TRLGNLTARKLVVLV | ASSLWDIYAEQF | 0.7227727 |
| WSQQRRLIYFPSAGP | ASSLWDIYAEQF | 0.6732772 |
| GPNGNPDPMAAAVDIR | ASSLWDIYAEQF | 0.7163476 |
| TQRCVSVADLRAAASR | ASSLWDIYAEQF | 0.6910409 |
| QFVLDVAEHPVEVLTETV | ASSLWDIYAEQF | 0.5842598 |
| GGAGGTGGAGGNGGTQAGDG | ASSLWDIYAEQF | 0.70969784 |
| PLLATLRAIAVDVDAKPAQV | ASSLWDIYAEQF | 0.72532696 |
| APKFRTGGYMIDFWGVGYQVAK | ASSLWDIYAEQF | 0.69561917 |
| GAGGAGGAGGAAGGPGGTGGTG | ASSLWDIYAEQF | 0.73371875 |
| IGALGLLATDVAPAGSVAFGVA | ASSLWDIYAEQF | 0.6314017 |
| AGGAGGIGGNGGAGGAGGVLMG | ASSLWDIYAEQF | 0.7203053 |
| ADELIGLL | ASSLWDIYAEQF | 0.72534215 |
| ADRALKA | ASSLWDIYAEQF | 0.66099495 |
| AGAVGQVGGAGGA | ASSLWDIYAEQF | 0.66639 |
| AGGGGGLGGAP | ASSLWDIYAEQF | 0.71172357 |
| AGGLLYGNGGAG | ASSLWDIYAEQF | 0.6983824 |

| | | |
|---|---|---|
| AGLESIGL | ASSLWDIYAEQF | 0.7156202 |
| AGLKTGII | ASSLWDIYAEQF | 0.7301244 |
| ASLAALLL | ASSLWDIYAEQF | 0.55024105 |
| ASVINISSI | ASSLWDIYAEQF | 0.7903883 |
| ATPPGPA | ASSLWDIYAEQF | 0.75190854 |
| ATVASALV | ASSLWDIYAEQF | 0.6944604 |
| ATVASLIL | ASSLWDIYAEQF | 0.5326575 |
| AVNGPVLIRH | ASSLWDIYAEQF | 0.2575228 |
| DGAAGAVGI | ASSLWDIYAEQF | 0.63906 |
| DPLLHGI | ASSLWDIYAEQF | 0.5232736 |
| ELGAALGAL | ASSLWDIYAEQF | 0.7731176 |
| ELGRTQAV | ASSLWDIYAEQF | 0.7709769 |
| FVAGGYLS | ASSLWDIYAEQF | 0.540472 |
| GASGGAGGAGGW | ASSLWDIYAEQF | 0.7764394 |
| GGAGAGGDGGAGGI | ASSLWDIYAEQF | 0.72101307 |
| GTGHNGSQPGMGGNG | ASSLWDIYAEQF | 0.45813268 |
| GYAAASASAS | ASSLWDIYAEQF | 0.71096796 |
| HQVDVAAVQQG | ASSLWDIYAEQF | 0.48066473 |
| IGAAIAHT | ASSLWDIYAEQF | 0.72580576 |
| IGMIEVAQL | ASSLWDIYAEQF | 0.4363442 |
| IRLAPEHTSAMD | ASSLWDIYAEQF | 0.7680896 |
| IYNVSVL | ASSLWDIYAEQF | 0.68207073 |
| KQGFLGM | ASSLWDIYAEQF | 0.21594739 |
| LESVITQL | ASSLWDIYAEQF | 0.75636816 |
| LGLGALVVV | ASSLWDIYAEQF | 0.7134761 |
| LIGNGGNGGSGGI | ASSLWDIYAEQF | 0.71974355 |
| LNAINAPTL | ASSLWDIYAEQF | 0.7829584 |
| LQLLNTGV | ASSLWDIYAEQF | 0.6049857 |
| LQTSIEF | ASSLWDIYAEQF | 0.5501114 |
| MSFAVSEL | ASSLWDIYAEQF | 0.76103896 |
| NTGALGL | ASSLWDIYAEQF | 0.7749446 |
| PPGAGGAGGAGTT | ASSLWDIYAEQF | 0.49757487 |
| QLVGLGI | ASSLWDIYAEQF | 0.75717354 |
| RFLEQVHQL | ASSLWDIYAEQF | 0.7807004 |
| RGVGYVLR | ASSLWDIYAEQF | 0.69571525 |
| SATGVFAGVI | ASSLWDIYAEQF | 0.77856493 |
| SLPAITLP | ASSLWDIYAEQF | 0.7828615 |
| STALAFTV | ASSLWDIYAEQF | 0.47580737 |
| SVLTVGLGK | ASSLWDIYAEQF | 0.78030664 |
| TALAGAGSGI | ASSLWDIYAEQF | 0.7850695 |
| TGLPFTGI | ASSLWDIYAEQF | 0.62402844 |
| VAEHVTV | ASSLWDIYAEQF | 0.7188159 |
| VDIVLNSL | ASSLWDIYAEQF | 0.46846792 |
| VLTGGGAVL | ASSLWDIYAEQF | 0.68474054 |
| VNVGLAYGI | ASSLWDIYAEQF | 0.3068532 |
| VSSAAVGAV | ASSLWDIYAEQF | 0.76382875 |
| VTSVYEAGV | ASSLWDIYAEQF | 0.7759056 |
| WVALDRGA | ASSLWDIYAEQF | 0.70385593 |
| YLVGSVVI | ASSLWDIYAEQF | 0.58522856 |
| GPPGTGKTLL | ASSLWDIYAEQF | 0.39237082 |
| GVGMTKF | ASSLWDIYAEQF | 0.7607046 |

| | | |
|---|---|---|
| IGIGGSDL | ASSLWDIYAEQF | 0.5693371 |
| IGTLLAEL | ASSLWDIYAEQF | 0.3642106 |
| IYTSGTT | ASSLWDIYAEQF | 0.63593286 |
| LSFPAGV | ASSLWDIYAEQF | 0.73868847 |
| LYTSGTT | ASSLWDIYAEQF | 0.7465985 |
| VGAFPFGF | ASSLWDIYAEQF | 0.61852556 |
| VGAGTVEF | ASSLWDIYAEQF | 0.79400676 |
| VGKSTLLL | ASSLWDIYAEQF | 0.77478313 |
| AALAHAGGGIV | ASSLWDIYAEQF | 0.76062393 |
| AALVAVSLL | ASSLWDIYAEQF | 0.6349197 |
| AAVSRAVAAL | ASSLWDIYAEQF | 0.57302916 |
| AGGSSGIGGF | ASSLWDIYAEQF | 0.7863547 |
| AGLTVANANG | ASSLWDIYAEQF | 0.67771804 |
| AGLVGAVL | ASSLWDIYAEQF | 0.6988268 |
| ALSQAFTV | ASSLWDIYAEQF | 0.75990635 |
| ALTFSVSL | ASSLWDIYAEQF | 0.69951713 |
| ALTGAALVAL | ASSLWDIYAEQF | 0.62831724 |
| ASFSIGAGVL | ASSLWDIYAEQF | 0.73660505 |
| DGSVYKI | ASSLWDIYAEQF | 0.7917853 |
| EQLAEIV | ASSLWDIYAEQF | 0.5506993 |
| FAVSELTVM | ASSLWDIYAEQF | 0.4136709 |
| GAAPLAAGL | ASSLWDIYAEQF | 0.6434829 |
| GADGGDGGSGGAGG | ASSLWDIYAEQF | 0.7364603 |
| GFGLDTGGAGG | ASSLWDIYAEQF | 0.7744584 |
| GLASGGLL | ASSLWDIYAEQF | 0.7408818 |
| GRIAGAAVLL | ASSLWDIYAEQF | 0.71936834 |
| GSAGSLNTSF | ASSLWDIYAEQF | 0.7631991 |
| GTGQIGI | ASSLWDIYAEQF | 0.75333095 |
| GTLIALIL | ASSLWDIYAEQF | 0.7353741 |
| IAATMGFVVG | ASSLWDIYAEQF | 0.7540934 |
| IFAQALL | ASSLWDIYAEQF | 0.6960423 |
| IGAAVALGF | ASSLWDIYAEQF | 0.7885183 |
| IGLTPLAL | ASSLWDIYAEQF | 0.7667382 |
| ILGSNTSTL | ASSLWDIYAEQF | 0.75568277 |
| IVFSLTVL | ASSLWDIYAEQF | 0.77664495 |
| LASKATGF | ASSLWDIYAEQF | 0.72999865 |
| LASVALIL | ASSLWDIYAEQF | 0.71982265 |
| LFAVGLTL | ASSLWDIYAEQF | 0.25805426 |
| LFNGGIGTYI | ASSLWDIYAEQF | 0.38172865 |
| LTLGAGII | ASSLWDIYAEQF | 0.7597847 |
| LVAAALGDI | ASSLWDIYAEQF | 0.7589004 |
| NTAFVQLGI | ASSLWDIYAEQF | 0.7773931 |
| QLFSGAEL | ASSLWDIYAEQF | 0.71536005 |
| SAATFITL | ASSLWDIYAEQF | 0.60705304 |
| SNYGLANLGV | ASSLWDIYAEQF | 0.7638167 |
| SYGGAYAVM | ASSLWDIYAEQF | 0.780218 |
| TTGVGGLTLG | ASSLWDIYAEQF | 0.41666508 |
| TTQTIYI | ASSLWDIYAEQF | 0.63166314 |
| VAAGAGLGAVI | ASSLWDIYAEQF | 0.79962134 |
| VAAVVTLL | ASSLWDIYAEQF | 0.7571702 |
| VGFLGGLI | ASSLWDIYAEQF | 0.618268 |

| | | |
|---|---|---|
| VGVIPILL | ASSLWDIYAEQF | 0.6750754 |
| VIFASSLI | ASSLWDIYAEQF | 0.7913698 |
| VKGGDGGAAAG | ASSLWDIYAEQF | 0.6978281 |
| VTKTFTNI | ASSLWDIYAEQF | 0.7948747 |
| VVAAAILL | ASSLWDIYAEQF | 0.25085035 |
| YAALDVEL | ASSLWDIYAEQF | 0.67631614 |
| LASVALLL | ASSLWDIYAEQF | 0.4952699 |
| LDAVHRV | ASSLWDIYAEQF | 0.2843619 |
| QGLSSSIF | ASSLWDIYAEQF | 0.75069696 |
| RGTFANIRL | ASSLWDIYAEQF | 0.557961 |
| RQLWWGHRI | ASSLWDIYAEQF | 0.7377499 |
| YGPPGTGKTLL | ASSLWDIYAEQF | 0.6401534 |
| AALVGAPANVA | ASSLWDIYAEQF | 0.77244395 |
| AATFRRTE | ASSLWDIYAEQF | 0.75180584 |
| AGALGGGI | ASSLWDIYAEQF | 0.69479954 |
| AGGAGGAGGIGAG | ASSLWDIYAEQF | 0.57760024 |
| AGSGALGL | ASSLWDIYAEQF | 0.7083657 |
| AQIADQI | ASSLWDIYAEQF | 0.77547216 |
| ASAVAVTYL | ASSLWDIYAEQF | 0.7741652 |
| ASTVLGGTGG | ASSLWDIYAEQF | 0.27708936 |
| AVGGEPAK | ASSLWDIYAEQF | 0.7534671 |
| CEQGTFGGTC | ASSLWDIYAEQF | 0.58898807 |
| DEGKAPAG | ASSLWDIYAEQF | 0.4204442 |
| DTVVFPAL | ASSLWDIYAEQF | 0.28486842 |
| FEGITMAG | ASSLWDIYAEQF | 0.6815057 |
| FPLVVQD | ASSLWDIYAEQF | 0.74512327 |
| GAFTTPQ | ASSLWDIYAEQF | 0.7670114 |
| GFGGFGTAGGVGGA | ASSLWDIYAEQF | 0.7699567 |
| GFLAQFL | ASSLWDIYAEQF | 0.7130525 |
| GGAGGAGGNAGVA | ASSLWDIYAEQF | 0.77383816 |
| GGVGTAGAGGAGGAGG | ASSLWDIYAEQF | 0.7421961 |
| GNGGGGDAGNAGSGGN | ASSLWDIYAEQF | 0.1835793 |
| GTGGAGGAGNDAGS | ASSLWDIYAEQF | 0.53752863 |
| GTTNTGWFNS | ASSLWDIYAEQF | 0.434057 |
| IFMGPLFTL | ASSLWDIYAEQF | 0.77055997 |
| ILAHAIEAI | ASSLWDIYAEQF | 0.5985702 |
| IRSAIREAL | ASSLWDIYAEQF | 0.7025161 |
| ITFAGGG | ASSLWDIYAEQF | 0.73633426 |
| ITGMSGHQEV | ASSLWDIYAEQF | 0.7625846 |
| IVAGLACTL | ASSLWDIYAEQF | 0.7013234 |
| IVGGGLAGA | ASSLWDIYAEQF | 0.6293328 |
| IVTGVLNI | ASSLWDIYAEQF | 0.30262363 |
| LAQQGCI | ASSLWDIYAEQF | 0.71788263 |
| LCLHVEN | ASSLWDIYAEQF | 0.47044033 |
| LDAISDGS | ASSLWDIYAEQF | 0.48145545 |
| LGGFAAML | ASSLWDIYAEQF | 0.4825704 |
| LGGKNPMI | ASSLWDIYAEQF | 0.73099864 |
| LGPKGVAAIL | ASSLWDIYAEQF | 0.2581966 |
| LMVGVSI | ASSLWDIYAEQF | 0.7760614 |
| LVGAAFVL | ASSLWDIYAEQF | 0.5739041 |
| LVLAVAPL | ASSLWDIYAEQF | 0.708022 |

| | | |
|---|---|---|
| MAAIAVIDAA | ASSLWDIYAEQF | 0.7920038 |
| NNGINAELQ | ASSLWDIYAEQF | 0.7578171 |
| RGEPLVLVHG | ASSLWDIYAEQF | 0.668876 |
| RGGAGGAAGLI | ASSLWDIYAEQF | 0.61322725 |
| RSYRMHPAV | ASSLWDIYAEQF | 0.5133643 |
| SAAHLLAGQLA | ASSLWDIYAEQF | 0.6255757 |
| SAAVGSY | ASSLWDIYAEQF | 0.61365044 |
| SPHAVLHRI | ASSLWDIYAEQF | 0.7824953 |
| SQIAKVL | ASSLWDIYAEQF | 0.7147062 |
| TALGHLVP | ASSLWDIYAEQF | 0.40985483 |
| TAPTTSGAA | ASSLWDIYAEQF | 0.72663057 |
| TDGDDGNGGSPGT | ASSLWDIYAEQF | 0.79414046 |
| TDGLGEN | ASSLWDIYAEQF | 0.79038686 |
| TGGPPLPGQA | ASSLWDIYAEQF | 0.6418351 |
| TIPAITI | ASSLWDIYAEQF | 0.48164487 |
| TIPGFTI | ASSLWDIYAEQF | 0.557875 |
| TSIASGALA | ASSLWDIYAEQF | 0.7164245 |
| TVTLPGAAI | ASSLWDIYAEQF | 0.60803485 |
| VASGLSAALP | ASSLWDIYAEQF | 0.5348212 |
| VGAGVIGI | ASSLWDIYAEQF | 0.7817073 |
| VGPKRFGV | ASSLWDIYAEQF | 0.79493463 |
| VNVAFPD | ASSLWDIYAEQF | 0.700351 |
| VSMTNSGVS | ASSLWDIYAEQF | 0.27748048 |
| VTDVLGDLSV | ASSLWDIYAEQF | 0.72044873 |
| VVVVGAGFAGL | ASSLWDIYAEQF | 0.55678314 |
| VYTVGTVV | ASSLWDIYAEQF | 0.68839216 |
| WTTFSGP | ASSLWDIYAEQF | 0.53527784 |
| YDELVEAF | ASSLWDIYAEQF | 0.47790748 |
| YGTGGAGGNGG | ASSLWDIYAEQF | 0.7067994 |
| YTNVMAV | ASSLWDIYAEQF | 0.7867879 |
| FLEQVHQL | ASSLWDIYAEQF | 0.5789822 |
| GLSLLAVAV | ASSLWDIYAEQF | 0.737044 |
| LEQVHQL | ASSLWDIYAEQF | 0.7906837 |
| LNAAGAI | ASSLWDIYAEQF | 0.7702377 |
| PAVHASL | ASSLWDIYAEQF | 0.7896452 |
| TELEDDA | ASSLWDIYAEQF | 0.7756289 |
| TPPGPPN | ASSLWDIYAEQF | 0.701142 |
| TPPPPPPP | ASSLWDIYAEQF | 0.7180995 |
| TQSPALI | ASSQERDREDEQY | 0.7800124 |
| EFAAAFPQ | ASSQERDREDEQY | 0.7130207 |
| LLSGNLDVL | ASSQERDREDEQY | 0.70995307 |
| GPANLGNYNI | ASSQERDREDEQY | 0.6125525 |
| TLGPTLP | ASSQERDREDEQY | 0.7733318 |
| VYAYSSCCK | ASSQERDREDEQY | 0.69949865 |
| APAPAPAPNI | ASSQERDREDEQY | 0.7852496 |
| PGGPVPIP | ASSQERDREDEQY | 0.31097645 |
| FLTVALEP | ASSQERDREDEQY | 0.5594245 |
| SQYFGANL | ASSQERDREDEQY | 0.7816628 |
| GGVKPAA | ASSQERDREDEQY | 0.19335686 |
| GGTGGAGGAGGW | ASSQERDREDEQY | 0.2713369 |
| GVGGVAGLGGAGG | ASSQERDREDEQY | 0.7468035 |

| | | |
|---|---|---|
| VLLVLATTI | ASSQERDREDEQY | 0.7855552 |
| IGIFAAML | ASSQERDREDEQY | 0.22118829 |
| AVGDLID | ASSQERDREDEQY | 0.5480568 |
| NISPLPL | ASSQERDREDEQY | 0.34502828 |
| VTAVFVAI | ASSQERDREDEQY | 0.7590105 |
| SGDDNDAAL | ASSQERDREDEQY | 0.70327777 |
| AVAALAHS | ASSQERDREDEQY | 0.26313093 |
| GGVTVMATP | ASSQERDREDEQY | 0.77193916 |
| AGLAANVRD | ASSQERDREDEQY | 0.35260963 |
| TNRPDLIDPALLRPG | ASSQERDREDEQY | 0.73659605 |
| LANLGVTAL | ASSQERDREDEQY | 0.7646222 |
| TGERIEL | ASSQERDREDEQY | 0.79078627 |
| NGEELAA | ASSQERDREDEQY | 0.5058062 |
| VLSLVWG | ASSQERDREDEQY | 0.75967133 |
| IGYALWL | ASSQERDREDEQY | 0.66362524 |
| LSLLLAPL | ASSQERDREDEQY | 0.796239 |
| TLDEALK | ASSQERDREDEQY | 0.4967718 |
| TDGAGTAEGG | ASSQERDREDEQY | 0.42633367 |
| NAPGGAGGNGG | ASSQERDREDEQY | 0.77739906 |
| VSVFMGMGGNF | ASSQERDREDEQY | 0.7723845 |
| LASYLAAL | ASSQERDREDEQY | 0.60162294 |
| LVGGTPAL | ASSQERDREDEQY | 0.77644813 |
| VRNADVL | ASSQERDREDEQY | 0.6775451 |
| TCSSPFLVSAM | ASSQERDREDEQY | 0.7206632 |
| VHITDAE | ASSQERDREDEQY | 0.7819899 |
| EYAAPSF | ASSQERDREDEQY | 0.78294194 |
| VVDAFHGGL | ASSQERDREDEQY | 0.54032326 |
| ASVVDPQ | ASSQERDREDEQY | 0.77818936 |
| SYNEFGGLS | ASSQERDREDEQY | 0.7355954 |
| DPDGSVY | ASSQERDREDEQY | 0.47480592 |
| MVSTIHATE | ASSQERDREDEQY | 0.78522754 |
| LAAGVADPV | ASSQERDREDEQY | 0.7118131 |
| TKEQIAA | ASSQERDREDEQY | 0.77974343 |
| SLASPAAL | ASSQERDREDEQY | 0.7181446 |
| IGLVFYGV | ASSQERDREDEQY | 0.7826853 |
| FLLDASNL | ASSQERDREDEQY | 0.28532618 |
| YQVGNFP | ASSQERDREDEQY | 0.72214615 |
| ESLLSQST | ASSQERDREDEQY | 0.7018469 |
| LFTVGFAAVG | ASSQERDREDEQY | 0.75288147 |
| ATPPTPT | ASSQERDREDEQY | 0.692073 |
| TDLATLG | ASSQERDREDEQY | 0.5983441 |
| SAGGTGR | ASSQERDREDEQY | 0.75574756 |
| GFSNSGS | ASSQERDREDEQY | 0.7873338 |
| GGNGGSGGTSV | ASSQERDREDEQY | 0.74418426 |
| GTSVATGGAGN | ASSQERDREDEQY | 0.7475174 |
| NGGNGGSGGTS | ASSQERDREDEQY | 0.7945174 |
| AVVKFPH | ASSQERDREDEQY | 0.5682552 |
| TPPPPPP | ASSQERDREDEQY | 0.7142323 |
| AAPAAAAAPA | ASSQERDREDEQY | 0.76779664 |
| IALNEDL | ASSQERDREDEQY | 0.775332 |
| IVNVSSM | ASSQERDREDEQY | 0.7041069 |

| | | |
|---|---|---|
| TGIPPAPR | ASSQERDREDEQY | 0.7917254 |
| FDLGGGTF | ASSQERDREDEQY | 0.53317714 |
| FDLGGGTFDVS | ASSQERDREDEQY | 0.7779552 |
| NEPTAAA | ASSQERDREDEQY | 0.6925568 |
| NEPTAAALA | ASSQERDREDEQY | 0.62471735 |
| GSIINISSI | ASSQERDREDEQY | 0.6465832 |
| VAPGNAGTA | ASSQERDREDEQY | 0.74193525 |
| TGGGASLE | ASSQERDREDEQY | 0.7787795 |
| VAVYNHYKRI | ASSQERDREDEQY | 0.76777697 |
| TDPAPATT | ASSQERDREDEQY | 0.25131536 |
| TNRPDILDPALLRPG | ASSQERDREDEQY | 0.77785265 |
| LTTTTTS | ASSQERDREDEQY | 0.76016563 |
| PTGSGKTL | ASSQERDREDEQY | 0.7952771 |
| TGSGKTL | ASSQERDREDEQY | 0.67092896 |
| TIGHVDHGK | ASSQERDREDEQY | 0.71785223 |
| TIGHVDHGKTT | ASSQERDREDEQY | 0.65732086 |
| GHVDHGKTT | ASSQERDREDEQY | 0.7192295 |
| AQFEHTLL | ASSQERDREDEQY | 0.5136917 |
| IVNQDEL | ASSQERDREDEQY | 0.5754097 |
| TNGSQFF | ASSQERDREDEQY | 0.72281075 |
| TGKGIVE | ASSQERDREDEQY | 0.6806202 |
| LISNASDA | ASSQERDREDEQY | 0.77881765 |
| AELRTLGELTKIAKAHGAQ | ASSQERDREDEQY | 0.78499436 |
| APNAVAAAEPSW | ASSQERDREDEQY | 0.76769584 |
| APQVAGAATAVL | ASSQERDREDEQY | 0.7860087 |
| ARRDPAAAAAILPTDA | ASSQERDREDEQY | 0.793792 |
| DAAARDVLAEAGLAEYF | ASSQERDREDEQY | 0.7758496 |
| DDQIVLSAKANLAKVEQ | ASSQERDREDEQY | 0.63333243 |
| DELLPDTGEDSDEDGH | ASSQERDREDEQY | 0.56541914 |
| ELPEAPANRATVPQD | ASSQERDREDEQY | 0.5312873 |
| FGQSKIERVTGIG | ASSQERDREDEQY | 0.71401733 |
| GAGGTGSTAGGAGGAGGAGGLY | ASSQERDREDEQY | 0.6403233 |
| GHGGTGAAGGNGGTGGAGGSGI | ASSQERDREDEQY | 0.7496284 |
| GNGGNGGAGGEGGPGGAGGAGA | ASSQERDREDEQY | 0.18908729 |
| GSTAGNGGNGGNATNGTIAGQP | ASSQERDREDEQY | 0.76074034 |
| GTGGDGGNGGNGGTGG | ASSQERDREDEQY | 0.7902554 |
| GVRRPAGIPADAPV | ASSQERDREDEQY | 0.67096376 |
| GYRQVTDAHLLTLARR | ASSQERDREDEQY | 0.7478471 |
| IAKMGQDGHDRG | ASSQERDREDEQY | 0.77528363 |
| KAEPSIEIQYT | ASSQERDREDEQY | 0.6763576 |
| KPVRIRMAADEDTVAVT | ASSQERDREDEQY | 0.76886046 |
| MPPRIAGMRLLVIKPEP | ASSQERDREDEQY | 0.6022131 |
| MKFYAVALFRSRGVPPTR | ASSQERDREDEQY | 0.77229774 |
| NAGGFGILGAVAHSP | ASSQERDREDEQY | 0.7707719 |
| NEGREITLAMAEM | ASSQERDREDEQY | 0.74540436 |
| NELVEWSHPLGHRWRWE | ASSQERDREDEQY | 0.6581101 |
| NLGAGGTGGNGGNPG | ASSQERDREDEQY | 0.6998267 |
| NTEWPKWETVMAADPVD | ASSQERDREDEQY | 0.24441862 |
| PASAAPAETQP | ASSQERDREDEQY | 0.6965759 |
| QQSSDGKNGVRAAEITGE | ASSQERDREDEQY | 0.5664071 |
| RAPDAVTTSWWKE | ASSQERDREDEQY | 0.735278 |

| | | |
|---|---|---|
| SGFGNVGAHTSGWFNQ | ASSQERDREDEQY | 0.43928558 |
| SGVAGAVGQAASVGGLKV | ASSQERDREDEQY | 0.73467165 |
| SNGGAGTGSAGGNGGT | ASSQERDREDEQY | 0.7392989 |
| TGGAGGEGGAAG | ASSQERDREDEQY | 0.75025165 |
| TGGDGGAAGAGGAPGHG | ASSQERDREDEQY | 0.36172426 |
| TMPPLLAAIANAGRHGV | ASSQERDREDEQY | 0.75279427 |
| TSSTAGNSGGA | ASSQERDREDEQY | 0.7254051 |
| VPATVPVGILC | ASSQERDREDEQY | 0.38700712 |
| VQRPDSGGVVLTGRIS | ASSQERDREDEQY | 0.77992153 |
| WTPNPVPATKGVPAGLVQAT | ASSQERDREDEQY | 0.70130646 |
| YALKRALAKFSDRFDVVII | ASSQERDREDEQY | 0.72112197 |
| YLIGSVIV | ASSQERDREDEQY | 0.7530049 |
| YGQLVAVID | ASSQERDREDEQY | 0.722219 |
| VTVGGMTLPA | ASSQERDREDEQY | 0.28571647 |
| VLALPAV | ASSQERDREDEQY | 0.7133703 |
| VIPPTLNYET | ASSQERDREDEQY | 0.702191 |
| VGMGGMHPSSG | ASSQERDREDEQY | 0.54002 |
| VGLAGGAATA | ASSQERDREDEQY | 0.2519712 |
| VGGAGGAGGSGP | ASSQERDREDEQY | 0.77812636 |
| VFAAFMV | ASSQERDREDEQY | 0.4638448 |
| TLYDKLR | ASSQERDREDEQY | 0.5595333 |
| TLAVFTVL | ASSQERDREDEQY | 0.7066323 |
| TGSVQPARV | ASSQERDREDEQY | 0.7792436 |
| TGGGAGSGGT | ASSQERDREDEQY | 0.66860795 |
| SHLETSL | ASSQERDREDEQY | 0.64385545 |
| SFTTGMT | ASSQERDREDEQY | 0.7722579 |
| QVEVAKRMGVRQPTVS | ASSQERDREDEQY | 0.728544 |
| QGSGGPALAM | ASSQERDREDEQY | 0.77410185 |
| MLAPSNP | ASSQERDREDEQY | 0.73537743 |
| LVNNAAI | ASSQERDREDEQY | 0.6527236 |
| LTGEINTLA | ASSQERDREDEQY | 0.69135845 |
| LSIFGPV | ASSQERDREDEQY | 0.5618749 |
| LLALLANL | ASSQERDREDEQY | 0.53544456 |
| LISNASDALDKLR | ASSQERDREDEQY | 0.47037843 |
| LINEAVP | ASSQERDREDEQY | 0.4673092 |
| LGYAFSGL | ASSQERDREDEQY | 0.37489933 |
| LDFVAVAL | ASSQERDREDEQY | 0.63649905 |
| LAQPGAE | ASSQERDREDEQY | 0.78658617 |
| ISNLGAVTGF | ASSQERDREDEQY | 0.647611 |
| HQEDPGRI | ASSQERDREDEQY | 0.37551576 |
| GTPGEPTP | ASSQERDREDEQY | 0.66947424 |
| GFFNSGS | ASSQERDREDEQY | 0.41811746 |
| GAGGGGLTAGAN | ASSQERDREDEQY | 0.76788306 |
| FVDALVNM | ASSQERDREDEQY | 0.6869835 |
| FLTVAPEL | ASSQERDREDEQY | 0.7964946 |
| FETLTAL | ASSQERDREDEQY | 0.77561206 |
| DVINPIEL | ASSQERDREDEQY | 0.69332623 |
| APVSGAPV | ASSQERDREDEQY | 0.7811874 |
| ALVGGLAW | ASSQERDREDEQY | 0.63610935 |
| ALAAAAATAI | ASSQERDREDEQY | 0.73180574 |
| AFGLGLLL | ASSQERDREDEQY | 0.7293756 |

| | | |
|---|---|---|
| AFFAAASL | ASSQERDREDEQY | 0.7636974 |
| AAAALPAFSPP | ASSQERDREDEQY | 0.6807597 |
| VVLAPAL | ASSQERDREDEQY | 0.6679521 |
| TIGHVDHGKTTL | ASSQERDREDEQY | 0.24660107 |
| LISNASDALDK | ASSQERDREDEQY | 0.7770128 |
| LISNASDAL | ASSQERDREDEQY | 0.40020677 |
| ISNASDALDK | ASSQERDREDEQY | 0.43919823 |
| NQAETLVYQTE | ASSQERDREDEQY | 0.5654752 |
| TGGTGGTAGAG | ASSQERDREDEQY | 0.7896423 |
| TGSTAGGAGGA | ASSQERDREDEQY | 0.7892637 |
| DPLAVPVAAGPV | ASSQERDREDEQY | 0.6664388 |
| GGTGGTPGNGGN | ASSQERDREDEQY | 0.5072619 |
| PAGLAGVDPATL | ASSQERDREDEQY | 0.7702634 |
| GNNLVVEAPHAGASV | ASSQERDREDEQY | 0.5081055 |
| KVTRIMSASTLEQPA | ASSQERDREDEQY | 0.76676476 |
| LGFALPIALPRRTVV | ASSQERDREDEQY | 0.7290994 |
| TRLGNLTARKLVVLV | ASSQERDREDEQY | 0.73980594 |
| WSQQRRLIYFPSAGP | ASSQERDREDEQY | 0.46935353 |
| GPNGNPDPMAAAVDIR | ASSQERDREDEQY | 0.7630286 |
| TQRCVSVADLRAAASR | ASSQERDREDEQY | 0.6807293 |
| QFVLDVAEHPVEVLTETV | ASSQERDREDEQY | 0.7537042 |
| GGAGGTGGAGGNGGTQAGDG | ASSQERDREDEQY | 0.2388418 |
| PLLATLRAIAVDVDAKPAQV | ASSQERDREDEQY | 0.7456623 |
| APKFRTGGYMIDFWGVGYQVAK | ASSQERDREDEQY | 0.6278138 |
| GAGGAGGAGGAAGGPGGTGGTG | ASSQERDREDEQY | 0.5671691 |
| IGALGLLATDVAPAGSVAFGVA | ASSQERDREDEQY | 0.33698606 |
| AGGAGGIGGNGGAGGAGGVLMG | ASSQERDREDEQY | 0.6351402 |
| ADELIGLL | ASSQERDREDEQY | 0.75641966 |
| ADRALKA | ASSQERDREDEQY | 0.7861626 |
| AGAVGQVGGAGGA | ASSQERDREDEQY | 0.68653834 |
| AGGGGGLGGAP | ASSQERDREDEQY | 0.7569511 |
| AGGLLYGNGGAG | ASSQERDREDEQY | 0.7763077 |
| AGLESIGL | ASSQERDREDEQY | 0.7032869 |
| AGLKTGII | ASSQERDREDEQY | 0.2445471 |
| ASLAALLL | ASSQERDREDEQY | 0.70135987 |
| ASVINISSI | ASSQERDREDEQY | 0.73478615 |
| ATPPGPA | ASSQERDREDEQY | 0.7757783 |
| ATVASALV | ASSQERDREDEQY | 0.74792916 |
| ATVASLIL | ASSQERDREDEQY | 0.5712359 |
| AVNGPVLIRH | ASSQERDREDEQY | 0.70278555 |
| DGAAGAVGI | ASSQERDREDEQY | 0.74154055 |
| DPLLHGI | ASSQERDREDEQY | 0.5114071 |
| ELGAALGAL | ASSQERDREDEQY | 0.57697916 |
| ELGRTQAV | ASSQERDREDEQY | 0.5195943 |
| FVAGGYLS | ASSQERDREDEQY | 0.443936 |
| GASGGAGGAGGW | ASSQERDREDEQY | 0.74412096 |
| GGAGAGGDGGAGGI | ASSQERDREDEQY | 0.7051069 |
| GTGHNGSQPGMGGNG | ASSQERDREDEQY | 0.64026076 |
| GYAAASASAS | ASSQERDREDEQY | 0.72751594 |
| HQVDVAAVQQG | ASSQERDREDEQY | 0.6177676 |
| IGAAIAHT | ASSQERDREDEQY | 0.6040963 |

| | | |
|---|---|---|
| IGMIEVAQL | ASSQERDREDEQY | 0.65411055 |
| IRLAPEHTSAMD | ASSQERDREDEQY | 0.6591076 |
| IYNVSVL | ASSQERDREDEQY | 0.7777573 |
| KQGFLGM | ASSQERDREDEQY | 0.6930304 |
| LESVITQL | ASSQERDREDEQY | 0.70092845 |
| LGLGALVVV | ASSQERDREDEQY | 0.79176605 |
| LIGNGGNGGSGGI | ASSQERDREDEQY | 0.40999767 |
| LNAINAPTL | ASSQERDREDEQY | 0.59093153 |
| LQLLNTGV | ASSQERDREDEQY | 0.7755641 |
| LQTSIEF | ASSQERDREDEQY | 0.55498886 |
| MSFAVSEL | ASSQERDREDEQY | 0.78194624 |
| NTGALGL | ASSQERDREDEQY | 0.7499299 |
| PPGAGGAGGAGTT | ASSQERDREDEQY | 0.767949 |
| QLVGLGI | ASSQERDREDEQY | 0.7266882 |
| RFLEQVHQL | ASSQERDREDEQY | 0.79595923 |
| RGVGYVLR | ASSQERDREDEQY | 0.7627702 |
| SATGVFAGVI | ASSQERDREDEQY | 0.6817391 |
| SLPAITLP | ASSQERDREDEQY | 0.7842971 |
| STALAFTV | ASSQERDREDEQY | 0.41820872 |
| SVLTVGLGK | ASSQERDREDEQY | 0.7649605 |
| TALAGAGSGI | ASSQERDREDEQY | 0.7579138 |
| TGLPFTGI | ASSQERDREDEQY | 0.52039754 |
| VAEHVTV | ASSQERDREDEQY | 0.7220841 |
| VDIVLNSL | ASSQERDREDEQY | 0.78917146 |
| VLTGGGAVL | ASSQERDREDEQY | 0.7871703 |
| VNVGLAYGI | ASSQERDREDEQY | 0.36447525 |
| VSSAAVGAV | ASSQERDREDEQY | 0.69706297 |
| VTSVYEAGV | ASSQERDREDEQY | 0.5072925 |
| WVALDRGA | ASSQERDREDEQY | 0.5865921 |
| YLVGSVVI | ASSQERDREDEQY | 0.34280533 |
| GPPGTGKTLL | ASSQERDREDEQY | 0.76672864 |
| GVGMTKF | ASSQERDREDEQY | 0.60064936 |
| IGIGGSDL | ASSQERDREDEQY | 0.7741798 |
| IGTLLAEL | ASSQERDREDEQY | 0.69402635 |
| IYTSGTT | ASSQERDREDEQY | 0.7638426 |
| LSFPAGV | ASSQERDREDEQY | 0.58897644 |
| LYTSGTT | ASSQERDREDEQY | 0.6882045 |
| VGAFPFGF | ASSQERDREDEQY | 0.36187637 |
| VGAGTVEF | ASSQERDREDEQY | 0.2780891 |
| VGKSTLLL | ASSQERDREDEQY | 0.6271608 |
| AALAHAGGGIV | ASSQERDREDEQY | 0.6483339 |
| AALVAVSLL | ASSQERDREDEQY | 0.2554736 |
| AAVSRAVAAL | ASSQERDREDEQY | 0.7939485 |
| AGGSSGIGGF | ASSQERDREDEQY | 0.7065962 |
| AGLTVANANG | ASSQERDREDEQY | 0.74336886 |
| AGLVGAVL | ASSQERDREDEQY | 0.5837353 |
| ALSQAFTV | ASSQERDREDEQY | 0.60677916 |
| ALTFSVSL | ASSQERDREDEQY | 0.6574211 |
| ALTGAALVAL | ASSQERDREDEQY | 0.76355594 |
| ASFSIGAGVL | ASSQERDREDEQY | 0.47807565 |
| DGSVYKI | ASSQERDREDEQY | 0.25317472 |

| | | |
|---|---|---|
| EQLAEIV | ASSQERDREDEQY | 0.51929045 |
| FAVSELTVM | ASSQERDREDEQY | 0.45590127 |
| GAAPLAAGL | ASSQERDREDEQY | 0.7935867 |
| GADGGDGGSGGAGG | ASSQERDREDEQY | 0.7808652 |
| GFGLDTGGAGG | ASSQERDREDEQY | 0.7387828 |
| GLASGGLL | ASSQERDREDEQY | 0.7919474 |
| GRIAGAAVLL | ASSQERDREDEQY | 0.20559895 |
| GSAGSLNTSF | ASSQERDREDEQY | 0.644271 |
| GTGQIGI | ASSQERDREDEQY | 0.775953 |
| GTLIALIL | ASSQERDREDEQY | 0.72775054 |
| IAATMGFVVG | ASSQERDREDEQY | 0.63061094 |
| IFAQALL | ASSQERDREDEQY | 0.7842765 |
| IGAAVALGF | ASSQERDREDEQY | 0.6219779 |
| IGLTPLAL | ASSQERDREDEQY | 0.48263127 |
| ILGSNTSTL | ASSQERDREDEQY | 0.74265313 |
| IVFSLTVL | ASSQERDREDEQY | 0.7618791 |
| LASKATGF | ASSQERDREDEQY | 0.76156294 |
| LASVALIL | ASSQERDREDEQY | 0.47011366 |
| LFAVGLTL | ASSQERDREDEQY | 0.71279407 |
| LFNGGIGTYI | ASSQERDREDEQY | 0.80195284 |
| LTLGAGII | ASSQERDREDEQY | 0.69881296 |
| LVAAALGDI | ASSQERDREDEQY | 0.7491109 |
| NTAFVQLGI | ASSQERDREDEQY | 0.7372507 |
| QLFSGAEL | ASSQERDREDEQY | 0.6005265 |
| SAATFITL | ASSQERDREDEQY | 0.54074466 |
| SNYGLANLGV | ASSQERDREDEQY | 0.6285819 |
| SYGGAYAVM | ASSQERDREDEQY | 0.7419559 |
| TTGVGGLTLG | ASSQERDREDEQY | 0.54968244 |
| TTQTIYI | ASSQERDREDEQY | 0.65811455 |
| VAAGAGLGAVI | ASSQERDREDEQY | 0.6833569 |
| VAAVVTLL | ASSQERDREDEQY | 0.80122817 |
| VGFLGGLI | ASSQERDREDEQY | 0.76764333 |
| VGVIPILL | ASSQERDREDEQY | 0.76347935 |
| VIFASSLI | ASSQERDREDEQY | 0.60266304 |
| VKGGDGGAAAG | ASSQERDREDEQY | 0.75248694 |
| VTKTFTNI | ASSQERDREDEQY | 0.76347476 |
| VVAAAILL | ASSQERDREDEQY | 0.263724 |
| YAALDVEL | ASSQERDREDEQY | 0.7872104 |
| LASVALLL | ASSQERDREDEQY | 0.5247772 |
| LDAVHRV | ASSQERDREDEQY | 0.716482 |
| QGLSSSIF | ASSQERDREDEQY | 0.73217964 |
| RGTFANIRL | ASSQERDREDEQY | 0.78835905 |
| RQLWWGHRI | ASSQERDREDEQY | 0.69540465 |
| YGPPGTGKTLL | ASSQERDREDEQY | 0.74939585 |
| AALVGAPANVA | ASSQERDREDEQY | 0.62050027 |
| AATFRRTE | ASSQERDREDEQY | 0.7745687 |
| AGALGGGI | ASSQERDREDEQY | 0.7846167 |
| AGGAGGAGGIGAG | ASSQERDREDEQY | 0.68579066 |
| AGSGALGL | ASSQERDREDEQY | 0.7954321 |
| AQIADQI | ASSQERDREDEQY | 0.7010882 |
| ASAVAVTYL | ASSQERDREDEQY | 0.28977722 |

| | | |
|---|---|---|
| ASTVLGGTGG | ASSQERDREDEQY | 0.68643326 |
| AVGGEPAK | ASSQERDREDEQY | 0.768421 |
| CEQGTFGGTC | ASSQERDREDEQY | 0.7895482 |
| DEGKAPAG | ASSQERDREDEQY | 0.73837715 |
| DTVVFPAL | ASSQERDREDEQY | 0.77401507 |
| FEGITMAG | ASSQERDREDEQY | 0.5920643 |
| FPLVVQD | ASSQERDREDEQY | 0.7885784 |
| GAFTTPQ | ASSQERDREDEQY | 0.795231 |
| GFGGFGTAGGVGGA | ASSQERDREDEQY | 0.7355329 |
| GFLAQFL | ASSQERDREDEQY | 0.6135614 |
| GGAGGAGGNAGVA | ASSQERDREDEQY | 0.5336098 |
| GGVGTAGAGGAGGAGG | ASSQERDREDEQY | 0.7497593 |
| GNGGGGDAGNAGSGGN | ASSQERDREDEQY | 0.7807616 |
| GTGGAGGAGNDAGS | ASSQERDREDEQY | 0.43800277 |
| GTTNTGWFNS | ASSQERDREDEQY | 0.7378863 |
| IFMGPLFTL | ASSQERDREDEQY | 0.7516682 |
| ILAHAIEAI | ASSQERDREDEQY | 0.28664893 |
| IRSAIREAL | ASSQERDREDEQY | 0.7509201 |
| ITFAGGG | ASSQERDREDEQY | 0.7920232 |
| ITGMSGHQEV | ASSQERDREDEQY | 0.7522501 |
| IVAGLACTL | ASSQERDREDEQY | 0.54276735 |
| IVGGGLAGA | ASSQERDREDEQY | 0.7902397 |
| IVTGVLNI | ASSQERDREDEQY | 0.7962674 |
| LAQQGCI | ASSQERDREDEQY | 0.78847325 |
| LCLHVEN | ASSQERDREDEQY | 0.3406675 |
| LDAISDGS | ASSQERDREDEQY | 0.246736 |
| LGGFAAML | ASSQERDREDEQY | 0.72548044 |
| LGGKNPMI | ASSQERDREDEQY | 0.7754496 |
| LGPKGVAAIL | ASSQERDREDEQY | 0.6618556 |
| LMVGVSI | ASSQERDREDEQY | 0.62206876 |
| LVGAAFVL | ASSQERDREDEQY | 0.7664258 |
| LVLAVAPL | ASSQERDREDEQY | 0.7806442 |
| MAAIAVIDAA | ASSQERDREDEQY | 0.48424792 |
| NNGINAELQ | ASSQERDREDEQY | 0.42728263 |
| RGEPLVLVHG | ASSQERDREDEQY | 0.7733394 |
| RGGAGGAAGLI | ASSQERDREDEQY | 0.76820207 |
| RSYRMHPAV | ASSQERDREDEQY | 0.6497866 |
| SAAHLLAGQLA | ASSQERDREDEQY | 0.7068058 |
| SAAVGSY | ASSQERDREDEQY | 0.7032381 |
| SPHAVLHRI | ASSQERDREDEQY | 0.6954305 |
| SQIAKVL | ASSQERDREDEQY | 0.74165463 |
| TALGHLVP | ASSQERDREDEQY | 0.53048766 |
| TAPTTSGAA | ASSQERDREDEQY | 0.7887181 |
| TDGDDGNGGSPGT | ASSQERDREDEQY | 0.52991295 |
| TDGLGEN | ASSQERDREDEQY | 0.7792419 |
| TGGPPLPGQA | ASSQERDREDEQY | 0.74684846 |
| TIPAITI | ASSQERDREDEQY | 0.7414734 |
| TIPGFTI | ASSQERDREDEQY | 0.6724771 |
| TSIASGALA | ASSQERDREDEQY | 0.49884143 |
| TVTLPGAAI | ASSQERDREDEQY | 0.40532094 |
| VASGLSAALP | ASSQERDREDEQY | 0.52932966 |

| | | |
|---|---|---|
| VGAGVIGI | ASSQERDREDEQY | 0.78228843 |
| VGPKRFGV | ASSQERDREDEQY | 0.6932779 |
| VNVAFPD | ASSQERDREDEQY | 0.78592175 |
| VSMTNSGVS | ASSQERDREDEQY | 0.6791346 |
| VTDVLGDLSV | ASSQERDREDEQY | 0.58965504 |
| VVVVGAGFAGL | ASSQERDREDEQY | 0.7719811 |
| VYTVGTVV | ASSQERDREDEQY | 0.5354529 |
| WTTFSGP | ASSQERDREDEQY | 0.6650561 |
| YDELVEAF | ASSQERDREDEQY | 0.6924691 |
| YGTGGAGGNGG | ASSQERDREDEQY | 0.7656164 |
| YTNVMAV | ASSQERDREDEQY | 0.48932034 |
| FLEQVHQL | ASSQERDREDEQY | 0.7597127 |
| GLSLLAVAV | ASSQERDREDEQY | 0.75393283 |
| LEQVHQL | ASSQERDREDEQY | 0.41128218 |
| LNAAGAI | ASSQERDREDEQY | 0.7337755 |
| PAVHASL | ASSQERDREDEQY | 0.78400946 |
| TELEDDA | ASSQERDREDEQY | 0.7447412 |
| TPPGPPN | ASSQERDREDEQY | 0.38624585 |
| TPPPPPPP | ASSQERDREDEQY | 0.7311715 |
| TQSPALI | ASSPDREGSGNTLY | 0.6349349 |
| EFAAAFPQ | ASSPDREGSGNTLY | 0.76370126 |
| LLSGNLDVL | ASSPDREGSGNTLY | 0.752731 |
| GPANLGNYNI | ASSPDREGSGNTLY | 0.5877346 |
| TLGPTLP | ASSPDREGSGNTLY | 0.7510061 |
| VYAYSSCCK | ASSPDREGSGNTLY | 0.32463336 |
| APAPAPAPNI | ASSPDREGSGNTLY | 0.6016943 |
| PGGPVPIP | ASSPDREGSGNTLY | 0.65312046 |
| FLTVALEP | ASSPDREGSGNTLY | 0.7153585 |
| SQYFGANL | ASSPDREGSGNTLY | 0.75503707 |
| GGVKPAA | ASSPDREGSGNTLY | 0.54828846 |
| GGTGGAGGAGGW | ASSPDREGSGNTLY | 0.7409667 |
| GVGGVAGLGGAGG | ASSPDREGSGNTLY | 0.47271228 |
| VLLVLATTI | ASSPDREGSGNTLY | 0.35840324 |
| IGIFAAML | ASSPDREGSGNTLY | 0.7252898 |
| AVGDLID | ASSPDREGSGNTLY | 0.6621423 |
| NISPLPL | ASSPDREGSGNTLY | 0.4402003 |
| VTAVFVAI | ASSPDREGSGNTLY | 0.7666638 |
| SGDDNDAAL | ASSPDREGSGNTLY | 0.5676588 |
| AVAALAHS | ASSPDREGSGNTLY | 0.7053592 |
| GGVTVMATP | ASSPDREGSGNTLY | 0.74127865 |
| AGLAANVRD | ASSPDREGSGNTLY | 0.48802948 |
| TNRPDLIDPALLRPG | ASSPDREGSGNTLY | 0.7723006 |
| LANLGVTAL | ASSPDREGSGNTLY | 0.72347367 |
| TGERIEL | ASSPDREGSGNTLY | 0.76757234 |
| NGEELAA | ASSPDREGSGNTLY | 0.67971957 |
| VLSLVWG | ASSPDREGSGNTLY | 0.47509152 |
| IGYALWL | ASSPDREGSGNTLY | 0.77996254 |
| LSLLLAPL | ASSPDREGSGNTLY | 0.7438793 |
| TLDEALK | ASSPDREGSGNTLY | 0.65440166 |
| TDGAGTAEGG | ASSPDREGSGNTLY | 0.77104455 |
| NAPGGAGGNGG | ASSPDREGSGNTLY | 0.77186525 |

| | | |
|---|---|---|
| VSVFMGMGGNF | ASSPDREGSGNTLY | 0.77523994 |
| LASYLAAL | ASSPDREGSGNTLY | 0.6812885 |
| LVGGTPAL | ASSPDREGSGNTLY | 0.34592792 |
| VRNADVL | ASSPDREGSGNTLY | 0.69040775 |
| TCSSPFLVSAM | ASSPDREGSGNTLY | 0.70417655 |
| VHITDAE | ASSPDREGSGNTLY | 0.5852913 |
| EYAAPSF | ASSPDREGSGNTLY | 0.38825908 |
| VVDAFHGGL | ASSPDREGSGNTLY | 0.7729232 |
| ASVVDPQ | ASSPDREGSGNTLY | 0.79934573 |
| SYNEFGGLS | ASSPDREGSGNTLY | 0.64095044 |
| DPDGSVY | ASSPDREGSGNTLY | 0.79730797 |
| MVSTIHATE | ASSPDREGSGNTLY | 0.42799413 |
| LAAGVADPV | ASSPDREGSGNTLY | 0.739908 |
| TKEQIAA | ASSPDREGSGNTLY | 0.77747256 |
| SLASPAAL | ASSPDREGSGNTLY | 0.5820483 |
| IGLVFYGV | ASSPDREGSGNTLY | 0.7011281 |
| FLLDASNL | ASSPDREGSGNTLY | 0.40477824 |
| YQVGNFP | ASSPDREGSGNTLY | 0.6009651 |
| ESLLSQST | ASSPDREGSGNTLY | 0.78121555 |
| LFTVGFAAVG | ASSPDREGSGNTLY | 0.57060075 |
| ATPPTPT | ASSPDREGSGNTLY | 0.73556507 |
| TDLATLG | ASSPDREGSGNTLY | 0.7037607 |
| SAGGTGR | ASSPDREGSGNTLY | 0.5503826 |
| GFSNSGS | ASSPDREGSGNTLY | 0.75864124 |
| GGNGGSGGTSV | ASSPDREGSGNTLY | 0.7607143 |
| GTSVATGGAGN | ASSPDREGSGNTLY | 0.6765684 |
| NGGNGGSGGTS | ASSPDREGSGNTLY | 0.7955514 |
| AVVKFPH | ASSPDREGSGNTLY | 0.5721137 |
| TPPPPPP | ASSPDREGSGNTLY | 0.7939615 |
| AAPAAAAAPA | ASSPDREGSGNTLY | 0.78990847 |
| IALNEDL | ASSPDREGSGNTLY | 0.74670964 |
| IVNVSSM | ASSPDREGSGNTLY | 0.64790976 |
| TGIPPAPR | ASSPDREGSGNTLY | 0.77711654 |
| FDLGGGTF | ASSPDREGSGNTLY | 0.61914957 |
| FDLGGGTFDVS | ASSPDREGSGNTLY | 0.75358796 |
| NEPTAAA | ASSPDREGSGNTLY | 0.7841679 |
| NEPTAAALA | ASSPDREGSGNTLY | 0.5526479 |
| GSIINISSI | ASSPDREGSGNTLY | 0.7986444 |
| VAPGNAGTA | ASSPDREGSGNTLY | 0.7738811 |
| TGGGASLE | ASSPDREGSGNTLY | 0.7391802 |
| VAVYNHYKRI | ASSPDREGSGNTLY | 0.59936297 |
| TDPAPATT | ASSPDREGSGNTLY | 0.33795074 |
| TNRPDILDPALLRPG | ASSPDREGSGNTLY | 0.7057184 |
| LTTTTTS | ASSPDREGSGNTLY | 0.50063634 |
| PTGSGKTL | ASSPDREGSGNTLY | 0.79532224 |
| TGSGKTL | ASSPDREGSGNTLY | 0.7960419 |
| TIGHVDHGK | ASSPDREGSGNTLY | 0.629719 |
| TIGHVDHGKTT | ASSPDREGSGNTLY | 0.779178 |
| GHVDHGKTT | ASSPDREGSGNTLY | 0.7251344 |
| AQFEHTLL | ASSPDREGSGNTLY | 0.46183354 |
| IVNQDEL | ASSPDREGSGNTLY | 0.7331251 |

| | | |
|---|---|---|
| TNGSQFF | ASSPDREGSGNTLY | 0.6792556 |
| TGKGIVE | ASSPDREGSGNTLY | 0.73483753 |
| LISNASDA | ASSPDREGSGNTLY | 0.4691158 |
| AELRTLGELTKIAKAHGAQ | ASSPDREGSGNTLY | 0.76960146 |
| APNAVAAAEPSW | ASSPDREGSGNTLY | 0.7184662 |
| APQVAGAATAVL | ASSPDREGSGNTLY | 0.28996056 |
| ARRDPAAAAAILPTDA | ASSPDREGSGNTLY | 0.7824968 |
| DAAARDVLAEAGLAEYF | ASSPDREGSGNTLY | 0.6779545 |
| DDQIVLSAKANLAKVEQ | ASSPDREGSGNTLY | 0.6355015 |
| DELLPDTGEDSDEDGH | ASSPDREGSGNTLY | 0.7996576 |
| ELPEAPANRATVPQD | ASSPDREGSGNTLY | 0.5510005 |
| FGQSKIERVTGIG | ASSPDREGSGNTLY | 0.50974137 |
| GAGGTGSTAGGAGGAGGAGGLY | ASSPDREGSGNTLY | 0.7741822 |
| GHGGTGAAGGNGGTGGAGGSGI | ASSPDREGSGNTLY | 0.79786587 |
| GNGGNGGAGGEGGPGGAGGAGA | ASSPDREGSGNTLY | 0.35743982 |
| GSTAGNGGNGGNATNGTIAGQP | ASSPDREGSGNTLY | 0.76679754 |
| GTGGDGGNGGNGGTGG | ASSPDREGSGNTLY | 0.7866467 |
| GVRRPAGIPADAPV | ASSPDREGSGNTLY | 0.71047044 |
| GYRQVTDAHLLTLARR | ASSPDREGSGNTLY | 0.8115684 |
| IAKMGQDGHDRG | ASSPDREGSGNTLY | 0.81280303 |
| KAEPSIEIQYT | ASSPDREGSGNTLY | 0.70739067 |
| KPVRIRMAADEDTVAVT | ASSPDREGSGNTLY | 0.8031154 |
| MPPRIAGMRLLVIKPEP | ASSPDREGSGNTLY | 0.8033532 |
| MKFYAVALFRSRGVPPTR | ASSPDREGSGNTLY | 0.62257445 |
| NAGGFGILGAVAHSP | ASSPDREGSGNTLY | 0.72524744 |
| NEGREITLAMAEM | ASSPDREGSGNTLY | 0.54449326 |
| NELVEWSHPLGHRWRWE | ASSPDREGSGNTLY | 0.72974783 |
| NLGAGGTGGNGGNPG | ASSPDREGSGNTLY | 0.40321732 |
| NTEWPKWETVMAADPVD | ASSPDREGSGNTLY | 0.780728 |
| PASAAPAETQP | ASSPDREGSGNTLY | 0.79179907 |
| QQSSDGKNGVRAAEITGE | ASSPDREGSGNTLY | 0.72451496 |
| RAPDAVTTSWWKE | ASSPDREGSGNTLY | 0.54581803 |
| SGFGNVGAHTSGWFNQ | ASSPDREGSGNTLY | 0.5022352 |
| SGVAGAVGQAASVGGLKV | ASSPDREGSGNTLY | 0.76127326 |
| SNGGAGTGSAGGNGGT | ASSPDREGSGNTLY | 0.51533836 |
| TGGAGGEGGAAG | ASSPDREGSGNTLY | 0.48468655 |
| TGGDGGAAGAGGAPGHG | ASSPDREGSGNTLY | 0.7424877 |
| TMPPLLAAIANAGRHGV | ASSPDREGSGNTLY | 0.7428274 |
| TSSTAGNSGGA | ASSPDREGSGNTLY | 0.76538455 |
| VPATVPVGILC | ASSPDREGSGNTLY | 0.5466087 |
| VQRPDSGGVVLTGRIS | ASSPDREGSGNTLY | 0.81216955 |
| WTPNPVPATKGVPAGLVQAT | ASSPDREGSGNTLY | 0.78470266 |
| YALKRALAKFSDRFDVVII | ASSPDREGSGNTLY | 0.80068886 |
| YLIGSVIV | ASSPDREGSGNTLY | 0.7166517 |
| YGQLVAVID | ASSPDREGSGNTLY | 0.5230388 |
| VTVGGMTLPA | ASSPDREGSGNTLY | 0.8003172 |
| VLALPAV | ASSPDREGSGNTLY | 0.6894567 |
| VIPPTLNYET | ASSPDREGSGNTLY | 0.7277589 |
| VGMGGMHPSSG | ASSPDREGSGNTLY | 0.79235935 |
| VGLAGGAATA | ASSPDREGSGNTLY | 0.66480744 |
| VGGAGGAGGSGP | ASSPDREGSGNTLY | 0.5660645 |

| | | |
|---|---|---|
| VFAAFMV | ASSPDREGSGNTLY | 0.7294709 |
| TLYDKLR | ASSPDREGSGNTLY | 0.8100246 |
| TLAVFTVL | ASSPDREGSGNTLY | 0.58480537 |
| TGSVQPARV | ASSPDREGSGNTLY | 0.44743523 |
| TGGGAGSGGT | ASSPDREGSGNTLY | 0.6661644 |
| SHLETSL | ASSPDREGSGNTLY | 0.70135343 |
| SFTTGMT | ASSPDREGSGNTLY | 0.79724187 |
| QVEVAKRMGVRQPTVS | ASSPDREGSGNTLY | 0.7642298 |
| QGSGGPALAM | ASSPDREGSGNTLY | 0.7286815 |
| MLAPSNP | ASSPDREGSGNTLY | 0.78171116 |
| LVNNAAI | ASSPDREGSGNTLY | 0.7628019 |
| LTGEINTLA | ASSPDREGSGNTLY | 0.7483603 |
| LSIFGPV | ASSPDREGSGNTLY | 0.7603719 |
| LLALLANL | ASSPDREGSGNTLY | 0.6585519 |
| LISNASDALDKLR | ASSPDREGSGNTLY | 0.804067 |
| LINEAVP | ASSPDREGSGNTLY | 0.78409004 |
| LGYAFSGL | ASSPDREGSGNTLY | 0.78974336 |
| LDFVAVAL | ASSPDREGSGNTLY | 0.8067699 |
| LAQPGAE | ASSPDREGSGNTLY | 0.67426646 |
| ISNLGAVTGF | ASSPDREGSGNTLY | 0.76592493 |
| HQEDPGRI | ASSPDREGSGNTLY | 0.3793015 |
| GTPGEPTP | ASSPDREGSGNTLY | 0.6894407 |
| GFFNSGS | ASSPDREGSGNTLY | 0.7727718 |
| GAGGGGLTAGAN | ASSPDREGSGNTLY | 0.77230304 |
| FVDALVNM | ASSPDREGSGNTLY | 0.7987883 |
| FLTVAPEL | ASSPDREGSGNTLY | 0.7419412 |
| FETLTAL | ASSPDREGSGNTLY | 0.65659916 |
| DVINPIEL | ASSPDREGSGNTLY | 0.7960635 |
| APVSGAPV | ASSPDREGSGNTLY | 0.79660153 |
| ALVGGLAW | ASSPDREGSGNTLY | 0.51776385 |
| ALAAAAATAI | ASSPDREGSGNTLY | 0.70861536 |
| AFGLGLLL | ASSPDREGSGNTLY | 0.812342 |
| AFFAAASL | ASSPDREGSGNTLY | 0.774734 |
| AAAALPAFSPP | ASSPDREGSGNTLY | 0.72360396 |
| VVLAPAL | ASSPDREGSGNTLY | 0.671701 |
| TIGHVDHGKTTL | ASSPDREGSGNTLY | 0.8042092 |
| LISNASDALDK | ASSPDREGSGNTLY | 0.726597 |
| LISNASDAL | ASSPDREGSGNTLY | 0.81099534 |
| ISNASDALDK | ASSPDREGSGNTLY | 0.25918046 |
| NQAETLVYQTE | ASSPDREGSGNTLY | 0.73068446 |
| TGGTGGTAGAG | ASSPDREGSGNTLY | 0.5483018 |
| TGSTAGGAGGA | ASSPDREGSGNTLY | 0.2555081 |
| DPLAVPVAAGPV | ASSPDREGSGNTLY | 0.780249 |
| GGTGGTPGNGGN | ASSPDREGSGNTLY | 0.6711593 |
| PAGLAGVDPATL | ASSPDREGSGNTLY | 0.7543859 |
| GNNLVVEAPHAGASV | ASSPDREGSGNTLY | 0.75769484 |
| KVTRIMSASTLEQPA | ASSPDREGSGNTLY | 0.7920575 |
| LGFALPIALPRRTVV | ASSPDREGSGNTLY | 0.7747586 |
| TRLGNLTARKLVVLV | ASSPDREGSGNTLY | 0.67242986 |
| WSQQRRLIYFPSAGP | ASSPDREGSGNTLY | 0.5983639 |
| GPNGNPDPMAAAVDIR | ASSPDREGSGNTLY | 0.7418653 |

| | | |
|---|---|---|
| TQRCVSVADLRAAASR | ASSPDREGSGNTLY | 0.80606407 |
| QFVLDVAEHPVEVLTETV | ASSPDREGSGNTLY | 0.7967204 |
| GGAGGTGGAGGNGGTQAGDG | ASSPDREGSGNTLY | 0.26705652 |
| PLLATLRAIAVDVDAKPAQV | ASSPDREGSGNTLY | 0.78395844 |
| APKFRTGGYMIDFWGVGYQVAK | ASSPDREGSGNTLY | 0.6374343 |
| GAGGAGGAGGAAGGPGGTGGTG | ASSPDREGSGNTLY | 0.3404566 |
| IGALGLLATDVAPAGSVAFGVA | ASSPDREGSGNTLY | 0.4035321 |
| AGGAGGIGGNGGAGGAGGVLMG | ASSPDREGSGNTLY | 0.67986226 |
| ADELIGLL | ASSPDREGSGNTLY | 0.74100333 |
| ADRALKA | ASSPDREGSGNTLY | 0.7889323 |
| AGAVGQVGGAGGA | ASSPDREGSGNTLY | 0.78271574 |
| AGGGGGLGGAP | ASSPDREGSGNTLY | 0.6755558 |
| AGGLLYGNGGAG | ASSPDREGSGNTLY | 0.78672236 |
| AGLESIGL | ASSPDREGSGNTLY | 0.68364894 |
| AGLKTGII | ASSPDREGSGNTLY | 0.16940378 |
| ASLAALLL | ASSPDREGSGNTLY | 0.6712327 |
| ASVINISSI | ASSPDREGSGNTLY | 0.4981699 |
| ATPPGPA | ASSPDREGSGNTLY | 0.8022016 |
| ATVASALV | ASSPDREGSGNTLY | 0.6389186 |
| ATVASLIL | ASSPDREGSGNTLY | 0.7194854 |
| AVNGPVLIRH | ASSPDREGSGNTLY | 0.71555793 |
| DGAAGAVGI | ASSPDREGSGNTLY | 0.7712861 |
| DPLLHGI | ASSPDREGSGNTLY | 0.74757713 |
| ELGAALGAL | ASSPDREGSGNTLY | 0.5567417 |
| ELGRTQAV | ASSPDREGSGNTLY | 0.45604885 |
| FVAGGYLS | ASSPDREGSGNTLY | 0.7316947 |
| GASGGAGGAGGW | ASSPDREGSGNTLY | 0.5796871 |
| GGAGAGGDGGAGGI | ASSPDREGSGNTLY | 0.42449698 |
| GTGHNGSQPGMGGNG | ASSPDREGSGNTLY | 0.64251924 |
| GYAAASASAS | ASSPDREGSGNTLY | 0.7866059 |
| HQVDVAAVQQG | ASSPDREGSGNTLY | 0.31991756 |
| IGAAIAHT | ASSPDREGSGNTLY | 0.8047178 |
| IGMIEVAQL | ASSPDREGSGNTLY | 0.702819 |
| IRLAPEHTSAMD | ASSPDREGSGNTLY | 0.70172775 |
| IYNVSVL | ASSPDREGSGNTLY | 0.8066685 |
| KQGFLGM | ASSPDREGSGNTLY | 0.7494158 |
| LESVITQL | ASSPDREGSGNTLY | 0.73149717 |
| LGLGALVVV | ASSPDREGSGNTLY | 0.5056194 |
| LIGNGGNGGSGGI | ASSPDREGSGNTLY | 0.64251894 |
| LNAINAPTL | ASSPDREGSGNTLY | 0.5590749 |
| LQLLNTGV | ASSPDREGSGNTLY | 0.51527727 |
| LQTSIEF | ASSPDREGSGNTLY | 0.8016801 |
| MSFAVSEL | ASSPDREGSGNTLY | 0.7423142 |
| NTGALGL | ASSPDREGSGNTLY | 0.5052371 |
| PPGAGGAGGAGTT | ASSPDREGSGNTLY | 0.7580162 |
| QLVGLGI | ASSPDREGSGNTLY | 0.81066835 |
| RFLEQVHQL | ASSPDREGSGNTLY | 0.81407815 |
| RGVGYVLR | ASSPDREGSGNTLY | 0.66822565 |
| SATGVFAGVI | ASSPDREGSGNTLY | 0.5022823 |
| SLPAITLP | ASSPDREGSGNTLY | 0.6683973 |
| STALAFTV | ASSPDREGSGNTLY | 0.7317896 |

| | | |
|---|---|---|
| SVLTVGLGK | ASSPDREGSGNTLY | 0.59910274 |
| TALAGAGSGI | ASSPDREGSGNTLY | 0.4568308 |
| TGLPFTGI | ASSPDREGSGNTLY | 0.7983631 |
| VAEHVTV | ASSPDREGSGNTLY | 0.8071959 |
| VDIVLNSL | ASSPDREGSGNTLY | 0.70495445 |
| VLTGGGAVL | ASSPDREGSGNTLY | 0.47907162 |
| VNVGLAYGI | ASSPDREGSGNTLY | 0.70755935 |
| VSSAAVGAV | ASSPDREGSGNTLY | 0.5652658 |
| VTSVYEAGV | ASSPDREGSGNTLY | 0.6231697 |
| WVALDRGA | ASSPDREGSGNTLY | 0.59216285 |
| YLVGSVVI | ASSPDREGSGNTLY | 0.46649373 |
| GPPGTGKTLL | ASSPDREGSGNTLY | 0.6997423 |
| GVGMTKF | ASSPDREGSGNTLY | 0.8025774 |
| IGIGGSDL | ASSPDREGSGNTLY | 0.6932721 |
| IGTLLAEL | ASSPDREGSGNTLY | 0.7788849 |
| IYTSGTT | ASSPDREGSGNTLY | 0.8097679 |
| LSFPAGV | ASSPDREGSGNTLY | 0.790227 |
| LYTSGTT | ASSPDREGSGNTLY | 0.80681777 |
| VGAFPFGF | ASSPDREGSGNTLY | 0.79096633 |
| VGAGTVEF | ASSPDREGSGNTLY | 0.71847224 |
| VGKSTLLL | ASSPDREGSGNTLY | 0.7277591 |
| AALAHAGGGIV | ASSPDREGSGNTLY | 0.8047688 |
| AALVAVSLL | ASSPDREGSGNTLY | 0.7084178 |
| AAVSRAVAAL | ASSPDREGSGNTLY | 0.7547799 |
| AGGSSGIGGF | ASSPDREGSGNTLY | 0.7410969 |
| AGLTVANANG | ASSPDREGSGNTLY | 0.7808851 |
| AGLVGAVL | ASSPDREGSGNTLY | 0.7025376 |
| ALSQAFTV | ASSPDREGSGNTLY | 0.80658567 |
| ALTFSVSL | ASSPDREGSGNTLY | 0.30706602 |
| ALTGAALVAL | ASSPDREGSGNTLY | 0.4295895 |
| ASFSIGAGVL | ASSPDREGSGNTLY | 0.80336225 |
| DGSVYKI | ASSPDREGSGNTLY | 0.18611343 |
| EQLAEIV | ASSPDREGSGNTLY | 0.37143505 |
| FAVSELTVM | ASSPDREGSGNTLY | 0.7677076 |
| GAAPLAAGL | ASSPDREGSGNTLY | 0.80214906 |
| GADGGDGGSGGAGG | ASSPDREGSGNTLY | 0.21460965 |
| GFGLDTGGAGG | ASSPDREGSGNTLY | 0.55588686 |
| GLASGGLL | ASSPDREGSGNTLY | 0.5264608 |
| GRIAGAAVLL | ASSPDREGSGNTLY | 0.7672487 |
| GSAGSLNTSF | ASSPDREGSGNTLY | 0.60577226 |
| GTGQIGI | ASSPDREGSGNTLY | 0.37610185 |
| GTLIALIL | ASSPDREGSGNTLY | 0.7847438 |
| IAATMGFVVG | ASSPDREGSGNTLY | 0.5215776 |
| IFAQALL | ASSPDREGSGNTLY | 0.72165585 |
| IGAAVALGF | ASSPDREGSGNTLY | 0.80082965 |
| IGLTPLAL | ASSPDREGSGNTLY | 0.8085595 |
| ILGSNTSTL | ASSPDREGSGNTLY | 0.6084618 |
| IVFSLTVL | ASSPDREGSGNTLY | 0.7778032 |
| LASKATGF | ASSPDREGSGNTLY | 0.7016438 |
| LASVALIL | ASSPDREGSGNTLY | 0.81793946 |
| LFAVGLTL | ASSPDREGSGNTLY | 0.6657777 |

| | | |
|---|---|---|
| LFNGGIGTYI | ASSPDREGSGNTLY | 0.4656511 |
| LTLGAGII | ASSPDREGSGNTLY | 0.7915015 |
| LVAAALGDI | ASSPDREGSGNTLY | 0.7945474 |
| NTAFVQLGI | ASSPDREGSGNTLY | 0.5982138 |
| QLFSGAEL | ASSPDREGSGNTLY | 0.78937495 |
| SAATFITL | ASSPDREGSGNTLY | 0.6880039 |
| SNYGLANLGV | ASSPDREGSGNTLY | 0.69578564 |
| SYGGAYAVM | ASSPDREGSGNTLY | 0.79971457 |
| TTGVGGLTLG | ASSPDREGSGNTLY | 0.7947254 |
| TTQTIYI | ASSPDREGSGNTLY | 0.498137 |
| VAAGAGLGAVI | ASSPDREGSGNTLY | 0.79752547 |
| VAAVVTLL | ASSPDREGSGNTLY | 0.69879186 |
| VGFLGGLI | ASSPDREGSGNTLY | 0.5293837 |
| VGVIPILL | ASSPDREGSGNTLY | 0.81177104 |
| VIFASSLI | ASSPDREGSGNTLY | 0.7350143 |
| VKGGDGGAAAG | ASSPDREGSGNTLY | 0.79805255 |
| VTKTFTNI | ASSPDREGSGNTLY | 0.7542771 |
| VVAAAILL | ASSPDREGSGNTLY | 0.7959842 |
| YAALDVEL | ASSPDREGSGNTLY | 0.5167322 |
| LASVALLL | ASSPDREGSGNTLY | 0.7691029 |
| LDAVHRV | ASSPDREGSGNTLY | 0.7359874 |
| QGLSSSIF | ASSPDREGSGNTLY | 0.79172873 |
| RGTFANIRL | ASSPDREGSGNTLY | 0.74979496 |
| RQLWWGHRI | ASSPDREGSGNTLY | 0.7516873 |
| YGPPGTGKTLL | ASSPDREGSGNTLY | 0.7923485 |
| AALVGAPANVA | ASSPDREGSGNTLY | 0.8018654 |
| AATFRRTE | ASSPDREGSGNTLY | 0.7548805 |
| AGALGGGI | ASSPDREGSGNTLY | 0.7433138 |
| AGGAGGAGGIGAG | ASSPDREGSGNTLY | 0.8102685 |
| AGSGALGL | ASSPDREGSGNTLY | 0.62793374 |
| AQIADQI | ASSPDREGSGNTLY | 0.69740593 |
| ASAVAVTYL | ASSPDREGSGNTLY | 0.78672254 |
| ASTVLGGTGG | ASSPDREGSGNTLY | 0.799281 |
| AVGGEPAK | ASSPDREGSGNTLY | 0.7422496 |
| CEQGTFGGTC | ASSPDREGSGNTLY | 0.809389 |
| DEGKAPAG | ASSPDREGSGNTLY | 0.6955923 |
| DTVVFPAL | ASSPDREGSGNTLY | 0.7792165 |
| FEGITMAG | ASSPDREGSGNTLY | 0.75219285 |
| FPLVVQD | ASSPDREGSGNTLY | 0.6858231 |
| GAFTTPQ | ASSPDREGSGNTLY | 0.5727652 |
| GFGGFGTAGGVGGA | ASSPDREGSGNTLY | 0.76076555 |
| GFLAQFL | ASSPDREGSGNTLY | 0.7961582 |
| GGAGGAGGNAGVA | ASSPDREGSGNTLY | 0.75675476 |
| GGVGTAGAGGAGGAGG | ASSPDREGSGNTLY | 0.3988974 |
| GNGGGGDAGNAGSGGN | ASSPDREGSGNTLY | 0.7750977 |
| GTGGAGGAGNDAGS | ASSPDREGSGNTLY | 0.75038046 |
| GTTNTGWFNS | ASSPDREGSGNTLY | 0.81324327 |
| IFMGPLFTL | ASSPDREGSGNTLY | 0.70456076 |
| ILAHAIEAI | ASSPDREGSGNTLY | 0.7669184 |
| IRSAIREAL | ASSPDREGSGNTLY | 0.6274072 |
| ITFAGGG | ASSPDREGSGNTLY | 0.74304795 |

| | | |
|---|---|---|
| ITGMSGHQEV | ASSPDREGSGNTLY | 0.48610133 |
| IVAGLACTL | ASSPDREGSGNTLY | 0.6668581 |
| IVGGGLAGA | ASSPDREGSGNTLY | 0.7258842 |
| IVTGVLNI | ASSPDREGSGNTLY | 0.6374829 |
| LAQQGCI | ASSPDREGSGNTLY | 0.78399605 |
| LCLHVEN | ASSPDREGSGNTLY | 0.80417585 |
| LDAISDGS | ASSPDREGSGNTLY | 0.7606538 |
| LGGFAAML | ASSPDREGSGNTLY | 0.79936564 |
| LGGKNPMI | ASSPDREGSGNTLY | 0.81256884 |
| LGPKGVAAIL | ASSPDREGSGNTLY | 0.79931295 |
| LMVGVSI | ASSPDREGSGNTLY | 0.5982742 |
| LVGAAFVL | ASSPDREGSGNTLY | 0.7383813 |
| LVLAVAPL | ASSPDREGSGNTLY | 0.47349888 |
| MAAIAVIDAA | ASSPDREGSGNTLY | 0.6994821 |
| NNGINAELQ | ASSPDREGSGNTLY | 0.6200429 |
| RGEPLVLVHG | ASSPDREGSGNTLY | 0.7483858 |
| RGGAGGAAGLI | ASSPDREGSGNTLY | 0.19508488 |
| RSYRMHPAV | ASSPDREGSGNTLY | 0.76494914 |
| SAAHLLAGQLA | ASSPDREGSGNTLY | 0.8096341 |
| SAAVGSY | ASSPDREGSGNTLY | 0.671252 |
| SPHAVLHRI | ASSPDREGSGNTLY | 0.7586357 |
| SQIAKVL | ASSPDREGSGNTLY | 0.79696065 |
| TALGHLVP | ASSPDREGSGNTLY | 0.70459545 |
| TAPTTSGAA | ASSPDREGSGNTLY | 0.7777505 |
| TDGDDGNGGSPGT | ASSPDREGSGNTLY | 0.6501195 |
| TDGLGEN | ASSPDREGSGNTLY | 0.7994287 |
| TGGPPLPGQA | ASSPDREGSGNTLY | 0.7986405 |
| TIPAITI | ASSPDREGSGNTLY | 0.7637377 |
| TIPGFTI | ASSPDREGSGNTLY | 0.70628554 |
| TSIASGALA | ASSPDREGSGNTLY | 0.72289264 |
| TVTLPGAAI | ASSPDREGSGNTLY | 0.39188558 |
| VASGLSAALP | ASSPDREGSGNTLY | 0.733152 |
| VGAGVIGI | ASSPDREGSGNTLY | 0.6594275 |
| VGPKRFGV | ASSPDREGSGNTLY | 0.75864816 |
| VNVAFPD | ASSPDREGSGNTLY | 0.5431637 |
| VSMTNSGVS | ASSPDREGSGNTLY | 0.7538026 |
| VTDVLGDLSV | ASSPDREGSGNTLY | 0.7025615 |
| VVVVGAGFAGL | ASSPDREGSGNTLY | 0.76611125 |
| VYTVGTVV | ASSPDREGSGNTLY | 0.6010232 |
| WTTFSGP | ASSPDREGSGNTLY | 0.77239263 |
| YDELVEAF | ASSPDREGSGNTLY | 0.7498963 |
| YGTGGAGGNGG | ASSPDREGSGNTLY | 0.36880115 |
| YTNVMAV | ASSPDREGSGNTLY | 0.80768293 |
| FLEQVHQL | ASSPDREGSGNTLY | 0.7266692 |
| GLSLLAVAV | ASSPDREGSGNTLY | 0.73512113 |
| LEQVHQL | ASSPDREGSGNTLY | 0.7646203 |
| LNAAGAI | ASSPDREGSGNTLY | 0.69875383 |
| PAVHASL | ASSPDREGSGNTLY | 0.49470416 |
| TELEDDA | ASSPDREGSGNTLY | 0.7061331 |
| TPPGPPN | ASSPDREGSGNTLY | 0.7647853 |
| TPPPPPPP | ASSPDREGSGNTLY | 0.65380174 |

| | | |
|---|---|---|
| TQSPALI | ASSSGTGVNSPLY | 0.2775675 |
| EFAAAFPQ | ASSSGTGVNSPLY | 0.79314834 |
| LLSGNLDVL | ASSSGTGVNSPLY | 0.5248506 |
| GPANLGNYNI | ASSSGTGVNSPLY | 0.68557274 |
| TLGPTLP | ASSSGTGVNSPLY | 0.68666244 |
| VYAYSSCCK | ASSSGTGVNSPLY | 0.794711 |
| APAPAPAPNI | ASSSGTGVNSPLY | 0.68996584 |
| PGGPVPIP | ASSSGTGVNSPLY | 0.54998726 |
| FLTVALEP | ASSSGTGVNSPLY | 0.7902287 |
| SQYFGANL | ASSSGTGVNSPLY | 0.6636306 |
| GGVKPAA | ASSSGTGVNSPLY | 0.7831416 |
| GGTGGAGGAGGW | ASSSGTGVNSPLY | 0.69853187 |
| GVGGVAGLGGAGG | ASSSGTGVNSPLY | 0.70851606 |
| VLLVLATTI | ASSSGTGVNSPLY | 0.7088796 |
| IGIFAAML | ASSSGTGVNSPLY | 0.64648724 |
| AVGDLID | ASSSGTGVNSPLY | 0.67313826 |
| NISPLPL | ASSSGTGVNSPLY | 0.5704622 |
| VTAVFVAI | ASSSGTGVNSPLY | 0.6711354 |
| SGDDNDAAL | ASSSGTGVNSPLY | 0.57888526 |
| AVAALAHS | ASSSGTGVNSPLY | 0.72857666 |
| GGVTVMATP | ASSSGTGVNSPLY | 0.8040159 |
| AGLAANVRD | ASSSGTGVNSPLY | 0.7537973 |
| TNRPDLIDPALLRPG | ASSSGTGVNSPLY | 0.46266204 |
| LANLGVTAL | ASSSGTGVNSPLY | 0.7131773 |
| TGERIEL | ASSSGTGVNSPLY | 0.6968428 |
| NGEELAA | ASSSGTGVNSPLY | 0.7992612 |
| VLSLVWG | ASSSGTGVNSPLY | 0.7735511 |
| IGYALWL | ASSSGTGVNSPLY | 0.8064642 |
| LSLLLAPL | ASSSGTGVNSPLY | 0.77088505 |
| TLDEALK | ASSSGTGVNSPLY | 0.62056786 |
| TDGAGTAEGG | ASSSGTGVNSPLY | 0.79885256 |
| NAPGGAGGNGG | ASSSGTGVNSPLY | 0.69775796 |
| VSVFMGMGGNF | ASSSGTGVNSPLY | 0.78597164 |
| LASYLAAL | ASSSGTGVNSPLY | 0.7790089 |
| LVGGTPAL | ASSSGTGVNSPLY | 0.774081 |
| VRNADVL | ASSSGTGVNSPLY | 0.63837135 |
| TCSSPFLVSAM | ASSSGTGVNSPLY | 0.7455065 |
| VHITDAE | ASSSGTGVNSPLY | 0.35613915 |
| EYAAPSF | ASSSGTGVNSPLY | 0.7844373 |
| VVDAFHGGL | ASSSGTGVNSPLY | 0.5600988 |
| ASVVDPQ | ASSSGTGVNSPLY | 0.578051 |
| SYNEFGGLS | ASSSGTGVNSPLY | 0.70265836 |
| DPDGSVY | ASSSGTGVNSPLY | 0.81278443 |
| MVSTIHATE | ASSSGTGVNSPLY | 0.80173665 |
| LAAGVADPV | ASSSGTGVNSPLY | 0.63118154 |
| TKEQIAA | ASSSGTGVNSPLY | 0.5529386 |
| SLASPAAL | ASSSGTGVNSPLY | 0.778613 |
| IGLVFYGV | ASSSGTGVNSPLY | 0.5381165 |
| FLLDASNL | ASSSGTGVNSPLY | 0.7864584 |
| YQVGNFP | ASSSGTGVNSPLY | 0.7255818 |
| ESLLSQST | ASSSGTGVNSPLY | 0.7176236 |

| | | |
|---|---|---|
| LFTVGFAAVG | ASSSGTGVNSPLY | 0.5742589 |
| ATPPTPT | ASSSGTGVNSPLY | 0.7872202 |
| TDLATLG | ASSSGTGVNSPLY | 0.6687578 |
| SAGGTGR | ASSSGTGVNSPLY | 0.7752705 |
| GFSNSGS | ASSSGTGVNSPLY | 0.2891773 |
| GGNGGSGGTSV | ASSSGTGVNSPLY | 0.77953684 |
| GTSVATGGAGN | ASSSGTGVNSPLY | 0.59987706 |
| NGGNGGSGGTS | ASSSGTGVNSPLY | 0.55875325 |
| AVVKFPH | ASSSGTGVNSPLY | 0.35089105 |
| TPPPPPP | ASSSGTGVNSPLY | 0.70528245 |
| AAPAAAAAPA | ASSSGTGVNSPLY | 0.789137 |
| IALNEDL | ASSSGTGVNSPLY | 0.8053719 |
| IVNVSSM | ASSSGTGVNSPLY | 0.7651612 |
| TGIPPAPR | ASSSGTGVNSPLY | 0.7747579 |
| FDLGGGTF | ASSSGTGVNSPLY | 0.806388 |
| FDLGGGTFDVS | ASSSGTGVNSPLY | 0.71338016 |
| NEPTAAA | ASSSGTGVNSPLY | 0.33897752 |
| NEPTAAALA | ASSSGTGVNSPLY | 0.6950314 |
| GSIINISSI | ASSSGTGVNSPLY | 0.75988597 |
| VAPGNAGTA | ASSSGTGVNSPLY | 0.79555094 |
| TGGGASLE | ASSSGTGVNSPLY | 0.774722 |
| VAVYNHYKRI | ASSSGTGVNSPLY | 0.5296769 |
| TDPAPATT | ASSSGTGVNSPLY | 0.70482105 |
| TNRPDILDPALLRPG | ASSSGTGVNSPLY | 0.78223085 |
| LTTTTTS | ASSSGTGVNSPLY | 0.50907815 |
| PTGSGKTL | ASSSGTGVNSPLY | 0.6988163 |
| TGSGKTL | ASSSGTGVNSPLY | 0.59563065 |
| TIGHVDHGK | ASSSGTGVNSPLY | 0.61419386 |
| TIGHVDHGKTT | ASSSGTGVNSPLY | 0.6981265 |
| GHVDHGKTT | ASSSGTGVNSPLY | 0.6915332 |
| AQFEHTLL | ASSSGTGVNSPLY | 0.53055876 |
| IVNQDEL | ASSSGTGVNSPLY | 0.6962999 |
| TNGSQFF | ASSSGTGVNSPLY | 0.4855333 |
| TGKGIVE | ASSSGTGVNSPLY | 0.5179399 |
| LISNASDA | ASSSGTGVNSPLY | 0.60037494 |
| AELRTLGELTKIAKAHGAQ | ASSSGTGVNSPLY | 0.7383283 |
| APNAVAAAEPSW | ASSSGTGVNSPLY | 0.7918538 |
| APQVAGAATAVL | ASSSGTGVNSPLY | 0.6915772 |
| ARRDPAAAAAILPTDA | ASSSGTGVNSPLY | 0.7546486 |
| DAAARDVLAEAGLAEYF | ASSSGTGVNSPLY | 0.8098768 |
| DDQIVLSAKANLAKVEQ | ASSSGTGVNSPLY | 0.32875347 |
| DELLPDTGEDSDEDGH | ASSSGTGVNSPLY | 0.521845 |
| ELPEAPANRATVPQD | ASSSGTGVNSPLY | 0.7744031 |
| FGQSKIERVTGIG | ASSSGTGVNSPLY | 0.62540585 |
| GAGGTGSTAGGAGGAGGAGGLY | ASSSGTGVNSPLY | 0.8020674 |
| GHGGTGAAGGNGGTGGAGGSGI | ASSSGTGVNSPLY | 0.71859944 |
| GNGGNGGAGGEGGPGGAGGAGA | ASSSGTGVNSPLY | 0.7674891 |
| GSTAGNGGNGGNATNGTIAGQP | ASSSGTGVNSPLY | 0.7595172 |
| GTGGDGGNGGNGGTGG | ASSSGTGVNSPLY | 0.81720924 |
| GVRRPAGIPADAPV | ASSSGTGVNSPLY | 0.7759657 |
| GYRQVTDAHLLTLARR | ASSSGTGVNSPLY | 0.71487767 |

| | | |
|---|---|---|
| IAKMGQDGHDRG | ASSSGTGVNSPLY | 0.7974975 |
| KAEPSIEIQYT | ASSSGTGVNSPLY | 0.41831383 |
| KPVRIRMAADEDTVAVT | ASSSGTGVNSPLY | 0.79087555 |
| MPPRIAGMRLLVIKPEP | ASSSGTGVNSPLY | 0.7934741 |
| MKFYAVALFRSRGVPPTR | ASSSGTGVNSPLY | 0.5765977 |
| NAGGFGILGAVAHSP | ASSSGTGVNSPLY | 0.7032546 |
| NEGREITLAMAEM | ASSSGTGVNSPLY | 0.8138468 |
| NELVEWSHPLGHRWRWE | ASSSGTGVNSPLY | 0.80658257 |
| NLGAGGTGGNGGNPG | ASSSGTGVNSPLY | 0.5197836 |
| NTEWPKWETVMAADPVD | ASSSGTGVNSPLY | 0.65016335 |
| PASAAPAETQP | ASSSGTGVNSPLY | 0.4669265 |
| QQSSDGKNGVRAAEITGE | ASSSGTGVNSPLY | 0.60936165 |
| RAPDAVTTSWWKE | ASSSGTGVNSPLY | 0.3447523 |
| SGFGNVGAHTSGWFNQ | ASSSGTGVNSPLY | 0.78632796 |
| SGVAGAVGQAASVGGLKV | ASSSGTGVNSPLY | 0.7102146 |
| SNGGAGTGSAGGNGGT | ASSSGTGVNSPLY | 0.7817466 |
| TGGAGGEGGAAG | ASSSGTGVNSPLY | 0.7406415 |
| TGGDGGAAGAGGAPGHG | ASSSGTGVNSPLY | 0.78488815 |
| TMPPLLAAIANAGRHGV | ASSSGTGVNSPLY | 0.7033348 |
| TSSTAGNSGGA | ASSSGTGVNSPLY | 0.6377357 |
| VPATVPVGILC | ASSSGTGVNSPLY | 0.4391802 |
| VQRPDSGGVVLTGRIS | ASSSGTGVNSPLY | 0.42527235 |
| WTPNPVPATKGVPAGLVQAT | ASSSGTGVNSPLY | 0.6946304 |
| YALKRALAKFSDRFDVVII | ASSSGTGVNSPLY | 0.5968803 |
| YLIGSVIV | ASSSGTGVNSPLY | 0.33149406 |
| YGQLVAVID | ASSSGTGVNSPLY | 0.8056422 |
| VTVGGMTLPA | ASSSGTGVNSPLY | 0.7205225 |
| VLALPAV | ASSSGTGVNSPLY | 0.75302714 |
| VIPPTLNYET | ASSSGTGVNSPLY | 0.5755567 |
| VGMGGMHPSSG | ASSSGTGVNSPLY | 0.61476475 |
| VGLAGGAATA | ASSSGTGVNSPLY | 0.6141658 |
| VGGAGGAGGSGP | ASSSGTGVNSPLY | 0.78689593 |
| VFAAFMV | ASSSGTGVNSPLY | 0.6096237 |
| TLYDKLR | ASSSGTGVNSPLY | 0.38594663 |
| TLAVFTVL | ASSSGTGVNSPLY | 0.5671042 |
| TGSVQPARV | ASSSGTGVNSPLY | 0.6837891 |
| TGGGAGSGGT | ASSSGTGVNSPLY | 0.8154831 |
| SHLETSL | ASSSGTGVNSPLY | 0.80033886 |
| SFTTGMT | ASSSGTGVNSPLY | 0.7581984 |
| QVEVAKRMGVRQPTVS | ASSSGTGVNSPLY | 0.8128532 |
| QGSGGPALAM | ASSSGTGVNSPLY | 0.21566424 |
| MLAPSNP | ASSSGTGVNSPLY | 0.59172976 |
| LVNNAAI | ASSSGTGVNSPLY | 0.7823006 |
| LTGEINTLA | ASSSGTGVNSPLY | 0.75366914 |
| LSIFGPV | ASSSGTGVNSPLY | 0.7263864 |
| LLALLANL | ASSSGTGVNSPLY | 0.8045124 |
| LISNASDALDKLR | ASSSGTGVNSPLY | 0.52577144 |
| LINEAVP | ASSSGTGVNSPLY | 0.63939965 |
| LGYAFSGL | ASSSGTGVNSPLY | 0.7223793 |
| LDFVAVAL | ASSSGTGVNSPLY | 0.801929 |
| LAQPGAE | ASSSGTGVNSPLY | 0.7925764 |

| | | |
|---|---|---|
| ISNLGAVTGF | ASSSGTGVNSPLY | 0.52592885 |
| HQEDPGRI | ASSSGTGVNSPLY | 0.7639619 |
| GTPGEPTP | ASSSGTGVNSPLY | 0.82053214 |
| GFFNSGS | ASSSGTGVNSPLY | 0.67556894 |
| GAGGGGLTAGAN | ASSSGTGVNSPLY | 0.7826338 |
| FVDALVNM | ASSSGTGVNSPLY | 0.7293732 |
| FLTVAPEL | ASSSGTGVNSPLY | 0.5608804 |
| FETLTAL | ASSSGTGVNSPLY | 0.6424704 |
| DVINPIEL | ASSSGTGVNSPLY | 0.6303221 |
| APVSGAPV | ASSSGTGVNSPLY | 0.7726375 |
| ALVGGLAW | ASSSGTGVNSPLY | 0.6165296 |
| ALAAAAATAI | ASSSGTGVNSPLY | 0.7475913 |
| AFGLGLLL | ASSSGTGVNSPLY | 0.71310496 |
| AFFAAASL | ASSSGTGVNSPLY | 0.81973314 |
| AAAALPAFSPP | ASSSGTGVNSPLY | 0.7769685 |
| VVLAPAL | ASSSGTGVNSPLY | 0.76089406 |
| TIGHVDHGKTTL | ASSSGTGVNSPLY | 0.5789345 |
| LISNASDALDK | ASSSGTGVNSPLY | 0.77922815 |
| LISNASDAL | ASSSGTGVNSPLY | 0.7796624 |
| ISNASDALDK | ASSSGTGVNSPLY | 0.3635813 |
| NQAETLVYQTE | ASSSGTGVNSPLY | 0.80102384 |
| TGGTGGTAGAG | ASSSGTGVNSPLY | 0.51227987 |
| TGSTAGGAGGA | ASSSGTGVNSPLY | 0.7507688 |
| DPLAVPVAAGPV | ASSSGTGVNSPLY | 0.7153863 |
| GGTGGTPGNGGN | ASSSGTGVNSPLY | 0.7898963 |
| PAGLAGVDPATL | ASSSGTGVNSPLY | 0.7596093 |
| GNNLVVEAPHAGASV | ASSSGTGVNSPLY | 0.76701176 |
| KVTRIMSASTLEQPA | ASSSGTGVNSPLY | 0.505888 |
| LGFALPIALPRRTVV | ASSSGTGVNSPLY | 0.7925781 |
| TRLGNLTARKLVVLV | ASSSGTGVNSPLY | 0.80011845 |
| WSQQRRLIYFPSAGP | ASSSGTGVNSPLY | 0.76507735 |
| GPNGNPDPMAAAVDIR | ASSSGTGVNSPLY | 0.8159058 |
| TQRCVSVADLRAAASR | ASSSGTGVNSPLY | 0.73833525 |
| QFVLDVAEHPVEVLTETV | ASSSGTGVNSPLY | 0.29756784 |
| GGAGGTGGAGGNGGTQAGDG | ASSSGTGVNSPLY | 0.7499479 |
| PLLATLRAIAVDVDAKPAQV | ASSSGTGVNSPLY | 0.79877335 |
| APKFRTGGYMIDFWGVGYQVAK | ASSSGTGVNSPLY | 0.8080676 |
| GAGGAGGAGGAAGGPGGTGGTG | ASSSGTGVNSPLY | 0.7654336 |
| IGALGLLATDVAPAGSVAFGVA | ASSSGTGVNSPLY | 0.79537165 |
| AGGAGGIGGNGGAGGAGGVLMG | ASSSGTGVNSPLY | 0.7289038 |
| ADELIGLL | ASSSGTGVNSPLY | 0.807498 |
| ADRALKA | ASSSGTGVNSPLY | 0.81810284 |
| AGAVGQVGGAGGA | ASSSGTGVNSPLY | 0.7565023 |
| AGGGGGLGGAP | ASSSGTGVNSPLY | 0.59303606 |
| AGGLLYGNGGAG | ASSSGTGVNSPLY | 0.48740408 |
| AGLESIGL | ASSSGTGVNSPLY | 0.78296137 |
| AGLKTGII | ASSSGTGVNSPLY | 0.7896052 |
| ASLAALLL | ASSSGTGVNSPLY | 0.5801686 |
| ASVINISSI | ASSSGTGVNSPLY | 0.7399641 |
| ATPPGPA | ASSSGTGVNSPLY | 0.73259693 |
| ATVASALV | ASSSGTGVNSPLY | 0.31988138 |

| | | |
|---|---|---|
| ATVASLIL | ASSSGTGVNSPLY | 0.78104323 |
| AVNGPVLIRH | ASSSGTGVNSPLY | 0.8116737 |
| DGAAGAVGI | ASSSGTGVNSPLY | 0.77105105 |
| DPLLHGI | ASSSGTGVNSPLY | 0.5541014 |
| ELGAALGAL | ASSSGTGVNSPLY | 0.8043352 |
| ELGRTQAV | ASSSGTGVNSPLY | 0.8154638 |
| FVAGGYLS | ASSSGTGVNSPLY | 0.8116152 |
| GASGGAGGAGGW | ASSSGTGVNSPLY | 0.30510077 |
| GGAGAGGDGGAGGI | ASSSGTGVNSPLY | 0.23512366 |
| GTGHNGSQPGMGGNG | ASSSGTGVNSPLY | 0.75635743 |
| GYAAASASAS | ASSSGTGVNSPLY | 0.77248347 |
| HQVDVAAVQQG | ASSSGTGVNSPLY | 0.55582106 |
| IGAAIAHT | ASSSGTGVNSPLY | 0.5252141 |
| IGMIEVAQL | ASSSGTGVNSPLY | 0.78418255 |
| IRLAPEHTSAMD | ASSSGTGVNSPLY | 0.7953779 |
| IYNVSVL | ASSSGTGVNSPLY | 0.59366685 |
| KQGFLGM | ASSSGTGVNSPLY | 0.43924025 |
| LESVITQL | ASSSGTGVNSPLY | 0.8058034 |
| LGLGALVVV | ASSSGTGVNSPLY | 0.79484355 |
| LIGNGGNGGSGGI | ASSSGTGVNSPLY | 0.6073643 |
| LNAINAPTL | ASSSGTGVNSPLY | 0.70883423 |
| LQLLNTGV | ASSSGTGVNSPLY | 0.719625 |
| LQTSIEF | ASSSGTGVNSPLY | 0.7372639 |
| MSFAVSEL | ASSSGTGVNSPLY | 0.7515378 |
| NTGALGL | ASSSGTGVNSPLY | 0.45376536 |
| PPGAGGAGGAGTT | ASSSGTGVNSPLY | 0.8005636 |
| QLVGLGI | ASSSGTGVNSPLY | 0.535359 |
| RFLEQVHQL | ASSSGTGVNSPLY | 0.79993373 |
| RGVGYVLR | ASSSGTGVNSPLY | 0.75035655 |
| SATGVFAGVI | ASSSGTGVNSPLY | 0.7773722 |
| SLPAITLP | ASSSGTGVNSPLY | 0.737849 |
| STALAFTV | ASSSGTGVNSPLY | 0.5303491 |
| SVLTVGLGK | ASSSGTGVNSPLY | 0.53256655 |
| TALAGAGSGI | ASSSGTGVNSPLY | 0.5181564 |
| TGLPFTGI | ASSSGTGVNSPLY | 0.80461633 |
| VAEHVTV | ASSSGTGVNSPLY | 0.75374115 |
| VDIVLNSL | ASSSGTGVNSPLY | 0.80395484 |
| VLTGGGAVL | ASSSGTGVNSPLY | 0.70617115 |
| VNVGLAYGI | ASSSGTGVNSPLY | 0.57621825 |
| VSSAAVGAV | ASSSGTGVNSPLY | 0.78886175 |
| VTSVYEAGV | ASSSGTGVNSPLY | 0.40516102 |
| WVALDRGA | ASSSGTGVNSPLY | 0.76476663 |
| YLVGSVVI | ASSSGTGVNSPLY | 0.7422959 |
| GPPGTGKTLL | ASSSGTGVNSPLY | 0.77075976 |
| GVGMTKF | ASSSGTGVNSPLY | 0.6856376 |
| IGIGGSDL | ASSSGTGVNSPLY | 0.790835 |
| IGTLLAEL | ASSSGTGVNSPLY | 0.7764623 |
| IYTSGTT | ASSSGTGVNSPLY | 0.6204763 |
| LSFPAGV | ASSSGTGVNSPLY | 0.73967254 |
| LYTSGTT | ASSSGTGVNSPLY | 0.7967199 |
| VGAFPFGF | ASSSGTGVNSPLY | 0.7693132 |

| | | |
|---|---|---|
| VGAGTVEF | ASSSGTGVNSPLY | 0.59267986 |
| VGKSTLLL | ASSSGTGVNSPLY | 0.71959174 |
| AALAHAGGGIV | ASSSGTGVNSPLY | 0.6740078 |
| AALVAVSLL | ASSSGTGVNSPLY | 0.7640736 |
| AAVSRAVAAL | ASSSGTGVNSPLY | 0.76166093 |
| AGGSSGIGGF | ASSSGTGVNSPLY | 0.57060665 |
| AGLTVANANG | ASSSGTGVNSPLY | 0.76491237 |
| AGLVGAVL | ASSSGTGVNSPLY | 0.4572668 |
| ALSQAFTV | ASSSGTGVNSPLY | 0.6838907 |
| ALTFSVSL | ASSSGTGVNSPLY | 0.7179985 |
| ALTGAALVAL | ASSSGTGVNSPLY | 0.70019007 |
| ASFSIGAGVL | ASSSGTGVNSPLY | 0.7645273 |
| DGSVYKI | ASSSGTGVNSPLY | 0.49726593 |
| EQLAEIV | ASSSGTGVNSPLY | 0.7654004 |
| FAVSELTVM | ASSSGTGVNSPLY | 0.6356633 |
| GAAPLAAGL | ASSSGTGVNSPLY | 0.48966855 |
| GADGGDGGSGGAGG | ASSSGTGVNSPLY | 0.76016754 |
| GFGLDTGGAGG | ASSSGTGVNSPLY | 0.7023843 |
| GLASGGLL | ASSSGTGVNSPLY | 0.49199778 |
| GRIAGAAVLL | ASSSGTGVNSPLY | 0.7983445 |
| GSAGSLNTSF | ASSSGTGVNSPLY | 0.5287359 |
| GTGQIGI | ASSSGTGVNSPLY | 0.67711836 |
| GTLIALIL | ASSSGTGVNSPLY | 0.79967815 |
| IAATMGFVVG | ASSSGTGVNSPLY | 0.49744862 |
| IFAQALL | ASSSGTGVNSPLY | 0.79517317 |
| IGAAVALGF | ASSSGTGVNSPLY | 0.7443733 |
| IGLTPLAL | ASSSGTGVNSPLY | 0.7803548 |
| ILGSNTSTL | ASSSGTGVNSPLY | 0.6284174 |
| IVFSLTVL | ASSSGTGVNSPLY | 0.64968884 |
| LASKATGF | ASSSGTGVNSPLY | 0.8029082 |
| LASVALIL | ASSSGTGVNSPLY | 0.71203274 |
| LFAVGLTL | ASSSGTGVNSPLY | 0.6684587 |
| LFNGGIGTYI | ASSSGTGVNSPLY | 0.77555114 |
| LTLGAGII | ASSSGTGVNSPLY | 0.79578483 |
| LVAAALGDI | ASSSGTGVNSPLY | 0.7934988 |
| NTAFVQLGI | ASSSGTGVNSPLY | 0.63281983 |
| QLFSGAEL | ASSSGTGVNSPLY | 0.42373294 |
| SAATFITL | ASSSGTGVNSPLY | 0.6428911 |
| SNYGLANLGV | ASSSGTGVNSPLY | 0.7097561 |
| SYGGAYAVM | ASSSGTGVNSPLY | 0.6701406 |
| TTGVGGLTLG | ASSSGTGVNSPLY | 0.35870382 |
| TTQTIYI | ASSSGTGVNSPLY | 0.77813345 |
| VAAGAGLGAVI | ASSSGTGVNSPLY | 0.819486 |
| VAAVVTLL | ASSSGTGVNSPLY | 0.5482348 |
| VGFLGGLI | ASSSGTGVNSPLY | 0.8191989 |
| VGVIPILL | ASSSGTGVNSPLY | 0.51253986 |
| VIFASSLI | ASSSGTGVNSPLY | 0.7227951 |
| VKGGDGGAAAG | ASSSGTGVNSPLY | 0.79298234 |
| VTKTFTNI | ASSSGTGVNSPLY | 0.49890062 |
| VVAAAILL | ASSSGTGVNSPLY | 0.6627322 |
| YAALDVEL | ASSSGTGVNSPLY | 0.7602134 |

| | | |
|---|---|---|
| LASVALLL | ASSSGTGVNSPLY | 0.62128496 |
| LDAVHRV | ASSSGTGVNSPLY | 0.8035705 |
| QGLSSSIF | ASSSGTGVNSPLY | 0.5015233 |
| RGTFANIRL | ASSSGTGVNSPLY | 0.7726711 |
| RQLWWGHRI | ASSSGTGVNSPLY | 0.60864943 |
| YGPPGTGKTLL | ASSSGTGVNSPLY | 0.52431166 |
| AALVGAPANVA | ASSSGTGVNSPLY | 0.79019994 |
| AATFRRTE | ASSSGTGVNSPLY | 0.78262216 |
| AGALGGGI | ASSSGTGVNSPLY | 0.74854636 |
| AGGAGGAGGIGAG | ASSSGTGVNSPLY | 0.81739473 |
| AGSGALGL | ASSSGTGVNSPLY | 0.69865847 |
| AQIADQI | ASSSGTGVNSPLY | 0.81746686 |
| ASAVAVTYL | ASSSGTGVNSPLY | 0.8081463 |
| ASTVLGGTGG | ASSSGTGVNSPLY | 0.74441576 |
| AVGGEPAK | ASSSGTGVNSPLY | 0.7181972 |
| CEQGTFGGTC | ASSSGTGVNSPLY | 0.78962374 |
| DEGKAPAG | ASSSGTGVNSPLY | 0.66131514 |
| DTVVFPAL | ASSSGTGVNSPLY | 0.76815397 |
| FEGITMAG | ASSSGTGVNSPLY | 0.8084854 |
| FPLVVQD | ASSSGTGVNSPLY | 0.39443627 |
| GAFTTPQ | ASSSGTGVNSPLY | 0.801849 |
| GFGGFGTAGGVGGA | ASSSGTGVNSPLY | 0.72719574 |
| GFLAQFL | ASSSGTGVNSPLY | 0.7956785 |
| GGAGGAGGNAGVA | ASSSGTGVNSPLY | 0.6238329 |
| GGVGTAGAGGAGGAGG | ASSSGTGVNSPLY | 0.36184084 |
| GNGGGGDAGNAGSGGN | ASSSGTGVNSPLY | 0.6905121 |
| GTGGAGGAGNDAGS | ASSSGTGVNSPLY | 0.56052136 |
| GTTNTGWFNS | ASSSGTGVNSPLY | 0.77745724 |
| IFMGPLFTL | ASSSGTGVNSPLY | 0.7878176 |
| ILAHAIEAI | ASSSGTGVNSPLY | 0.57085025 |
| IRSAIREAL | ASSSGTGVNSPLY | 0.7897148 |
| ITFAGGG | ASSSGTGVNSPLY | 0.7640958 |
| ITGMSGHQEV | ASSSGTGVNSPLY | 0.3988437 |
| IVAGLACTL | ASSSGTGVNSPLY | 0.6639508 |
| IVGGGLAGA | ASSSGTGVNSPLY | 0.69835776 |
| IVTGVLNI | ASSSGTGVNSPLY | 0.6899756 |
| LAQQGCI | ASSSGTGVNSPLY | 0.2966166 |
| LCLHVEN | ASSSGTGVNSPLY | 0.75817585 |
| LDAISDGS | ASSSGTGVNSPLY | 0.57079273 |
| LGGFAAML | ASSSGTGVNSPLY | 0.20889072 |
| LGGKNPMI | ASSSGTGVNSPLY | 0.76577747 |
| LGPKGVAAIL | ASSSGTGVNSPLY | 0.56794214 |
| LMVGVSI | ASSSGTGVNSPLY | 0.6241317 |
| LVGAAFVL | ASSSGTGVNSPLY | 0.7950851 |
| LVLAVAPL | ASSSGTGVNSPLY | 0.6322571 |
| MAAIAVIDAA | ASSSGTGVNSPLY | 0.5401646 |
| NNGINAELQ | ASSSGTGVNSPLY | 0.74983776 |
| RGEPLVLVHG | ASSSGTGVNSPLY | 0.8125982 |
| RGGAGGAAGLI | ASSSGTGVNSPLY | 0.47095224 |
| RSYRMHPAV | ASSSGTGVNSPLY | 0.6784638 |
| SAAHLLAGQLA | ASSSGTGVNSPLY | 0.8277577 |

| | | |
|---|---|---|
| SAAVGSY | ASSSGTGVNSPLY | 0.7446397 |
| SPHAVLHRI | ASSSGTGVNSPLY | 0.8458084 |
| SQIAKVL | ASSSGTGVNSPLY | 0.8463775 |
| TALGHLVP | ASSSGTGVNSPLY | 0.55063146 |
| TAPTTSGAA | ASSSGTGVNSPLY | 0.8110216 |
| TDGDDGNGGSPGT | ASSSGTGVNSPLY | 0.80434996 |
| TDGLGEN | ASSSGTGVNSPLY | 0.6573336 |
| TGGPPLPGQA | ASSSGTGVNSPLY | 0.7196918 |
| TIPAITI | ASSSGTGVNSPLY | 0.45102283 |
| TIPGFTI | ASSSGTGVNSPLY | 0.63182616 |
| TSIASGALA | ASSSGTGVNSPLY | 0.46787363 |
| TVTLPGAAI | ASSSGTGVNSPLY | 0.7261639 |
| VASGLSAALP | ASSSGTGVNSPLY | 0.8180277 |
| VGAGVIGI | ASSSGTGVNSPLY | 0.71091306 |
| VGPKRFGV | ASSSGTGVNSPLY | 0.38066262 |
| VNVAFPD | ASSSGTGVNSPLY | 0.39382777 |
| VSMTNSGVS | ASSSGTGVNSPLY | 0.7350309 |
| VTDVLGDLSV | ASSSGTGVNSPLY | 0.45190266 |
| VVVVGAGFAGL | ASSSGTGVNSPLY | 0.52226925 |
| VYTVGTVV | ASSSGTGVNSPLY | 0.5499169 |
| WTTFSGP | ASSSGTGVNSPLY | 0.80779266 |
| YDELVEAF | ASSSGTGVNSPLY | 0.8099208 |
| YGTGGAGGNGG | ASSSGTGVNSPLY | 0.5772848 |
| YTNVMAV | ASSSGTGVNSPLY | 0.82323813 |
| FLEQVHQL | ASSSGTGVNSPLY | 0.75082815 |
| GLSLLAVAV | ASSSGTGVNSPLY | 0.8215213 |
| LEQVHQL | ASSSGTGVNSPLY | 0.73991704 |
| LNAAGAI | ASSSGTGVNSPLY | 0.42031398 |
| PAVHASL | ASSSGTGVNSPLY | 0.7883679 |
| TELEDDA | ASSSGTGVNSPLY | 0.65922475 |
| TPPGPPN | ASSSGTGVNSPLY | 0.7830292 |
| TPPPPPPP | ASSSGTGVNSPLY | 0.8055649 |
| TQSPALI | AWSLGLGEDQNTLY | 0.68164814 |
| EFAAAFPQ | AWSLGLGEDQNTLY | 0.5519515 |
| LLSGNLDVL | AWSLGLGEDQNTLY | 0.65164524 |
| GPANLGNYNI | AWSLGLGEDQNTLY | 0.82885337 |
| TLGPTLP | AWSLGLGEDQNTLY | 0.47180775 |
| VYAYSSCCK | AWSLGLGEDQNTLY | 0.42723072 |
| APAPAPAPNI | AWSLGLGEDQNTLY | 0.6694795 |
| PGGPVPIP | AWSLGLGEDQNTLY | 0.68150157 |
| FLTVALEP | AWSLGLGEDQNTLY | 0.8271657 |
| SQYFGANL | AWSLGLGEDQNTLY | 0.75478464 |
| GGVKPAA | AWSLGLGEDQNTLY | 0.65032744 |
| GGTGGAGGAGGW | AWSLGLGEDQNTLY | 0.7652105 |
| GVGGVAGLGGAGG | AWSLGLGEDQNTLY | 0.7964809 |
| VLLVLATTI | AWSLGLGEDQNTLY | 0.7366944 |
| IGIFAAML | AWSLGLGEDQNTLY | 0.71281624 |
| AVGDLID | AWSLGLGEDQNTLY | 0.5843174 |
| NISPLPL | AWSLGLGEDQNTLY | 0.8197955 |
| VTAVFVAI | AWSLGLGEDQNTLY | 0.8135036 |
| SGDDNDAAL | AWSLGLGEDQNTLY | 0.7989173 |

| | | |
|---|---|---|
| AVAALAHS | AWSLGLGEDQNTLY | 0.8087114 |
| GGVTVMATP | AWSLGLGEDQNTLY | 0.6705631 |
| AGLAANVRD | AWSLGLGEDQNTLY | 0.7376557 |
| TNRPDLIDPALLRPG | AWSLGLGEDQNTLY | 0.50443876 |
| LANLGVTAL | AWSLGLGEDQNTLY | 0.60388434 |
| TGERIEL | AWSLGLGEDQNTLY | 0.6489079 |
| NGEELAA | AWSLGLGEDQNTLY | 0.7263263 |
| VLSLVWG | AWSLGLGEDQNTLY | 0.763122 |
| IGYALWL | AWSLGLGEDQNTLY | 0.75155556 |
| LSLLLAPL | AWSLGLGEDQNTLY | 0.54107654 |
| TLDEALK | AWSLGLGEDQNTLY | 0.8102925 |
| TDGAGTAEGG | AWSLGLGEDQNTLY | 0.80453974 |
| NAPGGAGGNGG | AWSLGLGEDQNTLY | 0.5278255 |
| VSVFMGMGGNF | AWSLGLGEDQNTLY | 0.6454276 |
| LASYLAAL | AWSLGLGEDQNTLY | 0.8024782 |
| LVGGTPAL | AWSLGLGEDQNTLY | 0.75151217 |
| VRNADVL | AWSLGLGEDQNTLY | 0.66768146 |
| TCSSPFLVSAM | AWSLGLGEDQNTLY | 0.7376628 |
| VHITDAE | AWSLGLGEDQNTLY | 0.7954749 |
| EYAAPSF | AWSLGLGEDQNTLY | 0.71474373 |
| VVDAFHGGL | AWSLGLGEDQNTLY | 0.821611 |
| ASVVDPQ | AWSLGLGEDQNTLY | 0.3115076 |
| SYNEFGGLS | AWSLGLGEDQNTLY | 0.69423044 |
| DPDGSVY | AWSLGLGEDQNTLY | 0.61725384 |
| MVSTIHATE | AWSLGLGEDQNTLY | 0.22416109 |
| LAAGVADPV | AWSLGLGEDQNTLY | 0.81722295 |
| TKEQIAA | AWSLGLGEDQNTLY | 0.5537782 |
| SLASPAAL | AWSLGLGEDQNTLY | 0.7601328 |
| IGLVFYGV | AWSLGLGEDQNTLY | 0.5825031 |
| FLLDASNL | AWSLGLGEDQNTLY | 0.8199816 |
| YQVGNFP | AWSLGLGEDQNTLY | 0.73874146 |
| ESLLSQST | AWSLGLGEDQNTLY | 0.5894219 |
| LFTVGFAAVG | AWSLGLGEDQNTLY | 0.5596815 |
| ATPPTPT | AWSLGLGEDQNTLY | 0.7859044 |
| TDLATLG | AWSLGLGEDQNTLY | 0.8226905 |
| SAGGTGR | AWSLGLGEDQNTLY | 0.83784556 |
| GFSNSGS | AWSLGLGEDQNTLY | 0.34648883 |
| GGNGGSGGTSV | AWSLGLGEDQNTLY | 0.7698368 |
| GTSVATGGAGN | AWSLGLGEDQNTLY | 0.7116738 |
| NGGNGGSGGTS | AWSLGLGEDQNTLY | 0.48511988 |
| AVVKFPH | AWSLGLGEDQNTLY | 0.351699 |
| TPPPPPP | AWSLGLGEDQNTLY | 0.57947814 |
| AAPAAAAAPA | AWSLGLGEDQNTLY | 0.6709627 |
| IALNEDL | AWSLGLGEDQNTLY | 0.7624645 |
| IVNVSSM | AWSLGLGEDQNTLY | 0.80654854 |
| TGIPPAPR | AWSLGLGEDQNTLY | 0.64452183 |
| FDLGGGTF | AWSLGLGEDQNTLY | 0.7524235 |
| FDLGGGTFDVS | AWSLGLGEDQNTLY | 0.698905 |
| NEPTAAA | AWSLGLGEDQNTLY | 0.11506076 |
| NEPTAAALA | AWSLGLGEDQNTLY | 0.60125554 |
| GSIINISSI | AWSLGLGEDQNTLY | 0.41104603 |

| | | |
|---|---|---|
| VAPGNAGTA | AWSLGLGEDQNTLY | 0.81902367 |
| TGGGASLE | AWSLGLGEDQNTLY | 0.43217647 |
| VAVYNHYKRI | AWSLGLGEDQNTLY | 0.6498871 |
| TDPAPATT | AWSLGLGEDQNTLY | 0.7123502 |
| TNRPDILDPALLRPG | AWSLGLGEDQNTLY | 0.7307913 |
| LTTTTTS | AWSLGLGEDQNTLY | 0.7220929 |
| PTGSGKTL | AWSLGLGEDQNTLY | 0.53499407 |
| TGSGKTL | AWSLGLGEDQNTLY | 0.4623831 |
| TIGHVDHGK | AWSLGLGEDQNTLY | 0.79610795 |
| TIGHVDHGKTT | AWSLGLGEDQNTLY | 0.6796258 |
| GHVDHGKTT | AWSLGLGEDQNTLY | 0.47556418 |
| AQFEHTLL | AWSLGLGEDQNTLY | 0.6048234 |
| IVNQDEL | AWSLGLGEDQNTLY | 0.81564367 |
| TNGSQFF | AWSLGLGEDQNTLY | 0.32566506 |
| TGKGIVE | AWSLGLGEDQNTLY | 0.83449936 |
| LISNASDA | AWSLGLGEDQNTLY | 0.51988125 |
| AELRTLGELTKIAKAHGAQ | AWSLGLGEDQNTLY | 0.6504775 |
| APNAVAAAEPSW | AWSLGLGEDQNTLY | 0.8231293 |
| APQVAGAATAVL | AWSLGLGEDQNTLY | 0.69580674 |
| ARRDPAAAAAILPTDA | AWSLGLGEDQNTLY | 0.7130214 |
| DAAARDVLAEAGLAEYF | AWSLGLGEDQNTLY | 0.58524257 |
| DDQIVLSAKANLAKVEQ | AWSLGLGEDQNTLY | 0.5304004 |
| DELLPDTGEDSDEDGH | AWSLGLGEDQNTLY | 0.5796329 |
| ELPEAPANRATVPQD | AWSLGLGEDQNTLY | 0.52503955 |
| FGQSKIERVTGIG | AWSLGLGEDQNTLY | 0.8189329 |
| GAGGTGSTAGGAGGAGGAGGLY | AWSLGLGEDQNTLY | 0.7568165 |
| GHGGTGAAGGNGGTGGAGGSGI | AWSLGLGEDQNTLY | 0.45071694 |
| GNGGNGGAGGEGGPGGAGGAGA | AWSLGLGEDQNTLY | 0.79221404 |
| GSTAGNGGNGGNATNGTIAGQP | AWSLGLGEDQNTLY | 0.81996846 |
| GTGGDGGNGGNGGTGG | AWSLGLGEDQNTLY | 0.8255724 |
| GVRRPAGIPADAPV | AWSLGLGEDQNTLY | 0.5063141 |
| GYRQVTDAHLLTLARR | AWSLGLGEDQNTLY | 0.5984316 |
| IAKMGQDGHDRG | AWSLGLGEDQNTLY | 0.7002048 |
| KAEPSIEIQYT | AWSLGLGEDQNTLY | 0.66115296 |
| KPVRIRMAADEDTVAVT | AWSLGLGEDQNTLY | 0.516584 |
| MPPRIAGMRLLVIKPEP | AWSLGLGEDQNTLY | 0.52352935 |
| MKFYAVALFRSRGVPPTR | AWSLGLGEDQNTLY | 0.8091035 |
| NAGGFGILGAVAHSP | AWSLGLGEDQNTLY | 0.8337001 |
| NEGREITLAMAEM | AWSLGLGEDQNTLY | 0.67027164 |
| NELVEWSHPLGHRWRWE | AWSLGLGEDQNTLY | 0.35100994 |
| NLGAGGTGGNGGNPG | AWSLGLGEDQNTLY | 0.6664307 |
| NTEWPKWETVMAADPVD | AWSLGLGEDQNTLY | 0.48939753 |
| PASAAPAETQP | AWSLGLGEDQNTLY | 0.5667296 |
| QQSSDGKNGVRAAEITGE | AWSLGLGEDQNTLY | 0.6317801 |
| RAPDAVTTSWWKE | AWSLGLGEDQNTLY | 0.3734353 |
| SGFGNVGAHTSGWFNQ | AWSLGLGEDQNTLY | 0.6561326 |
| SGVAGAVGQAASVGGLKV | AWSLGLGEDQNTLY | 0.83791935 |
| SNGGAGTGSAGGNGGT | AWSLGLGEDQNTLY | 0.6190459 |
| TGGAGGEGGAAG | AWSLGLGEDQNTLY | 0.7750629 |
| TGGDGGAAGAGGAPGHG | AWSLGLGEDQNTLY | 0.8233466 |
| TMPPLLAAIANAGRHGV | AWSLGLGEDQNTLY | 0.7728138 |

| | | |
|---|---|---|
| TSSTAGNSGGA | AWSLGLGEDQNTLY | 0.8142868 |
| VPATVPVGILC | AWSLGLGEDQNTLY | 0.7632245 |
| VQRPDSGGVVLTGRIS | AWSLGLGEDQNTLY | 0.7238667 |
| WTPNPVPATKGVPAGLVQAT | AWSLGLGEDQNTLY | 0.7240066 |
| YALKRALAKFSDRFDVVII | AWSLGLGEDQNTLY | 0.77638626 |
| YLIGSVIV | AWSLGLGEDQNTLY | 0.6643363 |
| YGQLVAVID | AWSLGLGEDQNTLY | 0.74008596 |
| VTVGGMTLPA | AWSLGLGEDQNTLY | 0.7586973 |
| VLALPAV | AWSLGLGEDQNTLY | 0.76205444 |
| VIPPTLNYET | AWSLGLGEDQNTLY | 0.7199489 |
| VGMGGMHPSSG | AWSLGLGEDQNTLY | 0.81954217 |
| VGLAGGAATA | AWSLGLGEDQNTLY | 0.24910483 |
| VGGAGGAGGSGP | AWSLGLGEDQNTLY | 0.50480163 |
| VFAAFMV | AWSLGLGEDQNTLY | 0.8097183 |
| TLYDKLR | AWSLGLGEDQNTLY | 0.20032533 |
| TLAVFTVL | AWSLGLGEDQNTLY | 0.41046214 |
| TGSVQPARV | AWSLGLGEDQNTLY | 0.7123856 |
| TGGGAGSGGT | AWSLGLGEDQNTLY | 0.80951476 |
| SHLETSL | AWSLGLGEDQNTLY | 0.21690041 |
| SFTTGMT | AWSLGLGEDQNTLY | 0.6536006 |
| QVEVAKRMGVRQPTVS | AWSLGLGEDQNTLY | 0.40311024 |
| QGSGGPALAM | AWSLGLGEDQNTLY | 0.8088846 |
| MLAPSNP | AWSLGLGEDQNTLY | 0.62923 |
| LVNNAAI | AWSLGLGEDQNTLY | 0.34675196 |
| LTGEINTLA | AWSLGLGEDQNTLY | 0.8058058 |
| LSIFGPV | AWSLGLGEDQNTLY | 0.5268401 |
| LLALLANL | AWSLGLGEDQNTLY | 0.75457764 |
| LISNASDALDKLR | AWSLGLGEDQNTLY | 0.822859 |
| LINEAVP | AWSLGLGEDQNTLY | 0.8331566 |
| LGYAFSGL | AWSLGLGEDQNTLY | 0.41407156 |
| LDFVAVAL | AWSLGLGEDQNTLY | 0.7822781 |
| LAQPGAE | AWSLGLGEDQNTLY | 0.54628867 |
| ISNLGAVTGF | AWSLGLGEDQNTLY | 0.8432587 |
| HQEDPGRI | AWSLGLGEDQNTLY | 0.67488205 |
| GTPGEPTP | AWSLGLGEDQNTLY | 0.38050896 |
| GFFNSGS | AWSLGLGEDQNTLY | 0.7722858 |
| GAGGGGLTAGAN | AWSLGLGEDQNTLY | 0.787388 |
| FVDALVNM | AWSLGLGEDQNTLY | 0.6535733 |
| FLTVAPEL | AWSLGLGEDQNTLY | 0.7614719 |
| FETLTAL | AWSLGLGEDQNTLY | 0.71716523 |
| DVINPIEL | AWSLGLGEDQNTLY | 0.63732135 |
| APVSGAPV | AWSLGLGEDQNTLY | 0.7728132 |
| ALVGGLAW | AWSLGLGEDQNTLY | 0.7305302 |
| ALAAAAATAI | AWSLGLGEDQNTLY | 0.6164182 |
| AFGLGLLL | AWSLGLGEDQNTLY | 0.8323649 |
| AFFAAASL | AWSLGLGEDQNTLY | 0.70917654 |
| AAAALPAFSPP | AWSLGLGEDQNTLY | 0.48974842 |
| VVLAPAL | AWSLGLGEDQNTLY | 0.8380512 |
| TIGHVDHGKTTL | AWSLGLGEDQNTLY | 0.7225158 |
| LISNASDALDK | AWSLGLGEDQNTLY | 0.7962159 |
| LISNASDAL | AWSLGLGEDQNTLY | 0.7894454 |

| | | |
|---|---|---|
| ISNASDALDK | AWSLGLGEDQNTLY | 0.8115828 |
| NQAETLVYQTE | AWSLGLGEDQNTLY | 0.52173656 |
| TGGTGGTAGAG | AWSLGLGEDQNTLY | 0.70361996 |
| TGSTAGGAGGA | AWSLGLGEDQNTLY | 0.72892 |
| DPLAVPVAAGPV | AWSLGLGEDQNTLY | 0.7929622 |
| GGTGGTPGNGGN | AWSLGLGEDQNTLY | 0.66454107 |
| PAGLAGVDPATL | AWSLGLGEDQNTLY | 0.7625721 |
| GNNLVVEAPHAGASV | AWSLGLGEDQNTLY | 0.7724967 |
| KVTRIMSASTLEQPA | AWSLGLGEDQNTLY | 0.8280163 |
| LGFALPIALPRRTVV | AWSLGLGEDQNTLY | 0.70762867 |
| TRLGNLTARKLVVLV | AWSLGLGEDQNTLY | 0.700987 |
| WSQQRRLIYFPSAGP | AWSLGLGEDQNTLY | 0.833718 |
| GPNGNPDPMAAAVDIR | AWSLGLGEDQNTLY | 0.47859898 |
| TQRCVSVADLRAAASR | AWSLGLGEDQNTLY | 0.7412454 |
| QFVLDVAEHPVEVLTETV | AWSLGLGEDQNTLY | 0.8123188 |
| GGAGGTGGAGGNGGTQAGDG | AWSLGLGEDQNTLY | 0.8227334 |
| PLLATLRAIAVDVDAKPAQV | AWSLGLGEDQNTLY | 0.7247957 |
| APKFRTGGYMIDFWGVGYQVAK | AWSLGLGEDQNTLY | 0.7904967 |
| GAGGAGGAGGAAGGPGGTGGTG | AWSLGLGEDQNTLY | 0.6215329 |
| IGALGLLATDVAPAGSVAFGVA | AWSLGLGEDQNTLY | 0.73243964 |
| AGGAGGIGGNGGAGGAGGVLMG | AWSLGLGEDQNTLY | 0.58428556 |
| ADELIGLL | AWSLGLGEDQNTLY | 0.64301556 |
| ADRALKA | AWSLGLGEDQNTLY | 0.52862155 |
| AGAVGQVGGAGGA | AWSLGLGEDQNTLY | 0.72660905 |
| AGGGGGLGGAP | AWSLGLGEDQNTLY | 0.7808466 |
| AGGLLYGNGGAG | AWSLGLGEDQNTLY | 0.7295736 |
| AGLESIGL | AWSLGLGEDQNTLY | 0.39385575 |
| AGLKTGII | AWSLGLGEDQNTLY | 0.7584152 |
| ASLAALLL | AWSLGLGEDQNTLY | 0.7488754 |
| ASVINISSI | AWSLGLGEDQNTLY | 0.83622074 |
| ATPPGPA | AWSLGLGEDQNTLY | 0.73302674 |
| ATVASALV | AWSLGLGEDQNTLY | 0.8074856 |
| ATVASLIL | AWSLGLGEDQNTLY | 0.6877979 |
| AVNGPVLIRH | AWSLGLGEDQNTLY | 0.7676426 |
| DGAAGAVGI | AWSLGLGEDQNTLY | 0.39424855 |
| DPLLHGI | AWSLGLGEDQNTLY | 0.5906714 |
| ELGAALGAL | AWSLGLGEDQNTLY | 0.67385477 |
| ELGRTQAV | AWSLGLGEDQNTLY | 0.5680752 |
| FVAGGYLS | AWSLGLGEDQNTLY | 0.7681258 |
| GASGGAGGAGGW | AWSLGLGEDQNTLY | 0.807471 |
| GGAGAGGDGGAGGI | AWSLGLGEDQNTLY | 0.79724926 |
| GTGHNGSQPGMGGNG | AWSLGLGEDQNTLY | 0.808697 |
| GYAAASASAS | AWSLGLGEDQNTLY | 0.81471545 |
| HQVDVAAVQQG | AWSLGLGEDQNTLY | 0.82065713 |
| IGAAIAHT | AWSLGLGEDQNTLY | 0.6400821 |
| IGMIEVAQL | AWSLGLGEDQNTLY | 0.60829115 |
| IRLAPEHTSAMD | AWSLGLGEDQNTLY | 0.40371907 |
| IYNVSVL | AWSLGLGEDQNTLY | 0.53896827 |
| KQGFLGM | AWSLGLGEDQNTLY | 0.6331458 |
| LESVITQL | AWSLGLGEDQNTLY | 0.7223364 |
| LGLGALVVV | AWSLGLGEDQNTLY | 0.17802426 |

| | | |
|---|---|---|
| LIGNGGNGGSGGI | AWSLGLGEDQNTLY | 0.68738097 |
| LNAINAPTL | AWSLGLGEDQNTLY | 0.8296297 |
| LQLLNTGV | AWSLGLGEDQNTLY | 0.6696662 |
| LQTSIEF | AWSLGLGEDQNTLY | 0.7584748 |
| MSFAVSEL | AWSLGLGEDQNTLY | 0.7578547 |
| NTGALGL | AWSLGLGEDQNTLY | 0.59330523 |
| PPGAGGAGGAGTT | AWSLGLGEDQNTLY | 0.7775383 |
| QLVGLGI | AWSLGLGEDQNTLY | 0.64708644 |
| RFLEQVHQL | AWSLGLGEDQNTLY | 0.82696164 |
| RGVGYVLR | AWSLGLGEDQNTLY | 0.8383355 |
| SATGVFAGVI | AWSLGLGEDQNTLY | 0.74612975 |
| SLPAITLP | AWSLGLGEDQNTLY | 0.608901 |
| STALAFTV | AWSLGLGEDQNTLY | 0.7805823 |
| SVLTVGLGK | AWSLGLGEDQNTLY | 0.52873594 |
| TALAGAGSGI | AWSLGLGEDQNTLY | 0.67112553 |
| TGLPFTGI | AWSLGLGEDQNTLY | 0.6568142 |
| VAEHVTV | AWSLGLGEDQNTLY | 0.7043657 |
| VDIVLNSL | AWSLGLGEDQNTLY | 0.54683316 |
| VLTGGGAVL | AWSLGLGEDQNTLY | 0.7591361 |
| VNVGLAYGI | AWSLGLGEDQNTLY | 0.7341879 |
| VSSAAVGAV | AWSLGLGEDQNTLY | 0.76079285 |
| VTSVYEAGV | AWSLGLGEDQNTLY | 0.52905226 |
| WVALDRGA | AWSLGLGEDQNTLY | 0.7826655 |
| YLVGSVVI | AWSLGLGEDQNTLY | 0.71883136 |
| GPPGTGKTLL | AWSLGLGEDQNTLY | 0.39345044 |
| GVGMTKF | AWSLGLGEDQNTLY | 0.79502195 |
| IGIGGSDL | AWSLGLGEDQNTLY | 0.672847 |
| IGTLLAEL | AWSLGLGEDQNTLY | 0.6633194 |
| IYTSGTT | AWSLGLGEDQNTLY | 0.796956 |
| LSFPAGV | AWSLGLGEDQNTLY | 0.7210419 |
| LYTSGTT | AWSLGLGEDQNTLY | 0.5796724 |
| VGAFPFGF | AWSLGLGEDQNTLY | 0.55746204 |
| VGAGTVEF | AWSLGLGEDQNTLY | 0.714132 |
| VGKSTLLL | AWSLGLGEDQNTLY | 0.6926797 |
| AALAHAGGGIV | AWSLGLGEDQNTLY | 0.2710221 |
| AALVAVSLL | AWSLGLGEDQNTLY | 0.74757814 |
| AAVSRAVAAL | AWSLGLGEDQNTLY | 0.49547702 |
| AGGSSGIGGF | AWSLGLGEDQNTLY | 0.5697143 |
| AGLTVANANG | AWSLGLGEDQNTLY | 0.6377059 |
| AGLVGAVL | AWSLGLGEDQNTLY | 0.8043651 |
| ALSQAFTV | AWSLGLGEDQNTLY | 0.7424069 |
| ALTFSVSL | AWSLGLGEDQNTLY | 0.6376349 |
| ALTGAALVAL | AWSLGLGEDQNTLY | 0.7507092 |
| ASFSIGAGVL | AWSLGLGEDQNTLY | 0.72685236 |
| DGSVYKI | AWSLGLGEDQNTLY | 0.80311453 |
| EQLAEIV | AWSLGLGEDQNTLY | 0.7056022 |
| FAVSELTVM | AWSLGLGEDQNTLY | 0.722036 |
| GAAPLAAGL | AWSLGLGEDQNTLY | 0.73731077 |
| GADGGDGGSGGAGG | AWSLGLGEDQNTLY | 0.6451333 |
| GFGLDTGGAGG | AWSLGLGEDQNTLY | 0.5952321 |
| GLASGGLL | AWSLGLGEDQNTLY | 0.58775234 |

| | | |
|---|---|---|
| GRIAGAAVLL | AWSLGLGEDQNTLY | 0.5844884 |
| GSAGSLNTSF | AWSLGLGEDQNTLY | 0.53598964 |
| GTGQIGI | AWSLGLGEDQNTLY | 0.7507192 |
| GTLIALIL | AWSLGLGEDQNTLY | 0.83296764 |
| IAATMGFVVG | AWSLGLGEDQNTLY | 0.6476452 |
| IFAQALL | AWSLGLGEDQNTLY | 0.33511746 |
| IGAAVALGF | AWSLGLGEDQNTLY | 0.6171973 |
| IGLTPLAL | AWSLGLGEDQNTLY | 0.6704067 |
| ILGSNTSTL | AWSLGLGEDQNTLY | 0.7925837 |
| IVFSLTVL | AWSLGLGEDQNTLY | 0.76967967 |
| LASKATGF | AWSLGLGEDQNTLY | 0.79462594 |
| LASVALIL | AWSLGLGEDQNTLY | 0.76398754 |
| LFAVGLTL | AWSLGLGEDQNTLY | 0.62390935 |
| LFNGGIGTYI | AWSLGLGEDQNTLY | 0.7997297 |
| LTLGAGII | AWSLGLGEDQNTLY | 0.66718566 |
| LVAAALGDI | AWSLGLGEDQNTLY | 0.7453401 |
| NTAFVQLGI | AWSLGLGEDQNTLY | 0.7302377 |
| QLFSGAEL | AWSLGLGEDQNTLY | 0.75664663 |
| SAATFITL | AWSLGLGEDQNTLY | 0.5127652 |
| SNYGLANLGV | AWSLGLGEDQNTLY | 0.6977396 |
| SYGGAYAVM | AWSLGLGEDQNTLY | 0.48745564 |
| TTGVGGLTLG | AWSLGLGEDQNTLY | 0.7606034 |
| TTQTIYI | AWSLGLGEDQNTLY | 0.59467185 |
| VAAGAGLGAVI | AWSLGLGEDQNTLY | 0.61787415 |
| VAAVVTLL | AWSLGLGEDQNTLY | 0.68069535 |
| VGFLGGLI | AWSLGLGEDQNTLY | 0.8205887 |
| VGVIPILL | AWSLGLGEDQNTLY | 0.8278823 |
| VIFASSLI | AWSLGLGEDQNTLY | 0.6104618 |
| VKGGDGGAAAG | AWSLGLGEDQNTLY | 0.59332883 |
| VTKTFTNI | AWSLGLGEDQNTLY | 0.73856807 |
| VVAAAILL | AWSLGLGEDQNTLY | 0.6140243 |
| YAALDVEL | AWSLGLGEDQNTLY | 0.7839818 |
| LASVALLL | AWSLGLGEDQNTLY | 0.6961106 |
| LDAVHRV | AWSLGLGEDQNTLY | 0.6401528 |
| QGLSSSIF | AWSLGLGEDQNTLY | 0.60041565 |
| RGTFANIRL | AWSLGLGEDQNTLY | 0.77635664 |
| RQLWWGHRI | AWSLGLGEDQNTLY | 0.6500714 |
| YGPPGTGKTLL | AWSLGLGEDQNTLY | 0.7787981 |
| AALVGAPANVA | AWSLGLGEDQNTLY | 0.42620307 |
| AATFRRTE | AWSLGLGEDQNTLY | 0.796803 |
| AGALGGGI | AWSLGLGEDQNTLY | 0.47174013 |
| AGGAGGAGGIGAG | AWSLGLGEDQNTLY | 0.55174786 |
| AGSGALGL | AWSLGLGEDQNTLY | 0.2550412 |
| AQIADQI | AWSLGLGEDQNTLY | 0.7230555 |
| ASAVAVTYL | AWSLGLGEDQNTLY | 0.7501038 |
| ASTVLGGTGG | AWSLGLGEDQNTLY | 0.82593834 |
| AVGGEPAK | AWSLGLGEDQNTLY | 0.74180365 |
| CEQGTFGGTC | AWSLGLGEDQNTLY | 0.71499974 |
| DEGKAPAG | AWSLGLGEDQNTLY | 0.81620896 |
| DTVVFPAL | AWSLGLGEDQNTLY | 0.64080715 |
| FEGITMAG | AWSLGLGEDQNTLY | 0.31724325 |

| | | |
|---|---|---|
| FPLVVQD | AWSLGLGEDQNTLY | 0.73221135 |
| GAFTTPQ | AWSLGLGEDQNTLY | 0.6228324 |
| GFGGFGTAGGVGGA | AWSLGLGEDQNTLY | 0.82737005 |
| GFLAQFL | AWSLGLGEDQNTLY | 0.7515461 |
| GGAGGAGGNAGVA | AWSLGLGEDQNTLY | 0.3809172 |
| GGVGTAGAGGAGGAGG | AWSLGLGEDQNTLY | 0.74604964 |
| GNGGGGDAGNAGSGGN | AWSLGLGEDQNTLY | 0.7811569 |
| GTGGAGGAGNDAGS | AWSLGLGEDQNTLY | 0.391401 |
| GTTNTGWFNS | AWSLGLGEDQNTLY | 0.63633937 |
| IFMGPLFTL | AWSLGLGEDQNTLY | 0.5786292 |
| ILAHAIEAI | AWSLGLGEDQNTLY | 0.5136161 |
| IRSAIREAL | AWSLGLGEDQNTLY | 0.7193527 |
| ITFAGGG | AWSLGLGEDQNTLY | 0.598703 |
| ITGMSGHQEV | AWSLGLGEDQNTLY | 0.60527295 |
| IVAGLACTL | AWSLGLGEDQNTLY | 0.65508807 |
| IVGGGLAGA | AWSLGLGEDQNTLY | 0.52409613 |
| IVTGVLNI | AWSLGLGEDQNTLY | 0.64287454 |
| LAQQGCI | AWSLGLGEDQNTLY | 0.6661634 |
| LCLHVEN | AWSLGLGEDQNTLY | 0.74413013 |
| LDAISDGS | AWSLGLGEDQNTLY | 0.7802331 |
| LGGFAAML | AWSLGLGEDQNTLY | 0.5997813 |
| LGGKNPMI | AWSLGLGEDQNTLY | 0.7304561 |
| LGPKGVAAIL | AWSLGLGEDQNTLY | 0.8337393 |
| LMVGVSI | AWSLGLGEDQNTLY | 0.4187924 |
| LVGAAFVL | AWSLGLGEDQNTLY | 0.5516165 |
| LVLAVAPL | AWSLGLGEDQNTLY | 0.7512711 |
| MAAIAVIDAA | AWSLGLGEDQNTLY | 0.669088 |
| NNGINAELQ | AWSLGLGEDQNTLY | 0.79296106 |
| RGEPLVLVHG | AWSLGLGEDQNTLY | 0.73758173 |
| RGGAGGAAGLI | AWSLGLGEDQNTLY | 0.7488258 |
| RSYRMHPAV | AWSLGLGEDQNTLY | 0.6605586 |
| SAAHLLAGQLA | AWSLGLGEDQNTLY | 0.83103347 |
| SAAVGSY | AWSLGLGEDQNTLY | 0.7722405 |
| SPHAVLHRI | AWSLGLGEDQNTLY | 0.6473642 |
| SQIAKVL | AWSLGLGEDQNTLY | 0.73004544 |
| TALGHLVP | AWSLGLGEDQNTLY | 0.43873608 |
| TAPTTSGAA | AWSLGLGEDQNTLY | 0.7421998 |
| TDGDDGNGGSPGT | AWSLGLGEDQNTLY | 0.7868558 |
| TDGLGEN | AWSLGLGEDQNTLY | 0.54486203 |
| TGGPPLPGQA | AWSLGLGEDQNTLY | 0.6359602 |
| TIPAITI | AWSLGLGEDQNTLY | 0.84223455 |
| TIPGFTI | AWSLGLGEDQNTLY | 0.8012202 |
| TSIASGALA | AWSLGLGEDQNTLY | 0.62812054 |
| TVTLPGAAI | AWSLGLGEDQNTLY | 0.5586797 |
| VASGLSAALP | AWSLGLGEDQNTLY | 0.6195475 |
| VGAGVIGI | AWSLGLGEDQNTLY | 0.62661666 |
| VGPKRFGV | AWSLGLGEDQNTLY | 0.37620306 |
| VNVAFPD | AWSLGLGEDQNTLY | 0.7729069 |
| VSMTNSGVS | AWSLGLGEDQNTLY | 0.6808352 |
| VTDVLGDLSV | AWSLGLGEDQNTLY | 0.783108 |
| VVVVGAGFAGL | AWSLGLGEDQNTLY | 0.7645371 |

| | | |
|---|---|---|
| VYTVGTVV | AWSLGLGEDQNTLY | 0.8301654 |
| WTTFSGP | AWSLGLGEDQNTLY | 0.5958343 |
| YDELVEAF | AWSLGLGEDQNTLY | 0.6589962 |
| YGTGGAGGNGG | AWSLGLGEDQNTLY | 0.3504253 |
| YTNVMAV | AWSLGLGEDQNTLY | 0.5929724 |
| FLEQVHQL | AWSLGLGEDQNTLY | 0.5181216 |
| GLSLLAVAV | AWSLGLGEDQNTLY | 0.6605403 |
| LEQVHQL | AWSLGLGEDQNTLY | 0.47489262 |
| LNAAGAI | AWSLGLGEDQNTLY | 0.8211608 |
| PAVHASL | AWSLGLGEDQNTLY | 0.66125023 |
| TELEDDA | AWSLGLGEDQNTLY | 0.80201256 |
| TPPGPPN | AWSLGLGEDQNTLY | 0.6385009 |
| TPPPPPPP | AWSLGLGEDQNTLY | 0.50492084 |
| TQSPALI | ASSPLGAINTGQLY | 0.64493364 |
| EFAAAFPQ | ASSPLGAINTGQLY | 0.78013575 |
| LLSGNLDVL | ASSPLGAINTGQLY | 0.40152377 |
| GPANLGNYNI | ASSPLGAINTGQLY | 0.397379 |
| TLGPTLP | ASSPLGAINTGQLY | 0.55750465 |
| VYAYSSCCK | ASSPLGAINTGQLY | 0.7197642 |
| APAPAPAPNI | ASSPLGAINTGQLY | 0.8234273 |
| PGGPVPIP | ASSPLGAINTGQLY | 0.8086559 |
| FLTVALEP | ASSPLGAINTGQLY | 0.7316185 |
| SQYFGANL | ASSPLGAINTGQLY | 0.8469331 |
| GGVKPAA | ASSPLGAINTGQLY | 0.1743055 |
| GGTGGAGGAGGW | ASSPLGAINTGQLY | 0.6174224 |
| GVGGVAGLGGAGG | ASSPLGAINTGQLY | 0.8153324 |
| VLLVLATTI | ASSPLGAINTGQLY | 0.75048614 |
| IGIFAAML | ASSPLGAINTGQLY | 0.70860386 |
| AVGDLID | ASSPLGAINTGQLY | 0.8194212 |
| NISPLPL | ASSPLGAINTGQLY | 0.5452 |
| VTAVFVAI | ASSPLGAINTGQLY | 0.69618434 |
| SGDDNDAAL | ASSPLGAINTGQLY | 0.69712013 |
| AVAALAHS | ASSPLGAINTGQLY | 0.79880345 |
| GGVTVMATP | ASSPLGAINTGQLY | 0.78766936 |
| AGLAANVRD | ASSPLGAINTGQLY | 0.33924773 |
| TNRPDLIDPALLRPG | ASSPLGAINTGQLY | 0.79727864 |
| LANLGVTAL | ASSPLGAINTGQLY | 0.8429942 |
| TGERIEL | ASSPLGAINTGQLY | 0.6310887 |
| NGEELAA | ASSPLGAINTGQLY | 0.8066529 |
| VLSLVWG | ASSPLGAINTGQLY | 0.73222005 |
| IGYALWL | ASSPLGAINTGQLY | 0.70318747 |
| LSLLLAPL | ASSPLGAINTGQLY | 0.5881322 |
| TLDEALK | ASSPLGAINTGQLY | 0.4757114 |
| TDGAGTAEGG | ASSPLGAINTGQLY | 0.7705368 |
| NAPGGAGGNGG | ASSPLGAINTGQLY | 0.5119122 |
| VSVFMGMGGNF | ASSPLGAINTGQLY | 0.73986804 |
| LASYLAAL | ASSPLGAINTGQLY | 0.7068325 |
| LVGGTPAL | ASSPLGAINTGQLY | 0.8541438 |
| VRNADVL | ASSPLGAINTGQLY | 0.7503009 |
| TCSSPFLVSAM | ASSPLGAINTGQLY | 0.77024245 |
| VHITDAE | ASSPLGAINTGQLY | 0.5552182 |

| | | |
|---|---|---|
| EYAAPSF | ASSPLGAINTGQLY | 0.74028665 |
| VVDAFHGGL | ASSPLGAINTGQLY | 0.7606345 |
| ASVVDPQ | ASSPLGAINTGQLY | 0.5320613 |
| SYNEFGGLS | ASSPLGAINTGQLY | 0.8097198 |
| DPDGSVY | ASSPLGAINTGQLY | 0.54167986 |
| MVSTIHATE | ASSPLGAINTGQLY | 0.7151284 |
| LAAGVADPV | ASSPLGAINTGQLY | 0.7134242 |
| TKEQIAA | ASSPLGAINTGQLY | 0.771549 |
| SLASPAAL | ASSPLGAINTGQLY | 0.7385938 |
| IGLVFYGV | ASSPLGAINTGQLY | 0.8098242 |
| FLLDASNL | ASSPLGAINTGQLY | 0.5226909 |
| YQVGNFP | ASSPLGAINTGQLY | 0.7874963 |
| ESLLSQST | ASSPLGAINTGQLY | 0.8186246 |
| LFTVGFAAVG | ASSPLGAINTGQLY | 0.71285903 |
| ATPPTPT | ASSPLGAINTGQLY | 0.82459587 |
| TDLATLG | ASSPLGAINTGQLY | 0.7532476 |
| SAGGTGR | ASSPLGAINTGQLY | 0.3302093 |
| GFSNSGS | ASSPLGAINTGQLY | 0.72325766 |
| GGNGGSGGTSV | ASSPLGAINTGQLY | 0.81251776 |
| GTSVATGGAGN | ASSPLGAINTGQLY | 0.81170416 |
| NGGNGGSGGTS | ASSPLGAINTGQLY | 0.79327106 |
| AVVKFPH | ASSPLGAINTGQLY | 0.7407012 |
| TPPPPPP | ASSPLGAINTGQLY | 0.62681013 |
| AAPAAAAAPA | ASSPLGAINTGQLY | 0.82857203 |
| IALNEDL | ASSPLGAINTGQLY | 0.86670434 |
| IVNVSSM | ASSPLGAINTGQLY | 0.79227 |
| TGIPPAPR | ASSPLGAINTGQLY | 0.5620196 |
| FDLGGGTF | ASSPLGAINTGQLY | 0.52791905 |
| FDLGGGTFDVS | ASSPLGAINTGQLY | 0.7803154 |
| NEPTAAA | ASSPLGAINTGQLY | 0.7863903 |
| NEPTAAALA | ASSPLGAINTGQLY | 0.5071515 |
| GSIINISSI | ASSPLGAINTGQLY | 0.68984747 |
| VAPGNAGTA | ASSPLGAINTGQLY | 0.7260436 |
| TGGGASLE | ASSPLGAINTGQLY | 0.27700505 |
| VAVYNHYKRI | ASSPLGAINTGQLY | 0.7362286 |
| TDPAPATT | ASSPLGAINTGQLY | 0.8022412 |
| TNRPDILDPALLRPG | ASSPLGAINTGQLY | 0.79242796 |
| LTTTTTS | ASSPLGAINTGQLY | 0.37132245 |
| PTGSGKTL | ASSPLGAINTGQLY | 0.8063477 |
| TGSGKTL | ASSPLGAINTGQLY | 0.8152186 |
| TIGHVDHGK | ASSPLGAINTGQLY | 0.83979964 |
| TIGHVDHGKTT | ASSPLGAINTGQLY | 0.39167115 |
| GHVDHGKTT | ASSPLGAINTGQLY | 0.20999882 |
| AQFEHTLL | ASSPLGAINTGQLY | 0.7559016 |
| IVNQDEL | ASSPLGAINTGQLY | 0.74816334 |
| TNGSQFF | ASSPLGAINTGQLY | 0.5743496 |
| TGKGIVE | ASSPLGAINTGQLY | 0.4728403 |
| LISNASDA | ASSPLGAINTGQLY | 0.69544953 |
| AELRTLGELTKIAKAHGAQ | ASSPLGAINTGQLY | 0.8142327 |
| APNAVAAAEPSW | ASSPLGAINTGQLY | 0.42539713 |
| APQVAGAATAVL | ASSPLGAINTGQLY | 0.4613994 |

| | | |
|---|---|---|
| ARRDPAAAAAILPTDA | ASSPLGAINTGQLY | 0.76313883 |
| DAAARDVLAEAGLAEYF | ASSPLGAINTGQLY | 0.8201085 |
| DDQIVLSAKANLAKVEQ | ASSPLGAINTGQLY | 0.48828125 |
| DELLPDTGEDSDEDGH | ASSPLGAINTGQLY | 0.61812645 |
| ELPEAPANRATVPQD | ASSPLGAINTGQLY | 0.766743 |
| FGQSKIERVTGIG | ASSPLGAINTGQLY | 0.7779895 |
| GAGGTGSTAGGAGGAGGAGGLY | ASSPLGAINTGQLY | 0.7285876 |
| GHGGTGAAGGNGGTGGAGGSGI | ASSPLGAINTGQLY | 0.46005657 |
| GNGGNGGAGGEGGPGGAGGAGA | ASSPLGAINTGQLY | 0.7724414 |
| GSTAGNGGNGGNATNGTIAGQP | ASSPLGAINTGQLY | 0.5439544 |
| GTGGDGGNGGNGGTGG | ASSPLGAINTGQLY | 0.7911767 |
| GVRRPAGIPADAPV | ASSPLGAINTGQLY | 0.6990823 |
| GYRQVTDAHLLTLARR | ASSPLGAINTGQLY | 0.7535363 |
| IAKMGQDGHDRG | ASSPLGAINTGQLY | 0.65803516 |
| KAEPSIEIQYT | ASSPLGAINTGQLY | 0.42615604 |
| KPVRIRMAADEDTVAVT | ASSPLGAINTGQLY | 0.47560984 |
| MPPRIAGMRLLVIKPEP | ASSPLGAINTGQLY | 0.530313 |
| MKFYAVALFRSRGVPPTR | ASSPLGAINTGQLY | 0.8147664 |
| NAGGFGILGAVAHSP | ASSPLGAINTGQLY | 0.73671305 |
| NEGREITLAMAEM | ASSPLGAINTGQLY | 0.80466187 |
| NELVEWSHPLGHRWRWE | ASSPLGAINTGQLY | 0.77850246 |
| NLGAGGTGGNGGNPG | ASSPLGAINTGQLY | 0.54347533 |
| NTEWPKWETVMAADPVD | ASSPLGAINTGQLY | 0.77915454 |
| PASAAPAETQP | ASSPLGAINTGQLY | 0.35503507 |
| QQSSDGKNGVRAAEITGE | ASSPLGAINTGQLY | 0.7640637 |
| RAPDAVTTSWWKE | ASSPLGAINTGQLY | 0.6441134 |
| SGFGNVGAHTSGWFNQ | ASSPLGAINTGQLY | 0.78808683 |
| SGVAGAVGQAASVGGLKV | ASSPLGAINTGQLY | 0.64759004 |
| SNGGAGTGSAGGNGGT | ASSPLGAINTGQLY | 0.80822104 |
| TGGAGGEGGAAG | ASSPLGAINTGQLY | 0.7568014 |
| TGGDGGAAGAGGAPGHG | ASSPLGAINTGQLY | 0.46193177 |
| TMPPLLAAIANAGRHGV | ASSPLGAINTGQLY | 0.7630597 |
| TSSTAGNSGGA | ASSPLGAINTGQLY | 0.78746456 |
| VPATVPVGILC | ASSPLGAINTGQLY | 0.80773544 |
| VQRPDSGGVVLTGRIS | ASSPLGAINTGQLY | 0.3525094 |
| WTPNPVPATKGVPAGLVQAT | ASSPLGAINTGQLY | 0.7330081 |
| YALKRALAKFSDRFDVVII | ASSPLGAINTGQLY | 0.5377233 |
| YLIGSVIV | ASSPLGAINTGQLY | 0.7280172 |
| YGQLVAVID | ASSPLGAINTGQLY | 0.7198443 |
| VTVGGMTLPA | ASSPLGAINTGQLY | 0.6446091 |
| VLALPAV | ASSPLGAINTGQLY | 0.77803755 |
| VIPPTLNYET | ASSPLGAINTGQLY | 0.4115922 |
| VGMGGMHPSSG | ASSPLGAINTGQLY | 0.5839 |
| VGLAGGAATA | ASSPLGAINTGQLY | 0.6729988 |
| VGGAGGAGGSGP | ASSPLGAINTGQLY | 0.74937797 |
| VFAAFMV | ASSPLGAINTGQLY | 0.7773379 |
| TLYDKLR | ASSPLGAINTGQLY | 0.48123646 |
| TLAVFTVL | ASSPLGAINTGQLY | 0.72885364 |
| TGSVQPARV | ASSPLGAINTGQLY | 0.67851627 |
| TGGGAGSGGT | ASSPLGAINTGQLY | 0.49505642 |
| SHLETSL | ASSPLGAINTGQLY | 0.7619708 |

| | | |
|---|---|---|
| SFTTGMT | ASSPLGAINTGQLY | 0.6094544 |
| QVEVAKRMGVRQPTVS | ASSPLGAINTGQLY | 0.36887515 |
| QGSGGPALAM | ASSPLGAINTGQLY | 0.78927886 |
| MLAPSNP | ASSPLGAINTGQLY | 0.559929 |
| LVNNAAI | ASSPLGAINTGQLY | 0.72694194 |
| LTGEINTLA | ASSPLGAINTGQLY | 0.79712826 |
| LSIFGPV | ASSPLGAINTGQLY | 0.37444034 |
| LLALLANL | ASSPLGAINTGQLY | 0.8005763 |
| LISNASDALDKLR | ASSPLGAINTGQLY | 0.6863854 |
| LINEAVP | ASSPLGAINTGQLY | 0.6697188 |
| LGYAFSGL | ASSPLGAINTGQLY | 0.6939485 |
| LDFVAVAL | ASSPLGAINTGQLY | 0.5170392 |
| LAQPGAE | ASSPLGAINTGQLY | 0.7941962 |
| ISNLGAVTGF | ASSPLGAINTGQLY | 0.77884555 |
| HQEDPGRI | ASSPLGAINTGQLY | 0.72524023 |
| GTPGEPTP | ASSPLGAINTGQLY | 0.71280897 |
| GFFNSGS | ASSPLGAINTGQLY | 0.7718376 |
| GAGGGGLTAGAN | ASSPLGAINTGQLY | 0.808306 |
| FVDALVNM | ASSPLGAINTGQLY | 0.55347914 |
| FLTVAPEL | ASSPLGAINTGQLY | 0.36917508 |
| FETLTAL | ASSPLGAINTGQLY | 0.6799383 |
| DVINPIEL | ASSPLGAINTGQLY | 0.68263644 |
| APVSGAPV | ASSPLGAINTGQLY | 0.60144055 |
| ALVGGLAW | ASSPLGAINTGQLY | 0.41595066 |
| ALAAAAATAI | ASSPLGAINTGQLY | 0.75690955 |
| AFGLGLLL | ASSPLGAINTGQLY | 0.8452679 |
| AFFAAASL | ASSPLGAINTGQLY | 0.5701168 |
| AAAALPAFSPP | ASSPLGAINTGQLY | 0.83927864 |
| VVLAPAL | ASSPLGAINTGQLY | 0.44513023 |
| TIGHVDHGKTTL | ASSPLGAINTGQLY | 0.788484 |
| LISNASDALDK | ASSPLGAINTGQLY | 0.81306887 |
| LISNASDAL | ASSPLGAINTGQLY | 0.5273458 |
| ISNASDALDK | ASSPLGAINTGQLY | 0.68512905 |
| NQAETLVYQTE | ASSPLGAINTGQLY | 0.66076577 |
| TGGTGGTAGAG | ASSPLGAINTGQLY | 0.52883816 |
| TGSTAGGAGGA | ASSPLGAINTGQLY | 0.8082511 |
| DPLAVPVAAGPV | ASSPLGAINTGQLY | 0.5974871 |
| GGTGGTPGNGGN | ASSPLGAINTGQLY | 0.7698824 |
| PAGLAGVDPATL | ASSPLGAINTGQLY | 0.6184367 |
| GNNLVVEAPHAGASV | ASSPLGAINTGQLY | 0.5724046 |
| KVTRIMSASTLEQPA | ASSPLGAINTGQLY | 0.7787745 |
| LGFALPIALPRRTVV | ASSPLGAINTGQLY | 0.75845265 |
| TRLGNLTARKLVVLV | ASSPLGAINTGQLY | 0.732103 |
| WSQQRRLIYFPSAGP | ASSPLGAINTGQLY | 0.8648995 |
| GPNGNPDPMAAAVDIR | ASSPLGAINTGQLY | 0.611032 |
| TQRCVSVADLRAAASR | ASSPLGAINTGQLY | 0.82424283 |
| QFVLDVAEHPVEVLTETV | ASSPLGAINTGQLY | 0.820876 |
| GGAGGTGGAGGNGGTQAGDG | ASSPLGAINTGQLY | 0.7057878 |
| PLLATLRAIAVDVDAKPAQV | ASSPLGAINTGQLY | 0.673264 |
| APKFRTGGYMIDFWGVGYQVAK | ASSPLGAINTGQLY | 0.7849424 |
| GAGGAGGAGGAAGGPGGTGGTG | ASSPLGAINTGQLY | 0.6487467 |

| | | |
|---|---|---|
| IGALGLLATDVAPAGSVAFGVA | ASSPLGAINTGQLY | 0.7565452 |
| AGGAGGIGGNGGAGGAGGVLMG | ASSPLGAINTGQLY | 0.82503366 |
| ADELIGLL | ASSPLGAINTGQLY | 0.48781222 |
| ADRALKA | ASSPLGAINTGQLY | 0.8376833 |
| AGAVGQVGGAGGA | ASSPLGAINTGQLY | 0.7676196 |
| AGGGGGLGGAP | ASSPLGAINTGQLY | 0.8033025 |
| AGGLLYGNGGAG | ASSPLGAINTGQLY | 0.6491037 |
| AGLESIGL | ASSPLGAINTGQLY | 0.33779845 |
| AGLKTGII | ASSPLGAINTGQLY | 0.695156 |
| ASLAALLL | ASSPLGAINTGQLY | 0.5961338 |
| ASVINISSI | ASSPLGAINTGQLY | 0.83711696 |
| ATPPGPA | ASSPLGAINTGQLY | 0.8210551 |
| ATVASALV | ASSPLGAINTGQLY | 0.66800195 |
| ATVASLIL | ASSPLGAINTGQLY | 0.7948783 |
| AVNGPVLIRH | ASSPLGAINTGQLY | 0.8185501 |
| DGAAGAVGI | ASSPLGAINTGQLY | 0.47377288 |
| DPLLHGI | ASSPLGAINTGQLY | 0.6762196 |
| ELGAALGAL | ASSPLGAINTGQLY | 0.6168681 |
| ELGRTQAV | ASSPLGAINTGQLY | 0.75881577 |
| FVAGGYLS | ASSPLGAINTGQLY | 0.37948996 |
| GASGGAGGAGGW | ASSPLGAINTGQLY | 0.7944139 |
| GGAGAGGDGGAGGI | ASSPLGAINTGQLY | 0.6159693 |
| GTGHNGSQPGMGGNG | ASSPLGAINTGQLY | 0.23735124 |
| GYAAASASAS | ASSPLGAINTGQLY | 0.7922851 |
| HQVDVAAVQQG | ASSPLGAINTGQLY | 0.6664145 |
| IGAAIAHT | ASSPLGAINTGQLY | 0.6991029 |
| IGMIEVAQL | ASSPLGAINTGQLY | 0.8079827 |
| IRLAPEHTSAMD | ASSPLGAINTGQLY | 0.5791782 |
| IYNVSVL | ASSPLGAINTGQLY | 0.45031452 |
| KQGFLGM | ASSPLGAINTGQLY | 0.7707305 |
| LESVITQL | ASSPLGAINTGQLY | 0.8221642 |
| LGLGALVVV | ASSPLGAINTGQLY | 0.39873466 |
| LIGNGGNGGSGGI | ASSPLGAINTGQLY | 0.7787458 |
| LNAINAPTL | ASSPLGAINTGQLY | 0.84238255 |
| LQLLNTGV | ASSPLGAINTGQLY | 0.7553639 |
| LQTSIEF | ASSPLGAINTGQLY | 0.8392679 |
| MSFAVSEL | ASSPLGAINTGQLY | 0.84801304 |
| NTGALGL | ASSPLGAINTGQLY | 0.69685197 |
| PPGAGGAGGAGTT | ASSPLGAINTGQLY | 0.83377004 |
| QLVGLGI | ASSPLGAINTGQLY | 0.8280977 |
| RFLEQVHQL | ASSPLGAINTGQLY | 0.674422 |
| RGVGYVLR | ASSPLGAINTGQLY | 0.75661755 |
| SATGVFAGVI | ASSPLGAINTGQLY | 0.49311528 |
| SLPAITLP | ASSPLGAINTGQLY | 0.7282568 |
| STALAFTV | ASSPLGAINTGQLY | 0.55865955 |
| SVLTVGLGK | ASSPLGAINTGQLY | 0.75553274 |
| TALAGAGSGI | ASSPLGAINTGQLY | 0.83901393 |
| TGLPFTGI | ASSPLGAINTGQLY | 0.7094662 |
| VAEHVTV | ASSPLGAINTGQLY | 0.44126803 |
| VDIVLNSL | ASSPLGAINTGQLY | 0.44944376 |
| VLTGGGAVL | ASSPLGAINTGQLY | 0.82566714 |

| | | |
|---|---|---|
| VNVGLAYGI | ASSPLGAINTGQLY | 0.4450159 |
| VSSAAVGAV | ASSPLGAINTGQLY | 0.49760747 |
| VTSVYEAGV | ASSPLGAINTGQLY | 0.70515215 |
| WVALDRGA | ASSPLGAINTGQLY | 0.7937094 |
| YLVGSVVI | ASSPLGAINTGQLY | 0.8190457 |
| GPPGTGKTLL | ASSPLGAINTGQLY | 0.5464317 |
| GVGMTKF | ASSPLGAINTGQLY | 0.8414767 |
| IGIGGSDL | ASSPLGAINTGQLY | 0.8012329 |
| IGTLLAEL | ASSPLGAINTGQLY | 0.8487039 |
| IYTSGTT | ASSPLGAINTGQLY | 0.744959 |
| LSFPAGV | ASSPLGAINTGQLY | 0.5165717 |
| LYTSGTT | ASSPLGAINTGQLY | 0.83465075 |
| VGAFPFGF | ASSPLGAINTGQLY | 0.6353396 |
| VGAGTVEF | ASSPLGAINTGQLY | 0.75327027 |
| VGKSTLLL | ASSPLGAINTGQLY | 0.8045336 |
| AALAHAGGGIV | ASSPLGAINTGQLY | 0.5692094 |
| AALVAVSLL | ASSPLGAINTGQLY | 0.5550811 |
| AAVSRAVAAL | ASSPLGAINTGQLY | 0.72031593 |
| AGGSSGIGGF | ASSPLGAINTGQLY | 0.85487986 |
| AGLTVANANG | ASSPLGAINTGQLY | 0.51225924 |
| AGLVGAVL | ASSPLGAINTGQLY | 0.489622 |
| ALSQAFTV | ASSPLGAINTGQLY | 0.6776973 |
| ALTFSVSL | ASSPLGAINTGQLY | 0.64158475 |
| ALTGAALVAL | ASSPLGAINTGQLY | 0.82788527 |
| ASFSIGAGVL | ASSPLGAINTGQLY | 0.78614885 |
| DGSVYKI | ASSPLGAINTGQLY | 0.72428584 |
| EQLAEIV | ASSPLGAINTGQLY | 0.78513575 |
| FAVSELTVM | ASSPLGAINTGQLY | 0.8019683 |
| GAAPLAAGL | ASSPLGAINTGQLY | 0.765107 |
| GADGGDGGSGGAGG | ASSPLGAINTGQLY | 0.71931875 |
| GFGLDTGGAGG | ASSPLGAINTGQLY | 0.64121974 |
| GLASGGLL | ASSPLGAINTGQLY | 0.8406743 |
| GRIAGAAVLL | ASSPLGAINTGQLY | 0.80848277 |
| GSAGSLNTSF | ASSPLGAINTGQLY | 0.8170204 |
| GTGQIGI | ASSPLGAINTGQLY | 0.8267439 |
| GTLIALIL | ASSPLGAINTGQLY | 0.7351036 |
| IAATMGFVVG | ASSPLGAINTGQLY | 0.78922665 |
| IFAQALL | ASSPLGAINTGQLY | 0.39994037 |
| IGAAVALGF | ASSPLGAINTGQLY | 0.63616884 |
| IGLTPLAL | ASSPLGAINTGQLY | 0.7433319 |
| ILGSNTSTL | ASSPLGAINTGQLY | 0.7458404 |
| IVFSLTVL | ASSPLGAINTGQLY | 0.80350447 |
| LASKATGF | ASSPLGAINTGQLY | 0.8042332 |
| LASVALIL | ASSPLGAINTGQLY | 0.65222615 |
| LFAVGLTL | ASSPLGAINTGQLY | 0.7945262 |
| LFNGGIGTYI | ASSPLGAINTGQLY | 0.84249353 |
| LTLGAGII | ASSPLGAINTGQLY | 0.5571816 |
| LVAAALGDI | ASSPLGAINTGQLY | 0.7478628 |
| NTAFVQLGI | ASSPLGAINTGQLY | 0.86342114 |
| QLFSGAEL | ASSPLGAINTGQLY | 0.79166806 |
| SAATFITL | ASSPLGAINTGQLY | 0.66421163 |

| | | |
|---|---|---|
| SNYGLANLGV | ASSPLGAINTGQLY | 0.7608744 |
| SYGGAYAVM | ASSPLGAINTGQLY | 0.84354305 |
| TTGVGGLTLG | ASSPLGAINTGQLY | 0.70499074 |
| TTQTIYI | ASSPLGAINTGQLY | 0.8446429 |
| VAAGAGLGAVI | ASSPLGAINTGQLY | 0.36051857 |
| VAAVVTLL | ASSPLGAINTGQLY | 0.77705204 |
| VGFLGGLI | ASSPLGAINTGQLY | 0.6514628 |
| VGVIPILL | ASSPLGAINTGQLY | 0.24408329 |
| VIFASSLI | ASSPLGAINTGQLY | 0.8310598 |
| VKGGDGGAAAG | ASSPLGAINTGQLY | 0.7001356 |
| VTKTFTNI | ASSPLGAINTGQLY | 0.78735906 |
| VVAAAILL | ASSPLGAINTGQLY | 0.69461393 |
| YAALDVEL | ASSPLGAINTGQLY | 0.8342113 |
| LASVALLL | ASSPLGAINTGQLY | 0.79009795 |
| LDAVHRV | ASSPLGAINTGQLY | 0.68674207 |
| QGLSSSIF | ASSPLGAINTGQLY | 0.6340764 |
| RGTFANIRL | ASSPLGAINTGQLY | 0.7358265 |
| RQLWWGHRI | ASSPLGAINTGQLY | 0.8100829 |
| YGPPGTGKTLL | ASSPLGAINTGQLY | 0.8505654 |
| AALVGAPANVA | ASSPLGAINTGQLY | 0.2510799 |
| AATFRRTE | ASSPLGAINTGQLY | 0.8029951 |
| AGALGGGI | ASSPLGAINTGQLY | 0.70878464 |
| AGGAGGAGGIGAG | ASSPLGAINTGQLY | 0.31983978 |
| AGSGALGL | ASSPLGAINTGQLY | 0.2706247 |
| AQIADQI | ASSPLGAINTGQLY | 0.6278111 |
| ASAVAVTYL | ASSPLGAINTGQLY | 0.74126333 |
| ASTVLGGTGG | ASSPLGAINTGQLY | 0.7604157 |
| AVGGEPAK | ASSPLGAINTGQLY | 0.7774025 |
| CEQGTFGGTC | ASSPLGAINTGQLY | 0.66900945 |
| DEGKAPAG | ASSPLGAINTGQLY | 0.7999761 |
| DTVVFPAL | ASSPLGAINTGQLY | 0.6686694 |
| FEGITMAG | ASSPLGAINTGQLY | 0.12418563 |
| FPLVVQD | ASSPLGAINTGQLY | 0.6479664 |
| GAFTTPQ | ASSPLGAINTGQLY | 0.48525208 |
| GFGGFGTAGGVGGA | ASSPLGAINTGQLY | 0.8387598 |
| GFLAQFL | ASSPLGAINTGQLY | 0.63566744 |
| GGAGGAGGNAGVA | ASSPLGAINTGQLY | 0.68097925 |
| GGVGTAGAGGAGGAGG | ASSPLGAINTGQLY | 0.7371695 |
| GNGGGGDAGNAGSGGN | ASSPLGAINTGQLY | 0.7854738 |
| GTGGAGGAGNDAGS | ASSPLGAINTGQLY | 0.72219 |
| GTTNTGWFNS | ASSPLGAINTGQLY | 0.5964794 |
| IFMGPLFTL | ASSPLGAINTGQLY | 0.32614788 |
| ILAHAIEAI | ASSPLGAINTGQLY | 0.75498635 |
| IRSAIREAL | ASSPLGAINTGQLY | 0.65217745 |
| ITFAGGG | ASSPLGAINTGQLY | 0.43965822 |
| ITGMSGHQEV | ASSPLGAINTGQLY | 0.55792665 |
| IVAGLACTL | ASSPLGAINTGQLY | 0.7624935 |
| IVGGGLAGA | ASSPLGAINTGQLY | 0.39389992 |
| IVTGVLNI | ASSPLGAINTGQLY | 0.83883905 |
| LAQQGCI | ASSPLGAINTGQLY | 0.48804122 |
| LCLHVEN | ASSPLGAINTGQLY | 0.595358 |

| | | |
|---|---|---|
| LDAISDGS | ASSPLGAINTGQLY | 0.84477794 |
| LGGFAAML | ASSPLGAINTGQLY | 0.7414174 |
| LGGKNPMI | ASSPLGAINTGQLY | 0.70070875 |
| LGPKGVAAIL | ASSPLGAINTGQLY | 0.6119025 |
| LMVGVSI | ASSPLGAINTGQLY | 0.64954567 |
| LVGAAFVL | ASSPLGAINTGQLY | 0.59007436 |
| LVLAVAPL | ASSPLGAINTGQLY | 0.5661031 |
| MAAIAVIDAA | ASSPLGAINTGQLY | 0.8209359 |
| NNGINAELQ | ASSPLGAINTGQLY | 0.8213949 |
| RGEPLVLVHG | ASSPLGAINTGQLY | 0.60351 |
| RGGAGGAAGLI | ASSPLGAINTGQLY | 0.85883754 |
| RSYRMHPAV | ASSPLGAINTGQLY | 0.8476781 |
| SAAHLLAGQLA | ASSPLGAINTGQLY | 0.85815144 |
| SAAVGSY | ASSPLGAINTGQLY | 0.55339545 |
| SPHAVLHRI | ASSPLGAINTGQLY | 0.7107374 |
| SQIAKVL | ASSPLGAINTGQLY | 0.7364011 |
| TALGHLVP | ASSPLGAINTGQLY | 0.687303 |
| TAPTTSGAA | ASSPLGAINTGQLY | 0.5339575 |
| TDGDDGNGGSPGT | ASSPLGAINTGQLY | 0.5801104 |
| TDGLGEN | ASSPLGAINTGQLY | 0.8376331 |
| TGGPPLPGQA | ASSPLGAINTGQLY | 0.8648473 |
| TIPAITI | ASSPLGAINTGQLY | 0.73552597 |
| TIPGFTI | ASSPLGAINTGQLY | 0.40054715 |
| TSIASGALA | ASSPLGAINTGQLY | 0.65375936 |
| TVTLPGAAI | ASSPLGAINTGQLY | 0.55587864 |
| VASGLSAALP | ASSPLGAINTGQLY | 0.6376399 |
| VGAGVIGI | ASSPLGAINTGQLY | 0.68482053 |
| VGPKRFGV | ASSPLGAINTGQLY | 0.4988951 |
| VNVAFPD | ASSPLGAINTGQLY | 0.7442279 |
| VSMTNSGVS | ASSPLGAINTGQLY | 0.8388971 |
| VTDVLGDLSV | ASSPLGAINTGQLY | 0.70425576 |
| VVVVGAGFAGL | ASSPLGAINTGQLY | 0.782899 |
| VYTVGTVV | ASSPLGAINTGQLY | 0.84311765 |
| WTTFSGP | ASSPLGAINTGQLY | 0.8030336 |
| YDELVEAF | ASSPLGAINTGQLY | 0.84861994 |
| YGTGGAGGNGG | ASSPLGAINTGQLY | 0.8149116 |
| YTNVMAV | ASSPLGAINTGQLY | 0.74252397 |
| FLEQVHQL | ASSPLGAINTGQLY | 0.7175708 |
| GLSLLAVAV | ASSPLGAINTGQLY | 0.83583814 |
| LEQVHQL | ASSPLGAINTGQLY | 0.704394 |
| LNAAGAI | ASSPLGAINTGQLY | 0.7484267 |
| PAVHASL | ASSPLGAINTGQLY | 0.77666444 |
| TELEDDA | ASSPLGAINTGQLY | 0.80521566 |
| TPPGPPN | ASSPLGAINTGQLY | 0.6866249 |
| TPPPPPPP | ASSPLGAINTGQLY | 0.85356486 |
| TQSPALI | ASSQDLGQGASAETLY | 0.23735228 |
| EFAAAFPQ | ASSQDLGQGASAETLY | 0.45280433 |
| LLSGNLDVL | ASSQDLGQGASAETLY | 0.8347217 |
| GPANLGNYNI | ASSQDLGQGASAETLY | 0.1429896 |
| TLGPTLP | ASSQDLGQGASAETLY | 0.42269006 |
| VYAYSSCCK | ASSQDLGQGASAETLY | 0.77987576 |

| | | |
|---|---|---|
| APAPAPAPNI | ASSQDLGQGASAETLY | 0.8249874 |
| PGGPVPIP | ASSQDLGQGASAETLY | 0.18832941 |
| FLTVALEP | ASSQDLGQGASAETLY | 0.57213396 |
| SQYFGANL | ASSQDLGQGASAETLY | 0.3077054 |
| GGVKPAA | ASSQDLGQGASAETLY | 0.83325005 |
| GGTGGAGGAGGW | ASSQDLGQGASAETLY | 0.7272358 |
| GVGGVAGLGGAGG | ASSQDLGQGASAETLY | 0.36840475 |
| VLLVLATTI | ASSQDLGQGASAETLY | 0.8293818 |
| IGIFAAML | ASSQDLGQGASAETLY | 0.58403206 |
| AVGDLID | ASSQDLGQGASAETLY | 0.77580285 |
| NISPLPL | ASSQDLGQGASAETLY | 0.8339927 |
| VTAVFVAI | ASSQDLGQGASAETLY | 0.85549355 |
| SGDDNDAAL | ASSQDLGQGASAETLY | 0.5176155 |
| AVAALAHS | ASSQDLGQGASAETLY | 0.78676736 |
| GGVTVMATP | ASSQDLGQGASAETLY | 0.6814475 |
| AGLAANVRD | ASSQDLGQGASAETLY | 0.86282986 |
| TNRPDLIDPALLRPG | ASSQDLGQGASAETLY | 0.63029295 |
| LANLGVTAL | ASSQDLGQGASAETLY | 0.44037032 |
| TGERIEL | ASSQDLGQGASAETLY | 0.82563484 |
| NGEELAA | ASSQDLGQGASAETLY | 0.8253141 |
| VLSLVWG | ASSQDLGQGASAETLY | 0.6673677 |
| IGYALWL | ASSQDLGQGASAETLY | 0.8196734 |
| LSLLLAPL | ASSQDLGQGASAETLY | 0.7790904 |
| TLDEALK | ASSQDLGQGASAETLY | 0.72086895 |
| TDGAGTAEGG | ASSQDLGQGASAETLY | 0.82715786 |
| NAPGGAGGNGG | ASSQDLGQGASAETLY | 0.79749894 |
| VSVFMGMGGNF | ASSQDLGQGASAETLY | 0.60599023 |
| LASYLAAL | ASSQDLGQGASAETLY | 0.85068893 |
| LVGGTPAL | ASSQDLGQGASAETLY | 0.71836406 |
| VRNADVL | ASSQDLGQGASAETLY | 0.460288 |
| TCSSPFLVSAM | ASSQDLGQGASAETLY | 0.8641921 |
| VHITDAE | ASSQDLGQGASAETLY | 0.75927347 |
| EYAAPSF | ASSQDLGQGASAETLY | 0.820159 |
| VVDAFHGGL | ASSQDLGQGASAETLY | 0.8093852 |
| ASVVDPQ | ASSQDLGQGASAETLY | 0.8557433 |
| SYNEFGGLS | ASSQDLGQGASAETLY | 0.49844587 |
| DPDGSVY | ASSQDLGQGASAETLY | 0.7530571 |
| MVSTIHATE | ASSQDLGQGASAETLY | 0.7282897 |
| LAAGVADPV | ASSQDLGQGASAETLY | 0.8026278 |
| TKEQIAA | ASSQDLGQGASAETLY | 0.7629925 |
| SLASPAAL | ASSQDLGQGASAETLY | 0.64886534 |
| IGLVFYGV | ASSQDLGQGASAETLY | 0.8200698 |
| FLLDASNL | ASSQDLGQGASAETLY | 0.84385693 |
| YQVGNFP | ASSQDLGQGASAETLY | 0.71457595 |
| ESLLSQST | ASSQDLGQGASAETLY | 0.6947798 |
| LFTVGFAAVG | ASSQDLGQGASAETLY | 0.8602515 |
| ATPPTPT | ASSQDLGQGASAETLY | 0.5278338 |
| TDLATLG | ASSQDLGQGASAETLY | 0.7558433 |
| SAGGTGR | ASSQDLGQGASAETLY | 0.8257017 |
| GFSNSGS | ASSQDLGQGASAETLY | 0.8023384 |
| GGNGGSGGTSV | ASSQDLGQGASAETLY | 0.7825409 |

| | | |
|---|---|---|
| GTSVATGGAGN | ASSQDLGQGASAETLY | 0.84152794 |
| NGGNGGSGGTS | ASSQDLGQGASAETLY | 0.607178 |
| AVVKFPH | ASSQDLGQGASAETLY | 0.7872827 |
| TPPPPPP | ASSQDLGQGASAETLY | 0.7319837 |
| AAPAAAAAPA | ASSQDLGQGASAETLY | 0.6743545 |
| IALNEDL | ASSQDLGQGASAETLY | 0.48091108 |
| IVNVSSM | ASSQDLGQGASAETLY | 0.7774037 |
| TGIPPAPR | ASSQDLGQGASAETLY | 0.81461126 |
| FDLGGGTF | ASSQDLGQGASAETLY | 0.7707967 |
| FDLGGGTFDVS | ASSQDLGQGASAETLY | 0.3490414 |
| NEPTAAA | ASSQDLGQGASAETLY | 0.80127347 |
| NEPTAAALA | ASSQDLGQGASAETLY | 0.7885373 |
| GSIINISSI | ASSQDLGQGASAETLY | 0.8652421 |
| VAPGNAGTA | ASSQDLGQGASAETLY | 0.6860887 |
| TGGGASLE | ASSQDLGQGASAETLY | 0.82431936 |
| VAVYNHYKRI | ASSQDLGQGASAETLY | 0.62066483 |
| TDPAPATT | ASSQDLGQGASAETLY | 0.79971504 |
| TNRPDILDPALLRPG | ASSQDLGQGASAETLY | 0.42876762 |
| LTTTTTS | ASSQDLGQGASAETLY | 0.4619881 |
| PTGSGKTL | ASSQDLGQGASAETLY | 0.68491507 |
| TGSGKTL | ASSQDLGQGASAETLY | 0.5895108 |
| TIGHVDHGK | ASSQDLGQGASAETLY | 0.8060386 |
| TIGHVDHGKTT | ASSQDLGQGASAETLY | 0.8135238 |
| GHVDHGKTT | ASSQDLGQGASAETLY | 0.7727357 |
| AQFEHTLL | ASSQDLGQGASAETLY | 0.82606876 |
| IVNQDEL | ASSQDLGQGASAETLY | 0.8508234 |
| TNGSQFF | ASSQDLGQGASAETLY | 0.8349513 |
| TGKGIVE | ASSQDLGQGASAETLY | 0.67387974 |
| LISNASDA | ASSQDLGQGASAETLY | 0.71946347 |
| AELRTLGELTKIAKAHGAQ | ASSQDLGQGASAETLY | 0.40835232 |
| APNAVAAAEPSW | ASSQDLGQGASAETLY | 0.6952944 |
| APQVAGAATAVL | ASSQDLGQGASAETLY | 0.6907588 |
| ARRDPAAAAAILPTDA | ASSQDLGQGASAETLY | 0.75455177 |
| DAAARDVLAEAGLAEYF | ASSQDLGQGASAETLY | 0.14690998 |
| DDQIVLSAKANLAKVEQ | ASSQDLGQGASAETLY | 0.77291024 |
| DELLPDTGEDSDEDGH | ASSQDLGQGASAETLY | 0.82302153 |
| ELPEAPANRATVPQD | ASSQDLGQGASAETLY | 0.6128702 |
| FGQSKIERVTGIG | ASSQDLGQGASAETLY | 0.7817587 |
| GAGGTGSTAGGAGGAGGAGGLY | ASSQDLGQGASAETLY | 0.78703684 |
| GHGGTGAAGGNGGTGGAGGSGI | ASSQDLGQGASAETLY | 0.62467843 |
| GNGGNGGAGGEGGPGGAGGAGA | ASSQDLGQGASAETLY | 0.7704036 |
| GSTAGNGGNGGNATNGTIAGQP | ASSQDLGQGASAETLY | 0.6889125 |
| GTGGDGGNGGNGGTGG | ASSQDLGQGASAETLY | 0.83654356 |
| GVRRPAGIPADAPV | ASSQDLGQGASAETLY | 0.82801753 |
| GYRQVTDAHLLTLARR | ASSQDLGQGASAETLY | 0.79305315 |
| IAKMGQDGHDRG | ASSQDLGQGASAETLY | 0.6578878 |
| KAEPSIEIQYT | ASSQDLGQGASAETLY | 0.79054534 |
| KPVRIRMAADEDTVAVT | ASSQDLGQGASAETLY | 0.40576816 |
| MPPRIAGMRLLVIKPEP | ASSQDLGQGASAETLY | 0.64884984 |
| MKFYAVALFRSRGVPPTR | ASSQDLGQGASAETLY | 0.3310844 |
| NAGGFGILGAVAHSP | ASSQDLGQGASAETLY | 0.7581911 |

| | | |
|---|---|---|
| NEGREITLAMAEM | ASSQDLGQGASAETLY | 0.6726397 |
| NELVEWSHPLGHRWRWE | ASSQDLGQGASAETLY | 0.7400742 |
| NLGAGGTGGNGGNPG | ASSQDLGQGASAETLY | 0.69234455 |
| NTEWPKWETVMAADPVD | ASSQDLGQGASAETLY | 0.75086784 |
| PASAAPAETQP | ASSQDLGQGASAETLY | 0.61045194 |
| QQSSDGKNGVRAAEITGE | ASSQDLGQGASAETLY | 0.8113694 |
| RAPDAVTTSWWKE | ASSQDLGQGASAETLY | 0.7726609 |
| SGFGNVGAHTSGWFNQ | ASSQDLGQGASAETLY | 0.38923073 |
| SGVAGAVGQAASVGGLKV | ASSQDLGQGASAETLY | 0.83280504 |
| SNGGAGTGSAGGNGGT | ASSQDLGQGASAETLY | 0.7677498 |
| TGGAGGEGGAAG | ASSQDLGQGASAETLY | 0.74617326 |
| TGGDGGAAGAGGAPGHG | ASSQDLGQGASAETLY | 0.8152889 |
| TMPPLLAAIANAGRHGV | ASSQDLGQGASAETLY | 0.7622247 |
| TSSTAGNSGGA | ASSQDLGQGASAETLY | 0.5193132 |
| VPATVPVGILC | ASSQDLGQGASAETLY | 0.62007195 |
| VQRPDSGGVVLTGRIS | ASSQDLGQGASAETLY | 0.77682126 |
| WTPNPVPATKGVPAGLVQAT | ASSQDLGQGASAETLY | 0.662583 |
| YALKRALAKFSDRFDVVII | ASSQDLGQGASAETLY | 0.18917069 |
| YLIGSVIV | ASSQDLGQGASAETLY | 0.82949984 |
| YGQLVAVID | ASSQDLGQGASAETLY | 0.44341105 |
| VTVGGMTLPA | ASSQDLGQGASAETLY | 0.6428444 |
| VLALPAV | ASSQDLGQGASAETLY | 0.758596 |
| VIPPTLNYET | ASSQDLGQGASAETLY | 0.8228155 |
| VGMGGMHPSSG | ASSQDLGQGASAETLY | 0.58355206 |
| VGLAGGAATA | ASSQDLGQGASAETLY | 0.6799882 |
| VGGAGGAGGSGP | ASSQDLGQGASAETLY | 0.8026041 |
| VFAAFMV | ASSQDLGQGASAETLY | 0.7141283 |
| TLYDKLR | ASSQDLGQGASAETLY | 0.80745065 |
| TLAVFTVL | ASSQDLGQGASAETLY | 0.74299264 |
| TGSVQPARV | ASSQDLGQGASAETLY | 0.732217 |
| TGGGAGSGGT | ASSQDLGQGASAETLY | 0.7806628 |
| SHLETSL | ASSQDLGQGASAETLY | 0.5886687 |
| SFTTGMT | ASSQDLGQGASAETLY | 0.68286705 |
| QVEVAKRMGVRQPTVS | ASSQDLGQGASAETLY | 0.61005765 |
| QGSGGPALAM | ASSQDLGQGASAETLY | 0.6135806 |
| MLAPSNP | ASSQDLGQGASAETLY | 0.5579531 |
| LVNNAAI | ASSQDLGQGASAETLY | 0.7680191 |
| LTGEINTLA | ASSQDLGQGASAETLY | 0.86442274 |
| LSIFGPV | ASSQDLGQGASAETLY | 0.7752938 |
| LLALLANL | ASSQDLGQGASAETLY | 0.36600673 |
| LISNASDALDKLR | ASSQDLGQGASAETLY | 0.6298845 |
| LINEAVP | ASSQDLGQGASAETLY | 0.72211504 |
| LGYAFSGL | ASSQDLGQGASAETLY | 0.80144596 |
| LDFVAVAL | ASSQDLGQGASAETLY | 0.8381193 |
| LAQPGAE | ASSQDLGQGASAETLY | 0.8502413 |
| ISNLGAVTGF | ASSQDLGQGASAETLY | 0.7993039 |
| HQEDPGRI | ASSQDLGQGASAETLY | 0.63998497 |
| GTPGEPTP | ASSQDLGQGASAETLY | 0.8320391 |
| GFFNSGS | ASSQDLGQGASAETLY | 0.7155732 |
| GAGGGGLTAGAN | ASSQDLGQGASAETLY | 0.70736325 |
| FVDALVNM | ASSQDLGQGASAETLY | 0.766178 |

| | | |
|---|---|---|
| FLTVAPEL | ASSQDLGQGASAETLY | 0.810536 |
| FETLTAL | ASSQDLGQGASAETLY | 0.66203773 |
| DVINPIEL | ASSQDLGQGASAETLY | 0.6961593 |
| APVSGAPV | ASSQDLGQGASAETLY | 0.5040543 |
| ALVGGLAW | ASSQDLGQGASAETLY | 0.81669223 |
| ALAAAAATAI | ASSQDLGQGASAETLY | 0.34624112 |
| AFGLGLLL | ASSQDLGQGASAETLY | 0.59314156 |
| AFFAAASL | ASSQDLGQGASAETLY | 0.6424389 |
| AAAALPAFSPP | ASSQDLGQGASAETLY | 0.8486776 |
| VVLAPAL | ASSQDLGQGASAETLY | 0.8282736 |
| TIGHVDHGKTTL | ASSQDLGQGASAETLY | 0.6606454 |
| LISNASDALDK | ASSQDLGQGASAETLY | 0.64502865 |
| LISNASDAL | ASSQDLGQGASAETLY | 0.7834997 |
| ISNASDALDK | ASSQDLGQGASAETLY | 0.5797801 |
| NQAETLVYQTE | ASSQDLGQGASAETLY | 0.8410578 |
| TGGTGGTAGAG | ASSQDLGQGASAETLY | 0.7181119 |
| TGSTAGGAGGA | ASSQDLGQGASAETLY | 0.7295097 |
| DPLAVPVAAGPV | ASSQDLGQGASAETLY | 0.6779356 |
| GGTGGTPGNGGN | ASSQDLGQGASAETLY | 0.7736877 |
| PAGLAGVDPATL | ASSQDLGQGASAETLY | 0.6886903 |
| GNNLVVEAPHAGASV | ASSQDLGQGASAETLY | 0.8054544 |
| KVTRIMSASTLEQPA | ASSQDLGQGASAETLY | 0.33113465 |
| LGFALPIALPRRTVV | ASSQDLGQGASAETLY | 0.834149 |
| TRLGNLTARKLVVLV | ASSQDLGQGASAETLY | 0.47464395 |
| WSQQRRLIYFPSAGP | ASSQDLGQGASAETLY | 0.5933826 |
| GPNGNPDPMAAAVDIR | ASSQDLGQGASAETLY | 0.2837981 |
| TQRCVSVADLRAAASR | ASSQDLGQGASAETLY | 0.70020604 |
| QFVLDVAEHPVEVLTETV | ASSQDLGQGASAETLY | 0.8015939 |
| GGAGGTGGAGGNGGTQAGDG | ASSQDLGQGASAETLY | 0.82867 |
| PLLATLRAIAVDVDAKPAQV | ASSQDLGQGASAETLY | 0.7561729 |
| APKFRTGGYMIDFWGVGYQVAK | ASSQDLGQGASAETLY | 0.79008305 |
| GAGGAGGAGGAAGGPGGTGGTG | ASSQDLGQGASAETLY | 0.8066338 |
| IGALGLLATDVAPAGSVAFGVA | ASSQDLGQGASAETLY | 0.6584208 |
| AGGAGGIGGNGGAGGAGGVLMG | ASSQDLGQGASAETLY | 0.2587497 |
| ADELIGLL | ASSQDLGQGASAETLY | 0.7863158 |
| ADRALKA | ASSQDLGQGASAETLY | 0.7400798 |
| AGAVGQVGGAGGA | ASSQDLGQGASAETLY | 0.8380674 |
| AGGGGGLGGAP | ASSQDLGQGASAETLY | 0.79717624 |
| AGGLLYGNGGAG | ASSQDLGQGASAETLY | 0.48578689 |
| AGLESIGL | ASSQDLGQGASAETLY | 0.79937685 |
| AGLKTGII | ASSQDLGQGASAETLY | 0.75986445 |
| ASLAALLL | ASSQDLGQGASAETLY | 0.42489713 |
| ASVINISSI | ASSQDLGQGASAETLY | 0.63618946 |
| ATPPGPA | ASSQDLGQGASAETLY | 0.64480716 |
| ATVASALV | ASSQDLGQGASAETLY | 0.5409602 |
| ATVASLIL | ASSQDLGQGASAETLY | 0.7552384 |
| AVNGPVLIRH | ASSQDLGQGASAETLY | 0.65005314 |
| DGAAGAVGI | ASSQDLGQGASAETLY | 0.5889413 |
| DPLLHGI | ASSQDLGQGASAETLY | 0.7384007 |
| ELGAALGAL | ASSQDLGQGASAETLY | 0.57754153 |
| ELGRTQAV | ASSQDLGQGASAETLY | 0.60663414 |

| | | |
|---|---|---|
| FVAGGYLS | ASSQDLGQGASAETLY | 0.5890694 |
| GASGGAGGAGGW | ASSQDLGQGASAETLY | 0.8145541 |
| GGAGAGGDGGAGGI | ASSQDLGQGASAETLY | 0.79851115 |
| GTGHNGSQPGMGGNG | ASSQDLGQGASAETLY | 0.7282579 |
| GYAAASASAS | ASSQDLGQGASAETLY | 0.80125254 |
| HQVDVAAVQQG | ASSQDLGQGASAETLY | 0.82315207 |
| IGAAIAHT | ASSQDLGQGASAETLY | 0.3454042 |
| IGMIEVAQL | ASSQDLGQGASAETLY | 0.5161205 |
| IRLAPEHTSAMD | ASSQDLGQGASAETLY | 0.8092837 |
| IYNVSVL | ASSQDLGQGASAETLY | 0.69501144 |
| KQGFLGM | ASSQDLGQGASAETLY | 0.83804107 |
| LESVITQL | ASSQDLGQGASAETLY | 0.67641556 |
| LGLGALVVV | ASSQDLGQGASAETLY | 0.76899415 |
| LIGNGGNGGSGGI | ASSQDLGQGASAETLY | 0.7170626 |
| LNAINAPTL | ASSQDLGQGASAETLY | 0.8574432 |
| LQLLNTGV | ASSQDLGQGASAETLY | 0.7993486 |
| LQTSIEF | ASSQDLGQGASAETLY | 0.7321116 |
| MSFAVSEL | ASSQDLGQGASAETLY | 0.79703844 |
| NTGALGL | ASSQDLGQGASAETLY | 0.4179042 |
| PPGAGGAGGAGTT | ASSQDLGQGASAETLY | 0.8010647 |
| QLVGLGI | ASSQDLGQGASAETLY | 0.82966 |
| RFLEQVHQL | ASSQDLGQGASAETLY | 0.55625206 |
| RGVGYVLR | ASSQDLGQGASAETLY | 0.71632564 |
| SATGVFAGVI | ASSQDLGQGASAETLY | 0.83844435 |
| SLPAITLP | ASSQDLGQGASAETLY | 0.8412007 |
| STALAFTV | ASSQDLGQGASAETLY | 0.63803554 |
| SVLTVGLGK | ASSQDLGQGASAETLY | 0.65135753 |
| TALAGAGSGI | ASSQDLGQGASAETLY | 0.37852618 |
| TGLPFTGI | ASSQDLGQGASAETLY | 0.5775629 |
| VAEHVTV | ASSQDLGQGASAETLY | 0.33385178 |
| VDIVLNSL | ASSQDLGQGASAETLY | 0.78451973 |
| VLTGGGAVL | ASSQDLGQGASAETLY | 0.6795705 |
| VNVGLAYGI | ASSQDLGQGASAETLY | 0.8254049 |
| VSSAAVGAV | ASSQDLGQGASAETLY | 0.7778063 |
| VTSVYEAGV | ASSQDLGQGASAETLY | 0.8270284 |
| WVALDRGA | ASSQDLGQGASAETLY | 0.7174245 |
| YLVGSVVI | ASSQDLGQGASAETLY | 0.64637387 |
| GPPGTGKTLL | ASSQDLGQGASAETLY | 0.46190277 |
| GVGMTKF | ASSQDLGQGASAETLY | 0.32221758 |
| IGIGGSDL | ASSQDLGQGASAETLY | 0.5919859 |
| IGTLLAEL | ASSQDLGQGASAETLY | 0.5817312 |
| IYTSGTT | ASSQDLGQGASAETLY | 0.29976636 |
| LSFPAGV | ASSQDLGQGASAETLY | 0.86023724 |
| LYTSGTT | ASSQDLGQGASAETLY | 0.7217326 |
| VGAFPFGF | ASSQDLGQGASAETLY | 0.78551173 |
| VGAGTVEF | ASSQDLGQGASAETLY | 0.65709555 |
| VGKSTLLL | ASSQDLGQGASAETLY | 0.47631767 |
| AALAHAGGGIV | ASSQDLGQGASAETLY | 0.69799566 |
| AALVAVSLL | ASSQDLGQGASAETLY | 0.7925719 |
| AAVSRAVAAL | ASSQDLGQGASAETLY | 0.5740734 |
| AGGSSGIGGF | ASSQDLGQGASAETLY | 0.4135629 |

| | | |
|---|---|---|
| AGLTVANANG | ASSQDLGQGASAETLY | 0.53754115 |
| AGLVGAVL | ASSQDLGQGASAETLY | 0.6448245 |
| ALSQAFTV | ASSQDLGQGASAETLY | 0.8617779 |
| ALTFSVSL | ASSQDLGQGASAETLY | 0.83297384 |
| ALTGAALVAL | ASSQDLGQGASAETLY | 0.78170264 |
| ASFSIGAGVL | ASSQDLGQGASAETLY | 0.87143 |
| DGSVYKI | ASSQDLGQGASAETLY | 0.18115255 |
| EQLAEIV | ASSQDLGQGASAETLY | 0.688501 |
| FAVSELTVM | ASSQDLGQGASAETLY | 0.8384715 |
| GAAPLAAGL | ASSQDLGQGASAETLY | 0.806986 |
| GADGGDGGSGGAGG | ASSQDLGQGASAETLY | 0.6773512 |
| GFGLDTGGAGG | ASSQDLGQGASAETLY | 0.8417635 |
| GLASGGLL | ASSQDLGQGASAETLY | 0.5669253 |
| GRIAGAAVLL | ASSQDLGQGASAETLY | 0.66788185 |
| GSAGSLNTSF | ASSQDLGQGASAETLY | 0.7392374 |
| GTGQIGI | ASSQDLGQGASAETLY | 0.8172035 |
| GTLIALIL | ASSQDLGQGASAETLY | 0.7947222 |
| IAATMGFVVG | ASSQDLGQGASAETLY | 0.40891284 |
| IFAQALL | ASSQDLGQGASAETLY | 0.81762284 |
| IGAAVALGF | ASSQDLGQGASAETLY | 0.85377634 |
| IGLTPLAL | ASSQDLGQGASAETLY | 0.66245383 |
| ILGSNTSTL | ASSQDLGQGASAETLY | 0.8152149 |
| IVFSLTVL | ASSQDLGQGASAETLY | 0.7609085 |
| LASKATGF | ASSQDLGQGASAETLY | 0.5516962 |
| LASVALIL | ASSQDLGQGASAETLY | 0.6657034 |
| LFAVGLTL | ASSQDLGQGASAETLY | 0.4448989 |
| LFNGGIGTYI | ASSQDLGQGASAETLY | 0.8070178 |
| LTLGAGII | ASSQDLGQGASAETLY | 0.5377425 |
| LVAAALGDI | ASSQDLGQGASAETLY | 0.76493806 |
| NTAFVQLGI | ASSQDLGQGASAETLY | 0.7650109 |
| QLFSGAEL | ASSQDLGQGASAETLY | 0.86336446 |
| SAATFITL | ASSQDLGQGASAETLY | 0.7980577 |
| SNYGLANLGV | ASSQDLGQGASAETLY | 0.82009155 |
| SYGGAYAVM | ASSQDLGQGASAETLY | 0.56520593 |
| TTGVGGLTLG | ASSQDLGQGASAETLY | 0.77843326 |
| TTQTIYI | ASSQDLGQGASAETLY | 0.7845825 |
| VAAGAGLGAVI | ASSQDLGQGASAETLY | 0.50349194 |
| VAAVVTLL | ASSQDLGQGASAETLY | 0.8208749 |
| VGFLGGLI | ASSQDLGQGASAETLY | 0.48473227 |
| VGVIPILL | ASSQDLGQGASAETLY | 0.7882438 |
| VIFASSLI | ASSQDLGQGASAETLY | 0.76763934 |
| VKGGDGGAAAG | ASSQDLGQGASAETLY | 0.82662463 |
| VTKTFTNI | ASSQDLGQGASAETLY | 0.7719432 |
| VVAAAILL | ASSQDLGQGASAETLY | 0.7709841 |
| YAALDVEL | ASSQDLGQGASAETLY | 0.554788 |
| LASVALLL | ASSQDLGQGASAETLY | 0.8236163 |
| LDAVHRV | ASSQDLGQGASAETLY | 0.8381761 |
| QGLSSSIF | ASSQDLGQGASAETLY | 0.7474929 |
| RGTFANIRL | ASSQDLGQGASAETLY | 0.85991967 |
| RQLWWGHRI | ASSQDLGQGASAETLY | 0.8169282 |
| YGPPGTGKTLL | ASSQDLGQGASAETLY | 0.30262077 |

| | | |
|---|---|---|
| AALVGAPANVA | ASSQDLGQGASAETLY | 0.6859472 |
| AATFRRTE | ASSQDLGQGASAETLY | 0.8359457 |
| AGALGGGI | ASSQDLGQGASAETLY | 0.8201796 |
| AGGAGGAGGIGAG | ASSQDLGQGASAETLY | 0.80899316 |
| AGSGALGL | ASSQDLGQGASAETLY | 0.81264114 |
| AQIADQI | ASSQDLGQGASAETLY | 0.675434 |
| ASAVAVTYL | ASSQDLGQGASAETLY | 0.84023833 |
| ASTVLGGTGG | ASSQDLGQGASAETLY | 0.8757081 |
| AVGGEPAK | ASSQDLGQGASAETLY | 0.83273435 |
| CEQGTFGGTC | ASSQDLGQGASAETLY | 0.5662216 |
| DEGKAPAG | ASSQDLGQGASAETLY | 0.51184857 |
| DTVVFPAL | ASSQDLGQGASAETLY | 0.8275627 |
| FEGITMAG | ASSQDLGQGASAETLY | 0.84550947 |
| FPLVVQD | ASSQDLGQGASAETLY | 0.59285724 |
| GAFTTPQ | ASSQDLGQGASAETLY | 0.71650296 |
| GFGGFGTAGGVGGA | ASSQDLGQGASAETLY | 0.6545614 |
| GFLAQFL | ASSQDLGQGASAETLY | 0.24138556 |
| GGAGGAGGNAGVA | ASSQDLGQGASAETLY | 0.82601297 |
| GGVGTAGAGGAGGAGG | ASSQDLGQGASAETLY | 0.8406795 |
| GNGGGGDAGNAGSGGN | ASSQDLGQGASAETLY | 0.82987463 |
| GTGGAGGAGNDAGS | ASSQDLGQGASAETLY | 0.3967908 |
| GTTNTGWFNS | ASSQDLGQGASAETLY | 0.8270074 |
| IFMGPLFTL | ASSQDLGQGASAETLY | 0.8206469 |
| ILAHAIEAI | ASSQDLGQGASAETLY | 0.8620477 |
| IRSAIREAL | ASSQDLGQGASAETLY | 0.3549534 |
| ITFAGGG | ASSQDLGQGASAETLY | 0.20276088 |
| ITGMSGHQEV | ASSQDLGQGASAETLY | 0.72192 |
| IVAGLACTL | ASSQDLGQGASAETLY | 0.79516166 |
| IVGGGLAGA | ASSQDLGQGASAETLY | 0.62156284 |
| IVTGVLNI | ASSQDLGQGASAETLY | 0.624718 |
| LAQQGCI | ASSQDLGQGASAETLY | 0.71639276 |
| LCLHVEN | ASSQDLGQGASAETLY | 0.8277781 |
| LDAISDGS | ASSQDLGQGASAETLY | 0.50105906 |
| LGGFAAML | ASSQDLGQGASAETLY | 0.43207777 |
| LGGKNPMI | ASSQDLGQGASAETLY | 0.773476 |
| LGPKGVAAIL | ASSQDLGQGASAETLY | 0.8494533 |
| LMVGVSI | ASSQDLGQGASAETLY | 0.5523999 |
| LVGAAFVL | ASSQDLGQGASAETLY | 0.69476825 |
| LVLAVAPL | ASSQDLGQGASAETLY | 0.71245563 |
| MAAIAVIDAA | ASSQDLGQGASAETLY | 0.7981488 |
| NNGINAELQ | ASSQDLGQGASAETLY | 0.7147465 |
| RGEPLVLVHG | ASSQDLGQGASAETLY | 0.4662078 |
| RGGAGGAAGLI | ASSQDLGQGASAETLY | 0.8327831 |
| RSYRMHPAV | ASSQDLGQGASAETLY | 0.5572256 |
| SAAHLLAGQLA | ASSQDLGQGASAETLY | 0.81973857 |
| SAAVGSY | ASSQDLGQGASAETLY | 0.70138794 |
| SPHAVLHRI | ASSQDLGQGASAETLY | 0.78921837 |
| SQIAKVL | ASSQDLGQGASAETLY | 0.6944243 |
| TALGHLVP | ASSQDLGQGASAETLY | 0.5142994 |
| TAPTTSGAA | ASSQDLGQGASAETLY | 0.5832354 |
| TDGDDGNGGSPGT | ASSQDLGQGASAETLY | 0.49586815 |

| | | |
|---|---|---|
| TDGLGEN | ASSQDLGQGASAETLY | 0.80764127 |
| TGGPPLPGQA | ASSQDLGQGASAETLY | 0.64053977 |
| TIPAITI | ASSQDLGQGASAETLY | 0.8361605 |
| TIPGFTI | ASSQDLGQGASAETLY | 0.7707933 |
| TSIASGALA | ASSQDLGQGASAETLY | 0.5751209 |
| TVTLPGAAI | ASSQDLGQGASAETLY | 0.7987187 |
| VASGLSAALP | ASSQDLGQGASAETLY | 0.42453682 |
| VGAGVIGI | ASSQDLGQGASAETLY | 0.7949285 |
| VGPKRFGV | ASSQDLGQGASAETLY | 0.72805625 |
| VNVAFPD | ASSQDLGQGASAETLY | 0.82496995 |
| VSMTNSGVS | ASSQDLGQGASAETLY | 0.6952638 |
| VTDVLGDLSV | ASSQDLGQGASAETLY | 0.8214487 |
| VVVVGAGFAGL | ASSQDLGQGASAETLY | 0.7891409 |
| VYTVGTVV | ASSQDLGQGASAETLY | 0.4777829 |
| WTTFSGP | ASSQDLGQGASAETLY | 0.8229413 |
| YDELVEAF | ASSQDLGQGASAETLY | 0.81887716 |
| YGTGGAGGNGG | ASSQDLGQGASAETLY | 0.82528275 |
| YTNVMAV | ASSQDLGQGASAETLY | 0.42365575 |
| FLEQVHQL | ASSQDLGQGASAETLY | 0.7634412 |
| GLSLLAVAV | ASSQDLGQGASAETLY | 0.62133765 |
| LEQVHQL | ASSQDLGQGASAETLY | 0.7796764 |
| LNAAGAI | ASSQDLGQGASAETLY | 0.7712462 |
| PAVHASL | ASSQDLGQGASAETLY | 0.6448696 |
| TELEDDA | ASSQDLGQGASAETLY | 0.80658185 |
| TPPGPPN | ASSQDLGQGASAETLY | 0.49069902 |
| TPPPPPPP | ASSQDLGQGASAETLY | 0.6458616 |
| TQSPALI | GARVGGASNTLY | 0.72356945 |
| EFAAAFPQ | GARVGGASNTLY | 0.74886954 |
| LLSGNLDVL | GARVGGASNTLY | 0.8244188 |
| GPANLGNYNI | GARVGGASNTLY | 0.52274203 |
| TLGPTLP | GARVGGASNTLY | 0.7608516 |
| VYAYSSCCK | GARVGGASNTLY | 0.7045402 |
| APAPAPAPNI | GARVGGASNTLY | 0.5243466 |
| PGGPVPIP | GARVGGASNTLY | 0.8272877 |
| FLTVALEP | GARVGGASNTLY | 0.68590975 |
| SQYFGANL | GARVGGASNTLY | 0.37379593 |
| GGVKPAA | GARVGGASNTLY | 0.8073845 |
| GGTGGAGGAGGW | GARVGGASNTLY | 0.5073984 |
| GVGGVAGLGGAGG | GARVGGASNTLY | 0.77076113 |
| VLLVLATTI | GARVGGASNTLY | 0.81749535 |
| IGIFAAML | GARVGGASNTLY | 0.36291468 |
| AVGDLID | GARVGGASNTLY | 0.808285 |
| NISPLPL | GARVGGASNTLY | 0.7868016 |
| VTAVFVAI | GARVGGASNTLY | 0.75506485 |
| SGDDNDAAL | GARVGGASNTLY | 0.5648959 |
| AVAALAHS | GARVGGASNTLY | 0.6204593 |
| GGVTVMATP | GARVGGASNTLY | 0.8520325 |
| AGLAANVRD | GARVGGASNTLY | 0.7901875 |
| TNRPDLIDPALLRPG | GARVGGASNTLY | 0.77302265 |
| LANLGVTAL | GARVGGASNTLY | 0.81345654 |
| TGERIEL | GARVGGASNTLY | 0.8108611 |

| | | |
|---|---|---|
| NGEELAA | GARVGGASNTLY | 0.8298555 |
| VLSLVWG | GARVGGASNTLY | 0.59193766 |
| IGYALWL | GARVGGASNTLY | 0.3586926 |
| LSLLLAPL | GARVGGASNTLY | 0.69561183 |
| TLDEALK | GARVGGASNTLY | 0.63895726 |
| TDGAGTAEGG | GARVGGASNTLY | 0.6516484 |
| NAPGGAGGNGG | GARVGGASNTLY | 0.3565767 |
| VSVFMGMGGNF | GARVGGASNTLY | 0.81460845 |
| LASYLAAL | GARVGGASNTLY | 0.8651192 |
| LVGGTPAL | GARVGGASNTLY | 0.44745952 |
| VRNADVL | GARVGGASNTLY | 0.8655796 |
| TCSSPFLVSAM | GARVGGASNTLY | 0.5773095 |
| VHITDAE | GARVGGASNTLY | 0.7862729 |
| EYAAPSF | GARVGGASNTLY | 0.8222902 |
| VVDAFHGGL | GARVGGASNTLY | 0.45298627 |
| ASVVDPQ | GARVGGASNTLY | 0.76104677 |
| SYNEFGGLS | GARVGGASNTLY | 0.6015361 |
| DPDGSVY | GARVGGASNTLY | 0.6401403 |
| MVSTIHATE | GARVGGASNTLY | 0.82920194 |
| LAAGVADPV | GARVGGASNTLY | 0.49262232 |
| TKEQIAA | GARVGGASNTLY | 0.761983 |
| SLASPAAL | GARVGGASNTLY | 0.70426726 |
| IGLVFYGV | GARVGGASNTLY | 0.55779046 |
| FLLDASNL | GARVGGASNTLY | 0.80014235 |
| YQVGNFP | GARVGGASNTLY | 0.8023979 |
| ESLLSQST | GARVGGASNTLY | 0.7688006 |
| LFTVGFAAVG | GARVGGASNTLY | 0.871645 |
| ATPPTPT | GARVGGASNTLY | 0.5916332 |
| TDLATLG | GARVGGASNTLY | 0.8655444 |
| SAGGTGR | GARVGGASNTLY | 0.857808 |
| GFSNSGS | GARVGGASNTLY | 0.7516537 |
| GGNGGSGGTSV | GARVGGASNTLY | 0.7251303 |
| GTSVATGGAGN | GARVGGASNTLY | 0.81519496 |
| NGGNGGSGGTS | GARVGGASNTLY | 0.6767983 |
| AVVKFPH | GARVGGASNTLY | 0.71594703 |
| TPPPPPP | GARVGGASNTLY | 0.8411515 |
| AAPAAAAAPA | GARVGGASNTLY | 0.42498708 |
| IALNEDL | GARVGGASNTLY | 0.77366376 |
| IVNVSSM | GARVGGASNTLY | 0.704346 |
| TGIPPAPR | GARVGGASNTLY | 0.6569551 |
| FDLGGGTF | GARVGGASNTLY | 0.57389283 |
| FDLGGGTFDVS | GARVGGASNTLY | 0.3957388 |
| NEPTAAA | GARVGGASNTLY | 0.5618666 |
| NEPTAAALA | GARVGGASNTLY | 0.47579306 |
| GSIINISSI | GARVGGASNTLY | 0.7599191 |
| VAPGNAGTA | GARVGGASNTLY | 0.75558895 |
| TGGGASLE | GARVGGASNTLY | 0.5434865 |
| VAVYNHYKRI | GARVGGASNTLY | 0.7737906 |
| TDPAPATT | GARVGGASNTLY | 0.6930444 |
| TNRPDILDPALLRPG | GARVGGASNTLY | 0.38342932 |
| LTTTTS | GARVGGASNTLY | 0.6337347 |

| | | |
|---|---|---|
| PTGSGKTL | GARVGGASNTLY | 0.5069999 |
| TGSGKTL | GARVGGASNTLY | 0.62330854 |
| TIGHVDHGK | GARVGGASNTLY | 0.29612458 |
| TIGHVDHGKTT | GARVGGASNTLY | 0.73275423 |
| GHVDHGKTT | GARVGGASNTLY | 0.5570826 |
| AQFEHTLL | GARVGGASNTLY | 0.26189166 |
| IVNQDEL | GARVGGASNTLY | 0.7284392 |
| TNGSQFF | GARVGGASNTLY | 0.6620015 |
| TGKGIVE | GARVGGASNTLY | 0.63597065 |
| LISNASDA | GARVGGASNTLY | 0.7674601 |
| AELRTLGELTKIAKAHGAQ | GARVGGASNTLY | 0.592263 |
| APNAVAAAEPSW | GARVGGASNTLY | 0.51511073 |
| APQVAGAATAVL | GARVGGASNTLY | 0.751083 |
| ARRDPAAAAAILPTDA | GARVGGASNTLY | 0.7739887 |
| DAAARDVLAEAGLAEYF | GARVGGASNTLY | 0.4597237 |
| DDQIVLSAKANLAKVEQ | GARVGGASNTLY | 0.70975155 |
| DELLPDTGEDSDEDGH | GARVGGASNTLY | 0.75932586 |
| ELPEAPANRATVPQD | GARVGGASNTLY | 0.7215054 |
| FGQSKIERVTGIG | GARVGGASNTLY | 0.7693342 |
| GAGGTGSTAGGAGGAGGAGGLY | GARVGGASNTLY | 0.7774974 |
| GHGGTGAAGGNGGTGGAGGSGI | GARVGGASNTLY | 0.5273832 |
| GNGGNGGAGGEGGPGGAGGAGA | GARVGGASNTLY | 0.77294433 |
| GSTAGNGGNGGNATNGTIAGQP | GARVGGASNTLY | 0.7766673 |
| GTGGDGGNGGNGGTGG | GARVGGASNTLY | 0.61274016 |
| GVRRPAGIPADAPV | GARVGGASNTLY | 0.66646457 |
| GYRQVTDAHLLTLARR | GARVGGASNTLY | 0.40250188 |
| IAKMGQDGHDRG | GARVGGASNTLY | 0.63464797 |
| KAEPSIEIQYT | GARVGGASNTLY | 0.46610653 |
| KPVRIRMAADEDTVAVT | GARVGGASNTLY | 0.7317006 |
| MPPRIAGMRLLVIKPEP | GARVGGASNTLY | 0.78029066 |
| MKFYAVALFRSRGVPPTR | GARVGGASNTLY | 0.6679375 |
| NAGGFGILGAVAHSP | GARVGGASNTLY | 0.42086706 |
| NEGREITLAMAEM | GARVGGASNTLY | 0.43197173 |
| NELVEWSHPLGHRWRWE | GARVGGASNTLY | 0.681172 |
| NLGAGGTGGNGGNPG | GARVGGASNTLY | 0.47919393 |
| NTEWPKWETVMAADPVD | GARVGGASNTLY | 0.35293746 |
| PASAAPAETQP | GARVGGASNTLY | 0.6214492 |
| QQSSDGKNGVRAAEITGE | GARVGGASNTLY | 0.76053965 |
| RAPDAVTTSWWKE | GARVGGASNTLY | 0.760418 |
| SGFGNVGAHTSGWFNQ | GARVGGASNTLY | 0.49594218 |
| SGVAGAVGQAASVGGLKV | GARVGGASNTLY | 0.7801337 |
| SNGGAGTGSAGGNGGT | GARVGGASNTLY | 0.7337595 |
| TGGAGGEGGAAG | GARVGGASNTLY | 0.7472667 |
| TGGDGGAAGAGGAPGHG | GARVGGASNTLY | 0.618335 |
| TMPPLLAAIANAGRHGV | GARVGGASNTLY | 0.4298191 |
| TSSTAGNSGGA | GARVGGASNTLY | 0.7621812 |
| VPATVPVGILC | GARVGGASNTLY | 0.6801295 |
| VQRPDSGGVVLTGRIS | GARVGGASNTLY | 0.7156935 |
| WTPNPVPATKGVPAGLVQAT | GARVGGASNTLY | 0.75215656 |
| YALKRALAKFSDRFDVVII | GARVGGASNTLY | 0.6659913 |
| YLIGSVIV | GARVGGASNTLY | 0.5791453 |

| | | |
|---|---|---|
| YGQLVAVID | GARVGGASNTLY | 0.6779276 |
| VTVGGMTLPA | GARVGGASNTLY | 0.7549362 |
| VLALPAV | GARVGGASNTLY | 0.53876394 |
| VIPPTLNYET | GARVGGASNTLY | 0.41971835 |
| VGMGGMHPSSG | GARVGGASNTLY | 0.6309256 |
| VGLAGGAATA | GARVGGASNTLY | 0.68062305 |
| VGGAGGAGGSGP | GARVGGASNTLY | 0.7709768 |
| VFAAFMV | GARVGGASNTLY | 0.69964755 |
| TLYDKLR | GARVGGASNTLY | 0.6004709 |
| TLAVFTVL | GARVGGASNTLY | 0.73930824 |
| TGSVQPARV | GARVGGASNTLY | 0.72013223 |
| TGGGAGSGGT | GARVGGASNTLY | 0.67495555 |
| SHLETSL | GARVGGASNTLY | 0.66683507 |
| SFTTGMT | GARVGGASNTLY | 0.5909877 |
| QVEVAKRMGVRQPTVS | GARVGGASNTLY | 0.7404089 |
| QGSGGPALAM | GARVGGASNTLY | 0.7402709 |
| MLAPSNP | GARVGGASNTLY | 0.7273693 |
| LVNNAAI | GARVGGASNTLY | 0.7613215 |
| LTGEINTLA | GARVGGASNTLY | 0.70378363 |
| LSIFGPV | GARVGGASNTLY | 0.67095697 |
| LLALLANL | GARVGGASNTLY | 0.38544363 |
| LISNASDALDKLR | GARVGGASNTLY | 0.5565282 |
| LINEAVP | GARVGGASNTLY | 0.7126961 |
| LGYAFSGL | GARVGGASNTLY | 0.72033364 |
| LDFVAVAL | GARVGGASNTLY | 0.760247 |
| LAQPGAE | GARVGGASNTLY | 0.67622054 |
| ISNLGAVTGF | GARVGGASNTLY | 0.6024294 |
| HQEDPGRI | GARVGGASNTLY | 0.7493721 |
| GTPGEPTP | GARVGGASNTLY | 0.7571792 |
| GFFNSGS | GARVGGASNTLY | 0.4043042 |
| GAGGGGLTAGAN | GARVGGASNTLY | 0.62177664 |
| FVDALVNM | GARVGGASNTLY | 0.7936404 |
| FLTVAPEL | GARVGGASNTLY | 0.72087634 |
| FETLTAL | GARVGGASNTLY | 0.6633688 |
| DVINPIEL | GARVGGASNTLY | 0.59539455 |
| APVSGAPV | GARVGGASNTLY | 0.7764801 |
| ALVGGLAW | GARVGGASNTLY | 0.66235864 |
| ALAAAAATAI | GARVGGASNTLY | 0.7854487 |
| AFGLGLLL | GARVGGASNTLY | 0.2954554 |
| AFFAAASL | GARVGGASNTLY | 0.5491972 |
| AAAALPAFSPP | GARVGGASNTLY | 0.42691272 |
| VVLAPAL | GARVGGASNTLY | 0.2573771 |
| TIGHVDHGKTTL | GARVGGASNTLY | 0.76947093 |
| LISNASDALDK | GARVGGASNTLY | 0.49306646 |
| LISNASDAL | GARVGGASNTLY | 0.7187911 |
| ISNASDALDK | GARVGGASNTLY | 0.641539 |
| NQAETLVYQTE | GARVGGASNTLY | 0.7690711 |
| TGGTGGTAGAG | GARVGGASNTLY | 0.6866745 |
| TGSTAGGAGGA | GARVGGASNTLY | 0.63792735 |
| DPLAVPVAAGPV | GARVGGASNTLY | 0.5288019 |
| GGTGGTPGNGGN | GARVGGASNTLY | 0.7479789 |

| | | |
|---|---|---|
| PAGLAGVDPATL | GARVGGASNTLY | 0.7689947 |
| GNNLVVEAPHAGASV | GARVGGASNTLY | 0.72309965 |
| KVTRIMSASTLEQPA | GARVGGASNTLY | 0.34068522 |
| LGFALPIALPRRTVV | GARVGGASNTLY | 0.7162213 |
| TRLGNLTARKLVVLV | GARVGGASNTLY | 0.66366976 |
| WSQQRRLIYFPSAGP | GARVGGASNTLY | 0.36887074 |
| GPNGNPDPMAAAVDIR | GARVGGASNTLY | 0.30498132 |
| TQRCVSVADLRAAASR | GARVGGASNTLY | 0.5703014 |
| QFVLDVAEHPVEVLTETV | GARVGGASNTLY | 0.6409122 |
| GGAGGTGGAGGNGGTQAGDG | GARVGGASNTLY | 0.75917864 |
| PLLATLRAIAVDVDAKPAQV | GARVGGASNTLY | 0.75121576 |
| APKFRTGGYMIDFWGVGYQVAK | GARVGGASNTLY | 0.60758233 |
| GAGGAGGAGGAAGGPGGTGGTG | GARVGGASNTLY | 0.73082983 |
| IGALGLLATDVAPAGSVAFGVA | GARVGGASNTLY | 0.702533 |
| AGGAGGIGGNGGAGGAGGVLMG | GARVGGASNTLY | 0.18630245 |
| ADELIGLL | GARVGGASNTLY | 0.48558974 |
| ADRALKA | GARVGGASNTLY | 0.40121016 |
| AGAVGQVGGAGGA | GARVGGASNTLY | 0.76899153 |
| AGGGGGLGGAP | GARVGGASNTLY | 0.51997006 |
| AGGLLYGNGGAG | GARVGGASNTLY | 0.6114764 |
| AGLESIGL | GARVGGASNTLY | 0.6708509 |
| AGLKTGII | GARVGGASNTLY | 0.71977365 |
| ASLAALLL | GARVGGASNTLY | 0.69661164 |
| ASVINISSI | GARVGGASNTLY | 0.45886418 |
| ATPPGPA | GARVGGASNTLY | 0.37597182 |
| ATVASALV | GARVGGASNTLY | 0.74438936 |
| ATVASLIL | GARVGGASNTLY | 0.58296883 |
| AVNGPVLIRH | GARVGGASNTLY | 0.4900635 |
| DGAAGAVGI | GARVGGASNTLY | 0.38394272 |
| DPLLHGI | GARVGGASNTLY | 0.7440983 |
| ELGAALGAL | GARVGGASNTLY | 0.2922003 |
| ELGRTQAV | GARVGGASNTLY | 0.7405224 |
| FVAGGYLS | GARVGGASNTLY | 0.57731736 |
| GASGGAGGAGGW | GARVGGASNTLY | 0.67756295 |
| GGAGAGGDGGAGGI | GARVGGASNTLY | 0.78842425 |
| GTGHNGSQPGMGGNG | GARVGGASNTLY | 0.7115002 |
| GYAAASASAS | GARVGGASNTLY | 0.66284466 |
| HQVDVAAVQQG | GARVGGASNTLY | 0.5415666 |
| IGAAIAHT | GARVGGASNTLY | 0.46706453 |
| IGMIEVAQL | GARVGGASNTLY | 0.5409146 |
| IRLAPEHTSAMD | GARVGGASNTLY | 0.5747925 |
| IYNVSVL | GARVGGASNTLY | 0.7807679 |
| KQGFLGM | GARVGGASNTLY | 0.6425405 |
| LESVITQL | GARVGGASNTLY | 0.36214483 |
| LGLGALVVV | GARVGGASNTLY | 0.6100246 |
| LIGNGGNGGSGGI | GARVGGASNTLY | 0.78105795 |
| LNAINAPTL | GARVGGASNTLY | 0.786378 |
| LQLLNTGV | GARVGGASNTLY | 0.5478265 |
| LQTSIEF | GARVGGASNTLY | 0.47894287 |
| MSFAVSEL | GARVGGASNTLY | 0.50719094 |
| NTGALGL | GARVGGASNTLY | 0.6740861 |

| | | |
|---|---|---|
| PPGAGGAGGAGTT | GARVGGASNTLY | 0.45187414 |
| QLVGLGI | GARVGGASNTLY | 0.47649103 |
| RFLEQVHQL | GARVGGASNTLY | 0.72288024 |
| RGVGYVLR | GARVGGASNTLY | 0.78778756 |
| SATGVFAGVI | GARVGGASNTLY | 0.6135441 |
| SLPAITLP | GARVGGASNTLY | 0.2898304 |
| STALAFTV | GARVGGASNTLY | 0.67075837 |
| SVLTVGLGK | GARVGGASNTLY | 0.5498117 |
| TALAGAGSGI | GARVGGASNTLY | 0.56564593 |
| TGLPFTGI | GARVGGASNTLY | 0.5859475 |
| VAEHVTV | GARVGGASNTLY | 0.3562681 |
| VDIVLNSL | GARVGGASNTLY | 0.67666674 |
| VLTGGGAVL | GARVGGASNTLY | 0.7857622 |
| VNVGLAYGI | GARVGGASNTLY | 0.65455204 |
| VSSAAVGAV | GARVGGASNTLY | 0.688076 |
| VTSVYEAGV | GARVGGASNTLY | 0.7819895 |
| WVALDRGA | GARVGGASNTLY | 0.745646 |
| YLVGSVVI | GARVGGASNTLY | 0.7760224 |
| GPPGTGKTLL | GARVGGASNTLY | 0.7521989 |
| GVGMTKF | GARVGGASNTLY | 0.6753421 |
| IGIGGSDL | GARVGGASNTLY | 0.6329603 |
| IGTLLAEL | GARVGGASNTLY | 0.7684009 |
| IYTSGTT | GARVGGASNTLY | 0.66731054 |
| LSFPAGV | GARVGGASNTLY | 0.68802303 |
| LYTSGTT | GARVGGASNTLY | 0.57892716 |
| VGAFPFGF | GARVGGASNTLY | 0.75564206 |
| VGAGTVEF | GARVGGASNTLY | 0.5558876 |
| VGKSTLLL | GARVGGASNTLY | 0.76728 |
| AALAHAGGGIV | GARVGGASNTLY | 0.2849302 |
| AALVAVSLL | GARVGGASNTLY | 0.48806357 |
| AAVSRAVAAL | GARVGGASNTLY | 0.76418954 |
| AGGSSGIGGF | GARVGGASNTLY | 0.20141904 |
| AGLTVANANG | GARVGGASNTLY | 0.30767336 |
| AGLVGAVL | GARVGGASNTLY | 0.6247472 |
| ALSQAFTV | GARVGGASNTLY | 0.76238704 |
| ALTFSVSL | GARVGGASNTLY | 0.22029899 |
| ALTGAALVAL | GARVGGASNTLY | 0.5292076 |
| ASFSIGAGVL | GARVGGASNTLY | 0.38784766 |
| DGSVYKI | GARVGGASNTLY | 0.71006006 |
| EQLAEIV | GARVGGASNTLY | 0.6677182 |
| FAVSELTVM | GARVGGASNTLY | 0.35430607 |
| GAAPLAAGL | GARVGGASNTLY | 0.7460212 |
| GADGGDGGSGGAGG | GARVGGASNTLY | 0.28573146 |
| GFGLDTGGAGG | GARVGGASNTLY | 0.7055594 |
| GLASGGLL | GARVGGASNTLY | 0.75861967 |
| GRIAGAAVLL | GARVGGASNTLY | 0.7799675 |
| GSAGSLNTSF | GARVGGASNTLY | 0.45701635 |
| GTGQIGI | GARVGGASNTLY | 0.74452734 |
| GTLIALIL | GARVGGASNTLY | 0.55927336 |
| IAATMGFVVG | GARVGGASNTLY | 0.7962036 |
| IFAQALL | GARVGGASNTLY | 0.6432094 |

| | | |
|---|---|---|
| IGAAVALGF | GARVGGASNTLY | 0.34933895 |
| IGLTPLAL | GARVGGASNTLY | 0.7591264 |
| ILGSNTSTL | GARVGGASNTLY | 0.7160105 |
| IVFSLTVL | GARVGGASNTLY | 0.6118947 |
| LASKATGF | GARVGGASNTLY | 0.7386221 |
| LASVALIL | GARVGGASNTLY | 0.53128594 |
| LFAVGLTL | GARVGGASNTLY | 0.6387543 |
| LFNGGIGTYI | GARVGGASNTLY | 0.7658903 |
| LTLGAGII | GARVGGASNTLY | 0.7485863 |
| LVAAALGDI | GARVGGASNTLY | 0.53297806 |
| NTAFVQLGI | GARVGGASNTLY | 0.7585726 |
| QLFSGAEL | GARVGGASNTLY | 0.68075883 |
| SAATFITL | GARVGGASNTLY | 0.44547576 |
| SNYGLANLGV | GARVGGASNTLY | 0.78853786 |
| SYGGAYAVM | GARVGGASNTLY | 0.5848985 |
| TTGVGGLTLG | GARVGGASNTLY | 0.7499954 |
| TTQTIYI | GARVGGASNTLY | 0.73632133 |
| VAAGAGLGAVI | GARVGGASNTLY | 0.7619453 |
| VAAVVTLL | GARVGGASNTLY | 0.51777273 |
| VGFLGGLI | GARVGGASNTLY | 0.65750474 |
| VGVIPILL | GARVGGASNTLY | 0.6692426 |
| VIFASSLI | GARVGGASNTLY | 0.7378659 |
| VKGGDGGAAAG | GARVGGASNTLY | 0.6177927 |
| VTKTFTNI | GARVGGASNTLY | 0.7037022 |
| VVAAAILL | GARVGGASNTLY | 0.7319875 |
| YAALDVEL | GARVGGASNTLY | 0.7758875 |
| LASVALLL | GARVGGASNTLY | 0.6164091 |
| LDAVHRV | GARVGGASNTLY | 0.66049737 |
| QGLSSSIF | GARVGGASNTLY | 0.7846229 |
| RGTFANIRL | GARVGGASNTLY | 0.5069692 |
| RQLWWGHRI | GARVGGASNTLY | 0.65884066 |
| YGPPGTGKTLL | GARVGGASNTLY | 0.7219262 |
| AALVGAPANVA | GARVGGASNTLY | 0.7659668 |
| AATFRRTE | GARVGGASNTLY | 0.6021137 |
| AGALGGGI | GARVGGASNTLY | 0.76824117 |
| AGGAGGAGGIGAG | GARVGGASNTLY | 0.6115066 |
| AGSGALGL | GARVGGASNTLY | 0.7518035 |
| AQIADQI | GARVGGASNTLY | 0.63478494 |
| ASAVAVTYL | GARVGGASNTLY | 0.6526378 |
| ASTVLGGTGG | GARVGGASNTLY | 0.5569774 |
| AVGGEPAK | GARVGGASNTLY | 0.74153143 |
| CEQGTFGGTC | GARVGGASNTLY | 0.7552482 |
| DEGKAPAG | GARVGGASNTLY | 0.75264746 |
| DTVVFPAL | GARVGGASNTLY | 0.2738632 |
| FEGITMAG | GARVGGASNTLY | 0.749889 |
| FPLVVQD | GARVGGASNTLY | 0.75037867 |
| GAFTTPQ | GARVGGASNTLY | 0.78697306 |
| GFGGFGTAGGVGGA | GARVGGASNTLY | 0.61198056 |
| GFLAQFL | GARVGGASNTLY | 0.686357 |
| GGAGGAGGNAGVA | GARVGGASNTLY | 0.58516157 |
| GGVGTAGAGGAGGAGG | GARVGGASNTLY | 0.64694864 |

| | | |
|---|---|---|
| GNGGGGDAGNAGSGGN | GARVGGASNTLY | 0.39579493 |
| GTGGAGGAGNDAGS | GARVGGASNTLY | 0.50298935 |
| GTTNTGWFNS | GARVGGASNTLY | 0.66206276 |
| IFMGPLFTL | GARVGGASNTLY | 0.5769649 |
| ILAHAIEAI | GARVGGASNTLY | 0.75473493 |
| IRSAIREAL | GARVGGASNTLY | 0.7695001 |
| ITFAGGG | GARVGGASNTLY | 0.7686329 |
| ITGMSGHQEV | GARVGGASNTLY | 0.76877797 |
| IVAGLACTL | GARVGGASNTLY | 0.7747871 |
| IVGGGLAGA | GARVGGASNTLY | 0.7536859 |
| IVTGVLNI | GARVGGASNTLY | 0.60165787 |
| LAQQGCI | GARVGGASNTLY | 0.6709376 |
| LCLHVEN | GARVGGASNTLY | 0.38354838 |
| LDAISDGS | GARVGGASNTLY | 0.6367463 |
| LGGFAAML | GARVGGASNTLY | 0.5960302 |
| LGGKNPMI | GARVGGASNTLY | 0.71254385 |
| LGPKGVAAIL | GARVGGASNTLY | 0.20610565 |
| LMVGVSI | GARVGGASNTLY | 0.71224844 |
| LVGAAFVL | GARVGGASNTLY | 0.7858306 |
| LVLAVAPL | GARVGGASNTLY | 0.6718127 |
| MAAIAVIDAA | GARVGGASNTLY | 0.7194189 |
| NNGINAELQ | GARVGGASNTLY | 0.7425709 |
| RGEPLVLVHG | GARVGGASNTLY | 0.56631243 |
| RGGAGGAAGLI | GARVGGASNTLY | 0.7379167 |
| RSYRMHPAV | GARVGGASNTLY | 0.58725965 |
| SAAHLLAGQLA | GARVGGASNTLY | 0.7544458 |
| SAAVGSY | GARVGGASNTLY | 0.7812954 |
| SPHAVLHRI | GARVGGASNTLY | 0.7111421 |
| SQIAKVL | GARVGGASNTLY | 0.5824886 |
| TALGHLVP | GARVGGASNTLY | 0.65019774 |
| TAPTTSGAA | GARVGGASNTLY | 0.41147014 |
| TDGDDGNGGSPGT | GARVGGASNTLY | 0.6673033 |
| TDGLGEN | GARVGGASNTLY | 0.619864 |
| TGGPPLPGQA | GARVGGASNTLY | 0.6889932 |
| TIPAITI | GARVGGASNTLY | 0.55200434 |
| TIPGFTI | GARVGGASNTLY | 0.73565733 |
| TSIASGALA | GARVGGASNTLY | 0.70344734 |
| TVTLPGAAI | GARVGGASNTLY | 0.6636632 |
| VASGLSAALP | GARVGGASNTLY | 0.5185138 |
| VGAGVIGI | GARVGGASNTLY | 0.74207747 |
| VGPKRFGV | GARVGGASNTLY | 0.7026883 |
| VNVAFPD | GARVGGASNTLY | 0.41962296 |
| VSMTNSGVS | GARVGGASNTLY | 0.73728406 |
| VTDVLGDLSV | GARVGGASNTLY | 0.56747705 |
| VVVVGAGFAGL | GARVGGASNTLY | 0.64979434 |
| VYTVGTVV | GARVGGASNTLY | 0.7649722 |
| WTTFSGP | GARVGGASNTLY | 0.6957364 |
| YDELVEAF | GARVGGASNTLY | 0.30218303 |
| YGTGGAGGNGG | GARVGGASNTLY | 0.6394434 |
| YTNVMAV | GARVGGASNTLY | 0.6892023 |
| FLEQVHQL | GARVGGASNTLY | 0.66734743 |

| | | |
|---|---|---|
| GLSLLAVAV | GARVGGASNTLY | 0.23768309 |
| LEQVHQL | GARVGGASNTLY | 0.73524296 |
| LNAAGAI | GARVGGASNTLY | 0.45024478 |
| PAVHASL | GARVGGASNTLY | 0.509306 |
| TELEDDA | GARVGGASNTLY | 0.57408416 |
| TPPGPPN | GARVGGASNTLY | 0.7639718 |
| TPPPPPPP | GARVGGASNTLY | 0.71596897 |
| TQSPALI | ASSSGTNEQY | 0.5707787 |
| EFAAAFPQ | ASSSGTNEQY | 0.72904444 |
| LLSGNLDVL | ASSSGTNEQY | 0.71508837 |
| GPANLGNYNI | ASSSGTNEQY | 0.7736261 |
| TLGPTLP | ASSSGTNEQY | 0.6985016 |
| VYAYSSCCK | ASSSGTNEQY | 0.5955255 |
| APAPAPAPNI | ASSSGTNEQY | 0.6751666 |
| PGGPVPIP | ASSSGTNEQY | 0.50958145 |
| FLTVALEP | ASSSGTNEQY | 0.55921865 |
| SQYFGANL | ASSSGTNEQY | 0.4667007 |
| GGVKPAA | ASSSGTNEQY | 0.5536164 |
| GGTGGAGGAGGW | ASSSGTNEQY | 0.5103167 |
| GVGGVAGLGGAGG | ASSSGTNEQY | 0.71286714 |
| VLLVLATTI | ASSSGTNEQY | 0.7536813 |
| IGIFAAML | ASSSGTNEQY | 0.59268665 |
| AVGDLID | ASSSGTNEQY | 0.34089252 |
| NISPLPL | ASSSGTNEQY | 0.59854865 |
| VTAVFVAI | ASSSGTNEQY | 0.6208997 |
| SGDDNDAAL | ASSSGTNEQY | 0.7593117 |
| AVAALAHS | ASSSGTNEQY | 0.71401405 |
| GGVTVMATP | ASSSGTNEQY | 0.7876184 |
| AGLAANVRD | ASSSGTNEQY | 0.76085544 |
| TNRPDLIDPALLRPG | ASSSGTNEQY | 0.6169342 |
| LANLGVTAL | ASSSGTNEQY | 0.75042 |
| TGERIEL | ASSSGTNEQY | 0.58081496 |
| NGEELAA | ASSSGTNEQY | 0.63838315 |
| VLSLVWG | ASSSGTNEQY | 0.68749154 |
| IGYALWL | ASSSGTNEQY | 0.6659571 |
| LSLLLAPL | ASSSGTNEQY | 0.59832287 |
| TLDEALK | ASSSGTNEQY | 0.6537986 |
| TDGAGTAEGG | ASSSGTNEQY | 0.32026938 |
| NAPGGAGGNGG | ASSSGTNEQY | 0.7280827 |
| VSVFMGMGGNF | ASSSGTNEQY | 0.4326346 |
| LASYLAAL | ASSSGTNEQY | 0.3800118 |
| LVGGTPAL | ASSSGTNEQY | 0.64521146 |
| VRNADVL | ASSSGTNEQY | 0.7875811 |
| TCSSPFLVSAM | ASSSGTNEQY | 0.7762672 |
| VHITDAE | ASSSGTNEQY | 0.59824824 |
| EYAAPSF | ASSSGTNEQY | 0.53117085 |
| VVDAFHGGL | ASSSGTNEQY | 0.724682 |
| ASVVDPQ | ASSSGTNEQY | 0.49249047 |
| SYNEFGGLS | ASSSGTNEQY | 0.70600903 |
| DPDGSVY | ASSSGTNEQY | 0.6899774 |
| MVSTIHATE | ASSSGTNEQY | 0.66585517 |

| | | |
|---|---|---|
| LAAGVADPV | ASSSGTNEQY | 0.43309176 |
| TKEQIAA | ASSSGTNEQY | 0.73425657 |
| SLASPAAL | ASSSGTNEQY | 0.613255 |
| IGLVFYGV | ASSSGTNEQY | 0.68765444 |
| FLLDASNL | ASSSGTNEQY | 0.34874767 |
| YQVGNFP | ASSSGTNEQY | 0.7476492 |
| ESLLSQST | ASSSGTNEQY | 0.5413958 |
| LFTVGFAAVG | ASSSGTNEQY | 0.43759954 |
| ATPPTPT | ASSSGTNEQY | 0.25477493 |
| TDLATLG | ASSSGTNEQY | 0.57949185 |
| SAGGTGR | ASSSGTNEQY | 0.73183477 |
| GFSNSGS | ASSSGTNEQY | 0.7701105 |
| GGNGGSGGTSV | ASSSGTNEQY | 0.6804908 |
| GTSVATGGAGN | ASSSGTNEQY | 0.75430036 |
| NGGNGGSGGTS | ASSSGTNEQY | 0.76686263 |
| AVVKFPH | ASSSGTNEQY | 0.66560835 |
| TPPPPPP | ASSSGTNEQY | 0.35779962 |
| AAPAAAAAPA | ASSSGTNEQY | 0.6913948 |
| IALNEDL | ASSSGTNEQY | 0.63361657 |
| IVNVSSM | ASSSGTNEQY | 0.7642585 |
| TGIPPAPR | ASSSGTNEQY | 0.7380254 |
| FDLGGGTF | ASSSGTNEQY | 0.4202817 |
| FDLGGGTFDVS | ASSSGTNEQY | 0.68152237 |
| NEPTAAA | ASSSGTNEQY | 0.738913 |
| NEPTAAALA | ASSSGTNEQY | 0.4494949 |
| GSIINISSI | ASSSGTNEQY | 0.43024868 |
| VAPGNAGTA | ASSSGTNEQY | 0.45680618 |
| TGGGASLE | ASSSGTNEQY | 0.4388227 |
| VAVYNHYKRI | ASSSGTNEQY | 0.7152502 |
| TDPAPATT | ASSSGTNEQY | 0.6475026 |
| TNRPDILDPALLRPG | ASSSGTNEQY | 0.6285599 |
| LTTTTTS | ASSSGTNEQY | 0.6456311 |
| PTGSGKTL | ASSSGTNEQY | 0.55374825 |
| TGSGKTL | ASSSGTNEQY | 0.58294785 |
| TIGHVDHGK | ASSSGTNEQY | 0.61958224 |
| TIGHVDHGKTT | ASSSGTNEQY | 0.68643 |
| GHVDHGKTT | ASSSGTNEQY | 0.7485266 |
| AQFEHTLL | ASSSGTNEQY | 0.60389376 |
| IVNQDEL | ASSSGTNEQY | 0.5649399 |
| TNGSQFF | ASSSGTNEQY | 0.78488445 |
| TGKGIVE | ASSSGTNEQY | 0.41858792 |
| LISNASDA | ASSSGTNEQY | 0.5127831 |
| AELRTLGELTKIAKAHGAQ | ASSSGTNEQY | 0.72428906 |
| APNAVAAAEPSW | ASSSGTNEQY | 0.55826116 |
| APQVAGAATAVL | ASSSGTNEQY | 0.75895596 |
| ARRDPAAAAAILPTDA | ASSSGTNEQY | 0.7153075 |
| DAAARDVLAEAGLAEYF | ASSSGTNEQY | 0.7241345 |
| DDQIVLSAKANLAKVEQ | ASSSGTNEQY | 0.6451371 |
| DELLPDTGEDSDEDGH | ASSSGTNEQY | 0.79062855 |
| ELPEAPANRATVPQD | ASSSGTNEQY | 0.7531389 |
| FGQSKIERVTGIG | ASSSGTNEQY | 0.54732996 |

| | | |
|---|---|---|
| GAGGTGSTAGGAGGAGGAGGLY | ASSSGTNEQY | 0.7515396 |
| GHGGTGAAGGNGGTGGAGGSGI | ASSSGTNEQY | 0.344955 |
| GNGGNGGAGGEGGPGGAGGAGA | ASSSGTNEQY | 0.7585979 |
| GSTAGNGGNGGNATNGTIAGQP | ASSSGTNEQY | 0.7322165 |
| GTGGDGGNGGNGGTGG | ASSSGTNEQY | 0.51880026 |
| GVRRPAGIPADAPV | ASSSGTNEQY | 0.6561638 |
| GYRQVTDAHLLTLARR | ASSSGTNEQY | 0.7680304 |
| IAKMGQDGHDRG | ASSSGTNEQY | 0.77585554 |
| KAEPSIEIQYT | ASSSGTNEQY | 0.55210453 |
| KPVRIRMAADEDTVAVT | ASSSGTNEQY | 0.5795901 |
| MPPRIAGMRLLVIKPEP | ASSSGTNEQY | 0.42306593 |
| MKFYAVALFRSRGVPPTR | ASSSGTNEQY | 0.40365314 |
| NAGGFGILGAVAHSP | ASSSGTNEQY | 0.33732557 |
| NEGREITLAMAEM | ASSSGTNEQY | 0.7312705 |
| NELVEWSHPLGHRWRWE | ASSSGTNEQY | 0.64066005 |
| NLGAGGTGGNGGNPG | ASSSGTNEQY | 0.7411065 |
| NTEWPKWETVMAADPVD | ASSSGTNEQY | 0.71436346 |
| PASAAPAETQP | ASSSGTNEQY | 0.7729523 |
| QQSSDGKNGVRAAEITGE | ASSSGTNEQY | 0.58548605 |
| RAPDAVTTSWWKE | ASSSGTNEQY | 0.65450364 |
| SGFGNVGAHTSGWFNQ | ASSSGTNEQY | 0.32519495 |
| SGVAGAVGQAASVGGLKV | ASSSGTNEQY | 0.5607933 |
| SNGGAGTGSAGGNGGT | ASSSGTNEQY | 0.46176443 |
| TGGAGGEGGAAG | ASSSGTNEQY | 0.6423032 |
| TGGDGGAAGAGGAPGHG | ASSSGTNEQY | 0.40778387 |
| TMPPLLAAIANAGRHGV | ASSSGTNEQY | 0.77669764 |
| TSSTAGNSGGA | ASSSGTNEQY | 0.6426564 |
| VPATVPVGILC | ASSSGTNEQY | 0.74248695 |
| VQRPDSGGVVLTGRIS | ASSSGTNEQY | 0.56814057 |
| WTPNPVPATKGVPAGLVQAT | ASSSGTNEQY | 0.4863627 |
| YALKRALAKFSDRFDVVII | ASSSGTNEQY | 0.57581925 |
| YLIGSVIV | ASSSGTNEQY | 0.7421628 |
| YGQLVAVID | ASSSGTNEQY | 0.4675763 |
| VTVGGMTLPA | ASSSGTNEQY | 0.24441265 |
| VLALPAV | ASSSGTNEQY | 0.43062794 |
| VIPPTLNYET | ASSSGTNEQY | 0.3842354 |
| VGMGGMHPSSG | ASSSGTNEQY | 0.7789217 |
| VGLAGGAATA | ASSSGTNEQY | 0.77198267 |
| VGGAGGAGGSGP | ASSSGTNEQY | 0.697507 |
| VFAAFMV | ASSSGTNEQY | 0.7712508 |
| TLYDKLR | ASSSGTNEQY | 0.2144495 |
| TLAVFTVL | ASSSGTNEQY | 0.5948589 |
| TGSVQPARV | ASSSGTNEQY | 0.7701179 |
| TGGGAGSGGT | ASSSGTNEQY | 0.72317386 |
| SHLETSL | ASSSGTNEQY | 0.6826426 |
| SFTTGMT | ASSSGTNEQY | 0.7713548 |
| QVEVAKRMGVRQPTVS | ASSSGTNEQY | 0.54470944 |
| QGSGGPALAM | ASSSGTNEQY | 0.66519636 |
| MLAPSNP | ASSSGTNEQY | 0.7011782 |
| LVNNAAI | ASSSGTNEQY | 0.750363 |
| LTGEINTLA | ASSSGTNEQY | 0.69067633 |

| | | |
|---|---|---|
| LSIFGPV | ASSSGTNEQY | 0.40506753 |
| LLALLANL | ASSSGTNEQY | 0.6798297 |
| LISNASDALDKLR | ASSSGTNEQY | 0.8000181 |
| LINEAVP | ASSSGTNEQY | 0.64128506 |
| LGYAFSGL | ASSSGTNEQY | 0.73865116 |
| LDFVAVAL | ASSSGTNEQY | 0.6967629 |
| LAQPGAE | ASSSGTNEQY | 0.6912237 |
| ISNLGAVTGF | ASSSGTNEQY | 0.55343795 |
| HQEDPGRI | ASSSGTNEQY | 0.5405813 |
| GTPGEPTP | ASSSGTNEQY | 0.7381562 |
| GFFNSGS | ASSSGTNEQY | 0.53968334 |
| GAGGGGLTAGAN | ASSSGTNEQY | 0.6056495 |
| FVDALVNM | ASSSGTNEQY | 0.5957906 |
| FLTVAPEL | ASSSGTNEQY | 0.7986882 |
| FETLTAL | ASSSGTNEQY | 0.72418755 |
| DVINPIEL | ASSSGTNEQY | 0.74747956 |
| APVSGAPV | ASSSGTNEQY | 0.5450438 |
| ALVGGLAW | ASSSGTNEQY | 0.6707966 |
| ALAAAAATAI | ASSSGTNEQY | 0.74075615 |
| AFGLGLLL | ASSSGTNEQY | 0.34221193 |
| AFFAAASL | ASSSGTNEQY | 0.7665305 |
| AAAALPAFSPP | ASSSGTNEQY | 0.53069925 |
| VVLAPAL | ASSSGTNEQY | 0.6679088 |
| TIGHVDHGKTTL | ASSSGTNEQY | 0.7130657 |
| LISNASDALDK | ASSSGTNEQY | 0.7722292 |
| LISNASDAL | ASSSGTNEQY | 0.66156816 |
| ISNASDALDK | ASSSGTNEQY | 0.7292059 |
| NQAETLVYQTE | ASSSGTNEQY | 0.51209867 |
| TGGTGGTAGAG | ASSSGTNEQY | 0.7240233 |
| TGSTAGGAGGA | ASSSGTNEQY | 0.77379525 |
| DPLAVPVAAGPV | ASSSGTNEQY | 0.6891953 |
| GGTGGTPGNGGN | ASSSGTNEQY | 0.7897289 |
| PAGLAGVDPATL | ASSSGTNEQY | 0.67564356 |
| GNNLVVEAPHAGASV | ASSSGTNEQY | 0.29224488 |
| KVTRIMSASTLEQPA | ASSSGTNEQY | 0.6607763 |
| LGFALPIALPRRTVV | ASSSGTNEQY | 0.7657455 |
| TRLGNLTARKLVVLV | ASSSGTNEQY | 0.77742577 |
| WSQQRRLIYFPSAGP | ASSSGTNEQY | 0.7586062 |
| GPNGNPDPMAAAVDIR | ASSSGTNEQY | 0.72475743 |
| TQRCVSVADLRAAASR | ASSSGTNEQY | 0.6320451 |
| QFVLDVAEHPVEVLTETV | ASSSGTNEQY | 0.77717865 |
| GGAGGTGGAGGNGGTQAGDG | ASSSGTNEQY | 0.7899449 |
| PLLATLRAIAVDVDAKPAQV | ASSSGTNEQY | 0.725316 |
| APKFRTGGYMIDFWGVGYQVAK | ASSSGTNEQY | 0.49064887 |
| GAGGAGGAGGAAGGPGGTGGTG | ASSSGTNEQY | 0.4297276 |
| IGALGLLATDVAPAGSVAFGVA | ASSSGTNEQY | 0.7248578 |
| AGGAGGIGGNGGAGGAGGVLMG | ASSSGTNEQY | 0.7562442 |
| ADELIGLL | ASSSGTNEQY | 0.3873622 |
| ADRALKA | ASSSGTNEQY | 0.71936345 |
| AGAVGQVGGAGGA | ASSSGTNEQY | 0.70991474 |
| AGGGGGLGGAP | ASSSGTNEQY | 0.28670582 |

| | | |
|---|---|---|
| AGGLLYGNGGAG | ASSSGTNEQY | 0.71397924 |
| AGLESIGL | ASSSGTNEQY | 0.77519155 |
| AGLKTGII | ASSSGTNEQY | 0.69094765 |
| ASLAALLL | ASSSGTNEQY | 0.45223135 |
| ASVINISSI | ASSSGTNEQY | 0.769536 |
| ATPPGPA | ASSSGTNEQY | 0.7920103 |
| ATVASALV | ASSSGTNEQY | 0.7844007 |
| ATVASLIL | ASSSGTNEQY | 0.34970963 |
| AVNGPVLIRH | ASSSGTNEQY | 0.21913503 |
| DGAAGAVGI | ASSSGTNEQY | 0.7103285 |
| DPLLHGI | ASSSGTNEQY | 0.739594 |
| ELGAALGAL | ASSSGTNEQY | 0.6097907 |
| ELGRTQAV | ASSSGTNEQY | 0.53250784 |
| FVAGGYLS | ASSSGTNEQY | 0.72025895 |
| GASGGAGGAGGW | ASSSGTNEQY | 0.75882053 |
| GGAGAGGDGGAGGI | ASSSGTNEQY | 0.45911908 |
| GTGHNGSQPGMGGNG | ASSSGTNEQY | 0.35046118 |
| GYAAASASAS | ASSSGTNEQY | 0.727253 |
| HQVDVAAVQQG | ASSSGTNEQY | 0.7322012 |
| IGAAIAHT | ASSSGTNEQY | 0.55240023 |
| IGMIEVAQL | ASSSGTNEQY | 0.6412164 |
| IRLAPEHTSAMD | ASSSGTNEQY | 0.7265452 |
| IYNVSVL | ASSSGTNEQY | 0.6982877 |
| KQGFLGM | ASSSGTNEQY | 0.71708107 |
| LESVITQL | ASSSGTNEQY | 0.40568697 |
| LGLGALVVV | ASSSGTNEQY | 0.77326274 |
| LIGNGGNGGSGGI | ASSSGTNEQY | 0.40272796 |
| LNAINAPTL | ASSSGTNEQY | 0.72135496 |
| LQLLNTGV | ASSSGTNEQY | 0.7144704 |
| LQTSIEF | ASSSGTNEQY | 0.7126249 |
| MSFAVSEL | ASSSGTNEQY | 0.6540705 |
| NTGALGL | ASSSGTNEQY | 0.5203732 |
| PPGAGGAGGAGTT | ASSSGTNEQY | 0.43420243 |
| QLVGLGI | ASSSGTNEQY | 0.4152457 |
| RFLEQVHQL | ASSSGTNEQY | 0.7719084 |
| RGVGYVLR | ASSSGTNEQY | 0.7271702 |
| SATGVFAGVI | ASSSGTNEQY | 0.7637907 |
| SLPAITLP | ASSSGTNEQY | 0.7260877 |
| STALAFTV | ASSSGTNEQY | 0.5720844 |
| SVLTVGLGK | ASSSGTNEQY | 0.70526254 |
| TALAGAGSGI | ASSSGTNEQY | 0.46419102 |
| TGLPFTGI | ASSSGTNEQY | 0.69998 |
| VAEHVTV | ASSSGTNEQY | 0.5989474 |
| VDIVLNSL | ASSSGTNEQY | 0.7122022 |
| VLTGGGAVL | ASSSGTNEQY | 0.53701884 |
| VNVGLAYGI | ASSSGTNEQY | 0.7077117 |
| VSSAAVGAV | ASSSGTNEQY | 0.7201264 |
| VTSVYEAGV | ASSSGTNEQY | 0.42679715 |
| WVALDRGA | ASSSGTNEQY | 0.66411877 |
| YLVGSVVI | ASSSGTNEQY | 0.77064526 |
| GPPGTGKTLL | ASSSGTNEQY | 0.7000113 |

| | | |
|---|---|---|
| GVGMTKF | ASSSGTNEQY | 0.40692705 |
| IGIGGSDL | ASSSGTNEQY | 0.737301 |
| IGTLLAEL | ASSSGTNEQY | 0.5613214 |
| IYTSGTT | ASSSGTNEQY | 0.7130409 |
| LSFPAGV | ASSSGTNEQY | 0.7112392 |
| LYTSGTT | ASSSGTNEQY | 0.53582263 |
| VGAFPFGF | ASSSGTNEQY | 0.7432989 |
| VGAGTVEF | ASSSGTNEQY | 0.33978432 |
| VGKSTLLL | ASSSGTNEQY | 0.55232215 |
| AALAHAGGGIV | ASSSGTNEQY | 0.645674 |
| AALVAVSLL | ASSSGTNEQY | 0.6963734 |
| AAVSRAVAAL | ASSSGTNEQY | 0.7441528 |
| AGGSSGIGGF | ASSSGTNEQY | 0.4815738 |
| AGLTVANANG | ASSSGTNEQY | 0.70051706 |
| AGLVGAVL | ASSSGTNEQY | 0.6265764 |
| ALSQAFTV | ASSSGTNEQY | 0.54649967 |
| ALTFSVSL | ASSSGTNEQY | 0.62171525 |
| ALTGAALVAL | ASSSGTNEQY | 0.5576163 |
| ASFSIGAGVL | ASSSGTNEQY | 0.38421723 |
| DGSVYKI | ASSSGTNEQY | 0.7475796 |
| EQLAEIV | ASSSGTNEQY | 0.5295111 |
| FAVSELTVM | ASSSGTNEQY | 0.68940616 |
| GAAPLAAGL | ASSSGTNEQY | 0.7671276 |
| GADGGDGGSGGAGG | ASSSGTNEQY | 0.37637258 |
| GFGLDTGGAGG | ASSSGTNEQY | 0.6964824 |
| GLASGGLL | ASSSGTNEQY | 0.69327444 |
| GRIAGAAVLL | ASSSGTNEQY | 0.7255321 |
| GSAGSLNTSF | ASSSGTNEQY | 0.65750146 |
| GTGQIGI | ASSSGTNEQY | 0.51807374 |
| GTLIALIL | ASSSGTNEQY | 0.76835775 |
| IAATMGFVVG | ASSSGTNEQY | 0.73946846 |
| IFAQALL | ASSSGTNEQY | 0.6039519 |
| IGAAVALGF | ASSSGTNEQY | 0.72173524 |
| IGLTPLAL | ASSSGTNEQY | 0.7336271 |
| ILGSNTSTL | ASSSGTNEQY | 0.75583947 |
| IVFSLTVL | ASSSGTNEQY | 0.613143 |
| LASKATGF | ASSSGTNEQY | 0.33739734 |
| LASVALIL | ASSSGTNEQY | 0.6578294 |
| LFAVGLTL | ASSSGTNEQY | 0.6461834 |
| LFNGGIGTYI | ASSSGTNEQY | 0.49660477 |
| LTLGAGII | ASSSGTNEQY | 0.3153183 |
| LVAAALGDI | ASSSGTNEQY | 0.7581279 |
| NTAFVQLGI | ASSSGTNEQY | 0.79414046 |
| QLFSGAEL | ASSSGTNEQY | 0.602631 |
| SAATFITL | ASSSGTNEQY | 0.7937269 |
| SNYGLANLGV | ASSSGTNEQY | 0.36675212 |
| SYGGAYAVM | ASSSGTNEQY | 0.75468457 |
| TTGVGGLTLG | ASSSGTNEQY | 0.76285577 |
| TTQTIYI | ASSSGTNEQY | 0.52391404 |
| VAAGAGLGAVI | ASSSGTNEQY | 0.72413266 |
| VAAVVTLL | ASSSGTNEQY | 0.5510536 |

| | | |
|---|---|---|
| VGFLGGLI | ASSSGTNEQY | 0.55767983 |
| VGVIPILL | ASSSGTNEQY | 0.76758325 |
| VIFASSLI | ASSSGTNEQY | 0.5604509 |
| VKGGDGGAAAG | ASSSGTNEQY | 0.71679926 |
| VTKTFTNI | ASSSGTNEQY | 0.62357306 |
| VVAAAILL | ASSSGTNEQY | 0.514959 |
| YAALDVEL | ASSSGTNEQY | 0.72616994 |
| LASVALLL | ASSSGTNEQY | 0.73601687 |
| LDAVHRV | ASSSGTNEQY | 0.63655514 |
| QGLSSSIF | ASSSGTNEQY | 0.7860167 |
| RGTFANIRL | ASSSGTNEQY | 0.5550385 |
| RQLWWGHRI | ASSSGTNEQY | 0.7814623 |
| YGPPGTGKTLL | ASSSGTNEQY | 0.76021194 |
| AALVGAPANVA | ASSSGTNEQY | 0.70299345 |
| AATFRRTE | ASSSGTNEQY | 0.6054265 |
| AGALGGGI | ASSSGTNEQY | 0.7720313 |
| AGGAGGAGGIGAG | ASSSGTNEQY | 0.6023586 |
| AGSGALGL | ASSSGTNEQY | 0.68502355 |
| AQIADQI | ASSSGTNEQY | 0.7738489 |
| ASAVAVTYL | ASSSGTNEQY | 0.06268455 |
| ASTVLGGTGG | ASSSGTNEQY | 0.07188533 |
| AVGGEPAK | ASSSGTNEQY | 0.06290066 |
| CEQGTFGGTC | ASSSGTNEQY | 0.06361795 |
| DEGKAPAG | ASSSGTNEQY | 0.06558221 |
| DTVVFPAL | ASSSGTNEQY | 0.06306495 |
| FEGITMAG | ASSSGTNEQY | 0.06377811 |
| FPLVVQD | ASSSGTNEQY | 0.06707116 |
| GAFTTPQ | ASSSGTNEQY | 0.06319673 |
| GFGGFGTAGGVGGA | ASSSGTNEQY | 0.063756496 |
| GFLAQFL | ASSSGTNEQY | 0.06605272 |
| GGAGGAGGNAGVA | ASSSGTNEQY | 0.07007638 |
| GGVGTAGAGGAGGAGG | ASSSGTNEQY | 0.067511015 |
| GNGGGGDAGNAGSGGN | ASSSGTNEQY | 0.06314343 |
| GTGGAGGAGNDAGS | ASSSGTNEQY | 0.06294357 |
| GTTNTGWFNS | ASSSGTNEQY | 0.06412106 |
| IFMGPLFTL | ASSSGTNEQY | 0.0640026 |
| ILAHAIEAI | ASSSGTNEQY | 0.06261218 |
| IRSAIREAL | ASSSGTNEQY | 0.06527205 |
| ITFAGGG | ASSSGTNEQY | 0.06227818 |
| ITGMSGHQEV | ASSSGTNEQY | 0.06367339 |
| IVAGLACTL | ASSSGTNEQY | 0.0643957 |
| IVGGGLAGA | ASSSGTNEQY | 0.06580405 |
| IVTGVLNI | ASSSGTNEQY | 0.0628709 |
| LAQQGCI | ASSSGTNEQY | 0.063543126 |
| LCLHVEN | ASSSGTNEQY | 0.06292603 |
| LDAISDGS | ASSSGTNEQY | 0.06538697 |
| LGGFAAML | ASSSGTNEQY | 0.06659019 |
| LGGKNPMI | ASSSGTNEQY | 0.06597062 |
| LGPKGVAAIL | ASSSGTNEQY | 0.06448957 |
| LMVGVSI | ASSSGTNEQY | 0.06273466 |
| LVGAAFVL | ASSSGTNEQY | 0.06805779 |

| | | |
|---|---|---|
| LVLAVAPL | ASSSGTNEQY | 0.065449186 |
| MAAIAVIDAA | ASSSGTNEQY | 0.06342412 |
| NNGINAELQ | ASSSGTNEQY | 0.063715965 |
| RGEPLVLVHG | ASSSGTNEQY | 0.06308783 |
| RGGAGGAAGLI | ASSSGTNEQY | 0.06469016 |
| RSYRMHPAV | ASSSGTNEQY | 0.0742204 |
| SAAHLLAGQLA | ASSSGTNEQY | 0.066212 |
| SAAVGSY | ASSSGTNEQY | 0.0667124 |
| SPHAVLHRI | ASSSGTNEQY | 0.06269037 |
| SQIAKVL | ASSSGTNEQY | 0.062865615 |
| TALGHLVP | ASSSGTNEQY | 0.06726673 |
| TAPTTSGAA | ASSSGTNEQY | 0.064697556 |
| TDGDDGNGGSPGT | ASSSGTNEQY | 0.067449786 |
| TDGLGEN | ASSSGTNEQY | 0.06443293 |
| TGGPPLPGQA | ASSSGTNEQY | 0.062693805 |
| TIPAITI | ASSSGTNEQY | 0.063291855 |
| TIPGFTI | ASSSGTNEQY | 0.06339694 |
| TSIASGALA | ASSSGTNEQY | 0.062581345 |
| TVTLPGAAI | ASSSGTNEQY | 0.06950483 |
| VASGLSAALP | ASSSGTNEQY | 0.06323967 |
| VGAGVIGI | ASSSGTNEQY | 0.07296259 |
| VGPKRFGV | ASSSGTNEQY | 0.06987104 |
| VNVAFPD | ASSSGTNEQY | 0.0797059 |
| VSMTNSGVS | ASSSGTNEQY | 0.06589031 |
| VTDVLGDLSV | ASSSGTNEQY | 0.06258119 |
| VVVVGAGFAGL | ASSSGTNEQY | 0.06625042 |
| VYTVGTVV | ASSSGTNEQY | 0.06466534 |
| WTTFSGP | ASSSGTNEQY | 0.062862314 |
| YDELVEAF | ASSSGTNEQY | 0.065303266 |
| YGTGGAGGNGG | ASSSGTNEQY | 0.06379622 |
| YTNVMAV | ASSSGTNEQY | 0.08275149 |
| FLEQVHQL | ASSSGTNEQY | 0.06598633 |
| GLSLLAVAV | ASSSGTNEQY | 0.065913476 |
| LEQVHQL | ASSSGTNEQY | 0.06469991 |
| LNAAGAI | ASSSGTNEQY | 0.062430058 |
| PAVHASL | ASSSGTNEQY | 0.06399354 |
| TELEDDA | ASSSGTNEQY | 0.06286058 |
| TPPGPPN | ASSSGTNEQY | 0.063672476 |
| TPPPPPPP | ASSSGTNEQY | 0.072027504 |
| TQSPALI | GARQGAQDTQY | 0.06429114 |
| EFAAAFPQ | GARQGAQDTQY | 0.066578075 |
| LLSGNLDVL | GARQGAQDTQY | 0.06487086 |
| GPANLGNYNI | GARQGAQDTQY | 0.06299754 |
| TLGPTLP | GARQGAQDTQY | 0.06310816 |
| VYAYSSCCK | GARQGAQDTQY | 0.0664688 |
| APAPAPAPNI | GARQGAQDTQY | 0.06560361 |
| PGGPVPIP | GARQGAQDTQY | 0.06295751 |
| FLTVALEP | GARQGAQDTQY | 0.06423981 |
| SQYFGANL | GARQGAQDTQY | 0.06401654 |
| GGVKPAA | GARQGAQDTQY | 0.06676545 |
| GGTGGAGGAGGW | GARQGAQDTQY | 0.06336573 |

| | | |
|---|---|---|
| GVGGVAGLGGAGG | GARQGAQDTQY | 0.06325965 |
| VLLVLATTI | GARQGAQDTQY | 0.06641535 |
| IGIFAAML | GARQGAQDTQY | 0.06364656 |
| AVGDLID | GARQGAQDTQY | 0.074318886 |
| NISPLPL | GARQGAQDTQY | 0.063362814 |
| VTAVFVAI | GARQGAQDTQY | 0.0626436 |
| SGDDNDAAL | GARQGAQDTQY | 0.06290921 |
| AVAALAHS | GARQGAQDTQY | 0.06285866 |
| GGVTVMATP | GARQGAQDTQY | 0.063709065 |
| AGLAANVRD | GARQGAQDTQY | 0.06363189 |
| TNRPDLIDPALLRPG | GARQGAQDTQY | 0.06352694 |
| LANLGVTAL | GARQGAQDTQY | 0.06564846 |
| TGERIEL | GARQGAQDTQY | 0.06344014 |
| NGEELAA | GARQGAQDTQY | 0.06365917 |
| VLSLVWG | GARQGAQDTQY | 0.07368104 |
| IGYALWL | GARQGAQDTQY | 0.06277702 |
| LSLLLAPL | GARQGAQDTQY | 0.06505725 |
| TLDEALK | GARQGAQDTQY | 0.062835425 |
| TDGAGTAEGG | GARQGAQDTQY | 0.06551703 |
| NAPGGAGGNGG | GARQGAQDTQY | 0.071416356 |
| VSVFMGMGGNF | GARQGAQDTQY | 0.06971756 |
| LASYLAAL | GARQGAQDTQY | 0.07321228 |
| LVGGTPAL | GARQGAQDTQY | 0.062497523 |
| VRNADVL | GARQGAQDTQY | 0.06297242 |
| TCSSPFLVSAM | GARQGAQDTQY | 0.062300835 |
| VHITDAE | GARQGAQDTQY | 0.06757474 |
| EYAAPSF | GARQGAQDTQY | 0.06328289 |
| VVDAFHGGL | GARQGAQDTQY | 0.06367424 |
| ASVVDPQ | GARQGAQDTQY | 0.06401213 |
| SYNEFGGLS | GARQGAQDTQY | 0.06488005 |
| DPDGSVY | GARQGAQDTQY | 0.065292895 |
| MVSTIHATE | GARQGAQDTQY | 0.0633582 |
| LAAGVADPV | GARQGAQDTQY | 0.06338058 |
| TKEQIAA | GARQGAQDTQY | 0.066301726 |
| SLASPAAL | GARQGAQDTQY | 0.06379344 |
| IGLVFYGV | GARQGAQDTQY | 0.063781954 |
| FLLDASNL | GARQGAQDTQY | 0.064000264 |
| YQVGNFP | GARQGAQDTQY | 0.07792718 |
| ESLLSQST | GARQGAQDTQY | 0.06575061 |
| LFTVGFAAVG | GARQGAQDTQY | 0.06567543 |
| ATPPTPT | GARQGAQDTQY | 0.06384102 |
| TDLATLG | GARQGAQDTQY | 0.06257557 |
| SAGGTGR | GARQGAQDTQY | 0.063516274 |
| GFSNSGS | GARQGAQDTQY | 0.067509815 |
| GGNGGSGGTSV | GARQGAQDTQY | 0.07057532 |
| GTSVATGGAGN | GARQGAQDTQY | 0.06473709 |
| NGGNGGSGGTS | GARQGAQDTQY | 0.065153636 |
| AVVKFPH | GARQGAQDTQY | 0.06658303 |
| TPPPPPP | GARQGAQDTQY | 0.062154975 |
| AAPAAAAAPA | GARQGAQDTQY | 0.06404831 |
| IALNEDL | GARQGAQDTQY | 0.063130625 |

| | | |
|---|---|---|
| IVNVSSM | GARQGAQDTQY | 0.06378538 |
| TGIPPAPR | GARQGAQDTQY | 0.0626442 |
| FDLGGGTF | GARQGAQDTQY | 0.063726455 |
| FDLGGGTFDVS | GARQGAQDTQY | 0.066696316 |
| NEPTAAA | GARQGAQDTQY | 0.06321912 |
| NEPTAAALA | GARQGAQDTQY | 0.06404225 |
| GSIINISSI | GARQGAQDTQY | 0.06269695 |
| VAPGNAGTA | GARQGAQDTQY | 0.06472899 |
| TGGGASLE | GARQGAQDTQY | 0.06871138 |
| VAVYNHYKRI | GARQGAQDTQY | 0.07354017 |
| TDPAPATT | GARQGAQDTQY | 0.06323483 |
| TNRPDILDPALLRPG | GARQGAQDTQY | 0.06706301 |
| LTTTTTS | GARQGAQDTQY | 0.062430333 |
| PTGSGKTL | GARQGAQDTQY | 0.06356497 |
| TGSGKTL | GARQGAQDTQY | 0.06364568 |
| TIGHVDHGK | GARQGAQDTQY | 0.062513135 |
| TIGHVDHGKTT | GARQGAQDTQY | 0.064429305 |
| GHVDHGKTT | GARQGAQDTQY | 0.07236907 |
| AQFEHTLL | GARQGAQDTQY | 0.06401838 |
| IVNQDEL | GARQGAQDTQY | 0.06482711 |
| TNGSQFF | GARQGAQDTQY | 0.06613655 |
| TGKGIVE | GARQGAQDTQY | 0.063500784 |
| LISNASDA | GARQGAQDTQY | 0.0645708 |
| AELRTLGELTKIAKAHGAQ | GARQGAQDTQY | 0.064244434 |
| APNAVAAAEPSW | GARQGAQDTQY | 0.06677341 |
| APQVAGAATAVL | GARQGAQDTQY | 0.062521264 |
| ARRDPAAAAAILPTDA | GARQGAQDTQY | 0.06327543 |
| DAAARDVLAEAGLAEYF | GARQGAQDTQY | 0.062964186 |
| DDQIVLSAKANLAKVEQ | GARQGAQDTQY | 0.06676461 |
| DELLPDTGEDSDEDGH | GARQGAQDTQY | 0.06879155 |
| ELPEAPANRATVPQD | GARQGAQDTQY | 0.062860265 |
| FGQSKIERVTGIG | GARQGAQDTQY | 0.06425768 |
| GAGGTGSTAGGAGGAGGAGGLY | GARQGAQDTQY | 0.06314155 |
| GHGGTGAAGGNGGTGGAGGSGI | GARQGAQDTQY | 0.06490709 |
| GNGGNGGAGGEGGPGGAGGAGA | GARQGAQDTQY | 0.062674776 |
| GSTAGNGGNGGNATNGTIAGQP | GARQGAQDTQY | 0.06451302 |
| GTGGDGGNGGNGGTGG | GARQGAQDTQY | 0.06347477 |
| GVRRPAGIPADAPV | GARQGAQDTQY | 0.065923735 |
| GYRQVTDAHLLTLARR | GARQGAQDTQY | 0.06476486 |
| IAKMGQDGHDRG | GARQGAQDTQY | 0.063355885 |
| KAEPSIEIQYT | GARQGAQDTQY | 0.06538015 |
| KPVRIRMAADEDTVAVT | GARQGAQDTQY | 0.063203074 |
| MPPRIAGMRLLVIKPEP | GARQGAQDTQY | 0.06470189 |
| MKFYAVALFRSRGVPPTR | GARQGAQDTQY | 0.064074524 |
| NAGGFGILGAVAHSP | GARQGAQDTQY | 0.06267602 |
| NEGREITLAMAEM | GARQGAQDTQY | 0.06442764 |
| NELVEWSHPLGHRWRWE | GARQGAQDTQY | 0.07305208 |
| NLGAGGTGGNGGNPG | GARQGAQDTQY | 0.06325385 |
| NTEWPKWETVMAADPVD | GARQGAQDTQY | 0.06391685 |
| PASAAPAETQP | GARQGAQDTQY | 0.06818476 |
| QQSSDGKNGVRAAEITGE | GARQGAQDTQY | 0.06429573 |

| | | |
|---|---|---|
| RAPDAVTTSWWKE | GARQGAQDTQY | 0.063896514 |
| SGFGNVGAHTSGWFNQ | GARQGAQDTQY | 0.062473655 |
| SGVAGAVGQAASVGGLKV | GARQGAQDTQY | 0.06534936 |
| SNGGAGTGSAGGNGGT | GARQGAQDTQY | 0.06381782 |
| TGGAGGEGGAAG | GARQGAQDTQY | 0.06418077 |
| TGGDGGAAGAGGAPGHG | GARQGAQDTQY | 0.0666642 |
| TMPPLLAAIANAGRHGV | GARQGAQDTQY | 0.06581541 |
| TSSTAGNSGGA | GARQGAQDTQY | 0.063287556 |
| VPATVPVGILC | GARQGAQDTQY | 0.06899833 |
| VQRPDSGGVVLTGRIS | GARQGAQDTQY | 0.06484285 |
| WTPNPVPATKGVPAGLVQAT | GARQGAQDTQY | 0.0634213 |
| YALKRALAKFSDRFDVVII | GARQGAQDTQY | 0.06237882 |
| YLIGSVIV | GARQGAQDTQY | 0.06439707 |
| YGQLVAVID | GARQGAQDTQY | 0.06299116 |
| VTVGGMTLPA | GARQGAQDTQY | 0.06544163 |
| VLALPAV | GARQGAQDTQY | 0.06562448 |
| VIPPTLNYET | GARQGAQDTQY | 0.06310569 |
| VGMGGMHPSSG | GARQGAQDTQY | 0.06755396 |
| VGLAGGAATA | GARQGAQDTQY | 0.06337222 |
| VGGAGGAGGSGP | GARQGAQDTQY | 0.06604451 |
| VFAAFMV | GARQGAQDTQY | 0.0623042 |
| TLYDKLR | GARQGAQDTQY | 0.063456975 |
| TLAVFTVL | GARQGAQDTQY | 0.06662963 |
| TGSVQPARV | GARQGAQDTQY | 0.0631496 |
| TGGGAGSGGT | GARQGAQDTQY | 0.06308526 |
| SHLETSL | GARQGAQDTQY | 0.0630004 |
| SFTTGMT | GARQGAQDTQY | 0.066120595 |
| QVEVAKRMGVRQPTVS | GARQGAQDTQY | 0.06470258 |
| QGSGGPALAM | GARQGAQDTQY | 0.062813155 |
| MLAPSNP | GARQGAQDTQY | 0.06266581 |
| LVNNAAI | GARQGAQDTQY | 0.064283326 |
| LTGEINTLA | GARQGAQDTQY | 0.062571466 |
| LSIFGPV | GARQGAQDTQY | 0.067646794 |
| LLALLANL | GARQGAQDTQY | 0.06636605 |
| LISNASDALDKLR | GARQGAQDTQY | 0.062544875 |
| LINEAVP | GARQGAQDTQY | 0.063534684 |
| LGYAFSGL | GARQGAQDTQY | 0.06312151 |
| LDFVAVAL | GARQGAQDTQY | 0.063625164 |
| LAQPGAE | GARQGAQDTQY | 0.06282029 |
| ISNLGAVTGF | GARQGAQDTQY | 0.06250381 |
| HQEDPGRI | GARQGAQDTQY | 0.06446153 |
| GTPGEPTP | GARQGAQDTQY | 0.06905274 |
| GFFNSGS | GARQGAQDTQY | 0.06337318 |
| GAGGGGLTAGAN | GARQGAQDTQY | 0.0645129 |
| FVDALVNM | GARQGAQDTQY | 0.06463168 |
| FLTVAPEL | GARQGAQDTQY | 0.06725835 |
| FETLTAL | GARQGAQDTQY | 0.06276862 |
| DVINPIEL | GARQGAQDTQY | 0.068047464 |
| APVSGAPV | GARQGAQDTQY | 0.062931865 |
| ALVGGLAW | GARQGAQDTQY | 0.06555577 |
| ALAAAAATAI | GARQGAQDTQY | 0.06517568 |

| | | |
|---|---|---|
| AFGLGLLL | GARQGAQDTQY | 0.063628726 |
| AFFAAASL | GARQGAQDTQY | 0.06319888 |
| AAAALPAFSPP | GARQGAQDTQY | 0.0632186 |
| VVLAPAL | GARQGAQDTQY | 0.06429337 |
| TIGHVDHGKTTL | GARQGAQDTQY | 0.06353562 |
| LISNASDALDK | GARQGAQDTQY | 0.06273518 |
| LISNASDAL | GARQGAQDTQY | 0.0735407 |
| ISNASDALDK | GARQGAQDTQY | 0.063719235 |
| NQAETLVYQTE | GARQGAQDTQY | 0.06965046 |
| TGGTGGTAGAG | GARQGAQDTQY | 0.063038066 |
| TGSTAGGAGGA | GARQGAQDTQY | 0.062984265 |
| DPLAVPVAAGPV | GARQGAQDTQY | 0.06498703 |
| GGTGGTPGNGGN | GARQGAQDTQY | 0.06252319 |
| PAGLAGVDPATL | GARQGAQDTQY | 0.06729888 |
| GNNLVVEAPHAGASV | GARQGAQDTQY | 0.064438984 |
| KVTRIMSASTLEQPA | GARQGAQDTQY | 0.06362848 |
| LGFALPIALPRRTVV | GARQGAQDTQY | 0.06276509 |
| TRLGNLTARKLVVLV | GARQGAQDTQY | 0.06492906 |
| WSQQRRLIYFPSAGP | GARQGAQDTQY | 0.06346842 |
| GPNGNPDPMAAAVDIR | GARQGAQDTQY | 0.06662395 |
| TQRCVSVADLRAAASR | GARQGAQDTQY | 0.062560245 |
| QFVLDVAEHPVEVLTETV | GARQGAQDTQY | 0.06583465 |
| GGAGGTGGAGGNGGTQAGDG | GARQGAQDTQY | 0.0630021 |
| PLLATLRAIAVDVDAKPAQV | GARQGAQDTQY | 0.06839105 |
| APKFRTGGYMIDFWGVGYQVAK | GARQGAQDTQY | 0.06644598 |
| GAGGAGGAGGAAGGPGGTGGTG | GARQGAQDTQY | 0.06837254 |
| IGALGLLATDVAPAGSVAFGVA | GARQGAQDTQY | 0.06547486 |
| AGGAGGIGGNGGAGGAGGVLMG | GARQGAQDTQY | 0.069916435 |
| ADELIGLL | GARQGAQDTQY | 0.06870479 |
| ADRALKA | GARQGAQDTQY | 0.063872434 |
| AGAVGQVGGAGGA | GARQGAQDTQY | 0.06365123 |
| AGGGGGLGGAP | GARQGAQDTQY | 0.064277105 |
| AGGLLYGNGGAG | GARQGAQDTQY | 0.066218235 |
| AGLESIGL | GARQGAQDTQY | 0.063589096 |
| AGLKTGII | GARQGAQDTQY | 0.062802754 |
| ASLAALLL | GARQGAQDTQY | 0.06262485 |
| ASVINISSI | GARQGAQDTQY | 0.06389012 |
| ATPPGPA | GARQGAQDTQY | 0.06345279 |
| ATVASALV | GARQGAQDTQY | 0.06619735 |
| ATVASLIL | GARQGAQDTQY | 0.063309304 |
| AVNGPVLIRH | GARQGAQDTQY | 0.07018822 |
| DGAAGAVGI | GARQGAQDTQY | 0.06623482 |
| DPLLHGI | GARQGAQDTQY | 0.067348816 |
| ELGAALGAL | GARQGAQDTQY | 0.062740475 |
| ELGRTQAV | GARQGAQDTQY | 0.06551635 |
| FVAGGYLS | GARQGAQDTQY | 0.06398533 |
| GASGGAGGAGGW | GARQGAQDTQY | 0.06325494 |
| GGAGAGGDGGAGGI | GARQGAQDTQY | 0.064703465 |
| GTGHNGSQPGMGGNG | GARQGAQDTQY | 0.06492532 |
| GYAAASASAS | GARQGAQDTQY | 0.06765801 |
| HQVDVAAVQQG | GARQGAQDTQY | 0.0620855 |

| | | |
|---|---|---|
| IGAAIAHT | GARQGAQDTQY | 0.0627999 |
| IGMIEVAQL | GARQGAQDTQY | 0.066468254 |
| IRLAPEHTSAMD | GARQGAQDTQY | 0.07051614 |
| IYNVSVL | GARQGAQDTQY | 0.06510085 |
| KQGFLGM | GARQGAQDTQY | 0.06492147 |
| LESVITQL | GARQGAQDTQY | 0.06253386 |
| LGLGALVVV | GARQGAQDTQY | 0.06603433 |
| LIGNGGNGGSGGI | GARQGAQDTQY | 0.06299573 |
| LNAINAPTL | GARQGAQDTQY | 0.063076206 |
| LQLLNTGV | GARQGAQDTQY | 0.06881288 |
| LQTSIEF | GARQGAQDTQY | 0.064675726 |
| MSFAVSEL | GARQGAQDTQY | 0.06260918 |
| NTGALGL | GARQGAQDTQY | 0.063088186 |
| PPGAGGAGGAGTT | GARQGAQDTQY | 0.063449554 |
| QLVGLGI | GARQGAQDTQY | 0.06275854 |
| RFLEQVHQL | GARQGAQDTQY | 0.062227223 |
| RGVGYVLR | GARQGAQDTQY | 0.063768536 |
| SATGVFAGVI | GARQGAQDTQY | 0.06370129 |
| SLPAITLP | GARQGAQDTQY | 0.06733643 |
| STALAFTV | GARQGAQDTQY | 0.06789694 |
| SVLTVGLGK | GARQGAQDTQY | 0.06356365 |
| TALAGAGSGI | GARQGAQDTQY | 0.06402006 |
| TGLPFTGI | GARQGAQDTQY | 0.07261318 |
| VAEHVTV | GARQGAQDTQY | 0.07744795 |
| VDIVLNSL | GARQGAQDTQY | 0.06364426 |
| VLTGGGAVL | GARQGAQDTQY | 0.063780725 |
| VNVGLAYGI | GARQGAQDTQY | 0.06334269 |
| VSSAAVGAV | GARQGAQDTQY | 0.06408312 |
| VTSVYEAGV | GARQGAQDTQY | 0.068864435 |
| WVALDRGA | GARQGAQDTQY | 0.06342295 |
| YLVGSVVI | GARQGAQDTQY | 0.0637619 |
| GPPGTGKTLL | GARQGAQDTQY | 0.06349074 |
| GVGMTKF | GARQGAQDTQY | 0.06438934 |
| IGIGGSDL | GARQGAQDTQY | 0.064484976 |
| IGTLLAEL | GARQGAQDTQY | 0.062141705 |
| IYTSGTT | GARQGAQDTQY | 0.06417176 |
| LSFPAGV | GARQGAQDTQY | 0.062759705 |
| LYTSGTT | GARQGAQDTQY | 0.06377235 |
| VGAFPFGF | GARQGAQDTQY | 0.06313697 |
| VGAGTVEF | GARQGAQDTQY | 0.06384861 |
| VGKSTLLL | GARQGAQDTQY | 0.062470783 |
| AALAHAGGGIV | GARQGAQDTQY | 0.063145116 |
| AALVAVSLL | GARQGAQDTQY | 0.063223675 |
| AAVSRAVAAL | GARQGAQDTQY | 0.07193602 |
| AGGSSGIGGF | GARQGAQDTQY | 0.068264045 |
| AGLTVANANG | GARQGAQDTQY | 0.06816853 |
| AGLVGAVL | GARQGAQDTQY | 0.06377173 |
| ALSQAFTV | GARQGAQDTQY | 0.0655301 |
| ALTFSVSL | GARQGAQDTQY | 0.064997286 |
| ALTGAALVAL | GARQGAQDTQY | 0.066942126 |
| ASFSIGAGVL | GARQGAQDTQY | 0.06840094 |

| | | |
|---|---|---|
| DGSVYKI | GARQGAQDTQY | 0.064810276 |
| EQLAEIV | GARQGAQDTQY | 0.069437616 |
| FAVSELTVM | GARQGAQDTQY | 0.067704685 |
| GAAPLAAGL | GARQGAQDTQY | 0.063937984 |
| GADGGDGGSGGAGG | GARQGAQDTQY | 0.06337675 |
| GFGLDTGGAGG | GARQGAQDTQY | 0.06242598 |
| GLASGGLL | GARQGAQDTQY | 0.062821604 |
| GRIAGAAVLL | GARQGAQDTQY | 0.06457039 |
| GSAGSLNTSF | GARQGAQDTQY | 0.0630326 |
| GTGQIGI | GARQGAQDTQY | 0.06365958 |
| GTLIALIL | GARQGAQDTQY | 0.06566361 |
| IAATMGFVVG | GARQGAQDTQY | 0.07210442 |
| IFAQALL | GARQGAQDTQY | 0.06485303 |
| IGAAVALGF | GARQGAQDTQY | 0.06346367 |
| IGLTPLAL | GARQGAQDTQY | 0.06327195 |
| ILGSNTSTL | GARQGAQDTQY | 0.06321761 |
| IVFSLTVL | GARQGAQDTQY | 0.063338995 |
| LASKATGF | GARQGAQDTQY | 0.06353956 |
| LASVALIL | GARQGAQDTQY | 0.06279227 |
| LFAVGLTL | GARQGAQDTQY | 0.06254258 |
| LFNGGIGTYI | GARQGAQDTQY | 0.06336183 |
| LTLGAGII | GARQGAQDTQY | 0.06279417 |
| LVAAALGDI | GARQGAQDTQY | 0.063536845 |
| NTAFVQLGI | GARQGAQDTQY | 0.06921715 |
| QLFSGAEL | GARQGAQDTQY | 0.06365351 |
| SAATFITL | GARQGAQDTQY | 0.06620954 |
| SNYGLANLGV | GARQGAQDTQY | 0.06961549 |
| SYGGAYAVM | GARQGAQDTQY | 0.062939204 |
| TTGVGGLTLG | GARQGAQDTQY | 0.06403873 |
| TTQTIYI | GARQGAQDTQY | 0.063542694 |
| VAAGAGLGAVI | GARQGAQDTQY | 0.086666256 |
| VAAVVTLL | GARQGAQDTQY | 0.06282735 |
| VGFLGGLI | GARQGAQDTQY | 0.0656047 |
| VGVIPILL | GARQGAQDTQY | 0.06592762 |
| VIFASSLI | GARQGAQDTQY | 0.062675044 |
| VKGGDGGAAAG | GARQGAQDTQY | 0.06280481 |
| VTKTFTNI | GARQGAQDTQY | 0.06463179 |
| VVAAAILL | GARQGAQDTQY | 0.06314896 |
| YAALDVEL | GARQGAQDTQY | 0.06374783 |
| LASVALLL | GARQGAQDTQY | 0.06408133 |
| LDAVHRV | GARQGAQDTQY | 0.06358433 |
| QGLSSSIF | GARQGAQDTQY | 0.06382737 |
| RGTFANIRL | GARQGAQDTQY | 0.062345196 |
| RQLWWGHRI | GARQGAQDTQY | 0.06401173 |
| YGPPGTGKTLL | GARQGAQDTQY | 0.0655911 |
| AALVGAPANVA | GARQGAQDTQY | 0.06401676 |
| AATFRRTE | GARQGAQDTQY | 0.0692569 |
| AGALGGGI | GARQGAQDTQY | 0.064567536 |
| AGGAGGAGGIGAG | GARQGAQDTQY | 0.066799484 |
| AGSGALGL | GARQGAQDTQY | 0.0637537 |
| AQIADQI | GARQGAQDTQY | 0.06284153 |

| | | |
|---|---|---|
| ASAVAVTYL | GARQGAQDTQY | 0.06462289 |
| ASTVLGGTGG | GARQGAQDTQY | 0.06261201 |
| AVGGEPAK | GARQGAQDTQY | 0.06323251 |
| CEQGTFGGTC | GARQGAQDTQY | 0.06510781 |
| DEGKAPAG | GARQGAQDTQY | 0.062941454 |
| DTVVFPAL | GARQGAQDTQY | 0.06439603 |
| FEGITMAG | GARQGAQDTQY | 0.068944186 |
| FPLVVQD | GARQGAQDTQY | 0.06342344 |
| GAFTTPQ | GARQGAQDTQY | 0.06318257 |
| GFGGFGTAGGVGGA | GARQGAQDTQY | 0.06379794 |
| GFLAQFL | GARQGAQDTQY | 0.06440619 |
| GGAGGAGGNAGVA | GARQGAQDTQY | 0.068477996 |
| GGVGTAGAGGAGGAGG | GARQGAQDTQY | 0.06314717 |
| GNGGGGDAGNAGSGGN | GARQGAQDTQY | 0.06507257 |
| GTGGAGGAGNDAGS | GARQGAQDTQY | 0.06481759 |
| GTTNTGWFNS | GARQGAQDTQY | 0.06515303 |
| IFMGPLFTL | GARQGAQDTQY | 0.06566968 |
| ILAHAIEAI | GARQGAQDTQY | 0.06712917 |
| IRSAIREAL | GARQGAQDTQY | 0.0655212 |
| ITFAGGG | GARQGAQDTQY | 0.0629279 |
| ITGMSGHQEV | GARQGAQDTQY | 0.06506982 |
| IVAGLACTL | GARQGAQDTQY | 0.063486814 |
| IVGGGLAGA | GARQGAQDTQY | 0.07255555 |
| IVTGVLNI | GARQGAQDTQY | 0.06401091 |
| LAQQGCI | GARQGAQDTQY | 0.06424269 |
| LCLHVEN | GARQGAQDTQY | 0.06311526 |
| LDAISDGS | GARQGAQDTQY | 0.0764179 |
| LGGFAAML | GARQGAQDTQY | 0.0630845 |
| LGGKNPMI | GARQGAQDTQY | 0.06718934 |
| LGPKGVAAIL | GARQGAQDTQY | 0.066727355 |
| LMVGVSI | GARQGAQDTQY | 0.06278775 |
| LVGAAFVL | GARQGAQDTQY | 0.06823585 |
| LVLAVAPL | GARQGAQDTQY | 0.06393058 |
| MAAIAVIDAA | GARQGAQDTQY | 0.06352518 |
| NNGINAELQ | GARQGAQDTQY | 0.062428094 |
| RGEPLVLVHG | GARQGAQDTQY | 0.06313086 |
| RGGAGGAAGLI | GARQGAQDTQY | 0.06418997 |
| RSYRMHPAV | GARQGAQDTQY | 0.06315945 |
| SAAHLLAGQLA | GARQGAQDTQY | 0.06361928 |
| SAAVGSY | GARQGAQDTQY | 0.06443204 |
| SPHAVLHRI | GARQGAQDTQY | 0.06395426 |
| SQIAKVL | GARQGAQDTQY | 0.0640578 |
| TALGHLVP | GARQGAQDTQY | 0.072074294 |
| TAPTTSGAA | GARQGAQDTQY | 0.0640197 |
| TDGDDGNGGSPGT | GARQGAQDTQY | 0.07568363 |
| TDGLGEN | GARQGAQDTQY | 0.06338287 |
| TGGPPLPGQA | GARQGAQDTQY | 0.06283566 |
| TIPAITI | GARQGAQDTQY | 0.06441387 |
| TIPGFTI | GARQGAQDTQY | 0.065428376 |
| TSIASGALA | GARQGAQDTQY | 0.06310204 |
| TVTLPGAAI | GARQGAQDTQY | 0.06909877 |

| | | |
|---|---|---|
| VASGLSAALP | GARQGAQDTQY | 0.07055578 |
| VGAGVIGI | GARQGAQDTQY | 0.06365683 |
| VGPKRFGV | GARQGAQDTQY | 0.06782614 |
| VNVAFPD | GARQGAQDTQY | 0.062307607 |
| VSMTNSGVS | GARQGAQDTQY | 0.062935926 |
| VTDVLGDLSV | GARQGAQDTQY | 0.0625533 |
| VVVVGAGFAGL | GARQGAQDTQY | 0.06505881 |
| VYTVGTVV | GARQGAQDTQY | 0.06611863 |
| WTTFSGP | GARQGAQDTQY | 0.070920765 |
| YDELVEAF | GARQGAQDTQY | 0.06296449 |
| YGTGGAGGNGG | GARQGAQDTQY | 0.06765575 |
| YTNVMAV | GARQGAQDTQY | 0.0661568 |
| FLEQVHQL | GARQGAQDTQY | 0.06272319 |
| GLSLLAVAV | GARQGAQDTQY | 0.0655196 |
| LEQVHQL | GARQGAQDTQY | 0.06326655 |
| LNAAGAI | GARQGAQDTQY | 0.0629559 |
| PAVHASL | GARQGAQDTQY | 0.06327389 |
| TELEDDA | GARQGAQDTQY | 0.06416391 |
| TPPGPPN | GARQGAQDTQY | 0.063774735 |
| TPPPPPPP | GARQGAQDTQY | 0.06645683 |
| TQSPALI | ASSEPSDQDTQY | 0.066130675 |
| EFAAAFPQ | ASSEPSDQDTQY | 0.06432341 |
| LLSGNLDVL | ASSEPSDQDTQY | 0.063804284 |
| GPANLGNYNI | ASSEPSDQDTQY | 0.062233064 |
| TLGPTLP | ASSEPSDQDTQY | 0.066282205 |
| VYAYSSCCK | ASSEPSDQDTQY | 0.07443882 |
| APAPAPAPNI | ASSEPSDQDTQY | 0.06533367 |
| PGGPVPIP | ASSEPSDQDTQY | 0.06587926 |
| FLTVALEP | ASSEPSDQDTQY | 0.064954646 |
| SQYFGANL | ASSEPSDQDTQY | 0.06538731 |
| GGVKPAA | ASSEPSDQDTQY | 0.06480599 |
| GGTGGAGGAGGW | ASSEPSDQDTQY | 0.06307206 |
| GVGGVAGLGGAGG | ASSEPSDQDTQY | 0.064258896 |
| VLLVLATTI | ASSEPSDQDTQY | 0.06317567 |
| IGIFAAML | ASSEPSDQDTQY | 0.06296835 |
| AVGDLID | ASSEPSDQDTQY | 0.063665815 |
| NISPLPL | ASSEPSDQDTQY | 0.07481947 |
| VTAVFVAI | ASSEPSDQDTQY | 0.06464212 |
| SGDDNDAAL | ASSEPSDQDTQY | 0.0666123 |
| AVAALAHS | ASSEPSDQDTQY | 0.063836776 |
| GGVTVMATP | ASSEPSDQDTQY | 0.06350506 |
| AGLAANVRD | ASSEPSDQDTQY | 0.064095624 |
| TNRPDLIDPALLRPG | ASSEPSDQDTQY | 0.06317645 |
| LANLGVTAL | ASSEPSDQDTQY | 0.072267756 |
| TGERIEL | ASSEPSDQDTQY | 0.063227475 |
| NGEELAA | ASSEPSDQDTQY | 0.06280698 |
| VLSLVWG | ASSEPSDQDTQY | 0.06520437 |
| IGYALWL | ASSEPSDQDTQY | 0.07586084 |
| LSLLLAPL | ASSEPSDQDTQY | 0.06456493 |
| TLDEALK | ASSEPSDQDTQY | 0.06462362 |
| TDGAGTAEGG | ASSEPSDQDTQY | 0.06526341 |

| | | |
|---|---|---|
| NAPGGAGGNGG | ASSEPSDQDTQY | 0.06297317 |
| VSVFMGMGGNF | ASSEPSDQDTQY | 0.064121656 |
| LASYLAAL | ASSEPSDQDTQY | 0.06386748 |
| LVGGTPAL | ASSEPSDQDTQY | 0.06782623 |
| VRNADVL | ASSEPSDQDTQY | 0.06380289 |
| TCSSPFLVSAM | ASSEPSDQDTQY | 0.064099364 |
| VHITDAE | ASSEPSDQDTQY | 0.0671023 |
| EYAAPSF | ASSEPSDQDTQY | 0.064756244 |
| VVDAFHGGL | ASSEPSDQDTQY | 0.064506784 |
| ASVVDPQ | ASSEPSDQDTQY | 0.08916074 |
| SYNEFGGLS | ASSEPSDQDTQY | 0.06564894 |
| DPDGSVY | ASSEPSDQDTQY | 0.064606264 |
| MVSTIHATE | ASSEPSDQDTQY | 0.06866438 |
| LAAGVADPV | ASSEPSDQDTQY | 0.06259526 |
| TKEQIAA | ASSEPSDQDTQY | 0.06393476 |
| SLASPAAL | ASSEPSDQDTQY | 0.06338259 |
| IGLVFYGV | ASSEPSDQDTQY | 0.06254422 |
| FLLDASNL | ASSEPSDQDTQY | 0.062698536 |
| YQVGNFP | ASSEPSDQDTQY | 0.06430169 |
| ESLLSQST | ASSEPSDQDTQY | 0.06341228 |
| LFTVGFAAVG | ASSEPSDQDTQY | 0.071535386 |
| ATPPTPT | ASSEPSDQDTQY | 0.062471017 |
| TDLATLG | ASSEPSDQDTQY | 0.06321553 |
| SAGGTGR | ASSEPSDQDTQY | 0.06760995 |
| GFSNSGS | ASSEPSDQDTQY | 0.067832984 |
| GGNGGSGGTSV | ASSEPSDQDTQY | 0.06538733 |
| GTSVATGGAGN | ASSEPSDQDTQY | 0.06331327 |
| NGGNGGSGGTS | ASSEPSDQDTQY | 0.06332534 |
| AVVKFPH | ASSEPSDQDTQY | 0.063777156 |
| TPPPPPP | ASSEPSDQDTQY | 0.06441751 |
| AAPAAAAAPA | ASSEPSDQDTQY | 0.06244388 |
| IALNEDL | ASSEPSDQDTQY | 0.06497717 |
| IVNVSSM | ASSEPSDQDTQY | 0.065065145 |
| TGIPPAPR | ASSEPSDQDTQY | 0.062421944 |
| FDLGGGTF | ASSEPSDQDTQY | 0.06499937 |
| FDLGGGTFDVS | ASSEPSDQDTQY | 0.06581154 |
| NEPTAAA | ASSEPSDQDTQY | 0.06295177 |
| NEPTAAALA | ASSEPSDQDTQY | 0.062423542 |
| GSIINISSI | ASSEPSDQDTQY | 0.06712669 |
| VAPGNAGTA | ASSEPSDQDTQY | 0.07337667 |
| TGGGASLE | ASSEPSDQDTQY | 0.063175276 |
| VAVYNHYKRI | ASSEPSDQDTQY | 0.06356133 |
| TDPAPATT | ASSEPSDQDTQY | 0.062485766 |
| TNRPDILDPALLRPG | ASSEPSDQDTQY | 0.08039135 |
| LTTTTTS | ASSEPSDQDTQY | 0.06442302 |
| PTGSGKTL | ASSEPSDQDTQY | 0.06502494 |
| TGSGKTL | ASSEPSDQDTQY | 0.06266436 |
| TIGHVDHGK | ASSEPSDQDTQY | 0.06264679 |
| TIGHVDHGKTT | ASSEPSDQDTQY | 0.07004036 |
| GHVDHGKTT | ASSEPSDQDTQY | 0.062554784 |
| AQFEHTLL | ASSEPSDQDTQY | 0.0628049 |

| | | |
|---|---|---|
| IVNQDEL | ASSEPSDQDTQY | 0.063934766 |
| TNGSQFF | ASSEPSDQDTQY | 0.06353944 |
| TGKGIVE | ASSEPSDQDTQY | 0.062921815 |
| LISNASDA | ASSEPSDQDTQY | 0.0633395 |
| AELRTLGELTKIAKAHGAQ | ASSEPSDQDTQY | 0.107782975 |
| APNAVAAAEPSW | ASSEPSDQDTQY | 0.062780075 |
| APQVAGAATAVL | ASSEPSDQDTQY | 0.06441981 |
| ARRDPAAAAAILPTDA | ASSEPSDQDTQY | 0.06402887 |
| DAAARDVLAEAGLAEYF | ASSEPSDQDTQY | 0.073252626 |
| DDQIVLSAKANLAKVEQ | ASSEPSDQDTQY | 0.06434459 |
| DELLPDTGEDSDEDGH | ASSEPSDQDTQY | 0.06323721 |
| ELPEAPANRATVPQD | ASSEPSDQDTQY | 0.06305088 |
| FGQSKIERVTGIG | ASSEPSDQDTQY | 0.06441527 |
| GAGGTGSTAGGAGGAGGAGGLY | ASSEPSDQDTQY | 0.06432504 |
| GHGGTGAAGGNGGTGGAGGSGI | ASSEPSDQDTQY | 0.06413581 |
| GNGGNGGAGGEGGPGGAGGAGA | ASSEPSDQDTQY | 0.06353518 |
| GSTAGNGGNGGNATNGTIAGQP | ASSEPSDQDTQY | 0.06811187 |
| GTGGDGGNGGNGGTGG | ASSEPSDQDTQY | 0.0634756 |
| GVRRPAGIPADAPV | ASSEPSDQDTQY | 0.062375117 |
| GYRQVTDAHLLTLARR | ASSEPSDQDTQY | 0.06394768 |
| IAKMGQDGHDRG | ASSEPSDQDTQY | 0.06634468 |
| KAEPSIEIQYT | ASSEPSDQDTQY | 0.06383546 |
| KPVRIRMAADEDTVAVT | ASSEPSDQDTQY | 0.063432306 |
| MPPRIAGMRLLVIKPEP | ASSEPSDQDTQY | 0.06249711 |
| MKFYAVALFRSRGVPPTR | ASSEPSDQDTQY | 0.07040928 |
| NAGGFGILGAVAHSP | ASSEPSDQDTQY | 0.064565234 |
| NEGREITLAMAEM | ASSEPSDQDTQY | 0.06289801 |
| NELVEWSHPLGHRWRWE | ASSEPSDQDTQY | 0.06465886 |
| NLGAGGTGGNGGNPG | ASSEPSDQDTQY | 0.0629189 |
| NTEWPKWETVMAADPVD | ASSEPSDQDTQY | 0.06424496 |
| PASAAPAETQP | ASSEPSDQDTQY | 0.06320962 |
| QQSSDGKNGVRAAEITGE | ASSEPSDQDTQY | 0.06312215 |
| RAPDAVTTSWWKE | ASSEPSDQDTQY | 0.06964682 |
| SGFGNVGAHTSGWFNQ | ASSEPSDQDTQY | 0.06343413 |
| SGVAGAVGQAASVGGLKV | ASSEPSDQDTQY | 0.06287274 |
| SNGGAGTGSAGGNGGT | ASSEPSDQDTQY | 0.06881824 |
| TGGAGGEGGAAG | ASSEPSDQDTQY | 0.062802896 |
| TGGDGGAAGAGGAPGHG | ASSEPSDQDTQY | 0.065856844 |
| TMPPLLAAIANAGRHGV | ASSEPSDQDTQY | 0.06463573 |
| TSSTAGNSGGA | ASSEPSDQDTQY | 0.06451287 |
| VPATVPVGILC | ASSEPSDQDTQY | 0.06392189 |
| VQRPDSGGVVLTGRIS | ASSEPSDQDTQY | 0.0632785 |
| WTPNPVPATKGVPAGLVQAT | ASSEPSDQDTQY | 0.063333906 |
| YALKRALAKFSDRFDVVII | ASSEPSDQDTQY | 0.06399385 |
| YLIGSVIV | ASSEPSDQDTQY | 0.07581562 |
| YGQLVAVID | ASSEPSDQDTQY | 0.0654383 |
| VTVGGMTLPA | ASSEPSDQDTQY | 0.06416219 |
| VLALPAV | ASSEPSDQDTQY | 0.06374676 |
| VIPPTLNYET | ASSEPSDQDTQY | 0.06369024 |
| VGMGGMHPSSG | ASSEPSDQDTQY | 0.06388583 |
| VGLAGGAATA | ASSEPSDQDTQY | 0.06280568 |

| | | |
|---|---|---|
| VGGAGGAGGSGP | ASSEPSDQDTQY | 0.063646555 |
| VFAAFMV | ASSEPSDQDTQY | 0.062539116 |
| TLYDKLR | ASSEPSDQDTQY | 0.063964486 |
| TLAVFTVL | ASSEPSDQDTQY | 0.06294438 |
| TGSVQPARV | ASSEPSDQDTQY | 0.0666019 |
| TGGGAGSGGT | ASSEPSDQDTQY | 0.06456974 |
| SHLETSL | ASSEPSDQDTQY | 0.06256523 |
| SFTTGMT | ASSEPSDQDTQY | 0.063402444 |
| QVEVAKRMGVRQPTVS | ASSEPSDQDTQY | 0.06413059 |
| QGSGGPALAM | ASSEPSDQDTQY | 0.06274995 |
| MLAPSNP | ASSEPSDQDTQY | 0.06610295 |
| LVNNAAI | ASSEPSDQDTQY | 0.06355604 |
| LTGEINTLA | ASSEPSDQDTQY | 0.066022485 |
| LSIFGPV | ASSEPSDQDTQY | 0.072546355 |
| LLALLANL | ASSEPSDQDTQY | 0.064732924 |
| LISNASDALDKLR | ASSEPSDQDTQY | 0.06342408 |
| LINEAVP | ASSEPSDQDTQY | 0.06301949 |
| LGYAFSGL | ASSEPSDQDTQY | 0.06339388 |
| LDFVAVAL | ASSEPSDQDTQY | 0.06429073 |
| LAQPGAE | ASSEPSDQDTQY | 0.062305763 |
| ISNLGAVTGF | ASSEPSDQDTQY | 0.06590988 |
| HQEDPGRI | ASSEPSDQDTQY | 0.06530192 |
| GTPGEPTP | ASSEPSDQDTQY | 0.0634314 |
| GFFNSGS | ASSEPSDQDTQY | 0.06258253 |
| GAGGGGLTAGAN | ASSEPSDQDTQY | 0.069365636 |
| FVDALVNM | ASSEPSDQDTQY | 0.06802765 |
| FLTVAPEL | ASSEPSDQDTQY | 0.06466156 |
| FETLTAL | ASSEPSDQDTQY | 0.064555906 |
| DVINPIEL | ASSEPSDQDTQY | 0.06254259 |
| APVSGAPV | ASSEPSDQDTQY | 0.06752955 |
| ALVGGLAW | ASSEPSDQDTQY | 0.068805486 |
| ALAAAAATAI | ASSEPSDQDTQY | 0.06354913 |
| AFGLGLLL | ASSEPSDQDTQY | 0.067595355 |
| AFFAAASL | ASSEPSDQDTQY | 0.06308022 |
| AAAALPAFSPP | ASSEPSDQDTQY | 0.06332908 |
| VVLAPAL | ASSEPSDQDTQY | 0.06535892 |
| TIGHVDHGKTTL | ASSEPSDQDTQY | 0.06698504 |
| LISNASDALDK | ASSEPSDQDTQY | 0.06407437 |
| LISNASDAL | ASSEPSDQDTQY | 0.062850274 |
| ISNASDALDK | ASSEPSDQDTQY | 0.06432927 |
| NQAETLVYQTE | ASSEPSDQDTQY | 0.06373053 |
| TGGTGGTAGAG | ASSEPSDQDTQY | 0.065707065 |
| TGSTAGGAGGA | ASSEPSDQDTQY | 0.0729425 |
| DPLAVPVAAGPV | ASSEPSDQDTQY | 0.06751485 |
| GGTGGTPGNGGN | ASSEPSDQDTQY | 0.0644207 |
| PAGLAGVDPATL | ASSEPSDQDTQY | 0.07463211 |
| GNNLVVEAPHAGASV | ASSEPSDQDTQY | 0.063366674 |
| KVTRIMSASTLEQPA | ASSEPSDQDTQY | 0.06447709 |
| LGFALPIALPRRTVV | ASSEPSDQDTQY | 0.06365782 |
| TRLGNLTARKLVVLV | ASSEPSDQDTQY | 0.0696475 |
| WSQQRRLIYFPSAGP | ASSEPSDQDTQY | 0.06279855 |

| | | |
|---|---|---|
| GPNGNPDPMAAAVDIR | ASSEPSDQDTQY | 0.063084066 |
| TQRCVSVADLRAAASR | ASSEPSDQDTQY | 0.06359625 |
| QFVLDVAEHPVEVLTETV | ASSEPSDQDTQY | 0.52075136 |
| GGAGGTGGAGGNGGTQAGDG | ASSEPSDQDTQY | 0.78083795 |
| PLLATLRAIAVDVDAKPAQV | ASSEPSDQDTQY | 0.64066756 |
| APKFRTGGYMIDFWGVGYQVAK | ASSEPSDQDTQY | 0.6088167 |
| GAGGAGGAGGAAGGPGGTGGTG | ASSEPSDQDTQY | 0.38873827 |
| IGALGLLATDVAPAGSVAFGVA | ASSEPSDQDTQY | 0.22755376 |
| AGGAGGIGGNGGAGGAGGVLMG | ASSEPSDQDTQY | 0.58576936 |
| ADELIGLL | ASSEPSDQDTQY | 0.3380544 |
| ADRALKA | ASSEPSDQDTQY | 0.75900984 |
| AGAVGQVGGAGGA | ASSEPSDQDTQY | 0.7266762 |
| AGGGGGLGGAP | ASSEPSDQDTQY | 0.4070269 |
| AGGLLYGNGGAG | ASSEPSDQDTQY | 0.80977947 |
| AGLESIGL | ASSEPSDQDTQY | 0.7915541 |
| AGLKTGII | ASSEPSDQDTQY | 0.26725513 |
| ASLAALLL | ASSEPSDQDTQY | 0.56707084 |
| ASVINISSI | ASSEPSDQDTQY | 0.42603716 |
| ATPPGPA | ASSEPSDQDTQY | 0.59574246 |
| ATVASALV | ASSEPSDQDTQY | 0.20397055 |
| ATVASLIL | ASSEPSDQDTQY | 0.7454282 |
| AVNGPVLIRH | ASSEPSDQDTQY | 0.41826203 |
| DGAAGAVGI | ASSEPSDQDTQY | 0.13992976 |
| DPLLHGI | ASSEPSDQDTQY | 0.63457924 |
| ELGAALGAL | ASSEPSDQDTQY | 0.61656433 |
| ELGRTQAV | ASSEPSDQDTQY | 0.5616149 |
| FVAGGYLS | ASSEPSDQDTQY | 0.7806252 |
| GASGGAGGAGGW | ASSEPSDQDTQY | 0.27832907 |
| GGAGAGGDGGAGGI | ASSEPSDQDTQY | 0.3750953 |
| GTGHNGSQPGMGGNG | ASSEPSDQDTQY | 0.76283884 |
| GYAAASASAS | ASSEPSDQDTQY | 0.8103455 |
| HQVDVAAVQQG | ASSEPSDQDTQY | 0.30826956 |
| IGAAIAHT | ASSEPSDQDTQY | 0.7036394 |
| IGMIEVAQL | ASSEPSDQDTQY | 0.75709724 |
| IRLAPEHTSAMD | ASSEPSDQDTQY | 0.6164031 |
| IYNVSVL | ASSEPSDQDTQY | 0.7676556 |
| KQGFLGM | ASSEPSDQDTQY | 0.7936653 |
| LESVITQL | ASSEPSDQDTQY | 0.3223759 |
| LGLGALVVV | ASSEPSDQDTQY | 0.802209 |
| LIGNGGNGGSGGI | ASSEPSDQDTQY | 0.80816317 |
| LNAINAPTL | ASSEPSDQDTQY | 0.4938898 |
| LQLLNTGV | ASSEPSDQDTQY | 0.6313601 |
| LQTSIEF | ASSEPSDQDTQY | 0.36410517 |
| MSFAVSEL | ASSEPSDQDTQY | 0.49381953 |
| NTGALGL | ASSEPSDQDTQY | 0.1994551 |
| PPGAGGAGGAGTT | ASSEPSDQDTQY | 0.7056905 |
| QLVGLGI | ASSEPSDQDTQY | 0.69774795 |
| RFLEQVHQL | ASSEPSDQDTQY | 0.5691172 |
| RGVGYVLR | ASSEPSDQDTQY | 0.38550425 |
| SATGVFAGVI | ASSEPSDQDTQY | 0.39180037 |
| SLPAITLP | ASSEPSDQDTQY | 0.39033356 |

| | | |
|---|---|---|
| STALAFTV | ASSEPSDQDTQY | 0.5661603 |
| SVLTVGLGK | ASSEPSDQDTQY | 0.21757016 |
| TALAGAGSGI | ASSEPSDQDTQY | 0.3759908 |
| TGLPFTGI | ASSEPSDQDTQY | 0.7413709 |
| VAEHVTV | ASSEPSDQDTQY | 0.72216016 |
| VDIVLNSL | ASSEPSDQDTQY | 0.22594915 |
| VLTGGGAVL | ASSEPSDQDTQY | 0.8027031 |
| VNVGLAYGI | ASSEPSDQDTQY | 0.7531514 |
| VSSAAVGAV | ASSEPSDQDTQY | 0.77278864 |
| VTSVYEAGV | ASSEPSDQDTQY | 0.5267925 |
| WVALDRGA | ASSEPSDQDTQY | 0.39770964 |
| YLVGSVVI | ASSEPSDQDTQY | 0.7520646 |
| GPPGTGKTLL | ASSEPSDQDTQY | 0.52510726 |
| GVGMTKF | ASSEPSDQDTQY | 0.31782234 |
| IGIGGSDL | ASSEPSDQDTQY | 0.7263836 |
| IGTLLAEL | ASSEPSDQDTQY | 0.48584405 |
| IYTSGTT | ASSEPSDQDTQY | 0.3882183 |
| LSFPAGV | ASSEPSDQDTQY | 0.55562854 |
| LYTSGTT | ASSEPSDQDTQY | 0.687 |
| VGAFPFGF | ASSEPSDQDTQY | 0.3519525 |
| VGAGTVEF | ASSEPSDQDTQY | 0.33319718 |
| VGKSTLLL | ASSEPSDQDTQY | 0.3559769 |
| AALAHAGGGIV | ASSEPSDQDTQY | 0.6261389 |
| AALVAVSLL | ASSEPSDQDTQY | 0.7936814 |
| AAVSRAVAAL | ASSEPSDQDTQY | 0.6774792 |
| AGGSSGIGGF | ASSEPSDQDTQY | 0.3787923 |
| AGLTVANANG | ASSEPSDQDTQY | 0.7405559 |
| AGLVGAVL | ASSEPSDQDTQY | 0.63317764 |
| ALSQAFTV | ASSEPSDQDTQY | 0.54481626 |
| ALTFSVSL | ASSEPSDQDTQY | 0.76984626 |
| ALTGAALVAL | ASSEPSDQDTQY | 0.4464289 |
| ASFSIGAGVL | ASSEPSDQDTQY | 0.7325119 |
| DGSVYKI | ASSEPSDQDTQY | 0.6711843 |
| EQLAEIV | ASSEPSDQDTQY | 0.74105257 |
| FAVSELTVM | ASSEPSDQDTQY | 0.7748399 |
| GAAPLAAGL | ASSEPSDQDTQY | 0.50081897 |
| GADGGDGGSGGAGG | ASSEPSDQDTQY | 0.5030722 |
| GFGLDTGGAGG | ASSEPSDQDTQY | 0.2023149 |
| GLASGGLL | ASSEPSDQDTQY | 0.35615253 |
| GRIAGAAVLL | ASSEPSDQDTQY | 0.71874726 |
| GSAGSLNTSF | ASSEPSDQDTQY | 0.65031576 |
| GTGQIGI | ASSEPSDQDTQY | 0.7603228 |
| GTLIALIL | ASSEPSDQDTQY | 0.66584504 |
| IAATMGFVVG | ASSEPSDQDTQY | 0.39192194 |
| IFAQALL | ASSEPSDQDTQY | 0.7609722 |
| IGAAVALGF | ASSEPSDQDTQY | 0.76653075 |
| IGLTPLAL | ASSEPSDQDTQY | 0.24869294 |
| ILGSNTSTL | ASSEPSDQDTQY | 0.42487586 |
| IVFSLTVL | ASSEPSDQDTQY | 0.8275182 |
| LASKATGF | ASSEPSDQDTQY | 0.71498466 |
| LASVALIL | ASSEPSDQDTQY | 0.5464761 |

| | | |
|---|---|---|
| LFAVGLTL | ASSEPSDQDTQY | 0.4668763 |
| LFNGGIGTYI | ASSEPSDQDTQY | 0.83178484 |
| LTLGAGII | ASSEPSDQDTQY | 0.50961256 |
| LVAAALGDI | ASSEPSDQDTQY | 0.80848235 |
| NTAFVQLGI | ASSEPSDQDTQY | 0.16690408 |
| QLFSGAEL | ASSEPSDQDTQY | 0.49893248 |
| SAATFITL | ASSEPSDQDTQY | 0.33180618 |
| SNYGLANLGV | ASSEPSDQDTQY | 0.24448797 |
| SYGGAYAVM | ASSEPSDQDTQY | 0.7350299 |
| TTGVGGLTLG | ASSEPSDQDTQY | 0.5769156 |
| TTQTIYI | ASSEPSDQDTQY | 0.76042545 |
| VAAGAGLGAVI | ASSEPSDQDTQY | 0.5804782 |
| VAAVVTLL | ASSEPSDQDTQY | 0.7760303 |
| VGFLGGLI | ASSEPSDQDTQY | 0.6660898 |
| VGVIPILL | ASSEPSDQDTQY | 0.6440023 |
| VIFASSLI | ASSEPSDQDTQY | 0.39769384 |
| VKGGDGGAAAG | ASSEPSDQDTQY | 0.7194873 |
| VTKTFTNI | ASSEPSDQDTQY | 0.7934821 |
| VVAAAILL | ASSEPSDQDTQY | 0.66794246 |
| YAALDVEL | ASSEPSDQDTQY | 0.25999835 |
| LASVALLL | ASSEPSDQDTQY | 0.5983355 |
| LDAVHRV | ASSEPSDQDTQY | 0.59113216 |
| QGLSSSIF | ASSEPSDQDTQY | 0.18032925 |
| RGTFANIRL | ASSEPSDQDTQY | 0.17438515 |
| RQLWWGHRI | ASSEPSDQDTQY | 0.3924787 |
| YGPPGTGKTLL | ASSEPSDQDTQY | 0.35238266 |
| AALVGAPANVA | ASSEPSDQDTQY | 0.7949041 |
| AATFRRTE | ASSEPSDQDTQY | 0.7541103 |
| AGALGGGI | ASSEPSDQDTQY | 0.60571575 |
| AGGAGGAGGIGAG | ASSEPSDQDTQY | 0.6279402 |
| AGSGALGL | ASSEPSDQDTQY | 0.6148571 |
| AQIADQI | ASSEPSDQDTQY | 0.1268036 |
| ASAVAVTYL | ASSEPSDQDTQY | 0.31633723 |
| ASTVLGGTGG | ASSEPSDQDTQY | 0.29263994 |
| AVGGEPAK | ASSEPSDQDTQY | 0.8000052 |
| CEQGTFGGTC | ASSEPSDQDTQY | 0.37964997 |
| DEGKAPAG | ASSEPSDQDTQY | 0.3625151 |
| DTVVFPAL | ASSEPSDQDTQY | 0.62271535 |
| FEGITMAG | ASSEPSDQDTQY | 0.7006406 |
| FPLVVQD | ASSEPSDQDTQY | 0.58861804 |
| GAFTTPQ | ASSEPSDQDTQY | 0.27889773 |
| GFGGFGTAGGVGGA | ASSEPSDQDTQY | 0.14192076 |
| GFLAQFL | ASSEPSDQDTQY | 0.7018415 |
| GGAGGAGGNAGVA | ASSEPSDQDTQY | 0.23765652 |
| GGVGTAGAGGAGGAGG | ASSEPSDQDTQY | 0.38102746 |
| GNGGGGDAGNAGSGGN | ASSEPSDQDTQY | 0.18676922 |
| GTGGAGGAGNDAGS | ASSEPSDQDTQY | 0.5808637 |
| GTTNTGWFNS | ASSEPSDQDTQY | 0.26370105 |
| IFMGPLFTL | ASSEPSDQDTQY | 0.75163317 |
| ILAHAIEAI | ASSEPSDQDTQY | 0.55421257 |
| IRSAIREAL | ASSEPSDQDTQY | 0.57727766 |

| | | |
|---|---|---|
| ITFAGGG | ASSEPSDQDTQY | 0.81745493 |
| ITGMSGHQEV | ASSEPSDQDTQY | 0.6927475 |
| IVAGLACTL | ASSEPSDQDTQY | 0.51023424 |
| IVGGGLAGA | ASSEPSDQDTQY | 0.37250584 |
| IVTGVLNI | ASSEPSDQDTQY | 0.38895288 |
| LAQQGCI | ASSEPSDQDTQY | 0.31777918 |
| LCLHVEN | ASSEPSDQDTQY | 0.37576863 |
| LDAISDGS | ASSEPSDQDTQY | 0.82001895 |
| LGGFAAML | ASSEPSDQDTQY | 0.4900593 |
| LGGKNPMI | ASSEPSDQDTQY | 0.3372528 |
| LGPKGVAAIL | ASSEPSDQDTQY | 0.5450072 |
| LMVGVSI | ASSEPSDQDTQY | 0.78927743 |
| LVGAAFVL | ASSEPSDQDTQY | 0.79796934 |
| LVLAVAPL | ASSEPSDQDTQY | 0.38780144 |
| MAAIAVIDAA | ASSEPSDQDTQY | 0.4537158 |
| NNGINAELQ | ASSEPSDQDTQY | 0.50474113 |
| RGEPLVLVHG | ASSEPSDQDTQY | 0.661813 |
| RGGAGGAAGLI | ASSEPSDQDTQY | 0.45949343 |
| RSYRMHPAV | ASSEPSDQDTQY | 0.321205 |
| SAAHLLAGQLA | ASSEPSDQDTQY | 0.64313865 |
| SAAVGSY | ASSEPSDQDTQY | 0.83682287 |
| SPHAVLHRI | ASSEPSDQDTQY | 0.38336974 |
| SQIAKVL | ASSEPSDQDTQY | 0.26636297 |
| TALGHLVP | ASSEPSDQDTQY | 0.61744404 |
| TAPTTSGAA | ASSEPSDQDTQY | 0.37705994 |
| TDGDDGNGGSPGT | ASSEPSDQDTQY | 0.46789345 |
| TDGLGEN | ASSEPSDQDTQY | 0.52621835 |
| TGGPPLPGQA | ASSEPSDQDTQY | 0.28395808 |
| TIPAITI | ASSEPSDQDTQY | 0.51565355 |
| TIPGFTI | ASSEPSDQDTQY | 0.81138283 |
| TSIASGALA | ASSEPSDQDTQY | 0.49598718 |
| TVTLPGAAI | ASSEPSDQDTQY | 0.7167679 |
| VASGLSAALP | ASSEPSDQDTQY | 0.8030801 |
| VGAGVIGI | ASSEPSDQDTQY | 0.73693895 |
| VGPKRFGV | ASSEPSDQDTQY | 0.80048513 |
| VNVAFPD | ASSEPSDQDTQY | 0.7866645 |
| VSMTNSGVS | ASSEPSDQDTQY | 0.54502803 |
| VTDVLGDLSV | ASSEPSDQDTQY | 0.52353865 |
| VVVVGAGFAGL | ASSEPSDQDTQY | 0.6907153 |
| VYTVGTVV | ASSEPSDQDTQY | 0.6595619 |
| WTTFSGP | ASSEPSDQDTQY | 0.5678458 |
| YDELVEAF | ASSEPSDQDTQY | 0.41244248 |
| YGTGGAGGNGG | ASSEPSDQDTQY | 0.7948857 |
| YTNVMAV | ASSEPSDQDTQY | 0.4592204 |
| FLEQVHQL | ASSEPSDQDTQY | 0.761182 |
| GLSLLAVAV | ASSEPSDQDTQY | 0.32237226 |
| LEQVHQL | ASSEPSDQDTQY | 0.30296475 |
| LNAAGAI | ASSEPSDQDTQY | 0.7512938 |
| PAVHASL | ASSEPSDQDTQY | 0.13004458 |
| TELEDDA | ASSEPSDQDTQY | 0.19218259 |
| TPPGPPN | ASSEPSDQDTQY | 0.5860979 |

| | | |
|---|---|---|
| TPPPPPPP | ASSEPSDQDTQY | 0.7683519 |
| TQSPALI | GASGTISNERLF | 0.14070682 |
| EFAAAFPQ | GASGTISNERLF | 0.32235625 |
| LLSGNLDVL | GASGTISNERLF | 0.32564962 |
| GPANLGNYNI | GASGTISNERLF | 0.6419428 |
| TLGPTLP | GASGTISNERLF | 0.5647222 |
| VYAYSSCCK | GASGTISNERLF | 0.22876678 |
| APAPAPAPNI | GASGTISNERLF | 0.7567707 |
| PGGPVPIP | GASGTISNERLF | 0.21150944 |
| FLTVALEP | GASGTISNERLF | 0.29120108 |
| SQYFGANL | GASGTISNERLF | 0.75449383 |
| GGVKPAA | GASGTISNERLF | 0.8000022 |
| GGTGGAGGAGGW | GASGTISNERLF | 0.3287166 |
| GVGGVAGLGGAGG | GASGTISNERLF | 0.75829697 |
| VLLVLATTI | GASGTISNERLF | 0.39470923 |
| IGIFAAML | GASGTISNERLF | 0.807646 |
| AVGDLID | GASGTISNERLF | 0.45311928 |
| NISPLPL | GASGTISNERLF | 0.29679537 |
| VTAVFVAI | GASGTISNERLF | 0.777789 |
| SGDDNDAAL | GASGTISNERLF | 0.6945497 |
| AVAALAHS | GASGTISNERLF | 0.5989107 |
| GGVTVMATP | GASGTISNERLF | 0.76571655 |
| AGLAANVRD | GASGTISNERLF | 0.35884696 |
| TNRPDLIDPALLRPG | GASGTISNERLF | 0.6357479 |
| LANLGVTAL | GASGTISNERLF | 0.8161747 |
| TGERIEL | GASGTISNERLF | 0.7702433 |
| NGEELAA | GASGTISNERLF | 0.32878262 |
| VLSLVWG | GASGTISNERLF | 0.74845505 |
| IGYALWL | GASGTISNERLF | 0.52710044 |
| LSLLLAPL | GASGTISNERLF | 0.41951758 |
| TLDEALK | GASGTISNERLF | 0.7887639 |
| TDGAGTAEGG | GASGTISNERLF | 0.5449597 |
| NAPGGAGGNGG | GASGTISNERLF | 0.73602444 |
| VSVFMGMGGNF | GASGTISNERLF | 0.6866641 |
| LASYLAAL | GASGTISNERLF | 0.7796384 |
| LVGGTPAL | GASGTISNERLF | 0.24590878 |
| VRNADVL | GASGTISNERLF | 0.5886435 |
| TCSSPFLVSAM | GASGTISNERLF | 0.56007177 |
| VHITDAE | GASGTISNERLF | 0.7636716 |
| EYAAPSF | GASGTISNERLF | 0.5897579 |
| VVDAFHGGL | GASGTISNERLF | 0.29658905 |
| ASVVDPQ | GASGTISNERLF | 0.6060701 |
| SYNEFGGLS | GASGTISNERLF | 0.7740723 |
| DPDGSVY | GASGTISNERLF | 0.5446423 |
| MVSTIHATE | GASGTISNERLF | 0.6745696 |
| LAAGVADPV | GASGTISNERLF | 0.8133179 |
| TKEQIAA | GASGTISNERLF | 0.46046883 |
| SLASPAAL | GASGTISNERLF | 0.4804216 |
| IGLVFYGV | GASGTISNERLF | 0.7150692 |
| FLLDASNL | GASGTISNERLF | 0.79118645 |
| YQVGNFP | GASGTISNERLF | 0.5503657 |

| | | |
|---|---|---|
| ESLLSQST | GASGTISNERLF | 0.797235 |
| LFTVGFAAVG | GASGTISNERLF | 0.3005237 |
| ATPPTPT | GASGTISNERLF | 0.7546159 |
| TDLATLG | GASGTISNERLF | 0.44683772 |
| SAGGTGR | GASGTISNERLF | 0.71098423 |
| GFSNSGS | GASGTISNERLF | 0.34908667 |
| GGNGGSGGTSV | GASGTISNERLF | 0.7679554 |
| GTSVATGGAGN | GASGTISNERLF | 0.7432525 |
| NGGNGGSGGTS | GASGTISNERLF | 0.77287304 |
| AVVKFPH | GASGTISNERLF | 0.17136742 |
| TPPPPPP | GASGTISNERLF | 0.76535624 |
| AAPAAAAAPA | GASGTISNERLF | 0.76508224 |
| IALNEDL | GASGTISNERLF | 0.83822805 |
| IVNVSSM | GASGTISNERLF | 0.50288934 |
| TGIPPAPR | GASGTISNERLF | 0.7138419 |
| FDLGGGTF | GASGTISNERLF | 0.38304007 |
| FDLGGGTFDVS | GASGTISNERLF | 0.4703083 |
| NEPTAAA | GASGTISNERLF | 0.28931412 |
| NEPTAAALA | GASGTISNERLF | 0.4693302 |
| GSIINISSI | GASGTISNERLF | 0.45980835 |
| VAPGNAGTA | GASGTISNERLF | 0.50448644 |
| TGGGASLE | GASGTISNERLF | 0.7526874 |
| VAVYNHYKRI | GASGTISNERLF | 0.79121006 |
| TDPAPATT | GASGTISNERLF | 0.74923 |
| TNRPDILDPALLRPG | GASGTISNERLF | 0.789112 |
| LTTTTTS | GASGTISNERLF | 0.77107394 |
| PTGSGKTL | GASGTISNERLF | 0.76263046 |
| TGSGKTL | GASGTISNERLF | 0.37581626 |
| TIGHVDHGK | GASGTISNERLF | 0.4651075 |
| TIGHVDHGKTT | GASGTISNERLF | 0.3138131 |
| GHVDHGKTT | GASGTISNERLF | 0.52200073 |
| AQFEHTLL | GASGTISNERLF | 0.399782 |
| IVNQDEL | GASGTISNERLF | 0.6916557 |
| TNGSQFF | GASGTISNERLF | 0.13953166 |
| TGKGIVE | GASGTISNERLF | 0.6679114 |
| LISNASDA | GASGTISNERLF | 0.81993765 |
| AELRTLGELTKIAKAHGAQ | GASGTISNERLF | 0.30867374 |
| APNAVAAAEPSW | GASGTISNERLF | 0.628001 |
| APQVAGAATAVL | GASGTISNERLF | 0.6379505 |
| ARRDPAAAAAILPTDA | GASGTISNERLF | 0.39368966 |
| DAAARDVLAEAGLAEYF | GASGTISNERLF | 0.7434102 |
| DDQIVLSAKANLAKVEQ | GASGTISNERLF | 0.36717257 |
| DELLPDTGEDSDEDGH | GASGTISNERLF | 0.72658426 |
| ELPEAPANRATVPQD | GASGTISNERLF | 0.7853466 |
| FGQSKIERVTGIG | GASGTISNERLF | 0.6481685 |
| GAGGTGSTAGGAGGAGGAGGLY | GASGTISNERLF | 0.65305156 |
| GHGGTGAAGGNGGTGGAGGSGI | GASGTISNERLF | 0.58732617 |
| GNGGNGGAGGEGGPGGAGGAGA | GASGTISNERLF | 0.22645122 |
| GSTAGNGGNGGNATNGTIAGQP | GASGTISNERLF | 0.70834243 |
| GTGGDGGNGGNGGTGG | GASGTISNERLF | 0.22474651 |
| GVRRPAGIPADAPV | GASGTISNERLF | 0.5871779 |

| | | |
|---|---|---|
| GYRQVTDAHLLTLARR | GASGTISNERLF | 0.32336003 |
| IAKMGQDGHDRG | GASGTISNERLF | 0.7086458 |
| KAEPSIEIQYT | GASGTISNERLF | 0.62165844 |
| KPVRIRMAADEDTVAVT | GASGTISNERLF | 0.6329168 |
| MPPRIAGMRLLVIKPEP | GASGTISNERLF | 0.38931155 |
| MKFYAVALFRSRGVPPTR | GASGTISNERLF | 0.6244607 |
| NAGGFGILGAVAHSP | GASGTISNERLF | 0.4028623 |
| NEGREITLAMAEM | GASGTISNERLF | 0.30395114 |
| NELVEWSHPLGHRWRWE | GASGTISNERLF | 0.7350637 |
| NLGAGGTGGNGGNPG | GASGTISNERLF | 0.46437362 |
| NTEWPKWETVMAADPVD | GASGTISNERLF | 0.47825187 |
| PASAAPAETQP | GASGTISNERLF | 0.7615008 |
| QQSSDGKNGVRAAEITGE | GASGTISNERLF | 0.6419778 |
| RAPDAVTTSWWKE | GASGTISNERLF | 0.2510891 |
| SGFGNVGAHTSGWFNQ | GASGTISNERLF | 0.58557975 |
| SGVAGAVGQAASVGGLKV | GASGTISNERLF | 0.62877595 |
| SNGGAGTGSAGGNGGT | GASGTISNERLF | 0.4506061 |
| TGGAGGEGGAAG | GASGTISNERLF | 0.14599076 |
| TGGDGGAAGAGGAPGHG | GASGTISNERLF | 0.56421006 |
| TMPPLLAAIANAGRHGV | GASGTISNERLF | 0.41731378 |
| TSSTAGNSGGA | GASGTISNERLF | 0.31974256 |
| VPATVPVGILC | GASGTISNERLF | 0.42293873 |
| VQRPDSGGVVLTGRIS | GASGTISNERLF | 0.778303 |
| WTPNPVPATKGVPAGLVQAT | GASGTISNERLF | 0.41560525 |
| YALKRALAKFSDRFDVVII | GASGTISNERLF | 0.61078 |
| YLIGSVIV | GASGTISNERLF | 0.7578076 |
| YGQLVAVID | GASGTISNERLF | 0.6861473 |
| VTVGGMTLPA | GASGTISNERLF | 0.81286204 |
| VLALPAV | GASGTISNERLF | 0.5687803 |
| VIPPTLNYET | GASGTISNERLF | 0.41079667 |
| VGMGGMHPSSG | GASGTISNERLF | 0.7071625 |
| VGLAGGAATA | GASGTISNERLF | 0.50011265 |
| VGGAGGAGGSGP | GASGTISNERLF | 0.2800749 |
| VFAAFMV | GASGTISNERLF | 0.17905556 |
| TLYDKLR | GASGTISNERLF | 0.6085312 |
| TLAVFTVL | GASGTISNERLF | 0.226764 |
| TGSVQPARV | GASGTISNERLF | 0.53574955 |
| TGGGAGSGGT | GASGTISNERLF | 0.7708244 |
| SHLETSL | GASGTISNERLF | 0.39097336 |
| SFTTGMT | GASGTISNERLF | 0.2948104 |
| QVEVAKRMGVRQPTVS | GASGTISNERLF | 0.50027746 |
| QGSGGPALAM | GASGTISNERLF | 0.39216787 |
| MLAPSNP | GASGTISNERLF | 0.73312294 |
| LVNNAAI | GASGTISNERLF | 0.58087134 |
| LTGEINTLA | GASGTISNERLF | 0.83737326 |
| LSIFGPV | GASGTISNERLF | 0.7721144 |
| LLALLANL | GASGTISNERLF | 0.4788422 |
| LISNASDALDKLR | GASGTISNERLF | 0.73939645 |
| LINEAVP | GASGTISNERLF | 0.5362068 |
| LGYAFSGL | GASGTISNERLF | 0.54732275 |
| LDFVAVAL | GASGTISNERLF | 0.5437555 |

| | | |
|---|---|---|
| LAQPGAE | GASGTISNERLF | 0.5644705 |
| ISNLGAVTGF | GASGTISNERLF | 0.5521328 |
| HQEDPGRI | GASGTISNERLF | 0.5319358 |
| GTPGEPTP | GASGTISNERLF | 0.2395013 |
| GFFNSGS | GASGTISNERLF | 0.6961572 |
| GAGGGGLTAGAN | GASGTISNERLF | 0.5730631 |
| FVDALVNM | GASGTISNERLF | 0.2280475 |
| FLTVAPEL | GASGTISNERLF | 0.47167557 |
| FETLTAL | GASGTISNERLF | 0.82480067 |
| DVINPIEL | GASGTISNERLF | 0.79263407 |
| APVSGAPV | GASGTISNERLF | 0.40053135 |
| ALVGGLAW | GASGTISNERLF | 0.35682583 |
| ALAAAAATAI | GASGTISNERLF | 0.6751071 |
| AFGLGLLL | GASGTISNERLF | 0.200712 |
| AFFAAASL | GASGTISNERLF | 0.6290097 |
| AAAALPAFSPP | GASGTISNERLF | 0.64225495 |
| VVLAPAL | GASGTISNERLF | 0.56994784 |
| TIGHVDHGKTTL | GASGTISNERLF | 0.4197185 |
| LISNASDALDK | GASGTISNERLF | 0.6371188 |
| LISNASDAL | GASGTISNERLF | 0.4432721 |
| ISNASDALDK | GASGTISNERLF | 0.6735135 |
| NQAETLVYQTE | GASGTISNERLF | 0.13254873 |
| TGGTGGTAGAG | GASGTISNERLF | 0.64213264 |
| TGSTAGGAGGA | GASGTISNERLF | 0.3389941 |
| DPLAVPVAAGPV | GASGTISNERLF | 0.32644516 |
| GGTGGTPGNGGN | GASGTISNERLF | 0.23431724 |
| PAGLAGVDPATL | GASGTISNERLF | 0.5718842 |
| GNNLVVEAPHAGASV | GASGTISNERLF | 0.74650896 |
| KVTRIMSASTLEQPA | GASGTISNERLF | 0.74561495 |
| LGFALPIALPRRTVV | GASGTISNERLF | 0.27444533 |
| TRLGNLTARKLVVLV | GASGTISNERLF | 0.7910327 |
| WSQQRRLIYFPSAGP | GASGTISNERLF | 0.77347016 |
| GPNGNPDPMAAAVDIR | GASGTISNERLF | 0.5099133 |
| TQRCVSVADLRAAASR | GASGTISNERLF | 0.23745582 |
| QFVLDVAEHPVEVLTETV | GASGTISNERLF | 0.6824262 |
| GGAGGTGGAGGNGGTQAGDG | GASGTISNERLF | 0.60341394 |
| PLLATLRAIAVDVDAKPAQV | GASGTISNERLF | 0.7312664 |
| APKFRTGGYMIDFWGVGYQVAK | GASGTISNERLF | 0.65076816 |
| GAGGAGGAGGAAGGPGGTGGTG | GASGTISNERLF | 0.3217968 |
| IGALGLLATDVAPAGSVAFGVA | GASGTISNERLF | 0.7536439 |
| AGGAGGIGGNGGAGGAGGVLMG | GASGTISNERLF | 0.77639997 |
| ADELIGLL | GASGTISNERLF | 0.3738773 |
| ADRALKA | GASGTISNERLF | 0.3930914 |
| AGAVGQVGGAGGA | GASGTISNERLF | 0.40375167 |
| AGGGGGLGGAP | GASGTISNERLF | 0.22897772 |
| AGGLLYGNGGAG | GASGTISNERLF | 0.7041044 |
| AGLESIGL | GASGTISNERLF | 0.4963569 |
| AGLKTGII | GASGTISNERLF | 0.3938676 |
| ASLAALLL | GASGTISNERLF | 0.66338176 |
| ASVINISSI | GASGTISNERLF | 0.41868016 |
| ATPPGPA | GASGTISNERLF | 0.4318722 |

| | | |
|---|---|---|
| ATVASALV | GASGTISNERLF | 0.4830646 |
| ATVASLIL | GASGTISNERLF | 0.5283573 |
| AVNGPVLIRH | GASGTISNERLF | 0.6501229 |
| DGAAGAVGI | GASGTISNERLF | 0.509672 |
| DPLLHGI | GASGTISNERLF | 0.5933168 |
| ELGAALGAL | GASGTISNERLF | 0.8119384 |
| ELGRTQAV | GASGTISNERLF | 0.27167 |
| FVAGGYLS | GASGTISNERLF | 0.40402985 |
| GASGGAGGAGGW | GASGTISNERLF | 0.6737224 |
| GGAGAGGDGGAGGI | GASGTISNERLF | 0.24506235 |
| GTGHNGSQPGMGGNG | GASGTISNERLF | 0.7353499 |
| GYAAASASAS | GASGTISNERLF | 0.6110839 |
| HQVDVAAVQQG | GASGTISNERLF | 0.6817258 |
| IGAAIAHT | GASGTISNERLF | 0.5086926 |
| IGMIEVAQL | GASGTISNERLF | 0.8296998 |
| IRLAPEHTSAMD | GASGTISNERLF | 0.78753394 |
| IYNVSVL | GASGTISNERLF | 0.5781486 |
| KQGFLGM | GASGTISNERLF | 0.760573 |
| LESVITQL | GASGTISNERLF | 0.34099156 |
| LGLGALVVV | GASGTISNERLF | 0.77820647 |
| LIGNGGNGGSGGI | GASGTISNERLF | 0.714844 |
| LNAINAPTL | GASGTISNERLF | 0.3278764 |
| LQLLNTGV | GASGTISNERLF | 0.6521915 |
| LQTSIEF | GASGTISNERLF | 0.77200335 |
| MSFAVSEL | GASGTISNERLF | 0.8086739 |
| NTGALGL | GASGTISNERLF | 0.217476 |
| PPGAGGAGGAGTT | GASGTISNERLF | 0.51957685 |
| QLVGLGI | GASGTISNERLF | 0.21570444 |
| RFLEQVHQL | GASGTISNERLF | 0.3317809 |
| RGVGYVLR | GASGTISNERLF | 0.32408184 |
| SATGVFAGVI | GASGTISNERLF | 0.6437423 |
| SLPAITLP | GASGTISNERLF | 0.5472754 |
| STALAFTV | GASGTISNERLF | 0.7225033 |
| SVLTVGLGK | GASGTISNERLF | 0.52082103 |
| TALAGAGSGI | GASGTISNERLF | 0.77358913 |
| TGLPFTGI | GASGTISNERLF | 0.4529361 |
| VAEHVTV | GASGTISNERLF | 0.48408034 |
| VDIVLNSL | GASGTISNERLF | 0.27632228 |
| VLTGGGAVL | GASGTISNERLF | 0.26230782 |
| VNVGLAYGI | GASGTISNERLF | 0.36367083 |
| VSSAAVGAV | GASGTISNERLF | 0.4756655 |
| VTSVYEAGV | GASGTISNERLF | 0.16320911 |
| WVALDRGA | GASGTISNERLF | 0.7502384 |
| YLVGSVVI | GASGTISNERLF | 0.71829855 |
| GPPGTGKTLL | GASGTISNERLF | 0.6852139 |
| GVGMTKF | GASGTISNERLF | 0.38837594 |
| IGIGGSDL | GASGTISNERLF | 0.64209926 |
| IGTLLAEL | GASGTISNERLF | 0.357381 |
| IYTSGTT | GASGTISNERLF | 0.7521528 |
| LSFPAGV | GASGTISNERLF | 0.3304668 |
| LYTSGTT | GASGTISNERLF | 0.25298017 |

| | | |
|---|---|---|
| VGAFPFGF | GASGTISNERLF | 0.41970298 |
| VGAGTVEF | GASGTISNERLF | 0.25827575 |
| VGKSTLLL | GASGTISNERLF | 0.7610009 |
| AALAHAGGGIV | GASGTISNERLF | 0.75947726 |
| AALVAVSLL | GASGTISNERLF | 0.7358506 |
| AAVSRAVAAL | GASGTISNERLF | 0.74663055 |
| AGGSSGIGGF | GASGTISNERLF | 0.18438797 |
| AGLTVANANG | GASGTISNERLF | 0.39188638 |
| AGLVGAVL | GASGTISNERLF | 0.8185128 |
| ALSQAFTV | GASGTISNERLF | 0.52188265 |
| ALTFSVSL | GASGTISNERLF | 0.58553493 |
| ALTGAALVAL | GASGTISNERLF | 0.79435575 |
| ASFSIGAGVL | GASGTISNERLF | 0.38668478 |
| DGSVYKI | GASGTISNERLF | 0.45070782 |
| EQLAEIV | GASGTISNERLF | 0.57071656 |
| FAVSELTVM | GASGTISNERLF | 0.67961156 |
| GAAPLAAGL | GASGTISNERLF | 0.6269818 |
| GADGGDGGSGGAGG | GASGTISNERLF | 0.3841763 |
| GFGLDTGGAGG | GASGTISNERLF | 0.6878562 |
| GLASGGLL | GASGTISNERLF | 0.8302479 |
| GRIAGAAVLL | GASGTISNERLF | 0.63400716 |
| GSAGSLNTSF | GASGTISNERLF | 0.69327 |
| GTGQIGI | GASGTISNERLF | 0.7045564 |
| GTLIALIL | GASGTISNERLF | 0.69568336 |
| IAATMGFVVG | GASGTISNERLF | 0.44240993 |
| IFAQALL | GASGTISNERLF | 0.44766915 |
| IGAAVALGF | GASGTISNERLF | 0.6327338 |
| IGLTPLAL | GASGTISNERLF | 0.32433844 |
| ILGSNTSTL | GASGTISNERLF | 0.5987356 |
| IVFSLTVL | GASGTISNERLF | 0.45772016 |
| LASKATGF | GASGTISNERLF | 0.8382548 |
| LASVALIL | GASGTISNERLF | 0.6781732 |
| LFAVGLTL | GASGTISNERLF | 0.66510254 |
| LFNGGIGTYI | GASGTISNERLF | 0.32475406 |
| LTLGAGII | GASGTISNERLF | 0.61159414 |
| LVAAALGDI | GASGTISNERLF | 0.69507843 |
| NTAFVQLGI | GASGTISNERLF | 0.2799608 |
| QLFSGAEL | GASGTISNERLF | 0.76922303 |
| SAATFITL | GASGTISNERLF | 0.4158277 |
| SNYGLANLGV | GASGTISNERLF | 0.49864382 |
| SYGGAYAVM | GASGTISNERLF | 0.71810627 |
| TTGVGGLTLG | GASGTISNERLF | 0.8175639 |
| TTQTIYI | GASGTISNERLF | 0.52419233 |
| VAAGAGLGAVI | GASGTISNERLF | 0.50933677 |
| VAAVVTLL | GASGTISNERLF | 0.3509569 |
| VGFLGGLI | GASGTISNERLF | 0.60702175 |
| VGVIPILL | GASGTISNERLF | 0.79321975 |
| VIFASSLI | GASGTISNERLF | 0.6392368 |
| VKGGDGGAAAG | GASGTISNERLF | 0.7958297 |
| VTKTFTNI | GASGTISNERLF | 0.59420943 |
| VVAAAILL | GASGTISNERLF | 0.2522371 |

| | | |
|---|---|---|
| YAALDVEL | GASGTISNERLF | 0.49572167 |
| LASVALLL | GASGTISNERLF | 0.7591733 |
| LDAVHRV | GASGTISNERLF | 0.7429703 |
| QGLSSSIF | GASGTISNERLF | 0.7946768 |
| RGTFANIRL | GASGTISNERLF | 0.7100475 |
| RQLWWGHRI | GASGTISNERLF | 0.43525165 |
| YGPPGTGKTLL | GASGTISNERLF | 0.79911613 |
| AALVGAPANVA | GASGTISNERLF | 0.8102492 |
| AATFRRTE | GASGTISNERLF | 0.7391051 |
| AGALGGGI | GASGTISNERLF | 0.39285713 |
| AGGAGGAGGIGAG | GASGTISNERLF | 0.32416427 |
| AGSGALGL | GASGTISNERLF | 0.5947922 |
| AQIADQI | GASGTISNERLF | 0.7470925 |
| ASAVAVTYL | GASGTISNERLF | 0.3588736 |
| ASTVLGGTGG | GASGTISNERLF | 0.6460962 |
| AVGGEPAK | GASGTISNERLF | 0.6711899 |
| CEQGTFGGTC | GASGTISNERLF | 0.24012461 |
| DEGKAPAG | GASGTISNERLF | 0.5066529 |
| DTVVFPAL | GASGTISNERLF | 0.7948767 |
| FEGITMAG | GASGTISNERLF | 0.60369754 |
| FPLVVQD | GASGTISNERLF | 0.52615064 |
| GAFTTPQ | GASGTISNERLF | 0.77335876 |
| GFGGFGTAGGVGGA | GASGTISNERLF | 0.8353787 |
| GFLAQFL | GASGTISNERLF | 0.7982272 |
| GGAGGAGGNAGVA | GASGTISNERLF | 0.26011524 |
| GGVGTAGAGGAGGAGG | GASGTISNERLF | 0.16980158 |
| GNGGGGDAGNAGSGGN | GASGTISNERLF | 0.59024 |
| GTGGAGGAGNDAGS | GASGTISNERLF | 0.7490016 |
| GTTNTGWFNS | GASGTISNERLF | 0.4819747 |
| IFMGPLFTL | GASGTISNERLF | 0.45424902 |
| ILAHAIEAI | GASGTISNERLF | 0.71028066 |
| IRSAIREAL | GASGTISNERLF | 0.7808598 |
| ITFAGGG | GASGTISNERLF | 0.43948036 |
| ITGMSGHQEV | GASGTISNERLF | 0.25276765 |
| IVAGLACTL | GASGTISNERLF | 0.69000447 |
| IVGGGLAGA | GASGTISNERLF | 0.7241818 |
| IVTGVLNI | GASGTISNERLF | 0.44828555 |
| LAQQGCI | GASGTISNERLF | 0.6057328 |
| LCLHVEN | GASGTISNERLF | 0.40159118 |
| LDAISDGS | GASGTISNERLF | 0.52828103 |
| LGGFAAML | GASGTISNERLF | 0.6605526 |
| LGGKNPMI | GASGTISNERLF | 0.3443516 |
| LGPKGVAAIL | GASGTISNERLF | 0.7911684 |
| LMVGVSI | GASGTISNERLF | 0.3978355 |
| LVGAAFVL | GASGTISNERLF | 0.65044403 |
| LVLAVAPL | GASGTISNERLF | 0.71298075 |
| MAAIAVIDAA | GASGTISNERLF | 0.67801833 |
| NNGINAELQ | GASGTISNERLF | 0.546366 |
| RGEPLVLVHG | GASGTISNERLF | 0.46747702 |
| RGGAGGAAGLI | GASGTISNERLF | 0.2732089 |
| RSYRMHPAV | GASGTISNERLF | 0.16609652 |

| | | |
|---|---|---|
| SAAHLLAGQLA | GASGTISNERLF | 0.79089093 |
| SAAVGSY | GASGTISNERLF | 0.47847855 |
| SPHAVLHRI | GASGTISNERLF | 0.8087819 |
| SQIAKVL | GASGTISNERLF | 0.6926246 |
| TALGHLVP | GASGTISNERLF | 0.35897753 |
| TAPTTSGAA | GASGTISNERLF | 0.65015274 |
| TDGDDGNGGSPGT | GASGTISNERLF | 0.30936438 |
| TDGLGEN | GASGTISNERLF | 0.31182182 |
| TGGPPLPGQA | GASGTISNERLF | 0.62464285 |
| TIPAITI | GASGTISNERLF | 0.73058784 |
| TIPGFTI | GASGTISNERLF | 0.369007 |
| TSIASGALA | GASGTISNERLF | 0.6326642 |
| TVTLPGAAI | GASGTISNERLF | 0.61539257 |
| VASGLSAALP | GASGTISNERLF | 0.31494433 |
| VGAGVIGI | GASGTISNERLF | 0.60384995 |
| VGPKRFGV | GASGTISNERLF | 0.8009541 |
| VNVAFPD | GASGTISNERLF | 0.6367874 |
| VSMTNSGVS | GASGTISNERLF | 0.41004407 |
| VTDVLGDLSV | GASGTISNERLF | 0.7097722 |
| VVVVGAGFAGL | GASGTISNERLF | 0.33659977 |
| VYTVGTVV | GASGTISNERLF | 0.70714307 |
| WTTFSGP | GASGTISNERLF | 0.56506073 |
| YDELVEAF | GASGTISNERLF | 0.4691878 |
| YGTGGAGGNGG | GASGTISNERLF | 0.7306938 |
| YTNVMAV | GASGTISNERLF | 0.28908947 |
| FLEQVHQL | GASGTISNERLF | 0.36061156 |
| GLSLLAVAV | GASGTISNERLF | 0.5294979 |
| LEQVHQL | GASGTISNERLF | 0.5888635 |
| LNAAGAI | GASGTISNERLF | 0.76414025 |
| PAVHASL | GASGTISNERLF | 0.37535968 |
| TELEDDA | GASGTISNERLF | 0.54837716 |
| TPPGPPN | GASGTISNERLF | 0.40146345 |
| TPPPPPPP | GASGTISNERLF | 0.31761795 |
| TQSPALI | GAPGGNGVF | 0.60646164 |
| EFAAAFPQ | GAPGGNGVF | 0.52045023 |
| LLSGNLDVL | GAPGGNGVF | 0.3557909 |
| GPANLGNYNI | GAPGGNGVF | 0.6782654 |
| TLGPTLP | GAPGGNGVF | 0.40001166 |
| VYAYSSCCK | GAPGGNGVF | 0.5932981 |
| APAPAPAPNI | GAPGGNGVF | 0.8309024 |
| PGGPVPIP | GAPGGNGVF | 0.5285327 |
| FLTVALEP | GAPGGNGVF | 0.6561847 |
| SQYFGANL | GAPGGNGVF | 0.69567 |
| GGVKPAA | GAPGGNGVF | 0.7398294 |
| GGTGGAGGAGGW | GAPGGNGVF | 0.47781783 |
| GVGGVAGLGGAGG | GAPGGNGVF | 0.3834205 |
| VLLVLATTI | GAPGGNGVF | 0.79849446 |
| IGIFAAML | GAPGGNGVF | 0.6815701 |
| AVGDLID | GAPGGNGVF | 0.74522716 |
| NISPLPL | GAPGGNGVF | 0.70785975 |
| VTAVFVAI | GAPGGNGVF | 0.7578162 |

| | | |
|---|---|---|
| SGDDNDAAL | GAPGGNGVF | 0.75349927 |
| AVAALAHS | GAPGGNGVF | 0.4970308 |
| GGVTVMATP | GAPGGNGVF | 0.26430845 |
| AGLAANVRD | GAPGGNGVF | 0.4900819 |
| TNRPDLIDPALLRPG | GAPGGNGVF | 0.61584926 |
| LANLGVTAL | GAPGGNGVF | 0.53163964 |
| TGERIEL | GAPGGNGVF | 0.22734252 |
| NGEELAA | GAPGGNGVF | 0.79382145 |
| VLSLVWG | GAPGGNGVF | 0.8361287 |
| IGYALWL | GAPGGNGVF | 0.34490255 |
| LSLLLAPL | GAPGGNGVF | 0.81076765 |
| TLDEALK | GAPGGNGVF | 0.30420464 |
| TDGAGTAEGG | GAPGGNGVF | 0.6598342 |
| NAPGGAGGNGG | GAPGGNGVF | 0.79655325 |
| VSVFMGMGGNF | GAPGGNGVF | 0.3363076 |
| LASYLAAL | GAPGGNGVF | 0.7255423 |
| LVGGTPAL | GAPGGNGVF | 0.36260936 |
| VRNADVL | GAPGGNGVF | 0.4702695 |
| TCSSPFLVSAM | GAPGGNGVF | 0.8174623 |
| VHITDAE | GAPGGNGVF | 0.46377444 |
| EYAAPSF | GAPGGNGVF | 0.65326345 |
| VVDAFHGGL | GAPGGNGVF | 0.571243 |
| ASVVDPQ | GAPGGNGVF | 0.34925544 |
| SYNEFGGLS | GAPGGNGVF | 0.672758 |
| DPDGSVY | GAPGGNGVF | 0.77770364 |
| MVSTIHATE | GAPGGNGVF | 0.31796235 |
| LAAGVADPV | GAPGGNGVF | 0.8065441 |
| TKEQIAA | GAPGGNGVF | 0.35620412 |
| SLASPAAL | GAPGGNGVF | 0.8255162 |
| IGLVFYGV | GAPGGNGVF | 0.74688363 |
| FLLDASNL | GAPGGNGVF | 0.6478964 |
| YQVGNFP | GAPGGNGVF | 0.44191492 |
| ESLLSQST | GAPGGNGVF | 0.7927784 |
| LFTVGFAAVG | GAPGGNGVF | 0.623788 |
| ATPPTPT | GAPGGNGVF | 0.73356247 |
| TDLATLG | GAPGGNGVF | 0.79846513 |
| SAGGTGR | GAPGGNGVF | 0.28564805 |
| GFSNSGS | GAPGGNGVF | 0.56732357 |
| GGNGGSGGTSV | GAPGGNGVF | 0.44453937 |
| GTSVATGGAGN | GAPGGNGVF | 0.3748364 |
| NGGNGGSGGTS | GAPGGNGVF | 0.33180365 |
| AVVKFPH | GAPGGNGVF | 0.23708338 |
| TPPPPPP | GAPGGNGVF | 0.3173984 |
| AAPAAAAAPA | GAPGGNGVF | 0.3016966 |
| IALNEDL | GAPGGNGVF | 0.7192594 |
| IVNVSSM | GAPGGNGVF | 0.34473303 |
| TGIPPAPR | GAPGGNGVF | 0.3126135 |
| FDLGGGTF | GAPGGNGVF | 0.7059289 |
| FDLGGGTFDVS | GAPGGNGVF | 0.6753386 |
| NEPTAAA | GAPGGNGVF | 0.2548702 |
| NEPTAAALA | GAPGGNGVF | 0.34909055 |

| | | |
|---|---|---|
| GSIINISSI | GAPGGNGVF | 0.29488802 |
| VAPGNAGTA | GAPGGNGVF | 0.33646125 |
| TGGGASLE | GAPGGNGVF | 0.22119726 |
| VAVYNHYKRI | GAPGGNGVF | 0.4344161 |
| TDPAPATT | GAPGGNGVF | 0.32594836 |
| TNRPDILDPALLRPG | GAPGGNGVF | 0.20299587 |
| LTTTTTS | GAPGGNGVF | 0.37675625 |
| PTGSGKTL | GAPGGNGVF | 0.4039467 |
| TGSGKTL | GAPGGNGVF | 0.36356086 |
| TIGHVDHGK | GAPGGNGVF | 0.68245256 |
| TIGHVDHGKTT | GAPGGNGVF | 0.22935453 |
| GHVDHGKTT | GAPGGNGVF | 0.2749581 |
| AQFEHTLL | GAPGGNGVF | 0.5898973 |
| IVNQDEL | GAPGGNGVF | 0.6346408 |
| TNGSQFF | GAPGGNGVF | 0.26374978 |
| TGKGIVE | GAPGGNGVF | 0.48485917 |
| LISNASDA | GAPGGNGVF | 0.58616793 |
| AELRTLGELTKIAKAHGAQ | GAPGGNGVF | 0.41148555 |
| APNAVAAAEPSW | GAPGGNGVF | 0.44351292 |
| APQVAGAATAVL | GAPGGNGVF | 0.48572314 |
| ARRDPAAAAAILPTDA | GAPGGNGVF | 0.26036474 |
| DAAARDVLAEAGLAEYF | GAPGGNGVF | 0.5501657 |
| DDQIVLSAKANLAKVEQ | GAPGGNGVF | 0.7073238 |
| DELLPDTGEDSDEDGH | GAPGGNGVF | 0.30654162 |
| ELPEAPANRATVPQD | GAPGGNGVF | 0.46104032 |
| FGQSKIERVTGIG | GAPGGNGVF | 0.2867723 |
| GAGGTGSTAGGAGGAGGAGGLY | GAPGGNGVF | 0.29546708 |
| GHGGTGAAGGNGGTGGAGGSGI | GAPGGNGVF | 0.22008382 |
| GNGGNGGAGGEGGPGGAGGAGA | GAPGGNGVF | 0.6030593 |
| GSTAGNGGNGGNATNGTIAGQP | GAPGGNGVF | 0.46845195 |
| GTGGDGGNGGNGGTGG | GAPGGNGVF | 0.43351287 |
| GVRRPAGIPADAPV | GAPGGNGVF | 0.26817355 |
| GYRQVTDAHLLTLARR | GAPGGNGVF | 0.25875902 |
| IAKMGQDGHDRG | GAPGGNGVF | 0.3037115 |
| KAEPSIEIQYT | GAPGGNGVF | 0.31578857 |
| KPVRIRMAADEDTVAVT | GAPGGNGVF | 0.21758334 |
| MPPRIAGMRLLVIKPEP | GAPGGNGVF | 0.26936203 |
| MKFYAVALFRSRGVPPTR | GAPGGNGVF | 0.6330552 |
| NAGGFGILGAVAHSP | GAPGGNGVF | 0.55565983 |
| NEGREITLAMAEM | GAPGGNGVF | 0.2735936 |
| NELVEWSHPLGHRWRWE | GAPGGNGVF | 0.61531067 |
| NLGAGGTGGNGGNPG | GAPGGNGVF | 0.58318263 |
| NTEWPKWETVMAADPVD | GAPGGNGVF | 0.4089741 |
| PASAAPAETQP | GAPGGNGVF | 0.29745346 |
| QQSSDGKNGVRAAEITGE | GAPGGNGVF | 0.2755742 |
| RAPDAVTTSWWKE | GAPGGNGVF | 0.73461765 |
| SGFGNVGAHTSGWFNQ | GAPGGNGVF | 0.5256956 |
| SGVAGAVGQAASVGGLKV | GAPGGNGVF | 0.27883524 |
| SNGGAGTGSAGGNGGT | GAPGGNGVF | 0.5774754 |
| TGGAGGEGGAAG | GAPGGNGVF | 0.3663883 |
| TGGDGGAAGAGGAPGHG | GAPGGNGVF | 0.38237238 |

| | | |
|---|---|---|
| TMPPLLAAIANAGRHGV | GAPGGNGVF | 0.56506217 |
| TSSTAGNSGGA | GAPGGNGVF | 0.36730188 |
| VPATVPVGILC | GAPGGNGVF | 0.2990712 |
| VQRPDSGGVVLTGRIS | GAPGGNGVF | 0.27013522 |
| WTPNPVPATKGVPAGLVQAT | GAPGGNGVF | 0.2597826 |
| YALKRALAKFSDRFDVVII | GAPGGNGVF | 0.4582342 |
| YLIGSVIV | GAPGGNGVF | 0.6076755 |
| YGQLVAVID | GAPGGNGVF | 0.3281842 |
| VTVGGMTLPA | GAPGGNGVF | 0.28973734 |
| VLALPAV | GAPGGNGVF | 0.6345993 |
| VIPPTLNYET | GAPGGNGVF | 0.34570333 |
| VGMGGMHPSSG | GAPGGNGVF | 0.36226004 |
| VGLAGGAATA | GAPGGNGVF | 0.60020185 |
| VGGAGGAGGSGP | GAPGGNGVF | 0.41832787 |
| VFAAFMV | GAPGGNGVF | 0.3709965 |
| TLYDKLR | GAPGGNGVF | 0.47295088 |
| TLAVFTVL | GAPGGNGVF | 0.37162176 |
| TGSVQPARV | GAPGGNGVF | 0.42191768 |
| TGGGAGSGGT | GAPGGNGVF | 0.32643032 |
| SHLETSL | GAPGGNGVF | 0.30643317 |
| SFTTGMT | GAPGGNGVF | 0.21737051 |
| QVEVAKRMGVRQPTVS | GAPGGNGVF | 0.29288563 |
| QGSGGPALAM | GAPGGNGVF | 0.592041 |
| MLAPSNP | GAPGGNGVF | 0.5108217 |
| LVNNAAI | GAPGGNGVF | 0.5625519 |
| LTGEINTLA | GAPGGNGVF | 0.34343898 |
| LSIFGPV | GAPGGNGVF | 0.29326007 |
| LLALLANL | GAPGGNGVF | 0.5034787 |
| LISNASDALDKLR | GAPGGNGVF | 0.5412602 |
| LINEAVP | GAPGGNGVF | 0.25722647 |
| LGYAFSGL | GAPGGNGVF | 0.31870154 |
| LDFVAVAL | GAPGGNGVF | 0.6693582 |
| LAQPGAE | GAPGGNGVF | 0.60345316 |
| ISNLGAVTGF | GAPGGNGVF | 0.40587384 |
| HQEDPGRI | GAPGGNGVF | 0.29178137 |
| GTPGEPTP | GAPGGNGVF | 0.7536596 |
| GFFNSGS | GAPGGNGVF | 0.29011074 |
| GAGGGGLTAGAN | GAPGGNGVF | 0.60304743 |
| FVDALVNM | GAPGGNGVF | 0.21320674 |
| FLTVAPEL | GAPGGNGVF | 0.2779449 |
| FETLTAL | GAPGGNGVF | 0.24961004 |
| DVINPIEL | GAPGGNGVF | 0.22496228 |
| APVSGAPV | GAPGGNGVF | 0.48127404 |
| ALVGGLAW | GAPGGNGVF | 0.29046816 |
| ALAAAAATAI | GAPGGNGVF | 0.5808288 |
| AFGLGLLL | GAPGGNGVF | 0.39032558 |
| AFFAAASL | GAPGGNGVF | 0.4639851 |
| AAAALPAFSPP | GAPGGNGVF | 0.4437288 |
| VVLAPAL | GAPGGNGVF | 0.40453893 |
| TIGHVDHGKTTL | GAPGGNGVF | 0.27333403 |
| LISNASDALDK | GAPGGNGVF | 0.50013775 |

| | | |
|---|---|---|
| LISNASDAL | GAPGGNGVF | 0.52270865 |
| ISNASDALDK | GAPGGNGVF | 0.32609692 |
| NQAETLVYQTE | GAPGGNGVF | 0.2441649 |
| TGGTGGTAGAG | GAPGGNGVF | 0.336761 |
| TGSTAGGAGGA | GAPGGNGVF | 0.37097296 |
| DPLAVPVAAGPV | GAPGGNGVF | 0.22140993 |
| GGTGGTPGNGGN | GAPGGNGVF | 0.21218847 |
| PAGLAGVDPATL | GAPGGNGVF | 0.30352515 |
| GNNLVVEAPHAGASV | GAPGGNGVF | 0.26974314 |
| KVTRIMSASTLEQPA | GAPGGNGVF | 0.64886653 |
| LGFALPIALPRRTVV | GAPGGNGVF | 0.6607456 |
| TRLGNLTARKLVVLV | GAPGGNGVF | 0.45083874 |
| WSQQRRLIYFPSAGP | GAPGGNGVF | 0.47149348 |
| GPNGNPDPMAAAVDIR | GAPGGNGVF | 0.5676247 |
| TQRCVSVADLRAAASR | GAPGGNGVF | 0.19897579 |
| QFVLDVAEHPVEVLTETV | GAPGGNGVF | 0.2887101 |
| GGAGGTGGAGGNGGTQAGDG | GAPGGNGVF | 0.26961505 |
| PLLATLRAIAVDVDAKPAQV | GAPGGNGVF | 0.4346383 |
| APKFRTGGYMIDFWGVGYQVAK | GAPGGNGVF | 0.2740402 |
| GAGGAGGAGGAAGGPGGTGGTG | GAPGGNGVF | 0.37139595 |
| IGALGLLATDVAPAGSVAFGVA | GAPGGNGVF | 0.54877985 |
| AGGAGGIGGNGGAGGAGGVLMG | GAPGGNGVF | 0.6640962 |
| ADELIGLL | GAPGGNGVF | 0.40728098 |
| ADRALKA | GAPGGNGVF | 0.25274453 |
| AGAVGQVGGAGGA | GAPGGNGVF | 0.20403707 |
| AGGGGGLGGAP | GAPGGNGVF | 0.5693914 |
| AGGLLYGNGGAG | GAPGGNGVF | 0.2377917 |
| AGLESIGL | GAPGGNGVF | 0.26207566 |
| AGLKTGII | GAPGGNGVF | 0.21285544 |
| ASLAALLL | GAPGGNGVF | 0.26633227 |
| ASVINISSI | GAPGGNGVF | 0.23783977 |
| ATPPGPA | GAPGGNGVF | 0.42149284 |
| ATVASALV | GAPGGNGVF | 0.36152843 |
| ATVASLIL | GAPGGNGVF | 0.4616976 |
| AVNGPVLIRH | GAPGGNGVF | 0.67767656 |
| DGAAGAVGI | GAPGGNGVF | 0.71412194 |
| DPLLHGI | GAPGGNGVF | 0.45235252 |
| ELGAALGAL | GAPGGNGVF | 0.31004468 |
| ELGRTQAV | GAPGGNGVF | 0.2834196 |
| FVAGGYLS | GAPGGNGVF | 0.27554736 |
| GASGGAGGAGGW | GAPGGNGVF | 0.2790735 |
| GGAGAGGDGGAGGI | GAPGGNGVF | 0.71941036 |
| GTGHNGSQPGMGGNG | GAPGGNGVF | 0.283751 |
| GYAAASASAS | GAPGGNGVF | 0.25470313 |
| HQVDVAAVQQG | GAPGGNGVF | 0.29741874 |
| IGAAIAHT | GAPGGNGVF | 0.63480896 |
| IGMIEVAQL | GAPGGNGVF | 0.6802442 |
| IRLAPEHTSAMD | GAPGGNGVF | 0.32726762 |
| IYNVSVL | GAPGGNGVF | 0.28454596 |
| KQGFLGM | GAPGGNGVF | 0.285132 |
| LESVITQL | GAPGGNGVF | 0.5561812 |

| | | |
|---|---|---|
| LGLGALVVV | GAPGGNGVF | 0.27014932 |
| LIGNGGNGGSGGI | GAPGGNGVF | 0.26565725 |
| LNAINAPTL | GAPGGNGVF | 0.35218388 |
| LQLLNTGV | GAPGGNGVF | 0.7508801 |
| LQTSIEF | GAPGGNGVF | 0.28825834 |
| MSFAVSEL | GAPGGNGVF | 0.2346639 |
| NTGALGL | GAPGGNGVF | 0.4915517 |
| PPGAGGAGGAGTT | GAPGGNGVF | 0.30144492 |
| QLVGLGI | GAPGGNGVF | 0.39358264 |
| RFLEQVHQL | GAPGGNGVF | 0.35139257 |
| RGVGYVLR | GAPGGNGVF | 0.2553699 |
| SATGVFAGVI | GAPGGNGVF | 0.3432803 |
| SLPAITLP | GAPGGNGVF | 0.59475976 |
| STALAFTV | GAPGGNGVF | 0.4943608 |
| SVLTVGLGK | GAPGGNGVF | 0.40550363 |
| TALAGAGSGI | GAPGGNGVF | 0.6802454 |
| TGLPFTGI | GAPGGNGVF | 0.5880074 |
| VAEHVTV | GAPGGNGVF | 0.70941997 |
| VDIVLNSL | GAPGGNGVF | 0.6389406 |
| VLTGGGAVL | GAPGGNGVF | 0.32023525 |
| VNVGLAYGI | GAPGGNGVF | 0.3081662 |
| VSSAAVGAV | GAPGGNGVF | 0.5569559 |
| VTSVYEAGV | GAPGGNGVF | 0.5604328 |
| WVALDRGA | GAPGGNGVF | 0.48315293 |
| YLVGSVVI | GAPGGNGVF | 0.26211527 |
| GPPGTGKTLL | GAPGGNGVF | 0.6822071 |
| GVGMTKF | GAPGGNGVF | 0.28746334 |
| IGIGGSDL | GAPGGNGVF | 0.60140735 |
| IGTLLAEL | GAPGGNGVF | 0.26009163 |
| IYTSGTT | GAPGGNGVF | 0.2716096 |
| LSFPAGV | GAPGGNGVF | 0.5859635 |
| LYTSGTT | GAPGGNGVF | 0.19868636 |
| VGAFPFGF | GAPGGNGVF | 0.20936735 |
| VGAGTVEF | GAPGGNGVF | 0.3230759 |
| VGKSTLLL | GAPGGNGVF | 0.55513847 |
| AALAHAGGGIV | GAPGGNGVF | 0.2018847 |
| AALVAVSLL | GAPGGNGVF | 0.27001297 |
| AAVSRAVAAL | GAPGGNGVF | 0.23695004 |
| AGGSSGIGGF | GAPGGNGVF | 0.32491642 |
| AGLTVANANG | GAPGGNGVF | 0.43147516 |
| AGLVGAVL | GAPGGNGVF | 0.23627503 |
| ALSQAFTV | GAPGGNGVF | 0.43405348 |
| ALTFSVSL | GAPGGNGVF | 0.25180086 |
| ALTGAALVAL | GAPGGNGVF | 0.27433085 |
| ASFSIGAGVL | GAPGGNGVF | 0.39067417 |
| DGSVYKI | GAPGGNGVF | 0.69191414 |
| EQLAEIV | GAPGGNGVF | 0.26196653 |
| FAVSELTVM | GAPGGNGVF | 0.4624383 |
| GAAPLAAGL | GAPGGNGVF | 0.2722881 |
| GADGGDGGSGGAGG | GAPGGNGVF | 0.73347795 |
| GFGLDTGGAGG | GAPGGNGVF | 0.33600962 |

| | | |
|---|---|---|
| GLASGGLL | GAPGGNGVF | 0.23867606 |
| GRIAGAAVLL | GAPGGNGVF | 0.52488124 |
| GSAGSLNTSF | GAPGGNGVF | 0.3500416 |
| GTGQIGI | GAPGGNGVF | 0.32553178 |
| GTLIALIL | GAPGGNGVF | 0.64651465 |
| IAATMGFVVG | GAPGGNGVF | 0.26142126 |
| IFAQALL | GAPGGNGVF | 0.49672395 |
| IGAAVALGF | GAPGGNGVF | 0.67935526 |
| IGLTPLAL | GAPGGNGVF | 0.64455795 |
| ILGSNTSTL | GAPGGNGVF | 0.28131476 |
| IVFSLTVL | GAPGGNGVF | 0.37833747 |
| LASKATGF | GAPGGNGVF | 0.57998693 |
| LASVALIL | GAPGGNGVF | 0.3039558 |
| LFAVGLTL | GAPGGNGVF | 0.46565133 |
| LFNGGIGTYI | GAPGGNGVF | 0.32052523 |
| LTLGAGII | GAPGGNGVF | 0.5477818 |
| LVAAALGDI | GAPGGNGVF | 0.37818635 |
| NTAFVQLGI | GAPGGNGVF | 0.73384225 |
| QLFSGAEL | GAPGGNGVF | 0.24871552 |
| SAATFITL | GAPGGNGVF | 0.3657489 |
| SNYGLANLGV | GAPGGNGVF | 0.4202749 |
| SYGGAYAVM | GAPGGNGVF | 0.4585476 |
| TTGVGGLTLG | GAPGGNGVF | 0.29942635 |
| TTQTIYI | GAPGGNGVF | 0.26726118 |
| VAAGAGLGAVI | GAPGGNGVF | 0.35869628 |
| VAAVVTLL | GAPGGNGVF | 0.63286877 |
| VGFLGGLI | GAPGGNGVF | 0.32471615 |
| VGVIPILL | GAPGGNGVF | 0.45556885 |
| VIFASSLI | GAPGGNGVF | 0.7437562 |
| VKGGDGGAAAG | GAPGGNGVF | 0.2964856 |
| VTKTFTNI | GAPGGNGVF | 0.43919995 |
| VVAAAILL | GAPGGNGVF | 0.36190426 |
| YAALDVEL | GAPGGNGVF | 0.6074787 |
| LASVALLL | GAPGGNGVF | 0.30662006 |
| LDAVHRV | GAPGGNGVF | 0.6525001 |
| QGLSSSIF | GAPGGNGVF | 0.25535876 |
| RGTFANIRL | GAPGGNGVF | 0.62965715 |
| RQLWWGHRI | GAPGGNGVF | 0.26620018 |
| YGPPGTGKTLL | GAPGGNGVF | 0.4835881 |
| AALVGAPANVA | GAPGGNGVF | 0.31981328 |
| AATFRRTE | GAPGGNGVF | 0.62804955 |
| AGALGGGI | GAPGGNGVF | 0.6449134 |
| AGGAGGAGGIGAG | GAPGGNGVF | 0.6850537 |
| AGSGALGL | GAPGGNGVF | 0.20847625 |
| AQIADQI | GAPGGNGVF | 0.64106566 |
| ASAVAVTYL | GAPGGNGVF | 0.60245395 |
| ASTVLGGTGG | GAPGGNGVF | 0.76309335 |
| AVGGEPAK | GAPGGNGVF | 0.30433637 |
| CEQGTFGGTC | GAPGGNGVF | 0.3845384 |
| DEGKAPAG | GAPGGNGVF | 0.37168607 |
| DTVVFPAL | GAPGGNGVF | 0.29377508 |

| | | |
|---|---|---|
| FEGITMAG | GAPGGNGVF | 0.29632136 |
| FPLVVQD | GAPGGNGVF | 0.32921255 |
| GAFTTPQ | GAPGGNGVF | 0.5130507 |
| GFGGFGTAGGVGGA | GAPGGNGVF | 0.38307437 |
| GFLAQFL | GAPGGNGVF | 0.7068728 |
| GGAGGAGGNAGVA | GAPGGNGVF | 0.68324494 |
| GGVGTAGAGGAGGAGG | GAPGGNGVF | 0.5272516 |
| GNGGGGDAGNAGSGGN | GAPGGNGVF | 0.6112241 |
| GTGGAGGAGNDAGS | GAPGGNGVF | 0.5718993 |
| GTTNTGWFNS | GAPGGNGVF | 0.3749354 |
| IFMGPLFTL | GAPGGNGVF | 0.2980507 |
| ILAHAIEAI | GAPGGNGVF | 0.32549897 |
| IRSAIREAL | GAPGGNGVF | 0.29623666 |
| ITFAGGG | GAPGGNGVF | 0.33069068 |
| ITGMSGHQEV | GAPGGNGVF | 0.32379484 |
| IVAGLACTL | GAPGGNGVF | 0.54179966 |
| IVGGGLAGA | GAPGGNGVF | 0.19846857 |
| IVTGVLNI | GAPGGNGVF | 0.6602489 |
| LAQQGCI | GAPGGNGVF | 0.7236241 |
| LCLHVEN | GAPGGNGVF | 0.4004333 |
| LDAISDGS | GAPGGNGVF | 0.4181615 |
| LGGFAAML | GAPGGNGVF | 0.4446215 |
| LGGKNPMI | GAPGGNGVF | 0.2836285 |
| LGPKGVAAIL | GAPGGNGVF | 0.5352434 |
| LMVGVSI | GAPGGNGVF | 0.27766714 |
| LVGAAFVL | GAPGGNGVF | 0.41183037 |
| LVLAVAPL | GAPGGNGVF | 0.53974986 |
| MAAIAVIDAA | GAPGGNGVF | 0.4051541 |
| NNGINAELQ | GAPGGNGVF | 0.41246656 |
| RGEPLVLVHG | GAPGGNGVF | 0.3106499 |
| RGGAGGAAGLI | GAPGGNGVF | 0.23997457 |
| RSYRMHPAV | GAPGGNGVF | 0.4431725 |
| SAAHLLAGQLA | GAPGGNGVF | 0.22457945 |
| SAAVGSY | GAPGGNGVF | 0.45441622 |
| SPHAVLHRI | GAPGGNGVF | 0.26022217 |
| SQIAKVL | GAPGGNGVF | 0.53470224 |
| TALGHLVP | GAPGGNGVF | 0.57884276 |
| TAPTTSGAA | GAPGGNGVF | 0.35473794 |
| TDGDDGNGGSPGT | GAPGGNGVF | 0.32021666 |
| TDGLGEN | GAPGGNGVF | 0.40362328 |
| TGGPPLPGQA | GAPGGNGVF | 0.3196596 |
| TIPAITI | GAPGGNGVF | 0.2459362 |
| TIPGFTI | GAPGGNGVF | 0.4626253 |
| TSIASGALA | GAPGGNGVF | 0.2943035 |
| TVTLPGAAI | GAPGGNGVF | 0.3302674 |
| VASGLSAALP | GAPGGNGVF | 0.55837893 |
| VGAGVIGI | GAPGGNGVF | 0.38936505 |
| VGPKRFGV | GAPGGNGVF | 0.23228803 |
| VNVAFPD | GAPGGNGVF | 0.49532935 |
| VSMTNSGVS | GAPGGNGVF | 0.44785228 |
| VTDVLGDLSV | GAPGGNGVF | 0.35409203 |

| | | |
|---|---|---|
| VVVVGAGFAGL | GAPGGNGVF | 0.20202485 |
| VYTVGTVV | GAPGGNGVF | 0.34760547 |
| WTTFSGP | GAPGGNGVF | 0.3116846 |
| YDELVEAF | GAPGGNGVF | 0.27793756 |
| YGTGGAGGNGG | GAPGGNGVF | 0.29796746 |
| YTNVMAV | GAPGGNGVF | 0.7018976 |
| FLEQVHQL | GAPGGNGVF | 0.2609469 |
| GLSLLAVAV | GAPGGNGVF | 0.33315313 |
| LEQVHQL | GAPGGNGVF | 0.6394148 |
| LNAAGAI | GAPGGNGVF | 0.5024168 |
| PAVHASL | GAPGGNGVF | 0.7095847 |
| TELEDDA | GAPGGNGVF | 0.38612315 |
| TPPGPPN | GAPGGNGVF | 0.26956505 |
| TPPPPPPP | GAPGGNGVF | 0.35179177 |
| TQSPALI | ASSLGMMDKDTQY | 0.2996251 |
| EFAAAFPQ | ASSLGMMDKDTQY | 0.22922489 |
| LLSGNLDVL | ASSLGMMDKDTQY | 0.21659958 |
| GPANLGNYNI | ASSLGMMDKDTQY | 0.37958446 |
| TLGPTLP | ASSLGMMDKDTQY | 0.22712699 |
| VYAYSSCCK | ASSLGMMDKDTQY | 0.39360985 |
| APAPAPAPNI | ASSLGMMDKDTQY | 0.37994742 |
| PGGPVPIP | ASSLGMMDKDTQY | 0.26245397 |
| FLTVALEP | ASSLGMMDKDTQY | 0.2565267 |
| SQYFGANL | ASSLGMMDKDTQY | 0.30248412 |
| GGVKPAA | ASSLGMMDKDTQY | 0.322863 |
| GGTGGAGGAGGW | ASSLGMMDKDTQY | 0.62952745 |
| GVGGVAGLGGAGG | ASSLGMMDKDTQY | 0.31215078 |
| VLLVLATTI | ASSLGMMDKDTQY | 0.7563745 |
| IGIFAAML | ASSLGMMDKDTQY | 0.6416898 |
| AVGDLID | ASSLGMMDKDTQY | 0.44261125 |
| NISPLPL | ASSLGMMDKDTQY | 0.62651634 |
| VTAVFVAI | ASSLGMMDKDTQY | 0.30330518 |
| SGDDNDAAL | ASSLGMMDKDTQY | 0.30806145 |
| AVAALAHS | ASSLGMMDKDTQY | 0.31412604 |
| GGVTVMATP | ASSLGMMDKDTQY | 0.31108862 |
| AGLAANVRD | ASSLGMMDKDTQY | 0.3666073 |
| TNRPDLIDPALLRPG | ASSLGMMDKDTQY | 0.29826385 |
| LANLGVTAL | ASSLGMMDKDTQY | 0.23217447 |
| TGERIEL | ASSLGMMDKDTQY | 0.38725805 |
| NGEELAA | ASSLGMMDKDTQY | 0.45949328 |
| VLSLVWG | ASSLGMMDKDTQY | 0.2225991 |
| IGYALWL | ASSLGMMDKDTQY | 0.28351477 |
| LSLLLAPL | ASSLGMMDKDTQY | 0.6931349 |
| TLDEALK | ASSLGMMDKDTQY | 0.6212034 |
| TDGAGTAEGG | ASSLGMMDKDTQY | 0.31075028 |
| NAPGGAGGNGG | ASSLGMMDKDTQY | 0.28101808 |
| VSVFMGMGGNF | ASSLGMMDKDTQY | 0.6537051 |
| LASYLAAL | ASSLGMMDKDTQY | 0.23427297 |
| LVGGTPAL | ASSLGMMDKDTQY | 0.32760566 |
| VRNADVL | ASSLGMMDKDTQY | 0.5853034 |
| TCSSPFLVSAM | ASSLGMMDKDTQY | 0.511505 |

| | | |
|---|---|---|
| VHITDAE | ASSLGMMDKDTQY | 0.26261765 |
| EYAAPSF | ASSLGMMDKDTQY | 0.4217124 |
| VVDAFHGGL | ASSLGMMDKDTQY | 0.37667772 |
| ASVVDPQ | ASSLGMMDKDTQY | 0.34666264 |
| SYNEFGGLS | ASSLGMMDKDTQY | 0.20271711 |
| DPDGSVY | ASSLGMMDKDTQY | 0.30525059 |
| MVSTIHATE | ASSLGMMDKDTQY | 0.36864218 |
| LAAGVADPV | ASSLGMMDKDTQY | 0.2625303 |
| TKEQIAA | ASSLGMMDKDTQY | 0.23273708 |
| SLASPAAL | ASSLGMMDKDTQY | 0.34090567 |
| IGLVFYGV | ASSLGMMDKDTQY | 0.6050843 |
| FLLDASNL | ASSLGMMDKDTQY | 0.6348164 |
| YQVGNFP | ASSLGMMDKDTQY | 0.25758338 |
| ESLLSQST | ASSLGMMDKDTQY | 0.64683014 |
| LFTVGFAAVG | ASSLGMMDKDTQY | 0.49463433 |
| ATPPTPT | ASSLGMMDKDTQY | 0.42497623 |
| TDLATLG | ASSLGMMDKDTQY | 0.23369044 |
| SAGGTGR | ASSLGMMDKDTQY | 0.4631691 |
| GFSNSGS | ASSLGMMDKDTQY | 0.3326723 |
| GGNGGSGGTSV | ASSLGMMDKDTQY | 0.35859826 |
| GTSVATGGAGN | ASSLGMMDKDTQY | 0.6223397 |
| NGGNGGSGGTS | ASSLGMMDKDTQY | 0.3356393 |
| AVVKFPH | ASSLGMMDKDTQY | 0.37753943 |
| TPPPPPP | ASSLGMMDKDTQY | 0.39609122 |
| AAPAAAAAPA | ASSLGMMDKDTQY | 0.27897012 |
| IALNEDL | ASSLGMMDKDTQY | 0.2706231 |
| IVNVSSM | ASSLGMMDKDTQY | 0.28184545 |
| TGIPPAPR | ASSLGMMDKDTQY | 0.21367459 |
| FDLGGGTF | ASSLGMMDKDTQY | 0.45330188 |
| FDLGGGTFDVS | ASSLGMMDKDTQY | 0.4342174 |
| NEPTAAA | ASSLGMMDKDTQY | 0.29609308 |
| NEPTAAALA | ASSLGMMDKDTQY | 0.50544417 |
| GSIINISSI | ASSLGMMDKDTQY | 0.28173012 |
| VAPGNAGTA | ASSLGMMDKDTQY | 0.351124 |
| TGGGASLE | ASSLGMMDKDTQY | 0.3443157 |
| VAVYNHYKRI | ASSLGMMDKDTQY | 0.30959633 |
| TDPAPATT | ASSLGMMDKDTQY | 0.46615505 |
| TNRPDILDPALLRPG | ASSLGMMDKDTQY | 0.34475347 |
| LTTTTTS | ASSLGMMDKDTQY | 0.3124171 |
| PTGSGKTL | ASSLGMMDKDTQY | 0.7405994 |
| TGSGKTL | ASSLGMMDKDTQY | 0.2496741 |
| TIGHVDHGK | ASSLGMMDKDTQY | 0.32500124 |
| TIGHVDHGKTT | ASSLGMMDKDTQY | 0.39576435 |
| GHVDHGKTT | ASSLGMMDKDTQY | 0.24906127 |
| AQFEHTLL | ASSLGMMDKDTQY | 0.47051066 |
| IVNQDEL | ASSLGMMDKDTQY | 0.5422696 |
| TNGSQFF | ASSLGMMDKDTQY | 0.61580634 |
| TGKGIVE | ASSLGMMDKDTQY | 0.30340567 |
| LISNASDA | ASSLGMMDKDTQY | 0.73427635 |
| AELRTLGELTKIAKAHGAQ | ASSLGMMDKDTQY | 0.58373344 |
| APNAVAAAEPSW | ASSLGMMDKDTQY | 0.3091316 |

| | | |
|---|---|---|
| APQVAGAATAVL | ASSLGMMDKDTQY | 0.70699245 |
| ARRDPAAAAAILPTDA | ASSLGMMDKDTQY | 0.2577301 |
| DAAARDVLAEAGLAEYF | ASSLGMMDKDTQY | 0.5502341 |
| DDQIVLSAKANLAKVEQ | ASSLGMMDKDTQY | 0.4233463 |
| DELLPDTGEDSDEDGH | ASSLGMMDKDTQY | 0.310819 |
| ELPEAPANRATVPQD | ASSLGMMDKDTQY | 0.47937593 |
| FGQSKIERVTGIG | ASSLGMMDKDTQY | 0.44943392 |
| GAGGTGSTAGGAGGAGGAGGLY | ASSLGMMDKDTQY | 0.6687573 |
| GHGGTGAAGGNGGTGGAGGSGI | ASSLGMMDKDTQY | 0.2262355 |
| GNGGNGGAGGEGGPGGAGGAGA | ASSLGMMDKDTQY | 0.45022124 |
| GSTAGNGGNGGNATNGTIAGQP | ASSLGMMDKDTQY | 0.22196895 |
| GTGGDGGNGGNGGTGG | ASSLGMMDKDTQY | 0.25364766 |
| GVRRPAGIPADAPV | ASSLGMMDKDTQY | 0.28383416 |
| GYRQVTDAHLLTLARR | ASSLGMMDKDTQY | 0.45010167 |
| IAKMGQDGHDRG | ASSLGMMDKDTQY | 0.40359837 |
| KAEPSIEIQYT | ASSLGMMDKDTQY | 0.68627185 |
| KPVRIRMAADEDTVAVT | ASSLGMMDKDTQY | 0.4611165 |
| MPPRIAGMRLLVIKPEP | ASSLGMMDKDTQY | 0.597944 |
| MKFYAVALFRSRGVPPTR | ASSLGMMDKDTQY | 0.2793926 |
| NAGGFGILGAVAHSP | ASSLGMMDKDTQY | 0.41421354 |
| NEGREITLAMAEM | ASSLGMMDKDTQY | 0.23493846 |
| NELVEWSHPLGHRWRWE | ASSLGMMDKDTQY | 0.24177286 |
| NLGAGGTGGNGGNPG | ASSLGMMDKDTQY | 0.2677339 |
| NTEWPKWETVMAADPVD | ASSLGMMDKDTQY | 0.3616224 |
| PASAAPAETQP | ASSLGMMDKDTQY | 0.21393599 |
| QQSSDGKNGVRAAEITGE | ASSLGMMDKDTQY | 0.47793937 |
| RAPDAVTTSWWKE | ASSLGMMDKDTQY | 0.43526906 |
| SGFGNVGAHTSGWFNQ | ASSLGMMDKDTQY | 0.58951277 |
| SGVAGAVGQAASVGGLKV | ASSLGMMDKDTQY | 0.28321344 |
| SNGGAGTGSAGGNGGT | ASSLGMMDKDTQY | 0.336616 |
| TGGAGGEGGAAG | ASSLGMMDKDTQY | 0.34294868 |
| TGGDGGAAGAGGAPGHG | ASSLGMMDKDTQY | 0.5738534 |
| TMPPLLAAIANAGRHGV | ASSLGMMDKDTQY | 0.27784258 |
| TSSTAGNSGGA | ASSLGMMDKDTQY | 0.22774085 |
| VPATVPVGILC | ASSLGMMDKDTQY | 0.27566385 |
| VQRPDSGGVVLTGRIS | ASSLGMMDKDTQY | 0.26581132 |
| WTPNPVPATKGVPAGLVQAT | ASSLGMMDKDTQY | 0.58087564 |
| YALKRALAKFSDRFDVVII | ASSLGMMDKDTQY | 0.52437985 |
| YLIGSVIV | ASSLGMMDKDTQY | 0.4696977 |
| YGQLVAVID | ASSLGMMDKDTQY | 0.5474926 |
| VTVGGMTLPA | ASSLGMMDKDTQY | 0.21602722 |
| VLALPAV | ASSLGMMDKDTQY | 0.29820716 |
| VIPPTLNYET | ASSLGMMDKDTQY | 0.73002744 |
| VGMGGMHPSSG | ASSLGMMDKDTQY | 0.35650435 |
| VGLAGGAATA | ASSLGMMDKDTQY | 0.45124638 |
| VGGAGGAGGSGP | ASSLGMMDKDTQY | 0.62500775 |
| VFAAFMV | ASSLGMMDKDTQY | 0.34931344 |
| TLYDKLR | ASSLGMMDKDTQY | 0.34795564 |
| TLAVFTVL | ASSLGMMDKDTQY | 0.39283293 |
| TGSVQPARV | ASSLGMMDKDTQY | 0.33784324 |
| TGGGAGSGGT | ASSLGMMDKDTQY | 0.30379084 |

| | | |
|---|---|---|
| SHLETSL | ASSLGMMDKDTQY | 0.26681247 |
| SFTTGMT | ASSLGMMDKDTQY | 0.35957718 |
| QVEVAKRMGVRQPTVS | ASSLGMMDKDTQY | 0.7636399 |
| QGSGGPALAM | ASSLGMMDKDTQY | 0.4560248 |
| MLAPSNP | ASSLGMMDKDTQY | 0.5649771 |
| LVNNAAI | ASSLGMMDKDTQY | 0.4268601 |
| LTGEINTLA | ASSLGMMDKDTQY | 0.69743145 |
| LSIFGPV | ASSLGMMDKDTQY | 0.35118943 |
| LLALLANL | ASSLGMMDKDTQY | 0.34153798 |
| LISNASDALDKLR | ASSLGMMDKDTQY | 0.37908527 |
| LINEAVP | ASSLGMMDKDTQY | 0.34864494 |
| LGYAFSGL | ASSLGMMDKDTQY | 0.32972527 |
| LDFVAVAL | ASSLGMMDKDTQY | 0.30740166 |
| LAQPGAE | ASSLGMMDKDTQY | 0.7613223 |
| ISNLGAVTGF | ASSLGMMDKDTQY | 0.4864989 |
| HQEDPGRI | ASSLGMMDKDTQY | 0.4663903 |
| GTPGEPTP | ASSLGMMDKDTQY | 0.30800956 |
| GFFNSGS | ASSLGMMDKDTQY | 0.31606895 |
| GAGGGGLTAGAN | ASSLGMMDKDTQY | 0.47760433 |
| FVDALVNM | ASSLGMMDKDTQY | 0.24665251 |
| FLTVAPEL | ASSLGMMDKDTQY | 0.676177 |
| FETLTAL | ASSLGMMDKDTQY | 0.3368185 |
| DVINPIEL | ASSLGMMDKDTQY | 0.2670766 |
| APVSGAPV | ASSLGMMDKDTQY | 0.45455366 |
| ALVGGLAW | ASSLGMMDKDTQY | 0.74750674 |
| ALAAAAATAI | ASSLGMMDKDTQY | 0.30915838 |
| AFGLGLLL | ASSLGMMDKDTQY | 0.39588618 |
| AFFAAASL | ASSLGMMDKDTQY | 0.36467373 |
| AAAALPAFSPP | ASSLGMMDKDTQY | 0.38810873 |
| VVLAPAL | ASSLGMMDKDTQY | 0.7044753 |
| TIGHVDHGKTTL | ASSLGMMDKDTQY | 0.39787996 |
| LISNASDALDK | ASSLGMMDKDTQY | 0.7369616 |
| LISNASDAL | ASSLGMMDKDTQY | 0.30191323 |
| ISNASDALDK | ASSLGMMDKDTQY | 0.24838892 |
| NQAETLVYQTE | ASSLGMMDKDTQY | 0.3210017 |
| TGGTGGTAGAG | ASSLGMMDKDTQY | 0.37967697 |
| TGSTAGGAGGA | ASSLGMMDKDTQY | 0.6697539 |
| DPLAVPVAAGPV | ASSLGMMDKDTQY | 0.529322 |
| GGTGGTPGNGGN | ASSLGMMDKDTQY | 0.43109757 |
| PAGLAGVDPATL | ASSLGMMDKDTQY | 0.26289305 |
| GNNLVVEAPHAGASV | ASSLGMMDKDTQY | 0.57054037 |
| KVTRIMSASTLEQPA | ASSLGMMDKDTQY | 0.4506021 |
| LGFALPIALPRRTVV | ASSLGMMDKDTQY | 0.50867546 |
| TRLGNLTARKLVVLV | ASSLGMMDKDTQY | 0.28224432 |
| WSQQRRLIYFPSAGP | ASSLGMMDKDTQY | 0.25475445 |
| GPNGNPDPMAAAVDIR | ASSLGMMDKDTQY | 0.3404146 |
| TQRCVSVADLRAAASR | ASSLGMMDKDTQY | 0.56043935 |
| QFVLDVAEHPVEVLTETV | ASSLGMMDKDTQY | 0.26129335 |
| GGAGGTGGAGGNGGTQAGDG | ASSLGMMDKDTQY | 0.5221836 |
| PLLATLRAIAVDVDAKPAQV | ASSLGMMDKDTQY | 0.54271066 |
| APKFRTGGYMIDFWGVGYQVAK | ASSLGMMDKDTQY | 0.25040224 |

| | | |
|---|---|---|
| GAGGAGGAGGAAGGPGGTGGTG | ASSLGMMDKDTQY | 0.3193585 |
| IGALGLLATDVAPAGSVAFGVA | ASSLGMMDKDTQY | 0.684867 |
| AGGAGGIGGNGGAGGAGGVLMG | ASSLGMMDKDTQY | 0.31203866 |
| ADELIGLL | ASSLGMMDKDTQY | 0.29192528 |
| ADRALKA | ASSLGMMDKDTQY | 0.64349866 |
| AGAVGQVGGAGGA | ASSLGMMDKDTQY | 0.75796306 |
| AGGGGGLGGAP | ASSLGMMDKDTQY | 0.49504554 |
| AGGLLYGNGGAG | ASSLGMMDKDTQY | 0.2407273 |
| AGLESIGL | ASSLGMMDKDTQY | 0.21502249 |
| AGLKTGII | ASSLGMMDKDTQY | 0.35030046 |
| ASLAALLL | ASSLGMMDKDTQY | 0.6372106 |
| ASVINISSI | ASSLGMMDKDTQY | 0.31791657 |
| ATPPGPA | ASSLGMMDKDTQY | 0.28380346 |
| ATVASALV | ASSLGMMDKDTQY | 0.5058456 |
| ATVASLIL | ASSLGMMDKDTQY | 0.55975604 |
| AVNGPVLIRH | ASSLGMMDKDTQY | 0.28594285 |
| DGAAGAVGI | ASSLGMMDKDTQY | 0.23813145 |
| DPLLHGI | ASSLGMMDKDTQY | 0.38721526 |
| ELGAALGAL | ASSLGMMDKDTQY | 0.48289526 |
| ELGRTQAV | ASSLGMMDKDTQY | 0.31918025 |
| FVAGGYLS | ASSLGMMDKDTQY | 0.4323163 |
| GASGGAGGAGGW | ASSLGMMDKDTQY | 0.3208065 |
| GGAGAGGDGGAGGI | ASSLGMMDKDTQY | 0.24077693 |
| GTGHNGSQPGMGGNG | ASSLGMMDKDTQY | 0.5365701 |
| GYAAASASAS | ASSLGMMDKDTQY | 0.26240492 |
| HQVDVAAVQQG | ASSLGMMDKDTQY | 0.60768306 |
| IGAAIAHT | ASSLGMMDKDTQY | 0.28192765 |
| IGMIEVAQL | ASSLGMMDKDTQY | 0.41400003 |
| IRLAPEHTSAMD | ASSLGMMDKDTQY | 0.5856677 |
| IYNVSVL | ASSLGMMDKDTQY | 0.3757373 |
| KQGFLGM | ASSLGMMDKDTQY | 0.40044636 |
| LESVITQL | ASSLGMMDKDTQY | 0.30947 |
| LGLGALVVV | ASSLGMMDKDTQY | 0.26612157 |
| LIGNGGNGGSGGI | ASSLGMMDKDTQY | 0.21495423 |
| LNAINAPTL | ASSLGMMDKDTQY | 0.6722725 |
| LQLLNTGV | ASSLGMMDKDTQY | 0.274002 |
| LQTSIEF | ASSLGMMDKDTQY | 0.6383385 |
| MSFAVSEL | ASSLGMMDKDTQY | 0.44681498 |
| NTGALGL | ASSLGMMDKDTQY | 0.30325985 |
| PPGAGGAGGAGTT | ASSLGMMDKDTQY | 0.38783768 |
| QLVGLGI | ASSLGMMDKDTQY | 0.2816438 |
| RFLEQVHQL | ASSLGMMDKDTQY | 0.27063712 |
| RGVGYVLR | ASSLGMMDKDTQY | 0.4265536 |
| SATGVFAGVI | ASSLGMMDKDTQY | 0.43827534 |
| SLPAITLP | ASSLGMMDKDTQY | 0.3021306 |
| STALAFTV | ASSLGMMDKDTQY | 0.35737795 |
| SVLTVGLGK | ASSLGMMDKDTQY | 0.41215968 |
| TALAGAGSGI | ASSLGMMDKDTQY | 0.25991756 |
| TGLPFTGI | ASSLGMMDKDTQY | 0.31216848 |
| VAEHVTV | ASSLGMMDKDTQY | 0.6094549 |
| VDIVLNSL | ASSLGMMDKDTQY | 0.32397202 |

| | | |
|---|---|---|
| VLTGGGAVL | ASSLGMMDKDTQY | 0.28721425 |
| VNVGLAYGI | ASSLGMMDKDTQY | 0.4241228 |
| VSSAAVGAV | ASSLGMMDKDTQY | 0.31210858 |
| VTSVYEAGV | ASSLGMMDKDTQY | 0.6410218 |
| WVALDRGA | ASSLGMMDKDTQY | 0.38211682 |
| YLVGSVVI | ASSLGMMDKDTQY | 0.32472923 |
| GPPGTGKTLL | ASSLGMMDKDTQY | 0.37105057 |
| GVGMTKF | ASSLGMMDKDTQY | 0.23946957 |
| IGIGGSDL | ASSLGMMDKDTQY | 0.26254466 |
| IGTLLAEL | ASSLGMMDKDTQY | 0.306274 |
| IYTSGTT | ASSLGMMDKDTQY | 0.4345099 |
| LSFPAGV | ASSLGMMDKDTQY | 0.5522978 |
| LYTSGTT | ASSLGMMDKDTQY | 0.32258445 |
| VGAFPFGF | ASSLGMMDKDTQY | 0.3634367 |
| VGAGTVEF | ASSLGMMDKDTQY | 0.32104954 |
| VGKSTLLL | ASSLGMMDKDTQY | 0.27902678 |
| AALAHAGGGIV | ASSLGMMDKDTQY | 0.30274668 |
| AALVAVSLL | ASSLGMMDKDTQY | 0.28523687 |
| AAVSRAVAAL | ASSLGMMDKDTQY | 0.26492763 |
| AGGSSGIGGF | ASSLGMMDKDTQY | 0.38488168 |
| AGLTVANANG | ASSLGMMDKDTQY | 0.2778754 |
| AGLVGAVL | ASSLGMMDKDTQY | 0.31222886 |
| ALSQAFTV | ASSLGMMDKDTQY | 0.57624507 |
| ALTFSVSL | ASSLGMMDKDTQY | 0.36861914 |
| ALTGAALVAL | ASSLGMMDKDTQY | 0.35823652 |
| ASFSIGAGVL | ASSLGMMDKDTQY | 0.33876517 |
| DGSVYKI | ASSLGMMDKDTQY | 0.6155796 |
| EQLAEIV | ASSLGMMDKDTQY | 0.37292892 |
| FAVSELTVM | ASSLGMMDKDTQY | 0.3171305 |
| GAAPLAAGL | ASSLGMMDKDTQY | 0.5400274 |
| GADGGDGGSGGAGG | ASSLGMMDKDTQY | 0.5563419 |
| GFGLDTGGAGG | ASSLGMMDKDTQY | 0.5786853 |
| GLASGGLL | ASSLGMMDKDTQY | 0.436032 |
| GRIAGAAVLL | ASSLGMMDKDTQY | 0.5987104 |
| GSAGSLNTSF | ASSLGMMDKDTQY | 0.5120293 |
| GTGQIGI | ASSLGMMDKDTQY | 0.4618871 |
| GTLIALIL | ASSLGMMDKDTQY | 0.23313838 |
| IAATMGFVVG | ASSLGMMDKDTQY | 0.35540617 |
| IFAQALL | ASSLGMMDKDTQY | 0.49746946 |
| IGAAVALGF | ASSLGMMDKDTQY | 0.29436246 |
| IGLTPLAL | ASSLGMMDKDTQY | 0.22816822 |
| ILGSNTSTL | ASSLGMMDKDTQY | 0.7016411 |
| IVFSLTVL | ASSLGMMDKDTQY | 0.7317574 |
| LASKATGF | ASSLGMMDKDTQY | 0.26398948 |
| LASVALIL | ASSLGMMDKDTQY | 0.5983064 |
| LFAVGLTL | ASSLGMMDKDTQY | 0.23906855 |
| LFNGGIGTYI | ASSLGMMDKDTQY | 0.4510694 |
| LTLGAGII | ASSLGMMDKDTQY | 0.6761265 |
| LVAAALGDI | ASSLGMMDKDTQY | 0.31575954 |
| NTAFVQLGI | ASSLGMMDKDTQY | 0.4812388 |
| QLFSGAEL | ASSLGMMDKDTQY | 0.2828645 |

| | | |
|---|---|---|
| SAATFITL | ASSLGMMDKDTQY | 0.31574556 |
| SNYGLANLGV | ASSLGMMDKDTQY | 0.690867 |
| SYGGAYAVM | ASSLGMMDKDTQY | 0.32609025 |
| TTGVGGLTLG | ASSLGMMDKDTQY | 0.3604738 |
| TTQTIYI | ASSLGMMDKDTQY | 0.33926722 |
| VAAGAGLGAVI | ASSLGMMDKDTQY | 0.29250124 |
| VAAVVTLL | ASSLGMMDKDTQY | 0.3561362 |
| VGFLGGLI | ASSLGMMDKDTQY | 0.6004927 |
| VGVIPILL | ASSLGMMDKDTQY | 0.25996116 |
| VIFASSLI | ASSLGMMDKDTQY | 0.42220885 |
| VKGGDGGAAAG | ASSLGMMDKDTQY | 0.28019217 |
| VTKTFTNI | ASSLGMMDKDTQY | 0.7421723 |
| VVAAAILL | ASSLGMMDKDTQY | 0.68256295 |
| YAALDVEL | ASSLGMMDKDTQY | 0.5828784 |
| LASVALLL | ASSLGMMDKDTQY | 0.29575583 |
| LDAVHRV | ASSLGMMDKDTQY | 0.6961756 |
| QGLSSSIF | ASSLGMMDKDTQY | 0.36909387 |
| RGTFANIRL | ASSLGMMDKDTQY | 0.5469589 |
| RQLWWGHRI | ASSLGMMDKDTQY | 0.40067932 |
| YGPPGTGKTLL | ASSLGMMDKDTQY | 0.11355106 |
| AALVGAPANVA | ASSLGMMDKDTQY | 0.12922612 |
| AATFRRTE | ASSLGMMDKDTQY | 0.115928024 |
| AGALGGGI | ASSLGMMDKDTQY | 0.13980958 |
| AGGAGGAGGIGAG | ASSLGMMDKDTQY | 0.123935014 |
| AGSGALGL | ASSLGMMDKDTQY | 0.10336307 |
| AQIADQI | ASSLGMMDKDTQY | 0.13679123 |
| ASAVAVTYL | ASSLGMMDKDTQY | 0.11181807 |
| ASTVLGGTGG | ASSLGMMDKDTQY | 0.11544231 |
| AVGGEPAK | ASSLGMMDKDTQY | 0.14014345 |
| CEQGTFGGTC | ASSLGMMDKDTQY | 0.12486804 |
| DEGKAPAG | ASSLGMMDKDTQY | 0.21645916 |
| DTVVFPAL | ASSLGMMDKDTQY | 0.17058006 |
| FEGITMAG | ASSLGMMDKDTQY | 0.1124565 |
| FPLVVQD | ASSLGMMDKDTQY | 0.1258866 |
| GAFTTPQ | ASSLGMMDKDTQY | 0.11154721 |
| GFGGFGTAGGVGGA | ASSLGMMDKDTQY | 0.12205914 |
| GFLAQFL | ASSLGMMDKDTQY | 0.09866862 |
| GGAGGAGGNAGVA | ASSLGMMDKDTQY | 0.1333099 |
| GGVGTAGAGGAGGAGG | ASSLGMMDKDTQY | 0.112959765 |
| GNGGGGDAGNAGSGGN | ASSLGMMDKDTQY | 0.09592864 |
| GTGGAGGAGNDAGS | ASSLGMMDKDTQY | 0.14502548 |
| GTTNTGWFNS | ASSLGMMDKDTQY | 0.11678777 |
| IFMGPLFTL | ASSLGMMDKDTQY | 0.12435063 |
| ILAHAIEAI | ASSLGMMDKDTQY | 0.17600402 |
| IRSAIREAL | ASSLGMMDKDTQY | 0.0979138 |
| ITFAGGG | ASSLGMMDKDTQY | 0.112253554 |
| ITGMSGHQEV | ASSLGMMDKDTQY | 0.16015394 |
| IVAGLACTL | ASSLGMMDKDTQY | 0.20850104 |
| IVGGGLAGA | ASSLGMMDKDTQY | 0.11004819 |
| IVTGVLNI | ASSLGMMDKDTQY | 0.13146399 |
| LAQQGCI | ASSLGMMDKDTQY | 0.18586066 |

| | | |
|---|---|---|
| LCLHVEN | ASSLGMMDKDTQY | 0.14178713 |
| LDAISDGS | ASSLGMMDKDTQY | 0.1563944 |
| LGGFAAML | ASSLGMMDKDTQY | 0.16162336 |
| LGGKNPMI | ASSLGMMDKDTQY | 0.10746056 |
| LGPKGVAAIL | ASSLGMMDKDTQY | 0.17955598 |
| LMVGVSI | ASSLGMMDKDTQY | 0.12653562 |
| LVGAAFVL | ASSLGMMDKDTQY | 0.12645914 |
| LVLAVAPL | ASSLGMMDKDTQY | 0.14091285 |
| MAAIAVIDAA | ASSLGMMDKDTQY | 0.12087138 |
| NNGINAELQ | ASSLGMMDKDTQY | 0.11439622 |
| RGEPLVLVHG | ASSLGMMDKDTQY | 0.097107284 |
| RGGAGGAAGLI | ASSLGMMDKDTQY | 0.12362351 |
| RSYRMHPAV | ASSLGMMDKDTQY | 0.16090523 |
| SAAHLLAGQLA | ASSLGMMDKDTQY | 0.13526574 |
| SAAVGSY | ASSLGMMDKDTQY | 0.10650422 |
| SPHAVLHRI | ASSLGMMDKDTQY | 0.11674278 |
| SQIAKVL | ASSLGMMDKDTQY | 0.099882215 |
| TALGHLVP | ASSLGMMDKDTQY | 0.12013638 |
| TAPTTSGAA | ASSLGMMDKDTQY | 0.09909605 |
| TDGDDGNGGSPGT | ASSLGMMDKDTQY | 0.10127 |
| TDGLGEN | ASSLGMMDKDTQY | 0.1956055 |
| TGGPPLPGQA | ASSLGMMDKDTQY | 0.17560546 |
| TIPAITI | ASSLGMMDKDTQY | 0.10359477 |
| TIPGFTI | ASSLGMMDKDTQY | 0.14186299 |
| TSIASGALA | ASSLGMMDKDTQY | 0.13191107 |
| TVTLPGAAI | ASSLGMMDKDTQY | 0.12603107 |
| VASGLSAALP | ASSLGMMDKDTQY | 0.12659259 |
| VGAGVIGI | ASSLGMMDKDTQY | 0.111993566 |
| VGPKRFGV | ASSLGMMDKDTQY | 0.22166252 |
| VNVAFPD | ASSLGMMDKDTQY | 0.12929037 |
| VSMTNSGVS | ASSLGMMDKDTQY | 0.10997343 |
| VTDVLGDLSV | ASSLGMMDKDTQY | 0.17546916 |
| VVVVGAGFAGL | ASSLGMMDKDTQY | 0.1156668 |
| VYTVGTVV | ASSLGMMDKDTQY | 0.11465727 |
| WTTFSGP | ASSLGMMDKDTQY | 0.15459186 |
| YDELVEAF | ASSLGMMDKDTQY | 0.13958105 |
| YGTGGAGGNGG | ASSLGMMDKDTQY | 0.1055246 |
| YTNVMAV | ASSLGMMDKDTQY | 0.10740519 |
| FLEQVHQL | ASSLGMMDKDTQY | 0.10453984 |
| GLSLLAVAV | ASSLGMMDKDTQY | 0.12854278 |
| LEQVHQL | ASSLGMMDKDTQY | 0.21352269 |
| LNAAGAI | ASSLGMMDKDTQY | 0.137846 |
| PAVHASL | ASSLGMMDKDTQY | 0.11172374 |
| TELEDDA | ASSLGMMDKDTQY | 0.1913253 |
| TPPGPPN | ASSLGMMDKDTQY | 0.12337975 |
| TPPPPPPP | ASSLGMMDKDTQY | 0.12048322 |
| TQSPALI | ASSRTGGIAEQF | 0.13666591 |
| EFAAAFPQ | ASSRTGGIAEQF | 0.10756415 |
| LLSGNLDVL | ASSRTGGIAEQF | 0.1444918 |
| GPANLGNYNI | ASSRTGGIAEQF | 0.14697658 |
| TLGPTLP | ASSRTGGIAEQF | 0.15825176 |

| | | |
|---|---|---|
| VYAYSSCCK | ASSRTGGIAEQF | 0.1578879 |
| APAPAPAPNI | ASSRTGGIAEQF | 0.11398494 |
| PGGPVPIP | ASSRTGGIAEQF | 0.123875834 |
| FLTVALEP | ASSRTGGIAEQF | 0.10172068 |
| SQYFGANL | ASSRTGGIAEQF | 0.10923777 |
| GGVKPAA | ASSRTGGIAEQF | 0.14845353 |
| GGTGGAGGAGGW | ASSRTGGIAEQF | 0.13534752 |
| GVGGVAGLGGAGG | ASSRTGGIAEQF | 0.15441845 |
| VLLVLATTI | ASSRTGGIAEQF | 0.15772295 |
| IGIFAAML | ASSRTGGIAEQF | 0.111795306 |
| AVGDLID | ASSRTGGIAEQF | 0.15076181 |
| NISPLPL | ASSRTGGIAEQF | 0.18524821 |
| VTAVFVAI | ASSRTGGIAEQF | 0.108093835 |
| SGDDNDAAL | ASSRTGGIAEQF | 0.11906451 |
| AVAALAHS | ASSRTGGIAEQF | 0.13958415 |
| GGVTVMATP | ASSRTGGIAEQF | 0.16135632 |
| AGLAANVRD | ASSRTGGIAEQF | 0.14088081 |
| TNRPDLIDPALLRPG | ASSRTGGIAEQF | 0.1192909 |
| LANLGVTAL | ASSRTGGIAEQF | 0.18525784 |
| TGERIEL | ASSRTGGIAEQF | 0.10633532 |
| NGEELAA | ASSRTGGIAEQF | 0.16721103 |
| VLSLVWG | ASSRTGGIAEQF | 0.10014321 |
| IGYALWL | ASSRTGGIAEQF | 0.12856661 |
| LSLLLAPL | ASSRTGGIAEQF | 0.10694069 |
| TLDEALK | ASSRTGGIAEQF | 0.1006826 |
| TDGAGTAEGG | ASSRTGGIAEQF | 0.15250862 |
| NAPGGAGGNGG | ASSRTGGIAEQF | 0.13401556 |
| VSVFMGMGGNF | ASSRTGGIAEQF | 0.16456953 |
| LASYLAAL | ASSRTGGIAEQF | 0.124877356 |
| LVGGTPAL | ASSRTGGIAEQF | 0.14473993 |
| VRNADVL | ASSRTGGIAEQF | 0.1164967 |
| TCSSPFLVSAM | ASSRTGGIAEQF | 0.12797196 |
| VHITDAE | ASSRTGGIAEQF | 0.11717339 |
| EYAAPSF | ASSRTGGIAEQF | 0.14197889 |
| VVDAFHGGL | ASSRTGGIAEQF | 0.17074326 |
| ASVVDPQ | ASSRTGGIAEQF | 0.14162958 |
| SYNEFGGLS | ASSRTGGIAEQF | 0.10487178 |
| DPDGSVY | ASSRTGGIAEQF | 0.10352411 |
| MVSTIHATE | ASSRTGGIAEQF | 0.13651942 |
| LAAGVADPV | ASSRTGGIAEQF | 0.0962928 |
| TKEQIAA | ASSRTGGIAEQF | 0.09752594 |
| SLASPAAL | ASSRTGGIAEQF | 0.10180818 |
| IGLVFYGV | ASSRTGGIAEQF | 0.098270334 |
| FLLDASNL | ASSRTGGIAEQF | 0.17133814 |
| YQVGNFP | ASSRTGGIAEQF | 0.190556 |
| ESLLSQST | ASSRTGGIAEQF | 0.1280101 |
| LFTVGFAAVG | ASSRTGGIAEQF | 0.13798387 |
| ATPPTPT | ASSRTGGIAEQF | 0.13259609 |
| TDLATLG | ASSRTGGIAEQF | 0.09458135 |
| SAGGTGR | ASSRTGGIAEQF | 0.11432312 |
| GFSNSGS | ASSRTGGIAEQF | 0.1193544 |

| | | |
|---|---|---|
| GGNGGSGGTSV | ASSRTGGIAEQF | 0.17861937 |
| GTSVATGGAGN | ASSRTGGIAEQF | 0.10860655 |
| NGGNGGSGGTS | ASSRTGGIAEQF | 0.11459138 |
| AVVKFPH | ASSRTGGIAEQF | 0.14321505 |
| TPPPPPP | ASSRTGGIAEQF | 0.13442062 |
| AAPAAAAAPA | ASSRTGGIAEQF | 0.114838704 |
| IALNEDL | ASSRTGGIAEQF | 0.10309984 |
| IVNVSSM | ASSRTGGIAEQF | 0.09493038 |
| TGIPPAPR | ASSRTGGIAEQF | 0.17164144 |
| FDLGGGTF | ASSRTGGIAEQF | 0.100471385 |
| FDLGGGTFDVS | ASSRTGGIAEQF | 0.11947085 |
| NEPTAAA | ASSRTGGIAEQF | 0.09753461 |
| NEPTAAALA | ASSRTGGIAEQF | 0.0976901 |
| GSIINISSI | ASSRTGGIAEQF | 0.100026816 |
| VAPGNAGTA | ASSRTGGIAEQF | 0.15453823 |
| TGGGASLE | ASSRTGGIAEQF | 0.12520124 |
| VAVYNHYKRI | ASSRTGGIAEQF | 0.12650149 |
| TDPAPATT | ASSRTGGIAEQF | 0.16715302 |
| TNRPDILDPALLRPG | ASSRTGGIAEQF | 0.2619966 |
| LTTTTTS | ASSRTGGIAEQF | 0.12795475 |
| PTGSGKTL | ASSRTGGIAEQF | 0.11519161 |
| TGSGKTL | ASSRTGGIAEQF | 0.108962744 |
| TIGHVDHGK | ASSRTGGIAEQF | 0.11057963 |
| TIGHVDHGKTT | ASSRTGGIAEQF | 0.11514493 |
| GHVDHGKTT | ASSRTGGIAEQF | 0.19996177 |
| AQFEHTLL | ASSRTGGIAEQF | 0.116468854 |
| IVNQDEL | ASSRTGGIAEQF | 0.10721152 |
| TNGSQFF | ASSRTGGIAEQF | 0.11659184 |
| TGKGIVE | ASSRTGGIAEQF | 0.16399874 |
| LISNASDA | ASSRTGGIAEQF | 0.15212 |
| AELRTLGELTKIAKAHGAQ | ASSRTGGIAEQF | 0.12692912 |
| APNAVAAAEPSW | ASSRTGGIAEQF | 0.12328336 |
| APQVAGAATAVL | ASSRTGGIAEQF | 0.123010874 |
| ARRDPAAAAAILPTDA | ASSRTGGIAEQF | 0.12457399 |
| DAAARDVLAEAGLAEYF | ASSRTGGIAEQF | 0.111717515 |
| DDQIVLSAKANLAKVEQ | ASSRTGGIAEQF | 0.1122394 |
| DELLPDTGEDSDEDGH | ASSRTGGIAEQF | 0.13617201 |
| ELPEAPANRATVPQD | ASSRTGGIAEQF | 0.4039383 |
| FGQSKIERVTGIG | ASSRTGGIAEQF | 0.11108365 |
| GAGGTGSTAGGAGGAGGAGGLY | ASSRTGGIAEQF | 0.10207844 |
| GHGGTGAAGGNGGTGGAGGSGI | ASSRTGGIAEQF | 0.13253386 |
| GNGGNGGAGGEGGPGGAGGAGA | ASSRTGGIAEQF | 0.10587775 |
| GSTAGNGGNGGNATNGTIAGQP | ASSRTGGIAEQF | 0.12562214 |
| GTGGDGGNGGNGGTGG | ASSRTGGIAEQF | 0.14436086 |
| GVRRPAGIPADAPV | ASSRTGGIAEQF | 0.10549296 |
| GYRQVTDAHLLTLARR | ASSRTGGIAEQF | 0.118218854 |
| IAKMGQDGHDRG | ASSRTGGIAEQF | 0.15608674 |
| KAEPSIEIQYT | ASSRTGGIAEQF | 0.14838657 |
| KPVRIRMAADEDTVAVT | ASSRTGGIAEQF | 0.14793326 |
| MPPRIAGMRLLVIKPEP | ASSRTGGIAEQF | 0.1862188 |
| MKFYAVALFRSRGVPPTR | ASSRTGGIAEQF | 0.14887452 |

| | | |
|---|---|---|
| NAGGFGILGAVAHSP | ASSRTGGIAEQF | 0.1951843 |
| NEGREITLAMAEM | ASSRTGGIAEQF | 0.15669532 |
| NELVEWSHPLGHRWRWE | ASSRTGGIAEQF | 0.12387705 |
| NLGAGGTGGNGGNPG | ASSRTGGIAEQF | 0.11232134 |
| NTEWPKWETVMAADPVD | ASSRTGGIAEQF | 0.12724616 |
| PASAAPAETQP | ASSRTGGIAEQF | 0.14204009 |
| QQSSDGKNGVRAAEITGE | ASSRTGGIAEQF | 0.15059447 |
| RAPDAVTTSWWKE | ASSRTGGIAEQF | 0.11020156 |
| SGFGNVGAHTSGWFNQ | ASSRTGGIAEQF | 0.19591017 |
| SGVAGAVGQAASVGGLKV | ASSRTGGIAEQF | 0.10941938 |
| SNGGAGTGSAGGNGGT | ASSRTGGIAEQF | 0.15344836 |
| TGGAGGEGGAAG | ASSRTGGIAEQF | 0.10656151 |
| TGGDGGAAGAGGAPGHG | ASSRTGGIAEQF | 0.110672385 |
| TMPPLLAAIANAGRHGV | ASSRTGGIAEQF | 0.14630422 |
| TSSTAGNSGGA | ASSRTGGIAEQF | 0.0956437 |
| VPATVPVGILC | ASSRTGGIAEQF | 0.097354665 |
| VQRPDSGGVVLTGRIS | ASSRTGGIAEQF | 0.14285359 |
| WTPNPVPATKGVPAGLVQAT | ASSRTGGIAEQF | 0.11511034 |
| YALKRALAKFSDRFDVVII | ASSRTGGIAEQF | 0.09490501 |
| YLIGSVIV | ASSRTGGIAEQF | 0.11497309 |
| YGQLVAVID | ASSRTGGIAEQF | 0.10768264 |
| VTVGGMTLPA | ASSRTGGIAEQF | 0.108966194 |
| VLALPAV | ASSRTGGIAEQF | 0.1345244 |
| VIPPTLNYET | ASSRTGGIAEQF | 0.10218754 |
| VGMGGMHPSSG | ASSRTGGIAEQF | 0.14255847 |
| VGLAGGAATA | ASSRTGGIAEQF | 0.10182853 |
| VGGAGGAGGSGP | ASSRTGGIAEQF | 0.09989877 |
| VFAAFMV | ASSRTGGIAEQF | 0.14002705 |
| TLYDKLR | ASSRTGGIAEQF | 0.212799 |
| TLAVFTVL | ASSRTGGIAEQF | 0.11714082 |
| TGSVQPARV | ASSRTGGIAEQF | 0.1477271 |
| TGGGAGSGGT | ASSRTGGIAEQF | 0.10480104 |
| SHLETSL | ASSRTGGIAEQF | 0.1540023 |
| SFTTGMT | ASSRTGGIAEQF | 0.1336046 |
| QVEVAKRMGVRQPTVS | ASSRTGGIAEQF | 0.10242595 |
| QGSGGPALAM | ASSRTGGIAEQF | 0.15443568 |
| MLAPSNP | ASSRTGGIAEQF | 0.12081432 |
| LVNNAAI | ASSRTGGIAEQF | 0.14973705 |
| LTGEINTLA | ASSRTGGIAEQF | 0.17407048 |
| LSIFGPV | ASSRTGGIAEQF | 0.102463916 |
| LLALLANL | ASSRTGGIAEQF | 0.13945292 |
| LISNASDALDKLR | ASSRTGGIAEQF | 0.17415236 |
| LINEAVP | ASSRTGGIAEQF | 0.14684172 |
| LGYAFSGL | ASSRTGGIAEQF | 0.107077524 |
| LDFVAVAL | ASSRTGGIAEQF | 0.15176368 |
| LAQPGAE | ASSRTGGIAEQF | 0.12903541 |
| ISNLGAVTGF | ASSRTGGIAEQF | 0.14109628 |
| HQEDPGRI | ASSRTGGIAEQF | 0.129373 |
| GTPGEPTP | ASSRTGGIAEQF | 0.13544714 |
| GFFNSGS | ASSRTGGIAEQF | 0.14922091 |
| GAGGGGLTAGAN | ASSRTGGIAEQF | 0.1397694 |

| | | |
|---|---|---|
| FVDALVNM | ASSRTGGIAEQF | 0.13173267 |
| FLTVAPEL | ASSRTGGIAEQF | 0.10574027 |
| FETLTAL | ASSRTGGIAEQF | 0.113767155 |
| DVINPIEL | ASSRTGGIAEQF | 0.12644868 |
| APVSGAPV | ASSRTGGIAEQF | 0.1684331 |
| ALVGGLAW | ASSRTGGIAEQF | 0.106435746 |
| ALAAAAATAI | ASSRTGGIAEQF | 0.10153909 |
| AFGLGLLL | ASSRTGGIAEQF | 0.13299927 |
| AFFAAASL | ASSRTGGIAEQF | 0.17808376 |
| AAAALPAFSPP | ASSRTGGIAEQF | 0.11895018 |
| VVLAPAL | ASSRTGGIAEQF | 0.12389297 |
| TIGHVDHGKTTL | ASSRTGGIAEQF | 0.2752958 |
| LISNASDALDK | ASSRTGGIAEQF | 0.13461305 |
| LISNASDAL | ASSRTGGIAEQF | 0.13483147 |
| ISNASDALDK | ASSRTGGIAEQF | 0.14305711 |
| NQAETLVYQTE | ASSRTGGIAEQF | 0.17650934 |
| TGGTGGTAGAG | ASSRTGGIAEQF | 0.12516977 |
| TGSTAGGAGGA | ASSRTGGIAEQF | 0.17632999 |
| DPLAVPVAAGPV | ASSRTGGIAEQF | 0.1046827 |
| GGTGGTPGNGGN | ASSRTGGIAEQF | 0.18441409 |
| PAGLAGVDPATL | ASSRTGGIAEQF | 0.1006862 |
| GNNLVVEAPHAGASV | ASSRTGGIAEQF | 0.14091189 |
| KVTRIMSASTLEQPA | ASSRTGGIAEQF | 0.102444366 |
| LGFALPIALPRRTVV | ASSRTGGIAEQF | 0.17485577 |
| TRLGNLTARKLVVLV | ASSRTGGIAEQF | 0.14970545 |
| WSQQRRLIYFPSAGP | ASSRTGGIAEQF | 0.17726108 |
| GPNGNPDPMAAAVDIR | ASSRTGGIAEQF | 0.09739308 |
| TQRCVSVADLRAAASR | ASSRTGGIAEQF | 0.15779741 |
| QFVLDVAEHPVEVLTETV | ASSRTGGIAEQF | 0.17788264 |
| GGAGGTGGAGGNGGTQAGDG | ASSRTGGIAEQF | 0.47554308 |
| PLLATLRAIAVDVDAKPAQV | ASSRTGGIAEQF | 0.10932374 |
| APKFRTGGYMIDFWGVGYQVAK | ASSRTGGIAEQF | 0.15614393 |
| GAGGAGGAGGAAGGPGGTGGTG | ASSRTGGIAEQF | 0.12991993 |
| IGALGLLATDVAPAGSVAFGVA | ASSRTGGIAEQF | 0.115934044 |
| AGGAGGIGGNGGAGGAGGVLMG | ASSRTGGIAEQF | 0.10559351 |
| ADELIGLL | ASSRTGGIAEQF | 0.12567241 |
| ADRALKA | ASSRTGGIAEQF | 0.12583552 |
| AGAVGQVGGAGGA | ASSRTGGIAEQF | 0.121281594 |
| AGGGGGLGGAP | ASSRTGGIAEQF | 0.1555243 |
| AGGLLYGNGGAG | ASSRTGGIAEQF | 0.16299658 |
| AGLESIGL | ASSRTGGIAEQF | 0.14577766 |
| AGLKTGII | ASSRTGGIAEQF | 0.20014372 |
| ASLAALLL | ASSRTGGIAEQF | 0.14161156 |
| ASVINISSI | ASSRTGGIAEQF | 0.15203267 |
| ATPPGPA | ASSRTGGIAEQF | 0.11231859 |
| ATVASALV | ASSRTGGIAEQF | 0.1223987 |
| ATVASLIL | ASSRTGGIAEQF | 0.10853964 |
| AVNGPVLIRH | ASSRTGGIAEQF | 0.11914462 |
| DGAAGAVGI | ASSRTGGIAEQF | 0.11304121 |
| DPLLHGI | ASSRTGGIAEQF | 0.1533606 |
| ELGAALGAL | ASSRTGGIAEQF | 0.09433121 |

| | | |
|---|---|---|
| ELGRTQAV | ASSRTGGIAEQF | 0.15219483 |
| FVAGGYLS | ASSRTGGIAEQF | 0.19866559 |
| GASGGAGGAGGW | ASSRTGGIAEQF | 0.100579806 |
| GGAGAGGDGGAGGI | ASSRTGGIAEQF | 0.12927236 |
| GTGHNGSQPGMGGNG | ASSRTGGIAEQF | 0.111940786 |
| GYAAASASAS | ASSRTGGIAEQF | 0.10275218 |
| HQVDVAAVQQG | ASSRTGGIAEQF | 0.17789552 |
| IGAAIAHT | ASSRTGGIAEQF | 0.11194069 |
| IGMIEVAQL | ASSRTGGIAEQF | 0.14402589 |
| IRLAPEHTSAMD | ASSRTGGIAEQF | 0.13700429 |
| IYNVSVL | ASSRTGGIAEQF | 0.1447062 |
| KQGFLGM | ASSRTGGIAEQF | 0.12445311 |
| LESVITQL | ASSRTGGIAEQF | 0.122750394 |
| LGLGALVVV | ASSRTGGIAEQF | 0.106500596 |
| LIGNGGNGGSGGI | ASSRTGGIAEQF | 0.14666347 |
| LNAINAPTL | ASSRTGGIAEQF | 0.09646729 |
| LQLLNTGV | ASSRTGGIAEQF | 0.13594529 |
| LQTSIEF | ASSRTGGIAEQF | 0.10986586 |
| MSFAVSEL | ASSRTGGIAEQF | 0.22190934 |
| NTGALGL | ASSRTGGIAEQF | 0.13787256 |
| PPGAGGAGGAGTT | ASSRTGGIAEQF | 0.12846579 |
| QLVGLGI | ASSRTGGIAEQF | 0.12498066 |
| RFLEQVHQL | ASSRTGGIAEQF | 0.11188952 |
| RGVGYVLR | ASSRTGGIAEQF | 0.10411073 |
| SATGVFAGVI | ASSRTGGIAEQF | 0.107537284 |
| SLPAITLP | ASSRTGGIAEQF | 0.11161555 |
| STALAFTV | ASSRTGGIAEQF | 0.123567075 |
| SVLTVGLGK | ASSRTGGIAEQF | 0.10220862 |
| TALAGAGSGI | ASSRTGGIAEQF | 0.17216913 |
| TGLPFTGI | ASSRTGGIAEQF | 0.14467026 |
| VAEHVTV | ASSRTGGIAEQF | 0.10101077 |
| VDIVLNSL | ASSRTGGIAEQF | 0.11421202 |
| VLTGGGAVL | ASSRTGGIAEQF | 0.15222363 |
| VNVGLAYGI | ASSRTGGIAEQF | 0.12838648 |
| VSSAAVGAV | ASSRTGGIAEQF | 0.0946032 |
| VTSVYEAGV | ASSRTGGIAEQF | 0.11016125 |
| WVALDRGA | ASSRTGGIAEQF | 0.14046551 |
| YLVGSVVI | ASSRTGGIAEQF | 0.103447974 |
| GPPGTGKTLL | ASSRTGGIAEQF | 0.11021337 |
| GVGMTKF | ASSRTGGIAEQF | 0.268112 |
| IGIGGSDL | ASSRTGGIAEQF | 0.09895098 |
| IGTLLAEL | ASSRTGGIAEQF | 0.12082508 |
| IYTSGTT | ASSRTGGIAEQF | 0.1620256 |
| LSFPAGV | ASSRTGGIAEQF | 0.17200905 |
| LYTSGTT | ASSRTGGIAEQF | 0.18050829 |
| VGAFPFGF | ASSRTGGIAEQF | 0.12662232 |
| VGAGTVEF | ASSRTGGIAEQF | 0.11367525 |
| VGKSTLLL | ASSRTGGIAEQF | 0.14173509 |
| AALAHAGGGIV | ASSRTGGIAEQF | 0.123455845 |
| AALVAVSLL | ASSRTGGIAEQF | 0.10126821 |
| AAVSRAVAAL | ASSRTGGIAEQF | 0.09723841 |

| | | |
|---|---|---|
| AGGSSGIGGF | ASSRTGGIAEQF | 0.12052807 |
| AGLTVANANG | ASSRTGGIAEQF | 0.0996314 |
| AGLVGAVL | ASSRTGGIAEQF | 0.14220387 |
| ALSQAFTV | ASSRTGGIAEQF | 0.16463919 |
| ALTFSVSL | ASSRTGGIAEQF | 0.10278818 |
| ALTGAALVAL | ASSRTGGIAEQF | 0.111106336 |
| ASFSIGAGVL | ASSRTGGIAEQF | 0.12027369 |
| DGSVYKI | ASSRTGGIAEQF | 0.118264675 |
| EQLAEIV | ASSRTGGIAEQF | 0.17193967 |
| FAVSELTVM | ASSRTGGIAEQF | 0.11008887 |
| GAAPLAAGL | ASSRTGGIAEQF | 0.57805526 |
| GADGGDGGSGGAGG | ASSRTGGIAEQF | 0.16472565 |
| GFGLDTGGAGG | ASSRTGGIAEQF | 0.11880556 |
| GLASGGLL | ASSRTGGIAEQF | 0.16851702 |
| GRIAGAAVLL | ASSRTGGIAEQF | 0.114173 |
| GSAGSLNTSF | ASSRTGGIAEQF | 0.10506391 |
| GTGQIGI | ASSRTGGIAEQF | 0.12437045 |
| GTLIALIL | ASSRTGGIAEQF | 0.10703453 |
| IAATMGFVVG | ASSRTGGIAEQF | 0.120356895 |
| IFAQALL | ASSRTGGIAEQF | 0.12549092 |
| IGAAVALGF | ASSRTGGIAEQF | 0.10130937 |
| IGLTPLAL | ASSRTGGIAEQF | 0.1293426 |
| ILGSNTSTL | ASSRTGGIAEQF | 0.11404182 |
| IVFSLTVL | ASSRTGGIAEQF | 0.097857535 |
| LASKATGF | ASSRTGGIAEQF | 0.10799157 |
| LASVALIL | ASSRTGGIAEQF | 0.17145205 |
| LFAVGLTL | ASSRTGGIAEQF | 0.22706348 |
| LFNGGIGTYI | ASSRTGGIAEQF | 0.11453799 |
| LTLGAGII | ASSRTGGIAEQF | 0.11259171 |
| LVAAALGDI | ASSRTGGIAEQF | 0.15589827 |
| NTAFVQLGI | ASSRTGGIAEQF | 0.10102497 |
| QLFSGAEL | ASSRTGGIAEQF | 0.12979949 |
| SAATFITL | ASSRTGGIAEQF | 0.116611876 |
| SNYGLANLGV | ASSRTGGIAEQF | 0.109972335 |
| SYGGAYAVM | ASSRTGGIAEQF | 0.11227108 |
| TTGVGGLTLG | ASSRTGGIAEQF | 0.1280209 |
| TTQTIYI | ASSRTGGIAEQF | 0.10520218 |
| VAAGAGLGAVI | ASSRTGGIAEQF | 0.1463468 |
| VAAVVTLL | ASSRTGGIAEQF | 0.0948795 |
| VGFLGGLI | ASSRTGGIAEQF | 0.109430805 |
| VGVIPILL | ASSRTGGIAEQF | 0.11514338 |
| VIFASSLI | ASSRTGGIAEQF | 0.10729786 |
| VKGGDGGAAAG | ASSRTGGIAEQF | 0.09996873 |
| VTKTFTNI | ASSRTGGIAEQF | 0.12676087 |
| VVAAAILL | ASSRTGGIAEQF | 0.14486848 |
| YAALDVEL | ASSRTGGIAEQF | 0.17001006 |
| LASVALLL | ASSRTGGIAEQF | 0.1025528 |
| LDAVHRV | ASSRTGGIAEQF | 0.16192259 |
| QGLSSSIF | ASSRTGGIAEQF | 0.12674391 |
| RGTFANIRL | ASSRTGGIAEQF | 0.1179598 |
| RQLWWGHRI | ASSRTGGIAEQF | 0.10656239 |

| | | |
|---|---|---|
| YGPPGTGKTLL | ASSRTGGIAEQF | 0.1532065 |
| AALVGAPANVA | ASSRTGGIAEQF | 0.09980439 |
| AATFRRTE | ASSRTGGIAEQF | 0.11706776 |
| AGALGGGI | ASSRTGGIAEQF | 0.13118601 |
| AGGAGGAGGIGAG | ASSRTGGIAEQF | 0.106080264 |
| AGSGALGL | ASSRTGGIAEQF | 0.15977208 |
| AQIADQI | ASSRTGGIAEQF | 0.17106378 |
| ASAVAVTYL | ASSRTGGIAEQF | 0.10724183 |
| ASTVLGGTGG | ASSRTGGIAEQF | 0.10845421 |
| AVGGEPAK | ASSRTGGIAEQF | 0.11089938 |
| CEQGTFGGTC | ASSRTGGIAEQF | 0.09568608 |
| DEGKAPAG | ASSRTGGIAEQF | 0.15787579 |
| DTVVFPAL | ASSRTGGIAEQF | 0.11361553 |
| FEGITMAG | ASSRTGGIAEQF | 0.11685932 |
| FPLVVQD | ASSRTGGIAEQF | 0.13540597 |
| GAFTTPQ | ASSRTGGIAEQF | 0.11463791 |
| GFGGFGTAGGVGGA | ASSRTGGIAEQF | 0.13174762 |
| GFLAQFL | ASSRTGGIAEQF | 0.12707937 |
| GGAGGAGGNAGVA | ASSRTGGIAEQF | 0.10610666 |
| GGVGTAGAGGAGGAGG | ASSRTGGIAEQF | 0.1303914 |
| GNGGGGDAGNAGSGGN | ASSRTGGIAEQF | 0.101239085 |
| GTGGAGGAGNDAGS | ASSRTGGIAEQF | 0.11278327 |
| GTTNTGWFNS | ASSRTGGIAEQF | 0.2453334 |
| IFMGPLFTL | ASSRTGGIAEQF | 0.10107977 |
| ILAHAIEAI | ASSRTGGIAEQF | 0.11009891 |
| IRSAIREAL | ASSRTGGIAEQF | 0.11835061 |
| ITFAGGG | ASSRTGGIAEQF | 0.101297416 |
| ITGMSGHQEV | ASSRTGGIAEQF | 0.14894444 |
| IVAGLACTL | ASSRTGGIAEQF | 0.12850001 |
| IVGGGLAGA | ASSRTGGIAEQF | 0.14313604 |
| IVTGVLNI | ASSRTGGIAEQF | 0.111847706 |
| LAQQGCI | ASSRTGGIAEQF | 0.15024179 |
| LCLHVEN | ASSRTGGIAEQF | 0.18256669 |
| LDAISDGS | ASSRTGGIAEQF | 0.11687894 |
| LGGFAAML | ASSRTGGIAEQF | 0.11274933 |
| LGGKNPMI | ASSRTGGIAEQF | 0.10676676 |
| LGPKGVAAIL | ASSRTGGIAEQF | 0.14044073 |
| LMVGVSI | ASSRTGGIAEQF | 0.16828308 |
| LVGAAFVL | ASSRTGGIAEQF | 0.10981383 |
| LVLAVAPL | ASSRTGGIAEQF | 0.13643946 |
| MAAIAVIDAA | ASSRTGGIAEQF | 0.13729277 |
| NNGINAELQ | ASSRTGGIAEQF | 0.19808199 |
| RGEPLVLVHG | ASSRTGGIAEQF | 0.10081863 |
| RGGAGGAAGLI | ASSRTGGIAEQF | 0.12397235 |
| RSYRMHPAV | ASSRTGGIAEQF | 0.10170593 |
| SAAHLLAGQLA | ASSRTGGIAEQF | 0.10790979 |
| SAAVGSY | ASSRTGGIAEQF | 0.118460186 |
| SPHAVLHRI | ASSRTGGIAEQF | 0.13202173 |
| SQIAKVL | ASSRTGGIAEQF | 0.1444601 |
| TALGHLVP | ASSRTGGIAEQF | 0.16507588 |
| TAPTTSGAA | ASSRTGGIAEQF | 0.155896 |

| | | |
|---|---|---|
| TDGDDGNGGSPGT | ASSRTGGIAEQF | 0.18058169 |
| TDGLGEN | ASSRTGGIAEQF | 0.12388744 |
| TGGPPLPGQA | ASSRTGGIAEQF | 0.12972353 |
| TIPAITI | ASSRTGGIAEQF | 0.105945155 |
| TIPGFTI | ASSRTGGIAEQF | 0.099704176 |
| TSIASGALA | ASSRTGGIAEQF | 0.10909199 |
| TVTLPGAAI | ASSRTGGIAEQF | 0.111260146 |
| VASGLSAALP | ASSRTGGIAEQF | 0.09598168 |
| VGAGVIGI | ASSRTGGIAEQF | 0.118147776 |
| VGPKRFGV | ASSRTGGIAEQF | 0.13922505 |
| VNVAFPD | ASSRTGGIAEQF | 0.16394825 |
| VSMTNSGVS | ASSRTGGIAEQF | 0.12015684 |
| VTDVLGDLSV | ASSRTGGIAEQF | 0.12281828 |
| VVVVGAGFAGL | ASSRTGGIAEQF | 0.112465486 |
| VYTVGTVV | ASSRTGGIAEQF | 0.20043668 |
| WTTFSGP | ASSRTGGIAEQF | 0.11052724 |
| YDELVEAF | ASSRTGGIAEQF | 0.100803815 |
| YGTGGAGGNGG | ASSRTGGIAEQF | 0.11365429 |
| YTNVMAV | ASSRTGGIAEQF | 0.10478962 |
| FLEQVHQL | ASSRTGGIAEQF | 0.12411385 |
| GLSLLAVAV | ASSRTGGIAEQF | 0.16057163 |
| LEQVHQL | ASSRTGGIAEQF | 0.15091679 |
| LNAAGAI | ASSRTGGIAEQF | 0.15509403 |
| PAVHASL | ASSRTGGIAEQF | 0.09998604 |
| TELEDDA | ASSRTGGIAEQF | 0.120084755 |
| TPPGPPN | ASSRTGGIAEQF | 0.24293949 |
| TPPPPPPP | ASSRTGGIAEQF | 0.13081999 |
| TQSPALI | ASGEWTNSYNSPLY | 0.14559782 |
| EFAAAFPQ | ASGEWTNSYNSPLY | 0.17108332 |
| LLSGNLDVL | ASGEWTNSYNSPLY | 0.11564908 |
| GPANLGNYNI | ASGEWTNSYNSPLY | 0.12681113 |
| TLGPTLP | ASGEWTNSYNSPLY | 0.13006431 |
| VYAYSSCCK | ASGEWTNSYNSPLY | 0.14009628 |
| APAPAPAPNI | ASGEWTNSYNSPLY | 0.1067342 |
| PGGPVPIP | ASGEWTNSYNSPLY | 0.109614775 |
| FLTVALEP | ASGEWTNSYNSPLY | 0.15462467 |
| SQYFGANL | ASGEWTNSYNSPLY | 0.15686034 |
| GGVKPAA | ASGEWTNSYNSPLY | 0.17673679 |
| GGTGGAGGAGGW | ASGEWTNSYNSPLY | 0.13413589 |
| GVGGVAGLGGAGG | ASGEWTNSYNSPLY | 0.14047748 |
| VLLVLATTI | ASGEWTNSYNSPLY | 0.11946716 |
| IGIFAAML | ASGEWTNSYNSPLY | 0.1231233 |
| AVGDLID | ASGEWTNSYNSPLY | 0.10953281 |
| NISPLPL | ASGEWTNSYNSPLY | 0.11037809 |
| VTAVFVAI | ASGEWTNSYNSPLY | 0.10981748 |
| SGDDNDAAL | ASGEWTNSYNSPLY | 0.1448635 |
| AVAALAHS | ASGEWTNSYNSPLY | 0.11642905 |
| GGVTVMATP | ASGEWTNSYNSPLY | 0.20701449 |
| AGLAANVRD | ASGEWTNSYNSPLY | 0.13797623 |
| TNRPDLIDPALLRPG | ASGEWTNSYNSPLY | 0.14027514 |
| LANLGVTAL | ASGEWTNSYNSPLY | 0.109993055 |

| | | |
|---|---|---|
| TGERIEL | ASGEWTNSYNSPLY | 0.11766492 |
| NGEELAA | ASGEWTNSYNSPLY | 0.1207275 |
| VLSLVWG | ASGEWTNSYNSPLY | 0.103565976 |
| IGYALWL | ASGEWTNSYNSPLY | 0.18678793 |
| LSLLLAPL | ASGEWTNSYNSPLY | 0.11786114 |
| TLDEALK | ASGEWTNSYNSPLY | 0.0996584 |
| TDGAGTAEGG | ASGEWTNSYNSPLY | 0.12793271 |
| NAPGGAGGNGG | ASGEWTNSYNSPLY | 0.528448 |
| VSVFMGMGGNF | ASGEWTNSYNSPLY | 0.11791798 |
| LASYLAAL | ASGEWTNSYNSPLY | 0.10745819 |
| LVGGTPAL | ASGEWTNSYNSPLY | 0.11645845 |
| VRNADVL | ASGEWTNSYNSPLY | 0.13142113 |
| TCSSPFLVSAM | ASGEWTNSYNSPLY | 0.30104065 |
| VHITDAE | ASGEWTNSYNSPLY | 0.10169555 |
| EYAAPSF | ASGEWTNSYNSPLY | 0.3243847 |
| VVDAFHGGL | ASGEWTNSYNSPLY | 0.11008762 |
| ASVVDPQ | ASGEWTNSYNSPLY | 0.108502425 |
| SYNEFGGLS | ASGEWTNSYNSPLY | 0.14293338 |
| DPDGSVY | ASGEWTNSYNSPLY | 0.123140536 |
| MVSTIHATE | ASGEWTNSYNSPLY | 0.13657472 |
| LAAGVADPV | ASGEWTNSYNSPLY | 0.1306213 |
| TKEQIAA | ASGEWTNSYNSPLY | 0.10901635 |
| SLASPAAL | ASGEWTNSYNSPLY | 0.11299102 |
| IGLVFYGV | ASGEWTNSYNSPLY | 0.17166078 |
| FLLDASNL | ASGEWTNSYNSPLY | 0.12732366 |
| YQVGNFP | ASGEWTNSYNSPLY | 0.16503546 |
| ESLLSQST | ASGEWTNSYNSPLY | 0.11447024 |
| LFTVGFAAVG | ASGEWTNSYNSPLY | 0.10409438 |
| ATPPTPT | ASGEWTNSYNSPLY | 0.124944106 |
| TDLATLG | ASGEWTNSYNSPLY | 0.12004078 |
| SAGGTGR | ASGEWTNSYNSPLY | 0.122095905 |
| GFSNSGS | ASGEWTNSYNSPLY | 0.12591267 |
| GGNGGSGGTSV | ASGEWTNSYNSPLY | 0.13749975 |
| GTSVATGGAGN | ASGEWTNSYNSPLY | 0.113935485 |
| NGGNGGSGGTS | ASGEWTNSYNSPLY | 0.11404452 |
| AVVKFPH | ASGEWTNSYNSPLY | 0.14246021 |
| TPPPPPP | ASGEWTNSYNSPLY | 0.12087531 |
| AAPAAAAAPA | ASGEWTNSYNSPLY | 0.11423868 |
| IALNEDL | ASGEWTNSYNSPLY | 0.17061384 |
| IVNVSSM | ASGEWTNSYNSPLY | 0.20660044 |
| TGIPPAPR | ASGEWTNSYNSPLY | 0.16627792 |
| FDLGGGTF | ASGEWTNSYNSPLY | 0.10519732 |
| FDLGGGTFDVS | ASGEWTNSYNSPLY | 0.096970946 |
| NEPTAAA | ASGEWTNSYNSPLY | 0.12463237 |
| NEPTAAALA | ASGEWTNSYNSPLY | 0.13210471 |
| GSIINISSI | ASGEWTNSYNSPLY | 0.12046577 |
| VAPGNAGTA | ASGEWTNSYNSPLY | 0.11612207 |
| TGGGASLE | ASGEWTNSYNSPLY | 0.11032315 |
| VAVYNHYKRI | ASGEWTNSYNSPLY | 0.15635021 |
| TDPAPATT | ASGEWTNSYNSPLY | 0.13535236 |
| TNRPDILDPALLRPG | ASGEWTNSYNSPLY | 0.10245674 |

| | | |
|---|---|---|
| LTTTTTS | ASGEWTNSYNSPLY | 0.10765166 |
| PTGSGKTL | ASGEWTNSYNSPLY | 0.19629915 |
| TGSGKTL | ASGEWTNSYNSPLY | 0.11373642 |
| TIGHVDHGK | ASGEWTNSYNSPLY | 0.14069718 |
| TIGHVDHGKTT | ASGEWTNSYNSPLY | 0.12531483 |
| GHVDHGKTT | ASGEWTNSYNSPLY | 0.09742975 |
| AQFEHTLL | ASGEWTNSYNSPLY | 0.116407715 |
| IVNQDEL | ASGEWTNSYNSPLY | 0.10849269 |
| TNGSQFF | ASGEWTNSYNSPLY | 0.11960701 |
| TGKGIVE | ASGEWTNSYNSPLY | 0.112856805 |
| LISNASDA | ASGEWTNSYNSPLY | 0.1313502 |
| AELRTLGELTKIAKAHGAQ | ASGEWTNSYNSPLY | 0.17140524 |
| APNAVAAAEPSW | ASGEWTNSYNSPLY | 0.1518443 |
| APQVAGAATAVL | ASGEWTNSYNSPLY | 0.13010904 |
| ARRDPAAAAAILPTDA | ASGEWTNSYNSPLY | 0.114765815 |
| DAAARDVLAEAGLAEYF | ASGEWTNSYNSPLY | 0.1092562 |
| DDQIVLSAKANLAKVEQ | ASGEWTNSYNSPLY | 0.09677956 |
| DELLPDTGEDSDEDGH | ASGEWTNSYNSPLY | 0.17365664 |
| ELPEAPANRATVPQD | ASGEWTNSYNSPLY | 0.09727662 |
| FGQSKIERVTGIG | ASGEWTNSYNSPLY | 0.17716353 |
| GAGGTGSTAGGAGGAGGAGGLY | ASGEWTNSYNSPLY | 0.14877157 |
| GHGGTGAAGGNGGTGGAGGSGI | ASGEWTNSYNSPLY | 0.115860246 |
| GNGGNGGAGGEGGPGGAGGAGA | ASGEWTNSYNSPLY | 0.12620436 |
| GSTAGNGGNGGNATNGTIAGQP | ASGEWTNSYNSPLY | 0.102927774 |
| GTGGDGGNGGNGGTGG | ASGEWTNSYNSPLY | 0.10492739 |
| GVRRPAGIPADAPV | ASGEWTNSYNSPLY | 0.11246992 |
| GYRQVTDAHLLTLARR | ASGEWTNSYNSPLY | 0.13370669 |
| IAKMGQDGHDRG | ASGEWTNSYNSPLY | 0.122275956 |
| KAEPSIEIQYT | ASGEWTNSYNSPLY | 0.13828011 |
| KPVRIRMAADEDTVAVT | ASGEWTNSYNSPLY | 0.14413269 |
| MPPRIAGMRLLVIKPEP | ASGEWTNSYNSPLY | 0.11397786 |
| MKFYAVALFRSRGVPPTR | ASGEWTNSYNSPLY | 0.12227144 |
| NAGGFGILGAVAHSP | ASGEWTNSYNSPLY | 0.15783815 |
| NEGREITLAMAEM | ASGEWTNSYNSPLY | 0.12924954 |
| NELVEWSHPLGHRWRWE | ASGEWTNSYNSPLY | 0.11820777 |
| NLGAGGTGGNGGNPG | ASGEWTNSYNSPLY | 0.13069965 |
| NTEWPKWETVMAADPVD | ASGEWTNSYNSPLY | 0.11032574 |
| PASAAPAETQP | ASGEWTNSYNSPLY | 0.14551258 |
| QQSSDGKNGVRAAEITGE | ASGEWTNSYNSPLY | 0.1231371 |
| RAPDAVTTSWWKE | ASGEWTNSYNSPLY | 0.12769951 |
| SGFGNVGAHTSGWFNQ | ASGEWTNSYNSPLY | 0.14122854 |
| SGVAGAVGQAASVGGLKV | ASGEWTNSYNSPLY | 0.09796381 |
| SNGGAGTGSAGGNGGT | ASGEWTNSYNSPLY | 0.11118939 |
| TGGAGGEGGAAG | ASGEWTNSYNSPLY | 0.11405377 |
| TGGDGGAAGAGGAPGHG | ASGEWTNSYNSPLY | 0.16222781 |
| TMPPLLAAIANAGRHGV | ASGEWTNSYNSPLY | 0.16620895 |
| TSSTAGNSGGA | ASGEWTNSYNSPLY | 0.10836129 |
| VPATVPVGILC | ASGEWTNSYNSPLY | 0.13520111 |
| VQRPDSGGVVLTGRIS | ASGEWTNSYNSPLY | 0.1237174 |
| WTPNPVPATKGVPAGLVQAT | ASGEWTNSYNSPLY | 0.11894183 |
| YALKRALAKFSDRFDVVII | ASGEWTNSYNSPLY | 0.13009822 |

| | | |
|---|---|---|
| YLIGSVIV | ASGEWTNSYNSPLY | 0.11541613 |
| YGQLVAVID | ASGEWTNSYNSPLY | 0.117173284 |
| VTVGGMTLPA | ASGEWTNSYNSPLY | 0.13863337 |
| VLALPAV | ASGEWTNSYNSPLY | 0.112053074 |
| VIPPTLNYET | ASGEWTNSYNSPLY | 0.13596429 |
| VGMGGMHPSSG | ASGEWTNSYNSPLY | 0.10777032 |
| VGLAGGAATA | ASGEWTNSYNSPLY | 0.121071264 |
| VGGAGGAGGSGP | ASGEWTNSYNSPLY | 0.12793697 |
| VFAAFMV | ASGEWTNSYNSPLY | 0.113448426 |
| TLYDKLR | ASGEWTNSYNSPLY | 0.15339741 |
| TLAVFTVL | ASGEWTNSYNSPLY | 0.123285994 |
| TGSVQPARV | ASGEWTNSYNSPLY | 0.11062222 |
| TGGGAGSGGT | ASGEWTNSYNSPLY | 0.12895936 |
| SHLETSL | ASGEWTNSYNSPLY | 0.17634705 |
| SFTTGMT | ASGEWTNSYNSPLY | 0.14057824 |
| QVEVAKRMGVRQPTVS | ASGEWTNSYNSPLY | 0.11270963 |
| QGSGGPALAM | ASGEWTNSYNSPLY | 0.14105263 |
| MLAPSNP | ASGEWTNSYNSPLY | 0.16353142 |
| LVNNAAI | ASGEWTNSYNSPLY | 0.11938455 |
| LTGEINTLA | ASGEWTNSYNSPLY | 0.10173561 |
| LSIFGPV | ASGEWTNSYNSPLY | 0.119952664 |
| LLALLANL | ASGEWTNSYNSPLY | 0.11210387 |
| LISNASDALDKLR | ASGEWTNSYNSPLY | 0.12380598 |
| LINEAVP | ASGEWTNSYNSPLY | 0.1000382 |
| LGYAFSGL | ASGEWTNSYNSPLY | 0.17209236 |
| LDFVAVAL | ASGEWTNSYNSPLY | 0.18886353 |
| LAQPGAE | ASGEWTNSYNSPLY | 0.11064619 |
| ISNLGAVTGF | ASGEWTNSYNSPLY | 0.13183998 |
| HQEDPGRI | ASGEWTNSYNSPLY | 0.10095662 |
| GTPGEPTP | ASGEWTNSYNSPLY | 0.13465938 |
| GFFNSGS | ASGEWTNSYNSPLY | 0.15962951 |
| GAGGGGLTAGAN | ASGEWTNSYNSPLY | 0.105970725 |
| FVDALVNM | ASGEWTNSYNSPLY | 0.13748471 |
| FLTVAPEL | ASGEWTNSYNSPLY | 0.109005496 |
| FETLTAL | ASGEWTNSYNSPLY | 0.11908497 |
| DVINPIEL | ASGEWTNSYNSPLY | 0.18933217 |
| APVSGAPV | ASGEWTNSYNSPLY | 0.12436935 |
| ALVGGLAW | ASGEWTNSYNSPLY | 0.12765802 |
| ALAAAAATAI | ASGEWTNSYNSPLY | 0.1281834 |
| AFGLGLLL | ASGEWTNSYNSPLY | 0.11322078 |
| AFFAAASL | ASGEWTNSYNSPLY | 0.13342847 |
| AAAALPAFSPP | ASGEWTNSYNSPLY | 0.15898973 |
| VVLAPAL | ASGEWTNSYNSPLY | 0.101371065 |
| TIGHVDHGKTTL | ASGEWTNSYNSPLY | 0.13217713 |
| LISNASDALDK | ASGEWTNSYNSPLY | 0.11400558 |
| LISNASDAL | ASGEWTNSYNSPLY | 0.14726673 |
| ISNASDALDK | ASGEWTNSYNSPLY | 0.19226013 |
| NQAETLVYQTE | ASGEWTNSYNSPLY | 0.1463157 |
| TGGTGGTAGAG | ASGEWTNSYNSPLY | 0.11533132 |
| TGSTAGGAGGA | ASGEWTNSYNSPLY | 0.20962977 |
| DPLAVPVAAGPV | ASGEWTNSYNSPLY | 0.16449519 |

| | | |
|---|---|---|
| GGTGGTPGNGGN | ASGEWTNSYNSPLY | 0.13358577 |
| PAGLAGVDPATL | ASGEWTNSYNSPLY | 0.16018683 |
| GNNLVVEAPHAGASV | ASGEWTNSYNSPLY | 0.29207495 |
| KVTRIMSASTLEQPA | ASGEWTNSYNSPLY | 0.30111194 |
| LGFALPIALPRRTVV | ASGEWTNSYNSPLY | 0.29114988 |
| TRLGNLTARKLVVLV | ASGEWTNSYNSPLY | 0.29163593 |
| WSQQRRLIYFPSAGP | ASGEWTNSYNSPLY | 0.29454738 |
| GPNGNPDPMAAAVDIR | ASGEWTNSYNSPLY | 0.28804392 |
| TQRCVSVADLRAAASR | ASGEWTNSYNSPLY | 0.2934349 |
| QFVLDVAEHPVEVLTETV | ASGEWTNSYNSPLY | 0.29430407 |
| GGAGGTGGAGGNGGTQAGDG | ASGEWTNSYNSPLY | 0.287743 |
| PLLATLRAIAVDVDAKPAQV | ASGEWTNSYNSPLY | 0.295035 |
| APKFRTGGYMIDFWGVGYQVAK | ASGEWTNSYNSPLY | 0.29368907 |
| GAGGAGGAGGAAGGPGGTGGTG | ASGEWTNSYNSPLY | 0.30496973 |
| IGALGLLATDVAPAGSVAFGVA | ASGEWTNSYNSPLY | 0.30089238 |
| AGGAGGIGGNGGAGGAGGVLMG | ASGEWTNSYNSPLY | 0.29004854 |
| ADELIGLL | ASGEWTNSYNSPLY | 0.29222077 |
| ADRALKA | ASGEWTNSYNSPLY | 0.2901904 |
| AGAVGQVGGAGGA | ASGEWTNSYNSPLY | 0.2915991 |
| AGGGGGLGGAP | ASGEWTNSYNSPLY | 0.28778628 |
| AGGLLYGNGGAG | ASGEWTNSYNSPLY | 0.2980077 |
| AGLESIGL | ASGEWTNSYNSPLY | 0.2875432 |
| AGLKTGII | ASGEWTNSYNSPLY | 0.28721946 |
| ASLAALLL | ASGEWTNSYNSPLY | 0.2968542 |
| ASVINISSI | ASGEWTNSYNSPLY | 0.29244006 |
| ATPPGPA | ASGEWTNSYNSPLY | 0.29256913 |
| ATVASALV | ASGEWTNSYNSPLY | 0.2999003 |
| ATVASLIL | ASGEWTNSYNSPLY | 0.2872272 |
| AVNGPVLIRH | ASGEWTNSYNSPLY | 0.29164666 |
| DGAAGAVGI | ASGEWTNSYNSPLY | 0.3019025 |
| DPLLHGI | ASGEWTNSYNSPLY | 0.3115878 |
| ELGAALGAL | ASGEWTNSYNSPLY | 0.2899724 |
| ELGRTQAV | ASGEWTNSYNSPLY | 0.29254234 |
| FVAGGYLS | ASGEWTNSYNSPLY | 0.2959664 |
| GASGGAGGAGGW | ASGEWTNSYNSPLY | 0.2971374 |
| GGAGAGGDGGAGGI | ASGEWTNSYNSPLY | 0.29581437 |
| GTGHNGSQPGMGGNG | ASGEWTNSYNSPLY | 0.29967535 |
| GYAAASASAS | ASGEWTNSYNSPLY | 0.2880555 |
| HQVDVAAVQQG | ASGEWTNSYNSPLY | 0.3010316 |
| IGAAIAHT | ASGEWTNSYNSPLY | 0.2945163 |
| IGMIEVAQL | ASGEWTNSYNSPLY | 0.29604918 |
| IRLAPEHTSAMD | ASGEWTNSYNSPLY | 0.29740393 |
| IYNVSVL | ASGEWTNSYNSPLY | 0.29035512 |
| KQGFLGM | ASGEWTNSYNSPLY | 0.28929397 |
| LESVITQL | ASGEWTNSYNSPLY | 0.28740656 |
| LGLGALVVV | ASGEWTNSYNSPLY | 0.29558244 |
| LIGNGGNGGSGGI | ASGEWTNSYNSPLY | 0.2990909 |
| LNAINAPTL | ASGEWTNSYNSPLY | 0.29311603 |
| LQLLNTGV | ASGEWTNSYNSPLY | 0.28913286 |
| LQTSIEF | ASGEWTNSYNSPLY | 0.29051632 |
| MSFAVSEL | ASGEWTNSYNSPLY | 0.28743163 |

| | | |
|---|---|---|
| NTGALGL | ASGEWTNSYNSPLY | 0.29085758 |
| PPGAGGAGGAGTT | ASGEWTNSYNSPLY | 0.2878668 |
| QLVGLGI | ASGEWTNSYNSPLY | 0.2885346 |
| RFLEQVHQL | ASGEWTNSYNSPLY | 0.3073263 |
| RGVGYVLR | ASGEWTNSYNSPLY | 0.3049152 |
| SATGVFAGVI | ASGEWTNSYNSPLY | 0.2935012 |
| SLPAITLP | ASGEWTNSYNSPLY | 0.29714602 |
| STALAFTV | ASGEWTNSYNSPLY | 0.2882641 |
| SVLTVGLGK | ASGEWTNSYNSPLY | 0.29523128 |
| TALAGAGSGI | ASGEWTNSYNSPLY | 0.29570362 |
| TGLPFTGI | ASGEWTNSYNSPLY | 0.29169518 |
| VAEHVTV | ASGEWTNSYNSPLY | 0.2893598 |
| VDIVLNSL | ASGEWTNSYNSPLY | 0.29111624 |
| VLTGGGAVL | ASGEWTNSYNSPLY | 0.29643765 |
| VNVGLAYGI | ASGEWTNSYNSPLY | 0.29879755 |
| VSSAAVGAV | ASGEWTNSYNSPLY | 0.29151526 |
| VTSVYEAGV | ASGEWTNSYNSPLY | 0.2930373 |
| WVALDRGA | ASGEWTNSYNSPLY | 0.2868033 |
| YLVGSVVI | ASGEWTNSYNSPLY | 0.29502708 |
| GPPGTGKTLL | ASGEWTNSYNSPLY | 0.28854722 |
| GVGMTKF | ASGEWTNSYNSPLY | 0.28941414 |
| IGIGGSDL | ASGEWTNSYNSPLY | 0.28930888 |
| IGTLLAEL | ASGEWTNSYNSPLY | 0.2971668 |
| IYTSGTT | ASGEWTNSYNSPLY | 0.30790544 |
| LSFPAGV | ASGEWTNSYNSPLY | 0.2982058 |
| LYTSGTT | ASGEWTNSYNSPLY | 0.29032752 |
| VGAFPFGF | ASGEWTNSYNSPLY | 0.29733592 |
| VGAGTVEF | ASGEWTNSYNSPLY | 0.29496643 |
| VGKSTLLL | ASGEWTNSYNSPLY | 0.29320464 |
| AALAHAGGGIV | ASGEWTNSYNSPLY | 0.30079293 |
| AALVAVSLL | ASGEWTNSYNSPLY | 0.2909886 |
| AAVSRAVAAL | ASGEWTNSYNSPLY | 0.29800323 |
| AGGSSGIGGF | ASGEWTNSYNSPLY | 0.29441983 |
| AGLTVANANG | ASGEWTNSYNSPLY | 0.297165 |
| AGLVGAVL | ASGEWTNSYNSPLY | 0.29558387 |
| ALSQAFTV | ASGEWTNSYNSPLY | 0.29340252 |
| ALTFSVSL | ASGEWTNSYNSPLY | 0.29047242 |
| ALTGAALVAL | ASGEWTNSYNSPLY | 0.2907109 |
| ASFSIGAGVL | ASGEWTNSYNSPLY | 0.2893742 |
| DGSVYKI | ASGEWTNSYNSPLY | 0.2944213 |
| EQLAEIV | ASGEWTNSYNSPLY | 0.2932461 |
| FAVSELTVM | ASGEWTNSYNSPLY | 0.29277045 |
| GAAPLAAGL | ASGEWTNSYNSPLY | 0.29884803 |
| GADGGDGGSGGAGG | ASGEWTNSYNSPLY | 0.2898459 |
| GFGLDTGGAGG | ASGEWTNSYNSPLY | 0.2956219 |
| GLASGGLL | ASGEWTNSYNSPLY | 0.29402575 |
| GRIAGAAVLL | ASGEWTNSYNSPLY | 0.29010248 |
| GSAGSLNTSF | ASGEWTNSYNSPLY | 0.2919088 |
| GTGQIGI | ASGEWTNSYNSPLY | 0.303151 |
| GTLIALIL | ASGEWTNSYNSPLY | 0.28892103 |
| IAATMGFVVG | ASGEWTNSYNSPLY | 0.2945416 |

| | | |
|---|---|---|
| IFAQALL | ASGEWTNSYNSPLY | 0.28847852 |
| IGAAVALGF | ASGEWTNSYNSPLY | 0.29244614 |
| IGLTPLAL | ASGEWTNSYNSPLY | 0.2903858 |
| ILGSNTSTL | ASGEWTNSYNSPLY | 0.29905573 |
| IVFSLTVL | ASGEWTNSYNSPLY | 0.29080442 |
| LASKATGF | ASGEWTNSYNSPLY | 0.2894889 |
| LASVALIL | ASGEWTNSYNSPLY | 0.28969258 |
| LFAVGLTL | ASGEWTNSYNSPLY | 0.2887059 |
| LFNGGIGTYI | ASGEWTNSYNSPLY | 0.30337638 |
| LTLGAGII | ASGEWTNSYNSPLY | 0.28947905 |
| LVAAALGDI | ASGEWTNSYNSPLY | 0.2914757 |
| NTAFVQLGI | ASGEWTNSYNSPLY | 0.29224283 |
| QLFSGAEL | ASGEWTNSYNSPLY | 0.2996792 |
| SAATFITL | ASGEWTNSYNSPLY | 0.2919425 |
| SNYGLANLGV | ASGEWTNSYNSPLY | 0.29258487 |
| SYGGAYAVM | ASGEWTNSYNSPLY | 0.29128987 |
| TTGVGGLTLG | ASGEWTNSYNSPLY | 0.29967088 |
| TTQTIYI | ASGEWTNSYNSPLY | 0.29849038 |
| VAAGAGLGAVI | ASGEWTNSYNSPLY | 0.2944615 |
| VAAVVTLL | ASGEWTNSYNSPLY | 0.2900484 |
| VGFLGGLI | ASGEWTNSYNSPLY | 0.2908329 |
| VGVIPILL | ASGEWTNSYNSPLY | 0.30000305 |
| VIFASSLI | ASGEWTNSYNSPLY | 0.28733093 |
| VKGGDGGAAAG | ASGEWTNSYNSPLY | 0.28746307 |
| VTKTFTNI | ASGEWTNSYNSPLY | 0.28736317 |
| VVAAAILL | ASGEWTNSYNSPLY | 0.28754637 |
| YAALDVEL | ASGEWTNSYNSPLY | 0.30937156 |
| LASVALLL | ASGEWTNSYNSPLY | 0.31180894 |
| LDAVHRV | ASGEWTNSYNSPLY | 0.2932226 |
| QGLSSSIF | ASGEWTNSYNSPLY | 0.2934939 |
| RGTFANIRL | ASGEWTNSYNSPLY | 0.29840252 |
| RQLWWGHRI | ASGEWTNSYNSPLY | 0.28685832 |
| YGPPGTGKTLL | ASGEWTNSYNSPLY | 0.29036665 |
| AALVGAPANVA | ASGEWTNSYNSPLY | 0.29005662 |
| AATFRRTE | ASGEWTNSYNSPLY | 0.3039016 |
| AGALGGGI | ASGEWTNSYNSPLY | 0.29026496 |
| AGGAGGAGGIGAG | ASGEWTNSYNSPLY | 0.28976375 |
| AGSGALGL | ASGEWTNSYNSPLY | 0.2968123 |
| AQIADQI | ASGEWTNSYNSPLY | 0.28807673 |
| ASAVAVTYL | ASGEWTNSYNSPLY | 0.28922287 |
| ASTVLGGTGG | ASGEWTNSYNSPLY | 0.28848162 |
| AVGGEPAK | ASGEWTNSYNSPLY | 0.28677413 |
| CEQGTFGGTC | ASGEWTNSYNSPLY | 0.30235708 |
| DEGKAPAG | ASGEWTNSYNSPLY | 0.28927413 |
| DTVVFPAL | ASGEWTNSYNSPLY | 0.2919644 |
| FEGITMAG | ASGEWTNSYNSPLY | 0.2877034 |
| FPLVVQD | ASGEWTNSYNSPLY | 0.28767914 |
| GAFTTPQ | ASGEWTNSYNSPLY | 0.2891895 |
| GFGGFGTAGGVGGA | ASGEWTNSYNSPLY | 0.29840735 |
| GFLAQFL | ASGEWTNSYNSPLY | 0.2888466 |
| GGAGGAGGNAGVA | ASGEWTNSYNSPLY | 0.29148188 |

| | | |
|---|---|---|
| GGVGTAGAGGAGGAGG | ASGEWTNSYNSPLY | 0.30145463 |
| GNGGGGDAGNAGSGGN | ASGEWTNSYNSPLY | 0.2931047 |
| GTGGAGGAGNDAGS | ASGEWTNSYNSPLY | 0.29095292 |
| GTTNTGWFNS | ASGEWTNSYNSPLY | 0.2944346 |
| IFMGPLFTL | ASGEWTNSYNSPLY | 0.29036638 |
| ILAHAIEAI | ASGEWTNSYNSPLY | 0.29069906 |
| IRSAIREAL | ASGEWTNSYNSPLY | 0.29093194 |
| ITFAGGG | ASGEWTNSYNSPLY | 0.31982258 |
| ITGMSGHQEV | ASGEWTNSYNSPLY | 0.28851718 |
| IVAGLACTL | ASGEWTNSYNSPLY | 0.29110885 |
| IVGGGLAGA | ASGEWTNSYNSPLY | 0.28918132 |
| IVTGVLNI | ASGEWTNSYNSPLY | 0.29904428 |
| LAQQGCI | ASGEWTNSYNSPLY | 0.29869688 |
| LCLHVEN | ASGEWTNSYNSPLY | 0.29151705 |
| LDAISDGS | ASGEWTNSYNSPLY | 0.29365176 |
| LGGFAAML | ASGEWTNSYNSPLY | 0.2901876 |
| LGGKNPMI | ASGEWTNSYNSPLY | 0.29730764 |
| LGPKGVAAIL | ASGEWTNSYNSPLY | 0.29083684 |
| LMVGVSI | ASGEWTNSYNSPLY | 0.29102716 |
| LVGAAFVL | ASGEWTNSYNSPLY | 0.29423296 |
| LVLAVAPL | ASGEWTNSYNSPLY | 0.29102862 |
| MAAIAVIDAA | ASGEWTNSYNSPLY | 0.2911368 |
| NNGINAELQ | ASGEWTNSYNSPLY | 0.28878823 |
| RGEPLVLVHG | ASGEWTNSYNSPLY | 0.29617143 |
| RGGAGGAAGLI | ASGEWTNSYNSPLY | 0.2893899 |
| RSYRMHPAV | ASGEWTNSYNSPLY | 0.2951868 |
| SAAHLLAGQLA | ASGEWTNSYNSPLY | 0.29693696 |
| SAAVGSY | ASGEWTNSYNSPLY | 0.28921628 |
| SPHAVLHRI | ASGEWTNSYNSPLY | 0.28974417 |
| SQIAKVL | ASGEWTNSYNSPLY | 0.30438372 |
| TALGHLVP | ASGEWTNSYNSPLY | 0.28894842 |
| TAPTTSGAA | ASGEWTNSYNSPLY | 0.29080114 |
| TDGDDGNGGSPGT | ASGEWTNSYNSPLY | 0.3050919 |
| TDGLGEN | ASGEWTNSYNSPLY | 0.2947051 |
| TGGPPLPGQA | ASGEWTNSYNSPLY | 0.3038759 |
| TIPAITI | ASGEWTNSYNSPLY | 0.2887566 |
| TIPGFTI | ASGEWTNSYNSPLY | 0.29380593 |
| TSIASGALA | ASGEWTNSYNSPLY | 0.2908933 |
| TVTLPGAAI | ASGEWTNSYNSPLY | 0.2898846 |
| VASGLSAALP | ASGEWTNSYNSPLY | 0.30257004 |
| VGAGVIGI | ASGEWTNSYNSPLY | 0.2944662 |
| VGPKRFGV | ASGEWTNSYNSPLY | 0.28826594 |
| VNVAFPD | ASGEWTNSYNSPLY | 0.31576252 |
| VSMTNSGVS | ASGEWTNSYNSPLY | 0.29287633 |
| VTDVLGDLSV | ASGEWTNSYNSPLY | 0.2880091 |
| VVVVGAGFAGL | ASGEWTNSYNSPLY | 0.28978392 |
| VYTVGTVV | ASGEWTNSYNSPLY | 0.29060292 |
| WTTFSGP | ASGEWTNSYNSPLY | 0.29403663 |
| YDELVEAF | ASGEWTNSYNSPLY | 0.288754 |
| YGTGGAGGNGG | ASGEWTNSYNSPLY | 0.2872558 |
| YTNVMAV | ASGEWTNSYNSPLY | 0.292468 |

| | | |
|---|---|---|
| FLEQVHQL | ASGEWTNSYNSPLY | 0.29490107 |
| GLSLLAVAV | ASGEWTNSYNSPLY | 0.28692278 |
| LEQVHQL | ASGEWTNSYNSPLY | 0.29120407 |
| LNAAGAI | ASGEWTNSYNSPLY | 0.2875813 |
| PAVHASL | ASGEWTNSYNSPLY | 0.2902962 |
| TELEDDA | ASGEWTNSYNSPLY | 0.29537874 |
| TPPGPPN | ASGEWTNSYNSPLY | 0.2885957 |
| TPPPPPPP | ASGEWTNSYNSPLY | 0.2946828 |
| TQSPALI | ASSPGQQNSGNTLY | 0.2882457 |
| EFAAAFPQ | ASSPGQQNSGNTLY | 0.28771928 |
| LLSGNLDVL | ASSPGQQNSGNTLY | 0.29803815 |
| GPANLGNYNI | ASSPGQQNSGNTLY | 0.28996623 |
| TLGPTLP | ASSPGQQNSGNTLY | 0.28954977 |
| VYAYSSCCK | ASSPGQQNSGNTLY | 0.29901254 |
| APAPAPAPNI | ASSPGQQNSGNTLY | 0.28865483 |
| PGGPVPIP | ASSPGQQNSGNTLY | 0.2966762 |
| FLTVALEP | ASSPGQQNSGNTLY | 0.29489985 |
| SQYFGANL | ASSPGQQNSGNTLY | 0.2882484 |
| GGVKPAA | ASSPGQQNSGNTLY | 0.29712155 |
| GGTGGAGGAGGW | ASSPGQQNSGNTLY | 0.28932804 |
| GVGGVAGLGGAGG | ASSPGQQNSGNTLY | 0.29366255 |
| VLLVLATTI | ASSPGQQNSGNTLY | 0.28931463 |
| IGIFAAML | ASSPGQQNSGNTLY | 0.28827187 |
| AVGDLID | ASSPGQQNSGNTLY | 0.29289106 |
| NISPLPL | ASSPGQQNSGNTLY | 0.30823135 |
| VTAVFVAI | ASSPGQQNSGNTLY | 0.30421594 |
| SGDDNDAAL | ASSPGQQNSGNTLY | 0.28946376 |
| AVAALAHS | ASSPGQQNSGNTLY | 0.29441386 |
| GGVTVMATP | ASSPGQQNSGNTLY | 0.29617965 |
| AGLAANVRD | ASSPGQQNSGNTLY | 0.29096848 |
| TNRPDLIDPALLRPG | ASSPGQQNSGNTLY | 0.29627308 |
| LANLGVTAL | ASSPGQQNSGNTLY | 0.29400918 |
| TGERIEL | ASSPGQQNSGNTLY | 0.29478055 |
| NGEELAA | ASSPGQQNSGNTLY | 0.29800287 |
| VLSLVWG | ASSPGQQNSGNTLY | 0.2885428 |
| IGYALWL | ASSPGQQNSGNTLY | 0.28925523 |
| LSLLLAPL | ASSPGQQNSGNTLY | 0.29162812 |
| TLDEALK | ASSPGQQNSGNTLY | 0.29269862 |
| TDGAGTAEGG | ASSPGQQNSGNTLY | 0.29811698 |
| NAPGGAGGNGG | ASSPGQQNSGNTLY | 0.28841132 |
| VSVFMGMGGNF | ASSPGQQNSGNTLY | 0.28820822 |
| LASYLAAL | ASSPGQQNSGNTLY | 0.29054514 |
| LVGGTPAL | ASSPGQQNSGNTLY | 0.30142236 |
| VRNADVL | ASSPGQQNSGNTLY | 0.2928846 |
| TCSSPFLVSAM | ASSPGQQNSGNTLY | 0.293399 |
| VHITDAE | ASSPGQQNSGNTLY | 0.31383008 |
| EYAAPSF | ASSPGQQNSGNTLY | 0.29084092 |
| VVDAFHGGL | ASSPGQQNSGNTLY | 0.295342 |
| ASVVDPQ | ASSPGQQNSGNTLY | 0.29867166 |
| SYNEFGGLS | ASSPGQQNSGNTLY | 0.2974934 |
| DPDGSVY | ASSPGQQNSGNTLY | 0.29175106 |

| | | |
|---|---|---|
| MVSTIHATE | ASSPGQQNSGNTLY | 0.30652648 |
| LAAGVADPV | ASSPGQQNSGNTLY | 0.2884406 |
| TKEQIAA | ASSPGQQNSGNTLY | 0.30028954 |
| SLASPAAL | ASSPGQQNSGNTLY | 0.28759792 |
| IGLVFYGV | ASSPGQQNSGNTLY | 0.29656067 |
| FLLDASNL | ASSPGQQNSGNTLY | 0.2881795 |
| YQVGNFP | ASSPGQQNSGNTLY | 0.2961254 |
| ESLLSQST | ASSPGQQNSGNTLY | 0.30028364 |
| LFTVGFAAVG | ASSPGQQNSGNTLY | 0.29565537 |
| ATPPTPT | ASSPGQQNSGNTLY | 0.2883 |
| TDLATLG | ASSPGQQNSGNTLY | 0.3052459 |
| SAGGTGR | ASSPGQQNSGNTLY | 0.2944919 |
| GFSNSGS | ASSPGQQNSGNTLY | 0.29783 |
| GGNGGSGGTSV | ASSPGQQNSGNTLY | 0.2911804 |
| GTSVATGGAGN | ASSPGQQNSGNTLY | 0.29406086 |
| NGGNGGSGGTS | ASSPGQQNSGNTLY | 0.29557854 |
| AVVKFPH | ASSPGQQNSGNTLY | 0.29061857 |
| TPPPPPP | ASSPGQQNSGNTLY | 0.28949493 |
| AAPAAAAAPA | ASSPGQQNSGNTLY | 0.29021376 |
| IALNEDL | ASSPGQQNSGNTLY | 0.29256758 |
| IVNVSSM | ASSPGQQNSGNTLY | 0.29137972 |
| TGIPPAPR | ASSPGQQNSGNTLY | 0.2930372 |
| FDLGGGTF | ASSPGQQNSGNTLY | 0.30111286 |
| FDLGGGTFDVS | ASSPGQQNSGNTLY | 0.29575422 |
| NEPTAAA | ASSPGQQNSGNTLY | 0.31182817 |
| NEPTAAALA | ASSPGQQNSGNTLY | 0.29436237 |
| GSIINISSI | ASSPGQQNSGNTLY | 0.294984 |
| VAPGNAGTA | ASSPGQQNSGNTLY | 0.29135847 |
| TGGGASLE | ASSPGQQNSGNTLY | 0.29152203 |
| VAVYNHYKRI | ASSPGQQNSGNTLY | 0.29063985 |
| TDPAPATT | ASSPGQQNSGNTLY | 0.29098237 |
| TNRPDILDPALLRPG | ASSPGQQNSGNTLY | 0.29297456 |
| LTTTTTS | ASSPGQQNSGNTLY | 0.2957842 |
| PTGSGKTL | ASSPGQQNSGNTLY | 0.28637826 |
| TGSGKTL | ASSPGQQNSGNTLY | 0.2878056 |
| TIGHVDHGK | ASSPGQQNSGNTLY | 0.30797327 |
| TIGHVDHGKTT | ASSPGQQNSGNTLY | 0.28808635 |
| GHVDHGKTT | ASSPGQQNSGNTLY | 0.29238173 |
| AQFEHTLL | ASSPGQQNSGNTLY | 0.28872135 |
| IVNQDEL | ASSPGQQNSGNTLY | 0.28856537 |
| TNGSQFF | ASSPGQQNSGNTLY | 0.30950114 |
| TGKGIVE | ASSPGQQNSGNTLY | 0.28923288 |
| LISNASDA | ASSPGQQNSGNTLY | 0.2925823 |
| AELRTLGELTKIAKAHGAQ | ASSPGQQNSGNTLY | 0.2954068 |
| APNAVAAAEPSW | ASSPGQQNSGNTLY | 0.29160413 |
| APQVAGAATAVL | ASSPGQQNSGNTLY | 0.29200804 |
| ARRDPAAAAAILPTDA | ASSPGQQNSGNTLY | 0.28986165 |
| DAAARDVLAEAGLAEYF | ASSPGQQNSGNTLY | 0.2890018 |
| DDQIVLSAKANLAKVEQ | ASSPGQQNSGNTLY | 0.294833 |
| DELLPDTGEDSDEDGH | ASSPGQQNSGNTLY | 0.28692415 |
| ELPEAPANRATVPQD | ASSPGQQNSGNTLY | 0.29060155 |

| | | |
|---|---|---|
| FGQSKIERVTGIG | ASSPGQQNSGNTLY | 0.2899816 |
| GAGGTGSTAGGAGGAGGAGGLY | ASSPGQQNSGNTLY | 0.30668476 |
| GHGGTGAAGGNGGTGGAGGSGI | ASSPGQQNSGNTLY | 0.2978543 |
| GNGGNGGAGGEGGPGGAGGAGA | ASSPGQQNSGNTLY | 0.29445058 |
| GSTAGNGGNGGNATNGTIAGQP | ASSPGQQNSGNTLY | 0.29093173 |
| GTGGDGGNGGNGGTGG | ASSPGQQNSGNTLY | 0.2896694 |
| GVRRPAGIPADAPV | ASSPGQQNSGNTLY | 0.2896912 |
| GYRQVTDAHLLTLARR | ASSPGQQNSGNTLY | 0.290347 |
| IAKMGQDGHDRG | ASSPGQQNSGNTLY | 0.29268634 |
| KAEPSIEIQYT | ASSPGQQNSGNTLY | 0.29280978 |
| KPVRIRMAADEDTVAVT | ASSPGQQNSGNTLY | 0.28834185 |
| MPPRIAGMRLLVIKPEP | ASSPGQQNSGNTLY | 0.3020243 |
| MKFYAVALFRSRGVPPTR | ASSPGQQNSGNTLY | 0.29290223 |
| NAGGFGILGAVAHSP | ASSPGQQNSGNTLY | 0.28982127 |
| NEGREITLAMAEM | ASSPGQQNSGNTLY | 0.290702 |
| NELVEWSHPLGHRWRWE | ASSPGQQNSGNTLY | 0.29346585 |
| NLGAGGTGGNGGNPG | ASSPGQQNSGNTLY | 0.29335448 |
| NTEWPKWETVMAADPVD | ASSPGQQNSGNTLY | 0.28646854 |
| PASAAPAETQP | ASSPGQQNSGNTLY | 0.2908009 |
| QQSSDGKNGVRAAEITGE | ASSPGQQNSGNTLY | 0.29111934 |
| RAPDAVTTSWWKE | ASSPGQQNSGNTLY | 0.2886337 |
| SGFGNVGAHTSGWFNQ | ASSPGQQNSGNTLY | 0.29139116 |
| SGVAGAVGQAASVGGLKV | ASSPGQQNSGNTLY | 0.3044522 |
| SNGGAGTGSAGGNGGT | ASSPGQQNSGNTLY | 0.28752822 |
| TGGAGGEGGAAG | ASSPGQQNSGNTLY | 0.28943756 |
| TGGDGGAAGAGGAPGHG | ASSPGQQNSGNTLY | 0.29725853 |
| TMPPLLAAIANAGRHGV | ASSPGQQNSGNTLY | 0.3093132 |
| TSSTAGNSGGA | ASSPGQQNSGNTLY | 0.3043688 |
| VPATVPVGILC | ASSPGQQNSGNTLY | 0.2975588 |
| VQRPDSGGVVLTGRIS | ASSPGQQNSGNTLY | 0.29086196 |
| WTPNPVPATKGVPAGLVQAT | ASSPGQQNSGNTLY | 0.29595393 |
| YALKRALAKFSDRFDVVII | ASSPGQQNSGNTLY | 0.29770583 |
| YLIGSVIV | ASSPGQQNSGNTLY | 0.2876514 |
| YGQLVAVID | ASSPGQQNSGNTLY | 0.28680122 |
| VTVGGMTLPA | ASSPGQQNSGNTLY | 0.29104686 |
| VLALPAV | ASSPGQQNSGNTLY | 0.28805786 |
| VIPPTLNYET | ASSPGQQNSGNTLY | 0.297195 |
| VGMGGMHPSSG | ASSPGQQNSGNTLY | 0.29909596 |
| VGLAGGAATA | ASSPGQQNSGNTLY | 0.28879863 |
| VGGAGGAGGSGP | ASSPGQQNSGNTLY | 0.2890855 |
| VFAAFMV | ASSPGQQNSGNTLY | 0.29176247 |
| TLYDKLR | ASSPGQQNSGNTLY | 0.29338163 |
| TLAVFTVL | ASSPGQQNSGNTLY | 0.295921 |
| TGSVQPARV | ASSPGQQNSGNTLY | 0.28949153 |
| TGGGAGSGGT | ASSPGQQNSGNTLY | 0.29446512 |
| SHLETSL | ASSPGQQNSGNTLY | 0.31289086 |
| SFTTGMT | ASSPGQQNSGNTLY | 0.29145426 |
| QVEVAKRMGVRQPTVS | ASSPGQQNSGNTLY | 0.29645464 |
| QGSGGPALAM | ASSPGQQNSGNTLY | 0.28913695 |
| MLAPSNP | ASSPGQQNSGNTLY | 0.28841597 |
| LVNNAAI | ASSPGQQNSGNTLY | 0.2913241 |

| | | |
|---|---|---|
| LTGEINTLA | ASSPGQQNSGNTLY | 0.29084757 |
| LSIFGPV | ASSPGQQNSGNTLY | 0.29361477 |
| LLALLANL | ASSPGQQNSGNTLY | 0.28962374 |
| LISNASDALDKLR | ASSPGQQNSGNTLY | 0.28862765 |
| LINEAVP | ASSPGQQNSGNTLY | 0.28921115 |
| LGYAFSGL | ASSPGQQNSGNTLY | 0.28745195 |
| LDFVAVAL | ASSPGQQNSGNTLY | 0.28733292 |
| LAQPGAE | ASSPGQQNSGNTLY | 0.28887975 |
| ISNLGAVTGF | ASSPGQQNSGNTLY | 0.30691752 |
| HQEDPGRI | ASSPGQQNSGNTLY | 0.31740662 |
| GTPGEPTP | ASSPGQQNSGNTLY | 0.2899611 |
| GFFNSGS | ASSPGQQNSGNTLY | 0.29139206 |
| GAGGGGLTAGAN | ASSPGQQNSGNTLY | 0.28830245 |
| FVDALVNM | ASSPGQQNSGNTLY | 0.29145575 |
| FLTVAPEL | ASSPGQQNSGNTLY | 0.29172322 |
| FETLTAL | ASSPGQQNSGNTLY | 0.29244158 |
| DVINPIEL | ASSPGQQNSGNTLY | 0.2920118 |
| APVSGAPV | ASSPGQQNSGNTLY | 0.28866422 |
| ALVGGLAW | ASSPGQQNSGNTLY | 0.28997442 |
| ALAAAAATAI | ASSPGQQNSGNTLY | 0.28982654 |
| AFGLGLLL | ASSPGQQNSGNTLY | 0.29857752 |
| AFFAAASL | ASSPGQQNSGNTLY | 0.2867817 |
| AAAALPAFSPP | ASSPGQQNSGNTLY | 0.29284737 |
| VVLAPAL | ASSPGQQNSGNTLY | 0.29006767 |
| TIGHVDHGKTTL | ASSPGQQNSGNTLY | 0.29069325 |
| LISNASDALDK | ASSPGQQNSGNTLY | 0.28910646 |
| LISNASDAL | ASSPGQQNSGNTLY | 0.29265597 |
| ISNASDALDK | ASSPGQQNSGNTLY | 0.29536098 |
| NQAETLVYQTE | ASSPGQQNSGNTLY | 0.29793164 |
| TGGTGGTAGAG | ASSPGQQNSGNTLY | 0.28750044 |
| TGSTAGGAGGA | ASSPGQQNSGNTLY | 0.30154523 |
| DPLAVPVAAGPV | ASSPGQQNSGNTLY | 0.29810268 |
| GGTGGTPGNGGN | ASSPGQQNSGNTLY | 0.28996375 |
| PAGLAGVDPATL | ASSPGQQNSGNTLY | 0.2894958 |
| GNNLVVEAPHAGASV | ASSPGQQNSGNTLY | 0.30018586 |
| KVTRIMSASTLEQPA | ASSPGQQNSGNTLY | 0.2877558 |
| LGFALPIALPRRTVV | ASSPGQQNSGNTLY | 0.2918188 |
| TRLGNLTARKLVVLV | ASSPGQQNSGNTLY | 0.29028612 |
| WSQQRRLIYFPSAGP | ASSPGQQNSGNTLY | 0.2889461 |
| GPNGNPDPMAAAVDIR | ASSPGQQNSGNTLY | 0.29325128 |
| TQRCVSVADLRAAASR | ASSPGQQNSGNTLY | 0.29719812 |
| QFVLDVAEHPVEVLTETV | ASSPGQQNSGNTLY | 0.2895777 |
| GGAGGTGGAGGNGGTQAGDG | ASSPGQQNSGNTLY | 0.29042736 |
| PLLATLRAIAVDVDAKPAQV | ASSPGQQNSGNTLY | 0.2917957 |
| APKFRTGGYMIDFWGVGYQVAK | ASSPGQQNSGNTLY | 0.28741497 |
| GAGGAGGAGGAAGGPGGTGGTG | ASSPGQQNSGNTLY | 0.30158123 |
| IGALGLLATDVAPAGSVAFGVA | ASSPGQQNSGNTLY | 0.28926185 |
| AGGAGGIGGNGGAGGAGGVLMG | ASSPGQQNSGNTLY | 0.29207152 |
| ADELIGLL | ASSPGQQNSGNTLY | 0.29774645 |
| ADRALKA | ASSPGQQNSGNTLY | 0.29289705 |
| AGAVGQVGGAGGA | ASSPGQQNSGNTLY | 0.29328668 |

| | | |
|---|---|---|
| AGGGGGLGGAP | ASSPGQQNSGNTLY | 0.29295912 |
| AGGLLYGNGGAG | ASSPGQQNSGNTLY | 0.29185438 |
| AGLESIGL | ASSPGQQNSGNTLY | 0.29094407 |
| AGLKTGII | ASSPGQQNSGNTLY | 0.2913508 |
| ASLAALLL | ASSPGQQNSGNTLY | 0.28889328 |
| ASVINISSI | ASSPGQQNSGNTLY | 0.31136906 |
| ATPPGPA | ASSPGQQNSGNTLY | 0.28883994 |
| ATVASALV | ASSPGQQNSGNTLY | 0.2919389 |
| ATVASLIL | ASSPGQQNSGNTLY | 0.29278386 |
| AVNGPVLIRH | ASSPGQQNSGNTLY | 0.2889687 |
| DGAAGAVGI | ASSPGQQNSGNTLY | 0.2940023 |
| DPLLHGI | ASSPGQQNSGNTLY | 0.2967919 |
| ELGAALGAL | ASSPGQQNSGNTLY | 0.29316154 |
| ELGRTQAV | ASSPGQQNSGNTLY | 0.28829432 |
| FVAGGYLS | ASSPGQQNSGNTLY | 0.30159205 |
| GASGGAGGAGGW | ASSPGQQNSGNTLY | 0.30369788 |
| GGAGAGGDGGAGGI | ASSPGQQNSGNTLY | 0.29160464 |
| GTGHNGSQPGMGGNG | ASSPGQQNSGNTLY | 0.28805837 |
| GYAAASASAS | ASSPGQQNSGNTLY | 0.29154712 |
| HQVDVAAVQQG | ASSPGQQNSGNTLY | 0.2986015 |
| IGAAIAHT | ASSPGQQNSGNTLY | 0.29602644 |
| IGMIEVAQL | ASSPGQQNSGNTLY | 0.2894017 |
| IRLAPEHTSAMD | ASSPGQQNSGNTLY | 0.29326037 |
| IYNVSVL | ASSPGQQNSGNTLY | 0.29616162 |
| KQGFLGM | ASSPGQQNSGNTLY | 0.2986274 |
| LESVITQL | ASSPGQQNSGNTLY | 0.28807393 |
| LGLGALVVV | ASSPGQQNSGNTLY | 0.29456118 |
| LIGNGGNGGSGGI | ASSPGQQNSGNTLY | 0.29149404 |
| LNAINAPTL | ASSPGQQNSGNTLY | 0.29004154 |
| LQLLNTGV | ASSPGQQNSGNTLY | 0.2914903 |
| LQTSIEF | ASSPGQQNSGNTLY | 0.2923538 |
| MSFAVSEL | ASSPGQQNSGNTLY | 0.29354286 |
| NTGALGL | ASSPGQQNSGNTLY | 0.2881101 |
| PPGAGGAGGAGTT | ASSPGQQNSGNTLY | 0.2997486 |
| QLVGLGI | ASSPGQQNSGNTLY | 0.3081639 |
| RFLEQVHQL | ASSPGQQNSGNTLY | 0.2906543 |
| RGVGYVLR | ASSPGQQNSGNTLY | 0.29771876 |
| SATGVFAGVI | ASSPGQQNSGNTLY | 0.28797022 |
| SLPAITLP | ASSPGQQNSGNTLY | 0.28724506 |
| STALAFTV | ASSPGQQNSGNTLY | 0.28872055 |
| SVLTVGLGK | ASSPGQQNSGNTLY | 0.2932045 |
| TALAGAGSGI | ASSPGQQNSGNTLY | 0.2877508 |
| TGLPFTGI | ASSPGQQNSGNTLY | 0.2940931 |
| VAEHVTV | ASSPGQQNSGNTLY | 0.29377022 |
| VDIVLNSL | ASSPGQQNSGNTLY | 0.3002427 |
| VLTGGGAVL | ASSPGQQNSGNTLY | 0.2922255 |
| VNVGLAYGI | ASSPGQQNSGNTLY | 0.2930166 |
| VSSAAVGAV | ASSPGQQNSGNTLY | 0.29322046 |
| VTSVYEAGV | ASSPGQQNSGNTLY | 0.29980454 |
| WVALDRGA | ASSPGQQNSGNTLY | 0.29067293 |
| YLVGSVVI | ASSPGQQNSGNTLY | 0.2885602 |

| | | |
|---|---|---|
| GPPGTGKTLL | ASSPGQQNSGNTLY | 0.29061115 |
| GVGMTKF | ASSPGQQNSGNTLY | 0.28966653 |
| IGIGGSDL | ASSPGQQNSGNTLY | 0.29092827 |
| IGTLLAEL | ASSPGQQNSGNTLY | 0.2982213 |
| IYTSGTT | ASSPGQQNSGNTLY | 0.29622126 |
| LSFPAGV | ASSPGQQNSGNTLY | 0.28866503 |
| LYTSGTT | ASSPGQQNSGNTLY | 0.28786546 |
| VGAFPFGF | ASSPGQQNSGNTLY | 0.2901451 |
| VGAGTVEF | ASSPGQQNSGNTLY | 0.30294365 |
| VGKSTLLL | ASSPGQQNSGNTLY | 0.2963597 |
| AALAHAGGGIV | ASSPGQQNSGNTLY | 0.2931789 |
| AALVAVSLL | ASSPGQQNSGNTLY | 0.3033132 |
| AAVSRAVAAL | ASSPGQQNSGNTLY | 0.29104793 |
| AGGSSGIGGF | ASSPGQQNSGNTLY | 0.29337135 |
| AGLTVANANG | ASSPGQQNSGNTLY | 0.2944382 |
| AGLVGAVL | ASSPGQQNSGNTLY | 0.2941157 |
| ALSQAFTV | ASSPGQQNSGNTLY | 0.29027495 |
| ALTFSVSL | ASSPGQQNSGNTLY | 0.29042557 |
| ALTGAALVAL | ASSPGQQNSGNTLY | 0.29243356 |
| ASFSIGAGVL | ASSPGQQNSGNTLY | 0.3049744 |
| DGSVYKI | ASSPGQQNSGNTLY | 0.2966466 |
| EQLAEIV | ASSPGQQNSGNTLY | 0.29394183 |
| FAVSELTVM | ASSPGQQNSGNTLY | 0.29445264 |
| GAAPLAAGL | ASSPGQQNSGNTLY | 0.2882119 |
| GADGGDGGSGGAGG | ASSPGQQNSGNTLY | 0.29135504 |
| GFGLDTGGAGG | ASSPGQQNSGNTLY | 0.29227498 |
| GLASGGLL | ASSPGQQNSGNTLY | 0.28993016 |
| GRIAGAAVLL | ASSPGQQNSGNTLY | 0.29022986 |
| GSAGSLNTSF | ASSPGQQNSGNTLY | 0.29193127 |
| GTGQIGI | ASSPGQQNSGNTLY | 0.2913782 |
| GTLIALIL | ASSPGQQNSGNTLY | 0.31012312 |
| IAATMGFVVG | ASSPGQQNSGNTLY | 0.2919991 |
| IFAQALL | ASSPGQQNSGNTLY | 0.29703516 |
| IGAAVALGF | ASSPGQQNSGNTLY | 0.29203603 |
| IGLTPLAL | ASSPGQQNSGNTLY | 0.2908874 |
| ILGSNTSTL | ASSPGQQNSGNTLY | 0.2931752 |
| IVFSLTVL | ASSPGQQNSGNTLY | 0.2894038 |
| LASKATGF | ASSPGQQNSGNTLY | 0.30625635 |
| LASVALIL | ASSPGQQNSGNTLY | 0.29114446 |
| LFAVGLTL | ASSPGQQNSGNTLY | 0.28786254 |
| LFNGGIGTYI | ASSPGQQNSGNTLY | 0.2971226 |
| LTLGAGII | ASSPGQQNSGNTLY | 0.2915657 |
| LVAAALGDI | ASSPGQQNSGNTLY | 0.29087365 |
| NTAFVQLGI | ASSPGQQNSGNTLY | 0.29027292 |
| QLFSGAEL | ASSPGQQNSGNTLY | 0.29408422 |
| SAATFITL | ASSPGQQNSGNTLY | 0.29413962 |
| SNYGLANLGV | ASSPGQQNSGNTLY | 0.31540692 |
| SYGGAYAVM | ASSPGQQNSGNTLY | 0.2870693 |
| TTGVGGLTLG | ASSPGQQNSGNTLY | 0.29159802 |
| TTQTIYI | ASSPGQQNSGNTLY | 0.29013112 |
| VAAGAGLGAVI | ASSPGQQNSGNTLY | 0.2895523 |

| | | |
|---|---|---|
| VAAVVTLL | ASSPGQQNSGNTLY | 0.29092544 |
| VGFLGGLI | ASSPGQQNSGNTLY | 0.29378927 |
| VGVIPILL | ASSPGQQNSGNTLY | 0.29276353 |
| VIFASSLI | ASSPGQQNSGNTLY | 0.30087057 |
| VKGGDGGAAAG | ASSPGQQNSGNTLY | 0.28843966 |
| VTKTFTNI | ASSPGQQNSGNTLY | 0.28860062 |
| VVAAAILL | ASSPGQQNSGNTLY | 0.30322802 |
| YAALDVEL | ASSPGQQNSGNTLY | 0.29581502 |
| LASVALLL | ASSPGQQNSGNTLY | 0.3010848 |
| LDAVHRV | ASSPGQQNSGNTLY | 0.28990847 |
| QGLSSSIF | ASSPGQQNSGNTLY | 0.28982225 |
| RGTFANIRL | ASSPGQQNSGNTLY | 0.28976682 |
| RQLWWGHRI | ASSPGQQNSGNTLY | 0.29501688 |
| YGPPGTGKTLL | ASSPGQQNSGNTLY | 0.28935975 |
| AALVGAPANVA | ASSPGQQNSGNTLY | 0.29142928 |
| AATFRRTE | ASSPGQQNSGNTLY | 0.29758283 |
| AGALGGGI | ASSPGQQNSGNTLY | 0.2889706 |
| AGGAGGAGGIGAG | ASSPGQQNSGNTLY | 0.2922488 |
| AGSGALGL | ASSPGQQNSGNTLY | 0.29834723 |
| AQIADQI | ASSPGQQNSGNTLY | 0.28925657 |
| ASAVAVTYL | ASSPGQQNSGNTLY | 0.28988138 |
| ASTVLGGTGG | ASSPGQQNSGNTLY | 0.299086 |
| AVGGEPAK | ASSPGQQNSGNTLY | 0.32448083 |
| CEQGTFGGTC | ASSPGQQNSGNTLY | 0.29531673 |
| DEGKAPAG | ASSPGQQNSGNTLY | 0.29018864 |
| DTVVFPAL | ASSPGQQNSGNTLY | 0.28743753 |
| FEGITMAG | ASSPGQQNSGNTLY | 0.30555698 |
| FPLVVQD | ASSPGQQNSGNTLY | 0.29006135 |
| GAFTTPQ | ASSPGQQNSGNTLY | 0.29210636 |
| GFGGFGTAGGVGGA | ASSPGQQNSGNTLY | 0.2908211 |
| GFLAQFL | ASSPGQQNSGNTLY | 0.29086095 |
| GGAGGAGGNAGVA | ASSPGQQNSGNTLY | 0.30280635 |
| GGVGTAGAGGAGGAGG | ASSPGQQNSGNTLY | 0.29087046 |
| GNGGGGDAGNAGSGGN | ASSPGQQNSGNTLY | 0.28911617 |
| GTGGAGGAGNDAGS | ASSPGQQNSGNTLY | 0.28922033 |
| GTTNTGWFNS | ASSPGQQNSGNTLY | 0.29665422 |
| IFMGPLFTL | ASSPGQQNSGNTLY | 0.28996488 |
| ILAHAIEAI | ASSPGQQNSGNTLY | 0.29363257 |
| IRSAIREAL | ASSPGQQNSGNTLY | 0.30992746 |
| ITFAGGG | ASSPGQQNSGNTLY | 0.28745967 |
| ITGMSGHQEV | ASSPGQQNSGNTLY | 0.29429212 |
| IVAGLACTL | ASSPGQQNSGNTLY | 0.29208177 |
| IVGGGLAGA | ASSPGQQNSGNTLY | 0.29923424 |
| IVTGVLNI | ASSPGQQNSGNTLY | 0.29201198 |
| LAQQGCI | ASSPGQQNSGNTLY | 0.29407695 |
| LCLHVEN | ASSPGQQNSGNTLY | 0.29914865 |
| LDAISDGS | ASSPGQQNSGNTLY | 0.2968252 |
| LGGFAAML | ASSPGQQNSGNTLY | 0.2922581 |
| LGGKNPMI | ASSPGQQNSGNTLY | 0.28961524 |
| LGPKGVAAIL | ASSPGQQNSGNTLY | 0.28932196 |
| LMVGVSI | ASSPGQQNSGNTLY | 0.28686994 |

| | | |
|---|---|---|
| LVGAAFVL | ASSPGQQNSGNTLY | 0.29951656 |
| LVLAVAPL | ASSPGQQNSGNTLY | 0.28759682 |
| MAAIAVIDAA | ASSPGQQNSGNTLY | 0.30389458 |
| NNGINAELQ | ASSPGQQNSGNTLY | 0.29990417 |
| RGEPLVLVHG | ASSPGQQNSGNTLY | 0.29083562 |
| RGGAGGAAGLI | ASSPGQQNSGNTLY | 0.29352102 |
| RSYRMHPAV | ASSPGQQNSGNTLY | 0.288743 |
| SAAHLLAGQLA | ASSPGQQNSGNTLY | 0.28799453 |
| SAAVGSY | ASSPGQQNSGNTLY | 0.29177803 |
| SPHAVLHRI | ASSPGQQNSGNTLY | 0.29378805 |
| SQIAKVL | ASSPGQQNSGNTLY | 0.2912344 |
| TALGHLVP | ASSPGQQNSGNTLY | 0.2918359 |
| TAPTTSGAA | ASSPGQQNSGNTLY | 0.29120824 |
| TDGDDGNGGSPGT | ASSPGQQNSGNTLY | 0.289613 |
| TDGLGEN | ASSPGQQNSGNTLY | 0.28926638 |
| TGGPPLPGQA | ASSPGQQNSGNTLY | 0.30779245 |
| TIPAITI | ASSPGQQNSGNTLY | 0.29107678 |
| TIPGFTI | ASSPGQQNSGNTLY | 0.29095986 |
| TSIASGALA | ASSPGQQNSGNTLY | 0.29720566 |
| TVTLPGAAI | ASSPGQQNSGNTLY | 0.28832796 |
| VASGLSAALP | ASSPGQQNSGNTLY | 0.29956293 |
| VGAGVIGI | ASSPGQQNSGNTLY | 0.293747 |
| VGPKRFGV | ASSPGQQNSGNTLY | 0.2923728 |
| VNVAFPD | ASSPGQQNSGNTLY | 0.29303947 |
| VSMTNSGVS | ASSPGQQNSGNTLY | 0.28954303 |
| VTDVLGDLSV | ASSPGQQNSGNTLY | 0.28972974 |
| VVVVGAGFAGL | ASSPGQQNSGNTLY | 0.29090327 |
| VYTVGTVV | ASSPGQQNSGNTLY | 0.30484077 |
| WTTFSGP | ASSPGQQNSGNTLY | 0.30217066 |
| YDELVEAF | ASSPGQQNSGNTLY | 0.2908146 |
| YGTGGAGGNGG | ASSPGQQNSGNTLY | 0.29201883 |
| YTNVMAV | ASSPGQQNSGNTLY | 0.29183146 |
| FLEQVHQL | ASSPGQQNSGNTLY | 0.2911312 |
| GLSLLAVAV | ASSPGQQNSGNTLY | 0.2910579 |
| LEQVHQL | ASSPGQQNSGNTLY | 0.29187793 |
| LNAAGAI | ASSPGQQNSGNTLY | 0.29097393 |
| PAVHASL | ASSPGQQNSGNTLY | 0.29491466 |
| TELEDDA | ASSPGQQNSGNTLY | 0.2893884 |
| TPPGPPN | ASSPGQQNSGNTLY | 0.29660177 |
| TPPPPPPP | ASSPGQQNSGNTLY | 0.2945886 |
| TQSPALI | ASGLGQDTQY | 0.29214337 |
| EFAAAFPQ | ASGLGQDTQY | 0.29415515 |
| LLSGNLDVL | ASGLGQDTQY | 0.2928967 |
| GPANLGNYNI | ASGLGQDTQY | 0.29382867 |
| TLGPTLP | ASGLGQDTQY | 0.29356948 |
| VYAYSSCCK | ASGLGQDTQY | 0.2905237 |
| APAPAPAPNI | ASGLGQDTQY | 0.29993293 |
| PGGPVPIP | ASGLGQDTQY | 0.30368435 |
| FLTVALEP | ASGLGQDTQY | 0.29193512 |
| SQYFGANL | ASGLGQDTQY | 0.29096222 |
| GGVKPAA | ASGLGQDTQY | 0.28995094 |

| | | |
|---|---|---|
| GGTGGAGGAGGW | ASGLGQDTQY | 0.29563648 |
| GVGGVAGLGGAGG | ASGLGQDTQY | 0.29409283 |
| VLLVLATTI | ASGLGQDTQY | 0.28738388 |
| IGIFAAML | ASGLGQDTQY | 0.29285958 |
| AVGDLID | ASGLGQDTQY | 0.28839648 |
| NISPLPL | ASGLGQDTQY | 0.29312027 |
| VTAVFVAI | ASGLGQDTQY | 0.28785 |
| SGDDNDAAL | ASGLGQDTQY | 0.29884645 |
| AVAALAHS | ASGLGQDTQY | 0.31491962 |
| GGVTVMATP | ASGLGQDTQY | 0.2926788 |
| AGLAANVRD | ASGLGQDTQY | 0.29227495 |
| TNRPDLIDPALLRPG | ASGLGQDTQY | 0.28789276 |
| LANLGVTAL | ASGLGQDTQY | 0.29611784 |
| TGERIEL | ASGLGQDTQY | 0.29699266 |
| NGEELAA | ASGLGQDTQY | 0.29052117 |
| VLSLVWG | ASGLGQDTQY | 0.29798266 |
| IGYALWL | ASGLGQDTQY | 0.28948817 |
| LSLLLAPL | ASGLGQDTQY | 0.2931672 |
| TLDEALK | ASGLGQDTQY | 0.30717513 |
| TDGAGTAEGG | ASGLGQDTQY | 0.29373398 |
| NAPGGAGGNGG | ASGLGQDTQY | 0.29062814 |
| VSVFMGMGGNF | ASGLGQDTQY | 0.29287392 |
| LASYLAAL | ASGLGQDTQY | 0.29254884 |
| LVGGTPAL | ASGLGQDTQY | 0.29335165 |
| VRNADVL | ASGLGQDTQY | 0.30604842 |
| TCSSPFLVSAM | ASGLGQDTQY | 0.28775233 |
| VHITDAE | ASGLGQDTQY | 0.29583344 |
| EYAAPSF | ASGLGQDTQY | 0.29113287 |
| VVDAFHGGL | ASGLGQDTQY | 0.30961072 |
| ASVVDPQ | ASGLGQDTQY | 0.29014504 |
| SYNEFGGLS | ASGLGQDTQY | 0.29761285 |
| DPDGSVY | ASGLGQDTQY | 0.2906486 |
| MVSTIHATE | ASGLGQDTQY | 0.31811714 |
| LAAGVADPV | ASGLGQDTQY | 0.29297325 |
| TKEQIAA | ASGLGQDTQY | 0.2966488 |
| SLASPAAL | ASGLGQDTQY | 0.29845306 |
| IGLVFYGV | ASGLGQDTQY | 0.21952955 |
| FLLDASNL | ASGLGQDTQY | 0.22025326 |
| YQVGNFP | ASGLGQDTQY | 0.21949051 |
| ESLLSQST | ASGLGQDTQY | 0.21949244 |
| LFTVGFAAVG | ASGLGQDTQY | 0.21983834 |
| ATPPTPT | ASGLGQDTQY | 0.21949328 |
| TDLATLG | ASGLGQDTQY | 0.21960548 |
| SAGGTGR | ASGLGQDTQY | 0.21967277 |
| GFSNSGS | ASGLGQDTQY | 0.21949801 |
| GGNGGSGGTSV | ASGLGQDTQY | 0.21962532 |
| GTSVATGGAGN | ASGLGQDTQY | 0.21988423 |
| NGGNGGSGGTS | ASGLGQDTQY | 0.22033341 |
| AVVKFPH | ASGLGQDTQY | 0.21988617 |
| TPPPPPP | ASGLGQDTQY | 0.2192849 |
| AAPAAAAAPA | ASGLGQDTQY | 0.219291 |

| | | |
|---|---|---|
| IALNEDL | ASGLGQDTQY | 0.2197318 |
| IVNVSSM | ASGLGQDTQY | 0.21952008 |
| TGIPPAPR | ASGLGQDTQY | 0.21919514 |
| FDLGGGTF | ASGLGQDTQY | 0.21944831 |
| FDLGGGTFDVS | ASGLGQDTQY | 0.21894361 |
| NEPTAAA | ASGLGQDTQY | 0.21918574 |
| NEPTAAALA | ASGLGQDTQY | 0.21956399 |
| GSIINISSI | ASGLGQDTQY | 0.2197023 |
| VAPGNAGTA | ASGLGQDTQY | 0.21956483 |
| TGGGASLE | ASGLGQDTQY | 0.21964155 |
| VAVYNHYKRI | ASGLGQDTQY | 0.21913771 |
| TDPAPATT | ASGLGQDTQY | 0.21965122 |
| TNRPDILDPALLRPG | ASGLGQDTQY | 0.22005996 |
| LTTTTTS | ASGLGQDTQY | 0.22002926 |
| PTGSGKTL | ASGLGQDTQY | 0.21971759 |
| TGSGKTL | ASGLGQDTQY | 0.21937886 |
| TIGHVDHGK | ASGLGQDTQY | 0.2200357 |
| TIGHVDHGKTT | ASGLGQDTQY | 0.22010842 |
| GHVDHGKTT | ASGLGQDTQY | 0.21939884 |
| AQFEHTLL | ASGLGQDTQY | 0.21933083 |
| IVNQDEL | ASGLGQDTQY | 0.2190547 |
| TNGSQFF | ASGLGQDTQY | 0.22007506 |
| TGKGIVE | ASGLGQDTQY | 0.22025017 |
| LISNASDA | ASGLGQDTQY | 0.22017361 |
| AELRTLGELTKIAKAHGAQ | ASGLGQDTQY | 0.21971199 |
| APNAVAAAEPSW | ASGLGQDTQY | 0.21950652 |
| APQVAGAATAVL | ASGLGQDTQY | 0.21934764 |
| ARRDPAAAAAILPTDA | ASGLGQDTQY | 0.2192301 |
| DAAARDVLAEAGLAEYF | ASGLGQDTQY | 0.21951933 |
| DDQIVLSAKANLAKVEQ | ASGLGQDTQY | 0.2199162 |
| DELLPDTGEDSDEDGH | ASGLGQDTQY | 0.21946143 |
| ELPEAPANRATVPQD | ASGLGQDTQY | 0.2193889 |
| FGQSKIERVTGIG | ASGLGQDTQY | 0.21945825 |
| GAGGTGSTAGGAGGAGGAGGLY | ASGLGQDTQY | 0.21894345 |
| GHGGTGAAGGNGGTGGAGGSGI | ASGLGQDTQY | 0.21952651 |
| GNGGNGGAGGEGGPGGAGGAGA | ASGLGQDTQY | 0.21923259 |
| GSTAGNGGNGGNATNGTIAGQP | ASGLGQDTQY | 0.21949354 |
| GTGGDGGNGGNGGTGG | ASGLGQDTQY | 0.2203449 |
| GVRRPAGIPADAPV | ASGLGQDTQY | 0.22024772 |
| GYRQVTDAHLLTLARR | ASGLGQDTQY | 0.21972623 |
| IAKMGQDGHDRG | ASGLGQDTQY | 0.22002341 |
| KAEPSIEIQYT | ASGLGQDTQY | 0.21900426 |
| KPVRIRMAADEDTVAVT | ASGLGQDTQY | 0.21982235 |
| MPPRIAGMRLLVIKPEP | ASGLGQDTQY | 0.21977027 |
| MKFYAVALFRSRGVPPTR | ASGLGQDTQY | 0.21950896 |
| NAGGFGILGAVAHSP | ASGLGQDTQY | 0.21958089 |
| NEGREITLAMAEM | ASGLGQDTQY | 0.21953453 |
| NELVEWSHPLGHRWRWE | ASGLGQDTQY | 0.21983358 |
| NLGAGGTGGNGGNPG | ASGLGQDTQY | 0.21963829 |
| NTEWPKWETVMAADPVD | ASGLGQDTQY | 0.21991761 |
| PASAAPAETQP | ASGLGQDTQY | 0.21956198 |

| | | |
|---|---|---|
| QQSSDGKNGVRAAEITGE | ASGLGQDTQY | 0.21862383 |
| RAPDAVTTSWWKE | ASGLGQDTQY | 0.21920036 |
| SGFGNVGAHTSGWFNQ | ASGLGQDTQY | 0.21945311 |
| SGVAGAVGQAASVGGLKV | ASGLGQDTQY | 0.2195605 |
| SNGGAGTGSAGGNGGT | ASGLGQDTQY | 0.21920468 |
| TGGAGGEGGAAG | ASGLGQDTQY | 0.21959278 |
| TGGDGGAAGAGGAPGHG | ASGLGQDTQY | 0.22003283 |
| TMPPLLAAIANAGRHGV | ASGLGQDTQY | 0.21976075 |
| TSSTAGNSGGA | ASGLGQDTQY | 0.21912962 |
| VPATVPVGILC | ASGLGQDTQY | 0.21965095 |
| VQRPDSGGVVLTGRIS | ASGLGQDTQY | 0.22004434 |
| WTPNPVPATKGVPAGLVQAT | ASGLGQDTQY | 0.21967533 |
| YALKRALAKFSDRFDVVII | ASGLGQDTQY | 0.2193033 |
| YLIGSVIV | ASGLGQDTQY | 0.21945049 |
| YGQLVAVID | ASGLGQDTQY | 0.21947445 |
| VTVGGMTLPA | ASGLGQDTQY | 0.21989715 |
| VLALPAV | ASGLGQDTQY | 0.21950883 |
| VIPPTLNYET | ASGLGQDTQY | 0.21929154 |
| VGMGGMHPSSG | ASGLGQDTQY | 0.2200403 |
| VGLAGGAATA | ASGLGQDTQY | 0.21949852 |
| VGGAGGAGGSGP | ASGLGQDTQY | 0.22000724 |
| VFAAFMV | ASGLGQDTQY | 0.21986672 |
| TLYDKLR | ASGLGQDTQY | 0.21936293 |
| TLAVFTVL | ASGLGQDTQY | 0.2194764 |
| TGSVQPARV | ASGLGQDTQY | 0.21928617 |
| TGGGAGSGGT | ASGLGQDTQY | 0.21937002 |
| SHLETSL | ASGLGQDTQY | 0.2193453 |
| SFTTGMT | ASGLGQDTQY | 0.2190496 |
| QVEVAKRMGVRQPTVS | ASGLGQDTQY | 0.22009633 |
| QGSGGPALAM | ASGLGQDTQY | 0.21937433 |
| MLAPSNP | ASGLGQDTQY | 0.21950543 |
| LVNNAAI | ASGLGQDTQY | 0.22003074 |
| LTGEINTLA | ASGLGQDTQY | 0.21903197 |
| LSIFGPV | ASGLGQDTQY | 0.21984448 |
| LLALLANL | ASGLGQDTQY | 0.21930505 |
| LISNASDALDKLR | ASGLGQDTQY | 0.21987033 |
| LINEAVP | ASGLGQDTQY | 0.21926157 |
| LGYAFSGL | ASGLGQDTQY | 0.2196894 |
| LDFVAVAL | ASGLGQDTQY | 0.21961871 |
| LAQPGAE | ASGLGQDTQY | 0.21925575 |
| ISNLGAVTGF | ASGLGQDTQY | 0.21935566 |
| HQEDPGRI | ASGLGQDTQY | 0.21946299 |
| GTPGEPTP | ASGLGQDTQY | 0.2203242 |
| GFFNSGS | ASGLGQDTQY | 0.21927096 |
| GAGGGGLTAGAN | ASGLGQDTQY | 0.21977742 |
| FVDALVNM | ASGLGQDTQY | 0.21985348 |
| FLTVAPEL | ASGLGQDTQY | 0.21992952 |
| FETLTAL | ASGLGQDTQY | 0.21972173 |
| DVINPIEL | ASGLGQDTQY | 0.21934095 |
| APVSGAPV | ASGLGQDTQY | 0.21919042 |
| ALVGGLAW | ASGLGQDTQY | 0.2202834 |

| | | |
|---|---|---|
| ALAAAAATAI | ASGLGQDTQY | 0.21909043 |
| AFGLGLLL | ASGLGQDTQY | 0.21918844 |
| AFFAAASL | ASGLGQDTQY | 0.21969129 |
| AAAALPAFSPP | ASGLGQDTQY | 0.21909943 |
| VVLAPAL | ASGLGQDTQY | 0.22017068 |
| TIGHVDHGKTTL | ASGLGQDTQY | 0.21934156 |
| LISNASDALDK | ASGLGQDTQY | 0.21894604 |
| LISNASDAL | ASGLGQDTQY | 0.2190701 |
| ISNASDALDK | ASGLGQDTQY | 0.21933302 |
| NQAETLVYQTE | ASGLGQDTQY | 0.22016841 |
| TGGTGGTAGAG | ASGLGQDTQY | 0.22015516 |
| TGSTAGGAGGA | ASGLGQDTQY | 0.21973635 |
| DPLAVPVAAGPV | ASGLGQDTQY | 0.21984102 |
| GGTGGTPGNGGN | ASGLGQDTQY | 0.21978283 |
| PAGLAGVDPATL | ASGLGQDTQY | 0.21905553 |
| GNNLVVEAPHAGASV | ASGLGQDTQY | 0.2198385 |
| KVTRIMSASTLEQPA | ASGLGQDTQY | 0.21940985 |
| LGFALPIALPRRTVV | ASGLGQDTQY | 0.21935736 |
| TRLGNLTARKLVVLV | ASGLGQDTQY | 0.21942537 |
| WSQQRRLIYFPSAGP | ASGLGQDTQY | 0.21947597 |
| GPNGNPDPMAAAVDIR | ASGLGQDTQY | 0.2197724 |
| TQRCVSVADLRAAASR | ASGLGQDTQY | 0.21908589 |
| QFVLDVAEHPVEVLTETV | ASGLGQDTQY | 0.21936671 |
| GGAGGTGGAGGNGGTQAGDG | ASGLGQDTQY | 0.21891706 |
| PLLATLRAIAVDVDAKPAQV | ASGLGQDTQY | 0.21893667 |
| APKFRTGGYMIDFWGVGYQVAK | ASGLGQDTQY | 0.22029163 |
| GAGGAGGAGGAAGGPGGTGGTG | ASGLGQDTQY | 0.21961775 |
| IGALGLLATDVAPAGSVAFGVA | ASGLGQDTQY | 0.21949406 |
| AGGAGGIGGNGGAGGAGGVLMG | ASGLGQDTQY | 0.21908623 |
| ADELIGLL | ASGLGQDTQY | 0.2196154 |
| ADRALKA | ASGLGQDTQY | 0.21962483 |
| AGAVGQVGGAGGA | ASGLGQDTQY | 0.21947993 |
| AGGGGGLGGAP | ASGLGQDTQY | 0.21952859 |
| AGGLLYGNGGAG | ASGLGQDTQY | 0.21956222 |
| AGLESIGL | ASGLGQDTQY | 0.2204861 |
| AGLKTGII | ASGLGQDTQY | 0.21944705 |
| ASLAALLL | ASGLGQDTQY | 0.2195843 |
| ASVINISSI | ASGLGQDTQY | 0.22005732 |
| ATPPGPA | ASGLGQDTQY | 0.21966138 |
| ATVASALV | ASGLGQDTQY | 0.21960978 |
| ATVASLIL | ASGLGQDTQY | 0.21971658 |
| AVNGPVLIRH | ASGLGQDTQY | 0.21990381 |
| DGAAGAVGI | ASGLGQDTQY | 0.2192704 |
| DPLLHGI | ASGLGQDTQY | 0.21951889 |
| ELGAALGAL | ASGLGQDTQY | 0.21941641 |
| ELGRTQAV | ASGLGQDTQY | 0.22000863 |
| FVAGGYLS | ASGLGQDTQY | 0.22022434 |
| GASGGAGGAGGW | ASGLGQDTQY | 0.21926291 |
| GGAGAGGDGGAGGI | ASGLGQDTQY | 0.21980894 |
| GTGHNGSQPGMGGNG | ASGLGQDTQY | 0.2194943 |
| GYAAASASAS | ASGLGQDTQY | 0.21953435 |

| | | |
|---|---|---|
| HQVDVAAVQQG | ASGLGQDTQY | 0.2197174 |
| IGAAIAHT | ASGLGQDTQY | 0.2197241 |
| IGMIEVAQL | ASGLGQDTQY | 0.21931913 |
| IRLAPEHTSAMD | ASGLGQDTQY | 0.21976769 |
| IYNVSVL | ASGLGQDTQY | 0.21955776 |
| KQGFLGM | ASGLGQDTQY | 0.2195557 |
| LESVITQL | ASGLGQDTQY | 0.21978037 |
| LGLGALVVV | ASGLGQDTQY | 0.21941751 |
| LIGNGGNGGSGGI | ASGLGQDTQY | 0.21963614 |
| LNAINAPTL | ASGLGQDTQY | 0.21980011 |
| LQLLNTGV | ASGLGQDTQY | 0.21936715 |
| LQTSIEF | ASGLGQDTQY | 0.21956198 |
| MSFAVSEL | ASGLGQDTQY | 0.22054447 |
| NTGALGL | ASGLGQDTQY | 0.2193866 |
| PPGAGGAGGAGTT | ASGLGQDTQY | 0.2196682 |
| QLVGLGI | ASGLGQDTQY | 0.22014895 |
| RFLEQVHQL | ASGLGQDTQY | 0.21964702 |
| RGVGYVLR | ASGLGQDTQY | 0.21969533 |
| SATGVFAGVI | ASGLGQDTQY | 0.2188671 |
| SLPAITLP | ASGLGQDTQY | 0.21938646 |
| STALAFTV | ASGLGQDTQY | 0.21944986 |
| SVLTVGLGK | ASGLGQDTQY | 0.2197309 |
| TALAGAGSGI | ASGLGQDTQY | 0.21985854 |
| TGLPFTGI | ASGLGQDTQY | 0.21983902 |
| VAEHVTV | ASGLGQDTQY | 0.21931456 |
| VDIVLNSL | ASGLGQDTQY | 0.22028895 |
| VLTGGGAVL | ASGLGQDTQY | 0.21954927 |
| VNVGLAYGI | ASGLGQDTQY | 0.21908739 |
| VSSAAVGAV | ASGLGQDTQY | 0.21948299 |
| VTSVYEAGV | ASGLGQDTQY | 0.21979873 |
| WVALDRGA | ASGLGQDTQY | 0.21935785 |
| YLVGSVVI | ASGLGQDTQY | 0.2192373 |
| GPPGTGKTLL | ASGLGQDTQY | 0.21862148 |
| GVGMTKF | ASGLGQDTQY | 0.21948464 |
| IGIGGSDL | ASGLGQDTQY | 0.21966608 |
| IGTLLAEL | ASGLGQDTQY | 0.21900749 |
| IYTSGTT | ASGLGQDTQY | 0.21955624 |
| LSFPAGV | ASGLGQDTQY | 0.2193126 |
| LYTSGTT | ASGLGQDTQY | 0.2190945 |
| VGAFPFGF | ASGLGQDTQY | 0.22016421 |
| VGAGTVEF | ASGLGQDTQY | 0.21947049 |
| VGKSTLLL | ASGLGQDTQY | 0.21948828 |
| AALAHAGGGIV | ASGLGQDTQY | 0.21943171 |
| AALVAVSLL | ASGLGQDTQY | 0.21958701 |
| AAVSRAVAAL | ASGLGQDTQY | 0.21988311 |
| AGGSSGIGGF | ASGLGQDTQY | 0.21917588 |
| AGLTVANANG | ASGLGQDTQY | 0.21939793 |
| AGLVGAVL | ASGLGQDTQY | 0.21994613 |
| ALSQAFTV | ASGLGQDTQY | 0.21920514 |
| ALTFSVSL | ASGLGQDTQY | 0.2203787 |
| ALTGAALVAL | ASGLGQDTQY | 0.21981302 |

| | | |
|---|---|---|
| ASFSIGAGVL | ASGLGQDTQY | 0.21939252 |
| DGSVYKI | ASGLGQDTQY | 0.21946134 |
| EQLAEIV | ASGLGQDTQY | 0.21950024 |
| FAVSELTVM | ASGLGQDTQY | 0.21985981 |
| GAAPLAAGL | ASGLGQDTQY | 0.21927597 |
| GADGGDGGSGGAGG | ASGLGQDTQY | 0.21922407 |
| GFGLDTGGAGG | ASGLGQDTQY | 0.21968044 |
| GLASGGLL | ASGLGQDTQY | 0.22032931 |
| GRIAGAAVLL | ASGLGQDTQY | 0.21945383 |
| GSAGSLNTSF | ASGLGQDTQY | 0.21979408 |
| GTGQIGI | ASGLGQDTQY | 0.21969357 |
| GTLIALIL | ASGLGQDTQY | 0.2197526 |
| IAATMGFVVG | ASGLGQDTQY | 0.21957153 |
| IFAQALL | ASGLGQDTQY | 0.2202456 |
| IGAAVALGF | ASGLGQDTQY | 0.21941929 |
| IGLTPLAL | ASGLGQDTQY | 0.22008723 |
| ILGSNTSTL | ASGLGQDTQY | 0.22006127 |
| IVFSLTVL | ASGLGQDTQY | 0.2193587 |
| LASKATGF | ASGLGQDTQY | 0.21960184 |
| LASVALIL | ASGLGQDTQY | 0.21962143 |
| LFAVGLTL | ASGLGQDTQY | 0.21964295 |
| LFNGGIGTYI | ASGLGQDTQY | 0.21958856 |
| LTLGAGII | ASGLGQDTQY | 0.21933396 |
| LVAAALGDI | ASGLGQDTQY | 0.21982247 |
| NTAFVQLGI | ASGLGQDTQY | 0.21931675 |
| QLFSGAEL | ASGLGQDTQY | 0.22021964 |
| SAATFITL | ASGLGQDTQY | 0.21944533 |
| SNYGLANLGV | ASGLGQDTQY | 0.21927708 |
| SYGGAYAVM | ASGLGQDTQY | 0.21967708 |
| TTGVGGLTLG | ASGLGQDTQY | 0.21943523 |
| TTQTIYI | ASGLGQDTQY | 0.21975511 |
| VAAGAGLGAVI | ASGLGQDTQY | 0.21971378 |
| VAAVVTLL | ASGLGQDTQY | 0.21937785 |
| VGFLGGLI | ASGLGQDTQY | 0.21946025 |
| VGVIPILL | ASGLGQDTQY | 0.21950783 |
| VIFASSLI | ASGLGQDTQY | 0.21920577 |
| VKGGDGGAAAG | ASGLGQDTQY | 0.22004004 |
| VTKTFTNI | ASGLGQDTQY | 0.21920164 |
| VVAAAILL | ASGLGQDTQY | 0.22004569 |
| YAALDVEL | ASGLGQDTQY | 0.219372 |
| LASVALLL | ASGLGQDTQY | 0.21996932 |
| LDAVHRV | ASGLGQDTQY | 0.21999575 |
| QGLSSSIF | ASGLGQDTQY | 0.22017227 |
| RGTFANIRL | ASGLGQDTQY | 0.21911253 |
| RQLWWGHRI | ASGLGQDTQY | 0.2203465 |
| YGPPGTGKTLL | ASGLGQDTQY | 0.22012116 |
| AALVGAPANVA | ASGLGQDTQY | 0.21935253 |
| AATFRRTE | ASGLGQDTQY | 0.2195909 |
| AGALGGGI | ASGLGQDTQY | 0.21958253 |
| AGGAGGAGGIGAG | ASGLGQDTQY | 0.21989869 |
| AGSGALGL | ASGLGQDTQY | 0.21950497 |

| | | |
|---|---|---|
| AQIADQI | ASGLGQDTQY | 0.21931034 |
| ASAVAVTYL | ASGLGQDTQY | 0.21954732 |
| ASTVLGGTGG | ASGLGQDTQY | 0.21938159 |
| AVGGEPAK | ASGLGQDTQY | 0.21943955 |
| CEQGTFGGTC | ASGLGQDTQY | 0.21966277 |
| DEGKAPAG | ASGLGQDTQY | 0.2194017 |
| DTVVFPAL | ASGLGQDTQY | 0.22039382 |
| FEGITMAG | ASGLGQDTQY | 0.22001334 |
| FPLVVQD | ASGLGQDTQY | 0.2201722 |
| GAFTTPQ | ASGLGQDTQY | 0.21922374 |
| GFGGFGTAGGVGGA | ASGLGQDTQY | 0.2197129 |
| GFLAQFL | ASGLGQDTQY | 0.21996123 |
| GGAGGAGGNAGVA | ASGLGQDTQY | 0.21954322 |
| GGVGTAGAGGAGGAGG | ASGLGQDTQY | 0.21959259 |
| GNGGGGDAGNAGSGGN | ASGLGQDTQY | 0.21969278 |
| GTGGAGGAGNDAGS | ASGLGQDTQY | 0.21981065 |
| GTTNTGWFNS | ASGLGQDTQY | 0.21884754 |
| IFMGPLFTL | ASGLGQDTQY | 0.21906641 |
| ILAHAIEAI | ASGLGQDTQY | 0.21990944 |
| IRSAIREAL | ASGLGQDTQY | 0.2193682 |
| ITFAGGG | ASGLGQDTQY | 0.21957515 |
| ITGMSGHQEV | ASGLGQDTQY | 0.21970099 |
| IVAGLACTL | ASGLGQDTQY | 0.21926686 |
| IVGGGLAGA | ASGLGQDTQY | 0.22004816 |
| IVTGVLNI | ASGLGQDTQY | 0.2193973 |
| LAQQGCI | ASGLGQDTQY | 0.21942101 |
| LCLHVEN | ASGLGQDTQY | 0.22036214 |
| LDAISDGS | ASGLGQDTQY | 0.21969676 |
| LGGFAAML | ASGLGQDTQY | 0.21930358 |
| LGGKNPMI | ASGLGQDTQY | 0.21926981 |
| LGPKGVAAIL | ASGLGQDTQY | 0.21972916 |
| LMVGVSI | ASGLGQDTQY | 0.21941607 |
| LVGAAFVL | ASGLGQDTQY | 0.21906208 |
| LVLAVAPL | ASGLGQDTQY | 0.21959537 |
| MAAIAVIDAA | ASGLGQDTQY | 0.21945253 |
| NNGINAELQ | ASGLGQDTQY | 0.21959904 |
| RGEPLVLVHG | ASGLGQDTQY | 0.21984613 |
| RGGAGGAAGLI | ASGLGQDTQY | 0.21939887 |
| RSYRMHPAV | ASGLGQDTQY | 0.21976277 |
| SAAHLLAGQLA | ASGLGQDTQY | 0.21954826 |
| SAAVGSY | ASGLGQDTQY | 0.21952926 |
| SPHAVLHRI | ASGLGQDTQY | 0.21953487 |
| SQIAKVL | ASGLGQDTQY | 0.21946286 |
| TALGHLVP | ASGLGQDTQY | 0.2195308 |
| TAPTTSGAA | ASGLGQDTQY | 0.21962577 |
| TDGDDGNGGSPGT | ASGLGQDTQY | 0.22024791 |
| TDGLGEN | ASGLGQDTQY | 0.21990491 |
| TGGPPLPGQA | ASGLGQDTQY | 0.21980093 |
| TIPAITI | ASGLGQDTQY | 0.21956463 |
| TIPGFTI | ASGLGQDTQY | 0.21969101 |
| TSIASGALA | ASGLGQDTQY | 0.21974082 |

| | | |
|---|---|---|
| TVTLPGAAI | ASGLGQDTQY | 0.21876168 |
| VASGLSAALP | ASGLGQDTQY | 0.21937454 |
| VGAGVIGI | ASGLGQDTQY | 0.21957216 |
| VGPKRFGV | ASGLGQDTQY | 0.21951443 |
| VNVAFPD | ASGLGQDTQY | 0.2195762 |
| VSMTNSGVS | ASGLGQDTQY | 0.21960348 |
| VTDVLGDLSV | ASGLGQDTQY | 0.21953431 |
| VVVVGAGFAGL | ASGLGQDTQY | 0.21933506 |
| VYTVGTVV | ASGLGQDTQY | 0.2194255 |
| WTTFSGP | ASGLGQDTQY | 0.2203081 |
| YDELVEAF | ASGLGQDTQY | 0.2201915 |
| YGTGGAGGNGG | ASGLGQDTQY | 0.22006068 |
| YTNVMAV | ASGLGQDTQY | 0.21937904 |
| FLEQVHQL | ASGLGQDTQY | 0.21942335 |
| GLSLLAVAV | ASGLGQDTQY | 0.21990466 |
| LEQVHQL | ASGLGQDTQY | 0.2192559 |
| LNAAGAI | ASGLGQDTQY | 0.21891205 |
| PAVHASL | ASGLGQDTQY | 0.21977232 |
| TELEDDA | ASGLGQDTQY | 0.21910757 |
| TPPGPPN | ASGLGQDTQY | 0.21987304 |
| TPPPPPPP | ASGLGQDTQY | 0.21928044 |
| TQSPALI | AWSQGGANSDYT | 0.21921557 |
| EFAAAFPQ | AWSQGGANSDYT | 0.21943775 |
| LLSGNLDVL | AWSQGGANSDYT | 0.21935561 |
| GPANLGNYNI | AWSQGGANSDYT | 0.21991254 |
| TLGPTLP | AWSQGGANSDYT | 0.21967012 |
| VYAYSSCCK | AWSQGGANSDYT | 0.2193472 |
| APAPAPAPNI | AWSQGGANSDYT | 0.21948029 |
| PGGPVPIP | AWSQGGANSDYT | 0.22039106 |
| FLTVALEP | AWSQGGANSDYT | 0.21967216 |
| SQYFGANL | AWSQGGANSDYT | 0.21947548 |
| GGVKPAA | AWSQGGANSDYT | 0.21942548 |
| GGTGGAGGAGGW | AWSQGGANSDYT | 0.2193518 |
| GVGGVAGLGGAGG | AWSQGGANSDYT | 0.21932109 |
| VLLVLATTI | AWSQGGANSDYT | 0.21940446 |
| IGIFAAML | AWSQGGANSDYT | 0.21929632 |
| AVGDLID | AWSQGGANSDYT | 0.21938373 |
| NISPLPL | AWSQGGANSDYT | 0.21966931 |
| VTAVFVAI | AWSQGGANSDYT | 0.21882212 |
| SGDDNDAAL | AWSQGGANSDYT | 0.21962577 |
| AVAALAHS | AWSQGGANSDYT | 0.21893199 |
| GGVTVMATP | AWSQGGANSDYT | 0.2193334 |
| AGLAANVRD | AWSQGGANSDYT | 0.22013877 |
| TNRPDLIDPALLRPG | AWSQGGANSDYT | 0.22007105 |
| LANLGVTAL | AWSQGGANSDYT | 0.21900502 |
| TGERIEL | AWSQGGANSDYT | 0.21964541 |
| NGEELAA | AWSQGGANSDYT | 0.219081 |
| VLSLVWG | AWSQGGANSDYT | 0.21962957 |
| IGYALWL | AWSQGGANSDYT | 0.21922514 |
| LSLLLAPL | AWSQGGANSDYT | 0.2196793 |
| TLDEALK | AWSQGGANSDYT | 0.2198314 |

| | | |
|---|---|---|
| TDGAGTAEGG | AWSQGGANSDYT | 0.21926336 |
| NAPGGAGGNGG | AWSQGGANSDYT | 0.21912968 |
| VSVFMGMGGNF | AWSQGGANSDYT | 0.21972336 |
| LASYLAAL | AWSQGGANSDYT | 0.21929635 |
| LVGGTPAL | AWSQGGANSDYT | 0.21898213 |
| VRNADVL | AWSQGGANSDYT | 0.21979018 |
| TCSSPFLVSAM | AWSQGGANSDYT | 0.21957996 |
| VHITDAE | AWSQGGANSDYT | 0.21937485 |
| EYAAPSF | AWSQGGANSDYT | 0.21950494 |
| VVDAFHGGL | AWSQGGANSDYT | 0.21956079 |
| ASVVDPQ | AWSQGGANSDYT | 0.22000526 |
| SYNEFGGLS | AWSQGGANSDYT | 0.21945712 |
| DPDGSVY | AWSQGGANSDYT | 0.21912782 |
| MVSTIHATE | AWSQGGANSDYT | 0.21977378 |
| LAAGVADPV | AWSQGGANSDYT | 0.22018467 |
| TKEQIAA | AWSQGGANSDYT | 0.2196573 |
| SLASPAAL | AWSQGGANSDYT | 0.21948409 |
| IGLVFYGV | AWSQGGANSDYT | 0.22000141 |
| FLLDASNL | AWSQGGANSDYT | 0.21902521 |
| YQVGNFP | AWSQGGANSDYT | 0.21947205 |
| ESLLSQST | AWSQGGANSDYT | 0.21980909 |
| LFTVGFAAVG | AWSQGGANSDYT | 0.21939929 |
| ATPPTPT | AWSQGGANSDYT | 0.21969396 |
| TDLATLG | AWSQGGANSDYT | 0.22037071 |
| SAGGTGR | AWSQGGANSDYT | 0.21935987 |
| GFSNSGS | AWSQGGANSDYT | 0.21951641 |
| GGNGGSGGTSV | AWSQGGANSDYT | 0.21958041 |
| GTSVATGGAGN | AWSQGGANSDYT | 0.21912271 |
| NGGNGGSGGTS | AWSQGGANSDYT | 0.2201894 |
| AVVKFPH | AWSQGGANSDYT | 0.21956919 |
| TPPPPPP | AWSQGGANSDYT | 0.2197063 |
| AAPAAAAAPA | AWSQGGANSDYT | 0.21957885 |
| IALNEDL | AWSQGGANSDYT | 0.2196192 |
| IVNVSSM | AWSQGGANSDYT | 0.2198395 |
| TGIPPAPR | AWSQGGANSDYT | 0.21985719 |
| FDLGGGTF | AWSQGGANSDYT | 0.22008379 |
| FDLGGGTFDVS | AWSQGGANSDYT | 0.21919706 |
| NEPTAAA | AWSQGGANSDYT | 0.2195334 |
| NEPTAAALA | AWSQGGANSDYT | 0.2192935 |
| GSIINISSI | AWSQGGANSDYT | 0.22044733 |
| VAPGNAGTA | AWSQGGANSDYT | 0.21970095 |
| TGGGASLE | AWSQGGANSDYT | 0.21962541 |
| VAVYNHYKRI | AWSQGGANSDYT | 0.21934333 |
| TDPAPATT | AWSQGGANSDYT | 0.21937142 |
| TNRPDILDPALLRPG | AWSQGGANSDYT | 0.21928364 |
| LTTTTTS | AWSQGGANSDYT | 0.21975952 |
| PTGSGKTL | AWSQGGANSDYT | 0.21986637 |
| TGSGKTL | AWSQGGANSDYT | 0.21903174 |
| TIGHVDHGK | AWSQGGANSDYT | 0.21980543 |
| TIGHVDHGKTT | AWSQGGANSDYT | 0.21967287 |
| GHVDHGKTT | AWSQGGANSDYT | 0.21937907 |

| | | |
|---|---|---|
| AQFEHTLL | AWSQGGANSDYT | 0.2192131 |
| IVNQDEL | AWSQGGANSDYT | 0.21945624 |
| TNGSQFF | AWSQGGANSDYT | 0.21959117 |
| TGKGIVE | AWSQGGANSDYT | 0.21939409 |
| LISNASDA | AWSQGGANSDYT | 0.21951029 |
| AELRTLGELTKIAKAHGAQ | AWSQGGANSDYT | 0.21970254 |
| APNAVAAAEPSW | AWSQGGANSDYT | 0.21944682 |
| APQVAGAATAVL | AWSQGGANSDYT | 0.21984082 |
| ARRDPAAAAAILPTDA | AWSQGGANSDYT | 0.21918148 |
| DAAARDVLAEAGLAEYF | AWSQGGANSDYT | 0.21935306 |
| DDQIVLSAKANLAKVEQ | AWSQGGANSDYT | 0.21950863 |
| DELLPDTGEDSDEDGH | AWSQGGANSDYT | 0.21935518 |
| ELPEAPANRATVPQD | AWSQGGANSDYT | 0.21957578 |
| FGQSKIERVTGIG | AWSQGGANSDYT | 0.21949703 |
| GAGGTGSTAGGAGGAGGAGGLY | AWSQGGANSDYT | 0.21986808 |
| GHGGTGAAGGNGGTGGAGGSGI | AWSQGGANSDYT | 0.21939607 |
| GNGGNGGAGGEGGPGGAGGAGA | AWSQGGANSDYT | 0.21989739 |
| GSTAGNGGNGGNATNGTIAGQP | AWSQGGANSDYT | 0.22033592 |
| GTGGDGGNGGNGGTGG | AWSQGGANSDYT | 0.21958013 |
| GVRRPAGIPADAPV | AWSQGGANSDYT | 0.22015998 |
| GYRQVTDAHLLTLARR | AWSQGGANSDYT | 0.21922882 |
| IAKMGQDGHDRG | AWSQGGANSDYT | 0.21943301 |
| KAEPSIEIQYT | AWSQGGANSDYT | 0.21926601 |
| KPVRIRMAADEDTVAVT | AWSQGGANSDYT | 0.21987136 |
| MPPRIAGMRLLVIKPEP | AWSQGGANSDYT | 0.21944071 |
| MKFYAVALFRSRGVPPTR | AWSQGGANSDYT | 0.22015522 |
| NAGGFGILGAVAHSP | AWSQGGANSDYT | 0.21967484 |
| NEGREITLAMAEM | AWSQGGANSDYT | 0.2202674 |
| NELVEWSHPLGHRWRWE | AWSQGGANSDYT | 0.2199911 |
| NLGAGGTGGNGGNPG | AWSQGGANSDYT | 0.2194881 |
| NTEWPKWETVMAADPVD | AWSQGGANSDYT | 0.2198097 |
| PASAAPAETQP | AWSQGGANSDYT | 0.2195388 |
| QQSSDGKNGVRAAEITGE | AWSQGGANSDYT | 0.21947166 |
| RAPDAVTTSWWKE | AWSQGGANSDYT | 0.21968648 |
| SGFGNVGAHTSGWFNQ | AWSQGGANSDYT | 0.21975832 |
| SGVAGAVGQAASVGGLKV | AWSQGGANSDYT | 0.2194863 |
| SNGGAGTGSAGGNGGT | AWSQGGANSDYT | 0.21964538 |
| TGGAGGEGGAAG | AWSQGGANSDYT | 0.22016965 |
| TGGDGGAAGAGGAPGHG | AWSQGGANSDYT | 0.2196823 |
| TMPPLLAAIANAGRHGV | AWSQGGANSDYT | 0.21916582 |
| TSSTAGNSGGA | AWSQGGANSDYT | 0.2193472 |
| VPATVPVGILC | AWSQGGANSDYT | 0.21989419 |
| VQRPDSGGVVLTGRIS | AWSQGGANSDYT | 0.2204158 |
| WTPNPVPATKGVPAGLVQAT | AWSQGGANSDYT | 0.21983024 |
| YALKRALAKFSDRFDVVII | AWSQGGANSDYT | 0.21989252 |
| YLIGSVIV | AWSQGGANSDYT | 0.22007586 |
| YGQLVAVID | AWSQGGANSDYT | 0.21967551 |
| VTVGGMTLPA | AWSQGGANSDYT | 0.21976756 |
| VLALPAV | AWSQGGANSDYT | 0.21955739 |
| VIPPTLNYET | AWSQGGANSDYT | 0.21926084 |
| VGMGGMHPSSG | AWSQGGANSDYT | 0.21944445 |

| | | |
|---|---|---|
| VGLAGGAATA | AWSQGGANSDYT | 0.21947683 |
| VGGAGGAGGSGP | AWSQGGANSDYT | 0.21953937 |
| VFAAFMV | AWSQGGANSDYT | 0.22036016 |
| TLYDKLR | AWSQGGANSDYT | 0.21966274 |
| TLAVFTVL | AWSQGGANSDYT | 0.22017291 |
| TGSVQPARV | AWSQGGANSDYT | 0.21994801 |
| TGGGAGSGGT | AWSQGGANSDYT | 0.21969117 |
| SHLETSL | AWSQGGANSDYT | 0.21972564 |
| SFTTGMT | AWSQGGANSDYT | 0.21963426 |
| QVEVAKRMGVRQPTVS | AWSQGGANSDYT | 0.21956044 |
| QGSGGPALAM | AWSQGGANSDYT | 0.21939458 |
| MLAPSNP | AWSQGGANSDYT | 0.2195888 |
| LVNNAAI | AWSQGGANSDYT | 0.21970525 |
| LTGEINTLA | AWSQGGANSDYT | 0.22045228 |
| LSIFGPV | AWSQGGANSDYT | 0.21990821 |
| LLALLANL | AWSQGGANSDYT | 0.21992956 |
| LISNASDALDKLR | AWSQGGANSDYT | 0.21942015 |
| LINEAVP | AWSQGGANSDYT | 0.21936513 |
| LGYAFSGL | AWSQGGANSDYT | 0.21949023 |
| LDFVAVAL | AWSQGGANSDYT | 0.2197822 |
| LAQPGAE | AWSQGGANSDYT | 0.2200392 |
| ISNLGAVTGF | AWSQGGANSDYT | 0.21960114 |
| HQEDPGRI | AWSQGGANSDYT | 0.21928279 |
| GTPGEPTP | AWSQGGANSDYT | 0.22000208 |
| GFFNSGS | AWSQGGANSDYT | 0.21956626 |
| GAGGGGLTAGAN | AWSQGGANSDYT | 0.21974291 |
| FVDALVNM | AWSQGGANSDYT | 0.21947071 |
| FLTVAPEL | AWSQGGANSDYT | 0.2196152 |
| FETLTAL | AWSQGGANSDYT | 0.21945694 |
| DVINPIEL | AWSQGGANSDYT | 0.21978392 |
| APVSGAPV | AWSQGGANSDYT | 0.2190061 |
| ALVGGLAW | AWSQGGANSDYT | 0.21919622 |
| ALAAAAATAI | AWSQGGANSDYT | 0.21930851 |
| AFGLGLLL | AWSQGGANSDYT | 0.2194627 |
| AFFAAASL | AWSQGGANSDYT | 0.2195535 |
| AAAALPAFSPP | AWSQGGANSDYT | 0.21980242 |
| VVLAPAL | AWSQGGANSDYT | 0.21944617 |
| TIGHVDHGKTTL | AWSQGGANSDYT | 0.22014876 |
| LISNASDALDK | AWSQGGANSDYT | 0.21907113 |
| LISNASDAL | AWSQGGANSDYT | 0.21936722 |
| ISNASDALDK | AWSQGGANSDYT | 0.22015458 |
| NQAETLVYQTE | AWSQGGANSDYT | 0.21985698 |
| TGGTGGTAGAG | AWSQGGANSDYT | 0.21998166 |
| TGSTAGGAGGA | AWSQGGANSDYT | 0.21935289 |
| DPLAVPVAAGPV | AWSQGGANSDYT | 0.21943289 |
| GGTGGTPGNGGN | AWSQGGANSDYT | 0.2194555 |
| PAGLAGVDPATL | AWSQGGANSDYT | 0.21971296 |
| GNNLVVEAPHAGASV | AWSQGGANSDYT | 0.2194874 |
| KVTRIMSASTLEQPA | AWSQGGANSDYT | 0.21983515 |
| LGFALPIALPRRTVV | AWSQGGANSDYT | 0.21966971 |
| TRLGNLTARKLVVLV | AWSQGGANSDYT | 0.21961807 |

| | | |
|---|---|---|
| WSQQRRLIYFPSAGP | AWSQGGANSDYT | 0.21990544 |
| GPNGNPDPMAAAVDIR | AWSQGGANSDYT | 0.21970509 |
| TQRCVSVADLRAAASR | AWSQGGANSDYT | 0.21929893 |
| QFVLDVAEHPVEVLTETV | AWSQGGANSDYT | 0.21931098 |
| GGAGGTGGAGGNGGTQAGDG | AWSQGGANSDYT | 0.22004664 |
| PLLATLRAIAVDVDAKPAQV | AWSQGGANSDYT | 0.22048734 |
| APKFRTGGYMIDFWGVGYQVAK | AWSQGGANSDYT | 0.21944818 |
| GAGGAGGAGGAAGGPGGTGGTG | AWSQGGANSDYT | 0.21962109 |
| IGALGLLATDVAPAGSVAFGVA | AWSQGGANSDYT | 0.21923774 |
| AGGAGGIGGNGGAGGAGGVLMG | AWSQGGANSDYT | 0.219765 |
| ADELIGLL | AWSQGGANSDYT | 0.2198019 |
| ADRALKA | AWSQGGANSDYT | 0.21978979 |
| AGAVGQVGGAGGA | AWSQGGANSDYT | 0.21948238 |
| AGGGGGLGGAP | AWSQGGANSDYT | 0.2195616 |
| AGGLLYGNGGAG | AWSQGGANSDYT | 0.22033958 |
| AGLESIGL | AWSQGGANSDYT | 0.21945456 |
| AGLKTGII | AWSQGGANSDYT | 0.21938507 |
| ASLAALLL | AWSQGGANSDYT | 0.2195199 |
| ASVINISSI | AWSQGGANSDYT | 0.2195239 |
| ATPPGPA | AWSQGGANSDYT | 0.21913457 |
| ATVASALV | AWSQGGANSDYT | 0.21958335 |
| ATVASLIL | AWSQGGANSDYT | 0.22022764 |
| AVNGPVLIRH | AWSQGGANSDYT | 0.2194261 |
| DGAAGAVGI | AWSQGGANSDYT | 0.21955 |
| DPLLHGI | AWSQGGANSDYT | 0.21953851 |
| ELGAALGAL | AWSQGGANSDYT | 0.22005029 |
| ELGRTQAV | AWSQGGANSDYT | 0.21967512 |
| FVAGGYLS | AWSQGGANSDYT | 0.21942574 |
| GASGGAGGAGGW | AWSQGGANSDYT | 0.21951045 |
| GGAGAGGDGGAGGI | AWSQGGANSDYT | 0.2195275 |
| GTGHNGSQPGMGGNG | AWSQGGANSDYT | 0.21982956 |
| GYAAASASAS | AWSQGGANSDYT | 0.21968195 |
| HQVDVAAVQQG | AWSQGGANSDYT | 0.21949074 |
| IGAAIAHT | AWSQGGANSDYT | 0.21868934 |
| IGMIEVAQL | AWSQGGANSDYT | 0.21952207 |
| IRLAPEHTSAMD | AWSQGGANSDYT | 0.21957366 |
| IYNVSVL | AWSQGGANSDYT | 0.21945693 |
| KQGFLGM | AWSQGGANSDYT | 0.22032508 |
| LESVITQL | AWSQGGANSDYT | 0.21939456 |
| LGLGALVVV | AWSQGGANSDYT | 0.21962042 |
| LIGNGGNGGSGGI | AWSQGGANSDYT | 0.21941565 |
| LNAINAPTL | AWSQGGANSDYT | 0.2191422 |
| LQLLNTGV | AWSQGGANSDYT | 0.2194907 |
| LQTSIEF | AWSQGGANSDYT | 0.21939053 |
| MSFAVSEL | AWSQGGANSDYT | 0.21968636 |
| NTGALGL | AWSQGGANSDYT | 0.2194538 |
| PPGAGGAGGAGTT | AWSQGGANSDYT | 0.21965192 |
| QLVGLGI | AWSQGGANSDYT | 0.21942388 |
| RFLEQVHQL | AWSQGGANSDYT | 0.2191612 |
| RGVGYVLR | AWSQGGANSDYT | 0.22018479 |
| SATGVFAGVI | AWSQGGANSDYT | 0.21938135 |

| | | |
|---|---|---|
| SLPAITLP | AWSQGGANSDYT | 0.21959873 |
| STALAFTV | AWSQGGANSDYT | 0.22022898 |
| SVLTVGLGK | AWSQGGANSDYT | 0.21932647 |
| TALAGAGSGI | AWSQGGANSDYT | 0.21966513 |
| TGLPFTGI | AWSQGGANSDYT | 0.2196121 |
| VAEHVTV | AWSQGGANSDYT | 0.21971491 |
| VDIVLNSL | AWSQGGANSDYT | 0.21995176 |
| VLTGGGAVL | AWSQGGANSDYT | 0.21976589 |
| VNVGLAYGI | AWSQGGANSDYT | 0.21924874 |
| VSSAAVGAV | AWSQGGANSDYT | 0.21945482 |
| VTSVYEAGV | AWSQGGANSDYT | 0.22023164 |
| WVALDRGA | AWSQGGANSDYT | 0.21994984 |
| YLVGSVVI | AWSQGGANSDYT | 0.21939588 |
| GPPGTGKTLL | AWSQGGANSDYT | 0.219579 |
| GVGMTKF | AWSQGGANSDYT | 0.21968709 |
| IGIGGSDL | AWSQGGANSDYT | 0.21972391 |
| IGTLLAEL | AWSQGGANSDYT | 0.2193771 |
| IYTSGTT | AWSQGGANSDYT | 0.21977028 |
| LSFPAGV | AWSQGGANSDYT | 0.21947677 |
| LYTSGTT | AWSQGGANSDYT | 0.21934323 |
| VGAFPFGF | AWSQGGANSDYT | 0.21961054 |
| VGAGTVEF | AWSQGGANSDYT | 0.21995415 |
| VGKSTLLL | AWSQGGANSDYT | 0.21984361 |
| AALAHAGGGIV | AWSQGGANSDYT | 0.21939166 |
| AALVAVSLL | AWSQGGANSDYT | 0.21958366 |
| AAVSRAVAAL | AWSQGGANSDYT | 0.21997617 |
| AGGSSGIGGF | AWSQGGANSDYT | 0.21940465 |
| AGLTVANANG | AWSQGGANSDYT | 0.21990333 |
| AGLVGAVL | AWSQGGANSDYT | 0.21954599 |
| ALSQAFTV | AWSQGGANSDYT | 0.22003174 |
| ALTFSVSL | AWSQGGANSDYT | 0.22032695 |
| ALTGAALVAL | AWSQGGANSDYT | 0.21967396 |
| ASFSIGAGVL | AWSQGGANSDYT | 0.21938536 |
| DGSVYKI | AWSQGGANSDYT | 0.21918635 |
| EQLAEIV | AWSQGGANSDYT | 0.21949749 |
| FAVSELTVM | AWSQGGANSDYT | 0.21944083 |
| GAAPLAAGL | AWSQGGANSDYT | 0.21922956 |
| GADGGDGGSGGAGG | AWSQGGANSDYT | 0.22009638 |
| GFGLDTGGAGG | AWSQGGANSDYT | 0.21984221 |
| GLASGGLL | AWSQGGANSDYT | 0.21961422 |
| GRIAGAAVLL | AWSQGGANSDYT | 0.21890946 |
| GSAGSLNTSF | AWSQGGANSDYT | 0.22024837 |
| GTGQIGI | AWSQGGANSDYT | 0.22021697 |
| GTLIALIL | AWSQGGANSDYT | 0.21968879 |
| IAATMGFVVG | AWSQGGANSDYT | 0.21947122 |
| IFAQALL | AWSQGGANSDYT | 0.21941242 |
| IGAAVALGF | AWSQGGANSDYT | 0.22015738 |
| IGLTPLAL | AWSQGGANSDYT | 0.22006516 |
| ILGSNTSTL | AWSQGGANSDYT | 0.21963234 |
| IVFSLTVL | AWSQGGANSDYT | 0.22015928 |
| LASKATGF | AWSQGGANSDYT | 0.21967472 |

| | | |
|---|---|---|
| LASVALIL | AWSQGGANSDYT | 0.21990415 |
| LFAVGLTL | AWSQGGANSDYT | 0.21998946 |
| LFNGGIGTYI | AWSQGGANSDYT | 0.21975572 |
| LTLGAGII | AWSQGGANSDYT | 0.21925575 |
| LVAAALGDI | AWSQGGANSDYT | 0.21952274 |
| NTAFVQLGI | AWSQGGANSDYT | 0.2197074 |
| QLFSGAEL | AWSQGGANSDYT | 0.21942006 |
| SAATFITL | AWSQGGANSDYT | 0.21966337 |
| SNYGLANLGV | AWSQGGANSDYT | 0.21919385 |
| SYGGAYAVM | AWSQGGANSDYT | 0.21982847 |
| TTGVGGLTLG | AWSQGGANSDYT | 0.21954824 |
| TTQTIYI | AWSQGGANSDYT | 0.22061801 |
| VAAGAGLGAVI | AWSQGGANSDYT | 0.2189148 |
| VAAVVTLL | AWSQGGANSDYT | 0.21948895 |
| VGFLGGLI | AWSQGGANSDYT | 0.21933956 |
| VGVIPILL | AWSQGGANSDYT | 0.22029331 |
| VIFASSLI | AWSQGGANSDYT | 0.21951465 |
| VKGGDGGAAAG | AWSQGGANSDYT | 0.21939075 |
| VTKTFTNI | AWSQGGANSDYT | 0.21929842 |
| VVAAAILL | AWSQGGANSDYT | 0.26462513 |
| YAALDVEL | AWSQGGANSDYT | 0.28872415 |
| LASVALLL | AWSQGGANSDYT | 0.27108413 |
| LDAVHRV | AWSQGGANSDYT | 0.28628018 |
| QGLSSSIF | AWSQGGANSDYT | 0.27927855 |
| RGTFANIRL | AWSQGGANSDYT | 0.25941643 |
| RQLWWGHRI | AWSQGGANSDYT | 0.282266 |
| YGPPGTGKTLL | AWSQGGANSDYT | 0.26700258 |
| AALVGAPANVA | AWSQGGANSDYT | 0.26423657 |
| AATFRRTE | AWSQGGANSDYT | 0.29594386 |
| AGALGGGI | AWSQGGANSDYT | 0.28021368 |
| AGGAGGAGGIGAG | AWSQGGANSDYT | 0.32596365 |
| AGSGALGL | AWSQGGANSDYT | 0.296618 |
| AQIADQI | AWSQGGANSDYT | 0.26054332 |
| ASAVAVTYL | AWSQGGANSDYT | 0.27953407 |
| ASTVLGGTGG | AWSQGGANSDYT | 0.27100918 |
| AVGGEPAK | AWSQGGANSDYT | 0.2746735 |
| CEQGTFGGTC | AWSQGGANSDYT | 0.25559857 |
| DEGKAPAG | AWSQGGANSDYT | 0.28511354 |
| DTVVFPAL | AWSQGGANSDYT | 0.2596417 |
| FEGITMAG | AWSQGGANSDYT | 0.2538799 |
| FPLVVQD | AWSQGGANSDYT | 0.29509407 |
| GAFTTPQ | AWSQGGANSDYT | 0.2797579 |
| GFGGFGTAGGVGGA | AWSQGGANSDYT | 0.27151766 |
| GFLAQFL | AWSQGGANSDYT | 0.301783 |
| GGAGGAGGNAGVA | AWSQGGANSDYT | 0.258701 |
| GGVGTAGAGGAGGAGG | AWSQGGANSDYT | 0.2669322 |
| GNGGGGDAGNAGSGGN | AWSQGGANSDYT | 0.28416067 |
| GTGGAGGAGNDAGS | AWSQGGANSDYT | 0.32880157 |
| GTTNTGWFNS | AWSQGGANSDYT | 0.2657259 |
| IFMGPLFTL | AWSQGGANSDYT | 0.27118847 |
| ILAHAIEAI | AWSQGGANSDYT | 0.31294146 |

| | | |
|---|---|---|
| IRSAIREAL | AWSQGGANSDYT | 0.28794947 |
| ITFAGGG | AWSQGGANSDYT | 0.30277395 |
| ITGMSGHQEV | AWSQGGANSDYT | 0.314064 |
| IVAGLACTL | AWSQGGANSDYT | 0.26047313 |
| IVGGGLAGA | AWSQGGANSDYT | 0.30172276 |
| IVTGVLNI | AWSQGGANSDYT | 0.29278362 |
| LAQQGCI | AWSQGGANSDYT | 0.28085917 |
| LCLHVEN | AWSQGGANSDYT | 0.28101408 |
| LDAISDGS | AWSQGGANSDYT | 0.26507956 |
| LGGFAAML | AWSQGGANSDYT | 0.26889518 |
| LGGKNPMI | AWSQGGANSDYT | 0.25649545 |
| LGPKGVAAIL | AWSQGGANSDYT | 0.27803636 |
| LMVGVSI | AWSQGGANSDYT | 0.30361733 |
| LVGAAFVL | AWSQGGANSDYT | 0.282232 |
| LVLAVAPL | AWSQGGANSDYT | 0.26071742 |
| MAAIAVIDAA | AWSQGGANSDYT | 0.2645443 |
| NNGINAELQ | AWSQGGANSDYT | 0.2604817 |
| RGEPLVLVHG | AWSQGGANSDYT | 0.2683471 |
| RGGAGGAAGLI | AWSQGGANSDYT | 0.25958604 |
| RSYRMHPAV | AWSQGGANSDYT | 0.2588803 |
| SAAHLLAGQLA | AWSQGGANSDYT | 0.30803326 |
| SAAVGSY | AWSQGGANSDYT | 0.31052095 |
| SPHAVLHRI | AWSQGGANSDYT | 0.27618405 |
| SQIAKVL | AWSQGGANSDYT | 0.2927418 |
| TALGHLVP | AWSQGGANSDYT | 0.2639756 |
| TAPTTSGAA | AWSQGGANSDYT | 0.28265917 |
| TDGDDGNGGSPGT | AWSQGGANSDYT | 0.29324248 |
| TDGLGEN | AWSQGGANSDYT | 0.2607726 |
| TGGPPLPGQA | AWSQGGANSDYT | 0.29371735 |
| TIPAITI | AWSQGGANSDYT | 0.2711527 |
| TIPGFTI | AWSQGGANSDYT | 0.26789293 |
| TSIASGALA | AWSQGGANSDYT | 0.29965228 |
| TVTLPGAAI | AWSQGGANSDYT | 0.2704135 |
| VASGLSAALP | AWSQGGANSDYT | 0.268519 |
| VGAGVIGI | AWSQGGANSDYT | 0.26122898 |
| VGPKRFGV | AWSQGGANSDYT | 0.29971406 |
| VNVAFPD | AWSQGGANSDYT | 0.26090783 |
| VSMTNSGVS | AWSQGGANSDYT | 0.2652678 |
| VTDVLGDLSV | AWSQGGANSDYT | 0.26559693 |
| VVVVGAGFAGL | AWSQGGANSDYT | 0.2892752 |
| VYTVGTVV | AWSQGGANSDYT | 0.31861353 |
| WTTFSGP | AWSQGGANSDYT | 0.2910952 |
| YDELVEAF | AWSQGGANSDYT | 0.26374936 |
| YGTGGAGGNGG | AWSQGGANSDYT | 0.29716513 |
| YTNVMAV | AWSQGGANSDYT | 0.28719103 |
| FLEQVHQL | AWSQGGANSDYT | 0.28307736 |
| GLSLLAVAV | AWSQGGANSDYT | 0.27325115 |
| LEQVHQL | AWSQGGANSDYT | 0.26022425 |
| LNAAGAI | AWSQGGANSDYT | 0.2960809 |
| PAVHASL | AWSQGGANSDYT | 0.29573756 |
| TELEDDA | AWSQGGANSDYT | 0.29961202 |

| | | |
|---|---|---|
| TPPGPPN | AWSQGGANSDYT | 0.28509158 |
| TPPPPPPP | AWSQGGANSDYT | 0.27478698 |
| TQSPALI | ASSSGTGGINTGQLY | 0.28463274 |
| EFAAAFPQ | ASSSGTGGINTGQLY | 0.27872533 |
| LLSGNLDVL | ASSSGTGGINTGQLY | 0.2649583 |
| GPANLGNYNI | ASSSGTGGINTGQLY | 0.27539873 |
| TLGPTLP | ASSSGTGGINTGQLY | 0.28479278 |
| VYAYSSCCK | ASSSGTGGINTGQLY | 0.27647164 |
| APAPAPAPNI | ASSSGTGGINTGQLY | 0.28318265 |
| PGGPVPIP | ASSSGTGGINTGQLY | 0.26694068 |
| FLTVALEP | ASSSGTGGINTGQLY | 0.28478107 |
| SQYFGANL | ASSSGTGGINTGQLY | 0.30476364 |
| GGVKPAA | ASSSGTGGINTGQLY | 0.26072043 |
| GGTGGAGGAGGW | ASSSGTGGINTGQLY | 0.26667336 |
| GVGGVAGLGGAGG | ASSSGTGGINTGQLY | 0.28999418 |
| VLLVLATTI | ASSSGTGGINTGQLY | 0.2670761 |
| IGIFAAML | ASSSGTGGINTGQLY | 0.296324 |
| AVGDLID | ASSSGTGGINTGQLY | 0.27548638 |
| NISPLPL | ASSSGTGGINTGQLY | 0.29749063 |
| VTAVFVAI | ASSSGTGGINTGQLY | 0.25941283 |
| SGDDNDAAL | ASSSGTGGINTGQLY | 0.31275937 |
| AVAALAHS | ASSSGTGGINTGQLY | 0.26389533 |
| GGVTVMATP | ASSSGTGGINTGQLY | 0.27879825 |
| AGLAANVRD | ASSSGTGGINTGQLY | 0.26765794 |
| TNRPDLIDPALLRPG | ASSSGTGGINTGQLY | 0.25619647 |
| LANLGVTAL | ASSSGTGGINTGQLY | 0.31378186 |
| TGERIEL | ASSSGTGGINTGQLY | 0.2757854 |
| NGEELAA | ASSSGTGGINTGQLY | 0.29546496 |
| VLSLVWG | ASSSGTGGINTGQLY | 0.27608478 |
| IGYALWL | ASSSGTGGINTGQLY | 0.2972216 |
| LSLLLAPL | ASSSGTGGINTGQLY | 0.26907152 |
| TLDEALK | ASSSGTGGINTGQLY | 0.27256072 |
| TDGAGTAEGG | ASSSGTGGINTGQLY | 0.268004 |
| NAPGGAGGNGG | ASSSGTGGINTGQLY | 0.30869856 |
| VSVFMGMGGNF | ASSSGTGGINTGQLY | 0.2914911 |
| LASYLAAL | ASSSGTGGINTGQLY | 0.28792763 |
| LVGGTPAL | ASSSGTGGINTGQLY | 0.26053953 |
| VRNADVL | ASSSGTGGINTGQLY | 0.2634503 |
| TCSSPFLVSAM | ASSSGTGGINTGQLY | 0.2824923 |
| VHITDAE | ASSSGTGGINTGQLY | 0.25572497 |
| EYAAPSF | ASSSGTGGINTGQLY | 0.25504705 |
| VVDAFHGGL | ASSSGTGGINTGQLY | 0.25766674 |
| ASVVDPQ | ASSSGTGGINTGQLY | 0.25878075 |
| SYNEFGGLS | ASSSGTGGINTGQLY | 0.31531635 |
| DPDGSVY | ASSSGTGGINTGQLY | 0.2975985 |
| MVSTIHATE | ASSSGTGGINTGQLY | 0.28246647 |
| LAAGVADPV | ASSSGTGGINTGQLY | 0.29306343 |
| TKEQIAA | ASSSGTGGINTGQLY | 0.2747549 |
| SLASPAAL | ASSSGTGGINTGQLY | 0.2522549 |
| IGLVFYGV | ASSSGTGGINTGQLY | 0.26776853 |
| FLLDASNL | ASSSGTGGINTGQLY | 0.26223007 |

| | | |
|---|---|---|
| YQVGNFP | ASSSGTGGINTGQLY | 0.308856 |
| ESLLSQST | ASSSGTGGINTGQLY | 0.26100746 |
| LFTVGFAAVG | ASSSGTGGINTGQLY | 0.26899225 |
| ATPPTPT | ASSSGTGGINTGQLY | 0.289587 |
| TDLATLG | ASSSGTGGINTGQLY | 0.26699913 |
| SAGGTGR | ASSSGTGGINTGQLY | 0.27994582 |
| GFSNSGS | ASSSGTGGINTGQLY | 0.2600077 |
| GGNGGSGGTSV | ASSSGTGGINTGQLY | 0.25390652 |
| GTSVATGGAGN | ASSSGTGGINTGQLY | 0.30598143 |
| NGGNGGSGGTS | ASSSGTGGINTGQLY | 0.25785294 |
| AVVKFPH | ASSSGTGGINTGQLY | 0.26572576 |
| TPPPPPP | ASSSGTGGINTGQLY | 0.2565739 |
| AAPAAAAAPA | ASSSGTGGINTGQLY | 0.26167545 |
| IALNEDL | ASSSGTGGINTGQLY | 0.25968924 |
| IVNVSSM | ASSSGTGGINTGQLY | 0.29948986 |
| TGIPPAPR | ASSSGTGGINTGQLY | 0.2653545 |
| FDLGGGTF | ASSSGTGGINTGQLY | 0.2795771 |
| FDLGGGTFDVS | ASSSGTGGINTGQLY | 0.29283407 |
| NEPTAAA | ASSSGTGGINTGQLY | 0.27025822 |
| NEPTAAALA | ASSSGTGGINTGQLY | 0.27277967 |
| GSIINISSI | ASSSGTGGINTGQLY | 0.27264732 |
| VAPGNAGTA | ASSSGTGGINTGQLY | 0.26763356 |
| TGGGASLE | ASSSGTGGINTGQLY | 0.2676333 |
| VAVYNHYKRI | ASSSGTGGINTGQLY | 0.2684464 |
| TDPAPATT | ASSSGTGGINTGQLY | 0.32628047 |
| TNRPDILDPALLRPG | ASSSGTGGINTGQLY | 0.2742617 |
| LTTTTTS | ASSSGTGGINTGQLY | 0.26302904 |
| PTGSGKTL | ASSSGTGGINTGQLY | 0.27332318 |
| TGSGKTL | ASSSGTGGINTGQLY | 0.31008 |
| TIGHVDHGK | ASSSGTGGINTGQLY | 0.29273662 |
| TIGHVDHGKTT | ASSSGTGGINTGQLY | 0.26448452 |
| GHVDHGKTT | ASSSGTGGINTGQLY | 0.27312085 |
| AQFEHTLL | ASSSGTGGINTGQLY | 0.27496505 |
| IVNQDEL | ASSSGTGGINTGQLY | 0.2755515 |
| TNGSQFF | ASSSGTGGINTGQLY | 0.27187672 |
| TGKGIVE | ASSSGTGGINTGQLY | 0.2673584 |
| LISNASDA | ASSSGTGGINTGQLY | 0.28842172 |
| AELRTLGELTKIAKAHGAQ | ASSSGTGGINTGQLY | 0.31786686 |
| APNAVAAAEPSW | ASSSGTGGINTGQLY | 0.27338973 |
| APQVAGAATAVL | ASSSGTGGINTGQLY | 0.2602333 |
| ARRDPAAAAAILPTDA | ASSSGTGGINTGQLY | 0.28898817 |
| DAAARDVLAEAGLAEYF | ASSSGTGGINTGQLY | 0.26103702 |
| DDQIVLSAKANLAKVEQ | ASSSGTGGINTGQLY | 0.274197 |
| DELLPDTGEDSDEDGH | ASSSGTGGINTGQLY | 0.27626744 |
| ELPEAPANRATVPQD | ASSSGTGGINTGQLY | 0.25813937 |
| FGQSKIERVTGIG | ASSSGTGGINTGQLY | 0.27430588 |
| GAGGTGSTAGGAGGAGGAGGLY | ASSSGTGGINTGQLY | 0.29713085 |
| GHGGTGAAGGNGGTGGAGGSGI | ASSSGTGGINTGQLY | 0.2625626 |
| GNGGNGGAGGEGGPGGAGGAGA | ASSSGTGGINTGQLY | 0.2791412 |
| GSTAGNGGNGGNATNGTIAGQP | ASSSGTGGINTGQLY | 0.30612415 |
| GTGGDGGNGGNGGTGG | ASSSGTGGINTGQLY | 0.29166862 |

| | | |
|---|---|---|
| GVRRPAGIPADAPV | ASSSGTGGINTGQLY | 0.31293827 |
| GYRQVTDAHLLTLARR | ASSSGTGGINTGQLY | 0.26534075 |
| IAKMGQDGHDRG | ASSSGTGGINTGQLY | 0.28065982 |
| KAEPSIEIQYT | ASSSGTGGINTGQLY | 0.27481166 |
| KPVRIRMAADEDTVAVT | ASSSGTGGINTGQLY | 0.2766921 |
| MPPRIAGMRLLVIKPEP | ASSSGTGGINTGQLY | 0.29648998 |
| MKFYAVALFRSRGVPPTR | ASSSGTGGINTGQLY | 0.2958125 |
| NAGGFGILGAVAHSP | ASSSGTGGINTGQLY | 0.2678762 |
| NEGREITLAMAEM | ASSSGTGGINTGQLY | 0.3086977 |
| NELVEWSHPLGHRWRWE | ASSSGTGGINTGQLY | 0.278892 |
| NLGAGGTGGNGGNPG | ASSSGTGGINTGQLY | 0.27633572 |
| NTEWPKWETVMAADPVD | ASSSGTGGINTGQLY | 0.26385298 |
| PASAAPAETQP | ASSSGTGGINTGQLY | 0.2682384 |
| QQSSDGKNGVRAAEITGE | ASSSGTGGINTGQLY | 0.2908217 |
| RAPDAVTTSWWKE | ASSSGTGGINTGQLY | 0.25966972 |
| SGFGNVGAHTSGWFNQ | ASSSGTGGINTGQLY | 0.25857714 |
| SGVAGAVGQAASVGGLKV | ASSSGTGGINTGQLY | 0.2789987 |
| SNGGAGTGSAGGNGGT | ASSSGTGGINTGQLY | 0.28746894 |
| TGGAGGEGGAAG | ASSSGTGGINTGQLY | 0.25329506 |
| TGGDGGAAGAGGAPGHG | ASSSGTGGINTGQLY | 0.2684148 |
| TMPPLLAAIANAGRHGV | ASSSGTGGINTGQLY | 0.2617761 |
| TSSTAGNSGGA | ASSSGTGGINTGQLY | 0.26965868 |
| VPATVPVGILC | ASSSGTGGINTGQLY | 0.27430183 |
| VQRPDSGGVVLTGRIS | ASSSGTGGINTGQLY | 0.25901312 |
| WTPNPVPATKGVPAGLVQAT | ASSSGTGGINTGQLY | 0.28846353 |
| YALKRALAKFSDRFDVVII | ASSSGTGGINTGQLY | 0.25964427 |
| YLIGSVIV | ASSSGTGGINTGQLY | 0.25864863 |
| YGQLVAVID | ASSSGTGGINTGQLY | 0.29596436 |
| VTVGGMTLPA | ASSSGTGGINTGQLY | 0.28222185 |
| VLALPAV | ASSSGTGGINTGQLY | 0.26610118 |
| VIPPTLNYET | ASSSGTGGINTGQLY | 0.30026615 |
| VGMGGMHPSSG | ASSSGTGGINTGQLY | 0.26283035 |
| VGLAGGAATA | ASSSGTGGINTGQLY | 0.29363093 |
| VGGAGGAGGSGP | ASSSGTGGINTGQLY | 0.2947892 |
| VFAAFMV | ASSSGTGGINTGQLY | 0.26362595 |
| TLYDKLR | ASSSGTGGINTGQLY | 0.2831988 |
| TLAVFTVL | ASSSGTGGINTGQLY | 0.27016005 |
| TGSVQPARV | ASSSGTGGINTGQLY | 0.28425008 |
| TGGGAGSGGT | ASSSGTGGINTGQLY | 0.26543963 |
| SHLETSL | ASSSGTGGINTGQLY | 0.26403835 |
| SFTTGMT | ASSSGTGGINTGQLY | 0.27893746 |
| QVEVAKRMGVRQPTVS | ASSSGTGGINTGQLY | 0.3053761 |
| QGSGGPALAM | ASSSGTGGINTGQLY | 0.28317472 |
| MLAPSNP | ASSSGTGGINTGQLY | 0.2623364 |
| LVNNAAI | ASSSGTGGINTGQLY | 0.29074442 |
| LTGEINTLA | ASSSGTGGINTGQLY | 0.27330813 |
| LSIFGPV | ASSSGTGGINTGQLY | 0.2709327 |
| LLALLANL | ASSSGTGGINTGQLY | 0.29442427 |
| LISNASDALDKLR | ASSSGTGGINTGQLY | 0.27354363 |
| LINEAVP | ASSSGTGGINTGQLY | 0.3014523 |
| LGYAFSGL | ASSSGTGGINTGQLY | 0.30141765 |

| | | |
|---|---|---|
| LDFVAVAL | ASSSGTGGINTGQLY | 0.2745621 |
| LAQPGAE | ASSSGTGGINTGQLY | 0.2643582 |
| ISNLGAVTGF | ASSSGTGGINTGQLY | 0.2668674 |
| HQEDPGRI | ASSSGTGGINTGQLY | 0.2804891 |
| GTPGEPTP | ASSSGTGGINTGQLY | 0.28826222 |
| GFFNSGS | ASSSGTGGINTGQLY | 0.26192835 |
| GAGGGGLTAGAN | ASSSGTGGINTGQLY | 0.25987542 |
| FVDALVNM | ASSSGTGGINTGQLY | 0.28454876 |
| FLTVAPEL | ASSSGTGGINTGQLY | 0.31891122 |
| FETLTAL | ASSSGTGGINTGQLY | 0.27368358 |
| DVINPIEL | ASSSGTGGINTGQLY | 0.26916793 |
| APVSGAPV | ASSSGTGGINTGQLY | 0.3246364 |
| ALVGGLAW | ASSSGTGGINTGQLY | 0.27052224 |
| ALAAAAATAI | ASSSGTGGINTGQLY | 0.2766882 |
| AFGLGLLL | ASSSGTGGINTGQLY | 0.30177185 |
| AFFAAASL | ASSSGTGGINTGQLY | 0.2972805 |
| AAAALPAFSPP | ASSSGTGGINTGQLY | 0.2760552 |
| VVLAPAL | ASSSGTGGINTGQLY | 0.30301583 |
| TIGHVDHGKTTL | ASSSGTGGINTGQLY | 0.25939065 |
| LISNASDALDK | ASSSGTGGINTGQLY | 0.29605553 |
| LISNASDAL | ASSSGTGGINTGQLY | 0.2565821 |
| ISNASDALDK | ASSSGTGGINTGQLY | 0.28330582 |
| NQAETLVYQTE | ASSSGTGGINTGQLY | 0.25881922 |
| TGGTGGTAGAG | ASSSGTGGINTGQLY | 0.28239626 |
| TGSTAGGAGGA | ASSSGTGGINTGQLY | 0.29564688 |
| DPLAVPVAAGPV | ASSSGTGGINTGQLY | 0.30066592 |
| GGTGGTPGNGGN | ASSSGTGGINTGQLY | 0.25654563 |
| PAGLAGVDPATL | ASSSGTGGINTGQLY | 0.29047897 |
| GNNLVVEAPHAGASV | ASSSGTGGINTGQLY | 0.30104762 |
| KVTRIMSASTLEQPA | ASSSGTGGINTGQLY | 0.33930284 |
| LGFALPIALPRRTVV | ASSSGTGGINTGQLY | 0.27610028 |
| TRLGNLTARKLVVLV | ASSSGTGGINTGQLY | 0.29699707 |
| WSQQRRLIYFPSAGP | ASSSGTGGINTGQLY | 0.27084926 |
| GPNGNPDPMAAAVDIR | ASSSGTGGINTGQLY | 0.27057868 |
| TQRCVSVADLRAAASR | ASSSGTGGINTGQLY | 0.26079705 |
| QFVLDVAEHPVEVLTETV | ASSSGTGGINTGQLY | 0.26470724 |
| GGAGGTGGAGGNGGTQAGDG | ASSSGTGGINTGQLY | 0.26865754 |
| PLLATLRAIAVDVDAKPAQV | ASSSGTGGINTGQLY | 0.26540613 |
| APKFRTGGYMIDFWGVGYQVAK | ASSSGTGGINTGQLY | 0.285726 |
| GAGGAGGAGGAAGGPGGTGGTG | ASSSGTGGINTGQLY | 0.2964824 |
| IGALGLLATDVAPAGSVAFGVA | ASSSGTGGINTGQLY | 0.28224242 |
| AGGAGGIGGNGGAGGAGGVLMG | ASSSGTGGINTGQLY | 0.32034907 |
| ADELIGLL | ASSSGTGGINTGQLY | 0.28996608 |
| ADRALKA | ASSSGTGGINTGQLY | 0.29646757 |
| AGAVGQVGGAGGA | ASSSGTGGINTGQLY | 0.26955846 |
| AGGGGGLGGAP | ASSSGTGGINTGQLY | 0.27177998 |
| AGGLLYGNGGAG | ASSSGTGGINTGQLY | 0.26391467 |
| AGLESIGL | ASSSGTGGINTGQLY | 0.27551666 |
| AGLKTGII | ASSSGTGGINTGQLY | 0.27075872 |
| ASLAALLL | ASSSGTGGINTGQLY | 0.29317892 |
| ASVINISSI | ASSSGTGGINTGQLY | 0.25284246 |

| | | |
|---|---|---|
| ATPPGPA | ASSSGTGGINTGQLY | 0.2624746 |
| ATVASALV | ASSSGTGGINTGQLY | 0.33216485 |
| ATVASLIL | ASSSGTGGINTGQLY | 0.2553312 |
| AVNGPVLIRH | ASSSGTGGINTGQLY | 0.2921453 |
| DGAAGAVGI | ASSSGTGGINTGQLY | 0.2754631 |
| DPLLHGI | ASSSGTGGINTGQLY | 0.26232663 |
| ELGAALGAL | ASSSGTGGINTGQLY | 0.31442243 |
| ELGRTQAV | ASSSGTGGINTGQLY | 0.26805833 |
| FVAGGYLS | ASSSGTGGINTGQLY | 0.29102048 |
| GASGGAGGAGGW | ASSSGTGGINTGQLY | 0.30629253 |
| GGAGAGGDGGAGGI | ASSSGTGGINTGQLY | 0.2832812 |
| GTGHNGSQPGMGGNG | ASSSGTGGINTGQLY | 0.26426235 |
| GYAAASASAS | ASSSGTGGINTGQLY | 0.27942994 |
| HQVDVAAVQQG | ASSSGTGGINTGQLY | 0.2624934 |
| IGAAIAHT | ASSSGTGGINTGQLY | 0.26816562 |
| IGMIEVAQL | ASSSGTGGINTGQLY | 0.2557497 |
| IRLAPEHTSAMD | ASSSGTGGINTGQLY | 0.27763766 |
| IYNVSVL | ASSSGTGGINTGQLY | 0.2701914 |
| KQGFLGM | ASSSGTGGINTGQLY | 0.3123975 |
| LESVITQL | ASSSGTGGINTGQLY | 0.27608955 |
| LGLGALVVV | ASSSGTGGINTGQLY | 0.27949974 |
| LIGNGGNGGSGGI | ASSSGTGGINTGQLY | 0.26864892 |
| LNAINAPTL | ASSSGTGGINTGQLY | 0.26436222 |
| LQLLNTGV | ASSSGTGGINTGQLY | 0.2624077 |
| LQTSIEF | ASSSGTGGINTGQLY | 0.2612852 |
| MSFAVSEL | ASSSGTGGINTGQLY | 0.27365625 |
| NTGALGL | ASSSGTGGINTGQLY | 0.27496693 |
| PPGAGGAGGAGTT | ASSSGTGGINTGQLY | 0.26021245 |
| QLVGLGI | ASSSGTGGINTGQLY | 0.30347484 |
| RFLEQVHQL | ASSSGTGGINTGQLY | 0.2823706 |
| RGVGYVLR | ASSSGTGGINTGQLY | 0.26140967 |
| SATGVFAGVI | ASSSGTGGINTGQLY | 0.26573154 |
| SLPAITLP | ASSSGTGGINTGQLY | 0.2870513 |
| STALAFTV | ASSSGTGGINTGQLY | 0.28724447 |
| SVLTVGLGK | ASSSGTGGINTGQLY | 0.2537422 |
| TALAGAGSGI | ASSSGTGGINTGQLY | 0.26791108 |
| TGLPFTGI | ASSSGTGGINTGQLY | 0.27139878 |
| VAEHVTV | ASSSGTGGINTGQLY | 0.25810298 |
| VDIVLNSL | ASSSGTGGINTGQLY | 0.26586437 |
| VLTGGGAVL | ASSSGTGGINTGQLY | 0.30734769 |
| VNVGLAYGI | ASSSGTGGINTGQLY | 0.25900567 |
| VSSAAVGAV | ASSSGTGGINTGQLY | 0.2697897 |
| VTSVYEAGV | ASSSGTGGINTGQLY | 0.27993438 |
| WVALDRGA | ASSSGTGGINTGQLY | 0.29630703 |
| YLVGSVVI | ASSSGTGGINTGQLY | 0.32526726 |
| GPPGTGKTLL | ASSSGTGGINTGQLY | 0.27732545 |
| GVGMTKF | ASSSGTGGINTGQLY | 0.27529636 |
| IGIGGSDL | ASSSGTGGINTGQLY | 0.2876433 |
| IGTLLAEL | ASSSGTGGINTGQLY | 0.29104915 |
| IYTSGTT | ASSSGTGGINTGQLY | 0.26028353 |
| LSFPAGV | ASSSGTGGINTGQLY | 0.25513333 |

| | | |
|---|---|---|
| LYTSGTT | ASSSGTGGINTGQLY | 0.2743115 |
| VGAFPFGF | ASSSGTGGINTGQLY | 0.26266593 |
| VGAGTVEF | ASSSGTGGINTGQLY | 0.30177623 |
| VGKSTLLL | ASSSGTGGINTGQLY | 0.29854125 |
| AALAHAGGGIV | ASSSGTGGINTGQLY | 0.26335207 |
| AALVAVSLL | ASSSGTGGINTGQLY | 0.26096022 |
| AAVSRAVAAL | ASSSGTGGINTGQLY | 0.27275968 |
| AGGSSGIGGF | ASSSGTGGINTGQLY | 0.26454008 |
| AGLTVANANG | ASSSGTGGINTGQLY | 0.29804 |
| AGLVGAVL | ASSSGTGGINTGQLY | 0.2697754 |
| ALSQAFTV | ASSSGTGGINTGQLY | 0.3571607 |
| ALTFSVSL | ASSSGTGGINTGQLY | 0.2909155 |
| ALTGAALVAL | ASSSGTGGINTGQLY | 0.26547545 |
| ASFSIGAGVL | ASSSGTGGINTGQLY | 0.3070473 |
| DGSVYKI | ASSSGTGGINTGQLY | 0.26881298 |
| EQLAEIV | ASSSGTGGINTGQLY | 0.26648098 |
| FAVSELTVM | ASSSGTGGINTGQLY | 0.27429062 |
| GAAPLAAGL | ASSSGTGGINTGQLY | 0.26611 |
| GADGGDGGSGGAGG | ASSSGTGGINTGQLY | 0.26824355 |
| GFGLDTGGAGG | ASSSGTGGINTGQLY | 0.2713202 |
| GLASGGLL | ASSSGTGGINTGQLY | 0.26224503 |
| GRIAGAAVLL | ASSSGTGGINTGQLY | 0.2700439 |
| GSAGSLNTSF | ASSSGTGGINTGQLY | 0.26170433 |
| GTGQIGI | ASSSGTGGINTGQLY | 0.2568956 |
| GTLIALIL | ASSSGTGGINTGQLY | 0.27310732 |
| IAATMGFVVG | ASSSGTGGINTGQLY | 0.30261803 |
| IFAQALL | ASSSGTGGINTGQLY | 0.35239542 |
| IGAAVALGF | ASSSGTGGINTGQLY | 0.26304045 |
| IGLTPLAL | ASSSGTGGINTGQLY | 0.2667956 |
| ILGSNTSTL | ASSSGTGGINTGQLY | 0.276469 |
| IVFSLTVL | ASSSGTGGINTGQLY | 0.2832976 |
| LASKATGF | ASSSGTGGINTGQLY | 0.28256938 |
| LASVALIL | ASSSGTGGINTGQLY | 0.2722408 |
| LFAVGLTL | ASSSGTGGINTGQLY | 0.2641896 |
| LFNGGIGTYI | ASSSGTGGINTGQLY | 0.26673135 |
| LTLGAGII | ASSSGTGGINTGQLY | 0.28086856 |
| LVAAALGDI | ASSSGTGGINTGQLY | 0.26103407 |
| NTAFVQLGI | ASSSGTGGINTGQLY | 0.2883178 |
| QLFSGAEL | ASSSGTGGINTGQLY | 0.2525614 |
| SAATFITL | ASSSGTGGINTGQLY | 0.2721045 |
| SNYGLANLGV | ASSSGTGGINTGQLY | 0.26998076 |
| SYGGAYAVM | ASSSGTGGINTGQLY | 0.26779887 |
| TTGVGGLTLG | ASSSGTGGINTGQLY | 0.25845516 |
| TTQTIYI | ASSSGTGGINTGQLY | 0.2771701 |
| VAAGAGLGAVI | ASSSGTGGINTGQLY | 0.27604106 |
| VAAVVTLL | ASSSGTGGINTGQLY | 0.31985137 |
| VGFLGGLI | ASSSGTGGINTGQLY | 0.2588966 |
| VGVIPILL | ASSSGTGGINTGQLY | 0.28750092 |
| VIFASSLI | ASSSGTGGINTGQLY | 0.28762645 |
| VKGGDGGAAAG | ASSSGTGGINTGQLY | 0.27565318 |
| VTKTFTNI | ASSSGTGGINTGQLY | 0.26234126 |

| | | |
|---|---|---|
| VVAAAILL | ASSSGTGGINTGQLY | 0.28364837 |
| YAALDVEL | ASSSGTGGINTGQLY | 0.25720775 |
| LASVALLL | ASSSGTGGINTGQLY | 0.27198696 |
| LDAVHRV | ASSSGTGGINTGQLY | 0.27402407 |
| QGLSSSIF | ASSSGTGGINTGQLY | 0.25863138 |
| RGTFANIRL | ASSSGTGGINTGQLY | 0.29031312 |
| RQLWWGHRI | ASSSGTGGINTGQLY | 0.29623196 |
| YGPPGTGKTLL | ASSSGTGGINTGQLY | 0.26273572 |
| AALVGAPANVA | ASSSGTGGINTGQLY | 0.26807776 |
| AATFRRTE | ASSSGTGGINTGQLY | 0.26832622 |
| AGALGGGI | ASSSGTGGINTGQLY | 0.2549599 |
| AGGAGGAGGIGAG | ASSSGTGGINTGQLY | 0.2869906 |
| AGSGALGL | ASSSGTGGINTGQLY | 0.26914033 |
| AQIADQI | ASSSGTGGINTGQLY | 0.27106103 |
| ASAVAVTYL | ASSSGTGGINTGQLY | 0.2764592 |
| ASTVLGGTGG | ASSSGTGGINTGQLY | 0.27235493 |
| AVGGEPAK | ASSSGTGGINTGQLY | 0.2749491 |
| CEQGTFGGTC | ASSSGTGGINTGQLY | 0.2773182 |
| DEGKAPAG | ASSSGTGGINTGQLY | 0.2693213 |
| DTVVFPAL | ASSSGTGGINTGQLY | 0.28183925 |
| FEGITMAG | ASSSGTGGINTGQLY | 0.26155823 |
| FPLVVQD | ASSSGTGGINTGQLY | 0.2699346 |
| GAFTTPQ | ASSSGTGGINTGQLY | 0.33572045 |
| GFGGFGTAGGVGGA | ASSSGTGGINTGQLY | 0.2582956 |
| GFLAQFL | ASSSGTGGINTGQLY | 0.26611492 |
| GGAGGAGGNAGVA | ASSSGTGGINTGQLY | 0.2743874 |
| GGVGTAGAGGAGGAGG | ASSSGTGGINTGQLY | 0.2586464 |
| GNGGGGDAGNAGSGGN | ASSSGTGGINTGQLY | 0.28381997 |
| GTGGAGGAGNDAGS | ASSSGTGGINTGQLY | 0.2772582 |
| GTTNTGWFNS | ASSSGTGGINTGQLY | 0.2926617 |
| IFMGPLFTL | ASSSGTGGINTGQLY | 0.26126352 |
| ILAHAIEAI | ASSSGTGGINTGQLY | 0.29482287 |
| IRSAIREAL | ASSSGTGGINTGQLY | 0.29627064 |
| ITFAGGG | ASSSGTGGINTGQLY | 0.27283385 |
| ITGMSGHQEV | ASSSGTGGINTGQLY | 0.26747307 |
| IVAGLACTL | ASSSGTGGINTGQLY | 0.26498154 |
| IVGGGLAGA | ASSSGTGGINTGQLY | 0.2849955 |
| IVTGVLNI | ASSSGTGGINTGQLY | 0.31031573 |
| LAQQGCI | ASSSGTGGINTGQLY | 0.2603578 |
| LCLHVEN | ASSSGTGGINTGQLY | 0.28158718 |
| LDAISDGS | ASSSGTGGINTGQLY | 0.30449817 |
| LGGFAAML | ASSSGTGGINTGQLY | 0.30428877 |
| LGGKNPMI | ASSSGTGGINTGQLY | 0.2613025 |
| LGPKGVAAIL | ASSSGTGGINTGQLY | 0.26796198 |
| LMVGVSI | ASSSGTGGINTGQLY | 0.2913609 |
| LVGAAFVL | ASSSGTGGINTGQLY | 0.27242514 |
| LVLAVAPL | ASSSGTGGINTGQLY | 0.26415905 |
| MAAIAVIDAA | ASSSGTGGINTGQLY | 0.28847772 |
| NNGINAELQ | ASSSGTGGINTGQLY | 0.29337168 |
| RGEPLVLVHG | ASSSGTGGINTGQLY | 0.27858844 |
| RGGAGGAAGLI | ASSSGTGGINTGQLY | 0.2996078 |

| | | |
|---|---|---|
| RSYRMHPAV | ASSSGTGGINTGQLY | 0.32415333 |
| SAAHLLAGQLA | ASSSGTGGINTGQLY | 0.27779132 |
| SAAVGSY | ASSSGTGGINTGQLY | 0.2769374 |
| SPHAVLHRI | ASSSGTGGINTGQLY | 0.2592965 |
| SQIAKVL | ASSSGTGGINTGQLY | 0.2583558 |
| TALGHLVP | ASSSGTGGINTGQLY | 0.26339698 |
| TAPTTSGAA | ASSSGTGGINTGQLY | 0.27172926 |
| TDGDDGNGGSPGT | ASSSGTGGINTGQLY | 0.25474074 |
| TDGLGEN | ASSSGTGGINTGQLY | 0.27918723 |
| TGGPPLPGQA | ASSSGTGGINTGQLY | 0.2814146 |
| TIPAITI | ASSSGTGGINTGQLY | 0.29931223 |
| TIPGFTI | ASSSGTGGINTGQLY | 0.27342397 |
| TSIASGALA | ASSSGTGGINTGQLY | 0.27082136 |
| TVTLPGAAI | ASSSGTGGINTGQLY | 0.26950172 |
| VASGLSAALP | ASSSGTGGINTGQLY | 0.30244887 |
| VGAGVIGI | ASSSGTGGINTGQLY | 0.26015335 |
| VGPKRFGV | ASSSGTGGINTGQLY | 0.26047394 |
| VNVAFPD | ASSSGTGGINTGQLY | 0.27086785 |
| VSMTNSGVS | ASSSGTGGINTGQLY | 0.26368538 |
| VTDVLGDLSV | ASSSGTGGINTGQLY | 0.2849889 |
| VVVVGAGFAGL | ASSSGTGGINTGQLY | 0.29648447 |
| VYTVGTVV | ASSSGTGGINTGQLY | 0.28739464 |
| WTTFSGP | ASSSGTGGINTGQLY | 0.28411952 |
| YDELVEAF | ASSSGTGGINTGQLY | 0.2555451 |
| YGTGGAGGNGG | ASSSGTGGINTGQLY | 0.2702273 |
| YTNVMAV | ASSSGTGGINTGQLY | 0.3259501 |
| FLEQVHQL | ASSSGTGGINTGQLY | 0.27860576 |
| GLSLLAVAV | ASSSGTGGINTGQLY | 0.29634252 |
| LEQVHQL | ASSSGTGGINTGQLY | 0.31676695 |
| LNAAGAI | ASSSGTGGINTGQLY | 0.26528615 |
| PAVHASL | ASSSGTGGINTGQLY | 0.28973186 |
| TELEDDA | ASSSGTGGINTGQLY | 0.27616975 |
| TPPGPPN | ASSSGTGGINTGQLY | 0.28919438 |
| TPPPPPPP | ASSSGTGGINTGQLY | 0.27204233 |
| TQSPALI | ASSDAGISDYT | 0.26103818 |
| EFAAAFPQ | ASSDAGISDYT | 0.29477313 |
| LLSGNLDVL | ASSDAGISDYT | 0.31345958 |
| GPANLGNYNI | ASSDAGISDYT | 0.28417647 |
| TLGPTLP | ASSDAGISDYT | 0.27868232 |
| VYAYSSCCK | ASSDAGISDYT | 0.28073835 |
| APAPAPAPNI | ASSDAGISDYT | 0.26048607 |
| PGGPVPIP | ASSDAGISDYT | 0.27352402 |
| FLTVALEP | ASSDAGISDYT | 0.26530248 |
| SQYFGANL | ASSDAGISDYT | 0.26527843 |
| GGVKPAA | ASSDAGISDYT | 0.26223934 |
| GGTGGAGGAGGW | ASSDAGISDYT | 0.29332376 |
| GVGGVAGLGGAGG | ASSDAGISDYT | 0.27259502 |
| VLLVLATTI | ASSDAGISDYT | 0.3307837 |
| IGIFAAML | ASSDAGISDYT | 0.28771836 |
| AVGDLID | ASSDAGISDYT | 0.28130978 |
| NISPLPL | ASSDAGISDYT | 0.2648013 |

| | | |
|---|---|---|
| VTAVFVAI | ASSDAGISDYT | 0.2716672 |
| SGDDNDAAL | ASSDAGISDYT | 0.27560377 |
| AVAALAHS | ASSDAGISDYT | 0.2628062 |
| GGVTVMATP | ASSDAGISDYT | 0.32673898 |
| AGLAANVRD | ASSDAGISDYT | 0.2737715 |
| TNRPDLIDPALLRPG | ASSDAGISDYT | 0.25997674 |
| LANLGVTAL | ASSDAGISDYT | 0.28956464 |
| TGERIEL | ASSDAGISDYT | 0.3218649 |
| NGEELAA | ASSDAGISDYT | 0.28618646 |
| VLSLVWG | ASSDAGISDYT | 0.26399672 |
| IGYALWL | ASSDAGISDYT | 0.26435906 |
| LSLLLAPL | ASSDAGISDYT | 0.2917882 |
| TLDEALK | ASSDAGISDYT | 0.34055474 |
| TDGAGTAEGG | ASSDAGISDYT | 0.25882038 |
| NAPGGAGGNGG | ASSDAGISDYT | 0.29213804 |
| VSVFMGMGGNF | ASSDAGISDYT | 0.26767766 |
| LASYLAAL | ASSDAGISDYT | 0.2614151 |
| LVGGTPAL | ASSDAGISDYT | 0.28428203 |
| VRNADVL | ASSDAGISDYT | 0.28433764 |
| TCSSPFLVSAM | ASSDAGISDYT | 0.29533765 |
| VHITDAE | ASSDAGISDYT | 0.29884517 |
| EYAAPSF | ASSDAGISDYT | 0.26279786 |
| VVDAFHGGL | ASSDAGISDYT | 0.26583135 |
| ASVVDPQ | ASSDAGISDYT | 0.30195925 |
| SYNEFGGLS | ASSDAGISDYT | 0.29500157 |
| DPDGSVY | ASSDAGISDYT | 0.29048732 |
| MVSTIHATE | ASSDAGISDYT | 0.26864228 |
| LAAGVADPV | ASSDAGISDYT | 0.26817954 |
| TKEQIAA | ASSDAGISDYT | 0.27974835 |
| SLASPAAL | ASSDAGISDYT | 0.27654108 |
| IGLVFYGV | ASSDAGISDYT | 0.27354112 |
| FLLDASNL | ASSDAGISDYT | 0.28783694 |
| YQVGNFP | ASSDAGISDYT | 0.2983454 |
| ESLLSQST | ASSDAGISDYT | 0.25996754 |
| LFTVGFAAVG | ASSDAGISDYT | 0.27189997 |
| ATPPTPT | ASSDAGISDYT | 0.2841536 |
| TDLATLG | ASSDAGISDYT | 0.2747223 |
| SAGGTGR | ASSDAGISDYT | 0.26298234 |
| GFSNSGS | ASSDAGISDYT | 0.32186833 |
| GGNGGSGGTSV | ASSDAGISDYT | 0.33079994 |
| GTSVATGGAGN | ASSDAGISDYT | 0.29563272 |
| NGGNGGSGGTS | ASSDAGISDYT | 0.26195768 |
| AVVKFPH | ASSDAGISDYT | 0.25642827 |
| TPPPPPP | ASSDAGISDYT | 0.32068866 |
| AAPAAAAAPA | ASSDAGISDYT | 0.27340057 |
| IALNEDL | ASSDAGISDYT | 0.27221915 |
| IVNVSSM | ASSDAGISDYT | 0.26431724 |
| TGIPPAPR | ASSDAGISDYT | 0.26860067 |
| FDLGGGTF | ASSDAGISDYT | 0.295884 |
| FDLGGGTFDVS | ASSDAGISDYT | 0.26883978 |
| NEPTAAA | ASSDAGISDYT | 0.2618159 |

| | | |
|---|---|---|
| NEPTAAALA | ASSDAGISDYT | 0.2713392 |
| GSIINISSI | ASSDAGISDYT | 0.31303793 |
| VAPGNAGTA | ASSDAGISDYT | 0.26092538 |
| TGGGASLE | ASSDAGISDYT | 0.27164617 |
| VAVYNHYKRI | ASSDAGISDYT | 0.3230024 |
| TDPAPATT | ASSDAGISDYT | 0.26666853 |
| TNRPDILDPALLRPG | ASSDAGISDYT | 0.27396655 |
| LTTTTTS | ASSDAGISDYT | 0.2683692 |
| PTGSGKTL | ASSDAGISDYT | 0.2742391 |
| TGSGKTL | ASSDAGISDYT | 0.2713062 |
| TIGHVDHGK | ASSDAGISDYT | 0.28403586 |
| TIGHVDHGKTT | ASSDAGISDYT | 0.28377092 |
| GHVDHGKTT | ASSDAGISDYT | 0.28531653 |
| AQFEHTLL | ASSDAGISDYT | 0.29031643 |
| IVNQDEL | ASSDAGISDYT | 0.26876733 |
| TNGSQFF | ASSDAGISDYT | 0.26205704 |
| TGKGIVE | ASSDAGISDYT | 0.25487638 |
| LISNASDA | ASSDAGISDYT | 0.29546136 |
| AELRTLGELTKIAKAHGAQ | ASSDAGISDYT | 0.25658286 |
| APNAVAAAEPSW | ASSDAGISDYT | 0.30031568 |
| APQVAGAATAVL | ASSDAGISDYT | 0.30643117 |
| ARRDPAAAAAILPTDA | ASSDAGISDYT | 0.26603952 |
| DAAARDVLAEAGLAEYF | ASSDAGISDYT | 0.28013843 |
| DDQIVLSAKANLAKVEQ | ASSDAGISDYT | 0.25732207 |
| DELLPDTGEDSDEDGH | ASSDAGISDYT | 0.26311892 |
| ELPEAPANRATVPQD | ASSDAGISDYT | 0.27029055 |
| FGQSKIERVTGIG | ASSDAGISDYT | 0.27761132 |
| GAGGTGSTAGGAGGAGGAGGLY | ASSDAGISDYT | 0.2796664 |
| GHGGTGAAGGNGGTGGAGGSGI | ASSDAGISDYT | 0.29233122 |
| GNGGNGGAGGEGGPGGAGGAGA | ASSDAGISDYT | 0.28393966 |
| GSTAGNGGNGGNATNGTIAGQP | ASSDAGISDYT | 0.2677165 |
| GTGGDGGNGGNGGTGG | ASSDAGISDYT | 0.27534142 |
| GVRRPAGIPADAPV | ASSDAGISDYT | 0.30944777 |
| GYRQVTDAHLLTLARR | ASSDAGISDYT | 0.292136 |
| IAKMGQDGHDRG | ASSDAGISDYT | 0.26453277 |
| KAEPSIEIQYT | ASSDAGISDYT | 0.2824026 |
| KPVRIRMAADEDTVAVT | ASSDAGISDYT | 0.2630271 |
| MPPRIAGMRLLVIKPEP | ASSDAGISDYT | 0.27846715 |
| MKFYAVALFRSRGVPPTR | ASSDAGISDYT | 0.27541727 |
| NAGGFGILGAVAHSP | ASSDAGISDYT | 0.2887241 |
| NEGREITLAMAEM | ASSDAGISDYT | 0.2758721 |
| NELVEWSHPLGHRWRWE | ASSDAGISDYT | 0.25880063 |
| NLGAGGTGGNGGNPG | ASSDAGISDYT | 0.26314795 |
| NTEWPKWETVMAADPVD | ASSDAGISDYT | 0.27443382 |
| PASAAPAETQP | ASSDAGISDYT | 0.2901935 |
| QQSSDGKNGVRAAEITGE | ASSDAGISDYT | 0.28762674 |
| RAPDAVTTSWWKE | ASSDAGISDYT | 0.26237363 |
| SGFGNVGAHTSGWFNQ | ASSDAGISDYT | 0.28526253 |
| SGVAGAVGQAASVGGLKV | ASSDAGISDYT | 0.2824625 |
| SNGGAGTGSAGGNGGT | ASSDAGISDYT | 0.27523723 |
| TGGAGGEGGAAG | ASSDAGISDYT | 0.287688 |

| | | |
|---|---|---|
| TGGDGGAAGAGGAPGHG | ASSDAGISDYT | 0.27344218 |
| TMPPLLAAIANAGRHGV | ASSDAGISDYT | 0.2658935 |
| TSSTAGNSGGA | ASSDAGISDYT | 0.28371185 |
| VPATVPVGILC | ASSDAGISDYT | 0.26452604 |
| VQRPDSGGVVLTGRIS | ASSDAGISDYT | 0.28578925 |
| WTPNPVPATKGVPAGLVQAT | ASSDAGISDYT | 0.27090377 |
| YALKRALAKFSDRFDVVII | ASSDAGISDYT | 0.26278543 |
| YLIGSVIV | ASSDAGISDYT | 0.2852021 |
| YGQLVAVID | ASSDAGISDYT | 0.26689163 |
| VTVGGMTLPA | ASSDAGISDYT | 0.28010488 |
| VLALPAV | ASSDAGISDYT | 0.28236315 |
| VIPPTLNYET | ASSDAGISDYT | 0.2680771 |
| VGMGGMHPSSG | ASSDAGISDYT | 0.2821807 |
| VGLAGGAATA | ASSDAGISDYT | 0.2948511 |
| VGGAGGAGGSGP | ASSDAGISDYT | 0.28628698 |
| VFAAFMV | ASSDAGISDYT | 0.26566777 |
| TLYDKLR | ASSDAGISDYT | 0.2738801 |
| TLAVFTVL | ASSDAGISDYT | 0.2913674 |
| TGSVQPARV | ASSDAGISDYT | 0.26743808 |
| TGGGAGSGGT | ASSDAGISDYT | 0.25810418 |
| SHLETSL | ASSDAGISDYT | 0.27338114 |
| SFTTGMT | ASSDAGISDYT | 0.26534557 |
| QVEVAKRMGVRQPTVS | ASSDAGISDYT | 0.27494276 |
| QGSGGPALAM | ASSDAGISDYT | 0.26059556 |
| MLAPSNP | ASSDAGISDYT | 0.29849923 |
| LVNNAAI | ASSDAGISDYT | 0.32282832 |
| LTGEINTLA | ASSDAGISDYT | 0.2719003 |
| LSIFGPV | ASSDAGISDYT | 0.2859083 |
| LLALLANL | ASSDAGISDYT | 0.25990057 |
| LISNASDALDKLR | ASSDAGISDYT | 0.29099998 |
| LINEAVP | ASSDAGISDYT | 0.3233137 |
| LGYAFSGL | ASSDAGISDYT | 0.2640346 |
| LDFVAVAL | ASSDAGISDYT | 0.2924785 |
| LAQPGAE | ASSDAGISDYT | 0.27001065 |
| ISNLGAVTGF | ASSDAGISDYT | 0.26762888 |
| HQEDPGRI | ASSDAGISDYT | 0.31996194 |
| GTPGEPTP | ASSDAGISDYT | 0.27595818 |
| GFFNSGS | ASSDAGISDYT | 0.2818785 |
| GAGGGGLTAGAN | ASSDAGISDYT | 0.27140048 |
| FVDALVNM | ASSDAGISDYT | 0.27131614 |
| FLTVAPEL | ASSDAGISDYT | 0.2794395 |
| FETLTAL | ASSDAGISDYT | 0.312603 |
| DVINPIEL | ASSDAGISDYT | 0.2625491 |
| APVSGAPV | ASSDAGISDYT | 0.29843706 |
| ALVGGLAW | ASSDAGISDYT | 0.27239197 |
| ALAAAAATAI | ASSDAGISDYT | 0.29048145 |
| AFGLGLLL | ASSDAGISDYT | 0.29977795 |
| AFFAAASL | ASSDAGISDYT | 0.28028017 |
| AAAALPAFSPP | ASSDAGISDYT | 0.2698121 |
| VVLAPAL | ASSDAGISDYT | 0.32348344 |
| TIGHVDHGKTTL | ASSDAGISDYT | 0.28719792 |

| | | |
|---|---|---|
| LISNASDALDK | ASSDAGISDYT | 0.27762404 |
| LISNASDAL | ASSDAGISDYT | 0.3033837 |
| ISNASDALDK | ASSDAGISDYT | 0.14028865 |
| NQAETLVYQTE | ASSDAGISDYT | 0.1604367 |
| TGGTGGTAGAG | ASSDAGISDYT | 0.14013039 |
| TGSTAGGAGGA | ASSDAGISDYT | 0.15593933 |
| DPLAVPVAAGPV | ASSDAGISDYT | 0.1440945 |
| GGTGGTPGNGGN | ASSDAGISDYT | 0.12968953 |
| PAGLAGVDPATL | ASSDAGISDYT | 0.16011682 |
| GNNLVVEAPHAGASV | ASSDAGISDYT | 0.13492613 |
| KVTRIMSASTLEQPA | ASSDAGISDYT | 0.1483589 |
| LGFALPIALPRRTVV | ASSDAGISDYT | 0.15784861 |
| TRLGNLTARKLVVLV | ASSDAGISDYT | 0.14895095 |
| WSQQRRLIYFPSAGP | ASSDAGISDYT | 0.21606767 |
| GPNGNPDPMAAAVDIR | ASSDAGISDYT | 0.18224172 |
| TQRCVSVADLRAAASR | ASSDAGISDYT | 0.13255066 |
| QFVLDVAEHPVEVLTETV | ASSDAGISDYT | 0.14537618 |
| GGAGGTGGAGGNGGTQAGDG | ASSDAGISDYT | 0.1376259 |
| PLLATLRAIAVDVDAKPAQV | ASSDAGISDYT | 0.14652497 |
| APKFRTGGYMIDFWGVGYQVAK | ASSDAGISDYT | 0.12897132 |
| GAGGAGGAGGAAGGPGGTGGTG | ASSDAGISDYT | 0.1594725 |
| IGALGLLATDVAPAGSVAFGVA | ASSDAGISDYT | 0.13264835 |
| AGGAGGIGGNGGAGGAGGVLMG | ASSDAGISDYT | 0.12688692 |
| ADELIGLL | ASSDAGISDYT | 0.15592319 |
| ADRALKA | ASSDAGISDYT | 0.14695129 |
| AGAVGQVGGAGGA | ASSDAGISDYT | 0.14003111 |
| AGGGGGLGGAP | ASSDAGISDYT | 0.17207159 |
| AGGLLYGNGGAG | ASSDAGISDYT | 0.12960841 |
| AGLESIGL | ASSDAGISDYT | 0.1346452 |
| AGLKTGII | ASSDAGISDYT | 0.16979107 |
| ASLAALLL | ASSDAGISDYT | 0.20440288 |
| ASVINISSI | ASSDAGISDYT | 0.1343874 |
| ATPPGPA | ASSDAGISDYT | 0.1490934 |
| ATVASALV | ASSDAGISDYT | 0.18423665 |
| ATVASLIL | ASSDAGISDYT | 0.1490791 |
| AVNGPVLIRH | ASSDAGISDYT | 0.1609097 |
| DGAAGAVGI | ASSDAGISDYT | 0.17026441 |
| DPLLHGI | ASSDAGISDYT | 0.13693619 |
| ELGAALGAL | ASSDAGISDYT | 0.18762071 |
| ELGRTQAV | ASSDAGISDYT | 0.17928544 |
| FVAGGYLS | ASSDAGISDYT | 0.14668353 |
| GASGGAGGAGGW | ASSDAGISDYT | 0.14986989 |
| GGAGAGGDGGAGGI | ASSDAGISDYT | 0.13785225 |
| GTGHNGSQPGMGGNG | ASSDAGISDYT | 0.14359565 |
| GYAAASASAS | ASSDAGISDYT | 0.12809514 |
| HQVDVAAVQQG | ASSDAGISDYT | 0.14857215 |
| IGAAIAHT | ASSDAGISDYT | 0.1695717 |
| IGMIEVAQL | ASSDAGISDYT | 0.1449018 |
| IRLAPEHTSAMD | ASSDAGISDYT | 0.13275848 |
| IYNVSVL | ASSDAGISDYT | 0.14063016 |
| KQGFLGM | ASSDAGISDYT | 0.1332832 |

| | | |
|---|---|---|
| LESVITQL | ASSDAGISDYT | 0.1405809 |
| LGLGALVVV | ASSDAGISDYT | 0.1283158 |
| LIGNGGNGGSGGI | ASSDAGISDYT | 0.1321908 |
| LNAINAPTL | ASSDAGISDYT | 0.18382625 |
| LQLLNTGV | ASSDAGISDYT | 0.17222092 |
| LQTSIEF | ASSDAGISDYT | 0.13439941 |
| MSFAVSEL | ASSDAGISDYT | 0.17178224 |
| NTGALGL | ASSDAGISDYT | 0.17560393 |
| PPGAGGAGGAGTT | ASSDAGISDYT | 0.1530177 |
| QLVGLGI | ASSDAGISDYT | 0.14533809 |
| RFLEQVHQL | ASSDAGISDYT | 0.13621211 |
| RGVGYVLR | ASSDAGISDYT | 0.23951986 |
| SATGVFAGVI | ASSDAGISDYT | 0.14234649 |
| SLPAITLP | ASSDAGISDYT | 0.13473757 |
| STALAFTV | ASSDAGISDYT | 0.16965772 |
| SVLTVGLGK | ASSDAGISDYT | 0.13782297 |
| TALAGAGSGI | ASSDAGISDYT | 0.13660811 |
| TGLPFTGI | ASSDAGISDYT | 0.1543103 |
| VAEHVTV | ASSDAGISDYT | 0.16562541 |
| VDIVLNSL | ASSDAGISDYT | 0.13252962 |
| VLTGGGAVL | ASSDAGISDYT | 0.13617799 |
| VNVGLAYGI | ASSDAGISDYT | 0.13456455 |
| VSSAAVGAV | ASSDAGISDYT | 0.15579058 |
| VTSVYEAGV | ASSDAGISDYT | 0.19667949 |
| WVALDRGA | ASSDAGISDYT | 0.15321411 |
| YLVGSVVI | ASSDAGISDYT | 0.13996494 |
| GPPGTGKTLL | ASSDAGISDYT | 0.1862224 |
| GVGMTKF | ASSDAGISDYT | 0.15020047 |
| IGIGGSDL | ASSDAGISDYT | 0.1515207 |
| IGTLLAEL | ASSDAGISDYT | 0.15809454 |
| IYTSGTT | ASSDAGISDYT | 0.13393821 |
| LSFPAGV | ASSDAGISDYT | 0.16329579 |
| LYTSGTT | ASSDAGISDYT | 0.15759474 |
| VGAFPFGF | ASSDAGISDYT | 0.16558811 |
| VGAGTVEF | ASSDAGISDYT | 0.16658998 |
| VGKSTLLL | ASSDAGISDYT | 0.14442883 |
| AALAHAGGGIV | ASSDAGISDYT | 0.15682688 |
| AALVAVSLL | ASSDAGISDYT | 0.13089442 |
| AAVSRAVAAL | ASSDAGISDYT | 0.13770938 |
| AGGSSGIGGF | ASSDAGISDYT | 0.16212963 |
| AGLTVANANG | ASSDAGISDYT | 0.14582841 |
| AGLVGAVL | ASSDAGISDYT | 0.16836748 |
| ALSQAFTV | ASSDAGISDYT | 0.16031432 |
| ALTFSVSL | ASSDAGISDYT | 0.13604146 |
| ALTGAALVAL | ASSDAGISDYT | 0.169447 |
| ASFSIGAGVL | ASSDAGISDYT | 0.19345589 |
| DGSVYKI | ASSDAGISDYT | 0.132008 |
| EQLAEIV | ASSDAGISDYT | 0.13786075 |
| FAVSELTVM | ASSDAGISDYT | 0.17884809 |
| GAAPLAAGL | ASSDAGISDYT | 0.16013166 |
| GADGGDGGSGGAGG | ASSDAGISDYT | 0.15657921 |

| | | |
|---|---|---|
| GFGLDTGGAGG | ASSDAGISDYT | 0.14394583 |
| GLASGGLL | ASSDAGISDYT | 0.20236807 |
| GRIAGAAVLL | ASSDAGISDYT | 0.13350253 |
| GSAGSLNTSF | ASSDAGISDYT | 0.19232301 |
| GTGQIGI | ASSDAGISDYT | 0.12879068 |
| GTLIALIL | ASSDAGISDYT | 0.15103394 |
| IAATMGFVVG | ASSDAGISDYT | 0.13512756 |
| IFAQALL | ASSDAGISDYT | 0.1290329 |
| IGAAVALGF | ASSDAGISDYT | 0.15445675 |
| IGLTPLAL | ASSDAGISDYT | 0.15674399 |
| ILGSNTSTL | ASSDAGISDYT | 0.17192616 |
| IVFSLTVL | ASSDAGISDYT | 0.14540604 |
| LASKATGF | ASSDAGISDYT | 0.15832222 |
| LASVALIL | ASSDAGISDYT | 0.14189383 |
| LFAVGLTL | ASSDAGISDYT | 0.15240617 |
| LFNGGIGTYI | ASSDAGISDYT | 0.13888381 |
| LTLGAGII | ASSDAGISDYT | 0.15439369 |
| LVAAALGDI | ASSDAGISDYT | 0.17849354 |
| NTAFVQLGI | ASSDAGISDYT | 0.16190377 |
| QLFSGAEL | ASSDAGISDYT | 0.13174017 |
| SAATFITL | ASSDAGISDYT | 0.13322587 |
| SNYGLANLGV | ASSDAGISDYT | 0.14803217 |
| SYGGAYAVM | ASSDAGISDYT | 0.12855372 |
| TTGVGGLTLG | ASSDAGISDYT | 0.12749016 |
| TTQTIYI | ASSDAGISDYT | 0.13002157 |
| VAAGAGLGAVI | ASSDAGISDYT | 0.1307099 |
| VAAVVTLL | ASSDAGISDYT | 0.19302619 |
| VGFLGGLI | ASSDAGISDYT | 0.18386334 |
| VGVIPILL | ASSDAGISDYT | 0.14742467 |
| VIFASSLI | ASSDAGISDYT | 0.16116926 |
| VKGGDGGAAAG | ASSDAGISDYT | 0.14940283 |
| VTKTFTNI | ASSDAGISDYT | 0.12580036 |
| VVAAAILL | ASSDAGISDYT | 0.13837187 |
| YAALDVEL | ASSDAGISDYT | 0.1356444 |
| LASVALLL | ASSDAGISDYT | 0.18194526 |
| LDAVHRV | ASSDAGISDYT | 0.134039 |
| QGLSSSIF | ASSDAGISDYT | 0.13518071 |
| RGTFANIRL | ASSDAGISDYT | 0.15657179 |
| RQLWWGHRI | ASSDAGISDYT | 0.14341666 |
| YGPPGTGKTLL | ASSDAGISDYT | 0.14191426 |
| AALVGAPANVA | ASSDAGISDYT | 0.13192862 |
| AATFRRTE | ASSDAGISDYT | 0.12648751 |
| AGALGGGI | ASSDAGISDYT | 0.16478072 |
| AGGAGGAGGIGAG | ASSDAGISDYT | 0.1295546 |
| AGSGALGL | ASSDAGISDYT | 0.13720332 |
| AQIADQI | ASSDAGISDYT | 0.1291639 |
| ASAVAVTYL | ASSDAGISDYT | 0.12776802 |
| ASTVLGGTGG | ASSDAGISDYT | 0.13026138 |
| AVGGEPAK | ASSDAGISDYT | 0.162094 |
| CEQGTFGGTC | ASSDAGISDYT | 0.1459274 |
| DEGKAPAG | ASSDAGISDYT | 0.14536572 |

| | | |
|---|---|---|
| DTVVFPAL | ASSDAGISDYT | 0.18615797 |
| FEGITMAG | ASSDAGISDYT | 0.27495795 |
| FPLVVQD | ASSDAGISDYT | 0.14335461 |
| GAFTTPQ | ASSDAGISDYT | 0.14063591 |
| GFGGFGTAGGVGGA | ASSDAGISDYT | 0.13755281 |
| GFLAQFL | ASSDAGISDYT | 0.13440165 |
| GGAGGAGGNAGVA | ASSDAGISDYT | 0.13734245 |
| GGVGTAGAGGAGGAGG | ASSDAGISDYT | 0.20151943 |
| GNGGGGDAGNAGSGGN | ASSDAGISDYT | 0.14296907 |
| GTGGAGGAGNDAGS | ASSDAGISDYT | 0.13479091 |
| GTTNTGWFNS | ASSDAGISDYT | 0.15537632 |
| IFMGPLFTL | ASSDAGISDYT | 0.18728517 |
| ILAHAIEAI | ASSDAGISDYT | 0.17653777 |
| IRSAIREAL | ASSDAGISDYT | 0.1387417 |
| ITFAGGG | ASSDAGISDYT | 0.14199737 |
| ITGMSGHQEV | ASSDAGISDYT | 0.14897773 |
| IVAGLACTL | ASSDAGISDYT | 0.14877501 |
| IVGGGLAGA | ASSDAGISDYT | 0.14044312 |
| IVTGVLNI | ASSDAGISDYT | 0.13532172 |
| LAQQGCI | ASSDAGISDYT | 0.15904307 |
| LCLHVEN | ASSDAGISDYT | 0.4694063 |
| LDAISDGS | ASSDAGISDYT | 0.14361262 |
| LGGFAAML | ASSDAGISDYT | 0.12998669 |
| LGGKNPMI | ASSDAGISDYT | 0.15727529 |
| LGPKGVAAIL | ASSDAGISDYT | 0.13325718 |
| LMVGVSI | ASSDAGISDYT | 0.1446742 |
| LVGAAFVL | ASSDAGISDYT | 0.15293398 |
| LVLAVAPL | ASSDAGISDYT | 0.13046284 |
| MAAIAVIDAA | ASSDAGISDYT | 0.14072691 |
| NNGINAELQ | ASSDAGISDYT | 0.1785637 |
| RGEPLVLVHG | ASSDAGISDYT | 0.14216606 |
| RGGAGGAAGLI | ASSDAGISDYT | 0.1725387 |
| RSYRMHPAV | ASSDAGISDYT | 0.1873068 |
| SAAHLLAGQLA | ASSDAGISDYT | 0.15757084 |
| SAAVGSY | ASSDAGISDYT | 0.18380508 |
| SPHAVLHRI | ASSDAGISDYT | 0.17274442 |
| SQIAKVL | ASSDAGISDYT | 0.14808017 |
| TALGHLVP | ASSDAGISDYT | 0.14443412 |
| TAPTTSGAA | ASSDAGISDYT | 0.16401003 |
| TDGDDGNGGSPGT | ASSDAGISDYT | 0.1619033 |
| TDGLGEN | ASSDAGISDYT | 0.15732948 |
| TGGPPLPGQA | ASSDAGISDYT | 0.13592084 |
| TIPAITI | ASSDAGISDYT | 0.20600937 |
| TIPGFTI | ASSDAGISDYT | 0.13844393 |
| TSIASGALA | ASSDAGISDYT | 0.177171 |
| TVTLPGAAI | ASSDAGISDYT | 0.13420022 |
| VASGLSAALP | ASSDAGISDYT | 0.13455643 |
| VGAGVIGI | ASSDAGISDYT | 0.1551236 |
| VGPKRFGV | ASSDAGISDYT | 0.12676738 |
| VNVAFPD | ASSDAGISDYT | 0.12881298 |
| VSMTNSGVS | ASSDAGISDYT | 0.15838544 |

| | | |
|---|---|---|
| VTDVLGDLSV | ASSDAGISDYT | 0.157773 |
| VVVVGAGFAGL | ASSDAGISDYT | 0.1261003 |
| VYTVGTVV | ASSDAGISDYT | 0.13683738 |
| WTTFSGP | ASSDAGISDYT | 0.13325989 |
| YDELVEAF | ASSDAGISDYT | 0.14262922 |
| YGTGGAGGNGG | ASSDAGISDYT | 0.14951125 |
| YTNVMAV | ASSDAGISDYT | 0.13065125 |
| FLEQVHQL | ASSDAGISDYT | 0.15579158 |
| GLSLLAVAV | ASSDAGISDYT | 0.12994836 |
| LEQVHQL | ASSDAGISDYT | 0.1317846 |
| LNAAGAI | ASSDAGISDYT | 0.15292215 |
| PAVHASL | ASSDAGISDYT | 0.21367972 |
| TELEDDA | ASSDAGISDYT | 0.13676909 |
| TPPGPPN | ASSDAGISDYT | 0.1605235 |
| TPPPPPPP | ASSDAGISDYT | 0.13481644 |
| TQSPALI | ASSQERGPGNTLY | 0.19082712 |
| EFAAAFPQ | ASSQERGPGNTLY | 0.15171084 |
| LLSGNLDVL | ASSQERGPGNTLY | 0.13141237 |
| GPANLGNYNI | ASSQERGPGNTLY | 0.16253474 |
| TLGPTLP | ASSQERGPGNTLY | 0.14698291 |
| VYAYSSCCK | ASSQERGPGNTLY | 0.15682185 |
| APAPAPAPNI | ASSQERGPGNTLY | 0.16666226 |
| PGGPVPIP | ASSQERGPGNTLY | 0.13531162 |
| FLTVALEP | ASSQERGPGNTLY | 0.15876408 |
| SQYFGANL | ASSQERGPGNTLY | 0.19001414 |
| GGVKPAA | ASSQERGPGNTLY | 0.16888672 |
| GGTGGAGGAGGW | ASSQERGPGNTLY | 0.13332012 |
| GVGGVAGLGGAGG | ASSQERGPGNTLY | 0.16665867 |
| VLLVLATTI | ASSQERGPGNTLY | 0.14592825 |
| IGIFAAML | ASSQERGPGNTLY | 0.14809361 |
| AVGDLID | ASSQERGPGNTLY | 0.1537779 |
| NISPLPL | ASSQERGPGNTLY | 0.14877915 |
| VTAVFVAI | ASSQERGPGNTLY | 0.16605368 |
| SGDDNDAAL | ASSQERGPGNTLY | 0.1547524 |
| AVAALAHS | ASSQERGPGNTLY | 0.20339438 |
| GGVTVMATP | ASSQERGPGNTLY | 0.13027278 |
| AGLAANVRD | ASSQERGPGNTLY | 0.15570244 |
| TNRPDLIDPALLRPG | ASSQERGPGNTLY | 0.15310143 |
| LANLGVTAL | ASSQERGPGNTLY | 0.17056507 |
| TGERIEL | ASSQERGPGNTLY | 0.13957705 |
| NGEELAA | ASSQERGPGNTLY | 0.13050988 |
| VLSLVWG | ASSQERGPGNTLY | 0.15628886 |
| IGYALWL | ASSQERGPGNTLY | 0.17583264 |
| LSLLLAPL | ASSQERGPGNTLY | 0.14618504 |
| TLDEALK | ASSQERGPGNTLY | 0.14722067 |
| TDGAGTAEGG | ASSQERGPGNTLY | 0.21609639 |
| NAPGGAGGNGG | ASSQERGPGNTLY | 0.14954121 |
| VSVFMGMGGNF | ASSQERGPGNTLY | 0.14718056 |
| LASYLAAL | ASSQERGPGNTLY | 0.15705651 |
| LVGGTPAL | ASSQERGPGNTLY | 0.18019417 |
| VRNADVL | ASSQERGPGNTLY | 0.14494444 |

| | | |
|---|---|---|
| TCSSPFLVSAM | ASSQERGPGNTLY | 0.1861922 |
| VHITDAE | ASSQERGPGNTLY | 0.13171189 |
| EYAAPSF | ASSQERGPGNTLY | 0.21346422 |
| VVDAFHGGL | ASSQERGPGNTLY | 0.13256764 |
| ASVVDPQ | ASSQERGPGNTLY | 0.15342745 |
| SYNEFGGLS | ASSQERGPGNTLY | 0.12949552 |
| DPDGSVY | ASSQERGPGNTLY | 0.17271379 |
| MVSTIHATE | ASSQERGPGNTLY | 0.16648178 |
| LAAGVADPV | ASSQERGPGNTLY | 0.19538534 |
| TKEQIAA | ASSQERGPGNTLY | 0.1273393 |
| SLASPAAL | ASSQERGPGNTLY | 0.17139408 |
| IGLVFYGV | ASSQERGPGNTLY | 0.1952787 |
| FLLDASNL | ASSQERGPGNTLY | 0.46437585 |
| YQVGNFP | ASSQERGPGNTLY | 0.14056532 |
| ESLLSQST | ASSQERGPGNTLY | 0.17374031 |
| LFTVGFAAVG | ASSQERGPGNTLY | 0.14606887 |
| ATPPTPT | ASSQERGPGNTLY | 0.1383211 |
| TDLATLG | ASSQERGPGNTLY | 0.132074 |
| SAGGTGR | ASSQERGPGNTLY | 0.14800332 |
| GFSNSGS | ASSQERGPGNTLY | 0.1417103 |
| GGNGGSGGTSV | ASSQERGPGNTLY | 0.14004321 |
| GTSVATGGAGN | ASSQERGPGNTLY | 0.16579577 |
| NGGNGGSGGTS | ASSQERGPGNTLY | 0.17359936 |
| AVVKFPH | ASSQERGPGNTLY | 0.1615716 |
| TPPPPPP | ASSQERGPGNTLY | 0.20893806 |
| AAPAAAAAPA | ASSQERGPGNTLY | 0.17959319 |
| IALNEDL | ASSQERGPGNTLY | 0.16260241 |
| IVNVSSM | ASSQERGPGNTLY | 0.135784 |
| TGIPPAPR | ASSQERGPGNTLY | 0.14322716 |
| FDLGGGTF | ASSQERGPGNTLY | 0.13258885 |
| FDLGGGTFDVS | ASSQERGPGNTLY | 0.14424025 |
| NEPTAAA | ASSQERGPGNTLY | 0.13652991 |
| NEPTAAALA | ASSQERGPGNTLY | 0.15916975 |
| GSIINISSI | ASSQERGPGNTLY | 0.12583958 |
| VAPGNAGTA | ASSQERGPGNTLY | 0.156882 |
| TGGGASLE | ASSQERGPGNTLY | 0.20609076 |
| VAVYNHYKRI | ASSQERGPGNTLY | 0.12964751 |
| TDPAPATT | ASSQERGPGNTLY | 0.15422048 |
| TNRPDILDPALLRPG | ASSQERGPGNTLY | 0.14222996 |
| LTTTTTS | ASSQERGPGNTLY | 0.13187248 |
| PTGSGKTL | ASSQERGPGNTLY | 0.1867977 |
| TGSGKTL | ASSQERGPGNTLY | 0.13519898 |
| TIGHVDHGK | ASSQERGPGNTLY | 0.15407601 |
| TIGHVDHGKTT | ASSQERGPGNTLY | 0.15696049 |
| GHVDHGKTT | ASSQERGPGNTLY | 0.1642594 |
| AQFEHTLL | ASSQERGPGNTLY | 0.14861509 |
| IVNQDEL | ASSQERGPGNTLY | 0.14651537 |
| TNGSQFF | ASSQERGPGNTLY | 0.12942621 |
| TGKGIVE | ASSQERGPGNTLY | 0.163072 |
| LISNASDA | ASSQERGPGNTLY | 0.12678955 |
| AELRTLGELTKIAKAHGAQ | ASSQERGPGNTLY | 0.15124756 |

| | | |
|---|---|---|
| APNAVAAAEPSW | ASSQERGPGNTLY | 0.13598701 |
| APQVAGAATAVL | ASSQERGPGNTLY | 0.18034236 |
| ARRDPAAAAAILPTDA | ASSQERGPGNTLY | 0.154111 |
| DAAARDVLAEAGLAEYF | ASSQERGPGNTLY | 0.15183465 |
| DDQIVLSAKANLAKVEQ | ASSQERGPGNTLY | 0.14284763 |
| DELLPDTGEDSDEDGH | ASSQERGPGNTLY | 0.13715991 |
| ELPEAPANRATVPQD | ASSQERGPGNTLY | 0.13296652 |
| FGQSKIERVTGIG | ASSQERGPGNTLY | 0.13285345 |
| GAGGTGSTAGGAGGAGGAGGLY | ASSQERGPGNTLY | 0.14940968 |
| GHGGTGAAGGNGGTGGAGGSGI | ASSQERGPGNTLY | 0.14544822 |
| GNGGNGGAGGEGGPGGAGGAGA | ASSQERGPGNTLY | 0.1335661 |
| GSTAGNGGNGGNATNGTIAGQP | ASSQERGPGNTLY | 0.17721584 |
| GTGGDGGNGGNGGTGG | ASSQERGPGNTLY | 0.15018767 |
| GVRRPAGIPADAPV | ASSQERGPGNTLY | 0.13047698 |
| GYRQVTDAHLLTLARR | ASSQERGPGNTLY | 0.14551085 |
| IAKMGQDGHDRG | ASSQERGPGNTLY | 0.16636607 |
| KAEPSIEIQYT | ASSQERGPGNTLY | 0.14683665 |
| KPVRIRMAADEDTVAVT | ASSQERGPGNTLY | 0.12599644 |
| MPPRIAGMRLLVIKPEP | ASSQERGPGNTLY | 0.1444519 |
| MKFYAVALFRSRGVPPTR | ASSQERGPGNTLY | 0.14861533 |
| NAGGFGILGAVAHSP | ASSQERGPGNTLY | 0.13309759 |
| NEGREITLAMAEM | ASSQERGPGNTLY | 0.13475375 |
| NELVEWSHPLGHRWRWE | ASSQERGPGNTLY | 0.20563224 |
| NLGAGGTGGNGGNPG | ASSQERGPGNTLY | 0.12954755 |
| NTEWPKWETVMAADPVD | ASSQERGPGNTLY | 0.1401947 |
| PASAAPAETQP | ASSQERGPGNTLY | 0.16049954 |
| QQSSDGKNGVRAAEITGE | ASSQERGPGNTLY | 0.17019013 |
| RAPDAVTTSWWKE | ASSQERGPGNTLY | 0.19689524 |
| SGFGNVGAHTSGWFNQ | ASSQERGPGNTLY | 0.15308605 |
| SGVAGAVGQAASVGGLKV | ASSQERGPGNTLY | 0.13946778 |
| SNGGAGTGSAGGNGGT | ASSQERGPGNTLY | 0.16474514 |
| TGGAGGEGGAAG | ASSQERGPGNTLY | 0.14824025 |
| TGGDGGAAGAGGAPGHG | ASSQERGPGNTLY | 0.13008426 |
| TMPPLLAAIANAGRHGV | ASSQERGPGNTLY | 0.12691574 |
| TSSTAGNSGGA | ASSQERGPGNTLY | 0.14428225 |
| VPATVPVGILC | ASSQERGPGNTLY | 0.13064162 |
| VQRPDSGGVVLTGRIS | ASSQERGPGNTLY | 0.15652125 |
| WTPNPVPATKGVPAGLVQAT | ASSQERGPGNTLY | 0.17060867 |
| YALKRALAKFSDRFDVVII | ASSQERGPGNTLY | 0.13427664 |
| YLIGSVIV | ASSQERGPGNTLY | 0.13573715 |
| YGQLVAVID | ASSQERGPGNTLY | 0.1402839 |
| VTVGGMTLPA | ASSQERGPGNTLY | 0.13803507 |
| VLALPAV | ASSQERGPGNTLY | 0.16368572 |
| VIPPTLNYET | ASSQERGPGNTLY | 0.14123788 |
| VGMGGMHPSSG | ASSQERGPGNTLY | 0.69970787 |
| VGLAGGAATA | ASSQERGPGNTLY | 0.17881781 |
| VGGAGGAGGSGP | ASSQERGPGNTLY | 0.1411116 |
| VFAAFMV | ASSQERGPGNTLY | 0.16484112 |
| TLYDKLR | ASSQERGPGNTLY | 0.13824068 |
| TLAVFTVL | ASSQERGPGNTLY | 0.13830525 |
| TGSVQPARV | ASSQERGPGNTLY | 0.14730783 |

| | | |
|---|---|---|
| TGGGAGSGGT | ASSQERGPGNTLY | 0.13897935 |
| SHLETSL | ASSQERGPGNTLY | 0.14223203 |
| SFTTGMT | ASSQERGPGNTLY | 0.14614876 |
| QVEVAKRMGVRQPTVS | ASSQERGPGNTLY | 0.13051283 |
| QGSGGPALAM | ASSQERGPGNTLY | 0.15424323 |
| MLAPSNP | ASSQERGPGNTLY | 0.13478601 |
| LVNNAAI | ASSQERGPGNTLY | 0.12921153 |
| LTGEINTLA | ASSQERGPGNTLY | 0.1387151 |
| LSIFGPV | ASSQERGPGNTLY | 0.20716047 |
| LLALLANL | ASSQERGPGNTLY | 0.22843157 |
| LISNASDALDKLR | ASSQERGPGNTLY | 0.13928291 |
| LINEAVP | ASSQERGPGNTLY | 0.13504924 |
| LGYAFSGL | ASSQERGPGNTLY | 0.1664831 |
| LDFVAVAL | ASSQERGPGNTLY | 0.13135156 |
| LAQPGAE | ASSQERGPGNTLY | 0.15075214 |
| ISNLGAVTGF | ASSQERGPGNTLY | 0.14107719 |
| HQEDPGRI | ASSQERGPGNTLY | 0.14098473 |
| GTPGEPTP | ASSQERGPGNTLY | 0.14468949 |
| GFFNSGS | ASSQERGPGNTLY | 0.14370875 |
| GAGGGGLTAGAN | ASSQERGPGNTLY | 0.13529937 |
| FVDALVNM | ASSQERGPGNTLY | 0.16344151 |
| FLTVAPEL | ASSQERGPGNTLY | 0.12598518 |
| FETLTAL | ASSQERGPGNTLY | 0.14086775 |
| DVINPIEL | ASSQERGPGNTLY | 0.13892208 |
| APVSGAPV | ASSQERGPGNTLY | 0.13609062 |
| ALVGGLAW | ASSQERGPGNTLY | 0.13041072 |
| ALAAAAATAI | ASSQERGPGNTLY | 0.14885002 |
| AFGLGLLL | ASSQERGPGNTLY | 0.16506651 |
| AFFAAASL | ASSQERGPGNTLY | 0.16106632 |
| AAAALPAFSPP | ASSQERGPGNTLY | 0.13071123 |
| VVLAPAL | ASSQERGPGNTLY | 0.17126885 |
| TIGHVDHGKTTL | ASSQERGPGNTLY | 0.15868299 |
| LISNASDALDK | ASSQERGPGNTLY | 0.13856763 |
| LISNASDAL | ASSQERGPGNTLY | 0.13058077 |
| ISNASDALDK | ASSQERGPGNTLY | 0.15750742 |
| NQAETLVYQTE | ASSQERGPGNTLY | 0.13543402 |
| TGGTGGTAGAG | ASSQERGPGNTLY | 0.14401728 |
| TGSTAGGAGGA | ASSQERGPGNTLY | 0.16706327 |
| DPLAVPVAAGPV | ASSQERGPGNTLY | 0.13260837 |
| GGTGGTPGNGGN | ASSQERGPGNTLY | 0.17881408 |
| PAGLAGVDPATL | ASSQERGPGNTLY | 0.18153486 |
| GNNLVVEAPHAGASV | ASSQERGPGNTLY | 0.13471366 |
| KVTRIMSASTLEQPA | ASSQERGPGNTLY | 0.13644284 |
| LGFALPIALPRRTVV | ASSQERGPGNTLY | 0.13985263 |
| TRLGNLTARKLVVLV | ASSQERGPGNTLY | 0.12731005 |
| WSQQRRLIYFPSAGP | ASSQERGPGNTLY | 0.16879572 |
| GPNGNPDPMAAAVDIR | ASSQERGPGNTLY | 0.13767742 |
| TQRCVSVADLRAAASR | ASSQERGPGNTLY | 0.13668424 |
| QFVLDVAEHPVEVLTETV | ASSQERGPGNTLY | 0.15161635 |
| GGAGGTGGAGGNGGTQAGDG | ASSQERGPGNTLY | 0.14005136 |
| PLLATLRAIAVDVDAKPAQV | ASSQERGPGNTLY | 0.1421608 |

| | | |
|---|---|---|
| APKFRTGGYMIDFWGVGYQVAK | ASSQERGPGNTLY | 0.14187247 |
| GAGGAGGAGGAAGGPGGTGGTG | ASSQERGPGNTLY | 0.1380148 |
| IGALGLLATDVAPAGSVAFGVA | ASSQERGPGNTLY | 0.14615247 |
| AGGAGGIGGNGGAGGAGGVLMG | ASSQERGPGNTLY | 0.1361339 |
| ADELIGLL | ASSQERGPGNTLY | 0.14406487 |
| ADRALKA | ASSQERGPGNTLY | 0.23915519 |
| AGAVGQVGGAGGA | ASSQERGPGNTLY | 0.13000154 |
| AGGGGGLGGAP | ASSQERGPGNTLY | 0.13764036 |
| AGGLLYGNGGAG | ASSQERGPGNTLY | 0.14476614 |
| AGLESIGL | ASSQERGPGNTLY | 0.12892863 |
| AGLKTGII | ASSQERGPGNTLY | 0.16451092 |
| ASLAALLL | ASSQERGPGNTLY | 0.15090929 |
| ASVINISSI | ASSQERGPGNTLY | 0.15452424 |
| ATPPGPA | ASSQERGPGNTLY | 0.1358488 |
| ATVASALV | ASSQERGPGNTLY | 0.1887424 |
| ATVASLIL | ASSQERGPGNTLY | 0.16540353 |
| AVNGPVLIRH | ASSQERGPGNTLY | 0.14718746 |
| DGAAGAVGI | ASSQERGPGNTLY | 0.14101398 |
| DPLLHGI | ASSQERGPGNTLY | 0.13888252 |
| ELGAALGAL | ASSQERGPGNTLY | 0.16592331 |
| ELGRTQAV | ASSQERGPGNTLY | 0.17205401 |
| FVAGGYLS | ASSQERGPGNTLY | 0.13391371 |
| GASGGAGGAGGW | ASSQERGPGNTLY | 0.1592445 |
| GGAGAGGDGGAGGI | ASSQERGPGNTLY | 0.16592276 |
| GTGHNGSQPGMGGNG | ASSQERGPGNTLY | 0.18859845 |
| GYAAASASAS | ASSQERGPGNTLY | 0.13043429 |
| HQVDVAAVQQG | ASSQERGPGNTLY | 0.14026403 |
| IGAAIAHT | ASSQERGPGNTLY | 0.1340218 |
| IGMIEVAQL | ASSQERGPGNTLY | 0.139709 |
| IRLAPEHTSAMD | ASSQERGPGNTLY | 0.13914947 |
| IYNVSVL | ASSQERGPGNTLY | 0.1537837 |
| KQGFLGM | ASSQERGPGNTLY | 0.15796086 |
| LESVITQL | ASSQERGPGNTLY | 0.1882329 |
| LGLGALVVV | ASSQERGPGNTLY | 0.1562418 |
| LIGNGGNGGSGGI | ASSQERGPGNTLY | 0.16774523 |
| LNAINAPTL | ASSQERGPGNTLY | 0.14827502 |
| LQLLNTGV | ASSQERGPGNTLY | 0.14839065 |
| LQTSIEF | ASSQERGPGNTLY | 0.13237254 |
| MSFAVSEL | ASSQERGPGNTLY | 0.12917624 |
| NTGALGL | ASSQERGPGNTLY | 0.13683741 |
| PPGAGGAGGAGTT | ASSQERGPGNTLY | 0.13664176 |
| QLVGLGI | ASSQERGPGNTLY | 0.1266484 |
| RFLEQVHQL | ASSQERGPGNTLY | 0.1571747 |
| RGVGYVLR | ASSQERGPGNTLY | 0.1484947 |
| SATGVFAGVI | ASSQERGPGNTLY | 0.1592815 |
| SLPAITLP | ASSQERGPGNTLY | 0.14527449 |
| STALAFTV | ASSQERGPGNTLY | 0.14415163 |
| SVLTVGLGK | ASSQERGPGNTLY | 0.13503952 |
| TALAGAGSGI | ASSQERGPGNTLY | 0.20720133 |
| TGLPFTGI | ASSQERGPGNTLY | 0.13342759 |
| VAEHVTV | ASSQERGPGNTLY | 0.13117217 |

| | | |
|---|---|---|
| VDIVLNSL | ASSQERGPGNTLY | 0.13905081 |
| VLTGGGAVL | ASSQERGPGNTLY | 0.13117535 |
| VNVGLAYGI | ASSQERGPGNTLY | 0.17427443 |
| VSSAAVGAV | ASSQERGPGNTLY | 0.17035969 |
| VTSVYEAGV | ASSQERGPGNTLY | 0.15991354 |
| WVALDRGA | ASSQERGPGNTLY | 0.17484711 |
| YLVGSVVI | ASSQERGPGNTLY | 0.12863566 |
| GPPGTGKTLL | ASSQERGPGNTLY | 0.14238763 |
| GVGMTKF | ASSQERGPGNTLY | 0.22610262 |
| IGIGGSDL | ASSQERGPGNTLY | 0.14669928 |
| IGTLLAEL | ASSQERGPGNTLY | 0.16035785 |
| IYTSGTT | ASSQERGPGNTLY | 0.16811448 |
| LSFPAGV | ASSQERGPGNTLY | 0.13875787 |
| LYTSGTT | ASSQERGPGNTLY | 0.14924489 |
| VGAFPFGF | ASSQERGPGNTLY | 0.14219564 |
| VGAGTVEF | ASSQERGPGNTLY | 0.16075474 |
| VGKSTLLL | ASSQERGPGNTLY | 0.1410498 |
| AALAHAGGGIV | ASSQERGPGNTLY | 0.13452545 |
| AALVAVSLL | ASSQERGPGNTLY | 0.17076235 |
| AAVSRAVAAL | ASSQERGPGNTLY | 0.26159993 |
| AGGSSGIGGF | ASSQERGPGNTLY | 0.15524527 |
| AGLTVANANG | ASSQERGPGNTLY | 0.15224877 |
| AGLVGAVL | ASSQERGPGNTLY | 0.1560294 |
| ALSQAFTV | ASSQERGPGNTLY | 0.15091144 |
| ALTFSVSL | ASSQERGPGNTLY | 0.1459841 |
| ALTGAALVAL | ASSQERGPGNTLY | 0.13602929 |
| ASFSIGAGVL | ASSQERGPGNTLY | 0.13890302 |
| DGSVYKI | ASSQERGPGNTLY | 0.13435392 |
| EQLAEIV | ASSQERGPGNTLY | 0.1571763 |
| FAVSELTVM | ASSQERGPGNTLY | 0.14247757 |
| GAAPLAAGL | ASSQERGPGNTLY | 0.3252247 |
| GADGGDGGSGGAGG | ASSQERGPGNTLY | 0.16236643 |
| GFGLDTGGAGG | ASSQERGPGNTLY | 0.15066119 |
| GLASGGLL | ASSQERGPGNTLY | 0.1331233 |
| GRIAGAAVLL | ASSQERGPGNTLY | 0.14753792 |
| GSAGSLNTSF | ASSQERGPGNTLY | 0.14336249 |
| GTGQIGI | ASSQERGPGNTLY | 0.13173899 |
| GTLIALIL | ASSQERGPGNTLY | 0.19062044 |
| IAATMGFVVG | ASSQERGPGNTLY | 0.13743757 |
| IFAQALL | ASSQERGPGNTLY | 0.13391018 |
| IGAAVALGF | ASSQERGPGNTLY | 0.15441275 |
| IGLTPLAL | ASSQERGPGNTLY | 0.600502 |
| ILGSNTSTL | ASSQERGPGNTLY | 0.14368948 |
| IVFSLTVL | ASSQERGPGNTLY | 0.13497691 |
| LASKATGF | ASSQERGPGNTLY | 0.13531904 |
| LASVALIL | ASSQERGPGNTLY | 0.1575338 |
| LFAVGLTL | ASSQERGPGNTLY | 0.21040612 |
| LFNGGIGTYI | ASSQERGPGNTLY | 0.13236368 |
| LTLGAGII | ASSQERGPGNTLY | 0.2981293 |
| LVAAALGDI | ASSQERGPGNTLY | 0.14292492 |
| NTAFVQLGI | ASSQERGPGNTLY | 0.13506633 |

| | | |
|---|---|---|
| QLFSGAEL | ASSQERGPGNTLY | 0.15815566 |
| SAATFITL | ASSQERGPGNTLY | 0.15029547 |
| SNYGLANLGV | ASSQERGPGNTLY | 0.15528238 |
| SYGGAYAVM | ASSQERGPGNTLY | 0.15868668 |
| TTGVGGLTLG | ASSQERGPGNTLY | 0.15182172 |
| TTQTIYI | ASSQERGPGNTLY | 0.13929632 |
| VAAGAGLGAVI | ASSQERGPGNTLY | 0.18175286 |
| VAAVVTLL | ASSQERGPGNTLY | 0.163107 |
| VGFLGGLI | ASSQERGPGNTLY | 0.16411689 |
| VGVIPILL | ASSQERGPGNTLY | 0.1394742 |
| VIFASSLI | ASSQERGPGNTLY | 0.13667883 |
| VKGGDGGAAAG | ASSQERGPGNTLY | 0.14814784 |
| VTKTFTNI | ASSQERGPGNTLY | 0.14518048 |
| VVAAAILL | ASSQERGPGNTLY | 0.14023522 |
| YAALDVEL | ASSQERGPGNTLY | 0.14798082 |
| LASVALLL | ASSQERGPGNTLY | 0.15759191 |
| LDAVHRV | ASSQERGPGNTLY | 0.13519493 |
| QGLSSSIF | ASSQERGPGNTLY | 0.15385558 |
| RGTFANIRL | ASSQERGPGNTLY | 0.16858713 |
| RQLWWGHRI | ASSQERGPGNTLY | 0.14700441 |
| YGPPGTGKTLL | ASSQERGPGNTLY | 0.13784666 |
| AALVGAPANVA | ASSQERGPGNTLY | 0.18763202 |
| AATFRRTE | ASSQERGPGNTLY | 0.28909513 |
| AGALGGGI | ASSQERGPGNTLY | 0.16857988 |
| AGGAGGAGGIGAG | ASSQERGPGNTLY | 0.13219444 |
| AGSGALGL | ASSQERGPGNTLY | 0.1280892 |
| AQIADQI | ASSQERGPGNTLY | 0.15535952 |
| ASAVAVTYL | ASSQERGPGNTLY | 0.15776885 |
| ASTVLGGTGG | ASSQERGPGNTLY | 0.14142318 |
| AVGGEPAK | ASSQERGPGNTLY | 0.1361078 |
| CEQGTFGGTC | ASSQERGPGNTLY | 0.14439343 |
| DEGKAPAG | ASSQERGPGNTLY | 0.1691904 |
| DTVVFPAL | ASSQERGPGNTLY | 0.14891276 |
| FEGITMAG | ASSQERGPGNTLY | 0.13093752 |
| FPLVVQD | ASSQERGPGNTLY | 0.14561903 |
| GAFTTPQ | ASSQERGPGNTLY | 0.17242943 |
| GFGGFGTAGGVGGA | ASSQERGPGNTLY | 0.13715589 |
| GFLAQFL | ASSQERGPGNTLY | 0.14540814 |
| GGAGGAGGNAGVA | ASSQERGPGNTLY | 0.14823306 |
| GGVGTAGAGGAGGAGG | ASSQERGPGNTLY | 0.13345048 |
| GNGGGGDAGNAGSGGN | ASSQERGPGNTLY | 0.14146474 |
| GTGGAGGAGNDAGS | ASSQERGPGNTLY | 0.13805954 |
| GTTNTGWFNS | ASSQERGPGNTLY | 0.15460692 |
| IFMGPLFTL | ASSQERGPGNTLY | 0.13819808 |
| ILAHAIEAI | ASSQERGPGNTLY | 0.15317518 |
| IRSAIREAL | ASSQERGPGNTLY | 0.17069589 |
| ITFAGGG | ASSQERGPGNTLY | 0.1616743 |
| ITGMSGHQEV | ASSQERGPGNTLY | 0.15396999 |
| IVAGLACTL | ASSQERGPGNTLY | 0.13799149 |
| IVGGGLAGA | ASSQERGPGNTLY | 0.13234828 |
| IVTGVLNI | ASSQERGPGNTLY | 0.1268548 |

| | | |
|---|---|---|
| LAQQGCI | ASSQERGPGNTLY | 0.1820509 |
| LCLHVEN | ASSQERGPGNTLY | 0.12739627 |
| LDAISDGS | ASSQERGPGNTLY | 0.17927308 |
| LGGFAAML | ASSQERGPGNTLY | 0.1521645 |
| LGGKNPMI | ASSQERGPGNTLY | 0.13565402 |
| LGPKGVAAIL | ASSQERGPGNTLY | 0.15027136 |
| LMVGVSI | ASSQERGPGNTLY | 0.13001299 |
| LVGAAFVL | ASSQERGPGNTLY | 0.13526972 |
| LVLAVAPL | ASSQERGPGNTLY | 0.148398 |
| MAAIAVIDAA | ASSQERGPGNTLY | 0.14956386 |
| NNGINAELQ | ASSQERGPGNTLY | 0.14440197 |
| RGEPLVLVHG | ASSQERGPGNTLY | 0.1514345 |
| RGGAGGAAGLI | ASSQERGPGNTLY | 0.17055842 |
| RSYRMHPAV | ASSQERGPGNTLY | 0.14219631 |
| SAAHLLAGQLA | ASSQERGPGNTLY | 0.15079685 |
| SAAVGSY | ASSQERGPGNTLY | 0.18676446 |
| SPHAVLHRI | ASSQERGPGNTLY | 0.15152653 |
| SQIAKVL | ASSQERGPGNTLY | 0.13702591 |
| TALGHLVP | ASSQERGPGNTLY | 0.15094559 |
| TAPTTSGAA | ASSQERGPGNTLY | 0.13705036 |
| TDGDDGNGGSPGT | ASSQERGPGNTLY | 0.16580307 |
| TDGLGEN | ASSQERGPGNTLY | 0.14340225 |
| TGGPPLPGQA | ASSQERGPGNTLY | 0.15397851 |
| TIPAITI | ASSQERGPGNTLY | 0.15318269 |
| TIPGFTI | ASSQERGPGNTLY | 0.13138929 |
| TSIASGALA | ASSQERGPGNTLY | 0.134226 |
| TVTLPGAAI | ASSQERGPGNTLY | 0.14144887 |
| VASGLSAALP | ASSQERGPGNTLY | 0.16234943 |
| VGAGVIGI | ASSQERGPGNTLY | 0.17019853 |
| VGPKRFGV | ASSQERGPGNTLY | 0.13478513 |
| VNVAFPD | ASSQERGPGNTLY | 0.16304159 |
| VSMTNSGVS | ASSQERGPGNTLY | 0.1458993 |
| VTDVLGDLSV | ASSQERGPGNTLY | 0.1407466 |
| VVVVGAGFAGL | ASSQERGPGNTLY | 0.14782318 |
| VYTVGTVV | ASSQERGPGNTLY | 0.14270063 |
| WTTFSGP | ASSQERGPGNTLY | 0.1365976 |
| YDELVEAF | ASSQERGPGNTLY | 0.15854575 |
| YGTGGAGGNGG | ASSQERGPGNTLY | 0.13477327 |
| YTNVMAV | ASSQERGPGNTLY | 0.15528306 |
| FLEQVHQL | ASSQERGPGNTLY | 0.14049749 |
| GLSLLAVAV | ASSQERGPGNTLY | 0.14001508 |
| LEQVHQL | ASSQERGPGNTLY | 0.15055679 |
| LNAAGAI | ASSQERGPGNTLY | 0.14417987 |
| PAVHASL | ASSQERGPGNTLY | 0.1647497 |
| TELEDDA | ASSQERGPGNTLY | 0.1397518 |
| TPPGPPN | ASSQERGPGNTLY | 0.13556668 |
| TPPPPPPP | ASSQERGPGNTLY | 0.160457 |
| TQSPALI | ASSPRLGGDTLY | 0.16759114 |
| EFAAAFPQ | ASSPRLGGDTLY | 0.1646508 |
| LLSGNLDVL | ASSPRLGGDTLY | 0.14324217 |
| GPANLGNYNI | ASSPRLGGDTLY | 0.18062893 |

| | | |
|---|---|---|
| TLGPTLP | ASSPRLGGDTLY | 0.1612309 |
| VYAYSSCCK | ASSPRLGGDTLY | 0.14325128 |
| APAPAPAPNI | ASSPRLGGDTLY | 0.12995566 |
| PGGPVPIP | ASSPRLGGDTLY | 0.14248352 |
| FLTVALEP | ASSPRLGGDTLY | 0.13816229 |
| SQYFGANL | ASSPRLGGDTLY | 0.14493534 |
| GGVKPAA | ASSPRLGGDTLY | 0.13014993 |
| GGTGGAGGAGGW | ASSPRLGGDTLY | 0.18649031 |
| GVGGVAGLGGAGG | ASSPRLGGDTLY | 0.22974405 |
| VLLVLATTI | ASSPRLGGDTLY | 0.13411354 |
| IGIFAAML | ASSPRLGGDTLY | 0.17156653 |
| AVGDLID | ASSPRLGGDTLY | 0.13029715 |
| NISPLPL | ASSPRLGGDTLY | 0.1542957 |
| VTAVFVAI | ASSPRLGGDTLY | 0.18473633 |
| SGDDNDAAL | ASSPRLGGDTLY | 0.13218592 |
| AVAALAHS | ASSPRLGGDTLY | 0.15572959 |
| GGVTVMATP | ASSPRLGGDTLY | 0.13470314 |
| AGLAANVRD | ASSPRLGGDTLY | 0.13883892 |
| TNRPDLIDPALLRPG | ASSPRLGGDTLY | 0.24291524 |
| LANLGVTAL | ASSPRLGGDTLY | 0.1437619 |
| TGERIEL | ASSPRLGGDTLY | 0.15446955 |
| NGEELAA | ASSPRLGGDTLY | 0.14121644 |
| VLSLVWG | ASSPRLGGDTLY | 0.13612075 |
| IGYALWL | ASSPRLGGDTLY | 0.15174308 |
| LSLLLAPL | ASSPRLGGDTLY | 0.17899518 |
| TLDEALK | ASSPRLGGDTLY | 0.13168341 |
| TDGAGTAEGG | ASSPRLGGDTLY | 0.16684738 |
| NAPGGAGGNGG | ASSPRLGGDTLY | 0.13912098 |
| VSVFMGMGGNF | ASSPRLGGDTLY | 0.21450536 |
| LASYLAAL | ASSPRLGGDTLY | 0.18823424 |
| LVGGTPAL | ASSPRLGGDTLY | 0.1523101 |
| VRNADVL | ASSPRLGGDTLY | 0.14076054 |
| TCSSPFLVSAM | ASSPRLGGDTLY | 0.22349212 |
| VHITDAE | ASSPRLGGDTLY | 0.15822701 |
| EYAAPSF | ASSPRLGGDTLY | 0.15325768 |
| VVDAFHGGL | ASSPRLGGDTLY | 0.17195213 |
| ASVVDPQ | ASSPRLGGDTLY | 0.16665323 |
| SYNEFGGLS | ASSPRLGGDTLY | 0.18053198 |
| DPDGSVY | ASSPRLGGDTLY | 0.17259574 |
| MVSTIHATE | ASSPRLGGDTLY | 0.19559515 |
| LAAGVADPV | ASSPRLGGDTLY | 0.17235991 |
| TKEQIAA | ASSPRLGGDTLY | 0.16412953 |
| SLASPAAL | ASSPRLGGDTLY | 0.18131106 |
| IGLVFYGV | ASSPRLGGDTLY | 0.16749749 |
| FLLDASNL | ASSPRLGGDTLY | 0.1887683 |
| YQVGNFP | ASSPRLGGDTLY | 0.18123735 |
| ESLLSQST | ASSPRLGGDTLY | 0.17515813 |
| LFTVGFAAVG | ASSPRLGGDTLY | 0.21837375 |
| ATPPTPT | ASSPRLGGDTLY | 0.19045421 |
| TDLATLG | ASSPRLGGDTLY | 0.16490205 |
| SAGGTGR | ASSPRLGGDTLY | 0.17076729 |

| | | |
|---|---|---|
| GFSNSGS | ASSPRLGGDTLY | 0.17564204 |
| GGNGGSGGTSV | ASSPRLGGDTLY | 0.17361856 |
| GTSVATGGAGN | ASSPRLGGDTLY | 0.16231868 |
| NGGNGGSGGTS | ASSPRLGGDTLY | 0.17541745 |
| AVVKFPH | ASSPRLGGDTLY | 0.16363682 |
| TPPPPPP | ASSPRLGGDTLY | 0.16075796 |
| AAPAAAAAPA | ASSPRLGGDTLY | 0.18764007 |
| IALNEDL | ASSPRLGGDTLY | 0.1689583 |
| IVNVSSM | ASSPRLGGDTLY | 0.16829488 |
| TGIPPAPR | ASSPRLGGDTLY | 0.185387 |
| FDLGGGTF | ASSPRLGGDTLY | 0.16450886 |
| FDLGGGTFDVS | ASSPRLGGDTLY | 0.16944575 |
| NEPTAAA | ASSPRLGGDTLY | 0.18565601 |
| NEPTAAALA | ASSPRLGGDTLY | 0.20055324 |
| GSIINISSI | ASSPRLGGDTLY | 0.16806427 |
| VAPGNAGTA | ASSPRLGGDTLY | 0.16993393 |
| TGGGASLE | ASSPRLGGDTLY | 0.21140526 |
| VAVYNHYKRI | ASSPRLGGDTLY | 0.18239643 |
| TDPAPATT | ASSPRLGGDTLY | 0.19475722 |
| TNRPDILDPALLRPG | ASSPRLGGDTLY | 0.19580536 |
| LTTTTTS | ASSPRLGGDTLY | 0.16691725 |
| PTGSGKTL | ASSPRLGGDTLY | 0.19024701 |
| TGSGKTL | ASSPRLGGDTLY | 0.23624665 |
| TIGHVDHGK | ASSPRLGGDTLY | 0.17442338 |
| TIGHVDHGKTT | ASSPRLGGDTLY | 0.18002577 |
| GHVDHGKTT | ASSPRLGGDTLY | 0.16962007 |
| AQFEHTLL | ASSPRLGGDTLY | 0.17468752 |
| IVNQDEL | ASSPRLGGDTLY | 0.1622299 |
| TNGSQFF | ASSPRLGGDTLY | 0.17235468 |
| TGKGIVE | ASSPRLGGDTLY | 0.18300407 |
| LISNASDA | ASSPRLGGDTLY | 0.17735519 |
| AELRTLGELTKIAKAHGAQ | ASSPRLGGDTLY | 0.1634751 |
| APNAVAAAEPSW | ASSPRLGGDTLY | 0.16488555 |
| APQVAGAATAVL | ASSPRLGGDTLY | 0.16515379 |
| ARRDPAAAAAILPTDA | ASSPRLGGDTLY | 0.16780192 |
| DAAARDVLAEAGLAEYF | ASSPRLGGDTLY | 0.16345963 |
| DDQIVLSAKANLAKVEQ | ASSPRLGGDTLY | 0.16523 |
| DELLPDTGEDSDEDGH | ASSPRLGGDTLY | 0.20903614 |
| ELPEAPANRATVPQD | ASSPRLGGDTLY | 0.19721918 |
| FGQSKIERVTGIG | ASSPRLGGDTLY | 0.16798042 |
| GAGGTGSTAGGAGGAGGAGGLY | ASSPRLGGDTLY | 0.18692887 |
| GHGGTGAAGGNGGTGGAGGSGI | ASSPRLGGDTLY | 0.18576449 |
| GNGGNGGAGGEGGPGGAGGAGA | ASSPRLGGDTLY | 0.18193991 |
| GSTAGNGGNGGNATNGTIAGQP | ASSPRLGGDTLY | 0.18034874 |
| GTGGDGGNGGNGGTGG | ASSPRLGGDTLY | 0.16640751 |
| GVRRPAGIPADAPV | ASSPRLGGDTLY | 0.26212966 |
| GYRQVTDAHLLTLARR | ASSPRLGGDTLY | 0.17464013 |
| IAKMGQDGHDRG | ASSPRLGGDTLY | 0.18805347 |
| KAEPSIEIQYT | ASSPRLGGDTLY | 0.19352797 |
| KPVRIRMAADEDTVAVT | ASSPRLGGDTLY | 0.17269981 |
| MPPRIAGMRLLVIKPEP | ASSPRLGGDTLY | 0.16617443 |

| | | |
|---|---|---|
| MKFYAVALFRSRGVPPTR | ASSPRLGGDTLY | 0.18720624 |
| NAGGFGILGAVAHSP | ASSPRLGGDTLY | 0.18381903 |
| NEGREITLAMAEM | ASSPRLGGDTLY | 0.16426381 |
| NELVEWSHPLGHRWRWE | ASSPRLGGDTLY | 0.16531903 |
| NLGAGGTGGNGGNPG | ASSPRLGGDTLY | 0.16649605 |
| NTEWPKWETVMAADPVD | ASSPRLGGDTLY | 0.17525591 |
| PASAAPAETQP | ASSPRLGGDTLY | 0.20502664 |
| QQSSDGKNGVRAAEITGE | ASSPRLGGDTLY | 0.17722534 |
| RAPDAVTTSWWKE | ASSPRLGGDTLY | 0.16501614 |
| SGFGNVGAHTSGWFNQ | ASSPRLGGDTLY | 0.18859 |
| SGVAGAVGQAASVGGLKV | ASSPRLGGDTLY | 0.17622668 |
| SNGGAGTGSAGGNGGT | ASSPRLGGDTLY | 0.17369494 |
| TGGAGGEGGAAG | ASSPRLGGDTLY | 0.17642142 |
| TGGDGGAAGAGGAPGHG | ASSPRLGGDTLY | 0.16627243 |
| TMPPLLAAIANAGRHGV | ASSPRLGGDTLY | 0.18096773 |
| TSSTAGNSGGA | ASSPRLGGDTLY | 0.19255324 |
| VPATVPVGILC | ASSPRLGGDTLY | 0.18435267 |
| VQRPDSGGVVLTGRIS | ASSPRLGGDTLY | 0.18298446 |
| WTPNPVPATKGVPAGLVQAT | ASSPRLGGDTLY | 0.1694859 |
| YALKRALAKFSDRFDVVII | ASSPRLGGDTLY | 0.17565084 |
| YLIGSVIV | ASSPRLGGDTLY | 0.16764118 |
| YGQLVAVID | ASSPRLGGDTLY | 0.17537457 |
| VTVGGMTLPA | ASSPRLGGDTLY | 0.17227682 |
| VLALPAV | ASSPRLGGDTLY | 0.17140278 |
| VIPPTLNYET | ASSPRLGGDTLY | 0.17910172 |
| VGMGGMHPSSG | ASSPRLGGDTLY | 0.18103965 |
| VGLAGGAATA | ASSPRLGGDTLY | 0.16681236 |
| VGGAGGAGGSGP | ASSPRLGGDTLY | 0.18941592 |
| VFAAFMV | ASSPRLGGDTLY | 0.20039263 |
| TLYDKLR | ASSPRLGGDTLY | 0.16576838 |
| TLAVFTVL | ASSPRLGGDTLY | 0.17500696 |
| TGSVQPARV | ASSPRLGGDTLY | 0.18733518 |
| TGGGAGSGGT | ASSPRLGGDTLY | 0.1958913 |
| SHLETSL | ASSPRLGGDTLY | 0.185967 |
| SFTTGMT | ASSPRLGGDTLY | 0.18410799 |
| QVEVAKRMGVRQPTVS | ASSPRLGGDTLY | 0.24300481 |
| QGSGGPALAM | ASSPRLGGDTLY | 0.1669866 |
| MLAPSNP | ASSPRLGGDTLY | 0.19914934 |
| LVNNAAI | ASSPRLGGDTLY | 0.16445787 |
| LTGEINTLA | ASSPRLGGDTLY | 0.17423667 |
| LSIFGPV | ASSPRLGGDTLY | 0.16745922 |
| LLALLANL | ASSPRLGGDTLY | 0.16128154 |
| LISNASDALDKLR | ASSPRLGGDTLY | 0.21497072 |
| LINEAVP | ASSPRLGGDTLY | 0.17572707 |
| LGYAFSGL | ASSPRLGGDTLY | 0.18594989 |
| LDFVAVAL | ASSPRLGGDTLY | 0.169723 |
| LAQPGAE | ASSPRLGGDTLY | 0.18607076 |
| ISNLGAVTGF | ASSPRLGGDTLY | 0.17943585 |
| HQEDPGRI | ASSPRLGGDTLY | 0.17239352 |
| GTPGEPTP | ASSPRLGGDTLY | 0.16937591 |
| GFFNSGS | ASSPRLGGDTLY | 0.19123636 |

| | | |
|---|---|---|
| GAGGGGLTAGAN | ASSPRLGGDTLY | 0.18932103 |
| FVDALVNM | ASSPRLGGDTLY | 0.18791987 |
| FLTVAPEL | ASSPRLGGDTLY | 0.16457011 |
| FETLTAL | ASSPRLGGDTLY | 0.17280369 |
| DVINPIEL | ASSPRLGGDTLY | 0.17721942 |
| APVSGAPV | ASSPRLGGDTLY | 0.16296808 |
| ALVGGLAW | ASSPRLGGDTLY | 0.16223569 |
| ALAAAAATAI | ASSPRLGGDTLY | 0.1621822 |
| AFGLGLLL | ASSPRLGGDTLY | 0.16364667 |
| AFFAAASL | ASSPRLGGDTLY | 0.19655867 |
| AAAALPAFSPP | ASSPRLGGDTLY | 0.2341679 |
| VVLAPAL | ASSPRLGGDTLY | 0.17316164 |
| TIGHVDHGKTTL | ASSPRLGGDTLY | 0.18326274 |
| LISNASDALDK | ASSPRLGGDTLY | 0.17749357 |
| LISNASDAL | ASSPRLGGDTLY | 0.15952997 |
| ISNASDALDK | ASSPRLGGDTLY | 0.16825616 |
| NQAETLVYQTE | ASSPRLGGDTLY | 0.16601647 |
| TGGTGGTAGAG | ASSPRLGGDTLY | 0.18917708 |
| TGSTAGGAGGA | ASSPRLGGDTLY | 0.16513795 |
| DPLAVPVAAGPV | ASSPRLGGDTLY | 0.18074574 |
| GGTGGTPGNGGN | ASSPRLGGDTLY | 0.17879194 |
| PAGLAGVDPATL | ASSPRLGGDTLY | 0.1942487 |
| GNNLVVEAPHAGASV | ASSPRLGGDTLY | 0.17771395 |
| KVTRIMSASTLEQPA | ASSPRLGGDTLY | 0.16300474 |
| LGFALPIALPRRTVV | ASSPRLGGDTLY | 0.1621402 |
| TRLGNLTARKLVVLV | ASSPRLGGDTLY | 0.19242343 |
| WSQQRRLIYFPSAGP | ASSPRLGGDTLY | 0.1662451 |
| GPNGNPDPMAAAVDIR | ASSPRLGGDTLY | 0.16908307 |
| TQRCVSVADLRAAASR | ASSPRLGGDTLY | 0.16673708 |
| QFVLDVAEHPVEVLTETV | ASSPRLGGDTLY | 0.1834937 |
| GGAGGTGGAGGNGGTQAGDG | ASSPRLGGDTLY | 0.16388604 |
| PLLATLRAIAVDVDAKPAQV | ASSPRLGGDTLY | 0.19076163 |
| APKFRTGGYMIDFWGVGYQVAK | ASSPRLGGDTLY | 0.16727132 |
| GAGGAGGAGGAAGGPGGTGGTG | ASSPRLGGDTLY | 0.17758025 |
| IGALGLLATDVAPAGSVAFGVA | ASSPRLGGDTLY | 0.18954927 |
| AGGAGGIGGNGGAGGAGGVLMG | ASSPRLGGDTLY | 0.25606412 |
| ADELIGLL | ASSPRLGGDTLY | 0.17688523 |
| ADRALKA | ASSPRLGGDTLY | 0.17019185 |
| AGAVGQVGGAGGA | ASSPRLGGDTLY | 0.16786861 |
| AGGGGGLGGAP | ASSPRLGGDTLY | 0.16728987 |
| AGGLLYGNGGAG | ASSPRLGGDTLY | 0.16736789 |
| AGLESIGL | ASSPRLGGDTLY | 0.2038291 |
| AGLKTGII | ASSPRLGGDTLY | 0.17789526 |
| ASLAALLL | ASSPRLGGDTLY | 0.16779327 |
| ASVINISSI | ASSPRLGGDTLY | 0.18684818 |
| ATPPGPA | ASSPRLGGDTLY | 0.19374947 |
| ATVASALV | ASSPRLGGDTLY | 0.19900313 |
| ATVASLIL | ASSPRLGGDTLY | 0.16690657 |
| AVNGPVLIRH | ASSPRLGGDTLY | 0.17305535 |
| DGAAGAVGI | ASSPRLGGDTLY | 0.18412389 |
| DPLLHGI | ASSPRLGGDTLY | 0.17579263 |

| | | |
|---|---|---|
| ELGAALGAL | ASSPRLGGDTLY | 0.16862236 |
| ELGRTQAV | ASSPRLGGDTLY | 0.16762999 |
| FVAGGYLS | ASSPRLGGDTLY | 0.18022801 |
| GASGGAGGAGGW | ASSPRLGGDTLY | 0.5101869 |
| GGAGAGGDGGAGGI | ASSPRLGGDTLY | 0.1703834 |
| GTGHNGSQPGMGGNG | ASSPRLGGDTLY | 0.1660627 |
| GYAAASASAS | ASSPRLGGDTLY | 0.17717499 |
| HQVDVAAVQQG | ASSPRLGGDTLY | 0.16362603 |
| IGAAIAHT | ASSPRLGGDTLY | 0.16832462 |
| IGMIEVAQL | ASSPRLGGDTLY | 0.17554301 |
| IRLAPEHTSAMD | ASSPRLGGDTLY | 0.1629966 |
| IYNVSVL | ASSPRLGGDTLY | 0.17084363 |
| KQGFLGM | ASSPRLGGDTLY | 0.19713986 |
| LESVITQL | ASSPRLGGDTLY | 0.18276657 |
| LGLGALVVV | ASSPRLGGDTLY | 0.1898964 |
| LIGNGGNGGSGGI | ASSPRLGGDTLY | 0.20187947 |
| LNAINAPTL | ASSPRLGGDTLY | 0.18098243 |
| LQLLNTGV | ASSPRLGGDTLY | 0.19866955 |
| LQTSIEF | ASSPRLGGDTLY | 0.18770328 |
| MSFAVSEL | ASSPRLGGDTLY | 0.17691968 |
| NTGALGL | ASSPRLGGDTLY | 0.16791372 |
| PPGAGGAGGAGTT | ASSPRLGGDTLY | 0.18686917 |
| QLVGLGI | ASSPRLGGDTLY | 0.17968921 |
| RFLEQVHQL | ASSPRLGGDTLY | 0.18840563 |
| RGVGYVLR | ASSPRLGGDTLY | 0.17783347 |
| SATGVFAGVI | ASSPRLGGDTLY | 0.2167705 |
| SLPAITLP | ASSPRLGGDTLY | 0.17018388 |
| STALAFTV | ASSPRLGGDTLY | 0.23519522 |
| SVLTVGLGK | ASSPRLGGDTLY | 0.16238055 |
| TALAGAGSGI | ASSPRLGGDTLY | 0.1667492 |
| TGLPFTGI | ASSPRLGGDTLY | 0.18754546 |
| VAEHVTV | ASSPRLGGDTLY | 0.16054761 |
| VDIVLNSL | ASSPRLGGDTLY | 0.16388465 |
| VLTGGGAVL | ASSPRLGGDTLY | 0.17376915 |
| VNVGLAYGI | ASSPRLGGDTLY | 0.18151087 |
| VSSAAVGAV | ASSPRLGGDTLY | 0.16016997 |
| VTSVYEAGV | ASSPRLGGDTLY | 0.17419527 |
| WVALDRGA | ASSPRLGGDTLY | 0.16565932 |
| YLVGSVVI | ASSPRLGGDTLY | 0.17407197 |
| GPPGTGKTLL | ASSPRLGGDTLY | 0.17421682 |
| GVGMTKF | ASSPRLGGDTLY | 0.16337939 |
| IGIGGSDL | ASSPRLGGDTLY | 0.18226674 |
| IGTLLAEL | ASSPRLGGDTLY | 0.16674684 |
| IYTSGTT | ASSPRLGGDTLY | 0.16597882 |
| LSFPAGV | ASSPRLGGDTLY | 0.19530621 |
| LYTSGTT | ASSPRLGGDTLY | 0.24029976 |
| VGAFPFGF | ASSPRLGGDTLY | 0.16759819 |
| VGAGTVEF | ASSPRLGGDTLY | 0.18022938 |
| VGKSTLLL | ASSPRLGGDTLY | 0.16458057 |
| AALAHAGGGIV | ASSPRLGGDTLY | 0.2471912 |
| AALVAVSLL | ASSPRLGGDTLY | 0.18390945 |

| | | |
|---|---|---|
| AAVSRAVAAL | ASSPRLGGDTLY | 0.16394143 |
| AGGSSGIGGF | ASSPRLGGDTLY | 0.18061411 |
| AGLTVANANG | ASSPRLGGDTLY | 0.18072587 |
| AGLVGAVL | ASSPRLGGDTLY | 0.17985746 |
| ALSQAFTV | ASSPRLGGDTLY | 0.19065239 |
| ALTFSVSL | ASSPRLGGDTLY | 0.1663848 |
| ALTGAALVAL | ASSPRLGGDTLY | 0.17081104 |
| ASFSIGAGVL | ASSPRLGGDTLY | 0.19494878 |
| DGSVYKI | ASSPRLGGDTLY | 0.17518075 |
| EQLAEIV | ASSPRLGGDTLY | 0.16722773 |
| FAVSELTVM | ASSPRLGGDTLY | 0.18810093 |
| GAAPLAAGL | ASSPRLGGDTLY | 0.17793675 |
| GADGGDGGSGGAGG | ASSPRLGGDTLY | 0.17416123 |
| GFGLDTGGAGG | ASSPRLGGDTLY | 0.21323779 |
| GLASGGLL | ASSPRLGGDTLY | 0.173457 |
| GRIAGAAVLL | ASSPRLGGDTLY | 0.1838674 |
| GSAGSLNTSF | ASSPRLGGDTLY | 0.18117972 |
| GTGQIGI | ASSPRLGGDTLY | 0.22205487 |
| GTLIALIL | ASSPRLGGDTLY | 0.16690372 |
| IAATMGFVVG | ASSPRLGGDTLY | 0.18722613 |
| IFAQALL | ASSPRLGGDTLY | 0.18688823 |
| IGAAVALGF | ASSPRLGGDTLY | 0.18451197 |
| IGLTPLAL | ASSPRLGGDTLY | 0.17050268 |
| ILGSNTSTL | ASSPRLGGDTLY | 0.17731468 |
| IVFSLTVL | ASSPRLGGDTLY | 0.1813104 |
| LASKATGF | ASSPRLGGDTLY | 0.2011306 |
| LASVALIL | ASSPRLGGDTLY | 0.17051925 |
| LFAVGLTL | ASSPRLGGDTLY | 0.17332666 |
| LFNGGIGTYI | ASSPRLGGDTLY | 0.21524093 |
| LTLGAGII | ASSPRLGGDTLY | 0.16898943 |
| LVAAALGDI | ASSPRLGGDTLY | 0.1798045 |
| NTAFVQLGI | ASSPRLGGDTLY | 0.19023953 |
| QLFSGAEL | ASSPRLGGDTLY | 0.19826669 |
| SAATFITL | ASSPRLGGDTLY | 0.17234218 |
| SNYGLANLGV | ASSPRLGGDTLY | 0.19033462 |
| SYGGAYAVM | ASSPRLGGDTLY | 0.16678175 |
| TTGVGGLTLG | ASSPRLGGDTLY | 0.19679001 |
| TTQTIYI | ASSPRLGGDTLY | 0.16428328 |
| VAAGAGLGAVI | ASSPRLGGDTLY | 0.17733668 |
| VAAVVTLL | ASSPRLGGDTLY | 0.16272895 |
| VGFLGGLI | ASSPRLGGDTLY | 0.19638036 |
| VGVIPILL | ASSPRLGGDTLY | 0.1859694 |
| VIFASSLI | ASSPRLGGDTLY | 0.19549713 |
| VKGGDGGAAAG | ASSPRLGGDTLY | 0.16517827 |
| VTKTFTNI | ASSPRLGGDTLY | 0.1836806 |
| VVAAAILL | ASSPRLGGDTLY | 0.19611913 |
| YAALDVEL | ASSPRLGGDTLY | 0.3163856 |
| LASVALLL | ASSPRLGGDTLY | 0.16954327 |
| LDAVHRV | ASSPRLGGDTLY | 0.20831424 |
| QGLSSSIF | ASSPRLGGDTLY | 0.18729167 |
| RGTFANIRL | ASSPRLGGDTLY | 0.17387822 |

| | | |
|---|---|---|
| RQLWWGHRI | ASSPRLGGDTLY | 0.16258611 |
| YGPPGTGKTLL | ASSPRLGGDTLY | 0.1737676 |
| AALVGAPANVA | ASSPRLGGDTLY | 0.17415866 |
| AATFRRTE | ASSPRLGGDTLY | 0.16627808 |
| AGALGGGI | ASSPRLGGDTLY | 0.26075605 |
| AGGAGGAGGIGAG | ASSPRLGGDTLY | 0.19707097 |
| AGSGALGL | ASSPRLGGDTLY | 0.1958599 |
| AQIADQI | ASSPRLGGDTLY | 0.19633122 |
| ASAVAVTYL | ASSPRLGGDTLY | 0.18985002 |
| ASTVLGGTGG | ASSPRLGGDTLY | 0.18374157 |
| AVGGEPAK | ASSPRLGGDTLY | 0.16926268 |
| CEQGTFGGTC | ASSPRLGGDTLY | 0.17720191 |
| DEGKAPAG | ASSPRLGGDTLY | 0.16396943 |
| DTVVFPAL | ASSPRLGGDTLY | 0.16872127 |
| FEGITMAG | ASSPRLGGDTLY | 0.16687325 |
| FPLVVQD | ASSPRLGGDTLY | 0.18110955 |
| GAFTTPQ | ASSPRLGGDTLY | 0.15992086 |
| GFGGFGTAGGVGGA | ASSPRLGGDTLY | 0.26284048 |
| GFLAQFL | ASSPRLGGDTLY | 0.206546 |
| GGAGGAGGNAGVA | ASSPRLGGDTLY | 0.16628242 |
| GGVGTAGAGGAGGAGG | ASSPRLGGDTLY | 0.1854954 |
| GNGGGDAGNAGSGGN | ASSPRLGGDTLY | 0.18409169 |
| GTGGAGGAGNDAGS | ASSPRLGGDTLY | 0.16361624 |
| GTTNTGWFNS | ASSPRLGGDTLY | 0.18597037 |
| IFMGPLFTL | ASSPRLGGDTLY | 0.16531219 |
| ILAHAIEAI | ASSPRLGGDTLY | 0.19170192 |
| IRSAIREAL | ASSPRLGGDTLY | 0.2069112 |
| ITFAGGG | ASSPRLGGDTLY | 0.18318729 |
| ITGMSGHQEV | ASSPRLGGDTLY | 0.17075197 |
| IVAGLACTL | ASSPRLGGDTLY | 0.18534468 |
| IVGGGLAGA | ASSPRLGGDTLY | 0.16758855 |
| IVTGVLNI | ASSPRLGGDTLY | 0.1722464 |
| LAQQGCI | ASSPRLGGDTLY | 0.16529727 |
| LCLHVEN | ASSPRLGGDTLY | 0.17916742 |
| LDAISDGS | ASSPRLGGDTLY | 0.16666734 |
| LGGFAAML | ASSPRLGGDTLY | 0.19806768 |
| LGGKNPMI | ASSPRLGGDTLY | 0.18195586 |
| LGPKGVAAIL | ASSPRLGGDTLY | 0.17595284 |
| LMVGVSI | ASSPRLGGDTLY | 0.175067 |
| LVGAAFVL | ASSPRLGGDTLY | 0.1712847 |
| LVLAVAPL | ASSPRLGGDTLY | 0.17311284 |
| MAAIAVIDAA | ASSPRLGGDTLY | 0.1643729 |
| NNGINAELQ | ASSPRLGGDTLY | 0.1883398 |
| RGEPLVLVHG | ASSPRLGGDTLY | 0.17138574 |
| RGGAGGAAGLI | ASSPRLGGDTLY | 0.16719367 |
| RSYRMHPAV | ASSPRLGGDTLY | 0.18825899 |
| SAAHLLAGQLA | ASSPRLGGDTLY | 0.17881441 |
| SAAVGSY | ASSPRLGGDTLY | 0.16613472 |
| SPHAVLHRI | ASSPRLGGDTLY | 0.16835374 |
| SQIAKVL | ASSPRLGGDTLY | 0.18240772 |
| TALGHLVP | ASSPRLGGDTLY | 0.1764331 |

| | | |
|---|---|---|
| TAPTTSGAA | ASSPRLGGDTLY | 0.1606327 |
| TDGDDGNGGSPGT | ASSPRLGGDTLY | 0.17322826 |
| TDGLGEN | ASSPRLGGDTLY | 0.17390965 |
| TGGPPLPGQA | ASSPRLGGDTLY | 0.16722421 |
| TIPAITI | ASSPRLGGDTLY | 0.16990693 |
| TIPGFTI | ASSPRLGGDTLY | 0.1960042 |
| TSIASGALA | ASSPRLGGDTLY | 0.1669313 |
| TVTLPGAAI | ASSPRLGGDTLY | 0.1744169 |
| VASGLSAALP | ASSPRLGGDTLY | 0.1823584 |
| VGAGVIGI | ASSPRLGGDTLY | 0.18929732 |
| VGPKRFGV | ASSPRLGGDTLY | 0.206381 |
| VNVAFPD | ASSPRLGGDTLY | 0.17900191 |
| VSMTNSGVS | ASSPRLGGDTLY | 0.16942754 |
| VTDVLGDLSV | ASSPRLGGDTLY | 0.18285821 |
| VVVVGAGFAGL | ASSPRLGGDTLY | 0.17711452 |
| VYTVGTVV | ASSPRLGGDTLY | 0.16568822 |
| WTTFSGP | ASSPRLGGDTLY | 0.16304469 |
| YDELVEAF | ASSPRLGGDTLY | 0.17474693 |
| YGTGGAGGNGG | ASSPRLGGDTLY | 0.16484882 |
| YTNVMAV | ASSPRLGGDTLY | 0.1991108 |
| FLEQVHQL | ASSPRLGGDTLY | 0.19317208 |
| GLSLLAVAV | ASSPRLGGDTLY | 0.16608772 |
| LEQVHQL | ASSPRLGGDTLY | 0.16466965 |
| LNAAGAI | ASSPRLGGDTLY | 0.16938256 |
| PAVHASL | ASSPRLGGDTLY | 0.16970947 |
| TELEDDA | ASSPRLGGDTLY | 0.18407866 |
| TPPGPPN | ASSPRLGGDTLY | 0.16816354 |
| TPPPPPPP | ASSPRLGGDTLY | 0.56163204 |
| TQSPALI | ASSLTGNQDTQY | 0.2022591 |
| EFAAAFPQ | ASSLTGNQDTQY | 0.17000383 |
| LLSGNLDVL | ASSLTGNQDTQY | 0.18903436 |
| GPANLGNYNI | ASSLTGNQDTQY | 0.16850899 |
| TLGPTLP | ASSLTGNQDTQY | 0.18352874 |
| VYAYSSCCK | ASSLTGNQDTQY | 0.17408645 |
| APAPAPAPNI | ASSLTGNQDTQY | 0.16753875 |
| PGGPVPIP | ASSLTGNQDTQY | 0.16723615 |
| FLTVALEP | ASSLTGNQDTQY | 0.17526436 |
| SQYFGANL | ASSLTGNQDTQY | 0.16648571 |
| GGVKPAA | ASSLTGNQDTQY | 0.17175359 |
| GGTGGAGGAGGW | ASSLTGNQDTQY | 0.1657602 |
| GVGGVAGLGGAGG | ASSLTGNQDTQY | 0.16239151 |
| VLLVLATTI | ASSLTGNQDTQY | 0.17274627 |
| IGIFAAML | ASSLTGNQDTQY | 0.2077133 |
| AVGDLID | ASSLTGNQDTQY | 0.20699945 |
| NISPLPL | ASSLTGNQDTQY | 0.16693445 |
| VTAVFVAI | ASSLTGNQDTQY | 0.16622289 |
| SGDDNDAAL | ASSLTGNQDTQY | 0.25740096 |
| AVAALAHS | ASSLTGNQDTQY | 0.16799346 |
| GGVTVMATP | ASSLTGNQDTQY | 0.17670614 |
| AGLAANVRD | ASSLTGNQDTQY | 0.17104837 |
| TNRPDLIDPALLRPG | ASSLTGNQDTQY | 0.16999322 |

| | | |
|---|---|---|
| LANLGVTAL | ASSLTGNQDTQY | 0.17357405 |
| TGERIEL | ASSLTGNQDTQY | 0.17665136 |
| NGEELAA | ASSLTGNQDTQY | 0.16857368 |
| VLSLVWG | ASSLTGNQDTQY | 0.18004382 |
| IGYALWL | ASSLTGNQDTQY | 0.16044185 |
| LSLLLAPL | ASSLTGNQDTQY | 0.17739339 |
| TLDEALK | ASSLTGNQDTQY | 0.16896917 |
| TDGAGTAEGG | ASSLTGNQDTQY | 0.16410759 |
| NAPGGAGGNGG | ASSLTGNQDTQY | 0.16180052 |
| VSVFMGMGGNF | ASSLTGNQDTQY | 0.18357144 |
| LASYLAAL | ASSLTGNQDTQY | 0.18336093 |
| LVGGTPAL | ASSLTGNQDTQY | 0.20093395 |
| VRNADVL | ASSLTGNQDTQY | 0.16905352 |
| TCSSPFLVSAM | ASSLTGNQDTQY | 0.18335834 |
| VHITDAE | ASSLTGNQDTQY | 0.19399317 |
| EYAAPSF | ASSLTGNQDTQY | 0.17525673 |
| VVDAFHGGL | ASSLTGNQDTQY | 0.16365273 |
| ASVVDPQ | ASSLTGNQDTQY | 0.17674461 |
| SYNEFGGLS | ASSLTGNQDTQY | 0.16688403 |
| DPDGSVY | ASSLTGNQDTQY | 0.17696206 |
| MVSTIHATE | ASSLTGNQDTQY | 0.23477045 |
| LAAGVADPV | ASSLTGNQDTQY | 0.16291647 |
| TKEQIAA | ASSLTGNQDTQY | 0.18446176 |
| SLASPAAL | ASSLTGNQDTQY | 0.18661167 |
| IGLVFYGV | ASSLTGNQDTQY | 0.16432002 |
| FLLDASNL | ASSLTGNQDTQY | 0.17429306 |
| YQVGNFP | ASSLTGNQDTQY | 0.17300484 |
| ESLLSQST | ASSLTGNQDTQY | 0.16327669 |
| LFTVGFAAVG | ASSLTGNQDTQY | 0.17739402 |
| ATPPTPT | ASSLTGNQDTQY | 0.16744351 |
| TDLATLG | ASSLTGNQDTQY | 0.17256887 |
| SAGGTGR | ASSLTGNQDTQY | 0.17126022 |
| GFSNSGS | ASSLTGNQDTQY | 0.16814315 |
| GGNGGSGGTSV | ASSLTGNQDTQY | 0.17497529 |
| GTSVATGGAGN | ASSLTGNQDTQY | 0.17605412 |
| NGGNGGSGGTS | ASSLTGNQDTQY | 0.1784051 |
| AVVKFPH | ASSLTGNQDTQY | 0.17582531 |
| TPPPPPP | ASSLTGNQDTQY | 0.16557053 |
| AAPAAAAAPA | ASSLTGNQDTQY | 0.18005472 |
| IALNEDL | ASSLTGNQDTQY | 0.27844036 |
| IVNVSSM | ASSLTGNQDTQY | 0.16386989 |
| TGIPPAPR | ASSLTGNQDTQY | 0.16617772 |
| FDLGGGTF | ASSLTGNQDTQY | 0.17096603 |
| FDLGGGTFDVS | ASSLTGNQDTQY | 0.1649801 |
| NEPTAAA | ASSLTGNQDTQY | 0.1797284 |
| NEPTAAALA | ASSLTGNQDTQY | 0.17900921 |
| GSIINISSI | ASSLTGNQDTQY | 0.18039675 |
| VAPGNAGTA | ASSLTGNQDTQY | 0.16638912 |
| TGGGASLE | ASSLTGNQDTQY | 0.2191691 |
| VAVYNHYKRI | ASSLTGNQDTQY | 0.18410899 |
| TDPAPATT | ASSLTGNQDTQY | 0.17253324 |

| | | |
|---|---|---|
| TNRPDILDPALLRPG | ASSLTGNQDTQY | 0.17055218 |
| LTTTTTS | ASSLTGNQDTQY | 0.16845435 |
| PTGSGKTL | ASSLTGNQDTQY | 0.17802028 |
| TGSGKTL | ASSLTGNQDTQY | 0.19286928 |
| TIGHVDHGK | ASSLTGNQDTQY | 0.16577001 |
| TIGHVDHGKTT | ASSLTGNQDTQY | 0.17096774 |
| GHVDHGKTT | ASSLTGNQDTQY | 0.19455343 |
| AQFEHTLL | ASSLTGNQDTQY | 0.19403179 |
| IVNQDEL | ASSLTGNQDTQY | 0.16493872 |
| TNGSQFF | ASSLTGNQDTQY | 0.1662744 |
| TGKGIVE | ASSLTGNQDTQY | 0.16702098 |
| LISNASDA | ASSLTGNQDTQY | 0.1688301 |
| AELRTLGELTKIAKAHGAQ | ASSLTGNQDTQY | 0.16849865 |
| APNAVAAAEPSW | ASSLTGNQDTQY | 0.17801513 |
| APQVAGAATAVL | ASSLTGNQDTQY | 0.19006559 |
| ARRDPAAAAAILPTDA | ASSLTGNQDTQY | 0.23107003 |
| DAAARDVLAEAGLAEYF | ASSLTGNQDTQY | 0.19166604 |
| DDQIVLSAKANLAKVEQ | ASSLTGNQDTQY | 0.20832135 |
| DELLPDTGEDSDEDGH | ASSLTGNQDTQY | 0.17242317 |
| ELPEAPANRATVPQD | ASSLTGNQDTQY | 0.17856702 |
| FGQSKIERVTGIG | ASSLTGNQDTQY | 0.1647579 |
| GAGGTGSTAGGAGGAGGAGGLY | ASSLTGNQDTQY | 0.16583349 |
| GHGGTGAAGGNGGTGGAGGSGI | ASSLTGNQDTQY | 0.16546543 |
| GNGGNGGAGGEGGPGGAGGAGA | ASSLTGNQDTQY | 0.17051323 |
| GSTAGNGGNGGNATNGTIAGQP | ASSLTGNQDTQY | 0.16155681 |
| GTGGDGGNGGNGGTGG | ASSLTGNQDTQY | 0.18275489 |
| GVRRPAGIPADAPV | ASSLTGNQDTQY | 0.17401673 |
| GYRQVTDAHLLTLARR | ASSLTGNQDTQY | 0.19118954 |
| IAKMGQDGHDRG | ASSLTGNQDTQY | 0.1745599 |
| KAEPSIEIQYT | ASSLTGNQDTQY | 0.17004912 |
| KPVRIRMAADEDTVAVT | ASSLTGNQDTQY | 0.16832139 |
| MPPRIAGMRLLVIKPEP | ASSLTGNQDTQY | 0.19579886 |
| MKFYAVALFRSRGVPPTR | ASSLTGNQDTQY | 0.16432726 |
| NAGGFGILGAVAHSP | ASSLTGNQDTQY | 0.1670137 |
| NEGREITLAMAEM | ASSLTGNQDTQY | 0.1708007 |
| NELVEWSHPLGHRWRWE | ASSLTGNQDTQY | 0.17383304 |
| NLGAGGTGGNGGNPG | ASSLTGNQDTQY | 0.21283121 |
| NTEWPKWETVMAADPVD | ASSLTGNQDTQY | 0.19055112 |
| PASAAPAETQP | ASSLTGNQDTQY | 0.17902331 |
| QQSSDGKNGVRAAEITGE | ASSLTGNQDTQY | 0.24329498 |
| RAPDAVTTSWWKE | ASSLTGNQDTQY | 0.16093263 |
| SGFGNVGAHTSGWFNQ | ASSLTGNQDTQY | 0.17234187 |
| SGVAGAVGQAASVGGLKV | ASSLTGNQDTQY | 0.23661901 |
| SNGGAGTGSAGGNGGT | ASSLTGNQDTQY | 0.17136055 |
| TGGAGGEGGAAG | ASSLTGNQDTQY | 0.19300352 |
| TGGDGGAAGAGGAPGHG | ASSLTGNQDTQY | 0.19218616 |
| TMPPLLAAIANAGRHGV | ASSLTGNQDTQY | 0.1698566 |
| TSSTAGNSGGA | ASSLTGNQDTQY | 0.17951354 |
| VPATVPVGILC | ASSLTGNQDTQY | 0.17267902 |
| VQRPDSGGVVLTGRIS | ASSLTGNQDTQY | 0.18168195 |
| WTPNPVPATKGVPAGLVQAT | ASSLTGNQDTQY | 0.17232756 |

| | | |
|---|---|---|
| YALKRALAKFSDRFDVVII | ASSLTGNQDTQY | 0.16408254 |
| YLIGSVIV | ASSLTGNQDTQY | 0.19743016 |
| YGQLVAVID | ASSLTGNQDTQY | 0.31534424 |
| VTVGGMTLPA | ASSLTGNQDTQY | 0.17842284 |
| VLALPAV | ASSLTGNQDTQY | 0.1851846 |
| VIPPTLNYET | ASSLTGNQDTQY | 0.17568164 |
| VGMGGMHPSSG | ASSLTGNQDTQY | 0.16811919 |
| VGLAGGAATA | ASSLTGNQDTQY | 0.17252819 |
| VGGAGGAGGSGP | ASSLTGNQDTQY | 0.1640248 |
| VFAAFMV | ASSLTGNQDTQY | 0.17118159 |
| TLYDKLR | ASSLTGNQDTQY | 0.1649064 |
| TLAVFTVL | ASSLTGNQDTQY | 0.18548347 |
| TGSVQPARV | ASSLTGNQDTQY | 0.1725724 |
| TGGGAGSGGT | ASSLTGNQDTQY | 0.32437974 |
| SHLETSL | ASSLTGNQDTQY | 0.18042262 |
| SFTTGMT | ASSLTGNQDTQY | 0.17679459 |
| QVEVAKRMGVRQPTVS | ASSLTGNQDTQY | 0.16620974 |
| QGSGGPALAM | ASSLTGNQDTQY | 0.17099872 |
| MLAPSNP | ASSLTGNQDTQY | 0.17020357 |
| LVNNAAI | ASSLTGNQDTQY | 0.16720755 |
| LTGEINTLA | ASSLTGNQDTQY | 0.19629402 |
| LSIFGPV | ASSLTGNQDTQY | 0.16780864 |
| LLALLANL | ASSLTGNQDTQY | 0.16788475 |
| LISNASDALDKLR | ASSLTGNQDTQY | 0.17626697 |
| LINEAVP | ASSLTGNQDTQY | 0.40426654 |
| LGYAFSGL | ASSLTGNQDTQY | 0.18407202 |
| LDFVAVAL | ASSLTGNQDTQY | 0.17864878 |
| LAQPGAE | ASSLTGNQDTQY | 0.16613837 |
| ISNLGAVTGF | ASSLTGNQDTQY | 0.17508581 |
| HQEDPGRI | ASSLTGNQDTQY | 0.24221617 |
| GTPGEPTP | ASSLTGNQDTQY | 0.16646801 |
| GFFNSGS | ASSLTGNQDTQY | 0.37455094 |
| GAGGGGLTAGAN | ASSLTGNQDTQY | 0.17194822 |
| FVDALVNM | ASSLTGNQDTQY | 0.16627681 |
| FLTVAPEL | ASSLTGNQDTQY | 0.17800295 |
| FETLTAL | ASSLTGNQDTQY | 0.18712842 |
| DVINPIEL | ASSLTGNQDTQY | 0.18944146 |
| APVSGAPV | ASSLTGNQDTQY | 0.18650137 |
| ALVGGLAW | ASSLTGNQDTQY | 0.17873886 |
| ALAAAAATAI | ASSLTGNQDTQY | 0.16793574 |
| AFGLGLLL | ASSLTGNQDTQY | 0.19877069 |
| AFFAAASL | ASSLTGNQDTQY | 0.21692146 |
| AAAALPAFSPP | ASSLTGNQDTQY | 0.18827608 |
| VVLAPAL | ASSLTGNQDTQY | 0.16651882 |
| TIGHVDHGKTTL | ASSLTGNQDTQY | 0.16560479 |
| LISNASDALDK | ASSLTGNQDTQY | 0.18564439 |
| LISNASDAL | ASSLTGNQDTQY | 0.1812836 |
| ISNASDALDK | ASSLTGNQDTQY | 0.16996695 |
| NQAETLVYQTE | ASSLTGNQDTQY | 0.1778197 |
| TGGTGGTAGAG | ASSLTGNQDTQY | 0.17673388 |
| TGSTAGGAGGA | ASSLTGNQDTQY | 0.16531725 |

| | | |
|---|---|---|
| DPLAVPVAAGPV | ASSLTGNQDTQY | 0.1761721 |
| GGTGGTPGNGGN | ASSLTGNQDTQY | 0.2107882 |
| PAGLAGVDPATL | ASSLTGNQDTQY | 0.18272963 |
| GNNLVVEAPHAGASV | ASSLTGNQDTQY | 0.16370775 |
| KVTRIMSASTLEQPA | ASSLTGNQDTQY | 0.20361574 |
| LGFALPIALPRRTVV | ASSLTGNQDTQY | 0.29430366 |
| TRLGNLTARKLVVLV | ASSLTGNQDTQY | 0.1915816 |
| WSQQRRLIYFPSAGP | ASSLTGNQDTQY | 0.16556694 |
| GPNGNPDPMAAAVDIR | ASSLTGNQDTQY | 0.16109021 |
| TQRCVSVADLRAAASR | ASSLTGNQDTQY | 0.17884952 |
| QFVLDVAEHPVEVLTETV | ASSLTGNQDTQY | 0.18144618 |
| GGAGGTGGAGGNGGTQAGDG | ASSLTGNQDTQY | 0.1692068 |
| PLLATLRAIAVDVDAKPAQV | ASSLTGNQDTQY | 0.16549087 |
| APKFRTGGYMIDFWGVGYQVAK | ASSLTGNQDTQY | 0.17200719 |
| GAGGAGGAGGAAGGPGGTGGTG | ASSLTGNQDTQY | 0.18982977 |
| IGALGLLATDVAPAGSVAFGVA | ASSLTGNQDTQY | 0.16971861 |
| AGGAGGIGGNGGAGGAGGVLMG | ASSLTGNQDTQY | 0.16333488 |
| ADELIGLL | ASSLTGNQDTQY | 0.1836289 |
| ADRALKA | ASSLTGNQDTQY | 0.20347324 |
| AGAVGQVGGAGGA | ASSLTGNQDTQY | 0.16362347 |
| AGGGGGLGGAP | ASSLTGNQDTQY | 0.17079885 |
| AGGLLYGNGGAG | ASSLTGNQDTQY | 0.20440567 |
| AGLESIGL | ASSLTGNQDTQY | 0.16771406 |
| AGLKTGII | ASSLTGNQDTQY | 0.16804086 |
| ASLAALLL | ASSLTGNQDTQY | 0.16466728 |
| ASVINISSI | ASSLTGNQDTQY | 0.18044762 |
| ATPPGPA | ASSLTGNQDTQY | 0.16820866 |
| ATVASALV | ASSLTGNQDTQY | 0.18293653 |
| ATVASLIL | ASSLTGNQDTQY | 0.1753807 |
| AVNGPVLIRH | ASSLTGNQDTQY | 0.17806505 |
| DGAAGAVGI | ASSLTGNQDTQY | 0.18713008 |
| DPLLHGI | ASSLTGNQDTQY | 0.16757615 |
| ELGAALGAL | ASSLTGNQDTQY | 0.16542938 |
| ELGRTQAV | ASSLTGNQDTQY | 0.16108687 |
| FVAGGYLS | ASSLTGNQDTQY | 0.19790903 |
| GASGGAGGAGGW | ASSLTGNQDTQY | 0.16829759 |
| GGAGAGGDGGAGGI | ASSLTGNQDTQY | 0.18609002 |
| GTGHNGSQPGMGGNG | ASSLTGNQDTQY | 0.18893725 |
| GYAAASASAS | ASSLTGNQDTQY | 0.16847433 |
| HQVDVAAVQQG | ASSLTGNQDTQY | 0.17276627 |
| IGAAIAHT | ASSLTGNQDTQY | 0.16243476 |
| IGMIEVAQL | ASSLTGNQDTQY | 0.17215699 |
| IRLAPEHTSAMD | ASSLTGNQDTQY | 0.1691854 |
| IYNVSVL | ASSLTGNQDTQY | 0.17914854 |
| KQGFLGM | ASSLTGNQDTQY | 0.18424036 |
| LESVITQL | ASSLTGNQDTQY | 0.18533088 |
| LGLGALVVV | ASSLTGNQDTQY | 0.18050687 |
| LIGNGGNGGSGGI | ASSLTGNQDTQY | 0.1673566 |
| LNAINAPTL | ASSLTGNQDTQY | 0.18390208 |
| LQLLNTGV | ASSLTGNQDTQY | 0.20395696 |
| LQTSIEF | ASSLTGNQDTQY | 0.19072627 |

| | | |
|---|---|---|
| MSFAVSEL | ASSLTGNQDTQY | 0.16447787 |
| NTGALGL | ASSLTGNQDTQY | 0.18352123 |
| PPGAGGAGGAGTT | ASSLTGNQDTQY | 0.1675155 |
| QLVGLGI | ASSLTGNQDTQY | 0.17584556 |
| RFLEQVHQL | ASSLTGNQDTQY | 0.17243002 |
| RGVGYVLR | ASSLTGNQDTQY | 0.1813501 |
| SATGVFAGVI | ASSLTGNQDTQY | 0.17957136 |
| SLPAITLP | ASSLTGNQDTQY | 0.1651331 |
| STALAFTV | ASSLTGNQDTQY | 0.16537257 |
| SVLTVGLGK | ASSLTGNQDTQY | 0.17420638 |
| TALAGAGSGI | ASSLTGNQDTQY | 0.20005685 |
| TGLPFTGI | ASSLTGNQDTQY | 0.18020104 |
| VAEHVTV | ASSLTGNQDTQY | 0.16570646 |
| VDIVLNSL | ASSLTGNQDTQY | 0.17515564 |
| VLTGGGAVL | ASSLTGNQDTQY | 0.17335989 |
| VNVGLAYGI | ASSLTGNQDTQY | 0.1714456 |
| VSSAAVGAV | ASSLTGNQDTQY | 0.17981207 |
| VTSVYEAGV | ASSLTGNQDTQY | 0.1699235 |
| WVALDRGA | ASSLTGNQDTQY | 0.1655385 |
| YLVGSVVI | ASSLTGNQDTQY | 0.18085302 |
| GPPGTGKTLL | ASSLTGNQDTQY | 0.16767491 |
| GVGMTKF | ASSLTGNQDTQY | 0.17698832 |
| IGIGGSDL | ASSLTGNQDTQY | 0.16806467 |
| IGTLLAEL | ASSLTGNQDTQY | 0.16514501 |
| IYTSGTT | ASSLTGNQDTQY | 0.17941251 |
| LSFPAGV | ASSLTGNQDTQY | 0.17033358 |
| LYTSGTT | ASSLTGNQDTQY | 0.17346054 |
| VGAFPFGF | ASSLTGNQDTQY | 0.17620696 |
| VGAGTVEF | ASSLTGNQDTQY | 0.16750029 |
| VGKSTLLL | ASSLTGNQDTQY | 0.18128055 |
| AALAHAGGGIV | ASSLTGNQDTQY | 0.19420606 |
| AALVAVSLL | ASSLTGNQDTQY | 0.18761306 |
| AAVSRAVAAL | ASSLTGNQDTQY | 0.1711263 |
| AGGSSGIGGF | ASSLTGNQDTQY | 0.21920155 |
| AGLTVANANG | ASSLTGNQDTQY | 0.18821123 |
| AGLVGAVL | ASSLTGNQDTQY | 0.16654932 |
| ALSQAFTV | ASSLTGNQDTQY | 0.16246203 |
| ALTFSVSL | ASSLTGNQDTQY | 0.1761123 |
| ALTGAALVAL | ASSLTGNQDTQY | 0.16895844 |
| ASFSIGAGVL | ASSLTGNQDTQY | 0.17291941 |
| DGSVYKI | ASSLTGNQDTQY | 0.16228466 |
| EQLAEIV | ASSLTGNQDTQY | 0.19759242 |
| FAVSELTVM | ASSLTGNQDTQY | 0.21735054 |
| GAAPLAAGL | ASSLTGNQDTQY | 0.16850413 |
| GADGGDGGSGGAGG | ASSLTGNQDTQY | 0.18566844 |
| GFGLDTGGAGG | ASSLTGNQDTQY | 0.1636923 |
| GLASGGLL | ASSLTGNQDTQY | 0.18663597 |
| GRIAGAAVLL | ASSLTGNQDTQY | 0.21345864 |
| GSAGSLNTSF | ASSLTGNQDTQY | 0.16425154 |
| GTGQIGI | ASSLTGNQDTQY | 0.17879999 |
| GTLIALIL | ASSLTGNQDTQY | 0.17257497 |

| | | |
|---|---|---|
| IAATMGFVVG | ASSLTGNQDTQY | 0.16738005 |
| IFAQALL | ASSLTGNQDTQY | 0.20331979 |
| IGAAVALGF | ASSLTGNQDTQY | 0.17594926 |
| IGLTPLAL | ASSLTGNQDTQY | 0.179589 |
| ILGSNTSTL | ASSLTGNQDTQY | 0.17035116 |
| IVFSLTVL | ASSLTGNQDTQY | 0.16640802 |
| LASKATGF | ASSLTGNQDTQY | 0.17754824 |
| LASVALIL | ASSLTGNQDTQY | 0.18936431 |
| LFAVGLTL | ASSLTGNQDTQY | 0.16532667 |
| LFNGGIGTYI | ASSLTGNQDTQY | 0.20476322 |
| LTLGAGII | ASSLTGNQDTQY | 0.17742196 |
| LVAAALGDI | ASSLTGNQDTQY | 0.3269689 |
| NTAFVQLGI | ASSLTGNQDTQY | 0.22610155 |
| QLFSGAEL | ASSLTGNQDTQY | 0.17433457 |
| SAATFITL | ASSLTGNQDTQY | 0.1695965 |
| SNYGLANLGV | ASSLTGNQDTQY | 0.2665785 |
| SYGGAYAVM | ASSLTGNQDTQY | 0.18025668 |
| TTGVGGLTLG | ASSLTGNQDTQY | 0.18161286 |
| TTQTIYI | ASSLTGNQDTQY | 0.18419775 |
| VAAGAGLGAVI | ASSLTGNQDTQY | 0.091655746 |
| VAAVVTLL | ASSLTGNQDTQY | 0.10839984 |
| VGFLGGLI | ASSLTGNQDTQY | 0.09261816 |
| VGVIPILL | ASSLTGNQDTQY | 0.09597018 |
| VIFASSLI | ASSLTGNQDTQY | 0.10338201 |
| VKGGDGGAAAG | ASSLTGNQDTQY | 0.09127919 |
| VTKTFTNI | ASSLTGNQDTQY | 0.09809988 |
| VVAAAILL | ASSLTGNQDTQY | 0.10458097 |
| YAALDVEL | ASSLTGNQDTQY | 0.09233139 |
| LASVALLL | ASSLTGNQDTQY | 0.094701976 |
| LDAVHRV | ASSLTGNQDTQY | 0.10216283 |
| QGLSSSIF | ASSLTGNQDTQY | 0.1177175 |
| RGTFANIRL | ASSLTGNQDTQY | 0.10196585 |
| RQLWWGHRI | ASSLTGNQDTQY | 0.09493739 |
| YGPPGTGKTLL | ASSLTGNQDTQY | 0.09373841 |
| AALVGAPANVA | ASSLTGNQDTQY | 0.09921656 |
| AATFRRTE | ASSLTGNQDTQY | 0.09353925 |
| AGALGGGI | ASSLTGNQDTQY | 0.08926873 |
| AGGAGGAGGIGAG | ASSLTGNQDTQY | 0.09944463 |
| AGSGALGL | ASSLTGNQDTQY | 0.09030589 |
| AQIADQI | ASSLTGNQDTQY | 0.093435094 |
| ASAVAVTYL | ASSLTGNQDTQY | 0.098180264 |
| ASTVLGGTGG | ASSLTGNQDTQY | 0.09744255 |
| AVGGEPAK | ASSLTGNQDTQY | 0.094834 |
| CEQGTFGGTC | ASSLTGNQDTQY | 0.108243935 |
| DEGKAPAG | ASSLTGNQDTQY | 0.100606754 |
| DTVVFPAL | ASSLTGNQDTQY | 0.09949638 |
| FEGITMAG | ASSLTGNQDTQY | 0.10296383 |
| FPLVVQD | ASSLTGNQDTQY | 0.11644872 |
| GAFTTPQ | ASSLTGNQDTQY | 0.09732855 |
| GFGGFGTAGGVGGA | ASSLTGNQDTQY | 0.09365245 |
| GFLAQFL | ASSLTGNQDTQY | 0.11177593 |

| | | |
|---|---|---|
| GGAGGAGGNAGVA | ASSLTGNQDTQY | 0.10372016 |
| GGVGTAGAGGAGGAGG | ASSLTGNQDTQY | 0.100211255 |
| GNGGGGDAGNAGSGGN | ASSLTGNQDTQY | 0.102465145 |
| GTGGAGGAGNDAGS | ASSLTGNQDTQY | 0.09074645 |
| GTTNTGWFNS | ASSLTGNQDTQY | 0.09977296 |
| IFMGPLFTL | ASSLTGNQDTQY | 0.10643317 |
| ILAHAIEAI | ASSLTGNQDTQY | 0.10222606 |
| IRSAIREAL | ASSLTGNQDTQY | 0.10276976 |
| ITFAGGG | ASSLTGNQDTQY | 0.09375903 |
| ITGMSGHQEV | ASSLTGNQDTQY | 0.09175542 |
| IVAGLACTL | ASSLTGNQDTQY | 0.1013388 |
| IVGGGLAGA | ASSLTGNQDTQY | 0.09885937 |
| IVTGVLNI | ASSLTGNQDTQY | 0.10123763 |
| LAQQGCI | ASSLTGNQDTQY | 0.10635783 |
| LCLHVEN | ASSLTGNQDTQY | 0.0906553 |
| LDAISDGS | ASSLTGNQDTQY | 0.09325823 |
| LGGFAAML | ASSLTGNQDTQY | 0.08985578 |
| LGGKNPMI | ASSLTGNQDTQY | 0.091460556 |
| LGPKGVAAIL | ASSLTGNQDTQY | 0.117926456 |
| LMVGVSI | ASSLTGNQDTQY | 0.090278596 |
| LVGAAFVL | ASSLTGNQDTQY | 0.13565062 |
| LVLAVAPL | ASSLTGNQDTQY | 0.12691772 |
| MAAIAVIDAA | ASSLTGNQDTQY | 0.121426776 |
| NNGINAELQ | ASSLTGNQDTQY | 0.09832952 |
| RGEPLVLVHG | ASSLTGNQDTQY | 0.09313721 |
| RGGAGGAAGLI | ASSLTGNQDTQY | 0.112321086 |
| RSYRMHPAV | ASSLTGNQDTQY | 0.10111232 |
| SAAHLLAGQLA | ASSLTGNQDTQY | 0.092312194 |
| SAAVGSY | ASSLTGNQDTQY | 0.10530167 |
| SPHAVLHRI | ASSLTGNQDTQY | 0.09787379 |
| SQIAKVL | ASSLTGNQDTQY | 0.13756713 |
| TALGHLVP | ASSLTGNQDTQY | 0.107587755 |
| TAPTTSGAA | ASSLTGNQDTQY | 0.09923976 |
| TDGDDGNGGSPGT | ASSLTGNQDTQY | 0.09878165 |
| TDGLGEN | ASSLTGNQDTQY | 0.09078919 |
| TGGPPLPGQA | ASSLTGNQDTQY | 0.099394165 |
| TIPAITI | ASSLTGNQDTQY | 0.09105527 |
| TIPGFTI | ASSLTGNQDTQY | 0.093653776 |
| TSIASGALA | ASSLTGNQDTQY | 0.13606502 |
| TVTLPGAAI | ASSLTGNQDTQY | 0.10416456 |
| VASGLSAALP | ASSLTGNQDTQY | 0.13162227 |
| VGAGVIGI | ASSLTGNQDTQY | 0.09896218 |
| VGPKRFGV | ASSLTGNQDTQY | 0.091524124 |
| VNVAFPD | ASSLTGNQDTQY | 0.1005759 |
| VSMTNSGVS | ASSLTGNQDTQY | 0.103172444 |
| VTDVLGDLSV | ASSLTGNQDTQY | 0.10302462 |
| VVVVGAGFAGL | ASSLTGNQDTQY | 0.09373638 |
| VYTVGTVV | ASSLTGNQDTQY | 0.09410445 |
| WTTFSGP | ASSLTGNQDTQY | 0.09893146 |
| YDELVEAF | ASSLTGNQDTQY | 0.109277576 |
| YGTGGAGGNGG | ASSLTGNQDTQY | 0.099563815 |

| | | |
|---|---|---|
| YTNVMAV | ASSLTGNQDTQY | 0.09468287 |
| FLEQVHQL | ASSLTGNQDTQY | 0.09922266 |
| GLSLLAVAV | ASSLTGNQDTQY | 0.097389504 |
| LEQVHQL | ASSLTGNQDTQY | 0.12022921 |
| LNAAGAI | ASSLTGNQDTQY | 0.093166985 |
| PAVHASL | ASSLTGNQDTQY | 0.09315831 |
| TELEDDA | ASSLTGNQDTQY | 0.095404714 |
| TPPGPPN | ASSLTGNQDTQY | 0.092018396 |
| TPPPPPPP | ASSLTGNQDTQY | 0.09885839 |
| TQSPALI | ASSPSTNTEVF | 0.09404663 |
| EFAAAFPQ | ASSPSTNTEVF | 0.095321596 |
| LLSGNLDVL | ASSPSTNTEVF | 0.09787881 |
| GPANLGNYNI | ASSPSTNTEVF | 0.09305434 |
| TLGPTLP | ASSPSTNTEVF | 0.09681998 |
| VYAYSSCCK | ASSPSTNTEVF | 0.09941233 |
| APAPAPAPNI | ASSPSTNTEVF | 0.09556644 |
| PGGPVPIP | ASSPSTNTEVF | 0.10602942 |
| FLTVALEP | ASSPSTNTEVF | 0.09274772 |
| SQYFGANL | ASSPSTNTEVF | 0.097995006 |
| GGVKPAA | ASSPSTNTEVF | 0.12036852 |
| GGTGGAGGAGGW | ASSPSTNTEVF | 0.10231177 |
| GVGGVAGLGGAGG | ASSPSTNTEVF | 0.108683154 |
| VLLVLATTI | ASSPSTNTEVF | 0.091808885 |
| IGIFAAML | ASSPSTNTEVF | 0.09217214 |
| AVGDLID | ASSPSTNTEVF | 0.08882862 |
| NISPLPL | ASSPSTNTEVF | 0.11470829 |
| VTAVFVAI | ASSPSTNTEVF | 0.09106732 |
| SGDDNDAAL | ASSPSTNTEVF | 0.09507982 |
| AVAALAHS | ASSPSTNTEVF | 0.09547928 |
| GGVTVMATP | ASSPSTNTEVF | 0.0992383 |
| AGLAANVRD | ASSPSTNTEVF | 0.098534904 |
| TNRPDLIDPALLRPG | ASSPSTNTEVF | 0.09141453 |
| LANLGVTAL | ASSPSTNTEVF | 0.09186837 |
| TGERIEL | ASSPSTNTEVF | 0.10947666 |
| NGEELAA | ASSPSTNTEVF | 0.0951021 |
| VLSLVWG | ASSPSTNTEVF | 0.09708101 |
| IGYALWL | ASSPSTNTEVF | 0.09588412 |
| LSLLLAPL | ASSPSTNTEVF | 0.13987824 |
| TLDEALK | ASSPSTNTEVF | 0.11002949 |
| TDGAGTAEGG | ASSPSTNTEVF | 0.099780634 |
| NAPGGAGGNGG | ASSPSTNTEVF | 0.10152572 |
| VSVFMGMGGNF | ASSPSTNTEVF | 0.08922538 |
| LASYLAAL | ASSPSTNTEVF | 0.09004246 |
| LVGGTPAL | ASSPSTNTEVF | 0.11235573 |
| VRNADVL | ASSPSTNTEVF | 0.13162029 |
| TCSSPFLVSAM | ASSPSTNTEVF | 0.09981873 |
| VHITDAE | ASSPSTNTEVF | 0.10338348 |
| EYAAPSF | ASSPSTNTEVF | 0.105618864 |
| VVDAFHGGL | ASSPSTNTEVF | 0.08836519 |
| ASVVDPQ | ASSPSTNTEVF | 0.095703885 |
| SYNEFGGLS | ASSPSTNTEVF | 0.09406455 |

| | | |
|---|---|---|
| DPDGSVY | ASSPSTNTEVF | 0.1001042 |
| MVSTIHATE | ASSPSTNTEVF | 0.089578144 |
| LAAGVADPV | ASSPSTNTEVF | 0.096424796 |
| TKEQIAA | ASSPSTNTEVF | 0.108676165 |
| SLASPAAL | ASSPSTNTEVF | 0.09179213 |
| IGLVFYGV | ASSPSTNTEVF | 0.09939369 |
| FLLDASNL | ASSPSTNTEVF | 0.09075377 |
| YQVGNFP | ASSPSTNTEVF | 0.09759091 |
| ESLLSQST | ASSPSTNTEVF | 0.12179617 |
| LFTVGFAAVG | ASSPSTNTEVF | 0.10884304 |
| ATPPTPT | ASSPSTNTEVF | 0.0993664 |
| TDLATLG | ASSPSTNTEVF | 0.11239055 |
| SAGGTGR | ASSPSTNTEVF | 0.112821326 |
| GFSNSGS | ASSPSTNTEVF | 0.093065314 |
| GGNGGSGGTSV | ASSPSTNTEVF | 0.10075028 |
| GTSVATGGAGN | ASSPSTNTEVF | 0.09281429 |
| NGGNGGSGGTS | ASSPSTNTEVF | 0.1053784 |
| AVVKFPH | ASSPSTNTEVF | 0.10732013 |
| TPPPPPP | ASSPSTNTEVF | 0.09930513 |
| AAPAAAAAPA | ASSPSTNTEVF | 0.09792817 |
| IALNEDL | ASSPSTNTEVF | 0.10361017 |
| IVNVSSM | ASSPSTNTEVF | 0.09359285 |
| TGIPPAPR | ASSPSTNTEVF | 0.096284315 |
| FDLGGGTF | ASSPSTNTEVF | 0.09521605 |
| FDLGGGTFDVS | ASSPSTNTEVF | 0.108975574 |
| NEPTAAA | ASSPSTNTEVF | 0.09129041 |
| NEPTAAALA | ASSPSTNTEVF | 0.09216395 |
| GSIINISSI | ASSPSTNTEVF | 0.09235158 |
| VAPGNAGTA | ASSPSTNTEVF | 0.10847238 |
| TGGGASLE | ASSPSTNTEVF | 0.12343922 |
| VAVYNHYKRI | ASSPSTNTEVF | 0.09376463 |
| TDPAPATT | ASSPSTNTEVF | 0.09696356 |
| TNRPDILDPALLRPG | ASSPSTNTEVF | 0.09293506 |
| LTTTTTS | ASSPSTNTEVF | 0.10051359 |
| PTGSGKTL | ASSPSTNTEVF | 0.0910432 |
| TGSGKTL | ASSPSTNTEVF | 0.09677622 |
| TIGHVDHGK | ASSPSTNTEVF | 0.097536705 |
| TIGHVDHGKTT | ASSPSTNTEVF | 0.10877238 |
| GHVDHGKTT | ASSPSTNTEVF | 0.0955074 |
| AQFEHTLL | ASSPSTNTEVF | 0.09355303 |
| IVNQDEL | ASSPSTNTEVF | 0.10864404 |
| TNGSQFF | ASSPSTNTEVF | 0.092367105 |
| TGKGIVE | ASSPSTNTEVF | 0.102672376 |
| LISNASDA | ASSPSTNTEVF | 0.09985327 |
| AELRTLGELTKIAKAHGAQ | ASSPSTNTEVF | 0.090653926 |
| APNAVAAAEPSW | ASSPSTNTEVF | 0.09448912 |
| APQVAGAATAVL | ASSPSTNTEVF | 0.12371998 |
| ARRDPAAAAAILPTDA | ASSPSTNTEVF | 0.09428791 |
| DAAARDVLAEAGLAEYF | ASSPSTNTEVF | 0.09448586 |
| DDQIVLSAKANLAKVEQ | ASSPSTNTEVF | 0.109672025 |
| DELLPDTGEDSDEDGH | ASSPSTNTEVF | 0.09716427 |

| | | |
|---|---|---|
| ELPEAPANRATVPQD | ASSPSTNTEVF | 0.10736334 |
| FGQSKIERVTGIG | ASSPSTNTEVF | 0.09316691 |
| GAGGTGSTAGGAGGAGGAGGLY | ASSPSTNTEVF | 0.094630785 |
| GHGGTGAAGGNGGTGGAGGSGI | ASSPSTNTEVF | 0.09443911 |
| GNGGNGGAGGEGGPGGAGGAGA | ASSPSTNTEVF | 0.09526292 |
| GSTAGNGGNGGNATNGTIAGQP | ASSPSTNTEVF | 0.10953482 |
| GTGGDGGNGGNGGTGG | ASSPSTNTEVF | 0.11393509 |
| GVRRPAGIPADAPV | ASSPSTNTEVF | 0.09238867 |
| GYRQVTDAHLLTLARR | ASSPSTNTEVF | 0.114703655 |
| IAKMGQDGHDRG | ASSPSTNTEVF | 0.10241386 |
| KAEPSIEIQYT | ASSPSTNTEVF | 0.097856745 |
| KPVRIRMAADEDTVAVT | ASSPSTNTEVF | 0.09055483 |
| MPPRIAGMRLLVIKPEP | ASSPSTNTEVF | 0.09868391 |
| MKFYAVALFRSRGVPPTR | ASSPSTNTEVF | 0.09681052 |
| NAGGFGILGAVAHSP | ASSPSTNTEVF | 0.1024809 |
| NEGREITLAMAEM | ASSPSTNTEVF | 0.107564144 |
| NELVEWSHPLGHRWRWE | ASSPSTNTEVF | 0.09447509 |
| NLGAGGTGGNGGNPG | ASSPSTNTEVF | 0.10787727 |
| NTEWPKWETVMAADPVD | ASSPSTNTEVF | 0.091807306 |
| PASAAPAETQP | ASSPSTNTEVF | 0.10491811 |
| QQSSDGKNGVRAAEITGE | ASSPSTNTEVF | 0.09052821 |
| RAPDAVTTSWWKE | ASSPSTNTEVF | 0.09261325 |
| SGFGNVGAHTSGWFNQ | ASSPSTNTEVF | 0.10439629 |
| SGVAGAVGQAASVGGLKV | ASSPSTNTEVF | 0.092725165 |
| SNGGAGTGSAGGNGGT | ASSPSTNTEVF | 0.0945365 |
| TGGAGGEGGAAG | ASSPSTNTEVF | 0.09136794 |
| TGGDGGAAGAGGAPGHG | ASSPSTNTEVF | 0.104435876 |
| TMPPLLAAIANAGRHGV | ASSPSTNTEVF | 0.09990821 |
| TSSTAGNSGGA | ASSPSTNTEVF | 0.097262524 |
| VPATVPVGILC | ASSPSTNTEVF | 0.09065672 |
| VQRPDSGGVVLTGRIS | ASSPSTNTEVF | 0.10411225 |
| WTPNPVPATKGVPAGLVQAT | ASSPSTNTEVF | 0.09124008 |
| YALKRALAKFSDRFDVVII | ASSPSTNTEVF | 0.1232094 |
| YLIGSVIV | ASSPSTNTEVF | 0.10906612 |
| YGQLVAVID | ASSPSTNTEVF | 0.08995546 |
| VTVGGMTLPA | ASSPSTNTEVF | 0.098658934 |
| VLALPAV | ASSPSTNTEVF | 0.09334434 |
| VIPPTLNYET | ASSPSTNTEVF | 0.10017275 |
| VGMGGMHPSSG | ASSPSTNTEVF | 0.09544088 |
| VGLAGGAATA | ASSPSTNTEVF | 0.09066383 |
| VGGAGGAGGSGP | ASSPSTNTEVF | 0.09744787 |
| VFAAFMV | ASSPSTNTEVF | 0.10793541 |
| TLYDKLR | ASSPSTNTEVF | 0.09583353 |
| TLAVFTVL | ASSPSTNTEVF | 0.09313035 |
| TGSVQPARV | ASSPSTNTEVF | 0.095455565 |
| TGGGAGSGGT | ASSPSTNTEVF | 0.11183822 |
| SHLETSL | ASSPSTNTEVF | 0.098899275 |
| SFTTGMT | ASSPSTNTEVF | 0.10719826 |
| QVEVAKRMGVRQPTVS | ASSPSTNTEVF | 0.09595436 |
| QGSGGPALAM | ASSPSTNTEVF | 0.10589121 |
| MLAPSNP | ASSPSTNTEVF | 0.1019547 |

| | | |
|---|---|---|
| LVNNAAI | ASSPSTNTEVF | 0.106329195 |
| LTGEINTLA | ASSPSTNTEVF | 0.093528554 |
| LSIFGPV | ASSPSTNTEVF | 0.09389897 |
| LLALLANL | ASSPSTNTEVF | 0.09862113 |
| LISNASDALDKLR | ASSPSTNTEVF | 0.09633397 |
| LINEAVP | ASSPSTNTEVF | 0.092155606 |
| LGYAFSGL | ASSPSTNTEVF | 0.114775755 |
| LDFVAVAL | ASSPSTNTEVF | 0.09548862 |
| LAQPGAE | ASSPSTNTEVF | 0.11642007 |
| ISNLGAVTGF | ASSPSTNTEVF | 0.09254214 |
| HQEDPGRI | ASSPSTNTEVF | 0.09318422 |
| GTPGEPTP | ASSPSTNTEVF | 0.12639779 |
| GFFNSGS | ASSPSTNTEVF | 0.091332056 |
| GAGGGGLTAGAN | ASSPSTNTEVF | 0.107836924 |
| FVDALVNM | ASSPSTNTEVF | 0.10000818 |
| FLTVAPEL | ASSPSTNTEVF | 0.09863583 |
| FETLTAL | ASSPSTNTEVF | 0.09398323 |
| DVINPIEL | ASSPSTNTEVF | 0.100027524 |
| APVSGAPV | ASSPSTNTEVF | 0.09256933 |
| ALVGGLAW | ASSPSTNTEVF | 0.10161527 |
| ALAAAAATAI | ASSPSTNTEVF | 0.09127753 |
| AFGLGLLL | ASSPSTNTEVF | 0.09821336 |
| AFFAAASL | ASSPSTNTEVF | 0.09015065 |
| AAAALPAFSPP | ASSPSTNTEVF | 0.10158287 |
| VVLAPAL | ASSPSTNTEVF | 0.11269851 |
| TIGHVDHGKTTL | ASSPSTNTEVF | 0.105126254 |
| LISNASDALDK | ASSPSTNTEVF | 0.10100064 |
| LISNASDAL | ASSPSTNTEVF | 0.1180432 |
| ISNASDALDK | ASSPSTNTEVF | 0.10819314 |
| NQAETLVYQTE | ASSPSTNTEVF | 0.11682613 |
| TGGTGGTAGAG | ASSPSTNTEVF | 0.097357854 |
| TGSTAGGAGGA | ASSPSTNTEVF | 0.09885539 |
| DPLAVPVAAGPV | ASSPSTNTEVF | 0.10848703 |
| GGTGGTPGNGGN | ASSPSTNTEVF | 0.09337539 |
| PAGLAGVDPATL | ASSPSTNTEVF | 0.0906858 |
| GNNLVVEAPHAGASV | ASSPSTNTEVF | 0.093497045 |
| KVTRIMSASTLEQPA | ASSPSTNTEVF | 0.09774891 |
| LGFALPIALPRRTVV | ASSPSTNTEVF | 0.09758789 |
| TRLGNLTARKLVVLV | ASSPSTNTEVF | 0.10333919 |
| WSQQRRLIYFPSAGP | ASSPSTNTEVF | 0.09974943 |
| GPNGNPDPMAAAVDIR | ASSPSTNTEVF | 0.1061629 |
| TQRCVSVADLRAAASR | ASSPSTNTEVF | 0.11494677 |
| QFVLDVAEHPVEVLTETV | ASSPSTNTEVF | 0.11393462 |
| GGAGGTGGAGGNGGTQAGDG | ASSPSTNTEVF | 0.09455336 |
| PLLATLRAIAVDVDAKPAQV | ASSPSTNTEVF | 0.09668919 |
| APKFRTGGYMIDFWGVGYQVAK | ASSPSTNTEVF | 0.09484781 |
| GAGGAGGAGGAAGGPGGTGGTG | ASSPSTNTEVF | 0.09429157 |
| IGALGLLATDVAPAGSVAFGVA | ASSPSTNTEVF | 0.09644655 |
| AGGAGGIGGNGGAGGAGGVLMG | ASSPSTNTEVF | 0.09597618 |
| ADELIGLL | ASSPSTNTEVF | 0.108351186 |
| ADRALKA | ASSPSTNTEVF | 0.08796323 |

| | | |
|---|---|---|
| AGAVGQVGGAGGA | ASSPSTNTEVF | 0.09430093 |
| AGGGGGLGGAP | ASSPSTNTEVF | 0.10487093 |
| AGGLLYGNGGAG | ASSPSTNTEVF | 0.11324169 |
| AGLESIGL | ASSPSTNTEVF | 0.10345898 |
| AGLKTGII | ASSPSTNTEVF | 0.09789136 |
| ASLAALLL | ASSPSTNTEVF | 0.08988491 |
| ASVINISSI | ASSPSTNTEVF | 0.124558166 |
| ATPPGPA | ASSPSTNTEVF | 0.09131581 |
| ATVASALV | ASSPSTNTEVF | 0.09878773 |
| ATVASLIL | ASSPSTNTEVF | 0.10946409 |
| AVNGPVLIRH | ASSPSTNTEVF | 0.09792558 |
| DGAAGAVGI | ASSPSTNTEVF | 0.091332115 |
| DPLLHGI | ASSPSTNTEVF | 0.09421779 |
| ELGAALGAL | ASSPSTNTEVF | 0.09527098 |
| ELGRTQAV | ASSPSTNTEVF | 0.09190004 |
| FVAGGYLS | ASSPSTNTEVF | 0.0893236 |
| GASGGAGGAGGW | ASSPSTNTEVF | 0.09836161 |
| GGAGAGGDGGAGGI | ASSPSTNTEVF | 0.092112385 |
| GTGHNGSQPGMGGNG | ASSPSTNTEVF | 0.12952806 |
| GYAAASASAS | ASSPSTNTEVF | 0.11259929 |
| HQVDVAAVQQG | ASSPSTNTEVF | 0.09631228 |
| IGAAIAHT | ASSPSTNTEVF | 0.09613642 |
| IGMIEVAQL | ASSPSTNTEVF | 0.13792287 |
| IRLAPEHTSAMD | ASSPSTNTEVF | 0.12343701 |
| IYNVSVL | ASSPSTNTEVF | 0.09368286 |
| KQGFLGM | ASSPSTNTEVF | 0.09976694 |
| LESVITQL | ASSPSTNTEVF | 0.09444314 |
| LGLGALVVV | ASSPSTNTEVF | 0.09337846 |
| LIGNGGNGGSGGI | ASSPSTNTEVF | 0.105752796 |
| LNAINAPTL | ASSPSTNTEVF | 0.09764368 |
| LQLLNTGV | ASSPSTNTEVF | 0.09430634 |
| LQTSIEF | ASSPSTNTEVF | 0.09604346 |
| MSFAVSEL | ASSPSTNTEVF | 0.09804074 |
| NTGALGL | ASSPSTNTEVF | 0.10024956 |
| PPGAGGAGGAGTT | ASSPSTNTEVF | 0.08896879 |
| QLVGLGI | ASSPSTNTEVF | 0.09208844 |
| RFLEQVHQL | ASSPSTNTEVF | 0.099330716 |
| RGVGYVLR | ASSPSTNTEVF | 0.091132924 |
| SATGVFAGVI | ASSPSTNTEVF | 0.09197368 |
| SLPAITLP | ASSPSTNTEVF | 0.12553078 |
| STALAFTV | ASSPSTNTEVF | 0.104943015 |
| SVLTVGLGK | ASSPSTNTEVF | 0.0934173 |
| TALAGAGSGI | ASSPSTNTEVF | 0.09930454 |
| TGLPFTGI | ASSPSTNTEVF | 0.13350402 |
| VAEHVTV | ASSPSTNTEVF | 0.10959729 |
| VDIVLNSL | ASSPSTNTEVF | 0.09894183 |
| VLTGGGAVL | ASSPSTNTEVF | 0.094040595 |
| VNVGLAYGI | ASSPSTNTEVF | 0.096420735 |
| VSSAAVGAV | ASSPSTNTEVF | 0.10164352 |
| VTSVYEAGV | ASSPSTNTEVF | 0.10712348 |
| WVALDRGA | ASSPSTNTEVF | 0.10359592 |

| | | |
|---|---|---|
| YLVGSVVI | ASSPSTNTEVF | 0.09951299 |
| GPPGTGKTLL | ASSPSTNTEVF | 0.116482176 |
| GVGMTKF | ASSPSTNTEVF | 0.11149046 |
| IGIGGSDL | ASSPSTNTEVF | 0.09851029 |
| IGTLLAEL | ASSPSTNTEVF | 0.0910889 |
| IYTSGTT | ASSPSTNTEVF | 0.09075318 |
| LSFPAGV | ASSPSTNTEVF | 0.09239023 |
| LYTSGTT | ASSPSTNTEVF | 0.100118786 |
| VGAFPFGF | ASSPSTNTEVF | 0.10354662 |
| VGAGTVEF | ASSPSTNTEVF | 0.094727 |
| VGKSTLLL | ASSPSTNTEVF | 0.11054427 |
| AALAHAGGGIV | ASSPSTNTEVF | 0.12668376 |
| AALVAVSLL | ASSPSTNTEVF | 0.09959898 |
| AAVSRAVAAL | ASSPSTNTEVF | 0.10336326 |
| AGGSSGIGGF | ASSPSTNTEVF | 0.09437376 |
| AGLTVANANG | ASSPSTNTEVF | 0.093383625 |
| AGLVGAVL | ASSPSTNTEVF | 0.09328088 |
| ALSQAFTV | ASSPSTNTEVF | 0.091070354 |
| ALTFSVSL | ASSPSTNTEVF | 0.0921345 |
| ALTGAALVAL | ASSPSTNTEVF | 0.09046885 |
| ASFSIGAGVL | ASSPSTNTEVF | 0.09327335 |
| DGSVYKI | ASSPSTNTEVF | 0.09344338 |
| EQLAEIV | ASSPSTNTEVF | 0.09191901 |
| FAVSELTVM | ASSPSTNTEVF | 0.12101276 |
| GAAPLAAGL | ASSPSTNTEVF | 0.093880884 |
| GADGGDGGSGGAGG | ASSPSTNTEVF | 0.10646544 |
| GFGLDTGGAGG | ASSPSTNTEVF | 0.15884858 |
| GLASGGLL | ASSPSTNTEVF | 0.09491062 |
| GRIAGAAVLL | ASSPSTNTEVF | 0.09674146 |
| GSAGSLNTSF | ASSPSTNTEVF | 0.097849235 |
| GTGQIGI | ASSPSTNTEVF | 0.11846574 |
| GTLIALIL | ASSPSTNTEVF | 0.09367508 |
| IAATMGFVVG | ASSPSTNTEVF | 0.10013799 |
| IFAQALL | ASSPSTNTEVF | 0.09608579 |
| IGAAVALGF | ASSPSTNTEVF | 0.0904091 |
| IGLTPLAL | ASSPSTNTEVF | 0.09139512 |
| ILGSNTSTL | ASSPSTNTEVF | 0.09369078 |
| IVFSLTVL | ASSPSTNTEVF | 0.09895347 |
| LASKATGF | ASSPSTNTEVF | 0.09037859 |
| LASVALIL | ASSPSTNTEVF | 0.09410607 |
| LFAVGLTL | ASSPSTNTEVF | 0.09556443 |
| LFNGGIGTYI | ASSPSTNTEVF | 0.09621057 |
| LTLGAGII | ASSPSTNTEVF | 0.08946372 |
| LVAAALGDI | ASSPSTNTEVF | 0.09558625 |
| NTAFVQLGI | ASSPSTNTEVF | 0.108434014 |
| QLFSGAEL | ASSPSTNTEVF | 0.10180973 |
| SAATFITL | ASSPSTNTEVF | 0.11889877 |
| SNYGLANLGV | ASSPSTNTEVF | 0.09499495 |
| SYGGAYAVM | ASSPSTNTEVF | 0.12994066 |
| TTGVGGLTLG | ASSPSTNTEVF | 0.097360924 |
| TTQTIYI | ASSPSTNTEVF | 0.09222292 |

| | | |
|---|---|---|
| VAAGAGLGAVI | ASSPSTNTEVF | 0.10572567 |
| VAAVVTLL | ASSPSTNTEVF | 0.09020443 |
| VGFLGGLI | ASSPSTNTEVF | 0.0929886 |
| VGVIPILL | ASSPSTNTEVF | 0.10157178 |
| VIFASSLI | ASSPSTNTEVF | 0.09074995 |
| VKGGDGGAAAG | ASSPSTNTEVF | 0.097654186 |
| VTKTFTNI | ASSPSTNTEVF | 0.106139 |
| VVAAAILL | ASSPSTNTEVF | 0.09262873 |
| YAALDVEL | ASSPSTNTEVF | 0.09301844 |
| LASVALLL | ASSPSTNTEVF | 0.09277656 |
| LDAVHRV | ASSPSTNTEVF | 0.09914383 |
| QGLSSSIF | ASSPSTNTEVF | 0.10833225 |
| RGTFANIRL | ASSPSTNTEVF | 0.093745604 |
| RQLWWGHRI | ASSPSTNTEVF | 0.10252188 |
| YGPPGTGKTLL | ASSPSTNTEVF | 0.09733157 |
| AALVGAPANVA | ASSPSTNTEVF | 0.09705597 |
| AATFRRTE | ASSPSTNTEVF | 0.10566962 |
| AGALGGGI | ASSPSTNTEVF | 0.111849524 |
| AGGAGGAGGIGAG | ASSPSTNTEVF | 0.09562487 |
| AGSGALGL | ASSPSTNTEVF | 0.09312254 |
| AQIADQI | ASSPSTNTEVF | 0.09629137 |
| ASAVAVTYL | ASSPSTNTEVF | 0.093903735 |
| ASTVLGGTGG | ASSPSTNTEVF | 0.12932445 |
| AVGGEPAK | ASSPSTNTEVF | 0.0918731 |
| CEQGTFGGTC | ASSPSTNTEVF | 0.095454164 |
| DEGKAPAG | ASSPSTNTEVF | 0.095273584 |
| DTVVFPAL | ASSPSTNTEVF | 0.11124822 |
| FEGITMAG | ASSPSTNTEVF | 0.09581154 |
| FPLVVQD | ASSPSTNTEVF | 0.10928275 |
| GAFTTPQ | ASSPSTNTEVF | 0.106476404 |
| GFGGFGTAGGVGGA | ASSPSTNTEVF | 0.09455055 |
| GFLAQFL | ASSPSTNTEVF | 0.1036265 |
| GGAGGAGGNAGVA | ASSPSTNTEVF | 0.10903583 |
| GGVGTAGAGGAGGAGG | ASSPSTNTEVF | 0.094166316 |
| GNGGGGDAGNAGSGGN | ASSPSTNTEVF | 0.09049434 |
| GTGGAGGAGNDAGS | ASSPSTNTEVF | 0.09126723 |
| GTTNTGWFNS | ASSPSTNTEVF | 0.09386782 |
| IFMGPLFTL | ASSPSTNTEVF | 0.09839507 |
| ILAHAIEAI | ASSPSTNTEVF | 0.09545553 |
| IRSAIREAL | ASSPSTNTEVF | 0.097633936 |
| ITFAGGG | ASSPSTNTEVF | 0.102034226 |
| ITGMSGHQEV | ASSPSTNTEVF | 0.09639286 |
| IVAGLACTL | ASSPSTNTEVF | 0.11322545 |
| IVGGGLAGA | ASSPSTNTEVF | 0.096589185 |
| IVTGVLNI | ASSPSTNTEVF | 0.13264102 |
| LAQQGCI | ASSPSTNTEVF | 0.09429638 |
| LCLHVEN | ASSPSTNTEVF | 0.09276359 |
| LDAISDGS | ASSPSTNTEVF | 0.10266198 |
| LGGFAAML | ASSPSTNTEVF | 0.10437406 |
| LGGKNPMI | ASSPSTNTEVF | 0.094953895 |
| LGPKGVAAIL | ASSPSTNTEVF | 0.12433154 |

| | | |
|---|---|---|
| LMVGVSI | ASSPSTNTEVF | 0.13093357 |
| LVGAAFVL | ASSPSTNTEVF | 0.09408085 |
| LVLAVAPL | ASSPSTNTEVF | 0.11106419 |
| MAAIAVIDAA | ASSPSTNTEVF | 0.08958501 |
| NNGINAELQ | ASSPSTNTEVF | 0.0910012 |
| RGEPLVLVHG | ASSPSTNTEVF | 0.0906122 |
| RGGAGGAAGLI | ASSPSTNTEVF | 0.10174854 |
| RSYRMHPAV | ASSPSTNTEVF | 0.09573656 |
| SAAHLLAGQLA | ASSPSTNTEVF | 0.100608416 |
| SAAVGSY | ASSPSTNTEVF | 0.09584168 |
| SPHAVLHRI | ASSPSTNTEVF | 0.12187589 |
| SQIAKVL | ASSPSTNTEVF | 0.09976815 |
| TALGHLVP | ASSPSTNTEVF | 0.09188629 |
| TAPTTSGAA | ASSPSTNTEVF | 0.09987542 |
| TDGDDGNGGSPGT | ASSPSTNTEVF | 0.10385976 |
| TDGLGEN | ASSPSTNTEVF | 0.09097439 |
| TGGPPLPGQA | ASSPSTNTEVF | 0.09156941 |
| TIPAITI | ASSPSTNTEVF | 0.09561754 |
| TIPGFTI | ASSPSTNTEVF | 0.094233334 |
| TSIASGALA | ASSPSTNTEVF | 0.1089712 |
| TVTLPGAAI | ASSPSTNTEVF | 0.10583445 |
| VASGLSAALP | ASSPSTNTEVF | 0.09764278 |
| VGAGVIGI | ASSPSTNTEVF | 0.09583734 |
| VGPKRFGV | ASSPSTNTEVF | 0.08903664 |
| VNVAFPD | ASSPSTNTEVF | 0.09772952 |
| VSMTNSGVS | ASSPSTNTEVF | 0.13936248 |
| VTDVLGDLSV | ASSPSTNTEVF | 0.0991536 |
| VVVVGAGFAGL | ASSPSTNTEVF | 0.10552622 |
| VYTVGTVV | ASSPSTNTEVF | 0.10659728 |
| WTTFSGP | ASSPSTNTEVF | 0.102276236 |
| YDELVEAF | ASSPSTNTEVF | 0.099707656 |
| YGTGGAGGNGG | ASSPSTNTEVF | 0.098943636 |
| YTNVMAV | ASSPSTNTEVF | 0.09454025 |
| FLEQVHQL | ASSPSTNTEVF | 0.09659798 |
| GLSLLAVAV | ASSPSTNTEVF | 0.09081246 |
| LEQVHQL | ASSPSTNTEVF | 0.099632725 |
| LNAAGAI | ASSPSTNTEVF | 0.16015448 |
| PAVHASL | ASSPSTNTEVF | 0.10831751 |
| TELEDDA | ASSPSTNTEVF | 0.107892446 |
| TPPGPPN | ASSPSTNTEVF | 0.09749166 |
| TPPPPPPP | ASSPSTNTEVF | 0.09886278 |
| TQSPALI | ASSLNWGEAETLY | 0.0990617 |
| EFAAAFPQ | ASSLNWGEAETLY | 0.09286585 |
| LLSGNLDVL | ASSLNWGEAETLY | 0.13797003 |
| GPANLGNYNI | ASSLNWGEAETLY | 0.09248304 |
| TLGPTLP | ASSLNWGEAETLY | 0.09416169 |
| VYAYSSCCK | ASSLNWGEAETLY | 0.09914961 |
| APAPAPAPNI | ASSLNWGEAETLY | 0.18352541 |
| PGGPVPIP | ASSLNWGEAETLY | 0.09793086 |
| FLTVALEP | ASSLNWGEAETLY | 0.102879584 |
| SQYFGANL | ASSLNWGEAETLY | 0.10053919 |

| | | |
|---|---|---|
| GGVKPAA | ASSLNWGEAETLY | 0.09329295 |
| GGTGGAGGAGGW | ASSLNWGEAETLY | 0.09532627 |
| GVGGVAGLGGAGG | ASSLNWGEAETLY | 0.09445488 |
| VLLVLATTI | ASSLNWGEAETLY | 0.1173282 |
| IGIFAAML | ASSLNWGEAETLY | 0.09705751 |
| AVGDLID | ASSLNWGEAETLY | 0.09277153 |
| NISPLPL | ASSLNWGEAETLY | 0.09864457 |
| VTAVFVAI | ASSLNWGEAETLY | 0.102774054 |
| SGDDNDAAL | ASSLNWGEAETLY | 0.09817137 |
| AVAALAHS | ASSLNWGEAETLY | 0.1637654 |
| GGVTVMATP | ASSLNWGEAETLY | 0.100021936 |
| AGLAANVRD | ASSLNWGEAETLY | 0.101516485 |
| TNRPDLIDPALLRPG | ASSLNWGEAETLY | 0.14786878 |
| LANLGVTAL | ASSLNWGEAETLY | 0.091011725 |
| TGERIEL | ASSLNWGEAETLY | 0.1017467 |
| NGEELAA | ASSLNWGEAETLY | 0.093945295 |
| VLSLVWG | ASSLNWGEAETLY | 0.09085583 |
| IGYALWL | ASSLNWGEAETLY | 0.09546633 |
| LSLLLAPL | ASSLNWGEAETLY | 0.095983684 |
| TLDEALK | ASSLNWGEAETLY | 0.10141912 |
| TDGAGTAEGG | ASSLNWGEAETLY | 0.10159863 |
| NAPGGAGGNGG | ASSLNWGEAETLY | 0.09041524 |
| VSVFMGMGGNF | ASSLNWGEAETLY | 0.092946485 |
| LASYLAAL | ASSLNWGEAETLY | 0.11114471 |
| LVGGTPAL | ASSLNWGEAETLY | 0.112627216 |
| VRNADVL | ASSLNWGEAETLY | 0.10600035 |
| TCSSPFLVSAM | ASSLNWGEAETLY | 0.09335299 |
| VHITDAE | ASSLNWGEAETLY | 0.09414039 |
| EYAAPSF | ASSLNWGEAETLY | 0.09500814 |
| VVDAFHGGL | ASSLNWGEAETLY | 0.09742235 |
| ASVVDPQ | ASSLNWGEAETLY | 0.09089609 |
| SYNEFGGLS | ASSLNWGEAETLY | 0.09978291 |
| DPDGSVY | ASSLNWGEAETLY | 0.11520569 |
| MVSTIHATE | ASSLNWGEAETLY | 0.09083437 |
| LAAGVADPV | ASSLNWGEAETLY | 0.10117266 |
| TKEQIAA | ASSLNWGEAETLY | 0.100280836 |
| SLASPAAL | ASSLNWGEAETLY | 0.09319701 |
| IGLVFYGV | ASSLNWGEAETLY | 0.09097383 |
| FLLDASNL | ASSLNWGEAETLY | 0.11033341 |
| YQVGNFP | ASSLNWGEAETLY | 0.15615343 |
| ESLLSQST | ASSLNWGEAETLY | 0.098564155 |
| LFTVGFAAVG | ASSLNWGEAETLY | 0.09248135 |
| ATPPTPT | ASSLNWGEAETLY | 0.08963692 |
| TDLATLG | ASSLNWGEAETLY | 0.1644013 |
| SAGGTGR | ASSLNWGEAETLY | 0.09744604 |
| GFSNSGS | ASSLNWGEAETLY | 0.097509354 |
| GGNGGSGGTSV | ASSLNWGEAETLY | 0.091386266 |
| GTSVATGGAGN | ASSLNWGEAETLY | 0.09528774 |
| NGGNGGSGGTS | ASSLNWGEAETLY | 0.11480771 |
| AVVKFPH | ASSLNWGEAETLY | 0.09107196 |
| TPPPPPP | ASSLNWGEAETLY | 0.09134962 |

| | | |
|---|---|---|
| AAPAAAAAPA | ASSLNWGEAETLY | 0.09639049 |
| IALNEDL | ASSLNWGEAETLY | 0.102060065 |
| IVNVSSM | ASSLNWGEAETLY | 0.0900706 |
| TGIPPAPR | ASSLNWGEAETLY | 0.095006384 |
| FDLGGGTF | ASSLNWGEAETLY | 0.1874949 |
| FDLGGGTFDVS | ASSLNWGEAETLY | 0.12302675 |
| NEPTAAA | ASSLNWGEAETLY | 0.09799201 |
| NEPTAAALA | ASSLNWGEAETLY | 0.09737457 |
| GSIINISSI | ASSLNWGEAETLY | 0.0995038 |
| VAPGNAGTA | ASSLNWGEAETLY | 0.09646008 |
| TGGGASLE | ASSLNWGEAETLY | 0.097002104 |
| VAVYNHYKRI | ASSLNWGEAETLY | 0.09944895 |
| TDPAPATT | ASSLNWGEAETLY | 0.09885694 |
| TNRPDILDPALLRPG | ASSLNWGEAETLY | 0.098553866 |
| LTTTTTS | ASSLNWGEAETLY | 0.09399031 |
| PTGSGKTL | ASSLNWGEAETLY | 0.09356939 |
| TGSGKTL | ASSLNWGEAETLY | 0.10070769 |
| TIGHVDHGK | ASSLNWGEAETLY | 0.09839574 |
| TIGHVDHGKTT | ASSLNWGEAETLY | 0.0900833 |
| GHVDHGKTT | ASSLNWGEAETLY | 0.09840947 |
| AQFEHTLL | ASSLNWGEAETLY | 0.10930206 |
| IVNQDEL | ASSLNWGEAETLY | 0.09458335 |
| TNGSQFF | ASSLNWGEAETLY | 0.09450369 |
| TGKGIVE | ASSLNWGEAETLY | 0.08976022 |
| LISNASDA | ASSLNWGEAETLY | 0.13474317 |
| AELRTLGELTKIAKAHGAQ | ASSLNWGEAETLY | 0.10119491 |
| APNAVAAAEPSW | ASSLNWGEAETLY | 0.09372604 |
| APQVAGAATAVL | ASSLNWGEAETLY | 0.09664096 |
| ARRDPAAAAAILPTDA | ASSLNWGEAETLY | 0.09782068 |
| DAAARDVLAEAGLAEYF | ASSLNWGEAETLY | 0.099023305 |
| DDQIVLSAKANLAKVEQ | ASSLNWGEAETLY | 0.09344854 |
| DELLPDTGEDSDEDGH | ASSLNWGEAETLY | 0.09261275 |
| ELPEAPANRATVPQD | ASSLNWGEAETLY | 0.11326038 |
| FGQSKIERVTGIG | ASSLNWGEAETLY | 0.096815534 |
| GAGGTGSTAGGAGGAGGAGGLY | ASSLNWGEAETLY | 0.09154422 |
| GHGGTGAAGGNGGTGGAGGSGI | ASSLNWGEAETLY | 0.10777199 |
| GNGGNGGAGGEGGPGGAGGAGA | ASSLNWGEAETLY | 0.09041046 |
| GSTAGNGGNGGNATNGTIAGQP | ASSLNWGEAETLY | 0.10338798 |
| GTGGDGGNGGNGGTGG | ASSLNWGEAETLY | 0.095382206 |
| GVRRPAGIPADAPV | ASSLNWGEAETLY | 0.09739505 |
| GYRQVTDAHLLTLARR | ASSLNWGEAETLY | 0.09748827 |
| IAKMGQDGHDRG | ASSLNWGEAETLY | 0.09154785 |
| KAEPSIEIQYT | ASSLNWGEAETLY | 0.09326935 |
| KPVRIRMAADEDTVAVT | ASSLNWGEAETLY | 0.095992565 |
| MPPRIAGMRLLVIKPEP | ASSLNWGEAETLY | 0.13416235 |
| MKFYAVALFRSRGVPPTR | ASSLNWGEAETLY | 0.10713338 |
| NAGGFGILGAVAHSP | ASSLNWGEAETLY | 0.09608825 |
| NEGREITLAMAEM | ASSLNWGEAETLY | 0.10034073 |
| NELVEWSHPLGHRWRWE | ASSLNWGEAETLY | 0.09389954 |
| NLGAGGTGGNGGNPG | ASSLNWGEAETLY | 0.09718987 |
| NTEWPKWETVMAADPVD | ASSLNWGEAETLY | 0.09475548 |

| | | |
|---|---|---|
| PASAAPAETQP | ASSLNWGEAETLY | 0.09420854 |
| QQSSDGKNGVRAAEITGE | ASSLNWGEAETLY | 0.093831986 |
| RAPDAVTTSWWKE | ASSLNWGEAETLY | 0.094739996 |
| SGFGNVGAHTSGWFNQ | ASSLNWGEAETLY | 0.0924247 |
| SGVAGAVGQAASVGGLKV | ASSLNWGEAETLY | 0.10202669 |
| SNGGAGTGSAGGNGGT | ASSLNWGEAETLY | 0.10182743 |
| TGGAGGEGGAAG | ASSLNWGEAETLY | 0.09248592 |
| TGGDGGAAGAGGAPGHG | ASSLNWGEAETLY | 0.09823195 |
| TMPPLLAAIANAGRHGV | ASSLNWGEAETLY | 0.09415575 |
| TSSTAGNSGGA | ASSLNWGEAETLY | 0.09377817 |
| VPATVPVGILC | ASSLNWGEAETLY | 0.11012075 |
| VQRPDSGGVVLTGRIS | ASSLNWGEAETLY | 0.0936182 |
| WTPNPVPATKGVPAGLVQAT | ASSLNWGEAETLY | 0.100002624 |
| YALKRALAKFSDRFDVVII | ASSLNWGEAETLY | 0.13551727 |
| YLIGSVIV | ASSLNWGEAETLY | 0.09515315 |
| YGQLVAVID | ASSLNWGEAETLY | 0.09128039 |
| VTVGGMTLPA | ASSLNWGEAETLY | 0.096407406 |
| VLALPAV | ASSLNWGEAETLY | 0.10164028 |
| VIPPTLNYET | ASSLNWGEAETLY | 0.09991068 |
| VGMGGMHPSSG | ASSLNWGEAETLY | 0.08932663 |
| VGLAGGAATA | ASSLNWGEAETLY | 0.103684135 |
| VGGAGGAGGSGP | ASSLNWGEAETLY | 0.09409735 |
| VFAAFMV | ASSLNWGEAETLY | 0.095316045 |
| TLYDKLR | ASSLNWGEAETLY | 0.089765266 |
| TLAVFTVL | ASSLNWGEAETLY | 0.106221765 |
| TGSVQPARV | ASSLNWGEAETLY | 0.11297774 |
| TGGGAGSGGT | ASSLNWGEAETLY | 0.10275201 |
| SHLETSL | ASSLNWGEAETLY | 0.100337215 |
| SFTTGMT | ASSLNWGEAETLY | 0.090171725 |
| QVEVAKRMGVRQPTVS | ASSLNWGEAETLY | 0.10677959 |
| QGSGGPALAM | ASSLNWGEAETLY | 0.105040394 |
| MLAPSNP | ASSLNWGEAETLY | 0.09527492 |
| LVNNAAI | ASSLNWGEAETLY | 0.10547539 |
| LTGEINTLA | ASSLNWGEAETLY | 0.09335508 |
| LSIFGPV | ASSLNWGEAETLY | 0.09507037 |
| LLALLANL | ASSLNWGEAETLY | 0.10151627 |
| LISNASDALDKLR | ASSLNWGEAETLY | 0.10859488 |
| LINEAVP | ASSLNWGEAETLY | 0.09327811 |
| LGYAFSGL | ASSLNWGEAETLY | 0.09317148 |
| LDFVAVAL | ASSLNWGEAETLY | 0.09662625 |
| LAQPGAE | ASSLNWGEAETLY | 0.0929476 |
| ISNLGAVTGF | ASSLNWGEAETLY | 0.09864228 |
| HQEDPGRI | ASSLNWGEAETLY | 0.13943568 |
| GTPGEPTP | ASSLNWGEAETLY | 0.11357538 |
| GFFNSGS | ASSLNWGEAETLY | 0.094855495 |
| GAGGGGLTAGAN | ASSLNWGEAETLY | 0.11503243 |
| FVDALVNM | ASSLNWGEAETLY | 0.09510261 |
| FLTVAPEL | ASSLNWGEAETLY | 0.108473696 |
| FETLTAL | ASSLNWGEAETLY | 0.09306587 |
| DVINPIEL | ASSLNWGEAETLY | 0.1294787 |
| APVSGAPV | ASSLNWGEAETLY | 0.096901625 |

| | | |
|---|---|---|
| ALVGGLAW | ASSLNWGEAETLY | 0.09564241 |
| ALAAAAATAI | ASSLNWGEAETLY | 0.098794974 |
| AFGLGLLL | ASSLNWGEAETLY | 0.12977745 |
| AFFAAASL | ASSLNWGEAETLY | 0.1889341 |
| AAAALPAFSPP | ASSLNWGEAETLY | 0.40332535 |
| VVLAPAL | ASSLNWGEAETLY | 0.482062 |
| TIGHVDHGKTTL | ASSLNWGEAETLY | 0.13350976 |
| LISNASDALDK | ASSLNWGEAETLY | 0.10383795 |
| LISNASDAL | ASSLNWGEAETLY | 0.30844706 |
| ISNASDALDK | ASSLNWGEAETLY | 0.10365193 |
| NQAETLVYQTE | ASSLNWGEAETLY | 0.1443696 |
| TGGTGGTAGAG | ASSLNWGEAETLY | 0.33091247 |
| TGSTAGGAGGA | ASSLNWGEAETLY | 0.13314384 |
| DPLAVPVAAGPV | ASSLNWGEAETLY | 0.53278524 |
| GGTGGTPGNGGN | ASSLNWGEAETLY | 0.4962841 |
| PAGLAGVDPATL | ASSLNWGEAETLY | 0.11562593 |
| GNNLVVEAPHAGASV | ASSLNWGEAETLY | 0.22987278 |
| KVTRIMSASTLEQPA | ASSLNWGEAETLY | 0.26142353 |
| LGFALPIALPRRTVV | ASSLNWGEAETLY | 0.20415163 |
| TRLGNLTARKLVVLV | ASSLNWGEAETLY | 0.09926474 |
| WSQQRRLIYFPSAGP | ASSLNWGEAETLY | 0.31306452 |
| GPNGNPDPMAAAVDIR | ASSLNWGEAETLY | 0.15275326 |
| TQRCVSVADLRAAASR | ASSLNWGEAETLY | 0.087108634 |
| QFVLDVAEHPVEVLTETV | ASSLNWGEAETLY | 0.41526228 |
| GGAGGTGGAGGNGGTQAGDG | ASSLNWGEAETLY | 0.16147578 |
| PLLATLRAIAVDVDAKPAQV | ASSLNWGEAETLY | 0.13216859 |
| APKFRTGGYMIDFWGVGYQVAK | ASSLNWGEAETLY | 0.6192065 |
| GAGGAGGAGGAAGGPGGTGGTG | ASSLNWGEAETLY | 0.09749715 |
| IGALGLLATDVAPAGSVAFGVA | ASSLNWGEAETLY | 0.12087051 |
| AGGAGGIGGNGGAGGAGGVLMG | ASSLNWGEAETLY | 0.417632 |
| ADELIGLL | ASSLNWGEAETLY | 0.48029864 |
| ADRALKA | ASSLNWGEAETLY | 0.11728494 |
| AGAVGQVGGAGGA | ASSLNWGEAETLY | 0.4306043 |
| AGGGGGLGGAP | ASSLNWGEAETLY | 0.7124494 |
| AGGLLYGNGGAG | ASSLNWGEAETLY | 0.19038133 |
| AGLESIGL | ASSLNWGEAETLY | 0.6801709 |
| AGLKTGII | ASSLNWGEAETLY | 0.7092249 |
| ASLAALLL | ASSLNWGEAETLY | 0.16634437 |
| ASVINISSI | ASSLNWGEAETLY | 0.42775464 |
| ATPPGPA | ASSLNWGEAETLY | 0.67175406 |
| ATVASALV | ASSLNWGEAETLY | 0.16249508 |
| ATVASLIL | ASSLNWGEAETLY | 0.20525414 |
| AVNGPVLIRH | ASSLNWGEAETLY | 0.15625718 |
| DGAAGAVGI | ASSLNWGEAETLY | 0.36658847 |
| DPLLHGI | ASSLNWGEAETLY | 0.08906367 |
| ELGAALGAL | ASSLNWGEAETLY | 0.3204978 |
| ELGRTQAV | ASSLNWGEAETLY | 0.38833973 |
| FVAGGYLS | ASSLNWGEAETLY | 0.22615096 |
| GASGGAGGAGGW | ASSLNWGEAETLY | 0.11901593 |
| GGAGAGGDGGAGGI | ASSLNWGEAETLY | 0.11946757 |
| GTGHNGSQPGMGGNG | ASSLNWGEAETLY | 0.1019798 |

| | | |
|---|---|---|
| GYAAASASAS | ASSLNWGEAETLY | 0.20796141 |
| HQVDVAAVQQG | ASSLNWGEAETLY | 0.09904302 |
| IGAAIAHT | ASSLNWGEAETLY | 0.1050655 |
| IGMIEVAQL | ASSLNWGEAETLY | 0.34564155 |
| IRLAPEHTSAMD | ASSLNWGEAETLY | 0.3866037 |
| IYNVSVL | ASSLNWGEAETLY | 0.15129171 |
| KQGFLGM | ASSLNWGEAETLY | 0.49463493 |
| LESVITQL | ASSLNWGEAETLY | 0.61306345 |
| LGLGALVVV | ASSLNWGEAETLY | 0.6216927 |
| LIGNGGNGGSGGI | ASSLNWGEAETLY | 0.28868806 |
| LNAINAPTL | ASSLNWGEAETLY | 0.1109071 |
| LQLLNTGV | ASSLNWGEAETLY | 0.7085182 |
| LQTSIEF | ASSLNWGEAETLY | 0.20852357 |
| MSFAVSEL | ASSLNWGEAETLY | 0.12786223 |
| NTGALGL | ASSLNWGEAETLY | 0.52507246 |
| PPGAGGAGGAGTT | ASSLNWGEAETLY | 0.12064346 |
| QLVGLGI | ASSLNWGEAETLY | 0.1178425 |
| RFLEQVHQL | ASSLNWGEAETLY | 0.56242436 |
| RGVGYVLR | ASSLNWGEAETLY | 0.60829383 |
| SATGVFAGVI | ASSLNWGEAETLY | 0.13898799 |
| SLPAITLP | ASSLNWGEAETLY | 0.10662844 |
| STALAFTV | ASSLNWGEAETLY | 0.13022016 |
| SVLTVGLGK | ASSLNWGEAETLY | 0.29175982 |
| TALAGAGSGI | ASSLNWGEAETLY | 0.44576633 |
| TGLPFTGI | ASSLNWGEAETLY | 0.26371443 |
| VAEHVTV | ASSLNWGEAETLY | 0.20571929 |
| VDIVLNSL | ASSLNWGEAETLY | 0.5814577 |
| VLTGGGAVL | ASSLNWGEAETLY | 0.22087511 |
| VNVGLAYGI | ASSLNWGEAETLY | 0.22168238 |
| VSSAAVGAV | ASSLNWGEAETLY | 0.54195404 |
| VTSVYEAGV | ASSLNWGEAETLY | 0.10234965 |
| WVALDRGA | ASSLNWGEAETLY | 0.5599597 |
| YLVGSVVI | ASSLNWGEAETLY | 0.503713 |
| GPPGTGKTLL | ASSLNWGEAETLY | 0.6051534 |
| GVGMTKF | ASSLNWGEAETLY | 0.66033435 |
| IGIGGSDL | ASSLNWGEAETLY | 0.12558272 |
| IGTLLAEL | ASSLNWGEAETLY | 0.40585193 |
| IYTSGTT | ASSLNWGEAETLY | 0.12759301 |
| LSFPAGV | ASSLNWGEAETLY | 0.23566058 |
| LYTSGTT | ASSLNWGEAETLY | 0.4881304 |
| VGAFPFGF | ASSLNWGEAETLY | 0.4276329 |
| VGAGTVEF | ASSLNWGEAETLY | 0.6200168 |
| VGKSTLLL | ASSLNWGEAETLY | 0.50014293 |
| AALAHAGGGIV | ASSLNWGEAETLY | 0.17942408 |
| AALVAVSLL | ASSLNWGEAETLY | 0.72912514 |
| AAVSRAVAAL | ASSLNWGEAETLY | 0.65754676 |
| AGGSSGIGGF | ASSLNWGEAETLY | 0.11531313 |
| AGLTVANANG | ASSLNWGEAETLY | 0.13111392 |
| AGLVGAVL | ASSLNWGEAETLY | 0.21077593 |
| ALSQAFTV | ASSLNWGEAETLY | 0.6060505 |
| ALTFSVSL | ASSLNWGEAETLY | 0.272954 |

| | | |
|---|---|---|
| ALTGAALVAL | ASSLNWGEAETLY | 0.45892733 |
| ASFSIGAGVL | ASSLNWGEAETLY | 0.3545164 |
| DGSVYKI | ASSLNWGEAETLY | 0.112966694 |
| EQLAEIV | ASSLNWGEAETLY | 0.6964965 |
| FAVSELTVM | ASSLNWGEAETLY | 0.11134038 |
| GAAPLAAGL | ASSLNWGEAETLY | 0.37312996 |
| GADGGDGGSGGAGG | ASSLNWGEAETLY | 0.13188732 |
| GFGLDTGGAGG | ASSLNWGEAETLY | 0.09136233 |
| GLASGGLL | ASSLNWGEAETLY | 0.3347296 |
| GRIAGAAVLL | ASSLNWGEAETLY | 0.22257374 |
| GSAGSLNTSF | ASSLNWGEAETLY | 0.6501628 |
| GTGQIGI | ASSLNWGEAETLY | 0.19822145 |
| GTLIALIL | ASSLNWGEAETLY | 0.33105898 |
| IAATMGFVVG | ASSLNWGEAETLY | 0.39360774 |
| IFAQALL | ASSLNWGEAETLY | 0.26534683 |
| IGAAVALGF | ASSLNWGEAETLY | 0.13052599 |
| IGLTPLAL | ASSLNWGEAETLY | 0.24456075 |
| ILGSNTSTL | ASSLNWGEAETLY | 0.71756804 |
| IVFSLTVL | ASSLNWGEAETLY | 0.54239184 |
| LASKATGF | ASSLNWGEAETLY | 0.103746265 |
| LASVALIL | ASSLNWGEAETLY | 0.20138484 |
| LFAVGLTL | ASSLNWGEAETLY | 0.15516958 |
| LFNGGIGTYI | ASSLNWGEAETLY | 0.08783438 |
| LTLGAGII | ASSLNWGEAETLY | 0.092378065 |
| LVAAALGDI | ASSLNWGEAETLY | 0.09345074 |
| NTAFVQLGI | ASSLNWGEAETLY | 0.103057295 |
| QLFSGAEL | ASSLNWGEAETLY | 0.48844245 |
| SAATFITL | ASSLNWGEAETLY | 0.46231818 |
| SNYGLANLGV | ASSLNWGEAETLY | 0.19515064 |
| SYGGAYAVM | ASSLNWGEAETLY | 0.5490631 |
| TTGVGGLTLG | ASSLNWGEAETLY | 0.14695293 |
| TTQTIYI | ASSLNWGEAETLY | 0.084594555 |
| VAAGAGLGAVI | ASSLNWGEAETLY | 0.14254564 |
| VAAVVTLL | ASSLNWGEAETLY | 0.13337673 |
| VGFLGGLI | ASSLNWGEAETLY | 0.6868328 |
| VGVIPILL | ASSLNWGEAETLY | 0.10589404 |
| VIFASSLI | ASSLNWGEAETLY | 0.21481173 |
| VKGGDGGAAAG | ASSLNWGEAETLY | 0.20886712 |
| VTKTFTNI | ASSLNWGEAETLY | 0.32281137 |
| VVAAAILL | ASSLNWGEAETLY | 0.4464477 |
| YAALDVEL | ASSLNWGEAETLY | 0.12691407 |
| LASVALLL | ASSLNWGEAETLY | 0.102898866 |
| LDAVHRV | ASSLNWGEAETLY | 0.25243744 |
| QGLSSSIF | ASSLNWGEAETLY | 0.095442034 |
| RGTFANIRL | ASSLNWGEAETLY | 0.1262787 |
| RQLWWGHRI | ASSLNWGEAETLY | 0.11703611 |
| YGPPGTGKTLL | ASSLNWGEAETLY | 0.09268807 |
| AALVGAPANVA | ASSLNWGEAETLY | 0.103443034 |
| AATFRRTE | ASSLNWGEAETLY | 0.65437055 |
| AGALGGGI | ASSLNWGEAETLY | 0.3091669 |
| AGGAGGAGGIGAG | ASSLNWGEAETLY | 0.20708069 |

| | | |
|---|---|---|
| AGSGALGL | ASSLNWGEAETLY | 0.39731562 |
| AQIADQI | ASSLNWGEAETLY | 0.2891751 |
| ASAVAVTYL | ASSLNWGEAETLY | 0.23889196 |
| ASTVLGGTGG | ASSLNWGEAETLY | 0.122524045 |
| AVGGEPAK | ASSLNWGEAETLY | 0.13857196 |
| CEQGTFGGTC | ASSLNWGEAETLY | 0.11503759 |
| DEGKAPAG | ASSLNWGEAETLY | 0.12240732 |
| DTVVFPAL | ASSLNWGEAETLY | 0.6973082 |
| FEGITMAG | ASSLNWGEAETLY | 0.4171337 |
| FPLVVQD | ASSLNWGEAETLY | 0.11869977 |
| GAFTTPQ | ASSLNWGEAETLY | 0.14294004 |
| GFGGFGTAGGVGGA | ASSLNWGEAETLY | 0.6418507 |
| GFLAQFL | ASSLNWGEAETLY | 0.66931033 |
| GGAGGAGGNAGVA | ASSLNWGEAETLY | 0.13935648 |
| GGVGTAGAGGAGGAGG | ASSLNWGEAETLY | 0.26438892 |
| GNGGGGDAGNAGSGGN | ASSLNWGEAETLY | 0.3891824 |
| GTGGAGGAGNDAGS | ASSLNWGEAETLY | 0.19056995 |
| GTTNTGWFNS | ASSLNWGEAETLY | 0.20506644 |
| IFMGPLFTL | ASSLNWGEAETLY | 0.111716904 |
| ILAHAIEAI | ASSLNWGEAETLY | 0.59412813 |
| IRSAIREAL | ASSLNWGEAETLY | 0.7763797 |
| ITFAGGG | ASSLNWGEAETLY | 0.1364477 |
| ITGMSGHQEV | ASSLNWGEAETLY | 0.13528518 |
| IVAGLACTL | ASSLNWGEAETLY | 0.22635852 |
| IVGGGLAGA | ASSLNWGEAETLY | 0.12816715 |
| IVTGVLNI | ASSLNWGEAETLY | 0.15751997 |
| LAQQGCI | ASSLNWGEAETLY | 0.20721504 |
| LCLHVEN | ASSLNWGEAETLY | 0.105947345 |
| LDAISDGS | ASSLNWGEAETLY | 0.2148022 |
| LGGFAAML | ASSLNWGEAETLY | 0.43739235 |
| LGGKNPMI | ASSLNWGEAETLY | 0.1763586 |
| LGPKGVAAIL | ASSLNWGEAETLY | 0.6569507 |
| LMVGVSI | ASSLNWGEAETLY | 0.63645655 |
| LVGAAFVL | ASSLNWGEAETLY | 0.5071026 |
| LVLAVAPL | ASSLNWGEAETLY | 0.6904147 |
| MAAIAVIDAA | ASSLNWGEAETLY | 0.64340425 |
| NNGINAELQ | ASSLNWGEAETLY | 0.18043591 |
| RGEPLVLVHG | ASSLNWGEAETLY | 0.24697745 |
| RGGAGGAAGLI | ASSLNWGEAETLY | 0.41515836 |
| RSYRMHPAV | ASSLNWGEAETLY | 0.209408 |
| SAAHLLAGQLA | ASSLNWGEAETLY | 0.3160461 |
| SAAVGSY | ASSLNWGEAETLY | 0.28876114 |
| SPHAVLHRI | ASSLNWGEAETLY | 0.5290563 |
| SQIAKVL | ASSLNWGEAETLY | 0.23510975 |
| TALGHLVP | ASSLNWGEAETLY | 0.6735964 |
| TAPTTSGAA | ASSLNWGEAETLY | 0.11938683 |
| TDGDDGNGGSPGT | ASSLNWGEAETLY | 0.11433269 |
| TDGLGEN | ASSLNWGEAETLY | 0.6554802 |
| TGGPPLPGQA | ASSLNWGEAETLY | 0.10083807 |
| TIPAITI | ASSLNWGEAETLY | 0.1152468 |
| TIPGFTI | ASSLNWGEAETLY | 0.3471892 |

| | | |
|---|---|---|
| TSIASGALA | ASSLNWGEAETLY | 0.5127337 |
| TVTLPGAAI | ASSLNWGEAETLY | 0.08525074 |
| VASGLSAALP | ASSLNWGEAETLY | 0.13380255 |
| VGAGVIGI | ASSLNWGEAETLY | 0.1712738 |
| VGPKRFGV | ASSLNWGEAETLY | 0.16990033 |
| VNVAFPD | ASSLNWGEAETLY | 0.1499539 |
| VSMTNSGVS | ASSLNWGEAETLY | 0.105751336 |
| VTDVLGDLSV | ASSLNWGEAETLY | 0.45637912 |
| VVVVGAGFAGL | ASSLNWGEAETLY | 0.13039738 |
| VYTVGTVV | ASSLNWGEAETLY | 0.09478814 |
| WTTFSGP | ASSLNWGEAETLY | 0.43380865 |
| YDELVEAF | ASSLNWGEAETLY | 0.73564994 |
| YGTGGAGGNGG | ASSLNWGEAETLY | 0.13757315 |
| YTNVMAV | ASSLNWGEAETLY | 0.3892016 |
| FLEQVHQL | ASSLNWGEAETLY | 0.19723947 |
| GLSLLAVAV | ASSLNWGEAETLY | 0.7345204 |
| LEQVHQL | ASSLNWGEAETLY | 0.1647971 |
| LNAAGAI | ASSLNWGEAETLY | 0.11685757 |
| PAVHASL | ASSLNWGEAETLY | 0.57013535 |
| TELEDDA | ASSLNWGEAETLY | 0.6779038 |
| TPPGPPN | ASSLNWGEAETLY | 0.25634176 |
| TPPPPPPP | ASSLNWGEAETLY | 0.61848354 |
| TQSPALI | ASSQDRGDERLF | 0.11620718 |
| EFAAAFPQ | ASSQDRGDERLF | 0.20598848 |
| LLSGNLDVL | ASSQDRGDERLF | 0.37397444 |
| GPANLGNYNI | ASSQDRGDERLF | 0.41999677 |
| TLGPTLP | ASSQDRGDERLF | 0.09968813 |
| VYAYSSCCK | ASSQDRGDERLF | 0.4318908 |
| APAPAPAPNI | ASSQDRGDERLF | 0.15667267 |
| PGGPVPIP | ASSQDRGDERLF | 0.1786431 |
| FLTVALEP | ASSQDRGDERLF | 0.49743316 |
| SQYFGANL | ASSQDRGDERLF | 0.3833775 |
| GGVKPAA | ASSQDRGDERLF | 0.5049976 |
| GGTGGAGGAGGW | ASSQDRGDERLF | 0.23553059 |
| GVGGVAGLGGAGG | ASSQDRGDERLF | 0.09109846 |
| VLLVLATTI | ASSQDRGDERLF | 0.109618224 |
| IGIFAAML | ASSQDRGDERLF | 0.57346225 |
| AVGDLID | ASSQDRGDERLF | 0.3669037 |
| NISPLPL | ASSQDRGDERLF | 0.5151249 |
| VTAVFVAI | ASSQDRGDERLF | 0.27498215 |
| SGDDNDAAL | ASSQDRGDERLF | 0.13274567 |
| AVAALAHS | ASSQDRGDERLF | 0.50322497 |
| GGVTVMATP | ASSQDRGDERLF | 0.52919984 |
| AGLAANVRD | ASSQDRGDERLF | 0.21163043 |
| TNRPDLIDPALLRPG | ASSQDRGDERLF | 0.3901947 |
| LANLGVTAL | ASSQDRGDERLF | 0.73944366 |
| TGERIEL | ASSQDRGDERLF | 0.16383277 |
| NGEELAA | ASSQDRGDERLF | 0.15222785 |
| VLSLVWG | ASSQDRGDERLF | 0.47005236 |
| IGYALWL | ASSQDRGDERLF | 0.729645 |
| LSLLLAPL | ASSQDRGDERLF | 0.2547719 |

| | | |
|---|---|---|
| TLDEALK | ASSQDRGDERLF | 0.69414717 |
| TDGAGTAEGG | ASSQDRGDERLF | 0.11487876 |
| NAPGGAGGNGG | ASSQDRGDERLF | 0.42617655 |
| VSVFMGMGGNF | ASSQDRGDERLF | 0.165018 |
| LASYLAAL | ASSQDRGDERLF | 0.34332734 |
| LVGGTPAL | ASSQDRGDERLF | 0.09263993 |
| VRNADVL | ASSQDRGDERLF | 0.4040436 |
| TCSSPFLVSAM | ASSQDRGDERLF | 0.56283724 |
| VHITDAE | ASSQDRGDERLF | 0.5000564 |
| EYAAPSF | ASSQDRGDERLF | 0.09316683 |
| VVDAFHGGL | ASSQDRGDERLF | 0.18292603 |
| ASVVDPQ | ASSQDRGDERLF | 0.5570241 |
| SYNEFGGLS | ASSQDRGDERLF | 0.7915128 |
| DPDGSVY | ASSQDRGDERLF | 0.23870489 |
| MVSTIHATE | ASSQDRGDERLF | 0.6160569 |
| LAAGVADPV | ASSQDRGDERLF | 0.13702722 |
| TKEQIAA | ASSQDRGDERLF | 0.18206389 |
| SLASPAAL | ASSQDRGDERLF | 0.13883565 |
| IGLVFYGV | ASSQDRGDERLF | 0.28900605 |
| FLLDASNL | ASSQDRGDERLF | 0.20791964 |
| YQVGNFP | ASSQDRGDERLF | 0.13500686 |
| ESLLSQST | ASSQDRGDERLF | 0.32076237 |
| LFTVGFAAVG | ASSQDRGDERLF | 0.7346699 |
| ATPPTPT | ASSQDRGDERLF | 0.2924338 |
| TDLATLG | ASSQDRGDERLF | 0.66795814 |
| SAGGTGR | ASSQDRGDERLF | 0.7069186 |
| GFSNSGS | ASSQDRGDERLF | 0.6760727 |
| GGNGGSGGTSV | ASSQDRGDERLF | 0.11425253 |
| GTSVATGGAGN | ASSQDRGDERLF | 0.25638688 |
| NGGNGGSGGTS | ASSQDRGDERLF | 0.10525622 |
| AVVKFPH | ASSQDRGDERLF | 0.1935249 |
| TPPPPPP | ASSQDRGDERLF | 0.11353927 |
| AAPAAAAAPA | ASSQDRGDERLF | 0.2634022 |
| IALNEDL | ASSQDRGDERLF | 0.08548871 |
| IVNVSSM | ASSQDRGDERLF | 0.5070321 |
| TGIPPAPR | ASSQDRGDERLF | 0.71421444 |
| FDLGGGTF | ASSQDRGDERLF | 0.0927475 |
| FDLGGGTFDVS | ASSQDRGDERLF | 0.57400405 |
| NEPTAAA | ASSQDRGDERLF | 0.65978175 |
| NEPTAAALA | ASSQDRGDERLF | 0.14007473 |
| GSIINISSI | ASSQDRGDERLF | 0.56429267 |
| VAPGNAGTA | ASSQDRGDERLF | 0.1336786 |
| TGGGASLE | ASSQDRGDERLF | 0.65210176 |
| VAVYNHYKRI | ASSQDRGDERLF | 0.38183522 |
| TDPAPATT | ASSQDRGDERLF | 0.42865515 |
| TNRPDILDPALLRPG | ASSQDRGDERLF | 0.42603952 |
| LTTTTTS | ASSQDRGDERLF | 0.525684 |
| PTGSGKTL | ASSQDRGDERLF | 0.11199799 |
| TGSGKTL | ASSQDRGDERLF | 0.46123177 |
| TIGHVDHGK | ASSQDRGDERLF | 0.08839192 |
| TIGHVDHGKTT | ASSQDRGDERLF | 0.40454113 |

| | | |
|---|---|---|
| GHVDHGKTT | ASSQDRGDERLF | 0.12346191 |
| AQFEHTLL | ASSQDRGDERLF | 0.4319017 |
| IVNQDEL | ASSQDRGDERLF | 0.14691544 |
| TNGSQFF | ASSQDRGDERLF | 0.42636928 |
| TGKGIVE | ASSQDRGDERLF | 0.19782612 |
| LISNASDA | ASSQDRGDERLF | 0.13124233 |
| AELRTLGELTKIAKAHGAQ | ASSQDRGDERLF | 0.10920708 |
| APNAVAAAEPSW | ASSQDRGDERLF | 0.10492012 |
| APQVAGAATAVL | ASSQDRGDERLF | 0.66246367 |
| ARRDPAAAAAILPTDA | ASSQDRGDERLF | 0.22175023 |
| DAAARDVLAEAGLAEYF | ASSQDRGDERLF | 0.11126231 |
| DDQIVLSAKANLAKVEQ | ASSQDRGDERLF | 0.37758225 |
| DELLPDTGEDSDEDGH | ASSQDRGDERLF | 0.23670565 |
| ELPEAPANRATVPQD | ASSQDRGDERLF | 0.112836875 |
| FGQSKIERVTGIG | ASSQDRGDERLF | 0.38511795 |
| GAGGTGSTAGGAGGAGGAGGLY | ASSQDRGDERLF | 0.30388144 |
| GHGGTGAAGGNGGTGGAGGSGI | ASSQDRGDERLF | 0.16542849 |
| GNGGNGGAGGEGGPGGAGGAGA | ASSQDRGDERLF | 0.085260294 |
| GSTAGNGGNGGNATNGTIAGQP | ASSQDRGDERLF | 0.13328837 |
| GTGGDGGNGGNGGTGG | ASSQDRGDERLF | 0.18229777 |
| GVRRPAGIPADAPV | ASSQDRGDERLF | 0.10178953 |
| GYRQVTDAHLLTLARR | ASSQDRGDERLF | 0.12273003 |
| IAKMGQDGHDRG | ASSQDRGDERLF | 0.61840063 |
| KAEPSIEIQYT | ASSQDRGDERLF | 0.09764311 |
| KPVRIRMAADEDTVAVT | ASSQDRGDERLF | 0.1478475 |
| MPPRIAGMRLLVIKPEP | ASSQDRGDERLF | 0.6049613 |
| MKFYAVALFRSRGVPPTR | ASSQDRGDERLF | 0.2260691 |
| NAGGFGILGAVAHSP | ASSQDRGDERLF | 0.6106087 |
| NEGREITLAMAEM | ASSQDRGDERLF | 0.13769017 |
| NELVEWSHPLGHRWRWE | ASSQDRGDERLF | 0.19673394 |
| NLGAGGTGGNGGNPG | ASSQDRGDERLF | 0.34367114 |
| NTEWPKWETVMAADPVD | ASSQDRGDERLF | 0.21763715 |
| PASAAPAETQP | ASSQDRGDERLF | 0.10342844 |
| QQSSDGKNGVRAAEITGE | ASSQDRGDERLF | 0.09381965 |
| RAPDAVTTSWWKE | ASSQDRGDERLF | 0.21866196 |
| SGFGNVGAHTSGWFNQ | ASSQDRGDERLF | 0.11461934 |
| SGVAGAVGQAASVGGLKV | ASSQDRGDERLF | 0.25764444 |
| SNGGAGTGSAGGNGGT | ASSQDRGDERLF | 0.6951586 |
| TGGAGGEGGAAG | ASSQDRGDERLF | 0.114420496 |
| TGGDGGAAGAGGAPGHG | ASSQDRGDERLF | 0.12443247 |
| TMPPLLAAIANAGRHGV | ASSQDRGDERLF | 0.23368636 |
| TSSTAGNSGGA | ASSQDRGDERLF | 0.13923356 |
| VPATVPVGILC | ASSQDRGDERLF | 0.5729587 |
| VQRPDSGGVVLTGRIS | ASSQDRGDERLF | 0.1412997 |
| WTPNPVPATKGVPAGLVQAT | ASSQDRGDERLF | 0.7578661 |
| YALKRALAKFSDRFDVVII | ASSQDRGDERLF | 0.2627023 |
| YLIGSVIV | ASSQDRGDERLF | 0.119995296 |
| YGQLVAVID | ASSQDRGDERLF | 0.58401954 |
| VTVGGMTLPA | ASSQDRGDERLF | 0.2360746 |
| VLALPAV | ASSQDRGDERLF | 0.54226923 |
| VIPPTLNYET | ASSQDRGDERLF | 0.3511218 |

| | | |
|---|---|---|
| VGMGGMHPSSG | ASSQDRGDERLF | 0.10946667 |
| VGLAGGAATA | ASSQDRGDERLF | 0.16087599 |
| VGGAGGAGGSGP | ASSQDRGDERLF | 0.34395155 |
| VFAAFMV | ASSQDRGDERLF | 0.12146573 |
| TLYDKLR | ASSQDRGDERLF | 0.31784272 |
| TLAVFTVL | ASSQDRGDERLF | 0.1005111 |
| TGSVQPARV | ASSQDRGDERLF | 0.10687514 |
| TGGGAGSGGT | ASSQDRGDERLF | 0.2951604 |
| SHLETSL | ASSQDRGDERLF | 0.48493662 |
| SFTTGMT | ASSQDRGDERLF | 0.739509 |
| QVEVAKRMGVRQPTVS | ASSQDRGDERLF | 0.16025288 |
| QGSGGPALAM | ASSQDRGDERLF | 0.109508246 |
| MLAPSNP | ASSQDRGDERLF | 0.668697 |
| LVNNAAI | ASSQDRGDERLF | 0.17448212 |
| LTGEINTLA | ASSQDRGDERLF | 0.54408056 |
| LSIFGPV | ASSQDRGDERLF | 0.16299349 |
| LLALLANL | ASSQDRGDERLF | 0.11407695 |
| LISNASDALDKLR | ASSQDRGDERLF | 0.26149696 |
| LINEAVP | ASSQDRGDERLF | 0.4652712 |
| LGYAFSGL | ASSQDRGDERLF | 0.10178144 |
| LDFVAVAL | ASSQDRGDERLF | 0.6041795 |
| LAQPGAE | ASSQDRGDERLF | 0.085404105 |
| ISNLGAVTGF | ASSQDRGDERLF | 0.13289122 |
| HQEDPGRI | ASSQDRGDERLF | 0.14832732 |
| GTPGEPTP | ASSQDRGDERLF | 0.13144676 |
| GFFNSGS | ASSQDRGDERLF | 0.09578481 |
| GAGGGGLTAGAN | ASSQDRGDERLF | 0.2681382 |
| FVDALVNM | ASSQDRGDERLF | 0.61148494 |
| FLTVAPEL | ASSQDRGDERLF | 0.67209256 |
| FETLTAL | ASSQDRGDERLF | 0.13213345 |
| DVINPIEL | ASSQDRGDERLF | 0.60586536 |
| APVSGAPV | ASSQDRGDERLF | 0.72929585 |
| ALVGGLAW | ASSQDRGDERLF | 0.20816733 |
| ALAAAAATAI | ASSQDRGDERLF | 0.100799926 |
| AFGLGLLL | ASSQDRGDERLF | 0.21196744 |
| AFFAAASL | ASSQDRGDERLF | 0.09294276 |
| AAAALPAFSPP | ASSQDRGDERLF | 0.15313289 |
| VVLAPAL | ASSQDRGDERLF | 0.7009268 |
| TIGHVDHGKTTL | ASSQDRGDERLF | 0.10396767 |
| LISNASDALDK | ASSQDRGDERLF | 0.4578193 |
| LISNASDAL | ASSQDRGDERLF | 0.5565029 |
| ISNASDALDK | ASSQDRGDERLF | 0.1668356 |
| NQAETLVYQTE | ASSQDRGDERLF | 0.19148873 |
| TGGTGGTAGAG | ASSQDRGDERLF | 0.12897514 |
| TGSTAGGAGGA | ASSQDRGDERLF | 0.1335469 |
| DPLAVPVAAGPV | ASSQDRGDERLF | 0.21929307 |
| GGTGGTPGNGGN | ASSQDRGDERLF | 0.20103258 |
| PAGLAGVDPATL | ASSQDRGDERLF | 0.14303063 |
| GNNLVVEAPHAGASV | ASSQDRGDERLF | 0.19895348 |
| KVTRIMSASTLEQPA | ASSQDRGDERLF | 0.13142464 |
| LGFALPIALPRRTVV | ASSQDRGDERLF | 0.14240953 |

| | | |
|---|---|---|
| TRLGNLTARKLVVLV | ASSQDRGDERLF | 0.13320205 |
| WSQQRRLIYFPSAGP | ASSQDRGDERLF | 0.1269459 |
| GPNGNPDPMAAAVDIR | ASSQDRGDERLF | 0.54022986 |
| TQRCVSVADLRAAASR | ASSQDRGDERLF | 0.1175807 |
| QFVLDVAEHPVEVLTETV | ASSQDRGDERLF | 0.51550555 |
| GGAGGTGGAGGNGGTQAGDG | ASSQDRGDERLF | 0.6156871 |
| PLLATLRAIAVDVDAKPAQV | ASSQDRGDERLF | 0.09549199 |
| APKFRTGGYMIDFWGVGYQVAK | ASSQDRGDERLF | 0.11855037 |
| GAGGAGGAGGAAGGPGGTGGTG | ASSQDRGDERLF | 0.24968275 |
| IGALGLLATDVAPAGSVAFGVA | ASSQDRGDERLF | 0.09624265 |
| AGGAGGIGGNGGAGGAGGVLMG | ASSQDRGDERLF | 0.6545365 |
| ADELIGLL | ASSQDRGDERLF | 0.16818838 |
| ADRALKA | ASSQDRGDERLF | 0.33251238 |
| AGAVGQVGGAGGA | ASSQDRGDERLF | 0.1949379 |
| AGGGGGLGGAP | ASSQDRGDERLF | 0.7632292 |
| AGGLLYGNGGAG | ASSQDRGDERLF | 0.4531267 |
| AGLESIGL | ASSQDRGDERLF | 0.20284972 |
| AGLKTGII | ASSQDRGDERLF | 0.09674896 |
| ASLAALLL | ASSQDRGDERLF | 0.11963406 |
| ASVINISSI | ASSQDRGDERLF | 0.590484 |
| ATPPGPA | ASSQDRGDERLF | 0.6642442 |
| ATVASALV | ASSQDRGDERLF | 0.1042732 |
| ATVASLIL | ASSQDRGDERLF | 0.21232313 |
| AVNGPVLIRH | ASSQDRGDERLF | 0.63755435 |
| DGAAGAVGI | ASSQDRGDERLF | 0.693579 |
| DPLLHGI | ASSQDRGDERLF | 0.10291905 |
| ELGAALGAL | ASSQDRGDERLF | 0.14026482 |
| ELGRTQAV | ASSQDRGDERLF | 0.2005578 |
| FVAGGYLS | ASSQDRGDERLF | 0.2189582 |
| GASGGAGGAGGW | ASSQDRGDERLF | 0.113459565 |
| GGAGAGGDGGAGGI | ASSQDRGDERLF | 0.600464 |
| GTGHNGSQPGMGGNG | ASSQDRGDERLF | 0.26822597 |
| GYAAASASAS | ASSQDRGDERLF | 0.7117041 |
| HQVDVAAVQQG | ASSQDRGDERLF | 0.23500103 |
| IGAAIAHT | ASSQDRGDERLF | 0.36181185 |
| IGMIEVAQL | ASSQDRGDERLF | 0.26947552 |
| IRLAPEHTSAMD | ASSQDRGDERLF | 0.13069518 |
| IYNVSVL | ASSQDRGDERLF | 0.13182178 |
| KQGFLGM | ASSQDRGDERLF | 0.09761946 |
| LESVITQL | ASSQDRGDERLF | 0.1509192 |
| LGLGALVVV | ASSQDRGDERLF | 0.13046628 |
| LIGNGGNGGSGGI | ASSQDRGDERLF | 0.08606582 |
| LNAINAPTL | ASSQDRGDERLF | 0.18478206 |
| LQLLNTGV | ASSQDRGDERLF | 0.30472565 |
| LQTSIEF | ASSQDRGDERLF | 0.27832758 |
| MSFAVSEL | ASSQDRGDERLF | 0.1610326 |
| NTGALGL | ASSQDRGDERLF | 0.19797815 |
| PPGAGGAGGAGTT | ASSQDRGDERLF | 0.1066966 |
| QLVGLGI | ASSQDRGDERLF | 0.67452294 |
| RFLEQVHQL | ASSQDRGDERLF | 0.100404225 |
| RGVGYVLR | ASSQDRGDERLF | 0.13499382 |

| | | |
|---|---|---|
| SATGVFAGVI | ASSQDRGDERLF | 0.1720714 |
| SLPAITLP | ASSQDRGDERLF | 0.18405245 |
| STALAFTV | ASSQDRGDERLF | 0.72123253 |
| SVLTVGLGK | ASSQDRGDERLF | 0.5610033 |
| TALAGAGSGI | ASSQDRGDERLF | 0.38306805 |
| TGLPFTGI | ASSQDRGDERLF | 0.67022485 |
| VAEHVTV | ASSQDRGDERLF | 0.09706358 |
| VDIVLNSL | ASSQDRGDERLF | 0.14044778 |
| VLTGGGAVL | ASSQDRGDERLF | 0.689325 |
| VNVGLAYGI | ASSQDRGDERLF | 0.1917561 |
| VSSAAVGAV | ASSQDRGDERLF | 0.30662128 |
| VTSVYEAGV | ASSQDRGDERLF | 0.46593884 |
| WVALDRGA | ASSQDRGDERLF | 0.118043914 |
| YLVGSVVI | ASSQDRGDERLF | 0.16856416 |
| GPPGTGKTLL | ASSQDRGDERLF | 0.15219022 |
| GVGMTKF | ASSQDRGDERLF | 0.4856763 |
| IGIGGSDL | ASSQDRGDERLF | 0.24212632 |
| IGTLLAEL | ASSQDRGDERLF | 0.11383117 |
| IYTSGTT | ASSQDRGDERLF | 0.6112359 |
| LSFPAGV | ASSQDRGDERLF | 0.8100805 |
| LYTSGTT | ASSQDRGDERLF | 0.2724253 |
| VGAFPFGF | ASSQDRGDERLF | 0.31387645 |
| VGAGTVEF | ASSQDRGDERLF | 0.27288586 |
| VGKSTLLL | ASSQDRGDERLF | 0.5280915 |
| AALAHAGGGIV | ASSQDRGDERLF | 0.14762762 |
| AALVAVSLL | ASSQDRGDERLF | 0.12526903 |
| AAVSRAVAAL | ASSQDRGDERLF | 0.13080154 |
| AGGSSGIGGF | ASSQDRGDERLF | 0.12496819 |
| AGLTVANANG | ASSQDRGDERLF | 0.4690044 |
| AGLVGAVL | ASSQDRGDERLF | 0.19770226 |
| ALSQAFTV | ASSQDRGDERLF | 0.7982932 |
| ALTFSVSL | ASSQDRGDERLF | 0.5152569 |
| ALTGAALVAL | ASSQDRGDERLF | 0.17446512 |
| ASFSIGAGVL | ASSQDRGDERLF | 0.10082786 |
| DGSVYKI | ASSQDRGDERLF | 0.3328057 |
| EQLAEIV | ASSQDRGDERLF | 0.26665673 |
| FAVSELTVM | ASSQDRGDERLF | 0.1198493 |
| GAAPLAAGL | ASSQDRGDERLF | 0.57176006 |
| GADGGDGGSGGAGG | ASSQDRGDERLF | 0.13238597 |
| GFGLDTGGAGG | ASSQDRGDERLF | 0.11624842 |
| GLASGGLL | ASSQDRGDERLF | 0.2309972 |
| GRIAGAAVLL | ASSQDRGDERLF | 0.7418767 |
| GSAGSLNTSF | ASSQDRGDERLF | 0.2857652 |
| GTGQIGI | ASSQDRGDERLF | 0.1358612 |
| GTLIALIL | ASSQDRGDERLF | 0.10211611 |
| IAATMGFVVG | ASSQDRGDERLF | 0.60150486 |
| IFAQALL | ASSQDRGDERLF | 0.6593345 |
| IGAAVALGF | ASSQDRGDERLF | 0.092087 |
| IGLTPLAL | ASSQDRGDERLF | 0.7613076 |
| ILGSNTSTL | ASSQDRGDERLF | 0.13193244 |
| IVFSLTVL | ASSQDRGDERLF | 0.11414094 |

| | | |
|---|---|---|
| LASKATGF | ASSQDRGDERLF | 0.36903834 |
| LASVALIL | ASSQDRGDERLF | 0.38276497 |
| LFAVGLTL | ASSQDRGDERLF | 0.71671927 |
| LFNGGIGTYI | ASSQDRGDERLF | 0.35066587 |
| LTLGAGII | ASSQDRGDERLF | 0.67077506 |
| LVAAALGDI | ASSQDRGDERLF | 0.24778435 |
| NTAFVQLGI | ASSQDRGDERLF | 0.6549623 |
| QLFSGAEL | ASSQDRGDERLF | 0.5182587 |
| SAATFITL | ASSQDRGDERLF | 0.36413974 |
| SNYGLANLGV | ASSQDRGDERLF | 0.24742146 |
| SYGGAYAVM | ASSQDRGDERLF | 0.1519405 |
| TTGVGGLTLG | ASSQDRGDERLF | 0.36560497 |
| TTQTIYI | ASSQDRGDERLF | 0.1401185 |
| VAAGAGLGAVI | ASSQDRGDERLF | 0.19804908 |
| VAAVVTLL | ASSQDRGDERLF | 0.30528367 |
| VGFLGGLI | ASSQDRGDERLF | 0.33597827 |
| VGVIPILL | ASSQDRGDERLF | 0.12623204 |
| VIFASSLI | ASSQDRGDERLF | 0.39378127 |
| VKGGDGGAAAG | ASSQDRGDERLF | 0.73790514 |
| VTKTFTNI | ASSQDRGDERLF | 0.44904348 |
| VVAAAILL | ASSQDRGDERLF | 0.14591292 |
| YAALDVEL | ASSQDRGDERLF | 0.6395749 |
| LASVALLL | ASSQDRGDERLF | 0.8012122 |
| LDAVHRV | ASSQDRGDERLF | 0.6250634 |
| QGLSSSIF | ASSQDRGDERLF | 0.108228214 |
| RGTFANIRL | ASSQDRGDERLF | 0.09862294 |
| RQLWWGHRI | ASSQDRGDERLF | 0.32242313 |
| YGPPGTGKTLL | ASSQDRGDERLF | 0.19326635 |
| AALVGAPANVA | ASSQDRGDERLF | 0.13734463 |
| AATFRRTE | ASSQDRGDERLF | 0.11459479 |
| AGALGGGI | ASSQDRGDERLF | 0.55908895 |
| AGGAGGAGGIGAG | ASSQDRGDERLF | 0.3646201 |
| AGSGALGL | ASSQDRGDERLF | 0.16316153 |
| AQIADQI | ASSQDRGDERLF | 0.10310058 |
| ASAVAVTYL | ASSQDRGDERLF | 0.6379463 |
| ASTVLGGTGG | ASSQDRGDERLF | 0.66466236 |
| AVGGEPAK | ASSQDRGDERLF | 0.13458881 |
| CEQGTFGGTC | ASSQDRGDERLF | 0.23407254 |
| DEGKAPAG | ASSQDRGDERLF | 0.26152736 |
| DTVVFPAL | ASSQDRGDERLF | 0.11323451 |
| FEGITMAG | ASSQDRGDERLF | 0.21595772 |
| FPLVVQD | ASSQDRGDERLF | 0.11632943 |
| GAFTTPQ | ASSQDRGDERLF | 0.15519609 |
| GFGGFGTAGGVGGA | ASSQDRGDERLF | 0.13345496 |
| GFLAQFL | ASSQDRGDERLF | 0.54524827 |
| GGAGGAGGNAGVA | ASSQDRGDERLF | 0.39456484 |
| GGVGTAGAGGAGGAGG | ASSQDRGDERLF | 0.38311028 |
| GNGGGGDAGNAGSGGN | ASSQDRGDERLF | 0.2618969 |
| GTGGAGGAGNDAGS | ASSQDRGDERLF | 0.18711704 |
| GTTNTGWFNS | ASSQDRGDERLF | 0.10913615 |
| IFMGPLFTL | ASSQDRGDERLF | 0.09209056 |

| | | |
|---|---|---|
| ILAHAIEAI | ASSQDRGDERLF | 0.72929704 |
| IRSAIREAL | ASSQDRGDERLF | 0.08549252 |
| ITFAGGG | ASSQDRGDERLF | 0.6100182 |
| ITGMSGHQEV | ASSQDRGDERLF | 0.2085984 |
| IVAGLACTL | ASSQDRGDERLF | 0.12268213 |
| IVGGGLAGA | ASSQDRGDERLF | 0.29559118 |
| IVTGVLNI | ASSQDRGDERLF | 0.091954544 |
| LAQQGCI | ASSQDRGDERLF | 0.15208219 |
| LCLHVEN | ASSQDRGDERLF | 0.63205785 |
| LDAISDGS | ASSQDRGDERLF | 0.4605948 |
| LGGFAAML | ASSQDRGDERLF | 0.23498204 |
| LGGKNPMI | ASSQDRGDERLF | 0.52683926 |
| LGPKGVAAIL | ASSQDRGDERLF | 0.47885376 |
| LMVGVSI | ASSQDRGDERLF | 0.18119672 |
| LVGAAFVL | ASSQDRGDERLF | 0.41900685 |
| LVLAVAPL | ASSQDRGDERLF | 0.4415472 |
| MAAIAVIDAA | ASSQDRGDERLF | 0.5433111 |
| NNGINAELQ | ASSQDRGDERLF | 0.111068435 |
| RGEPLVLVHG | ASSQDRGDERLF | 0.1887539 |
| RGGAGGAAGLI | ASSQDRGDERLF | 0.1502773 |
| RSYRMHPAV | ASSQDRGDERLF | 0.2987709 |
| SAAHLLAGQLA | ASSQDRGDERLF | 0.16388121 |
| SAAVGSY | ASSQDRGDERLF | 0.35510546 |
| SPHAVLHRI | ASSQDRGDERLF | 0.2605761 |
| SQIAKVL | ASSQDRGDERLF | 0.09625998 |
| TALGHLVP | ASSQDRGDERLF | 0.19254285 |
| TAPTTSGAA | ASSQDRGDERLF | 0.27649903 |
| TDGDDGNGGSPGT | ASSQDRGDERLF | 0.17264706 |
| TDGLGEN | ASSQDRGDERLF | 0.2788004 |
| TGGPPLPGQA | ASSQDRGDERLF | 0.11140183 |
| TIPAITI | ASSQDRGDERLF | 0.45151612 |
| TIPGFTI | ASSQDRGDERLF | 0.15548952 |
| TSIASGALA | ASSQDRGDERLF | 0.13207631 |
| TVTLPGAAI | ASSQDRGDERLF | 0.45193103 |
| VASGLSAALP | ASSQDRGDERLF | 0.17014864 |
| VGAGVIGI | ASSQDRGDERLF | 0.17202312 |
| VGPKRFGV | ASSQDRGDERLF | 0.5221883 |
| VNVAFPD | ASSQDRGDERLF | 0.11939748 |
| VSMTNSGVS | ASSQDRGDERLF | 0.2467381 |
| VTDVLGDLSV | ASSQDRGDERLF | 0.20181 |
| VVVVGAGFAGL | ASSQDRGDERLF | 0.15659814 |
| VYTVGTVV | ASSQDRGDERLF | 0.44999433 |
| WTTFSGP | ASSQDRGDERLF | 0.11526132 |
| YDELVEAF | ASSQDRGDERLF | 0.5414317 |
| YGTGGAGGNGG | ASSQDRGDERLF | 0.14073612 |
| YTNVMAV | ASSQDRGDERLF | 0.1415463 |
| FLEQVHQL | ASSQDRGDERLF | 0.2147822 |
| GLSLLAVAV | ASSQDRGDERLF | 0.24486285 |
| LEQVHQL | ASSQDRGDERLF | 0.63419306 |
| LNAAGAI | ASSQDRGDERLF | 0.13049905 |
| PAVHASL | ASSQDRGDERLF | 0.5507765 |

| | | |
|---|---|---|
| TELEDDA | ASSQDRGDERLF | 0.57933766 |
| TPPGPPN | ASSQDRGDERLF | 0.16841492 |
| TPPPPPPP | ASSQDRGDERLF | 0.11028705 |
| TQSPALI | ASADWGPSAETLY | 0.120528534 |
| EFAAAFPQ | ASADWGPSAETLY | 0.099945895 |
| LLSGNLDVL | ASADWGPSAETLY | 0.20454946 |
| GPANLGNYNI | ASADWGPSAETLY | 0.11208679 |
| TLGPTLP | ASADWGPSAETLY | 0.4343609 |
| VYAYSSCCK | ASADWGPSAETLY | 0.6491297 |
| APAPAPAPNI | ASADWGPSAETLY | 0.13130608 |
| PGGPVPIP | ASADWGPSAETLY | 0.58187735 |
| FLTVALEP | ASADWGPSAETLY | 0.10966323 |
| SQYFGANL | ASADWGPSAETLY | 0.25909007 |
| GGVKPAA | ASADWGPSAETLY | 0.67668855 |
| GGTGGAGGAGGW | ASADWGPSAETLY | 0.11198515 |
| GVGGVAGLGGAGG | ASADWGPSAETLY | 0.19523466 |
| VLLVLATTI | ASADWGPSAETLY | 0.1911645 |
| IGIFAAML | ASADWGPSAETLY | 0.120668426 |
| AVGDLID | ASADWGPSAETLY | 0.64526254 |
| NISPLPL | ASADWGPSAETLY | 0.15488519 |
| VTAVFVAI | ASADWGPSAETLY | 0.46724612 |
| SGDDNDAAL | ASADWGPSAETLY | 0.14956747 |
| AVAALAHS | ASADWGPSAETLY | 0.10694392 |
| GGVTVMATP | ASADWGPSAETLY | 0.28995663 |
| AGLAANVRD | ASADWGPSAETLY | 0.6950374 |
| TNRPDLIDPALLRPG | ASADWGPSAETLY | 0.14091945 |
| LANLGVTAL | ASADWGPSAETLY | 0.54901993 |
| TGERIEL | ASADWGPSAETLY | 0.20139259 |
| NGEELAA | ASADWGPSAETLY | 0.53303266 |
| VLSLVWG | ASADWGPSAETLY | 0.7544007 |
| IGYALWL | ASADWGPSAETLY | 0.28274024 |
| LSLLLAPL | ASADWGPSAETLY | 0.20737934 |
| TLDEALK | ASADWGPSAETLY | 0.65287113 |
| TDGAGTAEGG | ASADWGPSAETLY | 0.39002955 |
| NAPGGAGGNGG | ASADWGPSAETLY | 0.6086127 |
| VSVFMGMGGNF | ASADWGPSAETLY | 0.67196023 |
| LASYLAAL | ASADWGPSAETLY | 0.10720491 |
| LVGGTPAL | ASADWGPSAETLY | 0.13142698 |
| VRNADVL | ASADWGPSAETLY | 0.12435674 |
| TCSSPFLVSAM | ASADWGPSAETLY | 0.18291996 |
| VHITDAE | ASADWGPSAETLY | 0.11272924 |
| EYAAPSF | ASADWGPSAETLY | 0.09475955 |
| VVDAFHGGL | ASADWGPSAETLY | 0.13853997 |
| ASVVDPQ | ASADWGPSAETLY | 0.09897996 |
| SYNEFGGLS | ASADWGPSAETLY | 0.10304216 |
| DPDGSVY | ASADWGPSAETLY | 0.13632776 |
| MVSTIHATE | ASADWGPSAETLY | 0.11523476 |
| LAAGVADPV | ASADWGPSAETLY | 0.22958104 |
| TKEQIAA | ASADWGPSAETLY | 0.19293322 |
| SLASPAAL | ASADWGPSAETLY | 0.1027592 |
| IGLVFYGV | ASADWGPSAETLY | 0.11941336 |

| | | |
|---|---|---|
| FLLDASNL | ASADWGPSAETLY | 0.11403428 |
| YQVGNFP | ASADWGPSAETLY | 0.123238295 |
| ESLLSQST | ASADWGPSAETLY | 0.094340846 |
| LFTVGFAAVG | ASADWGPSAETLY | 0.13126044 |
| ATPPTPT | ASADWGPSAETLY | 0.10400711 |
| TDLATLG | ASADWGPSAETLY | 0.09208984 |
| SAGGTGR | ASADWGPSAETLY | 0.15732442 |
| GFSNSGS | ASADWGPSAETLY | 0.119124115 |
| GGNGGSGGTSV | ASADWGPSAETLY | 0.10514526 |
| GTSVATGGAGN | ASADWGPSAETLY | 0.23708162 |
| NGGNGGSGGTS | ASADWGPSAETLY | 0.094186686 |
| AVVKFPH | ASADWGPSAETLY | 0.101350635 |
| TPPPPPP | ASADWGPSAETLY | 0.14056322 |
| AAPAAAAAPA | ASADWGPSAETLY | 0.17558235 |
| IALNEDL | ASADWGPSAETLY | 0.10354379 |
| IVNVSSM | ASADWGPSAETLY | 0.13910532 |
| TGIPPAPR | ASADWGPSAETLY | 0.3870322 |
| FDLGGGTF | ASADWGPSAETLY | 0.1301329 |
| FDLGGGTFDVS | ASADWGPSAETLY | 0.2372835 |
| NEPTAAA | ASADWGPSAETLY | 0.26562873 |
| NEPTAAALA | ASADWGPSAETLY | 0.10674119 |
| GSIINISSI | ASADWGPSAETLY | 0.1401758 |
| VAPGNAGTA | ASADWGPSAETLY | 0.1687774 |
| TGGGASLE | ASADWGPSAETLY | 0.12896843 |
| VAVYNHYKRI | ASADWGPSAETLY | 0.1329754 |
| TDPAPATT | ASADWGPSAETLY | 0.11562916 |
| TNRPDILDPALLRPG | ASADWGPSAETLY | 0.13477889 |
| LTTTTTS | ASADWGPSAETLY | 0.091968134 |
| PTGSGKTL | ASADWGPSAETLY | 0.13572137 |
| TGSGKTL | ASADWGPSAETLY | 0.1489995 |
| TIGHVDHGK | ASADWGPSAETLY | 0.13449876 |
| TIGHVDHGKTT | ASADWGPSAETLY | 0.09912928 |
| GHVDHGKTT | ASADWGPSAETLY | 0.10234196 |
| AQFEHTLL | ASADWGPSAETLY | 0.096368894 |
| IVNQDEL | ASADWGPSAETLY | 0.11748168 |
| TNGSQFF | ASADWGPSAETLY | 0.0976122 |
| TGKGIVE | ASADWGPSAETLY | 0.0957884 |
| LISNASDA | ASADWGPSAETLY | 0.169825 |
| AELRTLGELTKIAKAHGAQ | ASADWGPSAETLY | 0.17567587 |
| APNAVAAAEPSW | ASADWGPSAETLY | 0.10889124 |
| APQVAGAATAVL | ASADWGPSAETLY | 0.1444829 |
| ARRDPAAAAAILPTDA | ASADWGPSAETLY | 0.25547194 |
| DAAARDVLAEAGLAEYF | ASADWGPSAETLY | 0.14923905 |
| DDQIVLSAKANLAKVEQ | ASADWGPSAETLY | 0.14174525 |
| DELLPDTGEDSDEDGH | ASADWGPSAETLY | 0.09892106 |
| ELPEAPANRATVPQD | ASADWGPSAETLY | 0.45716515 |
| FGQSKIERVTGIG | ASADWGPSAETLY | 0.12281909 |
| GAGGTGSTAGGAGGAGGAGGLY | ASADWGPSAETLY | 0.10699301 |
| GHGGTGAAGGNGGTGGAGGSGI | ASADWGPSAETLY | 0.19500758 |
| GNGGNGGAGGEGGPGGAGGAGA | ASADWGPSAETLY | 0.10501049 |
| GSTAGNGGNGGNATNGTIAGQP | ASADWGPSAETLY | 0.10444349 |

| | | |
|---|---|---|
| GTGGDGGNGGNGGTGG | ASADWGPSAETLY | 0.27100497 |
| GVRRPAGIPADAPV | ASADWGPSAETLY | 0.19972867 |
| GYRQVTDAHLLTLARR | ASADWGPSAETLY | 0.10232003 |
| IAKMGQDGHDRG | ASADWGPSAETLY | 0.10063116 |
| KAEPSIEIQYT | ASADWGPSAETLY | 0.10712068 |
| KPVRIRMAADEDTVAVT | ASADWGPSAETLY | 0.15642053 |
| MPPRIAGMRLLVIKPEP | ASADWGPSAETLY | 0.17380847 |
| MKFYAVALFRSRGVPPTR | ASADWGPSAETLY | 0.13856342 |
| NAGGFGILGAVAHSP | ASADWGPSAETLY | 0.11716182 |
| NEGREITLAMAEM | ASADWGPSAETLY | 0.20940587 |
| NELVEWSHPLGHRWRWE | ASADWGPSAETLY | 0.13779247 |
| NLGAGGTGGNGGNPG | ASADWGPSAETLY | 0.13001512 |
| NTEWPKWETVMAADPVD | ASADWGPSAETLY | 0.18908392 |
| PASAAPAETQP | ASADWGPSAETLY | 0.096222304 |
| QQSSDGKNGVRAAEITGE | ASADWGPSAETLY | 0.18080966 |
| RAPDAVTTSWWKE | ASADWGPSAETLY | 0.21079105 |
| SGFGNVGAHTSGWFNQ | ASADWGPSAETLY | 0.1793976 |
| SGVAGAVGQAASVGGLKV | ASADWGPSAETLY | 0.21905375 |
| SNGGAGTGSAGGNGGT | ASADWGPSAETLY | 0.10708319 |
| TGGAGGEGGAAG | ASADWGPSAETLY | 0.16312921 |
| TGGDGGAAGAGGAPGHG | ASADWGPSAETLY | 0.10178624 |
| TMPPLLAAIANAGRHGV | ASADWGPSAETLY | 0.12075658 |
| TSSTAGNSGGA | ASADWGPSAETLY | 0.16882032 |
| VPATVPVGILC | ASADWGPSAETLY | 0.14537312 |
| VQRPDSGGVVLTGRIS | ASADWGPSAETLY | 0.19742432 |
| WTPNPVPATKGVPAGLVQAT | ASADWGPSAETLY | 0.16646446 |
| YALKRALAKFSDRFDVVII | ASADWGPSAETLY | 0.11838934 |
| YLIGSVIV | ASADWGPSAETLY | 0.3582344 |
| YGQLVAVID | ASADWGPSAETLY | 0.28648758 |
| VTVGGMTLPA | ASADWGPSAETLY | 0.10247968 |
| VLALPAV | ASADWGPSAETLY | 0.103616945 |
| VIPPTLNYET | ASADWGPSAETLY | 0.12181461 |
| VGMGGMHPSSG | ASADWGPSAETLY | 0.22504056 |
| VGLAGGAATA | ASADWGPSAETLY | 0.14637813 |
| VGGAGGAGGSGP | ASADWGPSAETLY | 0.15904278 |
| VFAAFMV | ASADWGPSAETLY | 0.13853471 |
| TLYDKLR | ASADWGPSAETLY | 0.09719809 |
| TLAVFTVL | ASADWGPSAETLY | 0.3537459 |
| TGSVQPARV | ASADWGPSAETLY | 0.0986276 |
| TGGGAGSGGT | ASADWGPSAETLY | 0.13679145 |
| SHLETSL | ASADWGPSAETLY | 0.104532264 |
| SFTTGMT | ASADWGPSAETLY | 0.09234297 |
| QVEVAKRMGVRQPTVS | ASADWGPSAETLY | 0.15242857 |
| QGSGGPALAM | ASADWGPSAETLY | 0.11730804 |
| MLAPSNP | ASADWGPSAETLY | 0.23006174 |
| LVNNAAI | ASADWGPSAETLY | 0.11439671 |
| LTGEINTLA | ASADWGPSAETLY | 0.1371414 |
| LSIFGPV | ASADWGPSAETLY | 0.13740794 |
| LLALLANL | ASADWGPSAETLY | 0.12697756 |
| LISNASDALDKLR | ASADWGPSAETLY | 0.104356036 |
| LINEAVP | ASADWGPSAETLY | 0.14188622 |

| | | |
|---|---|---|
| LGYAFSGL | ASADWGPSAETLY | 0.31598842 |
| LDFVAVAL | ASADWGPSAETLY | 0.16757911 |
| LAQPGAE | ASADWGPSAETLY | 0.09722185 |
| ISNLGAVTGF | ASADWGPSAETLY | 0.10388229 |
| HQEDPGRI | ASADWGPSAETLY | 0.12108141 |
| GTPGEPTP | ASADWGPSAETLY | 0.09188276 |
| GFFNSGS | ASADWGPSAETLY | 0.09231832 |
| GAGGGGLTAGAN | ASADWGPSAETLY | 0.09222564 |
| FVDALVNM | ASADWGPSAETLY | 0.097186625 |
| FLTVAPEL | ASADWGPSAETLY | 0.19005704 |
| FETLTAL | ASADWGPSAETLY | 0.16152546 |
| DVINPIEL | ASADWGPSAETLY | 0.13752055 |
| APVSGAPV | ASADWGPSAETLY | 0.19533297 |
| ALVGGLAW | ASADWGPSAETLY | 0.11324637 |
| ALAAAAATAI | ASADWGPSAETLY | 0.09004582 |
| AFGLGLLL | ASADWGPSAETLY | 0.10520297 |
| AFFAAASL | ASADWGPSAETLY | 0.10706758 |
| AAAALPAFSPP | ASADWGPSAETLY | 0.24336231 |
| VVLAPAL | ASADWGPSAETLY | 0.09588459 |
| TIGHVDHGKTTL | ASADWGPSAETLY | 0.12527135 |
| LISNASDALDK | ASADWGPSAETLY | 0.14220576 |
| LISNASDAL | ASADWGPSAETLY | 0.12411196 |
| ISNASDALDK | ASADWGPSAETLY | 0.19177896 |
| NQAETLVYQTE | ASADWGPSAETLY | 0.105311826 |
| TGGTGGTAGAG | ASADWGPSAETLY | 0.09531351 |
| TGSTAGGAGGA | ASADWGPSAETLY | 0.15132615 |
| DPLAVPVAAGPV | ASADWGPSAETLY | 0.093249746 |
| GGTGGTPGNGGN | ASADWGPSAETLY | 0.108699165 |
| PAGLAGVDPATL | ASADWGPSAETLY | 0.10230569 |
| GNNLVVEAPHAGASV | ASADWGPSAETLY | 0.09010932 |
| KVTRIMSASTLEQPA | ASADWGPSAETLY | 0.09570534 |
| LGFALPIALPRRTVV | ASADWGPSAETLY | 0.22778156 |
| TRLGNLTARKLVVLV | ASADWGPSAETLY | 0.12536666 |
| WSQQRRLIYFPSAGP | ASADWGPSAETLY | 0.1385293 |
| GPNGNPDPMAAAVDIR | ASADWGPSAETLY | 0.14274363 |
| TQRCVSVADLRAAASR | ASADWGPSAETLY | 0.12949467 |
| QFVLDVAEHPVEVLTETV | ASADWGPSAETLY | 0.13125004 |
| GGAGGTGGAGGNGGTQAGDG | ASADWGPSAETLY | 0.10944412 |
| PLLATLRAIAVDVDAKPAQV | ASADWGPSAETLY | 0.10927639 |
| APKFRTGGYMIDFWGVGYQVAK | ASADWGPSAETLY | 0.101488605 |
| GAGGAGGAGGAAGGPGGTGGTG | ASADWGPSAETLY | 0.10611372 |
| IGALGLLATDVAPAGSVAFGVA | ASADWGPSAETLY | 0.28160524 |
| AGGAGGIGGNGGAGGAGGVLMG | ASADWGPSAETLY | 0.13166524 |
| ADELIGLL | ASADWGPSAETLY | 0.10200485 |
| ADRALKA | ASADWGPSAETLY | 0.109618016 |
| AGAVGQVGGAGGA | ASADWGPSAETLY | 0.28734535 |
| AGGGGGLGGAP | ASADWGPSAETLY | 0.27450332 |
| AGGLLYGNGGAG | ASADWGPSAETLY | 0.10884233 |
| AGLESIGL | ASADWGPSAETLY | 0.12665975 |
| AGLKTGII | ASADWGPSAETLY | 0.14257914 |
| ASLAALLL | ASADWGPSAETLY | 0.11931668 |

| | | |
|---|---|---|
| ASVINISSI | ASADWGPSAETLY | 0.12270747 |
| ATPPGPA | ASADWGPSAETLY | 0.10383586 |
| ATVASALV | ASADWGPSAETLY | 0.19768837 |
| ATVASLIL | ASADWGPSAETLY | 0.73370695 |
| AVNGPVLIRH | ASADWGPSAETLY | 0.11082098 |
| DGAAGAVGI | ASADWGPSAETLY | 0.096079655 |
| DPLLHGI | ASADWGPSAETLY | 0.14047016 |
| ELGAALGAL | ASADWGPSAETLY | 0.10077552 |
| ELGRTQAV | ASADWGPSAETLY | 0.12065349 |
| FVAGGYLS | ASADWGPSAETLY | 0.11785435 |
| GASGGAGGAGGW | ASADWGPSAETLY | 0.09926064 |
| GGAGAGGDGGAGGI | ASADWGPSAETLY | 0.12320091 |
| GTGHNGSQPGMGGNG | ASADWGPSAETLY | 0.15757343 |
| GYAAASASAS | ASADWGPSAETLY | 0.105907634 |
| HQVDVAAVQQG | ASADWGPSAETLY | 0.23966575 |
| IGAAIAHT | ASADWGPSAETLY | 0.2497356 |
| IGMIEVAQL | ASADWGPSAETLY | 0.17393047 |
| IRLAPEHTSAMD | ASADWGPSAETLY | 0.34638673 |
| IYNVSVL | ASADWGPSAETLY | 0.2530706 |
| KQGFLGM | ASADWGPSAETLY | 0.122331716 |
| LESVITQL | ASADWGPSAETLY | 0.12312377 |
| LGLGALVVV | ASADWGPSAETLY | 0.13471033 |
| LIGNGGNGGSGGI | ASADWGPSAETLY | 0.1330405 |
| LNAINAPTL | ASADWGPSAETLY | 0.15150456 |
| LQLLNTGV | ASADWGPSAETLY | 0.12888485 |
| LQTSIEF | ASADWGPSAETLY | 0.21287234 |
| MSFAVSEL | ASADWGPSAETLY | 0.13529497 |
| NTGALGL | ASADWGPSAETLY | 0.27750307 |
| PPGAGGAGGAGTT | ASADWGPSAETLY | 0.10236918 |
| QLVGLGI | ASADWGPSAETLY | 0.1041188 |
| RFLEQVHQL | ASADWGPSAETLY | 0.28179604 |
| RGVGYVLR | ASADWGPSAETLY | 0.09638729 |
| SATGVFAGVI | ASADWGPSAETLY | 0.09943451 |
| SLPAITLP | ASADWGPSAETLY | 0.12958996 |
| STALAFTV | ASADWGPSAETLY | 0.15301234 |
| SVLTVGLGK | ASADWGPSAETLY | 0.09036525 |
| TALAGAGSGI | ASADWGPSAETLY | 0.1097269 |
| TGLPFTGI | ASADWGPSAETLY | 0.11284611 |
| VAEHVTV | ASADWGPSAETLY | 0.11173456 |
| VDIVLNSL | ASADWGPSAETLY | 0.114699714 |
| VLTGGGAVL | ASADWGPSAETLY | 0.096751876 |
| VNVGLAYGI | ASADWGPSAETLY | 0.15742789 |
| VSSAAVGAV | ASADWGPSAETLY | 0.09888902 |
| VTSVYEAGV | ASADWGPSAETLY | 0.09595966 |
| WVALDRGA | ASADWGPSAETLY | 0.13869596 |
| YLVGSVVI | ASADWGPSAETLY | 0.44350976 |
| GPPGTGKTLL | ASADWGPSAETLY | 0.1102482 |
| GVGMTKF | ASADWGPSAETLY | 0.15581301 |
| IGIGGSDL | ASADWGPSAETLY | 0.109210975 |
| IGTLLAEL | ASADWGPSAETLY | 0.59309447 |
| IYTSGTT | ASADWGPSAETLY | 0.12518059 |

| | | |
|---|---|---|
| LSFPAGV | ASADWGPSAETLY | 0.10360154 |
| LYTSGTT | ASADWGPSAETLY | 0.18062937 |
| VGAFPFGF | ASADWGPSAETLY | 0.25050598 |
| VGAGTVEF | ASADWGPSAETLY | 0.14549288 |
| VGKSTLLL | ASADWGPSAETLY | 0.25661618 |
| AALAHAGGGIV | ASADWGPSAETLY | 0.10310386 |
| AALVAVSLL | ASADWGPSAETLY | 0.12350386 |
| AAVSRAVAAL | ASADWGPSAETLY | 0.16958149 |
| AGGSSGIGGF | ASADWGPSAETLY | 0.15669593 |
| AGLTVANANG | ASADWGPSAETLY | 0.09692564 |
| AGLVGAVL | ASADWGPSAETLY | 0.14300616 |
| ALSQAFTV | ASADWGPSAETLY | 0.1147175 |
| ALTFSVSL | ASADWGPSAETLY | 0.12272859 |
| ALTGAALVAL | ASADWGPSAETLY | 0.1452002 |
| ASFSIGAGVL | ASADWGPSAETLY | 0.14174224 |
| DGSVYKI | ASADWGPSAETLY | 0.20116407 |
| EQLAEIV | ASADWGPSAETLY | 0.14075917 |
| FAVSELTVM | ASADWGPSAETLY | 0.092155024 |
| GAAPLAAGL | ASADWGPSAETLY | 0.09724346 |
| GADGGDGGSGGAGG | ASADWGPSAETLY | 0.15347719 |
| GFGLDTGGAGG | ASADWGPSAETLY | 0.13852067 |
| GLASGGLL | ASADWGPSAETLY | 0.15209709 |
| GRIAGAAVLL | ASADWGPSAETLY | 0.103878856 |
| GSAGSLNTSF | ASADWGPSAETLY | 0.09667434 |
| GTGQIGI | ASADWGPSAETLY | 0.15855074 |
| GTLIALIL | ASADWGPSAETLY | 0.22387259 |
| IAATMGFVVG | ASADWGPSAETLY | 0.119548105 |
| IFAQALL | ASADWGPSAETLY | 0.14953001 |
| IGAAVALGF | ASADWGPSAETLY | 0.4075406 |
| IGLTPLAL | ASADWGPSAETLY | 0.110685326 |
| ILGSNTSTL | ASADWGPSAETLY | 0.11791275 |
| IVFSLTVL | ASADWGPSAETLY | 0.18740241 |
| LASKATGF | ASADWGPSAETLY | 0.3229558 |
| LASVALIL | ASADWGPSAETLY | 0.12740466 |
| LFAVGLTL | ASADWGPSAETLY | 0.29162827 |
| LFNGGIGTYI | ASADWGPSAETLY | 0.09611161 |
| LTLGAGII | ASADWGPSAETLY | 0.17443776 |
| LVAAALGDI | ASADWGPSAETLY | 0.09806517 |
| NTAFVQLGI | ASADWGPSAETLY | 0.13581863 |
| QLFSGAEL | ASADWGPSAETLY | 0.09356347 |
| SAATFITL | ASADWGPSAETLY | 0.13926692 |
| SNYGLANLGV | ASADWGPSAETLY | 0.20234114 |
| SYGGAYAVM | ASADWGPSAETLY | 0.19525202 |
| TTGVGGLTLG | ASADWGPSAETLY | 0.09325625 |
| TTQTIYI | ASADWGPSAETLY | 0.12397142 |
| VAAGAGLGAVI | ASADWGPSAETLY | 0.21747679 |
| VAAVVTLL | ASADWGPSAETLY | 0.67959213 |
| VGFLGGLI | ASADWGPSAETLY | 0.12464259 |
| VGVIPILL | ASADWGPSAETLY | 0.250226 |
| VIFASSLI | ASADWGPSAETLY | 0.10675839 |
| VKGGDGGAAAG | ASADWGPSAETLY | 0.112465054 |

| | | |
|---|---|---|
| VTKTFTNI | ASADWGPSAETLY | 0.10359097 |
| VVAAAILL | ASADWGPSAETLY | 0.122396514 |
| YAALDVEL | ASADWGPSAETLY | 0.13194516 |
| LASVALLL | ASADWGPSAETLY | 0.10643845 |
| LDAVHRV | ASADWGPSAETLY | 0.12894125 |
| QGLSSSIF | ASADWGPSAETLY | 0.39768597 |
| RGTFANIRL | ASADWGPSAETLY | 0.14172503 |
| RQLWWGHRI | ASADWGPSAETLY | 0.30312702 |
| YGPPGTGKTLL | ASADWGPSAETLY | 0.5098988 |
| AALVGAPANVA | ASADWGPSAETLY | 0.1811019 |
| AATFRRTE | ASADWGPSAETLY | 0.10257447 |
| AGALGGGI | ASADWGPSAETLY | 0.12209998 |
| AGGAGGAGGIGAG | ASADWGPSAETLY | 0.09777386 |
| AGSGALGL | ASADWGPSAETLY | 0.12352292 |
| AQIADQI | ASADWGPSAETLY | 0.10111624 |
| ASAVAVTYL | ASADWGPSAETLY | 0.14997765 |
| ASTVLGGTGG | ASADWGPSAETLY | 0.090309694 |
| AVGGEPAK | ASADWGPSAETLY | 0.2391425 |
| CEQGTFGGTC | ASADWGPSAETLY | 0.288272 |
| DEGKAPAG | ASADWGPSAETLY | 0.09153114 |
| DTVVFPAL | ASADWGPSAETLY | 0.19664544 |
| FEGITMAG | ASADWGPSAETLY | 0.3329974 |
| FPLVVQD | ASADWGPSAETLY | 0.09863979 |
| GAFTTPQ | ASADWGPSAETLY | 0.23133445 |
| GFGGFGTAGGVGGA | ASADWGPSAETLY | 0.10676433 |
| GFLAQFL | ASADWGPSAETLY | 0.2355048 |
| GGAGGAGGNAGVA | ASADWGPSAETLY | 0.16181807 |
| GGVGTAGAGGAGGAGG | ASADWGPSAETLY | 0.15093744 |
| GNGGGGDAGNAGSGGN | ASADWGPSAETLY | 0.14513625 |
| GTGGAGGAGNDAGS | ASADWGPSAETLY | 0.17459586 |
| GTTNTGWFNS | ASADWGPSAETLY | 0.09546729 |
| IFMGPLFTL | ASADWGPSAETLY | 0.14758688 |
| ILAHAIEAI | ASADWGPSAETLY | 0.08990333 |
| IRSAIREAL | ASADWGPSAETLY | 0.16317165 |
| ITFAGGG | ASADWGPSAETLY | 0.109479554 |
| ITGMSGHQEV | ASADWGPSAETLY | 0.16631863 |
| IVAGLACTL | ASADWGPSAETLY | 0.11315481 |
| IVGGGLAGA | ASADWGPSAETLY | 0.15710208 |
| IVTGVLNI | ASADWGPSAETLY | 0.11186019 |
| LAQQGCI | ASADWGPSAETLY | 0.10288027 |
| LCLHVEN | ASADWGPSAETLY | 0.09889643 |
| LDAISDGS | ASADWGPSAETLY | 0.097550385 |
| LGGFAAML | ASADWGPSAETLY | 0.21517625 |
| LGGKNPMI | ASADWGPSAETLY | 0.12345933 |
| LGPKGVAAIL | ASADWGPSAETLY | 0.10041112 |
| LMVGVSI | ASADWGPSAETLY | 0.15646546 |
| LVGAAFVL | ASADWGPSAETLY | 0.12715869 |
| LVLAVAPL | ASADWGPSAETLY | 0.09481543 |
| MAAIAVIDAA | ASADWGPSAETLY | 0.13564306 |
| NNGINAELQ | ASADWGPSAETLY | 0.13481273 |
| RGEPLVLVHG | ASADWGPSAETLY | 0.11962429 |

| | | |
|---|---|---|
| RGGAGGAAGLI | ASADWGPSAETLY | 0.09006318 |
| RSYRMHPAV | ASADWGPSAETLY | 0.10452761 |
| SAAHLLAGQLA | ASADWGPSAETLY | 0.12257887 |
| SAAVGSY | ASADWGPSAETLY | 0.094802186 |
| SPHAVLHRI | ASADWGPSAETLY | 0.10001919 |
| SQIAKVL | ASADWGPSAETLY | 0.23504642 |
| TALGHLVP | ASADWGPSAETLY | 0.091089696 |
| TAPTTSGAA | ASADWGPSAETLY | 0.11417314 |
| TDGDDGNGGSPGT | ASADWGPSAETLY | 0.2416139 |
| TDGLGEN | ASADWGPSAETLY | 0.14823326 |
| TGGPPLPGQA | ASADWGPSAETLY | 0.29952985 |
| TIPAITI | ASADWGPSAETLY | 0.11142995 |
| TIPGFTI | ASADWGPSAETLY | 0.116010085 |
| TSIASGALA | ASADWGPSAETLY | 0.1477582 |
| TVTLPGAAI | ASADWGPSAETLY | 0.13380389 |
| VASGLSAALP | ASADWGPSAETLY | 0.096966885 |
| VGAGVIGI | ASADWGPSAETLY | 0.0934108 |
| VGPKRFGV | ASADWGPSAETLY | 0.11651851 |
| VNVAFPD | ASADWGPSAETLY | 0.10209762 |
| VSMTNSGVS | ASADWGPSAETLY | 0.14436385 |
| VTDVLGDLSV | ASADWGPSAETLY | 0.23157501 |
| VVVVGAGFAGL | ASADWGPSAETLY | 0.09942783 |
| VYTVGTVV | ASADWGPSAETLY | 0.10390556 |
| WTTFSGP | ASADWGPSAETLY | 0.12021736 |
| YDELVEAF | ASADWGPSAETLY | 0.1006605 |
| YGTGGAGGNGG | ASADWGPSAETLY | 0.18782744 |
| YTNVMAV | ASADWGPSAETLY | 0.10054934 |
| FLEQVHQL | ASADWGPSAETLY | 0.7040111 |
| GLSLLAVAV | ASADWGPSAETLY | 0.14089048 |
| LEQVHQL | ASADWGPSAETLY | 0.103582345 |
| LNAAGAI | ASADWGPSAETLY | 0.22225556 |
| PAVHASL | ASADWGPSAETLY | 0.1309347 |
| TELEDDA | ASADWGPSAETLY | 0.13236293 |
| TPPGPPN | ASADWGPSAETLY | 0.13784078 |
| TPPPPPPP | ASADWGPSAETLY | 0.09735676 |
| TQSPALI | ASSLWGINTLY | 0.11289625 |
| EFAAAFPQ | ASSLWGINTLY | 0.13443616 |
| LLSGNLDVL | ASSLWGINTLY | 0.10015527 |
| GPANLGNYNI | ASSLWGINTLY | 0.12757486 |
| TLGPTLP | ASSLWGINTLY | 0.09563073 |
| VYAYSSCCK | ASSLWGINTLY | 0.0981894 |
| APAPAPAPNI | ASSLWGINTLY | 0.1448299 |
| PGGPVPIP | ASSLWGINTLY | 0.15779798 |
| FLTVALEP | ASSLWGINTLY | 0.47228968 |
| SQYFGANL | ASSLWGINTLY | 0.11090999 |
| GGVKPAA | ASSLWGINTLY | 0.101111636 |
| GGTGGAGGAGGW | ASSLWGINTLY | 0.4328749 |
| GVGGVAGLGGAGG | ASSLWGINTLY | 0.109430246 |
| VLLVLATTI | ASSLWGINTLY | 0.15611812 |
| IGIFAAML | ASSLWGINTLY | 0.110035256 |
| AVGDLID | ASSLWGINTLY | 0.1003981 |

| | | |
|---|---|---|
| NISPLPL | ASSLWGINTLY | 0.11715458 |
| VTAVFVAI | ASSLWGINTLY | 0.1447171 |
| SGDDNDAAL | ASSLWGINTLY | 0.094341345 |
| AVAALAHS | ASSLWGINTLY | 0.18312642 |
| GGVTVMATP | ASSLWGINTLY | 0.09013342 |
| AGLAANVRD | ASSLWGINTLY | 0.10488474 |
| TNRPDLIDPALLRPG | ASSLWGINTLY | 0.116270185 |
| LANLGVTAL | ASSLWGINTLY | 0.108971566 |
| TGERIEL | ASSLWGINTLY | 0.09602948 |
| NGEELAA | ASSLWGINTLY | 0.13229553 |
| VLSLVWG | ASSLWGINTLY | 0.14805672 |
| IGYALWL | ASSLWGINTLY | 0.2705568 |
| LSLLLAPL | ASSLWGINTLY | 0.0988446 |
| TLDEALK | ASSLWGINTLY | 0.17775162 |
| TDGAGTAEGG | ASSLWGINTLY | 0.36797926 |
| NAPGGAGGNGG | ASSLWGINTLY | 0.13493663 |
| VSVFMGMGGNF | ASSLWGINTLY | 0.095902376 |
| LASYLAAL | ASSLWGINTLY | 0.121698916 |
| LVGGTPAL | ASSLWGINTLY | 0.092782475 |
| VRNADVL | ASSLWGINTLY | 0.10447425 |
| TCSSPFLVSAM | ASSLWGINTLY | 0.2741319 |
| VHITDAE | ASSLWGINTLY | 0.09758838 |
| EYAAPSF | ASSLWGINTLY | 0.15332796 |
| VVDAFHGGL | ASSLWGINTLY | 0.20381932 |
| ASVVDPQ | ASSLWGINTLY | 0.11344864 |
| SYNEFGGLS | ASSLWGINTLY | 0.11314128 |
| DPDGSVY | ASSLWGINTLY | 0.10442272 |
| MVSTIHATE | ASSLWGINTLY | 0.09695335 |
| LAAGVADPV | ASSLWGINTLY | 0.13784464 |
| TKEQIAA | ASSLWGINTLY | 0.11936973 |
| SLASPAAL | ASSLWGINTLY | 0.11042654 |
| IGLVFYGV | ASSLWGINTLY | 0.12452153 |
| FLLDASNL | ASSLWGINTLY | 0.110104315 |
| YQVGNFP | ASSLWGINTLY | 0.11793925 |
| ESLLSQST | ASSLWGINTLY | 0.11193465 |
| LFTVGFAAVG | ASSLWGINTLY | 0.10267426 |
| ATPPTPT | ASSLWGINTLY | 0.16566601 |
| TDLATLG | ASSLWGINTLY | 0.097554356 |
| SAGGTGR | ASSLWGINTLY | 0.16175397 |
| GFSNSGS | ASSLWGINTLY | 0.3088203 |
| GGNGGSGGTSV | ASSLWGINTLY | 0.09356242 |
| GTSVATGGAGN | ASSLWGINTLY | 0.10699877 |
| NGGNGGSGGTS | ASSLWGINTLY | 0.12314596 |
| AVVKFPH | ASSLWGINTLY | 0.095979966 |
| TPPPPPP | ASSLWGINTLY | 0.27852207 |
| AAPAAAAAPA | ASSLWGINTLY | 0.11820384 |
| IALNEDL | ASSLWGINTLY | 0.16263628 |
| IVNVSSM | ASSLWGINTLY | 0.115852304 |
| TGIPPAPR | ASSLWGINTLY | 0.5542523 |
| FDLGGGTF | ASSLWGINTLY | 0.17467809 |
| FDLGGGTFDVS | ASSLWGINTLY | 0.1199429 |

| | | |
|---|---|---|
| NEPTAAA | ASSLWGINTLY | 0.09324235 |
| NEPTAAALA | ASSLWGINTLY | 0.104787104 |
| GSIINISSI | ASSLWGINTLY | 0.1718907 |
| VAPGNAGTA | ASSLWGINTLY | 0.21788839 |
| TGGGASLE | ASSLWGINTLY | 0.098467454 |
| VAVYNHYKRI | ASSLWGINTLY | 0.12535204 |
| TDPAPATT | ASSLWGINTLY | 0.23953557 |
| TNRPDILDPALLRPG | ASSLWGINTLY | 0.26133907 |
| LTTTTTS | ASSLWGINTLY | 0.09893529 |
| PTGSGKTL | ASSLWGINTLY | 0.11229102 |
| TGSGKTL | ASSLWGINTLY | 0.13329516 |
| TIGHVDHGK | ASSLWGINTLY | 0.13285065 |
| TIGHVDHGKTT | ASSLWGINTLY | 0.10106523 |
| GHVDHGKTT | ASSLWGINTLY | 0.23963389 |
| AQFEHTLL | ASSLWGINTLY | 0.14516753 |
| IVNQDEL | ASSLWGINTLY | 0.45378196 |
| TNGSQFF | ASSLWGINTLY | 0.1392526 |
| TGKGIVE | ASSLWGINTLY | 0.1719236 |
| LISNASDA | ASSLWGINTLY | 0.13478906 |
| AELRTLGELTKIAKAHGAQ | ASSLWGINTLY | 0.11306233 |
| APNAVAAAEPSW | ASSLWGINTLY | 0.10451826 |
| APQVAGAATAVL | ASSLWGINTLY | 0.09400496 |
| ARRDPAAAAAILPTDA | ASSLWGINTLY | 0.10154759 |
| DAAARDVLAEAGLAEYF | ASSLWGINTLY | 0.10857332 |
| DDQIVLSAKANLAKVEQ | ASSLWGINTLY | 0.09048845 |
| DELLPDTGEDSDEDGH | ASSLWGINTLY | 0.117951855 |
| ELPEAPANRATVPQD | ASSLWGINTLY | 0.13704848 |
| FGQSKIERVTGIG | ASSLWGINTLY | 0.15300667 |
| GAGGTGSTAGGAGGAGGAGGLY | ASSLWGINTLY | 0.12061219 |
| GHGGTGAAGGNGGTGGAGGSGI | ASSLWGINTLY | 0.122013 |
| GNGGNGGAGGEGGPGGAGGAGA | ASSLWGINTLY | 0.099235766 |
| GSTAGNGGNGGNATNGTIAGQP | ASSLWGINTLY | 0.27859342 |
| GTGGDGGNGGNGGTGG | ASSLWGINTLY | 0.09490362 |
| GVRRPAGIPADAPV | ASSLWGINTLY | 0.0999258 |
| GYRQVTDAHLLTLARR | ASSLWGINTLY | 0.11359967 |
| IAKMGQDGHDRG | ASSLWGINTLY | 0.0990317 |
| KAEPSIEIQYT | ASSLWGINTLY | 0.48945507 |
| KPVRIRMAADEDTVAVT | ASSLWGINTLY | 0.20765984 |
| MPPRIAGMRLLVIKPEP | ASSLWGINTLY | 0.1427954 |
| MKFYAVALFRSRGVPPTR | ASSLWGINTLY | 0.29208994 |
| NAGGFGILGAVAHSP | ASSLWGINTLY | 0.09469615 |
| NEGREITLAMAEM | ASSLWGINTLY | 0.10882688 |
| NELVEWSHPLGHRWRWE | ASSLWGINTLY | 0.36276385 |
| NLGAGGTGGNGGNPG | ASSLWGINTLY | 0.11462529 |
| NTEWPKWETVMAADPVD | ASSLWGINTLY | 0.15617062 |
| PASAAPAETQP | ASSLWGINTLY | 0.15548371 |
| QQSSDGKNGVRAAEITGE | ASSLWGINTLY | 0.10393464 |
| RAPDAVTTSWWKE | ASSLWGINTLY | 0.120445825 |
| SGFGNVGAHTSGWFNQ | ASSLWGINTLY | 0.10831194 |
| SGVAGAVGQAASVGGLKV | ASSLWGINTLY | 0.16295412 |
| SNGGAGTGSAGGNGGT | ASSLWGINTLY | 0.11296156 |

| | | |
|---|---|---|
| TGGAGGEGGAAG | ASSLWGINTLY | 0.100900166 |
| TGGDGGAAGAGGAPGHG | ASSLWGINTLY | 0.24138942 |
| TMPPLLAAIANAGRHGV | ASSLWGINTLY | 0.7572057 |
| TSSTAGNSGGA | ASSLWGINTLY | 0.12333244 |
| VPATVPVGILC | ASSLWGINTLY | 0.11687592 |
| VQRPDSGGVVLTGRIS | ASSLWGINTLY | 0.12833495 |
| WTPNPVPATKGVPAGLVQAT | ASSLWGINTLY | 0.1113645 |
| YALKRALAKFSDRFDVVII | ASSLWGINTLY | 0.110944666 |
| YLIGSVIV | ASSLWGINTLY | 0.103334844 |
| YGQLVAVID | ASSLWGINTLY | 0.10190423 |
| VTVGGMTLPA | ASSLWGINTLY | 0.104430966 |
| VLALPAV | ASSLWGINTLY | 0.15833445 |
| VIPPTLNYET | ASSLWGINTLY | 0.12334892 |
| VGMGGMHPSSG | ASSLWGINTLY | 0.74972296 |
| VGLAGGAATA | ASSLWGINTLY | 0.1968268 |
| VGGAGGAGGSGP | ASSLWGINTLY | 0.11087759 |
| VFAAFMV | ASSLWGINTLY | 0.097652376 |
| TLYDKLR | ASSLWGINTLY | 0.120914616 |
| TLAVFTVL | ASSLWGINTLY | 0.120535724 |
| TGSVQPARV | ASSLWGINTLY | 0.098631166 |
| TGGGAGSGGT | ASSLWGINTLY | 0.27510133 |
| SHLETSL | ASSLWGINTLY | 0.111002356 |
| SFTTGMT | ASSLWGINTLY | 0.103516474 |
| QVEVAKRMGVRQPTVS | ASSLWGINTLY | 0.14077865 |
| QGSGGPALAM | ASSLWGINTLY | 0.589138 |
| MLAPSNP | ASSLWGINTLY | 0.15502928 |
| LVNNAAI | ASSLWGINTLY | 0.10930365 |
| LTGEINTLA | ASSLWGINTLY | 0.098901935 |
| LSIFGPV | ASSLWGINTLY | 0.26059678 |
| LLALLANL | ASSLWGINTLY | 0.264646 |
| LISNASDALDKLR | ASSLWGINTLY | 0.09043574 |
| LINEAVP | ASSLWGINTLY | 0.46654528 |
| LGYAFSGL | ASSLWGINTLY | 0.10521578 |
| LDFVAVAL | ASSLWGINTLY | 0.10313461 |
| LAQPGAE | ASSLWGINTLY | 0.1445259 |
| ISNLGAVTGF | ASSLWGINTLY | 0.1290378 |
| HQEDPGRI | ASSLWGINTLY | 0.41977382 |
| GTPGEPTP | ASSLWGINTLY | 0.1217972 |
| GFFNSGS | ASSLWGINTLY | 0.32891017 |
| GAGGGGLTAGAN | ASSLWGINTLY | 0.10922378 |
| FVDALVNM | ASSLWGINTLY | 0.28578496 |
| FLTVAPEL | ASSLWGINTLY | 0.1469074 |
| FETLTAL | ASSLWGINTLY | 0.14248723 |
| DVINPIEL | ASSLWGINTLY | 0.13476634 |
| APVSGAPV | ASSLWGINTLY | 0.1209105 |
| ALVGGLAW | ASSLWGINTLY | 0.13897216 |
| ALAAAAATAI | ASSLWGINTLY | 0.10495802 |
| AFGLGLLL | ASSLWGINTLY | 0.11366896 |
| AFFAAASL | ASSLWGINTLY | 0.15478504 |
| AAAALPAFSPP | ASSLWGINTLY | 0.16060759 |
| VVLAPAL | ASSLWGINTLY | 0.10298764 |

| | | |
|---|---|---|
| TIGHVDHGKTTL | ASSLWGINTLY | 0.13708574 |
| LISNASDALDK | ASSLWGINTLY | 0.53841186 |
| LISNASDAL | ASSLWGINTLY | 0.14811872 |
| ISNASDALDK | ASSLWGINTLY | 0.10563264 |
| NQAETLVYQTE | ASSLWGINTLY | 0.33993095 |
| TGGTGGTAGAG | ASSLWGINTLY | 0.61432326 |
| TGSTAGGAGGA | ASSLWGINTLY | 0.2034536 |
| DPLAVPVAAGPV | ASSLWGINTLY | 0.099952616 |
| GGTGGTPGNGGN | ASSLWGINTLY | 0.098307714 |
| PAGLAGVDPATL | ASSLWGINTLY | 0.14333731 |
| GNNLVVEAPHAGASV | ASSLWGINTLY | 0.13092566 |
| KVTRIMSASTLEQPA | ASSLWGINTLY | 0.10850922 |
| LGFALPIALPRRTVV | ASSLWGINTLY | 0.09947827 |
| TRLGNLTARKLVVLV | ASSLWGINTLY | 0.18274662 |
| WSQQRRLIYFPSAGP | ASSLWGINTLY | 0.17668636 |
| GPNGNPDPMAAAVDIR | ASSLWGINTLY | 0.11098827 |
| TQRCVSVADLRAAASR | ASSLWGINTLY | 0.09728788 |
| QFVLDVAEHPVEVLTETV | ASSLWGINTLY | 0.17850694 |
| GGAGGTGGAGGNGGTQAGDG | ASSLWGINTLY | 0.23579387 |
| PLLATLRAIAVDVDAKPAQV | ASSLWGINTLY | 0.10477912 |
| APKFRTGGYMIDFWGVGYQVAK | ASSLWGINTLY | 0.11292872 |
| GAGGAGGAGGAAGGPGGTGGTG | ASSLWGINTLY | 0.14199752 |
| IGALGLLATDVAPAGSVAFGVA | ASSLWGINTLY | 0.09729151 |
| AGGAGGIGGNGGAGGAGGVLMG | ASSLWGINTLY | 0.11733061 |
| ADELIGLL | ASSLWGINTLY | 0.109781966 |
| ADRALKA | ASSLWGINTLY | 0.10538912 |
| AGAVGQVGGAGGA | ASSLWGINTLY | 0.1124623 |
| AGGGGGLGGAP | ASSLWGINTLY | 0.20948935 |
| AGGLLYGNGGAG | ASSLWGINTLY | 0.14808464 |
| AGLESIGL | ASSLWGINTLY | 0.15352505 |
| AGLKTGII | ASSLWGINTLY | 0.14359878 |
| ASLAALLL | ASSLWGINTLY | 0.11632904 |
| ASVINISSI | ASSLWGINTLY | 0.09826817 |
| ATPPGPA | ASSLWGINTLY | 0.09303492 |
| ATVASALV | ASSLWGINTLY | 0.36600995 |
| ATVASLIL | ASSLWGINTLY | 0.0899553 |
| AVNGPVLIRH | ASSLWGINTLY | 0.22120556 |
| DGAAGAVGI | ASSLWGINTLY | 0.13238673 |
| DPLLHGI | ASSLWGINTLY | 0.106165856 |
| ELGAALGAL | ASSLWGINTLY | 0.13070843 |
| ELGRTQAV | ASSLWGINTLY | 0.092604704 |
| FVAGGYLS | ASSLWGINTLY | 0.104795136 |
| GASGGAGGAGGW | ASSLWGINTLY | 0.2785054 |
| GGAGAGGDGGAGGI | ASSLWGINTLY | 0.14951652 |
| GTGHNGSQPGMGGNG | ASSLWGINTLY | 0.13012017 |
| GYAAASASAS | ASSLWGINTLY | 0.1709741 |
| HQVDVAAVQQG | ASSLWGINTLY | 0.16613005 |
| IGAAIAHT | ASSLWGINTLY | 0.10948325 |
| IGMIEVAQL | ASSLWGINTLY | 0.14849669 |
| IRLAPEHTSAMD | ASSLWGINTLY | 0.1972591 |
| IYNVSVL | ASSLWGINTLY | 0.18220657 |

| | | |
|---|---|---|
| KQGFLGM | ASSLWGINTLY | 0.099203154 |
| LESVITQL | ASSLWGINTLY | 0.12126951 |
| LGLGALVVV | ASSLWGINTLY | 0.107909575 |
| LIGNGGNGGSGGI | ASSLWGINTLY | 0.14030151 |
| LNAINAPTL | ASSLWGINTLY | 0.113360934 |
| LQLLNTGV | ASSLWGINTLY | 0.15966597 |
| LQTSIEF | ASSLWGINTLY | 0.12609296 |
| MSFAVSEL | ASSLWGINTLY | 0.09444889 |
| NTGALGL | ASSLWGINTLY | 0.10704734 |
| PPGAGGAGGAGTT | ASSLWGINTLY | 0.14687356 |
| QLVGLGI | ASSLWGINTLY | 0.13396342 |
| RFLEQVHQL | ASSLWGINTLY | 0.13038522 |
| RGVGYVLR | ASSLWGINTLY | 0.10424645 |
| SATGVFAGVI | ASSLWGINTLY | 0.16614388 |
| SLPAITLP | ASSLWGINTLY | 0.11613401 |
| STALAFTV | ASSLWGINTLY | 0.11191075 |
| SVLTVGLGK | ASSLWGINTLY | 0.15427543 |
| TALAGAGSGI | ASSLWGINTLY | 0.114541434 |
| TGLPFTGI | ASSLWGINTLY | 0.10955944 |
| VAEHVTV | ASSLWGINTLY | 0.17790112 |
| VDIVLNSL | ASSLWGINTLY | 0.1019504 |
| VLTGGGAVL | ASSLWGINTLY | 0.1501577 |
| VNVGLAYGI | ASSLWGINTLY | 0.0991485 |
| VSSAAVGAV | ASSLWGINTLY | 0.113607295 |
| VTSVYEAGV | ASSLWGINTLY | 0.17454553 |
| WVALDRGA | ASSLWGINTLY | 0.09843482 |
| YLVGSVVI | ASSLWGINTLY | 0.18319836 |
| GPPGTGKTLL | ASSLWGINTLY | 0.11262453 |
| GVGMTKF | ASSLWGINTLY | 0.10721765 |
| IGIGGSDL | ASSLWGINTLY | 0.11752269 |
| IGTLLAEL | ASSLWGINTLY | 0.1464307 |
| IYTSGTT | ASSLWGINTLY | 0.21350014 |
| LSFPAGV | ASSLWGINTLY | 0.10047228 |
| LYTSGTT | ASSLWGINTLY | 0.18697676 |
| VGAFPFGF | ASSLWGINTLY | 0.20439915 |
| VGAGTVEF | ASSLWGINTLY | 0.11472823 |
| VGKSTLLL | ASSLWGINTLY | 0.09788896 |
| AALAHAGGGIV | ASSLWGINTLY | 0.10706009 |
| AALVAVSLL | ASSLWGINTLY | 0.096884266 |
| AAVSRAVAAL | ASSLWGINTLY | 0.12032388 |
| AGGSSGIGGF | ASSLWGINTLY | 0.10362583 |
| AGLTVANANG | ASSLWGINTLY | 0.18250895 |
| AGLVGAVL | ASSLWGINTLY | 0.22798464 |
| ALSQAFTV | ASSLWGINTLY | 0.10964574 |
| ALTFSVSL | ASSLWGINTLY | 0.17026266 |
| ALTGAALVAL | ASSLWGINTLY | 0.10245021 |
| ASFSIGAGVL | ASSLWGINTLY | 0.13919674 |
| DGSVYKI | ASSLWGINTLY | 0.3658914 |
| EQLAEIV | ASSLWGINTLY | 0.1028415 |
| FAVSELTVM | ASSLWGINTLY | 0.12785047 |
| GAAPLAAGL | ASSLWGINTLY | 0.10636275 |

| | | |
|---|---|---|
| GADGGDGGSGGAGG | ASSLWGINTLY | 0.1027529 |
| GFGLDTGGAGG | ASSLWGINTLY | 0.2608899 |
| GLASGGLL | ASSLWGINTLY | 0.118107 |
| GRIAGAAVLL | ASSLWGINTLY | 0.16239747 |
| GSAGSLNTSF | ASSLWGINTLY | 0.11057535 |
| GTGQIGI | ASSLWGINTLY | 0.09926988 |
| GTLIALIL | ASSLWGINTLY | 0.13817713 |
| IAATMGFVVG | ASSLWGINTLY | 0.25979966 |
| IFAQALL | ASSLWGINTLY | 0.10560043 |
| IGAAVALGF | ASSLWGINTLY | 0.16142495 |
| IGLTPLAL | ASSLWGINTLY | 0.12141116 |
| ILGSNTSTL | ASSLWGINTLY | 0.14374083 |
| IVFSLTVL | ASSLWGINTLY | 0.62871957 |
| LASKATGF | ASSLWGINTLY | 0.1346405 |
| LASVALIL | ASSLWGINTLY | 0.11791826 |
| LFAVGLTL | ASSLWGINTLY | 0.32575002 |
| LFNGGIGTYI | ASSLWGINTLY | 0.14884079 |
| LTLGAGII | ASSLWGINTLY | 0.25220972 |
| LVAAALGDI | ASSLWGINTLY | 0.19987899 |
| NTAFVQLGI | ASSLWGINTLY | 0.103641346 |
| QLFSGAEL | ASSLWGINTLY | 0.27814677 |
| SAATFITL | ASSLWGINTLY | 0.14148691 |
| SNYGLANLGV | ASSLWGINTLY | 0.12860717 |
| SYGGAYAVM | ASSLWGINTLY | 0.13878101 |
| TTGVGGLTLG | ASSLWGINTLY | 0.110439144 |
| TTQTIYI | ASSLWGINTLY | 0.10702373 |
| VAAGAGLGAVI | ASSLWGINTLY | 0.12512667 |
| VAAVVTLL | ASSLWGINTLY | 0.11906257 |
| VGFLGGLI | ASSLWGINTLY | 0.11739493 |
| VGVIPILL | ASSLWGINTLY | 0.13676186 |
| VIFASSLI | ASSLWGINTLY | 0.29324573 |
| VKGGDGGAAAG | ASSLWGINTLY | 0.1678307 |
| VTKTFTNI | ASSLWGINTLY | 0.108411305 |
| VVAAAILL | ASSLWGINTLY | 0.13653573 |
| YAALDVEL | ASSLWGINTLY | 0.11957388 |
| LASVALLL | ASSLWGINTLY | 0.12114401 |
| LDAVHRV | ASSLWGINTLY | 0.09808085 |
| QGLSSSIF | ASSLWGINTLY | 0.16777258 |
| RGTFANIRL | ASSLWGINTLY | 0.1042523 |
| RQLWWGHRI | ASSLWGINTLY | 0.097653866 |
| YGPPGTGKTLL | ASSLWGINTLY | 0.14206721 |
| AALVGAPANVA | ASSLWGINTLY | 0.13726808 |
| AATFRRTE | ASSLWGINTLY | 0.119149044 |
| AGALGGGI | ASSLWGINTLY | 0.17429383 |
| AGGAGGAGGIGAG | ASSLWGINTLY | 0.1311651 |
| AGSGALGL | ASSLWGINTLY | 0.12780185 |
| AQIADQI | ASSLWGINTLY | 0.20735615 |
| ASAVAVTYL | ASSLWGINTLY | 0.2074404 |
| ASTVLGGTGG | ASSLWGINTLY | 0.12960964 |
| AVGGEPAK | ASSLWGINTLY | 0.116854146 |
| CEQGTFGGTC | ASSLWGINTLY | 0.40223318 |

| | | |
|---|---|---|
| DEGKAPAG | ASSLWGINTLY | 0.14463133 |
| DTVVFPAL | ASSLWGINTLY | 0.12397746 |
| FEGITMAG | ASSLWGINTLY | 0.13888068 |
| FPLVVQD | ASSLWGINTLY | 0.1052362 |
| GAFTTPQ | ASSLWGINTLY | 0.1437639 |
| GFGGFGTAGGVGGA | ASSLWGINTLY | 0.16195157 |
| GFLAQFL | ASSLWGINTLY | 0.1574873 |
| GGAGGAGGNAGVA | ASSLWGINTLY | 0.15128385 |
| GGVGTAGAGGAGGAGG | ASSLWGINTLY | 0.10308705 |
| GNGGGGDAGNAGSGGN | ASSLWGINTLY | 0.1200444 |
| GTGGAGGAGNDAGS | ASSLWGINTLY | 0.10574039 |
| GTTNTGWFNS | ASSLWGINTLY | 0.21212584 |
| IFMGPLFTL | ASSLWGINTLY | 0.222092 |
| ILAHAIEAI | ASSLWGINTLY | 0.16835144 |
| IRSAIREAL | ASSLWGINTLY | 0.102271624 |
| ITFAGGG | ASSLWGINTLY | 0.111136295 |
| ITGMSGHQEV | ASSLWGINTLY | 0.11664778 |
| IVAGLACTL | ASSLWGINTLY | 0.102666296 |
| IVGGGLAGA | ASSLWGINTLY | 0.108877204 |
| IVTGVLNI | ASSLWGINTLY | 0.10550711 |
| LAQQGCI | ASSLWGINTLY | 0.2658841 |
| LCLHVEN | ASSLWGINTLY | 0.24558619 |
| LDAISDGS | ASSLWGINTLY | 0.150467 |
| LGGFAAML | ASSLWGINTLY | 0.14441192 |
| LGGKNPMI | ASSLWGINTLY | 0.14124884 |
| LGPKGVAAIL | ASSLWGINTLY | 0.18703252 |
| LMVGVSI | ASSLWGINTLY | 0.13955894 |
| LVGAAFVL | ASSLWGINTLY | 0.10554998 |
| LVLAVAPL | ASSLWGINTLY | 0.34315294 |
| MAAIAVIDAA | ASSLWGINTLY | 0.1682155 |
| NNGINAELQ | ASSLWGINTLY | 0.13089433 |
| RGEPLVLVHG | ASSLWGINTLY | 0.24754721 |
| RGGAGGAAGLI | ASSLWGINTLY | 0.14947778 |
| RSYRMHPAV | ASSLWGINTLY | 0.12711467 |
| SAAHLLAGQLA | ASSLWGINTLY | 0.19593191 |
| SAAVGSY | ASSLWGINTLY | 0.15745655 |
| SPHAVLHRI | ASSLWGINTLY | 0.11099287 |
| SQIAKVL | ASSLWGINTLY | 0.13065147 |
| TALGHLVP | ASSLWGINTLY | 0.12148054 |
| TAPTTSGAA | ASSLWGINTLY | 0.18052319 |
| TDGDDGNGGSPGT | ASSLWGINTLY | 0.19879298 |
| TDGLGEN | ASSLWGINTLY | 0.15145795 |
| TGGPPLPGQA | ASSLWGINTLY | 0.12860309 |
| TIPAITI | ASSLWGINTLY | 0.15831286 |
| TIPGFTI | ASSLWGINTLY | 0.16787174 |
| TSIASGALA | ASSLWGINTLY | 0.1559633 |
| TVTLPGAAI | ASSLWGINTLY | 0.13634674 |
| VASGLSAALP | ASSLWGINTLY | 0.116427585 |
| VGAGVIGI | ASSLWGINTLY | 0.15868865 |
| VGPKRFGV | ASSLWGINTLY | 0.24110705 |
| VNVAFPD | ASSLWGINTLY | 0.124788664 |

| | | |
|---|---|---|
| VSMTNSGVS | ASSLWGINTLY | 0.12311282 |
| VTDVLGDLSV | ASSLWGINTLY | 0.18366992 |
| VVVVGAGFAGL | ASSLWGINTLY | 0.14718464 |
| VYTVGTVV | ASSLWGINTLY | 0.12994736 |
| WTTFSGP | ASSLWGINTLY | 0.10904772 |
| YDELVEAF | ASSLWGINTLY | 0.12116625 |
| YGTGGAGGNGG | ASSLWGINTLY | 0.17814842 |
| YTNVMAV | ASSLWGINTLY | 0.11329226 |
| FLEQVHQL | ASSLWGINTLY | 0.13672614 |
| GLSLLAVAV | ASSLWGINTLY | 0.12044206 |
| LEQVHQL | ASSLWGINTLY | 0.13064136 |
| LNAAGAI | ASSLWGINTLY | 0.23216265 |
| PAVHASL | ASSLWGINTLY | 0.11032358 |
| TELEDDA | ASSLWGINTLY | 0.11688047 |
| TPPGPPN | ASSLWGINTLY | 0.22740313 |
| TPPPPPPP | ASSLWGINTLY | 0.11422371 |
| TQSPALI | ASRTDSGNTLY | 0.15930074 |
| EFAAAFPQ | ASRTDSGNTLY | 0.104785666 |
| LLSGNLDVL | ASRTDSGNTLY | 0.24625729 |
| GPANLGNYNI | ASRTDSGNTLY | 0.1140382 |
| TLGPTLP | ASRTDSGNTLY | 0.270279 |
| VYAYSSCCK | ASRTDSGNTLY | 0.117354855 |
| APAPAPAPNI | ASRTDSGNTLY | 0.10701224 |
| PGGPVPIP | ASRTDSGNTLY | 0.116001084 |
| FLTVALEP | ASRTDSGNTLY | 0.09826254 |
| SQYFGANL | ASRTDSGNTLY | 0.17842573 |
| GGVKPAA | ASRTDSGNTLY | 0.10083867 |
| GGTGGAGGAGGW | ASRTDSGNTLY | 0.12833741 |
| GVGGVAGLGGAGG | ASRTDSGNTLY | 0.1166464 |
| VLLVLATTI | ASRTDSGNTLY | 0.14067657 |
| IGIFAAML | ASRTDSGNTLY | 0.14723043 |
| AVGDLID | ASRTDSGNTLY | 0.1239584 |
| NISPLPL | ASRTDSGNTLY | 0.110830136 |
| VTAVFVAI | ASRTDSGNTLY | 0.1667176 |
| SGDDNDAAL | ASRTDSGNTLY | 0.1296484 |
| AVAALAHS | ASRTDSGNTLY | 0.13834147 |
| GGVTVMATP | ASRTDSGNTLY | 0.11519728 |
| AGLAANVRD | ASRTDSGNTLY | 0.12791751 |
| TNRPDLIDPALLRPG | ASRTDSGNTLY | 0.14753221 |
| LANLGVTAL | ASRTDSGNTLY | 0.099467054 |
| TGERIEL | ASRTDSGNTLY | 0.10077608 |
| NGEELAA | ASRTDSGNTLY | 0.10823305 |
| VLSLVWG | ASRTDSGNTLY | 0.11092593 |
| IGYALWL | ASRTDSGNTLY | 0.28421432 |
| LSLLLAPL | ASRTDSGNTLY | 0.30016607 |
| TLDEALK | ASRTDSGNTLY | 0.13740835 |
| TDGAGTAEGG | ASRTDSGNTLY | 0.23906937 |
| NAPGGAGGNGG | ASRTDSGNTLY | 0.15231192 |
| VSVFMGMGGNF | ASRTDSGNTLY | 0.09705036 |
| LASYLAAL | ASRTDSGNTLY | 0.118402295 |
| LVGGTPAL | ASRTDSGNTLY | 0.10737356 |

| | | |
|---|---|---|
| VRNADVL | ASRTDSGNTLY | 0.13515753 |
| TCSSPFLVSAM | ASRTDSGNTLY | 0.101985104 |
| VHITDAE | ASRTDSGNTLY | 0.12284792 |
| EYAAPSF | ASRTDSGNTLY | 0.18842463 |
| VVDAFHGGL | ASRTDSGNTLY | 0.13211907 |
| ASVVDPQ | ASRTDSGNTLY | 0.1779382 |
| SYNEFGGLS | ASRTDSGNTLY | 0.11233223 |
| DPDGSVY | ASRTDSGNTLY | 0.104630806 |
| MVSTIHATE | ASRTDSGNTLY | 0.18650593 |
| LAAGVADPV | ASRTDSGNTLY | 0.10544498 |
| TKEQIAA | ASRTDSGNTLY | 0.12859572 |
| SLASPAAL | ASRTDSGNTLY | 0.110790424 |
| IGLVFYGV | ASRTDSGNTLY | 0.15442175 |
| FLLDASNL | ASRTDSGNTLY | 0.11100357 |
| YQVGNFP | ASRTDSGNTLY | 0.12819342 |
| ESLLSQST | ASRTDSGNTLY | 0.10737355 |
| LFTVGFAAVG | ASRTDSGNTLY | 0.1629368 |
| ATPPTPT | ASRTDSGNTLY | 0.37062782 |
| TDLATLG | ASRTDSGNTLY | 0.16190225 |
| SAGGTGR | ASRTDSGNTLY | 0.17927653 |
| GFSNSGS | ASRTDSGNTLY | 0.14150698 |
| GGNGGSGGTSV | ASRTDSGNTLY | 0.11770736 |
| GTSVATGGAGN | ASRTDSGNTLY | 0.12623945 |
| NGGNGGSGGTS | ASRTDSGNTLY | 0.13217166 |
| AVVKFPH | ASRTDSGNTLY | 0.15700617 |
| TPPPPPP | ASRTDSGNTLY | 0.10213014 |
| AAPAAAAAPA | ASRTDSGNTLY | 0.10306174 |
| IALNEDL | ASRTDSGNTLY | 0.122329585 |
| IVNVSSM | ASRTDSGNTLY | 0.28949076 |
| TGIPPAPR | ASRTDSGNTLY | 0.2680289 |
| FDLGGGTF | ASRTDSGNTLY | 0.1128728 |
| FDLGGGTFDVS | ASRTDSGNTLY | 0.12949714 |
| NEPTAAA | ASRTDSGNTLY | 0.108460724 |
| NEPTAAALA | ASRTDSGNTLY | 0.1590699 |
| GSIINISSI | ASRTDSGNTLY | 0.10139014 |
| VAPGNAGTA | ASRTDSGNTLY | 0.129497 |
| TGGGASLE | ASRTDSGNTLY | 0.15892793 |
| VAVYNHYKRI | ASRTDSGNTLY | 0.5156152 |
| TDPAPATT | ASRTDSGNTLY | 0.12390453 |
| TNRPDILDPALLRPG | ASRTDSGNTLY | 0.101384275 |
| LTTTTTS | ASRTDSGNTLY | 0.1825425 |
| PTGSGKTL | ASRTDSGNTLY | 0.11906297 |
| TGSGKTL | ASRTDSGNTLY | 0.1472955 |
| TIGHVDHGK | ASRTDSGNTLY | 0.13015768 |
| TIGHVDHGKTT | ASRTDSGNTLY | 0.10140295 |
| GHVDHGKTT | ASRTDSGNTLY | 0.12472258 |
| AQFEHTLL | ASRTDSGNTLY | 0.51218027 |
| IVNQDEL | ASRTDSGNTLY | 0.1269398 |
| TNGSQFF | ASRTDSGNTLY | 0.114624724 |
| TGKGIVE | ASRTDSGNTLY | 0.32976645 |
| LISNASDA | ASRTDSGNTLY | 0.18641251 |

| | | |
|---|---|---|
| AELRTLGELTKIAKAHGAQ | ASRTDSGNTLY | 0.23090711 |
| APNAVAAAEPSW | ASRTDSGNTLY | 0.114001356 |
| APQVAGAATAVL | ASRTDSGNTLY | 0.13149504 |
| ARRDPAAAAAILPTDA | ASRTDSGNTLY | 0.14055738 |
| DAAARDVLAEAGLAEYF | ASRTDSGNTLY | 0.14942908 |
| DDQIVLSAKANLAKVEQ | ASRTDSGNTLY | 0.17096958 |
| DELLPDTGEDSDEDGH | ASRTDSGNTLY | 0.20420307 |
| ELPEAPANRATVPQD | ASRTDSGNTLY | 0.11179788 |
| FGQSKIERVTGIG | ASRTDSGNTLY | 0.27928036 |
| GAGGTGSTAGGAGGAGGAGGLY | ASRTDSGNTLY | 0.15921769 |
| GHGGTGAAGGNGGTGGAGGSGI | ASRTDSGNTLY | 0.14749706 |
| GNGGNGGAGGEGGPGGAGGAGA | ASRTDSGNTLY | 0.100302555 |
| GSTAGNGGNGGNATNGTIAGQP | ASRTDSGNTLY | 0.12853533 |
| GTGGDGGNGGNGGTGG | ASRTDSGNTLY | 0.15269358 |
| GVRRPAGIPADAPV | ASRTDSGNTLY | 0.110232495 |
| GYRQVTDAHLLTLARR | ASRTDSGNTLY | 0.10906108 |
| IAKMGQDGHDRG | ASRTDSGNTLY | 0.12471217 |
| KAEPSIEIQYT | ASRTDSGNTLY | 0.35479724 |
| KPVRIRMAADEDTVAVT | ASRTDSGNTLY | 0.097479284 |
| MPPRIAGMRLLVIKPEP | ASRTDSGNTLY | 0.13097967 |
| MKFYAVALFRSRGVPPTR | ASRTDSGNTLY | 0.10310946 |
| NAGGFGILGAVAHSP | ASRTDSGNTLY | 0.123926744 |
| NEGREITLAMAEM | ASRTDSGNTLY | 0.15876366 |
| NELVEWSHPLGHRWRWE | ASRTDSGNTLY | 0.108022 |
| NLGAGGTGGNGGNPG | ASRTDSGNTLY | 0.13399231 |
| NTEWPKWETVMAADPVD | ASRTDSGNTLY | 0.1027592 |
| PASAAPAETQP | ASRTDSGNTLY | 0.11763297 |
| QQSSDGKNGVRAAEITGE | ASRTDSGNTLY | 0.12549587 |
| RAPDAVTTSWWKE | ASRTDSGNTLY | 0.15438753 |
| SGFGNVGAHTSGWFNQ | ASRTDSGNTLY | 0.10064218 |
| SGVAGAVGQAASVGGLKV | ASRTDSGNTLY | 0.14783844 |
| SNGGAGTGSAGGNGGT | ASRTDSGNTLY | 0.10953272 |
| TGGAGGEGGAAG | ASRTDSGNTLY | 0.4361245 |
| TGGDGGAAGAGGAPGHG | ASRTDSGNTLY | 0.14872888 |
| TMPPLLAAIANAGRHGV | ASRTDSGNTLY | 0.10046411 |
| TSSTAGNSGGA | ASRTDSGNTLY | 0.16258948 |
| VPATVPVGILC | ASRTDSGNTLY | 0.11112988 |
| VQRPDSGGVVLTGRIS | ASRTDSGNTLY | 0.15499797 |
| WTPNPVPATKGVPAGLVQAT | ASRTDSGNTLY | 0.192971 |
| YALKRALAKFSDRFDVVII | ASRTDSGNTLY | 0.11066275 |
| YLIGSVIV | ASRTDSGNTLY | 0.1543017 |
| YGQLVAVID | ASRTDSGNTLY | 0.21251427 |
| VTVGGMTLPA | ASRTDSGNTLY | 0.13610835 |
| VLALPAV | ASRTDSGNTLY | 0.12575704 |
| VIPPTLNYET | ASRTDSGNTLY | 0.12636547 |
| VGMGGMHPSSG | ASRTDSGNTLY | 0.14771709 |
| VGLAGGAATA | ASRTDSGNTLY | 0.109975845 |
| VGGAGGAGGSGP | ASRTDSGNTLY | 0.27004188 |
| VFAAFMV | ASRTDSGNTLY | 0.11370566 |
| TLYDKLR | ASRTDSGNTLY | 0.24963863 |
| TLAVFTVL | ASRTDSGNTLY | 0.15208234 |

| | | |
|---|---|---|
| TGSVQPARV | ASRTDSGNTLY | 0.15181893 |
| TGGGAGSGGT | ASRTDSGNTLY | 0.111954436 |
| SHLETSL | ASRTDSGNTLY | 0.10079742 |
| SFTTGMT | ASRTDSGNTLY | 0.15165807 |
| QVEVAKRMGVRQPTVS | ASRTDSGNTLY | 0.1394345 |
| QGSGGPALAM | ASRTDSGNTLY | 0.10024743 |
| MLAPSNP | ASRTDSGNTLY | 0.1492914 |
| LVNNAAI | ASRTDSGNTLY | 0.14328523 |
| LTGEINTLA | ASRTDSGNTLY | 0.2651435 |
| LSIFGPV | ASRTDSGNTLY | 0.11980656 |
| LLALLANL | ASRTDSGNTLY | 0.12225456 |
| LISNASDALDKLR | ASRTDSGNTLY | 0.3913707 |
| LINEAVP | ASRTDSGNTLY | 0.10133072 |
| LGYAFSGL | ASRTDSGNTLY | 0.1470816 |
| LDFVAVAL | ASRTDSGNTLY | 0.17219117 |
| LAQPGAE | ASRTDSGNTLY | 0.16013609 |
| ISNLGAVTGF | ASRTDSGNTLY | 0.118205115 |
| HQEDPGRI | ASRTDSGNTLY | 0.1322159 |
| GTPGEPTP | ASRTDSGNTLY | 0.1068683 |
| GFFNSGS | ASRTDSGNTLY | 0.2378635 |
| GAGGGGLTAGAN | ASRTDSGNTLY | 0.10891309 |
| FVDALVNM | ASRTDSGNTLY | 0.13426566 |
| FLTVAPEL | ASRTDSGNTLY | 0.10930497 |
| FETLTAL | ASRTDSGNTLY | 0.19870807 |
| DVINPIEL | ASRTDSGNTLY | 0.29036152 |
| APVSGAPV | ASRTDSGNTLY | 0.2530614 |
| ALVGGLAW | ASRTDSGNTLY | 0.09976152 |
| ALAAAAATAI | ASRTDSGNTLY | 0.19020948 |
| AFGLGLLL | ASRTDSGNTLY | 0.27409893 |
| AFFAAASL | ASRTDSGNTLY | 0.30575532 |
| AAAALPAFSPP | ASRTDSGNTLY | 0.15581582 |
| VVLAPAL | ASRTDSGNTLY | 0.13448305 |
| TIGHVDHGKTTL | ASRTDSGNTLY | 0.13054343 |
| LISNASDALDK | ASRTDSGNTLY | 0.12429084 |
| LISNASDAL | ASRTDSGNTLY | 0.11415764 |
| ISNASDALDK | ASRTDSGNTLY | 0.10391783 |
| NQAETLVYQTE | ASRTDSGNTLY | 0.15167615 |
| TGGTGGTAGAG | ASRTDSGNTLY | 0.12576854 |
| TGSTAGGAGGA | ASRTDSGNTLY | 0.19679411 |
| DPLAVPVAAGPV | ASRTDSGNTLY | 0.16202421 |
| GGTGGTPGNGGN | ASRTDSGNTLY | 0.27222556 |
| PAGLAGVDPATL | ASRTDSGNTLY | 0.38180253 |
| GNNLVVEAPHAGASV | ASRTDSGNTLY | 0.36526608 |
| KVTRIMSASTLEQPA | ASRTDSGNTLY | 0.10777913 |
| LGFALPIALPRRTVV | ASRTDSGNTLY | 0.1317908 |
| TRLGNLTARKLVVLV | ASRTDSGNTLY | 0.11519969 |
| WSQQRRLIYFPSAGP | ASRTDSGNTLY | 0.1071051 |
| GPNGNPDPMAAAVDIR | ASRTDSGNTLY | 0.12019539 |
| TQRCVSVADLRAAASR | ASRTDSGNTLY | 0.1329045 |
| QFVLDVAEHPVEVLTETV | ASRTDSGNTLY | 0.19362915 |
| GGAGGTGGAGGNGGTQAGDG | ASRTDSGNTLY | 0.09700314 |

| | | |
|---|---|---|
| PLLATLRAIAVDVDAKPAQV | ASRTDSGNTLY | 0.17869982 |
| APKFRTGGYMIDFWGVGYQVAK | ASRTDSGNTLY | 0.15122372 |
| GAGGAGGAGGAAGGPGGTGGTG | ASRTDSGNTLY | 0.112328395 |
| IGALGLLATDVAPAGSVAFGVA | ASRTDSGNTLY | 0.19048478 |
| AGGAGGIGGNGGAGGAGGVLMG | ASRTDSGNTLY | 0.14602259 |
| ADELIGLL | ASRTDSGNTLY | 0.107266836 |
| ADRALKA | ASRTDSGNTLY | 0.33011043 |
| AGAVGQVGGAGGA | ASRTDSGNTLY | 0.117499396 |
| AGGGGGLGGAP | ASRTDSGNTLY | 0.12762354 |
| AGGLLYGNGGAG | ASRTDSGNTLY | 0.24073407 |
| AGLESIGL | ASRTDSGNTLY | 0.15948883 |
| AGLKTGII | ASRTDSGNTLY | 0.10818965 |
| ASLAALLL | ASRTDSGNTLY | 0.11107073 |
| ASVINISSI | ASRTDSGNTLY | 0.11181474 |
| ATPPGPA | ASRTDSGNTLY | 0.111793794 |
| ATVASALV | ASRTDSGNTLY | 0.1133247 |
| ATVASLIL | ASRTDSGNTLY | 0.16578338 |
| AVNGPVLIRH | ASRTDSGNTLY | 0.11552441 |
| DGAAGAVGI | ASRTDSGNTLY | 0.22889873 |
| DPLLHGI | ASRTDSGNTLY | 0.14744864 |
| ELGAALGAL | ASRTDSGNTLY | 0.15611196 |
| ELGRTQAV | ASRTDSGNTLY | 0.12130531 |
| FVAGGYLS | ASRTDSGNTLY | 0.1276152 |
| GASGGAGGAGGW | ASRTDSGNTLY | 0.11659682 |
| GGAGAGGDGGAGGI | ASRTDSGNTLY | 0.11785747 |
| GTGHNGSQPGMGGNG | ASRTDSGNTLY | 0.10187469 |
| GYAAASASAS | ASRTDSGNTLY | 0.12266257 |
| HQVDVAAVQQG | ASRTDSGNTLY | 0.12922871 |
| IGAAIAHT | ASRTDSGNTLY | 0.23279177 |
| IGMIEVAQL | ASRTDSGNTLY | 0.14734092 |
| IRLAPEHTSAMD | ASRTDSGNTLY | 0.10755271 |
| IYNVSVL | ASRTDSGNTLY | 0.154912 |
| KQGFLGM | ASRTDSGNTLY | 0.14676915 |
| LESVITQL | ASRTDSGNTLY | 0.14613108 |
| LGLGALVVV | ASRTDSGNTLY | 0.10006843 |
| LIGNGGNGGSGGI | ASRTDSGNTLY | 0.1250528 |
| LNAINAPTL | ASRTDSGNTLY | 0.11076601 |
| LQLLNTGV | ASRTDSGNTLY | 0.108782046 |
| LQTSIEF | ASRTDSGNTLY | 0.11026474 |
| MSFAVSEL | ASRTDSGNTLY | 0.26057887 |
| NTGALGL | ASRTDSGNTLY | 0.25343692 |
| PPGAGGAGGAGTT | ASRTDSGNTLY | 0.11689916 |
| QLVGLGI | ASRTDSGNTLY | 0.13673222 |
| RFLEQVHQL | ASRTDSGNTLY | 0.21985511 |
| RGVGYVLR | ASRTDSGNTLY | 0.30484957 |
| SATGVFAGVI | ASRTDSGNTLY | 0.1630531 |
| SLPAITLP | ASRTDSGNTLY | 0.13067028 |
| STALAFTV | ASRTDSGNTLY | 0.15476055 |
| SVLTVGLGK | ASRTDSGNTLY | 0.12269119 |
| TALAGAGSGI | ASRTDSGNTLY | 0.10474296 |
| TGLPFTGI | ASRTDSGNTLY | 0.10333755 |

| | | |
|---|---|---|
| VAEHVTV | ASRTDSGNTLY | 0.123158336 |
| VDIVLNSL | ASRTDSGNTLY | 0.11227152 |
| VLTGGGAVL | ASRTDSGNTLY | 0.1833537 |
| VNVGLAYGI | ASRTDSGNTLY | 0.12521763 |
| VSSAAVGAV | ASRTDSGNTLY | 0.11959991 |
| VTSVYEAGV | ASRTDSGNTLY | 0.098967314 |
| WVALDRGA | ASRTDSGNTLY | 0.12190645 |
| YLVGSVVI | ASRTDSGNTLY | 0.1213445 |
| GPPGTGKTLL | ASRTDSGNTLY | 0.15255056 |
| GVGMTKF | ASRTDSGNTLY | 0.13609214 |
| IGIGGSDL | ASRTDSGNTLY | 0.5984743 |
| IGTLLAEL | ASRTDSGNTLY | 0.22202541 |
| IYTSGTT | ASRTDSGNTLY | 0.1254389 |
| LSFPAGV | ASRTDSGNTLY | 0.19025393 |
| LYTSGTT | ASRTDSGNTLY | 0.11643042 |
| VGAFPFGF | ASRTDSGNTLY | 0.0979986 |
| VGAGTVEF | ASRTDSGNTLY | 0.11539491 |
| VGKSTLLL | ASRTDSGNTLY | 0.122270025 |
| AALAHAGGGIV | ASRTDSGNTLY | 0.119953156 |
| AALVAVSLL | ASRTDSGNTLY | 0.10530164 |
| AAVSRAVAAL | ASRTDSGNTLY | 0.11041934 |
| AGGSSGIGGF | ASRTDSGNTLY | 0.11830689 |
| AGLTVANANG | ASRTDSGNTLY | 0.11446798 |
| AGLVGAVL | ASRTDSGNTLY | 0.11305908 |
| ALSQAFTV | ASRTDSGNTLY | 0.12920949 |
| ALTFSVSL | ASRTDSGNTLY | 0.16401197 |
| ALTGAALVAL | ASRTDSGNTLY | 0.56108975 |
| ASFSIGAGVL | ASRTDSGNTLY | 0.13123187 |
| DGSVYKI | ASRTDSGNTLY | 0.12787944 |
| EQLAEIV | ASRTDSGNTLY | 0.2602094 |
| FAVSELTVM | ASRTDSGNTLY | 0.16209327 |
| GAAPLAAGL | ASRTDSGNTLY | 0.114116006 |
| GADGGDGGSGGAGG | ASRTDSGNTLY | 0.15694654 |
| GFGLDTGGAGG | ASRTDSGNTLY | 0.1358599 |
| GLASGGLL | ASRTDSGNTLY | 0.10163041 |
| GRIAGAAVLL | ASRTDSGNTLY | 0.1457518 |
| GSAGSLNTSF | ASRTDSGNTLY | 0.11983691 |
| GTGQIGI | ASRTDSGNTLY | 0.12438046 |
| GTLIALIL | ASRTDSGNTLY | 0.096772656 |
| IAATMGFVVG | ASRTDSGNTLY | 0.11439052 |
| IFAQALL | ASRTDSGNTLY | 0.12575029 |
| IGAAVALGF | ASRTDSGNTLY | 0.12655894 |
| IGLTPLAL | ASRTDSGNTLY | 0.099014744 |
| ILGSNTSTL | ASRTDSGNTLY | 0.10695498 |
| IVFSLTVL | ASRTDSGNTLY | 0.16666694 |
| LASKATGF | ASRTDSGNTLY | 0.31407273 |
| LASVALIL | ASRTDSGNTLY | 0.11359516 |
| LFAVGLTL | ASRTDSGNTLY | 0.12514174 |
| LFNGGIGTYI | ASRTDSGNTLY | 0.3436135 |
| LTLGAGII | ASRTDSGNTLY | 0.13818498 |
| LVAAALGDI | ASRTDSGNTLY | 0.10565571 |

| | | |
|---|---|---|
| NTAFVQLGI | ASRTDSGNTLY | 0.15025376 |
| QLFSGAEL | ASRTDSGNTLY | 0.09758621 |
| SAATFITL | ASRTDSGNTLY | 0.11977363 |
| SNYGLANLGV | ASRTDSGNTLY | 0.23708785 |
| SYGGAYAVM | ASRTDSGNTLY | 0.10625181 |
| TTGVGGLTLG | ASRTDSGNTLY | 0.13973494 |
| TTQTIYI | ASRTDSGNTLY | 0.27129424 |
| VAAGAGLGAVI | ASRTDSGNTLY | 0.119504556 |
| VAAVVTLL | ASRTDSGNTLY | 0.09999315 |
| VGFLGGLI | ASRTDSGNTLY | 0.10397913 |
| VGVIPILL | ASRTDSGNTLY | 0.110521466 |
| VIFASSLI | ASRTDSGNTLY | 0.18968344 |
| VKGGDGGAAAG | ASRTDSGNTLY | 0.13242005 |
| VTKTFTNI | ASRTDSGNTLY | 0.14434431 |
| VVAAAILL | ASRTDSGNTLY | 0.1400432 |
| YAALDVEL | ASRTDSGNTLY | 0.14102422 |
| LASVALLL | ASRTDSGNTLY | 0.14399219 |
| LDAVHRV | ASRTDSGNTLY | 0.15089358 |
| QGLSSSIF | ASRTDSGNTLY | 0.12293233 |
| RGTFANIRL | ASRTDSGNTLY | 0.179013 |
| RQLWWGHRI | ASRTDSGNTLY | 0.11404333 |
| YGPPGTGKTLL | ASRTDSGNTLY | 0.10232963 |
| AALVGAPANVA | ASRTDSGNTLY | 0.46335638 |
| AATFRRTE | ASRTDSGNTLY | 0.11809563 |
| AGALGGGI | ASRTDSGNTLY | 0.1227306 |
| AGGAGGAGGIGAG | ASRTDSGNTLY | 0.13767534 |
| AGSGALGL | ASRTDSGNTLY | 0.10840105 |
| AQIADQI | ASRTDSGNTLY | 0.14677763 |
| ASAVAVTYL | ASRTDSGNTLY | 0.15273812 |
| ASTVLGGTGG | ASRTDSGNTLY | 0.22782093 |
| AVGGEPAK | ASRTDSGNTLY | 0.120092206 |
| CEQGTFGGTC | ASRTDSGNTLY | 0.12293948 |
| DEGKAPAG | ASRTDSGNTLY | 0.16314399 |
| DTVVFPAL | ASRTDSGNTLY | 0.10184896 |
| FEGITMAG | ASRTDSGNTLY | 0.09846847 |
| FPLVVQD | ASRTDSGNTLY | 0.10278312 |
| GAFTTPQ | ASRTDSGNTLY | 0.111959964 |
| GFGGFGTAGGVGGA | ASRTDSGNTLY | 0.1329521 |
| GFLAQFL | ASRTDSGNTLY | 0.11444721 |
| GGAGGAGGNAGVA | ASRTDSGNTLY | 0.15616992 |
| GGVGTAGAGGAGGAGG | ASRTDSGNTLY | 0.19453992 |
| GNGGGGDAGNAGSGGN | ASRTDSGNTLY | 0.1448594 |
| GTGGAGGAGNDAGS | ASRTDSGNTLY | 0.11061401 |
| GTTNTGWFNS | ASRTDSGNTLY | 0.121642604 |
| IFMGPLFTL | ASRTDSGNTLY | 0.15686047 |
| ILAHAIEAI | ASRTDSGNTLY | 0.1386347 |
| IRSAIREAL | ASRTDSGNTLY | 0.10598097 |
| ITFAGGG | ASRTDSGNTLY | 0.289141 |
| ITGMSGHQEV | ASRTDSGNTLY | 0.15026148 |
| IVAGLACTL | ASRTDSGNTLY | 0.21963963 |
| IVGGGLAGA | ASRTDSGNTLY | 0.19503127 |

| | | |
|---|---|---|
| IVTGVLNI | ASRTDSGNTLY | 0.24000846 |
| LAQQGCI | ASRTDSGNTLY | 0.12859094 |
| LCLHVEN | ASRTDSGNTLY | 0.14587532 |
| LDAISDGS | ASRTDSGNTLY | 0.100871935 |
| LGGFAAML | ASRTDSGNTLY | 0.11055809 |
| LGGKNPMI | ASRTDSGNTLY | 0.10313947 |
| LGPKGVAAIL | ASRTDSGNTLY | 0.15247007 |
| LMVGVSI | ASRTDSGNTLY | 0.09811302 |
| LVGAAFVL | ASRTDSGNTLY | 0.2215781 |
| LVLAVAPL | ASRTDSGNTLY | 0.11822259 |
| MAAIAVIDAA | ASRTDSGNTLY | 0.18891285 |
| NNGINAELQ | ASRTDSGNTLY | 0.1388604 |
| RGEPLVLVHG | ASRTDSGNTLY | 0.10066494 |
| RGGAGGAAGLI | ASRTDSGNTLY | 0.12785578 |
| RSYRMHPAV | ASRTDSGNTLY | 0.19254902 |
| SAAHLLAGQLA | ASRTDSGNTLY | 0.10624926 |
| SAAVGSY | ASRTDSGNTLY | 0.1017789 |
| SPHAVLHRI | ASRTDSGNTLY | 0.109447055 |
| SQIAKVL | ASRTDSGNTLY | 0.10406804 |
| TALGHLVP | ASRTDSGNTLY | 0.33345193 |
| TAPTTSGAA | ASRTDSGNTLY | 0.23888567 |
| TDGDDGNGGSPGT | ASRTDSGNTLY | 0.16083658 |
| TDGLGEN | ASRTDSGNTLY | 0.14424568 |
| TGGPPLPGQA | ASRTDSGNTLY | 0.09728087 |
| TIPAITI | ASRTDSGNTLY | 0.13105088 |
| TIPGFTI | ASRTDSGNTLY | 0.45382315 |
| TSIASGALA | ASRTDSGNTLY | 0.14290544 |
| TVTLPGAAI | ASRTDSGNTLY | 0.16839705 |
| VASGLSAALP | ASRTDSGNTLY | 0.15515922 |
| VGAGVIGI | ASRTDSGNTLY | 0.12854537 |
| VGPKRFGV | ASRTDSGNTLY | 0.1449304 |
| VNVAFPD | ASRTDSGNTLY | 0.11942575 |
| VSMTNSGVS | ASRTDSGNTLY | 0.13148966 |
| VTDVLGDLSV | ASRTDSGNTLY | 0.10884552 |
| VVVVGAGFAGL | ASRTDSGNTLY | 0.10698597 |
| VYTVGTVV | ASRTDSGNTLY | 0.12891284 |
| WTTFSGP | ASRTDSGNTLY | 0.5819507 |
| YDELVEAF | ASRTDSGNTLY | 0.16489711 |
| YGTGGAGGNGG | ASRTDSGNTLY | 0.20263232 |
| YTNVMAV | ASRTDSGNTLY | 0.123039044 |
| FLEQVHQL | ASRTDSGNTLY | 0.21585439 |
| GLSLLAVAV | ASRTDSGNTLY | 0.13435194 |
| LEQVHQL | ASRTDSGNTLY | 0.11670874 |
| LNAAGAI | ASRTDSGNTLY | 0.12637874 |
| PAVHASL | ASRTDSGNTLY | 0.11306223 |
| TELEDDA | ASRTDSGNTLY | 0.1146771 |
| TPPGPPN | ASRTDSGNTLY | 0.13432825 |
| TPPPPPPP | ASRTDSGNTLY | 0.6561562 |
| TQSPALI | ASSQEDRGSNERLF | 0.21228552 |
| EFAAAFPQ | ASSQEDRGSNERLF | 0.12647417 |
| LLSGNLDVL | ASSQEDRGSNERLF | 0.11659067 |

| | | |
|---|---|---|
| GPANLGNYNI | ASSQEDRGSNERLF | 0.11721337 |
| TLGPTLP | ASSQEDRGSNERLF | 0.15647334 |
| VYAYSSCCK | ASSQEDRGSNERLF | 0.11040579 |
| APAPAPAPNI | ASSQEDRGSNERLF | 0.39745706 |
| PGGPVPIP | ASSQEDRGSNERLF | 0.12287469 |
| FLTVALEP | ASSQEDRGSNERLF | 0.13684809 |
| SQYFGANL | ASSQEDRGSNERLF | 0.14567025 |
| GGVKPAA | ASSQEDRGSNERLF | 0.4412742 |
| GGTGGAGGAGGW | ASSQEDRGSNERLF | 0.1337173 |
| GVGGVAGLGGAGG | ASSQEDRGSNERLF | 0.13322362 |
| VLLVLATTI | ASSQEDRGSNERLF | 0.118720934 |
| IGIFAAML | ASSQEDRGSNERLF | 0.25098294 |
| AVGDLID | ASSQEDRGSNERLF | 0.42188346 |
| NISPLPL | ASSQEDRGSNERLF | 0.10467991 |
| VTAVFVAI | ASSQEDRGSNERLF | 0.34228593 |
| SGDDNDAAL | ASSQEDRGSNERLF | 0.113834515 |
| AVAALAHS | ASSQEDRGSNERLF | 0.104359575 |
| GGVTVMATP | ASSQEDRGSNERLF | 0.12696171 |
| AGLAANVRD | ASSQEDRGSNERLF | 0.116539575 |
| TNRPDLIDPALLRPG | ASSQEDRGSNERLF | 0.37025666 |
| LANLGVTAL | ASSQEDRGSNERLF | 0.15618502 |
| TGERIEL | ASSQEDRGSNERLF | 0.096777365 |
| NGEELAA | ASSQEDRGSNERLF | 0.12322897 |
| VLSLVWG | ASSQEDRGSNERLF | 0.31225348 |
| IGYALWL | ASSQEDRGSNERLF | 0.2315759 |
| LSLLLAPL | ASSQEDRGSNERLF | 0.15728492 |
| TLDEALK | ASSQEDRGSNERLF | 0.13337065 |
| TDGAGTAEGG | ASSQEDRGSNERLF | 0.13343033 |
| NAPGGAGGNGG | ASSQEDRGSNERLF | 0.13616201 |
| VSVFMGMGGNF | ASSQEDRGSNERLF | 0.12963408 |
| LASYLAAL | ASSQEDRGSNERLF | 0.10266509 |
| LVGGTPAL | ASSQEDRGSNERLF | 0.17007504 |
| VRNADVL | ASSQEDRGSNERLF | 0.22011831 |
| TCSSPFLVSAM | ASSQEDRGSNERLF | 0.09943917 |
| VHITDAE | ASSQEDRGSNERLF | 0.11936653 |
| EYAAPSF | ASSQEDRGSNERLF | 0.11365916 |
| VVDAFHGGL | ASSQEDRGSNERLF | 0.104736544 |
| ASVVDPQ | ASSQEDRGSNERLF | 0.101783596 |
| SYNEFGGLS | ASSQEDRGSNERLF | 0.37208134 |
| DPDGSVY | ASSQEDRGSNERLF | 0.65667886 |
| MVSTIHATE | ASSQEDRGSNERLF | 0.12702818 |
| LAAGVADPV | ASSQEDRGSNERLF | 0.11177721 |
| TKEQIAA | ASSQEDRGSNERLF | 0.099577144 |
| SLASPAAL | ASSQEDRGSNERLF | 0.21560906 |
| IGLVFYGV | ASSQEDRGSNERLF | 0.25493377 |
| FLLDASNL | ASSQEDRGSNERLF | 0.14239764 |
| YQVGNFP | ASSQEDRGSNERLF | 0.10993807 |
| ESLLSQST | ASSQEDRGSNERLF | 0.10291035 |
| LFTVGFAAVG | ASSQEDRGSNERLF | 0.24343225 |
| ATPPTPT | ASSQEDRGSNERLF | 0.10219459 |
| TDLATLG | ASSQEDRGSNERLF | 0.1130129 |

| | | |
|---|---|---|
| SAGGTGR | ASSQEDRGSNERLF | 0.098946944 |
| GFSNSGS | ASSQEDRGSNERLF | 0.17242657 |
| GGNGGSGGTSV | ASSQEDRGSNERLF | 0.10649299 |
| GTSVATGGAGN | ASSQEDRGSNERLF | 0.13803019 |
| NGGNGGSGGTS | ASSQEDRGSNERLF | 0.23731014 |
| AVVKFPH | ASSQEDRGSNERLF | 0.16711763 |
| TPPPPPP | ASSQEDRGSNERLF | 0.16124555 |
| AAPAAAAAPA | ASSQEDRGSNERLF | 0.1217415 |
| IALNEDL | ASSQEDRGSNERLF | 0.19204918 |
| IVNVSSM | ASSQEDRGSNERLF | 0.116840646 |
| TGIPPAPR | ASSQEDRGSNERLF | 0.16478063 |
| FDLGGGTF | ASSQEDRGSNERLF | 0.13713707 |
| FDLGGGTFDVS | ASSQEDRGSNERLF | 0.15486632 |
| NEPTAAA | ASSQEDRGSNERLF | 0.14696382 |
| NEPTAAALA | ASSQEDRGSNERLF | 0.11516524 |
| GSIINISSI | ASSQEDRGSNERLF | 0.11088058 |
| VAPGNAGTA | ASSQEDRGSNERLF | 0.10228854 |
| TGGGASLE | ASSQEDRGSNERLF | 0.15672585 |
| VAVYNHYKRI | ASSQEDRGSNERLF | 0.1525507 |
| TDPAPATT | ASSQEDRGSNERLF | 0.14345549 |
| TNRPDILDPALLRPG | ASSQEDRGSNERLF | 0.1531961 |
| LTTTTTS | ASSQEDRGSNERLF | 0.11420618 |
| PTGSGKTL | ASSQEDRGSNERLF | 0.14465506 |
| TGSGKTL | ASSQEDRGSNERLF | 0.10364232 |
| TIGHVDHGK | ASSQEDRGSNERLF | 0.12417838 |
| TIGHVDHGKTT | ASSQEDRGSNERLF | 0.19843641 |
| GHVDHGKTT | ASSQEDRGSNERLF | 0.11335131 |
| AQFEHTLL | ASSQEDRGSNERLF | 0.12224589 |
| IVNQDEL | ASSQEDRGSNERLF | 0.11577183 |
| TNGSQFF | ASSQEDRGSNERLF | 0.18768051 |
| TGKGIVE | ASSQEDRGSNERLF | 0.113653176 |
| LISNASDA | ASSQEDRGSNERLF | 0.10516419 |
| AELRTLGELTKIAKAHGAQ | ASSQEDRGSNERLF | 0.31096327 |
| APNAVAAAEPSW | ASSQEDRGSNERLF | 0.1111621 |
| APQVAGAATAVL | ASSQEDRGSNERLF | 0.10670707 |
| ARRDPAAAAAILPTDA | ASSQEDRGSNERLF | 0.1880305 |
| DAAARDVLAEAGLAEYF | ASSQEDRGSNERLF | 0.11304554 |
| DDQIVLSAKANLAKVEQ | ASSQEDRGSNERLF | 0.19843975 |
| DELLPDTGEDSDEDGH | ASSQEDRGSNERLF | 0.12866026 |
| ELPEAPANRATVPQD | ASSQEDRGSNERLF | 0.15051821 |
| FGQSKIERVTGIG | ASSQEDRGSNERLF | 0.18142612 |
| GAGGTGSTAGGAGGAGGAGGLY | ASSQEDRGSNERLF | 0.1006979 |
| GHGGTGAAGGNGGTGGAGGSGI | ASSQEDRGSNERLF | 0.11871566 |
| GNGGNGGAGGEGGPGGAGGAGA | ASSQEDRGSNERLF | 0.13514325 |
| GSTAGNGGNGGNATNGTIAGQP | ASSQEDRGSNERLF | 0.16703154 |
| GTGGDGGNGGNGGTGG | ASSQEDRGSNERLF | 0.15263255 |
| GVRRPAGIPADAPV | ASSQEDRGSNERLF | 0.11741312 |
| GYRQVTDAHLLTLARR | ASSQEDRGSNERLF | 0.18782331 |
| IAKMGQDGHDRG | ASSQEDRGSNERLF | 0.13118075 |
| KAEPSIEIQYT | ASSQEDRGSNERLF | 0.12664497 |
| KPVRIRMAADEDTVAVT | ASSQEDRGSNERLF | 0.11096466 |

| | | |
|---|---|---|
| MPPRIAGMRLLVIKPEP | ASSQEDRGSNERLF | 0.1047752 |
| MKFYAVALFRSRGVPPTR | ASSQEDRGSNERLF | 0.10677251 |
| NAGGFGILGAVAHSP | ASSQEDRGSNERLF | 0.14716676 |
| NEGREITLAMAEM | ASSQEDRGSNERLF | 0.12367825 |
| NELVEWSHPLGHRWRWE | ASSQEDRGSNERLF | 0.18043913 |
| NLGAGGTGGNGGNPG | ASSQEDRGSNERLF | 0.101910084 |
| NTEWPKWETVMAADPVD | ASSQEDRGSNERLF | 0.10766289 |
| PASAAPAETQP | ASSQEDRGSNERLF | 0.14518127 |
| QQSSDGKNGVRAAEITGE | ASSQEDRGSNERLF | 0.11392522 |
| RAPDAVTTSWWKE | ASSQEDRGSNERLF | 0.12746438 |
| SGFGNVGAHTSGWFNQ | ASSQEDRGSNERLF | 0.15008627 |
| SGVAGAVGQAASVGGLKV | ASSQEDRGSNERLF | 0.11215706 |
| SNGGAGTGSAGGNGGT | ASSQEDRGSNERLF | 0.13206366 |
| TGGAGGEGGAAG | ASSQEDRGSNERLF | 0.2434111 |
| TGGDGGAAGAGGAPGHG | ASSQEDRGSNERLF | 0.11491331 |
| TMPPLLAAIANAGRHGV | ASSQEDRGSNERLF | 0.109144256 |
| TSSTAGNSGGA | ASSQEDRGSNERLF | 0.10124793 |
| VPATVPVGILC | ASSQEDRGSNERLF | 0.16837172 |
| VQRPDSGGVVLTGRIS | ASSQEDRGSNERLF | 0.16135177 |
| WTPNPVPATKGVPAGLVQAT | ASSQEDRGSNERLF | 0.10191107 |
| YALKRALAKFSDRFDVVII | ASSQEDRGSNERLF | 0.1436453 |
| YLIGSVIV | ASSQEDRGSNERLF | 0.11437036 |
| YGQLVAVID | ASSQEDRGSNERLF | 0.10242339 |
| VTVGGMTLPA | ASSQEDRGSNERLF | 0.11094845 |
| VLALPAV | ASSQEDRGSNERLF | 0.23257166 |
| VIPPTLNYET | ASSQEDRGSNERLF | 0.27830285 |
| VGMGGMHPSSG | ASSQEDRGSNERLF | 0.1410179 |
| VGLAGGAATA | ASSQEDRGSNERLF | 0.16881841 |
| VGGAGGAGGSGP | ASSQEDRGSNERLF | 0.10101602 |
| VFAAFMV | ASSQEDRGSNERLF | 0.20843448 |
| TLYDKLR | ASSQEDRGSNERLF | 0.22822231 |
| TLAVFTVL | ASSQEDRGSNERLF | 0.12399849 |
| TGSVQPARV | ASSQEDRGSNERLF | 0.16289094 |
| TGGGAGSGGT | ASSQEDRGSNERLF | 0.1087196 |
| SHLETSL | ASSQEDRGSNERLF | 0.10920383 |
| SFTTGMT | ASSQEDRGSNERLF | 0.15540184 |
| QVEVAKRMGVRQPTVS | ASSQEDRGSNERLF | 0.15514868 |
| QGSGGPALAM | ASSQEDRGSNERLF | 0.12680174 |
| MLAPSNP | ASSQEDRGSNERLF | 0.118579224 |
| LVNNAAI | ASSQEDRGSNERLF | 0.12744446 |
| LTGEINTLA | ASSQEDRGSNERLF | 0.13390195 |
| LSIFGPV | ASSQEDRGSNERLF | 0.12937875 |
| LLALLANL | ASSQEDRGSNERLF | 0.12417423 |
| LISNASDALDKLR | ASSQEDRGSNERLF | 0.23534602 |
| LINEAVP | ASSQEDRGSNERLF | 0.11553228 |
| LGYAFSGL | ASSQEDRGSNERLF | 0.23312156 |
| LDFVAVAL | ASSQEDRGSNERLF | 0.14529645 |
| LAQPGAE | ASSQEDRGSNERLF | 0.2121481 |
| ISNLGAVTGF | ASSQEDRGSNERLF | 0.1279767 |
| HQEDPGRI | ASSQEDRGSNERLF | 0.36684835 |
| GTPGEPTP | ASSQEDRGSNERLF | 0.13388054 |

| | | |
|---|---|---|
| GFFNSGS | ASSQEDRGSNERLF | 0.12187126 |
| GAGGGGLTAGAN | ASSQEDRGSNERLF | 0.12562169 |
| FVDALVNM | ASSQEDRGSNERLF | 0.11728665 |
| FLTVAPEL | ASSQEDRGSNERLF | 0.15597916 |
| FETLTAL | ASSQEDRGSNERLF | 0.28006968 |
| DVINPIEL | ASSQEDRGSNERLF | 0.40018165 |
| APVSGAPV | ASSQEDRGSNERLF | 0.12923779 |
| ALVGGLAW | ASSQEDRGSNERLF | 0.10376735 |
| ALAAAAATAI | ASSQEDRGSNERLF | 0.21955287 |
| AFGLGLLL | ASSQEDRGSNERLF | 0.10438775 |
| AFFAAASL | ASSQEDRGSNERLF | 0.1365807 |
| AAAALPAFSPP | ASSQEDRGSNERLF | 0.2339895 |
| VVLAPAL | ASSQEDRGSNERLF | 0.13390104 |
| TIGHVDHGKTTL | ASSQEDRGSNERLF | 0.33484465 |
| LISNASDALDK | ASSQEDRGSNERLF | 0.27947128 |
| LISNASDAL | ASSQEDRGSNERLF | 0.10853026 |
| ISNASDALDK | ASSQEDRGSNERLF | 0.17354201 |
| NQAETLVYQTE | ASSQEDRGSNERLF | 0.2260636 |
| TGGTGGTAGAG | ASSQEDRGSNERLF | 0.15415618 |
| TGSTAGGAGGA | ASSQEDRGSNERLF | 0.09669484 |
| DPLAVPVAAGPV | ASSQEDRGSNERLF | 0.19660325 |
| GGTGGTPGNGGN | ASSQEDRGSNERLF | 0.13343562 |
| PAGLAGVDPATL | ASSQEDRGSNERLF | 0.08946897 |
| GNNLVVEAPHAGASV | ASSQEDRGSNERLF | 0.30745763 |
| KVTRIMSASTLEQPA | ASSQEDRGSNERLF | 0.1338935 |
| LGFALPIALPRRTVV | ASSQEDRGSNERLF | 0.114159 |
| TRLGNLTARKLVVLV | ASSQEDRGSNERLF | 0.41947138 |
| WSQQRRLIYFPSAGP | ASSQEDRGSNERLF | 0.09220328 |
| GPNGNPDPMAAAVDIR | ASSQEDRGSNERLF | 0.11979327 |
| TQRCVSVADLRAAASR | ASSQEDRGSNERLF | 0.25706303 |
| QFVLDVAEHPVEVLTETV | ASSQEDRGSNERLF | 0.33604604 |
| GGAGGTGGAGGNGGTQAGDG | ASSQEDRGSNERLF | 0.11444256 |
| PLLATLRAIAVDVDAKPAQV | ASSQEDRGSNERLF | 0.23773228 |
| APKFRTGGYMIDFWGVGYQVAK | ASSQEDRGSNERLF | 0.5991799 |
| GAGGAGGAGGAAGGPGGTGGTG | ASSQEDRGSNERLF | 0.16617882 |
| IGALGLLATDVAPAGSVAFGVA | ASSQEDRGSNERLF | 0.59198487 |
| AGGAGGIGGNGGAGGAGGVLMG | ASSQEDRGSNERLF | 0.63263357 |
| ADELIGLL | ASSQEDRGSNERLF | 0.14258711 |
| ADRALKA | ASSQEDRGSNERLF | 0.24290287 |
| AGAVGQVGGAGGA | ASSQEDRGSNERLF | 0.3355502 |
| AGGGGGLGGAP | ASSQEDRGSNERLF | 0.15593435 |
| AGGLLYGNGGAG | ASSQEDRGSNERLF | 0.17696318 |
| AGLESIGL | ASSQEDRGSNERLF | 0.13881572 |
| AGLKTGII | ASSQEDRGSNERLF | 0.22853985 |
| ASLAALLL | ASSQEDRGSNERLF | 0.092127495 |
| ASVINISSI | ASSQEDRGSNERLF | 0.20980766 |
| ATPPGPA | ASSQEDRGSNERLF | 0.2548178 |
| ATVASALV | ASSQEDRGSNERLF | 0.18789916 |
| ATVASLIL | ASSQEDRGSNERLF | 0.10548373 |
| AVNGPVLIRH | ASSQEDRGSNERLF | 0.11443512 |
| DGAAGAVGI | ASSQEDRGSNERLF | 0.105397105 |

| | | |
|---|---|---|
| DPLLHGI | ASSQEDRGSNERLF | 0.14875914 |
| ELGAALGAL | ASSQEDRGSNERLF | 0.10169339 |
| ELGRTQAV | ASSQEDRGSNERLF | 0.100698926 |
| FVAGGYLS | ASSQEDRGSNERLF | 0.23533458 |
| GASGGAGGAGGW | ASSQEDRGSNERLF | 0.26014408 |
| GGAGAGGDGGAGGI | ASSQEDRGSNERLF | 0.14432317 |
| GTGHNGSQPGMGGNG | ASSQEDRGSNERLF | 0.28103268 |
| GYAAASASAS | ASSQEDRGSNERLF | 0.51206714 |
| HQVDVAAVQQG | ASSQEDRGSNERLF | 0.56018996 |
| IGAAIAHT | ASSQEDRGSNERLF | 0.24930936 |
| IGMIEVAQL | ASSQEDRGSNERLF | 0.10401623 |
| IRLAPEHTSAMD | ASSQEDRGSNERLF | 0.54895556 |
| IYNVSVL | ASSQEDRGSNERLF | 0.17722988 |
| KQGFLGM | ASSQEDRGSNERLF | 0.13423721 |
| LESVITQL | ASSQEDRGSNERLF | 0.3519627 |
| LGLGALVVV | ASSQEDRGSNERLF | 0.12280043 |
| LIGNGGNGGSGGI | ASSQEDRGSNERLF | 0.11205693 |
| LNAINAPTL | ASSQEDRGSNERLF | 0.4970982 |
| LQLLNTGV | ASSQEDRGSNERLF | 0.4746948 |
| LQTSIEF | ASSQEDRGSNERLF | 0.12054376 |
| MSFAVSEL | ASSQEDRGSNERLF | 0.101838656 |
| NTGALGL | ASSQEDRGSNERLF | 0.12644374 |
| PPGAGGAGGAGTT | ASSQEDRGSNERLF | 0.22164261 |
| QLVGLGI | ASSQEDRGSNERLF | 0.2985038 |
| RFLEQVHQL | ASSQEDRGSNERLF | 0.18926236 |
| RGVGYVLR | ASSQEDRGSNERLF | 0.17104715 |
| SATGVFAGVI | ASSQEDRGSNERLF | 0.4281837 |
| SLPAITLP | ASSQEDRGSNERLF | 0.19248581 |
| STALAFTV | ASSQEDRGSNERLF | 0.16823764 |
| SVLTVGLGK | ASSQEDRGSNERLF | 0.28634784 |
| TALAGAGSGI | ASSQEDRGSNERLF | 0.1024578 |
| TGLPFTGI | ASSQEDRGSNERLF | 0.41033936 |
| VAEHVTV | ASSQEDRGSNERLF | 0.45244816 |
| VDIVLNSL | ASSQEDRGSNERLF | 0.41679788 |
| VLTGGGAVL | ASSQEDRGSNERLF | 0.51970196 |
| VNVGLAYGI | ASSQEDRGSNERLF | 0.11438985 |
| VSSAAVGAV | ASSQEDRGSNERLF | 0.32060683 |
| VTSVYEAGV | ASSQEDRGSNERLF | 0.12985675 |
| WVALDRGA | ASSQEDRGSNERLF | 0.21873175 |
| YLVGSVVI | ASSQEDRGSNERLF | 0.27261215 |
| GPPGTGKTLL | ASSQEDRGSNERLF | 0.29406887 |
| GVGMTKF | ASSQEDRGSNERLF | 0.3815329 |
| IGIGGSDL | ASSQEDRGSNERLF | 0.34800935 |
| IGTLLAEL | ASSQEDRGSNERLF | 0.15410236 |
| IYTSGTT | ASSQEDRGSNERLF | 0.5797814 |
| LSFPAGV | ASSQEDRGSNERLF | 0.5227832 |
| LYTSGTT | ASSQEDRGSNERLF | 0.11610989 |
| VGAFPFGF | ASSQEDRGSNERLF | 0.11847962 |
| VGAGTVEF | ASSQEDRGSNERLF | 0.14656012 |
| VGKSTLLL | ASSQEDRGSNERLF | 0.43015766 |
| AALAHAGGGIV | ASSQEDRGSNERLF | 0.23689763 |

| | | |
|---|---|---|
| AALVAVSLL | ASSQEDRGSNERLF | 0.36772943 |
| AAVSRAVAAL | ASSQEDRGSNERLF | 0.21552116 |
| AGGSSGIGGF | ASSQEDRGSNERLF | 0.10531252 |
| AGLTVANANG | ASSQEDRGSNERLF | 0.5586933 |
| AGLVGAVL | ASSQEDRGSNERLF | 0.11411482 |
| ALSQAFTV | ASSQEDRGSNERLF | 0.32094985 |
| ALTFSVSL | ASSQEDRGSNERLF | 0.12546958 |
| ALTGAALVAL | ASSQEDRGSNERLF | 0.09086305 |
| ASFSIGAGVL | ASSQEDRGSNERLF | 0.24272051 |
| DGSVYKI | ASSQEDRGSNERLF | 0.15760295 |
| EQLAEIV | ASSQEDRGSNERLF | 0.40088382 |
| FAVSELTVM | ASSQEDRGSNERLF | 0.16411915 |
| GAAPLAAGL | ASSQEDRGSNERLF | 0.23341133 |
| GADGGDGGSGGAGG | ASSQEDRGSNERLF | 0.27504268 |
| GFGLDTGGAGG | ASSQEDRGSNERLF | 0.16654587 |
| GLASGGLL | ASSQEDRGSNERLF | 0.11667961 |
| GRIAGAAVLL | ASSQEDRGSNERLF | 0.194398 |
| GSAGSLNTSF | ASSQEDRGSNERLF | 0.5396308 |
| GTGQIGI | ASSQEDRGSNERLF | 0.46857464 |
| GTLIALIL | ASSQEDRGSNERLF | 0.105163574 |
| IAATMGFVVG | ASSQEDRGSNERLF | 0.15616077 |
| IFAQALL | ASSQEDRGSNERLF | 0.14038162 |
| IGAAVALGF | ASSQEDRGSNERLF | 0.08909453 |
| IGLTPLAL | ASSQEDRGSNERLF | 0.0939151 |
| ILGSNTSTL | ASSQEDRGSNERLF | 0.091783874 |
| IVFSLTVL | ASSQEDRGSNERLF | 0.10316699 |
| LASKATGF | ASSQEDRGSNERLF | 0.26177114 |
| LASVALIL | ASSQEDRGSNERLF | 0.38026154 |
| LFAVGLTL | ASSQEDRGSNERLF | 0.16950211 |
| LFNGGIGTYI | ASSQEDRGSNERLF | 0.4259871 |
| LTLGAGII | ASSQEDRGSNERLF | 0.13393909 |
| LVAAALGDI | ASSQEDRGSNERLF | 0.0864519 |
| NTAFVQLGI | ASSQEDRGSNERLF | 0.12878937 |
| QLFSGAEL | ASSQEDRGSNERLF | 0.11654036 |
| SAATFITL | ASSQEDRGSNERLF | 0.56733245 |
| SNYGLANLGV | ASSQEDRGSNERLF | 0.09909685 |
| SYGGAYAVM | ASSQEDRGSNERLF | 0.18719423 |
| TTGVGGLTLG | ASSQEDRGSNERLF | 0.17505088 |
| TTQTIYI | ASSQEDRGSNERLF | 0.20824403 |
| VAAGAGLGAVI | ASSQEDRGSNERLF | 0.3676844 |
| VAAVVTLL | ASSQEDRGSNERLF | 0.12552652 |
| VGFLGGLI | ASSQEDRGSNERLF | 0.103207946 |
| VGVIPILL | ASSQEDRGSNERLF | 0.19057871 |
| VIFASSLI | ASSQEDRGSNERLF | 0.09674285 |
| VKGGDGGAAAG | ASSQEDRGSNERLF | 0.124845825 |
| VTKTFTNI | ASSQEDRGSNERLF | 0.11525928 |
| VVAAAILL | ASSQEDRGSNERLF | 0.09644148 |
| YAALDVEL | ASSQEDRGSNERLF | 0.10348836 |
| LASVALLL | ASSQEDRGSNERLF | 0.577299 |
| LDAVHRV | ASSQEDRGSNERLF | 0.21720292 |
| QGLSSSIF | ASSQEDRGSNERLF | 0.17640403 |

| | | |
|---|---|---|
| RGTFANIRL | ASSQEDRGSNERLF | 0.22371864 |
| RQLWWGHRI | ASSQEDRGSNERLF | 0.19435076 |
| YGPPGTGKTLL | ASSQEDRGSNERLF | 0.19496457 |
| AALVGAPANVA | ASSQEDRGSNERLF | 0.12312893 |
| AATFRRTE | ASSQEDRGSNERLF | 0.12649553 |
| AGALGGGI | ASSQEDRGSNERLF | 0.11202338 |
| AGGAGGAGGIGAG | ASSQEDRGSNERLF | 0.11639082 |
| AGSGALGL | ASSQEDRGSNERLF | 0.49860156 |
| AQIADQI | ASSQEDRGSNERLF | 0.33262485 |
| ASAVAVTYL | ASSQEDRGSNERLF | 0.10751914 |
| ASTVLGGTGG | ASSQEDRGSNERLF | 0.15576549 |
| AVGGEPAK | ASSQEDRGSNERLF | 0.53494275 |
| CEQGTFGGTC | ASSQEDRGSNERLF | 0.52800936 |
| DEGKAPAG | ASSQEDRGSNERLF | 0.11911569 |
| DTVVFPAL | ASSQEDRGSNERLF | 0.19506404 |
| FEGITMAG | ASSQEDRGSNERLF | 0.28983676 |
| FPLVVQD | ASSQEDRGSNERLF | 0.13992764 |
| GAFTTPQ | ASSQEDRGSNERLF | 0.16590002 |
| GFGGFGTAGGVGGA | ASSQEDRGSNERLF | 0.1128794 |
| GFLAQFL | ASSQEDRGSNERLF | 0.4619583 |
| GGAGGAGGNAGVA | ASSQEDRGSNERLF | 0.73066974 |
| GGVGTAGAGGAGGAGG | ASSQEDRGSNERLF | 0.12103624 |
| GNGGGGDAGNAGSGGN | ASSQEDRGSNERLF | 0.10827553 |
| GTGGAGGAGNDAGS | ASSQEDRGSNERLF | 0.17895547 |
| GTTNTGWFNS | ASSQEDRGSNERLF | 0.11558367 |
| IFMGPLFTL | ASSQEDRGSNERLF | 0.13350429 |
| ILAHAIEAI | ASSQEDRGSNERLF | 0.15057887 |
| IRSAIREAL | ASSQEDRGSNERLF | 0.10211569 |
| ITFAGGG | ASSQEDRGSNERLF | 0.16697043 |
| ITGMSGHQEV | ASSQEDRGSNERLF | 0.3505674 |
| IVAGLACTL | ASSQEDRGSNERLF | 0.13390373 |
| IVGGGLAGA | ASSQEDRGSNERLF | 0.57070553 |
| IVTGVLNI | ASSQEDRGSNERLF | 0.43919677 |
| LAQQGCI | ASSQEDRGSNERLF | 0.3037064 |
| LCLHVEN | ASSQEDRGSNERLF | 0.5566461 |
| LDAISDGS | ASSQEDRGSNERLF | 0.5437372 |
| LGGFAAML | ASSQEDRGSNERLF | 0.14943811 |
| LGGKNPMI | ASSQEDRGSNERLF | 0.18203622 |
| LGPKGVAAIL | ASSQEDRGSNERLF | 0.25322032 |
| LMVGVSI | ASSQEDRGSNERLF | 0.15952824 |
| LVGAAFVL | ASSQEDRGSNERLF | 0.26356378 |
| LVLAVAPL | ASSQEDRGSNERLF | 0.2998612 |
| MAAIAVIDAA | ASSQEDRGSNERLF | 0.28486723 |
| NNGINAELQ | ASSQEDRGSNERLF | 0.2025496 |
| RGEPLVLVHG | ASSQEDRGSNERLF | 0.55779815 |
| RGGAGGAAGLI | ASSQEDRGSNERLF | 0.108645916 |
| RSYRMHPAV | ASSQEDRGSNERLF | 0.113147065 |
| SAAHLLAGQLA | ASSQEDRGSNERLF | 0.5356947 |
| SAAVGSY | ASSQEDRGSNERLF | 0.10511104 |
| SPHAVLHRI | ASSQEDRGSNERLF | 0.12365446 |
| SQIAKVL | ASSQEDRGSNERLF | 0.26377422 |

| | | |
|---|---|---|
| TALGHLVP | ASSQEDRGSNERLF | 0.41671866 |
| TAPTTSGAA | ASSQEDRGSNERLF | 0.087404385 |
| TDGDDGNGGSPGT | ASSQEDRGSNERLF | 0.14008234 |
| TDGLGEN | ASSQEDRGSNERLF | 0.14599925 |
| TGGPPLPGQA | ASSQEDRGSNERLF | 0.16654901 |
| TIPAITI | ASSQEDRGSNERLF | 0.12975962 |
| TIPGFTI | ASSQEDRGSNERLF | 0.1037363 |
| TSIASGALA | ASSQEDRGSNERLF | 0.2944786 |
| TVTLPGAAI | ASSQEDRGSNERLF | 0.12237917 |
| VASGLSAALP | ASSQEDRGSNERLF | 0.09701313 |
| VGAGVIGI | ASSQEDRGSNERLF | 0.3287981 |
| VGPKRFGV | ASSQEDRGSNERLF | 0.6396346 |
| VNVAFPD | ASSQEDRGSNERLF | 0.12794682 |
| VSMTNSGVS | ASSQEDRGSNERLF | 0.27887708 |
| VTDVLGDLSV | ASSQEDRGSNERLF | 0.14566824 |
| VVVVGAGFAGL | ASSQEDRGSNERLF | 0.6970389 |
| VYTVGTVV | ASSQEDRGSNERLF | 0.15916775 |
| WTTFSGP | ASSQEDRGSNERLF | 0.11691266 |
| YDELVEAF | ASSQEDRGSNERLF | 0.32934403 |
| YGTGGAGGNGG | ASSQEDRGSNERLF | 0.4951824 |
| YTNVMAV | ASSQEDRGSNERLF | 0.1884169 |
| FLEQVHQL | ASSQEDRGSNERLF | 0.48953167 |
| GLSLLAVAV | ASSQEDRGSNERLF | 0.12484686 |
| LEQVHQL | ASSQEDRGSNERLF | 0.15464073 |
| LNAAGAI | ASSQEDRGSNERLF | 0.23515512 |
| PAVHASL | ASSQEDRGSNERLF | 0.22956027 |
| TELEDDA | ASSQEDRGSNERLF | 0.09895326 |
| TPPGPPN | ASSQEDRGSNERLF | 0.29495215 |
| TPPPPPPP | ASSQEDRGSNERLF | 0.13577335 |
| TQSPALI | ASSHRDFQAPL | 0.14943415 |
| EFAAAFPQ | ASSHRDFQAPL | 0.33369705 |
| LLSGNLDVL | ASSHRDFQAPL | 0.24676542 |
| GPANLGNYNI | ASSHRDFQAPL | 0.36816916 |
| TLGPTLP | ASSHRDFQAPL | 0.1888891 |
| VYAYSSCCK | ASSHRDFQAPL | 0.19327565 |
| APAPAPAPNI | ASSHRDFQAPL | 0.110898994 |
| PGGPVPIP | ASSHRDFQAPL | 0.3205705 |
| FLTVALEP | ASSHRDFQAPL | 0.2576614 |
| SQYFGANL | ASSHRDFQAPL | 0.29752126 |
| GGVKPAA | ASSHRDFQAPL | 0.20583454 |
| GGTGGAGGAGGW | ASSHRDFQAPL | 0.111184 |
| GVGGVAGLGGAGG | ASSHRDFQAPL | 0.32259816 |
| VLLVLATTI | ASSHRDFQAPL | 0.40325996 |
| IGIFAAML | ASSHRDFQAPL | 0.15643448 |
| AVGDLID | ASSHRDFQAPL | 0.20475109 |
| NISPLPL | ASSHRDFQAPL | 0.5980251 |
| VTAVFVAI | ASSHRDFQAPL | 0.14035723 |
| SGDDNDAAL | ASSHRDFQAPL | 0.1409576 |
| AVAALAHS | ASSHRDFQAPL | 0.43733168 |
| GGVTVMATP | ASSHRDFQAPL | 0.5550268 |
| AGLAANVRD | ASSHRDFQAPL | 0.20130332 |

| | | |
|---|---|---|
| TNRPDLIDPALLRPG | ASSHRDFQAPL | 0.4893245 |
| LANLGVTAL | ASSHRDFQAPL | 0.10611831 |
| TGERIEL | ASSHRDFQAPL | 0.2650265 |
| NGEELAA | ASSHRDFQAPL | 0.13195458 |
| VLSLVWG | ASSHRDFQAPL | 0.2120671 |
| IGYALWL | ASSHRDFQAPL | 0.092075676 |
| LSLLLAPL | ASSHRDFQAPL | 0.19901028 |
| TLDEALK | ASSHRDFQAPL | 0.41010964 |
| TDGAGTAEGG | ASSHRDFQAPL | 0.26304227 |
| NAPGGAGGNGG | ASSHRDFQAPL | 0.0980453 |
| VSVFMGMGGNF | ASSHRDFQAPL | 0.14944778 |
| LASYLAAL | ASSHRDFQAPL | 0.3414535 |
| LVGGTPAL | ASSHRDFQAPL | 0.7094966 |
| VRNADVL | ASSHRDFQAPL | 0.22511275 |
| TCSSPFLVSAM | ASSHRDFQAPL | 0.5303345 |
| VHITDAE | ASSHRDFQAPL | 0.14449899 |
| EYAAPSF | ASSHRDFQAPL | 0.14206423 |
| VVDAFHGGL | ASSHRDFQAPL | 0.11637007 |
| ASVVDPQ | ASSHRDFQAPL | 0.19299778 |
| SYNEFGGLS | ASSHRDFQAPL | 0.18836284 |
| DPDGSVY | ASSHRDFQAPL | 0.117915176 |
| MVSTIHATE | ASSHRDFQAPL | 0.20056516 |
| LAAGVADPV | ASSHRDFQAPL | 0.600488 |
| TKEQIAA | ASSHRDFQAPL | 0.19374329 |
| SLASPAAL | ASSHRDFQAPL | 0.5601488 |
| IGLVFYGV | ASSHRDFQAPL | 0.5962002 |
| FLLDASNL | ASSHRDFQAPL | 0.47638714 |
| YQVGNFP | ASSHRDFQAPL | 0.10940283 |
| ESLLSQST | ASSHRDFQAPL | 0.2103405 |
| LFTVGFAAVG | ASSHRDFQAPL | 0.10107468 |
| ATPPTPT | ASSHRDFQAPL | 0.15087657 |
| TDLATLG | ASSHRDFQAPL | 0.1075365 |
| SAGGTGR | ASSHRDFQAPL | 0.20634422 |
| GFSNSGS | ASSHRDFQAPL | 0.08769586 |
| GGNGGSGGTSV | ASSHRDFQAPL | 0.3874056 |
| GTSVATGGAGN | ASSHRDFQAPL | 0.5114545 |
| NGGNGGSGGTS | ASSHRDFQAPL | 0.09297965 |
| AVVKFPH | ASSHRDFQAPL | 0.3985752 |
| TPPPPPP | ASSHRDFQAPL | 0.5148773 |
| AAPAAAAAPA | ASSHRDFQAPL | 0.1132134 |
| IALNEDL | ASSHRDFQAPL | 0.4516152 |
| IVNVSSM | ASSHRDFQAPL | 0.13335611 |
| TGIPPAPR | ASSHRDFQAPL | 0.5621387 |
| FDLGGGTF | ASSHRDFQAPL | 0.27861097 |
| FDLGGGTFDVS | ASSHRDFQAPL | 0.28068426 |
| NEPTAAA | ASSHRDFQAPL | 0.22372544 |
| NEPTAAALA | ASSHRDFQAPL | 0.44129032 |
| GSIINISSI | ASSHRDFQAPL | 0.115452014 |
| VAPGNAGTA | ASSHRDFQAPL | 0.23008999 |
| TGGGASLE | ASSHRDFQAPL | 0.090047054 |
| VAVYNHYKRI | ASSHRDFQAPL | 0.33459502 |

| | | |
|---|---|---|
| TDPAPATT | ASSHRDFQAPL | 0.11976028 |
| TNRPDILDPALLRPG | ASSHRDFQAPL | 0.28137124 |
| LTTTTTS | ASSHRDFQAPL | 0.13580832 |
| PTGSGKTL | ASSHRDFQAPL | 0.31689754 |
| TGSGKTL | ASSHRDFQAPL | 0.16079554 |
| TIGHVDHGK | ASSHRDFQAPL | 0.11320768 |
| TIGHVDHGKTT | ASSHRDFQAPL | 0.10600595 |
| GHVDHGKTT | ASSHRDFQAPL | 0.1032536 |
| AQFEHTLL | ASSHRDFQAPL | 0.48943207 |
| IVNQDEL | ASSHRDFQAPL | 0.18145405 |
| TNGSQFF | ASSHRDFQAPL | 0.10593028 |
| TGKGIVE | ASSHRDFQAPL | 0.23205012 |
| LISNASDA | ASSHRDFQAPL | 0.17326586 |
| AELRTLGELTKIAKAHGAQ | ASSHRDFQAPL | 0.11159449 |
| APNAVAAAEPSW | ASSHRDFQAPL | 0.22909114 |
| APQVAGAATAVL | ASSHRDFQAPL | 0.2967139 |
| ARRDPAAAAAILPTDA | ASSHRDFQAPL | 0.16247655 |
| DAAARDVLAEAGLAEYF | ASSHRDFQAPL | 0.08703536 |
| DDQIVLSAKANLAKVEQ | ASSHRDFQAPL | 0.11722614 |
| DELLPDTGEDSDEDGH | ASSHRDFQAPL | 0.15021427 |
| ELPEAPANRATVPQD | ASSHRDFQAPL | 0.10250522 |
| FGQSKIERVTGIG | ASSHRDFQAPL | 0.10746389 |
| GAGGTGSTAGGAGGAGGAGGLY | ASSHRDFQAPL | 0.43393674 |
| GHGGTGAAGGNGGTGGAGGSGI | ASSHRDFQAPL | 0.09270177 |
| GNGGNGGAGGEGGPGGAGGAGA | ASSHRDFQAPL | 0.1288805 |
| GSTAGNGGNGGNATNGTIAGQP | ASSHRDFQAPL | 0.39114627 |
| GTGGDGGNGGNGGTGG | ASSHRDFQAPL | 0.19891915 |
| GVRRPAGIPADAPV | ASSHRDFQAPL | 0.45277107 |
| GYRQVTDAHLLTLARR | ASSHRDFQAPL | 0.12206768 |
| IAKMGQDGHDRG | ASSHRDFQAPL | 0.15660566 |
| KAEPSIEIQYT | ASSHRDFQAPL | 0.233708 |
| KPVRIRMAADEDTVAVT | ASSHRDFQAPL | 0.19842806 |
| MPPRIAGMRLLVIKPEP | ASSHRDFQAPL | 0.10290088 |
| MKFYAVALFRSRGVPPTR | ASSHRDFQAPL | 0.094023295 |
| NAGGFGILGAVAHSP | ASSHRDFQAPL | 0.17128326 |
| NEGREITLAMAEM | ASSHRDFQAPL | 0.12007316 |
| NELVEWSHPLGHRWRWE | ASSHRDFQAPL | 0.22627942 |
| NLGAGGTGGNGGNPG | ASSHRDFQAPL | 0.60164684 |
| NTEWPKWETVMAADPVD | ASSHRDFQAPL | 0.11304022 |
| PASAAPAETQP | ASSHRDFQAPL | 0.115102425 |
| QQSSDGKNGVRAAEITGE | ASSHRDFQAPL | 0.1744997 |
| RAPDAVTTSWWKE | ASSHRDFQAPL | 0.13518897 |
| SGFGNVGAHTSGWFNQ | ASSHRDFQAPL | 0.39617854 |
| SGVAGAVGQAASVGGLKV | ASSHRDFQAPL | 0.13402173 |
| SNGGAGTGSAGGNGGT | ASSHRDFQAPL | 0.69300103 |
| TGGAGGEGGAAG | ASSHRDFQAPL | 0.18799552 |
| TGGDGGAAGAGGAPGHG | ASSHRDFQAPL | 0.11049469 |
| TMPPLLAAIANAGRHGV | ASSHRDFQAPL | 0.40923685 |
| TSSTAGNSGGA | ASSHRDFQAPL | 0.19690081 |
| VPATVPVGILC | ASSHRDFQAPL | 0.36238098 |
| VQRPDSGGVVLTGRIS | ASSHRDFQAPL | 0.24652287 |

| | | |
|---|---|---|
| WTPNPVPATKGVPAGLVQAT | ASSHRDFQAPL | 0.1087773 |
| YALKRALAKFSDRFDVVII | ASSHRDFQAPL | 0.13365419 |
| YLIGSVIV | ASSHRDFQAPL | 0.24120155 |
| YGQLVAVID | ASSHRDFQAPL | 0.12603585 |
| VTVGGMTLPA | ASSHRDFQAPL | 0.24690478 |
| VLALPAV | ASSHRDFQAPL | 0.09972237 |
| VIPPTLNYET | ASSHRDFQAPL | 0.10617379 |
| VGMGGMHPSSG | ASSHRDFQAPL | 0.2612391 |
| VGLAGGAATA | ASSHRDFQAPL | 0.36827338 |
| VGGAGGAGGSGP | ASSHRDFQAPL | 0.65771073 |
| VFAAFMV | ASSHRDFQAPL | 0.144111 |
| TLYDKLR | ASSHRDFQAPL | 0.10769187 |
| TLAVFTVL | ASSHRDFQAPL | 0.56537294 |
| TGSVQPARV | ASSHRDFQAPL | 0.1695769 |
| TGGGAGSGGT | ASSHRDFQAPL | 0.39148748 |
| SHLETSL | ASSHRDFQAPL | 0.15161356 |
| SFTTGMT | ASSHRDFQAPL | 0.1096464 |
| QVEVAKRMGVRQPTVS | ASSHRDFQAPL | 0.19970314 |
| QGSGGPALAM | ASSHRDFQAPL | 0.34684643 |
| MLAPSNP | ASSHRDFQAPL | 0.09820408 |
| LVNNAAI | ASSHRDFQAPL | 0.4141693 |
| LTGEINTLA | ASSHRDFQAPL | 0.08756317 |
| LSIFGPV | ASSHRDFQAPL | 0.12069352 |
| LLALLANL | ASSHRDFQAPL | 0.1379141 |
| LISNASDALDKLR | ASSHRDFQAPL | 0.11853744 |
| LINEAVP | ASSHRDFQAPL | 0.09838355 |
| LGYAFSGL | ASSHRDFQAPL | 0.18594494 |
| LDFVAVAL | ASSHRDFQAPL | 0.4706918 |
| LAQPGAE | ASSHRDFQAPL | 0.59943414 |
| ISNLGAVTGF | ASSHRDFQAPL | 0.11874311 |
| HQEDPGRI | ASSHRDFQAPL | 0.33688432 |
| GTPGEPTP | ASSHRDFQAPL | 0.6758225 |
| GFFNSGS | ASSHRDFQAPL | 0.18242633 |
| GAGGGGLTAGAN | ASSHRDFQAPL | 0.10060636 |
| FVDALVNM | ASSHRDFQAPL | 0.15270092 |
| FLTVAPEL | ASSHRDFQAPL | 0.093251646 |
| FETLTAL | ASSHRDFQAPL | 0.1335524 |
| DVINPIEL | ASSHRDFQAPL | 0.6137631 |
| APVSGAPV | ASSHRDFQAPL | 0.09899775 |
| ALVGGLAW | ASSHRDFQAPL | 0.29632056 |
| ALAAAAATAI | ASSHRDFQAPL | 0.38423502 |
| AFGLGLLL | ASSHRDFQAPL | 0.14593948 |
| AFFAAASL | ASSHRDFQAPL | 0.14987984 |
| AAAALPAFSPP | ASSHRDFQAPL | 0.11659097 |
| VVLAPAL | ASSHRDFQAPL | 0.10903762 |
| TIGHVDHGKTTL | ASSHRDFQAPL | 0.17653523 |
| LISNASDALDK | ASSHRDFQAPL | 0.15734169 |
| LISNASDAL | ASSHRDFQAPL | 0.14315829 |
| ISNASDALDK | ASSHRDFQAPL | 0.15513402 |
| NQAETLVYQTE | ASSHRDFQAPL | 0.12296357 |
| TGGTGGTAGAG | ASSHRDFQAPL | 0.13785669 |

| | | |
|---|---|---|
| TGSTAGGAGGA | ASSHRDFQAPL | 0.1364388 |
| DPLAVPVAAGPV | ASSHRDFQAPL | 0.12790836 |
| GGTGGTPGNGGN | ASSHRDFQAPL | 0.39354026 |
| PAGLAGVDPATL | ASSHRDFQAPL | 0.10286659 |
| GNNLVVEAPHAGASV | ASSHRDFQAPL | 0.3379478 |
| KVTRIMSASTLEQPA | ASSHRDFQAPL | 0.48922783 |
| LGFALPIALPRRTVV | ASSHRDFQAPL | 0.096329555 |
| TRLGNLTARKLVVLV | ASSHRDFQAPL | 0.11572365 |
| WSQQRRLIYFPSAGP | ASSHRDFQAPL | 0.17127131 |
| GPNGNPDPMAAAVDIR | ASSHRDFQAPL | 0.09801049 |
| TQRCVSVADLRAAASR | ASSHRDFQAPL | 0.49872297 |
| QFVLDVAEHPVEVLTETV | ASSHRDFQAPL | 0.1422134 |
| GGAGGTGGAGGNGGTQAGDG | ASSHRDFQAPL | 0.2380832 |
| PLLATLRAIAVDVDAKPAQV | ASSHRDFQAPL | 0.1733174 |
| APKFRTGGYMIDFWGVGYQVAK | ASSHRDFQAPL | 0.66647744 |
| GAGGAGGAGGAAGGPGGTGGTG | ASSHRDFQAPL | 0.27683443 |
| IGALGLLATDVAPAGSVAFGVA | ASSHRDFQAPL | 0.15426078 |
| AGGAGGIGGNGGAGGAGGVLMG | ASSHRDFQAPL | 0.09418427 |
| ADELIGLL | ASSHRDFQAPL | 0.115373276 |
| ADRALKA | ASSHRDFQAPL | 0.35910356 |
| AGAVGQVGGAGGA | ASSHRDFQAPL | 0.5909922 |
| AGGGGGLGGAP | ASSHRDFQAPL | 0.10412249 |
| AGGLLYGNGGAG | ASSHRDFQAPL | 0.15851973 |
| AGLESIGL | ASSHRDFQAPL | 0.5829079 |
| AGLKTGII | ASSHRDFQAPL | 0.45089734 |
| ASLAALLL | ASSHRDFQAPL | 0.10450563 |
| ASVINISSI | ASSHRDFQAPL | 0.11750057 |
| ATPPGPA | ASSHRDFQAPL | 0.22554839 |
| ATVASALV | ASSHRDFQAPL | 0.19524851 |
| ATVASLIL | ASSHRDFQAPL | 0.107868806 |
| AVNGPVLIRH | ASSHRDFQAPL | 0.4288575 |
| DGAAGAVGI | ASSHRDFQAPL | 0.2436132 |
| DPLLHGI | ASSHRDFQAPL | 0.61406726 |
| ELGAALGAL | ASSHRDFQAPL | 0.20075367 |
| ELGRTQAV | ASSHRDFQAPL | 0.25676012 |
| FVAGGYLS | ASSHRDFQAPL | 0.2518931 |
| GASGGAGGAGGW | ASSHRDFQAPL | 0.12797485 |
| GGAGAGGDGGAGGI | ASSHRDFQAPL | 0.12880184 |
| GTGHNGSQPGMGGNG | ASSHRDFQAPL | 0.09650446 |
| GYAAASASAS | ASSHRDFQAPL | 0.13289015 |
| HQVDVAAVQQG | ASSHRDFQAPL | 0.13042031 |
| IGAAIAHT | ASSHRDFQAPL | 0.08833784 |
| IGMIEVAQL | ASSHRDFQAPL | 0.14847995 |
| IRLAPEHTSAMD | ASSHRDFQAPL | 0.20180817 |
| IYNVSVL | ASSHRDFQAPL | 0.20129728 |
| KQGFLGM | ASSHRDFQAPL | 0.15097134 |
| LESVITQL | ASSHRDFQAPL | 0.1558153 |
| LGLGALVVV | ASSHRDFQAPL | 0.105191424 |
| LIGNGGNGGSGGI | ASSHRDFQAPL | 0.56347656 |
| LNAINAPTL | ASSHRDFQAPL | 0.096659966 |
| LQLLNTGV | ASSHRDFQAPL | 0.12992997 |

| | | |
|---|---|---|
| LQTSIEF | ASSHRDFQAPL | 0.15041512 |
| MSFAVSEL | ASSHRDFQAPL | 0.20043045 |
| NTGALGL | ASSHRDFQAPL | 0.65834314 |
| PPGAGGAGGAGTT | ASSHRDFQAPL | 0.39575464 |
| QLVGLGI | ASSHRDFQAPL | 0.2257367 |
| RFLEQVHQL | ASSHRDFQAPL | 0.63236654 |
| RGVGYVLR | ASSHRDFQAPL | 0.09912482 |
| SATGVFAGVI | ASSHRDFQAPL | 0.123706885 |
| SLPAITLP | ASSHRDFQAPL | 0.5166893 |
| STALAFTV | ASSHRDFQAPL | 0.14688778 |
| SVLTVGLGK | ASSHRDFQAPL | 0.26067618 |
| TALAGAGSGI | ASSHRDFQAPL | 0.32829767 |
| TGLPFTGI | ASSHRDFQAPL | 0.11263489 |
| VAEHVTV | ASSHRDFQAPL | 0.16441151 |
| VDIVLNSL | ASSHRDFQAPL | 0.12716144 |
| VLTGGGAVL | ASSHRDFQAPL | 0.32806894 |
| VNVGLAYGI | ASSHRDFQAPL | 0.24965483 |
| VSSAAVGAV | ASSHRDFQAPL | 0.10587131 |
| VTSVYEAGV | ASSHRDFQAPL | 0.5464266 |
| WVALDRGA | ASSHRDFQAPL | 0.76031375 |
| YLVGSVVI | ASSHRDFQAPL | 0.16514796 |
| GPPGTGKTLL | ASSHRDFQAPL | 0.28849736 |
| GVGMTKF | ASSHRDFQAPL | 0.18356302 |
| IGIGGSDL | ASSHRDFQAPL | 0.21854371 |
| IGTLLAEL | ASSHRDFQAPL | 0.13896935 |
| IYTSGTT | ASSHRDFQAPL | 0.11169475 |
| LSFPAGV | ASSHRDFQAPL | 0.113849685 |
| LYTSGTT | ASSHRDFQAPL | 0.12692654 |
| VGAFPFGF | ASSHRDFQAPL | 0.42332178 |
| VGAGTVEF | ASSHRDFQAPL | 0.17117037 |
| VGKSTLLL | ASSHRDFQAPL | 0.74731714 |
| AALAHAGGGIV | ASSHRDFQAPL | 0.34623367 |
| AALVAVSLL | ASSHRDFQAPL | 0.1297038 |
| AAVSRAVAAL | ASSHRDFQAPL | 0.10067971 |
| AGGSSGIGGF | ASSHRDFQAPL | 0.23544122 |
| AGLTVANANG | ASSHRDFQAPL | 0.19841674 |
| AGLVGAVL | ASSHRDFQAPL | 0.1240641 |
| ALSQAFTV | ASSHRDFQAPL | 0.47750458 |
| ALTFSVSL | ASSHRDFQAPL | 0.12723146 |
| ALTGAALVAL | ASSHRDFQAPL | 0.11951292 |
| ASFSIGAGVL | ASSHRDFQAPL | 0.18724667 |
| DGSVYKI | ASSHRDFQAPL | 0.6434194 |
| EQLAEIV | ASSHRDFQAPL | 0.2692229 |
| FAVSELTVM | ASSHRDFQAPL | 0.13278313 |
| GAAPLAAGL | ASSHRDFQAPL | 0.101735786 |
| GADGGDGGSGGAGG | ASSHRDFQAPL | 0.45531586 |
| GFGLDTGGAGG | ASSHRDFQAPL | 0.4624738 |
| GLASGGLL | ASSHRDFQAPL | 0.09769472 |
| GRIAGAAVLL | ASSHRDFQAPL | 0.6551707 |
| GSAGSLNTSF | ASSHRDFQAPL | 0.120403394 |
| GTGQIGI | ASSHRDFQAPL | 0.112204246 |

| | | |
|---|---|---|
| GTLIALIL | ASSHRDFQAPL | 0.2554621 |
| IAATMGFVVG | ASSHRDFQAPL | 0.27682668 |
| IFAQALL | ASSHRDFQAPL | 0.64466834 |
| IGAAVALGF | ASSHRDFQAPL | 0.20710935 |
| IGLTPLAL | ASSHRDFQAPL | 0.56918186 |
| ILGSNTSTL | ASSHRDFQAPL | 0.17306405 |
| IVFSLTVL | ASSHRDFQAPL | 0.5190887 |
| LASKATGF | ASSHRDFQAPL | 0.50045884 |
| LASVALIL | ASSHRDFQAPL | 0.25595537 |
| LFAVGLTL | ASSHRDFQAPL | 0.18317054 |
| LFNGGIGTYI | ASSHRDFQAPL | 0.14130776 |
| LTLGAGII | ASSHRDFQAPL | 0.27473474 |
| LVAAALGDI | ASSHRDFQAPL | 0.115861304 |
| NTAFVQLGI | ASSHRDFQAPL | 0.18917489 |
| QLFSGAEL | ASSHRDFQAPL | 0.2370931 |
| SAATFITL | ASSHRDFQAPL | 0.24284121 |
| SNYGLANLGV | ASSHRDFQAPL | 0.11691805 |
| SYGGAYAVM | ASSHRDFQAPL | 0.3109879 |
| TTGVGGLTLG | ASSHRDFQAPL | 0.6365932 |
| TTQTIYI | ASSHRDFQAPL | 0.33829725 |
| VAAGAGLGAVI | ASSHRDFQAPL | 0.122032225 |
| VAAVVTLL | ASSHRDFQAPL | 0.532371 |
| VGFLGGLI | ASSHRDFQAPL | 0.7173478 |
| VGVIPILL | ASSHRDFQAPL | 0.51163375 |
| VIFASSLI | ASSHRDFQAPL | 0.10951993 |
| VKGGDGGAAAG | ASSHRDFQAPL | 0.09818857 |
| VTKTFTNI | ASSHRDFQAPL | 0.3196414 |
| VVAAAILL | ASSHRDFQAPL | 0.1732162 |
| YAALDVEL | ASSHRDFQAPL | 0.123245865 |
| LASVALLL | ASSHRDFQAPL | 0.105362825 |
| LDAVHRV | ASSHRDFQAPL | 0.29435188 |
| QGLSSSIF | ASSHRDFQAPL | 0.2678734 |
| RGTFANIRL | ASSHRDFQAPL | 0.13901785 |
| RQLWWGHRI | ASSHRDFQAPL | 0.10158643 |
| YGPPGTGKTLL | ASSHRDFQAPL | 0.3914922 |
| AALVGAPANVA | ASSHRDFQAPL | 0.5836189 |
| AATFRRTE | ASSHRDFQAPL | 0.11410421 |
| AGALGGGI | ASSHRDFQAPL | 0.14737456 |
| AGGAGGAGGIGAG | ASSHRDFQAPL | 0.25194293 |
| AGSGALGL | ASSHRDFQAPL | 0.10556316 |
| AQIADQI | ASSHRDFQAPL | 0.1673148 |
| ASAVAVTYL | ASSHRDFQAPL | 0.11700474 |
| ASTVLGGTGG | ASSHRDFQAPL | 0.12214138 |
| AVGGEPAK | ASSHRDFQAPL | 0.13740206 |
| CEQGTFGGTC | ASSHRDFQAPL | 0.4978477 |
| DEGKAPAG | ASSHRDFQAPL | 0.22011441 |
| DTVVFPAL | ASSHRDFQAPL | 0.2443237 |
| FEGITMAG | ASSHRDFQAPL | 0.23817763 |
| FPLVVQD | ASSHRDFQAPL | 0.14650309 |
| GAFTTPQ | ASSHRDFQAPL | 0.1080617 |
| GFGGFGTAGGVGGA | ASSHRDFQAPL | 0.09325357 |

| | | |
|---|---|---|
| GFLAQFL | ASSHRDFQAPL | 0.5818906 |
| GGAGGAGGNAGVA | ASSHRDFQAPL | 0.087429486 |
| GGVGTAGAGGAGGAGG | ASSHRDFQAPL | 0.3505327 |
| GNGGGGDAGNAGSGGN | ASSHRDFQAPL | 0.16748184 |
| GTGGAGGAGNDAGS | ASSHRDFQAPL | 0.114312835 |
| GTTNTGWFNS | ASSHRDFQAPL | 0.19823933 |
| IFMGPLFTL | ASSHRDFQAPL | 0.09041435 |
| ILAHAIEAI | ASSHRDFQAPL | 0.13507044 |
| IRSAIREAL | ASSHRDFQAPL | 0.47705993 |
| ITFAGGG | ASSHRDFQAPL | 0.26945683 |
| ITGMSGHQEV | ASSHRDFQAPL | 0.22593893 |
| IVAGLACTL | ASSHRDFQAPL | 0.41744632 |
| IVGGGLAGA | ASSHRDFQAPL | 0.3208614 |
| IVTGVLNI | ASSHRDFQAPL | 0.15268475 |
| LAQQGCI | ASSHRDFQAPL | 0.32746062 |
| LCLHVEN | ASSHRDFQAPL | 0.3053373 |
| LDAISDGS | ASSHRDFQAPL | 0.46307725 |
| LGGFAAML | ASSHRDFQAPL | 0.10573784 |
| LGGKNPMI | ASSHRDFQAPL | 0.14568754 |
| LGPKGVAAIL | ASSHRDFQAPL | 0.12200168 |
| LMVGVSI | ASSHRDFQAPL | 0.19940177 |
| LVGAAFVL | ASSHRDFQAPL | 0.13161491 |
| LVLAVAPL | ASSHRDFQAPL | 0.27505347 |
| MAAIAVIDAA | ASSHRDFQAPL | 0.17887157 |
| NNGINAELQ | ASSHRDFQAPL | 0.09554834 |
| RGEPLVLVHG | ASSHRDFQAPL | 0.13700353 |
| RGGAGGAAGLI | ASSHRDFQAPL | 0.22890466 |
| RSYRMHPAV | ASSHRDFQAPL | 0.17900065 |
| SAAHLLAGQLA | ASSHRDFQAPL | 0.17920375 |
| SAAVGSY | ASSHRDFQAPL | 0.10902028 |
| SPHAVLHRI | ASSHRDFQAPL | 0.33268428 |
| SQIAKVL | ASSHRDFQAPL | 0.15100825 |
| TALGHLVP | ASSHRDFQAPL | 0.13066182 |
| TAPTTSGAA | ASSHRDFQAPL | 0.33873436 |
| TDGDDGNGGSPGT | ASSHRDFQAPL | 0.15055296 |
| TDGLGEN | ASSHRDFQAPL | 0.14111435 |
| TGGPPLPGQA | ASSHRDFQAPL | 0.38398933 |
| TIPAITI | ASSHRDFQAPL | 0.11158653 |
| TIPGFTI | ASSHRDFQAPL | 0.20927338 |
| TSIASGALA | ASSHRDFQAPL | 0.14212734 |
| TVTLPGAAI | ASSHRDFQAPL | 0.13024996 |
| VASGLSAALP | ASSHRDFQAPL | 0.41653413 |
| VGAGVIGI | ASSHRDFQAPL | 0.102053046 |
| VGPKRFGV | ASSHRDFQAPL | 0.31147367 |
| VNVAFPD | ASSHRDFQAPL | 0.13938306 |
| VSMTNSGVS | ASSHRDFQAPL | 0.13576524 |
| VTDVLGDLSV | ASSHRDFQAPL | 0.16037212 |
| VVVVGAGFAGL | ASSHRDFQAPL | 0.19281556 |
| VYTVGTVV | ASSHRDFQAPL | 0.41067645 |
| WTTFSGP | ASSHRDFQAPL | 0.111817166 |
| YDELVEAF | ASSHRDFQAPL | 0.39523014 |

| | | |
|---|---|---|
| YGTGGAGGNGG | ASSHRDFQAPL | 0.41640344 |
| YTNVMAV | ASSHRDFQAPL | 0.13301955 |
| FLEQVHQL | ASSHRDFQAPL | 0.10630228 |
| GLSLLAVAV | ASSHRDFQAPL | 0.11730857 |
| LEQVHQL | ASSHRDFQAPL | 0.101503946 |
| LNAAGAI | ASSHRDFQAPL | 0.15925121 |
| PAVHASL | ASSHRDFQAPL | 0.11286634 |
| TELEDDA | ASSHRDFQAPL | 0.25446787 |
| TPPGPPN | ASSHRDFQAPL | 0.48515406 |
| TPPPPPPP | ASSHRDFQAPL | 0.1420877 |
| TQSPALI | ASSPRDSAETLY | 0.48295557 |
| EFAAAFPQ | ASSPRDSAETLY | 0.11474468 |
| LLSGNLDVL | ASSPRDSAETLY | 0.21689779 |
| GPANLGNYNI | ASSPRDSAETLY | 0.5341049 |
| TLGPTLP | ASSPRDSAETLY | 0.1127457 |
| VYAYSSCCK | ASSPRDSAETLY | 0.14944965 |
| APAPAPAPNI | ASSPRDSAETLY | 0.20072463 |
| PGGPVPIP | ASSPRDSAETLY | 0.11144275 |
| FLTVALEP | ASSPRDSAETLY | 0.44451368 |
| SQYFGANL | ASSPRDSAETLY | 0.14810176 |
| GGVKPAA | ASSPRDSAETLY | 0.34876704 |
| GGTGGAGGAGGW | ASSPRDSAETLY | 0.12378872 |
| GVGGVAGLGGAGG | ASSPRDSAETLY | 0.102415085 |
| VLLVLATTI | ASSPRDSAETLY | 0.20296225 |
| IGIFAAML | ASSPRDSAETLY | 0.47090757 |
| AVGDLID | ASSPRDSAETLY | 0.12754714 |
| NISPLPL | ASSPRDSAETLY | 0.50009894 |
| VTAVFVAI | ASSPRDSAETLY | 0.17278606 |
| SGDDNDAAL | ASSPRDSAETLY | 0.48895603 |
| AVAALAHS | ASSPRDSAETLY | 0.69822407 |
| GGVTVMATP | ASSPRDSAETLY | 0.20206362 |
| AGLAANVRD | ASSPRDSAETLY | 0.15670341 |
| TNRPDLIDPALLRPG | ASSPRDSAETLY | 0.5142671 |
| LANLGVTAL | ASSPRDSAETLY | 0.2530033 |
| TGERIEL | ASSPRDSAETLY | 0.41558397 |
| NGEELAA | ASSPRDSAETLY | 0.49138474 |
| VLSLVWG | ASSPRDSAETLY | 0.34794763 |
| IGYALWL | ASSPRDSAETLY | 0.6465266 |
| LSLLLAPL | ASSPRDSAETLY | 0.40480655 |
| TLDEALK | ASSPRDSAETLY | 0.49401328 |
| TDGAGTAEGG | ASSPRDSAETLY | 0.37437242 |
| NAPGGAGGNGG | ASSPRDSAETLY | 0.19002615 |
| VSVFMGMGGNF | ASSPRDSAETLY | 0.4392217 |
| LASYLAAL | ASSPRDSAETLY | 0.25028205 |
| LVGGTPAL | ASSPRDSAETLY | 0.19150363 |
| VRNADVL | ASSPRDSAETLY | 0.5682163 |
| TCSSPFLVSAM | ASSPRDSAETLY | 0.3973168 |
| VHITDAE | ASSPRDSAETLY | 0.7230637 |
| EYAAPSF | ASSPRDSAETLY | 0.7312436 |
| VVDAFHGGL | ASSPRDSAETLY | 0.22519583 |
| ASVVDPQ | ASSPRDSAETLY | 0.3988812 |

| | | |
|---|---|---|
| SYNEFGGLS | ASSPRDSAETLY | 0.27847067 |
| DPDGSVY | ASSPRDSAETLY | 0.4937053 |
| MVSTIHATE | ASSPRDSAETLY | 0.15270562 |
| LAAGVADPV | ASSPRDSAETLY | 0.61121315 |
| TKEQIAA | ASSPRDSAETLY | 0.21533005 |
| SLASPAAL | ASSPRDSAETLY | 0.12424367 |
| IGLVFYGV | ASSPRDSAETLY | 0.49811822 |
| FLLDASNL | ASSPRDSAETLY | 0.3945999 |
| YQVGNFP | ASSPRDSAETLY | 0.39262104 |
| ESLLSQST | ASSPRDSAETLY | 0.7501296 |
| LFTVGFAAVG | ASSPRDSAETLY | 0.16035777 |
| ATPPTPT | ASSPRDSAETLY | 0.2258421 |
| TDLATLG | ASSPRDSAETLY | 0.5819362 |
| SAGGTGR | ASSPRDSAETLY | 0.7969284 |
| GFSNSGS | ASSPRDSAETLY | 0.22358905 |
| GGNGGSGGTSV | ASSPRDSAETLY | 0.6368804 |
| GTSVATGGAGN | ASSPRDSAETLY | 0.7826438 |
| NGGNGGSGGTS | ASSPRDSAETLY | 0.46333486 |
| AVVKFPH | ASSPRDSAETLY | 0.6942661 |
| TPPPPPP | ASSPRDSAETLY | 0.7399824 |
| AAPAAAAAPA | ASSPRDSAETLY | 0.26254565 |
| IALNEDL | ASSPRDSAETLY | 0.7404772 |
| IVNVSSM | ASSPRDSAETLY | 0.6773967 |
| TGIPPAPR | ASSPRDSAETLY | 0.40495563 |
| FDLGGGTF | ASSPRDSAETLY | 0.5274462 |
| FDLGGGTFDVS | ASSPRDSAETLY | 0.3515364 |
| NEPTAAA | ASSPRDSAETLY | 0.45169693 |
| NEPTAAALA | ASSPRDSAETLY | 0.15588939 |
| GSIINISSI | ASSPRDSAETLY | 0.61490977 |
| VAPGNAGTA | ASSPRDSAETLY | 0.6260918 |
| TGGGASLE | ASSPRDSAETLY | 0.45558423 |
| VAVYNHYKRI | ASSPRDSAETLY | 0.24532917 |
| TDPAPATT | ASSPRDSAETLY | 0.27063236 |
| TNRPDILDPALLRPG | ASSPRDSAETLY | 0.29634562 |
| LTTTTTS | ASSPRDSAETLY | 0.42192334 |
| PTGSGKTL | ASSPRDSAETLY | 0.16685703 |
| TGSGKTL | ASSPRDSAETLY | 0.2532339 |
| TIGHVDHGK | ASSPRDSAETLY | 0.64366 |
| TIGHVDHGKTT | ASSPRDSAETLY | 0.649294 |
| GHVDHGKTT | ASSPRDSAETLY | 0.20955281 |
| AQFEHTLL | ASSPRDSAETLY | 0.74742883 |
| IVNQDEL | ASSPRDSAETLY | 0.5203313 |
| TNGSQFF | ASSPRDSAETLY | 0.61332023 |
| TGKGIVE | ASSPRDSAETLY | 0.37652397 |
| LISNASDA | ASSPRDSAETLY | 0.331068 |
| AELRTLGELTKIAKAHGAQ | ASSPRDSAETLY | 0.5779336 |
| APNAVAAAEPSW | ASSPRDSAETLY | 0.42437124 |
| APQVAGAATAVL | ASSPRDSAETLY | 0.23943529 |
| ARRDPAAAAAILPTDA | ASSPRDSAETLY | 0.7175777 |
| DAAARDVLAEAGLAEYF | ASSPRDSAETLY | 0.27131978 |
| DDQIVLSAKANLAKVEQ | ASSPRDSAETLY | 0.3145181 |

| | | |
|---|---|---|
| DELLPDTGEDSDEDGH | ASSPRDSAETLY | 0.44048017 |
| ELPEAPANRATVPQD | ASSPRDSAETLY | 0.60161483 |
| FGQSKIERVTGIG | ASSPRDSAETLY | 0.20887686 |
| GAGGTGSTAGGAGGAGGAGGLY | ASSPRDSAETLY | 0.28540614 |
| GHGGTGAAGGNGGTGGAGGSGI | ASSPRDSAETLY | 0.22351116 |
| GNGGNGGAGGEGGPGGAGGAGA | ASSPRDSAETLY | 0.4835907 |
| GSTAGNGGNGGNATNGTIAGQP | ASSPRDSAETLY | 0.77742875 |
| GTGGDGGNGGNGGTGG | ASSPRDSAETLY | 0.46131337 |
| GVRRPAGIPADAPV | ASSPRDSAETLY | 0.34305274 |
| GYRQVTDAHLLTLARR | ASSPRDSAETLY | 0.7156087 |
| IAKMGQDGHDRG | ASSPRDSAETLY | 0.48642343 |
| KAEPSIEIQYT | ASSPRDSAETLY | 0.459573 |
| KPVRIRMAADEDTVAVT | ASSPRDSAETLY | 0.67524815 |
| MPPRIAGMRLLVIKPEP | ASSPRDSAETLY | 0.3140034 |
| MKFYAVALFRSRGVPPTR | ASSPRDSAETLY | 0.5917357 |
| NAGGFGILGAVAHSP | ASSPRDSAETLY | 0.53509766 |
| NEGREITLAMAEM | ASSPRDSAETLY | 0.65367484 |
| NELVEWSHPLGHRWRWE | ASSPRDSAETLY | 0.6744224 |
| NLGAGGTGGNGGNPG | ASSPRDSAETLY | 0.38698766 |
| NTEWPKWETVMAADPVD | ASSPRDSAETLY | 0.46823862 |
| PASAAPAETQP | ASSPRDSAETLY | 0.18590064 |
| QQSSDGKNGVRAAEITGE | ASSPRDSAETLY | 0.29467905 |
| RAPDAVTTSWWKE | ASSPRDSAETLY | 0.6306706 |
| SGFGNVGAHTSGWFNQ | ASSPRDSAETLY | 0.53541785 |
| SGVAGAVGQAASVGGLKV | ASSPRDSAETLY | 0.658707 |
| SNGGAGTGSAGGNGGT | ASSPRDSAETLY | 0.5633519 |
| TGGAGGEGGAAG | ASSPRDSAETLY | 0.2760007 |
| TGGDGGAAGAGGAPGHG | ASSPRDSAETLY | 0.7263298 |
| TMPPLLAAIANAGRHGV | ASSPRDSAETLY | 0.739888 |
| TSSTAGNSGGA | ASSPRDSAETLY | 0.21728902 |
| VPATVPVGILC | ASSPRDSAETLY | 0.33822477 |
| VQRPDSGGVVLTGRIS | ASSPRDSAETLY | 0.77427745 |
| WTPNPVPATKGVPAGLVQAT | ASSPRDSAETLY | 0.5740137 |
| YALKRALAKFSDRFDVVII | ASSPRDSAETLY | 0.4297163 |
| YLIGSVIV | ASSPRDSAETLY | 0.4418262 |
| YGQLVAVID | ASSPRDSAETLY | 0.3872757 |
| VTVGGMTLPA | ASSPRDSAETLY | 0.26823547 |
| VLALPAV | ASSPRDSAETLY | 0.73095334 |
| VIPPTLNYET | ASSPRDSAETLY | 0.1596954 |
| VGMGGMHPSSG | ASSPRDSAETLY | 0.4167214 |
| VGLAGGAATA | ASSPRDSAETLY | 0.26464817 |
| VGGAGGAGGSGP | ASSPRDSAETLY | 0.18394956 |
| VFAAFMV | ASSPRDSAETLY | 0.57319343 |
| TLYDKLR | ASSPRDSAETLY | 0.37091994 |
| TLAVFTVL | ASSPRDSAETLY | 0.7240798 |
| TGSVQPARV | ASSPRDSAETLY | 0.38298762 |
| TGGGAGSGGT | ASSPRDSAETLY | 0.6271999 |
| SHLETSL | ASSPRDSAETLY | 0.54739875 |
| SFTTGMT | ASSPRDSAETLY | 0.57001436 |
| QVEVAKRMGVRQPTVS | ASSPRDSAETLY | 0.35237223 |
| QGSGGPALAM | ASSPRDSAETLY | 0.4676643 |

| | | |
|---|---|---|
| MLAPSNP | ASSPRDSAETLY | 0.7259679 |
| LVNNAAI | ASSPRDSAETLY | 0.54620147 |
| LTGEINTLA | ASSPRDSAETLY | 0.18689336 |
| LSIFGPV | ASSPRDSAETLY | 0.27462748 |
| LLALLANL | ASSPRDSAETLY | 0.44889 |
| LISNASDALDKLR | ASSPRDSAETLY | 0.14729378 |
| LINEAVP | ASSPRDSAETLY | 0.15950395 |
| LGYAFSGL | ASSPRDSAETLY | 0.21311705 |
| LDFVAVAL | ASSPRDSAETLY | 0.2726534 |
| LAQPGAE | ASSPRDSAETLY | 0.68305755 |
| ISNLGAVTGF | ASSPRDSAETLY | 0.6248313 |
| HQEDPGRI | ASSPRDSAETLY | 0.39928353 |
| GTPGEPTP | ASSPRDSAETLY | 0.633096 |
| GFFNSGS | ASSPRDSAETLY | 0.4704643 |
| GAGGGGLTAGAN | ASSPRDSAETLY | 0.10749461 |
| FVDALVNM | ASSPRDSAETLY | 0.32693884 |
| FLTVAPEL | ASSPRDSAETLY | 0.26205295 |
| FETLTAL | ASSPRDSAETLY | 0.7151695 |
| DVINPIEL | ASSPRDSAETLY | 0.25492135 |
| APVSGAPV | ASSPRDSAETLY | 0.2748618 |
| ALVGGLAW | ASSPRDSAETLY | 0.5287787 |
| ALAAAAATAI | ASSPRDSAETLY | 0.32587495 |
| AFGLGLLL | ASSPRDSAETLY | 0.52230376 |
| AFFAAASL | ASSPRDSAETLY | 0.24546024 |
| AAAALPAFSPP | ASSPRDSAETLY | 0.128745 |
| VVLAPAL | ASSPRDSAETLY | 0.5746505 |
| TIGHVDHGKTTL | ASSPRDSAETLY | 0.14225397 |
| LISNASDALDK | ASSPRDSAETLY | 0.30227005 |
| LISNASDAL | ASSPRDSAETLY | 0.16979654 |
| ISNASDALDK | ASSPRDSAETLY | 0.13630046 |
| NQAETLVYQTE | ASSPRDSAETLY | 0.18559547 |
| TGGTGGTAGAG | ASSPRDSAETLY | 0.674504 |
| TGSTAGGAGGA | ASSPRDSAETLY | 0.47886258 |
| DPLAVPVAAGPV | ASSPRDSAETLY | 0.3797841 |
| GGTGGTPGNGGN | ASSPRDSAETLY | 0.7162727 |
| PAGLAGVDPATL | ASSPRDSAETLY | 0.5596087 |
| GNNLVVEAPHAGASV | ASSPRDSAETLY | 0.47063968 |
| KVTRIMSASTLEQPA | ASSPRDSAETLY | 0.31829628 |
| LGFALPIALPRRTVV | ASSPRDSAETLY | 0.32319432 |
| TRLGNLTARKLVVLV | ASSPRDSAETLY | 0.25097668 |
| WSQQRRLIYFPSAGP | ASSPRDSAETLY | 0.24692559 |
| GPNGNPDPMAAAVDIR | ASSPRDSAETLY | 0.757761 |
| TQRCVSVADLRAAASR | ASSPRDSAETLY | 0.4150021 |
| QFVLDVAEHPVEVLTETV | ASSPRDSAETLY | 0.221575 |
| GGAGGTGGAGGNGGTQAGDG | ASSPRDSAETLY | 0.5036683 |
| PLLATLRAIAVDVDAKPAQV | ASSPRDSAETLY | 0.7387912 |
| APKFRTGGYMIDFWGVGYQVAK | ASSPRDSAETLY | 0.790282 |
| GAGGAGGAGGAAGGPGGTGGTG | ASSPRDSAETLY | 0.32285452 |
| IGALGLLATDVAPAGSVAFGVA | ASSPRDSAETLY | 0.35276976 |
| AGGAGGIGGNGGAGGAGGVLMG | ASSPRDSAETLY | 0.41939628 |
| ADELIGLL | ASSPRDSAETLY | 0.5240243 |

| | | |
|---|---|---|
| ADRALKA | ASSPRDSAETLY | 0.3206962 |
| AGAVGQVGGAGGA | ASSPRDSAETLY | 0.25901383 |
| AGGGGGLGGAP | ASSPRDSAETLY | 0.5832764 |
| AGGLLYGNGGAG | ASSPRDSAETLY | 0.7764962 |
| AGLESIGL | ASSPRDSAETLY | 0.32468188 |
| AGLKTGII | ASSPRDSAETLY | 0.18454222 |
| ASLAALLL | ASSPRDSAETLY | 0.42819744 |
| ASVINISSI | ASSPRDSAETLY | 0.2559214 |
| ATPPGPA | ASSPRDSAETLY | 0.3964314 |
| ATVASALV | ASSPRDSAETLY | 0.47986603 |
| ATVASLIL | ASSPRDSAETLY | 0.21381797 |
| AVNGPVLIRH | ASSPRDSAETLY | 0.31714898 |
| DGAAGAVGI | ASSPRDSAETLY | 0.7578348 |
| DPLLHGI | ASSPRDSAETLY | 0.3835305 |
| ELGAALGAL | ASSPRDSAETLY | 0.5781672 |
| ELGRTQAV | ASSPRDSAETLY | 0.7359216 |
| FVAGGYLS | ASSPRDSAETLY | 0.676514 |
| GASGGAGGAGGW | ASSPRDSAETLY | 0.7900132 |
| GGAGAGGDGGAGGI | ASSPRDSAETLY | 0.6073352 |
| GTGHNGSQPGMGGNG | ASSPRDSAETLY | 0.46013892 |
| GYAAASASAS | ASSPRDSAETLY | 0.4366126 |
| HQVDVAAVQQG | ASSPRDSAETLY | 0.55803484 |
| IGAAIAHT | ASSPRDSAETLY | 0.575405 |
| IGMIEVAQL | ASSPRDSAETLY | 0.45187986 |
| IRLAPEHTSAMD | ASSPRDSAETLY | 0.3341119 |
| IYNVSVL | ASSPRDSAETLY | 0.69699734 |
| KQGFLGM | ASSPRDSAETLY | 0.3512713 |
| LESVITQL | ASSPRDSAETLY | 0.5407078 |
| LGLGALVVV | ASSPRDSAETLY | 0.22321269 |
| LIGNGGNGGSGGI | ASSPRDSAETLY | 0.23975688 |
| LNAINAPTL | ASSPRDSAETLY | 0.7225472 |
| LQLLNTGV | ASSPRDSAETLY | 0.13249102 |
| LQTSIEF | ASSPRDSAETLY | 0.2059939 |
| MSFAVSEL | ASSPRDSAETLY | 0.4805271 |
| NTGALGL | ASSPRDSAETLY | 0.7146765 |
| PPGAGGAGGAGTT | ASSPRDSAETLY | 0.11253237 |
| QLVGLGI | ASSPRDSAETLY | 0.26179403 |
| RFLEQVHQL | ASSPRDSAETLY | 0.29720753 |
| RGVGYVLR | ASSPRDSAETLY | 0.5169441 |
| SATGVFAGVI | ASSPRDSAETLY | 0.44106892 |
| SLPAITLP | ASSPRDSAETLY | 0.18389328 |
| STALAFTV | ASSPRDSAETLY | 0.6587821 |
| SVLTVGLGK | ASSPRDSAETLY | 0.17280512 |
| TALAGAGSGI | ASSPRDSAETLY | 0.17725764 |
| TGLPFTGI | ASSPRDSAETLY | 0.5890441 |
| VAEHVTV | ASSPRDSAETLY | 0.7059058 |
| VDIVLNSL | ASSPRDSAETLY | 0.2516996 |
| VLTGGGAVL | ASSPRDSAETLY | 0.7110977 |
| VNVGLAYGI | ASSPRDSAETLY | 0.30813235 |
| VSSAAVGAV | ASSPRDSAETLY | 0.7837375 |
| VTSVYEAGV | ASSPRDSAETLY | 0.39064747 |

| | | |
|---|---|---|
| WVALDRGA | ASSPRDSAETLY | 0.22144738 |
| YLVGSVVI | ASSPRDSAETLY | 0.63598675 |
| GPPGTGKTLL | ASSPRDSAETLY | 0.66151845 |
| GVGMTKF | ASSPRDSAETLY | 0.53776634 |
| IGIGGSDL | ASSPRDSAETLY | 0.5573722 |
| IGTLLAEL | ASSPRDSAETLY | 0.37536213 |
| IYTSGTT | ASSPRDSAETLY | 0.49349374 |
| LSFPAGV | ASSPRDSAETLY | 0.74649453 |
| LYTSGTT | ASSPRDSAETLY | 0.66847944 |
| VGAFPFGF | ASSPRDSAETLY | 0.23963186 |
| VGAGTVEF | ASSPRDSAETLY | 0.62747175 |
| VGKSTLLL | ASSPRDSAETLY | 0.4094763 |
| AALAHAGGGIV | ASSPRDSAETLY | 0.39156473 |
| AALVAVSLL | ASSPRDSAETLY | 0.6474461 |
| AAVSRAVAAL | ASSPRDSAETLY | 0.5214921 |
| AGGSSGIGGF | ASSPRDSAETLY | 0.68125355 |
| AGLTVANANG | ASSPRDSAETLY | 0.49636218 |
| AGLVGAVL | ASSPRDSAETLY | 0.7561811 |
| ALSQAFTV | ASSPRDSAETLY | 0.2064566 |
| ALTFSVSL | ASSPRDSAETLY | 0.4887538 |
| ALTGAALVAL | ASSPRDSAETLY | 0.4810487 |
| ASFSIGAGVL | ASSPRDSAETLY | 0.7293401 |
| DGSVYKI | ASSPRDSAETLY | 0.37917358 |
| EQLAEIV | ASSPRDSAETLY | 0.21162048 |
| FAVSELTVM | ASSPRDSAETLY | 0.53851503 |
| GAAPLAAGL | ASSPRDSAETLY | 0.69067395 |
| GADGGDGGSGGAGG | ASSPRDSAETLY | 0.4041443 |
| GFGLDTGGAGG | ASSPRDSAETLY | 0.56185555 |
| GLASGGLL | ASSPRDSAETLY | 0.80947673 |
| GRIAGAAVLL | ASSPRDSAETLY | 0.36700463 |
| GSAGSLNTSF | ASSPRDSAETLY | 0.44130662 |
| GTGQIGI | ASSPRDSAETLY | 0.5198321 |
| GTLIALIL | ASSPRDSAETLY | 0.7356268 |
| IAATMGFVVG | ASSPRDSAETLY | 0.4596905 |
| IFAQALL | ASSPRDSAETLY | 0.71579796 |
| IGAAVALGF | ASSPRDSAETLY | 0.22018357 |
| IGLTPLAL | ASSPRDSAETLY | 0.6262517 |
| ILGSNTSTL | ASSPRDSAETLY | 0.21543825 |
| IVFSLTVL | ASSPRDSAETLY | 0.6053482 |
| LASKATGF | ASSPRDSAETLY | 0.20370024 |
| LASVALIL | ASSPRDSAETLY | 0.6048962 |
| LFAVGLTL | ASSPRDSAETLY | 0.6762178 |
| LFNGGIGTYI | ASSPRDSAETLY | 0.6794665 |
| LTLGAGII | ASSPRDSAETLY | 0.13199918 |
| LVAAALGDI | ASSPRDSAETLY | 0.7082034 |
| NTAFVQLGI | ASSPRDSAETLY | 0.68242157 |
| QLFSGAEL | ASSPRDSAETLY | 0.80709076 |
| SAATFITL | ASSPRDSAETLY | 0.45123577 |
| SNYGLANLGV | ASSPRDSAETLY | 0.6456947 |
| SYGGAYAVM | ASSPRDSAETLY | 0.3587109 |
| TTGVGGLTLG | ASSPRDSAETLY | 0.4278582 |

| | | |
|---|---|---|
| TTQTIYI | ASSPRDSAETLY | 0.24730304 |
| VAAGAGLGAVI | ASSPRDSAETLY | 0.45074117 |
| VAAVVTLL | ASSPRDSAETLY | 0.4531963 |
| VGFLGGLI | ASSPRDSAETLY | 0.3711513 |
| VGVIPILL | ASSPRDSAETLY | 0.43147743 |
| VIFASSLI | ASSPRDSAETLY | 0.7443629 |
| VKGGDGGAAAG | ASSPRDSAETLY | 0.6288641 |
| VTKTFTNI | ASSPRDSAETLY | 0.7258552 |
| VVAAAILL | ASSPRDSAETLY | 0.75987244 |
| YAALDVEL | ASSPRDSAETLY | 0.6630632 |
| LASVALLL | ASSPRDSAETLY | 0.27752006 |
| LDAVHRV | ASSPRDSAETLY | 0.35840577 |
| QGLSSSIF | ASSPRDSAETLY | 0.24391833 |
| RGTFANIRL | ASSPRDSAETLY | 0.33769184 |
| RQLWWGHRI | ASSPRDSAETLY | 0.28253475 |
| YGPPGTGKTLL | ASSPRDSAETLY | 0.57451755 |
| AALVGAPANVA | ASSPRDSAETLY | 0.11353987 |
| AATFRRTE | ASSPRDSAETLY | 0.4880338 |
| AGALGGGI | ASSPRDSAETLY | 0.7552589 |
| AGGAGGAGGIGAG | ASSPRDSAETLY | 0.15220383 |
| AGSGALGL | ASSPRDSAETLY | 0.5540012 |
| AQIADQI | ASSPRDSAETLY | 0.46533388 |
| ASAVAVTYL | ASSPRDSAETLY | 0.27728254 |
| ASTVLGGTGG | ASSPRDSAETLY | 0.6716914 |
| AVGGEPAK | ASSPRDSAETLY | 0.2783613 |
| CEQGTFGGTC | ASSPRDSAETLY | 0.70134073 |
| DEGKAPAG | ASSPRDSAETLY | 0.6438951 |
| DTVVFPAL | ASSPRDSAETLY | 0.5076512 |
| FEGITMAG | ASSPRDSAETLY | 0.54700273 |
| FPLVVQD | ASSPRDSAETLY | 0.4796832 |
| GAFTTPQ | ASSPRDSAETLY | 0.20022207 |
| GFGGFGTAGGVGGA | ASSPRDSAETLY | 0.6258124 |
| GFLAQFL | ASSPRDSAETLY | 0.11938883 |
| GGAGGAGGNAGVA | ASSPRDSAETLY | 0.5489132 |
| GGVGTAGAGGAGGAGG | ASSPRDSAETLY | 0.24808101 |
| GNGGGGDAGNAGSGGN | ASSPRDSAETLY | 0.6485723 |
| GTGGAGGAGNDAGS | ASSPRDSAETLY | 0.46006268 |
| GTTNTGWFNS | ASSPRDSAETLY | 0.524498 |
| IFMGPLFTL | ASSPRDSAETLY | 0.29779547 |
| ILAHAIEAI | ASSPRDSAETLY | 0.32822487 |
| IRSAIREAL | ASSPRDSAETLY | 0.20260149 |
| ITFAGGG | ASSPRDSAETLY | 0.20786211 |
| ITGMSGHQEV | ASSPRDSAETLY | 0.56822336 |
| IVAGLACTL | ASSPRDSAETLY | 0.37739363 |
| IVGGGLAGA | ASSPRDSAETLY | 0.40969265 |
| IVTGVLNI | ASSPRDSAETLY | 0.62220937 |
| LAQQGCI | ASSPRDSAETLY | 0.55679244 |
| LCLHVEN | ASSPRDSAETLY | 0.18980828 |
| LDAISDGS | ASSPRDSAETLY | 0.43167925 |
| LGGFAAML | ASSPRDSAETLY | 0.51718 |
| LGGKNPMI | ASSPRDSAETLY | 0.4363231 |

| | | |
|---|---|---|
| LGPKGVAAIL | ASSPRDSAETLY | 0.10809124 |
| LMVGVSI | ASSPRDSAETLY | 0.5219705 |
| LVGAAFVL | ASSPRDSAETLY | 0.3814654 |
| LVLAVAPL | ASSPRDSAETLY | 0.28710812 |
| MAAIAVIDAA | ASSPRDSAETLY | 0.31314313 |
| NNGINAELQ | ASSPRDSAETLY | 0.76474226 |
| RGEPLVLVHG | ASSPRDSAETLY | 0.19711661 |
| RGGAGGAAGLI | ASSPRDSAETLY | 0.45358095 |
| RSYRMHPAV | ASSPRDSAETLY | 0.70511055 |
| SAAHLLAGQLA | ASSPRDSAETLY | 0.549 |
| SAAVGSY | ASSPRDSAETLY | 0.76606566 |
| SPHAVLHRI | ASSPRDSAETLY | 0.55411434 |
| SQIAKVL | ASSPRDSAETLY | 0.31443927 |
| TALGHLVP | ASSPRDSAETLY | 0.620544 |
| TAPTTSGAA | ASSPRDSAETLY | 0.3880729 |
| TDGDDGNGGSPGT | ASSPRDSAETLY | 0.20426773 |
| TDGLGEN | ASSPRDSAETLY | 0.14644617 |
| TGGPPLPGQA | ASSPRDSAETLY | 0.54683876 |
| TIPAITI | ASSPRDSAETLY | 0.20302553 |
| TIPGFTI | ASSPRDSAETLY | 0.5423956 |
| TSIASGALA | ASSPRDSAETLY | 0.69438624 |
| TVTLPGAAI | ASSPRDSAETLY | 0.31309208 |
| VASGLSAALP | ASSPRDSAETLY | 0.257312 |
| VGAGVIGI | ASSPRDSAETLY | 0.32147604 |
| VGPKRFGV | ASSPRDSAETLY | 0.32596523 |
| VNVAFPD | ASSPRDSAETLY | 0.74159366 |
| VSMTNSGVS | ASSPRDSAETLY | 0.37080994 |
| VTDVLGDLSV | ASSPRDSAETLY | 0.7329129 |
| VVVVGAGFAGL | ASSPRDSAETLY | 0.6511234 |
| VYTVGTVV | ASSPRDSAETLY | 0.3279416 |
| WTTFSGP | ASSPRDSAETLY | 0.71490324 |
| YDELVEAF | ASSPRDSAETLY | 0.40557134 |
| YGTGGAGGNGG | ASSPRDSAETLY | 0.3110981 |
| YTNVMAV | ASSPRDSAETLY | 0.46374893 |
| FLEQVHQL | ASSPRDSAETLY | 0.45831215 |
| GLSLLAVAV | ASSPRDSAETLY | 0.35361356 |
| LEQVHQL | ASSPRDSAETLY | 0.4211377 |
| LNAAGAI | ASSPRDSAETLY | 0.2109632 |
| PAVHASL | ASSPRDSAETLY | 0.42849168 |
| TELEDDA | ASSPRDSAETLY | 0.17929351 |
| TPPGPPN | ASSPRDSAETLY | 0.2010133 |
| TPPPPPPP | ASSPRDSAETLY | 0.3450895 |
| TQSPALI | ASSQDQQTGPYEQY | 0.80837524 |
| EFAAAFPQ | ASSQDQQTGPYEQY | 0.7722448 |
| LLSGNLDVL | ASSQDQQTGPYEQY | 0.31557032 |
| GPANLGNYNI | ASSQDQQTGPYEQY | 0.30969173 |
| TLGPTLP | ASSQDQQTGPYEQY | 0.47346258 |
| VYAYSSCCK | ASSQDQQTGPYEQY | 0.20733641 |
| APAPAPAPNI | ASSQDQQTGPYEQY | 0.5308716 |
| PGGPVPIP | ASSQDQQTGPYEQY | 0.5234317 |
| FLTVALEP | ASSQDQQTGPYEQY | 0.4671407 |

| | | |
|---|---|---|
| SQYFGANL | ASSQDQQTGPYEQY | 0.3516022 |
| GGVKPAA | ASSQDQQTGPYEQY | 0.47230175 |
| GGTGGAGGAGGW | ASSQDQQTGPYEQY | 0.3127376 |
| GVGGVAGLGGAGG | ASSQDQQTGPYEQY | 0.64796597 |
| VLLVLATTI | ASSQDQQTGPYEQY | 0.110034026 |
| IGIFAAML | ASSQDQQTGPYEQY | 0.36853164 |
| AVGDLID | ASSQDQQTGPYEQY | 0.26665062 |
| NISPLPL | ASSQDQQTGPYEQY | 0.28623578 |
| VTAVFVAI | ASSQDQQTGPYEQY | 0.17857943 |
| SGDDNDAAL | ASSQDQQTGPYEQY | 0.41323805 |
| AVAALAHS | ASSQDQQTGPYEQY | 0.6605126 |
| GGVTVMATP | ASSQDQQTGPYEQY | 0.6947036 |
| AGLAANVRD | ASSQDQQTGPYEQY | 0.23823887 |
| TNRPDLIDPALLRPG | ASSQDQQTGPYEQY | 0.66886616 |
| LANLGVTAL | ASSQDQQTGPYEQY | 0.7440059 |
| TGERIEL | ASSQDQQTGPYEQY | 0.39416778 |
| NGEELAA | ASSQDQQTGPYEQY | 0.1994189 |
| VLSLVWG | ASSQDQQTGPYEQY | 0.5872733 |
| IGYALWL | ASSQDQQTGPYEQY | 0.29038405 |
| LSLLLAPL | ASSQDQQTGPYEQY | 0.6371963 |
| TLDEALK | ASSQDQQTGPYEQY | 0.6904576 |
| TDGAGTAEGG | ASSQDQQTGPYEQY | 0.26024234 |
| NAPGGAGGNGG | ASSQDQQTGPYEQY | 0.72930586 |
| VSVFMGMGGNF | ASSQDQQTGPYEQY | 0.6958121 |
| LASYLAAL | ASSQDQQTGPYEQY | 0.24441397 |
| LVGGTPAL | ASSQDQQTGPYEQY | 0.27710405 |
| VRNADVL | ASSQDQQTGPYEQY | 0.28401232 |
| TCSSPFLVSAM | ASSQDQQTGPYEQY | 0.14398378 |
| VHITDAE | ASSQDQQTGPYEQY | 0.51647544 |
| EYAAPSF | ASSQDQQTGPYEQY | 0.3076381 |
| VVDAFHGGL | ASSQDQQTGPYEQY | 0.2769091 |
| ASVVDPQ | ASSQDQQTGPYEQY | 0.5638416 |
| SYNEFGGLS | ASSQDQQTGPYEQY | 0.27185315 |
| DPDGSVY | ASSQDQQTGPYEQY | 0.36633947 |
| MVSTIHATE | ASSQDQQTGPYEQY | 0.30919403 |
| LAAGVADPV | ASSQDQQTGPYEQY | 0.28388798 |
| TKEQIAA | ASSQDQQTGPYEQY | 0.51954895 |
| SLASPAAL | ASSQDQQTGPYEQY | 0.32454735 |
| IGLVFYGV | ASSQDQQTGPYEQY | 0.46964216 |
| FLLDASNL | ASSQDQQTGPYEQY | 0.76323044 |
| YQVGNFP | ASSQDQQTGPYEQY | 0.17885664 |
| ESLLSQST | ASSQDQQTGPYEQY | 0.28488562 |
| LFTVGFAAVG | ASSQDQQTGPYEQY | 0.59285784 |
| ATPPTPT | ASSQDQQTGPYEQY | 0.15888889 |
| TDLATLG | ASSQDQQTGPYEQY | 0.703392 |
| SAGGTGR | ASSQDQQTGPYEQY | 0.37869823 |
| GFSNSGS | ASSQDQQTGPYEQY | 0.5443355 |
| GGNGGSGGTSV | ASSQDQQTGPYEQY | 0.23612629 |
| GTSVATGGAGN | ASSQDQQTGPYEQY | 0.7754066 |
| NGGNGGSGGTS | ASSQDQQTGPYEQY | 0.6969252 |
| AVVKFPH | ASSQDQQTGPYEQY | 0.35222656 |

| | | |
|---|---|---|
| TPPPPPP | ASSQDQQTGPYEQY | 0.18062456 |
| AAPAAAAAPA | ASSQDQQTGPYEQY | 0.29430243 |
| IALNEDL | ASSQDQQTGPYEQY | 0.63286626 |
| IVNVSSM | ASSQDQQTGPYEQY | 0.6773101 |
| TGIPPAPR | ASSQDQQTGPYEQY | 0.300931 |
| FDLGGGTF | ASSQDQQTGPYEQY | 0.48821992 |
| FDLGGGTFDVS | ASSQDQQTGPYEQY | 0.6653276 |
| NEPTAAA | ASSQDQQTGPYEQY | 0.7547216 |
| NEPTAAALA | ASSQDQQTGPYEQY | 0.20286097 |
| GSIINISSI | ASSQDQQTGPYEQY | 0.44424424 |
| VAPGNAGTA | ASSQDQQTGPYEQY | 0.2024918 |
| TGGGASLE | ASSQDQQTGPYEQY | 0.32513058 |
| VAVYNHYKRI | ASSQDQQTGPYEQY | 0.29476574 |
| TDPAPATT | ASSQDQQTGPYEQY | 0.59499943 |
| TNRPDILDPALLRPG | ASSQDQQTGPYEQY | 0.4289149 |
| LTTTTTS | ASSQDQQTGPYEQY | 0.6050977 |
| PTGSGKTL | ASSQDQQTGPYEQY | 0.5554129 |
| TGSGKTL | ASSQDQQTGPYEQY | 0.660004 |
| TIGHVDHGK | ASSQDQQTGPYEQY | 0.35397893 |
| TIGHVDHGKTT | ASSQDQQTGPYEQY | 0.4103267 |
| GHVDHGKTT | ASSQDQQTGPYEQY | 0.2344192 |
| AQFEHTLL | ASSQDQQTGPYEQY | 0.13823599 |
| IVNQDEL | ASSQDQQTGPYEQY | 0.26744354 |
| TNGSQFF | ASSQDQQTGPYEQY | 0.28165305 |
| TGKGIVE | ASSQDQQTGPYEQY | 0.11365866 |
| LISNASDA | ASSQDQQTGPYEQY | 0.6879946 |
| AELRTLGELTKIAKAHGAQ | ASSQDQQTGPYEQY | 0.65360385 |
| APNAVAAAEPSW | ASSQDQQTGPYEQY | 0.5317658 |
| APQVAGAATAVL | ASSQDQQTGPYEQY | 0.3045963 |
| ARRDPAAAAAILPTDA | ASSQDQQTGPYEQY | 0.34743738 |
| DAAARDVLAEAGLAEYF | ASSQDQQTGPYEQY | 0.2740728 |
| DDQIVLSAKANLAKVEQ | ASSQDQQTGPYEQY | 0.72998476 |
| DELLPDTGEDSDEDGH | ASSQDQQTGPYEQY | 0.26324034 |
| ELPEAPANRATVPQD | ASSQDQQTGPYEQY | 0.17941085 |
| FGQSKIERVTGIG | ASSQDQQTGPYEQY | 0.29177046 |
| GAGGTGSTAGGAGGAGGAGGLY | ASSQDQQTGPYEQY | 0.19453432 |
| GHGGTGAAGGNGGTGGAGGSGI | ASSQDQQTGPYEQY | 0.730482 |
| GNGGNGGAGGEGGPGGAGGAGA | ASSQDQQTGPYEQY | 0.62232476 |
| GSTAGNGGNGGNATNGTIAGQP | ASSQDQQTGPYEQY | 0.6360838 |
| GTGGDGGNGGNGGTGG | ASSQDQQTGPYEQY | 0.5048287 |
| GVRRPAGIPADAPV | ASSQDQQTGPYEQY | 0.17608689 |
| GYRQVTDAHLLTLARR | ASSQDQQTGPYEQY | 0.31079873 |
| IAKMGQDGHDRG | ASSQDQQTGPYEQY | 0.8018373 |
| KAEPSIEIQYT | ASSQDQQTGPYEQY | 0.45094723 |
| KPVRIRMAADEDTVAVT | ASSQDQQTGPYEQY | 0.46163213 |
| MPPRIAGMRLLVIKPEP | ASSQDQQTGPYEQY | 0.7510962 |
| MKFYAVALFRSRGVPPTR | ASSQDQQTGPYEQY | 0.25746503 |
| NAGGFGILGAVAHSP | ASSQDQQTGPYEQY | 0.45075926 |
| NEGREITLAMAEM | ASSQDQQTGPYEQY | 0.4753055 |
| NELVEWSHPLGHRWRWE | ASSQDQQTGPYEQY | 0.582998 |
| NLGAGGTGGNGGNPG | ASSQDQQTGPYEQY | 0.3918339 |

| | | |
|---|---|---|
| NTEWPKWETVMAADPVD | ASSQDQQTGPYEQY | 0.2641012 |
| PASAAPAETQP | ASSQDQQTGPYEQY | 0.6259505 |
| QQSSDGKNGVRAAEITGE | ASSQDQQTGPYEQY | 0.7803805 |
| RAPDAVTTSWWKE | ASSQDQQTGPYEQY | 0.5367291 |
| SGFGNVGAHTSGWFNQ | ASSQDQQTGPYEQY | 0.6461858 |
| SGVAGAVGQAASVGGLKV | ASSQDQQTGPYEQY | 0.6272952 |
| SNGGAGTGSAGGNGGT | ASSQDQQTGPYEQY | 0.25724518 |
| TGGAGGEGGAAG | ASSQDQQTGPYEQY | 0.35193878 |
| TGGDGGAAGAGGAPGHG | ASSQDQQTGPYEQY | 0.37154397 |
| TMPPLLAAIANAGRHGV | ASSQDQQTGPYEQY | 0.3098805 |
| TSSTAGNSGGA | ASSQDQQTGPYEQY | 0.28768277 |
| VPATVPVGILC | ASSQDQQTGPYEQY | 0.5487199 |
| VQRPDSGGVVLTGRIS | ASSQDQQTGPYEQY | 0.36595023 |
| WTPNPVPATKGVPAGLVQAT | ASSQDQQTGPYEQY | 0.8082925 |
| YALKRALAKFSDRFDVVII | ASSQDQQTGPYEQY | 0.6343819 |
| YLIGSVIV | ASSQDQQTGPYEQY | 0.6068891 |
| YGQLVAVID | ASSQDQQTGPYEQY | 0.24215241 |
| VTVGGMTLPA | ASSQDQQTGPYEQY | 0.4205267 |
| VLALPAV | ASSQDQQTGPYEQY | 0.5293647 |
| VIPPTLNYET | ASSQDQQTGPYEQY | 0.2240519 |
| VGMGGMHPSSG | ASSQDQQTGPYEQY | 0.74700654 |
| VGLAGGAATA | ASSQDQQTGPYEQY | 0.30380192 |
| VGGAGGAGGSGP | ASSQDQQTGPYEQY | 0.33844337 |
| VFAAFMV | ASSQDQQTGPYEQY | 0.59207726 |
| TLYDKLR | ASSQDQQTGPYEQY | 0.71256864 |
| TLAVFTVL | ASSQDQQTGPYEQY | 0.41021734 |
| TGSVQPARV | ASSQDQQTGPYEQY | 0.27456135 |
| TGGGAGSGGT | ASSQDQQTGPYEQY | 0.28250635 |
| SHLETSL | ASSQDQQTGPYEQY | 0.615126 |
| SFTTGMT | ASSQDQQTGPYEQY | 0.77397 |
| QVEVAKRMGVRQPTVS | ASSQDQQTGPYEQY | 0.19155744 |
| QGSGGPALAM | ASSQDQQTGPYEQY | 0.7534512 |
| MLAPSNP | ASSQDQQTGPYEQY | 0.43148163 |
| LVNNAAI | ASSQDQQTGPYEQY | 0.26782998 |
| LTGEINTLA | ASSQDQQTGPYEQY | 0.55500704 |
| LSIFGPV | ASSQDQQTGPYEQY | 0.563178 |
| LLALLANL | ASSQDQQTGPYEQY | 0.6770853 |
| LISNASDALDKLR | ASSQDQQTGPYEQY | 0.5530592 |
| LINEAVP | ASSQDQQTGPYEQY | 0.47714624 |
| LGYAFSGL | ASSQDQQTGPYEQY | 0.31374335 |
| LDFVAVAL | ASSQDQQTGPYEQY | 0.7371697 |
| LAQPGAE | ASSQDQQTGPYEQY | 0.70847285 |
| ISNLGAVTGF | ASSQDQQTGPYEQY | 0.7114799 |
| HQEDPGRI | ASSQDQQTGPYEQY | 0.3736473 |
| GTPGEPTP | ASSQDQQTGPYEQY | 0.31798768 |
| GFFNSGS | ASSQDQQTGPYEQY | 0.41997498 |
| GAGGGGLTAGAN | ASSQDQQTGPYEQY | 0.6862701 |
| FVDALVNM | ASSQDQQTGPYEQY | 0.3345641 |
| FLTVAPEL | ASSQDQQTGPYEQY | 0.53408754 |
| FETLTAL | ASSQDQQTGPYEQY | 0.4423021 |
| DVINPIEL | ASSQDQQTGPYEQY | 0.22838923 |

| | | |
|---|---|---|
| APVSGAPV | ASSQDQQTGPYEQY | 0.39012533 |
| ALVGGLAW | ASSQDQQTGPYEQY | 0.71390396 |
| ALAAAAATAI | ASSQDQQTGPYEQY | 0.4410702 |
| AFGLGLLL | ASSQDQQTGPYEQY | 0.33345675 |
| AFFAAASL | ASSQDQQTGPYEQY | 0.7277423 |
| AAAALPAFSPP | ASSQDQQTGPYEQY | 0.8031546 |
| VVLAPAL | ASSQDQQTGPYEQY | 0.72832453 |
| TIGHVDHGKTTL | ASSQDQQTGPYEQY | 0.20923918 |
| LISNASDALDK | ASSQDQQTGPYEQY | 0.1482339 |
| LISNASDAL | ASSQDQQTGPYEQY | 0.40425923 |
| ISNASDALDK | ASSQDQQTGPYEQY | 0.36099893 |
| NQAETLVYQTE | ASSQDQQTGPYEQY | 0.28402847 |
| TGGTGGTAGAG | ASSQDQQTGPYEQY | 0.33068883 |
| TGSTAGGAGGA | ASSQDQQTGPYEQY | 0.57298124 |
| DPLAVPVAAGPV | ASSQDQQTGPYEQY | 0.6828824 |
| GGTGGTPGNGGN | ASSQDQQTGPYEQY | 0.3512599 |
| PAGLAGVDPATL | ASSQDQQTGPYEQY | 0.21184829 |
| GNNLVVEAPHAGASV | ASSQDQQTGPYEQY | 0.39177227 |
| KVTRIMSASTLEQPA | ASSQDQQTGPYEQY | 0.7180395 |
| LGFALPIALPRRTVV | ASSQDQQTGPYEQY | 0.30183977 |
| TRLGNLTARKLVVLV | ASSQDQQTGPYEQY | 0.49372768 |
| WSQQRRLIYFPSAGP | ASSQDQQTGPYEQY | 0.3498017 |
| GPNGNPDPMAAAVDIR | ASSQDQQTGPYEQY | 0.2699178 |
| TQRCVSVADLRAAASR | ASSQDQQTGPYEQY | 0.49836683 |
| QFVLDVAEHPVEVLTETV | ASSQDQQTGPYEQY | 0.2870906 |
| GGAGGTGGAGGNGGTQAGDG | ASSQDQQTGPYEQY | 0.67941535 |
| PLLATLRAIAVDVDAKPAQV | ASSQDQQTGPYEQY | 0.26744375 |
| APKFRTGGYMIDFWGVGYQVAK | ASSQDQQTGPYEQY | 0.6108354 |
| GAGGAGGAGGAAGGPGGTGGTG | ASSQDQQTGPYEQY | 0.5477177 |
| IGALGLLATDVAPAGSVAFGVA | ASSQDQQTGPYEQY | 0.56940293 |
| AGGAGGIGGNGGAGGAGGVLMG | ASSQDQQTGPYEQY | 0.44556516 |
| ADELIGLL | ASSQDQQTGPYEQY | 0.34427208 |
| ADRALKA | ASSQDQQTGPYEQY | 0.22718006 |
| AGAVGQVGGAGGA | ASSQDQQTGPYEQY | 0.13150278 |
| AGGGGGLGGAP | ASSQDQQTGPYEQY | 0.73715425 |
| AGGLLYGNGGAG | ASSQDQQTGPYEQY | 0.121334635 |
| AGLESIGL | ASSQDQQTGPYEQY | 0.7418975 |
| AGLKTGII | ASSQDQQTGPYEQY | 0.5272185 |
| ASLAALLL | ASSQDQQTGPYEQY | 0.29465222 |
| ASVINISSI | ASSQDQQTGPYEQY | 0.5328374 |
| ATPPGPA | ASSQDQQTGPYEQY | 0.22556627 |
| ATVASALV | ASSQDQQTGPYEQY | 0.3393007 |
| ATVASLIL | ASSQDQQTGPYEQY | 0.6256497 |
| AVNGPVLIRH | ASSQDQQTGPYEQY | 0.6382313 |
| DGAAGAVGI | ASSQDQQTGPYEQY | 0.322056 |
| DPLLHGI | ASSQDQQTGPYEQY | 0.5502967 |
| ELGAALGAL | ASSQDQQTGPYEQY | 0.5590936 |
| ELGRTQAV | ASSQDQQTGPYEQY | 0.30077106 |
| FVAGGYLS | ASSQDQQTGPYEQY | 0.4462856 |
| GASGGAGGAGGW | ASSQDQQTGPYEQY | 0.71312857 |
| GGAGAGGDGGAGGI | ASSQDQQTGPYEQY | 0.5137149 |

| | | |
|---|---|---|
| GTGHNGSQPGMGGNG | ASSQDQQTGPYEQY | 0.30523524 |
| GYAAASASAS | ASSQDQQTGPYEQY | 0.54236937 |
| HQVDVAAVQQG | ASSQDQQTGPYEQY | 0.3677189 |
| IGAAIAHT | ASSQDQQTGPYEQY | 0.6196544 |
| IGMIEVAQL | ASSQDQQTGPYEQY | 0.5117241 |
| IRLAPEHTSAMD | ASSQDQQTGPYEQY | 0.4812875 |
| IYNVSVL | ASSQDQQTGPYEQY | 0.65003806 |
| KQGFLGM | ASSQDQQTGPYEQY | 0.20413013 |
| LESVITQL | ASSQDQQTGPYEQY | 0.25475863 |
| LGLGALVVV | ASSQDQQTGPYEQY | 0.45120338 |
| LIGNGGNGGSGGI | ASSQDQQTGPYEQY | 0.48100787 |
| LNAINAPTL | ASSQDQQTGPYEQY | 0.70822686 |
| LQLLNTGV | ASSQDQQTGPYEQY | 0.24654481 |
| LQTSIEF | ASSQDQQTGPYEQY | 0.5334699 |
| MSFAVSEL | ASSQDQQTGPYEQY | 0.4028916 |
| NTGALGL | ASSQDQQTGPYEQY | 0.31080186 |
| PPGAGGAGGAGTT | ASSQDQQTGPYEQY | 0.48690522 |
| QLVGLGI | ASSQDQQTGPYEQY | 0.3094699 |
| RFLEQVHQL | ASSQDQQTGPYEQY | 0.37453702 |
| RGVGYVLR | ASSQDQQTGPYEQY | 0.6341959 |
| SATGVFAGVI | ASSQDQQTGPYEQY | 0.28814498 |
| SLPAITLP | ASSQDQQTGPYEQY | 0.440541 |
| STALAFTV | ASSQDQQTGPYEQY | 0.3260296 |
| SVLTVGLGK | ASSQDQQTGPYEQY | 0.35688525 |
| TALAGAGSGI | ASSQDQQTGPYEQY | 0.5450107 |
| TGLPFTGI | ASSQDQQTGPYEQY | 0.5957938 |
| VAEHVTV | ASSQDQQTGPYEQY | 0.64248383 |
| VDIVLNSL | ASSQDQQTGPYEQY | 0.2796153 |
| VLTGGGAVL | ASSQDQQTGPYEQY | 0.27680275 |
| VNVGLAYGI | ASSQDQQTGPYEQY | 0.62356573 |
| VSSAAVGAV | ASSQDQQTGPYEQY | 0.6106514 |
| VTSVYEAGV | ASSQDQQTGPYEQY | 0.6587537 |
| WVALDRGA | ASSQDQQTGPYEQY | 0.57487106 |
| YLVGSVVI | ASSQDQQTGPYEQY | 0.61888653 |
| GPPGTGKTLL | ASSQDQQTGPYEQY | 0.719085 |
| GVGMTKF | ASSQDQQTGPYEQY | 0.41303065 |
| IGIGGSDL | ASSQDQQTGPYEQY | 0.2293885 |
| IGTLLAEL | ASSQDQQTGPYEQY | 0.3246691 |
| IYTSGTT | ASSQDQQTGPYEQY | 0.27528638 |
| LSFPAGV | ASSQDQQTGPYEQY | 0.3588025 |
| LYTSGTT | ASSQDQQTGPYEQY | 0.21537936 |
| VGAFPFGF | ASSQDQQTGPYEQY | 0.74886954 |
| VGAGTVEF | ASSQDQQTGPYEQY | 0.82778347 |
| VGKSTLLL | ASSQDQQTGPYEQY | 0.24427122 |
| AALAHAGGGIV | ASSQDQQTGPYEQY | 0.77311087 |
| AALVAVSLL | ASSQDQQTGPYEQY | 0.21193673 |
| AAVSRAVAAL | ASSQDQQTGPYEQY | 0.5341213 |
| AGGSSGIGGF | ASSQDQQTGPYEQY | 0.7371937 |
| AGLTVANANG | ASSQDQQTGPYEQY | 0.26167884 |
| AGLVGAVL | ASSQDQQTGPYEQY | 0.5041289 |
| ALSQAFTV | ASSQDQQTGPYEQY | 0.26166472 |

| | | |
|---|---|---|
| ALTFSVSL | ASSQDQQTGPYEQY | 0.3347142 |
| ALTGAALVAL | ASSQDQQTGPYEQY | 0.74232256 |
| ASFSIGAGVL | ASSQDQQTGPYEQY | 0.29008985 |
| DGSVYKI | ASSQDQQTGPYEQY | 0.58344257 |
| EQLAEIV | ASSQDQQTGPYEQY | 0.42479315 |
| FAVSELTVM | ASSQDQQTGPYEQY | 0.2695249 |
| GAAPLAAGL | ASSQDQQTGPYEQY | 0.5484153 |
| GADGGDGGSGGAGG | ASSQDQQTGPYEQY | 0.6940687 |
| GFGLDTGGAGG | ASSQDQQTGPYEQY | 0.24564081 |
| GLASGGLL | ASSQDQQTGPYEQY | 0.7010542 |
| GRIAGAAVLL | ASSQDQQTGPYEQY | 0.24572125 |
| GSAGSLNTSF | ASSQDQQTGPYEQY | 0.7503121 |
| GTGQIGI | ASSQDQQTGPYEQY | 0.6310433 |
| GTLIALIL | ASSQDQQTGPYEQY | 0.5396416 |
| IAATMGFVVG | ASSQDQQTGPYEQY | 0.37145704 |
| IFAQALL | ASSQDQQTGPYEQY | 0.71827686 |
| IGAAVALGF | ASSQDQQTGPYEQY | 0.56261164 |
| IGLTPLAL | ASSQDQQTGPYEQY | 0.6355722 |
| ILGSNTSTL | ASSQDQQTGPYEQY | 0.68147 |
| IVFSLTVL | ASSQDQQTGPYEQY | 0.09567432 |
| LASKATGF | ASSQDQQTGPYEQY | 0.098707765 |
| LASVALIL | ASSQDQQTGPYEQY | 0.09576149 |
| LFAVGLTL | ASSQDQQTGPYEQY | 0.097638026 |
| LFNGGIGTYI | ASSQDQQTGPYEQY | 0.1046859 |
| LTLGAGII | ASSQDQQTGPYEQY | 0.097149074 |
| LVAAALGDI | ASSQDQQTGPYEQY | 0.09757387 |
| NTAFVQLGI | ASSQDQQTGPYEQY | 0.10870031 |
| QLFSGAEL | ASSQDQQTGPYEQY | 0.09567632 |
| SAATFITL | ASSQDQQTGPYEQY | 0.09655931 |
| SNYGLANLGV | ASSQDQQTGPYEQY | 0.10522096 |
| SYGGAYAVM | ASSQDQQTGPYEQY | 0.10773867 |
| TTGVGGLTLG | ASSQDQQTGPYEQY | 0.09947997 |
| TTQTIYI | ASSQDQQTGPYEQY | 0.09656369 |
| VAAGAGLGAVI | ASSQDQQTGPYEQY | 0.09618631 |
| VAAVVTLL | ASSQDQQTGPYEQY | 0.0990841 |
| VGFLGGLI | ASSQDQQTGPYEQY | 0.09666225 |
| VGVIPILL | ASSQDQQTGPYEQY | 0.09548973 |
| VIFASSLI | ASSQDQQTGPYEQY | 0.09686372 |
| VKGGDGGAAAG | ASSQDQQTGPYEQY | 0.09535417 |
| VTKTFTNI | ASSQDQQTGPYEQY | 0.096636795 |
| VVAAAILL | ASSQDQQTGPYEQY | 0.09778119 |
| YAALDVEL | ASSQDQQTGPYEQY | 0.098763086 |
| LASVALLL | ASSQDQQTGPYEQY | 0.096682884 |
| LDAVHRV | ASSQDQQTGPYEQY | 0.09739238 |
| QGLSSSIF | ASSQDQQTGPYEQY | 0.10331635 |
| RGTFANIRL | ASSQDQQTGPYEQY | 0.101407215 |
| RQLWWGHRI | ASSQDQQTGPYEQY | 0.09974255 |
| YGPPGTGKTLL | ASSQDQQTGPYEQY | 0.10434806 |
| AALVGAPANVA | ASSQDQQTGPYEQY | 0.10134002 |
| AATFRRTE | ASSQDQQTGPYEQY | 0.09570403 |
| AGALGGGI | ASSQDQQTGPYEQY | 0.107401505 |

| | | |
|---|---|---|
| AGGAGGAGGIGAG | ASSQDQQTGPYEQY | 0.105131336 |
| AGSGALGL | ASSQDQQTGPYEQY | 0.09590516 |
| AQIADQI | ASSQDQQTGPYEQY | 0.09600047 |
| ASAVAVTYL | ASSQDQQTGPYEQY | 0.09561663 |
| ASTVLGGTGG | ASSQDQQTGPYEQY | 0.099865384 |
| AVGGEPAK | ASSQDQQTGPYEQY | 0.09816035 |
| CEQGTFGGTC | ASSQDQQTGPYEQY | 0.10152842 |
| DEGKAPAG | ASSQDQQTGPYEQY | 0.10582984 |
| DTVVFPAL | ASSQDQQTGPYEQY | 0.09575056 |
| FEGITMAG | ASSQDQQTGPYEQY | 0.095734924 |
| FPLVVQD | ASSQDQQTGPYEQY | 0.10057919 |
| GAFTTPQ | ASSQDQQTGPYEQY | 0.09765326 |
| GFGGFGTAGGVGGA | ASSQDQQTGPYEQY | 0.099746175 |
| GFLAQFL | ASSQDQQTGPYEQY | 0.105150625 |
| GGAGGAGGNAGVA | ASSQDQQTGPYEQY | 0.095567435 |
| GGVGTAGAGGAGGAGG | ASSQDQQTGPYEQY | 0.09773181 |
| GNGGGGDAGNAGSGGN | ASSQDQQTGPYEQY | 0.09509711 |
| GTGGAGGAGNDAGS | ASSQDQQTGPYEQY | 0.095544465 |
| GTTNTGWFNS | ASSQDQQTGPYEQY | 0.103539154 |
| IFMGPLFTL | ASSQDQQTGPYEQY | 0.095617324 |
| ILAHAIEAI | ASSQDQQTGPYEQY | 0.12680236 |
| IRSAIREAL | ASSQDQQTGPYEQY | 0.11508118 |
| ITFAGGG | ASSQDQQTGPYEQY | 0.10826506 |
| ITGMSGHQEV | ASSQDQQTGPYEQY | 0.09726346 |
| IVAGLACTL | ASSQDQQTGPYEQY | 0.0954005 |
| IVGGGLAGA | ASSQDQQTGPYEQY | 0.09721758 |
| IVTGVLNI | ASSQDQQTGPYEQY | 0.09936821 |
| LAQQGCI | ASSQDQQTGPYEQY | 0.09581688 |
| LCLHVEN | ASSQDQQTGPYEQY | 0.09855562 |
| LDAISDGS | ASSQDQQTGPYEQY | 0.10141031 |
| LGGFAAML | ASSQDQQTGPYEQY | 0.12277754 |
| LGGKNPMI | ASSQDQQTGPYEQY | 0.10010556 |
| LGPKGVAAIL | ASSQDQQTGPYEQY | 0.10195737 |
| LMVGVSI | ASSQDQQTGPYEQY | 0.10155955 |
| LVGAAFVL | ASSQDQQTGPYEQY | 0.095738605 |
| LVLAVAPL | ASSQDQQTGPYEQY | 0.0957154 |
| MAAIAVIDAA | ASSQDQQTGPYEQY | 0.09629624 |
| NNGINAELQ | ASSQDQQTGPYEQY | 0.09674473 |
| RGEPLVLVHG | ASSQDQQTGPYEQY | 0.104635 |
| RGGAGGAAGLI | ASSQDQQTGPYEQY | 0.10153651 |
| RSYRMHPAV | ASSQDQQTGPYEQY | 0.109993726 |
| SAAHLLAGQLA | ASSQDQQTGPYEQY | 0.09778288 |
| SAAVGSY | ASSQDQQTGPYEQY | 0.09576551 |
| SPHAVLHRI | ASSQDQQTGPYEQY | 0.09707448 |
| SQIAKVL | ASSQDQQTGPYEQY | 0.09834337 |
| TALGHLVP | ASSQDQQTGPYEQY | 0.09919029 |
| TAPTTSGAA | ASSQDQQTGPYEQY | 0.09604761 |
| TDGDDGNGGSPGT | ASSQDQQTGPYEQY | 0.09995169 |
| TDGLGEN | ASSQDQQTGPYEQY | 0.096360534 |
| TGGPPLPGQA | ASSQDQQTGPYEQY | 0.10566029 |
| TIPAITI | ASSQDQQTGPYEQY | 0.09618431 |

| | | |
|---|---|---|
| TIPGFTI | ASSQDQQTGPYEQY | 0.095896795 |
| TSIASGALA | ASSQDQQTGPYEQY | 0.09872422 |
| TVTLPGAAI | ASSQDQQTGPYEQY | 0.0964041 |
| VASGLSAALP | ASSQDQQTGPYEQY | 0.105069436 |
| VGAGVIGI | ASSQDQQTGPYEQY | 0.09778304 |
| VGPKRFGV | ASSQDQQTGPYEQY | 0.09574886 |
| VNVAFPD | ASSQDQQTGPYEQY | 0.09637117 |
| VSMTNSGVS | ASSQDQQTGPYEQY | 0.09571414 |
| VTDVLGDLSV | ASSQDQQTGPYEQY | 0.09652564 |
| VVVVGAGFAGL | ASSQDQQTGPYEQY | 0.096464336 |
| VYTVGTVV | ASSQDQQTGPYEQY | 0.095595814 |
| WTTFSGP | ASSQDQQTGPYEQY | 0.100111276 |
| YDELVEAF | ASSQDQQTGPYEQY | 0.099069215 |
| YGTGGAGGNGG | ASSQDQQTGPYEQY | 0.098522626 |
| YTNVMAV | ASSQDQQTGPYEQY | 0.09779934 |
| FLEQVHQL | ASSQDQQTGPYEQY | 0.09660932 |
| GLSLLAVAV | ASSQDQQTGPYEQY | 0.103792384 |
| LEQVHQL | ASSQDQQTGPYEQY | 0.095786326 |
| LNAAGAI | ASSQDQQTGPYEQY | 0.09693767 |
| PAVHASL | ASSQDQQTGPYEQY | 0.107556954 |
| TELEDDA | ASSQDQQTGPYEQY | 0.097277805 |
| TPPGPPN | ASSQDQQTGPYEQY | 0.10258964 |
| TPPPPPPP | ASSQDQQTGPYEQY | 0.09544577 |
| TQSPALI | ASSEGDNERLF | 0.09557422 |
| EFAAAFPQ | ASSEGDNERLF | 0.095371686 |
| LLSGNLDVL | ASSEGDNERLF | 0.11115638 |
| GPANLGNYNI | ASSEGDNERLF | 0.095847055 |
| TLGPTLP | ASSEGDNERLF | 0.09619878 |
| VYAYSSCCK | ASSEGDNERLF | 0.09719602 |
| APAPAPAPNI | ASSEGDNERLF | 0.09802957 |
| PGGPVPIP | ASSEGDNERLF | 0.09892294 |
| FLTVALEP | ASSEGDNERLF | 0.096132584 |
| SQYFGANL | ASSEGDNERLF | 0.095822856 |
| GGVKPAA | ASSEGDNERLF | 0.10696808 |
| GGTGGAGGAGGW | ASSEGDNERLF | 0.09575917 |
| GVGGVAGLGGAGG | ASSEGDNERLF | 0.09599602 |
| VLLVLATTI | ASSEGDNERLF | 0.09996002 |
| IGIFAAML | ASSEGDNERLF | 0.10692273 |
| AVGDLID | ASSEGDNERLF | 0.11118554 |
| NISPLPL | ASSEGDNERLF | 0.10156294 |
| VTAVFVAI | ASSEGDNERLF | 0.09751548 |
| SGDDNDAAL | ASSEGDNERLF | 0.095136434 |
| AVAALAHS | ASSEGDNERLF | 0.09552624 |
| GGVTVMATP | ASSEGDNERLF | 0.10626398 |
| AGLAANVRD | ASSEGDNERLF | 0.11357699 |
| TNRPDLIDPALLRPG | ASSEGDNERLF | 0.10131464 |
| LANLGVTAL | ASSEGDNERLF | 0.09847056 |
| TGERIEL | ASSEGDNERLF | 0.10826037 |
| NGEELAA | ASSEGDNERLF | 0.09542899 |
| VLSLVWG | ASSEGDNERLF | 0.09753596 |
| IGYALWL | ASSEGDNERLF | 0.09650414 |

| | | |
|---|---|---|
| LSLLLAPL | ASSEGDNERLF | 0.096231125 |
| TLDEALK | ASSEGDNERLF | 0.09595814 |
| TDGAGTAEGG | ASSEGDNERLF | 0.099252276 |
| NAPGGAGGNGG | ASSEGDNERLF | 0.10794777 |
| VSVFMGMGGNF | ASSEGDNERLF | 0.09594104 |
| LASYLAAL | ASSEGDNERLF | 0.09714588 |
| LVGGTPAL | ASSEGDNERLF | 0.095387034 |
| VRNADVL | ASSEGDNERLF | 0.097848184 |
| TCSSPFLVSAM | ASSEGDNERLF | 0.11808669 |
| VHITDAE | ASSEGDNERLF | 0.111388415 |
| EYAAPSF | ASSEGDNERLF | 0.09788891 |
| VVDAFHGGL | ASSEGDNERLF | 0.100250475 |
| ASVVDPQ | ASSEGDNERLF | 0.1081075 |
| SYNEFGGLS | ASSEGDNERLF | 0.09695493 |
| DPDGSVY | ASSEGDNERLF | 0.09598728 |
| MVSTIHATE | ASSEGDNERLF | 0.09597735 |
| LAAGVADPV | ASSEGDNERLF | 0.10421442 |
| TKEQIAA | ASSEGDNERLF | 0.101145834 |
| SLASPAAL | ASSEGDNERLF | 0.10112088 |
| IGLVFYGV | ASSEGDNERLF | 0.101711914 |
| FLLDASNL | ASSEGDNERLF | 0.101957195 |
| YQVGNFP | ASSEGDNERLF | 0.09679029 |
| ESLLSQST | ASSEGDNERLF | 0.09807197 |
| LFTVGFAAVG | ASSEGDNERLF | 0.099582866 |
| ATPPTPT | ASSEGDNERLF | 0.100681596 |
| TDLATLG | ASSEGDNERLF | 0.09529708 |
| SAGGTGR | ASSEGDNERLF | 0.096165255 |
| GFSNSGS | ASSEGDNERLF | 0.0954318 |
| GGNGGSGGTSV | ASSEGDNERLF | 0.0990504 |
| GTSVATGGAGN | ASSEGDNERLF | 0.100291565 |
| NGGNGGSGGTS | ASSEGDNERLF | 0.09609921 |
| AVVKFPH | ASSEGDNERLF | 0.09780686 |
| TPPPPPP | ASSEGDNERLF | 0.09642111 |
| AAPAAAAAPA | ASSEGDNERLF | 0.101634264 |
| IALNEDL | ASSEGDNERLF | 0.095632255 |
| IVNVSSM | ASSEGDNERLF | 0.10255502 |
| TGIPPAPR | ASSEGDNERLF | 0.09594726 |
| FDLGGGTF | ASSEGDNERLF | 0.097156666 |
| FDLGGGTFDVS | ASSEGDNERLF | 0.0970031 |
| NEPTAAA | ASSEGDNERLF | 0.096493594 |
| NEPTAAALA | ASSEGDNERLF | 0.104271136 |
| GSIINISSI | ASSEGDNERLF | 0.09650175 |
| VAPGNAGTA | ASSEGDNERLF | 0.10284523 |
| TGGGASLE | ASSEGDNERLF | 0.09830657 |
| VAVYNHYKRI | ASSEGDNERLF | 0.096028745 |
| TDPAPATT | ASSEGDNERLF | 0.09722712 |
| TNRPDILDPALLRPG | ASSEGDNERLF | 0.106011756 |
| LTTTTTS | ASSEGDNERLF | 0.09828098 |
| PTGSGKTL | ASSEGDNERLF | 0.09687545 |
| TGSGKTL | ASSEGDNERLF | 0.10057944 |
| TIGHVDHGK | ASSEGDNERLF | 0.097074 |

| | | |
|---|---|---|
| TIGHVDHGKTT | ASSEGDNERLF | 0.09764387 |
| GHVDHGKTT | ASSEGDNERLF | 0.095356174 |
| AQFEHTLL | ASSEGDNERLF | 0.0965287 |
| IVNQDEL | ASSEGDNERLF | 0.09619265 |
| TNGSQFF | ASSEGDNERLF | 0.09651346 |
| TGKGIVE | ASSEGDNERLF | 0.10839903 |
| LISNASDA | ASSEGDNERLF | 0.10753152 |
| AELRTLGELTKIAKAHGAQ | ASSEGDNERLF | 0.09648242 |
| APNAVAAAEPSW | ASSEGDNERLF | 0.108128324 |
| APQVAGAATAVL | ASSEGDNERLF | 0.09739491 |
| ARRDPAAAAAILPTDA | ASSEGDNERLF | 0.095753655 |
| DAAARDVLAEAGLAEYF | ASSEGDNERLF | 0.09543537 |
| DDQIVLSAKANLAKVEQ | ASSEGDNERLF | 0.100753926 |
| DELLPDTGEDSDEDGH | ASSEGDNERLF | 0.09596956 |
| ELPEAPANRATVPQD | ASSEGDNERLF | 0.09722554 |
| FGQSKIERVTGIG | ASSEGDNERLF | 0.09911642 |
| GAGGTGSTAGGAGGAGGAGGLY | ASSEGDNERLF | 0.096234776 |
| GHGGTGAAGGNGGTGGAGGSGI | ASSEGDNERLF | 0.097104326 |
| GNGGNGGAGGEGGPGGAGGAGA | ASSEGDNERLF | 0.09620606 |
| GSTAGNGGNGGNATNGTIAGQP | ASSEGDNERLF | 0.10747928 |
| GTGGDGGNGGNGGTGG | ASSEGDNERLF | 0.09543861 |
| GVRRPAGIPADAPV | ASSEGDNERLF | 0.095204905 |
| GYRQVTDAHLLTLARR | ASSEGDNERLF | 0.10699996 |
| IAKMGQDGHDRG | ASSEGDNERLF | 0.098972484 |
| KAEPSIEIQYT | ASSEGDNERLF | 0.0963761 |
| KPVRIRMAADEDTVAVT | ASSEGDNERLF | 0.09699959 |
| MPPRIAGMRLLVIKPEP | ASSEGDNERLF | 0.11131131 |
| MKFYAVALFRSRGVPPTR | ASSEGDNERLF | 0.097241946 |
| NAGGFGILGAVAHSP | ASSEGDNERLF | 0.0956346 |
| NEGREITLAMAEM | ASSEGDNERLF | 0.09619078 |
| NELVEWSHPLGHRWRWE | ASSEGDNERLF | 0.09816414 |
| NLGAGGTGGNGGNPG | ASSEGDNERLF | 0.09540582 |
| NTEWPKWETVMAADPVD | ASSEGDNERLF | 0.10022874 |
| PASAAPAETQP | ASSEGDNERLF | 0.10857793 |
| QQSSDGKNGVRAAEITGE | ASSEGDNERLF | 0.09537358 |
| RAPDAVTTSWWKE | ASSEGDNERLF | 0.09683213 |
| SGFGNVGAHTSGWFNQ | ASSEGDNERLF | 0.09579486 |
| SGVAGAVGQAASVGGLKV | ASSEGDNERLF | 0.09873242 |
| SNGGAGTGSAGGNGGT | ASSEGDNERLF | 0.09577288 |
| TGGAGGEGGAAG | ASSEGDNERLF | 0.095246434 |
| TGGDGGAAGAGGAPGHG | ASSEGDNERLF | 0.10062562 |
| TMPPLLAAIANAGRHGV | ASSEGDNERLF | 0.10577577 |
| TSSTAGNSGGA | ASSEGDNERLF | 0.09630387 |
| VPATVPVGILC | ASSEGDNERLF | 0.09955443 |
| VQRPDSGGVVLTGRIS | ASSEGDNERLF | 0.09744608 |
| WTPNPVPATKGVPAGLVQAT | ASSEGDNERLF | 0.11187432 |
| YALKRALAKFSDRFDVVII | ASSEGDNERLF | 0.0967982 |
| YLIGSVIV | ASSEGDNERLF | 0.1002009 |
| YGQLVAVID | ASSEGDNERLF | 0.09584819 |
| VTVGGMTLPA | ASSEGDNERLF | 0.09992719 |
| VLALPAV | ASSEGDNERLF | 0.099090114 |

| | | |
|---|---|---|
| VIPPTLNYET | ASSEGDNERLF | 0.095723756 |
| VGMGGMHPSSG | ASSEGDNERLF | 0.099294476 |
| VGLAGGAATA | ASSEGDNERLF | 0.09585789 |
| VGGAGGAGGSGP | ASSEGDNERLF | 0.10116938 |
| VFAAFMV | ASSEGDNERLF | 0.09682533 |
| TLYDKLR | ASSEGDNERLF | 0.09520591 |
| TLAVFTVL | ASSEGDNERLF | 0.11240201 |
| TGSVQPARV | ASSEGDNERLF | 0.09618883 |
| TGGGAGSGGT | ASSEGDNERLF | 0.105925225 |
| SHLETSL | ASSEGDNERLF | 0.09572735 |
| SFTTGMT | ASSEGDNERLF | 0.09570848 |
| QVEVAKRMGVRQPTVS | ASSEGDNERLF | 0.10109914 |
| QGSGGPALAM | ASSEGDNERLF | 0.09572506 |
| MLAPSNP | ASSEGDNERLF | 0.11332678 |
| LVNNAAI | ASSEGDNERLF | 0.097547166 |
| LTGEINTLA | ASSEGDNERLF | 0.09657751 |
| LSIFGPV | ASSEGDNERLF | 0.095740855 |
| LLALLANL | ASSEGDNERLF | 0.0968542 |
| LISNASDALDKLR | ASSEGDNERLF | 0.09658007 |
| LINEAVP | ASSEGDNERLF | 0.1009349 |
| LGYAFSGL | ASSEGDNERLF | 0.095315956 |
| LDFVAVAL | ASSEGDNERLF | 0.09866854 |
| LAQPGAE | ASSEGDNERLF | 0.09580964 |
| ISNLGAVTGF | ASSEGDNERLF | 0.0996428 |
| HQEDPGRI | ASSEGDNERLF | 0.100677766 |
| GTPGEPTP | ASSEGDNERLF | 0.10214682 |
| GFFNSGS | ASSEGDNERLF | 0.09859909 |
| GAGGGGLTAGAN | ASSEGDNERLF | 0.113514796 |
| FVDALVNM | ASSEGDNERLF | 0.10162202 |
| FLTVAPEL | ASSEGDNERLF | 0.09707912 |
| FETLTAL | ASSEGDNERLF | 0.0968718 |
| DVINPIEL | ASSEGDNERLF | 0.099240765 |
| APVSGAPV | ASSEGDNERLF | 0.105843075 |
| ALVGGLAW | ASSEGDNERLF | 0.09642676 |
| ALAAAAATAI | ASSEGDNERLF | 0.09579046 |
| AFGLGLLL | ASSEGDNERLF | 0.09566051 |
| AFFAAASL | ASSEGDNERLF | 0.10185427 |
| AAAALPAFSPP | ASSEGDNERLF | 0.098667584 |
| VVLAPAL | ASSEGDNERLF | 0.10011435 |
| TIGHVDHGKTTL | ASSEGDNERLF | 0.09629767 |
| LISNASDALDK | ASSEGDNERLF | 0.10758937 |
| LISNASDAL | ASSEGDNERLF | 0.104369596 |
| ISNASDALDK | ASSEGDNERLF | 0.0998244 |
| NQAETLVYQTE | ASSEGDNERLF | 0.09554058 |
| TGGTGGTAGAG | ASSEGDNERLF | 0.101051934 |
| TGSTAGGAGGA | ASSEGDNERLF | 0.0987936 |
| DPLAVPVAAGPV | ASSEGDNERLF | 0.096951604 |
| GGTGGTPGNGGN | ASSEGDNERLF | 0.097307324 |
| PAGLAGVDPATL | ASSEGDNERLF | 0.09934351 |
| GNNLVVEAPHAGASV | ASSEGDNERLF | 0.10930704 |
| KVTRIMSASTLEQPA | ASSEGDNERLF | 0.095113605 |

| | | |
|---|---|---|
| LGFALPIALPRRTVV | ASSEGDNERLF | 0.09559113 |
| TRLGNLTARKLVVLV | ASSEGDNERLF | 0.09914425 |
| WSQQRRLIYFPSAGP | ASSEGDNERLF | 0.10842598 |
| GPNGNPDPMAAAVDIR | ASSEGDNERLF | 0.09880889 |
| TQRCVSVADLRAAASR | ASSEGDNERLF | 0.0972651 |
| QFVLDVAEHPVEVLTETV | ASSEGDNERLF | 0.09531039 |
| GGAGGTGGAGGNGGTQAGDG | ASSEGDNERLF | 0.10920169 |
| PLLATLRAIAVDVDAKPAQV | ASSEGDNERLF | 0.09610837 |
| APKFRTGGYMIDFWGVGYQVAK | ASSEGDNERLF | 0.09602035 |
| GAGGAGGAGGAAGGPGGTGGTG | ASSEGDNERLF | 0.100317195 |
| IGALGLLATDVAPAGSVAFGVA | ASSEGDNERLF | 0.099315 |
| AGGAGGIGGNGGAGGAGGVLMG | ASSEGDNERLF | 0.09557715 |
| ADELIGLL | ASSEGDNERLF | 0.09601749 |
| ADRALKA | ASSEGDNERLF | 0.10126167 |
| AGAVGQVGGAGGA | ASSEGDNERLF | 0.09587505 |
| AGGGGGLGGAP | ASSEGDNERLF | 0.09929749 |
| AGGLLYGNGGAG | ASSEGDNERLF | 0.09895687 |
| AGLESIGL | ASSEGDNERLF | 0.09608605 |
| AGLKTGII | ASSEGDNERLF | 0.10516466 |
| ASLAALLL | ASSEGDNERLF | 0.11551032 |
| ASVINISSI | ASSEGDNERLF | 0.096243836 |
| ATPPGPA | ASSEGDNERLF | 0.099913545 |
| ATVASALV | ASSEGDNERLF | 0.11151428 |
| ATVASLIL | ASSEGDNERLF | 0.10530877 |
| AVNGPVLIRH | ASSEGDNERLF | 0.098487526 |
| DGAAGAVGI | ASSEGDNERLF | 0.09579554 |
| DPLLHGI | ASSEGDNERLF | 0.09602298 |
| ELGAALGAL | ASSEGDNERLF | 0.096946 |
| ELGRTQAV | ASSEGDNERLF | 0.10397618 |
| FVAGGYLS | ASSEGDNERLF | 0.09812299 |
| GASGGAGGAGGW | ASSEGDNERLF | 0.09836936 |
| GGAGAGGDGGAGGI | ASSEGDNERLF | 0.097294785 |
| GTGHNGSQPGMGGNG | ASSEGDNERLF | 0.09873254 |
| GYAAASASAS | ASSEGDNERLF | 0.10120372 |
| HQVDVAAVQQG | ASSEGDNERLF | 0.09636074 |
| IGAAIAHT | ASSEGDNERLF | 0.095766686 |
| IGMIEVAQL | ASSEGDNERLF | 0.09675404 |
| IRLAPEHTSAMD | ASSEGDNERLF | 0.09733086 |
| IYNVSVL | ASSEGDNERLF | 0.09632992 |
| KQGFLGM | ASSEGDNERLF | 0.10174465 |
| LESVITQL | ASSEGDNERLF | 0.11126089 |
| LGLGALVVV | ASSEGDNERLF | 0.09563476 |
| LIGNGGNGGSGGI | ASSEGDNERLF | 0.0964936 |
| LNAINAPTL | ASSEGDNERLF | 0.12701459 |
| LQLLNTGV | ASSEGDNERLF | 0.103684105 |
| LQTSIEF | ASSEGDNERLF | 0.101200126 |
| MSFAVSEL | ASSEGDNERLF | 0.09605086 |
| NTGALGL | ASSEGDNERLF | 0.09735534 |
| PPGAGGAGGAGTT | ASSEGDNERLF | 0.09984741 |
| QLVGLGI | ASSEGDNERLF | 0.09956176 |
| RFLEQVHQL | ASSEGDNERLF | 0.09903384 |

| | | |
|---|---|---|
| RGVGYVLR | ASSEGDNERLF | 0.09896398 |
| SATGVFAGVI | ASSEGDNERLF | 0.10071098 |
| SLPAITLP | ASSEGDNERLF | 0.10857037 |
| STALAFTV | ASSEGDNERLF | 0.095998526 |
| SVLTVGLGK | ASSEGDNERLF | 0.095392406 |
| TALAGAGSGI | ASSEGDNERLF | 0.09539513 |
| TGLPFTGI | ASSEGDNERLF | 0.09598698 |
| VAEHVTV | ASSEGDNERLF | 0.104582906 |
| VDIVLNSL | ASSEGDNERLF | 0.097133406 |
| VLTGGGAVL | ASSEGDNERLF | 0.095461264 |
| VNVGLAYGI | ASSEGDNERLF | 0.09750501 |
| VSSAAVGAV | ASSEGDNERLF | 0.112376384 |
| VTSVYEAGV | ASSEGDNERLF | 0.10228077 |
| WVALDRGA | ASSEGDNERLF | 0.09723057 |
| YLVGSVVI | ASSEGDNERLF | 0.09676667 |
| GPPGTGKTLL | ASSEGDNERLF | 0.09542825 |
| GVGMTKF | ASSEGDNERLF | 0.09584923 |
| IGIGGSDL | ASSEGDNERLF | 0.09551566 |
| IGTLLAEL | ASSEGDNERLF | 0.09573047 |
| IYTSGTT | ASSEGDNERLF | 0.095489636 |
| LSFPAGV | ASSEGDNERLF | 0.096934505 |
| LYTSGTT | ASSEGDNERLF | 0.095247515 |
| VGAFPFGF | ASSEGDNERLF | 0.09792921 |
| VGAGTVEF | ASSEGDNERLF | 0.10169901 |
| VGKSTLLL | ASSEGDNERLF | 0.096606396 |
| AALAHAGGGIV | ASSEGDNERLF | 0.09972348 |
| AALVAVSLL | ASSEGDNERLF | 0.11782298 |
| AAVSRAVAAL | ASSEGDNERLF | 0.09853254 |
| AGGSSGIGGF | ASSEGDNERLF | 0.10035128 |
| AGLTVANANG | ASSEGDNERLF | 0.09579511 |
| AGLVGAVL | ASSEGDNERLF | 0.10786241 |
| ALSQAFTV | ASSEGDNERLF | 0.095465824 |
| ALTFSVSL | ASSEGDNERLF | 0.10213684 |
| ALTGAALVAL | ASSEGDNERLF | 0.10059937 |
| ASFSIGAGVL | ASSEGDNERLF | 0.09567258 |
| DGSVYKI | ASSEGDNERLF | 0.0955616 |
| EQLAEIV | ASSEGDNERLF | 0.09958038 |
| FAVSELTVM | ASSEGDNERLF | 0.09583316 |
| GAAPLAAGL | ASSEGDNERLF | 0.09759782 |
| GADGGDGGSGGAGG | ASSEGDNERLF | 0.09625329 |
| GFGLDTGGAGG | ASSEGDNERLF | 0.09909921 |
| GLASGGLL | ASSEGDNERLF | 0.096132874 |
| GRIAGAAVLL | ASSEGDNERLF | 0.09537572 |
| GSAGSLNTSF | ASSEGDNERLF | 0.09737682 |
| GTGQIGI | ASSEGDNERLF | 0.09876886 |
| GTLIALIL | ASSEGDNERLF | 0.09768518 |
| IAATMGFVVG | ASSEGDNERLF | 0.10281794 |
| IFAQALL | ASSEGDNERLF | 0.0969854 |
| IGAAVALGF | ASSEGDNERLF | 0.098385096 |
| IGLTPLAL | ASSEGDNERLF | 0.09933753 |
| ILGSNTSTL | ASSEGDNERLF | 0.09619332 |

| | | |
|---|---|---|
| IVFSLTVL | ASSEGDNERLF | 0.09983167 |
| LASKATGF | ASSEGDNERLF | 0.095344454 |
| LASVALIL | ASSEGDNERLF | 0.09568781 |
| LFAVGLTL | ASSEGDNERLF | 0.09895411 |
| LFNGGIGTYI | ASSEGDNERLF | 0.09648752 |
| LTLGAGII | ASSEGDNERLF | 0.09825107 |
| LVAAALGDI | ASSEGDNERLF | 0.102738306 |
| NTAFVQLGI | ASSEGDNERLF | 0.09615497 |
| QLFSGAEL | ASSEGDNERLF | 0.09596183 |
| SAATFITL | ASSEGDNERLF | 0.096277885 |
| SNYGLANLGV | ASSEGDNERLF | 0.098317064 |
| SYGGAYAVM | ASSEGDNERLF | 0.1048598 |
| TTGVGGLTLG | ASSEGDNERLF | 0.09713966 |
| TTQTIYI | ASSEGDNERLF | 0.104252696 |
| VAAGAGLGAVI | ASSEGDNERLF | 0.09805289 |
| VAAVVTLL | ASSEGDNERLF | 0.0990744 |
| VGFLGGLI | ASSEGDNERLF | 0.11022485 |
| VGVIPILL | ASSEGDNERLF | 0.11267969 |
| VIFASSLI | ASSEGDNERLF | 0.09798142 |
| VKGGDGGAAAG | ASSEGDNERLF | 0.095712826 |
| VTKTFTNI | ASSEGDNERLF | 0.096181005 |
| VVAAAILL | ASSEGDNERLF | 0.09577337 |
| YAALDVEL | ASSEGDNERLF | 0.11175453 |
| LASVALLL | ASSEGDNERLF | 0.09831201 |
| LDAVHRV | ASSEGDNERLF | 0.09976644 |
| QGLSSSIF | ASSEGDNERLF | 0.09589775 |
| RGTFANIRL | ASSEGDNERLF | 0.10450466 |
| RQLWWGHRI | ASSEGDNERLF | 0.09599054 |
| YGPPGTGKTLL | ASSEGDNERLF | 0.10606854 |
| AALVGAPANVA | ASSEGDNERLF | 0.10505515 |
| AATFRRTE | ASSEGDNERLF | 0.09557949 |
| AGALGGGI | ASSEGDNERLF | 0.0966613 |
| AGGAGGAGGIGAG | ASSEGDNERLF | 0.09862412 |
| AGSGALGL | ASSEGDNERLF | 0.09599722 |
| AQIADQI | ASSEGDNERLF | 0.09535618 |
| ASAVAVTYL | ASSEGDNERLF | 0.09589847 |
| ASTVLGGTGG | ASSEGDNERLF | 0.09620185 |
| AVGGEPAK | ASSEGDNERLF | 0.09624949 |
| CEQGTFGGTC | ASSEGDNERLF | 0.101149246 |
| DEGKAPAG | ASSEGDNERLF | 0.10031179 |
| DTVVFPAL | ASSEGDNERLF | 0.09628196 |
| FEGITMAG | ASSEGDNERLF | 0.09694183 |
| FPLVVQD | ASSEGDNERLF | 0.10301584 |
| GAFTTPQ | ASSEGDNERLF | 0.09695666 |
| GFGGFGTAGGVGGA | ASSEGDNERLF | 0.10291611 |
| GFLAQFL | ASSEGDNERLF | 0.09577472 |
| GGAGGAGGNAGVA | ASSEGDNERLF | 0.09582361 |
| GGVGTAGAGGAGGAGG | ASSEGDNERLF | 0.09750508 |
| GNGGGGDAGNAGSGGN | ASSEGDNERLF | 0.10363889 |
| GTGGAGGAGNDAGS | ASSEGDNERLF | 0.09610477 |
| GTTNTGWFNS | ASSEGDNERLF | 0.11739151 |

| | | |
|---|---|---|
| IFMGPLFTL | ASSEGDNERLF | 0.119066134 |
| ILAHAIEAI | ASSEGDNERLF | 0.09654126 |
| IRSAIREAL | ASSEGDNERLF | 0.11109025 |
| ITFAGGG | ASSEGDNERLF | 0.09524287 |
| ITGMSGHQEV | ASSEGDNERLF | 0.09832693 |
| IVAGLACTL | ASSEGDNERLF | 0.095356956 |
| IVGGGLAGA | ASSEGDNERLF | 0.10331988 |
| IVTGVLNI | ASSEGDNERLF | 0.09983618 |
| LAQQGCI | ASSEGDNERLF | 0.09759177 |
| LCLHVEN | ASSEGDNERLF | 0.096263364 |
| LDAISDGS | ASSEGDNERLF | 0.11402148 |
| LGGFAAML | ASSEGDNERLF | 0.10197916 |
| LGGKNPMI | ASSEGDNERLF | 0.09559187 |
| LGPKGVAAIL | ASSEGDNERLF | 0.1053651 |
| LMVGVSI | ASSEGDNERLF | 0.097756825 |
| LVGAAFVL | ASSEGDNERLF | 0.09630764 |
| LVLAVAPL | ASSEGDNERLF | 0.09604229 |
| MAAIAVIDAA | ASSEGDNERLF | 0.097208716 |
| NNGINAELQ | ASSEGDNERLF | 0.09788851 |
| RGEPLVLVHG | ASSEGDNERLF | 0.09679971 |
| RGGAGGAAGLI | ASSEGDNERLF | 0.099759325 |
| RSYRMHPAV | ASSEGDNERLF | 0.0977821 |
| SAAHLLAGQLA | ASSEGDNERLF | 0.09548105 |
| SAAVGSY | ASSEGDNERLF | 0.09522779 |
| SPHAVLHRI | ASSEGDNERLF | 0.10299355 |
| SQIAKVL | ASSEGDNERLF | 0.122975096 |
| TALGHLVP | ASSEGDNERLF | 0.097739056 |
| TAPTTSGAA | ASSEGDNERLF | 0.104037955 |
| TDGDDGNGGSPGT | ASSEGDNERLF | 0.09903028 |
| TDGLGEN | ASSEGDNERLF | 0.10485392 |
| TGGPPLPGQA | ASSEGDNERLF | 0.1012356 |
| TIPAITI | ASSEGDNERLF | 0.096929684 |
| TIPGFTI | ASSEGDNERLF | 0.09608392 |
| TSIASGALA | ASSEGDNERLF | 0.09546878 |
| TVTLPGAAI | ASSEGDNERLF | 0.09594233 |
| VASGLSAALP | ASSEGDNERLF | 0.0981128 |
| VGAGVIGI | ASSEGDNERLF | 0.098911546 |
| VGPKRFGV | ASSEGDNERLF | 0.09981088 |
| VNVAFPD | ASSEGDNERLF | 0.10259356 |
| VSMTNSGVS | ASSEGDNERLF | 0.09780024 |
| VTDVLGDLSV | ASSEGDNERLF | 0.09799121 |
| VVVVGAGFAGL | ASSEGDNERLF | 0.10147044 |
| VYTVGTVV | ASSEGDNERLF | 0.09656643 |
| WTTFSGP | ASSEGDNERLF | 0.11042675 |
| YDELVEAF | ASSEGDNERLF | 0.09788985 |
| YGTGGAGGNGG | ASSEGDNERLF | 0.095988296 |
| YTNVMAV | ASSEGDNERLF | 0.100423224 |
| FLEQVHQL | ASSEGDNERLF | 0.09949507 |
| GLSLLAVAV | ASSEGDNERLF | 0.0982114 |
| LEQVHQL | ASSEGDNERLF | 0.09909471 |
| LNAAGAI | ASSEGDNERLF | 0.1017227 |

| | | |
|---|---|---|
| PAVHASL | ASSEGDNERLF | 0.095549375 |
| TELEDDA | ASSEGDNERLF | 0.09645762 |
| TPPGPPN | ASSEGDNERLF | 0.0977354 |
| TPPPPPPP | ASSEGDNERLF | 0.108268484 |
| TQSPALI | ASSQDPGADTQY | 0.10105609 |
| EFAAAFPQ | ASSQDPGADTQY | 0.09570674 |
| LLSGNLDVL | ASSQDPGADTQY | 0.09791016 |
| GPANLGNYNI | ASSQDPGADTQY | 0.09700062 |
| TLGPTLP | ASSQDPGADTQY | 0.09833932 |
| VYAYSSCCK | ASSQDPGADTQY | 0.11610625 |
| APAPAPAPNI | ASSQDPGADTQY | 0.10034822 |
| PGGPVPIP | ASSQDPGADTQY | 0.09706618 |
| FLTVALEP | ASSQDPGADTQY | 0.106281474 |
| SQYFGANL | ASSQDPGADTQY | 0.095324315 |
| GGVKPAA | ASSQDPGADTQY | 0.096347585 |
| GGTGGAGGAGGW | ASSQDPGADTQY | 0.09538995 |
| GVGGVAGLGGAGG | ASSQDPGADTQY | 0.095508635 |
| VLLVLATTI | ASSQDPGADTQY | 0.09678367 |
| IGIFAAML | ASSQDPGADTQY | 0.09686922 |
| AVGDLID | ASSQDPGADTQY | 0.09602822 |
| NISPLPL | ASSQDPGADTQY | 0.09765324 |
| VTAVFVAI | ASSQDPGADTQY | 0.09504997 |
| SGDDNDAAL | ASSQDPGADTQY | 0.09642518 |
| AVAALAHS | ASSQDPGADTQY | 0.10101091 |
| GGVTVMATP | ASSQDPGADTQY | 0.09654561 |
| AGLAANVRD | ASSQDPGADTQY | 0.1003086 |
| TNRPDLIDPALLRPG | ASSQDPGADTQY | 0.0964571 |
| LANLGVTAL | ASSQDPGADTQY | 0.09617945 |
| TGERIEL | ASSQDPGADTQY | 0.09623462 |
| NGEELAA | ASSQDPGADTQY | 0.0967775 |
| VLSLVWG | ASSQDPGADTQY | 0.09545956 |
| IGYALWL | ASSQDPGADTQY | 0.09983673 |
| LSLLLAPL | ASSQDPGADTQY | 0.1029298 |
| TLDEALK | ASSQDPGADTQY | 0.095576435 |
| TDGAGTAEGG | ASSQDPGADTQY | 0.0965979 |
| NAPGGAGGNGG | ASSQDPGADTQY | 0.096455894 |
| VSVFMGMGGNF | ASSQDPGADTQY | 0.09589017 |
| LASYLAAL | ASSQDPGADTQY | 0.09554472 |
| LVGGTPAL | ASSQDPGADTQY | 0.103614524 |
| VRNADVL | ASSQDPGADTQY | 0.105312206 |
| TCSSPFLVSAM | ASSQDPGADTQY | 0.096401304 |
| VHITDAE | ASSQDPGADTQY | 0.09819681 |
| EYAAPSF | ASSQDPGADTQY | 0.09533793 |
| VVDAFHGGL | ASSQDPGADTQY | 0.11256687 |
| ASVVDPQ | ASSQDPGADTQY | 0.09791575 |
| SYNEFGGLS | ASSQDPGADTQY | 0.1013448 |
| DPDGSVY | ASSQDPGADTQY | 0.095967054 |
| MVSTIHATE | ASSQDPGADTQY | 0.0955751 |
| LAAGVADPV | ASSQDPGADTQY | 0.10576662 |
| TKEQIAA | ASSQDPGADTQY | 0.095772736 |
| SLASPAAL | ASSQDPGADTQY | 0.09581182 |

| | | |
|---|---|---|
| IGLVFYGV | ASSQDPGADTQY | 0.09551348 |
| FLLDASNL | ASSQDPGADTQY | 0.09772747 |
| YQVGNFP | ASSQDPGADTQY | 0.09615759 |
| ESLLSQST | ASSQDPGADTQY | 0.09699055 |
| LFTVGFAAVG | ASSQDPGADTQY | 0.12320713 |
| ATPPTPT | ASSQDPGADTQY | 0.11385919 |
| TDLATLG | ASSQDPGADTQY | 0.0980669 |
| SAGGTGR | ASSQDPGADTQY | 0.099143706 |
| GFSNSGS | ASSQDPGADTQY | 0.097989246 |
| GGNGGSGGTSV | ASSQDPGADTQY | 0.100029916 |
| GTSVATGGAGN | ASSQDPGADTQY | 0.09629506 |
| NGGNGGSGGTS | ASSQDPGADTQY | 0.096555285 |
| AVVKFPH | ASSQDPGADTQY | 0.09794648 |
| TPPPPPP | ASSQDPGADTQY | 0.100902446 |
| AAPAAAAAPA | ASSQDPGADTQY | 0.09767887 |
| IALNEDL | ASSQDPGADTQY | 0.099292174 |
| IVNVSSM | ASSQDPGADTQY | 0.09818646 |
| TGIPPAPR | ASSQDPGADTQY | 0.09651388 |
| FDLGGGTF | ASSQDPGADTQY | 0.10671013 |
| FDLGGGTFDVS | ASSQDPGADTQY | 0.09670632 |
| NEPTAAA | ASSQDPGADTQY | 0.108310215 |
| NEPTAAALA | ASSQDPGADTQY | 0.09666997 |
| GSIINISSI | ASSQDPGADTQY | 0.0964356 |
| VAPGNAGTA | ASSQDPGADTQY | 0.09547472 |
| TGGGASLE | ASSQDPGADTQY | 0.1050941 |
| VAVYNHYKRI | ASSQDPGADTQY | 0.09605462 |
| TDPAPATT | ASSQDPGADTQY | 0.095726326 |
| TNRPDILDPALLRPG | ASSQDPGADTQY | 0.09958108 |
| LTTTTTS | ASSQDPGADTQY | 0.09578937 |
| PTGSGKTL | ASSQDPGADTQY | 0.099417046 |
| TGSGKTL | ASSQDPGADTQY | 0.096585974 |
| TIGHVDHGK | ASSQDPGADTQY | 0.095684655 |
| TIGHVDHGKTT | ASSQDPGADTQY | 0.10315262 |
| GHVDHGKTT | ASSQDPGADTQY | 0.09630388 |
| AQFEHTLL | ASSQDPGADTQY | 0.09621771 |
| IVNQDEL | ASSQDPGADTQY | 0.10451289 |
| TNGSQFF | ASSQDPGADTQY | 0.09624081 |
| TGKGIVE | ASSQDPGADTQY | 0.10241153 |
| LISNASDA | ASSQDPGADTQY | 0.097075425 |
| AELRTLGELTKIAKAHGAQ | ASSQDPGADTQY | 0.099199206 |
| APNAVAAAEPSW | ASSQDPGADTQY | 0.097432084 |
| APQVAGAATAVL | ASSQDPGADTQY | 0.09579907 |
| ARRDPAAAAAILPTDA | ASSQDPGADTQY | 0.09639583 |
| DAAARDVLAEAGLAEYF | ASSQDPGADTQY | 0.09778661 |
| DDQIVLSAKANLAKVEQ | ASSQDPGADTQY | 0.12508121 |
| DELLPDTGEDSDEDGH | ASSQDPGADTQY | 0.10029301 |
| ELPEAPANRATVPQD | ASSQDPGADTQY | 0.099890456 |
| FGQSKIERVTGIG | ASSQDPGADTQY | 0.09871915 |
| GAGGTGSTAGGAGGAGGAGGLY | ASSQDPGADTQY | 0.0973314 |
| GHGGTGAAGGNGGTGGAGGSGI | ASSQDPGADTQY | 0.10060626 |
| GNGGNGGAGGEGGPGGAGGAGA | ASSQDPGADTQY | 0.09563652 |

| | | |
|---|---|---|
| GSTAGNGGNGGNATNGTIAGQP | ASSQDPGADTQY | 0.09619127 |
| GTGGDGGNGGNGGTGG | ASSQDPGADTQY | 0.09585816 |
| GVRRPAGIPADAPV | ASSQDPGADTQY | 0.09654689 |
| GYRQVTDAHLLTLARR | ASSQDPGADTQY | 0.09630952 |
| IAKMGQDGHDRG | ASSQDPGADTQY | 0.09955914 |
| KAEPSIEIQYT | ASSQDPGADTQY | 0.09580592 |
| KPVRIRMAADEDTVAVT | ASSQDPGADTQY | 0.095968075 |
| MPPRIAGMRLLVIKPEP | ASSQDPGADTQY | 0.09653415 |
| MKFYAVALFRSRGVPPTR | ASSQDPGADTQY | 0.09681017 |
| NAGGFGILGAVAHSP | ASSQDPGADTQY | 0.09599577 |
| NEGREITLAMAEM | ASSQDPGADTQY | 0.10801879 |
| NELVEWSHPLGHRWRWE | ASSQDPGADTQY | 0.097331144 |
| NLGAGGTGGNGGNPG | ASSQDPGADTQY | 0.09765278 |
| NTEWPKWETVMAADPVD | ASSQDPGADTQY | 0.13135585 |
| PASAAPAETQP | ASSQDPGADTQY | 0.09656505 |
| QQSSDGKNGVRAAEITGE | ASSQDPGADTQY | 0.09545636 |
| RAPDAVTTSWWKE | ASSQDPGADTQY | 0.09620872 |
| SGFGNVGAHTSGWFNQ | ASSQDPGADTQY | 0.0974981 |
| SGVAGAVGQAASVGGLKV | ASSQDPGADTQY | 0.100788206 |
| SNGGAGTGSAGGNGGT | ASSQDPGADTQY | 0.09521694 |
| TGGAGGEGGAAG | ASSQDPGADTQY | 0.10523747 |
| TGGDGGAAGAGGAPGHG | ASSQDPGADTQY | 0.102132685 |
| TMPPLLAAIANAGRHGV | ASSQDPGADTQY | 0.09651736 |
| TSSTAGNSGGA | ASSQDPGADTQY | 0.095261596 |
| VPATVPVGILC | ASSQDPGADTQY | 0.10456548 |
| VQRPDSGGVVLTGRIS | ASSQDPGADTQY | 0.099589124 |
| WTPNPVPATKGVPAGLVQAT | ASSQDPGADTQY | 0.10201963 |
| YALKRALAKFSDRFDVVII | ASSQDPGADTQY | 0.095790826 |
| YLIGSVIV | ASSQDPGADTQY | 0.09549692 |
| YGQLVAVID | ASSQDPGADTQY | 0.103491694 |
| VTVGGMTLPA | ASSQDPGADTQY | 0.099834666 |
| VLALPAV | ASSQDPGADTQY | 0.09941551 |
| VIPPTLNYET | ASSQDPGADTQY | 0.105202235 |
| VGMGGMHPSSG | ASSQDPGADTQY | 0.09716626 |
| VGLAGGAATA | ASSQDPGADTQY | 0.097341 |
| VGGAGGAGGSGP | ASSQDPGADTQY | 0.09915397 |
| VFAAFMV | ASSQDPGADTQY | 0.1085812 |
| TLYDKLR | ASSQDPGADTQY | 0.09581251 |
| TLAVFTVL | ASSQDPGADTQY | 0.09647875 |
| TGSVQPARV | ASSQDPGADTQY | 0.099118106 |
| TGGGAGSGGT | ASSQDPGADTQY | 0.096275024 |
| SHLETSL | ASSQDPGADTQY | 0.09687515 |
| SFTTGMT | ASSQDPGADTQY | 0.10477625 |
| QVEVAKRMGVRQPTVS | ASSQDPGADTQY | 0.09646022 |
| QGSGGPALAM | ASSQDPGADTQY | 0.09811833 |
| MLAPSNP | ASSQDPGADTQY | 0.10172386 |
| LVNNAAI | ASSQDPGADTQY | 0.09579691 |
| LTGEINTLA | ASSQDPGADTQY | 0.10179684 |
| LSIFGPV | ASSQDPGADTQY | 0.096222594 |
| LLALLANL | ASSQDPGADTQY | 0.113859706 |
| LISNASDALDKLR | ASSQDPGADTQY | 0.096756615 |

| | | |
|---|---|---|
| LINEAVP | ASSQDPGADTQY | 0.09588976 |
| LGYAFSGL | ASSQDPGADTQY | 0.09586958 |
| LDFVAVAL | ASSQDPGADTQY | 0.34583348 |
| LAQPGAE | ASSQDPGADTQY | 0.67630553 |
| ISNLGAVTGF | ASSQDPGADTQY | 0.57402915 |
| HQEDPGRI | ASSQDPGADTQY | 0.51435304 |
| GTPGEPTP | ASSQDPGADTQY | 0.22683623 |
| GFFNSGS | ASSQDPGADTQY | 0.18343899 |
| GAGGGGLTAGAN | ASSQDPGADTQY | 0.52351123 |
| FVDALVNM | ASSQDPGADTQY | 0.18439685 |
| FLTVAPEL | ASSQDPGADTQY | 0.7808875 |
| FETLTAL | ASSQDPGADTQY | 0.64618385 |
| DVINPIEL | ASSQDPGADTQY | 0.21815696 |
| APVSGAPV | ASSQDPGADTQY | 0.67350036 |
| ALVGGLAW | ASSQDPGADTQY | 0.7738589 |
| ALAAAAATAI | ASSQDPGADTQY | 0.31782356 |
| AFGLGLLL | ASSQDPGADTQY | 0.42289978 |
| AFFAAASL | ASSQDPGADTQY | 0.24144512 |
| AAAALPAFSPP | ASSQDPGADTQY | 0.5380614 |
| VVLAPAL | ASSQDPGADTQY | 0.17192623 |
| TIGHVDHGKTTL | ASSQDPGADTQY | 0.4580813 |
| LISNASDALDK | ASSQDPGADTQY | 0.5506567 |
| LISNASDAL | ASSQDPGADTQY | 0.14727113 |
| ISNASDALDK | ASSQDPGADTQY | 0.7597743 |
| NQAETLVYQTE | ASSQDPGADTQY | 0.31906223 |
| TGGTGGTAGAG | ASSQDPGADTQY | 0.5526283 |
| TGSTAGGAGGA | ASSQDPGADTQY | 0.77561164 |
| DPLAVPVAAGPV | ASSQDPGADTQY | 0.234626 |
| GGTGGTPGNGGN | ASSQDPGADTQY | 0.22125658 |
| PAGLAGVDPATL | ASSQDPGADTQY | 0.58081913 |
| GNNLVVEAPHAGASV | ASSQDPGADTQY | 0.7988026 |
| KVTRIMSASTLEQPA | ASSQDPGADTQY | 0.18506463 |
| LGFALPIALPRRTVV | ASSQDPGADTQY | 0.6266452 |
| TRLGNLTARKLVVLV | ASSQDPGADTQY | 0.7665311 |
| WSQQRRLIYFPSAGP | ASSQDPGADTQY | 0.38939852 |
| GPNGNPDPMAAAVDIR | ASSQDPGADTQY | 0.8022116 |
| TQRCVSVADLRAAASR | ASSQDPGADTQY | 0.8054656 |
| QFVLDVAEHPVEVLTETV | ASSQDPGADTQY | 0.31031668 |
| GGAGGTGGAGGNGGTQAGDG | ASSQDPGADTQY | 0.70337284 |
| PLLATLRAIAVDVDAKPAQV | ASSQDPGADTQY | 0.77263963 |
| APKFRTGGYMIDFWGVGYQVAK | ASSQDPGADTQY | 0.46952137 |
| GAGGAGGAGGAAGGPGGTGGTG | ASSQDPGADTQY | 0.39936596 |
| IGALGLLATDVAPAGSVAFGVA | ASSQDPGADTQY | 0.43107092 |
| AGGAGGIGGNGGAGGAGGVLMG | ASSQDPGADTQY | 0.63411176 |
| ADELIGLL | ASSQDPGADTQY | 0.16266935 |
| ADRALKA | ASSQDPGADTQY | 0.52090025 |
| AGAVGQVGGAGGA | ASSQDPGADTQY | 0.6839653 |
| AGGGGGLGGAP | ASSQDPGADTQY | 0.3259815 |
| AGGLLYGNGGAG | ASSQDPGADTQY | 0.2113534 |
| AGLESIGL | ASSQDPGADTQY | 0.20353961 |
| AGLKTGII | ASSQDPGADTQY | 0.23781364 |

| | | |
|---|---|---|
| ASLAALLL | ASSQDPGADTQY | 0.4286675 |
| ASVINISSI | ASSQDPGADTQY | 0.17678186 |
| ATPPGPA | ASSQDPGADTQY | 0.53455174 |
| ATVASALV | ASSQDPGADTQY | 0.5578613 |
| ATVASLIL | ASSQDPGADTQY | 0.6783707 |
| AVNGPVLIRH | ASSQDPGADTQY | 0.16886237 |
| DGAAGAVGI | ASSQDPGADTQY | 0.7611524 |
| DPLLHGI | ASSQDPGADTQY | 0.74767077 |
| ELGAALGAL | ASSQDPGADTQY | 0.66130686 |
| ELGRTQAV | ASSQDPGADTQY | 0.35967278 |
| FVAGGYLS | ASSQDPGADTQY | 0.3282705 |
| GASGGAGGAGGW | ASSQDPGADTQY | 0.7458163 |
| GGAGAGGDGGAGGI | ASSQDPGADTQY | 0.30573517 |
| GTGHNGSQPGMGGNG | ASSQDPGADTQY | 0.403601 |
| GYAAASASAS | ASSQDPGADTQY | 0.7730836 |
| HQVDVAAVQQG | ASSQDPGADTQY | 0.23614497 |
| IGAAIAHT | ASSQDPGADTQY | 0.19725856 |
| IGMIEVAQL | ASSQDPGADTQY | 0.6625804 |
| IRLAPEHTSAMD | ASSQDPGADTQY | 0.6260431 |
| IYNVSVL | ASSQDPGADTQY | 0.2161971 |
| KQGFLGM | ASSQDPGADTQY | 0.2543332 |
| LESVITQL | ASSQDPGADTQY | 0.30659696 |
| LGLGALVVV | ASSQDPGADTQY | 0.37158632 |
| LIGNGGNGGSGGI | ASSQDPGADTQY | 0.7790885 |
| LNAINAPTL | ASSQDPGADTQY | 0.4470359 |
| LQLLNTGV | ASSQDPGADTQY | 0.4228536 |
| LQTSIEF | ASSQDPGADTQY | 0.7958329 |
| MSFAVSEL | ASSQDPGADTQY | 0.51252097 |
| NTGALGL | ASSQDPGADTQY | 0.4061 |
| PPGAGGAGGAGTT | ASSQDPGADTQY | 0.80440843 |
| QLVGLGI | ASSQDPGADTQY | 0.21365216 |
| RFLEQVHQL | ASSQDPGADTQY | 0.62162274 |
| RGVGYVLR | ASSQDPGADTQY | 0.42043757 |
| SATGVFAGVI | ASSQDPGADTQY | 0.73174834 |
| SLPAITLP | ASSQDPGADTQY | 0.7510816 |
| STALAFTV | ASSQDPGADTQY | 0.48370457 |
| SVLTVGLGK | ASSQDPGADTQY | 0.44960874 |
| TALAGAGSGI | ASSQDPGADTQY | 0.18242002 |
| TGLPFTGI | ASSQDPGADTQY | 0.35447043 |
| VAEHVTV | ASSQDPGADTQY | 0.7764079 |
| VDIVLNSL | ASSQDPGADTQY | 0.55715257 |
| VLTGGGAVL | ASSQDPGADTQY | 0.79151833 |
| VNVGLAYGI | ASSQDPGADTQY | 0.660148 |
| VSSAAVGAV | ASSQDPGADTQY | 0.32352787 |
| VTSVYEAGV | ASSQDPGADTQY | 0.8114704 |
| WVALDRGA | ASSQDPGADTQY | 0.7568032 |
| YLVGSVVI | ASSQDPGADTQY | 0.17851202 |
| GPPGTGKTLL | ASSQDPGADTQY | 0.55003715 |
| GVGMTKF | ASSQDPGADTQY | 0.7886815 |
| IGIGGSDL | ASSQDPGADTQY | 0.73583156 |
| IGTLLAEL | ASSQDPGADTQY | 0.33248445 |

| | | |
|---|---|---|
| IYTSGTT | ASSQDPGADTQY | 0.52246124 |
| LSFPAGV | ASSQDPGADTQY | 0.7996764 |
| LYTSGTT | ASSQDPGADTQY | 0.41334176 |
| VGAFPFGF | ASSQDPGADTQY | 0.7787878 |
| VGAGTVEF | ASSQDPGADTQY | 0.1625213 |
| VGKSTLLL | ASSQDPGADTQY | 0.38591444 |
| AALAHAGGGIV | ASSQDPGADTQY | 0.25635028 |
| AALVAVSLL | ASSQDPGADTQY | 0.16873391 |
| AAVSRAVAAL | ASSQDPGADTQY | 0.5882597 |
| AGGSSGIGGF | ASSQDPGADTQY | 0.4689705 |
| AGLTVANANG | ASSQDPGADTQY | 0.8073797 |
| AGLVGAVL | ASSQDPGADTQY | 0.4678308 |
| ALSQAFTV | ASSQDPGADTQY | 0.7050398 |
| ALTFSVSL | ASSQDPGADTQY | 0.40112185 |
| ALTGAALVAL | ASSQDPGADTQY | 0.7492715 |
| ASFSIGAGVL | ASSQDPGADTQY | 0.4059342 |
| DGSVYKI | ASSQDPGADTQY | 0.4154796 |
| EQLAEIV | ASSQDPGADTQY | 0.81445533 |
| FAVSELTVM | ASSQDPGADTQY | 0.52241665 |
| GAAPLAAGL | ASSQDPGADTQY | 0.16726097 |
| GADGGDGGSGGAGG | ASSQDPGADTQY | 0.6259107 |
| GFGLDTGGAGG | ASSQDPGADTQY | 0.31331822 |
| GLASGGLL | ASSQDPGADTQY | 0.16649964 |
| GRIAGAAVLL | ASSQDPGADTQY | 0.18459195 |
| GSAGSLNTSF | ASSQDPGADTQY | 0.20776743 |
| GTGQIGI | ASSQDPGADTQY | 0.26156902 |
| GTLIALIL | ASSQDPGADTQY | 0.5457626 |
| IAATMGFVVG | ASSQDPGADTQY | 0.58676857 |
| IFAQALL | ASSQDPGADTQY | 0.23117599 |
| IGAAVALGF | ASSQDPGADTQY | 0.67750967 |
| IGLTPLAL | ASSQDPGADTQY | 0.3145159 |
| ILGSNTSTL | ASSQDPGADTQY | 0.13752463 |
| IVFSLTVL | ASSQDPGADTQY | 0.35049313 |
| LASKATGF | ASSQDPGADTQY | 0.3775943 |
| LASVALIL | ASSQDPGADTQY | 0.7455765 |
| LFAVGLTL | ASSQDPGADTQY | 0.4491349 |
| LFNGGIGTYI | ASSQDPGADTQY | 0.31265202 |
| LTLGAGII | ASSQDPGADTQY | 0.3079841 |
| LVAAALGDI | ASSQDPGADTQY | 0.7249968 |
| NTAFVQLGI | ASSQDPGADTQY | 0.5358513 |
| QLFSGAEL | ASSQDPGADTQY | 0.2408829 |
| SAATFITL | ASSQDPGADTQY | 0.1563288 |
| SNYGLANLGV | ASSQDPGADTQY | 0.4996071 |
| SYGGAYAVM | ASSQDPGADTQY | 0.22024013 |
| TTGVGGLTLG | ASSQDPGADTQY | 0.4071438 |
| TTQTIYI | ASSQDPGADTQY | 0.29497585 |
| VAAGAGLGAVI | ASSQDPGADTQY | 0.23862846 |
| VAAVVTLL | ASSQDPGADTQY | 0.1764963 |
| VGFLGGLI | ASSQDPGADTQY | 0.7916765 |
| VGVIPILL | ASSQDPGADTQY | 0.6199191 |
| VIFASSLI | ASSQDPGADTQY | 0.26193485 |

| | | |
|---|---|---|
| VKGGDGGAAAG | ASSQDPGADTQY | 0.6841238 |
| VTKTFTNI | ASSQDPGADTQY | 0.68609744 |
| VVAAAILL | ASSQDPGADTQY | 0.29887098 |
| YAALDVEL | ASSQDPGADTQY | 0.35285527 |
| LASVALLL | ASSQDPGADTQY | 0.3362115 |
| LDAVHRV | ASSQDPGADTQY | 0.25658387 |
| QGLSSSIF | ASSQDPGADTQY | 0.22797008 |
| RGTFANIRL | ASSQDPGADTQY | 0.7925879 |
| RQLWWGHRI | ASSQDPGADTQY | 0.40571442 |
| YGPPGTGKTLL | ASSQDPGADTQY | 0.26929688 |
| AALVGAPANVA | ASSQDPGADTQY | 0.41236714 |
| AATFRRTE | ASSQDPGADTQY | 0.71839476 |
| AGALGGGI | ASSQDPGADTQY | 0.7658074 |
| AGGAGGAGGIGAG | ASSQDPGADTQY | 0.55556965 |
| AGSGALGL | ASSQDPGADTQY | 0.3264572 |
| AQIADQI | ASSQDPGADTQY | 0.4633097 |
| ASAVAVTYL | ASSQDPGADTQY | 0.3513192 |
| ASTVLGGTGG | ASSQDPGADTQY | 0.5397582 |
| AVGGEPAK | ASSQDPGADTQY | 0.19902757 |
| CEQGTFGGTC | ASSQDPGADTQY | 0.6389225 |
| DEGKAPAG | ASSQDPGADTQY | 0.80760205 |
| DTVVFPAL | ASSQDPGADTQY | 0.36970493 |
| FEGITMAG | ASSQDPGADTQY | 0.22581813 |
| FPLVVQD | ASSQDPGADTQY | 0.28827685 |
| GAFTTPQ | ASSQDPGADTQY | 0.2331331 |
| GFGGFGTAGGVGGA | ASSQDPGADTQY | 0.20420808 |
| GFLAQFL | ASSQDPGADTQY | 0.6408677 |
| GGAGGAGGNAGVA | ASSQDPGADTQY | 0.22892505 |
| GGVGTAGAGGAGGAGG | ASSQDPGADTQY | 0.3157007 |
| GNGGGGDAGNAGSGGN | ASSQDPGADTQY | 0.7083757 |
| GTGGAGGAGNDAGS | ASSQDPGADTQY | 0.49869984 |
| GTTNTGWFNS | ASSQDPGADTQY | 0.6561558 |
| IFMGPLFTL | ASSQDPGADTQY | 0.67686146 |
| ILAHAIEAI | ASSQDPGADTQY | 0.641327 |
| IRSAIREAL | ASSQDPGADTQY | 0.80986303 |
| ITFAGGG | ASSQDPGADTQY | 0.7962698 |
| ITGMSGHQEV | ASSQDPGADTQY | 0.6495503 |
| IVAGLACTL | ASSQDPGADTQY | 0.48351115 |
| IVGGGLAGA | ASSQDPGADTQY | 0.6290786 |
| IVTGVLNI | ASSQDPGADTQY | 0.46797377 |
| LAQQGCI | ASSQDPGADTQY | 0.30659106 |
| LCLHVEN | ASSQDPGADTQY | 0.4223578 |
| LDAISDGS | ASSQDPGADTQY | 0.65357685 |
| LGGFAAML | ASSQDPGADTQY | 0.51992387 |
| LGGKNPMI | ASSQDPGADTQY | 0.7495886 |
| LGPKGVAAIL | ASSQDPGADTQY | 0.21632528 |
| LMVGVSI | ASSQDPGADTQY | 0.1983273 |
| LVGAAFVL | ASSQDPGADTQY | 0.7566841 |
| LVLAVAPL | ASSQDPGADTQY | 0.15120682 |
| MAAIAVIDAA | ASSQDPGADTQY | 0.20599245 |
| NNGINAELQ | ASSQDPGADTQY | 0.5650773 |

| | | |
|---|---|---|
| RGEPLVLVHG | ASSQDPGADTQY | 0.6384187 |
| RGGAGGAAGLI | ASSQDPGADTQY | 0.14992043 |
| RSYRMHPAV | ASSQDPGADTQY | 0.24456486 |
| SAAHLLAGQLA | ASSQDPGADTQY | 0.30711156 |
| SAAVGSY | ASSQDPGADTQY | 0.49259713 |
| SPHAVLHRI | ASSQDPGADTQY | 0.2737077 |
| SQIAKVL | ASSQDPGADTQY | 0.16401608 |
| TALGHLVP | ASSQDPGADTQY | 0.7356936 |
| TAPTTSGAA | ASSQDPGADTQY | 0.1721612 |
| TDGDDGNGGSPGT | ASSQDPGADTQY | 0.23633668 |
| TDGLGEN | ASSQDPGADTQY | 0.7358084 |
| TGGPPLPGQA | ASSQDPGADTQY | 0.8242223 |
| TIPAITI | ASSQDPGADTQY | 0.38016674 |
| TIPGFTI | ASSQDPGADTQY | 0.73275566 |
| TSIASGALA | ASSQDPGADTQY | 0.23278326 |
| TVTLPGAAI | ASSQDPGADTQY | 0.79408145 |
| VASGLSAALP | ASSQDPGADTQY | 0.27158776 |
| VGAGVIGI | ASSQDPGADTQY | 0.19087948 |
| VGPKRFGV | ASSQDPGADTQY | 0.7264204 |
| VNVAFPD | ASSQDPGADTQY | 0.72249687 |
| VSMTNSGVS | ASSQDPGADTQY | 0.6663133 |
| VTDVLGDLSV | ASSQDPGADTQY | 0.75024414 |
| VVVVGAGFAGL | ASSQDPGADTQY | 0.2551353 |
| VYTVGTVV | ASSQDPGADTQY | 0.3119827 |
| WTTFSGP | ASSQDPGADTQY | 0.67584527 |
| YDELVEAF | ASSQDPGADTQY | 0.7795738 |
| YGTGGAGGNGG | ASSQDPGADTQY | 0.18964425 |
| YTNVMAV | ASSQDPGADTQY | 0.59624696 |
| FLEQVHQL | ASSQDPGADTQY | 0.29592344 |
| GLSLLAVAV | ASSQDPGADTQY | 0.45822963 |
| LEQVHQL | ASSQDPGADTQY | 0.67562234 |
| LNAAGAI | ASSQDPGADTQY | 0.6628986 |
| PAVHASL | ASSQDPGADTQY | 0.6203231 |
| TELEDDA | ASSQDPGADTQY | 0.4689969 |
| TPPGPPN | ASSQDPGADTQY | 0.8095063 |
| TPPPPPPP | ASSQDPGADTQY | 0.21907815 |
| TQSPALI | ASSLSAGGYAEQF | 0.6950572 |
| EFAAAFPQ | ASSLSAGGYAEQF | 0.5503689 |
| LLSGNLDVL | ASSLSAGGYAEQF | 0.78751475 |
| GPANLGNYNI | ASSLSAGGYAEQF | 0.5176422 |
| TLGPTLP | ASSLSAGGYAEQF | 0.2837779 |
| VYAYSSCCK | ASSLSAGGYAEQF | 0.52067745 |
| APAPAPAPNI | ASSLSAGGYAEQF | 0.70356214 |
| PGGPVPIP | ASSLSAGGYAEQF | 0.46315297 |
| FLTVALEP | ASSLSAGGYAEQF | 0.7717304 |
| SQYFGANL | ASSLSAGGYAEQF | 0.8167466 |
| GGVKPAA | ASSLSAGGYAEQF | 0.37063438 |
| GGTGGAGGAGGW | ASSLSAGGYAEQF | 0.33963156 |
| GVGGVAGLGGAGG | ASSLSAGGYAEQF | 0.7312011 |
| VLLVLATTI | ASSLSAGGYAEQF | 0.8173811 |
| IGIFAAML | ASSLSAGGYAEQF | 0.46744132 |

| | | |
|---|---|---|
| AVGDLID | ASSLSAGGYAEQF | 0.7966283 |
| NISPLPL | ASSLSAGGYAEQF | 0.5412279 |
| VTAVFVAI | ASSLSAGGYAEQF | 0.59476006 |
| SGDDNDAAL | ASSLSAGGYAEQF | 0.42627543 |
| AVAALAHS | ASSLSAGGYAEQF | 0.55840904 |
| GGVTVMATP | ASSLSAGGYAEQF | 0.32859662 |
| AGLAANVRD | ASSLSAGGYAEQF | 0.56167567 |
| TNRPDLIDPALLRPG | ASSLSAGGYAEQF | 0.63831276 |
| LANLGVTAL | ASSLSAGGYAEQF | 0.60668534 |
| TGERIEL | ASSLSAGGYAEQF | 0.21433939 |
| NGEELAA | ASSLSAGGYAEQF | 0.43851435 |
| VLSLVWG | ASSLSAGGYAEQF | 0.6909524 |
| IGYALWL | ASSLSAGGYAEQF | 0.82802516 |
| LSLLLAPL | ASSLSAGGYAEQF | 0.51954365 |
| TLDEALK | ASSLSAGGYAEQF | 0.6743078 |
| TDGAGTAEGG | ASSLSAGGYAEQF | 0.2997429 |
| NAPGGAGGNGG | ASSLSAGGYAEQF | 0.3961287 |
| VSVFMGMGGNF | ASSLSAGGYAEQF | 0.24758926 |
| LASYLAAL | ASSLSAGGYAEQF | 0.650857 |
| LVGGTPAL | ASSLSAGGYAEQF | 0.3554315 |
| VRNADVL | ASSLSAGGYAEQF | 0.37784863 |
| TCSSPFLVSAM | ASSLSAGGYAEQF | 0.74233496 |
| VHITDAE | ASSLSAGGYAEQF | 0.82135713 |
| EYAAPSF | ASSLSAGGYAEQF | 0.4681881 |
| VVDAFHGGL | ASSLSAGGYAEQF | 0.70488954 |
| ASVVDPQ | ASSLSAGGYAEQF | 0.6842196 |
| SYNEFGGLS | ASSLSAGGYAEQF | 0.74613583 |
| DPDGSVY | ASSLSAGGYAEQF | 0.22551858 |
| MVSTIHATE | ASSLSAGGYAEQF | 0.45706904 |
| LAAGVADPV | ASSLSAGGYAEQF | 0.22414733 |
| TKEQIAA | ASSLSAGGYAEQF | 0.28359827 |
| SLASPAAL | ASSLSAGGYAEQF | 0.23217759 |
| IGLVFYGV | ASSLSAGGYAEQF | 0.33566105 |
| FLLDASNL | ASSLSAGGYAEQF | 0.14950661 |
| YQVGNFP | ASSLSAGGYAEQF | 0.709872 |
| ESLLSQST | ASSLSAGGYAEQF | 0.7914361 |
| LFTVGFAAVG | ASSLSAGGYAEQF | 0.25759858 |
| ATPPTPT | ASSLSAGGYAEQF | 0.6361526 |
| TDLATLG | ASSLSAGGYAEQF | 0.35165098 |
| SAGGTGR | ASSLSAGGYAEQF | 0.2764002 |
| GFSNSGS | ASSLSAGGYAEQF | 0.5856364 |
| GGNGGSGGTSV | ASSLSAGGYAEQF | 0.27882332 |
| GTSVATGGAGN | ASSLSAGGYAEQF | 0.77718174 |
| NGGNGGSGGTS | ASSLSAGGYAEQF | 0.57175004 |
| AVVKFPH | ASSLSAGGYAEQF | 0.48108202 |
| TPPPPPP | ASSLSAGGYAEQF | 0.7238112 |
| AAPAAAAAPA | ASSLSAGGYAEQF | 0.54480577 |
| IALNEDL | ASSLSAGGYAEQF | 0.2015868 |
| IVNVSSM | ASSLSAGGYAEQF | 0.7325066 |
| TGIPPAPR | ASSLSAGGYAEQF | 0.25129354 |
| FDLGGGTF | ASSLSAGGYAEQF | 0.45424587 |

| | | |
|---|---|---|
| FDLGGGTFDVS | ASSLSAGGYAEQF | 0.23003678 |
| NEPTAAA | ASSLSAGGYAEQF | 0.7474368 |
| NEPTAAALA | ASSLSAGGYAEQF | 0.2896781 |
| GSIINISSI | ASSLSAGGYAEQF | 0.6233834 |
| VAPGNAGTA | ASSLSAGGYAEQF | 0.26470387 |
| TGGGASLE | ASSLSAGGYAEQF | 0.513772 |
| VAVYNHYKRI | ASSLSAGGYAEQF | 0.57558495 |
| TDPAPATT | ASSLSAGGYAEQF | 0.18739271 |
| TNRPDILDPALLRPG | ASSLSAGGYAEQF | 0.7064216 |
| LTTTTTS | ASSLSAGGYAEQF | 0.4109154 |
| PTGSGKTL | ASSLSAGGYAEQF | 0.24798751 |
| TGSGKTL | ASSLSAGGYAEQF | 0.64272606 |
| TIGHVDHGK | ASSLSAGGYAEQF | 0.6825175 |
| TIGHVDHGKTT | ASSLSAGGYAEQF | 0.18078856 |
| GHVDHGKTT | ASSLSAGGYAEQF | 0.35700312 |
| AQFEHTLL | ASSLSAGGYAEQF | 0.4930942 |
| IVNQDEL | ASSLSAGGYAEQF | 0.3223357 |
| TNGSQFF | ASSLSAGGYAEQF | 0.15646693 |
| TGKGIVE | ASSLSAGGYAEQF | 0.6339716 |
| LISNASDA | ASSLSAGGYAEQF | 0.45984462 |
| AELRTLGELTKIAKAHGAQ | ASSLSAGGYAEQF | 0.21919537 |
| APNAVAAAEPSW | ASSLSAGGYAEQF | 0.3949436 |
| APQVAGAATAVL | ASSLSAGGYAEQF | 0.79339564 |
| ARRDPAAAAAILPTDA | ASSLSAGGYAEQF | 0.26230592 |
| DAAARDVLAEAGLAEYF | ASSLSAGGYAEQF | 0.681654 |
| DDQIVLSAKANLAKVEQ | ASSLSAGGYAEQF | 0.79808044 |
| DELLPDTGEDSDEDGH | ASSLSAGGYAEQF | 0.354713 |
| ELPEAPANRATVPQD | ASSLSAGGYAEQF | 0.7136284 |
| FGQSKIERVTGIG | ASSLSAGGYAEQF | 0.6922123 |
| GAGGTGSTAGGAGGAGGAGGLY | ASSLSAGGYAEQF | 0.37004995 |
| GHGGTGAAGGNGGTGGAGGSGI | ASSLSAGGYAEQF | 0.77016294 |
| GNGGNGGAGGEGGPGGAGGAGA | ASSLSAGGYAEQF | 0.42244005 |
| GSTAGNGGNGGNATNGTIAGQP | ASSLSAGGYAEQF | 0.25461647 |
| GTGGDGGNGGNGGTGG | ASSLSAGGYAEQF | 0.24622326 |
| GVRRPAGIPADAPV | ASSLSAGGYAEQF | 0.3413111 |
| GYRQVTDAHLLTLARR | ASSLSAGGYAEQF | 0.25710082 |
| IAKMGQDGHDRG | ASSLSAGGYAEQF | 0.46246457 |
| KAEPSIEIQYT | ASSLSAGGYAEQF | 0.7578884 |
| KPVRIRMAADEDTVAVT | ASSLSAGGYAEQF | 0.42908168 |
| MPPRIAGMRLLVIKPEP | ASSLSAGGYAEQF | 0.26688272 |
| MKFYAVALFRSRGVPPTR | ASSLSAGGYAEQF | 0.3702066 |
| NAGGFGILGAVAHSP | ASSLSAGGYAEQF | 0.23031689 |
| NEGREITLAMAEM | ASSLSAGGYAEQF | 0.78993917 |
| NELVEWSHPLGHRWRWE | ASSLSAGGYAEQF | 0.4520684 |
| NLGAGGTGGNGGNPG | ASSLSAGGYAEQF | 0.8007436 |
| NTEWPKWETVMAADPVD | ASSLSAGGYAEQF | 0.5779259 |
| PASAAPAETQP | ASSLSAGGYAEQF | 0.22458799 |
| QQSSDGKNGVRAAEITGE | ASSLSAGGYAEQF | 0.7624663 |
| RAPDAVTTSWWKE | ASSLSAGGYAEQF | 0.27459526 |
| SGFGNVGAHTSGWFNQ | ASSLSAGGYAEQF | 0.503391 |
| SGVAGAVGQAASVGGLKV | ASSLSAGGYAEQF | 0.59624815 |

| | | |
|---|---|---|
| SNGGAGTGSAGGNGGT | ASSLSAGGYAEQF | 0.41095054 |
| TGGAGGEGGAAG | ASSLSAGGYAEQF | 0.6877005 |
| TGGDGGAAGAGGAPGHG | ASSLSAGGYAEQF | 0.7234939 |
| TMPPLLAAIANAGRHGV | ASSLSAGGYAEQF | 0.19546317 |
| TSSTAGNSGGA | ASSLSAGGYAEQF | 0.7018509 |
| VPATVPVGILC | ASSLSAGGYAEQF | 0.19467974 |
| VQRPDSGGVVLTGRIS | ASSLSAGGYAEQF | 0.17683925 |
| WTPNPVPATKGVPAGLVQAT | ASSLSAGGYAEQF | 0.39562 |
| YALKRALAKFSDRFDVVII | ASSLSAGGYAEQF | 0.8165512 |
| YLIGSVIV | ASSLSAGGYAEQF | 0.76259494 |
| YGQLVAVID | ASSLSAGGYAEQF | 0.29386902 |
| VTVGGMTLPA | ASSLSAGGYAEQF | 0.2176985 |
| VLALPAV | ASSLSAGGYAEQF | 0.75690466 |
| VIPPTLNYET | ASSLSAGGYAEQF | 0.19289106 |
| VGMGGMHPSSG | ASSLSAGGYAEQF | 0.4698434 |
| VGLAGGAATA | ASSLSAGGYAEQF | 0.35147288 |
| VGGAGGAGGSGP | ASSLSAGGYAEQF | 0.24485016 |
| VFAAFMV | ASSLSAGGYAEQF | 0.31930652 |
| TLYDKLR | ASSLSAGGYAEQF | 0.5668936 |
| TLAVFTVL | ASSLSAGGYAEQF | 0.23771602 |
| TGSVQPARV | ASSLSAGGYAEQF | 0.6942206 |
| TGGGAGSGGT | ASSLSAGGYAEQF | 0.14283353 |
| SHLETSL | ASSLSAGGYAEQF | 0.5835096 |
| SFTTGMT | ASSLSAGGYAEQF | 0.20224822 |
| QVEVAKRMGVRQPTVS | ASSLSAGGYAEQF | 0.30339324 |
| QGSGGPALAM | ASSLSAGGYAEQF | 0.18294132 |
| MLAPSNP | ASSLSAGGYAEQF | 0.5116852 |
| LVNNAAI | ASSLSAGGYAEQF | 0.6844516 |
| LTGEINTLA | ASSLSAGGYAEQF | 0.7545481 |
| LSIFGPV | ASSLSAGGYAEQF | 0.4430132 |
| LLALLANL | ASSLSAGGYAEQF | 0.75148475 |
| LISNASDALDKLR | ASSLSAGGYAEQF | 0.7204595 |
| LINEAVP | ASSLSAGGYAEQF | 0.28433216 |
| LGYAFSGL | ASSLSAGGYAEQF | 0.2923882 |
| LDFVAVAL | ASSLSAGGYAEQF | 0.6347997 |
| LAQPGAE | ASSLSAGGYAEQF | 0.681635 |
| ISNLGAVTGF | ASSLSAGGYAEQF | 0.6810138 |
| HQEDPGRI | ASSLSAGGYAEQF | 0.6380993 |
| GTPGEPTP | ASSLSAGGYAEQF | 0.2547208 |
| GFFNSGS | ASSLSAGGYAEQF | 0.61412966 |
| GAGGGGLTAGAN | ASSLSAGGYAEQF | 0.70582783 |
| FVDALVNM | ASSLSAGGYAEQF | 0.2677071 |
| FLTVAPEL | ASSLSAGGYAEQF | 0.45076904 |
| FETLTAL | ASSLSAGGYAEQF | 0.37686825 |
| DVINPIEL | ASSLSAGGYAEQF | 0.17435929 |
| APVSGAPV | ASSLSAGGYAEQF | 0.4608348 |
| ALVGGLAW | ASSLSAGGYAEQF | 0.28523716 |
| ALAAAAATAI | ASSLSAGGYAEQF | 0.28071624 |
| AFGLGLLL | ASSLSAGGYAEQF | 0.35291052 |
| AFFAAASL | ASSLSAGGYAEQF | 0.24928713 |
| AAAALPAFSPP | ASSLSAGGYAEQF | 0.26625443 |

| | | |
|---|---|---|
| VVLAPAL | ASSLSAGGYAEQF | 0.23814872 |
| TIGHVDHGKTTL | ASSLSAGGYAEQF | 0.37395117 |
| LISNASDALDK | ASSLSAGGYAEQF | 0.6054747 |
| LISNASDAL | ASSLSAGGYAEQF | 0.28815538 |
| ISNASDALDK | ASSLSAGGYAEQF | 0.5648275 |
| NQAETLVYQTE | ASSLSAGGYAEQF | 0.6956878 |
| TGGTGGTAGAG | ASSLSAGGYAEQF | 0.17332073 |
| TGSTAGGAGGA | ASSLSAGGYAEQF | 0.21302713 |
| DPLAVPVAAGPV | ASSLSAGGYAEQF | 0.65105426 |
| GGTGGTPGNGGN | ASSLSAGGYAEQF | 0.22486092 |
| PAGLAGVDPATL | ASSLSAGGYAEQF | 0.7640841 |
| GNNLVVEAPHAGASV | ASSLSAGGYAEQF | 0.2989692 |
| KVTRIMSASTLEQPA | ASSLSAGGYAEQF | 0.33652356 |
| LGFALPIALPRRTVV | ASSLSAGGYAEQF | 0.7037542 |
| TRLGNLTARKLVVLV | ASSLSAGGYAEQF | 0.814085 |
| WSQQRRLIYFPSAGP | ASSLSAGGYAEQF | 0.7657096 |
| GPNGNPDPMAAAVDIR | ASSLSAGGYAEQF | 0.3807683 |
| TQRCVSVADLRAAASR | ASSLSAGGYAEQF | 0.7825091 |
| QFVLDVAEHPVEVLTETV | ASSLSAGGYAEQF | 0.320919 |
| GGAGGTGGAGGNGGTQAGDG | ASSLSAGGYAEQF | 0.73942494 |
| PLLATLRAIAVDVDAKPAQV | ASSLSAGGYAEQF | 0.77697766 |
| APKFRTGGYMIDFWGVGYQVAK | ASSLSAGGYAEQF | 0.17802908 |
| GAGGAGGAGGAAGGPGGTGGTG | ASSLSAGGYAEQF | 0.32742298 |
| IGALGLLATDVAPAGSVAFGVA | ASSLSAGGYAEQF | 0.7627591 |
| AGGAGGIGGNGGAGGAGGVLMG | ASSLSAGGYAEQF | 0.81472635 |
| ADELIGLL | ASSLSAGGYAEQF | 0.22267354 |
| ADRALKA | ASSLSAGGYAEQF | 0.46943325 |
| AGAVGQVGGAGGA | ASSLSAGGYAEQF | 0.2807041 |
| AGGGGGLGGAP | ASSLSAGGYAEQF | 0.41514522 |
| AGGLLYGNGGAG | ASSLSAGGYAEQF | 0.3055008 |
| AGLESIGL | ASSLSAGGYAEQF | 0.6375638 |
| AGLKTGII | ASSLSAGGYAEQF | 0.3712715 |
| ASLAALLL | ASSLSAGGYAEQF | 0.77073383 |
| ASVINISSI | ASSLSAGGYAEQF | 0.3878224 |
| ATPPGPA | ASSLSAGGYAEQF | 0.6248752 |
| ATVASALV | ASSLSAGGYAEQF | 0.35005346 |
| ATVASLIL | ASSLSAGGYAEQF | 0.27407226 |
| AVNGPVLIRH | ASSLSAGGYAEQF | 0.259872 |
| DGAAGAVGI | ASSLSAGGYAEQF | 0.19852118 |
| DPLLHGI | ASSLSAGGYAEQF | 0.28196308 |
| ELGAALGAL | ASSLSAGGYAEQF | 0.24865088 |
| ELGRTQAV | ASSLSAGGYAEQF | 0.15727144 |
| FVAGGYLS | ASSLSAGGYAEQF | 0.5282811 |
| GASGGAGGAGGW | ASSLSAGGYAEQF | 0.684144 |
| GGAGAGGDGGAGGI | ASSLSAGGYAEQF | 0.46544448 |
| GTGHNGSQPGMGGNG | ASSLSAGGYAEQF | 0.28211296 |
| GYAAASASAS | ASSLSAGGYAEQF | 0.56871265 |
| HQVDVAAVQQG | ASSLSAGGYAEQF | 0.20071712 |
| IGAAIAHT | ASSLSAGGYAEQF | 0.7965561 |
| IGMIEVAQL | ASSLSAGGYAEQF | 0.39612687 |
| IRLAPEHTSAMD | ASSLSAGGYAEQF | 0.17616948 |

| | | |
|---|---|---|
| IYNVSVL | ASSLSAGGYAEQF | 0.33544374 |
| KQGFLGM | ASSLSAGGYAEQF | 0.23205209 |
| LESVITQL | ASSLSAGGYAEQF | 0.70356715 |
| LGLGALVVV | ASSLSAGGYAEQF | 0.6339866 |
| LIGNGGNGGSGGI | ASSLSAGGYAEQF | 0.5530704 |
| LNAINAPTL | ASSLSAGGYAEQF | 0.779848 |
| LQLLNTGV | ASSLSAGGYAEQF | 0.15955703 |
| LQTSIEF | ASSLSAGGYAEQF | 0.25384954 |
| MSFAVSEL | ASSLSAGGYAEQF | 0.75548834 |
| NTGALGL | ASSLSAGGYAEQF | 0.51407456 |
| PPGAGGAGGAGTT | ASSLSAGGYAEQF | 0.37073576 |
| QLVGLGI | ASSLSAGGYAEQF | 0.78017664 |
| RFLEQVHQL | ASSLSAGGYAEQF | 0.21156184 |
| RGVGYVLR | ASSLSAGGYAEQF | 0.3896503 |
| SATGVFAGVI | ASSLSAGGYAEQF | 0.6669917 |
| SLPAITLP | ASSLSAGGYAEQF | 0.6853982 |
| STALAFTV | ASSLSAGGYAEQF | 0.42735547 |
| SVLTVGLGK | ASSLSAGGYAEQF | 0.4469983 |
| TALAGAGSGI | ASSLSAGGYAEQF | 0.5973922 |
| TGLPFTGI | ASSLSAGGYAEQF | 0.80144715 |
| VAEHVTV | ASSLSAGGYAEQF | 0.5982847 |
| VDIVLNSL | ASSLSAGGYAEQF | 0.40933758 |
| VLTGGGAVL | ASSLSAGGYAEQF | 0.6357351 |
| VNVGLAYGI | ASSLSAGGYAEQF | 0.578521 |
| VSSAAVGAV | ASSLSAGGYAEQF | 0.21526985 |
| VTSVYEAGV | ASSLSAGGYAEQF | 0.32013863 |
| WVALDRGA | ASSLSAGGYAEQF | 0.5572852 |
| YLVGSVVI | ASSLSAGGYAEQF | 0.2087724 |
| GPPGTGKTLL | ASSLSAGGYAEQF | 0.6736391 |
| GVGMTKF | ASSLSAGGYAEQF | 0.28352016 |
| IGIGGSDL | ASSLSAGGYAEQF | 0.8160633 |
| IGTLLAEL | ASSLSAGGYAEQF | 0.6216771 |
| IYTSGTT | ASSLSAGGYAEQF | 0.6365829 |
| LSFPAGV | ASSLSAGGYAEQF | 0.21215773 |
| LYTSGTT | ASSLSAGGYAEQF | 0.47904208 |
| VGAFPFGF | ASSLSAGGYAEQF | 0.5318238 |
| VGAGTVEF | ASSLSAGGYAEQF | 0.18438303 |
| VGKSTLLL | ASSLSAGGYAEQF | 0.71626973 |
| AALAHAGGGIV | ASSLSAGGYAEQF | 0.20849419 |
| AALVAVSLL | ASSLSAGGYAEQF | 0.35115004 |
| AAVSRAVAAL | ASSLSAGGYAEQF | 0.6606614 |
| AGGSSGIGGF | ASSLSAGGYAEQF | 0.7884871 |
| AGLTVANANG | ASSLSAGGYAEQF | 0.28554964 |
| AGLVGAVL | ASSLSAGGYAEQF | 0.42866817 |
| ALSQAFTV | ASSLSAGGYAEQF | 0.29988566 |
| ALTFSVSL | ASSLSAGGYAEQF | 0.63246834 |
| ALTGAALVAL | ASSLSAGGYAEQF | 0.8047529 |
| ASFSIGAGVL | ASSLSAGGYAEQF | 0.6881218 |
| DGSVYKI | ASSLSAGGYAEQF | 0.81943166 |
| EQLAEIV | ASSLSAGGYAEQF | 0.57941115 |
| FAVSELTVM | ASSLSAGGYAEQF | 0.24006991 |

| | | |
|---|---|---|
| GAAPLAAGL | ASSLSAGGYAEQF | 0.6244537 |
| GADGGDGGSGGAGG | ASSLSAGGYAEQF | 0.68804336 |
| GFGLDTGGAGG | ASSLSAGGYAEQF | 0.7452134 |
| GLASGGLL | ASSLSAGGYAEQF | 0.68931717 |
| GRIAGAAVLL | ASSLSAGGYAEQF | 0.79882103 |
| GSAGSLNTSF | ASSLSAGGYAEQF | 0.36598307 |
| GTGQIGI | ASSLSAGGYAEQF | 0.64663005 |
| GTLIALIL | ASSLSAGGYAEQF | 0.72612226 |
| IAATMGFVVG | ASSLSAGGYAEQF | 0.72791094 |
| IFAQALL | ASSLSAGGYAEQF | 0.3749103 |
| IGAAVALGF | ASSLSAGGYAEQF | 0.2748366 |
| IGLTPLAL | ASSLSAGGYAEQF | 0.51537883 |
| ILGSNTSTL | ASSLSAGGYAEQF | 0.6651856 |
| IVFSLTVL | ASSLSAGGYAEQF | 0.39324757 |
| LASKATGF | ASSLSAGGYAEQF | 0.32357675 |
| LASVALIL | ASSLSAGGYAEQF | 0.37395704 |
| LFAVGLTL | ASSLSAGGYAEQF | 0.2743447 |
| LFNGGIGTYI | ASSLSAGGYAEQF | 0.52916396 |
| LTLGAGII | ASSLSAGGYAEQF | 0.80337775 |
| LVAAALGDI | ASSLSAGGYAEQF | 0.3850922 |
| NTAFVQLGI | ASSLSAGGYAEQF | 0.30304992 |
| QLFSGAEL | ASSLSAGGYAEQF | 0.72555494 |
| SAATFITL | ASSLSAGGYAEQF | 0.78518724 |
| SNYGLANLGV | ASSLSAGGYAEQF | 0.6691333 |
| SYGGAYAVM | ASSLSAGGYAEQF | 0.18670729 |
| TTGVGGLTLG | ASSLSAGGYAEQF | 0.14773639 |
| TTQTIYI | ASSLSAGGYAEQF | 0.5897863 |
| VAAGAGLGAVI | ASSLSAGGYAEQF | 0.7224363 |
| VAAVVTLL | ASSLSAGGYAEQF | 0.2872687 |
| VGFLGGLI | ASSLSAGGYAEQF | 0.38383114 |
| VGVIPILL | ASSLSAGGYAEQF | 0.6087804 |
| VIFASSLI | ASSLSAGGYAEQF | 0.48062423 |
| VKGGDGGAAAG | ASSLSAGGYAEQF | 0.36713582 |
| VTKTFTNI | ASSLSAGGYAEQF | 0.19858736 |
| VVAAAILL | ASSLSAGGYAEQF | 0.4241728 |
| YAALDVEL | ASSLSAGGYAEQF | 0.7617081 |
| LASVALLL | ASSLSAGGYAEQF | 0.24181086 |
| LDAVHRV | ASSLSAGGYAEQF | 0.4256559 |
| QGLSSSIF | ASSLSAGGYAEQF | 0.35439605 |
| RGTFANIRL | ASSLSAGGYAEQF | 0.22695148 |
| RQLWWGHRI | ASSLSAGGYAEQF | 0.39494288 |
| YGPPGTGKTLL | ASSLSAGGYAEQF | 0.21097219 |
| AALVGAPANVA | ASSLSAGGYAEQF | 0.6176599 |
| AATFRRTE | ASSLSAGGYAEQF | 0.17695053 |
| AGALGGGI | ASSLSAGGYAEQF | 0.71920985 |
| AGGAGGAGGIGAG | ASSLSAGGYAEQF | 0.75124794 |
| AGSGALGL | ASSLSAGGYAEQF | 0.6546832 |
| AQIADQI | ASSLSAGGYAEQF | 0.27352917 |
| ASAVAVTYL | ASSLSAGGYAEQF | 0.25647756 |
| ASTVLGGTGG | ASSLSAGGYAEQF | 0.19144166 |
| AVGGEPAK | ASSLSAGGYAEQF | 0.2075915 |

| | | |
|---|---|---|
| CEQGTFGGTC | ASSLSAGGYAEQF | 0.8160789 |
| DEGKAPAG | ASSLSAGGYAEQF | 0.20316823 |
| DTVVFPAL | ASSLSAGGYAEQF | 0.8068472 |
| FEGITMAG | ASSLSAGGYAEQF | 0.4783622 |
| FPLVVQD | ASSLSAGGYAEQF | 0.46500784 |
| GAFTTPQ | ASSLSAGGYAEQF | 0.5767678 |
| GFGGFGTAGGVGGA | ASSLSAGGYAEQF | 0.26200372 |
| GFLAQFL | ASSLSAGGYAEQF | 0.41910923 |
| GGAGGAGGNAGVA | ASSLSAGGYAEQF | 0.6058108 |
| GGVGTAGAGGAGGAGG | ASSLSAGGYAEQF | 0.71981716 |
| GNGGGGDAGNAGSGGN | ASSLSAGGYAEQF | 0.3728739 |
| GTGGAGGAGNDAGS | ASSLSAGGYAEQF | 0.61537397 |
| GTTNTGWFNS | ASSLSAGGYAEQF | 0.5045158 |
| IFMGPLFTL | ASSLSAGGYAEQF | 0.36190063 |
| ILAHAIEAI | ASSLSAGGYAEQF | 0.38024658 |
| IRSAIREAL | ASSLSAGGYAEQF | 0.6692425 |
| ITFAGGG | ASSLSAGGYAEQF | 0.58142674 |
| ITGMSGHQEV | ASSLSAGGYAEQF | 0.4104746 |
| IVAGLACTL | ASSLSAGGYAEQF | 0.44486222 |
| IVGGGLAGA | ASSLSAGGYAEQF | 0.41776907 |
| IVTGVLNI | ASSLSAGGYAEQF | 0.44759262 |
| LAQQGCI | ASSLSAGGYAEQF | 0.6756097 |
| LCLHVEN | ASSLSAGGYAEQF | 0.560612 |
| LDAISDGS | ASSLSAGGYAEQF | 0.70649123 |
| LGGFAAML | ASSLSAGGYAEQF | 0.21352603 |
| LGGKNPMI | ASSLSAGGYAEQF | 0.33025974 |
| LGPKGVAAIL | ASSLSAGGYAEQF | 0.33491674 |
| LMVGVSI | ASSLSAGGYAEQF | 0.3291733 |
| LVGAAFVL | ASSLSAGGYAEQF | 0.70861346 |
| LVLAVAPL | ASSLSAGGYAEQF | 0.19014491 |
| MAAIAVIDAA | ASSLSAGGYAEQF | 0.48111397 |
| NNGINAELQ | ASSLSAGGYAEQF | 0.3480698 |
| RGEPLVLVHG | ASSLSAGGYAEQF | 0.23097005 |
| RGGAGGAAGLI | ASSLSAGGYAEQF | 0.44064134 |
| RSYRMHPAV | ASSLSAGGYAEQF | 0.36572513 |
| SAAHLLAGQLA | ASSLSAGGYAEQF | 0.349562 |
| SAAVGSY | ASSLSAGGYAEQF | 0.7474771 |
| SPHAVLHRI | ASSLSAGGYAEQF | 0.38651574 |
| SQIAKVL | ASSLSAGGYAEQF | 0.56276643 |
| TALGHLVP | ASSLSAGGYAEQF | 0.6534415 |
| TAPTTSGAA | ASSLSAGGYAEQF | 0.5298784 |
| TDGDDGNGGSPGT | ASSLSAGGYAEQF | 0.6692188 |
| TDGLGEN | ASSLSAGGYAEQF | 0.573352 |
| TGGPPLPGQA | ASSLSAGGYAEQF | 0.78055525 |
| TIPAITI | ASSLSAGGYAEQF | 0.23121682 |
| TIPGFTI | ASSLSAGGYAEQF | 0.27026325 |
| TSIASGALA | ASSLSAGGYAEQF | 0.7142513 |
| TVTLPGAAI | ASSLSAGGYAEQF | 0.4361213 |
| VASGLSAALP | ASSLSAGGYAEQF | 0.7748041 |
| VGAGVIGI | ASSLSAGGYAEQF | 0.68305755 |
| VGPKRFGV | ASSLSAGGYAEQF | 0.78670925 |

| | | |
|---|---|---|
| VNVAFPD | ASSLSAGGYAEQF | 0.75825393 |
| VSMTNSGVS | ASSLSAGGYAEQF | 0.22656336 |
| VTDVLGDLSV | ASSLSAGGYAEQF | 0.2546967 |
| VVVVGAGFAGL | ASSLSAGGYAEQF | 0.263094 |
| VYTVGTVV | ASSLSAGGYAEQF | 0.2528308 |
| WTTFSGP | ASSLSAGGYAEQF | 0.38407817 |
| YDELVEAF | ASSLSAGGYAEQF | 0.18177404 |
| YGTGGAGGNGG | ASSLSAGGYAEQF | 0.68698883 |
| YTNVMAV | ASSLSAGGYAEQF | 0.8282567 |
| FLEQVHQL | ASSLSAGGYAEQF | 0.2493722 |
| GLSLLAVAV | ASSLSAGGYAEQF | 0.6977736 |
| LEQVHQL | ASSLSAGGYAEQF | 0.195902 |
| LNAAGAI | ASSLSAGGYAEQF | 0.571782 |
| PAVHASL | ASSLSAGGYAEQF | 0.74550295 |
| TELEDDA | ASSLSAGGYAEQF | 0.18700728 |
| TPPGPPN | ASSLSAGGYAEQF | 0.46240473 |
| TPPPPPPP | ASSLSAGGYAEQF | 0.3453074 |
| TQSPALI | AWSLWEETLY | 0.4126294 |
| EFAAAFPQ | AWSLWEETLY | 0.7871506 |
| LLSGNLDVL | AWSLWEETLY | 0.26679537 |
| GPANLGNYNI | AWSLWEETLY | 0.7090723 |
| TLGPTLP | AWSLWEETLY | 0.63041115 |
| VYAYSSCCK | AWSLWEETLY | 0.3138792 |
| APAPAPAPNI | AWSLWEETLY | 0.7332073 |
| PGGPVPIP | AWSLWEETLY | 0.79388547 |
| FLTVALEP | AWSLWEETLY | 0.34814587 |
| SQYFGANL | AWSLWEETLY | 0.6899588 |
| GGVKPAA | AWSLWEETLY | 0.38035023 |
| GGTGGAGGAGGW | AWSLWEETLY | 0.7333436 |
| GVGGVAGLGGAGG | AWSLWEETLY | 0.79989123 |
| VLLVLATTI | AWSLWEETLY | 0.38216233 |
| IGIFAAML | AWSLWEETLY | 0.4560445 |
| AVGDLID | AWSLWEETLY | 0.6953305 |
| NISPLPL | AWSLWEETLY | 0.73543125 |
| VTAVFVAI | AWSLWEETLY | 0.8022643 |
| SGDDNDAAL | AWSLWEETLY | 0.6971949 |
| AVAALAHS | AWSLWEETLY | 0.328662 |
| GGVTVMATP | AWSLWEETLY | 0.691785 |
| AGLAANVRD | AWSLWEETLY | 0.62195724 |
| TNRPDLIDPALLRPG | AWSLWEETLY | 0.4978356 |
| LANLGVTAL | AWSLWEETLY | 0.35854542 |
| TGERIEL | AWSLWEETLY | 0.24130929 |
| NGEELAA | AWSLWEETLY | 0.39153054 |
| VLSLVWG | AWSLWEETLY | 0.34158117 |
| IGYALWL | AWSLWEETLY | 0.66129744 |
| LSLLLAPL | AWSLWEETLY | 0.58861715 |
| TLDEALK | AWSLWEETLY | 0.39382896 |
| TDGAGTAEGG | AWSLWEETLY | 0.6741067 |
| NAPGGAGGNGG | AWSLWEETLY | 0.43627203 |
| VSVFMGMGGNF | AWSLWEETLY | 0.29806027 |
| LASYLAAL | AWSLWEETLY | 0.52202797 |

| | | |
|---|---|---|
| LVGGTPAL | AWSLWEETLY | 0.29516354 |
| VRNADVL | AWSLWEETLY | 0.5186347 |
| TCSSPFLVSAM | AWSLWEETLY | 0.2531796 |
| VHITDAE | AWSLWEETLY | 0.6101279 |
| EYAAPSF | AWSLWEETLY | 0.50319636 |
| VVDAFHGGL | AWSLWEETLY | 0.23738457 |
| ASVVDPQ | AWSLWEETLY | 0.574003 |
| SYNEFGGLS | AWSLWEETLY | 0.4462884 |
| DPDGSVY | AWSLWEETLY | 0.46959227 |
| MVSTIHATE | AWSLWEETLY | 0.6986526 |
| LAAGVADPV | AWSLWEETLY | 0.28038886 |
| TKEQIAA | AWSLWEETLY | 0.34316945 |
| SLASPAAL | AWSLWEETLY | 0.68011916 |
| IGLVFYGV | AWSLWEETLY | 0.6391294 |
| FLLDASNL | AWSLWEETLY | 0.32587016 |
| YQVGNFP | AWSLWEETLY | 0.5512229 |
| ESLLSQST | AWSLWEETLY | 0.65920436 |
| LFTVGFAAVG | AWSLWEETLY | 0.4502517 |
| ATPPTPT | AWSLWEETLY | 0.6469344 |
| TDLATLG | AWSLWEETLY | 0.6352525 |
| SAGGTGR | AWSLWEETLY | 0.27933246 |
| GFSNSGS | AWSLWEETLY | 0.69197667 |
| GGNGGSGGTSV | AWSLWEETLY | 0.7205561 |
| GTSVATGGAGN | AWSLWEETLY | 0.39258298 |
| NGGNGGSGGTS | AWSLWEETLY | 0.5546888 |
| AVVKFPH | AWSLWEETLY | 0.31871602 |
| TPPPPPP | AWSLWEETLY | 0.39636952 |
| AAPAAAAAPA | AWSLWEETLY | 0.25072575 |
| IALNEDL | AWSLWEETLY | 0.6656912 |
| IVNVSSM | AWSLWEETLY | 0.60825765 |
| TGIPPAPR | AWSLWEETLY | 0.5804222 |
| FDLGGGTF | AWSLWEETLY | 0.39416355 |
| FDLGGGTFDVS | AWSLWEETLY | 0.29777846 |
| NEPTAAA | AWSLWEETLY | 0.49915832 |
| NEPTAAALA | AWSLWEETLY | 0.32520515 |
| GSIINISSI | AWSLWEETLY | 0.27488968 |
| VAPGNAGTA | AWSLWEETLY | 0.27618676 |
| TGGGASLE | AWSLWEETLY | 0.57102895 |
| VAVYNHYKRI | AWSLWEETLY | 0.5773902 |
| TDPAPATT | AWSLWEETLY | 0.37476623 |
| TNRPDILDPALLRPG | AWSLWEETLY | 0.719331 |
| LTTTTTS | AWSLWEETLY | 0.6849567 |
| PTGSGKTL | AWSLWEETLY | 0.4473129 |
| TGSGKTL | AWSLWEETLY | 0.38473567 |
| TIGHVDHGK | AWSLWEETLY | 0.33054703 |
| TIGHVDHGKTT | AWSLWEETLY | 0.7422106 |
| GHVDHGKTT | AWSLWEETLY | 0.48803076 |
| AQFEHTLL | AWSLWEETLY | 0.58981735 |
| IVNQDEL | AWSLWEETLY | 0.5961382 |
| TNGSQFF | AWSLWEETLY | 0.462497 |
| TGKGIVE | AWSLWEETLY | 0.4058983 |

| | | |
|---|---|---|
| LISNASDA | AWSLWEETLY | 0.626519 |
| AELRTLGELTKIAKAHGAQ | AWSLWEETLY | 0.68534863 |
| APNAVAAAEPSW | AWSLWEETLY | 0.37667376 |
| APQVAGAATAVL | AWSLWEETLY | 0.28080833 |
| ARRDPAAAAAILPTDA | AWSLWEETLY | 0.40210426 |
| DAAARDVLAEAGLAEYF | AWSLWEETLY | 0.47215897 |
| DDQIVLSAKANLAKVEQ | AWSLWEETLY | 0.66022146 |
| DELLPDTGEDSDEDGH | AWSLWEETLY | 0.44294024 |
| ELPEAPANRATVPQD | AWSLWEETLY | 0.42339468 |
| FGQSKIERVTGIG | AWSLWEETLY | 0.6575494 |
| GAGGTGSTAGGAGGAGGAGGLY | AWSLWEETLY | 0.5097951 |
| GHGGTGAAGGNGGTGGAGGSGI | AWSLWEETLY | 0.46742803 |
| GNGGNGGAGGEGGPGGAGGAGA | AWSLWEETLY | 0.41997212 |
| GSTAGNGGNGGNATNGTIAGQP | AWSLWEETLY | 0.54830694 |
| GTGGDGGNGGNGGTGG | AWSLWEETLY | 0.6789682 |
| GVRRPAGIPADAPV | AWSLWEETLY | 0.6219901 |
| GYRQVTDAHLLTLARR | AWSLWEETLY | 0.597404 |
| IAKMGQDGHDRG | AWSLWEETLY | 0.6162077 |
| KAEPSIEIQYT | AWSLWEETLY | 0.39612722 |
| KPVRIRMAADEDTVAVT | AWSLWEETLY | 0.48149282 |
| MPPRIAGMRLLVIKPEP | AWSLWEETLY | 0.2545915 |
| MKFYAVALFRSRGVPPTR | AWSLWEETLY | 0.25522766 |
| NAGGFGILGAVAHSP | AWSLWEETLY | 0.5405818 |
| NEGREITLAMAEM | AWSLWEETLY | 0.64722764 |
| NELVEWSHPLGHRWRWE | AWSLWEETLY | 0.6405482 |
| NLGAGGTGGNGGNPG | AWSLWEETLY | 0.51778674 |
| NTEWPKWETVMAADPVD | AWSLWEETLY | 0.44354713 |
| PASAAPAETQP | AWSLWEETLY | 0.5559194 |
| QQSSDGKNGVRAAEITGE | AWSLWEETLY | 0.6812203 |
| RAPDAVTTSWWKE | AWSLWEETLY | 0.28983516 |
| SGFGNVGAHTSGWFNQ | AWSLWEETLY | 0.38145846 |
| SGVAGAVGQAASVGGLKV | AWSLWEETLY | 0.71914977 |
| SNGGAGTGSAGGNGGT | AWSLWEETLY | 0.671618 |
| TGGAGGEGGAAG | AWSLWEETLY | 0.50081474 |
| TGGDGGAAGAGGAPGHG | AWSLWEETLY | 0.4474072 |
| TMPPLLAAIANAGRHGV | AWSLWEETLY | 0.7272343 |
| TSSTAGNSGGA | AWSLWEETLY | 0.45267698 |
| VPATVPVGILC | AWSLWEETLY | 0.7339742 |
| VQRPDSGGVVLTGRIS | AWSLWEETLY | 0.23523417 |
| WTPNPVPATKGVPAGLVQAT | AWSLWEETLY | 0.47056377 |
| YALKRALAKFSDRFDVVII | AWSLWEETLY | 0.32420114 |
| YLIGSVIV | AWSLWEETLY | 0.23908356 |
| YGQLVAVID | AWSLWEETLY | 0.6483383 |
| VTVGGMTLPA | AWSLWEETLY | 0.3339601 |
| VLALPAV | AWSLWEETLY | 0.5708202 |
| VIPPTLNYET | AWSLWEETLY | 0.32378432 |
| VGMGGMHPSSG | AWSLWEETLY | 0.6454122 |
| VGLAGGAATA | AWSLWEETLY | 0.6376026 |
| VGGAGGAGGSGP | AWSLWEETLY | 0.32094416 |
| VFAAFMV | AWSLWEETLY | 0.30846536 |
| TLYDKLR | AWSLWEETLY | 0.5477519 |

| | | |
|---|---|---|
| TLAVFTVL | AWSLWEETLY | 0.6389072 |
| TGSVQPARV | AWSLWEETLY | 0.60317373 |
| TGGGAGSGGT | AWSLWEETLY | 0.2538427 |
| SHLETSL | AWSLWEETLY | 0.6488906 |
| SFTTGMT | AWSLWEETLY | 0.42185748 |
| QVEVAKRMGVRQPTVS | AWSLWEETLY | 0.2578662 |
| QGSGGPALAM | AWSLWEETLY | 0.24468875 |
| MLAPSNP | AWSLWEETLY | 0.38314846 |
| LVNNAAI | AWSLWEETLY | 0.47956735 |
| LTGEINTLA | AWSLWEETLY | 0.70171654 |
| LSIFGPV | AWSLWEETLY | 0.71176493 |
| LLALLANL | AWSLWEETLY | 0.5028459 |
| LISNASDALDKLR | AWSLWEETLY | 0.60175467 |
| LINEAVP | AWSLWEETLY | 0.5356308 |
| LGYAFSGL | AWSLWEETLY | 0.20760551 |
| LDFVAVAL | AWSLWEETLY | 0.29949114 |
| LAQPGAE | AWSLWEETLY | 0.3030575 |
| ISNLGAVTGF | AWSLWEETLY | 0.6146642 |
| HQEDPGRI | AWSLWEETLY | 0.29812777 |
| GTPGEPTP | AWSLWEETLY | 0.553715 |
| GFFNSGS | AWSLWEETLY | 0.5824818 |
| GAGGGGLTAGAN | AWSLWEETLY | 0.70643985 |
| FVDALVNM | AWSLWEETLY | 0.5278131 |
| FLTVAPEL | AWSLWEETLY | 0.30398253 |
| FETLTAL | AWSLWEETLY | 0.23049447 |
| DVINPIEL | AWSLWEETLY | 0.53054297 |
| APVSGAPV | AWSLWEETLY | 0.28523412 |
| ALVGGLAW | AWSLWEETLY | 0.34155214 |
| ALAAAAATAI | AWSLWEETLY | 0.2866459 |
| AFGLGLLL | AWSLWEETLY | 0.6513865 |
| AFFAAASL | AWSLWEETLY | 0.2592661 |
| AAAALPAFSPP | AWSLWEETLY | 0.62088525 |
| VVLAPAL | AWSLWEETLY | 0.3767631 |
| TIGHVDHGKTTL | AWSLWEETLY | 0.59045845 |
| LISNASDALDK | AWSLWEETLY | 0.7033331 |
| LISNASDAL | AWSLWEETLY | 0.6922988 |
| ISNASDALDK | AWSLWEETLY | 0.5406674 |
| NQAETLVYQTE | AWSLWEETLY | 0.36257428 |
| TGGTGGTAGAG | AWSLWEETLY | 0.37674427 |
| TGSTAGGAGGA | AWSLWEETLY | 0.42056188 |
| DPLAVPVAAGPV | AWSLWEETLY | 0.39993244 |
| GGTGGTPGNGGN | AWSLWEETLY | 0.6013583 |
| PAGLAGVDPATL | AWSLWEETLY | 0.55901176 |
| GNNLVVEAPHAGASV | AWSLWEETLY | 0.29331696 |
| KVTRIMSASTLEQPA | AWSLWEETLY | 0.5633054 |
| LGFALPIALPRRTVV | AWSLWEETLY | 0.70686114 |
| TRLGNLTARKLVVLV | AWSLWEETLY | 0.7051569 |
| WSQQRRLIYFPSAGP | AWSLWEETLY | 0.36894134 |
| GPNGNPDPMAAAVDIR | AWSLWEETLY | 0.29684195 |
| TQRCVSVADLRAAASR | AWSLWEETLY | 0.35571668 |
| QFVLDVAEHPVEVLTETV | AWSLWEETLY | 0.61507916 |

| | | |
|---|---|---|
| GGAGGTGGAGGNGGTQAGDG | AWSLWEETLY | 0.30745536 |
| PLLATLRAIAVDVDAKPAQV | AWSLWEETLY | 0.32066822 |
| APKFRTGGYMIDFWGVGYQVAK | AWSLWEETLY | 0.6634308 |
| GAGGAGGAGGAAGGPGGTGGTG | AWSLWEETLY | 0.7453265 |
| IGALGLLATDVAPAGSVAFGVA | AWSLWEETLY | 0.40134275 |
| AGGAGGIGGNGGAGGAGGVLMG | AWSLWEETLY | 0.2525481 |
| ADELIGLL | AWSLWEETLY | 0.5421917 |
| ADRALKA | AWSLWEETLY | 0.36006558 |
| AGAVGQVGGAGGA | AWSLWEETLY | 0.47657898 |
| AGGGGGLGGAP | AWSLWEETLY | 0.34439304 |
| AGGLLYGNGGAG | AWSLWEETLY | 0.31637317 |
| AGLESIGL | AWSLWEETLY | 0.53471005 |
| AGLKTGII | AWSLWEETLY | 0.65084255 |
| ASLAALLL | AWSLWEETLY | 0.47133625 |
| ASVINISSI | AWSLWEETLY | 0.5936557 |
| ATPPGPA | AWSLWEETLY | 0.73813885 |
| ATVASALV | AWSLWEETLY | 0.7005366 |
| ATVASLIL | AWSLWEETLY | 0.7191169 |
| AVNGPVLIRH | AWSLWEETLY | 0.67660964 |
| DGAAGAVGI | AWSLWEETLY | 0.3765546 |
| DPLLHGI | AWSLWEETLY | 0.44022688 |
| ELGAALGAL | AWSLWEETLY | 0.7302683 |
| ELGRTQAV | AWSLWEETLY | 0.548396 |
| FVAGGYLS | AWSLWEETLY | 0.57768285 |
| GASGGAGGAGGW | AWSLWEETLY | 0.5242992 |
| GGAGAGGDGGAGGI | AWSLWEETLY | 0.70912266 |
| GTGHNGSQPGMGGNG | AWSLWEETLY | 0.4329158 |
| GYAAASASAS | AWSLWEETLY | 0.7208623 |
| HQVDVAAVQQG | AWSLWEETLY | 0.24755223 |
| IGAAIAHT | AWSLWEETLY | 0.33180112 |
| IGMIEVAQL | AWSLWEETLY | 0.7214121 |
| IRLAPEHTSAMD | AWSLWEETLY | 0.21629538 |
| IYNVSVL | AWSLWEETLY | 0.26755562 |
| KQGFLGM | AWSLWEETLY | 0.58362615 |
| LESVITQL | AWSLWEETLY | 0.66227645 |
| LGLGALVVV | AWSLWEETLY | 0.22098395 |
| LIGNGGNGGSGGI | AWSLWEETLY | 0.32569674 |
| LNAINAPTL | AWSLWEETLY | 0.27793062 |
| LQLLNTGV | AWSLWEETLY | 0.38996035 |
| LQTSIEF | AWSLWEETLY | 0.48911378 |
| MSFAVSEL | AWSLWEETLY | 0.23149385 |
| NTGALGL | AWSLWEETLY | 0.5897944 |
| PPGAGGAGGAGTT | AWSLWEETLY | 0.313538 |
| QLVGLGI | AWSLWEETLY | 0.47193986 |
| RFLEQVHQL | AWSLWEETLY | 0.64565337 |
| RGVGYVLR | AWSLWEETLY | 0.74480045 |
| SATGVFAGVI | AWSLWEETLY | 0.3052942 |
| SLPAITLP | AWSLWEETLY | 0.6262617 |
| STALAFTV | AWSLWEETLY | 0.4612416 |
| SVLTVGLGK | AWSLWEETLY | 0.7401539 |
| TALAGAGSGI | AWSLWEETLY | 0.37307185 |

| | | |
|---|---|---|
| TGLPFTGI | AWSLWEETLY | 0.2793144 |
| VAEHVTV | AWSLWEETLY | 0.68228406 |
| VDIVLNSL | AWSLWEETLY | 0.5922264 |
| VLTGGGAVL | AWSLWEETLY | 0.4248091 |
| VNVGLAYGI | AWSLWEETLY | 0.70656383 |
| VSSAAVGAV | AWSLWEETLY | 0.35811442 |
| VTSVYEAGV | AWSLWEETLY | 0.5859451 |
| WVALDRGA | AWSLWEETLY | 0.67038125 |
| YLVGSVVI | AWSLWEETLY | 0.61133456 |
| GPPGTGKTLL | AWSLWEETLY | 0.32666674 |
| GVGMTKF | AWSLWEETLY | 0.6292455 |
| IGIGGSDL | AWSLWEETLY | 0.5403843 |
| IGTLLAEL | AWSLWEETLY | 0.32655388 |
| IYTSGTT | AWSLWEETLY | 0.6774082 |
| LSFPAGV | AWSLWEETLY | 0.41148898 |
| LYTSGTT | AWSLWEETLY | 0.6868626 |
| VGAFPFGF | AWSLWEETLY | 0.5689157 |
| VGAGTVEF | AWSLWEETLY | 0.7100314 |
| VGKSTLLL | AWSLWEETLY | 0.2439251 |
| AALAHAGGGIV | AWSLWEETLY | 0.39714596 |
| AALVAVSLL | AWSLWEETLY | 0.3800615 |
| AAVSRAVAAL | AWSLWEETLY | 0.59079266 |
| AGGSSGIGGF | AWSLWEETLY | 0.3763485 |
| AGLTVANANG | AWSLWEETLY | 0.41574568 |
| AGLVGAVL | AWSLWEETLY | 0.58615994 |
| ALSQAFTV | AWSLWEETLY | 0.7139019 |
| ALTFSVSL | AWSLWEETLY | 0.4923486 |
| ALTGAALVAL | AWSLWEETLY | 0.45404828 |
| ASFSIGAGVL | AWSLWEETLY | 0.745684 |
| DGSVYKI | AWSLWEETLY | 0.32149202 |
| EQLAEIV | AWSLWEETLY | 0.5150019 |
| FAVSELTVM | AWSLWEETLY | 0.66079384 |
| GAAPLAAGL | AWSLWEETLY | 0.63222325 |
| GADGGDGGSGGAGG | AWSLWEETLY | 0.52060413 |
| GFGLDTGGAGG | AWSLWEETLY | 0.5963299 |
| GLASGGLL | AWSLWEETLY | 0.26519516 |
| GRIAGAAVLL | AWSLWEETLY | 0.6436592 |
| GSAGSLNTSF | AWSLWEETLY | 0.41292912 |
| GTGQIGI | AWSLWEETLY | 0.4065216 |
| GTLIALIL | AWSLWEETLY | 0.40281004 |
| IAATMGFVVG | AWSLWEETLY | 0.62930846 |
| IFAQALL | AWSLWEETLY | 0.7037666 |
| IGAAVALGF | AWSLWEETLY | 0.6515038 |
| IGLTPLAL | AWSLWEETLY | 0.24795905 |
| ILGSNTSTL | AWSLWEETLY | 0.68938833 |
| IVFSLTVL | AWSLWEETLY | 0.62301445 |
| LASKATGF | AWSLWEETLY | 0.7566869 |
| LASVALIL | AWSLWEETLY | 0.38021207 |
| LFAVGLTL | AWSLWEETLY | 0.5143292 |
| LFNGGIGTYI | AWSLWEETLY | 0.43085924 |
| LTLGAGII | AWSLWEETLY | 0.5457746 |

| | | |
|---|---|---|
| LVAAALGDI | AWSLWEETLY | 0.30362853 |
| NTAFVQLGI | AWSLWEETLY | 0.34188974 |
| QLFSGAEL | AWSLWEETLY | 0.5003341 |
| SAATFITL | AWSLWEETLY | 0.5068616 |
| SNYGLANLGV | AWSLWEETLY | 0.73117816 |
| SYGGAYAVM | AWSLWEETLY | 0.6683961 |
| TTGVGGLTLG | AWSLWEETLY | 0.6437516 |
| TTQTIYI | AWSLWEETLY | 0.6998069 |
| VAAGAGLGAVI | AWSLWEETLY | 0.7270073 |
| VAAVVTLL | AWSLWEETLY | 0.5578683 |
| VGFLGGLI | AWSLWEETLY | 0.46496576 |
| VGVIPILL | AWSLWEETLY | 0.28622127 |
| VIFASSLI | AWSLWEETLY | 0.38054925 |
| VKGGDGGAAAG | AWSLWEETLY | 0.45945477 |
| VTKTFTNI | AWSLWEETLY | 0.4430772 |
| VVAAAILL | AWSLWEETLY | 0.5942316 |
| YAALDVEL | AWSLWEETLY | 0.2188967 |
| LASVALLL | AWSLWEETLY | 0.7019197 |
| LDAVHRV | AWSLWEETLY | 0.65152633 |
| QGLSSSIF | AWSLWEETLY | 0.361147 |
| RGTFANIRL | AWSLWEETLY | 0.52909917 |
| RQLWWGHRI | AWSLWEETLY | 0.6656445 |
| YGPPGTGKTLL | AWSLWEETLY | 0.47003442 |
| AALVGAPANVA | AWSLWEETLY | 0.6356244 |
| AATFRRTE | AWSLWEETLY | 0.44157404 |
| AGALGGGI | AWSLWEETLY | 0.6305953 |
| AGGAGGAGGIGAG | AWSLWEETLY | 0.6597098 |
| AGSGALGL | AWSLWEETLY | 0.6071378 |
| AQIADQI | AWSLWEETLY | 0.41295996 |
| ASAVAVTYL | AWSLWEETLY | 0.479258 |
| ASTVLGGTGG | AWSLWEETLY | 0.24515548 |
| AVGGEPAK | AWSLWEETLY | 0.43558994 |
| CEQGTFGGTC | AWSLWEETLY | 0.2915457 |
| DEGKAPAG | AWSLWEETLY | 0.5834713 |
| DTVVFPAL | AWSLWEETLY | 0.31613278 |
| FEGITMAG | AWSLWEETLY | 0.6290003 |
| FPLVVQD | AWSLWEETLY | 0.5806657 |
| GAFTTPQ | AWSLWEETLY | 0.5507304 |
| GFGGFGTAGGVGGA | AWSLWEETLY | 0.33077407 |
| GFLAQFL | AWSLWEETLY | 0.66006935 |
| GGAGGAGGNAGVA | AWSLWEETLY | 0.6577431 |
| GGVGTAGAGGAGGAGG | AWSLWEETLY | 0.28052092 |
| GNGGGGDAGNAGSGGN | AWSLWEETLY | 0.5362607 |
| GTGGAGGAGNDAGS | AWSLWEETLY | 0.51313806 |
| GTTNTGWFNS | AWSLWEETLY | 0.50887656 |
| IFMGPLFTL | AWSLWEETLY | 0.6126027 |
| ILAHAIEAI | AWSLWEETLY | 0.5869154 |
| IRSAIREAL | AWSLWEETLY | 0.26188686 |
| ITFAGGG | AWSLWEETLY | 0.4292803 |
| ITGMSGHQEV | AWSLWEETLY | 0.617921 |
| IVAGLACTL | AWSLWEETLY | 0.33843017 |

| | | |
|---|---|---|
| IVGGGLAGA | AWSLWEETLY | 0.22424087 |
| IVTGVLNI | AWSLWEETLY | 0.48480916 |
| LAQQGCI | AWSLWEETLY | 0.3243167 |
| LCLHVEN | AWSLWEETLY | 0.33912933 |
| LDAISDGS | AWSLWEETLY | 0.4664885 |
| LGGFAAML | AWSLWEETLY | 0.7058764 |
| LGGKNPMI | AWSLWEETLY | 0.3437159 |
| LGPKGVAAIL | AWSLWEETLY | 0.37115094 |
| LMVGVSI | AWSLWEETLY | 0.6273847 |
| LVGAAFVL | AWSLWEETLY | 0.53180623 |
| LVLAVAPL | AWSLWEETLY | 0.5884212 |
| MAAIAVIDAA | AWSLWEETLY | 0.46786293 |
| NNGINAELQ | AWSLWEETLY | 0.40376776 |
| RGEPLVLVHG | AWSLWEETLY | 0.40974146 |
| RGGAGGAAGLI | AWSLWEETLY | 0.34175387 |
| RSYRMHPAV | AWSLWEETLY | 0.5062659 |
| SAAHLLAGQLA | AWSLWEETLY | 0.41412205 |
| SAAVGSY | AWSLWEETLY | 0.37359607 |
| SPHAVLHRI | AWSLWEETLY | 0.392245 |
| SQIAKVL | AWSLWEETLY | 0.44627285 |
| TALGHLVP | AWSLWEETLY | 0.6283824 |
| TAPTTSGAA | AWSLWEETLY | 0.27974966 |
| TDGDDGNGGSPGT | AWSLWEETLY | 0.28085575 |
| TDGLGEN | AWSLWEETLY | 0.49483395 |
| TGGPPLPGQA | AWSLWEETLY | 0.26009348 |
| TIPAITI | AWSLWEETLY | 0.64440274 |
| TIPGFTI | AWSLWEETLY | 0.38783252 |
| TSIASGALA | AWSLWEETLY | 0.73589873 |
| TVTLPGAAI | AWSLWEETLY | 0.67092824 |
| VASGLSAALP | AWSLWEETLY | 0.4852435 |
| VGAGVIGI | AWSLWEETLY | 0.7127664 |
| VGPKRFGV | AWSLWEETLY | 0.40907797 |
| VNVAFPD | AWSLWEETLY | 0.43427306 |
| VSMTNSGVS | AWSLWEETLY | 0.47343144 |
| VTDVLGDLSV | AWSLWEETLY | 0.54352856 |
| VVVVGAGFAGL | AWSLWEETLY | 0.33993372 |
| VYTVGTVV | AWSLWEETLY | 0.43111983 |
| WTTFSGP | AWSLWEETLY | 0.2543748 |
| YDELVEAF | AWSLWEETLY | 0.6739309 |
| YGTGGAGGNGG | AWSLWEETLY | 0.27362654 |
| YTNVMAV | AWSLWEETLY | 0.29392046 |
| FLEQVHQL | AWSLWEETLY | 0.41348988 |
| GLSLLAVAV | AWSLWEETLY | 0.70017713 |
| LEQVHQL | AWSLWEETLY | 0.6805122 |
| LNAAGAI | AWSLWEETLY | 0.44167706 |
| PAVHASL | AWSLWEETLY | 0.3652201 |
| TELEDDA | AWSLWEETLY | 0.7176238 |
| TPPGPPN | AWSLWEETLY | 0.33355922 |
| TPPPPPPP | AWSLWEETLY | 0.5992929 |
| TQSPALI | ASSDARDGNTLY | 0.5871793 |
| EFAAAFPQ | ASSDARDGNTLY | 0.5144779 |

| | | |
|---|---|---|
| LLSGNLDVL | ASSDARDGNTLY | 0.30209947 |
| GPANLGNYNI | ASSDARDGNTLY | 0.6713049 |
| TLGPTLP | ASSDARDGNTLY | 0.4177969 |
| VYAYSSCCK | ASSDARDGNTLY | 0.4797269 |
| APAPAPAPNI | ASSDARDGNTLY | 0.226666 |
| PGGPVPIP | ASSDARDGNTLY | 0.4773804 |
| FLTVALEP | ASSDARDGNTLY | 0.4500779 |
| SQYFGANL | ASSDARDGNTLY | 0.32236677 |
| GGVKPAA | ASSDARDGNTLY | 0.24376528 |
| GGTGGAGGAGGW | ASSDARDGNTLY | 0.4019317 |
| GVGGVAGLGGAGG | ASSDARDGNTLY | 0.6508273 |
| VLLVLATTI | ASSDARDGNTLY | 0.7340128 |
| IGIFAAML | ASSDARDGNTLY | 0.53496903 |
| AVGDLID | ASSDARDGNTLY | 0.46873438 |
| NISPLPL | ASSDARDGNTLY | 0.65006375 |
| VTAVFVAI | ASSDARDGNTLY | 0.6002015 |
| SGDDNDAAL | ASSDARDGNTLY | 0.24145973 |
| AVAALAHS | ASSDARDGNTLY | 0.4602816 |
| GGVTVMATP | ASSDARDGNTLY | 0.5019698 |
| AGLAANVRD | ASSDARDGNTLY | 0.6232485 |
| TNRPDLIDPALLRPG | ASSDARDGNTLY | 0.5904738 |
| LANLGVTAL | ASSDARDGNTLY | 0.36600405 |
| TGERIEL | ASSDARDGNTLY | 0.43210605 |
| NGEELAA | ASSDARDGNTLY | 0.6078168 |
| VLSLVWG | ASSDARDGNTLY | 0.3097201 |
| IGYALWL | ASSDARDGNTLY | 0.3661708 |
| LSLLLAPL | ASSDARDGNTLY | 0.32203725 |
| TLDEALK | ASSDARDGNTLY | 0.2420951 |
| TDGAGTAEGG | ASSDARDGNTLY | 0.54108423 |
| NAPGGAGGNGG | ASSDARDGNTLY | 0.55850554 |
| VSVFMGMGGNF | ASSDARDGNTLY | 0.39340702 |
| LASYLAAL | ASSDARDGNTLY | 0.5148699 |
| LVGGTPAL | ASSDARDGNTLY | 0.39456975 |
| VRNADVL | ASSDARDGNTLY | 0.37407416 |
| TCSSPFLVSAM | ASSDARDGNTLY | 0.43611902 |
| VHITDAE | ASSDARDGNTLY | 0.33627334 |
| EYAAPSF | ASSDARDGNTLY | 0.6933292 |
| VVDAFHGGL | ASSDARDGNTLY | 0.4088698 |
| ASVVDPQ | ASSDARDGNTLY | 0.395267 |
| SYNEFGGLS | ASSDARDGNTLY | 0.7524253 |
| DPDGSVY | ASSDARDGNTLY | 0.27526617 |
| MVSTIHATE | ASSDARDGNTLY | 0.3222464 |
| LAAGVADPV | ASSDARDGNTLY | 0.5399407 |
| TKEQIAA | ASSDARDGNTLY | 0.44571576 |
| SLASPAAL | ASSDARDGNTLY | 0.62213945 |
| IGLVFYGV | ASSDARDGNTLY | 0.54406756 |
| FLLDASNL | ASSDARDGNTLY | 0.63038456 |
| YQVGNFP | ASSDARDGNTLY | 0.42862996 |
| ESLLSQST | ASSDARDGNTLY | 0.7026653 |
| LFTVGFAAVG | ASSDARDGNTLY | 0.6677808 |
| ATPPTPT | ASSDARDGNTLY | 0.38695973 |

| | | |
|---|---|---|
| TDLATLG | ASSDARDGNTLY | 0.69801676 |
| SAGGTGR | ASSDARDGNTLY | 0.28130943 |
| GFSNSGS | ASSDARDGNTLY | 0.45582664 |
| GGNGGSGGTSV | ASSDARDGNTLY | 0.6560583 |
| GTSVATGGAGN | ASSDARDGNTLY | 0.3516249 |
| NGGNGGSGGTS | ASSDARDGNTLY | 0.5800624 |
| AVVKFPH | ASSDARDGNTLY | 0.6908637 |
| TPPPPPP | ASSDARDGNTLY | 0.71489584 |
| AAPAAAAAPA | ASSDARDGNTLY | 0.32514194 |
| IALNEDL | ASSDARDGNTLY | 0.46537763 |
| IVNVSSM | ASSDARDGNTLY | 0.28709003 |
| TGIPPAPR | ASSDARDGNTLY | 0.32661733 |
| FDLGGGTF | ASSDARDGNTLY | 0.2830031 |
| FDLGGGTFDVS | ASSDARDGNTLY | 0.58152217 |
| NEPTAAA | ASSDARDGNTLY | 0.5370985 |
| NEPTAAALA | ASSDARDGNTLY | 0.71071696 |
| GSIINISSI | ASSDARDGNTLY | 0.49784276 |
| VAPGNAGTA | ASSDARDGNTLY | 0.6588106 |
| TGGGASLE | ASSDARDGNTLY | 0.45249933 |
| VAVYNHYKRI | ASSDARDGNTLY | 0.47134864 |
| TDPAPATT | ASSDARDGNTLY | 0.27483788 |
| TNRPDILDPALLRPG | ASSDARDGNTLY | 0.25757602 |
| LTTTTTS | ASSDARDGNTLY | 0.31377286 |
| PTGSGKTL | ASSDARDGNTLY | 0.45075902 |
| TGSGKTL | ASSDARDGNTLY | 0.23580973 |
| TIGHVDHGK | ASSDARDGNTLY | 0.64878964 |
| TIGHVDHGKTT | ASSDARDGNTLY | 0.49127805 |
| GHVDHGKTT | ASSDARDGNTLY | 0.565637 |
| AQFEHTLL | ASSDARDGNTLY | 0.38044488 |
| IVNQDEL | ASSDARDGNTLY | 0.33540562 |
| TNGSQFF | ASSDARDGNTLY | 0.45409465 |
| TGKGIVE | ASSDARDGNTLY | 0.6022017 |
| LISNASDA | ASSDARDGNTLY | 0.2668059 |
| AELRTLGELTKIAKAHGAQ | ASSDARDGNTLY | 0.27718312 |
| APNAVAAAEPSW | ASSDARDGNTLY | 0.3252388 |
| APQVAGAATAVL | ASSDARDGNTLY | 0.36636943 |
| ARRDPAAAAAILPTDA | ASSDARDGNTLY | 0.74251854 |
| DAAARDVLAEAGLAEYF | ASSDARDGNTLY | 0.69629514 |
| DDQIVLSAKANLAKVEQ | ASSDARDGNTLY | 0.65925115 |
| DELLPDTGEDSDEDGH | ASSDARDGNTLY | 0.73737574 |
| ELPEAPANRATVPQD | ASSDARDGNTLY | 0.22469886 |
| FGQSKIERVTGIG | ASSDARDGNTLY | 0.37382066 |
| GAGGTGSTAGGAGGAGGAGGLY | ASSDARDGNTLY | 0.63454807 |
| GHGGTGAAGGNGGTGGAGGSGI | ASSDARDGNTLY | 0.4525923 |
| GNGGNGGAGGEGGPGGAGGAGA | ASSDARDGNTLY | 0.5310049 |
| GSTAGNGGNGGNATNGTIAGQP | ASSDARDGNTLY | 0.7032138 |
| GTGGDGGNGGNGGTGG | ASSDARDGNTLY | 0.42688426 |
| GVRRPAGIPADAPV | ASSDARDGNTLY | 0.32907817 |
| GYRQVTDAHLLTLARR | ASSDARDGNTLY | 0.5415035 |
| IAKMGQDGHDRG | ASSDARDGNTLY | 0.5525955 |
| KAEPSIEIQYT | ASSDARDGNTLY | 0.48158076 |

| | | |
|---|---|---|
| KPVRIRMAADEDTVAVT | ASSDARDGNTLY | 0.26436594 |
| MPPRIAGMRLLVIKPEP | ASSDARDGNTLY | 0.52831864 |
| MKFYAVALFRSRGVPPTR | ASSDARDGNTLY | 0.76283425 |
| NAGGFGILGAVAHSP | ASSDARDGNTLY | 0.6034937 |
| NEGREITLAMAEM | ASSDARDGNTLY | 0.5216047 |
| NELVEWSHPLGHRWRWE | ASSDARDGNTLY | 0.4755441 |
| NLGAGGTGGNGGNPG | ASSDARDGNTLY | 0.36797437 |
| NTEWPKWETVMAADPVD | ASSDARDGNTLY | 0.44083536 |
| PASAAPAETQP | ASSDARDGNTLY | 0.40827346 |
| QQSSDGKNGVRAAEITGE | ASSDARDGNTLY | 0.6597682 |
| RAPDAVTTSWWKE | ASSDARDGNTLY | 0.30390653 |
| SGFGNVGAHTSGWFNQ | ASSDARDGNTLY | 0.53639275 |
| SGVAGAVGQAASVGGLKV | ASSDARDGNTLY | 0.39270505 |
| SNGGAGTGSAGGNGGT | ASSDARDGNTLY | 0.75827014 |
| TGGAGGEGGAAG | ASSDARDGNTLY | 0.6312477 |
| TGGDGGAAGAGGAPGHG | ASSDARDGNTLY | 0.53352374 |
| TMPPLLAAIANAGRHGV | ASSDARDGNTLY | 0.4008238 |
| TSSTAGNSGGA | ASSDARDGNTLY | 0.5281462 |
| VPATVPVGILC | ASSDARDGNTLY | 0.6683661 |
| VQRPDSGGVVLTGRIS | ASSDARDGNTLY | 0.25921357 |
| WTPNPVPATKGVPAGLVQAT | ASSDARDGNTLY | 0.6694082 |
| YALKRALAKFSDRFDVVII | ASSDARDGNTLY | 0.36690792 |
| YLIGSVIV | ASSDARDGNTLY | 0.45184734 |
| YGQLVAVID | ASSDARDGNTLY | 0.3959944 |
| VTVGGMTLPA | ASSDARDGNTLY | 0.7316538 |
| VLALPAV | ASSDARDGNTLY | 0.54695636 |
| VIPPTLNYET | ASSDARDGNTLY | 0.6592002 |
| VGMGGMHPSSG | ASSDARDGNTLY | 0.3878631 |
| VGLAGGAATA | ASSDARDGNTLY | 0.58940816 |
| VGGAGGAGGSGP | ASSDARDGNTLY | 0.74177754 |
| VFAAFMV | ASSDARDGNTLY | 0.33367094 |
| TLYDKLR | ASSDARDGNTLY | 0.74435735 |
| TLAVFTVL | ASSDARDGNTLY | 0.36629477 |
| TGSVQPARV | ASSDARDGNTLY | 0.2645571 |
| TGGGAGSGGT | ASSDARDGNTLY | 0.41573894 |
| SHLETSL | ASSDARDGNTLY | 0.59373176 |
| SFTTGMT | ASSDARDGNTLY | 0.7312254 |
| QVEVAKRMGVRQPTVS | ASSDARDGNTLY | 0.47016934 |
| QGSGGPALAM | ASSDARDGNTLY | 0.5146109 |
| MLAPSNP | ASSDARDGNTLY | 0.40040255 |
| LVNNAAI | ASSDARDGNTLY | 0.71427155 |
| LTGEINTLA | ASSDARDGNTLY | 0.6592382 |
| LSIFGPV | ASSDARDGNTLY | 0.50307477 |
| LLALLANL | ASSDARDGNTLY | 0.313515 |
| LISNASDALDKLR | ASSDARDGNTLY | 0.2825681 |
| LINEAVP | ASSDARDGNTLY | 0.64746845 |
| LGYAFSGL | ASSDARDGNTLY | 0.69826984 |
| LDFVAVAL | ASSDARDGNTLY | 0.3355831 |
| LAQPGAE | ASSDARDGNTLY | 0.5698079 |
| ISNLGAVTGF | ASSDARDGNTLY | 0.5353528 |
| HQEDPGRI | ASSDARDGNTLY | 0.2590415 |

| | | |
|---|---|---|
| GTPGEPTP | ASSDARDGNTLY | 0.44083792 |
| GFFNSGS | ASSDARDGNTLY | 0.6868534 |
| GAGGGGLTAGAN | ASSDARDGNTLY | 0.55826104 |
| FVDALVNM | ASSDARDGNTLY | 0.26928857 |
| FLTVAPEL | ASSDARDGNTLY | 0.6964274 |
| FETLTAL | ASSDARDGNTLY | 0.7550943 |
| DVINPIEL | ASSDARDGNTLY | 0.69058716 |
| APVSGAPV | ASSDARDGNTLY | 0.27144033 |
| ALVGGLAW | ASSDARDGNTLY | 0.24003306 |
| ALAAAAATAI | ASSDARDGNTLY | 0.441507 |
| AFGLGLLL | ASSDARDGNTLY | 0.69351435 |
| AFFAAASL | ASSDARDGNTLY | 0.45007005 |
| AAAALPAFSPP | ASSDARDGNTLY | 0.35350478 |
| VVLAPAL | ASSDARDGNTLY | 0.53852105 |
| TIGHVDHGKTTL | ASSDARDGNTLY | 0.67044955 |
| LISNASDALDK | ASSDARDGNTLY | 0.30395 |
| LISNASDAL | ASSDARDGNTLY | 0.26991966 |
| ISNASDALDK | ASSDARDGNTLY | 0.55645776 |
| NQAETLVYQTE | ASSDARDGNTLY | 0.68982935 |
| TGGTGGTAGAG | ASSDARDGNTLY | 0.3731199 |
| TGSTAGGAGGA | ASSDARDGNTLY | 0.55618024 |
| DPLAVPVAAGPV | ASSDARDGNTLY | 0.52381414 |
| GGTGGTPGNGGN | ASSDARDGNTLY | 0.49183363 |
| PAGLAGVDPATL | ASSDARDGNTLY | 0.58572155 |
| GNNLVVEAPHAGASV | ASSDARDGNTLY | 0.31174952 |
| KVTRIMSASTLEQPA | ASSDARDGNTLY | 0.6797954 |
| LGFALPIALPRRTVV | ASSDARDGNTLY | 0.3666821 |
| TRLGNLTARKLVVLV | ASSDARDGNTLY | 0.5937513 |
| WSQQRRLIYFPSAGP | ASSDARDGNTLY | 0.56143445 |
| GPNGNPDPMAAAVDIR | ASSDARDGNTLY | 0.52665937 |
| TQRCVSVADLRAAASR | ASSDARDGNTLY | 0.5338639 |
| QFVLDVAEHPVEVLTETV | ASSDARDGNTLY | 0.33253732 |
| GGAGGTGGAGGNGGTQAGDG | ASSDARDGNTLY | 0.26926523 |
| PLLATLRAIAVDVDAKPAQV | ASSDARDGNTLY | 0.34292322 |
| APKFRTGGYMIDFWGVGYQVAK | ASSDARDGNTLY | 0.67089254 |
| GAGGAGGAGGAAGGPGGTGGTG | ASSDARDGNTLY | 0.3342727 |
| IGALGLLATDVAPAGSVAFGVA | ASSDARDGNTLY | 0.67250055 |
| AGGAGGIGGNGGAGGAGGVLMG | ASSDARDGNTLY | 0.56493324 |
| ADELIGLL | ASSDARDGNTLY | 0.34381425 |
| ADRALKA | ASSDARDGNTLY | 0.6334616 |
| AGAVGQVGGAGGA | ASSDARDGNTLY | 0.2785339 |
| AGGGGGLGGAP | ASSDARDGNTLY | 0.5727403 |
| AGGLLYGNGGAG | ASSDARDGNTLY | 0.42717886 |
| AGLESIGL | ASSDARDGNTLY | 0.51940286 |
| AGLKTGII | ASSDARDGNTLY | 0.38848776 |
| ASLAALLL | ASSDARDGNTLY | 0.6335679 |
| ASVINISSI | ASSDARDGNTLY | 0.5803865 |
| ATPPGPA | ASSDARDGNTLY | 0.26325428 |
| ATVASALV | ASSDARDGNTLY | 0.5458528 |
| ATVASLIL | ASSDARDGNTLY | 0.71068203 |
| AVNGPVLIRH | ASSDARDGNTLY | 0.55102855 |

| | | |
|---|---|---|
| DGAAGAVGI | ASSDARDGNTLY | 0.27528343 |
| DPLLHGI | ASSDARDGNTLY | 0.59043777 |
| ELGAALGAL | ASSDARDGNTLY | 0.39147526 |
| ELGRTQAV | ASSDARDGNTLY | 0.6410209 |
| FVAGGYLS | ASSDARDGNTLY | 0.5532645 |
| GASGGAGGAGGW | ASSDARDGNTLY | 0.36944154 |
| GGAGAGGDGGAGGI | ASSDARDGNTLY | 0.5976887 |
| GTGHNGSQPGMGGNG | ASSDARDGNTLY | 0.25595194 |
| GYAAASASAS | ASSDARDGNTLY | 0.46130383 |
| HQVDVAAVQQG | ASSDARDGNTLY | 0.4024642 |
| IGAAIAHT | ASSDARDGNTLY | 0.56944156 |
| IGMIEVAQL | ASSDARDGNTLY | 0.5410951 |
| IRLAPEHTSAMD | ASSDARDGNTLY | 0.34046277 |
| IYNVSVL | ASSDARDGNTLY | 0.5282012 |
| KQGFLGM | ASSDARDGNTLY | 0.38138133 |
| LESVITQL | ASSDARDGNTLY | 0.30251077 |
| LGLGALVVV | ASSDARDGNTLY | 0.56067836 |
| LIGNGGNGGSGGI | ASSDARDGNTLY | 0.38155526 |
| LNAINAPTL | ASSDARDGNTLY | 0.28499067 |
| LQLLNTGV | ASSDARDGNTLY | 0.5477575 |
| LQTSIEF | ASSDARDGNTLY | 0.36300403 |
| MSFAVSEL | ASSDARDGNTLY | 0.5051387 |
| NTGALGL | ASSDARDGNTLY | 0.36069858 |
| PPGAGGAGGAGTT | ASSDARDGNTLY | 0.26937425 |
| QLVGLGI | ASSDARDGNTLY | 0.61645603 |
| RFLEQVHQL | ASSDARDGNTLY | 0.46692067 |
| RGVGYVLR | ASSDARDGNTLY | 0.5618441 |
| SATGVFAGVI | ASSDARDGNTLY | 0.48011827 |
| SLPAITLP | ASSDARDGNTLY | 0.28770825 |
| STALAFTV | ASSDARDGNTLY | 0.6449769 |
| SVLTVGLGK | ASSDARDGNTLY | 0.61525667 |
| TALAGAGSGI | ASSDARDGNTLY | 0.35542148 |
| TGLPFTGI | ASSDARDGNTLY | 0.46363825 |
| VAEHVTV | ASSDARDGNTLY | 0.68285143 |
| VDIVLNSL | ASSDARDGNTLY | 0.62358296 |
| VLTGGGAVL | ASSDARDGNTLY | 0.50257593 |
| VNVGLAYGI | ASSDARDGNTLY | 0.26269406 |
| VSSAAVGAV | ASSDARDGNTLY | 0.5158504 |
| VTSVYEAGV | ASSDARDGNTLY | 0.47703585 |
| WVALDRGA | ASSDARDGNTLY | 0.34904426 |
| YLVGSVVI | ASSDARDGNTLY | 0.25097546 |
| GPPGTGKTLL | ASSDARDGNTLY | 0.6257514 |
| GVGMTKF | ASSDARDGNTLY | 0.7441455 |
| IGIGGSDL | ASSDARDGNTLY | 0.35664275 |
| IGTLLAEL | ASSDARDGNTLY | 0.73901975 |
| IYTSGTT | ASSDARDGNTLY | 0.31058887 |
| LSFPAGV | ASSDARDGNTLY | 0.5410702 |
| LYTSGTT | ASSDARDGNTLY | 0.64928925 |
| VGAFPFGF | ASSDARDGNTLY | 0.378907 |
| VGAGTVEF | ASSDARDGNTLY | 0.369004 |
| VGKSTLLL | ASSDARDGNTLY | 0.266383 |

| | | |
|---|---|---|
| AALAHAGGGIV | ASSDARDGNTLY | 0.3005736 |
| AALVAVSLL | ASSDARDGNTLY | 0.6702313 |
| AAVSRAVAAL | ASSDARDGNTLY | 0.3525648 |
| AGGSSGIGGF | ASSDARDGNTLY | 0.4618838 |
| AGLTVANANG | ASSDARDGNTLY | 0.44183826 |
| AGLVGAVL | ASSDARDGNTLY | 0.35575587 |
| ALSQAFTV | ASSDARDGNTLY | 0.5246074 |
| ALTFSVSL | ASSDARDGNTLY | 0.50555164 |
| ALTGAALVAL | ASSDARDGNTLY | 0.4853259 |
| ASFSIGAGVL | ASSDARDGNTLY | 0.6444279 |
| DGSVYKI | ASSDARDGNTLY | 0.42041236 |
| EQLAEIV | ASSDARDGNTLY | 0.73666775 |
| FAVSELTVM | ASSDARDGNTLY | 0.7417166 |
| GAAPLAAGL | ASSDARDGNTLY | 0.64566076 |
| GADGGDGGSGGAGG | ASSDARDGNTLY | 0.40348467 |
| GFGLDTGGAGG | ASSDARDGNTLY | 0.7305603 |
| GLASGGLL | ASSDARDGNTLY | 0.45398262 |
| GRIAGAAVLL | ASSDARDGNTLY | 0.50479287 |
| GSAGSLNTSF | ASSDARDGNTLY | 0.6247991 |
| GTGQIGI | ASSDARDGNTLY | 0.13652411 |
| GTLIALIL | ASSDARDGNTLY | 0.15001804 |
| IAATMGFVVG | ASSDARDGNTLY | 0.13225923 |
| IFAQALL | ASSDARDGNTLY | 0.14337388 |
| IGAAVALGF | ASSDARDGNTLY | 0.17323196 |
| IGLTPLAL | ASSDARDGNTLY | 0.13243459 |
| ILGSNTSTL | ASSDARDGNTLY | 0.14849547 |
| IVFSLTVL | ASSDARDGNTLY | 0.1722959 |
| LASKATGF | ASSDARDGNTLY | 0.12949014 |
| LASVALIL | ASSDARDGNTLY | 0.13783227 |
| LFAVGLTL | ASSDARDGNTLY | 0.16462639 |
| LFNGGIGTYI | ASSDARDGNTLY | 0.20290405 |
| LTLGAGII | ASSDARDGNTLY | 0.15437563 |
| LVAAALGDI | ASSDARDGNTLY | 0.14446141 |
| NTAFVQLGI | ASSDARDGNTLY | 0.13992324 |
| QLFSGAEL | ASSDARDGNTLY | 0.14287317 |
| SAATFITL | ASSDARDGNTLY | 0.13989231 |
| SNYGLANLGV | ASSDARDGNTLY | 0.12829055 |
| SYGGAYAVM | ASSDARDGNTLY | 0.14604943 |
| TTGVGGLTLG | ASSDARDGNTLY | 0.12882815 |
| TTQTIYI | ASSDARDGNTLY | 0.14030321 |
| VAAGAGLGAVI | ASSDARDGNTLY | 0.15845744 |
| VAAVVTLL | ASSDARDGNTLY | 0.14877278 |
| VGFLGGLI | ASSDARDGNTLY | 0.15281859 |
| VGVIPILL | ASSDARDGNTLY | 0.17090651 |
| VIFASSLI | ASSDARDGNTLY | 0.154935 |
| VKGGDGGAAAG | ASSDARDGNTLY | 0.15212634 |
| VTKTFTNI | ASSDARDGNTLY | 0.15212072 |
| VVAAAILL | ASSDARDGNTLY | 0.19881596 |
| YAALDVEL | ASSDARDGNTLY | 0.14836821 |
| LASVALLL | ASSDARDGNTLY | 0.13749963 |
| LDAVHRV | ASSDARDGNTLY | 0.16842093 |

| | | |
|---|---|---|
| QGLSSSIF | ASSDARDGNTLY | 0.16101247 |
| RGTFANIRL | ASSDARDGNTLY | 0.14889064 |
| RQLWWGHRI | ASSDARDGNTLY | 0.14994416 |
| YGPPGTGKTLL | ASSDARDGNTLY | 0.13004148 |
| AALVGAPANVA | ASSDARDGNTLY | 0.15902661 |
| AATFRRTE | ASSDARDGNTLY | 0.15098554 |
| AGALGGGI | ASSDARDGNTLY | 0.15871245 |
| AGGAGGAGGIGAG | ASSDARDGNTLY | 0.17082497 |
| AGSGALGL | ASSDARDGNTLY | 0.1377628 |
| AQIADQI | ASSDARDGNTLY | 0.13528767 |
| ASAVAVTYL | ASSDARDGNTLY | 0.16126609 |
| ASTVLGGTGG | ASSDARDGNTLY | 0.14485087 |
| AVGGEPAK | ASSDARDGNTLY | 0.15730716 |
| CEQGTFGGTC | ASSDARDGNTLY | 0.17257364 |
| DEGKAPAG | ASSDARDGNTLY | 0.13212228 |
| DTVVFPAL | ASSDARDGNTLY | 0.14202845 |
| FEGITMAG | ASSDARDGNTLY | 0.12808625 |
| FPLVVQD | ASSDARDGNTLY | 0.13406728 |
| GAFTTPQ | ASSDARDGNTLY | 0.18133943 |
| GFGGFGTAGGVGGA | ASSDARDGNTLY | 0.13011901 |
| GFLAQFL | ASSDARDGNTLY | 0.25734517 |
| GGAGGAGGNAGVA | ASSDARDGNTLY | 0.20784709 |
| GGVGTAGAGGAGGAGG | ASSDARDGNTLY | 0.2679577 |
| GNGGGGDAGNAGSGGN | ASSDARDGNTLY | 0.1504235 |
| GTGGAGGAGNDAGS | ASSDARDGNTLY | 0.12989725 |
| GTTNTGWFNS | ASSDARDGNTLY | 0.13904409 |
| IFMGPLFTL | ASSDARDGNTLY | 0.14625692 |
| ILAHAIEAI | ASSDARDGNTLY | 0.13885193 |
| IRSAIREAL | ASSDARDGNTLY | 0.15526639 |
| ITFAGGG | ASSDARDGNTLY | 0.15225118 |
| ITGMSGHQEV | ASSDARDGNTLY | 0.29780877 |
| IVAGLACTL | ASSDARDGNTLY | 0.18828703 |
| IVGGGLAGA | ASSDARDGNTLY | 0.1526457 |
| IVTGVLNI | ASSDARDGNTLY | 0.15519017 |
| LAQQGCI | ASSDARDGNTLY | 0.13044171 |
| LCLHVEN | ASSDARDGNTLY | 0.14279145 |
| LDAISDGS | ASSDARDGNTLY | 0.13213485 |
| LGGFAAML | ASSDARDGNTLY | 0.13535745 |
| LGGKNPMI | ASSDARDGNTLY | 0.23637605 |
| LGPKGVAAIL | ASSDARDGNTLY | 0.16330053 |
| LMVGVSI | ASSDARDGNTLY | 0.24095294 |
| LVGAAFVL | ASSDARDGNTLY | 0.14209808 |
| LVLAVAPL | ASSDARDGNTLY | 0.13279858 |
| MAAIAVIDAA | ASSDARDGNTLY | 0.16613713 |
| NNGINAELQ | ASSDARDGNTLY | 0.14412524 |
| RGEPLVLVHG | ASSDARDGNTLY | 0.14836216 |
| RGGAGGAAGLI | ASSDARDGNTLY | 0.14436118 |
| RSYRMHPAV | ASSDARDGNTLY | 0.14675044 |
| SAAHLLAGQLA | ASSDARDGNTLY | 0.14264603 |
| SAAVGSY | ASSDARDGNTLY | 0.16910832 |
| SPHAVLHRI | ASSDARDGNTLY | 0.14560705 |

| | | |
|---|---|---|
| SQIAKVL | ASSDARDGNTLY | 0.14198174 |
| TALGHLVP | ASSDARDGNTLY | 0.1401507 |
| TAPTTSGAA | ASSDARDGNTLY | 0.14382224 |
| TDGDDGNGGSPGT | ASSDARDGNTLY | 0.22477213 |
| TDGLGEN | ASSDARDGNTLY | 0.13404375 |
| TGGPPLPGQA | ASSDARDGNTLY | 0.14241453 |
| TIPAITI | ASSDARDGNTLY | 0.14125508 |
| TIPGFTI | ASSDARDGNTLY | 0.14116502 |
| TSIASGALA | ASSDARDGNTLY | 0.1385744 |
| TVTLPGAAI | ASSDARDGNTLY | 0.13475941 |
| VASGLSAALP | ASSDARDGNTLY | 0.14291827 |
| VGAGVIGI | ASSDARDGNTLY | 0.15881996 |
| VGPKRFGV | ASSDARDGNTLY | 0.14119811 |
| VNVAFPD | ASSDARDGNTLY | 0.15353368 |
| VSMTNSGVS | ASSDARDGNTLY | 0.14710854 |
| VTDVLGDLSV | ASSDARDGNTLY | 0.14 |
| VVVVGAGFAGL | ASSDARDGNTLY | 0.17855325 |
| VYTVGTVV | ASSDARDGNTLY | 0.13549955 |
| WTTFSGP | ASSDARDGNTLY | 0.142179 |
| YDELVEAF | ASSDARDGNTLY | 0.21588838 |
| YGTGGAGGNGG | ASSDARDGNTLY | 0.15614453 |
| YTNVMAV | ASSDARDGNTLY | 0.19609617 |
| FLEQVHQL | ASSDARDGNTLY | 0.1340412 |
| GLSLLAVAV | ASSDARDGNTLY | 0.1312623 |
| LEQVHQL | ASSDARDGNTLY | 0.12920113 |
| LNAAGAI | ASSDARDGNTLY | 0.17870916 |
| PAVHASL | ASSDARDGNTLY | 0.13388507 |
| TELEDDA | ASSDARDGNTLY | 0.14285743 |
| TPPGPPN | ASSDARDGNTLY | 0.14294699 |
| TPPPPPPP | ASSDARDGNTLY | 0.15000425 |
| TQSPALI | ASSPDLGGRRYEQY | 0.14365503 |
| EFAAAFPQ | ASSPDLGGRRYEQY | 0.1369036 |
| LLSGNLDVL | ASSPDLGGRRYEQY | 0.13600287 |
| GPANLGNYNI | ASSPDLGGRRYEQY | 0.17321117 |
| TLGPTLP | ASSPDLGGRRYEQY | 0.14876254 |
| VYAYSSCCK | ASSPDLGGRRYEQY | 0.14067677 |
| APAPAPAPNI | ASSPDLGGRRYEQY | 0.14136632 |
| PGGPVPIP | ASSPDLGGRRYEQY | 0.24400929 |
| FLTVALEP | ASSPDLGGRRYEQY | 0.17643984 |
| SQYFGANL | ASSPDLGGRRYEQY | 0.15198559 |
| GGVKPAA | ASSPDLGGRRYEQY | 0.16893251 |
| GGTGGAGGAGGW | ASSPDLGGRRYEQY | 0.12723951 |
| GVGGVAGLGGAGG | ASSPDLGGRRYEQY | 0.13022226 |
| VLLVLATTI | ASSPDLGGRRYEQY | 0.17921902 |
| IGIFAAML | ASSPDLGGRRYEQY | 0.21846999 |
| AVGDLID | ASSPDLGGRRYEQY | 0.16069937 |
| NISPLPL | ASSPDLGGRRYEQY | 0.15096694 |
| VTAVFVAI | ASSPDLGGRRYEQY | 0.18015413 |
| SGDDNDAAL | ASSPDLGGRRYEQY | 0.12757437 |
| AVAALAHS | ASSPDLGGRRYEQY | 0.1357504 |
| GGVTVMATP | ASSPDLGGRRYEQY | 0.14561467 |

| | | |
|---|---|---|
| AGLAANVRD | ASSPDLGGRRYEQY | 0.14484292 |
| TNRPDLIDPALLRPG | ASSPDLGGRRYEQY | 0.13417105 |
| LANLGVTAL | ASSPDLGGRRYEQY | 0.1492251 |
| TGERIEL | ASSPDLGGRRYEQY | 0.20600276 |
| NGEELAA | ASSPDLGGRRYEQY | 0.13230982 |
| VLSLVWG | ASSPDLGGRRYEQY | 0.14354767 |
| IGYALWL | ASSPDLGGRRYEQY | 0.12940983 |
| LSLLLAPL | ASSPDLGGRRYEQY | 0.14208202 |
| TLDEALK | ASSPDLGGRRYEQY | 0.21981195 |
| TDGAGTAEGG | ASSPDLGGRRYEQY | 0.17045404 |
| NAPGGAGGNGG | ASSPDLGGRRYEQY | 0.15241781 |
| VSVFMGMGGNF | ASSPDLGGRRYEQY | 0.18621866 |
| LASYLAAL | ASSPDLGGRRYEQY | 0.16686143 |
| LVGGTPAL | ASSPDLGGRRYEQY | 0.13217098 |
| VRNADVL | ASSPDLGGRRYEQY | 0.14702708 |
| TCSSPFLVSAM | ASSPDLGGRRYEQY | 0.13712119 |
| VHITDAE | ASSPDLGGRRYEQY | 0.17306039 |
| EYAAPSF | ASSPDLGGRRYEQY | 0.17095639 |
| VVDAFHGGL | ASSPDLGGRRYEQY | 0.15011232 |
| ASVVDPQ | ASSPDLGGRRYEQY | 0.16029923 |
| SYNEFGGLS | ASSPDLGGRRYEQY | 0.16209507 |
| DPDGSVY | ASSPDLGGRRYEQY | 0.1381913 |
| MVSTIHATE | ASSPDLGGRRYEQY | 0.1431621 |
| LAAGVADPV | ASSPDLGGRRYEQY | 0.14654087 |
| TKEQIAA | ASSPDLGGRRYEQY | 0.1756645 |
| SLASPAAL | ASSPDLGGRRYEQY | 0.13255613 |
| IGLVFYGV | ASSPDLGGRRYEQY | 0.13429578 |
| FLLDASNL | ASSPDLGGRRYEQY | 0.13447833 |
| YQVGNFP | ASSPDLGGRRYEQY | 0.16588624 |
| ESLLSQST | ASSPDLGGRRYEQY | 0.15756012 |
| LFTVGFAAVG | ASSPDLGGRRYEQY | 0.1472329 |
| ATPPTPT | ASSPDLGGRRYEQY | 0.13585491 |
| TDLATLG | ASSPDLGGRRYEQY | 0.13594276 |
| SAGGTGR | ASSPDLGGRRYEQY | 0.15549783 |
| GFSNSGS | ASSPDLGGRRYEQY | 0.13242127 |
| GGNGGSGGTSV | ASSPDLGGRRYEQY | 0.14503141 |
| GTSVATGGAGN | ASSPDLGGRRYEQY | 0.14189114 |
| NGGNGGSGGTS | ASSPDLGGRRYEQY | 0.14675398 |
| AVVKFPH | ASSPDLGGRRYEQY | 0.1393289 |
| TPPPPPP | ASSPDLGGRRYEQY | 0.13149893 |
| AAPAAAAAPA | ASSPDLGGRRYEQY | 0.18661518 |
| IALNEDL | ASSPDLGGRRYEQY | 0.13432328 |
| IVNVSSM | ASSPDLGGRRYEQY | 0.17189623 |
| TGIPPAPR | ASSPDLGGRRYEQY | 0.15233104 |
| FDLGGGTF | ASSPDLGGRRYEQY | 0.13413154 |
| FDLGGGTFDVS | ASSPDLGGRRYEQY | 0.14216898 |
| NEPTAAA | ASSPDLGGRRYEQY | 0.16345966 |
| NEPTAAALA | ASSPDLGGRRYEQY | 0.14183213 |
| GSIINISSI | ASSPDLGGRRYEQY | 0.14633074 |
| VAPGNAGTA | ASSPDLGGRRYEQY | 0.18111554 |
| TGGGASLE | ASSPDLGGRRYEQY | 0.15643933 |

| | | |
|---|---|---|
| VAVYNHYKRI | ASSPDLGGRRYEQY | 0.17075147 |
| TDPAPATT | ASSPDLGGRRYEQY | 0.1300123 |
| TNRPDILDPALLRPG | ASSPDLGGRRYEQY | 0.13999292 |
| LTTTTTS | ASSPDLGGRRYEQY | 0.13939269 |
| PTGSGKTL | ASSPDLGGRRYEQY | 0.14907916 |
| TGSGKTL | ASSPDLGGRRYEQY | 0.19157752 |
| TIGHVDHGK | ASSPDLGGRRYEQY | 0.19683805 |
| TIGHVDHGKTT | ASSPDLGGRRYEQY | 0.13582861 |
| GHVDHGKTT | ASSPDLGGRRYEQY | 0.18891373 |
| AQFEHTLL | ASSPDLGGRRYEQY | 0.14292118 |
| IVNQDEL | ASSPDLGGRRYEQY | 0.13507962 |
| TNGSQFF | ASSPDLGGRRYEQY | 0.1315195 |
| TGKGIVE | ASSPDLGGRRYEQY | 0.14935155 |
| LISNASDA | ASSPDLGGRRYEQY | 0.14126314 |
| AELRTLGELTKIAKAHGAQ | ASSPDLGGRRYEQY | 0.15001656 |
| APNAVAAAEPSW | ASSPDLGGRRYEQY | 0.1820037 |
| APQVAGAATAVL | ASSPDLGGRRYEQY | 0.14012022 |
| ARRDPAAAAAILPTDA | ASSPDLGGRRYEQY | 0.14606433 |
| DAAARDVLAEAGLAEYF | ASSPDLGGRRYEQY | 0.13218887 |
| DDQIVLSAKANLAKVEQ | ASSPDLGGRRYEQY | 0.1741921 |
| DELLPDTGEDSDEDGH | ASSPDLGGRRYEQY | 0.13107622 |
| ELPEAPANRATVPQD | ASSPDLGGRRYEQY | 0.13037743 |
| FGQSKIERVTGIG | ASSPDLGGRRYEQY | 0.16039555 |
| GAGGTGSTAGGAGGAGGAGGLY | ASSPDLGGRRYEQY | 0.13579454 |
| GHGGTGAAGGNGGTGGAGGSGI | ASSPDLGGRRYEQY | 0.14254989 |
| GNGGNGGAGGEGGPGGAGGAGA | ASSPDLGGRRYEQY | 0.13529238 |
| GSTAGNGGNGGNATNGTIAGQP | ASSPDLGGRRYEQY | 0.16619703 |
| GTGGDGGNGGNGGTGG | ASSPDLGGRRYEQY | 0.1483998 |
| GVRRPAGIPADAPV | ASSPDLGGRRYEQY | 0.13626936 |
| GYRQVTDAHLLTLARR | ASSPDLGGRRYEQY | 0.13703424 |
| IAKMGQDGHDRG | ASSPDLGGRRYEQY | 0.1511479 |
| KAEPSIEIQYT | ASSPDLGGRRYEQY | 0.12984976 |
| KPVRIRMAADEDTVAVT | ASSPDLGGRRYEQY | 0.1629732 |
| MPPRIAGMRLLVIKPEP | ASSPDLGGRRYEQY | 0.19877768 |
| MKFYAVALFRSRGVPPTR | ASSPDLGGRRYEQY | 0.12960984 |
| NAGGFGILGAVAHSP | ASSPDLGGRRYEQY | 0.1497723 |
| NEGREITLAMAEM | ASSPDLGGRRYEQY | 0.13607629 |
| NELVEWSHPLGHRWRWE | ASSPDLGGRRYEQY | 0.15206088 |
| NLGAGGTGGNGGNPG | ASSPDLGGRRYEQY | 0.13361709 |
| NTEWPKWETVMAADPVD | ASSPDLGGRRYEQY | 0.12876315 |
| PASAAPAETQP | ASSPDLGGRRYEQY | 0.16522765 |
| QQSSDGKNGVRAAEITGE | ASSPDLGGRRYEQY | 0.17300169 |
| RAPDAVTTSWWKE | ASSPDLGGRRYEQY | 0.15119135 |
| SGFGNVGAHTSGWFNQ | ASSPDLGGRRYEQY | 0.13704175 |
| SGVAGAVGQAASVGGLKV | ASSPDLGGRRYEQY | 0.1430044 |
| SNGGAGTGSAGGNGGT | ASSPDLGGRRYEQY | 0.19434561 |
| TGGAGGEGGAAG | ASSPDLGGRRYEQY | 0.15793324 |
| TGGDGGAAGAGGAPGHG | ASSPDLGGRRYEQY | 0.14638716 |
| TMPPLLAAIANAGRHGV | ASSPDLGGRRYEQY | 0.1376022 |
| TSSTAGNSGGA | ASSPDLGGRRYEQY | 0.16230083 |
| VPATVPVGILC | ASSPDLGGRRYEQY | 0.14823003 |

| | | |
|---|---|---|
| VQRPDSGGVVLTGRIS | ASSPDLGGRRYEQY | 0.13338968 |
| WTPNPVPATKGVPAGLVQAT | ASSPDLGGRRYEQY | 0.13797665 |
| YALKRALAKFSDRFDVVII | ASSPDLGGRRYEQY | 0.13885334 |
| YLIGSVIV | ASSPDLGGRRYEQY | 0.15370046 |
| YGQLVAVID | ASSPDLGGRRYEQY | 0.14961028 |
| VTVGGMTLPA | ASSPDLGGRRYEQY | 0.13107535 |
| VLALPAV | ASSPDLGGRRYEQY | 0.18011603 |
| VIPPTLNYET | ASSPDLGGRRYEQY | 0.14266004 |
| VGMGGMHPSSG | ASSPDLGGRRYEQY | 0.17900899 |
| VGLAGGAATA | ASSPDLGGRRYEQY | 0.13469906 |
| VGGAGGAGGSGP | ASSPDLGGRRYEQY | 0.13954552 |
| VFAAFMV | ASSPDLGGRRYEQY | 0.22358805 |
| TLYDKLR | ASSPDLGGRRYEQY | 0.13590255 |
| TLAVFTVL | ASSPDLGGRRYEQY | 0.1911527 |
| TGSVQPARV | ASSPDLGGRRYEQY | 0.15047945 |
| TGGGAGSGGT | ASSPDLGGRRYEQY | 0.16302952 |
| SHLETSL | ASSPDLGGRRYEQY | 0.13658072 |
| SFTTGMT | ASSPDLGGRRYEQY | 0.1472338 |
| QVEVAKRMGVRQPTVS | ASSPDLGGRRYEQY | 0.13554285 |
| QGSGGPALAM | ASSPDLGGRRYEQY | 0.15922613 |
| MLAPSNP | ASSPDLGGRRYEQY | 0.1298055 |
| LVNNAAI | ASSPDLGGRRYEQY | 0.14676344 |
| LTGEINTLA | ASSPDLGGRRYEQY | 0.13188818 |
| LSIFGPV | ASSPDLGGRRYEQY | 0.16521232 |
| LLALLANL | ASSPDLGGRRYEQY | 0.16365004 |
| LISNASDALDKLR | ASSPDLGGRRYEQY | 0.17362712 |
| LINEAVP | ASSPDLGGRRYEQY | 0.16782048 |
| LGYAFSGL | ASSPDLGGRRYEQY | 0.19471268 |
| LDFVAVAL | ASSPDLGGRRYEQY | 0.17321156 |
| LAQPGAE | ASSPDLGGRRYEQY | 0.17225905 |
| ISNLGAVTGF | ASSPDLGGRRYEQY | 0.13965912 |
| HQEDPGRI | ASSPDLGGRRYEQY | 0.1505901 |
| GTPGEPTP | ASSPDLGGRRYEQY | 0.17974773 |
| GFFNSGS | ASSPDLGGRRYEQY | 0.13537022 |
| GAGGGGLTAGAN | ASSPDLGGRRYEQY | 0.13143142 |
| FVDALVNM | ASSPDLGGRRYEQY | 0.13705955 |
| FLTVAPEL | ASSPDLGGRRYEQY | 0.15832262 |
| FETLTAL | ASSPDLGGRRYEQY | 0.1512891 |
| DVINPIEL | ASSPDLGGRRYEQY | 0.14642985 |
| APVSGAPV | ASSPDLGGRRYEQY | 0.15985222 |
| ALVGGLAW | ASSPDLGGRRYEQY | 0.15785803 |
| ALAAAAATAI | ASSPDLGGRRYEQY | 0.17926522 |
| AFGLGLLL | ASSPDLGGRRYEQY | 0.16709459 |
| AFFAAASL | ASSPDLGGRRYEQY | 0.13795811 |
| AAAALPAFSPP | ASSPDLGGRRYEQY | 0.15417835 |
| VVLAPAL | ASSPDLGGRRYEQY | 0.14005038 |
| TIGHVDHGKTTL | ASSPDLGGRRYEQY | 0.14064664 |
| LISNASDALDK | ASSPDLGGRRYEQY | 0.14250603 |
| LISNASDAL | ASSPDLGGRRYEQY | 0.14827462 |
| ISNASDALDK | ASSPDLGGRRYEQY | 0.19815207 |
| NQAETLVYQTE | ASSPDLGGRRYEQY | 0.12585813 |

| | | |
|---|---|---|
| TGGTGGTAGAG | ASSPDLGGRRYEQY | 0.13263106 |
| TGSTAGGAGGA | ASSPDLGGRRYEQY | 0.17249855 |
| DPLAVPVAAGPV | ASSPDLGGRRYEQY | 0.15740287 |
| GGTGGTPGNGGN | ASSPDLGGRRYEQY | 0.14996237 |
| PAGLAGVDPATL | ASSPDLGGRRYEQY | 0.14964743 |
| GNNLVVEAPHAGASV | ASSPDLGGRRYEQY | 0.129545 |
| KVTRIMSASTLEQPA | ASSPDLGGRRYEQY | 0.19783227 |
| LGFALPIALPRRTVV | ASSPDLGGRRYEQY | 0.13390763 |
| TRLGNLTARKLVVLV | ASSPDLGGRRYEQY | 0.14740689 |
| WSQQRRLIYFPSAGP | ASSPDLGGRRYEQY | 0.15589252 |
| GPNGNPDPMAAAVDIR | ASSPDLGGRRYEQY | 0.15684915 |
| TQRCVSVADLRAAASR | ASSPDLGGRRYEQY | 0.13571924 |
| QFVLDVAEHPVEVLTETV | ASSPDLGGRRYEQY | 0.13775788 |
| GGAGGTGGAGGNGGTQAGDG | ASSPDLGGRRYEQY | 0.14290726 |
| PLLATLRAIAVDVDAKPAQV | ASSPDLGGRRYEQY | 0.14034048 |
| APKFRTGGYMIDFWGVGYQVAK | ASSPDLGGRRYEQY | 0.14089933 |
| GAGGAGGAGGAAGGPGGTGGTG | ASSPDLGGRRYEQY | 0.14821266 |
| IGALGLLATDVAPAGSVAFGVA | ASSPDLGGRRYEQY | 0.13547496 |
| AGGAGGIGGNGGAGGAGGVLMG | ASSPDLGGRRYEQY | 0.2636271 |
| ADELIGLL | ASSPDLGGRRYEQY | 0.21027976 |
| ADRALKA | ASSPDLGGRRYEQY | 0.14119276 |
| AGAVGQVGGAGGA | ASSPDLGGRRYEQY | 0.14898983 |
| AGGGGGLGGAP | ASSPDLGGRRYEQY | 0.24641526 |
| AGGLLYGNGGAG | ASSPDLGGRRYEQY | 0.22604948 |
| AGLESIGL | ASSPDLGGRRYEQY | 0.13993055 |
| AGLKTGII | ASSPDLGGRRYEQY | 0.1423266 |
| ASLAALLL | ASSPDLGGRRYEQY | 0.13562095 |
| ASVINISSI | ASSPDLGGRRYEQY | 0.13404739 |
| ATPPGPA | ASSPDLGGRRYEQY | 0.17207322 |
| ATVASALV | ASSPDLGGRRYEQY | 0.15594518 |
| ATVASLIL | ASSPDLGGRRYEQY | 0.13764918 |
| AVNGPVLIRH | ASSPDLGGRRYEQY | 0.13924402 |
| DGAAGAVGI | ASSPDLGGRRYEQY | 0.15660766 |
| DPLLHGI | ASSPDLGGRRYEQY | 0.15839761 |
| ELGAALGAL | ASSPDLGGRRYEQY | 0.12986776 |
| ELGRTQAV | ASSPDLGGRRYEQY | 0.13532074 |
| FVAGGYLS | ASSPDLGGRRYEQY | 0.16031796 |
| GASGGAGGAGGW | ASSPDLGGRRYEQY | 0.13613857 |
| GGAGAGGDGGAGGI | ASSPDLGGRRYEQY | 0.13558517 |
| GTGHNGSQPGMGGNG | ASSPDLGGRRYEQY | 0.23098934 |
| GYAAASASAS | ASSPDLGGRRYEQY | 0.14818847 |
| HQVDVAAVQQG | ASSPDLGGRRYEQY | 0.13407658 |
| IGAAIAHT | ASSPDLGGRRYEQY | 0.15128236 |
| IGMIEVAQL | ASSPDLGGRRYEQY | 0.23739685 |
| IRLAPEHTSAMD | ASSPDLGGRRYEQY | 0.17010047 |
| IYNVSVL | ASSPDLGGRRYEQY | 0.15714495 |
| KQGFLGM | ASSPDLGGRRYEQY | 0.13514009 |
| LESVITQL | ASSPDLGGRRYEQY | 0.14762768 |
| LGLGALVVV | ASSPDLGGRRYEQY | 0.1482448 |
| LIGNGGNGGSGGI | ASSPDLGGRRYEQY | 0.20982909 |
| LNAINAPTL | ASSPDLGGRRYEQY | 0.17178276 |

| | | |
|---|---|---|
| LQLLNTGV | ASSPDLGGRRYEQY | 0.14416696 |
| LQTSIEF | ASSPDLGGRRYEQY | 0.2225736 |
| MSFAVSEL | ASSPDLGGRRYEQY | 0.19304353 |
| NTGALGL | ASSPDLGGRRYEQY | 0.14054866 |
| PPGAGGAGGAGTT | ASSPDLGGRRYEQY | 0.12957968 |
| QLVGLGI | ASSPDLGGRRYEQY | 0.13220342 |
| RFLEQVHQL | ASSPDLGGRRYEQY | 0.136457 |
| RGVGYVLR | ASSPDLGGRRYEQY | 0.15024748 |
| SATGVFAGVI | ASSPDLGGRRYEQY | 0.164957 |
| SLPAITLP | ASSPDLGGRRYEQY | 0.1308371 |
| STALAFTV | ASSPDLGGRRYEQY | 0.15133768 |
| SVLTVGLGK | ASSPDLGGRRYEQY | 0.20242059 |
| TALAGAGSGI | ASSPDLGGRRYEQY | 0.16009085 |
| TGLPFTGI | ASSPDLGGRRYEQY | 0.17170098 |
| VAEHVTV | ASSPDLGGRRYEQY | 0.135971 |
| VDIVLNSL | ASSPDLGGRRYEQY | 0.13578895 |
| VLTGGGAVL | ASSPDLGGRRYEQY | 0.13875104 |
| VNVGLAYGI | ASSPDLGGRRYEQY | 0.1318584 |
| VSSAAVGAV | ASSPDLGGRRYEQY | 0.1362078 |
| VTSVYEAGV | ASSPDLGGRRYEQY | 0.1336943 |
| WVALDRGA | ASSPDLGGRRYEQY | 0.13242994 |
| YLVGSVVI | ASSPDLGGRRYEQY | 0.1300669 |
| GPPGTGKTLL | ASSPDLGGRRYEQY | 0.13042043 |
| GVGMTKF | ASSPDLGGRRYEQY | 0.17342618 |
| IGIGGSDL | ASSPDLGGRRYEQY | 0.13653801 |
| IGTLLAEL | ASSPDLGGRRYEQY | 0.16604625 |
| IYTSGTT | ASSPDLGGRRYEQY | 0.23010044 |
| LSFPAGV | ASSPDLGGRRYEQY | 0.14334957 |
| LYTSGTT | ASSPDLGGRRYEQY | 0.1492845 |
| VGAFPFGF | ASSPDLGGRRYEQY | 0.13672866 |
| VGAGTVEF | ASSPDLGGRRYEQY | 0.24419333 |
| VGKSTLLL | ASSPDLGGRRYEQY | 0.13404843 |
| AALAHAGGGIV | ASSPDLGGRRYEQY | 0.15855464 |
| AALVAVSLL | ASSPDLGGRRYEQY | 0.14590195 |
| AAVSRAVAAL | ASSPDLGGRRYEQY | 0.1315208 |
| AGGSSGIGGF | ASSPDLGGRRYEQY | 0.13561888 |
| AGLTVANANG | ASSPDLGGRRYEQY | 0.13921985 |
| AGLVGAVL | ASSPDLGGRRYEQY | 0.14028417 |
| ALSQAFTV | ASSPDLGGRRYEQY | 0.13054918 |
| ALTFSVSL | ASSPDLGGRRYEQY | 0.13401262 |
| ALTGAALVAL | ASSPDLGGRRYEQY | 0.14888322 |
| ASFSIGAGVL | ASSPDLGGRRYEQY | 0.13599293 |
| DGSVYKI | ASSPDLGGRRYEQY | 0.12826714 |
| EQLAEIV | ASSPDLGGRRYEQY | 0.14540742 |
| FAVSELTVM | ASSPDLGGRRYEQY | 0.14563978 |
| GAAPLAAGL | ASSPDLGGRRYEQY | 0.15594211 |
| GADGGDGGSGGAGG | ASSPDLGGRRYEQY | 0.22994468 |
| GFGLDTGGAGG | ASSPDLGGRRYEQY | 0.14291473 |
| GLASGGLL | ASSPDLGGRRYEQY | 0.14791512 |
| GRIAGAAVLL | ASSPDLGGRRYEQY | 0.14975044 |
| GSAGSLNTSF | ASSPDLGGRRYEQY | 0.13981335 |

| | | |
|---|---|---|
| GTGQIGI | ASSPDLGGRRYEQY | 0.16223964 |
| GTLIALIL | ASSPDLGGRRYEQY | 0.12824216 |
| IAATMGFVVG | ASSPDLGGRRYEQY | 0.13379991 |
| IFAQALL | ASSPDLGGRRYEQY | 0.1443444 |
| IGAAVALGF | ASSPDLGGRRYEQY | 0.13630673 |
| IGLTPLAL | ASSPDLGGRRYEQY | 0.13931131 |
| ILGSNTSTL | ASSPDLGGRRYEQY | 0.16979823 |
| IVFSLTVL | ASSPDLGGRRYEQY | 0.1334782 |
| LASKATGF | ASSPDLGGRRYEQY | 0.13596381 |
| LASVALIL | ASSPDLGGRRYEQY | 0.13667108 |
| LFAVGLTL | ASSPDLGGRRYEQY | 0.14289853 |
| LFNGGIGTYI | ASSPDLGGRRYEQY | 0.17115252 |
| LTLGAGII | ASSPDLGGRRYEQY | 0.13857786 |
| LVAAALGDI | ASSPDLGGRRYEQY | 0.15728992 |
| NTAFVQLGI | ASSPDLGGRRYEQY | 0.15206574 |
| QLFSGAEL | ASSPDLGGRRYEQY | 0.14586201 |
| SAATFITL | ASSPDLGGRRYEQY | 0.17710625 |
| SNYGLANLGV | ASSPDLGGRRYEQY | 0.19356437 |
| SYGGAYAVM | ASSPDLGGRRYEQY | 0.1344791 |
| TTGVGGLTLG | ASSPDLGGRRYEQY | 0.13642828 |
| TTQTIYI | ASSPDLGGRRYEQY | 0.13431312 |
| VAAGAGLGAVI | ASSPDLGGRRYEQY | 0.13430078 |
| VAAVVTLL | ASSPDLGGRRYEQY | 0.23371041 |
| VGFLGGLI | ASSPDLGGRRYEQY | 0.13401505 |
| VGVIPILL | ASSPDLGGRRYEQY | 0.14160496 |
| VIFASSLI | ASSPDLGGRRYEQY | 0.1367088 |
| VKGGDGGAAAG | ASSPDLGGRRYEQY | 0.20418496 |
| VTKTFTNI | ASSPDLGGRRYEQY | 0.14413941 |
| VVAAAILL | ASSPDLGGRRYEQY | 0.1874985 |
| YAALDVEL | ASSPDLGGRRYEQY | 0.17880824 |
| LASVALLL | ASSPDLGGRRYEQY | 0.13096674 |
| LDAVHRV | ASSPDLGGRRYEQY | 0.15029994 |
| QGLSSSIF | ASSPDLGGRRYEQY | 0.18522258 |
| RGTFANIRL | ASSPDLGGRRYEQY | 0.13676162 |
| RQLWWGHRI | ASSPDLGGRRYEQY | 0.13033235 |
| YGPPGTGKTLL | ASSPDLGGRRYEQY | 0.13351797 |
| AALVGAPANVA | ASSPDLGGRRYEQY | 0.13903454 |
| AATFRRTE | ASSPDLGGRRYEQY | 0.14774586 |
| AGALGGGI | ASSPDLGGRRYEQY | 0.14953248 |
| AGGAGGAGGIGAG | ASSPDLGGRRYEQY | 0.1625399 |
| AGSGALGL | ASSPDLGGRRYEQY | 0.14879902 |
| AQIADQI | ASSPDLGGRRYEQY | 0.1553365 |
| ASAVAVTYL | ASSPDLGGRRYEQY | 0.223709 |
| ASTVLGGTGG | ASSPDLGGRRYEQY | 0.15504447 |
| AVGGEPAK | ASSPDLGGRRYEQY | 0.26943314 |
| CEQGTFGGTC | ASSPDLGGRRYEQY | 0.13559072 |
| DEGKAPAG | ASSPDLGGRRYEQY | 0.14343889 |
| DTVVFPAL | ASSPDLGGRRYEQY | 0.15310311 |
| FEGITMAG | ASSPDLGGRRYEQY | 0.17807557 |
| FPLVVQD | ASSPDLGGRRYEQY | 0.13376604 |
| GAFTTPQ | ASSPDLGGRRYEQY | 0.23569953 |

| | | |
|---|---|---|
| GFGGFGTAGGVGGA | ASSPDLGGRRYEQY | 0.2293967 |
| GFLAQFL | ASSPDLGGRRYEQY | 0.13915448 |
| GGAGGAGGNAGVA | ASSPDLGGRRYEQY | 0.19624561 |
| GGVGTAGAGGAGGAGG | ASSPDLGGRRYEQY | 0.128746 |
| GNGGGGDAGNAGSGGN | ASSPDLGGRRYEQY | 0.13189025 |
| GTGGAGGAGNDAGS | ASSPDLGGRRYEQY | 0.13118006 |
| GTTNTGWFNS | ASSPDLGGRRYEQY | 0.15496525 |
| IFMGPLFTL | ASSPDLGGRRYEQY | 0.13886148 |
| ILAHAIEAI | ASSPDLGGRRYEQY | 0.14370398 |
| IRSAIREAL | ASSPDLGGRRYEQY | 0.1459442 |
| ITFAGGG | ASSPDLGGRRYEQY | 0.20747279 |
| ITGMSGHQEV | ASSPDLGGRRYEQY | 0.15004556 |
| IVAGLACTL | ASSPDLGGRRYEQY | 0.13841484 |
| IVGGGLAGA | ASSPDLGGRRYEQY | 0.16258268 |
| IVTGVLNI | ASSPDLGGRRYEQY | 0.17213266 |
| LAQQGCI | ASSPDLGGRRYEQY | 0.13894579 |
| LCLHVEN | ASSPDLGGRRYEQY | 0.1320183 |
| LDAISDGS | ASSPDLGGRRYEQY | 0.13691477 |
| LGGFAAML | ASSPDLGGRRYEQY | 0.13980642 |
| LGGKNPMI | ASSPDLGGRRYEQY | 0.1524688 |
| LGPKGVAAIL | ASSPDLGGRRYEQY | 0.15803692 |
| LMVGVSI | ASSPDLGGRRYEQY | 0.15999807 |
| LVGAAFVL | ASSPDLGGRRYEQY | 0.13724558 |
| LVLAVAPL | ASSPDLGGRRYEQY | 0.12732708 |
| MAAIAVIDAA | ASSPDLGGRRYEQY | 0.14957668 |
| NNGINAELQ | ASSPDLGGRRYEQY | 0.22804013 |
| RGEPLVLVHG | ASSPDLGGRRYEQY | 0.14337806 |
| RGGAGGAAGLI | ASSPDLGGRRYEQY | 0.18258356 |
| RSYRMHPAV | ASSPDLGGRRYEQY | 0.16014288 |
| SAAHLLAGQLA | ASSPDLGGRRYEQY | 0.16673818 |
| SAAVGSY | ASSPDLGGRRYEQY | 0.1602693 |
| SPHAVLHRI | ASSPDLGGRRYEQY | 0.15837017 |
| SQIAKVL | ASSPDLGGRRYEQY | 0.14229974 |
| TALGHLVP | ASSPDLGGRRYEQY | 0.13442855 |
| TAPTTSGAA | ASSPDLGGRRYEQY | 0.13390628 |
| TDGDDGNGGSPGT | ASSPDLGGRRYEQY | 0.14968884 |
| TDGLGEN | ASSPDLGGRRYEQY | 0.1926083 |
| TGGPPLPGQA | ASSPDLGGRRYEQY | 0.2004673 |
| TIPAITI | ASSPDLGGRRYEQY | 0.15264265 |
| TIPGFTI | ASSPDLGGRRYEQY | 0.14420722 |
| TSIASGALA | ASSPDLGGRRYEQY | 0.13206862 |
| TVTLPGAAI | ASSPDLGGRRYEQY | 0.16052143 |
| VASGLSAALP | ASSPDLGGRRYEQY | 0.13748223 |
| VGAGVIGI | ASSPDLGGRRYEQY | 0.24850693 |
| VGPKRFGV | ASSPDLGGRRYEQY | 0.13808647 |
| VNVAFPD | ASSPDLGGRRYEQY | 0.1423984 |
| VSMTNSGVS | ASSPDLGGRRYEQY | 0.14115828 |
| VTDVLGDLSV | ASSPDLGGRRYEQY | 0.18985224 |
| VVVVGAGFAGL | ASSPDLGGRRYEQY | 0.1472622 |
| VYTVGTVV | ASSPDLGGRRYEQY | 0.15904039 |
| WTTFSGP | ASSPDLGGRRYEQY | 0.1552563 |

| | | |
|---|---|---|
| YDELVEAF | ASSPDLGGRRYEQY | 0.13471967 |
| YGTGGAGGNGG | ASSPDLGGRRYEQY | 0.1389753 |
| YTNVMAV | ASSPDLGGRRYEQY | 0.13330469 |
| FLEQVHQL | ASSPDLGGRRYEQY | 0.19279872 |
| GLSLLAVAV | ASSPDLGGRRYEQY | 0.15147974 |
| LEQVHQL | ASSPDLGGRRYEQY | 0.13117905 |
| LNAAGAI | ASSPDLGGRRYEQY | 0.14301229 |
| PAVHASL | ASSPDLGGRRYEQY | 0.14414631 |
| TELEDDA | ASSPDLGGRRYEQY | 0.14832003 |
| TPPGPPN | ASSPDLGGRRYEQY | 0.3247662 |
| TPPPPPPP | ASSPDLGGRRYEQY | 0.16543092 |
| TQSPALI | ASSPTVSYNSPLY | 0.14776565 |
| EFAAAFPQ | ASSPTVSYNSPLY | 0.286482 |
| LLSGNLDVL | ASSPTVSYNSPLY | 0.1272118 |
| GPANLGNYNI | ASSPTVSYNSPLY | 0.14623368 |
| TLGPTLP | ASSPTVSYNSPLY | 0.13143109 |
| VYAYSSCCK | ASSPTVSYNSPLY | 0.13264903 |
| APAPAPAPNI | ASSPTVSYNSPLY | 0.14448966 |
| PGGPVPIP | ASSPTVSYNSPLY | 0.14156519 |
| FLTVALEP | ASSPTVSYNSPLY | 0.14470266 |
| SQYFGANL | ASSPTVSYNSPLY | 0.14389977 |
| GGVKPAA | ASSPTVSYNSPLY | 0.13074914 |
| GGTGGAGGAGGW | ASSPTVSYNSPLY | 0.13635795 |
| GVGGVAGLGGAGG | ASSPTVSYNSPLY | 0.16903318 |
| VLLVLATTI | ASSPTVSYNSPLY | 0.14004222 |
| IGIFAAML | ASSPTVSYNSPLY | 0.16614278 |
| AVGDLID | ASSPTVSYNSPLY | 0.13461101 |
| NISPLPL | ASSPTVSYNSPLY | 0.13354573 |
| VTAVFVAI | ASSPTVSYNSPLY | 0.1430322 |
| SGDDNDAAL | ASSPTVSYNSPLY | 0.14024472 |
| AVAALAHS | ASSPTVSYNSPLY | 0.1327142 |
| GGVTVMATP | ASSPTVSYNSPLY | 0.15277566 |
| AGLAANVRD | ASSPTVSYNSPLY | 0.17828849 |
| TNRPDLIDPALLRPG | ASSPTVSYNSPLY | 0.13483703 |
| LANLGVTAL | ASSPTVSYNSPLY | 0.13874792 |
| TGERIEL | ASSPTVSYNSPLY | 0.14591765 |
| NGEELAA | ASSPTVSYNSPLY | 0.13766043 |
| VLSLVWG | ASSPTVSYNSPLY | 0.13227178 |
| IGYALWL | ASSPTVSYNSPLY | 0.17823714 |
| LSLLLAPL | ASSPTVSYNSPLY | 0.19783397 |
| TLDEALK | ASSPTVSYNSPLY | 0.15014026 |
| TDGAGTAEGG | ASSPTVSYNSPLY | 0.13786231 |
| NAPGGAGGNGG | ASSPTVSYNSPLY | 0.12851425 |
| VSVFMGMGGNF | ASSPTVSYNSPLY | 0.35657072 |
| LASYLAAL | ASSPTVSYNSPLY | 0.13794188 |
| LVGGTPAL | ASSPTVSYNSPLY | 0.15428662 |
| VRNADVL | ASSPTVSYNSPLY | 0.13643499 |
| TCSSPFLVSAM | ASSPTVSYNSPLY | 0.1382384 |
| VHITDAE | ASSPTVSYNSPLY | 0.16402894 |
| EYAAPSF | ASSPTVSYNSPLY | 0.13471335 |
| VVDAFHGGL | ASSPTVSYNSPLY | 0.13027297 |

| | | |
|---|---|---|
| ASVVDPQ | ASSPTVSYNSPLY | 0.13815649 |
| SYNEFGGLS | ASSPTVSYNSPLY | 0.16598485 |
| DPDGSVY | ASSPTVSYNSPLY | 0.13374087 |
| MVSTIHATE | ASSPTVSYNSPLY | 0.14661735 |
| LAAGVADPV | ASSPTVSYNSPLY | 0.41289172 |
| TKEQIAA | ASSPTVSYNSPLY | 0.1658646 |
| SLASPAAL | ASSPTVSYNSPLY | 0.14425136 |
| IGLVFYGV | ASSPTVSYNSPLY | 0.14948778 |
| FLLDASNL | ASSPTVSYNSPLY | 0.1448221 |
| YQVGNFP | ASSPTVSYNSPLY | 0.1593888 |
| ESLLSQST | ASSPTVSYNSPLY | 0.13927181 |
| LFTVGFAAVG | ASSPTVSYNSPLY | 0.16473448 |
| ATPPTPT | ASSPTVSYNSPLY | 0.1540046 |
| TDLATLG | ASSPTVSYNSPLY | 0.15849204 |
| SAGGTGR | ASSPTVSYNSPLY | 0.13752213 |
| GFSNSGS | ASSPTVSYNSPLY | 0.14285618 |
| GGNGGSGGTSV | ASSPTVSYNSPLY | 0.16081765 |
| GTSVATGGAGN | ASSPTVSYNSPLY | 0.16238832 |
| NGGNGGSGGTS | ASSPTVSYNSPLY | 0.13649455 |
| AVVKFPH | ASSPTVSYNSPLY | 0.1575065 |
| TPPPPPP | ASSPTVSYNSPLY | 0.17258702 |
| AAPAAAAAPA | ASSPTVSYNSPLY | 0.14926253 |
| IALNEDL | ASSPTVSYNSPLY | 0.14028436 |
| IVNVSSM | ASSPTVSYNSPLY | 0.13061255 |
| TGIPPAPR | ASSPTVSYNSPLY | 0.2551382 |
| FDLGGGTF | ASSPTVSYNSPLY | 0.1370377 |
| FDLGGGTFDVS | ASSPTVSYNSPLY | 0.13649575 |
| NEPTAAA | ASSPTVSYNSPLY | 0.15115437 |
| NEPTAAALA | ASSPTVSYNSPLY | 0.14400314 |
| GSIINISSI | ASSPTVSYNSPLY | 0.15730359 |
| VAPGNAGTA | ASSPTVSYNSPLY | 0.13596007 |
| TGGGASLE | ASSPTVSYNSPLY | 0.13578403 |
| VAVYNHYKRI | ASSPTVSYNSPLY | 0.17954373 |
| TDPAPATT | ASSPTVSYNSPLY | 0.14440566 |
| TNRPDILDPALLRPG | ASSPTVSYNSPLY | 0.13979104 |
| LTTTTTS | ASSPTVSYNSPLY | 0.1636756 |
| PTGSGKTL | ASSPTVSYNSPLY | 0.13376848 |
| TGSGKTL | ASSPTVSYNSPLY | 0.1659689 |
| TIGHVDHGK | ASSPTVSYNSPLY | 0.15030196 |
| TIGHVDHGKTT | ASSPTVSYNSPLY | 0.14929162 |
| GHVDHGKTT | ASSPTVSYNSPLY | 0.14359884 |
| AQFEHTLL | ASSPTVSYNSPLY | 0.12849873 |
| IVNQDEL | ASSPTVSYNSPLY | 0.13617544 |
| TNGSQFF | ASSPTVSYNSPLY | 0.13935228 |
| TGKGIVE | ASSPTVSYNSPLY | 0.26817527 |
| LISNASDA | ASSPTVSYNSPLY | 0.16638993 |
| AELRTLGELTKIAKAHGAQ | ASSPTVSYNSPLY | 0.14869574 |
| APNAVAAAEPSW | ASSPTVSYNSPLY | 0.14916642 |
| APQVAGAATAVL | ASSPTVSYNSPLY | 0.14572744 |
| ARRDPAAAAAILPTDA | ASSPTVSYNSPLY | 0.15361091 |
| DAAARDVLAEAGLAEYF | ASSPTVSYNSPLY | 0.13999544 |

| | | |
|---|---|---|
| DDQIVLSAKANLAKVEQ | ASSPTVSYNSPLY | 0.13603112 |
| DELLPDTGEDSDEDGH | ASSPTVSYNSPLY | 0.13640684 |
| ELPEAPANRATVPQD | ASSPTVSYNSPLY | 0.14296594 |
| FGQSKIERVTGIG | ASSPTVSYNSPLY | 0.13780458 |
| GAGGTGSTAGGAGGAGGAGGLY | ASSPTVSYNSPLY | 0.14905122 |
| GHGGTGAAGGNGGTGGAGGSGI | ASSPTVSYNSPLY | 0.13263206 |
| GNGGNGGAGGEGGPGGAGGAGA | ASSPTVSYNSPLY | 0.13983536 |
| GSTAGNGGNGGNATNGTIAGQP | ASSPTVSYNSPLY | 0.14102076 |
| GTGGDGGNGGNGGTGG | ASSPTVSYNSPLY | 0.1350044 |
| GVRRPAGIPADAPV | ASSPTVSYNSPLY | 0.14695974 |
| GYRQVTDAHLLTLARR | ASSPTVSYNSPLY | 0.18109679 |
| IAKMGQDGHDRG | ASSPTVSYNSPLY | 0.13690501 |
| KAEPSIEIQYT | ASSPTVSYNSPLY | 0.14471197 |
| KPVRIRMAADEDTVAVT | ASSPTVSYNSPLY | 0.24310347 |
| MPPRIAGMRLLVIKPEP | ASSPTVSYNSPLY | 0.1425692 |
| MKFYAVALFRSRGVPPTR | ASSPTVSYNSPLY | 0.1329464 |
| NAGGFGILGAVAHSP | ASSPTVSYNSPLY | 0.13668577 |
| NEGREITLAMAEM | ASSPTVSYNSPLY | 0.16061316 |
| NELVEWSHPLGHRWRWE | ASSPTVSYNSPLY | 0.15963788 |
| NLGAGGTGGNGGNPG | ASSPTVSYNSPLY | 0.12879108 |
| NTEWPKWETVMAADPVD | ASSPTVSYNSPLY | 0.16595894 |
| PASAAPAETQP | ASSPTVSYNSPLY | 0.14943287 |
| QQSSDGKNGVRAAEITGE | ASSPTVSYNSPLY | 0.14012288 |
| RAPDAVTTSWWKE | ASSPTVSYNSPLY | 0.12781711 |
| SGFGNVGAHTSGWFNQ | ASSPTVSYNSPLY | 0.16314623 |
| SGVAGAVGQAASVGGLKV | ASSPTVSYNSPLY | 0.17408143 |
| SNGGAGTGSAGGNGGT | ASSPTVSYNSPLY | 0.15474923 |
| TGGAGGEGGAAG | ASSPTVSYNSPLY | 0.14112572 |
| TGGDGGAAGAGGAPGHG | ASSPTVSYNSPLY | 0.12945776 |
| TMPPLLAAIANAGRHGV | ASSPTVSYNSPLY | 0.15668209 |
| TSSTAGNSGGA | ASSPTVSYNSPLY | 0.16498026 |
| VPATVPVGILC | ASSPTVSYNSPLY | 0.14537527 |
| VQRPDSGGVVLTGRIS | ASSPTVSYNSPLY | 0.16156234 |
| WTPNPVPATKGVPAGLVQAT | ASSPTVSYNSPLY | 0.13640936 |
| YALKRALAKFSDRFDVVII | ASSPTVSYNSPLY | 0.14074416 |
| YLIGSVIV | ASSPTVSYNSPLY | 0.16466132 |
| YGQLVAVID | ASSPTVSYNSPLY | 0.17082353 |
| VTVGGMTLPA | ASSPTVSYNSPLY | 0.13911018 |
| VLALPAV | ASSPTVSYNSPLY | 0.14694597 |
| VIPPTLNYET | ASSPTVSYNSPLY | 0.15134065 |
| VGMGGMHPSSG | ASSPTVSYNSPLY | 0.14026213 |
| VGLAGGAATA | ASSPTVSYNSPLY | 0.14741516 |
| VGGAGGAGGSGP | ASSPTVSYNSPLY | 0.22519235 |
| VFAAFMV | ASSPTVSYNSPLY | 0.14348994 |
| TLYDKLR | ASSPTVSYNSPLY | 0.147988 |
| TLAVFTVL | ASSPTVSYNSPLY | 0.16253747 |
| TGSVQPARV | ASSPTVSYNSPLY | 0.13970593 |
| TGGGAGSGGT | ASSPTVSYNSPLY | 0.16819881 |
| SHLETSL | ASSPTVSYNSPLY | 0.13622056 |
| SFTTGMT | ASSPTVSYNSPLY | 0.20749453 |
| QVEVAKRMGVRQPTVS | ASSPTVSYNSPLY | 0.15898725 |

| | | |
|---|---|---|
| QGSGGPALAM | ASSPTVSYNSPLY | 0.14735737 |
| MLAPSNP | ASSPTVSYNSPLY | 0.14414543 |
| LVNNAAI | ASSPTVSYNSPLY | 0.11284207 |
| LTGEINTLA | ASSPTVSYNSPLY | 0.11374827 |
| LSIFGPV | ASSPTVSYNSPLY | 0.11285846 |
| LLALLANL | ASSPTVSYNSPLY | 0.11291485 |
| LISNASDALDKLR | ASSPTVSYNSPLY | 0.113608696 |
| LINEAVP | ASSPTVSYNSPLY | 0.11292608 |
| LGYAFSGL | ASSPTVSYNSPLY | 0.112989105 |
| LDFVAVAL | ASSPTVSYNSPLY | 0.11370685 |
| LAQPGAE | ASSPTVSYNSPLY | 0.11286658 |
| ISNLGAVTGF | ASSPTVSYNSPLY | 0.11290784 |
| HQEDPGRI | ASSPTVSYNSPLY | 0.113634884 |
| GTPGEPTP | ASSPTVSYNSPLY | 0.11350616 |
| GFFNSGS | ASSPTVSYNSPLY | 0.1131773 |
| GAGGGGLTAGAN | ASSPTVSYNSPLY | 0.11282308 |
| FVDALVNM | ASSPTVSYNSPLY | 0.11274552 |
| FLTVAPEL | ASSPTVSYNSPLY | 0.11314075 |
| FETLTAL | ASSPTVSYNSPLY | 0.112875104 |
| DVINPIEL | ASSPTVSYNSPLY | 0.11271774 |
| APVSGAPV | ASSPTVSYNSPLY | 0.112817355 |
| ALVGGLAW | ASSPTVSYNSPLY | 0.11257408 |
| ALAAAAATAI | ASSPTVSYNSPLY | 0.11271841 |
| AFGLGLLL | ASSPTVSYNSPLY | 0.112905666 |
| AFFAAASL | ASSPTVSYNSPLY | 0.113119945 |
| AAAALPAFSPP | ASSPTVSYNSPLY | 0.11288064 |
| VVLAPAL | ASSPTVSYNSPLY | 0.112945385 |
| TIGHVDHGKTTL | ASSPTVSYNSPLY | 0.11269438 |
| LISNASDALDK | ASSPTVSYNSPLY | 0.11338156 |
| LISNASDAL | ASSPTVSYNSPLY | 0.11328281 |
| ISNASDALDK | ASSPTVSYNSPLY | 0.113358185 |
| NQAETLVYQTE | ASSPTVSYNSPLY | 0.11333269 |
| TGGTGGTAGAG | ASSPTVSYNSPLY | 0.112743825 |
| TGSTAGGAGGA | ASSPTVSYNSPLY | 0.113762416 |
| DPLAVPVAAGPV | ASSPTVSYNSPLY | 0.113701105 |
| GGTGGTPGNGGN | ASSPTVSYNSPLY | 0.11280007 |
| PAGLAGVDPATL | ASSPTVSYNSPLY | 0.1127683 |
| GNNLVVEAPHAGASV | ASSPTVSYNSPLY | 0.11264734 |
| KVTRIMSASTLEQPA | ASSPTVSYNSPLY | 0.113230646 |
| LGFALPIALPRRTVV | ASSPTVSYNSPLY | 0.113270715 |
| TRLGNLTARKLVVLV | ASSPTVSYNSPLY | 0.1136528 |
| WSQQRRLIYFPSAGP | ASSPTVSYNSPLY | 0.11347569 |
| GPNGNPDPMAAAVDIR | ASSPTVSYNSPLY | 0.1128435 |
| TQRCVSVADLRAAASR | ASSPTVSYNSPLY | 0.11273473 |
| QFVLDVAEHPVEVLTETV | ASSPTVSYNSPLY | 0.11275998 |
| GGAGGTGGAGGNGGTQAGDG | ASSPTVSYNSPLY | 0.11292273 |
| PLLATLRAIAVDVDAKPAQV | ASSPTVSYNSPLY | 0.11323566 |
| APKFRTGGYMIDFWGVGYQVAK | ASSPTVSYNSPLY | 0.11306076 |
| GAGGAGGAGGAAGGPGGTGGTG | ASSPTVSYNSPLY | 0.112808846 |
| IGALGLLATDVAPAGSVAFGVA | ASSPTVSYNSPLY | 0.11292347 |
| AGGAGGIGGNGGAGGAGGVLMG | ASSPTVSYNSPLY | 0.11251445 |

| | | |
|---|---|---|
| ADELIGLL | ASSPTVSYNSPLY | 0.11281 |
| ADRALKA | ASSPTVSYNSPLY | 0.11271171 |
| AGAVGQVGGAGGA | ASSPTVSYNSPLY | 0.11281228 |
| AGGGGGLGGAP | ASSPTVSYNSPLY | 0.114177145 |
| AGGLLYGNGGAG | ASSPTVSYNSPLY | 0.11399061 |
| AGLESIGL | ASSPTVSYNSPLY | 0.11308366 |
| AGLKTGII | ASSPTVSYNSPLY | 0.113159366 |
| ASLAALLL | ASSPTVSYNSPLY | 0.11261312 |
| ASVINISSI | ASSPTVSYNSPLY | 0.113368504 |
| ATPPGPA | ASSPTVSYNSPLY | 0.113170594 |
| ATVASALV | ASSPTVSYNSPLY | 0.11286735 |
| ATVASLIL | ASSPTVSYNSPLY | 0.11306062 |
| AVNGPVLIRH | ASSPTVSYNSPLY | 0.11316101 |
| DGAAGAVGI | ASSPTVSYNSPLY | 0.11312155 |
| DPLLHGI | ASSPTVSYNSPLY | 0.112938076 |
| ELGAALGAL | ASSPTVSYNSPLY | 0.11359879 |
| ELGRTQAV | ASSPTVSYNSPLY | 0.11330404 |
| FVAGGYLS | ASSPTVSYNSPLY | 0.112479456 |
| GASGGAGGAGGW | ASSPTVSYNSPLY | 0.11266331 |
| GGAGAGGDGGAGGI | ASSPTVSYNSPLY | 0.11288498 |
| GTGHNGSQPGMGGNG | ASSPTVSYNSPLY | 0.112938955 |
| GYAAASASAS | ASSPTVSYNSPLY | 0.11272768 |
| HQVDVAAVQQG | ASSPTVSYNSPLY | 0.113080986 |
| IGAAIAHT | ASSPTVSYNSPLY | 0.11335882 |
| IGMIEVAQL | ASSPTVSYNSPLY | 0.11309817 |
| IRLAPEHTSAMD | ASSPTVSYNSPLY | 0.112695396 |
| IYNVSVL | ASSPTVSYNSPLY | 0.1129437 |
| KQGFLGM | ASSPTVSYNSPLY | 0.11351065 |
| LESVITQL | ASSPTVSYNSPLY | 0.11317378 |
| LGLGALVVV | ASSPTVSYNSPLY | 0.112759635 |
| LIGNGGNGGSGGI | ASSPTVSYNSPLY | 0.11319484 |
| LNAINAPTL | ASSPTVSYNSPLY | 0.1128369 |
| LQLLNTGV | ASSPTVSYNSPLY | 0.113510706 |
| LQTSIEF | ASSPTVSYNSPLY | 0.112829186 |
| MSFAVSEL | ASSPTVSYNSPLY | 0.112758115 |
| NTGALGL | ASSPTVSYNSPLY | 0.11364525 |
| PPGAGGAGGAGTT | ASSPTVSYNSPLY | 0.112857655 |
| QLVGLGI | ASSPTVSYNSPLY | 0.11308426 |
| RFLEQVHQL | ASSPTVSYNSPLY | 0.11311016 |
| RGVGYVLR | ASSPTVSYNSPLY | 0.11272246 |
| SATGVFAGVI | ASSPTVSYNSPLY | 0.11282605 |
| SLPAITLP | ASSPTVSYNSPLY | 0.11268977 |
| STALAFTV | ASSPTVSYNSPLY | 0.11282693 |
| SVLTVGLGK | ASSPTVSYNSPLY | 0.1128252 |
| TALAGAGSGI | ASSPTVSYNSPLY | 0.11261912 |
| TGLPFTGI | ASSPTVSYNSPLY | 0.11327976 |
| VAEHVTV | ASSPTVSYNSPLY | 0.11308652 |
| VDIVLNSL | ASSPTVSYNSPLY | 0.113019764 |
| VLTGGGAVL | ASSPTVSYNSPLY | 0.11330883 |
| VNVGLAYGI | ASSPTVSYNSPLY | 0.11268253 |
| VSSAAVGAV | ASSPTVSYNSPLY | 0.11318862 |

| | | |
|---|---|---|
| VTSVYEAGV | ASSPTVSYNSPLY | 0.11273631 |
| WVALDRGA | ASSPTVSYNSPLY | 0.11337927 |
| YLVGSVVI | ASSPTVSYNSPLY | 0.11268101 |
| GPPGTGKTLL | ASSPTVSYNSPLY | 0.11293869 |
| GVGMTKF | ASSPTVSYNSPLY | 0.11294517 |
| IGIGGSDL | ASSPTVSYNSPLY | 0.11268047 |
| IGTLLAEL | ASSPTVSYNSPLY | 0.11280126 |
| IYTSGTT | ASSPTVSYNSPLY | 0.11280683 |
| LSFPAGV | ASSPTVSYNSPLY | 0.113982975 |
| LYTSGTT | ASSPTVSYNSPLY | 0.11271463 |
| VGAFPFGF | ASSPTVSYNSPLY | 0.11290126 |
| VGAGTVEF | ASSPTVSYNSPLY | 0.1129706 |
| VGKSTLLL | ASSPTVSYNSPLY | 0.11324456 |
| AALAHAGGGIV | ASSPTVSYNSPLY | 0.11336693 |
| AALVAVSLL | ASSPTVSYNSPLY | 0.11277635 |
| AAVSRAVAAL | ASSPTVSYNSPLY | 0.11270246 |
| AGGSSGIGGF | ASSPTVSYNSPLY | 0.1136641 |
| AGLTVANANG | ASSPTVSYNSPLY | 0.11263298 |
| AGLVGAVL | ASSPTVSYNSPLY | 0.11272587 |
| ALSQAFTV | ASSPTVSYNSPLY | 0.113226704 |
| ALTFSVSL | ASSPTVSYNSPLY | 0.11260387 |
| ALTGAALVAL | ASSPTVSYNSPLY | 0.11363241 |
| ASFSIGAGVL | ASSPTVSYNSPLY | 0.1128128 |
| DGSVYKI | ASSPTVSYNSPLY | 0.112552345 |
| EQLAEIV | ASSPTVSYNSPLY | 0.112536356 |
| FAVSELTVM | ASSPTVSYNSPLY | 0.11277131 |
| GAAPLAAGL | ASSPTVSYNSPLY | 0.11323458 |
| GADGGDGGSGGAGG | ASSPTVSYNSPLY | 0.11364484 |
| GFGLDTGGAGG | ASSPTVSYNSPLY | 0.11321234 |
| GLASGGLL | ASSPTVSYNSPLY | 0.11313331 |
| GRIAGAAVLL | ASSPTVSYNSPLY | 0.114104025 |
| GSAGSLNTSF | ASSPTVSYNSPLY | 0.11268893 |
| GTGQIGI | ASSPTVSYNSPLY | 0.11314948 |
| GTLIALIL | ASSPTVSYNSPLY | 0.11282828 |
| IAATMGFVVG | ASSPTVSYNSPLY | 0.112792626 |
| IFAQALL | ASSPTVSYNSPLY | 0.112833634 |
| IGAAVALGF | ASSPTVSYNSPLY | 0.11300497 |
| IGLTPLAL | ASSPTVSYNSPLY | 0.11344706 |
| ILGSNTSTL | ASSPTVSYNSPLY | 0.1128267 |
| IVFSLTVL | ASSPTVSYNSPLY | 0.11286455 |
| LASKATGF | ASSPTVSYNSPLY | 0.112608485 |
| LASVALIL | ASSPTVSYNSPLY | 0.11260282 |
| LFAVGLTL | ASSPTVSYNSPLY | 0.11404666 |
| LFNGGIGTYI | ASSPTVSYNSPLY | 0.11300695 |
| LTLGAGII | ASSPTVSYNSPLY | 0.11288239 |
| LVAAALGDI | ASSPTVSYNSPLY | 0.11261316 |
| NTAFVQLGI | ASSPTVSYNSPLY | 0.112915225 |
| QLFSGAEL | ASSPTVSYNSPLY | 0.11298946 |
| SAATFITL | ASSPTVSYNSPLY | 0.11284353 |
| SNYGLANLGV | ASSPTVSYNSPLY | 0.112848654 |
| SYGGAYAVM | ASSPTVSYNSPLY | 0.11315914 |

| | | |
|---|---|---|
| TTGVGGLTLG | ASSPTVSYNSPLY | 0.113721274 |
| TTQTIYI | ASSPTVSYNSPLY | 0.113397546 |
| VAAGAGLGAVI | ASSPTVSYNSPLY | 0.11309617 |
| VAAVVTLL | ASSPTVSYNSPLY | 0.11357325 |
| VGFLGGLI | ASSPTVSYNSPLY | 0.11294682 |
| VGVIPILL | ASSPTVSYNSPLY | 0.11307655 |
| VIFASSLI | ASSPTVSYNSPLY | 0.11320021 |
| VKGGDGGAAAG | ASSPTVSYNSPLY | 0.11309631 |
| VTKTFTNI | ASSPTVSYNSPLY | 0.11267045 |
| VVAAAILL | ASSPTVSYNSPLY | 0.112929374 |
| YAALDVEL | ASSPTVSYNSPLY | 0.112818874 |
| LASVALLL | ASSPTVSYNSPLY | 0.113140926 |
| LDAVHRV | ASSPTVSYNSPLY | 0.11363525 |
| QGLSSSIF | ASSPTVSYNSPLY | 0.11276225 |
| RGTFANIRL | ASSPTVSYNSPLY | 0.11317814 |
| RQLWWGHRI | ASSPTVSYNSPLY | 0.11288904 |
| YGPPGTGKTLL | ASSPTVSYNSPLY | 0.11307476 |
| AALVGAPANVA | ASSPTVSYNSPLY | 0.11288901 |
| AATFRRTE | ASSPTVSYNSPLY | 0.11359031 |
| AGALGGGI | ASSPTVSYNSPLY | 0.11273162 |
| AGGAGGAGGIGAG | ASSPTVSYNSPLY | 0.113073364 |
| AGSGALGL | ASSPTVSYNSPLY | 0.113035955 |
| AQIADQI | ASSPTVSYNSPLY | 0.11295654 |
| ASAVAVTYL | ASSPTVSYNSPLY | 0.11321412 |
| ASTVLGGTGG | ASSPTVSYNSPLY | 0.11290702 |
| AVGGEPAK | ASSPTVSYNSPLY | 0.113104574 |
| CEQGTFGGTC | ASSPTVSYNSPLY | 0.11314514 |
| DEGKAPAG | ASSPTVSYNSPLY | 0.11285219 |
| DTVVFPAL | ASSPTVSYNSPLY | 0.11299738 |
| FEGITMAG | ASSPTVSYNSPLY | 0.114674255 |
| FPLVVQD | ASSPTVSYNSPLY | 0.112965316 |
| GAFTTPQ | ASSPTVSYNSPLY | 0.11295424 |
| GFGGFGTAGGVGGA | ASSPTVSYNSPLY | 0.11332945 |
| GFLAQFL | ASSPTVSYNSPLY | 0.112882726 |
| GGAGGAGGNAGVA | ASSPTVSYNSPLY | 0.11297806 |
| GGVGTAGAGGAGGAGG | ASSPTVSYNSPLY | 0.1125771 |
| GNGGGGDAGNAGSGGN | ASSPTVSYNSPLY | 0.11282989 |
| GTGGAGGAGNDAGS | ASSPTVSYNSPLY | 0.11287191 |
| GTTNTGWFNS | ASSPTVSYNSPLY | 0.112945735 |
| IFMGPLFTL | ASSPTVSYNSPLY | 0.11343631 |
| ILAHAIEAI | ASSPTVSYNSPLY | 0.113260016 |
| IRSAIREAL | ASSPTVSYNSPLY | 0.112886354 |
| ITFAGGG | ASSPTVSYNSPLY | 0.11351982 |
| ITGMSGHQEV | ASSPTVSYNSPLY | 0.113017604 |
| IVAGLACTL | ASSPTVSYNSPLY | 0.11271826 |
| IVGGGLAGA | ASSPTVSYNSPLY | 0.11280015 |
| IVTGVLNI | ASSPTVSYNSPLY | 0.11328202 |
| LAQQGCI | ASSPTVSYNSPLY | 0.11275157 |
| LCLHVEN | ASSPTVSYNSPLY | 0.11269707 |
| LDAISDGS | ASSPTVSYNSPLY | 0.11239245 |
| LGGFAAML | ASSPTVSYNSPLY | 0.11282939 |

| | | |
|---|---|---|
| LGGKNPMI | ASSPTVSYNSPLY | 0.11314533 |
| LGPKGVAAIL | ASSPTVSYNSPLY | 0.1126434 |
| LMVGVSI | ASSPTVSYNSPLY | 0.113460936 |
| LVGAAFVL | ASSPTVSYNSPLY | 0.112727724 |
| LVLAVAPL | ASSPTVSYNSPLY | 0.112648785 |
| MAAIAVIDAA | ASSPTVSYNSPLY | 0.11384861 |
| NNGINAELQ | ASSPTVSYNSPLY | 0.11317188 |
| RGEPLVLVHG | ASSPTVSYNSPLY | 0.11282854 |
| RGGAGGAAGLI | ASSPTVSYNSPLY | 0.1129438 |
| RSYRMHPAV | ASSPTVSYNSPLY | 0.11292488 |
| SAAHLLAGQLA | ASSPTVSYNSPLY | 0.113133036 |
| SAAVGSY | ASSPTVSYNSPLY | 0.11271449 |
| SPHAVLHRI | ASSPTVSYNSPLY | 0.11282488 |
| SQIAKVL | ASSPTVSYNSPLY | 0.11311002 |
| TALGHLVP | ASSPTVSYNSPLY | 0.11269201 |
| TAPTTSGAA | ASSPTVSYNSPLY | 0.11355894 |
| TDGDDGNGGSPGT | ASSPTVSYNSPLY | 0.11343224 |
| TDGLGEN | ASSPTVSYNSPLY | 0.112762675 |
| TGGPPLPGQA | ASSPTVSYNSPLY | 0.11285771 |
| TIPAITI | ASSPTVSYNSPLY | 0.11282472 |
| TIPGFTI | ASSPTVSYNSPLY | 0.1130455 |
| TSIASGALA | ASSPTVSYNSPLY | 0.11276043 |
| TVTLPGAAI | ASSPTVSYNSPLY | 0.11264096 |
| VASGLSAALP | ASSPTVSYNSPLY | 0.11305376 |
| VGAGVIGI | ASSPTVSYNSPLY | 0.113704555 |
| VGPKRFGV | ASSPTVSYNSPLY | 0.11283137 |
| VNVAFPD | ASSPTVSYNSPLY | 0.1135105 |
| VSMTNSGVS | ASSPTVSYNSPLY | 0.11309412 |
| VTDVLGDLSV | ASSPTVSYNSPLY | 0.11359168 |
| VVVVGAGFAGL | ASSPTVSYNSPLY | 0.11290987 |
| VYTVGTVV | ASSPTVSYNSPLY | 0.11413633 |
| WTTFSGP | ASSPTVSYNSPLY | 0.11279872 |
| YDELVEAF | ASSPTVSYNSPLY | 0.11326677 |
| YGTGGAGGNGG | ASSPTVSYNSPLY | 0.11329971 |
| YTNVMAV | ASSPTVSYNSPLY | 0.11292477 |
| FLEQVHQL | ASSPTVSYNSPLY | 0.11313627 |
| GLSLLAVAV | ASSPTVSYNSPLY | 0.11297725 |
| LEQVHQL | ASSPTVSYNSPLY | 0.11311449 |
| LNAAGAI | ASSPTVSYNSPLY | 0.11294802 |
| PAVHASL | ASSPTVSYNSPLY | 0.11267627 |
| TELEDDA | ASSPTVSYNSPLY | 0.11300739 |
| TPPGPPN | ASSPTVSYNSPLY | 0.112728946 |
| TPPPPPPP | ASSPTVSYNSPLY | 0.11367155 |
| TQSPALI | ASSLRLGPSQNTLY | 0.11278579 |
| EFAAAFPQ | ASSLRLGPSQNTLY | 0.11274771 |
| LLSGNLDVL | ASSLRLGPSQNTLY | 0.11301155 |
| GPANLGNYNI | ASSLRLGPSQNTLY | 0.11278183 |
| TLGPTLP | ASSLRLGPSQNTLY | 0.11404132 |
| VYAYSSCCK | ASSLRLGPSQNTLY | 0.113003366 |
| APAPAPAPNI | ASSLRLGPSQNTLY | 0.11280398 |
| PGGPVPIP | ASSLRLGPSQNTLY | 0.112821564 |

| | | |
|---|---|---|
| FLTVALEP | ASSLRLGPSQNTLY | 0.11289034 |
| SQYFGANL | ASSLRLGPSQNTLY | 0.11292017 |
| GGVKPAA | ASSLRLGPSQNTLY | 0.113149844 |
| GGTGGAGGAGGW | ASSLRLGPSQNTLY | 0.11272755 |
| GVGGVAGLGGAGG | ASSLRLGPSQNTLY | 0.113142125 |
| VLLVLATTI | ASSLRLGPSQNTLY | 0.11278332 |
| IGIFAAML | ASSLRLGPSQNTLY | 0.11304555 |
| AVGDLID | ASSLRLGPSQNTLY | 0.11326218 |
| NISPLPL | ASSLRLGPSQNTLY | 0.1132258 |
| VTAVFVAI | ASSLRLGPSQNTLY | 0.11266463 |
| SGDDNDAAL | ASSLRLGPSQNTLY | 0.114466466 |
| AVAALAHS | ASSLRLGPSQNTLY | 0.11317757 |
| GGVTVMATP | ASSLRLGPSQNTLY | 0.11282493 |
| AGLAANVRD | ASSLRLGPSQNTLY | 0.11302304 |
| TNRPDLIDPALLRPG | ASSLRLGPSQNTLY | 0.11306507 |
| LANLGVTAL | ASSLRLGPSQNTLY | 0.113647886 |
| TGERIEL | ASSLRLGPSQNTLY | 0.112886675 |
| NGEELAA | ASSLRLGPSQNTLY | 0.11276381 |
| VLSLVWG | ASSLRLGPSQNTLY | 0.112861164 |
| IGYALWL | ASSLRLGPSQNTLY | 0.113044456 |
| LSLLLAPL | ASSLRLGPSQNTLY | 0.112896636 |
| TLDEALK | ASSLRLGPSQNTLY | 0.113704614 |
| TDGAGTAEGG | ASSLRLGPSQNTLY | 0.11278897 |
| NAPGGAGGNGG | ASSLRLGPSQNTLY | 0.114559196 |
| VSVFMGMGGNF | ASSLRLGPSQNTLY | 0.11320141 |
| LASYLAAL | ASSLRLGPSQNTLY | 0.11330765 |
| LVGGTPAL | ASSLRLGPSQNTLY | 0.1126561 |
| VRNADVL | ASSLRLGPSQNTLY | 0.11345986 |
| TCSSPFLVSAM | ASSLRLGPSQNTLY | 0.113127984 |
| VHITDAE | ASSLRLGPSQNTLY | 0.11294253 |
| EYAAPSF | ASSLRLGPSQNTLY | 0.113010615 |
| VVDAFHGGL | ASSLRLGPSQNTLY | 0.1132071 |
| ASVVDPQ | ASSLRLGPSQNTLY | 0.11349127 |
| SYNEFGGLS | ASSLRLGPSQNTLY | 0.11252171 |
| DPDGSVY | ASSLRLGPSQNTLY | 0.112682454 |
| MVSTIHATE | ASSLRLGPSQNTLY | 0.11309245 |
| LAAGVADPV | ASSLRLGPSQNTLY | 0.1128835 |
| TKEQIAA | ASSLRLGPSQNTLY | 0.112993106 |
| SLASPAAL | ASSLRLGPSQNTLY | 0.11307804 |
| IGLVFYGV | ASSLRLGPSQNTLY | 0.1126857 |
| FLLDASNL | ASSLRLGPSQNTLY | 0.11322837 |
| YQVGNFP | ASSLRLGPSQNTLY | 0.11284967 |
| ESLLSQST | ASSLRLGPSQNTLY | 0.112819314 |
| LFTVGFAAVG | ASSLRLGPSQNTLY | 0.11391174 |
| ATPPTPT | ASSLRLGPSQNTLY | 0.11304049 |
| TDLATLG | ASSLRLGPSQNTLY | 0.11273564 |
| SAGGTGR | ASSLRLGPSQNTLY | 0.11277173 |
| GFSNSGS | ASSLRLGPSQNTLY | 0.1132415 |
| GGNGGSGGTSV | ASSLRLGPSQNTLY | 0.11279925 |
| GTSVATGGAGN | ASSLRLGPSQNTLY | 0.11267537 |
| NGGNGGSGGTS | ASSLRLGPSQNTLY | 0.113027126 |

| | | |
|---|---|---|
| AVVKFPH | ASSLRLGPSQNTLY | 0.11285126 |
| TPPPPPP | ASSLRLGPSQNTLY | 0.11298204 |
| AAPAAAAAPA | ASSLRLGPSQNTLY | 0.11412382 |
| IALNEDL | ASSLRLGPSQNTLY | 0.11278495 |
| IVNVSSM | ASSLRLGPSQNTLY | 0.11309289 |
| TGIPPAPR | ASSLRLGPSQNTLY | 0.11284204 |
| FDLGGGTF | ASSLRLGPSQNTLY | 0.11287786 |
| FDLGGGTFDVS | ASSLRLGPSQNTLY | 0.11307787 |
| NEPTAAA | ASSLRLGPSQNTLY | 0.112866454 |
| NEPTAAALA | ASSLRLGPSQNTLY | 0.11284329 |
| GSIINISSI | ASSLRLGPSQNTLY | 0.113212585 |
| VAPGNAGTA | ASSLRLGPSQNTLY | 0.11352796 |
| TGGGASLE | ASSLRLGPSQNTLY | 0.11305806 |
| VAVYNHYKRI | ASSLRLGPSQNTLY | 0.11323564 |
| TDPAPATT | ASSLRLGPSQNTLY | 0.11299216 |
| TNRPDILDPALLRPG | ASSLRLGPSQNTLY | 0.113036804 |
| LTTTTTS | ASSLRLGPSQNTLY | 0.113233514 |
| PTGSGKTL | ASSLRLGPSQNTLY | 0.11253728 |
| TGSGKTL | ASSLRLGPSQNTLY | 0.11280548 |
| TIGHVDHGK | ASSLRLGPSQNTLY | 0.11290958 |
| TIGHVDHGKTT | ASSLRLGPSQNTLY | 0.1131126 |
| GHVDHGKTT | ASSLRLGPSQNTLY | 0.112965494 |
| AQFEHTLL | ASSLRLGPSQNTLY | 0.112981334 |
| IVNQDEL | ASSLRLGPSQNTLY | 0.11292337 |
| TNGSQFF | ASSLRLGPSQNTLY | 0.11275249 |
| TGKGIVE | ASSLRLGPSQNTLY | 0.11282167 |
| LISNASDA | ASSLRLGPSQNTLY | 0.11462273 |
| AELRTLGELTKIAKAHGAQ | ASSLRLGPSQNTLY | 0.11353855 |
| APNAVAAAEPSW | ASSLRLGPSQNTLY | 0.11343328 |
| APQVAGAATAVL | ASSLRLGPSQNTLY | 0.11285169 |
| ARRDPAAAAAILPTDA | ASSLRLGPSQNTLY | 0.1128545 |
| DAAARDVLAEAGLAEYF | ASSLRLGPSQNTLY | 0.11317681 |
| DDQIVLSAKANLAKVEQ | ASSLRLGPSQNTLY | 0.11272236 |
| DELLPDTGEDSDEDGH | ASSLRLGPSQNTLY | 0.11254385 |
| ELPEAPANRATVPQD | ASSLRLGPSQNTLY | 0.11314722 |
| FGQSKIERVTGIG | ASSLRLGPSQNTLY | 0.1126209 |
| GAGGTGSTAGGAGGAGGAGGLY | ASSLRLGPSQNTLY | 0.11386063 |
| GHGGTGAAGGNGGTGGAGGSGI | ASSLRLGPSQNTLY | 0.112794705 |
| GNGGNGGAGGEGGPGGAGGAGA | ASSLRLGPSQNTLY | 0.11269373 |
| GSTAGNGGNGGNATNGTIAGQP | ASSLRLGPSQNTLY | 0.112784386 |
| GTGGDGGNGGNGGTGG | ASSLRLGPSQNTLY | 0.11279771 |
| GVRRPAGIPADAPV | ASSLRLGPSQNTLY | 0.11350292 |
| GYRQVTDAHLLTLARR | ASSLRLGPSQNTLY | 0.11295141 |
| IAKMGQDGHDRG | ASSLRLGPSQNTLY | 0.11279653 |
| KAEPSIEIQYT | ASSLRLGPSQNTLY | 0.11291773 |
| KPVRIRMAADEDTVAVT | ASSLRLGPSQNTLY | 0.1145491 |
| MPPRIAGMRLLVIKPEP | ASSLRLGPSQNTLY | 0.11340613 |
| MKFYAVALFRSRGVPPTR | ASSLRLGPSQNTLY | 0.11284109 |
| NAGGFGILGAVAHSP | ASSLRLGPSQNTLY | 0.11291782 |
| NEGREITLAMAEM | ASSLRLGPSQNTLY | 0.11274697 |
| NELVEWSHPLGHRWRWE | ASSLRLGPSQNTLY | 0.112727545 |

| | | |
|---|---|---|
| NLGAGGTGGNGGNPG | ASSLRLGPSQNTLY | 0.11276614 |
| NTEWPKWETVMAADPVD | ASSLRLGPSQNTLY | 0.11274689 |
| PASAAPAETQP | ASSLRLGPSQNTLY | 0.112712465 |
| QQSSDGKNGVRAAEITGE | ASSLRLGPSQNTLY | 0.113023266 |
| RAPDAVTTSWWKE | ASSLRLGPSQNTLY | 0.112535045 |
| SGFGNVGAHTSGWFNQ | ASSLRLGPSQNTLY | 0.113033056 |
| SGVAGAVGQAASVGGLKV | ASSLRLGPSQNTLY | 0.11254412 |
| SNGGAGTGSAGGNGGT | ASSLRLGPSQNTLY | 0.11284813 |
| TGGAGGEGGAAG | ASSLRLGPSQNTLY | 0.11348586 |
| TGGDGGAAGAGGAPGHG | ASSLRLGPSQNTLY | 0.113588326 |
| TMPPLLAAIANAGRHGV | ASSLRLGPSQNTLY | 0.11275727 |
| TSSTAGNSGGA | ASSLRLGPSQNTLY | 0.113245964 |
| VPATVPVGILC | ASSLRLGPSQNTLY | 0.112684965 |
| VQRPDSGGVVLTGRIS | ASSLRLGPSQNTLY | 0.113070756 |
| WTPNPVPATKGVPAGLVQAT | ASSLRLGPSQNTLY | 0.11266062 |
| YALKRALAKFSDRFDVVII | ASSLRLGPSQNTLY | 0.11352677 |
| YLIGSVIV | ASSLRLGPSQNTLY | 0.11361844 |
| YGQLVAVID | ASSLRLGPSQNTLY | 0.11271788 |
| VTVGGMTLPA | ASSLRLGPSQNTLY | 0.112619445 |
| VLALPAV | ASSLRLGPSQNTLY | 0.11374721 |
| VIPPTLNYET | ASSLRLGPSQNTLY | 0.11274041 |
| VGMGGMHPSSG | ASSLRLGPSQNTLY | 0.11266868 |
| VGLAGGAATA | ASSLRLGPSQNTLY | 0.11313389 |
| VGGAGGAGGSGP | ASSLRLGPSQNTLY | 0.1129765 |
| VFAAFMV | ASSLRLGPSQNTLY | 0.11286665 |
| TLYDKLR | ASSLRLGPSQNTLY | 0.11280252 |
| TLAVFTVL | ASSLRLGPSQNTLY | 0.11293638 |
| TGSVQPARV | ASSLRLGPSQNTLY | 0.113204345 |
| TGGGAGSGGT | ASSLRLGPSQNTLY | 0.11286448 |
| SHLETSL | ASSLRLGPSQNTLY | 0.11264512 |
| SFTTGMT | ASSLRLGPSQNTLY | 0.11296744 |
| QVEVAKRMGVRQPTVS | ASSLRLGPSQNTLY | 0.11348337 |
| QGSGGPALAM | ASSLRLGPSQNTLY | 0.11308754 |
| MLAPSNP | ASSLRLGPSQNTLY | 0.11284021 |
| LVNNAAI | ASSLRLGPSQNTLY | 0.113267854 |
| LTGEINTLA | ASSLRLGPSQNTLY | 0.112679034 |
| LSIFGPV | ASSLRLGPSQNTLY | 0.11282089 |
| LLALLANL | ASSLRLGPSQNTLY | 0.11337351 |
| LISNASDALDKLR | ASSLRLGPSQNTLY | 0.11286333 |
| LINEAVP | ASSLRLGPSQNTLY | 0.11299648 |
| LGYAFSGL | ASSLRLGPSQNTLY | 0.11356906 |
| LDFVAVAL | ASSLRLGPSQNTLY | 0.11280976 |
| LAQPGAE | ASSLRLGPSQNTLY | 0.112880774 |
| ISNLGAVTGF | ASSLRLGPSQNTLY | 0.11295551 |
| HQEDPGRI | ASSLRLGPSQNTLY | 0.11272043 |
| GTPGEPTP | ASSLRLGPSQNTLY | 0.11373616 |
| GFFNSGS | ASSLRLGPSQNTLY | 0.112937145 |
| GAGGGGLTAGAN | ASSLRLGPSQNTLY | 0.11361355 |
| FVDALVNM | ASSLRLGPSQNTLY | 0.11289875 |
| FLTVAPEL | ASSLRLGPSQNTLY | 0.1131004 |
| FETLTAL | ASSLRLGPSQNTLY | 0.1136094 |

| | | |
|---|---|---|
| DVINPIEL | ASSLRLGPSQNTLY | 0.11397865 |
| APVSGAPV | ASSLRLGPSQNTLY | 0.11340286 |
| ALVGGLAW | ASSLRLGPSQNTLY | 0.11265221 |
| ALAAAAATAI | ASSLRLGPSQNTLY | 0.11293476 |
| AFGLGLLL | ASSLRLGPSQNTLY | 0.11274864 |
| AFFAAASL | ASSLRLGPSQNTLY | 0.1139188 |
| AAAALPAFSPP | ASSLRLGPSQNTLY | 0.11326987 |
| VVLAPAL | ASSLRLGPSQNTLY | 0.11333722 |
| TIGHVDHGKTTL | ASSLRLGPSQNTLY | 0.11276661 |
| LISNASDALDK | ASSLRLGPSQNTLY | 0.11285435 |
| LISNASDAL | ASSLRLGPSQNTLY | 0.11273391 |
| ISNASDALDK | ASSLRLGPSQNTLY | 0.113800846 |
| NQAETLVYQTE | ASSLRLGPSQNTLY | 0.11341308 |
| TGGTGGTAGAG | ASSLRLGPSQNTLY | 0.1126646 |
| TGSTAGGAGGA | ASSLRLGPSQNTLY | 0.11312618 |
| DPLAVPVAAGPV | ASSLRLGPSQNTLY | 0.11302105 |
| GGTGGTPGNGGN | ASSLRLGPSQNTLY | 0.112799674 |
| PAGLAGVDPATL | ASSLRLGPSQNTLY | 0.112724535 |
| GNNLVVEAPHAGASV | ASSLRLGPSQNTLY | 0.11286253 |
| KVTRIMSASTLEQPA | ASSLRLGPSQNTLY | 0.11285471 |
| LGFALPIALPRRTVV | ASSLRLGPSQNTLY | 0.112796724 |
| TRLGNLTARKLVVLV | ASSLRLGPSQNTLY | 0.11315049 |
| WSQQRRLIYFPSAGP | ASSLRLGPSQNTLY | 0.113058135 |
| GPNGNPDPMAAAVDIR | ASSLRLGPSQNTLY | 0.11280078 |
| TQRCVSVADLRAAASR | ASSLRLGPSQNTLY | 0.11295093 |
| QFVLDVAEHPVEVLTETV | ASSLRLGPSQNTLY | 0.112705335 |
| GGAGGTGGAGGNGGTQAGDG | ASSLRLGPSQNTLY | 0.112825505 |
| PLLATLRAIAVDVDAKPAQV | ASSLRLGPSQNTLY | 0.11284312 |
| APKFRTGGYMIDFWGVGYQVAK | ASSLRLGPSQNTLY | 0.11281449 |
| GAGGAGGAGGAAGGPGGTGGTG | ASSLRLGPSQNTLY | 0.11288577 |
| IGALGLLATDVAPAGSVAFGVA | ASSLRLGPSQNTLY | 0.11287962 |
| AGGAGGIGGNGGAGGAGGVLMG | ASSLRLGPSQNTLY | 0.11319672 |
| ADELIGLL | ASSLRLGPSQNTLY | 0.11285426 |
| ADRALKA | ASSLRLGPSQNTLY | 0.11396091 |
| AGAVGQVGGAGGA | ASSLRLGPSQNTLY | 0.11399193 |
| AGGGGGLGGAP | ASSLRLGPSQNTLY | 0.11290796 |
| AGGLLYGNGGAG | ASSLRLGPSQNTLY | 0.11400744 |
| AGLESIGL | ASSLRLGPSQNTLY | 0.11267249 |
| AGLKTGII | ASSLRLGPSQNTLY | 0.11288703 |
| ASLAALLL | ASSLRLGPSQNTLY | 0.11266564 |
| ASVINISSI | ASSLRLGPSQNTLY | 0.11344475 |
| ATPPGPA | ASSLRLGPSQNTLY | 0.11285443 |
| ATVASALV | ASSLRLGPSQNTLY | 0.11388727 |
| ATVASLIL | ASSLRLGPSQNTLY | 0.11294535 |
| AVNGPVLIRH | ASSLRLGPSQNTLY | 0.11388696 |
| DGAAGAVGI | ASSLRLGPSQNTLY | 0.113600984 |
| DPLLHGI | ASSLRLGPSQNTLY | 0.11282117 |
| ELGAALGAL | ASSLRLGPSQNTLY | 0.113713875 |
| ELGRTQAV | ASSLRLGPSQNTLY | 0.11292287 |
| FVAGGYLS | ASSLRLGPSQNTLY | 0.112896405 |
| GASGGAGGAGGW | ASSLRLGPSQNTLY | 0.11293305 |

| | | |
|---|---|---|
| GGAGAGGDGGAGGI | ASSLRLGPSQNTLY | 0.11301509 |
| GTGHNGSQPGMGGNG | ASSLRLGPSQNTLY | 0.11299454 |
| GYAAASASAS | ASSLRLGPSQNTLY | 0.11299865 |
| HQVDVAAVQQG | ASSLRLGPSQNTLY | 0.11335606 |
| IGAAIAHT | ASSLRLGPSQNTLY | 0.11293149 |
| IGMIEVAQL | ASSLRLGPSQNTLY | 0.11276145 |
| IRLAPEHTSAMD | ASSLRLGPSQNTLY | 0.11274796 |
| IYNVSVL | ASSLRLGPSQNTLY | 0.11383562 |
| KQGFLGM | ASSLRLGPSQNTLY | 0.1142461 |
| LESVITQL | ASSLRLGPSQNTLY | 0.11314111 |
| LGLGALVVV | ASSLRLGPSQNTLY | 0.11319991 |
| LIGNGGNGGSGGI | ASSLRLGPSQNTLY | 0.11325033 |
| LNAINAPTL | ASSLRLGPSQNTLY | 0.11343622 |
| LQLLNTGV | ASSLRLGPSQNTLY | 0.113220304 |
| LQTSIEF | ASSLRLGPSQNTLY | 0.1129488 |
| MSFAVSEL | ASSLRLGPSQNTLY | 0.11271527 |
| NTGALGL | ASSLRLGPSQNTLY | 0.11277835 |
| PPGAGGAGGAGTT | ASSLRLGPSQNTLY | 0.112830006 |
| QLVGLGI | ASSLRLGPSQNTLY | 0.11298762 |
| RFLEQVHQL | ASSLRLGPSQNTLY | 0.11347095 |
| RGVGYVLR | ASSLRLGPSQNTLY | 0.112988785 |
| SATGVFAGVI | ASSLRLGPSQNTLY | 0.11362996 |
| SLPAITLP | ASSLRLGPSQNTLY | 0.11308859 |
| STALAFTV | ASSLRLGPSQNTLY | 0.11304989 |
| SVLTVGLGK | ASSLRLGPSQNTLY | 0.1131559 |
| TALAGAGSGI | ASSLRLGPSQNTLY | 0.11291701 |
| TGLPFTGI | ASSLRLGPSQNTLY | 0.11283711 |
| VAEHVTV | ASSLRLGPSQNTLY | 0.11296065 |
| VDIVLNSL | ASSLRLGPSQNTLY | 0.11289371 |
| VLTGGGAVL | ASSLRLGPSQNTLY | 0.1135003 |
| VNVGLAYGI | ASSLRLGPSQNTLY | 0.11380796 |
| VSSAAVGAV | ASSLRLGPSQNTLY | 0.113060825 |
| VTSVYEAGV | ASSLRLGPSQNTLY | 0.11324476 |
| WVALDRGA | ASSLRLGPSQNTLY | 0.11306935 |
| YLVGSVVI | ASSLRLGPSQNTLY | 0.11272724 |
| GPPGTGKTLL | ASSLRLGPSQNTLY | 0.11294566 |
| GVGMTKF | ASSLRLGPSQNTLY | 0.11318307 |
| IGIGGSDL | ASSLRLGPSQNTLY | 0.11342904 |
| IGTLLAEL | ASSLRLGPSQNTLY | 0.113152616 |
| IYTSGTT | ASSLRLGPSQNTLY | 0.11295192 |
| LSFPAGV | ASSLRLGPSQNTLY | 0.11336189 |
| LYTSGTT | ASSLRLGPSQNTLY | 0.11291922 |
| VGAFPFGF | ASSLRLGPSQNTLY | 0.11315053 |
| VGAGTVEF | ASSLRLGPSQNTLY | 0.112863556 |
| VGKSTLLL | ASSLRLGPSQNTLY | 0.113031685 |
| AALAHAGGGIV | ASSLRLGPSQNTLY | 0.11286139 |
| AALVAVSLL | ASSLRLGPSQNTLY | 0.11311719 |
| AAVSRAVAAL | ASSLRLGPSQNTLY | 0.11262226 |
| AGGSSGIGGF | ASSLRLGPSQNTLY | 0.11273683 |
| AGLTVANANG | ASSLRLGPSQNTLY | 0.11274306 |
| AGLVGAVL | ASSLRLGPSQNTLY | 0.112820946 |

| | | |
|---|---|---|
| ALSQAFTV | ASSLRLGPSQNTLY | 0.11286714 |
| ALTFSVSL | ASSLRLGPSQNTLY | 0.11311628 |
| ALTGAALVAL | ASSLRLGPSQNTLY | 0.11279654 |
| ASFSIGAGVL | ASSLRLGPSQNTLY | 0.1133406 |
| DGSVYKI | ASSLRLGPSQNTLY | 0.112536505 |
| EQLAEIV | ASSLRLGPSQNTLY | 0.11289697 |
| FAVSELTVM | ASSLRLGPSQNTLY | 0.11336417 |
| GAAPLAAGL | ASSLRLGPSQNTLY | 0.11356336 |
| GADGGDGGSGGAGG | ASSLRLGPSQNTLY | 0.11324747 |
| GFGLDTGGAGG | ASSLRLGPSQNTLY | 0.11280282 |
| GLASGGLL | ASSLRLGPSQNTLY | 0.11286885 |
| GRIAGAAVLL | ASSLRLGPSQNTLY | 0.11283962 |
| GSAGSLNTSF | ASSLRLGPSQNTLY | 0.11315406 |
| GTGQIGI | ASSLRLGPSQNTLY | 0.112817615 |
| GTLIALIL | ASSLRLGPSQNTLY | 0.11323692 |
| IAATMGFVVG | ASSLRLGPSQNTLY | 0.113123804 |
| IFAQALL | ASSLRLGPSQNTLY | 0.11288881 |
| IGAAVALGF | ASSLRLGPSQNTLY | 0.11310041 |
| IGLTPLAL | ASSLRLGPSQNTLY | 0.11300192 |
| ILGSNTSTL | ASSLRLGPSQNTLY | 0.1127541 |
| IVFSLTVL | ASSLRLGPSQNTLY | 0.11272117 |
| LASKATGF | ASSLRLGPSQNTLY | 0.11340202 |
| LASVALIL | ASSLRLGPSQNTLY | 0.11357219 |
| LFAVGLTL | ASSLRLGPSQNTLY | 0.112820156 |
| LFNGGIGTYI | ASSLRLGPSQNTLY | 0.112988904 |
| LTLGAGII | ASSLRLGPSQNTLY | 0.11273182 |
| LVAAALGDI | ASSLRLGPSQNTLY | 0.11289238 |
| NTAFVQLGI | ASSLRLGPSQNTLY | 0.11333681 |
| QLFSGAEL | ASSLRLGPSQNTLY | 0.11328926 |
| SAATFITL | ASSLRLGPSQNTLY | 0.11284298 |
| SNYGLANLGV | ASSLRLGPSQNTLY | 0.11286587 |
| SYGGAYAVM | ASSLRLGPSQNTLY | 0.114255704 |
| TTGVGGLTLG | ASSLRLGPSQNTLY | 0.11279464 |
| TTQTIYI | ASSLRLGPSQNTLY | 0.11284436 |
| VAAGAGLGAVI | ASSLRLGPSQNTLY | 0.1128361 |
| VAAVVTLL | ASSLRLGPSQNTLY | 0.11290921 |
| VGFLGGLI | ASSLRLGPSQNTLY | 0.11266357 |
| VGVIPILL | ASSLRLGPSQNTLY | 0.112937786 |
| VIFASSLI | ASSLRLGPSQNTLY | 0.11334499 |
| VKGGDGGAAAG | ASSLRLGPSQNTLY | 0.11284999 |
| VTKTFTNI | ASSLRLGPSQNTLY | 0.11302293 |
| VVAAAILL | ASSLRLGPSQNTLY | 0.11308843 |
| YAALDVEL | ASSLRLGPSQNTLY | 0.11358044 |
| LASVALLL | ASSLRLGPSQNTLY | 0.113106765 |
| LDAVHRV | ASSLRLGPSQNTLY | 0.112808876 |
| QGLSSSIF | ASSLRLGPSQNTLY | 0.11281432 |
| RGTFANIRL | ASSLRLGPSQNTLY | 0.112903506 |
| RQLWWGHRI | ASSLRLGPSQNTLY | 0.11315363 |
| YGPPGTGKTLL | ASSLRLGPSQNTLY | 0.113056615 |
| AALVGAPANVA | ASSLRLGPSQNTLY | 0.1130448 |
| AATFRRTE | ASSLRLGPSQNTLY | 0.112440616 |

| | | |
|---|---|---|
| AGALGGGI | ASSLRLGPSQNTLY | 0.11285865 |
| AGGAGGAGGIGAG | ASSLRLGPSQNTLY | 0.11290648 |
| AGSGALGL | ASSLRLGPSQNTLY | 0.11282241 |
| AQIADQI | ASSLRLGPSQNTLY | 0.11367822 |
| ASAVAVTYL | ASSLRLGPSQNTLY | 0.11283445 |
| ASTVLGGTGG | ASSLRLGPSQNTLY | 0.11289864 |
| AVGGEPAK | ASSLRLGPSQNTLY | 0.112784274 |
| CEQGTFGGTC | ASSLRLGPSQNTLY | 0.112655185 |
| DEGKAPAG | ASSLRLGPSQNTLY | 0.112863675 |
| DTVVFPAL | ASSLRLGPSQNTLY | 0.11280328 |
| FEGITMAG | ASSLRLGPSQNTLY | 0.11308224 |
| FPLVVQD | ASSLRLGPSQNTLY | 0.11280039 |
| GAFTTPQ | ASSLRLGPSQNTLY | 0.11300998 |
| GFGGFGTAGGVGGA | ASSLRLGPSQNTLY | 0.11282047 |
| GFLAQFL | ASSLRLGPSQNTLY | 0.1126916 |
| GGAGGAGGNAGVA | ASSLRLGPSQNTLY | 0.11378302 |
| GGVGTAGAGGAGGAGG | ASSLRLGPSQNTLY | 0.112815864 |
| GNGGGGDAGNAGSGGN | ASSLRLGPSQNTLY | 0.112914525 |
| GTGGAGGAGNDAGS | ASSLRLGPSQNTLY | 0.11386358 |
| GTTNTGWFNS | ASSLRLGPSQNTLY | 0.11280166 |
| IFMGPLFTL | ASSLRLGPSQNTLY | 0.113122664 |
| ILAHAIEAI | ASSLRLGPSQNTLY | 0.11292234 |
| IRSAIREAL | ASSLRLGPSQNTLY | 0.113051 |
| ITFAGGG | ASSLRLGPSQNTLY | 0.11308679 |
| ITGMSGHQEV | ASSLRLGPSQNTLY | 0.11303587 |
| IVAGLACTL | ASSLRLGPSQNTLY | 0.112821415 |
| IVGGGLAGA | ASSLRLGPSQNTLY | 0.112955496 |
| IVTGVLNI | ASSLRLGPSQNTLY | 0.1147479 |
| LAQQGCI | ASSLRLGPSQNTLY | 0.11326018 |
| LCLHVEN | ASSLRLGPSQNTLY | 0.1131611 |
| LDAISDGS | ASSLRLGPSQNTLY | 0.11296699 |
| LGGFAAML | ASSLRLGPSQNTLY | 0.11299168 |
| LGGKNPMI | ASSLRLGPSQNTLY | 0.113179706 |
| LGPKGVAAIL | ASSLRLGPSQNTLY | 0.1127546 |
| LMVGVSI | ASSLRLGPSQNTLY | 0.11296792 |
| LVGAAFVL | ASSLRLGPSQNTLY | 0.112893306 |
| LVLAVAPL | ASSLRLGPSQNTLY | 0.11276048 |
| MAAIAVIDAA | ASSLRLGPSQNTLY | 0.1129059 |
| NNGINAELQ | ASSLRLGPSQNTLY | 0.11335423 |
| RGEPLVLVHG | ASSLRLGPSQNTLY | 0.11305534 |
| RGGAGGAAGLI | ASSLRLGPSQNTLY | 0.112800516 |
| RSYRMHPAV | ASSLRLGPSQNTLY | 0.11290804 |
| SAAHLLAGQLA | ASSLRLGPSQNTLY | 0.1132204 |
| SAAVGSY | ASSLRLGPSQNTLY | 0.112752005 |
| SPHAVLHRI | ASSLRLGPSQNTLY | 0.113688216 |
| SQIAKVL | ASSLRLGPSQNTLY | 0.11292405 |
| TALGHLVP | ASSLRLGPSQNTLY | 0.113336705 |
| TAPTTSGAA | ASSLRLGPSQNTLY | 0.114911355 |
| TDGDDGNGGSPGT | ASSLRLGPSQNTLY | 0.1129141 |
| TDGLGEN | ASSLRLGPSQNTLY | 0.1127749 |
| TGGPPLPGQA | ASSLRLGPSQNTLY | 0.112751395 |

| | | |
|---|---|---|
| TIPAITI | ASSLRLGPSQNTLY | 0.11289244 |
| TIPGFTI | ASSLRLGPSQNTLY | 0.11300453 |
| TSIASGALA | ASSLRLGPSQNTLY | 0.11266846 |
| TVTLPGAAI | ASSLRLGPSQNTLY | 0.11378324 |
| VASGLSAALP | ASSLRLGPSQNTLY | 0.11333953 |
| VGAGVIGI | ASSLRLGPSQNTLY | 0.11294135 |
| VGPKRFGV | ASSLRLGPSQNTLY | 0.11254228 |
| VNVAFPD | ASSLRLGPSQNTLY | 0.113625735 |
| VSMTNSGVS | ASSLRLGPSQNTLY | 0.113534465 |
| VTDVLGDLSV | ASSLRLGPSQNTLY | 0.113371685 |
| VVVVGAGFAGL | ASSLRLGPSQNTLY | 0.11277184 |
| VYTVGTVV | ASSLRLGPSQNTLY | 0.11280897 |
| WTTFSGP | ASSLRLGPSQNTLY | 0.113846056 |
| YDELVEAF | ASSLRLGPSQNTLY | 0.113284364 |
| YGTGGAGGNGG | ASSLRLGPSQNTLY | 0.11318935 |
| YTNVMAV | ASSLRLGPSQNTLY | 0.113848254 |
| FLEQVHQL | ASSLRLGPSQNTLY | 0.112970546 |
| GLSLLAVAV | ASSLRLGPSQNTLY | 0.11311579 |
| LEQVHQL | ASSLRLGPSQNTLY | 0.11315065 |
| LNAAGAI | ASSLRLGPSQNTLY | 0.113596894 |
| PAVHASL | ASSLRLGPSQNTLY | 0.112670906 |
| TELEDDA | ASSLRLGPSQNTLY | 0.112869754 |
| TPPGPPN | ASSLRLGPSQNTLY | 0.11310067 |
| TPPPPPPP | ASSLRLGPSQNTLY | 0.11279913 |
| TQSPALI | ASSFSQGGNSPLY | 0.11291125 |
| EFAAAFPQ | ASSFSQGGNSPLY | 0.112687364 |
| LLSGNLDVL | ASSFSQGGNSPLY | 0.11345633 |
| GPANLGNYNI | ASSFSQGGNSPLY | 0.11295459 |
| TLGPTLP | ASSFSQGGNSPLY | 0.11380532 |
| VYAYSSCCK | ASSFSQGGNSPLY | 0.11264408 |
| APAPAPAPNI | ASSFSQGGNSPLY | 0.11298201 |
| PGGPVPIP | ASSFSQGGNSPLY | 0.11279502 |
| FLTVALEP | ASSFSQGGNSPLY | 0.1136703 |
| SQYFGANL | ASSFSQGGNSPLY | 0.11283979 |
| GGVKPAA | ASSFSQGGNSPLY | 0.11278808 |
| GGTGGAGGAGGW | ASSFSQGGNSPLY | 0.11274178 |
| GVGGVAGLGGAGG | ASSFSQGGNSPLY | 0.19916517 |
| VLLVLATTI | ASSFSQGGNSPLY | 0.3689173 |
| IGIFAAML | ASSFSQGGNSPLY | 0.443056 |
| AVGDLID | ASSFSQGGNSPLY | 0.34313655 |
| NISPLPL | ASSFSQGGNSPLY | 0.16229825 |
| VTAVFVAI | ASSFSQGGNSPLY | 0.10673144 |
| SGDDNDAAL | ASSFSQGGNSPLY | 0.40745643 |
| AVAALAHS | ASSFSQGGNSPLY | 0.10995163 |
| GGVTVMATP | ASSFSQGGNSPLY | 0.15319456 |
| AGLAANVRD | ASSFSQGGNSPLY | 0.40726 |
| TNRPDLIDPALLRPG | ASSFSQGGNSPLY | 0.18573773 |
| LANLGVTAL | ASSFSQGGNSPLY | 0.6363006 |
| TGERIEL | ASSFSQGGNSPLY | 0.6523255 |
| NGEELAA | ASSFSQGGNSPLY | 0.13972007 |
| VLSLVWG | ASSFSQGGNSPLY | 0.27028856 |

| | | |
|---|---|---|
| IGYALWL | ASSFSQGGNSPLY | 0.16888623 |
| LSLLLAPL | ASSFSQGGNSPLY | 0.3237918 |
| TLDEALK | ASSFSQGGNSPLY | 0.10410128 |
| TDGAGTAEGG | ASSFSQGGNSPLY | 0.35816103 |
| NAPGGAGGNGG | ASSFSQGGNSPLY | 0.1328039 |
| VSVFMGMGGNF | ASSFSQGGNSPLY | 0.092150986 |
| LASYLAAL | ASSFSQGGNSPLY | 0.48839137 |
| LVGGTPAL | ASSFSQGGNSPLY | 0.22236699 |
| VRNADVL | ASSFSQGGNSPLY | 0.15372278 |
| TCSSPFLVSAM | ASSFSQGGNSPLY | 0.67039955 |
| VHITDAE | ASSFSQGGNSPLY | 0.12477696 |
| EYAAPSF | ASSFSQGGNSPLY | 0.11588138 |
| VVDAFHGGL | ASSFSQGGNSPLY | 0.45256466 |
| ASVVDPQ | ASSFSQGGNSPLY | 0.71170926 |
| SYNEFGGLS | ASSFSQGGNSPLY | 0.116557464 |
| DPDGSVY | ASSFSQGGNSPLY | 0.48683828 |
| MVSTIHATE | ASSFSQGGNSPLY | 0.67006445 |
| LAAGVADPV | ASSFSQGGNSPLY | 0.2937787 |
| TKEQIAA | ASSFSQGGNSPLY | 0.57477474 |
| SLASPAAL | ASSFSQGGNSPLY | 0.6448079 |
| IGLVFYGV | ASSFSQGGNSPLY | 0.22492187 |
| FLLDASNL | ASSFSQGGNSPLY | 0.64571273 |
| YQVGNFP | ASSFSQGGNSPLY | 0.68180424 |
| ESLLSQST | ASSFSQGGNSPLY | 0.19549696 |
| LFTVGFAAVG | ASSFSQGGNSPLY | 0.25493133 |
| ATPPTPT | ASSFSQGGNSPLY | 0.20726907 |
| TDLATLG | ASSFSQGGNSPLY | 0.5019467 |
| SAGGTGR | ASSFSQGGNSPLY | 0.097957194 |
| GFSNSGS | ASSFSQGGNSPLY | 0.48376745 |
| GGNGGSGGTSV | ASSFSQGGNSPLY | 0.4165445 |
| GTSVATGGAGN | ASSFSQGGNSPLY | 0.2498489 |
| NGGNGGSGGTS | ASSFSQGGNSPLY | 0.1574198 |
| AVVKFPH | ASSFSQGGNSPLY | 0.13968898 |
| TPPPPPP | ASSFSQGGNSPLY | 0.15825194 |
| AAPAAAAAPA | ASSFSQGGNSPLY | 0.20759416 |
| IALNEDL | ASSFSQGGNSPLY | 0.104090564 |
| IVNVSSM | ASSFSQGGNSPLY | 0.2567308 |
| TGIPPAPR | ASSFSQGGNSPLY | 0.4265815 |
| FDLGGGTF | ASSFSQGGNSPLY | 0.42549783 |
| FDLGGGTFDVS | ASSFSQGGNSPLY | 0.10758545 |
| NEPTAAA | ASSFSQGGNSPLY | 0.7169862 |
| NEPTAAALA | ASSFSQGGNSPLY | 0.3429432 |
| GSIINISSI | ASSFSQGGNSPLY | 0.55900824 |
| VAPGNAGTA | ASSFSQGGNSPLY | 0.21918952 |
| TGGGASLE | ASSFSQGGNSPLY | 0.15770762 |
| VAVYNHYKRI | ASSFSQGGNSPLY | 0.5164517 |
| TDPAPATT | ASSFSQGGNSPLY | 0.32472324 |
| TNRPDILDPALLRPG | ASSFSQGGNSPLY | 0.124625996 |
| LTTTTTS | ASSFSQGGNSPLY | 0.61355597 |
| PTGSGKTL | ASSFSQGGNSPLY | 0.12778097 |
| TGSGKTL | ASSFSQGGNSPLY | 0.12999219 |

| | | |
|---|---|---|
| TIGHVDHGK | ASSFSQGGNSPLY | 0.27982235 |
| TIGHVDHGKTT | ASSFSQGGNSPLY | 0.6353243 |
| GHVDHGKTT | ASSFSQGGNSPLY | 0.14628416 |
| AQFEHTLL | ASSFSQGGNSPLY | 0.13225026 |
| IVNQDEL | ASSFSQGGNSPLY | 0.11893574 |
| TNGSQFF | ASSFSQGGNSPLY | 0.3359139 |
| TGKGIVE | ASSFSQGGNSPLY | 0.6471339 |
| LISNASDA | ASSFSQGGNSPLY | 0.32538223 |
| AELRTLGELTKIAKAHGAQ | ASSFSQGGNSPLY | 0.20390563 |
| APNAVAAAEPSW | ASSFSQGGNSPLY | 0.64960897 |
| APQVAGAATAVL | ASSFSQGGNSPLY | 0.23067942 |
| ARRDPAAAAAILPTDA | ASSFSQGGNSPLY | 0.2916303 |
| DAAARDVLAEAGLAEYF | ASSFSQGGNSPLY | 0.6829406 |
| DDQIVLSAKANLAKVEQ | ASSFSQGGNSPLY | 0.12492767 |
| DELLPDTGEDSDEDGH | ASSFSQGGNSPLY | 0.58268416 |
| ELPEAPANRATVPQD | ASSFSQGGNSPLY | 0.29923913 |
| FGQSKIERVTGIG | ASSFSQGGNSPLY | 0.47394493 |
| GAGGTGSTAGGAGGAGGAGGLY | ASSFSQGGNSPLY | 0.58657634 |
| GHGGTGAAGGNGGTGGAGGSGI | ASSFSQGGNSPLY | 0.1375648 |
| GNGGNGGAGGEGGPGGAGGAGA | ASSFSQGGNSPLY | 0.43799177 |
| GSTAGNGGNGGNATNGTIAGQP | ASSFSQGGNSPLY | 0.11253456 |
| GTGGDGGNGGNGGTGG | ASSFSQGGNSPLY | 0.277968 |
| GVRRPAGIPADAPV | ASSFSQGGNSPLY | 0.6560178 |
| GYRQVTDAHLLTLARR | ASSFSQGGNSPLY | 0.42382556 |
| IAKMGQDGHDRG | ASSFSQGGNSPLY | 0.69652426 |
| KAEPSIEIQYT | ASSFSQGGNSPLY | 0.3532604 |
| KPVRIRMAADEDTVAVT | ASSFSQGGNSPLY | 0.18302336 |
| MPPRIAGMRLLVIKPEP | ASSFSQGGNSPLY | 0.7463139 |
| MKFYAVALFRSRGVPPTR | ASSFSQGGNSPLY | 0.7238938 |
| NAGGFGILGAVAHSP | ASSFSQGGNSPLY | 0.11269379 |
| NEGREITLAMAEM | ASSFSQGGNSPLY | 0.19693837 |
| NELVEWSHPLGHRWRWE | ASSFSQGGNSPLY | 0.68135023 |
| NLGAGGTGGNGGNPG | ASSFSQGGNSPLY | 0.39226705 |
| NTEWPKWETVMAADPVD | ASSFSQGGNSPLY | 0.36185157 |
| PASAAPAETQP | ASSFSQGGNSPLY | 0.31152263 |
| QQSSDGKNGVRAAEITGE | ASSFSQGGNSPLY | 0.31774712 |
| RAPDAVTTSWWKE | ASSFSQGGNSPLY | 0.13111772 |
| SGFGNVGAHTSGWFNQ | ASSFSQGGNSPLY | 0.70305777 |
| SGVAGAVGQAASVGGLKV | ASSFSQGGNSPLY | 0.10263747 |
| SNGGAGTGSAGGNGGT | ASSFSQGGNSPLY | 0.3321151 |
| TGGAGGEGGAAG | ASSFSQGGNSPLY | 0.14620788 |
| TGGDGGAAGAGGAPGHG | ASSFSQGGNSPLY | 0.106400855 |
| TMPPLLAAIANAGRHGV | ASSFSQGGNSPLY | 0.37889406 |
| TSSTAGNSGGA | ASSFSQGGNSPLY | 0.2956678 |
| VPATVPVGILC | ASSFSQGGNSPLY | 0.6739152 |
| VQRPDSGGVVLTGRIS | ASSFSQGGNSPLY | 0.28340846 |
| WTPNPVPATKGVPAGLVQAT | ASSFSQGGNSPLY | 0.419761 |
| YALKRALAKFSDRFDVVII | ASSFSQGGNSPLY | 0.35241908 |
| YLIGSVIV | ASSFSQGGNSPLY | 0.48328573 |
| YGQLVAVID | ASSFSQGGNSPLY | 0.19017267 |
| VTVGGMTLPA | ASSFSQGGNSPLY | 0.28058696 |

| | | |
|---|---|---|
| VLALPAV | ASSFSQGGNSPLY | 0.75858337 |
| VIPPTLNYET | ASSFSQGGNSPLY | 0.44478315 |
| VGMGGMHPSSG | ASSFSQGGNSPLY | 0.102550946 |
| VGLAGGAATA | ASSFSQGGNSPLY | 0.19577697 |
| VGGAGGAGGSGP | ASSFSQGGNSPLY | 0.19618843 |
| VFAAFMV | ASSFSQGGNSPLY | 0.09226011 |
| TLYDKLR | ASSFSQGGNSPLY | 0.10227808 |
| TLAVFTVL | ASSFSQGGNSPLY | 0.11458969 |
| TGSVQPARV | ASSFSQGGNSPLY | 0.18847981 |
| TGGGAGSGGT | ASSFSQGGNSPLY | 0.6161209 |
| SHLETSL | ASSFSQGGNSPLY | 0.41531605 |
| SFTTGMT | ASSFSQGGNSPLY | 0.22167753 |
| QVEVAKRMGVRQPTVS | ASSFSQGGNSPLY | 0.5836426 |
| QGSGGPALAM | ASSFSQGGNSPLY | 0.21413797 |
| MLAPSNP | ASSFSQGGNSPLY | 0.08854221 |
| LVNNAAI | ASSFSQGGNSPLY | 0.1708486 |
| LTGEINTLA | ASSFSQGGNSPLY | 0.17114663 |
| LSIFGPV | ASSFSQGGNSPLY | 0.573748 |
| LLALLANL | ASSFSQGGNSPLY | 0.1474246 |
| LISNASDALDKLR | ASSFSQGGNSPLY | 0.19844905 |
| LINEAVP | ASSFSQGGNSPLY | 0.25340372 |
| LGYAFSGL | ASSFSQGGNSPLY | 0.19872043 |
| LDFVAVAL | ASSFSQGGNSPLY | 0.41307485 |
| LAQPGAE | ASSFSQGGNSPLY | 0.12532344 |
| ISNLGAVTGF | ASSFSQGGNSPLY | 0.09833632 |
| HQEDPGRI | ASSFSQGGNSPLY | 0.29348844 |
| GTPGEPTP | ASSFSQGGNSPLY | 0.097082205 |
| GFFNSGS | ASSFSQGGNSPLY | 0.14399856 |
| GAGGGGLTAGAN | ASSFSQGGNSPLY | 0.114867166 |
| FVDALVNM | ASSFSQGGNSPLY | 0.11076374 |
| FLTVAPEL | ASSFSQGGNSPLY | 0.10167132 |
| FETLTAL | ASSFSQGGNSPLY | 0.5402386 |
| DVINPIEL | ASSFSQGGNSPLY | 0.46182424 |
| APVSGAPV | ASSFSQGGNSPLY | 0.27294472 |
| ALVGGLAW | ASSFSQGGNSPLY | 0.5810362 |
| ALAAAAATAI | ASSFSQGGNSPLY | 0.23695785 |
| AFGLGLLL | ASSFSQGGNSPLY | 0.32000768 |
| AFFAAASL | ASSFSQGGNSPLY | 0.123718075 |
| AAAALPAFSPP | ASSFSQGGNSPLY | 0.2122863 |
| VVLAPAL | ASSFSQGGNSPLY | 0.12512417 |
| TIGHVDHGKTTL | ASSFSQGGNSPLY | 0.123334125 |
| LISNASDALDK | ASSFSQGGNSPLY | 0.75475335 |
| LISNASDAL | ASSFSQGGNSPLY | 0.30607578 |
| ISNASDALDK | ASSFSQGGNSPLY | 0.13440631 |
| NQAETLVYQTE | ASSFSQGGNSPLY | 0.27904877 |
| TGGTGGTAGAG | ASSFSQGGNSPLY | 0.64891905 |
| TGSTAGGAGGA | ASSFSQGGNSPLY | 0.7355437 |
| DPLAVPVAAGPV | ASSFSQGGNSPLY | 0.26590657 |
| GGTGGTPGNGGN | ASSFSQGGNSPLY | 0.19928518 |
| PAGLAGVDPATL | ASSFSQGGNSPLY | 0.4162132 |
| GNNLVVEAPHAGASV | ASSFSQGGNSPLY | 0.29685065 |

| | | |
|---|---|---|
| KVTRIMSASTLEQPA | ASSFSQGGNSPLY | 0.22973612 |
| LGFALPIALPRRTVV | ASSFSQGGNSPLY | 0.112303495 |
| TRLGNLTARKLVVLV | ASSFSQGGNSPLY | 0.5445511 |
| WSQQRRLIYFPSAGP | ASSFSQGGNSPLY | 0.6788657 |
| GPNGNPDPMAAAVDIR | ASSFSQGGNSPLY | 0.18913323 |
| TQRCVSVADLRAAASR | ASSFSQGGNSPLY | 0.11489831 |
| QFVLDVAEHPVEVLTETV | ASSFSQGGNSPLY | 0.29947382 |
| GGAGGTGGAGGNGGTQAGDG | ASSFSQGGNSPLY | 0.13587955 |
| PLLATLRAIAVDVDAKPAQV | ASSFSQGGNSPLY | 0.19991273 |
| APKFRTGGYMIDFWGVGYQVAK | ASSFSQGGNSPLY | 0.22901085 |
| GAGGAGGAGGAAGGPGGTGGTG | ASSFSQGGNSPLY | 0.123518795 |
| IGALGLLATDVAPAGSVAFGVA | ASSFSQGGNSPLY | 0.20682588 |
| AGGAGGIGGNGGAGGAGGVLMG | ASSFSQGGNSPLY | 0.6024127 |
| ADELIGLL | ASSFSQGGNSPLY | 0.22526535 |
| ADRALKA | ASSFSQGGNSPLY | 0.6702888 |
| AGAVGQVGGAGGA | ASSFSQGGNSPLY | 0.6889543 |
| AGGGGGLGGAP | ASSFSQGGNSPLY | 0.5712184 |
| AGGLLYGNGGAG | ASSFSQGGNSPLY | 0.7274873 |
| AGLESIGL | ASSFSQGGNSPLY | 0.3910011 |
| AGLKTGII | ASSFSQGGNSPLY | 0.2513258 |
| ASLAALLL | ASSFSQGGNSPLY | 0.2720647 |
| ASVINISSI | ASSFSQGGNSPLY | 0.68198025 |
| ATPPGPA | ASSFSQGGNSPLY | 0.3610523 |
| ATVASALV | ASSFSQGGNSPLY | 0.34087837 |
| ATVASLIL | ASSFSQGGNSPLY | 0.25625306 |
| AVNGPVLIRH | ASSFSQGGNSPLY | 0.59318185 |
| DGAAGAVGI | ASSFSQGGNSPLY | 0.20478374 |
| DPLLHGI | ASSFSQGGNSPLY | 0.45472246 |
| ELGAALGAL | ASSFSQGGNSPLY | 0.12450753 |
| ELGRTQAV | ASSFSQGGNSPLY | 0.12107754 |
| FVAGGYLS | ASSFSQGGNSPLY | 0.6304915 |
| GASGGAGGAGGW | ASSFSQGGNSPLY | 0.0933969 |
| GGAGAGGDGGAGGI | ASSFSQGGNSPLY | 0.11846646 |
| GTGHNGSQPGMGGNG | ASSFSQGGNSPLY | 0.45990083 |
| GYAAASASAS | ASSFSQGGNSPLY | 0.5997876 |
| HQVDVAAVQQG | ASSFSQGGNSPLY | 0.089465894 |
| IGAAIAHT | ASSFSQGGNSPLY | 0.12044237 |
| IGMIEVAQL | ASSFSQGGNSPLY | 0.17937456 |
| IRLAPEHTSAMD | ASSFSQGGNSPLY | 0.24466363 |
| IYNVSVL | ASSFSQGGNSPLY | 0.17262301 |
| KQGFLGM | ASSFSQGGNSPLY | 0.10687331 |
| LESVITQL | ASSFSQGGNSPLY | 0.43146414 |
| LGLGALVVV | ASSFSQGGNSPLY | 0.10930005 |
| LIGNGGNGGSGGI | ASSFSQGGNSPLY | 0.105234966 |
| LNAINAPTL | ASSFSQGGNSPLY | 0.44369406 |
| LQLLNTGV | ASSFSQGGNSPLY | 0.57769513 |
| LQTSIEF | ASSFSQGGNSPLY | 0.15676163 |
| MSFAVSEL | ASSFSQGGNSPLY | 0.46716207 |
| NTGALGL | ASSFSQGGNSPLY | 0.21629336 |
| PPGAGGAGGAGTT | ASSFSQGGNSPLY | 0.77286613 |
| QLVGLGI | ASSFSQGGNSPLY | 0.24042812 |

| | | |
|---|---|---|
| RFLEQVHQL | ASSFSQGGNSPLY | 0.13592882 |
| RGVGYVLR | ASSFSQGGNSPLY | 0.5332319 |
| SATGVFAGVI | ASSFSQGGNSPLY | 0.68055594 |
| SLPAITLP | ASSFSQGGNSPLY | 0.34044945 |
| STALAFTV | ASSFSQGGNSPLY | 0.47594982 |
| SVLTVGLGK | ASSFSQGGNSPLY | 0.178201 |
| TALAGAGSGI | ASSFSQGGNSPLY | 0.33046216 |
| TGLPFTGI | ASSFSQGGNSPLY | 0.6568601 |
| VAEHVTV | ASSFSQGGNSPLY | 0.61599267 |
| VDIVLNSL | ASSFSQGGNSPLY | 0.1047378 |
| VLTGGGAVL | ASSFSQGGNSPLY | 0.5668008 |
| VNVGLAYGI | ASSFSQGGNSPLY | 0.16182892 |
| VSSAAVGAV | ASSFSQGGNSPLY | 0.24674919 |
| VTSVYEAGV | ASSFSQGGNSPLY | 0.54218173 |
| WVALDRGA | ASSFSQGGNSPLY | 0.30822748 |
| YLVGSVVI | ASSFSQGGNSPLY | 0.5916678 |
| GPPGTGKTLL | ASSFSQGGNSPLY | 0.28110856 |
| GVGMTKF | ASSFSQGGNSPLY | 0.5051163 |
| IGIGGSDL | ASSFSQGGNSPLY | 0.118855916 |
| IGTLLAEL | ASSFSQGGNSPLY | 0.62559927 |
| IYTSGTT | ASSFSQGGNSPLY | 0.48207998 |
| LSFPAGV | ASSFSQGGNSPLY | 0.55779576 |
| LYTSGTT | ASSFSQGGNSPLY | 0.2961744 |
| VGAFPFGF | ASSFSQGGNSPLY | 0.12475127 |
| VGAGTVEF | ASSFSQGGNSPLY | 0.5430852 |
| VGKSTLLL | ASSFSQGGNSPLY | 0.5546793 |
| AALAHAGGGIV | ASSFSQGGNSPLY | 0.28604108 |
| AALVAVSLL | ASSFSQGGNSPLY | 0.5373416 |
| AAVSRAVAAL | ASSFSQGGNSPLY | 0.7364658 |
| AGGSSGIGGF | ASSFSQGGNSPLY | 0.26067048 |
| AGLTVANANG | ASSFSQGGNSPLY | 0.16882516 |
| AGLVGAVL | ASSFSQGGNSPLY | 0.468871 |
| ALSQAFTV | ASSFSQGGNSPLY | 0.7727661 |
| ALTFSVSL | ASSFSQGGNSPLY | 0.27607512 |
| ALTGAALVAL | ASSFSQGGNSPLY | 0.7184196 |
| ASFSIGAGVL | ASSFSQGGNSPLY | 0.15175171 |
| DGSVYKI | ASSFSQGGNSPLY | 0.58909416 |
| EQLAEIV | ASSFSQGGNSPLY | 0.34044114 |
| FAVSELTVM | ASSFSQGGNSPLY | 0.4139823 |
| GAAPLAAGL | ASSFSQGGNSPLY | 0.10182551 |
| GADGGDGGSGGAGG | ASSFSQGGNSPLY | 0.54891264 |
| GFGLDTGGAGG | ASSFSQGGNSPLY | 0.60001516 |
| GLASGGLL | ASSFSQGGNSPLY | 0.57504594 |
| GRIAGAAVLL | ASSFSQGGNSPLY | 0.09697763 |
| GSAGSLNTSF | ASSFSQGGNSPLY | 0.30899298 |
| GTGQIGI | ASSFSQGGNSPLY | 0.55894166 |
| GTLIALIL | ASSFSQGGNSPLY | 0.7152784 |
| IAATMGFVVG | ASSFSQGGNSPLY | 0.3011495 |
| IFAQALL | ASSFSQGGNSPLY | 0.6714828 |
| IGAAVALGF | ASSFSQGGNSPLY | 0.14536685 |
| IGLTPLAL | ASSFSQGGNSPLY | 0.21826808 |

| | | |
|---|---|---|
| ILGSNTSTL | ASSFSQGGNSPLY | 0.1479539 |
| IVFSLTVL | ASSFSQGGNSPLY | 0.4138744 |
| LASKATGF | ASSFSQGGNSPLY | 0.25696248 |
| LASVALIL | ASSFSQGGNSPLY | 0.22707988 |
| LFAVGLTL | ASSFSQGGNSPLY | 0.26233983 |
| LFNGGIGTYI | ASSFSQGGNSPLY | 0.75844216 |
| LTLGAGII | ASSFSQGGNSPLY | 0.36096513 |
| LVAAALGDI | ASSFSQGGNSPLY | 0.6321759 |
| NTAFVQLGI | ASSFSQGGNSPLY | 0.7565615 |
| QLFSGAEL | ASSFSQGGNSPLY | 0.53702676 |
| SAATFITL | ASSFSQGGNSPLY | 0.1286288 |
| SNYGLANLGV | ASSFSQGGNSPLY | 0.30433074 |
| SYGGAYAVM | ASSFSQGGNSPLY | 0.12745784 |
| TTGVGGLTLG | ASSFSQGGNSPLY | 0.21444711 |
| TTQTIYI | ASSFSQGGNSPLY | 0.12630951 |
| VAAGAGLGAVI | ASSFSQGGNSPLY | 0.3422956 |
| VAAVVTLL | ASSFSQGGNSPLY | 0.08912979 |
| VGFLGGLI | ASSFSQGGNSPLY | 0.45541477 |
| VGVIPILL | ASSFSQGGNSPLY | 0.7731972 |
| VIFASSLI | ASSFSQGGNSPLY | 0.10017551 |
| VKGGDGGAAAG | ASSFSQGGNSPLY | 0.4595446 |
| VTKTFTNI | ASSFSQGGNSPLY | 0.61502296 |
| VVAAAILL | ASSFSQGGNSPLY | 0.17689681 |
| YAALDVEL | ASSFSQGGNSPLY | 0.5263678 |
| LASVALLL | ASSFSQGGNSPLY | 0.14622626 |
| LDAVHRV | ASSFSQGGNSPLY | 0.52015257 |
| QGLSSSIF | ASSFSQGGNSPLY | 0.4774282 |
| RGTFANIRL | ASSFSQGGNSPLY | 0.47198176 |
| RQLWWGHRI | ASSFSQGGNSPLY | 0.6104591 |
| YGPPGTGKTLL | ASSFSQGGNSPLY | 0.3653841 |
| AALVGAPANVA | ASSFSQGGNSPLY | 0.113788806 |
| AATFRRTE | ASSFSQGGNSPLY | 0.6768602 |
| AGALGGGI | ASSFSQGGNSPLY | 0.11665906 |
| AGGAGGAGGIGAG | ASSFSQGGNSPLY | 0.46988901 |
| AGSGALGL | ASSFSQGGNSPLY | 0.15137857 |
| AQIADQI | ASSFSQGGNSPLY | 0.5146668 |
| ASAVAVTYL | ASSFSQGGNSPLY | 0.20172338 |
| ASTVLGGTGG | ASSFSQGGNSPLY | 0.36829638 |
| AVGGEPAK | ASSFSQGGNSPLY | 0.22937137 |
| CEQGTFGGTC | ASSFSQGGNSPLY | 0.16260004 |
| DEGKAPAG | ASSFSQGGNSPLY | 0.1117443 |
| DTVVFPAL | ASSFSQGGNSPLY | 0.10900857 |
| FEGITMAG | ASSFSQGGNSPLY | 0.6763119 |
| FPLVVQD | ASSFSQGGNSPLY | 0.24371046 |
| GAFTTPQ | ASSFSQGGNSPLY | 0.1972045 |
| GFGGFGTAGGVGGA | ASSFSQGGNSPLY | 0.45660543 |
| GFLAQFL | ASSFSQGGNSPLY | 0.32786888 |
| GGAGGAGGNAGVA | ASSFSQGGNSPLY | 0.11201455 |
| GGVGTAGAGGAGGAGG | ASSFSQGGNSPLY | 0.38348818 |
| GNGGGGDAGNAGSGGN | ASSFSQGGNSPLY | 0.4148056 |
| GTGGAGGAGNDAGS | ASSFSQGGNSPLY | 0.2569545 |

| | | |
|---|---|---|
| GTTNTGWFNS | ASSFSQGGNSPLY | 0.08978498 |
| IFMGPLFTL | ASSFSQGGNSPLY | 0.47133952 |
| ILAHAIEAI | ASSFSQGGNSPLY | 0.24022526 |
| IRSAIREAL | ASSFSQGGNSPLY | 0.12875934 |
| ITFAGGG | ASSFSQGGNSPLY | 0.15723392 |
| ITGMSGHQEV | ASSFSQGGNSPLY | 0.6990194 |
| IVAGLACTL | ASSFSQGGNSPLY | 0.13705318 |
| IVGGGLAGA | ASSFSQGGNSPLY | 0.15524337 |
| IVTGVLNI | ASSFSQGGNSPLY | 0.69951385 |
| LAQQGCI | ASSFSQGGNSPLY | 0.2412265 |
| LCLHVEN | ASSFSQGGNSPLY | 0.70937115 |
| LDAISDGS | ASSFSQGGNSPLY | 0.18661396 |
| LGGFAAML | ASSFSQGGNSPLY | 0.19870394 |
| LGGKNPMI | ASSFSQGGNSPLY | 0.4818299 |
| LGPKGVAAIL | ASSFSQGGNSPLY | 0.22507763 |
| LMVGVSI | ASSFSQGGNSPLY | 0.12179013 |
| LVGAAFVL | ASSFSQGGNSPLY | 0.0986933 |
| LVLAVAPL | ASSFSQGGNSPLY | 0.20571284 |
| MAAIAVIDAA | ASSFSQGGNSPLY | 0.12156275 |
| NNGINAELQ | ASSFSQGGNSPLY | 0.32302266 |
| RGEPLVLVHG | ASSFSQGGNSPLY | 0.55849975 |
| RGGAGGAAGLI | ASSFSQGGNSPLY | 0.19357434 |
| RSYRMHPAV | ASSFSQGGNSPLY | 0.15615994 |
| SAAHLLAGQLA | ASSFSQGGNSPLY | 0.23827288 |
| SAAVGSY | ASSFSQGGNSPLY | 0.17063983 |
| SPHAVLHRI | ASSFSQGGNSPLY | 0.6228283 |
| SQIAKVL | ASSFSQGGNSPLY | 0.25823414 |
| TALGHLVP | ASSFSQGGNSPLY | 0.68959993 |
| TAPTTSGAA | ASSFSQGGNSPLY | 0.3514551 |
| TDGDDGNGGSPGT | ASSFSQGGNSPLY | 0.14382449 |
| TDGLGEN | ASSFSQGGNSPLY | 0.6174756 |
| TGGPPLPGQA | ASSFSQGGNSPLY | 0.26048172 |
| TIPAITI | ASSFSQGGNSPLY | 0.49993336 |
| TIPGFTI | ASSFSQGGNSPLY | 0.40480912 |
| TSIASGALA | ASSFSQGGNSPLY | 0.26358837 |
| TVTLPGAAI | ASSFSQGGNSPLY | 0.2378237 |
| VASGLSAALP | ASSFSQGGNSPLY | 0.40589842 |
| VGAGVIGI | ASSFSQGGNSPLY | 0.11812729 |
| VGPKRFGV | ASSFSQGGNSPLY | 0.25014257 |
| VNVAFPD | ASSFSQGGNSPLY | 0.099514425 |
| VSMTNSGVS | ASSFSQGGNSPLY | 0.10728875 |
| VTDVLGDLSV | ASSFSQGGNSPLY | 0.23990077 |
| VVVVGAGFAGL | ASSFSQGGNSPLY | 0.7602552 |
| VYTVGTVV | ASSFSQGGNSPLY | 0.6957678 |
| WTTFSGP | ASSFSQGGNSPLY | 0.16455325 |
| YDELVEAF | ASSFSQGGNSPLY | 0.12237146 |
| YGTGGAGGNGG | ASSFSQGGNSPLY | 0.50685656 |
| YTNVMAV | ASSFSQGGNSPLY | 0.11300807 |
| FLEQVHQL | ASSFSQGGNSPLY | 0.448187 |
| GLSLLAVAV | ASSFSQGGNSPLY | 0.21456578 |
| LEQVHQL | ASSFSQGGNSPLY | 0.16415992 |

| | | |
|---|---|---|
| LNAAGAI | ASSFSQGGNSPLY | 0.32402676 |
| PAVHASL | ASSFSQGGNSPLY | 0.38213953 |
| TELEDDA | ASSFSQGGNSPLY | 0.11776039 |
| TPPGPPN | ASSFSQGGNSPLY | 0.48484117 |
| TPPPPPPP | ASSFSQGGNSPLY | 0.08827266 |
| TQSPALI | ASSPRDRDEQY | 0.20303151 |
| EFAAAFPQ | ASSPRDRDEQY | 0.16559587 |
| LLSGNLDVL | ASSPRDRDEQY | 0.14279935 |
| GPANLGNYNI | ASSPRDRDEQY | 0.11293141 |
| TLGPTLP | ASSPRDRDEQY | 0.306009 |
| VYAYSSCCK | ASSPRDRDEQY | 0.62842757 |
| APAPAPAPNI | ASSPRDRDEQY | 0.62841254 |
| PGGPVPIP | ASSPRDRDEQY | 0.17518522 |
| FLTVALEP | ASSPRDRDEQY | 0.696393 |
| SQYFGANL | ASSPRDRDEQY | 0.59659004 |
| GGVKPAA | ASSPRDRDEQY | 0.30309713 |
| GGTGGAGGAGGW | ASSPRDRDEQY | 0.116063856 |
| GVGGVAGLGGAGG | ASSPRDRDEQY | 0.25018883 |
| VLLVLATTI | ASSPRDRDEQY | 0.10172535 |
| IGIFAAML | ASSPRDRDEQY | 0.34879556 |
| AVGDLID | ASSPRDRDEQY | 0.65452605 |
| NISPLPL | ASSPRDRDEQY | 0.14782003 |
| VTAVFVAI | ASSPRDRDEQY | 0.41826883 |
| SGDDNDAAL | ASSPRDRDEQY | 0.6070503 |
| AVAALAHS | ASSPRDRDEQY | 0.15832593 |
| GGVTVMATP | ASSPRDRDEQY | 0.23235464 |
| AGLAANVRD | ASSPRDRDEQY | 0.17903921 |
| TNRPDLIDPALLRPG | ASSPRDRDEQY | 0.12404522 |
| LANLGVTAL | ASSPRDRDEQY | 0.22750323 |
| TGERIEL | ASSPRDRDEQY | 0.21787584 |
| NGEELAA | ASSPRDRDEQY | 0.12476383 |
| VLSLVWG | ASSPRDRDEQY | 0.24898754 |
| IGYALWL | ASSPRDRDEQY | 0.12949564 |
| LSLLLAPL | ASSPRDRDEQY | 0.14419259 |
| TLDEALK | ASSPRDRDEQY | 0.14471647 |
| TDGAGTAEGG | ASSPRDRDEQY | 0.14168993 |
| NAPGGAGGNGG | ASSPRDRDEQY | 0.4813227 |
| VSVFMGMGGNF | ASSPRDRDEQY | 0.14249624 |
| LASYLAAL | ASSPRDRDEQY | 0.37128782 |
| LVGGTPAL | ASSPRDRDEQY | 0.69430095 |
| VRNADVL | ASSPRDRDEQY | 0.09694683 |
| TCSSPFLVSAM | ASSPRDRDEQY | 0.11846981 |
| VHITDAE | ASSPRDRDEQY | 0.37017184 |
| EYAAPSF | ASSPRDRDEQY | 0.09900013 |
| VVDAFHGGL | ASSPRDRDEQY | 0.597966 |
| ASVVDPQ | ASSPRDRDEQY | 0.18592148 |
| SYNEFGGLS | ASSPRDRDEQY | 0.32586575 |
| DPDGSVY | ASSPRDRDEQY | 0.13901687 |
| MVSTIHATE | ASSPRDRDEQY | 0.7921488 |
| LAAGVADPV | ASSPRDRDEQY | 0.51702666 |
| TKEQIAA | ASSPRDRDEQY | 0.25029004 |

| | | |
|---|---|---|
| SLASPAAL | ASSPRDRDEQY | 0.21935731 |
| IGLVFYGV | ASSPRDRDEQY | 0.16601762 |
| FLLDASNL | ASSPRDRDEQY | 0.6562357 |
| YQVGNFP | ASSPRDRDEQY | 0.63001853 |
| ESLLSQST | ASSPRDRDEQY | 0.13566646 |
| LFTVGFAAVG | ASSPRDRDEQY | 0.3347566 |
| ATPPTPT | ASSPRDRDEQY | 0.6879068 |
| TDLATLG | ASSPRDRDEQY | 0.7579562 |
| SAGGTGR | ASSPRDRDEQY | 0.11136535 |
| GFSNSGS | ASSPRDRDEQY | 0.21366882 |
| GGNGGSGGTSV | ASSPRDRDEQY | 0.12910488 |
| GTSVATGGAGN | ASSPRDRDEQY | 0.20271339 |
| NGGNGGSGGTS | ASSPRDRDEQY | 0.13980977 |
| AVVKFPH | ASSPRDRDEQY | 0.6388709 |
| TPPPPPP | ASSPRDRDEQY | 0.3462854 |
| AAPAAAAAPA | ASSPRDRDEQY | 0.4414978 |
| IALNEDL | ASSPRDRDEQY | 0.2792311 |
| IVNVSSM | ASSPRDRDEQY | 0.40399033 |
| TGIPPAPR | ASSPRDRDEQY | 0.29107508 |
| FDLGGGTF | ASSPRDRDEQY | 0.14416791 |
| FDLGGGTFDVS | ASSPRDRDEQY | 0.1557102 |
| NEPTAAA | ASSPRDRDEQY | 0.10036578 |
| NEPTAAALA | ASSPRDRDEQY | 0.16804819 |
| GSIINISSI | ASSPRDRDEQY | 0.11618624 |
| VAPGNAGTA | ASSPRDRDEQY | 0.089479804 |
| TGGGASLE | ASSPRDRDEQY | 0.3997205 |
| VAVYNHYKRI | ASSPRDRDEQY | 0.42984828 |
| TDPAPATT | ASSPRDRDEQY | 0.28039432 |
| TNRPDILDPALLRPG | ASSPRDRDEQY | 0.14640528 |
| LTTTTTS | ASSPRDRDEQY | 0.2922197 |
| PTGSGKTL | ASSPRDRDEQY | 0.11413877 |
| TGSGKTL | ASSPRDRDEQY | 0.6962968 |
| TIGHVDHGK | ASSPRDRDEQY | 0.1381042 |
| TIGHVDHGKTT | ASSPRDRDEQY | 0.11333095 |
| GHVDHGKTT | ASSPRDRDEQY | 0.15752426 |
| AQFEHTLL | ASSPRDRDEQY | 0.113862455 |
| IVNQDEL | ASSPRDRDEQY | 0.72757095 |
| TNGSQFF | ASSPRDRDEQY | 0.5106181 |
| TGKGIVE | ASSPRDRDEQY | 0.4481354 |
| LISNASDA | ASSPRDRDEQY | 0.49007076 |
| AELRTLGELTKIAKAHGAQ | ASSPRDRDEQY | 0.106225066 |
| APNAVAAAEPSW | ASSPRDRDEQY | 0.12936051 |
| APQVAGAATAVL | ASSPRDRDEQY | 0.72946507 |
| ARRDPAAAAAILPTDA | ASSPRDRDEQY | 0.27550277 |
| DAAARDVLAEAGLAEYF | ASSPRDRDEQY | 0.4033143 |
| DDQIVLSAKANLAKVEQ | ASSPRDRDEQY | 0.59451044 |
| DELLPDTGEDSDEDGH | ASSPRDRDEQY | 0.122008234 |
| ELPEAPANRATVPQD | ASSPRDRDEQY | 0.26499838 |
| FGQSKIERVTGIG | ASSPRDRDEQY | 0.19593605 |
| GAGGTGSTAGGAGGAGGAGGLY | ASSPRDRDEQY | 0.5717959 |
| GHGGTGAAGGNGGTGGAGGSGI | ASSPRDRDEQY | 0.40570694 |

| | | |
|---|---|---|
| GNGGNGGAGGEGGPGGAGGAGA | ASSPRDRDEQY | 0.14734824 |
| GSTAGNGGNGGNATNGTIAGQP | ASSPRDRDEQY | 0.5827508 |
| GTGGDGGNGGNGGTGG | ASSPRDRDEQY | 0.7589372 |
| GVRRPAGIPADAPV | ASSPRDRDEQY | 0.39207435 |
| GYRQVTDAHLLTLARR | ASSPRDRDEQY | 0.2901687 |
| IAKMGQDGHDRG | ASSPRDRDEQY | 0.35799363 |
| KAEPSIEIQYT | ASSPRDRDEQY | 0.14650513 |
| KPVRIRMAADEDTVAVT | ASSPRDRDEQY | 0.2350454 |
| MPPRIAGMRLLVIKPEP | ASSPRDRDEQY | 0.18555725 |
| MKFYAVALFRSRGVPPTR | ASSPRDRDEQY | 0.15509965 |
| NAGGFGILGAVAHSP | ASSPRDRDEQY | 0.14819992 |
| NEGREITLAMAEM | ASSPRDRDEQY | 0.4201786 |
| NELVEWSHPLGHRWRWE | ASSPRDRDEQY | 0.18090847 |
| NLGAGGTGGNGGNPG | ASSPRDRDEQY | 0.7855717 |
| NTEWPKWETVMAADPVD | ASSPRDRDEQY | 0.5205704 |
| PASAAPAETQP | ASSPRDRDEQY | 0.29499388 |
| QQSSDGKNGVRAAEITGE | ASSPRDRDEQY | 0.107629456 |
| RAPDAVTTSWWKE | ASSPRDRDEQY | 0.39787674 |
| SGFGNVGAHTSGWFNQ | ASSPRDRDEQY | 0.32775566 |
| SGVAGAVGQAASVGGLKV | ASSPRDRDEQY | 0.123328246 |
| SNGGAGTGSAGGNGGT | ASSPRDRDEQY | 0.6026962 |
| TGGAGGEGGAAG | ASSPRDRDEQY | 0.14779401 |
| TGGDGGAAGAGGAPGHG | ASSPRDRDEQY | 0.23319995 |
| TMPPLLAAIANAGRHGV | ASSPRDRDEQY | 0.3163453 |
| TSSTAGNSGGA | ASSPRDRDEQY | 0.63102317 |
| VPATVPVGILC | ASSPRDRDEQY | 0.35053152 |
| VQRPDSGGVVLTGRIS | ASSPRDRDEQY | 0.13111305 |
| WTPNPVPATKGVPAGLVQAT | ASSPRDRDEQY | 0.11252029 |
| YALKRALAKFSDRFDVVII | ASSPRDRDEQY | 0.57679236 |
| YLIGSVIV | ASSPRDRDEQY | 0.69364476 |
| YGQLVAVID | ASSPRDRDEQY | 0.15773264 |
| VTVGGMTLPA | ASSPRDRDEQY | 0.73275125 |
| VLALPAV | ASSPRDRDEQY | 0.2089128 |
| VIPPTLNYET | ASSPRDRDEQY | 0.1355703 |
| VGMGGMHPSSG | ASSPRDRDEQY | 0.48115778 |
| VGLAGGAATA | ASSPRDRDEQY | 0.48731178 |
| VGGAGGAGGSGP | ASSPRDRDEQY | 0.7156398 |
| VFAAFMV | ASSPRDRDEQY | 0.371302 |
| TLYDKLR | ASSPRDRDEQY | 0.69462085 |
| TLAVFTVL | ASSPRDRDEQY | 0.34473303 |
| TGSVQPARV | ASSPRDRDEQY | 0.6134513 |
| TGGGAGSGGT | ASSPRDRDEQY | 0.6828336 |
| SHLETSL | ASSPRDRDEQY | 0.4093644 |
| SFTTGMT | ASSPRDRDEQY | 0.25218192 |
| QVEVAKRMGVRQPTVS | ASSPRDRDEQY | 0.1688377 |
| QGSGGPALAM | ASSPRDRDEQY | 0.38202834 |
| MLAPSNP | ASSPRDRDEQY | 0.4765189 |
| LVNNAAI | ASSPRDRDEQY | 0.20135099 |
| LTGEINTLA | ASSPRDRDEQY | 0.43012872 |
| LSIFGPV | ASSPRDRDEQY | 0.2921831 |
| LLALLANL | ASSPRDRDEQY | 0.14877386 |

| | | |
|---|---|---|
| LISNASDALDKLR | ASSPRDRDEQY | 0.3461002 |
| LINEAVP | ASSPRDRDEQY | 0.7511883 |
| LGYAFSGL | ASSPRDRDEQY | 0.403849 |
| LDFVAVAL | ASSPRDRDEQY | 0.14172786 |
| LAQPGAE | ASSPRDRDEQY | 0.6892929 |
| ISNLGAVTGF | ASSPRDRDEQY | 0.7910038 |
| HQEDPGRI | ASSPRDRDEQY | 0.64903116 |
| GTPGEPTP | ASSPRDRDEQY | 0.110027745 |
| GFFNSGS | ASSPRDRDEQY | 0.098892845 |
| GAGGGGLTAGAN | ASSPRDRDEQY | 0.4108026 |
| FVDALVNM | ASSPRDRDEQY | 0.13665739 |
| FLTVAPEL | ASSPRDRDEQY | 0.131556 |
| FETLTAL | ASSPRDRDEQY | 0.13598646 |
| DVINPIEL | ASSPRDRDEQY | 0.5680616 |
| APVSGAPV | ASSPRDRDEQY | 0.3994227 |
| ALVGGLAW | ASSPRDRDEQY | 0.25708205 |
| ALAAAAATAI | ASSPRDRDEQY | 0.11581874 |
| AFGLGLLL | ASSPRDRDEQY | 0.5033798 |
| AFFAAASL | ASSPRDRDEQY | 0.57753146 |
| AAAALPAFSPP | ASSPRDRDEQY | 0.17172107 |
| VVLAPAL | ASSPRDRDEQY | 0.3338533 |
| TIGHVDHGKTTL | ASSPRDRDEQY | 0.26069367 |
| LISNASDALDK | ASSPRDRDEQY | 0.1860582 |
| LISNASDAL | ASSPRDRDEQY | 0.22552304 |
| ISNASDALDK | ASSPRDRDEQY | 0.13652423 |
| NQAETLVYQTE | ASSPRDRDEQY | 0.48860922 |
| TGGTGGTAGAG | ASSPRDRDEQY | 0.14253643 |
| TGSTAGGAGGA | ASSPRDRDEQY | 0.62758684 |
| DPLAVPVAAGPV | ASSPRDRDEQY | 0.5207359 |
| GGTGGTPGNGGN | ASSPRDRDEQY | 0.5214515 |
| PAGLAGVDPATL | ASSPRDRDEQY | 0.4066934 |
| GNNLVVEAPHAGASV | ASSPRDRDEQY | 0.1860876 |
| KVTRIMSASTLEQPA | ASSPRDRDEQY | 0.124624394 |
| LGFALPIALPRRTVV | ASSPRDRDEQY | 0.094243206 |
| TRLGNLTARKLVVLV | ASSPRDRDEQY | 0.7668693 |
| WSQQRRLIYFPSAGP | ASSPRDRDEQY | 0.11643118 |
| GPNGNPDPMAAAVDIR | ASSPRDRDEQY | 0.7501404 |
| TQRCVSVADLRAAASR | ASSPRDRDEQY | 0.2791307 |
| QFVLDVAEHPVEVLTETV | ASSPRDRDEQY | 0.16903691 |
| GGAGGTGGAGGNGGTQAGDG | ASSPRDRDEQY | 0.42224815 |
| PLLATLRAIAVDVDAKPAQV | ASSPRDRDEQY | 0.099837214 |
| APKFRTGGYMIDFWGVGYQVAK | ASSPRDRDEQY | 0.21145661 |
| GAGGAGGAGGAAGGPGGTGGTG | ASSPRDRDEQY | 0.5633147 |
| IGALGLLATDVAPAGSVAFGVA | ASSPRDRDEQY | 0.385917 |
| AGGAGGIGGNGGAGGAGGVLMG | ASSPRDRDEQY | 0.25039554 |
| ADELIGLL | ASSPRDRDEQY | 0.4642524 |
| ADRALKA | ASSPRDRDEQY | 0.5700574 |
| AGAVGQVGGAGGA | ASSPRDRDEQY | 0.18690062 |
| AGGGGGLGGAP | ASSPRDRDEQY | 0.4002211 |
| AGGLLYGNGGAG | ASSPRDRDEQY | 0.56185555 |
| AGLESIGL | ASSPRDRDEQY | 0.4685247 |

| | | |
|---|---|---|
| AGLKTGII | ASSPRDRDEQY | 0.15096472 |
| ASLAALLL | ASSPRDRDEQY | 0.23353511 |
| ASVINISSI | ASSPRDRDEQY | 0.28889787 |
| ATPPGPA | ASSPRDRDEQY | 0.36520335 |
| ATVASALV | ASSPRDRDEQY | 0.35885894 |
| ATVASLIL | ASSPRDRDEQY | 0.3828861 |
| AVNGPVLIRH | ASSPRDRDEQY | 0.4921004 |
| DGAAGAVGI | ASSPRDRDEQY | 0.11327847 |
| DPLLHGI | ASSPRDRDEQY | 0.1851141 |
| ELGAALGAL | ASSPRDRDEQY | 0.39938098 |
| ELGRTQAV | ASSPRDRDEQY | 0.1996685 |
| FVAGGYLS | ASSPRDRDEQY | 0.4342634 |
| GASGGAGGAGGW | ASSPRDRDEQY | 0.1241996 |
| GGAGAGGDGGAGGI | ASSPRDRDEQY | 0.47725004 |
| GTGHNGSQPGMGGNG | ASSPRDRDEQY | 0.19344857 |
| GYAAASASAS | ASSPRDRDEQY | 0.16362843 |
| HQVDVAAVQQG | ASSPRDRDEQY | 0.36433095 |
| IGAAIAHT | ASSPRDRDEQY | 0.21291395 |
| IGMIEVAQL | ASSPRDRDEQY | 0.19170178 |
| IRLAPEHTSAMD | ASSPRDRDEQY | 0.53665435 |
| IYNVSVL | ASSPRDRDEQY | 0.14351346 |
| KQGFLGM | ASSPRDRDEQY | 0.2232692 |
| LESVITQL | ASSPRDRDEQY | 0.4101851 |
| LGLGALVVV | ASSPRDRDEQY | 0.19404405 |
| LIGNGGNGGSGGI | ASSPRDRDEQY | 0.5424999 |
| LNAINAPTL | ASSPRDRDEQY | 0.22488943 |
| LQLLNTGV | ASSPRDRDEQY | 0.6772342 |
| LQTSIEF | ASSPRDRDEQY | 0.1333924 |
| MSFAVSEL | ASSPRDRDEQY | 0.13456242 |
| NTGALGL | ASSPRDRDEQY | 0.46936652 |
| PPGAGGAGGAGTT | ASSPRDRDEQY | 0.27208835 |
| QLVGLGI | ASSPRDRDEQY | 0.68372464 |
| RFLEQVHQL | ASSPRDRDEQY | 0.45285863 |
| RGVGYVLR | ASSPRDRDEQY | 0.57517564 |
| SATGVFAGVI | ASSPRDRDEQY | 0.56414706 |
| SLPAITLP | ASSPRDRDEQY | 0.20269501 |
| STALAFTV | ASSPRDRDEQY | 0.12667157 |
| SVLTVGLGK | ASSPRDRDEQY | 0.13281913 |
| TALAGAGSGI | ASSPRDRDEQY | 0.10893681 |
| TGLPFTGI | ASSPRDRDEQY | 0.22969563 |
| VAEHVTV | ASSPRDRDEQY | 0.114657834 |
| VDIVLNSL | ASSPRDRDEQY | 0.6251844 |
| VLTGGGAVL | ASSPRDRDEQY | 0.7870013 |
| VNVGLAYGI | ASSPRDRDEQY | 0.120358005 |
| VSSAAVGAV | ASSPRDRDEQY | 0.7720432 |
| VTSVYEAGV | ASSPRDRDEQY | 0.12930317 |
| WVALDRGA | ASSPRDRDEQY | 0.2933897 |
| YLVGSVVI | ASSPRDRDEQY | 0.68252015 |
| GPPGTGKTLL | ASSPRDRDEQY | 0.12169386 |
| GVGMTKF | ASSPRDRDEQY | 0.27241734 |
| IGIGGSDL | ASSPRDRDEQY | 0.18465893 |

| | | |
|---|---|---|
| IGTLLAEL | ASSPRDRDEQY | 0.16092911 |
| IYTSGTT | ASSPRDRDEQY | 0.7710456 |
| LSFPAGV | ASSPRDRDEQY | 0.14271952 |
| LYTSGTT | ASSPRDRDEQY | 0.47789997 |
| VGAFPFGF | ASSPRDRDEQY | 0.15873851 |
| VGAGTVEF | ASSPRDRDEQY | 0.12428792 |
| VGKSTLLL | ASSPRDRDEQY | 0.45522267 |
| AALAHAGGGIV | ASSPRDRDEQY | 0.7022457 |
| AALVAVSLL | ASSPRDRDEQY | 0.12867408 |
| AAVSRAVAAL | ASSPRDRDEQY | 0.6517875 |
| AGGSSGIGGF | ASSPRDRDEQY | 0.20798753 |
| AGLTVANANG | ASSPRDRDEQY | 0.6696615 |
| AGLVGAVL | ASSPRDRDEQY | 0.62750614 |
| ALSQAFTV | ASSPRDRDEQY | 0.3206321 |
| ALTFSVSL | ASSPRDRDEQY | 0.2320638 |
| ALTGAALVAL | ASSPRDRDEQY | 0.58148074 |
| ASFSIGAGVL | ASSPRDRDEQY | 0.43476743 |
| DGSVYKI | ASSPRDRDEQY | 0.6673432 |
| EQLAEIV | ASSPRDRDEQY | 0.5861404 |
| FAVSELTVM | ASSPRDRDEQY | 0.13484536 |
| GAAPLAAGL | ASSPRDRDEQY | 0.55243844 |
| GADGGDGGSGGAGG | ASSPRDRDEQY | 0.28190315 |
| GFGLDTGGAGG | ASSPRDRDEQY | 0.23539498 |
| GLASGGLL | ASSPRDRDEQY | 0.19134203 |
| GRIAGAAVLL | ASSPRDRDEQY | 0.12354391 |
| GSAGSLNTSF | ASSPRDRDEQY | 0.17275675 |
| GTGQIGI | ASSPRDRDEQY | 0.16397008 |
| GTLIALIL | ASSPRDRDEQY | 0.24353915 |
| IAATMGFVVG | ASSPRDRDEQY | 0.20929071 |
| IFAQALL | ASSPRDRDEQY | 0.17631596 |
| IGAAVALGF | ASSPRDRDEQY | 0.6282721 |
| IGLTPLAL | ASSPRDRDEQY | 0.5251955 |
| ILGSNTSTL | ASSPRDRDEQY | 0.119287595 |
| IVFSLTVL | ASSPRDRDEQY | 0.15865211 |
| LASKATGF | ASSPRDRDEQY | 0.16401522 |
| LASVALIL | ASSPRDRDEQY | 0.1940791 |
| LFAVGLTL | ASSPRDRDEQY | 0.0972208 |
| LFNGGIGTYI | ASSPRDRDEQY | 0.26887837 |
| LTLGAGII | ASSPRDRDEQY | 0.124076195 |
| LVAAALGDI | ASSPRDRDEQY | 0.09415726 |
| NTAFVQLGI | ASSPRDRDEQY | 0.18872224 |
| QLFSGAEL | ASSPRDRDEQY | 0.17246714 |
| SAATFITL | ASSPRDRDEQY | 0.1774439 |
| SNYGLANLGV | ASSPRDRDEQY | 0.60974634 |
| SYGGAYAVM | ASSPRDRDEQY | 0.1016881 |
| TTGVGGLTLG | ASSPRDRDEQY | 0.14682539 |
| TTQTIYI | ASSPRDRDEQY | 0.5225999 |
| VAAGAGLGAVI | ASSPRDRDEQY | 0.66649294 |
| VAAVVTLL | ASSPRDRDEQY | 0.14588684 |
| VGFLGGLI | ASSPRDRDEQY | 0.28717142 |
| VGVIPILL | ASSPRDRDEQY | 0.55999196 |

| | | |
|---|---|---|
| VIFASSLI | ASSPRDRDEQY | 0.29408383 |
| VKGGDGGAAAG | ASSPRDRDEQY | 0.29095718 |
| VTKTFTNI | ASSPRDRDEQY | 0.3134051 |
| VVAAAILL | ASSPRDRDEQY | 0.11353862 |
| YAALDVEL | ASSPRDRDEQY | 0.51224947 |
| LASVALLL | ASSPRDRDEQY | 0.6232573 |
| LDAVHRV | ASSPRDRDEQY | 0.18086599 |
| QGLSSSIF | ASSPRDRDEQY | 0.28043333 |
| RGTFANIRL | ASSPRDRDEQY | 0.13951507 |
| RQLWWGHRI | ASSPRDRDEQY | 0.14653306 |
| YGPPGTGKTLL | ASSPRDRDEQY | 0.101085305 |
| AALVGAPANVA | ASSPRDRDEQY | 0.43509075 |
| AATFRRTE | ASSPRDRDEQY | 0.34746078 |
| AGALGGGI | ASSPRDRDEQY | 0.24904916 |
| AGGAGGAGGIGAG | ASSPRDRDEQY | 0.12170321 |
| AGSGALGL | ASSPRDRDEQY | 0.12964812 |
| AQIADQI | ASSPRDRDEQY | 0.12048795 |
| ASAVAVTYL | ASSPRDRDEQY | 0.16930576 |
| ASTVLGGTGG | ASSPRDRDEQY | 0.10514695 |
| AVGGEPAK | ASSPRDRDEQY | 0.119349115 |
| CEQGTFGGTC | ASSPRDRDEQY | 0.52893126 |
| DEGKAPAG | ASSPRDRDEQY | 0.44101757 |
| DTVVFPAL | ASSPRDRDEQY | 0.12724474 |
| FEGITMAG | ASSPRDRDEQY | 0.55592734 |
| FPLVVQD | ASSPRDRDEQY | 0.32957703 |
| GAFTTPQ | ASSPRDRDEQY | 0.3934288 |
| GFGGFGTAGGVGGA | ASSPRDRDEQY | 0.19586018 |
| GFLAQFL | ASSPRDRDEQY | 0.13954425 |
| GGAGGAGGNAGVA | ASSPRDRDEQY | 0.6155782 |
| GGVGTAGAGGAGGAGG | ASSPRDRDEQY | 0.321238 |
| GNGGGGDAGNAGSGGN | ASSPRDRDEQY | 0.14342558 |
| GTGGAGGAGNDAGS | ASSPRDRDEQY | 0.30484197 |
| GTTNTGWFNS | ASSPRDRDEQY | 0.2003544 |
| IFMGPLFTL | ASSPRDRDEQY | 0.18200956 |
| ILAHAIEAI | ASSPRDRDEQY | 0.29753816 |
| IRSAIREAL | ASSPRDRDEQY | 0.20613553 |
| ITFAGGG | ASSPRDRDEQY | 0.14539701 |
| ITGMSGHQEV | ASSPRDRDEQY | 0.1519633 |
| IVAGLACTL | ASSPRDRDEQY | 0.13071898 |
| IVGGGLAGA | ASSPRDRDEQY | 0.322429 |
| IVTGVLNI | ASSPRDRDEQY | 0.56584257 |
| LAQQGCI | ASSPRDRDEQY | 0.21265292 |
| LCLHVEN | ASSPRDRDEQY | 0.15264842 |
| LDAISDGS | ASSPRDRDEQY | 0.42222846 |
| LGGFAAML | ASSPRDRDEQY | 0.2862217 |
| LGGKNPMI | ASSPRDRDEQY | 0.25287542 |
| LGPKGVAAIL | ASSPRDRDEQY | 0.27610713 |
| LMVGVSI | ASSPRDRDEQY | 0.23160854 |
| LVGAAFVL | ASSPRDRDEQY | 0.26404846 |
| LVLAVAPL | ASSPRDRDEQY | 0.35175607 |
| MAAIAVIDAA | ASSPRDRDEQY | 0.2499431 |

| | | |
|---|---|---|
| NNGINAELQ | ASSPRDRDEQY | 0.2667026 |
| RGEPLVLVHG | ASSPRDRDEQY | 0.18390527 |
| RGGAGGAAGLI | ASSPRDRDEQY | 0.19198044 |
| RSYRMHPAV | ASSPRDRDEQY | 0.11236169 |
| SAAHLLAGQLA | ASSPRDRDEQY | 0.1386014 |
| SAAVGSY | ASSPRDRDEQY | 0.21980241 |
| SPHAVLHRI | ASSPRDRDEQY | 0.33457074 |
| SQIAKVL | ASSPRDRDEQY | 0.2786212 |
| TALGHLVP | ASSPRDRDEQY | 0.20845456 |
| TAPTTSGAA | ASSPRDRDEQY | 0.13290441 |
| TDGDDGNGGSPGT | ASSPRDRDEQY | 0.27260855 |
| TDGLGEN | ASSPRDRDEQY | 0.49561375 |
| TGGPPLPGQA | ASSPRDRDEQY | 0.13074401 |
| TIPAITI | ASSPRDRDEQY | 0.18716095 |
| TIPGFTI | ASSPRDRDEQY | 0.52264404 |
| TSIASGALA | ASSPRDRDEQY | 0.38973704 |
| TVTLPGAAI | ASSPRDRDEQY | 0.2229532 |
| VASGLSAALP | ASSPRDRDEQY | 0.179064 |
| VGAGVIGI | ASSPRDRDEQY | 0.45864844 |
| VGPKRFGV | ASSPRDRDEQY | 0.12755413 |
| VNVAFPD | ASSPRDRDEQY | 0.45759356 |
| VSMTNSGVS | ASSPRDRDEQY | 0.10576162 |
| VTDVLGDLSV | ASSPRDRDEQY | 0.13959047 |
| VVVVGAGFAGL | ASSPRDRDEQY | 0.13699088 |
| VYTVGTVV | ASSPRDRDEQY | 0.10930038 |
| WTTFSGP | ASSPRDRDEQY | 0.47469905 |
| YDELVEAF | ASSPRDRDEQY | 0.12854739 |
| YGTGGAGGNGG | ASSPRDRDEQY | 0.43871891 |
| YTNVMAV | ASSPRDRDEQY | 0.15102607 |
| FLEQVHQL | ASSPRDRDEQY | 0.40161395 |
| GLSLLAVAV | ASSPRDRDEQY | 0.34239388 |
| LEQVHQL | ASSPRDRDEQY | 0.18591604 |
| LNAAGAI | ASSPRDRDEQY | 0.14730398 |
| PAVHASL | ASSPRDRDEQY | 0.47967404 |
| TELEDDA | ASSPRDRDEQY | 0.2592749 |
| TPPGPPN | ASSPRDRDEQY | 0.20912069 |
| TPPPPPPP | ASSPRDRDEQY | 0.12699386 |
| TQSPALI | ASSQDRGQEGSDYT | 0.12251254 |
| EFAAAFPQ | ASSQDRGQEGSDYT | 0.32408857 |
| LLSGNLDVL | ASSQDRGQEGSDYT | 0.105093256 |
| GPANLGNYNI | ASSQDRGQEGSDYT | 0.09638164 |
| TLGPTLP | ASSQDRGQEGSDYT | 0.11390524 |
| VYAYSSCCK | ASSQDRGQEGSDYT | 0.12134283 |
| APAPAPAPNI | ASSQDRGQEGSDYT | 0.51305115 |
| PGGPVPIP | ASSQDRGQEGSDYT | 0.537433 |
| FLTVALEP | ASSQDRGQEGSDYT | 0.23784935 |
| SQYFGANL | ASSQDRGQEGSDYT | 0.3394546 |
| GGVKPAA | ASSQDRGQEGSDYT | 0.33376443 |
| GGTGGAGGAGGW | ASSQDRGQEGSDYT | 0.09103736 |
| GVGGVAGLGGAGG | ASSQDRGQEGSDYT | 0.14477253 |
| VLLVLATTI | ASSQDRGQEGSDYT | 0.123399734 |

| | | |
|---|---|---|
| IGIFAAML | ASSQDRGQEGSDYT | 0.31806487 |
| AVGDLID | ASSQDRGQEGSDYT | 0.12907617 |
| NISPLPL | ASSQDRGQEGSDYT | 0.16379243 |
| VTAVFVAI | ASSQDRGQEGSDYT | 0.27995557 |
| SGDDNDAAL | ASSQDRGQEGSDYT | 0.21415305 |
| AVAALAHS | ASSQDRGQEGSDYT | 0.2627918 |
| GGVTVMATP | ASSQDRGQEGSDYT | 0.12222825 |
| AGLAANVRD | ASSQDRGQEGSDYT | 0.093566835 |
| TNRPDLIDPALLRPG | ASSQDRGQEGSDYT | 0.4087432 |
| LANLGVTAL | ASSQDRGQEGSDYT | 0.1038791 |
| TGERIEL | ASSQDRGQEGSDYT | 0.1547979 |
| NGEELAA | ASSQDRGQEGSDYT | 0.09541841 |
| VLSLVWG | ASSQDRGQEGSDYT | 0.107053176 |
| IGYALWL | ASSQDRGQEGSDYT | 0.11589305 |
| LSLLLAPL | ASSQDRGQEGSDYT | 0.25898474 |
| TLDEALK | ASSQDRGQEGSDYT | 0.19500345 |
| TDGAGTAEGG | ASSQDRGQEGSDYT | 0.29683745 |
| NAPGGAGGNGG | ASSQDRGQEGSDYT | 0.728829 |
| VSVFMGMGGNF | ASSQDRGQEGSDYT | 0.6442801 |
| LASYLAAL | ASSQDRGQEGSDYT | 0.22791582 |
| LVGGTPAL | ASSQDRGQEGSDYT | 0.16015048 |
| VRNADVL | ASSQDRGQEGSDYT | 0.13405113 |
| TCSSPFLVSAM | ASSQDRGQEGSDYT | 0.14862727 |
| VHITDAE | ASSQDRGQEGSDYT | 0.15856117 |
| EYAAPSF | ASSQDRGQEGSDYT | 0.6028396 |
| VVDAFHGGL | ASSQDRGQEGSDYT | 0.15214612 |
| ASVVDPQ | ASSQDRGQEGSDYT | 0.11993947 |
| SYNEFGGLS | ASSQDRGQEGSDYT | 0.17317396 |
| DPDGSVY | ASSQDRGQEGSDYT | 0.49509704 |
| MVSTIHATE | ASSQDRGQEGSDYT | 0.6771926 |
| LAAGVADPV | ASSQDRGQEGSDYT | 0.13345306 |
| TKEQIAA | ASSQDRGQEGSDYT | 0.20374918 |
| SLASPAAL | ASSQDRGQEGSDYT | 0.18635276 |
| IGLVFYGV | ASSQDRGQEGSDYT | 0.4072407 |
| FLLDASNL | ASSQDRGQEGSDYT | 0.13600327 |
| YQVGNFP | ASSQDRGQEGSDYT | 0.1501924 |
| ESLLSQST | ASSQDRGQEGSDYT | 0.22293706 |
| LFTVGFAAVG | ASSQDRGQEGSDYT | 0.73116255 |
| ATPPTPT | ASSQDRGQEGSDYT | 0.16518943 |
| TDLATLG | ASSQDRGQEGSDYT | 0.10921624 |
| SAGGTGR | ASSQDRGQEGSDYT | 0.2722777 |
| GFSNSGS | ASSQDRGQEGSDYT | 0.14171338 |
| GGNGGSGGTSV | ASSQDRGQEGSDYT | 0.19723934 |
| GTSVATGGAGN | ASSQDRGQEGSDYT | 0.21075949 |
| NGGNGGSGGTS | ASSQDRGQEGSDYT | 0.12109853 |
| AVVKFPH | ASSQDRGQEGSDYT | 0.16725156 |
| TPPPPPP | ASSQDRGQEGSDYT | 0.6514611 |
| AAPAAAAAPA | ASSQDRGQEGSDYT | 0.27996513 |
| IALNEDL | ASSQDRGQEGSDYT | 0.29388568 |
| IVNVSSM | ASSQDRGQEGSDYT | 0.5957409 |
| TGIPPAPR | ASSQDRGQEGSDYT | 0.42711666 |

| | | |
|---|---|---|
| FDLGGGTF | ASSQDRGQEGSDYT | 0.69603294 |
| FDLGGGTFDVS | ASSQDRGQEGSDYT | 0.28642207 |
| NEPTAAA | ASSQDRGQEGSDYT | 0.16417027 |
| NEPTAAALA | ASSQDRGQEGSDYT | 0.20988256 |
| GSIINISSI | ASSQDRGQEGSDYT | 0.27561614 |
| VAPGNAGTA | ASSQDRGQEGSDYT | 0.2990098 |
| TGGGASLE | ASSQDRGQEGSDYT | 0.22718213 |
| VAVYNHYKRI | ASSQDRGQEGSDYT | 0.14874354 |
| TDPAPATT | ASSQDRGQEGSDYT | 0.57673293 |
| TNRPDILDPALLRPG | ASSQDRGQEGSDYT | 0.22137982 |
| LTTTTTS | ASSQDRGQEGSDYT | 0.28081265 |
| PTGSGKTL | ASSQDRGQEGSDYT | 0.1137926 |
| TGSGKTL | ASSQDRGQEGSDYT | 0.15370071 |
| TIGHVDHGK | ASSQDRGQEGSDYT | 0.5186044 |
| TIGHVDHGKTT | ASSQDRGQEGSDYT | 0.09380735 |
| GHVDHGKTT | ASSQDRGQEGSDYT | 0.09742149 |
| AQFEHTLL | ASSQDRGQEGSDYT | 0.17440507 |
| IVNQDEL | ASSQDRGQEGSDYT | 0.48582911 |
| TNGSQFF | ASSQDRGQEGSDYT | 0.092524655 |
| TGKGIVE | ASSQDRGQEGSDYT | 0.1445826 |
| LISNASDA | ASSQDRGQEGSDYT | 0.11626343 |
| AELRTLGELTKIAKAHGAQ | ASSQDRGQEGSDYT | 0.16518104 |
| APNAVAAAEPSW | ASSQDRGQEGSDYT | 0.24039951 |
| APQVAGAATAVL | ASSQDRGQEGSDYT | 0.11633473 |
| ARRDPAAAAAILPTDA | ASSQDRGQEGSDYT | 0.29462105 |
| DAAARDVLAEAGLAEYF | ASSQDRGQEGSDYT | 0.10938223 |
| DDQIVLSAKANLAKVEQ | ASSQDRGQEGSDYT | 0.113981515 |
| DELLPDTGEDSDEDGH | ASSQDRGQEGSDYT | 0.3268372 |
| ELPEAPANRATVPQD | ASSQDRGQEGSDYT | 0.5375809 |
| FGQSKIERVTGIG | ASSQDRGQEGSDYT | 0.116695665 |
| GAGGTGSTAGGAGGAGGAGGLY | ASSQDRGQEGSDYT | 0.3677388 |
| GHGGTGAAGGNGGTGGAGGSGI | ASSQDRGQEGSDYT | 0.12783222 |
| GNGGNGGAGGEGGPGGAGGAGA | ASSQDRGQEGSDYT | 0.732106 |
| GSTAGNGGNGGNATNGTIAGQP | ASSQDRGQEGSDYT | 0.23158014 |
| GTGGDGGNGGNGGTGG | ASSQDRGQEGSDYT | 0.108116195 |
| GVRRPAGIPADAPV | ASSQDRGQEGSDYT | 0.38704175 |
| GYRQVTDAHLLTLARR | ASSQDRGQEGSDYT | 0.26656577 |
| IAKMGQDGHDRG | ASSQDRGQEGSDYT | 0.2046184 |
| KAEPSIEIQYT | ASSQDRGQEGSDYT | 0.39168778 |
| KPVRIRMAADEDTVAVT | ASSQDRGQEGSDYT | 0.12790963 |
| MPPRIAGMRLLVIKPEP | ASSQDRGQEGSDYT | 0.26690513 |
| MKFYAVALFRSRGVPPTR | ASSQDRGQEGSDYT | 0.60677385 |
| NAGGFGILGAVAHSP | ASSQDRGQEGSDYT | 0.30670834 |
| NEGREITLAMAEM | ASSQDRGQEGSDYT | 0.14455116 |
| NELVEWSHPLGHRWRWE | ASSQDRGQEGSDYT | 0.2429706 |
| NLGAGGTGGNGGNPG | ASSQDRGQEGSDYT | 0.2955262 |
| NTEWPKWETVMAADPVD | ASSQDRGQEGSDYT | 0.14104474 |
| PASAAPAETQP | ASSQDRGQEGSDYT | 0.61940885 |
| QQSSDGKNGVRAAEITGE | ASSQDRGQEGSDYT | 0.18025088 |
| RAPDAVTTSWWKE | ASSQDRGQEGSDYT | 0.48063082 |
| SGFGNVGAHTSGWFNQ | ASSQDRGQEGSDYT | 0.30364442 |

| | | |
|---|---|---|
| SGVAGAVGQAASVGGLKV | ASSQDRGQEGSDYT | 0.49837503 |
| SNGGAGTGSAGGNGGT | ASSQDRGQEGSDYT | 0.1330551 |
| TGGAGGEGGAAG | ASSQDRGQEGSDYT | 0.1709265 |
| TGGDGGAAGAGGAPGHG | ASSQDRGQEGSDYT | 0.2838685 |
| TMPPLLAAIANAGRHGV | ASSQDRGQEGSDYT | 0.38721627 |
| TSSTAGNSGGA | ASSQDRGQEGSDYT | 0.13606219 |
| VPATVPVGILC | ASSQDRGQEGSDYT | 0.11589124 |
| VQRPDSGGVVLTGRIS | ASSQDRGQEGSDYT | 0.20281592 |
| WTPNPVPATKGVPAGLVQAT | ASSQDRGQEGSDYT | 0.5200552 |
| YALKRALAKFSDRFDVVII | ASSQDRGQEGSDYT | 0.16172928 |
| YLIGSVIV | ASSQDRGQEGSDYT | 0.20300281 |
| YGQLVAVID | ASSQDRGQEGSDYT | 0.7232636 |
| VTVGGMTLPA | ASSQDRGQEGSDYT | 0.1300853 |
| VLALPAV | ASSQDRGQEGSDYT | 0.24888597 |
| VIPPTLNYET | ASSQDRGQEGSDYT | 0.3038852 |
| VGMGGMHPSSG | ASSQDRGQEGSDYT | 0.4122492 |
| VGLAGGAATA | ASSQDRGQEGSDYT | 0.19128686 |
| VGGAGGAGGSGP | ASSQDRGQEGSDYT | 0.39675248 |
| VFAAFMV | ASSQDRGQEGSDYT | 0.122969426 |
| TLYDKLR | ASSQDRGQEGSDYT | 0.39732468 |
| TLAVFTVL | ASSQDRGQEGSDYT | 0.13528381 |
| TGSVQPARV | ASSQDRGQEGSDYT | 0.2651925 |
| TGGGAGSGGT | ASSQDRGQEGSDYT | 0.11687882 |
| SHLETSL | ASSQDRGQEGSDYT | 0.35506523 |
| SFTTGMT | ASSQDRGQEGSDYT | 0.5758997 |
| QVEVAKRMGVRQPTVS | ASSQDRGQEGSDYT | 0.49168834 |
| QGSGGPALAM | ASSQDRGQEGSDYT | 0.09604559 |
| MLAPSNP | ASSQDRGQEGSDYT | 0.522778 |
| LVNNAAI | ASSQDRGQEGSDYT | 0.40731433 |
| LTGEINTLA | ASSQDRGQEGSDYT | 0.7888917 |
| LSIFGPV | ASSQDRGQEGSDYT | 0.23600262 |
| LLALLANL | ASSQDRGQEGSDYT | 0.36182898 |
| LISNASDALDKLR | ASSQDRGQEGSDYT | 0.17834632 |
| LINEAVP | ASSQDRGQEGSDYT | 0.17794605 |
| LGYAFSGL | ASSQDRGQEGSDYT | 0.12378599 |
| LDFVAVAL | ASSQDRGQEGSDYT | 0.14839873 |
| LAQPGAE | ASSQDRGQEGSDYT | 0.24560592 |
| ISNLGAVTGF | ASSQDRGQEGSDYT | 0.21494853 |
| HQEDPGRI | ASSQDRGQEGSDYT | 0.57539785 |
| GTPGEPTP | ASSQDRGQEGSDYT | 0.44263968 |
| GFFNSGS | ASSQDRGQEGSDYT | 0.50760424 |
| GAGGGGLTAGAN | ASSQDRGQEGSDYT | 0.5421306 |
| FVDALVNM | ASSQDRGQEGSDYT | 0.5820034 |
| FLTVAPEL | ASSQDRGQEGSDYT | 0.22328934 |
| FETLTAL | ASSQDRGQEGSDYT | 0.18869412 |
| DVINPIEL | ASSQDRGQEGSDYT | 0.1546536 |
| APVSGAPV | ASSQDRGQEGSDYT | 0.13183537 |
| ALVGGLAW | ASSQDRGQEGSDYT | 0.16157418 |
| ALAAAAATAI | ASSQDRGQEGSDYT | 0.1952188 |
| AFGLGLLL | ASSQDRGQEGSDYT | 0.32097796 |
| AFFAAASL | ASSQDRGQEGSDYT | 0.09123081 |

| | | |
|---|---|---|
| AAAALPAFSPP | ASSQDRGQEGSDYT | 0.41845477 |
| VVLAPAL | ASSQDRGQEGSDYT | 0.60387015 |
| TIGHVDHGKTTL | ASSQDRGQEGSDYT | 0.1312357 |
| LISNASDALDK | ASSQDRGQEGSDYT | 0.27963576 |
| LISNASDAL | ASSQDRGQEGSDYT | 0.24238051 |
| ISNASDALDK | ASSQDRGQEGSDYT | 0.13480058 |
| NQAETLVYQTE | ASSQDRGQEGSDYT | 0.41714174 |
| TGGTGGTAGAG | ASSQDRGQEGSDYT | 0.14779991 |
| TGSTAGGAGGA | ASSQDRGQEGSDYT | 0.34560823 |
| DPLAVPVAAGPV | ASSQDRGQEGSDYT | 0.68046886 |
| GGTGGTPGNGGN | ASSQDRGQEGSDYT | 0.2896056 |
| PAGLAGVDPATL | ASSQDRGQEGSDYT | 0.19521767 |
| GNNLVVEAPHAGASV | ASSQDRGQEGSDYT | 0.2048076 |
| KVTRIMSASTLEQPA | ASSQDRGQEGSDYT | 0.13394457 |
| LGFALPIALPRRTVV | ASSQDRGQEGSDYT | 0.23306584 |
| TRLGNLTARKLVVLV | ASSQDRGQEGSDYT | 0.09454339 |
| WSQQRRLIYFPSAGP | ASSQDRGQEGSDYT | 0.29873112 |
| GPNGNPDPMAAAVDIR | ASSQDRGQEGSDYT | 0.12963884 |
| TQRCVSVADLRAAASR | ASSQDRGQEGSDYT | 0.39811784 |
| QFVLDVAEHPVEVLTETV | ASSQDRGQEGSDYT | 0.36080712 |
| GGAGGTGGAGGNGGTQAGDG | ASSQDRGQEGSDYT | 0.19263318 |
| PLLATLRAIAVDVDAKPAQV | ASSQDRGQEGSDYT | 0.15296403 |
| APKFRTGGYMIDFWGVGYQVAK | ASSQDRGQEGSDYT | 0.17580125 |
| GAGGAGGAGGAAGGPGGTGGTG | ASSQDRGQEGSDYT | 0.13980594 |
| IGALGLLATDVAPAGSVAFGVA | ASSQDRGQEGSDYT | 0.13348296 |
| AGGAGGIGGNGGAGGAGGVLMG | ASSQDRGQEGSDYT | 0.23002589 |
| ADELIGLL | ASSQDRGQEGSDYT | 0.15076798 |
| ADRALKA | ASSQDRGQEGSDYT | 0.22780928 |
| AGAVGQVGGAGGA | ASSQDRGQEGSDYT | 0.519142 |
| AGGGGGLGGAP | ASSQDRGQEGSDYT | 0.2375137 |
| AGGLLYGNGGAG | ASSQDRGQEGSDYT | 0.122942016 |
| AGLESIGL | ASSQDRGQEGSDYT | 0.3280258 |
| AGLKTGII | ASSQDRGQEGSDYT | 0.24219182 |
| ASLAALLL | ASSQDRGQEGSDYT | 0.20746022 |
| ASVINISSI | ASSQDRGQEGSDYT | 0.09116187 |
| ATPPGPA | ASSQDRGQEGSDYT | 0.1634115 |
| ATVASALV | ASSQDRGQEGSDYT | 0.14110333 |
| ATVASLIL | ASSQDRGQEGSDYT | 0.1422721 |
| AVNGPVLIRH | ASSQDRGQEGSDYT | 0.17692612 |
| DGAAGAVGI | ASSQDRGQEGSDYT | 0.5949451 |
| DPLLHGI | ASSQDRGQEGSDYT | 0.13090867 |
| ELGAALGAL | ASSQDRGQEGSDYT | 0.18448466 |
| ELGRTQAV | ASSQDRGQEGSDYT | 0.306349 |
| FVAGGYLS | ASSQDRGQEGSDYT | 0.29341757 |
| GASGGAGGAGGW | ASSQDRGQEGSDYT | 0.6566285 |
| GGAGAGGDGGAGGI | ASSQDRGQEGSDYT | 0.20884345 |
| GTGHNGSQPGMGGNG | ASSQDRGQEGSDYT | 0.16953513 |
| GYAAASASAS | ASSQDRGQEGSDYT | 0.19319528 |
| HQVDVAAVQQG | ASSQDRGQEGSDYT | 0.17033596 |
| IGAAIAHT | ASSQDRGQEGSDYT | 0.10607807 |
| IGMIEVAQL | ASSQDRGQEGSDYT | 0.09891475 |

| | | |
|---|---|---|
| IRLAPEHTSAMD | ASSQDRGQEGSDYT | 0.21088617 |
| IYNVSVL | ASSQDRGQEGSDYT | 0.10117695 |
| KQGFLGM | ASSQDRGQEGSDYT | 0.31285423 |
| LESVITQL | ASSQDRGQEGSDYT | 0.24799523 |
| LGLGALVVV | ASSQDRGQEGSDYT | 0.12292875 |
| LIGNGGNGGSGGI | ASSQDRGQEGSDYT | 0.114219934 |
| LNAINAPTL | ASSQDRGQEGSDYT | 0.16194491 |
| LQLLNTGV | ASSQDRGQEGSDYT | 0.16590461 |
| LQTSIEF | ASSQDRGQEGSDYT | 0.49903828 |
| MSFAVSEL | ASSQDRGQEGSDYT | 0.21611753 |
| NTGALGL | ASSQDRGQEGSDYT | 0.7166962 |
| PPGAGGAGGAGTT | ASSQDRGQEGSDYT | 0.5167889 |
| QLVGLGI | ASSQDRGQEGSDYT | 0.2298851 |
| RFLEQVHQL | ASSQDRGQEGSDYT | 0.34582186 |
| RGVGYVLR | ASSQDRGQEGSDYT | 0.18577123 |
| SATGVFAGVI | ASSQDRGQEGSDYT | 0.13193333 |
| SLPAITLP | ASSQDRGQEGSDYT | 0.14970143 |
| STALAFTV | ASSQDRGQEGSDYT | 0.16229488 |
| SVLTVGLGK | ASSQDRGQEGSDYT | 0.14710914 |
| TALAGAGSGI | ASSQDRGQEGSDYT | 0.14170523 |
| TGLPFTGI | ASSQDRGQEGSDYT | 0.11656731 |
| VAEHVTV | ASSQDRGQEGSDYT | 0.14697447 |
| VDIVLNSL | ASSQDRGQEGSDYT | 0.14026374 |
| VLTGGGAVL | ASSQDRGQEGSDYT | 0.106738515 |
| VNVGLAYGI | ASSQDRGQEGSDYT | 0.14666897 |
| VSSAAVGAV | ASSQDRGQEGSDYT | 0.70906675 |
| VTSVYEAGV | ASSQDRGQEGSDYT | 0.63747907 |
| WVALDRGA | ASSQDRGQEGSDYT | 0.18126284 |
| YLVGSVVI | ASSQDRGQEGSDYT | 0.16627118 |
| GPPGTGKTLL | ASSQDRGQEGSDYT | 0.38234776 |
| GVGMTKF | ASSQDRGQEGSDYT | 0.12302485 |
| IGIGGSDL | ASSQDRGQEGSDYT | 0.17296562 |
| IGTLLAEL | ASSQDRGQEGSDYT | 0.43050647 |
| IYTSGTT | ASSQDRGQEGSDYT | 0.3178957 |
| LSFPAGV | ASSQDRGQEGSDYT | 0.13092282 |
| LYTSGTT | ASSQDRGQEGSDYT | 0.24232331 |
| VGAFPFGF | ASSQDRGQEGSDYT | 0.21752691 |
| VGAGTVEF | ASSQDRGQEGSDYT | 0.20221792 |
| VGKSTLLL | ASSQDRGQEGSDYT | 0.09215562 |
| AALAHAGGGIV | ASSQDRGQEGSDYT | 0.20311569 |
| AALVAVSLL | ASSQDRGQEGSDYT | 0.15327038 |
| AAVSRAVAAL | ASSQDRGQEGSDYT | 0.14018328 |
| AGGSSGIGGF | ASSQDRGQEGSDYT | 0.10563113 |
| AGLTVANANG | ASSQDRGQEGSDYT | 0.16538322 |
| AGLVGAVL | ASSQDRGQEGSDYT | 0.4916101 |
| ALSQAFTV | ASSQDRGQEGSDYT | 0.5596515 |
| ALTFSVSL | ASSQDRGQEGSDYT | 0.10802685 |
| ALTGAALVAL | ASSQDRGQEGSDYT | 0.4383928 |
| ASFSIGAGVL | ASSQDRGQEGSDYT | 0.6758947 |
| DGSVYKI | ASSQDRGQEGSDYT | 0.26849532 |
| EQLAEIV | ASSQDRGQEGSDYT | 0.11673839 |

| | | |
|---|---|---|
| FAVSELTVM | ASSQDRGQEGSDYT | 0.32420108 |
| GAAPLAAGL | ASSQDRGQEGSDYT | 0.11462537 |
| GADGGDGGSGGAGG | ASSQDRGQEGSDYT | 0.25132227 |
| GFGLDTGGAGG | ASSQDRGQEGSDYT | 0.3701153 |
| GLASGGLL | ASSQDRGQEGSDYT | 0.12508486 |
| GRIAGAAVLL | ASSQDRGQEGSDYT | 0.30691245 |
| GSAGSLNTSF | ASSQDRGQEGSDYT | 0.44031304 |
| GTGQIGI | ASSQDRGQEGSDYT | 0.1285136 |
| GTLIALIL | ASSQDRGQEGSDYT | 0.1271436 |
| IAATMGFVVG | ASSQDRGQEGSDYT | 0.13872746 |
| IFAQALL | ASSQDRGQEGSDYT | 0.09742461 |
| IGAAVALGF | ASSQDRGQEGSDYT | 0.2568859 |
| IGLTPLAL | ASSQDRGQEGSDYT | 0.20302683 |
| ILGSNTSTL | ASSQDRGQEGSDYT | 0.1662772 |
| IVFSLTVL | ASSQDRGQEGSDYT | 0.23580164 |
| LASKATGF | ASSQDRGQEGSDYT | 0.15130611 |
| LASVALIL | ASSQDRGQEGSDYT | 0.19052726 |
| LFAVGLTL | ASSQDRGQEGSDYT | 0.20608416 |
| LFNGGIGTYI | ASSQDRGQEGSDYT | 0.18414183 |
| LTLGAGII | ASSQDRGQEGSDYT | 0.26596892 |
| LVAAALGDI | ASSQDRGQEGSDYT | 0.15916151 |
| NTAFVQLGI | ASSQDRGQEGSDYT | 0.14069839 |
| QLFSGAEL | ASSQDRGQEGSDYT | 0.7318464 |
| SAATFITL | ASSQDRGQEGSDYT | 0.120310545 |
| SNYGLANLGV | ASSQDRGQEGSDYT | 0.14724535 |
| SYGGAYAVM | ASSQDRGQEGSDYT | 0.25025296 |
| TTGVGGLTLG | ASSQDRGQEGSDYT | 0.11874383 |
| TTQTIYI | ASSQDRGQEGSDYT | 0.39366204 |
| VAAGAGLGAVI | ASSQDRGQEGSDYT | 0.29224885 |
| VAAVVTLL | ASSQDRGQEGSDYT | 0.32873738 |
| VGFLGGLI | ASSQDRGQEGSDYT | 0.14342427 |
| VGVIPILL | ASSQDRGQEGSDYT | 0.62125504 |
| VIFASSLI | ASSQDRGQEGSDYT | 0.47429305 |
| VKGGDGGAAAG | ASSQDRGQEGSDYT | 0.14768228 |
| VTKTFTNI | ASSQDRGQEGSDYT | 0.19123909 |
| VVAAAILL | ASSQDRGQEGSDYT | 0.11767931 |
| YAALDVEL | ASSQDRGQEGSDYT | 0.2957338 |
| LASVALLL | ASSQDRGQEGSDYT | 0.25484923 |
| LDAVHRV | ASSQDRGQEGSDYT | 0.18331882 |
| QGLSSSIF | ASSQDRGQEGSDYT | 0.27201974 |
| RGTFANIRL | ASSQDRGQEGSDYT | 0.31676596 |
| RQLWWGHRI | ASSQDRGQEGSDYT | 0.53686315 |
| YGPPGTGKTLL | ASSQDRGQEGSDYT | 0.114069104 |
| AALVGAPANVA | ASSQDRGQEGSDYT | 0.18153782 |
| AATFRRTE | ASSQDRGQEGSDYT | 0.11027026 |
| AGALGGGI | ASSQDRGQEGSDYT | 0.15521397 |
| AGGAGGAGGIGAG | ASSQDRGQEGSDYT | 0.12897305 |
| AGSGALGL | ASSQDRGQEGSDYT | 0.29733068 |
| AQIADQI | ASSQDRGQEGSDYT | 0.20252791 |
| ASAVAVTYL | ASSQDRGQEGSDYT | 0.4832949 |
| ASTVLGGTGG | ASSQDRGQEGSDYT | 0.31348345 |

| | | |
|---|---|---|
| AVGGEPAK | ASSQDRGQEGSDYT | 0.43519616 |
| CEQGTFGGTC | ASSQDRGQEGSDYT | 0.13770309 |
| DEGKAPAG | ASSQDRGQEGSDYT | 0.2173694 |
| DTVVFPAL | ASSQDRGQEGSDYT | 0.11290546 |
| FEGITMAG | ASSQDRGQEGSDYT | 0.11775596 |
| FPLVVQD | ASSQDRGQEGSDYT | 0.116915174 |
| GAFTTPQ | ASSQDRGQEGSDYT | 0.20118104 |
| GFGGFGTAGGVGGA | ASSQDRGQEGSDYT | 0.094869174 |
| GFLAQFL | ASSQDRGQEGSDYT | 0.38389495 |
| GGAGGAGGNAGVA | ASSQDRGQEGSDYT | 0.30275252 |
| GGVGTAGAGGAGGAGG | ASSQDRGQEGSDYT | 0.40779898 |
| GNGGGGDAGNAGSGGN | ASSQDRGQEGSDYT | 0.14792603 |
| GTGGAGGAGNDAGS | ASSQDRGQEGSDYT | 0.13880342 |
| GTTNTGWFNS | ASSQDRGQEGSDYT | 0.18631314 |
| IFMGPLFTL | ASSQDRGQEGSDYT | 0.3856113 |
| ILAHAIEAI | ASSQDRGQEGSDYT | 0.12854378 |
| IRSAIREAL | ASSQDRGQEGSDYT | 0.108435854 |
| ITFAGGG | ASSQDRGQEGSDYT | 0.12915093 |
| ITGMSGHQEV | ASSQDRGQEGSDYT | 0.11943963 |
| IVAGLACTL | ASSQDRGQEGSDYT | 0.496737 |
| IVGGGLAGA | ASSQDRGQEGSDYT | 0.56270796 |
| IVTGVLNI | ASSQDRGQEGSDYT | 0.33629096 |
| LAQQGCI | ASSQDRGQEGSDYT | 0.25417742 |
| LCLHVEN | ASSQDRGQEGSDYT | 0.09701447 |
| LDAISDGS | ASSQDRGQEGSDYT | 0.15394107 |
| LGGFAAML | ASSQDRGQEGSDYT | 0.67166436 |
| LGGKNPMI | ASSQDRGQEGSDYT | 0.2812661 |
| LGPKGVAAIL | ASSQDRGQEGSDYT | 0.20789073 |
| LMVGVSI | ASSQDRGQEGSDYT | 0.54481745 |
| LVGAAFVL | ASSQDRGQEGSDYT | 0.1802809 |
| LVLAVAPL | ASSQDRGQEGSDYT | 0.19541192 |
| MAAIAVIDAA | ASSQDRGQEGSDYT | 0.21193294 |
| NNGINAELQ | ASSQDRGQEGSDYT | 0.16555268 |
| RGEPLVLVHG | ASSQDRGQEGSDYT | 0.15320377 |
| RGGAGGAAGLI | ASSQDRGQEGSDYT | 0.11882082 |
| RSYRMHPAV | ASSQDRGQEGSDYT | 0.2757704 |
| SAAHLLAGQLA | ASSQDRGQEGSDYT | 0.7568632 |
| SAAVGSY | ASSQDRGQEGSDYT | 0.30182835 |
| SPHAVLHRI | ASSQDRGQEGSDYT | 0.3992402 |
| SQIAKVL | ASSQDRGQEGSDYT | 0.3276161 |
| TALGHLVP | ASSQDRGQEGSDYT | 0.54022026 |
| TAPTTSGAA | ASSQDRGQEGSDYT | 0.17214534 |
| TDGDDGNGGSPGT | ASSQDRGQEGSDYT | 0.14066212 |
| TDGLGEN | ASSQDRGQEGSDYT | 0.17696339 |
| TGGPPLPGQA | ASSQDRGQEGSDYT | 0.13875721 |
| TIPAITI | ASSQDRGQEGSDYT | 0.20159411 |
| TIPGFTI | ASSQDRGQEGSDYT | 0.19429359 |
| TSIASGALA | ASSQDRGQEGSDYT | 0.78209674 |
| TVTLPGAAI | ASSQDRGQEGSDYT | 0.3551551 |
| VASGLSAALP | ASSQDRGQEGSDYT | 0.30842835 |
| VGAGVIGI | ASSQDRGQEGSDYT | 0.13397944 |

| | | |
|---|---|---|
| VGPKRFGV | ASSQDRGQEGSDYT | 0.16260833 |
| VNVAFPD | ASSQDRGQEGSDYT | 0.34305733 |
| VSMTNSGVS | ASSQDRGQEGSDYT | 0.12857452 |
| VTDVLGDLSV | ASSQDRGQEGSDYT | 0.5316781 |
| VVVVGAGFAGL | ASSQDRGQEGSDYT | 0.16880712 |
| VYTVGTVV | ASSQDRGQEGSDYT | 0.16074032 |
| WTTFSGP | ASSQDRGQEGSDYT | 0.27570403 |
| YDELVEAF | ASSQDRGQEGSDYT | 0.70560586 |
| YGTGGAGGNGG | ASSQDRGQEGSDYT | 0.18169749 |
| YTNVMAV | ASSQDRGQEGSDYT | 0.1754152 |
| FLEQVHQL | ASSQDRGQEGSDYT | 0.14026378 |
| GLSLLAVAV | ASSQDRGQEGSDYT | 0.290521 |
| LEQVHQL | ASSQDRGQEGSDYT | 0.6511632 |
| LNAAGAI | ASSQDRGQEGSDYT | 0.13235366 |
| PAVHASL | ASSQDRGQEGSDYT | 0.50444233 |
| TELEDDA | ASSQDRGQEGSDYT | 0.16826604 |
| TPPGPPN | ASSQDRGQEGSDYT | 0.116133794 |
| TPPPPPPP | ASSQDRGQEGSDYT | 0.2080706 |
| TQSPALI | ASSLGGADTGQLY | 0.26716655 |
| EFAAAFPQ | ASSLGGADTGQLY | 0.6211837 |
| LLSGNLDVL | ASSLGGADTGQLY | 0.39277697 |
| GPANLGNYNI | ASSLGGADTGQLY | 0.14890313 |
| TLGPTLP | ASSLGGADTGQLY | 0.20456025 |
| VYAYSSCCK | ASSLGGADTGQLY | 0.63387907 |
| APAPAPAPNI | ASSLGGADTGQLY | 0.4716334 |
| PGGPVPIP | ASSLGGADTGQLY | 0.37282404 |
| FLTVALEP | ASSLGGADTGQLY | 0.14225796 |
| SQYFGANL | ASSLGGADTGQLY | 0.14199418 |
| GGVKPAA | ASSLGGADTGQLY | 0.23412538 |
| GGTGGAGGAGGW | ASSLGGADTGQLY | 0.56919056 |
| GVGGVAGLGGAGG | ASSLGGADTGQLY | 0.12784965 |
| VLLVLATTI | ASSLGGADTGQLY | 0.30965805 |
| IGIFAAML | ASSLGGADTGQLY | 0.36329067 |
| AVGDLID | ASSLGGADTGQLY | 0.11018268 |
| NISPLPL | ASSLGGADTGQLY | 0.16733745 |
| VTAVFVAI | ASSLGGADTGQLY | 0.48189563 |
| SGDDNDAAL | ASSLGGADTGQLY | 0.1824964 |
| AVAALAHS | ASSLGGADTGQLY | 0.13430268 |
| GGVTVMATP | ASSLGGADTGQLY | 0.59650975 |
| AGLAANVRD | ASSLGGADTGQLY | 0.7859431 |
| TNRPDLIDPALLRPG | ASSLGGADTGQLY | 0.39210308 |
| LANLGVTAL | ASSLGGADTGQLY | 0.123859935 |
| TGERIEL | ASSLGGADTGQLY | 0.09931449 |
| NGEELAA | ASSLGGADTGQLY | 0.14183764 |
| VLSLVWG | ASSLGGADTGQLY | 0.2717084 |
| IGYALWL | ASSLGGADTGQLY | 0.17505749 |
| LSLLLAPL | ASSLGGADTGQLY | 0.15496331 |
| TLDEALK | ASSLGGADTGQLY | 0.34368235 |
| TDGAGTAEGG | ASSLGGADTGQLY | 0.59359217 |
| NAPGGAGGNGG | ASSLGGADTGQLY | 0.12606709 |
| VSVFMGMGGNF | ASSLGGADTGQLY | 0.12450166 |

| | | |
|---|---|---|
| LASYLAAL | ASSLGGADTGQLY | 0.18490352 |
| LVGGTPAL | ASSLGGADTGQLY | 0.34801787 |
| VRNADVL | ASSLGGADTGQLY | 0.12758377 |
| TCSSPFLVSAM | ASSLGGADTGQLY | 0.25035548 |
| VHITDAE | ASSLGGADTGQLY | 0.22801857 |
| EYAAPSF | ASSLGGADTGQLY | 0.11394166 |
| VVDAFHGGL | ASSLGGADTGQLY | 0.33652103 |
| ASVVDPQ | ASSLGGADTGQLY | 0.13647617 |
| SYNEFGGLS | ASSLGGADTGQLY | 0.44888473 |
| DPDGSVY | ASSLGGADTGQLY | 0.14815678 |
| MVSTIHATE | ASSLGGADTGQLY | 0.24704742 |
| LAAGVADPV | ASSLGGADTGQLY | 0.22508034 |
| TKEQIAA | ASSLGGADTGQLY | 0.2171134 |
| SLASPAAL | ASSLGGADTGQLY | 0.207095 |
| IGLVFYGV | ASSLGGADTGQLY | 0.15618606 |
| FLLDASNL | ASSLGGADTGQLY | 0.1345852 |
| YQVGNFP | ASSLGGADTGQLY | 0.09589798 |
| ESLLSQST | ASSLGGADTGQLY | 0.47161102 |
| LFTVGFAAVG | ASSLGGADTGQLY | 0.109051734 |
| ATPPTPT | ASSLGGADTGQLY | 0.32261246 |
| TDLATLG | ASSLGGADTGQLY | 0.38572225 |
| SAGGTGR | ASSLGGADTGQLY | 0.14434727 |
| GFSNSGS | ASSLGGADTGQLY | 0.24473989 |
| GGNGGSGGTSV | ASSLGGADTGQLY | 0.11859345 |
| GTSVATGGAGN | ASSLGGADTGQLY | 0.12218714 |
| NGGNGGSGGTS | ASSLGGADTGQLY | 0.31310892 |
| AVVKFPH | ASSLGGADTGQLY | 0.32760286 |
| TPPPPPP | ASSLGGADTGQLY | 0.13356751 |
| AAPAAAAAPA | ASSLGGADTGQLY | 0.23522624 |
| IALNEDL | ASSLGGADTGQLY | 0.31723678 |
| IVNVSSM | ASSLGGADTGQLY | 0.12572941 |
| TGIPPAPR | ASSLGGADTGQLY | 0.16975257 |
| FDLGGGTF | ASSLGGADTGQLY | 0.5934274 |
| FDLGGGTFDVS | ASSLGGADTGQLY | 0.20306848 |
| NEPTAAA | ASSLGGADTGQLY | 0.13054518 |
| NEPTAAALA | ASSLGGADTGQLY | 0.37373918 |
| GSIINISSI | ASSLGGADTGQLY | 0.15457621 |
| VAPGNAGTA | ASSLGGADTGQLY | 0.37939507 |
| TGGGASLE | ASSLGGADTGQLY | 0.1741414 |
| VAVYNHYKRI | ASSLGGADTGQLY | 0.18859364 |
| TDPAPATT | ASSLGGADTGQLY | 0.32432228 |
| TNRPDILDPALLRPG | ASSLGGADTGQLY | 0.118473604 |
| LTTTTTS | ASSLGGADTGQLY | 0.13524231 |
| PTGSGKTL | ASSLGGADTGQLY | 0.17666814 |
| TGSGKTL | ASSLGGADTGQLY | 0.2658071 |
| TIGHVDHGK | ASSLGGADTGQLY | 0.46517387 |
| TIGHVDHGKTT | ASSLGGADTGQLY | 0.18067299 |
| GHVDHGKTT | ASSLGGADTGQLY | 0.2411839 |
| AQFEHTLL | ASSLGGADTGQLY | 0.20141792 |
| IVNQDEL | ASSLGGADTGQLY | 0.155083 |
| TNGSQFF | ASSLGGADTGQLY | 0.17499688 |

| | | |
|---|---|---|
| TGKGIVE | ASSLGGADTGQLY | 0.13968724 |
| LISNASDA | ASSLGGADTGQLY | 0.13529077 |
| AELRTLGELTKIAKAHGAQ | ASSLGGADTGQLY | 0.18178897 |
| APNAVAAAEPSW | ASSLGGADTGQLY | 0.16166438 |
| APQVAGAATAVL | ASSLGGADTGQLY | 0.2124604 |
| ARRDPAAAAAILPTDA | ASSLGGADTGQLY | 0.33078516 |
| DAAARDVLAEAGLAEYF | ASSLGGADTGQLY | 0.12939753 |
| DDQIVLSAKANLAKVEQ | ASSLGGADTGQLY | 0.2463459 |
| DELLPDTGEDSDEDGH | ASSLGGADTGQLY | 0.28846616 |
| ELPEAPANRATVPQD | ASSLGGADTGQLY | 0.23861949 |
| FGQSKIERVTGIG | ASSLGGADTGQLY | 0.21100992 |
| GAGGTGSTAGGAGGAGGAGGLY | ASSLGGADTGQLY | 0.13395916 |
| GHGGTGAAGGNGGTGGAGGSGI | ASSLGGADTGQLY | 0.40661085 |
| GNGGNGGAGGEGGPGGAGGAGA | ASSLGGADTGQLY | 0.3404723 |
| GSTAGNGGNGGNATNGTIAGQP | ASSLGGADTGQLY | 0.3177234 |
| GTGGDGGNGGNGGTGG | ASSLGGADTGQLY | 0.20511004 |
| GVRRPAGIPADAPV | ASSLGGADTGQLY | 0.2705624 |
| GYRQVTDAHLLTLARR | ASSLGGADTGQLY | 0.3866216 |
| IAKMGQDGHDRG | ASSLGGADTGQLY | 0.18978317 |
| KAEPSIEIQYT | ASSLGGADTGQLY | 0.11150015 |
| KPVRIRMAADEDTVAVT | ASSLGGADTGQLY | 0.192407 |
| MPPRIAGMRLLVIKPEP | ASSLGGADTGQLY | 0.13674876 |
| MKFYAVALFRSRGVPPTR | ASSLGGADTGQLY | 0.149209 |
| NAGGFGILGAVAHSP | ASSLGGADTGQLY | 0.11121308 |
| NEGREITLAMAEM | ASSLGGADTGQLY | 0.5937383 |
| NELVEWSHPLGHRWRWE | ASSLGGADTGQLY | 0.76372904 |
| NLGAGGTGGNGGNPG | ASSLGGADTGQLY | 0.14588699 |
| NTEWPKWETVMAADPVD | ASSLGGADTGQLY | 0.4221421 |
| PASAAPAETQP | ASSLGGADTGQLY | 0.119472936 |
| QQSSDGKNGVRAAEITGE | ASSLGGADTGQLY | 0.27881014 |
| RAPDAVTTSWWKE | ASSLGGADTGQLY | 0.48728293 |
| SGFGNVGAHTSGWFNQ | ASSLGGADTGQLY | 0.15311433 |
| SGVAGAVGQAASVGGLKV | ASSLGGADTGQLY | 0.3166517 |
| SNGGAGTGSAGGNGGT | ASSLGGADTGQLY | 0.13844174 |
| TGGAGGEGGAAG | ASSLGGADTGQLY | 0.16324551 |
| TGGDGGAAGAGGAPGHG | ASSLGGADTGQLY | 0.60204303 |
| TMPPLLAAIANAGRHGV | ASSLGGADTGQLY | 0.17972697 |
| TSSTAGNSGGA | ASSLGGADTGQLY | 0.16643189 |
| VPATVPVGILC | ASSLGGADTGQLY | 0.18280944 |
| VQRPDSGGVVLTGRIS | ASSLGGADTGQLY | 0.15953177 |
| WTPNPVPATKGVPAGLVQAT | ASSLGGADTGQLY | 0.17770064 |
| YALKRALAKFSDRFDVVII | ASSLGGADTGQLY | 0.29283094 |
| YLIGSVIV | ASSLGGADTGQLY | 0.119991444 |
| YGQLVAVID | ASSLGGADTGQLY | 0.4418344 |
| VTVGGMTLPA | ASSLGGADTGQLY | 0.1250294 |
| VLALPAV | ASSLGGADTGQLY | 0.6235721 |
| VIPPTLNYET | ASSLGGADTGQLY | 0.38225883 |
| VGMGGMHPSSG | ASSLGGADTGQLY | 0.3418166 |
| VGLAGGAATA | ASSLGGADTGQLY | 0.15005873 |
| VGGAGGAGGSGP | ASSLGGADTGQLY | 0.6334992 |
| VFAAFMV | ASSLGGADTGQLY | 0.20844743 |

| | | |
|---|---|---|
| TLYDKLR | ASSLGGADTGQLY | 0.24359089 |
| TLAVFTVL | ASSLGGADTGQLY | 0.22822493 |
| TGSVQPARV | ASSLGGADTGQLY | 0.24739048 |
| TGGGAGSGGT | ASSLGGADTGQLY | 0.61842614 |
| SHLETSL | ASSLGGADTGQLY | 0.542172 |
| SFTTGMT | ASSLGGADTGQLY | 0.36740565 |
| QVEVAKRMGVRQPTVS | ASSLGGADTGQLY | 0.35937837 |
| QGSGGPALAM | ASSLGGADTGQLY | 0.18766695 |
| MLAPSNP | ASSLGGADTGQLY | 0.27050844 |
| LVNNAAI | ASSLGGADTGQLY | 0.320065 |
| LTGEINTLA | ASSLGGADTGQLY | 0.4526667 |
| LSIFGPV | ASSLGGADTGQLY | 0.41464972 |
| LLALLANL | ASSLGGADTGQLY | 0.3411278 |
| LISNASDALDKLR | ASSLGGADTGQLY | 0.51984566 |
| LINEAVP | ASSLGGADTGQLY | 0.3347473 |
| LGYAFSGL | ASSLGGADTGQLY | 0.20233811 |
| LDFVAVAL | ASSLGGADTGQLY | 0.34953147 |
| LAQPGAE | ASSLGGADTGQLY | 0.34674287 |
| ISNLGAVTGF | ASSLGGADTGQLY | 0.44805545 |
| HQEDPGRI | ASSLGGADTGQLY | 0.14254875 |
| GTPGEPTP | ASSLGGADTGQLY | 0.5342406 |
| GFFNSGS | ASSLGGADTGQLY | 0.37571135 |
| GAGGGGLTAGAN | ASSLGGADTGQLY | 0.13097285 |
| FVDALVNM | ASSLGGADTGQLY | 0.35677254 |
| FLTVAPEL | ASSLGGADTGQLY | 0.38789713 |
| FETLTAL | ASSLGGADTGQLY | 0.38963684 |
| DVINPIEL | ASSLGGADTGQLY | 0.7275622 |
| APVSGAPV | ASSLGGADTGQLY | 0.14837532 |
| ALVGGLAW | ASSLGGADTGQLY | 0.24048254 |
| ALAAAAATAI | ASSLGGADTGQLY | 0.4842564 |
| AFGLGLLL | ASSLGGADTGQLY | 0.63816416 |
| AFFAAASL | ASSLGGADTGQLY | 0.23372447 |
| AAAALPAFSPP | ASSLGGADTGQLY | 0.5768906 |
| VVLAPAL | ASSLGGADTGQLY | 0.46651453 |
| TIGHVDHGKTTL | ASSLGGADTGQLY | 0.43080157 |
| LISNASDALDK | ASSLGGADTGQLY | 0.44295773 |
| LISNASDAL | ASSLGGADTGQLY | 0.5004925 |
| ISNASDALDK | ASSLGGADTGQLY | 0.17116664 |
| NQAETLVYQTE | ASSLGGADTGQLY | 0.5526455 |
| TGGTGGTAGAG | ASSLGGADTGQLY | 0.5469221 |
| TGSTAGGAGGA | ASSLGGADTGQLY | 0.26393208 |
| DPLAVPVAAGPV | ASSLGGADTGQLY | 0.46232325 |
| GGTGGTPGNGGN | ASSLGGADTGQLY | 0.25960097 |
| PAGLAGVDPATL | ASSLGGADTGQLY | 0.18471104 |
| GNNLVVEAPHAGASV | ASSLGGADTGQLY | 0.16844203 |
| KVTRIMSASTLEQPA | ASSLGGADTGQLY | 0.63113296 |
| LGFALPIALPRRTVV | ASSLGGADTGQLY | 0.4228312 |
| TRLGNLTARKLVVLV | ASSLGGADTGQLY | 0.47305036 |
| WSQQRRLIYFPSAGP | ASSLGGADTGQLY | 0.27651355 |
| GPNGNPDPMAAAVDIR | ASSLGGADTGQLY | 0.21048675 |
| TQRCVSVADLRAAASR | ASSLGGADTGQLY | 0.35802886 |

| | | |
|---|---|---|
| QFVLDVAEHPVEVLTETV | ASSLGGADTGQLY | 0.31979755 |
| GGAGGTGGAGGNGGTQAGDG | ASSLGGADTGQLY | 0.15861046 |
| PLLATLRAIAVDVDAKPAQV | ASSLGGADTGQLY | 0.2782638 |
| APKFRTGGYMIDFWGVGYQVAK | ASSLGGADTGQLY | 0.4139916 |
| GAGGAGGAGGAAGGPGGTGGTG | ASSLGGADTGQLY | 0.4511805 |
| IGALGLLATDVAPAGSVAFGVA | ASSLGGADTGQLY | 0.16023815 |
| AGGAGGIGGNGGAGGAGGVLMG | ASSLGGADTGQLY | 0.6968727 |
| ADELIGLL | ASSLGGADTGQLY | 0.3247735 |
| ADRALKA | ASSLGGADTGQLY | 0.3779431 |
| AGAVGQVGGAGGA | ASSLGGADTGQLY | 0.36985326 |
| AGGGGGLGGAP | ASSLGGADTGQLY | 0.27003855 |
| AGGLLYGNGGAG | ASSLGGADTGQLY | 0.6784706 |
| AGLESIGL | ASSLGGADTGQLY | 0.40271652 |
| AGLKTGII | ASSLGGADTGQLY | 0.1881335 |
| ASLAALLL | ASSLGGADTGQLY | 0.45419872 |
| ASVINISSI | ASSLGGADTGQLY | 0.3689126 |
| ATPPGPA | ASSLGGADTGQLY | 0.28051263 |
| ATVASALV | ASSLGGADTGQLY | 0.3756302 |
| ATVASLIL | ASSLGGADTGQLY | 0.548704 |
| AVNGPVLIRH | ASSLGGADTGQLY | 0.2062101 |
| DGAAGAVGI | ASSLGGADTGQLY | 0.31872794 |
| DPLLHGI | ASSLGGADTGQLY | 0.17942196 |
| ELGAALGAL | ASSLGGADTGQLY | 0.4295874 |
| ELGRTQAV | ASSLGGADTGQLY | 0.5612009 |
| FVAGGYLS | ASSLGGADTGQLY | 0.3337231 |
| GASGGAGGAGGW | ASSLGGADTGQLY | 0.21144256 |
| GGAGAGGDGGAGGI | ASSLGGADTGQLY | 0.5721322 |
| GTGHNGSQPGMGGNG | ASSLGGADTGQLY | 0.38364342 |
| GYAAASASAS | ASSLGGADTGQLY | 0.42095757 |
| HQVDVAAVQQG | ASSLGGADTGQLY | 0.32481262 |
| IGAAIAHT | ASSLGGADTGQLY | 0.3910514 |
| IGMIEVAQL | ASSLGGADTGQLY | 0.51285905 |
| IRLAPEHTSAMD | ASSLGGADTGQLY | 0.5595099 |
| IYNVSVL | ASSLGGADTGQLY | 0.4042442 |
| KQGFLGM | ASSLGGADTGQLY | 0.37963355 |
| LESVITQL | ASSLGGADTGQLY | 0.26557118 |
| LGLGALVVV | ASSLGGADTGQLY | 0.37349814 |
| LIGNGGNGGSGGI | ASSLGGADTGQLY | 0.15327103 |
| LNAINAPTL | ASSLGGADTGQLY | 0.1756583 |
| LQLLNTGV | ASSLGGADTGQLY | 0.366001 |
| LQTSIEF | ASSLGGADTGQLY | 0.560465 |
| MSFAVSEL | ASSLGGADTGQLY | 0.36036316 |
| NTGALGL | ASSLGGADTGQLY | 0.30974418 |
| PPGAGGAGGAGTT | ASSLGGADTGQLY | 0.30400947 |
| QLVGLGI | ASSLGGADTGQLY | 0.27382326 |
| RFLEQVHQL | ASSLGGADTGQLY | 0.451861 |
| RGVGYVLR | ASSLGGADTGQLY | 0.2641367 |
| SATGVFAGVI | ASSLGGADTGQLY | 0.4652745 |
| SLPAITLP | ASSLGGADTGQLY | 0.7020786 |
| STALAFTV | ASSLGGADTGQLY | 0.4622231 |
| SVLTVGLGK | ASSLGGADTGQLY | 0.4244122 |

| | | |
|---|---|---|
| TALAGAGSGI | ASSLGGADTGQLY | 0.2773623 |
| TGLPFTGI | ASSLGGADTGQLY | 0.557068 |
| VAEHVTV | ASSLGGADTGQLY | 0.2116564 |
| VDIVLNSL | ASSLGGADTGQLY | 0.53255296 |
| VLTGGGAVL | ASSLGGADTGQLY | 0.13892342 |
| VNVGLAYGI | ASSLGGADTGQLY | 0.36206633 |
| VSSAAVGAV | ASSLGGADTGQLY | 0.23124264 |
| VTSVYEAGV | ASSLGGADTGQLY | 0.24578191 |
| WVALDRGA | ASSLGGADTGQLY | 0.61548346 |
| YLVGSVVI | ASSLGGADTGQLY | 0.21293288 |
| GPPGTGKTLL | ASSLGGADTGQLY | 0.32465327 |
| GVGMTKF | ASSLGGADTGQLY | 0.34529313 |
| IGIGGSDL | ASSLGGADTGQLY | 0.50354743 |
| IGTLLAEL | ASSLGGADTGQLY | 0.53161716 |
| IYTSGTT | ASSLGGADTGQLY | 0.2618128 |
| LSFPAGV | ASSLGGADTGQLY | 0.26301166 |
| LYTSGTT | ASSLGGADTGQLY | 0.49907494 |
| VGAFPFGF | ASSLGGADTGQLY | 0.30953503 |
| VGAGTVEF | ASSLGGADTGQLY | 0.41207352 |
| VGKSTLLL | ASSLGGADTGQLY | 0.1752381 |
| AALAHAGGGIV | ASSLGGADTGQLY | 0.26604193 |
| AALVAVSLL | ASSLGGADTGQLY | 0.42436558 |
| AAVSRAVAAL | ASSLGGADTGQLY | 0.16638985 |
| AGGSSGIGGF | ASSLGGADTGQLY | 0.13588035 |
| AGLTVANANG | ASSLGGADTGQLY | 0.30517504 |
| AGLVGAVL | ASSLGGADTGQLY | 0.38066357 |
| ALSQAFTV | ASSLGGADTGQLY | 0.6645279 |
| ALTFSVSL | ASSLGGADTGQLY | 0.54073036 |
| ALTGAALVAL | ASSLGGADTGQLY | 0.5355557 |
| ASFSIGAGVL | ASSLGGADTGQLY | 0.41393098 |
| DGSVYKI | ASSLGGADTGQLY | 0.27703768 |
| EQLAEIV | ASSLGGADTGQLY | 0.1232787 |
| FAVSELTVM | ASSLGGADTGQLY | 0.22186905 |
| GAAPLAAGL | ASSLGGADTGQLY | 0.20995264 |
| GADGGDGGSGGAGG | ASSLGGADTGQLY | 0.35610288 |
| GFGLDTGGAGG | ASSLGGADTGQLY | 0.36782968 |
| GLASGGLL | ASSLGGADTGQLY | 0.34806758 |
| GRIAGAAVLL | ASSLGGADTGQLY | 0.5761887 |
| GSAGSLNTSF | ASSLGGADTGQLY | 0.5426679 |
| GTGQIGI | ASSLGGADTGQLY | 0.36843872 |
| GTLIALIL | ASSLGGADTGQLY | 0.19461769 |
| IAATMGFVVG | ASSLGGADTGQLY | 0.12878509 |
| IFAQALL | ASSLGGADTGQLY | 0.3931332 |
| IGAAVALGF | ASSLGGADTGQLY | 0.1532413 |
| IGLTPLAL | ASSLGGADTGQLY | 0.23143226 |
| ILGSNTSTL | ASSLGGADTGQLY | 0.13668849 |
| IVFSLTVL | ASSLGGADTGQLY | 0.13619936 |
| LASKATGF | ASSLGGADTGQLY | 0.17652842 |
| LASVALIL | ASSLGGADTGQLY | 0.42423895 |
| LFAVGLTL | ASSLGGADTGQLY | 0.32609406 |
| LFNGGIGTYI | ASSLGGADTGQLY | 0.3430584 |

| | | |
|---|---|---|
| LTLGAGII | ASSLGGADTGQLY | 0.53988683 |
| LVAAALGDI | ASSLGGADTGQLY | 0.52607 |
| NTAFVQLGI | ASSLGGADTGQLY | 0.4416052 |
| QLFSGAEL | ASSLGGADTGQLY | 0.23150952 |
| SAATFITL | ASSLGGADTGQLY | 0.41501218 |
| SNYGLANLGV | ASSLGGADTGQLY | 0.3156982 |
| SYGGAYAVM | ASSLGGADTGQLY | 0.26586288 |
| TTGVGGLTLG | ASSLGGADTGQLY | 0.4631911 |
| TTQTIYI | ASSLGGADTGQLY | 0.35966468 |
| VAAGAGLGAVI | ASSLGGADTGQLY | 0.21291278 |
| VAAVVTLL | ASSLGGADTGQLY | 0.38618124 |
| VGFLGGLI | ASSLGGADTGQLY | 0.5536801 |
| VGVIPILL | ASSLGGADTGQLY | 0.724632 |
| VIFASSLI | ASSLGGADTGQLY | 0.23578003 |
| VKGGDGGAAAG | ASSLGGADTGQLY | 0.1909355 |
| VTKTFTNI | ASSLGGADTGQLY | 0.25731274 |
| VVAAAILL | ASSLGGADTGQLY | 0.5023757 |
| YAALDVEL | ASSLGGADTGQLY | 0.20714237 |
| LASVALLL | ASSLGGADTGQLY | 0.25783446 |
| LDAVHRV | ASSLGGADTGQLY | 0.5056981 |
| QGLSSSIF | ASSLGGADTGQLY | 0.61570644 |
| RGTFANIRL | ASSLGGADTGQLY | 0.34361622 |
| RQLWWGHRI | ASSLGGADTGQLY | 0.25166616 |
| YGPPGTGKTLL | ASSLGGADTGQLY | 0.45980746 |
| AALVGAPANVA | ASSLGGADTGQLY | 0.24479116 |
| AATFRRTE | ASSLGGADTGQLY | 0.53930753 |
| AGALGGGI | ASSLGGADTGQLY | 0.27776936 |
| AGGAGGAGGIGAG | ASSLGGADTGQLY | 0.33161578 |
| AGSGALGL | ASSLGGADTGQLY | 0.35159206 |
| AQIADQI | ASSLGGADTGQLY | 0.62816805 |
| ASAVAVTYL | ASSLGGADTGQLY | 0.36667395 |
| ASTVLGGTGG | ASSLGGADTGQLY | 0.5352899 |
| AVGGEPAK | ASSLGGADTGQLY | 0.72082686 |
| CEQGTFGGTC | ASSLGGADTGQLY | 0.70022357 |
| DEGKAPAG | ASSLGGADTGQLY | 0.7822225 |
| DTVVFPAL | ASSLGGADTGQLY | 0.36501777 |
| FEGITMAG | ASSLGGADTGQLY | 0.2907329 |
| FPLVVQD | ASSLGGADTGQLY | 0.23394337 |
| GAFTTPQ | ASSLGGADTGQLY | 0.6106553 |
| GFGGFGTAGGVGGA | ASSLGGADTGQLY | 0.46188146 |
| GFLAQFL | ASSLGGADTGQLY | 0.5321616 |
| GGAGGAGGNAGVA | ASSLGGADTGQLY | 0.24517609 |
| GGVGTAGAGGAGGAGG | ASSLGGADTGQLY | 0.7155628 |
| GNGGGGDAGNAGSGGN | ASSLGGADTGQLY | 0.4405419 |
| GTGGAGGAGNDAGS | ASSLGGADTGQLY | 0.6074676 |
| GTTNTGWFNS | ASSLGGADTGQLY | 0.2263221 |
| IFMGPLFTL | ASSLGGADTGQLY | 0.28857157 |
| ILAHAIEAI | ASSLGGADTGQLY | 0.6915961 |
| IRSAIREAL | ASSLGGADTGQLY | 0.12171162 |
| ITFAGGG | ASSLGGADTGQLY | 0.14258084 |
| ITGMSGHQEV | ASSLGGADTGQLY | 0.40786868 |

| | | |
|---|---|---|
| IVAGLACTL | ASSLGGADTGQLY | 0.6527442 |
| IVGGGLAGA | ASSLGGADTGQLY | 0.12866391 |
| IVTGVLNI | ASSLGGADTGQLY | 0.2418265 |
| LAQQGCI | ASSLGGADTGQLY | 0.18394338 |
| LCLHVEN | ASSLGGADTGQLY | 0.31841925 |
| LDAISDGS | ASSLGGADTGQLY | 0.3167209 |
| LGGFAAML | ASSLGGADTGQLY | 0.21092314 |
| LGGKNPMI | ASSLGGADTGQLY | 0.5435198 |
| LGPKGVAAIL | ASSLGGADTGQLY | 0.20831685 |
| LMVGVSI | ASSLGGADTGQLY | 0.19063328 |
| LVGAAFVL | ASSLGGADTGQLY | 0.49316806 |
| LVLAVAPL | ASSLGGADTGQLY | 0.5480401 |
| MAAIAVIDAA | ASSLGGADTGQLY | 0.34275228 |
| NNGINAELQ | ASSLGGADTGQLY | 0.45456833 |
| RGEPLVLVHG | ASSLGGADTGQLY | 0.22534847 |
| RGGAGGAAGLI | ASSLGGADTGQLY | 0.72481203 |
| RSYRMHPAV | ASSLGGADTGQLY | 0.32424837 |
| SAAHLLAGQLA | ASSLGGADTGQLY | 0.24412227 |
| SAAVGSY | ASSLGGADTGQLY | 0.59640193 |
| SPHAVLHRI | ASSLGGADTGQLY | 0.5956092 |
| SQIAKVL | ASSLGGADTGQLY | 0.27836743 |
| TALGHLVP | ASSLGGADTGQLY | 0.59379005 |
| TAPTTSGAA | ASSLGGADTGQLY | 0.36217493 |
| TDGDDGNGGSPGT | ASSLGGADTGQLY | 0.52854216 |
| TDGLGEN | ASSLGGADTGQLY | 0.5850086 |
| TGGPPLPGQA | ASSLGGADTGQLY | 0.62125456 |
| TIPAITI | ASSLGGADTGQLY | 0.29115206 |
| TIPGFTI | ASSLGGADTGQLY | 0.41533542 |
| TSIASGALA | ASSLGGADTGQLY | 0.3675541 |
| TVTLPGAAI | ASSLGGADTGQLY | 0.19484106 |
| VASGLSAALP | ASSLGGADTGQLY | 0.7283291 |
| VGAGVIGI | ASSLGGADTGQLY | 0.3223086 |
| VGPKRFGV | ASSLGGADTGQLY | 0.6354704 |
| VNVAFPD | ASSLGGADTGQLY | 0.5055644 |
| VSMTNSGVS | ASSLGGADTGQLY | 0.4685977 |
| VTDVLGDLSV | ASSLGGADTGQLY | 0.24979094 |
| VVVVGAGFAGL | ASSLGGADTGQLY | 0.23584974 |
| VYTVGTVV | ASSLGGADTGQLY | 0.39207858 |
| WTTFSGP | ASSLGGADTGQLY | 0.3479192 |
| YDELVEAF | ASSLGGADTGQLY | 0.29953393 |
| YGTGGAGGNGG | ASSLGGADTGQLY | 0.16238561 |
| YTNVMAV | ASSLGGADTGQLY | 0.4373639 |
| FLEQVHQL | ASSLGGADTGQLY | 0.5893155 |
| GLSLLAVAV | ASSLGGADTGQLY | 0.31990397 |
| LEQVHQL | ASSLGGADTGQLY | 0.2479422 |
| LNAAGAI | ASSLGGADTGQLY | 0.79835093 |
| PAVHASL | ASSLGGADTGQLY | 0.25998387 |
| TELEDDA | ASSLGGADTGQLY | 0.33595258 |
| TPPGPPN | ASSLGGADTGQLY | 0.4253195 |
| TPPPPPPP | ASSLGGADTGQLY | 0.34940284 |
| TQSPALI | ASGRLGQDTQY | 0.41892174 |

| | | |
|---|---|---|
| EFAAAFPQ | ASGRLGQDTQY | 0.48654753 |
| LLSGNLDVL | ASGRLGQDTQY | 0.1799458 |
| GPANLGNYNI | ASGRLGQDTQY | 0.47400373 |
| TLGPTLP | ASGRLGQDTQY | 0.21989031 |
| VYAYSSCCK | ASGRLGQDTQY | 0.3101037 |
| APAPAPAPNI | ASGRLGQDTQY | 0.2536546 |
| PGGPVPIP | ASGRLGQDTQY | 0.41983694 |
| FLTVALEP | ASGRLGQDTQY | 0.6545704 |
| SQYFGANL | ASGRLGQDTQY | 0.46651644 |
| GGVKPAA | ASGRLGQDTQY | 0.13331282 |
| GGTGGAGGAGGW | ASGRLGQDTQY | 0.6801579 |
| GVGGVAGLGGAGG | ASGRLGQDTQY | 0.35391504 |
| VLLVLATTI | ASGRLGQDTQY | 0.7981488 |
| IGIFAAML | ASGRLGQDTQY | 0.25388974 |
| AVGDLID | ASGRLGQDTQY | 0.43994117 |
| NISPLPL | ASGRLGQDTQY | 0.2384015 |
| VTAVFVAI | ASGRLGQDTQY | 0.43148074 |
| SGDDNDAAL | ASGRLGQDTQY | 0.19649185 |
| AVAALAHS | ASGRLGQDTQY | 0.20010151 |
| GGVTVMATP | ASGRLGQDTQY | 0.33976728 |
| AGLAANVRD | ASGRLGQDTQY | 0.39515725 |
| TNRPDLIDPALLRPG | ASGRLGQDTQY | 0.656974 |
| LANLGVTAL | ASGRLGQDTQY | 0.43942797 |
| TGERIEL | ASGRLGQDTQY | 0.35574928 |
| NGEELAA | ASGRLGQDTQY | 0.674032 |
| VLSLVWG | ASGRLGQDTQY | 0.71173394 |
| IGYALWL | ASGRLGQDTQY | 0.48275414 |
| LSLLLAPL | ASGRLGQDTQY | 0.34209287 |
| TLDEALK | ASGRLGQDTQY | 0.23006491 |
| TDGAGTAEGG | ASGRLGQDTQY | 0.3470297 |
| NAPGGAGGNGG | ASGRLGQDTQY | 0.3693279 |
| VSVFMGMGGNF | ASGRLGQDTQY | 0.35635626 |
| LASYLAAL | ASGRLGQDTQY | 0.574895 |
| LVGGTPAL | ASGRLGQDTQY | 0.12214142 |
| VRNADVL | ASGRLGQDTQY | 0.5834608 |
| TCSSPFLVSAM | ASGRLGQDTQY | 0.50575036 |
| VHITDAE | ASGRLGQDTQY | 0.1794278 |
| EYAAPSF | ASGRLGQDTQY | 0.41581142 |
| VVDAFHGGL | ASGRLGQDTQY | 0.51814044 |
| ASVVDPQ | ASGRLGQDTQY | 0.28292993 |
| SYNEFGGLS | ASGRLGQDTQY | 0.59970945 |
| DPDGSVY | ASGRLGQDTQY | 0.25877947 |
| MVSTIHATE | ASGRLGQDTQY | 0.50049776 |
| LAAGVADPV | ASGRLGQDTQY | 0.67643625 |
| TKEQIAA | ASGRLGQDTQY | 0.45148063 |
| SLASPAAL | ASGRLGQDTQY | 0.21244872 |
| IGLVFYGV | ASGRLGQDTQY | 0.3138405 |
| FLLDASNL | ASGRLGQDTQY | 0.2173637 |
| YQVGNFP | ASGRLGQDTQY | 0.22815564 |
| ESLLSQST | ASGRLGQDTQY | 0.123935096 |
| LFTVGFAAVG | ASGRLGQDTQY | 0.5046889 |

| | | |
|---|---|---|
| ATPPTPT | ASGRLGQDTQY | 0.3178312 |
| TDLATLG | ASGRLGQDTQY | 0.50710934 |
| SAGGTGR | ASGRLGQDTQY | 0.37364185 |
| GFSNSGS | ASGRLGQDTQY | 0.30860534 |
| GGNGGSGGTSV | ASGRLGQDTQY | 0.24275012 |
| GTSVATGGAGN | ASGRLGQDTQY | 0.39110047 |
| NGGNGGSGGTS | ASGRLGQDTQY | 0.2021153 |
| AVVKFPH | ASGRLGQDTQY | 0.20079081 |
| TPPPPPP | ASGRLGQDTQY | 0.5333544 |
| AAPAAAAAPA | ASGRLGQDTQY | 0.34324032 |
| IALNEDL | ASGRLGQDTQY | 0.4514776 |
| IVNVSSM | ASGRLGQDTQY | 0.38580775 |
| TGIPPAPR | ASGRLGQDTQY | 0.37961257 |
| FDLGGGTF | ASGRLGQDTQY | 0.27353615 |
| FDLGGGTFDVS | ASGRLGQDTQY | 0.4930567 |
| NEPTAAA | ASGRLGQDTQY | 0.4730333 |
| NEPTAAALA | ASGRLGQDTQY | 0.4106654 |
| GSIINISSI | ASGRLGQDTQY | 0.12264362 |
| VAPGNAGTA | ASGRLGQDTQY | 0.3346659 |
| TGGGASLE | ASGRLGQDTQY | 0.19179356 |
| VAVYNHYKRI | ASGRLGQDTQY | 0.264056 |
| TDPAPATT | ASGRLGQDTQY | 0.46228802 |
| TNRPDILDPALLRPG | ASGRLGQDTQY | 0.7186717 |
| LTTTTTS | ASGRLGQDTQY | 0.14982937 |
| PTGSGKTL | ASGRLGQDTQY | 0.31850663 |
| TGSGKTL | ASGRLGQDTQY | 0.57083046 |
| TIGHVDHGK | ASGRLGQDTQY | 0.30361855 |
| TIGHVDHGKTT | ASGRLGQDTQY | 0.5446874 |
| GHVDHGKTT | ASGRLGQDTQY | 0.31710345 |
| AQFEHTLL | ASGRLGQDTQY | 0.24965647 |
| IVNQDEL | ASGRLGQDTQY | 0.25228894 |
| TNGSQFF | ASGRLGQDTQY | 0.23803368 |
| TGKGIVE | ASGRLGQDTQY | 0.18104155 |
| LISNASDA | ASGRLGQDTQY | 0.16422713 |
| AELRTLGELTKIAKAHGAQ | ASGRLGQDTQY | 0.33326352 |
| APNAVAAAEPSW | ASGRLGQDTQY | 0.15223972 |
| APQVAGAATAVL | ASGRLGQDTQY | 0.39158773 |
| ARRDPAAAAAILPTDA | ASGRLGQDTQY | 0.41656435 |
| DAAARDVLAEAGLAEYF | ASGRLGQDTQY | 0.38826838 |
| DDQIVLSAKANLAKVEQ | ASGRLGQDTQY | 0.17979604 |
| DELLPDTGEDSDEDGH | ASGRLGQDTQY | 0.30693698 |
| ELPEAPANRATVPQD | ASGRLGQDTQY | 0.3230869 |
| FGQSKIERVTGIG | ASGRLGQDTQY | 0.64191306 |
| GAGGTGSTAGGAGGAGGAGGLY | ASGRLGQDTQY | 0.31821316 |
| GHGGTGAAGGNGGTGGAGGSGI | ASGRLGQDTQY | 0.65648746 |
| GNGGNGGAGGEGGPGGAGGAGA | ASGRLGQDTQY | 0.5090901 |
| GSTAGNGGNGGNATNGTIAGQP | ASGRLGQDTQY | 0.2949937 |
| GTGGDGGNGGNGGTGG | ASGRLGQDTQY | 0.7403565 |
| GVRRPAGIPADAPV | ASGRLGQDTQY | 0.37619495 |
| GYRQVTDAHLLTLARR | ASGRLGQDTQY | 0.2272136 |
| IAKMGQDGHDRG | ASGRLGQDTQY | 0.29733437 |

| | | |
|---|---|---|
| KAEPSIEIQYT | ASGRLGQDTQY | 0.37492433 |
| KPVRIRMAADEDTVAVT | ASGRLGQDTQY | 0.2346892 |
| MPPRIAGMRLLVIKPEP | ASGRLGQDTQY | 0.22356169 |
| MKFYAVALFRSRGVPPTR | ASGRLGQDTQY | 0.20575166 |
| NAGGFGILGAVAHSP | ASGRLGQDTQY | 0.2590946 |
| NEGREITLAMAEM | ASGRLGQDTQY | 0.19086671 |
| NELVEWSHPLGHRWRWE | ASGRLGQDTQY | 0.17132857 |
| NLGAGGTGGNGGNPG | ASGRLGQDTQY | 0.19896318 |
| NTEWPKWETVMAADPVD | ASGRLGQDTQY | 0.73484886 |
| PASAAPAETQP | ASGRLGQDTQY | 0.5830821 |
| QQSSDGKNGVRAAEITGE | ASGRLGQDTQY | 0.21833591 |
| RAPDAVTTSWWKE | ASGRLGQDTQY | 0.27482742 |
| SGFGNVGAHTSGWFNQ | ASGRLGQDTQY | 0.55081356 |
| SGVAGAVGQAASVGGLKV | ASGRLGQDTQY | 0.18848588 |
| SNGGAGTGSAGGNGGT | ASGRLGQDTQY | 0.5645998 |
| TGGAGGEGGAAG | ASGRLGQDTQY | 0.49072766 |
| TGGDGGAAGAGGAPGHG | ASGRLGQDTQY | 0.4730795 |
| TMPPLLAAIANAGRHGV | ASGRLGQDTQY | 0.20471554 |
| TSSTAGNSGGA | ASGRLGQDTQY | 0.44252214 |
| VPATVPVGILC | ASGRLGQDTQY | 0.22555558 |
| VQRPDSGGVVLTGRIS | ASGRLGQDTQY | 0.35003725 |
| WTPNPVPATKGVPAGLVQAT | ASGRLGQDTQY | 0.13803062 |
| YALKRALAKFSDRFDVVII | ASGRLGQDTQY | 0.42380267 |
| YLIGSVIV | ASGRLGQDTQY | 0.2648704 |
| YGQLVAVID | ASGRLGQDTQY | 0.26490286 |
| VTVGGMTLPA | ASGRLGQDTQY | 0.17227815 |
| VLALPAV | ASGRLGQDTQY | 0.25633097 |
| VIPPTLNYET | ASGRLGQDTQY | 0.55578864 |
| VGMGGMHPSSG | ASGRLGQDTQY | 0.6955601 |
| VGLAGGAATA | ASGRLGQDTQY | 0.18439999 |
| VGGAGGAGGSGP | ASGRLGQDTQY | 0.33767158 |
| VFAAFMV | ASGRLGQDTQY | 0.5419581 |
| TLYDKLR | ASGRLGQDTQY | 0.40084264 |
| TLAVFTVL | ASGRLGQDTQY | 0.18132932 |
| TGSVQPARV | ASGRLGQDTQY | 0.39988917 |
| TGGGAGSGGT | ASGRLGQDTQY | 0.17678334 |
| SHLETSL | ASGRLGQDTQY | 0.48401076 |
| SFTTGMT | ASGRLGQDTQY | 0.228324 |
| QVEVAKRMGVRQPTVS | ASGRLGQDTQY | 0.24779366 |
| QGSGGPALAM | ASGRLGQDTQY | 0.31074518 |
| MLAPSNP | ASGRLGQDTQY | 0.485318 |
| LVNNAAI | ASGRLGQDTQY | 0.2509351 |
| LTGEINTLA | ASGRLGQDTQY | 0.2359188 |
| LSIFGPV | ASGRLGQDTQY | 0.21651593 |
| LLALLANL | ASGRLGQDTQY | 0.14590752 |
| LISNASDALDKLR | ASGRLGQDTQY | 0.28308862 |
| LINEAVP | ASGRLGQDTQY | 0.36916304 |
| LGYAFSGL | ASGRLGQDTQY | 0.2684253 |
| LDFVAVAL | ASGRLGQDTQY | 0.5027871 |
| LAQPGAE | ASGRLGQDTQY | 0.27700976 |
| ISNLGAVTGF | ASGRLGQDTQY | 0.3337632 |

| | | |
|---|---|---|
| HQEDPGRI | ASGRLGQDTQY | 0.35462424 |
| GTPGEPTP | ASGRLGQDTQY | 0.29008535 |
| GFFNSGS | ASGRLGQDTQY | 0.587059 |
| GAGGGGLTAGAN | ASGRLGQDTQY | 0.3520257 |
| FVDALVNM | ASGRLGQDTQY | 0.2620169 |
| FLTVAPEL | ASGRLGQDTQY | 0.748629 |
| FETLTAL | ASGRLGQDTQY | 0.19632941 |
| DVINPIEL | ASGRLGQDTQY | 0.20058331 |
| APVSGAPV | ASGRLGQDTQY | 0.5717912 |
| ALVGGLAW | ASGRLGQDTQY | 0.2173607 |
| ALAAAAATAI | ASGRLGQDTQY | 0.5844133 |
| AFGLGLLL | ASGRLGQDTQY | 0.28044423 |
| AFFAAASL | ASGRLGQDTQY | 0.63096094 |
| AAAALPAFSPP | ASGRLGQDTQY | 0.25310934 |
| VVLAPAL | ASGRLGQDTQY | 0.6245339 |
| TIGHVDHGKTTL | ASGRLGQDTQY | 0.6844595 |
| LISNASDALDK | ASGRLGQDTQY | 0.28461602 |
| LISNASDAL | ASGRLGQDTQY | 0.5121443 |
| ISNASDALDK | ASGRLGQDTQY | 0.17995599 |
| NQAETLVYQTE | ASGRLGQDTQY | 0.2958506 |
| TGGTGGTAGAG | ASGRLGQDTQY | 0.5272489 |
| TGSTAGGAGGA | ASGRLGQDTQY | 0.2941834 |
| DPLAVPVAAGPV | ASGRLGQDTQY | 0.5153241 |
| GGTGGTPGNGGN | ASGRLGQDTQY | 0.5365876 |
| PAGLAGVDPATL | ASGRLGQDTQY | 0.6615646 |
| GNNLVVEAPHAGASV | ASGRLGQDTQY | 0.17227744 |
| KVTRIMSASTLEQPA | ASGRLGQDTQY | 0.38759273 |
| LGFALPIALPRRTVV | ASGRLGQDTQY | 0.14518912 |
| TRLGNLTARKLVVLV | ASGRLGQDTQY | 0.18970008 |
| WSQQRRLIYFPSAGP | ASGRLGQDTQY | 0.18949775 |
| GPNGNPDPMAAAVDIR | ASGRLGQDTQY | 0.48951215 |
| TQRCVSVADLRAAASR | ASGRLGQDTQY | 0.3813167 |
| QFVLDVAEHPVEVLTETV | ASGRLGQDTQY | 0.4727738 |
| GGAGGTGGAGGNGGTQAGDG | ASGRLGQDTQY | 0.5090409 |
| PLLATLRAIAVDVDAKPAQV | ASGRLGQDTQY | 0.7075858 |
| APKFRTGGYMIDFWGVGYQVAK | ASGRLGQDTQY | 0.35591745 |
| GAGGAGGAGGAAGGPGGTGGTG | ASGRLGQDTQY | 0.25537157 |
| IGALGLLATDVAPAGSVAFGVA | ASGRLGQDTQY | 0.18210268 |
| AGGAGGIGGNGGAGGAGGVLMG | ASGRLGQDTQY | 0.13628006 |
| ADELIGLL | ASGRLGQDTQY | 0.25505742 |
| ADRALKA | ASGRLGQDTQY | 0.3390397 |
| AGAVGQVGGAGGA | ASGRLGQDTQY | 0.13994873 |
| AGGGGGLGGAP | ASGRLGQDTQY | 0.50882745 |
| AGGLLYGNGGAG | ASGRLGQDTQY | 0.54202175 |
| AGLESIGL | ASGRLGQDTQY | 0.50333285 |
| AGLKTGII | ASGRLGQDTQY | 0.21409766 |
| ASLAALLL | ASGRLGQDTQY | 0.26603493 |
| ASVINISSI | ASGRLGQDTQY | 0.42903316 |
| ATPPGPA | ASGRLGQDTQY | 0.53695977 |
| ATVASALV | ASGRLGQDTQY | 0.35330698 |
| ATVASLIL | ASGRLGQDTQY | 0.19336188 |

| | | |
|---|---|---|
| AVNGPVLIRH | ASGRLGQDTQY | 0.19663775 |
| DGAAGAVGI | ASGRLGQDTQY | 0.24852633 |
| DPLLHGI | ASGRLGQDTQY | 0.7038947 |
| ELGAALGAL | ASGRLGQDTQY | 0.5248127 |
| ELGRTQAV | ASGRLGQDTQY | 0.6235486 |
| FVAGGYLS | ASGRLGQDTQY | 0.6050743 |
| GASGGAGGAGGW | ASGRLGQDTQY | 0.13891217 |
| GGAGAGGDGGAGGI | ASGRLGQDTQY | 0.20823392 |
| GTGHNGSQPGMGGNG | ASGRLGQDTQY | 0.536357 |
| GYAAASASAS | ASGRLGQDTQY | 0.3742891 |
| HQVDVAAVQQG | ASGRLGQDTQY | 0.4010667 |
| IGAAIAHT | ASGRLGQDTQY | 0.6966233 |
| IGMIEVAQL | ASGRLGQDTQY | 0.20472363 |
| IRLAPEHTSAMD | ASGRLGQDTQY | 0.33571288 |
| IYNVSVL | ASGRLGQDTQY | 0.46926647 |
| KQGFLGM | ASGRLGQDTQY | 0.34652054 |
| LESVITQL | ASGRLGQDTQY | 0.38579527 |
| LGLGALVVV | ASGRLGQDTQY | 0.21426576 |
| LIGNGGNGGSGGI | ASGRLGQDTQY | 0.4257394 |
| LNAINAPTL | ASGRLGQDTQY | 0.78470165 |
| LQLLNTGV | ASGRLGQDTQY | 0.52470744 |
| LQTSIEF | ASGRLGQDTQY | 0.51592684 |
| MSFAVSEL | ASGRLGQDTQY | 0.4079231 |
| NTGALGL | ASGRLGQDTQY | 0.2343281 |
| PPGAGGAGGAGTT | ASGRLGQDTQY | 0.3656492 |
| QLVGLGI | ASGRLGQDTQY | 0.36362666 |
| RFLEQVHQL | ASGRLGQDTQY | 0.276626 |
| RGVGYVLR | ASGRLGQDTQY | 0.23446003 |
| SATGVFAGVI | ASGRLGQDTQY | 0.34296453 |
| SLPAITLP | ASGRLGQDTQY | 0.37688574 |
| STALAFTV | ASGRLGQDTQY | 0.790444 |
| SVLTVGLGK | ASGRLGQDTQY | 0.40642428 |
| TALAGAGSGI | ASGRLGQDTQY | 0.5171712 |
| TGLPFTGI | ASGRLGQDTQY | 0.27986312 |
| VAEHVTV | ASGRLGQDTQY | 0.32000196 |
| VDIVLNSL | ASGRLGQDTQY | 0.52359354 |
| VLTGGGAVL | ASGRLGQDTQY | 0.23721065 |
| VNVGLAYGI | ASGRLGQDTQY | 0.56575584 |
| VSSAAVGAV | ASGRLGQDTQY | 0.27231953 |
| VTSVYEAGV | ASGRLGQDTQY | 0.263157 |
| WVALDRGA | ASGRLGQDTQY | 0.30002058 |
| YLVGSVVI | ASGRLGQDTQY | 0.5627885 |
| GPPGTGKTLL | ASGRLGQDTQY | 0.45234543 |
| GVGMTKF | ASGRLGQDTQY | 0.36086905 |
| IGIGGSDL | ASGRLGQDTQY | 0.32800776 |
| IGTLLAEL | ASGRLGQDTQY | 0.5283827 |
| IYTSGTT | ASGRLGQDTQY | 0.7705473 |
| LSFPAGV | ASGRLGQDTQY | 0.19646999 |
| LYTSGTT | ASGRLGQDTQY | 0.69198143 |
| VGAFPFGF | ASGRLGQDTQY | 0.3437691 |
| VGAGTVEF | ASGRLGQDTQY | 0.16448548 |

| | | |
|---|---|---|
| VGKSTLLL | ASGRLGQDTQY | 0.36223972 |
| AALAHAGGGIV | ASGRLGQDTQY | 0.53620195 |
| AALVAVSLL | ASGRLGQDTQY | 0.70381975 |
| AAVSRAVAAL | ASGRLGQDTQY | 0.4868405 |
| AGGSSGIGGF | ASGRLGQDTQY | 0.29012817 |
| AGLTVANANG | ASGRLGQDTQY | 0.22982283 |
| AGLVGAVL | ASGRLGQDTQY | 0.63032925 |
| ALSQAFTV | ASGRLGQDTQY | 0.48668393 |
| ALTFSVSL | ASGRLGQDTQY | 0.5313865 |
| ALTGAALVAL | ASGRLGQDTQY | 0.23732415 |
| ASFSIGAGVL | ASGRLGQDTQY | 0.21524648 |
| DGSVYKI | ASGRLGQDTQY | 0.39564258 |
| EQLAEIV | ASGRLGQDTQY | 0.7565812 |
| FAVSELTVM | ASGRLGQDTQY | 0.249426 |
| GAAPLAAGL | ASGRLGQDTQY | 0.5080656 |
| GADGGDGGSGGAGG | ASGRLGQDTQY | 0.47479677 |
| GFGLDTGGAGG | ASGRLGQDTQY | 0.15133767 |
| GLASGGLL | ASGRLGQDTQY | 0.2562883 |
| GRIAGAAVLL | ASGRLGQDTQY | 0.5884414 |
| GSAGSLNTSF | ASGRLGQDTQY | 0.3986234 |
| GTGQIGI | ASGRLGQDTQY | 0.29646444 |
| GTLIALIL | ASGRLGQDTQY | 0.57539475 |
| IAATMGFVVG | ASGRLGQDTQY | 0.786817 |
| IFAQALL | ASGRLGQDTQY | 0.65252465 |
| IGAAVALGF | ASGRLGQDTQY | 0.19495142 |
| IGLTPLAL | ASGRLGQDTQY | 0.15372626 |
| ILGSNTSTL | ASGRLGQDTQY | 0.20389792 |
| IVFSLTVL | ASGRLGQDTQY | 0.3600315 |
| LASKATGF | ASGRLGQDTQY | 0.2693897 |
| LASVALIL | ASGRLGQDTQY | 0.2886687 |
| LFAVGLTL | ASGRLGQDTQY | 0.59142315 |
| LFNGGIGTYI | ASGRLGQDTQY | 0.5882555 |
| LTLGAGII | ASGRLGQDTQY | 0.20501162 |
| LVAAALGDI | ASGRLGQDTQY | 0.18729736 |
| NTAFVQLGI | ASGRLGQDTQY | 0.39185336 |
| QLFSGAEL | ASGRLGQDTQY | 0.6488564 |
| SAATFITL | ASGRLGQDTQY | 0.22164018 |
| SNYGLANLGV | ASGRLGQDTQY | 0.5975035 |
| SYGGAYAVM | ASGRLGQDTQY | 0.19450718 |
| TTGVGGLTLG | ASGRLGQDTQY | 0.17925186 |
| TTQTIYI | ASGRLGQDTQY | 0.45742682 |
| VAAGAGLGAVI | ASGRLGQDTQY | 0.26772743 |
| VAAVVTLL | ASGRLGQDTQY | 0.55627596 |
| VGFLGGLI | ASGRLGQDTQY | 0.4398995 |
| VGVIPILL | ASGRLGQDTQY | 0.49259022 |
| VIFASSLI | ASGRLGQDTQY | 0.34612918 |
| VKGGDGGAAAG | ASGRLGQDTQY | 0.3632018 |
| VTKTFTNI | ASGRLGQDTQY | 0.47284132 |
| VVAAAILL | ASGRLGQDTQY | 0.2755945 |
| YAALDVEL | ASGRLGQDTQY | 0.3298438 |
| LASVALLL | ASGRLGQDTQY | 0.15240833 |

| | | |
|---|---|---|
| LDAVHRV | ASGRLGQDTQY | 0.46407574 |
| QGLSSSIF | ASGRLGQDTQY | 0.13309443 |
| RGTFANIRL | ASGRLGQDTQY | 0.5953121 |
| RQLWWGHRI | ASGRLGQDTQY | 0.5539565 |
| YGPPGTGKTLL | ASGRLGQDTQY | 0.24519865 |
| AALVGAPANVA | ASGRLGQDTQY | 0.47198638 |
| AATFRRTE | ASGRLGQDTQY | 0.2501003 |
| AGALGGGI | ASGRLGQDTQY | 0.19855602 |
| AGGAGGAGGIGAG | ASGRLGQDTQY | 0.2770162 |
| AGSGALGL | ASGRLGQDTQY | 0.46722582 |
| AQIADQI | ASGRLGQDTQY | 0.32278544 |
| ASAVAVTYL | ASGRLGQDTQY | 0.52653027 |
| ASTVLGGTGG | ASGRLGQDTQY | 0.475196 |
| AVGGEPAK | ASGRLGQDTQY | 0.22845837 |
| CEQGTFGGTC | ASGRLGQDTQY | 0.32462615 |
| DEGKAPAG | ASGRLGQDTQY | 0.5051035 |
| DTVVFPAL | ASGRLGQDTQY | 0.40724087 |
| FEGITMAG | ASGRLGQDTQY | 0.3862008 |
| FPLVVQD | ASGRLGQDTQY | 0.34939253 |
| GAFTTPQ | ASGRLGQDTQY | 0.20406559 |
| GFGGFGTAGGVGGA | ASGRLGQDTQY | 0.6052736 |
| GFLAQFL | ASGRLGQDTQY | 0.4115014 |
| GGAGGAGGNAGVA | ASGRLGQDTQY | 0.2533233 |
| GGVGTAGAGGAGGAGG | ASGRLGQDTQY | 0.41415215 |
| GNGGGGDAGNAGSGGN | ASGRLGQDTQY | 0.17014885 |
| GTGGAGGAGNDAGS | ASGRLGQDTQY | 0.27548173 |
| GTTNTGWFNS | ASGRLGQDTQY | 0.24600808 |
| IFMGPLFTL | ASGRLGQDTQY | 0.37337443 |
| ILAHAIEAI | ASGRLGQDTQY | 0.5156465 |
| IRSAIREAL | ASGRLGQDTQY | 0.22056909 |
| ITFAGGG | ASGRLGQDTQY | 0.40266177 |
| ITGMSGHQEV | ASGRLGQDTQY | 0.27564284 |
| IVAGLACTL | ASGRLGQDTQY | 0.23282976 |
| IVGGGLAGA | ASGRLGQDTQY | 0.47414786 |
| IVTGVLNI | ASGRLGQDTQY | 0.24663506 |
| LAQQGCI | ASGRLGQDTQY | 0.25525883 |
| LCLHVEN | ASGRLGQDTQY | 0.29586717 |
| LDAISDGS | ASGRLGQDTQY | 0.3002988 |
| LGGFAAML | ASGRLGQDTQY | 0.26795086 |
| LGGKNPMI | ASGRLGQDTQY | 0.42734784 |
| LGPKGVAAIL | ASGRLGQDTQY | 0.25343996 |
| LMVGVSI | ASGRLGQDTQY | 0.5202037 |
| LVGAAFVL | ASGRLGQDTQY | 0.380761 |
| LVLAVAPL | ASGRLGQDTQY | 0.4232441 |
| MAAIAVIDAA | ASGRLGQDTQY | 0.39513332 |
| NNGINAELQ | ASGRLGQDTQY | 0.3673496 |
| RGEPLVLVHG | ASGRLGQDTQY | 0.67771745 |
| RGGAGGAAGLI | ASGRLGQDTQY | 0.42801034 |
| RSYRMHPAV | ASGRLGQDTQY | 0.2793989 |
| SAAHLLAGQLA | ASGRLGQDTQY | 0.30242833 |
| SAAVGSY | ASGRLGQDTQY | 0.3771939 |

| | | |
|---|---|---|
| SPHAVLHRI | ASGRLGQDTQY | 0.5789165 |
| SQIAKVL | ASGRLGQDTQY | 0.3271373 |
| TALGHLVP | ASGRLGQDTQY | 0.19494554 |
| TAPTTSGAA | ASGRLGQDTQY | 0.27049053 |
| TDGDDGNGGSPGT | ASGRLGQDTQY | 0.2707339 |
| TDGLGEN | ASGRLGQDTQY | 0.3170564 |
| TGGPPLPGQA | ASGRLGQDTQY | 0.18180086 |
| TIPAITI | ASGRLGQDTQY | 0.47700092 |
| TIPGFTI | ASGRLGQDTQY | 0.7667198 |
| TSIASGALA | ASGRLGQDTQY | 0.23960193 |
| TVTLPGAAI | ASGRLGQDTQY | 0.7496928 |
| VASGLSAALP | ASGRLGQDTQY | 0.23991345 |
| VGAGVIGI | ASGRLGQDTQY | 0.35006225 |
| VGPKRFGV | ASGRLGQDTQY | 0.45452988 |
| VNVAFPD | ASGRLGQDTQY | 0.26303762 |
| VSMTNSGVS | ASGRLGQDTQY | 0.2590567 |
| VTDVLGDLSV | ASGRLGQDTQY | 0.34727642 |
| VVVVGAGFAGL | ASGRLGQDTQY | 0.395621 |
| VYTVGTVV | ASGRLGQDTQY | 0.53328127 |
| WTTFSGP | ASGRLGQDTQY | 0.26100686 |
| YDELVEAF | ASGRLGQDTQY | 0.24501583 |
| YGTGGAGGNGG | ASGRLGQDTQY | 0.37335315 |
| YTNVMAV | ASGRLGQDTQY | 0.31814617 |
| FLEQVHQL | ASGRLGQDTQY | 0.4038881 |
| GLSLLAVAV | ASGRLGQDTQY | 0.27772215 |
| LEQVHQL | ASGRLGQDTQY | 0.21041037 |
| LNAAGAI | ASGRLGQDTQY | 0.44217226 |
| PAVHASL | ASGRLGQDTQY | 0.20975015 |
| TELEDDA | ASGRLGQDTQY | 0.74086857 |
| TPPGPPN | ASGRLGQDTQY | 0.69005907 |
| TPPPPPPP | ASGRLGQDTQY | 0.61524165 |
| TQSPALI | AWSHRGKSQNTLY | 0.2923905 |
| EFAAAFPQ | AWSHRGKSQNTLY | 0.7144662 |
| LLSGNLDVL | AWSHRGKSQNTLY | 0.35179263 |
| GPANLGNYNI | AWSHRGKSQNTLY | 0.31035042 |
| TLGPTLP | AWSHRGKSQNTLY | 0.46257457 |
| VYAYSSCCK | AWSHRGKSQNTLY | 0.08292385 |
| APAPAPAPNI | AWSHRGKSQNTLY | 0.0837616 |
| PGGPVPIP | AWSHRGKSQNTLY | 0.08286225 |
| FLTVALEP | AWSHRGKSQNTLY | 0.08322122 |
| SQYFGANL | AWSHRGKSQNTLY | 0.08468411 |
| GGVKPAA | AWSHRGKSQNTLY | 0.08303395 |
| GGTGGAGGAGGW | AWSHRGKSQNTLY | 0.083624475 |
| GVGGVAGLGGAGG | AWSHRGKSQNTLY | 0.085442595 |
| VLLVLATTI | AWSHRGKSQNTLY | 0.08281548 |
| IGIFAAML | AWSHRGKSQNTLY | 0.08311493 |
| AVGDLID | AWSHRGKSQNTLY | 0.084734924 |
| NISPLPL | AWSHRGKSQNTLY | 0.08501342 |
| VTAVFVAI | AWSHRGKSQNTLY | 0.08443757 |
| SGDDNDAAL | AWSHRGKSQNTLY | 0.08312268 |
| AVAALAHS | AWSHRGKSQNTLY | 0.082884856 |

| | | |
|---|---|---|
| GGVTVMATP | AWSHRGKSQNTLY | 0.08348663 |
| AGLAANVRD | AWSHRGKSQNTLY | 0.08307859 |
| TNRPDLIDPALLRPG | AWSHRGKSQNTLY | 0.08282949 |
| LANLGVTAL | AWSHRGKSQNTLY | 0.083072126 |
| TGERIEL | AWSHRGKSQNTLY | 0.08272101 |
| NGEELAA | AWSHRGKSQNTLY | 0.08306405 |
| VLSLVWG | AWSHRGKSQNTLY | 0.083642095 |
| IGYALWL | AWSHRGKSQNTLY | 0.08350359 |
| LSLLLAPL | AWSHRGKSQNTLY | 0.083250456 |
| TLDEALK | AWSHRGKSQNTLY | 0.08340013 |
| TDGAGTAEGG | AWSHRGKSQNTLY | 0.08354256 |
| NAPGGAGGNGG | AWSHRGKSQNTLY | 0.08443107 |
| VSVFMGMGGNF | AWSHRGKSQNTLY | 0.08369977 |
| LASYLAAL | AWSHRGKSQNTLY | 0.08533528 |
| LVGGTPAL | AWSHRGKSQNTLY | 0.084272586 |
| VRNADVL | AWSHRGKSQNTLY | 0.08282847 |
| TCSSPFLVSAM | AWSHRGKSQNTLY | 0.085097544 |
| VHITDAE | AWSHRGKSQNTLY | 0.08506215 |
| EYAAPSF | AWSHRGKSQNTLY | 0.08291179 |
| VVDAFHGGL | AWSHRGKSQNTLY | 0.08294291 |
| ASVVDPQ | AWSHRGKSQNTLY | 0.0827198 |
| SYNEFGGLS | AWSHRGKSQNTLY | 0.08377638 |
| DPDGSVY | AWSHRGKSQNTLY | 0.083303094 |
| MVSTIHATE | AWSHRGKSQNTLY | 0.08442095 |
| LAAGVADPV | AWSHRGKSQNTLY | 0.08429851 |
| TKEQIAA | AWSHRGKSQNTLY | 0.08290395 |
| SLASPAAL | AWSHRGKSQNTLY | 0.082829125 |
| IGLVFYGV | AWSHRGKSQNTLY | 0.08371185 |
| FLLDASNL | AWSHRGKSQNTLY | 0.08322931 |
| YQVGNFP | AWSHRGKSQNTLY | 0.08348071 |
| ESLLSQST | AWSHRGKSQNTLY | 0.0840346 |
| LFTVGFAAVG | AWSHRGKSQNTLY | 0.08283902 |
| ATPPTPT | AWSHRGKSQNTLY | 0.08316824 |
| TDLATLG | AWSHRGKSQNTLY | 0.08257346 |
| SAGGTGR | AWSHRGKSQNTLY | 0.08284564 |
| GFSNSGS | AWSHRGKSQNTLY | 0.08397329 |
| GGNGGSGGTSV | AWSHRGKSQNTLY | 0.08284793 |
| GTSVATGGAGN | AWSHRGKSQNTLY | 0.08831783 |
| NGGNGGSGGTS | AWSHRGKSQNTLY | 0.087486684 |
| AVVKFPH | AWSHRGKSQNTLY | 0.084750585 |
| TPPPPPP | AWSHRGKSQNTLY | 0.08322717 |
| AAPAAAAAPA | AWSHRGKSQNTLY | 0.08273143 |
| IALNEDL | AWSHRGKSQNTLY | 0.08314903 |
| IVNVSSM | AWSHRGKSQNTLY | 0.08368485 |
| TGIPPAPR | AWSHRGKSQNTLY | 0.08293125 |
| FDLGGGTF | AWSHRGKSQNTLY | 0.083203524 |
| FDLGGGTFDVS | AWSHRGKSQNTLY | 0.08363489 |
| NEPTAAA | AWSHRGKSQNTLY | 0.08651277 |
| NEPTAAALA | AWSHRGKSQNTLY | 0.08393551 |
| GSIINISSI | AWSHRGKSQNTLY | 0.0842926 |
| VAPGNAGTA | AWSHRGKSQNTLY | 0.0840768 |

| | | |
|---|---|---|
| TGGGASLE | AWSHRGKSQNTLY | 0.082685694 |
| VAVYNHYKRI | AWSHRGKSQNTLY | 0.08275675 |
| TDPAPATT | AWSHRGKSQNTLY | 0.082907684 |
| TNRPDILDPALLRPG | AWSHRGKSQNTLY | 0.08320248 |
| LTTTTTS | AWSHRGKSQNTLY | 0.084118366 |
| PTGSGKTL | AWSHRGKSQNTLY | 0.084174916 |
| TGSGKTL | AWSHRGKSQNTLY | 0.08574752 |
| TIGHVDHGK | AWSHRGKSQNTLY | 0.08347964 |
| TIGHVDHGKTT | AWSHRGKSQNTLY | 0.082772054 |
| GHVDHGKTT | AWSHRGKSQNTLY | 0.08336324 |
| AQFEHTLL | AWSHRGKSQNTLY | 0.08345256 |
| IVNQDEL | AWSHRGKSQNTLY | 0.08374641 |
| TNGSQFF | AWSHRGKSQNTLY | 0.08339222 |
| TGKGIVE | AWSHRGKSQNTLY | 0.08363803 |
| LISNASDA | AWSHRGKSQNTLY | 0.082973264 |
| AELRTLGELTKIAKAHGAQ | AWSHRGKSQNTLY | 0.08462129 |
| APNAVAAAEPSW | AWSHRGKSQNTLY | 0.08298258 |
| APQVAGAATAVL | AWSHRGKSQNTLY | 0.08289954 |
| ARRDPAAAAAILPTDA | AWSHRGKSQNTLY | 0.083745986 |
| DAAARDVLAEAGLAEYF | AWSHRGKSQNTLY | 0.082940735 |
| DDQIVLSAKANLAKVEQ | AWSHRGKSQNTLY | 0.08434541 |
| DELLPDTGEDSDEDGH | AWSHRGKSQNTLY | 0.08341513 |
| ELPEAPANRATVPQD | AWSHRGKSQNTLY | 0.082949184 |
| FGQSKIERVTGIG | AWSHRGKSQNTLY | 0.082926676 |
| GAGGTGSTAGGAGGAGGAGGLY | AWSHRGKSQNTLY | 0.08281498 |
| GHGGTGAAGGNGGTGGAGGSGI | AWSHRGKSQNTLY | 0.08306898 |
| GNGGNGGAGGEGGPGGAGGAGA | AWSHRGKSQNTLY | 0.08308853 |
| GSTAGNGGNGGNATNGTIAGQP | AWSHRGKSQNTLY | 0.08285127 |
| GTGGDGGNGGNGGTGG | AWSHRGKSQNTLY | 0.083454594 |
| GVRRPAGIPADAPV | AWSHRGKSQNTLY | 0.083427 |
| GYRQVTDAHLLTLARR | AWSHRGKSQNTLY | 0.08359424 |
| IAKMGQDGHDRG | AWSHRGKSQNTLY | 0.08333948 |
| KAEPSIEIQYT | AWSHRGKSQNTLY | 0.083037585 |
| KPVRIRMAADEDTVAVT | AWSHRGKSQNTLY | 0.08419615 |
| MPPRIAGMRLLVIKPEP | AWSHRGKSQNTLY | 0.082795024 |
| MKFYAVALFRSRGVPPTR | AWSHRGKSQNTLY | 0.08318462 |
| NAGGFGILGAVAHSP | AWSHRGKSQNTLY | 0.08447805 |
| NEGREITLAMAEM | AWSHRGKSQNTLY | 0.08307586 |
| NELVEWSHPLGHRWRWE | AWSHRGKSQNTLY | 0.08396736 |
| NLGAGGTGGNGGNPG | AWSHRGKSQNTLY | 0.082704514 |
| NTEWPKWETVMAADPVD | AWSHRGKSQNTLY | 0.08277475 |
| PASAAPAETQP | AWSHRGKSQNTLY | 0.08284178 |
| QQSSDGKNGVRAAEITGE | AWSHRGKSQNTLY | 0.08590215 |
| RAPDAVTTSWWKE | AWSHRGKSQNTLY | 0.08288096 |
| SGFGNVGAHTSGWFNQ | AWSHRGKSQNTLY | 0.08307989 |
| SGVAGAVGQAASVGGLKV | AWSHRGKSQNTLY | 0.083082475 |
| SNGGAGTGSAGGNGGT | AWSHRGKSQNTLY | 0.08366669 |
| TGGAGGEGGAAG | AWSHRGKSQNTLY | 0.083332695 |
| TGGDGGAAGAGGAPGHG | AWSHRGKSQNTLY | 0.083121076 |
| TMPPLLAAIANAGRHGV | AWSHRGKSQNTLY | 0.08288591 |
| TSSTAGNSGGA | AWSHRGKSQNTLY | 0.08521444 |

| | | |
|---|---|---|
| VPATVPVGILC | AWSHRGKSQNTLY | 0.08295543 |
| VQRPDSGGVVLTGRIS | AWSHRGKSQNTLY | 0.082762815 |
| WTPNPVPATKGVPAGLVQAT | AWSHRGKSQNTLY | 0.084019125 |
| YALKRALAKFSDRFDVVII | AWSHRGKSQNTLY | 0.084674165 |
| YLIGSVIV | AWSHRGKSQNTLY | 0.08548695 |
| YGQLVAVID | AWSHRGKSQNTLY | 0.083751164 |
| VTVGGMTLPA | AWSHRGKSQNTLY | 0.082987286 |
| VLALPAV | AWSHRGKSQNTLY | 0.082603574 |
| VIPPTLNYET | AWSHRGKSQNTLY | 0.08276739 |
| VGMGGMHPSSG | AWSHRGKSQNTLY | 0.08514405 |
| VGLAGGAATA | AWSHRGKSQNTLY | 0.085515045 |
| VGGAGGAGGSGP | AWSHRGKSQNTLY | 0.08386555 |
| VFAAFMV | AWSHRGKSQNTLY | 0.08340971 |
| TLYDKLR | AWSHRGKSQNTLY | 0.08534316 |
| TLAVFTVL | AWSHRGKSQNTLY | 0.08276388 |
| TGSVQPARV | AWSHRGKSQNTLY | 0.083234206 |
| TGGGAGSGGT | AWSHRGKSQNTLY | 0.08307571 |
| SHLETSL | AWSHRGKSQNTLY | 0.08295297 |
| SFTTGMT | AWSHRGKSQNTLY | 0.083054975 |
| QVEVAKRMGVRQPTVS | AWSHRGKSQNTLY | 0.08347229 |
| QGSGGPALAM | AWSHRGKSQNTLY | 0.084794685 |
| MLAPSNP | AWSHRGKSQNTLY | 0.082824126 |
| LVNNAAI | AWSHRGKSQNTLY | 0.08302403 |
| LTGEINTLA | AWSHRGKSQNTLY | 0.08267028 |
| LSIFGPV | AWSHRGKSQNTLY | 0.08297171 |
| LLALLANL | AWSHRGKSQNTLY | 0.086993225 |
| LISNASDALDKLR | AWSHRGKSQNTLY | 0.08590078 |
| LINEAVP | AWSHRGKSQNTLY | 0.08352625 |
| LGYAFSGL | AWSHRGKSQNTLY | 0.0841509 |
| LDFVAVAL | AWSHRGKSQNTLY | 0.083190456 |
| LAQPGAE | AWSHRGKSQNTLY | 0.083212726 |
| ISNLGAVTGF | AWSHRGKSQNTLY | 0.082952134 |
| HQEDPGRI | AWSHRGKSQNTLY | 0.08291727 |
| GTPGEPTP | AWSHRGKSQNTLY | 0.08405272 |
| GFFNSGS | AWSHRGKSQNTLY | 0.08396186 |
| GAGGGGLTAGAN | AWSHRGKSQNTLY | 0.08392707 |
| FVDALVNM | AWSHRGKSQNTLY | 0.08359271 |
| FLTVAPEL | AWSHRGKSQNTLY | 0.08460719 |
| FETLTAL | AWSHRGKSQNTLY | 0.08313494 |
| DVINPIEL | AWSHRGKSQNTLY | 0.08336442 |
| APVSGAPV | AWSHRGKSQNTLY | 0.08383364 |
| ALVGGLAW | AWSHRGKSQNTLY | 0.0846063 |
| ALAAAAATAI | AWSHRGKSQNTLY | 0.08268038 |
| AFGLGLLL | AWSHRGKSQNTLY | 0.0831282 |
| AFFAAASL | AWSHRGKSQNTLY | 0.082749136 |
| AAAALPAFSPP | AWSHRGKSQNTLY | 0.08326048 |
| VVLAPAL | AWSHRGKSQNTLY | 0.08359942 |
| TIGHVDHGKTTL | AWSHRGKSQNTLY | 0.083132476 |
| LISNASDALDK | AWSHRGKSQNTLY | 0.08336067 |
| LISNASDAL | AWSHRGKSQNTLY | 0.08303057 |
| ISNASDALDK | AWSHRGKSQNTLY | 0.08436241 |

| | | |
|---|---|---|
| NQAETLVYQTE | AWSHRGKSQNTLY | 0.08291485 |
| TGGTGGTAGAG | AWSHRGKSQNTLY | 0.08447577 |
| TGSTAGGAGGA | AWSHRGKSQNTLY | 0.0828333 |
| DPLAVPVAAGPV | AWSHRGKSQNTLY | 0.083101444 |
| GGTGGTPGNGGN | AWSHRGKSQNTLY | 0.08317893 |
| PAGLAGVDPATL | AWSHRGKSQNTLY | 0.083103575 |
| GNNLVVEAPHAGASV | AWSHRGKSQNTLY | 0.08464769 |
| KVTRIMSASTLEQPA | AWSHRGKSQNTLY | 0.08298083 |
| LGFALPIALPRRTVV | AWSHRGKSQNTLY | 0.08402566 |
| TRLGNLTARKLVVLV | AWSHRGKSQNTLY | 0.0837367 |
| WSQQRRLIYFPSAGP | AWSHRGKSQNTLY | 0.082951695 |
| GPNGNPDPMAAAVDIR | AWSHRGKSQNTLY | 0.08298794 |
| TQRCVSVADLRAAASR | AWSHRGKSQNTLY | 0.08489345 |
| QFVLDVAEHPVEVLTETV | AWSHRGKSQNTLY | 0.0833847 |
| GGAGGTGGAGGNGGTQAGDG | AWSHRGKSQNTLY | 0.0830816 |
| PLLATLRAIAVDVDAKPAQV | AWSHRGKSQNTLY | 0.083499625 |
| APKFRTGGYMIDFWGVGYQVAK | AWSHRGKSQNTLY | 0.08301177 |
| GAGGAGGAGGAAGGPGGTGGTG | AWSHRGKSQNTLY | 0.08346048 |
| IGALGLLATDVAPAGSVAFGVA | AWSHRGKSQNTLY | 0.08274838 |
| AGGAGGIGGNGGAGGAGGVLMG | AWSHRGKSQNTLY | 0.08309768 |
| ADELIGLL | AWSHRGKSQNTLY | 0.08294812 |
| ADRALKA | AWSHRGKSQNTLY | 0.08291587 |
| AGAVGQVGGAGGA | AWSHRGKSQNTLY | 0.0852462 |
| AGGGGGLGGAP | AWSHRGKSQNTLY | 0.08451235 |
| AGGLLYGNGGAG | AWSHRGKSQNTLY | 0.08282112 |
| AGLESIGL | AWSHRGKSQNTLY | 0.08501966 |
| AGLKTGII | AWSHRGKSQNTLY | 0.083286874 |
| ASLAALLL | AWSHRGKSQNTLY | 0.082735896 |
| ASVINISSI | AWSHRGKSQNTLY | 0.08285222 |
| ATPPGPA | AWSHRGKSQNTLY | 0.084254816 |
| ATVASALV | AWSHRGKSQNTLY | 0.0828756 |
| ATVASLIL | AWSHRGKSQNTLY | 0.083101854 |
| AVNGPVLIRH | AWSHRGKSQNTLY | 0.083215445 |
| DGAAGAVGI | AWSHRGKSQNTLY | 0.08292662 |
| DPLLHGI | AWSHRGKSQNTLY | 0.083207205 |
| ELGAALGAL | AWSHRGKSQNTLY | 0.08304881 |
| ELGRTQAV | AWSHRGKSQNTLY | 0.08517464 |
| FVAGGYLS | AWSHRGKSQNTLY | 0.08275634 |
| GASGGAGGAGGW | AWSHRGKSQNTLY | 0.0826405 |
| GGAGAGGDGGAGGI | AWSHRGKSQNTLY | 0.085041985 |
| GTGHNGSQPGMGGNG | AWSHRGKSQNTLY | 0.08342122 |
| GYAAASASAS | AWSHRGKSQNTLY | 0.08303712 |
| HQVDVAAVQQG | AWSHRGKSQNTLY | 0.0830733 |
| IGAAIAHT | AWSHRGKSQNTLY | 0.08433307 |
| IGMIEVAQL | AWSHRGKSQNTLY | 0.08335349 |
| IRLAPEHTSAMD | AWSHRGKSQNTLY | 0.08284933 |
| IYNVSVL | AWSHRGKSQNTLY | 0.08300182 |
| KQGFLGM | AWSHRGKSQNTLY | 0.083557256 |
| LESVITQL | AWSHRGKSQNTLY | 0.082756616 |
| LGLGALVVV | AWSHRGKSQNTLY | 0.08354356 |
| LIGNGGNGGSGGI | AWSHRGKSQNTLY | 0.08492556 |

| | | |
|---|---|---|
| LNAINAPTL | AWSHRGKSQNTLY | 0.082778096 |
| LQLLNTGV | AWSHRGKSQNTLY | 0.0831539 |
| LQTSIEF | AWSHRGKSQNTLY | 0.082852684 |
| MSFAVSEL | AWSHRGKSQNTLY | 0.08395651 |
| NTGALGL | AWSHRGKSQNTLY | 0.08287089 |
| PPGAGGAGGAGTT | AWSHRGKSQNTLY | 0.082669966 |
| QLVGLGI | AWSHRGKSQNTLY | 0.083487734 |
| RFLEQVHQL | AWSHRGKSQNTLY | 0.08479899 |
| RGVGYVLR | AWSHRGKSQNTLY | 0.08339812 |
| SATGVFAGVI | AWSHRGKSQNTLY | 0.08361699 |
| SLPAITLP | AWSHRGKSQNTLY | 0.08323786 |
| STALAFTV | AWSHRGKSQNTLY | 0.08498136 |
| SVLTVGLGK | AWSHRGKSQNTLY | 0.08320526 |
| TALAGAGSGI | AWSHRGKSQNTLY | 0.083779305 |
| TGLPFTGI | AWSHRGKSQNTLY | 0.08293724 |
| VAEHVTV | AWSHRGKSQNTLY | 0.083503045 |
| VDIVLNSL | AWSHRGKSQNTLY | 0.08351576 |
| VLTGGGAVL | AWSHRGKSQNTLY | 0.0828137 |
| VNVGLAYGI | AWSHRGKSQNTLY | 0.083466955 |
| VSSAAVGAV | AWSHRGKSQNTLY | 0.0831061 |
| VTSVYEAGV | AWSHRGKSQNTLY | 0.08396622 |
| WVALDRGA | AWSHRGKSQNTLY | 0.08330945 |
| YLVGSVVI | AWSHRGKSQNTLY | 0.08270586 |
| GPPGTGKTLL | AWSHRGKSQNTLY | 0.085649624 |
| GVGMTKF | AWSHRGKSQNTLY | 0.08281579 |
| IGIGGSDL | AWSHRGKSQNTLY | 0.08447221 |
| IGTLLAEL | AWSHRGKSQNTLY | 0.08284812 |
| IYTSGTT | AWSHRGKSQNTLY | 0.08314022 |
| LSFPAGV | AWSHRGKSQNTLY | 0.08427492 |
| LYTSGTT | AWSHRGKSQNTLY | 0.082848825 |
| VGAFPFGF | AWSHRGKSQNTLY | 0.08520019 |
| VGAGTVEF | AWSHRGKSQNTLY | 0.08320676 |
| VGKSTLLL | AWSHRGKSQNTLY | 0.08334878 |
| AALAHAGGGIV | AWSHRGKSQNTLY | 0.08284144 |
| AALVAVSLL | AWSHRGKSQNTLY | 0.083313994 |
| AAVSRAVAAL | AWSHRGKSQNTLY | 0.083007485 |
| AGGSSGIGGF | AWSHRGKSQNTLY | 0.08399768 |
| AGLTVANANG | AWSHRGKSQNTLY | 0.0827512 |
| AGLVGAVL | AWSHRGKSQNTLY | 0.08343792 |
| ALSQAFTV | AWSHRGKSQNTLY | 0.08290544 |
| ALTFSVSL | AWSHRGKSQNTLY | 0.083649024 |
| ALTGAALVAL | AWSHRGKSQNTLY | 0.08378052 |
| ASFSIGAGVL | AWSHRGKSQNTLY | 0.084073156 |
| DGSVYKI | AWSHRGKSQNTLY | 0.08350676 |
| EQLAEIV | AWSHRGKSQNTLY | 0.086537726 |
| FAVSELTVM | AWSHRGKSQNTLY | 0.08408401 |
| GAAPLAAGL | AWSHRGKSQNTLY | 0.08320906 |
| GADGGDGGSGGAGG | AWSHRGKSQNTLY | 0.08316765 |
| GFGLDTGGAGG | AWSHRGKSQNTLY | 0.08381613 |
| GLASGGLL | AWSHRGKSQNTLY | 0.08475535 |
| GRIAGAAVLL | AWSHRGKSQNTLY | 0.082967974 |

| | | |
|---|---|---|
| GSAGSLNTSF | AWSHRGKSQNTLY | 0.08279073 |
| GTGQIGI | AWSHRGKSQNTLY | 0.08292847 |
| GTLIALIL | AWSHRGKSQNTLY | 0.08384141 |
| IAATMGFVVG | AWSHRGKSQNTLY | 0.08336312 |
| IFAQALL | AWSHRGKSQNTLY | 0.084058456 |
| IGAAVALGF | AWSHRGKSQNTLY | 0.083308615 |
| IGLTPLAL | AWSHRGKSQNTLY | 0.08510029 |
| ILGSNTSTL | AWSHRGKSQNTLY | 0.08433314 |
| IVFSLTVL | AWSHRGKSQNTLY | 0.0834265 |
| LASKATGF | AWSHRGKSQNTLY | 0.08277141 |
| LASVALIL | AWSHRGKSQNTLY | 0.084089465 |
| LFAVGLTL | AWSHRGKSQNTLY | 0.083455615 |
| LFNGGIGTYI | AWSHRGKSQNTLY | 0.083202 |
| LTLGAGII | AWSHRGKSQNTLY | 0.08310696 |
| LVAAALGDI | AWSHRGKSQNTLY | 0.08368495 |
| NTAFVQLGI | AWSHRGKSQNTLY | 0.08479362 |
| QLFSGAEL | AWSHRGKSQNTLY | 0.08265856 |
| SAATFITL | AWSHRGKSQNTLY | 0.08280225 |
| SNYGLANLGV | AWSHRGKSQNTLY | 0.08387071 |
| SYGGAYAVM | AWSHRGKSQNTLY | 0.08446177 |
| TTGVGGLTLG | AWSHRGKSQNTLY | 0.08344432 |
| TTQTIYI | AWSHRGKSQNTLY | 0.082992874 |
| VAAGAGLGAVI | AWSHRGKSQNTLY | 0.08269431 |
| VAAVVTLL | AWSHRGKSQNTLY | 0.08510892 |
| VGFLGGLI | AWSHRGKSQNTLY | 0.08284528 |
| VGVIPILL | AWSHRGKSQNTLY | 0.08295106 |
| VIFASSLI | AWSHRGKSQNTLY | 0.08365242 |
| VKGGDGGAAAG | AWSHRGKSQNTLY | 0.08341411 |
| VTKTFTNI | AWSHRGKSQNTLY | 0.08289606 |
| VVAAAILL | AWSHRGKSQNTLY | 0.082844816 |
| YAALDVEL | AWSHRGKSQNTLY | 0.083757475 |
| LASVALLL | AWSHRGKSQNTLY | 0.08304369 |
| LDAVHRV | AWSHRGKSQNTLY | 0.082797974 |
| QGLSSSIF | AWSHRGKSQNTLY | 0.08337831 |
| RGTFANIRL | AWSHRGKSQNTLY | 0.08287784 |
| RQLWWGHRI | AWSHRGKSQNTLY | 0.084646575 |
| YGPPGTGKTLL | AWSHRGKSQNTLY | 0.08527202 |
| AALVGAPANVA | AWSHRGKSQNTLY | 0.082912214 |
| AATFRRTE | AWSHRGKSQNTLY | 0.08348716 |
| AGALGGGI | AWSHRGKSQNTLY | 0.084957406 |
| AGGAGGAGGIGAG | AWSHRGKSQNTLY | 0.08449521 |
| AGSGALGL | AWSHRGKSQNTLY | 0.0835294 |
| AQIADQI | AWSHRGKSQNTLY | 0.08293596 |
| ASAVAVTYL | AWSHRGKSQNTLY | 0.08287677 |
| ASTVLGGTGG | AWSHRGKSQNTLY | 0.08309284 |
| AVGGEPAK | AWSHRGKSQNTLY | 0.08470758 |
| CEQGTFGGTC | AWSHRGKSQNTLY | 0.08335073 |
| DEGKAPAG | AWSHRGKSQNTLY | 0.083474405 |
| DTVVFPAL | AWSHRGKSQNTLY | 0.08320059 |
| FEGITMAG | AWSHRGKSQNTLY | 0.08347126 |
| FPLVVQD | AWSHRGKSQNTLY | 0.083935075 |

| | | |
|---|---|---|
| GAFTTPQ | AWSHRGKSQNTLY | 0.08271923 |
| GFGGFGTAGGVGGA | AWSHRGKSQNTLY | 0.08285582 |
| GFLAQFL | AWSHRGKSQNTLY | 0.08320645 |
| GGAGGAGGNAGVA | AWSHRGKSQNTLY | 0.08321465 |
| GGVGTAGAGGAGGAGG | AWSHRGKSQNTLY | 0.08306515 |
| GNGGGGDAGNAGSGGN | AWSHRGKSQNTLY | 0.08411688 |
| GTGGAGGAGNDAGS | AWSHRGKSQNTLY | 0.08394784 |
| GTTNTGWFNS | AWSHRGKSQNTLY | 0.08288656 |
| IFMGPLFTL | AWSHRGKSQNTLY | 0.08331495 |
| ILAHAIEAI | AWSHRGKSQNTLY | 0.089335375 |
| IRSAIREAL | AWSHRGKSQNTLY | 0.08458964 |
| ITFAGGG | AWSHRGKSQNTLY | 0.085264586 |
| ITGMSGHQEV | AWSHRGKSQNTLY | 0.08290146 |
| IVAGLACTL | AWSHRGKSQNTLY | 0.083485514 |
| IVGGGLAGA | AWSHRGKSQNTLY | 0.08391416 |
| IVTGVLNI | AWSHRGKSQNTLY | 0.08325638 |
| LAQQGCI | AWSHRGKSQNTLY | 0.08316299 |
| LCLHVEN | AWSHRGKSQNTLY | 0.083281696 |
| LDAISDGS | AWSHRGKSQNTLY | 0.083433755 |
| LGGFAAML | AWSHRGKSQNTLY | 0.08572874 |
| LGGKNPMI | AWSHRGKSQNTLY | 0.08290936 |
| LGPKGVAAIL | AWSHRGKSQNTLY | 0.08274458 |
| LMVGVSI | AWSHRGKSQNTLY | 0.082783125 |
| LVGAAFVL | AWSHRGKSQNTLY | 0.0828482 |
| LVLAVAPL | AWSHRGKSQNTLY | 0.084124714 |
| MAAIAVIDAA | AWSHRGKSQNTLY | 0.08323888 |
| NNGINAELQ | AWSHRGKSQNTLY | 0.08277211 |
| RGEPLVLVHG | AWSHRGKSQNTLY | 0.08311336 |
| RGGAGGAAGLI | AWSHRGKSQNTLY | 0.08745627 |
| RSYRMHPAV | AWSHRGKSQNTLY | 0.083990075 |
| SAAHLLAGQLA | AWSHRGKSQNTLY | 0.083233505 |
| SAAVGSY | AWSHRGKSQNTLY | 0.082936555 |
| SPHAVLHRI | AWSHRGKSQNTLY | 0.08277609 |
| SQIAKVL | AWSHRGKSQNTLY | 0.08281391 |
| TALGHLVP | AWSHRGKSQNTLY | 0.082764074 |
| TAPTTSGAA | AWSHRGKSQNTLY | 0.08296116 |
| TDGDDGNGGSPGT | AWSHRGKSQNTLY | 0.082748376 |
| TDGLGEN | AWSHRGKSQNTLY | 0.0830608 |
| TGGPPLPGQA | AWSHRGKSQNTLY | 0.082658134 |
| TIPAITI | AWSHRGKSQNTLY | 0.08330222 |
| TIPGFTI | AWSHRGKSQNTLY | 0.0836265 |
| TSIASGALA | AWSHRGKSQNTLY | 0.08292563 |
| TVTLPGAAI | AWSHRGKSQNTLY | 0.0843504 |
| VASGLSAALP | AWSHRGKSQNTLY | 0.087725565 |
| VGAGVIGI | AWSHRGKSQNTLY | 0.08338928 |
| VGPKRFGV | AWSHRGKSQNTLY | 0.08377067 |
| VNVAFPD | AWSHRGKSQNTLY | 0.08282591 |
| VSMTNSGVS | AWSHRGKSQNTLY | 0.0852023 |
| VTDVLGDLSV | AWSHRGKSQNTLY | 0.08272607 |
| VVVVGAGFAGL | AWSHRGKSQNTLY | 0.084431656 |
| VYTVGTVV | AWSHRGKSQNTLY | 0.08366439 |

| | | |
|---|---|---|
| WTTFSGP | AWSHRGKSQNTLY | 0.08273084 |
| YDELVEAF | AWSHRGKSQNTLY | 0.082654044 |
| YGTGGAGGNGG | AWSHRGKSQNTLY | 0.083533905 |
| YTNVMAV | AWSHRGKSQNTLY | 0.08297386 |
| FLEQVHQL | AWSHRGKSQNTLY | 0.083072565 |
| GLSLLAVAV | AWSHRGKSQNTLY | 0.083171636 |
| LEQVHQL | AWSHRGKSQNTLY | 0.083463594 |
| LNAAGAI | AWSHRGKSQNTLY | 0.08298804 |
| PAVHASL | AWSHRGKSQNTLY | 0.082797214 |
| TELEDDA | AWSHRGKSQNTLY | 0.08342326 |
| TPPGPPN | AWSHRGKSQNTLY | 0.083440565 |
| TPPPPPPP | AWSHRGKSQNTLY | 0.08312133 |
| TQSPALI | ASGIGGDSETLY | 0.08343293 |
| EFAAAFPQ | ASGIGGDSETLY | 0.08342133 |
| LLSGNLDVL | ASGIGGDSETLY | 0.083611645 |
| GPANLGNYNI | ASGIGGDSETLY | 0.08347569 |
| TLGPTLP | ASGIGGDSETLY | 0.08311629 |
| VYAYSSCCK | ASGIGGDSETLY | 0.084144786 |
| APAPAPAPNI | ASGIGGDSETLY | 0.08282206 |
| PGGPVPIP | ASGIGGDSETLY | 0.082833506 |
| FLTVALEP | ASGIGGDSETLY | 0.08360466 |
| SQYFGANL | ASGIGGDSETLY | 0.08301747 |
| GGVKPAA | ASGIGGDSETLY | 0.08354323 |
| GGTGGAGGAGGW | ASGIGGDSETLY | 0.08468336 |
| GVGGVAGLGGAGG | ASGIGGDSETLY | 0.08287479 |
| VLLVLATTI | ASGIGGDSETLY | 0.08304864 |
| IGIFAAML | ASGIGGDSETLY | 0.08308953 |
| AVGDLID | ASGIGGDSETLY | 0.08348096 |
| NISPLPL | ASGIGGDSETLY | 0.085728824 |
| VTAVFVAI | ASGIGGDSETLY | 0.08307111 |
| SGDDNDAAL | ASGIGGDSETLY | 0.085046425 |
| AVAALAHS | ASGIGGDSETLY | 0.08313384 |
| GGVTVMATP | ASGIGGDSETLY | 0.08391258 |
| AGLAANVRD | ASGIGGDSETLY | 0.08575433 |
| TNRPDLIDPALLRPG | ASGIGGDSETLY | 0.0859499 |
| LANLGVTAL | ASGIGGDSETLY | 0.08350336 |
| TGERIEL | ASGIGGDSETLY | 0.082717374 |
| NGEELAA | ASGIGGDSETLY | 0.08301422 |
| VLSLVWG | ASGIGGDSETLY | 0.08287071 |
| IGYALWL | ASGIGGDSETLY | 0.08541621 |
| LSLLLAPL | ASGIGGDSETLY | 0.083398126 |
| TLDEALK | ASGIGGDSETLY | 0.08350326 |
| TDGAGTAEGG | ASGIGGDSETLY | 0.08294374 |
| NAPGGAGGNGG | ASGIGGDSETLY | 0.084259465 |
| VSVFMGMGGNF | ASGIGGDSETLY | 0.0829153 |
| LASYLAAL | ASGIGGDSETLY | 0.08542262 |
| LVGGTPAL | ASGIGGDSETLY | 0.08422599 |
| VRNADVL | ASGIGGDSETLY | 0.08280975 |
| TCSSPFLVSAM | ASGIGGDSETLY | 0.08322407 |
| VHITDAE | ASGIGGDSETLY | 0.08369494 |
| EYAAPSF | ASGIGGDSETLY | 0.08290571 |

| | | |
|---|---|---|
| VVDAFHGGL | ASGIGGDSETLY | 0.08283667 |
| ASVVDPQ | ASGIGGDSETLY | 0.082954906 |
| SYNEFGGLS | ASGIGGDSETLY | 0.08300813 |
| DPDGSVY | ASGIGGDSETLY | 0.08301425 |
| MVSTIHATE | ASGIGGDSETLY | 0.08350859 |
| LAAGVADPV | ASGIGGDSETLY | 0.08351202 |
| TKEQIAA | ASGIGGDSETLY | 0.082916096 |
| SLASPAAL | ASGIGGDSETLY | 0.08315424 |
| IGLVFYGV | ASGIGGDSETLY | 0.08394251 |
| FLLDASNL | ASGIGGDSETLY | 0.08331326 |
| YQVGNFP | ASGIGGDSETLY | 0.084192716 |
| ESLLSQST | ASGIGGDSETLY | 0.082859494 |
| LFTVGFAAVG | ASGIGGDSETLY | 0.082965896 |
| ATPPTPT | ASGIGGDSETLY | 0.08323185 |
| TDLATLG | ASGIGGDSETLY | 0.08410799 |
| SAGGTGR | ASGIGGDSETLY | 0.08284579 |
| GFSNSGS | ASGIGGDSETLY | 0.08612222 |
| GGNGGSGGTSV | ASGIGGDSETLY | 0.08822192 |
| GTSVATGGAGN | ASGIGGDSETLY | 0.08300241 |
| NGGNGGSGGTS | ASGIGGDSETLY | 0.085815445 |
| AVVKFPH | ASGIGGDSETLY | 0.08269039 |
| TPPPPPP | ASGIGGDSETLY | 0.08354251 |
| AAPAAAAAPA | ASGIGGDSETLY | 0.082698725 |
| IALNEDL | ASGIGGDSETLY | 0.084620565 |
| IVNVSSM | ASGIGGDSETLY | 0.083614685 |
| TGIPPAPR | ASGIGGDSETLY | 0.08318896 |
| FDLGGGTF | ASGIGGDSETLY | 0.08307634 |
| FDLGGGTFDVS | ASGIGGDSETLY | 0.08632967 |
| NEPTAAA | ASGIGGDSETLY | 0.08463046 |
| NEPTAAALA | ASGIGGDSETLY | 0.08296002 |
| GSIINISSI | ASGIGGDSETLY | 0.08517453 |
| VAPGNAGTA | ASGIGGDSETLY | 0.0836101 |
| TGGGASLE | ASGIGGDSETLY | 0.0832644 |
| VAVYNHYKRI | ASGIGGDSETLY | 0.0829561 |
| TDPAPATT | ASGIGGDSETLY | 0.08319901 |
| TNRPDILDPALLRPG | ASGIGGDSETLY | 0.08335159 |
| LTTTTTS | ASGIGGDSETLY | 0.08296235 |
| PTGSGKTL | ASGIGGDSETLY | 0.08362662 |
| TGSGKTL | ASGIGGDSETLY | 0.08317547 |
| TIGHVDHGK | ASGIGGDSETLY | 0.082697466 |
| TIGHVDHGKTT | ASGIGGDSETLY | 0.08273158 |
| GHVDHGKTT | ASGIGGDSETLY | 0.08525957 |
| AQFEHTLL | ASGIGGDSETLY | 0.08952746 |
| IVNQDEL | ASGIGGDSETLY | 0.083278455 |
| TNGSQFF | ASGIGGDSETLY | 0.08419099 |
| TGKGIVE | ASGIGGDSETLY | 0.08367151 |
| LISNASDA | ASGIGGDSETLY | 0.084392816 |
| AELRTLGELTKIAKAHGAQ | ASGIGGDSETLY | 0.08388218 |
| APNAVAAAEPSW | ASGIGGDSETLY | 0.083420366 |
| APQVAGAATAVL | ASGIGGDSETLY | 0.08286635 |
| ARRDPAAAAAILPTDA | ASGIGGDSETLY | 0.082821764 |

| | | |
|---|---|---|
| DAAARDVLAEAGLAEYF | ASGIGGDSETLY | 0.08304263 |
| DDQIVLSAKANLAKVEQ | ASGIGGDSETLY | 0.08345115 |
| DELLPDTGEDSDEDGH | ASGIGGDSETLY | 0.08345937 |
| ELPEAPANRATVPQD | ASGIGGDSETLY | 0.0834586 |
| FGQSKIERVTGIG | ASGIGGDSETLY | 0.08404511 |
| GAGGTGSTAGGAGGAGGAGGLY | ASGIGGDSETLY | 0.083297946 |
| GHGGTGAAGGNGGTGGAGGSGI | ASGIGGDSETLY | 0.083407104 |
| GNGGNGGAGGEGGPGGAGGAGA | ASGIGGDSETLY | 0.08377926 |
| GSTAGNGGNGGNATNGTIAGQP | ASGIGGDSETLY | 0.08308851 |
| GTGGDGGNGGNGGTGG | ASGIGGDSETLY | 0.08501023 |
| GVRRPAGIPADAPV | ASGIGGDSETLY | 0.08324277 |
| GYRQVTDAHLLTLARR | ASGIGGDSETLY | 0.08292406 |
| IAKMGQDGHDRG | ASGIGGDSETLY | 0.08383909 |
| KAEPSIEIQYT | ASGIGGDSETLY | 0.08345744 |
| KPVRIRMAADEDTVAVT | ASGIGGDSETLY | 0.083189785 |
| MPPRIAGMRLLVIKPEP | ASGIGGDSETLY | 0.08361577 |
| MKFYAVALFRSRGVPPTR | ASGIGGDSETLY | 0.08431488 |
| NAGGFGILGAVAHSP | ASGIGGDSETLY | 0.08275667 |
| NEGREITLAMAEM | ASGIGGDSETLY | 0.083016805 |
| NELVEWSHPLGHRWRWE | ASGIGGDSETLY | 0.0832122 |
| NLGAGGTGGNGGNPG | ASGIGGDSETLY | 0.08485636 |
| NTEWPKWETVMAADPVD | ASGIGGDSETLY | 0.08373299 |
| PASAAPAETQP | ASGIGGDSETLY | 0.082792155 |
| QQSSDGKNGVRAAEITGE | ASGIGGDSETLY | 0.08338208 |
| RAPDAVTTSWWKE | ASGIGGDSETLY | 0.08304069 |
| SGFGNVGAHTSGWFNQ | ASGIGGDSETLY | 0.08328691 |
| SGVAGAVGQAASVGGLKV | ASGIGGDSETLY | 0.08518245 |
| SNGGAGTGSAGGNGGT | ASGIGGDSETLY | 0.08388037 |
| TGGAGGEGGAAG | ASGIGGDSETLY | 0.08309765 |
| TGGDGGAAGAGGAPGHG | ASGIGGDSETLY | 0.08493505 |
| TMPPLLAAIANAGRHGV | ASGIGGDSETLY | 0.08275253 |
| TSSTAGNSGGA | ASGIGGDSETLY | 0.08291899 |
| VPATVPVGILC | ASGIGGDSETLY | 0.0827765 |
| VQRPDSGGVVLTGRIS | ASGIGGDSETLY | 0.08281971 |
| WTPNPVPATKGVPAGLVQAT | ASGIGGDSETLY | 0.08308742 |
| YALKRALAKFSDRFDVVII | ASGIGGDSETLY | 0.08327201 |
| YLIGSVIV | ASGIGGDSETLY | 0.082887344 |
| YGQLVAVID | ASGIGGDSETLY | 0.08354193 |
| VTVGGMTLPA | ASGIGGDSETLY | 0.08262211 |
| VLALPAV | ASGIGGDSETLY | 0.08297413 |
| VIPPTLNYET | ASGIGGDSETLY | 0.08380516 |
| VGMGGMHPSSG | ASGIGGDSETLY | 0.083162114 |
| VGLAGGAATA | ASGIGGDSETLY | 0.08382424 |
| VGGAGGAGGSGP | ASGIGGDSETLY | 0.0829555 |
| VFAAFMV | ASGIGGDSETLY | 0.08297834 |
| TLYDKLR | ASGIGGDSETLY | 0.082851864 |
| TLAVFTVL | ASGIGGDSETLY | 0.08320033 |
| TGSVQPARV | ASGIGGDSETLY | 0.08277179 |
| TGGGAGSGGT | ASGIGGDSETLY | 0.08337976 |
| SHLETSL | ASGIGGDSETLY | 0.08444619 |
| SFTTGMT | ASGIGGDSETLY | 0.082849704 |

| | | |
|---|---|---|
| QVEVAKRMGVRQPTVS | ASGIGGDSETLY | 0.08313192 |
| QGSGGPALAM | ASGIGGDSETLY | 0.08318744 |
| MLAPSNP | ASGIGGDSETLY | 0.08276838 |
| LVNNAAI | ASGIGGDSETLY | 0.08279095 |
| LTGEINTLA | ASGIGGDSETLY | 0.084197305 |
| LSIFGPV | ASGIGGDSETLY | 0.084485285 |
| LLALLANL | ASGIGGDSETLY | 0.082895555 |
| LISNASDALDKLR | ASGIGGDSETLY | 0.08353206 |
| LINEAVP | ASGIGGDSETLY | 0.08273725 |
| LGYAFSGL | ASGIGGDSETLY | 0.08727094 |
| LDFVAVAL | ASGIGGDSETLY | 0.08344427 |
| LAQPGAE | ASGIGGDSETLY | 0.08426205 |
| ISNLGAVTGF | ASGIGGDSETLY | 0.08296181 |
| HQEDPGRI | ASGIGGDSETLY | 0.08290781 |
| GTPGEPTP | ASGIGGDSETLY | 0.08499652 |
| GFFNSGS | ASGIGGDSETLY | 0.08286389 |
| GAGGGGLTAGAN | ASGIGGDSETLY | 0.08289452 |
| FVDALVNM | ASGIGGDSETLY | 0.08286259 |
| FLTVAPEL | ASGIGGDSETLY | 0.083242275 |
| FETLTAL | ASGIGGDSETLY | 0.08277015 |
| DVINPIEL | ASGIGGDSETLY | 0.08311432 |
| APVSGAPV | ASGIGGDSETLY | 0.087958835 |
| ALVGGLAW | ASGIGGDSETLY | 0.08349341 |
| ALAAAAATAI | ASGIGGDSETLY | 0.083368406 |
| AFGLGLLL | ASGIGGDSETLY | 0.083567716 |
| AFFAAASL | ASGIGGDSETLY | 0.083373904 |
| AAAALPAFSPP | ASGIGGDSETLY | 0.08368759 |
| VVLAPAL | ASGIGGDSETLY | 0.0830199 |
| TIGHVDHGKTTL | ASGIGGDSETLY | 0.08337677 |
| LISNASDALDK | ASGIGGDSETLY | 0.08346173 |
| LISNASDAL | ASGIGGDSETLY | 0.08384408 |
| ISNASDALDK | ASGIGGDSETLY | 0.08309847 |
| NQAETLVYQTE | ASGIGGDSETLY | 0.08345487 |
| TGGTGGTAGAG | ASGIGGDSETLY | 0.08299971 |
| TGSTAGGAGGA | ASGIGGDSETLY | 0.08334538 |
| DPLAVPVAAGPV | ASGIGGDSETLY | 0.0829959 |
| GGTGGTPGNGGN | ASGIGGDSETLY | 0.08333353 |
| PAGLAGVDPATL | ASGIGGDSETLY | 0.0849998 |
| GNNLVVEAPHAGASV | ASGIGGDSETLY | 0.08311888 |
| KVTRIMSASTLEQPA | ASGIGGDSETLY | 0.08298884 |
| LGFALPIALPRRTVV | ASGIGGDSETLY | 0.082838535 |
| TRLGNLTARKLVVLV | ASGIGGDSETLY | 0.08374 |
| WSQQRRLIYFPSAGP | ASGIGGDSETLY | 0.082960315 |
| GPNGNPDPMAAAVDIR | ASGIGGDSETLY | 0.08287841 |
| TQRCVSVADLRAAASR | ASGIGGDSETLY | 0.08354228 |
| QFVLDVAEHPVEVLTETV | ASGIGGDSETLY | 0.08283391 |
| GGAGGTGGAGGNGGTQAGDG | ASGIGGDSETLY | 0.08340791 |
| PLLATLRAIAVDVDAKPAQV | ASGIGGDSETLY | 0.082957156 |
| APKFRTGGYMIDFWGVGYQVAK | ASGIGGDSETLY | 0.08268989 |
| GAGGAGGAGGAAGGPGGTGGTG | ASGIGGDSETLY | 0.084718235 |
| IGALGLLATDVAPAGSVAFGVA | ASGIGGDSETLY | 0.0829504 |

| | | |
|---|---|---|
| AGGAGGIGGNGGAGGAGGVLMG | ASGIGGDSETLY | 0.083051495 |
| ADELIGLL | ASGIGGDSETLY | 0.08477897 |
| ADRALKA | ASGIGGDSETLY | 0.08294183 |
| AGAVGQVGGAGGA | ASGIGGDSETLY | 0.083898455 |
| AGGGGGLGGAP | ASGIGGDSETLY | 0.083328284 |
| AGGLLYGNGGAG | ASGIGGDSETLY | 0.08357241 |
| AGLESIGL | ASGIGGDSETLY | 0.08323834 |
| AGLKTGII | ASGIGGDSETLY | 0.08301409 |
| ASLAALLL | ASGIGGDSETLY | 0.082984164 |
| ASVINISSI | ASGIGGDSETLY | 0.08331873 |
| ATPPGPA | ASGIGGDSETLY | 0.08762574 |
| ATVASALV | ASGIGGDSETLY | 0.08405854 |
| ATVASLIL | ASGIGGDSETLY | 0.08371919 |
| AVNGPVLIRH | ASGIGGDSETLY | 0.083410405 |
| DGAAGAVGI | ASGIGGDSETLY | 0.08313868 |
| DPLLHGI | ASGIGGDSETLY | 0.08391331 |
| ELGAALGAL | ASGIGGDSETLY | 0.08279848 |
| ELGRTQAV | ASGIGGDSETLY | 0.08299276 |
| FVAGGYLS | ASGIGGDSETLY | 0.083009385 |
| GASGGAGGAGGW | ASGIGGDSETLY | 0.082906485 |
| GGAGAGGDGGAGGI | ASGIGGDSETLY | 0.08303123 |
| GTGHNGSQPGMGGNG | ASGIGGDSETLY | 0.0837881 |
| GYAAASASAS | ASGIGGDSETLY | 0.08309376 |
| HQVDVAAVQQG | ASGIGGDSETLY | 0.08315663 |
| IGAAIAHT | ASGIGGDSETLY | 0.08305641 |
| IGMIEVAQL | ASGIGGDSETLY | 0.0832227 |
| IRLAPEHTSAMD | ASGIGGDSETLY | 0.083005324 |
| IYNVSVL | ASGIGGDSETLY | 0.08531705 |
| KQGFLGM | ASGIGGDSETLY | 0.0830361 |
| LESVITQL | ASGIGGDSETLY | 0.08347251 |
| LGLGALVVV | ASGIGGDSETLY | 0.08775118 |
| LIGNGGNGGSGGI | ASGIGGDSETLY | 0.08327124 |
| LNAINAPTL | ASGIGGDSETLY | 0.08279315 |
| LQLLNTGV | ASGIGGDSETLY | 0.08307053 |
| LQTSIEF | ASGIGGDSETLY | 0.08338768 |
| MSFAVSEL | ASGIGGDSETLY | 0.08393031 |
| NTGALGL | ASGIGGDSETLY | 0.08268174 |
| PPGAGGAGGAGTT | ASGIGGDSETLY | 0.08523288 |
| QLVGLGI | ASGIGGDSETLY | 0.08442771 |
| RFLEQVHQL | ASGIGGDSETLY | 0.08314293 |
| RGVGYVLR | ASGIGGDSETLY | 0.08256735 |
| SATGVFAGVI | ASGIGGDSETLY | 0.0849354 |
| SLPAITLP | ASGIGGDSETLY | 0.084078826 |
| STALAFTV | ASGIGGDSETLY | 0.08474744 |
| SVLTVGLGK | ASGIGGDSETLY | 0.08281621 |
| TALAGAGSGI | ASGIGGDSETLY | 0.08279062 |
| TGLPFTGI | ASGIGGDSETLY | 0.08443508 |
| VAEHVTV | ASGIGGDSETLY | 0.08381924 |
| VDIVLNSL | ASGIGGDSETLY | 0.08342245 |
| VLTGGGAVL | ASGIGGDSETLY | 0.08614127 |
| VNVGLAYGI | ASGIGGDSETLY | 0.083067894 |

| | | |
|---|---|---|
| VSSAAVGAV | ASGIGGDSETLY | 0.083332576 |
| VTSVYEAGV | ASGIGGDSETLY | 0.08402744 |
| WVALDRGA | ASGIGGDSETLY | 0.086232066 |
| YLVGSVVI | ASGIGGDSETLY | 0.08273732 |
| GPPGTGKTLL | ASGIGGDSETLY | 0.08324092 |
| GVGMTKF | ASGIGGDSETLY | 0.084022276 |
| IGIGGSDL | ASGIGGDSETLY | 0.082989514 |
| IGTLLAEL | ASGIGGDSETLY | 0.0832883 |
| IYTSGTT | ASGIGGDSETLY | 0.083871804 |
| LSFPAGV | ASGIGGDSETLY | 0.08317648 |
| LYTSGTT | ASGIGGDSETLY | 0.08329792 |
| VGAFPFGF | ASGIGGDSETLY | 0.08428001 |
| VGAGTVEF | ASGIGGDSETLY | 0.0827722 |
| VGKSTLLL | ASGIGGDSETLY | 0.08382821 |
| AALAHAGGGIV | ASGIGGDSETLY | 0.08299504 |
| AALVAVSLL | ASGIGGDSETLY | 0.08603565 |
| AAVSRAVAAL | ASGIGGDSETLY | 0.08324134 |
| AGGSSGIGGF | ASGIGGDSETLY | 0.083194666 |
| AGLTVANANG | ASGIGGDSETLY | 0.082911745 |
| AGLVGAVL | ASGIGGDSETLY | 0.1202419 |
| ALSQAFTV | ASGIGGDSETLY | 0.17652676 |
| ALTFSVSL | ASGIGGDSETLY | 0.30171478 |
| ALTGAALVAL | ASGIGGDSETLY | 0.2495875 |
| ASFSIGAGVL | ASGIGGDSETLY | 0.13910437 |
| DGSVYKI | ASGIGGDSETLY | 0.11299227 |
| EQLAEIV | ASGIGGDSETLY | 0.16337214 |
| FAVSELTVM | ASGIGGDSETLY | 0.11879665 |
| GAAPLAAGL | ASGIGGDSETLY | 0.13511547 |
| GADGGDGGSGGAGG | ASGIGGDSETLY | 0.18200059 |
| GFGLDTGGAGG | ASGIGGDSETLY | 0.13522512 |
| GLASGGLL | ASGIGGDSETLY | 0.61261666 |
| GRIAGAAVLL | ASGIGGDSETLY | 0.34551537 |
| GSAGSLNTSF | ASGIGGDSETLY | 0.119287625 |
| GTGQIGI | ASGIGGDSETLY | 0.17559803 |
| GTLIALIL | ASGIGGDSETLY | 0.1506498 |
| IAATMGFVVG | ASGIGGDSETLY | 0.15128782 |
| IFAQALL | ASGIGGDSETLY | 0.10913298 |
| IGAAVALGF | ASGIGGDSETLY | 0.24247968 |
| IGLTPLAL | ASGIGGDSETLY | 0.15474701 |
| ILGSNTSTL | ASGIGGDSETLY | 0.10715241 |
| IVFSLTVL | ASGIGGDSETLY | 0.2214756 |
| LASKATGF | ASGIGGDSETLY | 0.15165228 |
| LASVALIL | ASGIGGDSETLY | 0.12225523 |
| LFAVGLTL | ASGIGGDSETLY | 0.48970452 |
| LFNGGIGTYI | ASGIGGDSETLY | 0.106787086 |
| LTLGAGII | ASGIGGDSETLY | 0.12534353 |
| LVAAALGDI | ASGIGGDSETLY | 0.45304936 |
| NTAFVQLGI | ASGIGGDSETLY | 0.4279849 |
| QLFSGAEL | ASGIGGDSETLY | 0.12395381 |
| SAATFITL | ASGIGGDSETLY | 0.30965573 |
| SNYGLANLGV | ASGIGGDSETLY | 0.70782655 |

| | | |
|---|---|---|
| SYGGAYAVM | ASGIGGDSETLY | 0.19852355 |
| TTGVGGLTLG | ASGIGGDSETLY | 0.4137534 |
| TTQTIYI | ASGIGGDSETLY | 0.5373528 |
| VAAGAGLGAVI | ASGIGGDSETLY | 0.13078392 |
| VAAVVTLL | ASGIGGDSETLY | 0.25597334 |
| VGFLGGLI | ASGIGGDSETLY | 0.6278468 |
| VGVIPILL | ASGIGGDSETLY | 0.14620174 |
| VIFASSLI | ASGIGGDSETLY | 0.20696007 |
| VKGGDGGAAAG | ASGIGGDSETLY | 0.13078919 |
| VTKTFTNI | ASGIGGDSETLY | 0.20846161 |
| VVAAAILL | ASGIGGDSETLY | 0.10899741 |
| YAALDVEL | ASGIGGDSETLY | 0.30878907 |
| LASVALLL | ASGIGGDSETLY | 0.37769002 |
| LDAVHRV | ASGIGGDSETLY | 0.23734374 |
| QGLSSSIF | ASGIGGDSETLY | 0.12054183 |
| RGTFANIRL | ASGIGGDSETLY | 0.13389382 |
| RQLWWGHRI | ASGIGGDSETLY | 0.115521185 |
| YGPPGTGKTLL | ASGIGGDSETLY | 0.17526647 |
| AALVGAPANVA | ASGIGGDSETLY | 0.11459239 |
| AATFRRTE | ASGIGGDSETLY | 0.11107615 |
| AGALGGGI | ASGIGGDSETLY | 0.39064103 |
| AGGAGGAGGIGAG | ASGIGGDSETLY | 0.36485577 |
| AGSGALGL | ASGIGGDSETLY | 0.1367093 |
| AQIADQI | ASGIGGDSETLY | 0.35213572 |
| ASAVAVTYL | ASGIGGDSETLY | 0.6090474 |
| ASTVLGGTGG | ASGIGGDSETLY | 0.50275576 |
| AVGGEPAK | ASGIGGDSETLY | 0.19542062 |
| CEQGTFGGTC | ASGIGGDSETLY | 0.11399146 |
| DEGKAPAG | ASGIGGDSETLY | 0.6747259 |
| DTVVFPAL | ASGIGGDSETLY | 0.32320285 |
| FEGITMAG | ASGIGGDSETLY | 0.121456034 |
| FPLVVQD | ASGIGGDSETLY | 0.40625104 |
| GAFTTPQ | ASGIGGDSETLY | 0.123726234 |
| GFGGFGTAGGVGGA | ASGIGGDSETLY | 0.13409959 |
| GFLAQFL | ASGIGGDSETLY | 0.5799502 |
| GGAGGAGGNAGVA | ASGIGGDSETLY | 0.4662833 |
| GGVGTAGAGGAGGAGG | ASGIGGDSETLY | 0.14514175 |
| GNGGGGDAGNAGSGGN | ASGIGGDSETLY | 0.11742313 |
| GTGGAGGAGNDAGS | ASGIGGDSETLY | 0.12836683 |
| GTTNTGWFNS | ASGIGGDSETLY | 0.34076318 |
| IFMGPLFTL | ASGIGGDSETLY | 0.32436946 |
| ILAHAIEAI | ASGIGGDSETLY | 0.17153025 |
| IRSAIREAL | ASGIGGDSETLY | 0.18959299 |
| ITFAGGG | ASGIGGDSETLY | 0.4579504 |
| ITGMSGHQEV | ASGIGGDSETLY | 0.18660595 |
| IVAGLACTL | ASGIGGDSETLY | 0.18250209 |
| IVGGGLAGA | ASGIGGDSETLY | 0.29231146 |
| IVTGVLNI | ASGIGGDSETLY | 0.11684978 |
| LAQQGCI | ASGIGGDSETLY | 0.40577123 |
| LCLHVEN | ASGIGGDSETLY | 0.37940896 |
| LDAISDGS | ASGIGGDSETLY | 0.3168947 |

| | | |
|---|---|---|
| LGGFAAML | ASGIGGDSETLY | 0.406645 |
| LGGKNPMI | ASGIGGDSETLY | 0.12692414 |
| LGPKGVAAIL | ASGIGGDSETLY | 0.30738848 |
| LMVGVSI | ASGIGGDSETLY | 0.1285256 |
| LVGAAFVL | ASGIGGDSETLY | 0.16951837 |
| LVLAVAPL | ASGIGGDSETLY | 0.37633872 |
| MAAIAVIDAA | ASGIGGDSETLY | 0.34993756 |
| NNGINAELQ | ASGIGGDSETLY | 0.4111135 |
| RGEPLVLVHG | ASGIGGDSETLY | 0.21559906 |
| RGGAGGAAGLI | ASGIGGDSETLY | 0.15495346 |
| RSYRMHPAV | ASGIGGDSETLY | 0.5951744 |
| SAAHLLAGQLA | ASGIGGDSETLY | 0.61259687 |
| SAAVGSY | ASGIGGDSETLY | 0.12425938 |
| SPHAVLHRI | ASGIGGDSETLY | 0.1252212 |
| SQIAKVL | ASGIGGDSETLY | 0.15570831 |
| TALGHLVP | ASGIGGDSETLY | 0.40668184 |
| TAPTTSGAA | ASGIGGDSETLY | 0.26989552 |
| TDGDDGNGGSPGT | ASGIGGDSETLY | 0.22636332 |
| TDGLGEN | ASGIGGDSETLY | 0.3877054 |
| TGGPPLPGQA | ASGIGGDSETLY | 0.11618974 |
| TIPAITI | ASGIGGDSETLY | 0.67965466 |
| TIPGFTI | ASGIGGDSETLY | 0.11862523 |
| TSIASGALA | ASGIGGDSETLY | 0.17867662 |
| TVTLPGAAI | ASGIGGDSETLY | 0.1308615 |
| VASGLSAALP | ASGIGGDSETLY | 0.10880222 |
| VGAGVIGI | ASGIGGDSETLY | 0.25875965 |
| VGPKRFGV | ASGIGGDSETLY | 0.13363026 |
| VNVAFPD | ASGIGGDSETLY | 0.4732211 |
| VSMTNSGVS | ASGIGGDSETLY | 0.17286903 |
| VTDVLGDLSV | ASGIGGDSETLY | 0.18817651 |
| VVVVGAGFAGL | ASGIGGDSETLY | 0.25694138 |
| VYTVGTVV | ASGIGGDSETLY | 0.16998583 |
| WTTFSGP | ASGIGGDSETLY | 0.11992295 |
| YDELVEAF | ASGIGGDSETLY | 0.2622951 |
| YGTGGAGGNGG | ASGIGGDSETLY | 0.5557499 |
| YTNVMAV | ASGIGGDSETLY | 0.3172225 |
| FLEQVHQL | ASGIGGDSETLY | 0.1182346 |
| GLSLLAVAV | ASGIGGDSETLY | 0.15403868 |
| LEQVHQL | ASGIGGDSETLY | 0.16526276 |
| LNAAGAI | ASGIGGDSETLY | 0.108399175 |
| PAVHASL | ASGIGGDSETLY | 0.108425304 |
| TELEDDA | ASGIGGDSETLY | 0.11116326 |
| TPPGPPN | ASGIGGDSETLY | 0.11210368 |
| TPPPPPPP | ASGIGGDSETLY | 0.5358007 |
| TQSPALI | ASSLGGGLF | 0.45322788 |
| EFAAAFPQ | ASSLGGGLF | 0.20316324 |
| LLSGNLDVL | ASSLGGGLF | 0.5621998 |
| GPANLGNYNI | ASSLGGGLF | 0.17213735 |
| TLGPTLP | ASSLGGGLF | 0.10609659 |
| VYAYSSCCK | ASSLGGGLF | 0.13467278 |
| APAPAPAPNI | ASSLGGGLF | 0.12409265 |

| | | |
|---|---|---|
| PGGPVPIP | ASSLGGGLF | 0.40925658 |
| FLTVALEP | ASSLGGGLF | 0.111884035 |
| SQYFGANL | ASSLGGGLF | 0.16643333 |
| GGVKPAA | ASSLGGGLF | 0.24763751 |
| GGTGGAGGAGGW | ASSLGGGLF | 0.24036321 |
| GVGGVAGLGGAGG | ASSLGGGLF | 0.45635158 |
| VLLVLATTI | ASSLGGGLF | 0.13228112 |
| IGIFAAML | ASSLGGGLF | 0.10962735 |
| AVGDLID | ASSLGGGLF | 0.2554943 |
| NISPLPL | ASSLGGGLF | 0.10907184 |
| VTAVFVAI | ASSLGGGLF | 0.12678608 |
| SGDDNDAAL | ASSLGGGLF | 0.11373725 |
| AVAALAHS | ASSLGGGLF | 0.10629056 |
| GGVTVMATP | ASSLGGGLF | 0.112213984 |
| AGLAANVRD | ASSLGGGLF | 0.37576985 |
| TNRPDLIDPALLRPG | ASSLGGGLF | 0.20115152 |
| LANLGVTAL | ASSLGGGLF | 0.2526369 |
| TGERIEL | ASSLGGGLF | 0.4564476 |
| NGEELAA | ASSLGGGLF | 0.26249856 |
| VLSLVWG | ASSLGGGLF | 0.2556113 |
| IGYALWL | ASSLGGGLF | 0.12730311 |
| LSLLLAPL | ASSLGGGLF | 0.12903392 |
| TLDEALK | ASSLGGGLF | 0.123833954 |
| TDGAGTAEGG | ASSLGGGLF | 0.12486811 |
| NAPGGAGGNGG | ASSLGGGLF | 0.63666284 |
| VSVFMGMGGNF | ASSLGGGLF | 0.16862369 |
| LASYLAAL | ASSLGGGLF | 0.113812245 |
| LVGGTPAL | ASSLGGGLF | 0.12876818 |
| VRNADVL | ASSLGGGLF | 0.6204257 |
| TCSSPFLVSAM | ASSLGGGLF | 0.64176846 |
| VHITDAE | ASSLGGGLF | 0.12918329 |
| EYAAPSF | ASSLGGGLF | 0.19600439 |
| VVDAFHGGL | ASSLGGGLF | 0.19515677 |
| ASVVDPQ | ASSLGGGLF | 0.19236954 |
| SYNEFGGLS | ASSLGGGLF | 0.1548945 |
| DPDGSVY | ASSLGGGLF | 0.12140158 |
| MVSTIHATE | ASSLGGGLF | 0.47838414 |
| LAAGVADPV | ASSLGGGLF | 0.781405 |
| TKEQIAA | ASSLGGGLF | 0.12960576 |
| SLASPAAL | ASSLGGGLF | 0.116440244 |
| IGLVFYGV | ASSLGGGLF | 0.27378023 |
| FLLDASNL | ASSLGGGLF | 0.1399239 |
| YQVGNFP | ASSLGGGLF | 0.18469349 |
| ESLLSQST | ASSLGGGLF | 0.16317275 |
| LFTVGFAAVG | ASSLGGGLF | 0.11401557 |
| ATPPTPT | ASSLGGGLF | 0.14851665 |
| TDLATLG | ASSLGGGLF | 0.5058208 |
| SAGGTGR | ASSLGGGLF | 0.18060234 |
| GFSNSGS | ASSLGGGLF | 0.42314094 |
| GGNGGSGGTSV | ASSLGGGLF | 0.61180544 |
| GTSVATGGAGN | ASSLGGGLF | 0.41774306 |

| | | |
|---|---|---|
| NGGNGGSGGTS | ASSLGGGLF | 0.61018395 |
| AVVKFPH | ASSLGGGLF | 0.56181824 |
| TPPPPPP | ASSLGGGLF | 0.14176269 |
| AAPAAAAAPA | ASSLGGGLF | 0.22699004 |
| IALNEDL | ASSLGGGLF | 0.16008456 |
| IVNVSSM | ASSLGGGLF | 0.2519632 |
| TGIPPAPR | ASSLGGGLF | 0.30213457 |
| FDLGGGTF | ASSLGGGLF | 0.18043914 |
| FDLGGGTFDVS | ASSLGGGLF | 0.5164632 |
| NEPTAAA | ASSLGGGLF | 0.23207042 |
| NEPTAAALA | ASSLGGGLF | 0.52269036 |
| GSIINISSI | ASSLGGGLF | 0.12269587 |
| VAPGNAGTA | ASSLGGGLF | 0.12390034 |
| TGGGASLE | ASSLGGGLF | 0.4882948 |
| VAVYNHYKRI | ASSLGGGLF | 0.11652586 |
| TDPAPATT | ASSLGGGLF | 0.11558462 |
| TNRPDILDPALLRPG | ASSLGGGLF | 0.27521265 |
| LTTTTTS | ASSLGGGLF | 0.56237507 |
| PTGSGKTL | ASSLGGGLF | 0.107042745 |
| TGSGKTL | ASSLGGGLF | 0.1257745 |
| TIGHVDHGK | ASSLGGGLF | 0.13235554 |
| TIGHVDHGKTT | ASSLGGGLF | 0.12899265 |
| GHVDHGKTT | ASSLGGGLF | 0.14704335 |
| AQFEHTLL | ASSLGGGLF | 0.115256935 |
| IVNQDEL | ASSLGGGLF | 0.20534858 |
| TNGSQFF | ASSLGGGLF | 0.11484909 |
| TGKGIVE | ASSLGGGLF | 0.11368251 |
| LISNASDA | ASSLGGGLF | 0.22871599 |
| AELRTLGELTKIAKAHGAQ | ASSLGGGLF | 0.6542197 |
| APNAVAAAEPSW | ASSLGGGLF | 0.122911155 |
| APQVAGAATAVL | ASSLGGGLF | 0.3073789 |
| ARRDPAAAAAILPTDA | ASSLGGGLF | 0.17632103 |
| DAAARDVLAEAGLAEYF | ASSLGGGLF | 0.7136263 |
| DDQIVLSAKANLAKVEQ | ASSLGGGLF | 0.16890368 |
| DELLPDTGEDSDEDGH | ASSLGGGLF | 0.11646515 |
| ELPEAPANRATVPQD | ASSLGGGLF | 0.28592017 |
| FGQSKIERVTGIG | ASSLGGGLF | 0.22699192 |
| GAGGTGSTAGGAGGAGGAGGLY | ASSLGGGLF | 0.16822462 |
| GHGGTGAAGGNGGTGGAGGSGI | ASSLGGGLF | 0.5810673 |
| GNGGNGGAGGEGGPGGAGGAGA | ASSLGGGLF | 0.11924447 |
| GSTAGNGGNGGNATNGTIAGQP | ASSLGGGLF | 0.21429044 |
| GTGGDGGNGGNGGTGG | ASSLGGGLF | 0.35706407 |
| GVRRPAGIPADAPV | ASSLGGGLF | 0.26108482 |
| GYRQVTDAHLLTLARR | ASSLGGGLF | 0.112835534 |
| IAKMGQDGHDRG | ASSLGGGLF | 0.20900127 |
| KAEPSIEIQYT | ASSLGGGLF | 0.15381974 |
| KPVRIRMAADEDTVAVT | ASSLGGGLF | 0.13917919 |
| MPPRIAGMRLLVIKPEP | ASSLGGGLF | 0.37853837 |
| MKFYAVALFRSRGVPPTR | ASSLGGGLF | 0.19628978 |
| NAGGFGILGAVAHSP | ASSLGGGLF | 0.5342127 |
| NEGREITLAMAEM | ASSLGGGLF | 0.22129346 |

| | | |
|---|---|---|
| NELVEWSHPLGHRWRWE | ASSLGGGLF | 0.106714696 |
| NLGAGGTGGNGGNPG | ASSLGGGLF | 0.11596105 |
| NTEWPKWETVMAADPVD | ASSLGGGLF | 0.20703298 |
| PASAAPAETQP | ASSLGGGLF | 0.2860949 |
| QQSSDGKNGVRAAEITGE | ASSLGGGLF | 0.25166374 |
| RAPDAVTTSWWKE | ASSLGGGLF | 0.13028152 |
| SGFGNVGAHTSGWFNQ | ASSLGGGLF | 0.11309075 |
| SGVAGAVGQAASVGGLKV | ASSLGGGLF | 0.2953386 |
| SNGGAGTGSAGGNGGT | ASSLGGGLF | 0.47080863 |
| TGGAGGEGGAAG | ASSLGGGLF | 0.1525303 |
| TGGDGGAAGAGGAPGHG | ASSLGGGLF | 0.20053846 |
| TMPPLLAAIANAGRHGV | ASSLGGGLF | 0.6500687 |
| TSSTAGNSGGA | ASSLGGGLF | 0.12609634 |
| VPATVPVGILC | ASSLGGGLF | 0.14739725 |
| VQRPDSGGVVLTGRIS | ASSLGGGLF | 0.34974504 |
| WTPNPVPATKGVPAGLVQAT | ASSLGGGLF | 0.6067047 |
| YALKRALAKFSDRFDVVII | ASSLGGGLF | 0.14980146 |
| YLIGSVIV | ASSLGGGLF | 0.58062285 |
| YGQLVAVID | ASSLGGGLF | 0.11422632 |
| VTVGGMTLPA | ASSLGGGLF | 0.5267186 |
| VLALPAV | ASSLGGGLF | 0.12163216 |
| VIPPTLNYET | ASSLGGGLF | 0.33213556 |
| VGMGGMHPSSG | ASSLGGGLF | 0.10952885 |
| VGLAGGAATA | ASSLGGGLF | 0.44634873 |
| VGGAGGAGGSGP | ASSLGGGLF | 0.5796572 |
| VFAAFMV | ASSLGGGLF | 0.5568525 |
| TLYDKLR | ASSLGGGLF | 0.10773131 |
| TLAVFTVL | ASSLGGGLF | 0.18465151 |
| TGSVQPARV | ASSLGGGLF | 0.6171988 |
| TGGGAGSGGT | ASSLGGGLF | 0.76200557 |
| SHLETSL | ASSLGGGLF | 0.259628 |
| SFTTGMT | ASSLGGGLF | 0.4009388 |
| QVEVAKRMGVRQPTVS | ASSLGGGLF | 0.13467674 |
| QGSGGPALAM | ASSLGGGLF | 0.13461795 |
| MLAPSNP | ASSLGGGLF | 0.15053996 |
| LVNNAAI | ASSLGGGLF | 0.14965257 |
| LTGEINTLA | ASSLGGGLF | 0.2286316 |
| LSIFGPV | ASSLGGGLF | 0.14509822 |
| LLALLANL | ASSLGGGLF | 0.32764983 |
| LISNASDALDKLR | ASSLGGGLF | 0.62670016 |
| LINEAVP | ASSLGGGLF | 0.2649554 |
| LGYAFSGL | ASSLGGGLF | 0.6513182 |
| LDFVAVAL | ASSLGGGLF | 0.6460446 |
| LAQPGAE | ASSLGGGLF | 0.2870271 |
| ISNLGAVTGF | ASSLGGGLF | 0.1240922 |
| HQEDPGRI | ASSLGGGLF | 0.17077148 |
| GTPGEPTP | ASSLGGGLF | 0.11824664 |
| GFFNSGS | ASSLGGGLF | 0.15106386 |
| GAGGGGLTAGAN | ASSLGGGLF | 0.12276605 |
| FVDALVNM | ASSLGGGLF | 0.25890952 |
| FLTVAPEL | ASSLGGGLF | 0.10649462 |

| | | |
|---|---|---|
| FETLTAL | ASSLGGGLF | 0.51014775 |
| DVINPIEL | ASSLGGGLF | 0.64110947 |
| APVSGAPV | ASSLGGGLF | 0.112599425 |
| ALVGGLAW | ASSLGGGLF | 0.4305034 |
| ALAAAAATAI | ASSLGGGLF | 0.29047126 |
| AFGLGLLL | ASSLGGGLF | 0.1299923 |
| AFFAAASL | ASSLGGGLF | 0.5758488 |
| AAAALPAFSPP | ASSLGGGLF | 0.1354386 |
| VVLAPAL | ASSLGGGLF | 0.3994314 |
| TIGHVDHGKTTL | ASSLGGGLF | 0.4549207 |
| LISNASDALDK | ASSLGGGLF | 0.2873878 |
| LISNASDAL | ASSLGGGLF | 0.3141296 |
| ISNASDALDK | ASSLGGGLF | 0.2636132 |
| NQAETLVYQTE | ASSLGGGLF | 0.11620167 |
| TGGTGGTAGAG | ASSLGGGLF | 0.34247974 |
| TGSTAGGAGGA | ASSLGGGLF | 0.106092274 |
| DPLAVPVAAGPV | ASSLGGGLF | 0.36555526 |
| GGTGGTPGNGGN | ASSLGGGLF | 0.1257951 |
| PAGLAGVDPATL | ASSLGGGLF | 0.36643243 |
| GNNLVVEAPHAGASV | ASSLGGGLF | 0.14688101 |
| KVTRIMSASTLEQPA | ASSLGGGLF | 0.35749778 |
| LGFALPIALPRRTVV | ASSLGGGLF | 0.17957744 |
| TRLGNLTARKLVVLV | ASSLGGGLF | 0.122628994 |
| WSQQRRLIYFPSAGP | ASSLGGGLF | 0.115971826 |
| GPNGNPDPMAAAVDIR | ASSLGGGLF | 0.11687283 |
| TQRCVSVADLRAAASR | ASSLGGGLF | 0.29073447 |
| QFVLDVAEHPVEVLTETV | ASSLGGGLF | 0.15636894 |
| GGAGGTGGAGGNGGTQAGDG | ASSLGGGLF | 0.115883514 |
| PLLATLRAIAVDVDAKPAQV | ASSLGGGLF | 0.3712676 |
| APKFRTGGYMIDFWGVGYQVAK | ASSLGGGLF | 0.1592913 |
| GAGGAGGAGGAAGGPGGTGGTG | ASSLGGGLF | 0.11407122 |
| IGALGLLATDVAPAGSVAFGVA | ASSLGGGLF | 0.4301521 |
| AGGAGGIGGNGGAGGAGGVLMG | ASSLGGGLF | 0.2320292 |
| ADELIGLL | ASSLGGGLF | 0.1793828 |
| ADRALKA | ASSLGGGLF | 0.10597072 |
| AGAVGQVGGAGGA | ASSLGGGLF | 0.12113388 |
| AGGGGGLGGAP | ASSLGGGLF | 0.14029661 |
| AGGLLYGNGGAG | ASSLGGGLF | 0.11104797 |
| AGLESIGL | ASSLGGGLF | 0.11778569 |
| AGLKTGII | ASSLGGGLF | 0.47993994 |
| ASLAALLL | ASSLGGGLF | 0.10657693 |
| ASVINISSI | ASSLGGGLF | 0.12665124 |
| ATPPGPA | ASSLGGGLF | 0.449388 |
| ATVASALV | ASSLGGGLF | 0.2322036 |
| ATVASLIL | ASSLGGGLF | 0.70727015 |
| AVNGPVLIRH | ASSLGGGLF | 0.12619594 |
| DGAAGAVGI | ASSLGGGLF | 0.16487174 |
| DPLLHGI | ASSLGGGLF | 0.20121 |
| ELGAALGAL | ASSLGGGLF | 0.18783015 |
| ELGRTQAV | ASSLGGGLF | 0.111379 |
| FVAGGYLS | ASSLGGGLF | 0.10916531 |

| | | |
|---|---|---|
| GASGGAGGAGGW | ASSLGGGLF | 0.2135318 |
| GGAGAGGDGGAGGI | ASSLGGGLF | 0.118548945 |
| GTGHNGSQPGMGGNG | ASSLGGGLF | 0.23845184 |
| GYAAASASAS | ASSLGGGLF | 0.3322327 |
| HQVDVAAVQQG | ASSLGGGLF | 0.11806329 |
| IGAAIAHT | ASSLGGGLF | 0.1162413 |
| IGMIEVAQL | ASSLGGGLF | 0.18396433 |
| IRLAPEHTSAMD | ASSLGGGLF | 0.1254711 |
| IYNVSVL | ASSLGGGLF | 0.3975938 |
| KQGFLGM | ASSLGGGLF | 0.15177017 |
| LESVITQL | ASSLGGGLF | 0.7645377 |
| LGLGALVVV | ASSLGGGLF | 0.22438933 |
| LIGNGGNGGSGGI | ASSLGGGLF | 0.12918976 |
| LNAINAPTL | ASSLGGGLF | 0.4537158 |
| LQLLNTGV | ASSLGGGLF | 0.19644499 |
| LQTSIEF | ASSLGGGLF | 0.16064276 |
| MSFAVSEL | ASSLGGGLF | 0.19638461 |
| NTGALGL | ASSLGGGLF | 0.11985761 |
| PPGAGGAGGAGTT | ASSLGGGLF | 0.13031058 |
| QLVGLGI | ASSLGGGLF | 0.16571516 |
| RFLEQVHQL | ASSLGGGLF | 0.122421175 |
| RGVGYVLR | ASSLGGGLF | 0.2614104 |
| SATGVFAGVI | ASSLGGGLF | 0.12419152 |
| SLPAITLP | ASSLGGGLF | 0.12043442 |
| STALAFTV | ASSLGGGLF | 0.26077303 |
| SVLTVGLGK | ASSLGGGLF | 0.40330422 |
| TALAGAGSGI | ASSLGGGLF | 0.7555852 |
| TGLPFTGI | ASSLGGGLF | 0.17196871 |
| VAEHVTV | ASSLGGGLF | 0.12108961 |
| VDIVLNSL | ASSLGGGLF | 0.6297816 |
| VLTGGGAVL | ASSLGGGLF | 0.13876268 |
| VNVGLAYGI | ASSLGGGLF | 0.2190447 |
| VSSAAVGAV | ASSLGGGLF | 0.17434146 |
| VTSVYEAGV | ASSLGGGLF | 0.1208118 |
| WVALDRGA | ASSLGGGLF | 0.15657727 |
| YLVGSVVI | ASSLGGGLF | 0.3634838 |
| GPPGTGKTLL | ASSLGGGLF | 0.1128962 |
| GVGMTKF | ASSLGGGLF | 0.33953393 |
| IGIGGSDL | ASSLGGGLF | 0.106779434 |
| IGTLLAEL | ASSLGGGLF | 0.117435224 |
| IYTSGTT | ASSLGGGLF | 0.16531886 |
| LSFPAGV | ASSLGGGLF | 0.13441704 |
| LYTSGTT | ASSLGGGLF | 0.11418053 |
| VGAFPFGF | ASSLGGGLF | 0.15015993 |
| VGAGTVEF | ASSLGGGLF | 0.5631604 |
| VGKSTLLL | ASSLGGGLF | 0.6206604 |
| AALAHAGGGIV | ASSLGGGLF | 0.12096855 |
| AALVAVSLL | ASSLGGGLF | 0.52555364 |
| AAVSRAVAAL | ASSLGGGLF | 0.73088455 |
| AGGSSGIGGF | ASSLGGGLF | 0.21804698 |
| AGLTVANANG | ASSLGGGLF | 0.11372723 |

| | | |
|---|---|---|
| AGLVGAVL | ASSLGGGLF | 0.15220796 |
| ALSQAFTV | ASSLGGGLF | 0.10847662 |
| ALTFSVSL | ASSLGGGLF | 0.12376187 |
| ALTGAALVAL | ASSLGGGLF | 0.68335325 |
| ASFSIGAGVL | ASSLGGGLF | 0.11705985 |
| DGSVYKI | ASSLGGGLF | 0.26370627 |
| EQLAEIV | ASSLGGGLF | 0.53317225 |
| FAVSELTVM | ASSLGGGLF | 0.17152394 |
| GAAPLAAGL | ASSLGGGLF | 0.12471448 |
| GADGGDGGSGGAGG | ASSLGGGLF | 0.11586457 |
| GFGLDTGGAGG | ASSLGGGLF | 0.11112104 |
| GLASGGLL | ASSLGGGLF | 0.19479299 |
| GRIAGAAVLL | ASSLGGGLF | 0.19089678 |
| GSAGSLNTSF | ASSLGGGLF | 0.13460924 |
| GTGQIGI | ASSLGGGLF | 0.18022874 |
| GTLIALIL | ASSLGGGLF | 0.13050082 |
| IAATMGFVVG | ASSLGGGLF | 0.1405162 |
| IFAQALL | ASSLGGGLF | 0.13663627 |
| IGAAVALGF | ASSLGGGLF | 0.117394045 |
| IGLTPLAL | ASSLGGGLF | 0.44439662 |
| ILGSNTSTL | ASSLGGGLF | 0.11816806 |
| IVFSLTVL | ASSLGGGLF | 0.1881508 |
| LASKATGF | ASSLGGGLF | 0.70831954 |
| LASVALIL | ASSLGGGLF | 0.110987954 |
| LFAVGLTL | ASSLGGGLF | 0.12657113 |
| LFNGGIGTYI | ASSLGGGLF | 0.16131356 |
| LTLGAGII | ASSLGGGLF | 0.11177316 |
| LVAAALGDI | ASSLGGGLF | 0.5088256 |
| NTAFVQLGI | ASSLGGGLF | 0.1616863 |
| QLFSGAEL | ASSLGGGLF | 0.33108252 |
| SAATFITL | ASSLGGGLF | 0.2115534 |
| SNYGLANLGV | ASSLGGGLF | 0.6891557 |
| SYGGAYAVM | ASSLGGGLF | 0.2665975 |
| TTGVGGLTLG | ASSLGGGLF | 0.13455126 |
| TTQTIYI | ASSLGGGLF | 0.10651712 |
| VAAGAGLGAVI | ASSLGGGLF | 0.11655217 |
| VAAVVTLL | ASSLGGGLF | 0.48915228 |
| VGFLGGLI | ASSLGGGLF | 0.41933963 |
| VGVIPILL | ASSLGGGLF | 0.11832073 |
| VIFASSLI | ASSLGGGLF | 0.22209969 |
| VKGGDGGAAAG | ASSLGGGLF | 0.59295547 |
| VTKTFTNI | ASSLGGGLF | 0.52769434 |
| VVAAAILL | ASSLGGGLF | 0.113952085 |
| YAALDVEL | ASSLGGGLF | 0.13703011 |
| LASVALLL | ASSLGGGLF | 0.17799586 |
| LDAVHRV | ASSLGGGLF | 0.2203146 |
| QGLSSSIF | ASSLGGGLF | 0.1170865 |
| RGTFANIRL | ASSLGGGLF | 0.5956023 |
| RQLWWGHRI | ASSLGGGLF | 0.24738431 |
| YGPPGTGKTLL | ASSLGGGLF | 0.6648922 |
| AALVGAPANVA | ASSLGGGLF | 0.22372419 |

| | | |
|---|---|---|
| AATFRRTE | ASSLGGGLF | 0.3594222 |
| AGALGGGI | ASSLGGGLF | 0.20483786 |
| AGGAGGAGGIGAG | ASSLGGGLF | 0.13632725 |
| AGSGALGL | ASSLGGGLF | 0.11990294 |
| AQIADQI | ASSLGGGLF | 0.11422047 |
| ASAVAVTYL | ASSLGGGLF | 0.13420202 |
| ASTVLGGTGG | ASSLGGGLF | 0.12797716 |
| AVGGEPAK | ASSLGGGLF | 0.10723812 |
| CEQGTFGGTC | ASSLGGGLF | 0.15531182 |
| DEGKAPAG | ASSLGGGLF | 0.26696342 |
| DTVVFPAL | ASSLGGGLF | 0.27065164 |
| FEGITMAG | ASSLGGGLF | 0.14279285 |
| FPLVVQD | ASSLGGGLF | 0.14901747 |
| GAFTTPQ | ASSLGGGLF | 0.12043728 |
| GFGGFGTAGGVGGA | ASSLGGGLF | 0.5829298 |
| GFLAQFL | ASSLGGGLF | 0.112225346 |
| GGAGGAGGNAGVA | ASSLGGGLF | 0.1170619 |
| GGVGTAGAGGAGGAGG | ASSLGGGLF | 0.14686936 |
| GNGGGGDAGNAGSGGN | ASSLGGGLF | 0.117440775 |
| GTGGAGGAGNDAGS | ASSLGGGLF | 0.6691355 |
| GTTNTGWFNS | ASSLGGGLF | 0.5073143 |
| IFMGPLFTL | ASSLGGGLF | 0.29425383 |
| ILAHAIEAI | ASSLGGGLF | 0.5470754 |
| IRSAIREAL | ASSLGGGLF | 0.10997903 |
| ITFAGGG | ASSLGGGLF | 0.1263635 |
| ITGMSGHQEV | ASSLGGGLF | 0.71516865 |
| IVAGLACTL | ASSLGGGLF | 0.21461426 |
| IVGGGLAGA | ASSLGGGLF | 0.29642588 |
| IVTGVLNI | ASSLGGGLF | 0.3352857 |
| LAQQGCI | ASSLGGGLF | 0.12536468 |
| LCLHVEN | ASSLGGGLF | 0.1849837 |
| LDAISDGS | ASSLGGGLF | 0.12687798 |
| LGGFAAML | ASSLGGGLF | 0.2653965 |
| LGGKNPMI | ASSLGGGLF | 0.15760495 |
| LGPKGVAAIL | ASSLGGGLF | 0.121021464 |
| LMVGVSI | ASSLGGGLF | 0.37380815 |
| LVGAAFVL | ASSLGGGLF | 0.75435805 |
| LVLAVAPL | ASSLGGGLF | 0.23845553 |
| MAAIAVIDAA | ASSLGGGLF | 0.23974586 |
| NNGINAELQ | ASSLGGGLF | 0.1986815 |
| RGEPLVLVHG | ASSLGGGLF | 0.22032732 |
| RGGAGGAAGLI | ASSLGGGLF | 0.14710832 |
| RSYRMHPAV | ASSLGGGLF | 0.12962529 |
| SAAHLLAGQLA | ASSLGGGLF | 0.12466599 |
| SAAVGSY | ASSLGGGLF | 0.14955235 |
| SPHAVLHRI | ASSLGGGLF | 0.22908895 |
| SQIAKVL | ASSLGGGLF | 0.14653477 |
| TALGHLVP | ASSLGGGLF | 0.7692406 |
| TAPTTSGAA | ASSLGGGLF | 0.5103668 |
| TDGDDGNGGSPGT | ASSLGGGLF | 0.13290906 |
| TDGLGEN | ASSLGGGLF | 0.11673799 |

| | | |
|---|---|---|
| TGGPPLPGQA | ASSLGGGLF | 0.2515276 |
| TIPAITI | ASSLGGGLF | 0.237145 |
| TIPGFTI | ASSLGGGLF | 0.12018359 |
| TSIASGALA | ASSLGGGLF | 0.64365697 |
| TVTLPGAAI | ASSLGGGLF | 0.14834338 |
| VASGLSAALP | ASSLGGGLF | 0.11987474 |
| VGAGVIGI | ASSLGGGLF | 0.18494354 |
| VGPKRFGV | ASSLGGGLF | 0.7521547 |
| VNVAFPD | ASSLGGGLF | 0.23621766 |
| VSMTNSGVS | ASSLGGGLF | 0.14113536 |
| VTDVLGDLSV | ASSLGGGLF | 0.11714783 |
| VVVVGAGFAGL | ASSLGGGLF | 0.57067674 |
| VYTVGTVV | ASSLGGGLF | 0.6278306 |
| WTTFSGP | ASSLGGGLF | 0.10619806 |
| YDELVEAF | ASSLGGGLF | 0.69566303 |
| YGTGGAGGNGG | ASSLGGGLF | 0.12036738 |
| YTNVMAV | ASSLGGGLF | 0.11680573 |
| FLEQVHQL | ASSLGGGLF | 0.22030587 |
| GLSLLAVAV | ASSLGGGLF | 0.1667316 |
| LEQVHQL | ASSLGGGLF | 0.7170499 |
| LNAAGAI | ASSLGGGLF | 0.24724823 |
| PAVHASL | ASSLGGGLF | 0.38379276 |
| TELEDDA | ASSLGGGLF | 0.2174843 |
| TPPGPPN | ASSLGGGLF | 0.64812756 |
| TPPPPPPP | ASSLGGGLF | 0.392819 |
| TQSPALI | ASSLGTGGWYEQY | 0.27442545 |
| EFAAAFPQ | ASSLGTGGWYEQY | 0.25729665 |
| LLSGNLDVL | ASSLGTGGWYEQY | 0.14142449 |
| GPANLGNYNI | ASSLGTGGWYEQY | 0.24235079 |
| TLGPTLP | ASSLGTGGWYEQY | 0.12550813 |
| VYAYSSCCK | ASSLGTGGWYEQY | 0.1386601 |
| APAPAPAPNI | ASSLGTGGWYEQY | 0.34357533 |
| PGGPVPIP | ASSLGTGGWYEQY | 0.37256372 |
| FLTVALEP | ASSLGTGGWYEQY | 0.11807336 |
| SQYFGANL | ASSLGTGGWYEQY | 0.20947947 |
| GGVKPAA | ASSLGTGGWYEQY | 0.5966274 |
| GGTGGAGGAGGW | ASSLGTGGWYEQY | 0.19355819 |
| GVGGVAGLGGAGG | ASSLGTGGWYEQY | 0.13381617 |
| VLLVLATTI | ASSLGTGGWYEQY | 0.6635132 |
| IGIFAAML | ASSLGTGGWYEQY | 0.76629543 |
| AVGDLID | ASSLGTGGWYEQY | 0.34760347 |
| NISPLPL | ASSLGTGGWYEQY | 0.116923675 |
| VTAVFVAI | ASSLGTGGWYEQY | 0.113559656 |
| SGDDNDAAL | ASSLGTGGWYEQY | 0.24194437 |
| AVAALAHS | ASSLGTGGWYEQY | 0.21614721 |
| GGVTVMATP | ASSLGTGGWYEQY | 0.12851399 |
| AGLAANVRD | ASSLGTGGWYEQY | 0.117967755 |
| TNRPDLIDPALLRPG | ASSLGTGGWYEQY | 0.2844476 |
| LANLGVTAL | ASSLGTGGWYEQY | 0.3570922 |
| TGERIEL | ASSLGTGGWYEQY | 0.12582567 |
| NGEELAA | ASSLGTGGWYEQY | 0.11549596 |

| | | |
|---|---|---|
| VLSLVWG | ASSLGTGGWYEQY | 0.20144427 |
| IGYALWL | ASSLGTGGWYEQY | 0.40210807 |
| LSLLLAPL | ASSLGTGGWYEQY | 0.13739398 |
| TLDEALK | ASSLGTGGWYEQY | 0.217594 |
| TDGAGTAEGG | ASSLGTGGWYEQY | 0.24304631 |
| NAPGGAGGNGG | ASSLGTGGWYEQY | 0.10779838 |
| VSVFMGMGGNF | ASSLGTGGWYEQY | 0.20169514 |
| LASYLAAL | ASSLGTGGWYEQY | 0.12925275 |
| LVGGTPAL | ASSLGTGGWYEQY | 0.13689211 |
| VRNADVL | ASSLGTGGWYEQY | 0.1368721 |
| TCSSPFLVSAM | ASSLGTGGWYEQY | 0.51016825 |
| VHITDAE | ASSLGTGGWYEQY | 0.3183355 |
| EYAAPSF | ASSLGTGGWYEQY | 0.2438961 |
| VVDAFHGGL | ASSLGTGGWYEQY | 0.2752205 |
| ASVVDPQ | ASSLGTGGWYEQY | 0.16934806 |
| SYNEFGGLS | ASSLGTGGWYEQY | 0.11670523 |
| DPDGSVY | ASSLGTGGWYEQY | 0.10916988 |
| MVSTIHATE | ASSLGTGGWYEQY | 0.6057269 |
| LAAGVADPV | ASSLGTGGWYEQY | 0.10616908 |
| TKEQIAA | ASSLGTGGWYEQY | 0.44118294 |
| SLASPAAL | ASSLGTGGWYEQY | 0.21245392 |
| IGLVFYGV | ASSLGTGGWYEQY | 0.12142404 |
| FLLDASNL | ASSLGTGGWYEQY | 0.2082142 |
| YQVGNFP | ASSLGTGGWYEQY | 0.10861926 |
| ESLLSQST | ASSLGTGGWYEQY | 0.13206612 |
| LFTVGFAAVG | ASSLGTGGWYEQY | 0.71017575 |
| ATPPTPT | ASSLGTGGWYEQY | 0.17375398 |
| TDLATLG | ASSLGTGGWYEQY | 0.16190559 |
| SAGGTGR | ASSLGTGGWYEQY | 0.2742563 |
| GFSNSGS | ASSLGTGGWYEQY | 0.37924182 |
| GGNGGSGGTSV | ASSLGTGGWYEQY | 0.15112175 |
| GTSVATGGAGN | ASSLGTGGWYEQY | 0.18921731 |
| NGGNGGSGGTS | ASSLGTGGWYEQY | 0.4621765 |
| AVVKFPH | ASSLGTGGWYEQY | 0.27088135 |
| TPPPPPP | ASSLGTGGWYEQY | 0.12115848 |
| AAPAAAAAPA | ASSLGTGGWYEQY | 0.17179312 |
| IALNEDL | ASSLGTGGWYEQY | 0.13199943 |
| IVNVSSM | ASSLGTGGWYEQY | 0.29312718 |
| TGIPPAPR | ASSLGTGGWYEQY | 0.13456343 |
| FDLGGGTF | ASSLGTGGWYEQY | 0.27118087 |
| FDLGGGTFDVS | ASSLGTGGWYEQY | 0.16367708 |
| NEPTAAA | ASSLGTGGWYEQY | 0.10948783 |
| NEPTAAALA | ASSLGTGGWYEQY | 0.15396512 |
| GSIINISSI | ASSLGTGGWYEQY | 0.25003698 |
| VAPGNAGTA | ASSLGTGGWYEQY | 0.16140275 |
| TGGGASLE | ASSLGTGGWYEQY | 0.18236494 |
| VAVYNHYKRI | ASSLGTGGWYEQY | 0.12859619 |
| TDPAPATT | ASSLGTGGWYEQY | 0.3807183 |
| TNRPDILDPALLRPG | ASSLGTGGWYEQY | 0.14662062 |
| LTTTTTS | ASSLGTGGWYEQY | 0.14157437 |
| PTGSGKTL | ASSLGTGGWYEQY | 0.1966756 |

| | | |
|---|---|---|
| TGSGKTL | ASSLGTGGWYEQY | 0.1482501 |
| TIGHVDHGK | ASSLGTGGWYEQY | 0.14000042 |
| TIGHVDHGKTT | ASSLGTGGWYEQY | 0.31916016 |
| GHVDHGKTT | ASSLGTGGWYEQY | 0.11550858 |
| AQFEHTLL | ASSLGTGGWYEQY | 0.21326086 |
| IVNQDEL | ASSLGTGGWYEQY | 0.106777586 |
| TNGSQFF | ASSLGTGGWYEQY | 0.16136782 |
| TGKGIVE | ASSLGTGGWYEQY | 0.4180287 |
| LISNASDA | ASSLGTGGWYEQY | 0.11652146 |
| AELRTLGELTKIAKAHGAQ | ASSLGTGGWYEQY | 0.4138844 |
| APNAVAAAEPSW | ASSLGTGGWYEQY | 0.13855378 |
| APQVAGAATAVL | ASSLGTGGWYEQY | 0.13661018 |
| ARRDPAAAAAILPTDA | ASSLGTGGWYEQY | 0.13636558 |
| DAAARDVLAEAGLAEYF | ASSLGTGGWYEQY | 0.23643935 |
| DDQIVLSAKANLAKVEQ | ASSLGTGGWYEQY | 0.5148163 |
| DELLPDTGEDSDEDGH | ASSLGTGGWYEQY | 0.11849065 |
| ELPEAPANRATVPQD | ASSLGTGGWYEQY | 0.28063768 |
| FGQSKIERVTGIG | ASSLGTGGWYEQY | 0.34096703 |
| GAGGTGSTAGGAGGAGGAGGLY | ASSLGTGGWYEQY | 0.22003666 |
| GHGGTGAAGGNGGTGGAGGSGI | ASSLGTGGWYEQY | 0.115969695 |
| GNGGNGGAGGEGGPGGAGGAGA | ASSLGTGGWYEQY | 0.12905362 |
| GSTAGNGGNGGNATNGTIAGQP | ASSLGTGGWYEQY | 0.11402753 |
| GTGGDGGNGGNGGTGG | ASSLGTGGWYEQY | 0.13486804 |
| GVRRPAGIPADAPV | ASSLGTGGWYEQY | 0.11957939 |
| GYRQVTDAHLLTLARR | ASSLGTGGWYEQY | 0.39203405 |
| IAKMGQDGHDRG | ASSLGTGGWYEQY | 0.6295506 |
| KAEPSIEIQYT | ASSLGTGGWYEQY | 0.13443701 |
| KPVRIRMAADEDTVAVT | ASSLGTGGWYEQY | 0.3579962 |
| MPPRIAGMRLLVIKPEP | ASSLGTGGWYEQY | 0.119971 |
| MKFYAVALFRSRGVPPTR | ASSLGTGGWYEQY | 0.2944708 |
| NAGGFGILGAVAHSP | ASSLGTGGWYEQY | 0.65954363 |
| NEGREITLAMAEM | ASSLGTGGWYEQY | 0.13280159 |
| NELVEWSHPLGHRWRWE | ASSLGTGGWYEQY | 0.1815545 |
| NLGAGGTGGNGGNPG | ASSLGTGGWYEQY | 0.14578788 |
| NTEWPKWETVMAADPVD | ASSLGTGGWYEQY | 0.12174994 |
| PASAAPAETQP | ASSLGTGGWYEQY | 0.6179496 |
| QQSSDGKNGVRAAEITGE | ASSLGTGGWYEQY | 0.1466032 |
| RAPDAVTTSWWKE | ASSLGTGGWYEQY | 0.27322102 |
| SGFGNVGAHTSGWFNQ | ASSLGTGGWYEQY | 0.12828647 |
| SGVAGAVGQAASVGGLKV | ASSLGTGGWYEQY | 0.119492374 |
| SNGGAGTGSAGGNGGT | ASSLGTGGWYEQY | 0.16999885 |
| TGGAGGEGGAAG | ASSLGTGGWYEQY | 0.60882294 |
| TGGDGGAAGAGGAPGHG | ASSLGTGGWYEQY | 0.14555672 |
| TMPPLLAAIANAGRHGV | ASSLGTGGWYEQY | 0.3994093 |
| TSSTAGNSGGA | ASSLGTGGWYEQY | 0.15385647 |
| VPATVPVGILC | ASSLGTGGWYEQY | 0.3923759 |
| VQRPDSGGVVLTGRIS | ASSLGTGGWYEQY | 0.6526624 |
| WTPNPVPATKGVPAGLVQAT | ASSLGTGGWYEQY | 0.30486706 |
| YALKRALAKFSDRFDVVII | ASSLGTGGWYEQY | 0.15416808 |
| YLIGSVIV | ASSLGTGGWYEQY | 0.61140734 |
| YGQLVAVID | ASSLGTGGWYEQY | 0.254252 |

| | | |
|---|---|---|
| VTVGGMTLPA | ASSLGTGGWYEQY | 0.42527184 |
| VLALPAV | ASSLGTGGWYEQY | 0.35871762 |
| VIPPTLNYET | ASSLGTGGWYEQY | 0.17501456 |
| VGMGGMHPSSG | ASSLGTGGWYEQY | 0.34211567 |
| VGLAGGAATA | ASSLGTGGWYEQY | 0.16923283 |
| VGGAGGAGGSGP | ASSLGTGGWYEQY | 0.22566992 |
| VFAAFMV | ASSLGTGGWYEQY | 0.17413317 |
| TLYDKLR | ASSLGTGGWYEQY | 0.14899495 |
| TLAVFTVL | ASSLGTGGWYEQY | 0.320721 |
| TGSVQPARV | ASSLGTGGWYEQY | 0.16058883 |
| TGGGAGSGGT | ASSLGTGGWYEQY | 0.19949189 |
| SHLETSL | ASSLGTGGWYEQY | 0.2437824 |
| SFTTGMT | ASSLGTGGWYEQY | 0.18051466 |
| QVEVAKRMGVRQPTVS | ASSLGTGGWYEQY | 0.6018477 |
| QGSGGPALAM | ASSLGTGGWYEQY | 0.6263554 |
| MLAPSNP | ASSLGTGGWYEQY | 0.1563794 |
| LVNNAAI | ASSLGTGGWYEQY | 0.19371946 |
| LTGEINTLA | ASSLGTGGWYEQY | 0.17087883 |
| LSIFGPV | ASSLGTGGWYEQY | 0.19165647 |
| LLALLANL | ASSLGTGGWYEQY | 0.15824705 |
| LISNASDALDKLR | ASSLGTGGWYEQY | 0.23105259 |
| LINEAVP | ASSLGTGGWYEQY | 0.15469335 |
| LGYAFSGL | ASSLGTGGWYEQY | 0.143724 |
| LDFVAVAL | ASSLGTGGWYEQY | 0.29052675 |
| LAQPGAE | ASSLGTGGWYEQY | 0.19804095 |
| ISNLGAVTGF | ASSLGTGGWYEQY | 0.17448106 |
| HQEDPGRI | ASSLGTGGWYEQY | 0.4617617 |
| GTPGEPTP | ASSLGTGGWYEQY | 0.14663409 |
| GFFNSGS | ASSLGTGGWYEQY | 0.16310109 |
| GAGGGGLTAGAN | ASSLGTGGWYEQY | 0.3127714 |
| FVDALVNM | ASSLGTGGWYEQY | 0.4848082 |
| FLTVAPEL | ASSLGTGGWYEQY | 0.16154552 |
| FETLTAL | ASSLGTGGWYEQY | 0.20805985 |
| DVINPIEL | ASSLGTGGWYEQY | 0.50729656 |
| APVSGAPV | ASSLGTGGWYEQY | 0.18655416 |
| ALVGGLAW | ASSLGTGGWYEQY | 0.3722062 |
| ALAAAAATAI | ASSLGTGGWYEQY | 0.3787685 |
| AFGLGLLL | ASSLGTGGWYEQY | 0.16174243 |
| AFFAAASL | ASSLGTGGWYEQY | 0.4164096 |
| AAAALPAFSPP | ASSLGTGGWYEQY | 0.6491033 |
| VVLAPAL | ASSLGTGGWYEQY | 0.18347016 |
| TIGHVDHGKTTL | ASSLGTGGWYEQY | 0.22724155 |
| LISNASDALDK | ASSLGTGGWYEQY | 0.17156693 |
| LISNASDAL | ASSLGTGGWYEQY | 0.20778224 |
| ISNASDALDK | ASSLGTGGWYEQY | 0.14520313 |
| NQAETLVYQTE | ASSLGTGGWYEQY | 0.25196505 |
| TGGTGGTAGAG | ASSLGTGGWYEQY | 0.23638514 |
| TGSTAGGAGGA | ASSLGTGGWYEQY | 0.21218145 |
| DPLAVPVAAGPV | ASSLGTGGWYEQY | 0.15351355 |
| GGTGGTPGNGGN | ASSLGTGGWYEQY | 0.15595627 |
| PAGLAGVDPATL | ASSLGTGGWYEQY | 0.15217225 |

| | | |
|---|---|---|
| GNNLVVEAPHAGASV | ASSLGTGGWYEQY | 0.18659563 |
| KVTRIMSASTLEQPA | ASSLGTGGWYEQY | 0.1449613 |
| LGFALPIALPRRTVV | ASSLGTGGWYEQY | 0.16336197 |
| TRLGNLTARKLVVLV | ASSLGTGGWYEQY | 0.34190303 |
| WSQQRRLIYFPSAGP | ASSLGTGGWYEQY | 0.23139907 |
| GPNGNPDPMAAAVDIR | ASSLGTGGWYEQY | 0.20272066 |
| TQRCVSVADLRAAASR | ASSLGTGGWYEQY | 0.5683155 |
| QFVLDVAEHPVEVLTETV | ASSLGTGGWYEQY | 0.32732248 |
| GGAGGTGGAGGNGGTQAGDG | ASSLGTGGWYEQY | 0.18869701 |
| PLLATLRAIAVDVDAKPAQV | ASSLGTGGWYEQY | 0.17114173 |
| APKFRTGGYMIDFWGVGYQVAK | ASSLGTGGWYEQY | 0.17040914 |
| GAGGAGGAGGAAGGPGGTGGTG | ASSLGTGGWYEQY | 0.51940894 |
| IGALGLLATDVAPAGSVAFGVA | ASSLGTGGWYEQY | 0.18678719 |
| AGGAGGIGGNGGAGGAGGVLMG | ASSLGTGGWYEQY | 0.16072802 |
| ADELIGLL | ASSLGTGGWYEQY | 0.3542115 |
| ADRALKA | ASSLGTGGWYEQY | 0.17515244 |
| AGAVGQVGGAGGA | ASSLGTGGWYEQY | 0.18929638 |
| AGGGGGLGGAP | ASSLGTGGWYEQY | 0.22216697 |
| AGGLLYGNGGAG | ASSLGTGGWYEQY | 0.2200727 |
| AGLESIGL | ASSLGTGGWYEQY | 0.16447018 |
| AGLKTGII | ASSLGTGGWYEQY | 0.15958382 |
| ASLAALLL | ASSLGTGGWYEQY | 0.15347356 |
| ASVINISSI | ASSLGTGGWYEQY | 0.24196064 |
| ATPPGPA | ASSLGTGGWYEQY | 0.5626384 |
| ATVASALV | ASSLGTGGWYEQY | 0.18167429 |
| ATVASLIL | ASSLGTGGWYEQY | 0.18183169 |
| AVNGPVLIRH | ASSLGTGGWYEQY | 0.41667503 |
| DGAAGAVGI | ASSLGTGGWYEQY | 0.18380529 |
| DPLLHGI | ASSLGTGGWYEQY | 0.19625276 |
| ELGAALGAL | ASSLGTGGWYEQY | 0.6216167 |
| ELGRTQAV | ASSLGTGGWYEQY | 0.17394444 |
| FVAGGYLS | ASSLGTGGWYEQY | 0.25369626 |
| GASGGAGGAGGW | ASSLGTGGWYEQY | 0.27175072 |
| GGAGAGGDGGAGGI | ASSLGTGGWYEQY | 0.3388136 |
| GTGHNGSQPGMGGNG | ASSLGTGGWYEQY | 0.4421633 |
| GYAAASASAS | ASSLGTGGWYEQY | 0.17199822 |
| HQVDVAAVQQG | ASSLGTGGWYEQY | 0.17303981 |
| IGAAIAHT | ASSLGTGGWYEQY | 0.1500025 |
| IGMIEVAQL | ASSLGTGGWYEQY | 0.16884339 |
| IRLAPEHTSAMD | ASSLGTGGWYEQY | 0.45781025 |
| IYNVSVL | ASSLGTGGWYEQY | 0.23359339 |
| KQGFLGM | ASSLGTGGWYEQY | 0.48587787 |
| LESVITQL | ASSLGTGGWYEQY | 0.27129236 |
| LGLGALVVV | ASSLGTGGWYEQY | 0.15783711 |
| LIGNGGNGGSGGI | ASSLGTGGWYEQY | 0.5139766 |
| LNAINAPTL | ASSLGTGGWYEQY | 0.38891402 |
| LQLLNTGV | ASSLGTGGWYEQY | 0.155014 |
| LQTSIEF | ASSLGTGGWYEQY | 0.1645116 |
| MSFAVSEL | ASSLGTGGWYEQY | 0.3933957 |
| NTGALGL | ASSLGTGGWYEQY | 0.26489344 |
| PPGAGGAGGAGTT | ASSLGTGGWYEQY | 0.20067598 |

| | | |
|---|---|---|
| QLVGLGI | ASSLGTGGWYEQY | 0.18460731 |
| RFLEQVHQL | ASSLGTGGWYEQY | 0.33243066 |
| RGVGYVLR | ASSLGTGGWYEQY | 0.14999487 |
| SATGVFAGVI | ASSLGTGGWYEQY | 0.54391867 |
| SLPAITLP | ASSLGTGGWYEQY | 0.15028632 |
| STALAFTV | ASSLGTGGWYEQY | 0.16548368 |
| SVLTVGLGK | ASSLGTGGWYEQY | 0.15111034 |
| TALAGAGSGI | ASSLGTGGWYEQY | 0.1490698 |
| TGLPFTGI | ASSLGTGGWYEQY | 0.22750084 |
| VAEHVTV | ASSLGTGGWYEQY | 0.23626447 |
| VDIVLNSL | ASSLGTGGWYEQY | 0.4188677 |
| VLTGGGAVL | ASSLGTGGWYEQY | 0.17170516 |
| VNVGLAYGI | ASSLGTGGWYEQY | 0.28071198 |
| VSSAAVGAV | ASSLGTGGWYEQY | 0.22596684 |
| VTSVYEAGV | ASSLGTGGWYEQY | 0.28258604 |
| WVALDRGA | ASSLGTGGWYEQY | 0.16561206 |
| YLVGSVVI | ASSLGTGGWYEQY | 0.21433415 |
| GPPGTGKTLL | ASSLGTGGWYEQY | 0.61678445 |
| GVGMTKF | ASSLGTGGWYEQY | 0.19816372 |
| IGIGGSDL | ASSLGTGGWYEQY | 0.15948085 |
| IGTLLAEL | ASSLGTGGWYEQY | 0.1536167 |
| IYTSGTT | ASSLGTGGWYEQY | 0.18905832 |
| LSFPAGV | ASSLGTGGWYEQY | 0.14792013 |
| LYTSGTT | ASSLGTGGWYEQY | 0.1440262 |
| VGAFPFGF | ASSLGTGGWYEQY | 0.14787297 |
| VGAGTVEF | ASSLGTGGWYEQY | 0.15817049 |
| VGKSTLLL | ASSLGTGGWYEQY | 0.50658363 |
| AALAHAGGGIV | ASSLGTGGWYEQY | 0.44780582 |
| AALVAVSLL | ASSLGTGGWYEQY | 0.19956136 |
| AAVSRAVAAL | ASSLGTGGWYEQY | 0.24799006 |
| AGGSSGIGGF | ASSLGTGGWYEQY | 0.4606436 |
| AGLTVANANG | ASSLGTGGWYEQY | 0.1425612 |
| AGLVGAVL | ASSLGTGGWYEQY | 0.16492848 |
| ALSQAFTV | ASSLGTGGWYEQY | 0.16505843 |
| ALTFSVSL | ASSLGTGGWYEQY | 0.4464658 |
| ALTGAALVAL | ASSLGTGGWYEQY | 0.15469502 |
| ASFSIGAGVL | ASSLGTGGWYEQY | 0.22082216 |
| DGSVYKI | ASSLGTGGWYEQY | 0.22248009 |
| EQLAEIV | ASSLGTGGWYEQY | 0.24978314 |
| FAVSELTVM | ASSLGTGGWYEQY | 0.1868698 |
| GAAPLAAGL | ASSLGTGGWYEQY | 0.15052277 |
| GADGGDGGSGGAGG | ASSLGTGGWYEQY | 0.14311284 |
| GFGLDTGGAGG | ASSLGTGGWYEQY | 0.2751219 |
| GLASGGLL | ASSLGTGGWYEQY | 0.14749402 |
| GRIAGAAVLL | ASSLGTGGWYEQY | 0.16823344 |
| GSAGSLNTSF | ASSLGTGGWYEQY | 0.15421052 |
| GTGQIGI | ASSLGTGGWYEQY | 0.14847472 |
| GTLIALIL | ASSLGTGGWYEQY | 0.15143715 |
| IAATMGFVVG | ASSLGTGGWYEQY | 0.3467131 |
| IFAQALL | ASSLGTGGWYEQY | 0.18532372 |
| IGAAVALGF | ASSLGTGGWYEQY | 0.29048103 |

| | | |
|---|---|---|
| IGLTPLAL | ASSLGTGGWYEQY | 0.3895403 |
| ILGSNTSTL | ASSLGTGGWYEQY | 0.5997261 |
| IVFSLTVL | ASSLGTGGWYEQY | 0.19827612 |
| LASKATGF | ASSLGTGGWYEQY | 0.17185318 |
| LASVALIL | ASSLGTGGWYEQY | 0.16194476 |
| LFAVGLTL | ASSLGTGGWYEQY | 0.15690163 |
| LFNGGIGTYI | ASSLGTGGWYEQY | 0.1629386 |
| LTLGAGII | ASSLGTGGWYEQY | 0.68402874 |
| LVAAALGDI | ASSLGTGGWYEQY | 0.16800296 |
| NTAFVQLGI | ASSLGTGGWYEQY | 0.17985779 |
| QLFSGAEL | ASSLGTGGWYEQY | 0.1789793 |
| SAATFITL | ASSLGTGGWYEQY | 0.43183094 |
| SNYGLANLGV | ASSLGTGGWYEQY | 0.4419946 |
| SYGGAYAVM | ASSLGTGGWYEQY | 0.18755282 |
| TTGVGGLTLG | ASSLGTGGWYEQY | 0.17783506 |
| TTQTIYI | ASSLGTGGWYEQY | 0.1894606 |
| VAAGAGLGAVI | ASSLGTGGWYEQY | 0.29155892 |
| VAAVVTLL | ASSLGTGGWYEQY | 0.20132677 |
| VGFLGGLI | ASSLGTGGWYEQY | 0.1614272 |
| VGVIPILL | ASSLGTGGWYEQY | 0.22259268 |
| VIFASSLI | ASSLGTGGWYEQY | 0.67941815 |
| VKGGDGGAAAG | ASSLGTGGWYEQY | 0.16715854 |
| VTKTFTNI | ASSLGTGGWYEQY | 0.15229233 |
| VVAAAILL | ASSLGTGGWYEQY | 0.23325491 |
| YAALDVEL | ASSLGTGGWYEQY | 0.1587462 |
| LASVALLL | ASSLGTGGWYEQY | 0.17742662 |
| LDAVHRV | ASSLGTGGWYEQY | 0.21090879 |
| QGLSSSIF | ASSLGTGGWYEQY | 0.15554656 |
| RGTFANIRL | ASSLGTGGWYEQY | 0.17235522 |
| RQLWWGHRI | ASSLGTGGWYEQY | 0.28915864 |
| YGPPGTGKTLL | ASSLGTGGWYEQY | 0.1630627 |
| AALVGAPANVA | ASSLGTGGWYEQY | 0.35499412 |
| AATFRRTE | ASSLGTGGWYEQY | 0.35144448 |
| AGALGGGI | ASSLGTGGWYEQY | 0.25130913 |
| AGGAGGAGGIGAG | ASSLGTGGWYEQY | 0.44980007 |
| AGSGALGL | ASSLGTGGWYEQY | 0.4981435 |
| AQIADQI | ASSLGTGGWYEQY | 0.20618145 |
| ASAVAVTYL | ASSLGTGGWYEQY | 0.17321856 |
| ASTVLGGTGG | ASSLGTGGWYEQY | 0.24804261 |
| AVGGEPAK | ASSLGTGGWYEQY | 0.33329242 |
| CEQGTFGGTC | ASSLGTGGWYEQY | 0.2111099 |
| DEGKAPAG | ASSLGTGGWYEQY | 0.16731665 |
| DTVVFPAL | ASSLGTGGWYEQY | 0.42234102 |
| FEGITMAG | ASSLGTGGWYEQY | 0.16386323 |
| FPLVVQD | ASSLGTGGWYEQY | 0.23191161 |
| GAFTTPQ | ASSLGTGGWYEQY | 0.15871368 |
| GFGGFGTAGGVGGA | ASSLGTGGWYEQY | 0.16298935 |
| GFLAQFL | ASSLGTGGWYEQY | 0.33892667 |
| GGAGGAGGNAGVA | ASSLGTGGWYEQY | 0.14423439 |
| GGVGTAGAGGAGGAGG | ASSLGTGGWYEQY | 0.14451964 |
| GNGGGGDAGNAGSGGN | ASSLGTGGWYEQY | 0.21713156 |

| | | |
|---|---|---|
| GTGGAGGAGNDAGS | ASSLGTGGWYEQY | 0.33047593 |
| GTTNTGWFNS | ASSLGTGGWYEQY | 0.14322509 |
| IFMGPLFTL | ASSLGTGGWYEQY | 0.1686468 |
| ILAHAIEAI | ASSLGTGGWYEQY | 0.15703613 |
| IRSAIREAL | ASSLGTGGWYEQY | 0.19060737 |
| ITFAGGG | ASSLGTGGWYEQY | 0.19194083 |
| ITGMSGHQEV | ASSLGTGGWYEQY | 0.14904453 |
| IVAGLACTL | ASSLGTGGWYEQY | 0.25911885 |
| IVGGGLAGA | ASSLGTGGWYEQY | 0.1543897 |
| IVTGVLNI | ASSLGTGGWYEQY | 0.14868128 |
| LAQQGCI | ASSLGTGGWYEQY | 0.23954152 |
| LCLHVEN | ASSLGTGGWYEQY | 0.6562825 |
| LDAISDGS | ASSLGTGGWYEQY | 0.15684424 |
| LGGFAAML | ASSLGTGGWYEQY | 0.3054631 |
| LGGKNPMI | ASSLGTGGWYEQY | 0.16173483 |
| LGPKGVAAIL | ASSLGTGGWYEQY | 0.5104382 |
| LMVGVSI | ASSLGTGGWYEQY | 0.18374214 |
| LVGAAFVL | ASSLGTGGWYEQY | 0.1508158 |
| LVLAVAPL | ASSLGTGGWYEQY | 0.27563673 |
| MAAIAVIDAA | ASSLGTGGWYEQY | 0.25236434 |
| NNGINAELQ | ASSLGTGGWYEQY | 0.21702589 |
| RGEPLVLVHG | ASSLGTGGWYEQY | 0.28723377 |
| RGGAGGAAGLI | ASSLGTGGWYEQY | 0.15758818 |
| RSYRMHPAV | ASSLGTGGWYEQY | 0.20671141 |
| SAAHLLAGQLA | ASSLGTGGWYEQY | 0.38276652 |
| SAAVGSY | ASSLGTGGWYEQY | 0.45108557 |
| SPHAVLHRI | ASSLGTGGWYEQY | 0.15735698 |
| SQIAKVL | ASSLGTGGWYEQY | 0.28966582 |
| TALGHLVP | ASSLGTGGWYEQY | 0.3133784 |
| TAPTTSGAA | ASSLGTGGWYEQY | 0.20219088 |
| TDGDDGNGGSPGT | ASSLGTGGWYEQY | 0.2149904 |
| TDGLGEN | ASSLGTGGWYEQY | 0.23985498 |
| TGGPPLPGQA | ASSLGTGGWYEQY | 0.2546916 |
| TIPAITI | ASSLGTGGWYEQY | 0.19315329 |
| TIPGFTI | ASSLGTGGWYEQY | 0.31733483 |
| TSIASGALA | ASSLGTGGWYEQY | 0.15014566 |
| TVTLPGAAI | ASSLGTGGWYEQY | 0.4312442 |
| VASGLSAALP | ASSLGTGGWYEQY | 0.19140866 |
| VGAGVIGI | ASSLGTGGWYEQY | 0.5442007 |
| VGPKRFGV | ASSLGTGGWYEQY | 0.19862568 |
| VNVAFPD | ASSLGTGGWYEQY | 0.14836442 |
| VSMTNSGVS | ASSLGTGGWYEQY | 0.18382981 |
| VTDVLGDLSV | ASSLGTGGWYEQY | 0.3187365 |
| VVVVGAGFAGL | ASSLGTGGWYEQY | 0.20265391 |
| VYTVGTVV | ASSLGTGGWYEQY | 0.49885213 |
| WTTFSGP | ASSLGTGGWYEQY | 0.617795 |
| YDELVEAF | ASSLGTGGWYEQY | 0.1668017 |
| YGTGGAGGNGG | ASSLGTGGWYEQY | 0.25999382 |
| YTNVMAV | ASSLGTGGWYEQY | 0.2659847 |
| FLEQVHQL | ASSLGTGGWYEQY | 0.67884266 |
| GLSLLAVAV | ASSLGTGGWYEQY | 0.1749509 |

| | | |
|---|---|---|
| LEQVHQL | ASSLGTGGWYEQY | 0.5767386 |
| LNAAGAI | ASSLGTGGWYEQY | 0.1533698 |
| PAVHASL | ASSLGTGGWYEQY | 0.56087184 |
| TELEDDA | ASSLGTGGWYEQY | 0.15917419 |
| TPPGPPN | ASSLGTGGWYEQY | 0.23708966 |
| TPPPPPPP | ASSLGTGGWYEQY | 0.15005858 |
| TQSPALI | ASSRDWGGYEQY | 0.5083803 |
| EFAAAFPQ | ASSRDWGGYEQY | 0.3754061 |
| LLSGNLDVL | ASSRDWGGYEQY | 0.54792356 |
| GPANLGNYNI | ASSRDWGGYEQY | 0.1465049 |
| TLGPTLP | ASSRDWGGYEQY | 0.27592567 |
| VYAYSSCCK | ASSRDWGGYEQY | 0.5738025 |
| APAPAPAPNI | ASSRDWGGYEQY | 0.71173185 |
| PGGPVPIP | ASSRDWGGYEQY | 0.16324426 |
| FLTVALEP | ASSRDWGGYEQY | 0.3308971 |
| SQYFGANL | ASSRDWGGYEQY | 0.21269527 |
| GGVKPAA | ASSRDWGGYEQY | 0.17072697 |
| GGTGGAGGAGGW | ASSRDWGGYEQY | 0.15805332 |
| GVGGVAGLGGAGG | ASSRDWGGYEQY | 0.20417805 |
| VLLVLATTI | ASSRDWGGYEQY | 0.20883335 |
| IGIFAAML | ASSRDWGGYEQY | 0.17789207 |
| AVGDLID | ASSRDWGGYEQY | 0.29733437 |
| NISPLPL | ASSRDWGGYEQY | 0.68414855 |
| VTAVFVAI | ASSRDWGGYEQY | 0.30913273 |
| SGDDNDAAL | ASSRDWGGYEQY | 0.45276332 |
| AVAALAHS | ASSRDWGGYEQY | 0.5287652 |
| GGVTVMATP | ASSRDWGGYEQY | 0.26850438 |
| AGLAANVRD | ASSRDWGGYEQY | 0.16545427 |
| TNRPDLIDPALLRPG | ASSRDWGGYEQY | 0.18058509 |
| LANLGVTAL | ASSRDWGGYEQY | 0.1639248 |
| TGERIEL | ASSRDWGGYEQY | 0.1709463 |
| NGEELAA | ASSRDWGGYEQY | 0.16355996 |
| VLSLVWG | ASSRDWGGYEQY | 0.23681355 |
| IGYALWL | ASSRDWGGYEQY | 0.1426599 |
| LSLLLAPL | ASSRDWGGYEQY | 0.20522171 |
| TLDEALK | ASSRDWGGYEQY | 0.6838939 |
| TDGAGTAEGG | ASSRDWGGYEQY | 0.14805613 |
| NAPGGAGGNGG | ASSRDWGGYEQY | 0.20879135 |
| VSVFMGMGGNF | ASSRDWGGYEQY | 0.27974647 |
| LASYLAAL | ASSRDWGGYEQY | 0.157172 |
| LVGGTPAL | ASSRDWGGYEQY | 0.3891995 |
| VRNADVL | ASSRDWGGYEQY | 0.15862545 |
| TCSSPFLVSAM | ASSRDWGGYEQY | 0.25494456 |
| VHITDAE | ASSRDWGGYEQY | 0.24122861 |
| EYAAPSF | ASSRDWGGYEQY | 0.21675558 |
| VVDAFHGGL | ASSRDWGGYEQY | 0.3918123 |
| ASVVDPQ | ASSRDWGGYEQY | 0.18920134 |
| SYNEFGGLS | ASSRDWGGYEQY | 0.15121962 |
| DPDGSVY | ASSRDWGGYEQY | 0.48176387 |
| MVSTIHATE | ASSRDWGGYEQY | 0.14568287 |
| LAAGVADPV | ASSRDWGGYEQY | 0.18988244 |

| | | |
|---|---|---|
| TKEQIAA | ASSRDWGGYEQY | 0.15612173 |
| SLASPAAL | ASSRDWGGYEQY | 0.34223217 |
| IGLVFYGV | ASSRDWGGYEQY | 0.3724749 |
| FLLDASNL | ASSRDWGGYEQY | 0.24462573 |
| YQVGNFP | ASSRDWGGYEQY | 0.17274456 |
| ESLLSQST | ASSRDWGGYEQY | 0.15646227 |
| LFTVGFAAVG | ASSRDWGGYEQY | 0.1523134 |
| ATPPTPT | ASSRDWGGYEQY | 0.15374838 |
| TDLATLG | ASSRDWGGYEQY | 0.48794228 |
| SAGGTGR | ASSRDWGGYEQY | 0.17710608 |
| GFSNSGS | ASSRDWGGYEQY | 0.17136553 |
| GGNGGSGGTSV | ASSRDWGGYEQY | 0.27893966 |
| GTSVATGGAGN | ASSRDWGGYEQY | 0.19070922 |
| NGGNGGSGGTS | ASSRDWGGYEQY | 0.15485497 |
| AVVKFPH | ASSRDWGGYEQY | 0.18001775 |
| TPPPPPP | ASSRDWGGYEQY | 0.21248397 |
| AAPAAAAAPA | ASSRDWGGYEQY | 0.16918157 |
| IALNEDL | ASSRDWGGYEQY | 0.14394562 |
| IVNVSSM | ASSRDWGGYEQY | 0.21909809 |
| TGIPPAPR | ASSRDWGGYEQY | 0.21024679 |
| FDLGGGTF | ASSRDWGGYEQY | 0.15650111 |
| FDLGGGTFDVS | ASSRDWGGYEQY | 0.17105792 |
| NEPTAAA | ASSRDWGGYEQY | 0.5375932 |
| NEPTAAALA | ASSRDWGGYEQY | 0.14703046 |
| GSIINISSI | ASSRDWGGYEQY | 0.17482628 |
| VAPGNAGTA | ASSRDWGGYEQY | 0.5851656 |
| TGGGASLE | ASSRDWGGYEQY | 0.4956298 |
| VAVYNHYKRI | ASSRDWGGYEQY | 0.45833635 |
| TDPAPATT | ASSRDWGGYEQY | 0.22642049 |
| TNRPDILDPALLRPG | ASSRDWGGYEQY | 0.16463514 |
| LTTTTTS | ASSRDWGGYEQY | 0.32758278 |
| PTGSGKTL | ASSRDWGGYEQY | 0.2972342 |
| TGSGKTL | ASSRDWGGYEQY | 0.14750275 |
| TIGHVDHGK | ASSRDWGGYEQY | 0.1439245 |
| TIGHVDHGKTT | ASSRDWGGYEQY | 0.17404929 |
| GHVDHGKTT | ASSRDWGGYEQY | 0.14882109 |
| AQFEHTLL | ASSRDWGGYEQY | 0.20054364 |
| IVNQDEL | ASSRDWGGYEQY | 0.4039993 |
| TNGSQFF | ASSRDWGGYEQY | 0.15230285 |
| TGKGIVE | ASSRDWGGYEQY | 0.1574247 |
| LISNASDA | ASSRDWGGYEQY | 0.17437069 |
| AELRTLGELTKIAKAHGAQ | ASSRDWGGYEQY | 0.15867716 |
| APNAVAAAEPSW | ASSRDWGGYEQY | 0.28520527 |
| APQVAGAATAVL | ASSRDWGGYEQY | 0.15964858 |
| ARRDPAAAAAILPTDA | ASSRDWGGYEQY | 0.727026 |
| DAAARDVLAEAGLAEYF | ASSRDWGGYEQY | 0.52076983 |
| DDQIVLSAKANLAKVEQ | ASSRDWGGYEQY | 0.23327202 |
| DELLPDTGEDSDEDGH | ASSRDWGGYEQY | 0.29164433 |
| ELPEAPANRATVPQD | ASSRDWGGYEQY | 0.16837686 |
| FGQSKIERVTGIG | ASSRDWGGYEQY | 0.2960311 |
| GAGGTGSTAGGAGGAGGAGGLY | ASSRDWGGYEQY | 0.19347945 |

| | | |
|---|---|---|
| GHGGTGAAGGNGGTGGAGGSGI | ASSRDWGGYEQY | 0.19689715 |
| GNGGNGGAGGEGGPGGAGGAGA | ASSRDWGGYEQY | 0.18938443 |
| GSTAGNGGNGGNATNGTIAGQP | ASSRDWGGYEQY | 0.18981057 |
| GTGGDGGNGGNGGTGG | ASSRDWGGYEQY | 0.1511708 |
| GVRRPAGIPADAPV | ASSRDWGGYEQY | 0.28948513 |
| GYRQVTDAHLLTLARR | ASSRDWGGYEQY | 0.15438129 |
| IAKMGQDGHDRG | ASSRDWGGYEQY | 0.14578667 |
| KAEPSIEIQYT | ASSRDWGGYEQY | 0.22916636 |
| KPVRIRMAADEDTVAVT | ASSRDWGGYEQY | 0.5720454 |
| MPPRIAGMRLLVIKPEP | ASSRDWGGYEQY | 0.6139953 |
| MKFYAVALFRSRGVPPTR | ASSRDWGGYEQY | 0.18024118 |
| NAGGFGILGAVAHSP | ASSRDWGGYEQY | 0.15816715 |
| NEGREITLAMAEM | ASSRDWGGYEQY | 0.3549407 |
| NELVEWSHPLGHRWRWE | ASSRDWGGYEQY | 0.15787195 |
| NLGAGGTGGNGGNPG | ASSRDWGGYEQY | 0.18114768 |
| NTEWPKWETVMAADPVD | ASSRDWGGYEQY | 0.24496296 |
| PASAAPAETQP | ASSRDWGGYEQY | 0.20036305 |
| QQSSDGKNGVRAAEITGE | ASSRDWGGYEQY | 0.16789389 |
| RAPDAVTTSWWKE | ASSRDWGGYEQY | 0.20981923 |
| SGFGNVGAHTSGWFNQ | ASSRDWGGYEQY | 0.19342725 |
| SGVAGAVGQAASVGGLKV | ASSRDWGGYEQY | 0.32614452 |
| SNGGAGTGSAGGNGGT | ASSRDWGGYEQY | 0.14277546 |
| TGGAGGEGGAAG | ASSRDWGGYEQY | 0.1897513 |
| TGGDGGAAGAGGAPGHG | ASSRDWGGYEQY | 0.21014856 |
| TMPPLLAAIANAGRHGV | ASSRDWGGYEQY | 0.15440209 |
| TSSTAGNSGGA | ASSRDWGGYEQY | 0.15034306 |
| VPATVPVGILC | ASSRDWGGYEQY | 0.33827373 |
| VQRPDSGGVVLTGRIS | ASSRDWGGYEQY | 0.32747483 |
| WTPNPVPATKGVPAGLVQAT | ASSRDWGGYEQY | 0.37545455 |
| YALKRALAKFSDRFDVVII | ASSRDWGGYEQY | 0.14825493 |
| YLIGSVIV | ASSRDWGGYEQY | 0.62347907 |
| YGQLVAVID | ASSRDWGGYEQY | 0.33594364 |
| VTVGGMTLPA | ASSRDWGGYEQY | 0.19986075 |
| VLALPAV | ASSRDWGGYEQY | 0.15064842 |
| VIPPTLNYET | ASSRDWGGYEQY | 0.23313005 |
| VGMGGMHPSSG | ASSRDWGGYEQY | 0.24658242 |
| VGLAGGAATA | ASSRDWGGYEQY | 0.25899148 |
| VGGAGGAGGSGP | ASSRDWGGYEQY | 0.50451696 |
| VFAAFMV | ASSRDWGGYEQY | 0.15646914 |
| TLYDKLR | ASSRDWGGYEQY | 0.38952458 |
| TLAVFTVL | ASSRDWGGYEQY | 0.3359146 |
| TGSVQPARV | ASSRDWGGYEQY | 0.15659502 |
| TGGGAGSGGT | ASSRDWGGYEQY | 0.24567297 |
| SHLETSL | ASSRDWGGYEQY | 0.20382935 |
| SFTTGMT | ASSRDWGGYEQY | 0.14548859 |
| QVEVAKRMGVRQPTVS | ASSRDWGGYEQY | 0.44449568 |
| QGSGGPALAM | ASSRDWGGYEQY | 0.20196077 |
| MLAPSNP | ASSRDWGGYEQY | 0.17123072 |
| LVNNAAI | ASSRDWGGYEQY | 0.22764 |
| LTGEINTLA | ASSRDWGGYEQY | 0.16927257 |
| LSIFGPV | ASSRDWGGYEQY | 0.18570238 |

| | | |
|---|---|---|
| LLALLANL | ASSRDWGGYEQY | 0.18238075 |
| LISNASDALDKLR | ASSRDWGGYEQY | 0.1714155 |
| LINEAVP | ASSRDWGGYEQY | 0.2130141 |
| LGYAFSGL | ASSRDWGGYEQY | 0.17753701 |
| LDFVAVAL | ASSRDWGGYEQY | 0.3774973 |
| LAQPGAE | ASSRDWGGYEQY | 0.55456257 |
| ISNLGAVTGF | ASSRDWGGYEQY | 0.15092765 |
| HQEDPGRI | ASSRDWGGYEQY | 0.18019481 |
| GTPGEPTP | ASSRDWGGYEQY | 0.2116496 |
| GFFNSGS | ASSRDWGGYEQY | 0.14786832 |
| GAGGGGLTAGAN | ASSRDWGGYEQY | 0.28569677 |
| FVDALVNM | ASSRDWGGYEQY | 0.38892552 |
| FLTVAPEL | ASSRDWGGYEQY | 0.25225428 |
| FETLTAL | ASSRDWGGYEQY | 0.1528102 |
| DVINPIEL | ASSRDWGGYEQY | 0.7441451 |
| APVSGAPV | ASSRDWGGYEQY | 0.27258366 |
| ALVGGLAW | ASSRDWGGYEQY | 0.22528006 |
| ALAAAAATAI | ASSRDWGGYEQY | 0.24055775 |
| AFGLGLLL | ASSRDWGGYEQY | 0.19053985 |
| AFFAAASL | ASSRDWGGYEQY | 0.51577336 |
| AAAALPAFSPP | ASSRDWGGYEQY | 0.30057484 |
| VVLAPAL | ASSRDWGGYEQY | 0.1598804 |
| TIGHVDHGKTTL | ASSRDWGGYEQY | 0.20880501 |
| LISNASDALDK | ASSRDWGGYEQY | 0.27846125 |
| LISNASDAL | ASSRDWGGYEQY | 0.42902806 |
| ISNASDALDK | ASSRDWGGYEQY | 0.14805475 |
| NQAETLVYQTE | ASSRDWGGYEQY | 0.18964615 |
| TGGTGGTAGAG | ASSRDWGGYEQY | 0.15379702 |
| TGSTAGGAGGA | ASSRDWGGYEQY | 0.16096309 |
| DPLAVPVAAGPV | ASSRDWGGYEQY | 0.18035139 |
| GGTGGTPGNGGN | ASSRDWGGYEQY | 0.20148425 |
| PAGLAGVDPATL | ASSRDWGGYEQY | 0.20005691 |
| GNNLVVEAPHAGASV | ASSRDWGGYEQY | 0.33442402 |
| KVTRIMSASTLEQPA | ASSRDWGGYEQY | 0.1997963 |
| LGFALPIALPRRTVV | ASSRDWGGYEQY | 0.26726252 |
| TRLGNLTARKLVVLV | ASSRDWGGYEQY | 0.16391556 |
| WSQQRRLIYFPSAGP | ASSRDWGGYEQY | 0.22002539 |
| GPNGNPDPMAAAVDIR | ASSRDWGGYEQY | 0.15311491 |
| TQRCVSVADLRAAASR | ASSRDWGGYEQY | 0.14696124 |
| QFVLDVAEHPVEVLTETV | ASSRDWGGYEQY | 0.15796486 |
| GGAGGTGGAGGNGGTQAGDG | ASSRDWGGYEQY | 0.17543605 |
| PLLATLRAIAVDVDAKPAQV | ASSRDWGGYEQY | 0.14348097 |
| APKFRTGGYMIDFWGVGYQVAK | ASSRDWGGYEQY | 0.20655608 |
| GAGGAGGAGGAAGGPGGTGGTG | ASSRDWGGYEQY | 0.21714787 |
| IGALGLLATDVAPAGSVAFGVA | ASSRDWGGYEQY | 0.2206349 |
| AGGAGGIGGNGGAGGAGGVLMG | ASSRDWGGYEQY | 0.20058484 |
| ADELIGLL | ASSRDWGGYEQY | 0.26156342 |
| ADRALKA | ASSRDWGGYEQY | 0.16446726 |
| AGAVGQVGGAGGA | ASSRDWGGYEQY | 0.4202963 |
| AGGGGGLGGAP | ASSRDWGGYEQY | 0.15174359 |
| AGGLLYGNGGAG | ASSRDWGGYEQY | 0.15619653 |

| | | |
|---|---|---|
| AGLESIGL | ASSRDWGGYEQY | 0.17976455 |
| AGLKTGII | ASSRDWGGYEQY | 0.16626094 |
| ASLAALLL | ASSRDWGGYEQY | 0.3860766 |
| ASVINISSI | ASSRDWGGYEQY | 0.26709384 |
| ATPPGPA | ASSRDWGGYEQY | 0.22295317 |
| ATVASALV | ASSRDWGGYEQY | 0.20415409 |
| ATVASLIL | ASSRDWGGYEQY | 0.14849353 |
| AVNGPVLIRH | ASSRDWGGYEQY | 0.19951078 |
| DGAAGAVGI | ASSRDWGGYEQY | 0.69878983 |
| DPLLHGI | ASSRDWGGYEQY | 0.16928683 |
| ELGAALGAL | ASSRDWGGYEQY | 0.21776824 |
| ELGRTQAV | ASSRDWGGYEQY | 0.31559697 |
| FVAGGYLS | ASSRDWGGYEQY | 0.20339388 |
| GASGGAGGAGGW | ASSRDWGGYEQY | 0.16942789 |
| GGAGAGGDGGAGGI | ASSRDWGGYEQY | 0.18348673 |
| GTGHNGSQPGMGGNG | ASSRDWGGYEQY | 0.23787118 |
| GYAAASASAS | ASSRDWGGYEQY | 0.18452188 |
| HQVDVAAVQQG | ASSRDWGGYEQY | 0.17113037 |
| IGAAIAHT | ASSRDWGGYEQY | 0.249926 |
| IGMIEVAQL | ASSRDWGGYEQY | 0.74349505 |
| IRLAPEHTSAMD | ASSRDWGGYEQY | 0.23452194 |
| IYNVSVL | ASSRDWGGYEQY | 0.23637559 |
| KQGFLGM | ASSRDWGGYEQY | 0.25980216 |
| LESVITQL | ASSRDWGGYEQY | 0.16893499 |
| LGLGALVVV | ASSRDWGGYEQY | 0.17352465 |
| LIGNGGNGGSGGI | ASSRDWGGYEQY | 0.18357486 |
| LNAINAPTL | ASSRDWGGYEQY | 0.15993634 |
| LQLLNTGV | ASSRDWGGYEQY | 0.16459802 |
| LQTSIEF | ASSRDWGGYEQY | 0.18341894 |
| MSFAVSEL | ASSRDWGGYEQY | 0.18070637 |
| NTGALGL | ASSRDWGGYEQY | 0.69370645 |
| PPGAGGAGGAGTT | ASSRDWGGYEQY | 0.265714 |
| QLVGLGI | ASSRDWGGYEQY | 0.21897882 |
| RFLEQVHQL | ASSRDWGGYEQY | 0.16034283 |
| RGVGYVLR | ASSRDWGGYEQY | 0.18229416 |
| SATGVFAGVI | ASSRDWGGYEQY | 0.2016405 |
| SLPAITLP | ASSRDWGGYEQY | 0.15518065 |
| STALAFTV | ASSRDWGGYEQY | 0.4039958 |
| SVLTVGLGK | ASSRDWGGYEQY | 0.18421897 |
| TALAGAGSGI | ASSRDWGGYEQY | 0.15815583 |
| TGLPFTGI | ASSRDWGGYEQY | 0.23418029 |
| VAEHVTV | ASSRDWGGYEQY | 0.7223048 |
| VDIVLNSL | ASSRDWGGYEQY | 0.17429931 |
| VLTGGGAVL | ASSRDWGGYEQY | 0.16299944 |
| VNVGLAYGI | ASSRDWGGYEQY | 0.16257916 |
| VSSAAVGAV | ASSRDWGGYEQY | 0.19579971 |
| VTSVYEAGV | ASSRDWGGYEQY | 0.48705485 |
| WVALDRGA | ASSRDWGGYEQY | 0.1631824 |
| YLVGSVVI | ASSRDWGGYEQY | 0.5152616 |
| GPPGTGKTLL | ASSRDWGGYEQY | 0.20369667 |
| GVGMTKF | ASSRDWGGYEQY | 0.1574966 |

| | | |
|---|---|---|
| IGIGGSDL | ASSRDWGGYEQY | 0.18761672 |
| IGTLLAEL | ASSRDWGGYEQY | 0.24787538 |
| IYTSGTT | ASSRDWGGYEQY | 0.48337233 |
| LSFPAGV | ASSRDWGGYEQY | 0.24966937 |
| LYTSGTT | ASSRDWGGYEQY | 0.46121243 |
| VGAFPFGF | ASSRDWGGYEQY | 0.15764728 |
| VGAGTVEF | ASSRDWGGYEQY | 0.29526263 |
| VGKSTLLL | ASSRDWGGYEQY | 0.26033753 |
| AALAHAGGGIV | ASSRDWGGYEQY | 0.30712414 |
| AALVAVSLL | ASSRDWGGYEQY | 0.16514471 |
| AAVSRAVAAL | ASSRDWGGYEQY | 0.15643911 |
| AGGSSGIGGF | ASSRDWGGYEQY | 0.1849029 |
| AGLTVANANG | ASSRDWGGYEQY | 0.23711094 |
| AGLVGAVL | ASSRDWGGYEQY | 0.15613437 |
| ALSQAFTV | ASSRDWGGYEQY | 0.20331034 |
| ALTFSVSL | ASSRDWGGYEQY | 0.28728062 |
| ALTGAALVAL | ASSRDWGGYEQY | 0.16113131 |
| ASFSIGAGVL | ASSRDWGGYEQY | 0.20742795 |
| DGSVYKI | ASSRDWGGYEQY | 0.73376 |
| EQLAEIV | ASSRDWGGYEQY | 0.18587355 |
| FAVSELTVM | ASSRDWGGYEQY | 0.15858905 |
| GAAPLAAGL | ASSRDWGGYEQY | 0.4336024 |
| GADGGDGGSGGAGG | ASSRDWGGYEQY | 0.7071109 |
| GFGLDTGGAGG | ASSRDWGGYEQY | 0.26087147 |
| GLASGGLL | ASSRDWGGYEQY | 0.1605019 |
| GRIAGAAVLL | ASSRDWGGYEQY | 0.14637963 |
| GSAGSLNTSF | ASSRDWGGYEQY | 0.2560245 |
| GTGQIGI | ASSRDWGGYEQY | 0.29206097 |
| GTLIALIL | ASSRDWGGYEQY | 0.18921076 |
| IAATMGFVVG | ASSRDWGGYEQY | 0.15856093 |
| IFAQALL | ASSRDWGGYEQY | 0.2586128 |
| IGAAVALGF | ASSRDWGGYEQY | 0.27732277 |
| IGLTPLAL | ASSRDWGGYEQY | 0.16867366 |
| ILGSNTSTL | ASSRDWGGYEQY | 0.14824337 |
| IVFSLTVL | ASSRDWGGYEQY | 0.36993158 |
| LASKATGF | ASSRDWGGYEQY | 0.2646805 |
| LASVALIL | ASSRDWGGYEQY | 0.16913874 |
| LFAVGLTL | ASSRDWGGYEQY | 0.17902103 |
| LFNGGIGTYI | ASSRDWGGYEQY | 0.2411693 |
| LTLGAGII | ASSRDWGGYEQY | 0.15328643 |
| LVAAALGDI | ASSRDWGGYEQY | 0.22827902 |
| NTAFVQLGI | ASSRDWGGYEQY | 0.16104184 |
| QLFSGAEL | ASSRDWGGYEQY | 0.43159252 |
| SAATFITL | ASSRDWGGYEQY | 0.16727631 |
| SNYGLANLGV | ASSRDWGGYEQY | 0.27943474 |
| SYGGAYAVM | ASSRDWGGYEQY | 0.5440008 |
| TTGVGGLTLG | ASSRDWGGYEQY | 0.26703689 |
| TTQTIYI | ASSRDWGGYEQY | 0.18629047 |
| VAAGAGLGAVI | ASSRDWGGYEQY | 0.16733742 |
| VAAVVTLL | ASSRDWGGYEQY | 0.1524613 |
| VGFLGGLI | ASSRDWGGYEQY | 0.14347112 |

| | | |
|---|---|---|
| VGVIPILL | ASSRDWGGYEQY | 0.6714916 |
| VIFASSLI | ASSRDWGGYEQY | 0.14502475 |
| VKGGDGGAAAG | ASSRDWGGYEQY | 0.62629664 |
| VTKTFTNI | ASSRDWGGYEQY | 0.20497918 |
| VVAAAILL | ASSRDWGGYEQY | 0.17258514 |
| YAALDVEL | ASSRDWGGYEQY | 0.218316 |
| LASVALLL | ASSRDWGGYEQY | 0.1500661 |
| LDAVHRV | ASSRDWGGYEQY | 0.14983845 |
| QGLSSSIF | ASSRDWGGYEQY | 0.19425337 |
| RGTFANIRL | ASSRDWGGYEQY | 0.24635066 |
| RQLWWGHRI | ASSRDWGGYEQY | 0.1682284 |
| YGPPGTGKTLL | ASSRDWGGYEQY | 0.19270985 |
| AALVGAPANVA | ASSRDWGGYEQY | 0.2222439 |
| AATFRRTE | ASSRDWGGYEQY | 0.15581487 |
| AGALGGGI | ASSRDWGGYEQY | 0.1860664 |
| AGGAGGAGGIGAG | ASSRDWGGYEQY | 0.43618488 |
| AGSGALGL | ASSRDWGGYEQY | 0.20382519 |
| AQIADQI | ASSRDWGGYEQY | 0.15503052 |
| ASAVAVTYL | ASSRDWGGYEQY | 0.25842685 |
| ASTVLGGTGG | ASSRDWGGYEQY | 0.17056607 |
| AVGGEPAK | ASSRDWGGYEQY | 0.27094457 |
| CEQGTFGGTC | ASSRDWGGYEQY | 0.24885684 |
| DEGKAPAG | ASSRDWGGYEQY | 0.18897684 |
| DTVVFPAL | ASSRDWGGYEQY | 0.22057524 |
| FEGITMAG | ASSRDWGGYEQY | 0.17044117 |
| FPLVVQD | ASSRDWGGYEQY | 0.15424597 |
| GAFTTPQ | ASSRDWGGYEQY | 0.18269973 |
| GFGGFGTAGGVGGA | ASSRDWGGYEQY | 0.2199913 |
| GFLAQFL | ASSRDWGGYEQY | 0.28624302 |
| GGAGGAGGNAGVA | ASSRDWGGYEQY | 0.16226038 |
| GGVGTAGAGGAGGAGG | ASSRDWGGYEQY | 0.19895478 |
| GNGGGGDAGNAGSGGN | ASSRDWGGYEQY | 0.16866204 |
| GTGGAGGAGNDAGS | ASSRDWGGYEQY | 0.1637124 |
| GTTNTGWFNS | ASSRDWGGYEQY | 0.18331213 |
| IFMGPLFTL | ASSRDWGGYEQY | 0.17813972 |
| ILAHAIEAI | ASSRDWGGYEQY | 0.16672707 |
| IRSAIREAL | ASSRDWGGYEQY | 0.40626806 |
| ITFAGGG | ASSRDWGGYEQY | 0.16454613 |
| ITGMSGHQEV | ASSRDWGGYEQY | 0.19769122 |
| IVAGLACTL | ASSRDWGGYEQY | 0.26967573 |
| IVGGGLAGA | ASSRDWGGYEQY | 0.23178443 |
| IVTGVLNI | ASSRDWGGYEQY | 0.22122315 |
| LAQQGCI | ASSRDWGGYEQY | 0.21660563 |
| LCLHVEN | ASSRDWGGYEQY | 0.49443927 |
| LDAISDGS | ASSRDWGGYEQY | 0.17902248 |
| LGGFAAML | ASSRDWGGYEQY | 0.155894 |
| LGGKNPMI | ASSRDWGGYEQY | 0.2752514 |
| LGPKGVAAIL | ASSRDWGGYEQY | 0.2527157 |
| LMVGVSI | ASSRDWGGYEQY | 0.524168 |
| LVGAAFVL | ASSRDWGGYEQY | 0.317303 |
| LVLAVAPL | ASSRDWGGYEQY | 0.53216785 |

| | | |
|---|---|---|
| MAAIAVIDAA | ASSRDWGGYEQY | 0.31756866 |
| NNGINAELQ | ASSRDWGGYEQY | 0.41901886 |
| RGEPLVLVHG | ASSRDWGGYEQY | 0.15068437 |
| RGGAGGAAGLI | ASSRDWGGYEQY | 0.16984388 |
| RSYRMHPAV | ASSRDWGGYEQY | 0.16502146 |
| SAAHLLAGQLA | ASSRDWGGYEQY | 0.21762341 |
| SAAVGSY | ASSRDWGGYEQY | 0.14749072 |
| SPHAVLHRI | ASSRDWGGYEQY | 0.4871025 |
| SQIAKVL | ASSRDWGGYEQY | 0.679036 |
| TALGHLVP | ASSRDWGGYEQY | 0.16668612 |
| TAPTTSGAA | ASSRDWGGYEQY | 0.3035761 |
| TDGDDGNGGSPGT | ASSRDWGGYEQY | 0.1513024 |
| TDGLGEN | ASSRDWGGYEQY | 0.19315013 |
| TGGPPLPGQA | ASSRDWGGYEQY | 0.4154446 |
| TIPAITI | ASSRDWGGYEQY | 0.16623007 |
| TIPGFTI | ASSRDWGGYEQY | 0.30938354 |
| TSIASGALA | ASSRDWGGYEQY | 0.15490025 |
| TVTLPGAAI | ASSRDWGGYEQY | 0.1672059 |
| VASGLSAALP | ASSRDWGGYEQY | 0.559538 |
| VGAGVIGI | ASSRDWGGYEQY | 0.19104473 |
| VGPKRFGV | ASSRDWGGYEQY | 0.25313112 |
| VNVAFPD | ASSRDWGGYEQY | 0.17921863 |
| VSMTNSGVS | ASSRDWGGYEQY | 0.16233516 |
| VTDVLGDLSV | ASSRDWGGYEQY | 0.21899214 |
| VVVVGAGFAGL | ASSRDWGGYEQY | 0.7092587 |
| VYTVGTVV | ASSRDWGGYEQY | 0.14876036 |
| WTTFSGP | ASSRDWGGYEQY | 0.2722531 |
| YDELVEAF | ASSRDWGGYEQY | 0.23289457 |
| YGTGGAGGNGG | ASSRDWGGYEQY | 0.6105003 |
| YTNVMAV | ASSRDWGGYEQY | 0.34435576 |
| FLEQVHQL | ASSRDWGGYEQY | 0.26957744 |
| GLSLLAVAV | ASSRDWGGYEQY | 0.17485811 |
| LEQVHQL | ASSRDWGGYEQY | 0.4875388 |
| LNAAGAI | ASSRDWGGYEQY | 0.22962302 |
| PAVHASL | ASSRDWGGYEQY | 0.5611545 |
| TELEDDA | ASSRDWGGYEQY | 0.33420298 |
| TPPGPPN | ASSRDWGGYEQY | 0.24261963 |
| TPPPPPPP | ASSRDWGGYEQY | 0.7634673 |
| TQSPALI | AWSLRDKYAEQF | 0.5692973 |
| EFAAAFPQ | AWSLRDKYAEQF | 0.36242914 |
| LLSGNLDVL | AWSLRDKYAEQF | 0.24373007 |
| GPANLGNYNI | AWSLRDKYAEQF | 0.1928515 |
| TLGPTLP | AWSLRDKYAEQF | 0.31534708 |
| VYAYSSCCK | AWSLRDKYAEQF | 0.22723716 |
| APAPAPAPNI | AWSLRDKYAEQF | 0.7809797 |
| PGGPVPIP | AWSLRDKYAEQF | 0.54903495 |
| FLTVALEP | AWSLRDKYAEQF | 0.28375313 |
| SQYFGANL | AWSLRDKYAEQF | 0.7994263 |
| GGVKPAA | AWSLRDKYAEQF | 0.60876495 |
| GGTGGAGGAGGW | AWSLRDKYAEQF | 0.19057584 |
| GVGGVAGLGGAGG | AWSLRDKYAEQF | 0.31609523 |

| | | |
|---|---|---|
| VLLVLATTI | AWSLRDKYAEQF | 0.2644362 |
| IGIFAAML | AWSLRDKYAEQF | 0.31150842 |
| AVGDLID | AWSLRDKYAEQF | 0.15246983 |
| NISPLPL | AWSLRDKYAEQF | 0.55477554 |
| VTAVFVAI | AWSLRDKYAEQF | 0.31552398 |
| SGDDNDAAL | AWSLRDKYAEQF | 0.13405992 |
| AVAALAHS | AWSLRDKYAEQF | 0.41379791 |
| GGVTVMATP | AWSLRDKYAEQF | 0.41848296 |
| AGLAANVRD | AWSLRDKYAEQF | 0.3760838 |
| TNRPDLIDPALLRPG | AWSLRDKYAEQF | 0.7474497 |
| LANLGVTAL | AWSLRDKYAEQF | 0.23055224 |
| TGERIEL | AWSLRDKYAEQF | 0.30947894 |
| NGEELAA | AWSLRDKYAEQF | 0.73168015 |
| VLSLVWG | AWSLRDKYAEQF | 0.7871225 |
| IGYALWL | AWSLRDKYAEQF | 0.23673 |
| LSLLLAPL | AWSLRDKYAEQF | 0.5923331 |
| TLDEALK | AWSLRDKYAEQF | 0.6914081 |
| TDGAGTAEGG | AWSLRDKYAEQF | 0.59661674 |
| NAPGGAGGNGG | AWSLRDKYAEQF | 0.604425 |
| VSVFMGMGGNF | AWSLRDKYAEQF | 0.67321193 |
| LASYLAAL | AWSLRDKYAEQF | 0.22911003 |
| LVGGTPAL | AWSLRDKYAEQF | 0.76067525 |
| VRNADVL | AWSLRDKYAEQF | 0.82238173 |
| TCSSPFLVSAM | AWSLRDKYAEQF | 0.3414464 |
| VHITDAE | AWSLRDKYAEQF | 0.4473235 |
| EYAAPSF | AWSLRDKYAEQF | 0.23974542 |
| VVDAFHGGL | AWSLRDKYAEQF | 0.3503461 |
| ASVVDPQ | AWSLRDKYAEQF | 0.16898449 |
| SYNEFGGLS | AWSLRDKYAEQF | 0.675366 |
| DPDGSVY | AWSLRDKYAEQF | 0.6749079 |
| MVSTIHATE | AWSLRDKYAEQF | 0.3886427 |
| LAAGVADPV | AWSLRDKYAEQF | 0.21905243 |
| TKEQIAA | AWSLRDKYAEQF | 0.22513042 |
| SLASPAAL | AWSLRDKYAEQF | 0.2959369 |
| IGLVFYGV | AWSLRDKYAEQF | 0.2745557 |
| FLLDASNL | AWSLRDKYAEQF | 0.17352585 |
| YQVGNFP | AWSLRDKYAEQF | 0.27126262 |
| ESLLSQST | AWSLRDKYAEQF | 0.6874629 |
| LFTVGFAAVG | AWSLRDKYAEQF | 0.6870763 |
| ATPPTPT | AWSLRDKYAEQF | 0.17878674 |
| TDLATLG | AWSLRDKYAEQF | 0.7902398 |
| SAGGTGR | AWSLRDKYAEQF | 0.7314584 |
| GFSNSGS | AWSLRDKYAEQF | 0.7121264 |
| GGNGGSGGTSV | AWSLRDKYAEQF | 0.34882164 |
| GTSVATGGAGN | AWSLRDKYAEQF | 0.23158568 |
| NGGNGGSGGTS | AWSLRDKYAEQF | 0.7969096 |
| AVVKFPH | AWSLRDKYAEQF | 0.49496108 |
| TPPPPPP | AWSLRDKYAEQF | 0.32858938 |
| AAPAAAAAPA | AWSLRDKYAEQF | 0.56284505 |
| IALNEDL | AWSLRDKYAEQF | 0.5022655 |
| IVNVSSM | AWSLRDKYAEQF | 0.32386443 |

| | | |
|---|---|---|
| TGIPPAPR | AWSLRDKYAEQF | 0.58711416 |
| FDLGGGTF | AWSLRDKYAEQF | 0.42640716 |
| FDLGGGTFDVS | AWSLRDKYAEQF | 0.24755847 |
| NEPTAAA | AWSLRDKYAEQF | 0.22117376 |
| NEPTAAALA | AWSLRDKYAEQF | 0.2882984 |
| GSIINISSI | AWSLRDKYAEQF | 0.5441921 |
| VAPGNAGTA | AWSLRDKYAEQF | 0.7357874 |
| TGGGASLE | AWSLRDKYAEQF | 0.436448 |
| VAVYNHYKRI | AWSLRDKYAEQF | 0.30092096 |
| TDPAPATT | AWSLRDKYAEQF | 0.6774276 |
| TNRPDILDPALLRPG | AWSLRDKYAEQF | 0.43301857 |
| LTTTTTS | AWSLRDKYAEQF | 0.3825084 |
| PTGSGKTL | AWSLRDKYAEQF | 0.45093775 |
| TGSGKTL | AWSLRDKYAEQF | 0.360107 |
| TIGHVDHGK | AWSLRDKYAEQF | 0.4572552 |
| TIGHVDHGKTT | AWSLRDKYAEQF | 0.43937778 |
| GHVDHGKTT | AWSLRDKYAEQF | 0.45307893 |
| AQFEHTLL | AWSLRDKYAEQF | 0.6336036 |
| IVNQDEL | AWSLRDKYAEQF | 0.58974355 |
| TNGSQFF | AWSLRDKYAEQF | 0.37003347 |
| TGKGIVE | AWSLRDKYAEQF | 0.14967813 |
| LISNASDA | AWSLRDKYAEQF | 0.3431965 |
| AELRTLGELTKIAKAHGAQ | AWSLRDKYAEQF | 0.60022485 |
| APNAVAAAEPSW | AWSLRDKYAEQF | 0.52511495 |
| APQVAGAATAVL | AWSLRDKYAEQF | 0.6704832 |
| ARRDPAAAAAILPTDA | AWSLRDKYAEQF | 0.44538885 |
| DAAARDVLAEAGLAEYF | AWSLRDKYAEQF | 0.3415643 |
| DDQIVLSAKANLAKVEQ | AWSLRDKYAEQF | 0.5610671 |
| DELLPDTGEDSDEDGH | AWSLRDKYAEQF | 0.703143 |
| ELPEAPANRATVPQD | AWSLRDKYAEQF | 0.1705915 |
| FGQSKIERVTGIG | AWSLRDKYAEQF | 0.34164265 |
| GAGGTGSTAGGAGGAGGAGGLY | AWSLRDKYAEQF | 0.7974403 |
| GHGGTGAAGGNGGTGGAGGSGI | AWSLRDKYAEQF | 0.7209564 |
| GNGGNGGAGGEGGPGGAGGAGA | AWSLRDKYAEQF | 0.39174896 |
| GSTAGNGGNGGNATNGTIAGQP | AWSLRDKYAEQF | 0.2526611 |
| GTGGDGGNGGNGGTGG | AWSLRDKYAEQF | 0.83284616 |
| GVRRPAGIPADAPV | AWSLRDKYAEQF | 0.2943151 |
| GYRQVTDAHLLTLARR | AWSLRDKYAEQF | 0.778284 |
| IAKMGQDGHDRG | AWSLRDKYAEQF | 0.13905095 |
| KAEPSIEIQYT | AWSLRDKYAEQF | 0.22471334 |
| KPVRIRMAADEDTVAVT | AWSLRDKYAEQF | 0.20344244 |
| MPPRIAGMRLLVIKPEP | AWSLRDKYAEQF | 0.15881024 |
| MKFYAVALFRSRGVPPTR | AWSLRDKYAEQF | 0.6369415 |
| NAGGFGILGAVAHSP | AWSLRDKYAEQF | 0.27408767 |
| NEGREITLAMAEM | AWSLRDKYAEQF | 0.6449808 |
| NELVEWSHPLGHRWRWE | AWSLRDKYAEQF | 0.48416248 |
| NLGAGGTGGNGGNPG | AWSLRDKYAEQF | 0.59648323 |
| NTEWPKWETVMAADPVD | AWSLRDKYAEQF | 0.54059505 |
| PASAAPAETQP | AWSLRDKYAEQF | 0.4272834 |
| QQSSDGKNGVRAAEITGE | AWSLRDKYAEQF | 0.23219249 |
| RAPDAVTTSWWKE | AWSLRDKYAEQF | 0.6018004 |

| | | |
|---|---|---|
| SGFGNVGAHTSGWFNQ | AWSLRDKYAEQF | 0.6417892 |
| SGVAGAVGQAASVGGLKV | AWSLRDKYAEQF | 0.36582348 |
| SNGGAGTGSAGGNGGT | AWSLRDKYAEQF | 0.20399256 |
| TGGAGGEGGAAG | AWSLRDKYAEQF | 0.39028722 |
| TGGDGGAAGAGGAPGHG | AWSLRDKYAEQF | 0.38316876 |
| TMPPLLAAIANAGRHGV | AWSLRDKYAEQF | 0.16164838 |
| TSSTAGNSGGA | AWSLRDKYAEQF | 0.1438889 |
| VPATVPVGILC | AWSLRDKYAEQF | 0.24736318 |
| VQRPDSGGVVLTGRIS | AWSLRDKYAEQF | 0.30962056 |
| WTPNPVPATKGVPAGLVQAT | AWSLRDKYAEQF | 0.7333087 |
| YALKRALAKFSDRFDVVII | AWSLRDKYAEQF | 0.7401874 |
| YLIGSVIV | AWSLRDKYAEQF | 0.39678586 |
| YGQLVAVID | AWSLRDKYAEQF | 0.6714966 |
| VTVGGMTLPA | AWSLRDKYAEQF | 0.54280156 |
| VLALPAV | AWSLRDKYAEQF | 0.12596151 |
| VIPPTLNYET | AWSLRDKYAEQF | 0.2488494 |
| VGMGGMHPSSG | AWSLRDKYAEQF | 0.2015519 |
| VGLAGGAATA | AWSLRDKYAEQF | 0.5794796 |
| VGGAGGAGGSGP | AWSLRDKYAEQF | 0.24855283 |
| VFAAFMV | AWSLRDKYAEQF | 0.31573313 |
| TLYDKLR | AWSLRDKYAEQF | 0.4783344 |
| TLAVFTVL | AWSLRDKYAEQF | 0.757287 |
| TGSVQPARV | AWSLRDKYAEQF | 0.4824718 |
| TGGGAGSGGT | AWSLRDKYAEQF | 0.20103732 |
| SHLETSL | AWSLRDKYAEQF | 0.14134231 |
| SFTTGMT | AWSLRDKYAEQF | 0.5930509 |
| QVEVAKRMGVRQPTVS | AWSLRDKYAEQF | 0.29334286 |
| QGSGGPALAM | AWSLRDKYAEQF | 0.27395263 |
| MLAPSNP | AWSLRDKYAEQF | 0.14270991 |
| LVNNAAI | AWSLRDKYAEQF | 0.54427564 |
| LTGEINTLA | AWSLRDKYAEQF | 0.17626798 |
| LSIFGPV | AWSLRDKYAEQF | 0.5484195 |
| LLALLANL | AWSLRDKYAEQF | 0.40573156 |
| LISNASDALDKLR | AWSLRDKYAEQF | 0.46753657 |
| LINEAVP | AWSLRDKYAEQF | 0.8212611 |
| LGYAFSGL | AWSLRDKYAEQF | 0.7535884 |
| LDFVAVAL | AWSLRDKYAEQF | 0.3620019 |
| LAQPGAE | AWSLRDKYAEQF | 0.28093034 |
| ISNLGAVTGF | AWSLRDKYAEQF | 0.21513991 |
| HQEDPGRI | AWSLRDKYAEQF | 0.27862358 |
| GTPGEPTP | AWSLRDKYAEQF | 0.2858395 |
| GFFNSGS | AWSLRDKYAEQF | 0.76175296 |
| GAGGGGLTAGAN | AWSLRDKYAEQF | 0.22538227 |
| FVDALVNM | AWSLRDKYAEQF | 0.1927308 |
| FLTVAPEL | AWSLRDKYAEQF | 0.2575397 |
| FETLTAL | AWSLRDKYAEQF | 0.72055537 |
| DVINPIEL | AWSLRDKYAEQF | 0.81354994 |
| APVSGAPV | AWSLRDKYAEQF | 0.29369426 |
| ALVGGLAW | AWSLRDKYAEQF | 0.31497404 |
| ALAAAAATAI | AWSLRDKYAEQF | 0.27992582 |
| AFGLGLLL | AWSLRDKYAEQF | 0.52246577 |

| | | |
|---|---|---|
| AFFAAASL | AWSLRDKYAEQF | 0.25812173 |
| AAAALPAFSPP | AWSLRDKYAEQF | 0.2157805 |
| VVLAPAL | AWSLRDKYAEQF | 0.4020338 |
| TIGHVDHGKTTL | AWSLRDKYAEQF | 0.83117706 |
| LISNASDALDK | AWSLRDKYAEQF | 0.31316116 |
| LISNASDAL | AWSLRDKYAEQF | 0.1650889 |
| ISNASDALDK | AWSLRDKYAEQF | 0.49155164 |
| NQAETLVYQTE | AWSLRDKYAEQF | 0.30458513 |
| TGGTGGTAGAG | AWSLRDKYAEQF | 0.34100494 |
| TGSTAGGAGGA | AWSLRDKYAEQF | 0.5500827 |
| DPLAVPVAAGPV | AWSLRDKYAEQF | 0.17814441 |
| GGTGGTPGNGGN | AWSLRDKYAEQF | 0.43352628 |
| PAGLAGVDPATL | AWSLRDKYAEQF | 0.82321113 |
| GNNLVVEAPHAGASV | AWSLRDKYAEQF | 0.50179106 |
| KVTRIMSASTLEQPA | AWSLRDKYAEQF | 0.57404417 |
| LGFALPIALPRRTVV | AWSLRDKYAEQF | 0.79895765 |
| TRLGNLTARKLVVLV | AWSLRDKYAEQF | 0.6193088 |
| WSQQRRLIYFPSAGP | AWSLRDKYAEQF | 0.7905791 |
| GPNGNPDPMAAAVDIR | AWSLRDKYAEQF | 0.77414334 |
| TQRCVSVADLRAAASR | AWSLRDKYAEQF | 0.31402826 |
| QFVLDVAEHPVEVLTETV | AWSLRDKYAEQF | 0.34521133 |
| GGAGGTGGAGGNGGTQAGDG | AWSLRDKYAEQF | 0.73886085 |
| PLLATLRAIAVDVDAKPAQV | AWSLRDKYAEQF | 0.5163623 |
| APKFRTGGYMIDFWGVGYQVAK | AWSLRDKYAEQF | 0.4085323 |
| GAGGAGGAGGAAGGPGGTGGTG | AWSLRDKYAEQF | 0.21884535 |
| IGALGLLATDVAPAGSVAFGVA | AWSLRDKYAEQF | 0.7645867 |
| AGGAGGIGGNGGAGGAGGVLMG | AWSLRDKYAEQF | 0.31199074 |
| ADELIGLL | AWSLRDKYAEQF | 0.6963614 |
| ADRALKA | AWSLRDKYAEQF | 0.19423954 |
| AGAVGQVGGAGGA | AWSLRDKYAEQF | 0.22390506 |
| AGGGGGLGGAP | AWSLRDKYAEQF | 0.68486285 |
| AGGLLYGNGGAG | AWSLRDKYAEQF | 0.12925008 |
| AGLESIGL | AWSLRDKYAEQF | 0.1409421 |
| AGLKTGII | AWSLRDKYAEQF | 0.35147125 |
| ASLAALLL | AWSLRDKYAEQF | 0.7660998 |
| ASVINISSI | AWSLRDKYAEQF | 0.13354972 |
| ATPPGPA | AWSLRDKYAEQF | 0.25198415 |
| ATVASALV | AWSLRDKYAEQF | 0.19350025 |
| ATVASLIL | AWSLRDKYAEQF | 0.35185137 |
| AVNGPVLIRH | AWSLRDKYAEQF | 0.53784484 |
| DGAAGAVGI | AWSLRDKYAEQF | 0.17387685 |
| DPLLHGI | AWSLRDKYAEQF | 0.48450038 |
| ELGAALGAL | AWSLRDKYAEQF | 0.15800819 |
| ELGRTQAV | AWSLRDKYAEQF | 0.30137604 |
| FVAGGYLS | AWSLRDKYAEQF | 0.5824181 |
| GASGGAGGAGGW | AWSLRDKYAEQF | 0.8023089 |
| GGAGAGGDGGAGGI | AWSLRDKYAEQF | 0.21683447 |
| GTGHNGSQPGMGGNG | AWSLRDKYAEQF | 0.6874341 |
| GYAAASASAS | AWSLRDKYAEQF | 0.22993955 |
| HQVDVAAVQQG | AWSLRDKYAEQF | 0.83727205 |
| IGAAIAHT | AWSLRDKYAEQF | 0.31462264 |

| | | |
|---|---|---|
| IGMIEVAQL | AWSLRDKYAEQF | 0.17211384 |
| IRLAPEHTSAMD | AWSLRDKYAEQF | 0.72088826 |
| IYNVSVL | AWSLRDKYAEQF | 0.4395232 |
| KQGFLGM | AWSLRDKYAEQF | 0.41806453 |
| LESVITQL | AWSLRDKYAEQF | 0.7088908 |
| LGLGALVVV | AWSLRDKYAEQF | 0.2075985 |
| LIGNGGNGGSGGI | AWSLRDKYAEQF | 0.48065832 |
| LNAINAPTL | AWSLRDKYAEQF | 0.79153395 |
| LQLLNTGV | AWSLRDKYAEQF | 0.68838274 |
| LQTSIEF | AWSLRDKYAEQF | 0.22544198 |
| MSFAVSEL | AWSLRDKYAEQF | 0.5494934 |
| NTGALGL | AWSLRDKYAEQF | 0.50336486 |
| PPGAGGAGGAGTT | AWSLRDKYAEQF | 0.24570048 |
| QLVGLGI | AWSLRDKYAEQF | 0.74317646 |
| RFLEQVHQL | AWSLRDKYAEQF | 0.34224224 |
| RGVGYVLR | AWSLRDKYAEQF | 0.6653844 |
| SATGVFAGVI | AWSLRDKYAEQF | 0.4986444 |
| SLPAITLP | AWSLRDKYAEQF | 0.7946463 |
| STALAFTV | AWSLRDKYAEQF | 0.23082934 |
| SVLTVGLGK | AWSLRDKYAEQF | 0.39293784 |
| TALAGAGSGI | AWSLRDKYAEQF | 0.51850843 |
| TGLPFTGI | AWSLRDKYAEQF | 0.5854782 |
| VAEHVTV | AWSLRDKYAEQF | 0.29405752 |
| VDIVLNSL | AWSLRDKYAEQF | 0.46786183 |
| VLTGGGAVL | AWSLRDKYAEQF | 0.4207934 |
| VNVGLAYGI | AWSLRDKYAEQF | 0.7647292 |
| VSSAAVGAV | AWSLRDKYAEQF | 0.27574727 |
| VTSVYEAGV | AWSLRDKYAEQF | 0.37586674 |
| WVALDRGA | AWSLRDKYAEQF | 0.8033953 |
| YLVGSVVI | AWSLRDKYAEQF | 0.2546685 |
| GPPGTGKTLL | AWSLRDKYAEQF | 0.37725395 |
| GVGMTKF | AWSLRDKYAEQF | 0.520185 |
| IGIGGSDL | AWSLRDKYAEQF | 0.6909749 |
| IGTLLAEL | AWSLRDKYAEQF | 0.26447415 |
| IYTSGTT | AWSLRDKYAEQF | 0.7391678 |
| LSFPAGV | AWSLRDKYAEQF | 0.40917856 |
| LYTSGTT | AWSLRDKYAEQF | 0.7828059 |
| VGAFPFGF | AWSLRDKYAEQF | 0.43651736 |
| VGAGTVEF | AWSLRDKYAEQF | 0.55353165 |
| VGKSTLLL | AWSLRDKYAEQF | 0.2612297 |
| AALAHAGGGIV | AWSLRDKYAEQF | 0.7278781 |
| AALVAVSLL | AWSLRDKYAEQF | 0.7570251 |
| AAVSRAVAAL | AWSLRDKYAEQF | 0.7553632 |
| AGGSSGIGGF | AWSLRDKYAEQF | 0.14119631 |
| AGLTVANANG | AWSLRDKYAEQF | 0.7401556 |
| AGLVGAVL | AWSLRDKYAEQF | 0.7565938 |
| ALSQAFTV | AWSLRDKYAEQF | 0.8326689 |
| ALTFSVSL | AWSLRDKYAEQF | 0.37672532 |
| ALTGAALVAL | AWSLRDKYAEQF | 0.48771387 |
| ASFSIGAGVL | AWSLRDKYAEQF | 0.47892618 |
| DGSVYKI | AWSLRDKYAEQF | 0.27584213 |

| | | |
|---|---|---|
| EQLAEIV | AWSLRDKYAEQF | 0.19678015 |
| FAVSELTVM | AWSLRDKYAEQF | 0.44764692 |
| GAAPLAAGL | AWSLRDKYAEQF | 0.5318417 |
| GADGGDGGSGGAGG | AWSLRDKYAEQF | 0.32385665 |
| GFGLDTGGAGG | AWSLRDKYAEQF | 0.77703893 |
| GLASGGLL | AWSLRDKYAEQF | 0.7458664 |
| GRIAGAAVLL | AWSLRDKYAEQF | 0.76972485 |
| GSAGSLNTSF | AWSLRDKYAEQF | 0.75216097 |
| GTGQIGI | AWSLRDKYAEQF | 0.77553463 |
| GTLIALIL | AWSLRDKYAEQF | 0.46793026 |
| IAATMGFVVG | AWSLRDKYAEQF | 0.2640942 |
| IFAQALL | AWSLRDKYAEQF | 0.4540194 |
| IGAAVALGF | AWSLRDKYAEQF | 0.19326085 |
| IGLTPLAL | AWSLRDKYAEQF | 0.39201108 |
| ILGSNTSTL | AWSLRDKYAEQF | 0.29696906 |
| IVFSLTVL | AWSLRDKYAEQF | 0.583112 |
| LASKATGF | AWSLRDKYAEQF | 0.1336857 |
| LASVALIL | AWSLRDKYAEQF | 0.71545255 |
| LFAVGLTL | AWSLRDKYAEQF | 0.7885745 |
| LFNGGIGTYI | AWSLRDKYAEQF | 0.49615085 |
| LTLGAGII | AWSLRDKYAEQF | 0.5366707 |
| LVAAALGDI | AWSLRDKYAEQF | 0.6370872 |
| NTAFVQLGI | AWSLRDKYAEQF | 0.22425598 |
| QLFSGAEL | AWSLRDKYAEQF | 0.6379576 |
| SAATFITL | AWSLRDKYAEQF | 0.24988542 |
| SNYGLANLGV | AWSLRDKYAEQF | 0.50536335 |
| SYGGAYAVM | AWSLRDKYAEQF | 0.74416363 |
| TTGVGGLTLG | AWSLRDKYAEQF | 0.5202499 |
| TTQTIYI | AWSLRDKYAEQF | 0.4173988 |
| VAAGAGLGAVI | AWSLRDKYAEQF | 0.3445094 |
| VAAVVTLL | AWSLRDKYAEQF | 0.18917316 |
| VGFLGGLI | AWSLRDKYAEQF | 0.4596603 |
| VGVIPILL | AWSLRDKYAEQF | 0.18639559 |
| VIFASSLI | AWSLRDKYAEQF | 0.50615895 |
| VKGGDGGAAAG | AWSLRDKYAEQF | 0.20576803 |
| VTKTFTNI | AWSLRDKYAEQF | 0.55920166 |
| VVAAAILL | AWSLRDKYAEQF | 0.5388229 |
| YAALDVEL | AWSLRDKYAEQF | 0.38782734 |
| LASVALLL | AWSLRDKYAEQF | 0.2550038 |
| LDAVHRV | AWSLRDKYAEQF | 0.5108476 |
| QGLSSSIF | AWSLRDKYAEQF | 0.42205527 |
| RGTFANIRL | AWSLRDKYAEQF | 0.21094039 |
| RQLWWGHRI | AWSLRDKYAEQF | 0.62132204 |
| YGPPGTGKTLL | AWSLRDKYAEQF | 0.26155642 |
| AALVGAPANVA | AWSLRDKYAEQF | 0.40699476 |
| AATFRRTE | AWSLRDKYAEQF | 0.6979964 |
| AGALGGGI | AWSLRDKYAEQF | 0.54109204 |
| AGGAGGAGGIGAG | AWSLRDKYAEQF | 0.15987004 |
| AGSGALGL | AWSLRDKYAEQF | 0.60819566 |
| AQIADQI | AWSLRDKYAEQF | 0.4777103 |
| ASAVAVTYL | AWSLRDKYAEQF | 0.3327595 |

| | | |
|---|---|---|
| ASTVLGGTGG | AWSLRDKYAEQF | 0.1326706 |
| AVGGEPAK | AWSLRDKYAEQF | 0.4583928 |
| CEQGTFGGTC | AWSLRDKYAEQF | 0.2415756 |
| DEGKAPAG | AWSLRDKYAEQF | 0.24908921 |
| DTVVFPAL | AWSLRDKYAEQF | 0.2497352 |
| FEGITMAG | AWSLRDKYAEQF | 0.7321135 |
| FPLVVQD | AWSLRDKYAEQF | 0.44165382 |
| GAFTTPQ | AWSLRDKYAEQF | 0.4114067 |
| GFGGFGTAGGVGGA | AWSLRDKYAEQF | 0.70123994 |
| GFLAQFL | AWSLRDKYAEQF | 0.6299288 |
| GGAGGAGGNAGVA | AWSLRDKYAEQF | 0.80854404 |
| GGVGTAGAGGAGGAGG | AWSLRDKYAEQF | 0.4501866 |
| GNGGGGDAGNAGSGGN | AWSLRDKYAEQF | 0.25469676 |
| GTGGAGGAGNDAGS | AWSLRDKYAEQF | 0.41393334 |
| GTTNTGWFNS | AWSLRDKYAEQF | 0.29832816 |
| IFMGPLFTL | AWSLRDKYAEQF | 0.18453169 |
| ILAHAIEAI | AWSLRDKYAEQF | 0.1596807 |
| IRSAIREAL | AWSLRDKYAEQF | 0.48026946 |
| ITFAGGG | AWSLRDKYAEQF | 0.16376199 |
| ITGMSGHQEV | AWSLRDKYAEQF | 0.42253548 |
| IVAGLACTL | AWSLRDKYAEQF | 0.54388523 |
| IVGGGLAGA | AWSLRDKYAEQF | 0.28995758 |
| IVTGVLNI | AWSLRDKYAEQF | 0.19409434 |
| LAQQGCI | AWSLRDKYAEQF | 0.30013758 |
| LCLHVEN | AWSLRDKYAEQF | 0.38736552 |
| LDAISDGS | AWSLRDKYAEQF | 0.7056479 |
| LGGFAAML | AWSLRDKYAEQF | 0.38573977 |
| LGGKNPMI | AWSLRDKYAEQF | 0.8449098 |
| LGPKGVAAIL | AWSLRDKYAEQF | 0.74237037 |
| LMVGVSI | AWSLRDKYAEQF | 0.38438332 |
| LVGAAFVL | AWSLRDKYAEQF | 0.62261474 |
| LVLAVAPL | AWSLRDKYAEQF | 0.29080877 |
| MAAIAVIDAA | AWSLRDKYAEQF | 0.30714047 |
| NNGINAELQ | AWSLRDKYAEQF | 0.32403317 |
| RGEPLVLVHG | AWSLRDKYAEQF | 0.37598592 |
| RGGAGGAAGLI | AWSLRDKYAEQF | 0.39017415 |
| RSYRMHPAV | AWSLRDKYAEQF | 0.30473733 |
| SAAHLLAGQLA | AWSLRDKYAEQF | 0.16696884 |
| SAAVGSY | AWSLRDKYAEQF | 0.5171387 |
| SPHAVLHRI | AWSLRDKYAEQF | 0.37944716 |
| SQIAKVL | AWSLRDKYAEQF | 0.16480947 |
| TALGHLVP | AWSLRDKYAEQF | 0.26166648 |
| TAPTTSGAA | AWSLRDKYAEQF | 0.8241443 |
| TDGDDGNGGSPGT | AWSLRDKYAEQF | 0.8007907 |
| TDGLGEN | AWSLRDKYAEQF | 0.29182994 |
| TGGPPLPGQA | AWSLRDKYAEQF | 0.23016477 |
| TIPAITI | AWSLRDKYAEQF | 0.74144167 |
| TIPGFTI | AWSLRDKYAEQF | 0.15558203 |
| TSIASGALA | AWSLRDKYAEQF | 0.30419418 |
| TVTLPGAAI | AWSLRDKYAEQF | 0.62434685 |
| VASGLSAALP | AWSLRDKYAEQF | 0.5613266 |

| | | |
|---|---|---|
| VGAGVIGI | AWSLRDKYAEQF | 0.21876301 |
| VGPKRFGV | AWSLRDKYAEQF | 0.4749388 |
| VNVAFPD | AWSLRDKYAEQF | 0.40794927 |
| VSMTNSGVS | AWSLRDKYAEQF | 0.4191136 |
| VTDVLGDLSV | AWSLRDKYAEQF | 0.14494696 |
| VVVVGAGFAGL | AWSLRDKYAEQF | 0.41779214 |
| VYTVGTVV | AWSLRDKYAEQF | 0.29330203 |
| WTTFSGP | AWSLRDKYAEQF | 0.24038035 |
| YDELVEAF | AWSLRDKYAEQF | 0.17188457 |
| YGTGGAGGNGG | AWSLRDKYAEQF | 0.27483806 |
| YTNVMAV | AWSLRDKYAEQF | 0.76940584 |
| FLEQVHQL | AWSLRDKYAEQF | 0.71596277 |
| GLSLLAVAV | AWSLRDKYAEQF | 0.22534585 |
| LEQVHQL | AWSLRDKYAEQF | 0.7390319 |
| LNAAGAI | AWSLRDKYAEQF | 0.75138295 |
| PAVHASL | AWSLRDKYAEQF | 0.4329443 |
| TELEDDA | AWSLRDKYAEQF | 0.18528041 |
| TPPGPPN | AWSLRDKYAEQF | 0.5837639 |
| TPPPPPPP | AWSLRDKYAEQF | 0.3584514 |
| TQSPALI | ASSDVGTANTEVF | 0.5468271 |
| EFAAAFPQ | ASSDVGTANTEVF | 0.7588973 |
| LLSGNLDVL | ASSDVGTANTEVF | 0.21717447 |
| GPANLGNYNI | ASSDVGTANTEVF | 0.49234954 |
| TLGPTLP | ASSDVGTANTEVF | 0.70719135 |
| VYAYSSCCK | ASSDVGTANTEVF | 0.24624059 |
| APAPAPAPNI | ASSDVGTANTEVF | 0.20193438 |
| PGGPVPIP | ASSDVGTANTEVF | 0.20933875 |
| FLTVALEP | ASSDVGTANTEVF | 0.17406896 |
| SQYFGANL | ASSDVGTANTEVF | 0.6384531 |
| GGVKPAA | ASSDVGTANTEVF | 0.36829716 |
| GGTGGAGGAGGW | ASSDVGTANTEVF | 0.39590597 |
| GVGGVAGLGGAGG | ASSDVGTANTEVF | 0.38905102 |
| VLLVLATTI | ASSDVGTANTEVF | 0.30982023 |
| IGIFAAML | ASSDVGTANTEVF | 0.33859783 |
| AVGDLID | ASSDVGTANTEVF | 0.4189158 |
| NISPLPL | ASSDVGTANTEVF | 0.45232242 |
| VTAVFVAI | ASSDVGTANTEVF | 0.54257745 |
| SGDDNDAAL | ASSDVGTANTEVF | 0.41736954 |
| AVAALAHS | ASSDVGTANTEVF | 0.32622474 |
| GGVTVMATP | ASSDVGTANTEVF | 0.8128114 |
| AGLAANVRD | ASSDVGTANTEVF | 0.21673727 |
| TNRPDLIDPALLRPG | ASSDVGTANTEVF | 0.23584355 |
| LANLGVTAL | ASSDVGTANTEVF | 0.6066261 |
| TGERIEL | ASSDVGTANTEVF | 0.19402891 |
| NGEELAA | ASSDVGTANTEVF | 0.71048665 |
| VLSLVWG | ASSDVGTANTEVF | 0.5179268 |
| IGYALWL | ASSDVGTANTEVF | 0.5957445 |
| LSLLLAPL | ASSDVGTANTEVF | 0.4756233 |
| TLDEALK | ASSDVGTANTEVF | 0.8232559 |
| TDGAGTAEGG | ASSDVGTANTEVF | 0.70229113 |
| NAPGGAGGNGG | ASSDVGTANTEVF | 0.26703686 |

| | | |
|---|---|---|
| VSVFMGMGGNF | ASSDVGTANTEVF | 0.78463423 |
| LASYLAAL | ASSDVGTANTEVF | 0.18739605 |
| LVGGTPAL | ASSDVGTANTEVF | 0.66328585 |
| VRNADVL | ASSDVGTANTEVF | 0.4935515 |
| TCSSPFLVSAM | ASSDVGTANTEVF | 0.23951456 |
| VHITDAE | ASSDVGTANTEVF | 0.5047324 |
| EYAAPSF | ASSDVGTANTEVF | 0.58803236 |
| VVDAFHGGL | ASSDVGTANTEVF | 0.7856369 |
| ASVVDPQ | ASSDVGTANTEVF | 0.16669333 |
| SYNEFGGLS | ASSDVGTANTEVF | 0.35147852 |
| DPDGSVY | ASSDVGTANTEVF | 0.17218302 |
| MVSTIHATE | ASSDVGTANTEVF | 0.21836099 |
| LAAGVADPV | ASSDVGTANTEVF | 0.20503865 |
| TKEQIAA | ASSDVGTANTEVF | 0.5934634 |
| SLASPAAL | ASSDVGTANTEVF | 0.39999354 |
| IGLVFYGV | ASSDVGTANTEVF | 0.73867786 |
| FLLDASNL | ASSDVGTANTEVF | 0.4107462 |
| YQVGNFP | ASSDVGTANTEVF | 0.6895988 |
| ESLLSQST | ASSDVGTANTEVF | 0.273332 |
| LFTVGFAAVG | ASSDVGTANTEVF | 0.40172946 |
| ATPPTPT | ASSDVGTANTEVF | 0.17615546 |
| TDLATLG | ASSDVGTANTEVF | 0.24308689 |
| SAGGTGR | ASSDVGTANTEVF | 0.2227436 |
| GFSNSGS | ASSDVGTANTEVF | 0.38166505 |
| GGNGGSGGTSV | ASSDVGTANTEVF | 0.16819258 |
| GTSVATGGAGN | ASSDVGTANTEVF | 0.5951437 |
| NGGNGGSGGTS | ASSDVGTANTEVF | 0.558533 |
| AVVKFPH | ASSDVGTANTEVF | 0.605593 |
| TPPPPPP | ASSDVGTANTEVF | 0.33931226 |
| AAPAAAAAPA | ASSDVGTANTEVF | 0.32062763 |
| IALNEDL | ASSDVGTANTEVF | 0.29161683 |
| IVNVSSM | ASSDVGTANTEVF | 0.7180767 |
| TGIPPAPR | ASSDVGTANTEVF | 0.24970052 |
| FDLGGGTF | ASSDVGTANTEVF | 0.15392072 |
| FDLGGGTFDVS | ASSDVGTANTEVF | 0.25590926 |
| NEPTAAA | ASSDVGTANTEVF | 0.17912923 |
| NEPTAAALA | ASSDVGTANTEVF | 0.75452816 |
| GSIINISSI | ASSDVGTANTEVF | 0.71022844 |
| VAPGNAGTA | ASSDVGTANTEVF | 0.5383097 |
| TGGGASLE | ASSDVGTANTEVF | 0.65733147 |
| VAVYNHYKRI | ASSDVGTANTEVF | 0.14585817 |
| TDPAPATT | ASSDVGTANTEVF | 0.36885023 |
| TNRPDILDPALLRPG | ASSDVGTANTEVF | 0.7940202 |
| LTTTTTS | ASSDVGTANTEVF | 0.54229355 |
| PTGSGKTL | ASSDVGTANTEVF | 0.4163075 |
| TGSGKTL | ASSDVGTANTEVF | 0.7602322 |
| TIGHVDHGK | ASSDVGTANTEVF | 0.3391415 |
| TIGHVDHGKTT | ASSDVGTANTEVF | 0.3378905 |
| GHVDHGKTT | ASSDVGTANTEVF | 0.5731286 |
| AQFEHTLL | ASSDVGTANTEVF | 0.41078424 |
| IVNQDEL | ASSDVGTANTEVF | 0.3890852 |

| | | |
|---|---|---|
| TNGSQFF | ASSDVGTANTEVF | 0.22687164 |
| TGKGIVE | ASSDVGTANTEVF | 0.48524064 |
| LISNASDA | ASSDVGTANTEVF | 0.84142506 |
| AELRTLGELTKIAKAHGAQ | ASSDVGTANTEVF | 0.46328127 |
| APNAVAAAEPSW | ASSDVGTANTEVF | 0.71599644 |
| APQVAGAATAVL | ASSDVGTANTEVF | 0.5594391 |
| ARRDPAAAAAILPTDA | ASSDVGTANTEVF | 0.74642646 |
| DAAARDVLAEAGLAEYF | ASSDVGTANTEVF | 0.276838 |
| DDQIVLSAKANLAKVEQ | ASSDVGTANTEVF | 0.28844854 |
| DELLPDTGEDSDEDGH | ASSDVGTANTEVF | 0.50358623 |
| ELPEAPANRATVPQD | ASSDVGTANTEVF | 0.3270492 |
| FGQSKIERVTGIG | ASSDVGTANTEVF | 0.32545346 |
| GAGGTGSTAGGAGGAGGAGGLY | ASSDVGTANTEVF | 0.26315743 |
| GHGGTGAAGGNGGTGGAGGSGI | ASSDVGTANTEVF | 0.8456483 |
| GNGGNGGAGGEGGPGGAGGAGA | ASSDVGTANTEVF | 0.55555415 |
| GSTAGNGGNGGNATNGTIAGQP | ASSDVGTANTEVF | 0.63620734 |
| GTGGDGGNGGNGGTGG | ASSDVGTANTEVF | 0.2688334 |
| GVRRPAGIPADAPV | ASSDVGTANTEVF | 0.3095778 |
| GYRQVTDAHLLTLARR | ASSDVGTANTEVF | 0.61998796 |
| IAKMGQDGHDRG | ASSDVGTANTEVF | 0.1798809 |
| KAEPSIEIQYT | ASSDVGTANTEVF | 0.74699926 |
| KPVRIRMAADEDTVAVT | ASSDVGTANTEVF | 0.2802245 |
| MPPRIAGMRLLVIKPEP | ASSDVGTANTEVF | 0.36262906 |
| MKFYAVALFRSRGVPPTR | ASSDVGTANTEVF | 0.5898396 |
| NAGGFGILGAVAHSP | ASSDVGTANTEVF | 0.82430124 |
| NEGREITLAMAEM | ASSDVGTANTEVF | 0.32614788 |
| NELVEWSHPLGHRWRWE | ASSDVGTANTEVF | 0.42952633 |
| NLGAGGTGGNGGNPG | ASSDVGTANTEVF | 0.2591314 |
| NTEWPKWETVMAADPVD | ASSDVGTANTEVF | 0.5247958 |
| PASAAPAETQP | ASSDVGTANTEVF | 0.77363825 |
| QQSSDGKNGVRAAEITGE | ASSDVGTANTEVF | 0.5795393 |
| RAPDAVTTSWWKE | ASSDVGTANTEVF | 0.8179429 |
| SGFGNVGAHTSGWFNQ | ASSDVGTANTEVF | 0.31322446 |
| SGVAGAVGQAASVGGLKV | ASSDVGTANTEVF | 0.17543441 |
| SNGGAGTGSAGGNGGT | ASSDVGTANTEVF | 0.37029606 |
| TGGAGGEGGAAG | ASSDVGTANTEVF | 0.55355954 |
| TGGDGGAAGAGGAPGHG | ASSDVGTANTEVF | 0.7495504 |
| TMPPLLAAIANAGRHGV | ASSDVGTANTEVF | 0.76614505 |
| TSSTAGNSGGA | ASSDVGTANTEVF | 0.58292186 |
| VPATVPVGILC | ASSDVGTANTEVF | 0.31440532 |
| VQRPDSGGVVLTGRIS | ASSDVGTANTEVF | 0.7186006 |
| WTPNPVPATKGVPAGLVQAT | ASSDVGTANTEVF | 0.7561024 |
| YALKRALAKFSDRFDVVII | ASSDVGTANTEVF | 0.647097 |
| YLIGSVIV | ASSDVGTANTEVF | 0.24896592 |
| YGQLVAVID | ASSDVGTANTEVF | 0.21826023 |
| VTVGGMTLPA | ASSDVGTANTEVF | 0.4478736 |
| VLALPAV | ASSDVGTANTEVF | 0.71326816 |
| VIPPTLNYET | ASSDVGTANTEVF | 0.18991615 |
| VGMGGMHPSSG | ASSDVGTANTEVF | 0.5812162 |
| VGLAGGAATA | ASSDVGTANTEVF | 0.560105 |
| VGGAGGAGGSGP | ASSDVGTANTEVF | 0.17322905 |

| | | |
|---|---|---|
| VFAAFMV | ASSDVGTANTEVF | 0.5149163 |
| TLYDKLR | ASSDVGTANTEVF | 0.80324084 |
| TLAVFTVL | ASSDVGTANTEVF | 0.30046988 |
| TGSVQPARV | ASSDVGTANTEVF | 0.29165116 |
| TGGGAGSGGT | ASSDVGTANTEVF | 0.77296114 |
| SHLETSL | ASSDVGTANTEVF | 0.84351456 |
| SFTTGMT | ASSDVGTANTEVF | 0.64244205 |
| QVEVAKRMGVRQPTVS | ASSDVGTANTEVF | 0.18968965 |
| QGSGGPALAM | ASSDVGTANTEVF | 0.13568427 |
| MLAPSNP | ASSDVGTANTEVF | 0.48206514 |
| LVNNAAI | ASSDVGTANTEVF | 0.753775 |
| LTGEINTLA | ASSDVGTANTEVF | 0.38167828 |
| LSIFGPV | ASSDVGTANTEVF | 0.33636394 |
| LLALLANL | ASSDVGTANTEVF | 0.6337058 |
| LISNASDALDKLR | ASSDVGTANTEVF | 0.6499496 |
| LINEAVP | ASSDVGTANTEVF | 0.24659386 |
| LGYAFSGL | ASSDVGTANTEVF | 0.1800766 |
| LDFVAVAL | ASSDVGTANTEVF | 0.46339628 |
| LAQPGAE | ASSDVGTANTEVF | 0.5191431 |
| ISNLGAVTGF | ASSDVGTANTEVF | 0.26025748 |
| HQEDPGRI | ASSDVGTANTEVF | 0.43730697 |
| GTPGEPTP | ASSDVGTANTEVF | 0.39593112 |
| GFFNSGS | ASSDVGTANTEVF | 0.28485632 |
| GAGGGGLTAGAN | ASSDVGTANTEVF | 0.5870354 |
| FVDALVNM | ASSDVGTANTEVF | 0.2241059 |
| FLTVAPEL | ASSDVGTANTEVF | 0.75759494 |
| FETLTAL | ASSDVGTANTEVF | 0.22230819 |
| DVINPIEL | ASSDVGTANTEVF | 0.5017153 |
| APVSGAPV | ASSDVGTANTEVF | 0.54133815 |
| ALVGGLAW | ASSDVGTANTEVF | 0.46274662 |
| ALAAAAATAI | ASSDVGTANTEVF | 0.42672473 |
| AFGLGLLL | ASSDVGTANTEVF | 0.33844194 |
| AFFAAASL | ASSDVGTANTEVF | 0.20244245 |
| AAAALPAFSPP | ASSDVGTANTEVF | 0.14918435 |
| VVLAPAL | ASSDVGTANTEVF | 0.7479594 |
| TIGHVDHGKTTL | ASSDVGTANTEVF | 0.5112022 |
| LISNASDALDK | ASSDVGTANTEVF | 0.70923424 |
| LISNASDAL | ASSDVGTANTEVF | 0.570882 |
| ISNASDALDK | ASSDVGTANTEVF | 0.2479171 |
| NQAETLVYQTE | ASSDVGTANTEVF | 0.38421535 |
| TGGTGGTAGAG | ASSDVGTANTEVF | 0.26863492 |
| TGSTAGGAGGA | ASSDVGTANTEVF | 0.2587026 |
| DPLAVPVAAGPV | ASSDVGTANTEVF | 0.53773034 |
| GGTGGTPGNGGN | ASSDVGTANTEVF | 0.4470364 |
| PAGLAGVDPATL | ASSDVGTANTEVF | 0.26015055 |
| GNNLVVEAPHAGASV | ASSDVGTANTEVF | 0.34527963 |
| KVTRIMSASTLEQPA | ASSDVGTANTEVF | 0.519572 |
| LGFALPIALPRRTVV | ASSDVGTANTEVF | 0.23869416 |
| TRLGNLTARKLVVLV | ASSDVGTANTEVF | 0.29829746 |
| WSQQRRLIYFPSAGP | ASSDVGTANTEVF | 0.7882067 |
| GPNGNPDPMAAAVDIR | ASSDVGTANTEVF | 0.3420902 |

| | | |
|---|---|---|
| TQRCVSVADLRAAASR | ASSDVGTANTEVF | 0.2264173 |
| QFVLDVAEHPVEVLTETV | ASSDVGTANTEVF | 0.71695226 |
| GGAGGTGGAGGNGGTQAGDG | ASSDVGTANTEVF | 0.23788363 |
| PLLATLRAIAVDVDAKPAQV | ASSDVGTANTEVF | 0.6099717 |
| APKFRTGGYMIDFWGVGYQVAK | ASSDVGTANTEVF | 0.40077284 |
| GAGGAGGAGGAAGGPGGTGGTG | ASSDVGTANTEVF | 0.3297364 |
| IGALGLLATDVAPAGSVAFGVA | ASSDVGTANTEVF | 0.6862782 |
| AGGAGGIGGNGGAGGAGGVLMG | ASSDVGTANTEVF | 0.18675593 |
| ADELIGLL | ASSDVGTANTEVF | 0.30568075 |
| ADRALKA | ASSDVGTANTEVF | 0.36628914 |
| AGAVGQVGGAGGA | ASSDVGTANTEVF | 0.5248118 |
| AGGGGGLGGAP | ASSDVGTANTEVF | 0.70047075 |
| AGGLLYGNGGAG | ASSDVGTANTEVF | 0.26597267 |
| AGLESIGL | ASSDVGTANTEVF | 0.49498314 |
| AGLKTGII | ASSDVGTANTEVF | 0.26704264 |
| ASLAALLL | ASSDVGTANTEVF | 0.21612358 |
| ASVINISSI | ASSDVGTANTEVF | 0.26194692 |
| ATPPGPA | ASSDVGTANTEVF | 0.28653103 |
| ATVASALV | ASSDVGTANTEVF | 0.29202253 |
| ATVASLIL | ASSDVGTANTEVF | 0.4366467 |
| AVNGPVLIRH | ASSDVGTANTEVF | 0.2822584 |
| DGAAGAVGI | ASSDVGTANTEVF | 0.5320226 |
| DPLLHGI | ASSDVGTANTEVF | 0.7229259 |
| ELGAALGAL | ASSDVGTANTEVF | 0.29754934 |
| ELGRTQAV | ASSDVGTANTEVF | 0.43179417 |
| FVAGGYLS | ASSDVGTANTEVF | 0.5616668 |
| GASGGAGGAGGW | ASSDVGTANTEVF | 0.66133034 |
| GGAGAGGDGGAGGI | ASSDVGTANTEVF | 0.37078798 |
| GTGHNGSQPGMGGNG | ASSDVGTANTEVF | 0.258836 |
| GYAAASASAS | ASSDVGTANTEVF | 0.7077328 |
| HQVDVAAVQQG | ASSDVGTANTEVF | 0.70495474 |
| IGAAIAHT | ASSDVGTANTEVF | 0.5295487 |
| IGMIEVAQL | ASSDVGTANTEVF | 0.5025785 |
| IRLAPEHTSAMD | ASSDVGTANTEVF | 0.674549 |
| IYNVSVL | ASSDVGTANTEVF | 0.6139517 |
| KQGFLGM | ASSDVGTANTEVF | 0.43170834 |
| LESVITQL | ASSDVGTANTEVF | 0.17061232 |
| LGLGALVVV | ASSDVGTANTEVF | 0.45509878 |
| LIGNGGNGGSGGI | ASSDVGTANTEVF | 0.4152111 |
| LNAINAPTL | ASSDVGTANTEVF | 0.25663164 |
| LQLLNTGV | ASSDVGTANTEVF | 0.1578195 |
| LQTSIEF | ASSDVGTANTEVF | 0.76550794 |
| MSFAVSEL | ASSDVGTANTEVF | 0.83991766 |
| NTGALGL | ASSDVGTANTEVF | 0.33154774 |
| PPGAGGAGGAGTT | ASSDVGTANTEVF | 0.75473475 |
| QLVGLGI | ASSDVGTANTEVF | 0.18017448 |
| RFLEQVHQL | ASSDVGTANTEVF | 0.7112505 |
| RGVGYVLR | ASSDVGTANTEVF | 0.77952874 |
| SATGVFAGVI | ASSDVGTANTEVF | 0.25614753 |
| SLPAITLP | ASSDVGTANTEVF | 0.6507136 |
| STALAFTV | ASSDVGTANTEVF | 0.27956462 |

| | | |
|---|---|---|
| SVLTVGLGK | ASSDVGTANTEVF | 0.30976963 |
| TALAGAGSGI | ASSDVGTANTEVF | 0.7972456 |
| TGLPFTGI | ASSDVGTANTEVF | 0.41008127 |
| VAEHVTV | ASSDVGTANTEVF | 0.39482477 |
| VDIVLNSL | ASSDVGTANTEVF | 0.3795161 |
| VLTGGGAVL | ASSDVGTANTEVF | 0.2217143 |
| VNVGLAYGI | ASSDVGTANTEVF | 0.3815823 |
| VSSAAVGAV | ASSDVGTANTEVF | 0.5853456 |
| VTSVYEAGV | ASSDVGTANTEVF | 0.24539675 |
| WVALDRGA | ASSDVGTANTEVF | 0.71049166 |
| YLVGSVVI | ASSDVGTANTEVF | 0.26468366 |
| GPPGTGKTLL | ASSDVGTANTEVF | 0.8268976 |
| GVGMTKF | ASSDVGTANTEVF | 0.7703761 |
| IGIGGSDL | ASSDVGTANTEVF | 0.57254726 |
| IGTLLAEL | ASSDVGTANTEVF | 0.281562 |
| IYTSGTT | ASSDVGTANTEVF | 0.7921299 |
| LSFPAGV | ASSDVGTANTEVF | 0.37161744 |
| LYTSGTT | ASSDVGTANTEVF | 0.50541794 |
| VGAFPFGF | ASSDVGTANTEVF | 0.5768393 |
| VGAGTVEF | ASSDVGTANTEVF | 0.24491242 |
| VGKSTLLL | ASSDVGTANTEVF | 0.36666444 |
| AALAHAGGGIV | ASSDVGTANTEVF | 0.26067632 |
| AALVAVSLL | ASSDVGTANTEVF | 0.33796358 |
| AAVSRAVAAL | ASSDVGTANTEVF | 0.2580342 |
| AGGSSGIGGF | ASSDVGTANTEVF | 0.21613961 |
| AGLTVANANG | ASSDVGTANTEVF | 0.24033111 |
| AGLVGAVL | ASSDVGTANTEVF | 0.24635315 |
| ALSQAFTV | ASSDVGTANTEVF | 0.46736848 |
| ALTFSVSL | ASSDVGTANTEVF | 0.37040275 |
| ALTGAALVAL | ASSDVGTANTEVF | 0.2745544 |
| ASFSIGAGVL | ASSDVGTANTEVF | 0.42656437 |
| DGSVYKI | ASSDVGTANTEVF | 0.4560653 |
| EQLAEIV | ASSDVGTANTEVF | 0.2704547 |
| FAVSELTVM | ASSDVGTANTEVF | 0.30391097 |
| GAAPLAAGL | ASSDVGTANTEVF | 0.23275778 |
| GADGGDGGSGGAGG | ASSDVGTANTEVF | 0.34217906 |
| GFGLDTGGAGG | ASSDVGTANTEVF | 0.21105786 |
| GLASGGLL | ASSDVGTANTEVF | 0.35607988 |
| GRIAGAAVLL | ASSDVGTANTEVF | 0.2640881 |
| GSAGSLNTSF | ASSDVGTANTEVF | 0.21239188 |
| GTGQIGI | ASSDVGTANTEVF | 0.44819498 |
| GTLIALIL | ASSDVGTANTEVF | 0.26584485 |
| IAATMGFVVG | ASSDVGTANTEVF | 0.5058595 |
| IFAQALL | ASSDVGTANTEVF | 0.63953125 |
| IGAAVALGF | ASSDVGTANTEVF | 0.34114847 |
| IGLTPLAL | ASSDVGTANTEVF | 0.23247205 |
| ILGSNTSTL | ASSDVGTANTEVF | 0.30787194 |
| IVFSLTVL | ASSDVGTANTEVF | 0.42000812 |
| LASKATGF | ASSDVGTANTEVF | 0.23231864 |
| LASVALIL | ASSDVGTANTEVF | 0.44986734 |
| LFAVGLTL | ASSDVGTANTEVF | 0.369003 |

| | | |
|---|---|---|
| LFNGGIGTYI | ASSDVGTANTEVF | 0.26379338 |
| LTLGAGII | ASSDVGTANTEVF | 0.43377656 |
| LVAAALGDI | ASSDVGTANTEVF | 0.43302184 |
| NTAFVQLGI | ASSDVGTANTEVF | 0.21847434 |
| QLFSGAEL | ASSDVGTANTEVF | 0.44640753 |
| SAATFITL | ASSDVGTANTEVF | 0.36306798 |
| SNYGLANLGV | ASSDVGTANTEVF | 0.25504684 |
| SYGGAYAVM | ASSDVGTANTEVF | 0.28551617 |
| TTGVGGLTLG | ASSDVGTANTEVF | 0.2394422 |
| TTQTIYI | ASSDVGTANTEVF | 0.331204 |
| VAAGAGLGAVI | ASSDVGTANTEVF | 0.22403681 |
| VAAVVTLL | ASSDVGTANTEVF | 0.41833842 |
| VGFLGGLI | ASSDVGTANTEVF | 0.35961342 |
| VGVIPILL | ASSDVGTANTEVF | 0.37310338 |
| VIFASSLI | ASSDVGTANTEVF | 0.2282649 |
| VKGGDGGAAAG | ASSDVGTANTEVF | 0.21889599 |
| VTKTFTNI | ASSDVGTANTEVF | 0.24962278 |
| VVAAAILL | ASSDVGTANTEVF | 0.26065361 |
| YAALDVEL | ASSDVGTANTEVF | 0.21992448 |
| LASVALLL | ASSDVGTANTEVF | 0.31530276 |
| LDAVHRV | ASSDVGTANTEVF | 0.3795187 |
| QGLSSSIF | ASSDVGTANTEVF | 0.36819488 |
| RGTFANIRL | ASSDVGTANTEVF | 0.31170794 |
| RQLWWGHRI | ASSDVGTANTEVF | 0.5116053 |
| YGPPGTGKTLL | ASSDVGTANTEVF | 0.4355299 |
| AALVGAPANVA | ASSDVGTANTEVF | 0.23327823 |
| AATFRRTE | ASSDVGTANTEVF | 0.2485717 |
| AGALGGGI | ASSDVGTANTEVF | 0.24378261 |
| AGGAGGAGGIGAG | ASSDVGTANTEVF | 0.63017774 |
| AGSGALGL | ASSDVGTANTEVF | 0.27469522 |
| AQIADQI | ASSDVGTANTEVF | 0.32114416 |
| ASAVAVTYL | ASSDVGTANTEVF | 0.54872984 |
| ASTVLGGTGG | ASSDVGTANTEVF | 0.30619478 |
| AVGGEPAK | ASSDVGTANTEVF | 0.24409269 |
| CEQGTFGGTC | ASSDVGTANTEVF | 0.455618 |
| DEGKAPAG | ASSDVGTANTEVF | 0.3435283 |
| DTVVFPAL | ASSDVGTANTEVF | 0.2235969 |
| FEGITMAG | ASSDVGTANTEVF | 0.24296972 |
| FPLVVQD | ASSDVGTANTEVF | 0.2800062 |
| GAFTTPQ | ASSDVGTANTEVF | 0.28849182 |
| GFGGFGTAGGVGGA | ASSDVGTANTEVF | 0.53272426 |
| GFLAQFL | ASSDVGTANTEVF | 0.27625865 |
| GGAGGAGGNAGVA | ASSDVGTANTEVF | 0.2223032 |
| GGVGTAGAGGAGGAGG | ASSDVGTANTEVF | 0.55225426 |
| GNGGGGDAGNAGSGGN | ASSDVGTANTEVF | 0.28326607 |
| GTGGAGGAGNDAGS | ASSDVGTANTEVF | 0.28172162 |
| GTTNTGWFNS | ASSDVGTANTEVF | 0.49707544 |
| IFMGPLFTL | ASSDVGTANTEVF | 0.27320367 |
| ILAHAIEAI | ASSDVGTANTEVF | 0.31243855 |
| IRSAIREAL | ASSDVGTANTEVF | 0.4052887 |
| ITFAGGG | ASSDVGTANTEVF | 0.40068698 |

| | | |
|---|---|---|
| ITGMSGHQEV | ASSDVGTANTEVF | 0.41090173 |
| IVAGLACTL | ASSDVGTANTEVF | 0.23863477 |
| IVGGGLAGA | ASSDVGTANTEVF | 0.27039286 |
| IVTGVLNI | ASSDVGTANTEVF | 0.26491314 |
| LAQQGCI | ASSDVGTANTEVF | 0.23221624 |
| LCLHVEN | ASSDVGTANTEVF | 0.47173178 |
| LDAISDGS | ASSDVGTANTEVF | 0.45030138 |
| LGGFAAML | ASSDVGTANTEVF | 0.5352013 |
| LGGKNPMI | ASSDVGTANTEVF | 0.30011547 |
| LGPKGVAAIL | ASSDVGTANTEVF | 0.23350947 |
| LMVGVSI | ASSDVGTANTEVF | 0.4865498 |
| LVGAAFVL | ASSDVGTANTEVF | 0.357587 |
| LVLAVAPL | ASSDVGTANTEVF | 0.22960953 |
| MAAIAVIDAA | ASSDVGTANTEVF | 0.2671255 |
| NNGINAELQ | ASSDVGTANTEVF | 0.5926484 |
| RGEPLVLVHG | ASSDVGTANTEVF | 0.53534293 |
| RGGAGGAAGLI | ASSDVGTANTEVF | 0.31988847 |
| RSYRMHPAV | ASSDVGTANTEVF | 0.28733495 |
| SAAHLLAGQLA | ASSDVGTANTEVF | 0.59807223 |
| SAAVGSY | ASSDVGTANTEVF | 0.30123514 |
| SPHAVLHRI | ASSDVGTANTEVF | 0.41036728 |
| SQIAKVL | ASSDVGTANTEVF | 0.23879398 |
| TALGHLVP | ASSDVGTANTEVF | 0.28378418 |
| TAPTTSGAA | ASSDVGTANTEVF | 0.2153516 |
| TDGDDGNGGSPGT | ASSDVGTANTEVF | 0.21570125 |
| TDGLGEN | ASSDVGTANTEVF | 0.45892173 |
| TGGPPLPGQA | ASSDVGTANTEVF | 0.2359419 |
| TIPAITI | ASSDVGTANTEVF | 0.490648 |
| TIPGFTI | ASSDVGTANTEVF | 0.25294846 |
| TSIASGALA | ASSDVGTANTEVF | 0.43053472 |
| TVTLPGAAI | ASSDVGTANTEVF | 0.2787667 |
| VASGLSAALP | ASSDVGTANTEVF | 0.36114824 |
| VGAGVIGI | ASSDVGTANTEVF | 0.27654067 |
| VGPKRFGV | ASSDVGTANTEVF | 0.34160024 |
| VNVAFPD | ASSDVGTANTEVF | 0.5018752 |
| VSMTNSGVS | ASSDVGTANTEVF | 0.30192754 |
| VTDVLGDLSV | ASSDVGTANTEVF | 0.21524245 |
| VVVVGAGFAGL | ASSDVGTANTEVF | 0.4181726 |
| VYTVGTVV | ASSDVGTANTEVF | 0.28119314 |
| WTTFSGP | ASSDVGTANTEVF | 0.22780575 |
| YDELVEAF | ASSDVGTANTEVF | 0.23111625 |
| YGTGGAGGNGG | ASSDVGTANTEVF | 0.23488057 |
| YTNVMAV | ASSDVGTANTEVF | 0.28488725 |
| FLEQVHQL | ASSDVGTANTEVF | 0.46885824 |
| GLSLLAVAV | ASSDVGTANTEVF | 0.3921072 |
| LEQVHQL | ASSDVGTANTEVF | 0.27620125 |
| LNAAGAI | ASSDVGTANTEVF | 0.2873829 |
| PAVHASL | ASSDVGTANTEVF | 0.26069218 |
| TELEDDA | ASSDVGTANTEVF | 0.21016788 |
| TPPGPPN | ASSDVGTANTEVF | 0.22600323 |
| TPPPPPPP | ASSDVGTANTEVF | 0.29656333 |

| | | |
|---|---|---|
| TQSPALI | ASSLGTHNNQAPL | 0.41853562 |
| EFAAAFPQ | ASSLGTHNNQAPL | 0.26192138 |
| LLSGNLDVL | ASSLGTHNNQAPL | 0.2884839 |
| GPANLGNYNI | ASSLGTHNNQAPL | 0.3452576 |
| TLGPTLP | ASSLGTHNNQAPL | 0.5133995 |
| VYAYSSCCK | ASSLGTHNNQAPL | 0.32863736 |
| APAPAPAPNI | ASSLGTHNNQAPL | 0.21504073 |
| PGGPVPIP | ASSLGTHNNQAPL | 0.24595524 |
| FLTVALEP | ASSLGTHNNQAPL | 0.32984352 |
| SQYFGANL | ASSLGTHNNQAPL | 0.22989674 |
| GGVKPAA | ASSLGTHNNQAPL | 0.28348258 |
| GGTGGAGGAGGW | ASSLGTHNNQAPL | 0.25354353 |
| GVGGVAGLGGAGG | ASSLGTHNNQAPL | 0.50971353 |
| VLLVLATTI | ASSLGTHNNQAPL | 0.21725388 |
| IGIFAAML | ASSLGTHNNQAPL | 0.37695062 |
| AVGDLID | ASSLGTHNNQAPL | 0.3817845 |
| NISPLPL | ASSLGTHNNQAPL | 0.29213974 |
| VTAVFVAI | ASSLGTHNNQAPL | 0.3491408 |
| SGDDNDAAL | ASSLGTHNNQAPL | 0.5555291 |
| AVAALAHS | ASSLGTHNNQAPL | 0.32794613 |
| GGVTVMATP | ASSLGTHNNQAPL | 0.24196708 |
| AGLAANVRD | ASSLGTHNNQAPL | 0.30823684 |
| TNRPDLIDPALLRPG | ASSLGTHNNQAPL | 0.2412228 |
| LANLGVTAL | ASSLGTHNNQAPL | 0.24036406 |
| TGERIEL | ASSLGTHNNQAPL | 0.5647372 |
| NGEELAA | ASSLGTHNNQAPL | 0.3161459 |
| VLSLVWG | ASSLGTHNNQAPL | 0.21995278 |
| IGYALWL | ASSLGTHNNQAPL | 0.30042383 |
| LSLLLAPL | ASSLGTHNNQAPL | 0.38663942 |
| TLDEALK | ASSLGTHNNQAPL | 0.3277543 |
| TDGAGTAEGG | ASSLGTHNNQAPL | 0.3034563 |
| NAPGGAGGNGG | ASSLGTHNNQAPL | 0.2247555 |
| VSVFMGMGGNF | ASSLGTHNNQAPL | 0.2597994 |
| LASYLAAL | ASSLGTHNNQAPL | 0.31597254 |
| LVGGTPAL | ASSLGTHNNQAPL | 0.22967649 |
| VRNADVL | ASSLGTHNNQAPL | 0.23429151 |
| TCSSPFLVSAM | ASSLGTHNNQAPL | 0.33304346 |
| VHITDAE | ASSLGTHNNQAPL | 0.65336955 |
| EYAAPSF | ASSLGTHNNQAPL | 0.24577178 |
| VVDAFHGGL | ASSLGTHNNQAPL | 0.2146189 |
| ASVVDPQ | ASSLGTHNNQAPL | 0.3099863 |
| SYNEFGGLS | ASSLGTHNNQAPL | 0.22476664 |
| DPDGSVY | ASSLGTHNNQAPL | 0.26950893 |
| MVSTIHATE | ASSLGTHNNQAPL | 0.24859953 |
| LAAGVADPV | ASSLGTHNNQAPL | 0.23279348 |
| TKEQIAA | ASSLGTHNNQAPL | 0.2784443 |
| SLASPAAL | ASSLGTHNNQAPL | 0.35562378 |
| IGLVFYGV | ASSLGTHNNQAPL | 0.30776346 |
| FLLDASNL | ASSLGTHNNQAPL | 0.40226823 |
| YQVGNFP | ASSLGTHNNQAPL | 0.5225562 |
| ESLLSQST | ASSLGTHNNQAPL | 0.5865141 |

| | | |
|---|---|---|
| LFTVGFAAVG | ASSLGTHNNQAPL | 0.65692526 |
| ATPPTPT | ASSLGTHNNQAPL | 0.52941555 |
| TDLATLG | ASSLGTHNNQAPL | 0.32469684 |
| SAGGTGR | ASSLGTHNNQAPL | 0.23457003 |
| GFSNSGS | ASSLGTHNNQAPL | 0.49884042 |
| GGNGGSGGTSV | ASSLGTHNNQAPL | 0.3037462 |
| GTSVATGGAGN | ASSLGTHNNQAPL | 0.39833555 |
| NGGNGGSGGTS | ASSLGTHNNQAPL | 0.24253644 |
| AVVKFPH | ASSLGTHNNQAPL | 0.5092041 |
| TPPPPPP | ASSLGTHNNQAPL | 0.24566032 |
| AAPAAAAAPA | ASSLGTHNNQAPL | 0.56532866 |
| IALNEDL | ASSLGTHNNQAPL | 0.22098501 |
| IVNVSSM | ASSLGTHNNQAPL | 0.25234142 |
| TGIPPAPR | ASSLGTHNNQAPL | 0.61052185 |
| FDLGGGTF | ASSLGTHNNQAPL | 0.21493535 |
| FDLGGGTFDVS | ASSLGTHNNQAPL | 0.25545466 |
| NEPTAAA | ASSLGTHNNQAPL | 0.2753068 |
| NEPTAAALA | ASSLGTHNNQAPL | 0.33552966 |
| GSIINISSI | ASSLGTHNNQAPL | 0.21006556 |
| VAPGNAGTA | ASSLGTHNNQAPL | 0.25393745 |
| TGGGASLE | ASSLGTHNNQAPL | 0.24965475 |
| VAVYNHYKRI | ASSLGTHNNQAPL | 0.2861013 |
| TDPAPATT | ASSLGTHNNQAPL | 0.25885114 |
| TNRPDILDPALLRPG | ASSLGTHNNQAPL | 0.21683094 |
| LTTTTTS | ASSLGTHNNQAPL | 0.59128547 |
| PTGSGKTL | ASSLGTHNNQAPL | 0.21579008 |
| TGSGKTL | ASSLGTHNNQAPL | 0.31146955 |
| TIGHVDHGK | ASSLGTHNNQAPL | 0.38314754 |
| TIGHVDHGKTT | ASSLGTHNNQAPL | 0.6557938 |
| GHVDHGKTT | ASSLGTHNNQAPL | 0.23981875 |
| AQFEHTLL | ASSLGTHNNQAPL | 0.4025107 |
| IVNQDEL | ASSLGTHNNQAPL | 0.22787188 |
| TNGSQFF | ASSLGTHNNQAPL | 0.44700834 |
| TGKGIVE | ASSLGTHNNQAPL | 0.28047633 |
| LISNASDA | ASSLGTHNNQAPL | 0.21908155 |
| AELRTLGELTKIAKAHGAQ | ASSLGTHNNQAPL | 0.5529915 |
| APNAVAAAEPSW | ASSLGTHNNQAPL | 0.48466888 |
| APQVAGAATAVL | ASSLGTHNNQAPL | 0.3390553 |
| ARRDPAAAAAILPTDA | ASSLGTHNNQAPL | 0.52069664 |
| DAAARDVLAEAGLAEYF | ASSLGTHNNQAPL | 0.2605617 |
| DDQIVLSAKANLAKVEQ | ASSLGTHNNQAPL | 0.3067599 |
| DELLPDTGEDSDEDGH | ASSLGTHNNQAPL | 0.46867388 |
| ELPEAPANRATVPQD | ASSLGTHNNQAPL | 0.53264165 |
| FGQSKIERVTGIG | ASSLGTHNNQAPL | 0.24384874 |
| GAGGTGSTAGGAGGAGGAGGLY | ASSLGTHNNQAPL | 0.3498252 |
| GHGGTGAAGGNGGTGGAGGSGI | ASSLGTHNNQAPL | 0.34204945 |
| GNGGNGGAGGEGGPGGAGGAGA | ASSLGTHNNQAPL | 0.25258783 |
| GSTAGNGGNGGNATNGTIAGQP | ASSLGTHNNQAPL | 0.37260556 |
| GTGGDGGNGGNGGTGG | ASSLGTHNNQAPL | 0.28576365 |
| GVRRPAGIPADAPV | ASSLGTHNNQAPL | 0.36939487 |
| GYRQVTDAHLLTLARR | ASSLGTHNNQAPL | 0.30288944 |

| | | |
|---|---|---|
| IAKMGQDGHDRG | ASSLGTHNNQAPL | 0.570538 |
| KAEPSIEIQYT | ASSLGTHNNQAPL | 0.22606236 |
| KPVRIRMAADEDTVAVT | ASSLGTHNNQAPL | 0.23726274 |
| MPPRIAGMRLLVIKPEP | ASSLGTHNNQAPL | 0.2670921 |
| MKFYAVALFRSRGVPPTR | ASSLGTHNNQAPL | 0.44815552 |
| NAGGFGILGAVAHSP | ASSLGTHNNQAPL | 0.23244825 |
| NEGREITLAMAEM | ASSLGTHNNQAPL | 0.34218812 |
| NELVEWSHPLGHRWRWE | ASSLGTHNNQAPL | 0.3774249 |
| NLGAGGTGGNGGNPG | ASSLGTHNNQAPL | 0.4212849 |
| NTEWPKWETVMAADPVD | ASSLGTHNNQAPL | 0.25870064 |
| PASAAPAETQP | ASSLGTHNNQAPL | 0.43365967 |
| QQSSDGKNGVRAAEITGE | ASSLGTHNNQAPL | 0.69465506 |
| RAPDAVTTSWWKE | ASSLGTHNNQAPL | 0.25782132 |
| SGFGNVGAHTSGWFNQ | ASSLGTHNNQAPL | 0.30994648 |
| SGVAGAVGQAASVGGLKV | ASSLGTHNNQAPL | 0.33463642 |
| SNGGAGTGSAGGNGGT | ASSLGTHNNQAPL | 0.5889291 |
| TGGAGGEGGAAG | ASSLGTHNNQAPL | 0.32014602 |
| TGGDGGAAGAGGAPGHG | ASSLGTHNNQAPL | 0.42052674 |
| TMPPLLAAIANAGRHGV | ASSLGTHNNQAPL | 0.23857483 |
| TSSTAGNSGGA | ASSLGTHNNQAPL | 0.40339017 |
| VPATVPVGILC | ASSLGTHNNQAPL | 0.2212358 |
| VQRPDSGGVVLTGRIS | ASSLGTHNNQAPL | 0.273886 |
| WTPNPVPATKGVPAGLVQAT | ASSLGTHNNQAPL | 0.3293579 |
| YALKRALAKFSDRFDVVII | ASSLGTHNNQAPL | 0.30310556 |
| YLIGSVIV | ASSLGTHNNQAPL | 0.39165932 |
| YGQLVAVID | ASSLGTHNNQAPL | 0.38421416 |
| VTVGGMTLPA | ASSLGTHNNQAPL | 0.2231049 |
| VLALPAV | ASSLGTHNNQAPL | 0.4540501 |
| VIPPTLNYET | ASSLGTHNNQAPL | 0.37419873 |
| VGMGGMHPSSG | ASSLGTHNNQAPL | 0.69828224 |
| VGLAGGAATA | ASSLGTHNNQAPL | 0.23275904 |
| VGGAGGAGGSGP | ASSLGTHNNQAPL | 0.38770163 |
| VFAAFMV | ASSLGTHNNQAPL | 0.23899594 |
| TLYDKLR | ASSLGTHNNQAPL | 0.34620628 |
| TLAVFTVL | ASSLGTHNNQAPL | 0.22290587 |
| TGSVQPARV | ASSLGTHNNQAPL | 0.24836321 |
| TGGGAGSGGT | ASSLGTHNNQAPL | 0.2779257 |
| SHLETSL | ASSLGTHNNQAPL | 0.3837586 |
| SFTTGMT | ASSLGTHNNQAPL | 0.5458387 |
| QVEVAKRMGVRQPTVS | ASSLGTHNNQAPL | 0.59270215 |
| QGSGGPALAM | ASSLGTHNNQAPL | 0.32305858 |
| MLAPSNP | ASSLGTHNNQAPL | 0.5052502 |
| LVNNAAI | ASSLGTHNNQAPL | 0.45349044 |
| LTGEINTLA | ASSLGTHNNQAPL | 0.48157692 |
| LSIFGPV | ASSLGTHNNQAPL | 0.2661988 |
| LLALLANL | ASSLGTHNNQAPL | 0.25318873 |
| LISNASDALDKLR | ASSLGTHNNQAPL | 0.24448153 |
| LINEAVP | ASSLGTHNNQAPL | 0.2398926 |
| LGYAFSGL | ASSLGTHNNQAPL | 0.25368556 |
| LDFVAVAL | ASSLGTHNNQAPL | 0.3367179 |
| LAQPGAE | ASSLGTHNNQAPL | 0.211268 |

| | | |
|---|---|---|
| ISNLGAVTGF | ASSLGTHNNQAPL | 0.49631292 |
| HQEDPGRI | ASSLGTHNNQAPL | 0.5897957 |
| GTPGEPTP | ASSLGTHNNQAPL | 0.25792873 |
| GFFNSGS | ASSLGTHNNQAPL | 0.3474234 |
| GAGGGGLTAGAN | ASSLGTHNNQAPL | 0.3780695 |
| FVDALVNM | ASSLGTHNNQAPL | 0.23849632 |
| FLTVAPEL | ASSLGTHNNQAPL | 0.4266677 |
| FETLTAL | ASSLGTHNNQAPL | 0.23572913 |
| DVINPIEL | ASSLGTHNNQAPL | 0.57094514 |
| APVSGAPV | ASSLGTHNNQAPL | 0.40064332 |
| ALVGGLAW | ASSLGTHNNQAPL | 0.3124926 |
| ALAAAAATAI | ASSLGTHNNQAPL | 0.32740217 |
| AFGLGLLL | ASSLGTHNNQAPL | 0.31841555 |
| AFFAAASL | ASSLGTHNNQAPL | 0.22314592 |
| AAAALPAFSPP | ASSLGTHNNQAPL | 0.33621985 |
| VVLAPAL | ASSLGTHNNQAPL | 0.4749511 |
| TIGHVDHGKTTL | ASSLGTHNNQAPL | 0.3444486 |
| LISNASDALDK | ASSLGTHNNQAPL | 0.22830328 |
| LISNASDAL | ASSLGTHNNQAPL | 0.61479783 |
| ISNASDALDK | ASSLGTHNNQAPL | 0.30083835 |
| NQAETLVYQTE | ASSLGTHNNQAPL | 0.2692495 |
| TGGTGGTAGAG | ASSLGTHNNQAPL | 0.2435615 |
| TGSTAGGAGGA | ASSLGTHNNQAPL | 0.40634555 |
| DPLAVPVAAGPV | ASSLGTHNNQAPL | 0.2912346 |
| GGTGGTPGNGGN | ASSLGTHNNQAPL | 0.21827091 |
| PAGLAGVDPATL | ASSLGTHNNQAPL | 0.27075684 |
| GNNLVVEAPHAGASV | ASSLGTHNNQAPL | 0.2530104 |
| KVTRIMSASTLEQPA | ASSLGTHNNQAPL | 0.29122382 |
| LGFALPIALPRRTVV | ASSLGTHNNQAPL | 0.33444348 |
| TRLGNLTARKLVVLV | ASSLGTHNNQAPL | 0.48421046 |
| WSQQRRLIYFPSAGP | ASSLGTHNNQAPL | 0.21837308 |
| GPNGNPDPMAAAVDIR | ASSLGTHNNQAPL | 0.25399098 |
| TQRCVSVADLRAAASR | ASSLGTHNNQAPL | 0.3176862 |
| QFVLDVAEHPVEVLTETV | ASSLGTHNNQAPL | 0.25748914 |
| GGAGGTGGAGGNGGTQAGDG | ASSLGTHNNQAPL | 0.21623808 |
| PLLATLRAIAVDVDAKPAQV | ASSLGTHNNQAPL | 0.40397248 |
| APKFRTGGYMIDFWGVGYQVAK | ASSLGTHNNQAPL | 0.25191233 |
| GAGGAGGAGGAAGGPGGTGGTG | ASSLGTHNNQAPL | 0.2435211 |
| IGALGLLATDVAPAGSVAFGVA | ASSLGTHNNQAPL | 0.29456204 |
| AGGAGGIGGNGGAGGAGGVLMG | ASSLGTHNNQAPL | 0.6679734 |
| ADELIGLL | ASSLGTHNNQAPL | 0.30766737 |
| ADRALKA | ASSLGTHNNQAPL | 0.336293 |
| AGAVGQVGGAGGA | ASSLGTHNNQAPL | 0.57601386 |
| AGGGGGLGGAP | ASSLGTHNNQAPL | 0.27229187 |
| AGGLLYGNGGAG | ASSLGTHNNQAPL | 0.39193735 |
| AGLESIGL | ASSLGTHNNQAPL | 0.42097077 |
| AGLKTGII | ASSLGTHNNQAPL | 0.22421144 |
| ASLAALLL | ASSLGTHNNQAPL | 0.38492113 |
| ASVINISSI | ASSLGTHNNQAPL | 0.22829244 |
| ATPPGPA | ASSLGTHNNQAPL | 0.33291134 |
| ATVASALV | ASSLGTHNNQAPL | 0.24035162 |

| | | |
|---|---|---|
| ATVASLIL | ASSLGTHNNQAPL | 0.22174999 |
| AVNGPVLIRH | ASSLGTHNNQAPL | 0.3197014 |
| DGAAGAVGI | ASSLGTHNNQAPL | 0.33922896 |
| DPLLHGI | ASSLGTHNNQAPL | 0.3917036 |
| ELGAALGAL | ASSLGTHNNQAPL | 0.24578804 |
| ELGRTQAV | ASSLGTHNNQAPL | 0.22027889 |
| FVAGGYLS | ASSLGTHNNQAPL | 0.26217753 |
| GASGGAGGAGGW | ASSLGTHNNQAPL | 0.25198993 |
| GGAGAGGDGGAGGI | ASSLGTHNNQAPL | 0.6290691 |
| GTGHNGSQPGMGGNG | ASSLGTHNNQAPL | 0.22667053 |
| GYAAASASAS | ASSLGTHNNQAPL | 0.65463364 |
| HQVDVAAVQQG | ASSLGTHNNQAPL | 0.37881035 |
| IGAAIAHT | ASSLGTHNNQAPL | 0.256106 |
| IGMIEVAQL | ASSLGTHNNQAPL | 0.6305784 |
| IRLAPEHTSAMD | ASSLGTHNNQAPL | 0.24287926 |
| IYNVSVL | ASSLGTHNNQAPL | 0.24538215 |
| KQGFLGM | ASSLGTHNNQAPL | 0.3238902 |
| LESVITQL | ASSLGTHNNQAPL | 0.27473405 |
| LGLGALVVV | ASSLGTHNNQAPL | 0.28978932 |
| LIGNGGNGGSGGI | ASSLGTHNNQAPL | 0.34663576 |
| LNAINAPTL | ASSLGTHNNQAPL | 0.21838546 |
| LQLLNTGV | ASSLGTHNNQAPL | 0.33761045 |
| LQTSIEF | ASSLGTHNNQAPL | 0.2153087 |
| MSFAVSEL | ASSLGTHNNQAPL | 0.22417513 |
| NTGALGL | ASSLGTHNNQAPL | 0.25476116 |
| PPGAGGAGGAGTT | ASSLGTHNNQAPL | 0.39377236 |
| QLVGLGI | ASSLGTHNNQAPL | 0.50715744 |
| RFLEQVHQL | ASSLGTHNNQAPL | 0.24756262 |
| RGVGYVLR | ASSLGTHNNQAPL | 0.24362926 |
| SATGVFAGVI | ASSLGTHNNQAPL | 0.5936692 |
| SLPAITLP | ASSLGTHNNQAPL | 0.34758604 |
| STALAFTV | ASSLGTHNNQAPL | 0.34398013 |
| SVLTVGLGK | ASSLGTHNNQAPL | 0.3474639 |
| TALAGAGSGI | ASSLGTHNNQAPL | 0.2730788 |
| TGLPFTGI | ASSLGTHNNQAPL | 0.22992079 |
| VAEHVTV | ASSLGTHNNQAPL | 0.3711776 |
| VDIVLNSL | ASSLGTHNNQAPL | 0.23804471 |
| VLTGGGAVL | ASSLGTHNNQAPL | 0.30630085 |
| VNVGLAYGI | ASSLGTHNNQAPL | 0.21101943 |
| VSSAAVGAV | ASSLGTHNNQAPL | 0.29113254 |
| VTSVYEAGV | ASSLGTHNNQAPL | 0.23957746 |
| WVALDRGA | ASSLGTHNNQAPL | 0.2233769 |
| YLVGSVVI | ASSLGTHNNQAPL | 0.21241485 |
| GPPGTGKTLL | ASSLGTHNNQAPL | 0.23921195 |
| GVGMTKF | ASSLGTHNNQAPL | 0.25862125 |
| IGIGGSDL | ASSLGTHNNQAPL | 0.5779439 |
| IGTLLAEL | ASSLGTHNNQAPL | 0.40314826 |
| IYTSGTT | ASSLGTHNNQAPL | 0.46991676 |
| LSFPAGV | ASSLGTHNNQAPL | 0.25609887 |
| LYTSGTT | ASSLGTHNNQAPL | 0.28628886 |
| VGAFPFGF | ASSLGTHNNQAPL | 0.22935806 |

| | | |
|---|---|---|
| VGAGTVEF | ASSLGTHNNQAPL | 0.36266512 |
| VGKSTLLL | ASSLGTHNNQAPL | 0.23283078 |
| AALAHAGGGIV | ASSLGTHNNQAPL | 0.51762456 |
| AALVAVSLL | ASSLGTHNNQAPL | 0.26576045 |
| AAVSRAVAAL | ASSLGTHNNQAPL | 0.2559287 |
| AGGSSGIGGF | ASSLGTHNNQAPL | 0.347537 |
| AGLTVANANG | ASSLGTHNNQAPL | 0.36100358 |
| AGLVGAVL | ASSLGTHNNQAPL | 0.22099924 |
| ALSQAFTV | ASSLGTHNNQAPL | 0.22711502 |
| ALTFSVSL | ASSLGTHNNQAPL | 0.22405112 |
| ALTGAALVAL | ASSLGTHNNQAPL | 0.25951862 |
| ASFSIGAGVL | ASSLGTHNNQAPL | 0.2524353 |
| DGSVYKI | ASSLGTHNNQAPL | 0.2774351 |
| EQLAEIV | ASSLGTHNNQAPL | 0.2676656 |
| FAVSELTVM | ASSLGTHNNQAPL | 0.35741934 |
| GAAPLAAGL | ASSLGTHNNQAPL | 0.23807591 |
| GADGGDGGSGGAGG | ASSLGTHNNQAPL | 0.27420205 |
| GFGLDTGGAGG | ASSLGTHNNQAPL | 0.29994425 |
| GLASGGLL | ASSLGTHNNQAPL | 0.24358076 |
| GRIAGAAVLL | ASSLGTHNNQAPL | 0.43987325 |
| GSAGSLNTSF | ASSLGTHNNQAPL | 0.22843787 |
| GTGQIGI | ASSLGTHNNQAPL | 0.24152564 |
| GTLIALIL | ASSLGTHNNQAPL | 0.57190883 |
| IAATMGFVVG | ASSLGTHNNQAPL | 0.21942519 |
| IFAQALL | ASSLGTHNNQAPL | 0.2283921 |
| IGAAVALGF | ASSLGTHNNQAPL | 0.47284356 |
| IGLTPLAL | ASSLGTHNNQAPL | 0.24568997 |
| ILGSNTSTL | ASSLGTHNNQAPL | 0.4984269 |
| IVFSLTVL | ASSLGTHNNQAPL | 0.28153616 |
| LASKATGF | ASSLGTHNNQAPL | 0.37459576 |
| LASVALIL | ASSLGTHNNQAPL | 0.2468472 |
| LFAVGLTL | ASSLGTHNNQAPL | 0.6035311 |
| LFNGGIGTYI | ASSLGTHNNQAPL | 0.6088618 |
| LTLGAGII | ASSLGTHNNQAPL | 0.2274756 |
| LVAAALGDI | ASSLGTHNNQAPL | 0.56416523 |
| NTAFVQLGI | ASSLGTHNNQAPL | 0.21683559 |
| QLFSGAEL | ASSLGTHNNQAPL | 0.367241 |
| SAATFITL | ASSLGTHNNQAPL | 0.51295197 |
| SNYGLANLGV | ASSLGTHNNQAPL | 0.26686308 |
| SYGGAYAVM | ASSLGTHNNQAPL | 0.307033 |
| TTGVGGLTLG | ASSLGTHNNQAPL | 0.3627373 |
| TTQTIYI | ASSLGTHNNQAPL | 0.61621976 |
| VAAGAGLGAVI | ASSLGTHNNQAPL | 0.23391958 |
| VAAVVTLL | ASSLGTHNNQAPL | 0.35595578 |
| VGFLGGLI | ASSLGTHNNQAPL | 0.2744166 |
| VGVIPILL | ASSLGTHNNQAPL | 0.22127709 |
| VIFASSLI | ASSLGTHNNQAPL | 0.22536539 |
| VKGGDGGAAAG | ASSLGTHNNQAPL | 0.41065282 |
| VTKTFTNI | ASSLGTHNNQAPL | 0.30852106 |
| VVAAAILL | ASSLGTHNNQAPL | 0.52894366 |
| YAALDVEL | ASSLGTHNNQAPL | 0.384852 |

| | | |
|---|---|---|
| LASVALLL | ASSLGTHNNQAPL | 0.49662474 |
| LDAVHRV | ASSLGTHNNQAPL | 0.25743118 |
| QGLSSSIF | ASSLGTHNNQAPL | 0.28276163 |
| RGTFANIRL | ASSLGTHNNQAPL | 0.22005294 |
| RQLWWGHRI | ASSLGTHNNQAPL | 0.22337978 |
| YGPPGTGKTLL | ASSLGTHNNQAPL | 0.22252904 |
| AALVGAPANVA | ASSLGTHNNQAPL | 0.31147203 |
| AATFRRTE | ASSLGTHNNQAPL | 0.21257792 |
| AGALGGGI | ASSLGTHNNQAPL | 0.35161385 |
| AGGAGGAGGIGAG | ASSLGTHNNQAPL | 0.4320477 |
| AGSGALGL | ASSLGTHNNQAPL | 0.35773638 |
| AQIADQI | ASSLGTHNNQAPL | 0.23671691 |
| ASAVAVTYL | ASSLGTHNNQAPL | 0.25150698 |
| ASTVLGGTGG | ASSLGTHNNQAPL | 0.2792954 |
| AVGGEPAK | ASSLGTHNNQAPL | 0.53381884 |
| CEQGTFGGTC | ASSLGTHNNQAPL | 0.27602273 |
| DEGKAPAG | ASSLGTHNNQAPL | 0.21346004 |
| DTVVFPAL | ASSLGTHNNQAPL | 0.21768613 |
| FEGITMAG | ASSLGTHNNQAPL | 0.2244634 |
| FPLVVQD | ASSLGTHNNQAPL | 0.49337345 |
| GAFTTPQ | ASSLGTHNNQAPL | 0.36393526 |
| GFGGFGTAGGVGGA | ASSLGTHNNQAPL | 0.512407 |
| GFLAQFL | ASSLGTHNNQAPL | 0.5623576 |
| GGAGGAGGNAGVA | ASSLGTHNNQAPL | 0.21038498 |
| GGVGTAGAGGAGGAGG | ASSLGTHNNQAPL | 0.2331379 |
| GNGGGGDAGNAGSGGN | ASSLGTHNNQAPL | 0.48477352 |
| GTGGAGGAGNDAGS | ASSLGTHNNQAPL | 0.25473705 |
| GTTNTGWFNS | ASSLGTHNNQAPL | 0.34826297 |
| IFMGPLFTL | ASSLGTHNNQAPL | 0.43624026 |
| ILAHAIEAI | ASSLGTHNNQAPL | 0.24405561 |
| IRSAIREAL | ASSLGTHNNQAPL | 0.25279912 |
| ITFAGGG | ASSLGTHNNQAPL | 0.4956532 |
| ITGMSGHQEV | ASSLGTHNNQAPL | 0.37244838 |
| IVAGLACTL | ASSLGTHNNQAPL | 0.2415265 |
| IVGGGLAGA | ASSLGTHNNQAPL | 0.2346237 |
| IVTGVLNI | ASSLGTHNNQAPL | 0.3706118 |
| LAQQGCI | ASSLGTHNNQAPL | 0.6323279 |
| LCLHVEN | ASSLGTHNNQAPL | 0.5394981 |
| LDAISDGS | ASSLGTHNNQAPL | 0.26932085 |
| LGGFAAML | ASSLGTHNNQAPL | 0.31626993 |
| LGGKNPMI | ASSLGTHNNQAPL | 0.22753252 |
| LGPKGVAAIL | ASSLGTHNNQAPL | 0.28500414 |
| LMVGVSI | ASSLGTHNNQAPL | 0.29518482 |
| LVGAAFVL | ASSLGTHNNQAPL | 0.36064708 |
| LVLAVAPL | ASSLGTHNNQAPL | 0.22482447 |
| MAAIAVIDAA | ASSLGTHNNQAPL | 0.31399786 |
| NNGINAELQ | ASSLGTHNNQAPL | 0.26179835 |
| RGEPLVLVHG | ASSLGTHNNQAPL | 0.6109637 |
| RGGAGGAAGLI | ASSLGTHNNQAPL | 0.289733 |
| RSYRMHPAV | ASSLGTHNNQAPL | 0.41549933 |
| SAAHLLAGQLA | ASSLGTHNNQAPL | 0.26109275 |

| | | |
|---|---|---|
| SAAVGSY | ASSLGTHNNQAPL | 0.2484594 |
| SPHAVLHRI | ASSLGTHNNQAPL | 0.32003438 |
| SQIAKVL | ASSLGTHNNQAPL | 0.21877894 |
| TALGHLVP | ASSLGTHNNQAPL | 0.35606408 |
| TAPTTSGAA | ASSLGTHNNQAPL | 0.24127442 |
| TDGDDGNGGSPGT | ASSLGTHNNQAPL | 0.25976366 |
| TDGLGEN | ASSLGTHNNQAPL | 0.28622028 |
| TGGPPLPGQA | ASSLGTHNNQAPL | 0.6346656 |
| TIPAITI | ASSLGTHNNQAPL | 0.2813871 |
| TIPGFTI | ASSLGTHNNQAPL | 0.40411317 |
| TSIASGALA | ASSLGTHNNQAPL | 0.3059922 |
| TVTLPGAAI | ASSLGTHNNQAPL | 0.38642162 |
| VASGLSAALP | ASSLGTHNNQAPL | 0.6949916 |
| VGAGVIGI | ASSLGTHNNQAPL | 0.26064223 |
| VGPKRFGV | ASSLGTHNNQAPL | 0.68245083 |
| VNVAFPD | ASSLGTHNNQAPL | 0.27596572 |
| VSMTNSGVS | ASSLGTHNNQAPL | 0.21769528 |
| VTDVLGDLSV | ASSLGTHNNQAPL | 0.40012404 |
| VVVVGAGFAGL | ASSLGTHNNQAPL | 0.3233607 |
| VYTVGTVV | ASSLGTHNNQAPL | 0.567574 |
| WTTFSGP | ASSLGTHNNQAPL | 0.26816154 |
| YDELVEAF | ASSLGTHNNQAPL | 0.44393674 |
| YGTGGAGGNGG | ASSLGTHNNQAPL | 0.23901169 |
| YTNVMAV | ASSLGTHNNQAPL | 0.40998062 |
| FLEQVHQL | ASSLGTHNNQAPL | 0.3006573 |
| GLSLLAVAV | ASSLGTHNNQAPL | 0.450083 |
| LEQVHQL | ASSLGTHNNQAPL | 0.23321575 |
| LNAAGAI | ASSLGTHNNQAPL | 0.2300741 |
| PAVHASL | ASSLGTHNNQAPL | 0.2897792 |
| TELEDDA | ASSLGTHNNQAPL | 0.5126297 |
| TPPGPPN | ASSLGTHNNQAPL | 0.2390071 |
| TPPPPPPP | ASSLGTHNNQAPL | 0.28509295 |
| TQSPALI | ASGAGGSNTGQLY | 0.33866918 |
| EFAAAFPQ | ASGAGGSNTGQLY | 0.21905597 |
| LLSGNLDVL | ASGAGGSNTGQLY | 0.23406252 |
| GPANLGNYNI | ASGAGGSNTGQLY | 0.56594753 |
| TLGPTLP | ASGAGGSNTGQLY | 0.30256033 |
| VYAYSSCCK | ASGAGGSNTGQLY | 0.23036894 |
| APAPAPAPNI | ASGAGGSNTGQLY | 0.40554786 |
| PGGPVPIP | ASGAGGSNTGQLY | 0.6635881 |
| FLTVALEP | ASGAGGSNTGQLY | 0.56485295 |
| SQYFGANL | ASGAGGSNTGQLY | 0.21892822 |
| GGVKPAA | ASGAGGSNTGQLY | 0.20960714 |
| GGTGGAGGAGGW | ASGAGGSNTGQLY | 0.64134014 |
| GVGGVAGLGGAGG | ASGAGGSNTGQLY | 0.3828339 |
| VLLVLATTI | ASGAGGSNTGQLY | 0.2528147 |
| IGIFAAML | ASGAGGSNTGQLY | 0.30293906 |
| AVGDLID | ASGAGGSNTGQLY | 0.4407647 |
| NISPLPL | ASGAGGSNTGQLY | 0.36025298 |
| VTAVFVAI | ASGAGGSNTGQLY | 0.24309266 |
| SGDDNDAAL | ASGAGGSNTGQLY | 0.21357352 |

| | | |
|---|---|---|
| AVAALAHS | ASGAGGSNTGQLY | 0.25227064 |
| GGVTVMATP | ASGAGGSNTGQLY | 0.57047194 |
| AGLAANVRD | ASGAGGSNTGQLY | 0.23046254 |
| TNRPDLIDPALLRPG | ASGAGGSNTGQLY | 0.41369906 |
| LANLGVTAL | ASGAGGSNTGQLY | 0.4652514 |
| TGERIEL | ASGAGGSNTGQLY | 0.31505448 |
| NGEELAA | ASGAGGSNTGQLY | 0.2963401 |
| VLSLVWG | ASGAGGSNTGQLY | 0.22917542 |
| IGYALWL | ASGAGGSNTGQLY | 0.32535186 |
| LSLLLAPL | ASGAGGSNTGQLY | 0.23859304 |
| TLDEALK | ASGAGGSNTGQLY | 0.3022033 |
| TDGAGTAEGG | ASGAGGSNTGQLY | 0.50326526 |
| NAPGGAGGNGG | ASGAGGSNTGQLY | 0.3908344 |
| VSVFMGMGGNF | ASGAGGSNTGQLY | 0.30290788 |
| LASYLAAL | ASGAGGSNTGQLY | 0.22896923 |
| LVGGTPAL | ASGAGGSNTGQLY | 0.22322728 |
| VRNADVL | ASGAGGSNTGQLY | 0.2216763 |
| TCSSPFLVSAM | ASGAGGSNTGQLY | 0.58516765 |
| VHITDAE | ASGAGGSNTGQLY | 0.4154333 |
| EYAAPSF | ASGAGGSNTGQLY | 0.6314299 |
| VVDAFHGGL | ASGAGGSNTGQLY | 0.3331934 |
| ASVVDPQ | ASGAGGSNTGQLY | 0.31274307 |
| SYNEFGGLS | ASGAGGSNTGQLY | 0.3219676 |
| DPDGSVY | ASGAGGSNTGQLY | 0.25742278 |
| MVSTIHATE | ASGAGGSNTGQLY | 0.3611614 |
| LAAGVADPV | ASGAGGSNTGQLY | 0.22582775 |
| TKEQIAA | ASGAGGSNTGQLY | 0.48549163 |
| SLASPAAL | ASGAGGSNTGQLY | 0.26118812 |
| IGLVFYGV | ASGAGGSNTGQLY | 0.42108837 |
| FLLDASNL | ASGAGGSNTGQLY | 0.32413915 |
| YQVGNFP | ASGAGGSNTGQLY | 0.22311956 |
| ESLLSQST | ASGAGGSNTGQLY | 0.28419155 |
| LFTVGFAAVG | ASGAGGSNTGQLY | 0.44695956 |
| ATPPTPT | ASGAGGSNTGQLY | 0.37746996 |
| TDLATLG | ASGAGGSNTGQLY | 0.27712652 |
| SAGGTGR | ASGAGGSNTGQLY | 0.27919745 |
| GFSNSGS | ASGAGGSNTGQLY | 0.30883506 |
| GGNGGSGGTSV | ASGAGGSNTGQLY | 0.36081368 |
| GTSVATGGAGN | ASGAGGSNTGQLY | 0.49313253 |
| NGGNGGSGGTS | ASGAGGSNTGQLY | 0.24852467 |
| AVVKFPH | ASGAGGSNTGQLY | 0.44417298 |
| TPPPPPP | ASGAGGSNTGQLY | 0.21421553 |
| AAPAAAAAPA | ASGAGGSNTGQLY | 0.2373743 |
| IALNEDL | ASGAGGSNTGQLY | 0.25651124 |
| IVNVSSM | ASGAGGSNTGQLY | 0.32202184 |
| TGIPPAPR | ASGAGGSNTGQLY | 0.40609342 |
| FDLGGGTF | ASGAGGSNTGQLY | 0.23438074 |
| FDLGGGTFDVS | ASGAGGSNTGQLY | 0.3050043 |
| NEPTAAA | ASGAGGSNTGQLY | 0.245968 |
| NEPTAAALA | ASGAGGSNTGQLY | 0.24269886 |
| GSIINISSI | ASGAGGSNTGQLY | 0.36223644 |

| | | |
|---|---|---|
| VAPGNAGTA | ASGAGGSNTGQLY | 0.22237939 |
| TGGGASLE | ASGAGGSNTGQLY | 0.24484229 |
| VAVYNHYKRI | ASGAGGSNTGQLY | 0.41409433 |
| TDPAPATT | ASGAGGSNTGQLY | 0.33388254 |
| TNRPDILDPALLRPG | ASGAGGSNTGQLY | 0.24346547 |
| LTTTTTS | ASGAGGSNTGQLY | 0.3395775 |
| PTGSGKTL | ASGAGGSNTGQLY | 0.24902423 |
| TGSGKTL | ASGAGGSNTGQLY | 0.28714725 |
| TIGHVDHGK | ASGAGGSNTGQLY | 0.28296125 |
| TIGHVDHGKTT | ASGAGGSNTGQLY | 0.5257368 |
| GHVDHGKTT | ASGAGGSNTGQLY | 0.3098909 |
| AQFEHTLL | ASGAGGSNTGQLY | 0.22531554 |
| IVNQDEL | ASGAGGSNTGQLY | 0.33267218 |
| TNGSQFF | ASGAGGSNTGQLY | 0.32972044 |
| TGKGIVE | ASGAGGSNTGQLY | 0.321641 |
| LISNASDA | ASGAGGSNTGQLY | 0.42245984 |
| AELRTLGELTKIAKAHGAQ | ASGAGGSNTGQLY | 0.3416376 |
| APNAVAAAEPSW | ASGAGGSNTGQLY | 0.6055907 |
| APQVAGAATAVL | ASGAGGSNTGQLY | 0.2820037 |
| ARRDPAAAAAILPTDA | ASGAGGSNTGQLY | 0.22822338 |
| DAAARDVLAEAGLAEYF | ASGAGGSNTGQLY | 0.2858687 |
| DDQIVLSAKANLAKVEQ | ASGAGGSNTGQLY | 0.2996196 |
| DELLPDTGEDSDEDGH | ASGAGGSNTGQLY | 0.23192483 |
| ELPEAPANRATVPQD | ASGAGGSNTGQLY | 0.21349573 |
| FGQSKIERVTGIG | ASGAGGSNTGQLY | 0.41500264 |
| GAGGTGSTAGGAGGAGGAGGLY | ASGAGGSNTGQLY | 0.56116366 |
| GHGGTGAAGGNGGTGGAGGSGI | ASGAGGSNTGQLY | 0.25612277 |
| GNGGNGGAGGEGGPGGAGGAGA | ASGAGGSNTGQLY | 0.6087744 |
| GSTAGNGGNGGNATNGTIAGQP | ASGAGGSNTGQLY | 0.21354632 |
| GTGGDGGNGGNGGTGG | ASGAGGSNTGQLY | 0.27543217 |
| GVRRPAGIPADAPV | ASGAGGSNTGQLY | 0.3820647 |
| GYRQVTDAHLLTLARR | ASGAGGSNTGQLY | 0.23613748 |
| IAKMGQDGHDRG | ASGAGGSNTGQLY | 0.29489815 |
| KAEPSIEIQYT | ASGAGGSNTGQLY | 0.24665919 |
| KPVRIRMAADEDTVAVT | ASGAGGSNTGQLY | 0.23482613 |
| MPPRIAGMRLLVIKPEP | ASGAGGSNTGQLY | 0.4961729 |
| MKFYAVALFRSRGVPPTR | ASGAGGSNTGQLY | 0.26284274 |
| NAGGFGILGAVAHSP | ASGAGGSNTGQLY | 0.35331315 |
| NEGREITLAMAEM | ASGAGGSNTGQLY | 0.46351665 |
| NELVEWSHPLGHRWRWE | ASGAGGSNTGQLY | 0.27330068 |
| NLGAGGTGGNGGNPG | ASGAGGSNTGQLY | 0.4991297 |
| NTEWPKWETVMAADPVD | ASGAGGSNTGQLY | 0.4278509 |
| PASAAPAETQP | ASGAGGSNTGQLY | 0.31465825 |
| QQSSDGKNGVRAAEITGE | ASGAGGSNTGQLY | 0.28181908 |
| RAPDAVTTSWWKE | ASGAGGSNTGQLY | 0.2567153 |
| SGFGNVGAHTSGWFNQ | ASGAGGSNTGQLY | 0.44841427 |
| SGVAGAVGQAASVGGLKV | ASGAGGSNTGQLY | 0.6194733 |
| SNGGAGTGSAGGNGGT | ASGAGGSNTGQLY | 0.4262991 |
| TGGAGGEGGAAG | ASGAGGSNTGQLY | 0.29223028 |
| TGGDGGAAGAGGAPGHG | ASGAGGSNTGQLY | 0.55278194 |
| TMPPLLAAIANAGRHGV | ASGAGGSNTGQLY | 0.5160713 |

| | | |
|---|---|---|
| TSSTAGNSGGA | ASGAGGSNTGQLY | 0.45430952 |
| VPATVPVGILC | ASGAGGSNTGQLY | 0.4445415 |
| VQRPDSGGVVLTGRIS | ASGAGGSNTGQLY | 0.45130336 |
| WTPNPVPATKGVPAGLVQAT | ASGAGGSNTGQLY | 0.6624449 |
| YALKRALAKFSDRFDVVII | ASGAGGSNTGQLY | 0.6783276 |
| YLIGSVIV | ASGAGGSNTGQLY | 0.6236769 |
| YGQLVAVID | ASGAGGSNTGQLY | 0.5280288 |
| VTVGGMTLPA | ASGAGGSNTGQLY | 0.35556865 |
| VLALPAV | ASGAGGSNTGQLY | 0.5137756 |
| VIPPTLNYET | ASGAGGSNTGQLY | 0.46395257 |
| VGMGGMHPSSG | ASGAGGSNTGQLY | 0.68358725 |
| VGLAGGAATA | ASGAGGSNTGQLY | 0.68308437 |
| VGGAGGAGGSGP | ASGAGGSNTGQLY | 0.5009848 |
| VFAAFMV | ASGAGGSNTGQLY | 0.6769493 |
| TLYDKLR | ASGAGGSNTGQLY | 0.6267681 |
| TLAVFTVL | ASGAGGSNTGQLY | 0.48658234 |
| TGSVQPARV | ASGAGGSNTGQLY | 0.6397521 |
| TGGGAGSGGT | ASGAGGSNTGQLY | 0.4908262 |
| SHLETSL | ASGAGGSNTGQLY | 0.62523115 |
| SFTTGMT | ASGAGGSNTGQLY | 0.29062298 |
| QVEVAKRMGVRQPTVS | ASGAGGSNTGQLY | 0.6654452 |
| QGSGGPALAM | ASGAGGSNTGQLY | 0.6457189 |
| MLAPSNP | ASGAGGSNTGQLY | 0.29372242 |
| LVNNAAI | ASGAGGSNTGQLY | 0.67622185 |
| LTGEINTLA | ASGAGGSNTGQLY | 0.5391443 |
| LSIFGPV | ASGAGGSNTGQLY | 0.6643214 |
| LLALLANL | ASGAGGSNTGQLY | 0.69865936 |
| LISNASDALDKLR | ASGAGGSNTGQLY | 0.59927976 |
| LINEAVP | ASGAGGSNTGQLY | 0.39974073 |
| LGYAFSGL | ASGAGGSNTGQLY | 0.6351082 |
| LDFVAVAL | ASGAGGSNTGQLY | 0.69458663 |
| LAQPGAE | ASGAGGSNTGQLY | 0.38893533 |
| ISNLGAVTGF | ASGAGGSNTGQLY | 0.65309924 |
| HQEDPGRI | ASGAGGSNTGQLY | 0.673815 |
| GTPGEPTP | ASGAGGSNTGQLY | 0.6382652 |
| GFFNSGS | ASGAGGSNTGQLY | 0.69703496 |
| GAGGGGLTAGAN | ASGAGGSNTGQLY | 0.7033245 |
| FVDALVNM | ASGAGGSNTGQLY | 0.5553731 |
| FLTVAPEL | ASGAGGSNTGQLY | 0.6884223 |
| FETLTAL | ASGAGGSNTGQLY | 0.70007235 |
| DVINPIEL | ASGAGGSNTGQLY | 0.47350067 |
| APVSGAPV | ASGAGGSNTGQLY | 0.53816664 |
| ALVGGLAW | ASGAGGSNTGQLY | 0.43993556 |
| ALAAAAATAI | ASGAGGSNTGQLY | 0.619734 |
| AFGLGLLL | ASGAGGSNTGQLY | 0.47564638 |
| AFFAAASL | ASGAGGSNTGQLY | 0.6536551 |
| AAAALPAFSPP | ASGAGGSNTGQLY | 0.6908688 |
| VVLAPAL | ASGAGGSNTGQLY | 0.62017953 |
| TIGHVDHGKTTL | ASGAGGSNTGQLY | 0.462367 |
| LISNASDALDK | ASGAGGSNTGQLY | 0.34940886 |
| LISNASDAL | ASGAGGSNTGQLY | 0.5942143 |

| | | |
|---|---|---|
| ISNASDALDK | ASGAGGSNTGQLY | 0.51333326 |
| NQAETLVYQTE | ASGAGGSNTGQLY | 0.375074 |
| TGGTGGTAGAG | ASGAGGSNTGQLY | 0.6245911 |
| TGSTAGGAGGA | ASGAGGSNTGQLY | 0.6557051 |
| DPLAVPVAAGPV | ASGAGGSNTGQLY | 0.6703966 |
| GGTGGTPGNGGN | ASGAGGSNTGQLY | 0.41984677 |
| PAGLAGVDPATL | ASGAGGSNTGQLY | 0.7037021 |
| GNNLVVEAPHAGASV | ASGAGGSNTGQLY | 0.6653923 |
| KVTRIMSASTLEQPA | ASGAGGSNTGQLY | 0.6618104 |
| LGFALPIALPRRTVV | ASGAGGSNTGQLY | 0.56353617 |
| TRLGNLTARKLVVLV | ASGAGGSNTGQLY | 0.45326036 |
| WSQQRRLIYFPSAGP | ASGAGGSNTGQLY | 0.6892868 |
| GPNGNPDPMAAAVDIR | ASGAGGSNTGQLY | 0.5988333 |
| TQRCVSVADLRAAASR | ASGAGGSNTGQLY | 0.558583 |
| QFVLDVAEHPVEVLTETV | ASGAGGSNTGQLY | 0.6733086 |
| GGAGGTGGAGGNGGTQAGDG | ASGAGGSNTGQLY | 0.5973178 |
| PLLATLRAIAVDVDAKPAQV | ASGAGGSNTGQLY | 0.45450658 |
| APKFRTGGYMIDFWGVGYQVAK | ASGAGGSNTGQLY | 0.63082457 |
| GAGGAGGAGGAAGGPGGTGGTG | ASGAGGSNTGQLY | 0.6957052 |
| IGALGLLATDVAPAGSVAFGVA | ASGAGGSNTGQLY | 0.45787165 |
| AGGAGGIGGNGGAGGAGGVLMG | ASGAGGSNTGQLY | 0.52067447 |
| ADELIGLL | ASGAGGSNTGQLY | 0.52921903 |
| ADRALKA | ASGAGGSNTGQLY | 0.5938386 |
| AGAVGQVGGAGGA | ASGAGGSNTGQLY | 0.69019836 |
| AGGGGGLGGAP | ASGAGGSNTGQLY | 0.6508398 |
| AGGLLYGNGGAG | ASGAGGSNTGQLY | 0.58849967 |
| AGLESIGL | ASGAGGSNTGQLY | 0.6776108 |
| AGLKTGII | ASGAGGSNTGQLY | 0.6105038 |
| ASLAALLL | ASGAGGSNTGQLY | 0.57305175 |
| ASVINISSI | ASGAGGSNTGQLY | 0.6424013 |
| ATPPGPA | ASGAGGSNTGQLY | 0.48934132 |
| ATVASALV | ASGAGGSNTGQLY | 0.6863727 |
| ATVASLIL | ASGAGGSNTGQLY | 0.6373728 |
| AVNGPVLIRH | ASGAGGSNTGQLY | 0.67597497 |
| DGAAGAVGI | ASGAGGSNTGQLY | 0.6874602 |
| DPLLHGI | ASGAGGSNTGQLY | 0.5437388 |
| ELGAALGAL | ASGAGGSNTGQLY | 0.6152061 |
| ELGRTQAV | ASGAGGSNTGQLY | 0.37798744 |
| FVAGGYLS | ASGAGGSNTGQLY | 0.58066905 |
| GASGGAGGAGGW | ASGAGGSNTGQLY | 0.66623557 |
| GGAGAGGDGGAGGI | ASGAGGSNTGQLY | 0.67601866 |
| GTGHNGSQPGMGGNG | ASGAGGSNTGQLY | 0.68649703 |
| GYAAASASAS | ASGAGGSNTGQLY | 0.6377874 |
| HQVDVAAVQQG | ASGAGGSNTGQLY | 0.5745875 |
| IGAAIAHT | ASGAGGSNTGQLY | 0.6650944 |
| IGMIEVAQL | ASGAGGSNTGQLY | 0.68006295 |
| IRLAPEHTSAMD | ASGAGGSNTGQLY | 0.36666387 |
| IYNVSVL | ASGAGGSNTGQLY | 0.6166197 |
| KQGFLGM | ASGAGGSNTGQLY | 0.70196104 |
| LESVITQL | ASGAGGSNTGQLY | 0.6623337 |
| LGLGALVVV | ASGAGGSNTGQLY | 0.60482776 |

| | | |
|---|---|---|
| LIGNGGNGGSGGI | ASGAGGSNTGQLY | 0.5938269 |
| LNAINAPTL | ASGAGGSNTGQLY | 0.70127594 |
| LQLLNTGV | ASGAGGSNTGQLY | 0.6310359 |
| LQTSIEF | ASGAGGSNTGQLY | 0.69831926 |
| MSFAVSEL | ASGAGGSNTGQLY | 0.30564234 |
| NTGALGL | ASGAGGSNTGQLY | 0.63781756 |
| PPGAGGAGGAGTT | ASGAGGSNTGQLY | 0.4772258 |
| QLVGLGI | ASGAGGSNTGQLY | 0.35356262 |
| RFLEQVHQL | ASGAGGSNTGQLY | 0.68333036 |
| RGVGYVLR | ASGAGGSNTGQLY | 0.5334367 |
| SATGVFAGVI | ASGAGGSNTGQLY | 0.677366 |
| SLPAITLP | ASGAGGSNTGQLY | 0.6644366 |
| STALAFTV | ASGAGGSNTGQLY | 0.6774229 |
| SVLTVGLGK | ASGAGGSNTGQLY | 0.5576829 |
| TALAGAGSGI | ASGAGGSNTGQLY | 0.6072493 |
| TGLPFTGI | ASGAGGSNTGQLY | 0.57515347 |
| VAEHVTV | ASGAGGSNTGQLY | 0.6663345 |
| VDIVLNSL | ASGAGGSNTGQLY | 0.69182664 |
| VLTGGGAVL | ASGAGGSNTGQLY | 0.6908102 |
| VNVGLAYGI | ASGAGGSNTGQLY | 0.2940198 |
| VSSAAVGAV | ASGAGGSNTGQLY | 0.6565138 |
| VTSVYEAGV | ASGAGGSNTGQLY | 0.5865791 |
| WVALDRGA | ASGAGGSNTGQLY | 0.3869562 |
| YLVGSVVI | ASGAGGSNTGQLY | 0.32485425 |
| GPPGTGKTLL | ASGAGGSNTGQLY | 0.5911207 |
| GVGMTKF | ASGAGGSNTGQLY | 0.596817 |
| IGIGGSDL | ASGAGGSNTGQLY | 0.68387556 |
| IGTLLAEL | ASGAGGSNTGQLY | 0.6407238 |
| IYTSGTT | ASGAGGSNTGQLY | 0.55324614 |
| LSFPAGV | ASGAGGSNTGQLY | 0.65163434 |
| LYTSGTT | ASGAGGSNTGQLY | 0.5693871 |
| VGAFPFGF | ASGAGGSNTGQLY | 0.25341275 |
| VGAGTVEF | ASGAGGSNTGQLY | 0.46197963 |
| VGKSTLLL | ASGAGGSNTGQLY | 0.51837134 |
| AALAHAGGGIV | ASGAGGSNTGQLY | 0.68800527 |
| AALVAVSLL | ASGAGGSNTGQLY | 0.6126144 |
| AAVSRAVAAL | ASGAGGSNTGQLY | 0.56547886 |
| AGGSSGIGGF | ASGAGGSNTGQLY | 0.58887863 |
| AGLTVANANG | ASGAGGSNTGQLY | 0.66062284 |
| AGLVGAVL | ASGAGGSNTGQLY | 0.60324967 |
| ALSQAFTV | ASGAGGSNTGQLY | 0.3890414 |
| ALTFSVSL | ASGAGGSNTGQLY | 0.37945947 |
| ALTGAALVAL | ASGAGGSNTGQLY | 0.6395247 |
| ASFSIGAGVL | ASGAGGSNTGQLY | 0.47841394 |
| DGSVYKI | ASGAGGSNTGQLY | 0.48864937 |
| EQLAEIV | ASGAGGSNTGQLY | 0.38019973 |
| FAVSELTVM | ASGAGGSNTGQLY | 0.6625485 |
| GAAPLAAGL | ASGAGGSNTGQLY | 0.35454735 |
| GADGGDGGSGGAGG | ASGAGGSNTGQLY | 0.6926785 |
| GFGLDTGGAGG | ASGAGGSNTGQLY | 0.61668843 |
| GLASGGLL | ASGAGGSNTGQLY | 0.5761757 |

| | | |
|---|---|---|
| GRIAGAAVLL | ASGAGGSNTGQLY | 0.69184005 |
| GSAGSLNTSF | ASGAGGSNTGQLY | 0.6403705 |
| GTGQIGI | ASGAGGSNTGQLY | 0.5933213 |
| GTLIALIL | ASGAGGSNTGQLY | 0.4524666 |
| IAATMGFVVG | ASGAGGSNTGQLY | 0.54128337 |
| IFAQALL | ASGAGGSNTGQLY | 0.5167778 |
| IGAAVALGF | ASGAGGSNTGQLY | 0.502071 |
| IGLTPLAL | ASGAGGSNTGQLY | 0.6923972 |
| ILGSNTSTL | ASGAGGSNTGQLY | 0.66467 |
| IVFSLTVL | ASGAGGSNTGQLY | 0.34757015 |
| LASKATGF | ASGAGGSNTGQLY | 0.6553515 |
| LASVALIL | ASGAGGSNTGQLY | 0.69025433 |
| LFAVGLTL | ASGAGGSNTGQLY | 0.7043568 |
| LFNGGIGTYI | ASGAGGSNTGQLY | 0.6022878 |
| LTLGAGII | ASGAGGSNTGQLY | 0.4038158 |
| LVAAALGDI | ASGAGGSNTGQLY | 0.4985752 |
| NTAFVQLGI | ASGAGGSNTGQLY | 0.5990995 |
| QLFSGAEL | ASGAGGSNTGQLY | 0.5110972 |
| SAATFITL | ASGAGGSNTGQLY | 0.37581915 |
| SNYGLANLGV | ASGAGGSNTGQLY | 0.6859129 |
| SYGGAYAVM | ASGAGGSNTGQLY | 0.7103616 |
| TTGVGGLTLG | ASGAGGSNTGQLY | 0.54680115 |
| TTQTIYI | ASGAGGSNTGQLY | 0.3466748 |
| VAAGAGLGAVI | ASGAGGSNTGQLY | 0.546491 |
| VAAVVTLL | ASGAGGSNTGQLY | 0.4406017 |
| VGFLGGLI | ASGAGGSNTGQLY | 0.5081099 |
| VGVIPILL | ASGAGGSNTGQLY | 0.5556864 |
| VIFASSLI | ASGAGGSNTGQLY | 0.39245474 |
| VKGGDGGAAAG | ASGAGGSNTGQLY | 0.611245 |
| VTKTFTNI | ASGAGGSNTGQLY | 0.6895976 |
| VVAAAILL | ASGAGGSNTGQLY | 0.6264671 |
| YAALDVEL | ASGAGGSNTGQLY | 0.68550694 |
| LASVALLL | ASGAGGSNTGQLY | 0.70092654 |
| LDAVHRV | ASGAGGSNTGQLY | 0.6950635 |
| QGLSSSIF | ASGAGGSNTGQLY | 0.7049413 |
| RGTFANIRL | ASGAGGSNTGQLY | 0.6732255 |
| RQLWWGHRI | ASGAGGSNTGQLY | 0.5808608 |
| YGPPGTGKTLL | ASGAGGSNTGQLY | 0.5751237 |
| AALVGAPANVA | ASGAGGSNTGQLY | 0.6951548 |
| AATFRRTE | ASGAGGSNTGQLY | 0.5600789 |
| AGALGGGI | ASGAGGSNTGQLY | 0.63913226 |
| AGGAGGAGGIGAG | ASGAGGSNTGQLY | 0.5502672 |
| AGSGALGL | ASGAGGSNTGQLY | 0.68347645 |
| AQIADQI | ASGAGGSNTGQLY | 0.53373384 |
| ASAVAVTYL | ASGAGGSNTGQLY | 0.6977749 |
| ASTVLGGTGG | ASGAGGSNTGQLY | 0.37130108 |
| AVGGEPAK | ASGAGGSNTGQLY | 0.48655325 |
| CEQGTFGGTC | ASGAGGSNTGQLY | 0.69508386 |
| DEGKAPAG | ASGAGGSNTGQLY | 0.24469176 |
| DTVVFPAL | ASGAGGSNTGQLY | 0.4703784 |
| FEGITMAG | ASGAGGSNTGQLY | 0.6469418 |

| | | |
|---|---|---|
| FPLVVQD | ASGAGGSNTGQLY | 0.6787586 |
| GAFTTPQ | ASGAGGSNTGQLY | 0.28053585 |
| GFGGFGTAGGVGGA | ASGAGGSNTGQLY | 0.38538587 |
| GFLAQFL | ASGAGGSNTGQLY | 0.4815472 |
| GGAGGAGGNAGVA | ASGAGGSNTGQLY | 0.5937177 |
| GGVGTAGAGGAGGAGG | ASGAGGSNTGQLY | 0.5231006 |
| GNGGGGDAGNAGSGGN | ASGAGGSNTGQLY | 0.30608624 |
| GTGGAGGAGNDAGS | ASGAGGSNTGQLY | 0.6931593 |
| GTTNTGWFNS | ASGAGGSNTGQLY | 0.30579364 |
| IFMGPLFTL | ASGAGGSNTGQLY | 0.6192528 |
| ILAHAIEAI | ASGAGGSNTGQLY | 0.6765815 |
| IRSAIREAL | ASGAGGSNTGQLY | 0.70270896 |
| ITFAGGG | ASGAGGSNTGQLY | 0.58752453 |
| ITGMSGHQEV | ASGAGGSNTGQLY | 0.6525644 |
| IVAGLACTL | ASGAGGSNTGQLY | 0.5560064 |
| IVGGGLAGA | ASGAGGSNTGQLY | 0.7134548 |
| IVTGVLNI | ASGAGGSNTGQLY | 0.5707885 |
| LAQQGCI | ASGAGGSNTGQLY | 0.40930417 |
| LCLHVEN | ASGAGGSNTGQLY | 0.6890918 |
| LDAISDGS | ASGAGGSNTGQLY | 0.6835942 |
| LGGFAAML | ASGAGGSNTGQLY | 0.6021732 |
| LGGKNPMI | ASGAGGSNTGQLY | 0.68166125 |
| LGPKGVAAIL | ASGAGGSNTGQLY | 0.56929886 |
| LMVGVSI | ASGAGGSNTGQLY | 0.572131 |
| LVGAAFVL | ASGAGGSNTGQLY | 0.6836486 |
| LVLAVAPL | ASGAGGSNTGQLY | 0.6807137 |
| MAAIAVIDAA | ASGAGGSNTGQLY | 0.44069344 |
| NNGINAELQ | ASGAGGSNTGQLY | 0.6962583 |
| RGEPLVLVHG | ASGAGGSNTGQLY | 0.5953971 |
| RGGAGGAAGLI | ASGAGGSNTGQLY | 0.39188635 |
| RSYRMHPAV | ASGAGGSNTGQLY | 0.70409 |
| SAAHLLAGQLA | ASGAGGSNTGQLY | 0.605835 |
| SAAVGSY | ASGAGGSNTGQLY | 0.67979157 |
| SPHAVLHRI | ASGAGGSNTGQLY | 0.6711853 |
| SQIAKVL | ASGAGGSNTGQLY | 0.69378835 |
| TALGHLVP | ASGAGGSNTGQLY | 0.49536794 |
| TAPTTSGAA | ASGAGGSNTGQLY | 0.5797479 |
| TDGDDGNGGSPGT | ASGAGGSNTGQLY | 0.60985047 |
| TDGLGEN | ASGAGGSNTGQLY | 0.6664115 |
| TGGPPLPGQA | ASGAGGSNTGQLY | 0.5869418 |
| TIPAITI | ASGAGGSNTGQLY | 0.6320205 |
| TIPGFTI | ASGAGGSNTGQLY | 0.6817957 |
| TSIASGALA | ASGAGGSNTGQLY | 0.6924276 |
| TVTLPGAAI | ASGAGGSNTGQLY | 0.6134486 |
| VASGLSAALP | ASGAGGSNTGQLY | 0.5668682 |
| VGAGVIGI | ASGAGGSNTGQLY | 0.71042943 |
| VGPKRFGV | ASGAGGSNTGQLY | 0.5520779 |
| VNVAFPD | ASGAGGSNTGQLY | 0.57015586 |
| VSMTNSGVS | ASGAGGSNTGQLY | 0.68847704 |
| VTDVLGDLSV | ASGAGGSNTGQLY | 0.69123244 |
| VVVVGAGFAGL | ASGAGGSNTGQLY | 0.63317275 |

| | | |
|---|---|---|
| VYTVGTVV | ASGAGGSNTGQLY | 0.6964359 |
| WTTFSGP | ASGAGGSNTGQLY | 0.56047577 |
| YDELVEAF | ASGAGGSNTGQLY | 0.66849506 |
| YGTGGAGGNGG | ASGAGGSNTGQLY | 0.61679363 |
| YTNVMAV | ASGAGGSNTGQLY | 0.57494533 |
| FLEQVHQL | ASGAGGSNTGQLY | 0.5823951 |
| GLSLLAVAV | ASGAGGSNTGQLY | 0.6435865 |
| LEQVHQL | ASGAGGSNTGQLY | 0.6788813 |
| LNAAGAI | ASGAGGSNTGQLY | 0.65987265 |
| PAVHASL | ASGAGGSNTGQLY | 0.30355912 |
| TELEDDA | ASGAGGSNTGQLY | 0.6470935 |
| TPPGPPN | ASGAGGSNTGQLY | 0.6571146 |
| TPPPPPPP | ASGAGGSNTGQLY | 0.7150671 |
| TQSPALI | ASSTDISYNSPLY | 0.58619213 |
| EFAAAFPQ | ASSTDISYNSPLY | 0.655064 |
| LLSGNLDVL | ASSTDISYNSPLY | 0.5405889 |
| GPANLGNYNI | ASSTDISYNSPLY | 0.6491535 |
| TLGPTLP | ASSTDISYNSPLY | 0.4114801 |
| VYAYSSCCK | ASSTDISYNSPLY | 0.49917218 |
| APAPAPAPNI | ASSTDISYNSPLY | 0.53852457 |
| PGGPVPIP | ASSTDISYNSPLY | 0.5449474 |
| FLTVALEP | ASSTDISYNSPLY | 0.6832673 |
| SQYFGANL | ASSTDISYNSPLY | 0.6929391 |
| GGVKPAA | ASSTDISYNSPLY | 0.6353694 |
| GGTGGAGGAGGW | ASSTDISYNSPLY | 0.68884957 |
| GVGGVAGLGGAGG | ASSTDISYNSPLY | 0.69796133 |
| VLLVLATTI | ASSTDISYNSPLY | 0.70023847 |
| IGIFAAML | ASSTDISYNSPLY | 0.5560479 |
| AVGDLID | ASSTDISYNSPLY | 0.621729 |
| NISPLPL | ASSTDISYNSPLY | 0.4618672 |
| VTAVFVAI | ASSTDISYNSPLY | 0.52575135 |
| SGDDNDAAL | ASSTDISYNSPLY | 0.5747342 |
| AVAALAHS | ASSTDISYNSPLY | 0.60166633 |
| GGVTVMATP | ASSTDISYNSPLY | 0.2521912 |
| AGLAANVRD | ASSTDISYNSPLY | 0.65245765 |
| TNRPDLIDPALLRPG | ASSTDISYNSPLY | 0.7027433 |
| LANLGVTAL | ASSTDISYNSPLY | 0.59918034 |
| TGERIEL | ASSTDISYNSPLY | 0.6432451 |
| NGEELAA | ASSTDISYNSPLY | 0.6590469 |
| VLSLVWG | ASSTDISYNSPLY | 0.59383655 |
| IGYALWL | ASSTDISYNSPLY | 0.66724676 |
| LSLLLAPL | ASSTDISYNSPLY | 0.53790736 |
| TLDEALK | ASSTDISYNSPLY | 0.6957759 |
| TDGAGTAEGG | ASSTDISYNSPLY | 0.7055781 |
| NAPGGAGGNGG | ASSTDISYNSPLY | 0.63742054 |
| VSVFMGMGGNF | ASSTDISYNSPLY | 0.5924306 |
| LASYLAAL | ASSTDISYNSPLY | 0.60063505 |
| LVGGTPAL | ASSTDISYNSPLY | 0.38187706 |
| VRNADVL | ASSTDISYNSPLY | 0.58410627 |
| TCSSPFLVSAM | ASSTDISYNSPLY | 0.6043706 |
| VHITDAE | ASSTDISYNSPLY | 0.6352644 |

| | | |
|---|---|---|
| EYAAPSF | ASSTDISYNSPLY | 0.5006168 |
| VVDAFHGGL | ASSTDISYNSPLY | 0.6910557 |
| ASVVDPQ | ASSTDISYNSPLY | 0.5725776 |
| SYNEFGGLS | ASSTDISYNSPLY | 0.6092334 |
| DPDGSVY | ASSTDISYNSPLY | 0.50131726 |
| MVSTIHATE | ASSTDISYNSPLY | 0.662112 |
| LAAGVADPV | ASSTDISYNSPLY | 0.6332512 |
| TKEQIAA | ASSTDISYNSPLY | 0.353992 |
| SLASPAAL | ASSTDISYNSPLY | 0.6764123 |
| IGLVFYGV | ASSTDISYNSPLY | 0.60213065 |
| FLLDASNL | ASSTDISYNSPLY | 0.58157575 |
| YQVGNFP | ASSTDISYNSPLY | 0.6666496 |
| ESLLSQST | ASSTDISYNSPLY | 0.678637 |
| LFTVGFAAVG | ASSTDISYNSPLY | 0.34258717 |
| ATPPTPT | ASSTDISYNSPLY | 0.5985966 |
| TDLATLG | ASSTDISYNSPLY | 0.6515987 |
| SAGGTGR | ASSTDISYNSPLY | 0.5993692 |
| GFSNSGS | ASSTDISYNSPLY | 0.2884579 |
| GGNGGSGGTSV | ASSTDISYNSPLY | 0.6600109 |
| GTSVATGGAGN | ASSTDISYNSPLY | 0.39111343 |
| NGGNGGSGGTS | ASSTDISYNSPLY | 0.4135239 |
| AVVKFPH | ASSTDISYNSPLY | 0.5688992 |
| TPPPPPP | ASSTDISYNSPLY | 0.7002406 |
| AAPAAAAAPA | ASSTDISYNSPLY | 0.6314659 |
| IALNEDL | ASSTDISYNSPLY | 0.6384876 |
| IVNVSSM | ASSTDISYNSPLY | 0.6871685 |
| TGIPPAPR | ASSTDISYNSPLY | 0.5118149 |
| FDLGGGTF | ASSTDISYNSPLY | 0.68705547 |
| FDLGGGTFDVS | ASSTDISYNSPLY | 0.6594419 |
| NEPTAAA | ASSTDISYNSPLY | 0.5794629 |
| NEPTAAALA | ASSTDISYNSPLY | 0.63034153 |
| GSIINISSI | ASSTDISYNSPLY | 0.5079124 |
| VAPGNAGTA | ASSTDISYNSPLY | 0.6252154 |
| TGGGASLE | ASSTDISYNSPLY | 0.5299257 |
| VAVYNHYKRI | ASSTDISYNSPLY | 0.50611734 |
| TDPAPATT | ASSTDISYNSPLY | 0.5600655 |
| TNRPDILDPALLRPG | ASSTDISYNSPLY | 0.64931834 |
| LTTTTTS | ASSTDISYNSPLY | 0.6945169 |
| PTGSGKTL | ASSTDISYNSPLY | 0.6370766 |
| TGSGKTL | ASSTDISYNSPLY | 0.33518666 |
| TIGHVDHGK | ASSTDISYNSPLY | 0.62490547 |
| TIGHVDHGKTT | ASSTDISYNSPLY | 0.55052996 |
| GHVDHGKTT | ASSTDISYNSPLY | 0.69817996 |
| AQFEHTLL | ASSTDISYNSPLY | 0.65341043 |
| IVNQDEL | ASSTDISYNSPLY | 0.699418 |
| TNGSQFF | ASSTDISYNSPLY | 0.64189893 |
| TGKGIVE | ASSTDISYNSPLY | 0.50985014 |
| LISNASDA | ASSTDISYNSPLY | 0.6973939 |
| AELRTLGELTKIAKAHGAQ | ASSTDISYNSPLY | 0.51934266 |
| APNAVAAAEPSW | ASSTDISYNSPLY | 0.5612807 |
| APQVAGAATAVL | ASSTDISYNSPLY | 0.6551019 |

| | | |
|---|---|---|
| ARRDPAAAAAILPTDA | ASSTDISYNSPLY | 0.65970445 |
| DAAARDVLAEAGLAEYF | ASSTDISYNSPLY | 0.6061559 |
| DDQIVLSAKANLAKVEQ | ASSTDISYNSPLY | 0.62699735 |
| DELLPDTGEDSDEDGH | ASSTDISYNSPLY | 0.3445929 |
| ELPEAPANRATVPQD | ASSTDISYNSPLY | 0.6826339 |
| FGQSKIERVTGIG | ASSTDISYNSPLY | 0.35997868 |
| GAGGTGSTAGGAGGAGGAGGLY | ASSTDISYNSPLY | 0.42600518 |
| GHGGTGAAGGNGGTGGAGGSGI | ASSTDISYNSPLY | 0.5356263 |
| GNGGNGGAGGEGGPGGAGGAGA | ASSTDISYNSPLY | 0.70708203 |
| GSTAGNGGNGGNATNGTIAGQP | ASSTDISYNSPLY | 0.6926706 |
| GTGGDGGNGGNGGTGG | ASSTDISYNSPLY | 0.44700742 |
| GVRRPAGIPADAPV | ASSTDISYNSPLY | 0.5271652 |
| GYRQVTDAHLLTLARR | ASSTDISYNSPLY | 0.66074395 |
| IAKMGQDGHDRG | ASSTDISYNSPLY | 0.4956243 |
| KAEPSIEIQYT | ASSTDISYNSPLY | 0.68019986 |
| KPVRIRMAADEDTVAVT | ASSTDISYNSPLY | 0.5894848 |
| MPPRIAGMRLLVIKPEP | ASSTDISYNSPLY | 0.5429095 |
| MKFYAVALFRSRGVPPTR | ASSTDISYNSPLY | 0.44959944 |
| NAGGFGILGAVAHSP | ASSTDISYNSPLY | 0.6780946 |
| NEGREITLAMAEM | ASSTDISYNSPLY | 0.50703347 |
| NELVEWSHPLGHRWRWE | ASSTDISYNSPLY | 0.6607862 |
| NLGAGGTGGNGGNPG | ASSTDISYNSPLY | 0.31409165 |
| NTEWPKWETVMAADPVD | ASSTDISYNSPLY | 0.65042317 |
| PASAAPAETQP | ASSTDISYNSPLY | 0.5033239 |
| QQSSDGKNGVRAAEITGE | ASSTDISYNSPLY | 0.4219038 |
| RAPDAVTTSWWKE | ASSTDISYNSPLY | 0.37243882 |
| SGFGNVGAHTSGWFNQ | ASSTDISYNSPLY | 0.47613314 |
| SGVAGAVGQAASVGGLKV | ASSTDISYNSPLY | 0.670894 |
| SNGGAGTGSAGGNGGT | ASSTDISYNSPLY | 0.7075181 |
| TGGAGGEGGAAG | ASSTDISYNSPLY | 0.6493748 |
| TGGDGGAAGAGGAPGHG | ASSTDISYNSPLY | 0.6705505 |
| TMPPLLAAIANAGRHGV | ASSTDISYNSPLY | 0.66916203 |
| TSSTAGNSGGA | ASSTDISYNSPLY | 0.5859395 |
| VPATVPVGILC | ASSTDISYNSPLY | 0.47338986 |
| VQRPDSGGVVLTGRIS | ASSTDISYNSPLY | 0.629567 |
| WTPNPVPATKGVPAGLVQAT | ASSTDISYNSPLY | 0.5590439 |
| YALKRALAKFSDRFDVVII | ASSTDISYNSPLY | 0.6847192 |
| YLIGSVIV | ASSTDISYNSPLY | 0.645537 |
| YGQLVAVID | ASSTDISYNSPLY | 0.4377826 |
| VTVGGMTLPA | ASSTDISYNSPLY | 0.61849535 |
| VLALPAV | ASSTDISYNSPLY | 0.67752844 |
| VIPPTLNYET | ASSTDISYNSPLY | 0.41211104 |
| VGMGGMHPSSG | ASSTDISYNSPLY | 0.40855563 |
| VGLAGGAATA | ASSTDISYNSPLY | 0.3872408 |
| VGGAGGAGGSGP | ASSTDISYNSPLY | 0.498101 |
| VFAAFMV | ASSTDISYNSPLY | 0.5364357 |
| TLYDKLR | ASSTDISYNSPLY | 0.5780462 |
| TLAVFTVL | ASSTDISYNSPLY | 0.49101323 |
| TGSVQPARV | ASSTDISYNSPLY | 0.6045227 |
| TGGGAGSGGT | ASSTDISYNSPLY | 0.5162169 |
| SHLETSL | ASSTDISYNSPLY | 0.5046098 |

| | | |
|---|---|---|
| SFTTGMT | ASSTDISYNSPLY | 0.5240397 |
| QVEVAKRMGVRQPTVS | ASSTDISYNSPLY | 0.60450804 |
| QGSGGPALAM | ASSTDISYNSPLY | 0.68339634 |
| MLAPSNP | ASSTDISYNSPLY | 0.49820316 |
| LVNNAAI | ASSTDISYNSPLY | 0.523549 |
| LTGEINTLA | ASSTDISYNSPLY | 0.6968658 |
| LSIFGPV | ASSTDISYNSPLY | 0.3350334 |
| LLALLANL | ASSTDISYNSPLY | 0.32424575 |
| LISNASDALDKLR | ASSTDISYNSPLY | 0.67054105 |
| LINEAVP | ASSTDISYNSPLY | 0.57175934 |
| LGYAFSGL | ASSTDISYNSPLY | 0.69258344 |
| LDFVAVAL | ASSTDISYNSPLY | 0.57243955 |
| LAQPGAE | ASSTDISYNSPLY | 0.6301016 |
| ISNLGAVTGF | ASSTDISYNSPLY | 0.642877 |
| HQEDPGRI | ASSTDISYNSPLY | 0.70798266 |
| GTPGEPTP | ASSTDISYNSPLY | 0.6902298 |
| GFFNSGS | ASSTDISYNSPLY | 0.49269438 |
| GAGGGGLTAGAN | ASSTDISYNSPLY | 0.6928617 |
| FVDALVNM | ASSTDISYNSPLY | 0.3005381 |
| FLTVAPEL | ASSTDISYNSPLY | 0.6676544 |
| FETLTAL | ASSTDISYNSPLY | 0.7016378 |
| DVINPIEL | ASSTDISYNSPLY | 0.4129526 |
| APVSGAPV | ASSTDISYNSPLY | 0.566682 |
| ALVGGLAW | ASSTDISYNSPLY | 0.69667256 |
| ALAAAAATAI | ASSTDISYNSPLY | 0.70638454 |
| AFGLGLLL | ASSTDISYNSPLY | 0.4662057 |
| AFFAAASL | ASSTDISYNSPLY | 0.59797996 |
| AAAALPAFSPP | ASSTDISYNSPLY | 0.36987826 |
| VVLAPAL | ASSTDISYNSPLY | 0.40446034 |
| TIGHVDHGKTTL | ASSTDISYNSPLY | 0.2996206 |
| LISNASDALDK | ASSTDISYNSPLY | 0.67408407 |
| LISNASDAL | ASSTDISYNSPLY | 0.6071178 |
| ISNASDALDK | ASSTDISYNSPLY | 0.68101156 |
| NQAETLVYQTE | ASSTDISYNSPLY | 0.65002024 |
| TGGTGGTAGAG | ASSTDISYNSPLY | 0.69996333 |
| TGSTAGGAGGA | ASSTDISYNSPLY | 0.55259997 |
| DPLAVPVAAGPV | ASSTDISYNSPLY | 0.54766417 |
| GGTGGTPGNGGN | ASSTDISYNSPLY | 0.45832226 |
| PAGLAGVDPATL | ASSTDISYNSPLY | 0.40630364 |
| GNNLVVEAPHAGASV | ASSTDISYNSPLY | 0.5872896 |
| KVTRIMSASTLEQPA | ASSTDISYNSPLY | 0.59639406 |
| LGFALPIALPRRTVV | ASSTDISYNSPLY | 0.34906414 |
| TRLGNLTARKLVVLV | ASSTDISYNSPLY | 0.6657043 |
| WSQQRRLIYFPSAGP | ASSTDISYNSPLY | 0.6465909 |
| GPNGNPDPMAAAVDIR | ASSTDISYNSPLY | 0.6643169 |
| TQRCVSVADLRAAASR | ASSTDISYNSPLY | 0.40505463 |
| QFVLDVAEHPVEVLTETV | ASSTDISYNSPLY | 0.56649435 |
| GGAGGTGGAGGNGGTQAGDG | ASSTDISYNSPLY | 0.55883926 |
| PLLATLRAIAVDVDAKPAQV | ASSTDISYNSPLY | 0.68060225 |
| APKFRTGGYMIDFWGVGYQVAK | ASSTDISYNSPLY | 0.60176665 |
| GAGGAGGAGGAAGGPGGTGGTG | ASSTDISYNSPLY | 0.27670318 |

| | | |
|---|---|---|
| IGALGLLATDVAPAGSVAFGVA | ASSTDISYNSPLY | 0.43930888 |
| AGGAGGIGGNGGAGGAGGVLMG | ASSTDISYNSPLY | 0.3448038 |
| ADELIGLL | ASSTDISYNSPLY | 0.70408213 |
| ADRALKA | ASSTDISYNSPLY | 0.68277276 |
| AGAVGQVGGAGGA | ASSTDISYNSPLY | 0.68527246 |
| AGGGGGLGGAP | ASSTDISYNSPLY | 0.7091357 |
| AGGLLYGNGGAG | ASSTDISYNSPLY | 0.2571742 |
| AGLESIGL | ASSTDISYNSPLY | 0.4094593 |
| AGLKTGII | ASSTDISYNSPLY | 0.6906969 |
| ASLAALLL | ASSTDISYNSPLY | 0.6422897 |
| ASVINISSI | ASSTDISYNSPLY | 0.63160384 |
| ATPPGPA | ASSTDISYNSPLY | 0.69169176 |
| ATVASALV | ASSTDISYNSPLY | 0.42777103 |
| ATVASLIL | ASSTDISYNSPLY | 0.5992106 |
| AVNGPVLIRH | ASSTDISYNSPLY | 0.6614844 |
| DGAAGAVGI | ASSTDISYNSPLY | 0.6891614 |
| DPLLHGI | ASSTDISYNSPLY | 0.66874623 |
| ELGAALGAL | ASSTDISYNSPLY | 0.5129659 |
| ELGRTQAV | ASSTDISYNSPLY | 0.6486218 |
| FVAGGYLS | ASSTDISYNSPLY | 0.70861506 |
| GASGGAGGAGGW | ASSTDISYNSPLY | 0.6716873 |
| GGAGAGGDGGAGGI | ASSTDISYNSPLY | 0.64207375 |
| GTGHNGSQPGMGGNG | ASSTDISYNSPLY | 0.62364435 |
| GYAAASASAS | ASSTDISYNSPLY | 0.6057159 |
| HQVDVAAVQQG | ASSTDISYNSPLY | 0.5427023 |
| IGAAIAHT | ASSTDISYNSPLY | 0.55173063 |
| IGMIEVAQL | ASSTDISYNSPLY | 0.64966047 |
| IRLAPEHTSAMD | ASSTDISYNSPLY | 0.42611074 |
| IYNVSVL | ASSTDISYNSPLY | 0.6614585 |
| KQGFLGM | ASSTDISYNSPLY | 0.47471082 |
| LESVITQL | ASSTDISYNSPLY | 0.71249527 |
| LGLGALVVV | ASSTDISYNSPLY | 0.6529267 |
| LIGNGGNGGSGGI | ASSTDISYNSPLY | 0.66407824 |
| LNAINAPTL | ASSTDISYNSPLY | 0.48985237 |
| LQLLNTGV | ASSTDISYNSPLY | 0.62599343 |
| LQTSIEF | ASSTDISYNSPLY | 0.6522267 |
| MSFAVSEL | ASSTDISYNSPLY | 0.33315718 |
| NTGALGL | ASSTDISYNSPLY | 0.6741325 |
| PPGAGGAGGAGTT | ASSTDISYNSPLY | 0.45318425 |
| QLVGLGI | ASSTDISYNSPLY | 0.55807394 |
| RFLEQVHQL | ASSTDISYNSPLY | 0.6137604 |
| RGVGYVLR | ASSTDISYNSPLY | 0.6901634 |
| SATGVFAGVI | ASSTDISYNSPLY | 0.60681003 |
| SLPAITLP | ASSTDISYNSPLY | 0.62873095 |
| STALAFTV | ASSTDISYNSPLY | 0.5120859 |
| SVLTVGLGK | ASSTDISYNSPLY | 0.679327 |
| TALAGAGSGI | ASSTDISYNSPLY | 0.71439743 |
| TGLPFTGI | ASSTDISYNSPLY | 0.6582294 |
| VAEHVTV | ASSTDISYNSPLY | 0.7120857 |
| VDIVLNSL | ASSTDISYNSPLY | 0.6080301 |
| VLTGGGAVL | ASSTDISYNSPLY | 0.2804926 |

| | | |
|---|---|---|
| VNVGLAYGI | ASSTDISYNSPLY | 0.6335921 |
| VSSAAVGAV | ASSTDISYNSPLY | 0.67996186 |
| VTSVYEAGV | ASSTDISYNSPLY | 0.7045634 |
| WVALDRGA | ASSTDISYNSPLY | 0.67832357 |
| YLVGSVVI | ASSTDISYNSPLY | 0.67518693 |
| GPPGTGKTLL | ASSTDISYNSPLY | 0.52294636 |
| GVGMTKF | ASSTDISYNSPLY | 0.6874232 |
| IGIGGSDL | ASSTDISYNSPLY | 0.70432985 |
| IGTLLAEL | ASSTDISYNSPLY | 0.66703194 |
| IYTSGTT | ASSTDISYNSPLY | 0.48948145 |
| LSFPAGV | ASSTDISYNSPLY | 0.40758106 |
| LYTSGTT | ASSTDISYNSPLY | 0.6588509 |
| VGAFPFGF | ASSTDISYNSPLY | 0.6856297 |
| VGAGTVEF | ASSTDISYNSPLY | 0.5355737 |
| VGKSTLLL | ASSTDISYNSPLY | 0.63353294 |
| AALAHAGGGIV | ASSTDISYNSPLY | 0.61795354 |
| AALVAVSLL | ASSTDISYNSPLY | 0.2893911 |
| AAVSRAVAAL | ASSTDISYNSPLY | 0.59136003 |
| AGGSSGIGGF | ASSTDISYNSPLY | 0.6947098 |
| AGLTVANANG | ASSTDISYNSPLY | 0.6475523 |
| AGLVGAVL | ASSTDISYNSPLY | 0.5036046 |
| ALSQAFTV | ASSTDISYNSPLY | 0.6837494 |
| ALTFSVSL | ASSTDISYNSPLY | 0.7159356 |
| ALTGAALVAL | ASSTDISYNSPLY | 0.71300876 |
| ASFSIGAGVL | ASSTDISYNSPLY | 0.31095827 |
| DGSVYKI | ASSTDISYNSPLY | 0.253744 |
| EQLAEIV | ASSTDISYNSPLY | 0.6467705 |
| FAVSELTVM | ASSTDISYNSPLY | 0.6755452 |
| GAAPLAAGL | ASSTDISYNSPLY | 0.5243087 |
| GADGGDGGSGGAGG | ASSTDISYNSPLY | 0.5934373 |
| GFGLDTGGAGG | ASSTDISYNSPLY | 0.6734646 |
| GLASGGLL | ASSTDISYNSPLY | 0.6536226 |
| GRIAGAAVLL | ASSTDISYNSPLY | 0.47441828 |
| GSAGSLNTSF | ASSTDISYNSPLY | 0.40923658 |
| GTGQIGI | ASSTDISYNSPLY | 0.6434612 |
| GTLIALIL | ASSTDISYNSPLY | 0.7074965 |
| IAATMGFVVG | ASSTDISYNSPLY | 0.5382954 |
| IFAQALL | ASSTDISYNSPLY | 0.64508843 |
| IGAAVALGF | ASSTDISYNSPLY | 0.5825378 |
| IGLTPLAL | ASSTDISYNSPLY | 0.6082653 |
| ILGSNTSTL | ASSTDISYNSPLY | 0.60860825 |
| IVFSLTVL | ASSTDISYNSPLY | 0.34783447 |
| LASKATGF | ASSTDISYNSPLY | 0.6646031 |
| LASVALIL | ASSTDISYNSPLY | 0.39668703 |
| LFAVGLTL | ASSTDISYNSPLY | 0.68397766 |
| LFNGGIGTYI | ASSTDISYNSPLY | 0.66333246 |
| LTLGAGII | ASSTDISYNSPLY | 0.66911995 |
| LVAAALGDI | ASSTDISYNSPLY | 0.6104641 |
| NTAFVQLGI | ASSTDISYNSPLY | 0.4202085 |
| QLFSGAEL | ASSTDISYNSPLY | 0.4968729 |
| SAATFITL | ASSTDISYNSPLY | 0.44057202 |

| | | |
|---|---|---|
| SNYGLANLGV | ASSTDISYNSPLY | 0.6907174 |
| SYGGAYAVM | ASSTDISYNSPLY | 0.6470514 |
| TTGVGGLTLG | ASSTDISYNSPLY | 0.7067218 |
| TTQTIYI | ASSTDISYNSPLY | 0.6666504 |
| VAAGAGLGAVI | ASSTDISYNSPLY | 0.56115353 |
| VAAVVTLL | ASSTDISYNSPLY | 0.6692975 |
| VGFLGGLI | ASSTDISYNSPLY | 0.53490704 |
| VGVIPILL | ASSTDISYNSPLY | 0.6267345 |
| VIFASSLI | ASSTDISYNSPLY | 0.57384694 |
| VKGGDGGAAAG | ASSTDISYNSPLY | 0.6666359 |
| VTKTFTNI | ASSTDISYNSPLY | 0.5071808 |
| VVAAAILL | ASSTDISYNSPLY | 0.68754315 |
| YAALDVEL | ASSTDISYNSPLY | 0.64754415 |
| LASVALLL | ASSTDISYNSPLY | 0.54112625 |
| LDAVHRV | ASSTDISYNSPLY | 0.62914574 |
| QGLSSSIF | ASSTDISYNSPLY | 0.669405 |
| RGTFANIRL | ASSTDISYNSPLY | 0.65391064 |
| RQLWWGHRI | ASSTDISYNSPLY | 0.6132755 |
| YGPPGTGKTLL | ASSTDISYNSPLY | 0.5861148 |
| AALVGAPANVA | ASSTDISYNSPLY | 0.558059 |
| AATFRRTE | ASSTDISYNSPLY | 0.63500655 |
| AGALGGGI | ASSTDISYNSPLY | 0.66061205 |
| AGGAGGAGGIGAG | ASSTDISYNSPLY | 0.5055453 |
| AGSGALGL | ASSTDISYNSPLY | 0.68207026 |
| AQIADQI | ASSTDISYNSPLY | 0.3402312 |
| ASAVAVTYL | ASSTDISYNSPLY | 0.540907 |
| ASTVLGGTGG | ASSTDISYNSPLY | 0.6084211 |
| AVGGEPAK | ASSTDISYNSPLY | 0.5963098 |
| CEQGTFGGTC | ASSTDISYNSPLY | 0.65917754 |
| DEGKAPAG | ASSTDISYNSPLY | 0.41762096 |
| DTVVFPAL | ASSTDISYNSPLY | 0.6299254 |
| FEGITMAG | ASSTDISYNSPLY | 0.6093509 |
| FPLVVQD | ASSTDISYNSPLY | 0.48610517 |
| GAFTTPQ | ASSTDISYNSPLY | 0.68013227 |
| GFGGFGTAGGVGGA | ASSTDISYNSPLY | 0.5681397 |
| GFLAQFL | ASSTDISYNSPLY | 0.5134288 |
| GGAGGAGGNAGVA | ASSTDISYNSPLY | 0.6725638 |
| GGVGTAGAGGAGGAGG | ASSTDISYNSPLY | 0.60408604 |
| GNGGGGDAGNAGSGGN | ASSTDISYNSPLY | 0.5434642 |
| GTGGAGGAGNDAGS | ASSTDISYNSPLY | 0.665262 |
| GTTNTGWFNS | ASSTDISYNSPLY | 0.5372399 |
| IFMGPLFTL | ASSTDISYNSPLY | 0.6762748 |
| ILAHAIEAI | ASSTDISYNSPLY | 0.5925834 |
| IRSAIREAL | ASSTDISYNSPLY | 0.6766839 |
| ITFAGGG | ASSTDISYNSPLY | 0.60658133 |
| ITGMSGHQEV | ASSTDISYNSPLY | 0.5412685 |
| IVAGLACTL | ASSTDISYNSPLY | 0.6806189 |
| IVGGGLAGA | ASSTDISYNSPLY | 0.60888344 |
| IVTGVLNI | ASSTDISYNSPLY | 0.62450266 |
| LAQQGCI | ASSTDISYNSPLY | 0.64258784 |
| LCLHVEN | ASSTDISYNSPLY | 0.66857976 |

| | | |
|---|---|---|
| LDAISDGS | ASSTDISYNSPLY | 0.6856446 |
| LGGFAAML | ASSTDISYNSPLY | 0.5023947 |
| LGGKNPMI | ASSTDISYNSPLY | 0.41528845 |
| LGPKGVAAIL | ASSTDISYNSPLY | 0.55890846 |
| LMVGVSI | ASSTDISYNSPLY | 0.5575904 |
| LVGAAFVL | ASSTDISYNSPLY | 0.46164095 |
| LVLAVAPL | ASSTDISYNSPLY | 0.2977794 |
| MAAIAVIDAA | ASSTDISYNSPLY | 0.6650919 |
| NNGINAELQ | ASSTDISYNSPLY | 0.71386075 |
| RGEPLVLVHG | ASSTDISYNSPLY | 0.44478798 |
| RGGAGGAAGLI | ASSTDISYNSPLY | 0.7228622 |
| RSYRMHPAV | ASSTDISYNSPLY | 0.40729597 |
| SAAHLLAGQLA | ASSTDISYNSPLY | 0.6213919 |
| SAAVGSY | ASSTDISYNSPLY | 0.6790736 |
| SPHAVLHRI | ASSTDISYNSPLY | 0.42542505 |
| SQIAKVL | ASSTDISYNSPLY | 0.6065928 |
| TALGHLVP | ASSTDISYNSPLY | 0.6269055 |
| TAPTTSGAA | ASSTDISYNSPLY | 0.5633465 |
| TDGDDGNGGSPGT | ASSTDISYNSPLY | 0.7019561 |
| TDGLGEN | ASSTDISYNSPLY | 0.5037636 |
| TGGPPLPGQA | ASSTDISYNSPLY | 0.63432443 |
| TIPAITI | ASSTDISYNSPLY | 0.66350853 |
| TIPGFTI | ASSTDISYNSPLY | 0.557963 |
| TSIASGALA | ASSTDISYNSPLY | 0.67838645 |
| TVTLPGAAI | ASSTDISYNSPLY | 0.65747404 |
| VASGLSAALP | ASSTDISYNSPLY | 0.6127535 |
| VGAGVIGI | ASSTDISYNSPLY | 0.6978102 |
| VGPKRFGV | ASSTDISYNSPLY | 0.49901795 |
| VNVAFPD | ASSTDISYNSPLY | 0.69372547 |
| VSMTNSGVS | ASSTDISYNSPLY | 0.69582665 |
| VTDVLGDLSV | ASSTDISYNSPLY | 0.64027506 |
| VVVVGAGFAGL | ASSTDISYNSPLY | 0.59914756 |
| VYTVGTVV | ASSTDISYNSPLY | 0.6846303 |
| WTTFSGP | ASSTDISYNSPLY | 0.6767179 |
| YDELVEAF | ASSTDISYNSPLY | 0.618483 |
| YGTGGAGGNGG | ASSTDISYNSPLY | 0.69968617 |
| YTNVMAV | ASSTDISYNSPLY | 0.1497146 |
| FLEQVHQL | ASSTDISYNSPLY | 0.1851322 |
| GLSLLAVAV | ASSTDISYNSPLY | 0.16242512 |
| LEQVHQL | ASSTDISYNSPLY | 0.16590229 |
| LNAAGAI | ASSTDISYNSPLY | 0.16501756 |
| PAVHASL | ASSTDISYNSPLY | 0.15897185 |
| TELEDDA | ASSTDISYNSPLY | 0.16000383 |
| TPPGPPN | ASSTDISYNSPLY | 0.1575701 |
| TPPPPPPP | ASSTDISYNSPLY | 0.16511574 |
| TQSPALI | ASSSTGGSGNTLY | 0.15597524 |
| EFAAAFPQ | ASSSTGGSGNTLY | 0.16536362 |
| LLSGNLDVL | ASSSTGGSGNTLY | 0.20012051 |
| GPANLGNYNI | ASSSTGGSGNTLY | 0.1685251 |
| TLGPTLP | ASSSTGGSGNTLY | 0.15127523 |
| VYAYSSCCK | ASSSTGGSGNTLY | 0.15544422 |

| | | |
|---|---|---|
| APAPAPAPNI | ASSSTGGSGNTLY | 0.18743217 |
| PGGPVPIP | ASSSTGGSGNTLY | 0.15689962 |
| FLTVALEP | ASSSTGGSGNTLY | 0.1479966 |
| SQYFGANL | ASSSTGGSGNTLY | 0.16798592 |
| GGVKPAA | ASSSTGGSGNTLY | 0.14983252 |
| GGTGGAGGAGGW | ASSSTGGSGNTLY | 0.14806525 |
| GVGGVAGLGGAGG | ASSSTGGSGNTLY | 0.17071709 |
| VLLVLATTI | ASSSTGGSGNTLY | 0.1618407 |
| IGIFAAML | ASSSTGGSGNTLY | 0.15256797 |
| AVGDLID | ASSSTGGSGNTLY | 0.17026514 |
| NISPLPL | ASSSTGGSGNTLY | 0.18144879 |
| VTAVFVAI | ASSSTGGSGNTLY | 0.16357076 |
| SGDDNDAAL | ASSSTGGSGNTLY | 0.17737956 |
| AVAALAHS | ASSSTGGSGNTLY | 0.17459154 |
| GGVTVMATP | ASSSTGGSGNTLY | 0.16153997 |
| AGLAANVRD | ASSSTGGSGNTLY | 0.15208332 |
| TNRPDLIDPALLRPG | ASSSTGGSGNTLY | 0.19741972 |
| LANLGVTAL | ASSSTGGSGNTLY | 0.17522487 |
| TGERIEL | ASSSTGGSGNTLY | 0.16627455 |
| NGEELAA | ASSSTGGSGNTLY | 0.1691571 |
| VLSLVWG | ASSSTGGSGNTLY | 0.15583804 |
| IGYALWL | ASSSTGGSGNTLY | 0.16361335 |
| LSLLLAPL | ASSSTGGSGNTLY | 0.19065146 |
| TLDEALK | ASSSTGGSGNTLY | 0.16951723 |
| TDGAGTAEGG | ASSSTGGSGNTLY | 0.16794635 |
| NAPGGAGGNGG | ASSSTGGSGNTLY | 0.1525209 |
| VSVFMGMGGNF | ASSSTGGSGNTLY | 0.16075009 |
| LASYLAAL | ASSSTGGSGNTLY | 0.15261424 |
| LVGGTPAL | ASSSTGGSGNTLY | 0.16067459 |
| VRNADVL | ASSSTGGSGNTLY | 0.18563655 |
| TCSSPFLVSAM | ASSSTGGSGNTLY | 0.16423471 |
| VHITDAE | ASSSTGGSGNTLY | 0.15061472 |
| EYAAPSF | ASSSTGGSGNTLY | 0.15375616 |
| VVDAFHGGL | ASSSTGGSGNTLY | 0.15189567 |
| ASVVDPQ | ASSSTGGSGNTLY | 0.14980485 |
| SYNEFGGLS | ASSSTGGSGNTLY | 0.1584313 |
| DPDGSVY | ASSSTGGSGNTLY | 0.15161674 |
| MVSTIHATE | ASSSTGGSGNTLY | 0.19699024 |
| LAAGVADPV | ASSSTGGSGNTLY | 0.19466574 |
| TKEQIAA | ASSSTGGSGNTLY | 0.16400285 |
| SLASPAAL | ASSSTGGSGNTLY | 0.1674283 |
| IGLVFYGV | ASSSTGGSGNTLY | 0.16501534 |
| FLLDASNL | ASSSTGGSGNTLY | 0.21364063 |
| YQVGNFP | ASSSTGGSGNTLY | 0.176864 |
| ESLLSQST | ASSSTGGSGNTLY | 0.14972216 |
| LFTVGFAAVG | ASSSTGGSGNTLY | 0.20073432 |
| ATPPTPT | ASSSTGGSGNTLY | 0.16259338 |
| TDLATLG | ASSSTGGSGNTLY | 0.18748668 |
| SAGGTGR | ASSSTGGSGNTLY | 0.18449871 |
| GFSNSGS | ASSSTGGSGNTLY | 0.18340465 |
| GGNGGSGGTSV | ASSSTGGSGNTLY | 0.15638988 |

| | | |
|---|---|---|
| GTSVATGGAGN | ASSSTGGSGNTLY | 0.15895012 |
| NGGNGGSGGTS | ASSSTGGSGNTLY | 0.17735143 |
| AVVKFPH | ASSSTGGSGNTLY | 0.15293543 |
| TPPPPPP | ASSSTGGSGNTLY | 0.15887599 |
| AAPAAAAAPA | ASSSTGGSGNTLY | 0.16874126 |
| IALNEDL | ASSSTGGSGNTLY | 0.17680506 |
| IVNVSSM | ASSSTGGSGNTLY | 0.17717144 |
| TGIPPAPR | ASSSTGGSGNTLY | 0.17133643 |
| FDLGGGTF | ASSSTGGSGNTLY | 0.15358639 |
| FDLGGGTFDVS | ASSSTGGSGNTLY | 0.16341084 |
| NEPTAAA | ASSSTGGSGNTLY | 0.1714886 |
| NEPTAAALA | ASSSTGGSGNTLY | 0.1718827 |
| GSIINISSI | ASSSTGGSGNTLY | 0.15744555 |
| VAPGNAGTA | ASSSTGGSGNTLY | 0.15224364 |
| TGGGASLE | ASSSTGGSGNTLY | 0.18094836 |
| VAVYNHYKRI | ASSSTGGSGNTLY | 0.19353828 |
| TDPAPATT | ASSSTGGSGNTLY | 0.16284145 |
| TNRPDILDPALLRPG | ASSSTGGSGNTLY | 0.16240358 |
| LTTTTTS | ASSSTGGSGNTLY | 0.17151016 |
| PTGSGKTL | ASSSTGGSGNTLY | 0.17502545 |
| TGSGKTL | ASSSTGGSGNTLY | 0.16873305 |
| TIGHVDHGK | ASSSTGGSGNTLY | 0.16263914 |
| TIGHVDHGKTT | ASSSTGGSGNTLY | 0.15350102 |
| GHVDHGKTT | ASSSTGGSGNTLY | 0.16078347 |
| AQFEHTLL | ASSSTGGSGNTLY | 0.15511517 |
| IVNQDEL | ASSSTGGSGNTLY | 0.16853128 |
| TNGSQFF | ASSSTGGSGNTLY | 0.15864098 |
| TGKGIVE | ASSSTGGSGNTLY | 0.16734098 |
| LISNASDA | ASSSTGGSGNTLY | 0.18642814 |
| AELRTLGELTKIAKAHGAQ | ASSSTGGSGNTLY | 0.15237196 |
| APNAVAAAEPSW | ASSSTGGSGNTLY | 0.15426141 |
| APQVAGAATAVL | ASSSTGGSGNTLY | 0.1675075 |
| ARRDPAAAAAILPTDA | ASSSTGGSGNTLY | 0.16650367 |
| DAAARDVLAEAGLAEYF | ASSSTGGSGNTLY | 0.18067104 |
| DDQIVLSAKANLAKVEQ | ASSSTGGSGNTLY | 0.1547316 |
| DELLPDTGEDSDEDGH | ASSSTGGSGNTLY | 0.18467137 |
| ELPEAPANRATVPQD | ASSSTGGSGNTLY | 0.16095844 |
| FGQSKIERVTGIG | ASSSTGGSGNTLY | 0.19146186 |
| GAGGTGSTAGGAGGAGGAGGLY | ASSSTGGSGNTLY | 0.16173328 |
| GHGGTGAAGGNGGTGGAGGSGI | ASSSTGGSGNTLY | 0.15269388 |
| GNGGNGGAGGEGGPGGAGGAGA | ASSSTGGSGNTLY | 0.16179255 |
| GSTAGNGGNGGNATNGTIAGQP | ASSSTGGSGNTLY | 0.14644112 |
| GTGGDGGNGGNGGTGG | ASSSTGGSGNTLY | 0.19158795 |
| GVRRPAGIPADAPV | ASSSTGGSGNTLY | 0.15129508 |
| GYRQVTDAHLLTLARR | ASSSTGGSGNTLY | 0.16051811 |
| IAKMGQDGHDRG | ASSSTGGSGNTLY | 0.16334093 |
| KAEPSIEIQYT | ASSSTGGSGNTLY | 0.1620196 |
| KPVRIRMAADEDTVAVT | ASSSTGGSGNTLY | 0.16623726 |
| MPPRIAGMRLLVIKPEP | ASSSTGGSGNTLY | 0.15684785 |
| MKFYAVALFRSRGVPPTR | ASSSTGGSGNTLY | 0.15346646 |
| NAGGFGILGAVAHSP | ASSSTGGSGNTLY | 0.18300314 |

| | | |
|---|---|---|
| NEGREITLAMAEM | ASSSTGGSGNTLY | 0.1666978 |
| NELVEWSHPLGHRWRWE | ASSSTGGSGNTLY | 0.17183995 |
| NLGAGGTGGNGGNPG | ASSSTGGSGNTLY | 0.15485701 |
| NTEWPKWETVMAADPVD | ASSSTGGSGNTLY | 0.18045692 |
| PASAAPAETQP | ASSSTGGSGNTLY | 0.16557242 |
| QQSSDGKNGVRAAEITGE | ASSSTGGSGNTLY | 0.1512876 |
| RAPDAVTTSWWKE | ASSSTGGSGNTLY | 0.15535642 |
| SGFGNVGAHTSGWFNQ | ASSSTGGSGNTLY | 0.15104626 |
| SGVAGAVGQAASVGGLKV | ASSSTGGSGNTLY | 0.1527436 |
| SNGGAGTGSAGGNGGT | ASSSTGGSGNTLY | 0.18716171 |
| TGGAGGEGGAAG | ASSSTGGSGNTLY | 0.19132379 |
| TGGDGGAAGAGGAPGHG | ASSSTGGSGNTLY | 0.16511025 |
| TMPPLLAAIANAGRHGV | ASSSTGGSGNTLY | 0.17797458 |
| TSSTAGNSGGA | ASSSTGGSGNTLY | 0.16194019 |
| VPATVPVGILC | ASSSTGGSGNTLY | 0.14588933 |
| VQRPDSGGVVLTGRIS | ASSSTGGSGNTLY | 0.15725279 |
| WTPNPVPATKGVPAGLVQAT | ASSSTGGSGNTLY | 0.15239704 |
| YALKRALAKFSDRFDVVII | ASSSTGGSGNTLY | 0.16851085 |
| YLIGSVIV | ASSSTGGSGNTLY | 0.1479439 |
| YGQLVAVID | ASSSTGGSGNTLY | 0.16046052 |
| VTVGGMTLPA | ASSSTGGSGNTLY | 0.17138846 |
| VLALPAV | ASSSTGGSGNTLY | 0.15637156 |
| VIPPTLNYET | ASSSTGGSGNTLY | 0.17201816 |
| VGMGGMHPSSG | ASSSTGGSGNTLY | 0.15379046 |
| VGLAGGAATA | ASSSTGGSGNTLY | 0.16754104 |
| VGGAGGAGGSGP | ASSSTGGSGNTLY | 0.18473715 |
| VFAAFMV | ASSSTGGSGNTLY | 0.15847525 |
| TLYDKLR | ASSSTGGSGNTLY | 0.15840364 |
| TLAVFTVL | ASSSTGGSGNTLY | 0.17013034 |
| TGSVQPARV | ASSSTGGSGNTLY | 0.2506979 |
| TGGGAGSGGT | ASSSTGGSGNTLY | 0.15525709 |
| SHLETSL | ASSSTGGSGNTLY | 0.1651446 |
| SFTTGMT | ASSSTGGSGNTLY | 0.15483975 |
| QVEVAKRMGVRQPTVS | ASSSTGGSGNTLY | 0.17349471 |
| QGSGGPALAM | ASSSTGGSGNTLY | 0.21669556 |
| MLAPSNP | ASSSTGGSGNTLY | 0.15886417 |
| LVNNAAI | ASSSTGGSGNTLY | 0.16855957 |
| LTGEINTLA | ASSSTGGSGNTLY | 0.15960403 |
| LSIFGPV | ASSSTGGSGNTLY | 0.15556002 |
| LLALLANL | ASSSTGGSGNTLY | 0.16060844 |
| LISNASDALDKLR | ASSSTGGSGNTLY | 0.16399562 |
| LINEAVP | ASSSTGGSGNTLY | 0.17448807 |
| LGYAFSGL | ASSSTGGSGNTLY | 0.15095983 |
| LDFVAVAL | ASSSTGGSGNTLY | 0.15178467 |
| LAQPGAE | ASSSTGGSGNTLY | 0.15686461 |
| ISNLGAVTGF | ASSSTGGSGNTLY | 0.19823815 |
| HQEDPGRI | ASSSTGGSGNTLY | 0.2330256 |
| GTPGEPTP | ASSSTGGSGNTLY | 0.1519758 |
| GFFNSGS | ASSSTGGSGNTLY | 0.17184585 |
| GAGGGGLTAGAN | ASSSTGGSGNTLY | 0.1571259 |
| FVDALVNM | ASSSTGGSGNTLY | 0.16140848 |

| | | |
|---|---|---|
| FLTVAPEL | ASSSTGGSGNTLY | 0.15173995 |
| FETLTAL | ASSSTGGSGNTLY | 0.16051444 |
| DVINPIEL | ASSSTGGSGNTLY | 0.1708804 |
| APVSGAPV | ASSSTGGSGNTLY | 0.21949483 |
| ALVGGLAW | ASSSTGGSGNTLY | 0.15972617 |
| ALAAAAATAI | ASSSTGGSGNTLY | 0.15509367 |
| AFGLGLLL | ASSSTGGSGNTLY | 0.16944456 |
| AFFAAASL | ASSSTGGSGNTLY | 0.15246472 |
| AAAALPAFSPP | ASSSTGGSGNTLY | 0.16029409 |
| VVLAPAL | ASSSTGGSGNTLY | 0.161938 |
| TIGHVDHGKTTL | ASSSTGGSGNTLY | 0.14858535 |
| LISNASDALDK | ASSSTGGSGNTLY | 0.16805092 |
| LISNASDAL | ASSSTGGSGNTLY | 0.23009825 |
| ISNASDALDK | ASSSTGGSGNTLY | 0.16050276 |
| NQAETLVYQTE | ASSSTGGSGNTLY | 0.16684814 |
| TGGTGGTAGAG | ASSSTGGSGNTLY | 0.19802217 |
| TGSTAGGAGGA | ASSSTGGSGNTLY | 0.1688206 |
| DPLAVPVAAGPV | ASSSTGGSGNTLY | 0.16973251 |
| GGTGGTPGNGGN | ASSSTGGSGNTLY | 0.15527867 |
| PAGLAGVDPATL | ASSSTGGSGNTLY | 0.16325222 |
| GNNLVVEAPHAGASV | ASSSTGGSGNTLY | 0.16507539 |
| KVTRIMSASTLEQPA | ASSSTGGSGNTLY | 0.19193207 |
| LGFALPIALPRRTVV | ASSSTGGSGNTLY | 0.1657508 |
| TRLGNLTARKLVVLV | ASSSTGGSGNTLY | 0.18120939 |
| WSQQRRLIYFPSAGP | ASSSTGGSGNTLY | 0.15976174 |
| GPNGNPDPMAAAVDIR | ASSSTGGSGNTLY | 0.1914137 |
| TQRCVSVADLRAAASR | ASSSTGGSGNTLY | 0.16572379 |
| QFVLDVAEHPVEVLTETV | ASSSTGGSGNTLY | 0.1792068 |
| GGAGGTGGAGGNGGTQAGDG | ASSSTGGSGNTLY | 0.1484207 |
| PLLATLRAIAVDVDAKPAQV | ASSSTGGSGNTLY | 0.1612195 |
| APKFRTGGYMIDFWGVGYQVAK | ASSSTGGSGNTLY | 0.15871662 |
| GAGGAGGAGGAAGGPGGTGGTG | ASSSTGGSGNTLY | 0.15287937 |
| IGALGLLATDVAPAGSVAFGVA | ASSSTGGSGNTLY | 0.15985712 |
| AGGAGGIGGNGGAGGAGGVLMG | ASSSTGGSGNTLY | 0.16168861 |
| ADELIGLL | ASSSTGGSGNTLY | 0.18766542 |
| ADRALKA | ASSSTGGSGNTLY | 0.14620093 |
| AGAVGQVGGAGGA | ASSSTGGSGNTLY | 0.16576669 |
| AGGGGGLGGAP | ASSSTGGSGNTLY | 0.15245795 |
| AGGLLYGNGGAG | ASSSTGGSGNTLY | 0.1555032 |
| AGLESIGL | ASSSTGGSGNTLY | 0.16817902 |
| AGLKTGII | ASSSTGGSGNTLY | 0.15326007 |
| ASLAALLL | ASSSTGGSGNTLY | 0.15729451 |
| ASVINISSI | ASSSTGGSGNTLY | 0.15685374 |
| ATPPGPA | ASSSTGGSGNTLY | 0.15976298 |
| ATVASALV | ASSSTGGSGNTLY | 0.16218358 |
| ATVASLIL | ASSSTGGSGNTLY | 0.16753449 |
| AVNGPVLIRH | ASSSTGGSGNTLY | 0.14865196 |
| DGAAGAVGI | ASSSTGGSGNTLY | 0.16281848 |
| DPLLHGI | ASSSTGGSGNTLY | 0.1529341 |
| ELGAALGAL | ASSSTGGSGNTLY | 0.3370588 |
| ELGRTQAV | ASSSTGGSGNTLY | 0.1762961 |

| | | |
|---|---|---|
| FVAGGYLS | ASSSTGGSGNTLY | 0.15005815 |
| GASGGAGGAGGW | ASSSTGGSGNTLY | 0.16974653 |
| GGAGAGGDGGAGGI | ASSSTGGSGNTLY | 0.15977156 |
| GTGHNGSQPGMGGNG | ASSSTGGSGNTLY | 0.17281985 |
| GYAAASASAS | ASSSTGGSGNTLY | 0.16510244 |
| HQVDVAAVQQG | ASSSTGGSGNTLY | 0.1500368 |
| IGAAIAHT | ASSSTGGSGNTLY | 0.16301756 |
| IGMIEVAQL | ASSSTGGSGNTLY | 0.17423396 |
| IRLAPEHTSAMD | ASSSTGGSGNTLY | 0.1575843 |
| IYNVSVL | ASSSTGGSGNTLY | 0.1611848 |
| KQGFLGM | ASSSTGGSGNTLY | 0.16056335 |
| LESVITQL | ASSSTGGSGNTLY | 0.1608801 |
| LGLGALVVV | ASSSTGGSGNTLY | 0.15543109 |
| LIGNGGNGGSGGI | ASSSTGGSGNTLY | 0.18663566 |
| LNAINAPTL | ASSSTGGSGNTLY | 0.156705 |
| LQLLNTGV | ASSSTGGSGNTLY | 0.18879452 |
| LQTSIEF | ASSSTGGSGNTLY | 0.17533296 |
| MSFAVSEL | ASSSTGGSGNTLY | 0.19441399 |
| NTGALGL | ASSSTGGSGNTLY | 0.1635486 |
| PPGAGGAGGAGTT | ASSSTGGSGNTLY | 0.15523914 |
| QLVGLGI | ASSSTGGSGNTLY | 0.17703237 |
| RFLEQVHQL | ASSSTGGSGNTLY | 0.16262138 |
| RGVGYVLR | ASSSTGGSGNTLY | 0.14969948 |
| SATGVFAGVI | ASSSTGGSGNTLY | 0.21036547 |
| SLPAITLP | ASSSTGGSGNTLY | 0.17275384 |
| STALAFTV | ASSSTGGSGNTLY | 0.22239661 |
| SVLTVGLGK | ASSSTGGSGNTLY | 0.15618022 |
| TALAGAGSGI | ASSSTGGSGNTLY | 0.15567183 |
| TGLPFTGI | ASSSTGGSGNTLY | 0.17480974 |
| VAEHVTV | ASSSTGGSGNTLY | 0.15081389 |
| VDIVLNSL | ASSSTGGSGNTLY | 0.16369425 |
| VLTGGGAVL | ASSSTGGSGNTLY | 0.1835938 |
| VNVGLAYGI | ASSSTGGSGNTLY | 0.17363252 |
| VSSAAVGAV | ASSSTGGSGNTLY | 0.15468383 |
| VTSVYEAGV | ASSSTGGSGNTLY | 0.16159166 |
| WVALDRGA | ASSSTGGSGNTLY | 0.15686232 |
| YLVGSVVI | ASSSTGGSGNTLY | 0.17324913 |
| GPPGTGKTLL | ASSSTGGSGNTLY | 0.15566948 |
| GVGMTKF | ASSSTGGSGNTLY | 0.16356249 |
| IGIGGSDL | ASSSTGGSGNTLY | 0.15027651 |
| IGTLLAEL | ASSSTGGSGNTLY | 0.16818443 |
| IYTSGTT | ASSSTGGSGNTLY | 0.19311139 |
| LSFPAGV | ASSSTGGSGNTLY | 0.17812501 |
| LYTSGTT | ASSSTGGSGNTLY | 0.15652515 |
| VGAFPFGF | ASSSTGGSGNTLY | 0.17580158 |
| VGAGTVEF | ASSSTGGSGNTLY | 0.1792171 |
| VGKSTLLL | ASSSTGGSGNTLY | 0.1799653 |
| AALAHAGGGIV | ASSSTGGSGNTLY | 0.16699752 |
| AALVAVSLL | ASSSTGGSGNTLY | 0.17080773 |
| AAVSRAVAAL | ASSSTGGSGNTLY | 0.16761467 |
| AGGSSGIGGF | ASSSTGGSGNTLY | 0.1584133 |

| | | |
|---|---|---|
| AGLTVANANG | ASSSTGGSGNTLY | 0.14970508 |
| AGLVGAVL | ASSSTGGSGNTLY | 0.1553143 |
| ALSQAFTV | ASSSTGGSGNTLY | 0.16012093 |
| ALTFSVSL | ASSSTGGSGNTLY | 0.15362456 |
| ALTGAALVAL | ASSSTGGSGNTLY | 0.17275773 |
| ASFSIGAGVL | ASSSTGGSGNTLY | 0.16887532 |
| DGSVYKI | ASSSTGGSGNTLY | 0.21976872 |
| EQLAEIV | ASSSTGGSGNTLY | 0.19899087 |
| FAVSELTVM | ASSSTGGSGNTLY | 0.2300307 |
| GAAPLAAGL | ASSSTGGSGNTLY | 0.15583985 |
| GADGGDGGSGGAGG | ASSSTGGSGNTLY | 0.16043438 |
| GFGLDTGGAGG | ASSSTGGSGNTLY | 0.1654988 |
| GLASGGLL | ASSSTGGSGNTLY | 0.15055107 |
| GRIAGAAVLL | ASSSTGGSGNTLY | 0.16103798 |
| GSAGSLNTSF | ASSSTGGSGNTLY | 0.15957487 |
| GTGQIGI | ASSSTGGSGNTLY | 0.18208542 |
| GTLIALIL | ASSSTGGSGNTLY | 0.14577179 |
| IAATMGFVVG | ASSSTGGSGNTLY | 0.16499609 |
| IFAQALL | ASSSTGGSGNTLY | 0.17567673 |
| IGAAVALGF | ASSSTGGSGNTLY | 0.15070638 |
| IGLTPLAL | ASSSTGGSGNTLY | 0.18918726 |
| ILGSNTSTL | ASSSTGGSGNTLY | 0.19359104 |
| IVFSLTVL | ASSSTGGSGNTLY | 0.14965735 |
| LASKATGF | ASSSTGGSGNTLY | 0.19060628 |
| LASVALIL | ASSSTGGSGNTLY | 0.15465501 |
| LFAVGLTL | ASSSTGGSGNTLY | 0.16166805 |
| LFNGGIGTYI | ASSSTGGSGNTLY | 0.19991648 |
| LTLGAGII | ASSSTGGSGNTLY | 0.18159433 |
| LVAAALGDI | ASSSTGGSGNTLY | 0.15259473 |
| NTAFVQLGI | ASSSTGGSGNTLY | 0.15984485 |
| QLFSGAEL | ASSSTGGSGNTLY | 0.16539629 |
| SAATFITL | ASSSTGGSGNTLY | 0.15416151 |
| SNYGLANLGV | ASSSTGGSGNTLY | 0.1986066 |
| SYGGAYAVM | ASSSTGGSGNTLY | 0.1676626 |
| TTGVGGLTLG | ASSSTGGSGNTLY | 0.15692429 |
| TTQTIYI | ASSSTGGSGNTLY | 0.1776733 |
| VAAGAGLGAVI | ASSSTGGSGNTLY | 0.17128398 |
| VAAVVTLL | ASSSTGGSGNTLY | 0.16385546 |
| VGFLGGLI | ASSSTGGSGNTLY | 0.16066536 |
| VGVIPILL | ASSSTGGSGNTLY | 0.16659214 |
| VIFASSLI | ASSSTGGSGNTLY | 0.16281751 |
| VKGGDGGAAAG | ASSSTGGSGNTLY | 0.15582632 |
| VTKTFTNI | ASSSTGGSGNTLY | 0.15958504 |
| VVAAAILL | ASSSTGGSGNTLY | 0.16406323 |
| YAALDVEL | ASSSTGGSGNTLY | 0.1582467 |
| LASVALLL | ASSSTGGSGNTLY | 0.18788584 |
| LDAVHRV | ASSSTGGSGNTLY | 0.16433147 |
| QGLSSSIF | ASSSTGGSGNTLY | 0.15942112 |
| RGTFANIRL | ASSSTGGSGNTLY | 0.16501047 |
| RQLWWGHRI | ASSSTGGSGNTLY | 0.1724822 |
| YGPPGTGKTLL | ASSSTGGSGNTLY | 0.16714245 |

| | | |
|---|---|---|
| AALVGAPANVA | ASSSTGGSGNTLY | 0.15304011 |
| AATFRRTE | ASSSTGGSGNTLY | 0.1582897 |
| AGALGGGI | ASSSTGGSGNTLY | 0.1556463 |
| AGGAGGAGGIGAG | ASSSTGGSGNTLY | 0.15213901 |
| AGSGALGL | ASSSTGGSGNTLY | 0.15118268 |
| AQIADQI | ASSSTGGSGNTLY | 0.17221448 |
| ASAVAVTYL | ASSSTGGSGNTLY | 0.20317118 |
| ASTVLGGTGG | ASSSTGGSGNTLY | 0.16164365 |
| AVGGEPAK | ASSSTGGSGNTLY | 0.16006108 |
| CEQGTFGGTC | ASSSTGGSGNTLY | 0.19248712 |
| DEGKAPAG | ASSSTGGSGNTLY | 0.19369276 |
| DTVVFPAL | ASSSTGGSGNTLY | 0.16932185 |
| FEGITMAG | ASSSTGGSGNTLY | 0.17002197 |
| FPLVVQD | ASSSTGGSGNTLY | 0.17092249 |
| GAFTTPQ | ASSSTGGSGNTLY | 0.16826564 |
| GFGGFGTAGGVGGA | ASSSTGGSGNTLY | 0.15868445 |
| GFLAQFL | ASSSTGGSGNTLY | 0.15070258 |
| GGAGGAGGNAGVA | ASSSTGGSGNTLY | 0.172912 |
| GGVGTAGAGGAGGAGG | ASSSTGGSGNTLY | 0.16581322 |
| GNGGGGDAGNAGSGGN | ASSSTGGSGNTLY | 0.18291356 |
| GTGGAGGAGNDAGS | ASSSTGGSGNTLY | 0.16548975 |
| GTTNTGWFNS | ASSSTGGSGNTLY | 0.15452293 |
| IFMGPLFTL | ASSSTGGSGNTLY | 0.14952065 |
| ILAHAIEAI | ASSSTGGSGNTLY | 0.15642029 |
| IRSAIREAL | ASSSTGGSGNTLY | 0.16498984 |
| ITFAGGG | ASSSTGGSGNTLY | 0.16565503 |
| ITGMSGHQEV | ASSSTGGSGNTLY | 0.16893065 |
| IVAGLACTL | ASSSTGGSGNTLY | 0.23024152 |
| IVGGGLAGA | ASSSTGGSGNTLY | 0.1752453 |
| IVTGVLNI | ASSSTGGSGNTLY | 0.1565745 |
| LAQQGCI | ASSSTGGSGNTLY | 0.1628709 |
| LCLHVEN | ASSSTGGSGNTLY | 0.16069649 |
| LDAISDGS | ASSSTGGSGNTLY | 0.15505543 |
| LGGFAAML | ASSSTGGSGNTLY | 0.16055883 |
| LGGKNPMI | ASSSTGGSGNTLY | 0.15426372 |
| LGPKGVAAIL | ASSSTGGSGNTLY | 0.15311807 |
| LMVGVSI | ASSSTGGSGNTLY | 0.15441118 |
| LVGAAFVL | ASSSTGGSGNTLY | 0.15526082 |
| LVLAVAPL | ASSSTGGSGNTLY | 0.16164331 |
| MAAIAVIDAA | ASSSTGGSGNTLY | 0.15990126 |
| NNGINAELQ | ASSSTGGSGNTLY | 0.15786906 |
| RGEPLVLVHG | ASSSTGGSGNTLY | 0.16100134 |
| RGGAGGAAGLI | ASSSTGGSGNTLY | 0.16904029 |
| RSYRMHPAV | ASSSTGGSGNTLY | 0.24326582 |
| SAAHLLAGQLA | ASSSTGGSGNTLY | 0.15645093 |
| SAAVGSY | ASSSTGGSGNTLY | 0.15811375 |
| SPHAVLHRI | ASSSTGGSGNTLY | 0.17962322 |
| SQIAKVL | ASSSTGGSGNTLY | 0.16514987 |
| TALGHLVP | ASSSTGGSGNTLY | 0.15605338 |
| TAPTTSGAA | ASSSTGGSGNTLY | 0.15949696 |
| TDGDDGNGGSPGT | ASSSTGGSGNTLY | 0.15813644 |

| | | |
|---|---|---|
| TDGLGEN | ASSSTGGSGNTLY | 0.15091708 |
| TGGPPLPGQA | ASSSTGGSGNTLY | 0.15983275 |
| TIPAITI | ASSSTGGSGNTLY | 0.15381946 |
| TIPGFTI | ASSSTGGSGNTLY | 0.1627692 |
| TSIASGALA | ASSSTGGSGNTLY | 0.14598241 |
| TVTLPGAAI | ASSSTGGSGNTLY | 0.1568445 |
| VASGLSAALP | ASSSTGGSGNTLY | 0.15827441 |
| VGAGVIGI | ASSSTGGSGNTLY | 0.1555863 |
| VGPKRFGV | ASSSTGGSGNTLY | 0.14809565 |
| VNVAFPD | ASSSTGGSGNTLY | 0.15977983 |
| VSMTNSGVS | ASSSTGGSGNTLY | 0.1970812 |
| VTDVLGDLSV | ASSSTGGSGNTLY | 0.17767362 |
| VVVVGAGFAGL | ASSSTGGSGNTLY | 0.1734007 |
| VYTVGTVV | ASSSTGGSGNTLY | 0.15718412 |
| WTTFSGP | ASSSTGGSGNTLY | 0.22169644 |
| YDELVEAF | ASSSTGGSGNTLY | 0.17210016 |
| YGTGGAGGNGG | ASSSTGGSGNTLY | 0.15328737 |
| YTNVMAV | ASSSTGGSGNTLY | 0.1651687 |
| FLEQVHQL | ASSSTGGSGNTLY | 0.1492213 |
| GLSLLAVAV | ASSSTGGSGNTLY | 0.15688643 |
| LEQVHQL | ASSSTGGSGNTLY | 0.19265759 |
| LNAAGAI | ASSSTGGSGNTLY | 0.14900835 |
| PAVHASL | ASSSTGGSGNTLY | 0.16060157 |
| TELEDDA | ASSSTGGSGNTLY | 0.20955035 |
| TPPGPPN | ASSSTGGSGNTLY | 0.15519066 |
| TPPPPPPP | ASSSTGGSGNTLY | 0.15338123 |
| TQSPALI | ASSGDVWEANTGQLY | 0.15304066 |
| EFAAAFPQ | ASSGDVWEANTGQLY | 0.17849554 |
| LLSGNLDVL | ASSGDVWEANTGQLY | 0.18635063 |
| GPANLGNYNI | ASSGDVWEANTGQLY | 0.15907398 |
| TLGPTLP | ASSGDVWEANTGQLY | 0.17096896 |
| VYAYSSCCK | ASSGDVWEANTGQLY | 0.1596257 |
| APAPAPAPNI | ASSGDVWEANTGQLY | 0.16537818 |
| PGGPVPIP | ASSGDVWEANTGQLY | 0.16607814 |
| FLTVALEP | ASSGDVWEANTGQLY | 0.18000431 |
| SQYFGANL | ASSGDVWEANTGQLY | 0.17294559 |
| GGVKPAA | ASSGDVWEANTGQLY | 0.16187084 |
| GGTGGAGGAGGW | ASSGDVWEANTGQLY | 0.15630952 |
| GVGGVAGLGGAGG | ASSGDVWEANTGQLY | 0.15314834 |
| VLLVLATTI | ASSGDVWEANTGQLY | 0.22965401 |
| IGIFAAML | ASSGDVWEANTGQLY | 0.15574814 |
| AVGDLID | ASSGDVWEANTGQLY | 0.154243 |
| NISPLPL | ASSGDVWEANTGQLY | 0.15682314 |
| VTAVFVAI | ASSGDVWEANTGQLY | 0.1562556 |
| SGDDNDAAL | ASSGDVWEANTGQLY | 0.1606873 |
| AVAALAHS | ASSGDVWEANTGQLY | 0.16722427 |
| GGVTVMATP | ASSGDVWEANTGQLY | 0.18091556 |
| AGLAANVRD | ASSGDVWEANTGQLY | 0.15848617 |
| TNRPDLIDPALLRPG | ASSGDVWEANTGQLY | 0.17012013 |
| LANLGVTAL | ASSGDVWEANTGQLY | 0.17375042 |
| TGERIEL | ASSGDVWEANTGQLY | 0.15396304 |

| | | |
|---|---|---|
| NGEELAA | ASSGDVWEANTGQLY | 0.15564418 |
| VLSLVWG | ASSGDVWEANTGQLY | 0.14944111 |
| IGYALWL | ASSGDVWEANTGQLY | 0.15484302 |
| LSLLLAPL | ASSGDVWEANTGQLY | 0.1620126 |
| TLDEALK | ASSGDVWEANTGQLY | 0.1537568 |
| TDGAGTAEGG | ASSGDVWEANTGQLY | 0.16336375 |
| NAPGGAGGNGG | ASSGDVWEANTGQLY | 0.18553606 |
| VSVFMGMGGNF | ASSGDVWEANTGQLY | 0.1620522 |
| LASYLAAL | ASSGDVWEANTGQLY | 0.15921432 |
| LVGGTPAL | ASSGDVWEANTGQLY | 0.15344058 |
| VRNADVL | ASSGDVWEANTGQLY | 0.17910632 |
| TCSSPFLVSAM | ASSGDVWEANTGQLY | 0.16272183 |
| VHITDAE | ASSGDVWEANTGQLY | 0.15046333 |
| EYAAPSF | ASSGDVWEANTGQLY | 0.18477535 |
| VVDAFHGGL | ASSGDVWEANTGQLY | 0.17822285 |
| ASVVDPQ | ASSGDVWEANTGQLY | 0.17267202 |
| SYNEFGGLS | ASSGDVWEANTGQLY | 0.18044001 |
| DPDGSVY | ASSGDVWEANTGQLY | 0.19055636 |
| MVSTIHATE | ASSGDVWEANTGQLY | 0.16452305 |
| LAAGVADPV | ASSGDVWEANTGQLY | 0.16559029 |
| TKEQIAA | ASSGDVWEANTGQLY | 0.14933275 |
| SLASPAAL | ASSGDVWEANTGQLY | 0.15835749 |
| IGLVFYGV | ASSGDVWEANTGQLY | 0.15286869 |
| FLLDASNL | ASSGDVWEANTGQLY | 0.16869068 |
| YQVGNFP | ASSGDVWEANTGQLY | 0.14791268 |
| ESLLSQST | ASSGDVWEANTGQLY | 0.17035618 |
| LFTVGFAAVG | ASSGDVWEANTGQLY | 0.15539742 |
| ATPPTPT | ASSGDVWEANTGQLY | 0.18539521 |
| TDLATLG | ASSGDVWEANTGQLY | 0.167974 |
| SAGGTGR | ASSGDVWEANTGQLY | 0.15182291 |
| GFSNSGS | ASSGDVWEANTGQLY | 0.1577889 |
| GGNGGSGGTSV | ASSGDVWEANTGQLY | 0.16593483 |
| GTSVATGGAGN | ASSGDVWEANTGQLY | 0.14941384 |
| NGGNGGSGGTS | ASSGDVWEANTGQLY | 0.15663399 |
| AVVKFPH | ASSGDVWEANTGQLY | 0.15642424 |
| TPPPPPP | ASSGDVWEANTGQLY | 0.16095999 |
| AAPAAAAAPA | ASSGDVWEANTGQLY | 0.23315755 |
| IALNEDL | ASSGDVWEANTGQLY | 0.1946396 |
| IVNVSSM | ASSGDVWEANTGQLY | 0.164976 |
| TGIPPAPR | ASSGDVWEANTGQLY | 0.17429194 |
| FDLGGGTF | ASSGDVWEANTGQLY | 0.14657524 |
| FDLGGGTFDVS | ASSGDVWEANTGQLY | 0.1635693 |
| NEPTAAA | ASSGDVWEANTGQLY | 0.20246843 |
| NEPTAAALA | ASSGDVWEANTGQLY | 0.16641764 |
| GSIINISSI | ASSGDVWEANTGQLY | 0.18438026 |
| VAPGNAGTA | ASSGDVWEANTGQLY | 0.16794516 |
| TGGGASLE | ASSGDVWEANTGQLY | 0.15889652 |
| VAVYNHYKRI | ASSGDVWEANTGQLY | 0.16714641 |
| TDPAPATT | ASSGDVWEANTGQLY | 0.16096179 |
| TNRPDILDPALLRPG | ASSGDVWEANTGQLY | 0.17203978 |
| LTTTTTS | ASSGDVWEANTGQLY | 0.16173886 |

| | | |
|---|---|---|
| PTGSGKTL | ASSGDVWEANTGQLY | 0.15181503 |
| TGSGKTL | ASSGDVWEANTGQLY | 0.1682044 |
| TIGHVDHGK | ASSGDVWEANTGQLY | 0.28044522 |
| TIGHVDHGKTT | ASSGDVWEANTGQLY | 0.15985018 |
| GHVDHGKTT | ASSGDVWEANTGQLY | 0.18116309 |
| AQFEHTLL | ASSGDVWEANTGQLY | 0.15820068 |
| IVNQDEL | ASSGDVWEANTGQLY | 0.18124801 |
| TNGSQFF | ASSGDVWEANTGQLY | 0.1620115 |
| TGKGIVE | ASSGDVWEANTGQLY | 0.15228556 |
| LISNASDA | ASSGDVWEANTGQLY | 0.16939867 |
| AELRTLGELTKIAKAHGAQ | ASSGDVWEANTGQLY | 0.15310906 |
| APNAVAAAEPSW | ASSGDVWEANTGQLY | 0.15933561 |
| APQVAGAATAVL | ASSGDVWEANTGQLY | 0.16460478 |
| ARRDPAAAAAILPTDA | ASSGDVWEANTGQLY | 0.35416654 |
| DAAARDVLAEAGLAEYF | ASSGDVWEANTGQLY | 0.17121898 |
| DDQIVLSAKANLAKVEQ | ASSGDVWEANTGQLY | 0.15933096 |
| DELLPDTGEDSDEDGH | ASSGDVWEANTGQLY | 0.15245324 |
| ELPEAPANRATVPQD | ASSGDVWEANTGQLY | 0.15946837 |
| FGQSKIERVTGIG | ASSGDVWEANTGQLY | 0.16183679 |
| GAGGTGSTAGGAGGAGGAGGLY | ASSGDVWEANTGQLY | 0.15998594 |
| GHGGTGAAGGNGGTGGAGGSGI | ASSGDVWEANTGQLY | 0.19385634 |
| GNGGNGGAGGEGGPGGAGGAGA | ASSGDVWEANTGQLY | 0.15804555 |
| GSTAGNGGNGGNATNGTIAGQP | ASSGDVWEANTGQLY | 0.1593205 |
| GTGGDGGNGGNGGTGG | ASSGDVWEANTGQLY | 0.16418451 |
| GVRRPAGIPADAPV | ASSGDVWEANTGQLY | 0.2092434 |
| GYRQVTDAHLLTLARR | ASSGDVWEANTGQLY | 0.17257978 |
| IAKMGQDGHDRG | ASSGDVWEANTGQLY | 0.18094833 |
| KAEPSIEIQYT | ASSGDVWEANTGQLY | 0.15225217 |
| KPVRIRMAADEDTVAVT | ASSGDVWEANTGQLY | 0.18161155 |
| MPPRIAGMRLLVIKPEP | ASSGDVWEANTGQLY | 0.19159143 |
| MKFYAVALFRSRGVPPTR | ASSGDVWEANTGQLY | 0.17048451 |
| NAGGFGILGAVAHSP | ASSGDVWEANTGQLY | 0.18677129 |
| NEGREITLAMAEM | ASSGDVWEANTGQLY | 0.15585454 |
| NELVEWSHPLGHRWRWE | ASSGDVWEANTGQLY | 0.14938276 |
| NLGAGGTGGNGGNPG | ASSGDVWEANTGQLY | 0.16365407 |
| NTEWPKWETVMAADPVD | ASSGDVWEANTGQLY | 0.15807986 |
| PASAAPAETQP | ASSGDVWEANTGQLY | 0.18412194 |
| QQSSDGKNGVRAAEITGE | ASSGDVWEANTGQLY | 0.1828899 |
| RAPDAVTTSWWKE | ASSGDVWEANTGQLY | 0.14840929 |
| SGFGNVGAHTSGWFNQ | ASSGDVWEANTGQLY | 0.15783747 |
| SGVAGAVGQAASVGGLKV | ASSGDVWEANTGQLY | 0.19789964 |
| SNGGAGTGSAGGNGGT | ASSGDVWEANTGQLY | 0.24527049 |
| TGGAGGEGGAAG | ASSGDVWEANTGQLY | 0.16221242 |
| TGGDGGAAGAGGAPGHG | ASSGDVWEANTGQLY | 0.15576193 |
| TMPPLLAAIANAGRHGV | ASSGDVWEANTGQLY | 0.16034433 |
| TSSTAGNSGGA | ASSGDVWEANTGQLY | 0.17984694 |
| VPATVPVGILC | ASSGDVWEANTGQLY | 0.1607165 |
| VQRPDSGGVVLTGRIS | ASSGDVWEANTGQLY | 0.15065408 |
| WTPNPVPATKGVPAGLVQAT | ASSGDVWEANTGQLY | 0.17466052 |
| YALKRALAKFSDRFDVVII | ASSGDVWEANTGQLY | 0.1882779 |
| YLIGSVIV | ASSGDVWEANTGQLY | 0.1491596 |

| | | |
|---|---|---|
| YGQLVAVID | ASSGDVWEANTGQLY | 0.16948773 |
| VTVGGMTLPA | ASSGDVWEANTGQLY | 0.16443661 |
| VLALPAV | ASSGDVWEANTGQLY | 0.15701938 |
| VIPPTLNYET | ASSGDVWEANTGQLY | 0.1490386 |
| VGMGGMHPSSG | ASSGDVWEANTGQLY | 0.2075769 |
| VGLAGGAATA | ASSGDVWEANTGQLY | 0.26231393 |
| VGGAGGAGGSGP | ASSGDVWEANTGQLY | 0.16545162 |
| VFAAFMV | ASSGDVWEANTGQLY | 0.15428804 |
| TLYDKLR | ASSGDVWEANTGQLY | 0.14807813 |
| TLAVFTVL | ASSGDVWEANTGQLY | 0.21163361 |
| TGSVQPARV | ASSGDVWEANTGQLY | 0.20404515 |
| TGGGAGSGGT | ASSGDVWEANTGQLY | 0.16630293 |
| SHLETSL | ASSGDVWEANTGQLY | 0.15089765 |
| SFTTGMT | ASSGDVWEANTGQLY | 0.15549585 |
| QVEVAKRMGVRQPTVS | ASSGDVWEANTGQLY | 0.18359762 |
| QGSGGPALAM | ASSGDVWEANTGQLY | 0.15128942 |
| MLAPSNP | ASSGDVWEANTGQLY | 0.15278272 |
| LVNNAAI | ASSGDVWEANTGQLY | 0.15772568 |
| LTGEINTLA | ASSGDVWEANTGQLY | 0.16577314 |
| LSIFGPV | ASSGDVWEANTGQLY | 0.14861792 |
| LLALLANL | ASSGDVWEANTGQLY | 0.1550775 |
| LISNASDALDKLR | ASSGDVWEANTGQLY | 0.20993894 |
| LINEAVP | ASSGDVWEANTGQLY | 0.20532621 |
| LGYAFSGL | ASSGDVWEANTGQLY | 0.16072008 |
| LDFVAVAL | ASSGDVWEANTGQLY | 0.15387805 |
| LAQPGAE | ASSGDVWEANTGQLY | 0.16307057 |
| ISNLGAVTGF | ASSGDVWEANTGQLY | 0.16168542 |
| HQEDPGRI | ASSGDVWEANTGQLY | 0.16959788 |
| GTPGEPTP | ASSGDVWEANTGQLY | 0.15794839 |
| GFFNSGS | ASSGDVWEANTGQLY | 0.17375167 |
| GAGGGGLTAGAN | ASSGDVWEANTGQLY | 0.17267217 |
| FVDALVNM | ASSGDVWEANTGQLY | 0.15665182 |
| FLTVAPEL | ASSGDVWEANTGQLY | 0.15629035 |
| FETLTAL | ASSGDVWEANTGQLY | 0.14767686 |
| DVINPIEL | ASSGDVWEANTGQLY | 0.17014 |
| APVSGAPV | ASSGDVWEANTGQLY | 0.22498046 |
| ALVGGLAW | ASSGDVWEANTGQLY | 0.16359267 |
| ALAAAAATAI | ASSGDVWEANTGQLY | 0.17781475 |
| AFGLGLLL | ASSGDVWEANTGQLY | 0.15488398 |
| AFFAAASL | ASSGDVWEANTGQLY | 0.16361858 |
| AAAALPAFSPP | ASSGDVWEANTGQLY | 0.14877509 |
| VVLAPAL | ASSGDVWEANTGQLY | 0.18571766 |
| TIGHVDHGKTTL | ASSGDVWEANTGQLY | 0.17377982 |
| LISNASDALDK | ASSGDVWEANTGQLY | 0.153974 |
| LISNASDAL | ASSGDVWEANTGQLY | 0.16675605 |
| ISNASDALDK | ASSGDVWEANTGQLY | 0.16093592 |
| NQAETLVYQTE | ASSGDVWEANTGQLY | 0.17975172 |
| TGGTGGTAGAG | ASSGDVWEANTGQLY | 0.15955637 |
| TGSTAGGAGGA | ASSGDVWEANTGQLY | 0.15560599 |
| DPLAVPVAAGPV | ASSGDVWEANTGQLY | 0.18551809 |
| GGTGGTPGNGGN | ASSGDVWEANTGQLY | 0.16110155 |

| | | |
|---|---|---|
| PAGLAGVDPATL | ASSGDVWEANTGQLY | 0.14968002 |
| GNNLVVEAPHAGASV | ASSGDVWEANTGQLY | 0.19733916 |
| KVTRIMSASTLEQPA | ASSGDVWEANTGQLY | 0.15181163 |
| LGFALPIALPRRTVV | ASSGDVWEANTGQLY | 0.16346025 |
| TRLGNLTARKLVVLV | ASSGDVWEANTGQLY | 0.15901147 |
| WSQQRRLIYFPSAGP | ASSGDVWEANTGQLY | 0.17913999 |
| GPNGNPDPMAAAVDIR | ASSGDVWEANTGQLY | 0.17713055 |
| TQRCVSVADLRAAASR | ASSGDVWEANTGQLY | 0.15064599 |
| QFVLDVAEHPVEVLTETV | ASSGDVWEANTGQLY | 0.15655906 |
| GGAGGTGGAGGNGGTQAGDG | ASSGDVWEANTGQLY | 0.16697511 |
| PLLATLRAIAVDVDAKPAQV | ASSGDVWEANTGQLY | 0.18481919 |
| APKFRTGGYMIDFWGVGYQVAK | ASSGDVWEANTGQLY | 0.16383697 |
| GAGGAGGAGGAAGGPGGTGGTG | ASSGDVWEANTGQLY | 0.15711452 |
| IGALGLLATDVAPAGSVAFGVA | ASSGDVWEANTGQLY | 0.17335814 |
| AGGAGGIGGNGGAGGAGGVLMG | ASSGDVWEANTGQLY | 0.16430329 |
| ADELIGLL | ASSGDVWEANTGQLY | 0.1599305 |
| ADRALKA | ASSGDVWEANTGQLY | 0.15594995 |
| AGAVGQVGGAGGA | ASSGDVWEANTGQLY | 0.15665883 |
| AGGGGGLGGAP | ASSGDVWEANTGQLY | 0.1523743 |
| AGGLLYGNGGAG | ASSGDVWEANTGQLY | 0.16678426 |
| AGLESIGL | ASSGDVWEANTGQLY | 0.16282363 |
| AGLKTGII | ASSGDVWEANTGQLY | 0.18674548 |
| ASLAALLL | ASSGDVWEANTGQLY | 0.16518041 |
| ASVINISSI | ASSGDVWEANTGQLY | 0.14988028 |
| ATPPGPA | ASSGDVWEANTGQLY | 0.1716205 |
| ATVASALV | ASSGDVWEANTGQLY | 0.15330611 |
| ATVASLIL | ASSGDVWEANTGQLY | 0.15497741 |
| AVNGPVLIRH | ASSGDVWEANTGQLY | 0.17614804 |
| DGAAGAVGI | ASSGDVWEANTGQLY | 0.15631053 |
| DPLLHGI | ASSGDVWEANTGQLY | 0.16497451 |
| ELGAALGAL | ASSGDVWEANTGQLY | 0.1982675 |
| ELGRTQAV | ASSGDVWEANTGQLY | 0.15924071 |
| FVAGGYLS | ASSGDVWEANTGQLY | 0.15193239 |
| GASGGAGGAGGW | ASSGDVWEANTGQLY | 0.15849622 |
| GGAGAGGDGGAGGI | ASSGDVWEANTGQLY | 0.16569811 |
| GTGHNGSQPGMGGNG | ASSGDVWEANTGQLY | 0.15642254 |
| GYAAASASAS | ASSGDVWEANTGQLY | 0.14770569 |
| HQVDVAAVQQG | ASSGDVWEANTGQLY | 0.16631787 |
| IGAAIAHT | ASSGDVWEANTGQLY | 0.1604573 |
| IGMIEVAQL | ASSGDVWEANTGQLY | 0.15503035 |
| IRLAPEHTSAMD | ASSGDVWEANTGQLY | 0.14925154 |
| IYNVSVL | ASSGDVWEANTGQLY | 0.17521298 |
| KQGFLGM | ASSGDVWEANTGQLY | 0.18084541 |
| LESVITQL | ASSGDVWEANTGQLY | 0.16433986 |
| LGLGALVVV | ASSGDVWEANTGQLY | 0.18498239 |
| LIGNGGNGGSGGI | ASSGDVWEANTGQLY | 0.15158768 |
| LNAINAPTL | ASSGDVWEANTGQLY | 0.18539429 |
| LQLLNTGV | ASSGDVWEANTGQLY | 0.18777336 |
| LQTSIEF | ASSGDVWEANTGQLY | 0.15385033 |
| MSFAVSEL | ASSGDVWEANTGQLY | 0.1751398 |
| NTGALGL | ASSGDVWEANTGQLY | 0.16631302 |

| | | |
|---|---|---|
| PPGAGGAGGAGTT | ASSGDVWEANTGQLY | 0.15291435 |
| QLVGLGI | ASSGDVWEANTGQLY | 0.17040026 |
| RFLEQVHQL | ASSGDVWEANTGQLY | 0.17165175 |
| RGVGYVLR | ASSGDVWEANTGQLY | 0.16826071 |
| SATGVFAGVI | ASSGDVWEANTGQLY | 0.15359959 |
| SLPAITLP | ASSGDVWEANTGQLY | 0.15540503 |
| STALAFTV | ASSGDVWEANTGQLY | 0.1605515 |
| SVLTVGLGK | ASSGDVWEANTGQLY | 0.16127022 |
| TALAGAGSGI | ASSGDVWEANTGQLY | 0.16957755 |
| TGLPFTGI | ASSGDVWEANTGQLY | 0.24478886 |
| VAEHVTV | ASSGDVWEANTGQLY | 0.16186492 |
| VDIVLNSL | ASSGDVWEANTGQLY | 0.20284574 |
| VLTGGGAVL | ASSGDVWEANTGQLY | 0.1583405 |
| VNVGLAYGI | ASSGDVWEANTGQLY | 0.15988834 |
| VSSAAVGAV | ASSGDVWEANTGQLY | 0.15850985 |
| VTSVYEAGV | ASSGDVWEANTGQLY | 0.20208615 |
| WVALDRGA | ASSGDVWEANTGQLY | 0.15528323 |
| YLVGSVVI | ASSGDVWEANTGQLY | 0.16181059 |
| GPPGTGKTLL | ASSGDVWEANTGQLY | 0.1632538 |
| GVGMTKF | ASSGDVWEANTGQLY | 0.16630909 |
| IGIGGSDL | ASSGDVWEANTGQLY | 0.26585373 |
| IGTLLAEL | ASSGDVWEANTGQLY | 0.22564858 |
| IYTSGTT | ASSGDVWEANTGQLY | 0.28647113 |
| LSFPAGV | ASSGDVWEANTGQLY | 0.19837748 |
| LYTSGTT | ASSGDVWEANTGQLY | 0.1357864 |
| VGAFPFGF | ASSGDVWEANTGQLY | 0.19758339 |
| VGAGTVEF | ASSGDVWEANTGQLY | 0.16215192 |
| VGKSTLLL | ASSGDVWEANTGQLY | 0.16961464 |
| AALAHAGGGIV | ASSGDVWEANTGQLY | 0.25766382 |
| AALVAVSLL | ASSGDVWEANTGQLY | 0.19156545 |
| AAVSRAVAAL | ASSGDVWEANTGQLY | 0.5776324 |
| AGGSSGIGGF | ASSGDVWEANTGQLY | 0.4508776 |
| AGLTVANANG | ASSGDVWEANTGQLY | 0.14921682 |
| AGLVGAVL | ASSGDVWEANTGQLY | 0.21049906 |
| ALSQAFTV | ASSGDVWEANTGQLY | 0.17593639 |
| ALTFSVSL | ASSGDVWEANTGQLY | 0.22052495 |
| ALTGAALVAL | ASSGDVWEANTGQLY | 0.11866861 |
| ASFSIGAGVL | ASSGDVWEANTGQLY | 0.29940474 |
| DGSVYKI | ASSGDVWEANTGQLY | 0.1470275 |
| EQLAEIV | ASSGDVWEANTGQLY | 0.113640465 |
| FAVSELTVM | ASSGDVWEANTGQLY | 0.2522236 |
| GAAPLAAGL | ASSGDVWEANTGQLY | 0.22433169 |
| GADGGDGGSGGAGG | ASSGDVWEANTGQLY | 0.19478117 |
| GFGLDTGGAGG | ASSGDVWEANTGQLY | 0.445363 |
| GLASGGLL | ASSGDVWEANTGQLY | 0.13376269 |
| GRIAGAAVLL | ASSGDVWEANTGQLY | 0.16803731 |
| GSAGSLNTSF | ASSGDVWEANTGQLY | 0.39532292 |
| GTGQIGI | ASSGDVWEANTGQLY | 0.45862812 |
| GTLIALIL | ASSGDVWEANTGQLY | 0.16160747 |
| IAATMGFVVG | ASSGDVWEANTGQLY | 0.28005734 |
| IFAQALL | ASSGDVWEANTGQLY | 0.55733854 |

| | | |
|---|---|---|
| IGAAVALGF | ASSGDVWEANTGQLY | 0.2608118 |
| IGLTPLAL | ASSGDVWEANTGQLY | 0.26430023 |
| ILGSNTSTL | ASSGDVWEANTGQLY | 0.331738 |
| IVFSLTVL | ASSGDVWEANTGQLY | 0.15116867 |
| LASKATGF | ASSGDVWEANTGQLY | 0.35357842 |
| LASVALIL | ASSGDVWEANTGQLY | 0.4102226 |
| LFAVGLTL | ASSGDVWEANTGQLY | 0.2231399 |
| LFNGGIGTYI | ASSGDVWEANTGQLY | 0.27636835 |
| LTLGAGII | ASSGDVWEANTGQLY | 0.16592127 |
| LVAAALGDI | ASSGDVWEANTGQLY | 0.19834672 |
| NTAFVQLGI | ASSGDVWEANTGQLY | 0.11805881 |
| QLFSGAEL | ASSGDVWEANTGQLY | 0.3050698 |
| SAATFITL | ASSGDVWEANTGQLY | 0.3166274 |
| SNYGLANLGV | ASSGDVWEANTGQLY | 0.23748828 |
| SYGGAYAVM | ASSGDVWEANTGQLY | 0.14858297 |
| TTGVGGLTLG | ASSGDVWEANTGQLY | 0.16783702 |
| TTQTIYI | ASSGDVWEANTGQLY | 0.1349923 |
| VAAGAGLGAVI | ASSGDVWEANTGQLY | 0.21006623 |
| VAAVVTLL | ASSGDVWEANTGQLY | 0.13287713 |
| VGFLGGLI | ASSGDVWEANTGQLY | 0.12631124 |
| VGVIPILL | ASSGDVWEANTGQLY | 0.40966666 |
| VIFASSLI | ASSGDVWEANTGQLY | 0.42654 |
| VKGGDGGAAAG | ASSGDVWEANTGQLY | 0.20489039 |
| VTKTFTNI | ASSGDVWEANTGQLY | 0.257149 |
| VVAAAILL | ASSGDVWEANTGQLY | 0.2620324 |
| YAALDVEL | ASSGDVWEANTGQLY | 0.25528806 |
| LASVALLL | ASSGDVWEANTGQLY | 0.23178066 |
| LDAVHRV | ASSGDVWEANTGQLY | 0.1433477 |
| QGLSSSIF | ASSGDVWEANTGQLY | 0.40732968 |
| RGTFANIRL | ASSGDVWEANTGQLY | 0.25990483 |
| RQLWWGHRI | ASSGDVWEANTGQLY | 0.18225697 |
| YGPPGTGKTLL | ASSGDVWEANTGQLY | 0.3642579 |
| AALVGAPANVA | ASSGDVWEANTGQLY | 0.17834052 |
| AATFRRTE | ASSGDVWEANTGQLY | 0.19077796 |
| AGALGGGI | ASSGDVWEANTGQLY | 0.25943702 |
| AGGAGGAGGIGAG | ASSGDVWEANTGQLY | 0.28438067 |
| AGSGALGL | ASSGDVWEANTGQLY | 0.14899752 |
| AQIADQI | ASSGDVWEANTGQLY | 0.1519489 |
| ASAVAVTYL | ASSGDVWEANTGQLY | 0.18649521 |
| ASTVLGGTGG | ASSGDVWEANTGQLY | 0.34131104 |
| AVGGEPAK | ASSGDVWEANTGQLY | 0.39856574 |
| CEQGTFGGTC | ASSGDVWEANTGQLY | 0.2558442 |
| DEGKAPAG | ASSGDVWEANTGQLY | 0.16707766 |
| DTVVFPAL | ASSGDVWEANTGQLY | 0.44517207 |
| FEGITMAG | ASSGDVWEANTGQLY | 0.25967434 |
| FPLVVQD | ASSGDVWEANTGQLY | 0.2643628 |
| GAFTTPQ | ASSGDVWEANTGQLY | 0.28868428 |
| GFGGFGTAGGVGGA | ASSGDVWEANTGQLY | 0.15110463 |
| GFLAQFL | ASSGDVWEANTGQLY | 0.29056072 |
| GGAGGAGGNAGVA | ASSGDVWEANTGQLY | 0.31911656 |
| GGVGTAGAGGAGGAGG | ASSGDVWEANTGQLY | 0.27222574 |

| | | |
|---|---|---|
| GNGGGGDAGNAGSGGN | ASSGDVWEANTGQLY | 0.23668805 |
| GTGGAGGAGNDAGS | ASSGDVWEANTGQLY | 0.19112018 |
| GTTNTGWFNS | ASSGDVWEANTGQLY | 0.24486054 |
| IFMGPLFTL | ASSGDVWEANTGQLY | 0.17949484 |
| ILAHAIEAI | ASSGDVWEANTGQLY | 0.15852228 |
| IRSAIREAL | ASSGDVWEANTGQLY | 0.31050628 |
| ITFAGGG | ASSGDVWEANTGQLY | 0.29302776 |
| ITGMSGHQEV | ASSGDVWEANTGQLY | 0.28901643 |
| IVAGLACTL | ASSGDVWEANTGQLY | 0.2789323 |
| IVGGGLAGA | ASSGDVWEANTGQLY | 0.1876947 |
| IVTGVLNI | ASSGDVWEANTGQLY | 0.3379287 |
| LAQQGCI | ASSGDVWEANTGQLY | 0.43219388 |
| LCLHVEN | ASSGDVWEANTGQLY | 0.14903176 |
| LDAISDGS | ASSGDVWEANTGQLY | 0.17523201 |
| LGGFAAML | ASSGDVWEANTGQLY | 0.27321774 |
| LGGKNPMI | ASSGDVWEANTGQLY | 0.26388013 |
| LGPKGVAAIL | ASSGDVWEANTGQLY | 0.26291102 |
| LMVGVSI | ASSGDVWEANTGQLY | 0.22119427 |
| LVGAAFVL | ASSGDVWEANTGQLY | 0.2664276 |
| LVLAVAPL | ASSGDVWEANTGQLY | 0.15676959 |
| MAAIAVIDAA | ASSGDVWEANTGQLY | 0.5198631 |
| NNGINAELQ | ASSGDVWEANTGQLY | 0.14961551 |
| RGEPLVLVHG | ASSGDVWEANTGQLY | 0.18810795 |
| RGGAGGAAGLI | ASSGDVWEANTGQLY | 0.16861725 |
| RSYRMHPAV | ASSGDVWEANTGQLY | 0.123202235 |
| SAAHLLAGQLA | ASSGDVWEANTGQLY | 0.34910896 |
| SAAVGSY | ASSGDVWEANTGQLY | 0.17344695 |
| SPHAVLHRI | ASSGDVWEANTGQLY | 0.42458284 |
| SQIAKVL | ASSGDVWEANTGQLY | 0.18922341 |
| TALGHLVP | ASSGDVWEANTGQLY | 0.26765692 |
| TAPTTSGAA | ASSGDVWEANTGQLY | 0.2055522 |
| TDGDDGNGGSPGT | ASSGDVWEANTGQLY | 0.2224009 |
| TDGLGEN | ASSGDVWEANTGQLY | 0.17192566 |
| TGGPPLPGQA | ASSGDVWEANTGQLY | 0.34867233 |
| TIPAITI | ASSGDVWEANTGQLY | 0.3833523 |
| TIPGFTI | ASSGDVWEANTGQLY | 0.2530428 |
| TSIASGALA | ASSGDVWEANTGQLY | 0.14708978 |
| TVTLPGAAI | ASSGDVWEANTGQLY | 0.16453731 |
| VASGLSAALP | ASSGDVWEANTGQLY | 0.26746362 |
| VGAGVIGI | ASSGDVWEANTGQLY | 0.11409615 |
| VGPKRFGV | ASSGDVWEANTGQLY | 0.12248944 |
| VNVAFPD | ASSGDVWEANTGQLY | 0.13112196 |
| VSMTNSGVS | ASSGDVWEANTGQLY | 0.12295648 |
| VTDVLGDLSV | ASSGDVWEANTGQLY | 0.5002928 |
| VVVVGAGFAGL | ASSGDVWEANTGQLY | 0.47523394 |
| VYTVGTVV | ASSGDVWEANTGQLY | 0.26375365 |
| WTTFSGP | ASSGDVWEANTGQLY | 0.40147883 |
| YDELVEAF | ASSGDVWEANTGQLY | 0.2873214 |
| YGTGGAGGNGG | ASSGDVWEANTGQLY | 0.1092075 |
| YTNVMAV | ASSGDVWEANTGQLY | 0.18451594 |
| FLEQVHQL | ASSGDVWEANTGQLY | 0.15820305 |

| | | |
|---|---|---|
| GLSLLAVAV | ASSGDVWEANTGQLY | 0.3366157 |
| LEQVHQL | ASSGDVWEANTGQLY | 0.13708009 |
| LNAAGAI | ASSGDVWEANTGQLY | 0.16954449 |
| PAVHASL | ASSGDVWEANTGQLY | 0.30964732 |
| TELEDDA | ASSGDVWEANTGQLY | 0.17719859 |
| TPPGPPN | ASSGDVWEANTGQLY | 0.2521424 |
| TPPPPPPP | ASSGDVWEANTGQLY | 0.15382162 |
| TQSPALI | ASSHLYEQY | 0.1146191 |
| EFAAAFPQ | ASSHLYEQY | 0.35604542 |
| LLSGNLDVL | ASSHLYEQY | 0.12068582 |
| GPANLGNYNI | ASSHLYEQY | 0.17692788 |
| TLGPTLP | ASSHLYEQY | 0.121684484 |
| VYAYSSCCK | ASSHLYEQY | 0.12639855 |
| APAPAPAPNI | ASSHLYEQY | 0.13662684 |
| PGGPVPIP | ASSHLYEQY | 0.28698438 |
| FLTVALEP | ASSHLYEQY | 0.16240509 |
| SQYFGANL | ASSHLYEQY | 0.22212926 |
| GGVKPAA | ASSHLYEQY | 0.31483692 |
| GGTGGAGGAGGW | ASSHLYEQY | 0.35550737 |
| GVGGVAGLGGAGG | ASSHLYEQY | 0.21312818 |
| VLLVLATTI | ASSHLYEQY | 0.18515253 |
| IGIFAAML | ASSHLYEQY | 0.16808167 |
| AVGDLID | ASSHLYEQY | 0.170999 |
| NISPLPL | ASSHLYEQY | 0.17878425 |
| VTAVFVAI | ASSHLYEQY | 0.52795446 |
| SGDDNDAAL | ASSHLYEQY | 0.18528305 |
| AVAALAHS | ASSHLYEQY | 0.14212303 |
| GGVTVMATP | ASSHLYEQY | 0.16397373 |
| AGLAANVRD | ASSHLYEQY | 0.5000597 |
| TNRPDLIDPALLRPG | ASSHLYEQY | 0.46650702 |
| LANLGVTAL | ASSHLYEQY | 0.16741672 |
| TGERIEL | ASSHLYEQY | 0.21082145 |
| NGEELAA | ASSHLYEQY | 0.20431434 |
| VLSLVWG | ASSHLYEQY | 0.27847207 |
| IGYALWL | ASSHLYEQY | 0.17078385 |
| LSLLLAPL | ASSHLYEQY | 0.16922723 |
| TLDEALK | ASSHLYEQY | 0.26246685 |
| TDGAGTAEGG | ASSHLYEQY | 0.60267526 |
| NAPGGAGGNGG | ASSHLYEQY | 0.19317015 |
| VSVFMGMGGNF | ASSHLYEQY | 0.1293014 |
| LASYLAAL | ASSHLYEQY | 0.34457853 |
| LVGGTPAL | ASSHLYEQY | 0.16049877 |
| VRNADVL | ASSHLYEQY | 0.23831074 |
| TCSSPFLVSAM | ASSHLYEQY | 0.23697375 |
| VHITDAE | ASSHLYEQY | 0.13336131 |
| EYAAPSF | ASSHLYEQY | 0.19616531 |
| VVDAFHGGL | ASSHLYEQY | 0.37297475 |
| ASVVDPQ | ASSHLYEQY | 0.18329267 |
| SYNEFGGLS | ASSHLYEQY | 0.19463953 |
| DPDGSVY | ASSHLYEQY | 0.44317293 |
| MVSTIHATE | ASSHLYEQY | 0.33608037 |

| | | |
|---|---|---|
| LAAGVADPV | ASSHLYEQY | 0.54417014 |
| TKEQIAA | ASSHLYEQY | 0.25243074 |
| SLASPAAL | ASSHLYEQY | 0.21499272 |
| IGLVFYGV | ASSHLYEQY | 0.2168275 |
| FLLDASNL | ASSHLYEQY | 0.18601193 |
| YQVGNFP | ASSHLYEQY | 0.3805393 |
| ESLLSQST | ASSHLYEQY | 0.28401726 |
| LFTVGFAAVG | ASSHLYEQY | 0.16273105 |
| ATPPTPT | ASSHLYEQY | 0.551666 |
| TDLATLG | ASSHLYEQY | 0.2213746 |
| SAGGTGR | ASSHLYEQY | 0.2899471 |
| GFSNSGS | ASSHLYEQY | 0.16657136 |
| GGNGGSGGTSV | ASSHLYEQY | 0.17147462 |
| GTSVATGGAGN | ASSHLYEQY | 0.3590982 |
| NGGNGGSGGTS | ASSHLYEQY | 0.1368251 |
| AVVKFPH | ASSHLYEQY | 0.13253249 |
| TPPPPPP | ASSHLYEQY | 0.20528689 |
| AAPAAAAAPA | ASSHLYEQY | 0.30960068 |
| IALNEDL | ASSHLYEQY | 0.10984366 |
| IVNVSSM | ASSHLYEQY | 0.17624116 |
| TGIPPAPR | ASSHLYEQY | 0.1402069 |
| FDLGGGTF | ASSHLYEQY | 0.17092516 |
| FDLGGGTFDVS | ASSHLYEQY | 0.24977048 |
| NEPTAAA | ASSHLYEQY | 0.13869059 |
| NEPTAAALA | ASSHLYEQY | 0.29228306 |
| GSIINISSI | ASSHLYEQY | 0.1326943 |
| VAPGNAGTA | ASSHLYEQY | 0.13122688 |
| TGGGASLE | ASSHLYEQY | 0.25383526 |
| VAVYNHYKRI | ASSHLYEQY | 0.31395173 |
| TDPAPATT | ASSHLYEQY | 0.1546353 |
| TNRPDILDPALLRPG | ASSHLYEQY | 0.30676705 |
| LTTTTTS | ASSHLYEQY | 0.15685204 |
| PTGSGKTL | ASSHLYEQY | 0.49821204 |
| TGSGKTL | ASSHLYEQY | 0.21097124 |
| TIGHVDHGK | ASSHLYEQY | 0.13466278 |
| TIGHVDHGKTT | ASSHLYEQY | 0.34205362 |
| GHVDHGKTT | ASSHLYEQY | 0.19382066 |
| AQFEHTLL | ASSHLYEQY | 0.29569456 |
| IVNQDEL | ASSHLYEQY | 0.2514375 |
| TNGSQFF | ASSHLYEQY | 0.14091747 |
| TGKGIVE | ASSHLYEQY | 0.26173833 |
| LISNASDA | ASSHLYEQY | 0.4406749 |
| AELRTLGELTKIAKAHGAQ | ASSHLYEQY | 0.3054958 |
| APNAVAAAEPSW | ASSHLYEQY | 0.14147997 |
| APQVAGAATAVL | ASSHLYEQY | 0.26122642 |
| ARRDPAAAAAILPTDA | ASSHLYEQY | 0.25094634 |
| DAAARDVLAEAGLAEYF | ASSHLYEQY | 0.1920354 |
| DDQIVLSAKANLAKVEQ | ASSHLYEQY | 0.2762829 |
| DELLPDTGEDSDEDGH | ASSHLYEQY | 0.2068138 |
| ELPEAPANRATVPQD | ASSHLYEQY | 0.3930645 |
| FGQSKIERVTGIG | ASSHLYEQY | 0.30928877 |

| | | |
|---|---|---|
| GAGGTGSTAGGAGGAGGAGGLY | ASSHLYEQY | 0.32042757 |
| GHGGTGAAGGNGGTGGAGGSGI | ASSHLYEQY | 0.14725107 |
| GNGGNGGAGGEGGPGGAGGAGA | ASSHLYEQY | 0.17025241 |
| GSTAGNGGNGGNATNGTIAGQP | ASSHLYEQY | 0.2566867 |
| GTGGDGGNGGNGGTGG | ASSHLYEQY | 0.2507737 |
| GVRRPAGIPADAPV | ASSHLYEQY | 0.1372083 |
| GYRQVTDAHLLTLARR | ASSHLYEQY | 0.13506372 |
| IAKMGQDGHDRG | ASSHLYEQY | 0.27360812 |
| KAEPSIEIQYT | ASSHLYEQY | 0.46192712 |
| KPVRIRMAADEDTVAVT | ASSHLYEQY | 0.20351066 |
| MPPRIAGMRLLVIKPEP | ASSHLYEQY | 0.21825507 |
| MKFYAVALFRSRGVPPTR | ASSHLYEQY | 0.6234715 |
| NAGGFGILGAVAHSP | ASSHLYEQY | 0.17379548 |
| NEGREITLAMAEM | ASSHLYEQY | 0.22719991 |
| NELVEWSHPLGHRWRWE | ASSHLYEQY | 0.288456 |
| NLGAGGTGGNGGNPG | ASSHLYEQY | 0.38118306 |
| NTEWPKWETVMAADPVD | ASSHLYEQY | 0.21295421 |
| PASAAPAETQP | ASSHLYEQY | 0.38450053 |
| QQSSDGKNGVRAAEITGE | ASSHLYEQY | 0.1388679 |
| RAPDAVTTSWWKE | ASSHLYEQY | 0.44528353 |
| SGFGNVGAHTSGWFNQ | ASSHLYEQY | 0.12441853 |
| SGVAGAVGQAASVGGLKV | ASSHLYEQY | 0.3171056 |
| SNGGAGTGSAGGNGGT | ASSHLYEQY | 0.12825176 |
| TGGAGGEGGAAG | ASSHLYEQY | 0.4344617 |
| TGGDGGAAGAGGAPGHG | ASSHLYEQY | 0.44872376 |
| TMPPLLAAIANAGRHGV | ASSHLYEQY | 0.47195667 |
| TSSTAGNSGGA | ASSHLYEQY | 0.118233316 |
| VPATVPVGILC | ASSHLYEQY | 0.337855 |
| VQRPDSGGVVLTGRIS | ASSHLYEQY | 0.48558247 |
| WTPNPVPATKGVPAGLVQAT | ASSHLYEQY | 0.59711826 |
| YALKRALAKFSDRFDVVII | ASSHLYEQY | 0.23174696 |
| YLIGSVIV | ASSHLYEQY | 0.29946852 |
| YGQLVAVID | ASSHLYEQY | 0.19412641 |
| VTVGGMTLPA | ASSHLYEQY | 0.1799529 |
| VLALPAV | ASSHLYEQY | 0.17093554 |
| VIPPTLNYET | ASSHLYEQY | 0.16463119 |
| VGMGGMHPSSG | ASSHLYEQY | 0.2401363 |
| VGLAGGAATA | ASSHLYEQY | 0.18479931 |
| VGGAGGAGGSGP | ASSHLYEQY | 0.3029185 |
| VFAAFMV | ASSHLYEQY | 0.39949167 |
| TLYDKLR | ASSHLYEQY | 0.2716241 |
| TLAVFTVL | ASSHLYEQY | 0.5512979 |
| TGSVQPARV | ASSHLYEQY | 0.33866474 |
| TGGGAGSGGT | ASSHLYEQY | 0.25934544 |
| SHLETSL | ASSHLYEQY | 0.16480953 |
| SFTTGMT | ASSHLYEQY | 0.19474271 |
| QVEVAKRMGVRQPTVS | ASSHLYEQY | 0.15173027 |
| QGSGGPALAM | ASSHLYEQY | 0.19825903 |
| MLAPSNP | ASSHLYEQY | 0.17202178 |
| LVNNAAI | ASSHLYEQY | 0.3094086 |
| LTGEINTLA | ASSHLYEQY | 0.1094867 |

| | | |
|---|---|---|
| LSIFGPV | ASSHLYEQY | 0.23529166 |
| LLALLANL | ASSHLYEQY | 0.49525118 |
| LISNASDALDKLR | ASSHLYEQY | 0.12338949 |
| LINEAVP | ASSHLYEQY | 0.34564584 |
| LGYAFSGL | ASSHLYEQY | 0.17228916 |
| LDFVAVAL | ASSHLYEQY | 0.16072474 |
| LAQPGAE | ASSHLYEQY | 0.49330258 |
| ISNLGAVTGF | ASSHLYEQY | 0.17372057 |
| HQEDPGRI | ASSHLYEQY | 0.2891392 |
| GTPGEPTP | ASSHLYEQY | 0.2959859 |
| GFFNSGS | ASSHLYEQY | 0.25270888 |
| GAGGGGLTAGAN | ASSHLYEQY | 0.21131125 |
| FVDALVNM | ASSHLYEQY | 0.23872532 |
| FLTVAPEL | ASSHLYEQY | 0.14561938 |
| FETLTAL | ASSHLYEQY | 0.23700671 |
| DVINPIEL | ASSHLYEQY | 0.114269584 |
| APVSGAPV | ASSHLYEQY | 0.24160358 |
| ALVGGLAW | ASSHLYEQY | 0.16637194 |
| ALAAAAATAI | ASSHLYEQY | 0.41345397 |
| AFGLGLLL | ASSHLYEQY | 0.24607666 |
| AFFAAASL | ASSHLYEQY | 0.25459403 |
| AAAALPAFSPP | ASSHLYEQY | 0.18441972 |
| VVLAPAL | ASSHLYEQY | 0.17879951 |
| TIGHVDHGKTTL | ASSHLYEQY | 0.15412654 |
| LISNASDALDK | ASSHLYEQY | 0.15144277 |
| LISNASDAL | ASSHLYEQY | 0.17402722 |
| ISNASDALDK | ASSHLYEQY | 0.20836245 |
| NQAETLVYQTE | ASSHLYEQY | 0.13732997 |
| TGGTGGTAGAG | ASSHLYEQY | 0.36333138 |
| TGSTAGGAGGA | ASSHLYEQY | 0.24562223 |
| DPLAVPVAAGPV | ASSHLYEQY | 0.13211638 |
| GGTGGTPGNGGN | ASSHLYEQY | 0.23632705 |
| PAGLAGVDPATL | ASSHLYEQY | 0.27377576 |
| GNNLVVEAPHAGASV | ASSHLYEQY | 0.21378487 |
| KVTRIMSASTLEQPA | ASSHLYEQY | 0.11021288 |
| LGFALPIALPRRTVV | ASSHLYEQY | 0.1638728 |
| TRLGNLTARKLVVLV | ASSHLYEQY | 0.19677712 |
| WSQQRRLIYFPSAGP | ASSHLYEQY | 0.12591666 |
| GPNGNPDPMAAAVDIR | ASSHLYEQY | 0.15716277 |
| TQRCVSVADLRAAASR | ASSHLYEQY | 0.47217065 |
| QFVLDVAEHPVEVLTETV | ASSHLYEQY | 0.14830543 |
| GGAGGTGGAGGNGGTQAGDG | ASSHLYEQY | 0.18391143 |
| PLLATLRAIAVDVDAKPAQV | ASSHLYEQY | 0.3041863 |
| APKFRTGGYMIDFWGVGYQVAK | ASSHLYEQY | 0.37636223 |
| GAGGAGGAGGAAGGPGGTGGTG | ASSHLYEQY | 0.5830527 |
| IGALGLLATDVAPAGSVAFGVA | ASSHLYEQY | 0.19206132 |
| AGGAGGIGGNGGAGGAGGVLMG | ASSHLYEQY | 0.1961555 |
| ADELIGLL | ASSHLYEQY | 0.27790856 |
| ADRALKA | ASSHLYEQY | 0.24266613 |
| AGAVGQVGGAGGA | ASSHLYEQY | 0.13206197 |
| AGGGGGLGGAP | ASSHLYEQY | 0.11976829 |

| | | |
|---|---|---|
| AGGLLYGNGGAG | ASSHLYEQY | 0.24018696 |
| AGLESIGL | ASSHLYEQY | 0.14681955 |
| AGLKTGII | ASSHLYEQY | 0.2595097 |
| ASLAALLL | ASSHLYEQY | 0.30407435 |
| ASVINISSI | ASSHLYEQY | 0.13763992 |
| ATPPGPA | ASSHLYEQY | 0.14964832 |
| ATVASALV | ASSHLYEQY | 0.20219985 |
| ATVASLIL | ASSHLYEQY | 0.18251644 |
| AVNGPVLIRH | ASSHLYEQY | 0.42169213 |
| DGAAGAVGI | ASSHLYEQY | 0.191398 |
| DPLLHGI | ASSHLYEQY | 0.5574424 |
| ELGAALGAL | ASSHLYEQY | 0.38811648 |
| ELGRTQAV | ASSHLYEQY | 0.19442298 |
| FVAGGYLS | ASSHLYEQY | 0.39370602 |
| GASGGAGGAGGW | ASSHLYEQY | 0.19049685 |
| GGAGAGGDGGAGGI | ASSHLYEQY | 0.13427955 |
| GTGHNGSQPGMGGNG | ASSHLYEQY | 0.20532133 |
| GYAAASASAS | ASSHLYEQY | 0.14574559 |
| HQVDVAAVQQG | ASSHLYEQY | 0.18638182 |
| IGAAIAHT | ASSHLYEQY | 0.18491778 |
| IGMIEVAQL | ASSHLYEQY | 0.14293316 |
| IRLAPEHTSAMD | ASSHLYEQY | 0.21279803 |
| IYNVSVL | ASSHLYEQY | 0.17807665 |
| KQGFLGM | ASSHLYEQY | 0.13261847 |
| LESVITQL | ASSHLYEQY | 0.2071358 |
| LGLGALVVV | ASSHLYEQY | 0.38466233 |
| LIGNGGNGGSGGI | ASSHLYEQY | 0.6176461 |
| LNAINAPTL | ASSHLYEQY | 0.197525 |
| LQLLNTGV | ASSHLYEQY | 0.17374884 |
| LQTSIEF | ASSHLYEQY | 0.2869351 |
| MSFAVSEL | ASSHLYEQY | 0.18375768 |
| NTGALGL | ASSHLYEQY | 0.22688836 |
| PPGAGGAGGAGTT | ASSHLYEQY | 0.23470142 |
| QLVGLGI | ASSHLYEQY | 0.16260287 |
| RFLEQVHQL | ASSHLYEQY | 0.17503694 |
| RGVGYVLR | ASSHLYEQY | 0.23139976 |
| SATGVFAGVI | ASSHLYEQY | 0.14051035 |
| SLPAITLP | ASSHLYEQY | 0.26461002 |
| STALAFTV | ASSHLYEQY | 0.1083351 |
| SVLTVGLGK | ASSHLYEQY | 0.16316216 |
| TALAGAGSGI | ASSHLYEQY | 0.1789999 |
| TGLPFTGI | ASSHLYEQY | 0.18099444 |
| VAEHVTV | ASSHLYEQY | 0.13033295 |
| VDIVLNSL | ASSHLYEQY | 0.18878753 |
| VLTGGGAVL | ASSHLYEQY | 0.29319155 |
| VNVGLAYGI | ASSHLYEQY | 0.43759495 |
| VSSAAVGAV | ASSHLYEQY | 0.13961907 |
| VTSVYEAGV | ASSHLYEQY | 0.36605358 |
| WVALDRGA | ASSHLYEQY | 0.40557867 |
| YLVGSVVI | ASSHLYEQY | 0.1968346 |
| GPPGTGKTLL | ASSHLYEQY | 0.13737662 |

| | | |
|---|---|---|
| GVGMTKF | ASSHLYEQY | 0.2862229 |
| IGIGGSDL | ASSHLYEQY | 0.115999155 |
| IGTLLAEL | ASSHLYEQY | 0.16268139 |
| IYTSGTT | ASSHLYEQY | 0.34398973 |
| LSFPAGV | ASSHLYEQY | 0.14623347 |
| LYTSGTT | ASSHLYEQY | 0.3523627 |
| VGAFPFGF | ASSHLYEQY | 0.2920452 |
| VGAGTVEF | ASSHLYEQY | 0.16959067 |
| VGKSTLLL | ASSHLYEQY | 0.15058398 |
| AALAHAGGGIV | ASSHLYEQY | 0.15221617 |
| AALVAVSLL | ASSHLYEQY | 0.1158066 |
| AAVSRAVAAL | ASSHLYEQY | 0.3319182 |
| AGGSSGIGGF | ASSHLYEQY | 0.18563326 |
| AGLTVANANG | ASSHLYEQY | 0.19010416 |
| AGLVGAVL | ASSHLYEQY | 0.2694927 |
| ALSQAFTV | ASSHLYEQY | 0.19320396 |
| ALTFSVSL | ASSHLYEQY | 0.20968433 |
| ALTGAALVAL | ASSHLYEQY | 0.20252988 |
| ASFSIGAGVL | ASSHLYEQY | 0.16669355 |
| DGSVYKI | ASSHLYEQY | 0.2611103 |
| EQLAEIV | ASSHLYEQY | 0.16039431 |
| FAVSELTVM | ASSHLYEQY | 0.1689986 |
| GAAPLAAGL | ASSHLYEQY | 0.5697338 |
| GADGGDGGSGGAGG | ASSHLYEQY | 0.13079123 |
| GFGLDTGGAGG | ASSHLYEQY | 0.18352099 |
| GLASGGLL | ASSHLYEQY | 0.21483248 |
| GRIAGAAVLL | ASSHLYEQY | 0.13526325 |
| GSAGSLNTSF | ASSHLYEQY | 0.31188738 |
| GTGQIGI | ASSHLYEQY | 0.27599266 |
| GTLIALIL | ASSHLYEQY | 0.32075858 |
| IAATMGFVVG | ASSHLYEQY | 0.15530866 |
| IFAQALL | ASSHLYEQY | 0.35480973 |
| IGAAVALGF | ASSHLYEQY | 0.378143 |
| IGLTPLAL | ASSHLYEQY | 0.1814332 |
| ILGSNTSTL | ASSHLYEQY | 0.14905402 |
| IVFSLTVL | ASSHLYEQY | 0.14451802 |
| LASKATGF | ASSHLYEQY | 0.29788008 |
| LASVALIL | ASSHLYEQY | 0.33899316 |
| LFAVGLTL | ASSHLYEQY | 0.1436912 |
| LFNGGIGTYI | ASSHLYEQY | 0.27600464 |
| LTLGAGII | ASSHLYEQY | 0.32836357 |
| LVAAALGDI | ASSHLYEQY | 0.4039808 |
| NTAFVQLGI | ASSHLYEQY | 0.13959661 |
| QLFSGAEL | ASSHLYEQY | 0.20070007 |
| SAATFITL | ASSHLYEQY | 0.2188907 |
| SNYGLANLGV | ASSHLYEQY | 0.19978423 |
| SYGGAYAVM | ASSHLYEQY | 0.15740347 |
| TTGVGGLTLG | ASSHLYEQY | 0.3492775 |
| TTQTIYI | ASSHLYEQY | 0.23726423 |
| VAAGAGLGAVI | ASSHLYEQY | 0.4017437 |
| VAAVVTLL | ASSHLYEQY | 0.27939567 |

| | | |
|---|---|---|
| VGFLGGLI | ASSHLYEQY | 0.43405658 |
| VGVIPILL | ASSHLYEQY | 0.20132011 |
| VIFASSLI | ASSHLYEQY | 0.22427323 |
| VKGGDGGAAAG | ASSHLYEQY | 0.13656424 |
| VTKTFTNI | ASSHLYEQY | 0.13049431 |
| VVAAAILL | ASSHLYEQY | 0.14823422 |
| YAALDVEL | ASSHLYEQY | 0.20035872 |
| LASVALLL | ASSHLYEQY | 0.11187054 |
| LDAVHRV | ASSHLYEQY | 0.23553771 |
| QGLSSSIF | ASSHLYEQY | 0.27853635 |
| RGTFANIRL | ASSHLYEQY | 0.35888848 |
| RQLWWGHRI | ASSHLYEQY | 0.21665275 |
| YGPPGTGKTLL | ASSHLYEQY | 0.1730333 |
| AALVGAPANVA | ASSHLYEQY | 0.16329785 |
| AATFRRTE | ASSHLYEQY | 0.48403108 |
| AGALGGGI | ASSHLYEQY | 0.13776593 |
| AGGAGGAGGIGAG | ASSHLYEQY | 0.13152015 |
| AGSGALGL | ASSHLYEQY | 0.18045759 |
| AQIADQI | ASSHLYEQY | 0.1435785 |
| ASAVAVTYL | ASSHLYEQY | 0.30604884 |
| ASTVLGGTGG | ASSHLYEQY | 0.40057904 |
| AVGGEPAK | ASSHLYEQY | 0.30984607 |
| CEQGTFGGTC | ASSHLYEQY | 0.28625545 |
| DEGKAPAG | ASSHLYEQY | 0.12056799 |
| DTVVFPAL | ASSHLYEQY | 0.19022742 |
| FEGITMAG | ASSHLYEQY | 0.62541103 |
| FPLVVQD | ASSHLYEQY | 0.23076831 |
| GAFTTPQ | ASSHLYEQY | 0.2587191 |
| GFGGFGTAGGVGGA | ASSHLYEQY | 0.38287473 |
| GFLAQFL | ASSHLYEQY | 0.18095453 |
| GGAGGAGGNAGVA | ASSHLYEQY | 0.21928264 |
| GGVGTAGAGGAGGAGG | ASSHLYEQY | 0.20446064 |
| GNGGGGDAGNAGSGGN | ASSHLYEQY | 0.25983566 |
| GTGGAGGAGNDAGS | ASSHLYEQY | 0.14904073 |
| GTTNTGWFNS | ASSHLYEQY | 0.14879496 |
| IFMGPLFTL | ASSHLYEQY | 0.29196233 |
| ILAHAIEAI | ASSHLYEQY | 0.5435906 |
| IRSAIREAL | ASSHLYEQY | 0.26580083 |
| ITFAGGG | ASSHLYEQY | 0.26627815 |
| ITGMSGHQEV | ASSHLYEQY | 0.24855249 |
| IVAGLACTL | ASSHLYEQY | 0.25235152 |
| IVGGGLAGA | ASSHLYEQY | 0.18541715 |
| IVTGVLNI | ASSHLYEQY | 0.17489353 |
| LAQQGCI | ASSHLYEQY | 0.19053307 |
| LCLHVEN | ASSHLYEQY | 0.16679008 |
| LDAISDGS | ASSHLYEQY | 0.22313832 |
| LGGFAAML | ASSHLYEQY | 0.2048566 |
| LGGKNPMI | ASSHLYEQY | 0.61736965 |
| LGPKGVAAIL | ASSHLYEQY | 0.36680365 |
| LMVGVSI | ASSHLYEQY | 0.22389363 |
| LVGAAFVL | ASSHLYEQY | 0.16374494 |

| | | |
|---|---|---|
| LVLAVAPL | ASSHLYEQY | 0.19879833 |
| MAAIAVIDAA | ASSHLYEQY | 0.2503175 |
| NNGINAELQ | ASSHLYEQY | 0.14853242 |
| RGEPLVLVHG | ASSHLYEQY | 0.59371907 |
| RGGAGGAAGLI | ASSHLYEQY | 0.18614773 |
| RSYRMHPAV | ASSHLYEQY | 0.14041777 |
| SAAHLLAGQLA | ASSHLYEQY | 0.2867626 |
| SAAVGSY | ASSHLYEQY | 0.56168413 |
| SPHAVLHRI | ASSHLYEQY | 0.21294273 |
| SQIAKVL | ASSHLYEQY | 0.17407951 |
| TALGHLVP | ASSHLYEQY | 0.16956411 |
| TAPTTSGAA | ASSHLYEQY | 0.3522292 |
| TDGDDGNGGSPGT | ASSHLYEQY | 0.5721983 |
| TDGLGEN | ASSHLYEQY | 0.13311437 |
| TGGPPLPGQA | ASSHLYEQY | 0.46534234 |
| TIPAITI | ASSHLYEQY | 0.16363665 |
| TIPGFTI | ASSHLYEQY | 0.1432681 |
| TSIASGALA | ASSHLYEQY | 0.25188783 |
| TVTLPGAAI | ASSHLYEQY | 0.21692592 |
| VASGLSAALP | ASSHLYEQY | 0.35669446 |
| VGAGVIGI | ASSHLYEQY | 0.31549877 |
| VGPKRFGV | ASSHLYEQY | 0.14915653 |
| VNVAFPD | ASSHLYEQY | 0.17556025 |
| VSMTNSGVS | ASSHLYEQY | 0.46544963 |
| VTDVLGDLSV | ASSHLYEQY | 0.2696869 |
| VVVVGAGFAGL | ASSHLYEQY | 0.31118408 |
| VYTVGTVV | ASSHLYEQY | 0.21169236 |
| WTTFSGP | ASSHLYEQY | 0.18729465 |
| YDELVEAF | ASSHLYEQY | 0.23577079 |
| YGTGGAGGNGG | ASSHLYEQY | 0.19562086 |
| YTNVMAV | ASSHLYEQY | 0.16340148 |
| FLEQVHQL | ASSHLYEQY | 0.26743168 |
| GLSLLAVAV | ASSHLYEQY | 0.40652522 |
| LEQVHQL | ASSHLYEQY | 0.14382623 |
| LNAAGAI | ASSHLYEQY | 0.17598899 |
| PAVHASL | ASSHLYEQY | 0.25165224 |
| TELEDDA | ASSHLYEQY | 0.20750299 |
| TPPGPPN | ASSHLYEQY | 0.18470402 |
| TPPPPPPP | ASSHLYEQY | 0.51946807 |
| TQSPALI | ASSPDRYNNQAPL | 0.6153796 |
| EFAAAFPQ | ASSPDRYNNQAPL | 0.34702536 |
| LLSGNLDVL | ASSPDRYNNQAPL | 0.14621213 |
| GPANLGNYNI | ASSPDRYNNQAPL | 0.12392812 |
| TLGPTLP | ASSPDRYNNQAPL | 0.29388097 |
| VYAYSSCCK | ASSPDRYNNQAPL | 0.2020702 |
| APAPAPAPNI | ASSPDRYNNQAPL | 0.19026524 |
| PGGPVPIP | ASSPDRYNNQAPL | 0.16742401 |
| FLTVALEP | ASSPDRYNNQAPL | 0.19096044 |
| SQYFGANL | ASSPDRYNNQAPL | 0.39078873 |
| GGVKPAA | ASSPDRYNNQAPL | 0.17588338 |
| GGTGGAGGAGGW | ASSPDRYNNQAPL | 0.144002 |

| | | |
|---|---|---|
| GVGGVAGLGGAGG | ASSPDRYNNQAPL | 0.14881389 |
| VLLVLATTI | ASSPDRYNNQAPL | 0.36078674 |
| IGIFAAML | ASSPDRYNNQAPL | 0.16377437 |
| AVGDLID | ASSPDRYNNQAPL | 0.2454696 |
| NISPLPL | ASSPDRYNNQAPL | 0.29650044 |
| VTAVFVAI | ASSPDRYNNQAPL | 0.16174215 |
| SGDDNDAAL | ASSPDRYNNQAPL | 0.26086769 |
| AVAALAHS | ASSPDRYNNQAPL | 0.16671202 |
| GGVTVMATP | ASSPDRYNNQAPL | 0.21601355 |
| AGLAANVRD | ASSPDRYNNQAPL | 0.17811136 |
| TNRPDLIDPALLRPG | ASSPDRYNNQAPL | 0.26079375 |
| LANLGVTAL | ASSPDRYNNQAPL | 0.29990563 |
| TGERIEL | ASSPDRYNNQAPL | 0.27263027 |
| NGEELAA | ASSPDRYNNQAPL | 0.23655409 |
| VLSLVWG | ASSPDRYNNQAPL | 0.17760073 |
| IGYALWL | ASSPDRYNNQAPL | 0.14341499 |
| LSLLLAPL | ASSPDRYNNQAPL | 0.119880654 |
| TLDEALK | ASSPDRYNNQAPL | 0.38107786 |
| TDGAGTAEGG | ASSPDRYNNQAPL | 0.1273671 |
| NAPGGAGGNGG | ASSPDRYNNQAPL | 0.36618888 |
| VSVFMGMGGNF | ASSPDRYNNQAPL | 0.30596524 |
| LASYLAAL | ASSPDRYNNQAPL | 0.1627335 |
| LVGGTPAL | ASSPDRYNNQAPL | 0.22787794 |
| VRNADVL | ASSPDRYNNQAPL | 0.12794754 |
| TCSSPFLVSAM | ASSPDRYNNQAPL | 0.17294738 |
| VHITDAE | ASSPDRYNNQAPL | 0.2931437 |
| EYAAPSF | ASSPDRYNNQAPL | 0.2738565 |
| VVDAFHGGL | ASSPDRYNNQAPL | 0.18215507 |
| ASVVDPQ | ASSPDRYNNQAPL | 0.23870602 |
| SYNEFGGLS | ASSPDRYNNQAPL | 0.26107156 |
| DPDGSVY | ASSPDRYNNQAPL | 0.17509393 |
| MVSTIHATE | ASSPDRYNNQAPL | 0.22657652 |
| LAAGVADPV | ASSPDRYNNQAPL | 0.4301239 |
| TKEQIAA | ASSPDRYNNQAPL | 0.25569147 |
| SLASPAAL | ASSPDRYNNQAPL | 0.16890927 |
| IGLVFYGV | ASSPDRYNNQAPL | 0.23445977 |
| FLLDASNL | ASSPDRYNNQAPL | 0.15302268 |
| YQVGNFP | ASSPDRYNNQAPL | 0.36598516 |
| ESLLSQST | ASSPDRYNNQAPL | 0.19241537 |
| LFTVGFAAVG | ASSPDRYNNQAPL | 0.26714313 |
| ATPPTPT | ASSPDRYNNQAPL | 0.21912953 |
| TDLATLG | ASSPDRYNNQAPL | 0.1249877 |
| SAGGTGR | ASSPDRYNNQAPL | 0.16344187 |
| GFSNSGS | ASSPDRYNNQAPL | 0.21425015 |
| GGNGGSGGTSV | ASSPDRYNNQAPL | 0.27038002 |
| GTSVATGGAGN | ASSPDRYNNQAPL | 0.3041079 |
| NGGNGGSGGTS | ASSPDRYNNQAPL | 0.1759709 |
| AVVKFPH | ASSPDRYNNQAPL | 0.2614986 |
| TPPPPPP | ASSPDRYNNQAPL | 0.19700578 |
| AAPAAAAAPA | ASSPDRYNNQAPL | 0.17756397 |
| IALNEDL | ASSPDRYNNQAPL | 0.22166991 |

| | | |
|---|---|---|
| IVNVSSM | ASSPDRYNNQAPL | 0.18465519 |
| TGIPPAPR | ASSPDRYNNQAPL | 0.18377693 |
| FDLGGGTF | ASSPDRYNNQAPL | 0.27255931 |
| FDLGGGTFDVS | ASSPDRYNNQAPL | 0.15042627 |
| NEPTAAA | ASSPDRYNNQAPL | 0.2838186 |
| NEPTAAALA | ASSPDRYNNQAPL | 0.1452355 |
| GSIINISSI | ASSPDRYNNQAPL | 0.17708477 |
| VAPGNAGTA | ASSPDRYNNQAPL | 0.24251534 |
| TGGGASLE | ASSPDRYNNQAPL | 0.16441275 |
| VAVYNHYKRI | ASSPDRYNNQAPL | 0.295448 |
| TDPAPATT | ASSPDRYNNQAPL | 0.21585886 |
| TNRPDILDPALLRPG | ASSPDRYNNQAPL | 0.18554167 |
| LTTTTTS | ASSPDRYNNQAPL | 0.23092031 |
| PTGSGKTL | ASSPDRYNNQAPL | 0.34974888 |
| TGSGKTL | ASSPDRYNNQAPL | 0.35267863 |
| TIGHVDHGK | ASSPDRYNNQAPL | 0.1587908 |
| TIGHVDHGKTT | ASSPDRYNNQAPL | 0.20802996 |
| GHVDHGKTT | ASSPDRYNNQAPL | 0.34185576 |
| AQFEHTLL | ASSPDRYNNQAPL | 0.2562885 |
| IVNQDEL | ASSPDRYNNQAPL | 0.13012701 |
| TNGSQFF | ASSPDRYNNQAPL | 0.1992883 |
| TGKGIVE | ASSPDRYNNQAPL | 0.14387393 |
| LISNASDA | ASSPDRYNNQAPL | 0.17995627 |
| AELRTLGELTKIAKAHGAQ | ASSPDRYNNQAPL | 0.14404336 |
| APNAVAAAEPSW | ASSPDRYNNQAPL | 0.47935373 |
| APQVAGAATAVL | ASSPDRYNNQAPL | 0.48038125 |
| ARRDPAAAAAILPTDA | ASSPDRYNNQAPL | 0.18090059 |
| DAAARDVLAEAGLAEYF | ASSPDRYNNQAPL | 0.27526298 |
| DDQIVLSAKANLAKVEQ | ASSPDRYNNQAPL | 0.13605584 |
| DELLPDTGEDSDEDGH | ASSPDRYNNQAPL | 0.31291866 |
| ELPEAPANRATVPQD | ASSPDRYNNQAPL | 0.5086052 |
| FGQSKIERVTGIG | ASSPDRYNNQAPL | 0.1614623 |
| GAGGTGSTAGGAGGAGGAGGLY | ASSPDRYNNQAPL | 0.2952321 |
| GHGGTGAAGGNGGTGGAGGSGI | ASSPDRYNNQAPL | 0.17390107 |
| GNGGNGGAGGEGGPGGAGGAGA | ASSPDRYNNQAPL | 0.18311529 |
| GSTAGNGGNGGNATNGTIAGQP | ASSPDRYNNQAPL | 0.51667154 |
| GTGGDGGNGGNGGTGG | ASSPDRYNNQAPL | 0.21063417 |
| GVRRPAGIPADAPV | ASSPDRYNNQAPL | 0.23381473 |
| GYRQVTDAHLLTLARR | ASSPDRYNNQAPL | 0.19954945 |
| IAKMGQDGHDRG | ASSPDRYNNQAPL | 0.16668382 |
| KAEPSIEIQYT | ASSPDRYNNQAPL | 0.22275299 |
| KPVRIRMAADEDTVAVT | ASSPDRYNNQAPL | 0.47584444 |
| MPPRIAGMRLLVIKPEP | ASSPDRYNNQAPL | 0.1640214 |
| MKFYAVALFRSRGVPPTR | ASSPDRYNNQAPL | 0.262529 |
| NAGGFGILGAVAHSP | ASSPDRYNNQAPL | 0.1778635 |
| NEGREITLAMAEM | ASSPDRYNNQAPL | 0.4029046 |
| NELVEWSHPLGHRWRWE | ASSPDRYNNQAPL | 0.32837856 |
| NLGAGGTGGNGGNPG | ASSPDRYNNQAPL | 0.32458162 |
| NTEWPKWETVMAADPVD | ASSPDRYNNQAPL | 0.19042237 |
| PASAAPAETQP | ASSPDRYNNQAPL | 0.57661945 |
| QQSSDGKNGVRAAEITGE | ASSPDRYNNQAPL | 0.3024654 |

| | | |
|---|---|---|
| RAPDAVTTSWWKE | ASSPDRYNNQAPL | 0.28867316 |
| SGFGNVGAHTSGWFNQ | ASSPDRYNNQAPL | 0.28775513 |
| SGVAGAVGQAASVGGLKV | ASSPDRYNNQAPL | 0.11924246 |
| SNGGAGTGSAGGNGGT | ASSPDRYNNQAPL | 0.29148078 |
| TGGAGGEGGAAG | ASSPDRYNNQAPL | 0.33950922 |
| TGGDGGAAGAGGAPGHG | ASSPDRYNNQAPL | 0.25140423 |
| TMPPLLAAIANAGRHGV | ASSPDRYNNQAPL | 0.13090006 |
| TSSTAGNSGGA | ASSPDRYNNQAPL | 0.103373945 |
| VPATVPVGILC | ASSPDRYNNQAPL | 0.16732588 |
| VQRPDSGGVVLTGRIS | ASSPDRYNNQAPL | 0.1099611 |
| WTPNPVPATKGVPAGLVQAT | ASSPDRYNNQAPL | 0.41964024 |
| YALKRALAKFSDRFDVVII | ASSPDRYNNQAPL | 0.22031008 |
| YLIGSVIV | ASSPDRYNNQAPL | 0.13309222 |
| YGQLVAVID | ASSPDRYNNQAPL | 0.5152056 |
| VTVGGMTLPA | ASSPDRYNNQAPL | 0.36486864 |
| VLALPAV | ASSPDRYNNQAPL | 0.10466055 |
| VIPPTLNYET | ASSPDRYNNQAPL | 0.15483478 |
| VGMGGMHPSSG | ASSPDRYNNQAPL | 0.15435001 |
| VGLAGGAATA | ASSPDRYNNQAPL | 0.16095498 |
| VGGAGGAGGSGP | ASSPDRYNNQAPL | 0.086428024 |
| VFAAFMV | ASSPDRYNNQAPL | 0.26675525 |
| TLYDKLR | ASSPDRYNNQAPL | 0.13143528 |
| TLAVFTVL | ASSPDRYNNQAPL | 0.084300496 |
| TGSVQPARV | ASSPDRYNNQAPL | 0.20279069 |
| TGGGAGSGGT | ASSPDRYNNQAPL | 0.1482617 |
| SHLETSL | ASSPDRYNNQAPL | 0.1337348 |
| SFTTGMT | ASSPDRYNNQAPL | 0.5436312 |
| QVEVAKRMGVRQPTVS | ASSPDRYNNQAPL | 0.18046388 |
| QGSGGPALAM | ASSPDRYNNQAPL | 0.1244427 |
| MLAPSNP | ASSPDRYNNQAPL | 0.37604055 |
| LVNNAAI | ASSPDRYNNQAPL | 0.51249105 |
| LTGEINTLA | ASSPDRYNNQAPL | 0.12777027 |
| LSIFGPV | ASSPDRYNNQAPL | 0.3244968 |
| LLALLANL | ASSPDRYNNQAPL | 0.6338335 |
| LISNASDALDKLR | ASSPDRYNNQAPL | 0.2273628 |
| LINEAVP | ASSPDRYNNQAPL | 0.4654451 |
| LGYAFSGL | ASSPDRYNNQAPL | 0.55178523 |
| LDFVAVAL | ASSPDRYNNQAPL | 0.1150408 |
| LAQPGAE | ASSPDRYNNQAPL | 0.33014488 |
| ISNLGAVTGF | ASSPDRYNNQAPL | 0.61718756 |
| HQEDPGRI | ASSPDRYNNQAPL | 0.15565658 |
| GTPGEPTP | ASSPDRYNNQAPL | 0.20217128 |
| GFFNSGS | ASSPDRYNNQAPL | 0.13898422 |
| GAGGGGLTAGAN | ASSPDRYNNQAPL | 0.17636082 |
| FVDALVNM | ASSPDRYNNQAPL | 0.1011809 |
| FLTVAPEL | ASSPDRYNNQAPL | 0.31483236 |
| FETLTAL | ASSPDRYNNQAPL | 0.4025669 |
| DVINPIEL | ASSPDRYNNQAPL | 0.22297017 |
| APVSGAPV | ASSPDRYNNQAPL | 0.11180045 |
| ALVGGLAW | ASSPDRYNNQAPL | 0.10875677 |
| ALAAAAATAI | ASSPDRYNNQAPL | 0.13104016 |

| | | |
|---|---|---|
| AFGLGLLL | ASSPDRYNNQAPL | 0.16155247 |
| AFFAAASL | ASSPDRYNNQAPL | 0.12208188 |
| AAAALPAFSPP | ASSPDRYNNQAPL | 0.10322893 |
| VVLAPAL | ASSPDRYNNQAPL | 0.38287902 |
| TIGHVDHGKTTL | ASSPDRYNNQAPL | 0.37430614 |
| LISNASDALDK | ASSPDRYNNQAPL | 0.14070545 |
| LISNASDAL | ASSPDRYNNQAPL | 0.45880353 |
| ISNASDALDK | ASSPDRYNNQAPL | 0.47308445 |
| NQAETLVYQTE | ASSPDRYNNQAPL | 0.5654527 |
| TGGTGGTAGAG | ASSPDRYNNQAPL | 0.19537409 |
| TGSTAGGAGGA | ASSPDRYNNQAPL | 0.10567334 |
| DPLAVPVAAGPV | ASSPDRYNNQAPL | 0.598392 |
| GGTGGTPGNGGN | ASSPDRYNNQAPL | 0.24665646 |
| PAGLAGVDPATL | ASSPDRYNNQAPL | 0.14180423 |
| GNNLVVEAPHAGASV | ASSPDRYNNQAPL | 0.31459808 |
| KVTRIMSASTLEQPA | ASSPDRYNNQAPL | 0.14747493 |
| LGFALPIALPRRTVV | ASSPDRYNNQAPL | 0.1262675 |
| TRLGNLTARKLVVLV | ASSPDRYNNQAPL | 0.3763159 |
| WSQQRRLIYFPSAGP | ASSPDRYNNQAPL | 0.3719481 |
| GPNGNPDPMAAAVDIR | ASSPDRYNNQAPL | 0.11620961 |
| TQRCVSVADLRAAASR | ASSPDRYNNQAPL | 0.12383243 |
| QFVLDVAEHPVEVLTETV | ASSPDRYNNQAPL | 0.14959985 |
| GGAGGTGGAGGNGGTQAGDG | ASSPDRYNNQAPL | 0.30435967 |
| PLLATLRAIAVDVDAKPAQV | ASSPDRYNNQAPL | 0.39252388 |
| APKFRTGGYMIDFWGVGYQVAK | ASSPDRYNNQAPL | 0.20450906 |
| GAGGAGGAGGAAGGPGGTGGTG | ASSPDRYNNQAPL | 0.15796472 |
| IGALGLLATDVAPAGSVAFGVA | ASSPDRYNNQAPL | 0.4808051 |
| AGGAGGIGGNGGAGGAGGVLMG | ASSPDRYNNQAPL | 0.23298842 |
| ADELIGLL | ASSPDRYNNQAPL | 0.19416744 |
| ADRALKA | ASSPDRYNNQAPL | 0.2859867 |
| AGAVGQVGGAGGA | ASSPDRYNNQAPL | 0.107189715 |
| AGGGGGLGGAP | ASSPDRYNNQAPL | 0.33897752 |
| AGGLLYGNGGAG | ASSPDRYNNQAPL | 0.3085269 |
| AGLESIGL | ASSPDRYNNQAPL | 0.33712167 |
| AGLKTGII | ASSPDRYNNQAPL | 0.400757 |
| ASLAALLL | ASSPDRYNNQAPL | 0.16856658 |
| ASVINISSI | ASSPDRYNNQAPL | 0.22199042 |
| ATPPGPA | ASSPDRYNNQAPL | 0.12052153 |
| ATVASALV | ASSPDRYNNQAPL | 0.15092725 |
| ATVASLIL | ASSPDRYNNQAPL | 0.30107045 |
| AVNGPVLIRH | ASSPDRYNNQAPL | 0.30171284 |
| DGAAGAVGI | ASSPDRYNNQAPL | 0.37335366 |
| DPLLHGI | ASSPDRYNNQAPL | 0.27580518 |
| ELGAALGAL | ASSPDRYNNQAPL | 0.13911368 |
| ELGRTQAV | ASSPDRYNNQAPL | 0.4754588 |
| FVAGGYLS | ASSPDRYNNQAPL | 0.54689944 |
| GASGGAGGAGGW | ASSPDRYNNQAPL | 0.10712527 |
| GGAGAGGDGGAGGI | ASSPDRYNNQAPL | 0.124089785 |
| GTGHNGSQPGMGGNG | ASSPDRYNNQAPL | 0.33034027 |
| GYAAASASAS | ASSPDRYNNQAPL | 0.35455632 |
| HQVDVAAVQQG | ASSPDRYNNQAPL | 0.21026964 |

| | | |
|---|---|---|
| IGAAIAHT | ASSPDRYNNQAPL | 0.22477932 |
| IGMIEVAQL | ASSPDRYNNQAPL | 0.4159483 |
| IRLAPEHTSAMD | ASSPDRYNNQAPL | 0.13334522 |
| IYNVSVL | ASSPDRYNNQAPL | 0.5232071 |
| KQGFLGM | ASSPDRYNNQAPL | 0.10193835 |
| LESVITQL | ASSPDRYNNQAPL | 0.181309 |
| LGLGALVVV | ASSPDRYNNQAPL | 0.13483611 |
| LIGNGGNGGSGGI | ASSPDRYNNQAPL | 0.08960416 |
| LNAINAPTL | ASSPDRYNNQAPL | 0.31230974 |
| LQLLNTGV | ASSPDRYNNQAPL | 0.14041291 |
| LQTSIEF | ASSPDRYNNQAPL | 0.40401512 |
| MSFAVSEL | ASSPDRYNNQAPL | 0.15872504 |
| NTGALGL | ASSPDRYNNQAPL | 0.24798635 |
| PPGAGGAGGAGTT | ASSPDRYNNQAPL | 0.27546504 |
| QLVGLGI | ASSPDRYNNQAPL | 0.17041317 |
| RFLEQVHQL | ASSPDRYNNQAPL | 0.11874051 |
| RGVGYVLR | ASSPDRYNNQAPL | 0.26701993 |
| SATGVFAGVI | ASSPDRYNNQAPL | 0.42462718 |
| SLPAITLP | ASSPDRYNNQAPL | 0.311702 |
| STALAFTV | ASSPDRYNNQAPL | 0.10572814 |
| SVLTVGLGK | ASSPDRYNNQAPL | 0.17495441 |
| TALAGAGSGI | ASSPDRYNNQAPL | 0.17456877 |
| TGLPFTGI | ASSPDRYNNQAPL | 0.08837307 |
| VAEHVTV | ASSPDRYNNQAPL | 0.10184331 |
| VDIVLNSL | ASSPDRYNNQAPL | 0.10734199 |
| VLTGGGAVL | ASSPDRYNNQAPL | 0.11326059 |
| VNVGLAYGI | ASSPDRYNNQAPL | 0.42174393 |
| VSSAAVGAV | ASSPDRYNNQAPL | 0.42314786 |
| VTSVYEAGV | ASSPDRYNNQAPL | 0.18596393 |
| WVALDRGA | ASSPDRYNNQAPL | 0.46487844 |
| YLVGSVVI | ASSPDRYNNQAPL | 0.1723141 |
| GPPGTGKTLL | ASSPDRYNNQAPL | 0.08230601 |
| GVGMTKF | ASSPDRYNNQAPL | 0.12879714 |
| IGIGGSDL | ASSPDRYNNQAPL | 0.11173125 |
| IGTLLAEL | ASSPDRYNNQAPL | 0.4038452 |
| IYTSGTT | ASSPDRYNNQAPL | 0.09813518 |
| LSFPAGV | ASSPDRYNNQAPL | 0.14455493 |
| LYTSGTT | ASSPDRYNNQAPL | 0.20789608 |
| VGAFPFGF | ASSPDRYNNQAPL | 0.19851021 |
| VGAGTVEF | ASSPDRYNNQAPL | 0.3012218 |
| VGKSTLLL | ASSPDRYNNQAPL | 0.119227484 |
| AALAHAGGGIV | ASSPDRYNNQAPL | 0.11161449 |
| AALVAVSLL | ASSPDRYNNQAPL | 0.27112615 |
| AAVSRAVAAL | ASSPDRYNNQAPL | 0.0944325 |
| AGGSSGIGGF | ASSPDRYNNQAPL | 0.13381888 |
| AGLTVANANG | ASSPDRYNNQAPL | 0.105959706 |
| AGLVGAVL | ASSPDRYNNQAPL | 0.29194123 |
| ALSQAFTV | ASSPDRYNNQAPL | 0.10033219 |
| ALTFSVSL | ASSPDRYNNQAPL | 0.41766512 |
| ALTGAALVAL | ASSPDRYNNQAPL | 0.18817209 |
| ASFSIGAGVL | ASSPDRYNNQAPL | 0.20429742 |

| | | |
|---|---|---|
| DGSVYKI | ASSPDRYNNQAPL | 0.4865334 |
| EQLAEIV | ASSPDRYNNQAPL | 0.2762096 |
| FAVSELTVM | ASSPDRYNNQAPL | 0.18721783 |
| GAAPLAAGL | ASSPDRYNNQAPL | 0.12786609 |
| GADGGDGGSGGAGG | ASSPDRYNNQAPL | 0.12184556 |
| GFGLDTGGAGG | ASSPDRYNNQAPL | 0.12708382 |
| GLASGGLL | ASSPDRYNNQAPL | 0.13100497 |
| GRIAGAAVLL | ASSPDRYNNQAPL | 0.5379607 |
| GSAGSLNTSF | ASSPDRYNNQAPL | 0.190905 |
| GTGQIGI | ASSPDRYNNQAPL | 0.10213856 |
| GTLIALIL | ASSPDRYNNQAPL | 0.16686295 |
| IAATMGFVVG | ASSPDRYNNQAPL | 0.5713314 |
| IFAQALL | ASSPDRYNNQAPL | 0.7228314 |
| IGAAVALGF | ASSPDRYNNQAPL | 0.11937973 |
| IGLTPLAL | ASSPDRYNNQAPL | 0.17966367 |
| ILGSNTSTL | ASSPDRYNNQAPL | 0.19086537 |
| IVFSLTVL | ASSPDRYNNQAPL | 0.20316802 |
| LASKATGF | ASSPDRYNNQAPL | 0.16226265 |
| LASVALIL | ASSPDRYNNQAPL | 0.11975831 |
| LFAVGLTL | ASSPDRYNNQAPL | 0.33382443 |
| LFNGGIGTYI | ASSPDRYNNQAPL | 0.7779155 |
| LTLGAGII | ASSPDRYNNQAPL | 0.130559 |
| LVAAALGDI | ASSPDRYNNQAPL | 0.10054758 |
| NTAFVQLGI | ASSPDRYNNQAPL | 0.1898132 |
| QLFSGAEL | ASSPDRYNNQAPL | 0.11737177 |
| SAATFITL | ASSPDRYNNQAPL | 0.15845378 |
| SNYGLANLGV | ASSPDRYNNQAPL | 0.19875342 |
| SYGGAYAVM | ASSPDRYNNQAPL | 0.09747658 |
| TTGVGGLTLG | ASSPDRYNNQAPL | 0.16645664 |
| TTQTIYI | ASSPDRYNNQAPL | 0.63257056 |
| VAAGAGLGAVI | ASSPDRYNNQAPL | 0.14306411 |
| VAAVVTLL | ASSPDRYNNQAPL | 0.32509512 |
| VGFLGGLI | ASSPDRYNNQAPL | 0.5938239 |
| VGVIPILL | ASSPDRYNNQAPL | 0.3787666 |
| VIFASSLI | ASSPDRYNNQAPL | 0.67877436 |
| VKGGDGGAAAG | ASSPDRYNNQAPL | 0.5111976 |
| VTKTFTNI | ASSPDRYNNQAPL | 0.15271795 |
| VVAAAILL | ASSPDRYNNQAPL | 0.18116957 |
| YAALDVEL | ASSPDRYNNQAPL | 0.49636352 |
| LASVALLL | ASSPDRYNNQAPL | 0.18242726 |
| LDAVHRV | ASSPDRYNNQAPL | 0.22247201 |
| QGLSSSIF | ASSPDRYNNQAPL | 0.15429987 |
| RGTFANIRL | ASSPDRYNNQAPL | 0.4834559 |
| RQLWWGHRI | ASSPDRYNNQAPL | 0.2135208 |
| YGPPGTGKTLL | ASSPDRYNNQAPL | 0.45014548 |
| AALVGAPANVA | ASSPDRYNNQAPL | 0.113959126 |
| AATFRRTE | ASSPDRYNNQAPL | 0.12482485 |
| AGALGGGI | ASSPDRYNNQAPL | 0.5251285 |
| AGGAGGAGGIGAG | ASSPDRYNNQAPL | 0.10020565 |
| AGSGALGL | ASSPDRYNNQAPL | 0.122364484 |
| AQIADQI | ASSPDRYNNQAPL | 0.20622402 |

| | | |
|---|---|---|
| ASAVAVTYL | ASSPDRYNNQAPL | 0.60596615 |
| ASTVLGGTGG | ASSPDRYNNQAPL | 0.083232224 |
| AVGGEPAK | ASSPDRYNNQAPL | 0.12580153 |
| CEQGTFGGTC | ASSPDRYNNQAPL | 0.12777752 |
| DEGKAPAG | ASSPDRYNNQAPL | 0.20460758 |
| DTVVFPAL | ASSPDRYNNQAPL | 0.17789686 |
| FEGITMAG | ASSPDRYNNQAPL | 0.09833373 |
| FPLVVQD | ASSPDRYNNQAPL | 0.27968597 |
| GAFTTPQ | ASSPDRYNNQAPL | 0.09844734 |
| GFGGFGTAGGVGGA | ASSPDRYNNQAPL | 0.14228019 |
| GFLAQFL | ASSPDRYNNQAPL | 0.26912928 |
| GGAGGAGGNAGVA | ASSPDRYNNQAPL | 0.6589196 |
| GGVGTAGAGGAGGAGG | ASSPDRYNNQAPL | 0.117324464 |
| GNGGGGDAGNAGSGGN | ASSPDRYNNQAPL | 0.35373014 |
| GTGGAGGAGNDAGS | ASSPDRYNNQAPL | 0.14313464 |
| GTTNTGWFNS | ASSPDRYNNQAPL | 0.7825217 |
| IFMGPLFTL | ASSPDRYNNQAPL | 0.15486453 |
| ILAHAIEAI | ASSPDRYNNQAPL | 0.111540034 |
| IRSAIREAL | ASSPDRYNNQAPL | 0.3903297 |
| ITFAGGG | ASSPDRYNNQAPL | 0.42006654 |
| ITGMSGHQEV | ASSPDRYNNQAPL | 0.21851705 |
| IVAGLACTL | ASSPDRYNNQAPL | 0.527666 |
| IVGGGLAGA | ASSPDRYNNQAPL | 0.13875715 |
| IVTGVLNI | ASSPDRYNNQAPL | 0.18583533 |
| LAQQGCI | ASSPDRYNNQAPL | 0.36131188 |
| LCLHVEN | ASSPDRYNNQAPL | 0.2621372 |
| LDAISDGS | ASSPDRYNNQAPL | 0.11123348 |
| LGGFAAML | ASSPDRYNNQAPL | 0.2808215 |
| LGGKNPMI | ASSPDRYNNQAPL | 0.15492786 |
| LGPKGVAAIL | ASSPDRYNNQAPL | 0.14068232 |
| LMVGVSI | ASSPDRYNNQAPL | 0.55673164 |
| LVGAAFVL | ASSPDRYNNQAPL | 0.23187974 |
| LVLAVAPL | ASSPDRYNNQAPL | 0.46494 |
| MAAIAVIDAA | ASSPDRYNNQAPL | 0.24919534 |
| NNGINAELQ | ASSPDRYNNQAPL | 0.720539 |
| RGEPLVLVHG | ASSPDRYNNQAPL | 0.10869068 |
| RGGAGGAAGLI | ASSPDRYNNQAPL | 0.21439368 |
| RSYRMHPAV | ASSPDRYNNQAPL | 0.23373917 |
| SAAHLLAGQLA | ASSPDRYNNQAPL | 0.27547345 |
| SAAVGSY | ASSPDRYNNQAPL | 0.24557188 |
| SPHAVLHRI | ASSPDRYNNQAPL | 0.17710154 |
| SQIAKVL | ASSPDRYNNQAPL | 0.27588448 |
| TALGHLVP | ASSPDRYNNQAPL | 0.48191032 |
| TAPTTSGAA | ASSPDRYNNQAPL | 0.16274095 |
| TDGDDGNGGSPGT | ASSPDRYNNQAPL | 0.21402425 |
| TDGLGEN | ASSPDRYNNQAPL | 0.70296097 |
| TGGPPLPGQA | ASSPDRYNNQAPL | 0.14227305 |
| TIPAITI | ASSPDRYNNQAPL | 0.15102777 |
| TIPGFTI | ASSPDRYNNQAPL | 0.2988706 |
| TSIASGALA | ASSPDRYNNQAPL | 0.5147668 |
| TVTLPGAAI | ASSPDRYNNQAPL | 0.17699781 |

| | | |
|---|---|---|
| VASGLSAALP | ASSPDRYNNQAPL | 0.4545118 |
| VGAGVIGI | ASSPDRYNNQAPL | 0.09785844 |
| VGPKRFGV | ASSPDRYNNQAPL | 0.40817666 |
| VNVAFPD | ASSPDRYNNQAPL | 0.26407674 |
| VSMTNSGVS | ASSPDRYNNQAPL | 0.27022517 |
| VTDVLGDLSV | ASSPDRYNNQAPL | 0.09667853 |
| VVVVGAGFAGL | ASSPDRYNNQAPL | 0.32445917 |
| VYTVGTVV | ASSPDRYNNQAPL | 0.52863574 |
| WTTFSGP | ASSPDRYNNQAPL | 0.4347542 |
| YDELVEAF | ASSPDRYNNQAPL | 0.08947484 |
| YGTGGAGGNGG | ASSPDRYNNQAPL | 0.27939183 |
| YTNVMAV | ASSPDRYNNQAPL | 0.40621543 |
| FLEQVHQL | ASSPDRYNNQAPL | 0.7627972 |
| GLSLLAVAV | ASSPDRYNNQAPL | 0.21191251 |
| LEQVHQL | ASSPDRYNNQAPL | 0.33644825 |
| LNAAGAI | ASSPDRYNNQAPL | 0.13091685 |
| PAVHASL | ASSPDRYNNQAPL | 0.1591501 |
| TELEDDA | ASSPDRYNNQAPL | 0.12110862 |
| TPPGPPN | ASSPDRYNNQAPL | 0.20748857 |
| TPPPPPPP | ASSPDRYNNQAPL | 0.2150175 |
| TQSPALI | AWSLLGGENTLY | 0.13597485 |
| EFAAAFPQ | AWSLLGGENTLY | 0.31498766 |
| LLSGNLDVL | AWSLLGGENTLY | 0.5657278 |
| GPANLGNYNI | AWSLLGGENTLY | 0.34209508 |
| TLGPTLP | AWSLLGGENTLY | 0.59264827 |
| VYAYSSCCK | AWSLLGGENTLY | 0.70051146 |
| APAPAPAPNI | AWSLLGGENTLY | 0.34780568 |
| PGGPVPIP | AWSLLGGENTLY | 0.12308027 |
| FLTVALEP | AWSLLGGENTLY | 0.16827475 |
| SQYFGANL | AWSLLGGENTLY | 0.10383403 |
| GGVKPAA | AWSLLGGENTLY | 0.1516155 |
| GGTGGAGGAGGW | AWSLLGGENTLY | 0.12552495 |
| GVGGVAGLGGAGG | AWSLLGGENTLY | 0.25211048 |
| VLLVLATTI | AWSLLGGENTLY | 0.082894325 |
| IGIFAAML | AWSLLGGENTLY | 0.43501225 |
| AVGDLID | AWSLLGGENTLY | 0.5733282 |
| NISPLPL | AWSLLGGENTLY | 0.12240046 |
| VTAVFVAI | AWSLLGGENTLY | 0.36146855 |
| SGDDNDAAL | AWSLLGGENTLY | 0.40332454 |
| AVAALAHS | AWSLLGGENTLY | 0.12069111 |
| GGVTVMATP | AWSLLGGENTLY | 0.48731095 |
| AGLAANVRD | AWSLLGGENTLY | 0.12697996 |
| TNRPDLIDPALLRPG | AWSLLGGENTLY | 0.4292703 |
| LANLGVTAL | AWSLLGGENTLY | 0.4828265 |
| TGERIEL | AWSLLGGENTLY | 0.24984263 |
| NGEELAA | AWSLLGGENTLY | 0.22897196 |
| VLSLVWG | AWSLLGGENTLY | 0.26268622 |
| IGYALWL | AWSLLGGENTLY | 0.10660585 |
| LSLLLAPL | AWSLLGGENTLY | 0.24330178 |
| TLDEALK | AWSLLGGENTLY | 0.12716042 |
| TDGAGTAEGG | AWSLLGGENTLY | 0.26266894 |

| | | |
|---|---|---|
| NAPGGAGGNGG | AWSLLGGENTLY | 0.11897787 |
| VSVFMGMGGNF | AWSLLGGENTLY | 0.30696282 |
| LASYLAAL | AWSLLGGENTLY | 0.15937227 |
| LVGGTPAL | AWSLLGGENTLY | 0.246205 |
| VRNADVL | AWSLLGGENTLY | 0.13199542 |
| TCSSPFLVSAM | AWSLLGGENTLY | 0.1613682 |
| VHITDAE | AWSLLGGENTLY | 0.117007405 |
| EYAAPSF | AWSLLGGENTLY | 0.108154275 |
| VVDAFHGGL | AWSLLGGENTLY | 0.3744126 |
| ASVVDPQ | AWSLLGGENTLY | 0.172261 |
| SYNEFGGLS | AWSLLGGENTLY | 0.13102227 |
| DPDGSVY | AWSLLGGENTLY | 0.38642392 |
| MVSTIHATE | AWSLLGGENTLY | 0.21766791 |
| LAAGVADPV | AWSLLGGENTLY | 0.099140696 |
| TKEQIAA | AWSLLGGENTLY | 0.32738936 |
| SLASPAAL | AWSLLGGENTLY | 0.19713849 |
| IGLVFYGV | AWSLLGGENTLY | 0.17104682 |
| FLLDASNL | AWSLLGGENTLY | 0.090162605 |
| YQVGNFP | AWSLLGGENTLY | 0.13513033 |
| ESLLSQST | AWSLLGGENTLY | 0.14214566 |
| LFTVGFAAVG | AWSLLGGENTLY | 0.0968644 |
| ATPPTPT | AWSLLGGENTLY | 0.11683231 |
| TDLATLG | AWSLLGGENTLY | 0.5407196 |
| SAGGTGR | AWSLLGGENTLY | 0.5411005 |
| GFSNSGS | AWSLLGGENTLY | 0.17129274 |
| GGNGGSGGTSV | AWSLLGGENTLY | 0.38701293 |
| GTSVATGGAGN | AWSLLGGENTLY | 0.23651536 |
| NGGNGGSGGTS | AWSLLGGENTLY | 0.6546054 |
| AVVKFPH | AWSLLGGENTLY | 0.14694591 |
| TPPPPPP | AWSLLGGENTLY | 0.16088086 |
| AAPAAAAAPA | AWSLLGGENTLY | 0.17639868 |
| IALNEDL | AWSLLGGENTLY | 0.1874469 |
| IVNVSSM | AWSLLGGENTLY | 0.11266751 |
| TGIPPAPR | AWSLLGGENTLY | 0.09383844 |
| FDLGGGTF | AWSLLGGENTLY | 0.18121187 |
| FDLGGGTFDVS | AWSLLGGENTLY | 0.120342694 |
| NEPTAAA | AWSLLGGENTLY | 0.20018563 |
| NEPTAAALA | AWSLLGGENTLY | 0.4067635 |
| GSIINISSI | AWSLLGGENTLY | 0.12717165 |
| VAPGNAGTA | AWSLLGGENTLY | 0.110656746 |
| TGGGASLE | AWSLLGGENTLY | 0.15567812 |
| VAVYNHYKRI | AWSLLGGENTLY | 0.12170157 |
| TDPAPATT | AWSLLGGENTLY | 0.49080685 |
| TNRPDILDPALLRPG | AWSLLGGENTLY | 0.20854026 |
| LTTTTTS | AWSLLGGENTLY | 0.74757904 |
| PTGSGKTL | AWSLLGGENTLY | 0.28906998 |
| TGSGKTL | AWSLLGGENTLY | 0.12833782 |
| TIGHVDHGK | AWSLLGGENTLY | 0.4034933 |
| TIGHVDHGKTT | AWSLLGGENTLY | 0.179541 |
| GHVDHGKTT | AWSLLGGENTLY | 0.16798732 |
| AQFEHTLL | AWSLLGGENTLY | 0.17751905 |

| | | |
|---|---|---|
| IVNQDEL | AWSLLGGENTLY | 0.14102337 |
| TNGSQFF | AWSLLGGENTLY | 0.14488575 |
| TGKGIVE | AWSLLGGENTLY | 0.16580625 |
| LISNASDA | AWSLLGGENTLY | 0.1059444 |
| AELRTLGELTKIAKAHGAQ | AWSLLGGENTLY | 0.19825947 |
| APNAVAAAEPSW | AWSLLGGENTLY | 0.12517607 |
| APQVAGAATAVL | AWSLLGGENTLY | 0.1343711 |
| ARRDPAAAAAILPTDA | AWSLLGGENTLY | 0.15959409 |
| DAAARDVLAEAGLAEYF | AWSLLGGENTLY | 0.6175617 |
| DDQIVLSAKANLAKVEQ | AWSLLGGENTLY | 0.7298821 |
| DELLPDTGEDSDEDGH | AWSLLGGENTLY | 0.15015519 |
| ELPEAPANRATVPQD | AWSLLGGENTLY | 0.11509065 |
| FGQSKIERVTGIG | AWSLLGGENTLY | 0.43595862 |
| GAGGTGSTAGGAGGAGGAGGLY | AWSLLGGENTLY | 0.13783997 |
| GHGGTGAAGGNGGTGGAGGSGI | AWSLLGGENTLY | 0.25410765 |
| GNGGNGGAGGEGGPGGAGGAGA | AWSLLGGENTLY | 0.1971715 |
| GSTAGNGGNGGNATNGTIAGQP | AWSLLGGENTLY | 0.1479739 |
| GTGGDGGNGGNGGTGG | AWSLLGGENTLY | 0.14490706 |
| GVRRPAGIPADAPV | AWSLLGGENTLY | 0.28552744 |
| GYRQVTDAHLLTLARR | AWSLLGGENTLY | 0.10417247 |
| IAKMGQDGHDRG | AWSLLGGENTLY | 0.33196372 |
| KAEPSIEIQYT | AWSLLGGENTLY | 0.080072306 |
| KPVRIRMAADEDTVAVT | AWSLLGGENTLY | 0.14231953 |
| MPPRIAGMRLLVIKPEP | AWSLLGGENTLY | 0.14029777 |
| MKFYAVALFRSRGVPPTR | AWSLLGGENTLY | 0.13056815 |
| NAGGFGILGAVAHSP | AWSLLGGENTLY | 0.10092127 |
| NEGREITLAMAEM | AWSLLGGENTLY | 0.15090889 |
| NELVEWSHPLGHRWRWE | AWSLLGGENTLY | 0.5505707 |
| NLGAGGTGGNGGNPG | AWSLLGGENTLY | 0.5444761 |
| NTEWPKWETVMAADPVD | AWSLLGGENTLY | 0.13958575 |
| PASAAPAETQP | AWSLLGGENTLY | 0.42726094 |
| QQSSDGKNGVRAAEITGE | AWSLLGGENTLY | 0.75422406 |
| RAPDAVTTSWWKE | AWSLLGGENTLY | 0.20072216 |
| SGFGNVGAHTSGWFNQ | AWSLLGGENTLY | 0.100420676 |
| SGVAGAVGQAASVGGLKV | AWSLLGGENTLY | 0.20932999 |
| SNGGAGTGSAGGNGGT | AWSLLGGENTLY | 0.089859515 |
| TGGAGGEGGAAG | AWSLLGGENTLY | 0.21678114 |
| TGGDGGAAGAGGAPGHG | AWSLLGGENTLY | 0.59303206 |
| TMPPLLAAIANAGRHGV | AWSLLGGENTLY | 0.09839851 |
| TSSTAGNSGGA | AWSLLGGENTLY | 0.30506518 |
| VPATVPVGILC | AWSLLGGENTLY | 0.36222547 |
| VQRPDSGGVVLTGRIS | AWSLLGGENTLY | 0.13448793 |
| WTPNPVPATKGVPAGLVQAT | AWSLLGGENTLY | 0.1201067 |
| YALKRALAKFSDRFDVVII | AWSLLGGENTLY | 0.107247554 |
| YLIGSVIV | AWSLLGGENTLY | 0.12310653 |
| YGQLVAVID | AWSLLGGENTLY | 0.20938195 |
| VTVGGMTLPA | AWSLLGGENTLY | 0.16209951 |
| VLALPAV | AWSLLGGENTLY | 0.15013625 |
| VIPPTLNYET | AWSLLGGENTLY | 0.18665153 |
| VGMGGMHPSSG | AWSLLGGENTLY | 0.142049 |
| VGLAGGAATA | AWSLLGGENTLY | 0.15384349 |

| | | |
|---|---|---|
| VGGAGGAGGSGP | AWSLLGGENTLY | 0.1568102 |
| VFAAFMV | AWSLLGGENTLY | 0.12981895 |
| TLYDKLR | AWSLLGGENTLY | 0.33962256 |
| TLAVFTVL | AWSLLGGENTLY | 0.12393609 |
| TGSVQPARV | AWSLLGGENTLY | 0.20321473 |
| TGGGAGSGGT | AWSLLGGENTLY | 0.6823983 |
| SHLETSL | AWSLLGGENTLY | 0.099815935 |
| SFTTGMT | AWSLLGGENTLY | 0.12345665 |
| QVEVAKRMGVRQPTVS | AWSLLGGENTLY | 0.23540384 |
| QGSGGPALAM | AWSLLGGENTLY | 0.09920555 |
| MLAPSNP | AWSLLGGENTLY | 0.523133 |
| LVNNAAI | AWSLLGGENTLY | 0.16280933 |
| LTGEINTLA | AWSLLGGENTLY | 0.28510863 |
| LSIFGPV | AWSLLGGENTLY | 0.16150829 |
| LLALLANL | AWSLLGGENTLY | 0.7091862 |
| LISNASDALDKLR | AWSLLGGENTLY | 0.3831156 |
| LINEAVP | AWSLLGGENTLY | 0.14275558 |
| LGYAFSGL | AWSLLGGENTLY | 0.47030932 |
| LDFVAVAL | AWSLLGGENTLY | 0.104766615 |
| LAQPGAE | AWSLLGGENTLY | 0.3295037 |
| ISNLGAVTGF | AWSLLGGENTLY | 0.31649005 |
| HQEDPGRI | AWSLLGGENTLY | 0.1094488 |
| GTPGEPTP | AWSLLGGENTLY | 0.18549332 |
| GFFNSGS | AWSLLGGENTLY | 0.4963378 |
| GAGGGGLTAGAN | AWSLLGGENTLY | 0.486402 |
| FVDALVNM | AWSLLGGENTLY | 0.10561359 |
| FLTVAPEL | AWSLLGGENTLY | 0.13461787 |
| FETLTAL | AWSLLGGENTLY | 0.16849293 |
| DVINPIEL | AWSLLGGENTLY | 0.16363493 |
| APVSGAPV | AWSLLGGENTLY | 0.10548844 |
| ALVGGLAW | AWSLLGGENTLY | 0.45331752 |
| ALAAAAATAI | AWSLLGGENTLY | 0.1926822 |
| AFGLGLLL | AWSLLGGENTLY | 0.64510024 |
| AFFAAASL | AWSLLGGENTLY | 0.1962204 |
| AAAALPAFSPP | AWSLLGGENTLY | 0.35169446 |
| VVLAPAL | AWSLLGGENTLY | 0.15855414 |
| TIGHVDHGKTTL | AWSLLGGENTLY | 0.13641356 |
| LISNASDALDK | AWSLLGGENTLY | 0.11669725 |
| LISNASDAL | AWSLLGGENTLY | 0.1406729 |
| ISNASDALDK | AWSLLGGENTLY | 0.12721846 |
| NQAETLVYQTE | AWSLLGGENTLY | 0.1423454 |
| TGGTGGTAGAG | AWSLLGGENTLY | 0.08384928 |
| TGSTAGGAGGA | AWSLLGGENTLY | 0.36836648 |
| DPLAVPVAAGPV | AWSLLGGENTLY | 0.2722205 |
| GGTGGTPGNGGN | AWSLLGGENTLY | 0.2801745 |
| PAGLAGVDPATL | AWSLLGGENTLY | 0.14517984 |
| GNNLVVEAPHAGASV | AWSLLGGENTLY | 0.15721405 |
| KVTRIMSASTLEQPA | AWSLLGGENTLY | 0.11608363 |
| LGFALPIALPRRTVV | AWSLLGGENTLY | 0.53835094 |
| TRLGNLTARKLVVLV | AWSLLGGENTLY | 0.09664983 |
| WSQQRRLIYFPSAGP | AWSLLGGENTLY | 0.09793633 |

| | | |
|---|---|---|
| GPNGNPDPMAAAVDIR | AWSLLGGENTLY | 0.1425884 |
| TQRCVSVADLRAAASR | AWSLLGGENTLY | 0.10096825 |
| QFVLDVAEHPVEVLTETV | AWSLLGGENTLY | 0.6911807 |
| GGAGGTGGAGGNGGTQAGDG | AWSLLGGENTLY | 0.4946264 |
| PLLATLRAIAVDVDAKPAQV | AWSLLGGENTLY | 0.26703343 |
| APKFRTGGYMIDFWGVGYQVAK | AWSLLGGENTLY | 0.43200526 |
| GAGGAGGAGGAAGGPGGTGGTG | AWSLLGGENTLY | 0.0912674 |
| IGALGLLATDVAPAGSVAFGVA | AWSLLGGENTLY | 0.1324074 |
| AGGAGGIGGNGGAGGAGGVLMG | AWSLLGGENTLY | 0.6294153 |
| ADELIGLL | AWSLLGGENTLY | 0.20228505 |
| ADRALKA | AWSLLGGENTLY | 0.22227523 |
| AGAVGQVGGAGGA | AWSLLGGENTLY | 0.4336863 |
| AGGGGGLGGAP | AWSLLGGENTLY | 0.12380709 |
| AGGLLYGNGGAG | AWSLLGGENTLY | 0.1671684 |
| AGLESIGL | AWSLLGGENTLY | 0.15094052 |
| AGLKTGII | AWSLLGGENTLY | 0.21055943 |
| ASLAALLL | AWSLLGGENTLY | 0.33827418 |
| ASVINISSI | AWSLLGGENTLY | 0.10698097 |
| ATPPGPA | AWSLLGGENTLY | 0.345595 |
| ATVASALV | AWSLLGGENTLY | 0.7932555 |
| ATVASLIL | AWSLLGGENTLY | 0.22199577 |
| AVNGPVLIRH | AWSLLGGENTLY | 0.35863286 |
| DGAAGAVGI | AWSLLGGENTLY | 0.26440144 |
| DPLLHGI | AWSLLGGENTLY | 0.61671925 |
| ELGAALGAL | AWSLLGGENTLY | 0.13330732 |
| ELGRTQAV | AWSLLGGENTLY | 0.12144649 |
| FVAGGYLS | AWSLLGGENTLY | 0.16058207 |
| GASGGAGGAGGW | AWSLLGGENTLY | 0.1287348 |
| GGAGAGGDGGAGGI | AWSLLGGENTLY | 0.2245447 |
| GTGHNGSQPGMGGNG | AWSLLGGENTLY | 0.15400396 |
| GYAAASASAS | AWSLLGGENTLY | 0.82473326 |
| HQVDVAAVQQG | AWSLLGGENTLY | 0.41265297 |
| IGAAIAHT | AWSLLGGENTLY | 0.19927289 |
| IGMIEVAQL | AWSLLGGENTLY | 0.105440624 |
| IRLAPEHTSAMD | AWSLLGGENTLY | 0.20644933 |
| IYNVSVL | AWSLLGGENTLY | 0.24748194 |
| KQGFLGM | AWSLLGGENTLY | 0.106925525 |
| LESVITQL | AWSLLGGENTLY | 0.55414593 |
| LGLGALVVV | AWSLLGGENTLY | 0.13899985 |
| LIGNGGNGGSGGI | AWSLLGGENTLY | 0.14012338 |
| LNAINAPTL | AWSLLGGENTLY | 0.21380818 |
| LQLLNTGV | AWSLLGGENTLY | 0.7104276 |
| LQTSIEF | AWSLLGGENTLY | 0.20072278 |
| MSFAVSEL | AWSLLGGENTLY | 0.17636918 |
| NTGALGL | AWSLLGGENTLY | 0.1099509 |
| PPGAGGAGGAGTT | AWSLLGGENTLY | 0.4542634 |
| QLVGLGI | AWSLLGGENTLY | 0.59858215 |
| RFLEQVHQL | AWSLLGGENTLY | 0.41867226 |
| RGVGYVLR | AWSLLGGENTLY | 0.6321598 |
| SATGVFAGVI | AWSLLGGENTLY | 0.16732281 |
| SLPAITLP | AWSLLGGENTLY | 0.1064411 |

| | | |
|---|---|---|
| STALAFTV | AWSLLGGENTLY | 0.2358262 |
| SVLTVGLGK | AWSLLGGENTLY | 0.24550329 |
| TALAGAGSGI | AWSLLGGENTLY | 0.6834203 |
| TGLPFTGI | AWSLLGGENTLY | 0.29891056 |
| VAEHVTV | AWSLLGGENTLY | 0.41587734 |
| VDIVLNSL | AWSLLGGENTLY | 0.14874803 |
| VLTGGGAVL | AWSLLGGENTLY | 0.5978174 |
| VNVGLAYGI | AWSLLGGENTLY | 0.68597525 |
| VSSAAVGAV | AWSLLGGENTLY | 0.3258556 |
| VTSVYEAGV | AWSLLGGENTLY | 0.18475217 |
| WVALDRGA | AWSLLGGENTLY | 0.14736548 |
| YLVGSVVI | AWSLLGGENTLY | 0.23966967 |
| GPPGTGKTLL | AWSLLGGENTLY | 0.2636094 |
| GVGMTKF | AWSLLGGENTLY | 0.13947746 |
| IGIGGSDL | AWSLLGGENTLY | 0.30812836 |
| IGTLLAEL | AWSLLGGENTLY | 0.3164678 |
| IYTSGTT | AWSLLGGENTLY | 0.107443266 |
| LSFPAGV | AWSLLGGENTLY | 0.31165734 |
| LYTSGTT | AWSLLGGENTLY | 0.6389092 |
| VGAFPFGF | AWSLLGGENTLY | 0.22651467 |
| VGAGTVEF | AWSLLGGENTLY | 0.13829125 |
| VGKSTLLL | AWSLLGGENTLY | 0.6206721 |
| AALAHAGGGIV | AWSLLGGENTLY | 0.80247307 |
| AALVAVSLL | AWSLLGGENTLY | 0.47248897 |
| AAVSRAVAAL | AWSLLGGENTLY | 0.10819487 |
| AGGSSGIGGF | AWSLLGGENTLY | 0.09240858 |
| AGLTVANANG | AWSLLGGENTLY | 0.28982872 |
| AGLVGAVL | AWSLLGGENTLY | 0.31211567 |
| ALSQAFTV | AWSLLGGENTLY | 0.14381354 |
| ALTFSVSL | AWSLLGGENTLY | 0.11854853 |
| ALTGAALVAL | AWSLLGGENTLY | 0.4428069 |
| ASFSIGAGVL | AWSLLGGENTLY | 0.41810927 |
| DGSVYKI | AWSLLGGENTLY | 0.13689703 |
| EQLAEIV | AWSLLGGENTLY | 0.10899005 |
| FAVSELTVM | AWSLLGGENTLY | 0.2733171 |
| GAAPLAAGL | AWSLLGGENTLY | 0.3711741 |
| GADGGDGGSGGAGG | AWSLLGGENTLY | 0.11560243 |
| GFGLDTGGAGG | AWSLLGGENTLY | 0.1975432 |
| GLASGGLL | AWSLLGGENTLY | 0.28082 |
| GRIAGAAVLL | AWSLLGGENTLY | 0.34824637 |
| GSAGSLNTSF | AWSLLGGENTLY | 0.20177913 |
| GTGQIGI | AWSLLGGENTLY | 0.11400462 |
| GTLIALIL | AWSLLGGENTLY | 0.23106807 |
| IAATMGFVVG | AWSLLGGENTLY | 0.123234615 |
| IFAQALL | AWSLLGGENTLY | 0.4036861 |
| IGAAVALGF | AWSLLGGENTLY | 0.25428575 |
| IGLTPLAL | AWSLLGGENTLY | 0.21488129 |
| ILGSNTSTL | AWSLLGGENTLY | 0.21958967 |
| IVFSLTVL | AWSLLGGENTLY | 0.14264461 |
| LASKATGF | AWSLLGGENTLY | 0.101347424 |
| LASVALIL | AWSLLGGENTLY | 0.089779586 |

| | | |
|---|---|---|
| LFAVGLTL | AWSLLGGENTLY | 0.5501203 |
| LFNGGIGTYI | AWSLLGGENTLY | 0.3279388 |
| LTLGAGII | AWSLLGGENTLY | 0.35328314 |
| LVAAALGDI | AWSLLGGENTLY | 0.2579965 |
| NTAFVQLGI | AWSLLGGENTLY | 0.11389004 |
| QLFSGAEL | AWSLLGGENTLY | 0.20709641 |
| SAATFITL | AWSLLGGENTLY | 0.09533797 |
| SNYGLANLGV | AWSLLGGENTLY | 0.16575931 |
| SYGGAYAVM | AWSLLGGENTLY | 0.55559576 |
| TTGVGGLTLG | AWSLLGGENTLY | 0.25295153 |
| TTQTIYI | AWSLLGGENTLY | 0.13960227 |
| VAAGAGLGAVI | AWSLLGGENTLY | 0.2796348 |
| VAAVVTLL | AWSLLGGENTLY | 0.277815 |
| VGFLGGLI | AWSLLGGENTLY | 0.13021365 |
| VGVIPILL | AWSLLGGENTLY | 0.21690992 |
| VIFASSLI | AWSLLGGENTLY | 0.41778618 |
| VKGGDGGAAAG | AWSLLGGENTLY | 0.26231182 |
| VTKTFTNI | AWSLLGGENTLY | 0.11219833 |
| VVAAAILL | AWSLLGGENTLY | 0.24698667 |
| YAALDVEL | AWSLLGGENTLY | 0.12737308 |
| LASVALLL | AWSLLGGENTLY | 0.27908245 |
| LDAVHRV | AWSLLGGENTLY | 0.14528456 |
| QGLSSSIF | AWSLLGGENTLY | 0.20502323 |
| RGTFANIRL | AWSLLGGENTLY | 0.28587544 |
| RQLWWGHRI | AWSLLGGENTLY | 0.09486194 |
| YGPPGTGKTLL | AWSLLGGENTLY | 0.13469082 |
| AALVGAPANVA | AWSLLGGENTLY | 0.20510457 |
| AATFRRTE | AWSLLGGENTLY | 0.17486037 |
| AGALGGGI | AWSLLGGENTLY | 0.28645626 |
| AGGAGGAGGIGAG | AWSLLGGENTLY | 0.11597105 |
| AGSGALGL | AWSLLGGENTLY | 0.25643837 |
| AQIADQI | AWSLLGGENTLY | 0.14364597 |
| ASAVAVTYL | AWSLLGGENTLY | 0.12723415 |
| ASTVLGGTGG | AWSLLGGENTLY | 0.21331616 |
| AVGGEPAK | AWSLLGGENTLY | 0.15904412 |
| CEQGTFGGTC | AWSLLGGENTLY | 0.14591 |
| DEGKAPAG | AWSLLGGENTLY | 0.2251697 |
| DTVVFPAL | AWSLLGGENTLY | 0.13405351 |
| FEGITMAG | AWSLLGGENTLY | 0.21570523 |
| FPLVVQD | AWSLLGGENTLY | 0.5197481 |
| GAFTTPQ | AWSLLGGENTLY | 0.14525875 |
| GFGGFGTAGGVGGA | AWSLLGGENTLY | 0.32855874 |
| GFLAQFL | AWSLLGGENTLY | 0.1446623 |
| GGAGGAGGNAGVA | AWSLLGGENTLY | 0.35756302 |
| GGVGTAGAGGAGGAGG | AWSLLGGENTLY | 0.15638363 |
| GNGGGGDAGNAGSGGN | AWSLLGGENTLY | 0.11996063 |
| GTGGAGGAGNDAGS | AWSLLGGENTLY | 0.24276087 |
| GTTNTGWFNS | AWSLLGGENTLY | 0.29427823 |
| IFMGPLFTL | AWSLLGGENTLY | 0.42254323 |
| ILAHAIEAI | AWSLLGGENTLY | 0.1731948 |
| IRSAIREAL | AWSLLGGENTLY | 0.2307358 |

| | | |
|---|---|---|
| ITFAGGG | AWSLLGGENTLY | 0.39684057 |
| ITGMSGHQEV | AWSLLGGENTLY | 0.1578632 |
| IVAGLACTL | AWSLLGGENTLY | 0.10249909 |
| IVGGGLAGA | AWSLLGGENTLY | 0.14161253 |
| IVTGVLNI | AWSLLGGENTLY | 0.10312507 |
| LAQQGCI | AWSLLGGENTLY | 0.14415096 |
| LCLHVEN | AWSLLGGENTLY | 0.11329377 |
| LDAISDGS | AWSLLGGENTLY | 0.39578012 |
| LGGFAAML | AWSLLGGENTLY | 0.72331864 |
| LGGKNPMI | AWSLLGGENTLY | 0.14370018 |
| LGPKGVAAIL | AWSLLGGENTLY | 0.70923036 |
| LMVGVSI | AWSLLGGENTLY | 0.11144839 |
| LVGAAFVL | AWSLLGGENTLY | 0.34388852 |
| LVLAVAPL | AWSLLGGENTLY | 0.5723747 |
| MAAIAVIDAA | AWSLLGGENTLY | 0.11944759 |
| NNGINAELQ | AWSLLGGENTLY | 0.20344508 |
| RGEPLVLVHG | AWSLLGGENTLY | 0.13655144 |
| RGGAGGAAGLI | AWSLLGGENTLY | 0.12282953 |
| RSYRMHPAV | AWSLLGGENTLY | 0.54781026 |
| SAAHLLAGQLA | AWSLLGGENTLY | 0.15890275 |
| SAAVGSY | AWSLLGGENTLY | 0.22248861 |
| SPHAVLHRI | AWSLLGGENTLY | 0.14450626 |
| SQIAKVL | AWSLLGGENTLY | 0.115220346 |
| TALGHLVP | AWSLLGGENTLY | 0.18276365 |
| TAPTTSGAA | AWSLLGGENTLY | 0.39930737 |
| TDGDDGNGGSPGT | AWSLLGGENTLY | 0.16586107 |
| TDGLGEN | AWSLLGGENTLY | 0.67289954 |
| TGGPPLPGQA | AWSLLGGENTLY | 0.12547827 |
| TIPAITI | AWSLLGGENTLY | 0.5043781 |
| TIPGFTI | AWSLLGGENTLY | 0.6008855 |
| TSIASGALA | AWSLLGGENTLY | 0.2745748 |
| TVTLPGAAI | AWSLLGGENTLY | 0.15164548 |
| VASGLSAALP | AWSLLGGENTLY | 0.59812415 |
| VGAGVIGI | AWSLLGGENTLY | 0.21282138 |
| VGPKRFGV | AWSLLGGENTLY | 0.372602 |
| VNVAFPD | AWSLLGGENTLY | 0.35396558 |
| VSMTNSGVS | AWSLLGGENTLY | 0.19297902 |
| VTDVLGDLSV | AWSLLGGENTLY | 0.7672669 |
| VVVVGAGFAGL | AWSLLGGENTLY | 0.5796274 |
| VYTVGTVV | AWSLLGGENTLY | 0.36634535 |
| WTTFSGP | AWSLLGGENTLY | 0.22948757 |
| YDELVEAF | AWSLLGGENTLY | 0.15149733 |
| YGTGGAGGNGG | AWSLLGGENTLY | 0.26995468 |
| YTNVMAV | AWSLLGGENTLY | 0.20913833 |
| FLEQVHQL | AWSLLGGENTLY | 0.6162123 |
| GLSLLAVAV | AWSLLGGENTLY | 0.3689568 |
| LEQVHQL | AWSLLGGENTLY | 0.2595669 |
| LNAAGAI | AWSLLGGENTLY | 0.7743357 |
| PAVHASL | AWSLLGGENTLY | 0.76996064 |
| TELEDDA | AWSLLGGENTLY | 0.16113308 |
| TPPGPPN | AWSLLGGENTLY | 0.35471237 |

| | | |
|---|---|---|
| TPPPPPPP | AWSLLGGENTLY | 0.19722779 |
| TQSPALI | ASSIVTGKNEQY | 0.26649022 |
| EFAAAFPQ | ASSIVTGKNEQY | 0.13306172 |
| LLSGNLDVL | ASSIVTGKNEQY | 0.49604136 |
| GPANLGNYNI | ASSIVTGKNEQY | 0.2263068 |
| TLGPTLP | ASSIVTGKNEQY | 0.12444308 |
| VYAYSSCCK | ASSIVTGKNEQY | 0.31587213 |
| APAPAPAPNI | ASSIVTGKNEQY | 0.24962547 |
| PGGPVPIP | ASSIVTGKNEQY | 0.25791264 |
| FLTVALEP | ASSIVTGKNEQY | 0.7323211 |
| SQYFGANL | ASSIVTGKNEQY | 0.1382618 |
| GGVKPAA | ASSIVTGKNEQY | 0.20391554 |
| GGTGGAGGAGGW | ASSIVTGKNEQY | 0.65179706 |
| GVGGVAGLGGAGG | ASSIVTGKNEQY | 0.7785201 |
| VLLVLATTI | ASSIVTGKNEQY | 0.18703853 |
| IGIFAAML | ASSIVTGKNEQY | 0.5564171 |
| AVGDLID | ASSIVTGKNEQY | 0.7943402 |
| NISPLPL | ASSIVTGKNEQY | 0.41614065 |
| VTAVFVAI | ASSIVTGKNEQY | 0.5333483 |
| SGDDNDAAL | ASSIVTGKNEQY | 0.6731901 |
| AVAALAHS | ASSIVTGKNEQY | 0.18301518 |
| GGVTVMATP | ASSIVTGKNEQY | 0.6409689 |
| AGLAANVRD | ASSIVTGKNEQY | 0.77392983 |
| TNRPDLIDPALLRPG | ASSIVTGKNEQY | 0.28571004 |
| LANLGVTAL | ASSIVTGKNEQY | 0.4004801 |
| TGERIEL | ASSIVTGKNEQY | 0.18738599 |
| NGEELAA | ASSIVTGKNEQY | 0.2306502 |
| VLSLVWG | ASSIVTGKNEQY | 0.13858104 |
| IGYALWL | ASSIVTGKNEQY | 0.62295854 |
| LSLLLAPL | ASSIVTGKNEQY | 0.5705546 |
| TLDEALK | ASSIVTGKNEQY | 0.3237461 |
| TDGAGTAEGG | ASSIVTGKNEQY | 0.1728747 |
| NAPGGAGGNGG | ASSIVTGKNEQY | 0.17586714 |
| VSVFMGMGGNF | ASSIVTGKNEQY | 0.30248564 |
| LASYLAAL | ASSIVTGKNEQY | 0.3833723 |
| LVGGTPAL | ASSIVTGKNEQY | 0.14086069 |
| VRNADVL | ASSIVTGKNEQY | 0.18623962 |
| TCSSPFLVSAM | ASSIVTGKNEQY | 0.6785813 |
| VHITDAE | ASSIVTGKNEQY | 0.70000154 |
| EYAAPSF | ASSIVTGKNEQY | 0.15186492 |
| VVDAFHGGL | ASSIVTGKNEQY | 0.76357263 |
| ASVVDPQ | ASSIVTGKNEQY | 0.5830848 |
| SYNEFGGLS | ASSIVTGKNEQY | 0.6899123 |
| DPDGSVY | ASSIVTGKNEQY | 0.26505938 |
| MVSTIHATE | ASSIVTGKNEQY | 0.20316043 |
| LAAGVADPV | ASSIVTGKNEQY | 0.67392063 |
| TKEQIAA | ASSIVTGKNEQY | 0.44370365 |
| SLASPAAL | ASSIVTGKNEQY | 0.1961447 |
| IGLVFYGV | ASSIVTGKNEQY | 0.5623857 |
| FLLDASNL | ASSIVTGKNEQY | 0.3133869 |
| YQVGNFP | ASSIVTGKNEQY | 0.28016102 |

| | | |
|---|---|---|
| ESLLSQST | ASSIVTGKNEQY | 0.5392439 |
| LFTVGFAAVG | ASSIVTGKNEQY | 0.48165676 |
| ATPPTPT | ASSIVTGKNEQY | 0.18440756 |
| TDLATLG | ASSIVTGKNEQY | 0.20471933 |
| SAGGTGR | ASSIVTGKNEQY | 0.18897286 |
| GFSNSGS | ASSIVTGKNEQY | 0.5292554 |
| GGNGGSGGTSV | ASSIVTGKNEQY | 0.7173013 |
| GTSVATGGAGN | ASSIVTGKNEQY | 0.301192 |
| NGGNGGSGGTS | ASSIVTGKNEQY | 0.32786697 |
| AVVKFPH | ASSIVTGKNEQY | 0.72565764 |
| TPPPPPP | ASSIVTGKNEQY | 0.49093586 |
| AAPAAAAAPA | ASSIVTGKNEQY | 0.3021371 |
| IALNEDL | ASSIVTGKNEQY | 0.6663359 |
| IVNVSSM | ASSIVTGKNEQY | 0.28343493 |
| TGIPPAPR | ASSIVTGKNEQY | 0.44837132 |
| FDLGGGTF | ASSIVTGKNEQY | 0.43190926 |
| FDLGGGTFDVS | ASSIVTGKNEQY | 0.4624315 |
| NEPTAAA | ASSIVTGKNEQY | 0.54766655 |
| NEPTAAALA | ASSIVTGKNEQY | 0.5434592 |
| GSIINISSI | ASSIVTGKNEQY | 0.3411485 |
| VAPGNAGTA | ASSIVTGKNEQY | 0.14050364 |
| TGGGASLE | ASSIVTGKNEQY | 0.23217131 |
| VAVYNHYKRI | ASSIVTGKNEQY | 0.6094255 |
| TDPAPATT | ASSIVTGKNEQY | 0.5507241 |
| TNRPDILDPALLRPG | ASSIVTGKNEQY | 0.5759088 |
| LTTTTTS | ASSIVTGKNEQY | 0.42514986 |
| PTGSGKTL | ASSIVTGKNEQY | 0.21934882 |
| TGSGKTL | ASSIVTGKNEQY | 0.694817 |
| TIGHVDHGK | ASSIVTGKNEQY | 0.7223549 |
| TIGHVDHGKTT | ASSIVTGKNEQY | 0.16214192 |
| GHVDHGKTT | ASSIVTGKNEQY | 0.26423597 |
| AQFEHTLL | ASSIVTGKNEQY | 0.77275425 |
| IVNQDEL | ASSIVTGKNEQY | 0.6026815 |
| TNGSQFF | ASSIVTGKNEQY | 0.34516585 |
| TGKGIVE | ASSIVTGKNEQY | 0.26311904 |
| LISNASDA | ASSIVTGKNEQY | 0.78081095 |
| AELRTLGELTKIAKAHGAQ | ASSIVTGKNEQY | 0.20601591 |
| APNAVAAAEPSW | ASSIVTGKNEQY | 0.772734 |
| APQVAGAATAVL | ASSIVTGKNEQY | 0.1335486 |
| ARRDPAAAAAILPTDA | ASSIVTGKNEQY | 0.25774598 |
| DAAARDVLAEAGLAEYF | ASSIVTGKNEQY | 0.20954551 |
| DDQIVLSAKANLAKVEQ | ASSIVTGKNEQY | 0.14832398 |
| DELLPDTGEDSDEDGH | ASSIVTGKNEQY | 0.5557376 |
| ELPEAPANRATVPQD | ASSIVTGKNEQY | 0.22546865 |
| FGQSKIERVTGIG | ASSIVTGKNEQY | 0.7100026 |
| GAGGTGSTAGGAGGAGGAGGLY | ASSIVTGKNEQY | 0.29881203 |
| GHGGTGAAGGNGGTGGAGGSGI | ASSIVTGKNEQY | 0.4501983 |
| GNGGNGGAGGEGGPGGAGGAGA | ASSIVTGKNEQY | 0.45900324 |
| GSTAGNGGNGGNATNGTIAGQP | ASSIVTGKNEQY | 0.3783721 |
| GTGGDGGNGGNGGTGG | ASSIVTGKNEQY | 0.19174764 |
| GVRRPAGIPADAPV | ASSIVTGKNEQY | 0.55270034 |

| | | |
|---|---|---|
| GYRQVTDAHLLTLARR | ASSIVTGKNEQY | 0.6884938 |
| IAKMGQDGHDRG | ASSIVTGKNEQY | 0.34528676 |
| KAEPSIEIQYT | ASSIVTGKNEQY | 0.17543565 |
| KPVRIRMAADEDTVAVT | ASSIVTGKNEQY | 0.28550506 |
| MPPRIAGMRLLVIKPEP | ASSIVTGKNEQY | 0.37654287 |
| MKFYAVALFRSRGVPPTR | ASSIVTGKNEQY | 0.12880291 |
| NAGGFGILGAVAHSP | ASSIVTGKNEQY | 0.1271247 |
| NEGREITLAMAEM | ASSIVTGKNEQY | 0.31737894 |
| NELVEWSHPLGHRWRWE | ASSIVTGKNEQY | 0.17662883 |
| NLGAGGTGGNGGNPG | ASSIVTGKNEQY | 0.7371558 |
| NTEWPKWETVMAADPVD | ASSIVTGKNEQY | 0.7120936 |
| PASAAPAETQP | ASSIVTGKNEQY | 0.37118703 |
| QQSSDGKNGVRAAEITGE | ASSIVTGKNEQY | 0.64740694 |
| RAPDAVTTSWWKE | ASSIVTGKNEQY | 0.6445001 |
| SGFGNVGAHTSGWFNQ | ASSIVTGKNEQY | 0.121877976 |
| SGVAGAVGQAASVGGLKV | ASSIVTGKNEQY | 0.22496554 |
| SNGGAGTGSAGGNGGT | ASSIVTGKNEQY | 0.16521844 |
| TGGAGGEGGAAG | ASSIVTGKNEQY | 0.5908336 |
| TGGDGGAAGAGGAPGHG | ASSIVTGKNEQY | 0.16871263 |
| TMPPLLAAIANAGRHGV | ASSIVTGKNEQY | 0.21632048 |
| TSSTAGNSGGA | ASSIVTGKNEQY | 0.44295037 |
| VPATVPVGILC | ASSIVTGKNEQY | 0.5948137 |
| VQRPDSGGVVLTGRIS | ASSIVTGKNEQY | 0.40995604 |
| WTPNPVPATKGVPAGLVQAT | ASSIVTGKNEQY | 0.19998205 |
| YALKRALAKFSDRFDVVII | ASSIVTGKNEQY | 0.12347314 |
| YLIGSVIV | ASSIVTGKNEQY | 0.59933364 |
| YGQLVAVID | ASSIVTGKNEQY | 0.14793126 |
| VTVGGMTLPA | ASSIVTGKNEQY | 0.20936583 |
| VLALPAV | ASSIVTGKNEQY | 0.1314803 |
| VIPPTLNYET | ASSIVTGKNEQY | 0.12569134 |
| VGMGGMHPSSG | ASSIVTGKNEQY | 0.16957174 |
| VGLAGGAATA | ASSIVTGKNEQY | 0.5498247 |
| VGGAGGAGGSGP | ASSIVTGKNEQY | 0.2715311 |
| VFAAFMV | ASSIVTGKNEQY | 0.3035076 |
| TLYDKLR | ASSIVTGKNEQY | 0.76090384 |
| TLAVFTVL | ASSIVTGKNEQY | 0.7090887 |
| TGSVQPARV | ASSIVTGKNEQY | 0.3518246 |
| TGGGAGSGGT | ASSIVTGKNEQY | 0.29178262 |
| SHLETSL | ASSIVTGKNEQY | 0.18625842 |
| SFTTGMT | ASSIVTGKNEQY | 0.19643608 |
| QVEVAKRMGVRQPTVS | ASSIVTGKNEQY | 0.2049901 |
| QGSGGPALAM | ASSIVTGKNEQY | 0.7979804 |
| MLAPSNP | ASSIVTGKNEQY | 0.24098013 |
| LVNNAAI | ASSIVTGKNEQY | 0.16195884 |
| LTGEINTLA | ASSIVTGKNEQY | 0.31830186 |
| LSIFGPV | ASSIVTGKNEQY | 0.71953106 |
| LLALLANL | ASSIVTGKNEQY | 0.7858834 |
| LISNASDALDKLR | ASSIVTGKNEQY | 0.19296601 |
| LINEAVP | ASSIVTGKNEQY | 0.2630418 |
| LGYAFSGL | ASSIVTGKNEQY | 0.25739264 |
| LDFVAVAL | ASSIVTGKNEQY | 0.50770926 |

| | | |
|---|---|---|
| LAQPGAE | ASSIVTGKNEQY | 0.2450186 |
| ISNLGAVTGF | ASSIVTGKNEQY | 0.195215 |
| HQEDPGRI | ASSIVTGKNEQY | 0.46165308 |
| GTPGEPTP | ASSIVTGKNEQY | 0.8197475 |
| GFFNSGS | ASSIVTGKNEQY | 0.2238906 |
| GAGGGGLTAGAN | ASSIVTGKNEQY | 0.1468117 |
| FVDALVNM | ASSIVTGKNEQY | 0.4830423 |
| FLTVAPEL | ASSIVTGKNEQY | 0.23426867 |
| FETLTAL | ASSIVTGKNEQY | 0.30761385 |
| DVINPIEL | ASSIVTGKNEQY | 0.47942808 |
| APVSGAPV | ASSIVTGKNEQY | 0.15316351 |
| ALVGGLAW | ASSIVTGKNEQY | 0.2020725 |
| ALAAAAATAI | ASSIVTGKNEQY | 0.81355566 |
| AFGLGLLL | ASSIVTGKNEQY | 0.30313066 |
| AFFAAASL | ASSIVTGKNEQY | 0.45365286 |
| AAAALPAFSPP | ASSIVTGKNEQY | 0.7298456 |
| VVLAPAL | ASSIVTGKNEQY | 0.5958645 |
| TIGHVDHGKTTL | ASSIVTGKNEQY | 0.7858559 |
| LISNASDALDK | ASSIVTGKNEQY | 0.6675322 |
| LISNASDAL | ASSIVTGKNEQY | 0.2579711 |
| ISNASDALDK | ASSIVTGKNEQY | 0.39435992 |
| NQAETLVYQTE | ASSIVTGKNEQY | 0.22074194 |
| TGGTGGTAGAG | ASSIVTGKNEQY | 0.5770228 |
| TGSTAGGAGGA | ASSIVTGKNEQY | 0.38913378 |
| DPLAVPVAAGPV | ASSIVTGKNEQY | 0.2070414 |
| GGTGGTPGNGGN | ASSIVTGKNEQY | 0.7418527 |
| PAGLAGVDPATL | ASSIVTGKNEQY | 0.4035995 |
| GNNLVVEAPHAGASV | ASSIVTGKNEQY | 0.49921328 |
| KVTRIMSASTLEQPA | ASSIVTGKNEQY | 0.20289908 |
| LGFALPIALPRRTVV | ASSIVTGKNEQY | 0.19456722 |
| TRLGNLTARKLVVLV | ASSIVTGKNEQY | 0.7086457 |
| WSQQRRLIYFPSAGP | ASSIVTGKNEQY | 0.12705164 |
| GPNGNPDPMAAAVDIR | ASSIVTGKNEQY | 0.13830473 |
| TQRCVSVADLRAAASR | ASSIVTGKNEQY | 0.33915082 |
| QFVLDVAEHPVEVLTETV | ASSIVTGKNEQY | 0.7783762 |
| GGAGGTGGAGGNGGTQAGDG | ASSIVTGKNEQY | 0.12654085 |
| PLLATLRAIAVDVDAKPAQV | ASSIVTGKNEQY | 0.20341787 |
| APKFRTGGYMIDFWGVGYQVAK | ASSIVTGKNEQY | 0.16100203 |
| GAGGAGGAGGAAGGPGGTGGTG | ASSIVTGKNEQY | 0.47032917 |
| IGALGLLATDVAPAGSVAFGVA | ASSIVTGKNEQY | 0.3840819 |
| AGGAGGIGGNGGAGGAGGVLMG | ASSIVTGKNEQY | 0.14908788 |
| ADELIGLL | ASSIVTGKNEQY | 0.46505678 |
| ADRALKA | ASSIVTGKNEQY | 0.14729904 |
| AGAVGQVGGAGGA | ASSIVTGKNEQY | 0.13695131 |
| AGGGGGLGGAP | ASSIVTGKNEQY | 0.4395014 |
| AGGLLYGNGGAG | ASSIVTGKNEQY | 0.72912097 |
| AGLESIGL | ASSIVTGKNEQY | 0.16013323 |
| AGLKTGII | ASSIVTGKNEQY | 0.64383423 |
| ASLAALLL | ASSIVTGKNEQY | 0.22646567 |
| ASVINISSI | ASSIVTGKNEQY | 0.78313553 |
| ATPPGPA | ASSIVTGKNEQY | 0.2530322 |

| | | |
|---|---|---|
| ATVASALV | ASSIVTGKNEQY | 0.15157202 |
| ATVASLIL | ASSIVTGKNEQY | 0.6033174 |
| AVNGPVLIRH | ASSIVTGKNEQY | 0.356077 |
| DGAAGAVGI | ASSIVTGKNEQY | 0.36719334 |
| DPLLHGI | ASSIVTGKNEQY | 0.6321455 |
| ELGAALGAL | ASSIVTGKNEQY | 0.23239593 |
| ELGRTQAV | ASSIVTGKNEQY | 0.367823 |
| FVAGGYLS | ASSIVTGKNEQY | 0.72809243 |
| GASGGAGGAGGW | ASSIVTGKNEQY | 0.661996 |
| GGAGAGGDGGAGGI | ASSIVTGKNEQY | 0.20092358 |
| GTGHNGSQPGMGGNG | ASSIVTGKNEQY | 0.4177957 |
| GYAAASASAS | ASSIVTGKNEQY | 0.5472476 |
| HQVDVAAVQQG | ASSIVTGKNEQY | 0.21849881 |
| IGAAIAHT | ASSIVTGKNEQY | 0.7333276 |
| IGMIEVAQL | ASSIVTGKNEQY | 0.30836764 |
| IRLAPEHTSAMD | ASSIVTGKNEQY | 0.6456063 |
| IYNVSVL | ASSIVTGKNEQY | 0.41056308 |
| KQGFLGM | ASSIVTGKNEQY | 0.7932311 |
| LESVITQL | ASSIVTGKNEQY | 0.16485241 |
| LGLGALVVV | ASSIVTGKNEQY | 0.5274592 |
| LIGNGGNGGSGGI | ASSIVTGKNEQY | 0.42949682 |
| LNAINAPTL | ASSIVTGKNEQY | 0.5278691 |
| LQLLNTGV | ASSIVTGKNEQY | 0.24707854 |
| LQTSIEF | ASSIVTGKNEQY | 0.14319092 |
| MSFAVSEL | ASSIVTGKNEQY | 0.31758687 |
| NTGALGL | ASSIVTGKNEQY | 0.7217485 |
| PPGAGGAGGAGTT | ASSIVTGKNEQY | 0.27138823 |
| QLVGLGI | ASSIVTGKNEQY | 0.4797949 |
| RFLEQVHQL | ASSIVTGKNEQY | 0.80597126 |
| RGVGYVLR | ASSIVTGKNEQY | 0.19399211 |
| SATGVFAGVI | ASSIVTGKNEQY | 0.37185308 |
| SLPAITLP | ASSIVTGKNEQY | 0.4539821 |
| STALAFTV | ASSIVTGKNEQY | 0.7024933 |
| SVLTVGLGK | ASSIVTGKNEQY | 0.2636471 |
| TALAGAGSGI | ASSIVTGKNEQY | 0.74855137 |
| TGLPFTGI | ASSIVTGKNEQY | 0.18749848 |
| VAEHVTV | ASSIVTGKNEQY | 0.697872 |
| VDIVLNSL | ASSIVTGKNEQY | 0.15751514 |
| VLTGGGAVL | ASSIVTGKNEQY | 0.5787518 |
| VNVGLAYGI | ASSIVTGKNEQY | 0.250014 |
| VSSAAVGAV | ASSIVTGKNEQY | 0.67414176 |
| VTSVYEAGV | ASSIVTGKNEQY | 0.72660303 |
| WVALDRGA | ASSIVTGKNEQY | 0.72775495 |
| YLVGSVVI | ASSIVTGKNEQY | 0.1294743 |
| GPPGTGKTLL | ASSIVTGKNEQY | 0.6753933 |
| GVGMTKF | ASSIVTGKNEQY | 0.75710297 |
| IGIGGSDL | ASSIVTGKNEQY | 0.8238026 |
| IGTLLAEL | ASSIVTGKNEQY | 0.42702043 |
| IYTSGTT | ASSIVTGKNEQY | 0.47777003 |
| LSFPAGV | ASSIVTGKNEQY | 0.38831016 |
| LYTSGTT | ASSIVTGKNEQY | 0.25206167 |

| | | |
|---|---|---|
| VGAFPFGF | ASSIVTGKNEQY | 0.2534485 |
| VGAGTVEF | ASSIVTGKNEQY | 0.29377472 |
| VGKSTLLL | ASSIVTGKNEQY | 0.4209666 |
| AALAHAGGGIV | ASSIVTGKNEQY | 0.24549356 |
| AALVAVSLL | ASSIVTGKNEQY | 0.741541 |
| AAVSRAVAAL | ASSIVTGKNEQY | 0.7218436 |
| AGGSSGIGGF | ASSIVTGKNEQY | 0.7233112 |
| AGLTVANANG | ASSIVTGKNEQY | 0.78333604 |
| AGLVGAVL | ASSIVTGKNEQY | 0.63426125 |
| ALSQAFTV | ASSIVTGKNEQY | 0.444179 |
| ALTFSVSL | ASSIVTGKNEQY | 0.22559988 |
| ALTGAALVAL | ASSIVTGKNEQY | 0.26618418 |
| ASFSIGAGVL | ASSIVTGKNEQY | 0.17391051 |
| DGSVYKI | ASSIVTGKNEQY | 0.2070876 |
| EQLAEIV | ASSIVTGKNEQY | 0.25297546 |
| FAVSELTVM | ASSIVTGKNEQY | 0.39950442 |
| GAAPLAAGL | ASSIVTGKNEQY | 0.12401414 |
| GADGGDGGSGGAGG | ASSIVTGKNEQY | 0.4974943 |
| GFGLDTGGAGG | ASSIVTGKNEQY | 0.7829641 |
| GLASGGLL | ASSIVTGKNEQY | 0.2602157 |
| GRIAGAAVLL | ASSIVTGKNEQY | 0.48000932 |
| GSAGSLNTSF | ASSIVTGKNEQY | 0.20346612 |
| GTGQIGI | ASSIVTGKNEQY | 0.2326301 |
| GTLIALIL | ASSIVTGKNEQY | 0.75255775 |
| IAATMGFVVG | ASSIVTGKNEQY | 0.20034757 |
| IFAQALL | ASSIVTGKNEQY | 0.50804234 |
| IGAAVALGF | ASSIVTGKNEQY | 0.71998346 |
| IGLTPLAL | ASSIVTGKNEQY | 0.36234066 |
| ILGSNTSTL | ASSIVTGKNEQY | 0.5311213 |
| IVFSLTVL | ASSIVTGKNEQY | 0.30098575 |
| LASKATGF | ASSIVTGKNEQY | 0.16053629 |
| LASVALIL | ASSIVTGKNEQY | 0.57944447 |
| LFAVGLTL | ASSIVTGKNEQY | 0.12206114 |
| LFNGGIGTYI | ASSIVTGKNEQY | 0.46418664 |
| LTLGAGII | ASSIVTGKNEQY | 0.18073206 |
| LVAAALGDI | ASSIVTGKNEQY | 0.55921483 |
| NTAFVQLGI | ASSIVTGKNEQY | 0.5120839 |
| QLFSGAEL | ASSIVTGKNEQY | 0.43742362 |
| SAATFITL | ASSIVTGKNEQY | 0.22745731 |
| SNYGLANLGV | ASSIVTGKNEQY | 0.29238334 |
| SYGGAYAVM | ASSIVTGKNEQY | 0.21607536 |
| TTGVGGLTLG | ASSIVTGKNEQY | 0.17641214 |
| TTQTIYI | ASSIVTGKNEQY | 0.4684168 |
| VAAGAGLGAVI | ASSIVTGKNEQY | 0.2557752 |
| VAAVVTLL | ASSIVTGKNEQY | 0.26167452 |
| VGFLGGLI | ASSIVTGKNEQY | 0.59392476 |
| VGVIPILL | ASSIVTGKNEQY | 0.3641796 |
| VIFASSLI | ASSIVTGKNEQY | 0.162992 |
| VKGGDGGAAAG | ASSIVTGKNEQY | 0.54803944 |
| VTKTFTNI | ASSIVTGKNEQY | 0.4355337 |
| VVAAAILL | ASSIVTGKNEQY | 0.27117354 |

| | | |
|---|---|---|
| YAALDVEL | ASSIVTGKNEQY | 0.122190066 |
| LASVALLL | ASSIVTGKNEQY | 0.32422912 |
| LDAVHRV | ASSIVTGKNEQY | 0.23154372 |
| QGLSSSIF | ASSIVTGKNEQY | 0.22377738 |
| RGTFANIRL | ASSIVTGKNEQY | 0.21563888 |
| RQLWWGHRI | ASSIVTGKNEQY | 0.73941475 |
| YGPPGTGKTLL | ASSIVTGKNEQY | 0.17086099 |
| AALVGAPANVA | ASSIVTGKNEQY | 0.5659008 |
| AATFRRTE | ASSIVTGKNEQY | 0.6314827 |
| AGALGGGI | ASSIVTGKNEQY | 0.64637905 |
| AGGAGGAGGIGAG | ASSIVTGKNEQY | 0.7987434 |
| AGSGALGL | ASSIVTGKNEQY | 0.5257449 |
| AQIADQI | ASSIVTGKNEQY | 0.24535598 |
| ASAVAVTYL | ASSIVTGKNEQY | 0.38678825 |
| ASTVLGGTGG | ASSIVTGKNEQY | 0.35031265 |
| AVGGEPAK | ASSIVTGKNEQY | 0.1331695 |
| CEQGTFGGTC | ASSIVTGKNEQY | 0.13611282 |
| DEGKAPAG | ASSIVTGKNEQY | 0.47213113 |
| DTVVFPAL | ASSIVTGKNEQY | 0.13565058 |
| FEGITMAG | ASSIVTGKNEQY | 0.3858522 |
| FPLVVQD | ASSIVTGKNEQY | 0.52799916 |
| GAFTTPQ | ASSIVTGKNEQY | 0.33553568 |
| GFGGFGTAGGVGGA | ASSIVTGKNEQY | 0.1601473 |
| GFLAQFL | ASSIVTGKNEQY | 0.28778222 |
| GGAGGAGGNAGVA | ASSIVTGKNEQY | 0.22868603 |
| GGVGTAGAGGAGGAGG | ASSIVTGKNEQY | 0.6484463 |
| GNGGGGDAGNAGSGGN | ASSIVTGKNEQY | 0.381073 |
| GTGGAGGAGNDAGS | ASSIVTGKNEQY | 0.79515225 |
| GTTNTGWFNS | ASSIVTGKNEQY | 0.728932 |
| IFMGPLFTL | ASSIVTGKNEQY | 0.35417366 |
| ILAHAIEAI | ASSIVTGKNEQY | 0.66829944 |
| IRSAIREAL | ASSIVTGKNEQY | 0.2387129 |
| ITFAGGG | ASSIVTGKNEQY | 0.19852714 |
| ITGMSGHQEV | ASSIVTGKNEQY | 0.24961498 |
| IVAGLACTL | ASSIVTGKNEQY | 0.37655365 |
| IVGGGLAGA | ASSIVTGKNEQY | 0.38362232 |
| IVTGVLNI | ASSIVTGKNEQY | 0.2228235 |
| LAQQGCI | ASSIVTGKNEQY | 0.1627379 |
| LCLHVEN | ASSIVTGKNEQY | 0.424501 |
| LDAISDGS | ASSIVTGKNEQY | 0.652675 |
| LGGFAAML | ASSIVTGKNEQY | 0.14200865 |
| LGGKNPMI | ASSIVTGKNEQY | 0.2826341 |
| LGPKGVAAIL | ASSIVTGKNEQY | 0.8169212 |
| LMVGVSI | ASSIVTGKNEQY | 0.8127625 |
| LVGAAFVL | ASSIVTGKNEQY | 0.2354931 |
| LVLAVAPL | ASSIVTGKNEQY | 0.21870147 |
| MAAIAVIDAA | ASSIVTGKNEQY | 0.60018 |
| NNGINAELQ | ASSIVTGKNEQY | 0.15055814 |
| RGEPLVLVHG | ASSIVTGKNEQY | 0.34812713 |
| RGGAGGAAGLI | ASSIVTGKNEQY | 0.5270622 |
| RSYRMHPAV | ASSIVTGKNEQY | 0.3940985 |

| | | |
|---|---|---|
| SAAHLLAGQLA | ASSIVTGKNEQY | 0.21280047 |
| SAAVGSY | ASSIVTGKNEQY | 0.44763362 |
| SPHAVLHRI | ASSIVTGKNEQY | 0.322339 |
| SQIAKVL | ASSIVTGKNEQY | 0.48572424 |
| TALGHLVP | ASSIVTGKNEQY | 0.12454795 |
| TAPTTSGAA | ASSIVTGKNEQY | 0.32150686 |
| TDGDDGNGGSPGT | ASSIVTGKNEQY | 0.22963898 |
| TDGLGEN | ASSIVTGKNEQY | 0.22431177 |
| TGGPPLPGQA | ASSIVTGKNEQY | 0.16009186 |
| TIPAITI | ASSIVTGKNEQY | 0.26527345 |
| TIPGFTI | ASSIVTGKNEQY | 0.71485394 |
| TSIASGALA | ASSIVTGKNEQY | 0.7285957 |
| TVTLPGAAI | ASSIVTGKNEQY | 0.14286698 |
| VASGLSAALP | ASSIVTGKNEQY | 0.7216222 |
| VGAGVIGI | ASSIVTGKNEQY | 0.805644 |
| VGPKRFGV | ASSIVTGKNEQY | 0.2450012 |
| VNVAFPD | ASSIVTGKNEQY | 0.15219891 |
| VSMTNSGVS | ASSIVTGKNEQY | 0.5250114 |
| VTDVLGDLSV | ASSIVTGKNEQY | 0.21821618 |
| VVVVGAGFAGL | ASSIVTGKNEQY | 0.49319506 |
| VYTVGTVV | ASSIVTGKNEQY | 0.7225392 |
| WTTFSGP | ASSIVTGKNEQY | 0.16652244 |
| YDELVEAF | ASSIVTGKNEQY | 0.6274728 |
| YGTGGAGGNGG | ASSIVTGKNEQY | 0.61543566 |
| YTNVMAV | ASSIVTGKNEQY | 0.2404705 |
| FLEQVHQL | ASSIVTGKNEQY | 0.18296213 |
| GLSLLAVAV | ASSIVTGKNEQY | 0.18785094 |
| LEQVHQL | ASSIVTGKNEQY | 0.13197078 |
| LNAAGAI | ASSIVTGKNEQY | 0.5537578 |
| PAVHASL | ASSIVTGKNEQY | 0.26891902 |
| TELEDDA | ASSIVTGKNEQY | 0.24377602 |
| TPPGPPN | ASSIVTGKNEQY | 0.40384585 |
| TPPPPPPP | ASSIVTGKNEQY | 0.2246818 |
| TQSPALI | ASSITGGGDTQY | 0.2516077 |
| EFAAAFPQ | ASSITGGGDTQY | 0.24802683 |
| LLSGNLDVL | ASSITGGGDTQY | 0.3396819 |
| GPANLGNYNI | ASSITGGGDTQY | 0.5191938 |
| TLGPTLP | ASSITGGGDTQY | 0.3845026 |
| VYAYSSCCK | ASSITGGGDTQY | 0.348966 |
| APAPAPAPNI | ASSITGGGDTQY | 0.7837088 |
| PGGPVPIP | ASSITGGGDTQY | 0.18844172 |
| FLTVALEP | ASSITGGGDTQY | 0.2366105 |
| SQYFGANL | ASSITGGGDTQY | 0.59947735 |
| GGVKPAA | ASSITGGGDTQY | 0.1626642 |
| GGTGGAGGAGGW | ASSITGGGDTQY | 0.6345104 |
| GVGGVAGLGGAGG | ASSITGGGDTQY | 0.5901071 |
| VLLVLATTI | ASSITGGGDTQY | 0.4689823 |
| IGIFAAML | ASSITGGGDTQY | 0.3644156 |
| AVGDLID | ASSITGGGDTQY | 0.783007 |
| NISPLPL | ASSITGGGDTQY | 0.6197845 |
| VTAVFVAI | ASSITGGGDTQY | 0.22891308 |

| | | |
|---|---|---|
| SGDDNDAAL | ASSITGGGDTQY | 0.5175115 |
| AVAALAHS | ASSITGGGDTQY | 0.1886382 |
| GGVTVMATP | ASSITGGGDTQY | 0.68970716 |
| AGLAANVRD | ASSITGGGDTQY | 0.6010799 |
| TNRPDLIDPALLRPG | ASSITGGGDTQY | 0.18855767 |
| LANLGVTAL | ASSITGGGDTQY | 0.3760006 |
| TGERIEL | ASSITGGGDTQY | 0.62637365 |
| NGEELAA | ASSITGGGDTQY | 0.7061499 |
| VLSLVWG | ASSITGGGDTQY | 0.14681785 |
| IGYALWL | ASSITGGGDTQY | 0.29107988 |
| LSLLLAPL | ASSITGGGDTQY | 0.15215428 |
| TLDEALK | ASSITGGGDTQY | 0.330153 |
| TDGAGTAEGG | ASSITGGGDTQY | 0.18928298 |
| NAPGGAGGNGG | ASSITGGGDTQY | 0.63524914 |
| VSVFMGMGGNF | ASSITGGGDTQY | 0.32747662 |
| LASYLAAL | ASSITGGGDTQY | 0.7191821 |
| LVGGTPAL | ASSITGGGDTQY | 0.3721068 |
| VRNADVL | ASSITGGGDTQY | 0.67173016 |
| TCSSPFLVSAM | ASSITGGGDTQY | 0.2560825 |
| VHITDAE | ASSITGGGDTQY | 0.4120521 |
| EYAAPSF | ASSITGGGDTQY | 0.16201803 |
| VVDAFHGGL | ASSITGGGDTQY | 0.14857362 |
| ASVVDPQ | ASSITGGGDTQY | 0.1770636 |
| SYNEFGGLS | ASSITGGGDTQY | 0.29660943 |
| DPDGSVY | ASSITGGGDTQY | 0.13139519 |
| MVSTIHATE | ASSITGGGDTQY | 0.73590666 |
| LAAGVADPV | ASSITGGGDTQY | 0.5205728 |
| TKEQIAA | ASSITGGGDTQY | 0.6128464 |
| SLASPAAL | ASSITGGGDTQY | 0.27105814 |
| IGLVFYGV | ASSITGGGDTQY | 0.27688712 |
| FLLDASNL | ASSITGGGDTQY | 0.23485014 |
| YQVGNFP | ASSITGGGDTQY | 0.75064886 |
| ESLLSQST | ASSITGGGDTQY | 0.1675579 |
| LFTVGFAAVG | ASSITGGGDTQY | 0.1455769 |
| ATPPTPT | ASSITGGGDTQY | 0.23565963 |
| TDLATLG | ASSITGGGDTQY | 0.15787292 |
| SAGGTGR | ASSITGGGDTQY | 0.607658 |
| GFSNSGS | ASSITGGGDTQY | 0.6631949 |
| GGNGGSGGTSV | ASSITGGGDTQY | 0.5527036 |
| GTSVATGGAGN | ASSITGGGDTQY | 0.51934844 |
| NGGNGGSGGTS | ASSITGGGDTQY | 0.135666 |
| AVVKFPH | ASSITGGGDTQY | 0.25290614 |
| TPPPPPP | ASSITGGGDTQY | 0.8113524 |
| AAPAAAAAPA | ASSITGGGDTQY | 0.3848148 |
| IALNEDL | ASSITGGGDTQY | 0.37052608 |
| IVNVSSM | ASSITGGGDTQY | 0.7153991 |
| TGIPPAPR | ASSITGGGDTQY | 0.3144432 |
| FDLGGGTF | ASSITGGGDTQY | 0.29902762 |
| FDLGGGTFDVS | ASSITGGGDTQY | 0.30548257 |
| NEPTAAA | ASSITGGGDTQY | 0.3279636 |
| NEPTAAALA | ASSITGGGDTQY | 0.25787032 |

| | | |
|---|---|---|
| GSIINISSI | ASSITGGGDTQY | 0.17499004 |
| VAPGNAGTA | ASSITGGGDTQY | 0.40144598 |
| TGGGASLE | ASSITGGGDTQY | 0.79429466 |
| VAVYNHYKRI | ASSITGGGDTQY | 0.41327834 |
| TDPAPATT | ASSITGGGDTQY | 0.7009365 |
| TNRPDILDPALLRPG | ASSITGGGDTQY | 0.517472 |
| LTTTTTS | ASSITGGGDTQY | 0.7389054 |
| PTGSGKTL | ASSITGGGDTQY | 0.2291312 |
| TGSGKTL | ASSITGGGDTQY | 0.2513988 |
| TIGHVDHGK | ASSITGGGDTQY | 0.28544974 |
| TIGHVDHGKTT | ASSITGGGDTQY | 0.27082902 |
| GHVDHGKTT | ASSITGGGDTQY | 0.35021263 |
| AQFEHTLL | ASSITGGGDTQY | 0.2260545 |
| IVNQDEL | ASSITGGGDTQY | 0.8240031 |
| TNGSQFF | ASSITGGGDTQY | 0.63056076 |
| TGKGIVE | ASSITGGGDTQY | 0.47523952 |
| LISNASDA | ASSITGGGDTQY | 0.2026894 |
| AELRTLGELTKIAKAHGAQ | ASSITGGGDTQY | 0.30278942 |
| APNAVAAAEPSW | ASSITGGGDTQY | 0.5261484 |
| APQVAGAATAVL | ASSITGGGDTQY | 0.16620448 |
| ARRDPAAAAAILPTDA | ASSITGGGDTQY | 0.7811853 |
| DAAARDVLAEAGLAEYF | ASSITGGGDTQY | 0.2655559 |
| DDQIVLSAKANLAKVEQ | ASSITGGGDTQY | 0.2634399 |
| DELLPDTGEDSDEDGH | ASSITGGGDTQY | 0.6058455 |
| ELPEAPANRATVPQD | ASSITGGGDTQY | 0.7851 |
| FGQSKIERVTGIG | ASSITGGGDTQY | 0.22993706 |
| GAGGTGSTAGGAGGAGGAGGLY | ASSITGGGDTQY | 0.27728397 |
| GHGGTGAAGGNGGTGGAGGSGI | ASSITGGGDTQY | 0.21136114 |
| GNGGNGGAGGEGGPGGAGGAGA | ASSITGGGDTQY | 0.65531087 |
| GSTAGNGGNGGNATNGTIAGQP | ASSITGGGDTQY | 0.7655623 |
| GTGGDGGNGGNGGTGG | ASSITGGGDTQY | 0.4694627 |
| GVRRPAGIPADAPV | ASSITGGGDTQY | 0.7328157 |
| GYRQVTDAHLLTLARR | ASSITGGGDTQY | 0.3250063 |
| IAKMGQDGHDRG | ASSITGGGDTQY | 0.17273194 |
| KAEPSIEIQYT | ASSITGGGDTQY | 0.31269187 |
| KPVRIRMAADEDTVAVT | ASSITGGGDTQY | 0.5539625 |
| MPPRIAGMRLLVIKPEP | ASSITGGGDTQY | 0.7516683 |
| MKFYAVALFRSRGVPPTR | ASSITGGGDTQY | 0.76216185 |
| NAGGFGILGAVAHSP | ASSITGGGDTQY | 0.46474484 |
| NEGREITLAMAEM | ASSITGGGDTQY | 0.23431903 |
| NELVEWSHPLGHRWRWE | ASSITGGGDTQY | 0.7558646 |
| NLGAGGTGGNGGNPG | ASSITGGGDTQY | 0.70580316 |
| NTEWPKWETVMAADPVD | ASSITGGGDTQY | 0.67665917 |
| PASAAPAETQP | ASSITGGGDTQY | 0.37298238 |
| QQSSDGKNGVRAAEITGE | ASSITGGGDTQY | 0.2897484 |
| RAPDAVTTSWWKE | ASSITGGGDTQY | 0.29541498 |
| SGFGNVGAHTSGWFNQ | ASSITGGGDTQY | 0.7028605 |
| SGVAGAVGQAASVGGLKV | ASSITGGGDTQY | 0.20385835 |
| SNGGAGTGSAGGNGGT | ASSITGGGDTQY | 0.3710266 |
| TGGAGGEGGAAG | ASSITGGGDTQY | 0.6037371 |
| TGGDGGAAGAGGAPGHG | ASSITGGGDTQY | 0.16063446 |

| | | |
|---|---|---|
| TMPPLLAAIANAGRHGV | ASSITGGGDTQY | 0.39146572 |
| TSSTAGNSGGA | ASSITGGGDTQY | 0.72218686 |
| VPATVPVGILC | ASSITGGGDTQY | 0.2415947 |
| VQRPDSGGVVLTGRIS | ASSITGGGDTQY | 0.25013912 |
| WTPNPVPATKGVPAGLVQAT | ASSITGGGDTQY | 0.7488098 |
| YALKRALAKFSDRFDVVII | ASSITGGGDTQY | 0.822022 |
| YLIGSVIV | ASSITGGGDTQY | 0.58706903 |
| YGQLVAVID | ASSITGGGDTQY | 0.17209028 |
| VTVGGMTLPA | ASSITGGGDTQY | 0.13771638 |
| VLALPAV | ASSITGGGDTQY | 0.2503038 |
| VIPPTLNYET | ASSITGGGDTQY | 0.7038809 |
| VGMGGMHPSSG | ASSITGGGDTQY | 0.2301088 |
| VGLAGGAATA | ASSITGGGDTQY | 0.2280573 |
| VGGAGGAGGSGP | ASSITGGGDTQY | 0.5058651 |
| VFAAFMV | ASSITGGGDTQY | 0.67026293 |
| TLYDKLR | ASSITGGGDTQY | 0.19466466 |
| TLAVFTVL | ASSITGGGDTQY | 0.18002093 |
| TGSVQPARV | ASSITGGGDTQY | 0.27603817 |
| TGGGAGSGGT | ASSITGGGDTQY | 0.5931838 |
| SHLETSL | ASSITGGGDTQY | 0.22506452 |
| SFTTGMT | ASSITGGGDTQY | 0.40140885 |
| QVEVAKRMGVRQPTVS | ASSITGGGDTQY | 0.20845613 |
| QGSGGPALAM | ASSITGGGDTQY | 0.15609914 |
| MLAPSNP | ASSITGGGDTQY | 0.5753772 |
| LVNNAAI | ASSITGGGDTQY | 0.21049647 |
| LTGEINTLA | ASSITGGGDTQY | 0.7316876 |
| LSIFGPV | ASSITGGGDTQY | 0.19603221 |
| LLALLANL | ASSITGGGDTQY | 0.5523385 |
| LISNASDALDKLR | ASSITGGGDTQY | 0.64823496 |
| LINEAVP | ASSITGGGDTQY | 0.3687955 |
| LGYAFSGL | ASSITGGGDTQY | 0.3270851 |
| LDFVAVAL | ASSITGGGDTQY | 0.19606581 |
| LAQPGAE | ASSITGGGDTQY | 0.16488643 |
| ISNLGAVTGF | ASSITGGGDTQY | 0.14106227 |
| HQEDPGRI | ASSITGGGDTQY | 0.7195319 |
| GTPGEPTP | ASSITGGGDTQY | 0.12539019 |
| GFFNSGS | ASSITGGGDTQY | 0.6898265 |
| GAGGGGLTAGAN | ASSITGGGDTQY | 0.4586712 |
| FVDALVNM | ASSITGGGDTQY | 0.18821147 |
| FLTVAPEL | ASSITGGGDTQY | 0.3856713 |
| FETLTAL | ASSITGGGDTQY | 0.24977884 |
| DVINPIEL | ASSITGGGDTQY | 0.16163838 |
| APVSGAPV | ASSITGGGDTQY | 0.734016 |
| ALVGGLAW | ASSITGGGDTQY | 0.4314139 |
| ALAAAAATAI | ASSITGGGDTQY | 0.23277894 |
| AFGLGLLL | ASSITGGGDTQY | 0.37053025 |
| AFFAAASL | ASSITGGGDTQY | 0.41859448 |
| AAAALPAFSPP | ASSITGGGDTQY | 0.20887786 |
| VVLAPAL | ASSITGGGDTQY | 0.24951905 |
| TIGHVDHGKTTL | ASSITGGGDTQY | 0.7078177 |
| LISNASDALDK | ASSITGGGDTQY | 0.31487894 |

| | | |
|---|---|---|
| LISNASDAL | ASSITGGGDTQY | 0.19612516 |
| ISNASDALDK | ASSITGGGDTQY | 0.59658295 |
| NQAETLVYQTE | ASSITGGGDTQY | 0.18692164 |
| TGGTGGTAGAG | ASSITGGGDTQY | 0.6049236 |
| TGSTAGGAGGA | ASSITGGGDTQY | 0.27063578 |
| DPLAVPVAAGPV | ASSITGGGDTQY | 0.4077516 |
| GGTGGTPGNGGN | ASSITGGGDTQY | 0.5776991 |
| PAGLAGVDPATL | ASSITGGGDTQY | 0.17720507 |
| GNNLVVEAPHAGASV | ASSITGGGDTQY | 0.20991956 |
| KVTRIMSASTLEQPA | ASSITGGGDTQY | 0.29658228 |
| LGFALPIALPRRTVV | ASSITGGGDTQY | 0.44754887 |
| TRLGNLTARKLVVLV | ASSITGGGDTQY | 0.6234676 |
| WSQQRRLIYFPSAGP | ASSITGGGDTQY | 0.22866471 |
| GPNGNPDPMAAAVDIR | ASSITGGGDTQY | 0.4327976 |
| TQRCVSVADLRAAASR | ASSITGGGDTQY | 0.23855571 |
| QFVLDVAEHPVEVLTETV | ASSITGGGDTQY | 0.19973752 |
| GGAGGTGGAGGNGGTQAGDG | ASSITGGGDTQY | 0.2727317 |
| PLLATLRAIAVDVDAKPAQV | ASSITGGGDTQY | 0.24256 |
| APKFRTGGYMIDFWGVGYQVAK | ASSITGGGDTQY | 0.24456646 |
| GAGGAGGAGGAAGGPGGTGGTG | ASSITGGGDTQY | 0.4433911 |
| IGALGLLATDVAPAGSVAFGVA | ASSITGGGDTQY | 0.21419042 |
| AGGAGGIGGNGGAGGAGGVLMG | ASSITGGGDTQY | 0.42611128 |
| ADELIGLL | ASSITGGGDTQY | 0.7606553 |
| ADRALKA | ASSITGGGDTQY | 0.34477258 |
| AGAVGQVGGAGGA | ASSITGGGDTQY | 0.4371498 |
| AGGGGGLGGAP | ASSITGGGDTQY | 0.5336566 |
| AGGLLYGNGGAG | ASSITGGGDTQY | 0.5862349 |
| AGLESIGL | ASSITGGGDTQY | 0.2604745 |
| AGLKTGII | ASSITGGGDTQY | 0.21209526 |
| ASLAALLL | ASSITGGGDTQY | 0.6557592 |
| ASVINISSI | ASSITGGGDTQY | 0.63044035 |
| ATPPGPA | ASSITGGGDTQY | 0.7138516 |
| ATVASALV | ASSITGGGDTQY | 0.52948654 |
| ATVASLIL | ASSITGGGDTQY | 0.53521824 |
| AVNGPVLIRH | ASSITGGGDTQY | 0.5815468 |
| DGAAGAVGI | ASSITGGGDTQY | 0.45744103 |
| DPLLHGI | ASSITGGGDTQY | 0.1695395 |
| ELGAALGAL | ASSITGGGDTQY | 0.2790673 |
| ELGRTQAV | ASSITGGGDTQY | 0.49161053 |
| FVAGGYLS | ASSITGGGDTQY | 0.20458573 |
| GASGGAGGAGGW | ASSITGGGDTQY | 0.16616915 |
| GGAGAGGDGGAGGI | ASSITGGGDTQY | 0.745568 |
| GTGHNGSQPGMGGNG | ASSITGGGDTQY | 0.8242304 |
| GYAAASASAS | ASSITGGGDTQY | 0.24339044 |
| HQVDVAAVQQG | ASSITGGGDTQY | 0.627825 |
| IGAAIAHT | ASSITGGGDTQY | 0.15821543 |
| IGMIEVAQL | ASSITGGGDTQY | 0.57426435 |
| IRLAPEHTSAMD | ASSITGGGDTQY | 0.76080596 |
| IYNVSVL | ASSITGGGDTQY | 0.24913123 |
| KQGFLGM | ASSITGGGDTQY | 0.4839147 |
| LESVITQL | ASSITGGGDTQY | 0.19369872 |

| | | |
|---|---|---|
| LGLGALVVV | ASSITGGGDTQY | 0.22343898 |
| LIGNGGNGGSGGI | ASSITGGGDTQY | 0.78237486 |
| LNAINAPTL | ASSITGGGDTQY | 0.30720237 |
| LQLLNTGV | ASSITGGGDTQY | 0.32573533 |
| LQTSIEF | ASSITGGGDTQY | 0.27805084 |
| MSFAVSEL | ASSITGGGDTQY | 0.20874551 |
| NTGALGL | ASSITGGGDTQY | 0.27176553 |
| PPGAGGAGGAGTT | ASSITGGGDTQY | 0.7731496 |
| QLVGLGI | ASSITGGGDTQY | 0.1842546 |
| RFLEQVHQL | ASSITGGGDTQY | 0.66928524 |
| RGVGYVLR | ASSITGGGDTQY | 0.2256361 |
| SATGVFAGVI | ASSITGGGDTQY | 0.7973243 |
| SLPAITLP | ASSITGGGDTQY | 0.7243067 |
| STALAFTV | ASSITGGGDTQY | 0.5457062 |
| SVLTVGLGK | ASSITGGGDTQY | 0.2627173 |
| TALAGAGSGI | ASSITGGGDTQY | 0.7793195 |
| TGLPFTGI | ASSITGGGDTQY | 0.46677896 |
| VAEHVTV | ASSITGGGDTQY | 0.64030266 |
| VDIVLNSL | ASSITGGGDTQY | 0.5471277 |
| VLTGGGAVL | ASSITGGGDTQY | 0.10440781 |
| VNVGLAYGI | ASSITGGGDTQY | 0.11444393 |
| VSSAAVGAV | ASSITGGGDTQY | 0.10735938 |
| VTSVYEAGV | ASSITGGGDTQY | 0.12152189 |
| WVALDRGA | ASSITGGGDTQY | 0.11016129 |
| YLVGSVVI | ASSITGGGDTQY | 0.1014001 |
| GPPGTGKTLL | ASSITGGGDTQY | 0.11281007 |
| GVGMTKF | ASSITGGGDTQY | 0.10511279 |
| IGIGGSDL | ASSITGGGDTQY | 0.11578262 |
| IGTLLAEL | ASSITGGGDTQY | 0.11383764 |
| IYTSGTT | ASSITGGGDTQY | 0.10981904 |
| LSFPAGV | ASSITGGGDTQY | 0.14935184 |
| LYTSGTT | ASSITGGGDTQY | 0.13421975 |
| VGAFPFGF | ASSITGGGDTQY | 0.10311209 |
| VGAGTVEF | ASSITGGGDTQY | 0.110979415 |
| VGKSTLLL | ASSITGGGDTQY | 0.10727565 |
| AALAHAGGGIV | ASSITGGGDTQY | 0.10996784 |
| AALVAVSLL | ASSITGGGDTQY | 0.09881381 |
| AAVSRAVAAL | ASSITGGGDTQY | 0.11637833 |
| AGGSSGIGGF | ASSITGGGDTQY | 0.103795566 |
| AGLTVANANG | ASSITGGGDTQY | 0.09830607 |
| AGLVGAVL | ASSITGGGDTQY | 0.11858881 |
| ALSQAFTV | ASSITGGGDTQY | 0.10797593 |
| ALTFSVSL | ASSITGGGDTQY | 0.10750138 |
| ALTGAALVAL | ASSITGGGDTQY | 0.13415523 |
| ASFSIGAGVL | ASSITGGGDTQY | 0.10005221 |
| DGSVYKI | ASSITGGGDTQY | 0.10552212 |
| EQLAEIV | ASSITGGGDTQY | 0.1228738 |
| FAVSELTVM | ASSITGGGDTQY | 0.17256892 |
| GAAPLAAGL | ASSITGGGDTQY | 0.10476607 |
| GADGGDGGSGGAGG | ASSITGGGDTQY | 0.109040596 |
| GFGLDTGGAGG | ASSITGGGDTQY | 0.17710875 |

| | | |
|---|---|---|
| GLASGGLL | ASSITGGGDTQY | 0.12167562 |
| GRIAGAAVLL | ASSITGGGDTQY | 0.12497274 |
| GSAGSLNTSF | ASSITGGGDTQY | 0.1323341 |
| GTGQIGI | ASSITGGGDTQY | 0.102199785 |
| GTLIALIL | ASSITGGGDTQY | 0.12852153 |
| IAATMGFVVG | ASSITGGGDTQY | 0.1221321 |
| IFAQALL | ASSITGGGDTQY | 0.114391215 |
| IGAAVALGF | ASSITGGGDTQY | 0.11669601 |
| IGLTPLAL | ASSITGGGDTQY | 0.10608925 |
| ILGSNTSTL | ASSITGGGDTQY | 0.10607512 |
| IVFSLTVL | ASSITGGGDTQY | 0.0992651 |
| LASKATGF | ASSITGGGDTQY | 0.11482745 |
| LASVALIL | ASSITGGGDTQY | 0.119608454 |
| LFAVGLTL | ASSITGGGDTQY | 0.1157036 |
| LFNGGIGTYI | ASSITGGGDTQY | 0.100976035 |
| LTLGAGII | ASSITGGGDTQY | 0.1043858 |
| LVAAALGDI | ASSITGGGDTQY | 0.10035945 |
| NTAFVQLGI | ASSITGGGDTQY | 0.10654558 |
| QLFSGAEL | ASSITGGGDTQY | 0.10088924 |
| SAATFITL | ASSITGGGDTQY | 0.09975778 |
| SNYGLANLGV | ASSITGGGDTQY | 0.15009305 |
| SYGGAYAVM | ASSITGGGDTQY | 0.15149507 |
| TTGVGGLTLG | ASSITGGGDTQY | 0.113574356 |
| TTQTIYI | ASSITGGGDTQY | 0.112257846 |
| VAAGAGLGAVI | ASSITGGGDTQY | 0.11892832 |
| VAAVVTLL | ASSITGGGDTQY | 0.11928673 |
| VGFLGGLI | ASSITGGGDTQY | 0.11592123 |
| VGVIPILL | ASSITGGGDTQY | 0.10169797 |
| VIFASSLI | ASSITGGGDTQY | 0.15376483 |
| VKGGDGGAAAG | ASSITGGGDTQY | 0.11130542 |
| VTKTFTNI | ASSITGGGDTQY | 0.10436228 |
| VVAAAILL | ASSITGGGDTQY | 0.13528322 |
| YAALDVEL | ASSITGGGDTQY | 0.10718964 |
| LASVALLL | ASSITGGGDTQY | 0.105413854 |
| LDAVHRV | ASSITGGGDTQY | 0.112944536 |
| QGLSSSIF | ASSITGGGDTQY | 0.1219594 |
| RGTFANIRL | ASSITGGGDTQY | 0.10157613 |
| RQLWWGHRI | ASSITGGGDTQY | 0.10251324 |
| YGPPGTGKTLL | ASSITGGGDTQY | 0.10398522 |
| AALVGAPANVA | ASSITGGGDTQY | 0.11816016 |
| AATFRRTE | ASSITGGGDTQY | 0.15768856 |
| AGALGGGI | ASSITGGGDTQY | 0.11747146 |
| AGGAGGAGGIGAG | ASSITGGGDTQY | 0.103945464 |
| AGSGALGL | ASSITGGGDTQY | 0.12750487 |
| AQIADQI | ASSITGGGDTQY | 0.11434095 |
| ASAVAVTYL | ASSITGGGDTQY | 0.11266165 |
| ASTVLGGTGG | ASSITGGGDTQY | 0.121196315 |
| AVGGEPAK | ASSITGGGDTQY | 0.10166579 |
| CEQGTFGGTC | ASSITGGGDTQY | 0.120153815 |
| DEGKAPAG | ASSITGGGDTQY | 0.13025068 |
| DTVVFPAL | ASSITGGGDTQY | 0.12213402 |

| | | |
|---|---|---|
| FEGITMAG | ASSITGGGDTQY | 0.11682187 |
| FPLVVQD | ASSITGGGDTQY | 0.10834656 |
| GAFTTPQ | ASSITGGGDTQY | 0.114388876 |
| GFGGFGTAGGVGGA | ASSITGGGDTQY | 0.111277156 |
| GFLAQFL | ASSITGGGDTQY | 0.10678813 |
| GGAGGAGGNAGVA | ASSITGGGDTQY | 0.11561538 |
| GGVGTAGAGGAGGAGG | ASSITGGGDTQY | 0.1137873 |
| GNGGGGDAGNAGSGGN | ASSITGGGDTQY | 0.112483375 |
| GTGGAGGAGNDAGS | ASSITGGGDTQY | 0.11924402 |
| GTTNTGWFNS | ASSITGGGDTQY | 0.10398701 |
| IFMGPLFTL | ASSITGGGDTQY | 0.12352745 |
| ILAHAIEAI | ASSITGGGDTQY | 0.12967758 |
| IRSAIREAL | ASSITGGGDTQY | 0.10283511 |
| ITFAGGG | ASSITGGGDTQY | 0.10610199 |
| ITGMSGHQEV | ASSITGGGDTQY | 0.111190274 |
| IVAGLACTL | ASSITGGGDTQY | 0.12478022 |
| IVGGGLAGA | ASSITGGGDTQY | 0.11443944 |
| IVTGVLNI | ASSITGGGDTQY | 0.11120256 |
| LAQQGCI | ASSITGGGDTQY | 0.14050701 |
| LCLHVEN | ASSITGGGDTQY | 0.10203705 |
| LDAISDGS | ASSITGGGDTQY | 0.1333736 |
| LGGFAAML | ASSITGGGDTQY | 0.10578161 |
| LGGKNPMI | ASSITGGGDTQY | 0.10879995 |
| LGPKGVAAIL | ASSITGGGDTQY | 0.104876526 |
| LMVGVSI | ASSITGGGDTQY | 0.09890296 |
| LVGAAFVL | ASSITGGGDTQY | 0.1310191 |
| LVLAVAPL | ASSITGGGDTQY | 0.10616923 |
| MAAIAVIDAA | ASSITGGGDTQY | 0.12499029 |
| NNGINAELQ | ASSITGGGDTQY | 0.10789005 |
| RGEPLVLVHG | ASSITGGGDTQY | 0.117739804 |
| RGGAGGAAGLI | ASSITGGGDTQY | 0.10594952 |
| RSYRMHPAV | ASSITGGGDTQY | 0.110973276 |
| SAAHLLAGQLA | ASSITGGGDTQY | 0.10325244 |
| SAAVGSY | ASSITGGGDTQY | 0.12791045 |
| SPHAVLHRI | ASSITGGGDTQY | 0.12273471 |
| SQIAKVL | ASSITGGGDTQY | 0.12088132 |
| TALGHLVP | ASSITGGGDTQY | 0.10233943 |
| TAPTTSGAA | ASSITGGGDTQY | 0.101789095 |
| TDGDDGNGGSPGT | ASSITGGGDTQY | 0.11918284 |
| TDGLGEN | ASSITGGGDTQY | 0.09901813 |
| TGGPPLPGQA | ASSITGGGDTQY | 0.09862776 |
| TIPAITI | ASSITGGGDTQY | 0.0997528 |
| TIPGFTI | ASSITGGGDTQY | 0.099347636 |
| TSIASGALA | ASSITGGGDTQY | 0.13417472 |
| TVTLPGAAI | ASSITGGGDTQY | 0.14120452 |
| VASGLSAALP | ASSITGGGDTQY | 0.113653585 |
| VGAGVIGI | ASSITGGGDTQY | 0.12950607 |
| VGPKRFGV | ASSITGGGDTQY | 0.11197036 |
| VNVAFPD | ASSITGGGDTQY | 0.097743005 |
| VSMTNSGVS | ASSITGGGDTQY | 0.1067896 |
| VTDVLGDLSV | ASSITGGGDTQY | 0.10390098 |

| | | |
|---|---|---|
| VVVVGAGFAGL | ASSITGGGDTQY | 0.12608105 |
| VYTVGTVV | ASSITGGGDTQY | 0.1010643 |
| WTTFSGP | ASSITGGGDTQY | 0.10558398 |
| YDELVEAF | ASSITGGGDTQY | 0.11831194 |
| YGTGGAGGNGG | ASSITGGGDTQY | 0.112197906 |
| YTNVMAV | ASSITGGGDTQY | 0.11137056 |
| FLEQVHQL | ASSITGGGDTQY | 0.102462605 |
| GLSLLAVAV | ASSITGGGDTQY | 0.098240465 |
| LEQVHQL | ASSITGGGDTQY | 0.13644056 |
| LNAAGAI | ASSITGGGDTQY | 0.10009886 |
| PAVHASL | ASSITGGGDTQY | 0.10712114 |
| TELEDDA | ASSITGGGDTQY | 0.09928298 |
| TPPGPPN | ASSITGGGDTQY | 0.100039005 |
| TPPPPPPP | ASSITGGGDTQY | 0.101206996 |
| TQSPALI | ASSLYWGGLEDTQY | 0.1255763 |
| EFAAAFPQ | ASSLYWGGLEDTQY | 0.102951355 |
| LLSGNLDVL | ASSLYWGGLEDTQY | 0.111192726 |
| GPANLGNYNI | ASSLYWGGLEDTQY | 0.122193955 |
| TLGPTLP | ASSLYWGGLEDTQY | 0.13714683 |
| VYAYSSCCK | ASSLYWGGLEDTQY | 0.10990973 |
| APAPAPAPNI | ASSLYWGGLEDTQY | 0.108933516 |
| PGGPVPIP | ASSLYWGGLEDTQY | 0.104386784 |
| FLTVALEP | ASSLYWGGLEDTQY | 0.104397535 |
| SQYFGANL | ASSLYWGGLEDTQY | 0.105822966 |
| GGVKPAA | ASSLYWGGLEDTQY | 0.15232967 |
| GGTGGAGGAGGW | ASSLYWGGLEDTQY | 0.10719376 |
| GVGGVAGLGGAGG | ASSLYWGGLEDTQY | 0.101713605 |
| VLLVLATTI | ASSLYWGGLEDTQY | 0.10827919 |
| IGIFAAML | ASSLYWGGLEDTQY | 0.1282459 |
| AVGDLID | ASSLYWGGLEDTQY | 0.12881798 |
| NISPLPL | ASSLYWGGLEDTQY | 0.105023086 |
| VTAVFVAI | ASSLYWGGLEDTQY | 0.11070055 |
| SGDDNDAAL | ASSLYWGGLEDTQY | 0.11030274 |
| AVAALAHS | ASSLYWGGLEDTQY | 0.109942436 |
| GGVTVMATP | ASSLYWGGLEDTQY | 0.10566855 |
| AGLAANVRD | ASSLYWGGLEDTQY | 0.10596796 |
| TNRPDLIDPALLRPG | ASSLYWGGLEDTQY | 0.11963155 |
| LANLGVTAL | ASSLYWGGLEDTQY | 0.30900526 |
| TGERIEL | ASSLYWGGLEDTQY | 0.10554412 |
| NGEELAA | ASSLYWGGLEDTQY | 0.10088955 |
| VLSLVWG | ASSLYWGGLEDTQY | 0.116494454 |
| IGYALWL | ASSLYWGGLEDTQY | 0.10263086 |
| LSLLLAPL | ASSLYWGGLEDTQY | 0.11026529 |
| TLDEALK | ASSLYWGGLEDTQY | 0.115589626 |
| TDGAGTAEGG | ASSLYWGGLEDTQY | 0.10070728 |
| NAPGGAGGNGG | ASSLYWGGLEDTQY | 0.104583085 |
| VSVFMGMGGNF | ASSLYWGGLEDTQY | 0.13228242 |
| LASYLAAL | ASSLYWGGLEDTQY | 0.10792443 |
| LVGGTPAL | ASSLYWGGLEDTQY | 0.10842725 |
| VRNADVL | ASSLYWGGLEDTQY | 0.13404877 |
| TCSSPFLVSAM | ASSLYWGGLEDTQY | 0.11380025 |

| | | |
|---|---|---|
| VHITDAE | ASSLYWGGLEDTQY | 0.15522905 |
| EYAAPSF | ASSLYWGGLEDTQY | 0.12957327 |
| VVDAFHGGL | ASSLYWGGLEDTQY | 0.10962472 |
| ASVVDPQ | ASSLYWGGLEDTQY | 0.10701959 |
| SYNEFGGLS | ASSLYWGGLEDTQY | 0.103128545 |
| DPDGSVY | ASSLYWGGLEDTQY | 0.119461775 |
| MVSTIHATE | ASSLYWGGLEDTQY | 0.12031563 |
| LAAGVADPV | ASSLYWGGLEDTQY | 0.105514616 |
| TKEQIAA | ASSLYWGGLEDTQY | 0.13918857 |
| SLASPAAL | ASSLYWGGLEDTQY | 0.112518564 |
| IGLVFYGV | ASSLYWGGLEDTQY | 0.116408885 |
| FLLDASNL | ASSLYWGGLEDTQY | 0.10285089 |
| YQVGNFP | ASSLYWGGLEDTQY | 0.10491559 |
| ESLLSQST | ASSLYWGGLEDTQY | 0.11501758 |
| LFTVGFAAVG | ASSLYWGGLEDTQY | 0.10277151 |
| ATPPTPT | ASSLYWGGLEDTQY | 0.09948289 |
| TDLATLG | ASSLYWGGLEDTQY | 0.11182527 |
| SAGGTGR | ASSLYWGGLEDTQY | 0.11632008 |
| GFSNSGS | ASSLYWGGLEDTQY | 0.09829324 |
| GGNGGSGGTSV | ASSLYWGGLEDTQY | 0.107584625 |
| GTSVATGGAGN | ASSLYWGGLEDTQY | 0.10177742 |
| NGGNGGSGGTS | ASSLYWGGLEDTQY | 0.10587323 |
| AVVKFPH | ASSLYWGGLEDTQY | 0.11271131 |
| TPPPPPP | ASSLYWGGLEDTQY | 0.10098398 |
| AAPAAAAAPA | ASSLYWGGLEDTQY | 0.117534384 |
| IALNEDL | ASSLYWGGLEDTQY | 0.100463405 |
| IVNVSSM | ASSLYWGGLEDTQY | 0.10100151 |
| TGIPPAPR | ASSLYWGGLEDTQY | 0.11791317 |
| FDLGGGTF | ASSLYWGGLEDTQY | 0.14757049 |
| FDLGGGTFDVS | ASSLYWGGLEDTQY | 0.10426975 |
| NEPTAAA | ASSLYWGGLEDTQY | 0.12665236 |
| NEPTAAALA | ASSLYWGGLEDTQY | 0.10255594 |
| GSIINISSI | ASSLYWGGLEDTQY | 0.14625171 |
| VAPGNAGTA | ASSLYWGGLEDTQY | 0.11467728 |
| TGGGASLE | ASSLYWGGLEDTQY | 0.10124256 |
| VAVYNHYKRI | ASSLYWGGLEDTQY | 0.11610072 |
| TDPAPATT | ASSLYWGGLEDTQY | 0.10629607 |
| TNRPDILDPALLRPG | ASSLYWGGLEDTQY | 0.12025387 |
| LTTTTTS | ASSLYWGGLEDTQY | 0.12568794 |
| PTGSGKTL | ASSLYWGGLEDTQY | 0.10228659 |
| TGSGKTL | ASSLYWGGLEDTQY | 0.11128224 |
| TIGHVDHGK | ASSLYWGGLEDTQY | 0.13729747 |
| TIGHVDHGKTT | ASSLYWGGLEDTQY | 0.11845781 |
| GHVDHGKTT | ASSLYWGGLEDTQY | 0.10206135 |
| AQFEHTLL | ASSLYWGGLEDTQY | 0.11451365 |
| IVNQDEL | ASSLYWGGLEDTQY | 0.11211609 |
| TNGSQFF | ASSLYWGGLEDTQY | 0.10992943 |
| TGKGIVE | ASSLYWGGLEDTQY | 0.119664446 |
| LISNASDA | ASSLYWGGLEDTQY | 0.11281636 |
| AELRTLGELTKIAKAHGAQ | ASSLYWGGLEDTQY | 0.12366006 |
| APNAVAAAEPSW | ASSLYWGGLEDTQY | 0.11959854 |

| | | |
|---|---|---|
| APQVAGAATAVL | ASSLYWGGLEDTQY | 0.15644407 |
| ARRDPAAAAAILPTDA | ASSLYWGGLEDTQY | 0.1028928 |
| DAAARDVLAEAGLAEYF | ASSLYWGGLEDTQY | 0.10434884 |
| DDQIVLSAKANLAKVEQ | ASSLYWGGLEDTQY | 0.11574189 |
| DELLPDTGEDSDEDGH | ASSLYWGGLEDTQY | 0.122369 |
| ELPEAPANRATVPQD | ASSLYWGGLEDTQY | 0.101718135 |
| FGQSKIERVTGIG | ASSLYWGGLEDTQY | 0.10124564 |
| GAGGTGSTAGGAGGAGGAGGLY | ASSLYWGGLEDTQY | 0.115444615 |
| GHGGTGAAGGNGGTGGAGGSGI | ASSLYWGGLEDTQY | 0.13361515 |
| GNGGNGGAGGEGGPGGAGGAGA | ASSLYWGGLEDTQY | 0.108819656 |
| GSTAGNGGNGGNATNGTIAGQP | ASSLYWGGLEDTQY | 0.11226271 |
| GTGGDGGNGGNGGTGG | ASSLYWGGLEDTQY | 0.19644272 |
| GVRRPAGIPADAPV | ASSLYWGGLEDTQY | 0.105818145 |
| GYRQVTDAHLLTLARR | ASSLYWGGLEDTQY | 0.11232407 |
| IAKMGQDGHDRG | ASSLYWGGLEDTQY | 0.122072875 |
| KAEPSIEIQYT | ASSLYWGGLEDTQY | 0.13640307 |
| KPVRIRMAADEDTVAVT | ASSLYWGGLEDTQY | 0.109521955 |
| MPPRIAGMRLLVIKPEP | ASSLYWGGLEDTQY | 0.12725836 |
| MKFYAVALFRSRGVPPTR | ASSLYWGGLEDTQY | 0.10095986 |
| NAGGFGILGAVAHSP | ASSLYWGGLEDTQY | 0.12806885 |
| NEGREITLAMAEM | ASSLYWGGLEDTQY | 0.09935363 |
| NELVEWSHPLGHRWRWE | ASSLYWGGLEDTQY | 0.11760974 |
| NLGAGGTGGNGGNPG | ASSLYWGGLEDTQY | 0.10001826 |
| NTEWPKWETVMAADPVD | ASSLYWGGLEDTQY | 0.12012683 |
| PASAAPAETQP | ASSLYWGGLEDTQY | 0.12690035 |
| QQSSDGKNGVRAAEITGE | ASSLYWGGLEDTQY | 0.12903467 |
| RAPDAVTTSWWKE | ASSLYWGGLEDTQY | 0.098556936 |
| SGFGNVGAHTSGWFNQ | ASSLYWGGLEDTQY | 0.119857594 |
| SGVAGAVGQAASVGGLKV | ASSLYWGGLEDTQY | 0.14001335 |
| SNGGAGTGSAGGNGGT | ASSLYWGGLEDTQY | 0.3929178 |
| TGGAGGEGGAAG | ASSLYWGGLEDTQY | 0.111179926 |
| TGGDGGAAGAGGAPGHG | ASSLYWGGLEDTQY | 0.12850618 |
| TMPPLLAAIANAGRHGV | ASSLYWGGLEDTQY | 0.109911494 |
| TSSTAGNSGGA | ASSLYWGGLEDTQY | 0.10547168 |
| VPATVPVGILC | ASSLYWGGLEDTQY | 0.10206893 |
| VQRPDSGGVVLTGRIS | ASSLYWGGLEDTQY | 0.10461268 |
| WTPNPVPATKGVPAGLVQAT | ASSLYWGGLEDTQY | 0.110964544 |
| YALKRALAKFSDRFDVVII | ASSLYWGGLEDTQY | 0.10573645 |
| YLIGSVIV | ASSLYWGGLEDTQY | 0.13808663 |
| YGQLVAVID | ASSLYWGGLEDTQY | 0.13844019 |
| VTVGGMTLPA | ASSLYWGGLEDTQY | 0.114956886 |
| VLALPAV | ASSLYWGGLEDTQY | 0.15082112 |
| VIPPTLNYET | ASSLYWGGLEDTQY | 0.11500217 |
| VGMGGMHPSSG | ASSLYWGGLEDTQY | 0.12013552 |
| VGLAGGAATA | ASSLYWGGLEDTQY | 0.104512855 |
| VGGAGGAGGSGP | ASSLYWGGLEDTQY | 0.10891519 |
| VFAAFMV | ASSLYWGGLEDTQY | 0.10182853 |
| TLYDKLR | ASSLYWGGLEDTQY | 0.10705479 |
| TLAVFTVL | ASSLYWGGLEDTQY | 0.10455324 |
| TGSVQPARV | ASSLYWGGLEDTQY | 0.1226356 |
| TGGGAGSGGT | ASSLYWGGLEDTQY | 0.09790525 |

| | | |
|---|---|---|
| SHLETSL | ASSLYWGGLEDTQY | 0.11850614 |
| SFTTGMT | ASSLYWGGLEDTQY | 0.14524572 |
| QVEVAKRMGVRQPTVS | ASSLYWGGLEDTQY | 0.100576945 |
| QGSGGPALAM | ASSLYWGGLEDTQY | 0.12135024 |
| MLAPSNP | ASSLYWGGLEDTQY | 0.10495752 |
| LVNNAAI | ASSLYWGGLEDTQY | 0.10130873 |
| LTGEINTLA | ASSLYWGGLEDTQY | 0.13758135 |
| LSIFGPV | ASSLYWGGLEDTQY | 0.10327723 |
| LLALLANL | ASSLYWGGLEDTQY | 0.11960142 |
| LISNASDALDKLR | ASSLYWGGLEDTQY | 0.1250704 |
| LINEAVP | ASSLYWGGLEDTQY | 0.11405991 |
| LGYAFSGL | ASSLYWGGLEDTQY | 0.10839755 |
| LDFVAVAL | ASSLYWGGLEDTQY | 0.11307388 |
| LAQPGAE | ASSLYWGGLEDTQY | 0.10293212 |
| ISNLGAVTGF | ASSLYWGGLEDTQY | 0.111370504 |
| HQEDPGRI | ASSLYWGGLEDTQY | 0.09835407 |
| GTPGEPTP | ASSLYWGGLEDTQY | 0.111862674 |
| GFFNSGS | ASSLYWGGLEDTQY | 0.10307319 |
| GAGGGGLTAGAN | ASSLYWGGLEDTQY | 0.13404456 |
| FVDALVNM | ASSLYWGGLEDTQY | 0.11065378 |
| FLTVAPEL | ASSLYWGGLEDTQY | 0.11764179 |
| FETLTAL | ASSLYWGGLEDTQY | 0.1053901 |
| DVINPIEL | ASSLYWGGLEDTQY | 0.10415085 |
| APVSGAPV | ASSLYWGGLEDTQY | 0.10220706 |
| ALVGGLAW | ASSLYWGGLEDTQY | 0.10280778 |
| ALAAAAATAI | ASSLYWGGLEDTQY | 0.10568786 |
| AFGLGLLL | ASSLYWGGLEDTQY | 0.10905989 |
| AFFAAASL | ASSLYWGGLEDTQY | 0.10055912 |
| AAAALPAFSPP | ASSLYWGGLEDTQY | 0.13053207 |
| VVLAPAL | ASSLYWGGLEDTQY | 0.112156115 |
| TIGHVDHGKTTL | ASSLYWGGLEDTQY | 0.10103039 |
| LISNASDALDK | ASSLYWGGLEDTQY | 0.107218884 |
| LISNASDAL | ASSLYWGGLEDTQY | 0.113078415 |
| ISNASDALDK | ASSLYWGGLEDTQY | 0.11411894 |
| NQAETLVYQTE | ASSLYWGGLEDTQY | 0.09782682 |
| TGGTGGTAGAG | ASSLYWGGLEDTQY | 0.10318457 |
| TGSTAGGAGGA | ASSLYWGGLEDTQY | 0.110104896 |
| DPLAVPVAAGPV | ASSLYWGGLEDTQY | 0.10043477 |
| GGTGGTPGNGGN | ASSLYWGGLEDTQY | 0.10358167 |
| PAGLAGVDPATL | ASSLYWGGLEDTQY | 0.15344575 |
| GNNLVVEAPHAGASV | ASSLYWGGLEDTQY | 0.10301081 |
| KVTRIMSASTLEQPA | ASSLYWGGLEDTQY | 0.10853156 |
| LGFALPIALPRRTVV | ASSLYWGGLEDTQY | 0.12643501 |
| TRLGNLTARKLVVLV | ASSLYWGGLEDTQY | 0.12726696 |
| WSQQRRLIYFPSAGP | ASSLYWGGLEDTQY | 0.17030896 |
| GPNGNPDPMAAAVDIR | ASSLYWGGLEDTQY | 0.10977463 |
| TQRCVSVADLRAAASR | ASSLYWGGLEDTQY | 0.10669878 |
| QFVLDVAEHPVEVLTETV | ASSLYWGGLEDTQY | 0.116037756 |
| GGAGGTGGAGGNGGTQAGDG | ASSLYWGGLEDTQY | 0.11693635 |
| PLLATLRAIAVDVDAKPAQV | ASSLYWGGLEDTQY | 0.09963918 |
| APKFRTGGYMIDFWGVGYQVAK | ASSLYWGGLEDTQY | 0.099168465 |

| | | |
|---|---|---|
| GAGGAGGAGGAAGGPGGTGGTG | ASSLYWGGLEDTQY | 0.107800476 |
| IGALGLLATDVAPAGSVAFGVA | ASSLYWGGLEDTQY | 0.10018976 |
| AGGAGGIGGNGGAGGAGGVLMG | ASSLYWGGLEDTQY | 0.12328102 |
| ADELIGLL | ASSLYWGGLEDTQY | 0.12590012 |
| ADRALKA | ASSLYWGGLEDTQY | 0.102030925 |
| AGAVGQVGGAGGA | ASSLYWGGLEDTQY | 0.10199256 |
| AGGGGGLGGAP | ASSLYWGGLEDTQY | 0.108473994 |
| AGGLLYGNGGAG | ASSLYWGGLEDTQY | 0.10695093 |
| AGLESIGL | ASSLYWGGLEDTQY | 0.12979129 |
| AGLKTGII | ASSLYWGGLEDTQY | 0.10596897 |
| ASLAALLL | ASSLYWGGLEDTQY | 0.4013198 |
| ASVINISSI | ASSLYWGGLEDTQY | 0.12738395 |
| ATPPGPA | ASSLYWGGLEDTQY | 0.105041705 |
| ATVASALV | ASSLYWGGLEDTQY | 0.12960076 |
| ATVASLIL | ASSLYWGGLEDTQY | 0.10659404 |
| AVNGPVLIRH | ASSLYWGGLEDTQY | 0.101461105 |
| DGAAGAVGI | ASSLYWGGLEDTQY | 0.1090064 |
| DPLLHGI | ASSLYWGGLEDTQY | 0.10223641 |
| ELGAALGAL | ASSLYWGGLEDTQY | 0.10712754 |
| ELGRTQAV | ASSLYWGGLEDTQY | 0.1075457 |
| FVAGGYLS | ASSLYWGGLEDTQY | 0.1034387 |
| GASGGAGGAGGW | ASSLYWGGLEDTQY | 0.110397376 |
| GGAGAGGDGGAGGI | ASSLYWGGLEDTQY | 0.10742102 |
| GTGHNGSQPGMGGNG | ASSLYWGGLEDTQY | 0.100521915 |
| GYAAASASAS | ASSLYWGGLEDTQY | 0.10630862 |
| HQVDVAAVQQG | ASSLYWGGLEDTQY | 0.14059122 |
| IGAAIAHT | ASSLYWGGLEDTQY | 0.21207735 |
| IGMIEVAQL | ASSLYWGGLEDTQY | 0.10763061 |
| IRLAPEHTSAMD | ASSLYWGGLEDTQY | 0.10471649 |
| IYNVSVL | ASSLYWGGLEDTQY | 0.124311335 |
| KQGFLGM | ASSLYWGGLEDTQY | 0.11505564 |
| LESVITQL | ASSLYWGGLEDTQY | 0.11029037 |
| LGLGALVVV | ASSLYWGGLEDTQY | 0.10972845 |
| LIGNGGNGGSGGI | ASSLYWGGLEDTQY | 0.10242446 |
| LNAINAPTL | ASSLYWGGLEDTQY | 0.10486731 |
| LQLLNTGV | ASSLYWGGLEDTQY | 0.10811404 |
| LQTSIEF | ASSLYWGGLEDTQY | 0.101373576 |
| MSFAVSEL | ASSLYWGGLEDTQY | 0.121219754 |
| NTGALGL | ASSLYWGGLEDTQY | 0.097932346 |
| PPGAGGAGGAGTT | ASSLYWGGLEDTQY | 0.103974395 |
| QLVGLGI | ASSLYWGGLEDTQY | 0.10534881 |
| RFLEQVHQL | ASSLYWGGLEDTQY | 0.10555816 |
| RGVGYVLR | ASSLYWGGLEDTQY | 0.10128584 |
| SATGVFAGVI | ASSLYWGGLEDTQY | 0.1099162 |
| SLPAITLP | ASSLYWGGLEDTQY | 0.11609409 |
| STALAFTV | ASSLYWGGLEDTQY | 0.13806465 |
| SVLTVGLGK | ASSLYWGGLEDTQY | 0.10029135 |
| TALAGAGSGI | ASSLYWGGLEDTQY | 0.12121335 |
| TGLPFTGI | ASSLYWGGLEDTQY | 0.13200203 |
| VAEHVTV | ASSLYWGGLEDTQY | 0.108481064 |
| VDIVLNSL | ASSLYWGGLEDTQY | 0.10193391 |

| | | |
|---|---|---|
| VLTGGGAVL | ASSLYWGGLEDTQY | 0.118320145 |
| VNVGLAYGI | ASSLYWGGLEDTQY | 0.100405954 |
| VSSAAVGAV | ASSLYWGGLEDTQY | 0.10518946 |
| VTSVYEAGV | ASSLYWGGLEDTQY | 0.11834442 |
| WVALDRGA | ASSLYWGGLEDTQY | 0.10104875 |
| YLVGSVVI | ASSLYWGGLEDTQY | 0.12371845 |
| GPPGTGKTLL | ASSLYWGGLEDTQY | 0.12204929 |
| GVGMTKF | ASSLYWGGLEDTQY | 0.10436382 |
| IGIGGSDL | ASSLYWGGLEDTQY | 0.10380952 |
| IGTLLAEL | ASSLYWGGLEDTQY | 0.102429755 |
| IYTSGTT | ASSLYWGGLEDTQY | 0.09870133 |
| LSFPAGV | ASSLYWGGLEDTQY | 0.118420646 |
| LYTSGTT | ASSLYWGGLEDTQY | 0.104462855 |
| VGAFPFGF | ASSLYWGGLEDTQY | 0.10889238 |
| VGAGTVEF | ASSLYWGGLEDTQY | 0.10986154 |
| VGKSTLLL | ASSLYWGGLEDTQY | 0.1075323 |
| AALAHAGGGIV | ASSLYWGGLEDTQY | 0.11368983 |
| AALVAVSLL | ASSLYWGGLEDTQY | 0.11183174 |
| AAVSRAVAAL | ASSLYWGGLEDTQY | 0.10413019 |
| AGGSSGIGGF | ASSLYWGGLEDTQY | 0.11025743 |
| AGLTVANANG | ASSLYWGGLEDTQY | 0.10148015 |
| AGLVGAVL | ASSLYWGGLEDTQY | 0.10543509 |
| ALSQAFTV | ASSLYWGGLEDTQY | 0.17552796 |
| ALTFSVSL | ASSLYWGGLEDTQY | 0.10069196 |
| ALTGAALVAL | ASSLYWGGLEDTQY | 0.10499059 |
| ASFSIGAGVL | ASSLYWGGLEDTQY | 0.10879408 |
| DGSVYKI | ASSLYWGGLEDTQY | 0.10069891 |
| EQLAEIV | ASSLYWGGLEDTQY | 0.118272066 |
| FAVSELTVM | ASSLYWGGLEDTQY | 0.11173778 |
| GAAPLAAGL | ASSLYWGGLEDTQY | 0.1220742 |
| GADGGDGGSGGAGG | ASSLYWGGLEDTQY | 0.1108605 |
| GFGLDTGGAGG | ASSLYWGGLEDTQY | 0.12717867 |
| GLASGGLL | ASSLYWGGLEDTQY | 0.12707204 |
| GRIAGAAVLL | ASSLYWGGLEDTQY | 0.10571338 |
| GSAGSLNTSF | ASSLYWGGLEDTQY | 0.104533985 |
| GTGQIGI | ASSLYWGGLEDTQY | 0.1032273 |
| GTLIALIL | ASSLYWGGLEDTQY | 0.1127356 |
| IAATMGFVVG | ASSLYWGGLEDTQY | 0.11918098 |
| IFAQALL | ASSLYWGGLEDTQY | 0.102731295 |
| IGAAVALGF | ASSLYWGGLEDTQY | 0.1119023 |
| IGLTPLAL | ASSLYWGGLEDTQY | 0.12836875 |
| ILGSNTSTL | ASSLYWGGLEDTQY | 0.13183981 |
| IVFSLTVL | ASSLYWGGLEDTQY | 0.100493036 |
| LASKATGF | ASSLYWGGLEDTQY | 0.10717847 |
| LASVALIL | ASSLYWGGLEDTQY | 0.12016283 |
| LFAVGLTL | ASSLYWGGLEDTQY | 0.11171894 |
| LFNGGIGTYI | ASSLYWGGLEDTQY | 0.10441153 |
| LTLGAGII | ASSLYWGGLEDTQY | 0.12226191 |
| LVAAALGDI | ASSLYWGGLEDTQY | 0.114362374 |
| NTAFVQLGI | ASSLYWGGLEDTQY | 0.12521504 |
| QLFSGAEL | ASSLYWGGLEDTQY | 0.123778395 |

| | | |
|---|---|---|
| SAATFITL | ASSLYWGGLEDTQY | 0.14720641 |
| SNYGLANLGV | ASSLYWGGLEDTQY | 0.108507775 |
| SYGGAYAVM | ASSLYWGGLEDTQY | 0.11129085 |
| TTGVGGLTLG | ASSLYWGGLEDTQY | 0.10155387 |
| TTQTIYI | ASSLYWGGLEDTQY | 0.10295102 |
| VAAGAGLGAVI | ASSLYWGGLEDTQY | 0.1033946 |
| VAAVVTLL | ASSLYWGGLEDTQY | 0.10661437 |
| VGFLGGLI | ASSLYWGGLEDTQY | 0.098721966 |
| VGVIPILL | ASSLYWGGLEDTQY | 0.10849731 |
| VIFASSLI | ASSLYWGGLEDTQY | 0.11563849 |
| VKGGDGGAAAG | ASSLYWGGLEDTQY | 0.1382255 |
| VTKTFTNI | ASSLYWGGLEDTQY | 0.10880172 |
| VVAAAILL | ASSLYWGGLEDTQY | 0.10779764 |
| YAALDVEL | ASSLYWGGLEDTQY | 0.1045261 |
| LASVALLL | ASSLYWGGLEDTQY | 0.14286965 |
| LDAVHRV | ASSLYWGGLEDTQY | 0.101039544 |
| QGLSSSIF | ASSLYWGGLEDTQY | 0.10071982 |
| RGTFANIRL | ASSLYWGGLEDTQY | 0.10801 |
| RQLWWGHRI | ASSLYWGGLEDTQY | 0.10134473 |
| YGPPGTGKTLL | ASSLYWGGLEDTQY | 0.11339826 |
| AALVGAPANVA | ASSLYWGGLEDTQY | 0.12285787 |
| AATFRRTE | ASSLYWGGLEDTQY | 0.11627066 |
| AGALGGGI | ASSLYWGGLEDTQY | 0.12085379 |
| AGGAGGAGGIGAG | ASSLYWGGLEDTQY | 0.09936671 |
| AGSGALGL | ASSLYWGGLEDTQY | 0.108081095 |
| AQIADQI | ASSLYWGGLEDTQY | 0.21109414 |
| ASAVAVTYL | ASSLYWGGLEDTQY | 0.110736236 |
| ASTVLGGTGG | ASSLYWGGLEDTQY | 0.11596419 |
| AVGGEPAK | ASSLYWGGLEDTQY | 0.13481008 |
| CEQGTFGGTC | ASSLYWGGLEDTQY | 0.10499964 |
| DEGKAPAG | ASSLYWGGLEDTQY | 0.11550791 |
| DTVVFPAL | ASSLYWGGLEDTQY | 0.10879853 |
| FEGITMAG | ASSLYWGGLEDTQY | 0.1141861 |
| FPLVVQD | ASSLYWGGLEDTQY | 0.104425706 |
| GAFTTPQ | ASSLYWGGLEDTQY | 0.103200585 |
| GFGGFGTAGGVGGA | ASSLYWGGLEDTQY | 0.12821692 |
| GFLAQFL | ASSLYWGGLEDTQY | 0.16098075 |
| GGAGGAGGNAGVA | ASSLYWGGLEDTQY | 0.116044395 |
| GGVGTAGAGGAGGAGG | ASSLYWGGLEDTQY | 0.11515255 |
| GNGGGGDAGNAGSGGN | ASSLYWGGLEDTQY | 0.11490091 |
| GTGGAGGAGNDAGS | ASSLYWGGLEDTQY | 0.11476628 |
| GTTNTGWFNS | ASSLYWGGLEDTQY | 0.10782479 |
| IFMGPLFTL | ASSLYWGGLEDTQY | 0.10424472 |
| ILAHAIEAI | ASSLYWGGLEDTQY | 0.104108095 |
| IRSAIREAL | ASSLYWGGLEDTQY | 0.10288793 |
| ITFAGGG | ASSLYWGGLEDTQY | 0.11402287 |
| ITGMSGHQEV | ASSLYWGGLEDTQY | 0.10910161 |
| IVAGLACTL | ASSLYWGGLEDTQY | 0.19847393 |
| IVGGGLAGA | ASSLYWGGLEDTQY | 0.117222436 |
| IVTGVLNI | ASSLYWGGLEDTQY | 0.111608736 |
| LAQQGCI | ASSLYWGGLEDTQY | 0.103303 |

| | | |
|---|---|---|
| LCLHVEN | ASSLYWGGLEDTQY | 0.107995324 |
| LDAISDGS | ASSLYWGGLEDTQY | 0.10738175 |
| LGGFAAML | ASSLYWGGLEDTQY | 0.10322644 |
| LGGKNPMI | ASSLYWGGLEDTQY | 0.16665508 |
| LGPKGVAAIL | ASSLYWGGLEDTQY | 0.10726452 |
| LMVGVSI | ASSLYWGGLEDTQY | 0.10140886 |
| LVGAAFVL | ASSLYWGGLEDTQY | 0.11640631 |
| LVLAVAPL | ASSLYWGGLEDTQY | 0.22804943 |
| MAAIAVIDAA | ASSLYWGGLEDTQY | 0.11029258 |
| NNGINAELQ | ASSLYWGGLEDTQY | 0.10598143 |
| RGEPLVLVHG | ASSLYWGGLEDTQY | 0.10393257 |
| RGGAGGAAGLI | ASSLYWGGLEDTQY | 0.120786905 |
| RSYRMHPAV | ASSLYWGGLEDTQY | 0.17131224 |
| SAAHLLAGQLA | ASSLYWGGLEDTQY | 0.1028929 |
| SAAVGSY | ASSLYWGGLEDTQY | 0.14458671 |
| SPHAVLHRI | ASSLYWGGLEDTQY | 0.10414637 |
| SQIAKVL | ASSLYWGGLEDTQY | 0.10373212 |
| TALGHLVP | ASSLYWGGLEDTQY | 0.11698163 |
| TAPTTSGAA | ASSLYWGGLEDTQY | 0.10985128 |
| TDGDDGNGGSPGT | ASSLYWGGLEDTQY | 0.117966585 |
| TDGLGEN | ASSLYWGGLEDTQY | 0.120052256 |
| TGGPPLPGQA | ASSLYWGGLEDTQY | 0.10363663 |
| TIPAITI | ASSLYWGGLEDTQY | 0.104392156 |
| TIPGFTI | ASSLYWGGLEDTQY | 0.13956344 |
| TSIASGALA | ASSLYWGGLEDTQY | 0.11845814 |
| TVTLPGAAI | ASSLYWGGLEDTQY | 0.13147005 |
| VASGLSAALP | ASSLYWGGLEDTQY | 0.110678926 |
| VGAGVIGI | ASSLYWGGLEDTQY | 0.10694827 |
| VGPKRFGV | ASSLYWGGLEDTQY | 0.10968027 |
| VNVAFPD | ASSLYWGGLEDTQY | 0.1057418 |
| VSMTNSGVS | ASSLYWGGLEDTQY | 0.10480487 |
| VTDVLGDLSV | ASSLYWGGLEDTQY | 0.113809295 |
| VVVVGAGFAGL | ASSLYWGGLEDTQY | 0.12428611 |
| VYTVGTVV | ASSLYWGGLEDTQY | 0.10203314 |
| WTTFSGP | ASSLYWGGLEDTQY | 0.10656694 |
| YDELVEAF | ASSLYWGGLEDTQY | 0.11634402 |
| YGTGGAGGNGG | ASSLYWGGLEDTQY | 0.1091944 |
| YTNVMAV | ASSLYWGGLEDTQY | 0.1048556 |
| FLEQVHQL | ASSLYWGGLEDTQY | 0.15621401 |
| GLSLLAVAV | ASSLYWGGLEDTQY | 0.18250984 |
| LEQVHQL | ASSLYWGGLEDTQY | 0.12375862 |
| LNAAGAI | ASSLYWGGLEDTQY | 0.102441065 |
| PAVHASL | ASSLYWGGLEDTQY | 0.099405795 |
| TELEDDA | ASSLYWGGLEDTQY | 0.15307428 |
| TPPGPPN | ASSLYWGGLEDTQY | 0.11340017 |
| TPPPPPPP | ASSLYWGGLEDTQY | 0.10723461 |
| TQSPALI | SSRSDREDNQAPL | 0.103808634 |
| EFAAAFPQ | SSRSDREDNQAPL | 0.103681765 |
| LLSGNLDVL | SSRSDREDNQAPL | 0.1301073 |
| GPANLGNYNI | SSRSDREDNQAPL | 0.106437355 |
| TLGPTLP | SSRSDREDNQAPL | 0.101705 |

| | | |
|---|---|---|
| VYAYSSCCK | SSRSDREDNQAPL | 0.10335092 |
| APAPAPAPNI | SSRSDREDNQAPL | 0.13451062 |
| PGGPVPIP | SSRSDREDNQAPL | 0.10289901 |
| FLTVALEP | SSRSDREDNQAPL | 0.1086457 |
| SQYFGANL | SSRSDREDNQAPL | 0.16324517 |
| GGVKPAA | SSRSDREDNQAPL | 0.10120648 |
| GGTGGAGGAGGW | SSRSDREDNQAPL | 0.10916977 |
| GVGGVAGLGGAGG | SSRSDREDNQAPL | 0.10676577 |
| VLLVLATTI | SSRSDREDNQAPL | 0.10634145 |
| IGIFAAML | SSRSDREDNQAPL | 0.10550984 |
| AVGDLID | SSRSDREDNQAPL | 0.12122032 |
| NISPLPL | SSRSDREDNQAPL | 0.118002765 |
| VTAVFVAI | SSRSDREDNQAPL | 0.11907474 |
| SGDDNDAAL | SSRSDREDNQAPL | 0.11170503 |
| AVAALAHS | SSRSDREDNQAPL | 0.10578872 |
| GGVTVMATP | SSRSDREDNQAPL | 0.10235137 |
| AGLAANVRD | SSRSDREDNQAPL | 0.098697044 |
| TNRPDLIDPALLRPG | SSRSDREDNQAPL | 0.13349345 |
| LANLGVTAL | SSRSDREDNQAPL | 0.09948143 |
| TGERIEL | SSRSDREDNQAPL | 0.12622629 |
| NGEELAA | SSRSDREDNQAPL | 0.12634228 |
| VLSLVWG | SSRSDREDNQAPL | 0.103776306 |
| IGYALWL | SSRSDREDNQAPL | 0.111133434 |
| LSLLLAPL | SSRSDREDNQAPL | 0.099802 |
| TLDEALK | SSRSDREDNQAPL | 0.10277627 |
| TDGAGTAEGG | SSRSDREDNQAPL | 0.11365221 |
| NAPGGAGGNGG | SSRSDREDNQAPL | 0.114551485 |
| VSVFMGMGGNF | SSRSDREDNQAPL | 0.10844963 |
| LASYLAAL | SSRSDREDNQAPL | 0.11416264 |
| LVGGTPAL | SSRSDREDNQAPL | 0.11371975 |
| VRNADVL | SSRSDREDNQAPL | 0.107280724 |
| TCSSPFLVSAM | SSRSDREDNQAPL | 0.10980883 |
| VHITDAE | SSRSDREDNQAPL | 0.13647006 |
| EYAAPSF | SSRSDREDNQAPL | 0.116223104 |
| VVDAFHGGL | SSRSDREDNQAPL | 0.104364425 |
| ASVVDPQ | SSRSDREDNQAPL | 0.11175361 |
| SYNEFGGLS | SSRSDREDNQAPL | 0.10384668 |
| DPDGSVY | SSRSDREDNQAPL | 0.11747414 |
| MVSTIHATE | SSRSDREDNQAPL | 0.10645415 |
| LAAGVADPV | SSRSDREDNQAPL | 0.116076015 |
| TKEQIAA | SSRSDREDNQAPL | 0.10942387 |
| SLASPAAL | SSRSDREDNQAPL | 0.10010525 |
| IGLVFYGV | SSRSDREDNQAPL | 0.10387212 |
| FLLDASNL | SSRSDREDNQAPL | 0.11100795 |
| YQVGNFP | SSRSDREDNQAPL | 0.12434916 |
| ESLLSQST | SSRSDREDNQAPL | 0.12603949 |
| LFTVGFAAVG | SSRSDREDNQAPL | 0.104138695 |
| ATPPTPT | SSRSDREDNQAPL | 0.11817133 |
| TDLATLG | SSRSDREDNQAPL | 0.108958006 |
| SAGGTGR | SSRSDREDNQAPL | 0.10803282 |
| GFSNSGS | SSRSDREDNQAPL | 0.11225074 |

| | | |
|---|---|---|
| GGNGGSGGTSV | SSRSDREDNQAPL | 0.10615085 |
| GTSVATGGAGN | SSRSDREDNQAPL | 0.10747279 |
| NGGNGGSGGTS | SSRSDREDNQAPL | 0.114634715 |
| AVVKFPH | SSRSDREDNQAPL | 0.10342144 |
| TPPPPPP | SSRSDREDNQAPL | 0.118351184 |
| AAPAAAAAPA | SSRSDREDNQAPL | 0.109306514 |
| IALNEDL | SSRSDREDNQAPL | 0.108644806 |
| IVNVSSM | SSRSDREDNQAPL | 0.11789258 |
| TGIPPAPR | SSRSDREDNQAPL | 0.10353383 |
| FDLGGGTF | SSRSDREDNQAPL | 0.11833122 |
| FDLGGGTFDVS | SSRSDREDNQAPL | 0.11247972 |
| NEPTAAA | SSRSDREDNQAPL | 0.10487975 |
| NEPTAAALA | SSRSDREDNQAPL | 0.10892424 |
| GSIINISSI | SSRSDREDNQAPL | 0.13367574 |
| VAPGNAGTA | SSRSDREDNQAPL | 0.122753054 |
| TGGGASLE | SSRSDREDNQAPL | 0.102792844 |
| VAVYNHYKRI | SSRSDREDNQAPL | 0.113213666 |
| TDPAPATT | SSRSDREDNQAPL | 0.12995806 |
| TNRPDILDPALLRPG | SSRSDREDNQAPL | 0.106382705 |
| LTTTTTS | SSRSDREDNQAPL | 0.099774964 |
| PTGSGKTL | SSRSDREDNQAPL | 0.10945962 |
| TGSGKTL | SSRSDREDNQAPL | 0.10432797 |
| TIGHVDHGK | SSRSDREDNQAPL | 0.106792696 |
| TIGHVDHGKTT | SSRSDREDNQAPL | 0.10105171 |
| GHVDHGKTT | SSRSDREDNQAPL | 0.13713372 |
| AQFEHTLL | SSRSDREDNQAPL | 0.16989984 |
| IVNQDEL | SSRSDREDNQAPL | 0.1089129 |
| TNGSQFF | SSRSDREDNQAPL | 0.109435245 |
| TGKGIVE | SSRSDREDNQAPL | 0.10079718 |
| LISNASDA | SSRSDREDNQAPL | 0.1209473 |
| AELRTLGELTKIAKAHGAQ | SSRSDREDNQAPL | 0.14464183 |
| APNAVAAAEPSW | SSRSDREDNQAPL | 0.1040978 |
| APQVAGAATAVL | SSRSDREDNQAPL | 0.122707985 |
| ARRDPAAAAAILPTDA | SSRSDREDNQAPL | 0.10631164 |
| DAAARDVLAEAGLAEYF | SSRSDREDNQAPL | 0.1057135 |
| DDQIVLSAKANLAKVEQ | SSRSDREDNQAPL | 0.14703031 |
| DELLPDTGEDSDEDGH | SSRSDREDNQAPL | 0.11301206 |
| ELPEAPANRATVPQD | SSRSDREDNQAPL | 0.11051129 |
| FGQSKIERVTGIG | SSRSDREDNQAPL | 0.10996272 |
| GAGGTGSTAGGAGGAGGAGGLY | SSRSDREDNQAPL | 0.104508124 |
| GHGGTGAAGGNGGTGGAGGSGI | SSRSDREDNQAPL | 0.11294867 |
| GNGGNGGAGGEGGPGGAGGAGA | SSRSDREDNQAPL | 0.1256486 |
| GSTAGNGGNGGNATNGTIAGQP | SSRSDREDNQAPL | 0.10254063 |
| GTGGDGGNGGNGGTGG | SSRSDREDNQAPL | 0.11883026 |
| GVRRPAGIPADAPV | SSRSDREDNQAPL | 0.10539774 |
| GYRQVTDAHLLTLARR | SSRSDREDNQAPL | 0.120499596 |
| IAKMGQDGHDRG | SSRSDREDNQAPL | 0.15252829 |
| KAEPSIEIQYT | SSRSDREDNQAPL | 0.11687897 |
| KPVRIRMAADEDTVAVT | SSRSDREDNQAPL | 0.10614008 |
| MPPRIAGMRLLVIKPEP | SSRSDREDNQAPL | 0.16074347 |
| MKFYAVALFRSRGVPPTR | SSRSDREDNQAPL | 0.11640976 |

| | | |
|---|---|---|
| NAGGFGILGAVAHSP | SSRSDREDNQAPL | 0.12107028 |
| NEGREITLAMAEM | SSRSDREDNQAPL | 0.123914264 |
| NELVEWSHPLGHRWRWE | SSRSDREDNQAPL | 0.090761766 |
| NLGAGGTGGNGGNPG | SSRSDREDNQAPL | 0.091402665 |
| NTEWPKWETVMAADPVD | SSRSDREDNQAPL | 0.0907572 |
| PASAAPAETQP | SSRSDREDNQAPL | 0.090871155 |
| QQSSDGKNGVRAAEITGE | SSRSDREDNQAPL | 0.09184097 |
| RAPDAVTTSWWKE | SSRSDREDNQAPL | 0.09081052 |
| SGFGNVGAHTSGWFNQ | SSRSDREDNQAPL | 0.0909472 |
| SGVAGAVGQAASVGGLKV | SSRSDREDNQAPL | 0.093433335 |
| SNGGAGTGSAGGNGGT | SSRSDREDNQAPL | 0.09073325 |
| TGGAGGEGGAAG | SSRSDREDNQAPL | 0.09080532 |
| TGGDGGAAGAGGAPGHG | SSRSDREDNQAPL | 0.09172756 |
| TMPPLLAAIANAGRHGV | SSRSDREDNQAPL | 0.09120711 |
| TSSTAGNSGGA | SSRSDREDNQAPL | 0.09137555 |
| VPATVPVGILC | SSRSDREDNQAPL | 0.090960756 |
| VQRPDSGGVVLTGRIS | SSRSDREDNQAPL | 0.09070565 |
| WTPNPVPATKGVPAGLVQAT | SSRSDREDNQAPL | 0.09095775 |
| YALKRALAKFSDRFDVVII | SSRSDREDNQAPL | 0.09079989 |
| YLIGSVIV | SSRSDREDNQAPL | 0.0906352 |
| YGQLVAVID | SSRSDREDNQAPL | 0.09075542 |
| VTVGGMTLPA | SSRSDREDNQAPL | 0.09054453 |
| VLALPAV | SSRSDREDNQAPL | 0.090934984 |
| VIPPTLNYET | SSRSDREDNQAPL | 0.09094085 |
| VGMGGMHPSSG | SSRSDREDNQAPL | 0.090888835 |
| VGLAGGAATA | SSRSDREDNQAPL | 0.09101325 |
| VGGAGGAGGSGP | SSRSDREDNQAPL | 0.090991564 |
| VFAAFMV | SSRSDREDNQAPL | 0.09105098 |
| TLYDKLR | SSRSDREDNQAPL | 0.09152788 |
| TLAVFTVL | SSRSDREDNQAPL | 0.0911208 |
| TGSVQPARV | SSRSDREDNQAPL | 0.09259808 |
| TGGGAGSGGT | SSRSDREDNQAPL | 0.091260105 |
| SHLETSL | SSRSDREDNQAPL | 0.090644605 |
| SFTTGMT | SSRSDREDNQAPL | 0.09146575 |
| QVEVAKRMGVRQPTVS | SSRSDREDNQAPL | 0.091724604 |
| QGSGGPALAM | SSRSDREDNQAPL | 0.0907199 |
| MLAPSNP | SSRSDREDNQAPL | 0.09068783 |
| LVNNAAI | SSRSDREDNQAPL | 0.090547286 |
| LTGEINTLA | SSRSDREDNQAPL | 0.09129336 |
| LSIFGPV | SSRSDREDNQAPL | 0.09095227 |
| LLALLANL | SSRSDREDNQAPL | 0.09161587 |
| LISNASDALDKLR | SSRSDREDNQAPL | 0.09134692 |
| LINEAVP | SSRSDREDNQAPL | 0.090748124 |
| LGYAFSGL | SSRSDREDNQAPL | 0.09064395 |
| LDFVAVAL | SSRSDREDNQAPL | 0.091554835 |
| LAQPGAE | SSRSDREDNQAPL | 0.09088291 |
| ISNLGAVTGF | SSRSDREDNQAPL | 0.09095676 |
| HQEDPGRI | SSRSDREDNQAPL | 0.09154556 |
| GTPGEPTP | SSRSDREDNQAPL | 0.09070616 |
| GFFNSGS | SSRSDREDNQAPL | 0.09091909 |
| GAGGGGLTAGAN | SSRSDREDNQAPL | 0.090425014 |

| | | |
|---|---|---|
| FVDALVNM | SSRSDREDNQAPL | 0.09071967 |
| FLTVAPEL | SSRSDREDNQAPL | 0.09148011 |
| FETLTAL | SSRSDREDNQAPL | 0.09069111 |
| DVINPIEL | SSRSDREDNQAPL | 0.0934502 |
| APVSGAPV | SSRSDREDNQAPL | 0.0927054 |
| ALVGGLAW | SSRSDREDNQAPL | 0.09461892 |
| ALAAAAATAI | SSRSDREDNQAPL | 0.09101828 |
| AFGLGLLL | SSRSDREDNQAPL | 0.09056674 |
| AFFAAASL | SSRSDREDNQAPL | 0.09104862 |
| AAAALPAFSPP | SSRSDREDNQAPL | 0.09105388 |
| VVLAPAL | SSRSDREDNQAPL | 0.09082074 |
| TIGHVDHGKTTL | SSRSDREDNQAPL | 0.09085341 |
| LISNASDALDK | SSRSDREDNQAPL | 0.09119456 |
| LISNASDAL | SSRSDREDNQAPL | 0.09445266 |
| ISNASDALDK | SSRSDREDNQAPL | 0.09112358 |
| NQAETLVYQTE | SSRSDREDNQAPL | 0.09147938 |
| TGGTGGTAGAG | SSRSDREDNQAPL | 0.09157967 |
| TGSTAGGAGGA | SSRSDREDNQAPL | 0.09045562 |
| DPLAVPVAAGPV | SSRSDREDNQAPL | 0.09054953 |
| GGTGGTPGNGGN | SSRSDREDNQAPL | 0.09077496 |
| PAGLAGVDPATL | SSRSDREDNQAPL | 0.09092755 |
| GNNLVVEAPHAGASV | SSRSDREDNQAPL | 0.09186813 |
| KVTRIMSASTLEQPA | SSRSDREDNQAPL | 0.0912453 |
| LGFALPIALPRRTVV | SSRSDREDNQAPL | 0.0937751 |
| TRLGNLTARKLVVLV | SSRSDREDNQAPL | 0.091032386 |
| WSQQRRLIYFPSAGP | SSRSDREDNQAPL | 0.090599276 |
| GPNGNPDPMAAAVDIR | SSRSDREDNQAPL | 0.09094302 |
| TQRCVSVADLRAAASR | SSRSDREDNQAPL | 0.09121964 |
| QFVLDVAEHPVEVLTETV | SSRSDREDNQAPL | 0.09099238 |
| GGAGGTGGAGGNGGTQAGDG | SSRSDREDNQAPL | 0.09073246 |
| PLLATLRAIAVDVDAKPAQV | SSRSDREDNQAPL | 0.09132531 |
| APKFRTGGYMIDFWGVGYQVAK | SSRSDREDNQAPL | 0.09072118 |
| GAGGAGGAGGAAGGPGGTGGTG | SSRSDREDNQAPL | 0.09134057 |
| IGALGLLATDVAPAGSVAFGVA | SSRSDREDNQAPL | 0.09074771 |
| AGGAGGIGGNGGAGGAGGVLMG | SSRSDREDNQAPL | 0.090659805 |
| ADELIGLL | SSRSDREDNQAPL | 0.09135086 |
| ADRALKA | SSRSDREDNQAPL | 0.09071251 |
| AGAVGQVGGAGGA | SSRSDREDNQAPL | 0.09298465 |
| AGGGGGLGGAP | SSRSDREDNQAPL | 0.09091836 |
| AGGLLYGNGGAG | SSRSDREDNQAPL | 0.09064904 |
| AGLESIGL | SSRSDREDNQAPL | 0.09076331 |
| AGLKTGII | SSRSDREDNQAPL | 0.09059363 |
| ASLAALLL | SSRSDREDNQAPL | 0.09077669 |
| ASVINISSI | SSRSDREDNQAPL | 0.090717115 |
| ATPPGPA | SSRSDREDNQAPL | 0.09052175 |
| ATVASALV | SSRSDREDNQAPL | 0.09100742 |
| ATVASLIL | SSRSDREDNQAPL | 0.091119796 |
| AVNGPVLIRH | SSRSDREDNQAPL | 0.09157375 |
| DGAAGAVGI | SSRSDREDNQAPL | 0.09107168 |
| DPLLHGI | SSRSDREDNQAPL | 0.09073834 |
| ELGAALGAL | SSRSDREDNQAPL | 0.09121099 |

| | | |
|---|---|---|
| ELGRTQAV | SSRSDREDNQAPL | 0.09065186 |
| FVAGGYLS | SSRSDREDNQAPL | 0.090846375 |
| GASGGAGGAGGW | SSRSDREDNQAPL | 0.09334917 |
| GGAGAGGDGGAGGI | SSRSDREDNQAPL | 0.09071653 |
| GTGHNGSQPGMGGNG | SSRSDREDNQAPL | 0.0925233 |
| GYAAASASAS | SSRSDREDNQAPL | 0.090580165 |
| HQVDVAAVQQG | SSRSDREDNQAPL | 0.09067342 |
| IGAAIAHT | SSRSDREDNQAPL | 0.09071741 |
| IGMIEVAQL | SSRSDREDNQAPL | 0.09201636 |
| IRLAPEHTSAMD | SSRSDREDNQAPL | 0.09062711 |
| IYNVSVL | SSRSDREDNQAPL | 0.09080869 |
| KQGFLGM | SSRSDREDNQAPL | 0.09080407 |
| LESVITQL | SSRSDREDNQAPL | 0.09129726 |
| LGLGALVVV | SSRSDREDNQAPL | 0.0910425 |
| LIGNGGNGGSGGI | SSRSDREDNQAPL | 0.09076693 |
| LNAINAPTL | SSRSDREDNQAPL | 0.090696104 |
| LQLLNTGV | SSRSDREDNQAPL | 0.09165726 |
| LQTSIEF | SSRSDREDNQAPL | 0.09054096 |
| MSFAVSEL | SSRSDREDNQAPL | 0.09059075 |
| NTGALGL | SSRSDREDNQAPL | 0.09120483 |
| PPGAGGAGGAGTT | SSRSDREDNQAPL | 0.09196553 |
| QLVGLGI | SSRSDREDNQAPL | 0.092467666 |
| RFLEQVHQL | SSRSDREDNQAPL | 0.09223897 |
| RGVGYVLR | SSRSDREDNQAPL | 0.090825036 |
| SATGVFAGVI | SSRSDREDNQAPL | 0.0904483 |
| SLPAITLP | SSRSDREDNQAPL | 0.09061387 |
| STALAFTV | SSRSDREDNQAPL | 0.09120021 |
| SVLTVGLGK | SSRSDREDNQAPL | 0.092200406 |
| TALAGAGSGI | SSRSDREDNQAPL | 0.09115969 |
| TGLPFTGI | SSRSDREDNQAPL | 0.09094854 |
| VAEHVTV | SSRSDREDNQAPL | 0.09210211 |
| VDIVLNSL | SSRSDREDNQAPL | 0.090691574 |
| VLTGGGAVL | SSRSDREDNQAPL | 0.09097352 |
| VNVGLAYGI | SSRSDREDNQAPL | 0.090817526 |
| VSSAAVGAV | SSRSDREDNQAPL | 0.09071152 |
| VTSVYEAGV | SSRSDREDNQAPL | 0.09086977 |
| WVALDRGA | SSRSDREDNQAPL | 0.090979114 |
| YLVGSVVI | SSRSDREDNQAPL | 0.09152074 |
| GPPGTGKTLL | SSRSDREDNQAPL | 0.09065993 |
| GVGMTKF | SSRSDREDNQAPL | 0.090696454 |
| IGIGGSDL | SSRSDREDNQAPL | 0.09053276 |
| IGTLLAEL | SSRSDREDNQAPL | 0.09089653 |
| IYTSGTT | SSRSDREDNQAPL | 0.09272855 |
| LSFPAGV | SSRSDREDNQAPL | 0.09427995 |
| LYTSGTT | SSRSDREDNQAPL | 0.09115034 |
| VGAFPFGF | SSRSDREDNQAPL | 0.09140233 |
| VGAGTVEF | SSRSDREDNQAPL | 0.090878785 |
| VGKSTLLL | SSRSDREDNQAPL | 0.09095124 |
| AALAHAGGGIV | SSRSDREDNQAPL | 0.09075866 |
| AALVAVSLL | SSRSDREDNQAPL | 0.09074377 |
| AAVSRAVAAL | SSRSDREDNQAPL | 0.091295175 |

| | | |
|---|---|---|
| AGGSSGIGGF | SSRSDREDNQAPL | 0.09128654 |
| AGLTVANANG | SSRSDREDNQAPL | 0.09133464 |
| AGLVGAVL | SSRSDREDNQAPL | 0.09099343 |
| ALSQAFTV | SSRSDREDNQAPL | 0.09184224 |
| ALTFSVSL | SSRSDREDNQAPL | 0.0908924 |
| ALTGAALVAL | SSRSDREDNQAPL | 0.09098426 |
| ASFSIGAGVL | SSRSDREDNQAPL | 0.09114539 |
| DGSVYKI | SSRSDREDNQAPL | 0.09126682 |
| EQLAEIV | SSRSDREDNQAPL | 0.09058186 |
| FAVSELTVM | SSRSDREDNQAPL | 0.09084452 |
| GAAPLAAGL | SSRSDREDNQAPL | 0.09070433 |
| GADGGDGGSGGAGG | SSRSDREDNQAPL | 0.090955034 |
| GFGLDTGGAGG | SSRSDREDNQAPL | 0.09130774 |
| GLASGGLL | SSRSDREDNQAPL | 0.09081455 |
| GRIAGAAVLL | SSRSDREDNQAPL | 0.090922296 |
| GSAGSLNTSF | SSRSDREDNQAPL | 0.09079488 |
| GTGQIGI | SSRSDREDNQAPL | 0.091363795 |
| GTLIALIL | SSRSDREDNQAPL | 0.09077737 |
| IAATMGFVVG | SSRSDREDNQAPL | 0.09166048 |
| IFAQALL | SSRSDREDNQAPL | 0.090622224 |
| IGAAVALGF | SSRSDREDNQAPL | 0.09080711 |
| IGLTPLAL | SSRSDREDNQAPL | 0.090862855 |
| ILGSNTSTL | SSRSDREDNQAPL | 0.09085404 |
| IVFSLTVL | SSRSDREDNQAPL | 0.09125536 |
| LASKATGF | SSRSDREDNQAPL | 0.090806276 |
| LASVALIL | SSRSDREDNQAPL | 0.09117743 |
| LFAVGLTL | SSRSDREDNQAPL | 0.09147047 |
| LFNGGIGTYI | SSRSDREDNQAPL | 0.090883344 |
| LTLGAGII | SSRSDREDNQAPL | 0.09077534 |
| LVAAALGDI | SSRSDREDNQAPL | 0.09176514 |
| NTAFVQLGI | SSRSDREDNQAPL | 0.090903744 |
| QLFSGAEL | SSRSDREDNQAPL | 0.090775385 |
| SAATFITL | SSRSDREDNQAPL | 0.09099355 |
| SNYGLANLGV | SSRSDREDNQAPL | 0.09079345 |
| SYGGAYAVM | SSRSDREDNQAPL | 0.091054566 |
| TTGVGGLTLG | SSRSDREDNQAPL | 0.090548135 |
| TTQTIYI | SSRSDREDNQAPL | 0.09075897 |
| VAAGAGLGAVI | SSRSDREDNQAPL | 0.09072904 |
| VAAVVTLL | SSRSDREDNQAPL | 0.09076044 |
| VGFLGGLI | SSRSDREDNQAPL | 0.09140552 |
| VGVIPILL | SSRSDREDNQAPL | 0.091501124 |
| VIFASSLI | SSRSDREDNQAPL | 0.09067802 |
| VKGGDGGAAAG | SSRSDREDNQAPL | 0.091315225 |
| VTKTFTNI | SSRSDREDNQAPL | 0.090915 |
| VVAAAILL | SSRSDREDNQAPL | 0.09056552 |
| YAALDVEL | SSRSDREDNQAPL | 0.090699896 |
| LASVALLL | SSRSDREDNQAPL | 0.09122998 |
| LDAVHRV | SSRSDREDNQAPL | 0.09069012 |
| QGLSSSIF | SSRSDREDNQAPL | 0.09130793 |
| RGTFANIRL | SSRSDREDNQAPL | 0.090976655 |
| RQLWWGHRI | SSRSDREDNQAPL | 0.09072827 |

| | | |
|---|---|---|
| YGPPGTGKTLL | SSRSDREDNQAPL | 0.090906605 |
| AALVGAPANVA | SSRSDREDNQAPL | 0.09099027 |
| AATFRRTE | SSRSDREDNQAPL | 0.09207823 |
| AGALGGGI | SSRSDREDNQAPL | 0.09063373 |
| AGGAGGAGGIGAG | SSRSDREDNQAPL | 0.0905642 |
| AGSGALGL | SSRSDREDNQAPL | 0.09195649 |
| AQIADQI | SSRSDREDNQAPL | 0.091033615 |
| ASAVAVTYL | SSRSDREDNQAPL | 0.090798035 |
| ASTVLGGTGG | SSRSDREDNQAPL | 0.090861015 |
| AVGGEPAK | SSRSDREDNQAPL | 0.09252726 |
| CEQGTFGGTC | SSRSDREDNQAPL | 0.09108493 |
| DEGKAPAG | SSRSDREDNQAPL | 0.090665326 |
| DTVVFPAL | SSRSDREDNQAPL | 0.090786435 |
| FEGITMAG | SSRSDREDNQAPL | 0.09115818 |
| FPLVVQD | SSRSDREDNQAPL | 0.090590544 |
| GAFTTPQ | SSRSDREDNQAPL | 0.09131284 |
| GFGGFGTAGGVGGA | SSRSDREDNQAPL | 0.0916425 |
| GFLAQFL | SSRSDREDNQAPL | 0.090638325 |
| GGAGGAGGNAGVA | SSRSDREDNQAPL | 0.09080496 |
| GGVGTAGAGGAGGAGG | SSRSDREDNQAPL | 0.09070949 |
| GNGGGGDAGNAGSGGN | SSRSDREDNQAPL | 0.091198646 |
| GTGGAGGAGNDAGS | SSRSDREDNQAPL | 0.090681635 |
| GTTNTGWFNS | SSRSDREDNQAPL | 0.09054204 |
| IFMGPLFTL | SSRSDREDNQAPL | 0.09101482 |
| ILAHAIEAI | SSRSDREDNQAPL | 0.09156593 |
| IRSAIREAL | SSRSDREDNQAPL | 0.09081865 |
| ITFAGGG | SSRSDREDNQAPL | 0.091264986 |
| ITGMSGHQEV | SSRSDREDNQAPL | 0.09089122 |
| IVAGLACTL | SSRSDREDNQAPL | 0.092985146 |
| IVGGGLAGA | SSRSDREDNQAPL | 0.09091217 |
| IVTGVLNI | SSRSDREDNQAPL | 0.09158371 |
| LAQQGCI | SSRSDREDNQAPL | 0.09073047 |
| LCLHVEN | SSRSDREDNQAPL | 0.09101606 |
| LDAISDGS | SSRSDREDNQAPL | 0.091062814 |
| LGGFAAML | SSRSDREDNQAPL | 0.09075168 |
| LGGKNPMI | SSRSDREDNQAPL | 0.09108149 |
| LGPKGVAAIL | SSRSDREDNQAPL | 0.090786345 |
| LMVGVSI | SSRSDREDNQAPL | 0.09112375 |
| LVGAAFVL | SSRSDREDNQAPL | 0.09094897 |
| LVLAVAPL | SSRSDREDNQAPL | 0.090582594 |
| MAAIAVIDAA | SSRSDREDNQAPL | 0.09297422 |
| NNGINAELQ | SSRSDREDNQAPL | 0.0906155 |
| RGEPLVLVHG | SSRSDREDNQAPL | 0.091345415 |
| RGGAGGAAGLI | SSRSDREDNQAPL | 0.09069428 |
| RSYRMHPAV | SSRSDREDNQAPL | 0.09067149 |
| SAAHLLAGQLA | SSRSDREDNQAPL | 0.09158477 |
| SAAVGSY | SSRSDREDNQAPL | 0.090692505 |
| SPHAVLHRI | SSRSDREDNQAPL | 0.09271646 |
| SQIAKVL | SSRSDREDNQAPL | 0.09089402 |
| TALGHLVP | SSRSDREDNQAPL | 0.09072957 |
| TAPTTSGAA | SSRSDREDNQAPL | 0.09073855 |

| | | |
|---|---|---|
| TDGDDGNGGSPGT | SSRSDREDNQAPL | 0.090722755 |
| TDGLGEN | SSRSDREDNQAPL | 0.09076747 |
| TGGPPLPGQA | SSRSDREDNQAPL | 0.09098855 |
| TIPAITI | SSRSDREDNQAPL | 0.09063093 |
| TIPGFTI | SSRSDREDNQAPL | 0.09096734 |
| TSIASGALA | SSRSDREDNQAPL | 0.0907802 |
| TVTLPGAAI | SSRSDREDNQAPL | 0.09087207 |
| VASGLSAALP | SSRSDREDNQAPL | 0.09112537 |
| VGAGVIGI | SSRSDREDNQAPL | 0.09096295 |
| VGPKRFGV | SSRSDREDNQAPL | 0.091166064 |
| VNVAFPD | SSRSDREDNQAPL | 0.09257727 |
| VSMTNSGVS | SSRSDREDNQAPL | 0.09096018 |
| VTDVLGDLSV | SSRSDREDNQAPL | 0.09083447 |
| VVVVGAGFAGL | SSRSDREDNQAPL | 0.09084604 |
| VYTVGTVV | SSRSDREDNQAPL | 0.09105615 |
| WTTFSGP | SSRSDREDNQAPL | 0.09220294 |
| YDELVEAF | SSRSDREDNQAPL | 0.09080079 |
| YGTGGAGGNGG | SSRSDREDNQAPL | 0.09068043 |
| YTNVMAV | SSRSDREDNQAPL | 0.09076111 |
| FLEQVHQL | SSRSDREDNQAPL | 0.09119944 |
| GLSLLAVAV | SSRSDREDNQAPL | 0.09116294 |
| LEQVHQL | SSRSDREDNQAPL | 0.091201894 |
| LNAAGAI | SSRSDREDNQAPL | 0.090715356 |
| PAVHASL | SSRSDREDNQAPL | 0.09159195 |
| TELEDDA | SSRSDREDNQAPL | 0.091369115 |
| TPPGPPN | SSRSDREDNQAPL | 0.09103787 |
| TPPPPPPP | SSRSDREDNQAPL | 0.09057777 |
| TQSPALI | TCSAEEASAETLY | 0.09131341 |
| EFAAAFPQ | TCSAEEASAETLY | 0.09092096 |
| LLSGNLDVL | TCSAEEASAETLY | 0.091097675 |
| GPANLGNYNI | TCSAEEASAETLY | 0.090831555 |
| TLGPTLP | TCSAEEASAETLY | 0.09125479 |
| VYAYSSCCK | TCSAEEASAETLY | 0.091516204 |
| APAPAPAPNI | TCSAEEASAETLY | 0.09050581 |
| PGGPVPIP | TCSAEEASAETLY | 0.090617046 |
| FLTVALEP | TCSAEEASAETLY | 0.09105859 |
| SQYFGANL | TCSAEEASAETLY | 0.09603905 |
| GGVKPAA | TCSAEEASAETLY | 0.09083175 |
| GGTGGAGGAGGW | TCSAEEASAETLY | 0.09078783 |
| GVGGVAGLGGAGG | TCSAEEASAETLY | 0.09058948 |
| VLLVLATTI | TCSAEEASAETLY | 0.09159724 |
| IGIFAAML | TCSAEEASAETLY | 0.09070631 |
| AVGDLID | TCSAEEASAETLY | 0.0907246 |
| NISPLPL | TCSAEEASAETLY | 0.09119899 |
| VTAVFVAI | TCSAEEASAETLY | 0.09091751 |
| SGDDNDAAL | TCSAEEASAETLY | 0.09064433 |
| AVAALAHS | TCSAEEASAETLY | 0.090676166 |
| GGVTVMATP | TCSAEEASAETLY | 0.09125224 |
| AGLAANVRD | TCSAEEASAETLY | 0.090707265 |
| TNRPDLIDPALLRPG | TCSAEEASAETLY | 0.09065605 |
| LANLGVTAL | TCSAEEASAETLY | 0.09093418 |

| | | |
|---|---|---|
| TGERIEL | TCSAEEASAETLY | 0.090688854 |
| NGEELAA | TCSAEEASAETLY | 0.0918291 |
| VLSLVWG | TCSAEEASAETLY | 0.09299121 |
| IGYALWL | TCSAEEASAETLY | 0.0907014 |
| LSLLLAPL | TCSAEEASAETLY | 0.090993986 |
| TLDEALK | TCSAEEASAETLY | 0.09455535 |
| TDGAGTAEGG | TCSAEEASAETLY | 0.0923601 |
| NAPGGAGGNGG | TCSAEEASAETLY | 0.09112785 |
| VSVFMGMGGNF | TCSAEEASAETLY | 0.09069298 |
| LASYLAAL | TCSAEEASAETLY | 0.09072327 |
| LVGGTPAL | TCSAEEASAETLY | 0.09091893 |
| VRNADVL | TCSAEEASAETLY | 0.09137827 |
| TCSSPFLVSAM | TCSAEEASAETLY | 0.09106966 |
| VHITDAE | TCSAEEASAETLY | 0.091094464 |
| EYAAPSF | TCSAEEASAETLY | 0.09088671 |
| VVDAFHGGL | TCSAEEASAETLY | 0.09096654 |
| ASVVDPQ | TCSAEEASAETLY | 0.09119124 |
| SYNEFGGLS | TCSAEEASAETLY | 0.09056002 |
| DPDGSVY | TCSAEEASAETLY | 0.09066577 |
| MVSTIHATE | TCSAEEASAETLY | 0.09091403 |
| LAAGVADPV | TCSAEEASAETLY | 0.0909088 |
| TKEQIAA | TCSAEEASAETLY | 0.09105361 |
| SLASPAAL | TCSAEEASAETLY | 0.09166195 |
| IGLVFYGV | TCSAEEASAETLY | 0.09158992 |
| FLLDASNL | TCSAEEASAETLY | 0.09065933 |
| YQVGNFP | TCSAEEASAETLY | 0.09077529 |
| ESLLSQST | TCSAEEASAETLY | 0.09318763 |
| LFTVGFAAVG | TCSAEEASAETLY | 0.09123779 |
| ATPPTPT | TCSAEEASAETLY | 0.091938905 |
| TDLATLG | TCSAEEASAETLY | 0.09069744 |
| SAGGTGR | TCSAEEASAETLY | 0.09083331 |
| GFSNSGS | TCSAEEASAETLY | 0.09111282 |
| GGNGGSGGTSV | TCSAEEASAETLY | 0.091140494 |
| GTSVATGGAGN | TCSAEEASAETLY | 0.09108511 |
| NGGNGGSGGTS | TCSAEEASAETLY | 0.090975076 |
| AVVKFPH | TCSAEEASAETLY | 0.09118341 |
| TPPPPPP | TCSAEEASAETLY | 0.09162345 |
| AAPAAAAAPA | TCSAEEASAETLY | 0.09073249 |
| IALNEDL | TCSAEEASAETLY | 0.09060552 |
| IVNVSSM | TCSAEEASAETLY | 0.09067645 |
| TGIPPAPR | TCSAEEASAETLY | 0.090705685 |
| FDLGGGTF | TCSAEEASAETLY | 0.0916132 |
| FDLGGGTFDVS | TCSAEEASAETLY | 0.09094416 |
| NEPTAAA | TCSAEEASAETLY | 0.090647474 |
| NEPTAAALA | TCSAEEASAETLY | 0.090722054 |
| GSIINISSI | TCSAEEASAETLY | 0.09264221 |
| VAPGNAGTA | TCSAEEASAETLY | 0.091418885 |
| TGGGASLE | TCSAEEASAETLY | 0.090846665 |
| VAVYNHYKRI | TCSAEEASAETLY | 0.09075086 |
| TDPAPATT | TCSAEEASAETLY | 0.09061854 |
| TNRPDILDPALLRPG | TCSAEEASAETLY | 0.09062491 |

| | | |
|---|---|---|
| LTTTTTS | TCSAEEASAETLY | 0.09064515 |
| PTGSGKTL | TCSAEEASAETLY | 0.09070192 |
| TGSGKTL | TCSAEEASAETLY | 0.09062952 |
| TIGHVDHGK | TCSAEEASAETLY | 0.09086665 |
| TIGHVDHGKTT | TCSAEEASAETLY | 0.090493284 |
| GHVDHGKTT | TCSAEEASAETLY | 0.09108464 |
| AQFEHTLL | TCSAEEASAETLY | 0.09125811 |
| IVNQDEL | TCSAEEASAETLY | 0.090625174 |
| TNGSQFF | TCSAEEASAETLY | 0.091784276 |
| TGKGIVE | TCSAEEASAETLY | 0.0923499 |
| LISNASDA | TCSAEEASAETLY | 0.090910465 |
| AELRTLGELTKIAKAHGAQ | TCSAEEASAETLY | 0.09135986 |
| APNAVAAAEPSW | TCSAEEASAETLY | 0.090613775 |
| APQVAGAATAVL | TCSAEEASAETLY | 0.09552644 |
| ARRDPAAAAAILPTDA | TCSAEEASAETLY | 0.090581656 |
| DAAARDVLAEAGLAEYF | TCSAEEASAETLY | 0.09155828 |
| DDQIVLSAKANLAKVEQ | TCSAEEASAETLY | 0.09133564 |
| DELLPDTGEDSDEDGH | TCSAEEASAETLY | 0.09060197 |
| ELPEAPANRATVPQD | TCSAEEASAETLY | 0.090534255 |
| FGQSKIERVTGIG | TCSAEEASAETLY | 0.0913444 |
| GAGGTGSTAGGAGGAGGAGGLY | TCSAEEASAETLY | 0.090662375 |
| GHGGTGAAGGNGGTGGAGGSGI | TCSAEEASAETLY | 0.09150734 |
| GNGGNGGAGGEGGPGGAGGAGA | TCSAEEASAETLY | 0.09102676 |
| GSTAGNGGNGGNATNGTIAGQP | TCSAEEASAETLY | 0.09092853 |
| GTGGDGGNGGNGGTGG | TCSAEEASAETLY | 0.090744495 |
| GVRRPAGIPADAPV | TCSAEEASAETLY | 0.090687156 |
| GYRQVTDAHLLTLARR | TCSAEEASAETLY | 0.09089063 |
| IAKMGQDGHDRG | TCSAEEASAETLY | 0.09113796 |
| KAEPSIEIQYT | TCSAEEASAETLY | 0.090857565 |
| KPVRIRMAADEDTVAVT | TCSAEEASAETLY | 0.09136298 |
| MPPRIAGMRLLVIKPEP | TCSAEEASAETLY | 0.090881325 |
| MKFYAVALFRSRGVPPTR | TCSAEEASAETLY | 0.09133345 |
| NAGGFGILGAVAHSP | TCSAEEASAETLY | 0.090941675 |
| NEGREITLAMAEM | TCSAEEASAETLY | 0.090809986 |
| NELVEWSHPLGHRWRWE | TCSAEEASAETLY | 0.09189425 |
| NLGAGGTGGNGGNPG | TCSAEEASAETLY | 0.090608954 |
| NTEWPKWETVMAADPVD | TCSAEEASAETLY | 0.090723604 |
| PASAAPAETQP | TCSAEEASAETLY | 0.09093895 |
| QQSSDGKNGVRAAEITGE | TCSAEEASAETLY | 0.09088683 |
| RAPDAVTTSWWKE | TCSAEEASAETLY | 0.09096759 |
| SGFGNVGAHTSGWFNQ | TCSAEEASAETLY | 0.0912448 |
| SGVAGAVGQAASVGGLKV | TCSAEEASAETLY | 0.0906916 |
| SNGGAGTGSAGGNGGT | TCSAEEASAETLY | 0.0907549 |
| TGGAGGEGGAAG | TCSAEEASAETLY | 0.0908617 |
| TGGDGGAAGAGGAPGHG | TCSAEEASAETLY | 0.09142885 |
| TMPPLLAAIANAGRHGV | TCSAEEASAETLY | 0.0916319 |
| TSSTAGNSGGA | TCSAEEASAETLY | 0.090819195 |
| VPATVPVGILC | TCSAEEASAETLY | 0.091643244 |
| VQRPDSGGVVLTGRIS | TCSAEEASAETLY | 0.09091344 |
| WTPNPVPATKGVPAGLVQAT | TCSAEEASAETLY | 0.091026895 |
| YALKRALAKFSDRFDVVII | TCSAEEASAETLY | 0.09230553 |

| | | |
|---|---|---|
| YLIGSVIV | TCSAEEASAETLY | 0.0927187 |
| YGQLVAVID | TCSAEEASAETLY | 0.091113314 |
| VTVGGMTLPA | TCSAEEASAETLY | 0.09056208 |
| VLALPAV | TCSAEEASAETLY | 0.090801634 |
| VIPPTLNYET | TCSAEEASAETLY | 0.090645015 |
| VGMGGMHPSSG | TCSAEEASAETLY | 0.091416456 |
| VGLAGGAATA | TCSAEEASAETLY | 0.09125575 |
| VGGAGGAGGSGP | TCSAEEASAETLY | 0.091157496 |
| VFAAFMV | TCSAEEASAETLY | 0.09069597 |
| TLYDKLR | TCSAEEASAETLY | 0.09325021 |
| TLAVFTVL | TCSAEEASAETLY | 0.0906571 |
| TGSVQPARV | TCSAEEASAETLY | 0.091814436 |
| TGGGAGSGGT | TCSAEEASAETLY | 0.091295145 |
| SHLETSL | TCSAEEASAETLY | 0.090614684 |
| SFTTGMT | TCSAEEASAETLY | 0.09087965 |
| QVEVAKRMGVRQPTVS | TCSAEEASAETLY | 0.091199055 |
| QGSGGPALAM | TCSAEEASAETLY | 0.090677135 |
| MLAPSNP | TCSAEEASAETLY | 0.09065008 |
| LVNNAAI | TCSAEEASAETLY | 0.09074079 |
| LTGEINTLA | TCSAEEASAETLY | 0.09071769 |
| LSIFGPV | TCSAEEASAETLY | 0.09073441 |
| LLALLANL | TCSAEEASAETLY | 0.09128869 |
| LISNASDALDKLR | TCSAEEASAETLY | 0.091021076 |
| LINEAVP | TCSAEEASAETLY | 0.0906792 |
| LGYAFSGL | TCSAEEASAETLY | 0.090860166 |
| LDFVAVAL | TCSAEEASAETLY | 0.091595694 |
| LAQPGAE | TCSAEEASAETLY | 0.090988085 |
| ISNLGAVTGF | TCSAEEASAETLY | 0.09217943 |
| HQEDPGRI | TCSAEEASAETLY | 0.09069037 |
| GTPGEPTP | TCSAEEASAETLY | 0.09086078 |
| GFFNSGS | TCSAEEASAETLY | 0.09077433 |
| GAGGGGLTAGAN | TCSAEEASAETLY | 0.09119751 |
| FVDALVNM | TCSAEEASAETLY | 0.09070152 |
| FLTVAPEL | TCSAEEASAETLY | 0.0928382 |
| FETLTAL | TCSAEEASAETLY | 0.09271298 |
| DVINPIEL | TCSAEEASAETLY | 0.0907645 |
| APVSGAPV | TCSAEEASAETLY | 0.0925628 |
| ALVGGLAW | TCSAEEASAETLY | 0.090591855 |
| ALAAAAATAI | TCSAEEASAETLY | 0.09099914 |
| AFGLGLLL | TCSAEEASAETLY | 0.09056368 |
| AFFAAASL | TCSAEEASAETLY | 0.09151055 |
| AAAALPAFSPP | TCSAEEASAETLY | 0.092351474 |
| VVLAPAL | TCSAEEASAETLY | 0.09126194 |
| TIGHVDHGKTTL | TCSAEEASAETLY | 0.0908719 |
| LISNASDALDK | TCSAEEASAETLY | 0.092626624 |
| LISNASDAL | TCSAEEASAETLY | 0.09128298 |
| ISNASDALDK | TCSAEEASAETLY | 0.090724766 |
| NQAETLVYQTE | TCSAEEASAETLY | 0.09244576 |
| TGGTGGTAGAG | TCSAEEASAETLY | 0.090940684 |
| TGSTAGGAGGA | TCSAEEASAETLY | 0.09100988 |
| DPLAVPVAAGPV | TCSAEEASAETLY | 0.09077801 |

| | | |
|---|---|---|
| GGTGGTPGNGGN | TCSAEEASAETLY | 0.09087922 |
| PAGLAGVDPATL | TCSAEEASAETLY | 0.09095488 |
| GNNLVVEAPHAGASV | TCSAEEASAETLY | 0.09077329 |
| KVTRIMSASTLEQPA | TCSAEEASAETLY | 0.09110353 |
| LGFALPIALPRRTVV | TCSAEEASAETLY | 0.09092442 |
| TRLGNLTARKLVVLV | TCSAEEASAETLY | 0.09058079 |
| WSQQRRLIYFPSAGP | TCSAEEASAETLY | 0.090636574 |
| GPNGNPDPMAAAVDIR | TCSAEEASAETLY | 0.09211462 |
| TQRCVSVADLRAAASR | TCSAEEASAETLY | 0.09270386 |
| QFVLDVAEHPVEVLTETV | TCSAEEASAETLY | 0.09094334 |
| GGAGGTGGAGGNGGTQAGDG | TCSAEEASAETLY | 0.091113776 |
| PLLATLRAIAVDVDAKPAQV | TCSAEEASAETLY | 0.091457814 |
| APKFRTGGYMIDFWGVGYQVAK | TCSAEEASAETLY | 0.091846995 |
| GAGGAGGAGGAAGGPGGTGGTG | TCSAEEASAETLY | 0.09109148 |
| IGALGLLATDVAPAGSVAFGVA | TCSAEEASAETLY | 0.09111652 |
| AGGAGGIGGNGGAGGAGGVLMG | TCSAEEASAETLY | 0.09061119 |
| ADELIGLL | TCSAEEASAETLY | 0.09065179 |
| ADRALKA | TCSAEEASAETLY | 0.09077127 |
| AGAVGQVGGAGGA | TCSAEEASAETLY | 0.09092815 |
| AGGGGGLGGAP | TCSAEEASAETLY | 0.09103425 |
| AGGLLYGNGGAG | TCSAEEASAETLY | 0.09142501 |
| AGLESIGL | TCSAEEASAETLY | 0.091429055 |
| AGLKTGII | TCSAEEASAETLY | 0.0911094 |
| ASLAALLL | TCSAEEASAETLY | 0.09101293 |
| ASVINISSI | TCSAEEASAETLY | 0.09121162 |
| ATPPGPA | TCSAEEASAETLY | 0.0909033 |
| ATVASALV | TCSAEEASAETLY | 0.09369011 |
| ATVASLIL | TCSAEEASAETLY | 0.09088883 |
| AVNGPVLIRH | TCSAEEASAETLY | 0.0908106 |
| DGAAGAVGI | TCSAEEASAETLY | 0.091116846 |
| DPLLHGI | TCSAEEASAETLY | 0.09109017 |
| ELGAALGAL | TCSAEEASAETLY | 0.090848565 |
| ELGRTQAV | TCSAEEASAETLY | 0.091839775 |
| FVAGGYLS | TCSAEEASAETLY | 0.09206961 |
| GASGGAGGAGGW | TCSAEEASAETLY | 0.09061593 |
| GGAGAGGDGGAGGI | TCSAEEASAETLY | 0.09082023 |
| GTGHNGSQPGMGGNG | TCSAEEASAETLY | 0.0909924 |
| GYAAASASAS | TCSAEEASAETLY | 0.091449514 |
| HQVDVAAVQQG | TCSAEEASAETLY | 0.091185 |
| IGAAIAHT | TCSAEEASAETLY | 0.0907058 |
| IGMIEVAQL | TCSAEEASAETLY | 0.09107548 |
| IRLAPEHTSAMD | TCSAEEASAETLY | 0.09072013 |
| IYNVSVL | TCSAEEASAETLY | 0.0908765 |
| KQGFLGM | TCSAEEASAETLY | 0.09441243 |
| LESVITQL | TCSAEEASAETLY | 0.09205212 |
| LGLGALVVV | TCSAEEASAETLY | 0.09074191 |
| LIGNGGNGGSGGI | TCSAEEASAETLY | 0.091909304 |
| LNAINAPTL | TCSAEEASAETLY | 0.090570815 |
| LQLLNTGV | TCSAEEASAETLY | 0.09065442 |
| LQTSIEF | TCSAEEASAETLY | 0.090666674 |
| MSFAVSEL | TCSAEEASAETLY | 0.09073346 |

| | | |
|---|---|---|
| NTGALGL | TCSAEEASAETLY | 0.09080295 |
| PPGAGGAGGAGTT | TCSAEEASAETLY | 0.09101756 |
| QLVGLGI | TCSAEEASAETLY | 0.0906773 |
| RFLEQVHQL | TCSAEEASAETLY | 0.09098847 |
| RGVGYVLR | TCSAEEASAETLY | 0.09045782 |
| SATGVFAGVI | TCSAEEASAETLY | 0.090743534 |
| SLPAITLP | TCSAEEASAETLY | 0.091200225 |
| STALAFTV | TCSAEEASAETLY | 0.091231264 |
| SVLTVGLGK | TCSAEEASAETLY | 0.091679536 |
| TALAGAGSGI | TCSAEEASAETLY | 0.090729624 |
| TGLPFTGI | TCSAEEASAETLY | 0.09077403 |
| VAEHVTV | TCSAEEASAETLY | 0.0906883 |
| VDIVLNSL | TCSAEEASAETLY | 0.09117493 |
| VLTGGGAVL | TCSAEEASAETLY | 0.09070937 |
| VNVGLAYGI | TCSAEEASAETLY | 0.090947844 |
| VSSAAVGAV | TCSAEEASAETLY | 0.091306314 |
| VTSVYEAGV | TCSAEEASAETLY | 0.090777144 |
| WVALDRGA | TCSAEEASAETLY | 0.09088845 |
| YLVGSVVI | TCSAEEASAETLY | 0.09081296 |
| GPPGTGKTLL | TCSAEEASAETLY | 0.09062147 |
| GVGMTKF | TCSAEEASAETLY | 0.09064445 |
| IGIGGSDL | TCSAEEASAETLY | 0.09117144 |
| IGTLLAEL | TCSAEEASAETLY | 0.091165826 |
| IYTSGTT | TCSAEEASAETLY | 0.0907225 |
| LSFPAGV | TCSAEEASAETLY | 0.09093058 |
| LYTSGTT | TCSAEEASAETLY | 0.0906238 |
| VGAFPFGF | TCSAEEASAETLY | 0.09377397 |
| VGAGTVEF | TCSAEEASAETLY | 0.0909083 |
| VGKSTLLL | TCSAEEASAETLY | 0.0912851 |
| AALAHAGGGIV | TCSAEEASAETLY | 0.09078954 |
| AALVAVSLL | TCSAEEASAETLY | 0.090744324 |
| AAVSRAVAAL | TCSAEEASAETLY | 0.09178655 |
| AGGSSGIGGF | TCSAEEASAETLY | 0.090711944 |
| AGLTVANANG | TCSAEEASAETLY | 0.09074039 |
| AGLVGAVL | TCSAEEASAETLY | 0.090689905 |
| ALSQAFTV | TCSAEEASAETLY | 0.09101435 |
| ALTFSVSL | TCSAEEASAETLY | 0.09061505 |
| ALTGAALVAL | TCSAEEASAETLY | 0.09092129 |
| ASFSIGAGVL | TCSAEEASAETLY | 0.095237285 |
| DGSVYKI | TCSAEEASAETLY | 0.091048546 |
| EQLAEIV | TCSAEEASAETLY | 0.090955205 |
| FAVSELTVM | TCSAEEASAETLY | 0.091189794 |
| GAAPLAAGL | TCSAEEASAETLY | 0.091029905 |
| GADGGDGGSGGAGG | TCSAEEASAETLY | 0.09105451 |
| GFGLDTGGAGG | TCSAEEASAETLY | 0.0907215 |
| GLASGGLL | TCSAEEASAETLY | 0.090792604 |
| GRIAGAAVLL | TCSAEEASAETLY | 0.09089766 |
| GSAGSLNTSF | TCSAEEASAETLY | 0.091132484 |
| GTGQIGI | TCSAEEASAETLY | 0.09084276 |
| GTLIALIL | TCSAEEASAETLY | 0.091167994 |
| IAATMGFVVG | TCSAEEASAETLY | 0.091020785 |

| | | |
|---|---|---|
| IFAQALL | TCSAEEASAETLY | 0.09078065 |
| IGAAVALGF | TCSAEEASAETLY | 0.09093669 |
| IGLTPLAL | TCSAEEASAETLY | 0.090761974 |
| ILGSNTSTL | TCSAEEASAETLY | 0.091589734 |
| IVFSLTVL | TCSAEEASAETLY | 0.090970576 |
| LASKATGF | TCSAEEASAETLY | 0.0907813 |
| LASVALIL | TCSAEEASAETLY | 0.09072876 |
| LFAVGLTL | TCSAEEASAETLY | 0.09175347 |
| LFNGGIGTYI | TCSAEEASAETLY | 0.09074403 |
| LTLGAGII | TCSAEEASAETLY | 0.090774074 |
| LVAAALGDI | TCSAEEASAETLY | 0.090929106 |
| NTAFVQLGI | TCSAEEASAETLY | 0.09069865 |
| QLFSGAEL | TCSAEEASAETLY | 0.09090245 |
| SAATFITL | TCSAEEASAETLY | 0.09071142 |
| SNYGLANLGV | TCSAEEASAETLY | 0.09056432 |
| SYGGAYAVM | TCSAEEASAETLY | 0.09125856 |
| TTGVGGLTLG | TCSAEEASAETLY | 0.09069544 |
| TTQTIYI | TCSAEEASAETLY | 0.09090258 |
| VAAGAGLGAVI | TCSAEEASAETLY | 0.09148092 |
| VAAVVTLL | TCSAEEASAETLY | 0.09077688 |
| VGFLGGLI | TCSAEEASAETLY | 0.09128907 |
| VGVIPILL | TCSAEEASAETLY | 0.090939336 |
| VIFASSLI | TCSAEEASAETLY | 0.09087977 |
| VKGGDGGAAAG | TCSAEEASAETLY | 0.09096688 |
| VTKTFTNI | TCSAEEASAETLY | 0.09086382 |
| VVAAAILL | TCSAEEASAETLY | 0.090715356 |
| YAALDVEL | TCSAEEASAETLY | 0.090806834 |
| LASVALLL | TCSAEEASAETLY | 0.09476458 |
| LDAVHRV | TCSAEEASAETLY | 0.0918525 |
| QGLSSSIF | TCSAEEASAETLY | 0.091188125 |
| RGTFANIRL | TCSAEEASAETLY | 0.0908943 |
| RQLWWGHRI | TCSAEEASAETLY | 0.09086085 |
| YGPPGTGKTLL | TCSAEEASAETLY | 0.091048345 |
| AALVGAPANVA | TCSAEEASAETLY | 0.09063968 |
| AATFRRTE | TCSAEEASAETLY | 0.09079508 |
| AGALGGGI | TCSAEEASAETLY | 0.09085022 |
| AGGAGGAGGIGAG | TCSAEEASAETLY | 0.090667345 |
| AGSGALGL | TCSAEEASAETLY | 0.090833634 |
| AQIADQI | TCSAEEASAETLY | 0.09110126 |
| ASAVAVTYL | TCSAEEASAETLY | 0.090852976 |
| ASTVLGGTGG | TCSAEEASAETLY | 0.090756334 |
| AVGGEPAK | TCSAEEASAETLY | 0.09079813 |
| CEQGTFGGTC | TCSAEEASAETLY | 0.09119334 |
| DEGKAPAG | TCSAEEASAETLY | 0.09067527 |
| DTVVFPAL | TCSAEEASAETLY | 0.09221165 |
| FEGITMAG | TCSAEEASAETLY | 0.090793885 |
| FPLVVQD | TCSAEEASAETLY | 0.091451615 |
| GAFTTPQ | TCSAEEASAETLY | 0.09411061 |
| GFGGFGTAGGVGGA | TCSAEEASAETLY | 0.090828404 |
| GFLAQFL | TCSAEEASAETLY | 0.090666905 |
| GGAGGAGGNAGVA | TCSAEEASAETLY | 0.090705015 |

| | | |
|---|---|---|
| GGVGTAGAGGAGGAGG | TCSAEEASAETLY | 0.09100443 |
| GNGGGGDAGNAGSGGN | TCSAEEASAETLY | 0.09103249 |
| GTGGAGGAGNDAGS | TCSAEEASAETLY | 0.09057544 |
| GTTNTGWFNS | TCSAEEASAETLY | 0.09192072 |
| IFMGPLFTL | TCSAEEASAETLY | 0.09128565 |
| ILAHAIEAI | TCSAEEASAETLY | 0.090815045 |
| IRSAIREAL | TCSAEEASAETLY | 0.090467244 |
| ITFAGGG | TCSAEEASAETLY | 0.09140172 |
| ITGMSGHQEV | TCSAEEASAETLY | 0.09161924 |
| IVAGLACTL | TCSAEEASAETLY | 0.09152771 |
| IVGGGLAGA | TCSAEEASAETLY | 0.09065117 |
| IVTGVLNI | TCSAEEASAETLY | 0.090677835 |
| LAQQGCI | TCSAEEASAETLY | 0.091345556 |
| LCLHVEN | TCSAEEASAETLY | 0.09100553 |
| LDAISDGS | TCSAEEASAETLY | 0.09118969 |
| LGGFAAML | TCSAEEASAETLY | 0.09186417 |
| LGGKNPMI | TCSAEEASAETLY | 0.09081305 |
| LGPKGVAAIL | TCSAEEASAETLY | 0.091274425 |
| LMVGVSI | TCSAEEASAETLY | 0.09118599 |
| LVGAAFVL | TCSAEEASAETLY | 0.092230305 |
| LVLAVAPL | TCSAEEASAETLY | 0.090579025 |
| MAAIAVIDAA | TCSAEEASAETLY | 0.09098851 |
| NNGINAELQ | TCSAEEASAETLY | 0.09167703 |
| RGEPLVLVHG | TCSAEEASAETLY | 0.09073964 |
| RGGAGGAAGLI | TCSAEEASAETLY | 0.09077999 |
| RSYRMHPAV | TCSAEEASAETLY | 0.0916512 |
| SAAHLLAGQLA | TCSAEEASAETLY | 0.091123484 |
| SAAVGSY | TCSAEEASAETLY | 0.09083498 |
| SPHAVLHRI | TCSAEEASAETLY | 0.09133796 |
| SQIAKVL | TCSAEEASAETLY | 0.09058066 |
| TALGHLVP | TCSAEEASAETLY | 0.091334246 |
| TAPTTSGAA | TCSAEEASAETLY | 0.09074028 |
| TDGDDGNGGSPGT | TCSAEEASAETLY | 0.09166569 |
| TDGLGEN | TCSAEEASAETLY | 0.09083037 |
| TGGPPLPGQA | TCSAEEASAETLY | 0.09070166 |
| TIPAITI | TCSAEEASAETLY | 0.09063812 |
| TIPGFTI | TCSAEEASAETLY | 0.2648022 |
| TSIASGALA | TCSAEEASAETLY | 0.6711463 |
| TVTLPGAAI | TCSAEEASAETLY | 0.5700975 |
| VASGLSAALP | TCSAEEASAETLY | 0.5533025 |
| VGAGVIGI | TCSAEEASAETLY | 0.38431448 |
| VGPKRFGV | TCSAEEASAETLY | 0.2754904 |
| VNVAFPD | TCSAEEASAETLY | 0.34641793 |
| VSMTNSGVS | TCSAEEASAETLY | 0.34871703 |
| VTDVLGDLSV | TCSAEEASAETLY | 0.63446784 |
| VVVVGAGFAGL | TCSAEEASAETLY | 0.590138 |
| VYTVGTVV | TCSAEEASAETLY | 0.35973215 |
| WTTFSGP | TCSAEEASAETLY | 0.6422156 |
| YDELVEAF | TCSAEEASAETLY | 0.67996716 |
| YGTGGAGGNGG | TCSAEEASAETLY | 0.24090354 |
| YTNVMAV | TCSAEEASAETLY | 0.46321622 |

| | | |
|---|---|---|
| FLEQVHQL | TCSAEEASAETLY | 0.27785748 |
| GLSLLAVAV | TCSAEEASAETLY | 0.3921988 |
| LEQVHQL | TCSAEEASAETLY | 0.19947965 |
| LNAAGAI | TCSAEEASAETLY | 0.568125 |
| PAVHASL | TCSAEEASAETLY | 0.2988981 |
| TELEDDA | TCSAEEASAETLY | 0.20004743 |
| TPPGPPN | TCSAEEASAETLY | 0.5132885 |
| TPPPPPPP | TCSAEEASAETLY | 0.4413678 |
| TQSPALI | ASSLPRGSYAEQF | 0.48401934 |
| EFAAAFPQ | ASSLPRGSYAEQF | 0.6156218 |
| LLSGNLDVL | ASSLPRGSYAEQF | 0.45244727 |
| GPANLGNYNI | ASSLPRGSYAEQF | 0.405066 |
| TLGPTLP | ASSLPRGSYAEQF | 0.57178247 |
| VYAYSSCCK | ASSLPRGSYAEQF | 0.6794155 |
| APAPAPAPNI | ASSLPRGSYAEQF | 0.39319417 |
| PGGPVPIP | ASSLPRGSYAEQF | 0.50057817 |
| FLTVALEP | ASSLPRGSYAEQF | 0.6724781 |
| SQYFGANL | ASSLPRGSYAEQF | 0.5075714 |
| GGVKPAA | ASSLPRGSYAEQF | 0.6265466 |
| GGTGGAGGAGGW | ASSLPRGSYAEQF | 0.67486465 |
| GVGGVAGLGGAGG | ASSLPRGSYAEQF | 0.3214583 |
| VLLVLATTI | ASSLPRGSYAEQF | 0.6521014 |
| IGIFAAML | ASSLPRGSYAEQF | 0.65080816 |
| AVGDLID | ASSLPRGSYAEQF | 0.46301728 |
| NISPLPL | ASSLPRGSYAEQF | 0.39264652 |
| VTAVFVAI | ASSLPRGSYAEQF | 0.24963026 |
| SGDDNDAAL | ASSLPRGSYAEQF | 0.4919824 |
| AVAALAHS | ASSLPRGSYAEQF | 0.312752 |
| GGVTVMATP | ASSLPRGSYAEQF | 0.5420474 |
| AGLAANVRD | ASSLPRGSYAEQF | 0.67516947 |
| TNRPDLIDPALLRPG | ASSLPRGSYAEQF | 0.39448863 |
| LANLGVTAL | ASSLPRGSYAEQF | 0.24270931 |
| TGERIEL | ASSLPRGSYAEQF | 0.2372046 |
| NGEELAA | ASSLPRGSYAEQF | 0.48212942 |
| VLSLVWG | ASSLPRGSYAEQF | 0.32772854 |
| IGYALWL | ASSLPRGSYAEQF | 0.22671433 |
| LSLLLAPL | ASSLPRGSYAEQF | 0.3605701 |
| TLDEALK | ASSLPRGSYAEQF | 0.6595473 |
| TDGAGTAEGG | ASSLPRGSYAEQF | 0.6630957 |
| NAPGGAGGNGG | ASSLPRGSYAEQF | 0.29831454 |
| VSVFMGMGGNF | ASSLPRGSYAEQF | 0.69100666 |
| LASYLAAL | ASSLPRGSYAEQF | 0.5859668 |
| LVGGTPAL | ASSLPRGSYAEQF | 0.55422026 |
| VRNADVL | ASSLPRGSYAEQF | 0.39125004 |
| TCSSPFLVSAM | ASSLPRGSYAEQF | 0.25943244 |
| VHITDAE | ASSLPRGSYAEQF | 0.6479851 |
| EYAAPSF | ASSLPRGSYAEQF | 0.47686136 |
| VVDAFHGGL | ASSLPRGSYAEQF | 0.50991 |
| ASVVDPQ | ASSLPRGSYAEQF | 0.5785801 |
| SYNEFGGLS | ASSLPRGSYAEQF | 0.549093 |
| DPDGSVY | ASSLPRGSYAEQF | 0.3526256 |

| | | |
|---|---|---|
| MVSTIHATE | ASSLPRGSYAEQF | 0.54955614 |
| LAAGVADPV | ASSLPRGSYAEQF | 0.5803082 |
| TKEQIAA | ASSLPRGSYAEQF | 0.29627547 |
| SLASPAAL | ASSLPRGSYAEQF | 0.33097503 |
| IGLVFYGV | ASSLPRGSYAEQF | 0.44005346 |
| FLLDASNL | ASSLPRGSYAEQF | 0.4973035 |
| YQVGNFP | ASSLPRGSYAEQF | 0.66325796 |
| ESLLSQST | ASSLPRGSYAEQF | 0.4729108 |
| LFTVGFAAVG | ASSLPRGSYAEQF | 0.3754216 |
| ATPPTPT | ASSLPRGSYAEQF | 0.59217644 |
| TDLATLG | ASSLPRGSYAEQF | 0.5563526 |
| SAGGTGR | ASSLPRGSYAEQF | 0.48576298 |
| GFSNSGS | ASSLPRGSYAEQF | 0.6385014 |
| GGNGGSGGTSV | ASSLPRGSYAEQF | 0.33971763 |
| GTSVATGGAGN | ASSLPRGSYAEQF | 0.539828 |
| NGGNGGSGGTS | ASSLPRGSYAEQF | 0.56960905 |
| AVVKFPH | ASSLPRGSYAEQF | 0.5628807 |
| TPPPPPP | ASSLPRGSYAEQF | 0.6222045 |
| AAPAAAAAPA | ASSLPRGSYAEQF | 0.60208094 |
| IALNEDL | ASSLPRGSYAEQF | 0.48774964 |
| IVNVSSM | ASSLPRGSYAEQF | 0.24982104 |
| TGIPPAPR | ASSLPRGSYAEQF | 0.34491116 |
| FDLGGGTF | ASSLPRGSYAEQF | 0.58128476 |
| FDLGGGTFDVS | ASSLPRGSYAEQF | 0.53008986 |
| NEPTAAA | ASSLPRGSYAEQF | 0.63239324 |
| NEPTAAALA | ASSLPRGSYAEQF | 0.5597015 |
| GSIINISSI | ASSLPRGSYAEQF | 0.37434137 |
| VAPGNAGTA | ASSLPRGSYAEQF | 0.68861365 |
| TGGGASLE | ASSLPRGSYAEQF | 0.631764 |
| VAVYNHYKRI | ASSLPRGSYAEQF | 0.23214325 |
| TDPAPATT | ASSLPRGSYAEQF | 0.4266706 |
| TNRPDILDPALLRPG | ASSLPRGSYAEQF | 0.68605304 |
| LTTTTTS | ASSLPRGSYAEQF | 0.5542736 |
| PTGSGKTL | ASSLPRGSYAEQF | 0.43938547 |
| TGSGKTL | ASSLPRGSYAEQF | 0.4478665 |
| TIGHVDHGK | ASSLPRGSYAEQF | 0.69751847 |
| TIGHVDHGKTT | ASSLPRGSYAEQF | 0.43372 |
| GHVDHGKTT | ASSLPRGSYAEQF | 0.67212975 |
| AQFEHTLL | ASSLPRGSYAEQF | 0.22498907 |
| IVNQDEL | ASSLPRGSYAEQF | 0.35400593 |
| TNGSQFF | ASSLPRGSYAEQF | 0.29643643 |
| TGKGIVE | ASSLPRGSYAEQF | 0.21110612 |
| LISNASDA | ASSLPRGSYAEQF | 0.63184875 |
| AELRTLGELTKIAKAHGAQ | ASSLPRGSYAEQF | 0.33660635 |
| APNAVAAAEPSW | ASSLPRGSYAEQF | 0.6190531 |
| APQVAGAATAVL | ASSLPRGSYAEQF | 0.3429342 |
| ARRDPAAAAAILPTDA | ASSLPRGSYAEQF | 0.6411071 |
| DAAARDVLAEAGLAEYF | ASSLPRGSYAEQF | 0.40671584 |
| DDQIVLSAKANLAKVEQ | ASSLPRGSYAEQF | 0.5307727 |
| DELLPDTGEDSDEDGH | ASSLPRGSYAEQF | 0.30529186 |
| ELPEAPANRATVPQD | ASSLPRGSYAEQF | 0.61794126 |

| | | |
|---|---|---|
| FGQSKIERVTGIG | ASSLPRGSYAEQF | 0.6982273 |
| GAGGTGSTAGGAGGAGGAGGLY | ASSLPRGSYAEQF | 0.53406 |
| GHGGTGAAGGNGGTGGAGGSGI | ASSLPRGSYAEQF | 0.30102572 |
| GNGGNGGAGGEGGPGGAGGAGA | ASSLPRGSYAEQF | 0.5224371 |
| GSTAGNGGNGGNATNGTIAGQP | ASSLPRGSYAEQF | 0.48625085 |
| GTGGDGGNGGNGGTGG | ASSLPRGSYAEQF | 0.23865603 |
| GVRRPAGIPADAPV | ASSLPRGSYAEQF | 0.21621048 |
| GYRQVTDAHLLTLARR | ASSLPRGSYAEQF | 0.4275722 |
| IAKMGQDGHDRG | ASSLPRGSYAEQF | 0.4097203 |
| KAEPSIEIQYT | ASSLPRGSYAEQF | 0.62435055 |
| KPVRIRMAADEDTVAVT | ASSLPRGSYAEQF | 0.5927484 |
| MPPRIAGMRLLVIKPEP | ASSLPRGSYAEQF | 0.34403044 |
| MKFYAVALFRSRGVPPTR | ASSLPRGSYAEQF | 0.58850133 |
| NAGGFGILGAVAHSP | ASSLPRGSYAEQF | 0.6067277 |
| NEGREITLAMAEM | ASSLPRGSYAEQF | 0.18585326 |
| NELVEWSHPLGHRWRWE | ASSLPRGSYAEQF | 0.31964552 |
| NLGAGGTGGNGGNPG | ASSLPRGSYAEQF | 0.24333338 |
| NTEWPKWETVMAADPVD | ASSLPRGSYAEQF | 0.6599112 |
| PASAAPAETQP | ASSLPRGSYAEQF | 0.30583832 |
| QQSSDGKNGVRAAEITGE | ASSLPRGSYAEQF | 0.33265805 |
| RAPDAVTTSWWKE | ASSLPRGSYAEQF | 0.42256618 |
| SGFGNVGAHTSGWFNQ | ASSLPRGSYAEQF | 0.60849386 |
| SGVAGAVGQAASVGGLKV | ASSLPRGSYAEQF | 0.5063884 |
| SNGGAGTGSAGGNGGT | ASSLPRGSYAEQF | 0.2860263 |
| TGGAGGEGGAAG | ASSLPRGSYAEQF | 0.263132 |
| TGGDGGAAGAGGAPGHG | ASSLPRGSYAEQF | 0.5960151 |
| TMPPLLAAIANAGRHGV | ASSLPRGSYAEQF | 0.37251887 |
| TSSTAGNSGGA | ASSLPRGSYAEQF | 0.41416857 |
| VPATVPVGILC | ASSLPRGSYAEQF | 0.27045587 |
| VQRPDSGGVVLTGRIS | ASSLPRGSYAEQF | 0.5577605 |
| WTPNPVPATKGVPAGLVQAT | ASSLPRGSYAEQF | 0.24671453 |
| YALKRALAKFSDRFDVVII | ASSLPRGSYAEQF | 0.5895065 |
| YLIGSVIV | ASSLPRGSYAEQF | 0.31732792 |
| YGQLVAVID | ASSLPRGSYAEQF | 0.4352666 |
| VTVGGMTLPA | ASSLPRGSYAEQF | 0.6959782 |
| VLALPAV | ASSLPRGSYAEQF | 0.60347044 |
| VIPPTLNYET | ASSLPRGSYAEQF | 0.4074245 |
| VGMGGMHPSSG | ASSLPRGSYAEQF | 0.4510972 |
| VGLAGGAATA | ASSLPRGSYAEQF | 0.27272448 |
| VGGAGGAGGSGP | ASSLPRGSYAEQF | 0.3733577 |
| VFAAFMV | ASSLPRGSYAEQF | 0.4596067 |
| TLYDKLR | ASSLPRGSYAEQF | 0.6927643 |
| TLAVFTVL | ASSLPRGSYAEQF | 0.41531354 |
| TGSVQPARV | ASSLPRGSYAEQF | 0.24413061 |
| TGGGAGSGGT | ASSLPRGSYAEQF | 0.44235563 |
| SHLETSL | ASSLPRGSYAEQF | 0.6652311 |
| SFTTGMT | ASSLPRGSYAEQF | 0.67819643 |
| QVEVAKRMGVRQPTVS | ASSLPRGSYAEQF | 0.3104084 |
| QGSGGPALAM | ASSLPRGSYAEQF | 0.36772823 |
| MLAPSNP | ASSLPRGSYAEQF | 0.42189553 |
| LVNNAAI | ASSLPRGSYAEQF | 0.43834817 |

| | | |
|---|---|---|
| LTGEINTLA | ASSLPRGSYAEQF | 0.33780664 |
| LSIFGPV | ASSLPRGSYAEQF | 0.35191143 |
| LLALLANL | ASSLPRGSYAEQF | 0.5370631 |
| LISNASDALDKLR | ASSLPRGSYAEQF | 0.7200089 |
| LINEAVP | ASSLPRGSYAEQF | 0.43437833 |
| LGYAFSGL | ASSLPRGSYAEQF | 0.20358984 |
| LDFVAVAL | ASSLPRGSYAEQF | 0.4845978 |
| LAQPGAE | ASSLPRGSYAEQF | 0.34452546 |
| ISNLGAVTGF | ASSLPRGSYAEQF | 0.31734827 |
| HQEDPGRI | ASSLPRGSYAEQF | 0.49740887 |
| GTPGEPTP | ASSLPRGSYAEQF | 0.22422962 |
| GFFNSGS | ASSLPRGSYAEQF | 0.45300663 |
| GAGGGGLTAGAN | ASSLPRGSYAEQF | 0.71413815 |
| FVDALVNM | ASSLPRGSYAEQF | 0.4408247 |
| FLTVAPEL | ASSLPRGSYAEQF | 0.5386145 |
| FETLTAL | ASSLPRGSYAEQF | 0.64967084 |
| DVINPIEL | ASSLPRGSYAEQF | 0.54004836 |
| APVSGAPV | ASSLPRGSYAEQF | 0.6514889 |
| ALVGGLAW | ASSLPRGSYAEQF | 0.623497 |
| ALAAAAATAI | ASSLPRGSYAEQF | 0.4784785 |
| AFGLGLLL | ASSLPRGSYAEQF | 0.46039706 |
| AFFAAASL | ASSLPRGSYAEQF | 0.6154667 |
| AAAALPAFSPP | ASSLPRGSYAEQF | 0.4849357 |
| VVLAPAL | ASSLPRGSYAEQF | 0.47397786 |
| TIGHVDHGKTTL | ASSLPRGSYAEQF | 0.31514758 |
| LISNASDALDK | ASSLPRGSYAEQF | 0.64629054 |
| LISNASDAL | ASSLPRGSYAEQF | 0.42702842 |
| ISNASDALDK | ASSLPRGSYAEQF | 0.6439339 |
| NQAETLVYQTE | ASSLPRGSYAEQF | 0.24306491 |
| TGGTGGTAGAG | ASSLPRGSYAEQF | 0.39337394 |
| TGSTAGGAGGA | ASSLPRGSYAEQF | 0.6362398 |
| DPLAVPVAAGPV | ASSLPRGSYAEQF | 0.19964388 |
| GGTGGTPGNGGN | ASSLPRGSYAEQF | 0.23481905 |
| PAGLAGVDPATL | ASSLPRGSYAEQF | 0.37852913 |
| GNNLVVEAPHAGASV | ASSLPRGSYAEQF | 0.66998494 |
| KVTRIMSASTLEQPA | ASSLPRGSYAEQF | 0.19718583 |
| LGFALPIALPRRTVV | ASSLPRGSYAEQF | 0.40146506 |
| TRLGNLTARKLVVLV | ASSLPRGSYAEQF | 0.2618737 |
| WSQQRRLIYFPSAGP | ASSLPRGSYAEQF | 0.5990328 |
| GPNGNPDPMAAAVDIR | ASSLPRGSYAEQF | 0.4762217 |
| TQRCVSVADLRAAASR | ASSLPRGSYAEQF | 0.23811553 |
| QFVLDVAEHPVEVLTETV | ASSLPRGSYAEQF | 0.5913694 |
| GGAGGTGGAGGNGGTQAGDG | ASSLPRGSYAEQF | 0.20860007 |
| PLLATLRAIAVDVDAKPAQV | ASSLPRGSYAEQF | 0.48265678 |
| APKFRTGGYMIDFWGVGYQVAK | ASSLPRGSYAEQF | 0.6102042 |
| GAGGAGGAGGAAGGPGGTGGTG | ASSLPRGSYAEQF | 0.6907675 |
| IGALGLLATDVAPAGSVAFGVA | ASSLPRGSYAEQF | 0.2525071 |
| AGGAGGIGGNGGAGGAGGVLMG | ASSLPRGSYAEQF | 0.5958767 |
| ADELIGLL | ASSLPRGSYAEQF | 0.3255839 |
| ADRALKA | ASSLPRGSYAEQF | 0.710889 |
| AGAVGQVGGAGGA | ASSLPRGSYAEQF | 0.45306942 |

| | | |
|---|---|---|
| AGGGGGLGGAP | ASSLPRGSYAEQF | 0.23012343 |
| AGGLLYGNGGAG | ASSLPRGSYAEQF | 0.6360111 |
| AGLESIGL | ASSLPRGSYAEQF | 0.5182005 |
| AGLKTGII | ASSLPRGSYAEQF | 0.538503 |
| ASLAALLL | ASSLPRGSYAEQF | 0.57867205 |
| ASVINISSI | ASSLPRGSYAEQF | 0.37050506 |
| ATPPGPA | ASSLPRGSYAEQF | 0.38748816 |
| ATVASALV | ASSLPRGSYAEQF | 0.6186358 |
| ATVASLIL | ASSLPRGSYAEQF | 0.61672366 |
| AVNGPVLIRH | ASSLPRGSYAEQF | 0.38770017 |
| DGAAGAVGI | ASSLPRGSYAEQF | 0.6426927 |
| DPLLHGI | ASSLPRGSYAEQF | 0.51928335 |
| ELGAALGAL | ASSLPRGSYAEQF | 0.28992712 |
| ELGRTQAV | ASSLPRGSYAEQF | 0.70737 |
| FVAGGYLS | ASSLPRGSYAEQF | 0.3889705 |
| GASGGAGGAGGW | ASSLPRGSYAEQF | 0.59451485 |
| GGAGAGGDGGAGGI | ASSLPRGSYAEQF | 0.58722925 |
| GTGHNGSQPGMGGNG | ASSLPRGSYAEQF | 0.6866497 |
| GYAAASASAS | ASSLPRGSYAEQF | 0.31061852 |
| HQVDVAAVQQG | ASSLPRGSYAEQF | 0.5237391 |
| IGAAIAHT | ASSLPRGSYAEQF | 0.5081735 |
| IGMIEVAQL | ASSLPRGSYAEQF | 0.58730054 |
| IRLAPEHTSAMD | ASSLPRGSYAEQF | 0.37575436 |
| IYNVSVL | ASSLPRGSYAEQF | 0.52512765 |
| KQGFLGM | ASSLPRGSYAEQF | 0.54144454 |
| LESVITQL | ASSLPRGSYAEQF | 0.6645223 |
| LGLGALVVV | ASSLPRGSYAEQF | 0.36910146 |
| LIGNGGNGGSGGI | ASSLPRGSYAEQF | 0.5966236 |
| LNAINAPTL | ASSLPRGSYAEQF | 0.68445736 |
| LQLLNTGV | ASSLPRGSYAEQF | 0.28260928 |
| LQTSIEF | ASSLPRGSYAEQF | 0.4745841 |
| MSFAVSEL | ASSLPRGSYAEQF | 0.56559694 |
| NTGALGL | ASSLPRGSYAEQF | 0.6892481 |
| PPGAGGAGGAGTT | ASSLPRGSYAEQF | 0.37912878 |
| QLVGLGI | ASSLPRGSYAEQF | 0.66166604 |
| RFLEQVHQL | ASSLPRGSYAEQF | 0.41031587 |
| RGVGYVLR | ASSLPRGSYAEQF | 0.6308589 |
| SATGVFAGVI | ASSLPRGSYAEQF | 0.32024235 |
| SLPAITLP | ASSLPRGSYAEQF | 0.43701905 |
| STALAFTV | ASSLPRGSYAEQF | 0.37782836 |
| SVLTVGLGK | ASSLPRGSYAEQF | 0.54768574 |
| TALAGAGSGI | ASSLPRGSYAEQF | 0.5960196 |
| TGLPFTGI | ASSLPRGSYAEQF | 0.6078279 |
| VAEHVTV | ASSLPRGSYAEQF | 0.21017693 |
| VDIVLNSL | ASSLPRGSYAEQF | 0.5610491 |
| VLTGGGAVL | ASSLPRGSYAEQF | 0.63314664 |
| VNVGLAYGI | ASSLPRGSYAEQF | 0.7040981 |
| VSSAAVGAV | ASSLPRGSYAEQF | 0.46215048 |
| VTSVYEAGV | ASSLPRGSYAEQF | 0.60863245 |
| WVALDRGA | ASSLPRGSYAEQF | 0.44298232 |
| YLVGSVVI | ASSLPRGSYAEQF | 0.38273537 |

| | | |
|---|---|---|
| GPPGTGKTLL | ASSLPRGSYAEQF | 0.2652763 |
| GVGMTKF | ASSLPRGSYAEQF | 0.32236418 |
| IGIGGSDL | ASSLPRGSYAEQF | 0.39424628 |
| IGTLLAEL | ASSLPRGSYAEQF | 0.3180694 |
| IYTSGTT | ASSLPRGSYAEQF | 0.656336 |
| LSFPAGV | ASSLPRGSYAEQF | 0.659953 |
| LYTSGTT | ASSLPRGSYAEQF | 0.66304135 |
| VGAFPFGF | ASSLPRGSYAEQF | 0.6540881 |
| VGAGTVEF | ASSLPRGSYAEQF | 0.6431413 |
| VGKSTLLL | ASSLPRGSYAEQF | 0.5926605 |
| AALAHAGGGIV | ASSLPRGSYAEQF | 0.42208517 |
| AALVAVSLL | ASSLPRGSYAEQF | 0.4062402 |
| AAVSRAVAAL | ASSLPRGSYAEQF | 0.24092479 |
| AGGSSGIGGF | ASSLPRGSYAEQF | 0.35109955 |
| AGLTVANANG | ASSLPRGSYAEQF | 0.4160692 |
| AGLVGAVL | ASSLPRGSYAEQF | 0.45659995 |
| ALSQAFTV | ASSLPRGSYAEQF | 0.2038437 |
| ALTFSVSL | ASSLPRGSYAEQF | 0.54910874 |
| ALTGAALVAL | ASSLPRGSYAEQF | 0.70439065 |
| ASFSIGAGVL | ASSLPRGSYAEQF | 0.5205623 |
| DGSVYKI | ASSLPRGSYAEQF | 0.5972707 |
| EQLAEIV | ASSLPRGSYAEQF | 0.54189646 |
| FAVSELTVM | ASSLPRGSYAEQF | 0.32284048 |
| GAAPLAAGL | ASSLPRGSYAEQF | 0.61369383 |
| GADGGDGGSGGAGG | ASSLPRGSYAEQF | 0.38743258 |
| GFGLDTGGAGG | ASSLPRGSYAEQF | 0.5968194 |
| GLASGGLL | ASSLPRGSYAEQF | 0.69753563 |
| GRIAGAAVLL | ASSLPRGSYAEQF | 0.51938426 |
| GSAGSLNTSF | ASSLPRGSYAEQF | 0.5533711 |
| GTGQIGI | ASSLPRGSYAEQF | 0.5084355 |
| GTLIALIL | ASSLPRGSYAEQF | 0.2720111 |
| IAATMGFVVG | ASSLPRGSYAEQF | 0.5733427 |
| IFAQALL | ASSLPRGSYAEQF | 0.40632436 |
| IGAAVALGF | ASSLPRGSYAEQF | 0.46015495 |
| IGLTPLAL | ASSLPRGSYAEQF | 0.33877772 |
| ILGSNTSTL | ASSLPRGSYAEQF | 0.57571036 |
| IVFSLTVL | ASSLPRGSYAEQF | 0.51344895 |
| LASKATGF | ASSLPRGSYAEQF | 0.48131728 |
| LASVALIL | ASSLPRGSYAEQF | 0.31465578 |
| LFAVGLTL | ASSLPRGSYAEQF | 0.5236732 |
| LFNGGIGTYI | ASSLPRGSYAEQF | 0.531761 |
| LTLGAGII | ASSLPRGSYAEQF | 0.32455665 |
| LVAAALGDI | ASSLPRGSYAEQF | 0.47233894 |
| NTAFVQLGI | ASSLPRGSYAEQF | 0.35545743 |
| QLFSGAEL | ASSLPRGSYAEQF | 0.41616017 |
| SAATFITL | ASSLPRGSYAEQF | 0.66191137 |
| SNYGLANLGV | ASSLPRGSYAEQF | 0.5170338 |
| SYGGAYAVM | ASSLPRGSYAEQF | 0.22178634 |
| TTGVGGLTLG | ASSLPRGSYAEQF | 0.42495522 |
| TTQTIYI | ASSLPRGSYAEQF | 0.49002057 |
| VAAGAGLGAVI | ASSLPRGSYAEQF | 0.46866938 |

| | | |
|---|---|---|
| VAAVVTLL | ASSLPRGSYAEQF | 0.20212826 |
| VGFLGGLI | ASSLPRGSYAEQF | 0.57429045 |
| VGVIPILL | ASSLPRGSYAEQF | 0.2986978 |
| VIFASSLI | ASSLPRGSYAEQF | 0.2748348 |
| VKGGDGGAAAG | ASSLPRGSYAEQF | 0.34450418 |
| VTKTFTNI | ASSLPRGSYAEQF | 0.60735255 |
| VVAAAILL | ASSLPRGSYAEQF | 0.58667254 |
| YAALDVEL | ASSLPRGSYAEQF | 0.5100844 |
| LASVALLL | ASSLPRGSYAEQF | 0.5828587 |
| LDAVHRV | ASSLPRGSYAEQF | 0.6229013 |
| QGLSSSIF | ASSLPRGSYAEQF | 0.6711684 |
| RGTFANIRL | ASSLPRGSYAEQF | 0.6272663 |
| RQLWWGHRI | ASSLPRGSYAEQF | 0.41589808 |
| YGPPGTGKTLL | ASSLPRGSYAEQF | 0.6089579 |
| AALVGAPANVA | ASSLPRGSYAEQF | 0.35652032 |
| AATFRRTE | ASSLPRGSYAEQF | 0.2861829 |
| AGALGGGI | ASSLPRGSYAEQF | 0.26782697 |
| AGGAGGAGGIGAG | ASSLPRGSYAEQF | 0.29898646 |
| AGSGALGL | ASSLPRGSYAEQF | 0.2769568 |
| AQIADQI | ASSLPRGSYAEQF | 0.5831121 |
| ASAVAVTYL | ASSLPRGSYAEQF | 0.62499034 |
| ASTVLGGTGG | ASSLPRGSYAEQF | 0.3861941 |
| AVGGEPAK | ASSLPRGSYAEQF | 0.23071028 |
| CEQGTFGGTC | ASSLPRGSYAEQF | 0.39561832 |
| DEGKAPAG | ASSLPRGSYAEQF | 0.3133544 |
| DTVVFPAL | ASSLPRGSYAEQF | 0.59711194 |
| FEGITMAG | ASSLPRGSYAEQF | 0.5855093 |
| FPLVVQD | ASSLPRGSYAEQF | 0.7053242 |
| GAFTTPQ | ASSLPRGSYAEQF | 0.637702 |
| GFGGFGTAGGVGGA | ASSLPRGSYAEQF | 0.42187124 |
| GFLAQFL | ASSLPRGSYAEQF | 0.5861107 |
| GGAGGAGGNAGVA | ASSLPRGSYAEQF | 0.31576446 |
| GGVGTAGAGGAGGAGG | ASSLPRGSYAEQF | 0.38635966 |
| GNGGGGDAGNAGSGGN | ASSLPRGSYAEQF | 0.433419 |
| GTGGAGGAGNDAGS | ASSLPRGSYAEQF | 0.4263955 |
| GTTNTGWFNS | ASSLPRGSYAEQF | 0.51589954 |
| IFMGPLFTL | ASSLPRGSYAEQF | 0.44768238 |
| ILAHAIEAI | ASSLPRGSYAEQF | 0.23175634 |
| IRSAIREAL | ASSLPRGSYAEQF | 0.55058616 |
| ITFAGGG | ASSLPRGSYAEQF | 0.3124616 |
| ITGMSGHQEV | ASSLPRGSYAEQF | 0.23934484 |
| IVAGLACTL | ASSLPRGSYAEQF | 0.4999964 |
| IVGGGLAGA | ASSLPRGSYAEQF | 0.6950582 |
| IVTGVLNI | ASSLPRGSYAEQF | 0.6950451 |
| LAQQGCI | ASSLPRGSYAEQF | 0.33690426 |
| LCLHVEN | ASSLPRGSYAEQF | 0.36523336 |
| LDAISDGS | ASSLPRGSYAEQF | 0.6071172 |
| LGGFAAML | ASSLPRGSYAEQF | 0.30929175 |
| LGGKNPMI | ASSLPRGSYAEQF | 0.50062525 |
| LGPKGVAAIL | ASSLPRGSYAEQF | 0.4941342 |
| LMVGVSI | ASSLPRGSYAEQF | 0.43286774 |

| | | |
|---|---|---|
| LVGAAFVL | ASSLPRGSYAEQF | 0.37742895 |
| LVLAVAPL | ASSLPRGSYAEQF | 0.5547154 |
| MAAIAVIDAA | ASSLPRGSYAEQF | 0.4249966 |
| NNGINAELQ | ASSLPRGSYAEQF | 0.5449604 |
| RGEPLVLVHG | ASSLPRGSYAEQF | 0.19659497 |
| RGGAGGAAGLI | ASSLPRGSYAEQF | 0.5946809 |
| RSYRMHPAV | ASSLPRGSYAEQF | 0.32895315 |
| SAAHLLAGQLA | ASSLPRGSYAEQF | 0.30303362 |
| SAAVGSY | ASSLPRGSYAEQF | 0.20863573 |
| SPHAVLHRI | ASSLPRGSYAEQF | 0.34200883 |
| SQIAKVL | ASSLPRGSYAEQF | 0.5135227 |
| TALGHLVP | ASSLPRGSYAEQF | 0.6549301 |
| TAPTTSGAA | ASSLPRGSYAEQF | 0.3830632 |
| TDGDDGNGGSPGT | ASSLPRGSYAEQF | 0.6310199 |
| TDGLGEN | ASSLPRGSYAEQF | 0.5961598 |
| TGGPPLPGQA | ASSLPRGSYAEQF | 0.37141883 |
| TIPAITI | ASSLPRGSYAEQF | 0.25696915 |
| TIPGFTI | ASSLPRGSYAEQF | 0.5792 |
| TSIASGALA | ASSLPRGSYAEQF | 0.3461465 |
| TVTLPGAAI | ASSLPRGSYAEQF | 0.63119984 |
| VASGLSAALP | ASSLPRGSYAEQF | 0.637277 |
| VGAGVIGI | ASSLPRGSYAEQF | 0.23355082 |
| VGPKRFGV | ASSLPRGSYAEQF | 0.6476191 |
| VNVAFPD | ASSLPRGSYAEQF | 0.6188295 |
| VSMTNSGVS | ASSLPRGSYAEQF | 0.32796866 |
| VTDVLGDLSV | ASSLPRGSYAEQF | 0.25145918 |
| VVVVGAGFAGL | ASSLPRGSYAEQF | 0.2940342 |
| VYTVGTVV | ASSLPRGSYAEQF | 0.24975158 |
| WTTFSGP | ASSLPRGSYAEQF | 0.6329936 |
| YDELVEAF | ASSLPRGSYAEQF | 0.40442562 |
| YGTGGAGGNGG | ASSLPRGSYAEQF | 0.5167984 |
| YTNVMAV | ASSLPRGSYAEQF | 0.3834507 |
| FLEQVHQL | ASSLPRGSYAEQF | 0.4842623 |
| GLSLLAVAV | ASSLPRGSYAEQF | 0.42181188 |
| LEQVHQL | ASSLPRGSYAEQF | 0.47401473 |
| LNAAGAI | ASSLPRGSYAEQF | 0.49329913 |
| PAVHASL | ASSLPRGSYAEQF | 0.5779774 |
| TELEDDA | ASSLPRGSYAEQF | 0.3574838 |
| TPPGPPN | ASSLPRGSYAEQF | 0.37897405 |
| TPPPPPPP | ASSLPRGSYAEQF | 0.6689651 |
| TQSPALI | ASGRTANTGQLY | 0.31893563 |
| EFAAAFPQ | ASGRTANTGQLY | 0.33805954 |
| LLSGNLDVL | ASGRTANTGQLY | 0.5809222 |
| GPANLGNYNI | ASGRTANTGQLY | 0.32933438 |
| TLGPTLP | ASGRTANTGQLY | 0.62197423 |
| VYAYSSCCK | ASGRTANTGQLY | 0.60143787 |
| APAPAPAPNI | ASGRTANTGQLY | 0.5108362 |
| PGGPVPIP | ASGRTANTGQLY | 0.37188047 |
| FLTVALEP | ASGRTANTGQLY | 0.6944732 |
| SQYFGANL | ASGRTANTGQLY | 0.5700303 |
| GGVKPAA | ASGRTANTGQLY | 0.32039595 |

| | | |
|---|---|---|
| GGTGGAGGAGGW | ASGRTANTGQLY | 0.5979786 |
| GVGGVAGLGGAGG | ASGRTANTGQLY | 0.2812428 |
| VLLVLATTI | ASGRTANTGQLY | 0.6213846 |
| IGIFAAML | ASGRTANTGQLY | 0.61798334 |
| AVGDLID | ASGRTANTGQLY | 0.2720027 |
| NISPLPL | ASGRTANTGQLY | 0.4106949 |
| VTAVFVAI | ASGRTANTGQLY | 0.62996864 |
| SGDDNDAAL | ASGRTANTGQLY | 0.6301472 |
| AVAALAHS | ASGRTANTGQLY | 0.32179686 |
| GGVTVMATP | ASGRTANTGQLY | 0.3054657 |
| AGLAANVRD | ASGRTANTGQLY | 0.31405586 |
| TNRPDLIDPALLRPG | ASGRTANTGQLY | 0.37433827 |
| LANLGVTAL | ASGRTANTGQLY | 0.25525132 |
| TGERIEL | ASGRTANTGQLY | 0.56880355 |
| NGEELAA | ASGRTANTGQLY | 0.40204605 |
| VLSLVWG | ASGRTANTGQLY | 0.6369009 |
| IGYALWL | ASGRTANTGQLY | 0.5592321 |
| LSLLLAPL | ASGRTANTGQLY | 0.66950655 |
| TLDEALK | ASGRTANTGQLY | 0.37497655 |
| TDGAGTAEGG | ASGRTANTGQLY | 0.552968 |
| NAPGGAGGNGG | ASGRTANTGQLY | 0.23536143 |
| VSVFMGMGGNF | ASGRTANTGQLY | 0.5142019 |
| LASYLAAL | ASGRTANTGQLY | 0.26718006 |
| LVGGTPAL | ASGRTANTGQLY | 0.5699533 |
| VRNADVL | ASGRTANTGQLY | 0.23451862 |
| TCSSPFLVSAM | ASGRTANTGQLY | 0.6704868 |
| VHITDAE | ASGRTANTGQLY | 0.40175557 |
| EYAAPSF | ASGRTANTGQLY | 0.59506774 |
| VVDAFHGGL | ASGRTANTGQLY | 0.43320388 |
| ASVVDPQ | ASGRTANTGQLY | 0.33893007 |
| SYNEFGGLS | ASGRTANTGQLY | 0.39972878 |
| DPDGSVY | ASGRTANTGQLY | 0.6328466 |
| MVSTIHATE | ASGRTANTGQLY | 0.26795802 |
| LAAGVADPV | ASGRTANTGQLY | 0.19857298 |
| TKEQIAA | ASGRTANTGQLY | 0.28044727 |
| SLASPAAL | ASGRTANTGQLY | 0.21531916 |
| IGLVFYGV | ASGRTANTGQLY | 0.6659199 |
| FLLDASNL | ASGRTANTGQLY | 0.64382 |
| YQVGNFP | ASGRTANTGQLY | 0.47093236 |
| ESLLSQST | ASGRTANTGQLY | 0.6557052 |
| LFTVGFAAVG | ASGRTANTGQLY | 0.1932271 |
| ATPPTPT | ASGRTANTGQLY | 0.40169334 |
| TDLATLG | ASGRTANTGQLY | 0.682927 |
| SAGGTGR | ASGRTANTGQLY | 0.5133256 |
| GFSNSGS | ASGRTANTGQLY | 0.46205264 |
| GGNGGSGGTSV | ASGRTANTGQLY | 0.64800835 |
| GTSVATGGAGN | ASGRTANTGQLY | 0.43110847 |
| NGGNGGSGGTS | ASGRTANTGQLY | 0.4966094 |
| AVVKFPH | ASGRTANTGQLY | 0.52549887 |
| TPPPPPP | ASGRTANTGQLY | 0.5707744 |
| AAPAAAAAPA | ASGRTANTGQLY | 0.3919037 |

| | | |
|---|---|---|
| IALNEDL | ASGRTANTGQLY | 0.25386462 |
| IVNVSSM | ASGRTANTGQLY | 0.5761306 |
| TGIPPAPR | ASGRTANTGQLY | 0.6995538 |
| FDLGGGTF | ASGRTANTGQLY | 0.39707905 |
| FDLGGGTFDVS | ASGRTANTGQLY | 0.5547829 |
| NEPTAAA | ASGRTANTGQLY | 0.52162606 |
| NEPTAAALA | ASGRTANTGQLY | 0.47996122 |
| GSIINISSI | ASGRTANTGQLY | 0.32552123 |
| VAPGNAGTA | ASGRTANTGQLY | 0.29283717 |
| TGGGASLE | ASGRTANTGQLY | 0.54645777 |
| VAVYNHYKRI | ASGRTANTGQLY | 0.27215806 |
| TDPAPATT | ASGRTANTGQLY | 0.42381108 |
| TNRPDILDPALLRPG | ASGRTANTGQLY | 0.3538135 |
| LTTTTTS | ASGRTANTGQLY | 0.7305232 |
| PTGSGKTL | ASGRTANTGQLY | 0.57478523 |
| TGSGKTL | ASGRTANTGQLY | 0.53672266 |
| TIGHVDHGK | ASGRTANTGQLY | 0.35289025 |
| TIGHVDHGKTT | ASGRTANTGQLY | 0.40612274 |
| GHVDHGKTT | ASGRTANTGQLY | 0.58126605 |
| AQFEHTLL | ASGRTANTGQLY | 0.23370515 |
| IVNQDEL | ASGRTANTGQLY | 0.64754546 |
| TNGSQFF | ASGRTANTGQLY | 0.32881525 |
| TGKGIVE | ASGRTANTGQLY | 0.47169286 |
| LISNASDA | ASGRTANTGQLY | 0.57781875 |
| AELRTLGELTKIAKAHGAQ | ASGRTANTGQLY | 0.6741296 |
| APNAVAAAEPSW | ASGRTANTGQLY | 0.3930152 |
| APQVAGAATAVL | ASGRTANTGQLY | 0.4938776 |
| ARRDPAAAAAILPTDA | ASGRTANTGQLY | 0.3262175 |
| DAAARDVLAEAGLAEYF | ASGRTANTGQLY | 0.58928674 |
| DDQIVLSAKANLAKVEQ | ASGRTANTGQLY | 0.6629107 |
| DELLPDTGEDSDEDGH | ASGRTANTGQLY | 0.49983534 |
| ELPEAPANRATVPQD | ASGRTANTGQLY | 0.70135427 |
| FGQSKIERVTGIG | ASGRTANTGQLY | 0.5216252 |
| GAGGTGSTAGGAGGAGGAGGLY | ASGRTANTGQLY | 0.23608924 |
| GHGGTGAAGGNGGTGGAGGSGI | ASGRTANTGQLY | 0.48326403 |
| GNGGNGGAGGEGGPGGAGGAGA | ASGRTANTGQLY | 0.624159 |
| GSTAGNGGNGGNATNGTIAGQP | ASGRTANTGQLY | 0.64146423 |
| GTGGDGGNGGNGGTGG | ASGRTANTGQLY | 0.6806758 |
| GVRRPAGIPADAPV | ASGRTANTGQLY | 0.62930524 |
| GYRQVTDAHLLTLARR | ASGRTANTGQLY | 0.39561674 |
| IAKMGQDGHDRG | ASGRTANTGQLY | 0.6538042 |
| KAEPSIEIQYT | ASGRTANTGQLY | 0.70544183 |
| KPVRIRMAADEDTVAVT | ASGRTANTGQLY | 0.5858675 |
| MPPRIAGMRLLVIKPEP | ASGRTANTGQLY | 0.3327409 |
| MKFYAVALFRSRGVPPTR | ASGRTANTGQLY | 0.352192 |
| NAGGFGILGAVAHSP | ASGRTANTGQLY | 0.5328976 |
| NEGREITLAMAEM | ASGRTANTGQLY | 0.64346015 |
| NELVEWSHPLGHRWRWE | ASGRTANTGQLY | 0.26245075 |
| NLGAGGTGGNGGNPG | ASGRTANTGQLY | 0.5083214 |
| NTEWPKWETVMAADPVD | ASGRTANTGQLY | 0.5521784 |
| PASAAPAETQP | ASGRTANTGQLY | 0.22256055 |

| | | |
|---|---|---|
| QQSSDGKNGVRAAEITGE | ASGRTANTGQLY | 0.47343874 |
| RAPDAVTTSWWKE | ASGRTANTGQLY | 0.6544956 |
| SGFGNVGAHTSGWFNQ | ASGRTANTGQLY | 0.4784186 |
| SGVAGAVGQAASVGGLKV | ASGRTANTGQLY | 0.28368136 |
| SNGGAGTGSAGGNGGT | ASGRTANTGQLY | 0.65454304 |
| TGGAGGEGGAAG | ASGRTANTGQLY | 0.7005531 |
| TGGDGGAAGAGGAPGHG | ASGRTANTGQLY | 0.64808553 |
| TMPPLLAAIANAGRHGV | ASGRTANTGQLY | 0.2720475 |
| TSSTAGNSGGA | ASGRTANTGQLY | 0.19441925 |
| VPATVPVGILC | ASGRTANTGQLY | 0.54814446 |
| VQRPDSGGVVLTGRIS | ASGRTANTGQLY | 0.6243695 |
| WTPNPVPATKGVPAGLVQAT | ASGRTANTGQLY | 0.54696906 |
| YALKRALAKFSDRFDVVII | ASGRTANTGQLY | 0.3990956 |
| YLIGSVIV | ASGRTANTGQLY | 0.49119473 |
| YGQLVAVID | ASGRTANTGQLY | 0.6040679 |
| VTVGGMTLPA | ASGRTANTGQLY | 0.2849328 |
| VLALPAV | ASGRTANTGQLY | 0.26842713 |
| VIPPTLNYET | ASGRTANTGQLY | 0.44643152 |
| VGMGGMHPSSG | ASGRTANTGQLY | 0.59185576 |
| VGLAGGAATA | ASGRTANTGQLY | 0.30286872 |
| VGGAGGAGGSGP | ASGRTANTGQLY | 0.38000578 |
| VFAAFMV | ASGRTANTGQLY | 0.53088194 |
| TLYDKLR | ASGRTANTGQLY | 0.5467316 |
| TLAVFTVL | ASGRTANTGQLY | 0.54128265 |
| TGSVQPARV | ASGRTANTGQLY | 0.28059325 |
| TGGGAGSGGT | ASGRTANTGQLY | 0.6100954 |
| SHLETSL | ASGRTANTGQLY | 0.23118237 |
| SFTTGMT | ASGRTANTGQLY | 0.5415754 |
| QVEVAKRMGVRQPTVS | ASGRTANTGQLY | 0.59968853 |
| QGSGGPALAM | ASGRTANTGQLY | 0.5397858 |
| MLAPSNP | ASGRTANTGQLY | 0.41431767 |
| LVNNAAI | ASGRTANTGQLY | 0.28346714 |
| LTGEINTLA | ASGRTANTGQLY | 0.264497 |
| LSIFGPV | ASGRTANTGQLY | 0.26066846 |
| LLALLANL | ASGRTANTGQLY | 0.66312456 |
| LISNASDALDKLR | ASGRTANTGQLY | 0.52817976 |
| LINEAVP | ASGRTANTGQLY | 0.6462431 |
| LGYAFSGL | ASGRTANTGQLY | 0.5193014 |
| LDFVAVAL | ASGRTANTGQLY | 0.3018341 |
| LAQPGAE | ASGRTANTGQLY | 0.48134327 |
| ISNLGAVTGF | ASGRTANTGQLY | 0.3381301 |
| HQEDPGRI | ASGRTANTGQLY | 0.40235573 |
| GTPGEPTP | ASGRTANTGQLY | 0.46314812 |
| GFFNSGS | ASGRTANTGQLY | 0.56662315 |
| GAGGGGLTAGAN | ASGRTANTGQLY | 0.3020533 |
| FVDALVNM | ASGRTANTGQLY | 0.4757954 |
| FLTVAPEL | ASGRTANTGQLY | 0.53147376 |
| FETLTAL | ASGRTANTGQLY | 0.30578598 |
| DVINPIEL | ASGRTANTGQLY | 0.5011506 |
| APVSGAPV | ASGRTANTGQLY | 0.64483714 |
| ALVGGLAW | ASGRTANTGQLY | 0.5308521 |

| | | |
|---|---|---|
| ALAAAAATAI | ASGRTANTGQLY | 0.23376574 |
| AFGLGLLL | ASGRTANTGQLY | 0.5796764 |
| AFFAAASL | ASGRTANTGQLY | 0.32892296 |
| AAAALPAFSPP | ASGRTANTGQLY | 0.5203502 |
| VVLAPAL | ASGRTANTGQLY | 0.46358526 |
| TIGHVDHGKTTL | ASGRTANTGQLY | 0.502614 |
| LISNASDALDK | ASGRTANTGQLY | 0.6086912 |
| LISNASDAL | ASGRTANTGQLY | 0.25089923 |
| ISNASDALDK | ASGRTANTGQLY | 0.33940464 |
| NQAETLVYQTE | ASGRTANTGQLY | 0.43874282 |
| TGGTGGTAGAG | ASGRTANTGQLY | 0.55412906 |
| TGSTAGGAGGA | ASGRTANTGQLY | 0.5809672 |
| DPLAVPVAAGPV | ASGRTANTGQLY | 0.30096787 |
| GGTGGTPGNGGN | ASGRTANTGQLY | 0.52155256 |
| PAGLAGVDPATL | ASGRTANTGQLY | 0.438127 |
| GNNLVVEAPHAGASV | ASGRTANTGQLY | 0.37218562 |
| KVTRIMSASTLEQPA | ASGRTANTGQLY | 0.40650854 |
| LGFALPIALPRRTVV | ASGRTANTGQLY | 0.34254032 |
| TRLGNLTARKLVVLV | ASGRTANTGQLY | 0.26058015 |
| WSQQRRLIYFPSAGP | ASGRTANTGQLY | 0.6061411 |
| GPNGNPDPMAAAVDIR | ASGRTANTGQLY | 0.4223696 |
| TQRCVSVADLRAAASR | ASGRTANTGQLY | 0.5906228 |
| QFVLDVAEHPVEVLTETV | ASGRTANTGQLY | 0.6478025 |
| GGAGGTGGAGGNGGTQAGDG | ASGRTANTGQLY | 0.30092704 |
| PLLATLRAIAVDVDAKPAQV | ASGRTANTGQLY | 0.4666112 |
| APKFRTGGYMIDFWGVGYQVAK | ASGRTANTGQLY | 0.4965598 |
| GAGGAGGAGGAAGGPGGTGGTG | ASGRTANTGQLY | 0.58991337 |
| IGALGLLATDVAPAGSVAFGVA | ASGRTANTGQLY | 0.53533083 |
| AGGAGGIGGNGGAGGAGGVLMG | ASGRTANTGQLY | 0.2865186 |
| ADELIGLL | ASGRTANTGQLY | 0.6050844 |
| ADRALKA | ASGRTANTGQLY | 0.60409886 |
| AGAVGQVGGAGGA | ASGRTANTGQLY | 0.6415158 |
| AGGGGGLGGAP | ASGRTANTGQLY | 0.59134495 |
| AGGLLYGNGGAG | ASGRTANTGQLY | 0.53492594 |
| AGLESIGL | ASGRTANTGQLY | 0.58493364 |
| AGLKTGII | ASGRTANTGQLY | 0.4495445 |
| ASLAALLL | ASGRTANTGQLY | 0.25585073 |
| ASVINISSI | ASGRTANTGQLY | 0.52262914 |
| ATPPGPA | ASGRTANTGQLY | 0.49267727 |
| ATVASALV | ASGRTANTGQLY | 0.28456524 |
| ATVASLIL | ASGRTANTGQLY | 0.24606752 |
| AVNGPVLIRH | ASGRTANTGQLY | 0.6317956 |
| DGAAGAVGI | ASGRTANTGQLY | 0.7231492 |
| DPLLHGI | ASGRTANTGQLY | 0.48705754 |
| ELGAALGAL | ASGRTANTGQLY | 0.6845891 |
| ELGRTQAV | ASGRTANTGQLY | 0.23420915 |
| FVAGGYLS | ASGRTANTGQLY | 0.65323806 |
| GASGGAGGAGGW | ASGRTANTGQLY | 0.65154237 |
| GGAGAGGDGGAGGI | ASGRTANTGQLY | 0.31743777 |
| GTGHNGSQPGMGGNG | ASGRTANTGQLY | 0.6288337 |
| GYAAASASAS | ASGRTANTGQLY | 0.26105458 |

| | | |
|---|---|---|
| HQVDVAAVQQG | ASGRTANTGQLY | 0.32951707 |
| IGAAIAHT | ASGRTANTGQLY | 0.68159807 |
| IGMIEVAQL | ASGRTANTGQLY | 0.57663655 |
| IRLAPEHTSAMD | ASGRTANTGQLY | 0.5602938 |
| IYNVSVL | ASGRTANTGQLY | 0.48673576 |
| KQGFLGM | ASGRTANTGQLY | 0.3712896 |
| LESVITQL | ASGRTANTGQLY | 0.51730454 |
| LGLGALVVV | ASGRTANTGQLY | 0.6979908 |
| LIGNGGNGGSGGI | ASGRTANTGQLY | 0.3928579 |
| LNAINAPTL | ASGRTANTGQLY | 0.68044686 |
| LQLLNTGV | ASGRTANTGQLY | 0.35668573 |
| LQTSIEF | ASGRTANTGQLY | 0.64420956 |
| MSFAVSEL | ASGRTANTGQLY | 0.6480137 |
| NTGALGL | ASGRTANTGQLY | 0.51357955 |
| PPGAGGAGGAGTT | ASGRTANTGQLY | 0.38884017 |
| QLVGLGI | ASGRTANTGQLY | 0.6534295 |
| RFLEQVHQL | ASGRTANTGQLY | 0.4975221 |
| RGVGYVLR | ASGRTANTGQLY | 0.6052489 |
| SATGVFAGVI | ASGRTANTGQLY | 0.6465657 |
| SLPAITLP | ASGRTANTGQLY | 0.65376556 |
| STALAFTV | ASGRTANTGQLY | 0.76686746 |
| SVLTVGLGK | ASGRTANTGQLY | 0.74958766 |
| TALAGAGSGI | ASGRTANTGQLY | 0.73717785 |
| TGLPFTGI | ASGRTANTGQLY | 0.70912987 |
| VAEHVTV | ASGRTANTGQLY | 0.4203695 |
| VDIVLNSL | ASGRTANTGQLY | 0.7233572 |
| VLTGGGAVL | ASGRTANTGQLY | 0.6635548 |
| VNVGLAYGI | ASGRTANTGQLY | 0.74910617 |
| VSSAAVGAV | ASGRTANTGQLY | 0.7469847 |
| VTSVYEAGV | ASGRTANTGQLY | 0.64815545 |
| WVALDRGA | ASGRTANTGQLY | 0.7684481 |
| YLVGSVVI | ASGRTANTGQLY | 0.769102 |
| GPPGTGKTLL | ASGRTANTGQLY | 0.50903064 |
| GVGMTKF | ASGRTANTGQLY | 0.7234156 |
| IGIGGSDL | ASGRTANTGQLY | 0.68863297 |
| IGTLLAEL | ASGRTANTGQLY | 0.7344794 |
| IYTSGTT | ASGRTANTGQLY | 0.31222236 |
| LSFPAGV | ASGRTANTGQLY | 0.7658535 |
| LYTSGTT | ASGRTANTGQLY | 0.6972487 |
| VGAFPFGF | ASGRTANTGQLY | 0.23190199 |
| VGAGTVEF | ASGRTANTGQLY | 0.7258122 |
| VGKSTLLL | ASGRTANTGQLY | 0.71656644 |
| AALAHAGGGIV | ASGRTANTGQLY | 0.73494756 |
| AALVAVSLL | ASGRTANTGQLY | 0.76555204 |
| AAVSRAVAAL | ASGRTANTGQLY | 0.2914522 |
| AGGSSGIGGF | ASGRTANTGQLY | 0.6329607 |
| AGLTVANANG | ASGRTANTGQLY | 0.7592481 |
| AGLVGAVL | ASGRTANTGQLY | 0.76885974 |
| ALSQAFTV | ASGRTANTGQLY | 0.57876253 |
| ALTFSVSL | ASGRTANTGQLY | 0.75401187 |
| ALTGAALVAL | ASGRTANTGQLY | 0.7672628 |

| | | |
|---|---|---|
| ASFSIGAGVL | ASGRTANTGQLY | 0.7489288 |
| DGSVYKI | ASGRTANTGQLY | 0.7647031 |
| EQLAEIV | ASGRTANTGQLY | 0.7695478 |
| FAVSELTVM | ASGRTANTGQLY | 0.5635396 |
| GAAPLAAGL | ASGRTANTGQLY | 0.7675555 |
| GADGGDGGSGGAGG | ASGRTANTGQLY | 0.77040994 |
| GFGLDTGGAGG | ASGRTANTGQLY | 0.6525791 |
| GLASGGLL | ASGRTANTGQLY | 0.7412149 |
| GRIAGAAVLL | ASGRTANTGQLY | 0.6282613 |
| GSAGSLNTSF | ASGRTANTGQLY | 0.67668545 |
| GTGQIGI | ASGRTANTGQLY | 0.39044136 |
| GTLIALIL | ASGRTANTGQLY | 0.75137484 |
| IAATMGFVVG | ASGRTANTGQLY | 0.7676842 |
| IFAQALL | ASGRTANTGQLY | 0.7314526 |
| IGAAVALGF | ASGRTANTGQLY | 0.6207254 |
| IGLTPLAL | ASGRTANTGQLY | 0.59280086 |
| ILGSNTSTL | ASGRTANTGQLY | 0.67388606 |
| IVFSLTVL | ASGRTANTGQLY | 0.72002757 |
| LASKATGF | ASGRTANTGQLY | 0.3490097 |
| LASVALIL | ASGRTANTGQLY | 0.58480763 |
| LFAVGLTL | ASGRTANTGQLY | 0.7675637 |
| LFNGGIGTYI | ASGRTANTGQLY | 0.76806945 |
| LTLGAGII | ASGRTANTGQLY | 0.5436952 |
| LVAAALGDI | ASGRTANTGQLY | 0.7680288 |
| NTAFVQLGI | ASGRTANTGQLY | 0.76113087 |
| QLFSGAEL | ASGRTANTGQLY | 0.76992387 |
| SAATFITL | ASGRTANTGQLY | 0.7133899 |
| SNYGLANLGV | ASGRTANTGQLY | 0.65135556 |
| SYGGAYAVM | ASGRTANTGQLY | 0.7581192 |
| TTGVGGLTLG | ASGRTANTGQLY | 0.7102789 |
| TTQTIYI | ASGRTANTGQLY | 0.56933224 |
| VAAGAGLGAVI | ASGRTANTGQLY | 0.76343703 |
| VAAVVTLL | ASGRTANTGQLY | 0.7125125 |
| VGFLGGLI | ASGRTANTGQLY | 0.6907294 |
| VGVIPILL | ASGRTANTGQLY | 0.7170601 |
| VIFASSLI | ASGRTANTGQLY | 0.7660363 |
| VKGGDGGAAAG | ASGRTANTGQLY | 0.6494729 |
| VTKTFTNI | ASGRTANTGQLY | 0.53778815 |
| VVAAAILL | ASGRTANTGQLY | 0.64368606 |
| YAALDVEL | ASGRTANTGQLY | 0.7399461 |
| LASVALLL | ASGRTANTGQLY | 0.76615393 |
| LDAVHRV | ASGRTANTGQLY | 0.73851955 |
| QGLSSSIF | ASGRTANTGQLY | 0.6451561 |
| RGTFANIRL | ASGRTANTGQLY | 0.75552833 |
| RQLWWGHRI | ASGRTANTGQLY | 0.7460728 |
| YGPPGTGKTLL | ASGRTANTGQLY | 0.73826206 |
| AALVGAPANVA | ASGRTANTGQLY | 0.73509073 |
| AATFRRTE | ASGRTANTGQLY | 0.685568 |
| AGALGGGI | ASGRTANTGQLY | 0.7667258 |
| AGGAGGAGGIGAG | ASGRTANTGQLY | 0.7575794 |
| AGSGALGL | ASGRTANTGQLY | 0.7632499 |

| | | |
|---|---|---|
| AQIADQI | ASGRTANTGQLY | 0.7616922 |
| ASAVAVTYL | ASGRTANTGQLY | 0.6867656 |
| ASTVLGGTGG | ASGRTANTGQLY | 0.7286449 |
| AVGGEPAK | ASGRTANTGQLY | 0.45262524 |
| CEQGTFGGTC | ASGRTANTGQLY | 0.6064527 |
| DEGKAPAG | ASGRTANTGQLY | 0.73710823 |
| DTVVFPAL | ASGRTANTGQLY | 0.75130534 |
| FEGITMAG | ASGRTANTGQLY | 0.7559329 |
| FPLVVQD | ASGRTANTGQLY | 0.7496695 |
| GAFTTPQ | ASGRTANTGQLY | 0.6626552 |
| GFGGFGTAGGVGGA | ASGRTANTGQLY | 0.7543609 |
| GFLAQFL | ASGRTANTGQLY | 0.7630205 |
| GGAGGAGGNAGVA | ASGRTANTGQLY | 0.5851736 |
| GGVGTAGAGGAGGAGG | ASGRTANTGQLY | 0.6728072 |
| GNGGGGDAGNAGSGGN | ASGRTANTGQLY | 0.77262366 |
| GTGGAGGAGNDAGS | ASGRTANTGQLY | 0.74844694 |
| GTTNTGWFNS | ASGRTANTGQLY | 0.7138928 |
| IFMGPLFTL | ASGRTANTGQLY | 0.6987913 |
| ILAHAIEAI | ASGRTANTGQLY | 0.7712152 |
| IRSAIREAL | ASGRTANTGQLY | 0.65949404 |
| ITFAGGG | ASGRTANTGQLY | 0.7703443 |
| ITGMSGHQEV | ASGRTANTGQLY | 0.34250152 |
| IVAGLACTL | ASGRTANTGQLY | 0.6991295 |
| IVGGGLAGA | ASGRTANTGQLY | 0.6070714 |
| IVTGVLNI | ASGRTANTGQLY | 0.39511168 |
| LAQQGCI | ASGRTANTGQLY | 0.76834464 |
| LCLHVEN | ASGRTANTGQLY | 0.6919215 |
| LDAISDGS | ASGRTANTGQLY | 0.7604588 |
| LGGFAAML | ASGRTANTGQLY | 0.73014677 |
| LGGKNPMI | ASGRTANTGQLY | 0.7644721 |
| LGPKGVAAIL | ASGRTANTGQLY | 0.7433698 |
| LMVGVSI | ASGRTANTGQLY | 0.7207382 |
| LVGAAFVL | ASGRTANTGQLY | 0.53267586 |
| LVLAVAPL | ASGRTANTGQLY | 0.76515454 |
| MAAIAVIDAA | ASGRTANTGQLY | 0.76123637 |
| NNGINAELQ | ASGRTANTGQLY | 0.7566606 |
| RGEPLVLVHG | ASGRTANTGQLY | 0.51343215 |
| RGGAGGAAGLI | ASGRTANTGQLY | 0.6974284 |
| RSYRMHPAV | ASGRTANTGQLY | 0.7529489 |
| SAAHLLAGQLA | ASGRTANTGQLY | 0.26164123 |
| SAAVGSY | ASGRTANTGQLY | 0.27741134 |
| SPHAVLHRI | ASGRTANTGQLY | 0.70222056 |
| SQIAKVL | ASGRTANTGQLY | 0.6135261 |
| TALGHLVP | ASGRTANTGQLY | 0.7673378 |
| TAPTTSGAA | ASGRTANTGQLY | 0.75761056 |
| TDGDDGNGGSPGT | ASGRTANTGQLY | 0.7347839 |
| TDGLGEN | ASGRTANTGQLY | 0.74362683 |
| TGGPPLPGQA | ASGRTANTGQLY | 0.7381046 |
| TIPAITI | ASGRTANTGQLY | 0.20384604 |
| TIPGFTI | ASGRTANTGQLY | 0.6029779 |
| TSIASGALA | ASGRTANTGQLY | 0.5471035 |

| | | |
|---|---|---|
| TVTLPGAAI | ASGRTANTGQLY | 0.774534 |
| VASGLSAALP | ASGRTANTGQLY | 0.59280676 |
| VGAGVIGI | ASGRTANTGQLY | 0.66298676 |
| VGPKRFGV | ASGRTANTGQLY | 0.7416985 |
| VNVAFPD | ASGRTANTGQLY | 0.74523866 |
| VSMTNSGVS | ASGRTANTGQLY | 0.73196584 |
| VTDVLGDLSV | ASGRTANTGQLY | 0.6030469 |
| VVVVGAGFAGL | ASGRTANTGQLY | 0.24413492 |
| VYTVGTVV | ASGRTANTGQLY | 0.76421165 |
| WTTFSGP | ASGRTANTGQLY | 0.35444987 |
| YDELVEAF | ASGRTANTGQLY | 0.5832054 |
| YGTGGAGGNGG | ASGRTANTGQLY | 0.31606197 |
| YTNVMAV | ASGRTANTGQLY | 0.26772404 |
| FLEQVHQL | ASGRTANTGQLY | 0.46082926 |
| GLSLLAVAV | ASGRTANTGQLY | 0.758034 |
| LEQVHQL | ASGRTANTGQLY | 0.7203616 |
| LNAAGAI | ASGRTANTGQLY | 0.70688474 |
| PAVHASL | ASGRTANTGQLY | 0.7727088 |
| TELEDDA | ASGRTANTGQLY | 0.7483845 |
| TPPGPPN | ASGRTANTGQLY | 0.7070283 |
| TPPPPPPP | ASGRTANTGQLY | 0.6639581 |
| TQSPALI | GARDRDRDYAEQF | 0.6502929 |
| EFAAAFPQ | GARDRDRDYAEQF | 0.6490599 |
| LLSGNLDVL | GARDRDRDYAEQF | 0.6463319 |
| GPANLGNYNI | GARDRDRDYAEQF | 0.77316016 |
| TLGPTLP | GARDRDRDYAEQF | 0.70289135 |
| VYAYSSCCK | GARDRDRDYAEQF | 0.52723575 |
| APAPAPAPNI | GARDRDRDYAEQF | 0.7208898 |
| PGGPVPIP | GARDRDRDYAEQF | 0.7728257 |
| FLTVALEP | GARDRDRDYAEQF | 0.7755206 |
| SQYFGANL | GARDRDRDYAEQF | 0.58626497 |
| GGVKPAA | GARDRDRDYAEQF | 0.6750094 |
| GGTGGAGGAGGW | GARDRDRDYAEQF | 0.67985404 |
| GVGGVAGLGGAGG | GARDRDRDYAEQF | 0.74014664 |
| VLLVLATTI | GARDRDRDYAEQF | 0.59701514 |
| IGIFAAML | GARDRDRDYAEQF | 0.52261585 |
| AVGDLID | GARDRDRDYAEQF | 0.7600268 |
| NISPLPL | GARDRDRDYAEQF | 0.77746373 |
| VTAVFVAI | GARDRDRDYAEQF | 0.6467756 |
| SGDDNDAAL | GARDRDRDYAEQF | 0.61615396 |
| AVAALAHS | GARDRDRDYAEQF | 0.72551113 |
| GGVTVMATP | GARDRDRDYAEQF | 0.67297864 |
| AGLAANVRD | GARDRDRDYAEQF | 0.7127119 |
| TNRPDLIDPALLRPG | GARDRDRDYAEQF | 0.7137656 |
| LANLGVTAL | GARDRDRDYAEQF | 0.58291113 |
| TGERIEL | GARDRDRDYAEQF | 0.71257424 |
| NGEELAA | GARDRDRDYAEQF | 0.7743572 |
| VLSLVWG | GARDRDRDYAEQF | 0.732335 |
| IGYALWL | GARDRDRDYAEQF | 0.7482221 |
| LSLLLAPL | GARDRDRDYAEQF | 0.7688168 |
| TLDEALK | GARDRDRDYAEQF | 0.7589183 |

| | | |
|---|---|---|
| TDGAGTAEGG | GARDRDRDYAEQF | 0.771321 |
| NAPGGAGGNGG | GARDRDRDYAEQF | 0.75946164 |
| VSVFMGMGGNF | GARDRDRDYAEQF | 0.6875199 |
| LASYLAAL | GARDRDRDYAEQF | 0.70899546 |
| LVGGTPAL | GARDRDRDYAEQF | 0.7430178 |
| VRNADVL | GARDRDRDYAEQF | 0.73883724 |
| TCSSPFLVSAM | GARDRDRDYAEQF | 0.72880673 |
| VHITDAE | GARDRDRDYAEQF | 0.64037025 |
| EYAAPSF | GARDRDRDYAEQF | 0.7707997 |
| VVDAFHGGL | GARDRDRDYAEQF | 0.6992103 |
| ASVVDPQ | GARDRDRDYAEQF | 0.76571095 |
| SYNEFGGLS | GARDRDRDYAEQF | 0.51425683 |
| DPDGSVY | GARDRDRDYAEQF | 0.60783863 |
| MVSTIHATE | GARDRDRDYAEQF | 0.7646799 |
| LAAGVADPV | GARDRDRDYAEQF | 0.21598321 |
| TKEQIAA | GARDRDRDYAEQF | 0.3013239 |
| SLASPAAL | GARDRDRDYAEQF | 0.7398945 |
| IGLVFYGV | GARDRDRDYAEQF | 0.76773524 |
| FLLDASNL | GARDRDRDYAEQF | 0.2205618 |
| YQVGNFP | GARDRDRDYAEQF | 0.5435605 |
| ESLLSQST | GARDRDRDYAEQF | 0.5479021 |
| LFTVGFAAVG | GARDRDRDYAEQF | 0.72854996 |
| ATPPTPT | GARDRDRDYAEQF | 0.735638 |
| TDLATLG | GARDRDRDYAEQF | 0.37148792 |
| SAGGTGR | GARDRDRDYAEQF | 0.7638954 |
| GFSNSGS | GARDRDRDYAEQF | 0.52110386 |
| GGNGGSGGTSV | GARDRDRDYAEQF | 0.40766945 |
| GTSVATGGAGN | GARDRDRDYAEQF | 0.7580297 |
| NGGNGGSGGTS | GARDRDRDYAEQF | 0.7689176 |
| AVVKFPH | GARDRDRDYAEQF | 0.52284324 |
| TPPPPPP | GARDRDRDYAEQF | 0.7605655 |
| AAPAAAAAPA | GARDRDRDYAEQF | 0.6846089 |
| IALNEDL | GARDRDRDYAEQF | 0.7745439 |
| IVNVSSM | GARDRDRDYAEQF | 0.7119417 |
| TGIPPAPR | GARDRDRDYAEQF | 0.47008002 |
| FDLGGGTF | GARDRDRDYAEQF | 0.7659316 |
| FDLGGGTFDVS | GARDRDRDYAEQF | 0.7513493 |
| NEPTAAA | GARDRDRDYAEQF | 0.70272017 |
| NEPTAAALA | GARDRDRDYAEQF | 0.76271164 |
| GSIINISSI | GARDRDRDYAEQF | 0.60875356 |
| VAPGNAGTA | GARDRDRDYAEQF | 0.7356517 |
| TGGGASLE | GARDRDRDYAEQF | 0.7684388 |
| VAVYNHYKRI | GARDRDRDYAEQF | 0.76106864 |
| TDPAPATT | GARDRDRDYAEQF | 0.55023247 |
| TNRPDILDPALLRPG | GARDRDRDYAEQF | 0.755677 |
| LTTTTTS | GARDRDRDYAEQF | 0.73359215 |
| PTGSGKTL | GARDRDRDYAEQF | 0.65968835 |
| TGSGKTL | GARDRDRDYAEQF | 0.7753179 |
| TIGHVDHGK | GARDRDRDYAEQF | 0.71228266 |
| TIGHVDHGKTT | GARDRDRDYAEQF | 0.76092124 |
| GHVDHGKTT | GARDRDRDYAEQF | 0.7683852 |

| | | |
|---|---|---|
| AQFEHTLL | GARDRDRDYAEQF | 0.76756024 |
| IVNQDEL | GARDRDRDYAEQF | 0.44547617 |
| TNGSQFF | GARDRDRDYAEQF | 0.73783445 |
| TGKGIVE | GARDRDRDYAEQF | 0.7201309 |
| LISNASDA | GARDRDRDYAEQF | 0.7602093 |
| AELRTLGELTKIAKAHGAQ | GARDRDRDYAEQF | 0.7347108 |
| APNAVAAAEPSW | GARDRDRDYAEQF | 0.57582206 |
| APQVAGAATAVL | GARDRDRDYAEQF | 0.74965245 |
| ARRDPAAAAAILPTDA | GARDRDRDYAEQF | 0.7689549 |
| DAAARDVLAEAGLAEYF | GARDRDRDYAEQF | 0.7261703 |
| DDQIVLSAKANLAKVEQ | GARDRDRDYAEQF | 0.7135669 |
| DELLPDTGEDSDEDGH | GARDRDRDYAEQF | 0.7750114 |
| ELPEAPANRATVPQD | GARDRDRDYAEQF | 0.6793592 |
| FGQSKIERVTGIG | GARDRDRDYAEQF | 0.7258227 |
| GAGGTGSTAGGAGGAGGAGGLY | GARDRDRDYAEQF | 0.7579664 |
| GHGGTGAAGGNGGTGGAGGSGI | GARDRDRDYAEQF | 0.76089805 |
| GNGGNGGAGGEGGPGGAGGAGA | GARDRDRDYAEQF | 0.71851385 |
| GSTAGNGGNGGNATNGTIAGQP | GARDRDRDYAEQF | 0.7636047 |
| GTGGDGGNGGNGGTGG | GARDRDRDYAEQF | 0.47505045 |
| GVRRPAGIPADAPV | GARDRDRDYAEQF | 0.7530377 |
| GYRQVTDAHLLTLARR | GARDRDRDYAEQF | 0.6387624 |
| IAKMGQDGHDRG | GARDRDRDYAEQF | 0.75179374 |
| KAEPSIEIQYT | GARDRDRDYAEQF | 0.6280813 |
| KPVRIRMAADEDTVAVT | GARDRDRDYAEQF | 0.75449467 |
| MPPRIAGMRLLVIKPEP | GARDRDRDYAEQF | 0.76188654 |
| MKFYAVALFRSRGVPPTR | GARDRDRDYAEQF | 0.7630155 |
| NAGGFGILGAVAHSP | GARDRDRDYAEQF | 0.2858225 |
| NEGREITLAMAEM | GARDRDRDYAEQF | 0.7610588 |
| NELVEWSHPLGHRWRWE | GARDRDRDYAEQF | 0.759308 |
| NLGAGGTGGNGGNPG | GARDRDRDYAEQF | 0.77719915 |
| NTEWPKWETVMAADPVD | GARDRDRDYAEQF | 0.6912964 |
| PASAAPAETQP | GARDRDRDYAEQF | 0.7502905 |
| QQSSDGKNGVRAAEITGE | GARDRDRDYAEQF | 0.6679946 |
| RAPDAVTTSWWKE | GARDRDRDYAEQF | 0.7203581 |
| SGFGNVGAHTSGWFNQ | GARDRDRDYAEQF | 0.64019036 |
| SGVAGAVGQAASVGGLKV | GARDRDRDYAEQF | 0.6821579 |
| SNGGAGTGSAGGNGGT | GARDRDRDYAEQF | 0.680729 |
| TGGAGGEGGAAG | GARDRDRDYAEQF | 0.6958675 |
| TGGDGGAAGAGGAPGHG | GARDRDRDYAEQF | 0.7593432 |
| TMPPLLAAIANAGRHGV | GARDRDRDYAEQF | 0.76054597 |
| TSSTAGNSGGA | GARDRDRDYAEQF | 0.75947666 |
| VPATVPVGILC | GARDRDRDYAEQF | 0.7702873 |
| VQRPDSGGVVLTGRIS | GARDRDRDYAEQF | 0.7673298 |
| WTPNPVPATKGVPAGLVQAT | GARDRDRDYAEQF | 0.7648283 |
| YALKRALAKFSDRFDVVII | GARDRDRDYAEQF | 0.7065574 |
| YLIGSVIV | GARDRDRDYAEQF | 0.6920973 |
| YGQLVAVID | GARDRDRDYAEQF | 0.646364 |
| VTVGGMTLPA | GARDRDRDYAEQF | 0.69722784 |
| VLALPAV | GARDRDRDYAEQF | 0.7082286 |
| VIPPTLNYET | GARDRDRDYAEQF | 0.7461575 |
| VGMGGMHPSSG | GARDRDRDYAEQF | 0.21321055 |

| | | |
|---|---|---|
| VGLAGGAATA | GARDRDRDYAEQF | 0.7328127 |
| VGGAGGAGGSGP | GARDRDRDYAEQF | 0.7715858 |
| VFAAFMV | GARDRDRDYAEQF | 0.6446397 |
| TLYDKLR | GARDRDRDYAEQF | 0.7543236 |
| TLAVFTVL | GARDRDRDYAEQF | 0.7239624 |
| TGSVQPARV | GARDRDRDYAEQF | 0.6860087 |
| TGGGAGSGGT | GARDRDRDYAEQF | 0.762841 |
| SHLETSL | GARDRDRDYAEQF | 0.65495884 |
| SFTTGMT | GARDRDRDYAEQF | 0.7599765 |
| QVEVAKRMGVRQPTVS | GARDRDRDYAEQF | 0.77470756 |
| QGSGGPALAM | GARDRDRDYAEQF | 0.7481221 |
| MLAPSNP | GARDRDRDYAEQF | 0.69572186 |
| LVNNAAI | GARDRDRDYAEQF | 0.7268432 |
| LTGEINTLA | GARDRDRDYAEQF | 0.43100923 |
| LSIFGPV | GARDRDRDYAEQF | 0.7214612 |
| LLALLANL | GARDRDRDYAEQF | 0.20809518 |
| LISNASDALDKLR | GARDRDRDYAEQF | 0.7369057 |
| LINEAVP | GARDRDRDYAEQF | 0.6238711 |
| LGYAFSGL | GARDRDRDYAEQF | 0.753924 |
| LDFVAVAL | GARDRDRDYAEQF | 0.73780006 |
| LAQPGAE | GARDRDRDYAEQF | 0.7411556 |
| ISNLGAVTGF | GARDRDRDYAEQF | 0.66632485 |
| HQEDPGRI | GARDRDRDYAEQF | 0.7313554 |
| GTPGEPTP | GARDRDRDYAEQF | 0.67245966 |
| GFFNSGS | GARDRDRDYAEQF | 0.5232873 |
| GAGGGGLTAGAN | GARDRDRDYAEQF | 0.7574791 |
| FVDALVNM | GARDRDRDYAEQF | 0.7142725 |
| FLTVAPEL | GARDRDRDYAEQF | 0.6933982 |
| FETLTAL | GARDRDRDYAEQF | 0.77005255 |
| DVINPIEL | GARDRDRDYAEQF | 0.7387241 |
| APVSGAPV | GARDRDRDYAEQF | 0.43962783 |
| ALVGGLAW | GARDRDRDYAEQF | 0.7276337 |
| ALAAAAATAI | GARDRDRDYAEQF | 0.73717844 |
| AFGLGLLL | GARDRDRDYAEQF | 0.7308539 |
| AFFAAASL | GARDRDRDYAEQF | 0.20769164 |
| AAAALPAFSPP | GARDRDRDYAEQF | 0.7042291 |
| VVLAPAL | GARDRDRDYAEQF | 0.6674631 |
| TIGHVDHGKTTL | GARDRDRDYAEQF | 0.6334559 |
| LISNASDALDK | GARDRDRDYAEQF | 0.6826425 |
| LISNASDAL | GARDRDRDYAEQF | 0.7659308 |
| ISNASDALDK | GARDRDRDYAEQF | 0.3577902 |
| NQAETLVYQTE | GARDRDRDYAEQF | 0.73800826 |
| TGGTGGTAGAG | GARDRDRDYAEQF | 0.7543062 |
| TGSTAGGAGGA | GARDRDRDYAEQF | 0.75033677 |
| DPLAVPVAAGPV | GARDRDRDYAEQF | 0.7710612 |
| GGTGGTPGNGGN | GARDRDRDYAEQF | 0.7281054 |
| PAGLAGVDPATL | GARDRDRDYAEQF | 0.69601333 |
| GNNLVVEAPHAGASV | GARDRDRDYAEQF | 0.74126387 |
| KVTRIMSASTLEQPA | GARDRDRDYAEQF | 0.6374732 |
| LGFALPIALPRRTVV | GARDRDRDYAEQF | 0.39342728 |
| TRLGNLTARKLVVLV | GARDRDRDYAEQF | 0.4428934 |

| | | |
|---|---|---|
| WSQQRRLIYFPSAGP | GARDRDRDYAEQF | 0.74202645 |
| GPNGNPDPMAAAVDIR | GARDRDRDYAEQF | 0.40029174 |
| TQRCVSVADLRAAASR | GARDRDRDYAEQF | 0.74505234 |
| QFVLDVAEHPVEVLTETV | GARDRDRDYAEQF | 0.76584876 |
| GGAGGTGGAGGNGGTQAGDG | GARDRDRDYAEQF | 0.6243441 |
| PLLATLRAIAVDVDAKPAQV | GARDRDRDYAEQF | 0.46578255 |
| APKFRTGGYMIDFWGVGYQVAK | GARDRDRDYAEQF | 0.69876856 |
| GAGGAGGAGGAAGGPGGTGGTG | GARDRDRDYAEQF | 0.65867114 |
| IGALGLLATDVAPAGSVAFGVA | GARDRDRDYAEQF | 0.76192915 |
| AGGAGGIGGNGGAGGAGGVLMG | GARDRDRDYAEQF | 0.7539556 |
| ADELIGLL | GARDRDRDYAEQF | 0.77201957 |
| ADRALKA | GARDRDRDYAEQF | 0.76293546 |
| AGAVGQVGGAGGA | GARDRDRDYAEQF | 0.71362007 |
| AGGGGGLGGAP | GARDRDRDYAEQF | 0.760088 |
| AGGLLYGNGGAG | GARDRDRDYAEQF | 0.7177485 |
| AGLESIGL | GARDRDRDYAEQF | 0.71263397 |
| AGLKTGII | GARDRDRDYAEQF | 0.70556414 |
| ASLAALLL | GARDRDRDYAEQF | 0.72614986 |
| ASVINISSI | GARDRDRDYAEQF | 0.6599258 |
| ATPPGPA | GARDRDRDYAEQF | 0.66157997 |
| ATVASALV | GARDRDRDYAEQF | 0.47549754 |
| ATVASLIL | GARDRDRDYAEQF | 0.75044614 |
| AVNGPVLIRH | GARDRDRDYAEQF | 0.73121315 |
| DGAAGAVGI | GARDRDRDYAEQF | 0.43793917 |
| DPLLHGI | GARDRDRDYAEQF | 0.66282153 |
| ELGAALGAL | GARDRDRDYAEQF | 0.7731845 |
| ELGRTQAV | GARDRDRDYAEQF | 0.76877964 |
| FVAGGYLS | GARDRDRDYAEQF | 0.680027 |
| GASGGAGGAGGW | GARDRDRDYAEQF | 0.6582804 |
| GGAGAGGDGGAGGI | GARDRDRDYAEQF | 0.72858465 |
| GTGHNGSQPGMGGNG | GARDRDRDYAEQF | 0.49866784 |
| GYAAASASAS | GARDRDRDYAEQF | 0.7475338 |
| HQVDVAAVQQG | GARDRDRDYAEQF | 0.73356605 |
| IGAAIAHT | GARDRDRDYAEQF | 0.71491873 |
| IGMIEVAQL | GARDRDRDYAEQF | 0.5832979 |
| IRLAPEHTSAMD | GARDRDRDYAEQF | 0.7452544 |
| IYNVSVL | GARDRDRDYAEQF | 0.6829685 |
| KQGFLGM | GARDRDRDYAEQF | 0.7505393 |
| LESVITQL | GARDRDRDYAEQF | 0.22231865 |
| LGLGALVVV | GARDRDRDYAEQF | 0.7120838 |
| LIGNGGNGGSGGI | GARDRDRDYAEQF | 0.64588976 |
| LNAINAPTL | GARDRDRDYAEQF | 0.586564 |
| LQLLNTGV | GARDRDRDYAEQF | 0.3352176 |
| LQTSIEF | GARDRDRDYAEQF | 0.72952294 |
| MSFAVSEL | GARDRDRDYAEQF | 0.7635523 |
| NTGALGL | GARDRDRDYAEQF | 0.76630116 |
| PPGAGGAGGAGTT | GARDRDRDYAEQF | 0.58849275 |
| QLVGLGI | GARDRDRDYAEQF | 0.76163614 |
| RFLEQVHQL | GARDRDRDYAEQF | 0.77073956 |
| RGVGYVLR | GARDRDRDYAEQF | 0.69897974 |
| SATGVFAGVI | GARDRDRDYAEQF | 0.31898355 |

| | | |
|---|---|---|
| SLPAITLP | GARDRDRDYAEQF | 0.7566098 |
| STALAFTV | GARDRDRDYAEQF | 0.6736208 |
| SVLTVGLGK | GARDRDRDYAEQF | 0.75558174 |
| TALAGAGSGI | GARDRDRDYAEQF | 0.7461185 |
| TGLPFTGI | GARDRDRDYAEQF | 0.62137496 |
| VAEHVTV | GARDRDRDYAEQF | 0.76024234 |
| VDIVLNSL | GARDRDRDYAEQF | 0.76342547 |
| VLTGGGAVL | GARDRDRDYAEQF | 0.6227375 |
| VNVGLAYGI | GARDRDRDYAEQF | 0.5985724 |
| VSSAAVGAV | GARDRDRDYAEQF | 0.6193751 |
| VTSVYEAGV | GARDRDRDYAEQF | 0.38910902 |
| WVALDRGA | GARDRDRDYAEQF | 0.7543063 |
| YLVGSVVI | GARDRDRDYAEQF | 0.7025398 |
| GPPGTGKTLL | GARDRDRDYAEQF | 0.6718099 |
| GVGMTKF | GARDRDRDYAEQF | 0.7545303 |
| IGIGGSDL | GARDRDRDYAEQF | 0.6397097 |
| IGTLLAEL | GARDRDRDYAEQF | 0.7095769 |
| IYTSGTT | GARDRDRDYAEQF | 0.70576894 |
| LSFPAGV | GARDRDRDYAEQF | 0.72623056 |
| LYTSGTT | GARDRDRDYAEQF | 0.7514962 |
| VGAFPFGF | GARDRDRDYAEQF | 0.7196026 |
| VGAGTVEF | GARDRDRDYAEQF | 0.73233134 |
| VGKSTLLL | GARDRDRDYAEQF | 0.77086663 |
| AALAHAGGGIV | GARDRDRDYAEQF | 0.47403544 |
| AALVAVSLL | GARDRDRDYAEQF | 0.65425503 |
| AAVSRAVAAL | GARDRDRDYAEQF | 0.7521836 |
| AGGSSGIGGF | GARDRDRDYAEQF | 0.57803863 |
| AGLTVANANG | GARDRDRDYAEQF | 0.7586039 |
| AGLVGAVL | GARDRDRDYAEQF | 0.73217285 |
| ALSQAFTV | GARDRDRDYAEQF | 0.75019526 |
| ALTFSVSL | GARDRDRDYAEQF | 0.68991685 |
| ALTGAALVAL | GARDRDRDYAEQF | 0.7728164 |
| ASFSIGAGVL | GARDRDRDYAEQF | 0.76817703 |
| DGSVYKI | GARDRDRDYAEQF | 0.7273501 |
| EQLAEIV | GARDRDRDYAEQF | 0.7284857 |
| FAVSELTVM | GARDRDRDYAEQF | 0.55698454 |
| GAAPLAAGL | GARDRDRDYAEQF | 0.7606328 |
| GADGGDGGSGGAGG | GARDRDRDYAEQF | 0.7575951 |
| GFGLDTGGAGG | GARDRDRDYAEQF | 0.6501622 |
| GLASGGLL | GARDRDRDYAEQF | 0.73606294 |
| GRIAGAAVLL | GARDRDRDYAEQF | 0.7686305 |
| GSAGSLNTSF | GARDRDRDYAEQF | 0.76772976 |
| GTGQIGI | GARDRDRDYAEQF | 0.4859576 |
| GTLIALIL | GARDRDRDYAEQF | 0.71562266 |
| IAATMGFVVG | GARDRDRDYAEQF | 0.33764014 |
| IFAQALL | GARDRDRDYAEQF | 0.53570735 |
| IGAAVALGF | GARDRDRDYAEQF | 0.5760945 |
| IGLTPLAL | GARDRDRDYAEQF | 0.75304145 |
| ILGSNTSTL | GARDRDRDYAEQF | 0.703948 |
| IVFSLTVL | GARDRDRDYAEQF | 0.7528814 |
| LASKATGF | GARDRDRDYAEQF | 0.7486971 |

| | | |
|---|---|---|
| LASVALIL | GARDRDRDYAEQF | 0.7738423 |
| LFAVGLTL | GARDRDRDYAEQF | 0.6716813 |
| LFNGGIGTYI | GARDRDRDYAEQF | 0.7099086 |
| LTLGAGII | GARDRDRDYAEQF | 0.42455614 |
| LVAAALGDI | GARDRDRDYAEQF | 0.38836253 |
| NTAFVQLGI | GARDRDRDYAEQF | 0.6457431 |
| QLFSGAEL | GARDRDRDYAEQF | 0.7259102 |
| SAATFITL | GARDRDRDYAEQF | 0.24908996 |
| SNYGLANLGV | GARDRDRDYAEQF | 0.7656147 |
| SYGGAYAVM | GARDRDRDYAEQF | 0.74920607 |
| TTGVGGLTLG | GARDRDRDYAEQF | 0.76578546 |
| TTQTIYI | GARDRDRDYAEQF | 0.58364856 |
| VAAGAGLGAVI | GARDRDRDYAEQF | 0.67900574 |
| VAAVVTLL | GARDRDRDYAEQF | 0.69792247 |
| VGFLGGLI | GARDRDRDYAEQF | 0.7562356 |
| VGVIPILL | GARDRDRDYAEQF | 0.5789159 |
| VIFASSLI | GARDRDRDYAEQF | 0.506757 |
| VKGGDGGAAAG | GARDRDRDYAEQF | 0.6124759 |
| VTKTFTNI | GARDRDRDYAEQF | 0.55432725 |
| VVAAAILL | GARDRDRDYAEQF | 0.7695743 |
| YAALDVEL | GARDRDRDYAEQF | 0.76702815 |
| LASVALLL | GARDRDRDYAEQF | 0.7543227 |
| LDAVHRV | GARDRDRDYAEQF | 0.7633402 |
| QGLSSSIF | GARDRDRDYAEQF | 0.25924098 |
| RGTFANIRL | GARDRDRDYAEQF | 0.5823532 |
| RQLWWGHRI | GARDRDRDYAEQF | 0.7732828 |
| YGPPGTGKTLL | GARDRDRDYAEQF | 0.7542136 |
| AALVGAPANVA | GARDRDRDYAEQF | 0.71853936 |
| AATFRRTE | GARDRDRDYAEQF | 0.7691265 |
| AGALGGGI | GARDRDRDYAEQF | 0.67657363 |
| AGGAGGAGGIGAG | GARDRDRDYAEQF | 0.72240263 |
| AGSGALGL | GARDRDRDYAEQF | 0.7408885 |
| AQIADQI | GARDRDRDYAEQF | 0.73664355 |
| ASAVAVTYL | GARDRDRDYAEQF | 0.7384708 |
| ASTVLGGTGG | GARDRDRDYAEQF | 0.58049846 |
| AVGGEPAK | GARDRDRDYAEQF | 0.74841964 |
| CEQGTFGGTC | GARDRDRDYAEQF | 0.7743479 |
| DEGKAPAG | GARDRDRDYAEQF | 0.73212135 |
| DTVVFPAL | GARDRDRDYAEQF | 0.76122606 |
| FEGITMAG | GARDRDRDYAEQF | 0.75067234 |
| FPLVVQD | GARDRDRDYAEQF | 0.7570288 |
| GAFTTPQ | GARDRDRDYAEQF | 0.67999125 |
| GFGGFGTAGGVGGA | GARDRDRDYAEQF | 0.7046821 |
| GFLAQFL | GARDRDRDYAEQF | 0.72846234 |
| GGAGGAGGNAGVA | GARDRDRDYAEQF | 0.5924529 |
| GGVGTAGAGGAGGAGG | GARDRDRDYAEQF | 0.7214973 |
| GNGGGGDAGNAGSGGN | GARDRDRDYAEQF | 0.70041835 |
| GTGGAGGAGNDAGS | GARDRDRDYAEQF | 0.77807224 |
| GTTNTGWFNS | GARDRDRDYAEQF | 0.75279605 |
| IFMGPLFTL | GARDRDRDYAEQF | 0.7483176 |
| ILAHAIEAI | GARDRDRDYAEQF | 0.6735631 |

| | | |
|---|---|---|
| IRSAIREAL | GARDRDRDYAEQF | 0.7389929 |
| ITFAGGG | GARDRDRDYAEQF | 0.751606 |
| ITGMSGHQEV | GARDRDRDYAEQF | 0.53793013 |
| IVAGLACTL | GARDRDRDYAEQF | 0.7679516 |
| IVGGGLAGA | GARDRDRDYAEQF | 0.67198694 |
| IVTGVLNI | GARDRDRDYAEQF | 0.65007734 |
| LAQQGCI | GARDRDRDYAEQF | 0.75637555 |
| LCLHVEN | GARDRDRDYAEQF | 0.77060854 |
| LDAISDGS | GARDRDRDYAEQF | 0.6992421 |
| LGGFAAML | GARDRDRDYAEQF | 0.7180849 |
| LGGKNPMI | GARDRDRDYAEQF | 0.69345576 |
| LGPKGVAAIL | GARDRDRDYAEQF | 0.76241827 |
| LMVGVSI | GARDRDRDYAEQF | 0.77088827 |
| LVGAAFVL | GARDRDRDYAEQF | 0.69085276 |
| LVLAVAPL | GARDRDRDYAEQF | 0.7712497 |
| MAAIAVIDAA | GARDRDRDYAEQF | 0.71275926 |
| NNGINAELQ | GARDRDRDYAEQF | 0.40620643 |
| RGEPLVLVHG | GARDRDRDYAEQF | 0.72390515 |
| RGGAGGAAGLI | GARDRDRDYAEQF | 0.7594038 |
| RSYRMHPAV | GARDRDRDYAEQF | 0.7639054 |
| SAAHLLAGQLA | GARDRDRDYAEQF | 0.77406794 |
| SAAVGSY | GARDRDRDYAEQF | 0.73488235 |
| SPHAVLHRI | GARDRDRDYAEQF | 0.6728809 |
| SQIAKVL | GARDRDRDYAEQF | 0.7709598 |
| TALGHLVP | GARDRDRDYAEQF | 0.7741735 |
| TAPTTSGAA | GARDRDRDYAEQF | 0.76393557 |
| TDGDDGNGGSPGT | GARDRDRDYAEQF | 0.6759536 |
| TDGLGEN | GARDRDRDYAEQF | 0.5649106 |
| TGGPPLPGQA | GARDRDRDYAEQF | 0.74163306 |
| TIPAITI | GARDRDRDYAEQF | 0.7614029 |
| TIPGFTI | GARDRDRDYAEQF | 0.6063601 |
| TSIASGALA | GARDRDRDYAEQF | 0.73825026 |
| TVTLPGAAI | GARDRDRDYAEQF | 0.7515619 |
| VASGLSAALP | GARDRDRDYAEQF | 0.5001191 |
| VGAGVIGI | GARDRDRDYAEQF | 0.7229638 |
| VGPKRFGV | GARDRDRDYAEQF | 0.7625127 |
| VNVAFPD | GARDRDRDYAEQF | 0.7320732 |
| VSMTNSGVS | GARDRDRDYAEQF | 0.6998059 |
| VTDVLGDLSV | GARDRDRDYAEQF | 0.7620002 |
| VVVVGAGFAGL | GARDRDRDYAEQF | 0.7773925 |
| VYTVGTVV | GARDRDRDYAEQF | 0.77348423 |
| WTTFSGP | GARDRDRDYAEQF | 0.42341542 |
| YDELVEAF | GARDRDRDYAEQF | 0.3543037 |
| YGTGGAGGNGG | GARDRDRDYAEQF | 0.67128086 |
| YTNVMAV | GARDRDRDYAEQF | 0.756629 |
| FLEQVHQL | GARDRDRDYAEQF | 0.7128005 |
| GLSLLAVAV | GARDRDRDYAEQF | 0.6945611 |
| LEQVHQL | GARDRDRDYAEQF | 0.7472459 |
| LNAAGAI | GARDRDRDYAEQF | 0.76825565 |
| PAVHASL | GARDRDRDYAEQF | 0.6342895 |
| TELEDDA | GARDRDRDYAEQF | 0.4650023 |

| | | |
|---|---|---|
| TPPGPPN | GARDRDRDYAEQF | 0.75075626 |
| TPPPPPPP | GARDRDRDYAEQF | 0.7627489 |
| TQSPALI | ASSSSRAPDTQY | 0.66520727 |
| EFAAAFPQ | ASSSSRAPDTQY | 0.7396351 |
| LLSGNLDVL | ASSSSRAPDTQY | 0.6713091 |
| GPANLGNYNI | ASSSSRAPDTQY | 0.417323 |
| TLGPTLP | ASSSSRAPDTQY | 0.74561536 |
| VYAYSSCCK | ASSSSRAPDTQY | 0.5764575 |
| APAPAPAPNI | ASSSSRAPDTQY | 0.7612179 |
| PGGPVPIP | ASSSSRAPDTQY | 0.6451436 |
| FLTVALEP | ASSSSRAPDTQY | 0.74867344 |
| SQYFGANL | ASSSSRAPDTQY | 0.75158155 |
| GGVKPAA | ASSSSRAPDTQY | 0.7463348 |
| GGTGGAGGAGGW | ASSSSRAPDTQY | 0.70335364 |
| GVGGVAGLGGAGG | ASSSSRAPDTQY | 0.6814007 |
| VLLVLATTI | ASSSSRAPDTQY | 0.6020366 |
| IGIFAAML | ASSSSRAPDTQY | 0.43211204 |
| AVGDLID | ASSSSRAPDTQY | 0.7632942 |
| NISPLPL | ASSSSRAPDTQY | 0.2475842 |
| VTAVFVAI | ASSSSRAPDTQY | 0.76564217 |
| SGDDNDAAL | ASSSSRAPDTQY | 0.7652938 |
| AVAALAHS | ASSSSRAPDTQY | 0.62223214 |
| GGVTVMATP | ASSSSRAPDTQY | 0.7535987 |
| AGLAANVRD | ASSSSRAPDTQY | 0.61193526 |
| TNRPDLIDPALLRPG | ASSSSRAPDTQY | 0.6509402 |
| LANLGVTAL | ASSSSRAPDTQY | 0.7538279 |
| TGERIEL | ASSSSRAPDTQY | 0.7605752 |
| NGEELAA | ASSSSRAPDTQY | 0.62620085 |
| VLSLVWG | ASSSSRAPDTQY | 0.7460078 |
| IGYALWL | ASSSSRAPDTQY | 0.747023 |
| LSLLLAPL | ASSSSRAPDTQY | 0.6239386 |
| TLDEALK | ASSSSRAPDTQY | 0.7225884 |
| TDGAGTAEGG | ASSSSRAPDTQY | 0.76398087 |
| NAPGGAGGNGG | ASSSSRAPDTQY | 0.7379732 |
| VSVFMGMGGNF | ASSSSRAPDTQY | 0.67789745 |
| LASYLAAL | ASSSSRAPDTQY | 0.7478565 |
| LVGGTPAL | ASSSSRAPDTQY | 0.6388693 |
| VRNADVL | ASSSSRAPDTQY | 0.75449073 |
| TCSSPFLVSAM | ASSSSRAPDTQY | 0.7142687 |
| VHITDAE | ASSSSRAPDTQY | 0.6870111 |
| EYAAPSF | ASSSSRAPDTQY | 0.75417894 |
| VVDAFHGGL | ASSSSRAPDTQY | 0.36499405 |
| ASVVDPQ | ASSSSRAPDTQY | 0.6378064 |
| SYNEFGGLS | ASSSSRAPDTQY | 0.67797 |
| DPDGSVY | ASSSSRAPDTQY | 0.7325505 |
| MVSTIHATE | ASSSSRAPDTQY | 0.7638236 |
| LAAGVADPV | ASSSSRAPDTQY | 0.6695106 |
| TKEQIAA | ASSSSRAPDTQY | 0.73857117 |
| SLASPAAL | ASSSSRAPDTQY | 0.7241428 |
| IGLVFYGV | ASSSSRAPDTQY | 0.6485071 |
| FLLDASNL | ASSSSRAPDTQY | 0.73512673 |

| | | |
|---|---|---|
| YQVGNFP | ASSSSRAPDTQY | 0.66393256 |
| ESLLSQST | ASSSSRAPDTQY | 0.6210867 |
| LFTVGFAAVG | ASSSSRAPDTQY | 0.74648845 |
| ATPPTPT | ASSSSRAPDTQY | 0.614229 |
| TDLATLG | ASSSSRAPDTQY | 0.6966289 |
| SAGGTGR | ASSSSRAPDTQY | 0.75820386 |
| GFSNSGS | ASSSSRAPDTQY | 0.63705057 |
| GGNGGSGGTSV | ASSSSRAPDTQY | 0.75139785 |
| GTSVATGGAGN | ASSSSRAPDTQY | 0.7385068 |
| NGGNGGSGGTS | ASSSSRAPDTQY | 0.7427505 |
| AVVKFPH | ASSSSRAPDTQY | 0.727904 |
| TPPPPPP | ASSSSRAPDTQY | 0.69438607 |
| AAPAAAAAPA | ASSSSRAPDTQY | 0.76873434 |
| IALNEDL | ASSSSRAPDTQY | 0.7530433 |
| IVNVSSM | ASSSSRAPDTQY | 0.75228316 |
| TGIPPAPR | ASSSSRAPDTQY | 0.74012697 |
| FDLGGGTF | ASSSSRAPDTQY | 0.7559528 |
| FDLGGGTFDVS | ASSSSRAPDTQY | 0.7614052 |
| NEPTAAA | ASSSSRAPDTQY | 0.69067955 |
| NEPTAAALA | ASSSSRAPDTQY | 0.5083573 |
| GSIINISSI | ASSSSRAPDTQY | 0.6985239 |
| VAPGNAGTA | ASSSSRAPDTQY | 0.70555747 |
| TGGGASLE | ASSSSRAPDTQY | 0.6919402 |
| VAVYNHYKRI | ASSSSRAPDTQY | 0.3526914 |
| TDPAPATT | ASSSSRAPDTQY | 0.7681824 |
| TNRPDILDPALLRPG | ASSSSRAPDTQY | 0.7770811 |
| LTTTTTS | ASSSSRAPDTQY | 0.5974734 |
| PTGSGKTL | ASSSSRAPDTQY | 0.7750827 |
| TGSGKTL | ASSSSRAPDTQY | 0.5633557 |
| TIGHVDHGK | ASSSSRAPDTQY | 0.7466372 |
| TIGHVDHGKTT | ASSSSRAPDTQY | 0.7667674 |
| GHVDHGKTT | ASSSSRAPDTQY | 0.65470666 |
| AQFEHTLL | ASSSSRAPDTQY | 0.7519022 |
| IVNQDEL | ASSSSRAPDTQY | 0.6493665 |
| TNGSQFF | ASSSSRAPDTQY | 0.69450283 |
| TGKGIVE | ASSSSRAPDTQY | 0.7676493 |
| LISNASDA | ASSSSRAPDTQY | 0.6871678 |
| AELRTLGELTKIAKAHGAQ | ASSSSRAPDTQY | 0.7351193 |
| APNAVAAAEPSW | ASSSSRAPDTQY | 0.73787534 |
| APQVAGAATAVL | ASSSSRAPDTQY | 0.65488 |
| ARRDPAAAAAILPTDA | ASSSSRAPDTQY | 0.73505604 |
| DAAARDVLAEAGLAEYF | ASSSSRAPDTQY | 0.76361006 |
| DDQIVLSAKANLAKVEQ | ASSSSRAPDTQY | 0.6367973 |
| DELLPDTGEDSDEDGH | ASSSSRAPDTQY | 0.7736248 |
| ELPEAPANRATVPQD | ASSSSRAPDTQY | 0.57606345 |
| FGQSKIERVTGIG | ASSSSRAPDTQY | 0.7751225 |
| GAGGTGSTAGGAGGAGGAGGLY | ASSSSRAPDTQY | 0.75260705 |
| GHGGTGAAGGNGGTGGAGGSGI | ASSSSRAPDTQY | 0.7464329 |
| GNGGNGGAGGEGGPGGAGGAGA | ASSSSRAPDTQY | 0.6867708 |
| GSTAGNGGNGGNATNGTIAGQP | ASSSSRAPDTQY | 0.76875484 |
| GTGGDGGNGGNGGTGG | ASSSSRAPDTQY | 0.7280389 |

| | | |
|---|---|---|
| GVRRPAGIPADAPV | ASSSSRAPDTQY | 0.7265014 |
| GYRQVTDAHLLTLARR | ASSSSRAPDTQY | 0.76852 |
| IAKMGQDGHDRG | ASSSSRAPDTQY | 0.20908421 |
| KAEPSIEIQYT | ASSSSRAPDTQY | 0.2105171 |
| KPVRIRMAADEDTVAVT | ASSSSRAPDTQY | 0.20907505 |
| MPPRIAGMRLLVIKPEP | ASSSSRAPDTQY | 0.20952754 |
| MKFYAVALFRSRGVPPTR | ASSSSRAPDTQY | 0.21169457 |
| NAGGFGILGAVAHSP | ASSSSRAPDTQY | 0.20920652 |
| NEGREITLAMAEM | ASSSSRAPDTQY | 0.20931599 |
| NELVEWSHPLGHRWRWE | ASSSSRAPDTQY | 0.21154009 |
| NLGAGGTGGNGGNPG | ASSSSRAPDTQY | 0.20929222 |
| NTEWPKWETVMAADPVD | ASSSSRAPDTQY | 0.2093371 |
| PASAAPAETQP | ASSSSRAPDTQY | 0.21133725 |
| QQSSDGKNGVRAAEITGE | ASSSSRAPDTQY | 0.21079452 |
| RAPDAVTTSWWKE | ASSSSRAPDTQY | 0.20981707 |
| SGFGNVGAHTSGWFNQ | ASSSSRAPDTQY | 0.20901035 |
| SGVAGAVGQAASVGGLKV | ASSSSRAPDTQY | 0.20891114 |
| SNGGAGTGSAGGNGGT | ASSSSRAPDTQY | 0.20988823 |
| TGGAGGEGGAAG | ASSSSRAPDTQY | 0.20927802 |
| TGGDGGAAGAGGAPGHG | ASSSSRAPDTQY | 0.2089541 |
| TMPPLLAAIANAGRHGV | ASSSSRAPDTQY | 0.2091079 |
| TSSTAGNSGGA | ASSSSRAPDTQY | 0.20864797 |
| VPATVPVGILC | ASSSSRAPDTQY | 0.20903683 |
| VQRPDSGGVVLTGRIS | ASSSSRAPDTQY | 0.20930977 |
| WTPNPVPATKGVPAGLVQAT | ASSSSRAPDTQY | 0.2102312 |
| YALKRALAKFSDRFDVVII | ASSSSRAPDTQY | 0.20913945 |
| YLIGSVIV | ASSSSRAPDTQY | 0.20924157 |
| YGQLVAVID | ASSSSRAPDTQY | 0.20889468 |
| VTVGGMTLPA | ASSSSRAPDTQY | 0.2101784 |
| VLALPAV | ASSSSRAPDTQY | 0.20953603 |
| VIPPTLNYET | ASSSSRAPDTQY | 0.21009496 |
| VGMGGMHPSSG | ASSSSRAPDTQY | 0.21082817 |
| VGLAGGAATA | ASSSSRAPDTQY | 0.20889448 |
| VGGAGGAGGSGP | ASSSSRAPDTQY | 0.21031079 |
| VFAAFMV | ASSSSRAPDTQY | 0.21108305 |
| TLYDKLR | ASSSSRAPDTQY | 0.2091322 |
| TLAVFTVL | ASSSSRAPDTQY | 0.20910628 |
| TGSVQPARV | ASSSSRAPDTQY | 0.20873569 |
| TGGGAGSGGT | ASSSSRAPDTQY | 0.20972483 |
| SHLETSL | ASSSSRAPDTQY | 0.21013175 |
| SFTTGMT | ASSSSRAPDTQY | 0.21040449 |
| QVEVAKRMGVRQPTVS | ASSSSRAPDTQY | 0.21099229 |
| QGSGGPALAM | ASSSSRAPDTQY | 0.20911197 |
| MLAPSNP | ASSSSRAPDTQY | 0.20894876 |
| LVNNAAI | ASSSSRAPDTQY | 0.20898765 |
| LTGEINTLA | ASSSSRAPDTQY | 0.20922534 |
| LSIFGPV | ASSSSRAPDTQY | 0.20953369 |
| LLALLANL | ASSSSRAPDTQY | 0.21011807 |
| LISNASDALDKLR | ASSSSRAPDTQY | 0.208991 |
| LINEAVP | ASSSSRAPDTQY | 0.20906393 |
| LGYAFSGL | ASSSSRAPDTQY | 0.2084405 |

| | | |
|---|---|---|
| LDFVAVAL | ASSSSRAPDTQY | 0.20903476 |
| LAQPGAE | ASSSSRAPDTQY | 0.20894635 |
| ISNLGAVTGF | ASSSSRAPDTQY | 0.20916754 |
| HQEDPGRI | ASSSSRAPDTQY | 0.21330073 |
| GTPGEPTP | ASSSSRAPDTQY | 0.21217877 |
| GFFNSGS | ASSSSRAPDTQY | 0.21197261 |
| GAGGGGLTAGAN | ASSSSRAPDTQY | 0.20993179 |
| FVDALVNM | ASSSSRAPDTQY | 0.20872183 |
| FLTVAPEL | ASSSSRAPDTQY | 0.20961832 |
| FETLTAL | ASSSSRAPDTQY | 0.20954134 |
| DVINPIEL | ASSSSRAPDTQY | 0.2090508 |
| APVSGAPV | ASSSSRAPDTQY | 0.2108844 |
| ALVGGLAW | ASSSSRAPDTQY | 0.20988458 |
| ALAAAAATAI | ASSSSRAPDTQY | 0.21153879 |
| AFGLGLLL | ASSSSRAPDTQY | 0.20943755 |
| AFFAAASL | ASSSSRAPDTQY | 0.21075934 |
| AAAALPAFSPP | ASSSSRAPDTQY | 0.21045905 |
| VVLAPAL | ASSSSRAPDTQY | 0.2085087 |
| TIGHVDHGKTTL | ASSSSRAPDTQY | 0.20873523 |
| LISNASDALDK | ASSSSRAPDTQY | 0.20900646 |
| LISNASDAL | ASSSSRAPDTQY | 0.20961185 |
| ISNASDALDK | ASSSSRAPDTQY | 0.20901094 |
| NQAETLVYQTE | ASSSSRAPDTQY | 0.20954482 |
| TGGTGGTAGAG | ASSSSRAPDTQY | 0.21025996 |
| TGSTAGGAGGA | ASSSSRAPDTQY | 0.2096404 |
| DPLAVPVAAGPV | ASSSSRAPDTQY | 0.2087443 |
| GGTGGTPGNGGN | ASSSSRAPDTQY | 0.20921013 |
| PAGLAGVDPATL | ASSSSRAPDTQY | 0.21051313 |
| GNNLVVEAPHAGASV | ASSSSRAPDTQY | 0.2099744 |
| KVTRIMSASTLEQPA | ASSSSRAPDTQY | 0.20892972 |
| LGFALPIALPRRTVV | ASSSSRAPDTQY | 0.20994991 |
| TRLGNLTARKLVVLV | ASSSSRAPDTQY | 0.20903385 |
| WSQQRRLIYFPSAGP | ASSSSRAPDTQY | 0.21177864 |
| GPNGNPDPMAAAVDIR | ASSSSRAPDTQY | 0.20914114 |
| TQRCVSVADLRAAASR | ASSSSRAPDTQY | 0.20896667 |
| QFVLDVAEHPVEVLTETV | ASSSSRAPDTQY | 0.21032342 |
| GGAGGTGGAGGNGGTQAGDG | ASSSSRAPDTQY | 0.20904106 |
| PLLATLRAIAVDVDAKPAQV | ASSSSRAPDTQY | 0.21109013 |
| APKFRTGGYMIDFWGVGYQVAK | ASSSSRAPDTQY | 0.20976989 |
| GAGGAGGAGGAAGGPGGTGGTG | ASSSSRAPDTQY | 0.20888467 |
| IGALGLLATDVAPAGSVAFGVA | ASSSSRAPDTQY | 0.20901509 |
| AGGAGGIGGNGGAGGAGGVLMG | ASSSSRAPDTQY | 0.20882735 |
| ADELIGLL | ASSSSRAPDTQY | 0.20906848 |
| ADRALKA | ASSSSRAPDTQY | 0.20898752 |
| AGAVGQVGGAGGA | ASSSSRAPDTQY | 0.20870535 |
| AGGGGGLGGAP | ASSSSRAPDTQY | 0.20957306 |
| AGGLLYGNGGAG | ASSSSRAPDTQY | 0.20966068 |
| AGLESIGL | ASSSSRAPDTQY | 0.20944868 |
| AGLKTGII | ASSSSRAPDTQY | 0.21067527 |
| ASLAALLL | ASSSSRAPDTQY | 0.2090381 |
| ASVINISSI | ASSSSRAPDTQY | 0.2098591 |

| | | |
|---|---|---|
| ATPPGPA | ASSSSRAPDTQY | 0.20912145 |
| ATVASALV | ASSSSRAPDTQY | 0.21273462 |
| ATVASLIL | ASSSSRAPDTQY | 0.20895071 |
| AVNGPVLIRH | ASSSSRAPDTQY | 0.20922983 |
| DGAAGAVGI | ASSSSRAPDTQY | 0.21055663 |
| DPLLHGI | ASSSSRAPDTQY | 0.20881215 |
| ELGAALGAL | ASSSSRAPDTQY | 0.20890394 |
| ELGRTQAV | ASSSSRAPDTQY | 0.20899178 |
| FVAGGYLS | ASSSSRAPDTQY | 0.21243308 |
| GASGGAGGAGGW | ASSSSRAPDTQY | 0.20889226 |
| GGAGAGGDGGAGGI | ASSSSRAPDTQY | 0.20921142 |
| GTGHNGSQPGMGGNG | ASSSSRAPDTQY | 0.20932604 |
| GYAAASASAS | ASSSSRAPDTQY | 0.21011987 |
| HQVDVAAVQQG | ASSSSRAPDTQY | 0.20990409 |
| IGAAIAHT | ASSSSRAPDTQY | 0.20901579 |
| IGMIEVAQL | ASSSSRAPDTQY | 0.20895119 |
| IRLAPEHTSAMD | ASSSSRAPDTQY | 0.21124823 |
| IYNVSVL | ASSSSRAPDTQY | 0.20873164 |
| KQGFLGM | ASSSSRAPDTQY | 0.20886521 |
| LESVITQL | ASSSSRAPDTQY | 0.2099327 |
| LGLGALVVV | ASSSSRAPDTQY | 0.20878208 |
| LIGNGGNGGSGGI | ASSSSRAPDTQY | 0.21147615 |
| LNAINAPTL | ASSSSRAPDTQY | 0.20923005 |
| LQLLNTGV | ASSSSRAPDTQY | 0.20869306 |
| LQTSIEF | ASSSSRAPDTQY | 0.20850922 |
| MSFAVSEL | ASSSSRAPDTQY | 0.20895077 |
| NTGALGL | ASSSSRAPDTQY | 0.2102122 |
| PPGAGGAGGAGTT | ASSSSRAPDTQY | 0.21104461 |
| QLVGLGI | ASSSSRAPDTQY | 0.21001261 |
| RFLEQVHQL | ASSSSRAPDTQY | 0.20950207 |
| RGVGYVLR | ASSSSRAPDTQY | 0.21151605 |
| SATGVFAGVI | ASSSSRAPDTQY | 0.20893517 |
| SLPAITLP | ASSSSRAPDTQY | 0.20950386 |
| STALAFTV | ASSSSRAPDTQY | 0.20905276 |
| SVLTVGLGK | ASSSSRAPDTQY | 0.20907092 |
| TALAGAGSGI | ASSSSRAPDTQY | 0.20907772 |
| TGLPFTGI | ASSSSRAPDTQY | 0.20963958 |
| VAEHVTV | ASSSSRAPDTQY | 0.2109808 |
| VDIVLNSL | ASSSSRAPDTQY | 0.20928071 |
| VLTGGGAVL | ASSSSRAPDTQY | 0.20936406 |
| VNVGLAYGI | ASSSSRAPDTQY | 0.20859149 |
| VSSAAVGAV | ASSSSRAPDTQY | 0.20876537 |
| VTSVYEAGV | ASSSSRAPDTQY | 0.21212132 |
| WVALDRGA | ASSSSRAPDTQY | 0.20975366 |
| YLVGSVVI | ASSSSRAPDTQY | 0.20903179 |
| GPPGTGKTLL | ASSSSRAPDTQY | 0.20904009 |
| GVGMTKF | ASSSSRAPDTQY | 0.20950265 |
| IGIGGSDL | ASSSSRAPDTQY | 0.2094293 |
| IGTLLAEL | ASSSSRAPDTQY | 0.20918232 |
| IYTSGTT | ASSSSRAPDTQY | 0.20904799 |
| LSFPAGV | ASSSSRAPDTQY | 0.2098671 |

| | | |
|---|---|---|
| LYTSGTT | ASSSSRAPDTQY | 0.20996158 |
| VGAFPFGF | ASSSSRAPDTQY | 0.21308212 |
| VGAGTVEF | ASSSSRAPDTQY | 0.2098426 |
| VGKSTLLL | ASSSSRAPDTQY | 0.21067467 |
| AALAHAGGGIV | ASSSSRAPDTQY | 0.20939465 |
| AALVAVSLL | ASSSSRAPDTQY | 0.20961368 |
| AAVSRAVAAL | ASSSSRAPDTQY | 0.2094675 |
| AGGSSGIGGF | ASSSSRAPDTQY | 0.20963359 |
| AGLTVANANG | ASSSSRAPDTQY | 0.20883586 |
| AGLVGAVL | ASSSSRAPDTQY | 0.20923372 |
| ALSQAFTV | ASSSSRAPDTQY | 0.20912594 |
| ALTFSVSL | ASSSSRAPDTQY | 0.20948167 |
| ALTGAALVAL | ASSSSRAPDTQY | 0.20978 |
| ASFSIGAGVL | ASSSSRAPDTQY | 0.20893425 |
| DGSVYKI | ASSSSRAPDTQY | 0.20943908 |
| EQLAEIV | ASSSSRAPDTQY | 0.20921083 |
| FAVSELTVM | ASSSSRAPDTQY | 0.21001226 |
| GAAPLAAGL | ASSSSRAPDTQY | 0.20916326 |
| GADGGDGGSGGAGG | ASSSSRAPDTQY | 0.21131837 |
| GFGLDTGGAGG | ASSSSRAPDTQY | 0.20885807 |
| GLASGGLL | ASSSSRAPDTQY | 0.20973252 |
| GRIAGAAVLL | ASSSSRAPDTQY | 0.20941857 |
| GSAGSLNTSF | ASSSSRAPDTQY | 0.20930767 |
| GTGQIGI | ASSSSRAPDTQY | 0.20993814 |
| GTLIALIL | ASSSSRAPDTQY | 0.20900975 |
| IAATMGFVVG | ASSSSRAPDTQY | 0.20968957 |
| IFAQALL | ASSSSRAPDTQY | 0.20946677 |
| IGAAVALGF | ASSSSRAPDTQY | 0.20899914 |
| IGLTPLAL | ASSSSRAPDTQY | 0.20923932 |
| ILGSNTSTL | ASSSSRAPDTQY | 0.21134162 |
| IVFSLTVL | ASSSSRAPDTQY | 0.2097288 |
| LASKATGF | ASSSSRAPDTQY | 0.20929427 |
| LASVALIL | ASSSSRAPDTQY | 0.2097519 |
| LFAVGLTL | ASSSSRAPDTQY | 0.20917019 |
| LFNGGIGTYI | ASSSSRAPDTQY | 0.20936652 |
| LTLGAGII | ASSSSRAPDTQY | 0.20870896 |
| LVAAALGDI | ASSSSRAPDTQY | 0.20949827 |
| NTAFVQLGI | ASSSSRAPDTQY | 0.20904656 |
| QLFSGAEL | ASSSSRAPDTQY | 0.20914133 |
| SAATFITL | ASSSSRAPDTQY | 0.21062003 |
| SNYGLANLGV | ASSSSRAPDTQY | 0.21034604 |
| SYGGAYAVM | ASSSSRAPDTQY | 0.20923412 |
| TTGVGGLTLG | ASSSSRAPDTQY | 0.21033531 |
| TTQTIYI | ASSSSRAPDTQY | 0.20932807 |
| VAAGAGLGAVI | ASSSSRAPDTQY | 0.20890303 |
| VAAVVTLL | ASSSSRAPDTQY | 0.2090191 |
| VGFLGGLI | ASSSSRAPDTQY | 0.21055132 |
| VGVIPILL | ASSSSRAPDTQY | 0.20890923 |
| VIFASSLI | ASSSSRAPDTQY | 0.20918219 |
| VKGGDGGAAAG | ASSSSRAPDTQY | 0.2084385 |
| VTKTFTNI | ASSSSRAPDTQY | 0.2090113 |

| | | |
|---|---|---|
| VVAAAILL | ASSSSRAPDTQY | 0.20967378 |
| YAALDVEL | ASSSSRAPDTQY | 0.20881005 |
| LASVALLL | ASSSSRAPDTQY | 0.21110767 |
| LDAVHRV | ASSSSRAPDTQY | 0.20886871 |
| QGLSSSIF | ASSSSRAPDTQY | 0.20881532 |
| RGTFANIRL | ASSSSRAPDTQY | 0.21181515 |
| RQLWWGHRI | ASSSSRAPDTQY | 0.21022877 |
| YGPPGTGKTLL | ASSSSRAPDTQY | 0.20913245 |
| AALVGAPANVA | ASSSSRAPDTQY | 0.20922568 |
| AATFRRTE | ASSSSRAPDTQY | 0.20932825 |
| AGALGGGI | ASSSSRAPDTQY | 0.2097993 |
| AGGAGGAGGIGAG | ASSSSRAPDTQY | 0.20901692 |
| AGSGALGL | ASSSSRAPDTQY | 0.2090277 |
| AQIADQI | ASSSSRAPDTQY | 0.20988233 |
| ASAVAVTYL | ASSSSRAPDTQY | 0.20879632 |
| ASTVLGGTGG | ASSSSRAPDTQY | 0.20993121 |
| AVGGEPAK | ASSSSRAPDTQY | 0.21085858 |
| CEQGTFGGTC | ASSSSRAPDTQY | 0.20894435 |
| DEGKAPAG | ASSSSRAPDTQY | 0.2090581 |
| DTVVFPAL | ASSSSRAPDTQY | 0.20905039 |
| FEGITMAG | ASSSSRAPDTQY | 0.20934384 |
| FPLVVQD | ASSSSRAPDTQY | 0.20898657 |
| GAFTTPQ | ASSSSRAPDTQY | 0.20873326 |
| GFGGFGTAGGVGGA | ASSSSRAPDTQY | 0.20920663 |
| GFLAQFL | ASSSSRAPDTQY | 0.21167381 |
| GGAGGAGGNAGVA | ASSSSRAPDTQY | 0.20906408 |
| GGVGTAGAGGAGGAGG | ASSSSRAPDTQY | 0.21082225 |
| GNGGGGDAGNAGSGGN | ASSSSRAPDTQY | 0.20965692 |
| GTGGAGGAGNDAGS | ASSSSRAPDTQY | 0.21142176 |
| GTTNTGWFNS | ASSSSRAPDTQY | 0.2091324 |
| IFMGPLFTL | ASSSSRAPDTQY | 0.21103206 |
| ILAHAIEAI | ASSSSRAPDTQY | 0.20904447 |
| IRSAIREAL | ASSSSRAPDTQY | 0.20967762 |
| ITFAGGG | ASSSSRAPDTQY | 0.2097654 |
| ITGMSGHQEV | ASSSSRAPDTQY | 0.20915505 |
| IVAGLACTL | ASSSSRAPDTQY | 0.21010205 |
| IVGGGLAGA | ASSSSRAPDTQY | 0.2092797 |
| IVTGVLNI | ASSSSRAPDTQY | 0.20947078 |
| LAQQGCI | ASSSSRAPDTQY | 0.20925497 |
| LCLHVEN | ASSSSRAPDTQY | 0.20886378 |
| LDAISDGS | ASSSSRAPDTQY | 0.20976727 |
| LGGFAAML | ASSSSRAPDTQY | 0.2089227 |
| LGGKNPMI | ASSSSRAPDTQY | 0.21001586 |
| LGPKGVAAIL | ASSSSRAPDTQY | 0.20898597 |
| LMVGVSI | ASSSSRAPDTQY | 0.20893218 |
| LVGAAFVL | ASSSSRAPDTQY | 0.20936991 |
| LVLAVAPL | ASSSSRAPDTQY | 0.20896769 |
| MAAIAVIDAA | ASSSSRAPDTQY | 0.21194671 |
| NNGINAELQ | ASSSSRAPDTQY | 0.20931089 |
| RGEPLVLVHG | ASSSSRAPDTQY | 0.20903225 |
| RGGAGGAAGLI | ASSSSRAPDTQY | 0.20906904 |

| | | |
|---|---|---|
| RSYRMHPAV | ASSSSRAPDTQY | 0.20908801 |
| SAAHLLAGQLA | ASSSSRAPDTQY | 0.2092523 |
| SAAVGSY | ASSSSRAPDTQY | 0.20999125 |
| SPHAVLHRI | ASSSSRAPDTQY | 0.208787 |
| SQIAKVL | ASSSSRAPDTQY | 0.20991774 |
| TALGHLVP | ASSSSRAPDTQY | 0.2090896 |
| TAPTTSGAA | ASSSSRAPDTQY | 0.20967937 |
| TDGDDGNGGSPGT | ASSSSRAPDTQY | 0.20957796 |
| TDGLGEN | ASSSSRAPDTQY | 0.21034612 |
| TGGPPLPGQA | ASSSSRAPDTQY | 0.20904554 |
| TIPAITI | ASSSSRAPDTQY | 0.21324927 |
| TIPGFTI | ASSSSRAPDTQY | 0.21021733 |
| TSIASGALA | ASSSSRAPDTQY | 0.20924999 |
| TVTLPGAAI | ASSSSRAPDTQY | 0.20927213 |
| VASGLSAALP | ASSSSRAPDTQY | 0.20951784 |
| VGAGVIGI | ASSSSRAPDTQY | 0.21065111 |
| VGPKRFGV | ASSSSRAPDTQY | 0.20930783 |
| VNVAFPD | ASSSSRAPDTQY | 0.20892629 |
| VSMTNSGVS | ASSSSRAPDTQY | 0.20912227 |
| VTDVLGDLSV | ASSSSRAPDTQY | 0.20972209 |
| VVVVGAGFAGL | ASSSSRAPDTQY | 0.209252 |
| VYTVGTVV | ASSSSRAPDTQY | 0.21523623 |
| WTTFSGP | ASSSSRAPDTQY | 0.20903447 |
| YDELVEAF | ASSSSRAPDTQY | 0.21201879 |
| YGTGGAGGNGG | ASSSSRAPDTQY | 0.20942901 |
| YTNVMAV | ASSSSRAPDTQY | 0.20964858 |
| FLEQVHQL | ASSSSRAPDTQY | 0.20879312 |
| GLSLLAVAV | ASSSSRAPDTQY | 0.21123622 |
| LEQVHQL | ASSSSRAPDTQY | 0.20989631 |
| LNAAGAI | ASSSSRAPDTQY | 0.20927891 |
| PAVHASL | ASSSSRAPDTQY | 0.20924902 |
| TELEDDA | ASSSSRAPDTQY | 0.20956488 |
| TPPGPPN | ASSSSRAPDTQY | 0.21101405 |
| TPPPPPPP | ASSSSRAPDTQY | 0.20862523 |
| TQSPALI | ASGEQGAHSGNTLY | 0.20878805 |
| EFAAAFPQ | ASGEQGAHSGNTLY | 0.20966306 |
| LLSGNLDVL | ASGEQGAHSGNTLY | 0.20953962 |
| GPANLGNYNI | ASGEQGAHSGNTLY | 0.20946051 |
| TLGPTLP | ASGEQGAHSGNTLY | 0.20973139 |
| VYAYSSCCK | ASGEQGAHSGNTLY | 0.20880935 |
| APAPAPAPNI | ASGEQGAHSGNTLY | 0.20957698 |
| PGGPVPIP | ASGEQGAHSGNTLY | 0.20897017 |
| FLTVALEP | ASGEQGAHSGNTLY | 0.20913307 |
| SQYFGANL | ASGEQGAHSGNTLY | 0.21172668 |
| GGVKPAA | ASGEQGAHSGNTLY | 0.20921575 |
| GGTGGAGGAGGW | ASGEQGAHSGNTLY | 0.2088973 |
| GVGGVAGLGGAGG | ASGEQGAHSGNTLY | 0.20922199 |
| VLLVLATTI | ASGEQGAHSGNTLY | 0.21043098 |
| IGIFAAML | ASGEQGAHSGNTLY | 0.20899275 |
| AVGDLID | ASGEQGAHSGNTLY | 0.20890245 |
| NISPLPL | ASGEQGAHSGNTLY | 0.20922068 |

| | | |
|---|---|---|
| VTAVFVAI | ASGEQGAHSGNTLY | 0.2090028 |
| SGDDNDAAL | ASGEQGAHSGNTLY | 0.2094026 |
| AVAALAHS | ASGEQGAHSGNTLY | 0.21188341 |
| GGVTVMATP | ASGEQGAHSGNTLY | 0.20898697 |
| AGLAANVRD | ASGEQGAHSGNTLY | 0.20949614 |
| TNRPDLIDPALLRPG | ASGEQGAHSGNTLY | 0.20921595 |
| LANLGVTAL | ASGEQGAHSGNTLY | 0.20931329 |
| TGERIEL | ASGEQGAHSGNTLY | 0.20918669 |
| NGEELAA | ASGEQGAHSGNTLY | 0.20913996 |
| VLSLVWG | ASGEQGAHSGNTLY | 0.20905134 |
| IGYALWL | ASGEQGAHSGNTLY | 0.2099576 |
| LSLLLAPL | ASGEQGAHSGNTLY | 0.20990665 |
| TLDEALK | ASGEQGAHSGNTLY | 0.20938426 |
| TDGAGTAEGG | ASGEQGAHSGNTLY | 0.21002343 |
| NAPGGAGGNGG | ASGEQGAHSGNTLY | 0.20922181 |
| VSVFMGMGGNF | ASGEQGAHSGNTLY | 0.20915094 |
| LASYLAAL | ASGEQGAHSGNTLY | 0.20996965 |
| LVGGTPAL | ASGEQGAHSGNTLY | 0.20862135 |
| VRNADVL | ASGEQGAHSGNTLY | 0.20907405 |
| TCSSPFLVSAM | ASGEQGAHSGNTLY | 0.20912965 |
| VHITDAE | ASGEQGAHSGNTLY | 0.2095483 |
| EYAAPSF | ASGEQGAHSGNTLY | 0.20945391 |
| VVDAFHGGL | ASGEQGAHSGNTLY | 0.2092578 |
| ASVVDPQ | ASGEQGAHSGNTLY | 0.20938946 |
| SYNEFGGLS | ASGEQGAHSGNTLY | 0.208944 |
| DPDGSVY | ASGEQGAHSGNTLY | 0.20905977 |
| MVSTIHATE | ASGEQGAHSGNTLY | 0.21296377 |
| LAAGVADPV | ASGEQGAHSGNTLY | 0.21094842 |
| TKEQIAA | ASGEQGAHSGNTLY | 0.21077652 |
| SLASPAAL | ASGEQGAHSGNTLY | 0.20898019 |
| IGLVFYGV | ASGEQGAHSGNTLY | 0.20931849 |
| FLLDASNL | ASGEQGAHSGNTLY | 0.20958753 |
| YQVGNFP | ASGEQGAHSGNTLY | 0.20896573 |
| ESLLSQST | ASGEQGAHSGNTLY | 0.20866221 |
| LFTVGFAAVG | ASGEQGAHSGNTLY | 0.2095521 |
| ATPPTPT | ASGEQGAHSGNTLY | 0.20887361 |
| TDLATLG | ASGEQGAHSGNTLY | 0.21241555 |
| SAGGTGR | ASGEQGAHSGNTLY | 0.2090992 |
| GFSNSGS | ASGEQGAHSGNTLY | 0.20883696 |
| GGNGGSGGTSV | ASGEQGAHSGNTLY | 0.2089867 |
| GTSVATGGAGN | ASGEQGAHSGNTLY | 0.20891024 |
| NGGNGGSGGTS | ASGEQGAHSGNTLY | 0.21134715 |
| AVVKFPH | ASGEQGAHSGNTLY | 0.20928077 |
| TPPPPPP | ASGEQGAHSGNTLY | 0.20886773 |
| AAPAAAAAPA | ASGEQGAHSGNTLY | 0.20936765 |
| IALNEDL | ASGEQGAHSGNTLY | 0.21277706 |
| IVNVSSM | ASGEQGAHSGNTLY | 0.21053468 |
| TGIPPAPR | ASGEQGAHSGNTLY | 0.20911779 |
| FDLGGGTF | ASGEQGAHSGNTLY | 0.20910491 |
| FDLGGGTFDVS | ASGEQGAHSGNTLY | 0.20902202 |
| NEPTAAA | ASGEQGAHSGNTLY | 0.20893098 |

| | | |
|---|---|---|
| NEPTAAALA | ASGEQGAHSGNTLY | 0.20892558 |
| GSIINISSI | ASGEQGAHSGNTLY | 0.20892479 |
| VAPGNAGTA | ASGEQGAHSGNTLY | 0.20889398 |
| TGGGASLE | ASGEQGAHSGNTLY | 0.20970097 |
| VAVYNHYKRI | ASGEQGAHSGNTLY | 0.20863706 |
| TDPAPATT | ASGEQGAHSGNTLY | 0.20951071 |
| TNRPDILDPALLRPG | ASGEQGAHSGNTLY | 0.20861322 |
| LTTTTTS | ASGEQGAHSGNTLY | 0.20946243 |
| PTGSGKTL | ASGEQGAHSGNTLY | 0.21012776 |
| TGSGKTL | ASGEQGAHSGNTLY | 0.21000093 |
| TIGHVDHGK | ASGEQGAHSGNTLY | 0.20879433 |
| TIGHVDHGKTT | ASGEQGAHSGNTLY | 0.20957193 |
| GHVDHGKTT | ASGEQGAHSGNTLY | 0.20895064 |
| AQFEHTLL | ASGEQGAHSGNTLY | 0.21320334 |
| IVNQDEL | ASGEQGAHSGNTLY | 0.2087943 |
| TNGSQFF | ASGEQGAHSGNTLY | 0.21056674 |
| TGKGIVE | ASGEQGAHSGNTLY | 0.21049657 |
| LISNASDA | ASGEQGAHSGNTLY | 0.20890503 |
| AELRTLGELTKIAKAHGAQ | ASGEQGAHSGNTLY | 0.20867671 |
| APNAVAAAEPSW | ASGEQGAHSGNTLY | 0.21020335 |
| APQVAGAATAVL | ASGEQGAHSGNTLY | 0.20895764 |
| ARRDPAAAAAILPTDA | ASGEQGAHSGNTLY | 0.20887859 |
| DAAARDVLAEAGLAEYF | ASGEQGAHSGNTLY | 0.2098468 |
| DDQIVLSAKANLAKVEQ | ASGEQGAHSGNTLY | 0.20944436 |
| DELLPDTGEDSDEDGH | ASGEQGAHSGNTLY | 0.2090988 |
| ELPEAPANRATVPQD | ASGEQGAHSGNTLY | 0.20900467 |
| FGQSKIERVTGIG | ASGEQGAHSGNTLY | 0.20933482 |
| GAGGTGSTAGGAGGAGGAGGLY | ASGEQGAHSGNTLY | 0.20945044 |
| GHGGTGAAGGNGGTGGAGGSGI | ASGEQGAHSGNTLY | 0.20911731 |
| GNGGNGGAGGEGGPGGAGGAGA | ASGEQGAHSGNTLY | 0.2088572 |
| GSTAGNGGNGGNATNGTIAGQP | ASGEQGAHSGNTLY | 0.20932844 |
| GTGGDGGNGGNGGTGG | ASGEQGAHSGNTLY | 0.20999047 |
| GVRRPAGIPADAPV | ASGEQGAHSGNTLY | 0.209623 |
| GYRQVTDAHLLTLARR | ASGEQGAHSGNTLY | 0.20915161 |
| IAKMGQDGHDRG | ASGEQGAHSGNTLY | 0.20993482 |
| KAEPSIEIQYT | ASGEQGAHSGNTLY | 0.20890845 |
| KPVRIRMAADEDTVAVT | ASGEQGAHSGNTLY | 0.20907936 |
| MPPRIAGMRLLVIKPEP | ASGEQGAHSGNTLY | 0.20975961 |
| MKFYAVALFRSRGVPPTR | ASGEQGAHSGNTLY | 0.20908266 |
| NAGGFGILGAVAHSP | ASGEQGAHSGNTLY | 0.20954257 |
| NEGREITLAMAEM | ASGEQGAHSGNTLY | 0.21010575 |
| NELVEWSHPLGHRWRWE | ASGEQGAHSGNTLY | 0.20900603 |
| NLGAGGTGGNGGNPG | ASGEQGAHSGNTLY | 0.20922185 |
| NTEWPKWETVMAADPVD | ASGEQGAHSGNTLY | 0.20938174 |
| PASAAPAETQP | ASGEQGAHSGNTLY | 0.2089716 |
| QQSSDGKNGVRAAEITGE | ASGEQGAHSGNTLY | 0.20997268 |
| RAPDAVTTSWWKE | ASGEQGAHSGNTLY | 0.20911102 |
| SGFGNVGAHTSGWFNQ | ASGEQGAHSGNTLY | 0.21172573 |
| SGVAGAVGQAASVGGLKV | ASGEQGAHSGNTLY | 0.20922333 |
| SNGGAGTGSAGGNGGT | ASGEQGAHSGNTLY | 0.20949997 |
| TGGAGGEGGAAG | ASGEQGAHSGNTLY | 0.21183702 |

| | | |
|---|---|---|
| TGGDGGAAGAGGAPGHG | ASGEQGAHSGNTLY | 0.21202558 |
| TMPPLLAAIANAGRHGV | ASGEQGAHSGNTLY | 0.21083647 |
| TSSTAGNSGGA | ASGEQGAHSGNTLY | 0.20872015 |
| VPATVPVGILC | ASGEQGAHSGNTLY | 0.20938127 |
| VQRPDSGGVVLTGRIS | ASGEQGAHSGNTLY | 0.20903684 |
| WTPNPVPATKGVPAGLVQAT | ASGEQGAHSGNTLY | 0.21179967 |
| YALKRALAKFSDRFDVVII | ASGEQGAHSGNTLY | 0.21013093 |
| YLIGSVIV | ASGEQGAHSGNTLY | 0.21086445 |
| YGQLVAVID | ASGEQGAHSGNTLY | 0.20895064 |
| VTVGGMTLPA | ASGEQGAHSGNTLY | 0.20921798 |
| VLALPAV | ASGEQGAHSGNTLY | 0.20887566 |
| VIPPTLNYET | ASGEQGAHSGNTLY | 0.21159844 |
| VGMGGMHPSSG | ASGEQGAHSGNTLY | 0.21098784 |
| VGLAGGAATA | ASGEQGAHSGNTLY | 0.20882179 |
| VGGAGGAGGSGP | ASGEQGAHSGNTLY | 0.20952705 |
| VFAAFMV | ASGEQGAHSGNTLY | 0.20928796 |
| TLYDKLR | ASGEQGAHSGNTLY | 0.20902704 |
| TLAVFTVL | ASGEQGAHSGNTLY | 0.20894198 |
| TGSVQPARV | ASGEQGAHSGNTLY | 0.2091781 |
| TGGGAGSGGT | ASGEQGAHSGNTLY | 0.20909484 |
| SHLETSL | ASGEQGAHSGNTLY | 0.20907548 |
| SFTTGMT | ASGEQGAHSGNTLY | 0.20994699 |
| QVEVAKRMGVRQPTVS | ASGEQGAHSGNTLY | 0.2092157 |
| QGSGGPALAM | ASGEQGAHSGNTLY | 0.20904316 |
| MLAPSNP | ASGEQGAHSGNTLY | 0.20927578 |
| LVNNAAI | ASGEQGAHSGNTLY | 0.20899768 |
| LTGEINTLA | ASGEQGAHSGNTLY | 0.20911488 |
| LSIFGPV | ASGEQGAHSGNTLY | 0.20952447 |
| LLALLANL | ASGEQGAHSGNTLY | 0.20895186 |
| LISNASDALDKLR | ASGEQGAHSGNTLY | 0.20917352 |
| LINEAVP | ASGEQGAHSGNTLY | 0.20917208 |
| LGYAFSGL | ASGEQGAHSGNTLY | 0.20991352 |
| LDFVAVAL | ASGEQGAHSGNTLY | 0.20933194 |
| LAQPGAE | ASGEQGAHSGNTLY | 0.21346718 |
| ISNLGAVTGF | ASGEQGAHSGNTLY | 0.21321814 |
| HQEDPGRI | ASGEQGAHSGNTLY | 0.2091341 |
| GTPGEPTP | ASGEQGAHSGNTLY | 0.21201535 |
| GFFNSGS | ASGEQGAHSGNTLY | 0.20878625 |
| GAGGGGLTAGAN | ASGEQGAHSGNTLY | 0.20936467 |
| FVDALVNM | ASGEQGAHSGNTLY | 0.2087526 |
| FLTVAPEL | ASGEQGAHSGNTLY | 0.21113433 |
| FETLTAL | ASGEQGAHSGNTLY | 0.20921318 |
| DVINPIEL | ASGEQGAHSGNTLY | 0.21112083 |
| APVSGAPV | ASGEQGAHSGNTLY | 0.20925424 |
| ALVGGLAW | ASGEQGAHSGNTLY | 0.21213108 |
| ALAAAAATAI | ASGEQGAHSGNTLY | 0.21105815 |
| AFGLGLLL | ASGEQGAHSGNTLY | 0.20909731 |
| AFFAAASL | ASGEQGAHSGNTLY | 0.21150064 |
| AAAALPAFSPP | ASGEQGAHSGNTLY | 0.2091059 |
| VVLAPAL | ASGEQGAHSGNTLY | 0.20915814 |
| TIGHVDHGKTTL | ASGEQGAHSGNTLY | 0.20931421 |

| | | |
|---|---|---|
| LISNASDALDK | ASGEQGAHSGNTLY | 0.2093789 |
| LISNASDAL | ASGEQGAHSGNTLY | 0.20954846 |
| ISNASDALDK | ASGEQGAHSGNTLY | 0.20913059 |
| NQAETLVYQTE | ASGEQGAHSGNTLY | 0.20977743 |
| TGGTGGTAGAG | ASGEQGAHSGNTLY | 0.20921472 |
| TGSTAGGAGGA | ASGEQGAHSGNTLY | 0.20893143 |
| DPLAVPVAAGPV | ASGEQGAHSGNTLY | 0.20892054 |
| GGTGGTPGNGGN | ASGEQGAHSGNTLY | 0.21126492 |
| PAGLAGVDPATL | ASGEQGAHSGNTLY | 0.21286291 |
| GNNLVVEAPHAGASV | ASGEQGAHSGNTLY | 0.20933789 |
| KVTRIMSASTLEQPA | ASGEQGAHSGNTLY | 0.2100586 |
| LGFALPIALPRRTVV | ASGEQGAHSGNTLY | 0.21014693 |
| TRLGNLTARKLVVLV | ASGEQGAHSGNTLY | 0.2106729 |
| WSQQRRLIYFPSAGP | ASGEQGAHSGNTLY | 0.21008947 |
| GPNGNPDPMAAAVDIR | ASGEQGAHSGNTLY | 0.20913218 |
| TQRCVSVADLRAAASR | ASGEQGAHSGNTLY | 0.20894411 |
| QFVLDVAEHPVEVLTETV | ASGEQGAHSGNTLY | 0.2089949 |
| GGAGGTGGAGGNGGTQAGDG | ASGEQGAHSGNTLY | 0.20913118 |
| PLLATLRAIAVDVDAKPAQV | ASGEQGAHSGNTLY | 0.20954503 |
| APKFRTGGYMIDFWGVGYQVAK | ASGEQGAHSGNTLY | 0.21041992 |
| GAGGAGGAGGAAGGPGGTGGTG | ASGEQGAHSGNTLY | 0.20935766 |
| IGALGLLATDVAPAGSVAFGVA | ASGEQGAHSGNTLY | 0.2108825 |
| AGGAGGIGGNGGAGGAGGVLMG | ASGEQGAHSGNTLY | 0.20973723 |
| ADELIGLL | ASGEQGAHSGNTLY | 0.20948324 |
| ADRALKA | ASGEQGAHSGNTLY | 0.20995843 |
| AGAVGQVGGAGGA | ASGEQGAHSGNTLY | 0.2092802 |
| AGGGGGLGGAP | ASGEQGAHSGNTLY | 0.20920932 |
| AGGLLYGNGGAG | ASGEQGAHSGNTLY | 0.20936604 |
| AGLESIGL | ASGEQGAHSGNTLY | 0.20917107 |
| AGLKTGII | ASGEQGAHSGNTLY | 0.21086991 |
| ASLAALLL | ASGEQGAHSGNTLY | 0.21041119 |
| ASVINISSI | ASGEQGAHSGNTLY | 0.20942958 |
| ATPPGPA | ASGEQGAHSGNTLY | 0.21003084 |
| ATVASALV | ASGEQGAHSGNTLY | 0.21023342 |
| ATVASLIL | ASGEQGAHSGNTLY | 0.20886682 |
| AVNGPVLIRH | ASGEQGAHSGNTLY | 0.20904808 |
| DGAAGAVGI | ASGEQGAHSGNTLY | 0.21009581 |
| DPLLHGI | ASGEQGAHSGNTLY | 0.20984463 |
| ELGAALGAL | ASGEQGAHSGNTLY | 0.21004356 |
| ELGRTQAV | ASGEQGAHSGNTLY | 0.20920941 |
| FVAGGYLS | ASGEQGAHSGNTLY | 0.21049622 |
| GASGGAGGAGGW | ASGEQGAHSGNTLY | 0.20930724 |
| GGAGAGGDGGAGGI | ASGEQGAHSGNTLY | 0.21000463 |
| GTGHNGSQPGMGGNG | ASGEQGAHSGNTLY | 0.20980813 |
| GYAAASASAS | ASGEQGAHSGNTLY | 0.2097898 |
| HQVDVAAVQQG | ASGEQGAHSGNTLY | 0.20906943 |
| IGAAIAHT | ASGEQGAHSGNTLY | 0.20987006 |
| IGMIEVAQL | ASGEQGAHSGNTLY | 0.20877403 |
| IRLAPEHTSAMD | ASGEQGAHSGNTLY | 0.20904009 |
| IYNVSVL | ASGEQGAHSGNTLY | 0.2090185 |
| KQGFLGM | ASGEQGAHSGNTLY | 0.2090467 |

| | | |
|---|---|---|
| LESVITQL | ASGEQGAHSGNTLY | 0.20905401 |
| LGLGALVVV | ASGEQGAHSGNTLY | 0.20983824 |
| LIGNGGNGGSGGI | ASGEQGAHSGNTLY | 0.20899518 |
| LNAINAPTL | ASGEQGAHSGNTLY | 0.2102514 |
| LQLLNTGV | ASGEQGAHSGNTLY | 0.20859326 |
| LQTSIEF | ASGEQGAHSGNTLY | 0.208994 |
| MSFAVSEL | ASGEQGAHSGNTLY | 0.20980401 |
| NTGALGL | ASGEQGAHSGNTLY | 0.2098212 |
| PPGAGGAGGAGTT | ASGEQGAHSGNTLY | 0.21002851 |
| QLVGLGI | ASGEQGAHSGNTLY | 0.20902999 |
| RFLEQVHQL | ASGEQGAHSGNTLY | 0.20916343 |
| RGVGYVLR | ASGEQGAHSGNTLY | 0.20901465 |
| SATGVFAGVI | ASGEQGAHSGNTLY | 0.20980357 |
| SLPAITLP | ASGEQGAHSGNTLY | 0.209054 |
| STALAFTV | ASGEQGAHSGNTLY | 0.21031581 |
| SVLTVGLGK | ASGEQGAHSGNTLY | 0.20955853 |
| TALAGAGSGI | ASGEQGAHSGNTLY | 0.20916739 |
| TGLPFTGI | ASGEQGAHSGNTLY | 0.2094039 |
| VAEHVTV | ASGEQGAHSGNTLY | 0.20933504 |
| VDIVLNSL | ASGEQGAHSGNTLY | 0.2090274 |
| VLTGGGAVL | ASGEQGAHSGNTLY | 0.20884869 |
| VNVGLAYGI | ASGEQGAHSGNTLY | 0.20976 |
| VSSAAVGAV | ASGEQGAHSGNTLY | 0.21027645 |
| VTSVYEAGV | ASGEQGAHSGNTLY | 0.20904961 |
| WVALDRGA | ASGEQGAHSGNTLY | 0.20986025 |
| YLVGSVVI | ASGEQGAHSGNTLY | 0.20886682 |
| GPPGTGKTLL | ASGEQGAHSGNTLY | 0.20982504 |
| GVGMTKF | ASGEQGAHSGNTLY | 0.21091601 |
| IGIGGSDL | ASGEQGAHSGNTLY | 0.20959662 |
| IGTLLAEL | ASGEQGAHSGNTLY | 0.20902926 |
| IYTSGTT | ASGEQGAHSGNTLY | 0.20915109 |
| LSFPAGV | ASGEQGAHSGNTLY | 0.21197861 |
| LYTSGTT | ASGEQGAHSGNTLY | 0.2089931 |
| VGAFPFGF | ASGEQGAHSGNTLY | 0.20901762 |
| VGAGTVEF | ASGEQGAHSGNTLY | 0.20916592 |
| VGKSTLLL | ASGEQGAHSGNTLY | 0.20926 |
| AALAHAGGGIV | ASGEQGAHSGNTLY | 0.2087615 |
| AALVAVSLL | ASGEQGAHSGNTLY | 0.2091197 |
| AAVSRAVAAL | ASGEQGAHSGNTLY | 0.21380334 |
| AGGSSGIGGF | ASGEQGAHSGNTLY | 0.20914878 |
| AGLTVANANG | ASGEQGAHSGNTLY | 0.20927532 |
| AGLVGAVL | ASGEQGAHSGNTLY | 0.20956878 |
| ALSQAFTV | ASGEQGAHSGNTLY | 0.21074136 |
| ALTFSVSL | ASGEQGAHSGNTLY | 0.20970987 |
| ALTGAALVAL | ASGEQGAHSGNTLY | 0.20903601 |
| ASFSIGAGVL | ASGEQGAHSGNTLY | 0.20903777 |
| DGSVYKI | ASGEQGAHSGNTLY | 0.20917502 |
| EQLAEIV | ASGEQGAHSGNTLY | 0.20979832 |
| FAVSELTVM | ASGEQGAHSGNTLY | 0.20963292 |
| GAAPLAAGL | ASGEQGAHSGNTLY | 0.20933956 |
| GADGGDGGSGGAGG | ASGEQGAHSGNTLY | 0.20846345 |

| | | |
|---|---|---|
| GFGLDTGGAGG | ASGEQGAHSGNTLY | 0.209138 |
| GLASGGLL | ASGEQGAHSGNTLY | 0.20933117 |
| GRIAGAAVLL | ASGEQGAHSGNTLY | 0.20904955 |
| GSAGSLNTSF | ASGEQGAHSGNTLY | 0.21163702 |
| GTGQIGI | ASGEQGAHSGNTLY | 0.20925303 |
| GTLIALIL | ASGEQGAHSGNTLY | 0.20917225 |
| IAATMGFVVG | ASGEQGAHSGNTLY | 0.2089773 |
| IFAQALL | ASGEQGAHSGNTLY | 0.20885508 |
| IGAAVALGF | ASGEQGAHSGNTLY | 0.20909368 |
| IGLTPLAL | ASGEQGAHSGNTLY | 0.2090429 |
| ILGSNTSTL | ASGEQGAHSGNTLY | 0.20983543 |
| IVFSLTVL | ASGEQGAHSGNTLY | 0.20907333 |
| LASKATGF | ASGEQGAHSGNTLY | 0.20914091 |
| LASVALIL | ASGEQGAHSGNTLY | 0.20899677 |
| LFAVGLTL | ASGEQGAHSGNTLY | 0.2089723 |
| LFNGGIGTYI | ASGEQGAHSGNTLY | 0.21195062 |
| LTLGAGII | ASGEQGAHSGNTLY | 0.20928939 |
| LVAAALGDI | ASGEQGAHSGNTLY | 0.20916496 |
| NTAFVQLGI | ASGEQGAHSGNTLY | 0.21084902 |
| QLFSGAEL | ASGEQGAHSGNTLY | 0.20906872 |
| SAATFITL | ASGEQGAHSGNTLY | 0.20939586 |
| SNYGLANLGV | ASGEQGAHSGNTLY | 0.20946972 |
| SYGGAYAVM | ASGEQGAHSGNTLY | 0.20954281 |
| TTGVGGLTLG | ASGEQGAHSGNTLY | 0.2093885 |
| TTQTIYI | ASGEQGAHSGNTLY | 0.20939569 |
| VAAGAGLGAVI | ASGEQGAHSGNTLY | 0.20890148 |
| VAAVVTLL | ASGEQGAHSGNTLY | 0.20961887 |
| VGFLGGLI | ASGEQGAHSGNTLY | 0.21348344 |
| VGVIPILL | ASGEQGAHSGNTLY | 0.2099729 |
| VIFASSLI | ASGEQGAHSGNTLY | 0.20987003 |
| VKGGDGGAAAG | ASGEQGAHSGNTLY | 0.20910844 |
| VTKTFTNI | ASGEQGAHSGNTLY | 0.20918614 |
| VVAAAILL | ASGEQGAHSGNTLY | 0.21011432 |
| YAALDVEL | ASGEQGAHSGNTLY | 0.20900127 |
| LASVALLL | ASGEQGAHSGNTLY | 0.20924178 |
| LDAVHRV | ASGEQGAHSGNTLY | 0.20906588 |
| QGLSSSIF | ASGEQGAHSGNTLY | 0.20899437 |
| RGTFANIRL | ASGEQGAHSGNTLY | 0.20911887 |
| RQLWWGHRI | ASGEQGAHSGNTLY | 0.20955735 |
| YGPPGTGKTLL | ASGEQGAHSGNTLY | 0.2093076 |
| AALVGAPANVA | ASGEQGAHSGNTLY | 0.20902446 |
| AATFRRTE | ASGEQGAHSGNTLY | 0.20920104 |
| AGALGGGI | ASGEQGAHSGNTLY | 0.2097245 |
| AGGAGGAGGIGAG | ASGEQGAHSGNTLY | 0.20894037 |
| AGSGALGL | ASGEQGAHSGNTLY | 0.21177964 |
| AQIADQI | ASGEQGAHSGNTLY | 0.2092018 |
| ASAVAVTYL | ASGEQGAHSGNTLY | 0.21023385 |
| ASTVLGGTGG | ASGEQGAHSGNTLY | 0.21422718 |
| AVGGEPAK | ASGEQGAHSGNTLY | 0.2093101 |
| CEQGTFGGTC | ASGEQGAHSGNTLY | 0.2089945 |
| DEGKAPAG | ASGEQGAHSGNTLY | 0.20913495 |

| | | |
|---|---|---|
| DTVVFPAL | ASGEQGAHSGNTLY | 0.20914735 |
| FEGITMAG | ASGEQGAHSGNTLY | 0.20942725 |
| FPLVVQD | ASGEQGAHSGNTLY | 0.208764 |
| GAFTTPQ | ASGEQGAHSGNTLY | 0.21181585 |
| GFGGFGTAGGVGGA | ASGEQGAHSGNTLY | 0.21036452 |
| GFLAQFL | ASGEQGAHSGNTLY | 0.20933703 |
| GGAGGAGGNAGVA | ASGEQGAHSGNTLY | 0.20858017 |
| GGVGTAGAGGAGGAGG | ASGEQGAHSGNTLY | 0.2116482 |
| GNGGGGDAGNAGSGGN | ASGEQGAHSGNTLY | 0.21035114 |
| GTGGAGGAGNDAGS | ASGEQGAHSGNTLY | 0.21057884 |
| GTTNTGWFNS | ASGEQGAHSGNTLY | 0.20895748 |
| IFMGPLFTL | ASGEQGAHSGNTLY | 0.20896797 |
| ILAHAIEAI | ASGEQGAHSGNTLY | 0.21149005 |
| IRSAIREAL | ASGEQGAHSGNTLY | 0.21055706 |
| ITFAGGG | ASGEQGAHSGNTLY | 0.21007201 |
| ITGMSGHQEV | ASGEQGAHSGNTLY | 0.21213573 |
| IVAGLACTL | ASGEQGAHSGNTLY | 0.20938565 |
| IVGGGLAGA | ASGEQGAHSGNTLY | 0.20965041 |
| IVTGVLNI | ASGEQGAHSGNTLY | 0.20964456 |
| LAQQGCI | ASGEQGAHSGNTLY | 0.21129602 |
| LCLHVEN | ASGEQGAHSGNTLY | 0.20889513 |
| LDAISDGS | ASGEQGAHSGNTLY | 0.20912698 |
| LGGFAAML | ASGEQGAHSGNTLY | 0.20993271 |
| LGGKNPMI | ASGEQGAHSGNTLY | 0.2091463 |
| LGPKGVAAIL | ASGEQGAHSGNTLY | 0.20937873 |
| LMVGVSI | ASGEQGAHSGNTLY | 0.20889458 |
| LVGAAFVL | ASGEQGAHSGNTLY | 0.20974976 |
| LVLAVAPL | ASGEQGAHSGNTLY | 0.20937681 |
| MAAIAVIDAA | ASGEQGAHSGNTLY | 0.21136877 |
| NNGINAELQ | ASGEQGAHSGNTLY | 0.20895761 |
| RGEPLVLVHG | ASGEQGAHSGNTLY | 0.2092138 |
| RGGAGGAAGLI | ASGEQGAHSGNTLY | 0.20915397 |
| RSYRMHPAV | ASGEQGAHSGNTLY | 0.21148887 |
| SAAHLLAGQLA | ASGEQGAHSGNTLY | 0.20905554 |
| SAAVGSY | ASGEQGAHSGNTLY | 0.20902447 |
| SPHAVLHRI | ASGEQGAHSGNTLY | 0.20896417 |
| SQIAKVL | ASGEQGAHSGNTLY | 0.08397329 |
| TALGHLVP | ASGEQGAHSGNTLY | 0.08887082 |
| TAPTTSGAA | ASGEQGAHSGNTLY | 0.083159074 |
| TDGDDGNGGSPGT | ASGEQGAHSGNTLY | 0.08797478 |
| TDGLGEN | ASGEQGAHSGNTLY | 0.08794549 |
| TGGPPLPGQA | ASGEQGAHSGNTLY | 0.08363968 |
| TIPAITI | ASGEQGAHSGNTLY | 0.09093436 |
| TIPGFTI | ASGEQGAHSGNTLY | 0.085074276 |
| TSIASGALA | ASGEQGAHSGNTLY | 0.08825487 |
| TVTLPGAAI | ASGEQGAHSGNTLY | 0.08625135 |
| VASGLSAALP | ASGEQGAHSGNTLY | 0.08877629 |
| VGAGVIGI | ASGEQGAHSGNTLY | 0.09753033 |
| VGPKRFGV | ASGEQGAHSGNTLY | 0.09474488 |
| VNVAFPD | ASGEQGAHSGNTLY | 0.08396617 |
| VSMTNSGVS | ASGEQGAHSGNTLY | 0.08372739 |

| | | |
|---|---|---|
| VTDVLGDLSV | ASGEQGAHSGNTLY | 0.09064963 |
| VVVVGAGFAGL | ASGEQGAHSGNTLY | 0.08514266 |
| VYTVGTVV | ASGEQGAHSGNTLY | 0.082710706 |
| WTTFSGP | ASGEQGAHSGNTLY | 0.08589474 |
| YDELVEAF | ASGEQGAHSGNTLY | 0.08208877 |
| YGTGGAGGNGG | ASGEQGAHSGNTLY | 0.08165043 |
| YTNVMAV | ASGEQGAHSGNTLY | 0.08816681 |
| FLEQVHQL | ASGEQGAHSGNTLY | 0.08669761 |
| GLSLLAVAV | ASGEQGAHSGNTLY | 0.084200144 |
| LEQVHQL | ASGEQGAHSGNTLY | 0.09250605 |
| LNAAGAI | ASGEQGAHSGNTLY | 0.08239265 |
| PAVHASL | ASGEQGAHSGNTLY | 0.085702315 |
| TELEDDA | ASGEQGAHSGNTLY | 0.089109406 |
| TPPGPPN | ASGEQGAHSGNTLY | 0.1040831 |
| TPPPPPPP | ASGEQGAHSGNTLY | 0.08462768 |
| TQSPALI | ASGDAGTAAEQF | 0.08433083 |
| EFAAAFPQ | ASGDAGTAAEQF | 0.10109667 |
| LLSGNLDVL | ASGDAGTAAEQF | 0.09244072 |
| GPANLGNYNI | ASGDAGTAAEQF | 0.08772007 |
| TLGPTLP | ASGDAGTAAEQF | 0.08828472 |
| VYAYSSCCK | ASGDAGTAAEQF | 0.083579086 |
| APAPAPAPNI | ASGDAGTAAEQF | 0.091793604 |
| PGGPVPIP | ASGDAGTAAEQF | 0.091216035 |
| FLTVALEP | ASGDAGTAAEQF | 0.088194095 |
| SQYFGANL | ASGDAGTAAEQF | 0.08928984 |
| GGVKPAA | ASGDAGTAAEQF | 0.08539655 |
| GGTGGAGGAGGW | ASGDAGTAAEQF | 0.084622234 |
| GVGGVAGLGGAGG | ASGDAGTAAEQF | 0.08159924 |
| VLLVLATTI | ASGDAGTAAEQF | 0.08520057 |
| IGIFAAML | ASGDAGTAAEQF | 0.089812316 |
| AVGDLID | ASGDAGTAAEQF | 0.086476296 |
| NISPLPL | ASGDAGTAAEQF | 0.082272515 |
| VTAVFVAI | ASGDAGTAAEQF | 0.08449485 |
| SGDDNDAAL | ASGDAGTAAEQF | 0.08202013 |
| AVAALAHS | ASGDAGTAAEQF | 0.08366089 |
| GGVTVMATP | ASGDAGTAAEQF | 0.08203104 |
| AGLAANVRD | ASGDAGTAAEQF | 0.08338211 |
| TNRPDLIDPALLRPG | ASGDAGTAAEQF | 0.10240749 |
| LANLGVTAL | ASGDAGTAAEQF | 0.09891165 |
| TGERIEL | ASGDAGTAAEQF | 0.08821192 |
| NGEELAA | ASGDAGTAAEQF | 0.09322104 |
| VLSLVWG | ASGDAGTAAEQF | 0.08538057 |
| IGYALWL | ASGDAGTAAEQF | 0.088730626 |
| LSLLLAPL | ASGDAGTAAEQF | 0.08791209 |
| TLDEALK | ASGDAGTAAEQF | 0.08410978 |
| TDGAGTAEGG | ASGDAGTAAEQF | 0.0971005 |
| NAPGGAGGNGG | ASGDAGTAAEQF | 0.08579143 |
| VSVFMGMGGNF | ASGDAGTAAEQF | 0.08753531 |
| LASYLAAL | ASGDAGTAAEQF | 0.09322535 |
| LVGGTPAL | ASGDAGTAAEQF | 0.086263016 |
| VRNADVL | ASGDAGTAAEQF | 0.08545719 |

| | | |
|---|---|---|
| TCSSPFLVSAM | ASGDAGTAAEQF | 0.0846914 |
| VHITDAE | ASGDAGTAAEQF | 0.08661103 |
| EYAAPSF | ASGDAGTAAEQF | 0.0830469 |
| VVDAFHGGL | ASGDAGTAAEQF | 0.083789885 |
| ASVVDPQ | ASGDAGTAAEQF | 0.08323527 |
| SYNEFGGLS | ASGDAGTAAEQF | 0.088242345 |
| DPDGSVY | ASGDAGTAAEQF | 0.10624167 |
| MVSTIHATE | ASGDAGTAAEQF | 0.08726421 |
| LAAGVADPV | ASGDAGTAAEQF | 0.0835084 |
| TKEQIAA | ASGDAGTAAEQF | 0.088460706 |
| SLASPAAL | ASGDAGTAAEQF | 0.086315826 |
| IGLVFYGV | ASGDAGTAAEQF | 0.088713 |
| FLLDASNL | ASGDAGTAAEQF | 0.08937082 |
| YQVGNFP | ASGDAGTAAEQF | 0.08322221 |
| ESLLSQST | ASGDAGTAAEQF | 0.0885891 |
| LFTVGFAAVG | ASGDAGTAAEQF | 0.09217678 |
| ATPPTPT | ASGDAGTAAEQF | 0.08781589 |
| TDLATLG | ASGDAGTAAEQF | 0.08768123 |
| SAGGTGR | ASGDAGTAAEQF | 0.08523379 |
| GFSNSGS | ASGDAGTAAEQF | 0.08598596 |
| GGNGGSGGTSV | ASGDAGTAAEQF | 0.08743177 |
| GTSVATGGAGN | ASGDAGTAAEQF | 0.087352574 |
| NGGNGGSGGTS | ASGDAGTAAEQF | 0.086551994 |
| AVVKFPH | ASGDAGTAAEQF | 0.08420351 |
| TPPPPPP | ASGDAGTAAEQF | 0.08548358 |
| AAPAAAAAPA | ASGDAGTAAEQF | 0.08940661 |
| IALNEDL | ASGDAGTAAEQF | 0.0848716 |
| IVNVSSM | ASGDAGTAAEQF | 0.088456914 |
| TGIPPAPR | ASGDAGTAAEQF | 0.09311572 |
| FDLGGGTF | ASGDAGTAAEQF | 0.08422319 |
| FDLGGGTFDVS | ASGDAGTAAEQF | 0.08634306 |
| NEPTAAA | ASGDAGTAAEQF | 0.08871346 |
| NEPTAAALA | ASGDAGTAAEQF | 0.09193604 |
| GSIINISSI | ASGDAGTAAEQF | 0.090294644 |
| VAPGNAGTA | ASGDAGTAAEQF | 0.08438656 |
| TGGGASLE | ASGDAGTAAEQF | 0.09428457 |
| VAVYNHYKRI | ASGDAGTAAEQF | 0.082126856 |
| TDPAPATT | ASGDAGTAAEQF | 0.08859381 |
| TNRPDILDPALLRPG | ASGDAGTAAEQF | 0.08622927 |
| LTTTTTS | ASGDAGTAAEQF | 0.08327878 |
| PTGSGKTL | ASGDAGTAAEQF | 0.0830294 |
| TGSGKTL | ASGDAGTAAEQF | 0.081844784 |
| TIGHVDHGK | ASGDAGTAAEQF | 0.09742798 |
| TIGHVDHGKTT | ASGDAGTAAEQF | 0.083984375 |
| GHVDHGKTT | ASGDAGTAAEQF | 0.08775679 |
| AQFEHTLL | ASGDAGTAAEQF | 0.085868336 |
| IVNQDEL | ASGDAGTAAEQF | 0.09079175 |
| TNGSQFF | ASGDAGTAAEQF | 0.088024996 |
| TGKGIVE | ASGDAGTAAEQF | 0.08698623 |
| LISNASDA | ASGDAGTAAEQF | 0.08402183 |
| AELRTLGELTKIAKAHGAQ | ASGDAGTAAEQF | 0.093570165 |

| | | |
|---|---|---|
| APNAVAAAEPSW | ASGDAGTAAEQF | 0.089789964 |
| APQVAGAATAVL | ASGDAGTAAEQF | 0.08659095 |
| ARRDPAAAAAILPTDA | ASGDAGTAAEQF | 0.08424975 |
| DAAARDVLAEAGLAEYF | ASGDAGTAAEQF | 0.082099654 |
| DDQIVLSAKANLAKVEQ | ASGDAGTAAEQF | 0.09268282 |
| DELLPDTGEDSDEDGH | ASGDAGTAAEQF | 0.08205272 |
| ELPEAPANRATVPQD | ASGDAGTAAEQF | 0.081326336 |
| FGQSKIERVTGIG | ASGDAGTAAEQF | 0.08151984 |
| GAGGTGSTAGGAGGAGGAGGLY | ASGDAGTAAEQF | 0.08249369 |
| GHGGTGAAGGNGGTGGAGGSGI | ASGDAGTAAEQF | 0.09344932 |
| GNGGNGGAGGEGGPGGAGGAGA | ASGDAGTAAEQF | 0.09794162 |
| GSTAGNGGNGGNATNGTIAGQP | ASGDAGTAAEQF | 0.08770801 |
| GTGGDGGNGGNGGTGG | ASGDAGTAAEQF | 0.08902347 |
| GVRRPAGIPADAPV | ASGDAGTAAEQF | 0.08749078 |
| GYRQVTDAHLLTLARR | ASGDAGTAAEQF | 0.08117838 |
| IAKMGQDGHDRG | ASGDAGTAAEQF | 0.086023994 |
| KAEPSIEIQYT | ASGDAGTAAEQF | 0.08437076 |
| KPVRIRMAADEDTVAVT | ASGDAGTAAEQF | 0.08999719 |
| MPPRIAGMRLLVIKPEP | ASGDAGTAAEQF | 0.082970925 |
| MKFYAVALFRSRGVPPTR | ASGDAGTAAEQF | 0.08628704 |
| NAGGFGILGAVAHSP | ASGDAGTAAEQF | 0.08987797 |
| NEGREITLAMAEM | ASGDAGTAAEQF | 0.08652402 |
| NELVEWSHPLGHRWRWE | ASGDAGTAAEQF | 0.0863966 |
| NLGAGGTGGNGGNPG | ASGDAGTAAEQF | 0.082980596 |
| NTEWPKWETVMAADPVD | ASGDAGTAAEQF | 0.081760146 |
| PASAAPAETQP | ASGDAGTAAEQF | 0.097139955 |
| QQSSDGKNGVRAAEITGE | ASGDAGTAAEQF | 0.083102286 |
| RAPDAVTTSWWKE | ASGDAGTAAEQF | 0.08567166 |
| SGFGNVGAHTSGWFNQ | ASGDAGTAAEQF | 0.08238744 |
| SGVAGAVGQAASVGGLKV | ASGDAGTAAEQF | 0.08158094 |
| SNGGAGTGSAGGNGGT | ASGDAGTAAEQF | 0.0829532 |
| TGGAGGEGGAAG | ASGDAGTAAEQF | 0.088277966 |
| TGGDGGAAGAGGAPGHG | ASGDAGTAAEQF | 0.08561361 |
| TMPPLLAAIANAGRHGV | ASGDAGTAAEQF | 0.08974762 |
| TSSTAGNSGGA | ASGDAGTAAEQF | 0.09821129 |
| VPATVPVGILC | ASGDAGTAAEQF | 0.08962204 |
| VQRPDSGGVVLTGRIS | ASGDAGTAAEQF | 0.08668315 |
| WTPNPVPATKGVPAGLVQAT | ASGDAGTAAEQF | 0.0846366 |
| YALKRALAKFSDRFDVVII | ASGDAGTAAEQF | 0.08492472 |
| YLIGSVIV | ASGDAGTAAEQF | 0.0842563 |
| YGQLVAVID | ASGDAGTAAEQF | 0.084479354 |
| VTVGGMTLPA | ASGDAGTAAEQF | 0.096558206 |
| VLALPAV | ASGDAGTAAEQF | 0.08334215 |
| VIPPTLNYET | ASGDAGTAAEQF | 0.084724225 |
| VGMGGMHPSSG | ASGDAGTAAEQF | 0.08448368 |
| VGLAGGAATA | ASGDAGTAAEQF | 0.08936755 |
| VGGAGGAGGSGP | ASGDAGTAAEQF | 0.097488545 |
| VFAAFMV | ASGDAGTAAEQF | 0.08492423 |
| TLYDKLR | ASGDAGTAAEQF | 0.087871216 |
| TLAVFTVL | ASGDAGTAAEQF | 0.08661448 |
| TGSVQPARV | ASGDAGTAAEQF | 0.08757634 |

| | | |
|---|---|---|
| TGGGAGSGGT | ASGDAGTAAEQF | 0.083922975 |
| SHLETSL | ASGDAGTAAEQF | 0.084854685 |
| SFTTGMT | ASGDAGTAAEQF | 0.08635615 |
| QVEVAKRMGVRQPTVS | ASGDAGTAAEQF | 0.095810354 |
| QGSGGPALAM | ASGDAGTAAEQF | 0.08500879 |
| MLAPSNP | ASGDAGTAAEQF | 0.08446222 |
| LVNNAAI | ASGDAGTAAEQF | 0.08921496 |
| LTGEINTLA | ASGDAGTAAEQF | 0.083249606 |
| LSIFGPV | ASGDAGTAAEQF | 0.08695455 |
| LLALLANL | ASGDAGTAAEQF | 0.090361245 |
| LISNASDALDKLR | ASGDAGTAAEQF | 0.083249465 |
| LINEAVP | ASGDAGTAAEQF | 0.08553179 |
| LGYAFSGL | ASGDAGTAAEQF | 0.097982645 |
| LDFVAVAL | ASGDAGTAAEQF | 0.08719227 |
| LAQPGAE | ASGDAGTAAEQF | 0.090275176 |
| ISNLGAVTGF | ASGDAGTAAEQF | 0.09180491 |
| HQEDPGRI | ASGDAGTAAEQF | 0.085990004 |
| GTPGEPTP | ASGDAGTAAEQF | 0.09452513 |
| GFFNSGS | ASGDAGTAAEQF | 0.085455075 |
| GAGGGGLTAGAN | ASGDAGTAAEQF | 0.08606441 |
| FVDALVNM | ASGDAGTAAEQF | 0.08509101 |
| FLTVAPEL | ASGDAGTAAEQF | 0.086001754 |
| FETLTAL | ASGDAGTAAEQF | 0.09003357 |
| DVINPIEL | ASGDAGTAAEQF | 0.09029473 |
| APVSGAPV | ASGDAGTAAEQF | 0.084362864 |
| ALVGGLAW | ASGDAGTAAEQF | 0.09546272 |
| ALAAAAATAI | ASGDAGTAAEQF | 0.08637212 |
| AFGLGLLL | ASGDAGTAAEQF | 0.08796066 |
| AFFAAASL | ASGDAGTAAEQF | 0.08275015 |
| AAAALPAFSPP | ASGDAGTAAEQF | 0.08432906 |
| VVLAPAL | ASGDAGTAAEQF | 0.085570455 |
| TIGHVDHGKTTL | ASGDAGTAAEQF | 0.08288624 |
| LISNASDALDK | ASGDAGTAAEQF | 0.08137258 |
| LISNASDAL | ASGDAGTAAEQF | 0.084974766 |
| ISNASDALDK | ASGDAGTAAEQF | 0.0878469 |
| NQAETLVYQTE | ASGDAGTAAEQF | 0.081300005 |
| TGGTGGTAGAG | ASGDAGTAAEQF | 0.08721227 |
| TGSTAGGAGGA | ASGDAGTAAEQF | 0.08355233 |
| DPLAVPVAAGPV | ASGDAGTAAEQF | 0.083870046 |
| GGTGGTPGNGGN | ASGDAGTAAEQF | 0.08900503 |
| PAGLAGVDPATL | ASGDAGTAAEQF | 0.08436772 |
| GNNLVVEAPHAGASV | ASGDAGTAAEQF | 0.087169565 |
| KVTRIMSASTLEQPA | ASGDAGTAAEQF | 0.08378078 |
| LGFALPIALPRRTVV | ASGDAGTAAEQF | 0.083453104 |
| TRLGNLTARKLVVLV | ASGDAGTAAEQF | 0.08995615 |
| WSQQRRLIYFPSAGP | ASGDAGTAAEQF | 0.08884559 |
| GPNGNPDPMAAAVDIR | ASGDAGTAAEQF | 0.08359928 |
| TQRCVSVADLRAAASR | ASGDAGTAAEQF | 0.09142703 |
| QFVLDVAEHPVEVLTETV | ASGDAGTAAEQF | 0.08247967 |
| GGAGGTGGAGGNGGTQAGDG | ASGDAGTAAEQF | 0.099262856 |
| PLLATLRAIAVDVDAKPAQV | ASGDAGTAAEQF | 0.09148287 |

| | | |
|---|---|---|
| APKFRTGGYMIDFWGVGYQVAK | ASGDAGTAAEQF | 0.082003154 |
| GAGGAGGAGGAAGGPGGTGGTG | ASGDAGTAAEQF | 0.088979885 |
| IGALGLLATDVAPAGSVAFGVA | ASGDAGTAAEQF | 0.08732896 |
| AGGAGGIGGNGGAGGAGGVLMG | ASGDAGTAAEQF | 0.090704344 |
| ADELIGLL | ASGDAGTAAEQF | 0.086132616 |
| ADRALKA | ASGDAGTAAEQF | 0.08267033 |
| AGAVGQVGGAGGA | ASGDAGTAAEQF | 0.08708868 |
| AGGGGGLGGAP | ASGDAGTAAEQF | 0.09433032 |
| AGGLLYGNGGAG | ASGDAGTAAEQF | 0.08956486 |
| AGLESIGL | ASGDAGTAAEQF | 0.083356835 |
| AGLKTGII | ASGDAGTAAEQF | 0.08884736 |
| ASLAALLL | ASGDAGTAAEQF | 0.087185845 |
| ASVINISSI | ASGDAGTAAEQF | 0.0869455 |
| ATPPGPA | ASGDAGTAAEQF | 0.09210259 |
| ATVASALV | ASGDAGTAAEQF | 0.086645186 |
| ATVASLIL | ASGDAGTAAEQF | 0.08930123 |
| AVNGPVLIRH | ASGDAGTAAEQF | 0.088627286 |
| DGAAGAVGI | ASGDAGTAAEQF | 0.08499588 |
| DPLLHGI | ASGDAGTAAEQF | 0.08477467 |
| ELGAALGAL | ASGDAGTAAEQF | 0.08742429 |
| ELGRTQAV | ASGDAGTAAEQF | 0.08918372 |
| FVAGGYLS | ASGDAGTAAEQF | 0.09103536 |
| GASGGAGGAGGW | ASGDAGTAAEQF | 0.083058536 |
| GGAGAGGDGGAGGI | ASGDAGTAAEQF | 0.08416997 |
| GTGHNGSQPGMGGNG | ASGDAGTAAEQF | 0.085657895 |
| GYAAASASAS | ASGDAGTAAEQF | 0.096224084 |
| HQVDVAAVQQG | ASGDAGTAAEQF | 0.084227405 |
| IGAAIAHT | ASGDAGTAAEQF | 0.087298326 |
| IGMIEVAQL | ASGDAGTAAEQF | 0.0998671 |
| IRLAPEHTSAMD | ASGDAGTAAEQF | 0.08362475 |
| IYNVSVL | ASGDAGTAAEQF | 0.0879295 |
| KQGFLGM | ASGDAGTAAEQF | 0.089205705 |
| LESVITQL | ASGDAGTAAEQF | 0.09431324 |
| LGLGALVVV | ASGDAGTAAEQF | 0.08489316 |
| LIGNGGNGGSGGI | ASGDAGTAAEQF | 0.08964427 |
| LNAINAPTL | ASGDAGTAAEQF | 0.08435491 |
| LQLLNTGV | ASGDAGTAAEQF | 0.09035859 |
| LQTSIEF | ASGDAGTAAEQF | 0.08211882 |
| MSFAVSEL | ASGDAGTAAEQF | 0.08736622 |
| NTGALGL | ASGDAGTAAEQF | 0.08189253 |
| PPGAGGAGGAGTT | ASGDAGTAAEQF | 0.08965232 |
| QLVGLGI | ASGDAGTAAEQF | 0.09155431 |
| RFLEQVHQL | ASGDAGTAAEQF | 0.09125577 |
| RGVGYVLR | ASGDAGTAAEQF | 0.08188575 |
| SATGVFAGVI | ASGDAGTAAEQF | 0.09387912 |
| SLPAITLP | ASGDAGTAAEQF | 0.09192297 |
| STALAFTV | ASGDAGTAAEQF | 0.104346834 |
| SVLTVGLGK | ASGDAGTAAEQF | 0.08546745 |
| TALAGAGSGI | ASGDAGTAAEQF | 0.09603261 |
| TGLPFTGI | ASGDAGTAAEQF | 0.09078318 |
| VAEHVTV | ASGDAGTAAEQF | 0.08430348 |

| | | |
|---|---|---|
| VDIVLNSL | ASGDAGTAAEQF | 0.08242072 |
| VLTGGGAVL | ASGDAGTAAEQF | 0.08591761 |
| VNVGLAYGI | ASGDAGTAAEQF | 0.087186925 |
| VSSAAVGAV | ASGDAGTAAEQF | 0.084139705 |
| VTSVYEAGV | ASGDAGTAAEQF | 0.08998158 |
| WVALDRGA | ASGDAGTAAEQF | 0.092019565 |
| YLVGSVVI | ASGDAGTAAEQF | 0.10032805 |
| GPPGTGKTLL | ASGDAGTAAEQF | 0.09352495 |
| GVGMTKF | ASGDAGTAAEQF | 0.09331172 |
| IGIGGSDL | ASGDAGTAAEQF | 0.08625318 |
| IGTLLAEL | ASGDAGTAAEQF | 0.084367804 |
| IYTSGTT | ASGDAGTAAEQF | 0.08767701 |
| LSFPAGV | ASGDAGTAAEQF | 0.08312766 |
| LYTSGTT | ASGDAGTAAEQF | 0.08586573 |
| VGAFPFGF | ASGDAGTAAEQF | 0.08388342 |
| VGAGTVEF | ASGDAGTAAEQF | 0.09281527 |
| VGKSTLLL | ASGDAGTAAEQF | 0.08118152 |
| AALAHAGGGIV | ASGDAGTAAEQF | 0.08463501 |
| AALVAVSLL | ASGDAGTAAEQF | 0.09464797 |
| AAVSRAVAAL | ASGDAGTAAEQF | 0.08201181 |
| AGGSSGIGGF | ASGDAGTAAEQF | 0.08881184 |
| AGLTVANANG | ASGDAGTAAEQF | 0.08870681 |
| AGLVGAVL | ASGDAGTAAEQF | 0.08191129 |
| ALSQAFTV | ASGDAGTAAEQF | 0.09283207 |
| ALTFSVSL | ASGDAGTAAEQF | 0.08289934 |
| ALTGAALVAL | ASGDAGTAAEQF | 0.08634326 |
| ASFSIGAGVL | ASGDAGTAAEQF | 0.09531337 |
| DGSVYKI | ASGDAGTAAEQF | 0.089060456 |
| EQLAEIV | ASGDAGTAAEQF | 0.084872626 |
| FAVSELTVM | ASGDAGTAAEQF | 0.085511394 |
| GAAPLAAGL | ASGDAGTAAEQF | 0.085854664 |
| GADGGDGGSGGAGG | ASGDAGTAAEQF | 0.086249396 |
| GFGLDTGGAGG | ASGDAGTAAEQF | 0.08123061 |
| GLASGGLL | ASGDAGTAAEQF | 0.0875273 |
| GRIAGAAVLL | ASGDAGTAAEQF | 0.08356129 |
| GSAGSLNTSF | ASGDAGTAAEQF | 0.0922514 |
| GTGQIGI | ASGDAGTAAEQF | 0.09023423 |
| GTLIALIL | ASGDAGTAAEQF | 0.08627544 |
| IAATMGFVVG | ASGDAGTAAEQF | 0.087529935 |
| IFAQALL | ASGDAGTAAEQF | 0.08339997 |
| IGAAVALGF | ASGDAGTAAEQF | 0.08397172 |
| IGLTPLAL | ASGDAGTAAEQF | 0.083762094 |
| ILGSNTSTL | ASGDAGTAAEQF | 0.08715425 |
| IVFSLTVL | ASGDAGTAAEQF | 0.08474554 |
| LASKATGF | ASGDAGTAAEQF | 0.08482539 |
| LASVALIL | ASGDAGTAAEQF | 0.09401742 |
| LFAVGLTL | ASGDAGTAAEQF | 0.08764614 |
| LFNGGIGTYI | ASGDAGTAAEQF | 0.08506098 |
| LTLGAGII | ASGDAGTAAEQF | 0.085768715 |
| LVAAALGDI | ASGDAGTAAEQF | 0.08819484 |
| NTAFVQLGI | ASGDAGTAAEQF | 0.086652055 |

| | | |
|---|---|---|
| QLFSGAEL | ASGDAGTAAEQF | 0.08117349 |
| SAATFITL | ASGDAGTAAEQF | 0.085229196 |
| SNYGLANLGV | ASGDAGTAAEQF | 0.0868942 |
| SYGGAYAVM | ASGDAGTAAEQF | 0.083751716 |
| TTGVGGLTLG | ASGDAGTAAEQF | 0.083966926 |
| TTQTIYI | ASGDAGTAAEQF | 0.095560625 |
| VAAGAGLGAVI | ASGDAGTAAEQF | 0.082087904 |
| VAAVVTLL | ASGDAGTAAEQF | 0.085052714 |
| VGFLGGLI | ASGDAGTAAEQF | 0.08999183 |
| VGVIPILL | ASGDAGTAAEQF | 0.1026571 |
| VIFASSLI | ASGDAGTAAEQF | 0.09462378 |
| VKGGDGGAAAG | ASGDAGTAAEQF | 0.08695174 |
| VTKTFTNI | ASGDAGTAAEQF | 0.085147 |
| VVAAAILL | ASGDAGTAAEQF | 0.09003192 |
| YAALDVEL | ASGDAGTAAEQF | 0.08910425 |
| LASVALLL | ASGDAGTAAEQF | 0.0819634 |
| LDAVHRV | ASGDAGTAAEQF | 0.081665315 |
| QGLSSSIF | ASGDAGTAAEQF | 0.08825637 |
| RGTFANIRL | ASGDAGTAAEQF | 0.08187542 |
| RQLWWGHRI | ASGDAGTAAEQF | 0.09378559 |
| YGPPGTGKTLL | ASGDAGTAAEQF | 0.08944577 |
| AALVGAPANVA | ASGDAGTAAEQF | 0.08285021 |
| AATFRRTE | ASGDAGTAAEQF | 0.08304328 |
| AGALGGGI | ASGDAGTAAEQF | 0.08342907 |
| AGGAGGAGGIGAG | ASGDAGTAAEQF | 0.09146569 |
| AGSGALGL | ASGDAGTAAEQF | 0.09022456 |
| AQIADQI | ASGDAGTAAEQF | 0.08392235 |
| ASAVAVTYL | ASGDAGTAAEQF | 0.10481917 |
| ASTVLGGTGG | ASGDAGTAAEQF | 0.09471734 |
| AVGGEPAK | ASGDAGTAAEQF | 0.08498313 |
| CEQGTFGGTC | ASGDAGTAAEQF | 0.0904424 |
| DEGKAPAG | ASGDAGTAAEQF | 0.08544825 |
| DTVVFPAL | ASGDAGTAAEQF | 0.084103435 |
| FEGITMAG | ASGDAGTAAEQF | 0.084841125 |
| FPLVVQD | ASGDAGTAAEQF | 0.08340463 |
| GAFTTPQ | ASGDAGTAAEQF | 0.08365088 |
| GFGGFGTAGGVGGA | ASGDAGTAAEQF | 0.08414536 |
| GFLAQFL | ASGDAGTAAEQF | 0.08435577 |
| GGAGGAGGNAGVA | ASGDAGTAAEQF | 0.08387835 |
| GGVGTAGAGGAGGAGG | ASGDAGTAAEQF | 0.08397963 |
| GNGGGGDAGNAGSGGN | ASGDAGTAAEQF | 0.08187931 |
| GTGGAGGAGNDAGS | ASGDAGTAAEQF | 0.08404345 |
| GTTNTGWFNS | ASGDAGTAAEQF | 0.10385896 |
| IFMGPLFTL | ASGDAGTAAEQF | 0.10819167 |
| ILAHAIEAI | ASGDAGTAAEQF | 0.084490255 |
| IRSAIREAL | ASGDAGTAAEQF | 0.0842167 |
| ITFAGGG | ASGDAGTAAEQF | 0.08672305 |
| ITGMSGHQEV | ASGDAGTAAEQF | 0.08732419 |
| IVAGLACTL | ASGDAGTAAEQF | 0.08505103 |
| IVGGGLAGA | ASGDAGTAAEQF | 0.08704346 |
| IVTGVLNI | ASGDAGTAAEQF | 0.085253276 |

| | | |
|---|---|---|
| LAQQGCI | ASGDAGTAAEQF | 0.08364292 |
| LCLHVEN | ASGDAGTAAEQF | 0.083204836 |
| LDAISDGS | ASGDAGTAAEQF | 0.08365725 |
| LGGFAAML | ASGDAGTAAEQF | 0.08866092 |
| LGGKNPMI | ASGDAGTAAEQF | 0.081434034 |
| LGPKGVAAIL | ASGDAGTAAEQF | 0.086429715 |
| LMVGVSI | ASGDAGTAAEQF | 0.08506628 |
| LVGAAFVL | ASGDAGTAAEQF | 0.08407782 |
| LVLAVAPL | ASGDAGTAAEQF | 0.08204996 |
| MAAIAVIDAA | ASGDAGTAAEQF | 0.08876045 |
| NNGINAELQ | ASGDAGTAAEQF | 0.08992601 |
| RGEPLVLVHG | ASGDAGTAAEQF | 0.08732645 |
| RGGAGGAAGLI | ASGDAGTAAEQF | 0.08239021 |
| RSYRMHPAV | ASGDAGTAAEQF | 0.08947402 |
| SAAHLLAGQLA | ASGDAGTAAEQF | 0.090254396 |
| SAAVGSY | ASGDAGTAAEQF | 0.08734701 |
| SPHAVLHRI | ASGDAGTAAEQF | 0.08325021 |
| SQIAKVL | ASGDAGTAAEQF | 0.08952667 |
| TALGHLVP | ASGDAGTAAEQF | 0.08364645 |
| TAPTTSGAA | ASGDAGTAAEQF | 0.085029975 |
| TDGDDGNGGSPGT | ASGDAGTAAEQF | 0.09323791 |
| TDGLGEN | ASGDAGTAAEQF | 0.082761124 |
| TGGPPLPGQA | ASGDAGTAAEQF | 0.08981367 |
| TIPAITI | ASGDAGTAAEQF | 0.09934097 |
| TIPGFTI | ASGDAGTAAEQF | 0.08312128 |
| TSIASGALA | ASGDAGTAAEQF | 0.085160114 |
| TVTLPGAAI | ASGDAGTAAEQF | 0.08500082 |
| VASGLSAALP | ASGDAGTAAEQF | 0.08237711 |
| VGAGVIGI | ASGDAGTAAEQF | 0.09372504 |
| VGPKRFGV | ASGDAGTAAEQF | 0.08399036 |
| VNVAFPD | ASGDAGTAAEQF | 0.08716629 |
| VSMTNSGVS | ASGDAGTAAEQF | 0.08536887 |
| VTDVLGDLSV | ASGDAGTAAEQF | 0.08491583 |
| VVVVGAGFAGL | ASGDAGTAAEQF | 0.08823396 |
| VYTVGTVV | ASGDAGTAAEQF | 0.088783175 |
| WTTFSGP | ASGDAGTAAEQF | 0.08693615 |
| YDELVEAF | ASGDAGTAAEQF | 0.08389334 |
| YGTGGAGGNGG | ASGDAGTAAEQF | 0.084502354 |
| YTNVMAV | ASGDAGTAAEQF | 0.08638285 |
| FLEQVHQL | ASGDAGTAAEQF | 0.1013345 |
| GLSLLAVAV | ASGDAGTAAEQF | 0.08266524 |
| LEQVHQL | ASGDAGTAAEQF | 0.08386701 |
| LNAAGAI | ASGDAGTAAEQF | 0.08558876 |
| PAVHASL | ASGDAGTAAEQF | 0.082591735 |
| TELEDDA | ASGDAGTAAEQF | 0.08748663 |
| TPPGPPN | ASGDAGTAAEQF | 0.088052414 |
| TPPPPPPP | ASGDAGTAAEQF | 0.0909405 |
| TQSPALI | ASSQGGEDTQY | 0.08394771 |
| EFAAAFPQ | ASSQGGEDTQY | 0.09483469 |
| LLSGNLDVL | ASSQGGEDTQY | 0.09293112 |
| GPANLGNYNI | ASSQGGEDTQY | 0.08462235 |

| | | |
|---|---|---|
| TLGPTLP | ASSQGGEDTQY | 0.08348672 |
| VYAYSSCCK | ASSQGGEDTQY | 0.083496615 |
| APAPAPAPNI | ASSQGGEDTQY | 0.08579567 |
| PGGPVPIP | ASSQGGEDTQY | 0.08760115 |
| FLTVALEP | ASSQGGEDTQY | 0.08413125 |
| SQYFGANL | ASSQGGEDTQY | 0.08709555 |
| GGVKPAA | ASSQGGEDTQY | 0.08728597 |
| GGTGGAGGAGGW | ASSQGGEDTQY | 0.090470396 |
| GVGGVAGLGGAGG | ASSQGGEDTQY | 0.08264635 |
| VLLVLATTI | ASSQGGEDTQY | 0.08531312 |
| IGIFAAML | ASSQGGEDTQY | 0.08738835 |
| AVGDLID | ASSQGGEDTQY | 0.08479566 |
| NISPLPL | ASSQGGEDTQY | 0.08413819 |
| VTAVFVAI | ASSQGGEDTQY | 0.08897697 |
| SGDDNDAAL | ASSQGGEDTQY | 0.09000483 |
| AVAALAHS | ASSQGGEDTQY | 0.08746362 |
| GGVTVMATP | ASSQGGEDTQY | 0.09362169 |
| AGLAANVRD | ASSQGGEDTQY | 0.09764635 |
| TNRPDLIDPALLRPG | ASSQGGEDTQY | 0.08491762 |
| LANLGVTAL | ASSQGGEDTQY | 0.08836657 |
| TGERIEL | ASSQGGEDTQY | 0.08242414 |
| NGEELAA | ASSQGGEDTQY | 0.08294387 |
| VLSLVWG | ASSQGGEDTQY | 0.08257348 |
| IGYALWL | ASSQGGEDTQY | 0.08546156 |
| LSLLLAPL | ASSQGGEDTQY | 0.0814839 |
| TLDEALK | ASSQGGEDTQY | 0.086479865 |
| TDGAGTAEGG | ASSQGGEDTQY | 0.08693213 |
| NAPGGAGGNGG | ASSQGGEDTQY | 0.09553445 |
| VSVFMGMGGNF | ASSQGGEDTQY | 0.08783309 |
| LASYLAAL | ASSQGGEDTQY | 0.08476128 |
| LVGGTPAL | ASSQGGEDTQY | 0.08455527 |
| VRNADVL | ASSQGGEDTQY | 0.090337664 |
| TCSSPFLVSAM | ASSQGGEDTQY | 0.08315097 |
| VHITDAE | ASSQGGEDTQY | 0.08425413 |
| EYAAPSF | ASSQGGEDTQY | 0.085300736 |
| VVDAFHGGL | ASSQGGEDTQY | 0.08649074 |
| ASVVDPQ | ASSQGGEDTQY | 0.09042368 |
| SYNEFGGLS | ASSQGGEDTQY | 0.08955652 |
| DPDGSVY | ASSQGGEDTQY | 0.08626072 |
| MVSTIHATE | ASSQGGEDTQY | 0.08599504 |
| LAAGVADPV | ASSQGGEDTQY | 0.08185335 |
| TKEQIAA | ASSQGGEDTQY | 0.08874123 |
| SLASPAAL | ASSQGGEDTQY | 0.110929504 |
| IGLVFYGV | ASSQGGEDTQY | 0.08650255 |
| FLLDASNL | ASSQGGEDTQY | 0.09079308 |
| YQVGNFP | ASSQGGEDTQY | 0.09296739 |
| ESLLSQST | ASSQGGEDTQY | 0.08494422 |
| LFTVGFAAVG | ASSQGGEDTQY | 0.08687657 |
| ATPPTPT | ASSQGGEDTQY | 0.08617114 |
| TDLATLG | ASSQGGEDTQY | 0.08755042 |
| SAGGTGR | ASSQGGEDTQY | 0.083282694 |

| | | |
|---|---|---|
| GFSNSGS | ASSQGGEDTQY | 0.084357 |
| GGNGGSGGTSV | ASSQGGEDTQY | 0.092456765 |
| GTSVATGGAGN | ASSQGGEDTQY | 0.10043219 |
| NGGNGGSGGTS | ASSQGGEDTQY | 0.08628793 |
| AVVKFPH | ASSQGGEDTQY | 0.09426604 |
| TPPPPPP | ASSQGGEDTQY | 0.08852499 |
| AAPAAAAAPA | ASSQGGEDTQY | 0.08471273 |
| IALNEDL | ASSQGGEDTQY | 0.087185465 |
| IVNVSSM | ASSQGGEDTQY | 0.08351691 |
| TGIPPAPR | ASSQGGEDTQY | 0.08345116 |
| FDLGGGTF | ASSQGGEDTQY | 0.08449312 |
| FDLGGGTFDVS | ASSQGGEDTQY | 0.08540757 |
| NEPTAAA | ASSQGGEDTQY | 0.08820217 |
| NEPTAAALA | ASSQGGEDTQY | 0.096678734 |
| GSIINISSI | ASSQGGEDTQY | 0.08673876 |
| VAPGNAGTA | ASSQGGEDTQY | 0.08784343 |
| TGGGASLE | ASSQGGEDTQY | 0.083711624 |
| VAVYNHYKRI | ASSQGGEDTQY | 0.08366747 |
| TDPAPATT | ASSQGGEDTQY | 0.08459428 |
| TNRPDILDPALLRPG | ASSQGGEDTQY | 0.08566818 |
| LTTTTTS | ASSQGGEDTQY | 0.09550044 |
| PTGSGKTL | ASSQGGEDTQY | 0.08493818 |
| TGSGKTL | ASSQGGEDTQY | 0.083275855 |
| TIGHVDHGK | ASSQGGEDTQY | 0.08639153 |
| TIGHVDHGKTT | ASSQGGEDTQY | 0.095858 |
| GHVDHGKTT | ASSQGGEDTQY | 0.08689127 |
| AQFEHTLL | ASSQGGEDTQY | 0.08561884 |
| IVNQDEL | ASSQGGEDTQY | 0.08326115 |
| TNGSQFF | ASSQGGEDTQY | 0.08846382 |
| TGKGIVE | ASSQGGEDTQY | 0.09594449 |
| LISNASDA | ASSQGGEDTQY | 0.08276543 |
| AELRTLGELTKIAKAHGAQ | ASSQGGEDTQY | 0.0933443 |
| APNAVAAAEPSW | ASSQGGEDTQY | 0.08468735 |
| APQVAGAATAVL | ASSQGGEDTQY | 0.08291429 |
| ARRDPAAAAAILPTDA | ASSQGGEDTQY | 0.08727203 |
| DAAARDVLAEAGLAEYF | ASSQGGEDTQY | 0.08896066 |
| DDQIVLSAKANLAKVEQ | ASSQGGEDTQY | 0.085357875 |
| DELLPDTGEDSDEDGH | ASSQGGEDTQY | 0.09037336 |
| ELPEAPANRATVPQD | ASSQGGEDTQY | 0.08398028 |
| FGQSKIERVTGIG | ASSQGGEDTQY | 0.08438695 |
| GAGGTGSTAGGAGGAGGAGGLY | ASSQGGEDTQY | 0.091737054 |
| GHGGTGAAGGNGGTGGAGGSGI | ASSQGGEDTQY | 0.08924186 |
| GNGGNGGAGGEGGPGGAGGAGA | ASSQGGEDTQY | 0.092637144 |
| GSTAGNGGNGGNATNGTIAGQP | ASSQGGEDTQY | 0.08446697 |
| GTGGDGGNGGNGGTGG | ASSQGGEDTQY | 0.08402745 |
| GVRRPAGIPADAPV | ASSQGGEDTQY | 0.08503438 |
| GYRQVTDAHLLTLARR | ASSQGGEDTQY | 0.08745912 |
| IAKMGQDGHDRG | ASSQGGEDTQY | 0.082902774 |
| KAEPSIEIQYT | ASSQGGEDTQY | 0.08776093 |
| KPVRIRMAADEDTVAVT | ASSQGGEDTQY | 0.089562334 |
| MPPRIAGMRLLVIKPEP | ASSQGGEDTQY | 0.083209015 |

| | | |
|---|---|---|
| MKFYAVALFRSRGVPPTR | ASSQGGEDTQY | 0.08654608 |
| NAGGFGILGAVAHSP | ASSQGGEDTQY | 0.09255267 |
| NEGREITLAMAEM | ASSQGGEDTQY | 0.08483204 |
| NELVEWSHPLGHRWRWE | ASSQGGEDTQY | 0.08285448 |
| NLGAGGTGGNGGNPG | ASSQGGEDTQY | 0.09545556 |
| NTEWPKWETVMAADPVD | ASSQGGEDTQY | 0.104531065 |
| PASAAPAETQP | ASSQGGEDTQY | 0.08782976 |
| QQSSDGKNGVRAAEITGE | ASSQGGEDTQY | 0.08409284 |
| RAPDAVTTSWWKE | ASSQGGEDTQY | 0.08174899 |
| SGFGNVGAHTSGWFNQ | ASSQGGEDTQY | 0.08747682 |
| SGVAGAVGQAASVGGLKV | ASSQGGEDTQY | 0.09364726 |
| SNGGAGTGSAGGNGGT | ASSQGGEDTQY | 0.08552862 |
| TGGAGGEGGAAG | ASSQGGEDTQY | 0.083153345 |
| TGGDGGAAGAGGAPGHG | ASSQGGEDTQY | 0.08455199 |
| TMPPLLAAIANAGRHGV | ASSQGGEDTQY | 0.09382711 |
| TSSTAGNSGGA | ASSQGGEDTQY | 0.083756305 |
| VPATVPVGILC | ASSQGGEDTQY | 0.08245778 |
| VQRPDSGGVVLTGRIS | ASSQGGEDTQY | 0.08451913 |
| WTPNPVPATKGVPAGLVQAT | ASSQGGEDTQY | 0.09063537 |
| YALKRALAKFSDRFDVVII | ASSQGGEDTQY | 0.08275368 |
| YLIGSVIV | ASSQGGEDTQY | 0.0842574 |
| YGQLVAVID | ASSQGGEDTQY | 0.10103537 |
| VTVGGMTLPA | ASSQGGEDTQY | 0.08237021 |
| VLALPAV | ASSQGGEDTQY | 0.085425645 |
| VIPPTLNYET | ASSQGGEDTQY | 0.08480903 |
| VGMGGMHPSSG | ASSQGGEDTQY | 0.08729891 |
| VGLAGGAATA | ASSQGGEDTQY | 0.08627076 |
| VGGAGGAGGSGP | ASSQGGEDTQY | 0.08721994 |
| VFAAFMV | ASSQGGEDTQY | 0.08701949 |
| TLYDKLR | ASSQGGEDTQY | 0.09053512 |
| TLAVFTVL | ASSQGGEDTQY | 0.088086806 |
| TGSVQPARV | ASSQGGEDTQY | 0.08596371 |
| TGGGAGSGGT | ASSQGGEDTQY | 0.084927656 |
| SHLETSL | ASSQGGEDTQY | 0.08136786 |
| SFTTGMT | ASSQGGEDTQY | 0.092438646 |
| QVEVAKRMGVRQPTVS | ASSQGGEDTQY | 0.08137705 |
| QGSGGPALAM | ASSQGGEDTQY | 0.09082505 |
| MLAPSNP | ASSQGGEDTQY | 0.093105674 |
| LVNNAAI | ASSQGGEDTQY | 0.08416066 |
| LTGEINTLA | ASSQGGEDTQY | 0.085566275 |
| LSIFGPV | ASSQGGEDTQY | 0.082061365 |
| LLALLANL | ASSQGGEDTQY | 0.08300979 |
| LISNASDALDKLR | ASSQGGEDTQY | 0.08564478 |
| LINEAVP | ASSQGGEDTQY | 0.08625226 |
| LGYAFSGL | ASSQGGEDTQY | 0.08754695 |
| LDFVAVAL | ASSQGGEDTQY | 0.086223006 |
| LAQPGAE | ASSQGGEDTQY | 0.09064438 |
| ISNLGAVTGF | ASSQGGEDTQY | 0.08483947 |
| HQEDPGRI | ASSQGGEDTQY | 0.08450561 |
| GTPGEPTP | ASSQGGEDTQY | 0.09341316 |
| GFFNSGS | ASSQGGEDTQY | 0.08663298 |

| | | |
|---|---|---|
| GAGGGGLTAGAN | ASSQGGEDTQY | 0.08335443 |
| FVDALVNM | ASSQGGEDTQY | 0.095264144 |
| FLTVAPEL | ASSQGGEDTQY | 0.083364144 |
| FETLTAL | ASSQGGEDTQY | 0.08829856 |
| DVINPIEL | ASSQGGEDTQY | 0.08582531 |
| APVSGAPV | ASSQGGEDTQY | 0.087872215 |
| ALVGGLAW | ASSQGGEDTQY | 0.09078888 |
| ALAAAAATAI | ASSQGGEDTQY | 0.08332239 |
| AFGLGLLL | ASSQGGEDTQY | 0.083965786 |
| AFFAAASL | ASSQGGEDTQY | 0.08641954 |
| AAAALPAFSPP | ASSQGGEDTQY | 0.09813315 |
| VVLAPAL | ASSQGGEDTQY | 0.09283609 |
| TIGHVDHGKTTL | ASSQGGEDTQY | 0.0848392 |
| LISNASDALDK | ASSQGGEDTQY | 0.08872812 |
| LISNASDAL | ASSQGGEDTQY | 0.08512873 |
| ISNASDALDK | ASSQGGEDTQY | 0.08624865 |
| NQAETLVYQTE | ASSQGGEDTQY | 0.0850791 |
| TGGTGGTAGAG | ASSQGGEDTQY | 0.08395951 |
| TGSTAGGAGGA | ASSQGGEDTQY | 0.085336216 |
| DPLAVPVAAGPV | ASSQGGEDTQY | 0.08679923 |
| GGTGGTPGNGGN | ASSQGGEDTQY | 0.08387272 |
| PAGLAGVDPATL | ASSQGGEDTQY | 0.08968372 |
| GNNLVVEAPHAGASV | ASSQGGEDTQY | 0.085819766 |
| KVTRIMSASTLEQPA | ASSQGGEDTQY | 0.08438784 |
| LGFALPIALPRRTVV | ASSQGGEDTQY | 0.08713318 |
| TRLGNLTARKLVVLV | ASSQGGEDTQY | 0.08563772 |
| WSQQRRLIYFPSAGP | ASSQGGEDTQY | 0.08772398 |
| GPNGNPDPMAAAVDIR | ASSQGGEDTQY | 0.08754177 |
| TQRCVSVADLRAAASR | ASSQGGEDTQY | 0.085168645 |
| QFVLDVAEHPVEVLTETV | ASSQGGEDTQY | 0.09226968 |
| GGAGGTGGAGGNGGTQAGDG | ASSQGGEDTQY | 0.100186296 |
| PLLATLRAIAVDVDAKPAQV | ASSQGGEDTQY | 0.08877707 |
| APKFRTGGYMIDFWGVGYQVAK | ASSQGGEDTQY | 0.08435646 |
| GAGGAGGAGGAAGGPGGTGGTG | ASSQGGEDTQY | 0.08844309 |
| IGALGLLATDVAPAGSVAFGVA | ASSQGGEDTQY | 0.09026122 |
| AGGAGGIGGNGGAGGAGGVLMG | ASSQGGEDTQY | 0.08523211 |
| ADELIGLL | ASSQGGEDTQY | 0.08205941 |
| ADRALKA | ASSQGGEDTQY | 0.08623089 |
| AGAVGQVGGAGGA | ASSQGGEDTQY | 0.08386756 |
| AGGGGGLGGAP | ASSQGGEDTQY | 0.0864542 |
| AGGLLYGNGGAG | ASSQGGEDTQY | 0.08174995 |
| AGLESIGL | ASSQGGEDTQY | 0.09453317 |
| AGLKTGII | ASSQGGEDTQY | 0.10730229 |
| ASLAALLL | ASSQGGEDTQY | 0.085656874 |
| ASVINISSI | ASSQGGEDTQY | 0.08497602 |
| ATPPGPA | ASSQGGEDTQY | 0.082524545 |
| ATVASALV | ASSQGGEDTQY | 0.08964967 |
| ATVASLIL | ASSQGGEDTQY | 0.091267705 |
| AVNGPVLIRH | ASSQGGEDTQY | 0.083630875 |
| DGAAGAVGI | ASSQGGEDTQY | 0.09228231 |
| DPLLHGI | ASSQGGEDTQY | 0.08553315 |

| | | |
|---|---|---|
| ELGAALGAL | ASSQGGEDTQY | 0.08617928 |
| ELGRTQAV | ASSQGGEDTQY | 0.093027435 |
| FVAGGYLS | ASSQGGEDTQY | 0.088291936 |
| GASGGAGGAGGW | ASSQGGEDTQY | 0.085788384 |
| GGAGAGGDGGAGGI | ASSQGGEDTQY | 0.08445777 |
| GTGHNGSQPGMGGNG | ASSQGGEDTQY | 0.08364085 |
| GYAAASASAS | ASSQGGEDTQY | 0.08642563 |
| HQVDVAAVQQG | ASSQGGEDTQY | 0.08834002 |
| IGAAIAHT | ASSQGGEDTQY | 0.08264434 |
| IGMIEVAQL | ASSQGGEDTQY | 0.08919822 |
| IRLAPEHTSAMD | ASSQGGEDTQY | 0.08574139 |
| IYNVSVL | ASSQGGEDTQY | 0.1066843 |
| KQGFLGM | ASSQGGEDTQY | 0.08732968 |
| LESVITQL | ASSQGGEDTQY | 0.086682625 |
| LGLGALVVV | ASSQGGEDTQY | 0.08437388 |
| LIGNGGNGGSGGI | ASSQGGEDTQY | 0.104380324 |
| LNAINAPTL | ASSQGGEDTQY | 0.085717894 |
| LQLLNTGV | ASSQGGEDTQY | 0.08994199 |
| LQTSIEF | ASSQGGEDTQY | 0.08729716 |
| MSFAVSEL | ASSQGGEDTQY | 0.212185 |
| NTGALGL | ASSQGGEDTQY | 0.22259371 |
| PPGAGGAGGAGTT | ASSQGGEDTQY | 0.21266113 |
| QLVGLGI | ASSQGGEDTQY | 0.21895812 |
| RFLEQVHQL | ASSQGGEDTQY | 0.22306491 |
| RGVGYVLR | ASSQGGEDTQY | 0.21425706 |
| SATGVFAGVI | ASSQGGEDTQY | 0.21959658 |
| SLPAITLP | ASSQGGEDTQY | 0.21855995 |
| STALAFTV | ASSQGGEDTQY | 0.22205006 |
| SVLTVGLGK | ASSQGGEDTQY | 0.21756288 |
| TALAGAGSGI | ASSQGGEDTQY | 0.2231975 |
| TGLPFTGI | ASSQGGEDTQY | 0.23402292 |
| VAEHVTV | ASSQGGEDTQY | 0.22271048 |
| VDIVLNSL | ASSQGGEDTQY | 0.2123888 |
| VLTGGGAVL | ASSQGGEDTQY | 0.2129197 |
| VNVGLAYGI | ASSQGGEDTQY | 0.2204538 |
| VSSAAVGAV | ASSQGGEDTQY | 0.21582708 |
| VTSVYEAGV | ASSQGGEDTQY | 0.21151431 |
| WVALDRGA | ASSQGGEDTQY | 0.2168903 |
| YLVGSVVI | ASSQGGEDTQY | 0.21155432 |
| GPPGTGKTLL | ASSQGGEDTQY | 0.21104519 |
| GVGMTKF | ASSQGGEDTQY | 0.21674752 |
| IGIGGSDL | ASSQGGEDTQY | 0.22055751 |
| IGTLLAEL | ASSQGGEDTQY | 0.21286811 |
| IYTSGTT | ASSQGGEDTQY | 0.21589455 |
| LSFPAGV | ASSQGGEDTQY | 0.21226197 |
| LYTSGTT | ASSQGGEDTQY | 0.2188654 |
| VGAFPFGF | ASSQGGEDTQY | 0.22016422 |
| VGAGTVEF | ASSQGGEDTQY | 0.23010525 |
| VGKSTLLL | ASSQGGEDTQY | 0.21760806 |
| AALAHAGGGIV | ASSQGGEDTQY | 0.21326348 |
| AALVAVSLL | ASSQGGEDTQY | 0.23040538 |

| | | |
|---|---|---|
| AAVSRAVAAL | ASSQGGEDTQY | 0.22342362 |
| AGGSSGIGGF | ASSQGGEDTQY | 0.21734624 |
| AGLTVANANG | ASSQGGEDTQY | 0.21852078 |
| AGLVGAVL | ASSQGGEDTQY | 0.2132843 |
| ALSQAFTV | ASSQGGEDTQY | 0.22429153 |
| ALTFSVSL | ASSQGGEDTQY | 0.23398212 |
| ALTGAALVAL | ASSQGGEDTQY | 0.22073647 |
| ASFSIGAGVL | ASSQGGEDTQY | 0.22335933 |
| DGSVYKI | ASSQGGEDTQY | 0.2125519 |
| EQLAEIV | ASSQGGEDTQY | 0.21393692 |
| FAVSELTVM | ASSQGGEDTQY | 0.2118704 |
| GAAPLAAGL | ASSQGGEDTQY | 0.21591777 |
| GADGGDGGSGGAGG | ASSQGGEDTQY | 0.22185397 |
| GFGLDTGGAGG | ASSQGGEDTQY | 0.21699734 |
| GLASGGLL | ASSQGGEDTQY | 0.21186265 |
| GRIAGAAVLL | ASSQGGEDTQY | 0.21578383 |
| GSAGSLNTSF | ASSQGGEDTQY | 0.21171804 |
| GTGQIGI | ASSQGGEDTQY | 0.21212718 |
| GTLIALIL | ASSQGGEDTQY | 0.21207738 |
| IAATMGFVVG | ASSQGGEDTQY | 0.21248022 |
| IFAQALL | ASSQGGEDTQY | 0.23431009 |
| IGAAVALGF | ASSQGGEDTQY | 0.23577446 |
| IGLTPLAL | ASSQGGEDTQY | 0.2140547 |
| ILGSNTSTL | ASSQGGEDTQY | 0.22264837 |
| IVFSLTVL | ASSQGGEDTQY | 0.21326782 |
| LASKATGF | ASSQGGEDTQY | 0.22477578 |
| LASVALIL | ASSQGGEDTQY | 0.22092164 |
| LFAVGLTL | ASSQGGEDTQY | 0.21204424 |
| LFNGGIGTYI | ASSQGGEDTQY | 0.23785275 |
| LTLGAGII | ASSQGGEDTQY | 0.21726874 |
| LVAAALGDI | ASSQGGEDTQY | 0.21955521 |
| NTAFVQLGI | ASSQGGEDTQY | 0.21835466 |
| QLFSGAEL | ASSQGGEDTQY | 0.22093241 |
| SAATFITL | ASSQGGEDTQY | 0.21669546 |
| SNYGLANLGV | ASSQGGEDTQY | 0.21310899 |
| SYGGAYAVM | ASSQGGEDTQY | 0.21690351 |
| TTGVGGLTLG | ASSQGGEDTQY | 0.21239634 |
| TTQTIYI | ASSQGGEDTQY | 0.21457273 |
| VAAGAGLGAVI | ASSQGGEDTQY | 0.21269965 |
| VAAVVTLL | ASSQGGEDTQY | 0.21867207 |
| VGFLGGLI | ASSQGGEDTQY | 0.22856429 |
| VGVIPILL | ASSQGGEDTQY | 0.21922195 |
| VIFASSLI | ASSQGGEDTQY | 0.21239212 |
| VKGGDGGAAAG | ASSQGGEDTQY | 0.21573648 |
| VTKTFTNI | ASSQGGEDTQY | 0.22049381 |
| VVAAAILL | ASSQGGEDTQY | 0.2206373 |
| YAALDVEL | ASSQGGEDTQY | 0.21349458 |
| LASVALLL | ASSQGGEDTQY | 0.21471438 |
| LDAVHRV | ASSQGGEDTQY | 0.21681659 |
| QGLSSSIF | ASSQGGEDTQY | 0.22557285 |
| RGTFANIRL | ASSQGGEDTQY | 0.21755865 |

| | | |
|---|---|---|
| RQLWWGHRI | ASSQGGEDTQY | 0.21664944 |
| YGPPGTGKTLL | ASSQGGEDTQY | 0.21920225 |
| AALVGAPANVA | ASSQGGEDTQY | 0.21730237 |
| AATFRRTE | ASSQGGEDTQY | 0.2145838 |
| AGALGGGI | ASSQGGEDTQY | 0.21673101 |
| AGGAGGAGGIGAG | ASSQGGEDTQY | 0.21303399 |
| AGSGALGL | ASSQGGEDTQY | 0.21300355 |
| AQIADQI | ASSQGGEDTQY | 0.21490556 |
| ASAVAVTYL | ASSQGGEDTQY | 0.2171447 |
| ASTVLGGTGG | ASSQGGEDTQY | 0.2141746 |
| AVGGEPAK | ASSQGGEDTQY | 0.21610071 |
| CEQGTFGGTC | ASSQGGEDTQY | 0.22958727 |
| DEGKAPAG | ASSQGGEDTQY | 0.21396862 |
| DTVVFPAL | ASSQGGEDTQY | 0.21541941 |
| FEGITMAG | ASSQGGEDTQY | 0.2228226 |
| FPLVVQD | ASSQGGEDTQY | 0.21550274 |
| GAFTTPQ | ASSQGGEDTQY | 0.22367759 |
| GFGGFGTAGGVGGA | ASSQGGEDTQY | 0.21553221 |
| GFLAQFL | ASSQGGEDTQY | 0.25589377 |
| GGAGGAGGNAGVA | ASSQGGEDTQY | 0.21271409 |
| GGVGTAGAGGAGGAGG | ASSQGGEDTQY | 0.22518308 |
| GNGGGGDAGNAGSGGN | ASSQGGEDTQY | 0.2126879 |
| GTGGAGGAGNDAGS | ASSQGGEDTQY | 0.21487667 |
| GTTNTGWFNS | ASSQGGEDTQY | 0.21232612 |
| IFMGPLFTL | ASSQGGEDTQY | 0.21111503 |
| ILAHAIEAI | ASSQGGEDTQY | 0.22900896 |
| IRSAIREAL | ASSQGGEDTQY | 0.21548788 |
| ITFAGGG | ASSQGGEDTQY | 0.21434398 |
| ITGMSGHQEV | ASSQGGEDTQY | 0.21766907 |
| IVAGLACTL | ASSQGGEDTQY | 0.22083381 |
| IVGGGLAGA | ASSQGGEDTQY | 0.21925208 |
| IVTGVLNI | ASSQGGEDTQY | 0.21380818 |
| LAQQGCI | ASSQGGEDTQY | 0.2131409 |
| LCLHVEN | ASSQGGEDTQY | 0.22643626 |
| LDAISDGS | ASSQGGEDTQY | 0.21748786 |
| LGGFAAML | ASSQGGEDTQY | 0.21773022 |
| LGGKNPMI | ASSQGGEDTQY | 0.21511903 |
| LGPKGVAAIL | ASSQGGEDTQY | 0.2122674 |
| LMVGVSI | ASSQGGEDTQY | 0.22259705 |
| LVGAAFVL | ASSQGGEDTQY | 0.21295683 |
| LVLAVAPL | ASSQGGEDTQY | 0.21090935 |
| MAAIAVIDAA | ASSQGGEDTQY | 0.21152726 |
| NNGINAELQ | ASSQGGEDTQY | 0.21265179 |
| RGEPLVLVHG | ASSQGGEDTQY | 0.2277917 |
| RGGAGGAAGLI | ASSQGGEDTQY | 0.23021303 |
| RSYRMHPAV | ASSQGGEDTQY | 0.2189067 |
| SAAHLLAGQLA | ASSQGGEDTQY | 0.22012474 |
| SAAVGSY | ASSQGGEDTQY | 0.21964167 |
| SPHAVLHRI | ASSQGGEDTQY | 0.21072277 |
| SQIAKVL | ASSQGGEDTQY | 0.21583636 |
| TALGHLVP | ASSQGGEDTQY | 0.21264441 |

| | | |
|---|---|---|
| TAPTTSGAA | ASSQGGEDTQY | 0.22009183 |
| TDGDDGNGGSPGT | ASSQGGEDTQY | 0.21215223 |
| TDGLGEN | ASSQGGEDTQY | 0.21548147 |
| TGGPPLPGQA | ASSQGGEDTQY | 0.22266789 |
| TIPAITI | ASSQGGEDTQY | 0.21879458 |
| TIPGFTI | ASSQGGEDTQY | 0.21863016 |
| TSIASGALA | ASSQGGEDTQY | 0.21169154 |
| TVTLPGAAI | ASSQGGEDTQY | 0.21125498 |
| VASGLSAALP | ASSQGGEDTQY | 0.22804035 |
| VGAGVIGI | ASSQGGEDTQY | 0.21646377 |
| VGPKRFGV | ASSQGGEDTQY | 0.21308573 |
| VNVAFPD | ASSQGGEDTQY | 0.21166085 |
| VSMTNSGVS | ASSQGGEDTQY | 0.21507001 |
| VTDVLGDLSV | ASSQGGEDTQY | 0.21419582 |
| VVVVGAGFAGL | ASSQGGEDTQY | 0.21735539 |
| VYTVGTVV | ASSQGGEDTQY | 0.21253845 |
| WTTFSGP | ASSQGGEDTQY | 0.21821825 |
| YDELVEAF | ASSQGGEDTQY | 0.23030514 |
| YGTGGAGGNGG | ASSQGGEDTQY | 0.22568765 |
| YTNVMAV | ASSQGGEDTQY | 0.21834065 |
| FLEQVHQL | ASSQGGEDTQY | 0.21735978 |
| GLSLLAVAV | ASSQGGEDTQY | 0.2143502 |
| LEQVHQL | ASSQGGEDTQY | 0.21685056 |
| LNAAGAI | ASSQGGEDTQY | 0.21639718 |
| PAVHASL | ASSQGGEDTQY | 0.22388056 |
| TELEDDA | ASSQGGEDTQY | 0.2138169 |
| TPPGPPN | ASSQGGEDTQY | 0.21371204 |
| TPPPPPPP | ASSQGGEDTQY | 0.21829313 |
| TQSPALI | ASSDQGARNSPLY | 0.22640558 |
| EFAAAFPQ | ASSDQGARNSPLY | 0.23588042 |
| LLSGNLDVL | ASSDQGARNSPLY | 0.21215343 |
| GPANLGNYNI | ASSDQGARNSPLY | 0.21746165 |
| TLGPTLP | ASSDQGARNSPLY | 0.21675287 |
| VYAYSSCCK | ASSDQGARNSPLY | 0.21682484 |
| APAPAPAPNI | ASSDQGARNSPLY | 0.21273535 |
| PGGPVPIP | ASSDQGARNSPLY | 0.21868885 |
| FLTVALEP | ASSDQGARNSPLY | 0.21540302 |
| SQYFGANL | ASSDQGARNSPLY | 0.24203876 |
| GGVKPAA | ASSDQGARNSPLY | 0.21593627 |
| GGTGGAGGAGGW | ASSDQGARNSPLY | 0.21478187 |
| GVGGVAGLGGAGG | ASSDQGARNSPLY | 0.2230833 |
| VLLVLATTI | ASSDQGARNSPLY | 0.21346559 |
| IGIFAAML | ASSDQGARNSPLY | 0.21822459 |
| AVGDLID | ASSDQGARNSPLY | 0.21781835 |
| NISPLPL | ASSDQGARNSPLY | 0.21207865 |
| VTAVFVAI | ASSDQGARNSPLY | 0.21774507 |
| SGDDNDAAL | ASSDQGARNSPLY | 0.24175644 |
| AVAALAHS | ASSDQGARNSPLY | 0.21898983 |
| GGVTVMATP | ASSDQGARNSPLY | 0.22113886 |
| AGLAANVRD | ASSDQGARNSPLY | 0.22835468 |
| TNRPDLIDPALLRPG | ASSDQGARNSPLY | 0.21622829 |

| | | |
|---|---|---|
| LANLGVTAL | ASSDQGARNSPLY | 0.21792896 |
| TGERIEL | ASSDQGARNSPLY | 0.21429564 |
| NGEELAA | ASSDQGARNSPLY | 0.21579437 |
| VLSLVWG | ASSDQGARNSPLY | 0.21593924 |
| IGYALWL | ASSDQGARNSPLY | 0.22064483 |
| LSLLLAPL | ASSDQGARNSPLY | 0.22547546 |
| TLDEALK | ASSDQGARNSPLY | 0.2251974 |
| TDGAGTAEGG | ASSDQGARNSPLY | 0.215849 |
| NAPGGAGGNGG | ASSDQGARNSPLY | 0.2329011 |
| VSVFMGMGGNF | ASSDQGARNSPLY | 0.21608941 |
| LASYLAAL | ASSDQGARNSPLY | 0.22429776 |
| LVGGTPAL | ASSDQGARNSPLY | 0.2116457 |
| VRNADVL | ASSDQGARNSPLY | 0.21642222 |
| TCSSPFLVSAM | ASSDQGARNSPLY | 0.21504702 |
| VHITDAE | ASSDQGARNSPLY | 0.21148066 |
| EYAAPSF | ASSDQGARNSPLY | 0.21086866 |
| VVDAFHGGL | ASSDQGARNSPLY | 0.21485724 |
| ASVVDPQ | ASSDQGARNSPLY | 0.22114982 |
| SYNEFGGLS | ASSDQGARNSPLY | 0.21081024 |
| DPDGSVY | ASSDQGARNSPLY | 0.2197093 |
| MVSTIHATE | ASSDQGARNSPLY | 0.2118566 |
| LAAGVADPV | ASSDQGARNSPLY | 0.21416537 |
| TKEQIAA | ASSDQGARNSPLY | 0.2226844 |
| SLASPAAL | ASSDQGARNSPLY | 0.21477601 |
| IGLVFYGV | ASSDQGARNSPLY | 0.21475102 |
| FLLDASNL | ASSDQGARNSPLY | 0.21317412 |
| YQVGNFP | ASSDQGARNSPLY | 0.21447504 |
| ESLLSQST | ASSDQGARNSPLY | 0.22111486 |
| LFTVGFAAVG | ASSDQGARNSPLY | 0.21813591 |
| ATPPTPT | ASSDQGARNSPLY | 0.21259741 |
| TDLATLG | ASSDQGARNSPLY | 0.22157365 |
| SAGGTGR | ASSDQGARNSPLY | 0.21205355 |
| GFSNSGS | ASSDQGARNSPLY | 0.23568094 |
| GGNGGSGGTSV | ASSDQGARNSPLY | 0.22785616 |
| GTSVATGGAGN | ASSDQGARNSPLY | 0.21207699 |
| NGGNGGSGGTS | ASSDQGARNSPLY | 0.21607086 |
| AVVKFPH | ASSDQGARNSPLY | 0.21473174 |
| TPPPPPP | ASSDQGARNSPLY | 0.21663652 |
| AAPAAAAAPA | ASSDQGARNSPLY | 0.21439031 |
| IALNEDL | ASSDQGARNSPLY | 0.21278487 |
| IVNVSSM | ASSDQGARNSPLY | 0.21857929 |
| TGIPPAPR | ASSDQGARNSPLY | 0.23172429 |
| FDLGGGTF | ASSDQGARNSPLY | 0.21408646 |
| FDLGGGTFDVS | ASSDQGARNSPLY | 0.21769536 |
| NEPTAAA | ASSDQGARNSPLY | 0.22312292 |
| NEPTAAALA | ASSDQGARNSPLY | 0.21822773 |
| GSIINISSI | ASSDQGARNSPLY | 0.21310097 |
| VAPGNAGTA | ASSDQGARNSPLY | 0.22685252 |
| TGGGASLE | ASSDQGARNSPLY | 0.21520855 |
| VAVYNHYKRI | ASSDQGARNSPLY | 0.2238335 |
| TDPAPATT | ASSDQGARNSPLY | 0.22312312 |

| | | |
|---|---|---|
| TNRPDILDPALLRPG | ASSDQGARNSPLY | 0.23250416 |
| LTTTTTS | ASSDQGARNSPLY | 0.21603633 |
| PTGSGKTL | ASSDQGARNSPLY | 0.21644099 |
| TGSGKTL | ASSDQGARNSPLY | 0.22141345 |
| TIGHVDHGK | ASSDQGARNSPLY | 0.21832764 |
| TIGHVDHGKTT | ASSDQGARNSPLY | 0.21244574 |
| GHVDHGKTT | ASSDQGARNSPLY | 0.21631835 |
| AQFEHTLL | ASSDQGARNSPLY | 0.21708466 |
| IVNQDEL | ASSDQGARNSPLY | 0.23020089 |
| TNGSQFF | ASSDQGARNSPLY | 0.21270035 |
| TGKGIVE | ASSDQGARNSPLY | 0.21329603 |
| LISNASDA | ASSDQGARNSPLY | 0.21941628 |
| AELRTLGELTKIAKAHGAQ | ASSDQGARNSPLY | 0.21321294 |
| APNAVAAAEPSW | ASSDQGARNSPLY | 0.22115608 |
| APQVAGAATAVL | ASSDQGARNSPLY | 0.21662046 |
| ARRDPAAAAAILPTDA | ASSDQGARNSPLY | 0.21722943 |
| DAAARDVLAEAGLAEYF | ASSDQGARNSPLY | 0.21389644 |
| DDQIVLSAKANLAKVEQ | ASSDQGARNSPLY | 0.21897556 |
| DELLPDTGEDSDEDGH | ASSDQGARNSPLY | 0.21458608 |
| ELPEAPANRATVPQD | ASSDQGARNSPLY | 0.22122982 |
| FGQSKIERVTGIG | ASSDQGARNSPLY | 0.21166414 |
| GAGGTGSTAGGAGGAGGAGGLY | ASSDQGARNSPLY | 0.22060767 |
| GHGGTGAAGGNGGTGGAGGSGI | ASSDQGARNSPLY | 0.21250372 |
| GNGGNGGAGGEGGPGGAGGAGA | ASSDQGARNSPLY | 0.22131893 |
| GSTAGNGGNGGNATNGTIAGQP | ASSDQGARNSPLY | 0.22249816 |
| GTGGDGGNGGNGGTGG | ASSDQGARNSPLY | 0.22523329 |
| GVRRPAGIPADAPV | ASSDQGARNSPLY | 0.2115309 |
| GYRQVTDAHLLTLARR | ASSDQGARNSPLY | 0.23411813 |
| IAKMGQDGHDRG | ASSDQGARNSPLY | 0.22606836 |
| KAEPSIEIQYT | ASSDQGARNSPLY | 0.22218077 |
| KPVRIRMAADEDTVAVT | ASSDQGARNSPLY | 0.21689825 |
| MPPRIAGMRLLVIKPEP | ASSDQGARNSPLY | 0.2250157 |
| MKFYAVALFRSRGVPPTR | ASSDQGARNSPLY | 0.21992722 |
| NAGGFGILGAVAHSP | ASSDQGARNSPLY | 0.21488102 |
| NEGREITLAMAEM | ASSDQGARNSPLY | 0.21179356 |
| NELVEWSHPLGHRWRWE | ASSDQGARNSPLY | 0.21315888 |
| NLGAGGTGGNGGNPG | ASSDQGARNSPLY | 0.21718131 |
| NTEWPKWETVMAADPVD | ASSDQGARNSPLY | 0.21391082 |
| PASAAPAETQP | ASSDQGARNSPLY | 0.23113725 |
| QQSSDGKNGVRAAEITGE | ASSDQGARNSPLY | 0.21590649 |
| RAPDAVTTSWWKE | ASSDQGARNSPLY | 0.2382576 |
| SGFGNVGAHTSGWFNQ | ASSDQGARNSPLY | 0.22740996 |
| SGVAGAVGQAASVGGLKV | ASSDQGARNSPLY | 0.22764923 |
| SNGGAGTGSAGGNGGT | ASSDQGARNSPLY | 0.2155017 |
| TGGAGGEGGAAG | ASSDQGARNSPLY | 0.21852207 |
| TGGDGGAAGAGGAPGHG | ASSDQGARNSPLY | 0.21939924 |
| TMPPLLAAIANAGRHGV | ASSDQGARNSPLY | 0.21317786 |
| TSSTAGNSGGA | ASSDQGARNSPLY | 0.21740419 |
| VPATVPVGILC | ASSDQGARNSPLY | 0.21597852 |
| VQRPDSGGVVLTGRIS | ASSDQGARNSPLY | 0.22808105 |
| WTPNPVPATKGVPAGLVQAT | ASSDQGARNSPLY | 0.21066362 |

| | | |
|---|---|---|
| YALKRALAKFSDRFDVVII | ASSDQGARNSPLY | 0.21342315 |
| YLIGSVIV | ASSDQGARNSPLY | 0.22463508 |
| YGQLVAVID | ASSDQGARNSPLY | 0.21380448 |
| VTVGGMTLPA | ASSDQGARNSPLY | 0.22261673 |
| VLALPAV | ASSDQGARNSPLY | 0.22331971 |
| VIPPTLNYET | ASSDQGARNSPLY | 0.21146397 |
| VGMGGMHPSSG | ASSDQGARNSPLY | 0.22740386 |
| VGLAGGAATA | ASSDQGARNSPLY | 0.21299969 |
| VGGAGGAGGSGP | ASSDQGARNSPLY | 0.21554604 |
| VFAAFMV | ASSDQGARNSPLY | 0.23428556 |
| TLYDKLR | ASSDQGARNSPLY | 0.22100106 |
| TLAVFTVL | ASSDQGARNSPLY | 0.21311428 |
| TGSVQPARV | ASSDQGARNSPLY | 0.21715558 |
| TGGGAGSGGT | ASSDQGARNSPLY | 0.21634693 |
| SHLETSL | ASSDQGARNSPLY | 0.21431711 |
| SFTTGMT | ASSDQGARNSPLY | 0.21181916 |
| QVEVAKRMGVRQPTVS | ASSDQGARNSPLY | 0.22024696 |
| QGSGGPALAM | ASSDQGARNSPLY | 0.21378371 |
| MLAPSNP | ASSDQGARNSPLY | 0.21913105 |
| LVNNAAI | ASSDQGARNSPLY | 0.22163542 |
| LTGEINTLA | ASSDQGARNSPLY | 0.21343824 |
| LSIFGPV | ASSDQGARNSPLY | 0.21840487 |
| LLALLANL | ASSDQGARNSPLY | 0.21389805 |
| LISNASDALDKLR | ASSDQGARNSPLY | 0.2133872 |
| LINEAVP | ASSDQGARNSPLY | 0.21437491 |
| LGYAFSGL | ASSDQGARNSPLY | 0.21673085 |
| LDFVAVAL | ASSDQGARNSPLY | 0.21371017 |
| LAQPGAE | ASSDQGARNSPLY | 0.21555069 |
| ISNLGAVTGF | ASSDQGARNSPLY | 0.22558697 |
| HQEDPGRI | ASSDQGARNSPLY | 0.21593468 |
| GTPGEPTP | ASSDQGARNSPLY | 0.21629338 |
| GFFNSGS | ASSDQGARNSPLY | 0.21576926 |
| GAGGGGLTAGAN | ASSDQGARNSPLY | 0.21807879 |
| FVDALVNM | ASSDQGARNSPLY | 0.22165427 |
| FLTVAPEL | ASSDQGARNSPLY | 0.21090338 |
| FETLTAL | ASSDQGARNSPLY | 0.21430622 |
| DVINPIEL | ASSDQGARNSPLY | 0.21301106 |
| APVSGAPV | ASSDQGARNSPLY | 0.21502876 |
| ALVGGLAW | ASSDQGARNSPLY | 0.21412894 |
| ALAAAAATAI | ASSDQGARNSPLY | 0.2179916 |
| AFGLGLLL | ASSDQGARNSPLY | 0.21449064 |
| AFFAAASL | ASSDQGARNSPLY | 0.2134376 |
| AAAALPAFSPP | ASSDQGARNSPLY | 0.21541962 |
| VVLAPAL | ASSDQGARNSPLY | 0.23626289 |
| TIGHVDHGKTTL | ASSDQGARNSPLY | 0.22862615 |
| LISNASDALDK | ASSDQGARNSPLY | 0.22024713 |
| LISNASDAL | ASSDQGARNSPLY | 0.21427071 |
| ISNASDALDK | ASSDQGARNSPLY | 0.21832673 |
| NQAETLVYQTE | ASSDQGARNSPLY | 0.22027217 |
| TGGTGGTAGAG | ASSDQGARNSPLY | 0.21206962 |
| TGSTAGGAGGA | ASSDQGARNSPLY | 0.21127124 |

| | | |
|---|---|---|
| DPLAVPVAAGPV | ASSDQGARNSPLY | 0.22387935 |
| GGTGGTPGNGGN | ASSDQGARNSPLY | 0.21141803 |
| PAGLAGVDPATL | ASSDQGARNSPLY | 0.23342094 |
| GNNLVVEAPHAGASV | ASSDQGARNSPLY | 0.22143905 |
| KVTRIMSASTLEQPA | ASSDQGARNSPLY | 0.21313332 |
| LGFALPIALPRRTVV | ASSDQGARNSPLY | 0.21189333 |
| TRLGNLTARKLVVLV | ASSDQGARNSPLY | 0.21267411 |
| WSQQRRLIYFPSAGP | ASSDQGARNSPLY | 0.21962324 |
| GPNGNPDPMAAAVDIR | ASSDQGARNSPLY | 0.21541297 |
| TQRCVSVADLRAAASR | ASSDQGARNSPLY | 0.21395627 |
| QFVLDVAEHPVEVLTETV | ASSDQGARNSPLY | 0.24114202 |
| GGAGGTGGAGGNGGTQAGDG | ASSDQGARNSPLY | 0.2297404 |
| PLLATLRAIAVDVDAKPAQV | ASSDQGARNSPLY | 0.21617296 |
| APKFRTGGYMIDFWGVGYQVAK | ASSDQGARNSPLY | 0.21478036 |
| GAGGAGGAGGAAGGPGGTGGTG | ASSDQGARNSPLY | 0.21573895 |
| IGALGLLATDVAPAGSVAFGVA | ASSDQGARNSPLY | 0.2148516 |
| AGGAGGIGGNGGAGGAGGVLMG | ASSDQGARNSPLY | 0.21458451 |
| ADELIGLL | ASSDQGARNSPLY | 0.21324694 |
| ADRALKA | ASSDQGARNSPLY | 0.21278496 |
| AGAVGQVGGAGGA | ASSDQGARNSPLY | 0.21259294 |
| AGGGGGLGGAP | ASSDQGARNSPLY | 0.21582158 |
| AGGLLYGNGGAG | ASSDQGARNSPLY | 0.21443063 |
| AGLESIGL | ASSDQGARNSPLY | 0.22077566 |
| AGLKTGII | ASSDQGARNSPLY | 0.21173176 |
| ASLAALLL | ASSDQGARNSPLY | 0.21684317 |
| ASVINISSI | ASSDQGARNSPLY | 0.23033376 |
| ATPPGPA | ASSDQGARNSPLY | 0.24322912 |
| ATVASALV | ASSDQGARNSPLY | 0.21409488 |
| ATVASLIL | ASSDQGARNSPLY | 0.2167526 |
| AVNGPVLIRH | ASSDQGARNSPLY | 0.22037865 |
| DGAAGAVGI | ASSDQGARNSPLY | 0.21423537 |
| DPLLHGI | ASSDQGARNSPLY | 0.21484838 |
| ELGAALGAL | ASSDQGARNSPLY | 0.21947175 |
| ELGRTQAV | ASSDQGARNSPLY | 0.21647139 |
| FVAGGYLS | ASSDQGARNSPLY | 0.21494953 |
| GASGGAGGAGGW | ASSDQGARNSPLY | 0.21272823 |
| GGAGAGGDGGAGGI | ASSDQGARNSPLY | 0.21621475 |
| GTGHNGSQPGMGGNG | ASSDQGARNSPLY | 0.21732366 |
| GYAAASASAS | ASSDQGARNSPLY | 0.21133503 |
| HQVDVAAVQQG | ASSDQGARNSPLY | 0.21679527 |
| IGAAIAHT | ASSDQGARNSPLY | 0.21478467 |
| IGMIEVAQL | ASSDQGARNSPLY | 0.21424137 |
| IRLAPEHTSAMD | ASSDQGARNSPLY | 0.21128523 |
| IYNVSVL | ASSDQGARNSPLY | 0.21628599 |
| KQGFLGM | ASSDQGARNSPLY | 0.22472897 |
| LESVITQL | ASSDQGARNSPLY | 0.21599454 |
| LGLGALVVV | ASSDQGARNSPLY | 0.21192905 |
| LIGNGGNGGSGGI | ASSDQGARNSPLY | 0.21746182 |
| LNAINAPTL | ASSDQGARNSPLY | 0.23249082 |
| LQLLNTGV | ASSDQGARNSPLY | 0.21801712 |
| LQTSIEF | ASSDQGARNSPLY | 0.21251878 |

| | | |
|---|---|---|
| MSFAVSEL | ASSDQGARNSPLY | 0.21901189 |
| NTGALGL | ASSDQGARNSPLY | 0.21428117 |
| PPGAGGAGGAGTT | ASSDQGARNSPLY | 0.21489161 |
| QLVGLGI | ASSDQGARNSPLY | 0.22630358 |
| RFLEQVHQL | ASSDQGARNSPLY | 0.21264707 |
| RGVGYVLR | ASSDQGARNSPLY | 0.22112356 |
| SATGVFAGVI | ASSDQGARNSPLY | 0.23333731 |
| SLPAITLP | ASSDQGARNSPLY | 0.21307418 |
| STALAFTV | ASSDQGARNSPLY | 0.21407352 |
| SVLTVGLGK | ASSDQGARNSPLY | 0.21459365 |
| TALAGAGSGI | ASSDQGARNSPLY | 0.21138698 |
| TGLPFTGI | ASSDQGARNSPLY | 0.22649112 |
| VAEHVTV | ASSDQGARNSPLY | 0.21352662 |
| VDIVLNSL | ASSDQGARNSPLY | 0.2214438 |
| VLTGGGAVL | ASSDQGARNSPLY | 0.21516997 |
| VNVGLAYGI | ASSDQGARNSPLY | 0.21794784 |
| VSSAAVGAV | ASSDQGARNSPLY | 0.21960838 |
| VTSVYEAGV | ASSDQGARNSPLY | 0.22446416 |
| WVALDRGA | ASSDQGARNSPLY | 0.2209565 |
| YLVGSVVI | ASSDQGARNSPLY | 0.21337007 |
| GPPGTGKTLL | ASSDQGARNSPLY | 0.21421105 |
| GVGMTKF | ASSDQGARNSPLY | 0.21542366 |
| IGIGGSDL | ASSDQGARNSPLY | 0.24203156 |
| IGTLLAEL | ASSDQGARNSPLY | 0.21529175 |
| IYTSGTT | ASSDQGARNSPLY | 0.216708 |
| LSFPAGV | ASSDQGARNSPLY | 0.21391441 |
| LYTSGTT | ASSDQGARNSPLY | 0.21278483 |
| VGAFPFGF | ASSDQGARNSPLY | 0.21490853 |
| VGAGTVEF | ASSDQGARNSPLY | 0.22012773 |
| VGKSTLLL | ASSDQGARNSPLY | 0.2248832 |
| AALAHAGGGIV | ASSDQGARNSPLY | 0.21504952 |
| AALVAVSLL | ASSDQGARNSPLY | 0.22455396 |
| AAVSRAVAAL | ASSDQGARNSPLY | 0.21657792 |
| AGGSSGIGGF | ASSDQGARNSPLY | 0.21486896 |
| AGLTVANANG | ASSDQGARNSPLY | 0.2139942 |
| AGLVGAVL | ASSDQGARNSPLY | 0.21378551 |
| ALSQAFTV | ASSDQGARNSPLY | 0.21691062 |
| ALTFSVSL | ASSDQGARNSPLY | 0.21882045 |
| ALTGAALVAL | ASSDQGARNSPLY | 0.21536589 |
| ASFSIGAGVL | ASSDQGARNSPLY | 0.21838573 |
| DGSVYKI | ASSDQGARNSPLY | 0.21621032 |
| EQLAEIV | ASSDQGARNSPLY | 0.21667218 |
| FAVSELTVM | ASSDQGARNSPLY | 0.21246535 |
| GAAPLAAGL | ASSDQGARNSPLY | 0.21306676 |
| GADGGDGGSGGAGG | ASSDQGARNSPLY | 0.21333733 |
| GFGLDTGGAGG | ASSDQGARNSPLY | 0.2143399 |
| GLASGGLL | ASSDQGARNSPLY | 0.2122037 |
| GRIAGAAVLL | ASSDQGARNSPLY | 0.21821044 |
| GSAGSLNTSF | ASSDQGARNSPLY | 0.22428405 |
| GTGQIGI | ASSDQGARNSPLY | 0.22205628 |
| GTLIALIL | ASSDQGARNSPLY | 0.23291765 |

| | | |
|---|---|---|
| IAATMGFVVG | ASSDQGARNSPLY | 0.23659116 |
| IFAQALL | ASSDQGARNSPLY | 0.21617833 |
| IGAAVALGF | ASSDQGARNSPLY | 0.22038862 |
| IGLTPLAL | ASSDQGARNSPLY | 0.2113632 |
| ILGSNTSTL | ASSDQGARNSPLY | 0.21548784 |
| IVFSLTVL | ASSDQGARNSPLY | 0.21210295 |
| LASKATGF | ASSDQGARNSPLY | 0.21881479 |
| LASVALIL | ASSDQGARNSPLY | 0.2120127 |
| LFAVGLTL | ASSDQGARNSPLY | 0.22384384 |
| LFNGGIGTYI | ASSDQGARNSPLY | 0.21378127 |
| LTLGAGII | ASSDQGARNSPLY | 0.22738352 |
| LVAAALGDI | ASSDQGARNSPLY | 0.2225743 |
| NTAFVQLGI | ASSDQGARNSPLY | 0.21281286 |
| QLFSGAEL | ASSDQGARNSPLY | 0.21866262 |
| SAATFITL | ASSDQGARNSPLY | 0.21738169 |
| SNYGLANLGV | ASSDQGARNSPLY | 0.21244147 |
| SYGGAYAVM | ASSDQGARNSPLY | 0.21445638 |
| TTGVGGLTLG | ASSDQGARNSPLY | 0.21605383 |
| TTQTIYI | ASSDQGARNSPLY | 0.21553202 |
| VAAGAGLGAVI | ASSDQGARNSPLY | 0.22702734 |
| VAAVVTLL | ASSDQGARNSPLY | 0.22370693 |
| VGFLGGLI | ASSDQGARNSPLY | 0.21662202 |
| VGVIPILL | ASSDQGARNSPLY | 0.22841218 |
| VIFASSLI | ASSDQGARNSPLY | 0.21111451 |
| VKGGDGGAAAG | ASSDQGARNSPLY | 0.22146477 |
| VTKTFTNI | ASSDQGARNSPLY | 0.23947598 |
| VVAAAILL | ASSDQGARNSPLY | 0.21698721 |
| YAALDVEL | ASSDQGARNSPLY | 0.22648787 |
| LASVALLL | ASSDQGARNSPLY | 0.22260219 |
| LDAVHRV | ASSDQGARNSPLY | 0.21708256 |
| QGLSSSIF | ASSDQGARNSPLY | 0.22236119 |
| RGTFANIRL | ASSDQGARNSPLY | 0.21376146 |
| RQLWWGHRI | ASSDQGARNSPLY | 0.21780519 |
| YGPPGTGKTLL | ASSDQGARNSPLY | 0.21344315 |
| AALVGAPANVA | ASSDQGARNSPLY | 0.2127586 |
| AATFRRTE | ASSDQGARNSPLY | 0.22356288 |
| AGALGGGI | ASSDQGARNSPLY | 0.23764437 |
| AGGAGGAGGIGAG | ASSDQGARNSPLY | 0.21579304 |
| AGSGALGL | ASSDQGARNSPLY | 0.22866005 |
| AQIADQI | ASSDQGARNSPLY | 0.2183603 |
| ASAVAVTYL | ASSDQGARNSPLY | 0.21730533 |
| ASTVLGGTGG | ASSDQGARNSPLY | 0.22138721 |
| AVGGEPAK | ASSDQGARNSPLY | 0.21320046 |
| CEQGTFGGTC | ASSDQGARNSPLY | 0.21360388 |
| DEGKAPAG | ASSDQGARNSPLY | 0.21381675 |
| DTVVFPAL | ASSDQGARNSPLY | 0.21552645 |
| FEGITMAG | ASSDQGARNSPLY | 0.21933524 |
| FPLVVQD | ASSDQGARNSPLY | 0.24667737 |
| GAFTTPQ | ASSDQGARNSPLY | 0.21984094 |
| GFGGFGTAGGVGGA | ASSDQGARNSPLY | 0.21934025 |
| GFLAQFL | ASSDQGARNSPLY | 0.214837 |

| | | |
|---|---|---|
| GGAGGAGGNAGVA | ASSDQGARNSPLY | 0.21308535 |
| GGVGTAGAGGAGGAGG | ASSDQGARNSPLY | 0.21463376 |
| GNGGGGDAGNAGSGGN | ASSDQGARNSPLY | 0.21674147 |
| GTGGAGGAGNDAGS | ASSDQGARNSPLY | 0.2259721 |
| GTTNTGWFNS | ASSDQGARNSPLY | 0.21605167 |
| IFMGPLFTL | ASSDQGARNSPLY | 0.21481803 |
| ILAHAIEAI | ASSDQGARNSPLY | 0.21962005 |
| IRSAIREAL | ASSDQGARNSPLY | 0.232635 |
| ITFAGGG | ASSDQGARNSPLY | 0.21929395 |
| ITGMSGHQEV | ASSDQGARNSPLY | 0.21661864 |
| IVAGLACTL | ASSDQGARNSPLY | 0.2144153 |
| IVGGGLAGA | ASSDQGARNSPLY | 0.2189318 |
| IVTGVLNI | ASSDQGARNSPLY | 0.22426179 |
| LAQQGCI | ASSDQGARNSPLY | 0.21523218 |
| LCLHVEN | ASSDQGARNSPLY | 0.22452481 |
| LDAISDGS | ASSDQGARNSPLY | 0.21437055 |
| LGGFAAML | ASSDQGARNSPLY | 0.21184175 |
| LGGKNPMI | ASSDQGARNSPLY | 0.21532398 |
| LGPKGVAAIL | ASSDQGARNSPLY | 0.21921322 |
| LMVGVSI | ASSDQGARNSPLY | 0.21446712 |
| LVGAAFVL | ASSDQGARNSPLY | 0.23138091 |
| LVLAVAPL | ASSDQGARNSPLY | 0.21180077 |
| MAAIAVIDAA | ASSDQGARNSPLY | 0.21536224 |
| NNGINAELQ | ASSDQGARNSPLY | 0.22562964 |
| RGEPLVLVHG | ASSDQGARNSPLY | 0.22688824 |
| RGGAGGAAGLI | ASSDQGARNSPLY | 0.22200102 |
| RSYRMHPAV | ASSDQGARNSPLY | 0.21402398 |
| SAAHLLAGQLA | ASSDQGARNSPLY | 0.21400616 |
| SAAVGSY | ASSDQGARNSPLY | 0.21566324 |
| SPHAVLHRI | ASSDQGARNSPLY | 0.21769817 |
| SQIAKVL | ASSDQGARNSPLY | 0.21299484 |
| TALGHLVP | ASSDQGARNSPLY | 0.2212857 |
| TAPTTSGAA | ASSDQGARNSPLY | 0.22091717 |
| TDGDDGNGGSPGT | ASSDQGARNSPLY | 0.21183081 |
| TDGLGEN | ASSDQGARNSPLY | 0.21882828 |
| TGGPPLPGQA | ASSDQGARNSPLY | 0.21994455 |
| TIPAITI | ASSDQGARNSPLY | 0.21648999 |
| TIPGFTI | ASSDQGARNSPLY | 0.21185578 |
| TSIASGALA | ASSDQGARNSPLY | 0.22688255 |
| TVTLPGAAI | ASSDQGARNSPLY | 0.24534115 |
| VASGLSAALP | ASSDQGARNSPLY | 0.2173538 |
| VGAGVIGI | ASSDQGARNSPLY | 0.21434835 |
| VGPKRFGV | ASSDQGARNSPLY | 0.21128753 |
| VNVAFPD | ASSDQGARNSPLY | 0.21380304 |
| VSMTNSGVS | ASSDQGARNSPLY | 0.2394075 |
| VTDVLGDLSV | ASSDQGARNSPLY | 0.21778949 |
| VVVVGAGFAGL | ASSDQGARNSPLY | 0.21248497 |
| VYTVGTVV | ASSDQGARNSPLY | 0.21304083 |
| WTTFSGP | ASSDQGARNSPLY | 0.2283874 |
| YDELVEAF | ASSDQGARNSPLY | 0.21313085 |
| YGTGGAGGNGG | ASSDQGARNSPLY | 0.21227862 |

| | | |
|---|---|---|
| YTNVMAV | ASSDQGARNSPLY | 0.21542384 |
| FLEQVHQL | ASSDQGARNSPLY | 0.22008428 |
| GLSLLAVAV | ASSDQGARNSPLY | 0.21220055 |
| LEQVHQL | ASSDQGARNSPLY | 0.21362083 |
| LNAAGAI | ASSDQGARNSPLY | 0.22176543 |
| PAVHASL | ASSDQGARNSPLY | 0.21528171 |
| TELEDDA | ASSDQGARNSPLY | 0.21571214 |
| TPPGPPN | ASSDQGARNSPLY | 0.21570675 |
| TPPPPPPP | ASSDQGARNSPLY | 0.22150506 |
| TQSPALI | ASSQEWGAYEQY | 0.21708286 |
| EFAAAFPQ | ASSQEWGAYEQY | 0.21407537 |
| LLSGNLDVL | ASSQEWGAYEQY | 0.2133658 |
| GPANLGNYNI | ASSQEWGAYEQY | 0.21715577 |
| TLGPTLP | ASSQEWGAYEQY | 0.22079507 |
| VYAYSSCCK | ASSQEWGAYEQY | 0.21767251 |
| APAPAPAPNI | ASSQEWGAYEQY | 0.21459275 |
| PGGPVPIP | ASSQEWGAYEQY | 0.21092783 |
| FLTVALEP | ASSQEWGAYEQY | 0.21591748 |
| SQYFGANL | ASSQEWGAYEQY | 0.21359676 |
| GGVKPAA | ASSQEWGAYEQY | 0.21808803 |
| GGTGGAGGAGGW | ASSQEWGAYEQY | 0.22714277 |
| GVGGVAGLGGAGG | ASSQEWGAYEQY | 0.21290937 |
| VLLVLATTI | ASSQEWGAYEQY | 0.21433079 |
| IGIFAAML | ASSQEWGAYEQY | 0.21178108 |
| AVGDLID | ASSQEWGAYEQY | 0.21289456 |
| NISPLPL | ASSQEWGAYEQY | 0.21595928 |
| VTAVFVAI | ASSQEWGAYEQY | 0.21365559 |
| SGDDNDAAL | ASSQEWGAYEQY | 0.22070965 |
| AVAALAHS | ASSQEWGAYEQY | 0.2153133 |
| GGVTVMATP | ASSQEWGAYEQY | 0.22117959 |
| AGLAANVRD | ASSQEWGAYEQY | 0.2155621 |
| TNRPDLIDPALLRPG | ASSQEWGAYEQY | 0.21757694 |
| LANLGVTAL | ASSQEWGAYEQY | 0.23353472 |
| TGERIEL | ASSQEWGAYEQY | 0.21901831 |
| NGEELAA | ASSQEWGAYEQY | 0.21272492 |
| VLSLVWG | ASSQEWGAYEQY | 0.22868216 |
| IGYALWL | ASSQEWGAYEQY | 0.21323727 |
| LSLLLAPL | ASSQEWGAYEQY | 0.21903801 |
| TLDEALK | ASSQEWGAYEQY | 0.21465328 |
| TDGAGTAEGG | ASSQEWGAYEQY | 0.22193535 |
| NAPGGAGGNGG | ASSQEWGAYEQY | 0.21642363 |
| VSVFMGMGGNF | ASSQEWGAYEQY | 0.21363187 |
| LASYLAAL | ASSQEWGAYEQY | 0.21466848 |
| LVGGTPAL | ASSQEWGAYEQY | 0.217132 |
| VRNADVL | ASSQEWGAYEQY | 0.23240423 |
| TCSSPFLVSAM | ASSQEWGAYEQY | 0.22158301 |
| VHITDAE | ASSQEWGAYEQY | 0.2150951 |
| EYAAPSF | ASSQEWGAYEQY | 0.22103998 |
| VVDAFHGGL | ASSQEWGAYEQY | 0.21503942 |
| ASVVDPQ | ASSQEWGAYEQY | 0.21931368 |
| SYNEFGGLS | ASSQEWGAYEQY | 0.2153928 |

| | | |
|---|---|---|
| DPDGSVY | ASSQEWGAYEQY | 0.21583813 |
| MVSTIHATE | ASSQEWGAYEQY | 0.21237391 |
| LAAGVADPV | ASSQEWGAYEQY | 0.21599844 |
| TKEQIAA | ASSQEWGAYEQY | 0.21302794 |
| SLASPAAL | ASSQEWGAYEQY | 0.22152837 |
| IGLVFYGV | ASSQEWGAYEQY | 0.21481088 |
| FLLDASNL | ASSQEWGAYEQY | 0.21219057 |
| YQVGNFP | ASSQEWGAYEQY | 0.21450621 |
| ESLLSQST | ASSQEWGAYEQY | 0.21698202 |
| LFTVGFAAVG | ASSQEWGAYEQY | 0.21357088 |
| ATPPTPT | ASSQEWGAYEQY | 0.22160074 |
| TDLATLG | ASSQEWGAYEQY | 0.2141298 |
| SAGGTGR | ASSQEWGAYEQY | 0.2208871 |
| GFSNSGS | ASSQEWGAYEQY | 0.2399341 |
| GGNGGSGGTSV | ASSQEWGAYEQY | 0.21714453 |
| GTSVATGGAGN | ASSQEWGAYEQY | 0.21326263 |
| NGGNGGSGGTS | ASSQEWGAYEQY | 0.21831101 |
| AVVKFPH | ASSQEWGAYEQY | 0.21567886 |
| TPPPPPP | ASSQEWGAYEQY | 0.21678455 |
| AAPAAAAAPA | ASSQEWGAYEQY | 0.21120232 |
| IALNEDL | ASSQEWGAYEQY | 0.22081505 |
| IVNVSSM | ASSQEWGAYEQY | 0.219111 |
| TGIPPAPR | ASSQEWGAYEQY | 0.21676025 |
| FDLGGGTF | ASSQEWGAYEQY | 0.21194327 |
| FDLGGGTFDVS | ASSQEWGAYEQY | 0.2260899 |
| NEPTAAA | ASSQEWGAYEQY | 0.23557624 |
| NEPTAAALA | ASSQEWGAYEQY | 0.22004655 |
| GSIINISSI | ASSQEWGAYEQY | 0.22006124 |
| VAPGNAGTA | ASSQEWGAYEQY | 0.2122909 |
| TGGGASLE | ASSQEWGAYEQY | 0.22894612 |
| VAVYNHYKRI | ASSQEWGAYEQY | 0.22630721 |
| TDPAPATT | ASSQEWGAYEQY | 0.21533392 |
| TNRPDILDPALLRPG | ASSQEWGAYEQY | 0.22773744 |
| LTTTTTS | ASSQEWGAYEQY | 0.21552376 |
| PTGSGKTL | ASSQEWGAYEQY | 0.21556962 |
| TGSGKTL | ASSQEWGAYEQY | 0.2253275 |
| TIGHVDHGK | ASSQEWGAYEQY | 0.22130442 |
| TIGHVDHGKTT | ASSQEWGAYEQY | 0.21551451 |
| GHVDHGKTT | ASSQEWGAYEQY | 0.21309546 |
| AQFEHTLL | ASSQEWGAYEQY | 0.21471933 |
| IVNQDEL | ASSQEWGAYEQY | 0.21535167 |
| TNGSQFF | ASSQEWGAYEQY | 0.22121346 |
| TGKGIVE | ASSQEWGAYEQY | 0.2125571 |
| LISNASDA | ASSQEWGAYEQY | 0.22726072 |
| AELRTLGELTKIAKAHGAQ | ASSQEWGAYEQY | 0.2174108 |
| APNAVAAAEPSW | ASSQEWGAYEQY | 0.23615977 |
| APQVAGAATAVL | ASSQEWGAYEQY | 0.22106239 |
| ARRDPAAAAAILPTDA | ASSQEWGAYEQY | 0.21631606 |
| DAAARDVLAEAGLAEYF | ASSQEWGAYEQY | 0.21436341 |
| DDQIVLSAKANLAKVEQ | ASSQEWGAYEQY | 0.23752603 |
| DELLPDTGEDSDEDGH | ASSQEWGAYEQY | 0.2130912 |

| | | |
|---|---|---|
| ELPEAPANRATVPQD | ASSQEWGAYEQY | 0.21415406 |
| FGQSKIERVTGIG | ASSQEWGAYEQY | 0.21834719 |
| GAGGTGSTAGGAGGAGGAGGLY | ASSQEWGAYEQY | 0.440362 |
| GHGGTGAAGGNGGTGGAGGSGI | ASSQEWGAYEQY | 0.7697276 |
| GNGGNGGAGGEGGPGGAGGAGA | ASSQEWGAYEQY | 0.70948946 |
| GSTAGNGGNGGNATNGTIAGQP | ASSQEWGAYEQY | 0.73454785 |
| GTGGDGGNGGNGGTGG | ASSQEWGAYEQY | 0.5609123 |
| GVRRPAGIPADAPV | ASSQEWGAYEQY | 0.41129047 |
| GYRQVTDAHLLTLARR | ASSQEWGAYEQY | 0.6828449 |
| IAKMGQDGHDRG | ASSQEWGAYEQY | 0.49702695 |
| KAEPSIEIQYT | ASSQEWGAYEQY | 0.77076125 |
| KPVRIRMAADEDTVAVT | ASSQEWGAYEQY | 0.76241827 |
| MPPRIAGMRLLVIKPEP | ASSQEWGAYEQY | 0.547717 |
| MKFYAVALFRSRGVPPTR | ASSQEWGAYEQY | 0.7519009 |
| NAGGFGILGAVAHSP | ASSQEWGAYEQY | 0.7781241 |
| NEGREITLAMAEM | ASSQEWGAYEQY | 0.37791044 |
| NELVEWSHPLGHRWRWE | ASSQEWGAYEQY | 0.642996 |
| NLGAGGTGGNGGNPG | ASSQEWGAYEQY | 0.6832334 |
| NTEWPKWETVMAADPVD | ASSQEWGAYEQY | 0.6181939 |
| PASAAPAETQP | ASSQEWGAYEQY | 0.32824647 |
| QQSSDGKNGVRAAEITGE | ASSQEWGAYEQY | 0.74505836 |
| RAPDAVTTSWWKE | ASSQEWGAYEQY | 0.5987634 |
| SGFGNVGAHTSGWFNQ | ASSQEWGAYEQY | 0.2682259 |
| SGVAGAVGQAASVGGLKV | ASSQEWGAYEQY | 0.7186879 |
| SNGGAGTGSAGGNGGT | ASSQEWGAYEQY | 0.5728846 |
| TGGAGGEGGAAG | ASSQEWGAYEQY | 0.533422 |
| TGGDGGAAGAGGAPGHG | ASSQEWGAYEQY | 0.761209 |
| TMPPLLAAIANAGRHGV | ASSQEWGAYEQY | 0.6671996 |
| TSSTAGNSGGA | ASSQEWGAYEQY | 0.5186809 |
| VPATVPVGILC | ASSQEWGAYEQY | 0.72835183 |
| VQRPDSGGVVLTGRIS | ASSQEWGAYEQY | 0.7769556 |
| WTPNPVPATKGVPAGLVQAT | ASSQEWGAYEQY | 0.42286336 |
| YALKRALAKFSDRFDVVII | ASSQEWGAYEQY | 0.69687486 |
| YLIGSVIV | ASSQEWGAYEQY | 0.7904675 |
| YGQLVAVID | ASSQEWGAYEQY | 0.69503665 |
| VTVGGMTLPA | ASSQEWGAYEQY | 0.79217136 |
| VLALPAV | ASSQEWGAYEQY | 0.8004359 |
| VIPPTLNYET | ASSQEWGAYEQY | 0.65282977 |
| VGMGGMHPSSG | ASSQEWGAYEQY | 0.7697808 |
| VGLAGGAATA | ASSQEWGAYEQY | 0.7934526 |
| VGGAGGAGGSGP | ASSQEWGAYEQY | 0.5854572 |
| VFAAFMV | ASSQEWGAYEQY | 0.668697 |
| TLYDKLR | ASSQEWGAYEQY | 0.4863164 |
| TLAVFTVL | ASSQEWGAYEQY | 0.69344157 |
| TGSVQPARV | ASSQEWGAYEQY | 0.41332963 |
| TGGGAGSGGT | ASSQEWGAYEQY | 0.70179605 |
| SHLETSL | ASSQEWGAYEQY | 0.7746029 |
| SFTTGMT | ASSQEWGAYEQY | 0.62949115 |
| QVEVAKRMGVRQPTVS | ASSQEWGAYEQY | 0.4423309 |
| QGSGGPALAM | ASSQEWGAYEQY | 0.45240104 |
| MLAPSNP | ASSQEWGAYEQY | 0.7254216 |

| | | |
|---|---|---|
| LVNNAAI | ASSQEWGAYEQY | 0.47067368 |
| LTGEINTLA | ASSQEWGAYEQY | 0.4600362 |
| LSIFGPV | ASSQEWGAYEQY | 0.5733753 |
| LLALLANL | ASSQEWGAYEQY | 0.7616258 |
| LISNASDALDKLR | ASSQEWGAYEQY | 0.7491759 |
| LINEAVP | ASSQEWGAYEQY | 0.49554378 |
| LGYAFSGL | ASSQEWGAYEQY | 0.7864494 |
| LDFVAVAL | ASSQEWGAYEQY | 0.76148653 |
| LAQPGAE | ASSQEWGAYEQY | 0.79480547 |
| ISNLGAVTGF | ASSQEWGAYEQY | 0.6838004 |
| HQEDPGRI | ASSQEWGAYEQY | 0.43947905 |
| GTPGEPTP | ASSQEWGAYEQY | 0.7639036 |
| GFFNSGS | ASSQEWGAYEQY | 0.60792583 |
| GAGGGGLTAGAN | ASSQEWGAYEQY | 0.7026745 |
| FVDALVNM | ASSQEWGAYEQY | 0.75775135 |
| FLTVAPEL | ASSQEWGAYEQY | 0.6207367 |
| FETLTAL | ASSQEWGAYEQY | 0.5139923 |
| DVINPIEL | ASSQEWGAYEQY | 0.6520684 |
| APVSGAPV | ASSQEWGAYEQY | 0.793218 |
| ALVGGLAW | ASSQEWGAYEQY | 0.5021529 |
| ALAAAAATAI | ASSQEWGAYEQY | 0.436255 |
| AFGLGLLL | ASSQEWGAYEQY | 0.61271083 |
| AFFAAASL | ASSQEWGAYEQY | 0.6334152 |
| AAAALPAFSPP | ASSQEWGAYEQY | 0.77533704 |
| VVLAPAL | ASSQEWGAYEQY | 0.71080554 |
| TIGHVDHGKTTL | ASSQEWGAYEQY | 0.62312025 |
| LISNASDALDK | ASSQEWGAYEQY | 0.7255248 |
| LISNASDAL | ASSQEWGAYEQY | 0.7258218 |
| ISNASDALDK | ASSQEWGAYEQY | 0.6860751 |
| NQAETLVYQTE | ASSQEWGAYEQY | 0.7689885 |
| TGGTGGTAGAG | ASSQEWGAYEQY | 0.545561 |
| TGSTAGGAGGA | ASSQEWGAYEQY | 0.7870674 |
| DPLAVPVAAGPV | ASSQEWGAYEQY | 0.7422462 |
| GGTGGTPGNGGN | ASSQEWGAYEQY | 0.7641131 |
| PAGLAGVDPATL | ASSQEWGAYEQY | 0.78297627 |
| GNNLVVEAPHAGASV | ASSQEWGAYEQY | 0.65782464 |
| KVTRIMSASTLEQPA | ASSQEWGAYEQY | 0.71840835 |
| LGFALPIALPRRTVV | ASSQEWGAYEQY | 0.4469577 |
| TRLGNLTARKLVVLV | ASSQEWGAYEQY | 0.684085 |
| WSQQRRLIYFPSAGP | ASSQEWGAYEQY | 0.7092497 |
| GPNGNPDPMAAAVDIR | ASSQEWGAYEQY | 0.7006922 |
| TQRCVSVADLRAAASR | ASSQEWGAYEQY | 0.76749295 |
| QFVLDVAEHPVEVLTETV | ASSQEWGAYEQY | 0.7661637 |
| GGAGGTGGAGGNGGTQAGDG | ASSQEWGAYEQY | 0.5734739 |
| PLLATLRAIAVDVDAKPAQV | ASSQEWGAYEQY | 0.79928625 |
| APKFRTGGYMIDFWGVGYQVAK | ASSQEWGAYEQY | 0.77861166 |
| GAGGAGGAGGAAGGPGGTGGTG | ASSQEWGAYEQY | 0.45095918 |
| IGALGLLATDVAPAGSVAFGVA | ASSQEWGAYEQY | 0.6501393 |
| AGGAGGIGGNGGAGGAGGVLMG | ASSQEWGAYEQY | 0.7838147 |
| ADELIGLL | ASSQEWGAYEQY | 0.7411244 |
| ADRALKA | ASSQEWGAYEQY | 0.60549927 |

| | | |
|---|---|---|
| AGAVGQVGGAGGA | ASSQEWGAYEQY | 0.68878394 |
| AGGGGGLGGAP | ASSQEWGAYEQY | 0.78517693 |
| AGGLLYGNGGAG | ASSQEWGAYEQY | 0.7016021 |
| AGLESIGL | ASSQEWGAYEQY | 0.79178596 |
| AGLKTGII | ASSQEWGAYEQY | 0.3118124 |
| ASLAALLL | ASSQEWGAYEQY | 0.6531971 |
| ASVINISSI | ASSQEWGAYEQY | 0.6415812 |
| ATPPGPA | ASSQEWGAYEQY | 0.264138 |
| ATVASALV | ASSQEWGAYEQY | 0.7622465 |
| ATVASLIL | ASSQEWGAYEQY | 0.6546441 |
| AVNGPVLIRH | ASSQEWGAYEQY | 0.74151766 |
| DGAAGAVGI | ASSQEWGAYEQY | 0.6432291 |
| DPLLHGI | ASSQEWGAYEQY | 0.76711816 |
| ELGAALGAL | ASSQEWGAYEQY | 0.70452625 |
| ELGRTQAV | ASSQEWGAYEQY | 0.65153813 |
| FVAGGYLS | ASSQEWGAYEQY | 0.48887718 |
| GASGGAGGAGGW | ASSQEWGAYEQY | 0.73366135 |
| GGAGAGGDGGAGGI | ASSQEWGAYEQY | 0.8060258 |
| GTGHNGSQPGMGGNG | ASSQEWGAYEQY | 0.78419733 |
| GYAAASASAS | ASSQEWGAYEQY | 0.3284314 |
| HQVDVAAVQQG | ASSQEWGAYEQY | 0.724364 |
| IGAAIAHT | ASSQEWGAYEQY | 0.67705214 |
| IGMIEVAQL | ASSQEWGAYEQY | 0.38626242 |
| IRLAPEHTSAMD | ASSQEWGAYEQY | 0.34269124 |
| IYNVSVL | ASSQEWGAYEQY | 0.6378056 |
| KQGFLGM | ASSQEWGAYEQY | 0.6416932 |
| LESVITQL | ASSQEWGAYEQY | 0.7233623 |
| LGLGALVVV | ASSQEWGAYEQY | 0.7345693 |
| LIGNGGNGGSGGI | ASSQEWGAYEQY | 0.5762712 |
| LNAINAPTL | ASSQEWGAYEQY | 0.74449885 |
| LQLLNTGV | ASSQEWGAYEQY | 0.6592601 |
| LQTSIEF | ASSQEWGAYEQY | 0.16103627 |
| MSFAVSEL | ASSQEWGAYEQY | 0.58403766 |
| NTGALGL | ASSQEWGAYEQY | 0.37850896 |
| PPGAGGAGGAGTT | ASSQEWGAYEQY | 0.79832006 |
| QLVGLGI | ASSQEWGAYEQY | 0.5187705 |
| RFLEQVHQL | ASSQEWGAYEQY | 0.60721564 |
| RGVGYVLR | ASSQEWGAYEQY | 0.6188736 |
| SATGVFAGVI | ASSQEWGAYEQY | 0.70916045 |
| SLPAITLP | ASSQEWGAYEQY | 0.6554705 |
| STALAFTV | ASSQEWGAYEQY | 0.5312948 |
| SVLTVGLGK | ASSQEWGAYEQY | 0.37937403 |
| TALAGAGSGI | ASSQEWGAYEQY | 0.73465836 |
| TGLPFTGI | ASSQEWGAYEQY | 0.5939842 |
| VAEHVTV | ASSQEWGAYEQY | 0.48290947 |
| VDIVLNSL | ASSQEWGAYEQY | 0.5401437 |
| VLTGGGAVL | ASSQEWGAYEQY | 0.74217165 |
| VNVGLAYGI | ASSQEWGAYEQY | 0.3588544 |
| VSSAAVGAV | ASSQEWGAYEQY | 0.7835839 |
| VTSVYEAGV | ASSQEWGAYEQY | 0.54018736 |
| WVALDRGA | ASSQEWGAYEQY | 0.61977327 |

| | | |
|---|---|---|
| YLVGSVVI | ASSQEWGAYEQY | 0.7912518 |
| GPPGTGKTLL | ASSQEWGAYEQY | 0.6711577 |
| GVGMTKF | ASSQEWGAYEQY | 0.62819505 |
| IGIGGSDL | ASSQEWGAYEQY | 0.47997862 |
| IGTLLAEL | ASSQEWGAYEQY | 0.4819013 |
| IYTSGTT | ASSQEWGAYEQY | 0.5258756 |
| LSFPAGV | ASSQEWGAYEQY | 0.5125347 |
| LYTSGTT | ASSQEWGAYEQY | 0.79473317 |
| VGAFPFGF | ASSQEWGAYEQY | 0.7063535 |
| VGAGTVEF | ASSQEWGAYEQY | 0.46929094 |
| VGKSTLLL | ASSQEWGAYEQY | 0.75307566 |
| AALAHAGGGIV | ASSQEWGAYEQY | 0.795352 |
| AALVAVSLL | ASSQEWGAYEQY | 0.81019354 |
| AAVSRAVAAL | ASSQEWGAYEQY | 0.49002403 |
| AGGSSGIGGF | ASSQEWGAYEQY | 0.6152132 |
| AGLTVANANG | ASSQEWGAYEQY | 0.6753176 |
| AGLVGAVL | ASSQEWGAYEQY | 0.6479418 |
| ALSQAFTV | ASSQEWGAYEQY | 0.55235213 |
| ALTFSVSL | ASSQEWGAYEQY | 0.45165727 |
| ALTGAALVAL | ASSQEWGAYEQY | 0.7698592 |
| ASFSIGAGVL | ASSQEWGAYEQY | 0.8090886 |
| DGSVYKI | ASSQEWGAYEQY | 0.6462364 |
| EQLAEIV | ASSQEWGAYEQY | 0.46261808 |
| FAVSELTVM | ASSQEWGAYEQY | 0.6144862 |
| GAAPLAAGL | ASSQEWGAYEQY | 0.49403867 |
| GADGGDGGSGGAGG | ASSQEWGAYEQY | 0.52776444 |
| GFGLDTGGAGG | ASSQEWGAYEQY | 0.66715753 |
| GLASGGLL | ASSQEWGAYEQY | 0.38444567 |
| GRIAGAAVLL | ASSQEWGAYEQY | 0.6441276 |
| GSAGSLNTSF | ASSQEWGAYEQY | 0.7946498 |
| GTGQIGI | ASSQEWGAYEQY | 0.6288202 |
| GTLIALIL | ASSQEWGAYEQY | 0.73647606 |
| IAATMGFVVG | ASSQEWGAYEQY | 0.7717812 |
| IFAQALL | ASSQEWGAYEQY | 0.7301819 |
| IGAAVALGF | ASSQEWGAYEQY | 0.77614945 |
| IGLTPLAL | ASSQEWGAYEQY | 0.76967764 |
| ILGSNTSTL | ASSQEWGAYEQY | 0.6795666 |
| IVFSLTVL | ASSQEWGAYEQY | 0.70292807 |
| LASKATGF | ASSQEWGAYEQY | 0.7574385 |
| LASVALIL | ASSQEWGAYEQY | 0.6028193 |
| LFAVGLTL | ASSQEWGAYEQY | 0.65505767 |
| LFNGGIGTYI | ASSQEWGAYEQY | 0.6672033 |
| LTLGAGII | ASSQEWGAYEQY | 0.74117327 |
| LVAAALGDI | ASSQEWGAYEQY | 0.6579172 |
| NTAFVQLGI | ASSQEWGAYEQY | 0.7945187 |
| QLFSGAEL | ASSQEWGAYEQY | 0.31515893 |
| SAATFITL | ASSQEWGAYEQY | 0.41709545 |
| SNYGLANLGV | ASSQEWGAYEQY | 0.76494163 |
| SYGGAYAVM | ASSQEWGAYEQY | 0.22862117 |
| TTGVGGLTLG | ASSQEWGAYEQY | 0.35545385 |
| TTQTIYI | ASSQEWGAYEQY | 0.6975719 |

| | | |
|---|---|---|
| VAAGAGLGAVI | ASSQEWGAYEQY | 0.77358186 |
| VAAVVTLL | ASSQEWGAYEQY | 0.22636984 |
| VGFLGGLI | ASSQEWGAYEQY | 0.5768293 |
| VGVIPILL | ASSQEWGAYEQY | 0.4018718 |
| VIFASSLI | ASSQEWGAYEQY | 0.77271426 |
| VKGGDGGAAAG | ASSQEWGAYEQY | 0.6273452 |
| VTKTFTNI | ASSQEWGAYEQY | 0.36847675 |
| VVAAAILL | ASSQEWGAYEQY | 0.74576646 |
| YAALDVEL | ASSQEWGAYEQY | 0.49609584 |
| LASVALLL | ASSQEWGAYEQY | 0.68768394 |
| LDAVHRV | ASSQEWGAYEQY | 0.7732686 |
| QGLSSSIF | ASSQEWGAYEQY | 0.79571694 |
| RGTFANIRL | ASSQEWGAYEQY | 0.4767604 |
| RQLWWGHRI | ASSQEWGAYEQY | 0.73331904 |
| YGPPGTGKTLL | ASSQEWGAYEQY | 0.6211989 |
| AALVGAPANVA | ASSQEWGAYEQY | 0.81394553 |
| AATFRRTE | ASSQEWGAYEQY | 0.6031395 |
| AGALGGGI | ASSQEWGAYEQY | 0.38636726 |
| AGGAGGAGGIGAG | ASSQEWGAYEQY | 0.7522347 |
| AGSGALGL | ASSQEWGAYEQY | 0.7491243 |
| AQIADQI | ASSQEWGAYEQY | 0.68196243 |
| ASAVAVTYL | ASSQEWGAYEQY | 0.75914454 |
| ASTVLGGTGG | ASSQEWGAYEQY | 0.6648307 |
| AVGGEPAK | ASSQEWGAYEQY | 0.60725856 |
| CEQGTFGGTC | ASSQEWGAYEQY | 0.7365765 |
| DEGKAPAG | ASSQEWGAYEQY | 0.75426877 |
| DTVVFPAL | ASSQEWGAYEQY | 0.4917084 |
| FEGITMAG | ASSQEWGAYEQY | 0.7867161 |
| FPLVVQD | ASSQEWGAYEQY | 0.6516144 |
| GAFTTPQ | ASSQEWGAYEQY | 0.4795398 |
| GFGGFGTAGGVGGA | ASSQEWGAYEQY | 0.80200946 |
| GFLAQFL | ASSQEWGAYEQY | 0.67699385 |
| GGAGGAGGNAGVA | ASSQEWGAYEQY | 0.75219154 |
| GGVGTAGAGGAGGAGG | ASSQEWGAYEQY | 0.7510785 |
| GNGGGGDAGNAGSGGN | ASSQEWGAYEQY | 0.7945161 |
| GTGGAGGAGNDAGS | ASSQEWGAYEQY | 0.50774765 |
| GTTNTGWFNS | ASSQEWGAYEQY | 0.7541853 |
| IFMGPLFTL | ASSQEWGAYEQY | 0.6887064 |
| ILAHAIEAI | ASSQEWGAYEQY | 0.75561607 |
| IRSAIREAL | ASSQEWGAYEQY | 0.68915534 |
| ITFAGGG | ASSQEWGAYEQY | 0.7037772 |
| ITGMSGHQEV | ASSQEWGAYEQY | 0.7487558 |
| IVAGLACTL | ASSQEWGAYEQY | 0.7803422 |
| IVGGGLAGA | ASSQEWGAYEQY | 0.69567096 |
| IVTGVLNI | ASSQEWGAYEQY | 0.7497244 |
| LAQQGCI | ASSQEWGAYEQY | 0.78635466 |
| LCLHVEN | ASSQEWGAYEQY | 0.494228 |
| LDAISDGS | ASSQEWGAYEQY | 0.652879 |
| LGGFAAML | ASSQEWGAYEQY | 0.76536024 |
| LGGKNPMI | ASSQEWGAYEQY | 0.79616886 |
| LGPKGVAAIL | ASSQEWGAYEQY | 0.6679717 |

| | | |
|---|---|---|
| LMVGVSI | ASSQEWGAYEQY | 0.78732336 |
| LVGAAFVL | ASSQEWGAYEQY | 0.63940895 |
| LVLAVAPL | ASSQEWGAYEQY | 0.7153307 |
| MAAIAVIDAA | ASSQEWGAYEQY | 0.61814827 |
| NNGINAELQ | ASSQEWGAYEQY | 0.6543905 |
| RGEPLVLVHG | ASSQEWGAYEQY | 0.46961933 |
| RGGAGGAAGLI | ASSQEWGAYEQY | 0.68462765 |
| RSYRMHPAV | ASSQEWGAYEQY | 0.75092286 |
| SAAHLLAGQLA | ASSQEWGAYEQY | 0.7038813 |
| SAAVGSY | ASSQEWGAYEQY | 0.3586712 |
| SPHAVLHRI | ASSQEWGAYEQY | 0.71136093 |
| SQIAKVL | ASSQEWGAYEQY | 0.7043363 |
| TALGHLVP | ASSQEWGAYEQY | 0.8075796 |
| TAPTTSGAA | ASSQEWGAYEQY | 0.6858361 |
| TDGDDGNGGSPGT | ASSQEWGAYEQY | 0.7732285 |
| TDGLGEN | ASSQEWGAYEQY | 0.6239387 |
| TGGPPLPGQA | ASSQEWGAYEQY | 0.686854 |
| TIPAITI | ASSQEWGAYEQY | 0.39949697 |
| TIPGFTI | ASSQEWGAYEQY | 0.63825464 |
| TSIASGALA | ASSQEWGAYEQY | 0.6457052 |
| TVTLPGAAI | ASSQEWGAYEQY | 0.5052299 |
| VASGLSAALP | ASSQEWGAYEQY | 0.74528074 |
| VGAGVIGI | ASSQEWGAYEQY | 0.78428805 |
| VGPKRFGV | ASSQEWGAYEQY | 0.76020753 |
| VNVAFPD | ASSQEWGAYEQY | 0.77012104 |
| VSMTNSGVS | ASSQEWGAYEQY | 0.7922348 |
| VTDVLGDLSV | ASSQEWGAYEQY | 0.7745824 |
| VVVVGAGFAGL | ASSQEWGAYEQY | 0.556141 |
| VYTVGTVV | ASSQEWGAYEQY | 0.66771364 |
| WTTFSGP | ASSQEWGAYEQY | 0.37136275 |
| YDELVEAF | ASSQEWGAYEQY | 0.6041474 |
| YGTGGAGGNGG | ASSQEWGAYEQY | 0.5813896 |
| YTNVMAV | ASSQEWGAYEQY | 0.6680145 |
| FLEQVHQL | ASSQEWGAYEQY | 0.20005186 |
| GLSLLAVAV | ASSQEWGAYEQY | 0.71020114 |
| LEQVHQL | ASSQEWGAYEQY | 0.7968826 |
| LNAAGAI | ASSQEWGAYEQY | 0.62436724 |
| PAVHASL | ASSQEWGAYEQY | 0.71709335 |
| TELEDDA | ASSQEWGAYEQY | 0.72960865 |
| TPPGPPN | ASSQEWGAYEQY | 0.5908515 |
| TPPPPPPP | ASSQEWGAYEQY | 0.73302424 |
| TQSPALI | ASSSGTGGGRNTGQLY | 0.5640652 |
| EFAAAFPQ | ASSSGTGGGRNTGQLY | 0.76041335 |
| LLSGNLDVL | ASSSGTGGGRNTGQLY | 0.7870233 |
| GPANLGNYNI | ASSSGTGGGRNTGQLY | 0.7209729 |
| TLGPTLP | ASSSGTGGGRNTGQLY | 0.7280717 |
| VYAYSSCCK | ASSSGTGGGRNTGQLY | 0.7270504 |
| APAPAPAPNI | ASSSGTGGGRNTGQLY | 0.46370173 |
| PGGPVPIP | ASSSGTGGGRNTGQLY | 0.74280167 |
| FLTVALEP | ASSSGTGGGRNTGQLY | 0.6003291 |
| SQYFGANL | ASSSGTGGGRNTGQLY | 0.6588696 |

| | | |
|---|---|---|
| GGVKPAA | ASSSGTGGGRNTGQLY | 0.52803636 |
| GGTGGAGGAGGW | ASSSGTGGGRNTGQLY | 0.69519424 |
| GVGGVAGLGGAGG | ASSSGTGGGRNTGQLY | 0.66687214 |
| VLLVLATTI | ASSSGTGGGRNTGQLY | 0.73448277 |
| IGIFAAML | ASSSGTGGGRNTGQLY | 0.4998805 |
| AVGDLID | ASSSGTGGGRNTGQLY | 0.7250101 |
| NISPLPL | ASSSGTGGGRNTGQLY | 0.700572 |
| VTAVFVAI | ASSSGTGGGRNTGQLY | 0.42231455 |
| SGDDNDAAL | ASSSGTGGGRNTGQLY | 0.7788063 |
| AVAALAHS | ASSSGTGGGRNTGQLY | 0.66523707 |
| GGVTVMATP | ASSSGTGGGRNTGQLY | 0.6466989 |
| AGLAANVRD | ASSSGTGGGRNTGQLY | 0.76696074 |
| TNRPDLIDPALLRPG | ASSSGTGGGRNTGQLY | 0.686491 |
| LANLGVTAL | ASSSGTGGGRNTGQLY | 0.4964148 |
| TGERIEL | ASSSGTGGGRNTGQLY | 0.59709525 |
| NGEELAA | ASSSGTGGGRNTGQLY | 0.69449055 |
| VLSLVWG | ASSSGTGGGRNTGQLY | 0.62715876 |
| IGYALWL | ASSSGTGGGRNTGQLY | 0.2997148 |
| LSLLLAPL | ASSSGTGGGRNTGQLY | 0.7421099 |
| TLDEALK | ASSSGTGGGRNTGQLY | 0.47777498 |
| TDGAGTAEGG | ASSSGTGGGRNTGQLY | 0.5764509 |
| NAPGGAGGNGG | ASSSGTGGGRNTGQLY | 0.6011684 |
| VSVFMGMGGNF | ASSSGTGGGRNTGQLY | 0.74943197 |
| LASYLAAL | ASSSGTGGGRNTGQLY | 0.69460577 |
| LVGGTPAL | ASSSGTGGGRNTGQLY | 0.5877242 |
| VRNADVL | ASSSGTGGGRNTGQLY | 0.75125253 |
| TCSSPFLVSAM | ASSSGTGGGRNTGQLY | 0.7092584 |
| VHITDAE | ASSSGTGGGRNTGQLY | 0.77053785 |
| EYAAPSF | ASSSGTGGGRNTGQLY | 0.6987497 |
| VVDAFHGGL | ASSSGTGGGRNTGQLY | 0.70541394 |
| ASVVDPQ | ASSSGTGGGRNTGQLY | 0.7353414 |
| SYNEFGGLS | ASSSGTGGGRNTGQLY | 0.6296949 |
| DPDGSVY | ASSSGTGGGRNTGQLY | 0.52499235 |
| MVSTIHATE | ASSSGTGGGRNTGQLY | 0.46050268 |
| LAAGVADPV | ASSSGTGGGRNTGQLY | 0.625008 |
| TKEQIAA | ASSSGTGGGRNTGQLY | 0.46340632 |
| SLASPAAL | ASSSGTGGGRNTGQLY | 0.6920991 |
| IGLVFYGV | ASSSGTGGGRNTGQLY | 0.7972365 |
| FLLDASNL | ASSSGTGGGRNTGQLY | 0.67514837 |
| YQVGNFP | ASSSGTGGGRNTGQLY | 0.40813026 |
| ESLLSQST | ASSSGTGGGRNTGQLY | 0.63193583 |
| LFTVGFAAVG | ASSSGTGGGRNTGQLY | 0.6491079 |
| ATPPTPT | ASSSGTGGGRNTGQLY | 0.74314845 |
| TDLATLG | ASSSGTGGGRNTGQLY | 0.77109075 |
| SAGGTGR | ASSSGTGGGRNTGQLY | 0.8005092 |
| GFSNSGS | ASSSGTGGGRNTGQLY | 0.74311936 |
| GGNGGSGGTSV | ASSSGTGGGRNTGQLY | 0.5557122 |
| GTSVATGGAGN | ASSSGTGGGRNTGQLY | 0.747063 |
| NGGNGGSGGTS | ASSSGTGGGRNTGQLY | 0.62843037 |
| AVVKFPH | ASSSGTGGGRNTGQLY | 0.7423755 |
| TPPPPPP | ASSSGTGGGRNTGQLY | 0.7075076 |

| | | |
|---|---|---|
| AAPAAAAAPA | ASSSGTGGGRNTGQLY | 0.72838485 |
| IALNEDL | ASSSGTGGGRNTGQLY | 0.63658726 |
| IVNVSSM | ASSSGTGGGRNTGQLY | 0.61668086 |
| TGIPPAPR | ASSSGTGGGRNTGQLY | 0.411743 |
| FDLGGGTF | ASSSGTGGGRNTGQLY | 0.76624334 |
| FDLGGGTFDVS | ASSSGTGGGRNTGQLY | 0.64349574 |
| NEPTAAA | ASSSGTGGGRNTGQLY | 0.51843077 |
| NEPTAAALA | ASSSGTGGGRNTGQLY | 0.58881444 |
| GSIINISSI | ASSSGTGGGRNTGQLY | 0.79963607 |
| VAPGNAGTA | ASSSGTGGGRNTGQLY | 0.77643955 |
| TGGGASLE | ASSSGTGGGRNTGQLY | 0.5779172 |
| VAVYNHYKRI | ASSSGTGGGRNTGQLY | 0.5674282 |
| TDPAPATT | ASSSGTGGGRNTGQLY | 0.72856593 |
| TNRPDILDPALLRPG | ASSSGTGGGRNTGQLY | 0.5018423 |
| LTTTTTS | ASSSGTGGGRNTGQLY | 0.75188035 |
| PTGSGKTL | ASSSGTGGGRNTGQLY | 0.6440693 |
| TGSGKTL | ASSSGTGGGRNTGQLY | 0.6500133 |
| TIGHVDHGK | ASSSGTGGGRNTGQLY | 0.6464401 |
| TIGHVDHGKTT | ASSSGTGGGRNTGQLY | 0.7307042 |
| GHVDHGKTT | ASSSGTGGGRNTGQLY | 0.59923834 |
| AQFEHTLL | ASSSGTGGGRNTGQLY | 0.75122726 |
| IVNQDEL | ASSSGTGGGRNTGQLY | 0.28231385 |
| TNGSQFF | ASSSGTGGGRNTGQLY | 0.77706766 |
| TGKGIVE | ASSSGTGGGRNTGQLY | 0.48259676 |
| LISNASDA | ASSSGTGGGRNTGQLY | 0.5337986 |
| AELRTLGELTKIAKAHGAQ | ASSSGTGGGRNTGQLY | 0.2966368 |
| APNAVAAAEPSW | ASSSGTGGGRNTGQLY | 0.70205194 |
| APQVAGAATAVL | ASSSGTGGGRNTGQLY | 0.74857026 |
| ARRDPAAAAAILPTDA | ASSSGTGGGRNTGQLY | 0.76869404 |
| DAAARDVLAEAGLAEYF | ASSSGTGGGRNTGQLY | 0.68813854 |
| DDQIVLSAKANLAKVEQ | ASSSGTGGGRNTGQLY | 0.75350356 |
| DELLPDTGEDSDEDGH | ASSSGTGGGRNTGQLY | 0.782295 |
| ELPEAPANRATVPQD | ASSSGTGGGRNTGQLY | 0.60297495 |
| FGQSKIERVTGIG | ASSSGTGGGRNTGQLY | 0.3342346 |
| GAGGTGSTAGGAGGAGGAGGLY | ASSSGTGGGRNTGQLY | 0.69948053 |
| GHGGTGAAGGNGGTGGAGGSGI | ASSSGTGGGRNTGQLY | 0.7096572 |
| GNGGNGGAGGEGGPGGAGGAGA | ASSSGTGGGRNTGQLY | 0.7770735 |
| GSTAGNGGNGGNATNGTIAGQP | ASSSGTGGGRNTGQLY | 0.7445031 |
| GTGGDGGNGGNGGTGG | ASSSGTGGGRNTGQLY | 0.3920138 |
| GVRRPAGIPADAPV | ASSSGTGGGRNTGQLY | 0.78627783 |
| GYRQVTDAHLLTLARR | ASSSGTGGGRNTGQLY | 0.72646594 |
| IAKMGQDGHDRG | ASSSGTGGGRNTGQLY | 0.4368003 |
| KAEPSIEIQYT | ASSSGTGGGRNTGQLY | 0.6439116 |
| KPVRIRMAADEDTVAVT | ASSSGTGGGRNTGQLY | 0.58232766 |
| MPPRIAGMRLLVIKPEP | ASSSGTGGGRNTGQLY | 0.47637776 |
| MKFYAVALFRSRGVPPTR | ASSSGTGGGRNTGQLY | 0.73562443 |
| NAGGFGILGAVAHSP | ASSSGTGGGRNTGQLY | 0.58965534 |
| NEGREITLAMAEM | ASSSGTGGGRNTGQLY | 0.557938 |
| NELVEWSHPLGHRWRWE | ASSSGTGGGRNTGQLY | 0.65734893 |
| NLGAGGTGGNGGNPG | ASSSGTGGGRNTGQLY | 0.56374013 |
| NTEWPKWETVMAADPVD | ASSSGTGGGRNTGQLY | 0.54428965 |

| | | |
|---|---|---|
| PASAAPAETQP | ASSSGTGGGRNTGQLY | 0.58808076 |
| QQSSDGKNGVRAAEITGE | ASSSGTGGGRNTGQLY | 0.73625207 |
| RAPDAVTTSWWKE | ASSSGTGGGRNTGQLY | 0.7419654 |
| SGFGNVGAHTSGWFNQ | ASSSGTGGGRNTGQLY | 0.65899444 |
| SGVAGAVGQAASVGGLKV | ASSSGTGGGRNTGQLY | 0.7263005 |
| SNGGAGTGSAGGNGGT | ASSSGTGGGRNTGQLY | 0.7750577 |
| TGGAGGEGGAAG | ASSSGTGGGRNTGQLY | 0.38084644 |
| TGGDGGAAGAGGAPGHG | ASSSGTGGGRNTGQLY | 0.5250634 |
| TMPPLLAAIANAGRHGV | ASSSGTGGGRNTGQLY | 0.71742034 |
| TSSTAGNSGGA | ASSSGTGGGRNTGQLY | 0.5794811 |
| VPATVPVGILC | ASSSGTGGGRNTGQLY | 0.7666004 |
| VQRPDSGGVVLTGRIS | ASSSGTGGGRNTGQLY | 0.67534745 |
| WTPNPVPATKGVPAGLVQAT | ASSSGTGGGRNTGQLY | 0.68687344 |
| YALKRALAKFSDRFDVVII | ASSSGTGGGRNTGQLY | 0.6998789 |
| YLIGSVIV | ASSSGTGGGRNTGQLY | 0.8046256 |
| YGQLVAVID | ASSSGTGGGRNTGQLY | 0.7272539 |
| VTVGGMTLPA | ASSSGTGGGRNTGQLY | 0.6908258 |
| VLALPAV | ASSSGTGGGRNTGQLY | 0.7769629 |
| VIPPTLNYET | ASSSGTGGGRNTGQLY | 0.49286866 |
| VGMGGMHPSSG | ASSSGTGGGRNTGQLY | 0.76635957 |
| VGLAGGAATA | ASSSGTGGGRNTGQLY | 0.74072754 |
| VGGAGGAGGSGP | ASSSGTGGGRNTGQLY | 0.475313 |
| VFAAFMV | ASSSGTGGGRNTGQLY | 0.6149246 |
| TLYDKLR | ASSSGTGGGRNTGQLY | 0.80090505 |
| TLAVFTVL | ASSSGTGGGRNTGQLY | 0.77086854 |
| TGSVQPARV | ASSSGTGGGRNTGQLY | 0.5368488 |
| TGGGAGSGGT | ASSSGTGGGRNTGQLY | 0.5507907 |
| SHLETSL | ASSSGTGGGRNTGQLY | 0.50509095 |
| SFTTGMT | ASSSGTGGGRNTGQLY | 0.6401502 |
| QVEVAKRMGVRQPTVS | ASSSGTGGGRNTGQLY | 0.3792234 |
| QGSGGPALAM | ASSSGTGGGRNTGQLY | 0.7366775 |
| MLAPSNP | ASSSGTGGGRNTGQLY | 0.58098924 |
| LVNNAAI | ASSSGTGGGRNTGQLY | 0.76108944 |
| LTGEINTLA | ASSSGTGGGRNTGQLY | 0.6922972 |
| LSIFGPV | ASSSGTGGGRNTGQLY | 0.7748914 |
| LLALLANL | ASSSGTGGGRNTGQLY | 0.59737504 |
| LISNASDALDKLR | ASSSGTGGGRNTGQLY | 0.5805172 |
| LINEAVP | ASSSGTGGGRNTGQLY | 0.35133404 |
| LGYAFSGL | ASSSGTGGGRNTGQLY | 0.5874935 |
| LDFVAVAL | ASSSGTGGGRNTGQLY | 0.59193146 |
| LAQPGAE | ASSSGTGGGRNTGQLY | 0.6116166 |
| ISNLGAVTGF | ASSSGTGGGRNTGQLY | 0.33455557 |
| HQEDPGRI | ASSSGTGGGRNTGQLY | 0.77828103 |
| GTPGEPTP | ASSSGTGGGRNTGQLY | 0.6240554 |
| GFFNSGS | ASSSGTGGGRNTGQLY | 0.7349274 |
| GAGGGGLTAGAN | ASSSGTGGGRNTGQLY | 0.58055824 |
| FVDALVNM | ASSSGTGGGRNTGQLY | 0.59300935 |
| FLTVAPEL | ASSSGTGGGRNTGQLY | 0.5562195 |
| FETLTAL | ASSSGTGGGRNTGQLY | 0.7439761 |
| DVINPIEL | ASSSGTGGGRNTGQLY | 0.4707638 |
| APVSGAPV | ASSSGTGGGRNTGQLY | 0.36207712 |

| | | |
|---|---|---|
| ALVGGLAW | ASSSGTGGGRNTGQLY | 0.5463009 |
| ALAAAAATAI | ASSSGTGGGRNTGQLY | 0.6602882 |
| AFGLGLLL | ASSSGTGGGRNTGQLY | 0.79706335 |
| AFFAAASL | ASSSGTGGGRNTGQLY | 0.765507 |
| AAAALPAFSPP | ASSSGTGGGRNTGQLY | 0.68537587 |
| VVLAPAL | ASSSGTGGGRNTGQLY | 0.80011487 |
| TIGHVDHGKTTL | ASSSGTGGGRNTGQLY | 0.24101757 |
| LISNASDALDK | ASSSGTGGGRNTGQLY | 0.6078065 |
| LISNASDAL | ASSSGTGGGRNTGQLY | 0.7875266 |
| ISNASDALDK | ASSSGTGGGRNTGQLY | 0.7308399 |
| NQAETLVYQTE | ASSSGTGGGRNTGQLY | 0.6234674 |
| TGGTGGTAGAG | ASSSGTGGGRNTGQLY | 0.7686876 |
| TGSTAGGAGGA | ASSSGTGGGRNTGQLY | 0.49458802 |
| DPLAVPVAAGPV | ASSSGTGGGRNTGQLY | 0.6072351 |
| GGTGGTPGNGGN | ASSSGTGGGRNTGQLY | 0.6477932 |
| PAGLAGVDPATL | ASSSGTGGGRNTGQLY | 0.7660616 |
| GNNLVVEAPHAGASV | ASSSGTGGGRNTGQLY | 0.7381362 |
| KVTRIMSASTLEQPA | ASSSGTGGGRNTGQLY | 0.41038758 |
| LGFALPIALPRRTVV | ASSSGTGGGRNTGQLY | 0.75338733 |
| TRLGNLTARKLVVLV | ASSSGTGGGRNTGQLY | 0.8043573 |
| WSQQRRLIYFPSAGP | ASSSGTGGGRNTGQLY | 0.559968 |
| GPNGNPDPMAAAVDIR | ASSSGTGGGRNTGQLY | 0.7732461 |
| TQRCVSVADLRAAASR | ASSSGTGGGRNTGQLY | 0.70398235 |
| QFVLDVAEHPVEVLTETV | ASSSGTGGGRNTGQLY | 0.6741836 |
| GGAGGTGGAGGNGGTQAGDG | ASSSGTGGGRNTGQLY | 0.5203219 |
| PLLATLRAIAVDVDAKPAQV | ASSSGTGGGRNTGQLY | 0.48480284 |
| APKFRTGGYMIDFWGVGYQVAK | ASSSGTGGGRNTGQLY | 0.7360302 |
| GAGGAGGAGGAAGGPGGTGGTG | ASSSGTGGGRNTGQLY | 0.5021169 |
| IGALGLLATDVAPAGSVAFGVA | ASSSGTGGGRNTGQLY | 0.6840143 |
| AGGAGGIGGNGGAGGAGGVLMG | ASSSGTGGGRNTGQLY | 0.64491916 |
| ADELIGLL | ASSSGTGGGRNTGQLY | 0.81085026 |
| ADRALKA | ASSSGTGGGRNTGQLY | 0.7286455 |
| AGAVGQVGGAGGA | ASSSGTGGGRNTGQLY | 0.7079197 |
| AGGGGGLGGAP | ASSSGTGGGRNTGQLY | 0.5164856 |
| AGGLLYGNGGAG | ASSSGTGGGRNTGQLY | 0.7331854 |
| AGLESIGL | ASSSGTGGGRNTGQLY | 0.72670037 |
| AGLKTGII | ASSSGTGGGRNTGQLY | 0.45708445 |
| ASLAALLL | ASSSGTGGGRNTGQLY | 0.7782201 |
| ASVINISSI | ASSSGTGGGRNTGQLY | 0.44984522 |
| ATPPGPA | ASSSGTGGGRNTGQLY | 0.6691767 |
| ATVASALV | ASSSGTGGGRNTGQLY | 0.70945764 |
| ATVASLIL | ASSSGTGGGRNTGQLY | 0.77594477 |
| AVNGPVLIRH | ASSSGTGGGRNTGQLY | 0.66919816 |
| DGAAGAVGI | ASSSGTGGGRNTGQLY | 0.7408529 |
| DPLLHGI | ASSSGTGGGRNTGQLY | 0.46063164 |
| ELGAALGAL | ASSSGTGGGRNTGQLY | 0.75507534 |
| ELGRTQAV | ASSSGTGGGRNTGQLY | 0.75214356 |
| FVAGGYLS | ASSSGTGGGRNTGQLY | 0.7439528 |
| GASGGAGGAGGW | ASSSGTGGGRNTGQLY | 0.80464566 |
| GGAGAGGDGGAGGI | ASSSGTGGGRNTGQLY | 0.7616081 |
| GTGHNGSQPGMGGNG | ASSSGTGGGRNTGQLY | 0.3268964 |

| | | |
|---|---|---|
| GYAAASASAS | ASSSGTGGGRNTGQLY | 0.66868114 |
| HQVDVAAVQQG | ASSSGTGGGRNTGQLY | 0.78010976 |
| IGAAIAHT | ASSSGTGGGRNTGQLY | 0.7788774 |
| IGMIEVAQL | ASSSGTGGGRNTGQLY | 0.78957164 |
| IRLAPEHTSAMD | ASSSGTGGGRNTGQLY | 0.7809061 |
| IYNVSVL | ASSSGTGGGRNTGQLY | 0.6401336 |
| KQGFLGM | ASSSGTGGGRNTGQLY | 0.78325796 |
| LESVITQL | ASSSGTGGGRNTGQLY | 0.8190602 |
| LGLGALVVV | ASSSGTGGGRNTGQLY | 0.7437619 |
| LIGNGGNGGSGGI | ASSSGTGGGRNTGQLY | 0.49887675 |
| LNAINAPTL | ASSSGTGGGRNTGQLY | 0.48674133 |
| LQLLNTGV | ASSSGTGGGRNTGQLY | 0.7490959 |
| LQTSIEF | ASSSGTGGGRNTGQLY | 0.7620936 |
| MSFAVSEL | ASSSGTGGGRNTGQLY | 0.46846378 |
| NTGALGL | ASSSGTGGGRNTGQLY | 0.63908184 |
| PPGAGGAGGAGTT | ASSSGTGGGRNTGQLY | 0.6666121 |
| QLVGLGI | ASSSGTGGGRNTGQLY | 0.29991937 |
| RFLEQVHQL | ASSSGTGGGRNTGQLY | 0.7391235 |
| RGVGYVLR | ASSSGTGGGRNTGQLY | 0.79309607 |
| SATGVFAGVI | ASSSGTGGGRNTGQLY | 0.7418533 |
| SLPAITLP | ASSSGTGGGRNTGQLY | 0.5044118 |
| STALAFTV | ASSSGTGGGRNTGQLY | 0.7781034 |
| SVLTVGLGK | ASSSGTGGGRNTGQLY | 0.7843782 |
| TALAGAGSGI | ASSSGTGGGRNTGQLY | 0.7958477 |
| TGLPFTGI | ASSSGTGGGRNTGQLY | 0.34136713 |
| VAEHVTV | ASSSGTGGGRNTGQLY | 0.23624168 |
| VDIVLNSL | ASSSGTGGGRNTGQLY | 0.7051523 |
| VLTGGGAVL | ASSSGTGGGRNTGQLY | 0.7386304 |
| VNVGLAYGI | ASSSGTGGGRNTGQLY | 0.5736317 |
| VSSAAVGAV | ASSSGTGGGRNTGQLY | 0.4705773 |
| VTSVYEAGV | ASSSGTGGGRNTGQLY | 0.7128031 |
| WVALDRGA | ASSSGTGGGRNTGQLY | 0.76084816 |
| YLVGSVVI | ASSSGTGGGRNTGQLY | 0.4903407 |
| GPPGTGKTLL | ASSSGTGGGRNTGQLY | 0.40224522 |
| GVGMTKF | ASSSGTGGGRNTGQLY | 0.7786957 |
| IGIGGSDL | ASSSGTGGGRNTGQLY | 0.75507736 |
| IGTLLAEL | ASSSGTGGGRNTGQLY | 0.5038882 |
| IYTSGTT | ASSSGTGGGRNTGQLY | 0.6023862 |
| LSFPAGV | ASSSGTGGGRNTGQLY | 0.7060311 |
| LYTSGTT | ASSSGTGGGRNTGQLY | 0.71803904 |
| VGAFPFGF | ASSSGTGGGRNTGQLY | 0.6953287 |
| VGAGTVEF | ASSSGTGGGRNTGQLY | 0.4119711 |
| VGKSTLLL | ASSSGTGGGRNTGQLY | 0.751614 |
| AALAHAGGGIV | ASSSGTGGGRNTGQLY | 0.43468767 |
| AALVAVSLL | ASSSGTGGGRNTGQLY | 0.77452135 |
| AAVSRAVAAL | ASSSGTGGGRNTGQLY | 0.7026874 |
| AGGSSGIGGF | ASSSGTGGGRNTGQLY | 0.7398186 |
| AGLTVANANG | ASSSGTGGGRNTGQLY | 0.61147106 |
| AGLVGAVL | ASSSGTGGGRNTGQLY | 0.44940227 |
| ALSQAFTV | ASSSGTGGGRNTGQLY | 0.49934965 |
| ALTFSVSL | ASSSGTGGGRNTGQLY | 0.41960514 |

| | | |
|---|---|---|
| ALTGAALVAL | ASSSGTGGGRNTGQLY | 0.790989 |
| ASFSIGAGVL | ASSSGTGGGRNTGQLY | 0.7012275 |
| DGSVYKI | ASSSGTGGGRNTGQLY | 0.7683368 |
| EQLAEIV | ASSSGTGGGRNTGQLY | 0.7022352 |
| FAVSELTVM | ASSSGTGGGRNTGQLY | 0.4558505 |
| GAAPLAAGL | ASSSGTGGGRNTGQLY | 0.732306 |
| GADGGDGGSGGAGG | ASSSGTGGGRNTGQLY | 0.37292016 |
| GFGLDTGGAGG | ASSSGTGGGRNTGQLY | 0.7260183 |
| GLASGGLL | ASSSGTGGGRNTGQLY | 0.6902001 |
| GRIAGAAVLL | ASSSGTGGGRNTGQLY | 0.7413393 |
| GSAGSLNTSF | ASSSGTGGGRNTGQLY | 0.56810904 |
| GTGQIGI | ASSSGTGGGRNTGQLY | 0.75128436 |
| GTLIALIL | ASSSGTGGGRNTGQLY | 0.7318483 |
| IAATMGFVVG | ASSSGTGGGRNTGQLY | 0.54867786 |
| IFAQALL | ASSSGTGGGRNTGQLY | 0.7526264 |
| IGAAVALGF | ASSSGTGGGRNTGQLY | 0.7319104 |
| IGLTPLAL | ASSSGTGGGRNTGQLY | 0.75133103 |
| ILGSNTSTL | ASSSGTGGGRNTGQLY | 0.37205428 |
| IVFSLTVL | ASSSGTGGGRNTGQLY | 0.70266676 |
| LASKATGF | ASSSGTGGGRNTGQLY | 0.5446062 |
| LASVALIL | ASSSGTGGGRNTGQLY | 0.70046866 |
| LFAVGLTL | ASSSGTGGGRNTGQLY | 0.6659583 |
| LFNGGIGTYI | ASSSGTGGGRNTGQLY | 0.5739631 |
| LTLGAGII | ASSSGTGGGRNTGQLY | 0.748447 |
| LVAAALGDI | ASSSGTGGGRNTGQLY | 0.4946499 |
| NTAFVQLGI | ASSSGTGGGRNTGQLY | 0.59271824 |
| QLFSGAEL | ASSSGTGGGRNTGQLY | 0.6111121 |
| SAATFITL | ASSSGTGGGRNTGQLY | 0.68696356 |
| SNYGLANLGV | ASSSGTGGGRNTGQLY | 0.73331285 |
| SYGGAYAVM | ASSSGTGGGRNTGQLY | 0.45136136 |
| TTGVGGLTLG | ASSSGTGGGRNTGQLY | 0.6971063 |
| TTQTIYI | ASSSGTGGGRNTGQLY | 0.6266594 |
| VAAGAGLGAVI | ASSSGTGGGRNTGQLY | 0.47076127 |
| VAAVVTLL | ASSSGTGGGRNTGQLY | 0.7178786 |
| VGFLGGLI | ASSSGTGGGRNTGQLY | 0.62844324 |
| VGVIPILL | ASSSGTGGGRNTGQLY | 0.44205916 |
| VIFASSLI | ASSSGTGGGRNTGQLY | 0.75676394 |
| VKGGDGGAAAG | ASSSGTGGGRNTGQLY | 0.50265026 |
| VTKTFTNI | ASSSGTGGGRNTGQLY | 0.72453713 |
| VVAAAILL | ASSSGTGGGRNTGQLY | 0.8134086 |
| YAALDVEL | ASSSGTGGGRNTGQLY | 0.54097223 |
| LASVALLL | ASSSGTGGGRNTGQLY | 0.76652926 |
| LDAVHRV | ASSSGTGGGRNTGQLY | 0.6937201 |
| QGLSSSIF | ASSSGTGGGRNTGQLY | 0.72018236 |
| RGTFANIRL | ASSSGTGGGRNTGQLY | 0.62442815 |
| RQLWWGHRI | ASSSGTGGGRNTGQLY | 0.55406725 |
| YGPPGTGKTLL | ASSSGTGGGRNTGQLY | 0.77905023 |
| AALVGAPANVA | ASSSGTGGGRNTGQLY | 0.72723556 |
| AATFRRTE | ASSSGTGGGRNTGQLY | 0.77432907 |
| AGALGGGI | ASSSGTGGGRNTGQLY | 0.7457799 |
| AGGAGGAGGIGAG | ASSSGTGGGRNTGQLY | 0.77340406 |

| | | |
|---|---|---|
| AGSGALGL | ASSSGTGGGRNTGQLY | 0.75357497 |
| AQIADQI | ASSSGTGGGRNTGQLY | 0.5147652 |
| ASAVAVTYL | ASSSGTGGGRNTGQLY | 0.338662 |
| ASTVLGGTGG | ASSSGTGGGRNTGQLY | 0.5869671 |
| AVGGEPAK | ASSSGTGGGRNTGQLY | 0.60948837 |
| CEQGTFGGTC | ASSSGTGGGRNTGQLY | 0.5911997 |
| DEGKAPAG | ASSSGTGGGRNTGQLY | 0.36702064 |
| DTVVFPAL | ASSSGTGGGRNTGQLY | 0.7418033 |
| FEGITMAG | ASSSGTGGGRNTGQLY | 0.811235 |
| FPLVVQD | ASSSGTGGGRNTGQLY | 0.53209037 |
| GAFTTPQ | ASSSGTGGGRNTGQLY | 0.8013053 |
| GFGGFGTAGGVGGA | ASSSGTGGGRNTGQLY | 0.46441913 |
| GFLAQFL | ASSSGTGGGRNTGQLY | 0.73803294 |
| GGAGGAGGNAGVA | ASSSGTGGGRNTGQLY | 0.76373327 |
| GGVGTAGAGGAGGAGG | ASSSGTGGGRNTGQLY | 0.42593533 |
| GNGGGGDAGNAGSGGN | ASSSGTGGGRNTGQLY | 0.7104795 |
| GTGGAGGAGNDAGS | ASSSGTGGGRNTGQLY | 0.6128149 |
| GTTNTGWFNS | ASSSGTGGGRNTGQLY | 0.5790193 |
| IFMGPLFTL | ASSSGTGGGRNTGQLY | 0.77888894 |
| ILAHAIEAI | ASSSGTGGGRNTGQLY | 0.57417774 |
| IRSAIREAL | ASSSGTGGGRNTGQLY | 0.7553543 |
| ITFAGGG | ASSSGTGGGRNTGQLY | 0.5627186 |
| ITGMSGHQEV | ASSSGTGGGRNTGQLY | 0.47205198 |
| IVAGLACTL | ASSSGTGGGRNTGQLY | 0.71197766 |
| IVGGGLAGA | ASSSGTGGGRNTGQLY | 0.76737845 |
| IVTGVLNI | ASSSGTGGGRNTGQLY | 0.6890226 |
| LAQQGCI | ASSSGTGGGRNTGQLY | 0.8141745 |
| LCLHVEN | ASSSGTGGGRNTGQLY | 0.5419426 |
| LDAISDGS | ASSSGTGGGRNTGQLY | 0.7958227 |
| LGGFAAML | ASSSGTGGGRNTGQLY | 0.774711 |
| LGGKNPMI | ASSSGTGGGRNTGQLY | 0.6605483 |
| LGPKGVAAIL | ASSSGTGGGRNTGQLY | 0.60971487 |
| LMVGVSI | ASSSGTGGGRNTGQLY | 0.7375163 |
| LVGAAFVL | ASSSGTGGGRNTGQLY | 0.5597632 |
| LVLAVAPL | ASSSGTGGGRNTGQLY | 0.7746557 |
| MAAIAVIDAA | ASSSGTGGGRNTGQLY | 0.7903829 |
| NNGINAELQ | ASSSGTGGGRNTGQLY | 0.28796056 |
| RGEPLVLVHG | ASSSGTGGGRNTGQLY | 0.6547904 |
| RGGAGGAAGLI | ASSSGTGGGRNTGQLY | 0.5480529 |
| RSYRMHPAV | ASSSGTGGGRNTGQLY | 0.6417885 |
| SAAHLLAGQLA | ASSSGTGGGRNTGQLY | 0.42136958 |
| SAAVGSY | ASSSGTGGGRNTGQLY | 0.32756576 |
| SPHAVLHRI | ASSSGTGGGRNTGQLY | 0.51175725 |
| SQIAKVL | ASSSGTGGGRNTGQLY | 0.38691062 |
| TALGHLVP | ASSSGTGGGRNTGQLY | 0.83627963 |
| TAPTTSGAA | ASSSGTGGGRNTGQLY | 0.65703297 |
| TDGDDGNGGSPGT | ASSSGTGGGRNTGQLY | 0.42451596 |
| TDGLGEN | ASSSGTGGGRNTGQLY | 0.73483074 |
| TGGPPLPGQA | ASSSGTGGGRNTGQLY | 0.734418 |
| TIPAITI | ASSSGTGGGRNTGQLY | 0.24392253 |
| TIPGFTI | ASSSGTGGGRNTGQLY | 0.5201773 |

| | | |
|---|---|---|
| TSIASGALA | ASSSGTGGGRNTGQLY | 0.29506359 |
| TVTLPGAAI | ASSSGTGGGRNTGQLY | 0.39329445 |
| VASGLSAALP | ASSSGTGGGRNTGQLY | 0.13719095 |
| VGAGVIGI | ASSSGTGGGRNTGQLY | 0.60633147 |
| VGPKRFGV | ASSSGTGGGRNTGQLY | 0.5982971 |
| VNVAFPD | ASSSGTGGGRNTGQLY | 0.19140126 |
| VSMTNSGVS | ASSSGTGGGRNTGQLY | 0.6015705 |
| VTDVLGDLSV | ASSSGTGGGRNTGQLY | 0.48390073 |
| VVVVGAGFAGL | ASSSGTGGGRNTGQLY | 0.51034355 |
| VYTVGTVV | ASSSGTGGGRNTGQLY | 0.66323316 |
| WTTFSGP | ASSSGTGGGRNTGQLY | 0.3518772 |
| YDELVEAF | ASSSGTGGGRNTGQLY | 0.464801 |
| YGTGGAGGNGG | ASSSGTGGGRNTGQLY | 0.6396308 |
| YTNVMAV | ASSSGTGGGRNTGQLY | 0.7522195 |
| FLEQVHQL | ASSSGTGGGRNTGQLY | 0.41273046 |
| GLSLLAVAV | ASSSGTGGGRNTGQLY | 0.5239055 |
| LEQVHQL | ASSSGTGGGRNTGQLY | 0.77389306 |
| LNAAGAI | ASSSGTGGGRNTGQLY | 0.5413355 |
| PAVHASL | ASSSGTGGGRNTGQLY | 0.6680294 |
| TELEDDA | ASSSGTGGGRNTGQLY | 0.7187052 |
| TPPGPPN | ASSSGTGGGRNTGQLY | 0.35461783 |
| TPPPPPPP | ASSSGTGGGRNTGQLY | 0.7463298 |
| TQSPALI | GAGQNSGNTLY | 0.72810996 |
| EFAAAFPQ | GAGQNSGNTLY | 0.45310718 |
| LLSGNLDVL | GAGQNSGNTLY | 0.47932577 |
| GPANLGNYNI | GAGQNSGNTLY | 0.30906174 |
| TLGPTLP | GAGQNSGNTLY | 0.5145439 |
| VYAYSSCCK | GAGQNSGNTLY | 0.278053 |
| APAPAPAPNI | GAGQNSGNTLY | 0.57311034 |
| PGGPVPIP | GAGQNSGNTLY | 0.7152877 |
| FLTVALEP | GAGQNSGNTLY | 0.52409995 |
| SQYFGANL | GAGQNSGNTLY | 0.20463319 |
| GGVKPAA | GAGQNSGNTLY | 0.29097784 |
| GGTGGAGGAGGW | GAGQNSGNTLY | 0.4285999 |
| GVGGVAGLGGAGG | GAGQNSGNTLY | 0.30569586 |
| VLLVLATTI | GAGQNSGNTLY | 0.28800216 |
| IGIFAAML | GAGQNSGNTLY | 0.251615 |
| AVGDLID | GAGQNSGNTLY | 0.7520611 |
| NISPLPL | GAGQNSGNTLY | 0.7438859 |
| VTAVFVAI | GAGQNSGNTLY | 0.46392548 |
| SGDDNDAAL | GAGQNSGNTLY | 0.68987167 |
| AVAALAHS | GAGQNSGNTLY | 0.78407454 |
| GGVTVMATP | GAGQNSGNTLY | 0.6257619 |
| AGLAANVRD | GAGQNSGNTLY | 0.48188823 |
| TNRPDLIDPALLRPG | GAGQNSGNTLY | 0.24110039 |
| LANLGVTAL | GAGQNSGNTLY | 0.8249972 |
| TGERIEL | GAGQNSGNTLY | 0.52767265 |
| NGEELAA | GAGQNSGNTLY | 0.6760911 |
| VLSLVWG | GAGQNSGNTLY | 0.6713896 |
| IGYALWL | GAGQNSGNTLY | 0.60066473 |
| LSLLLAPL | GAGQNSGNTLY | 0.31336695 |

| | | |
|---|---|---|
| TLDEALK | GAGQNSGNTLY | 0.673626 |
| TDGAGTAEGG | GAGQNSGNTLY | 0.5436393 |
| NAPGGAGGNGG | GAGQNSGNTLY | 0.3133523 |
| VSVFMGMGGNF | GAGQNSGNTLY | 0.2849167 |
| LASYLAAL | GAGQNSGNTLY | 0.610596 |
| LVGGTPAL | GAGQNSGNTLY | 0.54767704 |
| VRNADVL | GAGQNSGNTLY | 0.75507855 |
| TCSSPFLVSAM | GAGQNSGNTLY | 0.4914686 |
| VHITDAE | GAGQNSGNTLY | 0.2814269 |
| EYAAPSF | GAGQNSGNTLY | 0.7031714 |
| VVDAFHGGL | GAGQNSGNTLY | 0.56148 |
| ASVVDPQ | GAGQNSGNTLY | 0.49189347 |
| SYNEFGGLS | GAGQNSGNTLY | 0.67356217 |
| DPDGSVY | GAGQNSGNTLY | 0.41086257 |
| MVSTIHATE | GAGQNSGNTLY | 0.56425804 |
| LAAGVADPV | GAGQNSGNTLY | 0.6220604 |
| TKEQIAA | GAGQNSGNTLY | 0.5500008 |
| SLASPAAL | GAGQNSGNTLY | 0.6833279 |
| IGLVFYGV | GAGQNSGNTLY | 0.49013892 |
| FLLDASNL | GAGQNSGNTLY | 0.4990165 |
| YQVGNFP | GAGQNSGNTLY | 0.34630972 |
| ESLLSQST | GAGQNSGNTLY | 0.41492927 |
| LFTVGFAAVG | GAGQNSGNTLY | 0.64563805 |
| ATPPTPT | GAGQNSGNTLY | 0.54948604 |
| TDLATLG | GAGQNSGNTLY | 0.7380178 |
| SAGGTGR | GAGQNSGNTLY | 0.62833637 |
| GFSNSGS | GAGQNSGNTLY | 0.45550412 |
| GGNGGSGGTSV | GAGQNSGNTLY | 0.75173914 |
| GTSVATGGAGN | GAGQNSGNTLY | 0.74084806 |
| NGGNGGSGGTS | GAGQNSGNTLY | 0.22658399 |
| AVVKFPH | GAGQNSGNTLY | 0.5031966 |
| TPPPPPP | GAGQNSGNTLY | 0.70694697 |
| AAPAAAAAPA | GAGQNSGNTLY | 0.75699174 |
| IALNEDL | GAGQNSGNTLY | 0.52190185 |
| IVNVSSM | GAGQNSGNTLY | 0.532292 |
| TGIPPAPR | GAGQNSGNTLY | 0.85161287 |
| FDLGGGTF | GAGQNSGNTLY | 0.6573866 |
| FDLGGGTFDVS | GAGQNSGNTLY | 0.76930654 |
| NEPTAAA | GAGQNSGNTLY | 0.23269403 |
| NEPTAAALA | GAGQNSGNTLY | 0.37815797 |
| GSIINISSI | GAGQNSGNTLY | 0.31884223 |
| VAPGNAGTA | GAGQNSGNTLY | 0.14670387 |
| TGGGASLE | GAGQNSGNTLY | 0.6865675 |
| VAVYNHYKRI | GAGQNSGNTLY | 0.495498 |
| TDPAPATT | GAGQNSGNTLY | 0.6443993 |
| TNRPDILDPALLRPG | GAGQNSGNTLY | 0.5064586 |
| LTTTTTS | GAGQNSGNTLY | 0.6710568 |
| PTGSGKTL | GAGQNSGNTLY | 0.3974548 |
| TGSGKTL | GAGQNSGNTLY | 0.4679879 |
| TIGHVDHGK | GAGQNSGNTLY | 0.27636528 |
| TIGHVDHGKTT | GAGQNSGNTLY | 0.6907387 |

| | | |
|---|---|---|
| GHVDHGKTT | GAGQNSGNTLY | 0.7725094 |
| AQFEHTLL | GAGQNSGNTLY | 0.47056895 |
| IVNQDEL | GAGQNSGNTLY | 0.2748649 |
| TNGSQFF | GAGQNSGNTLY | 0.6336336 |
| TGKGIVE | GAGQNSGNTLY | 0.55533266 |
| LISNASDA | GAGQNSGNTLY | 0.23640634 |
| AELRTLGELTKIAKAHGAQ | GAGQNSGNTLY | 0.26622456 |
| APNAVAAAEPSW | GAGQNSGNTLY | 0.40404662 |
| APQVAGAATAVL | GAGQNSGNTLY | 0.3048234 |
| ARRDPAAAAAILPTDA | GAGQNSGNTLY | 0.69354445 |
| DAAARDVLAEAGLAEYF | GAGQNSGNTLY | 0.67461216 |
| DDQIVLSAKANLAKVEQ | GAGQNSGNTLY | 0.4229241 |
| DELLPDTGEDSDEDGH | GAGQNSGNTLY | 0.6318712 |
| ELPEAPANRATVPQD | GAGQNSGNTLY | 0.642069 |
| FGQSKIERVTGIG | GAGQNSGNTLY | 0.11509717 |
| GAGGTGSTAGGAGGAGGAGGLY | GAGQNSGNTLY | 0.34852484 |
| GHGGTGAAGGNGGTGGAGGSGI | GAGQNSGNTLY | 0.26256534 |
| GNGGNGGAGGEGGPGGAGGAGA | GAGQNSGNTLY | 0.6908587 |
| GSTAGNGGNGGNATNGTIAGQP | GAGQNSGNTLY | 0.32016665 |
| GTGGDGGNGGNGGTGG | GAGQNSGNTLY | 0.43415445 |
| GVRRPAGIPADAPV | GAGQNSGNTLY | 0.55381083 |
| GYRQVTDAHLLTLARR | GAGQNSGNTLY | 0.75971663 |
| IAKMGQDGHDRG | GAGQNSGNTLY | 0.588549 |
| KAEPSIEIQYT | GAGQNSGNTLY | 0.23195396 |
| KPVRIRMAADEDTVAVT | GAGQNSGNTLY | 0.19644713 |
| MPPRIAGMRLLVIKPEP | GAGQNSGNTLY | 0.6855184 |
| MKFYAVALFRSRGVPPTR | GAGQNSGNTLY | 0.39764598 |
| NAGGFGILGAVAHSP | GAGQNSGNTLY | 0.4519549 |
| NEGREITLAMAEM | GAGQNSGNTLY | 0.31812814 |
| NELVEWSHPLGHRWRWE | GAGQNSGNTLY | 0.5752436 |
| NLGAGGTGGNGGNPG | GAGQNSGNTLY | 0.2652311 |
| NTEWPKWETVMAADPVD | GAGQNSGNTLY | 0.589281 |
| PASAAPAETQP | GAGQNSGNTLY | 0.32002088 |
| QQSSDGKNGVRAAEITGE | GAGQNSGNTLY | 0.5255778 |
| RAPDAVTTSWWKE | GAGQNSGNTLY | 0.75907695 |
| SGFGNVGAHTSGWFNQ | GAGQNSGNTLY | 0.7615788 |
| SGVAGAVGQAASVGGLKV | GAGQNSGNTLY | 0.42775008 |
| SNGGAGTGSAGGNGGT | GAGQNSGNTLY | 0.4818902 |
| TGGAGGEGGAAG | GAGQNSGNTLY | 0.28977656 |
| TGGDGGAAGAGGAPGHG | GAGQNSGNTLY | 0.42860368 |
| TMPPLLAAIANAGRHGV | GAGQNSGNTLY | 0.4856022 |
| TSSTAGNSGGA | GAGQNSGNTLY | 0.74759424 |
| VPATVPVGILC | GAGQNSGNTLY | 0.39490676 |
| VQRPDSGGVVLTGRIS | GAGQNSGNTLY | 0.26955023 |
| WTPNPVPATKGVPAGLVQAT | GAGQNSGNTLY | 0.46481642 |
| YALKRALAKFSDRFDVVII | GAGQNSGNTLY | 0.76377606 |
| YLIGSVIV | GAGQNSGNTLY | 0.7467295 |
| YGQLVAVID | GAGQNSGNTLY | 0.30295414 |
| VTVGGMTLPA | GAGQNSGNTLY | 0.39029574 |
| VLALPAV | GAGQNSGNTLY | 0.46824765 |
| VIPPTLNYET | GAGQNSGNTLY | 0.5170688 |

| | | |
|---|---|---|
| VGMGGMHPSSG | GAGQNSGNTLY | 0.29024503 |
| VGLAGGAATA | GAGQNSGNTLY | 0.33197868 |
| VGGAGGAGGSGP | GAGQNSGNTLY | 0.51403785 |
| VFAAFMV | GAGQNSGNTLY | 0.86206883 |
| TLYDKLR | GAGQNSGNTLY | 0.44205093 |
| TLAVFTVL | GAGQNSGNTLY | 0.17206192 |
| TGSVQPARV | GAGQNSGNTLY | 0.58821857 |
| TGGGAGSGGT | GAGQNSGNTLY | 0.30741528 |
| SHLETSL | GAGQNSGNTLY | 0.38083872 |
| SFTTGMT | GAGQNSGNTLY | 0.5348358 |
| QVEVAKRMGVRQPTVS | GAGQNSGNTLY | 0.18709737 |
| QGSGGPALAM | GAGQNSGNTLY | 0.5520235 |
| MLAPSNP | GAGQNSGNTLY | 0.75257003 |
| LVNNAAI | GAGQNSGNTLY | 0.709278 |
| LTGEINTLA | GAGQNSGNTLY | 0.5582092 |
| LSIFGPV | GAGQNSGNTLY | 0.717407 |
| LLALLANL | GAGQNSGNTLY | 0.5842047 |
| LISNASDALDKLR | GAGQNSGNTLY | 0.7269097 |
| LINEAVP | GAGQNSGNTLY | 0.8049911 |
| LGYAFSGL | GAGQNSGNTLY | 0.46707004 |
| LDFVAVAL | GAGQNSGNTLY | 0.46241105 |
| LAQPGAE | GAGQNSGNTLY | 0.68820703 |
| ISNLGAVTGF | GAGQNSGNTLY | 0.57631737 |
| HQEDPGRI | GAGQNSGNTLY | 0.602209 |
| GTPGEPTP | GAGQNSGNTLY | 0.43284598 |
| GFFNSGS | GAGQNSGNTLY | 0.7605339 |
| GAGGGGLTAGAN | GAGQNSGNTLY | 0.3978455 |
| FVDALVNM | GAGQNSGNTLY | 0.828596 |
| FLTVAPEL | GAGQNSGNTLY | 0.21712065 |
| FETLTAL | GAGQNSGNTLY | 0.36565527 |
| DVINPIEL | GAGQNSGNTLY | 0.67124426 |
| APVSGAPV | GAGQNSGNTLY | 0.1552303 |
| ALVGGLAW | GAGQNSGNTLY | 0.2860536 |
| ALAAAAATAI | GAGQNSGNTLY | 0.47787756 |
| AFGLGLLL | GAGQNSGNTLY | 0.59458554 |
| AFFAAASL | GAGQNSGNTLY | 0.13852814 |
| AAAALPAFSPP | GAGQNSGNTLY | 0.5109291 |
| VVLAPAL | GAGQNSGNTLY | 0.260913 |
| TIGHVDHGKTTL | GAGQNSGNTLY | 0.48195666 |
| LISNASDALDK | GAGQNSGNTLY | 0.584951 |
| LISNASDAL | GAGQNSGNTLY | 0.23623538 |
| ISNASDALDK | GAGQNSGNTLY | 0.5871626 |
| NQAETLVYQTE | GAGQNSGNTLY | 0.18506622 |
| TGGTGGTAGAG | GAGQNSGNTLY | 0.4930566 |
| TGSTAGGAGGA | GAGQNSGNTLY | 0.66131365 |
| DPLAVPVAAGPV | GAGQNSGNTLY | 0.8325782 |
| GGTGGTPGNGGN | GAGQNSGNTLY | 0.30104458 |
| PAGLAGVDPATL | GAGQNSGNTLY | 0.6633148 |
| GNNLVVEAPHAGASV | GAGQNSGNTLY | 0.30620217 |
| KVTRIMSASTLEQPA | GAGQNSGNTLY | 0.795487 |
| LGFALPIALPRRTVV | GAGQNSGNTLY | 0.5180271 |

| | | |
|---|---|---|
| TRLGNLTARKLVVLV | GAGQNSGNTLY | 0.19881795 |
| WSQQRRLIYFPSAGP | GAGQNSGNTLY | 0.6916355 |
| GPNGNPDPMAAAVDIR | GAGQNSGNTLY | 0.43748412 |
| TQRCVSVADLRAAASR | GAGQNSGNTLY | 0.5906663 |
| QFVLDVAEHPVEVLTETV | GAGQNSGNTLY | 0.7786663 |
| GGAGGTGGAGGNGGTQAGDG | GAGQNSGNTLY | 0.3013025 |
| PLLATLRAIAVDVDAKPAQV | GAGQNSGNTLY | 0.54037476 |
| APKFRTGGYMIDFWGVGYQVAK | GAGQNSGNTLY | 0.7068519 |
| GAGGAGGAGGAAGGPGGTGGTG | GAGQNSGNTLY | 0.6630616 |
| IGALGLLATDVAPAGSVAFGVA | GAGQNSGNTLY | 0.32040003 |
| AGGAGGIGGNGGAGGAGGVLMG | GAGQNSGNTLY | 0.67274725 |
| ADELIGLL | GAGQNSGNTLY | 0.6452409 |
| ADRALKA | GAGQNSGNTLY | 0.4119527 |
| AGAVGQVGGAGGA | GAGQNSGNTLY | 0.7422805 |
| AGGGGGLGGAP | GAGQNSGNTLY | 0.36713392 |
| AGGLLYGNGGAG | GAGQNSGNTLY | 0.63068545 |
| AGLESIGL | GAGQNSGNTLY | 0.6729232 |
| AGLKTGII | GAGQNSGNTLY | 0.8557554 |
| ASLAALLL | GAGQNSGNTLY | 0.3332286 |
| ASVINISSI | GAGQNSGNTLY | 0.5133352 |
| ATPPGPA | GAGQNSGNTLY | 0.59626806 |
| ATVASALV | GAGQNSGNTLY | 0.6508532 |
| ATVASLIL | GAGQNSGNTLY | 0.31959584 |
| AVNGPVLIRH | GAGQNSGNTLY | 0.5272833 |
| DGAAGAVGI | GAGQNSGNTLY | 0.56356347 |
| DPLLHGI | GAGQNSGNTLY | 0.744437 |
| ELGAALGAL | GAGQNSGNTLY | 0.37907442 |
| ELGRTQAV | GAGQNSGNTLY | 0.63019514 |
| FVAGGYLS | GAGQNSGNTLY | 0.7781593 |
| GASGGAGGAGGW | GAGQNSGNTLY | 0.34207165 |
| GGAGAGGDGGAGGI | GAGQNSGNTLY | 0.57441413 |
| GTGHNGSQPGMGGNG | GAGQNSGNTLY | 0.66326046 |
| GYAAASASAS | GAGQNSGNTLY | 0.77490073 |
| HQVDVAAVQQG | GAGQNSGNTLY | 0.3561061 |
| IGAAIAHT | GAGQNSGNTLY | 0.6795224 |
| IGMIEVAQL | GAGQNSGNTLY | 0.42971075 |
| IRLAPEHTSAMD | GAGQNSGNTLY | 0.6388953 |
| IYNVSVL | GAGQNSGNTLY | 0.23120692 |
| KQGFLGM | GAGQNSGNTLY | 0.49381953 |
| LESVITQL | GAGQNSGNTLY | 0.34806293 |
| LGLGALVVV | GAGQNSGNTLY | 0.6331197 |
| LIGNGGNGGSGGI | GAGQNSGNTLY | 0.6717198 |
| LNAINAPTL | GAGQNSGNTLY | 0.67722285 |
| LQLLNTGV | GAGQNSGNTLY | 0.29752883 |
| LQTSIEF | GAGQNSGNTLY | 0.6716281 |
| MSFAVSEL | GAGQNSGNTLY | 0.6704914 |
| NTGALGL | GAGQNSGNTLY | 0.850981 |
| PPGAGGAGGAGTT | GAGQNSGNTLY | 0.4644779 |
| QLVGLGI | GAGQNSGNTLY | 0.7109518 |
| RFLEQVHQL | GAGQNSGNTLY | 0.53284156 |
| RGVGYVLR | GAGQNSGNTLY | 0.37858492 |

| | | |
|---|---|---|
| SATGVFAGVI | GAGQNSGNTLY | 0.22515522 |
| SLPAITLP | GAGQNSGNTLY | 0.3867901 |
| STALAFTV | GAGQNSGNTLY | 0.44750217 |
| SVLTVGLGK | GAGQNSGNTLY | 0.33312282 |
| TALAGAGSGI | GAGQNSGNTLY | 0.8145087 |
| TGLPFTGI | GAGQNSGNTLY | 0.72833633 |
| VAEHVTV | GAGQNSGNTLY | 0.70860136 |
| VDIVLNSL | GAGQNSGNTLY | 0.76256615 |
| VLTGGGAVL | GAGQNSGNTLY | 0.6646967 |
| VNVGLAYGI | GAGQNSGNTLY | 0.52095217 |
| VSSAAVGAV | GAGQNSGNTLY | 0.45874023 |
| VTSVYEAGV | GAGQNSGNTLY | 0.55223286 |
| WVALDRGA | GAGQNSGNTLY | 0.23802313 |
| YLVGSVVI | GAGQNSGNTLY | 0.4353839 |
| GPPGTGKTLL | GAGQNSGNTLY | 0.43623656 |
| GVGMTKF | GAGQNSGNTLY | 0.586367 |
| IGIGGSDL | GAGQNSGNTLY | 0.14240214 |
| IGTLLAEL | GAGQNSGNTLY | 0.75269127 |
| IYTSGTT | GAGQNSGNTLY | 0.76813877 |
| LSFPAGV | GAGQNSGNTLY | 0.59823334 |
| LYTSGTT | GAGQNSGNTLY | 0.695204 |
| VGAFPFGF | GAGQNSGNTLY | 0.5732248 |
| VGAGTVEF | GAGQNSGNTLY | 0.26923323 |
| VGKSTLLL | GAGQNSGNTLY | 0.67550236 |
| AALAHAGGGIV | GAGQNSGNTLY | 0.3450658 |
| AALVAVSLL | GAGQNSGNTLY | 0.59125674 |
| AAVSRAVAAL | GAGQNSGNTLY | 0.7295809 |
| AGGSSGIGGF | GAGQNSGNTLY | 0.64792264 |
| AGLTVANANG | GAGQNSGNTLY | 0.62310064 |
| AGLVGAVL | GAGQNSGNTLY | 0.54775935 |
| ALSQAFTV | GAGQNSGNTLY | 0.31130427 |
| ALTFSVSL | GAGQNSGNTLY | 0.6414534 |
| ALTGAALVAL | GAGQNSGNTLY | 0.3187456 |
| ASFSIGAGVL | GAGQNSGNTLY | 0.57991135 |
| DGSVYKI | GAGQNSGNTLY | 0.32016277 |
| EQLAEIV | GAGQNSGNTLY | 0.6981969 |
| FAVSELTVM | GAGQNSGNTLY | 0.63093495 |
| GAAPLAAGL | GAGQNSGNTLY | 0.5380813 |
| GADGGDGGSGGAGG | GAGQNSGNTLY | 0.40875536 |
| GFGLDTGGAGG | GAGQNSGNTLY | 0.67317295 |
| GLASGGLL | GAGQNSGNTLY | 0.6706933 |
| GRIAGAAVLL | GAGQNSGNTLY | 0.3331912 |
| GSAGSLNTSF | GAGQNSGNTLY | 0.44709164 |
| GTGQIGI | GAGQNSGNTLY | 0.3718341 |
| GTLIALIL | GAGQNSGNTLY | 0.36423758 |
| IAATMGFVVG | GAGQNSGNTLY | 0.73884755 |
| IFAQALL | GAGQNSGNTLY | 0.61358744 |
| IGAAVALGF | GAGQNSGNTLY | 0.1952839 |
| IGLTPLAL | GAGQNSGNTLY | 0.47911566 |
| ILGSNTSTL | GAGQNSGNTLY | 0.65335655 |
| IVFSLTVL | GAGQNSGNTLY | 0.5035876 |

| | | |
|---|---|---|
| LASKATGF | GAGQNSGNTLY | 0.15525079 |
| LASVALIL | GAGQNSGNTLY | 0.47970653 |
| LFAVGLTL | GAGQNSGNTLY | 0.41262344 |
| LFNGGIGTYI | GAGQNSGNTLY | 0.30171797 |
| LTLGAGII | GAGQNSGNTLY | 0.37679994 |
| LVAAALGDI | GAGQNSGNTLY | 0.7235654 |
| NTAFVQLGI | GAGQNSGNTLY | 0.59237206 |
| QLFSGAEL | GAGQNSGNTLY | 0.5180976 |
| SAATFITL | GAGQNSGNTLY | 0.67418236 |
| SNYGLANLGV | GAGQNSGNTLY | 0.6720617 |
| SYGGAYAVM | GAGQNSGNTLY | 0.7395824 |
| TTGVGGLTLG | GAGQNSGNTLY | 0.6910012 |
| TTQTIYI | GAGQNSGNTLY | 0.4249358 |
| VAAGAGLGAVI | GAGQNSGNTLY | 0.6143302 |
| VAAVVTLL | GAGQNSGNTLY | 0.3789025 |
| VGFLGGLI | GAGQNSGNTLY | 0.41319093 |
| VGVIPILL | GAGQNSGNTLY | 0.34550148 |
| VIFASSLI | GAGQNSGNTLY | 0.39596248 |
| VKGGDGGAAAG | GAGQNSGNTLY | 0.45475227 |
| VTKTFTNI | GAGQNSGNTLY | 0.68666327 |
| VVAAAILL | GAGQNSGNTLY | 0.65318173 |
| YAALDVEL | GAGQNSGNTLY | 0.35988995 |
| LASVALLL | GAGQNSGNTLY | 0.25824717 |
| LDAVHRV | GAGQNSGNTLY | 0.44553918 |
| QGLSSSIF | GAGQNSGNTLY | 0.3310202 |
| RGTFANIRL | GAGQNSGNTLY | 0.64654994 |
| RQLWWGHRI | GAGQNSGNTLY | 0.565504 |
| YGPPGTGKTLL | GAGQNSGNTLY | 0.85168946 |
| AALVGAPANVA | GAGQNSGNTLY | 0.7277314 |
| AATFRRTE | GAGQNSGNTLY | 0.5166926 |
| AGALGGGI | GAGQNSGNTLY | 0.65218365 |
| AGGAGGAGGIGAG | GAGQNSGNTLY | 0.3334 |
| AGSGALGL | GAGQNSGNTLY | 0.3974707 |
| AQIADQI | GAGQNSGNTLY | 0.424495 |
| ASAVAVTYL | GAGQNSGNTLY | 0.4077426 |
| ASTVLGGTGG | GAGQNSGNTLY | 0.5437379 |
| AVGGEPAK | GAGQNSGNTLY | 0.41376653 |
| CEQGTFGGTC | GAGQNSGNTLY | 0.21543403 |
| DEGKAPAG | GAGQNSGNTLY | 0.75958705 |
| DTVVFPAL | GAGQNSGNTLY | 0.5482597 |
| FEGITMAG | GAGQNSGNTLY | 0.2592294 |
| FPLVVQD | GAGQNSGNTLY | 0.4027506 |
| GAFTTPQ | GAGQNSGNTLY | 0.76773036 |
| GFGGFGTAGGVGGA | GAGQNSGNTLY | 0.77589166 |
| GFLAQFL | GAGQNSGNTLY | 0.38524938 |
| GGAGGAGGNAGVA | GAGQNSGNTLY | 0.4164381 |
| GGVGTAGAGGAGGAGG | GAGQNSGNTLY | 0.7566587 |
| GNGGGGDAGNAGSGGN | GAGQNSGNTLY | 0.43237838 |
| GTGGAGGAGNDAGS | GAGQNSGNTLY | 0.4274042 |
| GTTNTGWFNS | GAGQNSGNTLY | 0.530236 |
| IFMGPLFTL | GAGQNSGNTLY | 0.44387025 |

| | | |
|---|---|---|
| ILAHAIEAI | GAGQNSGNTLY | 0.38378388 |
| IRSAIREAL | GAGQNSGNTLY | 0.6233081 |
| ITFAGGG | GAGQNSGNTLY | 0.5035505 |
| ITGMSGHQEV | GAGQNSGNTLY | 0.46675608 |
| IVAGLACTL | GAGQNSGNTLY | 0.15043579 |
| IVGGGLAGA | GAGQNSGNTLY | 0.5549576 |
| IVTGVLNI | GAGQNSGNTLY | 0.35924068 |
| LAQQGCI | GAGQNSGNTLY | 0.26224804 |
| LCLHVEN | GAGQNSGNTLY | 0.15731098 |
| LDAISDGS | GAGQNSGNTLY | 0.48731083 |
| LGGFAAML | GAGQNSGNTLY | 0.6597555 |
| LGGKNPMI | GAGQNSGNTLY | 0.72909975 |
| LGPKGVAAIL | GAGQNSGNTLY | 0.5283222 |
| LMVGVSI | GAGQNSGNTLY | 0.6679791 |
| LVGAAFVL | GAGQNSGNTLY | 0.6178678 |
| LVLAVAPL | GAGQNSGNTLY | 0.40843767 |
| MAAIAVIDAA | GAGQNSGNTLY | 0.23712507 |
| NNGINAELQ | GAGQNSGNTLY | 0.6191847 |
| RGEPLVLVHG | GAGQNSGNTLY | 0.6919073 |
| RGGAGGAAGLI | GAGQNSGNTLY | 0.5420049 |
| RSYRMHPAV | GAGQNSGNTLY | 0.73340106 |
| SAAHLLAGQLA | GAGQNSGNTLY | 0.25968635 |
| SAAVGSY | GAGQNSGNTLY | 0.74369514 |
| SPHAVLHRI | GAGQNSGNTLY | 0.59636134 |
| SQIAKVL | GAGQNSGNTLY | 0.24553534 |
| TALGHLVP | GAGQNSGNTLY | 0.32655042 |
| TAPTTSGAA | GAGQNSGNTLY | 0.36159134 |
| TDGDDGNGGSPGT | GAGQNSGNTLY | 0.20205495 |
| TDGLGEN | GAGQNSGNTLY | 0.6771338 |
| TGGPPLPGQA | GAGQNSGNTLY | 0.44769865 |
| TIPAITI | GAGQNSGNTLY | 0.57750267 |
| TIPGFTI | GAGQNSGNTLY | 0.41596693 |
| TSIASGALA | GAGQNSGNTLY | 0.53023154 |
| TVTLPGAAI | GAGQNSGNTLY | 0.4883692 |
| VASGLSAALP | GAGQNSGNTLY | 0.60806054 |
| VGAGVIGI | GAGQNSGNTLY | 0.5016151 |
| VGPKRFGV | GAGQNSGNTLY | 0.6115993 |
| VNVAFPD | GAGQNSGNTLY | 0.34840375 |
| VSMTNSGVS | GAGQNSGNTLY | 0.49235037 |
| VTDVLGDLSV | GAGQNSGNTLY | 0.77190834 |
| VVVVGAGFAGL | GAGQNSGNTLY | 0.28411418 |
| VYTVGTVV | GAGQNSGNTLY | 0.34865856 |
| WTTFSGP | GAGQNSGNTLY | 0.53203243 |
| YDELVEAF | GAGQNSGNTLY | 0.47897458 |
| YGTGGAGGNGG | GAGQNSGNTLY | 0.6302959 |
| YTNVMAV | GAGQNSGNTLY | 0.64531493 |
| FLEQVHQL | GAGQNSGNTLY | 0.6108002 |
| GLSLLAVAV | GAGQNSGNTLY | 0.7207738 |
| LEQVHQL | GAGQNSGNTLY | 0.7500529 |
| LNAAGAI | GAGQNSGNTLY | 0.6499741 |
| PAVHASL | GAGQNSGNTLY | 0.41529238 |

| | | |
|---|---|---|
| TELEDDA | GAGQNSGNTLY | 0.79593647 |
| TPPGPPN | GAGQNSGNTLY | 0.3052606 |
| TPPPPPPP | GAGQNSGNTLY | 0.65031374 |
| TQSPALI | ASSPRNYAEQF | 0.70255053 |
| EFAAAFPQ | ASSPRNYAEQF | 0.31287664 |
| LLSGNLDVL | ASSPRNYAEQF | 0.54330385 |
| GPANLGNYNI | ASSPRNYAEQF | 0.68672717 |
| TLGPTLP | ASSPRNYAEQF | 0.68723506 |
| VYAYSSCCK | ASSPRNYAEQF | 0.4081611 |
| APAPAPAPNI | ASSPRNYAEQF | 0.30913514 |
| PGGPVPIP | ASSPRNYAEQF | 0.36955684 |
| FLTVALEP | ASSPRNYAEQF | 0.31310314 |
| SQYFGANL | ASSPRNYAEQF | 0.2928204 |
| GGVKPAA | ASSPRNYAEQF | 0.556106 |
| GGTGGAGGAGGW | ASSPRNYAEQF | 0.53431916 |
| GVGGVAGLGGAGG | ASSPRNYAEQF | 0.8146936 |
| VLLVLATTI | ASSPRNYAEQF | 0.6468851 |
| IGIFAAML | ASSPRNYAEQF | 0.76308006 |
| AVGDLID | ASSPRNYAEQF | 0.4485488 |
| NISPLPL | ASSPRNYAEQF | 0.5713017 |
| VTAVFVAI | ASSPRNYAEQF | 0.20405798 |
| SGDDNDAAL | ASSPRNYAEQF | 0.46644676 |
| AVAALAHS | ASSPRNYAEQF | 0.2841224 |
| GGVTVMATP | ASSPRNYAEQF | 0.5901885 |
| AGLAANVRD | ASSPRNYAEQF | 0.20586741 |
| TNRPDLIDPALLRPG | ASSPRNYAEQF | 0.6815536 |
| LANLGVTAL | ASSPRNYAEQF | 0.48084253 |
| TGERIEL | ASSPRNYAEQF | 0.69297373 |
| NGEELAA | ASSPRNYAEQF | 0.42880693 |
| VLSLVWG | ASSPRNYAEQF | 0.34887522 |
| IGYALWL | ASSPRNYAEQF | 0.43846172 |
| LSLLLAPL | ASSPRNYAEQF | 0.7549777 |
| TLDEALK | ASSPRNYAEQF | 0.26769722 |
| TDGAGTAEGG | ASSPRNYAEQF | 0.16994222 |
| NAPGGAGGNGG | ASSPRNYAEQF | 0.33649546 |
| VSVFMGMGGNF | ASSPRNYAEQF | 0.21385524 |
| LASYLAAL | ASSPRNYAEQF | 0.75044346 |
| LVGGTPAL | ASSPRNYAEQF | 0.7034509 |
| VRNADVL | ASSPRNYAEQF | 0.5601062 |
| TCSSPFLVSAM | ASSPRNYAEQF | 0.8388685 |
| VHITDAE | ASSPRNYAEQF | 0.13841692 |
| EYAAPSF | ASSPRNYAEQF | 0.42254865 |
| VVDAFHGGL | ASSPRNYAEQF | 0.78929996 |
| ASVVDPQ | ASSPRNYAEQF | 0.5775496 |
| SYNEFGGLS | ASSPRNYAEQF | 0.6084519 |
| DPDGSVY | ASSPRNYAEQF | 0.67909074 |
| MVSTIHATE | ASSPRNYAEQF | 0.47964084 |
| LAAGVADPV | ASSPRNYAEQF | 0.5778965 |
| TKEQIAA | ASSPRNYAEQF | 0.5517268 |
| SLASPAAL | ASSPRNYAEQF | 0.55747855 |
| IGLVFYGV | ASSPRNYAEQF | 0.29933923 |

| | | |
|---|---|---|
| FLLDASNL | ASSPRNYAEQF | 0.26559874 |
| YQVGNFP | ASSPRNYAEQF | 0.649593 |
| ESLLSQST | ASSPRNYAEQF | 0.7901216 |
| LFTVGFAAVG | ASSPRNYAEQF | 0.57331085 |
| ATPPTPT | ASSPRNYAEQF | 0.5681721 |
| TDLATLG | ASSPRNYAEQF | 0.5990007 |
| SAGGTGR | ASSPRNYAEQF | 0.715359 |
| GFSNSGS | ASSPRNYAEQF | 0.4286005 |
| GGNGGSGGTSV | ASSPRNYAEQF | 0.26462492 |
| GTSVATGGAGN | ASSPRNYAEQF | 0.7201104 |
| NGGNGGSGGTS | ASSPRNYAEQF | 0.23087987 |
| AVVKFPH | ASSPRNYAEQF | 0.542436 |
| TPPPPPP | ASSPRNYAEQF | 0.34517923 |
| AAPAAAAAPA | ASSPRNYAEQF | 0.8137238 |
| IALNEDL | ASSPRNYAEQF | 0.6523458 |
| IVNVSSM | ASSPRNYAEQF | 0.53537786 |
| TGIPPAPR | ASSPRNYAEQF | 0.45217457 |
| FDLGGGTF | ASSPRNYAEQF | 0.44224316 |
| FDLGGGTFDVS | ASSPRNYAEQF | 0.6031767 |
| NEPTAAA | ASSPRNYAEQF | 0.2160873 |
| NEPTAAALA | ASSPRNYAEQF | 0.7396195 |
| GSIINISSI | ASSPRNYAEQF | 0.32534316 |
| VAPGNAGTA | ASSPRNYAEQF | 0.41002184 |
| TGGGASLE | ASSPRNYAEQF | 0.5794226 |
| VAVYNHYKRI | ASSPRNYAEQF | 0.83162117 |
| TDPAPATT | ASSPRNYAEQF | 0.3917967 |
| TNRPDILDPALLRPG | ASSPRNYAEQF | 0.6714181 |
| LTTTTTS | ASSPRNYAEQF | 0.38371676 |
| PTGSGKTL | ASSPRNYAEQF | 0.5799706 |
| TGSGKTL | ASSPRNYAEQF | 0.7592963 |
| TIGHVDHGK | ASSPRNYAEQF | 0.74180067 |
| TIGHVDHGKTT | ASSPRNYAEQF | 0.84183073 |
| GHVDHGKTT | ASSPRNYAEQF | 0.45420867 |
| AQFEHTLL | ASSPRNYAEQF | 0.22281027 |
| IVNQDEL | ASSPRNYAEQF | 0.4386838 |
| TNGSQFF | ASSPRNYAEQF | 0.63314736 |
| TGKGIVE | ASSPRNYAEQF | 0.6519613 |
| LISNASDA | ASSPRNYAEQF | 0.6929133 |
| AELRTLGELTKIAKAHGAQ | ASSPRNYAEQF | 0.59864783 |
| APNAVAAAEPSW | ASSPRNYAEQF | 0.4305957 |
| APQVAGAATAVL | ASSPRNYAEQF | 0.72744405 |
| ARRDPAAAAAILPTDA | ASSPRNYAEQF | 0.7469065 |
| DAAARDVLAEAGLAEYF | ASSPRNYAEQF | 0.6800994 |
| DDQIVLSAKANLAKVEQ | ASSPRNYAEQF | 0.30381754 |
| DELLPDTGEDSDEDGH | ASSPRNYAEQF | 0.3174225 |
| ELPEAPANRATVPQD | ASSPRNYAEQF | 0.636488 |
| FGQSKIERVTGIG | ASSPRNYAEQF | 0.5757128 |
| GAGGTGSTAGGAGGAGGAGGLY | ASSPRNYAEQF | 0.29052603 |
| GHGGTGAAGGNGGTGGAGGSGI | ASSPRNYAEQF | 0.5630193 |
| GNGGNGGAGGEGGPGGAGGAGA | ASSPRNYAEQF | 0.63826543 |
| GSTAGNGGNGGNATNGTIAGQP | ASSPRNYAEQF | 0.24236709 |

| | | |
|---|---|---|
| GTGGDGGNGGNGGTGG | ASSPRNYAEQF | 0.3801883 |
| GVRRPAGIPADAPV | ASSPRNYAEQF | 0.70106506 |
| GYRQVTDAHLLTLARR | ASSPRNYAEQF | 0.43393245 |
| IAKMGQDGHDRG | ASSPRNYAEQF | 0.33569098 |
| KAEPSIEIQYT | ASSPRNYAEQF | 0.7462027 |
| KPVRIRMAADEDTVAVT | ASSPRNYAEQF | 0.79075783 |
| MPPRIAGMRLLVIKPEP | ASSPRNYAEQF | 0.6663382 |
| MKFYAVALFRSRGVPPTR | ASSPRNYAEQF | 0.24717668 |
| NAGGFGILGAVAHSP | ASSPRNYAEQF | 0.1337779 |
| NEGREITLAMAEM | ASSPRNYAEQF | 0.61762315 |
| NELVEWSHPLGHRWRWE | ASSPRNYAEQF | 0.8088138 |
| NLGAGGTGGNGGNPG | ASSPRNYAEQF | 0.56804746 |
| NTEWPKWETVMAADPVD | ASSPRNYAEQF | 0.44018352 |
| PASAAPAETQP | ASSPRNYAEQF | 0.37696373 |
| QQSSDGKNGVRAAEITGE | ASSPRNYAEQF | 0.66051376 |
| RAPDAVTTSWWKE | ASSPRNYAEQF | 0.37427306 |
| SGFGNVGAHTSGWFNQ | ASSPRNYAEQF | 0.2395533 |
| SGVAGAVGQAASVGGLKV | ASSPRNYAEQF | 0.42239848 |
| SNGGAGTGSAGGNGGT | ASSPRNYAEQF | 0.70796025 |
| TGGAGGEGGAAG | ASSPRNYAEQF | 0.26971465 |
| TGGDGGAAGAGGAPGHG | ASSPRNYAEQF | 0.42093194 |
| TMPPLLAAIANAGRHGV | ASSPRNYAEQF | 0.68461716 |
| TSSTAGNSGGA | ASSPRNYAEQF | 0.45175973 |
| VPATVPVGILC | ASSPRNYAEQF | 0.547025 |
| VQRPDSGGVVLTGRIS | ASSPRNYAEQF | 0.30027175 |
| WTPNPVPATKGVPAGLVQAT | ASSPRNYAEQF | 0.6364049 |
| YALKRALAKFSDRFDVVII | ASSPRNYAEQF | 0.25126714 |
| YLIGSVIV | ASSPRNYAEQF | 0.5256058 |
| YGQLVAVID | ASSPRNYAEQF | 0.64247054 |
| VTVGGMTLPA | ASSPRNYAEQF | 0.59062874 |
| VLALPAV | ASSPRNYAEQF | 0.5042415 |
| VIPPTLNYET | ASSPRNYAEQF | 0.35051453 |
| VGMGGMHPSSG | ASSPRNYAEQF | 0.33435884 |
| VGLAGGAATA | ASSPRNYAEQF | 0.31207067 |
| VGGAGGAGGSGP | ASSPRNYAEQF | 0.7441651 |
| VFAAFMV | ASSPRNYAEQF | 0.48463285 |
| TLYDKLR | ASSPRNYAEQF | 0.70172113 |
| TLAVFTVL | ASSPRNYAEQF | 0.62046516 |
| TGSVQPARV | ASSPRNYAEQF | 0.29984725 |
| TGGGAGSGGT | ASSPRNYAEQF | 0.47720623 |
| SHLETSL | ASSPRNYAEQF | 0.2704295 |
| SFTTGMT | ASSPRNYAEQF | 0.6116744 |
| QVEVAKRMGVRQPTVS | ASSPRNYAEQF | 0.40664446 |
| QGSGGPALAM | ASSPRNYAEQF | 0.5180508 |
| MLAPSNP | ASSPRNYAEQF | 0.41674852 |
| LVNNAAI | ASSPRNYAEQF | 0.46676975 |
| LTGEINTLA | ASSPRNYAEQF | 0.605042 |
| LSIFGPV | ASSPRNYAEQF | 0.37199038 |
| LLALLANL | ASSPRNYAEQF | 0.54170716 |
| LISNASDALDKLR | ASSPRNYAEQF | 0.70161647 |
| LINEAVP | ASSPRNYAEQF | 0.5344262 |

| | | |
|---|---|---|
| LGYAFSGL | ASSPRNYAEQF | 0.26914644 |
| LDFVAVAL | ASSPRNYAEQF | 0.61539924 |
| LAQPGAE | ASSPRNYAEQF | 0.30214062 |
| ISNLGAVTGF | ASSPRNYAEQF | 0.6200511 |
| HQEDPGRI | ASSPRNYAEQF | 0.4263274 |
| GTPGEPTP | ASSPRNYAEQF | 0.4763999 |
| GFFNSGS | ASSPRNYAEQF | 0.56882274 |
| GAGGGGLTAGAN | ASSPRNYAEQF | 0.19827278 |
| FVDALVNM | ASSPRNYAEQF | 0.38674614 |
| FLTVAPEL | ASSPRNYAEQF | 0.43476757 |
| FETLTAL | ASSPRNYAEQF | 0.6870942 |
| DVINPIEL | ASSPRNYAEQF | 0.6286499 |
| APVSGAPV | ASSPRNYAEQF | 0.2879221 |
| ALVGGLAW | ASSPRNYAEQF | 0.56854296 |
| ALAAAAATAI | ASSPRNYAEQF | 0.51597375 |
| AFGLGLLL | ASSPRNYAEQF | 0.41538715 |
| AFFAAASL | ASSPRNYAEQF | 0.4108405 |
| AAAALPAFSPP | ASSPRNYAEQF | 0.34534228 |
| VVLAPAL | ASSPRNYAEQF | 0.24046725 |
| TIGHVDHGKTTL | ASSPRNYAEQF | 0.602396 |
| LISNASDALDK | ASSPRNYAEQF | 0.38742638 |
| LISNASDAL | ASSPRNYAEQF | 0.5963206 |
| ISNASDALDK | ASSPRNYAEQF | 0.62341386 |
| NQAETLVYQTE | ASSPRNYAEQF | 0.35336608 |
| TGGTGGTAGAG | ASSPRNYAEQF | 0.4931701 |
| TGSTAGGAGGA | ASSPRNYAEQF | 0.4745232 |
| DPLAVPVAAGPV | ASSPRNYAEQF | 0.638329 |
| GGTGGTPGNGGN | ASSPRNYAEQF | 0.63638103 |
| PAGLAGVDPATL | ASSPRNYAEQF | 0.36598727 |
| GNNLVVEAPHAGASV | ASSPRNYAEQF | 0.6105865 |
| KVTRIMSASTLEQPA | ASSPRNYAEQF | 0.717129 |
| LGFALPIALPRRTVV | ASSPRNYAEQF | 0.6724533 |
| TRLGNLTARKLVVLV | ASSPRNYAEQF | 0.55695975 |
| WSQQRRLIYFPSAGP | ASSPRNYAEQF | 0.79313624 |
| GPNGNPDPMAAAVDIR | ASSPRNYAEQF | 0.61294013 |
| TQRCVSVADLRAAASR | ASSPRNYAEQF | 0.4123211 |
| QFVLDVAEHPVEVLTETV | ASSPRNYAEQF | 0.1879203 |
| GGAGGTGGAGGNGGTQAGDG | ASSPRNYAEQF | 0.5983392 |
| PLLATLRAIAVDVDAKPAQV | ASSPRNYAEQF | 0.58335024 |
| APKFRTGGYMIDFWGVGYQVAK | ASSPRNYAEQF | 0.3448763 |
| GAGGAGGAGGAAGGPGGTGGTG | ASSPRNYAEQF | 0.19080134 |
| IGALGLLATDVAPAGSVAFGVA | ASSPRNYAEQF | 0.70268667 |
| AGGAGGIGGNGGAGGAGGVLMG | ASSPRNYAEQF | 0.8046718 |
| ADELIGLL | ASSPRNYAEQF | 0.4909944 |
| ADRALKA | ASSPRNYAEQF | 0.6815349 |
| AGAVGQVGGAGGA | ASSPRNYAEQF | 0.22880551 |
| AGGGGGLGGAP | ASSPRNYAEQF | 0.7101109 |
| AGGLLYGNGGAG | ASSPRNYAEQF | 0.71256465 |
| AGLESIGL | ASSPRNYAEQF | 0.27826518 |
| AGLKTGII | ASSPRNYAEQF | 0.6575469 |
| ASLAALLL | ASSPRNYAEQF | 0.3333258 |

| | | |
|---|---|---|
| ASVINISSI | ASSPRNYAEQF | 0.36238602 |
| ATPPGPA | ASSPRNYAEQF | 0.75436556 |
| ATVASALV | ASSPRNYAEQF | 0.6124381 |
| ATVASLIL | ASSPRNYAEQF | 0.58333874 |
| AVNGPVLIRH | ASSPRNYAEQF | 0.5179914 |
| DGAAGAVGI | ASSPRNYAEQF | 0.37599748 |
| DPLLHGI | ASSPRNYAEQF | 0.47154942 |
| ELGAALGAL | ASSPRNYAEQF | 0.70036125 |
| ELGRTQAV | ASSPRNYAEQF | 0.5569516 |
| FVAGGYLS | ASSPRNYAEQF | 0.69038224 |
| GASGGAGGAGGW | ASSPRNYAEQF | 0.3660677 |
| GGAGAGGDGGAGGI | ASSPRNYAEQF | 0.76127446 |
| GTGHNGSQPGMGGNG | ASSPRNYAEQF | 0.81748414 |
| GYAAASASAS | ASSPRNYAEQF | 0.5793989 |
| HQVDVAAVQQG | ASSPRNYAEQF | 0.32936934 |
| IGAAIAHT | ASSPRNYAEQF | 0.75195754 |
| IGMIEVAQL | ASSPRNYAEQF | 0.5853376 |
| IRLAPEHTSAMD | ASSPRNYAEQF | 0.67944646 |
| IYNVSVL | ASSPRNYAEQF | 0.6148387 |
| KQGFLGM | ASSPRNYAEQF | 0.4048787 |
| LESVITQL | ASSPRNYAEQF | 0.8352523 |
| LGLGALVVV | ASSPRNYAEQF | 0.7239461 |
| LIGNGGNGGSGGI | ASSPRNYAEQF | 0.6374978 |
| LNAINAPTL | ASSPRNYAEQF | 0.50058615 |
| LQLLNTGV | ASSPRNYAEQF | 0.27466196 |
| LQTSIEF | ASSPRNYAEQF | 0.4957785 |
| MSFAVSEL | ASSPRNYAEQF | 0.42745048 |
| NTGALGL | ASSPRNYAEQF | 0.67550063 |
| PPGAGGAGGAGTT | ASSPRNYAEQF | 0.72874373 |
| QLVGLGI | ASSPRNYAEQF | 0.47819683 |
| RFLEQVHQL | ASSPRNYAEQF | 0.8019967 |
| RGVGYVLR | ASSPRNYAEQF | 0.7757379 |
| SATGVFAGVI | ASSPRNYAEQF | 0.292302 |
| SLPAITLP | ASSPRNYAEQF | 0.58504033 |
| STALAFTV | ASSPRNYAEQF | 0.47083908 |
| SVLTVGLGK | ASSPRNYAEQF | 0.5784153 |
| TALAGAGSGI | ASSPRNYAEQF | 0.18830392 |
| TGLPFTGI | ASSPRNYAEQF | 0.7423091 |
| VAEHVTV | ASSPRNYAEQF | 0.48264682 |
| VDIVLNSL | ASSPRNYAEQF | 0.115829684 |
| VLTGGGAVL | ASSPRNYAEQF | 0.5972068 |
| VNVGLAYGI | ASSPRNYAEQF | 0.56426084 |
| VSSAAVGAV | ASSPRNYAEQF | 0.55093914 |
| VTSVYEAGV | ASSPRNYAEQF | 0.800871 |
| WVALDRGA | ASSPRNYAEQF | 0.26203465 |
| YLVGSVVI | ASSPRNYAEQF | 0.4138872 |
| GPPGTGKTLL | ASSPRNYAEQF | 0.761717 |
| GVGMTKF | ASSPRNYAEQF | 0.8160938 |
| IGIGGSDL | ASSPRNYAEQF | 0.3485469 |
| IGTLLAEL | ASSPRNYAEQF | 0.70422846 |
| IYTSGTT | ASSPRNYAEQF | 0.8070775 |

| | | |
|---|---|---|
| LSFPAGV | ASSPRNYAEQF | 0.71562594 |
| LYTSGTT | ASSPRNYAEQF | 0.78461885 |
| VGAFPFGF | ASSPRNYAEQF | 0.81016684 |
| VGAGTVEF | ASSPRNYAEQF | 0.3815275 |
| VGKSTLLL | ASSPRNYAEQF | 0.7952275 |
| AALAHAGGGIV | ASSPRNYAEQF | 0.7641839 |
| AALVAVSLL | ASSPRNYAEQF | 0.5523363 |
| AAVSRAVAAL | ASSPRNYAEQF | 0.6377549 |
| AGGSSGIGGF | ASSPRNYAEQF | 0.40677258 |
| AGLTVANANG | ASSPRNYAEQF | 0.49373984 |
| AGLVGAVL | ASSPRNYAEQF | 0.2031255 |
| ALSQAFTV | ASSPRNYAEQF | 0.7228857 |
| ALTFSVSL | ASSPRNYAEQF | 0.80069077 |
| ALTGAALVAL | ASSPRNYAEQF | 0.5765922 |
| ASFSIGAGVL | ASSPRNYAEQF | 0.36009216 |
| DGSVYKI | ASSPRNYAEQF | 0.32239383 |
| EQLAEIV | ASSPRNYAEQF | 0.54446185 |
| FAVSELTVM | ASSPRNYAEQF | 0.51407933 |
| GAAPLAAGL | ASSPRNYAEQF | 0.23958683 |
| GADGGDGGSGGAGG | ASSPRNYAEQF | 0.49447173 |
| GFGLDTGGAGG | ASSPRNYAEQF | 0.7889405 |
| GLASGGLL | ASSPRNYAEQF | 0.7805946 |
| GRIAGAAVLL | ASSPRNYAEQF | 0.29904422 |
| GSAGSLNTSF | ASSPRNYAEQF | 0.8204626 |
| GTGQIGI | ASSPRNYAEQF | 0.708702 |
| GTLIALIL | ASSPRNYAEQF | 0.79759204 |
| IAATMGFVVG | ASSPRNYAEQF | 0.59178066 |
| IFAQALL | ASSPRNYAEQF | 0.45605335 |
| IGAAVALGF | ASSPRNYAEQF | 0.7447007 |
| IGLTPLAL | ASSPRNYAEQF | 0.598752 |
| ILGSNTSTL | ASSPRNYAEQF | 0.4549464 |
| IVFSLTVL | ASSPRNYAEQF | 0.77035576 |
| LASKATGF | ASSPRNYAEQF | 0.5626625 |
| LASVALIL | ASSPRNYAEQF | 0.5103179 |
| LFAVGLTL | ASSPRNYAEQF | 0.53608 |
| LFNGGIGTYI | ASSPRNYAEQF | 0.79830766 |
| LTLGAGII | ASSPRNYAEQF | 0.3632568 |
| LVAAALGDI | ASSPRNYAEQF | 0.44891083 |
| NTAFVQLGI | ASSPRNYAEQF | 0.34435228 |
| QLFSGAEL | ASSPRNYAEQF | 0.6287418 |
| SAATFITL | ASSPRNYAEQF | 0.8019694 |
| SNYGLANLGV | ASSPRNYAEQF | 0.69197345 |
| SYGGAYAVM | ASSPRNYAEQF | 0.58984125 |
| TTGVGGLTLG | ASSPRNYAEQF | 0.7224593 |
| TTQTIYI | ASSPRNYAEQF | 0.7402043 |
| VAAGAGLGAVI | ASSPRNYAEQF | 0.6534056 |
| VAAVVTLL | ASSPRNYAEQF | 0.7612809 |
| VGFLGGLI | ASSPRNYAEQF | 0.50577724 |
| VGVIPILL | ASSPRNYAEQF | 0.80271 |
| VIFASSLI | ASSPRNYAEQF | 0.744964 |
| VKGGDGGAAAG | ASSPRNYAEQF | 0.74893653 |

| | | |
|---|---|---|
| VTKTFTNI | ASSPRNYAEQF | 0.7703986 |
| VVAAAILL | ASSPRNYAEQF | 0.7139623 |
| YAALDVEL | ASSPRNYAEQF | 0.6239969 |
| LASVALLL | ASSPRNYAEQF | 0.2167337 |
| LDAVHRV | ASSPRNYAEQF | 0.42935717 |
| QGLSSSIF | ASSPRNYAEQF | 0.6998943 |
| RGTFANIRL | ASSPRNYAEQF | 0.73539704 |
| RQLWWGHRI | ASSPRNYAEQF | 0.6996766 |
| YGPPGTGKTLL | ASSPRNYAEQF | 0.67371786 |
| AALVGAPANVA | ASSPRNYAEQF | 0.44538152 |
| AATFRRTE | ASSPRNYAEQF | 0.7291349 |
| AGALGGGI | ASSPRNYAEQF | 0.78353167 |
| AGGAGGAGGIGAG | ASSPRNYAEQF | 0.28532767 |
| AGSGALGL | ASSPRNYAEQF | 0.51103765 |
| AQIADQI | ASSPRNYAEQF | 0.84527516 |
| ASAVAVTYL | ASSPRNYAEQF | 0.6846389 |
| ASTVLGGTGG | ASSPRNYAEQF | 0.5591966 |
| AVGGEPAK | ASSPRNYAEQF | 0.4827596 |
| CEQGTFGGTC | ASSPRNYAEQF | 0.79651284 |
| DEGKAPAG | ASSPRNYAEQF | 0.41785884 |
| DTVVFPAL | ASSPRNYAEQF | 0.8152224 |
| FEGITMAG | ASSPRNYAEQF | 0.16275325 |
| FPLVVQD | ASSPRNYAEQF | 0.48293293 |
| GAFTTPQ | ASSPRNYAEQF | 0.44109565 |
| GFGGFGTAGGVGGA | ASSPRNYAEQF | 0.24554347 |
| GFLAQFL | ASSPRNYAEQF | 0.7719405 |
| GGAGGAGGNAGVA | ASSPRNYAEQF | 0.3853273 |
| GGVGTAGAGGAGGAGG | ASSPRNYAEQF | 0.7329979 |
| GNGGGGDAGNAGSGGN | ASSPRNYAEQF | 0.5902271 |
| GTGGAGGAGNDAGS | ASSPRNYAEQF | 0.7913588 |
| GTTNTGWFNS | ASSPRNYAEQF | 0.699262 |
| IFMGPLFTL | ASSPRNYAEQF | 0.6053434 |
| ILAHAIEAI | ASSPRNYAEQF | 0.4577774 |
| IRSAIREAL | ASSPRNYAEQF | 0.73733187 |
| ITFAGGG | ASSPRNYAEQF | 0.75510573 |
| ITGMSGHQEV | ASSPRNYAEQF | 0.7272978 |
| IVAGLACTL | ASSPRNYAEQF | 0.2657695 |
| IVGGGLAGA | ASSPRNYAEQF | 0.5285487 |
| IVTGVLNI | ASSPRNYAEQF | 0.6096325 |
| LAQQGCI | ASSPRNYAEQF | 0.22473547 |
| LCLHVEN | ASSPRNYAEQF | 0.14939763 |
| LDAISDGS | ASSPRNYAEQF | 0.46024734 |
| LGGFAAML | ASSPRNYAEQF | 0.43943757 |
| LGGKNPMI | ASSPRNYAEQF | 0.7858275 |
| LGPKGVAAIL | ASSPRNYAEQF | 0.7756717 |
| LMVGVSI | ASSPRNYAEQF | 0.5897568 |
| LVGAAFVL | ASSPRNYAEQF | 0.7600292 |
| LVLAVAPL | ASSPRNYAEQF | 0.6666701 |
| MAAIAVIDAA | ASSPRNYAEQF | 0.0919092 |
| NNGINAELQ | ASSPRNYAEQF | 0.4929213 |
| RGEPLVLVHG | ASSPRNYAEQF | 0.28523675 |

| | | |
|---|---|---|
| RGGAGGAAGLI | ASSPRNYAEQF | 0.77170414 |
| RSYRMHPAV | ASSPRNYAEQF | 0.36933932 |
| SAAHLLAGQLA | ASSPRNYAEQF | 0.43738574 |
| SAAVGSY | ASSPRNYAEQF | 0.65216565 |
| SPHAVLHRI | ASSPRNYAEQF | 0.6603831 |
| SQIAKVL | ASSPRNYAEQF | 0.60922354 |
| TALGHLVP | ASSPRNYAEQF | 0.43251258 |
| TAPTTSGAA | ASSPRNYAEQF | 0.12314276 |
| TDGDDGNGGSPGT | ASSPRNYAEQF | 0.7739196 |
| TDGLGEN | ASSPRNYAEQF | 0.32093224 |
| TGGPPLPGQA | ASSPRNYAEQF | 0.36427358 |
| TIPAITI | ASSPRNYAEQF | 0.18937986 |
| TIPGFTI | ASSPRNYAEQF | 0.17745475 |
| TSIASGALA | ASSPRNYAEQF | 0.25100923 |
| TVTLPGAAI | ASSPRNYAEQF | 0.77161616 |
| VASGLSAALP | ASSPRNYAEQF | 0.5305947 |
| VGAGVIGI | ASSPRNYAEQF | 0.56725025 |
| VGPKRFGV | ASSPRNYAEQF | 0.8310369 |
| VNVAFPD | ASSPRNYAEQF | 0.63251275 |
| VSMTNSGVS | ASSPRNYAEQF | 0.6083106 |
| VTDVLGDLSV | ASSPRNYAEQF | 0.47040677 |
| VVVVGAGFAGL | ASSPRNYAEQF | 0.40405965 |
| VYTVGTVV | ASSPRNYAEQF | 0.43708682 |
| WTTFSGP | ASSPRNYAEQF | 0.4227677 |
| YDELVEAF | ASSPRNYAEQF | 0.8247193 |
| YGTGGAGGNGG | ASSPRNYAEQF | 0.48061532 |
| YTNVMAV | ASSPRNYAEQF | 0.32858613 |
| FLEQVHQL | ASSPRNYAEQF | 0.59578407 |
| GLSLLAVAV | ASSPRNYAEQF | 0.80829 |
| LEQVHQL | ASSPRNYAEQF | 0.8535539 |
| LNAAGAI | ASSPRNYAEQF | 0.42246604 |
| PAVHASL | ASSPRNYAEQF | 0.51236045 |
| TELEDDA | ASSPRNYAEQF | 0.5032307 |
| TPPGPPN | ASSPRNYAEQF | 0.63988596 |
| TPPPPPPP | ASSPRNYAEQF | 0.47919652 |
| TQSPALI | AWSWGDNYAEQF | 0.37438473 |
| EFAAAFPQ | AWSWGDNYAEQF | 0.7685611 |
| LLSGNLDVL | AWSWGDNYAEQF | 0.8156321 |
| GPANLGNYNI | AWSWGDNYAEQF | 0.51777476 |
| TLGPTLP | AWSWGDNYAEQF | 0.24577531 |
| VYAYSSCCK | AWSWGDNYAEQF | 0.5790789 |
| APAPAPAPNI | AWSWGDNYAEQF | 0.4459378 |
| PGGPVPIP | AWSWGDNYAEQF | 0.51915395 |
| FLTVALEP | AWSWGDNYAEQF | 0.65197945 |
| SQYFGANL | AWSWGDNYAEQF | 0.27660745 |
| GGVKPAA | AWSWGDNYAEQF | 0.52985823 |
| GGTGGAGGAGGW | AWSWGDNYAEQF | 0.85209966 |
| GVGGVAGLGGAGG | AWSWGDNYAEQF | 0.5253053 |
| VLLVLATTI | AWSWGDNYAEQF | 0.68016064 |
| IGIFAAML | AWSWGDNYAEQF | 0.80617636 |
| AVGDLID | AWSWGDNYAEQF | 0.7532743 |

| | | |
|---|---|---|
| NISPLPL | AWSWGDNYAEQF | 0.82345647 |
| VTAVFVAI | AWSWGDNYAEQF | 0.722358 |
| SGDDNDAAL | AWSWGDNYAEQF | 0.56047696 |
| AVAALAHS | AWSWGDNYAEQF | 0.6491899 |
| GGVTVMATP | AWSWGDNYAEQF | 0.61530316 |
| AGLAANVRD | AWSWGDNYAEQF | 0.5971466 |
| TNRPDLIDPALLRPG | AWSWGDNYAEQF | 0.6228305 |
| LANLGVTAL | AWSWGDNYAEQF | 0.46834657 |
| TGERIEL | AWSWGDNYAEQF | 0.7805014 |
| NGEELAA | AWSWGDNYAEQF | 0.64291114 |
| VLSLVWG | AWSWGDNYAEQF | 0.72154176 |
| IGYALWL | AWSWGDNYAEQF | 0.29846668 |
| LSLLLAPL | AWSWGDNYAEQF | 0.3823028 |
| TLDEALK | AWSWGDNYAEQF | 0.78995645 |
| TDGAGTAEGG | AWSWGDNYAEQF | 0.13089481 |
| NAPGGAGGNGG | AWSWGDNYAEQF | 0.17610158 |
| VSVFMGMGGNF | AWSWGDNYAEQF | 0.6779849 |
| LASYLAAL | AWSWGDNYAEQF | 0.82082504 |
| LVGGTPAL | AWSWGDNYAEQF | 0.15533113 |
| VRNADVL | AWSWGDNYAEQF | 0.37154117 |
| TCSSPFLVSAM | AWSWGDNYAEQF | 0.31109175 |
| VHITDAE | AWSWGDNYAEQF | 0.6819 |
| EYAAPSF | AWSWGDNYAEQF | 0.65888596 |
| VVDAFHGGL | AWSWGDNYAEQF | 0.2439593 |
| ASVVDPQ | AWSWGDNYAEQF | 0.7443037 |
| SYNEFGGLS | AWSWGDNYAEQF | 0.24384254 |
| DPDGSVY | AWSWGDNYAEQF | 0.34011883 |
| MVSTIHATE | AWSWGDNYAEQF | 0.7696756 |
| LAAGVADPV | AWSWGDNYAEQF | 0.7928714 |
| TKEQIAA | AWSWGDNYAEQF | 0.3268222 |
| SLASPAAL | AWSWGDNYAEQF | 0.7743521 |
| IGLVFYGV | AWSWGDNYAEQF | 0.41869187 |
| FLLDASNL | AWSWGDNYAEQF | 0.84757566 |
| YQVGNFP | AWSWGDNYAEQF | 0.5025586 |
| ESLLSQST | AWSWGDNYAEQF | 0.26106107 |
| LFTVGFAAVG | AWSWGDNYAEQF | 0.7698237 |
| ATPPTPT | AWSWGDNYAEQF | 0.71345 |
| TDLATLG | AWSWGDNYAEQF | 0.59117657 |
| SAGGTGR | AWSWGDNYAEQF | 0.74318886 |
| GFSNSGS | AWSWGDNYAEQF | 0.46789423 |
| GGNGGSGGTSV | AWSWGDNYAEQF | 0.5782492 |
| GTSVATGGAGN | AWSWGDNYAEQF | 0.7986385 |
| NGGNGGSGGTS | AWSWGDNYAEQF | 0.77690315 |
| AVVKFPH | AWSWGDNYAEQF | 0.39345908 |
| TPPPPPP | AWSWGDNYAEQF | 0.78686833 |
| AAPAAAAAPA | AWSWGDNYAEQF | 0.62879395 |
| IALNEDL | AWSWGDNYAEQF | 0.40666473 |
| IVNVSSM | AWSWGDNYAEQF | 0.838966 |
| TGIPPAPR | AWSWGDNYAEQF | 0.6417651 |
| FDLGGGTF | AWSWGDNYAEQF | 0.78863955 |
| FDLGGGTFDVS | AWSWGDNYAEQF | 0.71876997 |

| | | |
|---|---|---|
| NEPTAAA | AWSWGDNYAEQF | 0.78874457 |
| NEPTAAALA | AWSWGDNYAEQF | 0.32350326 |
| GSIINISSI | AWSWGDNYAEQF | 0.5425432 |
| VAPGNAGTA | AWSWGDNYAEQF | 0.62447464 |
| TGGGASLE | AWSWGDNYAEQF | 0.764062 |
| VAVYNHYKRI | AWSWGDNYAEQF | 0.53814304 |
| TDPAPATT | AWSWGDNYAEQF | 0.30159864 |
| TNRPDILDPALLRPG | AWSWGDNYAEQF | 0.65441346 |
| LTTTTTS | AWSWGDNYAEQF | 0.8237659 |
| PTGSGKTL | AWSWGDNYAEQF | 0.66742414 |
| TGSGKTL | AWSWGDNYAEQF | 0.6542476 |
| TIGHVDHGK | AWSWGDNYAEQF | 0.84345055 |
| TIGHVDHGKTT | AWSWGDNYAEQF | 0.4196701 |
| GHVDHGKTT | AWSWGDNYAEQF | 0.597788 |
| AQFEHTLL | AWSWGDNYAEQF | 0.7607957 |
| IVNQDEL | AWSWGDNYAEQF | 0.7671725 |
| TNGSQFF | AWSWGDNYAEQF | 0.5822086 |
| TGKGIVE | AWSWGDNYAEQF | 0.80196095 |
| LISNASDA | AWSWGDNYAEQF | 0.3597613 |
| AELRTLGELTKIAKAHGAQ | AWSWGDNYAEQF | 0.7463006 |
| APNAVAAAEPSW | AWSWGDNYAEQF | 0.4049314 |
| APQVAGAATAVL | AWSWGDNYAEQF | 0.67254114 |
| ARRDPAAAAAILPTDA | AWSWGDNYAEQF | 0.3894649 |
| DAAARDVLAEAGLAEYF | AWSWGDNYAEQF | 0.72918963 |
| DDQIVLSAKANLAKVEQ | AWSWGDNYAEQF | 0.79245996 |
| DELLPDTGEDSDEDGH | AWSWGDNYAEQF | 0.765826 |
| ELPEAPANRATVPQD | AWSWGDNYAEQF | 0.15442932 |
| FGQSKIERVTGIG | AWSWGDNYAEQF | 0.79118955 |
| GAGGTGSTAGGAGGAGGAGGLY | AWSWGDNYAEQF | 0.7324888 |
| GHGGTGAAGGNGGTGGAGGSGI | AWSWGDNYAEQF | 0.8397516 |
| GNGGNGGAGGEGGPGGAGGAGA | AWSWGDNYAEQF | 0.64348054 |
| GSTAGNGGNGGNATNGTIAGQP | AWSWGDNYAEQF | 0.70120585 |
| GTGGDGGNGGNGGTGG | AWSWGDNYAEQF | 0.53506136 |
| GVRRPAGIPADAPV | AWSWGDNYAEQF | 0.60596085 |
| GYRQVTDAHLLTLARR | AWSWGDNYAEQF | 0.368431 |
| IAKMGQDGHDRG | AWSWGDNYAEQF | 0.4647898 |
| KAEPSIEIQYT | AWSWGDNYAEQF | 0.55878645 |
| KPVRIRMAADEDTVAVT | AWSWGDNYAEQF | 0.4708757 |
| MPPRIAGMRLLVIKPEP | AWSWGDNYAEQF | 0.7493254 |
| MKFYAVALFRSRGVPPTR | AWSWGDNYAEQF | 0.78942776 |
| NAGGFGILGAVAHSP | AWSWGDNYAEQF | 0.79251695 |
| NEGREITLAMAEM | AWSWGDNYAEQF | 0.8067408 |
| NELVEWSHPLGHRWRWE | AWSWGDNYAEQF | 0.81422174 |
| NLGAGGTGGNGGNPG | AWSWGDNYAEQF | 0.76304513 |
| NTEWPKWETVMAADPVD | AWSWGDNYAEQF | 0.50954324 |
| PASAAPAETQP | AWSWGDNYAEQF | 0.5140584 |
| QQSSDGKNGVRAAEITGE | AWSWGDNYAEQF | 0.3496988 |
| RAPDAVTTSWWKE | AWSWGDNYAEQF | 0.5144077 |
| SGFGNVGAHTSGWFNQ | AWSWGDNYAEQF | 0.5507244 |
| SGVAGAVGQAASVGGLKV | AWSWGDNYAEQF | 0.6899663 |
| SNGGAGTGSAGGNGGT | AWSWGDNYAEQF | 0.12026561 |

| | | |
|---|---|---|
| TGGAGGEGGAAG | AWSWGDNYAEQF | 0.6180612 |
| TGGDGGAAGAGGAPGHG | AWSWGDNYAEQF | 0.824981 |
| TMPPLLAAIANAGRHGV | AWSWGDNYAEQF | 0.55042344 |
| TSSTAGNSGGA | AWSWGDNYAEQF | 0.66697055 |
| VPATVPVGILC | AWSWGDNYAEQF | 0.60080075 |
| VQRPDSGGVVLTGRIS | AWSWGDNYAEQF | 0.5300789 |
| WTPNPVPATKGVPAGLVQAT | AWSWGDNYAEQF | 0.76756877 |
| YALKRALAKFSDRFDVVII | AWSWGDNYAEQF | 0.44255772 |
| YLIGSVIV | AWSWGDNYAEQF | 0.7370553 |
| YGQLVAVID | AWSWGDNYAEQF | 0.8235755 |
| VTVGGMTLPA | AWSWGDNYAEQF | 0.6546114 |
| VLALPAV | AWSWGDNYAEQF | 0.6160942 |
| VIPPTLNYET | AWSWGDNYAEQF | 0.58303595 |
| VGMGGMHPSSG | AWSWGDNYAEQF | 0.28086087 |
| VGLAGGAATA | AWSWGDNYAEQF | 0.6442521 |
| VGGAGGAGGSGP | AWSWGDNYAEQF | 0.11333078 |
| VFAAFMV | AWSWGDNYAEQF | 0.6382295 |
| TLYDKLR | AWSWGDNYAEQF | 0.39282554 |
| TLAVFTVL | AWSWGDNYAEQF | 0.661963 |
| TGSVQPARV | AWSWGDNYAEQF | 0.67120564 |
| TGGGAGSGGT | AWSWGDNYAEQF | 0.7426617 |
| SHLETSL | AWSWGDNYAEQF | 0.3525111 |
| SFTTGMT | AWSWGDNYAEQF | 0.61471725 |
| QVEVAKRMGVRQPTVS | AWSWGDNYAEQF | 0.47816658 |
| QGSGGPALAM | AWSWGDNYAEQF | 0.31522936 |
| MLAPSNP | AWSWGDNYAEQF | 0.724257 |
| LVNNAAI | AWSWGDNYAEQF | 0.58812773 |
| LTGEINTLA | AWSWGDNYAEQF | 0.6104749 |
| LSIFGPV | AWSWGDNYAEQF | 0.7953678 |
| LLALLANL | AWSWGDNYAEQF | 0.6993144 |
| LISNASDALDKLR | AWSWGDNYAEQF | 0.3042587 |
| LINEAVP | AWSWGDNYAEQF | 0.6280283 |
| LGYAFSGL | AWSWGDNYAEQF | 0.6181264 |
| LDFVAVAL | AWSWGDNYAEQF | 0.5737625 |
| LAQPGAE | AWSWGDNYAEQF | 0.107448526 |
| ISNLGAVTGF | AWSWGDNYAEQF | 0.6765509 |
| HQEDPGRI | AWSWGDNYAEQF | 0.416439 |
| GTPGEPTP | AWSWGDNYAEQF | 0.42268646 |
| GFFNSGS | AWSWGDNYAEQF | 0.50068533 |
| GAGGGGLTAGAN | AWSWGDNYAEQF | 0.7932038 |
| FVDALVNM | AWSWGDNYAEQF | 0.4531001 |
| FLTVAPEL | AWSWGDNYAEQF | 0.49631935 |
| FETLTAL | AWSWGDNYAEQF | 0.75988126 |
| DVINPIEL | AWSWGDNYAEQF | 0.72332585 |
| APVSGAPV | AWSWGDNYAEQF | 0.818099 |
| ALVGGLAW | AWSWGDNYAEQF | 0.69543976 |
| ALAAAAATAI | AWSWGDNYAEQF | 0.58919054 |
| AFGLGLLL | AWSWGDNYAEQF | 0.6161913 |
| AFFAAASL | AWSWGDNYAEQF | 0.5719441 |
| AAAALPAFSPP | AWSWGDNYAEQF | 0.19119528 |
| VVLAPAL | AWSWGDNYAEQF | 0.20316963 |

| | | |
|---|---|---|
| TIGHVDHGKTTL | AWSWGDNYAEQF | 0.64845467 |
| LISNASDALDK | AWSWGDNYAEQF | 0.21217567 |
| LISNASDAL | AWSWGDNYAEQF | 0.611639 |
| ISNASDALDK | AWSWGDNYAEQF | 0.7626288 |
| NQAETLVYQTE | AWSWGDNYAEQF | 0.50383955 |
| TGGTGGTAGAG | AWSWGDNYAEQF | 0.31594214 |
| TGSTAGGAGGA | AWSWGDNYAEQF | 0.56042224 |
| DPLAVPVAAGPV | AWSWGDNYAEQF | 0.52638364 |
| GGTGGTPGNGGN | AWSWGDNYAEQF | 0.77394384 |
| PAGLAGVDPATL | AWSWGDNYAEQF | 0.6607505 |
| GNNLVVEAPHAGASV | AWSWGDNYAEQF | 0.78083074 |
| KVTRIMSASTLEQPA | AWSWGDNYAEQF | 0.77701813 |
| LGFALPIALPRRTVV | AWSWGDNYAEQF | 0.5331052 |
| TRLGNLTARKLVVLV | AWSWGDNYAEQF | 0.7737005 |
| WSQQRRLIYFPSAGP | AWSWGDNYAEQF | 0.5422362 |
| GPNGNPDPMAAAVDIR | AWSWGDNYAEQF | 0.4131981 |
| TQRCVSVADLRAAASR | AWSWGDNYAEQF | 0.5684582 |
| QFVLDVAEHPVEVLTETV | AWSWGDNYAEQF | 0.7008939 |
| GGAGGTGGAGGNGGTQAGDG | AWSWGDNYAEQF | 0.57125485 |
| PLLATLRAIAVDVDAKPAQV | AWSWGDNYAEQF | 0.46857244 |
| APKFRTGGYMIDFWGVGYQVAK | AWSWGDNYAEQF | 0.27661452 |
| GAGGAGGAGGAAGGPGGTGGTG | AWSWGDNYAEQF | 0.65189713 |
| IGALGLLATDVAPAGSVAFGVA | AWSWGDNYAEQF | 0.71605533 |
| AGGAGGIGGNGGAGGAGGVLMG | AWSWGDNYAEQF | 0.24893005 |
| ADELIGLL | AWSWGDNYAEQF | 0.42461056 |
| ADRALKA | AWSWGDNYAEQF | 0.8402482 |
| AGAVGQVGGAGGA | AWSWGDNYAEQF | 0.82747483 |
| AGGGGGLGGAP | AWSWGDNYAEQF | 0.40187633 |
| AGGLLYGNGGAG | AWSWGDNYAEQF | 0.45293844 |
| AGLESIGL | AWSWGDNYAEQF | 0.64144754 |
| AGLKTGII | AWSWGDNYAEQF | 0.27618968 |
| ASLAALLL | AWSWGDNYAEQF | 0.67433214 |
| ASVINISSI | AWSWGDNYAEQF | 0.6629 |
| ATPPGPA | AWSWGDNYAEQF | 0.6620364 |
| ATVASALV | AWSWGDNYAEQF | 0.39472887 |
| ATVASLIL | AWSWGDNYAEQF | 0.6868136 |
| AVNGPVLIRH | AWSWGDNYAEQF | 0.5792599 |
| DGAAGAVGI | AWSWGDNYAEQF | 0.7292423 |
| DPLLHGI | AWSWGDNYAEQF | 0.13531548 |
| ELGAALGAL | AWSWGDNYAEQF | 0.6922669 |
| ELGRTQAV | AWSWGDNYAEQF | 0.4043065 |
| FVAGGYLS | AWSWGDNYAEQF | 0.49229556 |
| GASGGAGGAGGW | AWSWGDNYAEQF | 0.26659575 |
| GGAGAGGDGGAGGI | AWSWGDNYAEQF | 0.5273008 |
| GTGHNGSQPGMGGNG | AWSWGDNYAEQF | 0.76480496 |
| GYAAASASAS | AWSWGDNYAEQF | 0.81051517 |
| HQVDVAAVQQG | AWSWGDNYAEQF | 0.3258811 |
| IGAAIAHT | AWSWGDNYAEQF | 0.7581033 |
| IGMIEVAQL | AWSWGDNYAEQF | 0.84598494 |
| IRLAPEHTSAMD | AWSWGDNYAEQF | 0.5537626 |
| IYNVSVL | AWSWGDNYAEQF | 0.2524536 |

| | | |
|---|---|---|
| KQGFLGM | AWSWGDNYAEQF | 0.7346867 |
| LESVITQL | AWSWGDNYAEQF | 0.44325787 |
| LGLGALVVV | AWSWGDNYAEQF | 0.76713204 |
| LIGNGGNGGSGGI | AWSWGDNYAEQF | 0.7380494 |
| LNAINAPTL | AWSWGDNYAEQF | 0.32209802 |
| LQLLNTGV | AWSWGDNYAEQF | 0.68665105 |
| LQTSIEF | AWSWGDNYAEQF | 0.79147094 |
| MSFAVSEL | AWSWGDNYAEQF | 0.43031514 |
| NTGALGL | AWSWGDNYAEQF | 0.4127245 |
| PPGAGGAGGAGTT | AWSWGDNYAEQF | 0.41625482 |
| QLVGLGI | AWSWGDNYAEQF | 0.1716552 |
| RFLEQVHQL | AWSWGDNYAEQF | 0.7402311 |
| RGVGYVLR | AWSWGDNYAEQF | 0.5252888 |
| SATGVFAGVI | AWSWGDNYAEQF | 0.43954262 |
| SLPAITLP | AWSWGDNYAEQF | 0.65800166 |
| STALAFTV | AWSWGDNYAEQF | 0.43051118 |
| SVLTVGLGK | AWSWGDNYAEQF | 0.5096952 |
| TALAGAGSGI | AWSWGDNYAEQF | 0.5365457 |
| TGLPFTGI | AWSWGDNYAEQF | 0.573761 |
| VAEHVTV | AWSWGDNYAEQF | 0.74261034 |
| VDIVLNSL | AWSWGDNYAEQF | 0.5949713 |
| VLTGGGAVL | AWSWGDNYAEQF | 0.5434361 |
| VNVGLAYGI | AWSWGDNYAEQF | 0.82757366 |
| VSSAAVGAV | AWSWGDNYAEQF | 0.32272235 |
| VTSVYEAGV | AWSWGDNYAEQF | 0.4063969 |
| WVALDRGA | AWSWGDNYAEQF | 0.7478845 |
| YLVGSVVI | AWSWGDNYAEQF | 0.32746696 |
| GPPGTGKTLL | AWSWGDNYAEQF | 0.74812067 |
| GVGMTKF | AWSWGDNYAEQF | 0.65086406 |
| IGIGGSDL | AWSWGDNYAEQF | 0.7247124 |
| IGTLLAEL | AWSWGDNYAEQF | 0.5458605 |
| IYTSGTT | AWSWGDNYAEQF | 0.8462281 |
| LSFPAGV | AWSWGDNYAEQF | 0.78113824 |
| LYTSGTT | AWSWGDNYAEQF | 0.48952812 |
| VGAFPFGF | AWSWGDNYAEQF | 0.57291776 |
| VGAGTVEF | AWSWGDNYAEQF | 0.35088342 |
| VGKSTLLL | AWSWGDNYAEQF | 0.745515 |
| AALAHAGGGIV | AWSWGDNYAEQF | 0.7050891 |
| AALVAVSLL | AWSWGDNYAEQF | 0.40927854 |
| AAVSRAVAAL | AWSWGDNYAEQF | 0.5844809 |
| AGGSSGIGGF | AWSWGDNYAEQF | 0.82312566 |
| AGLTVANANG | AWSWGDNYAEQF | 0.8067595 |
| AGLVGAVL | AWSWGDNYAEQF | 0.24994615 |
| ALSQAFTV | AWSWGDNYAEQF | 0.54219735 |
| ALTFSVSL | AWSWGDNYAEQF | 0.28984943 |
| ALTGAALVAL | AWSWGDNYAEQF | 0.49068666 |
| ASFSIGAGVL | AWSWGDNYAEQF | 0.36111236 |
| DGSVYKI | AWSWGDNYAEQF | 0.7423574 |
| EQLAEIV | AWSWGDNYAEQF | 0.5323955 |
| FAVSELTVM | AWSWGDNYAEQF | 0.67644024 |
| GAAPLAAGL | AWSWGDNYAEQF | 0.6627902 |

| | | |
|---|---|---|
| GADGGDGGSGGAGG | AWSWGDNYAEQF | 0.8029858 |
| GFGLDTGGAGG | AWSWGDNYAEQF | 0.43995237 |
| GLASGGLL | AWSWGDNYAEQF | 0.561613 |
| GRIAGAAVLL | AWSWGDNYAEQF | 0.29394916 |
| GSAGSLNTSF | AWSWGDNYAEQF | 0.32622206 |
| GTGQIGI | AWSWGDNYAEQF | 0.44553077 |
| GTLIALIL | AWSWGDNYAEQF | 0.5447539 |
| IAATMGFVVG | AWSWGDNYAEQF | 0.20136584 |
| IFAQALL | AWSWGDNYAEQF | 0.82257104 |
| IGAAVALGF | AWSWGDNYAEQF | 0.6765255 |
| IGLTPLAL | AWSWGDNYAEQF | 0.7697977 |
| ILGSNTSTL | AWSWGDNYAEQF | 0.4530344 |
| IVFSLTVL | AWSWGDNYAEQF | 0.5092665 |
| LASKATGF | AWSWGDNYAEQF | 0.5365899 |
| LASVALIL | AWSWGDNYAEQF | 0.72402275 |
| LFAVGLTL | AWSWGDNYAEQF | 0.36256057 |
| LFNGGIGTYI | AWSWGDNYAEQF | 0.22876067 |
| LTLGAGII | AWSWGDNYAEQF | 0.47124025 |
| LVAAALGDI | AWSWGDNYAEQF | 0.2884896 |
| NTAFVQLGI | AWSWGDNYAEQF | 0.8220628 |
| QLFSGAEL | AWSWGDNYAEQF | 0.7983168 |
| SAATFITL | AWSWGDNYAEQF | 0.6860652 |
| SNYGLANLGV | AWSWGDNYAEQF | 0.7465935 |
| SYGGAYAVM | AWSWGDNYAEQF | 0.18304864 |
| TTGVGGLTLG | AWSWGDNYAEQF | 0.5091566 |
| TTQTIYI | AWSWGDNYAEQF | 0.82190657 |
| VAAGAGLGAVI | AWSWGDNYAEQF | 0.715032 |
| VAAVVTLL | AWSWGDNYAEQF | 0.5714612 |
| VGFLGGLI | AWSWGDNYAEQF | 0.82442456 |
| VGVIPILL | AWSWGDNYAEQF | 0.42133522 |
| VIFASSLI | AWSWGDNYAEQF | 0.5337515 |
| VKGGDGGAAAG | AWSWGDNYAEQF | 0.6367154 |
| VTKTFTNI | AWSWGDNYAEQF | 0.658285 |
| VVAAAILL | AWSWGDNYAEQF | 0.60444164 |
| YAALDVEL | AWSWGDNYAEQF | 0.3764625 |
| LASVALLL | AWSWGDNYAEQF | 0.66595244 |
| LDAVHRV | AWSWGDNYAEQF | 0.85890806 |
| QGLSSSIF | AWSWGDNYAEQF | 0.55491674 |
| RGTFANIRL | AWSWGDNYAEQF | 0.8008193 |
| RQLWWGHRI | AWSWGDNYAEQF | 0.7280153 |
| YGPPGTGKTLL | AWSWGDNYAEQF | 0.62192905 |
| AALVGAPANVA | AWSWGDNYAEQF | 0.42572576 |
| AATFRRTE | AWSWGDNYAEQF | 0.51119566 |
| AGALGGGI | AWSWGDNYAEQF | 0.58684045 |
| AGGAGGAGGIGAG | AWSWGDNYAEQF | 0.4036452 |
| AGSGALGL | AWSWGDNYAEQF | 0.60309803 |
| AQIADQI | AWSWGDNYAEQF | 0.5303781 |
| ASAVAVTYL | AWSWGDNYAEQF | 0.8630923 |
| ASTVLGGTGG | AWSWGDNYAEQF | 0.7366613 |
| AVGGEPAK | AWSWGDNYAEQF | 0.7633997 |
| CEQGTFGGTC | AWSWGDNYAEQF | 0.38949665 |

| | | |
|---|---|---|
| DEGKAPAG | AWSWGDNYAEQF | 0.6762594 |
| DTVVFPAL | AWSWGDNYAEQF | 0.72904015 |
| FEGITMAG | AWSWGDNYAEQF | 0.32172993 |
| FPLVVQD | AWSWGDNYAEQF | 0.8294188 |
| GAFTTPQ | AWSWGDNYAEQF | 0.4644273 |
| GFGGFGTAGGVGGA | AWSWGDNYAEQF | 0.5729358 |
| GFLAQFL | AWSWGDNYAEQF | 0.7153223 |
| GGAGGAGGNAGVA | AWSWGDNYAEQF | 0.7422037 |
| GGVGTAGAGGAGGAGG | AWSWGDNYAEQF | 0.5530251 |
| GNGGGGDAGNAGSGGN | AWSWGDNYAEQF | 0.5976022 |
| GTGGAGGAGNDAGS | AWSWGDNYAEQF | 0.42457482 |
| GTTNTGWFNS | AWSWGDNYAEQF | 0.7689349 |
| IFMGPLFTL | AWSWGDNYAEQF | 0.8021573 |
| ILAHAIEAI | AWSWGDNYAEQF | 0.39349332 |
| IRSAIREAL | AWSWGDNYAEQF | 0.8081782 |
| ITFAGGG | AWSWGDNYAEQF | 0.61437887 |
| ITGMSGHQEV | AWSWGDNYAEQF | 0.30257803 |
| IVAGLACTL | AWSWGDNYAEQF | 0.58708143 |
| IVGGGLAGA | AWSWGDNYAEQF | 0.78114974 |
| IVTGVLNI | AWSWGDNYAEQF | 0.8140209 |
| LAQQGCI | AWSWGDNYAEQF | 0.770581 |
| LCLHVEN | AWSWGDNYAEQF | 0.5796644 |
| LDAISDGS | AWSWGDNYAEQF | 0.48480996 |
| LGGFAAML | AWSWGDNYAEQF | 0.8242184 |
| LGGKNPMI | AWSWGDNYAEQF | 0.8560034 |
| LGPKGVAAIL | AWSWGDNYAEQF | 0.7644559 |
| LMVGVSI | AWSWGDNYAEQF | 0.50886154 |
| LVGAAFVL | AWSWGDNYAEQF | 0.45423394 |
| LVLAVAPL | AWSWGDNYAEQF | 0.62024117 |
| MAAIAVIDAA | AWSWGDNYAEQF | 0.7887615 |
| NNGINAELQ | AWSWGDNYAEQF | 0.38839835 |
| RGEPLVLVHG | AWSWGDNYAEQF | 0.6757816 |
| RGGAGGAAGLI | AWSWGDNYAEQF | 0.68193793 |
| RSYRMHPAV | AWSWGDNYAEQF | 0.23987795 |
| SAAHLLAGQLA | AWSWGDNYAEQF | 0.67246103 |
| SAAVGSY | AWSWGDNYAEQF | 0.7993753 |
| SPHAVLHRI | AWSWGDNYAEQF | 0.5690781 |
| SQIAKVL | AWSWGDNYAEQF | 0.43233004 |
| TALGHLVP | AWSWGDNYAEQF | 0.8117895 |
| TAPTTSGAA | AWSWGDNYAEQF | 0.8609221 |
| TDGDDGNGGSPGT | AWSWGDNYAEQF | 0.79277956 |
| TDGLGEN | AWSWGDNYAEQF | 0.2605015 |
| TGGPPLPGQA | AWSWGDNYAEQF | 0.16726166 |
| TIPAITI | AWSWGDNYAEQF | 0.53362054 |
| TIPGFTI | AWSWGDNYAEQF | 0.7131572 |
| TSIASGALA | AWSWGDNYAEQF | 0.51236033 |
| TVTLPGAAI | AWSWGDNYAEQF | 0.4361866 |
| VASGLSAALP | AWSWGDNYAEQF | 0.69494843 |
| VGAGVIGI | AWSWGDNYAEQF | 0.8117273 |
| VGPKRFGV | AWSWGDNYAEQF | 0.35046715 |
| VNVAFPD | AWSWGDNYAEQF | 0.38758123 |

| | | |
|---|---|---|
| VSMTNSGVS | AWSWGDNYAEQF | 0.6035993 |
| VTDVLGDLSV | AWSWGDNYAEQF | 0.75613105 |
| VVVVGAGFAGL | AWSWGDNYAEQF | 0.41652974 |
| VYTVGTVV | AWSWGDNYAEQF | 0.597425 |
| WTTFSGP | AWSWGDNYAEQF | 0.54098666 |
| YDELVEAF | AWSWGDNYAEQF | 0.48838806 |
| YGTGGAGGNGG | AWSWGDNYAEQF | 0.7097436 |
| YTNVMAV | AWSWGDNYAEQF | 0.346898 |
| FLEQVHQL | AWSWGDNYAEQF | 0.8238627 |
| GLSLLAVAV | AWSWGDNYAEQF | 0.36257815 |
| LEQVHQL | AWSWGDNYAEQF | 0.7737108 |
| LNAAGAI | AWSWGDNYAEQF | 0.66731155 |
| PAVHASL | AWSWGDNYAEQF | 0.6669719 |
| TELEDDA | AWSWGDNYAEQF | 0.5567033 |
| TPPGPPN | AWSWGDNYAEQF | 0.43984258 |
| TPPPPPPP | AWSWGDNYAEQF | 0.35814434 |
| TQSPALI | ASSSLRDREGEQY | 0.20986398 |
| EFAAAFPQ | ASSSLRDREGEQY | 0.7868103 |
| LLSGNLDVL | ASSSLRDREGEQY | 0.14815295 |
| GPANLGNYNI | ASSSLRDREGEQY | 0.7874643 |
| TLGPTLP | ASSSLRDREGEQY | 0.7270215 |
| VYAYSSCCK | ASSSLRDREGEQY | 0.43065947 |
| APAPAPAPNI | ASSSLRDREGEQY | 0.74762416 |
| PGGPVPIP | ASSSLRDREGEQY | 0.3578151 |
| FLTVALEP | ASSSLRDREGEQY | 0.35326552 |
| SQYFGANL | ASSSLRDREGEQY | 0.742715 |
| GGVKPAA | ASSSLRDREGEQY | 0.73132485 |
| GGTGGAGGAGGW | ASSSLRDREGEQY | 0.39047253 |
| GVGGVAGLGGAGG | ASSSLRDREGEQY | 0.6974963 |
| VLLVLATTI | ASSSLRDREGEQY | 0.6837899 |
| IGIFAAML | ASSSLRDREGEQY | 0.36672243 |
| AVGDLID | ASSSLRDREGEQY | 0.5126562 |
| NISPLPL | ASSSLRDREGEQY | 0.7950948 |
| VTAVFVAI | ASSSLRDREGEQY | 0.6358465 |
| SGDDNDAAL | ASSSLRDREGEQY | 0.38686308 |
| AVAALAHS | ASSSLRDREGEQY | 0.75684476 |
| GGVTVMATP | ASSSLRDREGEQY | 0.39467645 |
| AGLAANVRD | ASSSLRDREGEQY | 0.71447957 |
| TNRPDLIDPALLRPG | ASSSLRDREGEQY | 0.6647805 |
| LANLGVTAL | ASSSLRDREGEQY | 0.48786265 |
| TGERIEL | ASSSLRDREGEQY | 0.7763742 |
| NGEELAA | ASSSLRDREGEQY | 0.33113834 |
| VLSLVWG | ASSSLRDREGEQY | 0.40731484 |
| IGYALWL | ASSSLRDREGEQY | 0.5197514 |
| LSLLLAPL | ASSSLRDREGEQY | 0.6264615 |
| TLDEALK | ASSSLRDREGEQY | 0.7900678 |
| TDGAGTAEGG | ASSSLRDREGEQY | 0.39281824 |
| NAPGGAGGNGG | ASSSLRDREGEQY | 0.6520948 |
| VSVFMGMGGNF | ASSSLRDREGEQY | 0.5113136 |
| LASYLAAL | ASSSLRDREGEQY | 0.34481877 |
| LVGGTPAL | ASSSLRDREGEQY | 0.5710474 |

| | | |
|---|---|---|
| VRNADVL | ASSSLRDREGEQY | 0.5109799 |
| TCSSPFLVSAM | ASSSLRDREGEQY | 0.43639517 |
| VHITDAE | ASSSLRDREGEQY | 0.68239737 |
| EYAAPSF | ASSSLRDREGEQY | 0.42483702 |
| VVDAFHGGL | ASSSLRDREGEQY | 0.66099775 |
| ASVVDPQ | ASSSLRDREGEQY | 0.83147967 |
| SYNEFGGLS | ASSSLRDREGEQY | 0.48781177 |
| DPDGSVY | ASSSLRDREGEQY | 0.7775221 |
| MVSTIHATE | ASSSLRDREGEQY | 0.72922623 |
| LAAGVADPV | ASSSLRDREGEQY | 0.72706354 |
| TKEQIAA | ASSSLRDREGEQY | 0.548486 |
| SLASPAAL | ASSSLRDREGEQY | 0.42990017 |
| IGLVFYGV | ASSSLRDREGEQY | 0.81540275 |
| FLLDASNL | ASSSLRDREGEQY | 0.7661314 |
| YQVGNFP | ASSSLRDREGEQY | 0.7217462 |
| ESLLSQST | ASSSLRDREGEQY | 0.7189723 |
| LFTVGFAAVG | ASSSLRDREGEQY | 0.6347141 |
| ATPPTPT | ASSSLRDREGEQY | 0.75900346 |
| TDLATLG | ASSSLRDREGEQY | 0.4870496 |
| SAGGTGR | ASSSLRDREGEQY | 0.30859756 |
| GFSNSGS | ASSSLRDREGEQY | 0.5546659 |
| GGNGGSGGTSV | ASSSLRDREGEQY | 0.5938567 |
| GTSVATGGAGN | ASSSLRDREGEQY | 0.46591553 |
| NGGNGGSGGTS | ASSSLRDREGEQY | 0.30777577 |
| AVVKFPH | ASSSLRDREGEQY | 0.78772783 |
| TPPPPPP | ASSSLRDREGEQY | 0.8653232 |
| AAPAAAAAPA | ASSSLRDREGEQY | 0.43510535 |
| IALNEDL | ASSSLRDREGEQY | 0.8343363 |
| IVNVSSM | ASSSLRDREGEQY | 0.28942788 |
| TGIPPAPR | ASSSLRDREGEQY | 0.7383257 |
| FDLGGGTF | ASSSLRDREGEQY | 0.79229665 |
| FDLGGGTFDVS | ASSSLRDREGEQY | 0.44225293 |
| NEPTAAA | ASSSLRDREGEQY | 0.68196225 |
| NEPTAAALA | ASSSLRDREGEQY | 0.42102125 |
| GSIINISSI | ASSSLRDREGEQY | 0.55336565 |
| VAPGNAGTA | ASSSLRDREGEQY | 0.80593276 |
| TGGGASLE | ASSSLRDREGEQY | 0.46903172 |
| VAVYNHYKRI | ASSSLRDREGEQY | 0.62077194 |
| TDPAPATT | ASSSLRDREGEQY | 0.52449816 |
| TNRPDILDPALLRPG | ASSSLRDREGEQY | 0.46402717 |
| LTTTTTS | ASSSLRDREGEQY | 0.66441333 |
| PTGSGKTL | ASSSLRDREGEQY | 0.76055455 |
| TGSGKTL | ASSSLRDREGEQY | 0.41328788 |
| TIGHVDHGK | ASSSLRDREGEQY | 0.84989 |
| TIGHVDHGKTT | ASSSLRDREGEQY | 0.39460772 |
| GHVDHGKTT | ASSSLRDREGEQY | 0.83494914 |
| AQFEHTLL | ASSSLRDREGEQY | 0.766873 |
| IVNQDEL | ASSSLRDREGEQY | 0.67252517 |
| TNGSQFF | ASSSLRDREGEQY | 0.5647604 |
| TGKGIVE | ASSSLRDREGEQY | 0.7955686 |
| LISNASDA | ASSSLRDREGEQY | 0.4824766 |

| | | |
|---|---|---|
| AELRTLGELTKIAKAHGAQ | ASSSLRDREGEQY | 0.7260958 |
| APNAVAAAEPSW | ASSSLRDREGEQY | 0.7860806 |
| APQVAGAATAVL | ASSSLRDREGEQY | 0.6444314 |
| ARRDPAAAAAILPTDA | ASSSLRDREGEQY | 0.67566484 |
| DAAARDVLAEAGLAEYF | ASSSLRDREGEQY | 0.66120327 |
| DDQIVLSAKANLAKVEQ | ASSSLRDREGEQY | 0.65639704 |
| DELLPDTGEDSDEDGH | ASSSLRDREGEQY | 0.6442606 |
| ELPEAPANRATVPQD | ASSSLRDREGEQY | 0.5267411 |
| FGQSKIERVTGIG | ASSSLRDREGEQY | 0.6592894 |
| GAGGTGSTAGGAGGAGGAGGLY | ASSSLRDREGEQY | 0.62141764 |
| GHGGTGAAGGNGGTGGAGGSGI | ASSSLRDREGEQY | 0.6626642 |
| GNGGNGGAGGEGGPGGAGGAGA | ASSSLRDREGEQY | 0.6722052 |
| GSTAGNGGNGGNATNGTIAGQP | ASSSLRDREGEQY | 0.64183664 |
| GTGGDGGNGGNGGTGG | ASSSLRDREGEQY | 0.67650527 |
| GVRRPAGIPADAPV | ASSSLRDREGEQY | 0.6722547 |
| GYRQVTDAHLLTLARR | ASSSLRDREGEQY | 0.5707481 |
| IAKMGQDGHDRG | ASSSLRDREGEQY | 0.6514751 |
| KAEPSIEIQYT | ASSSLRDREGEQY | 0.6449045 |
| KPVRIRMAADEDTVAVT | ASSSLRDREGEQY | 0.6548036 |
| MPPRIAGMRLLVIKPEP | ASSSLRDREGEQY | 0.5854988 |
| MKFYAVALFRSRGVPPTR | ASSSLRDREGEQY | 0.67083037 |
| NAGGFGILGAVAHSP | ASSSLRDREGEQY | 0.6408633 |
| NEGREITLAMAEM | ASSSLRDREGEQY | 0.35593128 |
| NELVEWSHPLGHRWRWE | ASSSLRDREGEQY | 0.6630522 |
| NLGAGGTGGNGGNPG | ASSSLRDREGEQY | 0.66043115 |
| NTEWPKWETVMAADPVD | ASSSLRDREGEQY | 0.6461545 |
| PASAAPAETQP | ASSSLRDREGEQY | 0.669606 |
| QQSSDGKNGVRAAEITGE | ASSSLRDREGEQY | 0.52418673 |
| RAPDAVTTSWWKE | ASSSLRDREGEQY | 0.63501734 |
| SGFGNVGAHTSGWFNQ | ASSSLRDREGEQY | 0.6725924 |
| SGVAGAVGQAASVGGLKV | ASSSLRDREGEQY | 0.6754717 |
| SNGGAGTGSAGGNGGT | ASSSLRDREGEQY | 0.5872225 |
| TGGAGGEGGAAG | ASSSLRDREGEQY | 0.6635707 |
| TGGDGGAAGAGGAPGHG | ASSSLRDREGEQY | 0.6721018 |
| TMPPLLAAIANAGRHGV | ASSSLRDREGEQY | 0.66429 |
| TSSTAGNSGGA | ASSSLRDREGEQY | 0.67337215 |
| VPATVPVGILC | ASSSLRDREGEQY | 0.6758746 |
| VQRPDSGGVVLTGRIS | ASSSLRDREGEQY | 0.6428186 |
| WTPNPVPATKGVPAGLVQAT | ASSSLRDREGEQY | 0.6745146 |
| YALKRALAKFSDRFDVVII | ASSSLRDREGEQY | 0.67613065 |
| YLIGSVIV | ASSSLRDREGEQY | 0.6548234 |
| YGQLVAVID | ASSSLRDREGEQY | 0.6601249 |
| VTVGGMTLPA | ASSSLRDREGEQY | 0.63644767 |
| VLALPAV | ASSSLRDREGEQY | 0.6521717 |
| VIPPTLNYET | ASSSLRDREGEQY | 0.47617483 |
| VGMGGMHPSSG | ASSSLRDREGEQY | 0.6629404 |
| VGLAGGAATA | ASSSLRDREGEQY | 0.6737844 |
| VGGAGGAGGSGP | ASSSLRDREGEQY | 0.64058995 |
| VFAAFMV | ASSSLRDREGEQY | 0.6183657 |
| TLYDKLR | ASSSLRDREGEQY | 0.63491654 |
| TLAVFTVL | ASSSLRDREGEQY | 0.6523352 |

| | | |
|---|---|---|
| TGSVQPARV | ASSSLRDREGEQY | 0.6489079 |
| TGGGAGSGGT | ASSSLRDREGEQY | 0.4896069 |
| SHLETSL | ASSSLRDREGEQY | 0.6422349 |
| SFTTGMT | ASSSLRDREGEQY | 0.6737937 |
| QVEVAKRMGVRQPTVS | ASSSLRDREGEQY | 0.6728201 |
| QGSGGPALAM | ASSSLRDREGEQY | 0.5386574 |
| MLAPSNP | ASSSLRDREGEQY | 0.6744091 |
| LVNNAAI | ASSSLRDREGEQY | 0.6705571 |
| LTGEINTLA | ASSSLRDREGEQY | 0.67411983 |
| LSIFGPV | ASSSLRDREGEQY | 0.6577932 |
| LLALLANL | ASSSLRDREGEQY | 0.6385629 |
| LISNASDALDKLR | ASSSLRDREGEQY | 0.6673238 |
| LINEAVP | ASSSLRDREGEQY | 0.6411963 |
| LGYAFSGL | ASSSLRDREGEQY | 0.60334945 |
| LDFVAVAL | ASSSLRDREGEQY | 0.6715711 |
| LAQPGAE | ASSSLRDREGEQY | 0.64515835 |
| ISNLGAVTGF | ASSSLRDREGEQY | 0.6375016 |
| HQEDPGRI | ASSSLRDREGEQY | 0.6222599 |
| GTPGEPTP | ASSSLRDREGEQY | 0.67436826 |
| GFFNSGS | ASSSLRDREGEQY | 0.61450803 |
| GAGGGGLTAGAN | ASSSLRDREGEQY | 0.603601 |
| FVDALVNM | ASSSLRDREGEQY | 0.60747266 |
| FLTVAPEL | ASSSLRDREGEQY | 0.6564516 |
| FETLTAL | ASSSLRDREGEQY | 0.67329234 |
| DVINPIEL | ASSSLRDREGEQY | 0.66649246 |
| APVSGAPV | ASSSLRDREGEQY | 0.6476854 |
| ALVGGLAW | ASSSLRDREGEQY | 0.6642618 |
| ALAAAAATAI | ASSSLRDREGEQY | 0.66681325 |
| AFGLGLLL | ASSSLRDREGEQY | 0.66397905 |
| AFFAAASL | ASSSLRDREGEQY | 0.6707041 |
| AAAALPAFSPP | ASSSLRDREGEQY | 0.62373614 |
| VVLAPAL | ASSSLRDREGEQY | 0.6745218 |
| TIGHVDHGKTTL | ASSSLRDREGEQY | 0.6652235 |
| LISNASDALDK | ASSSLRDREGEQY | 0.67303264 |
| LISNASDAL | ASSSLRDREGEQY | 0.6740101 |
| ISNASDALDK | ASSSLRDREGEQY | 0.6663616 |
| NQAETLVYQTE | ASSSLRDREGEQY | 0.6609658 |
| TGGTGGTAGAG | ASSSLRDREGEQY | 0.48289332 |
| TGSTAGGAGGA | ASSSLRDREGEQY | 0.64382976 |
| DPLAVPVAAGPV | ASSSLRDREGEQY | 0.66670954 |
| GGTGGTPGNGGN | ASSSLRDREGEQY | 0.66077745 |
| PAGLAGVDPATL | ASSSLRDREGEQY | 0.6697373 |
| GNNLVVEAPHAGASV | ASSSLRDREGEQY | 0.6663645 |
| KVTRIMSASTLEQPA | ASSSLRDREGEQY | 0.64086056 |
| LGFALPIALPRRTVV | ASSSLRDREGEQY | 0.67155236 |
| TRLGNLTARKLVVLV | ASSSLRDREGEQY | 0.6715691 |
| WSQQRRLIYFPSAGP | ASSSLRDREGEQY | 0.55338585 |
| GPNGNPDPMAAAVDIR | ASSSLRDREGEQY | 0.651739 |
| TQRCVSVADLRAAASR | ASSSLRDREGEQY | 0.6784208 |
| QFVLDVAEHPVEVLTETV | ASSSLRDREGEQY | 0.6568464 |
| GGAGGTGGAGGNGGTQAGDG | ASSSLRDREGEQY | 0.65248466 |

| | | |
|---|---|---|
| PLLATLRAIAVDVDAKPAQV | ASSSLRDREGEQY | 0.63769674 |
| APKFRTGGYMIDFWGVGYQVAK | ASSSLRDREGEQY | 0.67576516 |
| GAGGAGGAGGAAGGPGGTGGTG | ASSSLRDREGEQY | 0.63209796 |
| IGALGLLATDVAPAGSVAFGVA | ASSSLRDREGEQY | 0.6774379 |
| AGGAGGIGGNGGAGGAGGVLMG | ASSSLRDREGEQY | 0.38845354 |
| ADELIGLL | ASSSLRDREGEQY | 0.63375247 |
| ADRALKA | ASSSLRDREGEQY | 0.6010339 |
| AGAVGQVGGAGGA | ASSSLRDREGEQY | 0.5118327 |
| AGGGGGLGGAP | ASSSLRDREGEQY | 0.67289174 |
| AGGLLYGNGGAG | ASSSLRDREGEQY | 0.64527 |
| AGLESIGL | ASSSLRDREGEQY | 0.66867316 |
| AGLKTGII | ASSSLRDREGEQY | 0.664028 |
| ASLAALLL | ASSSLRDREGEQY | 0.67504096 |
| ASVINISSI | ASSSLRDREGEQY | 0.6579722 |
| ATPPGPA | ASSSLRDREGEQY | 0.6636713 |
| ATVASALV | ASSSLRDREGEQY | 0.62025625 |
| ATVASLIL | ASSSLRDREGEQY | 0.6719053 |
| AVNGPVLIRH | ASSSLRDREGEQY | 0.6742805 |
| DGAAGAVGI | ASSSLRDREGEQY | 0.6676618 |
| DPLLHGI | ASSSLRDREGEQY | 0.5753379 |
| ELGAALGAL | ASSSLRDREGEQY | 0.64330745 |
| ELGRTQAV | ASSSLRDREGEQY | 0.66465706 |
| FVAGGYLS | ASSSLRDREGEQY | 0.39526218 |
| GASGGAGGAGGW | ASSSLRDREGEQY | 0.439077 |
| GGAGAGGDGGAGGI | ASSSLRDREGEQY | 0.6473842 |
| GTGHNGSQPGMGGNG | ASSSLRDREGEQY | 0.6167654 |
| GYAAASASAS | ASSSLRDREGEQY | 0.6748254 |
| HQVDVAAVQQG | ASSSLRDREGEQY | 0.6718849 |
| IGAAIAHT | ASSSLRDREGEQY | 0.64939487 |
| IGMIEVAQL | ASSSLRDREGEQY | 0.6707375 |
| IRLAPEHTSAMD | ASSSLRDREGEQY | 0.6575734 |
| IYNVSVL | ASSSLRDREGEQY | 0.30478245 |
| KQGFLGM | ASSSLRDREGEQY | 0.64898187 |
| LESVITQL | ASSSLRDREGEQY | 0.5980952 |
| LGLGALVVV | ASSSLRDREGEQY | 0.67683184 |
| LIGNGGNGGSGGI | ASSSLRDREGEQY | 0.6400594 |
| LNAINAPTL | ASSSLRDREGEQY | 0.6238785 |
| LQLLNTGV | ASSSLRDREGEQY | 0.6544344 |
| LQTSIEF | ASSSLRDREGEQY | 0.6541405 |
| MSFAVSEL | ASSSLRDREGEQY | 0.65141755 |
| NTGALGL | ASSSLRDREGEQY | 0.62284136 |
| PPGAGGAGGAGTT | ASSSLRDREGEQY | 0.34643558 |
| QLVGLGI | ASSSLRDREGEQY | 0.672842 |
| RFLEQVHQL | ASSSLRDREGEQY | 0.5384414 |
| RGVGYVLR | ASSSLRDREGEQY | 0.61863416 |
| SATGVFAGVI | ASSSLRDREGEQY | 0.5338848 |
| SLPAITLP | ASSSLRDREGEQY | 0.3708385 |
| STALAFTV | ASSSLRDREGEQY | 0.53950465 |
| SVLTVGLGK | ASSSLRDREGEQY | 0.67221946 |
| TALAGAGSGI | ASSSLRDREGEQY | 0.6448629 |
| TGLPFTGI | ASSSLRDREGEQY | 0.6425916 |

| | | |
|---|---|---|
| VAEHVTV | ASSSLRDREGEQY | 0.67685705 |
| VDIVLNSL | ASSSLRDREGEQY | 0.65213215 |
| VLTGGGAVL | ASSSLRDREGEQY | 0.6410311 |
| VNVGLAYGI | ASSSLRDREGEQY | 0.63955885 |
| VSSAAVGAV | ASSSLRDREGEQY | 0.6294232 |
| VTSVYEAGV | ASSSLRDREGEQY | 0.6238046 |
| WVALDRGA | ASSSLRDREGEQY | 0.623595 |
| YLVGSVVI | ASSSLRDREGEQY | 0.6775214 |
| GPPGTGKTLL | ASSSLRDREGEQY | 0.64449537 |
| GVGMTKF | ASSSLRDREGEQY | 0.5992901 |
| IGIGGSDL | ASSSLRDREGEQY | 0.6515082 |
| IGTLLAEL | ASSSLRDREGEQY | 0.6763511 |
| IYTSGTT | ASSSLRDREGEQY | 0.67843467 |
| LSFPAGV | ASSSLRDREGEQY | 0.6352877 |
| LYTSGTT | ASSSLRDREGEQY | 0.64128685 |
| VGAFPFGF | ASSSLRDREGEQY | 0.6466731 |
| VGAGTVEF | ASSSLRDREGEQY | 0.6555817 |
| VGKSTLLL | ASSSLRDREGEQY | 0.64650786 |
| AALAHAGGGIV | ASSSLRDREGEQY | 0.5657916 |
| AALVAVSLL | ASSSLRDREGEQY | 0.67033947 |
| AAVSRAVAAL | ASSSLRDREGEQY | 0.6778028 |
| AGGSSGIGGF | ASSSLRDREGEQY | 0.649875 |
| AGLTVANANG | ASSSLRDREGEQY | 0.6094045 |
| AGLVGAVL | ASSSLRDREGEQY | 0.6547815 |
| ALSQAFTV | ASSSLRDREGEQY | 0.63104075 |
| ALTFSVSL | ASSSLRDREGEQY | 0.6410084 |
| ALTGAALVAL | ASSSLRDREGEQY | 0.66200584 |
| ASFSIGAGVL | ASSSLRDREGEQY | 0.55475616 |
| DGSVYKI | ASSSLRDREGEQY | 0.6410606 |
| EQLAEIV | ASSSLRDREGEQY | 0.67755204 |
| FAVSELTVM | ASSSLRDREGEQY | 0.64585423 |
| GAAPLAAGL | ASSSLRDREGEQY | 0.6580107 |
| GADGGDGGSGGAGG | ASSSLRDREGEQY | 0.6733324 |
| GFGLDTGGAGG | ASSSLRDREGEQY | 0.6649505 |
| GLASGGLL | ASSSLRDREGEQY | 0.67323905 |
| GRIAGAAVLL | ASSSLRDREGEQY | 0.67297137 |
| GSAGSLNTSF | ASSSLRDREGEQY | 0.6602842 |
| GTGQIGI | ASSSLRDREGEQY | 0.662826 |
| GTLIALIL | ASSSLRDREGEQY | 0.65570784 |
| IAATMGFVVG | ASSSLRDREGEQY | 0.65913033 |
| IFAQALL | ASSSLRDREGEQY | 0.64838934 |
| IGAAVALGF | ASSSLRDREGEQY | 0.6343565 |
| IGLTPLAL | ASSSLRDREGEQY | 0.67385507 |
| ILGSNTSTL | ASSSLRDREGEQY | 0.65667725 |
| IVFSLTVL | ASSSLRDREGEQY | 0.67047244 |
| LASKATGF | ASSSLRDREGEQY | 0.60476124 |
| LASVALIL | ASSSLRDREGEQY | 0.59548634 |
| LFAVGLTL | ASSSLRDREGEQY | 0.669034 |
| LFNGGIGTYI | ASSSLRDREGEQY | 0.32731575 |
| LTLGAGII | ASSSLRDREGEQY | 0.41735765 |
| LVAAALGDI | ASSSLRDREGEQY | 0.6609045 |

| | | |
|---|---|---|
| NTAFVQLGI | ASSSLRDREGEQY | 0.6743731 |
| QLFSGAEL | ASSSLRDREGEQY | 0.33699217 |
| SAATFITL | ASSSLRDREGEQY | 0.62533695 |
| SNYGLANLGV | ASSSLRDREGEQY | 0.5913616 |
| SYGGAYAVM | ASSSLRDREGEQY | 0.6686248 |
| TTGVGGLTLG | ASSSLRDREGEQY | 0.6595889 |
| TTQTIYI | ASSSLRDREGEQY | 0.54724824 |
| VAAGAGLGAVI | ASSSLRDREGEQY | 0.6678877 |
| VAAVVTLL | ASSSLRDREGEQY | 0.5645997 |
| VGFLGGLI | ASSSLRDREGEQY | 0.5610528 |
| VGVIPILL | ASSSLRDREGEQY | 0.67436165 |
| VIFASSLI | ASSSLRDREGEQY | 0.67442036 |
| VKGGDGGAAAG | ASSSLRDREGEQY | 0.6278612 |
| VTKTFTNI | ASSSLRDREGEQY | 0.67212856 |
| VVAAAILL | ASSSLRDREGEQY | 0.64024764 |
| YAALDVEL | ASSSLRDREGEQY | 0.6779576 |
| LASVALLL | ASSSLRDREGEQY | 0.6485015 |
| LDAVHRV | ASSSLRDREGEQY | 0.5437584 |
| QGLSSSIF | ASSSLRDREGEQY | 0.6687269 |
| RGTFANIRL | ASSSLRDREGEQY | 0.660744 |
| RQLWWGHRI | ASSSLRDREGEQY | 0.6598048 |
| YGPPGTGKTLL | ASSSLRDREGEQY | 0.6693017 |
| AALVGAPANVA | ASSSLRDREGEQY | 0.6238561 |
| AATFRRTE | ASSSLRDREGEQY | 0.64624804 |
| AGALGGGI | ASSSLRDREGEQY | 0.6753413 |
| AGGAGGAGGIGAG | ASSSLRDREGEQY | 0.674499 |
| AGSGALGL | ASSSLRDREGEQY | 0.5790379 |
| AQIADQI | ASSSLRDREGEQY | 0.6721885 |
| ASAVAVTYL | ASSSLRDREGEQY | 0.648973 |
| ASTVLGGTGG | ASSSLRDREGEQY | 0.6392802 |
| AVGGEPAK | ASSSLRDREGEQY | 0.6770085 |
| CEQGTFGGTC | ASSSLRDREGEQY | 0.6593813 |
| DEGKAPAG | ASSSLRDREGEQY | 0.67000437 |
| DTVVFPAL | ASSSLRDREGEQY | 0.6710994 |
| FEGITMAG | ASSSLRDREGEQY | 0.6675322 |
| FPLVVQD | ASSSLRDREGEQY | 0.53716505 |
| GAFTTPQ | ASSSLRDREGEQY | 0.6626796 |
| GFGGFGTAGGVGGA | ASSSLRDREGEQY | 0.6557833 |
| GFLAQFL | ASSSLRDREGEQY | 0.67254543 |
| GGAGGAGGNAGVA | ASSSLRDREGEQY | 0.66097176 |
| GGVGTAGAGGAGGAGG | ASSSLRDREGEQY | 0.5868252 |
| GNGGGGDAGNAGSGGN | ASSSLRDREGEQY | 0.66464305 |
| GTGGAGGAGNDAGS | ASSSLRDREGEQY | 0.6740012 |
| GTTNTGWFNS | ASSSLRDREGEQY | 0.6655942 |
| IFMGPLFTL | ASSSLRDREGEQY | 0.66500354 |
| ILAHAIEAI | ASSSLRDREGEQY | 0.67499024 |
| IRSAIREAL | ASSSLRDREGEQY | 0.6361192 |
| ITFAGGG | ASSSLRDREGEQY | 0.64939004 |
| ITGMSGHQEV | ASSSLRDREGEQY | 0.671544 |
| IVAGLACTL | ASSSLRDREGEQY | 0.67191607 |
| IVGGGLAGA | ASSSLRDREGEQY | 0.6534522 |

| | | |
|---|---|---|
| IVTGVLNI | ASSSLRDREGEQY | 0.6766728 |
| LAQQGCI | ASSSLRDREGEQY | 0.6247376 |
| LCLHVEN | ASSSLRDREGEQY | 0.67276514 |
| LDAISDGS | ASSSLRDREGEQY | 0.62378174 |
| LGGFAAML | ASSSLRDREGEQY | 0.66807246 |
| LGGKNPMI | ASSSLRDREGEQY | 0.6208898 |
| LGPKGVAAIL | ASSSLRDREGEQY | 0.67218477 |
| LMVGVSI | ASSSLRDREGEQY | 0.67145365 |
| LVGAAFVL | ASSSLRDREGEQY | 0.6729404 |
| LVLAVAPL | ASSSLRDREGEQY | 0.42732602 |
| MAAIAVIDAA | ASSSLRDREGEQY | 0.6711491 |
| NNGINAELQ | ASSSLRDREGEQY | 0.6716056 |
| RGEPLVLVHG | ASSSLRDREGEQY | 0.6774579 |
| RGGAGGAAGLI | ASSSLRDREGEQY | 0.6577281 |
| RSYRMHPAV | ASSSLRDREGEQY | 0.6640965 |
| SAAHLLAGQLA | ASSSLRDREGEQY | 0.6531208 |
| SAAVGSY | ASSSLRDREGEQY | 0.6479329 |
| SPHAVLHRI | ASSSLRDREGEQY | 0.6166289 |
| SQIAKVL | ASSSLRDREGEQY | 0.6438635 |
| TALGHLVP | ASSSLRDREGEQY | 0.6395112 |
| TAPTTSGAA | ASSSLRDREGEQY | 0.6227545 |
| TDGDDGNGGSPGT | ASSSLRDREGEQY | 0.6686131 |
| TDGLGEN | ASSSLRDREGEQY | 0.6728413 |
| TGGPPLPGQA | ASSSLRDREGEQY | 0.67193896 |
| TIPAITI | ASSSLRDREGEQY | 0.6748249 |
| TIPGFTI | ASSSLRDREGEQY | 0.6731236 |
| TSIASGALA | ASSSLRDREGEQY | 0.67241526 |
| TVTLPGAAI | ASSSLRDREGEQY | 0.63553613 |
| VASGLSAALP | ASSSLRDREGEQY | 0.6591357 |
| VGAGVIGI | ASSSLRDREGEQY | 0.6137906 |
| VGPKRFGV | ASSSLRDREGEQY | 0.6537364 |
| VNVAFPD | ASSSLRDREGEQY | 0.6376089 |
| VSMTNSGVS | ASSSLRDREGEQY | 0.6631034 |
| VTDVLGDLSV | ASSSLRDREGEQY | 0.31694824 |
| VVVVGAGFAGL | ASSSLRDREGEQY | 0.65366167 |
| VYTVGTVV | ASSSLRDREGEQY | 0.6763528 |
| WTTFSGP | ASSSLRDREGEQY | 0.62330693 |
| YDELVEAF | ASSSLRDREGEQY | 0.6620254 |
| YGTGGAGGNGG | ASSSLRDREGEQY | 0.64965415 |
| YTNVMAV | ASSSLRDREGEQY | 0.643291 |
| FLEQVHQL | ASSSLRDREGEQY | 0.6710431 |
| GLSLLAVAV | ASSSLRDREGEQY | 0.6227678 |
| LEQVHQL | ASSSLRDREGEQY | 0.6642258 |
| LNAAGAI | ASSSLRDREGEQY | 0.6758879 |
| PAVHASL | ASSSLRDREGEQY | 0.6591242 |
| TELEDDA | ASSSLRDREGEQY | 0.65963703 |
| TPPGPPN | ASSSLRDREGEQY | 0.6535324 |
| TPPPPPPP | ASSSLRDREGEQY | 0.5069816 |
| TQSPALI | ASSWGHLYEQY | 0.66584504 |
| EFAAAFPQ | ASSWGHLYEQY | 0.30816537 |
| LLSGNLDVL | ASSWGHLYEQY | 0.65246147 |

| | | |
|---|---|---|
| GPANLGNYNI | ASSWGHLYEQY | 0.620507 |
| TLGPTLP | ASSWGHLYEQY | 0.6633638 |
| VYAYSSCCK | ASSWGHLYEQY | 0.65695494 |
| APAPAPAPNI | ASSWGHLYEQY | 0.66892195 |
| PGGPVPIP | ASSWGHLYEQY | 0.6264389 |
| FLTVALEP | ASSWGHLYEQY | 0.643843 |
| SQYFGANL | ASSWGHLYEQY | 0.6281946 |
| GGVKPAA | ASSWGHLYEQY | 0.5976371 |
| GGTGGAGGAGGW | ASSWGHLYEQY | 0.6702095 |
| GVGGVAGLGGAGG | ASSWGHLYEQY | 0.65322477 |
| VLLVLATTI | ASSWGHLYEQY | 0.6427973 |
| IGIFAAML | ASSWGHLYEQY | 0.675001 |
| AVGDLID | ASSWGHLYEQY | 0.6596441 |
| NISPLPL | ASSWGHLYEQY | 0.5447453 |
| VTAVFVAI | ASSWGHLYEQY | 0.6487698 |
| SGDDNDAAL | ASSWGHLYEQY | 0.6579739 |
| AVAALAHS | ASSWGHLYEQY | 0.6491035 |
| GGVTVMATP | ASSWGHLYEQY | 0.3104556 |
| AGLAANVRD | ASSWGHLYEQY | 0.66749835 |
| TNRPDLIDPALLRPG | ASSWGHLYEQY | 0.6403972 |
| LANLGVTAL | ASSWGHLYEQY | 0.621153 |
| TGERIEL | ASSWGHLYEQY | 0.6377739 |
| NGEELAA | ASSWGHLYEQY | 0.6671878 |
| VLSLVWG | ASSWGHLYEQY | 0.5703321 |
| IGYALWL | ASSWGHLYEQY | 0.65001893 |
| LSLLLAPL | ASSWGHLYEQY | 0.6690378 |
| TLDEALK | ASSWGHLYEQY | 0.67141473 |
| TDGAGTAEGG | ASSWGHLYEQY | 0.6755246 |
| NAPGGAGGNGG | ASSWGHLYEQY | 0.66637295 |
| VSVFMGMGGNF | ASSWGHLYEQY | 0.65058255 |
| LASYLAAL | ASSWGHLYEQY | 0.66646725 |
| LVGGTPAL | ASSWGHLYEQY | 0.66305745 |
| VRNADVL | ASSWGHLYEQY | 0.5357888 |
| TCSSPFLVSAM | ASSWGHLYEQY | 0.4741084 |
| VHITDAE | ASSWGHLYEQY | 0.6643868 |
| EYAAPSF | ASSWGHLYEQY | 0.48455355 |
| VVDAFHGGL | ASSWGHLYEQY | 0.6607753 |
| ASVVDPQ | ASSWGHLYEQY | 0.67546123 |
| SYNEFGGLS | ASSWGHLYEQY | 0.654915 |
| DPDGSVY | ASSWGHLYEQY | 0.6006539 |
| MVSTIHATE | ASSWGHLYEQY | 0.6529504 |
| LAAGVADPV | ASSWGHLYEQY | 0.6455419 |
| TKEQIAA | ASSWGHLYEQY | 0.66716826 |
| SLASPAAL | ASSWGHLYEQY | 0.67021763 |
| IGLVFYGV | ASSWGHLYEQY | 0.67780185 |
| FLLDASNL | ASSWGHLYEQY | 0.67401636 |
| YQVGNFP | ASSWGHLYEQY | 0.6392005 |
| ESLLSQST | ASSWGHLYEQY | 0.6657115 |
| LFTVGFAAVG | ASSWGHLYEQY | 0.6425351 |
| ATPPTPT | ASSWGHLYEQY | 0.6541579 |
| TDLATLG | ASSWGHLYEQY | 0.65933466 |

| | | |
|---|---|---|
| SAGGTGR | ASSWGHLYEQY | 0.66427207 |
| GFSNSGS | ASSWGHLYEQY | 0.65780455 |
| GGNGGSGGTSV | ASSWGHLYEQY | 0.6483393 |
| GTSVATGGAGN | ASSWGHLYEQY | 0.587242 |
| NGGNGGSGGTS | ASSWGHLYEQY | 0.67202204 |
| AVVKFPH | ASSWGHLYEQY | 0.66159993 |
| TPPPPPP | ASSWGHLYEQY | 0.5378679 |
| AAPAAAAAPA | ASSWGHLYEQY | 0.6449324 |
| IALNEDL | ASSWGHLYEQY | 0.67751217 |
| IVNVSSM | ASSWGHLYEQY | 0.67550975 |
| TGIPPAPR | ASSWGHLYEQY | 0.6256257 |
| FDLGGGTF | ASSWGHLYEQY | 0.63659006 |
| FDLGGGTFDVS | ASSWGHLYEQY | 0.65633035 |
| NEPTAAA | ASSWGHLYEQY | 0.49764484 |
| NEPTAAALA | ASSWGHLYEQY | 0.6662524 |
| GSIINISSI | ASSWGHLYEQY | 0.65341353 |
| VAPGNAGTA | ASSWGHLYEQY | 0.65087 |
| TGGGASLE | ASSWGHLYEQY | 0.631619 |
| VAVYNHYKRI | ASSWGHLYEQY | 0.6570233 |
| TDPAPATT | ASSWGHLYEQY | 0.64363337 |
| TNRPDILDPALLRPG | ASSWGHLYEQY | 0.670281 |
| LTTTTTS | ASSWGHLYEQY | 0.3506885 |
| PTGSGKTL | ASSWGHLYEQY | 0.66964304 |
| TGSGKTL | ASSWGHLYEQY | 0.6214131 |
| TIGHVDHGK | ASSWGHLYEQY | 0.6222759 |
| TIGHVDHGKTT | ASSWGHLYEQY | 0.4995803 |
| GHVDHGKTT | ASSWGHLYEQY | 0.66029376 |
| AQFEHTLL | ASSWGHLYEQY | 0.67140955 |
| IVNQDEL | ASSWGHLYEQY | 0.6704081 |
| TNGSQFF | ASSWGHLYEQY | 0.60371387 |
| TGKGIVE | ASSWGHLYEQY | 0.6737045 |
| LISNASDA | ASSWGHLYEQY | 0.6778876 |
| AELRTLGELTKIAKAHGAQ | ASSWGHLYEQY | 0.64086574 |
| APNAVAAAEPSW | ASSWGHLYEQY | 0.49750164 |
| APQVAGAATAVL | ASSWGHLYEQY | 0.6662486 |
| ARRDPAAAAAILPTDA | ASSWGHLYEQY | 0.6556255 |
| DAAARDVLAEAGLAEYF | ASSWGHLYEQY | 0.6688037 |
| DDQIVLSAKANLAKVEQ | ASSWGHLYEQY | 0.6638789 |
| DELLPDTGEDSDEDGH | ASSWGHLYEQY | 0.60459596 |
| ELPEAPANRATVPQD | ASSWGHLYEQY | 0.66928476 |
| FGQSKIERVTGIG | ASSWGHLYEQY | 0.67360574 |
| GAGGTGSTAGGAGGAGGAGGLY | ASSWGHLYEQY | 0.6304481 |
| GHGGTGAAGGNGGTGGAGGSGI | ASSWGHLYEQY | 0.64642686 |
| GNGGNGGAGGEGGPGGAGGAGA | ASSWGHLYEQY | 0.6461807 |
| GSTAGNGGNGGNATNGTIAGQP | ASSWGHLYEQY | 0.48998028 |
| GTGGDGGNGGNGGTGG | ASSWGHLYEQY | 0.6709332 |
| GVRRPAGIPADAPV | ASSWGHLYEQY | 0.63146603 |
| GYRQVTDAHLLTLARR | ASSWGHLYEQY | 0.6287799 |
| IAKMGQDGHDRG | ASSWGHLYEQY | 0.6597796 |
| KAEPSIEIQYT | ASSWGHLYEQY | 0.63408923 |
| KPVRIRMAADEDTVAVT | ASSWGHLYEQY | 0.63608617 |

| | | |
|---|---|---|
| MPPRIAGMRLLVIKPEP | ASSWGHLYEQY | 0.63966876 |
| MKFYAVALFRSRGVPPTR | ASSWGHLYEQY | 0.6640127 |
| NAGGFGILGAVAHSP | ASSWGHLYEQY | 0.66122484 |
| NEGREITLAMAEM | ASSWGHLYEQY | 0.6600734 |
| NELVEWSHPLGHRWRWE | ASSWGHLYEQY | 0.657667 |
| NLGAGGTGGNGGNPG | ASSWGHLYEQY | 0.6749593 |
| NTEWPKWETVMAADPVD | ASSWGHLYEQY | 0.5896935 |
| PASAAPAETQP | ASSWGHLYEQY | 0.6332659 |
| QQSSDGKNGVRAAEITGE | ASSWGHLYEQY | 0.6621046 |
| RAPDAVTTSWWKE | ASSWGHLYEQY | 0.54009926 |
| SGFGNVGAHTSGWFNQ | ASSWGHLYEQY | 0.6672698 |
| SGVAGAVGQAASVGGLKV | ASSWGHLYEQY | 0.65745217 |
| SNGGAGTGSAGGNGGT | ASSWGHLYEQY | 0.6617155 |
| TGGAGGEGGAAG | ASSWGHLYEQY | 0.65956086 |
| TGGDGGAAGAGGAPGHG | ASSWGHLYEQY | 0.67819554 |
| TMPPLLAAIANAGRHGV | ASSWGHLYEQY | 0.66926444 |
| TSSTAGNSGGA | ASSWGHLYEQY | 0.660899 |
| VPATVPVGILC | ASSWGHLYEQY | 0.6586929 |
| VQRPDSGGVVLTGRIS | ASSWGHLYEQY | 0.62652177 |
| WTPNPVPATKGVPAGLVQAT | ASSWGHLYEQY | 0.6744641 |
| YALKRALAKFSDRFDVVII | ASSWGHLYEQY | 0.666928 |
| YLIGSVIV | ASSWGHLYEQY | 0.60280746 |
| YGQLVAVID | ASSWGHLYEQY | 0.6510804 |
| VTVGGMTLPA | ASSWGHLYEQY | 0.6753426 |
| VLALPAV | ASSWGHLYEQY | 0.67288804 |
| VIPPTLNYET | ASSWGHLYEQY | 0.5161105 |
| VGMGGMHPSSG | ASSWGHLYEQY | 0.6447544 |
| VGLAGGAATA | ASSWGHLYEQY | 0.48993668 |
| VGGAGGAGGSGP | ASSWGHLYEQY | 0.6301068 |
| VFAAFMV | ASSWGHLYEQY | 0.613212 |
| TLYDKLR | ASSWGHLYEQY | 0.6651755 |
| TLAVFTVL | ASSWGHLYEQY | 0.64449847 |
| TGSVQPARV | ASSWGHLYEQY | 0.66050196 |
| TGGGAGSGGT | ASSWGHLYEQY | 0.6607807 |
| SHLETSL | ASSWGHLYEQY | 0.6756025 |
| SFTTGMT | ASSWGHLYEQY | 0.6400896 |
| QVEVAKRMGVRQPTVS | ASSWGHLYEQY | 0.6562456 |
| QGSGGPALAM | ASSWGHLYEQY | 0.55403167 |
| MLAPSNP | ASSWGHLYEQY | 0.55107564 |
| LVNNAAI | ASSWGHLYEQY | 0.6419092 |
| LTGEINTLA | ASSWGHLYEQY | 0.647786 |
| LSIFGPV | ASSWGHLYEQY | 0.41490832 |
| LLALLANL | ASSWGHLYEQY | 0.67420155 |
| LISNASDALDKLR | ASSWGHLYEQY | 0.66186464 |
| LINEAVP | ASSWGHLYEQY | 0.66966057 |
| LGYAFSGL | ASSWGHLYEQY | 0.6435705 |
| LDFVAVAL | ASSWGHLYEQY | 0.65713173 |
| LAQPGAE | ASSWGHLYEQY | 0.63097996 |
| ISNLGAVTGF | ASSWGHLYEQY | 0.6661927 |
| HQEDPGRI | ASSWGHLYEQY | 0.6325392 |
| GTPGEPTP | ASSWGHLYEQY | 0.584736 |

| | | |
|---|---|---|
| GFFNSGS | ASSWGHLYEQY | 0.648957 |
| GAGGGGLTAGAN | ASSWGHLYEQY | 0.620501 |
| FVDALVNM | ASSWGHLYEQY | 0.67320806 |
| FLTVAPEL | ASSWGHLYEQY | 0.67302066 |
| FETLTAL | ASSWGHLYEQY | 0.6622095 |
| DVINPIEL | ASSWGHLYEQY | 0.6731433 |
| APVSGAPV | ASSWGHLYEQY | 0.43308073 |
| ALVGGLAW | ASSWGHLYEQY | 0.644112 |
| ALAAAAATAI | ASSWGHLYEQY | 0.6760386 |
| AFGLGLLL | ASSWGHLYEQY | 0.6656417 |
| AFFAAASL | ASSWGHLYEQY | 0.65019375 |
| AAAALPAFSPP | ASSWGHLYEQY | 0.6739724 |
| VVLAPAL | ASSWGHLYEQY | 0.6295183 |
| TIGHVDHGKTTL | ASSWGHLYEQY | 0.65021014 |
| LISNASDALDK | ASSWGHLYEQY | 0.65187174 |
| LISNASDAL | ASSWGHLYEQY | 0.67039895 |
| ISNASDALDK | ASSWGHLYEQY | 0.6640666 |
| NQAETLVYQTE | ASSWGHLYEQY | 0.6410767 |
| TGGTGGTAGAG | ASSWGHLYEQY | 0.66234344 |
| TGSTAGGAGGA | ASSWGHLYEQY | 0.6782899 |
| DPLAVPVAAGPV | ASSWGHLYEQY | 0.64420587 |
| GGTGGTPGNGGN | ASSWGHLYEQY | 0.6727801 |
| PAGLAGVDPATL | ASSWGHLYEQY | 0.6646192 |
| GNNLVVEAPHAGASV | ASSWGHLYEQY | 0.6609379 |
| KVTRIMSASTLEQPA | ASSWGHLYEQY | 0.63008285 |
| LGFALPIALPRRTVV | ASSWGHLYEQY | 0.64089096 |
| TRLGNLTARKLVVLV | ASSWGHLYEQY | 0.65002024 |
| WSQQRRLIYFPSAGP | ASSWGHLYEQY | 0.6202902 |
| GPNGNPDPMAAAVDIR | ASSWGHLYEQY | 0.65222627 |
| TQRCVSVADLRAAASR | ASSWGHLYEQY | 0.64775306 |
| QFVLDVAEHPVEVLTETV | ASSWGHLYEQY | 0.6792966 |
| GGAGGTGGAGGNGGTQAGDG | ASSWGHLYEQY | 0.66507715 |
| PLLATLRAIAVDVDAKPAQV | ASSWGHLYEQY | 0.664564 |
| APKFRTGGYMIDFWGVGYQVAK | ASSWGHLYEQY | 0.6248213 |
| GAGGAGGAGGAAGGPGGTGGTG | ASSWGHLYEQY | 0.66617286 |
| IGALGLLATDVAPAGSVAFGVA | ASSWGHLYEQY | 0.6655802 |
| AGGAGGIGGNGGAGGAGGVLMG | ASSWGHLYEQY | 0.5894232 |
| ADELIGLL | ASSWGHLYEQY | 0.676632 |
| ADRALKA | ASSWGHLYEQY | 0.63072944 |
| AGAVGQVGGAGGA | ASSWGHLYEQY | 0.6365769 |
| AGGGGGLGGAP | ASSWGHLYEQY | 0.6714097 |
| AGGLLYGNGGAG | ASSWGHLYEQY | 0.67468065 |
| AGLESIGL | ASSWGHLYEQY | 0.64563733 |
| AGLKTGII | ASSWGHLYEQY | 0.63613766 |
| ASLAALLL | ASSWGHLYEQY | 0.62066483 |
| ASVINISSI | ASSWGHLYEQY | 0.67060554 |
| ATPPGPA | ASSWGHLYEQY | 0.6715561 |
| ATVASALV | ASSWGHLYEQY | 0.64992225 |
| ATVASLIL | ASSWGHLYEQY | 0.6767498 |
| AVNGPVLIRH | ASSWGHLYEQY | 0.6658136 |
| DGAAGAVGI | ASSWGHLYEQY | 0.55740243 |

| | | |
|---|---|---|
| DPLLHGI | ASSWGHLYEQY | 0.6560987 |
| ELGAALGAL | ASSWGHLYEQY | 0.67335 |
| ELGRTQAV | ASSWGHLYEQY | 0.66985255 |
| FVAGGYLS | ASSWGHLYEQY | 0.6778629 |
| GASGGAGGAGGW | ASSWGHLYEQY | 0.6681539 |
| GGAGAGGDGGAGGI | ASSWGHLYEQY | 0.6465318 |
| GTGHNGSQPGMGGNG | ASSWGHLYEQY | 0.67535937 |
| GYAAASASAS | ASSWGHLYEQY | 0.67752 |
| HQVDVAAVQQG | ASSWGHLYEQY | 0.6709918 |
| IGAAIAHT | ASSWGHLYEQY | 0.6450709 |
| IGMIEVAQL | ASSWGHLYEQY | 0.61669683 |
| IRLAPEHTSAMD | ASSWGHLYEQY | 0.6577723 |
| IYNVSVL | ASSWGHLYEQY | 0.6684224 |
| KQGFLGM | ASSWGHLYEQY | 0.61462045 |
| LESVITQL | ASSWGHLYEQY | 0.6571392 |
| LGLGALVVV | ASSWGHLYEQY | 0.661598 |
| LIGNGGNGGSGGI | ASSWGHLYEQY | 0.58617806 |
| LNAINAPTL | ASSWGHLYEQY | 0.6721276 |
| LQLLNTGV | ASSWGHLYEQY | 0.6751695 |
| LQTSIEF | ASSWGHLYEQY | 0.64946467 |
| MSFAVSEL | ASSWGHLYEQY | 0.64513505 |
| NTGALGL | ASSWGHLYEQY | 0.6738618 |
| PPGAGGAGGAGTT | ASSWGHLYEQY | 0.6794317 |
| QLVGLGI | ASSWGHLYEQY | 0.6734191 |
| RFLEQVHQL | ASSWGHLYEQY | 0.54348654 |
| RGVGYVLR | ASSWGHLYEQY | 0.48375127 |
| SATGVFAGVI | ASSWGHLYEQY | 0.62699515 |
| SLPAITLP | ASSWGHLYEQY | 0.6659035 |
| STALAFTV | ASSWGHLYEQY | 0.6468786 |
| SVLTVGLGK | ASSWGHLYEQY | 0.6364033 |
| TALAGAGSGI | ASSWGHLYEQY | 0.6608751 |
| TGLPFTGI | ASSWGHLYEQY | 0.6737908 |
| VAEHVTV | ASSWGHLYEQY | 0.63544977 |
| VDIVLNSL | ASSWGHLYEQY | 0.57215047 |
| VLTGGGAVL | ASSWGHLYEQY | 0.66462374 |
| VNVGLAYGI | ASSWGHLYEQY | 0.6677089 |
| VSSAAVGAV | ASSWGHLYEQY | 0.63078296 |
| VTSVYEAGV | ASSWGHLYEQY | 0.6473774 |
| WVALDRGA | ASSWGHLYEQY | 0.6253292 |
| YLVGSVVI | ASSWGHLYEQY | 0.55294263 |
| GPPGTGKTLL | ASSWGHLYEQY | 0.661999 |
| GVGMTKF | ASSWGHLYEQY | 0.6192775 |
| IGIGGSDL | ASSWGHLYEQY | 0.6767781 |
| IGTLLAEL | ASSWGHLYEQY | 0.61989486 |
| IYTSGTT | ASSWGHLYEQY | 0.670393 |
| LSFPAGV | ASSWGHLYEQY | 0.66523284 |
| LYTSGTT | ASSWGHLYEQY | 0.66833216 |
| VGAFPFGF | ASSWGHLYEQY | 0.6442784 |
| VGAGTVEF | ASSWGHLYEQY | 0.6404499 |
| VGKSTLLL | ASSWGHLYEQY | 0.61064434 |
| AALAHAGGGIV | ASSWGHLYEQY | 0.5111555 |

| | | |
|---|---|---|
| AALVAVSLL | ASSWGHLYEQY | 0.6730012 |
| AAVSRAVAAL | ASSWGHLYEQY | 0.33072957 |
| AGGSSGIGGF | ASSWGHLYEQY | 0.6743316 |
| AGLTVANANG | ASSWGHLYEQY | 0.66923386 |
| AGLVGAVL | ASSWGHLYEQY | 0.63141423 |
| ALSQAFTV | ASSWGHLYEQY | 0.6670219 |
| ALTFSVSL | ASSWGHLYEQY | 0.6111324 |
| ALTGAALVAL | ASSWGHLYEQY | 0.6269263 |
| ASFSIGAGVL | ASSWGHLYEQY | 0.6698416 |
| DGSVYKI | ASSWGHLYEQY | 0.6671201 |
| EQLAEIV | ASSWGHLYEQY | 0.6334144 |
| FAVSELTVM | ASSWGHLYEQY | 0.66288817 |
| GAAPLAAGL | ASSWGHLYEQY | 0.661513 |
| GADGGDGGSGGAGG | ASSWGHLYEQY | 0.64407617 |
| GFGLDTGGAGG | ASSWGHLYEQY | 0.6561497 |
| GLASGGLL | ASSWGHLYEQY | 0.67468154 |
| GRIAGAAVLL | ASSWGHLYEQY | 0.66252714 |
| GSAGSLNTSF | ASSWGHLYEQY | 0.634645 |
| GTGQIGI | ASSWGHLYEQY | 0.6675085 |
| GTLIALIL | ASSWGHLYEQY | 0.61042213 |
| IAATMGFVVG | ASSWGHLYEQY | 0.66477036 |
| IFAQALL | ASSWGHLYEQY | 0.65885997 |
| IGAAVALGF | ASSWGHLYEQY | 0.6487838 |
| IGLTPLAL | ASSWGHLYEQY | 0.6665367 |
| ILGSNTSTL | ASSWGHLYEQY | 0.6069988 |
| IVFSLTVL | ASSWGHLYEQY | 0.6227102 |
| LASKATGF | ASSWGHLYEQY | 0.64817965 |
| LASVALIL | ASSWGHLYEQY | 0.66187453 |
| LFAVGLTL | ASSWGHLYEQY | 0.6711943 |
| LFNGGIGTYI | ASSWGHLYEQY | 0.62288797 |
| LTLGAGII | ASSWGHLYEQY | 0.66170275 |
| LVAAALGDI | ASSWGHLYEQY | 0.6444222 |
| NTAFVQLGI | ASSWGHLYEQY | 0.6219215 |
| QLFSGAEL | ASSWGHLYEQY | 0.6528264 |
| SAATFITL | ASSWGHLYEQY | 0.656355 |
| SNYGLANLGV | ASSWGHLYEQY | 0.6371216 |
| SYGGAYAVM | ASSWGHLYEQY | 0.6693883 |
| TTGVGGLTLG | ASSWGHLYEQY | 0.6059472 |
| TTQTIYI | ASSWGHLYEQY | 0.66686636 |
| VAAGAGLGAVI | ASSWGHLYEQY | 0.6760647 |
| VAAVVTLL | ASSWGHLYEQY | 0.65058595 |
| VGFLGGLI | ASSWGHLYEQY | 0.6692603 |
| VGVIPILL | ASSWGHLYEQY | 0.6667553 |
| VIFASSLI | ASSWGHLYEQY | 0.66659784 |
| VKGGDGGAAAG | ASSWGHLYEQY | 0.6456867 |
| VTKTFTNI | ASSWGHLYEQY | 0.648152 |
| VVAAAILL | ASSWGHLYEQY | 0.6747771 |
| YAALDVEL | ASSWGHLYEQY | 0.66917515 |
| LASVALLL | ASSWGHLYEQY | 0.6661944 |
| LDAVHRV | ASSWGHLYEQY | 0.6681237 |
| QGLSSSIF | ASSWGHLYEQY | 0.6694933 |

| | | |
|---|---|---|
| RGTFANIRL | ASSWGHLYEQY | 0.66600794 |
| RQLWWGHRI | ASSWGHLYEQY | 0.6345941 |
| YGPPGTGKTLL | ASSWGHLYEQY | 0.5798868 |
| AALVGAPANVA | ASSWGHLYEQY | 0.6476152 |
| AATFRRTE | ASSWGHLYEQY | 0.65722775 |
| AGALGGGI | ASSWGHLYEQY | 0.6520442 |
| AGGAGGAGGIGAG | ASSWGHLYEQY | 0.5218663 |
| AGSGALGL | ASSWGHLYEQY | 0.67330635 |
| AQIADQI | ASSWGHLYEQY | 0.67926586 |
| ASAVAVTYL | ASSWGHLYEQY | 0.631146 |
| ASTVLGGTGG | ASSWGHLYEQY | 0.67653656 |
| AVGGEPAK | ASSWGHLYEQY | 0.59527415 |
| CEQGTFGGTC | ASSWGHLYEQY | 0.66818804 |
| DEGKAPAG | ASSWGHLYEQY | 0.6735543 |
| DTVVFPAL | ASSWGHLYEQY | 0.6183697 |
| FEGITMAG | ASSWGHLYEQY | 0.6710404 |
| FPLVVQD | ASSWGHLYEQY | 0.6388961 |
| GAFTTPQ | ASSWGHLYEQY | 0.6543441 |
| GFGGFGTAGGVGGA | ASSWGHLYEQY | 0.6759621 |
| GFLAQFL | ASSWGHLYEQY | 0.6408949 |
| GGAGGAGGNAGVA | ASSWGHLYEQY | 0.66409105 |
| GGVGTAGAGGAGGAGG | ASSWGHLYEQY | 0.6469612 |
| GNGGGGDAGNAGSGGN | ASSWGHLYEQY | 0.6213197 |
| GTGGAGGAGNDAGS | ASSWGHLYEQY | 0.6646347 |
| GTTNTGWFNS | ASSWGHLYEQY | 0.67421544 |
| IFMGPLFTL | ASSWGHLYEQY | 0.62589216 |
| ILAHAIEAI | ASSWGHLYEQY | 0.6765863 |
| IRSAIREAL | ASSWGHLYEQY | 0.6400791 |
| ITFAGGG | ASSWGHLYEQY | 0.6796185 |
| ITGMSGHQEV | ASSWGHLYEQY | 0.66721165 |
| IVAGLACTL | ASSWGHLYEQY | 0.65382314 |
| IVGGGLAGA | ASSWGHLYEQY | 0.6485063 |
| IVTGVLNI | ASSWGHLYEQY | 0.6738411 |
| LAQQGCI | ASSWGHLYEQY | 0.6451164 |
| LCLHVEN | ASSWGHLYEQY | 0.6691506 |
| LDAISDGS | ASSWGHLYEQY | 0.6764119 |
| LGGFAAML | ASSWGHLYEQY | 0.15863362 |
| LGGKNPMI | ASSWGHLYEQY | 0.75177985 |
| LGPKGVAAIL | ASSWGHLYEQY | 0.39458063 |
| LMVGVSI | ASSWGHLYEQY | 0.41415054 |
| LVGAAFVL | ASSWGHLYEQY | 0.18850198 |
| LVLAVAPL | ASSWGHLYEQY | 0.13070664 |
| MAAIAVIDAA | ASSWGHLYEQY | 0.27980632 |
| NNGINAELQ | ASSWGHLYEQY | 0.17638066 |
| RGEPLVLVHG | ASSWGHLYEQY | 0.52849466 |
| RGGAGGAAGLI | ASSWGHLYEQY | 0.53431535 |
| RSYRMHPAV | ASSWGHLYEQY | 0.20561844 |
| SAAHLLAGQLA | ASSWGHLYEQY | 0.67380416 |
| SAAVGSY | ASSWGHLYEQY | 0.6875769 |
| SPHAVLHRI | ASSWGHLYEQY | 0.14284796 |
| SQIAKVL | ASSWGHLYEQY | 0.30263975 |

| | | |
|---|---|---|
| TALGHLVP | ASSWGHLYEQY | 0.16502737 |
| TAPTTSGAA | ASSWGHLYEQY | 0.22995658 |
| TDGDDGNGGSPGT | ASSWGHLYEQY | 0.12313163 |
| TDGLGEN | ASSWGHLYEQY | 0.48266047 |
| TGGPPLPGQA | ASSWGHLYEQY | 0.18590786 |
| TIPAITI | ASSWGHLYEQY | 0.11183388 |
| TIPGFTI | ASSWGHLYEQY | 0.31350985 |
| TSIASGALA | ASSWGHLYEQY | 0.28290978 |
| TVTLPGAAI | ASSWGHLYEQY | 0.21604079 |
| VASGLSAALP | ASSWGHLYEQY | 0.5545659 |
| VGAGVIGI | ASSWGHLYEQY | 0.19772114 |
| VGPKRFGV | ASSWGHLYEQY | 0.22458681 |
| VNVAFPD | ASSWGHLYEQY | 0.6375887 |
| VSMTNSGVS | ASSWGHLYEQY | 0.55855346 |
| VTDVLGDLSV | ASSWGHLYEQY | 0.18648677 |
| VVVVGAGFAGL | ASSWGHLYEQY | 0.39996552 |
| VYTVGTVV | ASSWGHLYEQY | 0.7100935 |
| WTTFSGP | ASSWGHLYEQY | 0.24814457 |
| YDELVEAF | ASSWGHLYEQY | 0.59015995 |
| YGTGGAGGNGG | ASSWGHLYEQY | 0.59851646 |
| YTNVMAV | ASSWGHLYEQY | 0.20359968 |
| FLEQVHQL | ASSWGHLYEQY | 0.65017796 |
| GLSLLAVAV | ASSWGHLYEQY | 0.750995 |
| LEQVHQL | ASSWGHLYEQY | 0.25392246 |
| LNAAGAI | ASSWGHLYEQY | 0.32099497 |
| PAVHASL | ASSWGHLYEQY | 0.16130112 |
| TELEDDA | ASSWGHLYEQY | 0.2414877 |
| TPPGPPN | ASSWGHLYEQY | 0.12709512 |
| TPPPPPPP | ASSWGHLYEQY | 0.5107594 |
| TQSPALI | ASSLSMAGGAREQY | 0.5685414 |
| EFAAAFPQ | ASSLSMAGGAREQY | 0.35209626 |
| LLSGNLDVL | ASSLSMAGGAREQY | 0.15649927 |
| GPANLGNYNI | ASSLSMAGGAREQY | 0.15157814 |
| TLGPTLP | ASSLSMAGGAREQY | 0.32112703 |
| VYAYSSCCK | ASSLSMAGGAREQY | 0.1818222 |
| APAPAPAPNI | ASSLSMAGGAREQY | 0.14203471 |
| PGGPVPIP | ASSLSMAGGAREQY | 0.16595924 |
| FLTVALEP | ASSLSMAGGAREQY | 0.5306152 |
| SQYFGANL | ASSLSMAGGAREQY | 0.461552 |
| GGVKPAA | ASSLSMAGGAREQY | 0.18917106 |
| GGTGGAGGAGGW | ASSLSMAGGAREQY | 0.7885587 |
| GVGGVAGLGGAGG | ASSLSMAGGAREQY | 0.6704978 |
| VLLVLATTI | ASSLSMAGGAREQY | 0.6706816 |
| IGIFAAML | ASSLSMAGGAREQY | 0.28441325 |
| AVGDLID | ASSLSMAGGAREQY | 0.16138922 |
| NISPLPL | ASSLSMAGGAREQY | 0.8107533 |
| VTAVFVAI | ASSLSMAGGAREQY | 0.20438999 |
| SGDDNDAAL | ASSLSMAGGAREQY | 0.4453384 |
| AVAALAHS | ASSLSMAGGAREQY | 0.38560903 |
| GGVTVMATP | ASSLSMAGGAREQY | 0.27936706 |
| AGLAANVRD | ASSLSMAGGAREQY | 0.20817442 |

| | | |
|---|---|---|
| TNRPDLIDPALLRPG | ASSLSMAGGAREQY | 0.52886665 |
| LANLGVTAL | ASSLSMAGGAREQY | 0.58115274 |
| TGERIEL | ASSLSMAGGAREQY | 0.19360474 |
| NGEELAA | ASSLSMAGGAREQY | 0.1725715 |
| VLSLVWG | ASSLSMAGGAREQY | 0.20832747 |
| IGYALWL | ASSLSMAGGAREQY | 0.2754723 |
| LSLLLAPL | ASSLSMAGGAREQY | 0.6271335 |
| TLDEALK | ASSLSMAGGAREQY | 0.32800364 |
| TDGAGTAEGG | ASSLSMAGGAREQY | 0.23589481 |
| NAPGGAGGNGG | ASSLSMAGGAREQY | 0.48781827 |
| VSVFMGMGGNF | ASSLSMAGGAREQY | 0.37405252 |
| LASYLAAL | ASSLSMAGGAREQY | 0.26953933 |
| LVGGTPAL | ASSLSMAGGAREQY | 0.3682924 |
| VRNADVL | ASSLSMAGGAREQY | 0.23529024 |
| TCSSPFLVSAM | ASSLSMAGGAREQY | 0.54266477 |
| VHITDAE | ASSLSMAGGAREQY | 0.50101334 |
| EYAAPSF | ASSLSMAGGAREQY | 0.45705646 |
| VVDAFHGGL | ASSLSMAGGAREQY | 0.63720274 |
| ASVVDPQ | ASSLSMAGGAREQY | 0.2989803 |
| SYNEFGGLS | ASSLSMAGGAREQY | 0.2844903 |
| DPDGSVY | ASSLSMAGGAREQY | 0.13802333 |
| MVSTIHATE | ASSLSMAGGAREQY | 0.18777153 |
| LAAGVADPV | ASSLSMAGGAREQY | 0.38396043 |
| TKEQIAA | ASSLSMAGGAREQY | 0.43430263 |
| SLASPAAL | ASSLSMAGGAREQY | 0.6553234 |
| IGLVFYGV | ASSLSMAGGAREQY | 0.4996318 |
| FLLDASNL | ASSLSMAGGAREQY | 0.24274376 |
| YQVGNFP | ASSLSMAGGAREQY | 0.6755189 |
| ESLLSQST | ASSLSMAGGAREQY | 0.66305494 |
| LFTVGFAAVG | ASSLSMAGGAREQY | 0.14189835 |
| ATPPTPT | ASSLSMAGGAREQY | 0.19093052 |
| TDLATLG | ASSLSMAGGAREQY | 0.764691 |
| SAGGTGR | ASSLSMAGGAREQY | 0.5582639 |
| GFSNSGS | ASSLSMAGGAREQY | 0.24483877 |
| GGNGGSGGTSV | ASSLSMAGGAREQY | 0.3014285 |
| GTSVATGGAGN | ASSLSMAGGAREQY | 0.83440226 |
| NGGNGGSGGTS | ASSLSMAGGAREQY | 0.24820395 |
| AVVKFPH | ASSLSMAGGAREQY | 0.7687874 |
| TPPPPPP | ASSLSMAGGAREQY | 0.12561642 |
| AAPAAAAAPA | ASSLSMAGGAREQY | 0.24823013 |
| IALNEDL | ASSLSMAGGAREQY | 0.18567175 |
| IVNVSSM | ASSLSMAGGAREQY | 0.11378673 |
| TGIPPAPR | ASSLSMAGGAREQY | 0.52952635 |
| FDLGGGTF | ASSLSMAGGAREQY | 0.25134057 |
| FDLGGGTFDVS | ASSLSMAGGAREQY | 0.49755967 |
| NEPTAAA | ASSLSMAGGAREQY | 0.23369734 |
| NEPTAAALA | ASSLSMAGGAREQY | 0.53792036 |
| GSIINISSI | ASSLSMAGGAREQY | 0.49327803 |
| VAPGNAGTA | ASSLSMAGGAREQY | 0.19512518 |
| TGGGASLE | ASSLSMAGGAREQY | 0.17689425 |
| VAVYNHYKRI | ASSLSMAGGAREQY | 0.3780823 |

| | | |
|---|---|---|
| TDPAPATT | ASSLSMAGGAREQY | 0.72857076 |
| TNRPDILDPALLRPG | ASSLSMAGGAREQY | 0.4608826 |
| LTTTTTS | ASSLSMAGGAREQY | 0.14563958 |
| PTGSGKTL | ASSLSMAGGAREQY | 0.35164404 |
| TGSGKTL | ASSLSMAGGAREQY | 0.26947832 |
| TIGHVDHGK | ASSLSMAGGAREQY | 0.14675996 |
| TIGHVDHGKTT | ASSLSMAGGAREQY | 0.11350294 |
| GHVDHGKTT | ASSLSMAGGAREQY | 0.21839184 |
| AQFEHTLL | ASSLSMAGGAREQY | 0.268622 |
| IVNQDEL | ASSLSMAGGAREQY | 0.60285795 |
| TNGSQFF | ASSLSMAGGAREQY | 0.6933632 |
| TGKGIVE | ASSLSMAGGAREQY | 0.2537841 |
| LISNASDA | ASSLSMAGGAREQY | 0.5062315 |
| AELRTLGELTKIAKAHGAQ | ASSLSMAGGAREQY | 0.31237882 |
| APNAVAAAEPSW | ASSLSMAGGAREQY | 0.0998613 |
| APQVAGAATAVL | ASSLSMAGGAREQY | 0.17665178 |
| ARRDPAAAAAILPTDA | ASSLSMAGGAREQY | 0.14147434 |
| DAAARDVLAEAGLAEYF | ASSLSMAGGAREQY | 0.665637 |
| DDQIVLSAKANLAKVEQ | ASSLSMAGGAREQY | 0.14174482 |
| DELLPDTGEDSDEDGH | ASSLSMAGGAREQY | 0.30228812 |
| ELPEAPANRATVPQD | ASSLSMAGGAREQY | 0.30442092 |
| FGQSKIERVTGIG | ASSLSMAGGAREQY | 0.78256 |
| GAGGTGSTAGGAGGAGGAGGLY | ASSLSMAGGAREQY | 0.35677952 |
| GHGGTGAAGGNGGTGGAGGSGI | ASSLSMAGGAREQY | 0.1743677 |
| GNGGNGGAGGEGGPGGAGGAGA | ASSLSMAGGAREQY | 0.11242069 |
| GSTAGNGGNGGNATNGTIAGQP | ASSLSMAGGAREQY | 0.38586986 |
| GTGGDGGNGGNGGTGG | ASSLSMAGGAREQY | 0.16043349 |
| GVRRPAGIPADAPV | ASSLSMAGGAREQY | 0.18156703 |
| GYRQVTDAHLLTLARR | ASSLSMAGGAREQY | 0.14254338 |
| IAKMGQDGHDRG | ASSLSMAGGAREQY | 0.19847639 |
| KAEPSIEIQYT | ASSLSMAGGAREQY | 0.14158137 |
| KPVRIRMAADEDTVAVT | ASSLSMAGGAREQY | 0.46736488 |
| MPPRIAGMRLLVIKPEP | ASSLSMAGGAREQY | 0.20861606 |
| MKFYAVALFRSRGVPPTR | ASSLSMAGGAREQY | 0.35349637 |
| NAGGFGILGAVAHSP | ASSLSMAGGAREQY | 0.77504313 |
| NEGREITLAMAEM | ASSLSMAGGAREQY | 0.71629673 |
| NELVEWSHPLGHRWRWE | ASSLSMAGGAREQY | 0.25455648 |
| NLGAGGTGGNGGNPG | ASSLSMAGGAREQY | 0.19064021 |
| NTEWPKWETVMAADPVD | ASSLSMAGGAREQY | 0.16405496 |
| PASAAPAETQP | ASSLSMAGGAREQY | 0.19506976 |
| QQSSDGKNGVRAAEITGE | ASSLSMAGGAREQY | 0.21786636 |
| RAPDAVTTSWWKE | ASSLSMAGGAREQY | 0.7312987 |
| SGFGNVGAHTSGWFNQ | ASSLSMAGGAREQY | 0.28064758 |
| SGVAGAVGQAASVGGLKV | ASSLSMAGGAREQY | 0.16607524 |
| SNGGAGTGSAGGNGGT | ASSLSMAGGAREQY | 0.3977003 |
| TGGAGGEGGAAG | ASSLSMAGGAREQY | 0.7220097 |
| TGGDGGAAGAGGAPGHG | ASSLSMAGGAREQY | 0.7402476 |
| TMPPLLAAIANAGRHGV | ASSLSMAGGAREQY | 0.15602109 |
| TSSTAGNSGGA | ASSLSMAGGAREQY | 0.20458223 |
| VPATVPVGILC | ASSLSMAGGAREQY | 0.27029306 |
| VQRPDSGGVVLTGRIS | ASSLSMAGGAREQY | 0.3843457 |

| | | |
|---|---|---|
| WTPNPVPATKGVPAGLVQAT | ASSLSMAGGAREQY | 0.18676446 |
| YALKRALAKFSDRFDVVII | ASSLSMAGGAREQY | 0.17228001 |
| YLIGSVIV | ASSLSMAGGAREQY | 0.49311736 |
| YGQLVAVID | ASSLSMAGGAREQY | 0.8508668 |
| VTVGGMTLPA | ASSLSMAGGAREQY | 0.22975692 |
| VLALPAV | ASSLSMAGGAREQY | 0.13046256 |
| VIPPTLNYET | ASSLSMAGGAREQY | 0.29493284 |
| VGMGGMHPSSG | ASSLSMAGGAREQY | 0.19348185 |
| VGLAGGAATA | ASSLSMAGGAREQY | 0.20811947 |
| VGGAGGAGGSGP | ASSLSMAGGAREQY | 0.29899466 |
| VFAAFMV | ASSLSMAGGAREQY | 0.1503916 |
| TLYDKLR | ASSLSMAGGAREQY | 0.37261283 |
| TLAVFTVL | ASSLSMAGGAREQY | 0.68609464 |
| TGSVQPARV | ASSLSMAGGAREQY | 0.19658038 |
| TGGGAGSGGT | ASSLSMAGGAREQY | 0.5240861 |
| SHLETSL | ASSLSMAGGAREQY | 0.7640292 |
| SFTTGMT | ASSLSMAGGAREQY | 0.5438597 |
| QVEVAKRMGVRQPTVS | ASSLSMAGGAREQY | 0.5981158 |
| QGSGGPALAM | ASSLSMAGGAREQY | 0.6843413 |
| MLAPSNP | ASSLSMAGGAREQY | 0.24179491 |
| LVNNAAI | ASSLSMAGGAREQY | 0.27451965 |
| LTGEINTLA | ASSLSMAGGAREQY | 0.6354207 |
| LSIFGPV | ASSLSMAGGAREQY | 0.27160823 |
| LLALLANL | ASSLSMAGGAREQY | 0.30447936 |
| LISNASDALDKLR | ASSLSMAGGAREQY | 0.29004806 |
| LINEAVP | ASSLSMAGGAREQY | 0.6893388 |
| LGYAFSGL | ASSLSMAGGAREQY | 0.233088 |
| LDFVAVAL | ASSLSMAGGAREQY | 0.68499666 |
| LAQPGAE | ASSLSMAGGAREQY | 0.13384312 |
| ISNLGAVTGF | ASSLSMAGGAREQY | 0.17687519 |
| HQEDPGRI | ASSLSMAGGAREQY | 0.71056557 |
| GTPGEPTP | ASSLSMAGGAREQY | 0.10698995 |
| GFFNSGS | ASSLSMAGGAREQY | 0.12316357 |
| GAGGGGLTAGAN | ASSLSMAGGAREQY | 0.35164672 |
| FVDALVNM | ASSLSMAGGAREQY | 0.7067329 |
| FLTVAPEL | ASSLSMAGGAREQY | 0.10703648 |
| FETLTAL | ASSLSMAGGAREQY | 0.22579633 |
| DVINPIEL | ASSLSMAGGAREQY | 0.16453743 |
| APVSGAPV | ASSLSMAGGAREQY | 0.4067987 |
| ALVGGLAW | ASSLSMAGGAREQY | 0.33541483 |
| ALAAAAATAI | ASSLSMAGGAREQY | 0.12202882 |
| AFGLGLLL | ASSLSMAGGAREQY | 0.44598103 |
| AFFAAASL | ASSLSMAGGAREQY | 0.14134145 |
| AAAALPAFSPP | ASSLSMAGGAREQY | 0.16433676 |
| VVLAPAL | ASSLSMAGGAREQY | 0.5113417 |
| TIGHVDHGKTTL | ASSLSMAGGAREQY | 0.7907958 |
| LISNASDALDK | ASSLSMAGGAREQY | 0.1544189 |
| LISNASDAL | ASSLSMAGGAREQY | 0.52544 |
| ISNASDALDK | ASSLSMAGGAREQY | 0.27605498 |
| NQAETLVYQTE | ASSLSMAGGAREQY | 0.8169724 |
| TGGTGGTAGAG | ASSLSMAGGAREQY | 0.20976165 |

| | | |
|---|---|---|
| TGSTAGGAGGA | ASSLSMAGGAREQY | 0.13890556 |
| DPLAVPVAAGPV | ASSLSMAGGAREQY | 0.6753082 |
| GGTGGTPGNGGN | ASSLSMAGGAREQY | 0.43497986 |
| PAGLAGVDPATL | ASSLSMAGGAREQY | 0.2873066 |
| GNNLVVEAPHAGASV | ASSLSMAGGAREQY | 0.6452609 |
| KVTRIMSASTLEQPA | ASSLSMAGGAREQY | 0.21344672 |
| LGFALPIALPRRTVV | ASSLSMAGGAREQY | 0.27406397 |
| TRLGNLTARKLVVLV | ASSLSMAGGAREQY | 0.6096039 |
| WSQQRRLIYFPSAGP | ASSLSMAGGAREQY | 0.43909043 |
| GPNGNPDPMAAAVDIR | ASSLSMAGGAREQY | 0.18574028 |
| TQRCVSVADLRAAASR | ASSLSMAGGAREQY | 0.6000319 |
| QFVLDVAEHPVEVLTETV | ASSLSMAGGAREQY | 0.3566687 |
| GGAGGTGGAGGNGGTQAGDG | ASSLSMAGGAREQY | 0.1645193 |
| PLLATLRAIAVDVDAKPAQV | ASSLSMAGGAREQY | 0.69433403 |
| APKFRTGGYMIDFWGVGYQVAK | ASSLSMAGGAREQY | 0.25276938 |
| GAGGAGGAGGAAGGPGGTGGTG | ASSLSMAGGAREQY | 0.52476 |
| IGALGLLATDVAPAGSVAFGVA | ASSLSMAGGAREQY | 0.40934116 |
| AGGAGGIGGNGGAGGAGGVLMG | ASSLSMAGGAREQY | 0.8140665 |
| ADELIGLL | ASSLSMAGGAREQY | 0.13031153 |
| ADRALKA | ASSLSMAGGAREQY | 0.3558269 |
| AGAVGQVGGAGGA | ASSLSMAGGAREQY | 0.24759313 |
| AGGGGGLGGAP | ASSLSMAGGAREQY | 0.4684241 |
| AGGLLYGNGGAG | ASSLSMAGGAREQY | 0.2620787 |
| AGLESIGL | ASSLSMAGGAREQY | 0.19212392 |
| AGLKTGII | ASSLSMAGGAREQY | 0.35110223 |
| ASLAALLL | ASSLSMAGGAREQY | 0.6815486 |
| ASVINISSI | ASSLSMAGGAREQY | 0.24245861 |
| ATPPGPA | ASSLSMAGGAREQY | 0.3947386 |
| ATVASALV | ASSLSMAGGAREQY | 0.76423407 |
| ATVASLIL | ASSLSMAGGAREQY | 0.16205548 |
| AVNGPVLIRH | ASSLSMAGGAREQY | 0.339678 |
| DGAAGAVGI | ASSLSMAGGAREQY | 0.49055433 |
| DPLLHGI | ASSLSMAGGAREQY | 0.7189159 |
| ELGAALGAL | ASSLSMAGGAREQY | 0.25263724 |
| ELGRTQAV | ASSLSMAGGAREQY | 0.6697307 |
| FVAGGYLS | ASSLSMAGGAREQY | 0.17037041 |
| GASGGAGGAGGW | ASSLSMAGGAREQY | 0.54991055 |
| GGAGAGGDGGAGGI | ASSLSMAGGAREQY | 0.41119933 |
| GTGHNGSQPGMGGNG | ASSLSMAGGAREQY | 0.29701298 |
| GYAAASASAS | ASSLSMAGGAREQY | 0.17314431 |
| HQVDVAAVQQG | ASSLSMAGGAREQY | 0.4196094 |
| IGAAIAHT | ASSLSMAGGAREQY | 0.59545135 |
| IGMIEVAQL | ASSLSMAGGAREQY | 0.59838593 |
| IRLAPEHTSAMD | ASSLSMAGGAREQY | 0.13063678 |
| IYNVSVL | ASSLSMAGGAREQY | 0.504579 |
| KQGFLGM | ASSLSMAGGAREQY | 0.5894735 |
| LESVITQL | ASSLSMAGGAREQY | 0.84084594 |
| LGLGALVVV | ASSLSMAGGAREQY | 0.20024711 |
| LIGNGGNGGSGGI | ASSLSMAGGAREQY | 0.55045545 |
| LNAINAPTL | ASSLSMAGGAREQY | 0.24143359 |
| LQLLNTGV | ASSLSMAGGAREQY | 0.2639392 |

| | | |
|---|---|---|
| LQTSIEF | ASSLSMAGGAREQY | 0.14730915 |
| MSFAVSEL | ASSLSMAGGAREQY | 0.18274932 |
| NTGALGL | ASSLSMAGGAREQY | 0.29607546 |
| PPGAGGAGGAGTT | ASSLSMAGGAREQY | 0.19286211 |
| QLVGLGI | ASSLSMAGGAREQY | 0.7692585 |
| RFLEQVHQL | ASSLSMAGGAREQY | 0.60129344 |
| RGVGYVLR | ASSLSMAGGAREQY | 0.6150861 |
| SATGVFAGVI | ASSLSMAGGAREQY | 0.71654475 |
| SLPAITLP | ASSLSMAGGAREQY | 0.79020536 |
| STALAFTV | ASSLSMAGGAREQY | 0.44329843 |
| SVLTVGLGK | ASSLSMAGGAREQY | 0.22529176 |
| TALAGAGSGI | ASSLSMAGGAREQY | 0.19128165 |
| TGLPFTGI | ASSLSMAGGAREQY | 0.14107788 |
| VAEHVTV | ASSLSMAGGAREQY | 0.3081252 |
| VDIVLNSL | ASSLSMAGGAREQY | 0.22051598 |
| VLTGGGAVL | ASSLSMAGGAREQY | 0.3976505 |
| VNVGLAYGI | ASSLSMAGGAREQY | 0.10948769 |
| VSSAAVGAV | ASSLSMAGGAREQY | 0.70714307 |
| VTSVYEAGV | ASSLSMAGGAREQY | 0.73707724 |
| WVALDRGA | ASSLSMAGGAREQY | 0.13962428 |
| YLVGSVVI | ASSLSMAGGAREQY | 0.4086157 |
| GPPGTGKTLL | ASSLSMAGGAREQY | 0.6505198 |
| GVGMTKF | ASSLSMAGGAREQY | 0.1941236 |
| IGIGGSDL | ASSLSMAGGAREQY | 0.5366093 |
| IGTLLAEL | ASSLSMAGGAREQY | 0.20395455 |
| IYTSGTT | ASSLSMAGGAREQY | 0.5461008 |
| LSFPAGV | ASSLSMAGGAREQY | 0.59202874 |
| LYTSGTT | ASSLSMAGGAREQY | 0.43254215 |
| VGAFPFGF | ASSLSMAGGAREQY | 0.3343044 |
| VGAGTVEF | ASSLSMAGGAREQY | 0.36989093 |
| VGKSTLLL | ASSLSMAGGAREQY | 0.1308232 |
| AALAHAGGGIV | ASSLSMAGGAREQY | 0.39818147 |
| AALVAVSLL | ASSLSMAGGAREQY | 0.11459618 |
| AAVSRAVAAL | ASSLSMAGGAREQY | 0.37644765 |
| AGGSSGIGGF | ASSLSMAGGAREQY | 0.1799928 |
| AGLTVANANG | ASSLSMAGGAREQY | 0.45636514 |
| AGLVGAVL | ASSLSMAGGAREQY | 0.3695922 |
| ALSQAFTV | ASSLSMAGGAREQY | 0.38093448 |
| ALTFSVSL | ASSLSMAGGAREQY | 0.15778029 |
| ALTGAALVAL | ASSLSMAGGAREQY | 0.35393563 |
| ASFSIGAGVL | ASSLSMAGGAREQY | 0.37774843 |
| DGSVYKI | ASSLSMAGGAREQY | 0.15797177 |
| EQLAEIV | ASSLSMAGGAREQY | 0.5871359 |
| FAVSELTVM | ASSLSMAGGAREQY | 0.29192674 |
| GAAPLAAGL | ASSLSMAGGAREQY | 0.2615723 |
| GADGGDGGSGGAGG | ASSLSMAGGAREQY | 0.58821756 |
| GFGLDTGGAGG | ASSLSMAGGAREQY | 0.33730718 |
| GLASGGLL | ASSLSMAGGAREQY | 0.13095742 |
| GRIAGAAVLL | ASSLSMAGGAREQY | 0.23909876 |
| GSAGSLNTSF | ASSLSMAGGAREQY | 0.3558796 |
| GTGQIGI | ASSLSMAGGAREQY | 0.18572597 |

| | | |
|---|---|---|
| GTLIALIL | ASSLSMAGGAREQY | 0.11048836 |
| IAATMGFVVG | ASSLSMAGGAREQY | 0.3448999 |
| IFAQALL | ASSLSMAGGAREQY | 0.1670153 |
| IGAAVALGF | ASSLSMAGGAREQY | 0.17971386 |
| IGLTPLAL | ASSLSMAGGAREQY | 0.20170122 |
| ILGSNTSTL | ASSLSMAGGAREQY | 0.6178372 |
| IVFSLTVL | ASSLSMAGGAREQY | 0.19771685 |
| LASKATGF | ASSLSMAGGAREQY | 0.28552675 |
| LASVALIL | ASSLSMAGGAREQY | 0.46360755 |
| LFAVGLTL | ASSLSMAGGAREQY | 0.47160697 |
| LFNGGIGTYI | ASSLSMAGGAREQY | 0.58283544 |
| LTLGAGII | ASSLSMAGGAREQY | 0.31782138 |
| LVAAALGDI | ASSLSMAGGAREQY | 0.27300924 |
| NTAFVQLGI | ASSLSMAGGAREQY | 0.37443665 |
| QLFSGAEL | ASSLSMAGGAREQY | 0.22614776 |
| SAATFITL | ASSLSMAGGAREQY | 0.14753899 |
| SNYGLANLGV | ASSLSMAGGAREQY | 0.15136072 |
| SYGGAYAVM | ASSLSMAGGAREQY | 0.24025986 |
| TTGVGGLTLG | ASSLSMAGGAREQY | 0.14585571 |
| TTQTIYI | ASSLSMAGGAREQY | 0.2996087 |
| VAAGAGLGAVI | ASSLSMAGGAREQY | 0.6187458 |
| VAAVVTLL | ASSLSMAGGAREQY | 0.20131974 |
| VGFLGGLI | ASSLSMAGGAREQY | 0.14455403 |
| VGVIPILL | ASSLSMAGGAREQY | 0.26681793 |
| VIFASSLI | ASSLSMAGGAREQY | 0.13817784 |
| VKGGDGGAAAG | ASSLSMAGGAREQY | 0.37575164 |
| VTKTFTNI | ASSLSMAGGAREQY | 0.3106494 |
| VVAAAILL | ASSLSMAGGAREQY | 0.84577584 |
| YAALDVEL | ASSLSMAGGAREQY | 0.68011343 |
| LASVALLL | ASSLSMAGGAREQY | 0.26192224 |
| LDAVHRV | ASSLSMAGGAREQY | 0.56817454 |
| QGLSSSIF | ASSLSMAGGAREQY | 0.22272979 |
| RGTFANIRL | ASSLSMAGGAREQY | 0.44467172 |
| RQLWWGHRI | ASSLSMAGGAREQY | 0.26695013 |
| YGPPGTGKTLL | ASSLSMAGGAREQY | 0.39155117 |
| AALVGAPANVA | ASSLSMAGGAREQY | 0.22679065 |
| AATFRRTE | ASSLSMAGGAREQY | 0.19293141 |
| AGALGGGI | ASSLSMAGGAREQY | 0.1325207 |
| AGGAGGAGGIGAG | ASSLSMAGGAREQY | 0.5239663 |
| AGSGALGL | ASSLSMAGGAREQY | 0.1856693 |
| AQIADQI | ASSLSMAGGAREQY | 0.1347195 |
| ASAVAVTYL | ASSLSMAGGAREQY | 0.2344923 |
| ASTVLGGTGG | ASSLSMAGGAREQY | 0.70384 |
| AVGGEPAK | ASSLSMAGGAREQY | 0.73074865 |
| CEQGTFGGTC | ASSLSMAGGAREQY | 0.24657917 |
| DEGKAPAG | ASSLSMAGGAREQY | 0.19302881 |
| DTVVFPAL | ASSLSMAGGAREQY | 0.7598284 |
| FEGITMAG | ASSLSMAGGAREQY | 0.17426337 |
| FPLVVQD | ASSLSMAGGAREQY | 0.38926303 |
| GAFTTPQ | ASSLSMAGGAREQY | 0.34795934 |
| GFGGFGTAGGVGGA | ASSLSMAGGAREQY | 0.25663012 |

| | | |
|---|---|---|
| GFLAQFL | ASSLSMAGGAREQY | 0.22062632 |
| GGAGGAGGNAGVA | ASSLSMAGGAREQY | 0.549208 |
| GGVGTAGAGGAGGAGG | ASSLSMAGGAREQY | 0.21516311 |
| GNGGGGDAGNAGSGGN | ASSLSMAGGAREQY | 0.37742507 |
| GTGGAGGAGNDAGS | ASSLSMAGGAREQY | 0.10733689 |
| GTTNTGWFNS | ASSLSMAGGAREQY | 0.33466554 |
| IFMGPLFTL | ASSLSMAGGAREQY | 0.20564571 |
| ILAHAIEAI | ASSLSMAGGAREQY | 0.16566864 |
| IRSAIREAL | ASSLSMAGGAREQY | 0.123581834 |
| ITFAGGG | ASSLSMAGGAREQY | 0.28721026 |
| ITGMSGHQEV | ASSLSMAGGAREQY | 0.53580225 |
| IVAGLACTL | ASSLSMAGGAREQY | 0.69972837 |
| IVGGGLAGA | ASSLSMAGGAREQY | 0.17106307 |
| IVTGVLNI | ASSLSMAGGAREQY | 0.5460491 |
| LAQQGCI | ASSLSMAGGAREQY | 0.68672645 |
| LCLHVEN | ASSLSMAGGAREQY | 0.31067836 |
| LDAISDGS | ASSLSMAGGAREQY | 0.12619072 |
| LGGFAAML | ASSLSMAGGAREQY | 0.41651428 |
| LGGKNPMI | ASSLSMAGGAREQY | 0.31920588 |
| LGPKGVAAIL | ASSLSMAGGAREQY | 0.55761075 |
| LMVGVSI | ASSLSMAGGAREQY | 0.4021912 |
| LVGAAFVL | ASSLSMAGGAREQY | 0.13780989 |
| LVLAVAPL | ASSLSMAGGAREQY | 0.63414896 |
| MAAIAVIDAA | ASSLSMAGGAREQY | 0.5403924 |
| NNGINAELQ | ASSLSMAGGAREQY | 0.1561857 |
| RGEPLVLVHG | ASSLSMAGGAREQY | 0.20298745 |
| RGGAGGAAGLI | ASSLSMAGGAREQY | 0.20646104 |
| RSYRMHPAV | ASSLSMAGGAREQY | 0.12996545 |
| SAAHLLAGQLA | ASSLSMAGGAREQY | 0.5313337 |
| SAAVGSY | ASSLSMAGGAREQY | 0.3140553 |
| SPHAVLHRI | ASSLSMAGGAREQY | 0.27573922 |
| SQIAKVL | ASSLSMAGGAREQY | 0.31851792 |
| TALGHLVP | ASSLSMAGGAREQY | 0.25255087 |
| TAPTTSGAA | ASSLSMAGGAREQY | 0.23135579 |
| TDGDDGNGGSPGT | ASSLSMAGGAREQY | 0.27000228 |
| TDGLGEN | ASSLSMAGGAREQY | 0.24298897 |
| TGGPPLPGQA | ASSLSMAGGAREQY | 0.58175945 |
| TIPAITI | ASSLSMAGGAREQY | 0.2663727 |
| TIPGFTI | ASSLSMAGGAREQY | 0.4355018 |
| TSIASGALA | ASSLSMAGGAREQY | 0.77616966 |
| TVTLPGAAI | ASSLSMAGGAREQY | 0.16061415 |
| VASGLSAALP | ASSLSMAGGAREQY | 0.19858986 |
| VGAGVIGI | ASSLSMAGGAREQY | 0.42418933 |
| VGPKRFGV | ASSLSMAGGAREQY | 0.19522455 |
| VNVAFPD | ASSLSMAGGAREQY | 0.6247328 |
| VSMTNSGVS | ASSLSMAGGAREQY | 0.43583435 |
| VTDVLGDLSV | ASSLSMAGGAREQY | 0.46138793 |
| VVVVGAGFAGL | ASSLSMAGGAREQY | 0.2934421 |
| VYTVGTVV | ASSLSMAGGAREQY | 0.8363527 |
| WTTFSGP | ASSLSMAGGAREQY | 0.5299081 |
| YDELVEAF | ASSLSMAGGAREQY | 0.29029882 |

| | | |
|---|---|---|
| YGTGGAGGNGG | ASSLSMAGGAREQY | 0.6322978 |
| YTNVMAV | ASSLSMAGGAREQY | 0.16535932 |
| FLEQVHQL | ASSLSMAGGAREQY | 0.62356097 |
| GLSLLAVAV | ASSLSMAGGAREQY | 0.5237754 |
| LEQVHQL | ASSLSMAGGAREQY | 0.15517445 |
| LNAAGAI | ASSLSMAGGAREQY | 0.2898858 |
| PAVHASL | ASSLSMAGGAREQY | 0.58906317 |
| TELEDDA | ASSLSMAGGAREQY | 0.6565503 |
| TPPGPPN | ASSLSMAGGAREQY | 0.15077195 |
| TPPPPPPP | ASSLSMAGGAREQY | 0.20642585 |
| TQSPALI | ASRRGGQNTLY | 0.16470295 |
| EFAAAFPQ | ASRRGGQNTLY | 0.18209146 |
| LLSGNLDVL | ASRRGGQNTLY | 0.14903213 |
| GPANLGNYNI | ASRRGGQNTLY | 0.36793935 |
| TLGPTLP | ASRRGGQNTLY | 0.2544749 |
| VYAYSSCCK | ASRRGGQNTLY | 0.7346718 |
| APAPAPAPNI | ASRRGGQNTLY | 0.29468676 |
| PGGPVPIP | ASRRGGQNTLY | 0.56466913 |
| FLTVALEP | ASRRGGQNTLY | 0.22797096 |
| SQYFGANL | ASRRGGQNTLY | 0.25317395 |
| GGVKPAA | ASRRGGQNTLY | 0.14221768 |
| GGTGGAGGAGGW | ASRRGGQNTLY | 0.15347336 |
| GVGGVAGLGGAGG | ASRRGGQNTLY | 0.17600754 |
| VLLVLATTI | ASRRGGQNTLY | 0.34332663 |
| IGIFAAML | ASRRGGQNTLY | 0.11910351 |
| AVGDLID | ASRRGGQNTLY | 0.68953526 |
| NISPLPL | ASRRGGQNTLY | 0.2674936 |
| VTAVFVAI | ASRRGGQNTLY | 0.38535565 |
| SGDDNDAAL | ASRRGGQNTLY | 0.28720427 |
| AVAALAHS | ASRRGGQNTLY | 0.19136713 |
| GGVTVMATP | ASRRGGQNTLY | 0.19899787 |
| AGLAANVRD | ASRRGGQNTLY | 0.48801273 |
| TNRPDLIDPALLRPG | ASRRGGQNTLY | 0.12911575 |
| LANLGVTAL | ASRRGGQNTLY | 0.14278622 |
| TGERIEL | ASRRGGQNTLY | 0.18855684 |
| NGEELAA | ASRRGGQNTLY | 0.15244307 |
| VLSLVWG | ASRRGGQNTLY | 0.77762437 |
| IGYALWL | ASRRGGQNTLY | 0.68543434 |
| LSLLLAPL | ASRRGGQNTLY | 0.42781067 |
| TLDEALK | ASRRGGQNTLY | 0.6396508 |
| TDGAGTAEGG | ASRRGGQNTLY | 0.11563748 |
| NAPGGAGGNGG | ASRRGGQNTLY | 0.26350814 |
| VSVFMGMGGNF | ASRRGGQNTLY | 0.7875353 |
| LASYLAAL | ASRRGGQNTLY | 0.30869517 |
| LVGGTPAL | ASRRGGQNTLY | 0.2655462 |
| VRNADVL | ASRRGGQNTLY | 0.55494475 |
| TCSSPFLVSAM | ASRRGGQNTLY | 0.24957842 |
| VHITDAE | ASRRGGQNTLY | 0.18417963 |
| EYAAPSF | ASRRGGQNTLY | 0.25632945 |
| VVDAFHGGL | ASRRGGQNTLY | 0.375973 |
| ASVVDPQ | ASRRGGQNTLY | 0.33834666 |

| | | |
|---|---|---|
| SYNEFGGLS | ASRRGGQNTLY | 0.13750802 |
| DPDGSVY | ASRRGGQNTLY | 0.4432777 |
| MVSTIHATE | ASRRGGQNTLY | 0.8689275 |
| LAAGVADPV | ASRRGGQNTLY | 0.27074385 |
| TKEQIAA | ASRRGGQNTLY | 0.3756143 |
| SLASPAAL | ASRRGGQNTLY | 0.38980094 |
| IGLVFYGV | ASRRGGQNTLY | 0.27865928 |
| FLLDASNL | ASRRGGQNTLY | 0.1788283 |
| YQVGNFP | ASRRGGQNTLY | 0.17668596 |
| ESLLSQST | ASRRGGQNTLY | 0.38774896 |
| LFTVGFAAVG | ASRRGGQNTLY | 0.15739119 |
| ATPPTPT | ASRRGGQNTLY | 0.2862826 |
| TDLATLG | ASRRGGQNTLY | 0.23652607 |
| SAGGTGR | ASRRGGQNTLY | 0.86128473 |
| GFSNSGS | ASRRGGQNTLY | 0.46766737 |
| GGNGGSGGTSV | ASRRGGQNTLY | 0.3533294 |
| GTSVATGGAGN | ASRRGGQNTLY | 0.21109757 |
| NGGNGGSGGTS | ASRRGGQNTLY | 0.35846317 |
| AVVKFPH | ASRRGGQNTLY | 0.5125634 |
| TPPPPPP | ASRRGGQNTLY | 0.14016171 |
| AAPAAAAAPA | ASRRGGQNTLY | 0.6077038 |
| IALNEDL | ASRRGGQNTLY | 0.19792739 |
| IVNVSSM | ASRRGGQNTLY | 0.3806325 |
| TGIPPAPR | ASRRGGQNTLY | 0.39150634 |
| FDLGGGTF | ASRRGGQNTLY | 0.8300296 |
| FDLGGGTFDVS | ASRRGGQNTLY | 0.26084393 |
| NEPTAAA | ASRRGGQNTLY | 0.42249918 |
| NEPTAAALA | ASRRGGQNTLY | 0.16967136 |
| GSIINISSI | ASRRGGQNTLY | 0.38798583 |
| VAPGNAGTA | ASRRGGQNTLY | 0.6856153 |
| TGGGASLE | ASRRGGQNTLY | 0.26102942 |
| VAVYNHYKRI | ASRRGGQNTLY | 0.7731991 |
| TDPAPATT | ASRRGGQNTLY | 0.29180783 |
| TNRPDILDPALLRPG | ASRRGGQNTLY | 0.13451193 |
| LTTTTTS | ASRRGGQNTLY | 0.2034795 |
| PTGSGKTL | ASRRGGQNTLY | 0.5157385 |
| TGSGKTL | ASRRGGQNTLY | 0.79231536 |
| TIGHVDHGK | ASRRGGQNTLY | 0.5019681 |
| TIGHVDHGKTT | ASRRGGQNTLY | 0.4919333 |
| GHVDHGKTT | ASRRGGQNTLY | 0.20869395 |
| AQFEHTLL | ASRRGGQNTLY | 0.73282415 |
| IVNQDEL | ASRRGGQNTLY | 0.8094828 |
| TNGSQFF | ASRRGGQNTLY | 0.46386015 |
| TGKGIVE | ASRRGGQNTLY | 0.18134913 |
| LISNASDA | ASRRGGQNTLY | 0.16626148 |
| AELRTLGELTKIAKAHGAQ | ASRRGGQNTLY | 0.45118064 |
| APNAVAAAEPSW | ASRRGGQNTLY | 0.5072592 |
| APQVAGAATAVL | ASRRGGQNTLY | 0.1649114 |
| ARRDPAAAAAILPTDA | ASRRGGQNTLY | 0.3555269 |
| DAAARDVLAEAGLAEYF | ASRRGGQNTLY | 0.3722758 |
| DDQIVLSAKANLAKVEQ | ASRRGGQNTLY | 0.12633984 |

| | | |
|---|---|---|
| DELLPDTGEDSDEDGH | ASRRGGQNTLY | 0.39727396 |
| ELPEAPANRATVPQD | ASRRGGQNTLY | 0.8082584 |
| FGQSKIERVTGIG | ASRRGGQNTLY | 0.3760237 |
| GAGGTGSTAGGAGGAGGAGGLY | ASRRGGQNTLY | 0.15807691 |
| GHGGTGAAGGNGGTGGAGGSGI | ASRRGGQNTLY | 0.6738943 |
| GNGGNGGAGGEGGPGGAGGAGA | ASRRGGQNTLY | 0.8388361 |
| GSTAGNGGNGGNATNGTIAGQP | ASRRGGQNTLY | 0.6938299 |
| GTGGDGGNGGNGGTGG | ASRRGGQNTLY | 0.1550023 |
| GVRRPAGIPADAPV | ASRRGGQNTLY | 0.11617275 |
| GYRQVTDAHLLTLARR | ASRRGGQNTLY | 0.2977034 |
| IAKMGQDGHDRG | ASRRGGQNTLY | 0.752261 |
| KAEPSIEIQYT | ASRRGGQNTLY | 0.29795113 |
| KPVRIRMAADEDTVAVT | ASRRGGQNTLY | 0.18778324 |
| MPPRIAGMRLLVIKPEP | ASRRGGQNTLY | 0.36869735 |
| MKFYAVALFRSRGVPPTR | ASRRGGQNTLY | 0.56493825 |
| NAGGFGILGAVAHSP | ASRRGGQNTLY | 0.15904176 |
| NEGREITLAMAEM | ASRRGGQNTLY | 0.14326558 |
| NELVEWSHPLGHRWRWE | ASRRGGQNTLY | 0.5944251 |
| NLGAGGTGGNGGNPG | ASRRGGQNTLY | 0.5345468 |
| NTEWPKWETVMAADPVD | ASRRGGQNTLY | 0.18106514 |
| PASAAPAETQP | ASRRGGQNTLY | 0.24372841 |
| QQSSDGKNGVRAAEITGE | ASRRGGQNTLY | 0.36486208 |
| RAPDAVTTSWWKE | ASRRGGQNTLY | 0.5012368 |
| SGFGNVGAHTSGWFNQ | ASRRGGQNTLY | 0.41851342 |
| SGVAGAVGQAASVGGLKV | ASRRGGQNTLY | 0.15053734 |
| SNGGAGTGSAGGNGGT | ASRRGGQNTLY | 0.61280596 |
| TGGAGGEGGAAG | ASRRGGQNTLY | 0.16343167 |
| TGGDGGAAGAGGAPGHG | ASRRGGQNTLY | 0.3870423 |
| TMPPLLAAIANAGRHGV | ASRRGGQNTLY | 0.43892735 |
| TSSTAGNSGGA | ASRRGGQNTLY | 0.3697808 |
| VPATVPVGILC | ASRRGGQNTLY | 0.23712213 |
| VQRPDSGGVVLTGRIS | ASRRGGQNTLY | 0.19237538 |
| WTPNPVPATKGVPAGLVQAT | ASRRGGQNTLY | 0.13660495 |
| YALKRALAKFSDRFDVVII | ASRRGGQNTLY | 0.13373058 |
| YLIGSVIV | ASRRGGQNTLY | 0.6770769 |
| YGQLVAVID | ASRRGGQNTLY | 0.11414766 |
| VTVGGMTLPA | ASRRGGQNTLY | 0.5542425 |
| VLALPAV | ASRRGGQNTLY | 0.32626265 |
| VIPPTLNYET | ASRRGGQNTLY | 0.16411093 |
| VGMGGMHPSSG | ASRRGGQNTLY | 0.34549153 |
| VGLAGGAATA | ASRRGGQNTLY | 0.14740774 |
| VGGAGGAGGSGP | ASRRGGQNTLY | 0.1772369 |
| VFAAFMV | ASRRGGQNTLY | 0.30156636 |
| TLYDKLR | ASRRGGQNTLY | 0.37860012 |
| TLAVFTVL | ASRRGGQNTLY | 0.19408965 |
| TGSVQPARV | ASRRGGQNTLY | 0.38158342 |
| TGGGAGSGGT | ASRRGGQNTLY | 0.39812872 |
| SHLETSL | ASRRGGQNTLY | 0.16265306 |
| SFTTGMT | ASRRGGQNTLY | 0.43349126 |
| QVEVAKRMGVRQPTVS | ASRRGGQNTLY | 0.6718241 |
| QGSGGPALAM | ASRRGGQNTLY | 0.39048856 |

| | | |
|---|---|---|
| MLAPSNP | ASRRGGQNTLY | 0.13215104 |
| LVNNAAI | ASRRGGQNTLY | 0.51097405 |
| LTGEINTLA | ASRRGGQNTLY | 0.17784892 |
| LSIFGPV | ASRRGGQNTLY | 0.42141604 |
| LLALLANL | ASRRGGQNTLY | 0.25542805 |
| LISNASDALDKLR | ASRRGGQNTLY | 0.23874885 |
| LINEAVP | ASRRGGQNTLY | 0.4745072 |
| LGYAFSGL | ASRRGGQNTLY | 0.16503684 |
| LDFVAVAL | ASRRGGQNTLY | 0.22937709 |
| LAQPGAE | ASRRGGQNTLY | 0.21329176 |
| ISNLGAVTGF | ASRRGGQNTLY | 0.39450753 |
| HQEDPGRI | ASRRGGQNTLY | 0.46207193 |
| GTPGEPTP | ASRRGGQNTLY | 0.17585874 |
| GFFNSGS | ASRRGGQNTLY | 0.3492198 |
| GAGGGGLTAGAN | ASRRGGQNTLY | 0.20637968 |
| FVDALVNM | ASRRGGQNTLY | 0.16674347 |
| FLTVAPEL | ASRRGGQNTLY | 0.33437032 |
| FETLTAL | ASRRGGQNTLY | 0.23209415 |
| DVINPIEL | ASRRGGQNTLY | 0.16661428 |
| APVSGAPV | ASRRGGQNTLY | 0.4119202 |
| ALVGGLAW | ASRRGGQNTLY | 0.19641621 |
| ALAAAAATAI | ASRRGGQNTLY | 0.4144633 |
| AFGLGLLL | ASRRGGQNTLY | 0.83278656 |
| AFFAAASL | ASRRGGQNTLY | 0.1584813 |
| AAAALPAFSPP | ASRRGGQNTLY | 0.33718592 |
| VVLAPAL | ASRRGGQNTLY | 0.30157036 |
| TIGHVDHGKTTL | ASRRGGQNTLY | 0.404768 |
| LISNASDALDK | ASRRGGQNTLY | 0.29908338 |
| LISNASDAL | ASRRGGQNTLY | 0.15710644 |
| ISNASDALDK | ASRRGGQNTLY | 0.5586206 |
| NQAETLVYQTE | ASRRGGQNTLY | 0.52460665 |
| TGGTGGTAGAG | ASRRGGQNTLY | 0.50124615 |
| TGSTAGGAGGA | ASRRGGQNTLY | 0.46289933 |
| DPLAVPVAAGPV | ASRRGGQNTLY | 0.623057 |
| GGTGGTPGNGGN | ASRRGGQNTLY | 0.5076395 |
| PAGLAGVDPATL | ASRRGGQNTLY | 0.26180783 |
| GNNLVVEAPHAGASV | ASRRGGQNTLY | 0.1398942 |
| KVTRIMSASTLEQPA | ASRRGGQNTLY | 0.26484627 |
| LGFALPIALPRRTVV | ASRRGGQNTLY | 0.2803491 |
| TRLGNLTARKLVVLV | ASRRGGQNTLY | 0.20252791 |
| WSQQRRLIYFPSAGP | ASRRGGQNTLY | 0.13407713 |
| GPNGNPDPMAAAVDIR | ASRRGGQNTLY | 0.65318346 |
| TQRCVSVADLRAAASR | ASRRGGQNTLY | 0.8210105 |
| QFVLDVAEHPVEVLTETV | ASRRGGQNTLY | 0.27395263 |
| GGAGGTGGAGGNGGTQAGDG | ASRRGGQNTLY | 0.81936204 |
| PLLATLRAIAVDVDAKPAQV | ASRRGGQNTLY | 0.15039003 |
| APKFRTGGYMIDFWGVGYQVAK | ASRRGGQNTLY | 0.42049074 |
| GAGGAGGAGGAAGGPGGTGGTG | ASRRGGQNTLY | 0.67066693 |
| IGALGLLATDVAPAGSVAFGVA | ASRRGGQNTLY | 0.177662 |
| AGGAGGIGGNGGAGGAGGVLMG | ASRRGGQNTLY | 0.39141545 |
| ADELIGLL | ASRRGGQNTLY | 0.14086461 |

| | | |
|---|---|---|
| ADRALKA | ASRRGGQNTLY | 0.15857787 |
| AGAVGQVGGAGGA | ASRRGGQNTLY | 0.66703093 |
| AGGGGGLGGAP | ASRRGGQNTLY | 0.28563324 |
| AGGLLYGNGGAG | ASRRGGQNTLY | 0.4052853 |
| AGLESIGL | ASRRGGQNTLY | 0.22932465 |
| AGLKTGII | ASRRGGQNTLY | 0.17389333 |
| ASLAALLL | ASRRGGQNTLY | 0.2619417 |
| ASVINISSI | ASRRGGQNTLY | 0.6507858 |
| ATPPGPA | ASRRGGQNTLY | 0.17697638 |
| ATVASALV | ASRRGGQNTLY | 0.82644284 |
| ATVASLIL | ASRRGGQNTLY | 0.19514906 |
| AVNGPVLIRH | ASRRGGQNTLY | 0.7602978 |
| DGAAGAVGI | ASRRGGQNTLY | 0.79391 |
| DPLLHGI | ASRRGGQNTLY | 0.4629717 |
| ELGAALGAL | ASRRGGQNTLY | 0.21603328 |
| ELGRTQAV | ASRRGGQNTLY | 0.74962926 |
| FVAGGYLS | ASRRGGQNTLY | 0.24794322 |
| GASGGAGGAGGW | ASRRGGQNTLY | 0.39711377 |
| GGAGAGGDGGAGGI | ASRRGGQNTLY | 0.6125162 |
| GTGHNGSQPGMGGNG | ASRRGGQNTLY | 0.14906655 |
| GYAAASASAS | ASRRGGQNTLY | 0.44446468 |
| HQVDVAAVQQG | ASRRGGQNTLY | 0.30835983 |
| IGAAIAHT | ASRRGGQNTLY | 0.34311867 |
| IGMIEVAQL | ASRRGGQNTLY | 0.1918122 |
| IRLAPEHTSAMD | ASRRGGQNTLY | 0.16463451 |
| IYNVSVL | ASRRGGQNTLY | 0.24228163 |
| KQGFLGM | ASRRGGQNTLY | 0.17335802 |
| LESVITQL | ASRRGGQNTLY | 0.65052927 |
| LGLGALVVV | ASRRGGQNTLY | 0.3607072 |
| LIGNGGNGGSGGI | ASRRGGQNTLY | 0.18156737 |
| LNAINAPTL | ASRRGGQNTLY | 0.38039237 |
| LQLLNTGV | ASRRGGQNTLY | 0.3935859 |
| LQTSIEF | ASRRGGQNTLY | 0.14391074 |
| MSFAVSEL | ASRRGGQNTLY | 0.2224408 |
| NTGALGL | ASRRGGQNTLY | 0.28957382 |
| PPGAGGAGGAGTT | ASRRGGQNTLY | 0.23718733 |
| QLVGLGI | ASRRGGQNTLY | 0.11551978 |
| RFLEQVHQL | ASRRGGQNTLY | 0.3107755 |
| RGVGYVLR | ASRRGGQNTLY | 0.17306922 |
| SATGVFAGVI | ASRRGGQNTLY | 0.09928226 |
| SLPAITLP | ASRRGGQNTLY | 0.309573 |
| STALAFTV | ASRRGGQNTLY | 0.18244357 |
| SVLTVGLGK | ASRRGGQNTLY | 0.23197591 |
| TALAGAGSGI | ASRRGGQNTLY | 0.38289845 |
| TGLPFTGI | ASRRGGQNTLY | 0.3185635 |
| VAEHVTV | ASRRGGQNTLY | 0.1949842 |
| VDIVLNSL | ASRRGGQNTLY | 0.29451954 |
| VLTGGGAVL | ASRRGGQNTLY | 0.4243294 |
| VNVGLAYGI | ASRRGGQNTLY | 0.17112301 |
| VSSAAVGAV | ASRRGGQNTLY | 0.22928508 |
| VTSVYEAGV | ASRRGGQNTLY | 0.5057357 |

| | | |
|---|---|---|
| WVALDRGA | ASRRGGQNTLY | 0.3135535 |
| YLVGSVVI | ASRRGGQNTLY | 0.50639546 |
| GPPGTGKTLL | ASRRGGQNTLY | 0.50430393 |
| GVGMTKF | ASRRGGQNTLY | 0.20988186 |
| IGIGGSDL | ASRRGGQNTLY | 0.3990114 |
| IGTLLAEL | ASRRGGQNTLY | 0.5287881 |
| IYTSGTT | ASRRGGQNTLY | 0.25748628 |
| LSFPAGV | ASRRGGQNTLY | 0.21877828 |
| LYTSGTT | ASRRGGQNTLY | 0.14824806 |
| VGAFPFGF | ASRRGGQNTLY | 0.21896482 |
| VGAGTVEF | ASRRGGQNTLY | 0.17739727 |
| VGKSTLLL | ASRRGGQNTLY | 0.28964716 |
| AALAHAGGGIV | ASRRGGQNTLY | 0.46584243 |
| AALVAVSLL | ASRRGGQNTLY | 0.21544985 |
| AAVSRAVAAL | ASRRGGQNTLY | 0.11520561 |
| AGGSSGIGGF | ASRRGGQNTLY | 0.113895155 |
| AGLTVANANG | ASRRGGQNTLY | 0.31348062 |
| AGLVGAVL | ASRRGGQNTLY | 0.15666525 |
| ALSQAFTV | ASRRGGQNTLY | 0.195029 |
| ALTFSVSL | ASRRGGQNTLY | 0.2048936 |
| ALTGAALVAL | ASRRGGQNTLY | 0.45368242 |
| ASFSIGAGVL | ASRRGGQNTLY | 0.44359058 |
| DGSVYKI | ASRRGGQNTLY | 0.17405963 |
| EQLAEIV | ASRRGGQNTLY | 0.45679384 |
| FAVSELTVM | ASRRGGQNTLY | 0.58341515 |
| GAAPLAAGL | ASRRGGQNTLY | 0.4479399 |
| GADGGDGGSGGAGG | ASRRGGQNTLY | 0.25626007 |
| GFGLDTGGAGG | ASRRGGQNTLY | 0.1555324 |
| GLASGGLL | ASRRGGQNTLY | 0.54954964 |
| GRIAGAAVLL | ASRRGGQNTLY | 0.18171732 |
| GSAGSLNTSF | ASRRGGQNTLY | 0.42558885 |
| GTGQIGI | ASRRGGQNTLY | 0.3739319 |
| GTLIALIL | ASRRGGQNTLY | 0.29270604 |
| IAATMGFVVG | ASRRGGQNTLY | 0.1574527 |
| IFAQALL | ASRRGGQNTLY | 0.39274633 |
| IGAAVALGF | ASRRGGQNTLY | 0.4643226 |
| IGLTPLAL | ASRRGGQNTLY | 0.1200557 |
| ILGSNTSTL | ASRRGGQNTLY | 0.18491574 |
| IVFSLTVL | ASRRGGQNTLY | 0.24167728 |
| LASKATGF | ASRRGGQNTLY | 0.21573052 |
| LASVALIL | ASRRGGQNTLY | 0.45078737 |
| LFAVGLTL | ASRRGGQNTLY | 0.34204903 |
| LFNGGIGTYI | ASRRGGQNTLY | 0.22966805 |
| LTLGAGII | ASRRGGQNTLY | 0.32719186 |
| LVAAALGDI | ASRRGGQNTLY | 0.3670593 |
| NTAFVQLGI | ASRRGGQNTLY | 0.26627302 |
| QLFSGAEL | ASRRGGQNTLY | 0.3155833 |
| SAATFITL | ASRRGGQNTLY | 0.1539519 |
| SNYGLANLGV | ASRRGGQNTLY | 0.44502202 |
| SYGGAYAVM | ASRRGGQNTLY | 0.40529263 |
| TTGVGGLTLG | ASRRGGQNTLY | 0.3587518 |

| | | |
|---|---|---|
| TTQTIYI | ASRRGGQNTLY | 0.44413018 |
| VAAGAGLGAVI | ASRRGGQNTLY | 0.2911052 |
| VAAVVTLL | ASRRGGQNTLY | 0.26558784 |
| VGFLGGLI | ASRRGGQNTLY | 0.16311604 |
| VGVIPILL | ASRRGGQNTLY | 0.23251821 |
| VIFASSLI | ASRRGGQNTLY | 0.21203387 |
| VKGGDGGAAAG | ASRRGGQNTLY | 0.27460632 |
| VTKTFTNI | ASRRGGQNTLY | 0.3526805 |
| VVAAAILL | ASRRGGQNTLY | 0.36433142 |
| YAALDVEL | ASRRGGQNTLY | 0.15017113 |
| LASVALLL | ASRRGGQNTLY | 0.42320928 |
| LDAVHRV | ASRRGGQNTLY | 0.43121594 |
| QGLSSSIF | ASRRGGQNTLY | 0.14691427 |
| RGTFANIRL | ASRRGGQNTLY | 0.21660545 |
| RQLWWGHRI | ASRRGGQNTLY | 0.50912684 |
| YGPPGTGKTLL | ASRRGGQNTLY | 0.43243966 |
| AALVGAPANVA | ASRRGGQNTLY | 0.20016214 |
| AATFRRTE | ASRRGGQNTLY | 0.28983414 |
| AGALGGGI | ASRRGGQNTLY | 0.6599489 |
| AGGAGGAGGIGAG | ASRRGGQNTLY | 0.34009448 |
| AGSGALGL | ASRRGGQNTLY | 0.46015614 |
| AQIADQI | ASRRGGQNTLY | 0.13256605 |
| ASAVAVTYL | ASRRGGQNTLY | 0.25991416 |
| ASTVLGGTGG | ASRRGGQNTLY | 0.21374974 |
| AVGGEPAK | ASRRGGQNTLY | 0.09695042 |
| CEQGTFGGTC | ASRRGGQNTLY | 0.4613701 |
| DEGKAPAG | ASRRGGQNTLY | 0.210678 |
| DTVVFPAL | ASRRGGQNTLY | 0.3481087 |
| FEGITMAG | ASRRGGQNTLY | 0.19720691 |
| FPLVVQD | ASRRGGQNTLY | 0.43373588 |
| GAFTTPQ | ASRRGGQNTLY | 0.262688 |
| GFGGFGTAGGVGGA | ASRRGGQNTLY | 0.2711282 |
| GFLAQFL | ASRRGGQNTLY | 0.23115674 |
| GGAGGAGGNAGVA | ASRRGGQNTLY | 0.358336 |
| GGVGTAGAGGAGGAGG | ASRRGGQNTLY | 0.42855927 |
| GNGGGGDAGNAGSGGN | ASRRGGQNTLY | 0.49613893 |
| GTGGAGGAGNDAGS | ASRRGGQNTLY | 0.14076509 |
| GTTNTGWFNS | ASRRGGQNTLY | 0.4657455 |
| IFMGPLFTL | ASRRGGQNTLY | 0.25125766 |
| ILAHAIEAI | ASRRGGQNTLY | 0.19978979 |
| IRSAIREAL | ASRRGGQNTLY | 0.14652745 |
| ITFAGGG | ASRRGGQNTLY | 0.19033682 |
| ITGMSGHQEV | ASRRGGQNTLY | 0.20491442 |
| IVAGLACTL | ASRRGGQNTLY | 0.2872416 |
| IVGGGLAGA | ASRRGGQNTLY | 0.40982383 |
| IVTGVLNI | ASRRGGQNTLY | 0.18670343 |
| LAQQGCI | ASRRGGQNTLY | 0.3066953 |
| LCLHVEN | ASRRGGQNTLY | 0.24382147 |
| LDAISDGS | ASRRGGQNTLY | 0.07267281 |
| LGGFAAML | ASRRGGQNTLY | 0.20456356 |
| LGGKNPMI | ASRRGGQNTLY | 0.13384445 |

| | | |
|---|---|---|
| LGPKGVAAIL | ASRRGGQNTLY | 0.45058212 |
| LMVGVSI | ASRRGGQNTLY | 0.16239163 |
| LVGAAFVL | ASRRGGQNTLY | 0.19293886 |
| LVLAVAPL | ASRRGGQNTLY | 0.21246551 |
| MAAIAVIDAA | ASRRGGQNTLY | 0.47042042 |
| NNGINAELQ | ASRRGGQNTLY | 0.2353452 |
| RGEPLVLVHG | ASRRGGQNTLY | 0.19104218 |
| RGGAGGAAGLI | ASRRGGQNTLY | 0.16949576 |
| RSYRMHPAV | ASRRGGQNTLY | 0.3678143 |
| SAAHLLAGQLA | ASRRGGQNTLY | 0.37352446 |
| SAAVGSY | ASRRGGQNTLY | 0.18498069 |
| SPHAVLHRI | ASRRGGQNTLY | 0.19864008 |
| SQIAKVL | ASRRGGQNTLY | 0.5805038 |
| TALGHLVP | ASRRGGQNTLY | 0.12654427 |
| TAPTTSGAA | ASRRGGQNTLY | 0.43607086 |
| TDGDDGNGGSPGT | ASRRGGQNTLY | 0.1582671 |
| TDGLGEN | ASRRGGQNTLY | 0.18065867 |
| TGGPPLPGQA | ASRRGGQNTLY | 0.5213889 |
| TIPAITI | ASRRGGQNTLY | 0.47273433 |
| TIPGFTI | ASRRGGQNTLY | 0.20440388 |
| TSIASGALA | ASRRGGQNTLY | 0.21460675 |
| TVTLPGAAI | ASRRGGQNTLY | 0.15613253 |
| VASGLSAALP | ASRRGGQNTLY | 0.17298943 |
| VGAGVIGI | ASRRGGQNTLY | 0.19386965 |
| VGPKRFGV | ASRRGGQNTLY | 0.42414227 |
| VNVAFPD | ASRRGGQNTLY | 0.3133891 |
| VSMTNSGVS | ASRRGGQNTLY | 0.16646673 |
| VTDVLGDLSV | ASRRGGQNTLY | 0.4083509 |
| VVVVGAGFAGL | ASRRGGQNTLY | 0.4066151 |
| VYTVGTVV | ASRRGGQNTLY | 0.54714096 |
| WTTFSGP | ASRRGGQNTLY | 0.16923037 |
| YDELVEAF | ASRRGGQNTLY | 0.22351123 |
| YGTGGAGGNGG | ASRRGGQNTLY | 0.23183368 |
| YTNVMAV | ASRRGGQNTLY | 0.18621634 |
| FLEQVHQL | ASRRGGQNTLY | 0.2152769 |
| GLSLLAVAV | ASRRGGQNTLY | 0.19292888 |
| LEQVHQL | ASRRGGQNTLY | 0.4345648 |
| LNAAGAI | ASRRGGQNTLY | 0.76548374 |
| PAVHASL | ASRRGGQNTLY | 0.21956654 |
| TELEDDA | ASRRGGQNTLY | 0.12573488 |
| TPPGPPN | ASRRGGQNTLY | 0.19337983 |
| TPPPPPPP | ASRRGGQNTLY | 0.13575907 |
| TQSPALI | ASSPGQPSQNTLY | 0.14264186 |
| EFAAAFPQ | ASSPGQPSQNTLY | 0.2327587 |
| LLSGNLDVL | ASSPGQPSQNTLY | 0.115579784 |
| GPANLGNYNI | ASSPGQPSQNTLY | 0.21076314 |
| TLGPTLP | ASSPGQPSQNTLY | 0.54003984 |
| VYAYSSCCK | ASSPGQPSQNTLY | 0.2216671 |
| APAPAPAPNI | ASSPGQPSQNTLY | 0.3077963 |
| PGGPVPIP | ASSPGQPSQNTLY | 0.46172458 |
| FLTVALEP | ASSPGQPSQNTLY | 0.29626766 |

| | | |
|---|---|---|
| SQYFGANL | ASSPGQPSQNTLY | 0.48422244 |
| GGVKPAA | ASSPGQPSQNTLY | 0.59126556 |
| GGTGGAGGAGGW | ASSPGQPSQNTLY | 0.30982238 |
| GVGGVAGLGGAGG | ASSPGQPSQNTLY | 0.24548155 |
| VLLVLATTI | ASSPGQPSQNTLY | 0.46449405 |
| IGIFAAML | ASSPGQPSQNTLY | 0.18656218 |
| AVGDLID | ASSPGQPSQNTLY | 0.24675564 |
| NISPLPL | ASSPGQPSQNTLY | 0.24537164 |
| VTAVFVAI | ASSPGQPSQNTLY | 0.30909213 |
| SGDDNDAAL | ASSPGQPSQNTLY | 0.24638943 |
| AVAALAHS | ASSPGQPSQNTLY | 0.6123395 |
| GGVTVMATP | ASSPGQPSQNTLY | 0.11092432 |
| AGLAANVRD | ASSPGQPSQNTLY | 0.17844446 |
| TNRPDLIDPALLRPG | ASSPGQPSQNTLY | 0.45492163 |
| LANLGVTAL | ASSPGQPSQNTLY | 0.0983417 |
| TGERIEL | ASSPGQPSQNTLY | 0.19105013 |
| NGEELAA | ASSPGQPSQNTLY | 0.2648772 |
| VLSLVWG | ASSPGQPSQNTLY | 0.47971344 |
| IGYALWL | ASSPGQPSQNTLY | 0.11290165 |
| LSLLLAPL | ASSPGQPSQNTLY | 0.23838927 |
| TLDEALK | ASSPGQPSQNTLY | 0.1449434 |
| TDGAGTAEGG | ASSPGQPSQNTLY | 0.45735335 |
| NAPGGAGGNGG | ASSPGQPSQNTLY | 0.23363496 |
| VSVFMGMGGNF | ASSPGQPSQNTLY | 0.13962948 |
| LASYLAAL | ASSPGQPSQNTLY | 0.41378152 |
| LVGGTPAL | ASSPGQPSQNTLY | 0.14688236 |
| VRNADVL | ASSPGQPSQNTLY | 0.44347453 |
| TCSSPFLVSAM | ASSPGQPSQNTLY | 0.41664308 |
| VHITDAE | ASSPGQPSQNTLY | 0.7033479 |
| EYAAPSF | ASSPGQPSQNTLY | 0.13957994 |
| VVDAFHGGL | ASSPGQPSQNTLY | 0.32156008 |
| ASVVDPQ | ASSPGQPSQNTLY | 0.16087697 |
| SYNEFGGLS | ASSPGQPSQNTLY | 0.69001937 |
| DPDGSVY | ASSPGQPSQNTLY | 0.20748667 |
| MVSTIHATE | ASSPGQPSQNTLY | 0.14379801 |
| LAAGVADPV | ASSPGQPSQNTLY | 0.35795438 |
| TKEQIAA | ASSPGQPSQNTLY | 0.36688718 |
| SLASPAAL | ASSPGQPSQNTLY | 0.2743098 |
| IGLVFYGV | ASSPGQPSQNTLY | 0.5164819 |
| FLLDASNL | ASSPGQPSQNTLY | 0.28387356 |
| YQVGNFP | ASSPGQPSQNTLY | 0.17348531 |
| ESLLSQST | ASSPGQPSQNTLY | 0.39014694 |
| LFTVGFAAVG | ASSPGQPSQNTLY | 0.2997673 |
| ATPPTPT | ASSPGQPSQNTLY | 0.2355484 |
| TDLATLG | ASSPGQPSQNTLY | 0.46008497 |
| SAGGTGR | ASSPGQPSQNTLY | 0.25746655 |
| GFSNSGS | ASSPGQPSQNTLY | 0.17056839 |
| GGNGGSGGTSV | ASSPGQPSQNTLY | 0.60952675 |
| GTSVATGGAGN | ASSPGQPSQNTLY | 0.28725997 |
| NGGNGGSGGTS | ASSPGQPSQNTLY | 0.32215023 |
| AVVKFPH | ASSPGQPSQNTLY | 0.32980368 |

| | | |
|---|---|---|
| TPPPPPP | ASSPGQPSQNTLY | 0.72405803 |
| AAPAAAAAPA | ASSPGQPSQNTLY | 0.2065304 |
| IALNEDL | ASSPGQPSQNTLY | 0.29779762 |
| IVNVSSM | ASSPGQPSQNTLY | 0.24355164 |
| TGIPPAPR | ASSPGQPSQNTLY | 0.3976684 |
| FDLGGGTF | ASSPGQPSQNTLY | 0.25038588 |
| FDLGGGTFDVS | ASSPGQPSQNTLY | 0.47146058 |
| NEPTAAA | ASSPGQPSQNTLY | 0.32321805 |
| NEPTAAALA | ASSPGQPSQNTLY | 0.4285547 |
| GSIINISSI | ASSPGQPSQNTLY | 0.26944166 |
| VAPGNAGTA | ASSPGQPSQNTLY | 0.28142837 |
| TGGGASLE | ASSPGQPSQNTLY | 0.57534003 |
| VAVYNHYKRI | ASSPGQPSQNTLY | 0.17728928 |
| TDPAPATT | ASSPGQPSQNTLY | 0.26794767 |
| TNRPDILDPALLRPG | ASSPGQPSQNTLY | 0.40162718 |
| LTTTTTS | ASSPGQPSQNTLY | 0.41106835 |
| PTGSGKTL | ASSPGQPSQNTLY | 0.2668957 |
| TGSGKTL | ASSPGQPSQNTLY | 0.464973 |
| TIGHVDHGK | ASSPGQPSQNTLY | 0.20933685 |
| TIGHVDHGKTT | ASSPGQPSQNTLY | 0.32451287 |
| GHVDHGKTT | ASSPGQPSQNTLY | 0.18310177 |
| AQFEHTLL | ASSPGQPSQNTLY | 0.20849153 |
| IVNQDEL | ASSPGQPSQNTLY | 0.14992318 |
| TNGSQFF | ASSPGQPSQNTLY | 0.24344851 |
| TGKGIVE | ASSPGQPSQNTLY | 0.34551498 |
| LISNASDA | ASSPGQPSQNTLY | 0.31211373 |
| AELRTLGELTKIAKAHGAQ | ASSPGQPSQNTLY | 0.14124946 |
| APNAVAAAEPSW | ASSPGQPSQNTLY | 0.33166617 |
| APQVAGAATAVL | ASSPGQPSQNTLY | 0.38701892 |
| ARRDPAAAAAILPTDA | ASSPGQPSQNTLY | 0.69761175 |
| DAAARDVLAEAGLAEYF | ASSPGQPSQNTLY | 0.24219517 |
| DDQIVLSAKANLAKVEQ | ASSPGQPSQNTLY | 0.3735325 |
| DELLPDTGEDSDEDGH | ASSPGQPSQNTLY | 0.26136702 |
| ELPEAPANRATVPQD | ASSPGQPSQNTLY | 0.31047672 |
| FGQSKIERVTGIG | ASSPGQPSQNTLY | 0.10544223 |
| GAGGTGSTAGGAGGAGGAGGLY | ASSPGQPSQNTLY | 0.1859549 |
| GHGGTGAAGGNGGTGGAGGSGI | ASSPGQPSQNTLY | 0.21335958 |
| GNGGNGGAGGEGGPGGAGGAGA | ASSPGQPSQNTLY | 0.15381509 |
| GSTAGNGGNGGNATNGTIAGQP | ASSPGQPSQNTLY | 0.5328001 |
| GTGGDGGNGGNGGTGG | ASSPGQPSQNTLY | 0.43279746 |
| GVRRPAGIPADAPV | ASSPGQPSQNTLY | 0.4072241 |
| GYRQVTDAHLLTLARR | ASSPGQPSQNTLY | 0.42011118 |
| IAKMGQDGHDRG | ASSPGQPSQNTLY | 0.45286474 |
| KAEPSIEIQYT | ASSPGQPSQNTLY | 0.4271193 |
| KPVRIRMAADEDTVAVT | ASSPGQPSQNTLY | 0.18862621 |
| MPPRIAGMRLLVIKPEP | ASSPGQPSQNTLY | 0.23514816 |
| MKFYAVALFRSRGVPPTR | ASSPGQPSQNTLY | 0.11962411 |
| NAGGFGILGAVAHSP | ASSPGQPSQNTLY | 0.17891222 |
| NEGREITLAMAEM | ASSPGQPSQNTLY | 0.19161539 |
| NELVEWSHPLGHRWRWE | ASSPGQPSQNTLY | 0.23785105 |
| NLGAGGTGGNGGNPG | ASSPGQPSQNTLY | 0.101403095 |

| | | |
|---|---|---|
| NTEWPKWETVMAADPVD | ASSPGQPSQNTLY | 0.4361822 |
| PASAAPAETQP | ASSPGQPSQNTLY | 0.43831977 |
| QQSSDGKNGVRAAEITGE | ASSPGQPSQNTLY | 0.3421296 |
| RAPDAVTTSWWKE | ASSPGQPSQNTLY | 0.31575242 |
| SGFGNVGAHTSGWFNQ | ASSPGQPSQNTLY | 0.29874006 |
| SGVAGAVGQAASVGGLKV | ASSPGQPSQNTLY | 0.16189818 |
| SNGGAGTGSAGGNGGT | ASSPGQPSQNTLY | 0.30506214 |
| TGGAGGEGGAAG | ASSPGQPSQNTLY | 0.18043181 |
| TGGDGGAAGAGGAPGHG | ASSPGQPSQNTLY | 0.46404803 |
| TMPPLLAAIANAGRHGV | ASSPGQPSQNTLY | 0.49078387 |
| TSSTAGNSGGA | ASSPGQPSQNTLY | 0.26998514 |
| VPATVPVGILC | ASSPGQPSQNTLY | 0.20432277 |
| VQRPDSGGVVLTGRIS | ASSPGQPSQNTLY | 0.32563776 |
| WTPNPVPATKGVPAGLVQAT | ASSPGQPSQNTLY | 0.20798188 |
| YALKRALAKFSDRFDVVII | ASSPGQPSQNTLY | 0.21921456 |
| YLIGSVIV | ASSPGQPSQNTLY | 0.3157988 |
| YGQLVAVID | ASSPGQPSQNTLY | 0.23613676 |
| VTVGGMTLPA | ASSPGQPSQNTLY | 0.15840977 |
| VLALPAV | ASSPGQPSQNTLY | 0.30558264 |
| VIPPTLNYET | ASSPGQPSQNTLY | 0.256449 |
| VGMGGMHPSSG | ASSPGQPSQNTLY | 0.2997939 |
| VGLAGGAATA | ASSPGQPSQNTLY | 0.13943389 |
| VGGAGGAGGSGP | ASSPGQPSQNTLY | 0.45619962 |
| VFAAFMV | ASSPGQPSQNTLY | 0.38303208 |
| TLYDKLR | ASSPGQPSQNTLY | 0.14594021 |
| TLAVFTVL | ASSPGQPSQNTLY | 0.3583964 |
| TGSVQPARV | ASSPGQPSQNTLY | 0.26771516 |
| TGGGAGSGGT | ASSPGQPSQNTLY | 0.2132372 |
| SHLETSL | ASSPGQPSQNTLY | 0.43945375 |
| SFTTGMT | ASSPGQPSQNTLY | 0.2538428 |
| QVEVAKRMGVRQPTVS | ASSPGQPSQNTLY | 0.1830105 |
| QGSGGPALAM | ASSPGQPSQNTLY | 0.16969024 |
| MLAPSNP | ASSPGQPSQNTLY | 0.23276281 |
| LVNNAAI | ASSPGQPSQNTLY | 0.21511346 |
| LTGEINTLA | ASSPGQPSQNTLY | 0.13435155 |
| LSIFGPV | ASSPGQPSQNTLY | 0.34417802 |
| LLALLANL | ASSPGQPSQNTLY | 0.17147398 |
| LISNASDALDKLR | ASSPGQPSQNTLY | 0.15471078 |
| LINEAVP | ASSPGQPSQNTLY | 0.20290038 |
| LGYAFSGL | ASSPGQPSQNTLY | 0.40431726 |
| LDFVAVAL | ASSPGQPSQNTLY | 0.51991117 |
| LAQPGAE | ASSPGQPSQNTLY | 0.32513756 |
| ISNLGAVTGF | ASSPGQPSQNTLY | 0.2894319 |
| HQEDPGRI | ASSPGQPSQNTLY | 0.3058206 |
| GTPGEPTP | ASSPGQPSQNTLY | 0.39200157 |
| GFFNSGS | ASSPGQPSQNTLY | 0.33769488 |
| GAGGGGLTAGAN | ASSPGQPSQNTLY | 0.27804112 |
| FVDALVNM | ASSPGQPSQNTLY | 0.32565936 |
| FLTVAPEL | ASSPGQPSQNTLY | 0.24871722 |
| FETLTAL | ASSPGQPSQNTLY | 0.21308218 |
| DVINPIEL | ASSPGQPSQNTLY | 0.19523332 |

| | | |
|---|---|---|
| APVSGAPV | ASSPGQPSQNTLY | 0.16899906 |
| ALVGGLAW | ASSPGQPSQNTLY | 0.19232628 |
| ALAAAAATAI | ASSPGQPSQNTLY | 0.2964735 |
| AFGLGLLL | ASSPGQPSQNTLY | 0.4860945 |
| AFFAAASL | ASSPGQPSQNTLY | 0.25410184 |
| AAAALPAFSPP | ASSPGQPSQNTLY | 0.13204654 |
| VVLAPAL | ASSPGQPSQNTLY | 0.18681498 |
| TIGHVDHGKTTL | ASSPGQPSQNTLY | 0.2748026 |
| LISNASDALDK | ASSPGQPSQNTLY | 0.3977139 |
| LISNASDAL | ASSPGQPSQNTLY | 0.38981646 |
| ISNASDALDK | ASSPGQPSQNTLY | 0.70885456 |
| NQAETLVYQTE | ASSPGQPSQNTLY | 0.3950022 |
| TGGTGGTAGAG | ASSPGQPSQNTLY | 0.17712767 |
| TGSTAGGAGGA | ASSPGQPSQNTLY | 0.34996846 |
| DPLAVPVAAGPV | ASSPGQPSQNTLY | 0.19126974 |
| GGTGGTPGNGGN | ASSPGQPSQNTLY | 0.28228787 |
| PAGLAGVDPATL | ASSPGQPSQNTLY | 0.26340023 |
| GNNLVVEAPHAGASV | ASSPGQPSQNTLY | 0.3270237 |
| KVTRIMSASTLEQPA | ASSPGQPSQNTLY | 0.21533665 |
| LGFALPIALPRRTVV | ASSPGQPSQNTLY | 0.26188794 |
| TRLGNLTARKLVVLV | ASSPGQPSQNTLY | 0.15748124 |
| WSQQRRLIYFPSAGP | ASSPGQPSQNTLY | 0.41455197 |
| GPNGNPDPMAAAVDIR | ASSPGQPSQNTLY | 0.22692695 |
| TQRCVSVADLRAAASR | ASSPGQPSQNTLY | 0.24156137 |
| QFVLDVAEHPVEVLTETV | ASSPGQPSQNTLY | 0.18351653 |
| GGAGGTGGAGGNGGTQAGDG | ASSPGQPSQNTLY | 0.5533819 |
| PLLATLRAIAVDVDAKPAQV | ASSPGQPSQNTLY | 0.5067669 |
| APKFRTGGYMIDFWGVGYQVAK | ASSPGQPSQNTLY | 0.16196203 |
| GAGGAGGAGGAAGGPGGTGGTG | ASSPGQPSQNTLY | 0.18433668 |
| IGALGLLATDVAPAGSVAFGVA | ASSPGQPSQNTLY | 0.44187123 |
| AGGAGGIGGNGGAGGAGGVLMG | ASSPGQPSQNTLY | 0.18837616 |
| ADELIGLL | ASSPGQPSQNTLY | 0.3620336 |
| ADRALKA | ASSPGQPSQNTLY | 0.2397952 |
| AGAVGQVGGAGGA | ASSPGQPSQNTLY | 0.20336476 |
| AGGGGGLGGAP | ASSPGQPSQNTLY | 0.20492958 |
| AGGLLYGNGGAG | ASSPGQPSQNTLY | 0.35937464 |
| AGLESIGL | ASSPGQPSQNTLY | 0.20786092 |
| AGLKTGII | ASSPGQPSQNTLY | 0.358963 |
| ASLAALLL | ASSPGQPSQNTLY | 0.16201122 |
| ASVINISSI | ASSPGQPSQNTLY | 0.43842494 |
| ATPPGPA | ASSPGQPSQNTLY | 0.13172755 |
| ATVASALV | ASSPGQPSQNTLY | 0.18153113 |
| ATVASLIL | ASSPGQPSQNTLY | 0.111813605 |
| AVNGPVLIRH | ASSPGQPSQNTLY | 0.24557608 |
| DGAAGAVGI | ASSPGQPSQNTLY | 0.32928923 |
| DPLLHGI | ASSPGQPSQNTLY | 0.5259015 |
| ELGAALGAL | ASSPGQPSQNTLY | 0.3615303 |
| ELGRTQAV | ASSPGQPSQNTLY | 0.27933574 |
| FVAGGYLS | ASSPGQPSQNTLY | 0.5007456 |
| GASGGAGGAGGW | ASSPGQPSQNTLY | 0.16728376 |
| GGAGAGGDGGAGGI | ASSPGQPSQNTLY | 0.14114682 |

| | | |
|---|---|---|
| GTGHNGSQPGMGGNG | ASSPGQPSQNTLY | 0.29621506 |
| GYAAASASAS | ASSPGQPSQNTLY | 0.24091767 |
| HQVDVAAVQQG | ASSPGQPSQNTLY | 0.50226414 |
| IGAAIAHT | ASSPGQPSQNTLY | 0.43061003 |
| IGMIEVAQL | ASSPGQPSQNTLY | 0.10657665 |
| IRLAPEHTSAMD | ASSPGQPSQNTLY | 0.34668446 |
| IYNVSVL | ASSPGQPSQNTLY | 0.3522264 |
| KQGFLGM | ASSPGQPSQNTLY | 0.13349755 |
| LESVITQL | ASSPGQPSQNTLY | 0.18164109 |
| LGLGALVVV | ASSPGQPSQNTLY | 0.2116045 |
| LIGNGGNGGSGGI | ASSPGQPSQNTLY | 0.17880547 |
| LNAINAPTL | ASSPGQPSQNTLY | 0.2962113 |
| LQLLNTGV | ASSPGQPSQNTLY | 0.15590933 |
| LQTSIEF | ASSPGQPSQNTLY | 0.25714397 |
| MSFAVSEL | ASSPGQPSQNTLY | 0.20060985 |
| NTGALGL | ASSPGQPSQNTLY | 0.19959301 |
| PPGAGGAGGAGTT | ASSPGQPSQNTLY | 0.25589085 |
| QLVGLGI | ASSPGQPSQNTLY | 0.25128078 |
| RFLEQVHQL | ASSPGQPSQNTLY | 0.3948715 |
| RGVGYVLR | ASSPGQPSQNTLY | 0.38415974 |
| SATGVFAGVI | ASSPGQPSQNTLY | 0.20288977 |
| SLPAITLP | ASSPGQPSQNTLY | 0.2750982 |
| STALAFTV | ASSPGQPSQNTLY | 0.4680025 |
| SVLTVGLGK | ASSPGQPSQNTLY | 0.16383182 |
| TALAGAGSGI | ASSPGQPSQNTLY | 0.15857378 |
| TGLPFTGI | ASSPGQPSQNTLY | 0.35683113 |
| VAEHVTV | ASSPGQPSQNTLY | 0.24047115 |
| VDIVLNSL | ASSPGQPSQNTLY | 0.46783084 |
| VLTGGGAVL | ASSPGQPSQNTLY | 0.28958067 |
| VNVGLAYGI | ASSPGQPSQNTLY | 0.26945907 |
| VSSAAVGAV | ASSPGQPSQNTLY | 0.28610972 |
| VTSVYEAGV | ASSPGQPSQNTLY | 0.56030226 |
| WVALDRGA | ASSPGQPSQNTLY | 0.3484189 |
| YLVGSVVI | ASSPGQPSQNTLY | 0.23095544 |
| GPPGTGKTLL | ASSPGQPSQNTLY | 0.4103523 |
| GVGMTKF | ASSPGQPSQNTLY | 0.17670487 |
| IGIGGSDL | ASSPGQPSQNTLY | 0.31675306 |
| IGTLLAEL | ASSPGQPSQNTLY | 0.34122548 |
| IYTSGTT | ASSPGQPSQNTLY | 0.13306402 |
| LSFPAGV | ASSPGQPSQNTLY | 0.17456977 |
| LYTSGTT | ASSPGQPSQNTLY | 0.5105094 |
| VGAFPFGF | ASSPGQPSQNTLY | 0.4436304 |
| VGAGTVEF | ASSPGQPSQNTLY | 0.21828413 |
| VGKSTLLL | ASSPGQPSQNTLY | 0.18061355 |
| AALAHAGGGIV | ASSPGQPSQNTLY | 0.21649638 |
| AALVAVSLL | ASSPGQPSQNTLY | 0.2324597 |
| AAVSRAVAAL | ASSPGQPSQNTLY | 0.15563542 |
| AGGSSGIGGF | ASSPGQPSQNTLY | 0.32158676 |
| AGLTVANANG | ASSPGQPSQNTLY | 0.20099525 |
| AGLVGAVL | ASSPGQPSQNTLY | 0.5663041 |
| ALSQAFTV | ASSPGQPSQNTLY | 0.28937897 |

| | | |
|---|---|---|
| ALTFSVSL | ASSPGQPSQNTLY | 0.4489003 |
| ALTGAALVAL | ASSPGQPSQNTLY | 0.172243 |
| ASFSIGAGVL | ASSPGQPSQNTLY | 0.23971 |
| DGSVYKI | ASSPGQPSQNTLY | 0.15330838 |
| EQLAEIV | ASSPGQPSQNTLY | 0.314394 |
| FAVSELTVM | ASSPGQPSQNTLY | 0.21509302 |
| GAAPLAAGL | ASSPGQPSQNTLY | 0.24046713 |
| GADGGDGGSGGAGG | ASSPGQPSQNTLY | 0.17680183 |
| GFGLDTGGAGG | ASSPGQPSQNTLY | 0.5339855 |
| GLASGGLL | ASSPGQPSQNTLY | 0.20746554 |
| GRIAGAAVLL | ASSPGQPSQNTLY | 0.39196512 |
| GSAGSLNTSF | ASSPGQPSQNTLY | 0.23428066 |
| GTGQIGI | ASSPGQPSQNTLY | 0.19424844 |
| GTLIALIL | ASSPGQPSQNTLY | 0.1871176 |
| IAATMGFVVG | ASSPGQPSQNTLY | 0.36302033 |
| IFAQALL | ASSPGQPSQNTLY | 0.1407642 |
| IGAAVALGF | ASSPGQPSQNTLY | 0.12640221 |
| IGLTPLAL | ASSPGQPSQNTLY | 0.19579175 |
| ILGSNTSTL | ASSPGQPSQNTLY | 0.24557105 |
| IVFSLTVL | ASSPGQPSQNTLY | 0.55461234 |
| LASKATGF | ASSPGQPSQNTLY | 0.41423506 |
| LASVALIL | ASSPGQPSQNTLY | 0.23052074 |
| LFAVGLTL | ASSPGQPSQNTLY | 0.61428 |
| LFNGGIGTYI | ASSPGQPSQNTLY | 0.088983715 |
| LTLGAGII | ASSPGQPSQNTLY | 0.26150572 |
| LVAAALGDI | ASSPGQPSQNTLY | 0.5179691 |
| NTAFVQLGI | ASSPGQPSQNTLY | 0.26947224 |
| QLFSGAEL | ASSPGQPSQNTLY | 0.21925613 |
| SAATFITL | ASSPGQPSQNTLY | 0.45117772 |
| SNYGLANLGV | ASSPGQPSQNTLY | 0.17131001 |
| SYGGAYAVM | ASSPGQPSQNTLY | 0.21362278 |
| TTGVGGLTLG | ASSPGQPSQNTLY | 0.29700708 |
| TTQTIYI | ASSPGQPSQNTLY | 0.35594714 |
| VAAGAGLGAVI | ASSPGQPSQNTLY | 0.37679467 |
| VAAVVTLL | ASSPGQPSQNTLY | 0.1410104 |
| VGFLGGLI | ASSPGQPSQNTLY | 0.35974842 |
| VGVIPILL | ASSPGQPSQNTLY | 0.5897081 |
| VIFASSLI | ASSPGQPSQNTLY | 0.17108737 |
| VKGGDGGAAAG | ASSPGQPSQNTLY | 0.48921055 |
| VTKTFTNI | ASSPGQPSQNTLY | 0.26766625 |
| VVAAAILL | ASSPGQPSQNTLY | 0.38983306 |
| YAALDVEL | ASSPGQPSQNTLY | 0.15582992 |
| LASVALLL | ASSPGQPSQNTLY | 0.13763326 |
| LDAVHRV | ASSPGQPSQNTLY | 0.43979448 |
| QGLSSSIF | ASSPGQPSQNTLY | 0.14397681 |
| RGTFANIRL | ASSPGQPSQNTLY | 0.2788808 |
| RQLWWGHRI | ASSPGQPSQNTLY | 0.26635063 |
| YGPPGTGKTLL | ASSPGQPSQNTLY | 0.6933702 |
| AALVGAPANVA | ASSPGQPSQNTLY | 0.2693717 |
| AATFRRTE | ASSPGQPSQNTLY | 0.33196282 |
| AGALGGGI | ASSPGQPSQNTLY | 0.14422598 |

| | | |
|---|---|---|
| AGGAGGAGGIGAG | ASSPGQPSQNTLY | 0.29002905 |
| AGSGALGL | ASSPGQPSQNTLY | 0.32127097 |
| AQIADQI | ASSPGQPSQNTLY | 0.18311693 |
| ASAVAVTYL | ASSPGQPSQNTLY | 0.40296322 |
| ASTVLGGTGG | ASSPGQPSQNTLY | 0.16700491 |
| AVGGEPAK | ASSPGQPSQNTLY | 0.2740649 |
| CEQGTFGGTC | ASSPGQPSQNTLY | 0.34030163 |
| DEGKAPAG | ASSPGQPSQNTLY | 0.65200615 |
| DTVVFPAL | ASSPGQPSQNTLY | 0.21966511 |
| FEGITMAG | ASSPGQPSQNTLY | 0.48549545 |
| FPLVVQD | ASSPGQPSQNTLY | 0.14450455 |
| GAFTTPQ | ASSPGQPSQNTLY | 0.36784336 |
| GFGGFGTAGGVGGA | ASSPGQPSQNTLY | 0.4810387 |
| GFLAQFL | ASSPGQPSQNTLY | 0.3956344 |
| GGAGGAGGNAGVA | ASSPGQPSQNTLY | 0.6823218 |
| GGVGTAGAGGAGGAGG | ASSPGQPSQNTLY | 0.38536644 |
| GNGGGGDAGNAGSGGN | ASSPGQPSQNTLY | 0.14138076 |
| GTGGAGGAGNDAGS | ASSPGQPSQNTLY | 0.2606746 |
| GTTNTGWFNS | ASSPGQPSQNTLY | 0.4451382 |
| IFMGPLFTL | ASSPGQPSQNTLY | 0.5342628 |
| ILAHAIEAI | ASSPGQPSQNTLY | 0.4458563 |
| IRSAIREAL | ASSPGQPSQNTLY | 0.43083233 |
| ITFAGGG | ASSPGQPSQNTLY | 0.16953713 |
| ITGMSGHQEV | ASSPGQPSQNTLY | 0.48016334 |
| IVAGLACTL | ASSPGQPSQNTLY | 0.75545764 |
| IVGGGLAGA | ASSPGQPSQNTLY | 0.33198372 |
| IVTGVLNI | ASSPGQPSQNTLY | 0.1658324 |
| LAQQGCI | ASSPGQPSQNTLY | 0.20874573 |
| LCLHVEN | ASSPGQPSQNTLY | 0.31128442 |
| LDAISDGS | ASSPGQPSQNTLY | 0.42884797 |
| LGGFAAML | ASSPGQPSQNTLY | 0.1669794 |
| LGGKNPMI | ASSPGQPSQNTLY | 0.21639596 |
| LGPKGVAAIL | ASSPGQPSQNTLY | 0.26150268 |
| LMVGVSI | ASSPGQPSQNTLY | 0.10785587 |
| LVGAAFVL | ASSPGQPSQNTLY | 0.3644227 |
| LVLAVAPL | ASSPGQPSQNTLY | 0.4990279 |
| MAAIAVIDAA | ASSPGQPSQNTLY | 0.33591688 |
| NNGINAELQ | ASSPGQPSQNTLY | 0.14816718 |
| RGEPLVLVHG | ASSPGQPSQNTLY | 0.42946172 |
| RGGAGGAAGLI | ASSPGQPSQNTLY | 0.57260364 |
| RSYRMHPAV | ASSPGQPSQNTLY | 0.51363266 |
| SAAHLLAGQLA | ASSPGQPSQNTLY | 0.15116638 |
| SAAVGSY | ASSPGQPSQNTLY | 0.08746049 |
| SPHAVLHRI | ASSPGQPSQNTLY | 0.3332355 |
| SQIAKVL | ASSPGQPSQNTLY | 0.4046418 |
| TALGHLVP | ASSPGQPSQNTLY | 0.21603191 |
| TAPTTSGAA | ASSPGQPSQNTLY | 0.16870373 |
| TDGDDGNGGSPGT | ASSPGQPSQNTLY | 0.28404078 |
| TDGLGEN | ASSPGQPSQNTLY | 0.41708735 |
| TGGPPLPGQA | ASSPGQPSQNTLY | 0.15991285 |
| TIPAITI | ASSPGQPSQNTLY | 0.15845887 |

| | | |
|---|---|---|
| TIPGFTI | ASSPGQPSQNTLY | 0.3139078 |
| TSIASGALA | ASSPGQPSQNTLY | 0.42141294 |
| TVTLPGAAI | ASSPGQPSQNTLY | 0.128667 |
| VASGLSAALP | ASSPGQPSQNTLY | 0.16357332 |
| VGAGVIGI | ASSPGQPSQNTLY | 0.38852358 |
| VGPKRFGV | ASSPGQPSQNTLY | 0.44686922 |
| VNVAFPD | ASSPGQPSQNTLY | 0.24583934 |
| VSMTNSGVS | ASSPGQPSQNTLY | 0.14102715 |
| VTDVLGDLSV | ASSPGQPSQNTLY | 0.36252093 |
| VVVVGAGFAGL | ASSPGQPSQNTLY | 0.14172326 |
| VYTVGTVV | ASSPGQPSQNTLY | 0.3968587 |
| WTTFSGP | ASSPGQPSQNTLY | 0.24686052 |
| YDELVEAF | ASSPGQPSQNTLY | 0.30351493 |
| YGTGGAGGNGG | ASSPGQPSQNTLY | 0.19092968 |
| YTNVMAV | ASSPGQPSQNTLY | 0.13689335 |
| FLEQVHQL | ASSPGQPSQNTLY | 0.16507131 |
| GLSLLAVAV | ASSPGQPSQNTLY | 0.14425004 |
| LEQVHQL | ASSPGQPSQNTLY | 0.4205861 |
| LNAAGAI | ASSPGQPSQNTLY | 0.51173055 |
| PAVHASL | ASSPGQPSQNTLY | 0.3413469 |
| TELEDDA | ASSPGQPSQNTLY | 0.32465172 |
| TPPGPPN | ASSPGQPSQNTLY | 0.18804845 |
| TPPPPPPP | ASSPGQPSQNTLY | 0.33212027 |
| TQSPALI | ASSLAGGARGEQY | 0.15048194 |
| EFAAAFPQ | ASSLAGGARGEQY | 0.38190106 |
| LLSGNLDVL | ASSLAGGARGEQY | 0.20414351 |
| GPANLGNYNI | ASSLAGGARGEQY | 0.38102013 |
| TLGPTLP | ASSLAGGARGEQY | 0.19166248 |
| VYAYSSCCK | ASSLAGGARGEQY | 0.36283007 |
| APAPAPAPNI | ASSLAGGARGEQY | 0.28333467 |
| PGGPVPIP | ASSLAGGARGEQY | 0.1716658 |
| FLTVALEP | ASSLAGGARGEQY | 0.3095173 |
| SQYFGANL | ASSLAGGARGEQY | 0.43518174 |
| GGVKPAA | ASSLAGGARGEQY | 0.3426711 |
| GGTGGAGGAGGW | ASSLAGGARGEQY | 0.12322678 |
| GVGGVAGLGGAGG | ASSLAGGARGEQY | 0.3109615 |
| VLLVLATTI | ASSLAGGARGEQY | 0.1474534 |
| IGIFAAML | ASSLAGGARGEQY | 0.24786231 |
| AVGDLID | ASSLAGGARGEQY | 0.27481052 |
| NISPLPL | ASSLAGGARGEQY | 0.21800351 |
| VTAVFVAI | ASSLAGGARGEQY | 0.3345772 |
| SGDDNDAAL | ASSLAGGARGEQY | 0.19359286 |
| AVAALAHS | ASSLAGGARGEQY | 0.1555879 |
| GGVTVMATP | ASSLAGGARGEQY | 0.21169522 |
| AGLAANVRD | ASSLAGGARGEQY | 0.31858742 |
| TNRPDLIDPALLRPG | ASSLAGGARGEQY | 0.35487503 |
| LANLGVTAL | ASSLAGGARGEQY | 0.13980784 |
| TGERIEL | ASSLAGGARGEQY | 0.24827002 |
| NGEELAA | ASSLAGGARGEQY | 0.19773224 |
| VLSLVWG | ASSLAGGARGEQY | 0.15743905 |
| IGYALWL | ASSLAGGARGEQY | 0.29344654 |

| | | |
|---|---|---|
| LSLLLAPL | ASSLAGGARGEQY | 0.22766379 |
| TLDEALK | ASSLAGGARGEQY | 0.15084274 |
| TDGAGTAEGG | ASSLAGGARGEQY | 0.33519307 |
| NAPGGAGGNGG | ASSLAGGARGEQY | 0.21970452 |
| VSVFMGMGGNF | ASSLAGGARGEQY | 0.29725492 |
| LASYLAAL | ASSLAGGARGEQY | 0.44184995 |
| LVGGTPAL | ASSLAGGARGEQY | 0.13394994 |
| VRNADVL | ASSLAGGARGEQY | 0.36512202 |
| TCSSPFLVSAM | ASSLAGGARGEQY | 0.23489395 |
| VHITDAE | ASSLAGGARGEQY | 0.24042796 |
| EYAAPSF | ASSLAGGARGEQY | 0.2777674 |
| VVDAFHGGL | ASSLAGGARGEQY | 0.18094727 |
| ASVVDPQ | ASSLAGGARGEQY | 0.41940677 |
| SYNEFGGLS | ASSLAGGARGEQY | 0.35044217 |
| DPDGSVY | ASSLAGGARGEQY | 0.312402 |
| MVSTIHATE | ASSLAGGARGEQY | 0.29307157 |
| LAAGVADPV | ASSLAGGARGEQY | 0.4115808 |
| TKEQIAA | ASSLAGGARGEQY | 0.39527515 |
| SLASPAAL | ASSLAGGARGEQY | 0.14543477 |
| IGLVFYGV | ASSLAGGARGEQY | 0.1252824 |
| FLLDASNL | ASSLAGGARGEQY | 0.25360078 |
| YQVGNFP | ASSLAGGARGEQY | 0.26028833 |
| ESLLSQST | ASSLAGGARGEQY | 0.20111322 |
| LFTVGFAAVG | ASSLAGGARGEQY | 0.16489267 |
| ATPPTPT | ASSLAGGARGEQY | 0.39544612 |
| TDLATLG | ASSLAGGARGEQY | 0.6214158 |
| SAGGTGR | ASSLAGGARGEQY | 0.20860732 |
| GFSNSGS | ASSLAGGARGEQY | 0.68203557 |
| GGNGGSGGTSV | ASSLAGGARGEQY | 0.14541398 |
| GTSVATGGAGN | ASSLAGGARGEQY | 0.3401033 |
| NGGNGGSGGTS | ASSLAGGARGEQY | 0.4490314 |
| AVVKFPH | ASSLAGGARGEQY | 0.13641801 |
| TPPPPPP | ASSLAGGARGEQY | 0.33188742 |
| AAPAAAAAPA | ASSLAGGARGEQY | 0.22780944 |
| IALNEDL | ASSLAGGARGEQY | 0.19395426 |
| IVNVSSM | ASSLAGGARGEQY | 0.3852154 |
| TGIPPAPR | ASSLAGGARGEQY | 0.25737408 |
| FDLGGGTF | ASSLAGGARGEQY | 0.32099175 |
| FDLGGGTFDVS | ASSLAGGARGEQY | 0.26691136 |
| NEPTAAA | ASSLAGGARGEQY | 0.17938882 |
| NEPTAAALA | ASSLAGGARGEQY | 0.33836964 |
| GSIINISSI | ASSLAGGARGEQY | 0.340613 |
| VAPGNAGTA | ASSLAGGARGEQY | 0.35368997 |
| TGGGASLE | ASSLAGGARGEQY | 0.7056402 |
| VAVYNHYKRI | ASSLAGGARGEQY | 0.15643816 |
| TDPAPATT | ASSLAGGARGEQY | 0.6139747 |
| TNRPDILDPALLRPG | ASSLAGGARGEQY | 0.57041204 |
| LTTTTTS | ASSLAGGARGEQY | 0.23581429 |
| PTGSGKTL | ASSLAGGARGEQY | 0.22907518 |
| TGSGKTL | ASSLAGGARGEQY | 0.44855994 |
| TIGHVDHGK | ASSLAGGARGEQY | 0.23439682 |

| | | |
|---|---|---|
| TIGHVDHGKTT | ASSLAGGARGEQY | 0.32071686 |
| GHVDHGKTT | ASSLAGGARGEQY | 0.45977455 |
| AQFEHTLL | ASSLAGGARGEQY | 0.5003276 |
| IVNQDEL | ASSLAGGARGEQY | 0.85349506 |
| TNGSQFF | ASSLAGGARGEQY | 0.8130981 |
| TGKGIVE | ASSLAGGARGEQY | 0.70669144 |
| LISNASDA | ASSLAGGARGEQY | 0.61114967 |
| AELRTLGELTKIAKAHGAQ | ASSLAGGARGEQY | 0.24539067 |
| APNAVAAAEPSW | ASSLAGGARGEQY | 0.6422956 |
| APQVAGAATAVL | ASSLAGGARGEQY | 0.53885573 |
| ARRDPAAAAAILPTDA | ASSLAGGARGEQY | 0.78253937 |
| DAAARDVLAEAGLAEYF | ASSLAGGARGEQY | 0.7334726 |
| DDQIVLSAKANLAKVEQ | ASSLAGGARGEQY | 0.5896369 |
| DELLPDTGEDSDEDGH | ASSLAGGARGEQY | 0.8548856 |
| ELPEAPANRATVPQD | ASSLAGGARGEQY | 0.8566299 |
| FGQSKIERVTGIG | ASSLAGGARGEQY | 0.37520078 |
| GAGGTGSTAGGAGGAGGAGGLY | ASSLAGGARGEQY | 0.7334188 |
| GHGGTGAAGGNGGTGGAGGSGI | ASSLAGGARGEQY | 0.53510034 |
| GNGGNGGAGGEGGPGGAGGAGA | ASSLAGGARGEQY | 0.63790226 |
| GSTAGNGGNGGNATNGTIAGQP | ASSLAGGARGEQY | 0.24103136 |
| GTGGDGGNGGNGGTGG | ASSLAGGARGEQY | 0.810251 |
| GVRRPAGIPADAPV | ASSLAGGARGEQY | 0.63007575 |
| GYRQVTDAHLLTLARR | ASSLAGGARGEQY | 0.13564366 |
| IAKMGQDGHDRG | ASSLAGGARGEQY | 0.7105837 |
| KAEPSIEIQYT | ASSLAGGARGEQY | 0.6624715 |
| KPVRIRMAADEDTVAVT | ASSLAGGARGEQY | 0.67182857 |
| MPPRIAGMRLLVIKPEP | ASSLAGGARGEQY | 0.8447077 |
| MKFYAVALFRSRGVPPTR | ASSLAGGARGEQY | 0.20333615 |
| NAGGFGILGAVAHSP | ASSLAGGARGEQY | 0.49234664 |
| NEGREITLAMAEM | ASSLAGGARGEQY | 0.83875936 |
| NELVEWSHPLGHRWRWE | ASSLAGGARGEQY | 0.85700357 |
| NLGAGGTGGNGGNPG | ASSLAGGARGEQY | 0.3886463 |
| NTEWPKWETVMAADPVD | ASSLAGGARGEQY | 0.81454384 |
| PASAAPAETQP | ASSLAGGARGEQY | 0.85375655 |
| QQSSDGKNGVRAAEITGE | ASSLAGGARGEQY | 0.7869089 |
| RAPDAVTTSWWKE | ASSLAGGARGEQY | 0.82224774 |
| SGFGNVGAHTSGWFNQ | ASSLAGGARGEQY | 0.8433952 |
| SGVAGAVGQAASVGGLKV | ASSLAGGARGEQY | 0.40225178 |
| SNGGAGTGSAGGNGGT | ASSLAGGARGEQY | 0.8400218 |
| TGGAGGEGGAAG | ASSLAGGARGEQY | 0.85177946 |
| TGGDGGAAGAGGAPGHG | ASSLAGGARGEQY | 0.58101463 |
| TMPPLLAAIANAGRHGV | ASSLAGGARGEQY | 0.7935934 |
| TSSTAGNSGGA | ASSLAGGARGEQY | 0.4490928 |
| VPATVPVGILC | ASSLAGGARGEQY | 0.48924226 |
| VQRPDSGGVVLTGRIS | ASSLAGGARGEQY | 0.18112649 |
| WTPNPVPATKGVPAGLVQAT | ASSLAGGARGEQY | 0.826668 |
| YALKRALAKFSDRFDVVII | ASSLAGGARGEQY | 0.8266358 |
| YLIGSVIV | ASSLAGGARGEQY | 0.769264 |
| YGQLVAVID | ASSLAGGARGEQY | 0.48897195 |
| VTVGGMTLPA | ASSLAGGARGEQY | 0.5351664 |
| VLALPAV | ASSLAGGARGEQY | 0.6762222 |

| | | |
|---|---|---|
| VIPPTLNYET | ASSLAGGARGEQY | 0.724355 |
| VGMGGMHPSSG | ASSLAGGARGEQY | 0.19746566 |
| VGLAGGAATA | ASSLAGGARGEQY | 0.5457348 |
| VGGAGGAGGSGP | ASSLAGGARGEQY | 0.85073507 |
| VFAAFMV | ASSLAGGARGEQY | 0.8474574 |
| TLYDKLR | ASSLAGGARGEQY | 0.32269895 |
| TLAVFTVL | ASSLAGGARGEQY | 0.8493944 |
| TGSVQPARV | ASSLAGGARGEQY | 0.8062854 |
| TGGGAGSGGT | ASSLAGGARGEQY | 0.84445024 |
| SHLETSL | ASSLAGGARGEQY | 0.6342162 |
| SFTTGMT | ASSLAGGARGEQY | 0.55103695 |
| QVEVAKRMGVRQPTVS | ASSLAGGARGEQY | 0.81863517 |
| QGSGGPALAM | ASSLAGGARGEQY | 0.77531916 |
| MLAPSNP | ASSLAGGARGEQY | 0.39853913 |
| LVNNAAI | ASSLAGGARGEQY | 0.8331833 |
| LTGEINTLA | ASSLAGGARGEQY | 0.68235445 |
| LSIFGPV | ASSLAGGARGEQY | 0.72529936 |
| LLALLANL | ASSLAGGARGEQY | 0.71631706 |
| LISNASDALDKLR | ASSLAGGARGEQY | 0.7868153 |
| LINEAVP | ASSLAGGARGEQY | 0.53497505 |
| LGYAFSGL | ASSLAGGARGEQY | 0.3998655 |
| LDFVAVAL | ASSLAGGARGEQY | 0.4000239 |
| LAQPGAE | ASSLAGGARGEQY | 0.8038443 |
| ISNLGAVTGF | ASSLAGGARGEQY | 0.85178566 |
| HQEDPGRI | ASSLAGGARGEQY | 0.67736083 |
| GTPGEPTP | ASSLAGGARGEQY | 0.6411004 |
| GFFNSGS | ASSLAGGARGEQY | 0.83217657 |
| GAGGGGLTAGAN | ASSLAGGARGEQY | 0.76031697 |
| FVDALVNM | ASSLAGGARGEQY | 0.690175 |
| FLTVAPEL | ASSLAGGARGEQY | 0.8360875 |
| FETLTAL | ASSLAGGARGEQY | 0.6982863 |
| DVINPIEL | ASSLAGGARGEQY | 0.7991394 |
| APVSGAPV | ASSLAGGARGEQY | 0.79710585 |
| ALVGGLAW | ASSLAGGARGEQY | 0.8019701 |
| ALAAAAATAI | ASSLAGGARGEQY | 0.8147851 |
| AFGLGLLL | ASSLAGGARGEQY | 0.79042876 |
| AFFAAASL | ASSLAGGARGEQY | 0.6347366 |
| AAAALPAFSPP | ASSLAGGARGEQY | 0.19753955 |
| VVLAPAL | ASSLAGGARGEQY | 0.60246915 |
| TIGHVDHGKTTL | ASSLAGGARGEQY | 0.8139458 |
| LISNASDALDK | ASSLAGGARGEQY | 0.8099454 |
| LISNASDAL | ASSLAGGARGEQY | 0.8098579 |
| ISNASDALDK | ASSLAGGARGEQY | 0.72820437 |
| NQAETLVYQTE | ASSLAGGARGEQY | 0.526724 |
| TGGTGGTAGAG | ASSLAGGARGEQY | 0.83007455 |
| TGSTAGGAGGA | ASSLAGGARGEQY | 0.83271694 |
| DPLAVPVAAGPV | ASSLAGGARGEQY | 0.3656444 |
| GGTGGTPGNGGN | ASSLAGGARGEQY | 0.6685884 |
| PAGLAGVDPATL | ASSLAGGARGEQY | 0.8590305 |
| GNNLVVEAPHAGASV | ASSLAGGARGEQY | 0.79598194 |
| KVTRIMSASTLEQPA | ASSLAGGARGEQY | 0.73791945 |

| | | |
|---|---|---|
| LGFALPIALPRRTVV | ASSLAGGARGEQY | 0.50176424 |
| TRLGNLTARKLVVLV | ASSLAGGARGEQY | 0.84467345 |
| WSQQRRLIYFPSAGP | ASSLAGGARGEQY | 0.5117663 |
| GPNGNPDPMAAAVDIR | ASSLAGGARGEQY | 0.85253716 |
| TQRCVSVADLRAAASR | ASSLAGGARGEQY | 0.16050237 |
| QFVLDVAEHPVEVLTETV | ASSLAGGARGEQY | 0.47538066 |
| GGAGGTGGAGGNGGTQAGDG | ASSLAGGARGEQY | 0.36559546 |
| PLLATLRAIAVDVDAKPAQV | ASSLAGGARGEQY | 0.29839543 |
| APKFRTGGYMIDFWGVGYQVAK | ASSLAGGARGEQY | 0.8292047 |
| GAGGAGGAGGAAGGPGGTGGTG | ASSLAGGARGEQY | 0.47548684 |
| IGALGLLATDVAPAGSVAFGVA | ASSLAGGARGEQY | 0.8360522 |
| AGGAGGIGGNGGAGGAGGVLMG | ASSLAGGARGEQY | 0.7347499 |
| ADELIGLL | ASSLAGGARGEQY | 0.81080115 |
| ADRALKA | ASSLAGGARGEQY | 0.793448 |
| AGAVGQVGGAGGA | ASSLAGGARGEQY | 0.7479237 |
| AGGGGGLGGAP | ASSLAGGARGEQY | 0.38269532 |
| AGGLLYGNGGAG | ASSLAGGARGEQY | 0.81984174 |
| AGLESIGL | ASSLAGGARGEQY | 0.8330595 |
| AGLKTGII | ASSLAGGARGEQY | 0.6958493 |
| ASLAALLL | ASSLAGGARGEQY | 0.3416815 |
| ASVINISSI | ASSLAGGARGEQY | 0.61062944 |
| ATPPGPA | ASSLAGGARGEQY | 0.7626427 |
| ATVASALV | ASSLAGGARGEQY | 0.16371897 |
| ATVASLIL | ASSLAGGARGEQY | 0.15570164 |
| AVNGPVLIRH | ASSLAGGARGEQY | 0.6762643 |
| DGAAGAVGI | ASSLAGGARGEQY | 0.504485 |
| DPLLHGI | ASSLAGGARGEQY | 0.8527674 |
| ELGAALGAL | ASSLAGGARGEQY | 0.8510687 |
| ELGRTQAV | ASSLAGGARGEQY | 0.7700741 |
| FVAGGYLS | ASSLAGGARGEQY | 0.82786655 |
| GASGGAGGAGGW | ASSLAGGARGEQY | 0.8099333 |
| GGAGAGGDGGAGGI | ASSLAGGARGEQY | 0.12529321 |
| GTGHNGSQPGMGGNG | ASSLAGGARGEQY | 0.599092 |
| GYAAASASAS | ASSLAGGARGEQY | 0.41587436 |
| HQVDVAAVQQG | ASSLAGGARGEQY | 0.8313425 |
| IGAAIAHT | ASSLAGGARGEQY | 0.47403955 |
| IGMIEVAQL | ASSLAGGARGEQY | 0.6510626 |
| IRLAPEHTSAMD | ASSLAGGARGEQY | 0.81410694 |
| IYNVSVL | ASSLAGGARGEQY | 0.7861672 |
| KQGFLGM | ASSLAGGARGEQY | 0.7424934 |
| LESVITQL | ASSLAGGARGEQY | 0.4650407 |
| LGLGALVVV | ASSLAGGARGEQY | 0.13143408 |
| LIGNGGNGGSGGI | ASSLAGGARGEQY | 0.83987117 |
| LNAINAPTL | ASSLAGGARGEQY | 0.24070114 |
| LQLLNTGV | ASSLAGGARGEQY | 0.4238245 |
| LQTSIEF | ASSLAGGARGEQY | 0.16737513 |
| MSFAVSEL | ASSLAGGARGEQY | 0.14268507 |
| NTGALGL | ASSLAGGARGEQY | 0.28813562 |
| PPGAGGAGGAGTT | ASSLAGGARGEQY | 0.80842733 |
| QLVGLGI | ASSLAGGARGEQY | 0.71426356 |
| RFLEQVHQL | ASSLAGGARGEQY | 0.73153615 |

| | | |
|---|---|---|
| RGVGYVLR | ASSLAGGARGEQY | 0.8522432 |
| SATGVFAGVI | ASSLAGGARGEQY | 0.8062682 |
| SLPAITLP | ASSLAGGARGEQY | 0.7531086 |
| STALAFTV | ASSLAGGARGEQY | 0.5654099 |
| SVLTVGLGK | ASSLAGGARGEQY | 0.45266598 |
| TALAGAGSGI | ASSLAGGARGEQY | 0.42842013 |
| TGLPFTGI | ASSLAGGARGEQY | 0.41273382 |
| VAEHVTV | ASSLAGGARGEQY | 0.86127603 |
| VDIVLNSL | ASSLAGGARGEQY | 0.4674691 |
| VLTGGGAVL | ASSLAGGARGEQY | 0.3135185 |
| VNVGLAYGI | ASSLAGGARGEQY | 0.602579 |
| VSSAAVGAV | ASSLAGGARGEQY | 0.8481761 |
| VTSVYEAGV | ASSLAGGARGEQY | 0.86147237 |
| WVALDRGA | ASSLAGGARGEQY | 0.6116954 |
| YLVGSVVI | ASSLAGGARGEQY | 0.5737233 |
| GPPGTGKTLL | ASSLAGGARGEQY | 0.5359386 |
| GVGMTKF | ASSLAGGARGEQY | 0.80838394 |
| IGIGGSDL | ASSLAGGARGEQY | 0.5558507 |
| IGTLLAEL | ASSLAGGARGEQY | 0.32386807 |
| IYTSGTT | ASSLAGGARGEQY | 0.78015 |
| LSFPAGV | ASSLAGGARGEQY | 0.85781103 |
| LYTSGTT | ASSLAGGARGEQY | 0.46571007 |
| VGAFPFGF | ASSLAGGARGEQY | 0.33144885 |
| VGAGTVEF | ASSLAGGARGEQY | 0.7907586 |
| VGKSTLLL | ASSLAGGARGEQY | 0.6238642 |
| AALAHAGGGIV | ASSLAGGARGEQY | 0.7655101 |
| AALVAVSLL | ASSLAGGARGEQY | 0.7894053 |
| AAVSRAVAAL | ASSLAGGARGEQY | 0.34612256 |
| AGGSSGIGGF | ASSLAGGARGEQY | 0.5071922 |
| AGLTVANANG | ASSLAGGARGEQY | 0.85994637 |
| AGLVGAVL | ASSLAGGARGEQY | 0.7134279 |
| ALSQAFTV | ASSLAGGARGEQY | 0.7419696 |
| ALTFSVSL | ASSLAGGARGEQY | 0.8470501 |
| ALTGAALVAL | ASSLAGGARGEQY | 0.82126933 |
| ASFSIGAGVL | ASSLAGGARGEQY | 0.85670173 |
| DGSVYKI | ASSLAGGARGEQY | 0.8222913 |
| EQLAEIV | ASSLAGGARGEQY | 0.53207725 |
| FAVSELTVM | ASSLAGGARGEQY | 0.69247806 |
| GAAPLAAGL | ASSLAGGARGEQY | 0.6807946 |
| GADGGDGGSGGAGG | ASSLAGGARGEQY | 0.8145964 |
| GFGLDTGGAGG | ASSLAGGARGEQY | 0.78366673 |
| GLASGGLL | ASSLAGGARGEQY | 0.48450124 |
| GRIAGAAVLL | ASSLAGGARGEQY | 0.85195816 |
| GSAGSLNTSF | ASSLAGGARGEQY | 0.7217369 |
| GTGQIGI | ASSLAGGARGEQY | 0.7837007 |
| GTLIALIL | ASSLAGGARGEQY | 0.44704217 |
| IAATMGFVVG | ASSLAGGARGEQY | 0.40341052 |
| IFAQALL | ASSLAGGARGEQY | 0.83618116 |
| IGAAVALGF | ASSLAGGARGEQY | 0.14063272 |
| IGLTPLAL | ASSLAGGARGEQY | 0.16608007 |
| ILGSNTSTL | ASSLAGGARGEQY | 0.7117481 |

| | | |
|---|---|---|
| IVFSLTVL | ASSLAGGARGEQY | 0.8484391 |
| LASKATGF | ASSLAGGARGEQY | 0.14366686 |
| LASVALIL | ASSLAGGARGEQY | 0.47834793 |
| LFAVGLTL | ASSLAGGARGEQY | 0.335831 |
| LFNGGIGTYI | ASSLAGGARGEQY | 0.7638644 |
| LTLGAGII | ASSLAGGARGEQY | 0.7760627 |
| LVAAALGDI | ASSLAGGARGEQY | 0.3193218 |
| NTAFVQLGI | ASSLAGGARGEQY | 0.803368 |
| QLFSGAEL | ASSLAGGARGEQY | 0.3578725 |
| SAATFITL | ASSLAGGARGEQY | 0.2357761 |
| SNYGLANLGV | ASSLAGGARGEQY | 0.79797816 |
| SYGGAYAVM | ASSLAGGARGEQY | 0.84144425 |
| TTGVGGLTLG | ASSLAGGARGEQY | 0.39024964 |
| TTQTIYI | ASSLAGGARGEQY | 0.837286 |
| VAAGAGLGAVI | ASSLAGGARGEQY | 0.5631199 |
| VAAVVTLL | ASSLAGGARGEQY | 0.8597641 |
| VGFLGGLI | ASSLAGGARGEQY | 0.68749696 |
| VGVIPILL | ASSLAGGARGEQY | 0.2609584 |
| VIFASSLI | ASSLAGGARGEQY | 0.8221801 |
| VKGGDGGAAAG | ASSLAGGARGEQY | 0.7435441 |
| VTKTFTNI | ASSLAGGARGEQY | 0.6966273 |
| VVAAAILL | ASSLAGGARGEQY | 0.8262494 |
| YAALDVEL | ASSLAGGARGEQY | 0.45482123 |
| LASVALLL | ASSLAGGARGEQY | 0.76153564 |
| LDAVHRV | ASSLAGGARGEQY | 0.85307974 |
| QGLSSSIF | ASSLAGGARGEQY | 0.8392631 |
| RGTFANIRL | ASSLAGGARGEQY | 0.40855408 |
| RQLWWGHRI | ASSLAGGARGEQY | 0.75135815 |
| YGPPGTGKTLL | ASSLAGGARGEQY | 0.808989 |
| AALVGAPANVA | ASSLAGGARGEQY | 0.620793 |
| AATFRRTE | ASSLAGGARGEQY | 0.8457688 |
| AGALGGGI | ASSLAGGARGEQY | 0.67501235 |
| AGGAGGAGGIGAG | ASSLAGGARGEQY | 0.829535 |
| AGSGALGL | ASSLAGGARGEQY | 0.75208765 |
| AQIADQI | ASSLAGGARGEQY | 0.8363572 |
| ASAVAVTYL | ASSLAGGARGEQY | 0.30085635 |
| ASTVLGGTGG | ASSLAGGARGEQY | 0.77509224 |
| AVGGEPAK | ASSLAGGARGEQY | 0.75343895 |
| CEQGTFGGTC | ASSLAGGARGEQY | 0.8188794 |
| DEGKAPAG | ASSLAGGARGEQY | 0.6791267 |
| DTVVFPAL | ASSLAGGARGEQY | 0.25343361 |
| FEGITMAG | ASSLAGGARGEQY | 0.68112063 |
| FPLVVQD | ASSLAGGARGEQY | 0.84618664 |
| GAFTTPQ | ASSLAGGARGEQY | 0.6666738 |
| GFGGFGTAGGVGGA | ASSLAGGARGEQY | 0.77673346 |
| GFLAQFL | ASSLAGGARGEQY | 0.86205864 |
| GGAGGAGGNAGVA | ASSLAGGARGEQY | 0.455566 |
| GGVGTAGAGGAGGAGG | ASSLAGGARGEQY | 0.75969684 |
| GNGGGGDAGNAGSGGN | ASSLAGGARGEQY | 0.7761582 |
| GTGGAGGAGNDAGS | ASSLAGGARGEQY | 0.8354348 |
| GTTNTGWFNS | ASSLAGGARGEQY | 0.5983741 |

| | | |
|---|---|---|
| IFMGPLFTL | ASSLAGGARGEQY | 0.8481046 |
| ILAHAIEAI | ASSLAGGARGEQY | 0.43474227 |
| IRSAIREAL | ASSLAGGARGEQY | 0.83850133 |
| ITFAGGG | ASSLAGGARGEQY | 0.41742942 |
| ITGMSGHQEV | ASSLAGGARGEQY | 0.82222587 |
| IVAGLACTL | ASSLAGGARGEQY | 0.6518773 |
| IVGGGLAGA | ASSLAGGARGEQY | 0.83580756 |
| IVTGVLNI | ASSLAGGARGEQY | 0.84345114 |
| LAQQGCI | ASSLAGGARGEQY | 0.8414897 |
| LCLHVEN | ASSLAGGARGEQY | 0.15504387 |
| LDAISDGS | ASSLAGGARGEQY | 0.8470243 |
| LGGFAAML | ASSLAGGARGEQY | 0.84758234 |
| LGGKNPMI | ASSLAGGARGEQY | 0.86293274 |
| LGPKGVAAIL | ASSLAGGARGEQY | 0.71522796 |
| LMVGVSI | ASSLAGGARGEQY | 0.7624155 |
| LVGAAFVL | ASSLAGGARGEQY | 0.7319574 |
| LVLAVAPL | ASSLAGGARGEQY | 0.61923224 |
| MAAIAVIDAA | ASSLAGGARGEQY | 0.65262395 |
| NNGINAELQ | ASSLAGGARGEQY | 0.65704393 |
| RGEPLVLVHG | ASSLAGGARGEQY | 0.78722525 |
| RGGAGGAAGLI | ASSLAGGARGEQY | 0.70735407 |
| RSYRMHPAV | ASSLAGGARGEQY | 0.836404 |
| SAAHLLAGQLA | ASSLAGGARGEQY | 0.83864164 |
| SAAVGSY | ASSLAGGARGEQY | 0.8354671 |
| SPHAVLHRI | ASSLAGGARGEQY | 0.8531467 |
| SQIAKVL | ASSLAGGARGEQY | 0.8380857 |
| TALGHLVP | ASSLAGGARGEQY | 0.79789376 |
| TAPTTSGAA | ASSLAGGARGEQY | 0.5714328 |
| TDGDDGNGGSPGT | ASSLAGGARGEQY | 0.68665 |
| TDGLGEN | ASSLAGGARGEQY | 0.55379415 |
| TGGPPLPGQA | ASSLAGGARGEQY | 0.55322105 |
| TIPAITI | ASSLAGGARGEQY | 0.61177 |
| TIPGFTI | ASSLAGGARGEQY | 0.8056632 |
| TSIASGALA | ASSLAGGARGEQY | 0.12565258 |
| TVTLPGAAI | ASSLAGGARGEQY | 0.763561 |
| VASGLSAALP | ASSLAGGARGEQY | 0.85671794 |
| VGAGVIGI | ASSLAGGARGEQY | 0.62563515 |
| VGPKRFGV | ASSLAGGARGEQY | 0.76086986 |
| VNVAFPD | ASSLAGGARGEQY | 0.6356679 |
| VSMTNSGVS | ASSLAGGARGEQY | 0.6078771 |
| VTDVLGDLSV | ASSLAGGARGEQY | 0.84597325 |
| VVVVGAGFAGL | ASSLAGGARGEQY | 0.43859428 |
| VYTVGTVV | ASSLAGGARGEQY | 0.7857506 |
| WTTFSGP | ASSLAGGARGEQY | 0.84303683 |
| YDELVEAF | ASSLAGGARGEQY | 0.71217895 |
| YGTGGAGGNGG | ASSLAGGARGEQY | 0.76866114 |
| YTNVMAV | ASSLAGGARGEQY | 0.6052456 |
| FLEQVHQL | ASSLAGGARGEQY | 0.27433944 |
| GLSLLAVAV | ASSLAGGARGEQY | 0.7997345 |
| LEQVHQL | ASSLAGGARGEQY | 0.12053273 |
| LNAAGAI | ASSLAGGARGEQY | 0.77309036 |

| | | |
|---|---|---|
| PAVHASL | ASSLAGGARGEQY | 0.33426535 |
| TELEDDA | ASSLAGGARGEQY | 0.8238565 |
| TPPGPPN | ASSLAGGARGEQY | 0.80780387 |
| TPPPPPPP | ASSLAGGARGEQY | 0.7927725 |
| TQSPALI | ASSLGLGDPEQY | 0.5970089 |
| EFAAAFPQ | ASSLGLGDPEQY | 0.6906065 |
| LLSGNLDVL | ASSLGLGDPEQY | 0.43311346 |
| GPANLGNYNI | ASSLGLGDPEQY | 0.33345 |
| TLGPTLP | ASSLGLGDPEQY | 0.80909646 |
| VYAYSSCCK | ASSLGLGDPEQY | 0.5197016 |
| APAPAPAPNI | ASSLGLGDPEQY | 0.6631667 |
| PGGPVPIP | ASSLGLGDPEQY | 0.8345679 |
| FLTVALEP | ASSLGLGDPEQY | 0.74907625 |
| SQYFGANL | ASSLGLGDPEQY | 0.29859704 |
| GGVKPAA | ASSLGLGDPEQY | 0.785149 |
| GGTGGAGGAGGW | ASSLGLGDPEQY | 0.7375591 |
| GVGGVAGLGGAGG | ASSLGLGDPEQY | 0.69227874 |
| VLLVLATTI | ASSLGLGDPEQY | 0.12062135 |
| IGIFAAML | ASSLGLGDPEQY | 0.6947247 |
| AVGDLID | ASSLGLGDPEQY | 0.63689655 |
| NISPLPL | ASSLGLGDPEQY | 0.6076187 |
| VTAVFVAI | ASSLGLGDPEQY | 0.5411962 |
| SGDDNDAAL | ASSLGLGDPEQY | 0.8468636 |
| AVAALAHS | ASSLGLGDPEQY | 0.3177292 |
| GGVTVMATP | ASSLGLGDPEQY | 0.76913166 |
| AGLAANVRD | ASSLGLGDPEQY | 0.8281846 |
| TNRPDLIDPALLRPG | ASSLGLGDPEQY | 0.824404 |
| LANLGVTAL | ASSLGLGDPEQY | 0.85897726 |
| TGERIEL | ASSLGLGDPEQY | 0.784182 |
| NGEELAA | ASSLGLGDPEQY | 0.5498091 |
| VLSLVWG | ASSLGLGDPEQY | 0.7083695 |
| IGYALWL | ASSLGLGDPEQY | 0.73460627 |
| LSLLLAPL | ASSLGLGDPEQY | 0.17655788 |
| TLDEALK | ASSLGLGDPEQY | 0.17587534 |
| TDGAGTAEGG | ASSLGLGDPEQY | 0.7999787 |
| NAPGGAGGNGG | ASSLGLGDPEQY | 0.1714488 |
| VSVFMGMGGNF | ASSLGLGDPEQY | 0.7552488 |
| LASYLAAL | ASSLGLGDPEQY | 0.80185676 |
| LVGGTPAL | ASSLGLGDPEQY | 0.6857103 |
| VRNADVL | ASSLGLGDPEQY | 0.31102636 |
| TCSSPFLVSAM | ASSLGLGDPEQY | 0.66520584 |
| VHITDAE | ASSLGLGDPEQY | 0.6976031 |
| EYAAPSF | ASSLGLGDPEQY | 0.8342652 |
| VVDAFHGGL | ASSLGLGDPEQY | 0.70769155 |
| ASVVDPQ | ASSLGLGDPEQY | 0.8531557 |
| SYNEFGGLS | ASSLGLGDPEQY | 0.84981483 |
| DPDGSVY | ASSLGLGDPEQY | 0.7399149 |
| MVSTIHATE | ASSLGLGDPEQY | 0.8308496 |
| LAAGVADPV | ASSLGLGDPEQY | 0.6161636 |
| TKEQIAA | ASSLGLGDPEQY | 0.5590658 |
| SLASPAAL | ASSLGLGDPEQY | 0.5600102 |

| | | |
|---|---|---|
| IGLVFYGV | ASSLGLGDPEQY | 0.7536112 |
| FLLDASNL | ASSLGLGDPEQY | 0.7091714 |
| YQVGNFP | ASSLGLGDPEQY | 0.4842167 |
| ESLLSQST | ASSLGLGDPEQY | 0.27896988 |
| LFTVGFAAVG | ASSLGLGDPEQY | 0.7371355 |
| ATPPTPT | ASSLGLGDPEQY | 0.7780827 |
| TDLATLG | ASSLGLGDPEQY | 0.22277352 |
| SAGGTGR | ASSLGLGDPEQY | 0.5267676 |
| GFSNSGS | ASSLGLGDPEQY | 0.86322623 |
| GGNGGSGGTSV | ASSLGLGDPEQY | 0.8584242 |
| GTSVATGGAGN | ASSLGLGDPEQY | 0.6573183 |
| NGGNGGSGGTS | ASSLGLGDPEQY | 0.47546571 |
| AVVKFPH | ASSLGLGDPEQY | 0.7811261 |
| TPPPPPP | ASSLGLGDPEQY | 0.24509269 |
| AAPAAAAAPA | ASSLGLGDPEQY | 0.6952498 |
| IALNEDL | ASSLGLGDPEQY | 0.79971635 |
| IVNVSSM | ASSLGLGDPEQY | 0.77969635 |
| TGIPPAPR | ASSLGLGDPEQY | 0.39569303 |
| FDLGGGTF | ASSLGLGDPEQY | 0.77178615 |
| FDLGGGTFDVS | ASSLGLGDPEQY | 0.6930363 |
| NEPTAAA | ASSLGLGDPEQY | 0.7797372 |
| NEPTAAALA | ASSLGLGDPEQY | 0.14170495 |
| GSIINISSI | ASSLGLGDPEQY | 0.72767425 |
| VAPGNAGTA | ASSLGLGDPEQY | 0.6506184 |
| TGGGASLE | ASSLGLGDPEQY | 0.47947755 |
| VAVYNHYKRI | ASSLGLGDPEQY | 0.29130155 |
| TDPAPATT | ASSLGLGDPEQY | 0.7092807 |
| TNRPDILDPALLRPG | ASSLGLGDPEQY | 0.84549594 |
| LTTTTTS | ASSLGLGDPEQY | 0.8442577 |
| PTGSGKTL | ASSLGLGDPEQY | 0.26575533 |
| TGSGKTL | ASSLGLGDPEQY | 0.8434993 |
| TIGHVDHGK | ASSLGLGDPEQY | 0.8628354 |
| TIGHVDHGKTT | ASSLGLGDPEQY | 0.6553961 |
| GHVDHGKTT | ASSLGLGDPEQY | 0.24825144 |
| AQFEHTLL | ASSLGLGDPEQY | 0.8181008 |
| IVNQDEL | ASSLGLGDPEQY | 0.63617975 |
| TNGSQFF | ASSLGLGDPEQY | 0.8077151 |
| TGKGIVE | ASSLGLGDPEQY | 0.81569046 |
| LISNASDA | ASSLGLGDPEQY | 0.55262303 |
| AELRTLGELTKIAKAHGAQ | ASSLGLGDPEQY | 0.81077945 |
| APNAVAAAEPSW | ASSLGLGDPEQY | 0.8210309 |
| APQVAGAATAVL | ASSLGLGDPEQY | 0.57883036 |
| ARRDPAAAAAILPTDA | ASSLGLGDPEQY | 0.4632231 |
| DAAARDVLAEAGLAEYF | ASSLGLGDPEQY | 0.43870783 |
| DDQIVLSAKANLAKVEQ | ASSLGLGDPEQY | 0.18958911 |
| DELLPDTGEDSDEDGH | ASSLGLGDPEQY | 0.7910625 |
| ELPEAPANRATVPQD | ASSLGLGDPEQY | 0.66394573 |
| FGQSKIERVTGIG | ASSLGLGDPEQY | 0.5774036 |
| GAGGTGSTAGGAGGAGGAGGLY | ASSLGLGDPEQY | 0.7992631 |
| GHGGTGAAGGNGGTGGAGGSGI | ASSLGLGDPEQY | 0.43846464 |
| GNGGNGGAGGEGGPGGAGGAGA | ASSLGLGDPEQY | 0.63214964 |

| | | |
|---|---|---|
| GSTAGNGGNGGNATNGTIAGQP | ASSLGLGDPEQY | 0.6000723 |
| GTGGDGGNGGNGGTGG | ASSLGLGDPEQY | 0.7141792 |
| GVRRPAGIPADAPV | ASSLGLGDPEQY | 0.80021197 |
| GYRQVTDAHLLTLARR | ASSLGLGDPEQY | 0.76786566 |
| IAKMGQDGHDRG | ASSLGLGDPEQY | 0.7215828 |
| KAEPSIEIQYT | ASSLGLGDPEQY | 0.85679644 |
| KPVRIRMAADEDTVAVT | ASSLGLGDPEQY | 0.4293817 |
| MPPRIAGMRLLVIKPEP | ASSLGLGDPEQY | 0.5726414 |
| MKFYAVALFRSRGVPPTR | ASSLGLGDPEQY | 0.8051684 |
| NAGGFGILGAVAHSP | ASSLGLGDPEQY | 0.27199775 |
| NEGREITLAMAEM | ASSLGLGDPEQY | 0.8146244 |
| NELVEWSHPLGHRWRWE | ASSLGLGDPEQY | 0.7976375 |
| NLGAGGTGGNGGNPG | ASSLGLGDPEQY | 0.80933887 |
| NTEWPKWETVMAADPVD | ASSLGLGDPEQY | 0.6969676 |
| PASAAPAETQP | ASSLGLGDPEQY | 0.8601835 |
| QQSSDGKNGVRAAEITGE | ASSLGLGDPEQY | 0.8377621 |
| RAPDAVTTSWWKE | ASSLGLGDPEQY | 0.56426454 |
| SGFGNVGAHTSGWFNQ | ASSLGLGDPEQY | 0.7784039 |
| SGVAGAVGQAASVGGLKV | ASSLGLGDPEQY | 0.36205882 |
| SNGGAGTGSAGGNGGT | ASSLGLGDPEQY | 0.8379725 |
| TGGAGGEGGAAG | ASSLGLGDPEQY | 0.78941035 |
| TGGDGGAAGAGGAPGHG | ASSLGLGDPEQY | 0.599102 |
| TMPPLLAAIANAGRHGV | ASSLGLGDPEQY | 0.76459706 |
| TSSTAGNSGGA | ASSLGLGDPEQY | 0.83116925 |
| VPATVPVGILC | ASSLGLGDPEQY | 0.8438381 |
| VQRPDSGGVVLTGRIS | ASSLGLGDPEQY | 0.1999706 |
| WTPNPVPATKGVPAGLVQAT | ASSLGLGDPEQY | 0.7502866 |
| YALKRALAKFSDRFDVVII | ASSLGLGDPEQY | 0.22699302 |
| YLIGSVIV | ASSLGLGDPEQY | 0.48698524 |
| YGQLVAVID | ASSLGLGDPEQY | 0.3940073 |
| VTVGGMTLPA | ASSLGLGDPEQY | 0.81096375 |
| VLALPAV | ASSLGLGDPEQY | 0.7209203 |
| VIPPTLNYET | ASSLGLGDPEQY | 0.80381846 |
| VGMGGMHPSSG | ASSLGLGDPEQY | 0.79989225 |
| VGLAGGAATA | ASSLGLGDPEQY | 0.8507352 |
| VGGAGGAGGSGP | ASSLGLGDPEQY | 0.5299165 |
| VFAAFMV | ASSLGLGDPEQY | 0.7274474 |
| TLYDKLR | ASSLGLGDPEQY | 0.28438482 |
| TLAVFTVL | ASSLGLGDPEQY | 0.2083346 |
| TGSVQPARV | ASSLGLGDPEQY | 0.4224428 |
| TGGGAGSGGT | ASSLGLGDPEQY | 0.70641214 |
| SHLETSL | ASSLGLGDPEQY | 0.17624143 |
| SFTTGMT | ASSLGLGDPEQY | 0.84447026 |
| QVEVAKRMGVRQPTVS | ASSLGLGDPEQY | 0.80927384 |
| QGSGGPALAM | ASSLGLGDPEQY | 0.83807296 |
| MLAPSNP | ASSLGLGDPEQY | 0.5114251 |
| LVNNAAI | ASSLGLGDPEQY | 0.6822168 |
| LTGEINTLA | ASSLGLGDPEQY | 0.6245428 |
| LSIFGPV | ASSLGLGDPEQY | 0.8337153 |
| LLALLANL | ASSLGLGDPEQY | 0.51877105 |
| LISNASDALDKLR | ASSLGLGDPEQY | 0.2635712 |

| | | |
|---|---|---|
| LINEAVP | ASSLGLGDPEQY | 0.622831 |
| LGYAFSGL | ASSLGLGDPEQY | 0.51102746 |
| LDFVAVAL | ASSLGLGDPEQY | 0.83046335 |
| LAQPGAE | ASSLGLGDPEQY | 0.82807577 |
| ISNLGAVTGF | ASSLGLGDPEQY | 0.81311274 |
| HQEDPGRI | ASSLGLGDPEQY | 0.77195823 |
| GTPGEPTP | ASSLGLGDPEQY | 0.20247836 |
| GFFNSGS | ASSLGLGDPEQY | 0.61887205 |
| GAGGGGLTAGAN | ASSLGLGDPEQY | 0.8634744 |
| FVDALVNM | ASSLGLGDPEQY | 0.7712749 |
| FLTVAPEL | ASSLGLGDPEQY | 0.7431512 |
| FETLTAL | ASSLGLGDPEQY | 0.8526674 |
| DVINPIEL | ASSLGLGDPEQY | 0.6653049 |
| APVSGAPV | ASSLGLGDPEQY | 0.6966883 |
| ALVGGLAW | ASSLGLGDPEQY | 0.7344978 |
| ALAAAAATAI | ASSLGLGDPEQY | 0.6637393 |
| AFGLGLLL | ASSLGLGDPEQY | 0.67721605 |
| AFFAAASL | ASSLGLGDPEQY | 0.46368948 |
| AAAALPAFSPP | ASSLGLGDPEQY | 0.70152044 |
| VVLAPAL | ASSLGLGDPEQY | 0.85912865 |
| TIGHVDHGKTTL | ASSLGLGDPEQY | 0.79099905 |
| LISNASDALDK | ASSLGLGDPEQY | 0.8466395 |
| LISNASDAL | ASSLGLGDPEQY | 0.8106153 |
| ISNASDALDK | ASSLGLGDPEQY | 0.8196263 |
| NQAETLVYQTE | ASSLGLGDPEQY | 0.62555623 |
| TGGTGGTAGAG | ASSLGLGDPEQY | 0.7264062 |
| TGSTAGGAGGA | ASSLGLGDPEQY | 0.6773848 |
| DPLAVPVAAGPV | ASSLGLGDPEQY | 0.7116938 |
| GGTGGTPGNGGN | ASSLGLGDPEQY | 0.6745459 |
| PAGLAGVDPATL | ASSLGLGDPEQY | 0.522387 |
| GNNLVVEAPHAGASV | ASSLGLGDPEQY | 0.8661393 |
| KVTRIMSASTLEQPA | ASSLGLGDPEQY | 0.80599546 |
| LGFALPIALPRRTVV | ASSLGLGDPEQY | 0.81033707 |
| TRLGNLTARKLVVLV | ASSLGLGDPEQY | 0.6148352 |
| WSQQRRLIYFPSAGP | ASSLGLGDPEQY | 0.6930943 |
| GPNGNPDPMAAAVDIR | ASSLGLGDPEQY | 0.8034575 |
| TQRCVSVADLRAAASR | ASSLGLGDPEQY | 0.3036191 |
| QFVLDVAEHPVEVLTETV | ASSLGLGDPEQY | 0.8569624 |
| GGAGGTGGAGGNGGTQAGDG | ASSLGLGDPEQY | 0.67831063 |
| PLLATLRAIAVDVDAKPAQV | ASSLGLGDPEQY | 0.5430717 |
| APKFRTGGYMIDFWGVGYQVAK | ASSLGLGDPEQY | 0.8227979 |
| GAGGAGGAGGAAGGPGGTGGTG | ASSLGLGDPEQY | 0.84352565 |
| IGALGLLATDVAPAGSVAFGVA | ASSLGLGDPEQY | 0.56477237 |
| AGGAGGIGGNGGAGGAGGVLMG | ASSLGLGDPEQY | 0.6638893 |
| ADELIGLL | ASSLGLGDPEQY | 0.6459042 |
| ADRALKA | ASSLGLGDPEQY | 0.81912136 |
| AGAVGQVGGAGGA | ASSLGLGDPEQY | 0.8545456 |
| AGGGGGLGGAP | ASSLGLGDPEQY | 0.71800494 |
| AGGLLYGNGGAG | ASSLGLGDPEQY | 0.84524584 |
| AGLESIGL | ASSLGLGDPEQY | 0.72461456 |
| AGLKTGII | ASSLGLGDPEQY | 0.3036528 |

| | | |
|---|---|---|
| ASLAALLL | ASSLGLGDPEQY | 0.66518927 |
| ASVINISSI | ASSLGLGDPEQY | 0.8230213 |
| ATPPGPA | ASSLGLGDPEQY | 0.8399847 |
| ATVASALV | ASSLGLGDPEQY | 0.85269606 |
| ATVASLIL | ASSLGLGDPEQY | 0.7660364 |
| AVNGPVLIRH | ASSLGLGDPEQY | 0.5432607 |
| DGAAGAVGI | ASSLGLGDPEQY | 0.85238767 |
| DPLLHGI | ASSLGLGDPEQY | 0.85240185 |
| ELGAALGAL | ASSLGLGDPEQY | 0.8456186 |
| ELGRTQAV | ASSLGLGDPEQY | 0.6977358 |
| FVAGGYLS | ASSLGLGDPEQY | 0.4778719 |
| GASGGAGGAGGW | ASSLGLGDPEQY | 0.6139027 |
| GGAGAGGDGGAGGI | ASSLGLGDPEQY | 0.83598554 |
| GTGHNGSQPGMGGNG | ASSLGLGDPEQY | 0.4232303 |
| GYAAASASAS | ASSLGLGDPEQY | 0.7570084 |
| HQVDVAAVQQG | ASSLGLGDPEQY | 0.82027733 |
| IGAAIAHT | ASSLGLGDPEQY | 0.3268666 |
| IGMIEVAQL | ASSLGLGDPEQY | 0.7622591 |
| IRLAPEHTSAMD | ASSLGLGDPEQY | 0.84690815 |
| IYNVSVL | ASSLGLGDPEQY | 0.5198145 |
| KQGFLGM | ASSLGLGDPEQY | 0.65971667 |
| LESVITQL | ASSLGLGDPEQY | 0.84624135 |
| LGLGALVVV | ASSLGLGDPEQY | 0.8672122 |
| LIGNGGNGGSGGI | ASSLGLGDPEQY | 0.82437104 |
| LNAINAPTL | ASSLGLGDPEQY | 0.30101547 |
| LQLLNTGV | ASSLGLGDPEQY | 0.2073078 |
| LQTSIEF | ASSLGLGDPEQY | 0.612392 |
| MSFAVSEL | ASSLGLGDPEQY | 0.81273043 |
| NTGALGL | ASSLGLGDPEQY | 0.5549575 |
| PPGAGGAGGAGTT | ASSLGLGDPEQY | 0.5530899 |
| QLVGLGI | ASSLGLGDPEQY | 0.8087999 |
| RFLEQVHQL | ASSLGLGDPEQY | 0.8439847 |
| RGVGYVLR | ASSLGLGDPEQY | 0.42500764 |
| SATGVFAGVI | ASSLGLGDPEQY | 0.33466017 |
| SLPAITLP | ASSLGLGDPEQY | 0.709846 |
| STALAFTV | ASSLGLGDPEQY | 0.82140994 |
| SVLTVGLGK | ASSLGLGDPEQY | 0.6433799 |
| TALAGAGSGI | ASSLGLGDPEQY | 0.7766837 |
| TGLPFTGI | ASSLGLGDPEQY | 0.52273035 |
| VAEHVTV | ASSLGLGDPEQY | 0.2455711 |
| VDIVLNSL | ASSLGLGDPEQY | 0.81574225 |
| VLTGGGAVL | ASSLGLGDPEQY | 0.51317644 |
| VNVGLAYGI | ASSLGLGDPEQY | 0.85135317 |
| VSSAAVGAV | ASSLGLGDPEQY | 0.58681357 |
| VTSVYEAGV | ASSLGLGDPEQY | 0.81437755 |
| WVALDRGA | ASSLGLGDPEQY | 0.8249899 |
| YLVGSVVI | ASSLGLGDPEQY | 0.71313256 |
| GPPGTGKTLL | ASSLGLGDPEQY | 0.7223674 |
| GVGMTKF | ASSLGLGDPEQY | 0.57536644 |
| IGIGGSDL | ASSLGLGDPEQY | 0.394568 |
| IGTLLAEL | ASSLGLGDPEQY | 0.22820508 |

| | | |
|---|---|---|
| IYTSGTT | ASSLGLGDPEQY | 0.84229857 |
| LSFPAGV | ASSLGLGDPEQY | 0.1451879 |
| LYTSGTT | ASSLGLGDPEQY | 0.8401742 |
| VGAFPFGF | ASSLGLGDPEQY | 0.8120729 |
| VGAGTVEF | ASSLGLGDPEQY | 0.52151453 |
| VGKSTLLL | ASSLGLGDPEQY | 0.765617 |
| AALAHAGGGIV | ASSLGLGDPEQY | 0.60573137 |
| AALVAVSLL | ASSLGLGDPEQY | 0.2998703 |
| AAVSRAVAAL | ASSLGLGDPEQY | 0.8342217 |
| AGGSSGIGGF | ASSLGLGDPEQY | 0.7849747 |
| AGLTVANANG | ASSLGLGDPEQY | 0.5964196 |
| AGLVGAVL | ASSLGLGDPEQY | 0.72456264 |
| ALSQAFTV | ASSLGLGDPEQY | 0.7519984 |
| ALTFSVSL | ASSLGLGDPEQY | 0.47585404 |
| ALTGAALVAL | ASSLGLGDPEQY | 0.49596912 |
| ASFSIGAGVL | ASSLGLGDPEQY | 0.8411531 |
| DGSVYKI | ASSLGLGDPEQY | 0.6298665 |
| EQLAEIV | ASSLGLGDPEQY | 0.59219265 |
| FAVSELTVM | ASSLGLGDPEQY | 0.8021654 |
| GAAPLAAGL | ASSLGLGDPEQY | 0.48811066 |
| GADGGDGGSGGAGG | ASSLGLGDPEQY | 0.83182514 |
| GFGLDTGGAGG | ASSLGLGDPEQY | 0.6661201 |
| GLASGGLL | ASSLGLGDPEQY | 0.6749695 |
| GRIAGAAVLL | ASSLGLGDPEQY | 0.7897668 |
| GSAGSLNTSF | ASSLGLGDPEQY | 0.4011472 |
| GTGQIGI | ASSLGLGDPEQY | 0.463791 |
| GTLIALIL | ASSLGLGDPEQY | 0.6206454 |
| IAATMGFVVG | ASSLGLGDPEQY | 0.78676057 |
| IFAQALL | ASSLGLGDPEQY | 0.8392881 |
| IGAAVALGF | ASSLGLGDPEQY | 0.60238415 |
| IGLTPLAL | ASSLGLGDPEQY | 0.75317353 |
| ILGSNTSTL | ASSLGLGDPEQY | 0.56354797 |
| IVFSLTVL | ASSLGLGDPEQY | 0.4852003 |
| LASKATGF | ASSLGLGDPEQY | 0.5420849 |
| LASVALIL | ASSLGLGDPEQY | 0.5711074 |
| LFAVGLTL | ASSLGLGDPEQY | 0.6310793 |
| LFNGGIGTYI | ASSLGLGDPEQY | 0.7397635 |
| LTLGAGII | ASSLGLGDPEQY | 0.4852308 |
| LVAAALGDI | ASSLGLGDPEQY | 0.6924932 |
| NTAFVQLGI | ASSLGLGDPEQY | 0.8495991 |
| QLFSGAEL | ASSLGLGDPEQY | 0.7669685 |
| SAATFITL | ASSLGLGDPEQY | 0.7856628 |
| SNYGLANLGV | ASSLGLGDPEQY | 0.8104037 |
| SYGGAYAVM | ASSLGLGDPEQY | 0.8119521 |
| TTGVGGLTLG | ASSLGLGDPEQY | 0.63230515 |
| TTQTIYI | ASSLGLGDPEQY | 0.60668707 |
| VAAGAGLGAVI | ASSLGLGDPEQY | 0.8492589 |
| VAAVVTLL | ASSLGLGDPEQY | 0.8393873 |
| VGFLGGLI | ASSLGLGDPEQY | 0.8411497 |
| VGVIPILL | ASSLGLGDPEQY | 0.8069719 |
| VIFASSLI | ASSLGLGDPEQY | 0.7956568 |

| | | |
|---|---|---|
| VKGGDGGAAAG | ASSLGLGDPEQY | 0.82495314 |
| VTKTFTNI | ASSLGLGDPEQY | 0.7706114 |
| VVAAAILL | ASSLGLGDPEQY | 0.3441276 |
| YAALDVEL | ASSLGLGDPEQY | 0.6400399 |
| LASVALLL | ASSLGLGDPEQY | 0.7714977 |
| LDAVHRV | ASSLGLGDPEQY | 0.5243386 |
| QGLSSSIF | ASSLGLGDPEQY | 0.2980984 |
| RGTFANIRL | ASSLGLGDPEQY | 0.8502248 |
| RQLWWGHRI | ASSLGLGDPEQY | 0.8656553 |
| YGPPGTGKTLL | ASSLGLGDPEQY | 0.5813348 |
| AALVGAPANVA | ASSLGLGDPEQY | 0.8448061 |
| AATFRRTE | ASSLGLGDPEQY | 0.31097457 |
| AGALGGGI | ASSLGLGDPEQY | 0.81458825 |
| AGGAGGAGGIGAG | ASSLGLGDPEQY | 0.85002226 |
| AGSGALGL | ASSLGLGDPEQY | 0.67279625 |
| AQIADQI | ASSLGLGDPEQY | 0.8169279 |
| ASAVAVTYL | ASSLGLGDPEQY | 0.51443344 |
| ASTVLGGTGG | ASSLGLGDPEQY | 0.6401491 |
| AVGGEPAK | ASSLGLGDPEQY | 0.8534542 |
| CEQGTFGGTC | ASSLGLGDPEQY | 0.643554 |
| DEGKAPAG | ASSLGLGDPEQY | 0.6662899 |
| DTVVFPAL | ASSLGLGDPEQY | 0.6572418 |
| FEGITMAG | ASSLGLGDPEQY | 0.4818344 |
| FPLVVQD | ASSLGLGDPEQY | 0.66298735 |
| GAFTTPQ | ASSLGLGDPEQY | 0.84823966 |
| GFGGFGTAGGVGGA | ASSLGLGDPEQY | 0.3909601 |
| GFLAQFL | ASSLGLGDPEQY | 0.8472847 |
| GGAGGAGGNAGVA | ASSLGLGDPEQY | 0.5129738 |
| GGVGTAGAGGAGGAGG | ASSLGLGDPEQY | 0.8641025 |
| GNGGGGDAGNAGSGGN | ASSLGLGDPEQY | 0.82730126 |
| GTGGAGGAGNDAGS | ASSLGLGDPEQY | 0.8191403 |
| GTTNTGWFNS | ASSLGLGDPEQY | 0.5927266 |
| IFMGPLFTL | ASSLGLGDPEQY | 0.8559099 |
| ILAHAIEAI | ASSLGLGDPEQY | 0.76653004 |
| IRSAIREAL | ASSLGLGDPEQY | 0.82336354 |
| ITFAGGG | ASSLGLGDPEQY | 0.8189629 |
| ITGMSGHQEV | ASSLGLGDPEQY | 0.130766 |
| IVAGLACTL | ASSLGLGDPEQY | 0.7221168 |
| IVGGGLAGA | ASSLGLGDPEQY | 0.48830426 |
| IVTGVLNI | ASSLGLGDPEQY | 0.29349053 |
| LAQQGCI | ASSLGLGDPEQY | 0.167064 |
| LCLHVEN | ASSLGLGDPEQY | 0.11630303 |
| LDAISDGS | ASSLGLGDPEQY | 0.16869669 |
| LGGFAAML | ASSLGLGDPEQY | 0.14829919 |
| LGGKNPMI | ASSLGLGDPEQY | 0.7593976 |
| LGPKGVAAIL | ASSLGLGDPEQY | 0.33964643 |
| LMVGVSI | ASSLGLGDPEQY | 0.1776268 |
| LVGAAFVL | ASSLGLGDPEQY | 0.5352927 |
| LVLAVAPL | ASSLGLGDPEQY | 0.60224056 |
| MAAIAVIDAA | ASSLGLGDPEQY | 0.11027224 |
| NNGINAELQ | ASSLGLGDPEQY | 0.22349186 |

| | | |
|---|---|---|
| RGEPLVLVHG | ASSLGLGDPEQY | 0.14558317 |
| RGGAGGAAGLI | ASSLGLGDPEQY | 0.19717652 |
| RSYRMHPAV | ASSLGLGDPEQY | 0.09190846 |
| SAAHLLAGQLA | ASSLGLGDPEQY | 0.3389899 |
| SAAVGSY | ASSLGLGDPEQY | 0.17717594 |
| SPHAVLHRI | ASSLGLGDPEQY | 0.08898285 |
| SQIAKVL | ASSLGLGDPEQY | 0.24275291 |
| TALGHLVP | ASSLGLGDPEQY | 0.18889523 |
| TAPTTSGAA | ASSLGLGDPEQY | 0.17883763 |
| TDGDDGNGGSPGT | ASSLGLGDPEQY | 0.52097905 |
| TDGLGEN | ASSLGLGDPEQY | 0.31223935 |
| TGGPPLPGQA | ASSLGLGDPEQY | 0.1428169 |
| TIPAITI | ASSLGLGDPEQY | 0.4233659 |
| TIPGFTI | ASSLGLGDPEQY | 0.6860366 |
| TSIASGALA | ASSLGLGDPEQY | 0.13686341 |
| TVTLPGAAI | ASSLGLGDPEQY | 0.31766838 |
| VASGLSAALP | ASSLGLGDPEQY | 0.70731604 |
| VGAGVIGI | ASSLGLGDPEQY | 0.30334437 |
| VGPKRFGV | ASSLGLGDPEQY | 0.569477 |
| VNVAFPD | ASSLGLGDPEQY | 0.6700422 |
| VSMTNSGVS | ASSLGLGDPEQY | 0.14761858 |
| VTDVLGDLSV | ASSLGLGDPEQY | 0.5092553 |
| VVVVGAGFAGL | ASSLGLGDPEQY | 0.7249741 |
| VYTVGTVV | ASSLGLGDPEQY | 0.23019165 |
| WTTFSGP | ASSLGLGDPEQY | 0.23636198 |
| YDELVEAF | ASSLGLGDPEQY | 0.13438426 |
| YGTGGAGGNGG | ASSLGLGDPEQY | 0.19106467 |
| YTNVMAV | ASSLGLGDPEQY | 0.12078317 |
| FLEQVHQL | ASSLGLGDPEQY | 0.42199907 |
| GLSLLAVAV | ASSLGLGDPEQY | 0.49408045 |
| LEQVHQL | ASSLGLGDPEQY | 0.22082338 |
| LNAAGAI | ASSLGLGDPEQY | 0.11168594 |
| PAVHASL | ASSLGLGDPEQY | 0.116265126 |
| TELEDDA | ASSLGLGDPEQY | 0.34490675 |
| TPPGPPN | ASSLGLGDPEQY | 0.17294036 |
| TPPPPPPP | ASSLGLGDPEQY | 0.1267691 |
| TQSPALI | ASGEDWGENTQY | 0.18333183 |
| EFAAAFPQ | ASGEDWGENTQY | 0.52020085 |
| LLSGNLDVL | ASGEDWGENTQY | 0.5374641 |
| GPANLGNYNI | ASGEDWGENTQY | 0.12489707 |
| TLGPTLP | ASGEDWGENTQY | 0.74694586 |
| VYAYSSCCK | ASGEDWGENTQY | 0.6763474 |
| APAPAPAPNI | ASGEDWGENTQY | 0.6910567 |
| PGGPVPIP | ASGEDWGENTQY | 0.21388248 |
| FLTVALEP | ASGEDWGENTQY | 0.12943897 |
| SQYFGANL | ASGEDWGENTQY | 0.722691 |
| GGVKPAA | ASGEDWGENTQY | 0.28975677 |
| GGTGGAGGAGGW | ASGEDWGENTQY | 0.21872528 |
| GVGGVAGLGGAGG | ASGEDWGENTQY | 0.40409064 |
| VLLVLATTI | ASGEDWGENTQY | 0.22155921 |
| IGIFAAML | ASGEDWGENTQY | 0.1696446 |

| | | |
|---|---|---|
| AVGDLID | ASGEDWGENTQY | 0.48831445 |
| NISPLPL | ASGEDWGENTQY | 0.42119986 |
| VTAVFVAI | ASGEDWGENTQY | 0.1173987 |
| SGDDNDAAL | ASGEDWGENTQY | 0.15516508 |
| AVAALAHS | ASGEDWGENTQY | 0.16544585 |
| GGVTVMATP | ASGEDWGENTQY | 0.29114115 |
| AGLAANVRD | ASGEDWGENTQY | 0.5733795 |
| TNRPDLIDPALLRPG | ASGEDWGENTQY | 0.28184706 |
| LANLGVTAL | ASGEDWGENTQY | 0.26692677 |
| TGERIEL | ASGEDWGENTQY | 0.5165251 |
| NGEELAA | ASGEDWGENTQY | 0.44397447 |
| VLSLVWG | ASGEDWGENTQY | 0.24214783 |
| IGYALWL | ASGEDWGENTQY | 0.51945186 |
| LSLLLAPL | ASGEDWGENTQY | 0.17676523 |
| TLDEALK | ASGEDWGENTQY | 0.45002568 |
| TDGAGTAEGG | ASGEDWGENTQY | 0.39822084 |
| NAPGGAGGNGG | ASGEDWGENTQY | 0.39446437 |
| VSVFMGMGGNF | ASGEDWGENTQY | 0.5377872 |
| LASYLAAL | ASGEDWGENTQY | 0.5145173 |
| LVGGTPAL | ASGEDWGENTQY | 0.2433216 |
| VRNADVL | ASGEDWGENTQY | 0.11067182 |
| TCSSPFLVSAM | ASGEDWGENTQY | 0.16548678 |
| VHITDAE | ASGEDWGENTQY | 0.37967938 |
| EYAAPSF | ASGEDWGENTQY | 0.34879 |
| VVDAFHGGL | ASGEDWGENTQY | 0.43549153 |
| ASVVDPQ | ASGEDWGENTQY | 0.36103883 |
| SYNEFGGLS | ASGEDWGENTQY | 0.16110472 |
| DPDGSVY | ASGEDWGENTQY | 0.57117414 |
| MVSTIHATE | ASGEDWGENTQY | 0.6285634 |
| LAAGVADPV | ASGEDWGENTQY | 0.113342956 |
| TKEQIAA | ASGEDWGENTQY | 0.19561084 |
| SLASPAAL | ASGEDWGENTQY | 0.7363807 |
| IGLVFYGV | ASGEDWGENTQY | 0.5435369 |
| FLLDASNL | ASGEDWGENTQY | 0.19057035 |
| YQVGNFP | ASGEDWGENTQY | 0.21621615 |
| ESLLSQST | ASGEDWGENTQY | 0.784979 |
| LFTVGFAAVG | ASGEDWGENTQY | 0.22283371 |
| ATPPTPT | ASGEDWGENTQY | 0.64602655 |
| TDLATLG | ASGEDWGENTQY | 0.098613285 |
| SAGGTGR | ASGEDWGENTQY | 0.21765086 |
| GFSNSGS | ASGEDWGENTQY | 0.16526704 |
| GGNGGSGGTSV | ASGEDWGENTQY | 0.099158555 |
| GTSVATGGAGN | ASGEDWGENTQY | 0.41652375 |
| NGGNGGSGGTS | ASGEDWGENTQY | 0.1581893 |
| AVVKFPH | ASGEDWGENTQY | 0.49850625 |
| TPPPPPP | ASGEDWGENTQY | 0.17277181 |
| AAPAAAAAPA | ASGEDWGENTQY | 0.4604652 |
| IALNEDL | ASGEDWGENTQY | 0.36594093 |
| IVNVSSM | ASGEDWGENTQY | 0.26367712 |
| TGIPPAPR | ASGEDWGENTQY | 0.15191966 |
| FDLGGGTF | ASGEDWGENTQY | 0.38364455 |

| | | |
|---|---|---|
| FDLGGGTFDVS | ASGEDWGENTQY | 0.5851685 |
| NEPTAAA | ASGEDWGENTQY | 0.3727294 |
| NEPTAAALA | ASGEDWGENTQY | 0.11401924 |
| GSIINISSI | ASGEDWGENTQY | 0.32941723 |
| VAPGNAGTA | ASGEDWGENTQY | 0.2421613 |
| TGGGASLE | ASGEDWGENTQY | 0.11816676 |
| VAVYNHYKRI | ASGEDWGENTQY | 0.09658836 |
| TDPAPATT | ASGEDWGENTQY | 0.16456787 |
| TNRPDILDPALLRPG | ASGEDWGENTQY | 0.22691646 |
| LTTTTTS | ASGEDWGENTQY | 0.45798707 |
| PTGSGKTL | ASGEDWGENTQY | 0.48605564 |
| TGSGKTL | ASGEDWGENTQY | 0.18117182 |
| TIGHVDHGK | ASGEDWGENTQY | 0.4910742 |
| TIGHVDHGKTT | ASGEDWGENTQY | 0.39644444 |
| GHVDHGKTT | ASGEDWGENTQY | 0.08361632 |
| AQFEHTLL | ASGEDWGENTQY | 0.16779427 |
| IVNQDEL | ASGEDWGENTQY | 0.10707924 |
| TNGSQFF | ASGEDWGENTQY | 0.491929 |
| TGKGIVE | ASGEDWGENTQY | 0.13416138 |
| LISNASDA | ASGEDWGENTQY | 0.13973011 |
| AELRTLGELTKIAKAHGAQ | ASGEDWGENTQY | 0.22750086 |
| APNAVAAAEPSW | ASGEDWGENTQY | 0.63513017 |
| APQVAGAATAVL | ASGEDWGENTQY | 0.29606766 |
| ARRDPAAAAAILPTDA | ASGEDWGENTQY | 0.15837571 |
| DAAARDVLAEAGLAEYF | ASGEDWGENTQY | 0.10544076 |
| DDQIVLSAKANLAKVEQ | ASGEDWGENTQY | 0.3958164 |
| DELLPDTGEDSDEDGH | ASGEDWGENTQY | 0.25063705 |
| ELPEAPANRATVPQD | ASGEDWGENTQY | 0.14483635 |
| FGQSKIERVTGIG | ASGEDWGENTQY | 0.10465861 |
| GAGGTGSTAGGAGGAGGAGGLY | ASGEDWGENTQY | 0.5312118 |
| GHGGTGAAGGNGGTGGAGGSGI | ASGEDWGENTQY | 0.12239489 |
| GNGGNGGAGGEGGPGGAGGAGA | ASGEDWGENTQY | 0.5039679 |
| GSTAGNGGNGGNATNGTIAGQP | ASGEDWGENTQY | 0.20571743 |
| GTGGDGGNGGNGGTGG | ASGEDWGENTQY | 0.200546 |
| GVRRPAGIPADAPV | ASGEDWGENTQY | 0.71638733 |
| GYRQVTDAHLLTLARR | ASGEDWGENTQY | 0.6668936 |
| IAKMGQDGHDRG | ASGEDWGENTQY | 0.21929377 |
| KAEPSIEIQYT | ASGEDWGENTQY | 0.18863167 |
| KPVRIRMAADEDTVAVT | ASGEDWGENTQY | 0.128807 |
| MPPRIAGMRLLVIKPEP | ASGEDWGENTQY | 0.13702062 |
| MKFYAVALFRSRGVPPTR | ASGEDWGENTQY | 0.15392049 |
| NAGGFGILGAVAHSP | ASGEDWGENTQY | 0.6351107 |
| NEGREITLAMAEM | ASGEDWGENTQY | 0.20792975 |
| NELVEWSHPLGHRWRWE | ASGEDWGENTQY | 0.115831695 |
| NLGAGGTGGNGGNPG | ASGEDWGENTQY | 0.40003154 |
| NTEWPKWETVMAADPVD | ASGEDWGENTQY | 0.58237046 |
| PASAAPAETQP | ASGEDWGENTQY | 0.8125421 |
| QQSSDGKNGVRAAEITGE | ASGEDWGENTQY | 0.13042697 |
| RAPDAVTTSWWKE | ASGEDWGENTQY | 0.20032556 |
| SGFGNVGAHTSGWFNQ | ASGEDWGENTQY | 0.18847564 |
| SGVAGAVGQAASVGGLKV | ASGEDWGENTQY | 0.25329262 |

| | | |
|---|---|---|
| SNGGAGTGSAGGNGGT | ASGEDWGENTQY | 0.18277827 |
| TGGAGGEGGAAG | ASGEDWGENTQY | 0.14275597 |
| TGGDGGAAGAGGAPGHG | ASGEDWGENTQY | 0.41640896 |
| TMPPLLAAIANAGRHGV | ASGEDWGENTQY | 0.84182066 |
| TSSTAGNSGGA | ASGEDWGENTQY | 0.17866492 |
| VPATVPVGILC | ASGEDWGENTQY | 0.09983446 |
| VQRPDSGGVVLTGRIS | ASGEDWGENTQY | 0.21326494 |
| WTPNPVPATKGVPAGLVQAT | ASGEDWGENTQY | 0.13907948 |
| YALKRALAKFSDRFDVVII | ASGEDWGENTQY | 0.14990483 |
| YLIGSVIV | ASGEDWGENTQY | 0.29916474 |
| YGQLVAVID | ASGEDWGENTQY | 0.10567246 |
| VTVGGMTLPA | ASGEDWGENTQY | 0.185118 |
| VLALPAV | ASGEDWGENTQY | 0.80615103 |
| VIPPTLNYET | ASGEDWGENTQY | 0.25990528 |
| VGMGGMHPSSG | ASGEDWGENTQY | 0.4851569 |
| VGLAGGAATA | ASGEDWGENTQY | 0.5908185 |
| VGGAGGAGGSGP | ASGEDWGENTQY | 0.37998587 |
| VFAAFMV | ASGEDWGENTQY | 0.70689535 |
| TLYDKLR | ASGEDWGENTQY | 0.71735454 |
| TLAVFTVL | ASGEDWGENTQY | 0.23397364 |
| TGSVQPARV | ASGEDWGENTQY | 0.27440035 |
| TGGGAGSGGT | ASGEDWGENTQY | 0.6025955 |
| SHLETSL | ASGEDWGENTQY | 0.23220676 |
| SFTTGMT | ASGEDWGENTQY | 0.25119758 |
| QVEVAKRMGVRQPTVS | ASGEDWGENTQY | 0.18471017 |
| QGSGGPALAM | ASGEDWGENTQY | 0.5022705 |
| MLAPSNP | ASGEDWGENTQY | 0.31877294 |
| LVNNAAI | ASGEDWGENTQY | 0.5688336 |
| LTGEINTLA | ASGEDWGENTQY | 0.12176578 |
| LSIFGPV | ASGEDWGENTQY | 0.1393892 |
| LLALLANL | ASGEDWGENTQY | 0.63403153 |
| LISNASDALDKLR | ASGEDWGENTQY | 0.093776666 |
| LINEAVP | ASGEDWGENTQY | 0.11708212 |
| LGYAFSGL | ASGEDWGENTQY | 0.25522512 |
| LDFVAVAL | ASGEDWGENTQY | 0.7879767 |
| LAQPGAE | ASGEDWGENTQY | 0.10456133 |
| ISNLGAVTGF | ASGEDWGENTQY | 0.14165764 |
| HQEDPGRI | ASGEDWGENTQY | 0.12288688 |
| GTPGEPTP | ASGEDWGENTQY | 0.5491585 |
| GFFNSGS | ASGEDWGENTQY | 0.25964952 |
| GAGGGGLTAGAN | ASGEDWGENTQY | 0.107646614 |
| FVDALVNM | ASGEDWGENTQY | 0.38392663 |
| FLTVAPEL | ASGEDWGENTQY | 0.09830929 |
| FETLTAL | ASGEDWGENTQY | 0.34995288 |
| DVINPIEL | ASGEDWGENTQY | 0.40903094 |
| APVSGAPV | ASGEDWGENTQY | 0.7701781 |
| ALVGGLAW | ASGEDWGENTQY | 0.12099129 |
| ALAAAAATAI | ASGEDWGENTQY | 0.47234994 |
| AFGLGLLL | ASGEDWGENTQY | 0.1656235 |
| AFFAAASL | ASGEDWGENTQY | 0.8440081 |
| AAAALPAFSPP | ASGEDWGENTQY | 0.1846259 |

| | | |
|---|---|---|
| VVLAPAL | ASGEDWGENTQY | 0.11761203 |
| TIGHVDHGKTTL | ASGEDWGENTQY | 0.44684428 |
| LISNASDALDK | ASGEDWGENTQY | 0.47963846 |
| LISNASDAL | ASGEDWGENTQY | 0.25373396 |
| ISNASDALDK | ASGEDWGENTQY | 0.6487441 |
| NQAETLVYQTE | ASGEDWGENTQY | 0.26289672 |
| TGGTGGTAGAG | ASGEDWGENTQY | 0.1869578 |
| TGSTAGGAGGA | ASGEDWGENTQY | 0.5526305 |
| DPLAVPVAAGPV | ASGEDWGENTQY | 0.46912295 |
| GGTGGTPGNGGN | ASGEDWGENTQY | 0.14653155 |
| PAGLAGVDPATL | ASGEDWGENTQY | 0.37859762 |
| GNNLVVEAPHAGASV | ASGEDWGENTQY | 0.29120737 |
| KVTRIMSASTLEQPA | ASGEDWGENTQY | 0.14549486 |
| LGFALPIALPRRTVV | ASGEDWGENTQY | 0.7386937 |
| TRLGNLTARKLVVLV | ASGEDWGENTQY | 0.2482146 |
| WSQQRRLIYFPSAGP | ASGEDWGENTQY | 0.46784368 |
| GPNGNPDPMAAAVDIR | ASGEDWGENTQY | 0.31884104 |
| TQRCVSVADLRAAASR | ASGEDWGENTQY | 0.8115134 |
| QFVLDVAEHPVEVLTETV | ASGEDWGENTQY | 0.12288492 |
| GGAGGTGGAGGNGGTQAGDG | ASGEDWGENTQY | 0.3545319 |
| PLLATLRAIAVDVDAKPAQV | ASGEDWGENTQY | 0.26567912 |
| APKFRTGGYMIDFWGVGYQVAK | ASGEDWGENTQY | 0.47342277 |
| GAGGAGGAGGAAGGPGGTGGTG | ASGEDWGENTQY | 0.3084755 |
| IGALGLLATDVAPAGSVAFGVA | ASGEDWGENTQY | 0.34408087 |
| AGGAGGIGGNGGAGGAGGVLMG | ASGEDWGENTQY | 0.27215356 |
| ADELIGLL | ASGEDWGENTQY | 0.5474064 |
| ADRALKA | ASGEDWGENTQY | 0.22841616 |
| AGAVGQVGGAGGA | ASGEDWGENTQY | 0.34987444 |
| AGGGGGLGGAP | ASGEDWGENTQY | 0.74656117 |
| AGGLLYGNGGAG | ASGEDWGENTQY | 0.14066575 |
| AGLESIGL | ASGEDWGENTQY | 0.24731104 |
| AGLKTGII | ASGEDWGENTQY | 0.43303502 |
| ASLAALLL | ASGEDWGENTQY | 0.6011129 |
| ASVINISSI | ASGEDWGENTQY | 0.2285772 |
| ATPPGPA | ASGEDWGENTQY | 0.58741796 |
| ATVASALV | ASGEDWGENTQY | 0.15030116 |
| ATVASLIL | ASGEDWGENTQY | 0.45872465 |
| AVNGPVLIRH | ASGEDWGENTQY | 0.34757268 |
| DGAAGAVGI | ASGEDWGENTQY | 0.2904474 |
| DPLLHGI | ASGEDWGENTQY | 0.13069251 |
| ELGAALGAL | ASGEDWGENTQY | 0.3783018 |
| ELGRTQAV | ASGEDWGENTQY | 0.5018794 |
| FVAGGYLS | ASGEDWGENTQY | 0.4770686 |
| GASGGAGGAGGW | ASGEDWGENTQY | 0.0963252 |
| GGAGAGGDGGAGGI | ASGEDWGENTQY | 0.46134776 |
| GTGHNGSQPGMGGNG | ASGEDWGENTQY | 0.53684056 |
| GYAAASASAS | ASGEDWGENTQY | 0.8238512 |
| HQVDVAAVQQG | ASGEDWGENTQY | 0.32939613 |
| IGAAIAHT | ASGEDWGENTQY | 0.35640848 |
| IGMIEVAQL | ASGEDWGENTQY | 0.18290704 |
| IRLAPEHTSAMD | ASGEDWGENTQY | 0.26180142 |

| | | |
|---|---|---|
| IYNVSVL | ASGEDWGENTQY | 0.12549953 |
| KQGFLGM | ASGEDWGENTQY | 0.23123908 |
| LESVITQL | ASGEDWGENTQY | 0.22095728 |
| LGLGALVVV | ASGEDWGENTQY | 0.14810103 |
| LIGNGGNGGSGGI | ASGEDWGENTQY | 0.7446807 |
| LNAINAPTL | ASGEDWGENTQY | 0.6506171 |
| LQLLNTGV | ASGEDWGENTQY | 0.6156357 |
| LQTSIEF | ASGEDWGENTQY | 0.6616999 |
| MSFAVSEL | ASGEDWGENTQY | 0.758715 |
| NTGALGL | ASGEDWGENTQY | 0.38679165 |
| PPGAGGAGGAGTT | ASGEDWGENTQY | 0.16948327 |
| QLVGLGI | ASGEDWGENTQY | 0.1719183 |
| RFLEQVHQL | ASGEDWGENTQY | 0.113545604 |
| RGVGYVLR | ASGEDWGENTQY | 0.16715132 |
| SATGVFAGVI | ASGEDWGENTQY | 0.18005441 |
| SLPAITLP | ASGEDWGENTQY | 0.2642308 |
| STALAFTV | ASGEDWGENTQY | 0.09286542 |
| SVLTVGLGK | ASGEDWGENTQY | 0.59502506 |
| TALAGAGSGI | ASGEDWGENTQY | 0.65985227 |
| TGLPFTGI | ASGEDWGENTQY | 0.36552858 |
| VAEHVTV | ASGEDWGENTQY | 0.3461787 |
| VDIVLNSL | ASGEDWGENTQY | 0.47282514 |
| VLTGGGAVL | ASGEDWGENTQY | 0.16621082 |
| VNVGLAYGI | ASGEDWGENTQY | 0.5198473 |
| VSSAAVGAV | ASGEDWGENTQY | 0.15791324 |
| VTSVYEAGV | ASGEDWGENTQY | 0.44939733 |
| WVALDRGA | ASGEDWGENTQY | 0.6519659 |
| YLVGSVVI | ASGEDWGENTQY | 0.2584828 |
| GPPGTGKTLL | ASGEDWGENTQY | 0.30596465 |
| GVGMTKF | ASGEDWGENTQY | 0.2551523 |
| IGIGGSDL | ASGEDWGENTQY | 0.117571056 |
| IGTLLAEL | ASGEDWGENTQY | 0.3357114 |
| IYTSGTT | ASGEDWGENTQY | 0.13730876 |
| LSFPAGV | ASGEDWGENTQY | 0.28832182 |
| LYTSGTT | ASGEDWGENTQY | 0.1285736 |
| VGAFPFGF | ASGEDWGENTQY | 0.33763155 |
| VGAGTVEF | ASGEDWGENTQY | 0.26861343 |
| VGKSTLLL | ASGEDWGENTQY | 0.32782865 |
| AALAHAGGGIV | ASGEDWGENTQY | 0.12795325 |
| AALVAVSLL | ASGEDWGENTQY | 0.34414476 |
| AAVSRAVAAL | ASGEDWGENTQY | 0.25406572 |
| AGGSSGIGGF | ASGEDWGENTQY | 0.12035076 |
| AGLTVANANG | ASGEDWGENTQY | 0.5293103 |
| AGLVGAVL | ASGEDWGENTQY | 0.21350873 |
| ALSQAFTV | ASGEDWGENTQY | 0.29546767 |
| ALTFSVSL | ASGEDWGENTQY | 0.52383214 |
| ALTGAALVAL | ASGEDWGENTQY | 0.26202172 |
| ASFSIGAGVL | ASGEDWGENTQY | 0.108331054 |
| DGSVYKI | ASGEDWGENTQY | 0.26668033 |
| EQLAEIV | ASGEDWGENTQY | 0.23741405 |
| FAVSELTVM | ASGEDWGENTQY | 0.21316567 |

| | | |
|---|---|---|
| GAAPLAAGL | ASGEDWGENTQY | 0.09245433 |
| GADGGDGGSGGAGG | ASGEDWGENTQY | 0.34897244 |
| GFGLDTGGAGG | ASGEDWGENTQY | 0.147757 |
| GLASGGLL | ASGEDWGENTQY | 0.14734107 |
| GRIAGAAVLL | ASGEDWGENTQY | 0.1586138 |
| GSAGSLNTSF | ASGEDWGENTQY | 0.5665966 |
| GTGQIGI | ASGEDWGENTQY | 0.6647947 |
| GTLIALIL | ASGEDWGENTQY | 0.26552075 |
| IAATMGFVVG | ASGEDWGENTQY | 0.43502277 |
| IFAQALL | ASGEDWGENTQY | 0.32891333 |
| IGAAVALGF | ASGEDWGENTQY | 0.722265 |
| IGLTPLAL | ASGEDWGENTQY | 0.3298821 |
| ILGSNTSTL | ASGEDWGENTQY | 0.23749964 |
| IVFSLTVL | ASGEDWGENTQY | 0.32043195 |
| LASKATGF | ASGEDWGENTQY | 0.21555883 |
| LASVALIL | ASGEDWGENTQY | 0.10914119 |
| LFAVGLTL | ASGEDWGENTQY | 0.113054425 |
| LFNGGIGTYI | ASGEDWGENTQY | 0.2506405 |
| LTLGAGII | ASGEDWGENTQY | 0.11038211 |
| LVAAALGDI | ASGEDWGENTQY | 0.30045024 |
| NTAFVQLGI | ASGEDWGENTQY | 0.4846608 |
| QLFSGAEL | ASGEDWGENTQY | 0.3785764 |
| SAATFITL | ASGEDWGENTQY | 0.11468695 |
| SNYGLANLGV | ASGEDWGENTQY | 0.17823832 |
| SYGGAYAVM | ASGEDWGENTQY | 0.1653096 |
| TTGVGGLTLG | ASGEDWGENTQY | 0.47730413 |
| TTQTIYI | ASGEDWGENTQY | 0.4414858 |
| VAAGAGLGAVI | ASGEDWGENTQY | 0.81844395 |
| VAAVVTLL | ASGEDWGENTQY | 0.56324387 |
| VGFLGGLI | ASGEDWGENTQY | 0.1813443 |
| VGVIPILL | ASGEDWGENTQY | 0.45372337 |
| VIFASSLI | ASGEDWGENTQY | 0.18551277 |
| VKGGDGGAAAG | ASGEDWGENTQY | 0.2641459 |
| VTKTFTNI | ASGEDWGENTQY | 0.19472863 |
| VVAAAILL | ASGEDWGENTQY | 0.425106 |
| YAALDVEL | ASGEDWGENTQY | 0.2781877 |
| LASVALLL | ASGEDWGENTQY | 0.18454143 |
| LDAVHRV | ASGEDWGENTQY | 0.11930331 |
| QGLSSSIF | ASGEDWGENTQY | 0.358378 |
| RGTFANIRL | ASGEDWGENTQY | 0.31378317 |
| RQLWWGHRI | ASGEDWGENTQY | 0.14356858 |
| YGPPGTGKTLL | ASGEDWGENTQY | 0.19263414 |
| AALVGAPANVA | ASGEDWGENTQY | 0.8082274 |
| AATFRRTE | ASGEDWGENTQY | 0.75134265 |
| AGALGGGI | ASGEDWGENTQY | 0.15981777 |
| AGGAGGAGGIGAG | ASGEDWGENTQY | 0.15739976 |
| AGSGALGL | ASGEDWGENTQY | 0.58177465 |
| AQIADQI | ASGEDWGENTQY | 0.12712768 |
| ASAVAVTYL | ASGEDWGENTQY | 0.3042966 |
| ASTVLGGTGG | ASGEDWGENTQY | 0.38917267 |
| AVGGEPAK | ASGEDWGENTQY | 0.31740516 |

| | | |
|---|---|---|
| CEQGTFGGTC | ASGEDWGENTQY | 0.16036375 |
| DEGKAPAG | ASGEDWGENTQY | 0.3446536 |
| DTVVFPAL | ASGEDWGENTQY | 0.2068753 |
| FEGITMAG | ASGEDWGENTQY | 0.39107147 |
| FPLVVQD | ASGEDWGENTQY | 0.094198965 |
| GAFTTPQ | ASGEDWGENTQY | 0.36913168 |
| GFGGFGTAGGVGGA | ASGEDWGENTQY | 0.13432105 |
| GFLAQFL | ASGEDWGENTQY | 0.18109915 |
| GGAGGAGGNAGVA | ASGEDWGENTQY | 0.1075003 |
| GGVGTAGAGGAGGAGG | ASGEDWGENTQY | 0.17066817 |
| GNGGGGDAGNAGSGGN | ASGEDWGENTQY | 0.6148851 |
| GTGGAGGAGNDAGS | ASGEDWGENTQY | 0.6423706 |
| GTTNTGWFNS | ASGEDWGENTQY | 0.15559343 |
| IFMGPLFTL | ASGEDWGENTQY | 0.49052745 |
| ILAHAIEAI | ASGEDWGENTQY | 0.80919313 |
| IRSAIREAL | ASGEDWGENTQY | 0.1725363 |
| ITFAGGG | ASGEDWGENTQY | 0.10600261 |
| ITGMSGHQEV | ASGEDWGENTQY | 0.366383 |
| IVAGLACTL | ASGEDWGENTQY | 0.24867558 |
| IVGGGLAGA | ASGEDWGENTQY | 0.5925894 |
| IVTGVLNI | ASGEDWGENTQY | 0.7134062 |
| LAQQGCI | ASGEDWGENTQY | 0.10760613 |
| LCLHVEN | ASGEDWGENTQY | 0.44075736 |
| LDAISDGS | ASGEDWGENTQY | 0.5713686 |
| LGGFAAML | ASGEDWGENTQY | 0.14765374 |
| LGGKNPMI | ASGEDWGENTQY | 0.1288826 |
| LGPKGVAAIL | ASGEDWGENTQY | 0.14165768 |
| LMVGVSI | ASGEDWGENTQY | 0.12097124 |
| LVGAAFVL | ASGEDWGENTQY | 0.3187917 |
| LVLAVAPL | ASGEDWGENTQY | 0.16137971 |
| MAAIAVIDAA | ASGEDWGENTQY | 0.18163225 |
| NNGINAELQ | ASGEDWGENTQY | 0.20795198 |
| RGEPLVLVHG | ASGEDWGENTQY | 0.16517763 |
| RGGAGGAAGLI | ASGEDWGENTQY | 0.18439677 |
| RSYRMHPAV | ASGEDWGENTQY | 0.18436006 |
| SAAHLLAGQLA | ASGEDWGENTQY | 0.26029798 |
| SAAVGSY | ASGEDWGENTQY | 0.40443617 |
| SPHAVLHRI | ASGEDWGENTQY | 0.23601341 |
| SQIAKVL | ASGEDWGENTQY | 0.2874577 |
| TALGHLVP | ASGEDWGENTQY | 0.65721345 |
| TAPTTSGAA | ASGEDWGENTQY | 0.12838869 |
| TDGDDGNGGSPGT | ASGEDWGENTQY | 0.15761201 |
| TDGLGEN | ASGEDWGENTQY | 0.46554756 |
| TGGPPLPGQA | ASGEDWGENTQY | 0.14710735 |
| TIPAITI | ASGEDWGENTQY | 0.58627623 |
| TIPGFTI | ASGEDWGENTQY | 0.30454227 |
| TSIASGALA | ASGEDWGENTQY | 0.32335842 |
| TVTLPGAAI | ASGEDWGENTQY | 0.36838442 |
| VASGLSAALP | ASGEDWGENTQY | 0.80597067 |
| VGAGVIGI | ASGEDWGENTQY | 0.38313353 |
| VGPKRFGV | ASGEDWGENTQY | 0.16553916 |

| | | |
|---|---|---|
| VNVAFPD | ASGEDWGENTQY | 0.6748607 |
| VSMTNSGVS | ASGEDWGENTQY | 0.1404314 |
| VTDVLGDLSV | ASGEDWGENTQY | 0.45954835 |
| VVVVGAGFAGL | ASGEDWGENTQY | 0.42516136 |
| VYTVGTVV | ASGEDWGENTQY | 0.13376838 |
| WTTFSGP | ASGEDWGENTQY | 0.19557053 |
| YDELVEAF | ASGEDWGENTQY | 0.61030805 |
| YGTGGAGGNGG | ASGEDWGENTQY | 0.5552171 |
| YTNVMAV | ASGEDWGENTQY | 0.122083865 |
| FLEQVHQL | ASGEDWGENTQY | 0.15406324 |
| GLSLLAVAV | ASGEDWGENTQY | 0.13778824 |
| LEQVHQL | ASGEDWGENTQY | 0.249326 |
| LNAAGAI | ASGEDWGENTQY | 0.1326972 |
| PAVHASL | ASGEDWGENTQY | 0.45479998 |
| TELEDDA | ASGEDWGENTQY | 0.18592025 |
| TPPGPPN | ASGEDWGENTQY | 0.7239648 |
| TPPPPPPP | ASGEDWGENTQY | 0.29602587 |
| TQSPALI | ASSEEDQDTQY | 0.532179 |
| EFAAAFPQ | ASSEEDQDTQY | 0.16615207 |
| LLSGNLDVL | ASSEEDQDTQY | 0.2089654 |
| GPANLGNYNI | ASSEEDQDTQY | 0.122991316 |
| TLGPTLP | ASSEEDQDTQY | 0.18428202 |
| VYAYSSCCK | ASSEEDQDTQY | 0.14841594 |
| APAPAPAPNI | ASSEEDQDTQY | 0.19975668 |
| PGGPVPIP | ASSEEDQDTQY | 0.11497202 |
| FLTVALEP | ASSEEDQDTQY | 0.7371939 |
| SQYFGANL | ASSEEDQDTQY | 0.27201426 |
| GGVKPAA | ASSEEDQDTQY | 0.4237466 |
| GGTGGAGGAGGW | ASSEEDQDTQY | 0.17721274 |
| GVGGVAGLGGAGG | ASSEEDQDTQY | 0.16654833 |
| VLLVLATTI | ASSEEDQDTQY | 0.17237708 |
| IGIFAAML | ASSEEDQDTQY | 0.6229687 |
| AVGDLID | ASSEEDQDTQY | 0.12686387 |
| NISPLPL | ASSEEDQDTQY | 0.09501978 |
| VTAVFVAI | ASSEEDQDTQY | 0.15474825 |
| SGDDNDAAL | ASSEEDQDTQY | 0.10382019 |
| AVAALAHS | ASSEEDQDTQY | 0.73619044 |
| GGVTVMATP | ASSEEDQDTQY | 0.54073787 |
| AGLAANVRD | ASSEEDQDTQY | 0.29318756 |
| TNRPDLIDPALLRPG | ASSEEDQDTQY | 0.5629931 |
| LANLGVTAL | ASSEEDQDTQY | 0.094900936 |
| TGERIEL | ASSEEDQDTQY | 0.19193643 |
| NGEELAA | ASSEEDQDTQY | 0.72184664 |
| VLSLVWG | ASSEEDQDTQY | 0.29893333 |
| IGYALWL | ASSEEDQDTQY | 0.21166457 |
| LSLLLAPL | ASSEEDQDTQY | 0.6236173 |
| TLDEALK | ASSEEDQDTQY | 0.16623485 |
| TDGAGTAEGG | ASSEEDQDTQY | 0.2305358 |
| NAPGGAGGNGG | ASSEEDQDTQY | 0.22863758 |
| VSVFMGMGGNF | ASSEEDQDTQY | 0.2772873 |
| LASYLAAL | ASSEEDQDTQY | 0.451567 |

| | | |
|---|---|---|
| LVGGTPAL | ASSEEDQDTQY | 0.12720968 |
| VRNADVL | ASSEEDQDTQY | 0.32980955 |
| TCSSPFLVSAM | ASSEEDQDTQY | 0.81397307 |
| VHITDAE | ASSEEDQDTQY | 0.20592937 |
| EYAAPSF | ASSEEDQDTQY | 0.59360766 |
| VVDAFHGGL | ASSEEDQDTQY | 0.3206532 |
| ASVVDPQ | ASSEEDQDTQY | 0.71293044 |
| SYNEFGGLS | ASSEEDQDTQY | 0.14819089 |
| DPDGSVY | ASSEEDQDTQY | 0.13498983 |
| MVSTIHATE | ASSEEDQDTQY | 0.34637004 |
| LAAGVADPV | ASSEEDQDTQY | 0.13631186 |
| TKEQIAA | ASSEEDQDTQY | 0.27823585 |
| SLASPAAL | ASSEEDQDTQY | 0.1800848 |
| IGLVFYGV | ASSEEDQDTQY | 0.85599554 |
| FLLDASNL | ASSEEDQDTQY | 0.41776568 |
| YQVGNFP | ASSEEDQDTQY | 0.37833458 |
| ESLLSQST | ASSEEDQDTQY | 0.13297825 |
| LFTVGFAAVG | ASSEEDQDTQY | 0.2582355 |
| ATPPTPT | ASSEEDQDTQY | 0.48564106 |
| TDLATLG | ASSEEDQDTQY | 0.12279245 |
| SAGGTGR | ASSEEDQDTQY | 0.65013313 |
| GFSNSGS | ASSEEDQDTQY | 0.15654811 |
| GGNGGSGGTSV | ASSEEDQDTQY | 0.34054196 |
| GTSVATGGAGN | ASSEEDQDTQY | 0.46068695 |
| NGGNGGSGGTS | ASSEEDQDTQY | 0.786103 |
| AVVKFPH | ASSEEDQDTQY | 0.20278394 |
| TPPPPPP | ASSEEDQDTQY | 0.29012623 |
| AAPAAAAAPA | ASSEEDQDTQY | 0.1315469 |
| IALNEDL | ASSEEDQDTQY | 0.506384 |
| IVNVSSM | ASSEEDQDTQY | 0.63008547 |
| TGIPPAPR | ASSEEDQDTQY | 0.663011 |
| FDLGGGTF | ASSEEDQDTQY | 0.7625711 |
| FDLGGGTFDVS | ASSEEDQDTQY | 0.3737532 |
| NEPTAAA | ASSEEDQDTQY | 0.12978134 |
| NEPTAAALA | ASSEEDQDTQY | 0.21429884 |
| GSIINISSI | ASSEEDQDTQY | 0.5345085 |
| VAPGNAGTA | ASSEEDQDTQY | 0.72897846 |
| TGGGASLE | ASSEEDQDTQY | 0.5506994 |
| VAVYNHYKRI | ASSEEDQDTQY | 0.55780613 |
| TDPAPATT | ASSEEDQDTQY | 0.16914722 |
| TNRPDILDPALLRPG | ASSEEDQDTQY | 0.6387746 |
| LTTTTTS | ASSEEDQDTQY | 0.83775574 |
| PTGSGKTL | ASSEEDQDTQY | 0.4747323 |
| TGSGKTL | ASSEEDQDTQY | 0.2109172 |
| TIGHVDHGK | ASSEEDQDTQY | 0.21717536 |
| TIGHVDHGKTT | ASSEEDQDTQY | 0.25946036 |
| GHVDHGKTT | ASSEEDQDTQY | 0.68664527 |
| AQFEHTLL | ASSEEDQDTQY | 0.14776306 |
| IVNQDEL | ASSEEDQDTQY | 0.32670987 |
| TNGSQFF | ASSEEDQDTQY | 0.31092036 |
| TGKGIVE | ASSEEDQDTQY | 0.10308306 |

| | | |
|---|---|---|
| LISNASDA | ASSEEDQDTQY | 0.44962493 |
| AELRTLGELTKIAKAHGAQ | ASSEEDQDTQY | 0.75278175 |
| APNAVAAAEPSW | ASSEEDQDTQY | 0.2241599 |
| APQVAGAATAVL | ASSEEDQDTQY | 0.15751098 |
| ARRDPAAAAAILPTDA | ASSEEDQDTQY | 0.65063536 |
| DAAARDVLAEAGLAEYF | ASSEEDQDTQY | 0.8347791 |
| DDQIVLSAKANLAKVEQ | ASSEEDQDTQY | 0.52180433 |
| DELLPDTGEDSDEDGH | ASSEEDQDTQY | 0.11133186 |
| ELPEAPANRATVPQD | ASSEEDQDTQY | 0.092773005 |
| FGQSKIERVTGIG | ASSEEDQDTQY | 0.260038 |
| GAGGTGSTAGGAGGAGGAGGLY | ASSEEDQDTQY | 0.72325826 |
| GHGGTGAAGGNGGTGGAGGSGI | ASSEEDQDTQY | 0.18662417 |
| GNGGNGGAGGEGGPGGAGGAGA | ASSEEDQDTQY | 0.15329267 |
| GSTAGNGGNGGNATNGTIAGQP | ASSEEDQDTQY | 0.51469815 |
| GTGGDGGNGGNGGTGG | ASSEEDQDTQY | 0.54282665 |
| GVRRPAGIPADAPV | ASSEEDQDTQY | 0.13998446 |
| GYRQVTDAHLLTLARR | ASSEEDQDTQY | 0.13836858 |
| IAKMGQDGHDRG | ASSEEDQDTQY | 0.35286373 |
| KAEPSIEIQYT | ASSEEDQDTQY | 0.50176924 |
| KPVRIRMAADEDTVAVT | ASSEEDQDTQY | 0.12972993 |
| MPPRIAGMRLLVIKPEP | ASSEEDQDTQY | 0.20063825 |
| MKFYAVALFRSRGVPPTR | ASSEEDQDTQY | 0.2398594 |
| NAGGFGILGAVAHSP | ASSEEDQDTQY | 0.51654375 |
| NEGREITLAMAEM | ASSEEDQDTQY | 0.37523317 |
| NELVEWSHPLGHRWRWE | ASSEEDQDTQY | 0.1402723 |
| NLGAGGTGGNGGNPG | ASSEEDQDTQY | 0.6679815 |
| NTEWPKWETVMAADPVD | ASSEEDQDTQY | 0.112868056 |
| PASAAPAETQP | ASSEEDQDTQY | 0.5315602 |
| QQSSDGKNGVRAAEITGE | ASSEEDQDTQY | 0.3329211 |
| RAPDAVTTSWWKE | ASSEEDQDTQY | 0.26878214 |
| SGFGNVGAHTSGWFNQ | ASSEEDQDTQY | 0.2118601 |
| SGVAGAVGQAASVGGLKV | ASSEEDQDTQY | 0.14187874 |
| SNGGAGTGSAGGNGGT | ASSEEDQDTQY | 0.12446143 |
| TGGAGGEGGAAG | ASSEEDQDTQY | 0.111388706 |
| TGGDGGAAGAGGAPGHG | ASSEEDQDTQY | 0.6414491 |
| TMPPLLAAIANAGRHGV | ASSEEDQDTQY | 0.48064935 |
| TSSTAGNSGGA | ASSEEDQDTQY | 0.48587427 |
| VPATVPVGILC | ASSEEDQDTQY | 0.31582862 |
| VQRPDSGGVVLTGRIS | ASSEEDQDTQY | 0.13318036 |
| WTPNPVPATKGVPAGLVQAT | ASSEEDQDTQY | 0.3324802 |
| YALKRALAKFSDRFDVVII | ASSEEDQDTQY | 0.13026588 |
| YLIGSVIV | ASSEEDQDTQY | 0.18714967 |
| YGQLVAVID | ASSEEDQDTQY | 0.6770617 |
| VTVGGMTLPA | ASSEEDQDTQY | 0.3804831 |
| VLALPAV | ASSEEDQDTQY | 0.13869536 |
| VIPPTLNYET | ASSEEDQDTQY | 0.31542838 |
| VGMGGMHPSSG | ASSEEDQDTQY | 0.2928576 |
| VGLAGGAATA | ASSEEDQDTQY | 0.14402461 |
| VGGAGGAGGSGP | ASSEEDQDTQY | 0.21792075 |
| VFAAFMV | ASSEEDQDTQY | 0.5910144 |
| TLYDKLR | ASSEEDQDTQY | 0.25962535 |

| | | |
|---|---|---|
| TLAVFTVL | ASSEEDQDTQY | 0.1268577 |
| TGSVQPARV | ASSEEDQDTQY | 0.39482117 |
| TGGGAGSGGT | ASSEEDQDTQY | 0.14580327 |
| SHLETSL | ASSEEDQDTQY | 0.3290912 |
| SFTTGMT | ASSEEDQDTQY | 0.22687128 |
| QVEVAKRMGVRQPTVS | ASSEEDQDTQY | 0.22057307 |
| QGSGGPALAM | ASSEEDQDTQY | 0.45943198 |
| MLAPSNP | ASSEEDQDTQY | 0.13958532 |
| LVNNAAI | ASSEEDQDTQY | 0.14230976 |
| LTGEINTLA | ASSEEDQDTQY | 0.22821999 |
| LSIFGPV | ASSEEDQDTQY | 0.24754444 |
| LLALLANL | ASSEEDQDTQY | 0.47274667 |
| LISNASDALDKLR | ASSEEDQDTQY | 0.14018208 |
| LINEAVP | ASSEEDQDTQY | 0.27818927 |
| LGYAFSGL | ASSEEDQDTQY | 0.177721 |
| LDFVAVAL | ASSEEDQDTQY | 0.14377601 |
| LAQPGAE | ASSEEDQDTQY | 0.23310983 |
| ISNLGAVTGF | ASSEEDQDTQY | 0.17733756 |
| HQEDPGRI | ASSEEDQDTQY | 0.14840339 |
| GTPGEPTP | ASSEEDQDTQY | 0.29683614 |
| GFFNSGS | ASSEEDQDTQY | 0.15406455 |
| GAGGGGLTAGAN | ASSEEDQDTQY | 0.33750862 |
| FVDALVNM | ASSEEDQDTQY | 0.65341127 |
| FLTVAPEL | ASSEEDQDTQY | 0.17166694 |
| FETLTAL | ASSEEDQDTQY | 0.42681164 |
| DVINPIEL | ASSEEDQDTQY | 0.40912092 |
| APVSGAPV | ASSEEDQDTQY | 0.41795328 |
| ALVGGLAW | ASSEEDQDTQY | 0.20046467 |
| ALAAAAATAI | ASSEEDQDTQY | 0.14395396 |
| AFGLGLLL | ASSEEDQDTQY | 0.57985586 |
| AFFAAASL | ASSEEDQDTQY | 0.4347474 |
| AAAALPAFSPP | ASSEEDQDTQY | 0.50603783 |
| VVLAPAL | ASSEEDQDTQY | 0.5137208 |
| TIGHVDHGKTTL | ASSEEDQDTQY | 0.426896 |
| LISNASDALDK | ASSEEDQDTQY | 0.45501626 |
| LISNASDAL | ASSEEDQDTQY | 0.18327087 |
| ISNASDALDK | ASSEEDQDTQY | 0.117176324 |
| NQAETLVYQTE | ASSEEDQDTQY | 0.19836828 |
| TGGTGGTAGAG | ASSEEDQDTQY | 0.22260731 |
| TGSTAGGAGGA | ASSEEDQDTQY | 0.14190947 |
| DPLAVPVAAGPV | ASSEEDQDTQY | 0.13637358 |
| GGTGGTPGNGGN | ASSEEDQDTQY | 0.60505295 |
| PAGLAGVDPATL | ASSEEDQDTQY | 0.8440516 |
| GNNLVVEAPHAGASV | ASSEEDQDTQY | 0.16755663 |
| KVTRIMSASTLEQPA | ASSEEDQDTQY | 0.81419957 |
| LGFALPIALPRRTVV | ASSEEDQDTQY | 0.121469684 |
| TRLGNLTARKLVVLV | ASSEEDQDTQY | 0.5137513 |
| WSQQRRLIYFPSAGP | ASSEEDQDTQY | 0.65557754 |
| GPNGNPDPMAAAVDIR | ASSEEDQDTQY | 0.14650136 |
| TQRCVSVADLRAAASR | ASSEEDQDTQY | 0.35752243 |
| QFVLDVAEHPVEVLTETV | ASSEEDQDTQY | 0.13853528 |

| | | |
|---|---|---|
| GGAGGTGGAGGNGGTQAGDG | ASSEEDQDTQY | 0.16544089 |
| PLLATLRAIAVDVDAKPAQV | ASSEEDQDTQY | 0.6754878 |
| APKFRTGGYMIDFWGVGYQVAK | ASSEEDQDTQY | 0.21436848 |
| GAGGAGGAGGAAGGPGGTGGTG | ASSEEDQDTQY | 0.26366726 |
| IGALGLLATDVAPAGSVAFGVA | ASSEEDQDTQY | 0.20584004 |
| AGGAGGIGGNGGAGGAGGVLMG | ASSEEDQDTQY | 0.15317662 |
| ADELIGLL | ASSEEDQDTQY | 0.2473611 |
| ADRALKA | ASSEEDQDTQY | 0.5769909 |
| AGAVGQVGGAGGA | ASSEEDQDTQY | 0.2586726 |
| AGGGGGLGGAP | ASSEEDQDTQY | 0.8184887 |
| AGGLLYGNGGAG | ASSEEDQDTQY | 0.13075441 |
| AGLESIGL | ASSEEDQDTQY | 0.7579678 |
| AGLKTGII | ASSEEDQDTQY | 0.7392783 |
| ASLAALLL | ASSEEDQDTQY | 0.31461298 |
| ASVINISSI | ASSEEDQDTQY | 0.21378328 |
| ATPPGPA | ASSEEDQDTQY | 0.6073236 |
| ATVASALV | ASSEEDQDTQY | 0.2524855 |
| ATVASLIL | ASSEEDQDTQY | 0.4607094 |
| AVNGPVLIRH | ASSEEDQDTQY | 0.47342172 |
| DGAAGAVGI | ASSEEDQDTQY | 0.8457875 |
| DPLLHGI | ASSEEDQDTQY | 0.913611 |
| ELGAALGAL | ASSEEDQDTQY | 0.8977034 |
| ELGRTQAV | ASSEEDQDTQY | 0.86731994 |
| FVAGGYLS | ASSEEDQDTQY | 0.8591652 |
| GASGGAGGAGGW | ASSEEDQDTQY | 0.46222305 |
| GGAGAGGDGGAGGI | ASSEEDQDTQY | 0.8409865 |
| GTGHNGSQPGMGGNG | ASSEEDQDTQY | 0.8193284 |
| GYAAASASAS | ASSEEDQDTQY | 0.86796874 |
| HQVDVAAVQQG | ASSEEDQDTQY | 0.90415525 |
| IGAAIAHT | ASSEEDQDTQY | 0.83716154 |
| IGMIEVAQL | ASSEEDQDTQY | 0.91128266 |
| IRLAPEHTSAMD | ASSEEDQDTQY | 0.88957065 |
| IYNVSVL | ASSEEDQDTQY | 0.7214953 |
| KQGFLGM | ASSEEDQDTQY | 0.86977017 |
| LESVITQL | ASSEEDQDTQY | 0.8247498 |
| LGLGALVVV | ASSEEDQDTQY | 0.8825188 |
| LIGNGGNGGSGGI | ASSEEDQDTQY | 0.54300183 |
| LNAINAPTL | ASSEEDQDTQY | 0.9074843 |
| LQLLNTGV | ASSEEDQDTQY | 0.8435533 |
| LQTSIEF | ASSEEDQDTQY | 0.1539062 |
| MSFAVSEL | ASSEEDQDTQY | 0.88919884 |
| NTGALGL | ASSEEDQDTQY | 0.87060964 |
| PPGAGGAGGAGTT | ASSEEDQDTQY | 0.87664807 |
| QLVGLGI | ASSEEDQDTQY | 0.90863276 |
| RFLEQVHQL | ASSEEDQDTQY | 0.3130636 |
| RGVGYVLR | ASSEEDQDTQY | 0.7895658 |
| SATGVFAGVI | ASSEEDQDTQY | 0.9044777 |
| SLPAITLP | ASSEEDQDTQY | 0.91303545 |
| STALAFTV | ASSEEDQDTQY | 0.7516389 |
| SVLTVGLGK | ASSEEDQDTQY | 0.9000791 |
| TALAGAGSGI | ASSEEDQDTQY | 0.90210795 |

| | | |
|---|---|---|
| TGLPFTGI | ASSEEDQDTQY | 0.899805 |
| VAEHVTV | ASSEEDQDTQY | 0.90579975 |
| VDIVLNSL | ASSEEDQDTQY | 0.9130655 |
| VLTGGGAVL | ASSEEDQDTQY | 0.72933984 |
| VNVGLAYGI | ASSEEDQDTQY | 0.9118973 |
| VSSAAVGAV | ASSEEDQDTQY | 0.9144255 |
| VTSVYEAGV | ASSEEDQDTQY | 0.8325218 |
| WVALDRGA | ASSEEDQDTQY | 0.88547367 |
| YLVGSVVI | ASSEEDQDTQY | 0.8151052 |
| GPPGTGKTLL | ASSEEDQDTQY | 0.8277113 |
| GVGMTKF | ASSEEDQDTQY | 0.37529182 |
| IGIGGSDL | ASSEEDQDTQY | 0.8961588 |
| IGTLLAEL | ASSEEDQDTQY | 0.9094342 |
| IYTSGTT | ASSEEDQDTQY | 0.86253774 |
| LSFPAGV | ASSEEDQDTQY | 0.7883674 |
| LYTSGTT | ASSEEDQDTQY | 0.80561125 |
| VGAFPFGF | ASSEEDQDTQY | 0.8681363 |
| VGAGTVEF | ASSEEDQDTQY | 0.87073755 |
| VGKSTLLL | ASSEEDQDTQY | 0.42033982 |
| AALAHAGGGIV | ASSEEDQDTQY | 0.80075294 |
| AALVAVSLL | ASSEEDQDTQY | 0.9085626 |
| AAVSRAVAAL | ASSEEDQDTQY | 0.9080442 |
| AGGSSGIGGF | ASSEEDQDTQY | 0.5885029 |
| AGLTVANANG | ASSEEDQDTQY | 0.91387415 |
| AGLVGAVL | ASSEEDQDTQY | 0.8963188 |
| ALSQAFTV | ASSEEDQDTQY | 0.91292155 |
| ALTFSVSL | ASSEEDQDTQY | 0.83981156 |
| ALTGAALVAL | ASSEEDQDTQY | 0.8270925 |
| ASFSIGAGVL | ASSEEDQDTQY | 0.8918005 |
| DGSVYKI | ASSEEDQDTQY | 0.85561097 |
| EQLAEIV | ASSEEDQDTQY | 0.73574734 |
| FAVSELTVM | ASSEEDQDTQY | 0.9070428 |
| GAAPLAAGL | ASSEEDQDTQY | 0.8625876 |
| GADGGDGGSGGAGG | ASSEEDQDTQY | 0.8338007 |
| GFGLDTGGAGG | ASSEEDQDTQY | 0.804124 |
| GLASGGLL | ASSEEDQDTQY | 0.9167919 |
| GRIAGAAVLL | ASSEEDQDTQY | 0.7612382 |
| GSAGSLNTSF | ASSEEDQDTQY | 0.7625543 |
| GTGQIGI | ASSEEDQDTQY | 0.69891554 |
| GTLIALIL | ASSEEDQDTQY | 0.87406623 |
| IAATMGFVVG | ASSEEDQDTQY | 0.908782 |
| IFAQALL | ASSEEDQDTQY | 0.8956804 |
| IGAAVALGF | ASSEEDQDTQY | 0.82286555 |
| IGLTPLAL | ASSEEDQDTQY | 0.8958052 |
| ILGSNTSTL | ASSEEDQDTQY | 0.8918134 |
| IVFSLTVL | ASSEEDQDTQY | 0.8846259 |
| LASKATGF | ASSEEDQDTQY | 0.8994193 |
| LASVALIL | ASSEEDQDTQY | 0.82199514 |
| LFAVGLTL | ASSEEDQDTQY | 0.9143469 |
| LFNGGIGTYI | ASSEEDQDTQY | 0.8981534 |
| LTLGAGII | ASSEEDQDTQY | 0.90524393 |

| | | |
|---|---|---|
| LVAAALGDI | ASSEEDQDTQY | 0.9026763 |
| NTAFVQLGI | ASSEEDQDTQY | 0.87320805 |
| QLFSGAEL | ASSEEDQDTQY | 0.8813271 |
| SAATFITL | ASSEEDQDTQY | 0.5210023 |
| SNYGLANLGV | ASSEEDQDTQY | 0.6593107 |
| SYGGAYAVM | ASSEEDQDTQY | 0.8965615 |
| TTGVGGLTLG | ASSEEDQDTQY | 0.89194524 |
| TTQTIYI | ASSEEDQDTQY | 0.9006075 |
| VAAGAGLGAVI | ASSEEDQDTQY | 0.8772316 |
| VAAVVTLL | ASSEEDQDTQY | 0.8302123 |
| VGFLGGLI | ASSEEDQDTQY | 0.8919285 |
| VGVIPILL | ASSEEDQDTQY | 0.89926094 |
| VIFASSLI | ASSEEDQDTQY | 0.67936444 |
| VKGGDGGAAAG | ASSEEDQDTQY | 0.8557173 |
| VTKTFTNI | ASSEEDQDTQY | 0.9196022 |
| VVAAAILL | ASSEEDQDTQY | 0.8830631 |
| YAALDVEL | ASSEEDQDTQY | 0.8671632 |
| LASVALLL | ASSEEDQDTQY | 0.8222357 |
| LDAVHRV | ASSEEDQDTQY | 0.9062613 |
| QGLSSSIF | ASSEEDQDTQY | 0.802144 |
| RGTFANIRL | ASSEEDQDTQY | 0.91961956 |
| RQLWWGHRI | ASSEEDQDTQY | 0.26899466 |
| YGPPGTGKTLL | ASSEEDQDTQY | 0.84549 |
| AALVGAPANVA | ASSEEDQDTQY | 0.7375966 |
| AATFRRTE | ASSEEDQDTQY | 0.68059945 |
| AGALGGGI | ASSEEDQDTQY | 0.9136816 |
| AGGAGGAGGIGAG | ASSEEDQDTQY | 0.7784326 |
| AGSGALGL | ASSEEDQDTQY | 0.89516515 |
| AQIADQI | ASSEEDQDTQY | 0.87848985 |
| ASAVAVTYL | ASSEEDQDTQY | 0.9110571 |
| ASTVLGGTGG | ASSEEDQDTQY | 0.88093 |
| AVGGEPAK | ASSEEDQDTQY | 0.85909575 |
| CEQGTFGGTC | ASSEEDQDTQY | 0.75046074 |
| DEGKAPAG | ASSEEDQDTQY | 0.90966547 |
| DTVVFPAL | ASSEEDQDTQY | 0.9021301 |
| FEGITMAG | ASSEEDQDTQY | 0.90085185 |
| FPLVVQD | ASSEEDQDTQY | 0.63060117 |
| GAFTTPQ | ASSEEDQDTQY | 0.8536719 |
| GFGGFGTAGGVGGA | ASSEEDQDTQY | 0.8943014 |
| GFLAQFL | ASSEEDQDTQY | 0.26380906 |
| GGAGGAGGNAGVA | ASSEEDQDTQY | 0.3045959 |
| GGVGTAGAGGAGGAGG | ASSEEDQDTQY | 0.87153244 |
| GNGGGGDAGNAGSGGN | ASSEEDQDTQY | 0.81526166 |
| GTGGAGGAGNDAGS | ASSEEDQDTQY | 0.91371876 |
| GTTNTGWFNS | ASSEEDQDTQY | 0.9027635 |
| IFMGPLFTL | ASSEEDQDTQY | 0.87580097 |
| ILAHAIEAI | ASSEEDQDTQY | 0.8952719 |
| IRSAIREAL | ASSEEDQDTQY | 0.8708677 |
| ITFAGGG | ASSEEDQDTQY | 0.08253676 |
| ITGMSGHQEV | ASSEEDQDTQY | 0.7538018 |
| IVAGLACTL | ASSEEDQDTQY | 0.7389606 |

| | | |
|---|---|---|
| IVGGGLAGA | ASSEEDQDTQY | 0.90757304 |
| IVTGVLNI | ASSEEDQDTQY | 0.83533084 |
| LAQQGCI | ASSEEDQDTQY | 0.7993772 |
| LCLHVEN | ASSEEDQDTQY | 0.8787786 |
| LDAISDGS | ASSEEDQDTQY | 0.87224555 |
| LGGFAAML | ASSEEDQDTQY | 0.861167 |
| LGGKNPMI | ASSEEDQDTQY | 0.7842097 |
| LGPKGVAAIL | ASSEEDQDTQY | 0.16268885 |
| LMVGVSI | ASSEEDQDTQY | 0.9081481 |
| LVGAAFVL | ASSEEDQDTQY | 0.37950778 |
| LVLAVAPL | ASSEEDQDTQY | 0.7797898 |
| MAAIAVIDAA | ASSEEDQDTQY | 0.30852598 |
| NNGINAELQ | ASSEEDQDTQY | 0.22823325 |
| RGEPLVLVHG | ASSEEDQDTQY | 0.45030355 |
| RGGAGGAAGLI | ASSEEDQDTQY | 0.9047272 |
| RSYRMHPAV | ASSEEDQDTQY | 0.84414756 |
| SAAHLLAGQLA | ASSEEDQDTQY | 0.8349081 |
| SAAVGSY | ASSEEDQDTQY | 0.9160396 |
| SPHAVLHRI | ASSEEDQDTQY | 0.8514888 |
| SQIAKVL | ASSEEDQDTQY | 0.8583286 |
| TALGHLVP | ASSEEDQDTQY | 0.8255206 |
| TAPTTSGAA | ASSEEDQDTQY | 0.8295022 |
| TDGDDGNGGSPGT | ASSEEDQDTQY | 0.8197967 |
| TDGLGEN | ASSEEDQDTQY | 0.79601586 |
| TGGPPLPGQA | ASSEEDQDTQY | 0.91491604 |
| TIPAITI | ASSEEDQDTQY | 0.85291994 |
| TIPGFTI | ASSEEDQDTQY | 0.5754767 |
| TSIASGALA | ASSEEDQDTQY | 0.8531321 |
| TVTLPGAAI | ASSEEDQDTQY | 0.91267353 |
| VASGLSAALP | ASSEEDQDTQY | 0.9208015 |
| VGAGVIGI | ASSEEDQDTQY | 0.8353392 |
| VGPKRFGV | ASSEEDQDTQY | 0.751884 |
| VNVAFPD | ASSEEDQDTQY | 0.8254887 |
| VSMTNSGVS | ASSEEDQDTQY | 0.87472034 |
| VTDVLGDLSV | ASSEEDQDTQY | 0.79383767 |
| VVVVGAGFAGL | ASSEEDQDTQY | 0.66349584 |
| VYTVGTVV | ASSEEDQDTQY | 0.90865445 |
| WTTFSGP | ASSEEDQDTQY | 0.9083898 |
| YDELVEAF | ASSEEDQDTQY | 0.8368223 |
| YGTGGAGGNGG | ASSEEDQDTQY | 0.7429372 |
| YTNVMAV | ASSEEDQDTQY | 0.87112343 |
| FLEQVHQL | ASSEEDQDTQY | 0.8220595 |
| GLSLLAVAV | ASSEEDQDTQY | 0.8644134 |
| LEQVHQL | ASSEEDQDTQY | 0.8552138 |
| LNAAGAI | ASSEEDQDTQY | 0.67863 |
| PAVHASL | ASSEEDQDTQY | 0.84152025 |
| TELEDDA | ASSEEDQDTQY | 0.9198688 |
| TPPGPPN | ASSEEDQDTQY | 0.8403775 |
| TPPPPPPP | ASSEEDQDTQY | 0.8776159 |
| TQSPALI | AWSHRGGNYAEQF | 0.9140095 |
| EFAAAFPQ | AWSHRGGNYAEQF | 0.90475124 |

| | | |
|---|---|---|
| LLSGNLDVL | AWSHRGGNYAEQF | 0.91619086 |
| GPANLGNYNI | AWSHRGGNYAEQF | 0.89510095 |
| TLGPTLP | AWSHRGGNYAEQF | 0.8656275 |
| VYAYSSCCK | AWSHRGGNYAEQF | 0.88005114 |
| APAPAPAPNI | AWSHRGGNYAEQF | 0.8702247 |
| PGGPVPIP | AWSHRGGNYAEQF | 0.8789801 |
| FLTVALEP | AWSHRGGNYAEQF | 0.8716155 |
| SQYFGANL | AWSHRGGNYAEQF | 0.8139412 |
| GGVKPAA | AWSHRGGNYAEQF | 0.91053724 |
| GGTGGAGGAGGW | AWSHRGGNYAEQF | 0.8733374 |
| GVGGVAGLGGAGG | AWSHRGGNYAEQF | 0.90297866 |
| VLLVLATTI | AWSHRGGNYAEQF | 0.70924056 |
| IGIFAAML | AWSHRGGNYAEQF | 0.77667165 |
| AVGDLID | AWSHRGGNYAEQF | 0.90854716 |
| NISPLPL | AWSHRGGNYAEQF | 0.13020092 |
| VTAVFVAI | AWSHRGGNYAEQF | 0.25549835 |
| SGDDNDAAL | AWSHRGGNYAEQF | 0.890277 |
| AVAALAHS | AWSHRGGNYAEQF | 0.9130757 |
| GGVTVMATP | AWSHRGGNYAEQF | 0.15171708 |
| AGLAANVRD | AWSHRGGNYAEQF | 0.73335695 |
| TNRPDLIDPALLRPG | AWSHRGGNYAEQF | 0.68540233 |
| LANLGVTAL | AWSHRGGNYAEQF | 0.8824806 |
| TGERIEL | AWSHRGGNYAEQF | 0.8833002 |
| NGEELAA | AWSHRGGNYAEQF | 0.53368205 |
| VLSLVWG | AWSHRGGNYAEQF | 0.9054467 |
| IGYALWL | AWSHRGGNYAEQF | 0.6542952 |
| LSLLLAPL | AWSHRGGNYAEQF | 0.6302938 |
| TLDEALK | AWSHRGGNYAEQF | 0.9090444 |
| TDGAGTAEGG | AWSHRGGNYAEQF | 0.91062593 |
| NAPGGAGGNGG | AWSHRGGNYAEQF | 0.7724225 |
| VSVFMGMGGNF | AWSHRGGNYAEQF | 0.90642524 |
| LASYLAAL | AWSHRGGNYAEQF | 0.8112764 |
| LVGGTPAL | AWSHRGGNYAEQF | 0.9196888 |
| VRNADVL | AWSHRGGNYAEQF | 0.85749495 |
| TCSSPFLVSAM | AWSHRGGNYAEQF | 0.603892 |
| VHITDAE | AWSHRGGNYAEQF | 0.9060865 |
| EYAAPSF | AWSHRGGNYAEQF | 0.8934144 |
| VVDAFHGGL | AWSHRGGNYAEQF | 0.8777815 |
| ASVVDPQ | AWSHRGGNYAEQF | 0.9058423 |
| SYNEFGGLS | AWSHRGGNYAEQF | 0.80862546 |
| DPDGSVY | AWSHRGGNYAEQF | 0.8649257 |
| MVSTIHATE | AWSHRGGNYAEQF | 0.91095304 |
| LAAGVADPV | AWSHRGGNYAEQF | 0.9098805 |
| TKEQIAA | AWSHRGGNYAEQF | 0.6894705 |
| SLASPAAL | AWSHRGGNYAEQF | 0.9105035 |
| IGLVFYGV | AWSHRGGNYAEQF | 0.866717 |
| FLLDASNL | AWSHRGGNYAEQF | 0.8077829 |
| YQVGNFP | AWSHRGGNYAEQF | 0.9201274 |
| ESLLSQST | AWSHRGGNYAEQF | 0.87026703 |
| LFTVGFAAVG | AWSHRGGNYAEQF | 0.90717113 |
| ATPPTPT | AWSHRGGNYAEQF | 0.9113887 |

| | | |
|---|---|---|
| TDLATLG | AWSHRGGNYAEQF | 0.8763552 |
| SAGGTGR | AWSHRGGNYAEQF | 0.60650396 |
| GFSNSGS | AWSHRGGNYAEQF | 0.85728717 |
| GGNGGSGGTSV | AWSHRGGNYAEQF | 0.86938953 |
| GTSVATGGAGN | AWSHRGGNYAEQF | 0.8982984 |
| NGGNGGSGGTS | AWSHRGGNYAEQF | 0.8623146 |
| AVVKFPH | AWSHRGGNYAEQF | 0.64061224 |
| TPPPPPP | AWSHRGGNYAEQF | 0.8943224 |
| AAPAAAAAPA | AWSHRGGNYAEQF | 0.9119401 |
| IALNEDL | AWSHRGGNYAEQF | 0.888458 |
| IVNVSSM | AWSHRGGNYAEQF | 0.8597155 |
| TGIPPAPR | AWSHRGGNYAEQF | 0.9184252 |
| FDLGGGTF | AWSHRGGNYAEQF | 0.84568715 |
| FDLGGGTFDVS | AWSHRGGNYAEQF | 0.86375785 |
| NEPTAAA | AWSHRGGNYAEQF | 0.9063505 |
| NEPTAAALA | AWSHRGGNYAEQF | 0.89836836 |
| GSIINISSI | AWSHRGGNYAEQF | 0.8675456 |
| VAPGNAGTA | AWSHRGGNYAEQF | 0.91448736 |
| TGGGASLE | AWSHRGGNYAEQF | 0.7212353 |
| VAVYNHYKRI | AWSHRGGNYAEQF | 0.9022967 |
| TDPAPATT | AWSHRGGNYAEQF | 0.7916143 |
| TNRPDILDPALLRPG | AWSHRGGNYAEQF | 0.891064 |
| LTTTTTS | AWSHRGGNYAEQF | 0.8155371 |
| PTGSGKTL | AWSHRGGNYAEQF | 0.90522325 |
| TGSGKTL | AWSHRGGNYAEQF | 0.9081131 |
| TIGHVDHGK | AWSHRGGNYAEQF | 0.90068555 |
| TIGHVDHGKTT | AWSHRGGNYAEQF | 0.20577928 |
| GHVDHGKTT | AWSHRGGNYAEQF | 0.90810406 |
| AQFEHTLL | AWSHRGGNYAEQF | 0.8849052 |
| IVNQDEL | AWSHRGGNYAEQF | 0.92113334 |
| TNGSQFF | AWSHRGGNYAEQF | 0.8768861 |
| TGKGIVE | AWSHRGGNYAEQF | 0.88047105 |
| LISNASDA | AWSHRGGNYAEQF | 0.8545846 |
| AELRTLGELTKIAKAHGAQ | AWSHRGGNYAEQF | 0.871508 |
| APNAVAAAEPSW | AWSHRGGNYAEQF | 0.7850332 |
| APQVAGAATAVL | AWSHRGGNYAEQF | 0.8334518 |
| ARRDPAAAAAILPTDA | AWSHRGGNYAEQF | 0.83183014 |
| DAAARDVLAEAGLAEYF | AWSHRGGNYAEQF | 0.8350768 |
| DDQIVLSAKANLAKVEQ | AWSHRGGNYAEQF | 0.90241534 |
| DELLPDTGEDSDEDGH | AWSHRGGNYAEQF | 0.9088883 |
| ELPEAPANRATVPQD | AWSHRGGNYAEQF | 0.9056592 |
| FGQSKIERVTGIG | AWSHRGGNYAEQF | 0.9081663 |
| GAGGTGSTAGGAGGAGGAGGLY | AWSHRGGNYAEQF | 0.9112172 |
| GHGGTGAAGGNGGTGGAGGSGI | AWSHRGGNYAEQF | 0.9109291 |
| GNGGNGGAGGEGGPGGAGGAGA | AWSHRGGNYAEQF | 0.8490914 |
| GSTAGNGGNGGNATNGTIAGQP | AWSHRGGNYAEQF | 0.8211366 |
| GTGGDGGNGGNGGTGG | AWSHRGGNYAEQF | 0.811429 |
| GVRRPAGIPADAPV | AWSHRGGNYAEQF | 0.8422359 |
| GYRQVTDAHLLTLARR | AWSHRGGNYAEQF | 0.8455421 |
| IAKMGQDGHDRG | AWSHRGGNYAEQF | 0.8908285 |
| KAEPSIEIQYT | AWSHRGGNYAEQF | 0.11107811 |

| | | |
|---|---|---|
| KPVRIRMAADEDTVAVT | AWSHRGGNYAEQF | 0.8688232 |
| MPPRIAGMRLLVIKPEP | AWSHRGGNYAEQF | 0.9145658 |
| MKFYAVALFRSRGVPPTR | AWSHRGGNYAEQF | 0.7828833 |
| NAGGFGILGAVAHSP | AWSHRGGNYAEQF | 0.8860125 |
| NEGREITLAMAEM | AWSHRGGNYAEQF | 0.8544878 |
| NELVEWSHPLGHRWRWE | AWSHRGGNYAEQF | 0.8539238 |
| NLGAGGTGGNGGNPG | AWSHRGGNYAEQF | 0.89980644 |
| NTEWPKWETVMAADPVD | AWSHRGGNYAEQF | 0.7966415 |
| PASAAPAETQP | AWSHRGGNYAEQF | 0.8990975 |
| QQSSDGKNGVRAAEITGE | AWSHRGGNYAEQF | 0.9191909 |
| RAPDAVTTSWWKE | AWSHRGGNYAEQF | 0.8815212 |
| SGFGNVGAHTSGWFNQ | AWSHRGGNYAEQF | 0.84807384 |
| SGVAGAVGQAASVGGLKV | AWSHRGGNYAEQF | 0.8643575 |
| SNGGAGTGSAGGNGGT | AWSHRGGNYAEQF | 0.49117294 |
| TGGAGGEGGAAG | AWSHRGGNYAEQF | 0.8705335 |
| TGGDGGAAGAGGAPGHG | AWSHRGGNYAEQF | 0.08765596 |
| TMPPLLAAIANAGRHGV | AWSHRGGNYAEQF | 0.87079537 |
| TSSTAGNSGGA | AWSHRGGNYAEQF | 0.78418994 |
| VPATVPVGILC | AWSHRGGNYAEQF | 0.88832116 |
| VQRPDSGGVVLTGRIS | AWSHRGGNYAEQF | 0.87185574 |
| WTPNPVPATKGVPAGLVQAT | AWSHRGGNYAEQF | 0.8980593 |
| YALKRALAKFSDRFDVVII | AWSHRGGNYAEQF | 0.79386246 |
| YLIGSVIV | AWSHRGGNYAEQF | 0.86167264 |
| YGQLVAVID | AWSHRGGNYAEQF | 0.7806885 |
| VTVGGMTLPA | AWSHRGGNYAEQF | 0.670247 |
| VLALPAV | AWSHRGGNYAEQF | 0.90342045 |
| VIPPTLNYET | AWSHRGGNYAEQF | 0.8694747 |
| VGMGGMHPSSG | AWSHRGGNYAEQF | 0.85333085 |
| VGLAGGAATA | AWSHRGGNYAEQF | 0.9098961 |
| VGGAGGAGGSGP | AWSHRGGNYAEQF | 0.8895055 |
| VFAAFMV | AWSHRGGNYAEQF | 0.58402145 |
| TLYDKLR | AWSHRGGNYAEQF | 0.85973334 |
| TLAVFTVL | AWSHRGGNYAEQF | 0.8747817 |
| TGSVQPARV | AWSHRGGNYAEQF | 0.8777383 |
| TGGGAGSGGT | AWSHRGGNYAEQF | 0.08583926 |
| SHLETSL | AWSHRGGNYAEQF | 0.88791627 |
| SFTTGMT | AWSHRGGNYAEQF | 0.8005333 |
| QVEVAKRMGVRQPTVS | AWSHRGGNYAEQF | 0.79279345 |
| QGSGGPALAM | AWSHRGGNYAEQF | 0.85600007 |
| MLAPSNP | AWSHRGGNYAEQF | 0.90431845 |
| LVNNAAI | AWSHRGGNYAEQF | 0.55391157 |
| LTGEINTLA | AWSHRGGNYAEQF | 0.85862124 |
| LSIFGPV | AWSHRGGNYAEQF | 0.9081956 |
| LLALLANL | AWSHRGGNYAEQF | 0.8970499 |
| LISNASDALDKLR | AWSHRGGNYAEQF | 0.91139615 |
| LINEAVP | AWSHRGGNYAEQF | 0.8859774 |
| LGYAFSGL | AWSHRGGNYAEQF | 0.8450619 |
| LDFVAVAL | AWSHRGGNYAEQF | 0.87074983 |
| LAQPGAE | AWSHRGGNYAEQF | 0.8123532 |
| ISNLGAVTGF | AWSHRGGNYAEQF | 0.5378177 |
| HQEDPGRI | AWSHRGGNYAEQF | 0.48377994 |

| | | |
|---|---|---|
| GTPGEPTP | AWSHRGGNYAEQF | 0.88742995 |
| GFFNSGS | AWSHRGGNYAEQF | 0.36731648 |
| GAGGGGLTAGAN | AWSHRGGNYAEQF | 0.88318276 |
| FVDALVNM | AWSHRGGNYAEQF | 0.90966994 |
| FLTVAPEL | AWSHRGGNYAEQF | 0.8477818 |
| FETLTAL | AWSHRGGNYAEQF | 0.64621913 |
| DVINPIEL | AWSHRGGNYAEQF | 0.86352205 |
| APVSGAPV | AWSHRGGNYAEQF | 0.83058727 |
| ALVGGLAW | AWSHRGGNYAEQF | 0.9064349 |
| ALAAAAATAI | AWSHRGGNYAEQF | 0.896966 |
| AFGLGLLL | AWSHRGGNYAEQF | 0.9148777 |
| AFFAAASL | AWSHRGGNYAEQF | 0.9062787 |
| AAAALPAFSPP | AWSHRGGNYAEQF | 0.84455436 |
| VVLAPAL | AWSHRGGNYAEQF | 0.9065409 |
| TIGHVDHGKTTL | AWSHRGGNYAEQF | 0.85041416 |
| LISNASDALDK | AWSHRGGNYAEQF | 0.8208057 |
| LISNASDAL | AWSHRGGNYAEQF | 0.8726666 |
| ISNASDALDK | AWSHRGGNYAEQF | 0.8935444 |
| NQAETLVYQTE | AWSHRGGNYAEQF | 0.85180306 |
| TGGTGGTAGAG | AWSHRGGNYAEQF | 0.8508915 |
| TGSTAGGAGGA | AWSHRGGNYAEQF | 0.61784226 |
| DPLAVPVAAGPV | AWSHRGGNYAEQF | 0.8974333 |
| GGTGGTPGNGGN | AWSHRGGNYAEQF | 0.8796775 |
| PAGLAGVDPATL | AWSHRGGNYAEQF | 0.60272074 |
| GNNLVVEAPHAGASV | AWSHRGGNYAEQF | 0.8285157 |
| KVTRIMSASTLEQPA | AWSHRGGNYAEQF | 0.9182062 |
| LGFALPIALPRRTVV | AWSHRGGNYAEQF | 0.9070587 |
| TRLGNLTARKLVVLV | AWSHRGGNYAEQF | 0.80416477 |
| WSQQRRLIYFPSAGP | AWSHRGGNYAEQF | 0.82443583 |
| GPNGNPDPMAAAVDIR | AWSHRGGNYAEQF | 0.8701613 |
| TQRCVSVADLRAAASR | AWSHRGGNYAEQF | 0.5681797 |
| QFVLDVAEHPVEVLTETV | AWSHRGGNYAEQF | 0.90003014 |
| GGAGGTGGAGGNGGTQAGDG | AWSHRGGNYAEQF | 0.870166 |
| PLLATLRAIAVDVDAKPAQV | AWSHRGGNYAEQF | 0.85715616 |
| APKFRTGGYMIDFWGVGYQVAK | AWSHRGGNYAEQF | 0.7687032 |
| GAGGAGGAGGAAGGPGGTGGTG | AWSHRGGNYAEQF | 0.88415515 |
| IGALGLLATDVAPAGSVAFGVA | AWSHRGGNYAEQF | 0.8328409 |
| AGGAGGIGGNGGAGGAGGVLMG | AWSHRGGNYAEQF | 0.90131986 |
| ADELIGLL | AWSHRGGNYAEQF | 0.17216095 |
| ADRALKA | AWSHRGGNYAEQF | 0.88350916 |
| AGAVGQVGGAGGA | AWSHRGGNYAEQF | 0.783977 |
| AGGGGGLGGAP | AWSHRGGNYAEQF | 0.8312038 |
| AGGLLYGNGGAG | AWSHRGGNYAEQF | 0.5373379 |
| AGLESIGL | AWSHRGGNYAEQF | 0.8410859 |
| AGLKTGII | AWSHRGGNYAEQF | 0.9068171 |
| ASLAALLL | AWSHRGGNYAEQF | 0.90982777 |
| ASVINISSI | AWSHRGGNYAEQF | 0.73962533 |
| ATPPGPA | AWSHRGGNYAEQF | 0.9045365 |
| ATVASALV | AWSHRGGNYAEQF | 0.9203196 |
| ATVASLIL | AWSHRGGNYAEQF | 0.84220344 |
| AVNGPVLIRH | AWSHRGGNYAEQF | 0.48402542 |

| | | |
|---|---|---|
| DGAAGAVGI | AWSHRGGNYAEQF | 0.89154696 |
| DPLLHGI | AWSHRGGNYAEQF | 0.82856905 |
| ELGAALGAL | AWSHRGGNYAEQF | 0.8992982 |
| ELGRTQAV | AWSHRGGNYAEQF | 0.8777673 |
| FVAGGYLS | AWSHRGGNYAEQF | 0.7765273 |
| GASGGAGGAGGW | AWSHRGGNYAEQF | 0.8873614 |
| GGAGAGGDGGAGGI | AWSHRGGNYAEQF | 0.9012209 |
| GTGHNGSQPGMGGNG | AWSHRGGNYAEQF | 0.81201124 |
| GYAAASASAS | AWSHRGGNYAEQF | 0.81371295 |
| HQVDVAAVQQG | AWSHRGGNYAEQF | 0.6644907 |
| IGAAIAHT | AWSHRGGNYAEQF | 0.41749534 |
| IGMIEVAQL | AWSHRGGNYAEQF | 0.89296496 |
| IRLAPEHTSAMD | AWSHRGGNYAEQF | 0.8238745 |
| IYNVSVL | AWSHRGGNYAEQF | 0.8134726 |
| KQGFLGM | AWSHRGGNYAEQF | 0.8861972 |
| LESVITQL | AWSHRGGNYAEQF | 0.82966125 |
| LGLGALVVV | AWSHRGGNYAEQF | 0.83918226 |
| LIGNGGNGGSGGI | AWSHRGGNYAEQF | 0.85036594 |
| LNAINAPTL | AWSHRGGNYAEQF | 0.86756885 |
| LQLLNTGV | AWSHRGGNYAEQF | 0.896564 |
| LQTSIEF | AWSHRGGNYAEQF | 0.8573224 |
| MSFAVSEL | AWSHRGGNYAEQF | 0.82761264 |
| NTGALGL | AWSHRGGNYAEQF | 0.91448337 |
| PPGAGGAGGAGTT | AWSHRGGNYAEQF | 0.68926555 |
| QLVGLGI | AWSHRGGNYAEQF | 0.7538253 |
| RFLEQVHQL | AWSHRGGNYAEQF | 0.8971547 |
| RGVGYVLR | AWSHRGGNYAEQF | 0.65652764 |
| SATGVFAGVI | AWSHRGGNYAEQF | 0.89795995 |
| SLPAITLP | AWSHRGGNYAEQF | 0.8717715 |
| STALAFTV | AWSHRGGNYAEQF | 0.8926812 |
| SVLTVGLGK | AWSHRGGNYAEQF | 0.85384625 |
| TALAGAGSGI | AWSHRGGNYAEQF | 0.918947 |
| TGLPFTGI | AWSHRGGNYAEQF | 0.90968096 |
| VAEHVTV | AWSHRGGNYAEQF | 0.83877575 |
| VDIVLNSL | AWSHRGGNYAEQF | 0.8701933 |
| VLTGGGAVL | AWSHRGGNYAEQF | 0.54863554 |
| VNVGLAYGI | AWSHRGGNYAEQF | 0.9018675 |
| VSSAAVGAV | AWSHRGGNYAEQF | 0.89353627 |
| VTSVYEAGV | AWSHRGGNYAEQF | 0.7999414 |
| WVALDRGA | AWSHRGGNYAEQF | 0.8678969 |
| YLVGSVVI | AWSHRGGNYAEQF | 0.9163981 |
| GPPGTGKTLL | AWSHRGGNYAEQF | 0.9119215 |
| GVGMTKF | AWSHRGGNYAEQF | 0.494381 |
| IGIGGSDL | AWSHRGGNYAEQF | 0.85894775 |
| IGTLLAEL | AWSHRGGNYAEQF | 0.3981546 |
| IYTSGTT | AWSHRGGNYAEQF | 0.7990011 |
| LSFPAGV | AWSHRGGNYAEQF | 0.65689534 |
| LYTSGTT | AWSHRGGNYAEQF | 0.9033305 |
| VGAFPFGF | AWSHRGGNYAEQF | 0.84947103 |
| VGAGTVEF | AWSHRGGNYAEQF | 0.86798304 |
| VGKSTLLL | AWSHRGGNYAEQF | 0.8896312 |

| | | |
|---|---|---|
| AALAHAGGGIV | AWSHRGGNYAEQF | 0.918521 |
| AALVAVSLL | AWSHRGGNYAEQF | 0.8403207 |
| AAVSRAVAAL | AWSHRGGNYAEQF | 0.8652811 |
| AGGSSGIGGF | AWSHRGGNYAEQF | 0.63571066 |
| AGLTVANANG | AWSHRGGNYAEQF | 0.42036957 |
| AGLVGAVL | AWSHRGGNYAEQF | 0.8286604 |
| ALSQAFTV | AWSHRGGNYAEQF | 0.8418435 |
| ALTFSVSL | AWSHRGGNYAEQF | 0.21085383 |
| ALTGAALVAL | AWSHRGGNYAEQF | 0.91085666 |
| ASFSIGAGVL | AWSHRGGNYAEQF | 0.8939332 |
| DGSVYKI | AWSHRGGNYAEQF | 0.9094075 |
| EQLAEIV | AWSHRGGNYAEQF | 0.81174684 |
| FAVSELTVM | AWSHRGGNYAEQF | 0.85513717 |
| GAAPLAAGL | AWSHRGGNYAEQF | 0.8517139 |
| GADGGDGGSGGAGG | AWSHRGGNYAEQF | 0.89706814 |
| GFGLDTGGAGG | AWSHRGGNYAEQF | 0.8224039 |
| GLASGGLL | AWSHRGGNYAEQF | 0.6372931 |
| GRIAGAAVLL | AWSHRGGNYAEQF | 0.79153824 |
| GSAGSLNTSF | AWSHRGGNYAEQF | 0.74679524 |
| GTGQIGI | AWSHRGGNYAEQF | 0.91423374 |
| GTLIALIL | AWSHRGGNYAEQF | 0.9093715 |
| IAATMGFVVG | AWSHRGGNYAEQF | 0.8970584 |
| IFAQALL | AWSHRGGNYAEQF | 0.91054535 |
| IGAAVALGF | AWSHRGGNYAEQF | 0.28350186 |
| IGLTPLAL | AWSHRGGNYAEQF | 0.7931465 |
| ILGSNTSTL | AWSHRGGNYAEQF | 0.908149 |
| IVFSLTVL | AWSHRGGNYAEQF | 0.89832336 |
| LASKATGF | AWSHRGGNYAEQF | 0.8583901 |
| LASVALIL | AWSHRGGNYAEQF | 0.9160302 |
| LFAVGLTL | AWSHRGGNYAEQF | 0.82017183 |
| LFNGGIGTYI | AWSHRGGNYAEQF | 0.8669592 |
| LTLGAGII | AWSHRGGNYAEQF | 0.8786183 |
| LVAAALGDI | AWSHRGGNYAEQF | 0.9009013 |
| NTAFVQLGI | AWSHRGGNYAEQF | 0.89168775 |
| QLFSGAEL | AWSHRGGNYAEQF | 0.8241991 |
| SAATFITL | AWSHRGGNYAEQF | 0.8775749 |
| SNYGLANLGV | AWSHRGGNYAEQF | 0.9215528 |
| SYGGAYAVM | AWSHRGGNYAEQF | 0.8684617 |
| TTGVGGLTLG | AWSHRGGNYAEQF | 0.9083515 |
| TTQTIYI | AWSHRGGNYAEQF | 0.894869 |
| VAAGAGLGAVI | AWSHRGGNYAEQF | 0.8690264 |
| VAAVVTLL | AWSHRGGNYAEQF | 0.82660186 |
| VGFLGGLI | AWSHRGGNYAEQF | 0.85854083 |
| VGVIPILL | AWSHRGGNYAEQF | 0.8606787 |
| VIFASSLI | AWSHRGGNYAEQF | 0.7707828 |
| VKGGDGGAAAG | AWSHRGGNYAEQF | 0.85681605 |
| VTKTFTNI | AWSHRGGNYAEQF | 0.8314273 |
| VVAAAILL | AWSHRGGNYAEQF | 0.92378604 |
| YAALDVEL | AWSHRGGNYAEQF | 0.89530873 |
| LASVALLL | AWSHRGGNYAEQF | 0.8996848 |
| LDAVHRV | AWSHRGGNYAEQF | 0.81715065 |

| | | |
|---|---|---|
| QGLSSSIF | AWSHRGGNYAEQF | 0.8963646 |
| RGTFANIRL | AWSHRGGNYAEQF | 0.8972214 |
| RQLWWGHRI | AWSHRGGNYAEQF | 0.65767515 |
| YGPPGTGKTLL | AWSHRGGNYAEQF | 0.91329086 |
| AALVGAPANVA | AWSHRGGNYAEQF | 0.8300319 |
| AATFRRTE | AWSHRGGNYAEQF | 0.83075607 |
| AGALGGGI | AWSHRGGNYAEQF | 0.8919538 |
| AGGAGGAGGIGAG | AWSHRGGNYAEQF | 0.8990721 |
| AGSGALGL | AWSHRGGNYAEQF | 0.85417044 |
| AQIADQI | AWSHRGGNYAEQF | 0.8425045 |
| ASAVAVTYL | AWSHRGGNYAEQF | 0.8244424 |
| ASTVLGGTGG | AWSHRGGNYAEQF | 0.9102138 |
| AVGGEPAK | AWSHRGGNYAEQF | 0.9125774 |
| CEQGTFGGTC | AWSHRGGNYAEQF | 0.8434659 |
| DEGKAPAG | AWSHRGGNYAEQF | 0.9172543 |
| DTVVFPAL | AWSHRGGNYAEQF | 0.84427756 |
| FEGITMAG | AWSHRGGNYAEQF | 0.5409277 |
| FPLVVQD | AWSHRGGNYAEQF | 0.8724741 |
| GAFTTPQ | AWSHRGGNYAEQF | 0.90497214 |
| GFGGFGTAGGVGGA | AWSHRGGNYAEQF | 0.91107225 |
| GFLAQFL | AWSHRGGNYAEQF | 0.91750646 |
| GGAGGAGGNAGVA | AWSHRGGNYAEQF | 0.88212824 |
| GGVGTAGAGGAGGAGG | AWSHRGGNYAEQF | 0.84770817 |
| GNGGGGDAGNAGSGGN | AWSHRGGNYAEQF | 0.91408634 |
| GTGGAGGAGNDAGS | AWSHRGGNYAEQF | 0.91689897 |
| GTTNTGWFNS | AWSHRGGNYAEQF | 0.9037142 |
| IFMGPLFTL | AWSHRGGNYAEQF | 0.8485315 |
| ILAHAIEAI | AWSHRGGNYAEQF | 0.7545561 |
| IRSAIREAL | AWSHRGGNYAEQF | 0.8815702 |
| ITFAGGG | AWSHRGGNYAEQF | 0.91329277 |
| ITGMSGHQEV | AWSHRGGNYAEQF | 0.78119075 |
| IVAGLACTL | AWSHRGGNYAEQF | 0.8753283 |
| IVGGGLAGA | AWSHRGGNYAEQF | 0.88860124 |
| IVTGVLNI | AWSHRGGNYAEQF | 0.63367313 |
| LAQQGCI | AWSHRGGNYAEQF | 0.87753016 |
| LCLHVEN | AWSHRGGNYAEQF | 0.91259855 |
| LDAISDGS | AWSHRGGNYAEQF | 0.8691492 |
| LGGFAAML | AWSHRGGNYAEQF | 0.82774884 |
| LGGKNPMI | AWSHRGGNYAEQF | 0.90817094 |
| LGPKGVAAIL | AWSHRGGNYAEQF | 0.9239687 |
| LMVGVSI | AWSHRGGNYAEQF | 0.9154205 |
| LVGAAFVL | AWSHRGGNYAEQF | 0.596743 |
| LVLAVAPL | AWSHRGGNYAEQF | 0.40478584 |
| MAAIAVIDAA | AWSHRGGNYAEQF | 0.8110033 |
| NNGINAELQ | AWSHRGGNYAEQF | 0.8854875 |
| RGEPLVLVHG | AWSHRGGNYAEQF | 0.8317406 |
| RGGAGGAAGLI | AWSHRGGNYAEQF | 0.83905244 |
| RSYRMHPAV | AWSHRGGNYAEQF | 0.8941073 |
| SAAHLLAGQLA | AWSHRGGNYAEQF | 0.9105996 |
| SAAVGSY | AWSHRGGNYAEQF | 0.8182485 |
| SPHAVLHRI | AWSHRGGNYAEQF | 0.66639256 |

| | | |
|---|---|---|
| SQIAKVL | AWSHRGGNYAEQF | 0.88205874 |
| TALGHLVP | AWSHRGGNYAEQF | 0.9017238 |
| TAPTTSGAA | AWSHRGGNYAEQF | 0.8209219 |
| TDGDDGNGGSPGT | AWSHRGGNYAEQF | 0.88019395 |
| TDGLGEN | AWSHRGGNYAEQF | 0.790259 |
| TGGPPLPGQA | AWSHRGGNYAEQF | 0.47063357 |
| TIPAITI | AWSHRGGNYAEQF | 0.8895661 |
| TIPGFTI | AWSHRGGNYAEQF | 0.7909446 |
| TSIASGALA | AWSHRGGNYAEQF | 0.91594917 |
| TVTLPGAAI | AWSHRGGNYAEQF | 0.82769847 |
| VASGLSAALP | AWSHRGGNYAEQF | 0.9060817 |
| VGAGVIGI | AWSHRGGNYAEQF | 0.89480805 |
| VGPKRFGV | AWSHRGGNYAEQF | 0.89772034 |
| VNVAFPD | AWSHRGGNYAEQF | 0.85833025 |
| VSMTNSGVS | AWSHRGGNYAEQF | 0.8148266 |
| VTDVLGDLSV | AWSHRGGNYAEQF | 0.76378304 |
| VVVVGAGFAGL | AWSHRGGNYAEQF | 0.58054256 |
| VYTVGTVV | AWSHRGGNYAEQF | 0.905522 |
| WTTFSGP | AWSHRGGNYAEQF | 0.120206475 |
| YDELVEAF | AWSHRGGNYAEQF | 0.9122648 |
| YGTGGAGGNGG | AWSHRGGNYAEQF | 0.91016424 |
| YTNVMAV | AWSHRGGNYAEQF | 0.8211421 |
| FLEQVHQL | AWSHRGGNYAEQF | 0.90381426 |
| GLSLLAVAV | AWSHRGGNYAEQF | 0.78018177 |
| LEQVHQL | AWSHRGGNYAEQF | 0.79749393 |
| LNAAGAI | AWSHRGGNYAEQF | 0.9036416 |
| PAVHASL | AWSHRGGNYAEQF | 0.89977366 |
| TELEDDA | AWSHRGGNYAEQF | 0.8214035 |
| TPPGPPN | AWSHRGGNYAEQF | 0.89373064 |
| TPPPPPPP | AWSHRGGNYAEQF | 0.88488376 |
| TQSPALI | ASGDASGGAETLY | 0.78205854 |
| EFAAAFPQ | ASGDASGGAETLY | 0.8671534 |
| LLSGNLDVL | ASGDASGGAETLY | 0.90853673 |
| GPANLGNYNI | ASGDASGGAETLY | 0.88917357 |
| TLGPTLP | ASGDASGGAETLY | 0.8405181 |
| VYAYSSCCK | ASGDASGGAETLY | 0.8991626 |
| APAPAPAPNI | ASGDASGGAETLY | 0.77621174 |
| PGGPVPIP | ASGDASGGAETLY | 0.892743 |
| FLTVALEP | ASGDASGGAETLY | 0.8818507 |
| SQYFGANL | ASGDASGGAETLY | 0.8489523 |
| GGVKPAA | ASGDASGGAETLY | 0.9002465 |
| GGTGGAGGAGGW | ASGDASGGAETLY | 0.49798256 |
| GVGGVAGLGGAGG | ASGDASGGAETLY | 0.8223534 |
| VLLVLATTI | ASGDASGGAETLY | 0.8545393 |
| IGIFAAML | ASGDASGGAETLY | 0.88763577 |
| AVGDLID | ASGDASGGAETLY | 0.90577364 |
| NISPLPL | ASGDASGGAETLY | 0.815066 |
| VTAVFVAI | ASGDASGGAETLY | 0.87964654 |
| SGDDNDAAL | ASGDASGGAETLY | 0.83785504 |
| AVAALAHS | ASGDASGGAETLY | 0.8023788 |
| GGVTVMATP | ASGDASGGAETLY | 0.8773214 |

| | | |
|---|---|---|
| AGLAANVRD | ASGDASGGAETLY | 0.83944535 |
| TNRPDLIDPALLRPG | ASGDASGGAETLY | 0.8007456 |
| LANLGVTAL | ASGDASGGAETLY | 0.89610565 |
| TGERIEL | ASGDASGGAETLY | 0.78104913 |
| NGEELAA | ASGDASGGAETLY | 0.8870871 |
| VLSLVWG | ASGDASGGAETLY | 0.9115672 |
| IGYALWL | ASGDASGGAETLY | 0.862187 |
| LSLLLAPL | ASGDASGGAETLY | 0.90641844 |
| TLDEALK | ASGDASGGAETLY | 0.8962454 |
| TDGAGTAEGG | ASGDASGGAETLY | 0.8971567 |
| NAPGGAGGNGG | ASGDASGGAETLY | 0.8601724 |
| VSVFMGMGGNF | ASGDASGGAETLY | 0.827186 |
| LASYLAAL | ASGDASGGAETLY | 0.91176283 |
| LVGGTPAL | ASGDASGGAETLY | 0.9008419 |
| VRNADVL | ASGDASGGAETLY | 0.8675108 |
| TCSSPFLVSAM | ASGDASGGAETLY | 0.8885353 |
| VHITDAE | ASGDASGGAETLY | 0.8865628 |
| EYAAPSF | ASGDASGGAETLY | 0.8996822 |
| VVDAFHGGL | ASGDASGGAETLY | 0.8420901 |
| ASVVDPQ | ASGDASGGAETLY | 0.69536614 |
| SYNEFGGLS | ASGDASGGAETLY | 0.86141604 |
| DPDGSVY | ASGDASGGAETLY | 0.86174613 |
| MVSTIHATE | ASGDASGGAETLY | 0.82845145 |
| LAAGVADPV | ASGDASGGAETLY | 0.48922467 |
| TKEQIAA | ASGDASGGAETLY | 0.90461457 |
| SLASPAAL | ASGDASGGAETLY | 0.9221825 |
| IGLVFYGV | ASGDASGGAETLY | 0.7923143 |
| FLLDASNL | ASGDASGGAETLY | 0.9172417 |
| YQVGNFP | ASGDASGGAETLY | 0.70638776 |
| ESLLSQST | ASGDASGGAETLY | 0.9018433 |
| LFTVGFAAVG | ASGDASGGAETLY | 0.9061048 |
| ATPPTPT | ASGDASGGAETLY | 0.8115083 |
| TDLATLG | ASGDASGGAETLY | 0.88163424 |
| SAGGTGR | ASGDASGGAETLY | 0.8263304 |
| GFSNSGS | ASGDASGGAETLY | 0.8573622 |
| GGNGGSGGTSV | ASGDASGGAETLY | 0.9132909 |
| GTSVATGGAGN | ASGDASGGAETLY | 0.8249611 |
| NGGNGGSGGTS | ASGDASGGAETLY | 0.87656033 |
| AVVKFPH | ASGDASGGAETLY | 0.8778905 |
| TPPPPPP | ASGDASGGAETLY | 0.81337476 |
| AAPAAAAAPA | ASGDASGGAETLY | 0.896015 |
| IALNEDL | ASGDASGGAETLY | 0.9064725 |
| IVNVSSM | ASGDASGGAETLY | 0.8117503 |
| TGIPPAPR | ASGDASGGAETLY | 0.916034 |
| FDLGGGTF | ASGDASGGAETLY | 0.8390305 |
| FDLGGGTFDVS | ASGDASGGAETLY | 0.9207948 |
| NEPTAAA | ASGDASGGAETLY | 0.9039078 |
| NEPTAAALA | ASGDASGGAETLY | 0.88131255 |
| GSIINISSI | ASGDASGGAETLY | 0.86221945 |
| VAPGNAGTA | ASGDASGGAETLY | 0.9116367 |
| TGGGASLE | ASGDASGGAETLY | 0.85626227 |

| | | |
|---|---|---|
| VAVYNHYKRI | ASGDASGGAETLY | 0.8881886 |
| TDPAPATT | ASGDASGGAETLY | 0.9145069 |
| TNRPDILDPALLRPG | ASGDASGGAETLY | 0.46583545 |
| LTTTTTS | ASGDASGGAETLY | 0.62635666 |
| PTGSGKTL | ASGDASGGAETLY | 0.58396494 |
| TGSGKTL | ASGDASGGAETLY | 0.6328439 |
| TIGHVDHGK | ASGDASGGAETLY | 0.52700865 |
| TIGHVDHGKTT | ASGDASGGAETLY | 0.44590515 |
| GHVDHGKTT | ASGDASGGAETLY | 0.6024008 |
| AQFEHTLL | ASGDASGGAETLY | 0.4859212 |
| IVNQDEL | ASGDASGGAETLY | 0.61019844 |
| TNGSQFF | ASGDASGGAETLY | 0.63489777 |
| TGKGIVE | ASGDASGGAETLY | 0.5123394 |
| LISNASDA | ASGDASGGAETLY | 0.632944 |
| AELRTLGELTKIAKAHGAQ | ASGDASGGAETLY | 0.63470304 |
| APNAVAAAEPSW | ASGDASGGAETLY | 0.4450559 |
| APQVAGAATAVL | ASGDASGGAETLY | 0.577322 |
| ARRDPAAAAAILPTDA | ASGDASGGAETLY | 0.5764748 |
| DAAARDVLAEAGLAEYF | ASGDASGGAETLY | 0.5544934 |
| DDQIVLSAKANLAKVEQ | ASGDASGGAETLY | 0.37920418 |
| DELLPDTGEDSDEDGH | ASGDASGGAETLY | 0.6092019 |
| ELPEAPANRATVPQD | ASGDASGGAETLY | 0.55223656 |
| FGQSKIERVTGIG | ASGDASGGAETLY | 0.3399884 |
| GAGGTGSTAGGAGGAGGAGGLY | ASGDASGGAETLY | 0.6169222 |
| GHGGTGAAGGNGGTGGAGGSGI | ASGDASGGAETLY | 0.54586565 |
| GNGGNGGAGGEGGPGGAGGAGA | ASGDASGGAETLY | 0.5650518 |
| GSTAGNGGNGGNATNGTIAGQP | ASGDASGGAETLY | 0.63998926 |
| GTGGDGGNGGNGGTGG | ASGDASGGAETLY | 0.4271805 |
| GVRRPAGIPADAPV | ASGDASGGAETLY | 0.5162678 |
| GYRQVTDAHLLTLARR | ASGDASGGAETLY | 0.6111667 |
| IAKMGQDGHDRG | ASGDASGGAETLY | 0.6482344 |
| KAEPSIEIQYT | ASGDASGGAETLY | 0.4828327 |
| KPVRIRMAADEDTVAVT | ASGDASGGAETLY | 0.59705067 |
| MPPRIAGMRLLVIKPEP | ASGDASGGAETLY | 0.642305 |
| MKFYAVALFRSRGVPPTR | ASGDASGGAETLY | 0.5858418 |
| NAGGFGILGAVAHSP | ASGDASGGAETLY | 0.6535584 |
| NEGREITLAMAEM | ASGDASGGAETLY | 0.6541474 |
| NELVEWSHPLGHRWRWE | ASGDASGGAETLY | 0.56051946 |
| NLGAGGTGGNGGNPG | ASGDASGGAETLY | 0.63799965 |
| NTEWPKWETVMAADPVD | ASGDASGGAETLY | 0.65089315 |
| PASAAPAETQP | ASGDASGGAETLY | 0.57031083 |
| QQSSDGKNGVRAAEITGE | ASGDASGGAETLY | 0.58646065 |
| RAPDAVTTSWWKE | ASGDASGGAETLY | 0.50612044 |
| SGFGNVGAHTSGWFNQ | ASGDASGGAETLY | 0.58398277 |
| SGVAGAVGQAASVGGLKV | ASGDASGGAETLY | 0.40943518 |
| SNGGAGTGSAGGNGGT | ASGDASGGAETLY | 0.580105 |
| TGGAGGEGGAAG | ASGDASGGAETLY | 0.6367253 |
| TGGDGGAAGAGGAPGHG | ASGDASGGAETLY | 0.5637632 |
| TMPPLLAAIANAGRHGV | ASGDASGGAETLY | 0.42147392 |
| TSSTAGNSGGA | ASGDASGGAETLY | 0.45336187 |
| VPATVPVGILC | ASGDASGGAETLY | 0.57605714 |

| | | |
|---|---|---|
| VQRPDSGGVVLTGRIS | ASGDASGGAETLY | 0.534784 |
| WTPNPVPATKGVPAGLVQAT | ASGDASGGAETLY | 0.4773001 |
| YALKRALAKFSDRFDVVII | ASGDASGGAETLY | 0.51643425 |
| YLIGSVIV | ASGDASGGAETLY | 0.6281527 |
| YGQLVAVID | ASGDASGGAETLY | 0.6330204 |
| VTVGGMTLPA | ASGDASGGAETLY | 0.4654308 |
| VLALPAV | ASGDASGGAETLY | 0.6459683 |
| VIPPTLNYET | ASGDASGGAETLY | 0.5952772 |
| VGMGGMHPSSG | ASGDASGGAETLY | 0.6431752 |
| VGLAGGAATA | ASGDASGGAETLY | 0.61269426 |
| VGGAGGAGGSGP | ASGDASGGAETLY | 0.44993556 |
| VFAAFMV | ASGDASGGAETLY | 0.6218904 |
| TLYDKLR | ASGDASGGAETLY | 0.5348506 |
| TLAVFTVL | ASGDASGGAETLY | 0.5857964 |
| TGSVQPARV | ASGDASGGAETLY | 0.6344293 |
| TGGGAGSGGT | ASGDASGGAETLY | 0.554654 |
| SHLETSL | ASGDASGGAETLY | 0.4857134 |
| SFTTGMT | ASGDASGGAETLY | 0.54132354 |
| QVEVAKRMGVRQPTVS | ASGDASGGAETLY | 0.6384233 |
| QGSGGPALAM | ASGDASGGAETLY | 0.46317646 |
| MLAPSNP | ASGDASGGAETLY | 0.43828848 |
| LVNNAAI | ASGDASGGAETLY | 0.5512947 |
| LTGEINTLA | ASGDASGGAETLY | 0.5647167 |
| LSIFGPV | ASGDASGGAETLY | 0.64707017 |
| LLALLANL | ASGDASGGAETLY | 0.6116964 |
| LISNASDALDKLR | ASGDASGGAETLY | 0.5662113 |
| LINEAVP | ASGDASGGAETLY | 0.6366677 |
| LGYAFSGL | ASGDASGGAETLY | 0.61380607 |
| LDFVAVAL | ASGDASGGAETLY | 0.5777812 |
| LAQPGAE | ASGDASGGAETLY | 0.5934632 |
| ISNLGAVTGF | ASGDASGGAETLY | 0.5022429 |
| HQEDPGRI | ASGDASGGAETLY | 0.62995094 |
| GTPGEPTP | ASGDASGGAETLY | 0.6243571 |
| GFFNSGS | ASGDASGGAETLY | 0.6403924 |
| GAGGGGLTAGAN | ASGDASGGAETLY | 0.63969964 |
| FVDALVNM | ASGDASGGAETLY | 0.5884604 |
| FLTVAPEL | ASGDASGGAETLY | 0.6036557 |
| FETLTAL | ASGDASGGAETLY | 0.4313772 |
| DVINPIEL | ASGDASGGAETLY | 0.56992066 |
| APVSGAPV | ASGDASGGAETLY | 0.6104656 |
| ALVGGLAW | ASGDASGGAETLY | 0.59440637 |
| ALAAAAATAI | ASGDASGGAETLY | 0.6291941 |
| AFGLGLLL | ASGDASGGAETLY | 0.63696986 |
| AFFAAASL | ASGDASGGAETLY | 0.5239004 |
| AAAALPAFSPP | ASGDASGGAETLY | 0.635255 |
| VVLAPAL | ASGDASGGAETLY | 0.64182764 |
| TIGHVDHGKTTL | ASGDASGGAETLY | 0.46559846 |
| LISNASDALDK | ASGDASGGAETLY | 0.57399553 |
| LISNASDAL | ASGDASGGAETLY | 0.63793474 |
| ISNASDALDK | ASGDASGGAETLY | 0.6024555 |
| NQAETLVYQTE | ASGDASGGAETLY | 0.55665445 |

| | | |
|---|---|---|
| TGGTGGTAGAG | ASGDASGGAETLY | 0.6162275 |
| TGSTAGGAGGA | ASGDASGGAETLY | 0.63335276 |
| DPLAVPVAAGPV | ASGDASGGAETLY | 0.58555573 |
| GGTGGTPGNGGN | ASGDASGGAETLY | 0.6478864 |
| PAGLAGVDPATL | ASGDASGGAETLY | 0.3780082 |
| GNNLVVEAPHAGASV | ASGDASGGAETLY | 0.6012504 |
| KVTRIMSASTLEQPA | ASGDASGGAETLY | 0.52294594 |
| LGFALPIALPRRTVV | ASGDASGGAETLY | 0.34981143 |
| TRLGNLTARKLVVLV | ASGDASGGAETLY | 0.621369 |
| WSQQRRLIYFPSAGP | ASGDASGGAETLY | 0.5855144 |
| GPNGNPDPMAAAVDIR | ASGDASGGAETLY | 0.6382151 |
| TQRCVSVADLRAAASR | ASGDASGGAETLY | 0.59852946 |
| QFVLDVAEHPVEVLTETV | ASGDASGGAETLY | 0.63250196 |
| GGAGGTGGAGGNGGTQAGDG | ASGDASGGAETLY | 0.57965475 |
| PLLATLRAIAVDVDAKPAQV | ASGDASGGAETLY | 0.57226974 |
| APKFRTGGYMIDFWGVGYQVAK | ASGDASGGAETLY | 0.50631744 |
| GAGGAGGAGGAAGGPGGTGGTG | ASGDASGGAETLY | 0.60646844 |
| IGALGLLATDVAPAGSVAFGVA | ASGDASGGAETLY | 0.6431204 |
| AGGAGGIGGNGGAGGAGGVLMG | ASGDASGGAETLY | 0.6423531 |
| ADELIGLL | ASGDASGGAETLY | 0.4164921 |
| ADRALKA | ASGDASGGAETLY | 0.5845814 |
| AGAVGQVGGAGGA | ASGDASGGAETLY | 0.5746066 |
| AGGGGGLGGAP | ASGDASGGAETLY | 0.36756375 |
| AGGLLYGNGGAG | ASGDASGGAETLY | 0.40179595 |
| AGLESIGL | ASGDASGGAETLY | 0.5163489 |
| AGLKTGII | ASGDASGGAETLY | 0.51193243 |
| ASLAALLL | ASGDASGGAETLY | 0.6143935 |
| ASVINISSI | ASGDASGGAETLY | 0.62032324 |
| ATPPGPA | ASGDASGGAETLY | 0.53523916 |
| ATVASALV | ASGDASGGAETLY | 0.618824 |
| ATVASLIL | ASGDASGGAETLY | 0.5679814 |
| AVNGPVLIRH | ASGDASGGAETLY | 0.30848432 |
| DGAAGAVGI | ASGDASGGAETLY | 0.5613024 |
| DPLLHGI | ASGDASGGAETLY | 0.47416934 |
| ELGAALGAL | ASGDASGGAETLY | 0.644842 |
| ELGRTQAV | ASGDASGGAETLY | 0.4980815 |
| FVAGGYLS | ASGDASGGAETLY | 0.53253525 |
| GASGGAGGAGGW | ASGDASGGAETLY | 0.5634004 |
| GGAGAGGDGGAGGI | ASGDASGGAETLY | 0.58162516 |
| GTGHNGSQPGMGGNG | ASGDASGGAETLY | 0.5808063 |
| GYAAASASAS | ASGDASGGAETLY | 0.47907883 |
| HQVDVAAVQQG | ASGDASGGAETLY | 0.35645622 |
| IGAAIAHT | ASGDASGGAETLY | 0.6120613 |
| IGMIEVAQL | ASGDASGGAETLY | 0.52580273 |
| IRLAPEHTSAMD | ASGDASGGAETLY | 0.5432005 |
| IYNVSVL | ASGDASGGAETLY | 0.50921625 |
| KQGFLGM | ASGDASGGAETLY | 0.56564564 |
| LESVITQL | ASGDASGGAETLY | 0.4186052 |
| LGLGALVVV | ASGDASGGAETLY | 0.65032536 |
| LIGNGGNGGSGGI | ASGDASGGAETLY | 0.570443 |
| LNAINAPTL | ASGDASGGAETLY | 0.53168863 |

| | | |
|---|---|---|
| LQLLNTGV | ASGDASGGAETLY | 0.6392041 |
| LQTSIEF | ASGDASGGAETLY | 0.56738347 |
| MSFAVSEL | ASGDASGGAETLY | 0.56151265 |
| NTGALGL | ASGDASGGAETLY | 0.53620845 |
| PPGAGGAGGAGTT | ASGDASGGAETLY | 0.47499794 |
| QLVGLGI | ASGDASGGAETLY | 0.5055521 |
| RFLEQVHQL | ASGDASGGAETLY | 0.51366866 |
| RGVGYVLR | ASGDASGGAETLY | 0.6493069 |
| SATGVFAGVI | ASGDASGGAETLY | 0.59825456 |
| SLPAITLP | ASGDASGGAETLY | 0.46173614 |
| STALAFTV | ASGDASGGAETLY | 0.6138705 |
| SVLTVGLGK | ASGDASGGAETLY | 0.6395696 |
| TALAGAGSGI | ASGDASGGAETLY | 0.6430759 |
| TGLPFTGI | ASGDASGGAETLY | 0.49286634 |
| VAEHVTV | ASGDASGGAETLY | 0.5666113 |
| VDIVLNSL | ASGDASGGAETLY | 0.6041466 |
| VLTGGGAVL | ASGDASGGAETLY | 0.5411118 |
| VNVGLAYGI | ASGDASGGAETLY | 0.5426496 |
| VSSAAVGAV | ASGDASGGAETLY | 0.4840982 |
| VTSVYEAGV | ASGDASGGAETLY | 0.63098276 |
| WVALDRGA | ASGDASGGAETLY | 0.64204544 |
| YLVGSVVI | ASGDASGGAETLY | 0.56208503 |
| GPPGTGKTLL | ASGDASGGAETLY | 0.43700612 |
| GVGMTKF | ASGDASGGAETLY | 0.54708505 |
| IGIGGSDL | ASGDASGGAETLY | 0.49201772 |
| IGTLLAEL | ASGDASGGAETLY | 0.49329013 |
| IYTSGTT | ASGDASGGAETLY | 0.59840983 |
| LSFPAGV | ASGDASGGAETLY | 0.43136036 |
| LYTSGTT | ASGDASGGAETLY | 0.5719027 |
| VGAFPFGF | ASGDASGGAETLY | 0.6344806 |
| VGAGTVEF | ASGDASGGAETLY | 0.55462956 |
| VGKSTLLL | ASGDASGGAETLY | 0.6336258 |
| AALAHAGGGIV | ASGDASGGAETLY | 0.63679016 |
| AALVAVSLL | ASGDASGGAETLY | 0.61399233 |
| AAVSRAVAAL | ASGDASGGAETLY | 0.645638 |
| AGGSSGIGGF | ASGDASGGAETLY | 0.6118151 |
| AGLTVANANG | ASGDASGGAETLY | 0.58847773 |
| AGLVGAVL | ASGDASGGAETLY | 0.5804016 |
| ALSQAFTV | ASGDASGGAETLY | 0.62040794 |
| ALTFSVSL | ASGDASGGAETLY | 0.56014276 |
| ALTGAALVAL | ASGDASGGAETLY | 0.5825584 |
| ASFSIGAGVL | ASGDASGGAETLY | 0.58891916 |
| DGSVYKI | ASGDASGGAETLY | 0.61834675 |
| EQLAEIV | ASGDASGGAETLY | 0.5765454 |
| FAVSELTVM | ASGDASGGAETLY | 0.62715113 |
| GAAPLAAGL | ASGDASGGAETLY | 0.4178692 |
| GADGGDGGSGGAGG | ASGDASGGAETLY | 0.47474605 |
| GFGLDTGGAGG | ASGDASGGAETLY | 0.6255106 |
| GLASGGLL | ASGDASGGAETLY | 0.33007348 |
| GRIAGAAVLL | ASGDASGGAETLY | 0.45176354 |
| GSAGSLNTSF | ASGDASGGAETLY | 0.6005717 |

| | | |
|---|---|---|
| GTGQIGI | ASGDASGGAETLY | 0.62189794 |
| GTLIALIL | ASGDASGGAETLY | 0.32516608 |
| IAATMGFVVG | ASGDASGGAETLY | 0.5550181 |
| IFAQALL | ASGDASGGAETLY | 0.5156901 |
| IGAAVALGF | ASGDASGGAETLY | 0.6246599 |
| IGLTPLAL | ASGDASGGAETLY | 0.5610341 |
| ILGSNTSTL | ASGDASGGAETLY | 0.42196438 |
| IVFSLTVL | ASGDASGGAETLY | 0.6375776 |
| LASKATGF | ASGDASGGAETLY | 0.4562622 |
| LASVALIL | ASGDASGGAETLY | 0.5063544 |
| LFAVGLTL | ASGDASGGAETLY | 0.6395516 |
| LFNGGIGTYI | ASGDASGGAETLY | 0.6409172 |
| LTLGAGII | ASGDASGGAETLY | 0.51140255 |
| LVAAALGDI | ASGDASGGAETLY | 0.6350254 |
| NTAFVQLGI | ASGDASGGAETLY | 0.52965415 |
| QLFSGAEL | ASGDASGGAETLY | 0.6510871 |
| SAATFITL | ASGDASGGAETLY | 0.56510204 |
| SNYGLANLGV | ASGDASGGAETLY | 0.42968658 |
| SYGGAYAVM | ASGDASGGAETLY | 0.62710875 |
| TTGVGGLTLG | ASGDASGGAETLY | 0.6271709 |
| TTQTIYI | ASGDASGGAETLY | 0.5998697 |
| VAAGAGLGAVI | ASGDASGGAETLY | 0.60242623 |
| VAAVVTLL | ASGDASGGAETLY | 0.54619175 |
| VGFLGGLI | ASGDASGGAETLY | 0.55360645 |
| VGVIPILL | ASGDASGGAETLY | 0.62675804 |
| VIFASSLI | ASGDASGGAETLY | 0.61374104 |
| VKGGDGGAAAG | ASGDASGGAETLY | 0.47957456 |
| VTKTFTNI | ASGDASGGAETLY | 0.6412905 |
| VVAAAILL | ASGDASGGAETLY | 0.5650923 |
| YAALDVEL | ASGDASGGAETLY | 0.53243077 |
| LASVALLL | ASGDASGGAETLY | 0.63617146 |
| LDAVHRV | ASGDASGGAETLY | 0.6050803 |
| QGLSSSIF | ASGDASGGAETLY | 0.6192857 |
| RGTFANIRL | ASGDASGGAETLY | 0.61568034 |
| RQLWWGHRI | ASGDASGGAETLY | 0.6260087 |
| YGPPGTGKTLL | ASGDASGGAETLY | 0.49323517 |
| AALVGAPANVA | ASGDASGGAETLY | 0.5943167 |
| AATFRRTE | ASGDASGGAETLY | 0.5958967 |
| AGALGGGI | ASGDASGGAETLY | 0.63607424 |
| AGGAGGAGGIGAG | ASGDASGGAETLY | 0.5880182 |
| AGSGALGL | ASGDASGGAETLY | 0.5688447 |
| AQIADQI | ASGDASGGAETLY | 0.62007666 |
| ASAVAVTYL | ASGDASGGAETLY | 0.6316972 |
| ASTVLGGTGG | ASGDASGGAETLY | 0.5716356 |
| AVGGEPAK | ASGDASGGAETLY | 0.5907992 |
| CEQGTFGGTC | ASGDASGGAETLY | 0.6503967 |
| DEGKAPAG | ASGDASGGAETLY | 0.5086951 |
| DTVVFPAL | ASGDASGGAETLY | 0.58423895 |
| FEGITMAG | ASGDASGGAETLY | 0.62862617 |
| FPLVVQD | ASGDASGGAETLY | 0.64698994 |
| GAFTTPQ | ASGDASGGAETLY | 0.5965709 |

| | | |
|---|---|---|
| GFGGFGTAGGVGGA | ASGDASGGAETLY | 0.63611877 |
| GFLAQFL | ASGDASGGAETLY | 0.5595056 |
| GGAGGAGGNAGVA | ASGDASGGAETLY | 0.61771816 |
| GGVGTAGAGGAGGAGG | ASGDASGGAETLY | 0.55975467 |
| GNGGGGDAGNAGSGGN | ASGDASGGAETLY | 0.5899238 |
| GTGGAGGAGNDAGS | ASGDASGGAETLY | 0.4659902 |
| GTTNTGWFNS | ASGDASGGAETLY | 0.607834 |
| IFMGPLFTL | ASGDASGGAETLY | 0.61309516 |
| ILAHAIEAI | ASGDASGGAETLY | 0.61774683 |
| IRSAIREAL | ASGDASGGAETLY | 0.4405997 |
| ITFAGGG | ASGDASGGAETLY | 0.59991145 |
| ITGMSGHQEV | ASGDASGGAETLY | 0.62071764 |
| IVAGLACTL | ASGDASGGAETLY | 0.6544055 |
| IVGGGLAGA | ASGDASGGAETLY | 0.5705657 |
| IVTGVLNI | ASGDASGGAETLY | 0.6410461 |
| LAQQGCI | ASGDASGGAETLY | 0.5588068 |
| LCLHVEN | ASGDASGGAETLY | 0.5869197 |
| LDAISDGS | ASGDASGGAETLY | 0.4379139 |
| LGGFAAML | ASGDASGGAETLY | 0.57003486 |
| LGGKNPMI | ASGDASGGAETLY | 0.55182487 |
| LGPKGVAAIL | ASGDASGGAETLY | 0.48594964 |
| LMVGVSI | ASGDASGGAETLY | 0.61590505 |
| LVGAAFVL | ASGDASGGAETLY | 0.64040893 |
| LVLAVAPL | ASGDASGGAETLY | 0.62033284 |
| MAAIAVIDAA | ASGDASGGAETLY | 0.6343688 |
| NNGINAELQ | ASGDASGGAETLY | 0.6318686 |
| RGEPLVLVHG | ASGDASGGAETLY | 0.64829606 |
| RGGAGGAAGLI | ASGDASGGAETLY | 0.5286526 |
| RSYRMHPAV | ASGDASGGAETLY | 0.5878779 |
| SAAHLLAGQLA | ASGDASGGAETLY | 0.3929976 |
| SAAVGSY | ASGDASGGAETLY | 0.5546967 |
| SPHAVLHRI | ASGDASGGAETLY | 0.53287005 |
| SQIAKVL | ASGDASGGAETLY | 0.5886047 |
| TALGHLVP | ASGDASGGAETLY | 0.32700902 |
| TAPTTSGAA | ASGDASGGAETLY | 0.575812 |
| TDGDDGNGGSPGT | ASGDASGGAETLY | 0.6521119 |
| TDGLGEN | ASGDASGGAETLY | 0.5267 |
| TGGPPLPGQA | ASGDASGGAETLY | 0.61365104 |
| TIPAITI | ASGDASGGAETLY | 0.60320586 |
| TIPGFTI | ASGDASGGAETLY | 0.5153078 |
| TSIASGALA | ASGDASGGAETLY | 0.61635214 |
| TVTLPGAAI | ASGDASGGAETLY | 0.54006433 |
| VASGLSAALP | ASGDASGGAETLY | 0.6445442 |
| VGAGVIGI | ASGDASGGAETLY | 0.64090717 |
| VGPKRFGV | ASGDASGGAETLY | 0.60705006 |
| VNVAFPD | ASGDASGGAETLY | 0.5563244 |
| VSMTNSGVS | ASGDASGGAETLY | 0.62711143 |
| VTDVLGDLSV | ASGDASGGAETLY | 0.47126663 |
| VVVVGAGFAGL | ASGDASGGAETLY | 0.5770138 |
| VYTVGTVV | ASGDASGGAETLY | 0.4106504 |
| WTTFSGP | ASGDASGGAETLY | 0.580583 |

| | | |
|---|---|---|
| YDELVEAF | ASGDASGGAETLY | 0.506916 |
| YGTGGAGGNGG | ASGDASGGAETLY | 0.6199546 |
| YTNVMAV | ASGDASGGAETLY | 0.56973755 |
| FLEQVHQL | ASGDASGGAETLY | 0.606777 |
| GLSLLAVAV | ASGDASGGAETLY | 0.5035096 |
| LEQVHQL | ASGDASGGAETLY | 0.59717053 |
| LNAAGAI | ASGDASGGAETLY | 0.5711807 |
| PAVHASL | ASGDASGGAETLY | 0.4515934 |
| TELEDDA | ASGDASGGAETLY | 0.62911946 |
| TPPGPPN | ASGDASGGAETLY | 0.5784321 |
| TPPPPPPP | ASGDASGGAETLY | 0.53135276 |
| TQSPALI | ASSLGGSYNSPLY | 0.62765473 |
| EFAAAFPQ | ASSLGGSYNSPLY | 0.61447245 |
| LLSGNLDVL | ASSLGGSYNSPLY | 0.4777308 |
| GPANLGNYNI | ASSLGGSYNSPLY | 0.5386145 |
| TLGPTLP | ASSLGGSYNSPLY | 0.60073364 |
| VYAYSSCCK | ASSLGGSYNSPLY | 0.56836224 |
| APAPAPAPNI | ASSLGGSYNSPLY | 0.33739996 |
| PGGPVPIP | ASSLGGSYNSPLY | 0.59127814 |
| FLTVALEP | ASSLGGSYNSPLY | 0.5344243 |
| SQYFGANL | ASSLGGSYNSPLY | 0.50580746 |
| GGVKPAA | ASSLGGSYNSPLY | 0.5444868 |
| GGTGGAGGAGGW | ASSLGGSYNSPLY | 0.64210093 |
| GVGGVAGLGGAGG | ASSLGGSYNSPLY | 0.52276033 |
| VLLVLATTI | ASSLGGSYNSPLY | 0.60278064 |
| IGIFAAML | ASSLGGSYNSPLY | 0.6237067 |
| AVGDLID | ASSLGGSYNSPLY | 0.5984305 |
| NISPLPL | ASSLGGSYNSPLY | 0.63617164 |
| VTAVFVAI | ASSLGGSYNSPLY | 0.58346796 |
| SGDDNDAAL | ASSLGGSYNSPLY | 0.58459795 |
| AVAALAHS | ASSLGGSYNSPLY | 0.62059927 |
| GGVTVMATP | ASSLGGSYNSPLY | 0.5704439 |
| AGLAANVRD | ASSLGGSYNSPLY | 0.5100922 |
| TNRPDLIDPALLRPG | ASSLGGSYNSPLY | 0.484083 |
| LANLGVTAL | ASSLGGSYNSPLY | 0.56607676 |
| TGERIEL | ASSLGGSYNSPLY | 0.47580087 |
| NGEELAA | ASSLGGSYNSPLY | 0.61169505 |
| VLSLVWG | ASSLGGSYNSPLY | 0.6473351 |
| IGYALWL | ASSLGGSYNSPLY | 0.57231915 |
| LSLLLAPL | ASSLGGSYNSPLY | 0.45237187 |
| TLDEALK | ASSLGGSYNSPLY | 0.56525224 |
| TDGAGTAEGG | ASSLGGSYNSPLY | 0.5562333 |
| NAPGGAGGNGG | ASSLGGSYNSPLY | 0.6393549 |
| VSVFMGMGGNF | ASSLGGSYNSPLY | 0.6069864 |
| LASYLAAL | ASSLGGSYNSPLY | 0.63245577 |
| LVGGTPAL | ASSLGGSYNSPLY | 0.6069221 |
| VRNADVL | ASSLGGSYNSPLY | 0.5230898 |
| TCSSPFLVSAM | ASSLGGSYNSPLY | 0.6389738 |
| VHITDAE | ASSLGGSYNSPLY | 0.53594255 |
| EYAAPSF | ASSLGGSYNSPLY | 0.60847324 |
| VVDAFHGGL | ASSLGGSYNSPLY | 0.599604 |

| | | |
|---|---|---|
| ASVVDPQ | ASSLGGSYNSPLY | 0.56075853 |
| SYNEFGGLS | ASSLGGSYNSPLY | 0.55793273 |
| DPDGSVY | ASSLGGSYNSPLY | 0.60829294 |
| MVSTIHATE | ASSLGGSYNSPLY | 0.44381505 |
| LAAGVADPV | ASSLGGSYNSPLY | 0.60150033 |
| TKEQIAA | ASSLGGSYNSPLY | 0.5460501 |
| SLASPAAL | ASSLGGSYNSPLY | 0.48843116 |
| IGLVFYGV | ASSLGGSYNSPLY | 0.5577843 |
| FLLDASNL | ASSLGGSYNSPLY | 0.6492233 |
| YQVGNFP | ASSLGGSYNSPLY | 0.64386785 |
| ESLLSQST | ASSLGGSYNSPLY | 0.5235417 |
| LFTVGFAAVG | ASSLGGSYNSPLY | 0.5111232 |
| ATPPTPT | ASSLGGSYNSPLY | 0.58332103 |
| TDLATLG | ASSLGGSYNSPLY | 0.49112484 |
| SAGGTGR | ASSLGGSYNSPLY | 0.61787486 |
| GFSNSGS | ASSLGGSYNSPLY | 0.55939156 |
| GGNGGSGGTSV | ASSLGGSYNSPLY | 0.5337753 |
| GTSVATGGAGN | ASSLGGSYNSPLY | 0.57809013 |
| NGGNGGSGGTS | ASSLGGSYNSPLY | 0.61572194 |
| AVVKFPH | ASSLGGSYNSPLY | 0.53342694 |
| TPPPPPP | ASSLGGSYNSPLY | 0.62654436 |
| AAPAAAAAPA | ASSLGGSYNSPLY | 0.35061228 |
| IALNEDL | ASSLGGSYNSPLY | 0.6152713 |
| IVNVSSM | ASSLGGSYNSPLY | 0.47244594 |
| TGIPPAPR | ASSLGGSYNSPLY | 0.5170455 |
| FDLGGGTF | ASSLGGSYNSPLY | 0.37706473 |
| FDLGGGTFDVS | ASSLGGSYNSPLY | 0.5898783 |
| NEPTAAA | ASSLGGSYNSPLY | 0.6157986 |
| NEPTAAALA | ASSLGGSYNSPLY | 0.63463366 |
| GSIINISSI | ASSLGGSYNSPLY | 0.560354 |
| VAPGNAGTA | ASSLGGSYNSPLY | 0.62055504 |
| TGGGASLE | ASSLGGSYNSPLY | 0.64051706 |
| VAVYNHYKRI | ASSLGGSYNSPLY | 0.5400625 |
| TDPAPATT | ASSLGGSYNSPLY | 0.42162144 |
| TNRPDILDPALLRPG | ASSLGGSYNSPLY | 0.61079985 |
| LTTTTTS | ASSLGGSYNSPLY | 0.5434984 |
| PTGSGKTL | ASSLGGSYNSPLY | 0.62727684 |
| TGSGKTL | ASSLGGSYNSPLY | 0.60459715 |
| TIGHVDHGK | ASSLGGSYNSPLY | 0.4135623 |
| TIGHVDHGKTT | ASSLGGSYNSPLY | 0.63645595 |
| GHVDHGKTT | ASSLGGSYNSPLY | 0.6340331 |
| AQFEHTLL | ASSLGGSYNSPLY | 0.46637323 |
| IVNQDEL | ASSLGGSYNSPLY | 0.5363617 |
| TNGSQFF | ASSLGGSYNSPLY | 0.52705985 |
| TGKGIVE | ASSLGGSYNSPLY | 0.3999759 |
| LISNASDA | ASSLGGSYNSPLY | 0.59806824 |
| AELRTLGELTKIAKAHGAQ | ASSLGGSYNSPLY | 0.51492697 |
| APNAVAAAEPSW | ASSLGGSYNSPLY | 0.5619675 |
| APQVAGAATAVL | ASSLGGSYNSPLY | 0.5458434 |
| ARRDPAAAAAILPTDA | ASSLGGSYNSPLY | 0.53849214 |
| DAAARDVLAEAGLAEYF | ASSLGGSYNSPLY | 0.54084337 |

| | | |
|---|---|---|
| DDQIVLSAKANLAKVEQ | ASSLGGSYNSPLY | 0.5529253 |
| DELLPDTGEDSDEDGH | ASSLGGSYNSPLY | 0.5998231 |
| ELPEAPANRATVPQD | ASSLGGSYNSPLY | 0.6124332 |
| FGQSKIERVTGIG | ASSLGGSYNSPLY | 0.53165364 |
| GAGGTGSTAGGAGGAGGAGGLY | ASSLGGSYNSPLY | 0.6123756 |
| GHGGTGAAGGNGGTGGAGGSGI | ASSLGGSYNSPLY | 0.6414422 |
| GNGGNGGAGGEGGPGGAGGAGA | ASSLGGSYNSPLY | 0.45753306 |
| GSTAGNGGNGGNATNGTIAGQP | ASSLGGSYNSPLY | 0.5034001 |
| GTGGDGGNGGNGGTGG | ASSLGGSYNSPLY | 0.6124026 |
| GVRRPAGIPADAPV | ASSLGGSYNSPLY | 0.52047354 |
| GYRQVTDAHLLTLARR | ASSLGGSYNSPLY | 0.63556826 |
| IAKMGQDGHDRG | ASSLGGSYNSPLY | 0.58387905 |
| KAEPSIEIQYT | ASSLGGSYNSPLY | 0.59467334 |
| KPVRIRMAADEDTVAVT | ASSLGGSYNSPLY | 0.5960205 |
| MPPRIAGMRLLVIKPEP | ASSLGGSYNSPLY | 0.6516105 |
| MKFYAVALFRSRGVPPTR | ASSLGGSYNSPLY | 0.633196 |
| NAGGFGILGAVAHSP | ASSLGGSYNSPLY | 0.57803553 |
| NEGREITLAMAEM | ASSLGGSYNSPLY | 0.612975 |
| NELVEWSHPLGHRWRWE | ASSLGGSYNSPLY | 0.45714447 |
| NLGAGGTGGNGGNPG | ASSLGGSYNSPLY | 0.6237583 |
| NTEWPKWETVMAADPVD | ASSLGGSYNSPLY | 0.64001393 |
| PASAAPAETQP | ASSLGGSYNSPLY | 0.4678194 |
| QQSSDGKNGVRAAEITGE | ASSLGGSYNSPLY | 0.55302024 |
| RAPDAVTTSWWKE | ASSLGGSYNSPLY | 0.6442329 |
| SGFGNVGAHTSGWFNQ | ASSLGGSYNSPLY | 0.6452304 |
| SGVAGAVGQAASVGGLKV | ASSLGGSYNSPLY | 0.4894938 |
| SNGGAGTGSAGGNGGT | ASSLGGSYNSPLY | 0.5155273 |
| TGGAGGEGGAAG | ASSLGGSYNSPLY | 0.4882803 |
| TGGDGGAAGAGGAPGHG | ASSLGGSYNSPLY | 0.53152525 |
| TMPPLLAAIANAGRHGV | ASSLGGSYNSPLY | 0.44011617 |
| TSSTAGNSGGA | ASSLGGSYNSPLY | 0.6119062 |
| VPATVPVGILC | ASSLGGSYNSPLY | 0.5651148 |
| VQRPDSGGVVLTGRIS | ASSLGGSYNSPLY | 0.60729367 |
| WTPNPVPATKGVPAGLVQAT | ASSLGGSYNSPLY | 0.6004004 |
| YALKRALAKFSDRFDVVII | ASSLGGSYNSPLY | 0.6355125 |
| YLIGSVIV | ASSLGGSYNSPLY | 0.5486248 |
| YGQLVAVID | ASSLGGSYNSPLY | 0.5480306 |
| VTVGGMTLPA | ASSLGGSYNSPLY | 0.436653 |
| VLALPAV | ASSLGGSYNSPLY | 0.47776484 |
| VIPPTLNYET | ASSLGGSYNSPLY | 0.5415147 |
| VGMGGMHPSSG | ASSLGGSYNSPLY | 0.55039936 |
| VGLAGGAATA | ASSLGGSYNSPLY | 0.3729938 |
| VGGAGGAGGSGP | ASSLGGSYNSPLY | 0.6257346 |
| VFAAFMV | ASSLGGSYNSPLY | 0.6016824 |
| TLYDKLR | ASSLGGSYNSPLY | 0.6114247 |
| TLAVFTVL | ASSLGGSYNSPLY | 0.56511456 |
| TGSVQPARV | ASSLGGSYNSPLY | 0.55262184 |
| TGGGAGSGGT | ASSLGGSYNSPLY | 0.518552 |
| SHLETSL | ASSLGGSYNSPLY | 0.6371002 |
| SFTTGMT | ASSLGGSYNSPLY | 0.47043583 |
| QVEVAKRMGVRQPTVS | ASSLGGSYNSPLY | 0.41353858 |

| | | |
|---|---|---|
| QGSGGPALAM | ASSLGGSYNSPLY | 0.55014455 |
| MLAPSNP | ASSLGGSYNSPLY | 0.5414435 |
| LVNNAAI | ASSLGGSYNSPLY | 0.6391219 |
| LTGEINTLA | ASSLGGSYNSPLY | 0.63027465 |
| LSIFGPV | ASSLGGSYNSPLY | 0.6056858 |
| LLALLANL | ASSLGGSYNSPLY | 0.64175296 |
| LISNASDALDKLR | ASSLGGSYNSPLY | 0.34520257 |
| LINEAVP | ASSLGGSYNSPLY | 0.5682316 |
| LGYAFSGL | ASSLGGSYNSPLY | 0.64244395 |
| LDFVAVAL | ASSLGGSYNSPLY | 0.5981022 |
| LAQPGAE | ASSLGGSYNSPLY | 0.5846045 |
| ISNLGAVTGF | ASSLGGSYNSPLY | 0.6428459 |
| HQEDPGRI | ASSLGGSYNSPLY | 0.51231134 |
| GTPGEPTP | ASSLGGSYNSPLY | 0.58498025 |
| GFFNSGS | ASSLGGSYNSPLY | 0.5958507 |
| GAGGGGLTAGAN | ASSLGGSYNSPLY | 0.6276147 |
| FVDALVNM | ASSLGGSYNSPLY | 0.6021751 |
| FLTVAPEL | ASSLGGSYNSPLY | 0.44497108 |
| FETLTAL | ASSLGGSYNSPLY | 0.6398572 |
| DVINPIEL | ASSLGGSYNSPLY | 0.65023595 |
| APVSGAPV | ASSLGGSYNSPLY | 0.56578445 |
| ALVGGLAW | ASSLGGSYNSPLY | 0.6173362 |
| ALAAAAATAI | ASSLGGSYNSPLY | 0.6184056 |
| AFGLGLLL | ASSLGGSYNSPLY | 0.5655761 |
| AFFAAASL | ASSLGGSYNSPLY | 0.50768673 |
| AAAALPAFSPP | ASSLGGSYNSPLY | 0.4454186 |
| VVLAPAL | ASSLGGSYNSPLY | 0.59392154 |
| TIGHVDHGKTTL | ASSLGGSYNSPLY | 0.47381485 |
| LISNASDALDK | ASSLGGSYNSPLY | 0.6225285 |
| LISNASDAL | ASSLGGSYNSPLY | 0.565815 |
| ISNASDALDK | ASSLGGSYNSPLY | 0.6551744 |
| NQAETLVYQTE | ASSLGGSYNSPLY | 0.6091781 |
| TGGTGGTAGAG | ASSLGGSYNSPLY | 0.6139715 |
| TGSTAGGAGGA | ASSLGGSYNSPLY | 0.47953382 |
| DPLAVPVAAGPV | ASSLGGSYNSPLY | 0.6006487 |
| GGTGGTPGNGGN | ASSLGGSYNSPLY | 0.5906974 |
| PAGLAGVDPATL | ASSLGGSYNSPLY | 0.46126503 |
| GNNLVVEAPHAGASV | ASSLGGSYNSPLY | 0.6328119 |
| KVTRIMSASTLEQPA | ASSLGGSYNSPLY | 0.49864903 |
| LGFALPIALPRRTVV | ASSLGGSYNSPLY | 0.55077255 |
| TRLGNLTARKLVVLV | ASSLGGSYNSPLY | 0.60244226 |
| WSQQRRLIYFPSAGP | ASSLGGSYNSPLY | 0.6111115 |
| GPNGNPDPMAAAVDIR | ASSLGGSYNSPLY | 0.5752784 |
| TQRCVSVADLRAAASR | ASSLGGSYNSPLY | 0.5997337 |
| QFVLDVAEHPVEVLTETV | ASSLGGSYNSPLY | 0.47724408 |
| GGAGGTGGAGGNGGTQAGDG | ASSLGGSYNSPLY | 0.6209166 |
| PLLATLRAIAVDVDAKPAQV | ASSLGGSYNSPLY | 0.64147925 |
| APKFRTGGYMIDFWGVGYQVAK | ASSLGGSYNSPLY | 0.5650378 |
| GAGGAGGAGGAAGGPGGTGGTG | ASSLGGSYNSPLY | 0.6409333 |
| IGALGLLATDVAPAGSVAFGVA | ASSLGGSYNSPLY | 0.6020291 |
| AGGAGGIGGNGGAGGAGGVLMG | ASSLGGSYNSPLY | 0.41217363 |

| | | |
|---|---|---|
| ADELIGLL | ASSLGGSYNSPLY | 0.6120596 |
| ADRALKA | ASSLGGSYNSPLY | 0.6338452 |
| AGAVGQVGGAGGA | ASSLGGSYNSPLY | 0.62791264 |
| AGGGGGLGGAP | ASSLGGSYNSPLY | 0.63931525 |
| AGGLLYGNGGAG | ASSLGGSYNSPLY | 0.634272 |
| AGLESIGL | ASSLGGSYNSPLY | 0.55191684 |
| AGLKTGII | ASSLGGSYNSPLY | 0.6413301 |
| ASLAALLL | ASSLGGSYNSPLY | 0.65596056 |
| ASVINISSI | ASSLGGSYNSPLY | 0.6347785 |
| ATPPGPA | ASSLGGSYNSPLY | 0.5174862 |
| ATVASALV | ASSLGGSYNSPLY | 0.49602056 |
| ATVASLIL | ASSLGGSYNSPLY | 0.6202008 |
| AVNGPVLIRH | ASSLGGSYNSPLY | 0.6096115 |
| DGAAGAVGI | ASSLGGSYNSPLY | 0.5293764 |
| DPLLHGI | ASSLGGSYNSPLY | 0.5850046 |
| ELGAALGAL | ASSLGGSYNSPLY | 0.5784313 |
| ELGRTQAV | ASSLGGSYNSPLY | 0.41560596 |
| FVAGGYLS | ASSLGGSYNSPLY | 0.61397815 |
| GASGGAGGAGGW | ASSLGGSYNSPLY | 0.63893294 |
| GGAGAGGDGGAGGI | ASSLGGSYNSPLY | 0.61760676 |
| GTGHNGSQPGMGGNG | ASSLGGSYNSPLY | 0.54371005 |
| GYAAASASAS | ASSLGGSYNSPLY | 0.63385546 |
| HQVDVAAVQQG | ASSLGGSYNSPLY | 0.64269865 |
| IGAAIAHT | ASSLGGSYNSPLY | 0.6484157 |
| IGMIEVAQL | ASSLGGSYNSPLY | 0.44141227 |
| IRLAPEHTSAMD | ASSLGGSYNSPLY | 0.33774716 |
| IYNVSVL | ASSLGGSYNSPLY | 0.5943604 |
| KQGFLGM | ASSLGGSYNSPLY | 0.6033992 |
| LESVITQL | ASSLGGSYNSPLY | 0.54211575 |
| LGLGALVVV | ASSLGGSYNSPLY | 0.51315176 |
| LIGNGGNGGSGGI | ASSLGGSYNSPLY | 0.60462964 |
| LNAINAPTL | ASSLGGSYNSPLY | 0.6248739 |
| LQLLNTGV | ASSLGGSYNSPLY | 0.5076223 |
| LQTSIEF | ASSLGGSYNSPLY | 0.44344252 |
| MSFAVSEL | ASSLGGSYNSPLY | 0.62759984 |
| NTGALGL | ASSLGGSYNSPLY | 0.64519835 |
| PPGAGGAGGAGTT | ASSLGGSYNSPLY | 0.5001168 |
| QLVGLGI | ASSLGGSYNSPLY | 0.52245754 |
| RFLEQVHQL | ASSLGGSYNSPLY | 0.59114873 |
| RGVGYVLR | ASSLGGSYNSPLY | 0.52882135 |
| SATGVFAGVI | ASSLGGSYNSPLY | 0.5685031 |
| SLPAITLP | ASSLGGSYNSPLY | 0.46223137 |
| STALAFTV | ASSLGGSYNSPLY | 0.61568177 |
| SVLTVGLGK | ASSLGGSYNSPLY | 0.46327978 |
| TALAGAGSGI | ASSLGGSYNSPLY | 0.618641 |
| TGLPFTGI | ASSLGGSYNSPLY | 0.61485595 |
| VAEHVTV | ASSLGGSYNSPLY | 0.60528535 |
| VDIVLNSL | ASSLGGSYNSPLY | 0.5496746 |
| VLTGGGAVL | ASSLGGSYNSPLY | 0.5004734 |
| VNVGLAYGI | ASSLGGSYNSPLY | 0.4740674 |
| VSSAAVGAV | ASSLGGSYNSPLY | 0.4673403 |

| | | |
|---|---|---|
| VTSVYEAGV | ASSLGGSYNSPLY | 0.6433247 |
| WVALDRGA | ASSLGGSYNSPLY | 0.4796952 |
| YLVGSVVI | ASSLGGSYNSPLY | 0.6418094 |
| GPPGTGKTLL | ASSLGGSYNSPLY | 0.608232 |
| GVGMTKF | ASSLGGSYNSPLY | 0.5065659 |
| IGIGGSDL | ASSLGGSYNSPLY | 0.6044241 |
| IGTLLAEL | ASSLGGSYNSPLY | 0.42106777 |
| IYTSGTT | ASSLGGSYNSPLY | 0.5902927 |
| LSFPAGV | ASSLGGSYNSPLY | 0.584545 |
| LYTSGTT | ASSLGGSYNSPLY | 0.6319119 |
| VGAFPFGF | ASSLGGSYNSPLY | 0.550593 |
| VGAGTVEF | ASSLGGSYNSPLY | 0.6339253 |
| VGKSTLLL | ASSLGGSYNSPLY | 0.6074177 |
| AALAHAGGGIV | ASSLGGSYNSPLY | 0.5744117 |
| AALVAVSLL | ASSLGGSYNSPLY | 0.62411726 |
| AAVSRAVAAL | ASSLGGSYNSPLY | 0.6296671 |
| AGGSSGIGGF | ASSLGGSYNSPLY | 0.6326704 |
| AGLTVANANG | ASSLGGSYNSPLY | 0.4900133 |
| AGLVGAVL | ASSLGGSYNSPLY | 0.59538776 |
| ALSQAFTV | ASSLGGSYNSPLY | 0.5081479 |
| ALTFSVSL | ASSLGGSYNSPLY | 0.5783005 |
| ALTGAALVAL | ASSLGGSYNSPLY | 0.56938034 |
| ASFSIGAGVL | ASSLGGSYNSPLY | 0.57018596 |
| DGSVYKI | ASSLGGSYNSPLY | 0.6322801 |
| EQLAEIV | ASSLGGSYNSPLY | 0.4204916 |
| FAVSELTVM | ASSLGGSYNSPLY | 0.54442143 |
| GAAPLAAGL | ASSLGGSYNSPLY | 0.55720437 |
| GADGGDGGSGGAGG | ASSLGGSYNSPLY | 0.601596 |
| GFGLDTGGAGG | ASSLGGSYNSPLY | 0.6262674 |
| GLASGGLL | ASSLGGSYNSPLY | 0.43905368 |
| GRIAGAAVLL | ASSLGGSYNSPLY | 0.6032387 |
| GSAGSLNTSF | ASSLGGSYNSPLY | 0.5840578 |
| GTGQIGI | ASSLGGSYNSPLY | 0.5270162 |
| GTLIALIL | ASSLGGSYNSPLY | 0.6246794 |
| IAATMGFVVG | ASSLGGSYNSPLY | 0.56768763 |
| IFAQALL | ASSLGGSYNSPLY | 0.50425005 |
| IGAAVALGF | ASSLGGSYNSPLY | 0.62418324 |
| IGLTPLAL | ASSLGGSYNSPLY | 0.54507214 |
| ILGSNTSTL | ASSLGGSYNSPLY | 0.61338264 |
| IVFSLTVL | ASSLGGSYNSPLY | 0.6214263 |
| LASKATGF | ASSLGGSYNSPLY | 0.49513942 |
| LASVALIL | ASSLGGSYNSPLY | 0.61937386 |
| LFAVGLTL | ASSLGGSYNSPLY | 0.58360016 |
| LFNGGIGTYI | ASSLGGSYNSPLY | 0.620285 |
| LTLGAGII | ASSLGGSYNSPLY | 0.5645955 |
| LVAAALGDI | ASSLGGSYNSPLY | 0.5298568 |
| NTAFVQLGI | ASSLGGSYNSPLY | 0.63494575 |
| QLFSGAEL | ASSLGGSYNSPLY | 0.61646533 |
| SAATFITL | ASSLGGSYNSPLY | 0.6363892 |
| SNYGLANLGV | ASSLGGSYNSPLY | 0.608253 |
| SYGGAYAVM | ASSLGGSYNSPLY | 0.62124425 |

| | | |
|---|---|---|
| TTGVGGLTLG | ASSLGGSYNSPLY | 0.64007896 |
| TTQTIYI | ASSLGGSYNSPLY | 0.50104594 |
| VAAGAGLGAVI | ASSLGGSYNSPLY | 0.44779134 |
| VAAVVTLL | ASSLGGSYNSPLY | 0.5528846 |
| VGFLGGLI | ASSLGGSYNSPLY | 0.5504527 |
| VGVIPILL | ASSLGGSYNSPLY | 0.5601909 |
| VIFASSLI | ASSLGGSYNSPLY | 0.4264215 |
| VKGGDGGAAAG | ASSLGGSYNSPLY | 0.6210538 |
| VTKTFTNI | ASSLGGSYNSPLY | 0.6571021 |
| VVAAAILL | ASSLGGSYNSPLY | 0.538214 |
| YAALDVEL | ASSLGGSYNSPLY | 0.64917374 |
| LASVALLL | ASSLGGSYNSPLY | 0.45030493 |
| LDAVHRV | ASSLGGSYNSPLY | 0.61873233 |
| QGLSSSIF | ASSLGGSYNSPLY | 0.63969046 |
| RGTFANIRL | ASSLGGSYNSPLY | 0.47588503 |
| RQLWWGHRI | ASSLGGSYNSPLY | 0.5999019 |
| YGPPGTGKTLL | ASSLGGSYNSPLY | 0.57430303 |
| AALVGAPANVA | ASSLGGSYNSPLY | 0.54064524 |
| AATFRRTE | ASSLGGSYNSPLY | 0.6473189 |
| AGALGGGI | ASSLGGSYNSPLY | 0.56073993 |
| AGGAGGAGGIGAG | ASSLGGSYNSPLY | 0.6275948 |
| AGSGALGL | ASSLGGSYNSPLY | 0.5767557 |
| AQIADQI | ASSLGGSYNSPLY | 0.48175758 |
| ASAVAVTYL | ASSLGGSYNSPLY | 0.6130563 |
| ASTVLGGTGG | ASSLGGSYNSPLY | 0.6363157 |
| AVGGEPAK | ASSLGGSYNSPLY | 0.56443495 |
| CEQGTFGGTC | ASSLGGSYNSPLY | 0.6560387 |
| DEGKAPAG | ASSLGGSYNSPLY | 0.527289 |
| DTVVFPAL | ASSLGGSYNSPLY | 0.644719 |
| FEGITMAG | ASSLGGSYNSPLY | 0.6404917 |
| FPLVVQD | ASSLGGSYNSPLY | 0.55619997 |
| GAFTTPQ | ASSLGGSYNSPLY | 0.53879714 |
| GFGGFGTAGGVGGA | ASSLGGSYNSPLY | 0.62528145 |
| GFLAQFL | ASSLGGSYNSPLY | 0.6034808 |
| GGAGGAGGNAGVA | ASSLGGSYNSPLY | 0.6151377 |
| GGVGTAGAGGAGGAGG | ASSLGGSYNSPLY | 0.6453329 |
| GNGGGGDAGNAGSGGN | ASSLGGSYNSPLY | 0.742804 |
| GTGGAGGAGNDAGS | ASSLGGSYNSPLY | 0.84055734 |
| GTTNTGWFNS | ASSLGGSYNSPLY | 0.812089 |
| IFMGPLFTL | ASSLGGSYNSPLY | 0.7995099 |
| ILAHAIEAI | ASSLGGSYNSPLY | 0.7591728 |
| IRSAIREAL | ASSLGGSYNSPLY | 0.4745307 |
| ITFAGGG | ASSLGGSYNSPLY | 0.79344213 |
| ITGMSGHQEV | ASSLGGSYNSPLY | 0.7056612 |
| IVAGLACTL | ASSLGGSYNSPLY | 0.79205215 |
| IVGGGLAGA | ASSLGGSYNSPLY | 0.8237982 |
| IVTGVLNI | ASSLGGSYNSPLY | 0.74297833 |
| LAQQGCI | ASSLGGSYNSPLY | 0.8401339 |
| LCLHVEN | ASSLGGSYNSPLY | 0.8267211 |
| LDAISDGS | ASSLGGSYNSPLY | 0.60039264 |
| LGGFAAML | ASSLGGSYNSPLY | 0.78765327 |

| | | |
|---|---|---|
| LGGKNPMI | ASSLGGSYNSPLY | 0.77010965 |
| LGPKGVAAIL | ASSLGGSYNSPLY | 0.79666203 |
| LMVGVSI | ASSLGGSYNSPLY | 0.5387546 |
| LVGAAFVL | ASSLGGSYNSPLY | 0.8265444 |
| LVLAVAPL | ASSLGGSYNSPLY | 0.75390255 |
| MAAIAVIDAA | ASSLGGSYNSPLY | 0.19140485 |
| NNGINAELQ | ASSLGGSYNSPLY | 0.80957437 |
| RGEPLVLVHG | ASSLGGSYNSPLY | 0.79798514 |
| RGGAGGAAGLI | ASSLGGSYNSPLY | 0.76439226 |
| RSYRMHPAV | ASSLGGSYNSPLY | 0.83381104 |
| SAAHLLAGQLA | ASSLGGSYNSPLY | 0.35243833 |
| SAAVGSY | ASSLGGSYNSPLY | 0.71415854 |
| SPHAVLHRI | ASSLGGSYNSPLY | 0.82651603 |
| SQIAKVL | ASSLGGSYNSPLY | 0.83597183 |
| TALGHLVP | ASSLGGSYNSPLY | 0.64493316 |
| TAPTTSGAA | ASSLGGSYNSPLY | 0.8122232 |
| TDGDDGNGGSPGT | ASSLGGSYNSPLY | 0.8317529 |
| TDGLGEN | ASSLGGSYNSPLY | 0.8140851 |
| TGGPPLPGQA | ASSLGGSYNSPLY | 0.83447254 |
| TIPAITI | ASSLGGSYNSPLY | 0.8392874 |
| TIPGFTI | ASSLGGSYNSPLY | 0.7106863 |
| TSIASGALA | ASSLGGSYNSPLY | 0.8348216 |
| TVTLPGAAI | ASSLGGSYNSPLY | 0.84575987 |
| VASGLSAALP | ASSLGGSYNSPLY | 0.76156235 |
| VGAGVIGI | ASSLGGSYNSPLY | 0.8021114 |
| VGPKRFGV | ASSLGGSYNSPLY | 0.74640363 |
| VNVAFPD | ASSLGGSYNSPLY | 0.7675709 |
| VSMTNSGVS | ASSLGGSYNSPLY | 0.34492224 |
| VTDVLGDLSV | ASSLGGSYNSPLY | 0.81216264 |
| VVVVGAGFAGL | ASSLGGSYNSPLY | 0.8359776 |
| VYTVGTVV | ASSLGGSYNSPLY | 0.77731 |
| WTTFSGP | ASSLGGSYNSPLY | 0.7205586 |
| YDELVEAF | ASSLGGSYNSPLY | 0.6888319 |
| YGTGGAGGNGG | ASSLGGSYNSPLY | 0.7809758 |
| YTNVMAV | ASSLGGSYNSPLY | 0.75816 |
| FLEQVHQL | ASSLGGSYNSPLY | 0.3835519 |
| GLSLLAVAV | ASSLGGSYNSPLY | 0.7370555 |
| LEQVHQL | ASSLGGSYNSPLY | 0.8324739 |
| LNAAGAI | ASSLGGSYNSPLY | 0.831468 |
| PAVHASL | ASSLGGSYNSPLY | 0.5355503 |
| TELEDDA | ASSLGGSYNSPLY | 0.8404146 |
| TPPGPPN | ASSLGGSYNSPLY | 0.81516635 |
| TPPPPPPP | ASSLGGSYNSPLY | 0.8386637 |
| TQSPALI | ASSQPGPEVF | 0.79647803 |
| EFAAAFPQ | ASSQPGPEVF | 0.7404434 |
| LLSGNLDVL | ASSQPGPEVF | 0.82443553 |
| GPANLGNYNI | ASSQPGPEVF | 0.7749562 |
| TLGPTLP | ASSQPGPEVF | 0.6755003 |
| VYAYSSCCK | ASSQPGPEVF | 0.8338472 |
| APAPAPAPNI | ASSQPGPEVF | 0.7726517 |
| PGGPVPIP | ASSQPGPEVF | 0.7368306 |

| | | |
|---|---|---|
| FLTVALEP | ASSQPGPEVF | 0.7303596 |
| SQYFGANL | ASSQPGPEVF | 0.84073484 |
| GGVKPAA | ASSQPGPEVF | 0.70745564 |
| GGTGGAGGAGGW | ASSQPGPEVF | 0.60743546 |
| GVGGVAGLGGAGG | ASSQPGPEVF | 0.6463838 |
| VLLVLATTI | ASSQPGPEVF | 0.79433 |
| IGIFAAML | ASSQPGPEVF | 0.83406866 |
| AVGDLID | ASSQPGPEVF | 0.8159166 |
| NISPLPL | ASSQPGPEVF | 0.73668957 |
| VTAVFVAI | ASSQPGPEVF | 0.8143653 |
| SGDDNDAAL | ASSQPGPEVF | 0.81595755 |
| AVAALAHS | ASSQPGPEVF | 0.80676794 |
| GGVTVMATP | ASSQPGPEVF | 0.8182528 |
| AGLAANVRD | ASSQPGPEVF | 0.73902786 |
| TNRPDLIDPALLRPG | ASSQPGPEVF | 0.8391795 |
| LANLGVTAL | ASSQPGPEVF | 0.8216398 |
| TGERIEL | ASSQPGPEVF | 0.8285254 |
| NGEELAA | ASSQPGPEVF | 0.82974195 |
| VLSLVWG | ASSQPGPEVF | 0.7878722 |
| IGYALWL | ASSQPGPEVF | 0.8063496 |
| LSLLLAPL | ASSQPGPEVF | 0.4182005 |
| TLDEALK | ASSQPGPEVF | 0.6773038 |
| TDGAGTAEGG | ASSQPGPEVF | 0.8137393 |
| NAPGGAGGNGG | ASSQPGPEVF | 0.8125487 |
| VSVFMGMGGNF | ASSQPGPEVF | 0.8232069 |
| LASYLAAL | ASSQPGPEVF | 0.80733156 |
| LVGGTPAL | ASSQPGPEVF | 0.7445419 |
| VRNADVL | ASSQPGPEVF | 0.82565933 |
| TCSSPFLVSAM | ASSQPGPEVF | 0.8286462 |
| VHITDAE | ASSQPGPEVF | 0.59980756 |
| EYAAPSF | ASSQPGPEVF | 0.7664758 |
| VVDAFHGGL | ASSQPGPEVF | 0.8455745 |
| ASVVDPQ | ASSQPGPEVF | 0.80114275 |
| SYNEFGGLS | ASSQPGPEVF | 0.78882897 |
| DPDGSVY | ASSQPGPEVF | 0.75504404 |
| MVSTIHATE | ASSQPGPEVF | 0.8301719 |
| LAAGVADPV | ASSQPGPEVF | 0.7164571 |
| TKEQIAA | ASSQPGPEVF | 0.8443481 |
| SLASPAAL | ASSQPGPEVF | 0.2648949 |
| IGLVFYGV | ASSQPGPEVF | 0.75635827 |
| FLLDASNL | ASSQPGPEVF | 0.7356873 |
| YQVGNFP | ASSQPGPEVF | 0.3968013 |
| ESLLSQST | ASSQPGPEVF | 0.8366711 |
| LFTVGFAAVG | ASSQPGPEVF | 0.7441755 |
| ATPPTPT | ASSQPGPEVF | 0.8197411 |
| TDLATLG | ASSQPGPEVF | 0.80578744 |
| SAGGTGR | ASSQPGPEVF | 0.83649296 |
| GFSNSGS | ASSQPGPEVF | 0.8008595 |
| GGNGGSGGTSV | ASSQPGPEVF | 0.7748912 |
| GTSVATGGAGN | ASSQPGPEVF | 0.651399 |
| NGGNGGSGGTS | ASSQPGPEVF | 0.8322245 |

| | | |
|---|---|---|
| AVVKFPH | ASSQPGPEVF | 0.83272505 |
| TPPPPPP | ASSQPGPEVF | 0.82581925 |
| AAPAAAAAPA | ASSQPGPEVF | 0.5306411 |
| IALNEDL | ASSQPGPEVF | 0.7685982 |
| IVNVSSM | ASSQPGPEVF | 0.80730575 |
| TGIPPAPR | ASSQPGPEVF | 0.2515892 |
| FDLGGGTF | ASSQPGPEVF | 0.30981678 |
| FDLGGGTFDVS | ASSQPGPEVF | 0.76223326 |
| NEPTAAA | ASSQPGPEVF | 0.7385103 |
| NEPTAAALA | ASSQPGPEVF | 0.8348539 |
| GSIINISSI | ASSQPGPEVF | 0.8281894 |
| VAPGNAGTA | ASSQPGPEVF | 0.788486 |
| TGGGASLE | ASSQPGPEVF | 0.82183284 |
| VAVYNHYKRI | ASSQPGPEVF | 0.7872524 |
| TDPAPATT | ASSQPGPEVF | 0.13936932 |
| TNRPDILDPALLRPG | ASSQPGPEVF | 0.73129845 |
| LTTTTTS | ASSQPGPEVF | 0.6476811 |
| PTGSGKTL | ASSQPGPEVF | 0.8334064 |
| TGSGKTL | ASSQPGPEVF | 0.7181215 |
| TIGHVDHGK | ASSQPGPEVF | 0.7416862 |
| TIGHVDHGKTT | ASSQPGPEVF | 0.7992086 |
| GHVDHGKTT | ASSQPGPEVF | 0.7839219 |
| AQFEHTLL | ASSQPGPEVF | 0.7955925 |
| IVNQDEL | ASSQPGPEVF | 0.70319664 |
| TNGSQFF | ASSQPGPEVF | 0.2397702 |
| TGKGIVE | ASSQPGPEVF | 0.83124185 |
| LISNASDA | ASSQPGPEVF | 0.42381498 |
| AELRTLGELTKIAKAHGAQ | ASSQPGPEVF | 0.67812526 |
| APNAVAAAEPSW | ASSQPGPEVF | 0.40422106 |
| APQVAGAATAVL | ASSQPGPEVF | 0.24629171 |
| ARRDPAAAAAILPTDA | ASSQPGPEVF | 0.4426678 |
| DAAARDVLAEAGLAEYF | ASSQPGPEVF | 0.82955164 |
| DDQIVLSAKANLAKVEQ | ASSQPGPEVF | 0.7726093 |
| DELLPDTGEDSDEDGH | ASSQPGPEVF | 0.77911097 |
| ELPEAPANRATVPQD | ASSQPGPEVF | 0.84415174 |
| FGQSKIERVTGIG | ASSQPGPEVF | 0.7711127 |
| GAGGTGSTAGGAGGAGGAGGLY | ASSQPGPEVF | 0.7829054 |
| GHGGTGAAGGNGGTGGAGGSGI | ASSQPGPEVF | 0.7094953 |
| GNGGNGGAGGEGGPGGAGGAGA | ASSQPGPEVF | 0.7310312 |
| GSTAGNGGNGGNATNGTIAGQP | ASSQPGPEVF | 0.7369467 |
| GTGGDGGNGGNGGTGG | ASSQPGPEVF | 0.6996839 |
| GVRRPAGIPADAPV | ASSQPGPEVF | 0.8435806 |
| GYRQVTDAHLLTLARR | ASSQPGPEVF | 0.7692986 |
| IAKMGQDGHDRG | ASSQPGPEVF | 0.6138711 |
| KAEPSIEIQYT | ASSQPGPEVF | 0.7894316 |
| KPVRIRMAADEDTVAVT | ASSQPGPEVF | 0.84087956 |
| MPPRIAGMRLLVIKPEP | ASSQPGPEVF | 0.8472175 |
| MKFYAVALFRSRGVPPTR | ASSQPGPEVF | 0.7203181 |
| NAGGFGILGAVAHSP | ASSQPGPEVF | 0.745929 |
| NEGREITLAMAEM | ASSQPGPEVF | 0.7686497 |
| NELVEWSHPLGHRWRWE | ASSQPGPEVF | 0.79081845 |

| | | |
|---|---|---|
| NLGAGGTGGNGGNPG | ASSQPGPEVF | 0.74425244 |
| NTEWPKWETVMAADPVD | ASSQPGPEVF | 0.5567965 |
| PASAAPAETQP | ASSQPGPEVF | 0.8310038 |
| QQSSDGKNGVRAAEITGE | ASSQPGPEVF | 0.83605504 |
| RAPDAVTTSWWKE | ASSQPGPEVF | 0.75430113 |
| SGFGNVGAHTSGWFNQ | ASSQPGPEVF | 0.70431006 |
| SGVAGAVGQAASVGGLKV | ASSQPGPEVF | 0.78702986 |
| SNGGAGTGSAGGNGGT | ASSQPGPEVF | 0.72378314 |
| TGGAGGEGGAAG | ASSQPGPEVF | 0.7669009 |
| TGGDGGAAGAGGAPGHG | ASSQPGPEVF | 0.77184534 |
| TMPPLLAAIANAGRHGV | ASSQPGPEVF | 0.612986 |
| TSSTAGNSGGA | ASSQPGPEVF | 0.78532124 |
| VPATVPVGILC | ASSQPGPEVF | 0.8466542 |
| VQRPDSGGVVLTGRIS | ASSQPGPEVF | 0.7594442 |
| WTPNPVPATKGVPAGLVQAT | ASSQPGPEVF | 0.80752224 |
| YALKRALAKFSDRFDVVII | ASSQPGPEVF | 0.8392621 |
| YLIGSVIV | ASSQPGPEVF | 0.82690394 |
| YGQLVAVID | ASSQPGPEVF | 0.83948016 |
| VTVGGMTLPA | ASSQPGPEVF | 0.81357414 |
| VLALPAV | ASSQPGPEVF | 0.7827373 |
| VIPPTLNYET | ASSQPGPEVF | 0.8031597 |
| VGMGGMHPSSG | ASSQPGPEVF | 0.8057833 |
| VGLAGGAATA | ASSQPGPEVF | 0.7930969 |
| VGGAGGAGGSGP | ASSQPGPEVF | 0.7898705 |
| VFAAFMV | ASSQPGPEVF | 0.754778 |
| TLYDKLR | ASSQPGPEVF | 0.8348386 |
| TLAVFTVL | ASSQPGPEVF | 0.80977154 |
| TGSVQPARV | ASSQPGPEVF | 0.823583 |
| TGGGAGSGGT | ASSQPGPEVF | 0.5463269 |
| SHLETSL | ASSQPGPEVF | 0.6491096 |
| SFTTGMT | ASSQPGPEVF | 0.82733405 |
| QVEVAKRMGVRQPTVS | ASSQPGPEVF | 0.16999795 |
| QGSGGPALAM | ASSQPGPEVF | 0.2732985 |
| MLAPSNP | ASSQPGPEVF | 0.812395 |
| LVNNAAI | ASSQPGPEVF | 0.83740306 |
| LTGEINTLA | ASSQPGPEVF | 0.17277929 |
| LSIFGPV | ASSQPGPEVF | 0.6440754 |
| LLALLANL | ASSQPGPEVF | 0.6284084 |
| LISNASDALDKLR | ASSQPGPEVF | 0.80373883 |
| LINEAVP | ASSQPGPEVF | 0.79651606 |
| LGYAFSGL | ASSQPGPEVF | 0.43283382 |
| LDFVAVAL | ASSQPGPEVF | 0.8220023 |
| LAQPGAE | ASSQPGPEVF | 0.59438896 |
| ISNLGAVTGF | ASSQPGPEVF | 0.5073989 |
| HQEDPGRI | ASSQPGPEVF | 0.835436 |
| GTPGEPTP | ASSQPGPEVF | 0.8313122 |
| GFFNSGS | ASSQPGPEVF | 0.69097936 |
| GAGGGGLTAGAN | ASSQPGPEVF | 0.82813287 |
| FVDALVNM | ASSQPGPEVF | 0.7382802 |
| FLTVAPEL | ASSQPGPEVF | 0.8487056 |
| FETLTAL | ASSQPGPEVF | 0.77936065 |

| | | |
|---|---|---|
| DVINPIEL | ASSQPGPEVF | 0.548779 |
| APVSGAPV | ASSQPGPEVF | 0.8289225 |
| ALVGGLAW | ASSQPGPEVF | 0.8148508 |
| ALAAAAATAI | ASSQPGPEVF | 0.7803904 |
| AFGLGLLL | ASSQPGPEVF | 0.8218218 |
| AFFAAASL | ASSQPGPEVF | 0.69347274 |
| AAAALPAFSPP | ASSQPGPEVF | 0.7853828 |
| VVLAPAL | ASSQPGPEVF | 0.83497065 |
| TIGHVDHGKTTL | ASSQPGPEVF | 0.83246887 |
| LISNASDALDK | ASSQPGPEVF | 0.5645219 |
| LISNASDAL | ASSQPGPEVF | 0.83411133 |
| ISNASDALDK | ASSQPGPEVF | 0.7794295 |
| NQAETLVYQTE | ASSQPGPEVF | 0.720951 |
| TGGTGGTAGAG | ASSQPGPEVF | 0.8464778 |
| TGSTAGGAGGA | ASSQPGPEVF | 0.7793465 |
| DPLAVPVAAGPV | ASSQPGPEVF | 0.83262384 |
| GGTGGTPGNGGN | ASSQPGPEVF | 0.831776 |
| PAGLAGVDPATL | ASSQPGPEVF | 0.81238025 |
| GNNLVVEAPHAGASV | ASSQPGPEVF | 0.47344324 |
| KVTRIMSASTLEQPA | ASSQPGPEVF | 0.79596484 |
| LGFALPIALPRRTVV | ASSQPGPEVF | 0.792771 |
| TRLGNLTARKLVVLV | ASSQPGPEVF | 0.8239428 |
| WSQQRRLIYFPSAGP | ASSQPGPEVF | 0.7950852 |
| GPNGNPDPMAAAVDIR | ASSQPGPEVF | 0.6403372 |
| TQRCVSVADLRAAASR | ASSQPGPEVF | 0.8170613 |
| QFVLDVAEHPVEVLTETV | ASSQPGPEVF | 0.8409364 |
| GGAGGTGGAGGNGGTQAGDG | ASSQPGPEVF | 0.8113967 |
| PLLATLRAIAVDVDAKPAQV | ASSQPGPEVF | 0.8057987 |
| APKFRTGGYMIDFWGVGYQVAK | ASSQPGPEVF | 0.84299874 |
| GAGGAGGAGGAAGGPGGTGGTG | ASSQPGPEVF | 0.752542 |
| IGALGLLATDVAPAGSVAFGVA | ASSQPGPEVF | 0.77342397 |
| AGGAGGIGGNGGAGGAGGVLMG | ASSQPGPEVF | 0.83202827 |
| ADELIGLL | ASSQPGPEVF | 0.83092445 |
| ADRALKA | ASSQPGPEVF | 0.78097534 |
| AGAVGQVGGAGGA | ASSQPGPEVF | 0.83484226 |
| AGGGGGLGGAP | ASSQPGPEVF | 0.6603708 |
| AGGLLYGNGGAG | ASSQPGPEVF | 0.8167141 |
| AGLESIGL | ASSQPGPEVF | 0.7337782 |
| AGLKTGII | ASSQPGPEVF | 0.8157431 |
| ASLAALLL | ASSQPGPEVF | 0.7024218 |
| ASVINISSI | ASSQPGPEVF | 0.82441175 |
| ATPPGPA | ASSQPGPEVF | 0.8330359 |
| ATVASALV | ASSQPGPEVF | 0.82976246 |
| ATVASLIL | ASSQPGPEVF | 0.28172395 |
| AVNGPVLIRH | ASSQPGPEVF | 0.83065414 |
| DGAAGAVGI | ASSQPGPEVF | 0.81082684 |
| DPLLHGI | ASSQPGPEVF | 0.84649205 |
| ELGAALGAL | ASSQPGPEVF | 0.7948609 |
| ELGRTQAV | ASSQPGPEVF | 0.8148129 |
| FVAGGYLS | ASSQPGPEVF | 0.759416 |
| GASGGAGGAGGW | ASSQPGPEVF | 0.7748333 |

| | | |
|---|---|---|
| GGAGAGGDGGAGGI | ASSQPGPEVF | 0.6916158 |
| GTGHNGSQPGMGGNG | ASSQPGPEVF | 0.7490308 |
| GYAAASASAS | ASSQPGPEVF | 0.7494815 |
| HQVDVAAVQQG | ASSQPGPEVF | 0.74105453 |
| IGAAIAHT | ASSQPGPEVF | 0.8209816 |
| IGMIEVAQL | ASSQPGPEVF | 0.8336424 |
| IRLAPEHTSAMD | ASSQPGPEVF | 0.8301132 |
| IYNVSVL | ASSQPGPEVF | 0.8343556 |
| KQGFLGM | ASSQPGPEVF | 0.838659 |
| LESVITQL | ASSQPGPEVF | 0.82948244 |
| LGLGALVVV | ASSQPGPEVF | 0.7524294 |
| LIGNGGNGGSGGI | ASSQPGPEVF | 0.7631408 |
| LNAINAPTL | ASSQPGPEVF | 0.6937305 |
| LQLLNTGV | ASSQPGPEVF | 0.7830516 |
| LQTSIEF | ASSQPGPEVF | 0.75338507 |
| MSFAVSEL | ASSQPGPEVF | 0.8154906 |
| NTGALGL | ASSQPGPEVF | 0.14746836 |
| PPGAGGAGGAGTT | ASSQPGPEVF | 0.782703 |
| QLVGLGI | ASSQPGPEVF | 0.8433764 |
| RFLEQVHQL | ASSQPGPEVF | 0.706475 |
| RGVGYVLR | ASSQPGPEVF | 0.81254673 |
| SATGVFAGVI | ASSQPGPEVF | 0.798375 |
| SLPAITLP | ASSQPGPEVF | 0.76479465 |
| STALAFTV | ASSQPGPEVF | 0.824468 |
| SVLTVGLGK | ASSQPGPEVF | 0.7034837 |
| TALAGAGSGI | ASSQPGPEVF | 0.8183212 |
| TGLPFTGI | ASSQPGPEVF | 0.8463471 |
| VAEHVTV | ASSQPGPEVF | 0.811036 |
| VDIVLNSL | ASSQPGPEVF | 0.78679484 |
| VLTGGGAVL | ASSQPGPEVF | 0.78930306 |
| VNVGLAYGI | ASSQPGPEVF | 0.44921243 |
| VSSAAVGAV | ASSQPGPEVF | 0.800133 |
| VTSVYEAGV | ASSQPGPEVF | 0.14704937 |
| WVALDRGA | ASSQPGPEVF | 0.7936946 |
| YLVGSVVI | ASSQPGPEVF | 0.6882414 |
| GPPGTGKTLL | ASSQPGPEVF | 0.8080678 |
| GVGMTKF | ASSQPGPEVF | 0.79833114 |
| IGIGGSDL | ASSQPGPEVF | 0.8227497 |
| IGTLLAEL | ASSQPGPEVF | 0.7223304 |
| IYTSGTT | ASSQPGPEVF | 0.7658977 |
| LSFPAGV | ASSQPGPEVF | 0.7112471 |
| LYTSGTT | ASSQPGPEVF | 0.600742 |
| VGAFPFGF | ASSQPGPEVF | 0.8288484 |
| VGAGTVEF | ASSQPGPEVF | 0.7965069 |
| VGKSTLLL | ASSQPGPEVF | 0.77729595 |
| AALAHAGGGIV | ASSQPGPEVF | 0.83444405 |
| AALVAVSLL | ASSQPGPEVF | 0.81035507 |
| AAVSRAVAAL | ASSQPGPEVF | 0.52334577 |
| AGGSSGIGGF | ASSQPGPEVF | 0.78429776 |
| AGLTVANANG | ASSQPGPEVF | 0.7948197 |
| AGLVGAVL | ASSQPGPEVF | 0.78292334 |

| | | |
|---|---|---|
| ALSQAFTV | ASSQPGPEVF | 0.1483989 |
| ALTFSVSL | ASSQPGPEVF | 0.8075291 |
| ALTGAALVAL | ASSQPGPEVF | 0.7168349 |
| ASFSIGAGVL | ASSQPGPEVF | 0.7288032 |
| DGSVYKI | ASSQPGPEVF | 0.7552748 |
| EQLAEIV | ASSQPGPEVF | 0.8274856 |
| FAVSELTVM | ASSQPGPEVF | 0.4821846 |
| GAAPLAAGL | ASSQPGPEVF | 0.74434614 |
| GADGGDGGSGGAGG | ASSQPGPEVF | 0.8264922 |
| GFGLDTGGAGG | ASSQPGPEVF | 0.81585145 |
| GLASGGLL | ASSQPGPEVF | 0.8367704 |
| GRIAGAAVLL | ASSQPGPEVF | 0.7933361 |
| GSAGSLNTSF | ASSQPGPEVF | 0.7931943 |
| GTGQIGI | ASSQPGPEVF | 0.8019824 |
| GTLIALIL | ASSQPGPEVF | 0.7847831 |
| IAATMGFVVG | ASSQPGPEVF | 0.5264964 |
| IFAQALL | ASSQPGPEVF | 0.4334107 |
| IGAAVALGF | ASSQPGPEVF | 0.8017698 |
| IGLTPLAL | ASSQPGPEVF | 0.40809897 |
| ILGSNTSTL | ASSQPGPEVF | 0.8055279 |
| IVFSLTVL | ASSQPGPEVF | 0.83410394 |
| LASKATGF | ASSQPGPEVF | 0.77196634 |
| LASVALIL | ASSQPGPEVF | 0.639246 |
| LFAVGLTL | ASSQPGPEVF | 0.7885425 |
| LFNGGIGTYI | ASSQPGPEVF | 0.72875476 |
| LTLGAGII | ASSQPGPEVF | 0.8274763 |
| LVAAALGDI | ASSQPGPEVF | 0.8301234 |
| NTAFVQLGI | ASSQPGPEVF | 0.8339247 |
| QLFSGAEL | ASSQPGPEVF | 0.8279404 |
| SAATFITL | ASSQPGPEVF | 0.7576448 |
| SNYGLANLGV | ASSQPGPEVF | 0.828584 |
| SYGGAYAVM | ASSQPGPEVF | 0.7717061 |
| TTGVGGLTLG | ASSQPGPEVF | 0.7891396 |
| TTQTIYI | ASSQPGPEVF | 0.8002243 |
| VAAGAGLGAVI | ASSQPGPEVF | 0.80524087 |
| VAAVVTLL | ASSQPGPEVF | 0.7620403 |
| VGFLGGLI | ASSQPGPEVF | 0.7656174 |
| VGVIPILL | ASSQPGPEVF | 0.588495 |
| VIFASSLI | ASSQPGPEVF | 0.8101099 |
| VKGGDGGAAAG | ASSQPGPEVF | 0.7984303 |
| VTKTFTNI | ASSQPGPEVF | 0.4957897 |
| VVAAAILL | ASSQPGPEVF | 0.7570827 |
| YAALDVEL | ASSQPGPEVF | 0.8433782 |
| LASVALLL | ASSQPGPEVF | 0.83605504 |
| LDAVHRV | ASSQPGPEVF | 0.7200134 |
| QGLSSSIF | ASSQPGPEVF | 0.70089126 |
| RGTFANIRL | ASSQPGPEVF | 0.79511315 |
| RQLWWGHRI | ASSQPGPEVF | 0.4578389 |
| YGPPGTGKTLL | ASSQPGPEVF | 0.81269443 |
| AALVGAPANVA | ASSQPGPEVF | 0.7874317 |
| AATFRRTE | ASSQPGPEVF | 0.78437185 |

| | | |
|---|---|---|
| AGALGGGI | ASSQPGPEVF | 0.7234222 |
| AGGAGGAGGIGAG | ASSQPGPEVF | 0.808984 |
| AGSGALGL | ASSQPGPEVF | 0.7489772 |
| AQIADQI | ASSQPGPEVF | 0.8156774 |
| ASAVAVTYL | ASSQPGPEVF | 0.17530124 |
| ASTVLGGTGG | ASSQPGPEVF | 0.8122413 |
| AVGGEPAK | ASSQPGPEVF | 0.71990365 |
| CEQGTFGGTC | ASSQPGPEVF | 0.73690236 |
| DEGKAPAG | ASSQPGPEVF | 0.4615176 |
| DTVVFPAL | ASSQPGPEVF | 0.7983192 |
| FEGITMAG | ASSQPGPEVF | 0.8338455 |
| FPLVVQD | ASSQPGPEVF | 0.83399904 |
| GAFTTPQ | ASSQPGPEVF | 0.69187856 |
| GFGGFGTAGGVGGA | ASSQPGPEVF | 0.8288874 |
| GFLAQFL | ASSQPGPEVF | 0.8427705 |
| GGAGGAGGNAGVA | ASSQPGPEVF | 0.78117263 |
| GGVGTAGAGGAGGAGG | ASSQPGPEVF | 0.37936112 |
| GNGGGGDAGNAGSGGN | ASSQPGPEVF | 0.81003964 |
| GTGGAGGAGNDAGS | ASSQPGPEVF | 0.75764567 |
| GTTNTGWFNS | ASSQPGPEVF | 0.82159024 |
| IFMGPLFTL | ASSQPGPEVF | 0.8057367 |
| ILAHAIEAI | ASSQPGPEVF | 0.6906777 |
| IRSAIREAL | ASSQPGPEVF | 0.8110218 |
| ITFAGGG | ASSQPGPEVF | 0.82948744 |
| ITGMSGHQEV | ASSQPGPEVF | 0.72406125 |
| IVAGLACTL | ASSQPGPEVF | 0.7630279 |
| IVGGGLAGA | ASSQPGPEVF | 0.7142117 |
| IVTGVLNI | ASSQPGPEVF | 0.4588918 |
| LAQQGCI | ASSQPGPEVF | 0.8132365 |
| LCLHVEN | ASSQPGPEVF | 0.7600062 |
| LDAISDGS | ASSQPGPEVF | 0.7152486 |
| LGGFAAML | ASSQPGPEVF | 0.8018895 |
| LGGKNPMI | ASSQPGPEVF | 0.7270463 |
| LGPKGVAAIL | ASSQPGPEVF | 0.7342344 |
| LMVGVSI | ASSQPGPEVF | 0.76358426 |
| LVGAAFVL | ASSQPGPEVF | 0.7977625 |
| LVLAVAPL | ASSQPGPEVF | 0.81699204 |
| MAAIAVIDAA | ASSQPGPEVF | 0.7901712 |
| NNGINAELQ | ASSQPGPEVF | 0.7854959 |
| RGEPLVLVHG | ASSQPGPEVF | 0.84310746 |
| RGGAGGAAGLI | ASSQPGPEVF | 0.5268627 |
| RSYRMHPAV | ASSQPGPEVF | 0.68705165 |
| SAAHLLAGQLA | ASSQPGPEVF | 0.8137791 |
| SAAVGSY | ASSQPGPEVF | 0.58935785 |
| SPHAVLHRI | ASSQPGPEVF | 0.81852496 |
| SQIAKVL | ASSQPGPEVF | 0.79420924 |
| TALGHLVP | ASSQPGPEVF | 0.8173962 |
| TAPTTSGAA | ASSQPGPEVF | 0.7852855 |
| TDGDDGNGGSPGT | ASSQPGPEVF | 0.84586954 |
| TDGLGEN | ASSQPGPEVF | 0.8314506 |
| TGGPPLPGQA | ASSQPGPEVF | 0.79251987 |

| | | |
|---|---|---|
| TIPAITI | ASSQPGPEVF | 0.7961192 |
| TIPGFTI | ASSQPGPEVF | 0.5864898 |
| TSIASGALA | ASSQPGPEVF | 0.828161 |
| TVTLPGAAI | ASSQPGPEVF | 0.81569123 |
| VASGLSAALP | ASSQPGPEVF | 0.71073365 |
| VGAGVIGI | ASSQPGPEVF | 0.78684413 |
| VGPKRFGV | ASSQPGPEVF | 0.8434612 |
| VNVAFPD | ASSQPGPEVF | 0.83724165 |
| VSMTNSGVS | ASSQPGPEVF | 0.43617332 |
| VTDVLGDLSV | ASSQPGPEVF | 0.77119863 |
| VVVVGAGFAGL | ASSQPGPEVF | 0.44504118 |
| VYTVGTVV | ASSQPGPEVF | 0.75408685 |
| WTTFSGP | ASSQPGPEVF | 0.5792796 |
| YDELVEAF | ASSQPGPEVF | 0.8244115 |
| YGTGGAGGNGG | ASSQPGPEVF | 0.7735928 |
| YTNVMAV | ASSQPGPEVF | 0.79795504 |
| FLEQVHQL | ASSQPGPEVF | 0.80678236 |
| GLSLLAVAV | ASSQPGPEVF | 0.8426187 |
| LEQVHQL | ASSQPGPEVF | 0.7611145 |
| LNAAGAI | ASSQPGPEVF | 0.7751802 |
| PAVHASL | ASSQPGPEVF | 0.505296 |
| TELEDDA | ASSQPGPEVF | 0.5646585 |
| TPPGPPN | ASSQPGPEVF | 0.7593981 |
| TPPPPPPP | ASSQPGPEVF | 0.76087546 |
| TQSPALI | ASSQDYRNTEVF | 0.23390546 |
| EFAAAFPQ | ASSQDYRNTEVF | 0.8361988 |
| LLSGNLDVL | ASSQDYRNTEVF | 0.8048401 |
| GPANLGNYNI | ASSQDYRNTEVF | 0.82939124 |
| TLGPTLP | ASSQDYRNTEVF | 0.7153079 |
| VYAYSSCCK | ASSQDYRNTEVF | 0.7823967 |
| APAPAPAPNI | ASSQDYRNTEVF | 0.7443441 |
| PGGPVPIP | ASSQDYRNTEVF | 0.8159055 |
| FLTVALEP | ASSQDYRNTEVF | 0.68956983 |
| SQYFGANL | ASSQDYRNTEVF | 0.5837176 |
| GGVKPAA | ASSQDYRNTEVF | 0.76361275 |
| GGTGGAGGAGGW | ASSQDYRNTEVF | 0.7015867 |
| GVGGVAGLGGAGG | ASSQDYRNTEVF | 0.83999956 |
| VLLVLATTI | ASSQDYRNTEVF | 0.83344537 |
| IGIFAAML | ASSQDYRNTEVF | 0.81237197 |
| AVGDLID | ASSQDYRNTEVF | 0.837376 |
| NISPLPL | ASSQDYRNTEVF | 0.24702626 |
| VTAVFVAI | ASSQDYRNTEVF | 0.72287834 |
| SGDDNDAAL | ASSQDYRNTEVF | 0.83657974 |
| AVAALAHS | ASSQDYRNTEVF | 0.81536734 |
| GGVTVMATP | ASSQDYRNTEVF | 0.78846014 |
| AGLAANVRD | ASSQDYRNTEVF | 0.83783895 |
| TNRPDLIDPALLRPG | ASSQDYRNTEVF | 0.7178698 |
| LANLGVTAL | ASSQDYRNTEVF | 0.7765807 |
| TGERIEL | ASSQDYRNTEVF | 0.7895849 |
| NGEELAA | ASSQDYRNTEVF | 0.8249716 |
| VLSLVWG | ASSQDYRNTEVF | 0.81708664 |

| | | |
|---|---|---|
| IGYALWL | ASSQDYRNTEVF | 0.7212083 |
| LSLLLAPL | ASSQDYRNTEVF | 0.8106657 |
| TLDEALK | ASSQDYRNTEVF | 0.85081875 |
| TDGAGTAEGG | ASSQDYRNTEVF | 0.7793286 |
| NAPGGAGGNGG | ASSQDYRNTEVF | 0.8301687 |
| VSVFMGMGGNF | ASSQDYRNTEVF | 0.81702 |
| LASYLAAL | ASSQDYRNTEVF | 0.785223 |
| LVGGTPAL | ASSQDYRNTEVF | 0.7319809 |
| VRNADVL | ASSQDYRNTEVF | 0.7539374 |
| TCSSPFLVSAM | ASSQDYRNTEVF | 0.7729248 |
| VHITDAE | ASSQDYRNTEVF | 0.66024673 |
| EYAAPSF | ASSQDYRNTEVF | 0.78606474 |
| VVDAFHGGL | ASSQDYRNTEVF | 0.7655386 |
| ASVVDPQ | ASSQDYRNTEVF | 0.85092735 |
| SYNEFGGLS | ASSQDYRNTEVF | 0.81967354 |
| DPDGSVY | ASSQDYRNTEVF | 0.81643736 |
| MVSTIHATE | ASSQDYRNTEVF | 0.7007756 |
| LAAGVADPV | ASSQDYRNTEVF | 0.8187915 |
| TKEQIAA | ASSQDYRNTEVF | 0.8129059 |
| SLASPAAL | ASSQDYRNTEVF | 0.63214815 |
| IGLVFYGV | ASSQDYRNTEVF | 0.8405763 |
| FLLDASNL | ASSQDYRNTEVF | 0.7297663 |
| YQVGNFP | ASSQDYRNTEVF | 0.767903 |
| ESLLSQST | ASSQDYRNTEVF | 0.81589293 |
| LFTVGFAAVG | ASSQDYRNTEVF | 0.82341146 |
| ATPPTPT | ASSQDYRNTEVF | 0.77890444 |
| TDLATLG | ASSQDYRNTEVF | 0.7716626 |
| SAGGTGR | ASSQDYRNTEVF | 0.7132634 |
| GFSNSGS | ASSQDYRNTEVF | 0.8303875 |
| GGNGGSGGTSV | ASSQDYRNTEVF | 0.8356066 |
| GTSVATGGAGN | ASSQDYRNTEVF | 0.7673603 |
| NGGNGGSGGTS | ASSQDYRNTEVF | 0.8397449 |
| AVVKFPH | ASSQDYRNTEVF | 0.795289 |
| TPPPPPP | ASSQDYRNTEVF | 0.47080874 |
| AAPAAAAAPA | ASSQDYRNTEVF | 0.78685427 |
| IALNEDL | ASSQDYRNTEVF | 0.83277076 |
| IVNVSSM | ASSQDYRNTEVF | 0.83536536 |
| TGIPPAPR | ASSQDYRNTEVF | 0.84229445 |
| FDLGGGTF | ASSQDYRNTEVF | 0.8215546 |
| FDLGGGTFDVS | ASSQDYRNTEVF | 0.7700341 |
| NEPTAAA | ASSQDYRNTEVF | 0.8399699 |
| NEPTAAALA | ASSQDYRNTEVF | 0.8458101 |
| GSIINISSI | ASSQDYRNTEVF | 0.8266417 |
| VAPGNAGTA | ASSQDYRNTEVF | 0.7558299 |
| TGGGASLE | ASSQDYRNTEVF | 0.6733376 |
| VAVYNHYKRI | ASSQDYRNTEVF | 0.81494164 |
| TDPAPATT | ASSQDYRNTEVF | 0.8266078 |
| TNRPDILDPALLRPG | ASSQDYRNTEVF | 0.6761446 |
| LTTTTTS | ASSQDYRNTEVF | 0.80573857 |
| PTGSGKTL | ASSQDYRNTEVF | 0.8092822 |
| TGSGKTL | ASSQDYRNTEVF | 0.5596254 |

| | | |
|---|---|---|
| TIGHVDHGK | ASSQDYRNTEVF | 0.8157296 |
| TIGHVDHGKTT | ASSQDYRNTEVF | 0.83624434 |
| GHVDHGKTT | ASSQDYRNTEVF | 0.797904 |
| AQFEHTLL | ASSQDYRNTEVF | 0.73346394 |
| IVNQDEL | ASSQDYRNTEVF | 0.8370002 |
| TNGSQFF | ASSQDYRNTEVF | 0.8503938 |
| TGKGIVE | ASSQDYRNTEVF | 0.8389393 |
| LISNASDA | ASSQDYRNTEVF | 0.49691963 |
| AELRTLGELTKIAKAHGAQ | ASSQDYRNTEVF | 0.39864045 |
| APNAVAAAEPSW | ASSQDYRNTEVF | 0.73041606 |
| APQVAGAATAVL | ASSQDYRNTEVF | 0.79989016 |
| ARRDPAAAAAILPTDA | ASSQDYRNTEVF | 0.74516726 |
| DAAARDVLAEAGLAEYF | ASSQDYRNTEVF | 0.7007446 |
| DDQIVLSAKANLAKVEQ | ASSQDYRNTEVF | 0.8112479 |
| DELLPDTGEDSDEDGH | ASSQDYRNTEVF | 0.8360217 |
| ELPEAPANRATVPQD | ASSQDYRNTEVF | 0.7127298 |
| FGQSKIERVTGIG | ASSQDYRNTEVF | 0.5596075 |
| GAGGTGSTAGGAGGAGGAGGLY | ASSQDYRNTEVF | 0.82428026 |
| GHGGTGAAGGNGGTGGAGGSGI | ASSQDYRNTEVF | 0.82291794 |
| GNGGNGGAGGEGGPGGAGGAGA | ASSQDYRNTEVF | 0.7309786 |
| GSTAGNGGNGGNATNGTIAGQP | ASSQDYRNTEVF | 0.7832404 |
| GTGGDGGNGGNGGTGG | ASSQDYRNTEVF | 0.73850054 |
| GVRRPAGIPADAPV | ASSQDYRNTEVF | 0.45344374 |
| GYRQVTDAHLLTLARR | ASSQDYRNTEVF | 0.8079031 |
| IAKMGQDGHDRG | ASSQDYRNTEVF | 0.66872907 |
| KAEPSIEIQYT | ASSQDYRNTEVF | 0.8407162 |
| KPVRIRMAADEDTVAVT | ASSQDYRNTEVF | 0.7247007 |
| MPPRIAGMRLLVIKPEP | ASSQDYRNTEVF | 0.8234649 |
| MKFYAVALFRSRGVPPTR | ASSQDYRNTEVF | 0.8137598 |
| NAGGFGILGAVAHSP | ASSQDYRNTEVF | 0.8185406 |
| NEGREITLAMAEM | ASSQDYRNTEVF | 0.779721 |
| NELVEWSHPLGHRWRWE | ASSQDYRNTEVF | 0.75030303 |
| NLGAGGTGGNGGNPG | ASSQDYRNTEVF | 0.65044403 |
| NTEWPKWETVMAADPVD | ASSQDYRNTEVF | 0.48092166 |
| PASAAPAETQP | ASSQDYRNTEVF | 0.83460724 |
| QQSSDGKNGVRAAEITGE | ASSQDYRNTEVF | 0.23890015 |
| RAPDAVTTSWWKE | ASSQDYRNTEVF | 0.83545756 |
| SGFGNVGAHTSGWFNQ | ASSQDYRNTEVF | 0.8317093 |
| SGVAGAVGQAASVGGLKV | ASSQDYRNTEVF | 0.72730064 |
| SNGGAGTGSAGGNGGT | ASSQDYRNTEVF | 0.8242241 |
| TGGAGGEGGAAG | ASSQDYRNTEVF | 0.5979619 |
| TGGDGGAAGAGGAPGHG | ASSQDYRNTEVF | 0.70115566 |
| TMPPLLAAIANAGRHGV | ASSQDYRNTEVF | 0.8225524 |
| TSSTAGNSGGA | ASSQDYRNTEVF | 0.8173437 |
| VPATVPVGILC | ASSQDYRNTEVF | 0.74253637 |
| VQRPDSGGVVLTGRIS | ASSQDYRNTEVF | 0.81491023 |
| WTPNPVPATKGVPAGLVQAT | ASSQDYRNTEVF | 0.81217307 |
| YALKRALAKFSDRFDVVII | ASSQDYRNTEVF | 0.7342359 |
| YLIGSVIV | ASSQDYRNTEVF | 0.7985539 |
| YGQLVAVID | ASSQDYRNTEVF | 0.8329625 |
| VTVGGMTLPA | ASSQDYRNTEVF | 0.8151541 |

| | | |
|---|---|---|
| VLALPAV | ASSQDYRNTEVF | 0.7277508 |
| VIPPTLNYET | ASSQDYRNTEVF | 0.8238312 |
| VGMGGMHPSSG | ASSQDYRNTEVF | 0.6995101 |
| VGLAGGAATA | ASSQDYRNTEVF | 0.81032735 |
| VGGAGGAGGSGP | ASSQDYRNTEVF | 0.80235994 |
| VFAAFMV | ASSQDYRNTEVF | 0.77219605 |
| TLYDKLR | ASSQDYRNTEVF | 0.82499933 |
| TLAVFTVL | ASSQDYRNTEVF | 0.57062817 |
| TGSVQPARV | ASSQDYRNTEVF | 0.74173534 |
| TGGGAGSGGT | ASSQDYRNTEVF | 0.7566632 |
| SHLETSL | ASSQDYRNTEVF | 0.80307233 |
| SFTTGMT | ASSQDYRNTEVF | 0.8290059 |
| QVEVAKRMGVRQPTVS | ASSQDYRNTEVF | 0.7140758 |
| QGSGGPALAM | ASSQDYRNTEVF | 0.7965753 |
| MLAPSNP | ASSQDYRNTEVF | 0.7636653 |
| LVNNAAI | ASSQDYRNTEVF | 0.69396925 |
| LTGEINTLA | ASSQDYRNTEVF | 0.7966653 |
| LSIFGPV | ASSQDYRNTEVF | 0.7897153 |
| LLALLANL | ASSQDYRNTEVF | 0.6648721 |
| LISNASDALDKLR | ASSQDYRNTEVF | 0.8182527 |
| LINEAVP | ASSQDYRNTEVF | 0.6725483 |
| LGYAFSGL | ASSQDYRNTEVF | 0.8061719 |
| LDFVAVAL | ASSQDYRNTEVF | 0.8382066 |
| LAQPGAE | ASSQDYRNTEVF | 0.7353313 |
| ISNLGAVTGF | ASSQDYRNTEVF | 0.82927 |
| HQEDPGRI | ASSQDYRNTEVF | 0.81219447 |
| GTPGEPTP | ASSQDYRNTEVF | 0.820042 |
| GFFNSGS | ASSQDYRNTEVF | 0.7750125 |
| GAGGGGLTAGAN | ASSQDYRNTEVF | 0.74430037 |
| FVDALVNM | ASSQDYRNTEVF | 0.83813214 |
| FLTVAPEL | ASSQDYRNTEVF | 0.8239367 |
| FETLTAL | ASSQDYRNTEVF | 0.8154557 |
| DVINPIEL | ASSQDYRNTEVF | 0.81265503 |
| APVSGAPV | ASSQDYRNTEVF | 0.8147205 |
| ALVGGLAW | ASSQDYRNTEVF | 0.82211703 |
| ALAAAAATAI | ASSQDYRNTEVF | 0.74453866 |
| AFGLGLLL | ASSQDYRNTEVF | 0.5617697 |
| AFFAAASL | ASSQDYRNTEVF | 0.7699951 |
| AAAALPAFSPP | ASSQDYRNTEVF | 0.77607495 |
| VVLAPAL | ASSQDYRNTEVF | 0.7751585 |
| TIGHVDHGKTTL | ASSQDYRNTEVF | 0.4164193 |
| LISNASDALDK | ASSQDYRNTEVF | 0.8319596 |
| LISNASDAL | ASSQDYRNTEVF | 0.8482949 |
| ISNASDALDK | ASSQDYRNTEVF | 0.70796704 |
| NQAETLVYQTE | ASSQDYRNTEVF | 0.8418845 |
| TGGTGGTAGAG | ASSQDYRNTEVF | 0.6493742 |
| TGSTAGGAGGA | ASSQDYRNTEVF | 0.82085896 |
| DPLAVPVAAGPV | ASSQDYRNTEVF | 0.83351725 |
| GGTGGTPGNGGN | ASSQDYRNTEVF | 0.695321 |
| PAGLAGVDPATL | ASSQDYRNTEVF | 0.80897915 |
| GNNLVVEAPHAGASV | ASSQDYRNTEVF | 0.7415925 |

| | | |
|---|---|---|
| KVTRIMSASTLEQPA | ASSQDYRNTEVF | 0.76312363 |
| LGFALPIALPRRTVV | ASSQDYRNTEVF | 0.83752173 |
| TRLGNLTARKLVVLV | ASSQDYRNTEVF | 0.7426151 |
| WSQQRRLIYFPSAGP | ASSQDYRNTEVF | 0.8082308 |
| GPNGNPDPMAAAVDIR | ASSQDYRNTEVF | 0.75621057 |
| TQRCVSVADLRAAASR | ASSQDYRNTEVF | 0.68111026 |
| QFVLDVAEHPVEVLTETV | ASSQDYRNTEVF | 0.8145304 |
| GGAGGTGGAGGNGGTQAGDG | ASSQDYRNTEVF | 0.83224237 |
| PLLATLRAIAVDVDAKPAQV | ASSQDYRNTEVF | 0.7241329 |
| APKFRTGGYMIDFWGVGYQVAK | ASSQDYRNTEVF | 0.84466124 |
| GAGGAGGAGGAAGGPGGTGGTG | ASSQDYRNTEVF | 0.7486869 |
| IGALGLLATDVAPAGSVAFGVA | ASSQDYRNTEVF | 0.8476178 |
| AGGAGGIGGNGGAGGAGGVLMG | ASSQDYRNTEVF | 0.8263118 |
| ADELIGLL | ASSQDYRNTEVF | 0.79216397 |
| ADRALKA | ASSQDYRNTEVF | 0.76739454 |
| AGAVGQVGGAGGA | ASSQDYRNTEVF | 0.83386827 |
| AGGGGGLGGAP | ASSQDYRNTEVF | 0.7561376 |
| AGGLLYGNGGAG | ASSQDYRNTEVF | 0.82112575 |
| AGLESIGL | ASSQDYRNTEVF | 0.83886623 |
| AGLKTGII | ASSQDYRNTEVF | 0.38622692 |
| ASLAALLL | ASSQDYRNTEVF | 0.6200958 |
| ASVINISSI | ASSQDYRNTEVF | 0.52740985 |
| ATPPGPA | ASSQDYRNTEVF | 0.58470994 |
| ATVASALV | ASSQDYRNTEVF | 0.4717297 |
| ATVASLIL | ASSQDYRNTEVF | 0.36256105 |
| AVNGPVLIRH | ASSQDYRNTEVF | 0.5010052 |
| DGAAGAVGI | ASSQDYRNTEVF | 0.441777 |
| DPLLHGI | ASSQDYRNTEVF | 0.6227472 |
| ELGAALGAL | ASSQDYRNTEVF | 0.59771997 |
| ELGRTQAV | ASSQDYRNTEVF | 0.46203342 |
| FVAGGYLS | ASSQDYRNTEVF | 0.6188998 |
| GASGGAGGAGGW | ASSQDYRNTEVF | 0.620007 |
| GGAGAGGDGGAGGI | ASSQDYRNTEVF | 0.39225882 |
| GTGHNGSQPGMGGNG | ASSQDYRNTEVF | 0.48501587 |
| GYAAASASAS | ASSQDYRNTEVF | 0.4188842 |
| HQVDVAAVQQG | ASSQDYRNTEVF | 0.52080476 |
| IGAAIAHT | ASSQDYRNTEVF | 0.33027172 |
| IGMIEVAQL | ASSQDYRNTEVF | 0.55162495 |
| IRLAPEHTSAMD | ASSQDYRNTEVF | 0.41898325 |
| IYNVSVL | ASSQDYRNTEVF | 0.31223613 |
| KQGFLGM | ASSQDYRNTEVF | 0.5723861 |
| LESVITQL | ASSQDYRNTEVF | 0.481997 |
| LGLGALVVV | ASSQDYRNTEVF | 0.49464136 |
| LIGNGGNGGSGGI | ASSQDYRNTEVF | 0.60871196 |
| LNAINAPTL | ASSQDYRNTEVF | 0.48407185 |
| LQLLNTGV | ASSQDYRNTEVF | 0.43710804 |
| LQTSIEF | ASSQDYRNTEVF | 0.5588432 |
| MSFAVSEL | ASSQDYRNTEVF | 0.63457525 |
| NTGALGL | ASSQDYRNTEVF | 0.42263493 |
| PPGAGGAGGAGTT | ASSQDYRNTEVF | 0.5358546 |
| QLVGLGI | ASSQDYRNTEVF | 0.6361412 |

| | | |
|---|---|---|
| RFLEQVHQL | ASSQDYRNTEVF | 0.5317265 |
| RGVGYVLR | ASSQDYRNTEVF | 0.63130915 |
| SATGVFAGVI | ASSQDYRNTEVF | 0.63437414 |
| SLPAITLP | ASSQDYRNTEVF | 0.4200006 |
| STALAFTV | ASSQDYRNTEVF | 0.6257365 |
| SVLTVGLGK | ASSQDYRNTEVF | 0.6256736 |
| TALAGAGSGI | ASSQDYRNTEVF | 0.5266738 |
| TGLPFTGI | ASSQDYRNTEVF | 0.53215516 |
| VAEHVTV | ASSQDYRNTEVF | 0.38931096 |
| VDIVLNSL | ASSQDYRNTEVF | 0.5183027 |
| VLTGGGAVL | ASSQDYRNTEVF | 0.3799744 |
| VNVGLAYGI | ASSQDYRNTEVF | 0.5482999 |
| VSSAAVGAV | ASSQDYRNTEVF | 0.60108536 |
| VTSVYEAGV | ASSQDYRNTEVF | 0.52750534 |
| WVALDRGA | ASSQDYRNTEVF | 0.3658191 |
| YLVGSVVI | ASSQDYRNTEVF | 0.34005514 |
| GPPGTGKTLL | ASSQDYRNTEVF | 0.50280124 |
| GVGMTKF | ASSQDYRNTEVF | 0.394605 |
| IGIGGSDL | ASSQDYRNTEVF | 0.3613043 |
| IGTLLAEL | ASSQDYRNTEVF | 0.45842052 |
| IYTSGTT | ASSQDYRNTEVF | 0.61463773 |
| LSFPAGV | ASSQDYRNTEVF | 0.62580943 |
| LYTSGTT | ASSQDYRNTEVF | 0.44818592 |
| VGAFPFGF | ASSQDYRNTEVF | 0.64116806 |
| VGAGTVEF | ASSQDYRNTEVF | 0.57653344 |
| VGKSTLLL | ASSQDYRNTEVF | 0.5928227 |
| AALAHAGGGIV | ASSQDYRNTEVF | 0.50111985 |
| AALVAVSLL | ASSQDYRNTEVF | 0.39545506 |
| AAVSRAVAAL | ASSQDYRNTEVF | 0.6278069 |
| AGGSSGIGGF | ASSQDYRNTEVF | 0.5184137 |
| AGLTVANANG | ASSQDYRNTEVF | 0.56856817 |
| AGLVGAVL | ASSQDYRNTEVF | 0.60814697 |
| ALSQAFTV | ASSQDYRNTEVF | 0.50770754 |
| ALTFSVSL | ASSQDYRNTEVF | 0.45493358 |
| ALTGAALVAL | ASSQDYRNTEVF | 0.55465204 |
| ASFSIGAGVL | ASSQDYRNTEVF | 0.5553318 |
| DGSVYKI | ASSQDYRNTEVF | 0.37678686 |
| EQLAEIV | ASSQDYRNTEVF | 0.40010244 |
| FAVSELTVM | ASSQDYRNTEVF | 0.47762322 |
| GAAPLAAGL | ASSQDYRNTEVF | 0.5270338 |
| GADGGDGGSGGAGG | ASSQDYRNTEVF | 0.6391455 |
| GFGLDTGGAGG | ASSQDYRNTEVF | 0.5449437 |
| GLASGGLL | ASSQDYRNTEVF | 0.44484213 |
| GRIAGAAVLL | ASSQDYRNTEVF | 0.59230983 |
| GSAGSLNTSF | ASSQDYRNTEVF | 0.5791124 |
| GTGQIGI | ASSQDYRNTEVF | 0.53903145 |
| GTLIALIL | ASSQDYRNTEVF | 0.5043201 |
| IAATMGFVVG | ASSQDYRNTEVF | 0.48764294 |
| IFAQALL | ASSQDYRNTEVF | 0.5514197 |
| IGAAVALGF | ASSQDYRNTEVF | 0.5994801 |
| IGLTPLAL | ASSQDYRNTEVF | 0.5808317 |

| | | |
|---|---|---|
| ILGSNTSTL | ASSQDYRNTEVF | 0.61335135 |
| IVFSLTVL | ASSQDYRNTEVF | 0.542103 |
| LASKATGF | ASSQDYRNTEVF | 0.50414276 |
| LASVALIL | ASSQDYRNTEVF | 0.36852205 |
| LFAVGLTL | ASSQDYRNTEVF | 0.49166816 |
| LFNGGIGTYI | ASSQDYRNTEVF | 0.50134224 |
| LTLGAGII | ASSQDYRNTEVF | 0.54066455 |
| LVAAALGDI | ASSQDYRNTEVF | 0.5882439 |
| NTAFVQLGI | ASSQDYRNTEVF | 0.5723437 |
| QLFSGAEL | ASSQDYRNTEVF | 0.4397832 |
| SAATFITL | ASSQDYRNTEVF | 0.56582135 |
| SNYGLANLGV | ASSQDYRNTEVF | 0.6266856 |
| SYGGAYAVM | ASSQDYRNTEVF | 0.39660224 |
| TTGVGGLTLG | ASSQDYRNTEVF | 0.52639496 |
| TTQTIYI | ASSQDYRNTEVF | 0.64724827 |
| VAAGAGLGAVI | ASSQDYRNTEVF | 0.60415196 |
| VAAVVTLL | ASSQDYRNTEVF | 0.4912759 |
| VGFLGGLI | ASSQDYRNTEVF | 0.5264473 |
| VGVIPILL | ASSQDYRNTEVF | 0.6360607 |
| VIFASSLI | ASSQDYRNTEVF | 0.514228 |
| VKGGDGGAAAG | ASSQDYRNTEVF | 0.61181927 |
| VTKTFTNI | ASSQDYRNTEVF | 0.35998517 |
| VVAAAILL | ASSQDYRNTEVF | 0.48832536 |
| YAALDVEL | ASSQDYRNTEVF | 0.39310595 |
| LASVALLL | ASSQDYRNTEVF | 0.3212332 |
| LDAVHRV | ASSQDYRNTEVF | 0.6158932 |
| QGLSSSIF | ASSQDYRNTEVF | 0.47704464 |
| RGTFANIRL | ASSQDYRNTEVF | 0.6009653 |
| RQLWWGHRI | ASSQDYRNTEVF | 0.4607369 |
| YGPPGTGKTLL | ASSQDYRNTEVF | 0.62784207 |
| AALVGAPANVA | ASSQDYRNTEVF | 0.5364349 |
| AATFRRTE | ASSQDYRNTEVF | 0.50316423 |
| AGALGGGI | ASSQDYRNTEVF | 0.45652002 |
| AGGAGGAGGIGAG | ASSQDYRNTEVF | 0.5637235 |
| AGSGALGL | ASSQDYRNTEVF | 0.59328085 |
| AQIADQI | ASSQDYRNTEVF | 0.56877655 |
| ASAVAVTYL | ASSQDYRNTEVF | 0.35230386 |
| ASTVLGGTGG | ASSQDYRNTEVF | 0.54951096 |
| AVGGEPAK | ASSQDYRNTEVF | 0.5140125 |
| CEQGTFGGTC | ASSQDYRNTEVF | 0.37527353 |
| DEGKAPAG | ASSQDYRNTEVF | 0.33586255 |
| DTVVFPAL | ASSQDYRNTEVF | 0.38823107 |
| FEGITMAG | ASSQDYRNTEVF | 0.49355507 |
| FPLVVQD | ASSQDYRNTEVF | 0.5865632 |
| GAFTTPQ | ASSQDYRNTEVF | 0.6042211 |
| GFGGFGTAGGVGGA | ASSQDYRNTEVF | 0.46607697 |
| GFLAQFL | ASSQDYRNTEVF | 0.56497073 |
| GGAGGAGGNAGVA | ASSQDYRNTEVF | 0.53926307 |
| GGVGTAGAGGAGGAGG | ASSQDYRNTEVF | 0.29539803 |
| GNGGGGDAGNAGSGGN | ASSQDYRNTEVF | 0.43450382 |
| GTGGAGGAGNDAGS | ASSQDYRNTEVF | 0.399054 |

| | | |
|---|---|---|
| GTTNTGWFNS | ASSQDYRNTEVF | 0.60612696 |
| IFMGPLFTL | ASSQDYRNTEVF | 0.4190343 |
| ILAHAIEAI | ASSQDYRNTEVF | 0.49172282 |
| IRSAIREAL | ASSQDYRNTEVF | 0.5151702 |
| ITFAGGG | ASSQDYRNTEVF | 0.5860747 |
| ITGMSGHQEV | ASSQDYRNTEVF | 0.5519225 |
| IVAGLACTL | ASSQDYRNTEVF | 0.37826347 |
| IVGGGLAGA | ASSQDYRNTEVF | 0.37287894 |
| IVTGVLNI | ASSQDYRNTEVF | 0.5811342 |
| LAQQGCI | ASSQDYRNTEVF | 0.48842454 |
| LCLHVEN | ASSQDYRNTEVF | 0.4248214 |
| LDAISDGS | ASSQDYRNTEVF | 0.42490605 |
| LGGFAAML | ASSQDYRNTEVF | 0.5985205 |
| LGGKNPMI | ASSQDYRNTEVF | 0.35759154 |
| LGPKGVAAIL | ASSQDYRNTEVF | 0.6009556 |
| LMVGVSI | ASSQDYRNTEVF | 0.4293488 |
| LVGAAFVL | ASSQDYRNTEVF | 0.5071799 |
| LVLAVAPL | ASSQDYRNTEVF | 0.6318625 |
| MAAIAVIDAA | ASSQDYRNTEVF | 0.5944841 |
| NNGINAELQ | ASSQDYRNTEVF | 0.51706666 |
| RGEPLVLVHG | ASSQDYRNTEVF | 0.46047616 |
| RGGAGGAAGLI | ASSQDYRNTEVF | 0.42443186 |
| RSYRMHPAV | ASSQDYRNTEVF | 0.42215055 |
| SAAHLLAGQLA | ASSQDYRNTEVF | 0.42458776 |
| SAAVGSY | ASSQDYRNTEVF | 0.6263177 |
| SPHAVLHRI | ASSQDYRNTEVF | 0.5214007 |
| SQIAKVL | ASSQDYRNTEVF | 0.3973159 |
| TALGHLVP | ASSQDYRNTEVF | 0.5771 |
| TAPTTSGAA | ASSQDYRNTEVF | 0.6103439 |
| TDGDDGNGGSPGT | ASSQDYRNTEVF | 0.62330955 |
| TDGLGEN | ASSQDYRNTEVF | 0.43042928 |
| TGGPPLPGQA | ASSQDYRNTEVF | 0.44437763 |
| TIPAITI | ASSQDYRNTEVF | 0.5254011 |
| TIPGFTI | ASSQDYRNTEVF | 0.52227664 |
| TSIASGALA | ASSQDYRNTEVF | 0.4292493 |
| TVTLPGAAI | ASSQDYRNTEVF | 0.4441707 |
| VASGLSAALP | ASSQDYRNTEVF | 0.54064727 |
| VGAGVIGI | ASSQDYRNTEVF | 0.6512086 |
| VGPKRFGV | ASSQDYRNTEVF | 0.492816 |
| VNVAFPD | ASSQDYRNTEVF | 0.33881897 |
| VSMTNSGVS | ASSQDYRNTEVF | 0.46929982 |
| VTDVLGDLSV | ASSQDYRNTEVF | 0.38949987 |
| VVVVGAGFAGL | ASSQDYRNTEVF | 0.43009442 |
| VYTVGTVV | ASSQDYRNTEVF | 0.51795596 |
| WTTFSGP | ASSQDYRNTEVF | 0.37236187 |
| YDELVEAF | ASSQDYRNTEVF | 0.48516244 |
| YGTGGAGGNGG | ASSQDYRNTEVF | 0.63512444 |
| YTNVMAV | ASSQDYRNTEVF | 0.5241537 |
| FLEQVHQL | ASSQDYRNTEVF | 0.586945 |
| GLSLLAVAV | ASSQDYRNTEVF | 0.62318116 |
| LEQVHQL | ASSQDYRNTEVF | 0.5688932 |

| | | |
|---|---|---|
| LNAAGAI | ASSQDYRNTEVF | 0.63408434 |
| PAVHASL | ASSQDYRNTEVF | 0.6123506 |
| TELEDDA | ASSQDYRNTEVF | 0.54522294 |
| TPPGPPN | ASSQDYRNTEVF | 0.48271233 |
| TPPPPPPP | ASSQDYRNTEVF | 0.60895026 |
| TQSPALI | ASGDPGGADTQY | 0.48218828 |
| EFAAAFPQ | ASGDPGGADTQY | 0.5349379 |
| LLSGNLDVL | ASGDPGGADTQY | 0.4841558 |
| GPANLGNYNI | ASGDPGGADTQY | 0.6101564 |
| TLGPTLP | ASGDPGGADTQY | 0.47825894 |
| VYAYSSCCK | ASGDPGGADTQY | 0.6271986 |
| APAPAPAPNI | ASGDPGGADTQY | 0.3401435 |
| PGGPVPIP | ASGDPGGADTQY | 0.43021092 |
| FLTVALEP | ASGDPGGADTQY | 0.6221728 |
| SQYFGANL | ASGDPGGADTQY | 0.31084976 |
| GGVKPAA | ASGDPGGADTQY | 0.3617213 |
| GGTGGAGGAGGW | ASGDPGGADTQY | 0.47109833 |
| GVGGVAGLGGAGG | ASGDPGGADTQY | 0.6116141 |
| VLLVLATTI | ASGDPGGADTQY | 0.3086344 |
| IGIFAAML | ASGDPGGADTQY | 0.5106977 |
| AVGDLID | ASGDPGGADTQY | 0.37599754 |
| NISPLPL | ASGDPGGADTQY | 0.5961858 |
| VTAVFVAI | ASGDPGGADTQY | 0.5303251 |
| SGDDNDAAL | ASGDPGGADTQY | 0.36956173 |
| AVAALAHS | ASGDPGGADTQY | 0.60521185 |
| GGVTVMATP | ASGDPGGADTQY | 0.3623914 |
| AGLAANVRD | ASGDPGGADTQY | 0.5393849 |
| TNRPDLIDPALLRPG | ASGDPGGADTQY | 0.61912066 |
| LANLGVTAL | ASGDPGGADTQY | 0.6461279 |
| TGERIEL | ASGDPGGADTQY | 0.39080068 |
| NGEELAA | ASGDPGGADTQY | 0.6023849 |
| VLSLVWG | ASGDPGGADTQY | 0.4117617 |
| IGYALWL | ASGDPGGADTQY | 0.6437857 |
| LSLLLAPL | ASGDPGGADTQY | 0.4960945 |
| TLDEALK | ASGDPGGADTQY | 0.35728434 |
| TDGAGTAEGG | ASGDPGGADTQY | 0.60378987 |
| NAPGGAGGNGG | ASGDPGGADTQY | 0.57364744 |
| VSVFMGMGGNF | ASGDPGGADTQY | 0.50399065 |
| LASYLAAL | ASGDPGGADTQY | 0.59185296 |
| LVGGTPAL | ASGDPGGADTQY | 0.5076965 |
| VRNADVL | ASGDPGGADTQY | 0.4628089 |
| TCSSPFLVSAM | ASGDPGGADTQY | 0.6300338 |
| VHITDAE | ASGDPGGADTQY | 0.55176294 |
| EYAAPSF | ASGDPGGADTQY | 0.46321398 |
| VVDAFHGGL | ASGDPGGADTQY | 0.6209183 |
| ASVVDPQ | ASGDPGGADTQY | 0.53766066 |
| SYNEFGGLS | ASGDPGGADTQY | 0.4042022 |
| DPDGSVY | ASGDPGGADTQY | 0.6384418 |
| MVSTIHATE | ASGDPGGADTQY | 0.5043649 |
| LAAGVADPV | ASGDPGGADTQY | 0.58159685 |
| TKEQIAA | ASGDPGGADTQY | 0.54113764 |

| | | |
|---|---|---|
| SLASPAAL | ASGDPGGADTQY | 0.6390747 |
| IGLVFYGV | ASGDPGGADTQY | 0.41589674 |
| FLLDASNL | ASGDPGGADTQY | 0.527516 |
| YQVGNFP | ASGDPGGADTQY | 0.5370779 |
| ESLLSQST | ASGDPGGADTQY | 0.6018552 |
| LFTVGFAAVG | ASGDPGGADTQY | 0.49292508 |
| ATPPTPT | ASGDPGGADTQY | 0.5317474 |
| TDLATLG | ASGDPGGADTQY | 0.5585035 |
| SAGGTGR | ASGDPGGADTQY | 0.61645055 |
| GFSNSGS | ASGDPGGADTQY | 0.45282596 |
| GGNGGSGGTSV | ASGDPGGADTQY | 0.48908412 |
| GTSVATGGAGN | ASGDPGGADTQY | 0.64907753 |
| NGGNGGSGGTS | ASGDPGGADTQY | 0.40707934 |
| AVVKFPH | ASGDPGGADTQY | 0.5502279 |
| TPPPPPP | ASGDPGGADTQY | 0.55205387 |
| AAPAAAAAPA | ASGDPGGADTQY | 0.5970328 |
| IALNEDL | ASGDPGGADTQY | 0.5221365 |
| IVNVSSM | ASGDPGGADTQY | 0.5990012 |
| TGIPPAPR | ASGDPGGADTQY | 0.50596285 |
| FDLGGGTF | ASGDPGGADTQY | 0.59718627 |
| FDLGGGTFDVS | ASGDPGGADTQY | 0.47275737 |
| NEPTAAA | ASGDPGGADTQY | 0.526289 |
| NEPTAAALA | ASGDPGGADTQY | 0.41092557 |
| GSIINISSI | ASGDPGGADTQY | 0.5216801 |
| VAPGNAGTA | ASGDPGGADTQY | 0.5846286 |
| TGGGASLE | ASGDPGGADTQY | 0.59251237 |
| VAVYNHYKRI | ASGDPGGADTQY | 0.36166456 |
| TDPAPATT | ASGDPGGADTQY | 0.597856 |
| TNRPDILDPALLRPG | ASGDPGGADTQY | 0.59835607 |
| LTTTTTS | ASGDPGGADTQY | 0.6541257 |
| PTGSGKTL | ASGDPGGADTQY | 0.48445606 |
| TGSGKTL | ASGDPGGADTQY | 0.620487 |
| TIGHVDHGK | ASGDPGGADTQY | 0.49604708 |
| TIGHVDHGKTT | ASGDPGGADTQY | 0.52895164 |
| GHVDHGKTT | ASGDPGGADTQY | 0.356824 |
| AQFEHTLL | ASGDPGGADTQY | 0.41382805 |
| IVNQDEL | ASGDPGGADTQY | 0.52959263 |
| TNGSQFF | ASGDPGGADTQY | 0.46154484 |
| TGKGIVE | ASGDPGGADTQY | 0.6191899 |
| LISNASDA | ASGDPGGADTQY | 0.59909827 |
| AELRTLGELTKIAKAHGAQ | ASGDPGGADTQY | 0.61973023 |
| APNAVAAAEPSW | ASGDPGGADTQY | 0.61280584 |
| APQVAGAATAVL | ASGDPGGADTQY | 0.6182943 |
| ARRDPAAAAAILPTDA | ASGDPGGADTQY | 0.5999274 |
| DAAARDVLAEAGLAEYF | ASGDPGGADTQY | 0.48117808 |
| DDQIVLSAKANLAKVEQ | ASGDPGGADTQY | 0.52077353 |
| DELLPDTGEDSDEDGH | ASGDPGGADTQY | 0.3767655 |
| ELPEAPANRATVPQD | ASGDPGGADTQY | 0.44516882 |
| FGQSKIERVTGIG | ASGDPGGADTQY | 0.46135202 |
| GAGGTGSTAGGAGGAGGAGGLY | ASGDPGGADTQY | 0.55016136 |
| GHGGTGAAGGNGGTGGAGGSGI | ASGDPGGADTQY | 0.31189948 |

| | | |
|---|---|---|
| GNGGNGGAGGEGGPGGAGGAGA | ASGDPGGADTQY | 0.5723175 |
| GSTAGNGGNGGNATNGTIAGQP | ASGDPGGADTQY | 0.6346355 |
| GTGGDGGNGGNGGTGG | ASGDPGGADTQY | 0.47233552 |
| GVRRPAGIPADAPV | ASGDPGGADTQY | 0.5824933 |
| GYRQVTDAHLLTLARR | ASGDPGGADTQY | 0.58147746 |
| IAKMGQDGHDRG | ASGDPGGADTQY | 0.4039997 |
| KAEPSIEIQYT | ASGDPGGADTQY | 0.57609516 |
| KPVRIRMAADEDTVAVT | ASGDPGGADTQY | 0.4331957 |
| MPPRIAGMRLLVIKPEP | ASGDPGGADTQY | 0.6178824 |
| MKFYAVALFRSRGVPPTR | ASGDPGGADTQY | 0.62851024 |
| NAGGFGILGAVAHSP | ASGDPGGADTQY | 0.54423964 |
| NEGREITLAMAEM | ASGDPGGADTQY | 0.42824373 |
| NELVEWSHPLGHRWRWE | ASGDPGGADTQY | 0.5675862 |
| NLGAGGTGGNGGNPG | ASGDPGGADTQY | 0.39811367 |
| NTEWPKWETVMAADPVD | ASGDPGGADTQY | 0.4544575 |
| PASAAPAETQP | ASGDPGGADTQY | 0.449125 |
| QQSSDGKNGVRAAEITGE | ASGDPGGADTQY | 0.54311293 |
| RAPDAVTTSWWKE | ASGDPGGADTQY | 0.38898176 |
| SGFGNVGAHTSGWFNQ | ASGDPGGADTQY | 0.5886314 |
| SGVAGAVGQAASVGGLKV | ASGDPGGADTQY | 0.54327565 |
| SNGGAGTGSAGGNGGT | ASGDPGGADTQY | 0.48399976 |
| TGGAGGEGGAAG | ASGDPGGADTQY | 0.3922939 |
| TGGDGGAAGAGGAPGHG | ASGDPGGADTQY | 0.580027 |
| TMPPLLAAIANAGRHGV | ASGDPGGADTQY | 0.5245038 |
| TSSTAGNSGGA | ASGDPGGADTQY | 0.36524054 |
| VPATVPVGILC | ASGDPGGADTQY | 0.57177114 |
| VQRPDSGGVVLTGRIS | ASGDPGGADTQY | 0.4408857 |
| WTPNPVPATKGVPAGLVQAT | ASGDPGGADTQY | 0.5309687 |
| YALKRALAKFSDRFDVVII | ASGDPGGADTQY | 0.6003216 |
| YLIGSVIV | ASGDPGGADTQY | 0.5402797 |
| YGQLVAVID | ASGDPGGADTQY | 0.35950124 |
| VTVGGMTLPA | ASGDPGGADTQY | 0.46465775 |
| VLALPAV | ASGDPGGADTQY | 0.49924287 |
| VIPPTLNYET | ASGDPGGADTQY | 0.49310198 |
| VGMGGMHPSSG | ASGDPGGADTQY | 0.32412812 |
| VGLAGGAATA | ASGDPGGADTQY | 0.5722511 |
| VGGAGGAGGSGP | ASGDPGGADTQY | 0.4064584 |
| VFAAFMV | ASGDPGGADTQY | 0.43847787 |
| TLYDKLR | ASGDPGGADTQY | 0.49787045 |
| TLAVFTVL | ASGDPGGADTQY | 0.6240692 |
| TGSVQPARV | ASGDPGGADTQY | 0.51922417 |
| TGGGAGSGGT | ASGDPGGADTQY | 0.49826467 |
| SHLETSL | ASGDPGGADTQY | 0.5676275 |
| SFTTGMT | ASGDPGGADTQY | 0.546896 |
| QVEVAKRMGVRQPTVS | ASGDPGGADTQY | 0.6337029 |
| QGSGGPALAM | ASGDPGGADTQY | 0.55281913 |
| MLAPSNP | ASGDPGGADTQY | 0.46054024 |
| LVNNAAI | ASGDPGGADTQY | 0.60056037 |
| LTGEINTLA | ASGDPGGADTQY | 0.47617364 |
| LSIFGPV | ASGDPGGADTQY | 0.40417075 |
| LLALLANL | ASGDPGGADTQY | 0.38523123 |

| | | |
|---|---|---|
| LISNASDALDKLR | ASGDPGGADTQY | 0.38495982 |
| LINEAVP | ASGDPGGADTQY | 0.36910638 |
| LGYAFSGL | ASGDPGGADTQY | 0.57782954 |
| LDFVAVAL | ASGDPGGADTQY | 0.62084866 |
| LAQPGAE | ASGDPGGADTQY | 0.47406507 |
| ISNLGAVTGF | ASGDPGGADTQY | 0.3598211 |
| HQEDPGRI | ASGDPGGADTQY | 0.43957886 |
| GTPGEPTP | ASGDPGGADTQY | 0.5005204 |
| GFFNSGS | ASGDPGGADTQY | 0.6065381 |
| GAGGGGLTAGAN | ASGDPGGADTQY | 0.53687257 |
| FVDALVNM | ASGDPGGADTQY | 0.6430244 |
| FLTVAPEL | ASGDPGGADTQY | 0.60913175 |
| FETLTAL | ASGDPGGADTQY | 0.4770173 |
| DVINPIEL | ASGDPGGADTQY | 0.60682106 |
| APVSGAPV | ASGDPGGADTQY | 0.4602336 |
| ALVGGLAW | ASGDPGGADTQY | 0.5056894 |
| ALAAAAATAI | ASGDPGGADTQY | 0.5177458 |
| AFGLGLLL | ASGDPGGADTQY | 0.48761976 |
| AFFAAASL | ASGDPGGADTQY | 0.45648113 |
| AAAALPAFSPP | ASGDPGGADTQY | 0.48004997 |
| VVLAPAL | ASGDPGGADTQY | 0.36058328 |
| TIGHVDHGKTTL | ASGDPGGADTQY | 0.5353887 |
| LISNASDALDK | ASGDPGGADTQY | 0.38083908 |
| LISNASDAL | ASGDPGGADTQY | 0.37392023 |
| ISNASDALDK | ASGDPGGADTQY | 0.4908323 |
| NQAETLVYQTE | ASGDPGGADTQY | 0.64750373 |
| TGGTGGTAGAG | ASGDPGGADTQY | 0.63735116 |
| TGSTAGGAGGA | ASGDPGGADTQY | 0.4438067 |
| DPLAVPVAAGPV | ASGDPGGADTQY | 0.41829973 |
| GGTGGTPGNGGN | ASGDPGGADTQY | 0.5915824 |
| PAGLAGVDPATL | ASGDPGGADTQY | 0.46355495 |
| GNNLVVEAPHAGASV | ASGDPGGADTQY | 0.5682451 |
| KVTRIMSASTLEQPA | ASGDPGGADTQY | 0.5553466 |
| LGFALPIALPRRTVV | ASGDPGGADTQY | 0.51679486 |
| TRLGNLTARKLVVLV | ASGDPGGADTQY | 0.45165005 |
| WSQQRRLIYFPSAGP | ASGDPGGADTQY | 0.5605161 |
| GPNGNPDPMAAAVDIR | ASGDPGGADTQY | 0.46727157 |
| TQRCVSVADLRAAASR | ASGDPGGADTQY | 0.5615664 |
| QFVLDVAEHPVEVLTETV | ASGDPGGADTQY | 0.3544909 |
| GGAGGTGGAGGNGGTQAGDG | ASGDPGGADTQY | 0.6230825 |
| PLLATLRAIAVDVDAKPAQV | ASGDPGGADTQY | 0.41498068 |
| APKFRTGGYMIDFWGVGYQVAK | ASGDPGGADTQY | 0.4104257 |
| GAGGAGGAGGAAGGPGGTGGTG | ASGDPGGADTQY | 0.32461038 |
| IGALGLLATDVAPAGSVAFGVA | ASGDPGGADTQY | 0.4999841 |
| AGGAGGIGGNGGAGGAGGVLMG | ASGDPGGADTQY | 0.5571718 |
| ADELIGLL | ASGDPGGADTQY | 0.61889005 |
| ADRALKA | ASGDPGGADTQY | 0.5146538 |
| AGAVGQVGGAGGA | ASGDPGGADTQY | 0.56463933 |
| AGGGGGLGGAP | ASGDPGGADTQY | 0.59503675 |
| AGGLLYGNGGAG | ASGDPGGADTQY | 0.48408097 |
| AGLESIGL | ASGDPGGADTQY | 0.35216686 |

| | | |
|---|---|---|
| AGLKTGII | ASGDPGGADTQY | 0.59216744 |
| ASLAALLL | ASGDPGGADTQY | 0.5235176 |
| ASVINISSI | ASGDPGGADTQY | 0.6273031 |
| ATPPGPA | ASGDPGGADTQY | 0.57669824 |
| ATVASALV | ASGDPGGADTQY | 0.38354605 |
| ATVASLIL | ASGDPGGADTQY | 0.5906567 |
| AVNGPVLIRH | ASGDPGGADTQY | 0.6152693 |
| DGAAGAVGI | ASGDPGGADTQY | 0.36445668 |
| DPLLHGI | ASGDPGGADTQY | 0.41622314 |
| ELGAALGAL | ASGDPGGADTQY | 0.44945446 |
| ELGRTQAV | ASGDPGGADTQY | 0.39670038 |
| FVAGGYLS | ASGDPGGADTQY | 0.5111709 |
| GASGGAGGAGGW | ASGDPGGADTQY | 0.44833618 |
| GGAGAGGDGGAGGI | ASGDPGGADTQY | 0.5326954 |
| GTGHNGSQPGMGGNG | ASGDPGGADTQY | 0.46487212 |
| GYAAASASAS | ASGDPGGADTQY | 0.4250584 |
| HQVDVAAVQQG | ASGDPGGADTQY | 0.5119627 |
| IGAAIAHT | ASGDPGGADTQY | 0.5173353 |
| IGMIEVAQL | ASGDPGGADTQY | 0.54731053 |
| IRLAPEHTSAMD | ASGDPGGADTQY | 0.5647868 |
| IYNVSVL | ASGDPGGADTQY | 0.49860793 |
| KQGFLGM | ASGDPGGADTQY | 0.5497899 |
| LESVITQL | ASGDPGGADTQY | 0.63807523 |
| LGLGALVVV | ASGDPGGADTQY | 0.37927288 |
| LIGNGGNGGSGGI | ASGDPGGADTQY | 0.43768016 |
| LNAINAPTL | ASGDPGGADTQY | 0.5878876 |
| LQLLNTGV | ASGDPGGADTQY | 0.43510926 |
| LQTSIEF | ASGDPGGADTQY | 0.61729825 |
| MSFAVSEL | ASGDPGGADTQY | 0.541342 |
| NTGALGL | ASGDPGGADTQY | 0.5593221 |
| PPGAGGAGGAGTT | ASGDPGGADTQY | 0.5153822 |
| QLVGLGI | ASGDPGGADTQY | 0.6421709 |
| RFLEQVHQL | ASGDPGGADTQY | 0.59311867 |
| RGVGYVLR | ASGDPGGADTQY | 0.46690673 |
| SATGVFAGVI | ASGDPGGADTQY | 0.5837139 |
| SLPAITLP | ASGDPGGADTQY | 0.39000732 |
| STALAFTV | ASGDPGGADTQY | 0.5652706 |
| SVLTVGLGK | ASGDPGGADTQY | 0.59764206 |
| TALAGAGSGI | ASGDPGGADTQY | 0.424722 |
| TGLPFTGI | ASGDPGGADTQY | 0.4619601 |
| VAEHVTV | ASGDPGGADTQY | 0.5956522 |
| VDIVLNSL | ASGDPGGADTQY | 0.62497073 |
| VLTGGGAVL | ASGDPGGADTQY | 0.38465765 |
| VNVGLAYGI | ASGDPGGADTQY | 0.48402107 |
| VSSAAVGAV | ASGDPGGADTQY | 0.4165812 |
| VTSVYEAGV | ASGDPGGADTQY | 0.40100005 |
| WVALDRGA | ASGDPGGADTQY | 0.37699068 |
| YLVGSVVI | ASGDPGGADTQY | 0.5739373 |
| GPPGTGKTLL | ASGDPGGADTQY | 0.47973973 |
| GVGMTKF | ASGDPGGADTQY | 0.6144768 |
| IGIGGSDL | ASGDPGGADTQY | 0.58198154 |

| | | |
|---|---|---|
| IGTLLAEL | ASGDPGGADTQY | 0.6250738 |
| IYTSGTT | ASGDPGGADTQY | 0.41775194 |
| LSFPAGV | ASGDPGGADTQY | 0.53787625 |
| LYTSGTT | ASGDPGGADTQY | 0.3549099 |
| VGAFPFGF | ASGDPGGADTQY | 0.4102053 |
| VGAGTVEF | ASGDPGGADTQY | 0.37651074 |
| VGKSTLLL | ASGDPGGADTQY | 0.524437 |
| AALAHAGGGIV | ASGDPGGADTQY | 0.34233198 |
| AALVAVSLL | ASGDPGGADTQY | 0.6312313 |
| AAVSRAVAAL | ASGDPGGADTQY | 0.50077444 |
| AGGSSGIGGF | ASGDPGGADTQY | 0.5818791 |
| AGLTVANANG | ASGDPGGADTQY | 0.47721076 |
| AGLVGAVL | ASGDPGGADTQY | 0.39182252 |
| ALSQAFTV | ASGDPGGADTQY | 0.48845476 |
| ALTFSVSL | ASGDPGGADTQY | 0.61048967 |
| ALTGAALVAL | ASGDPGGADTQY | 0.4026013 |
| ASFSIGAGVL | ASGDPGGADTQY | 0.32805434 |
| DGSVYKI | ASGDPGGADTQY | 0.4218967 |
| EQLAEIV | ASGDPGGADTQY | 0.4003834 |
| FAVSELTVM | ASGDPGGADTQY | 0.6260327 |
| GAAPLAAGL | ASGDPGGADTQY | 0.60233957 |
| GADGGDGGSGGAGG | ASGDPGGADTQY | 0.5391596 |
| GFGLDTGGAGG | ASGDPGGADTQY | 0.63082796 |
| GLASGGLL | ASGDPGGADTQY | 0.31165588 |
| GRIAGAAVLL | ASGDPGGADTQY | 0.51078534 |
| GSAGSLNTSF | ASGDPGGADTQY | 0.6501298 |
| GTGQIGI | ASGDPGGADTQY | 0.5229062 |
| GTLIALIL | ASGDPGGADTQY | 0.50412524 |
| IAATMGFVVG | ASGDPGGADTQY | 0.62304354 |
| IFAQALL | ASGDPGGADTQY | 0.45368603 |
| IGAAVALGF | ASGDPGGADTQY | 0.49397707 |
| IGLTPLAL | ASGDPGGADTQY | 0.5471833 |
| ILGSNTSTL | ASGDPGGADTQY | 0.58262557 |
| IVFSLTVL | ASGDPGGADTQY | 0.5270952 |
| LASKATGF | ASGDPGGADTQY | 0.36738315 |
| LASVALIL | ASGDPGGADTQY | 0.6074937 |
| LFAVGLTL | ASGDPGGADTQY | 0.6505728 |
| LFNGGIGTYI | ASGDPGGADTQY | 0.50814193 |
| LTLGAGII | ASGDPGGADTQY | 0.5791402 |
| LVAAALGDI | ASGDPGGADTQY | 0.562262 |
| NTAFVQLGI | ASGDPGGADTQY | 0.421176 |
| QLFSGAEL | ASGDPGGADTQY | 0.44697452 |
| SAATFITL | ASGDPGGADTQY | 0.38917795 |
| SNYGLANLGV | ASGDPGGADTQY | 0.5710371 |
| SYGGAYAVM | ASGDPGGADTQY | 0.4218583 |
| TTGVGGLTLG | ASGDPGGADTQY | 0.52777416 |
| TTQTIYI | ASGDPGGADTQY | 0.53118664 |
| VAAGAGLGAVI | ASGDPGGADTQY | 0.6541714 |
| VAAVVTLL | ASGDPGGADTQY | 0.5536348 |
| VGFLGGLI | ASGDPGGADTQY | 0.6007609 |
| VGVIPILL | ASGDPGGADTQY | 0.44536656 |

| | | |
|---|---|---|
| VIFASSLI | ASGDPGGADTQY | 0.46106488 |
| VKGGDGGAAAG | ASGDPGGADTQY | 0.5625262 |
| VTKTFTNI | ASGDPGGADTQY | 0.3712155 |
| VVAAAILL | ASGDPGGADTQY | 0.61357576 |
| YAALDVEL | ASGDPGGADTQY | 0.4271905 |
| LASVALLL | ASGDPGGADTQY | 0.53648686 |
| LDAVHRV | ASGDPGGADTQY | 0.5867182 |
| QGLSSSIF | ASGDPGGADTQY | 0.6291008 |
| RGTFANIRL | ASGDPGGADTQY | 0.5320942 |
| RQLWWGHRI | ASGDPGGADTQY | 0.5731339 |
| YGPPGTGKTLL | ASGDPGGADTQY | 0.43068963 |
| AALVGAPANVA | ASGDPGGADTQY | 0.5801344 |
| AATFRRTE | ASGDPGGADTQY | 0.6366154 |
| AGALGGGI | ASGDPGGADTQY | 0.5449257 |
| AGGAGGAGGIGAG | ASGDPGGADTQY | 0.64169455 |
| AGSGALGL | ASGDPGGADTQY | 0.58633643 |
| AQIADQI | ASGDPGGADTQY | 0.3619714 |
| ASAVAVTYL | ASGDPGGADTQY | 0.53478235 |
| ASTVLGGTGG | ASGDPGGADTQY | 0.6342739 |
| AVGGEPAK | ASGDPGGADTQY | 0.6031827 |
| CEQGTFGGTC | ASGDPGGADTQY | 0.5595664 |
| DEGKAPAG | ASGDPGGADTQY | 0.53671753 |
| DTVVFPAL | ASGDPGGADTQY | 0.44272 |
| FEGITMAG | ASGDPGGADTQY | 0.6217278 |
| FPLVVQD | ASGDPGGADTQY | 0.65189016 |
| GAFTTPQ | ASGDPGGADTQY | 0.62001526 |
| GFGGFGTAGGVGGA | ASGDPGGADTQY | 0.44138527 |
| GFLAQFL | ASGDPGGADTQY | 0.42426753 |
| GGAGGAGGNAGVA | ASGDPGGADTQY | 0.53950226 |
| GGVGTAGAGGAGGAGG | ASGDPGGADTQY | 0.62271434 |
| GNGGGGDAGNAGSGGN | ASGDPGGADTQY | 0.39826602 |
| GTGGAGGAGNDAGS | ASGDPGGADTQY | 0.5385983 |
| GTTNTGWFNS | ASGDPGGADTQY | 0.52311856 |
| IFMGPLFTL | ASGDPGGADTQY | 0.33398727 |
| ILAHAIEAI | ASGDPGGADTQY | 0.5484016 |
| IRSAIREAL | ASGDPGGADTQY | 0.6231857 |
| ITFAGGG | ASGDPGGADTQY | 0.5582413 |
| ITGMSGHQEV | ASGDPGGADTQY | 0.38740104 |
| IVAGLACTL | ASGDPGGADTQY | 0.62029386 |
| IVGGGLAGA | ASGDPGGADTQY | 0.6448322 |
| IVTGVLNI | ASGDPGGADTQY | 0.630785 |
| LAQQGCI | ASGDPGGADTQY | 0.38153073 |
| LCLHVEN | ASGDPGGADTQY | 0.3142994 |
| LDAISDGS | ASGDPGGADTQY | 0.56352365 |
| LGGFAAML | ASGDPGGADTQY | 0.6036598 |
| LGGKNPMI | ASGDPGGADTQY | 0.45248395 |
| LGPKGVAAIL | ASGDPGGADTQY | 0.42835864 |
| LMVGVSI | ASGDPGGADTQY | 0.54205203 |
| LVGAAFVL | ASGDPGGADTQY | 0.61593014 |
| LVLAVAPL | ASGDPGGADTQY | 0.40151387 |
| MAAIAVIDAA | ASGDPGGADTQY | 0.357779 |

| | | |
|---|---|---|
| NNGINAELQ | ASGDPGGADTQY | 0.57461154 |
| RGEPLVLVHG | ASGDPGGADTQY | 0.6230638 |
| RGGAGGAAGLI | ASGDPGGADTQY | 0.39931086 |
| RSYRMHPAV | ASGDPGGADTQY | 0.46739373 |
| SAAHLLAGQLA | ASGDPGGADTQY | 0.5790396 |
| SAAVGSY | ASGDPGGADTQY | 0.54029095 |
| SPHAVLHRI | ASGDPGGADTQY | 0.53106326 |
| SQIAKVL | ASGDPGGADTQY | 0.38193816 |
| TALGHLVP | ASGDPGGADTQY | 0.59297276 |
| TAPTTSGAA | ASGDPGGADTQY | 0.37539658 |
| TDGDDGNGGSPGT | ASGDPGGADTQY | 0.5445275 |
| TDGLGEN | ASGDPGGADTQY | 0.52752733 |
| TGGPPLPGQA | ASGDPGGADTQY | 0.54366434 |
| TIPAITI | ASGDPGGADTQY | 0.5048784 |
| TIPGFTI | ASGDPGGADTQY | 0.39990854 |
| TSIASGALA | ASGDPGGADTQY | 0.39306256 |
| TVTLPGAAI | ASGDPGGADTQY | 0.35333094 |
| VASGLSAALP | ASGDPGGADTQY | 0.60206413 |
| VGAGVIGI | ASGDPGGADTQY | 0.5359457 |
| VGPKRFGV | ASGDPGGADTQY | 0.5978956 |
| VNVAFPD | ASGDPGGADTQY | 0.5516799 |
| VSMTNSGVS | ASGDPGGADTQY | 0.4572023 |
| VTDVLGDLSV | ASGDPGGADTQY | 0.521497 |
| VVVVGAGFAGL | ASGDPGGADTQY | 0.38426656 |
| VYTVGTVV | ASGDPGGADTQY | 0.51929396 |
| WTTFSGP | ASGDPGGADTQY | 0.4755148 |
| YDELVEAF | ASGDPGGADTQY | 0.59174156 |
| YGTGGAGGNGG | ASGDPGGADTQY | 0.44315732 |
| YTNVMAV | ASGDPGGADTQY | 0.5721705 |
| FLEQVHQL | ASGDPGGADTQY | 0.5550982 |
| GLSLLAVAV | ASGDPGGADTQY | 0.39536378 |
| LEQVHQL | ASGDPGGADTQY | 0.5457189 |
| LNAAGAI | ASGDPGGADTQY | 0.62920165 |
| PAVHASL | ASGDPGGADTQY | 0.57535684 |
| TELEDDA | ASGDPGGADTQY | 0.37153068 |
| TPPGPPN | ASGDPGGADTQY | 0.5784043 |
| TPPPPPPP | ASGDPGGADTQY | 0.46694303 |
| TQSPALI | ASSLGEGAEQF | 0.53815806 |
| EFAAAFPQ | ASSLGEGAEQF | 0.5546302 |
| LLSGNLDVL | ASSLGEGAEQF | 0.4852742 |
| GPANLGNYNI | ASSLGEGAEQF | 0.58091974 |
| TLGPTLP | ASSLGEGAEQF | 0.39438975 |
| VYAYSSCCK | ASSLGEGAEQF | 0.42947847 |
| APAPAPAPNI | ASSLGEGAEQF | 0.51319313 |
| PGGPVPIP | ASSLGEGAEQF | 0.58492595 |
| FLTVALEP | ASSLGEGAEQF | 0.61482805 |
| SQYFGANL | ASSLGEGAEQF | 0.41165674 |
| GGVKPAA | ASSLGEGAEQF | 0.53482974 |
| GGTGGAGGAGGW | ASSLGEGAEQF | 0.45553347 |
| GVGGVAGLGGAGG | ASSLGEGAEQF | 0.43792754 |
| VLLVLATTI | ASSLGEGAEQF | 0.5359774 |

| | | |
|---|---|---|
| IGIFAAML | ASSLGEGAEQF | 0.4511426 |
| AVGDLID | ASSLGEGAEQF | 0.37298074 |
| NISPLPL | ASSLGEGAEQF | 0.5752638 |
| VTAVFVAI | ASSLGEGAEQF | 0.43083754 |
| SGDDNDAAL | ASSLGEGAEQF | 0.51666707 |
| AVAALAHS | ASSLGEGAEQF | 0.6122166 |
| GGVTVMATP | ASSLGEGAEQF | 0.36573094 |
| AGLAANVRD | ASSLGEGAEQF | 0.53059554 |
| TNRPDLIDPALLRPG | ASSLGEGAEQF | 0.5566013 |
| LANLGVTAL | ASSLGEGAEQF | 0.5338422 |
| TGERIEL | ASSLGEGAEQF | 0.5158508 |
| NGEELAA | ASSLGEGAEQF | 0.40980184 |
| VLSLVWG | ASSLGEGAEQF | 0.6343101 |
| IGYALWL | ASSLGEGAEQF | 0.59885377 |
| LSLLLAPL | ASSLGEGAEQF | 0.55049694 |
| TLDEALK | ASSLGEGAEQF | 0.57405454 |
| TDGAGTAEGG | ASSLGEGAEQF | 0.61047935 |
| NAPGGAGGNGG | ASSLGEGAEQF | 0.62056977 |
| VSVFMGMGGNF | ASSLGEGAEQF | 0.46480113 |
| LASYLAAL | ASSLGEGAEQF | 0.36013818 |
| LVGGTPAL | ASSLGEGAEQF | 0.5466821 |
| VRNADVL | ASSLGEGAEQF | 0.5367052 |
| TCSSPFLVSAM | ASSLGEGAEQF | 0.44049948 |
| VHITDAE | ASSLGEGAEQF | 0.35987785 |
| EYAAPSF | ASSLGEGAEQF | 0.6327913 |
| VVDAFHGGL | ASSLGEGAEQF | 0.65593046 |
| ASVVDPQ | ASSLGEGAEQF | 0.48675808 |
| SYNEFGGLS | ASSLGEGAEQF | 0.64451987 |
| DPDGSVY | ASSLGEGAEQF | 0.36161238 |
| MVSTIHATE | ASSLGEGAEQF | 0.5948119 |
| LAAGVADPV | ASSLGEGAEQF | 0.636989 |
| TKEQIAA | ASSLGEGAEQF | 0.4177361 |
| SLASPAAL | ASSLGEGAEQF | 0.5723243 |
| IGLVFYGV | ASSLGEGAEQF | 0.43558434 |
| FLLDASNL | ASSLGEGAEQF | 0.43625396 |
| YQVGNFP | ASSLGEGAEQF | 0.62443686 |
| ESLLSQST | ASSLGEGAEQF | 0.503092 |
| LFTVGFAAVG | ASSLGEGAEQF | 0.5488275 |
| ATPPTPT | ASSLGEGAEQF | 0.5032238 |
| TDLATLG | ASSLGEGAEQF | 0.43596143 |
| SAGGTGR | ASSLGEGAEQF | 0.5753831 |
| GFSNSGS | ASSLGEGAEQF | 0.5906793 |
| GGNGGSGGTSV | ASSLGEGAEQF | 0.49310824 |
| GTSVATGGAGN | ASSLGEGAEQF | 0.64628625 |
| NGGNGGSGGTS | ASSLGEGAEQF | 0.4182992 |
| AVVKFPH | ASSLGEGAEQF | 0.6425798 |
| TPPPPPP | ASSLGEGAEQF | 0.63826877 |
| AAPAAAAAPA | ASSLGEGAEQF | 0.53395325 |
| IALNEDL | ASSLGEGAEQF | 0.47701883 |
| IVNVSSM | ASSLGEGAEQF | 0.6277354 |
| TGIPPAPR | ASSLGEGAEQF | 0.5099081 |

| | | |
|---|---|---|
| FDLGGGTF | ASSLGEGAEQF | 0.4988872 |
| FDLGGGTFDVS | ASSLGEGAEQF | 0.61813295 |
| NEPTAAA | ASSLGEGAEQF | 0.22070932 |
| NEPTAAALA | ASSLGEGAEQF | 0.79563487 |
| GSIINISSI | ASSLGEGAEQF | 0.69582874 |
| VAPGNAGTA | ASSLGEGAEQF | 0.6787276 |
| TGGGASLE | ASSLGEGAEQF | 0.39581013 |
| VAVYNHYKRI | ASSLGEGAEQF | 0.228159 |
| TDPAPATT | ASSLGEGAEQF | 0.39027703 |
| TNRPDILDPALLRPG | ASSLGEGAEQF | 0.29052627 |
| LTTTTTS | ASSLGEGAEQF | 0.812332 |
| PTGSGKTL | ASSLGEGAEQF | 0.6973091 |
| TGSGKTL | ASSLGEGAEQF | 0.39394525 |
| TIGHVDHGK | ASSLGEGAEQF | 0.77279353 |
| TIGHVDHGKTT | ASSLGEGAEQF | 0.7844151 |
| GHVDHGKTT | ASSLGEGAEQF | 0.20635389 |
| AQFEHTLL | ASSLGEGAEQF | 0.46764332 |
| IVNQDEL | ASSLGEGAEQF | 0.3073579 |
| TNGSQFF | ASSLGEGAEQF | 0.500602 |
| TGKGIVE | ASSLGEGAEQF | 0.141376 |
| LISNASDA | ASSLGEGAEQF | 0.7153617 |
| AELRTLGELTKIAKAHGAQ | ASSLGEGAEQF | 0.44116524 |
| APNAVAAAEPSW | ASSLGEGAEQF | 0.1490933 |
| APQVAGAATAVL | ASSLGEGAEQF | 0.5978434 |
| ARRDPAAAAAILPTDA | ASSLGEGAEQF | 0.48618335 |
| DAAARDVLAEAGLAEYF | ASSLGEGAEQF | 0.43069896 |
| DDQIVLSAKANLAKVEQ | ASSLGEGAEQF | 0.7805661 |
| DELLPDTGEDSDEDGH | ASSLGEGAEQF | 0.56862843 |
| ELPEAPANRATVPQD | ASSLGEGAEQF | 0.34337226 |
| FGQSKIERVTGIG | ASSLGEGAEQF | 0.71434915 |
| GAGGTGSTAGGAGGAGGAGGLY | ASSLGEGAEQF | 0.796725 |
| GHGGTGAAGGNGGTGGAGGSGI | ASSLGEGAEQF | 0.34321314 |
| GNGGNGGAGGEGGPGGAGGAGA | ASSLGEGAEQF | 0.67761016 |
| GSTAGNGGNGGNATNGTIAGQP | ASSLGEGAEQF | 0.8316975 |
| GTGGDGGNGGNGGTGG | ASSLGEGAEQF | 0.5922319 |
| GVRRPAGIPADAPV | ASSLGEGAEQF | 0.803959 |
| GYRQVTDAHLLTLARR | ASSLGEGAEQF | 0.8203649 |
| IAKMGQDGHDRG | ASSLGEGAEQF | 0.38108146 |
| KAEPSIEIQYT | ASSLGEGAEQF | 0.78209937 |
| KPVRIRMAADEDTVAVT | ASSLGEGAEQF | 0.77390337 |
| MPPRIAGMRLLVIKPEP | ASSLGEGAEQF | 0.4840632 |
| MKFYAVALFRSRGVPPTR | ASSLGEGAEQF | 0.586087 |
| NAGGFGILGAVAHSP | ASSLGEGAEQF | 0.21826017 |
| NEGREITLAMAEM | ASSLGEGAEQF | 0.56335324 |
| NELVEWSHPLGHRWRWE | ASSLGEGAEQF | 0.2324745 |
| NLGAGGTGGNGGNPG | ASSLGEGAEQF | 0.7284963 |
| NTEWPKWETVMAADPVD | ASSLGEGAEQF | 0.80039895 |
| PASAAPAETQP | ASSLGEGAEQF | 0.57465833 |
| QQSSDGKNGVRAAEITGE | ASSLGEGAEQF | 0.23730934 |
| RAPDAVTTSWWKE | ASSLGEGAEQF | 0.21829735 |
| SGFGNVGAHTSGWFNQ | ASSLGEGAEQF | 0.50455415 |

| | | |
|---|---|---|
| SGVAGAVGQAASVGGLKV | ASSLGEGAEQF | 0.34084725 |
| SNGGAGTGSAGGNGGT | ASSLGEGAEQF | 0.2353305 |
| TGGAGGEGGAAG | ASSLGEGAEQF | 0.40303493 |
| TGGDGGAAGAGGAPGHG | ASSLGEGAEQF | 0.78041327 |
| TMPPLLAAIANAGRHGV | ASSLGEGAEQF | 0.7890436 |
| TSSTAGNSGGA | ASSLGEGAEQF | 0.35638994 |
| VPATVPVGILC | ASSLGEGAEQF | 0.8125257 |
| VQRPDSGGVVLTGRIS | ASSLGEGAEQF | 0.7923268 |
| WTPNPVPATKGVPAGLVQAT | ASSLGEGAEQF | 0.8190938 |
| YALKRALAKFSDRFDVVII | ASSLGEGAEQF | 0.51352954 |
| YLIGSVIV | ASSLGEGAEQF | 0.21935745 |
| YGQLVAVID | ASSLGEGAEQF | 0.8200611 |
| VTVGGMTLPA | ASSLGEGAEQF | 0.6082046 |
| VLALPAV | ASSLGEGAEQF | 0.774147 |
| VIPPTLNYET | ASSLGEGAEQF | 0.7484725 |
| VGMGGMHPSSG | ASSLGEGAEQF | 0.49198323 |
| VGLAGGAATA | ASSLGEGAEQF | 0.39015752 |
| VGGAGGAGGSGP | ASSLGEGAEQF | 0.7388228 |
| VFAAFMV | ASSLGEGAEQF | 0.7591851 |
| TLYDKLR | ASSLGEGAEQF | 0.31341368 |
| TLAVFTVL | ASSLGEGAEQF | 0.2798428 |
| TGSVQPARV | ASSLGEGAEQF | 0.59412754 |
| TGGGAGSGGT | ASSLGEGAEQF | 0.61327696 |
| SHLETSL | ASSLGEGAEQF | 0.7980621 |
| SFTTGMT | ASSLGEGAEQF | 0.5742121 |
| QVEVAKRMGVRQPTVS | ASSLGEGAEQF | 0.59647715 |
| QGSGGPALAM | ASSLGEGAEQF | 0.72157025 |
| MLAPSNP | ASSLGEGAEQF | 0.7206181 |
| LVNNAAI | ASSLGEGAEQF | 0.62483925 |
| LTGEINTLA | ASSLGEGAEQF | 0.7090516 |
| LSIFGPV | ASSLGEGAEQF | 0.42541784 |
| LLALLANL | ASSLGEGAEQF | 0.7524723 |
| LISNASDALDKLR | ASSLGEGAEQF | 0.7688111 |
| LINEAVP | ASSLGEGAEQF | 0.72678417 |
| LGYAFSGL | ASSLGEGAEQF | 0.7891566 |
| LDFVAVAL | ASSLGEGAEQF | 0.6410338 |
| LAQPGAE | ASSLGEGAEQF | 0.6256142 |
| ISNLGAVTGF | ASSLGEGAEQF | 0.23277129 |
| HQEDPGRI | ASSLGEGAEQF | 0.49273038 |
| GTPGEPTP | ASSLGEGAEQF | 0.6003374 |
| GFFNSGS | ASSLGEGAEQF | 0.68091565 |
| GAGGGGLTAGAN | ASSLGEGAEQF | 0.730667 |
| FVDALVNM | ASSLGEGAEQF | 0.67996407 |
| FLTVAPEL | ASSLGEGAEQF | 0.42103964 |
| FETLTAL | ASSLGEGAEQF | 0.7831295 |
| DVINPIEL | ASSLGEGAEQF | 0.81572473 |
| APVSGAPV | ASSLGEGAEQF | 0.2422137 |
| ALVGGLAW | ASSLGEGAEQF | 0.46912047 |
| ALAAAAATAI | ASSLGEGAEQF | 0.8170187 |
| AFGLGLLL | ASSLGEGAEQF | 0.7588116 |
| AFFAAASL | ASSLGEGAEQF | 0.51369655 |

| | | |
|---|---|---|
| AAAALPAFSPP | ASSLGEGAEQF | 0.57532793 |
| VVLAPAL | ASSLGEGAEQF | 0.8342886 |
| TIGHVDHGKTTL | ASSLGEGAEQF | 0.7289518 |
| LISNASDALDK | ASSLGEGAEQF | 0.80129457 |
| LISNASDAL | ASSLGEGAEQF | 0.19949192 |
| ISNASDALDK | ASSLGEGAEQF | 0.4537527 |
| NQAETLVYQTE | ASSLGEGAEQF | 0.35850462 |
| TGGTGGTAGAG | ASSLGEGAEQF | 0.14224234 |
| TGSTAGGAGGA | ASSLGEGAEQF | 0.7665422 |
| DPLAVPVAAGPV | ASSLGEGAEQF | 0.3306995 |
| GGTGGTPGNGGN | ASSLGEGAEQF | 0.75557864 |
| PAGLAGVDPATL | ASSLGEGAEQF | 0.46647793 |
| GNNLVVEAPHAGASV | ASSLGEGAEQF | 0.7478806 |
| KVTRIMSASTLEQPA | ASSLGEGAEQF | 0.6679566 |
| LGFALPIALPRRTVV | ASSLGEGAEQF | 0.5633999 |
| TRLGNLTARKLVVLV | ASSLGEGAEQF | 0.31176084 |
| WSQQRRLIYFPSAGP | ASSLGEGAEQF | 0.68113923 |
| GPNGNPDPMAAAVDIR | ASSLGEGAEQF | 0.7767521 |
| TQRCVSVADLRAAASR | ASSLGEGAEQF | 0.7312292 |
| QFVLDVAEHPVEVLTETV | ASSLGEGAEQF | 0.20862457 |
| GGAGGTGGAGGNGGTQAGDG | ASSLGEGAEQF | 0.7402233 |
| PLLATLRAIAVDVDAKPAQV | ASSLGEGAEQF | 0.5284785 |
| APKFRTGGYMIDFWGVGYQVAK | ASSLGEGAEQF | 0.18868914 |
| GAGGAGGAGGAAGGPGGTGGTG | ASSLGEGAEQF | 0.17225355 |
| IGALGLLATDVAPAGSVAFGVA | ASSLGEGAEQF | 0.33796316 |
| AGGAGGIGGNGGAGGAGGVLMG | ASSLGEGAEQF | 0.47475475 |
| ADELIGLL | ASSLGEGAEQF | 0.71607876 |
| ADRALKA | ASSLGEGAEQF | 0.73398703 |
| AGAVGQVGGAGGA | ASSLGEGAEQF | 0.46730745 |
| AGGGGGLGGAP | ASSLGEGAEQF | 0.75084376 |
| AGGLLYGNGGAG | ASSLGEGAEQF | 0.63741016 |
| AGLESIGL | ASSLGEGAEQF | 0.112186566 |
| AGLKTGII | ASSLGEGAEQF | 0.4305535 |
| ASLAALLL | ASSLGEGAEQF | 0.20555158 |
| ASVINISSI | ASSLGEGAEQF | 0.78405523 |
| ATPPGPA | ASSLGEGAEQF | 0.2688855 |
| ATVASALV | ASSLGEGAEQF | 0.42528856 |
| ATVASLIL | ASSLGEGAEQF | 0.5582705 |
| AVNGPVLIRH | ASSLGEGAEQF | 0.78480947 |
| DGAAGAVGI | ASSLGEGAEQF | 0.67692447 |
| DPLLHGI | ASSLGEGAEQF | 0.32200027 |
| ELGAALGAL | ASSLGEGAEQF | 0.18668124 |
| ELGRTQAV | ASSLGEGAEQF | 0.70486426 |
| FVAGGYLS | ASSLGEGAEQF | 0.38646686 |
| GASGGAGGAGGW | ASSLGEGAEQF | 0.3209653 |
| GGAGAGGDGGAGGI | ASSLGEGAEQF | 0.22447276 |
| GTGHNGSQPGMGGNG | ASSLGEGAEQF | 0.7038971 |
| GYAAASASAS | ASSLGEGAEQF | 0.2086346 |
| HQVDVAAVQQG | ASSLGEGAEQF | 0.77007395 |
| IGAAIAHT | ASSLGEGAEQF | 0.36723185 |
| IGMIEVAQL | ASSLGEGAEQF | 0.5325562 |

| | | |
|---|---|---|
| IRLAPEHTSAMD | ASSLGEGAEQF | 0.82554686 |
| IYNVSVL | ASSLGEGAEQF | 0.7679438 |
| KQGFLGM | ASSLGEGAEQF | 0.59491724 |
| LESVITQL | ASSLGEGAEQF | 0.3743925 |
| LGLGALVVV | ASSLGEGAEQF | 0.27616 |
| LIGNGGNGGSGGI | ASSLGEGAEQF | 0.31901723 |
| LNAINAPTL | ASSLGEGAEQF | 0.36248654 |
| LQLLNTGV | ASSLGEGAEQF | 0.7955219 |
| LQTSIEF | ASSLGEGAEQF | 0.58068395 |
| MSFAVSEL | ASSLGEGAEQF | 0.20664853 |
| NTGALGL | ASSLGEGAEQF | 0.64846504 |
| PPGAGGAGGAGTT | ASSLGEGAEQF | 0.8234242 |
| QLVGLGI | ASSLGEGAEQF | 0.8339002 |
| RFLEQVHQL | ASSLGEGAEQF | 0.30042443 |
| RGVGYVLR | ASSLGEGAEQF | 0.49192142 |
| SATGVFAGVI | ASSLGEGAEQF | 0.50924397 |
| SLPAITLP | ASSLGEGAEQF | 0.6064024 |
| STALAFTV | ASSLGEGAEQF | 0.44408855 |
| SVLTVGLGK | ASSLGEGAEQF | 0.29010016 |
| TALAGAGSGI | ASSLGEGAEQF | 0.73786896 |
| TGLPFTGI | ASSLGEGAEQF | 0.8637173 |
| VAEHVTV | ASSLGEGAEQF | 0.4541118 |
| VDIVLNSL | ASSLGEGAEQF | 0.14988483 |
| VLTGGGAVL | ASSLGEGAEQF | 0.5302311 |
| VNVGLAYGI | ASSLGEGAEQF | 0.34944242 |
| VSSAAVGAV | ASSLGEGAEQF | 0.39428717 |
| VTSVYEAGV | ASSLGEGAEQF | 0.550943 |
| WVALDRGA | ASSLGEGAEQF | 0.20904367 |
| YLVGSVVI | ASSLGEGAEQF | 0.5301789 |
| GPPGTGKTLL | ASSLGEGAEQF | 0.83532816 |
| GVGMTKF | ASSLGEGAEQF | 0.5899432 |
| IGIGGSDL | ASSLGEGAEQF | 0.69809085 |
| IGTLLAEL | ASSLGEGAEQF | 0.8146144 |
| IYTSGTT | ASSLGEGAEQF | 0.73010147 |
| LSFPAGV | ASSLGEGAEQF | 0.8163987 |
| LYTSGTT | ASSLGEGAEQF | 0.80383044 |
| VGAFPFGF | ASSLGEGAEQF | 0.487051 |
| VGAGTVEF | ASSLGEGAEQF | 0.63136804 |
| VGKSTLLL | ASSLGEGAEQF | 0.72336537 |
| AALAHAGGGIV | ASSLGEGAEQF | 0.5037321 |
| AALVAVSLL | ASSLGEGAEQF | 0.62793386 |
| AAVSRAVAAL | ASSLGEGAEQF | 0.49062485 |
| AGGSSGIGGF | ASSLGEGAEQF | 0.7583251 |
| AGLTVANANG | ASSLGEGAEQF | 0.6283253 |
| AGLVGAVL | ASSLGEGAEQF | 0.83951974 |
| ALSQAFTV | ASSLGEGAEQF | 0.17897403 |
| ALTFSVSL | ASSLGEGAEQF | 0.3536304 |
| ALTGAALVAL | ASSLGEGAEQF | 0.81372654 |
| ASFSIGAGVL | ASSLGEGAEQF | 0.1345145 |
| DGSVYKI | ASSLGEGAEQF | 0.21215057 |
| EQLAEIV | ASSLGEGAEQF | 0.5328207 |

| | | |
|---|---|---|
| FAVSELTVM | ASSLGEGAEQF | 0.81971514 |
| GAAPLAAGL | ASSLGEGAEQF | 0.13227972 |
| GADGGDGGSGGAGG | ASSLGEGAEQF | 0.4422961 |
| GFGLDTGGAGG | ASSLGEGAEQF | 0.22483301 |
| GLASGGLL | ASSLGEGAEQF | 0.7263888 |
| GRIAGAAVLL | ASSLGEGAEQF | 0.5786584 |
| GSAGSLNTSF | ASSLGEGAEQF | 0.22498009 |
| GTGQIGI | ASSLGEGAEQF | 0.7254023 |
| GTLIALIL | ASSLGEGAEQF | 0.17052849 |
| IAATMGFVVG | ASSLGEGAEQF | 0.5365896 |
| IFAQALL | ASSLGEGAEQF | 0.7250155 |
| IGAAVALGF | ASSLGEGAEQF | 0.84141517 |
| IGLTPLAL | ASSLGEGAEQF | 0.22237006 |
| ILGSNTSTL | ASSLGEGAEQF | 0.75638777 |
| IVFSLTVL | ASSLGEGAEQF | 0.47294566 |
| LASKATGF | ASSLGEGAEQF | 0.85792315 |
| LASVALIL | ASSLGEGAEQF | 0.46723714 |
| LFAVGLTL | ASSLGEGAEQF | 0.20958547 |
| LFNGGIGTYI | ASSLGEGAEQF | 0.7634727 |
| LTLGAGII | ASSLGEGAEQF | 0.75343525 |
| LVAAALGDI | ASSLGEGAEQF | 0.5253302 |
| NTAFVQLGI | ASSLGEGAEQF | 0.80360854 |
| QLFSGAEL | ASSLGEGAEQF | 0.49046114 |
| SAATFITL | ASSLGEGAEQF | 0.5186231 |
| SNYGLANLGV | ASSLGEGAEQF | 0.76895714 |
| SYGGAYAVM | ASSLGEGAEQF | 0.69216263 |
| TTGVGGLTLG | ASSLGEGAEQF | 0.3079394 |
| TTQTIYI | ASSLGEGAEQF | 0.75460124 |
| VAAGAGLGAVI | ASSLGEGAEQF | 0.61346745 |
| VAAVVTLL | ASSLGEGAEQF | 0.26528412 |
| VGFLGGLI | ASSLGEGAEQF | 0.8349098 |
| VGVIPILL | ASSLGEGAEQF | 0.5841948 |
| VIFASSLI | ASSLGEGAEQF | 0.77220863 |
| VKGGDGGAAAG | ASSLGEGAEQF | 0.67086756 |
| VTKTFTNI | ASSLGEGAEQF | 0.84673065 |
| VVAAAILL | ASSLGEGAEQF | 0.2743708 |
| YAALDVEL | ASSLGEGAEQF | 0.5754773 |
| LASVALLL | ASSLGEGAEQF | 0.61123073 |
| LDAVHRV | ASSLGEGAEQF | 0.7330135 |
| QGLSSSIF | ASSLGEGAEQF | 0.59150714 |
| RGTFANIRL | ASSLGEGAEQF | 0.4755646 |
| RQLWWGHRI | ASSLGEGAEQF | 0.7107599 |
| YGPPGTGKTLL | ASSLGEGAEQF | 0.7979213 |
| AALVGAPANVA | ASSLGEGAEQF | 0.486771 |
| AATFRRTE | ASSLGEGAEQF | 0.6480328 |
| AGALGGGI | ASSLGEGAEQF | 0.8366853 |
| AGGAGGAGGIGAG | ASSLGEGAEQF | 0.23801944 |
| AGSGALGL | ASSLGEGAEQF | 0.56085277 |
| AQIADQI | ASSLGEGAEQF | 0.7126764 |
| ASAVAVTYL | ASSLGEGAEQF | 0.76545626 |
| ASTVLGGTGG | ASSLGEGAEQF | 0.52673835 |

| | | |
|---|---|---|
| AVGGEPAK | ASSLGEGAEQF | 0.79853785 |
| CEQGTFGGTC | ASSLGEGAEQF | 0.42031193 |
| DEGKAPAG | ASSLGEGAEQF | 0.7413579 |
| DTVVFPAL | ASSLGEGAEQF | 0.5648647 |
| FEGITMAG | ASSLGEGAEQF | 0.59278214 |
| FPLVVQD | ASSLGEGAEQF | 0.27943394 |
| GAFTTPQ | ASSLGEGAEQF | 0.626516 |
| GFGGFGTAGGVGGA | ASSLGEGAEQF | 0.7740365 |
| GFLAQFL | ASSLGEGAEQF | 0.7440829 |
| GGAGGAGGNAGVA | ASSLGEGAEQF | 0.18035841 |
| GGVGTAGAGGAGGAGG | ASSLGEGAEQF | 0.7282256 |
| GNGGGGDAGNAGSGGN | ASSLGEGAEQF | 0.75535655 |
| GTGGAGGAGNDAGS | ASSLGEGAEQF | 0.8548833 |
| GTTNTGWFNS | ASSLGEGAEQF | 0.63573205 |
| IFMGPLFTL | ASSLGEGAEQF | 0.7342081 |
| ILAHAIEAI | ASSLGEGAEQF | 0.41140133 |
| IRSAIREAL | ASSLGEGAEQF | 0.5652206 |
| ITFAGGG | ASSLGEGAEQF | 0.2821359 |
| ITGMSGHQEV | ASSLGEGAEQF | 0.2960847 |
| IVAGLACTL | ASSLGEGAEQF | 0.56990814 |
| IVGGGLAGA | ASSLGEGAEQF | 0.38453147 |
| IVTGVLNI | ASSLGEGAEQF | 0.8164695 |
| LAQQGCI | ASSLGEGAEQF | 0.77782214 |
| LCLHVEN | ASSLGEGAEQF | 0.76887363 |
| LDAISDGS | ASSLGEGAEQF | 0.80690014 |
| LGGFAAML | ASSLGEGAEQF | 0.82820565 |
| LGGKNPMI | ASSLGEGAEQF | 0.74409866 |
| LGPKGVAAIL | ASSLGEGAEQF | 0.3923264 |
| LMVGVSI | ASSLGEGAEQF | 0.55094475 |
| LVGAAFVL | ASSLGEGAEQF | 0.19847803 |
| LVLAVAPL | ASSLGEGAEQF | 0.47304666 |
| MAAIAVIDAA | ASSLGEGAEQF | 0.4430694 |
| NNGINAELQ | ASSLGEGAEQF | 0.6343555 |
| RGEPLVLVHG | ASSLGEGAEQF | 0.1289467 |
| RGGAGGAAGLI | ASSLGEGAEQF | 0.77747226 |
| RSYRMHPAV | ASSLGEGAEQF | 0.8020909 |
| SAAHLLAGQLA | ASSLGEGAEQF | 0.5975673 |
| SAAVGSY | ASSLGEGAEQF | 0.7428615 |
| SPHAVLHRI | ASSLGEGAEQF | 0.7521065 |
| SQIAKVL | ASSLGEGAEQF | 0.40387303 |
| TALGHLVP | ASSLGEGAEQF | 0.75243795 |
| TAPTTSGAA | ASSLGEGAEQF | 0.40393585 |
| TDGDDGNGGSPGT | ASSLGEGAEQF | 0.7853867 |
| TDGLGEN | ASSLGEGAEQF | 0.79859704 |
| TGGPPLPGQA | ASSLGEGAEQF | 0.6862588 |
| TIPAITI | ASSLGEGAEQF | 0.5691898 |
| TIPGFTI | ASSLGEGAEQF | 0.65485156 |
| TSIASGALA | ASSLGEGAEQF | 0.25885117 |
| TVTLPGAAI | ASSLGEGAEQF | 0.5922694 |
| VASGLSAALP | ASSLGEGAEQF | 0.24859697 |
| VGAGVIGI | ASSLGEGAEQF | 0.66793746 |

| | | |
|---|---|---|
| VGPKRFGV | ASSLGEGAEQF | 0.29574513 |
| VNVAFPD | ASSLGEGAEQF | 0.6707865 |
| VSMTNSGVS | ASSLGEGAEQF | 0.5830319 |
| VTDVLGDLSV | ASSLGEGAEQF | 0.6131667 |
| VVVVGAGFAGL | ASSLGEGAEQF | 0.31591564 |
| VYTVGTVV | ASSLGEGAEQF | 0.80074525 |
| WTTFSGP | ASSLGEGAEQF | 0.7756239 |
| YDELVEAF | ASSLGEGAEQF | 0.2420125 |
| YGTGGAGGNGG | ASSLGEGAEQF | 0.72143924 |
| YTNVMAV | ASSLGEGAEQF | 0.4532649 |
| FLEQVHQL | ASSLGEGAEQF | 0.5511629 |
| GLSLLAVAV | ASSLGEGAEQF | 0.78264326 |
| LEQVHQL | ASSLGEGAEQF | 0.6136441 |
| LNAAGAI | ASSLGEGAEQF | 0.1769588 |
| PAVHASL | ASSLGEGAEQF | 0.56489134 |
| TELEDDA | ASSLGEGAEQF | 0.56073 |
| TPPGPPN | ASSLGEGAEQF | 0.50871086 |
| TPPPPPPP | ASSLGEGAEQF | 0.15121621 |
| TQSPALI | ASSSTGEGSDYT | 0.62699306 |
| EFAAAFPQ | ASSSTGEGSDYT | 0.2695487 |
| LLSGNLDVL | ASSSTGEGSDYT | 0.31913826 |
| GPANLGNYNI | ASSSTGEGSDYT | 0.3811139 |
| TLGPTLP | ASSSTGEGSDYT | 0.7878791 |
| VYAYSSCCK | ASSSTGEGSDYT | 0.8000587 |
| APAPAPAPNI | ASSSTGEGSDYT | 0.46547565 |
| PGGPVPIP | ASSSTGEGSDYT | 0.68218946 |
| FLTVALEP | ASSSTGEGSDYT | 0.64630127 |
| SQYFGANL | ASSSTGEGSDYT | 0.8149868 |
| GGVKPAA | ASSSTGEGSDYT | 0.577401 |
| GGTGGAGGAGGW | ASSSTGEGSDYT | 0.53068787 |
| GVGGVAGLGGAGG | ASSSTGEGSDYT | 0.6719495 |
| VLLVLATTI | ASSSTGEGSDYT | 0.49150634 |
| IGIFAAML | ASSSTGEGSDYT | 0.38477796 |
| AVGDLID | ASSSTGEGSDYT | 0.4374268 |
| NISPLPL | ASSSTGEGSDYT | 0.33665875 |
| VTAVFVAI | ASSSTGEGSDYT | 0.26864952 |
| SGDDNDAAL | ASSSTGEGSDYT | 0.67291 |
| AVAALAHS | ASSSTGEGSDYT | 0.7984427 |
| GGVTVMATP | ASSSTGEGSDYT | 0.61984926 |
| AGLAANVRD | ASSSTGEGSDYT | 0.19246528 |
| TNRPDLIDPALLRPG | ASSSTGEGSDYT | 0.4018861 |
| LANLGVTAL | ASSSTGEGSDYT | 0.4758759 |
| TGERIEL | ASSSTGEGSDYT | 0.74929285 |
| NGEELAA | ASSSTGEGSDYT | 0.7251322 |
| VLSLVWG | ASSSTGEGSDYT | 0.84848225 |
| IGYALWL | ASSSTGEGSDYT | 0.75474405 |
| LSLLLAPL | ASSSTGEGSDYT | 0.4472301 |
| TLDEALK | ASSSTGEGSDYT | 0.7561058 |
| TDGAGTAEGG | ASSSTGEGSDYT | 0.5445633 |
| NAPGGAGGNGG | ASSSTGEGSDYT | 0.50000715 |
| VSVFMGMGGNF | ASSSTGEGSDYT | 0.5669019 |

| | | |
|---|---|---|
| LASYLAAL | ASSSTGEGSDYT | 0.6272416 |
| LVGGTPAL | ASSSTGEGSDYT | 0.4428959 |
| VRNADVL | ASSSTGEGSDYT | 0.41826898 |
| TCSSPFLVSAM | ASSSTGEGSDYT | 0.20678064 |
| VHITDAE | ASSSTGEGSDYT | 0.6340898 |
| EYAAPSF | ASSSTGEGSDYT | 0.3576289 |
| VVDAFHGGL | ASSSTGEGSDYT | 0.3654309 |
| ASVVDPQ | ASSSTGEGSDYT | 0.5261624 |
| SYNEFGGLS | ASSSTGEGSDYT | 0.83143264 |
| DPDGSVY | ASSSTGEGSDYT | 0.82905895 |
| MVSTIHATE | ASSSTGEGSDYT | 0.40581083 |
| LAAGVADPV | ASSSTGEGSDYT | 0.3180002 |
| TKEQIAA | ASSSTGEGSDYT | 0.7738487 |
| SLASPAAL | ASSSTGEGSDYT | 0.35984248 |
| IGLVFYGV | ASSSTGEGSDYT | 0.7033285 |
| FLLDASNL | ASSSTGEGSDYT | 0.6560358 |
| YQVGNFP | ASSSTGEGSDYT | 0.58191085 |
| ESLLSQST | ASSSTGEGSDYT | 0.40567693 |
| LFTVGFAAVG | ASSSTGEGSDYT | 0.7004804 |
| ATPPTPT | ASSSTGEGSDYT | 0.38580433 |
| TDLATLG | ASSSTGEGSDYT | 0.70889163 |
| SAGGTGR | ASSSTGEGSDYT | 0.13243645 |
| GFSNSGS | ASSSTGEGSDYT | 0.6897046 |
| GGNGGSGGTSV | ASSSTGEGSDYT | 0.34937805 |
| GTSVATGGAGN | ASSSTGEGSDYT | 0.40205514 |
| NGGNGGSGGTS | ASSSTGEGSDYT | 0.1674979 |
| AVVKFPH | ASSSTGEGSDYT | 0.41031182 |
| TPPPPPP | ASSSTGEGSDYT | 0.7218163 |
| AAPAAAAAPA | ASSSTGEGSDYT | 0.80770564 |
| IALNEDL | ASSSTGEGSDYT | 0.6648137 |
| IVNVSSM | ASSSTGEGSDYT | 0.71330684 |
| TGIPPAPR | ASSSTGEGSDYT | 0.8252181 |
| FDLGGGTF | ASSSTGEGSDYT | 0.50559676 |
| FDLGGGTFDVS | ASSSTGEGSDYT | 0.17818245 |
| NEPTAAA | ASSSTGEGSDYT | 0.6703379 |
| NEPTAAALA | ASSSTGEGSDYT | 0.571335 |
| GSIINISSI | ASSSTGEGSDYT | 0.76763284 |
| VAPGNAGTA | ASSSTGEGSDYT | 0.76412934 |
| TGGGASLE | ASSSTGEGSDYT | 0.22332746 |
| VAVYNHYKRI | ASSSTGEGSDYT | 0.7498428 |
| TDPAPATT | ASSSTGEGSDYT | 0.748314 |
| TNRPDILDPALLRPG | ASSSTGEGSDYT | 0.3456967 |
| LTTTTTS | ASSSTGEGSDYT | 0.24319561 |
| PTGSGKTL | ASSSTGEGSDYT | 0.28660318 |
| TGSGKTL | ASSSTGEGSDYT | 0.20157546 |
| TIGHVDHGK | ASSSTGEGSDYT | 0.6214388 |
| TIGHVDHGKTT | ASSSTGEGSDYT | 0.49793637 |
| GHVDHGKTT | ASSSTGEGSDYT | 0.5786197 |
| AQFEHTLL | ASSSTGEGSDYT | 0.5198023 |
| IVNQDEL | ASSSTGEGSDYT | 0.3717538 |
| TNGSQFF | ASSSTGEGSDYT | 0.4878185 |

| | | |
|---|---|---|
| TGKGIVE | ASSSTGEGSDYT | 0.5258521 |
| LISNASDA | ASSSTGEGSDYT | 0.4930762 |
| AELRTLGELTKIAKAHGAQ | ASSSTGEGSDYT | 0.7328862 |
| APNAVAAAEPSW | ASSSTGEGSDYT | 0.49038348 |
| APQVAGAATAVL | ASSSTGEGSDYT | 0.5676074 |
| ARRDPAAAAAILPTDA | ASSSTGEGSDYT | 0.8174672 |
| DAAARDVLAEAGLAEYF | ASSSTGEGSDYT | 0.25607795 |
| DDQIVLSAKANLAKVEQ | ASSSTGEGSDYT | 0.35383397 |
| DELLPDTGEDSDEDGH | ASSSTGEGSDYT | 0.6921351 |
| ELPEAPANRATVPQD | ASSSTGEGSDYT | 0.50205594 |
| FGQSKIERVTGIG | ASSSTGEGSDYT | 0.8115536 |
| GAGGTGSTAGGAGGAGGAGGLY | ASSSTGEGSDYT | 0.6220788 |
| GHGGTGAAGGNGGTGGAGGSGI | ASSSTGEGSDYT | 0.6840366 |
| GNGGNGGAGGEGGPGGAGGAGA | ASSSTGEGSDYT | 0.7023798 |
| GSTAGNGGNGGNATNGTIAGQP | ASSSTGEGSDYT | 0.8238561 |
| GTGGDGGNGGNGGTGG | ASSSTGEGSDYT | 0.70673084 |
| GVRRPAGIPADAPV | ASSSTGEGSDYT | 0.4037196 |
| GYRQVTDAHLLTLARR | ASSSTGEGSDYT | 0.77005136 |
| IAKMGQDGHDRG | ASSSTGEGSDYT | 0.22582905 |
| KAEPSIEIQYT | ASSSTGEGSDYT | 0.71907747 |
| KPVRIRMAADEDTVAVT | ASSSTGEGSDYT | 0.6930634 |
| MPPRIAGMRLLVIKPEP | ASSSTGEGSDYT | 0.3459411 |
| MKFYAVALFRSRGVPPTR | ASSSTGEGSDYT | 0.5256795 |
| NAGGFGILGAVAHSP | ASSSTGEGSDYT | 0.80816764 |
| NEGREITLAMAEM | ASSSTGEGSDYT | 0.76872844 |
| NELVEWSHPLGHRWRWE | ASSSTGEGSDYT | 0.41388202 |
| NLGAGGTGGNGGNPG | ASSSTGEGSDYT | 0.4142518 |
| NTEWPKWETVMAADPVD | ASSSTGEGSDYT | 0.3754778 |
| PASAAPAETQP | ASSSTGEGSDYT | 0.48351198 |
| QQSSDGKNGVRAAEITGE | ASSSTGEGSDYT | 0.19261308 |
| RAPDAVTTSWWKE | ASSSTGEGSDYT | 0.7548144 |
| SGFGNVGAHTSGWFNQ | ASSSTGEGSDYT | 0.48533815 |
| SGVAGAVGQAASVGGLKV | ASSSTGEGSDYT | 0.82684606 |
| SNGGAGTGSAGGNGGT | ASSSTGEGSDYT | 0.6663669 |
| TGGAGGEGGAAG | ASSSTGEGSDYT | 0.77885485 |
| TGGDGGAAGAGGAPGHG | ASSSTGEGSDYT | 0.3976879 |
| TMPPLLAAIANAGRHGV | ASSSTGEGSDYT | 0.5008028 |
| TSSTAGNSGGA | ASSSTGEGSDYT | 0.2081253 |
| VPATVPVGILC | ASSSTGEGSDYT | 0.32347482 |
| VQRPDSGGVVLTGRIS | ASSSTGEGSDYT | 0.32977003 |
| WTPNPVPATKGVPAGLVQAT | ASSSTGEGSDYT | 0.5976038 |
| YALKRALAKFSDRFDVVII | ASSSTGEGSDYT | 0.17601982 |
| YLIGSVIV | ASSSTGEGSDYT | 0.82943285 |
| YGQLVAVID | ASSSTGEGSDYT | 0.54292643 |
| VTVGGMTLPA | ASSSTGEGSDYT | 0.7255961 |
| VLALPAV | ASSSTGEGSDYT | 0.4203934 |
| VIPPTLNYET | ASSSTGEGSDYT | 0.27181488 |
| VGMGGMHPSSG | ASSSTGEGSDYT | 0.39727378 |
| VGLAGGAATA | ASSSTGEGSDYT | 0.7552287 |
| VGGAGGAGGSGP | ASSSTGEGSDYT | 0.24335551 |
| VFAAFMV | ASSSTGEGSDYT | 0.13952748 |

| | | |
|---|---|---|
| TLYDKLR | ASSSTGEGSDYT | 0.34041744 |
| TLAVFTVL | ASSSTGEGSDYT | 0.18701732 |
| TGSVQPARV | ASSSTGEGSDYT | 0.8378422 |
| TGGGAGSGGT | ASSSTGEGSDYT | 0.79861677 |
| SHLETSL | ASSSTGEGSDYT | 0.6516034 |
| SFTTGMT | ASSSTGEGSDYT | 0.8404903 |
| QVEVAKRMGVRQPTVS | ASSSTGEGSDYT | 0.12940775 |
| QGSGGPALAM | ASSSTGEGSDYT | 0.4413327 |
| MLAPSNP | ASSSTGEGSDYT | 0.8290901 |
| LVNNAAI | ASSSTGEGSDYT | 0.671524 |
| LTGEINTLA | ASSSTGEGSDYT | 0.53435886 |
| LSIFGPV | ASSSTGEGSDYT | 0.78040934 |
| LLALLANL | ASSSTGEGSDYT | 0.39555404 |
| LISNASDALDKLR | ASSSTGEGSDYT | 0.51872814 |
| LINEAVP | ASSSTGEGSDYT | 0.53095317 |
| LGYAFSGL | ASSSTGEGSDYT | 0.66420805 |
| LDFVAVAL | ASSSTGEGSDYT | 0.61840427 |
| LAQPGAE | ASSSTGEGSDYT | 0.20686737 |
| ISNLGAVTGF | ASSSTGEGSDYT | 0.72678125 |
| HQEDPGRI | ASSSTGEGSDYT | 0.8393235 |
| GTPGEPTP | ASSSTGEGSDYT | 0.4913569 |
| GFFNSGS | ASSSTGEGSDYT | 0.71975493 |
| GAGGGGLTAGAN | ASSSTGEGSDYT | 0.6844752 |
| FVDALVNM | ASSSTGEGSDYT | 0.7142569 |
| FLTVAPEL | ASSSTGEGSDYT | 0.4046309 |
| FETLTAL | ASSSTGEGSDYT | 0.2886481 |
| DVINPIEL | ASSSTGEGSDYT | 0.8044617 |
| APVSGAPV | ASSSTGEGSDYT | 0.34852135 |
| ALVGGLAW | ASSSTGEGSDYT | 0.5559802 |
| ALAAAAATAI | ASSSTGEGSDYT | 0.5288074 |
| AFGLGLLL | ASSSTGEGSDYT | 0.86130846 |
| AFFAAASL | ASSSTGEGSDYT | 0.7504782 |
| AAAALPAFSPP | ASSSTGEGSDYT | 0.71913195 |
| VVLAPAL | ASSSTGEGSDYT | 0.3479168 |
| TIGHVDHGKTTL | ASSSTGEGSDYT | 0.59367645 |
| LISNASDALDK | ASSSTGEGSDYT | 0.7425037 |
| LISNASDAL | ASSSTGEGSDYT | 0.20925637 |
| ISNASDALDK | ASSSTGEGSDYT | 0.812446 |
| NQAETLVYQTE | ASSSTGEGSDYT | 0.3670634 |
| TGGTGGTAGAG | ASSSTGEGSDYT | 0.62628794 |
| TGSTAGGAGGA | ASSSTGEGSDYT | 0.69950736 |
| DPLAVPVAAGPV | ASSSTGEGSDYT | 0.8345523 |
| GGTGGTPGNGGN | ASSSTGEGSDYT | 0.5525036 |
| PAGLAGVDPATL | ASSSTGEGSDYT | 0.8164235 |
| GNNLVVEAPHAGASV | ASSSTGEGSDYT | 0.3090613 |
| KVTRIMSASTLEQPA | ASSSTGEGSDYT | 0.79230684 |
| LGFALPIALPRRTVV | ASSSTGEGSDYT | 0.8033277 |
| TRLGNLTARKLVVLV | ASSSTGEGSDYT | 0.7505553 |
| WSQQRRLIYFPSAGP | ASSSTGEGSDYT | 0.8464422 |
| GPNGNPDPMAAAVDIR | ASSSTGEGSDYT | 0.70013714 |
| TQRCVSVADLRAAASR | ASSSTGEGSDYT | 0.18317357 |

| | | |
|---|---|---|
| QFVLDVAEHPVEVLTETV | ASSSTGEGSDYT | 0.54185367 |
| GGAGGTGGAGGNGGTQAGDG | ASSSTGEGSDYT | 0.7611511 |
| PLLATLRAIAVDVDAKPAQV | ASSSTGEGSDYT | 0.81965184 |
| APKFRTGGYMIDFWGVGYQVAK | ASSSTGEGSDYT | 0.65012383 |
| GAGGAGGAGGAAGGPGGTGGTG | ASSSTGEGSDYT | 0.6606022 |
| IGALGLLATDVAPAGSVAFGVA | ASSSTGEGSDYT | 0.5211566 |
| AGGAGGIGGNGGAGGAGGVLMG | ASSSTGEGSDYT | 0.82530314 |
| ADELIGLL | ASSSTGEGSDYT | 0.85646236 |
| ADRALKA | ASSSTGEGSDYT | 0.74973524 |
| AGAVGQVGGAGGA | ASSSTGEGSDYT | 0.4716189 |
| AGGGGGLGGAP | ASSSTGEGSDYT | 0.42456612 |
| AGGLLYGNGGAG | ASSSTGEGSDYT | 0.6850865 |
| AGLESIGL | ASSSTGEGSDYT | 0.7732401 |
| AGLKTGII | ASSSTGEGSDYT | 0.265156 |
| ASLAALLL | ASSSTGEGSDYT | 0.6677971 |
| ASVINISSI | ASSSTGEGSDYT | 0.6561843 |
| ATPPGPA | ASSSTGEGSDYT | 0.15356824 |
| ATVASALV | ASSSTGEGSDYT | 0.691794 |
| ATVASLIL | ASSSTGEGSDYT | 0.7904256 |
| AVNGPVLIRH | ASSSTGEGSDYT | 0.66295856 |
| DGAAGAVGI | ASSSTGEGSDYT | 0.33043832 |
| DPLLHGI | ASSSTGEGSDYT | 0.8075672 |
| ELGAALGAL | ASSSTGEGSDYT | 0.8141286 |
| ELGRTQAV | ASSSTGEGSDYT | 0.8171725 |
| FVAGGYLS | ASSSTGEGSDYT | 0.21773297 |
| GASGGAGGAGGW | ASSSTGEGSDYT | 0.13651462 |
| GGAGAGGDGGAGGI | ASSSTGEGSDYT | 0.62122756 |
| GTGHNGSQPGMGGNG | ASSSTGEGSDYT | 0.79921055 |
| GYAAASASAS | ASSSTGEGSDYT | 0.48361167 |
| HQVDVAAVQQG | ASSSTGEGSDYT | 0.332102 |
| IGAAIAHT | ASSSTGEGSDYT | 0.6700428 |
| IGMIEVAQL | ASSSTGEGSDYT | 0.7949519 |
| IRLAPEHTSAMD | ASSSTGEGSDYT | 0.22435394 |
| IYNVSVL | ASSSTGEGSDYT | 0.255957 |
| KQGFLGM | ASSSTGEGSDYT | 0.6203828 |
| LESVITQL | ASSSTGEGSDYT | 0.750952 |
| LGLGALVVV | ASSSTGEGSDYT | 0.33200395 |
| LIGNGGNGGSGGI | ASSSTGEGSDYT | 0.47206303 |
| LNAINAPTL | ASSSTGEGSDYT | 0.6983277 |
| LQLLNTGV | ASSSTGEGSDYT | 0.68318 |
| LQTSIEF | ASSSTGEGSDYT | 0.6710421 |
| MSFAVSEL | ASSSTGEGSDYT | 0.24712707 |
| NTGALGL | ASSSTGEGSDYT | 0.7382191 |
| PPGAGGAGGAGTT | ASSSTGEGSDYT | 0.22470553 |
| QLVGLGI | ASSSTGEGSDYT | 0.7705297 |
| RFLEQVHQL | ASSSTGEGSDYT | 0.5829577 |
| RGVGYVLR | ASSSTGEGSDYT | 0.6270972 |
| SATGVFAGVI | ASSSTGEGSDYT | 0.5150608 |
| SLPAITLP | ASSSTGEGSDYT | 0.3309242 |
| STALAFTV | ASSSTGEGSDYT | 0.2755513 |
| SVLTVGLGK | ASSSTGEGSDYT | 0.31457642 |

| | | |
|---|---|---|
| TALAGAGSGI | ASSSTGEGSDYT | 0.7768235 |
| TGLPFTGI | ASSSTGEGSDYT | 0.763312 |
| VAEHVTV | ASSSTGEGSDYT | 0.7146152 |
| VDIVLNSL | ASSSTGEGSDYT | 0.62986684 |
| VLTGGGAVL | ASSSTGEGSDYT | 0.2884668 |
| VNVGLAYGI | ASSSTGEGSDYT | 0.6609913 |
| VSSAAVGAV | ASSSTGEGSDYT | 0.23543823 |
| VTSVYEAGV | ASSSTGEGSDYT | 0.605791 |
| WVALDRGA | ASSSTGEGSDYT | 0.7294114 |
| YLVGSVVI | ASSSTGEGSDYT | 0.7098394 |
| GPPGTGKTLL | ASSSTGEGSDYT | 0.314565 |
| GVGMTKF | ASSSTGEGSDYT | 0.6790384 |
| IGIGGSDL | ASSSTGEGSDYT | 0.7094355 |
| IGTLLAEL | ASSSTGEGSDYT | 0.34047717 |
| IYTSGTT | ASSSTGEGSDYT | 0.61927783 |
| LSFPAGV | ASSSTGEGSDYT | 0.7891073 |
| LYTSGTT | ASSSTGEGSDYT | 0.64696705 |
| VGAFPFGF | ASSSTGEGSDYT | 0.20818645 |
| VGAGTVEF | ASSSTGEGSDYT | 0.67286956 |
| VGKSTLLL | ASSSTGEGSDYT | 0.44819283 |
| AALAHAGGGIV | ASSSTGEGSDYT | 0.68810487 |
| AALVAVSLL | ASSSTGEGSDYT | 0.5457076 |
| AAVSRAVAAL | ASSSTGEGSDYT | 0.5656569 |
| AGGSSGIGGF | ASSSTGEGSDYT | 0.7230113 |
| AGLTVANANG | ASSSTGEGSDYT | 0.29498962 |
| AGLVGAVL | ASSSTGEGSDYT | 0.4223105 |
| ALSQAFTV | ASSSTGEGSDYT | 0.5703206 |
| ALTFSVSL | ASSSTGEGSDYT | 0.62205005 |
| ALTGAALVAL | ASSSTGEGSDYT | 0.7554544 |
| ASFSIGAGVL | ASSSTGEGSDYT | 0.32525134 |
| DGSVYKI | ASSSTGEGSDYT | 0.67797995 |
| EQLAEIV | ASSSTGEGSDYT | 0.4674148 |
| FAVSELTVM | ASSSTGEGSDYT | 0.32703182 |
| GAAPLAAGL | ASSSTGEGSDYT | 0.5719569 |
| GADGGDGGSGGAGG | ASSSTGEGSDYT | 0.48913112 |
| GFGLDTGGAGG | ASSSTGEGSDYT | 0.2611895 |
| GLASGGLL | ASSSTGEGSDYT | 0.698572 |
| GRIAGAAVLL | ASSSTGEGSDYT | 0.29710278 |
| GSAGSLNTSF | ASSSTGEGSDYT | 0.6331531 |
| GTGQIGI | ASSSTGEGSDYT | 0.8324437 |
| GTLIALIL | ASSSTGEGSDYT | 0.26386124 |
| IAATMGFVVG | ASSSTGEGSDYT | 0.730657 |
| IFAQALL | ASSSTGEGSDYT | 0.6028205 |
| IGAAVALGF | ASSSTGEGSDYT | 0.6668363 |
| IGLTPLAL | ASSSTGEGSDYT | 0.56554353 |
| ILGSNTSTL | ASSSTGEGSDYT | 0.3564748 |
| IVFSLTVL | ASSSTGEGSDYT | 0.77996767 |
| LASKATGF | ASSSTGEGSDYT | 0.7371026 |
| LASVALIL | ASSSTGEGSDYT | 0.71522325 |
| LFAVGLTL | ASSSTGEGSDYT | 0.69178355 |
| LFNGGIGTYI | ASSSTGEGSDYT | 0.72240317 |

| | | |
|---|---|---|
| LTLGAGII | ASSSTGEGSDYT | 0.77260077 |
| LVAAALGDI | ASSSTGEGSDYT | 0.47487947 |
| NTAFVQLGI | ASSSTGEGSDYT | 0.21174978 |
| QLFSGAEL | ASSSTGEGSDYT | 0.52156997 |
| SAATFITL | ASSSTGEGSDYT | 0.5391527 |
| SNYGLANLGV | ASSSTGEGSDYT | 0.29037878 |
| SYGGAYAVM | ASSSTGEGSDYT | 0.22908603 |
| TTGVGGLTLG | ASSSTGEGSDYT | 0.7868412 |
| TTQTIYI | ASSSTGEGSDYT | 0.8460747 |
| VAAGAGLGAVI | ASSSTGEGSDYT | 0.5082921 |
| VAAVVTLL | ASSSTGEGSDYT | 0.84056515 |
| VGFLGGLI | ASSSTGEGSDYT | 0.2585874 |
| VGVIPILL | ASSSTGEGSDYT | 0.75488925 |
| VIFASSLI | ASSSTGEGSDYT | 0.8074715 |
| VKGGDGGAAAG | ASSSTGEGSDYT | 0.34614375 |
| VTKTFTNI | ASSSTGEGSDYT | 0.6792723 |
| VVAAAILL | ASSSTGEGSDYT | 0.27752843 |
| YAALDVEL | ASSSTGEGSDYT | 0.35177788 |
| LASVALLL | ASSSTGEGSDYT | 0.7805141 |
| LDAVHRV | ASSSTGEGSDYT | 0.5690516 |
| QGLSSSIF | ASSSTGEGSDYT | 0.68154 |
| RGTFANIRL | ASSSTGEGSDYT | 0.45477164 |
| RQLWWGHRI | ASSSTGEGSDYT | 0.2990002 |
| YGPPGTGKTLL | ASSSTGEGSDYT | 0.58966875 |
| AALVGAPANVA | ASSSTGEGSDYT | 0.7553767 |
| AATFRRTE | ASSSTGEGSDYT | 0.7024063 |
| AGALGGGI | ASSSTGEGSDYT | 0.8537462 |
| AGGAGGAGGIGAG | ASSSTGEGSDYT | 0.28782573 |
| AGSGALGL | ASSSTGEGSDYT | 0.7925726 |
| AQIADQI | ASSSTGEGSDYT | 0.83098876 |
| ASAVAVTYL | ASSSTGEGSDYT | 0.67036617 |
| ASTVLGGTGG | ASSSTGEGSDYT | 0.5174811 |
| AVGGEPAK | ASSSTGEGSDYT | 0.80257744 |
| CEQGTFGGTC | ASSSTGEGSDYT | 0.51537395 |
| DEGKAPAG | ASSSTGEGSDYT | 0.6848217 |
| DTVVFPAL | ASSSTGEGSDYT | 0.78341895 |
| FEGITMAG | ASSSTGEGSDYT | 0.6336041 |
| FPLVVQD | ASSSTGEGSDYT | 0.70386565 |
| GAFTTPQ | ASSSTGEGSDYT | 0.6581362 |
| GFGGFGTAGGVGGA | ASSSTGEGSDYT | 0.6749804 |
| GFLAQFL | ASSSTGEGSDYT | 0.592206 |
| GGAGGAGGNAGVA | ASSSTGEGSDYT | 0.48888665 |
| GGVGTAGAGGAGGAGG | ASSSTGEGSDYT | 0.6648938 |
| GNGGGGDAGNAGSGGN | ASSSTGEGSDYT | 0.59127885 |
| GTGGAGGAGNDAGS | ASSSTGEGSDYT | 0.69148284 |
| GTTNTGWFNS | ASSSTGEGSDYT | 0.7066958 |
| IFMGPLFTL | ASSSTGEGSDYT | 0.5889623 |
| ILAHAIEAI | ASSSTGEGSDYT | 0.71798146 |
| IRSAIREAL | ASSSTGEGSDYT | 0.69925666 |
| ITFAGGG | ASSSTGEGSDYT | 0.5451491 |
| ITGMSGHQEV | ASSSTGEGSDYT | 0.6450361 |

| | | |
|---|---|---|
| IVAGLACTL | ASSSTGEGSDYT | 0.59673834 |
| IVGGGLAGA | ASSSTGEGSDYT | 0.6675811 |
| IVTGVLNI | ASSSTGEGSDYT | 0.5308887 |
| LAQQGCI | ASSSTGEGSDYT | 0.7033133 |
| LCLHVEN | ASSSTGEGSDYT | 0.5980555 |
| LDAISDGS | ASSSTGEGSDYT | 0.35466877 |
| LGGFAAML | ASSSTGEGSDYT | 0.68763274 |
| LGGKNPMI | ASSSTGEGSDYT | 0.6566357 |
| LGPKGVAAIL | ASSSTGEGSDYT | 0.68444 |
| LMVGVSI | ASSSTGEGSDYT | 0.70522505 |
| LVGAAFVL | ASSSTGEGSDYT | 0.5823078 |
| LVLAVAPL | ASSSTGEGSDYT | 0.6302424 |
| MAAIAVIDAA | ASSSTGEGSDYT | 0.67460716 |
| NNGINAELQ | ASSSTGEGSDYT | 0.7097784 |
| RGEPLVLVHG | ASSSTGEGSDYT | 0.63794976 |
| RGGAGGAAGLI | ASSSTGEGSDYT | 0.6676674 |
| RSYRMHPAV | ASSSTGEGSDYT | 0.71030605 |
| SAAHLLAGQLA | ASSSTGEGSDYT | 0.6787225 |
| SAAVGSY | ASSSTGEGSDYT | 0.71367836 |
| SPHAVLHRI | ASSSTGEGSDYT | 0.7164553 |
| SQIAKVL | ASSSTGEGSDYT | 0.6080058 |
| TALGHLVP | ASSSTGEGSDYT | 0.7052485 |
| TAPTTSGAA | ASSSTGEGSDYT | 0.71423054 |
| TDGDDGNGGSPGT | ASSSTGEGSDYT | 0.6511706 |
| TDGLGEN | ASSSTGEGSDYT | 0.66446424 |
| TGGPPLPGQA | ASSSTGEGSDYT | 0.5429495 |
| TIPAITI | ASSSTGEGSDYT | 0.65802217 |
| TIPGFTI | ASSSTGEGSDYT | 0.3855437 |
| TSIASGALA | ASSSTGEGSDYT | 0.6686746 |
| TVTLPGAAI | ASSSTGEGSDYT | 0.7171445 |
| VASGLSAALP | ASSSTGEGSDYT | 0.64284843 |
| VGAGVIGI | ASSSTGEGSDYT | 0.58570707 |
| VGPKRFGV | ASSSTGEGSDYT | 0.52270365 |
| VNVAFPD | ASSSTGEGSDYT | 0.65874004 |
| VSMTNSGVS | ASSSTGEGSDYT | 0.6331436 |
| VTDVLGDLSV | ASSSTGEGSDYT | 0.5339167 |
| VVVVGAGFAGL | ASSSTGEGSDYT | 0.61695504 |
| VYTVGTVV | ASSSTGEGSDYT | 0.70773554 |
| WTTFSGP | ASSSTGEGSDYT | 0.70749784 |
| YDELVEAF | ASSSTGEGSDYT | 0.59417856 |
| YGTGGAGGNGG | ASSSTGEGSDYT | 0.7131548 |
| YTNVMAV | ASSSTGEGSDYT | 0.68734217 |
| FLEQVHQL | ASSSTGEGSDYT | 0.68985724 |
| GLSLLAVAV | ASSSTGEGSDYT | 0.6511152 |
| LEQVHQL | ASSSTGEGSDYT | 0.61427546 |
| LNAAGAI | ASSSTGEGSDYT | 0.69969636 |
| PAVHASL | ASSSTGEGSDYT | 0.60663456 |
| TELEDDA | ASSSTGEGSDYT | 0.6879355 |
| TPPGPPN | ASSSTGEGSDYT | 0.69900525 |
| TPPPPPPP | ASSSTGEGSDYT | 0.6720449 |
| TQSPALI | ASSSGQFSNERLF | 0.6087243 |

| | | |
|---|---|---|
| EFAAAFPQ | ASSSGQFSNERLF | 0.6277482 |
| LLSGNLDVL | ASSSGQFSNERLF | 0.71315706 |
| GPANLGNYNI | ASSSGQFSNERLF | 0.5475137 |
| TLGPTLP | ASSSGQFSNERLF | 0.56345 |
| VYAYSSCCK | ASSSGQFSNERLF | 0.6309796 |
| APAPAPAPNI | ASSSGQFSNERLF | 0.6659051 |
| PGGPVPIP | ASSSGQFSNERLF | 0.70911837 |
| FLTVALEP | ASSSGQFSNERLF | 0.69674104 |
| SQYFGANL | ASSSGQFSNERLF | 0.6496211 |
| GGVKPAA | ASSSGQFSNERLF | 0.69170797 |
| GGTGGAGGAGGW | ASSSGQFSNERLF | 0.6967337 |
| GVGGVAGLGGAGG | ASSSGQFSNERLF | 0.68341756 |
| VLLVLATTI | ASSSGQFSNERLF | 0.69397175 |
| IGIFAAML | ASSSGQFSNERLF | 0.60649824 |
| AVGDLID | ASSSGQFSNERLF | 0.70578074 |
| NISPLPL | ASSSGQFSNERLF | 0.6951378 |
| VTAVFVAI | ASSSGQFSNERLF | 0.70364606 |
| SGDDNDAAL | ASSSGQFSNERLF | 0.705944 |
| AVAALAHS | ASSSGQFSNERLF | 0.6957028 |
| GGVTVMATP | ASSSGQFSNERLF | 0.6635859 |
| AGLAANVRD | ASSSGQFSNERLF | 0.41316122 |
| TNRPDLIDPALLRPG | ASSSGQFSNERLF | 0.5658958 |
| LANLGVTAL | ASSSGQFSNERLF | 0.67804575 |
| TGERIEL | ASSSGQFSNERLF | 0.6701293 |
| NGEELAA | ASSSGQFSNERLF | 0.6937305 |
| VLSLVWG | ASSSGQFSNERLF | 0.6948424 |
| IGYALWL | ASSSGQFSNERLF | 0.5967419 |
| LSLLLAPL | ASSSGQFSNERLF | 0.7056166 |
| TLDEALK | ASSSGQFSNERLF | 0.7067524 |
| TDGAGTAEGG | ASSSGQFSNERLF | 0.4843927 |
| NAPGGAGGNGG | ASSSGQFSNERLF | 0.6175644 |
| VSVFMGMGGNF | ASSSGQFSNERLF | 0.723644 |
| LASYLAAL | ASSSGQFSNERLF | 0.6741135 |
| LVGGTPAL | ASSSGQFSNERLF | 0.62008536 |
| VRNADVL | ASSSGQFSNERLF | 0.65939856 |
| TCSSPFLVSAM | ASSSGQFSNERLF | 0.72355866 |
| VHITDAE | ASSSGQFSNERLF | 0.68675256 |
| EYAAPSF | ASSSGQFSNERLF | 0.72167003 |
| VVDAFHGGL | ASSSGQFSNERLF | 0.42032978 |
| ASVVDPQ | ASSSGQFSNERLF | 0.6424588 |
| SYNEFGGLS | ASSSGQFSNERLF | 0.6080518 |
| DPDGSVY | ASSSGQFSNERLF | 0.40290508 |
| MVSTIHATE | ASSSGQFSNERLF | 0.69670296 |
| LAAGVADPV | ASSSGQFSNERLF | 0.64607805 |
| TKEQIAA | ASSSGQFSNERLF | 0.6945833 |
| SLASPAAL | ASSSGQFSNERLF | 0.6509657 |
| IGLVFYGV | ASSSGQFSNERLF | 0.7065538 |
| FLLDASNL | ASSSGQFSNERLF | 0.6394924 |
| YQVGNFP | ASSSGQFSNERLF | 0.67927825 |
| ESLLSQST | ASSSGQFSNERLF | 0.66061306 |
| LFTVGFAAVG | ASSSGQFSNERLF | 0.68955994 |

| | | |
|---|---|---|
| ATPPTPT | ASSSGQFSNERLF | 0.7118039 |
| TDLATLG | ASSSGQFSNERLF | 0.7075665 |
| SAGGTGR | ASSSGQFSNERLF | 0.5377449 |
| GFSNSGS | ASSSGQFSNERLF | 0.6660475 |
| GGNGGSGGTSV | ASSSGQFSNERLF | 0.6544728 |
| GTSVATGGAGN | ASSSGQFSNERLF | 0.41077468 |
| NGGNGGSGGTS | ASSSGQFSNERLF | 0.44512188 |
| AVVKFPH | ASSSGQFSNERLF | 0.6371019 |
| TPPPPPP | ASSSGQFSNERLF | 0.60969496 |
| AAPAAAAAPA | ASSSGQFSNERLF | 0.69094205 |
| IALNEDL | ASSSGQFSNERLF | 0.68380725 |
| IVNVSSM | ASSSGQFSNERLF | 0.6499609 |
| TGIPPAPR | ASSSGQFSNERLF | 0.69104046 |
| FDLGGGTF | ASSSGQFSNERLF | 0.6815152 |
| FDLGGGTFDVS | ASSSGQFSNERLF | 0.25296447 |
| NEPTAAA | ASSSGQFSNERLF | 0.612815 |
| NEPTAAALA | ASSSGQFSNERLF | 0.55921817 |
| GSIINISSI | ASSSGQFSNERLF | 0.7151171 |
| VAPGNAGTA | ASSSGQFSNERLF | 0.61334866 |
| TGGGASLE | ASSSGQFSNERLF | 0.5967469 |
| VAVYNHYKRI | ASSSGQFSNERLF | 0.66515017 |
| TDPAPATT | ASSSGQFSNERLF | 0.6631732 |
| TNRPDILDPALLRPG | ASSSGQFSNERLF | 0.6396735 |
| LTTTTTS | ASSSGQFSNERLF | 0.59579796 |
| PTGSGKTL | ASSSGQFSNERLF | 0.3364241 |
| TGSGKTL | ASSSGQFSNERLF | 0.7077534 |
| TIGHVDHGK | ASSSGQFSNERLF | 0.5523505 |
| TIGHVDHGKTT | ASSSGQFSNERLF | 0.63295156 |
| GHVDHGKTT | ASSSGQFSNERLF | 0.54638034 |
| AQFEHTLL | ASSSGQFSNERLF | 0.6383287 |
| IVNQDEL | ASSSGQFSNERLF | 0.4422542 |
| TNGSQFF | ASSSGQFSNERLF | 0.71231735 |
| TGKGIVE | ASSSGQFSNERLF | 0.58857554 |
| LISNASDA | ASSSGQFSNERLF | 0.63410807 |
| AELRTLGELTKIAKAHGAQ | ASSSGQFSNERLF | 0.71618724 |
| APNAVAAAEPSW | ASSSGQFSNERLF | 0.67432606 |
| APQVAGAATAVL | ASSSGQFSNERLF | 0.6313568 |
| ARRDPAAAAAILPTDA | ASSSGQFSNERLF | 0.67336375 |
| DAAARDVLAEAGLAEYF | ASSSGQFSNERLF | 0.571798 |
| DDQIVLSAKANLAKVEQ | ASSSGQFSNERLF | 0.63297343 |
| DELLPDTGEDSDEDGH | ASSSGQFSNERLF | 0.66061413 |
| ELPEAPANRATVPQD | ASSSGQFSNERLF | 0.7185099 |
| FGQSKIERVTGIG | ASSSGQFSNERLF | 0.6614762 |
| GAGGTGSTAGGAGGAGGAGGLY | ASSSGQFSNERLF | 0.5831624 |
| GHGGTGAAGGNGGTGGAGGSGI | ASSSGQFSNERLF | 0.67327 |
| GNGGNGGAGGEGGPGGAGGAGA | ASSSGQFSNERLF | 0.7170372 |
| GSTAGNGGNGGNATNGTIAGQP | ASSSGQFSNERLF | 0.70296186 |
| GTGGDGGNGGNGGTGG | ASSSGQFSNERLF | 0.6305044 |
| GVRRPAGIPADAPV | ASSSGQFSNERLF | 0.62780946 |
| GYRQVTDAHLLTLARR | ASSSGQFSNERLF | 0.6221221 |
| IAKMGQDGHDRG | ASSSGQFSNERLF | 0.6357218 |

| | | |
|---|---|---|
| KAEPSIEIQYT | ASSSGQFSNERLF | 0.65347767 |
| KPVRIRMAADEDTVAVT | ASSSGQFSNERLF | 0.6160835 |
| MPPRIAGMRLLVIKPEP | ASSSGQFSNERLF | 0.69974506 |
| MKFYAVALFRSRGVPPTR | ASSSGQFSNERLF | 0.7284168 |
| NAGGFGILGAVAHSP | ASSSGQFSNERLF | 0.66847885 |
| NEGREITLAMAEM | ASSSGQFSNERLF | 0.40133727 |
| NELVEWSHPLGHRWRWE | ASSSGQFSNERLF | 0.6433402 |
| NLGAGGTGGNGGNPG | ASSSGQFSNERLF | 0.5678082 |
| NTEWPKWETVMAADPVD | ASSSGQFSNERLF | 0.61938477 |
| PASAAPAETQP | ASSSGQFSNERLF | 0.6554085 |
| QQSSDGKNGVRAAEITGE | ASSSGQFSNERLF | 0.5218612 |
| RAPDAVTTSWWKE | ASSSGQFSNERLF | 0.66296977 |
| SGFGNVGAHTSGWFNQ | ASSSGQFSNERLF | 0.7152363 |
| SGVAGAVGQAASVGGLKV | ASSSGQFSNERLF | 0.56475717 |
| SNGGAGTGSAGGNGGT | ASSSGQFSNERLF | 0.6776695 |
| TGGAGGEGGAAG | ASSSGQFSNERLF | 0.71110797 |
| TGGDGGAAGAGGAPGHG | ASSSGQFSNERLF | 0.6914927 |
| TMPPLLAAIANAGRHGV | ASSSGQFSNERLF | 0.7084706 |
| TSSTAGNSGGA | ASSSGQFSNERLF | 0.69356513 |
| VPATVPVGILC | ASSSGQFSNERLF | 0.6921423 |
| VQRPDSGGVVLTGRIS | ASSSGQFSNERLF | 0.6769752 |
| WTPNPVPATKGVPAGLVQAT | ASSSGQFSNERLF | 0.6674197 |
| YALKRALAKFSDRFDVVII | ASSSGQFSNERLF | 0.65105015 |
| YLIGSVIV | ASSSGQFSNERLF | 0.6438387 |
| YGQLVAVID | ASSSGQFSNERLF | 0.5959648 |
| VTVGGMTLPA | ASSSGQFSNERLF | 0.69839394 |
| VLALPAV | ASSSGQFSNERLF | 0.64603853 |
| VIPPTLNYET | ASSSGQFSNERLF | 0.7044364 |
| VGMGGMHPSSG | ASSSGQFSNERLF | 0.525206 |
| VGLAGGAATA | ASSSGQFSNERLF | 0.63294923 |
| VGGAGGAGGSGP | ASSSGQFSNERLF | 0.69665545 |
| VFAAFMV | ASSSGQFSNERLF | 0.27613732 |
| TLYDKLR | ASSSGQFSNERLF | 0.41821787 |
| TLAVFTVL | ASSSGQFSNERLF | 0.66414255 |
| TGSVQPARV | ASSSGQFSNERLF | 0.70130885 |
| TGGGAGSGGT | ASSSGQFSNERLF | 0.29674944 |
| SHLETSL | ASSSGQFSNERLF | 0.6425903 |
| SFTTGMT | ASSSGQFSNERLF | 0.5048521 |
| QVEVAKRMGVRQPTVS | ASSSGQFSNERLF | 0.701417 |
| QGSGGPALAM | ASSSGQFSNERLF | 0.685878 |
| MLAPSNP | ASSSGQFSNERLF | 0.4101674 |
| LVNNAAI | ASSSGQFSNERLF | 0.7031517 |
| LTGEINTLA | ASSSGQFSNERLF | 0.48960742 |
| LSIFGPV | ASSSGQFSNERLF | 0.63760597 |
| LLALLANL | ASSSGQFSNERLF | 0.6993707 |
| LISNASDALDKLR | ASSSGQFSNERLF | 0.71296906 |
| LINEAVP | ASSSGQFSNERLF | 0.57170606 |
| LGYAFSGL | ASSSGQFSNERLF | 0.6948598 |
| LDFVAVAL | ASSSGQFSNERLF | 0.6139078 |
| LAQPGAE | ASSSGQFSNERLF | 0.71106964 |
| ISNLGAVTGF | ASSSGQFSNERLF | 0.6166109 |

| | | |
|---|---|---|
| HQEDPGRI | ASSSGQFSNERLF | 0.5231974 |
| GTPGEPTP | ASSSGQFSNERLF | 0.7060257 |
| GFFNSGS | ASSSGQFSNERLF | 0.6670667 |
| GAGGGGLTAGAN | ASSSGQFSNERLF | 0.67980176 |
| FVDALVNM | ASSSGQFSNERLF | 0.69002676 |
| FLTVAPEL | ASSSGQFSNERLF | 0.62838304 |
| FETLTAL | ASSSGQFSNERLF | 0.6362804 |
| DVINPIEL | ASSSGQFSNERLF | 0.70483387 |
| APVSGAPV | ASSSGQFSNERLF | 0.7019285 |
| ALVGGLAW | ASSSGQFSNERLF | 0.625467 |
| ALAAAAATAI | ASSSGQFSNERLF | 0.7105879 |
| AFGLGLLL | ASSSGQFSNERLF | 0.67322075 |
| AFFAAASL | ASSSGQFSNERLF | 0.6229965 |
| AAAALPAFSPP | ASSSGQFSNERLF | 0.7086203 |
| VVLAPAL | ASSSGQFSNERLF | 0.6634779 |
| TIGHVDHGKTTL | ASSSGQFSNERLF | 0.696226 |
| LISNASDALDK | ASSSGQFSNERLF | 0.6964274 |
| LISNASDAL | ASSSGQFSNERLF | 0.7057494 |
| ISNASDALDK | ASSSGQFSNERLF | 0.47802177 |
| NQAETLVYQTE | ASSSGQFSNERLF | 0.6566633 |
| TGGTGGTAGAG | ASSSGQFSNERLF | 0.650138 |
| TGSTAGGAGGA | ASSSGQFSNERLF | 0.6976637 |
| DPLAVPVAAGPV | ASSSGQFSNERLF | 0.6553713 |
| GGTGGTPGNGGN | ASSSGQFSNERLF | 0.5805266 |
| PAGLAGVDPATL | ASSSGQFSNERLF | 0.68816036 |
| GNNLVVEAPHAGASV | ASSSGQFSNERLF | 0.7128805 |
| KVTRIMSASTLEQPA | ASSSGQFSNERLF | 0.6829739 |
| LGFALPIALPRRTVV | ASSSGQFSNERLF | 0.6897486 |
| TRLGNLTARKLVVLV | ASSSGQFSNERLF | 0.71255654 |
| WSQQRRLIYFPSAGP | ASSSGQFSNERLF | 0.59962046 |
| GPNGNPDPMAAAVDIR | ASSSGQFSNERLF | 0.6761339 |
| TQRCVSVADLRAAASR | ASSSGQFSNERLF | 0.69789416 |
| QFVLDVAEHPVEVLTETV | ASSSGQFSNERLF | 0.7077403 |
| GGAGGTGGAGGNGGTQAGDG | ASSSGQFSNERLF | 0.6584943 |
| PLLATLRAIAVDVDAKPAQV | ASSSGQFSNERLF | 0.7128459 |
| APKFRTGGYMIDFWGVGYQVAK | ASSSGQFSNERLF | 0.58472586 |
| GAGGAGGAGGAAGGPGGTGGTG | ASSSGQFSNERLF | 0.7044141 |
| IGALGLLATDVAPAGSVAFGVA | ASSSGQFSNERLF | 0.5819967 |
| AGGAGGIGGNGGAGGAGGVLMG | ASSSGQFSNERLF | 0.662619 |
| ADELIGLL | ASSSGQFSNERLF | 0.6098335 |
| ADRALKA | ASSSGQFSNERLF | 0.68545544 |
| AGAVGQVGGAGGA | ASSSGQFSNERLF | 0.6904993 |
| AGGGGGLGGAP | ASSSGQFSNERLF | 0.70844185 |
| AGGLLYGNGGAG | ASSSGQFSNERLF | 0.50761133 |
| AGLESIGL | ASSSGQFSNERLF | 0.69404864 |
| AGLKTGII | ASSSGQFSNERLF | 0.7115507 |
| ASLAALLL | ASSSGQFSNERLF | 0.7236221 |
| ASVINISSI | ASSSGQFSNERLF | 0.65195423 |
| ATPPGPA | ASSSGQFSNERLF | 0.6768539 |
| ATVASALV | ASSSGQFSNERLF | 0.6419718 |
| ATVASLIL | ASSSGQFSNERLF | 0.666193 |

| | | |
|---|---|---|
| AVNGPVLIRH | ASSSGQFSNERLF | 0.5563669 |
| DGAAGAVGI | ASSSGQFSNERLF | 0.5789038 |
| DPLLHGI | ASSSGQFSNERLF | 0.6222686 |
| ELGAALGAL | ASSSGQFSNERLF | 0.602663 |
| ELGRTQAV | ASSSGQFSNERLF | 0.695327 |
| FVAGGYLS | ASSSGQFSNERLF | 0.70816934 |
| GASGGAGGAGGW | ASSSGQFSNERLF | 0.70048404 |
| GGAGAGGDGGAGGI | ASSSGQFSNERLF | 0.69934976 |
| GTGHNGSQPGMGGNG | ASSSGQFSNERLF | 0.7003807 |
| GYAAASASAS | ASSSGQFSNERLF | 0.711567 |
| HQVDVAAVQQG | ASSSGQFSNERLF | 0.63813514 |
| IGAAIAHT | ASSSGQFSNERLF | 0.6296859 |
| IGMIEVAQL | ASSSGQFSNERLF | 0.54715514 |
| IRLAPEHTSAMD | ASSSGQFSNERLF | 0.6405436 |
| IYNVSVL | ASSSGQFSNERLF | 0.6419677 |
| KQGFLGM | ASSSGQFSNERLF | 0.673863 |
| LESVITQL | ASSSGQFSNERLF | 0.27110833 |
| LGLGALVVV | ASSSGQFSNERLF | 0.65404564 |
| LIGNGGNGGSGGI | ASSSGQFSNERLF | 0.71865535 |
| LNAINAPTL | ASSSGQFSNERLF | 0.57304275 |
| LQLLNTGV | ASSSGQFSNERLF | 0.6865924 |
| LQTSIEF | ASSSGQFSNERLF | 0.65976334 |
| MSFAVSEL | ASSSGQFSNERLF | 0.61767626 |
| NTGALGL | ASSSGQFSNERLF | 0.6844598 |
| PPGAGGAGGAGTT | ASSSGQFSNERLF | 0.6179586 |
| QLVGLGI | ASSSGQFSNERLF | 0.70335466 |
| RFLEQVHQL | ASSSGQFSNERLF | 0.7085986 |
| RGVGYVLR | ASSSGQFSNERLF | 0.6813701 |
| SATGVFAGVI | ASSSGQFSNERLF | 0.6496947 |
| SLPAITLP | ASSSGQFSNERLF | 0.6741075 |
| STALAFTV | ASSSGQFSNERLF | 0.43191615 |
| SVLTVGLGK | ASSSGQFSNERLF | 0.6642648 |
| TALAGAGSGI | ASSSGQFSNERLF | 0.46208465 |
| TGLPFTGI | ASSSGQFSNERLF | 0.6300019 |
| VAEHVTV | ASSSGQFSNERLF | 0.57524157 |
| VDIVLNSL | ASSSGQFSNERLF | 0.68505734 |
| VLTGGGAVL | ASSSGQFSNERLF | 0.68177676 |
| VNVGLAYGI | ASSSGQFSNERLF | 0.67942 |
| VSSAAVGAV | ASSSGQFSNERLF | 0.48397046 |
| VTSVYEAGV | ASSSGQFSNERLF | 0.69768095 |
| WVALDRGA | ASSSGQFSNERLF | 0.67661583 |
| YLVGSVVI | ASSSGQFSNERLF | 0.58505845 |
| GPPGTGKTLL | ASSSGQFSNERLF | 0.6844242 |
| GVGMTKF | ASSSGQFSNERLF | 0.67391235 |
| IGIGGSDL | ASSSGQFSNERLF | 0.6149133 |
| IGTLLAEL | ASSSGQFSNERLF | 0.7072078 |
| IYTSGTT | ASSSGQFSNERLF | 0.68539923 |
| LSFPAGV | ASSSGQFSNERLF | 0.4014203 |
| LYTSGTT | ASSSGQFSNERLF | 0.609559 |
| VGAFPFGF | ASSSGQFSNERLF | 0.6367621 |
| VGAGTVEF | ASSSGQFSNERLF | 0.60957414 |

| | | |
|---|---|---|
| VGKSTLLL | ASSSGQFSNERLF | 0.32683063 |
| AALAHAGGGIV | ASSSGQFSNERLF | 0.6909565 |
| AALVAVSLL | ASSSGQFSNERLF | 0.6285604 |
| AAVSRAVAAL | ASSSGQFSNERLF | 0.5411756 |
| AGGSSGIGGF | ASSSGQFSNERLF | 0.6454934 |
| AGLTVANANG | ASSSGQFSNERLF | 0.70150656 |
| AGLVGAVL | ASSSGQFSNERLF | 0.5977103 |
| ALSQAFTV | ASSSGQFSNERLF | 0.7005086 |
| ALTFSVSL | ASSSGQFSNERLF | 0.676085 |
| ALTGAALVAL | ASSSGQFSNERLF | 0.70279944 |
| ASFSIGAGVL | ASSSGQFSNERLF | 0.7102073 |
| DGSVYKI | ASSSGQFSNERLF | 0.70267 |
| EQLAEIV | ASSSGQFSNERLF | 0.66235965 |
| FAVSELTVM | ASSSGQFSNERLF | 0.6885828 |
| GAAPLAAGL | ASSSGQFSNERLF | 0.65774155 |
| GADGGDGGSGGAGG | ASSSGQFSNERLF | 0.59259206 |
| GFGLDTGGAGG | ASSSGQFSNERLF | 0.58834314 |
| GLASGGLL | ASSSGQFSNERLF | 0.5762189 |
| GRIAGAAVLL | ASSSGQFSNERLF | 0.5347284 |
| GSAGSLNTSF | ASSSGQFSNERLF | 0.64809126 |
| GTGQIGI | ASSSGQFSNERLF | 0.7146689 |
| GTLIALIL | ASSSGQFSNERLF | 0.61542594 |
| IAATMGFVVG | ASSSGQFSNERLF | 0.53519917 |
| IFAQALL | ASSSGQFSNERLF | 0.63234717 |
| IGAAVALGF | ASSSGQFSNERLF | 0.54930764 |
| IGLTPLAL | ASSSGQFSNERLF | 0.69213116 |
| ILGSNTSTL | ASSSGQFSNERLF | 0.6810407 |
| IVFSLTVL | ASSSGQFSNERLF | 0.7294347 |
| LASKATGF | ASSSGQFSNERLF | 0.707623 |
| LASVALIL | ASSSGQFSNERLF | 0.64472675 |
| LFAVGLTL | ASSSGQFSNERLF | 0.6950054 |
| LFNGGIGTYI | ASSSGQFSNERLF | 0.61538535 |
| LTLGAGII | ASSSGQFSNERLF | 0.6544167 |
| LVAAALGDI | ASSSGQFSNERLF | 0.6689953 |
| NTAFVQLGI | ASSSGQFSNERLF | 0.66941005 |
| QLFSGAEL | ASSSGQFSNERLF | 0.68375903 |
| SAATFITL | ASSSGQFSNERLF | 0.65184367 |
| SNYGLANLGV | ASSSGQFSNERLF | 0.36514792 |
| SYGGAYAVM | ASSSGQFSNERLF | 0.69901806 |
| TTGVGGLTLG | ASSSGQFSNERLF | 0.56239676 |
| TTQTIYI | ASSSGQFSNERLF | 0.5490447 |
| VAAGAGLGAVI | ASSSGQFSNERLF | 0.6134976 |
| VAAVVTLL | ASSSGQFSNERLF | 0.70719373 |
| VGFLGGLI | ASSSGQFSNERLF | 0.70966804 |
| VGVIPILL | ASSSGQFSNERLF | 0.60646635 |
| VIFASSLI | ASSSGQFSNERLF | 0.6018996 |
| VKGGDGGAAAG | ASSSGQFSNERLF | 0.6500082 |
| VTKTFTNI | ASSSGQFSNERLF | 0.53594685 |
| VVAAAILL | ASSSGQFSNERLF | 0.6866583 |
| YAALDVEL | ASSSGQFSNERLF | 0.64169705 |
| LASVALLL | ASSSGQFSNERLF | 0.6535802 |

| | | |
|---|---|---|
| LDAVHRV | ASSSGQFSNERLF | 0.58892286 |
| QGLSSSIF | ASSSGQFSNERLF | 0.6741247 |
| RGTFANIRL | ASSSGQFSNERLF | 0.6461868 |
| RQLWWGHRI | ASSSGQFSNERLF | 0.6960321 |
| YGPPGTGKTLL | ASSSGQFSNERLF | 0.27263695 |
| AALVGAPANVA | ASSSGQFSNERLF | 0.69453585 |
| AATFRRTE | ASSSGQFSNERLF | 0.5838057 |
| AGALGGGI | ASSSGQFSNERLF | 0.6253219 |
| AGGAGGAGGIGAG | ASSSGQFSNERLF | 0.4729478 |
| AGSGALGL | ASSSGQFSNERLF | 0.6751323 |
| AQIADQI | ASSSGQFSNERLF | 0.6591762 |
| ASAVAVTYL | ASSSGQFSNERLF | 0.69664586 |
| ASTVLGGTGG | ASSSGQFSNERLF | 0.6132421 |
| AVGGEPAK | ASSSGQFSNERLF | 0.6920484 |
| CEQGTFGGTC | ASSSGQFSNERLF | 0.69118935 |
| DEGKAPAG | ASSSGQFSNERLF | 0.6238748 |
| DTVVFPAL | ASSSGQFSNERLF | 0.4537383 |
| FEGITMAG | ASSSGQFSNERLF | 0.6979754 |
| FPLVVQD | ASSSGQFSNERLF | 0.6529287 |
| GAFTTPQ | ASSSGQFSNERLF | 0.69230795 |
| GFGGFGTAGGVGGA | ASSSGQFSNERLF | 0.6586529 |
| GFLAQFL | ASSSGQFSNERLF | 0.5515 |
| GGAGGAGGNAGVA | ASSSGQFSNERLF | 0.6940729 |
| GGVGTAGAGGAGGAGG | ASSSGQFSNERLF | 0.6930102 |
| GNGGGGDAGNAGSGGN | ASSSGQFSNERLF | 0.5701356 |
| GTGGAGGAGNDAGS | ASSSGQFSNERLF | 0.61104035 |
| GTTNTGWFNS | ASSSGQFSNERLF | 0.6523832 |
| IFMGPLFTL | ASSSGQFSNERLF | 0.44538796 |
| ILAHAIEAI | ASSSGQFSNERLF | 0.70367444 |
| IRSAIREAL | ASSSGQFSNERLF | 0.6123423 |
| ITFAGGG | ASSSGQFSNERLF | 0.6655036 |
| ITGMSGHQEV | ASSSGQFSNERLF | 0.6709137 |
| IVAGLACTL | ASSSGQFSNERLF | 0.66583633 |
| IVGGGLAGA | ASSSGQFSNERLF | 0.6705587 |
| IVTGVLNI | ASSSGQFSNERLF | 0.6745585 |
| LAQQGCI | ASSSGQFSNERLF | 0.673277 |
| LCLHVEN | ASSSGQFSNERLF | 0.6825176 |
| LDAISDGS | ASSSGQFSNERLF | 0.64049953 |
| LGGFAAML | ASSSGQFSNERLF | 0.6593968 |
| LGGKNPMI | ASSSGQFSNERLF | 0.7106397 |
| LGPKGVAAIL | ASSSGQFSNERLF | 0.58028626 |
| LMVGVSI | ASSSGQFSNERLF | 0.647238 |
| LVGAAFVL | ASSSGQFSNERLF | 0.6878631 |
| LVLAVAPL | ASSSGQFSNERLF | 0.6266263 |
| MAAIAVIDAA | ASSSGQFSNERLF | 0.70292246 |
| NNGINAELQ | ASSSGQFSNERLF | 0.68505096 |
| RGEPLVLVHG | ASSSGQFSNERLF | 0.67791605 |
| RGGAGGAAGLI | ASSSGQFSNERLF | 0.6429343 |
| RSYRMHPAV | ASSSGQFSNERLF | 0.71116936 |
| SAAHLLAGQLA | ASSSGQFSNERLF | 0.69873524 |
| SAAVGSY | ASSSGQFSNERLF | 0.6710383 |

| | | |
|---|---|---|
| SPHAVLHRI | ASSSGQFSNERLF | 0.694343 |
| SQIAKVL | ASSSGQFSNERLF | 0.6075266 |
| TALGHLVP | ASSSGQFSNERLF | 0.6967183 |
| TAPTTSGAA | ASSSGQFSNERLF | 0.68261707 |
| TDGDDGNGGSPGT | ASSSGQFSNERLF | 0.53635323 |
| TDGLGEN | ASSSGQFSNERLF | 0.6376713 |
| TGGPPLPGQA | ASSSGQFSNERLF | 0.71109843 |
| TIPAITI | ASSSGQFSNERLF | 0.70825124 |
| TIPGFTI | ASSSGQFSNERLF | 0.58286464 |
| TSIASGALA | ASSSGQFSNERLF | 0.64543104 |
| TVTLPGAAI | ASSSGQFSNERLF | 0.58359396 |
| VASGLSAALP | ASSSGQFSNERLF | 0.6077484 |
| VGAGVIGI | ASSSGQFSNERLF | 0.5955604 |
| VGPKRFGV | ASSSGQFSNERLF | 0.68337244 |
| VNVAFPD | ASSSGQFSNERLF | 0.6115223 |
| VSMTNSGVS | ASSSGQFSNERLF | 0.6946937 |
| VTDVLGDLSV | ASSSGQFSNERLF | 0.6579398 |
| VVVVGAGFAGL | ASSSGQFSNERLF | 0.70842826 |
| VYTVGTVV | ASSSGQFSNERLF | 0.58561635 |
| WTTFSGP | ASSSGQFSNERLF | 0.6796053 |
| YDELVEAF | ASSSGQFSNERLF | 0.51526 |
| YGTGGAGGNGG | ASSSGQFSNERLF | 0.5307147 |
| YTNVMAV | ASSSGQFSNERLF | 0.6269374 |
| FLEQVHQL | ASSSGQFSNERLF | 0.66529447 |
| GLSLLAVAV | ASSSGQFSNERLF | 0.33746597 |
| LEQVHQL | ASSSGQFSNERLF | 0.71462893 |
| LNAAGAI | ASSSGQFSNERLF | 0.6560337 |
| PAVHASL | ASSSGQFSNERLF | 0.6984437 |
| TELEDDA | ASSSGQFSNERLF | 0.65829456 |
| TPPGPPN | ASSSGQFSNERLF | 0.6582169 |
| TPPPPPPP | ASSSGQFSNERLF | 0.62966454 |
| TQSPALI | ASLRDRYEQY | 0.69009376 |
| EFAAAFPQ | ASLRDRYEQY | 0.5709269 |
| LLSGNLDVL | ASLRDRYEQY | 0.42278007 |
| GPANLGNYNI | ASLRDRYEQY | 0.62294865 |
| TLGPTLP | ASLRDRYEQY | 0.5298463 |
| VYAYSSCCK | ASLRDRYEQY | 0.70083356 |
| APAPAPAPNI | ASLRDRYEQY | 0.7046418 |
| PGGPVPIP | ASLRDRYEQY | 0.66753787 |
| FLTVALEP | ASLRDRYEQY | 0.69602287 |
| SQYFGANL | ASLRDRYEQY | 0.30707714 |
| GGVKPAA | ASLRDRYEQY | 0.6704716 |
| GGTGGAGGAGGW | ASLRDRYEQY | 0.711614 |
| GVGGVAGLGGAGG | ASLRDRYEQY | 0.6753552 |
| VLLVLATTI | ASLRDRYEQY | 0.63468534 |
| IGIFAAML | ASLRDRYEQY | 0.71228516 |
| AVGDLID | ASLRDRYEQY | 0.63957024 |
| NISPLPL | ASLRDRYEQY | 0.59825414 |
| VTAVFVAI | ASLRDRYEQY | 0.6928136 |
| SGDDNDAAL | ASLRDRYEQY | 0.69378614 |
| AVAALAHS | ASLRDRYEQY | 0.6758733 |

| | | |
|---|---|---|
| GGVTVMATP | ASLRDRYEQY | 0.58530694 |
| AGLAANVRD | ASLRDRYEQY | 0.68186456 |
| TNRPDLIDPALLRPG | ASLRDRYEQY | 0.7221451 |
| LANLGVTAL | ASLRDRYEQY | 0.62852025 |
| TGERIEL | ASLRDRYEQY | 0.6777158 |
| NGEELAA | ASLRDRYEQY | 0.6885591 |
| VLSLVWG | ASLRDRYEQY | 0.6161674 |
| IGYALWL | ASLRDRYEQY | 0.57310957 |
| LSLLLAPL | ASLRDRYEQY | 0.60692215 |
| TLDEALK | ASLRDRYEQY | 0.6968775 |
| TDGAGTAEGG | ASLRDRYEQY | 0.5678934 |
| NAPGGAGGNGG | ASLRDRYEQY | 0.6537247 |
| VSVFMGMGGNF | ASLRDRYEQY | 0.65581274 |
| LASYLAAL | ASLRDRYEQY | 0.7216455 |
| LVGGTPAL | ASLRDRYEQY | 0.6872201 |
| VRNADVL | ASLRDRYEQY | 0.6991168 |
| TCSSPFLVSAM | ASLRDRYEQY | 0.6022402 |
| VHITDAE | ASLRDRYEQY | 0.6767618 |
| EYAAPSF | ASLRDRYEQY | 0.67676425 |
| VVDAFHGGL | ASLRDRYEQY | 0.38784575 |
| ASVVDPQ | ASLRDRYEQY | 0.71483624 |
| SYNEFGGLS | ASLRDRYEQY | 0.6103209 |
| DPDGSVY | ASLRDRYEQY | 0.6245922 |
| MVSTIHATE | ASLRDRYEQY | 0.70280904 |
| LAAGVADPV | ASLRDRYEQY | 0.72335756 |
| TKEQIAA | ASLRDRYEQY | 0.6436792 |
| SLASPAAL | ASLRDRYEQY | 0.69376004 |
| IGLVFYGV | ASLRDRYEQY | 0.62443197 |
| FLLDASNL | ASLRDRYEQY | 0.6923395 |
| YQVGNFP | ASLRDRYEQY | 0.70151055 |
| ESLLSQST | ASLRDRYEQY | 0.60393494 |
| LFTVGFAAVG | ASLRDRYEQY | 0.7104138 |
| ATPPTPT | ASLRDRYEQY | 0.6936196 |
| TDLATLG | ASLRDRYEQY | 0.48771876 |
| SAGGTGR | ASLRDRYEQY | 0.6564963 |
| GFSNSGS | ASLRDRYEQY | 0.6953227 |
| GGNGGSGGTSV | ASLRDRYEQY | 0.6970663 |
| GTSVATGGAGN | ASLRDRYEQY | 0.7074909 |
| NGGNGGSGGTS | ASLRDRYEQY | 0.6879322 |
| AVVKFPH | ASLRDRYEQY | 0.5857192 |
| TPPPPPP | ASLRDRYEQY | 0.7088301 |
| AAPAAAAAPA | ASLRDRYEQY | 0.71560895 |
| IALNEDL | ASLRDRYEQY | 0.68922997 |
| IVNVSSM | ASLRDRYEQY | 0.61703193 |
| TGIPPAPR | ASLRDRYEQY | 0.6136984 |
| FDLGGGTF | ASLRDRYEQY | 0.6885653 |
| FDLGGGTFDVS | ASLRDRYEQY | 0.6931421 |
| NEPTAAA | ASLRDRYEQY | 0.5569529 |
| NEPTAAALA | ASLRDRYEQY | 0.664345 |
| GSIINISSI | ASLRDRYEQY | 0.6775987 |
| VAPGNAGTA | ASLRDRYEQY | 0.46485662 |

| | | |
|---|---|---|
| TGGGASLE | ASLRDRYEQY | 0.6865475 |
| VAVYNHYKRI | ASLRDRYEQY | 0.6981552 |
| TDPAPATT | ASLRDRYEQY | 0.6867533 |
| TNRPDILDPALLRPG | ASLRDRYEQY | 0.6171139 |
| LTTTTTS | ASLRDRYEQY | 0.7101242 |
| PTGSGKTL | ASLRDRYEQY | 0.70947444 |
| TGSGKTL | ASLRDRYEQY | 0.71715367 |
| TIGHVDHGK | ASLRDRYEQY | 0.57678276 |
| TIGHVDHGKTT | ASLRDRYEQY | 0.38234973 |
| GHVDHGKTT | ASLRDRYEQY | 0.66394883 |
| AQFEHTLL | ASLRDRYEQY | 0.697801 |
| IVNQDEL | ASLRDRYEQY | 0.67860013 |
| TNGSQFF | ASLRDRYEQY | 0.6473768 |
| TGKGIVE | ASLRDRYEQY | 0.6673639 |
| LISNASDA | ASLRDRYEQY | 0.70307934 |
| AELRTLGELTKIAKAHGAQ | ASLRDRYEQY | 0.6007836 |
| APNAVAAAEPSW | ASLRDRYEQY | 0.51301855 |
| APQVAGAATAVL | ASLRDRYEQY | 0.680343 |
| ARRDPAAAAAILPTDA | ASLRDRYEQY | 0.6873623 |
| DAAARDVLAEAGLAEYF | ASLRDRYEQY | 0.6179543 |
| DDQIVLSAKANLAKVEQ | ASLRDRYEQY | 0.6309936 |
| DELLPDTGEDSDEDGH | ASLRDRYEQY | 0.67304367 |
| ELPEAPANRATVPQD | ASLRDRYEQY | 0.65381235 |
| FGQSKIERVTGIG | ASLRDRYEQY | 0.6589968 |
| GAGGTGSTAGGAGGAGGAGGLY | ASLRDRYEQY | 0.55093783 |
| GHGGTGAAGGNGGTGGAGGSGI | ASLRDRYEQY | 0.70454264 |
| GNGGNGGAGGEGGPGGAGGAGA | ASLRDRYEQY | 0.5787474 |
| GSTAGNGGNGGNATNGTIAGQP | ASLRDRYEQY | 0.6956929 |
| GTGGDGGNGGNGGTGG | ASLRDRYEQY | 0.68311936 |
| GVRRPAGIPADAPV | ASLRDRYEQY | 0.68325007 |
| GYRQVTDAHLLTLARR | ASLRDRYEQY | 0.6164222 |
| IAKMGQDGHDRG | ASLRDRYEQY | 0.58173674 |
| KAEPSIEIQYT | ASLRDRYEQY | 0.5079937 |
| KPVRIRMAADEDTVAVT | ASLRDRYEQY | 0.56654304 |
| MPPRIAGMRLLVIKPEP | ASLRDRYEQY | 0.7077728 |
| MKFYAVALFRSRGVPPTR | ASLRDRYEQY | 0.58289504 |
| NAGGFGILGAVAHSP | ASLRDRYEQY | 0.704918 |
| NEGREITLAMAEM | ASLRDRYEQY | 0.68582165 |
| NELVEWSHPLGHRWRWE | ASLRDRYEQY | 0.6367865 |
| NLGAGGTGGNGGNPG | ASLRDRYEQY | 0.69020057 |
| NTEWPKWETVMAADPVD | ASLRDRYEQY | 0.60513604 |
| PASAAPAETQP | ASLRDRYEQY | 0.6240173 |
| QQSSDGKNGVRAAEITGE | ASLRDRYEQY | 0.6321253 |
| RAPDAVTTSWWKE | ASLRDRYEQY | 0.701125 |
| SGFGNVGAHTSGWFNQ | ASLRDRYEQY | 0.55693585 |
| SGVAGAVGQAASVGGLKV | ASLRDRYEQY | 0.6882453 |
| SNGGAGTGSAGGNGGT | ASLRDRYEQY | 0.67557216 |
| TGGAGGEGGAAG | ASLRDRYEQY | 0.576493 |
| TGGDGGAAGAGGAPGHG | ASLRDRYEQY | 0.6828754 |
| TMPPLLAAIANAGRHGV | ASLRDRYEQY | 0.71084154 |
| TSSTAGNSGGA | ASLRDRYEQY | 0.69144964 |

| | | |
|---|---|---|
| VPATVPVGILC | ASLRDRYEQY | 0.5501953 |
| VQRPDSGGVVLTGRIS | ASLRDRYEQY | 0.68853664 |
| WTPNPVPATKGVPAGLVQAT | ASLRDRYEQY | 0.59927183 |
| YALKRALAKFSDRFDVVII | ASLRDRYEQY | 0.669363 |
| YLIGSVIV | ASLRDRYEQY | 0.6801806 |
| YGQLVAVID | ASLRDRYEQY | 0.6522263 |
| VTVGGMTLPA | ASLRDRYEQY | 0.6988915 |
| VLALPAV | ASLRDRYEQY | 0.48953974 |
| VIPPTLNYET | ASLRDRYEQY | 0.5856274 |
| VGMGGMHPSSG | ASLRDRYEQY | 0.6392559 |
| VGLAGGAATA | ASLRDRYEQY | 0.6878882 |
| VGGAGGAGGSGP | ASLRDRYEQY | 0.6996261 |
| VFAAFMV | ASLRDRYEQY | 0.587379 |
| TLYDKLR | ASLRDRYEQY | 0.6615689 |
| TLAVFTVL | ASLRDRYEQY | 0.62200594 |
| TGSVQPARV | ASLRDRYEQY | 0.5507353 |
| TGGGAGSGGT | ASLRDRYEQY | 0.6792097 |
| SHLETSL | ASLRDRYEQY | 0.6595543 |
| SFTTGMT | ASLRDRYEQY | 0.574684 |
| QVEVAKRMGVRQPTVS | ASLRDRYEQY | 0.690004 |
| QGSGGPALAM | ASLRDRYEQY | 0.6573537 |
| MLAPSNP | ASLRDRYEQY | 0.69690883 |
| LVNNAAI | ASLRDRYEQY | 0.7072712 |
| LTGEINTLA | ASLRDRYEQY | 0.58073765 |
| LSIFGPV | ASLRDRYEQY | 0.6908169 |
| LLALLANL | ASLRDRYEQY | 0.6607535 |
| LISNASDALDKLR | ASLRDRYEQY | 0.6858034 |
| LINEAVP | ASLRDRYEQY | 0.6700433 |
| LGYAFSGL | ASLRDRYEQY | 0.5770377 |
| LDFVAVAL | ASLRDRYEQY | 0.70054364 |
| LAQPGAE | ASLRDRYEQY | 0.70349085 |
| ISNLGAVTGF | ASLRDRYEQY | 0.67441744 |
| HQEDPGRI | ASLRDRYEQY | 0.6882402 |
| GTPGEPTP | ASLRDRYEQY | 0.69861424 |
| GFFNSGS | ASLRDRYEQY | 0.7000081 |
| GAGGGGLTAGAN | ASLRDRYEQY | 0.61006296 |
| FVDALVNM | ASLRDRYEQY | 0.5040767 |
| FLTVAPEL | ASLRDRYEQY | 0.6797455 |
| FETLTAL | ASLRDRYEQY | 0.64219683 |
| DVINPIEL | ASLRDRYEQY | 0.6523646 |
| APVSGAPV | ASLRDRYEQY | 0.497622 |
| ALVGGLAW | ASLRDRYEQY | 0.7046324 |
| ALAAAAATAI | ASLRDRYEQY | 0.72059655 |
| AFGLGLLL | ASLRDRYEQY | 0.6504362 |
| AFFAAASL | ASLRDRYEQY | 0.7170281 |
| AAAALPAFSPP | ASLRDRYEQY | 0.52947086 |
| VVLAPAL | ASLRDRYEQY | 0.7004168 |
| TIGHVDHGKTTL | ASLRDRYEQY | 0.7120007 |
| LISNASDALDK | ASLRDRYEQY | 0.5689695 |
| LISNASDAL | ASLRDRYEQY | 0.70661664 |
| ISNASDALDK | ASLRDRYEQY | 0.56444854 |

| | | |
|---|---|---|
| NQAETLVYQTE | ASLRDRYEQY | 0.62000406 |
| TGGTGGTAGAG | ASLRDRYEQY | 0.7124084 |
| TGSTAGGAGGA | ASLRDRYEQY | 0.6626092 |
| DPLAVPVAAGPV | ASLRDRYEQY | 0.6866565 |
| GGTGGTPGNGGN | ASLRDRYEQY | 0.6862528 |
| PAGLAGVDPATL | ASLRDRYEQY | 0.6260266 |
| GNNLVVEAPHAGASV | ASLRDRYEQY | 0.6871342 |
| KVTRIMSASTLEQPA | ASLRDRYEQY | 0.70330733 |
| LGFALPIALPRRTVV | ASLRDRYEQY | 0.62815917 |
| TRLGNLTARKLVVLV | ASLRDRYEQY | 0.71788585 |
| WSQQRRLIYFPSAGP | ASLRDRYEQY | 0.60717815 |
| GPNGNPDPMAAAVDIR | ASLRDRYEQY | 0.71227205 |
| TQRCVSVADLRAAASR | ASLRDRYEQY | 0.6837107 |
| QFVLDVAEHPVEVLTETV | ASLRDRYEQY | 0.64653146 |
| GGAGGTGGAGGNGGTQAGDG | ASLRDRYEQY | 0.65396506 |
| PLLATLRAIAVDVDAKPAQV | ASLRDRYEQY | 0.6989028 |
| APKFRTGGYMIDFWGVGYQVAK | ASLRDRYEQY | 0.64653695 |
| GAGGAGGAGGAAGGPGGTGGTG | ASLRDRYEQY | 0.69171447 |
| IGALGLLATDVAPAGSVAFGVA | ASLRDRYEQY | 0.7142267 |
| AGGAGGIGGNGGAGGAGGVLMG | ASLRDRYEQY | 0.40028203 |
| ADELIGLL | ASLRDRYEQY | 0.8339101 |
| ADRALKA | ASLRDRYEQY | 0.79606175 |
| AGAVGQVGGAGGA | ASLRDRYEQY | 0.77690727 |
| AGGGGGLGGAP | ASLRDRYEQY | 0.60381174 |
| AGGLLYGNGGAG | ASLRDRYEQY | 0.38586512 |
| AGLESIGL | ASLRDRYEQY | 0.6386558 |
| AGLKTGII | ASLRDRYEQY | 0.53479934 |
| ASLAALLL | ASLRDRYEQY | 0.81488174 |
| ASVINISSI | ASLRDRYEQY | 0.8308178 |
| ATPPGPA | ASLRDRYEQY | 0.53817844 |
| ATVASALV | ASLRDRYEQY | 0.7737496 |
| ATVASLIL | ASLRDRYEQY | 0.7789986 |
| AVNGPVLIRH | ASLRDRYEQY | 0.33359504 |
| DGAAGAVGI | ASLRDRYEQY | 0.710075 |
| DPLLHGI | ASLRDRYEQY | 0.67970884 |
| ELGAALGAL | ASLRDRYEQY | 0.68074334 |
| ELGRTQAV | ASLRDRYEQY | 0.3343014 |
| FVAGGYLS | ASLRDRYEQY | 0.78429115 |
| GASGGAGGAGGW | ASLRDRYEQY | 0.62670064 |
| GGAGAGGDGGAGGI | ASLRDRYEQY | 0.21390657 |
| GTGHNGSQPGMGGNG | ASLRDRYEQY | 0.77862823 |
| GYAAASASAS | ASLRDRYEQY | 0.57363963 |
| HQVDVAAVQQG | ASLRDRYEQY | 0.6650668 |
| IGAAIAHT | ASLRDRYEQY | 0.80385536 |
| IGMIEVAQL | ASLRDRYEQY | 0.6103243 |
| IRLAPEHTSAMD | ASLRDRYEQY | 0.5854333 |
| IYNVSVL | ASLRDRYEQY | 0.7646562 |
| KQGFLGM | ASLRDRYEQY | 0.8200561 |
| LESVITQL | ASLRDRYEQY | 0.5421639 |
| LGLGALVVV | ASLRDRYEQY | 0.7498889 |
| LIGNGGNGGSGGI | ASLRDRYEQY | 0.836331 |

| | | |
|---|---|---|
| LNAINAPTL | ASLRDRYEQY | 0.7101364 |
| LQLLNTGV | ASLRDRYEQY | 0.8580095 |
| LQTSIEF | ASLRDRYEQY | 0.86227536 |
| MSFAVSEL | ASLRDRYEQY | 0.58676606 |
| NTGALGL | ASLRDRYEQY | 0.82465744 |
| PPGAGGAGGAGTT | ASLRDRYEQY | 0.8154036 |
| QLVGLGI | ASLRDRYEQY | 0.6111282 |
| RFLEQVHQL | ASLRDRYEQY | 0.6798907 |
| RGVGYVLR | ASLRDRYEQY | 0.35660505 |
| SATGVFAGVI | ASLRDRYEQY | 0.68797183 |
| SLPAITLP | ASLRDRYEQY | 0.4802419 |
| STALAFTV | ASLRDRYEQY | 0.7485486 |
| SVLTVGLGK | ASLRDRYEQY | 0.83130693 |
| TALAGAGSGI | ASLRDRYEQY | 0.6688862 |
| TGLPFTGI | ASLRDRYEQY | 0.42861396 |
| VAEHVTV | ASLRDRYEQY | 0.38856712 |
| VDIVLNSL | ASLRDRYEQY | 0.7836513 |
| VLTGGGAVL | ASLRDRYEQY | 0.46405828 |
| VNVGLAYGI | ASLRDRYEQY | 0.39930022 |
| VSSAAVGAV | ASLRDRYEQY | 0.64115 |
| VTSVYEAGV | ASLRDRYEQY | 0.77808845 |
| WVALDRGA | ASLRDRYEQY | 0.80334044 |
| YLVGSVVI | ASLRDRYEQY | 0.5165159 |
| GPPGTGKTLL | ASLRDRYEQY | 0.8419768 |
| GVGMTKF | ASLRDRYEQY | 0.8034885 |
| IGIGGSDL | ASLRDRYEQY | 0.8577096 |
| IGTLLAEL | ASLRDRYEQY | 0.71742725 |
| IYTSGTT | ASLRDRYEQY | 0.4322443 |
| LSFPAGV | ASLRDRYEQY | 0.79889566 |
| LYTSGTT | ASLRDRYEQY | 0.614958 |
| VGAFPFGF | ASLRDRYEQY | 0.82442296 |
| VGAGTVEF | ASLRDRYEQY | 0.79188496 |
| VGKSTLLL | ASLRDRYEQY | 0.6860358 |
| AALAHAGGGIV | ASLRDRYEQY | 0.5162414 |
| AALVAVSLL | ASLRDRYEQY | 0.70584136 |
| AAVSRAVAAL | ASLRDRYEQY | 0.86232316 |
| AGGSSGIGGF | ASLRDRYEQY | 0.4792629 |
| AGLTVANANG | ASLRDRYEQY | 0.4482146 |
| AGLVGAVL | ASLRDRYEQY | 0.68403566 |
| ALSQAFTV | ASLRDRYEQY | 0.67457485 |
| ALTFSVSL | ASLRDRYEQY | 0.82900006 |
| ALTGAALVAL | ASLRDRYEQY | 0.7869061 |
| ASFSIGAGVL | ASLRDRYEQY | 0.6904439 |
| DGSVYKI | ASLRDRYEQY | 0.77193624 |
| EQLAEIV | ASLRDRYEQY | 0.7825799 |
| FAVSELTVM | ASLRDRYEQY | 0.72262716 |
| GAAPLAAGL | ASLRDRYEQY | 0.7928213 |
| GADGGDGGSGGAGG | ASLRDRYEQY | 0.5565764 |
| GFGLDTGGAGG | ASLRDRYEQY | 0.8534454 |
| GLASGGLL | ASLRDRYEQY | 0.80691373 |
| GRIAGAAVLL | ASLRDRYEQY | 0.82145154 |

| | | |
|---|---|---|
| GSAGSLNTSF | ASLRDRYEQY | 0.842172 |
| GTGQIGI | ASLRDRYEQY | 0.7556404 |
| GTLIALIL | ASLRDRYEQY | 0.71909744 |
| IAATMGFVVG | ASLRDRYEQY | 0.40792498 |
| IFAQALL | ASLRDRYEQY | 0.5316523 |
| IGAAVALGF | ASLRDRYEQY | 0.724206 |
| IGLTPLAL | ASLRDRYEQY | 0.76697713 |
| ILGSNTSTL | ASLRDRYEQY | 0.7956094 |
| IVFSLTVL | ASLRDRYEQY | 0.8072537 |
| LASKATGF | ASLRDRYEQY | 0.52730596 |
| LASVALIL | ASLRDRYEQY | 0.83322287 |
| LFAVGLTL | ASLRDRYEQY | 0.82145053 |
| LFNGGIGTYI | ASLRDRYEQY | 0.41371655 |
| LTLGAGII | ASLRDRYEQY | 0.6760355 |
| LVAAALGDI | ASLRDRYEQY | 0.8459583 |
| NTAFVQLGI | ASLRDRYEQY | 0.77042705 |
| QLFSGAEL | ASLRDRYEQY | 0.6180202 |
| SAATFITL | ASLRDRYEQY | 0.7135676 |
| SNYGLANLGV | ASLRDRYEQY | 0.8311219 |
| SYGGAYAVM | ASLRDRYEQY | 0.7838208 |
| TTGVGGLTLG | ASLRDRYEQY | 0.8408952 |
| TTQTIYI | ASLRDRYEQY | 0.25867096 |
| VAAGAGLGAVI | ASLRDRYEQY | 0.7097396 |
| VAAVVTLL | ASLRDRYEQY | 0.6938492 |
| VGFLGGLI | ASLRDRYEQY | 0.21093851 |
| VGVIPILL | ASLRDRYEQY | 0.8223685 |
| VIFASSLI | ASLRDRYEQY | 0.5473088 |
| VKGGDGGAAAG | ASLRDRYEQY | 0.7678889 |
| VTKTFTNI | ASLRDRYEQY | 0.5916095 |
| VVAAAILL | ASLRDRYEQY | 0.8381308 |
| YAALDVEL | ASLRDRYEQY | 0.7431405 |
| LASVALLL | ASLRDRYEQY | 0.6306386 |
| LDAVHRV | ASLRDRYEQY | 0.52776647 |
| QGLSSSIF | ASLRDRYEQY | 0.7653892 |
| RGTFANIRL | ASLRDRYEQY | 0.84056115 |
| RQLWWGHRI | ASLRDRYEQY | 0.85450196 |
| YGPPGTGKTLL | ASLRDRYEQY | 0.3669929 |
| AALVGAPANVA | ASLRDRYEQY | 0.8154546 |
| AATFRRTE | ASLRDRYEQY | 0.68532443 |
| AGALGGGI | ASLRDRYEQY | 0.34595767 |
| AGGAGGAGGIGAG | ASLRDRYEQY | 0.3247003 |
| AGSGALGL | ASLRDRYEQY | 0.70011723 |
| AQIADQI | ASLRDRYEQY | 0.6844907 |
| ASAVAVTYL | ASLRDRYEQY | 0.7445525 |
| ASTVLGGTGG | ASLRDRYEQY | 0.7939371 |
| AVGGEPAK | ASLRDRYEQY | 0.595657 |
| CEQGTFGGTC | ASLRDRYEQY | 0.77937806 |
| DEGKAPAG | ASLRDRYEQY | 0.6848203 |
| DTVVFPAL | ASLRDRYEQY | 0.10031159 |
| FEGITMAG | ASLRDRYEQY | 0.5111629 |
| FPLVVQD | ASLRDRYEQY | 0.3132519 |

| | | |
|---|---|---|
| GAFTTPQ | ASLRDRYEQY | 0.8469776 |
| GFGGFGTAGGVGGA | ASLRDRYEQY | 0.5380374 |
| GFLAQFL | ASLRDRYEQY | 0.62317604 |
| GGAGGAGGNAGVA | ASLRDRYEQY | 0.6573404 |
| GGVGTAGAGGAGGAGG | ASLRDRYEQY | 0.75858164 |
| GNGGGGDAGNAGSGGN | ASLRDRYEQY | 0.69515383 |
| GTGGAGGAGNDAGS | ASLRDRYEQY | 0.58260334 |
| GTTNTGWFNS | ASLRDRYEQY | 0.44993547 |
| IFMGPLFTL | ASLRDRYEQY | 0.76942086 |
| ILAHAIEAI | ASLRDRYEQY | 0.6779234 |
| IRSAIREAL | ASLRDRYEQY | 0.57860625 |
| ITFAGGG | ASLRDRYEQY | 0.5848329 |
| ITGMSGHQEV | ASLRDRYEQY | 0.79197204 |
| IVAGLACTL | ASLRDRYEQY | 0.33300573 |
| IVGGGLAGA | ASLRDRYEQY | 0.84143245 |
| IVTGVLNI | ASLRDRYEQY | 0.4415486 |
| LAQQGCI | ASLRDRYEQY | 0.6198621 |
| LCLHVEN | ASLRDRYEQY | 0.8420234 |
| LDAISDGS | ASLRDRYEQY | 0.7115104 |
| LGGFAAML | ASLRDRYEQY | 0.6679287 |
| LGGKNPMI | ASLRDRYEQY | 0.59270436 |
| LGPKGVAAIL | ASLRDRYEQY | 0.459809 |
| LMVGVSI | ASLRDRYEQY | 0.5695253 |
| LVGAAFVL | ASLRDRYEQY | 0.58930403 |
| LVLAVAPL | ASLRDRYEQY | 0.8309034 |
| MAAIAVIDAA | ASLRDRYEQY | 0.7822113 |
| NNGINAELQ | ASLRDRYEQY | 0.39309967 |
| RGEPLVLVHG | ASLRDRYEQY | 0.8284693 |
| RGGAGGAAGLI | ASLRDRYEQY | 0.8378521 |
| RSYRMHPAV | ASLRDRYEQY | 0.8527477 |
| SAAHLLAGQLA | ASLRDRYEQY | 0.4625042 |
| SAAVGSY | ASLRDRYEQY | 0.64147234 |
| SPHAVLHRI | ASLRDRYEQY | 0.67807746 |
| SQIAKVL | ASLRDRYEQY | 0.646193 |
| TALGHLVP | ASLRDRYEQY | 0.60524035 |
| TAPTTSGAA | ASLRDRYEQY | 0.5200958 |
| TDGDDGNGGSPGT | ASLRDRYEQY | 0.8393396 |
| TDGLGEN | ASLRDRYEQY | 0.843827 |
| TGGPPLPGQA | ASLRDRYEQY | 0.6610396 |
| TIPAITI | ASLRDRYEQY | 0.37797093 |
| TIPGFTI | ASLRDRYEQY | 0.64416635 |
| TSIASGALA | ASLRDRYEQY | 0.51625085 |
| TVTLPGAAI | ASLRDRYEQY | 0.51938933 |
| VASGLSAALP | ASLRDRYEQY | 0.5428095 |
| VGAGVIGI | ASLRDRYEQY | 0.32585046 |
| VGPKRFGV | ASLRDRYEQY | 0.6858438 |
| VNVAFPD | ASLRDRYEQY | 0.8538852 |
| VSMTNSGVS | ASLRDRYEQY | 0.6427111 |
| VTDVLGDLSV | ASLRDRYEQY | 0.7324498 |
| VVVVGAGFAGL | ASLRDRYEQY | 0.8311024 |
| VYTVGTVV | ASLRDRYEQY | 0.7858522 |

| | | |
|---|---|---|
| WTTFSGP | ASLRDRYEQY | 0.8318217 |
| YDELVEAF | ASLRDRYEQY | 0.81548613 |
| YGTGGAGGNGG | ASLRDRYEQY | 0.7180152 |
| YTNVMAV | ASLRDRYEQY | 0.7387266 |
| FLEQVHQL | ASLRDRYEQY | 0.79054606 |
| GLSLLAVAV | ASLRDRYEQY | 0.6147824 |
| LEQVHQL | ASLRDRYEQY | 0.70188594 |
| LNAAGAI | ASLRDRYEQY | 0.70623785 |
| PAVHASL | ASLRDRYEQY | 0.7574036 |
| TELEDDA | ASLRDRYEQY | 0.70209277 |
| TPPGPPN | ASLRDRYEQY | 0.842051 |
| TPPPPPPP | ASLRDRYEQY | 0.21903418 |
| TQSPALI | ASSWDRGGAEQF | 0.5010661 |
| EFAAAFPQ | ASSWDRGGAEQF | 0.82983375 |
| LLSGNLDVL | ASSWDRGGAEQF | 0.15939404 |
| GPANLGNYNI | ASSWDRGGAEQF | 0.36813837 |
| TLGPTLP | ASSWDRGGAEQF | 0.74297667 |
| VYAYSSCCK | ASSWDRGGAEQF | 0.8239794 |
| APAPAPAPNI | ASSWDRGGAEQF | 0.1847184 |
| PGGPVPIP | ASSWDRGGAEQF | 0.6437672 |
| FLTVALEP | ASSWDRGGAEQF | 0.32140225 |
| SQYFGANL | ASSWDRGGAEQF | 0.8531302 |
| GGVKPAA | ASSWDRGGAEQF | 0.68771696 |
| GGTGGAGGAGGW | ASSWDRGGAEQF | 0.31512362 |
| GVGGVAGLGGAGG | ASSWDRGGAEQF | 0.8220789 |
| VLLVLATTI | ASSWDRGGAEQF | 0.49171644 |
| IGIFAAML | ASSWDRGGAEQF | 0.7422881 |
| AVGDLID | ASSWDRGGAEQF | 0.8368781 |
| NISPLPL | ASSWDRGGAEQF | 0.855368 |
| VTAVFVAI | ASSWDRGGAEQF | 0.42745572 |
| SGDDNDAAL | ASSWDRGGAEQF | 0.7911882 |
| AVAALAHS | ASSWDRGGAEQF | 0.6101582 |
| GGVTVMATP | ASSWDRGGAEQF | 0.87320244 |
| AGLAANVRD | ASSWDRGGAEQF | 0.6249223 |
| TNRPDLIDPALLRPG | ASSWDRGGAEQF | 0.38318065 |
| LANLGVTAL | ASSWDRGGAEQF | 0.82039213 |
| TGERIEL | ASSWDRGGAEQF | 0.79960287 |
| NGEELAA | ASSWDRGGAEQF | 0.69029063 |
| VLSLVWG | ASSWDRGGAEQF | 0.8077805 |
| IGYALWL | ASSWDRGGAEQF | 0.75262475 |
| LSLLLAPL | ASSWDRGGAEQF | 0.6194558 |
| TLDEALK | ASSWDRGGAEQF | 0.77842146 |
| TDGAGTAEGG | ASSWDRGGAEQF | 0.79258347 |
| NAPGGAGGNGG | ASSWDRGGAEQF | 0.5843853 |
| VSVFMGMGGNF | ASSWDRGGAEQF | 0.851897 |
| LASYLAAL | ASSWDRGGAEQF | 0.68512976 |
| LVGGTPAL | ASSWDRGGAEQF | 0.42160156 |
| VRNADVL | ASSWDRGGAEQF | 0.86470854 |
| TCSSPFLVSAM | ASSWDRGGAEQF | 0.72939104 |
| VHITDAE | ASSWDRGGAEQF | 0.7938864 |
| EYAAPSF | ASSWDRGGAEQF | 0.7987563 |

| | | |
|---|---|---|
| VVDAFHGGL | ASSWDRGGAEQF | 0.82891405 |
| ASVVDPQ | ASSWDRGGAEQF | 0.5602535 |
| SYNEFGGLS | ASSWDRGGAEQF | 0.7375721 |
| DPDGSVY | ASSWDRGGAEQF | 0.7073631 |
| MVSTIHATE | ASSWDRGGAEQF | 0.8124175 |
| LAAGVADPV | ASSWDRGGAEQF | 0.7540262 |
| TKEQIAA | ASSWDRGGAEQF | 0.76864266 |
| SLASPAAL | ASSWDRGGAEQF | 0.7966188 |
| IGLVFYGV | ASSWDRGGAEQF | 0.8317915 |
| FLLDASNL | ASSWDRGGAEQF | 0.7605554 |
| YQVGNFP | ASSWDRGGAEQF | 0.7395146 |
| ESLLSQST | ASSWDRGGAEQF | 0.8438066 |
| LFTVGFAAVG | ASSWDRGGAEQF | 0.45023325 |
| ATPPTPT | ASSWDRGGAEQF | 0.70630014 |
| TDLATLG | ASSWDRGGAEQF | 0.82966983 |
| SAGGTGR | ASSWDRGGAEQF | 0.82584953 |
| GFSNSGS | ASSWDRGGAEQF | 0.73988104 |
| GGNGGSGGTSV | ASSWDRGGAEQF | 0.83736277 |
| GTSVATGGAGN | ASSWDRGGAEQF | 0.64963126 |
| NGGNGGSGGTS | ASSWDRGGAEQF | 0.72448146 |
| AVVKFPH | ASSWDRGGAEQF | 0.66636455 |
| TPPPPPP | ASSWDRGGAEQF | 0.625164 |
| AAPAAAAAPA | ASSWDRGGAEQF | 0.5361283 |
| IALNEDL | ASSWDRGGAEQF | 0.7066135 |
| IVNVSSM | ASSWDRGGAEQF | 0.7905838 |
| TGIPPAPR | ASSWDRGGAEQF | 0.7268758 |
| FDLGGGTF | ASSWDRGGAEQF | 0.34100914 |
| FDLGGGTFDVS | ASSWDRGGAEQF | 0.7505276 |
| NEPTAAA | ASSWDRGGAEQF | 0.7154024 |
| NEPTAAALA | ASSWDRGGAEQF | 0.8531511 |
| GSIINISSI | ASSWDRGGAEQF | 0.73189855 |
| VAPGNAGTA | ASSWDRGGAEQF | 0.8012003 |
| TGGGASLE | ASSWDRGGAEQF | 0.65406895 |
| VAVYNHYKRI | ASSWDRGGAEQF | 0.7565377 |
| TDPAPATT | ASSWDRGGAEQF | 0.350859 |
| TNRPDILDPALLRPG | ASSWDRGGAEQF | 0.4714152 |
| LTTTTTS | ASSWDRGGAEQF | 0.655483 |
| PTGSGKTL | ASSWDRGGAEQF | 0.5206177 |
| TGSGKTL | ASSWDRGGAEQF | 0.8013004 |
| TIGHVDHGK | ASSWDRGGAEQF | 0.83897734 |
| TIGHVDHGKTT | ASSWDRGGAEQF | 0.80450267 |
| GHVDHGKTT | ASSWDRGGAEQF | 0.81081045 |
| AQFEHTLL | ASSWDRGGAEQF | 0.82909703 |
| IVNQDEL | ASSWDRGGAEQF | 0.85277045 |
| TNGSQFF | ASSWDRGGAEQF | 0.6452596 |
| TGKGIVE | ASSWDRGGAEQF | 0.6016498 |
| LISNASDA | ASSWDRGGAEQF | 0.33803654 |
| AELRTLGELTKIAKAHGAQ | ASSWDRGGAEQF | 0.56145275 |
| APNAVAAAEPSW | ASSWDRGGAEQF | 0.6498152 |
| APQVAGAATAVL | ASSWDRGGAEQF | 0.68308854 |
| ARRDPAAAAAILPTDA | ASSWDRGGAEQF | 0.14197989 |

| | | |
|---|---|---|
| DAAARDVLAEAGLAEYF | ASSWDRGGAEQF | 0.7406094 |
| DDQIVLSAKANLAKVEQ | ASSWDRGGAEQF | 0.8368602 |
| DELLPDTGEDSDEDGH | ASSWDRGGAEQF | 0.697477 |
| ELPEAPANRATVPQD | ASSWDRGGAEQF | 0.7645729 |
| FGQSKIERVTGIG | ASSWDRGGAEQF | 0.75739676 |
| GAGGTGSTAGGAGGAGGAGGLY | ASSWDRGGAEQF | 0.64581597 |
| GHGGTGAAGGNGGTGGAGGSGI | ASSWDRGGAEQF | 0.759477 |
| GNGGNGGAGGEGGPGGAGGAGA | ASSWDRGGAEQF | 0.62445635 |
| GSTAGNGGNGGNATNGTIAGQP | ASSWDRGGAEQF | 0.82846534 |
| GTGGDGGNGGNGGTGG | ASSWDRGGAEQF | 0.84374857 |
| GVRRPAGIPADAPV | ASSWDRGGAEQF | 0.7545072 |
| GYRQVTDAHLLTLARR | ASSWDRGGAEQF | 0.6771575 |
| IAKMGQDGHDRG | ASSWDRGGAEQF | 0.76819265 |
| KAEPSIEIQYT | ASSWDRGGAEQF | 0.5686492 |
| KPVRIRMAADEDTVAVT | ASSWDRGGAEQF | 0.700599 |
| MPPRIAGMRLLVIKPEP | ASSWDRGGAEQF | 0.5356599 |
| MKFYAVALFRSRGVPPTR | ASSWDRGGAEQF | 0.684464 |
| NAGGFGILGAVAHSP | ASSWDRGGAEQF | 0.53295445 |
| NEGREITLAMAEM | ASSWDRGGAEQF | 0.74981695 |
| NELVEWSHPLGHRWRWE | ASSWDRGGAEQF | 0.69055665 |
| NLGAGGTGGNGGNPG | ASSWDRGGAEQF | 0.7890742 |
| NTEWPKWETVMAADPVD | ASSWDRGGAEQF | 0.4480937 |
| PASAAPAETQP | ASSWDRGGAEQF | 0.83231276 |
| QQSSDGKNGVRAAEITGE | ASSWDRGGAEQF | 0.79149294 |
| RAPDAVTTSWWKE | ASSWDRGGAEQF | 0.41726965 |
| SGFGNVGAHTSGWFNQ | ASSWDRGGAEQF | 0.8285682 |
| SGVAGAVGQAASVGGLKV | ASSWDRGGAEQF | 0.73153704 |
| SNGGAGTGSAGGNGGT | ASSWDRGGAEQF | 0.66475093 |
| TGGAGGEGGAAG | ASSWDRGGAEQF | 0.7944952 |
| TGGDGGAAGAGGAPGHG | ASSWDRGGAEQF | 0.74536955 |
| TMPPLLAAIANAGRHGV | ASSWDRGGAEQF | 0.54677254 |
| TSSTAGNSGGA | ASSWDRGGAEQF | 0.59440035 |
| VPATVPVGILC | ASSWDRGGAEQF | 0.6926835 |
| VQRPDSGGVVLTGRIS | ASSWDRGGAEQF | 0.6863979 |
| WTPNPVPATKGVPAGLVQAT | ASSWDRGGAEQF | 0.27033302 |
| YALKRALAKFSDRFDVVII | ASSWDRGGAEQF | 0.7986471 |
| YLIGSVIV | ASSWDRGGAEQF | 0.4192954 |
| YGQLVAVID | ASSWDRGGAEQF | 0.58453155 |
| VTVGGMTLPA | ASSWDRGGAEQF | 0.7088214 |
| VLALPAV | ASSWDRGGAEQF | 0.80608654 |
| VIPPTLNYET | ASSWDRGGAEQF | 0.76561964 |
| VGMGGMHPSSG | ASSWDRGGAEQF | 0.638402 |
| VGLAGGAATA | ASSWDRGGAEQF | 0.7723567 |
| VGGAGGAGGSGP | ASSWDRGGAEQF | 0.7493235 |
| VFAAFMV | ASSWDRGGAEQF | 0.79600435 |
| TLYDKLR | ASSWDRGGAEQF | 0.7408722 |
| TLAVFTVL | ASSWDRGGAEQF | 0.7717107 |
| TGSVQPARV | ASSWDRGGAEQF | 0.73998797 |
| TGGGAGSGGT | ASSWDRGGAEQF | 0.6188427 |
| SHLETSL | ASSWDRGGAEQF | 0.63783693 |
| SFTTGMT | ASSWDRGGAEQF | 0.64527404 |

| | | |
|---|---|---|
| QVEVAKRMGVRQPTVS | ASSWDRGGAEQF | 0.6029196 |
| QGSGGPALAM | ASSWDRGGAEQF | 0.5050751 |
| MLAPSNP | ASSWDRGGAEQF | 0.7398163 |
| LVNNAAI | ASSWDRGGAEQF | 0.85809636 |
| LTGEINTLA | ASSWDRGGAEQF | 0.7504041 |
| LSIFGPV | ASSWDRGGAEQF | 0.33466107 |
| LLALLANL | ASSWDRGGAEQF | 0.6675916 |
| LISNASDALDKLR | ASSWDRGGAEQF | 0.6759069 |
| LINEAVP | ASSWDRGGAEQF | 0.7887487 |
| LGYAFSGL | ASSWDRGGAEQF | 0.83667976 |
| LDFVAVAL | ASSWDRGGAEQF | 0.83320415 |
| LAQPGAE | ASSWDRGGAEQF | 0.7859317 |
| ISNLGAVTGF | ASSWDRGGAEQF | 0.59442836 |
| HQEDPGRI | ASSWDRGGAEQF | 0.81074965 |
| GTPGEPTP | ASSWDRGGAEQF | 0.6419929 |
| GFFNSGS | ASSWDRGGAEQF | 0.73791265 |
| GAGGGGLTAGAN | ASSWDRGGAEQF | 0.7492683 |
| FVDALVNM | ASSWDRGGAEQF | 0.7941712 |
| FLTVAPEL | ASSWDRGGAEQF | 0.64311004 |
| FETLTAL | ASSWDRGGAEQF | 0.69054693 |
| DVINPIEL | ASSWDRGGAEQF | 0.4257154 |
| APVSGAPV | ASSWDRGGAEQF | 0.81965053 |
| ALVGGLAW | ASSWDRGGAEQF | 0.5366883 |
| ALAAAAATAI | ASSWDRGGAEQF | 0.5727465 |
| AFGLGLLL | ASSWDRGGAEQF | 0.64520484 |
| AFFAAASL | ASSWDRGGAEQF | 0.8391293 |
| AAAALPAFSPP | ASSWDRGGAEQF | 0.822001 |
| VVLAPAL | ASSWDRGGAEQF | 0.5678855 |
| TIGHVDHGKTTL | ASSWDRGGAEQF | 0.57056177 |
| LISNASDALDK | ASSWDRGGAEQF | 0.7681768 |
| LISNASDAL | ASSWDRGGAEQF | 0.5665406 |
| ISNASDALDK | ASSWDRGGAEQF | 0.8263056 |
| NQAETLVYQTE | ASSWDRGGAEQF | 0.6764561 |
| TGGTGGTAGAG | ASSWDRGGAEQF | 0.68509114 |
| TGSTAGGAGGA | ASSWDRGGAEQF | 0.6093682 |
| DPLAVPVAAGPV | ASSWDRGGAEQF | 0.8054964 |
| GGTGGTPGNGGN | ASSWDRGGAEQF | 0.661971 |
| PAGLAGVDPATL | ASSWDRGGAEQF | 0.81180656 |
| GNNLVVEAPHAGASV | ASSWDRGGAEQF | 0.24052504 |
| KVTRIMSASTLEQPA | ASSWDRGGAEQF | 0.84695446 |
| LGFALPIALPRRTVV | ASSWDRGGAEQF | 0.45098978 |
| TRLGNLTARKLVVLV | ASSWDRGGAEQF | 0.54151165 |
| WSQQRRLIYFPSAGP | ASSWDRGGAEQF | 0.29176718 |
| GPNGNPDPMAAAVDIR | ASSWDRGGAEQF | 0.6960515 |
| TQRCVSVADLRAAASR | ASSWDRGGAEQF | 0.7879293 |
| QFVLDVAEHPVEVLTETV | ASSWDRGGAEQF | 0.84003246 |
| GGAGGTGGAGGNGGTQAGDG | ASSWDRGGAEQF | 0.76901424 |
| PLLATLRAIAVDVDAKPAQV | ASSWDRGGAEQF | 0.77285767 |
| APKFRTGGYMIDFWGVGYQVAK | ASSWDRGGAEQF | 0.8353276 |
| GAGGAGGAGGAAGGPGGTGGTG | ASSWDRGGAEQF | 0.60026366 |
| IGALGLLATDVAPAGSVAFGVA | ASSWDRGGAEQF | 0.29714713 |

| | | |
|---|---|---|
| AGGAGGIGGNGGAGGAGGVLMG | ASSWDRGGAEQF | 0.718685 |
| ADELIGLL | ASSWDRGGAEQF | 0.7276943 |
| ADRALKA | ASSWDRGGAEQF | 0.8492762 |
| AGAVGQVGGAGGA | ASSWDRGGAEQF | 0.77776694 |
| AGGGGGLGGAP | ASSWDRGGAEQF | 0.3288854 |
| AGGLLYGNGGAG | ASSWDRGGAEQF | 0.7747011 |
| AGLESIGL | ASSWDRGGAEQF | 0.7432022 |
| AGLKTGII | ASSWDRGGAEQF | 0.42431664 |
| ASLAALLL | ASSWDRGGAEQF | 0.62271094 |
| ASVINISSI | ASSWDRGGAEQF | 0.5497689 |
| ATPPGPA | ASSWDRGGAEQF | 0.473865 |
| ATVASALV | ASSWDRGGAEQF | 0.7664293 |
| ATVASLIL | ASSWDRGGAEQF | 0.61365664 |
| AVNGPVLIRH | ASSWDRGGAEQF | 0.6886451 |
| DGAAGAVGI | ASSWDRGGAEQF | 0.64417756 |
| DPLLHGI | ASSWDRGGAEQF | 0.6266525 |
| ELGAALGAL | ASSWDRGGAEQF | 0.6504893 |
| ELGRTQAV | ASSWDRGGAEQF | 0.6702306 |
| FVAGGYLS | ASSWDRGGAEQF | 0.8058828 |
| GASGGAGGAGGW | ASSWDRGGAEQF | 0.8195157 |
| GGAGAGGDGGAGGI | ASSWDRGGAEQF | 0.6774926 |
| GTGHNGSQPGMGGNG | ASSWDRGGAEQF | 0.708235 |
| GYAAASASAS | ASSWDRGGAEQF | 0.8168597 |
| HQVDVAAVQQG | ASSWDRGGAEQF | 0.47649536 |
| IGAAIAHT | ASSWDRGGAEQF | 0.56599116 |
| IGMIEVAQL | ASSWDRGGAEQF | 0.79781866 |
| IRLAPEHTSAMD | ASSWDRGGAEQF | 0.70481265 |
| IYNVSVL | ASSWDRGGAEQF | 0.825619 |
| KQGFLGM | ASSWDRGGAEQF | 0.7128657 |
| LESVITQL | ASSWDRGGAEQF | 0.72594476 |
| LGLGALVVV | ASSWDRGGAEQF | 0.7178186 |
| LIGNGGNGGSGGI | ASSWDRGGAEQF | 0.8501735 |
| LNAINAPTL | ASSWDRGGAEQF | 0.77672154 |
| LQLLNTGV | ASSWDRGGAEQF | 0.6795945 |
| LQTSIEF | ASSWDRGGAEQF | 0.8126216 |
| MSFAVSEL | ASSWDRGGAEQF | 0.39564192 |
| NTGALGL | ASSWDRGGAEQF | 0.8073251 |
| PPGAGGAGGAGTT | ASSWDRGGAEQF | 0.7954187 |
| QLVGLGI | ASSWDRGGAEQF | 0.48679182 |
| RFLEQVHQL | ASSWDRGGAEQF | 0.6311759 |
| RGVGYVLR | ASSWDRGGAEQF | 0.86267966 |
| SATGVFAGVI | ASSWDRGGAEQF | 0.82132816 |
| SLPAITLP | ASSWDRGGAEQF | 0.69235057 |
| STALAFTV | ASSWDRGGAEQF | 0.5305635 |
| SVLTVGLGK | ASSWDRGGAEQF | 0.535522 |
| TALAGAGSGI | ASSWDRGGAEQF | 0.65058506 |
| TGLPFTGI | ASSWDRGGAEQF | 0.30913752 |
| VAEHVTV | ASSWDRGGAEQF | 0.77093416 |
| VDIVLNSL | ASSWDRGGAEQF | 0.5904696 |
| VLTGGGAVL | ASSWDRGGAEQF | 0.7926149 |
| VNVGLAYGI | ASSWDRGGAEQF | 0.7562876 |

| | | |
|---|---|---|
| VSSAAVGAV | ASSWDRGGAEQF | 0.8286176 |
| VTSVYEAGV | ASSWDRGGAEQF | 0.5644899 |
| WVALDRGA | ASSWDRGGAEQF | 0.66781914 |
| YLVGSVVI | ASSWDRGGAEQF | 0.33924007 |
| GPPGTGKTLL | ASSWDRGGAEQF | 0.5451492 |
| GVGMTKF | ASSWDRGGAEQF | 0.6316968 |
| IGIGGSDL | ASSWDRGGAEQF | 0.72387767 |
| IGTLLAEL | ASSWDRGGAEQF | 0.32927775 |
| IYTSGTT | ASSWDRGGAEQF | 0.8486906 |
| LSFPAGV | ASSWDRGGAEQF | 0.6236069 |
| LYTSGTT | ASSWDRGGAEQF | 0.7797153 |
| VGAFPFGF | ASSWDRGGAEQF | 0.6201371 |
| VGAGTVEF | ASSWDRGGAEQF | 0.5179819 |
| VGKSTLLL | ASSWDRGGAEQF | 0.6285391 |
| AALAHAGGGIV | ASSWDRGGAEQF | 0.78564847 |
| AALVAVSLL | ASSWDRGGAEQF | 0.43370834 |
| AAVSRAVAAL | ASSWDRGGAEQF | 0.30332488 |
| AGGSSGIGGF | ASSWDRGGAEQF | 0.5727898 |
| AGLTVANANG | ASSWDRGGAEQF | 0.685851 |
| AGLVGAVL | ASSWDRGGAEQF | 0.85423523 |
| ALSQAFTV | ASSWDRGGAEQF | 0.82146466 |
| ALTFSVSL | ASSWDRGGAEQF | 0.7178787 |
| ALTGAALVAL | ASSWDRGGAEQF | 0.86005783 |
| ASFSIGAGVL | ASSWDRGGAEQF | 0.1506249 |
| DGSVYKI | ASSWDRGGAEQF | 0.6242388 |
| EQLAEIV | ASSWDRGGAEQF | 0.81609815 |
| FAVSELTVM | ASSWDRGGAEQF | 0.7722205 |
| GAAPLAAGL | ASSWDRGGAEQF | 0.62639844 |
| GADGGDGGSGGAGG | ASSWDRGGAEQF | 0.83356524 |
| GFGLDTGGAGG | ASSWDRGGAEQF | 0.5680282 |
| GLASGGLL | ASSWDRGGAEQF | 0.6694318 |
| GRIAGAAVLL | ASSWDRGGAEQF | 0.72902507 |
| GSAGSLNTSF | ASSWDRGGAEQF | 0.8145174 |
| GTGQIGI | ASSWDRGGAEQF | 0.80478686 |
| GTLIALIL | ASSWDRGGAEQF | 0.34772032 |
| IAATMGFVVG | ASSWDRGGAEQF | 0.7828162 |
| IFAQALL | ASSWDRGGAEQF | 0.8604171 |
| IGAAVALGF | ASSWDRGGAEQF | 0.6149539 |
| IGLTPLAL | ASSWDRGGAEQF | 0.81426823 |
| ILGSNTSTL | ASSWDRGGAEQF | 0.7188232 |
| IVFSLTVL | ASSWDRGGAEQF | 0.6560304 |
| LASKATGF | ASSWDRGGAEQF | 0.52156234 |
| LASVALIL | ASSWDRGGAEQF | 0.4720751 |
| LFAVGLTL | ASSWDRGGAEQF | 0.8341313 |
| LFNGGIGTYI | ASSWDRGGAEQF | 0.4455974 |
| LTLGAGII | ASSWDRGGAEQF | 0.73330283 |
| LVAAALGDI | ASSWDRGGAEQF | 0.67836857 |
| NTAFVQLGI | ASSWDRGGAEQF | 0.8701525 |
| QLFSGAEL | ASSWDRGGAEQF | 0.75899124 |
| SAATFITL | ASSWDRGGAEQF | 0.7784979 |
| SNYGLANLGV | ASSWDRGGAEQF | 0.5585783 |

| | | |
|---|---|---|
| SYGGAYAVM | ASSWDRGGAEQF | 0.79496944 |
| TTGVGGLTLG | ASSWDRGGAEQF | 0.7781528 |
| TTQTIYI | ASSWDRGGAEQF | 0.4716441 |
| VAAGAGLGAVI | ASSWDRGGAEQF | 0.80937845 |
| VAAVVTLL | ASSWDRGGAEQF | 0.45602274 |
| VGFLGGLI | ASSWDRGGAEQF | 0.7596451 |
| VGVIPILL | ASSWDRGGAEQF | 0.74495304 |
| VIFASSLI | ASSWDRGGAEQF | 0.8004414 |
| VKGGDGGAAAG | ASSWDRGGAEQF | 0.7118368 |
| VTKTFTNI | ASSWDRGGAEQF | 0.83319026 |
| VVAAAILL | ASSWDRGGAEQF | 0.49968475 |
| YAALDVEL | ASSWDRGGAEQF | 0.8012924 |
| LASVALLL | ASSWDRGGAEQF | 0.81360006 |
| LDAVHRV | ASSWDRGGAEQF | 0.7543644 |
| QGLSSSIF | ASSWDRGGAEQF | 0.8598443 |
| RGTFANIRL | ASSWDRGGAEQF | 0.8274527 |
| RQLWWGHRI | ASSWDRGGAEQF | 0.27994397 |
| YGPPGTGKTLL | ASSWDRGGAEQF | 0.67607725 |
| AALVGAPANVA | ASSWDRGGAEQF | 0.8427806 |
| AATFRRTE | ASSWDRGGAEQF | 0.844735 |
| AGALGGGI | ASSWDRGGAEQF | 0.83485293 |
| AGGAGGAGGIGAG | ASSWDRGGAEQF | 0.7956371 |
| AGSGALGL | ASSWDRGGAEQF | 0.65657276 |
| AQIADQI | ASSWDRGGAEQF | 0.8339838 |
| ASAVAVTYL | ASSWDRGGAEQF | 0.88205564 |
| ASTVLGGTGG | ASSWDRGGAEQF | 0.7566196 |
| AVGGEPAK | ASSWDRGGAEQF | 0.5396036 |
| CEQGTFGGTC | ASSWDRGGAEQF | 0.5125408 |
| DEGKAPAG | ASSWDRGGAEQF | 0.7964559 |
| DTVVFPAL | ASSWDRGGAEQF | 0.8319573 |
| FEGITMAG | ASSWDRGGAEQF | 0.4510567 |
| FPLVVQD | ASSWDRGGAEQF | 0.6763943 |
| GAFTTPQ | ASSWDRGGAEQF | 0.7040273 |
| GFGGFGTAGGVGGA | ASSWDRGGAEQF | 0.21861935 |
| GFLAQFL | ASSWDRGGAEQF | 0.78683084 |
| GGAGGAGGNAGVA | ASSWDRGGAEQF | 0.83160603 |
| GGVGTAGAGGAGGAGG | ASSWDRGGAEQF | 0.8075466 |
| GNGGGGDAGNAGSGGN | ASSWDRGGAEQF | 0.43647116 |
| GTGGAGGAGNDAGS | ASSWDRGGAEQF | 0.81368935 |
| GTTNTGWFNS | ASSWDRGGAEQF | 0.80145323 |
| IFMGPLFTL | ASSWDRGGAEQF | 0.85648775 |
| ILAHAIEAI | ASSWDRGGAEQF | 0.42576334 |
| IRSAIREAL | ASSWDRGGAEQF | 0.1713038 |
| ITFAGGG | ASSWDRGGAEQF | 0.7625896 |
| ITGMSGHQEV | ASSWDRGGAEQF | 0.7672405 |
| IVAGLACTL | ASSWDRGGAEQF | 0.64121765 |
| IVGGGLAGA | ASSWDRGGAEQF | 0.5389626 |
| IVTGVLNI | ASSWDRGGAEQF | 0.73624444 |
| LAQQGCI | ASSWDRGGAEQF | 0.8067437 |
| LCLHVEN | ASSWDRGGAEQF | 0.3820715 |
| LDAISDGS | ASSWDRGGAEQF | 0.5069171 |

| | | |
|---|---|---|
| LGGFAAML | ASSWDRGGAEQF | 0.79714054 |
| LGGKNPMI | ASSWDRGGAEQF | 0.82182634 |
| LGPKGVAAIL | ASSWDRGGAEQF | 0.5067529 |
| LMVGVSI | ASSWDRGGAEQF | 0.6211927 |
| LVGAAFVL | ASSWDRGGAEQF | 0.75692296 |
| LVLAVAPL | ASSWDRGGAEQF | 0.75130653 |
| MAAIAVIDAA | ASSWDRGGAEQF | 0.72926354 |
| NNGINAELQ | ASSWDRGGAEQF | 0.3685854 |
| RGEPLVLVHG | ASSWDRGGAEQF | 0.81332374 |
| RGGAGGAAGLI | ASSWDRGGAEQF | 0.4111144 |
| RSYRMHPAV | ASSWDRGGAEQF | 0.8189198 |
| SAAHLLAGQLA | ASSWDRGGAEQF | 0.7059466 |
| SAAVGSY | ASSWDRGGAEQF | 0.75835216 |
| SPHAVLHRI | ASSWDRGGAEQF | 0.63517416 |
| SQIAKVL | ASSWDRGGAEQF | 0.3884336 |
| TALGHLVP | ASSWDRGGAEQF | 0.44605935 |
| TAPTTSGAA | ASSWDRGGAEQF | 0.55172944 |
| TDGDDGNGGSPGT | ASSWDRGGAEQF | 0.8275168 |
| TDGLGEN | ASSWDRGGAEQF | 0.7567811 |
| TGGPPLPGQA | ASSWDRGGAEQF | 0.81257963 |
| TIPAITI | ASSWDRGGAEQF | 0.7475456 |
| TIPGFTI | ASSWDRGGAEQF | 0.517042 |
| TSIASGALA | ASSWDRGGAEQF | 0.80409336 |
| TVTLPGAAI | ASSWDRGGAEQF | 0.38933825 |
| VASGLSAALP | ASSWDRGGAEQF | 0.7487674 |
| VGAGVIGI | ASSWDRGGAEQF | 0.68553865 |
| VGPKRFGV | ASSWDRGGAEQF | 0.80119884 |
| VNVAFPD | ASSWDRGGAEQF | 0.52391994 |
| VSMTNSGVS | ASSWDRGGAEQF | 0.81357765 |
| VTDVLGDLSV | ASSWDRGGAEQF | 0.73991644 |
| VVVVGAGFAGL | ASSWDRGGAEQF | 0.4284932 |
| VYTVGTVV | ASSWDRGGAEQF | 0.7792688 |
| WTTFSGP | ASSWDRGGAEQF | 0.79518044 |
| YDELVEAF | ASSWDRGGAEQF | 0.80168897 |
| YGTGGAGGNGG | ASSWDRGGAEQF | 0.29879776 |
| YTNVMAV | ASSWDRGGAEQF | 0.7579262 |
| FLEQVHQL | ASSWDRGGAEQF | 0.5217491 |
| GLSLLAVAV | ASSWDRGGAEQF | 0.7185091 |
| LEQVHQL | ASSWDRGGAEQF | 0.70917994 |
| LNAAGAI | ASSWDRGGAEQF | 0.5983261 |
| PAVHASL | ASSWDRGGAEQF | 0.80933976 |
| TELEDDA | ASSWDRGGAEQF | 0.54145586 |
| TPPGPPN | ASSWDRGGAEQF | 0.600132 |
| TPPPPPPP | ASSWDRGGAEQF | 0.6290695 |
| TQSPALI | ASSWTGGNTLY | 0.7473099 |
| EFAAAFPQ | ASSWTGGNTLY | 0.7640741 |
| LLSGNLDVL | ASSWTGGNTLY | 0.43752354 |
| GPANLGNYNI | ASSWTGGNTLY | 0.70535475 |
| TLGPTLP | ASSWTGGNTLY | 0.66246355 |
| VYAYSSCCK | ASSWTGGNTLY | 0.50780725 |
| APAPAPAPNI | ASSWTGGNTLY | 0.7844765 |

| | | |
|---|---|---|
| PGGPVPIP | ASSWTGGNTLY | 0.68716013 |
| FLTVALEP | ASSWTGGNTLY | 0.32808447 |
| SQYFGANL | ASSWTGGNTLY | 0.8013927 |
| GGVKPAA | ASSWTGGNTLY | 0.5511705 |
| GGTGGAGGAGGW | ASSWTGGNTLY | 0.76156545 |
| GVGGVAGLGGAGG | ASSWTGGNTLY | 0.85440564 |
| VLLVLATTI | ASSWTGGNTLY | 0.33883694 |
| IGIFAAML | ASSWTGGNTLY | 0.8233016 |
| AVGDLID | ASSWTGGNTLY | 0.72961366 |
| NISPLPL | ASSWTGGNTLY | 0.7379735 |
| VTAVFVAI | ASSWTGGNTLY | 0.7251492 |
| SGDDNDAAL | ASSWTGGNTLY | 0.5341531 |
| AVAALAHS | ASSWTGGNTLY | 0.8362111 |
| GGVTVMATP | ASSWTGGNTLY | 0.7815983 |
| AGLAANVRD | ASSWTGGNTLY | 0.74442196 |
| TNRPDLIDPALLRPG | ASSWTGGNTLY | 0.7811107 |
| LANLGVTAL | ASSWTGGNTLY | 0.79776955 |
| TGERIEL | ASSWTGGNTLY | 0.8173878 |
| NGEELAA | ASSWTGGNTLY | 0.5344204 |
| VLSLVWG | ASSWTGGNTLY | 0.3557667 |
| IGYALWL | ASSWTGGNTLY | 0.70218635 |
| LSLLLAPL | ASSWTGGNTLY | 0.651478 |
| TLDEALK | ASSWTGGNTLY | 0.59568095 |
| TDGAGTAEGG | ASSWTGGNTLY | 0.3635475 |
| NAPGGAGGNGG | ASSWTGGNTLY | 0.7758054 |
| VSVFMGMGGNF | ASSWTGGNTLY | 0.86648023 |
| LASYLAAL | ASSWTGGNTLY | 0.65683323 |
| LVGGTPAL | ASSWTGGNTLY | 0.8634656 |
| VRNADVL | ASSWTGGNTLY | 0.45356584 |
| TCSSPFLVSAM | ASSWTGGNTLY | 0.80183226 |
| VHITDAE | ASSWTGGNTLY | 0.81163025 |
| EYAAPSF | ASSWTGGNTLY | 0.47149712 |
| VVDAFHGGL | ASSWTGGNTLY | 0.73243874 |
| ASVVDPQ | ASSWTGGNTLY | 0.5764445 |
| SYNEFGGLS | ASSWTGGNTLY | 0.5732485 |
| DPDGSVY | ASSWTGGNTLY | 0.820447 |
| MVSTIHATE | ASSWTGGNTLY | 0.6151894 |
| LAAGVADPV | ASSWTGGNTLY | 0.78427607 |
| TKEQIAA | ASSWTGGNTLY | 0.6768472 |
| SLASPAAL | ASSWTGGNTLY | 0.5430012 |
| IGLVFYGV | ASSWTGGNTLY | 0.7827072 |
| FLLDASNL | ASSWTGGNTLY | 0.80322707 |
| YQVGNFP | ASSWTGGNTLY | 0.77592516 |
| ESLLSQST | ASSWTGGNTLY | 0.8798611 |
| LFTVGFAAVG | ASSWTGGNTLY | 0.5722084 |
| ATPPTPT | ASSWTGGNTLY | 0.8489609 |
| TDLATLG | ASSWTGGNTLY | 0.84092826 |
| SAGGTGR | ASSWTGGNTLY | 0.7073704 |
| GFSNSGS | ASSWTGGNTLY | 0.64810395 |
| GGNGGSGGTSV | ASSWTGGNTLY | 0.7845325 |
| GTSVATGGAGN | ASSWTGGNTLY | 0.59589696 |

| | | |
|---|---|---|
| NGGNGGSGGTS | ASSWTGGNTLY | 0.7820653 |
| AVVKFPH | ASSWTGGNTLY | 0.85582733 |
| TPPPPPP | ASSWTGGNTLY | 0.32080543 |
| AAPAAAAAPA | ASSWTGGNTLY | 0.72819114 |
| IALNEDL | ASSWTGGNTLY | 0.55667144 |
| IVNVSSM | ASSWTGGNTLY | 0.58109075 |
| TGIPPAPR | ASSWTGGNTLY | 0.21419582 |
| FDLGGGTF | ASSWTGGNTLY | 0.12487975 |
| FDLGGGTFDVS | ASSWTGGNTLY | 0.549367 |
| NEPTAAA | ASSWTGGNTLY | 0.16615382 |
| NEPTAAALA | ASSWTGGNTLY | 0.23815782 |
| GSIINISSI | ASSWTGGNTLY | 0.6316853 |
| VAPGNAGTA | ASSWTGGNTLY | 0.28140524 |
| TGGGASLE | ASSWTGGNTLY | 0.7755537 |
| VAVYNHYKRI | ASSWTGGNTLY | 0.7243928 |
| TDPAPATT | ASSWTGGNTLY | 0.20265609 |
| TNRPDILDPALLRPG | ASSWTGGNTLY | 0.47107887 |
| LTTTTTS | ASSWTGGNTLY | 0.29255867 |
| PTGSGKTL | ASSWTGGNTLY | 0.51260483 |
| TGSGKTL | ASSWTGGNTLY | 0.14100838 |
| TIGHVDHGK | ASSWTGGNTLY | 0.5969629 |
| TIGHVDHGKTT | ASSWTGGNTLY | 0.32264698 |
| GHVDHGKTT | ASSWTGGNTLY | 0.10816239 |
| AQFEHTLL | ASSWTGGNTLY | 0.62619364 |
| IVNQDEL | ASSWTGGNTLY | 0.4834904 |
| TNGSQFF | ASSWTGGNTLY | 0.34075046 |
| TGKGIVE | ASSWTGGNTLY | 0.7692085 |
| LISNASDA | ASSWTGGNTLY | 0.12514664 |
| AELRTLGELTKIAKAHGAQ | ASSWTGGNTLY | 0.18694475 |
| APNAVAAAEPSW | ASSWTGGNTLY | 0.6949244 |
| APQVAGAATAVL | ASSWTGGNTLY | 0.7445437 |
| ARRDPAAAAAILPTDA | ASSWTGGNTLY | 0.183595 |
| DAAARDVLAEAGLAEYF | ASSWTGGNTLY | 0.6698816 |
| DDQIVLSAKANLAKVEQ | ASSWTGGNTLY | 0.7634709 |
| DELLPDTGEDSDEDGH | ASSWTGGNTLY | 0.40539396 |
| ELPEAPANRATVPQD | ASSWTGGNTLY | 0.7126973 |
| FGQSKIERVTGIG | ASSWTGGNTLY | 0.73614717 |
| GAGGTGSTAGGAGGAGGAGGLY | ASSWTGGNTLY | 0.18705851 |
| GHGGTGAAGGNGGTGGAGGSGI | ASSWTGGNTLY | 0.6986045 |
| GNGGNGGAGGEGGPGGAGGAGA | ASSWTGGNTLY | 0.72080356 |
| GSTAGNGGNGGNATNGTIAGQP | ASSWTGGNTLY | 0.38572463 |
| GTGGDGGNGGNGGTGG | ASSWTGGNTLY | 0.514112 |
| GVRRPAGIPADAPV | ASSWTGGNTLY | 0.2663909 |
| GYRQVTDAHLLTLARR | ASSWTGGNTLY | 0.45470554 |
| IAKMGQDGHDRG | ASSWTGGNTLY | 0.10950513 |
| KAEPSIEIQYT | ASSWTGGNTLY | 0.6268259 |
| KPVRIRMAADEDTVAVT | ASSWTGGNTLY | 0.58034605 |
| MPPRIAGMRLLVIKPEP | ASSWTGGNTLY | 0.5063648 |
| MKFYAVALFRSRGVPPTR | ASSWTGGNTLY | 0.3870781 |
| NAGGFGILGAVAHSP | ASSWTGGNTLY | 0.20259029 |
| NEGREITLAMAEM | ASSWTGGNTLY | 0.22860776 |

| | | |
|---|---|---|
| NELVEWSHPLGHRWRWE | ASSWTGGNTLY | 0.5163442 |
| NLGAGGTGGNGGNPG | ASSWTGGNTLY | 0.13203177 |
| NTEWPKWETVMAADPVD | ASSWTGGNTLY | 0.2452156 |
| PASAAPAETQP | ASSWTGGNTLY | 0.60572976 |
| QQSSDGKNGVRAAEITGE | ASSWTGGNTLY | 0.62191343 |
| RAPDAVTTSWWKE | ASSWTGGNTLY | 0.17718336 |
| SGFGNVGAHTSGWFNQ | ASSWTGGNTLY | 0.7545712 |
| SGVAGAVGQAASVGGLKV | ASSWTGGNTLY | 0.6096951 |
| SNGGAGTGSAGGNGGT | ASSWTGGNTLY | 0.6639898 |
| TGGAGGEGGAAG | ASSWTGGNTLY | 0.47218806 |
| TGGDGGAAGAGGAPGHG | ASSWTGGNTLY | 0.28816587 |
| TMPPLLAAIANAGRHGV | ASSWTGGNTLY | 0.7050974 |
| TSSTAGNSGGA | ASSWTGGNTLY | 0.46019286 |
| VPATVPVGILC | ASSWTGGNTLY | 0.19753596 |
| VQRPDSGGVVLTGRIS | ASSWTGGNTLY | 0.6594026 |
| WTPNPVPATKGVPAGLVQAT | ASSWTGGNTLY | 0.2321853 |
| YALKRALAKFSDRFDVVII | ASSWTGGNTLY | 0.31334156 |
| YLIGSVIV | ASSWTGGNTLY | 0.5962521 |
| YGQLVAVID | ASSWTGGNTLY | 0.6881268 |
| VTVGGMTLPA | ASSWTGGNTLY | 0.3346966 |
| VLALPAV | ASSWTGGNTLY | 0.15948719 |
| VIPPTLNYET | ASSWTGGNTLY | 0.28464353 |
| VGMGGMHPSSG | ASSWTGGNTLY | 0.6110989 |
| VGLAGGAATA | ASSWTGGNTLY | 0.7415613 |
| VGGAGGAGGSGP | ASSWTGGNTLY | 0.5033824 |
| VFAAFMV | ASSWTGGNTLY | 0.31036633 |
| TLYDKLR | ASSWTGGNTLY | 0.7267442 |
| TLAVFTVL | ASSWTGGNTLY | 0.49710244 |
| TGSVQPARV | ASSWTGGNTLY | 0.43625724 |
| TGGGAGSGGT | ASSWTGGNTLY | 0.61030525 |
| SHLETSL | ASSWTGGNTLY | 0.28387597 |
| SFTTGMT | ASSWTGGNTLY | 0.69091296 |
| QVEVAKRMGVRQPTVS | ASSWTGGNTLY | 0.6206419 |
| QGSGGPALAM | ASSWTGGNTLY | 0.6981107 |
| MLAPSNP | ASSWTGGNTLY | 0.706229 |
| LVNNAAI | ASSWTGGNTLY | 0.21881834 |
| LTGEINTLA | ASSWTGGNTLY | 0.5184952 |
| LSIFGPV | ASSWTGGNTLY | 0.13293695 |
| LLALLANL | ASSWTGGNTLY | 0.25005698 |
| LISNASDALDKLR | ASSWTGGNTLY | 0.66866374 |
| LINEAVP | ASSWTGGNTLY | 0.6540233 |
| LGYAFSGL | ASSWTGGNTLY | 0.6915697 |
| LDFVAVAL | ASSWTGGNTLY | 0.5891469 |
| LAQPGAE | ASSWTGGNTLY | 0.44256458 |
| ISNLGAVTGF | ASSWTGGNTLY | 0.7002924 |
| HQEDPGRI | ASSWTGGNTLY | 0.7568134 |
| GTPGEPTP | ASSWTGGNTLY | 0.17103098 |
| GFFNSGS | ASSWTGGNTLY | 0.25734815 |
| GAGGGGLTAGAN | ASSWTGGNTLY | 0.7674469 |
| FVDALVNM | ASSWTGGNTLY | 0.6367811 |
| FLTVAPEL | ASSWTGGNTLY | 0.5215137 |

| | | |
|---|---|---|
| FETLTAL | ASSWTGGNTLY | 0.5025861 |
| DVINPIEL | ASSWTGGNTLY | 0.5787783 |
| APVSGAPV | ASSWTGGNTLY | 0.1796793 |
| ALVGGLAW | ASSWTGGNTLY | 0.7698177 |
| ALAAAAATAI | ASSWTGGNTLY | 0.12265975 |
| AFGLGLLL | ASSWTGGNTLY | 0.5010668 |
| AFFAAASL | ASSWTGGNTLY | 0.17223865 |
| AAAALPAFSPP | ASSWTGGNTLY | 0.14236437 |
| VVLAPAL | ASSWTGGNTLY | 0.5708977 |
| TIGHVDHGKTTL | ASSWTGGNTLY | 0.5114266 |
| LISNASDALDK | ASSWTGGNTLY | 0.7744092 |
| LISNASDAL | ASSWTGGNTLY | 0.46295017 |
| ISNASDALDK | ASSWTGGNTLY | 0.5919539 |
| NQAETLVYQTE | ASSWTGGNTLY | 0.5801485 |
| TGGTGGTAGAG | ASSWTGGNTLY | 0.5130478 |
| TGSTAGGAGGA | ASSWTGGNTLY | 0.28050333 |
| DPLAVPVAAGPV | ASSWTGGNTLY | 0.48219988 |
| GGTGGTPGNGGN | ASSWTGGNTLY | 0.74036086 |
| PAGLAGVDPATL | ASSWTGGNTLY | 0.6190214 |
| GNNLVVEAPHAGASV | ASSWTGGNTLY | 0.1427583 |
| KVTRIMSASTLEQPA | ASSWTGGNTLY | 0.41290268 |
| LGFALPIALPRRTVV | ASSWTGGNTLY | 0.3359266 |
| TRLGNLTARKLVVLV | ASSWTGGNTLY | 0.10601168 |
| WSQQRRLIYFPSAGP | ASSWTGGNTLY | 0.114448115 |
| GPNGNPDPMAAAVDIR | ASSWTGGNTLY | 0.15203759 |
| TQRCVSVADLRAAASR | ASSWTGGNTLY | 0.30371547 |
| QFVLDVAEHPVEVLTETV | ASSWTGGNTLY | 0.7602544 |
| GGAGGTGGAGGNGGTQAGDG | ASSWTGGNTLY | 0.7226444 |
| PLLATLRAIAVDVDAKPAQV | ASSWTGGNTLY | 0.44695923 |
| APKFRTGGYMIDFWGVGYQVAK | ASSWTGGNTLY | 0.6648304 |
| GAGGAGGAGGAAGGPGGTGGTG | ASSWTGGNTLY | 0.42716214 |
| IGALGLLATDVAPAGSVAFGVA | ASSWTGGNTLY | 0.1007301 |
| AGGAGGIGGNGGAGGAGGVLMG | ASSWTGGNTLY | 0.21392716 |
| ADELIGLL | ASSWTGGNTLY | 0.26748917 |
| ADRALKA | ASSWTGGNTLY | 0.74192315 |
| AGAVGQVGGAGGA | ASSWTGGNTLY | 0.20625016 |
| AGGGGGLGGAP | ASSWTGGNTLY | 0.50729024 |
| AGGLLYGNGGAG | ASSWTGGNTLY | 0.520715 |
| AGLESIGL | ASSWTGGNTLY | 0.50889874 |
| AGLKTGII | ASSWTGGNTLY | 0.5827892 |
| ASLAALLL | ASSWTGGNTLY | 0.17478482 |
| ASVINISSI | ASSWTGGNTLY | 0.11322731 |
| ATPPGPA | ASSWTGGNTLY | 0.49698803 |
| ATVASALV | ASSWTGGNTLY | 0.11563543 |
| ATVASLIL | ASSWTGGNTLY | 0.25413585 |
| AVNGPVLIRH | ASSWTGGNTLY | 0.13805275 |
| DGAAGAVGI | ASSWTGGNTLY | 0.21900868 |
| DPLLHGI | ASSWTGGNTLY | 0.13019931 |
| ELGAALGAL | ASSWTGGNTLY | 0.6580672 |
| ELGRTQAV | ASSWTGGNTLY | 0.67078465 |
| FVAGGYLS | ASSWTGGNTLY | 0.61870337 |

| | | |
|---|---|---|
| GASGGAGGAGGW | ASSWTGGNTLY | 0.7470598 |
| GGAGAGGDGGAGGI | ASSWTGGNTLY | 0.61006635 |
| GTGHNGSQPGMGGNG | ASSWTGGNTLY | 0.49966317 |
| GYAAASASAS | ASSWTGGNTLY | 0.19624586 |
| HQVDVAAVQQG | ASSWTGGNTLY | 0.28334844 |
| IGAAIAHT | ASSWTGGNTLY | 0.18813558 |
| IGMIEVAQL | ASSWTGGNTLY | 0.19078751 |
| IRLAPEHTSAMD | ASSWTGGNTLY | 0.7961048 |
| IYNVSVL | ASSWTGGNTLY | 0.4663633 |
| KQGFLGM | ASSWTGGNTLY | 0.18342069 |
| LESVITQL | ASSWTGGNTLY | 0.32778597 |
| LGLGALVVV | ASSWTGGNTLY | 0.7355001 |
| LIGNGGNGGSGGI | ASSWTGGNTLY | 0.7465729 |
| LNAINAPTL | ASSWTGGNTLY | 0.35815555 |
| LQLLNTGV | ASSWTGGNTLY | 0.32498464 |
| LQTSIEF | ASSWTGGNTLY | 0.52872384 |
| MSFAVSEL | ASSWTGGNTLY | 0.5885097 |
| NTGALGL | ASSWTGGNTLY | 0.43281272 |
| PPGAGGAGGAGTT | ASSWTGGNTLY | 0.16279264 |
| QLVGLGI | ASSWTGGNTLY | 0.6844032 |
| RFLEQVHQL | ASSWTGGNTLY | 0.7588515 |
| RGVGYVLR | ASSWTGGNTLY | 0.31237245 |
| SATGVFAGVI | ASSWTGGNTLY | 0.17736463 |
| SLPAITLP | ASSWTGGNTLY | 0.49789008 |
| STALAFTV | ASSWTGGNTLY | 0.23540807 |
| SVLTVGLGK | ASSWTGGNTLY | 0.44016975 |
| TALAGAGSGI | ASSWTGGNTLY | 0.3837256 |
| TGLPFTGI | ASSWTGGNTLY | 0.21651575 |
| VAEHVTV | ASSWTGGNTLY | 0.45241126 |
| VDIVLNSL | ASSWTGGNTLY | 0.7055079 |
| VLTGGGAVL | ASSWTGGNTLY | 0.2418226 |
| VNVGLAYGI | ASSWTGGNTLY | 0.7513049 |
| VSSAAVGAV | ASSWTGGNTLY | 0.77056646 |
| VTSVYEAGV | ASSWTGGNTLY | 0.720913 |
| WVALDRGA | ASSWTGGNTLY | 0.77312416 |
| YLVGSVVI | ASSWTGGNTLY | 0.64411116 |
| GPPGTGKTLL | ASSWTGGNTLY | 0.40024346 |
| GVGMTKF | ASSWTGGNTLY | 0.36515445 |
| IGIGGSDL | ASSWTGGNTLY | 0.6642526 |
| IGTLLAEL | ASSWTGGNTLY | 0.5041815 |
| IYTSGTT | ASSWTGGNTLY | 0.5285722 |
| LSFPAGV | ASSWTGGNTLY | 0.31470004 |
| LYTSGTT | ASSWTGGNTLY | 0.75458914 |
| VGAFPFGF | ASSWTGGNTLY | 0.33523923 |
| VGAGTVEF | ASSWTGGNTLY | 0.64892715 |
| VGKSTLLL | ASSWTGGNTLY | 0.22363967 |
| AALAHAGGGIV | ASSWTGGNTLY | 0.18143019 |
| AALVAVSLL | ASSWTGGNTLY | 0.72851795 |
| AAVSRAVAAL | ASSWTGGNTLY | 0.10937802 |
| AGGSSGIGGF | ASSWTGGNTLY | 0.13003369 |
| AGLTVANANG | ASSWTGGNTLY | 0.58204824 |

| | | |
|---|---|---|
| AGLVGAVL | ASSWTGGNTLY | 0.678308 |
| ALSQAFTV | ASSWTGGNTLY | 0.10185926 |
| ALTFSVSL | ASSWTGGNTLY | 0.20608522 |
| ALTGAALVAL | ASSWTGGNTLY | 0.40316066 |
| ASFSIGAGVL | ASSWTGGNTLY | 0.30366826 |
| DGSVYKI | ASSWTGGNTLY | 0.31148946 |
| EQLAEIV | ASSWTGGNTLY | 0.12332697 |
| FAVSELTVM | ASSWTGGNTLY | 0.699263 |
| GAAPLAAGL | ASSWTGGNTLY | 0.1759272 |
| GADGGDGGSGGAGG | ASSWTGGNTLY | 0.11644799 |
| GFGLDTGGAGG | ASSWTGGNTLY | 0.60635877 |
| GLASGGLL | ASSWTGGNTLY | 0.71413183 |
| GRIAGAAVLL | ASSWTGGNTLY | 0.27713686 |
| GSAGSLNTSF | ASSWTGGNTLY | 0.6324692 |
| GTGQIGI | ASSWTGGNTLY | 0.4486677 |
| GTLIALIL | ASSWTGGNTLY | 0.7490245 |
| IAATMGFVVG | ASSWTGGNTLY | 0.28640157 |
| IFAQALL | ASSWTGGNTLY | 0.2075633 |
| IGAAVALGF | ASSWTGGNTLY | 0.72449857 |
| IGLTPLAL | ASSWTGGNTLY | 0.7131376 |
| ILGSNTSTL | ASSWTGGNTLY | 0.47834635 |
| IVFSLTVL | ASSWTGGNTLY | 0.66541004 |
| LASKATGF | ASSWTGGNTLY | 0.23576628 |
| LASVALIL | ASSWTGGNTLY | 0.4806942 |
| LFAVGLTL | ASSWTGGNTLY | 0.73658174 |
| LFNGGIGTYI | ASSWTGGNTLY | 0.6789878 |
| LTLGAGII | ASSWTGGNTLY | 0.13408656 |
| LVAAALGDI | ASSWTGGNTLY | 0.5931536 |
| NTAFVQLGI | ASSWTGGNTLY | 0.42497724 |
| QLFSGAEL | ASSWTGGNTLY | 0.3507262 |
| SAATFITL | ASSWTGGNTLY | 0.63159704 |
| SNYGLANLGV | ASSWTGGNTLY | 0.5244769 |
| SYGGAYAVM | ASSWTGGNTLY | 0.7047339 |
| TTGVGGLTLG | ASSWTGGNTLY | 0.41347635 |
| TTQTIYI | ASSWTGGNTLY | 0.6549673 |
| VAAGAGLGAVI | ASSWTGGNTLY | 0.13007396 |
| VAAVVTLL | ASSWTGGNTLY | 0.5494718 |
| VGFLGGLI | ASSWTGGNTLY | 0.5215251 |
| VGVIPILL | ASSWTGGNTLY | 0.6869128 |
| VIFASSLI | ASSWTGGNTLY | 0.44995084 |
| VKGGDGGAAAG | ASSWTGGNTLY | 0.17473565 |
| VTKTFTNI | ASSWTGGNTLY | 0.5996965 |
| VVAAAILL | ASSWTGGNTLY | 0.73276734 |
| YAALDVEL | ASSWTGGNTLY | 0.39365262 |
| LASVALLL | ASSWTGGNTLY | 0.5330236 |
| LDAVHRV | ASSWTGGNTLY | 0.79470026 |
| QGLSSSIF | ASSWTGGNTLY | 0.34195882 |
| RGTFANIRL | ASSWTGGNTLY | 0.37124252 |
| RQLWWGHRI | ASSWTGGNTLY | 0.6211493 |
| YGPPGTGKTLL | ASSWTGGNTLY | 0.7615284 |
| AALVGAPANVA | ASSWTGGNTLY | 0.4842366 |

| | | |
|---|---|---|
| AATFRRTE | ASSWTGGNTLY | 0.75973064 |
| AGALGGGI | ASSWTGGNTLY | 0.18926197 |
| AGGAGGAGGIGAG | ASSWTGGNTLY | 0.74553406 |
| AGSGALGL | ASSWTGGNTLY | 0.4692087 |
| AQIADQI | ASSWTGGNTLY | 0.6585426 |
| ASAVAVTYL | ASSWTGGNTLY | 0.14168935 |
| ASTVLGGTGG | ASSWTGGNTLY | 0.67916465 |
| AVGGEPAK | ASSWTGGNTLY | 0.71345717 |
| CEQGTFGGTC | ASSWTGGNTLY | 0.7472435 |
| DEGKAPAG | ASSWTGGNTLY | 0.11525046 |
| DTVVFPAL | ASSWTGGNTLY | 0.68439406 |
| FEGITMAG | ASSWTGGNTLY | 0.7561736 |
| FPLVVQD | ASSWTGGNTLY | 0.7814176 |
| GAFTTPQ | ASSWTGGNTLY | 0.31244126 |
| GFGGFGTAGGVGGA | ASSWTGGNTLY | 0.72122097 |
| GFLAQFL | ASSWTGGNTLY | 0.3030825 |
| GGAGGAGGNAGVA | ASSWTGGNTLY | 0.3703106 |
| GGVGTAGAGGAGGAGG | ASSWTGGNTLY | 0.26179785 |
| GNGGGGDAGNAGSGGN | ASSWTGGNTLY | 0.5222606 |
| GTGGAGGAGNDAGS | ASSWTGGNTLY | 0.50136274 |
| GTTNTGWFNS | ASSWTGGNTLY | 0.43469498 |
| IFMGPLFTL | ASSWTGGNTLY | 0.57115215 |
| ILAHAIEAI | ASSWTGGNTLY | 0.74219763 |
| IRSAIREAL | ASSWTGGNTLY | 0.6310733 |
| ITFAGGG | ASSWTGGNTLY | 0.7547686 |
| ITGMSGHQEV | ASSWTGGNTLY | 0.7630354 |
| IVAGLACTL | ASSWTGGNTLY | 0.73331386 |
| IVGGGLAGA | ASSWTGGNTLY | 0.1874846 |
| IVTGVLNI | ASSWTGGNTLY | 0.3752898 |
| LAQQGCI | ASSWTGGNTLY | 0.287327 |
| LCLHVEN | ASSWTGGNTLY | 0.46878108 |
| LDAISDGS | ASSWTGGNTLY | 0.18248397 |
| LGGFAAML | ASSWTGGNTLY | 0.6285168 |
| LGGKNPMI | ASSWTGGNTLY | 0.10176338 |
| LGPKGVAAIL | ASSWTGGNTLY | 0.64359677 |
| LMVGVSI | ASSWTGGNTLY | 0.79905 |
| LVGAAFVL | ASSWTGGNTLY | 0.119046204 |
| LVLAVAPL | ASSWTGGNTLY | 0.6033207 |
| MAAIAVIDAA | ASSWTGGNTLY | 0.68768287 |
| NNGINAELQ | ASSWTGGNTLY | 0.3186996 |
| RGEPLVLVHG | ASSWTGGNTLY | 0.7179425 |
| RGGAGGAAGLI | ASSWTGGNTLY | 0.23370668 |
| RSYRMHPAV | ASSWTGGNTLY | 0.71245354 |
| SAAHLLAGQLA | ASSWTGGNTLY | 0.69456846 |
| SAAVGSY | ASSWTGGNTLY | 0.5838921 |
| SPHAVLHRI | ASSWTGGNTLY | 0.49756688 |
| SQIAKVL | ASSWTGGNTLY | 0.5479435 |
| TALGHLVP | ASSWTGGNTLY | 0.12587593 |
| TAPTTSGAA | ASSWTGGNTLY | 0.5894401 |
| TDGDDGNGGSPGT | ASSWTGGNTLY | 0.10780699 |
| TDGLGEN | ASSWTGGNTLY | 0.60266227 |

| | | |
|---|---|---|
| TGGPPLPGQA | ASSWTGGNTLY | 0.18689574 |
| TIPAITI | ASSWTGGNTLY | 0.7317219 |
| TIPGFTI | ASSWTGGNTLY | 0.47533983 |
| TSIASGALA | ASSWTGGNTLY | 0.5259106 |
| TVTLPGAAI | ASSWTGGNTLY | 0.32595235 |
| VASGLSAALP | ASSWTGGNTLY | 0.2847743 |
| VGAGVIGI | ASSWTGGNTLY | 0.19067775 |
| VGPKRFGV | ASSWTGGNTLY | 0.14818959 |
| VNVAFPD | ASSWTGGNTLY | 0.6817942 |
| VSMTNSGVS | ASSWTGGNTLY | 0.36093706 |
| VTDVLGDLSV | ASSWTGGNTLY | 0.36853525 |
| VVVVGAGFAGL | ASSWTGGNTLY | 0.5942599 |
| VYTVGTVV | ASSWTGGNTLY | 0.49761438 |
| WTTFSGP | ASSWTGGNTLY | 0.13776161 |
| YDELVEAF | ASSWTGGNTLY | 0.55827993 |
| YGTGGAGGNGG | ASSWTGGNTLY | 0.5629915 |
| YTNVMAV | ASSWTGGNTLY | 0.25531325 |
| FLEQVHQL | ASSWTGGNTLY | 0.100853406 |
| GLSLLAVAV | ASSWTGGNTLY | 0.54704523 |
| LEQVHQL | ASSWTGGNTLY | 0.34887993 |
| LNAAGAI | ASSWTGGNTLY | 0.2634505 |
| PAVHASL | ASSWTGGNTLY | 0.24171048 |
| TELEDDA | ASSWTGGNTLY | 0.77103657 |
| TPPGPPN | ASSWTGGNTLY | 0.11153877 |
| TPPPPPPP | ASSWTGGNTLY | 0.29744968 |
| TQSPALI | ASSLLGYEQY | 0.71599805 |
| EFAAAFPQ | ASSLLGYEQY | 0.49871978 |
| LLSGNLDVL | ASSLLGYEQY | 0.77086294 |
| GPANLGNYNI | ASSLLGYEQY | 0.29309502 |
| TLGPTLP | ASSLLGYEQY | 0.24759056 |
| VYAYSSCCK | ASSLLGYEQY | 0.62177134 |
| APAPAPAPNI | ASSLLGYEQY | 0.4072973 |
| PGGPVPIP | ASSLLGYEQY | 0.18713361 |
| FLTVALEP | ASSLLGYEQY | 0.118393734 |
| SQYFGANL | ASSLLGYEQY | 0.40509298 |
| GGVKPAA | ASSLLGYEQY | 0.14988367 |
| GGTGGAGGAGGW | ASSLLGYEQY | 0.40559274 |
| GVGGVAGLGGAGG | ASSLLGYEQY | 0.71600235 |
| VLLVLATTI | ASSLLGYEQY | 0.2507606 |
| IGIFAAML | ASSLLGYEQY | 0.18943572 |
| AVGDLID | ASSLLGYEQY | 0.47133952 |
| NISPLPL | ASSLLGYEQY | 0.15690154 |
| VTAVFVAI | ASSLLGYEQY | 0.71335804 |
| SGDDNDAAL | ASSLLGYEQY | 0.20557907 |
| AVAALAHS | ASSLLGYEQY | 0.75670874 |
| GGVTVMATP | ASSLLGYEQY | 0.70529443 |
| AGLAANVRD | ASSLLGYEQY | 0.33706895 |
| TNRPDLIDPALLRPG | ASSLLGYEQY | 0.75723976 |
| LANLGVTAL | ASSLLGYEQY | 0.4081072 |
| TGERIEL | ASSLLGYEQY | 0.5977535 |
| NGEELAA | ASSLLGYEQY | 0.53361297 |

| | | |
|---|---|---|
| VLSLVWG | ASSLLGYEQY | 0.31494698 |
| IGYALWL | ASSLLGYEQY | 0.32381025 |
| LSLLLAPL | ASSLLGYEQY | 0.5394573 |
| TLDEALK | ASSLLGYEQY | 0.13342427 |
| TDGAGTAEGG | ASSLLGYEQY | 0.5531447 |
| NAPGGAGGNGG | ASSLLGYEQY | 0.13682084 |
| VSVFMGMGGNF | ASSLLGYEQY | 0.13067086 |
| LASYLAAL | ASSLLGYEQY | 0.46341008 |
| LVGGTPAL | ASSLLGYEQY | 0.7400906 |
| VRNADVL | ASSLLGYEQY | 0.79288805 |
| TCSSPFLVSAM | ASSLLGYEQY | 0.3423162 |
| VHITDAE | ASSLLGYEQY | 0.16032475 |
| EYAAPSF | ASSLLGYEQY | 0.67427015 |
| VVDAFHGGL | ASSLLGYEQY | 0.13833024 |
| ASVVDPQ | ASSLLGYEQY | 0.6388492 |
| SYNEFGGLS | ASSLLGYEQY | 0.45476425 |
| DPDGSVY | ASSLLGYEQY | 0.32377803 |
| MVSTIHATE | ASSLLGYEQY | 0.37524772 |
| LAAGVADPV | ASSLLGYEQY | 0.627468 |
| TKEQIAA | ASSLLGYEQY | 0.20186904 |
| SLASPAAL | ASSLLGYEQY | 0.66000473 |
| IGLVFYGV | ASSLLGYEQY | 0.10096156 |
| FLLDASNL | ASSLLGYEQY | 0.23816904 |
| YQVGNFP | ASSLLGYEQY | 0.47745013 |
| ESLLSQST | ASSLLGYEQY | 0.19250846 |
| LFTVGFAAVG | ASSLLGYEQY | 0.13573062 |
| ATPPTPT | ASSLLGYEQY | 0.46351 |
| TDLATLG | ASSLLGYEQY | 0.6982497 |
| SAGGTGR | ASSLLGYEQY | 0.734411 |
| GFSNSGS | ASSLLGYEQY | 0.20237859 |
| GGNGGSGGTSV | ASSLLGYEQY | 0.72214735 |
| GTSVATGGAGN | ASSLLGYEQY | 0.7262588 |
| NGGNGGSGGTS | ASSLLGYEQY | 0.5986893 |
| AVVKFPH | ASSLLGYEQY | 0.14196888 |
| TPPPPPP | ASSLLGYEQY | 0.44424102 |
| AAPAAAAAPA | ASSLLGYEQY | 0.12224178 |
| IALNEDL | ASSLLGYEQY | 0.5221468 |
| IVNVSSM | ASSLLGYEQY | 0.6347721 |
| TGIPPAPR | ASSLLGYEQY | 0.21064073 |
| FDLGGGTF | ASSLLGYEQY | 0.6268429 |
| FDLGGGTFDVS | ASSLLGYEQY | 0.8042742 |
| NEPTAAA | ASSLLGYEQY | 0.28581554 |
| NEPTAAALA | ASSLLGYEQY | 0.2824979 |
| GSIINISSI | ASSLLGYEQY | 0.23310015 |
| VAPGNAGTA | ASSLLGYEQY | 0.15782417 |
| TGGGASLE | ASSLLGYEQY | 0.43358803 |
| VAVYNHYKRI | ASSLLGYEQY | 0.50584894 |
| TDPAPATT | ASSLLGYEQY | 0.24822298 |
| TNRPDILDPALLRPG | ASSLLGYEQY | 0.5278574 |
| LTTTTTS | ASSLLGYEQY | 0.21484019 |
| PTGSGKTL | ASSLLGYEQY | 0.2507429 |

| | | |
|---|---|---|
| TGSGKTL | ASSLLGYEQY | 0.26385665 |
| TIGHVDHGK | ASSLLGYEQY | 0.18692511 |
| TIGHVDHGKTT | ASSLLGYEQY | 0.6677948 |
| GHVDHGKTT | ASSLLGYEQY | 0.22575456 |
| AQFEHTLL | ASSLLGYEQY | 0.5642278 |
| IVNQDEL | ASSLLGYEQY | 0.7948315 |
| TNGSQFF | ASSLLGYEQY | 0.12275384 |
| TGKGIVE | ASSLLGYEQY | 0.21611428 |
| LISNASDA | ASSLLGYEQY | 0.64677006 |
| AELRTLGELTKIAKAHGAQ | ASSLLGYEQY | 0.12099018 |
| APNAVAAAEPSW | ASSLLGYEQY | 0.7308227 |
| APQVAGAATAVL | ASSLLGYEQY | 0.4765525 |
| ARRDPAAAAAILPTDA | ASSLLGYEQY | 0.6373884 |
| DAAARDVLAEAGLAEYF | ASSLLGYEQY | 0.25300437 |
| DDQIVLSAKANLAKVEQ | ASSLLGYEQY | 0.7824087 |
| DELLPDTGEDSDEDGH | ASSLLGYEQY | 0.7367357 |
| ELPEAPANRATVPQD | ASSLLGYEQY | 0.45270845 |
| FGQSKIERVTGIG | ASSLLGYEQY | 0.27071607 |
| GAGGTGSTAGGAGGAGGAGGLY | ASSLLGYEQY | 0.17846954 |
| GHGGTGAAGGNGGTGGAGGSGI | ASSLLGYEQY | 0.7009636 |
| GNGGNGGAGGEGGPGGAGGAGA | ASSLLGYEQY | 0.7235749 |
| GSTAGNGGNGGNATNGTIAGQP | ASSLLGYEQY | 0.21480191 |
| GTGGDGGNGGNGGTGG | ASSLLGYEQY | 0.5002271 |
| GVRRPAGIPADAPV | ASSLLGYEQY | 0.69530666 |
| GYRQVTDAHLLTLARR | ASSLLGYEQY | 0.8015772 |
| IAKMGQDGHDRG | ASSLLGYEQY | 0.12892622 |
| KAEPSIEIQYT | ASSLLGYEQY | 0.47546127 |
| KPVRIRMAADEDTVAVT | ASSLLGYEQY | 0.17922139 |
| MPPRIAGMRLLVIKPEP | ASSLLGYEQY | 0.22911741 |
| MKFYAVALFRSRGVPPTR | ASSLLGYEQY | 0.18708421 |
| NAGGFGILGAVAHSP | ASSLLGYEQY | 0.64518946 |
| NEGREITLAMAEM | ASSLLGYEQY | 0.51625985 |
| NELVEWSHPLGHRWRWE | ASSLLGYEQY | 0.6661937 |
| NLGAGGTGGNGGNPG | ASSLLGYEQY | 0.39507502 |
| NTEWPKWETVMAADPVD | ASSLLGYEQY | 0.6080278 |
| PASAAPAETQP | ASSLLGYEQY | 0.4117289 |
| QQSSDGKNGVRAAEITGE | ASSLLGYEQY | 0.28980604 |
| RAPDAVTTSWWKE | ASSLLGYEQY | 0.18540327 |
| SGFGNVGAHTSGWFNQ | ASSLLGYEQY | 0.11565625 |
| SGVAGAVGQAASVGGLKV | ASSLLGYEQY | 0.24519 |
| SNGGAGTGSAGGNGGT | ASSLLGYEQY | 0.25629058 |
| TGGAGGEGGAAG | ASSLLGYEQY | 0.103089474 |
| TGGDGGAAGAGGAPGHG | ASSLLGYEQY | 0.64796317 |
| TMPPLLAAIANAGRHGV | ASSLLGYEQY | 0.6645757 |
| TSSTAGNSGGA | ASSLLGYEQY | 0.49421257 |
| VPATVPVGILC | ASSLLGYEQY | 0.2771663 |
| VQRPDSGGVVLTGRIS | ASSLLGYEQY | 0.4636235 |
| WTPNPVPATKGVPAGLVQAT | ASSLLGYEQY | 0.17494622 |
| YALKRALAKFSDRFDVVII | ASSLLGYEQY | 0.74832654 |
| YLIGSVIV | ASSLLGYEQY | 0.17179221 |
| YGQLVAVID | ASSLLGYEQY | 0.16117145 |

| | | |
|---|---|---|
| VTVGGMTLPA | ASSLLGYEQY | 0.25996208 |
| VLALPAV | ASSLLGYEQY | 0.2725099 |
| VIPPTLNYET | ASSLLGYEQY | 0.72745836 |
| VGMGGMHPSSG | ASSLLGYEQY | 0.70382035 |
| VGLAGGAATA | ASSLLGYEQY | 0.69098705 |
| VGGAGGAGGSGP | ASSLLGYEQY | 0.6487772 |
| VFAAFMV | ASSLLGYEQY | 0.12729605 |
| TLYDKLR | ASSLLGYEQY | 0.23141988 |
| TLAVFTVL | ASSLLGYEQY | 0.80922884 |
| TGSVQPARV | ASSLLGYEQY | 0.36742374 |
| TGGGAGSGGT | ASSLLGYEQY | 0.5670267 |
| SHLETSL | ASSLLGYEQY | 0.67811394 |
| SFTTGMT | ASSLLGYEQY | 0.273435 |
| QVEVAKRMGVRQPTVS | ASSLLGYEQY | 0.2574492 |
| QGSGGPALAM | ASSLLGYEQY | 0.4574778 |
| MLAPSNP | ASSLLGYEQY | 0.6681844 |
| LVNNAAI | ASSLLGYEQY | 0.5139594 |
| LTGEINTLA | ASSLLGYEQY | 0.20129359 |
| LSIFGPV | ASSLLGYEQY | 0.6734152 |
| LLALLANL | ASSLLGYEQY | 0.81560504 |
| LISNASDALDKLR | ASSLLGYEQY | 0.6326525 |
| LINEAVP | ASSLLGYEQY | 0.46584725 |
| LGYAFSGL | ASSLLGYEQY | 0.5541614 |
| LDFVAVAL | ASSLLGYEQY | 0.5696038 |
| LAQPGAE | ASSLLGYEQY | 0.30536366 |
| ISNLGAVTGF | ASSLLGYEQY | 0.34903675 |
| HQEDPGRI | ASSLLGYEQY | 0.2928443 |
| GTPGEPTP | ASSLLGYEQY | 0.34006223 |
| GFFNSGS | ASSLLGYEQY | 0.6074772 |
| GAGGGGLTAGAN | ASSLLGYEQY | 0.34331715 |
| FVDALVNM | ASSLLGYEQY | 0.81382495 |
| FLTVAPEL | ASSLLGYEQY | 0.64686686 |
| FETLTAL | ASSLLGYEQY | 0.40719888 |
| DVINPIEL | ASSLLGYEQY | 0.19150321 |
| APVSGAPV | ASSLLGYEQY | 0.48150846 |
| ALVGGLAW | ASSLLGYEQY | 0.5107076 |
| ALAAAAATAI | ASSLLGYEQY | 0.1377444 |
| AFGLGLLL | ASSLLGYEQY | 0.7259653 |
| AFFAAASL | ASSLLGYEQY | 0.27553537 |
| AAAALPAFSPP | ASSLLGYEQY | 0.369089 |
| VVLAPAL | ASSLLGYEQY | 0.61751467 |
| TIGHVDHGKTTL | ASSLLGYEQY | 0.7273484 |
| LISNASDALDK | ASSLLGYEQY | 0.46201253 |
| LISNASDAL | ASSLLGYEQY | 0.2995084 |
| ISNASDALDK | ASSLLGYEQY | 0.17847599 |
| NQAETLVYQTE | ASSLLGYEQY | 0.64522594 |
| TGGTGGTAGAG | ASSLLGYEQY | 0.78843796 |
| TGSTAGGAGGA | ASSLLGYEQY | 0.15431245 |
| DPLAVPVAAGPV | ASSLLGYEQY | 0.73797905 |
| GGTGGTPGNGGN | ASSLLGYEQY | 0.27138233 |
| PAGLAGVDPATL | ASSLLGYEQY | 0.15862769 |

| | | |
|---|---|---|
| GNNLVVEAPHAGASV | ASSLLGYEQY | 0.5607697 |
| KVTRIMSASTLEQPA | ASSLLGYEQY | 0.5783256 |
| LGFALPIALPRRTVV | ASSLLGYEQY | 0.73203516 |
| TRLGNLTARKLVVLV | ASSLLGYEQY | 0.5999192 |
| WSQQRRLIYFPSAGP | ASSLLGYEQY | 0.65881884 |
| GPNGNPDPMAAAVDIR | ASSLLGYEQY | 0.3522735 |
| TQRCVSVADLRAAASR | ASSLLGYEQY | 0.7583239 |
| QFVLDVAEHPVEVLTETV | ASSLLGYEQY | 0.7231792 |
| GGAGGTGGAGGNGGTQAGDG | ASSLLGYEQY | 0.6341946 |
| PLLATLRAIAVDVDAKPAQV | ASSLLGYEQY | 0.3042039 |
| APKFRTGGYMIDFWGVGYQVAK | ASSLLGYEQY | 0.19194052 |
| GAGGAGGAGGAAGGPGGTGGTG | ASSLLGYEQY | 0.5391138 |
| IGALGLLATDVAPAGSVAFGVA | ASSLLGYEQY | 0.56423914 |
| AGGAGGIGGNGGAGGAGGVLMG | ASSLLGYEQY | 0.32023853 |
| ADELIGLL | ASSLLGYEQY | 0.5969 |
| ADRALKA | ASSLLGYEQY | 0.6147417 |
| AGAVGQVGGAGGA | ASSLLGYEQY | 0.17939597 |
| AGGGGGLGGAP | ASSLLGYEQY | 0.5602331 |
| AGGLLYGNGGAG | ASSLLGYEQY | 0.7544534 |
| AGLESIGL | ASSLLGYEQY | 0.5661038 |
| AGLKTGII | ASSLLGYEQY | 0.36328393 |
| ASLAALLL | ASSLLGYEQY | 0.75660145 |
| ASVINISSI | ASSLLGYEQY | 0.8235786 |
| ATPPGPA | ASSLLGYEQY | 0.7455647 |
| ATVASALV | ASSLLGYEQY | 0.15165676 |
| ATVASLIL | ASSLLGYEQY | 0.13461307 |
| AVNGPVLIRH | ASSLLGYEQY | 0.5850688 |
| DGAAGAVGI | ASSLLGYEQY | 0.42333764 |
| DPLLHGI | ASSLLGYEQY | 0.24835305 |
| ELGAALGAL | ASSLLGYEQY | 0.24753329 |
| ELGRTQAV | ASSLLGYEQY | 0.6806483 |
| FVAGGYLS | ASSLLGYEQY | 0.6789822 |
| GASGGAGGAGGW | ASSLLGYEQY | 0.3233171 |
| GGAGAGGDGGAGGI | ASSLLGYEQY | 0.13411742 |
| GTGHNGSQPGMGGNG | ASSLLGYEQY | 0.655483 |
| GYAAASASAS | ASSLLGYEQY | 0.7237451 |
| HQVDVAAVQQG | ASSLLGYEQY | 0.35039896 |
| IGAAIAHT | ASSLLGYEQY | 0.5815851 |
| IGMIEVAQL | ASSLLGYEQY | 0.3894117 |
| IRLAPEHTSAMD | ASSLLGYEQY | 0.18647282 |
| IYNVSVL | ASSLLGYEQY | 0.48617867 |
| KQGFLGM | ASSLLGYEQY | 0.19117683 |
| LESVITQL | ASSLLGYEQY | 0.6628807 |
| LGLGALVVV | ASSLLGYEQY | 0.27506736 |
| LIGNGGNGGSGGI | ASSLLGYEQY | 0.6446907 |
| LNAINAPTL | ASSLLGYEQY | 0.68416715 |
| LQLLNTGV | ASSLLGYEQY | 0.61660904 |
| LQTSIEF | ASSLLGYEQY | 0.5298187 |
| MSFAVSEL | ASSLLGYEQY | 0.38209212 |
| NTGALGL | ASSLLGYEQY | 0.15045562 |
| PPGAGGAGGAGTT | ASSLLGYEQY | 0.11350185 |

| | | |
|---|---|---|
| QLVGLGI | ASSLLGYEQY | 0.74606526 |
| RFLEQVHQL | ASSLLGYEQY | 0.10475176 |
| RGVGYVLR | ASSLLGYEQY | 0.7678997 |
| SATGVFAGVI | ASSLLGYEQY | 0.43153793 |
| SLPAITLP | ASSLLGYEQY | 0.2641114 |
| STALAFTV | ASSLLGYEQY | 0.5181575 |
| SVLTVGLGK | ASSLLGYEQY | 0.12809646 |
| TALAGAGSGI | ASSLLGYEQY | 0.23192519 |
| TGLPFTGI | ASSLLGYEQY | 0.577563 |
| VAEHVTV | ASSLLGYEQY | 0.6556264 |
| VDIVLNSL | ASSLLGYEQY | 0.36455774 |
| VLTGGGAVL | ASSLLGYEQY | 0.6653863 |
| VNVGLAYGI | ASSLLGYEQY | 0.6232544 |
| VSSAAVGAV | ASSLLGYEQY | 0.29209167 |
| VTSVYEAGV | ASSLLGYEQY | 0.563931 |
| WVALDRGA | ASSLLGYEQY | 0.7392022 |
| YLVGSVVI | ASSLLGYEQY | 0.6181817 |
| GPPGTGKTLL | ASSLLGYEQY | 0.2454719 |
| GVGMTKF | ASSLLGYEQY | 0.5271393 |
| IGIGGSDL | ASSLLGYEQY | 0.3346693 |
| IGTLLAEL | ASSLLGYEQY | 0.5646748 |
| IYTSGTT | ASSLLGYEQY | 0.51674896 |
| LSFPAGV | ASSLLGYEQY | 0.48742786 |
| LYTSGTT | ASSLLGYEQY | 0.6568949 |
| VGAFPFGF | ASSLLGYEQY | 0.13708615 |
| VGAGTVEF | ASSLLGYEQY | 0.32500172 |
| VGKSTLLL | ASSLLGYEQY | 0.4048599 |
| AALAHAGGGIV | ASSLLGYEQY | 0.4709964 |
| AALVAVSLL | ASSLLGYEQY | 0.5859335 |
| AAVSRAVAAL | ASSLLGYEQY | 0.2135427 |
| AGGSSGIGGF | ASSLLGYEQY | 0.5724784 |
| AGLTVANANG | ASSLLGYEQY | 0.22782192 |
| AGLVGAVL | ASSLLGYEQY | 0.19690226 |
| ALSQAFTV | ASSLLGYEQY | 0.60955334 |
| ALTFSVSL | ASSLLGYEQY | 0.38622177 |
| ALTGAALVAL | ASSLLGYEQY | 0.38722405 |
| ASFSIGAGVL | ASSLLGYEQY | 0.64383 |
| DGSVYKI | ASSLLGYEQY | 0.21940833 |
| EQLAEIV | ASSLLGYEQY | 0.3831861 |
| FAVSELTVM | ASSLLGYEQY | 0.64542377 |
| GAAPLAAGL | ASSLLGYEQY | 0.29357216 |
| GADGGDGGSGGAGG | ASSLLGYEQY | 0.60008895 |
| GFGLDTGGAGG | ASSLLGYEQY | 0.285476 |
| GLASGGLL | ASSLLGYEQY | 0.7028725 |
| GRIAGAAVLL | ASSLLGYEQY | 0.2644604 |
| GSAGSLNTSF | ASSLLGYEQY | 0.19960971 |
| GTGQIGI | ASSLLGYEQY | 0.53020006 |
| GTLIALIL | ASSLLGYEQY | 0.5555335 |
| IAATMGFVVG | ASSLLGYEQY | 0.6760282 |
| IFAQALL | ASSLLGYEQY | 0.56654805 |
| IGAAVALGF | ASSLLGYEQY | 0.64605 |

| | | |
|---|---|---|
| IGLTPLAL | ASSLLGYEQY | 0.7390384 |
| ILGSNTSTL | ASSLLGYEQY | 0.49104893 |
| IVFSLTVL | ASSLLGYEQY | 0.22246213 |
| LASKATGF | ASSLLGYEQY | 0.24344715 |
| LASVALIL | ASSLLGYEQY | 0.17281404 |
| LFAVGLTL | ASSLLGYEQY | 0.4003084 |
| LFNGGIGTYI | ASSLLGYEQY | 0.13605197 |
| LTLGAGII | ASSLLGYEQY | 0.74191844 |
| LVAAALGDI | ASSLLGYEQY | 0.7703171 |
| NTAFVQLGI | ASSLLGYEQY | 0.21081764 |
| QLFSGAEL | ASSLLGYEQY | 0.74006313 |
| SAATFITL | ASSLLGYEQY | 0.22519147 |
| SNYGLANLGV | ASSLLGYEQY | 0.45636 |
| SYGGAYAVM | ASSLLGYEQY | 0.7687613 |
| TTGVGGLTLG | ASSLLGYEQY | 0.23007375 |
| TTQTIYI | ASSLLGYEQY | 0.44615105 |
| VAAGAGLGAVI | ASSLLGYEQY | 0.21656774 |
| VAAVVTLL | ASSLLGYEQY | 0.18115653 |
| VGFLGGLI | ASSLLGYEQY | 0.7665918 |
| VGVIPILL | ASSLLGYEQY | 0.29145712 |
| VIFASSLI | ASSLLGYEQY | 0.6084018 |
| VKGGDGGAAAG | ASSLLGYEQY | 0.35753655 |
| VTKTFTNI | ASSLLGYEQY | 0.1679259 |
| VVAAAILL | ASSLLGYEQY | 0.5783598 |
| YAALDVEL | ASSLLGYEQY | 0.7610297 |
| LASVALLL | ASSLLGYEQY | 0.18783683 |
| LDAVHRV | ASSLLGYEQY | 0.727563 |
| QGLSSSIF | ASSLLGYEQY | 0.27379134 |
| RGTFANIRL | ASSLLGYEQY | 0.75322247 |
| RQLWWGHRI | ASSLLGYEQY | 0.7206301 |
| YGPPGTGKTLL | ASSLLGYEQY | 0.6148859 |
| AALVGAPANVA | ASSLLGYEQY | 0.33258888 |
| AATFRRTE | ASSLLGYEQY | 0.75118876 |
| AGALGGGI | ASSLLGYEQY | 0.6751506 |
| AGGAGGAGGIGAG | ASSLLGYEQY | 0.6312009 |
| AGSGALGL | ASSLLGYEQY | 0.72486913 |
| AQIADQI | ASSLLGYEQY | 0.34790653 |
| ASAVAVTYL | ASSLLGYEQY | 0.6865766 |
| ASTVLGGTGG | ASSLLGYEQY | 0.59337354 |
| AVGGEPAK | ASSLLGYEQY | 0.52085245 |
| CEQGTFGGTC | ASSLLGYEQY | 0.43460494 |
| DEGKAPAG | ASSLLGYEQY | 0.33107424 |
| DTVVFPAL | ASSLLGYEQY | 0.45519862 |
| FEGITMAG | ASSLLGYEQY | 0.40064946 |
| FPLVVQD | ASSLLGYEQY | 0.70429945 |
| GAFTTPQ | ASSLLGYEQY | 0.5798016 |
| GFGGFGTAGGVGGA | ASSLLGYEQY | 0.4307188 |
| GFLAQFL | ASSLLGYEQY | 0.6715441 |
| GGAGGAGGNAGVA | ASSLLGYEQY | 0.64501333 |
| GGVGTAGAGGAGGAGG | ASSLLGYEQY | 0.32705128 |
| GNGGGGDAGNAGSGGN | ASSLLGYEQY | 0.45692763 |

| | | |
|---|---|---|
| GTGGAGGAGNDAGS | ASSLLGYEQY | 0.45374936 |
| GTTNTGWFNS | ASSLLGYEQY | 0.45176613 |
| IFMGPLFTL | ASSLLGYEQY | 0.3079222 |
| ILAHAIEAI | ASSLLGYEQY | 0.5880095 |
| IRSAIREAL | ASSLLGYEQY | 0.47211277 |
| ITFAGGG | ASSLLGYEQY | 0.2745889 |
| ITGMSGHQEV | ASSLLGYEQY | 0.530882 |
| IVAGLACTL | ASSLLGYEQY | 0.5137168 |
| IVGGGLAGA | ASSLLGYEQY | 0.46290627 |
| IVTGVLNI | ASSLLGYEQY | 0.66596425 |
| LAQQGCI | ASSLLGYEQY | 0.521397 |
| LCLHVEN | ASSLLGYEQY | 0.44413275 |
| LDAISDGS | ASSLLGYEQY | 0.64919055 |
| LGGFAAML | ASSLLGYEQY | 0.68643427 |
| LGGKNPMI | ASSLLGYEQY | 0.37377143 |
| LGPKGVAAIL | ASSLLGYEQY | 0.589488 |
| LMVGVSI | ASSLLGYEQY | 0.6961949 |
| LVGAAFVL | ASSLLGYEQY | 0.5777722 |
| LVLAVAPL | ASSLLGYEQY | 0.6701392 |
| MAAIAVIDAA | ASSLLGYEQY | 0.6696483 |
| NNGINAELQ | ASSLLGYEQY | 0.36593872 |
| RGEPLVLVHG | ASSLLGYEQY | 0.6774949 |
| RGGAGGAAGLI | ASSLLGYEQY | 0.702026 |
| RSYRMHPAV | ASSLLGYEQY | 0.44406998 |
| SAAHLLAGQLA | ASSLLGYEQY | 0.5505373 |
| SAAVGSY | ASSLLGYEQY | 0.35964453 |
| SPHAVLHRI | ASSLLGYEQY | 0.48961186 |
| SQIAKVL | ASSLLGYEQY | 0.34580654 |
| TALGHLVP | ASSLLGYEQY | 0.6509714 |
| TAPTTSGAA | ASSLLGYEQY | 0.67499197 |
| TDGDDGNGGSPGT | ASSLLGYEQY | 0.54199886 |
| TDGLGEN | ASSLLGYEQY | 0.34875864 |
| TGGPPLPGQA | ASSLLGYEQY | 0.36768392 |
| TIPAITI | ASSLLGYEQY | 0.5312152 |
| TIPGFTI | ASSLLGYEQY | 0.3769455 |
| TSIASGALA | ASSLLGYEQY | 0.3120224 |
| TVTLPGAAI | ASSLLGYEQY | 0.42848545 |
| VASGLSAALP | ASSLLGYEQY | 0.6666297 |
| VGAGVIGI | ASSLLGYEQY | 0.6691242 |
| VGPKRFGV | ASSLLGYEQY | 0.37874055 |
| VNVAFPD | ASSLLGYEQY | 0.69879484 |
| VSMTNSGVS | ASSLLGYEQY | 0.68541765 |
| VTDVLGDLSV | ASSLLGYEQY | 0.6668724 |
| VVVVGAGFAGL | ASSLLGYEQY | 0.5190203 |
| VYTVGTVV | ASSLLGYEQY | 0.3690701 |
| WTTFSGP | ASSLLGYEQY | 0.70246184 |
| YDELVEAF | ASSLLGYEQY | 0.5366253 |
| YGTGGAGGNGG | ASSLLGYEQY | 0.53830487 |
| YTNVMAV | ASSLLGYEQY | 0.6220238 |
| FLEQVHQL | ASSLLGYEQY | 0.5402839 |
| GLSLLAVAV | ASSLLGYEQY | 0.46658194 |

| | | |
|---|---|---|
| LEQVHQL | ASSLLGYEQY | 0.63315964 |
| LNAAGAI | ASSLLGYEQY | 0.6265061 |
| PAVHASL | ASSLLGYEQY | 0.40606803 |
| TELEDDA | ASSLLGYEQY | 0.36636478 |
| TPPGPPN | ASSLLGYEQY | 0.434058 |
| TPPPPPPP | ASSLLGYEQY | 0.55855584 |
| TQSPALI | AWTGLSAETLY | 0.68364054 |
| EFAAAFPQ | AWTGLSAETLY | 0.51507926 |
| LLSGNLDVL | AWTGLSAETLY | 0.507406 |
| GPANLGNYNI | AWTGLSAETLY | 0.6397928 |
| TLGPTLP | AWTGLSAETLY | 0.6127851 |
| VYAYSSCCK | AWTGLSAETLY | 0.5267377 |
| APAPAPAPNI | AWTGLSAETLY | 0.5446231 |
| PGGPVPIP | AWTGLSAETLY | 0.4890089 |
| FLTVALEP | AWTGLSAETLY | 0.62932456 |
| SQYFGANL | AWTGLSAETLY | 0.63849247 |
| GGVKPAA | AWTGLSAETLY | 0.5948833 |
| GGTGGAGGAGGW | AWTGLSAETLY | 0.649876 |
| GVGGVAGLGGAGG | AWTGLSAETLY | 0.6151304 |
| VLLVLATTI | AWTGLSAETLY | 0.50707924 |
| IGIFAAML | AWTGLSAETLY | 0.32775545 |
| AVGDLID | AWTGLSAETLY | 0.4091304 |
| NISPLPL | AWTGLSAETLY | 0.5448139 |
| VTAVFVAI | AWTGLSAETLY | 0.5868967 |
| SGDDNDAAL | AWTGLSAETLY | 0.64018023 |
| AVAALAHS | AWTGLSAETLY | 0.5562887 |
| GGVTVMATP | AWTGLSAETLY | 0.42376807 |
| AGLAANVRD | AWTGLSAETLY | 0.5926468 |
| TNRPDLIDPALLRPG | AWTGLSAETLY | 0.6836073 |
| LANLGVTAL | AWTGLSAETLY | 0.36125088 |
| TGERIEL | AWTGLSAETLY | 0.5031488 |
| NGEELAA | AWTGLSAETLY | 0.7007356 |
| VLSLVWG | AWTGLSAETLY | 0.6617354 |
| IGYALWL | AWTGLSAETLY | 0.49221194 |
| LSLLLAPL | AWTGLSAETLY | 0.45941728 |
| TLDEALK | AWTGLSAETLY | 0.7076655 |
| TDGAGTAEGG | AWTGLSAETLY | 0.43805152 |
| NAPGGAGGNGG | AWTGLSAETLY | 0.6690829 |
| VSVFMGMGGNF | AWTGLSAETLY | 0.29572 |
| LASYLAAL | AWTGLSAETLY | 0.42475846 |
| LVGGTPAL | AWTGLSAETLY | 0.35750088 |
| VRNADVL | AWTGLSAETLY | 0.29022524 |
| TCSSPFLVSAM | AWTGLSAETLY | 0.66331315 |
| VHITDAE | AWTGLSAETLY | 0.38395044 |
| EYAAPSF | AWTGLSAETLY | 0.62760144 |
| VVDAFHGGL | AWTGLSAETLY | 0.508411 |
| ASVVDPQ | AWTGLSAETLY | 0.65665597 |
| SYNEFGGLS | AWTGLSAETLY | 0.63144934 |
| DPDGSVY | AWTGLSAETLY | 0.50394285 |
| MVSTIHATE | AWTGLSAETLY | 0.37151167 |
| LAAGVADPV | AWTGLSAETLY | 0.61203396 |

| | | |
|---|---|---|
| TKEQIAA | AWTGLSAETLY | 0.6076664 |
| SLASPAAL | AWTGLSAETLY | 0.6080364 |
| IGLVFYGV | AWTGLSAETLY | 0.32352927 |
| FLLDASNL | AWTGLSAETLY | 0.5298141 |
| YQVGNFP | AWTGLSAETLY | 0.5337868 |
| ESLLSQST | AWTGLSAETLY | 0.39273858 |
| LFTVGFAAVG | AWTGLSAETLY | 0.28776276 |
| ATPPTPT | AWTGLSAETLY | 0.40287626 |
| TDLATLG | AWTGLSAETLY | 0.55354726 |
| SAGGTGR | AWTGLSAETLY | 0.6569123 |
| GFSNSGS | AWTGLSAETLY | 0.6791302 |
| GGNGGSGGTSV | AWTGLSAETLY | 0.48191768 |
| GTSVATGGAGN | AWTGLSAETLY | 0.65897304 |
| NGGNGGSGGTS | AWTGLSAETLY | 0.52469337 |
| AVVKFPH | AWTGLSAETLY | 0.25543505 |
| TPPPPPP | AWTGLSAETLY | 0.46401125 |
| AAPAAAAAPA | AWTGLSAETLY | 0.32703334 |
| IALNEDL | AWTGLSAETLY | 0.64809763 |
| IVNVSSM | AWTGLSAETLY | 0.3809923 |
| TGIPPAPR | AWTGLSAETLY | 0.48782578 |
| FDLGGGTF | AWTGLSAETLY | 0.56991315 |
| FDLGGGTFDVS | AWTGLSAETLY | 0.6788142 |
| NEPTAAA | AWTGLSAETLY | 0.5806154 |
| NEPTAAALA | AWTGLSAETLY | 0.3947932 |
| GSIINISSI | AWTGLSAETLY | 0.31642497 |
| VAPGNAGTA | AWTGLSAETLY | 0.62781894 |
| TGGGASLE | AWTGLSAETLY | 0.5146131 |
| VAVYNHYKRI | AWTGLSAETLY | 0.36615133 |
| TDPAPATT | AWTGLSAETLY | 0.31188264 |
| TNRPDILDPALLRPG | AWTGLSAETLY | 0.6373221 |
| LTTTTTS | AWTGLSAETLY | 0.33758283 |
| PTGSGKTL | AWTGLSAETLY | 0.62663966 |
| TGSGKTL | AWTGLSAETLY | 0.43002915 |
| TIGHVDHGK | AWTGLSAETLY | 0.53195846 |
| TIGHVDHGKTT | AWTGLSAETLY | 0.70420194 |
| GHVDHGKTT | AWTGLSAETLY | 0.6773011 |
| AQFEHTLL | AWTGLSAETLY | 0.51081926 |
| IVNQDEL | AWTGLSAETLY | 0.41661012 |
| TNGSQFF | AWTGLSAETLY | 0.3637424 |
| TGKGIVE | AWTGLSAETLY | 0.40546864 |
| LISNASDA | AWTGLSAETLY | 0.4182998 |
| AELRTLGELTKIAKAHGAQ | AWTGLSAETLY | 0.67107093 |
| APNAVAAAEPSW | AWTGLSAETLY | 0.45493856 |
| APQVAGAATAVL | AWTGLSAETLY | 0.3373872 |
| ARRDPAAAAAILPTDA | AWTGLSAETLY | 0.54506034 |
| DAAARDVLAEAGLAEYF | AWTGLSAETLY | 0.68621457 |
| DDQIVLSAKANLAKVEQ | AWTGLSAETLY | 0.7129563 |
| DELLPDTGEDSDEDGH | AWTGLSAETLY | 0.37092978 |
| ELPEAPANRATVPQD | AWTGLSAETLY | 0.46763885 |
| FGQSKIERVTGIG | AWTGLSAETLY | 0.450883 |
| GAGGTGSTAGGAGGAGGAGGLY | AWTGLSAETLY | 0.572181 |

| | | |
|---|---|---|
| GHGGTGAAGGNGGTGGAGGSGI | AWTGLSAETLY | 0.38886666 |
| GNGGNGGAGGEGGPGGAGGAGA | AWTGLSAETLY | 0.3751814 |
| GSTAGNGGNGGNATNGTIAGQP | AWTGLSAETLY | 0.61559725 |
| GTGGDGGNGGNGGTGG | AWTGLSAETLY | 0.7180173 |
| GVRRPAGIPADAPV | AWTGLSAETLY | 0.45101815 |
| GYRQVTDAHLLTLARR | AWTGLSAETLY | 0.38870353 |
| IAKMGQDGHDRG | AWTGLSAETLY | 0.5370916 |
| KAEPSIEIQYT | AWTGLSAETLY | 0.43390435 |
| KPVRIRMAADEDTVAVT | AWTGLSAETLY | 0.47138956 |
| MPPRIAGMRLLVIKPEP | AWTGLSAETLY | 0.55768585 |
| MKFYAVALFRSRGVPPTR | AWTGLSAETLY | 0.32744405 |
| NAGGFGILGAVAHSP | AWTGLSAETLY | 0.52124286 |
| NEGREITLAMAEM | AWTGLSAETLY | 0.7058605 |
| NELVEWSHPLGHRWRWE | AWTGLSAETLY | 0.5885703 |
| NLGAGGTGGNGGNPG | AWTGLSAETLY | 0.56620455 |
| NTEWPKWETVMAADPVD | AWTGLSAETLY | 0.6887232 |
| PASAAPAETQP | AWTGLSAETLY | 0.62542576 |
| QQSSDGKNGVRAAEITGE | AWTGLSAETLY | 0.6922523 |
| RAPDAVTTSWWKE | AWTGLSAETLY | 0.6899619 |
| SGFGNVGAHTSGWFNQ | AWTGLSAETLY | 0.4655112 |
| SGVAGAVGQAASVGGLKV | AWTGLSAETLY | 0.49118096 |
| SNGGAGTGSAGGNGGT | AWTGLSAETLY | 0.69396675 |
| TGGAGGEGGAAG | AWTGLSAETLY | 0.50636184 |
| TGGDGGAAGAGGAPGHG | AWTGLSAETLY | 0.55532587 |
| TMPPLLAAIANAGRHGV | AWTGLSAETLY | 0.43708253 |
| TSSTAGNSGGA | AWTGLSAETLY | 0.6607962 |
| VPATVPVGILC | AWTGLSAETLY | 0.55960053 |
| VQRPDSGGVVLTGRIS | AWTGLSAETLY | 0.6924644 |
| WTPNPVPATKGVPAGLVQAT | AWTGLSAETLY | 0.30345657 |
| YALKRALAKFSDRFDVVII | AWTGLSAETLY | 0.38444147 |
| YLIGSVIV | AWTGLSAETLY | 0.6795205 |
| YGQLVAVID | AWTGLSAETLY | 0.26951322 |
| VTVGGMTLPA | AWTGLSAETLY | 0.2960593 |
| VLALPAV | AWTGLSAETLY | 0.51300436 |
| VIPPTLNYET | AWTGLSAETLY | 0.6930404 |
| VGMGGMHPSSG | AWTGLSAETLY | 0.2824098 |
| VGLAGGAATA | AWTGLSAETLY | 0.43742216 |
| VGGAGGAGGSGP | AWTGLSAETLY | 0.32832068 |
| VFAAFMV | AWTGLSAETLY | 0.5846341 |
| TLYDKLR | AWTGLSAETLY | 0.55820006 |
| TLAVFTVL | AWTGLSAETLY | 0.3190545 |
| TGSVQPARV | AWTGLSAETLY | 0.59869564 |
| TGGGAGSGGT | AWTGLSAETLY | 0.34488803 |
| SHLETSL | AWTGLSAETLY | 0.59121096 |
| SFTTGMT | AWTGLSAETLY | 0.655871 |
| QVEVAKRMGVRQPTVS | AWTGLSAETLY | 0.7102459 |
| QGSGGPALAM | AWTGLSAETLY | 0.35294652 |
| MLAPSNP | AWTGLSAETLY | 0.6464412 |
| LVNNAAI | AWTGLSAETLY | 0.45011985 |
| LTGEINTLA | AWTGLSAETLY | 0.7169305 |
| LSIFGPV | AWTGLSAETLY | 0.4682759 |

| | | |
|---|---|---|
| LLALLANL | AWTGLSAETLY | 0.3271935 |
| LISNASDALDKLR | AWTGLSAETLY | 0.6421224 |
| LINEAVP | AWTGLSAETLY | 0.57924086 |
| LGYAFSGL | AWTGLSAETLY | 0.4368409 |
| LDFVAVAL | AWTGLSAETLY | 0.6849369 |
| LAQPGAE | AWTGLSAETLY | 0.44105452 |
| ISNLGAVTGF | AWTGLSAETLY | 0.542763 |
| HQEDPGRI | AWTGLSAETLY | 0.6739851 |
| GTPGEPTP | AWTGLSAETLY | 0.6265929 |
| GFFNSGS | AWTGLSAETLY | 0.41738713 |
| GAGGGGLTAGAN | AWTGLSAETLY | 0.56258905 |
| FVDALVNM | AWTGLSAETLY | 0.5799887 |
| FLTVAPEL | AWTGLSAETLY | 0.38666144 |
| FETLTAL | AWTGLSAETLY | 0.7066363 |
| DVINPIEL | AWTGLSAETLY | 0.4747221 |
| APVSGAPV | AWTGLSAETLY | 0.6565877 |
| ALVGGLAW | AWTGLSAETLY | 0.5797158 |
| ALAAAAATAI | AWTGLSAETLY | 0.7110617 |
| AFGLGLLL | AWTGLSAETLY | 0.34285825 |
| AFFAAASL | AWTGLSAETLY | 0.5359093 |
| AAAALPAFSPP | AWTGLSAETLY | 0.5563596 |
| VVLAPAL | AWTGLSAETLY | 0.63276273 |
| TIGHVDHGKTTL | AWTGLSAETLY | 0.52493924 |
| LISNASDALDK | AWTGLSAETLY | 0.59485483 |
| LISNASDAL | AWTGLSAETLY | 0.551368 |
| ISNASDALDK | AWTGLSAETLY | 0.6783006 |
| NQAETLVYQTE | AWTGLSAETLY | 0.46817023 |
| TGGTGGTAGAG | AWTGLSAETLY | 0.4989543 |
| TGSTAGGAGGA | AWTGLSAETLY | 0.7063074 |
| DPLAVPVAAGPV | AWTGLSAETLY | 0.36300498 |
| GGTGGTPGNGGN | AWTGLSAETLY | 0.5458189 |
| PAGLAGVDPATL | AWTGLSAETLY | 0.5991161 |
| GNNLVVEAPHAGASV | AWTGLSAETLY | 0.63583076 |
| KVTRIMSASTLEQPA | AWTGLSAETLY | 0.47148198 |
| LGFALPIALPRRTVV | AWTGLSAETLY | 0.62134945 |
| TRLGNLTARKLVVLV | AWTGLSAETLY | 0.41155934 |
| WSQQRRLIYFPSAGP | AWTGLSAETLY | 0.64776254 |
| GPNGNPDPMAAAVDIR | AWTGLSAETLY | 0.5255984 |
| TQRCVSVADLRAAASR | AWTGLSAETLY | 0.5768471 |
| QFVLDVAEHPVEVLTETV | AWTGLSAETLY | 0.39341667 |
| GGAGGTGGAGGNGGTQAGDG | AWTGLSAETLY | 0.613589 |
| PLLATLRAIAVDVDAKPAQV | AWTGLSAETLY | 0.68085957 |
| APKFRTGGYMIDFWGVGYQVAK | AWTGLSAETLY | 0.6524363 |
| GAGGAGGAGGAAGGPGGTGGTG | AWTGLSAETLY | 0.2870009 |
| IGALGLLATDVAPAGSVAFGVA | AWTGLSAETLY | 0.6659486 |
| AGGAGGIGGNGGAGGAGGVLMG | AWTGLSAETLY | 0.6558929 |
| ADELIGLL | AWTGLSAETLY | 0.71434975 |
| ADRALKA | AWTGLSAETLY | 0.48542994 |
| AGAVGQVGGAGGA | AWTGLSAETLY | 0.594018 |
| AGGGGGLGGAP | AWTGLSAETLY | 0.45745724 |
| AGGLLYGNGGAG | AWTGLSAETLY | 0.5381592 |

| | | |
|---|---|---|
| AGLESIGL | AWTGLSAETLY | 0.3495523 |
| AGLKTGII | AWTGLSAETLY | 0.4596756 |
| ASLAALLL | AWTGLSAETLY | 0.5784993 |
| ASVINISSI | AWTGLSAETLY | 0.42967212 |
| ATPPGPA | AWTGLSAETLY | 0.6941412 |
| ATVASALV | AWTGLSAETLY | 0.65398526 |
| ATVASLIL | AWTGLSAETLY | 0.67569256 |
| AVNGPVLIRH | AWTGLSAETLY | 0.69206583 |
| DGAAGAVGI | AWTGLSAETLY | 0.7007612 |
| DPLLHGI | AWTGLSAETLY | 0.59811956 |
| ELGAALGAL | AWTGLSAETLY | 0.46104372 |
| ELGRTQAV | AWTGLSAETLY | 0.5043591 |
| FVAGGYLS | AWTGLSAETLY | 0.36086306 |
| GASGGAGGAGGW | AWTGLSAETLY | 0.51224184 |
| GGAGAGGDGGAGGI | AWTGLSAETLY | 0.47495392 |
| GTGHNGSQPGMGGNG | AWTGLSAETLY | 0.6124495 |
| GYAAASASAS | AWTGLSAETLY | 0.27073452 |
| HQVDVAAVQQG | AWTGLSAETLY | 0.6711922 |
| IGAAIAHT | AWTGLSAETLY | 0.6767329 |
| IGMIEVAQL | AWTGLSAETLY | 0.56672364 |
| IRLAPEHTSAMD | AWTGLSAETLY | 0.59541166 |
| IYNVSVL | AWTGLSAETLY | 0.6497234 |
| KQGFLGM | AWTGLSAETLY | 0.4196915 |
| LESVITQL | AWTGLSAETLY | 0.65971154 |
| LGLGALVVV | AWTGLSAETLY | 0.40274578 |
| LIGNGGNGGSGGI | AWTGLSAETLY | 0.6351522 |
| LNAINAPTL | AWTGLSAETLY | 0.68125564 |
| LQLLNTGV | AWTGLSAETLY | 0.56384176 |
| LQTSIEF | AWTGLSAETLY | 0.42856258 |
| MSFAVSEL | AWTGLSAETLY | 0.51356757 |
| NTGALGL | AWTGLSAETLY | 0.3325397 |
| PPGAGGAGGAGTT | AWTGLSAETLY | 0.4781845 |
| QLVGLGI | AWTGLSAETLY | 0.36929178 |
| RFLEQVHQL | AWTGLSAETLY | 0.59473073 |
| RGVGYVLR | AWTGLSAETLY | 0.3645584 |
| SATGVFAGVI | AWTGLSAETLY | 0.59606254 |
| SLPAITLP | AWTGLSAETLY | 0.5719099 |
| STALAFTV | AWTGLSAETLY | 0.5597904 |
| SVLTVGLGK | AWTGLSAETLY | 0.3983937 |
| TALAGAGSGI | AWTGLSAETLY | 0.6058694 |
| TGLPFTGI | AWTGLSAETLY | 0.48771805 |
| VAEHVTV | AWTGLSAETLY | 0.35730356 |
| VDIVLNSL | AWTGLSAETLY | 0.63114333 |
| VLTGGGAVL | AWTGLSAETLY | 0.44265586 |
| VNVGLAYGI | AWTGLSAETLY | 0.5713215 |
| VSSAAVGAV | AWTGLSAETLY | 0.6547532 |
| VTSVYEAGV | AWTGLSAETLY | 0.5347562 |
| WVALDRGA | AWTGLSAETLY | 0.34596744 |
| YLVGSVVI | AWTGLSAETLY | 0.56866795 |
| GPPGTGKTLL | AWTGLSAETLY | 0.5243873 |
| GVGMTKF | AWTGLSAETLY | 0.4749545 |

| | | |
|---|---|---|
| IGIGGSDL | AWTGLSAETLY | 0.28170088 |
| IGTLLAEL | AWTGLSAETLY | 0.57426035 |
| IYTSGTT | AWTGLSAETLY | 0.38306826 |
| LSFPAGV | AWTGLSAETLY | 0.44972962 |
| LYTSGTT | AWTGLSAETLY | 0.4402942 |
| VGAFPFGF | AWTGLSAETLY | 0.6747934 |
| VGAGTVEF | AWTGLSAETLY | 0.6416414 |
| VGKSTLLL | AWTGLSAETLY | 0.51214886 |
| AALAHAGGGIV | AWTGLSAETLY | 0.6080822 |
| AALVAVSLL | AWTGLSAETLY | 0.5057356 |
| AAVSRAVAAL | AWTGLSAETLY | 0.68314314 |
| AGGSSGIGGF | AWTGLSAETLY | 0.5312844 |
| AGLTVANANG | AWTGLSAETLY | 0.44198778 |
| AGLVGAVL | AWTGLSAETLY | 0.54909563 |
| ALSQAFTV | AWTGLSAETLY | 0.43762767 |
| ALTFSVSL | AWTGLSAETLY | 0.3334986 |
| ALTGAALVAL | AWTGLSAETLY | 0.31476972 |
| ASFSIGAGVL | AWTGLSAETLY | 0.53414977 |
| DGSVYKI | AWTGLSAETLY | 0.31732726 |
| EQLAEIV | AWTGLSAETLY | 0.5843357 |
| FAVSELTVM | AWTGLSAETLY | 0.6339562 |
| GAAPLAAGL | AWTGLSAETLY | 0.52998453 |
| GADGGDGGSGGAGG | AWTGLSAETLY | 0.3286813 |
| GFGLDTGGAGG | AWTGLSAETLY | 0.41780013 |
| GLASGGLL | AWTGLSAETLY | 0.46125904 |
| GRIAGAAVLL | AWTGLSAETLY | 0.6635961 |
| GSAGSLNTSF | AWTGLSAETLY | 0.54360694 |
| GTGQIGI | AWTGLSAETLY | 0.7133542 |
| GTLIALIL | AWTGLSAETLY | 0.6668001 |
| IAATMGFVVG | AWTGLSAETLY | 0.48241684 |
| IFAQALL | AWTGLSAETLY | 0.64099944 |
| IGAAVALGF | AWTGLSAETLY | 0.47515118 |
| IGLTPLAL | AWTGLSAETLY | 0.50493205 |
| ILGSNTSTL | AWTGLSAETLY | 0.46584487 |
| IVFSLTVL | AWTGLSAETLY | 0.5876335 |
| LASKATGF | AWTGLSAETLY | 0.4483416 |
| LASVALIL | AWTGLSAETLY | 0.40482733 |
| LFAVGLTL | AWTGLSAETLY | 0.33800453 |
| LFNGGIGTYI | AWTGLSAETLY | 0.5657846 |
| LTLGAGII | AWTGLSAETLY | 0.42285824 |
| LVAAALGDI | AWTGLSAETLY | 0.33982056 |
| NTAFVQLGI | AWTGLSAETLY | 0.36125088 |
| QLFSGAEL | AWTGLSAETLY | 0.7087991 |
| SAATFITL | AWTGLSAETLY | 0.7019784 |
| SNYGLANLGV | AWTGLSAETLY | 0.47301722 |
| SYGGAYAVM | AWTGLSAETLY | 0.4102719 |
| TTGVGGLTLG | AWTGLSAETLY | 0.678759 |
| TTQTIYI | AWTGLSAETLY | 0.3702229 |
| VAAGAGLGAVI | AWTGLSAETLY | 0.5558951 |
| VAAVVTLL | AWTGLSAETLY | 0.6243707 |
| VGFLGGLI | AWTGLSAETLY | 0.6086185 |

| | | |
|---|---|---|
| VGVIPILL | AWTGLSAETLY | 0.37772498 |
| VIFASSLI | AWTGLSAETLY | 0.59989864 |
| VKGGDGGAAAG | AWTGLSAETLY | 0.5154591 |
| VTKTFTNI | AWTGLSAETLY | 0.5449543 |
| VVAAAILL | AWTGLSAETLY | 0.38857263 |
| YAALDVEL | AWTGLSAETLY | 0.6502182 |
| LASVALLL | AWTGLSAETLY | 0.41058892 |
| LDAVHRV | AWTGLSAETLY | 0.39941904 |
| QGLSSSIF | AWTGLSAETLY | 0.29670548 |
| RGTFANIRL | AWTGLSAETLY | 0.46903187 |
| RQLWWGHRI | AWTGLSAETLY | 0.6863034 |
| YGPPGTGKTLL | AWTGLSAETLY | 0.6812674 |
| AALVGAPANVA | AWTGLSAETLY | 0.4316874 |
| AATFRRTE | AWTGLSAETLY | 0.62352127 |
| AGALGGGI | AWTGLSAETLY | 0.7043339 |
| AGGAGGAGGIGAG | AWTGLSAETLY | 0.5279212 |
| AGSGALGL | AWTGLSAETLY | 0.29950744 |
| AQIADQI | AWTGLSAETLY | 0.59533286 |
| ASAVAVTYL | AWTGLSAETLY | 0.56664324 |
| ASTVLGGTGG | AWTGLSAETLY | 0.6777602 |
| AVGGEPAK | AWTGLSAETLY | 0.62066936 |
| CEQGTFGGTC | AWTGLSAETLY | 0.35097045 |
| DEGKAPAG | AWTGLSAETLY | 0.5444082 |
| DTVVFPAL | AWTGLSAETLY | 0.60696316 |
| FEGITMAG | AWTGLSAETLY | 0.35192224 |
| FPLVVQD | AWTGLSAETLY | 0.3532218 |
| GAFTTPQ | AWTGLSAETLY | 0.39153513 |
| GFGGFGTAGGVGGA | AWTGLSAETLY | 0.3330561 |
| GFLAQFL | AWTGLSAETLY | 0.51447505 |
| GGAGGAGGNAGVA | AWTGLSAETLY | 0.49328896 |
| GGVGTAGAGGAGGAGG | AWTGLSAETLY | 0.5159518 |
| GNGGGGDAGNAGSGGN | AWTGLSAETLY | 0.4931669 |
| GTGGAGGAGNDAGS | AWTGLSAETLY | 0.4145981 |
| GTTNTGWFNS | AWTGLSAETLY | 0.47484183 |
| IFMGPLFTL | AWTGLSAETLY | 0.4720206 |
| ILAHAIEAI | AWTGLSAETLY | 0.60849404 |
| IRSAIREAL | AWTGLSAETLY | 0.6248741 |
| ITFAGGG | AWTGLSAETLY | 0.60192835 |
| ITGMSGHQEV | AWTGLSAETLY | 0.5449652 |
| IVAGLACTL | AWTGLSAETLY | 0.6960833 |
| IVGGGLAGA | AWTGLSAETLY | 0.38075 |
| IVTGVLNI | AWTGLSAETLY | 0.41313872 |
| LAQQGCI | AWTGLSAETLY | 0.61923075 |
| LCLHVEN | AWTGLSAETLY | 0.378692 |
| LDAISDGS | AWTGLSAETLY | 0.67631704 |
| LGGFAAML | AWTGLSAETLY | 0.5589151 |
| LGGKNPMI | AWTGLSAETLY | 0.6333778 |
| LGPKGVAAIL | AWTGLSAETLY | 0.6107403 |
| LMVGVSI | AWTGLSAETLY | 0.69913137 |
| LVGAAFVL | AWTGLSAETLY | 0.6268588 |
| LVLAVAPL | AWTGLSAETLY | 0.42982408 |

| | | |
|---|---|---|
| MAAIAVIDAA | AWTGLSAETLY | 0.6822882 |
| NNGINAELQ | AWTGLSAETLY | 0.33903795 |
| RGEPLVLVHG | AWTGLSAETLY | 0.5949981 |
| RGGAGGAAGLI | AWTGLSAETLY | 0.58536184 |
| RSYRMHPAV | AWTGLSAETLY | 0.42562705 |
| SAAHLLAGQLA | AWTGLSAETLY | 0.54603565 |
| SAAVGSY | AWTGLSAETLY | 0.66036475 |
| SPHAVLHRI | AWTGLSAETLY | 0.6645675 |
| SQIAKVL | AWTGLSAETLY | 0.33842307 |
| TALGHLVP | AWTGLSAETLY | 0.45972833 |
| TAPTTSGAA | AWTGLSAETLY | 0.34353828 |
| TDGDDGNGGSPGT | AWTGLSAETLY | 0.36512417 |
| TDGLGEN | AWTGLSAETLY | 0.33703858 |
| TGGPPLPGQA | AWTGLSAETLY | 0.6225172 |
| TIPAITI | AWTGLSAETLY | 0.4800794 |
| TIPGFTI | AWTGLSAETLY | 0.69331014 |
| TSIASGALA | AWTGLSAETLY | 0.60324806 |
| TVTLPGAAI | AWTGLSAETLY | 0.6705548 |
| VASGLSAALP | AWTGLSAETLY | 0.40771237 |
| VGAGVIGI | AWTGLSAETLY | 0.47556716 |
| VGPKRFGV | AWTGLSAETLY | 0.30620402 |
| VNVAFPD | AWTGLSAETLY | 0.4131864 |
| VSMTNSGVS | AWTGLSAETLY | 0.3908652 |
| VTDVLGDLSV | AWTGLSAETLY | 0.5308564 |
| VVVVGAGFAGL | AWTGLSAETLY | 0.38847235 |
| VYTVGTVV | AWTGLSAETLY | 0.6967616 |
| WTTFSGP | AWTGLSAETLY | 0.53357506 |
| YDELVEAF | AWTGLSAETLY | 0.6265516 |
| YGTGGAGGNGG | AWTGLSAETLY | 0.40073964 |
| YTNVMAV | AWTGLSAETLY | 0.38085943 |
| FLEQVHQL | AWTGLSAETLY | 0.4838271 |
| GLSLLAVAV | AWTGLSAETLY | 0.6429509 |
| LEQVHQL | AWTGLSAETLY | 0.35781765 |
| LNAAGAI | AWTGLSAETLY | 0.3132991 |
| PAVHASL | AWTGLSAETLY | 0.4278927 |
| TELEDDA | AWTGLSAETLY | 0.431978 |
| TPPGPPN | AWTGLSAETLY | 0.70230496 |
| TPPPPPPP | AWTGLSAETLY | 0.67295814 |
| TQSPALI | ASSRLGGRDTQY | 0.5776611 |
| EFAAAFPQ | ASSRLGGRDTQY | 0.6891931 |
| LLSGNLDVL | ASSRLGGRDTQY | 0.27751997 |
| GPANLGNYNI | ASSRLGGRDTQY | 0.45372683 |
| TLGPTLP | ASSRLGGRDTQY | 0.6993315 |
| VYAYSSCCK | ASSRLGGRDTQY | 0.6158867 |
| APAPAPAPNI | ASSRLGGRDTQY | 0.51396847 |
| PGGPVPIP | ASSRLGGRDTQY | 0.6798174 |
| FLTVALEP | ASSRLGGRDTQY | 0.41627067 |
| SQYFGANL | ASSRLGGRDTQY | 0.47879282 |
| GGVKPAA | ASSRLGGRDTQY | 0.5746223 |
| GGTGGAGGAGGW | ASSRLGGRDTQY | 0.51649266 |
| GVGGVAGLGGAGG | ASSRLGGRDTQY | 0.5996234 |

| | | |
|---|---|---|
| VLLVLATTI | ASSRLGGRDTQY | 0.33937758 |
| IGIFAAML | ASSRLGGRDTQY | 0.5627877 |
| AVGDLID | ASSRLGGRDTQY | 0.7134769 |
| NISPLPL | ASSRLGGRDTQY | 0.52648836 |
| VTAVFVAI | ASSRLGGRDTQY | 0.6918044 |
| SGDDNDAAL | ASSRLGGRDTQY | 0.59549963 |
| AVAALAHS | ASSRLGGRDTQY | 0.6668861 |
| GGVTVMATP | ASSRLGGRDTQY | 0.42380732 |
| AGLAANVRD | ASSRLGGRDTQY | 0.387451 |
| TNRPDLIDPALLRPG | ASSRLGGRDTQY | 0.58254373 |
| LANLGVTAL | ASSRLGGRDTQY | 0.44371742 |
| TGERIEL | ASSRLGGRDTQY | 0.49732542 |
| NGEELAA | ASSRLGGRDTQY | 0.4781574 |
| VLSLVWG | ASSRLGGRDTQY | 0.71621656 |
| IGYALWL | ASSRLGGRDTQY | 0.6293754 |
| LSLLLAPL | ASSRLGGRDTQY | 0.6348538 |
| TLDEALK | ASSRLGGRDTQY | 0.40291968 |
| TDGAGTAEGG | ASSRLGGRDTQY | 0.53288877 |
| NAPGGAGGNGG | ASSRLGGRDTQY | 0.64653945 |
| VSVFMGMGGNF | ASSRLGGRDTQY | 0.3468345 |
| LASYLAAL | ASSRLGGRDTQY | 0.68502545 |
| LVGGTPAL | ASSRLGGRDTQY | 0.40567482 |
| VRNADVL | ASSRLGGRDTQY | 0.5479928 |
| TCSSPFLVSAM | ASSRLGGRDTQY | 0.6065293 |
| VHITDAE | ASSRLGGRDTQY | 0.70754516 |
| EYAAPSF | ASSRLGGRDTQY | 0.4850788 |
| VVDAFHGGL | ASSRLGGRDTQY | 0.60022616 |
| ASVVDPQ | ASSRLGGRDTQY | 0.39210021 |
| SYNEFGGLS | ASSRLGGRDTQY | 0.6360754 |
| DPDGSVY | ASSRLGGRDTQY | 0.6875648 |
| MVSTIHATE | ASSRLGGRDTQY | 0.64076304 |
| LAAGVADPV | ASSRLGGRDTQY | 0.7122568 |
| TKEQIAA | ASSRLGGRDTQY | 0.5767409 |
| SLASPAAL | ASSRLGGRDTQY | 0.31932995 |
| IGLVFYGV | ASSRLGGRDTQY | 0.4703669 |
| FLLDASNL | ASSRLGGRDTQY | 0.67371094 |
| YQVGNFP | ASSRLGGRDTQY | 0.69567955 |
| ESLLSQST | ASSRLGGRDTQY | 0.6723807 |
| LFTVGFAAVG | ASSRLGGRDTQY | 0.60468924 |
| ATPPTPT | ASSRLGGRDTQY | 0.46648836 |
| TDLATLG | ASSRLGGRDTQY | 0.6926502 |
| SAGGTGR | ASSRLGGRDTQY | 0.7032319 |
| GFSNSGS | ASSRLGGRDTQY | 0.656188 |
| GGNGGSGGTSV | ASSRLGGRDTQY | 0.42375413 |
| GTSVATGGAGN | ASSRLGGRDTQY | 0.39239633 |
| NGGNGGSGGTS | ASSRLGGRDTQY | 0.54913193 |
| AVVKFPH | ASSRLGGRDTQY | 0.6869075 |
| TPPPPPP | ASSRLGGRDTQY | 0.34512907 |
| AAPAAAAAPA | ASSRLGGRDTQY | 0.60196 |
| IALNEDL | ASSRLGGRDTQY | 0.5811745 |
| IVNVSSM | ASSRLGGRDTQY | 0.3056583 |

| | | |
|---|---|---|
| TGIPPAPR | ASSRLGGRDTQY | 0.62649643 |
| FDLGGGTF | ASSRLGGRDTQY | 0.6808648 |
| FDLGGGTFDVS | ASSRLGGRDTQY | 0.49138337 |
| NEPTAAA | ASSRLGGRDTQY | 0.3850308 |
| NEPTAAALA | ASSRLGGRDTQY | 0.6874778 |
| GSIINISSI | ASSRLGGRDTQY | 0.713382 |
| VAPGNAGTA | ASSRLGGRDTQY | 0.65832007 |
| TGGGASLE | ASSRLGGRDTQY | 0.33346906 |
| VAVYNHYKRI | ASSRLGGRDTQY | 0.28113657 |
| TDPAPATT | ASSRLGGRDTQY | 0.49691108 |
| TNRPDILDPALLRPG | ASSRLGGRDTQY | 0.6833183 |
| LTTTTTS | ASSRLGGRDTQY | 0.48124364 |
| PTGSGKTL | ASSRLGGRDTQY | 0.41506702 |
| TGSGKTL | ASSRLGGRDTQY | 0.63431394 |
| TIGHVDHGK | ASSRLGGRDTQY | 0.673645 |
| TIGHVDHGKTT | ASSRLGGRDTQY | 0.35087803 |
| GHVDHGKTT | ASSRLGGRDTQY | 0.34126344 |
| AQFEHTLL | ASSRLGGRDTQY | 0.638392 |
| IVNQDEL | ASSRLGGRDTQY | 0.64523196 |
| TNGSQFF | ASSRLGGRDTQY | 0.40031585 |
| TGKGIVE | ASSRLGGRDTQY | 0.50329083 |
| LISNASDA | ASSRLGGRDTQY | 0.5587887 |
| AELRTLGELTKIAKAHGAQ | ASSRLGGRDTQY | 0.59170955 |
| APNAVAAAEPSW | ASSRLGGRDTQY | 0.6228428 |
| APQVAGAATAVL | ASSRLGGRDTQY | 0.36968452 |
| ARRDPAAAAAILPTDA | ASSRLGGRDTQY | 0.6887214 |
| DAAARDVLAEAGLAEYF | ASSRLGGRDTQY | 0.36164212 |
| DDQIVLSAKANLAKVEQ | ASSRLGGRDTQY | 0.6218592 |
| DELLPDTGEDSDEDGH | ASSRLGGRDTQY | 0.5402367 |
| ELPEAPANRATVPQD | ASSRLGGRDTQY | 0.52017754 |
| FGQSKIERVTGIG | ASSRLGGRDTQY | 0.49140894 |
| GAGGTGSTAGGAGGAGGAGGLY | ASSRLGGRDTQY | 0.4729497 |
| GHGGTGAAGGNGGTGGAGGSGI | ASSRLGGRDTQY | 0.35334015 |
| GNGGNGGAGGEGGPGGAGGAGA | ASSRLGGRDTQY | 0.31416464 |
| GSTAGNGGNGGNATNGTIAGQP | ASSRLGGRDTQY | 0.6630933 |
| GTGGDGGNGGNGGTGG | ASSRLGGRDTQY | 0.6368231 |
| GVRRPAGIPADAPV | ASSRLGGRDTQY | 0.62519896 |
| GYRQVTDAHLLTLARR | ASSRLGGRDTQY | 0.60120535 |
| IAKMGQDGHDRG | ASSRLGGRDTQY | 0.38380012 |
| KAEPSIEIQYT | ASSRLGGRDTQY | 0.5771304 |
| KPVRIRMAADEDTVAVT | ASSRLGGRDTQY | 0.38728404 |
| MPPRIAGMRLLVIKPEP | ASSRLGGRDTQY | 0.450524 |
| MKFYAVALFRSRGVPPTR | ASSRLGGRDTQY | 0.59062624 |
| NAGGFGILGAVAHSP | ASSRLGGRDTQY | 0.5834514 |
| NEGREITLAMAEM | ASSRLGGRDTQY | 0.4033435 |
| NELVEWSHPLGHRWRWE | ASSRLGGRDTQY | 0.54452276 |
| NLGAGGTGGNGGNPG | ASSRLGGRDTQY | 0.59597456 |
| NTEWPKWETVMAADPVD | ASSRLGGRDTQY | 0.402308 |
| PASAAPAETQP | ASSRLGGRDTQY | 0.4617645 |
| QQSSDGKNGVRAAEITGE | ASSRLGGRDTQY | 0.6627252 |
| RAPDAVTTSWWKE | ASSRLGGRDTQY | 0.51362544 |

| | | |
|---|---|---|
| SGFGNVGAHTSGWFNQ | ASSRLGGRDTQY | 0.34622353 |
| SGVAGAVGQAASVGGLKV | ASSRLGGRDTQY | 0.6457113 |
| SNGGAGTGSAGGNGGT | ASSRLGGRDTQY | 0.41756633 |
| TGGAGGEGGAAG | ASSRLGGRDTQY | 0.6278248 |
| TGGDGGAAGAGGAPGHG | ASSRLGGRDTQY | 0.529478 |
| TMPPLLAAIANAGRHGV | ASSRLGGRDTQY | 0.45468318 |
| TSSTAGNSGGA | ASSRLGGRDTQY | 0.59939593 |
| VPATVPVGILC | ASSRLGGRDTQY | 0.3688267 |
| VQRPDSGGVVLTGRIS | ASSRLGGRDTQY | 0.40603596 |
| WTPNPVPATKGVPAGLVQAT | ASSRLGGRDTQY | 0.43278128 |
| YALKRALAKFSDRFDVVII | ASSRLGGRDTQY | 0.52676046 |
| YLIGSVIV | ASSRLGGRDTQY | 0.6541974 |
| YGQLVAVID | ASSRLGGRDTQY | 0.37697512 |
| VTVGGMTLPA | ASSRLGGRDTQY | 0.57733303 |
| VLALPAV | ASSRLGGRDTQY | 0.44023257 |
| VIPPTLNYET | ASSRLGGRDTQY | 0.38567644 |
| VGMGGMHPSSG | ASSRLGGRDTQY | 0.45931923 |
| VGLAGGAATA | ASSRLGGRDTQY | 0.45150945 |
| VGGAGGAGGSGP | ASSRLGGRDTQY | 0.3698488 |
| VFAAFMV | ASSRLGGRDTQY | 0.52793205 |
| TLYDKLR | ASSRLGGRDTQY | 0.39315766 |
| TLAVFTVL | ASSRLGGRDTQY | 0.5391476 |
| TGSVQPARV | ASSRLGGRDTQY | 0.67014205 |
| TGGGAGSGGT | ASSRLGGRDTQY | 0.35003886 |
| SHLETSL | ASSRLGGRDTQY | 0.59729517 |
| SFTTGMT | ASSRLGGRDTQY | 0.62405354 |
| QVEVAKRMGVRQPTVS | ASSRLGGRDTQY | 0.59278184 |
| QGSGGPALAM | ASSRLGGRDTQY | 0.492486 |
| MLAPSNP | ASSRLGGRDTQY | 0.41664496 |
| LVNNAAI | ASSRLGGRDTQY | 0.6827476 |
| LTGEINTLA | ASSRLGGRDTQY | 0.65270734 |
| LSIFGPV | ASSRLGGRDTQY | 0.5549169 |
| LLALLANL | ASSRLGGRDTQY | 0.56972986 |
| LISNASDALDKLR | ASSRLGGRDTQY | 0.66403663 |
| LINEAVP | ASSRLGGRDTQY | 0.6492728 |
| LGYAFSGL | ASSRLGGRDTQY | 0.42710948 |
| LDFVAVAL | ASSRLGGRDTQY | 0.3110101 |
| LAQPGAE | ASSRLGGRDTQY | 0.48801935 |
| ISNLGAVTGF | ASSRLGGRDTQY | 0.46955016 |
| HQEDPGRI | ASSRLGGRDTQY | 0.3989725 |
| GTPGEPTP | ASSRLGGRDTQY | 0.3303597 |
| GFFNSGS | ASSRLGGRDTQY | 0.67687225 |
| GAGGGGLTAGAN | ASSRLGGRDTQY | 0.7150011 |
| FVDALVNM | ASSRLGGRDTQY | 0.4867111 |
| FLTVAPEL | ASSRLGGRDTQY | 0.70991546 |
| FETLTAL | ASSRLGGRDTQY | 0.3430647 |
| DVINPIEL | ASSRLGGRDTQY | 0.6355485 |
| APVSGAPV | ASSRLGGRDTQY | 0.67348003 |
| ALVGGLAW | ASSRLGGRDTQY | 0.4268537 |
| ALAAAAATAI | ASSRLGGRDTQY | 0.5852081 |
| AFGLGLLL | ASSRLGGRDTQY | 0.37493572 |

| | | |
|---|---|---|
| AFFAAASL | ASSRLGGRDTQY | 0.44999832 |
| AAAALPAFSPP | ASSRLGGRDTQY | 0.64968264 |
| VVLAPAL | ASSRLGGRDTQY | 0.47655874 |
| TIGHVDHGKTTL | ASSRLGGRDTQY | 0.5212711 |
| LISNASDALDK | ASSRLGGRDTQY | 0.47095698 |
| LISNASDAL | ASSRLGGRDTQY | 0.38912848 |
| ISNASDALDK | ASSRLGGRDTQY | 0.5024631 |
| NQAETLVYQTE | ASSRLGGRDTQY | 0.5457656 |
| TGGTGGTAGAG | ASSRLGGRDTQY | 0.47706825 |
| TGSTAGGAGGA | ASSRLGGRDTQY | 0.69795936 |
| DPLAVPVAAGPV | ASSRLGGRDTQY | 0.3336992 |
| GGTGGTPGNGGN | ASSRLGGRDTQY | 0.70868874 |
| PAGLAGVDPATL | ASSRLGGRDTQY | 0.6971431 |
| GNNLVVEAPHAGASV | ASSRLGGRDTQY | 0.6067964 |
| KVTRIMSASTLEQPA | ASSRLGGRDTQY | 0.45190978 |
| LGFALPIALPRRTVV | ASSRLGGRDTQY | 0.6947856 |
| TRLGNLTARKLVVLV | ASSRLGGRDTQY | 0.44974232 |
| WSQQRRLIYFPSAGP | ASSRLGGRDTQY | 0.5286319 |
| GPNGNPDPMAAAVDIR | ASSRLGGRDTQY | 0.6457466 |
| TQRCVSVADLRAAASR | ASSRLGGRDTQY | 0.57368433 |
| QFVLDVAEHPVEVLTETV | ASSRLGGRDTQY | 0.7213354 |
| GGAGGTGGAGGNGGTQAGDG | ASSRLGGRDTQY | 0.6946218 |
| PLLATLRAIAVDVDAKPAQV | ASSRLGGRDTQY | 0.6821228 |
| APKFRTGGYMIDFWGVGYQVAK | ASSRLGGRDTQY | 0.57440376 |
| GAGGAGGAGGAAGGPGGTGGTG | ASSRLGGRDTQY | 0.37657586 |
| IGALGLLATDVAPAGSVAFGVA | ASSRLGGRDTQY | 0.6844425 |
| AGGAGGIGGNGGAGGAGGVLMG | ASSRLGGRDTQY | 0.51161885 |
| ADELIGLL | ASSRLGGRDTQY | 0.7181035 |
| ADRALKA | ASSRLGGRDTQY | 0.71283805 |
| AGAVGQVGGAGGA | ASSRLGGRDTQY | 0.5772396 |
| AGGGGGLGGAP | ASSRLGGRDTQY | 0.72629786 |
| AGGLLYGNGGAG | ASSRLGGRDTQY | 0.72435296 |
| AGLESIGL | ASSRLGGRDTQY | 0.42105755 |
| AGLKTGII | ASSRLGGRDTQY | 0.64422953 |
| ASLAALLL | ASSRLGGRDTQY | 0.6058796 |
| ASVINISSI | ASSRLGGRDTQY | 0.64226127 |
| ATPPGPA | ASSRLGGRDTQY | 0.45050615 |
| ATVASALV | ASSRLGGRDTQY | 0.70891356 |
| ATVASLIL | ASSRLGGRDTQY | 0.64039934 |
| AVNGPVLIRH | ASSRLGGRDTQY | 0.278788 |
| DGAAGAVGI | ASSRLGGRDTQY | 0.6866561 |
| DPLLHGI | ASSRLGGRDTQY | 0.6552857 |
| ELGAALGAL | ASSRLGGRDTQY | 0.62872016 |
| ELGRTQAV | ASSRLGGRDTQY | 0.71990955 |
| FVAGGYLS | ASSRLGGRDTQY | 0.6304797 |
| GASGGAGGAGGW | ASSRLGGRDTQY | 0.56740505 |
| GGAGAGGDGGAGGI | ASSRLGGRDTQY | 0.710931 |
| GTGHNGSQPGMGGNG | ASSRLGGRDTQY | 0.72688144 |
| GYAAASASAS | ASSRLGGRDTQY | 0.47952348 |
| HQVDVAAVQQG | ASSRLGGRDTQY | 0.69556653 |
| IGAAIAHT | ASSRLGGRDTQY | 0.72753155 |

| | | |
|---|---|---|
| IGMIEVAQL | ASSRLGGRDTQY | 0.702513 |
| IRLAPEHTSAMD | ASSRLGGRDTQY | 0.72623193 |
| IYNVSVL | ASSRLGGRDTQY | 0.72906613 |
| KQGFLGM | ASSRLGGRDTQY | 0.6240748 |
| LESVITQL | ASSRLGGRDTQY | 0.7207309 |
| LGLGALVVV | ASSRLGGRDTQY | 0.72901094 |
| LIGNGGNGGSGGI | ASSRLGGRDTQY | 0.5874748 |
| LNAINAPTL | ASSRLGGRDTQY | 0.6875901 |
| LQLLNTGV | ASSRLGGRDTQY | 0.5517123 |
| LQTSIEF | ASSRLGGRDTQY | 0.6856042 |
| MSFAVSEL | ASSRLGGRDTQY | 0.34084412 |
| NTGALGL | ASSRLGGRDTQY | 0.7054112 |
| PPGAGGAGGAGTT | ASSRLGGRDTQY | 0.7236582 |
| QLVGLGI | ASSRLGGRDTQY | 0.6648546 |
| RFLEQVHQL | ASSRLGGRDTQY | 0.53764147 |
| RGVGYVLR | ASSRLGGRDTQY | 0.5508027 |
| SATGVFAGVI | ASSRLGGRDTQY | 0.6028615 |
| SLPAITLP | ASSRLGGRDTQY | 0.6492635 |
| STALAFTV | ASSRLGGRDTQY | 0.3704055 |
| SVLTVGLGK | ASSRLGGRDTQY | 0.5992379 |
| TALAGAGSGI | ASSRLGGRDTQY | 0.72333515 |
| TGLPFTGI | ASSRLGGRDTQY | 0.72286963 |
| VAEHVTV | ASSRLGGRDTQY | 0.47307253 |
| VDIVLNSL | ASSRLGGRDTQY | 0.72389066 |
| VLTGGGAVL | ASSRLGGRDTQY | 0.71618986 |
| VNVGLAYGI | ASSRLGGRDTQY | 0.7275099 |
| VSSAAVGAV | ASSRLGGRDTQY | 0.6621946 |
| VTSVYEAGV | ASSRLGGRDTQY | 0.54253185 |
| WVALDRGA | ASSRLGGRDTQY | 0.71939063 |
| YLVGSVVI | ASSRLGGRDTQY | 0.6684936 |
| GPPGTGKTLL | ASSRLGGRDTQY | 0.66779125 |
| GVGMTKF | ASSRLGGRDTQY | 0.7203492 |
| IGIGGSDL | ASSRLGGRDTQY | 0.622707 |
| IGTLLAEL | ASSRLGGRDTQY | 0.6237223 |
| IYTSGTT | ASSRLGGRDTQY | 0.6823956 |
| LSFPAGV | ASSRLGGRDTQY | 0.7145177 |
| LYTSGTT | ASSRLGGRDTQY | 0.56199145 |
| VGAFPFGF | ASSRLGGRDTQY | 0.48489845 |
| VGAGTVEF | ASSRLGGRDTQY | 0.5865346 |
| VGKSTLLL | ASSRLGGRDTQY | 0.6830473 |
| AALAHAGGGIV | ASSRLGGRDTQY | 0.7231118 |
| AALVAVSLL | ASSRLGGRDTQY | 0.6960225 |
| AAVSRAVAAL | ASSRLGGRDTQY | 0.6623937 |
| AGGSSGIGGF | ASSRLGGRDTQY | 0.71082675 |
| AGLTVANANG | ASSRLGGRDTQY | 0.7022158 |
| AGLVGAVL | ASSRLGGRDTQY | 0.6742806 |
| ALSQAFTV | ASSRLGGRDTQY | 0.71727467 |
| ALTFSVSL | ASSRLGGRDTQY | 0.62189245 |
| ALTGAALVAL | ASSRLGGRDTQY | 0.71006525 |
| ASFSIGAGVL | ASSRLGGRDTQY | 0.7060573 |
| DGSVYKI | ASSRLGGRDTQY | 0.71686685 |

| | | |
|---|---|---|
| EQLAEIV | ASSRLGGRDTQY | 0.72117573 |
| FAVSELTVM | ASSRLGGRDTQY | 0.69301623 |
| GAAPLAAGL | ASSRLGGRDTQY | 0.6804345 |
| GADGGDGGSGGAGG | ASSRLGGRDTQY | 0.39338517 |
| GFGLDTGGAGG | ASSRLGGRDTQY | 0.66432995 |
| GLASGGLL | ASSRLGGRDTQY | 0.7062707 |
| GRIAGAAVLL | ASSRLGGRDTQY | 0.6962076 |
| GSAGSLNTSF | ASSRLGGRDTQY | 0.7162069 |
| GTGQIGI | ASSRLGGRDTQY | 0.7024621 |
| GTLIALIL | ASSRLGGRDTQY | 0.60913193 |
| IAATMGFVVG | ASSRLGGRDTQY | 0.7259761 |
| IFAQALL | ASSRLGGRDTQY | 0.7220768 |
| IGAAVALGF | ASSRLGGRDTQY | 0.43481913 |
| IGLTPLAL | ASSRLGGRDTQY | 0.638206 |
| ILGSNTSTL | ASSRLGGRDTQY | 0.7264905 |
| IVFSLTVL | ASSRLGGRDTQY | 0.70985246 |
| LASKATGF | ASSRLGGRDTQY | 0.65865123 |
| LASVALIL | ASSRLGGRDTQY | 0.64958423 |
| LFAVGLTL | ASSRLGGRDTQY | 0.7319029 |
| LFNGGIGTYI | ASSRLGGRDTQY | 0.6520282 |
| LTLGAGII | ASSRLGGRDTQY | 0.7294879 |
| LVAAALGDI | ASSRLGGRDTQY | 0.31378296 |
| NTAFVQLGI | ASSRLGGRDTQY | 0.585144 |
| QLFSGAEL | ASSRLGGRDTQY | 0.4571418 |
| SAATFITL | ASSRLGGRDTQY | 0.34204057 |
| SNYGLANLGV | ASSRLGGRDTQY | 0.71822464 |
| SYGGAYAVM | ASSRLGGRDTQY | 0.64185977 |
| TTGVGGLTLG | ASSRLGGRDTQY | 0.7173416 |
| TTQTIYI | ASSRLGGRDTQY | 0.68492186 |
| VAAGAGLGAVI | ASSRLGGRDTQY | 0.72125304 |
| VAAVVTLL | ASSRLGGRDTQY | 0.6975687 |
| VGFLGGLI | ASSRLGGRDTQY | 0.70225775 |
| VGVIPILL | ASSRLGGRDTQY | 0.5422128 |
| VIFASSLI | ASSRLGGRDTQY | 0.7115846 |
| VKGGDGGAAAG | ASSRLGGRDTQY | 0.72856236 |
| VTKTFTNI | ASSRLGGRDTQY | 0.696489 |
| VVAAAILL | ASSRLGGRDTQY | 0.45397726 |
| YAALDVEL | ASSRLGGRDTQY | 0.6805768 |
| LASVALLL | ASSRLGGRDTQY | 0.6835153 |
| LDAVHRV | ASSRLGGRDTQY | 0.34251255 |
| QGLSSSIF | ASSRLGGRDTQY | 0.31622583 |
| RGTFANIRL | ASSRLGGRDTQY | 0.52776515 |
| RQLWWGHRI | ASSRLGGRDTQY | 0.5851488 |
| YGPPGTGKTLL | ASSRLGGRDTQY | 0.7192762 |
| AALVGAPANVA | ASSRLGGRDTQY | 0.7149776 |
| AATFRRTE | ASSRLGGRDTQY | 0.65903986 |
| AGALGGGI | ASSRLGGRDTQY | 0.7140018 |
| AGGAGGAGGIGAG | ASSRLGGRDTQY | 0.69906986 |
| AGSGALGL | ASSRLGGRDTQY | 0.24900189 |
| AQIADQI | ASSRLGGRDTQY | 0.6199316 |
| ASAVAVTYL | ASSRLGGRDTQY | 0.45468742 |

| | | |
|---|---|---|
| ASTVLGGTGG | ASSRLGGRDTQY | 0.7260133 |
| AVGGEPAK | ASSRLGGRDTQY | 0.5525246 |
| CEQGTFGGTC | ASSRLGGRDTQY | 0.6146023 |
| DEGKAPAG | ASSRLGGRDTQY | 0.68219364 |
| DTVVFPAL | ASSRLGGRDTQY | 0.70387983 |
| FEGITMAG | ASSRLGGRDTQY | 0.6901994 |
| FPLVVQD | ASSRLGGRDTQY | 0.5376471 |
| GAFTTPQ | ASSRLGGRDTQY | 0.28257054 |
| GFGGFGTAGGVGGA | ASSRLGGRDTQY | 0.717222 |
| GFLAQFL | ASSRLGGRDTQY | 0.55896044 |
| GGAGGAGGNAGVA | ASSRLGGRDTQY | 0.4957059 |
| GGVGTAGAGGAGGAGG | ASSRLGGRDTQY | 0.4376645 |
| GNGGGGDAGNAGSGGN | ASSRLGGRDTQY | 0.6831075 |
| GTGGAGGAGNDAGS | ASSRLGGRDTQY | 0.41560692 |
| GTTNTGWFNS | ASSRLGGRDTQY | 0.72471344 |
| IFMGPLFTL | ASSRLGGRDTQY | 0.63547236 |
| ILAHAIEAI | ASSRLGGRDTQY | 0.6577094 |
| IRSAIREAL | ASSRLGGRDTQY | 0.72731876 |
| ITFAGGG | ASSRLGGRDTQY | 0.70339036 |
| ITGMSGHQEV | ASSRLGGRDTQY | 0.66733897 |
| IVAGLACTL | ASSRLGGRDTQY | 0.556405 |
| IVGGGLAGA | ASSRLGGRDTQY | 0.5134598 |
| IVTGVLNI | ASSRLGGRDTQY | 0.52866745 |
| LAQQGCI | ASSRLGGRDTQY | 0.5383673 |
| LCLHVEN | ASSRLGGRDTQY | 0.7297631 |
| LDAISDGS | ASSRLGGRDTQY | 0.59798515 |
| LGGFAAML | ASSRLGGRDTQY | 0.49068934 |
| LGGKNPMI | ASSRLGGRDTQY | 0.62160575 |
| LGPKGVAAIL | ASSRLGGRDTQY | 0.72608757 |
| LMVGVSI | ASSRLGGRDTQY | 0.733219 |
| LVGAAFVL | ASSRLGGRDTQY | 0.6029289 |
| LVLAVAPL | ASSRLGGRDTQY | 0.6424149 |
| MAAIAVIDAA | ASSRLGGRDTQY | 0.6476759 |
| NNGINAELQ | ASSRLGGRDTQY | 0.68726766 |
| RGEPLVLVHG | ASSRLGGRDTQY | 0.65228283 |
| RGGAGGAAGLI | ASSRLGGRDTQY | 0.42785203 |
| RSYRMHPAV | ASSRLGGRDTQY | 0.7054524 |
| SAAHLLAGQLA | ASSRLGGRDTQY | 0.7337248 |
| SAAVGSY | ASSRLGGRDTQY | 0.61011636 |
| SPHAVLHRI | ASSRLGGRDTQY | 0.53709304 |
| SQIAKVL | ASSRLGGRDTQY | 0.67571867 |
| TALGHLVP | ASSRLGGRDTQY | 0.60805786 |
| TAPTTSGAA | ASSRLGGRDTQY | 0.6290144 |
| TDGDDGNGGSPGT | ASSRLGGRDTQY | 0.6943786 |
| TDGLGEN | ASSRLGGRDTQY | 0.41027656 |
| TGGPPLPGQA | ASSRLGGRDTQY | 0.6326505 |
| TIPAITI | ASSRLGGRDTQY | 0.7319014 |
| TIPGFTI | ASSRLGGRDTQY | 0.65623385 |
| TSIASGALA | ASSRLGGRDTQY | 0.6904224 |
| TVTLPGAAI | ASSRLGGRDTQY | 0.72284365 |
| VASGLSAALP | ASSRLGGRDTQY | 0.708359 |

| | | |
|---|---|---|
| VGAGVIGI | ASSRLGGRDTQY | 0.7250465 |
| VGPKRFGV | ASSRLGGRDTQY | 0.7212912 |
| VNVAFPD | ASSRLGGRDTQY | 0.66888773 |
| VSMTNSGVS | ASSRLGGRDTQY | 0.6817847 |
| VTDVLGDLSV | ASSRLGGRDTQY | 0.6927736 |
| VVVVGAGFAGL | ASSRLGGRDTQY | 0.6902306 |
| VYTVGTVV | ASSRLGGRDTQY | 0.6758241 |
| WTTFSGP | ASSRLGGRDTQY | 0.6192224 |
| YDELVEAF | ASSRLGGRDTQY | 0.71520466 |
| YGTGGAGGNGG | ASSRLGGRDTQY | 0.6573781 |
| YTNVMAV | ASSRLGGRDTQY | 0.71924007 |
| FLEQVHQL | ASSRLGGRDTQY | 0.41867635 |
| GLSLLAVAV | ASSRLGGRDTQY | 0.48584026 |
| LEQVHQL | ASSRLGGRDTQY | 0.7163359 |
| LNAAGAI | ASSRLGGRDTQY | 0.2689324 |
| PAVHASL | ASSRLGGRDTQY | 0.31739467 |
| TELEDDA | ASSRLGGRDTQY | 0.6714475 |
| TPPGPPN | ASSRLGGRDTQY | 0.72612983 |
| TPPPPPPP | ASSRLGGRDTQY | 0.27372402 |
| TQSPALI | ASRRQISNERLF | 0.5230958 |
| EFAAAFPQ | ASRRQISNERLF | 0.50059724 |
| LLSGNLDVL | ASRRQISNERLF | 0.68917966 |
| GPANLGNYNI | ASRRQISNERLF | 0.68728524 |
| TLGPTLP | ASRRQISNERLF | 0.365371 |
| VYAYSSCCK | ASRRQISNERLF | 0.71033347 |
| APAPAPAPNI | ASRRQISNERLF | 0.4655163 |
| PGGPVPIP | ASRRQISNERLF | 0.55789554 |
| FLTVALEP | ASRRQISNERLF | 0.7204899 |
| SQYFGANL | ASRRQISNERLF | 0.72740567 |
| GGVKPAA | ASRRQISNERLF | 0.5017109 |
| GGTGGAGGAGGW | ASRRQISNERLF | 0.719956 |
| GVGGVAGLGGAGG | ASRRQISNERLF | 0.6296502 |
| VLLVLATTI | ASRRQISNERLF | 0.7336112 |
| IGIFAAML | ASRRQISNERLF | 0.62142825 |
| AVGDLID | ASRRQISNERLF | 0.3856311 |
| NISPLPL | ASRRQISNERLF | 0.71338856 |
| VTAVFVAI | ASRRQISNERLF | 0.68748975 |
| SGDDNDAAL | ASRRQISNERLF | 0.6703569 |
| AVAALAHS | ASRRQISNERLF | 0.71486294 |
| GGVTVMATP | ASRRQISNERLF | 0.5236513 |
| AGLAANVRD | ASRRQISNERLF | 0.6546092 |
| TNRPDLIDPALLRPG | ASRRQISNERLF | 0.72374225 |
| LANLGVTAL | ASRRQISNERLF | 0.717838 |
| TGERIEL | ASRRQISNERLF | 0.41966024 |
| NGEELAA | ASRRQISNERLF | 0.7200668 |
| VLSLVWG | ASRRQISNERLF | 0.685693 |
| IGYALWL | ASRRQISNERLF | 0.607373 |
| LSLLLAPL | ASRRQISNERLF | 0.7283641 |
| TLDEALK | ASRRQISNERLF | 0.6904433 |
| TDGAGTAEGG | ASRRQISNERLF | 0.71797967 |
| NAPGGAGGNGG | ASRRQISNERLF | 0.70478827 |

| | | |
|---|---|---|
| VSVFMGMGGNF | ASRRQISNERLF | 0.72759134 |
| LASYLAAL | ASRRQISNERLF | 0.39830863 |
| LVGGTPAL | ASRRQISNERLF | 0.70808053 |
| VRNADVL | ASRRQISNERLF | 0.6844444 |
| TCSSPFLVSAM | ASRRQISNERLF | 0.7191441 |
| VHITDAE | ASRRQISNERLF | 0.6745833 |
| EYAAPSF | ASRRQISNERLF | 0.6211045 |
| VVDAFHGGL | ASRRQISNERLF | 0.6921892 |
| ASVVDPQ | ASRRQISNERLF | 0.72401464 |
| SYNEFGGLS | ASRRQISNERLF | 0.66530144 |
| DPDGSVY | ASRRQISNERLF | 0.70840406 |
| MVSTIHATE | ASRRQISNERLF | 0.72719777 |
| LAAGVADPV | ASRRQISNERLF | 0.55652416 |
| TKEQIAA | ASRRQISNERLF | 0.6722749 |
| SLASPAAL | ASRRQISNERLF | 0.7032814 |
| IGLVFYGV | ASRRQISNERLF | 0.7240566 |
| FLLDASNL | ASRRQISNERLF | 0.63846034 |
| YQVGNFP | ASRRQISNERLF | 0.7268722 |
| ESLLSQST | ASRRQISNERLF | 0.60815346 |
| LFTVGFAAVG | ASRRQISNERLF | 0.7166317 |
| ATPPTPT | ASRRQISNERLF | 0.6358007 |
| TDLATLG | ASRRQISNERLF | 0.7043536 |
| SAGGTGR | ASRRQISNERLF | 0.51283973 |
| GFSNSGS | ASRRQISNERLF | 0.71206367 |
| GGNGGSGGTSV | ASRRQISNERLF | 0.71986467 |
| GTSVATGGAGN | ASRRQISNERLF | 0.71884084 |
| NGGNGGSGGTS | ASRRQISNERLF | 0.3323607 |
| AVVKFPH | ASRRQISNERLF | 0.71209407 |
| TPPPPPP | ASRRQISNERLF | 0.7189503 |
| AAPAAAAAPA | ASRRQISNERLF | 0.7321349 |
| IALNEDL | ASRRQISNERLF | 0.67804253 |
| IVNVSSM | ASRRQISNERLF | 0.70790815 |
| TGIPPAPR | ASRRQISNERLF | 0.65633464 |
| FDLGGGTF | ASRRQISNERLF | 0.65676975 |
| FDLGGGTFDVS | ASRRQISNERLF | 0.5556096 |
| NEPTAAA | ASRRQISNERLF | 0.62639093 |
| NEPTAAALA | ASRRQISNERLF | 0.6734538 |
| GSIINISSI | ASRRQISNERLF | 0.5893511 |
| VAPGNAGTA | ASRRQISNERLF | 0.7211523 |
| TGGGASLE | ASRRQISNERLF | 0.7266408 |
| VAVYNHYKRI | ASRRQISNERLF | 0.72155404 |
| TDPAPATT | ASRRQISNERLF | 0.7212689 |
| TNRPDILDPALLRPG | ASRRQISNERLF | 0.7243639 |
| LTTTTTS | ASRRQISNERLF | 0.71825993 |
| PTGSGKTL | ASRRQISNERLF | 0.60757875 |
| TGSGKTL | ASRRQISNERLF | 0.67223096 |
| TIGHVDHGK | ASRRQISNERLF | 0.5172447 |
| TIGHVDHGKTT | ASRRQISNERLF | 0.6177132 |
| GHVDHGKTT | ASRRQISNERLF | 0.59544307 |
| AQFEHTLL | ASRRQISNERLF | 0.6939961 |
| IVNQDEL | ASRRQISNERLF | 0.26619622 |

| | | |
|---|---|---|
| TNGSQFF | ASRRQISNERLF | 0.6996002 |
| TGKGIVE | ASRRQISNERLF | 0.7290718 |
| LISNASDA | ASRRQISNERLF | 0.6536368 |
| AELRTLGELTKIAKAHGAQ | ASRRQISNERLF | 0.7019814 |
| APNAVAAAEPSW | ASRRQISNERLF | 0.68177223 |
| APQVAGAATAVL | ASRRQISNERLF | 0.6138127 |
| ARRDPAAAAAILPTDA | ASRRQISNERLF | 0.714147 |
| DAAARDVLAEAGLAEYF | ASRRQISNERLF | 0.5858715 |
| DDQIVLSAKANLAKVEQ | ASRRQISNERLF | 0.71265763 |
| DELLPDTGEDSDEDGH | ASRRQISNERLF | 0.7280903 |
| ELPEAPANRATVPQD | ASRRQISNERLF | 0.6865405 |
| FGQSKIERVTGIG | ASRRQISNERLF | 0.7010857 |
| GAGGTGSTAGGAGGAGGAGGLY | ASRRQISNERLF | 0.68190515 |
| GHGGTGAAGGNGGTGGAGGSGI | ASRRQISNERLF | 0.37508014 |
| GNGGNGGAGGEGGPGGAGGAGA | ASRRQISNERLF | 0.7086923 |
| GSTAGNGGNGGNATNGTIAGQP | ASRRQISNERLF | 0.40026724 |
| GTGGDGGNGGNGGTGG | ASRRQISNERLF | 0.679541 |
| GVRRPAGIPADAPV | ASRRQISNERLF | 0.5099292 |
| GYRQVTDAHLLTLARR | ASRRQISNERLF | 0.7005799 |
| IAKMGQDGHDRG | ASRRQISNERLF | 0.6870859 |
| KAEPSIEIQYT | ASRRQISNERLF | 0.70123035 |
| KPVRIRMAADEDTVAVT | ASRRQISNERLF | 0.5884203 |
| MPPRIAGMRLLVIKPEP | ASRRQISNERLF | 0.69279253 |
| MKFYAVALFRSRGVPPTR | ASRRQISNERLF | 0.67097545 |
| NAGGFGILGAVAHSP | ASRRQISNERLF | 0.43183792 |
| NEGREITLAMAEM | ASRRQISNERLF | 0.7157252 |
| NELVEWSHPLGHRWRWE | ASRRQISNERLF | 0.58854353 |
| NLGAGGTGGNGGNPG | ASRRQISNERLF | 0.6591818 |
| NTEWPKWETVMAADPVD | ASRRQISNERLF | 0.7234899 |
| PASAAPAETQP | ASRRQISNERLF | 0.69210213 |
| QQSSDGKNGVRAAEITGE | ASRRQISNERLF | 0.41716564 |
| RAPDAVTTSWWKE | ASRRQISNERLF | 0.6776204 |
| SGFGNVGAHTSGWFNQ | ASRRQISNERLF | 0.67472124 |
| SGVAGAVGQAASVGGLKV | ASRRQISNERLF | 0.6327785 |
| SNGGAGTGSAGGNGGT | ASRRQISNERLF | 0.2807439 |
| TGGAGGEGGAAG | ASRRQISNERLF | 0.6897719 |
| TGGDGGAAGAGGAPGHG | ASRRQISNERLF | 0.6152242 |
| TMPPLLAAIANAGRHGV | ASRRQISNERLF | 0.5859327 |
| TSSTAGNSGGA | ASRRQISNERLF | 0.60832036 |
| VPATVPVGILC | ASRRQISNERLF | 0.7170639 |
| VQRPDSGGVVLTGRIS | ASRRQISNERLF | 0.69236255 |
| WTPNPVPATKGVPAGLVQAT | ASRRQISNERLF | 0.6511587 |
| YALKRALAKFSDRFDVVII | ASRRQISNERLF | 0.7142564 |
| YLIGSVIV | ASRRQISNERLF | 0.7153981 |
| YGQLVAVID | ASRRQISNERLF | 0.72828233 |
| VTVGGMTLPA | ASRRQISNERLF | 0.68369085 |
| VLALPAV | ASRRQISNERLF | 0.60264885 |
| VIPPTLNYET | ASRRQISNERLF | 0.711201 |
| VGMGGMHPSSG | ASRRQISNERLF | 0.6918764 |
| VGLAGGAATA | ASRRQISNERLF | 0.4163077 |
| VGGAGGAGGSGP | ASRRQISNERLF | 0.40578708 |

| | | |
|---|---|---|
| VFAAFMV | ASRRQISNERLF | 0.6989521 |
| TLYDKLR | ASRRQISNERLF | 0.423293 |
| TLAVFTVL | ASRRQISNERLF | 0.67945457 |
| TGSVQPARV | ASRRQISNERLF | 0.72301567 |
| TGGGAGSGGT | ASRRQISNERLF | 0.6444835 |
| SHLETSL | ASRRQISNERLF | 0.4656941 |
| SFTTGMT | ASRRQISNERLF | 0.6296571 |
| QVEVAKRMGVRQPTVS | ASRRQISNERLF | 0.62009835 |
| QGSGGPALAM | ASRRQISNERLF | 0.7168815 |
| MLAPSNP | ASRRQISNERLF | 0.69706476 |
| LVNNAAI | ASRRQISNERLF | 0.7309953 |
| LTGEINTLA | ASRRQISNERLF | 0.7176585 |
| LSIFGPV | ASRRQISNERLF | 0.64853024 |
| LLALLANL | ASRRQISNERLF | 0.7111769 |
| LISNASDALDKLR | ASRRQISNERLF | 0.6331153 |
| LINEAVP | ASRRQISNERLF | 0.67403626 |
| LGYAFSGL | ASRRQISNERLF | 0.67610896 |
| LDFVAVAL | ASRRQISNERLF | 0.6880824 |
| LAQPGAE | ASRRQISNERLF | 0.6510056 |
| ISNLGAVTGF | ASRRQISNERLF | 0.6645266 |
| HQEDPGRI | ASRRQISNERLF | 0.39335373 |
| GTPGEPTP | ASRRQISNERLF | 0.70361125 |
| GFFNSGS | ASRRQISNERLF | 0.70259535 |
| GAGGGGLTAGAN | ASRRQISNERLF | 0.46578968 |
| FVDALVNM | ASRRQISNERLF | 0.6588113 |
| FLTVAPEL | ASRRQISNERLF | 0.72964233 |
| FETLTAL | ASRRQISNERLF | 0.72776204 |
| DVINPIEL | ASRRQISNERLF | 0.59146726 |
| APVSGAPV | ASRRQISNERLF | 0.5495109 |
| ALVGGLAW | ASRRQISNERLF | 0.697268 |
| ALAAAAATAI | ASRRQISNERLF | 0.4453485 |
| AFGLGLLL | ASRRQISNERLF | 0.6757933 |
| AFFAAASL | ASRRQISNERLF | 0.6893407 |
| AAAALPAFSPP | ASRRQISNERLF | 0.6768447 |
| VVLAPAL | ASRRQISNERLF | 0.6132767 |
| TIGHVDHGKTTL | ASRRQISNERLF | 0.69578934 |
| LISNASDALDK | ASRRQISNERLF | 0.64298797 |
| LISNASDAL | ASRRQISNERLF | 0.71535003 |
| ISNASDALDK | ASRRQISNERLF | 0.29454428 |
| NQAETLVYQTE | ASRRQISNERLF | 0.7016665 |
| TGGTGGTAGAG | ASRRQISNERLF | 0.5962826 |
| TGSTAGGAGGA | ASRRQISNERLF | 0.60095346 |
| DPLAVPVAAGPV | ASRRQISNERLF | 0.3660589 |
| GGTGGTPGNGGN | ASRRQISNERLF | 0.68689907 |
| PAGLAGVDPATL | ASRRQISNERLF | 0.7196169 |
| GNNLVVEAPHAGASV | ASRRQISNERLF | 0.7191367 |
| KVTRIMSASTLEQPA | ASRRQISNERLF | 0.6038065 |
| LGFALPIALPRRTVV | ASRRQISNERLF | 0.7195374 |
| TRLGNLTARKLVVLV | ASRRQISNERLF | 0.7322509 |
| WSQQRRLIYFPSAGP | ASRRQISNERLF | 0.65708494 |
| GPNGNPDPMAAAVDIR | ASRRQISNERLF | 0.34354603 |

| | | |
|---|---|---|
| TQRCVSVADLRAAASR | ASRRQISNERLF | 0.7027023 |
| QFVLDVAEHPVEVLTETV | ASRRQISNERLF | 0.69403183 |
| GGAGGTGGAGGNGGTQAGDG | ASRRQISNERLF | 0.7131876 |
| PLLATLRAIAVDVDAKPAQV | ASRRQISNERLF | 0.71714246 |
| APKFRTGGYMIDFWGVGYQVAK | ASRRQISNERLF | 0.5157016 |
| GAGGAGGAGGAAGGPGGTGGTG | ASRRQISNERLF | 0.71226513 |
| IGALGLLATDVAPAGSVAFGVA | ASRRQISNERLF | 0.7162968 |
| AGGAGGIGGNGGAGGAGGVLMG | ASRRQISNERLF | 0.5687661 |
| ADELIGLL | ASRRQISNERLF | 0.62109625 |
| ADRALKA | ASRRQISNERLF | 0.6159309 |
| AGAVGQVGGAGGA | ASRRQISNERLF | 0.42556703 |
| AGGGGGLGGAP | ASRRQISNERLF | 0.71629333 |
| AGGLLYGNGGAG | ASRRQISNERLF | 0.62931406 |
| AGLESIGL | ASRRQISNERLF | 0.6182748 |
| AGLKTGII | ASRRQISNERLF | 0.666093 |
| ASLAALLL | ASRRQISNERLF | 0.56739926 |
| ASVINISSI | ASRRQISNERLF | 0.6131289 |
| ATPPGPA | ASRRQISNERLF | 0.58893955 |
| ATVASALV | ASRRQISNERLF | 0.6817199 |
| ATVASLIL | ASRRQISNERLF | 0.70264184 |
| AVNGPVLIRH | ASRRQISNERLF | 0.6854232 |
| DGAAGAVGI | ASRRQISNERLF | 0.70055723 |
| DPLLHGI | ASRRQISNERLF | 0.7257276 |
| ELGAALGAL | ASRRQISNERLF | 0.4403817 |
| ELGRTQAV | ASRRQISNERLF | 0.60078555 |
| FVAGGYLS | ASRRQISNERLF | 0.7019086 |
| GASGGAGGAGGW | ASRRQISNERLF | 0.45097798 |
| GGAGAGGDGGAGGI | ASRRQISNERLF | 0.7156323 |
| GTGHNGSQPGMGGNG | ASRRQISNERLF | 0.6937789 |
| GYAAASASAS | ASRRQISNERLF | 0.6996465 |
| HQVDVAAVQQG | ASRRQISNERLF | 0.69220763 |
| IGAAIAHT | ASRRQISNERLF | 0.72990227 |
| IGMIEVAQL | ASRRQISNERLF | 0.7120833 |
| IRLAPEHTSAMD | ASRRQISNERLF | 0.66055465 |
| IYNVSVL | ASRRQISNERLF | 0.71181965 |
| KQGFLGM | ASRRQISNERLF | 0.56251186 |
| LESVITQL | ASRRQISNERLF | 0.7211276 |
| LGLGALVVV | ASRRQISNERLF | 0.70991886 |
| LIGNGGNGGSGGI | ASRRQISNERLF | 0.5467212 |
| LNAINAPTL | ASRRQISNERLF | 0.65914994 |
| LQLLNTGV | ASRRQISNERLF | 0.72611177 |
| LQTSIEF | ASRRQISNERLF | 0.7228852 |
| MSFAVSEL | ASRRQISNERLF | 0.40999296 |
| NTGALGL | ASRRQISNERLF | 0.6434375 |
| PPGAGGAGGAGTT | ASRRQISNERLF | 0.41941622 |
| QLVGLGI | ASRRQISNERLF | 0.52364093 |
| RFLEQVHQL | ASRRQISNERLF | 0.48329085 |
| RGVGYVLR | ASRRQISNERLF | 0.7109088 |
| SATGVFAGVI | ASRRQISNERLF | 0.6564852 |
| SLPAITLP | ASRRQISNERLF | 0.71573555 |
| STALAFTV | ASRRQISNERLF | 0.67880094 |

| | | |
|---|---|---|
| SVLTVGLGK | ASRRQISNERLF | 0.723706 |
| TALAGAGSGI | ASRRQISNERLF | 0.5891815 |
| TGLPFTGI | ASRRQISNERLF | 0.6657605 |
| VAEHVTV | ASRRQISNERLF | 0.40581155 |
| VDIVLNSL | ASRRQISNERLF | 0.63347924 |
| VLTGGGAVL | ASRRQISNERLF | 0.57597405 |
| VNVGLAYGI | ASRRQISNERLF | 0.63920367 |
| VSSAAVGAV | ASRRQISNERLF | 0.36147973 |
| VTSVYEAGV | ASRRQISNERLF | 0.7218478 |
| WVALDRGA | ASRRQISNERLF | 0.69404507 |
| YLVGSVVI | ASRRQISNERLF | 0.71405804 |
| GPPGTGKTLL | ASRRQISNERLF | 0.64188325 |
| GVGMTKF | ASRRQISNERLF | 0.65832055 |
| IGIGGSDL | ASRRQISNERLF | 0.58891475 |
| IGTLLAEL | ASRRQISNERLF | 0.7105708 |
| IYTSGTT | ASRRQISNERLF | 0.5032272 |
| LSFPAGV | ASRRQISNERLF | 0.4129628 |
| LYTSGTT | ASRRQISNERLF | 0.6711823 |
| VGAFPFGF | ASRRQISNERLF | 0.55610037 |
| VGAGTVEF | ASRRQISNERLF | 0.7260762 |
| VGKSTLLL | ASRRQISNERLF | 0.72159076 |
| AALAHAGGGIV | ASRRQISNERLF | 0.6924765 |
| AALVAVSLL | ASRRQISNERLF | 0.7152219 |
| AAVSRAVAAL | ASRRQISNERLF | 0.28770292 |
| AGGSSGIGGF | ASRRQISNERLF | 0.65001273 |
| AGLTVANANG | ASRRQISNERLF | 0.7295521 |
| AGLVGAVL | ASRRQISNERLF | 0.70227474 |
| ALSQAFTV | ASRRQISNERLF | 0.66909766 |
| ALTFSVSL | ASRRQISNERLF | 0.7259101 |
| ALTGAALVAL | ASRRQISNERLF | 0.6258304 |
| ASFSIGAGVL | ASRRQISNERLF | 0.6453551 |
| DGSVYKI | ASRRQISNERLF | 0.67908967 |
| EQLAEIV | ASRRQISNERLF | 0.7087833 |
| FAVSELTVM | ASRRQISNERLF | 0.6957036 |
| GAAPLAAGL | ASRRQISNERLF | 0.5508944 |
| GADGGDGGSGGAGG | ASRRQISNERLF | 0.69869924 |
| GFGLDTGGAGG | ASRRQISNERLF | 0.7315286 |
| GLASGGLL | ASRRQISNERLF | 0.6613929 |
| GRIAGAAVLL | ASRRQISNERLF | 0.72323835 |
| GSAGSLNTSF | ASRRQISNERLF | 0.7058938 |
| GTGQIGI | ASRRQISNERLF | 0.71302223 |
| GTLIALIL | ASRRQISNERLF | 0.5912713 |
| IAATMGFVVG | ASRRQISNERLF | 0.62269366 |
| IFAQALL | ASRRQISNERLF | 0.6996691 |
| IGAAVALGF | ASRRQISNERLF | 0.6239894 |
| IGLTPLAL | ASRRQISNERLF | 0.6742266 |
| ILGSNTSTL | ASRRQISNERLF | 0.6028475 |
| IVFSLTVL | ASRRQISNERLF | 0.73398167 |
| LASKATGF | ASRRQISNERLF | 0.7121613 |
| LASVALIL | ASRRQISNERLF | 0.7049737 |
| LFAVGLTL | ASRRQISNERLF | 0.55781305 |

| | | |
|---|---|---|
| LFNGGIGTYI | ASRRQISNERLF | 0.6841753 |
| LTLGAGII | ASRRQISNERLF | 0.7077657 |
| LVAAALGDI | ASRRQISNERLF | 0.40906787 |
| NTAFVQLGI | ASRRQISNERLF | 0.72581244 |
| QLFSGAEL | ASRRQISNERLF | 0.608503 |
| SAATFITL | ASRRQISNERLF | 0.6389241 |
| SNYGLANLGV | ASRRQISNERLF | 0.707885 |
| SYGGAYAVM | ASRRQISNERLF | 0.7269409 |
| TTGVGGLTLG | ASRRQISNERLF | 0.6492528 |
| TTQTIYI | ASRRQISNERLF | 0.69667196 |
| VAAGAGLGAVI | ASRRQISNERLF | 0.5273379 |
| VAAVVTLL | ASRRQISNERLF | 0.7158096 |
| VGFLGGLI | ASRRQISNERLF | 0.72035825 |
| VGVIPILL | ASRRQISNERLF | 0.70242906 |
| VIFASSLI | ASRRQISNERLF | 0.7287182 |
| VKGGDGGAAAG | ASRRQISNERLF | 0.6824187 |
| VTKTFTNI | ASRRQISNERLF | 0.37522298 |
| VVAAAILL | ASRRQISNERLF | 0.6718936 |
| YAALDVEL | ASRRQISNERLF | 0.7204081 |
| LASVALLL | ASRRQISNERLF | 0.72197974 |
| LDAVHRV | ASRRQISNERLF | 0.72847176 |
| QGLSSSIF | ASRRQISNERLF | 0.7043673 |
| RGTFANIRL | ASRRQISNERLF | 0.6488001 |
| RQLWWGHRI | ASRRQISNERLF | 0.726234 |
| YGPPGTGKTLL | ASRRQISNERLF | 0.7328053 |
| AALVGAPANVA | ASRRQISNERLF | 0.7153446 |
| AATFRRTE | ASRRQISNERLF | 0.6240408 |
| AGALGGGI | ASRRQISNERLF | 0.51772034 |
| AGGAGGAGGIGAG | ASRRQISNERLF | 0.6828301 |
| AGSGALGL | ASRRQISNERLF | 0.7121822 |
| AQIADQI | ASRRQISNERLF | 0.5346753 |
| ASAVAVTYL | ASRRQISNERLF | 0.69384813 |
| ASTVLGGTGG | ASRRQISNERLF | 0.69527763 |
| AVGGEPAK | ASRRQISNERLF | 0.4430367 |
| CEQGTFGGTC | ASRRQISNERLF | 0.7034756 |
| DEGKAPAG | ASRRQISNERLF | 0.72327983 |
| DTVVFPAL | ASRRQISNERLF | 0.6404149 |
| FEGITMAG | ASRRQISNERLF | 0.601561 |
| FPLVVQD | ASRRQISNERLF | 0.72206926 |
| GAFTTPQ | ASRRQISNERLF | 0.73153406 |
| GFGGFGTAGGVGGA | ASRRQISNERLF | 0.7217796 |
| GFLAQFL | ASRRQISNERLF | 0.43139488 |
| GGAGGAGGNAGVA | ASRRQISNERLF | 0.33108068 |
| GGVGTAGAGGAGGAGG | ASRRQISNERLF | 0.66130203 |
| GNGGGGDAGNAGSGGN | ASRRQISNERLF | 0.71891916 |
| GTGGAGGAGNDAGS | ASRRQISNERLF | 0.631728 |
| GTTNTGWFNS | ASRRQISNERLF | 0.57557535 |
| IFMGPLFTL | ASRRQISNERLF | 0.7010156 |
| ILAHAIEAI | ASRRQISNERLF | 0.7216638 |
| IRSAIREAL | ASRRQISNERLF | 0.5384252 |
| ITFAGGG | ASRRQISNERLF | 0.4198535 |

| | | |
|---|---|---|
| ITGMSGHQEV | ASRRQISNERLF | 0.7094773 |
| IVAGLACTL | ASRRQISNERLF | 0.71023643 |
| IVGGGLAGA | ASRRQISNERLF | 0.6126907 |
| IVTGVLNI | ASRRQISNERLF | 0.65255237 |
| LAQQGCI | ASRRQISNERLF | 0.65843487 |
| LCLHVEN | ASRRQISNERLF | 0.6775737 |
| LDAISDGS | ASRRQISNERLF | 0.6973083 |
| LGGFAAML | ASRRQISNERLF | 0.48982474 |
| LGGKNPMI | ASRRQISNERLF | 0.7198769 |
| LGPKGVAAIL | ASRRQISNERLF | 0.49742037 |
| LMVGVSI | ASRRQISNERLF | 0.7144946 |
| LVGAAFVL | ASRRQISNERLF | 0.70440525 |
| LVLAVAPL | ASRRQISNERLF | 0.69530606 |
| MAAIAVIDAA | ASRRQISNERLF | 0.6546358 |
| NNGINAELQ | ASRRQISNERLF | 0.62993765 |
| RGEPLVLVHG | ASRRQISNERLF | 0.4366017 |
| RGGAGGAAGLI | ASRRQISNERLF | 0.39351717 |
| RSYRMHPAV | ASRRQISNERLF | 0.7265011 |
| SAAHLLAGQLA | ASRRQISNERLF | 0.6613882 |
| SAAVGSY | ASRRQISNERLF | 0.72195244 |
| SPHAVLHRI | ASRRQISNERLF | 0.7089191 |
| SQIAKVL | ASRRQISNERLF | 0.53604305 |
| TALGHLVP | ASRRQISNERLF | 0.7027422 |
| TAPTTSGAA | ASRRQISNERLF | 0.47342384 |
| TDGDDGNGGSPGT | ASRRQISNERLF | 0.6348237 |
| TDGLGEN | ASRRQISNERLF | 0.71076554 |
| TGGPPLPGQA | ASRRQISNERLF | 0.7079444 |
| TIPAITI | ASRRQISNERLF | 0.59430516 |
| TIPGFTI | ASRRQISNERLF | 0.6926634 |
| TSIASGALA | ASRRQISNERLF | 0.6982484 |
| TVTLPGAAI | ASRRQISNERLF | 0.6171212 |
| VASGLSAALP | ASRRQISNERLF | 0.64181846 |
| VGAGVIGI | ASRRQISNERLF | 0.7228211 |
| VGPKRFGV | ASRRQISNERLF | 0.6872815 |
| VNVAFPD | ASRRQISNERLF | 0.555091 |
| VSMTNSGVS | ASRRQISNERLF | 0.7149873 |
| VTDVLGDLSV | ASRRQISNERLF | 0.59683585 |
| VVVVGAGFAGL | ASRRQISNERLF | 0.6979139 |
| VYTVGTVV | ASRRQISNERLF | 0.66763616 |
| WTTFSGP | ASRRQISNERLF | 0.66276336 |
| YDELVEAF | ASRRQISNERLF | 0.71415555 |
| YGTGGAGGNGG | ASRRQISNERLF | 0.5801722 |
| YTNVMAV | ASRRQISNERLF | 0.5915483 |
| FLEQVHQL | ASRRQISNERLF | 0.6624248 |
| GLSLLAVAV | ASRRQISNERLF | 0.6715195 |
| LEQVHQL | ASRRQISNERLF | 0.7168591 |
| LNAAGAI | ASRRQISNERLF | 0.5822847 |
| PAVHASL | ASRRQISNERLF | 0.6941407 |
| TELEDDA | ASRRQISNERLF | 0.6243944 |
| TPPGPPN | ASRRQISNERLF | 0.564283 |
| TPPPPPPP | ASRRQISNERLF | 0.63870066 |

| | | |
|---|---|---|
| TQSPALI | ASSRQRSQNTLY | 0.67205334 |
| EFAAAFPQ | ASSRQRSQNTLY | 0.5572811 |
| LLSGNLDVL | ASSRQRSQNTLY | 0.71175456 |
| GPANLGNYNI | ASSRQRSQNTLY | 0.47342092 |
| TLGPTLP | ASSRQRSQNTLY | 0.6995877 |
| VYAYSSCCK | ASSRQRSQNTLY | 0.7316416 |
| APAPAPAPNI | ASSRQRSQNTLY | 0.6326839 |
| PGGPVPIP | ASSRQRSQNTLY | 0.7087427 |
| FLTVALEP | ASSRQRSQNTLY | 0.7022728 |
| SQYFGANL | ASSRQRSQNTLY | 0.70978475 |
| GGVKPAA | ASSRQRSQNTLY | 0.60873497 |
| GGTGGAGGAGGW | ASSRQRSQNTLY | 0.5942898 |
| GVGGVAGLGGAGG | ASSRQRSQNTLY | 0.7211354 |
| VLLVLATTI | ASSRQRSQNTLY | 0.7165963 |
| IGIFAAML | ASSRQRSQNTLY | 0.72010803 |
| AVGDLID | ASSRQRSQNTLY | 0.699256 |
| NISPLPL | ASSRQRSQNTLY | 0.7206132 |
| VTAVFVAI | ASSRQRSQNTLY | 0.7107849 |
| SGDDNDAAL | ASSRQRSQNTLY | 0.66240394 |
| AVAALAHS | ASSRQRSQNTLY | 0.44970286 |
| GGVTVMATP | ASSRQRSQNTLY | 0.63837993 |
| AGLAANVRD | ASSRQRSQNTLY | 0.6285025 |
| TNRPDLIDPALLRPG | ASSRQRSQNTLY | 0.64823914 |
| LANLGVTAL | ASSRQRSQNTLY | 0.37152708 |
| TGERIEL | ASSRQRSQNTLY | 0.7225311 |
| NGEELAA | ASSRQRSQNTLY | 0.73258555 |
| VLSLVWG | ASSRQRSQNTLY | 0.58908165 |
| IGYALWL | ASSRQRSQNTLY | 0.7286826 |
| LSLLLAPL | ASSRQRSQNTLY | 0.48707357 |
| TLDEALK | ASSRQRSQNTLY | 0.71581745 |
| TDGAGTAEGG | ASSRQRSQNTLY | 0.725543 |
| NAPGGAGGNGG | ASSRQRSQNTLY | 0.5958272 |
| VSVFMGMGGNF | ASSRQRSQNTLY | 0.7135854 |
| LASYLAAL | ASSRQRSQNTLY | 0.58082736 |
| LVGGTPAL | ASSRQRSQNTLY | 0.64427257 |
| VRNADVL | ASSRQRSQNTLY | 0.72654057 |
| TCSSPFLVSAM | ASSRQRSQNTLY | 0.6320559 |
| VHITDAE | ASSRQRSQNTLY | 0.7076379 |
| EYAAPSF | ASSRQRSQNTLY | 0.6464863 |
| VVDAFHGGL | ASSRQRSQNTLY | 0.5438955 |
| ASVVDPQ | ASSRQRSQNTLY | 0.6901815 |
| SYNEFGGLS | ASSRQRSQNTLY | 0.7247451 |
| DPDGSVY | ASSRQRSQNTLY | 0.62383986 |
| MVSTIHATE | ASSRQRSQNTLY | 0.7320322 |
| LAAGVADPV | ASSRQRSQNTLY | 0.6124641 |
| TKEQIAA | ASSRQRSQNTLY | 0.7262131 |
| SLASPAAL | ASSRQRSQNTLY | 0.7127774 |
| IGLVFYGV | ASSRQRSQNTLY | 0.6843535 |
| FLLDASNL | ASSRQRSQNTLY | 0.65309244 |
| YQVGNFP | ASSRQRSQNTLY | 0.7218549 |
| ESLLSQST | ASSRQRSQNTLY | 0.66573775 |

| | | |
|---|---|---|
| LFTVGFAAVG | ASSRQRSQNTLY | 0.717044 |
| ATPPTPT | ASSRQRSQNTLY | 0.7235689 |
| TDLATLG | ASSRQRSQNTLY | 0.74181795 |
| SAGGTGR | ASSRQRSQNTLY | 0.89752316 |
| GFSNSGS | ASSRQRSQNTLY | 0.85976356 |
| GGNGGSGGTSV | ASSRQRSQNTLY | 0.8567751 |
| GTSVATGGAGN | ASSRQRSQNTLY | 0.7865739 |
| NGGNGGSGGTS | ASSRQRSQNTLY | 0.452473 |
| AVVKFPH | ASSRQRSQNTLY | 0.82545733 |
| TPPPPPP | ASSRQRSQNTLY | 0.6892871 |
| AAPAAAAAPA | ASSRQRSQNTLY | 0.87677693 |
| IALNEDL | ASSRQRSQNTLY | 0.89119476 |
| IVNVSSM | ASSRQRSQNTLY | 0.77729034 |
| TGIPPAPR | ASSRQRSQNTLY | 0.8874175 |
| FDLGGGTF | ASSRQRSQNTLY | 0.87330043 |
| FDLGGGTFDVS | ASSRQRSQNTLY | 0.58082813 |
| NEPTAAA | ASSRQRSQNTLY | 0.83340997 |
| NEPTAAALA | ASSRQRSQNTLY | 0.68761426 |
| GSIINISSI | ASSRQRSQNTLY | 0.8434763 |
| VAPGNAGTA | ASSRQRSQNTLY | 0.41043815 |
| TGGGASLE | ASSRQRSQNTLY | 0.8861516 |
| VAVYNHYKRI | ASSRQRSQNTLY | 0.78044105 |
| TDPAPATT | ASSRQRSQNTLY | 0.23341915 |
| TNRPDILDPALLRPG | ASSRQRSQNTLY | 0.8662721 |
| LTTTTTS | ASSRQRSQNTLY | 0.83388793 |
| PTGSGKTL | ASSRQRSQNTLY | 0.806658 |
| TGSGKTL | ASSRQRSQNTLY | 0.89379036 |
| TIGHVDHGK | ASSRQRSQNTLY | 0.69302607 |
| TIGHVDHGKTT | ASSRQRSQNTLY | 0.66427577 |
| GHVDHGKTT | ASSRQRSQNTLY | 0.8819146 |
| AQFEHTLL | ASSRQRSQNTLY | 0.8923962 |
| IVNQDEL | ASSRQRSQNTLY | 0.53514254 |
| TNGSQFF | ASSRQRSQNTLY | 0.87517524 |
| TGKGIVE | ASSRQRSQNTLY | 0.8892747 |
| LISNASDA | ASSRQRSQNTLY | 0.8546617 |
| AELRTLGELTKIAKAHGAQ | ASSRQRSQNTLY | 0.8952466 |
| APNAVAAAEPSW | ASSRQRSQNTLY | 0.89997107 |
| APQVAGAATAVL | ASSRQRSQNTLY | 0.8162067 |
| ARRDPAAAAAILPTDA | ASSRQRSQNTLY | 0.89342916 |
| DAAARDVLAEAGLAEYF | ASSRQRSQNTLY | 0.8939715 |
| DDQIVLSAKANLAKVEQ | ASSRQRSQNTLY | 0.81621504 |
| DELLPDTGEDSDEDGH | ASSRQRSQNTLY | 0.8395351 |
| ELPEAPANRATVPQD | ASSRQRSQNTLY | 0.69761825 |
| FGQSKIERVTGIG | ASSRQRSQNTLY | 0.8441204 |
| GAGGTGSTAGGAGGAGGAGGLY | ASSRQRSQNTLY | 0.5764724 |
| GHGGTGAAGGNGGTGGAGGSGI | ASSRQRSQNTLY | 0.8632208 |
| GNGGNGGAGGEGGPGGAGGAGA | ASSRQRSQNTLY | 0.8949754 |
| GSTAGNGGNGGNATNGTIAGQP | ASSRQRSQNTLY | 0.8130084 |
| GTGGDGGNGGNGGTGG | ASSRQRSQNTLY | 0.7322843 |
| GVRRPAGIPADAPV | ASSRQRSQNTLY | 0.7325299 |
| GYRQVTDAHLLTLARR | ASSRQRSQNTLY | 0.8692433 |

| | | |
|---|---|---|
| IAKMGQDGHDRG | ASSRQRSQNTLY | 0.7856759 |
| KAEPSIEIQYT | ASSRQRSQNTLY | 0.54840887 |
| KPVRIRMAADEDTVAVT | ASSRQRSQNTLY | 0.819947 |
| MPPRIAGMRLLVIKPEP | ASSRQRSQNTLY | 0.8785825 |
| MKFYAVALFRSRGVPPTR | ASSRQRSQNTLY | 0.8784549 |
| NAGGFGILGAVAHSP | ASSRQRSQNTLY | 0.62091285 |
| NEGREITLAMAEM | ASSRQRSQNTLY | 0.90226305 |
| NELVEWSHPLGHRWRWE | ASSRQRSQNTLY | 0.87705374 |
| NLGAGGTGGNGGNPG | ASSRQRSQNTLY | 0.89027154 |
| NTEWPKWETVMAADPVD | ASSRQRSQNTLY | 0.81455374 |
| PASAAPAETQP | ASSRQRSQNTLY | 0.73544407 |
| QQSSDGKNGVRAAEITGE | ASSRQRSQNTLY | 0.88921475 |
| RAPDAVTTSWWKE | ASSRQRSQNTLY | 0.8192902 |
| SGFGNVGAHTSGWFNQ | ASSRQRSQNTLY | 0.78902036 |
| SGVAGAVGQAASVGGLKV | ASSRQRSQNTLY | 0.88531685 |
| SNGGAGTGSAGGNGGT | ASSRQRSQNTLY | 0.7877165 |
| TGGAGGEGGAAG | ASSRQRSQNTLY | 0.74947417 |
| TGGDGGAAGAGGAPGHG | ASSRQRSQNTLY | 0.8053428 |
| TMPPLLAAIANAGRHGV | ASSRQRSQNTLY | 0.9007347 |
| TSSTAGNSGGA | ASSRQRSQNTLY | 0.7479808 |
| VPATVPVGILC | ASSRQRSQNTLY | 0.6420842 |
| VQRPDSGGVVLTGRIS | ASSRQRSQNTLY | 0.791167 |
| WTPNPVPATKGVPAGLVQAT | ASSRQRSQNTLY | 0.82529426 |
| YALKRALAKFSDRFDVVII | ASSRQRSQNTLY | 0.8902002 |
| YLIGSVIV | ASSRQRSQNTLY | 0.8730348 |
| YGQLVAVID | ASSRQRSQNTLY | 0.82857764 |
| VTVGGMTLPA | ASSRQRSQNTLY | 0.86665493 |
| VLALPAV | ASSRQRSQNTLY | 0.8734242 |
| VIPPTLNYET | ASSRQRSQNTLY | 0.85950434 |
| VGMGGMHPSSG | ASSRQRSQNTLY | 0.8780089 |
| VGLAGGAATA | ASSRQRSQNTLY | 0.7604186 |
| VGGAGGAGGSGP | ASSRQRSQNTLY | 0.8978846 |
| VFAAFMV | ASSRQRSQNTLY | 0.8785066 |
| TLYDKLR | ASSRQRSQNTLY | 0.89134353 |
| TLAVFTVL | ASSRQRSQNTLY | 0.89552397 |
| TGSVQPARV | ASSRQRSQNTLY | 0.86567366 |
| TGGGAGSGGT | ASSRQRSQNTLY | 0.86801493 |
| SHLETSL | ASSRQRSQNTLY | 0.49534547 |
| SFTTGMT | ASSRQRSQNTLY | 0.74674106 |
| QVEVAKRMGVRQPTVS | ASSRQRSQNTLY | 0.86501676 |
| QGSGGPALAM | ASSRQRSQNTLY | 0.8682227 |
| MLAPSNP | ASSRQRSQNTLY | 0.88597196 |
| LVNNAAI | ASSRQRSQNTLY | 0.86944413 |
| LTGEINTLA | ASSRQRSQNTLY | 0.80135787 |
| LSIFGPV | ASSRQRSQNTLY | 0.8919903 |
| LLALLANL | ASSRQRSQNTLY | 0.89012957 |
| LISNASDALDKLR | ASSRQRSQNTLY | 0.59748524 |
| LINEAVP | ASSRQRSQNTLY | 0.8155663 |
| LGYAFSGL | ASSRQRSQNTLY | 0.90511215 |
| LDFVAVAL | ASSRQRSQNTLY | 0.8626852 |
| LAQPGAE | ASSRQRSQNTLY | 0.82846045 |

| | | |
|---|---|---|
| ISNLGAVTGF | ASSRQRSQNTLY | 0.79452175 |
| HQEDPGRI | ASSRQRSQNTLY | 0.8947675 |
| GTPGEPTP | ASSRQRSQNTLY | 0.8387172 |
| GFFNSGS | ASSRQRSQNTLY | 0.90320355 |
| GAGGGGLTAGAN | ASSRQRSQNTLY | 0.29938945 |
| FVDALVNM | ASSRQRSQNTLY | 0.8356676 |
| FLTVAPEL | ASSRQRSQNTLY | 0.732952 |
| FETLTAL | ASSRQRSQNTLY | 0.3448069 |
| DVINPIEL | ASSRQRSQNTLY | 0.8830604 |
| APVSGAPV | ASSRQRSQNTLY | 0.8080256 |
| ALVGGLAW | ASSRQRSQNTLY | 0.8702167 |
| ALAAAAATAI | ASSRQRSQNTLY | 0.8249169 |
| AFGLGLLL | ASSRQRSQNTLY | 0.89082205 |
| AFFAAASL | ASSRQRSQNTLY | 0.8544894 |
| AAAALPAFSPP | ASSRQRSQNTLY | 0.83532333 |
| VVLAPAL | ASSRQRSQNTLY | 0.7402971 |
| TIGHVDHGKTTL | ASSRQRSQNTLY | 0.8726645 |
| LISNASDALDK | ASSRQRSQNTLY | 0.89854777 |
| LISNASDAL | ASSRQRSQNTLY | 0.89053357 |
| ISNASDALDK | ASSRQRSQNTLY | 0.4099629 |
| NQAETLVYQTE | ASSRQRSQNTLY | 0.8602922 |
| TGGTGGTAGAG | ASSRQRSQNTLY | 0.82811177 |
| TGSTAGGAGGA | ASSRQRSQNTLY | 0.26151943 |
| DPLAVPVAAGPV | ASSRQRSQNTLY | 0.44332647 |
| GGTGGTPGNGGN | ASSRQRSQNTLY | 0.8382013 |
| PAGLAGVDPATL | ASSRQRSQNTLY | 0.8328916 |
| GNNLVVEAPHAGASV | ASSRQRSQNTLY | 0.88663507 |
| KVTRIMSASTLEQPA | ASSRQRSQNTLY | 0.8754424 |
| LGFALPIALPRRTVV | ASSRQRSQNTLY | 0.8120756 |
| TRLGNLTARKLVVLV | ASSRQRSQNTLY | 0.8775065 |
| WSQQRRLIYFPSAGP | ASSRQRSQNTLY | 0.850729 |
| GPNGNPDPMAAAVDIR | ASSRQRSQNTLY | 0.0939306 |
| TQRCVSVADLRAAASR | ASSRQRSQNTLY | 0.79136217 |
| QFVLDVAEHPVEVLTETV | ASSRQRSQNTLY | 0.583735 |
| GGAGGTGGAGGNGGTQAGDG | ASSRQRSQNTLY | 0.8979578 |
| PLLATLRAIAVDVDAKPAQV | ASSRQRSQNTLY | 0.76953375 |
| APKFRTGGYMIDFWGVGYQVAK | ASSRQRSQNTLY | 0.77744293 |
| GAGGAGGAGGAAGGPGGTGGTG | ASSRQRSQNTLY | 0.83852166 |
| IGALGLLATDVAPAGSVAFGVA | ASSRQRSQNTLY | 0.8513537 |
| AGGAGGIGGNGGAGGAGGVLMG | ASSRQRSQNTLY | 0.8382439 |
| ADELIGLL | ASSRQRSQNTLY | 0.73794675 |
| ADRALKA | ASSRQRSQNTLY | 0.33191097 |
| AGAVGQVGGAGGA | ASSRQRSQNTLY | 0.8725122 |
| AGGGGGLGGAP | ASSRQRSQNTLY | 0.57037944 |
| AGGLLYGNGGAG | ASSRQRSQNTLY | 0.65664494 |
| AGLESIGL | ASSRQRSQNTLY | 0.6216904 |
| AGLKTGII | ASSRQRSQNTLY | 0.5016147 |
| ASLAALLL | ASSRQRSQNTLY | 0.4039228 |
| ASVINISSI | ASSRQRSQNTLY | 0.8890668 |
| ATPPGPA | ASSRQRSQNTLY | 0.81441814 |
| ATVASALV | ASSRQRSQNTLY | 0.82274675 |

| | | |
|---|---|---|
| ATVASLIL | ASSRQRSQNTLY | 0.90203065 |
| AVNGPVLIRH | ASSRQRSQNTLY | 0.8282317 |
| DGAAGAVGI | ASSRQRSQNTLY | 0.83323145 |
| DPLLHGI | ASSRQRSQNTLY | 0.72227216 |
| ELGAALGAL | ASSRQRSQNTLY | 0.7498697 |
| ELGRTQAV | ASSRQRSQNTLY | 0.72979844 |
| FVAGGYLS | ASSRQRSQNTLY | 0.72525716 |
| GASGGAGGAGGW | ASSRQRSQNTLY | 0.89742154 |
| GGAGAGGDGGAGGI | ASSRQRSQNTLY | 0.8482706 |
| GTGHNGSQPGMGGNG | ASSRQRSQNTLY | 0.6151686 |
| GYAAASASAS | ASSRQRSQNTLY | 0.862885 |
| HQVDVAAVQQG | ASSRQRSQNTLY | 0.9002098 |
| IGAAIAHT | ASSRQRSQNTLY | 0.90386677 |
| IGMIEVAQL | ASSRQRSQNTLY | 0.7774118 |
| IRLAPEHTSAMD | ASSRQRSQNTLY | 0.7287203 |
| IYNVSVL | ASSRQRSQNTLY | 0.8062383 |
| KQGFLGM | ASSRQRSQNTLY | 0.82684445 |
| LESVITQL | ASSRQRSQNTLY | 0.756454 |
| LGLGALVVV | ASSRQRSQNTLY | 0.59849894 |
| LIGNGGNGGSGGI | ASSRQRSQNTLY | 0.8912361 |
| LNAINAPTL | ASSRQRSQNTLY | 0.9012958 |
| LQLLNTGV | ASSRQRSQNTLY | 0.8377916 |
| LQTSIEF | ASSRQRSQNTLY | 0.37964547 |
| MSFAVSEL | ASSRQRSQNTLY | 0.8314639 |
| NTGALGL | ASSRQRSQNTLY | 0.76043004 |
| PPGAGGAGGAGTT | ASSRQRSQNTLY | 0.789016 |
| QLVGLGI | ASSRQRSQNTLY | 0.80312836 |
| RFLEQVHQL | ASSRQRSQNTLY | 0.61506724 |
| RGVGYVLR | ASSRQRSQNTLY | 0.82904947 |
| SATGVFAGVI | ASSRQRSQNTLY | 0.9009688 |
| SLPAITLP | ASSRQRSQNTLY | 0.8002451 |
| STALAFTV | ASSRQRSQNTLY | 0.8751044 |
| SVLTVGLGK | ASSRQRSQNTLY | 0.89656144 |
| TALAGAGSGI | ASSRQRSQNTLY | 0.888188 |
| TGLPFTGI | ASSRQRSQNTLY | 0.8978802 |
| VAEHVTV | ASSRQRSQNTLY | 0.8811216 |
| VDIVLNSL | ASSRQRSQNTLY | 0.84463096 |
| VLTGGGAVL | ASSRQRSQNTLY | 0.8580238 |
| VNVGLAYGI | ASSRQRSQNTLY | 0.88242054 |
| VSSAAVGAV | ASSRQRSQNTLY | 0.8364016 |
| VTSVYEAGV | ASSRQRSQNTLY | 0.8390839 |
| WVALDRGA | ASSRQRSQNTLY | 0.81271535 |
| YLVGSVVI | ASSRQRSQNTLY | 0.8819798 |
| GPPGTGKTLL | ASSRQRSQNTLY | 0.818753 |
| GVGMTKF | ASSRQRSQNTLY | 0.8946827 |
| IGIGGSDL | ASSRQRSQNTLY | 0.40788573 |
| IGTLLAEL | ASSRQRSQNTLY | 0.56148386 |
| IYTSGTT | ASSRQRSQNTLY | 0.8935554 |
| LSFPAGV | ASSRQRSQNTLY | 0.13629223 |
| LYTSGTT | ASSRQRSQNTLY | 0.4017409 |
| VGAFPFGF | ASSRQRSQNTLY | 0.86067384 |

| | | |
|---|---|---|
| VGAGTVEF | ASSRQRSQNTLY | 0.89518434 |
| VGKSTLLL | ASSRQRSQNTLY | 0.14987399 |
| AALAHAGGGIV | ASSRQRSQNTLY | 0.66829884 |
| AALVAVSLL | ASSRQRSQNTLY | 0.6554738 |
| AAVSRAVAAL | ASSRQRSQNTLY | 0.88374436 |
| AGGSSGIGGF | ASSRQRSQNTLY | 0.79804146 |
| AGLTVANANG | ASSRQRSQNTLY | 0.50467145 |
| AGLVGAVL | ASSRQRSQNTLY | 0.8879654 |
| ALSQAFTV | ASSRQRSQNTLY | 0.43361732 |
| ALTFSVSL | ASSRQRSQNTLY | 0.79131114 |
| ALTGAALVAL | ASSRQRSQNTLY | 0.8886173 |
| ASFSIGAGVL | ASSRQRSQNTLY | 0.89761126 |
| DGSVYKI | ASSRQRSQNTLY | 0.746402 |
| EQLAEIV | ASSRQRSQNTLY | 0.87963736 |
| FAVSELTVM | ASSRQRSQNTLY | 0.80499005 |
| GAAPLAAGL | ASSRQRSQNTLY | 0.9036177 |
| GADGGDGGSGGAGG | ASSRQRSQNTLY | 0.8066628 |
| GFGLDTGGAGG | ASSRQRSQNTLY | 0.64733386 |
| GLASGGLL | ASSRQRSQNTLY | 0.88919824 |
| GRIAGAAVLL | ASSRQRSQNTLY | 0.89252675 |
| GSAGSLNTSF | ASSRQRSQNTLY | 0.8376426 |
| GTGQIGI | ASSRQRSQNTLY | 0.8821668 |
| GTLIALIL | ASSRQRSQNTLY | 0.82608134 |
| IAATMGFVVG | ASSRQRSQNTLY | 0.8256844 |
| IFAQALL | ASSRQRSQNTLY | 0.887526 |
| IGAAVALGF | ASSRQRSQNTLY | 0.8897139 |
| IGLTPLAL | ASSRQRSQNTLY | 0.638492 |
| ILGSNTSTL | ASSRQRSQNTLY | 0.89734954 |
| IVFSLTVL | ASSRQRSQNTLY | 0.81812227 |
| LASKATGF | ASSRQRSQNTLY | 0.7135778 |
| LASVALIL | ASSRQRSQNTLY | 0.9027135 |
| LFAVGLTL | ASSRQRSQNTLY | 0.8572352 |
| LFNGGIGTYI | ASSRQRSQNTLY | 0.8881823 |
| LTLGAGII | ASSRQRSQNTLY | 0.8785598 |
| LVAAALGDI | ASSRQRSQNTLY | 0.876385 |
| NTAFVQLGI | ASSRQRSQNTLY | 0.57494295 |
| QLFSGAEL | ASSRQRSQNTLY | 0.85068035 |
| SAATFITL | ASSRQRSQNTLY | 0.84028554 |
| SNYGLANLGV | ASSRQRSQNTLY | 0.8833581 |
| SYGGAYAVM | ASSRQRSQNTLY | 0.8475591 |
| TTGVGGLTLG | ASSRQRSQNTLY | 0.7320936 |
| TTQTIYI | ASSRQRSQNTLY | 0.88652205 |
| VAAGAGLGAVI | ASSRQRSQNTLY | 0.89494514 |
| VAAVVTLL | ASSRQRSQNTLY | 0.8713071 |
| VGFLGGLI | ASSRQRSQNTLY | 0.87030494 |
| VGVIPILL | ASSRQRSQNTLY | 0.90142643 |
| VIFASSLI | ASSRQRSQNTLY | 0.77430993 |
| VKGGDGGAAAG | ASSRQRSQNTLY | 0.8137598 |
| VTKTFTNI | ASSRQRSQNTLY | 0.88648534 |
| VVAAAILL | ASSRQRSQNTLY | 0.89221525 |
| YAALDVEL | ASSRQRSQNTLY | 0.8491155 |

| | | |
|---|---|---|
| LASVALLL | ASSRQRSQNTLY | 0.8980937 |
| LDAVHRV | ASSRQRSQNTLY | 0.7696792 |
| QGLSSSIF | ASSRQRSQNTLY | 0.87668073 |
| RGTFANIRL | ASSRQRSQNTLY | 0.7841724 |
| RQLWWGHRI | ASSRQRSQNTLY | 0.8338357 |
| YGPPGTGKTLL | ASSRQRSQNTLY | 0.743567 |
| AALVGAPANVA | ASSRQRSQNTLY | 0.87369615 |
| AATFRRTE | ASSRQRSQNTLY | 0.88361144 |
| AGALGGGI | ASSRQRSQNTLY | 0.87250715 |
| AGGAGGAGGIGAG | ASSRQRSQNTLY | 0.28320175 |
| AGSGALGL | ASSRQRSQNTLY | 0.87798357 |
| AQIADQI | ASSRQRSQNTLY | 0.8705306 |
| ASAVAVTYL | ASSRQRSQNTLY | 0.9054242 |
| ASTVLGGTGG | ASSRQRSQNTLY | 0.835392 |
| AVGGEPAK | ASSRQRSQNTLY | 0.8721323 |
| CEQGTFGGTC | ASSRQRSQNTLY | 0.75049174 |
| DEGKAPAG | ASSRQRSQNTLY | 0.85371584 |
| DTVVFPAL | ASSRQRSQNTLY | 0.66031563 |
| FEGITMAG | ASSRQRSQNTLY | 0.7887305 |
| FPLVVQD | ASSRQRSQNTLY | 0.7942058 |
| GAFTTPQ | ASSRQRSQNTLY | 0.7606435 |
| GFGGFGTAGGVGGA | ASSRQRSQNTLY | 0.878928 |
| GFLAQFL | ASSRQRSQNTLY | 0.8895129 |
| GGAGGAGGNAGVA | ASSRQRSQNTLY | 0.887813 |
| GGVGTAGAGGAGGAGG | ASSRQRSQNTLY | 0.89033556 |
| GNGGGGDAGNAGSGGN | ASSRQRSQNTLY | 0.9004129 |
| GTGGAGGAGNDAGS | ASSRQRSQNTLY | 0.89738417 |
| GTTNTGWFNS | ASSRQRSQNTLY | 0.7809172 |
| IFMGPLFTL | ASSRQRSQNTLY | 0.8131498 |
| ILAHAIEAI | ASSRQRSQNTLY | 0.6025347 |
| IRSAIREAL | ASSRQRSQNTLY | 0.83634514 |
| ITFAGGG | ASSRQRSQNTLY | 0.79314435 |
| ITGMSGHQEV | ASSRQRSQNTLY | 0.8569914 |
| IVAGLACTL | ASSRQRSQNTLY | 0.13895306 |
| IVGGGLAGA | ASSRQRSQNTLY | 0.85251 |
| IVTGVLNI | ASSRQRSQNTLY | 0.90003884 |
| LAQQGCI | ASSRQRSQNTLY | 0.77627563 |
| LCLHVEN | ASSRQRSQNTLY | 0.858334 |
| LDAISDGS | ASSRQRSQNTLY | 0.8770679 |
| LGGFAAML | ASSRQRSQNTLY | 0.81701636 |
| LGGKNPMI | ASSRQRSQNTLY | 0.8732132 |
| LGPKGVAAIL | ASSRQRSQNTLY | 0.77391744 |
| LMVGVSI | ASSRQRSQNTLY | 0.885181 |
| LVGAAFVL | ASSRQRSQNTLY | 0.8986834 |
| LVLAVAPL | ASSRQRSQNTLY | 0.86263347 |
| MAAIAVIDAA | ASSRQRSQNTLY | 0.8564545 |
| NNGINAELQ | ASSRQRSQNTLY | 0.8493674 |
| RGEPLVLVHG | ASSRQRSQNTLY | 0.4101195 |
| RGGAGGAAGLI | ASSRQRSQNTLY | 0.86884165 |
| RSYRMHPAV | ASSRQRSQNTLY | 0.16660632 |
| SAAHLLAGQLA | ASSRQRSQNTLY | 0.845378 |

| | | |
|---|---|---|
| SAAVGSY | ASSRQRSQNTLY | 0.7895607 |
| SPHAVLHRI | ASSRQRSQNTLY | 0.8569863 |
| SQIAKVL | ASSRQRSQNTLY | 0.83954984 |
| TALGHLVP | ASSRQRSQNTLY | 0.8763386 |
| TAPTTSGAA | ASSRQRSQNTLY | 0.7339424 |
| TDGDDGNGGSPGT | ASSRQRSQNTLY | 0.8716763 |
| TDGLGEN | ASSRQRSQNTLY | 0.8236641 |
| TGGPPLPGQA | ASSRQRSQNTLY | 0.60112333 |
| TIPAITI | ASSRQRSQNTLY | 0.8907959 |
| TIPGFTI | ASSRQRSQNTLY | 0.84777355 |
| TSIASGALA | ASSRQRSQNTLY | 0.841578 |
| TVTLPGAAI | ASSRQRSQNTLY | 0.88733023 |
| VASGLSAALP | ASSRQRSQNTLY | 0.8568969 |
| VGAGVIGI | ASSRQRSQNTLY | 0.39279053 |
| VGPKRFGV | ASSRQRSQNTLY | 0.8122082 |
| VNVAFPD | ASSRQRSQNTLY | 0.85724026 |
| VSMTNSGVS | ASSRQRSQNTLY | 0.82676697 |
| VTDVLGDLSV | ASSRQRSQNTLY | 0.16243455 |
| VVVVGAGFAGL | ASSRQRSQNTLY | 0.8885705 |
| VYTVGTVV | ASSRQRSQNTLY | 0.7156099 |
| WTTFSGP | ASSRQRSQNTLY | 0.75671387 |
| YDELVEAF | ASSRQRSQNTLY | 0.8001947 |
| YGTGGAGGNGG | ASSRQRSQNTLY | 0.88702905 |
| YTNVMAV | ASSRQRSQNTLY | 0.82185864 |
| FLEQVHQL | ASSRQRSQNTLY | 0.7568793 |
| GLSLLAVAV | ASSRQRSQNTLY | 0.8803159 |
| LEQVHQL | ASSRQRSQNTLY | 0.8512678 |
| LNAAGAI | ASSRQRSQNTLY | 0.8916111 |
| PAVHASL | ASSRQRSQNTLY | 0.8490137 |
| TELEDDA | ASSRQRSQNTLY | 0.84168464 |
| TPPGPPN | ASSRQRSQNTLY | 0.85635555 |
| TPPPPPPP | ASSRQRSQNTLY | 0.7670499 |
| TQSPALI | ASSQGDSYAEQF | 0.7538718 |
| EFAAAFPQ | ASSQGDSYAEQF | 0.5955217 |
| LLSGNLDVL | ASSQGDSYAEQF | 0.8052926 |
| GPANLGNYNI | ASSQGDSYAEQF | 0.64419436 |
| TLGPTLP | ASSQGDSYAEQF | 0.8572054 |
| VYAYSSCCK | ASSQGDSYAEQF | 0.8963691 |
| APAPAPAPNI | ASSQGDSYAEQF | 0.8357476 |
| PGGPVPIP | ASSQGDSYAEQF | 0.65268314 |
| FLTVALEP | ASSQGDSYAEQF | 0.83537734 |
| SQYFGANL | ASSQGDSYAEQF | 0.7833901 |
| GGVKPAA | ASSQGDSYAEQF | 0.88393855 |
| GGTGGAGGAGGW | ASSQGDSYAEQF | 0.878995 |
| GVGGVAGLGGAGG | ASSQGDSYAEQF | 0.89805996 |
| VLLVLATTI | ASSQGDSYAEQF | 0.87943476 |
| IGIFAAML | ASSQGDSYAEQF | 0.7798196 |
| AVGDLID | ASSQGDSYAEQF | 0.8896687 |
| NISPLPL | ASSQGDSYAEQF | 0.8280686 |
| VTAVFVAI | ASSQGDSYAEQF | 0.8530719 |
| SGDDNDAAL | ASSQGDSYAEQF | 0.86804545 |

| | | |
|---|---|---|
| AVAALAHS | ASSQGDSYAEQF | 0.8741629 |
| GGVTVMATP | ASSQGDSYAEQF | 0.8191508 |
| AGLAANVRD | ASSQGDSYAEQF | 0.83280766 |
| TNRPDLIDPALLRPG | ASSQGDSYAEQF | 0.4945574 |
| LANLGVTAL | ASSQGDSYAEQF | 0.8799459 |
| TGERIEL | ASSQGDSYAEQF | 0.7786066 |
| NGEELAA | ASSQGDSYAEQF | 0.6793654 |
| VLSLVWG | ASSQGDSYAEQF | 0.82609713 |
| IGYALWL | ASSQGDSYAEQF | 0.8996022 |
| LSLLLAPL | ASSQGDSYAEQF | 0.8916916 |
| TLDEALK | ASSQGDSYAEQF | 0.7737733 |
| TDGAGTAEGG | ASSQGDSYAEQF | 0.76930547 |
| NAPGGAGGNGG | ASSQGDSYAEQF | 0.86155903 |
| VSVFMGMGGNF | ASSQGDSYAEQF | 0.6081898 |
| LASYLAAL | ASSQGDSYAEQF | 0.88718027 |
| LVGGTPAL | ASSQGDSYAEQF | 0.83304715 |
| VRNADVL | ASSQGDSYAEQF | 0.8401678 |
| TCSSPFLVSAM | ASSQGDSYAEQF | 0.7973396 |
| VHITDAE | ASSQGDSYAEQF | 0.8715519 |
| EYAAPSF | ASSQGDSYAEQF | 0.8139485 |
| VVDAFHGGL | ASSQGDSYAEQF | 0.88235307 |
| ASVVDPQ | ASSQGDSYAEQF | 0.16565853 |
| SYNEFGGLS | ASSQGDSYAEQF | 0.8842814 |
| DPDGSVY | ASSQGDSYAEQF | 0.7436948 |
| MVSTIHATE | ASSQGDSYAEQF | 0.7422067 |
| LAAGVADPV | ASSQGDSYAEQF | 0.34632987 |
| TKEQIAA | ASSQGDSYAEQF | 0.83510447 |
| SLASPAAL | ASSQGDSYAEQF | 0.8720137 |
| IGLVFYGV | ASSQGDSYAEQF | 0.8916739 |
| FLLDASNL | ASSQGDSYAEQF | 0.8356683 |
| YQVGNFP | ASSQGDSYAEQF | 0.87956345 |
| ESLLSQST | ASSQGDSYAEQF | 0.89530766 |
| LFTVGFAAVG | ASSQGDSYAEQF | 0.8168133 |
| ATPPTPT | ASSQGDSYAEQF | 0.35511717 |
| TDLATLG | ASSQGDSYAEQF | 0.8501184 |
| SAGGTGR | ASSQGDSYAEQF | 0.82006294 |
| GFSNSGS | ASSQGDSYAEQF | 0.8926758 |
| GGNGGSGGTSV | ASSQGDSYAEQF | 0.8666581 |
| GTSVATGGAGN | ASSQGDSYAEQF | 0.67127883 |
| NGGNGGSGGTS | ASSQGDSYAEQF | 0.8770479 |
| AVVKFPH | ASSQGDSYAEQF | 0.8873776 |
| TPPPPPP | ASSQGDSYAEQF | 0.74469805 |
| AAPAAAAAPA | ASSQGDSYAEQF | 0.7665137 |
| IALNEDL | ASSQGDSYAEQF | 0.75063765 |
| IVNVSSM | ASSQGDSYAEQF | 0.52958536 |
| TGIPPAPR | ASSQGDSYAEQF | 0.8706831 |
| FDLGGGTF | ASSQGDSYAEQF | 0.80889666 |
| FDLGGGTFDVS | ASSQGDSYAEQF | 0.6919447 |
| NEPTAAA | ASSQGDSYAEQF | 0.8533465 |
| NEPTAAALA | ASSQGDSYAEQF | 0.7424927 |
| GSIINISSI | ASSQGDSYAEQF | 0.67800105 |

| | | |
|---|---|---|
| VAPGNAGTA | ASSQGDSYAEQF | 0.7435423 |
| TGGGASLE | ASSQGDSYAEQF | 0.82757354 |
| VAVYNHYKRI | ASSQGDSYAEQF | 0.8829264 |
| TDPAPATT | ASSQGDSYAEQF | 0.8166299 |
| TNRPDILDPALLRPG | ASSQGDSYAEQF | 0.84720135 |
| LTTTTTS | ASSQGDSYAEQF | 0.8956989 |
| PTGSGKTL | ASSQGDSYAEQF | 0.51357067 |
| TGSGKTL | ASSQGDSYAEQF | 0.72180223 |
| TIGHVDHGK | ASSQGDSYAEQF | 0.8790661 |
| TIGHVDHGKTT | ASSQGDSYAEQF | 0.7118487 |
| GHVDHGKTT | ASSQGDSYAEQF | 0.8907202 |
| AQFEHTLL | ASSQGDSYAEQF | 0.85241604 |
| IVNQDEL | ASSQGDSYAEQF | 0.8673748 |
| TNGSQFF | ASSQGDSYAEQF | 0.8165882 |
| TGKGIVE | ASSQGDSYAEQF | 0.90428126 |
| LISNASDA | ASSQGDSYAEQF | 0.8817738 |
| AELRTLGELTKIAKAHGAQ | ASSQGDSYAEQF | 0.8485293 |
| APNAVAAAEPSW | ASSQGDSYAEQF | 0.87521744 |
| APQVAGAATAVL | ASSQGDSYAEQF | 0.6585797 |
| ARRDPAAAAAILPTDA | ASSQGDSYAEQF | 0.88883996 |
| DAAARDVLAEAGLAEYF | ASSQGDSYAEQF | 0.87713325 |
| DDQIVLSAKANLAKVEQ | ASSQGDSYAEQF | 0.70457506 |
| DELLPDTGEDSDEDGH | ASSQGDSYAEQF | 0.83369815 |
| ELPEAPANRATVPQD | ASSQGDSYAEQF | 0.8999665 |
| FGQSKIERVTGIG | ASSQGDSYAEQF | 0.89520586 |
| GAGGTGSTAGGAGGAGGAGGLY | ASSQGDSYAEQF | 0.68055403 |
| GHGGTGAAGGNGGTGGAGGSGI | ASSQGDSYAEQF | 0.79073536 |
| GNGGNGGAGGEGGPGGAGGAGA | ASSQGDSYAEQF | 0.57909375 |
| GSTAGNGGNGGNATNGTIAGQP | ASSQGDSYAEQF | 0.8050382 |
| GTGGDGGNGGNGGTGG | ASSQGDSYAEQF | 0.5056211 |
| GVRRPAGIPADAPV | ASSQGDSYAEQF | 0.8772832 |
| GYRQVTDAHLLTLARR | ASSQGDSYAEQF | 0.80533403 |
| IAKMGQDGHDRG | ASSQGDSYAEQF | 0.8696747 |
| KAEPSIEIQYT | ASSQGDSYAEQF | 0.85745233 |
| KPVRIRMAADEDTVAVT | ASSQGDSYAEQF | 0.89023334 |
| MPPRIAGMRLLVIKPEP | ASSQGDSYAEQF | 0.81520134 |
| MKFYAVALFRSRGVPPTR | ASSQGDSYAEQF | 0.79632723 |
| NAGGFGILGAVAHSP | ASSQGDSYAEQF | 0.64312696 |
| NEGREITLAMAEM | ASSQGDSYAEQF | 0.51499474 |
| NELVEWSHPLGHRWRWE | ASSQGDSYAEQF | 0.7827293 |
| NLGAGGTGGNGGNPG | ASSQGDSYAEQF | 0.7773242 |
| NTEWPKWETVMAADPVD | ASSQGDSYAEQF | 0.26523948 |
| PASAAPAETQP | ASSQGDSYAEQF | 0.8989625 |
| QQSSDGKNGVRAAEITGE | ASSQGDSYAEQF | 0.8513104 |
| RAPDAVTTSWWKE | ASSQGDSYAEQF | 0.87082654 |
| SGFGNVGAHTSGWFNQ | ASSQGDSYAEQF | 0.7641568 |
| SGVAGAVGQAASVGGLKV | ASSQGDSYAEQF | 0.75013506 |
| SNGGAGTGSAGGNGGT | ASSQGDSYAEQF | 0.7685426 |
| TGGAGGEGGAAG | ASSQGDSYAEQF | 0.8730998 |
| TGGDGGAAGAGGAPGHG | ASSQGDSYAEQF | 0.71064997 |
| TMPPLLAAIANAGRHGV | ASSQGDSYAEQF | 0.378712 |

| | | |
|---|---|---|
| TSSTAGNSGGA | ASSQGDSYAEQF | 0.6893654 |
| VPATVPVGILC | ASSQGDSYAEQF | 0.59052134 |
| VQRPDSGGVVLTGRIS | ASSQGDSYAEQF | 0.90350974 |
| WTPNPVPATKGVPAGLVQAT | ASSQGDSYAEQF | 0.8915634 |
| YALKRALAKFSDRFDVVII | ASSQGDSYAEQF | 0.8658377 |
| YLIGSVIV | ASSQGDSYAEQF | 0.9025363 |
| YGQLVAVID | ASSQGDSYAEQF | 0.22534484 |
| VTVGGMTLPA | ASSQGDSYAEQF | 0.7695712 |
| VLALPAV | ASSQGDSYAEQF | 0.8901541 |
| VIPPTLNYET | ASSQGDSYAEQF | 0.8732886 |
| VGMGGMHPSSG | ASSQGDSYAEQF | 0.83413804 |
| VGLAGGAATA | ASSQGDSYAEQF | 0.89374125 |
| VGGAGGAGGSGP | ASSQGDSYAEQF | 0.7189088 |
| VFAAFMV | ASSQGDSYAEQF | 0.82309425 |
| TLYDKLR | ASSQGDSYAEQF | 0.83553267 |
| TLAVFTVL | ASSQGDSYAEQF | 0.8921011 |
| TGSVQPARV | ASSQGDSYAEQF | 0.860203 |
| TGGGAGSGGT | ASSQGDSYAEQF | 0.65347475 |
| SHLETSL | ASSQGDSYAEQF | 0.86985826 |
| SFTTGMT | ASSQGDSYAEQF | 0.9068899 |
| QVEVAKRMGVRQPTVS | ASSQGDSYAEQF | 0.7957528 |
| QGSGGPALAM | ASSQGDSYAEQF | 0.87246317 |
| MLAPSNP | ASSQGDSYAEQF | 0.8653192 |
| LVNNAAI | ASSQGDSYAEQF | 0.82055265 |
| LTGEINTLA | ASSQGDSYAEQF | 0.7550604 |
| LSIFGPV | ASSQGDSYAEQF | 0.77143085 |
| LLALLANL | ASSQGDSYAEQF | 0.86786765 |
| LISNASDALDKLR | ASSQGDSYAEQF | 0.706668 |
| LINEAVP | ASSQGDSYAEQF | 0.83547604 |
| LGYAFSGL | ASSQGDSYAEQF | 0.81444776 |
| LDFVAVAL | ASSQGDSYAEQF | 0.90833914 |
| LAQPGAE | ASSQGDSYAEQF | 0.8754101 |
| ISNLGAVTGF | ASSQGDSYAEQF | 0.8803334 |
| HQEDPGRI | ASSQGDSYAEQF | 0.704806 |
| GTPGEPTP | ASSQGDSYAEQF | 0.87799305 |
| GFFNSGS | ASSQGDSYAEQF | 0.8756151 |
| GAGGGGLTAGAN | ASSQGDSYAEQF | 0.51980495 |
| FVDALVNM | ASSQGDSYAEQF | 0.89481896 |
| FLTVAPEL | ASSQGDSYAEQF | 0.6777755 |
| FETLTAL | ASSQGDSYAEQF | 0.8490024 |
| DVINPIEL | ASSQGDSYAEQF | 0.86686367 |
| APVSGAPV | ASSQGDSYAEQF | 0.88664925 |
| ALVGGLAW | ASSQGDSYAEQF | 0.8371505 |
| ALAAAAATAI | ASSQGDSYAEQF | 0.8537129 |
| AFGLGLLL | ASSQGDSYAEQF | 0.6469815 |
| AFFAAASL | ASSQGDSYAEQF | 0.8865082 |
| AAAALPAFSPP | ASSQGDSYAEQF | 0.8895315 |
| VVLAPAL | ASSQGDSYAEQF | 0.7999468 |
| TIGHVDHGKTTL | ASSQGDSYAEQF | 0.9043074 |
| LISNASDALDK | ASSQGDSYAEQF | 0.85642254 |
| LISNASDAL | ASSQGDSYAEQF | 0.41860712 |

| | | |
|---|---|---|
| ISNASDALDK | ASSQGDSYAEQF | 0.85147774 |
| NQAETLVYQTE | ASSQGDSYAEQF | 0.89457595 |
| TGGTGGTAGAG | ASSQGDSYAEQF | 0.8959898 |
| TGSTAGGAGGA | ASSQGDSYAEQF | 0.8856947 |
| DPLAVPVAAGPV | ASSQGDSYAEQF | 0.8716982 |
| GGTGGTPGNGGN | ASSQGDSYAEQF | 0.82787716 |
| PAGLAGVDPATL | ASSQGDSYAEQF | 0.8973143 |
| GNNLVVEAPHAGASV | ASSQGDSYAEQF | 0.9048503 |
| KVTRIMSASTLEQPA | ASSQGDSYAEQF | 0.880731 |
| LGFALPIALPRRTVV | ASSQGDSYAEQF | 0.7740953 |
| TRLGNLTARKLVVLV | ASSQGDSYAEQF | 0.7124361 |
| WSQQRRLIYFPSAGP | ASSQGDSYAEQF | 0.87027097 |
| GPNGNPDPMAAAVDIR | ASSQGDSYAEQF | 0.8913126 |
| TQRCVSVADLRAAASR | ASSQGDSYAEQF | 0.68899083 |
| QFVLDVAEHPVEVLTETV | ASSQGDSYAEQF | 0.8555247 |
| GGAGGTGGAGGNGGTQAGDG | ASSQGDSYAEQF | 0.8529959 |
| PLLATLRAIAVDVDAKPAQV | ASSQGDSYAEQF | 0.42023417 |
| APKFRTGGYMIDFWGVGYQVAK | ASSQGDSYAEQF | 0.86979514 |
| GAGGAGGAGGAAGGPGGTGGTG | ASSQGDSYAEQF | 0.897889 |
| IGALGLLATDVAPAGSVAFGVA | ASSQGDSYAEQF | 0.8784377 |
| AGGAGGIGGNGGAGGAGGVLMG | ASSQGDSYAEQF | 0.7583121 |
| ADELIGLL | ASSQGDSYAEQF | 0.89370906 |
| ADRALKA | ASSQGDSYAEQF | 0.9038812 |
| AGAVGQVGGAGGA | ASSQGDSYAEQF | 0.90388536 |
| AGGGGGLGGAP | ASSQGDSYAEQF | 0.37513697 |
| AGGLLYGNGGAG | ASSQGDSYAEQF | 0.28445223 |
| AGLESIGL | ASSQGDSYAEQF | 0.82817644 |
| AGLKTGII | ASSQGDSYAEQF | 0.8678338 |
| ASLAALLL | ASSQGDSYAEQF | 0.77621216 |
| ASVINISSI | ASSQGDSYAEQF | 0.7759305 |
| ATPPGPA | ASSQGDSYAEQF | 0.8672309 |
| ATVASALV | ASSQGDSYAEQF | 0.88954926 |
| ATVASLIL | ASSQGDSYAEQF | 0.74791706 |
| AVNGPVLIRH | ASSQGDSYAEQF | 0.60212517 |
| DGAAGAVGI | ASSQGDSYAEQF | 0.8797101 |
| DPLLHGI | ASSQGDSYAEQF | 0.88124335 |
| ELGAALGAL | ASSQGDSYAEQF | 0.7634766 |
| ELGRTQAV | ASSQGDSYAEQF | 0.8304843 |
| FVAGGYLS | ASSQGDSYAEQF | 0.821742 |
| GASGGAGGAGGW | ASSQGDSYAEQF | 0.82686967 |
| GGAGAGGDGGAGGI | ASSQGDSYAEQF | 0.86256456 |
| GTGHNGSQPGMGGNG | ASSQGDSYAEQF | 0.6697265 |
| GYAAASASAS | ASSQGDSYAEQF | 0.8979366 |
| HQVDVAAVQQG | ASSQGDSYAEQF | 0.6613659 |
| IGAAIAHT | ASSQGDSYAEQF | 0.88411593 |
| IGMIEVAQL | ASSQGDSYAEQF | 0.8592138 |
| IRLAPEHTSAMD | ASSQGDSYAEQF | 0.8769074 |
| IYNVSVL | ASSQGDSYAEQF | 0.81992847 |
| KQGFLGM | ASSQGDSYAEQF | 0.7336421 |
| LESVITQL | ASSQGDSYAEQF | 0.6997036 |
| LGLGALVVV | ASSQGDSYAEQF | 0.5914552 |

| | | |
|---|---|---|
| LIGNGGNGGSGGI | ASSQGDSYAEQF | 0.8916222 |
| LNAINAPTL | ASSQGDSYAEQF | 0.36740837 |
| LQLLNTGV | ASSQGDSYAEQF | 0.895807 |
| LQTSIEF | ASSQGDSYAEQF | 0.8499334 |
| MSFAVSEL | ASSQGDSYAEQF | 0.718788 |
| NTGALGL | ASSQGDSYAEQF | 0.87934333 |
| PPGAGGAGGAGTT | ASSQGDSYAEQF | 0.66110873 |
| QLVGLGI | ASSQGDSYAEQF | 0.84683996 |
| RFLEQVHQL | ASSQGDSYAEQF | 0.8669059 |
| RGVGYVLR | ASSQGDSYAEQF | 0.8867159 |
| SATGVFAGVI | ASSQGDSYAEQF | 0.76382697 |
| SLPAITLP | ASSQGDSYAEQF | 0.8851905 |
| STALAFTV | ASSQGDSYAEQF | 0.87130564 |
| SVLTVGLGK | ASSQGDSYAEQF | 0.72231734 |
| TALAGAGSGI | ASSQGDSYAEQF | 0.873322 |
| TGLPFTGI | ASSQGDSYAEQF | 0.8860388 |
| VAEHVTV | ASSQGDSYAEQF | 0.8731985 |
| VDIVLNSL | ASSQGDSYAEQF | 0.6328323 |
| VLTGGGAVL | ASSQGDSYAEQF | 0.8663403 |
| VNVGLAYGI | ASSQGDSYAEQF | 0.76809466 |
| VSSAAVGAV | ASSQGDSYAEQF | 0.86568356 |
| VTSVYEAGV | ASSQGDSYAEQF | 0.8615658 |
| WVALDRGA | ASSQGDSYAEQF | 0.78057027 |
| YLVGSVVI | ASSQGDSYAEQF | 0.8886615 |
| GPPGTGKTLL | ASSQGDSYAEQF | 0.41936734 |
| GVGMTKF | ASSQGDSYAEQF | 0.7851815 |
| IGIGGSDL | ASSQGDSYAEQF | 0.8295635 |
| IGTLLAEL | ASSQGDSYAEQF | 0.82080674 |
| IYTSGTT | ASSQGDSYAEQF | 0.88190985 |
| LSFPAGV | ASSQGDSYAEQF | 0.68822986 |
| LYTSGTT | ASSQGDSYAEQF | 0.8575386 |
| VGAFPFGF | ASSQGDSYAEQF | 0.8102563 |
| VGAGTVEF | ASSQGDSYAEQF | 0.70423776 |
| VGKSTLLL | ASSQGDSYAEQF | 0.8642324 |
| AALAHAGGGIV | ASSQGDSYAEQF | 0.817837 |
| AALVAVSLL | ASSQGDSYAEQF | 0.6813618 |
| AAVSRAVAAL | ASSQGDSYAEQF | 0.8881843 |
| AGGSSGIGGF | ASSQGDSYAEQF | 0.70003307 |
| AGLTVANANG | ASSQGDSYAEQF | 0.87313235 |
| AGLVGAVL | ASSQGDSYAEQF | 0.8976399 |
| ALSQAFTV | ASSQGDSYAEQF | 0.64852864 |
| ALTFSVSL | ASSQGDSYAEQF | 0.8840802 |
| ALTGAALVAL | ASSQGDSYAEQF | 0.8723257 |
| ASFSIGAGVL | ASSQGDSYAEQF | 0.8728778 |
| DGSVYKI | ASSQGDSYAEQF | 0.7820403 |
| EQLAEIV | ASSQGDSYAEQF | 0.77148247 |
| FAVSELTVM | ASSQGDSYAEQF | 0.89723206 |
| GAAPLAAGL | ASSQGDSYAEQF | 0.86376536 |
| GADGGDGGSGGAGG | ASSQGDSYAEQF | 0.8512914 |
| GFGLDTGGAGG | ASSQGDSYAEQF | 0.88610077 |
| GLASGGLL | ASSQGDSYAEQF | 0.87194437 |

| | | |
|---|---|---|
| GRIAGAAVLL | ASSQGDSYAEQF | 0.8874036 |
| GSAGSLNTSF | ASSQGDSYAEQF | 0.77007353 |
| GTGQIGI | ASSQGDSYAEQF | 0.6091354 |
| GTLIALIL | ASSQGDSYAEQF | 0.7910764 |
| IAATMGFVVG | ASSQGDSYAEQF | 0.83572894 |
| IFAQALL | ASSQGDSYAEQF | 0.8023757 |
| IGAAVALGF | ASSQGDSYAEQF | 0.45337075 |
| IGLTPLAL | ASSQGDSYAEQF | 0.880268 |
| ILGSNTSTL | ASSQGDSYAEQF | 0.90586424 |
| IVFSLTVL | ASSQGDSYAEQF | 0.7113332 |
| LASKATGF | ASSQGDSYAEQF | 0.9084581 |
| LASVALIL | ASSQGDSYAEQF | 0.67577946 |
| LFAVGLTL | ASSQGDSYAEQF | 0.874874 |
| LFNGGIGTYI | ASSQGDSYAEQF | 0.8841883 |
| LTLGAGII | ASSQGDSYAEQF | 0.6896067 |
| LVAAALGDI | ASSQGDSYAEQF | 0.8462899 |
| NTAFVQLGI | ASSQGDSYAEQF | 0.683515 |
| QLFSGAEL | ASSQGDSYAEQF | 0.797892 |
| SAATFITL | ASSQGDSYAEQF | 0.8921987 |
| SNYGLANLGV | ASSQGDSYAEQF | 0.72711587 |
| SYGGAYAVM | ASSQGDSYAEQF | 0.8763593 |
| TTGVGGLTLG | ASSQGDSYAEQF | 0.7987118 |
| TTQTIYI | ASSQGDSYAEQF | 0.70510197 |
| VAAGAGLGAVI | ASSQGDSYAEQF | 0.87904936 |
| VAAVVTLL | ASSQGDSYAEQF | 0.885183 |
| VGFLGGLI | ASSQGDSYAEQF | 0.83192277 |
| VGVIPILL | ASSQGDSYAEQF | 0.9031161 |
| VIFASSLI | ASSQGDSYAEQF | 0.78910124 |
| VKGGDGGAAAG | ASSQGDSYAEQF | 0.9002522 |
| VTKTFTNI | ASSQGDSYAEQF | 0.8947621 |
| VVAAAILL | ASSQGDSYAEQF | 0.8470636 |
| YAALDVEL | ASSQGDSYAEQF | 0.8245598 |
| LASVALLL | ASSQGDSYAEQF | 0.8797413 |
| LDAVHRV | ASSQGDSYAEQF | 0.79592705 |
| QGLSSSIF | ASSQGDSYAEQF | 0.8768331 |
| RGTFANIRL | ASSQGDSYAEQF | 0.90157115 |
| RQLWWGHRI | ASSQGDSYAEQF | 0.56034243 |
| YGPPGTGKTLL | ASSQGDSYAEQF | 0.8779303 |
| AALVGAPANVA | ASSQGDSYAEQF | 0.73995125 |
| AATFRRTE | ASSQGDSYAEQF | 0.7757464 |
| AGALGGGI | ASSQGDSYAEQF | 0.5212461 |
| AGGAGGAGGIGAG | ASSQGDSYAEQF | 0.26136795 |
| AGSGALGL | ASSQGDSYAEQF | 0.732375 |
| AQIADQI | ASSQGDSYAEQF | 0.49083605 |
| ASAVAVTYL | ASSQGDSYAEQF | 0.75038016 |
| ASTVLGGTGG | ASSQGDSYAEQF | 0.8104247 |
| AVGGEPAK | ASSQGDSYAEQF | 0.5365132 |
| CEQGTFGGTC | ASSQGDSYAEQF | 0.87265456 |
| DEGKAPAG | ASSQGDSYAEQF | 0.8632004 |
| DTVVFPAL | ASSQGDSYAEQF | 0.34932157 |
| FEGITMAG | ASSQGDSYAEQF | 0.6782321 |

| | | |
|---|---|---|
| FPLVVQD | ASSQGDSYAEQF | 0.53956974 |
| GAFTTPQ | ASSQGDSYAEQF | 0.7274046 |
| GFGGFGTAGGVGGA | ASSQGDSYAEQF | 0.2977073 |
| GFLAQFL | ASSQGDSYAEQF | 0.8282656 |
| GGAGGAGGNAGVA | ASSQGDSYAEQF | 0.53680396 |
| GGVGTAGAGGAGGAGG | ASSQGDSYAEQF | 0.11792149 |
| GNGGGGDAGNAGSGGN | ASSQGDSYAEQF | 0.7705585 |
| GTGGAGGAGNDAGS | ASSQGDSYAEQF | 0.7112551 |
| GTTNTGWFNS | ASSQGDSYAEQF | 0.6419416 |
| IFMGPLFTL | ASSQGDSYAEQF | 0.8670943 |
| ILAHAIEAI | ASSQGDSYAEQF | 0.2904302 |
| IRSAIREAL | ASSQGDSYAEQF | 0.5210999 |
| ITFAGGG | ASSQGDSYAEQF | 0.821754 |
| ITGMSGHQEV | ASSQGDSYAEQF | 0.87096816 |
| IVAGLACTL | ASSQGDSYAEQF | 0.47605467 |
| IVGGGLAGA | ASSQGDSYAEQF | 0.7726846 |
| IVTGVLNI | ASSQGDSYAEQF | 0.87054837 |
| LAQQGCI | ASSQGDSYAEQF | 0.76619637 |
| LCLHVEN | ASSQGDSYAEQF | 0.84985846 |
| LDAISDGS | ASSQGDSYAEQF | 0.85756576 |
| LGGFAAML | ASSQGDSYAEQF | 0.42910886 |
| LGGKNPMI | ASSQGDSYAEQF | 0.85321784 |
| LGPKGVAAIL | ASSQGDSYAEQF | 0.8670335 |
| LMVGVSI | ASSQGDSYAEQF | 0.6204214 |
| LVGAAFVL | ASSQGDSYAEQF | 0.7845632 |
| LVLAVAPL | ASSQGDSYAEQF | 0.48307392 |
| MAAIAVIDAA | ASSQGDSYAEQF | 0.6134949 |
| NNGINAELQ | ASSQGDSYAEQF | 0.17272408 |
| RGEPLVLVHG | ASSQGDSYAEQF | 0.80456656 |
| RGGAGGAAGLI | ASSQGDSYAEQF | 0.84681165 |
| RSYRMHPAV | ASSQGDSYAEQF | 0.7232298 |
| SAAHLLAGQLA | ASSQGDSYAEQF | 0.5440554 |
| SAAVGSY | ASSQGDSYAEQF | 0.343218 |
| SPHAVLHRI | ASSQGDSYAEQF | 0.69806635 |
| SQIAKVL | ASSQGDSYAEQF | 0.5892015 |
| TALGHLVP | ASSQGDSYAEQF | 0.24439216 |
| TAPTTSGAA | ASSQGDSYAEQF | 0.53628 |
| TDGDDGNGGSPGT | ASSQGDSYAEQF | 0.84613717 |
| TDGLGEN | ASSQGDSYAEQF | 0.8433596 |
| TGGPPLPGQA | ASSQGDSYAEQF | 0.3923288 |
| TIPAITI | ASSQGDSYAEQF | 0.8819853 |
| TIPGFTI | ASSQGDSYAEQF | 0.76960766 |
| TSIASGALA | ASSQGDSYAEQF | 0.82706 |
| TVTLPGAAI | ASSQGDSYAEQF | 0.71083117 |
| VASGLSAALP | ASSQGDSYAEQF | 0.58457255 |
| VGAGVIGI | ASSQGDSYAEQF | 0.8250505 |
| VGPKRFGV | ASSQGDSYAEQF | 0.6830523 |
| VNVAFPD | ASSQGDSYAEQF | 0.5246538 |
| VSMTNSGVS | ASSQGDSYAEQF | 0.82004225 |
| VTDVLGDLSV | ASSQGDSYAEQF | 0.6430616 |
| VVVVGAGFAGL | ASSQGDSYAEQF | 0.62933433 |

| | | |
|---|---|---|
| VYTVGTVV | ASSQGDSYAEQF | 0.6605563 |
| WTTFSGP | ASSQGDSYAEQF | 0.8389043 |
| YDELVEAF | ASSQGDSYAEQF | 0.47209704 |
| YGTGGAGGNGG | ASSQGDSYAEQF | 0.4037249 |
| YTNVMAV | ASSQGDSYAEQF | 0.43499452 |
| FLEQVHQL | ASSQGDSYAEQF | 0.7622341 |
| GLSLLAVAV | ASSQGDSYAEQF | 0.86377656 |
| LEQVHQL | ASSQGDSYAEQF | 0.7670306 |
| LNAAGAI | ASSQGDSYAEQF | 0.6825851 |
| PAVHASL | ASSQGDSYAEQF | 0.8195684 |
| TELEDDA | ASSQGDSYAEQF | 0.8060087 |
| TPPGPPN | ASSQGDSYAEQF | 0.76497835 |
| TPPPPPPP | ASSQGDSYAEQF | 0.8034396 |
| TQSPALI | ASRMRGFYAEQF | 0.67571473 |
| EFAAAFPQ | ASRMRGFYAEQF | 0.8469058 |
| LLSGNLDVL | ASRMRGFYAEQF | 0.83494145 |
| GPANLGNYNI | ASRMRGFYAEQF | 0.8107401 |
| TLGPTLP | ASRMRGFYAEQF | 0.8358675 |
| VYAYSSCCK | ASRMRGFYAEQF | 0.6971934 |
| APAPAPAPNI | ASRMRGFYAEQF | 0.7084176 |
| PGGPVPIP | ASRMRGFYAEQF | 0.19853927 |
| FLTVALEP | ASRMRGFYAEQF | 0.48942518 |
| SQYFGANL | ASRMRGFYAEQF | 0.7932985 |
| GGVKPAA | ASRMRGFYAEQF | 0.7952464 |
| GGTGGAGGAGGW | ASRMRGFYAEQF | 0.8013933 |
| GVGGVAGLGGAGG | ASRMRGFYAEQF | 0.77482545 |
| VLLVLATTI | ASRMRGFYAEQF | 0.5958129 |
| IGIFAAML | ASRMRGFYAEQF | 0.8280057 |
| AVGDLID | ASRMRGFYAEQF | 0.85196203 |
| NISPLPL | ASRMRGFYAEQF | 0.35381442 |
| VTAVFVAI | ASRMRGFYAEQF | 0.62610894 |
| SGDDNDAAL | ASRMRGFYAEQF | 0.8922869 |
| AVAALAHS | ASRMRGFYAEQF | 0.78457916 |
| GGVTVMATP | ASRMRGFYAEQF | 0.63956857 |
| AGLAANVRD | ASRMRGFYAEQF | 0.62196267 |
| TNRPDLIDPALLRPG | ASRMRGFYAEQF | 0.8642708 |
| LANLGVTAL | ASRMRGFYAEQF | 0.5243938 |
| TGERIEL | ASRMRGFYAEQF | 0.877156 |
| NGEELAA | ASRMRGFYAEQF | 0.16575728 |
| VLSLVWG | ASRMRGFYAEQF | 0.59040797 |
| IGYALWL | ASRMRGFYAEQF | 0.46150026 |
| LSLLLAPL | ASRMRGFYAEQF | 0.24531375 |
| TLDEALK | ASRMRGFYAEQF | 0.8413906 |
| TDGAGTAEGG | ASRMRGFYAEQF | 0.63692105 |
| NAPGGAGGNGG | ASRMRGFYAEQF | 0.8192584 |
| VSVFMGMGGNF | ASRMRGFYAEQF | 0.7149417 |
| LASYLAAL | ASRMRGFYAEQF | 0.84587675 |
| LVGGTPAL | ASRMRGFYAEQF | 0.78886056 |
| VRNADVL | ASRMRGFYAEQF | 0.69531584 |
| TCSSPFLVSAM | ASRMRGFYAEQF | 0.5091255 |
| VHITDAE | ASRMRGFYAEQF | 0.80225104 |

| | | |
|---|---|---|
| EYAAPSF | ASRMRGFYAEQF | 0.851328 |
| VVDAFHGGL | ASRMRGFYAEQF | 0.8052949 |
| ASVVDPQ | ASRMRGFYAEQF | 0.30182013 |
| SYNEFGGLS | ASRMRGFYAEQF | 0.6851331 |
| DPDGSVY | ASRMRGFYAEQF | 0.70325756 |
| MVSTIHATE | ASRMRGFYAEQF | 0.20437944 |
| LAAGVADPV | ASRMRGFYAEQF | 0.15104167 |
| TKEQIAA | ASRMRGFYAEQF | 0.48618233 |
| SLASPAAL | ASRMRGFYAEQF | 0.60078955 |
| IGLVFYGV | ASRMRGFYAEQF | 0.8560506 |
| FLLDASNL | ASRMRGFYAEQF | 0.8458626 |
| YQVGNFP | ASRMRGFYAEQF | 0.7011998 |
| ESLLSQST | ASRMRGFYAEQF | 0.83356893 |
| LFTVGFAAVG | ASRMRGFYAEQF | 0.7830459 |
| ATPPTPT | ASRMRGFYAEQF | 0.08144349 |
| TDLATLG | ASRMRGFYAEQF | 0.55945694 |
| SAGGTGR | ASRMRGFYAEQF | 0.39035583 |
| GFSNSGS | ASRMRGFYAEQF | 0.83885187 |
| GGNGGSGGTSV | ASRMRGFYAEQF | 0.4724859 |
| GTSVATGGAGN | ASRMRGFYAEQF | 0.66554093 |
| NGGNGGSGGTS | ASRMRGFYAEQF | 0.7741895 |
| AVVKFPH | ASRMRGFYAEQF | 0.7615557 |
| TPPPPPP | ASRMRGFYAEQF | 0.7148342 |
| AAPAAAAAPA | ASRMRGFYAEQF | 0.46989828 |
| IALNEDL | ASRMRGFYAEQF | 0.10932052 |
| IVNVSSM | ASRMRGFYAEQF | 0.8236215 |
| TGIPPAPR | ASRMRGFYAEQF | 0.30069044 |
| FDLGGGTF | ASRMRGFYAEQF | 0.46491298 |
| FDLGGGTFDVS | ASRMRGFYAEQF | 0.2656678 |
| NEPTAAA | ASRMRGFYAEQF | 0.14797592 |
| NEPTAAALA | ASRMRGFYAEQF | 0.26374772 |
| GSIINISSI | ASRMRGFYAEQF | 0.8464436 |
| VAPGNAGTA | ASRMRGFYAEQF | 0.6154568 |
| TGGGASLE | ASRMRGFYAEQF | 0.7226818 |
| VAVYNHYKRI | ASRMRGFYAEQF | 0.8840921 |
| TDPAPATT | ASRMRGFYAEQF | 0.7487936 |
| TNRPDILDPALLRPG | ASRMRGFYAEQF | 0.71666706 |
| LTTTTTS | ASRMRGFYAEQF | 0.54946953 |
| PTGSGKTL | ASRMRGFYAEQF | 0.46682715 |
| TGSGKTL | ASRMRGFYAEQF | 0.52474976 |
| TIGHVDHGK | ASRMRGFYAEQF | 0.54406327 |
| TIGHVDHGKTT | ASRMRGFYAEQF | 0.8848618 |
| GHVDHGKTT | ASRMRGFYAEQF | 0.6089885 |
| AQFEHTLL | ASRMRGFYAEQF | 0.4494804 |
| IVNQDEL | ASRMRGFYAEQF | 0.7204708 |
| TNGSQFF | ASRMRGFYAEQF | 0.8671248 |
| TGKGIVE | ASRMRGFYAEQF | 0.87999237 |
| LISNASDA | ASRMRGFYAEQF | 0.50097764 |
| AELRTLGELTKIAKAHGAQ | ASRMRGFYAEQF | 0.6030332 |
| APNAVAAAEPSW | ASRMRGFYAEQF | 0.6676649 |
| APQVAGAATAVL | ASRMRGFYAEQF | 0.7597933 |

| | | |
|---|---|---|
| ARRDPAAAAAILPTDA | ASRMRGFYAEQF | 0.590654 |
| DAAARDVLAEAGLAEYF | ASRMRGFYAEQF | 0.44458586 |
| DDQIVLSAKANLAKVEQ | ASRMRGFYAEQF | 0.8244586 |
| DELLPDTGEDSDEDGH | ASRMRGFYAEQF | 0.8768959 |
| ELPEAPANRATVPQD | ASRMRGFYAEQF | 0.6587337 |
| FGQSKIERVTGIG | ASRMRGFYAEQF | 0.30695483 |
| GAGGTGSTAGGAGGAGGAGGLY | ASRMRGFYAEQF | 0.7098955 |
| GHGGTGAAGGNGGTGGAGGSGI | ASRMRGFYAEQF | 0.48730758 |
| GNGGNGGAGGEGGPGGAGGAGA | ASRMRGFYAEQF | 0.65271795 |
| GSTAGNGGNGGNATNGTIAGQP | ASRMRGFYAEQF | 0.71836436 |
| GTGGDGGNGGNGGTGG | ASRMRGFYAEQF | 0.41432625 |
| GVRRPAGIPADAPV | ASRMRGFYAEQF | 0.678572 |
| GYRQVTDAHLLTLARR | ASRMRGFYAEQF | 0.8829597 |
| IAKMGQDGHDRG | ASRMRGFYAEQF | 0.5760486 |
| KAEPSIEIQYT | ASRMRGFYAEQF | 0.80428433 |
| KPVRIRMAADEDTVAVT | ASRMRGFYAEQF | 0.87280595 |
| MPPRIAGMRLLVIKPEP | ASRMRGFYAEQF | 0.8295942 |
| MKFYAVALFRSRGVPPTR | ASRMRGFYAEQF | 0.88126504 |
| NAGGFGILGAVAHSP | ASRMRGFYAEQF | 0.7969604 |
| NEGREITLAMAEM | ASRMRGFYAEQF | 0.648077 |
| NELVEWSHPLGHRWRWE | ASRMRGFYAEQF | 0.7014264 |
| NLGAGGTGGNGGNPG | ASRMRGFYAEQF | 0.7477473 |
| NTEWPKWETVMAADPVD | ASRMRGFYAEQF | 0.7384612 |
| PASAAPAETQP | ASRMRGFYAEQF | 0.7091607 |
| QQSSDGKNGVRAAEITGE | ASRMRGFYAEQF | 0.59046274 |
| RAPDAVTTSWWKE | ASRMRGFYAEQF | 0.8600837 |
| SGFGNVGAHTSGWFNQ | ASRMRGFYAEQF | 0.7172433 |
| SGVAGAVGQAASVGGLKV | ASRMRGFYAEQF | 0.8014748 |
| SNGGAGTGSAGGNGGT | ASRMRGFYAEQF | 0.33462536 |
| TGGAGGEGGAAG | ASRMRGFYAEQF | 0.47621608 |
| TGGDGGAAGAGGAPGHG | ASRMRGFYAEQF | 0.84512484 |
| TMPPLLAAIANAGRHGV | ASRMRGFYAEQF | 0.11070275 |
| TSSTAGNSGGA | ASRMRGFYAEQF | 0.17434356 |
| VPATVPVGILC | ASRMRGFYAEQF | 0.7766291 |
| VQRPDSGGVVLTGRIS | ASRMRGFYAEQF | 0.8621695 |
| WTPNPVPATKGVPAGLVQAT | ASRMRGFYAEQF | 0.12446746 |
| YALKRALAKFSDRFDVVII | ASRMRGFYAEQF | 0.543103 |
| YLIGSVIV | ASRMRGFYAEQF | 0.3394743 |
| YGQLVAVID | ASRMRGFYAEQF | 0.7819308 |
| VTVGGMTLPA | ASRMRGFYAEQF | 0.74459374 |
| VLALPAV | ASRMRGFYAEQF | 0.24924432 |
| VIPPTLNYET | ASRMRGFYAEQF | 0.8144058 |
| VGMGGMHPSSG | ASRMRGFYAEQF | 0.3671985 |
| VGLAGGAATA | ASRMRGFYAEQF | 0.35476196 |
| VGGAGGAGGSGP | ASRMRGFYAEQF | 0.8339405 |
| VFAAFMV | ASRMRGFYAEQF | 0.8493792 |
| TLYDKLR | ASRMRGFYAEQF | 0.4565981 |
| TLAVFTVL | ASRMRGFYAEQF | 0.8370207 |
| TGSVQPARV | ASRMRGFYAEQF | 0.58763903 |
| TGGGAGSGGT | ASRMRGFYAEQF | 0.88096374 |
| SHLETSL | ASRMRGFYAEQF | 0.53866315 |

| | | |
|---|---|---|
| SFTTGMT | ASRMRGFYAEQF | 0.29323846 |
| QVEVAKRMGVRQPTVS | ASRMRGFYAEQF | 0.84884846 |
| QGSGGPALAM | ASRMRGFYAEQF | 0.79620343 |
| MLAPSNP | ASRMRGFYAEQF | 0.6475648 |
| LVNNAAI | ASRMRGFYAEQF | 0.8100552 |
| LTGEINTLA | ASRMRGFYAEQF | 0.5159769 |
| LSIFGPV | ASRMRGFYAEQF | 0.70287937 |
| LLALLANL | ASRMRGFYAEQF | 0.86742616 |
| LISNASDALDKLR | ASRMRGFYAEQF | 0.8473641 |
| LINEAVP | ASRMRGFYAEQF | 0.43053225 |
| LGYAFSGL | ASRMRGFYAEQF | 0.841688 |
| LDFVAVAL | ASRMRGFYAEQF | 0.7665595 |
| LAQPGAE | ASRMRGFYAEQF | 0.47318786 |
| ISNLGAVTGF | ASRMRGFYAEQF | 0.8764511 |
| HQEDPGRI | ASRMRGFYAEQF | 0.691743 |
| GTPGEPTP | ASRMRGFYAEQF | 0.8483592 |
| GFFNSGS | ASRMRGFYAEQF | 0.7792769 |
| GAGGGGLTAGAN | ASRMRGFYAEQF | 0.8280138 |
| FVDALVNM | ASRMRGFYAEQF | 0.29591656 |
| FLTVAPEL | ASRMRGFYAEQF | 0.7578341 |
| FETLTAL | ASRMRGFYAEQF | 0.71187323 |
| DVINPIEL | ASRMRGFYAEQF | 0.8230996 |
| APVSGAPV | ASRMRGFYAEQF | 0.7069432 |
| ALVGGLAW | ASRMRGFYAEQF | 0.31050575 |
| ALAAAAATAI | ASRMRGFYAEQF | 0.74567235 |
| AFGLGLLL | ASRMRGFYAEQF | 0.8773256 |
| AFFAAASL | ASRMRGFYAEQF | 0.7060287 |
| AAAALPAFSPP | ASRMRGFYAEQF | 0.78043514 |
| VVLAPAL | ASRMRGFYAEQF | 0.8902822 |
| TIGHVDHGKTTL | ASRMRGFYAEQF | 0.49516976 |
| LISNASDALDK | ASRMRGFYAEQF | 0.751299 |
| LISNASDAL | ASRMRGFYAEQF | 0.8200301 |
| ISNASDALDK | ASRMRGFYAEQF | 0.8538673 |
| NQAETLVYQTE | ASRMRGFYAEQF | 0.67229855 |
| TGGTGGTAGAG | ASRMRGFYAEQF | 0.8697864 |
| TGSTAGGAGGA | ASRMRGFYAEQF | 0.42559436 |
| DPLAVPVAAGPV | ASRMRGFYAEQF | 0.8368099 |
| GGTGGTPGNGGN | ASRMRGFYAEQF | 0.509337 |
| PAGLAGVDPATL | ASRMRGFYAEQF | 0.7698371 |
| GNNLVVEAPHAGASV | ASRMRGFYAEQF | 0.44378594 |
| KVTRIMSASTLEQPA | ASRMRGFYAEQF | 0.83299816 |
| LGFALPIALPRRTVV | ASRMRGFYAEQF | 0.85877025 |
| TRLGNLTARKLVVLV | ASRMRGFYAEQF | 0.8471212 |
| WSQQRRLIYFPSAGP | ASRMRGFYAEQF | 0.18961737 |
| GPNGNPDPMAAAVDIR | ASRMRGFYAEQF | 0.86475706 |
| TQRCVSVADLRAAASR | ASRMRGFYAEQF | 0.8345872 |
| QFVLDVAEHPVEVLTETV | ASRMRGFYAEQF | 0.8911985 |
| GGAGGTGGAGGNGGTQAGDG | ASRMRGFYAEQF | 0.6553863 |
| PLLATLRAIAVDVDAKPAQV | ASRMRGFYAEQF | 0.82251215 |
| APKFRTGGYMIDFWGVGYQVAK | ASRMRGFYAEQF | 0.6234923 |
| GAGGAGGAGGAAGGPGGTGGTG | ASRMRGFYAEQF | 0.6709181 |

| | | |
|---|---|---|
| IGALGLLATDVAPAGSVAFGVA | ASRMRGFYAEQF | 0.4663608 |
| AGGAGGIGGNGGAGGAGGVLMG | ASRMRGFYAEQF | 0.57471734 |
| ADELIGLL | ASRMRGFYAEQF | 0.7210915 |
| ADRALKA | ASRMRGFYAEQF | 0.61168545 |
| AGAVGQVGGAGGA | ASRMRGFYAEQF | 0.832829 |
| AGGGGGLGGAP | ASRMRGFYAEQF | 0.8518745 |
| AGGLLYGNGGAG | ASRMRGFYAEQF | 0.8412947 |
| AGLESIGL | ASRMRGFYAEQF | 0.86286664 |
| AGLKTGII | ASRMRGFYAEQF | 0.864012 |
| ASLAALLL | ASRMRGFYAEQF | 0.80198765 |
| ASVINISSI | ASRMRGFYAEQF | 0.6268028 |
| ATPPGPA | ASRMRGFYAEQF | 0.6025143 |
| ATVASALV | ASRMRGFYAEQF | 0.4669258 |
| ATVASLIL | ASRMRGFYAEQF | 0.68102837 |
| AVNGPVLIRH | ASRMRGFYAEQF | 0.5998935 |
| DGAAGAVGI | ASRMRGFYAEQF | 0.8131585 |
| DPLLHGI | ASRMRGFYAEQF | 0.09398549 |
| ELGAALGAL | ASRMRGFYAEQF | 0.73601425 |
| ELGRTQAV | ASRMRGFYAEQF | 0.8849697 |
| FVAGGYLS | ASRMRGFYAEQF | 0.4561725 |
| GASGGAGGAGGW | ASRMRGFYAEQF | 0.7685319 |
| GGAGAGGDGGAGGI | ASRMRGFYAEQF | 0.72655034 |
| GTGHNGSQPGMGGNG | ASRMRGFYAEQF | 0.58454907 |
| GYAAASASAS | ASRMRGFYAEQF | 0.8416903 |
| HQVDVAAVQQG | ASRMRGFYAEQF | 0.52554816 |
| IGAAIAHT | ASRMRGFYAEQF | 0.8144952 |
| IGMIEVAQL | ASRMRGFYAEQF | 0.8726219 |
| IRLAPEHTSAMD | ASRMRGFYAEQF | 0.77205986 |
| IYNVSVL | ASRMRGFYAEQF | 0.71004045 |
| KQGFLGM | ASRMRGFYAEQF | 0.70518935 |
| LESVITQL | ASRMRGFYAEQF | 0.25786853 |
| LGLGALVVV | ASRMRGFYAEQF | 0.74825716 |
| LIGNGGNGGSGGI | ASRMRGFYAEQF | 0.0910287 |
| LNAINAPTL | ASRMRGFYAEQF | 0.7379813 |
| LQLLNTGV | ASRMRGFYAEQF | 0.4224802 |
| LQTSIEF | ASRMRGFYAEQF | 0.78149116 |
| MSFAVSEL | ASRMRGFYAEQF | 0.7940626 |
| NTGALGL | ASRMRGFYAEQF | 0.7936255 |
| PPGAGGAGGAGTT | ASRMRGFYAEQF | 0.41422856 |
| QLVGLGI | ASRMRGFYAEQF | 0.6758299 |
| RFLEQVHQL | ASRMRGFYAEQF | 0.59415054 |
| RGVGYVLR | ASRMRGFYAEQF | 0.37568232 |
| SATGVFAGVI | ASRMRGFYAEQF | 0.8395396 |
| SLPAITLP | ASRMRGFYAEQF | 0.6739373 |
| STALAFTV | ASRMRGFYAEQF | 0.68866026 |
| SVLTVGLGK | ASRMRGFYAEQF | 0.85031736 |
| TALAGAGSGI | ASRMRGFYAEQF | 0.75265515 |
| TGLPFTGI | ASRMRGFYAEQF | 0.29248387 |
| VAEHVTV | ASRMRGFYAEQF | 0.6897899 |
| VDIVLNSL | ASRMRGFYAEQF | 0.71291375 |
| VLTGGGAVL | ASRMRGFYAEQF | 0.5277359 |

| | | |
|---|---|---|
| VNVGLAYGI | ASRMRGFYAEQF | 0.102255605 |
| VSSAAVGAV | ASRMRGFYAEQF | 0.7676203 |
| VTSVYEAGV | ASRMRGFYAEQF | 0.4803772 |
| WVALDRGA | ASRMRGFYAEQF | 0.5453037 |
| YLVGSVVI | ASRMRGFYAEQF | 0.65120447 |
| GPPGTGKTLL | ASRMRGFYAEQF | 0.85966724 |
| GVGMTKF | ASRMRGFYAEQF | 0.26630425 |
| IGIGGSDL | ASRMRGFYAEQF | 0.565251 |
| IGTLLAEL | ASRMRGFYAEQF | 0.8267267 |
| IYTSGTT | ASRMRGFYAEQF | 0.7994691 |
| LSFPAGV | ASRMRGFYAEQF | 0.874109 |
| LYTSGTT | ASRMRGFYAEQF | 0.72814983 |
| VGAFPFGF | ASRMRGFYAEQF | 0.60692704 |
| VGAGTVEF | ASRMRGFYAEQF | 0.7661572 |
| VGKSTLLL | ASRMRGFYAEQF | 0.7203078 |
| AALAHAGGGIV | ASRMRGFYAEQF | 0.23725453 |
| AALVAVSLL | ASRMRGFYAEQF | 0.21577656 |
| AAVSRAVAAL | ASRMRGFYAEQF | 0.62881774 |
| AGGSSGIGGF | ASRMRGFYAEQF | 0.20560588 |
| AGLTVANANG | ASRMRGFYAEQF | 0.68706065 |
| AGLVGAVL | ASRMRGFYAEQF | 0.85417455 |
| ALSQAFTV | ASRMRGFYAEQF | 0.53008544 |
| ALTFSVSL | ASRMRGFYAEQF | 0.3708231 |
| ALTGAALVAL | ASRMRGFYAEQF | 0.6796913 |
| ASFSIGAGVL | ASRMRGFYAEQF | 0.4647909 |
| DGSVYKI | ASRMRGFYAEQF | 0.8248482 |
| EQLAEIV | ASRMRGFYAEQF | 0.6375637 |
| FAVSELTVM | ASRMRGFYAEQF | 0.8648279 |
| GAAPLAAGL | ASRMRGFYAEQF | 0.8512032 |
| GADGGDGGSGGAGG | ASRMRGFYAEQF | 0.7189069 |
| GFGLDTGGAGG | ASRMRGFYAEQF | 0.8474058 |
| GLASGGLL | ASRMRGFYAEQF | 0.61772203 |
| GRIAGAAVLL | ASRMRGFYAEQF | 0.7028425 |
| GSAGSLNTSF | ASRMRGFYAEQF | 0.70852256 |
| GTGQIGI | ASRMRGFYAEQF | 0.7657802 |
| GTLIALIL | ASRMRGFYAEQF | 0.6192833 |
| IAATMGFVVG | ASRMRGFYAEQF | 0.56565154 |
| IFAQALL | ASRMRGFYAEQF | 0.2634724 |
| IGAAVALGF | ASRMRGFYAEQF | 0.7542677 |
| IGLTPLAL | ASRMRGFYAEQF | 0.52527374 |
| ILGSNTSTL | ASRMRGFYAEQF | 0.25931293 |
| IVFSLTVL | ASRMRGFYAEQF | 0.57741964 |
| LASKATGF | ASRMRGFYAEQF | 0.8861595 |
| LASVALIL | ASRMRGFYAEQF | 0.8808348 |
| LFAVGLTL | ASRMRGFYAEQF | 0.58609474 |
| LFNGGIGTYI | ASRMRGFYAEQF | 0.48551446 |
| LTLGAGII | ASRMRGFYAEQF | 0.75367785 |
| LVAAALGDI | ASRMRGFYAEQF | 0.2942839 |
| NTAFVQLGI | ASRMRGFYAEQF | 0.7534535 |
| QLFSGAEL | ASRMRGFYAEQF | 0.7630098 |
| SAATFITL | ASRMRGFYAEQF | 0.77529716 |

| | | |
|---|---|---|
| SNYGLANLGV | ASRMRGFYAEQF | 0.49728286 |
| SYGGAYAVM | ASRMRGFYAEQF | 0.7655987 |
| TTGVGGLTLG | ASRMRGFYAEQF | 0.65244526 |
| TTQTIYI | ASRMRGFYAEQF | 0.7701789 |
| VAAGAGLGAVI | ASRMRGFYAEQF | 0.112491734 |
| VAAVVTLL | ASRMRGFYAEQF | 0.75780714 |
| VGFLGGLI | ASRMRGFYAEQF | 0.5931883 |
| VGVIPILL | ASRMRGFYAEQF | 0.5234068 |
| VIFASSLI | ASRMRGFYAEQF | 0.28621536 |
| VKGGDGGAAAG | ASRMRGFYAEQF | 0.7391541 |
| VTKTFTNI | ASRMRGFYAEQF | 0.81768215 |
| VVAAAILL | ASRMRGFYAEQF | 0.8547935 |
| YAALDVEL | ASRMRGFYAEQF | 0.3564986 |
| LASVALLL | ASRMRGFYAEQF | 0.84585714 |
| LDAVHRV | ASRMRGFYAEQF | 0.8495662 |
| QGLSSSIF | ASRMRGFYAEQF | 0.69598335 |
| RGTFANIRL | ASRMRGFYAEQF | 0.2594956 |
| RQLWWGHRI | ASRMRGFYAEQF | 0.8122407 |
| YGPPGTGKTLL | ASRMRGFYAEQF | 0.65949154 |
| AALVGAPANVA | ASRMRGFYAEQF | 0.816352 |
| AATFRRTE | ASRMRGFYAEQF | 0.76650566 |
| AGALGGGI | ASRMRGFYAEQF | 0.446786 |
| AGGAGGAGGIGAG | ASRMRGFYAEQF | 0.81145763 |
| AGSGALGL | ASRMRGFYAEQF | 0.8581003 |
| AQIADQI | ASRMRGFYAEQF | 0.48362067 |
| ASAVAVTYL | ASRMRGFYAEQF | 0.59522605 |
| ASTVLGGTGG | ASRMRGFYAEQF | 0.57767975 |
| AVGGEPAK | ASRMRGFYAEQF | 0.2200549 |
| CEQGTFGGTC | ASRMRGFYAEQF | 0.8065051 |
| DEGKAPAG | ASRMRGFYAEQF | 0.670315 |
| DTVVFPAL | ASRMRGFYAEQF | 0.61608434 |
| FEGITMAG | ASRMRGFYAEQF | 0.7798313 |
| FPLVVQD | ASRMRGFYAEQF | 0.58194435 |
| GAFTTPQ | ASRMRGFYAEQF | 0.62769777 |
| GFGGFGTAGGVGGA | ASRMRGFYAEQF | 0.6631706 |
| GFLAQFL | ASRMRGFYAEQF | 0.6038374 |
| GGAGGAGGNAGVA | ASRMRGFYAEQF | 0.80056417 |
| GGVGTAGAGGAGGAGG | ASRMRGFYAEQF | 0.7174906 |
| GNGGGGDAGNAGSGGN | ASRMRGFYAEQF | 0.7679038 |
| GTGGAGGAGNDAGS | ASRMRGFYAEQF | 0.88793886 |
| GTTNTGWFNS | ASRMRGFYAEQF | 0.33045137 |
| IFMGPLFTL | ASRMRGFYAEQF | 0.52471817 |
| ILAHAIEAI | ASRMRGFYAEQF | 0.80608416 |
| IRSAIREAL | ASRMRGFYAEQF | 0.33006042 |
| ITFAGGG | ASRMRGFYAEQF | 0.8434299 |
| ITGMSGHQEV | ASRMRGFYAEQF | 0.7855464 |
| IVAGLACTL | ASRMRGFYAEQF | 0.81969965 |
| IVGGGLAGA | ASRMRGFYAEQF | 0.61724496 |
| IVTGVLNI | ASRMRGFYAEQF | 0.8949454 |
| LAQQGCI | ASRMRGFYAEQF | 0.8478657 |
| LCLHVEN | ASRMRGFYAEQF | 0.70126724 |

| | | |
|---|---|---|
| LDAISDGS | ASRMRGFYAEQF | 0.77348524 |
| LGGFAAML | ASRMRGFYAEQF | 0.457387 |
| LGGKNPMI | ASRMRGFYAEQF | 0.8280448 |
| LGPKGVAAIL | ASRMRGFYAEQF | 0.8022413 |
| LMVGVSI | ASRMRGFYAEQF | 0.45874587 |
| LVGAAFVL | ASRMRGFYAEQF | 0.7059182 |
| LVLAVAPL | ASRMRGFYAEQF | 0.85597837 |
| MAAIAVIDAA | ASRMRGFYAEQF | 0.874234 |
| NNGINAELQ | ASRMRGFYAEQF | 0.24986355 |
| RGEPLVLVHG | ASRMRGFYAEQF | 0.70891607 |
| RGGAGGAAGLI | ASRMRGFYAEQF | 0.29154083 |
| RSYRMHPAV | ASRMRGFYAEQF | 0.52930176 |
| SAAHLLAGQLA | ASRMRGFYAEQF | 0.3950129 |
| SAAVGSY | ASRMRGFYAEQF | 0.77853346 |
| SPHAVLHRI | ASRMRGFYAEQF | 0.6351413 |
| SQIAKVL | ASRMRGFYAEQF | 0.77710867 |
| TALGHLVP | ASRMRGFYAEQF | 0.714453 |
| TAPTTSGAA | ASRMRGFYAEQF | 0.86051226 |
| TDGDDGNGGSPGT | ASRMRGFYAEQF | 0.53257924 |
| TDGLGEN | ASRMRGFYAEQF | 0.68539375 |
| TGGPPLPGQA | ASRMRGFYAEQF | 0.31018895 |
| TIPAITI | ASRMRGFYAEQF | 0.21891424 |
| TIPGFTI | ASRMRGFYAEQF | 0.5048381 |
| TSIASGALA | ASRMRGFYAEQF | 0.66605115 |
| TVTLPGAAI | ASRMRGFYAEQF | 0.16650395 |
| VASGLSAALP | ASRMRGFYAEQF | 0.87718534 |
| VGAGVIGI | ASRMRGFYAEQF | 0.77224475 |
| VGPKRFGV | ASRMRGFYAEQF | 0.81867677 |
| VNVAFPD | ASRMRGFYAEQF | 0.60435003 |
| VSMTNSGVS | ASRMRGFYAEQF | 0.65300107 |
| VTDVLGDLSV | ASRMRGFYAEQF | 0.6087173 |
| VVVVGAGFAGL | ASRMRGFYAEQF | 0.81601965 |
| VYTVGTVV | ASRMRGFYAEQF | 0.40372193 |
| WTTFSGP | ASRMRGFYAEQF | 0.29823816 |
| YDELVEAF | ASRMRGFYAEQF | 0.6254275 |
| YGTGGAGGNGG | ASRMRGFYAEQF | 0.4918828 |
| YTNVMAV | ASRMRGFYAEQF | 0.85652673 |
| FLEQVHQL | ASRMRGFYAEQF | 0.85779905 |
| GLSLLAVAV | ASRMRGFYAEQF | 0.79017746 |
| LEQVHQL | ASRMRGFYAEQF | 0.82631814 |
| LNAAGAI | ASRMRGFYAEQF | 0.18321031 |
| PAVHASL | ASRMRGFYAEQF | 0.62369764 |
| TELEDDA | ASRMRGFYAEQF | 0.8831342 |
| TPPGPPN | ASRMRGFYAEQF | 0.73192024 |
| TPPPPPPP | ASRMRGFYAEQF | 0.6668583 |
| TQSPALI | GARTGNSDYT | 0.8743954 |
| EFAAAFPQ | GARTGNSDYT | 0.6041063 |
| LLSGNLDVL | GARTGNSDYT | 0.5137488 |
| GPANLGNYNI | GARTGNSDYT | 0.74231696 |
| TLGPTLP | GARTGNSDYT | 0.7845916 |
| VYAYSSCCK | GARTGNSDYT | 0.746945 |

| | | |
|---|---|---|
| APAPAPAPNI | GARTGNSDYT | 0.413396 |
| PGGPVPIP | GARTGNSDYT | 0.7969898 |
| FLTVALEP | GARTGNSDYT | 0.9020169 |
| SQYFGANL | GARTGNSDYT | 0.69409 |
| GGVKPAA | GARTGNSDYT | 0.8376628 |
| GGTGGAGGAGGW | GARTGNSDYT | 0.81742054 |
| GVGGVAGLGGAGG | GARTGNSDYT | 0.5550834 |
| VLLVLATTI | GARTGNSDYT | 0.50743157 |
| IGIFAAML | GARTGNSDYT | 0.5602823 |
| AVGDLID | GARTGNSDYT | 0.667362 |
| NISPLPL | GARTGNSDYT | 0.5455202 |
| VTAVFVAI | GARTGNSDYT | 0.70633703 |
| SGDDNDAAL | GARTGNSDYT | 0.70267665 |
| AVAALAHS | GARTGNSDYT | 0.8976673 |
| GGVTVMATP | GARTGNSDYT | 0.8209971 |
| AGLAANVRD | GARTGNSDYT | 0.82360095 |
| TNRPDLIDPALLRPG | GARTGNSDYT | 0.539277 |
| LANLGVTAL | GARTGNSDYT | 0.728982 |
| TGERIEL | GARTGNSDYT | 0.79694235 |
| NGEELAA | GARTGNSDYT | 0.29912227 |
| VLSLVWG | GARTGNSDYT | 0.88144195 |
| IGYALWL | GARTGNSDYT | 0.57250863 |
| LSLLLAPL | GARTGNSDYT | 0.6526736 |
| TLDEALK | GARTGNSDYT | 0.7917701 |
| TDGAGTAEGG | GARTGNSDYT | 0.8354245 |
| NAPGGAGGNGG | GARTGNSDYT | 0.68786186 |
| VSVFMGMGGNF | GARTGNSDYT | 0.69831765 |
| LASYLAAL | GARTGNSDYT | 0.5595602 |
| LVGGTPAL | GARTGNSDYT | 0.800256 |
| VRNADVL | GARTGNSDYT | 0.8667088 |
| TCSSPFLVSAM | GARTGNSDYT | 0.41656092 |
| VHITDAE | GARTGNSDYT | 0.8477373 |
| EYAAPSF | GARTGNSDYT | 0.7579211 |
| VVDAFHGGL | GARTGNSDYT | 0.3135588 |
| ASVVDPQ | GARTGNSDYT | 0.6477458 |
| SYNEFGGLS | GARTGNSDYT | 0.8311237 |
| DPDGSVY | GARTGNSDYT | 0.8509697 |
| MVSTIHATE | GARTGNSDYT | 0.81646365 |
| LAAGVADPV | GARTGNSDYT | 0.7978394 |
| TKEQIAA | GARTGNSDYT | 0.5271086 |
| SLASPAAL | GARTGNSDYT | 0.8789356 |
| IGLVFYGV | GARTGNSDYT | 0.8832399 |
| FLLDASNL | GARTGNSDYT | 0.840901 |
| YQVGNFP | GARTGNSDYT | 0.52904934 |
| ESLLSQST | GARTGNSDYT | 0.42720795 |
| LFTVGFAAVG | GARTGNSDYT | 0.7486932 |
| ATPPTPT | GARTGNSDYT | 0.84340656 |
| TDLATLG | GARTGNSDYT | 0.4436823 |
| SAGGTGR | GARTGNSDYT | 0.75559807 |
| GFSNSGS | GARTGNSDYT | 0.783724 |
| GGNGGSGGTSV | GARTGNSDYT | 0.26557103 |

| | | |
|---|---|---|
| GTSVATGGAGN | GARTGNSDYT | 0.76943433 |
| NGGNGGSGGTS | GARTGNSDYT | 0.87646186 |
| AVVKFPH | GARTGNSDYT | 0.71387804 |
| TPPPPPP | GARTGNSDYT | 0.5679823 |
| AAPAAAAAPA | GARTGNSDYT | 0.8749163 |
| IALNEDL | GARTGNSDYT | 0.89222264 |
| IVNVSSM | GARTGNSDYT | 0.85761607 |
| TGIPPAPR | GARTGNSDYT | 0.36268902 |
| FDLGGGTF | GARTGNSDYT | 0.22059333 |
| FDLGGGTFDVS | GARTGNSDYT | 0.63467586 |
| NEPTAAA | GARTGNSDYT | 0.7951269 |
| NEPTAAALA | GARTGNSDYT | 0.65195054 |
| GSIINISSI | GARTGNSDYT | 0.50404024 |
| VAPGNAGTA | GARTGNSDYT | 0.78570056 |
| TGGGASLE | GARTGNSDYT | 0.8737018 |
| VAVYNHYKRI | GARTGNSDYT | 0.44918975 |
| TDPAPATT | GARTGNSDYT | 0.34990343 |
| TNRPDILDPALLRPG | GARTGNSDYT | 0.8286183 |
| LTTTTTS | GARTGNSDYT | 0.8317478 |
| PTGSGKTL | GARTGNSDYT | 0.6090143 |
| TGSGKTL | GARTGNSDYT | 0.6805959 |
| TIGHVDHGK | GARTGNSDYT | 0.57799435 |
| TIGHVDHGKTT | GARTGNSDYT | 0.516073 |
| GHVDHGKTT | GARTGNSDYT | 0.7914293 |
| AQFEHTLL | GARTGNSDYT | 0.40882963 |
| IVNQDEL | GARTGNSDYT | 0.87374914 |
| TNGSQFF | GARTGNSDYT | 0.5122153 |
| TGKGIVE | GARTGNSDYT | 0.826062 |
| LISNASDA | GARTGNSDYT | 0.79179025 |
| AELRTLGELTKIAKAHGAQ | GARTGNSDYT | 0.77006793 |
| APNAVAAAEPSW | GARTGNSDYT | 0.65592194 |
| APQVAGAATAVL | GARTGNSDYT | 0.5142524 |
| ARRDPAAAAAILPTDA | GARTGNSDYT | 0.36037436 |
| DAAARDVLAEAGLAEYF | GARTGNSDYT | 0.22161138 |
| DDQIVLSAKANLAKVEQ | GARTGNSDYT | 0.86283815 |
| DELLPDTGEDSDEDGH | GARTGNSDYT | 0.10139225 |
| ELPEAPANRATVPQD | GARTGNSDYT | 0.85154194 |
| FGQSKIERVTGIG | GARTGNSDYT | 0.7952293 |
| GAGGTGSTAGGAGGAGGAGGLY | GARTGNSDYT | 0.5086259 |
| GHGGTGAAGGNGGTGGAGGSGI | GARTGNSDYT | 0.7977046 |
| GNGGNGGAGGEGGPGGAGGAGA | GARTGNSDYT | 0.3607705 |
| GSTAGNGGNGGNATNGTIAGQP | GARTGNSDYT | 0.39496696 |
| GTGGDGGNGGNGGTGG | GARTGNSDYT | 0.6901107 |
| GVRRPAGIPADAPV | GARTGNSDYT | 0.80614746 |
| GYRQVTDAHLLTLARR | GARTGNSDYT | 0.5181394 |
| IAKMGQDGHDRG | GARTGNSDYT | 0.7630336 |
| KAEPSIEIQYT | GARTGNSDYT | 0.7727562 |
| KPVRIRMAADEDTVAVT | GARTGNSDYT | 0.48949662 |
| MPPRIAGMRLLVIKPEP | GARTGNSDYT | 0.70618796 |
| MKFYAVALFRSRGVPPTR | GARTGNSDYT | 0.86277175 |
| NAGGFGILGAVAHSP | GARTGNSDYT | 0.76685274 |

| | | |
|---|---|---|
| NEGREITLAMAEM | GARTGNSDYT | 0.4100992 |
| NELVEWSHPLGHRWRWE | GARTGNSDYT | 0.8269963 |
| NLGAGGTGGNGGNPG | GARTGNSDYT | 0.4906562 |
| NTEWPKWETVMAADPVD | GARTGNSDYT | 0.81427366 |
| PASAAPAETQP | GARTGNSDYT | 0.76916134 |
| QQSSDGKNGVRAAEITGE | GARTGNSDYT | 0.64669335 |
| RAPDAVTTSWWKE | GARTGNSDYT | 0.8191426 |
| SGFGNVGAHTSGWFNQ | GARTGNSDYT | 0.38866076 |
| SGVAGAVGQAASVGGLKV | GARTGNSDYT | 0.56771016 |
| SNGGAGTGSAGGNGGT | GARTGNSDYT | 0.61797047 |
| TGGAGGEGGAAG | GARTGNSDYT | 0.7658059 |
| TGGDGGAAGAGGAPGHG | GARTGNSDYT | 0.85420036 |
| TMPPLLAAIANAGRHGV | GARTGNSDYT | 0.5293962 |
| TSSTAGNSGGA | GARTGNSDYT | 0.7383826 |
| VPATVPVGILC | GARTGNSDYT | 0.5212097 |
| VQRPDSGGVVLTGRIS | GARTGNSDYT | 0.3942521 |
| WTPNPVPATKGVPAGLVQAT | GARTGNSDYT | 0.7078718 |
| YALKRALAKFSDRFDVVII | GARTGNSDYT | 0.6455154 |
| YLIGSVIV | GARTGNSDYT | 0.4721946 |
| YGQLVAVID | GARTGNSDYT | 0.7940383 |
| VTVGGMTLPA | GARTGNSDYT | 0.4719894 |
| VLALPAV | GARTGNSDYT | 0.7270036 |
| VIPPTLNYET | GARTGNSDYT | 0.8941754 |
| VGMGGMHPSSG | GARTGNSDYT | 0.5691764 |
| VGLAGGAATA | GARTGNSDYT | 0.81755334 |
| VGGAGGAGGSGP | GARTGNSDYT | 0.77092946 |
| VFAAFMV | GARTGNSDYT | 0.80868423 |
| TLYDKLR | GARTGNSDYT | 0.65631294 |
| TLAVFTVL | GARTGNSDYT | 0.49807143 |
| TGSVQPARV | GARTGNSDYT | 0.86949 |
| TGGGAGSGGT | GARTGNSDYT | 0.8366811 |
| SHLETSL | GARTGNSDYT | 0.8275229 |
| SFTTGMT | GARTGNSDYT | 0.7994069 |
| QVEVAKRMGVRQPTVS | GARTGNSDYT | 0.8118843 |
| QGSGGPALAM | GARTGNSDYT | 0.844462 |
| MLAPSNP | GARTGNSDYT | 0.6834721 |
| LVNNAAI | GARTGNSDYT | 0.3174375 |
| LTGEINTLA | GARTGNSDYT | 0.69261515 |
| LSIFGPV | GARTGNSDYT | 0.6530408 |
| LLALLANL | GARTGNSDYT | 0.6623657 |
| LISNASDALDKLR | GARTGNSDYT | 0.31301466 |
| LINEAVP | GARTGNSDYT | 0.85721487 |
| LGYAFSGL | GARTGNSDYT | 0.8980212 |
| LDFVAVAL | GARTGNSDYT | 0.5231607 |
| LAQPGAE | GARTGNSDYT | 0.8713455 |
| ISNLGAVTGF | GARTGNSDYT | 0.40059942 |
| HQEDPGRI | GARTGNSDYT | 0.80228007 |
| GTPGEPTP | GARTGNSDYT | 0.87070596 |
| GFFNSGS | GARTGNSDYT | 0.4909958 |
| GAGGGGLTAGAN | GARTGNSDYT | 0.7772524 |
| FVDALVNM | GARTGNSDYT | 0.4491973 |

| | | |
|---|---|---|
| FLTVAPEL | GARTGNSDYT | 0.60932255 |
| FETLTAL | GARTGNSDYT | 0.8717225 |
| DVINPIEL | GARTGNSDYT | 0.61709034 |
| APVSGAPV | GARTGNSDYT | 0.7446236 |
| ALVGGLAW | GARTGNSDYT | 0.6080897 |
| ALAAAAATAI | GARTGNSDYT | 0.49617755 |
| AFGLGLLL | GARTGNSDYT | 0.75429857 |
| AFFAAASL | GARTGNSDYT | 0.8498199 |
| AAAALPAFSPP | GARTGNSDYT | 0.39115202 |
| VVLAPAL | GARTGNSDYT | 0.8831023 |
| TIGHVDHGKTTL | GARTGNSDYT | 0.5563844 |
| LISNASDALDK | GARTGNSDYT | 0.89133203 |
| LISNASDAL | GARTGNSDYT | 0.8533355 |
| ISNASDALDK | GARTGNSDYT | 0.7731547 |
| NQAETLVYQTE | GARTGNSDYT | 0.65597004 |
| TGGTGGTAGAG | GARTGNSDYT | 0.86339724 |
| TGSTAGGAGGA | GARTGNSDYT | 0.609877 |
| DPLAVPVAAGPV | GARTGNSDYT | 0.81611943 |
| GGTGGTPGNGGN | GARTGNSDYT | 0.84295106 |
| PAGLAGVDPATL | GARTGNSDYT | 0.42766863 |
| GNNLVVEAPHAGASV | GARTGNSDYT | 0.6860126 |
| KVTRIMSASTLEQPA | GARTGNSDYT | 0.6368017 |
| LGFALPIALPRRTVV | GARTGNSDYT | 0.6343783 |
| TRLGNLTARKLVVLV | GARTGNSDYT | 0.53029543 |
| WSQQRRLIYFPSAGP | GARTGNSDYT | 0.4170215 |
| GPNGNPDPMAAAVDIR | GARTGNSDYT | 0.55403435 |
| TQRCVSVADLRAAASR | GARTGNSDYT | 0.49112928 |
| QFVLDVAEHPVEVLTETV | GARTGNSDYT | 0.6881331 |
| GGAGGTGGAGGNGGTQAGDG | GARTGNSDYT | 0.66752714 |
| PLLATLRAIAVDVDAKPAQV | GARTGNSDYT | 0.5119082 |
| APKFRTGGYMIDFWGVGYQVAK | GARTGNSDYT | 0.6718724 |
| GAGGAGGAGGAAGGPGGTGGTG | GARTGNSDYT | 0.6676046 |
| IGALGLLATDVAPAGSVAFGVA | GARTGNSDYT | 0.40911546 |
| AGGAGGIGGNGGAGGAGGVLMG | GARTGNSDYT | 0.5998534 |
| ADELIGLL | GARTGNSDYT | 0.568648 |
| ADRALKA | GARTGNSDYT | 0.58598113 |
| AGAVGQVGGAGGA | GARTGNSDYT | 0.40779623 |
| AGGGGGLGGAP | GARTGNSDYT | 0.6544728 |
| AGGLLYGNGGAG | GARTGNSDYT | 0.4998657 |
| AGLESIGL | GARTGNSDYT | 0.30295816 |
| AGLKTGII | GARTGNSDYT | 0.65102094 |
| ASLAALLL | GARTGNSDYT | 0.55377156 |
| ASVINISSI | GARTGNSDYT | 0.5712545 |
| ATPPGPA | GARTGNSDYT | 0.6547754 |
| ATVASALV | GARTGNSDYT | 0.5494399 |
| ATVASLIL | GARTGNSDYT | 0.530615 |
| AVNGPVLIRH | GARTGNSDYT | 0.65868545 |
| DGAAGAVGI | GARTGNSDYT | 0.6721051 |
| DPLLHGI | GARTGNSDYT | 0.46205503 |
| ELGAALGAL | GARTGNSDYT | 0.62374544 |
| ELGRTQAV | GARTGNSDYT | 0.69535124 |

| | | |
|---|---|---|
| FVAGGYLS | GARTGNSDYT | 0.58533156 |
| GASGGAGGAGGW | GARTGNSDYT | 0.6806773 |
| GGAGAGGDGGAGGI | GARTGNSDYT | 0.685941 |
| GTGHNGSQPGMGGNG | GARTGNSDYT | 0.5734787 |
| GYAAASASAS | GARTGNSDYT | 0.6796609 |
| HQVDVAAVQQG | GARTGNSDYT | 0.68129617 |
| IGAAIAHT | GARTGNSDYT | 0.587704 |
| IGMIEVAQL | GARTGNSDYT | 0.6167218 |
| IRLAPEHTSAMD | GARTGNSDYT | 0.38752607 |
| IYNVSVL | GARTGNSDYT | 0.6027324 |
| KQGFLGM | GARTGNSDYT | 0.45822406 |
| LESVITQL | GARTGNSDYT | 0.64429724 |
| LGLGALVVV | GARTGNSDYT | 0.6868782 |
| LIGNGGNGGSGGI | GARTGNSDYT | 0.58749783 |
| LNAINAPTL | GARTGNSDYT | 0.41047898 |
| LQLLNTGV | GARTGNSDYT | 0.45857662 |
| LQTSIEF | GARTGNSDYT | 0.6508626 |
| MSFAVSEL | GARTGNSDYT | 0.39984614 |
| NTGALGL | GARTGNSDYT | 0.3626981 |
| PPGAGGAGGAGTT | GARTGNSDYT | 0.56793284 |
| QLVGLGI | GARTGNSDYT | 0.6754714 |
| RFLEQVHQL | GARTGNSDYT | 0.675484 |
| RGVGYVLR | GARTGNSDYT | 0.54931974 |
| SATGVFAGVI | GARTGNSDYT | 0.6844156 |
| SLPAITLP | GARTGNSDYT | 0.6788196 |
| STALAFTV | GARTGNSDYT | 0.6839372 |
| SVLTVGLGK | GARTGNSDYT | 0.5778039 |
| TALAGAGSGI | GARTGNSDYT | 0.4385113 |
| TGLPFTGI | GARTGNSDYT | 0.6801659 |
| VAEHVTV | GARTGNSDYT | 0.56670946 |
| VDIVLNSL | GARTGNSDYT | 0.6571161 |
| VLTGGGAVL | GARTGNSDYT | 0.6724597 |
| VNVGLAYGI | GARTGNSDYT | 0.57234585 |
| VSSAAVGAV | GARTGNSDYT | 0.52627325 |
| VTSVYEAGV | GARTGNSDYT | 0.6305722 |
| WVALDRGA | GARTGNSDYT | 0.68492496 |
| YLVGSVVI | GARTGNSDYT | 0.52576447 |
| GPPGTGKTLL | GARTGNSDYT | 0.42084482 |
| GVGMTKF | GARTGNSDYT | 0.5958284 |
| IGIGGSDL | GARTGNSDYT | 0.60078245 |
| IGTLLAEL | GARTGNSDYT | 0.6850351 |
| IYTSGTT | GARTGNSDYT | 0.62785685 |
| LSFPAGV | GARTGNSDYT | 0.6131409 |
| LYTSGTT | GARTGNSDYT | 0.637814 |
| VGAFPFGF | GARTGNSDYT | 0.65105915 |
| VGAGTVEF | GARTGNSDYT | 0.6371689 |
| VGKSTLLL | GARTGNSDYT | 0.65771073 |
| AALAHAGGGIV | GARTGNSDYT | 0.5521792 |
| AALVAVSLL | GARTGNSDYT | 0.68396413 |
| AAVSRAVAAL | GARTGNSDYT | 0.6739154 |
| AGGSSGIGGF | GARTGNSDYT | 0.6465068 |

| | | |
|---|---|---|
| AGLTVANANG | GARTGNSDYT | 0.68226314 |
| AGLVGAVL | GARTGNSDYT | 0.63973117 |
| ALSQAFTV | GARTGNSDYT | 0.61300164 |
| ALTFSVSL | GARTGNSDYT | 0.43317586 |
| ALTGAALVAL | GARTGNSDYT | 0.54050446 |
| ASFSIGAGVL | GARTGNSDYT | 0.6097825 |
| DGSVYKI | GARTGNSDYT | 0.63676083 |
| EQLAEIV | GARTGNSDYT | 0.6621134 |
| FAVSELTVM | GARTGNSDYT | 0.64010906 |
| GAAPLAAGL | GARTGNSDYT | 0.535934 |
| GADGGDGGSGGAGG | GARTGNSDYT | 0.68345535 |
| GFGLDTGGAGG | GARTGNSDYT | 0.6804018 |
| GLASGGLL | GARTGNSDYT | 0.45448083 |
| GRIAGAAVLL | GARTGNSDYT | 0.6028714 |
| GSAGSLNTSF | GARTGNSDYT | 0.69603264 |
| GTGQIGI | GARTGNSDYT | 0.64704084 |
| GTLIALIL | GARTGNSDYT | 0.55217516 |
| IAATMGFVVG | GARTGNSDYT | 0.5639123 |
| IFAQALL | GARTGNSDYT | 0.6945373 |
| IGAAVALGF | GARTGNSDYT | 0.57362163 |
| IGLTPLAL | GARTGNSDYT | 0.6955837 |
| ILGSNTSTL | GARTGNSDYT | 0.3531027 |
| IVFSLTVL | GARTGNSDYT | 0.5068157 |
| LASKATGF | GARTGNSDYT | 0.53819746 |
| LASVALIL | GARTGNSDYT | 0.2975184 |
| LFAVGLTL | GARTGNSDYT | 0.6692846 |
| LFNGGIGTYI | GARTGNSDYT | 0.4590829 |
| LTLGAGII | GARTGNSDYT | 0.6476836 |
| LVAAALGDI | GARTGNSDYT | 0.5423837 |
| NTAFVQLGI | GARTGNSDYT | 0.6812685 |
| QLFSGAEL | GARTGNSDYT | 0.6381308 |
| SAATFITL | GARTGNSDYT | 0.6055512 |
| SNYGLANLGV | GARTGNSDYT | 0.48342603 |
| SYGGAYAVM | GARTGNSDYT | 0.6361712 |
| TTGVGGLTLG | GARTGNSDYT | 0.6856828 |
| TTQTIYI | GARTGNSDYT | 0.67525387 |
| VAAGAGLGAVI | GARTGNSDYT | 0.39794162 |
| VAAVVTLL | GARTGNSDYT | 0.6552322 |
| VGFLGGLI | GARTGNSDYT | 0.5926584 |
| VGVIPILL | GARTGNSDYT | 0.4293838 |
| VIFASSLI | GARTGNSDYT | 0.32579282 |
| VKGGDGGAAAG | GARTGNSDYT | 0.5946362 |
| VTKTFTNI | GARTGNSDYT | 0.5900669 |
| VVAAAILL | GARTGNSDYT | 0.6487259 |
| YAALDVEL | GARTGNSDYT | 0.67363214 |
| LASVALLL | GARTGNSDYT | 0.5336025 |
| LDAVHRV | GARTGNSDYT | 0.6544713 |
| QGLSSSIF | GARTGNSDYT | 0.63244724 |
| RGTFANIRL | GARTGNSDYT | 0.2583705 |
| RQLWWGHRI | GARTGNSDYT | 0.5389606 |
| YGPPGTGKTLL | GARTGNSDYT | 0.40793303 |

| | | |
|---|---|---|
| AALVGAPANVA | GARTGNSDYT | 0.6927496 |
| AATFRRTE | GARTGNSDYT | 0.5005891 |
| AGALGGGI | GARTGNSDYT | 0.59130013 |
| AGGAGGAGGIGAG | GARTGNSDYT | 0.61233115 |
| AGSGALGL | GARTGNSDYT | 0.6522647 |
| AQIADQI | GARTGNSDYT | 0.60589594 |
| ASAVAVTYL | GARTGNSDYT | 0.52280784 |
| ASTVLGGTGG | GARTGNSDYT | 0.44254893 |
| AVGGEPAK | GARTGNSDYT | 0.6644701 |
| CEQGTFGGTC | GARTGNSDYT | 0.6118091 |
| DEGKAPAG | GARTGNSDYT | 0.50108075 |
| DTVVFPAL | GARTGNSDYT | 0.53194296 |
| FEGITMAG | GARTGNSDYT | 0.64975154 |
| FPLVVQD | GARTGNSDYT | 0.38668287 |
| GAFTTPQ | GARTGNSDYT | 0.66393673 |
| GFGGFGTAGGVGGA | GARTGNSDYT | 0.40446097 |
| GFLAQFL | GARTGNSDYT | 0.58753973 |
| GGAGGAGGNAGVA | GARTGNSDYT | 0.6950521 |
| GGVGTAGAGGAGGAGG | GARTGNSDYT | 0.6537775 |
| GNGGGGDAGNAGSGGN | GARTGNSDYT | 0.60540587 |
| GTGGAGGAGNDAGS | GARTGNSDYT | 0.5570273 |
| GTTNTGWFNS | GARTGNSDYT | 0.46805057 |
| IFMGPLFTL | GARTGNSDYT | 0.51740164 |
| ILAHAIEAI | GARTGNSDYT | 0.53552866 |
| IRSAIREAL | GARTGNSDYT | 0.68697417 |
| ITFAGGG | GARTGNSDYT | 0.5906679 |
| ITGMSGHQEV | GARTGNSDYT | 0.4312309 |
| IVAGLACTL | GARTGNSDYT | 0.63871145 |
| IVGGGLAGA | GARTGNSDYT | 0.6939516 |
| IVTGVLNI | GARTGNSDYT | 0.6935382 |
| LAQQGCI | GARTGNSDYT | 0.49327144 |
| LCLHVEN | GARTGNSDYT | 0.5837045 |
| LDAISDGS | GARTGNSDYT | 0.52649844 |
| LGGFAAML | GARTGNSDYT | 0.6065705 |
| LGGKNPMI | GARTGNSDYT | 0.5011928 |
| LGPKGVAAIL | GARTGNSDYT | 0.49509406 |
| LMVGVSI | GARTGNSDYT | 0.66852397 |
| LVGAAFVL | GARTGNSDYT | 0.7076602 |
| LVLAVAPL | GARTGNSDYT | 0.586131 |
| MAAIAVIDAA | GARTGNSDYT | 0.36180225 |
| NNGINAELQ | GARTGNSDYT | 0.60283494 |
| RGEPLVLVHG | GARTGNSDYT | 0.49543402 |
| RGGAGGAAGLI | GARTGNSDYT | 0.5355455 |
| RSYRMHPAV | GARTGNSDYT | 0.501758 |
| SAAHLLAGQLA | GARTGNSDYT | 0.37806344 |
| SAAVGSY | GARTGNSDYT | 0.58338267 |
| SPHAVLHRI | GARTGNSDYT | 0.6938913 |
| SQIAKVL | GARTGNSDYT | 0.5860528 |
| TALGHLVP | GARTGNSDYT | 0.5359742 |
| TAPTTSGAA | GARTGNSDYT | 0.6800149 |
| TDGDDGNGGSPGT | GARTGNSDYT | 0.6477047 |

| | | |
|---|---|---|
| TDGLGEN | GARTGNSDYT | 0.67597735 |
| TGGPPLPGQA | GARTGNSDYT | 0.684028 |
| TIPAITI | GARTGNSDYT | 0.63620496 |
| TIPGFTI | GARTGNSDYT | 0.62889767 |
| TSIASGALA | GARTGNSDYT | 0.6663904 |
| TVTLPGAAI | GARTGNSDYT | 0.60337806 |
| VASGLSAALP | GARTGNSDYT | 0.61864597 |
| VGAGVIGI | GARTGNSDYT | 0.56443006 |
| VGPKRFGV | GARTGNSDYT | 0.6642802 |
| VNVAFPD | GARTGNSDYT | 0.5623222 |
| VSMTNSGVS | GARTGNSDYT | 0.6937758 |
| VTDVLGDLSV | GARTGNSDYT | 0.30952743 |
| VVVVGAGFAGL | GARTGNSDYT | 0.46277106 |
| VYTVGTVV | GARTGNSDYT | 0.6764562 |
| WTTFSGP | GARTGNSDYT | 0.29631028 |
| YDELVEAF | GARTGNSDYT | 0.33223003 |
| YGTGGAGGNGG | GARTGNSDYT | 0.59025073 |
| YTNVMAV | GARTGNSDYT | 0.6769509 |
| FLEQVHQL | GARTGNSDYT | 0.29914865 |
| GLSLLAVAV | GARTGNSDYT | 0.58897066 |
| LEQVHQL | GARTGNSDYT | 0.35886627 |
| LNAAGAI | GARTGNSDYT | 0.6816257 |
| PAVHASL | GARTGNSDYT | 0.6067495 |
| TELEDDA | GARTGNSDYT | 0.39835912 |
| TPPGPPN | GARTGNSDYT | 0.66808367 |
| TPPPPPPP | GARTGNSDYT | 0.47919863 |
| TQSPALI | ASSLEGAGNTLY | 0.65313935 |
| EFAAAFPQ | ASSLEGAGNTLY | 0.6734559 |
| LLSGNLDVL | ASSLEGAGNTLY | 0.6997992 |
| GPANLGNYNI | ASSLEGAGNTLY | 0.4273482 |
| TLGPTLP | ASSLEGAGNTLY | 0.64016783 |
| VYAYSSCCK | ASSLEGAGNTLY | 0.54907084 |
| APAPAPAPNI | ASSLEGAGNTLY | 0.70118463 |
| PGGPVPIP | ASSLEGAGNTLY | 0.5496187 |
| FLTVALEP | ASSLEGAGNTLY | 0.36726832 |
| SQYFGANL | ASSLEGAGNTLY | 0.67049325 |
| GGVKPAA | ASSLEGAGNTLY | 0.63292354 |
| GGTGGAGGAGGW | ASSLEGAGNTLY | 0.5742744 |
| GVGGVAGLGGAGG | ASSLEGAGNTLY | 0.6686418 |
| VLLVLATTI | ASSLEGAGNTLY | 0.58219314 |
| IGIFAAML | ASSLEGAGNTLY | 0.57823944 |
| AVGDLID | ASSLEGAGNTLY | 0.66178644 |
| NISPLPL | ASSLEGAGNTLY | 0.6685286 |
| VTAVFVAI | ASSLEGAGNTLY | 0.52400196 |
| SGDDNDAAL | ASSLEGAGNTLY | 0.66387975 |
| AVAALAHS | ASSLEGAGNTLY | 0.6358118 |
| GGVTVMATP | ASSLEGAGNTLY | 0.47073328 |
| AGLAANVRD | ASSLEGAGNTLY | 0.6971117 |
| TNRPDLIDPALLRPG | ASSLEGAGNTLY | 0.54808813 |
| LANLGVTAL | ASSLEGAGNTLY | 0.65433455 |
| TGERIEL | ASSLEGAGNTLY | 0.64466155 |

| | | |
|---|---|---|
| NGEELAA | ASSLEGAGNTLY | 0.68939406 |
| VLSLVWG | ASSLEGAGNTLY | 0.49737597 |
| IGYALWL | ASSLEGAGNTLY | 0.6198704 |
| LSLLLAPL | ASSLEGAGNTLY | 0.6091151 |
| TLDEALK | ASSLEGAGNTLY | 0.66868466 |
| TDGAGTAEGG | ASSLEGAGNTLY | 0.5624457 |
| NAPGGAGGNGG | ASSLEGAGNTLY | 0.6357027 |
| VSVFMGMGGNF | ASSLEGAGNTLY | 0.6544008 |
| LASYLAAL | ASSLEGAGNTLY | 0.6838988 |
| LVGGTPAL | ASSLEGAGNTLY | 0.6289097 |
| VRNADVL | ASSLEGAGNTLY | 0.638893 |
| TCSSPFLVSAM | ASSLEGAGNTLY | 0.6928429 |
| VHITDAE | ASSLEGAGNTLY | 0.41827965 |
| EYAAPSF | ASSLEGAGNTLY | 0.63082075 |
| VVDAFHGGL | ASSLEGAGNTLY | 0.66730976 |
| ASVVDPQ | ASSLEGAGNTLY | 0.6746057 |
| SYNEFGGLS | ASSLEGAGNTLY | 0.57976186 |
| DPDGSVY | ASSLEGAGNTLY | 0.68740016 |
| MVSTIHATE | ASSLEGAGNTLY | 0.5637778 |
| LAAGVADPV | ASSLEGAGNTLY | 0.65714693 |
| TKEQIAA | ASSLEGAGNTLY | 0.5681949 |
| SLASPAAL | ASSLEGAGNTLY | 0.5493513 |
| IGLVFYGV | ASSLEGAGNTLY | 0.4687202 |
| FLLDASNL | ASSLEGAGNTLY | 0.63985324 |
| YQVGNFP | ASSLEGAGNTLY | 0.6606401 |
| ESLLSQST | ASSLEGAGNTLY | 0.65515673 |
| LFTVGFAAVG | ASSLEGAGNTLY | 0.34266365 |
| ATPPTPT | ASSLEGAGNTLY | 0.6407591 |
| TDLATLG | ASSLEGAGNTLY | 0.6745746 |
| SAGGTGR | ASSLEGAGNTLY | 0.69794166 |
| GFSNSGS | ASSLEGAGNTLY | 0.6004159 |
| GGNGGSGGTSV | ASSLEGAGNTLY | 0.63598573 |
| GTSVATGGAGN | ASSLEGAGNTLY | 0.60304224 |
| NGGNGGSGGTS | ASSLEGAGNTLY | 0.60275567 |
| AVVKFPH | ASSLEGAGNTLY | 0.44591334 |
| TPPPPPP | ASSLEGAGNTLY | 0.45238835 |
| AAPAAAAAPA | ASSLEGAGNTLY | 0.60699457 |
| IALNEDL | ASSLEGAGNTLY | 0.5095035 |
| IVNVSSM | ASSLEGAGNTLY | 0.6715737 |
| TGIPPAPR | ASSLEGAGNTLY | 0.6825094 |
| FDLGGGTF | ASSLEGAGNTLY | 0.66754353 |
| FDLGGGTFDVS | ASSLEGAGNTLY | 0.6832855 |
| NEPTAAA | ASSLEGAGNTLY | 0.67987907 |
| NEPTAAALA | ASSLEGAGNTLY | 0.65493846 |
| GSIINISSI | ASSLEGAGNTLY | 0.54895806 |
| VAPGNAGTA | ASSLEGAGNTLY | 0.5842821 |
| TGGGASLE | ASSLEGAGNTLY | 0.3877848 |
| VAVYNHYKRI | ASSLEGAGNTLY | 0.55258906 |
| TDPAPATT | ASSLEGAGNTLY | 0.56178176 |
| TNRPDILDPALLRPG | ASSLEGAGNTLY | 0.6141743 |
| LTTTTTS | ASSLEGAGNTLY | 0.27469587 |

| | | |
|---|---|---|
| PTGSGKTL | ASSLEGAGNTLY | 0.6464207 |
| TGSGKTL | ASSLEGAGNTLY | 0.6861875 |
| TIGHVDHGK | ASSLEGAGNTLY | 0.6131341 |
| TIGHVDHGKTT | ASSLEGAGNTLY | 0.64854383 |
| GHVDHGKTT | ASSLEGAGNTLY | 0.6147052 |
| AQFEHTLL | ASSLEGAGNTLY | 0.55025715 |
| IVNQDEL | ASSLEGAGNTLY | 0.65468836 |
| TNGSQFF | ASSLEGAGNTLY | 0.5273732 |
| TGKGIVE | ASSLEGAGNTLY | 0.6608672 |
| LISNASDA | ASSLEGAGNTLY | 0.6719771 |
| AELRTLGELTKIAKAHGAQ | ASSLEGAGNTLY | 0.6414081 |
| APNAVAAAEPSW | ASSLEGAGNTLY | 0.5554433 |
| APQVAGAATAVL | ASSLEGAGNTLY | 0.5904875 |
| ARRDPAAAAAILPTDA | ASSLEGAGNTLY | 0.48407176 |
| DAAARDVLAEAGLAEYF | ASSLEGAGNTLY | 0.5980947 |
| DDQIVLSAKANLAKVEQ | ASSLEGAGNTLY | 0.49459627 |
| DELLPDTGEDSDEDGH | ASSLEGAGNTLY | 0.6012418 |
| ELPEAPANRATVPQD | ASSLEGAGNTLY | 0.48566806 |
| FGQSKIERVTGIG | ASSLEGAGNTLY | 0.64512277 |
| GAGGTGSTAGGAGGAGGAGGLY | ASSLEGAGNTLY | 0.63384163 |
| GHGGTGAAGGNGGTGGAGGSGI | ASSLEGAGNTLY | 0.6549587 |
| GNGGNGGAGGEGGPGGAGGAGA | ASSLEGAGNTLY | 0.4929732 |
| GSTAGNGGNGGNATNGTIAGQP | ASSLEGAGNTLY | 0.655506 |
| GTGGDGGNGGNGGTGG | ASSLEGAGNTLY | 0.6482301 |
| GVRRPAGIPADAPV | ASSLEGAGNTLY | 0.45041648 |
| GYRQVTDAHLLTLARR | ASSLEGAGNTLY | 0.6474135 |
| IAKMGQDGHDRG | ASSLEGAGNTLY | 0.5741392 |
| KAEPSIEIQYT | ASSLEGAGNTLY | 0.5968961 |
| KPVRIRMAADEDTVAVT | ASSLEGAGNTLY | 0.67563677 |
| MPPRIAGMRLLVIKPEP | ASSLEGAGNTLY | 0.5953652 |
| MKFYAVALFRSRGVPPTR | ASSLEGAGNTLY | 0.4672874 |
| NAGGFGILGAVAHSP | ASSLEGAGNTLY | 0.5297296 |
| NEGREITLAMAEM | ASSLEGAGNTLY | 0.6078799 |
| NELVEWSHPLGHRWRWE | ASSLEGAGNTLY | 0.5726906 |
| NLGAGGTGGNGGNPG | ASSLEGAGNTLY | 0.30832806 |
| NTEWPKWETVMAADPVD | ASSLEGAGNTLY | 0.66261923 |
| PASAAPAETQP | ASSLEGAGNTLY | 0.4700574 |
| QQSSDGKNGVRAAEITGE | ASSLEGAGNTLY | 0.5468187 |
| RAPDAVTTSWWKE | ASSLEGAGNTLY | 0.5764582 |
| SGFGNVGAHTSGWFNQ | ASSLEGAGNTLY | 0.67038673 |
| SGVAGAVGQAASVGGLKV | ASSLEGAGNTLY | 0.6111599 |
| SNGGAGTGSAGGNGGT | ASSLEGAGNTLY | 0.5539217 |
| TGGAGGEGGAAG | ASSLEGAGNTLY | 0.63475937 |
| TGGDGGAAGAGGAPGHG | ASSLEGAGNTLY | 0.6499884 |
| TMPPLLAAIANAGRHGV | ASSLEGAGNTLY | 0.66670895 |
| TSSTAGNSGGA | ASSLEGAGNTLY | 0.641201 |
| VPATVPVGILC | ASSLEGAGNTLY | 0.6303768 |
| VQRPDSGGVVLTGRIS | ASSLEGAGNTLY | 0.6448925 |
| WTPNPVPATKGVPAGLVQAT | ASSLEGAGNTLY | 0.5613554 |
| YALKRALAKFSDRFDVVII | ASSLEGAGNTLY | 0.476824 |
| YLIGSVIV | ASSLEGAGNTLY | 0.41859648 |

| | | |
|---|---|---|
| YGQLVAVID | ASSLEGAGNTLY | 0.49943447 |
| VTVGGMTLPA | ASSLEGAGNTLY | 0.45085764 |
| VLALPAV | ASSLEGAGNTLY | 0.6253414 |
| VIPPTLNYET | ASSLEGAGNTLY | 0.6837163 |
| VGMGGMHPSSG | ASSLEGAGNTLY | 0.58896196 |
| VGLAGGAATA | ASSLEGAGNTLY | 0.37174985 |
| VGGAGGAGGSGP | ASSLEGAGNTLY | 0.56985426 |
| VFAAFMV | ASSLEGAGNTLY | 0.5947597 |
| TLYDKLR | ASSLEGAGNTLY | 0.6462966 |
| TLAVFTVL | ASSLEGAGNTLY | 0.68027896 |
| TGSVQPARV | ASSLEGAGNTLY | 0.7042254 |
| TGGGAGSGGT | ASSLEGAGNTLY | 0.68300486 |
| SHLETSL | ASSLEGAGNTLY | 0.56519705 |
| SFTTGMT | ASSLEGAGNTLY | 0.6634634 |
| QVEVAKRMGVRQPTVS | ASSLEGAGNTLY | 0.53850365 |
| QGSGGPALAM | ASSLEGAGNTLY | 0.58471656 |
| MLAPSNP | ASSLEGAGNTLY | 0.61543775 |
| LVNNAAI | ASSLEGAGNTLY | 0.6465049 |
| LTGEINTLA | ASSLEGAGNTLY | 0.55042934 |
| LSIFGPV | ASSLEGAGNTLY | 0.5876538 |
| LLALLANL | ASSLEGAGNTLY | 0.42313287 |
| LISNASDALDKLR | ASSLEGAGNTLY | 0.6738839 |
| LINEAVP | ASSLEGAGNTLY | 0.49282554 |
| LGYAFSGL | ASSLEGAGNTLY | 0.4037213 |
| LDFVAVAL | ASSLEGAGNTLY | 0.58605015 |
| LAQPGAE | ASSLEGAGNTLY | 0.6854863 |
| ISNLGAVTGF | ASSLEGAGNTLY | 0.68590736 |
| HQEDPGRI | ASSLEGAGNTLY | 0.5702552 |
| GTPGEPTP | ASSLEGAGNTLY | 0.5516382 |
| GFFNSGS | ASSLEGAGNTLY | 0.66169256 |
| GAGGGGLTAGAN | ASSLEGAGNTLY | 0.523025 |
| FVDALVNM | ASSLEGAGNTLY | 0.6501403 |
| FLTVAPEL | ASSLEGAGNTLY | 0.60991234 |
| FETLTAL | ASSLEGAGNTLY | 0.6267556 |
| DVINPIEL | ASSLEGAGNTLY | 0.46922415 |
| APVSGAPV | ASSLEGAGNTLY | 0.6594192 |
| ALVGGLAW | ASSLEGAGNTLY | 0.6161251 |
| ALAAAAATAI | ASSLEGAGNTLY | 0.6371307 |
| AFGLGLLL | ASSLEGAGNTLY | 0.39131173 |
| AFFAAASL | ASSLEGAGNTLY | 0.6828544 |
| AAAALPAFSPP | ASSLEGAGNTLY | 0.5109215 |
| VVLAPAL | ASSLEGAGNTLY | 0.46562484 |
| TIGHVDHGKTTL | ASSLEGAGNTLY | 0.31044775 |
| LISNASDALDK | ASSLEGAGNTLY | 0.6230922 |
| LISNASDAL | ASSLEGAGNTLY | 0.64030415 |
| ISNASDALDK | ASSLEGAGNTLY | 0.68260765 |
| NQAETLVYQTE | ASSLEGAGNTLY | 0.61672544 |
| TGGTGGTAGAG | ASSLEGAGNTLY | 0.65630233 |
| TGSTAGGAGGA | ASSLEGAGNTLY | 0.6781845 |
| DPLAVPVAAGPV | ASSLEGAGNTLY | 0.53564864 |
| GGTGGTPGNGGN | ASSLEGAGNTLY | 0.3570551 |

| | | |
|---|---|---|
| PAGLAGVDPATL | ASSLEGAGNTLY | 0.5979866 |
| GNNLVVEAPHAGASV | ASSLEGAGNTLY | 0.616966 |
| KVTRIMSASTLEQPA | ASSLEGAGNTLY | 0.6815292 |
| LGFALPIALPRRTVV | ASSLEGAGNTLY | 0.6676279 |
| TRLGNLTARKLVVLV | ASSLEGAGNTLY | 0.4275403 |
| WSQQRRLIYFPSAGP | ASSLEGAGNTLY | 0.6240065 |
| GPNGNPDPMAAAVDIR | ASSLEGAGNTLY | 0.6377255 |
| TQRCVSVADLRAAASR | ASSLEGAGNTLY | 0.4369828 |
| QFVLDVAEHPVEVLTETV | ASSLEGAGNTLY | 0.5183157 |
| GGAGGTGGAGGNGGTQAGDG | ASSLEGAGNTLY | 0.53162295 |
| PLLATLRAIAVDVDAKPAQV | ASSLEGAGNTLY | 0.40585247 |
| APKFRTGGYMIDFWGVGYQVAK | ASSLEGAGNTLY | 0.6628486 |
| GAGGAGGAGGAAGGPGGTGGTG | ASSLEGAGNTLY | 0.5823044 |
| IGALGLLATDVAPAGSVAFGVA | ASSLEGAGNTLY | 0.56830835 |
| AGGAGGIGGNGGAGGAGGVLMG | ASSLEGAGNTLY | 0.61268866 |
| ADELIGLL | ASSLEGAGNTLY | 0.52822304 |
| ADRALKA | ASSLEGAGNTLY | 0.54523385 |
| AGAVGQVGGAGGA | ASSLEGAGNTLY | 0.560407 |
| AGGGGGLGGAP | ASSLEGAGNTLY | 0.66657144 |
| AGGLLYGNGGAG | ASSLEGAGNTLY | 0.6650237 |
| AGLESIGL | ASSLEGAGNTLY | 0.5995587 |
| AGLKTGII | ASSLEGAGNTLY | 0.5862435 |
| ASLAALLL | ASSLEGAGNTLY | 0.677021 |
| ASVINISSI | ASSLEGAGNTLY | 0.40817118 |
| ATPPGPA | ASSLEGAGNTLY | 0.5773783 |
| ATVASALV | ASSLEGAGNTLY | 0.6474265 |
| ATVASLIL | ASSLEGAGNTLY | 0.5287024 |
| AVNGPVLIRH | ASSLEGAGNTLY | 0.67290395 |
| DGAAGAVGI | ASSLEGAGNTLY | 0.6503087 |
| DPLLHGI | ASSLEGAGNTLY | 0.63745815 |
| ELGAALGAL | ASSLEGAGNTLY | 0.6024872 |
| ELGRTQAV | ASSLEGAGNTLY | 0.6900791 |
| FVAGGYLS | ASSLEGAGNTLY | 0.6324484 |
| GASGGAGGAGGW | ASSLEGAGNTLY | 0.57639515 |
| GGAGAGGDGGAGGI | ASSLEGAGNTLY | 0.6594068 |
| GTGHNGSQPGMGGNG | ASSLEGAGNTLY | 0.4374441 |
| GYAAASASAS | ASSLEGAGNTLY | 0.672732 |
| HQVDVAAVQQG | ASSLEGAGNTLY | 0.6404652 |
| IGAAIAHT | ASSLEGAGNTLY | 0.4742571 |
| IGMIEVAQL | ASSLEGAGNTLY | 0.58991337 |
| IRLAPEHTSAMD | ASSLEGAGNTLY | 0.68894637 |
| IYNVSVL | ASSLEGAGNTLY | 0.66794443 |
| KQGFLGM | ASSLEGAGNTLY | 0.5216402 |
| LESVITQL | ASSLEGAGNTLY | 0.5483631 |
| LGLGALVVV | ASSLEGAGNTLY | 0.51026785 |
| LIGNGGNGGSGGI | ASSLEGAGNTLY | 0.553672 |
| LNAINAPTL | ASSLEGAGNTLY | 0.39261305 |
| LQLLNTGV | ASSLEGAGNTLY | 0.63654304 |
| LQTSIEF | ASSLEGAGNTLY | 0.53427374 |
| MSFAVSEL | ASSLEGAGNTLY | 0.6782948 |
| NTGALGL | ASSLEGAGNTLY | 0.6404903 |

| | | |
|---|---|---|
| PPGAGGAGGAGTT | ASSLEGAGNTLY | 0.6798079 |
| QLVGLGI | ASSLEGAGNTLY | 0.5252493 |
| RFLEQVHQL | ASSLEGAGNTLY | 0.62465835 |
| RGVGYVLR | ASSLEGAGNTLY | 0.35083744 |
| SATGVFAGVI | ASSLEGAGNTLY | 0.56703913 |
| SLPAITLP | ASSLEGAGNTLY | 0.48924804 |
| STALAFTV | ASSLEGAGNTLY | 0.6129986 |
| SVLTVGLGK | ASSLEGAGNTLY | 0.4003067 |
| TALAGAGSGI | ASSLEGAGNTLY | 0.6936849 |
| TGLPFTGI | ASSLEGAGNTLY | 0.52412534 |
| VAEHVTV | ASSLEGAGNTLY | 0.6552565 |
| VDIVLNSL | ASSLEGAGNTLY | 0.56890064 |
| VLTGGGAVL | ASSLEGAGNTLY | 0.43187812 |
| VNVGLAYGI | ASSLEGAGNTLY | 0.54800355 |
| VSSAAVGAV | ASSLEGAGNTLY | 0.6510596 |
| VTSVYEAGV | ASSLEGAGNTLY | 0.4621876 |
| WVALDRGA | ASSLEGAGNTLY | 0.3518722 |
| YLVGSVVI | ASSLEGAGNTLY | 0.5228186 |
| GPPGTGKTLL | ASSLEGAGNTLY | 0.6063841 |
| GVGMTKF | ASSLEGAGNTLY | 0.6909927 |
| IGIGGSDL | ASSLEGAGNTLY | 0.68132687 |
| IGTLLAEL | ASSLEGAGNTLY | 0.61236066 |
| IYTSGTT | ASSLEGAGNTLY | 0.69610065 |
| LSFPAGV | ASSLEGAGNTLY | 0.28107232 |
| LYTSGTT | ASSLEGAGNTLY | 0.6070252 |
| VGAFPFGF | ASSLEGAGNTLY | 0.69450325 |
| VGAGTVEF | ASSLEGAGNTLY | 0.6479789 |
| VGKSTLLL | ASSLEGAGNTLY | 0.56168795 |
| AALAHAGGGIV | ASSLEGAGNTLY | 0.6674502 |
| AALVAVSLL | ASSLEGAGNTLY | 0.55743694 |
| AAVSRAVAAL | ASSLEGAGNTLY | 0.57185405 |
| AGGSSGIGGF | ASSLEGAGNTLY | 0.6045481 |
| AGLTVANANG | ASSLEGAGNTLY | 0.6640586 |
| AGLVGAVL | ASSLEGAGNTLY | 0.62097716 |
| ALSQAFTV | ASSLEGAGNTLY | 0.40921196 |
| ALTFSVSL | ASSLEGAGNTLY | 0.6044065 |
| ALTGAALVAL | ASSLEGAGNTLY | 0.7001402 |
| ASFSIGAGVL | ASSLEGAGNTLY | 0.54186034 |
| DGSVYKI | ASSLEGAGNTLY | 0.6800952 |
| EQLAEIV | ASSLEGAGNTLY | 0.58430797 |
| FAVSELTVM | ASSLEGAGNTLY | 0.6193627 |
| GAAPLAAGL | ASSLEGAGNTLY | 0.50731623 |
| GADGGDGGSGGAGG | ASSLEGAGNTLY | 0.47375423 |
| GFGLDTGGAGG | ASSLEGAGNTLY | 0.66778183 |
| GLASGGLL | ASSLEGAGNTLY | 0.495983 |
| GRIAGAAVLL | ASSLEGAGNTLY | 0.5624245 |
| GSAGSLNTSF | ASSLEGAGNTLY | 0.6059968 |
| GTGQIGI | ASSLEGAGNTLY | 0.7009673 |
| GTLIALIL | ASSLEGAGNTLY | 0.6463889 |
| IAATMGFVVG | ASSLEGAGNTLY | 0.6242043 |
| IFAQALL | ASSLEGAGNTLY | 0.5461876 |

| | | |
|---|---|---|
| IGAAVALGF | ASSLEGAGNTLY | 0.6340283 |
| IGLTPLAL | ASSLEGAGNTLY | 0.6517807 |
| ILGSNTSTL | ASSLEGAGNTLY | 0.43022248 |
| IVFSLTVL | ASSLEGAGNTLY | 0.6854335 |
| LASKATGF | ASSLEGAGNTLY | 0.436489 |
| LASVALIL | ASSLEGAGNTLY | 0.6267545 |
| LFAVGLTL | ASSLEGAGNTLY | 0.64630055 |
| LFNGGIGTYI | ASSLEGAGNTLY | 0.69619894 |
| LTLGAGII | ASSLEGAGNTLY | 0.58278847 |
| LVAAALGDI | ASSLEGAGNTLY | 0.6601703 |
| NTAFVQLGI | ASSLEGAGNTLY | 0.52147895 |
| QLFSGAEL | ASSLEGAGNTLY | 0.6557385 |
| SAATFITL | ASSLEGAGNTLY | 0.67636025 |
| SNYGLANLGV | ASSLEGAGNTLY | 0.6279809 |
| SYGGAYAVM | ASSLEGAGNTLY | 0.7049617 |
| TTGVGGLTLG | ASSLEGAGNTLY | 0.6530261 |
| TTQTIYI | ASSLEGAGNTLY | 0.35725257 |
| VAAGAGLGAVI | ASSLEGAGNTLY | 0.577382 |
| VAAVVTLL | ASSLEGAGNTLY | 0.68406296 |
| VGFLGGLI | ASSLEGAGNTLY | 0.68196857 |
| VGVIPILL | ASSLEGAGNTLY | 0.6831996 |
| VIFASSLI | ASSLEGAGNTLY | 0.6295825 |
| VKGGDGGAAAG | ASSLEGAGNTLY | 0.5796306 |
| VTKTFTNI | ASSLEGAGNTLY | 0.68852746 |
| VVAAAILL | ASSLEGAGNTLY | 0.7071718 |
| YAALDVEL | ASSLEGAGNTLY | 0.62897027 |
| LASVALLL | ASSLEGAGNTLY | 0.50038093 |
| LDAVHRV | ASSLEGAGNTLY | 0.47422016 |
| QGLSSSIF | ASSLEGAGNTLY | 0.6531167 |
| RGTFANIRL | ASSLEGAGNTLY | 0.67290133 |
| RQLWWGHRI | ASSLEGAGNTLY | 0.3772644 |
| YGPPGTGKTLL | ASSLEGAGNTLY | 0.5730549 |
| AALVGAPANVA | ASSLEGAGNTLY | 0.6276063 |
| AATFRRTE | ASSLEGAGNTLY | 0.3283235 |
| AGALGGGI | ASSLEGAGNTLY | 0.6594023 |
| AGGAGGAGGIGAG | ASSLEGAGNTLY | 0.68307775 |
| AGSGALGL | ASSLEGAGNTLY | 0.6187873 |
| AQIADQI | ASSLEGAGNTLY | 0.43478 |
| ASAVAVTYL | ASSLEGAGNTLY | 0.67837065 |
| ASTVLGGTGG | ASSLEGAGNTLY | 0.6732631 |
| AVGGEPAK | ASSLEGAGNTLY | 0.68917346 |
| CEQGTFGGTC | ASSLEGAGNTLY | 0.40581226 |
| DEGKAPAG | ASSLEGAGNTLY | 0.29512358 |
| DTVVFPAL | ASSLEGAGNTLY | 0.63191634 |
| FEGITMAG | ASSLEGAGNTLY | 0.6602397 |
| FPLVVQD | ASSLEGAGNTLY | 0.58450395 |
| GAFTTPQ | ASSLEGAGNTLY | 0.4958959 |
| GFGGFGTAGGVGGA | ASSLEGAGNTLY | 0.57269806 |
| GFLAQFL | ASSLEGAGNTLY | 0.67936456 |
| GGAGGAGGNAGVA | ASSLEGAGNTLY | 0.41758752 |
| GGVGTAGAGGAGGAGG | ASSLEGAGNTLY | 0.4270417 |

| | | |
|---|---|---|
| GNGGGGDAGNAGSGGN | ASSLEGAGNTLY | 0.63433325 |
| GTGGAGGAGNDAGS | ASSLEGAGNTLY | 0.66028863 |
| GTTNTGWFNS | ASSLEGAGNTLY | 0.5393115 |
| IFMGPLFTL | ASSLEGAGNTLY | 0.5661702 |
| ILAHAIEAI | ASSLEGAGNTLY | 0.654649 |
| IRSAIREAL | ASSLEGAGNTLY | 0.6374979 |
| ITFAGGG | ASSLEGAGNTLY | 0.63979936 |
| ITGMSGHQEV | ASSLEGAGNTLY | 0.42919478 |
| IVAGLACTL | ASSLEGAGNTLY | 0.6816939 |
| IVGGGLAGA | ASSLEGAGNTLY | 0.42793015 |
| IVTGVLNI | ASSLEGAGNTLY | 0.658692 |
| LAQQGCI | ASSLEGAGNTLY | 0.61790824 |
| LCLHVEN | ASSLEGAGNTLY | 0.643214 |
| LDAISDGS | ASSLEGAGNTLY | 0.5448111 |
| LGGFAAML | ASSLEGAGNTLY | 0.42428362 |
| LGGKNPMI | ASSLEGAGNTLY | 0.4701175 |
| LGPKGVAAIL | ASSLEGAGNTLY | 0.42309427 |
| LMVGVSI | ASSLEGAGNTLY | 0.6769857 |
| LVGAAFVL | ASSLEGAGNTLY | 0.6435206 |
| LVLAVAPL | ASSLEGAGNTLY | 0.66536963 |
| MAAIAVIDAA | ASSLEGAGNTLY | 0.5871148 |
| NNGINAELQ | ASSLEGAGNTLY | 0.52288437 |
| RGEPLVLVHG | ASSLEGAGNTLY | 0.65408516 |
| RGGAGGAAGLI | ASSLEGAGNTLY | 0.39822522 |
| RSYRMHPAV | ASSLEGAGNTLY | 0.6143768 |
| SAAHLLAGQLA | ASSLEGAGNTLY | 0.5292274 |
| SAAVGSY | ASSLEGAGNTLY | 0.64258397 |
| SPHAVLHRI | ASSLEGAGNTLY | 0.5058154 |
| SQIAKVL | ASSLEGAGNTLY | 0.6239487 |
| TALGHLVP | ASSLEGAGNTLY | 0.6414131 |
| TAPTTSGAA | ASSLEGAGNTLY | 0.45952153 |
| TDGDDGNGGSPGT | ASSLEGAGNTLY | 0.60865796 |
| TDGLGEN | ASSLEGAGNTLY | 0.682159 |
| TGGPPLPGQA | ASSLEGAGNTLY | 0.61319107 |
| TIPAITI | ASSLEGAGNTLY | 0.3691839 |
| TIPGFTI | ASSLEGAGNTLY | 0.625154 |
| TSIASGALA | ASSLEGAGNTLY | 0.52175295 |
| TVTLPGAAI | ASSLEGAGNTLY | 0.64985704 |
| VASGLSAALP | ASSLEGAGNTLY | 0.6249099 |
| VGAGVIGI | ASSLEGAGNTLY | 0.53463966 |
| VGPKRFGV | ASSLEGAGNTLY | 0.64491796 |
| VNVAFPD | ASSLEGAGNTLY | 0.45448166 |
| VSMTNSGVS | ASSLEGAGNTLY | 0.53245866 |
| VTDVLGDLSV | ASSLEGAGNTLY | 0.55268395 |
| VVVVGAGFAGL | ASSLEGAGNTLY | 0.633172 |
| VYTVGTVV | ASSLEGAGNTLY | 0.63794434 |
| WTTFSGP | ASSLEGAGNTLY | 0.4718216 |
| YDELVEAF | ASSLEGAGNTLY | 0.6278511 |
| YGTGGAGGNGG | ASSLEGAGNTLY | 0.58019906 |
| YTNVMAV | ASSLEGAGNTLY | 0.47295234 |
| FLEQVHQL | ASSLEGAGNTLY | 0.5670375 |

| | | |
|---|---|---|
| GLSLLAVAV | ASSLEGAGNTLY | 0.54552865 |
| LEQVHQL | ASSLEGAGNTLY | 0.35666367 |
| LNAAGAI | ASSLEGAGNTLY | 0.6749696 |
| PAVHASL | ASSLEGAGNTLY | 0.47452027 |
| TELEDDA | ASSLEGAGNTLY | 0.6418776 |
| TPPGPPN | ASSLEGAGNTLY | 0.66955626 |
| TPPPPPPP | ASSLEGAGNTLY | 0.3981728 |
| TQSPALI | ASSPGTGDTLY | 0.64391875 |
| EFAAAFPQ | ASSPGTGDTLY | 0.59918696 |
| LLSGNLDVL | ASSPGTGDTLY | 0.61564356 |
| GPANLGNYNI | ASSPGTGDTLY | 0.5972787 |
| TLGPTLP | ASSPGTGDTLY | 0.5411017 |
| VYAYSSCCK | ASSPGTGDTLY | 0.6789752 |
| APAPAPAPNI | ASSPGTGDTLY | 0.65492463 |
| PGGPVPIP | ASSPGTGDTLY | 0.6174381 |
| FLTVALEP | ASSPGTGDTLY | 0.6423504 |
| SQYFGANL | ASSPGTGDTLY | 0.6659913 |
| GGVKPAA | ASSPGTGDTLY | 0.6748843 |
| GGTGGAGGAGGW | ASSPGTGDTLY | 0.55442405 |
| GVGGVAGLGGAGG | ASSPGTGDTLY | 0.3512313 |
| VLLVLATTI | ASSPGTGDTLY | 0.61279607 |
| IGIFAAML | ASSPGTGDTLY | 0.59885776 |
| AVGDLID | ASSPGTGDTLY | 0.46885544 |
| NISPLPL | ASSPGTGDTLY | 0.3776623 |
| VTAVFVAI | ASSPGTGDTLY | 0.67317504 |
| SGDDNDAAL | ASSPGTGDTLY | 0.70254296 |
| AVAALAHS | ASSPGTGDTLY | 0.5486148 |
| GGVTVMATP | ASSPGTGDTLY | 0.6987864 |
| AGLAANVRD | ASSPGTGDTLY | 0.39530388 |
| TNRPDLIDPALLRPG | ASSPGTGDTLY | 0.6531753 |
| LANLGVTAL | ASSPGTGDTLY | 0.6734065 |
| TGERIEL | ASSPGTGDTLY | 0.44508427 |
| NGEELAA | ASSPGTGDTLY | 0.65867555 |
| VLSLVWG | ASSPGTGDTLY | 0.48083496 |
| IGYALWL | ASSPGTGDTLY | 0.506144 |
| LSLLLAPL | ASSPGTGDTLY | 0.6645043 |
| TLDEALK | ASSPGTGDTLY | 0.5448285 |
| TDGAGTAEGG | ASSPGTGDTLY | 0.65085596 |
| NAPGGAGGNGG | ASSPGTGDTLY | 0.5590029 |
| VSVFMGMGGNF | ASSPGTGDTLY | 0.50349206 |
| LASYLAAL | ASSPGTGDTLY | 0.64842355 |
| LVGGTPAL | ASSPGTGDTLY | 0.6733986 |
| VRNADVL | ASSPGTGDTLY | 0.60137165 |
| TCSSPFLVSAM | ASSPGTGDTLY | 0.70448786 |
| VHITDAE | ASSPGTGDTLY | 0.5289167 |
| EYAAPSF | ASSPGTGDTLY | 0.698913 |
| VVDAFHGGL | ASSPGTGDTLY | 0.6818083 |
| ASVVDPQ | ASSPGTGDTLY | 0.6327263 |
| SYNEFGGLS | ASSPGTGDTLY | 0.5791992 |
| DPDGSVY | ASSPGTGDTLY | 0.6657499 |
| MVSTIHATE | ASSPGTGDTLY | 0.5293516 |

| | | |
|---|---|---|
| LAAGVADPV | ASSPGTGDTLY | 0.63786316 |
| TKEQIAA | ASSPGTGDTLY | 0.68441373 |
| SLASPAAL | ASSPGTGDTLY | 0.69453484 |
| IGLVFYGV | ASSPGTGDTLY | 0.76303035 |
| FLLDASNL | ASSPGTGDTLY | 0.72843534 |
| YQVGNFP | ASSPGTGDTLY | 0.72225356 |
| ESLLSQST | ASSPGTGDTLY | 0.68617624 |
| LFTVGFAAVG | ASSPGTGDTLY | 0.54179776 |
| ATPPTPT | ASSPGTGDTLY | 0.728317 |
| TDLATLG | ASSPGTGDTLY | 0.6405514 |
| SAGGTGR | ASSPGTGDTLY | 0.71312106 |
| GFSNSGS | ASSPGTGDTLY | 0.7582055 |
| GGNGGSGGTSV | ASSPGTGDTLY | 0.68749297 |
| GTSVATGGAGN | ASSPGTGDTLY | 0.76964533 |
| NGGNGGSGGTS | ASSPGTGDTLY | 0.7708608 |
| AVVKFPH | ASSPGTGDTLY | 0.5976521 |
| TPPPPPP | ASSPGTGDTLY | 0.69075906 |
| AAPAAAAAPA | ASSPGTGDTLY | 0.69497454 |
| IALNEDL | ASSPGTGDTLY | 0.7144217 |
| IVNVSSM | ASSPGTGDTLY | 0.5402293 |
| TGIPPAPR | ASSPGTGDTLY | 0.757209 |
| FDLGGGTF | ASSPGTGDTLY | 0.6593573 |
| FDLGGGTFDVS | ASSPGTGDTLY | 0.27405393 |
| NEPTAAA | ASSPGTGDTLY | 0.7313802 |
| NEPTAAALA | ASSPGTGDTLY | 0.721796 |
| GSIINISSI | ASSPGTGDTLY | 0.70107085 |
| VAPGNAGTA | ASSPGTGDTLY | 0.75445473 |
| TGGGASLE | ASSPGTGDTLY | 0.43178824 |
| VAVYNHYKRI | ASSPGTGDTLY | 0.67844015 |
| TDPAPATT | ASSPGTGDTLY | 0.7578182 |
| TNRPDILDPALLRPG | ASSPGTGDTLY | 0.7675357 |
| LTTTTTS | ASSPGTGDTLY | 0.6192371 |
| PTGSGKTL | ASSPGTGDTLY | 0.73388815 |
| TGSGKTL | ASSPGTGDTLY | 0.7620959 |
| TIGHVDHGK | ASSPGTGDTLY | 0.7408028 |
| TIGHVDHGKTT | ASSPGTGDTLY | 0.7599961 |
| GHVDHGKTT | ASSPGTGDTLY | 0.76796854 |
| AQFEHTLL | ASSPGTGDTLY | 0.6880282 |
| IVNQDEL | ASSPGTGDTLY | 0.76257086 |
| TNGSQFF | ASSPGTGDTLY | 0.7672088 |
| TGKGIVE | ASSPGTGDTLY | 0.7175202 |
| LISNASDA | ASSPGTGDTLY | 0.7310049 |
| AELRTLGELTKIAKAHGAQ | ASSPGTGDTLY | 0.68875057 |
| APNAVAAAEPSW | ASSPGTGDTLY | 0.7015033 |
| APQVAGAATAVL | ASSPGTGDTLY | 0.42413792 |
| ARRDPAAAAAILPTDA | ASSPGTGDTLY | 0.73138106 |
| DAAARDVLAEAGLAEYF | ASSPGTGDTLY | 0.7605076 |
| DDQIVLSAKANLAKVEQ | ASSPGTGDTLY | 0.68062663 |
| DELLPDTGEDSDEDGH | ASSPGTGDTLY | 0.63532764 |
| ELPEAPANRATVPQD | ASSPGTGDTLY | 0.648229 |
| FGQSKIERVTGIG | ASSPGTGDTLY | 0.6938173 |

| | | |
|---|---|---|
| GAGGTGSTAGGAGGAGGAGGLY | ASSPGTGDTLY | 0.70503026 |
| GHGGTGAAGGNGGTGGAGGSGI | ASSPGTGDTLY | 0.4375479 |
| GNGGNGGAGGEGGPGGAGGAGA | ASSPGTGDTLY | 0.6793978 |
| GSTAGNGGNGGNATNGTIAGQP | ASSPGTGDTLY | 0.76417047 |
| GTGGDGGNGGNGGTGG | ASSPGTGDTLY | 0.7610415 |
| GVRRPAGIPADAPV | ASSPGTGDTLY | 0.46818265 |
| GYRQVTDAHLLTLARR | ASSPGTGDTLY | 0.76417434 |
| IAKMGQDGHDRG | ASSPGTGDTLY | 0.7463182 |
| KAEPSIEIQYT | ASSPGTGDTLY | 0.7646192 |
| KPVRIRMAADEDTVAVT | ASSPGTGDTLY | 0.7287376 |
| MPPRIAGMRLLVIKPEP | ASSPGTGDTLY | 0.66068864 |
| MKFYAVALFRSRGVPPTR | ASSPGTGDTLY | 0.7403364 |
| NAGGFGILGAVAHSP | ASSPGTGDTLY | 0.67468494 |
| NEGREITLAMAEM | ASSPGTGDTLY | 0.58174723 |
| NELVEWSHPLGHRWRWE | ASSPGTGDTLY | 0.75756836 |
| NLGAGGTGGNGGNPG | ASSPGTGDTLY | 0.7068155 |
| NTEWPKWETVMAADPVD | ASSPGTGDTLY | 0.6578134 |
| PASAAPAETQP | ASSPGTGDTLY | 0.60038817 |
| QQSSDGKNGVRAAEITGE | ASSPGTGDTLY | 0.7617239 |
| RAPDAVTTSWWKE | ASSPGTGDTLY | 0.59983504 |
| SGFGNVGAHTSGWFNQ | ASSPGTGDTLY | 0.65332687 |
| SGVAGAVGQAASVGGLKV | ASSPGTGDTLY | 0.58449626 |
| SNGGAGTGSAGGNGGT | ASSPGTGDTLY | 0.71411926 |
| TGGAGGEGGAAG | ASSPGTGDTLY | 0.7593583 |
| TGGDGGAAGAGGAPGHG | ASSPGTGDTLY | 0.7480651 |
| TMPPLLAAIANAGRHGV | ASSPGTGDTLY | 0.6771914 |
| TSSTAGNSGGA | ASSPGTGDTLY | 0.73689854 |
| VPATVPVGILC | ASSPGTGDTLY | 0.74152505 |
| VQRPDSGGVVLTGRIS | ASSPGTGDTLY | 0.7359787 |
| WTPNPVPATKGVPAGLVQAT | ASSPGTGDTLY | 0.75139713 |
| YALKRALAKFSDRFDVVII | ASSPGTGDTLY | 0.6290114 |
| YLIGSVIV | ASSPGTGDTLY | 0.76249266 |
| YGQLVAVID | ASSPGTGDTLY | 0.7386221 |
| VTVGGMTLPA | ASSPGTGDTLY | 0.7623564 |
| VLALPAV | ASSPGTGDTLY | 0.7597781 |
| VIPPTLNYET | ASSPGTGDTLY | 0.7315566 |
| VGMGGMHPSSG | ASSPGTGDTLY | 0.72373366 |
| VGLAGGAATA | ASSPGTGDTLY | 0.40817368 |
| VGGAGGAGGSGP | ASSPGTGDTLY | 0.7062993 |
| VFAAFMV | ASSPGTGDTLY | 0.7362163 |
| TLYDKLR | ASSPGTGDTLY | 0.7220999 |
| TLAVFTVL | ASSPGTGDTLY | 0.7508384 |
| TGSVQPARV | ASSPGTGDTLY | 0.7570177 |
| TGGGAGSGGT | ASSPGTGDTLY | 0.67149556 |
| SHLETSL | ASSPGTGDTLY | 0.76153964 |
| SFTTGMT | ASSPGTGDTLY | 0.7532513 |
| QVEVAKRMGVRQPTVS | ASSPGTGDTLY | 0.5465143 |
| QGSGGPALAM | ASSPGTGDTLY | 0.7012249 |
| MLAPSNP | ASSPGTGDTLY | 0.7720853 |
| LVNNAAI | ASSPGTGDTLY | 0.70845675 |
| LTGEINTLA | ASSPGTGDTLY | 0.70789814 |

| | | |
|---|---|---|
| LSIFGPV | ASSPGTGDTLY | 0.67884964 |
| LLALLANL | ASSPGTGDTLY | 0.76519066 |
| LISNASDALDKLR | ASSPGTGDTLY | 0.655972 |
| LINEAVP | ASSPGTGDTLY | 0.7697549 |
| LGYAFSGL | ASSPGTGDTLY | 0.3271348 |
| LDFVAVAL | ASSPGTGDTLY | 0.67766374 |
| LAQPGAE | ASSPGTGDTLY | 0.62078047 |
| ISNLGAVTGF | ASSPGTGDTLY | 0.5167496 |
| HQEDPGRI | ASSPGTGDTLY | 0.7608924 |
| GTPGEPTP | ASSPGTGDTLY | 0.723691 |
| GFFNSGS | ASSPGTGDTLY | 0.751847 |
| GAGGGGLTAGAN | ASSPGTGDTLY | 0.72971416 |
| FVDALVNM | ASSPGTGDTLY | 0.7614692 |
| FLTVAPEL | ASSPGTGDTLY | 0.71579945 |
| FETLTAL | ASSPGTGDTLY | 0.74162585 |
| DVINPIEL | ASSPGTGDTLY | 0.6600986 |
| APVSGAPV | ASSPGTGDTLY | 0.7580016 |
| ALVGGLAW | ASSPGTGDTLY | 0.76955986 |
| ALAAAAATAI | ASSPGTGDTLY | 0.74602234 |
| AFGLGLLL | ASSPGTGDTLY | 0.6126238 |
| AFFAAASL | ASSPGTGDTLY | 0.6434908 |
| AAAALPAFSPP | ASSPGTGDTLY | 0.73872894 |
| VVLAPAL | ASSPGTGDTLY | 0.34062198 |
| TIGHVDHGKTTL | ASSPGTGDTLY | 0.35405 |
| LISNASDALDK | ASSPGTGDTLY | 0.68789554 |
| LISNASDAL | ASSPGTGDTLY | 0.6128566 |
| ISNASDALDK | ASSPGTGDTLY | 0.76419985 |
| NQAETLVYQTE | ASSPGTGDTLY | 0.7556005 |
| TGGTGGTAGAG | ASSPGTGDTLY | 0.70085406 |
| TGSTAGGAGGA | ASSPGTGDTLY | 0.74804825 |
| DPLAVPVAAGPV | ASSPGTGDTLY | 0.7205502 |
| GGTGGTPGNGGN | ASSPGTGDTLY | 0.22384866 |
| PAGLAGVDPATL | ASSPGTGDTLY | 0.7085143 |
| GNNLVVEAPHAGASV | ASSPGTGDTLY | 0.61997336 |
| KVTRIMSASTLEQPA | ASSPGTGDTLY | 0.7742339 |
| LGFALPIALPRRTVV | ASSPGTGDTLY | 0.6766771 |
| TRLGNLTARKLVVLV | ASSPGTGDTLY | 0.63093036 |
| WSQQRRLIYFPSAGP | ASSPGTGDTLY | 0.70800394 |
| GPNGNPDPMAAAVDIR | ASSPGTGDTLY | 0.69759625 |
| TQRCVSVADLRAAASR | ASSPGTGDTLY | 0.69816977 |
| QFVLDVAEHPVEVLTETV | ASSPGTGDTLY | 0.62616825 |
| GGAGGTGGAGGNGGTQAGDG | ASSPGTGDTLY | 0.2588285 |
| PLLATLRAIAVDVDAKPAQV | ASSPGTGDTLY | 0.76048654 |
| APKFRTGGYMIDFWGVGYQVAK | ASSPGTGDTLY | 0.51549256 |
| GAGGAGGAGGAAGGPGGTGGTG | ASSPGTGDTLY | 0.6604565 |
| IGALGLLATDVAPAGSVAFGVA | ASSPGTGDTLY | 0.42399794 |
| AGGAGGIGGNGGAGGAGGVLMG | ASSPGTGDTLY | 0.28311 |
| ADELIGLL | ASSPGTGDTLY | 0.5677948 |
| ADRALKA | ASSPGTGDTLY | 0.75999486 |
| AGAVGQVGGAGGA | ASSPGTGDTLY | 0.6861805 |
| AGGGGGLGGAP | ASSPGTGDTLY | 0.66907096 |

| | | |
|---|---|---|
| AGGLLYGNGGAG | ASSPGTGDTLY | 0.771594 |
| AGLESIGL | ASSPGTGDTLY | 0.69983965 |
| AGLKTGII | ASSPGTGDTLY | 0.66758406 |
| ASLAALLL | ASSPGTGDTLY | 0.675829 |
| ASVINISSI | ASSPGTGDTLY | 0.6528857 |
| ATPPGPA | ASSPGTGDTLY | 0.6624198 |
| ATVASALV | ASSPGTGDTLY | 0.66240996 |
| ATVASLIL | ASSPGTGDTLY | 0.7741074 |
| AVNGPVLIRH | ASSPGTGDTLY | 0.6920518 |
| DGAAGAVGI | ASSPGTGDTLY | 0.6510928 |
| DPLLHGI | ASSPGTGDTLY | 0.71422887 |
| ELGAALGAL | ASSPGTGDTLY | 0.76801085 |
| ELGRTQAV | ASSPGTGDTLY | 0.77491903 |
| FVAGGYLS | ASSPGTGDTLY | 0.64869803 |
| GASGGAGGAGGW | ASSPGTGDTLY | 0.7077114 |
| GGAGAGGDGGAGGI | ASSPGTGDTLY | 0.697348 |
| GTGHNGSQPGMGGNG | ASSPGTGDTLY | 0.71267194 |
| GYAAASASAS | ASSPGTGDTLY | 0.6900828 |
| HQVDVAAVQQG | ASSPGTGDTLY | 0.5968192 |
| IGAAIAHT | ASSPGTGDTLY | 0.7500977 |
| IGMIEVAQL | ASSPGTGDTLY | 0.77484155 |
| IRLAPEHTSAMD | ASSPGTGDTLY | 0.7023196 |
| IYNVSVL | ASSPGTGDTLY | 0.61447287 |
| KQGFLGM | ASSPGTGDTLY | 0.7013128 |
| LESVITQL | ASSPGTGDTLY | 0.6379467 |
| LGLGALVVV | ASSPGTGDTLY | 0.67215323 |
| LIGNGGNGGSGGI | ASSPGTGDTLY | 0.7401398 |
| LNAINAPTL | ASSPGTGDTLY | 0.5429865 |
| LQLLNTGV | ASSPGTGDTLY | 0.6857451 |
| LQTSIEF | ASSPGTGDTLY | 0.77307206 |
| MSFAVSEL | ASSPGTGDTLY | 0.677641 |
| NTGALGL | ASSPGTGDTLY | 0.721094 |
| PPGAGGAGGAGTT | ASSPGTGDTLY | 0.7580119 |
| QLVGLGI | ASSPGTGDTLY | 0.736721 |
| RFLEQVHQL | ASSPGTGDTLY | 0.762785 |
| RGVGYVLR | ASSPGTGDTLY | 0.7538177 |
| SATGVFAGVI | ASSPGTGDTLY | 0.73024714 |
| SLPAITLP | ASSPGTGDTLY | 0.71720344 |
| STALAFTV | ASSPGTGDTLY | 0.71111494 |
| SVLTVGLGK | ASSPGTGDTLY | 0.72015506 |
| TALAGAGSGI | ASSPGTGDTLY | 0.6885185 |
| TGLPFTGI | ASSPGTGDTLY | 0.6553046 |
| VAEHVTV | ASSPGTGDTLY | 0.76484436 |
| VDIVLNSL | ASSPGTGDTLY | 0.7126544 |
| VLTGGGAVL | ASSPGTGDTLY | 0.74913573 |
| VNVGLAYGI | ASSPGTGDTLY | 0.6470262 |
| VSSAAVGAV | ASSPGTGDTLY | 0.61812437 |
| VTSVYEAGV | ASSPGTGDTLY | 0.748365 |
| WVALDRGA | ASSPGTGDTLY | 0.24573591 |
| YLVGSVVI | ASSPGTGDTLY | 0.34244362 |
| GPPGTGKTLL | ASSPGTGDTLY | 0.72805107 |

| | | |
|---|---|---|
| GVGMTKF | ASSPGTGDTLY | 0.763093 |
| IGIGGSDL | ASSPGTGDTLY | 0.25804937 |
| IGTLLAEL | ASSPGTGDTLY | 0.6533191 |
| IYTSGTT | ASSPGTGDTLY | 0.5735521 |
| LSFPAGV | ASSPGTGDTLY | 0.7494683 |
| LYTSGTT | ASSPGTGDTLY | 0.7253041 |
| VGAFPFGF | ASSPGTGDTLY | 0.55074084 |
| VGAGTVEF | ASSPGTGDTLY | 0.74777603 |
| VGKSTLLL | ASSPGTGDTLY | 0.5212151 |
| AALAHAGGGIV | ASSPGTGDTLY | 0.45363265 |
| AALVAVSLL | ASSPGTGDTLY | 0.7620876 |
| AAVSRAVAAL | ASSPGTGDTLY | 0.7592986 |
| AGGSSGIGGF | ASSPGTGDTLY | 0.6576194 |
| AGLTVANANG | ASSPGTGDTLY | 0.7553601 |
| AGLVGAVL | ASSPGTGDTLY | 0.6659766 |
| ALSQAFTV | ASSPGTGDTLY | 0.7739792 |
| ALTFSVSL | ASSPGTGDTLY | 0.69260895 |
| ALTGAALVAL | ASSPGTGDTLY | 0.5547011 |
| ASFSIGAGVL | ASSPGTGDTLY | 0.7514252 |
| DGSVYKI | ASSPGTGDTLY | 0.7314831 |
| EQLAEIV | ASSPGTGDTLY | 0.74277467 |
| FAVSELTVM | ASSPGTGDTLY | 0.74494135 |
| GAAPLAAGL | ASSPGTGDTLY | 0.6536284 |
| GADGGDGGSGGAGG | ASSPGTGDTLY | 0.6853881 |
| GFGLDTGGAGG | ASSPGTGDTLY | 0.76699114 |
| GLASGGLL | ASSPGTGDTLY | 0.7590914 |
| GRIAGAAVLL | ASSPGTGDTLY | 0.63111776 |
| GSAGSLNTSF | ASSPGTGDTLY | 0.7565583 |
| GTGQIGI | ASSPGTGDTLY | 0.6938554 |
| GTLIALIL | ASSPGTGDTLY | 0.675435 |
| IAATMGFVVG | ASSPGTGDTLY | 0.77102727 |
| IFAQALL | ASSPGTGDTLY | 0.7402998 |
| IGAAVALGF | ASSPGTGDTLY | 0.7492806 |
| IGLTPLAL | ASSPGTGDTLY | 0.75850934 |
| ILGSNTSTL | ASSPGTGDTLY | 0.7418493 |
| IVFSLTVL | ASSPGTGDTLY | 0.58093905 |
| LASKATGF | ASSPGTGDTLY | 0.74001247 |
| LASVALIL | ASSPGTGDTLY | 0.7122442 |
| LFAVGLTL | ASSPGTGDTLY | 0.7541681 |
| LFNGGIGTYI | ASSPGTGDTLY | 0.7277323 |
| LTLGAGII | ASSPGTGDTLY | 0.5562113 |
| LVAAALGDI | ASSPGTGDTLY | 0.7368357 |
| NTAFVQLGI | ASSPGTGDTLY | 0.76285005 |
| QLFSGAEL | ASSPGTGDTLY | 0.7343216 |
| SAATFITL | ASSPGTGDTLY | 0.73811656 |
| SNYGLANLGV | ASSPGTGDTLY | 0.76752377 |
| SYGGAYAVM | ASSPGTGDTLY | 0.6932901 |
| TTGVGGLTLG | ASSPGTGDTLY | 0.70269036 |
| TTQTIYI | ASSPGTGDTLY | 0.756324 |
| VAAGAGLGAVI | ASSPGTGDTLY | 0.7622649 |
| VAAVVTLL | ASSPGTGDTLY | 0.71373594 |

| | | |
|---|---|---|
| VGFLGGLI | ASSPGTGDTLY | 0.76514035 |
| VGVIPILL | ASSPGTGDTLY | 0.62925184 |
| VIFASSLI | ASSPGTGDTLY | 0.7545948 |
| VKGGDGGAAAG | ASSPGTGDTLY | 0.62113446 |
| VTKTFTNI | ASSPGTGDTLY | 0.74728924 |
| VVAAAILL | ASSPGTGDTLY | 0.6327454 |
| YAALDVEL | ASSPGTGDTLY | 0.7559549 |
| LASVALLL | ASSPGTGDTLY | 0.7538278 |
| LDAVHRV | ASSPGTGDTLY | 0.75482976 |
| QGLSSSIF | ASSPGTGDTLY | 0.37183186 |
| RGTFANIRL | ASSPGTGDTLY | 0.75254625 |
| RQLWWGHRI | ASSPGTGDTLY | 0.75758266 |
| YGPPGTGKTLL | ASSPGTGDTLY | 0.774806 |
| AALVGAPANVA | ASSPGTGDTLY | 0.7082084 |
| AATFRRTE | ASSPGTGDTLY | 0.7431701 |
| AGALGGGI | ASSPGTGDTLY | 0.711618 |
| AGGAGGAGGIGAG | ASSPGTGDTLY | 0.6998385 |
| AGSGALGL | ASSPGTGDTLY | 0.6143523 |
| AQIADQI | ASSPGTGDTLY | 0.69261765 |
| ASAVAVTYL | ASSPGTGDTLY | 0.6697834 |
| ASTVLGGTGG | ASSPGTGDTLY | 0.6350361 |
| AVGGEPAK | ASSPGTGDTLY | 0.74352086 |
| CEQGTFGGTC | ASSPGTGDTLY | 0.755571 |
| DEGKAPAG | ASSPGTGDTLY | 0.75825393 |
| DTVVFPAL | ASSPGTGDTLY | 0.7642417 |
| FEGITMAG | ASSPGTGDTLY | 0.7595245 |
| FPLVVQD | ASSPGTGDTLY | 0.76085055 |
| GAFTTPQ | ASSPGTGDTLY | 0.66096944 |
| GFGGFGTAGGVGGA | ASSPGTGDTLY | 0.7241986 |
| GFLAQFL | ASSPGTGDTLY | 0.6192875 |
| GGAGGAGGNAGVA | ASSPGTGDTLY | 0.7105362 |
| GGVGTAGAGGAGGAGG | ASSPGTGDTLY | 0.6772495 |
| GNGGGGDAGNAGSGGN | ASSPGTGDTLY | 0.7328031 |
| GTGGAGGAGNDAGS | ASSPGTGDTLY | 0.23293665 |
| GTTNTGWFNS | ASSPGTGDTLY | 0.69816434 |
| IFMGPLFTL | ASSPGTGDTLY | 0.7707956 |
| ILAHAIEAI | ASSPGTGDTLY | 0.5951619 |
| IRSAIREAL | ASSPGTGDTLY | 0.73481154 |
| ITFAGGG | ASSPGTGDTLY | 0.69666886 |
| ITGMSGHQEV | ASSPGTGDTLY | 0.66812533 |
| IVAGLACTL | ASSPGTGDTLY | 0.7539334 |
| IVGGGLAGA | ASSPGTGDTLY | 0.64183503 |
| IVTGVLNI | ASSPGTGDTLY | 0.73607594 |
| LAQQGCI | ASSPGTGDTLY | 0.7674428 |
| LCLHVEN | ASSPGTGDTLY | 0.7210756 |
| LDAISDGS | ASSPGTGDTLY | 0.7356985 |
| LGGFAAML | ASSPGTGDTLY | 0.7122485 |
| LGGKNPMI | ASSPGTGDTLY | 0.5313589 |
| LGPKGVAAIL | ASSPGTGDTLY | 0.74427915 |
| LMVGVSI | ASSPGTGDTLY | 0.22056043 |
| LVGAAFVL | ASSPGTGDTLY | 0.70187384 |

| | | |
|---|---|---|
| LVLAVAPL | ASSPGTGDTLY | 0.6562819 |
| MAAIAVIDAA | ASSPGTGDTLY | 0.7330353 |
| NNGINAELQ | ASSPGTGDTLY | 0.7176286 |
| RGEPLVLVHG | ASSPGTGDTLY | 0.7410927 |
| RGGAGGAAGLI | ASSPGTGDTLY | 0.6459669 |
| RSYRMHPAV | ASSPGTGDTLY | 0.66332746 |
| SAAHLLAGQLA | ASSPGTGDTLY | 0.5978223 |
| SAAVGSY | ASSPGTGDTLY | 0.6240741 |
| SPHAVLHRI | ASSPGTGDTLY | 0.7467568 |
| SQIAKVL | ASSPGTGDTLY | 0.72534883 |
| TALGHLVP | ASSPGTGDTLY | 0.67994094 |
| TAPTTSGAA | ASSPGTGDTLY | 0.76714194 |
| TDGDDGNGGSPGT | ASSPGTGDTLY | 0.7294841 |
| TDGLGEN | ASSPGTGDTLY | 0.5615192 |
| TGGPPLPGQA | ASSPGTGDTLY | 0.69386446 |
| TIPAITI | ASSPGTGDTLY | 0.7105675 |
| TIPGFTI | ASSPGTGDTLY | 0.69794166 |
| TSIASGALA | ASSPGTGDTLY | 0.22191347 |
| TVTLPGAAI | ASSPGTGDTLY | 0.73680973 |
| VASGLSAALP | ASSPGTGDTLY | 0.67850804 |
| VGAGVIGI | ASSPGTGDTLY | 0.61929643 |
| VGPKRFGV | ASSPGTGDTLY | 0.6693126 |
| VNVAFPD | ASSPGTGDTLY | 0.7455984 |
| VSMTNSGVS | ASSPGTGDTLY | 0.4602563 |
| VTDVLGDLSV | ASSPGTGDTLY | 0.7249745 |
| VVVVGAGFAGL | ASSPGTGDTLY | 0.7506339 |
| VYTVGTVV | ASSPGTGDTLY | 0.7617123 |
| WTTFSGP | ASSPGTGDTLY | 0.76830673 |
| YDELVEAF | ASSPGTGDTLY | 0.74438816 |
| YGTGGAGGNGG | ASSPGTGDTLY | 0.689167 |
| YTNVMAV | ASSPGTGDTLY | 0.7547241 |
| FLEQVHQL | ASSPGTGDTLY | 0.7375939 |
| GLSLLAVAV | ASSPGTGDTLY | 0.46052676 |
| LEQVHQL | ASSPGTGDTLY | 0.38296336 |
| LNAAGAI | ASSPGTGDTLY | 0.73026615 |
| PAVHASL | ASSPGTGDTLY | 0.39608225 |
| TELEDDA | ASSPGTGDTLY | 0.72374135 |
| TPPGPPN | ASSPGTGDTLY | 0.77044636 |
| TPPPPPPP | ASSPGTGDTLY | 0.7102672 |
| TQSPALI | ASSQERTNYAEQF | 0.6340737 |
| EFAAAFPQ | ASSQERTNYAEQF | 0.6963499 |
| LLSGNLDVL | ASSQERTNYAEQF | 0.71273756 |
| GPANLGNYNI | ASSQERTNYAEQF | 0.74778926 |
| TLGPTLP | ASSQERTNYAEQF | 0.75264704 |
| VYAYSSCCK | ASSQERTNYAEQF | 0.76982516 |
| APAPAPAPNI | ASSQERTNYAEQF | 0.7581579 |
| PGGPVPIP | ASSQERTNYAEQF | 0.6662229 |
| FLTVALEP | ASSQERTNYAEQF | 0.73765385 |
| SQYFGANL | ASSQERTNYAEQF | 0.68826735 |
| GGVKPAA | ASSQERTNYAEQF | 0.70337987 |
| GGTGGAGGAGGW | ASSQERTNYAEQF | 0.7246173 |

| | | |
|---|---|---|
| GVGGVAGLGGAGG | ASSQERTNYAEQF | 0.72989535 |
| VLLVLATTI | ASSQERTNYAEQF | 0.71796703 |
| IGIFAAML | ASSQERTNYAEQF | 0.6895328 |
| AVGDLID | ASSQERTNYAEQF | 0.56341827 |
| NISPLPL | ASSQERTNYAEQF | 0.7446861 |
| VTAVFVAI | ASSQERTNYAEQF | 0.7230626 |
| SGDDNDAAL | ASSQERTNYAEQF | 0.4613352 |
| AVAALAHS | ASSQERTNYAEQF | 0.67615896 |
| GGVTVMATP | ASSQERTNYAEQF | 0.77292687 |
| AGLAANVRD | ASSQERTNYAEQF | 0.7667624 |
| TNRPDLIDPALLRPG | ASSQERTNYAEQF | 0.6241368 |
| LANLGVTAL | ASSQERTNYAEQF | 0.6665156 |
| TGERIEL | ASSQERTNYAEQF | 0.69997615 |
| NGEELAA | ASSQERTNYAEQF | 0.4116798 |
| VLSLVWG | ASSQERTNYAEQF | 0.73873174 |
| IGYALWL | ASSQERTNYAEQF | 0.69982505 |
| LSLLLAPL | ASSQERTNYAEQF | 0.6963886 |
| TLDEALK | ASSQERTNYAEQF | 0.6752225 |
| TDGAGTAEGG | ASSQERTNYAEQF | 0.70273125 |
| NAPGGAGGNGG | ASSQERTNYAEQF | 0.6617236 |
| VSVFMGMGGNF | ASSQERTNYAEQF | 0.7561445 |
| LASYLAAL | ASSQERTNYAEQF | 0.25265345 |
| LVGGTPAL | ASSQERTNYAEQF | 0.7494776 |
| VRNADVL | ASSQERTNYAEQF | 0.62610173 |
| TCSSPFLVSAM | ASSQERTNYAEQF | 0.66055876 |
| VHITDAE | ASSQERTNYAEQF | 0.51028675 |
| EYAAPSF | ASSQERTNYAEQF | 0.72153217 |
| VVDAFHGGL | ASSQERTNYAEQF | 0.7568011 |
| ASVVDPQ | ASSQERTNYAEQF | 0.74868953 |
| SYNEFGGLS | ASSQERTNYAEQF | 0.5467694 |
| DPDGSVY | ASSQERTNYAEQF | 0.76409316 |
| MVSTIHATE | ASSQERTNYAEQF | 0.77144 |
| LAAGVADPV | ASSQERTNYAEQF | 0.6747507 |
| TKEQIAA | ASSQERTNYAEQF | 0.5304834 |
| SLASPAAL | ASSQERTNYAEQF | 0.7480131 |
| IGLVFYGV | ASSQERTNYAEQF | 0.6870893 |
| FLLDASNL | ASSQERTNYAEQF | 0.74792135 |
| YQVGNFP | ASSQERTNYAEQF | 0.73796123 |
| ESLLSQST | ASSQERTNYAEQF | 0.5947397 |
| LFTVGFAAVG | ASSQERTNYAEQF | 0.7682433 |
| ATPPTPT | ASSQERTNYAEQF | 0.761148 |
| TDLATLG | ASSQERTNYAEQF | 0.638616 |
| SAGGTGR | ASSQERTNYAEQF | 0.69042367 |
| GFSNSGS | ASSQERTNYAEQF | 0.69974124 |
| GGNGGSGGTSV | ASSQERTNYAEQF | 0.41711366 |
| GTSVATGGAGN | ASSQERTNYAEQF | 0.7633179 |
| NGGNGGSGGTS | ASSQERTNYAEQF | 0.6362037 |
| AVVKFPH | ASSQERTNYAEQF | 0.67512447 |
| TPPPPPP | ASSQERTNYAEQF | 0.72542703 |
| AAPAAAAAPA | ASSQERTNYAEQF | 0.68917567 |
| IALNEDL | ASSQERTNYAEQF | 0.68691295 |

| | | |
|---|---|---|
| IVNVSSM | ASSQERTNYAEQF | 0.6911915 |
| TGIPPAPR | ASSQERTNYAEQF | 0.73601806 |
| FDLGGGTF | ASSQERTNYAEQF | 0.7181778 |
| FDLGGGTFDVS | ASSQERTNYAEQF | 0.72730106 |
| NEPTAAA | ASSQERTNYAEQF | 0.72381395 |
| NEPTAAALA | ASSQERTNYAEQF | 0.7652937 |
| GSIINISSI | ASSQERTNYAEQF | 0.6005748 |
| VAPGNAGTA | ASSQERTNYAEQF | 0.6644532 |
| TGGGASLE | ASSQERTNYAEQF | 0.7273697 |
| VAVYNHYKRI | ASSQERTNYAEQF | 0.4970309 |
| TDPAPATT | ASSQERTNYAEQF | 0.7457238 |
| TNRPDILDPALLRPG | ASSQERTNYAEQF | 0.71777207 |
| LTTTTTS | ASSQERTNYAEQF | 0.7282852 |
| PTGSGKTL | ASSQERTNYAEQF | 0.7020645 |
| TGSGKTL | ASSQERTNYAEQF | 0.77538186 |
| TIGHVDHGK | ASSQERTNYAEQF | 0.75584686 |
| TIGHVDHGKTT | ASSQERTNYAEQF | 0.7346381 |
| GHVDHGKTT | ASSQERTNYAEQF | 0.7101016 |
| AQFEHTLL | ASSQERTNYAEQF | 0.672807 |
| IVNQDEL | ASSQERTNYAEQF | 0.76474965 |
| TNGSQFF | ASSQERTNYAEQF | 0.7417586 |
| TGKGIVE | ASSQERTNYAEQF | 0.5890997 |
| LISNASDA | ASSQERTNYAEQF | 0.69285905 |
| AELRTLGELTKIAKAHGAQ | ASSQERTNYAEQF | 0.7640016 |
| APNAVAAAEPSW | ASSQERTNYAEQF | 0.75539935 |
| APQVAGAATAVL | ASSQERTNYAEQF | 0.43823704 |
| ARRDPAAAAAILPTDA | ASSQERTNYAEQF | 0.693025 |
| DAAARDVLAEAGLAEYF | ASSQERTNYAEQF | 0.4180366 |
| DDQIVLSAKANLAKVEQ | ASSQERTNYAEQF | 0.6308859 |
| DELLPDTGEDSDEDGH | ASSQERTNYAEQF | 0.6659166 |
| ELPEAPANRATVPQD | ASSQERTNYAEQF | 0.7382969 |
| FGQSKIERVTGIG | ASSQERTNYAEQF | 0.6848819 |
| GAGGTGSTAGGAGGAGGAGGLY | ASSQERTNYAEQF | 0.7230171 |
| GHGGTGAAGGNGGTGGAGGSGI | ASSQERTNYAEQF | 0.72369874 |
| GNGGNGGAGGEGGPGGAGGAGA | ASSQERTNYAEQF | 0.767369 |
| GSTAGNGGNGGNATNGTIAGQP | ASSQERTNYAEQF | 0.66295576 |
| GTGGDGGNGGNGGTGG | ASSQERTNYAEQF | 0.7198258 |
| GVRRPAGIPADAPV | ASSQERTNYAEQF | 0.55906165 |
| GYRQVTDAHLLTLARR | ASSQERTNYAEQF | 0.53541034 |
| IAKMGQDGHDRG | ASSQERTNYAEQF | 0.67637205 |
| KAEPSIEIQYT | ASSQERTNYAEQF | 0.69860125 |
| KPVRIRMAADEDTVAVT | ASSQERTNYAEQF | 0.3429155 |
| MPPRIAGMRLLVIKPEP | ASSQERTNYAEQF | 0.7608298 |
| MKFYAVALFRSRGVPPTR | ASSQERTNYAEQF | 0.7340442 |
| NAGGFGILGAVAHSP | ASSQERTNYAEQF | 0.7507074 |
| NEGREITLAMAEM | ASSQERTNYAEQF | 0.7103185 |
| NELVEWSHPLGHRWRWE | ASSQERTNYAEQF | 0.7211571 |
| NLGAGGTGGNGGNPG | ASSQERTNYAEQF | 0.65427047 |
| NTEWPKWETVMAADPVD | ASSQERTNYAEQF | 0.7422988 |
| PASAAPAETQP | ASSQERTNYAEQF | 0.6615319 |
| QQSSDGKNGVRAAEITGE | ASSQERTNYAEQF | 0.5764635 |

| | | |
|---|---|---|
| RAPDAVTTSWWKE | ASSQERTNYAEQF | 0.69378847 |
| SGFGNVGAHTSGWFNQ | ASSQERTNYAEQF | 0.6208987 |
| SGVAGAVGQAASVGGLKV | ASSQERTNYAEQF | 0.759271 |
| SNGGAGTGSAGGNGGT | ASSQERTNYAEQF | 0.7574968 |
| TGGAGGEGGAAG | ASSQERTNYAEQF | 0.73271495 |
| TGGDGGAAGAGGAPGHG | ASSQERTNYAEQF | 0.7531526 |
| TMPPLLAAIANAGRHGV | ASSQERTNYAEQF | 0.4203294 |
| TSSTAGNSGGA | ASSQERTNYAEQF | 0.7194208 |
| VPATVPVGILC | ASSQERTNYAEQF | 0.7717551 |
| VQRPDSGGVVLTGRIS | ASSQERTNYAEQF | 0.7379657 |
| WTPNPVPATKGVPAGLVQAT | ASSQERTNYAEQF | 0.69798183 |
| YALKRALAKFSDRFDVVII | ASSQERTNYAEQF | 0.76337105 |
| YLIGSVIV | ASSQERTNYAEQF | 0.6533319 |
| YGQLVAVID | ASSQERTNYAEQF | 0.7027398 |
| VTVGGMTLPA | ASSQERTNYAEQF | 0.7127428 |
| VLALPAV | ASSQERTNYAEQF | 0.75016886 |
| VIPPTLNYET | ASSQERTNYAEQF | 0.7332055 |
| VGMGGMHPSSG | ASSQERTNYAEQF | 0.6724843 |
| VGLAGGAATA | ASSQERTNYAEQF | 0.73937774 |
| VGGAGGAGGSGP | ASSQERTNYAEQF | 0.7734407 |
| VFAAFMV | ASSQERTNYAEQF | 0.688293 |
| TLYDKLR | ASSQERTNYAEQF | 0.7575717 |
| TLAVFTVL | ASSQERTNYAEQF | 0.7411405 |
| TGSVQPARV | ASSQERTNYAEQF | 0.73725647 |
| TGGGAGSGGT | ASSQERTNYAEQF | 0.6444466 |
| SHLETSL | ASSQERTNYAEQF | 0.67224634 |
| SFTTGMT | ASSQERTNYAEQF | 0.67896533 |
| QVEVAKRMGVRQPTVS | ASSQERTNYAEQF | 0.6248963 |
| QGSGGPALAM | ASSQERTNYAEQF | 0.70861745 |
| MLAPSNP | ASSQERTNYAEQF | 0.70670044 |
| LVNNAAI | ASSQERTNYAEQF | 0.7779152 |
| LTGEINTLA | ASSQERTNYAEQF | 0.73597026 |
| LSIFGPV | ASSQERTNYAEQF | 0.73164964 |
| LLALLANL | ASSQERTNYAEQF | 0.62847924 |
| LISNASDALDKLR | ASSQERTNYAEQF | 0.73751426 |
| LINEAVP | ASSQERTNYAEQF | 0.73688793 |
| LGYAFSGL | ASSQERTNYAEQF | 0.59794146 |
| LDFVAVAL | ASSQERTNYAEQF | 0.76967704 |
| LAQPGAE | ASSQERTNYAEQF | 0.65859884 |
| ISNLGAVTGF | ASSQERTNYAEQF | 0.6509366 |
| HQEDPGRI | ASSQERTNYAEQF | 0.7525755 |
| GTPGEPTP | ASSQERTNYAEQF | 0.7655714 |
| GFFNSGS | ASSQERTNYAEQF | 0.6888562 |
| GAGGGGLTAGAN | ASSQERTNYAEQF | 0.64507437 |
| FVDALVNM | ASSQERTNYAEQF | 0.6439365 |
| FLTVAPEL | ASSQERTNYAEQF | 0.75490785 |
| FETLTAL | ASSQERTNYAEQF | 0.7591928 |
| DVINPIEL | ASSQERTNYAEQF | 0.67091596 |
| APVSGAPV | ASSQERTNYAEQF | 0.7676382 |
| ALVGGLAW | ASSQERTNYAEQF | 0.75038123 |
| ALAAAAATAI | ASSQERTNYAEQF | 0.556684 |

| | | |
|---|---|---|
| AFGLGLLL | ASSQERTNYAEQF | 0.71894455 |
| AFFAAASL | ASSQERTNYAEQF | 0.75754964 |
| AAAALPAFSPP | ASSQERTNYAEQF | 0.74537545 |
| VVLAPAL | ASSQERTNYAEQF | 0.7771114 |
| TIGHVDHGKTTL | ASSQERTNYAEQF | 0.74820083 |
| LISNASDALDK | ASSQERTNYAEQF | 0.66836107 |
| LISNASDAL | ASSQERTNYAEQF | 0.7662629 |
| ISNASDALDK | ASSQERTNYAEQF | 0.7743236 |
| NQAETLVYQTE | ASSQERTNYAEQF | 0.75761443 |
| TGGTGGTAGAG | ASSQERTNYAEQF | 0.6863158 |
| TGSTAGGAGGA | ASSQERTNYAEQF | 0.6532872 |
| DPLAVPVAAGPV | ASSQERTNYAEQF | 0.7180811 |
| GGTGGTPGNGGN | ASSQERTNYAEQF | 0.74528056 |
| PAGLAGVDPATL | ASSQERTNYAEQF | 0.6412791 |
| GNNLVVEAPHAGASV | ASSQERTNYAEQF | 0.7181935 |
| KVTRIMSASTLEQPA | ASSQERTNYAEQF | 0.7286691 |
| LGFALPIALPRRTVV | ASSQERTNYAEQF | 0.59701055 |
| TRLGNLTARKLVVLV | ASSQERTNYAEQF | 0.7566875 |
| WSQQRRLIYFPSAGP | ASSQERTNYAEQF | 0.7640505 |
| GPNGNPDPMAAAVDIR | ASSQERTNYAEQF | 0.70430374 |
| TQRCVSVADLRAAASR | ASSQERTNYAEQF | 0.69224966 |
| QFVLDVAEHPVEVLTETV | ASSQERTNYAEQF | 0.75971043 |
| GGAGGTGGAGGNGGTQAGDG | ASSQERTNYAEQF | 0.7792819 |
| PLLATLRAIAVDVDAKPAQV | ASSQERTNYAEQF | 0.76104325 |
| APKFRTGGYMIDFWGVGYQVAK | ASSQERTNYAEQF | 0.5582205 |
| GAGGAGGAGGAAGGPGGTGGTG | ASSQERTNYAEQF | 0.4143659 |
| IGALGLLATDVAPAGSVAFGVA | ASSQERTNYAEQF | 0.62600017 |
| AGGAGGIGGNGGAGGAGGVLMG | ASSQERTNYAEQF | 0.735427 |
| ADELIGLL | ASSQERTNYAEQF | 0.7025517 |
| ADRALKA | ASSQERTNYAEQF | 0.69013464 |
| AGAVGQVGGAGGA | ASSQERTNYAEQF | 0.7313808 |
| AGGGGGLGGAP | ASSQERTNYAEQF | 0.764014 |
| AGGLLYGNGGAG | ASSQERTNYAEQF | 0.6905273 |
| AGLESIGL | ASSQERTNYAEQF | 0.5517901 |
| AGLKTGII | ASSQERTNYAEQF | 0.7350301 |
| ASLAALLL | ASSQERTNYAEQF | 0.744602 |
| ASVINISSI | ASSQERTNYAEQF | 0.639669 |
| ATPPGPA | ASSQERTNYAEQF | 0.68639576 |
| ATVASALV | ASSQERTNYAEQF | 0.6230961 |
| ATVASLIL | ASSQERTNYAEQF | 0.45927557 |
| AVNGPVLIRH | ASSQERTNYAEQF | 0.7243824 |
| DGAAGAVGI | ASSQERTNYAEQF | 0.62558997 |
| DPLLHGI | ASSQERTNYAEQF | 0.76609623 |
| ELGAALGAL | ASSQERTNYAEQF | 0.6399685 |
| ELGRTQAV | ASSQERTNYAEQF | 0.7487354 |
| FVAGGYLS | ASSQERTNYAEQF | 0.7358345 |
| GASGGAGGAGGW | ASSQERTNYAEQF | 0.7495314 |
| GGAGAGGDGGAGGI | ASSQERTNYAEQF | 0.6851455 |
| GTGHNGSQPGMGGNG | ASSQERTNYAEQF | 0.6805014 |
| GYAAASASAS | ASSQERTNYAEQF | 0.61904097 |
| HQVDVAAVQQG | ASSQERTNYAEQF | 0.4089367 |

| | | |
|---|---|---|
| IGAAIAHT | ASSQERTNYAEQF | 0.76217186 |
| IGMIEVAQL | ASSQERTNYAEQF | 0.26397946 |
| IRLAPEHTSAMD | ASSQERTNYAEQF | 0.7631097 |
| IYNVSVL | ASSQERTNYAEQF | 0.75390697 |
| KQGFLGM | ASSQERTNYAEQF | 0.650389 |
| LESVITQL | ASSQERTNYAEQF | 0.7364143 |
| LGLGALVVV | ASSQERTNYAEQF | 0.63197994 |
| LIGNGGNGGSGGI | ASSQERTNYAEQF | 0.6060883 |
| LNAINAPTL | ASSQERTNYAEQF | 0.7518614 |
| LQLLNTGV | ASSQERTNYAEQF | 0.7484829 |
| LQTSIEF | ASSQERTNYAEQF | 0.6543404 |
| MSFAVSEL | ASSQERTNYAEQF | 0.7340572 |
| NTGALGL | ASSQERTNYAEQF | 0.72356117 |
| PPGAGGAGGAGTT | ASSQERTNYAEQF | 0.67628527 |
| QLVGLGI | ASSQERTNYAEQF | 0.72115487 |
| RFLEQVHQL | ASSQERTNYAEQF | 0.75918293 |
| RGVGYVLR | ASSQERTNYAEQF | 0.7316281 |
| SATGVFAGVI | ASSQERTNYAEQF | 0.6773731 |
| SLPAITLP | ASSQERTNYAEQF | 0.7474546 |
| STALAFTV | ASSQERTNYAEQF | 0.6019276 |
| SVLTVGLGK | ASSQERTNYAEQF | 0.7369021 |
| TALAGAGSGI | ASSQERTNYAEQF | 0.7146206 |
| TGLPFTGI | ASSQERTNYAEQF | 0.7017381 |
| VAEHVTV | ASSQERTNYAEQF | 0.746391 |
| VDIVLNSL | ASSQERTNYAEQF | 0.65448344 |
| VLTGGGAVL | ASSQERTNYAEQF | 0.62639904 |
| VNVGLAYGI | ASSQERTNYAEQF | 0.6989023 |
| VSSAAVGAV | ASSQERTNYAEQF | 0.7376039 |
| VTSVYEAGV | ASSQERTNYAEQF | 0.7561358 |
| WVALDRGA | ASSQERTNYAEQF | 0.63923347 |
| YLVGSVVI | ASSQERTNYAEQF | 0.7264505 |
| GPPGTGKTLL | ASSQERTNYAEQF | 0.6881775 |
| GVGMTKF | ASSQERTNYAEQF | 0.64046776 |
| IGIGGSDL | ASSQERTNYAEQF | 0.71739024 |
| IGTLLAEL | ASSQERTNYAEQF | 0.7208135 |
| IYTSGTT | ASSQERTNYAEQF | 0.6747612 |
| LSFPAGV | ASSQERTNYAEQF | 0.74478686 |
| LYTSGTT | ASSQERTNYAEQF | 0.66870314 |
| VGAFPFGF | ASSQERTNYAEQF | 0.7450214 |
| VGAGTVEF | ASSQERTNYAEQF | 0.76974666 |
| VGKSTLLL | ASSQERTNYAEQF | 0.7008624 |
| AALAHAGGGIV | ASSQERTNYAEQF | 0.7481524 |
| AALVAVSLL | ASSQERTNYAEQF | 0.7388015 |
| AAVSRAVAAL | ASSQERTNYAEQF | 0.73647314 |
| AGGSSGIGGF | ASSQERTNYAEQF | 0.6993882 |
| AGLTVANANG | ASSQERTNYAEQF | 0.6871778 |
| AGLVGAVL | ASSQERTNYAEQF | 0.76615155 |
| ALSQAFTV | ASSQERTNYAEQF | 0.7485179 |
| ALTFSVSL | ASSQERTNYAEQF | 0.7615315 |
| ALTGAALVAL | ASSQERTNYAEQF | 0.7449622 |
| ASFSIGAGVL | ASSQERTNYAEQF | 0.7435676 |

| | | |
|---|---|---|
| DGSVYKI | ASSQERTNYAEQF | 0.74410033 |
| EQLAEIV | ASSQERTNYAEQF | 0.6595665 |
| FAVSELTVM | ASSQERTNYAEQF | 0.57262033 |
| GAAPLAAGL | ASSQERTNYAEQF | 0.6979061 |
| GADGGDGGSGGAGG | ASSQERTNYAEQF | 0.72573775 |
| GFGLDTGGAGG | ASSQERTNYAEQF | 0.7217144 |
| GLASGGLL | ASSQERTNYAEQF | 0.5104401 |
| GRIAGAAVLL | ASSQERTNYAEQF | 0.7612762 |
| GSAGSLNTSF | ASSQERTNYAEQF | 0.7776867 |
| GTGQIGI | ASSQERTNYAEQF | 0.6683167 |
| GTLIALIL | ASSQERTNYAEQF | 0.7699467 |
| IAATMGFVVG | ASSQERTNYAEQF | 0.59456027 |
| IFAQALL | ASSQERTNYAEQF | 0.74729097 |
| IGAAVALGF | ASSQERTNYAEQF | 0.7608354 |
| IGLTPLAL | ASSQERTNYAEQF | 0.6088575 |
| ILGSNTSTL | ASSQERTNYAEQF | 0.76156753 |
| IVFSLTVL | ASSQERTNYAEQF | 0.6868999 |
| LASKATGF | ASSQERTNYAEQF | 0.7208445 |
| LASVALIL | ASSQERTNYAEQF | 0.7684547 |
| LFAVGLTL | ASSQERTNYAEQF | 0.70588136 |
| LFNGGIGTYI | ASSQERTNYAEQF | 0.73775876 |
| LTLGAGII | ASSQERTNYAEQF | 0.71174264 |
| LVAAALGDI | ASSQERTNYAEQF | 0.649247 |
| NTAFVQLGI | ASSQERTNYAEQF | 0.73673356 |
| QLFSGAEL | ASSQERTNYAEQF | 0.7666648 |
| SAATFITL | ASSQERTNYAEQF | 0.6087575 |
| SNYGLANLGV | ASSQERTNYAEQF | 0.7722103 |
| SYGGAYAVM | ASSQERTNYAEQF | 0.671427 |
| TTGVGGLTLG | ASSQERTNYAEQF | 0.77865 |
| TTQTIYI | ASSQERTNYAEQF | 0.74301267 |
| VAAGAGLGAVI | ASSQERTNYAEQF | 0.7048277 |
| VAAVVTLL | ASSQERTNYAEQF | 0.69457364 |
| VGFLGGLI | ASSQERTNYAEQF | 0.7644691 |
| VGVIPILL | ASSQERTNYAEQF | 0.6943806 |
| VIFASSLI | ASSQERTNYAEQF | 0.74847454 |
| VKGGDGGAAAG | ASSQERTNYAEQF | 0.76957667 |
| VTKTFTNI | ASSQERTNYAEQF | 0.22396785 |
| VVAAAILL | ASSQERTNYAEQF | 0.3068363 |
| YAALDVEL | ASSQERTNYAEQF | 0.22982723 |
| LASVALLL | ASSQERTNYAEQF | 0.24785987 |
| LDAVHRV | ASSQERTNYAEQF | 0.2448149 |
| QGLSSSIF | ASSQERTNYAEQF | 0.22518057 |
| RGTFANIRL | ASSQERTNYAEQF | 0.237142 |
| RQLWWGHRI | ASSQERTNYAEQF | 0.23401335 |
| YGPPGTGKTLL | ASSQERTNYAEQF | 0.3479967 |
| AALVGAPANVA | ASSQERTNYAEQF | 0.28222057 |
| AATFRRTE | ASSQERTNYAEQF | 0.24407355 |
| AGALGGGI | ASSQERTNYAEQF | 0.39836594 |
| AGGAGGAGGIGAG | ASSQERTNYAEQF | 0.2965436 |
| AGSGALGL | ASSQERTNYAEQF | 0.23221987 |
| AQIADQI | ASSQERTNYAEQF | 0.23299246 |

| | | |
|---|---|---|
| ASAVAVTYL | ASSQERTNYAEQF | 0.24136746 |
| ASTVLGGTGG | ASSQERTNYAEQF | 0.25574192 |
| AVGGEPAK | ASSQERTNYAEQF | 0.2195955 |
| CEQGTFGGTC | ASSQERTNYAEQF | 0.27071306 |
| DEGKAPAG | ASSQERTNYAEQF | 0.22336464 |
| DTVVFPAL | ASSQERTNYAEQF | 0.21307 |
| FEGITMAG | ASSQERTNYAEQF | 0.32153445 |
| FPLVVQD | ASSQERTNYAEQF | 0.2694331 |
| GAFTTPQ | ASSQERTNYAEQF | 0.23939924 |
| GFGGFGTAGGVGGA | ASSQERTNYAEQF | 0.2637753 |
| GFLAQFL | ASSQERTNYAEQF | 0.22571532 |
| GGAGGAGGNAGVA | ASSQERTNYAEQF | 0.23435646 |
| GGVGTAGAGGAGGAGG | ASSQERTNYAEQF | 0.2703342 |
| GNGGGGDAGNAGSGGN | ASSQERTNYAEQF | 0.3187823 |
| GTGGAGGAGNDAGS | ASSQERTNYAEQF | 0.23268016 |
| GTTNTGWFNS | ASSQERTNYAEQF | 0.23113959 |
| IFMGPLFTL | ASSQERTNYAEQF | 0.29627872 |
| ILAHAIEAI | ASSQERTNYAEQF | 0.27325842 |
| IRSAIREAL | ASSQERTNYAEQF | 0.25972328 |
| ITFAGGG | ASSQERTNYAEQF | 0.2676579 |
| ITGMSGHQEV | ASSQERTNYAEQF | 0.23151474 |
| IVAGLACTL | ASSQERTNYAEQF | 0.28758043 |
| IVGGGLAGA | ASSQERTNYAEQF | 0.3728879 |
| IVTGVLNI | ASSQERTNYAEQF | 0.25227007 |
| LAQQGCI | ASSQERTNYAEQF | 0.2558971 |
| LCLHVEN | ASSQERTNYAEQF | 0.22181001 |
| LDAISDGS | ASSQERTNYAEQF | 0.25148338 |
| LGGFAAML | ASSQERTNYAEQF | 0.21490066 |
| LGGKNPMI | ASSQERTNYAEQF | 0.2874916 |
| LGPKGVAAIL | ASSQERTNYAEQF | 0.2845691 |
| LMVGVSI | ASSQERTNYAEQF | 0.27957067 |
| LVGAAFVL | ASSQERTNYAEQF | 0.2228603 |
| LVLAVAPL | ASSQERTNYAEQF | 0.22183459 |
| MAAIAVIDAA | ASSQERTNYAEQF | 0.2194153 |
| NNGINAELQ | ASSQERTNYAEQF | 0.2227613 |
| RGEPLVLVHG | ASSQERTNYAEQF | 0.2144368 |
| RGGAGGAAGLI | ASSQERTNYAEQF | 0.2539799 |
| RSYRMHPAV | ASSQERTNYAEQF | 0.4242652 |
| SAAHLLAGQLA | ASSQERTNYAEQF | 0.3317971 |
| SAAVGSY | ASSQERTNYAEQF | 0.2245244 |
| SPHAVLHRI | ASSQERTNYAEQF | 0.32440448 |
| SQIAKVL | ASSQERTNYAEQF | 0.24773279 |
| TALGHLVP | ASSQERTNYAEQF | 0.24599105 |
| TAPTTSGAA | ASSQERTNYAEQF | 0.23812158 |
| TDGDDGNGGSPGT | ASSQERTNYAEQF | 0.22223715 |
| TDGLGEN | ASSQERTNYAEQF | 0.49284655 |
| TGGPPLPGQA | ASSQERTNYAEQF | 0.25714484 |
| TIPAITI | ASSQERTNYAEQF | 0.26273265 |
| TIPGFTI | ASSQERTNYAEQF | 0.37929994 |
| TSIASGALA | ASSQERTNYAEQF | 0.24719311 |
| TVTLPGAAI | ASSQERTNYAEQF | 0.2376483 |

| | | |
|---|---|---|
| VASGLSAALP | ASSQERTNYAEQF | 0.26077354 |
| VGAGVIGI | ASSQERTNYAEQF | 0.24359383 |
| VGPKRFGV | ASSQERTNYAEQF | 0.21974526 |
| VNVAFPD | ASSQERTNYAEQF | 0.23644613 |
| VSMTNSGVS | ASSQERTNYAEQF | 0.2172177 |
| VTDVLGDLSV | ASSQERTNYAEQF | 0.24549459 |
| VVVVGAGFAGL | ASSQERTNYAEQF | 0.31724152 |
| VYTVGTVV | ASSQERTNYAEQF | 0.26575524 |
| WTTFSGP | ASSQERTNYAEQF | 0.22183728 |
| YDELVEAF | ASSQERTNYAEQF | 0.2559805 |
| YGTGGAGGNGG | ASSQERTNYAEQF | 0.2539901 |
| YTNVMAV | ASSQERTNYAEQF | 0.25018194 |
| FLEQVHQL | ASSQERTNYAEQF | 0.2393324 |
| GLSLLAVAV | ASSQERTNYAEQF | 0.23679882 |
| LEQVHQL | ASSQERTNYAEQF | 0.24475387 |
| LNAAGAI | ASSQERTNYAEQF | 0.280801 |
| PAVHASL | ASSQERTNYAEQF | 0.24722803 |
| TELEDDA | ASSQERTNYAEQF | 0.25010487 |
| TPPGPPN | ASSQERTNYAEQF | 0.25433895 |
| TPPPPPPP | ASSQERTNYAEQF | 0.22929884 |
| TQSPALI | ASSQDRANTEVF | 0.22047949 |
| EFAAAFPQ | ASSQDRANTEVF | 0.24558042 |
| LLSGNLDVL | ASSQDRANTEVF | 0.23777577 |
| GPANLGNYNI | ASSQDRANTEVF | 0.23632376 |
| TLGPTLP | ASSQDRANTEVF | 0.2618418 |
| VYAYSSCCK | ASSQDRANTEVF | 0.24231663 |
| APAPAPAPNI | ASSQDRANTEVF | 0.23042218 |
| PGGPVPIP | ASSQDRANTEVF | 0.25375816 |
| FLTVALEP | ASSQDRANTEVF | 0.2679132 |
| SQYFGANL | ASSQDRANTEVF | 0.22085892 |
| GGVKPAA | ASSQDRANTEVF | 0.24316004 |
| GGTGGAGGAGGW | ASSQDRANTEVF | 0.51019025 |
| GVGGVAGLGGAGG | ASSQDRANTEVF | 0.2630587 |
| VLLVLATTI | ASSQDRANTEVF | 0.25604972 |
| IGIFAAML | ASSQDRANTEVF | 0.2345344 |
| AVGDLID | ASSQDRANTEVF | 0.5403949 |
| NISPLPL | ASSQDRANTEVF | 0.22642928 |
| VTAVFVAI | ASSQDRANTEVF | 0.3423646 |
| SGDDNDAAL | ASSQDRANTEVF | 0.21713951 |
| AVAALAHS | ASSQDRANTEVF | 0.22447613 |
| GGVTVMATP | ASSQDRANTEVF | 0.21575844 |
| AGLAANVRD | ASSQDRANTEVF | 0.21494028 |
| TNRPDLIDPALLRPG | ASSQDRANTEVF | 0.31293276 |
| LANLGVTAL | ASSQDRANTEVF | 0.23889455 |
| TGERIEL | ASSQDRANTEVF | 0.26549798 |
| NGEELAA | ASSQDRANTEVF | 0.2522804 |
| VLSLVWG | ASSQDRANTEVF | 0.29380512 |
| IGYALWL | ASSQDRANTEVF | 0.23803444 |
| LSLLLAPL | ASSQDRANTEVF | 0.2764419 |
| TLDEALK | ASSQDRANTEVF | 0.25181678 |
| TDGAGTAEGG | ASSQDRANTEVF | 0.29273817 |

| | | |
|---|---|---|
| NAPGGAGGNGG | ASSQDRANTEVF | 0.27386454 |
| VSVFMGMGGNF | ASSQDRANTEVF | 0.25117743 |
| LASYLAAL | ASSQDRANTEVF | 0.22736348 |
| LVGGTPAL | ASSQDRANTEVF | 0.22138919 |
| VRNADVL | ASSQDRANTEVF | 0.2593452 |
| TCSSPFLVSAM | ASSQDRANTEVF | 0.22079279 |
| VHITDAE | ASSQDRANTEVF | 0.21388608 |
| EYAAPSF | ASSQDRANTEVF | 0.21848789 |
| VVDAFHGGL | ASSQDRANTEVF | 0.22717303 |
| ASVVDPQ | ASSQDRANTEVF | 0.38457024 |
| SYNEFGGLS | ASSQDRANTEVF | 0.3400537 |
| DPDGSVY | ASSQDRANTEVF | 0.24302305 |
| MVSTIHATE | ASSQDRANTEVF | 0.24931541 |
| LAAGVADPV | ASSQDRANTEVF | 0.28483817 |
| TKEQIAA | ASSQDRANTEVF | 0.21151768 |
| SLASPAAL | ASSQDRANTEVF | 0.2302175 |
| IGLVFYGV | ASSQDRANTEVF | 0.23554212 |
| FLLDASNL | ASSQDRANTEVF | 0.26778746 |
| YQVGNFP | ASSQDRANTEVF | 0.23601332 |
| ESLLSQST | ASSQDRANTEVF | 0.24184291 |
| LFTVGFAAVG | ASSQDRANTEVF | 0.2606157 |
| ATPPTPT | ASSQDRANTEVF | 0.3939637 |
| TDLATLG | ASSQDRANTEVF | 0.24851158 |
| SAGGTGR | ASSQDRANTEVF | 0.21627066 |
| GFSNSGS | ASSQDRANTEVF | 0.21498245 |
| GGNGGSGGTSV | ASSQDRANTEVF | 0.31344876 |
| GTSVATGGAGN | ASSQDRANTEVF | 0.22889338 |
| NGGNGGSGGTS | ASSQDRANTEVF | 0.22578406 |
| AVVKFPH | ASSQDRANTEVF | 0.21660891 |
| TPPPPPP | ASSQDRANTEVF | 0.26293656 |
| AAPAAAAAPA | ASSQDRANTEVF | 0.2200916 |
| IALNEDL | ASSQDRANTEVF | 0.24875021 |
| IVNVSSM | ASSQDRANTEVF | 0.22337228 |
| TGIPPAPR | ASSQDRANTEVF | 0.25042364 |
| FDLGGGTF | ASSQDRANTEVF | 0.3452623 |
| FDLGGGTFDVS | ASSQDRANTEVF | 0.4220392 |
| NEPTAAA | ASSQDRANTEVF | 0.25511444 |
| NEPTAAALA | ASSQDRANTEVF | 0.24608508 |
| GSIINISSI | ASSQDRANTEVF | 0.23907354 |
| VAPGNAGTA | ASSQDRANTEVF | 0.22989714 |
| TGGGASLE | ASSQDRANTEVF | 0.23234555 |
| VAVYNHYKRI | ASSQDRANTEVF | 0.3378615 |
| TDPAPATT | ASSQDRANTEVF | 0.22610117 |
| TNRPDILDPALLRPG | ASSQDRANTEVF | 0.22236353 |
| LTTTTTS | ASSQDRANTEVF | 0.22899248 |
| PTGSGKTL | ASSQDRANTEVF | 0.268041 |
| TGSGKTL | ASSQDRANTEVF | 0.29168686 |
| TIGHVDHGK | ASSQDRANTEVF | 0.25197762 |
| TIGHVDHGKTT | ASSQDRANTEVF | 0.23185241 |
| GHVDHGKTT | ASSQDRANTEVF | 0.23135667 |
| AQFEHTLL | ASSQDRANTEVF | 0.2971366 |

| | | |
|---|---|---|
| IVNQDEL | ASSQDRANTEVF | 0.22682065 |
| TNGSQFF | ASSQDRANTEVF | 0.23356299 |
| TGKGIVE | ASSQDRANTEVF | 0.2378189 |
| LISNASDA | ASSQDRANTEVF | 0.55005443 |
| AELRTLGELTKIAKAHGAQ | ASSQDRANTEVF | 0.23576424 |
| APNAVAAAEPSW | ASSQDRANTEVF | 0.22483674 |
| APQVAGAATAVL | ASSQDRANTEVF | 0.26654217 |
| ARRDPAAAAAILPTDA | ASSQDRANTEVF | 0.22106043 |
| DAAARDVLAEAGLAEYF | ASSQDRANTEVF | 0.23648319 |
| DDQIVLSAKANLAKVEQ | ASSQDRANTEVF | 0.23808846 |
| DELLPDTGEDSDEDGH | ASSQDRANTEVF | 0.21704783 |
| ELPEAPANRATVPQD | ASSQDRANTEVF | 0.23590188 |
| FGQSKIERVTGIG | ASSQDRANTEVF | 0.34100437 |
| GAGGTGSTAGGAGGAGGAGGLY | ASSQDRANTEVF | 0.27381992 |
| GHGGTGAAGGNGGTGGAGGSGI | ASSQDRANTEVF | 0.25279108 |
| GNGGNGGAGGEGGPGGAGGAGA | ASSQDRANTEVF | 0.3161757 |
| GSTAGNGGNGGNATNGTIAGQP | ASSQDRANTEVF | 0.2942099 |
| GTGGDGGNGGNGGTGG | ASSQDRANTEVF | 0.29899883 |
| GVRRPAGIPADAPV | ASSQDRANTEVF | 0.27107054 |
| GYRQVTDAHLLTLARR | ASSQDRANTEVF | 0.29398894 |
| IAKMGQDGHDRG | ASSQDRANTEVF | 0.22819437 |
| KAEPSIEIQYT | ASSQDRANTEVF | 0.2406584 |
| KPVRIRMAADEDTVAVT | ASSQDRANTEVF | 0.2739113 |
| MPPRIAGMRLLVIKPEP | ASSQDRANTEVF | 0.26729655 |
| MKFYAVALFRSRGVPPTR | ASSQDRANTEVF | 0.2279985 |
| NAGGFGILGAVAHSP | ASSQDRANTEVF | 0.33707267 |
| NEGREITLAMAEM | ASSQDRANTEVF | 0.23298398 |
| NELVEWSHPLGHRWRWE | ASSQDRANTEVF | 0.29296148 |
| NLGAGGTGGNGGNPG | ASSQDRANTEVF | 0.21950513 |
| NTEWPKWETVMAADPVD | ASSQDRANTEVF | 0.23931877 |
| PASAAPAETQP | ASSQDRANTEVF | 0.2580472 |
| QQSSDGKNGVRAAEITGE | ASSQDRANTEVF | 0.2128566 |
| RAPDAVTTSWWKE | ASSQDRANTEVF | 0.21479662 |
| SGFGNVGAHTSGWFNQ | ASSQDRANTEVF | 0.22846147 |
| SGVAGAVGQAASVGGLKV | ASSQDRANTEVF | 0.28469095 |
| SNGGAGTGSAGGNGGT | ASSQDRANTEVF | 0.21245167 |
| TGGAGGEGGAAG | ASSQDRANTEVF | 0.2398469 |
| TGGDGGAAGAGGAPGHG | ASSQDRANTEVF | 0.21824117 |
| TMPPLLAAIANAGRHGV | ASSQDRANTEVF | 0.23729256 |
| TSSTAGNSGGA | ASSQDRANTEVF | 0.26281157 |
| VPATVPVGILC | ASSQDRANTEVF | 0.22399464 |
| VQRPDSGGVVLTGRIS | ASSQDRANTEVF | 0.260208 |
| WTPNPVPATKGVPAGLVQAT | ASSQDRANTEVF | 0.21722883 |
| YALKRALAKFSDRFDVVII | ASSQDRANTEVF | 0.22754186 |
| YLIGSVIV | ASSQDRANTEVF | 0.27430525 |
| YGQLVAVID | ASSQDRANTEVF | 0.3563751 |
| VTVGGMTLPA | ASSQDRANTEVF | 0.2318204 |
| VLALPAV | ASSQDRANTEVF | 0.26051548 |
| VIPPTLNYET | ASSQDRANTEVF | 0.21779372 |
| VGMGGMHPSSG | ASSQDRANTEVF | 0.34770083 |
| VGLAGGAATA | ASSQDRANTEVF | 0.25894552 |

| | | |
|---|---|---|
| VGGAGGAGGSGP | ASSQDRANTEVF | 0.21978559 |
| VFAAFMV | ASSQDRANTEVF | 0.25185615 |
| TLYDKLR | ASSQDRANTEVF | 0.23696548 |
| TLAVFTVL | ASSQDRANTEVF | 0.24802673 |
| TGSVQPARV | ASSQDRANTEVF | 0.2643106 |
| TGGGAGSGGT | ASSQDRANTEVF | 0.21957427 |
| SHLETSL | ASSQDRANTEVF | 0.2351101 |
| SFTTGMT | ASSQDRANTEVF | 0.38113013 |
| QVEVAKRMGVRQPTVS | ASSQDRANTEVF | 0.2585135 |
| QGSGGPALAM | ASSQDRANTEVF | 0.23262908 |
| MLAPSNP | ASSQDRANTEVF | 0.2808893 |
| LVNNAAI | ASSQDRANTEVF | 0.27297154 |
| LTGEINTLA | ASSQDRANTEVF | 0.22712432 |
| LSIFGPV | ASSQDRANTEVF | 0.30829746 |
| LLALLANL | ASSQDRANTEVF | 0.22899175 |
| LISNASDALDKLR | ASSQDRANTEVF | 0.25675955 |
| LINEAVP | ASSQDRANTEVF | 0.2538644 |
| LGYAFSGL | ASSQDRANTEVF | 0.3638242 |
| LDFVAVAL | ASSQDRANTEVF | 0.23607287 |
| LAQPGAE | ASSQDRANTEVF | 0.23900524 |
| ISNLGAVTGF | ASSQDRANTEVF | 0.25510168 |
| HQEDPGRI | ASSQDRANTEVF | 0.2547494 |
| GTPGEPTP | ASSQDRANTEVF | 0.22325158 |
| GFFNSGS | ASSQDRANTEVF | 0.24652354 |
| GAGGGGLTAGAN | ASSQDRANTEVF | 0.26105574 |
| FVDALVNM | ASSQDRANTEVF | 0.28504992 |
| FLTVAPEL | ASSQDRANTEVF | 0.2361871 |
| FETLTAL | ASSQDRANTEVF | 0.23576015 |
| DVINPIEL | ASSQDRANTEVF | 0.36000842 |
| APVSGAPV | ASSQDRANTEVF | 0.22778578 |
| ALVGGLAW | ASSQDRANTEVF | 0.25685358 |
| ALAAAAATAI | ASSQDRANTEVF | 0.24981616 |
| AFGLGLLL | ASSQDRANTEVF | 0.275274 |
| AFFAAASL | ASSQDRANTEVF | 0.22785042 |
| AAAALPAFSPP | ASSQDRANTEVF | 0.2735542 |
| VVLAPAL | ASSQDRANTEVF | 0.23355974 |
| TIGHVDHGKTTL | ASSQDRANTEVF | 0.4142707 |
| LISNASDALDK | ASSQDRANTEVF | 0.21545172 |
| LISNASDAL | ASSQDRANTEVF | 0.27295566 |
| ISNASDALDK | ASSQDRANTEVF | 0.24701539 |
| NQAETLVYQTE | ASSQDRANTEVF | 0.34665608 |
| TGGTGGTAGAG | ASSQDRANTEVF | 0.25746238 |
| TGSTAGGAGGA | ASSQDRANTEVF | 0.36093217 |
| DPLAVPVAAGPV | ASSQDRANTEVF | 0.21548888 |
| GGTGGTPGNGGN | ASSQDRANTEVF | 0.30299854 |
| PAGLAGVDPATL | ASSQDRANTEVF | 0.3879748 |
| GNNLVVEAPHAGASV | ASSQDRANTEVF | 0.48322108 |
| KVTRIMSASTLEQPA | ASSQDRANTEVF | 0.23307341 |
| LGFALPIALPRRTVV | ASSQDRANTEVF | 0.25508162 |
| TRLGNLTARKLVVLV | ASSQDRANTEVF | 0.2473965 |
| WSQQRRLIYFPSAGP | ASSQDRANTEVF | 0.24270397 |

| | | |
|---|---|---|
| GPNGNPDPMAAAVDIR | ASSQDRANTEVF | 0.22152925 |
| TQRCVSVADLRAAASR | ASSQDRANTEVF | 0.22381385 |
| QFVLDVAEHPVEVLTETV | ASSQDRANTEVF | 0.2546911 |
| GGAGGTGGAGGNGGTQAGDG | ASSQDRANTEVF | 0.2544088 |
| PLLATLRAIAVDVDAKPAQV | ASSQDRANTEVF | 0.48475164 |
| APKFRTGGYMIDFWGVGYQVAK | ASSQDRANTEVF | 0.26543266 |
| GAGGAGGAGGAAGGPGGTGGTG | ASSQDRANTEVF | 0.30772486 |
| IGALGLLATDVAPAGSVAFGVA | ASSQDRANTEVF | 0.26141044 |
| AGGAGGIGGNGGAGGAGGVLMG | ASSQDRANTEVF | 0.26316288 |
| ADELIGLL | ASSQDRANTEVF | 0.2428033 |
| ADRALKA | ASSQDRANTEVF | 0.23845527 |
| AGAVGQVGGAGGA | ASSQDRANTEVF | 0.26562467 |
| AGGGGGLGGAP | ASSQDRANTEVF | 0.23114094 |
| AGGLLYGNGGAG | ASSQDRANTEVF | 0.23635487 |
| AGLESIGL | ASSQDRANTEVF | 0.23093948 |
| AGLKTGII | ASSQDRANTEVF | 0.27845857 |
| ASLAALLL | ASSQDRANTEVF | 0.21222565 |
| ASVINISSI | ASSQDRANTEVF | 0.25312304 |
| ATPPGPA | ASSQDRANTEVF | 0.3457992 |
| ATVASALV | ASSQDRANTEVF | 0.24268073 |
| ATVASLIL | ASSQDRANTEVF | 0.25549904 |
| AVNGPVLIRH | ASSQDRANTEVF | 0.25986496 |
| DGAAGAVGI | ASSQDRANTEVF | 0.22059135 |
| DPLLHGI | ASSQDRANTEVF | 0.26095128 |
| ELGAALGAL | ASSQDRANTEVF | 0.22054306 |
| ELGRTQAV | ASSQDRANTEVF | 0.25203753 |
| FVAGGYLS | ASSQDRANTEVF | 0.30785218 |
| GASGGAGGAGGW | ASSQDRANTEVF | 0.24194266 |
| GGAGAGGDGGAGGI | ASSQDRANTEVF | 0.22802238 |
| GTGHNGSQPGMGGNG | ASSQDRANTEVF | 0.23927593 |
| GYAAASASAS | ASSQDRANTEVF | 0.2335361 |
| HQVDVAAVQQG | ASSQDRANTEVF | 0.23496337 |
| IGAAIAHT | ASSQDRANTEVF | 0.23193216 |
| IGMIEVAQL | ASSQDRANTEVF | 0.23845026 |
| IRLAPEHTSAMD | ASSQDRANTEVF | 0.22204117 |
| IYNVSVL | ASSQDRANTEVF | 0.39171493 |
| KQGFLGM | ASSQDRANTEVF | 0.2691576 |
| LESVITQL | ASSQDRANTEVF | 0.2375756 |
| LGLGALVVV | ASSQDRANTEVF | 0.23369995 |
| LIGNGGNGGSGGI | ASSQDRANTEVF | 0.22887434 |
| LNAINAPTL | ASSQDRANTEVF | 0.22698398 |
| LQLLNTGV | ASSQDRANTEVF | 0.22295848 |
| LQTSIEF | ASSQDRANTEVF | 0.24637653 |
| MSFAVSEL | ASSQDRANTEVF | 0.22705323 |
| NTGALGL | ASSQDRANTEVF | 0.23045117 |
| PPGAGGAGGAGTT | ASSQDRANTEVF | 0.32888407 |
| QLVGLGI | ASSQDRANTEVF | 0.24379669 |
| RFLEQVHQL | ASSQDRANTEVF | 0.2235176 |
| RGVGYVLR | ASSQDRANTEVF | 0.22912838 |
| SATGVFAGVI | ASSQDRANTEVF | 0.23122707 |
| SLPAITLP | ASSQDRANTEVF | 0.2521423 |

| | | |
|---|---|---|
| STALAFTV | ASSQDRANTEVF | 0.21296713 |
| SVLTVGLGK | ASSQDRANTEVF | 0.2803129 |
| TALAGAGSGI | ASSQDRANTEVF | 0.22779001 |
| TGLPFTGI | ASSQDRANTEVF | 0.22456414 |
| VAEHVTV | ASSQDRANTEVF | 0.22933316 |
| VDIVLNSL | ASSQDRANTEVF | 0.28491437 |
| VLTGGGAVL | ASSQDRANTEVF | 0.23562095 |
| VNVGLAYGI | ASSQDRANTEVF | 0.23136452 |
| VSSAAVGAV | ASSQDRANTEVF | 0.2488338 |
| VTSVYEAGV | ASSQDRANTEVF | 0.30451357 |
| WVALDRGA | ASSQDRANTEVF | 0.41330534 |
| YLVGSVVI | ASSQDRANTEVF | 0.31755388 |
| GPPGTGKTLL | ASSQDRANTEVF | 0.22195618 |
| GVGMTKF | ASSQDRANTEVF | 0.3242371 |
| IGIGGSDL | ASSQDRANTEVF | 0.24580371 |
| IGTLLAEL | ASSQDRANTEVF | 0.22088338 |
| IYTSGTT | ASSQDRANTEVF | 0.21608213 |
| LSFPAGV | ASSQDRANTEVF | 0.24181478 |
| LYTSGTT | ASSQDRANTEVF | 0.21845967 |
| VGAFPFGF | ASSQDRANTEVF | 0.28571478 |
| VGAGTVEF | ASSQDRANTEVF | 0.2565343 |
| VGKSTLLL | ASSQDRANTEVF | 0.23010002 |
| AALAHAGGGIV | ASSQDRANTEVF | 0.21898128 |
| AALVAVSLL | ASSQDRANTEVF | 0.22561128 |
| AAVSRAVAAL | ASSQDRANTEVF | 0.24753664 |
| AGGSSGIGGF | ASSQDRANTEVF | 0.26022965 |
| AGLTVANANG | ASSQDRANTEVF | 0.22152425 |
| AGLVGAVL | ASSQDRANTEVF | 0.5568857 |
| ALSQAFTV | ASSQDRANTEVF | 0.35346583 |
| ALTFSVSL | ASSQDRANTEVF | 0.2381652 |
| ALTGAALVAL | ASSQDRANTEVF | 0.25744474 |
| ASFSIGAGVL | ASSQDRANTEVF | 0.22845319 |
| DGSVYKI | ASSQDRANTEVF | 0.23121078 |
| EQLAEIV | ASSQDRANTEVF | 0.26009628 |
| FAVSELTVM | ASSQDRANTEVF | 0.22812064 |
| GAAPLAAGL | ASSQDRANTEVF | 0.23116349 |
| GADGGDGGSGGAGG | ASSQDRANTEVF | 0.2382321 |
| GFGLDTGGAGG | ASSQDRANTEVF | 0.2258711 |
| GLASGGLL | ASSQDRANTEVF | 0.23300107 |
| GRIAGAAVLL | ASSQDRANTEVF | 0.2380735 |
| GSAGSLNTSF | ASSQDRANTEVF | 0.21460554 |
| GTGQIGI | ASSQDRANTEVF | 0.2568783 |
| GTLIALIL | ASSQDRANTEVF | 0.36438572 |
| IAATMGFVVG | ASSQDRANTEVF | 0.29180712 |
| IFAQALL | ASSQDRANTEVF | 0.22286448 |
| IGAAVALGF | ASSQDRANTEVF | 0.22878507 |
| IGLTPLAL | ASSQDRANTEVF | 0.39970464 |
| ILGSNTSTL | ASSQDRANTEVF | 0.2318804 |
| IVFSLTVL | ASSQDRANTEVF | 0.23396333 |
| LASKATGF | ASSQDRANTEVF | 0.26627398 |
| LASVALIL | ASSQDRANTEVF | 0.24425998 |

| | | |
|---|---|---|
| LFAVGLTL | ASSQDRANTEVF | 0.22793643 |
| LFNGGIGTYI | ASSQDRANTEVF | 0.2357491 |
| LTLGAGII | ASSQDRANTEVF | 0.23820767 |
| LVAAALGDI | ASSQDRANTEVF | 0.23860057 |
| NTAFVQLGI | ASSQDRANTEVF | 0.21316266 |
| QLFSGAEL | ASSQDRANTEVF | 0.25946826 |
| SAATFITL | ASSQDRANTEVF | 0.23301065 |
| SNYGLANLGV | ASSQDRANTEVF | 0.2258355 |
| SYGGAYAVM | ASSQDRANTEVF | 0.21545029 |
| TTGVGGLTLG | ASSQDRANTEVF | 0.23819417 |
| TTQTIYI | ASSQDRANTEVF | 0.2424551 |
| VAAGAGLGAVI | ASSQDRANTEVF | 0.2759402 |
| VAAVVTLL | ASSQDRANTEVF | 0.22334819 |
| VGFLGGLI | ASSQDRANTEVF | 0.28415483 |
| VGVIPILL | ASSQDRANTEVF | 0.2731439 |
| VIFASSLI | ASSQDRANTEVF | 0.24293399 |
| VKGGDGGAAAG | ASSQDRANTEVF | 0.23706649 |
| VTKTFTNI | ASSQDRANTEVF | 0.25757474 |
| VVAAAILL | ASSQDRANTEVF | 0.22750792 |
| YAALDVEL | ASSQDRANTEVF | 0.24266721 |
| LASVALLL | ASSQDRANTEVF | 0.3066765 |
| LDAVHRV | ASSQDRANTEVF | 0.23334417 |
| QGLSSSIF | ASSQDRANTEVF | 0.27375284 |
| RGTFANIRL | ASSQDRANTEVF | 0.3235753 |
| RQLWWGHRI | ASSQDRANTEVF | 0.22536385 |
| YGPPGTGKTLL | ASSQDRANTEVF | 0.22399116 |
| AALVGAPANVA | ASSQDRANTEVF | 0.23372263 |
| AATFRRTE | ASSQDRANTEVF | 0.21559083 |
| AGALGGGI | ASSQDRANTEVF | 0.27261674 |
| AGGAGGAGGIGAG | ASSQDRANTEVF | 0.22641398 |
| AGSGALGL | ASSQDRANTEVF | 0.24521416 |
| AQIADQI | ASSQDRANTEVF | 0.2548034 |
| ASAVAVTYL | ASSQDRANTEVF | 0.24010564 |
| ASTVLGGTGG | ASSQDRANTEVF | 0.2526681 |
| AVGGEPAK | ASSQDRANTEVF | 0.24961239 |
| CEQGTFGGTC | ASSQDRANTEVF | 0.2640052 |
| DEGKAPAG | ASSQDRANTEVF | 0.2361635 |
| DTVVFPAL | ASSQDRANTEVF | 0.23280606 |
| FEGITMAG | ASSQDRANTEVF | 0.23515028 |
| FPLVVQD | ASSQDRANTEVF | 0.40650353 |
| GAFTTPQ | ASSQDRANTEVF | 0.22582084 |
| GFGGFGTAGGVGGA | ASSQDRANTEVF | 0.23583281 |
| GFLAQFL | ASSQDRANTEVF | 0.2342863 |
| GGAGGAGGNAGVA | ASSQDRANTEVF | 0.2183174 |
| GGVGTAGAGGAGGAGG | ASSQDRANTEVF | 0.2543786 |
| GNGGGGDAGNAGSGGN | ASSQDRANTEVF | 0.28275535 |
| GTGGAGGAGNDAGS | ASSQDRANTEVF | 0.2818336 |
| GTTNTGWFNS | ASSQDRANTEVF | 0.2365281 |
| IFMGPLFTL | ASSQDRANTEVF | 0.35273457 |
| ILAHAIEAI | ASSQDRANTEVF | 0.2552318 |
| IRSAIREAL | ASSQDRANTEVF | 0.22949369 |

| | | |
|---|---|---|
| ITFAGGG | ASSQDRANTEVF | 0.270495 |
| ITGMSGHQEV | ASSQDRANTEVF | 0.22402069 |
| IVAGLACTL | ASSQDRANTEVF | 0.2709979 |
| IVGGGLAGA | ASSQDRANTEVF | 0.2477456 |
| IVTGVLNI | ASSQDRANTEVF | 0.22404341 |
| LAQQGCI | ASSQDRANTEVF | 0.23784599 |
| LCLHVEN | ASSQDRANTEVF | 0.26021677 |
| LDAISDGS | ASSQDRANTEVF | 0.3058279 |
| LGGFAAML | ASSQDRANTEVF | 0.21753278 |
| LGGKNPMI | ASSQDRANTEVF | 0.26725116 |
| LGPKGVAAIL | ASSQDRANTEVF | 0.22036766 |
| LMVGVSI | ASSQDRANTEVF | 0.22112314 |
| LVGAAFVL | ASSQDRANTEVF | 0.22681771 |
| LVLAVAPL | ASSQDRANTEVF | 0.2609166 |
| MAAIAVIDAA | ASSQDRANTEVF | 0.25726566 |
| NNGINAELQ | ASSQDRANTEVF | 0.3320266 |
| RGEPLVLVHG | ASSQDRANTEVF | 0.28732842 |
| RGGAGGAAGLI | ASSQDRANTEVF | 0.37672722 |
| RSYRMHPAV | ASSQDRANTEVF | 0.22483389 |
| SAAHLLAGQLA | ASSQDRANTEVF | 0.27601382 |
| SAAVGSY | ASSQDRANTEVF | 0.21665832 |
| SPHAVLHRI | ASSQDRANTEVF | 0.23961945 |
| SQIAKVL | ASSQDRANTEVF | 0.22446805 |
| TALGHLVP | ASSQDRANTEVF | 0.24946676 |
| TAPTTSGAA | ASSQDRANTEVF | 0.21590567 |
| TDGDDGNGGSPGT | ASSQDRANTEVF | 0.28762496 |
| TDGLGEN | ASSQDRANTEVF | 0.23143402 |
| TGGPPLPGQA | ASSQDRANTEVF | 0.3023304 |
| TIPAITI | ASSQDRANTEVF | 0.26232958 |
| TIPGFTI | ASSQDRANTEVF | 0.22861966 |
| TSIASGALA | ASSQDRANTEVF | 0.23915085 |
| TVTLPGAAI | ASSQDRANTEVF | 0.25645557 |
| VASGLSAALP | ASSQDRANTEVF | 0.24900025 |
| VGAGVIGI | ASSQDRANTEVF | 0.21793365 |
| VGPKRFGV | ASSQDRANTEVF | 0.23304743 |
| VNVAFPD | ASSQDRANTEVF | 0.22422847 |
| VSMTNSGVS | ASSQDRANTEVF | 0.26 |
| VTDVLGDLSV | ASSQDRANTEVF | 0.2699145 |
| VVVVGAGFAGL | ASSQDRANTEVF | 0.28436118 |
| VYTVGTVV | ASSQDRANTEVF | 0.28156188 |
| WTTFSGP | ASSQDRANTEVF | 0.21311106 |
| YDELVEAF | ASSQDRANTEVF | 0.25387013 |
| YGTGGAGGNGG | ASSQDRANTEVF | 0.4470501 |
| YTNVMAV | ASSQDRANTEVF | 0.23256642 |
| FLEQVHQL | ASSQDRANTEVF | 0.26707128 |
| GLSLLAVAV | ASSQDRANTEVF | 0.2878227 |
| LEQVHQL | ASSQDRANTEVF | 0.23860204 |
| LNAAGAI | ASSQDRANTEVF | 0.25918102 |
| PAVHASL | ASSQDRANTEVF | 0.24100997 |
| TELEDDA | ASSQDRANTEVF | 0.29556066 |
| TPPGPPN | ASSQDRANTEVF | 0.23190852 |

| | | |
|---|---|---|
| TPPPPPPP | ASSQDRANTEVF | 0.24544731 |
| TQSPALI | ASSSDWGGYEQY | 0.25170863 |
| EFAAAFPQ | ASSSDWGGYEQY | 0.5112765 |
| LLSGNLDVL | ASSSDWGGYEQY | 0.2996877 |
| GPANLGNYNI | ASSSDWGGYEQY | 0.2768396 |
| TLGPTLP | ASSSDWGGYEQY | 0.24831788 |
| VYAYSSCCK | ASSSDWGGYEQY | 0.25744832 |
| APAPAPAPNI | ASSSDWGGYEQY | 0.23743555 |
| PGGPVPIP | ASSSDWGGYEQY | 0.24609792 |
| FLTVALEP | ASSSDWGGYEQY | 0.2282321 |
| SQYFGANL | ASSSDWGGYEQY | 0.2286359 |
| GGVKPAA | ASSSDWGGYEQY | 0.231232 |
| GGTGGAGGAGGW | ASSSDWGGYEQY | 0.23903027 |
| GVGGVAGLGGAGG | ASSSDWGGYEQY | 0.51584184 |
| VLLVLATTI | ASSSDWGGYEQY | 0.254098 |
| IGIFAAML | ASSSDWGGYEQY | 0.26627392 |
| AVGDLID | ASSSDWGGYEQY | 0.25326037 |
| NISPLPL | ASSSDWGGYEQY | 0.2297085 |
| VTAVFVAI | ASSSDWGGYEQY | 0.23313197 |
| SGDDNDAAL | ASSSDWGGYEQY | 0.22874735 |
| AVAALAHS | ASSSDWGGYEQY | 0.2941908 |
| GGVTVMATP | ASSSDWGGYEQY | 0.2351051 |
| AGLAANVRD | ASSSDWGGYEQY | 0.22727986 |
| TNRPDLIDPALLRPG | ASSSDWGGYEQY | 0.2875883 |
| LANLGVTAL | ASSSDWGGYEQY | 0.5205455 |
| TGERIEL | ASSSDWGGYEQY | 0.24477935 |
| NGEELAA | ASSSDWGGYEQY | 0.25205675 |
| VLSLVWG | ASSSDWGGYEQY | 0.2587227 |
| IGYALWL | ASSSDWGGYEQY | 0.24457705 |
| LSLLLAPL | ASSSDWGGYEQY | 0.51115364 |
| TLDEALK | ASSSDWGGYEQY | 0.23331824 |
| TDGAGTAEGG | ASSSDWGGYEQY | 0.5279074 |
| NAPGGAGGNGG | ASSSDWGGYEQY | 0.233389 |
| VSVFMGMGGNF | ASSSDWGGYEQY | 0.2197056 |
| LASYLAAL | ASSSDWGGYEQY | 0.23593499 |
| LVGGTPAL | ASSSDWGGYEQY | 0.2768349 |
| VRNADVL | ASSSDWGGYEQY | 0.26557407 |
| TCSSPFLVSAM | ASSSDWGGYEQY | 0.3360432 |
| VHITDAE | ASSSDWGGYEQY | 0.22954375 |
| EYAAPSF | ASSSDWGGYEQY | 0.22609146 |
| VVDAFHGGL | ASSSDWGGYEQY | 0.2830147 |
| ASVVDPQ | ASSSDWGGYEQY | 0.25416487 |
| SYNEFGGLS | ASSSDWGGYEQY | 0.26905036 |
| DPDGSVY | ASSSDWGGYEQY | 0.22899878 |
| MVSTIHATE | ASSSDWGGYEQY | 0.2251671 |
| LAAGVADPV | ASSSDWGGYEQY | 0.22868143 |
| TKEQIAA | ASSSDWGGYEQY | 0.26397118 |
| SLASPAAL | ASSSDWGGYEQY | 0.22014427 |
| IGLVFYGV | ASSSDWGGYEQY | 0.27750498 |
| FLLDASNL | ASSSDWGGYEQY | 0.256133 |
| YQVGNFP | ASSSDWGGYEQY | 0.21756923 |

| | | |
|---|---|---|
| ESLLSQST | ASSSDWGGYEQY | 0.24507068 |
| LFTVGFAAVG | ASSSDWGGYEQY | 0.2949283 |
| ATPPTPT | ASSSDWGGYEQY | 0.2383843 |
| TDLATLG | ASSSDWGGYEQY | 0.2223896 |
| SAGGTGR | ASSSDWGGYEQY | 0.28299782 |
| GFSNSGS | ASSSDWGGYEQY | 0.5420554 |
| GGNGGSGGTSV | ASSSDWGGYEQY | 0.25446087 |
| GTSVATGGAGN | ASSSDWGGYEQY | 0.22714384 |
| NGGNGGSGGTS | ASSSDWGGYEQY | 0.21435049 |
| AVVKFPH | ASSSDWGGYEQY | 0.24423054 |
| TPPPPPP | ASSSDWGGYEQY | 0.36601847 |
| AAPAAAAAPA | ASSSDWGGYEQY | 0.23726727 |
| IALNEDL | ASSSDWGGYEQY | 0.22238915 |
| IVNVSSM | ASSSDWGGYEQY | 0.2326464 |
| TGIPPAPR | ASSSDWGGYEQY | 0.28325862 |
| FDLGGGTF | ASSSDWGGYEQY | 0.22662118 |
| FDLGGGTFDVS | ASSSDWGGYEQY | 0.21722029 |
| NEPTAAA | ASSSDWGGYEQY | 0.24240354 |
| NEPTAAALA | ASSSDWGGYEQY | 0.24939126 |
| GSIINISSI | ASSSDWGGYEQY | 0.23554027 |
| VAPGNAGTA | ASSSDWGGYEQY | 0.2502562 |
| TGGGASLE | ASSSDWGGYEQY | 0.27356213 |
| VAVYNHYKRI | ASSSDWGGYEQY | 0.2265767 |
| TDPAPATT | ASSSDWGGYEQY | 0.24774644 |
| TNRPDILDPALLRPG | ASSSDWGGYEQY | 0.22378443 |
| LTTTTTS | ASSSDWGGYEQY | 0.37596938 |
| PTGSGKTL | ASSSDWGGYEQY | 0.2255807 |
| TGSGKTL | ASSSDWGGYEQY | 0.24082547 |
| TIGHVDHGK | ASSSDWGGYEQY | 0.24715331 |
| TIGHVDHGKTT | ASSSDWGGYEQY | 0.27735743 |
| GHVDHGKTT | ASSSDWGGYEQY | 0.25036833 |
| AQFEHTLL | ASSSDWGGYEQY | 0.23640227 |
| IVNQDEL | ASSSDWGGYEQY | 0.22509345 |
| TNGSQFF | ASSSDWGGYEQY | 0.21339203 |
| TGKGIVE | ASSSDWGGYEQY | 0.27735236 |
| LISNASDA | ASSSDWGGYEQY | 0.22831799 |
| AELRTLGELTKIAKAHGAQ | ASSSDWGGYEQY | 0.27111664 |
| APNAVAAAEPSW | ASSSDWGGYEQY | 0.27449444 |
| APQVAGAATAVL | ASSSDWGGYEQY | 0.2595787 |
| ARRDPAAAAAILPTDA | ASSSDWGGYEQY | 0.23946717 |
| DAAARDVLAEAGLAEYF | ASSSDWGGYEQY | 0.2194423 |
| DDQIVLSAKANLAKVEQ | ASSSDWGGYEQY | 0.22450995 |
| DELLPDTGEDSDEDGH | ASSSDWGGYEQY | 0.23833174 |
| ELPEAPANRATVPQD | ASSSDWGGYEQY | 0.23774886 |
| FGQSKIERVTGIG | ASSSDWGGYEQY | 0.24367385 |
| GAGGTGSTAGGAGGAGGAGGLY | ASSSDWGGYEQY | 0.2358053 |
| GHGGTGAAGGNGGTGGAGGSGI | ASSSDWGGYEQY | 0.24273047 |
| GNGGNGGAGGEGGPGGAGGAGA | ASSSDWGGYEQY | 0.22946411 |
| GSTAGNGGNGGNATNGTIAGQP | ASSSDWGGYEQY | 0.2388708 |
| GTGGDGGNGGNGGTGG | ASSSDWGGYEQY | 0.35094494 |
| GVRRPAGIPADAPV | ASSSDWGGYEQY | 0.24895781 |

| | | |
|---|---|---|
| GYRQVTDAHLLTLARR | ASSSDWGGYEQY | 0.24759412 |
| IAKMGQDGHDRG | ASSSDWGGYEQY | 0.275876 |
| KAEPSIEIQYT | ASSSDWGGYEQY | 0.2467446 |
| KPVRIRMAADEDTVAVT | ASSSDWGGYEQY | 0.26737097 |
| MPPRIAGMRLLVIKPEP | ASSSDWGGYEQY | 0.30468577 |
| MKFYAVALFRSRGVPPTR | ASSSDWGGYEQY | 0.24968894 |
| NAGGFGILGAVAHSP | ASSSDWGGYEQY | 0.25017628 |
| NEGREITLAMAEM | ASSSDWGGYEQY | 0.22472516 |
| NELVEWSHPLGHRWRWE | ASSSDWGGYEQY | 0.22264163 |
| NLGAGGTGGNGGNPG | ASSSDWGGYEQY | 0.25835815 |
| NTEWPKWETVMAADPVD | ASSSDWGGYEQY | 0.28640875 |
| PASAAPAETQP | ASSSDWGGYEQY | 0.2741429 |
| QQSSDGKNGVRAAEITGE | ASSSDWGGYEQY | 0.23741037 |
| RAPDAVTTSWWKE | ASSSDWGGYEQY | 0.23730963 |
| SGFGNVGAHTSGWFNQ | ASSSDWGGYEQY | 0.22991286 |
| SGVAGAVGQAASVGGLKV | ASSSDWGGYEQY | 0.24129063 |
| SNGGAGTGSAGGNGGT | ASSSDWGGYEQY | 0.23049483 |
| TGGAGGEGGAAG | ASSSDWGGYEQY | 0.23166189 |
| TGGDGGAAGAGGAPGHG | ASSSDWGGYEQY | 0.21933985 |
| TMPPLLAAIANAGRHGV | ASSSDWGGYEQY | 0.29375225 |
| TSSTAGNSGGA | ASSSDWGGYEQY | 0.22511937 |
| VPATVPVGILC | ASSSDWGGYEQY | 0.25261354 |
| VQRPDSGGVVLTGRIS | ASSSDWGGYEQY | 0.2607 |
| WTPNPVPATKGVPAGLVQAT | ASSSDWGGYEQY | 0.22231625 |
| YALKRALAKFSDRFDVVII | ASSSDWGGYEQY | 0.24045582 |
| YLIGSVIV | ASSSDWGGYEQY | 0.2396477 |
| YGQLVAVID | ASSSDWGGYEQY | 0.24532773 |
| VTVGGMTLPA | ASSSDWGGYEQY | 0.25665775 |
| VLALPAV | ASSSDWGGYEQY | 0.23357615 |
| VIPPTLNYET | ASSSDWGGYEQY | 0.28243202 |
| VGMGGMHPSSG | ASSSDWGGYEQY | 0.3071942 |
| VGLAGGAATA | ASSSDWGGYEQY | 0.2634663 |
| VGGAGGAGGSGP | ASSSDWGGYEQY | 0.24201047 |
| VFAAFMV | ASSSDWGGYEQY | 0.29689085 |
| TLYDKLR | ASSSDWGGYEQY | 0.2567846 |
| TLAVFTVL | ASSSDWGGYEQY | 0.25160182 |
| TGSVQPARV | ASSSDWGGYEQY | 0.21586674 |
| TGGGAGSGGT | ASSSDWGGYEQY | 0.25974572 |
| SHLETSL | ASSSDWGGYEQY | 0.25174022 |
| SFTTGMT | ASSSDWGGYEQY | 0.23083854 |
| QVEVAKRMGVRQPTVS | ASSSDWGGYEQY | 0.21848264 |
| QGSGGPALAM | ASSSDWGGYEQY | 0.38321432 |
| MLAPSNP | ASSSDWGGYEQY | 0.39799136 |
| LVNNAAI | ASSSDWGGYEQY | 0.23894279 |
| LTGEINTLA | ASSSDWGGYEQY | 0.2805665 |
| LSIFGPV | ASSSDWGGYEQY | 0.21642347 |
| LLALLANL | ASSSDWGGYEQY | 0.29260182 |
| LISNASDALDKLR | ASSSDWGGYEQY | 0.36973834 |
| LINEAVP | ASSSDWGGYEQY | 0.22731534 |
| LGYAFSGL | ASSSDWGGYEQY | 0.31910628 |
| LDFVAVAL | ASSSDWGGYEQY | 0.24499026 |

| | | |
|---|---|---|
| LAQPGAE | ASSSDWGGYEQY | 0.23627676 |
| ISNLGAVTGF | ASSSDWGGYEQY | 0.32181236 |
| HQEDPGRI | ASSSDWGGYEQY | 0.26808423 |
| GTPGEPTP | ASSSDWGGYEQY | 0.286499 |
| GFFNSGS | ASSSDWGGYEQY | 0.23894808 |
| GAGGGGLTAGAN | ASSSDWGGYEQY | 0.25278383 |
| FVDALVNM | ASSSDWGGYEQY | 0.3052667 |
| FLTVAPEL | ASSSDWGGYEQY | 0.29355434 |
| FETLTAL | ASSSDWGGYEQY | 0.21811366 |
| DVINPIEL | ASSSDWGGYEQY | 0.25099382 |
| APVSGAPV | ASSSDWGGYEQY | 0.24890584 |
| ALVGGLAW | ASSSDWGGYEQY | 0.43343592 |
| ALAAAAATAI | ASSSDWGGYEQY | 0.38702333 |
| AFGLGLLL | ASSSDWGGYEQY | 0.27154544 |
| AFFAAASL | ASSSDWGGYEQY | 0.24741532 |
| AAAALPAFSPP | ASSSDWGGYEQY | 0.3928306 |
| VVLAPAL | ASSSDWGGYEQY | 0.23660214 |
| TIGHVDHGKTTL | ASSSDWGGYEQY | 0.23917446 |
| LISNASDALDK | ASSSDWGGYEQY | 0.2614378 |
| LISNASDAL | ASSSDWGGYEQY | 0.5656798 |
| ISNASDALDK | ASSSDWGGYEQY | 0.8786133 |
| NQAETLVYQTE | ASSSDWGGYEQY | 0.8000478 |
| TGGTGGTAGAG | ASSSDWGGYEQY | 0.79094005 |
| TGSTAGGAGGA | ASSSDWGGYEQY | 0.5239665 |
| DPLAVPVAAGPV | ASSSDWGGYEQY | 0.23824854 |
| GGTGGTPGNGGN | ASSSDWGGYEQY | 0.70500076 |
| PAGLAGVDPATL | ASSSDWGGYEQY | 0.50455034 |
| GNNLVVEAPHAGASV | ASSSDWGGYEQY | 0.8314165 |
| KVTRIMSASTLEQPA | ASSSDWGGYEQY | 0.8532794 |
| LGFALPIALPRRTVV | ASSSDWGGYEQY | 0.5209601 |
| TRLGNLTARKLVVLV | ASSSDWGGYEQY | 0.877524 |
| WSQQRRLIYFPSAGP | ASSSDWGGYEQY | 0.87226105 |
| GPNGNPDPMAAAVDIR | ASSSDWGGYEQY | 0.35614994 |
| TQRCVSVADLRAAASR | ASSSDWGGYEQY | 0.7470386 |
| QFVLDVAEHPVEVLTETV | ASSSDWGGYEQY | 0.5356573 |
| GGAGGTGGAGGNGGTQAGDG | ASSSDWGGYEQY | 0.74156797 |
| PLLATLRAIAVDVDAKPAQV | ASSSDWGGYEQY | 0.35477167 |
| APKFRTGGYMIDFWGVGYQVAK | ASSSDWGGYEQY | 0.8548753 |
| GAGGAGGAGGAAGGPGGTGGTG | ASSSDWGGYEQY | 0.6237329 |
| IGALGLLATDVAPAGSVAFGVA | ASSSDWGGYEQY | 0.16571344 |
| AGGAGGIGGNGGAGGAGGVLMG | ASSSDWGGYEQY | 0.8041347 |
| ADELIGLL | ASSSDWGGYEQY | 0.7539439 |
| ADRALKA | ASSSDWGGYEQY | 0.73723245 |
| AGAVGQVGGAGGA | ASSSDWGGYEQY | 0.8690993 |
| AGGGGGLGGAP | ASSSDWGGYEQY | 0.5284617 |
| AGGLLYGNGGAG | ASSSDWGGYEQY | 0.54066336 |
| AGLESIGL | ASSSDWGGYEQY | 0.84584653 |
| AGLKTGII | ASSSDWGGYEQY | 0.8734067 |
| ASLAALLL | ASSSDWGGYEQY | 0.51730835 |
| ASVINISSI | ASSSDWGGYEQY | 0.8235872 |
| ATPPGPA | ASSSDWGGYEQY | 0.8822025 |

| | | |
|---|---|---|
| ATVASALV | ASSSDWGGYEQY | 0.7571836 |
| ATVASLIL | ASSSDWGGYEQY | 0.85968363 |
| AVNGPVLIRH | ASSSDWGGYEQY | 0.8721118 |
| DGAAGAVGI | ASSSDWGGYEQY | 0.62232625 |
| DPLLHGI | ASSSDWGGYEQY | 0.86812997 |
| ELGAALGAL | ASSSDWGGYEQY | 0.87977517 |
| ELGRTQAV | ASSSDWGGYEQY | 0.66988695 |
| FVAGGYLS | ASSSDWGGYEQY | 0.8031462 |
| GASGGAGGAGGW | ASSSDWGGYEQY | 0.37918913 |
| GGAGAGGDGGAGGI | ASSSDWGGYEQY | 0.71800804 |
| GTGHNGSQPGMGGNG | ASSSDWGGYEQY | 0.23395512 |
| GYAAASASAS | ASSSDWGGYEQY | 0.822044 |
| HQVDVAAVQQG | ASSSDWGGYEQY | 0.87725234 |
| IGAAIAHT | ASSSDWGGYEQY | 0.7630148 |
| IGMIEVAQL | ASSSDWGGYEQY | 0.5707153 |
| IRLAPEHTSAMD | ASSSDWGGYEQY | 0.37031695 |
| IYNVSVL | ASSSDWGGYEQY | 0.76209706 |
| KQGFLGM | ASSSDWGGYEQY | 0.604663 |
| LESVITQL | ASSSDWGGYEQY | 0.24051185 |
| LGLGALVVV | ASSSDWGGYEQY | 0.65425146 |
| LIGNGGNGGSGGI | ASSSDWGGYEQY | 0.8650986 |
| LNAINAPTL | ASSSDWGGYEQY | 0.8508326 |
| LQLLNTGV | ASSSDWGGYEQY | 0.6264601 |
| LQTSIEF | ASSSDWGGYEQY | 0.88783264 |
| MSFAVSEL | ASSSDWGGYEQY | 0.848203 |
| NTGALGL | ASSSDWGGYEQY | 0.84275717 |
| PPGAGGAGGAGTT | ASSSDWGGYEQY | 0.6458466 |
| QLVGLGI | ASSSDWGGYEQY | 0.48297876 |
| RFLEQVHQL | ASSSDWGGYEQY | 0.85918075 |
| RGVGYVLR | ASSSDWGGYEQY | 0.6936748 |
| SATGVFAGVI | ASSSDWGGYEQY | 0.6818954 |
| SLPAITLP | ASSSDWGGYEQY | 0.841739 |
| STALAFTV | ASSSDWGGYEQY | 0.72243416 |
| SVLTVGLGK | ASSSDWGGYEQY | 0.6567342 |
| TALAGAGSGI | ASSSDWGGYEQY | 0.75871694 |
| TGLPFTGI | ASSSDWGGYEQY | 0.86408836 |
| VAEHVTV | ASSSDWGGYEQY | 0.5797807 |
| VDIVLNSL | ASSSDWGGYEQY | 0.41063285 |
| VLTGGGAVL | ASSSDWGGYEQY | 0.64137363 |
| VNVGLAYGI | ASSSDWGGYEQY | 0.7874604 |
| VSSAAVGAV | ASSSDWGGYEQY | 0.87031746 |
| VTSVYEAGV | ASSSDWGGYEQY | 0.797498 |
| WVALDRGA | ASSSDWGGYEQY | 0.759374 |
| YLVGSVVI | ASSSDWGGYEQY | 0.83083737 |
| GPPGTGKTLL | ASSSDWGGYEQY | 0.8181858 |
| GVGMTKF | ASSSDWGGYEQY | 0.78840387 |
| IGIGGSDL | ASSSDWGGYEQY | 0.8255521 |
| IGTLLAEL | ASSSDWGGYEQY | 0.67918366 |
| IYTSGTT | ASSSDWGGYEQY | 0.8631346 |
| LSFPAGV | ASSSDWGGYEQY | 0.84107125 |
| LYTSGTT | ASSSDWGGYEQY | 0.83491564 |

| | | |
|---|---|---|
| VGAFPFGF | ASSSDWGGYEQY | 0.8654103 |
| VGAGTVEF | ASSSDWGGYEQY | 0.80487955 |
| VGKSTLLL | ASSSDWGGYEQY | 0.7408019 |
| AALAHAGGGIV | ASSSDWGGYEQY | 0.23810339 |
| AALVAVSLL | ASSSDWGGYEQY | 0.58442354 |
| AAVSRAVAAL | ASSSDWGGYEQY | 0.80946326 |
| AGGSSGIGGF | ASSSDWGGYEQY | 0.8131408 |
| AGLTVANANG | ASSSDWGGYEQY | 0.84509885 |
| AGLVGAVL | ASSSDWGGYEQY | 0.83121884 |
| ALSQAFTV | ASSSDWGGYEQY | 0.6659738 |
| ALTFSVSL | ASSSDWGGYEQY | 0.8677051 |
| ALTGAALVAL | ASSSDWGGYEQY | 0.8745693 |
| ASFSIGAGVL | ASSSDWGGYEQY | 0.35405824 |
| DGSVYKI | ASSSDWGGYEQY | 0.6452869 |
| EQLAEIV | ASSSDWGGYEQY | 0.8921849 |
| FAVSELTVM | ASSSDWGGYEQY | 0.8194917 |
| GAAPLAAGL | ASSSDWGGYEQY | 0.63932407 |
| GADGGDGGSGGAGG | ASSSDWGGYEQY | 0.67679065 |
| GFGLDTGGAGG | ASSSDWGGYEQY | 0.8946248 |
| GLASGGLL | ASSSDWGGYEQY | 0.64857376 |
| GRIAGAAVLL | ASSSDWGGYEQY | 0.89146364 |
| GSAGSLNTSF | ASSSDWGGYEQY | 0.23722856 |
| GTGQIGI | ASSSDWGGYEQY | 0.6498556 |
| GTLIALIL | ASSSDWGGYEQY | 0.43828645 |
| IAATMGFVVG | ASSSDWGGYEQY | 0.21505207 |
| IFAQALL | ASSSDWGGYEQY | 0.83715653 |
| IGAAVALGF | ASSSDWGGYEQY | 0.70689857 |
| IGLTPLAL | ASSSDWGGYEQY | 0.8502984 |
| ILGSNTSTL | ASSSDWGGYEQY | 0.7296065 |
| IVFSLTVL | ASSSDWGGYEQY | 0.8620635 |
| LASKATGF | ASSSDWGGYEQY | 0.7925832 |
| LASVALIL | ASSSDWGGYEQY | 0.78866935 |
| LFAVGLTL | ASSSDWGGYEQY | 0.4725228 |
| LFNGGIGTYI | ASSSDWGGYEQY | 0.80435014 |
| LTLGAGII | ASSSDWGGYEQY | 0.87960654 |
| LVAAALGDI | ASSSDWGGYEQY | 0.8352486 |
| NTAFVQLGI | ASSSDWGGYEQY | 0.3449427 |
| QLFSGAEL | ASSSDWGGYEQY | 0.7498462 |
| SAATFITL | ASSSDWGGYEQY | 0.717521 |
| SNYGLANLGV | ASSSDWGGYEQY | 0.18449707 |
| SYGGAYAVM | ASSSDWGGYEQY | 0.15252163 |
| TTGVGGLTLG | ASSSDWGGYEQY | 0.6391109 |
| TTQTIYI | ASSSDWGGYEQY | 0.69802344 |
| VAAGAGLGAVI | ASSSDWGGYEQY | 0.857679 |
| VAAVVTLL | ASSSDWGGYEQY | 0.84839714 |
| VGFLGGLI | ASSSDWGGYEQY | 0.7310095 |
| VGVIPILL | ASSSDWGGYEQY | 0.84317327 |
| VIFASSLI | ASSSDWGGYEQY | 0.82111275 |
| VKGGDGGAAAG | ASSSDWGGYEQY | 0.08319222 |
| VTKTFTNI | ASSSDWGGYEQY | 0.576511 |
| VVAAAILL | ASSSDWGGYEQY | 0.35066158 |

| | | |
|---|---|---|
| YAALDVEL | ASSSDWGGYEQY | 0.8816766 |
| LASVALLL | ASSSDWGGYEQY | 0.57958937 |
| LDAVHRV | ASSSDWGGYEQY | 0.6839539 |
| QGLSSSIF | ASSSDWGGYEQY | 0.7881509 |
| RGTFANIRL | ASSSDWGGYEQY | 0.8295342 |
| RQLWWGHRI | ASSSDWGGYEQY | 0.74986535 |
| YGPPGTGKTLL | ASSSDWGGYEQY | 0.54056203 |
| AALVGAPANVA | ASSSDWGGYEQY | 0.11870155 |
| AATFRRTE | ASSSDWGGYEQY | 0.8404406 |
| AGALGGGI | ASSSDWGGYEQY | 0.42387193 |
| AGGAGGAGGIGAG | ASSSDWGGYEQY | 0.5436412 |
| AGSGALGL | ASSSDWGGYEQY | 0.2693987 |
| AQIADQI | ASSSDWGGYEQY | 0.1550156 |
| ASAVAVTYL | ASSSDWGGYEQY | 0.28092828 |
| ASTVLGGTGG | ASSSDWGGYEQY | 0.84898865 |
| AVGGEPAK | ASSSDWGGYEQY | 0.584789 |
| CEQGTFGGTC | ASSSDWGGYEQY | 0.7583176 |
| DEGKAPAG | ASSSDWGGYEQY | 0.8965433 |
| DTVVFPAL | ASSSDWGGYEQY | 0.82021344 |
| FEGITMAG | ASSSDWGGYEQY | 0.6921989 |
| FPLVVQD | ASSSDWGGYEQY | 0.65568745 |
| GAFTTPQ | ASSSDWGGYEQY | 0.44734317 |
| GFGGFGTAGGVGGA | ASSSDWGGYEQY | 0.51830596 |
| GFLAQFL | ASSSDWGGYEQY | 0.6449541 |
| GGAGGAGGNAGVA | ASSSDWGGYEQY | 0.8865209 |
| GGVGTAGAGGAGGAGG | ASSSDWGGYEQY | 0.6391996 |
| GNGGGGDAGNAGSGGN | ASSSDWGGYEQY | 0.43322933 |
| GTGGAGGAGNDAGS | ASSSDWGGYEQY | 0.7052732 |
| GTTNTGWFNS | ASSSDWGGYEQY | 0.88499725 |
| IFMGPLFTL | ASSSDWGGYEQY | 0.88372105 |
| ILAHAIEAI | ASSSDWGGYEQY | 0.61963284 |
| IRSAIREAL | ASSSDWGGYEQY | 0.5910014 |
| ITFAGGG | ASSSDWGGYEQY | 0.651345 |
| ITGMSGHQEV | ASSSDWGGYEQY | 0.78018856 |
| IVAGLACTL | ASSSDWGGYEQY | 0.60682195 |
| IVGGGLAGA | ASSSDWGGYEQY | 0.4906069 |
| IVTGVLNI | ASSSDWGGYEQY | 0.844895 |
| LAQQGCI | ASSSDWGGYEQY | 0.9008751 |
| LCLHVEN | ASSSDWGGYEQY | 0.7007954 |
| LDAISDGS | ASSSDWGGYEQY | 0.23580042 |
| LGGFAAML | ASSSDWGGYEQY | 0.7519477 |
| LGGKNPMI | ASSSDWGGYEQY | 0.5480233 |
| LGPKGVAAIL | ASSSDWGGYEQY | 0.6549598 |
| LMVGVSI | ASSSDWGGYEQY | 0.7307749 |
| LVGAAFVL | ASSSDWGGYEQY | 0.44287455 |
| LVLAVAPL | ASSSDWGGYEQY | 0.785705 |
| MAAIAVIDAA | ASSSDWGGYEQY | 0.88859487 |
| NNGINAELQ | ASSSDWGGYEQY | 0.5993308 |
| RGEPLVLVHG | ASSSDWGGYEQY | 0.81979096 |
| RGGAGGAAGLI | ASSSDWGGYEQY | 0.8794897 |
| RSYRMHPAV | ASSSDWGGYEQY | 0.85106474 |

| | | |
|---|---|---|
| SAAHLLAGQLA | ASSSDWGGYEQY | 0.88014257 |
| SAAVGSY | ASSSDWGGYEQY | 0.85001236 |
| SPHAVLHRI | ASSSDWGGYEQY | 0.7633295 |
| SQIAKVL | ASSSDWGGYEQY | 0.7628879 |
| TALGHLVP | ASSSDWGGYEQY | 0.81549966 |
| TAPTTSGAA | ASSSDWGGYEQY | 0.7727083 |
| TDGDDGNGGSPGT | ASSSDWGGYEQY | 0.7249638 |
| TDGLGEN | ASSSDWGGYEQY | 0.59778035 |
| TGGPPLPGQA | ASSSDWGGYEQY | 0.8712103 |
| TIPAITI | ASSSDWGGYEQY | 0.59414995 |
| TIPGFTI | ASSSDWGGYEQY | 0.8590942 |
| TSIASGALA | ASSSDWGGYEQY | 0.29616618 |
| TVTLPGAAI | ASSSDWGGYEQY | 0.48386353 |
| VASGLSAALP | ASSSDWGGYEQY | 0.86427 |
| VGAGVIGI | ASSSDWGGYEQY | 0.10055075 |
| VGPKRFGV | ASSSDWGGYEQY | 0.1745012 |
| VNVAFPD | ASSSDWGGYEQY | 0.7790806 |
| VSMTNSGVS | ASSSDWGGYEQY | 0.870538 |
| VTDVLGDLSV | ASSSDWGGYEQY | 0.12111634 |
| VVVVGAGFAGL | ASSSDWGGYEQY | 0.5634166 |
| VYTVGTVV | ASSSDWGGYEQY | 0.2986193 |
| WTTFSGP | ASSSDWGGYEQY | 0.797392 |
| YDELVEAF | ASSSDWGGYEQY | 0.79538745 |
| YGTGGAGGNGG | ASSSDWGGYEQY | 0.2134646 |
| YTNVMAV | ASSSDWGGYEQY | 0.8482038 |
| FLEQVHQL | ASSSDWGGYEQY | 0.344734 |
| GLSLLAVAV | ASSSDWGGYEQY | 0.43611333 |
| LEQVHQL | ASSSDWGGYEQY | 0.82851744 |
| LNAAGAI | ASSSDWGGYEQY | 0.87318766 |
| PAVHASL | ASSSDWGGYEQY | 0.43707305 |
| TELEDDA | ASSSDWGGYEQY | 0.8334035 |
| TPPGPPN | ASSSDWGGYEQY | 0.6672174 |
| TPPPPPPP | ASSSDWGGYEQY | 0.88907456 |
| TQSPALI | ASSLEGGSNERLF | 0.5506817 |
| EFAAAFPQ | ASSLEGGSNERLF | 0.3423267 |
| LLSGNLDVL | ASSLEGGSNERLF | 0.8666858 |
| GPANLGNYNI | ASSLEGGSNERLF | 0.82773054 |
| TLGPTLP | ASSLEGGSNERLF | 0.75162613 |
| VYAYSSCCK | ASSLEGGSNERLF | 0.84737587 |
| APAPAPAPNI | ASSLEGGSNERLF | 0.544886 |
| PGGPVPIP | ASSLEGGSNERLF | 0.7510394 |
| FLTVALEP | ASSLEGGSNERLF | 0.8704276 |
| SQYFGANL | ASSLEGGSNERLF | 0.86170805 |
| GGVKPAA | ASSLEGGSNERLF | 0.44573963 |
| GGTGGAGGAGGW | ASSLEGGSNERLF | 0.86073524 |
| GVGGVAGLGGAGG | ASSLEGGSNERLF | 0.8002597 |
| VLLVLATTI | ASSLEGGSNERLF | 0.5421844 |
| IGIFAAML | ASSLEGGSNERLF | 0.88510567 |
| AVGDLID | ASSLEGGSNERLF | 0.7305115 |
| NISPLPL | ASSLEGGSNERLF | 0.85297096 |
| VTAVFVAI | ASSLEGGSNERLF | 0.8254041 |

| | | |
|---|---|---|
| SGDDNDAAL | ASSLEGGSNERLF | 0.8776089 |
| AVAALAHS | ASSLEGGSNERLF | 0.26861295 |
| GGVTVMATP | ASSLEGGSNERLF | 0.80558103 |
| AGLAANVRD | ASSLEGGSNERLF | 0.760327 |
| TNRPDLIDPALLRPG | ASSLEGGSNERLF | 0.8454473 |
| LANLGVTAL | ASSLEGGSNERLF | 0.7204411 |
| TGERIEL | ASSLEGGSNERLF | 0.5163301 |
| NGEELAA | ASSLEGGSNERLF | 0.807505 |
| VLSLVWG | ASSLEGGSNERLF | 0.8843518 |
| IGYALWL | ASSLEGGSNERLF | 0.7843373 |
| LSLLLAPL | ASSLEGGSNERLF | 0.80922127 |
| TLDEALK | ASSLEGGSNERLF | 0.8897406 |
| TDGAGTAEGG | ASSLEGGSNERLF | 0.47878432 |
| NAPGGAGGNGG | ASSLEGGSNERLF | 0.7979483 |
| VSVFMGMGGNF | ASSLEGGSNERLF | 0.85839933 |
| LASYLAAL | ASSLEGGSNERLF | 0.86531794 |
| LVGGTPAL | ASSLEGGSNERLF | 0.7182269 |
| VRNADVL | ASSLEGGSNERLF | 0.8843008 |
| TCSSPFLVSAM | ASSLEGGSNERLF | 0.51127964 |
| VHITDAE | ASSLEGGSNERLF | 0.8615928 |
| EYAAPSF | ASSLEGGSNERLF | 0.68519413 |
| VVDAFHGGL | ASSLEGGSNERLF | 0.7770837 |
| ASVVDPQ | ASSLEGGSNERLF | 0.52442247 |
| SYNEFGGLS | ASSLEGGSNERLF | 0.8349115 |
| DPDGSVY | ASSLEGGSNERLF | 0.85192895 |
| MVSTIHATE | ASSLEGGSNERLF | 0.8606615 |
| LAAGVADPV | ASSLEGGSNERLF | 0.17426285 |
| TKEQIAA | ASSLEGGSNERLF | 0.8521607 |
| SLASPAAL | ASSLEGGSNERLF | 0.865171 |
| IGLVFYGV | ASSLEGGSNERLF | 0.89802504 |
| FLLDASNL | ASSLEGGSNERLF | 0.6583477 |
| YQVGNFP | ASSLEGGSNERLF | 0.8284091 |
| ESLLSQST | ASSLEGGSNERLF | 0.67349195 |
| LFTVGFAAVG | ASSLEGGSNERLF | 0.71076906 |
| ATPPTPT | ASSLEGGSNERLF | 0.50615466 |
| TDLATLG | ASSLEGGSNERLF | 0.5654668 |
| SAGGTGR | ASSLEGGSNERLF | 0.746858 |
| GFSNSGS | ASSLEGGSNERLF | 0.6459579 |
| GGNGGSGGTSV | ASSLEGGSNERLF | 0.8549412 |
| GTSVATGGAGN | ASSLEGGSNERLF | 0.8580664 |
| NGGNGGSGGTS | ASSLEGGSNERLF | 0.87038314 |
| AVVKFPH | ASSLEGGSNERLF | 0.8710053 |
| TPPPPPP | ASSLEGGSNERLF | 0.8738186 |
| AAPAAAAAPA | ASSLEGGSNERLF | 0.83333707 |
| IALNEDL | ASSLEGGSNERLF | 0.6099003 |
| IVNVSSM | ASSLEGGSNERLF | 0.68694884 |
| TGIPPAPR | ASSLEGGSNERLF | 0.45578524 |
| FDLGGGTF | ASSLEGGSNERLF | 0.75454956 |
| FDLGGGTFDVS | ASSLEGGSNERLF | 0.6514209 |
| NEPTAAA | ASSLEGGSNERLF | 0.8093885 |
| NEPTAAALA | ASSLEGGSNERLF | 0.09997088 |

| | | |
|---|---|---|
| GSIINISSI | ASSLEGGSNERLF | 0.80931973 |
| VAPGNAGTA | ASSLEGGSNERLF | 0.88914305 |
| TGGGASLE | ASSLEGGSNERLF | 0.62305653 |
| VAVYNHYKRI | ASSLEGGSNERLF | 0.8008231 |
| TDPAPATT | ASSLEGGSNERLF | 0.81175077 |
| TNRPDILDPALLRPG | ASSLEGGSNERLF | 0.6619998 |
| LTTTTTS | ASSLEGGSNERLF | 0.84166765 |
| PTGSGKTL | ASSLEGGSNERLF | 0.60173833 |
| TGSGKTL | ASSLEGGSNERLF | 0.8418188 |
| TIGHVDHGK | ASSLEGGSNERLF | 0.8775605 |
| TIGHVDHGKTT | ASSLEGGSNERLF | 0.81035495 |
| GHVDHGKTT | ASSLEGGSNERLF | 0.7670231 |
| AQFEHTLL | ASSLEGGSNERLF | 0.74642134 |
| IVNQDEL | ASSLEGGSNERLF | 0.19529237 |
| TNGSQFF | ASSLEGGSNERLF | 0.79453176 |
| TGKGIVE | ASSLEGGSNERLF | 0.08907727 |
| LISNASDA | ASSLEGGSNERLF | 0.76475596 |
| AELRTLGELTKIAKAHGAQ | ASSLEGGSNERLF | 0.429207 |
| APNAVAAAEPSW | ASSLEGGSNERLF | 0.82374763 |
| APQVAGAATAVL | ASSLEGGSNERLF | 0.81701636 |
| ARRDPAAAAAILPTDA | ASSLEGGSNERLF | 0.8121467 |
| DAAARDVLAEAGLAEYF | ASSLEGGSNERLF | 0.42652744 |
| DDQIVLSAKANLAKVEQ | ASSLEGGSNERLF | 0.7730234 |
| DELLPDTGEDSDEDGH | ASSLEGGSNERLF | 0.7701991 |
| ELPEAPANRATVPQD | ASSLEGGSNERLF | 0.43123394 |
| FGQSKIERVTGIG | ASSLEGGSNERLF | 0.85322416 |
| GAGGTGSTAGGAGGAGGAGGLY | ASSLEGGSNERLF | 0.72908735 |
| GHGGTGAAGGNGGTGGAGGSGI | ASSLEGGSNERLF | 0.71212435 |
| GNGGNGGAGGEGGPGGAGGAGA | ASSLEGGSNERLF | 0.87337995 |
| GSTAGNGGNGGNATNGTIAGQP | ASSLEGGSNERLF | 0.7887291 |
| GTGGDGGNGGNGGTGG | ASSLEGGSNERLF | 0.21432333 |
| GVRRPAGIPADAPV | ASSLEGGSNERLF | 0.7171784 |
| GYRQVTDAHLLTLARR | ASSLEGGSNERLF | 0.76687443 |
| IAKMGQDGHDRG | ASSLEGGSNERLF | 0.5523985 |
| KAEPSIEIQYT | ASSLEGGSNERLF | 0.09954916 |
| KPVRIRMAADEDTVAVT | ASSLEGGSNERLF | 0.8170333 |
| MPPRIAGMRLLVIKPEP | ASSLEGGSNERLF | 0.5710553 |
| MKFYAVALFRSRGVPPTR | ASSLEGGSNERLF | 0.46778297 |
| NAGGFGILGAVAHSP | ASSLEGGSNERLF | 0.66747963 |
| NEGREITLAMAEM | ASSLEGGSNERLF | 0.8657099 |
| NELVEWSHPLGHRWRWE | ASSLEGGSNERLF | 0.4687822 |
| NLGAGGTGGNGGNPG | ASSLEGGSNERLF | 0.70425224 |
| NTEWPKWETVMAADPVD | ASSLEGGSNERLF | 0.8278425 |
| PASAAPAETQP | ASSLEGGSNERLF | 0.8550387 |
| QQSSDGKNGVRAAEITGE | ASSLEGGSNERLF | 0.87706864 |
| RAPDAVTTSWWKE | ASSLEGGSNERLF | 0.79604954 |
| SGFGNVGAHTSGWFNQ | ASSLEGGSNERLF | 0.65101194 |
| SGVAGAVGQAASVGGLKV | ASSLEGGSNERLF | 0.8333086 |
| SNGGAGTGSAGGNGGT | ASSLEGGSNERLF | 0.60845685 |
| TGGAGGEGGAAG | ASSLEGGSNERLF | 0.35177904 |
| TGGDGGAAGAGGAPGHG | ASSLEGGSNERLF | 0.2713275 |

| | | |
|---|---|---|
| TMPPLLAAIANAGRHGV | ASSLEGGSNERLF | 0.6170642 |
| TSSTAGNSGGA | ASSLEGGSNERLF | 0.30140784 |
| VPATVPVGILC | ASSLEGGSNERLF | 0.72438467 |
| VQRPDSGGVVLTGRIS | ASSLEGGSNERLF | 0.864941 |
| WTPNPVPATKGVPAGLVQAT | ASSLEGGSNERLF | 0.6279454 |
| YALKRALAKFSDRFDVVII | ASSLEGGSNERLF | 0.32480276 |
| YLIGSVIV | ASSLEGGSNERLF | 0.6867775 |
| YGQLVAVID | ASSLEGGSNERLF | 0.45942146 |
| VTVGGMTLPA | ASSLEGGSNERLF | 0.8265097 |
| VLALPAV | ASSLEGGSNERLF | 0.7438797 |
| VIPPTLNYET | ASSLEGGSNERLF | 0.89981514 |
| VGMGGMHPSSG | ASSLEGGSNERLF | 0.86520386 |
| VGLAGGAATA | ASSLEGGSNERLF | 0.73542464 |
| VGGAGGAGGSGP | ASSLEGGSNERLF | 0.8542269 |
| VFAAFMV | ASSLEGGSNERLF | 0.6612568 |
| TLYDKLR | ASSLEGGSNERLF | 0.7752571 |
| TLAVFTVL | ASSLEGGSNERLF | 0.74977046 |
| TGSVQPARV | ASSLEGGSNERLF | 0.80542207 |
| TGGGAGSGGT | ASSLEGGSNERLF | 0.7326296 |
| SHLETSL | ASSLEGGSNERLF | 0.6672213 |
| SFTTGMT | ASSLEGGSNERLF | 0.19234167 |
| QVEVAKRMGVRQPTVS | ASSLEGGSNERLF | 0.83574986 |
| QGSGGPALAM | ASSLEGGSNERLF | 0.66850084 |
| MLAPSNP | ASSLEGGSNERLF | 0.2675864 |
| LVNNAAI | ASSLEGGSNERLF | 0.6793879 |
| LTGEINTLA | ASSLEGGSNERLF | 0.8872993 |
| LSIFGPV | ASSLEGGSNERLF | 0.88130856 |
| LLALLANL | ASSLEGGSNERLF | 0.6939791 |
| LISNASDALDKLR | ASSLEGGSNERLF | 0.49036008 |
| LINEAVP | ASSLEGGSNERLF | 0.8000413 |
| LGYAFSGL | ASSLEGGSNERLF | 0.44696122 |
| LDFVAVAL | ASSLEGGSNERLF | 0.8085475 |
| LAQPGAE | ASSLEGGSNERLF | 0.7810289 |
| ISNLGAVTGF | ASSLEGGSNERLF | 0.79007876 |
| HQEDPGRI | ASSLEGGSNERLF | 0.587636 |
| GTPGEPTP | ASSLEGGSNERLF | 0.8168529 |
| GFFNSGS | ASSLEGGSNERLF | 0.7105243 |
| GAGGGGLTAGAN | ASSLEGGSNERLF | 0.81322134 |
| FVDALVNM | ASSLEGGSNERLF | 0.11163728 |
| FLTVAPEL | ASSLEGGSNERLF | 0.8022282 |
| FETLTAL | ASSLEGGSNERLF | 0.67168725 |
| DVINPIEL | ASSLEGGSNERLF | 0.4301218 |
| APVSGAPV | ASSLEGGSNERLF | 0.28227782 |
| ALVGGLAW | ASSLEGGSNERLF | 0.74903786 |
| ALAAAAATAI | ASSLEGGSNERLF | 0.82114005 |
| AFGLGLLL | ASSLEGGSNERLF | 0.85862184 |
| AFFAAASL | ASSLEGGSNERLF | 0.47354594 |
| AAAALPAFSPP | ASSLEGGSNERLF | 0.84657985 |
| VVLAPAL | ASSLEGGSNERLF | 0.85867715 |
| TIGHVDHGKTTL | ASSLEGGSNERLF | 0.7142597 |
| LISNASDALDK | ASSLEGGSNERLF | 0.21496123 |

| | | |
|---|---|---|
| LISNASDAL | ASSLEGGSNERLF | 0.82179105 |
| ISNASDALDK | ASSLEGGSNERLF | 0.74969995 |
| NQAETLVYQTE | ASSLEGGSNERLF | 0.8394307 |
| TGGTGGTAGAG | ASSLEGGSNERLF | 0.8164114 |
| TGSTAGGAGGA | ASSLEGGSNERLF | 0.47466815 |
| DPLAVPVAAGPV | ASSLEGGSNERLF | 0.857628 |
| GGTGGTPGNGGN | ASSLEGGSNERLF | 0.8565099 |
| PAGLAGVDPATL | ASSLEGGSNERLF | 0.51246566 |
| GNNLVVEAPHAGASV | ASSLEGGSNERLF | 0.5016198 |
| KVTRIMSASTLEQPA | ASSLEGGSNERLF | 0.6084603 |
| LGFALPIALPRRTVV | ASSLEGGSNERLF | 0.30206138 |
| TRLGNLTARKLVVLV | ASSLEGGSNERLF | 0.8727816 |
| WSQQRRLIYFPSAGP | ASSLEGGSNERLF | 0.7177633 |
| GPNGNPDPMAAAVDIR | ASSLEGGSNERLF | 0.6805874 |
| TQRCVSVADLRAAASR | ASSLEGGSNERLF | 0.81067705 |
| QFVLDVAEHPVEVLTETV | ASSLEGGSNERLF | 0.6850125 |
| GGAGGTGGAGGNGGTQAGDG | ASSLEGGSNERLF | 0.66439867 |
| PLLATLRAIAVDVDAKPAQV | ASSLEGGSNERLF | 0.68631566 |
| APKFRTGGYMIDFWGVGYQVAK | ASSLEGGSNERLF | 0.7103373 |
| GAGGAGGAGGAAGGPGGTGGTG | ASSLEGGSNERLF | 0.83186483 |
| IGALGLLATDVAPAGSVAFGVA | ASSLEGGSNERLF | 0.7767695 |
| AGGAGGIGGNGGAGGAGGVLMG | ASSLEGGSNERLF | 0.77377766 |
| ADELIGLL | ASSLEGGSNERLF | 0.8847286 |
| ADRALKA | ASSLEGGSNERLF | 0.36172348 |
| AGAVGQVGGAGGA | ASSLEGGSNERLF | 0.7036095 |
| AGGGGGLGGAP | ASSLEGGSNERLF | 0.83042645 |
| AGGLLYGNGGAG | ASSLEGGSNERLF | 0.41645697 |
| AGLESIGL | ASSLEGGSNERLF | 0.86756444 |
| AGLKTGII | ASSLEGGSNERLF | 0.8247155 |
| ASLAALLL | ASSLEGGSNERLF | 0.8245437 |
| ASVINISSI | ASSLEGGSNERLF | 0.66733825 |
| ATPPGPA | ASSLEGGSNERLF | 0.8924445 |
| ATVASALV | ASSLEGGSNERLF | 0.8522721 |
| ATVASLIL | ASSLEGGSNERLF | 0.76666737 |
| AVNGPVLIRH | ASSLEGGSNERLF | 0.8492093 |
| DGAAGAVGI | ASSLEGGSNERLF | 0.49912828 |
| DPLLHGI | ASSLEGGSNERLF | 0.8596756 |
| ELGAALGAL | ASSLEGGSNERLF | 0.82131046 |
| ELGRTQAV | ASSLEGGSNERLF | 0.407495 |
| FVAGGYLS | ASSLEGGSNERLF | 0.7589532 |
| GASGGAGGAGGW | ASSLEGGSNERLF | 0.8714825 |
| GGAGAGGDGGAGGI | ASSLEGGSNERLF | 0.8755332 |
| GTGHNGSQPGMGGNG | ASSLEGGSNERLF | 0.3183015 |
| GYAAASASAS | ASSLEGGSNERLF | 0.73985094 |
| HQVDVAAVQQG | ASSLEGGSNERLF | 0.39289892 |
| IGAAIAHT | ASSLEGGSNERLF | 0.502018 |
| IGMIEVAQL | ASSLEGGSNERLF | 0.42515355 |
| IRLAPEHTSAMD | ASSLEGGSNERLF | 0.8105293 |
| IYNVSVL | ASSLEGGSNERLF | 0.6418482 |
| KQGFLGM | ASSLEGGSNERLF | 0.83920777 |
| LESVITQL | ASSLEGGSNERLF | 0.735878 |

| | | |
|---|---|---|
| LGLGALVVV | ASSLEGGSNERLF | 0.86164975 |
| LIGNGGNGGSGGI | ASSLEGGSNERLF | 0.5669118 |
| LNAINAPTL | ASSLEGGSNERLF | 0.7735034 |
| LQLLNTGV | ASSLEGGSNERLF | 0.31795782 |
| LQTSIEF | ASSLEGGSNERLF | 0.2745368 |
| MSFAVSEL | ASSLEGGSNERLF | 0.56368023 |
| NTGALGL | ASSLEGGSNERLF | 0.7200376 |
| PPGAGGAGGAGTT | ASSLEGGSNERLF | 0.16388026 |
| QLVGLGI | ASSLEGGSNERLF | 0.8784238 |
| RFLEQVHQL | ASSLEGGSNERLF | 0.7689233 |
| RGVGYVLR | ASSLEGGSNERLF | 0.8252176 |
| SATGVFAGVI | ASSLEGGSNERLF | 0.71974105 |
| SLPAITLP | ASSLEGGSNERLF | 0.6335919 |
| STALAFTV | ASSLEGGSNERLF | 0.6161399 |
| SVLTVGLGK | ASSLEGGSNERLF | 0.837641 |
| TALAGAGSGI | ASSLEGGSNERLF | 0.478096 |
| TGLPFTGI | ASSLEGGSNERLF | 0.20883578 |
| VAEHVTV | ASSLEGGSNERLF | 0.475438 |
| VDIVLNSL | ASSLEGGSNERLF | 0.41436106 |
| VLTGGGAVL | ASSLEGGSNERLF | 0.87149286 |
| VNVGLAYGI | ASSLEGGSNERLF | 0.87094784 |
| VSSAAVGAV | ASSLEGGSNERLF | 0.8104258 |
| VTSVYEAGV | ASSLEGGSNERLF | 0.8559749 |
| WVALDRGA | ASSLEGGSNERLF | 0.1422283 |
| YLVGSVVI | ASSLEGGSNERLF | 0.710238 |
| GPPGTGKTLL | ASSLEGGSNERLF | 0.8901545 |
| GVGMTKF | ASSLEGGSNERLF | 0.79387605 |
| IGIGGSDL | ASSLEGGSNERLF | 0.6800807 |
| IGTLLAEL | ASSLEGGSNERLF | 0.8658873 |
| IYTSGTT | ASSLEGGSNERLF | 0.7127645 |
| LSFPAGV | ASSLEGGSNERLF | 0.55506086 |
| LYTSGTT | ASSLEGGSNERLF | 0.7746812 |
| VGAFPFGF | ASSLEGGSNERLF | 0.8246056 |
| VGAGTVEF | ASSLEGGSNERLF | 0.7863352 |
| VGKSTLLL | ASSLEGGSNERLF | 0.43116343 |
| AALAHAGGGIV | ASSLEGGSNERLF | 0.8132578 |
| AALVAVSLL | ASSLEGGSNERLF | 0.89830106 |
| AAVSRAVAAL | ASSLEGGSNERLF | 0.7214122 |
| AGGSSGIGGF | ASSLEGGSNERLF | 0.84373105 |
| AGLTVANANG | ASSLEGGSNERLF | 0.81541145 |
| AGLVGAVL | ASSLEGGSNERLF | 0.7827512 |
| ALSQAFTV | ASSLEGGSNERLF | 0.51949465 |
| ALTFSVSL | ASSLEGGSNERLF | 0.6326623 |
| ALTGAALVAL | ASSLEGGSNERLF | 0.78534824 |
| ASFSIGAGVL | ASSLEGGSNERLF | 0.6259073 |
| DGSVYKI | ASSLEGGSNERLF | 0.72762835 |
| EQLAEIV | ASSLEGGSNERLF | 0.72907317 |
| FAVSELTVM | ASSLEGGSNERLF | 0.89928067 |
| GAAPLAAGL | ASSLEGGSNERLF | 0.8283021 |
| GADGGDGGSGGAGG | ASSLEGGSNERLF | 0.8231747 |
| GFGLDTGGAGG | ASSLEGGSNERLF | 0.59296775 |

| | | |
|---|---|---|
| GLASGGLL | ASSLEGGSNERLF | 0.7909601 |
| GRIAGAAVLL | ASSLEGGSNERLF | 0.83119905 |
| GSAGSLNTSF | ASSLEGGSNERLF | 0.22225487 |
| GTGQIGI | ASSLEGGSNERLF | 0.8844604 |
| GTLIALIL | ASSLEGGSNERLF | 0.5726929 |
| IAATMGFVVG | ASSLEGGSNERLF | 0.7018841 |
| IFAQALL | ASSLEGGSNERLF | 0.8462205 |
| IGAAVALGF | ASSLEGGSNERLF | 0.88495404 |
| IGLTPLAL | ASSLEGGSNERLF | 0.6899754 |
| ILGSNTSTL | ASSLEGGSNERLF | 0.75696576 |
| IVFSLTVL | ASSLEGGSNERLF | 0.6241087 |
| LASKATGF | ASSLEGGSNERLF | 0.8302901 |
| LASVALIL | ASSLEGGSNERLF | 0.86737764 |
| LFAVGLTL | ASSLEGGSNERLF | 0.6475285 |
| LFNGGIGTYI | ASSLEGGSNERLF | 0.87683237 |
| LTLGAGII | ASSLEGGSNERLF | 0.75900614 |
| LVAAALGDI | ASSLEGGSNERLF | 0.26155746 |
| NTAFVQLGI | ASSLEGGSNERLF | 0.6560668 |
| QLFSGAEL | ASSLEGGSNERLF | 0.8527559 |
| SAATFITL | ASSLEGGSNERLF | 0.86068094 |
| SNYGLANLGV | ASSLEGGSNERLF | 0.8636636 |
| SYGGAYAVM | ASSLEGGSNERLF | 0.8367987 |
| TTGVGGLTLG | ASSLEGGSNERLF | 0.5312701 |
| TTQTIYI | ASSLEGGSNERLF | 0.8825106 |
| VAAGAGLGAVI | ASSLEGGSNERLF | 0.88356465 |
| VAAVVTLL | ASSLEGGSNERLF | 0.82847464 |
| VGFLGGLI | ASSLEGGSNERLF | 0.53976345 |
| VGVIPILL | ASSLEGGSNERLF | 0.41738266 |
| VIFASSLI | ASSLEGGSNERLF | 0.81152105 |
| VKGGDGGAAAG | ASSLEGGSNERLF | 0.85841376 |
| VTKTFTNI | ASSLEGGSNERLF | 0.3997948 |
| VVAAAILL | ASSLEGGSNERLF | 0.79035556 |
| YAALDVEL | ASSLEGGSNERLF | 0.8106672 |
| LASVALLL | ASSLEGGSNERLF | 0.2220703 |
| LDAVHRV | ASSLEGGSNERLF | 0.815086 |
| QGLSSSIF | ASSLEGGSNERLF | 0.87897855 |
| RGTFANIRL | ASSLEGGSNERLF | 0.7908007 |
| RQLWWGHRI | ASSLEGGSNERLF | 0.61821616 |
| YGPPGTGKTLL | ASSLEGGSNERLF | 0.87802523 |
| AALVGAPANVA | ASSLEGGSNERLF | 0.88729227 |
| AATFRRTE | ASSLEGGSNERLF | 0.88593274 |
| AGALGGGI | ASSLEGGSNERLF | 0.3602724 |
| AGGAGGAGGIGAG | ASSLEGGSNERLF | 0.19032553 |
| AGSGALGL | ASSLEGGSNERLF | 0.76533103 |
| AQIADQI | ASSLEGGSNERLF | 0.86026055 |
| ASAVAVTYL | ASSLEGGSNERLF | 0.7603649 |
| ASTVLGGTGG | ASSLEGGSNERLF | 0.6432438 |
| AVGGEPAK | ASSLEGGSNERLF | 0.8231152 |
| CEQGTFGGTC | ASSLEGGSNERLF | 0.8740975 |
| DEGKAPAG | ASSLEGGSNERLF | 0.4716285 |
| DTVVFPAL | ASSLEGGSNERLF | 0.2622667 |

| | | |
|---|---|---|
| FEGITMAG | ASSLEGGSNERLF | 0.831884 |
| FPLVVQD | ASSLEGGSNERLF | 0.83198655 |
| GAFTTPQ | ASSLEGGSNERLF | 0.6792295 |
| GFGGFGTAGGVGGA | ASSLEGGSNERLF | 0.73759615 |
| GFLAQFL | ASSLEGGSNERLF | 0.77376926 |
| GGAGGAGGNAGVA | ASSLEGGSNERLF | 0.69290483 |
| GGVGTAGAGGAGGAGG | ASSLEGGSNERLF | 0.8049008 |
| GNGGGGDAGNAGSGGN | ASSLEGGSNERLF | 0.4137932 |
| GTGGAGGAGNDAGS | ASSLEGGSNERLF | 0.864068 |
| GTTNTGWFNS | ASSLEGGSNERLF | 0.44623926 |
| IFMGPLFTL | ASSLEGGSNERLF | 0.84846926 |
| ILAHAIEAI | ASSLEGGSNERLF | 0.81926 |
| IRSAIREAL | ASSLEGGSNERLF | 0.8055345 |
| ITFAGGG | ASSLEGGSNERLF | 0.6754982 |
| ITGMSGHQEV | ASSLEGGSNERLF | 0.5590455 |
| IVAGLACTL | ASSLEGGSNERLF | 0.38911495 |
| IVGGGLAGA | ASSLEGGSNERLF | 0.2991384 |
| IVTGVLNI | ASSLEGGSNERLF | 0.86617064 |
| LAQQGCI | ASSLEGGSNERLF | 0.105944805 |
| LCLHVEN | ASSLEGGSNERLF | 0.86171126 |
| LDAISDGS | ASSLEGGSNERLF | 0.77623427 |
| LGGFAAML | ASSLEGGSNERLF | 0.5103101 |
| LGGKNPMI | ASSLEGGSNERLF | 0.8258563 |
| LGPKGVAAIL | ASSLEGGSNERLF | 0.4273619 |
| LMVGVSI | ASSLEGGSNERLF | 0.6091752 |
| LVGAAFVL | ASSLEGGSNERLF | 0.7574353 |
| LVLAVAPL | ASSLEGGSNERLF | 0.8463199 |
| MAAIAVIDAA | ASSLEGGSNERLF | 0.46373144 |
| NNGINAELQ | ASSLEGGSNERLF | 0.81099313 |
| RGEPLVLVHG | ASSLEGGSNERLF | 0.8014957 |
| RGGAGGAAGLI | ASSLEGGSNERLF | 0.4375998 |
| RSYRMHPAV | ASSLEGGSNERLF | 0.7977836 |
| SAAHLLAGQLA | ASSLEGGSNERLF | 0.86932325 |
| SAAVGSY | ASSLEGGSNERLF | 0.7780827 |
| SPHAVLHRI | ASSLEGGSNERLF | 0.39520633 |
| SQIAKVL | ASSLEGGSNERLF | 0.831998 |
| TALGHLVP | ASSLEGGSNERLF | 0.6154736 |
| TAPTTSGAA | ASSLEGGSNERLF | 0.8291984 |
| TDGDDGNGGSPGT | ASSLEGGSNERLF | 0.77355933 |
| TDGLGEN | ASSLEGGSNERLF | 0.67886937 |
| TGGPPLPGQA | ASSLEGGSNERLF | 0.8502389 |
| TIPAITI | ASSLEGGSNERLF | 0.28610486 |
| TIPGFTI | ASSLEGGSNERLF | 0.49732643 |
| TSIASGALA | ASSLEGGSNERLF | 0.6582739 |
| TVTLPGAAI | ASSLEGGSNERLF | 0.79559404 |
| VASGLSAALP | ASSLEGGSNERLF | 0.85487103 |
| VGAGVIGI | ASSLEGGSNERLF | 0.61098385 |
| VGPKRFGV | ASSLEGGSNERLF | 0.7649098 |
| VNVAFPD | ASSLEGGSNERLF | 0.61404115 |
| VSMTNSGVS | ASSLEGGSNERLF | 0.39966637 |
| VTDVLGDLSV | ASSLEGGSNERLF | 0.7334939 |

| | | |
|---|---|---|
| VVVVGAGFAGL | ASSLEGGSNERLF | 0.6697912 |
| VYTVGTVV | ASSLEGGSNERLF | 0.45384812 |
| WTTFSGP | ASSLEGGSNERLF | 0.8326502 |
| YDELVEAF | ASSLEGGSNERLF | 0.5459126 |
| YGTGGAGGNGG | ASSLEGGSNERLF | 0.8349595 |
| YTNVMAV | ASSLEGGSNERLF | 0.89434904 |
| FLEQVHQL | ASSLEGGSNERLF | 0.5382318 |
| GLSLLAVAV | ASSLEGGSNERLF | 0.8242108 |
| LEQVHQL | ASSLEGGSNERLF | 0.78858197 |
| LNAAGAI | ASSLEGGSNERLF | 0.82461226 |
| PAVHASL | ASSLEGGSNERLF | 0.7074293 |
| TELEDDA | ASSLEGGSNERLF | 0.54895604 |
| TPPGPPN | ASSLEGGSNERLF | 0.8798901 |
| TPPPPPPP | ASSLEGGSNERLF | 0.8525828 |
| TQSPALI | ASSSGTGASDYT | 0.8118833 |
| EFAAAFPQ | ASSSGTGASDYT | 0.824595 |
| LLSGNLDVL | ASSSGTGASDYT | 0.84430623 |
| GPANLGNYNI | ASSSGTGASDYT | 0.851753 |
| TLGPTLP | ASSSGTGASDYT | 0.6855165 |
| VYAYSSCCK | ASSSGTGASDYT | 0.29480392 |
| APAPAPAPNI | ASSSGTGASDYT | 0.729784 |
| PGGPVPIP | ASSSGTGASDYT | 0.7582563 |
| FLTVALEP | ASSSGTGASDYT | 0.6257714 |
| SQYFGANL | ASSSGTGASDYT | 0.25538585 |
| GGVKPAA | ASSSGTGASDYT | 0.8717425 |
| GGTGGAGGAGGW | ASSSGTGASDYT | 0.8974503 |
| GVGGVAGLGGAGG | ASSSGTGASDYT | 0.6092012 |
| VLLVLATTI | ASSSGTGASDYT | 0.8944304 |
| IGIFAAML | ASSSGTGASDYT | 0.4686941 |
| AVGDLID | ASSSGTGASDYT | 0.84359026 |
| NISPLPL | ASSSGTGASDYT | 0.87342906 |
| VTAVFVAI | ASSSGTGASDYT | 0.52521515 |
| SGDDNDAAL | ASSSGTGASDYT | 0.8341838 |
| AVAALAHS | ASSSGTGASDYT | 0.43983284 |
| GGVTVMATP | ASSSGTGASDYT | 0.60792136 |
| AGLAANVRD | ASSSGTGASDYT | 0.8705088 |
| TNRPDLIDPALLRPG | ASSSGTGASDYT | 0.71931326 |
| LANLGVTAL | ASSSGTGASDYT | 0.82174677 |
| TGERIEL | ASSSGTGASDYT | 0.73717123 |
| NGEELAA | ASSSGTGASDYT | 0.52235395 |
| VLSLVWG | ASSSGTGASDYT | 0.79205954 |
| IGYALWL | ASSSGTGASDYT | 0.864342 |
| LSLLLAPL | ASSSGTGASDYT | 0.48758727 |
| TLDEALK | ASSSGTGASDYT | 0.8836865 |
| TDGAGTAEGG | ASSSGTGASDYT | 0.55478024 |
| NAPGGAGGNGG | ASSSGTGASDYT | 0.8913685 |
| VSVFMGMGGNF | ASSSGTGASDYT | 0.86135495 |
| LASYLAAL | ASSSGTGASDYT | 0.8038402 |
| LVGGTPAL | ASSSGTGASDYT | 0.68320084 |
| VRNADVL | ASSSGTGASDYT | 0.86384743 |
| TCSSPFLVSAM | ASSSGTGASDYT | 0.68885016 |

| | | |
|---|---|---|
| VHITDAE | ASSSGTGASDYT | 0.8127378 |
| EYAAPSF | ASSSGTGASDYT | 0.8699365 |
| VVDAFHGGL | ASSSGTGASDYT | 0.6757657 |
| ASVVDPQ | ASSSGTGASDYT | 0.84479445 |
| SYNEFGGLS | ASSSGTGASDYT | 0.795982 |
| DPDGSVY | ASSSGTGASDYT | 0.82719857 |
| MVSTIHATE | ASSSGTGASDYT | 0.6954224 |
| LAAGVADPV | ASSSGTGASDYT | 0.52098733 |
| TKEQIAA | ASSSGTGASDYT | 0.75841343 |
| SLASPAAL | ASSSGTGASDYT | 0.61771107 |
| IGLVFYGV | ASSSGTGASDYT | 0.8030696 |
| FLLDASNL | ASSSGTGASDYT | 0.8521643 |
| YQVGNFP | ASSSGTGASDYT | 0.7105806 |
| ESLLSQST | ASSSGTGASDYT | 0.8529716 |
| LFTVGFAAVG | ASSSGTGASDYT | 0.8602514 |
| ATPPTPT | ASSSGTGASDYT | 0.46467942 |
| TDLATLG | ASSSGTGASDYT | 0.73822796 |
| SAGGTGR | ASSSGTGASDYT | 0.68536496 |
| GFSNSGS | ASSSGTGASDYT | 0.779907 |
| GGNGGSGGTSV | ASSSGTGASDYT | 0.29992932 |
| GTSVATGGAGN | ASSSGTGASDYT | 0.8457167 |
| NGGNGGSGGTS | ASSSGTGASDYT | 0.67025423 |
| AVVKFPH | ASSSGTGASDYT | 0.23648077 |
| TPPPPPP | ASSSGTGASDYT | 0.80360144 |
| AAPAAAAAPA | ASSSGTGASDYT | 0.76346034 |
| IALNEDL | ASSSGTGASDYT | 0.7545413 |
| IVNVSSM | ASSSGTGASDYT | 0.8455043 |
| TGIPPAPR | ASSSGTGASDYT | 0.6249225 |
| FDLGGGTF | ASSSGTGASDYT | 0.6624851 |
| FDLGGGTFDVS | ASSSGTGASDYT | 0.79499197 |
| NEPTAAA | ASSSGTGASDYT | 0.86075914 |
| NEPTAAALA | ASSSGTGASDYT | 0.64075136 |
| GSIINISSI | ASSSGTGASDYT | 0.80118066 |
| VAPGNAGTA | ASSSGTGASDYT | 0.8657348 |
| TGGGASLE | ASSSGTGASDYT | 0.8130314 |
| VAVYNHYKRI | ASSSGTGASDYT | 0.8645259 |
| TDPAPATT | ASSSGTGASDYT | 0.8776893 |
| TNRPDILDPALLRPG | ASSSGTGASDYT | 0.70329005 |
| LTTTTTS | ASSSGTGASDYT | 0.8455532 |
| PTGSGKTL | ASSSGTGASDYT | 0.84955406 |
| TGSGKTL | ASSSGTGASDYT | 0.76246136 |
| TIGHVDHGK | ASSSGTGASDYT | 0.7829687 |
| TIGHVDHGKTT | ASSSGTGASDYT | 0.67201674 |
| GHVDHGKTT | ASSSGTGASDYT | 0.77293324 |
| AQFEHTLL | ASSSGTGASDYT | 0.44036877 |
| IVNQDEL | ASSSGTGASDYT | 0.77626944 |
| TNGSQFF | ASSSGTGASDYT | 0.85520333 |
| TGKGIVE | ASSSGTGASDYT | 0.73301965 |
| LISNASDA | ASSSGTGASDYT | 0.56974995 |
| AELRTLGELTKIAKAHGAQ | ASSSGTGASDYT | 0.5809883 |
| APNAVAAAEPSW | ASSSGTGASDYT | 0.75742114 |

| | | |
|---|---|---|
| APQVAGAATAVL | ASSSGTGASDYT | 0.75788546 |
| ARRDPAAAAAILPTDA | ASSSGTGASDYT | 0.5323325 |
| DAAARDVLAEAGLAEYF | ASSSGTGASDYT | 0.66103417 |
| DDQIVLSAKANLAKVEQ | ASSSGTGASDYT | 0.8498336 |
| DELLPDTGEDSDEDGH | ASSSGTGASDYT | 0.8586965 |
| ELPEAPANRATVPQD | ASSSGTGASDYT | 0.5069982 |
| FGQSKIERVTGIG | ASSSGTGASDYT | 0.8484496 |
| GAGGTGSTAGGAGGAGGAGGLY | ASSSGTGASDYT | 0.8226143 |
| GHGGTGAAGGNGGTGGAGGSGI | ASSSGTGASDYT | 0.86910224 |
| GNGGNGGAGGEGGPGGAGGAGA | ASSSGTGASDYT | 0.8105217 |
| GSTAGNGGNGGNATNGTIAGQP | ASSSGTGASDYT | 0.6208393 |
| GTGGDGGNGGNGGTGG | ASSSGTGASDYT | 0.8373418 |
| GVRRPAGIPADAPV | ASSSGTGASDYT | 0.7082854 |
| GYRQVTDAHLLTLARR | ASSSGTGASDYT | 0.7697983 |
| IAKMGQDGHDRG | ASSSGTGASDYT | 0.8540282 |
| KAEPSIEIQYT | ASSSGTGASDYT | 0.7517731 |
| KPVRIRMAADEDTVAVT | ASSSGTGASDYT | 0.65344185 |
| MPPRIAGMRLLVIKPEP | ASSSGTGASDYT | 0.6879297 |
| MKFYAVALFRSRGVPPTR | ASSSGTGASDYT | 0.8718898 |
| NAGGFGILGAVAHSP | ASSSGTGASDYT | 0.54666376 |
| NEGREITLAMAEM | ASSSGTGASDYT | 0.71590185 |
| NELVEWSHPLGHRWRWE | ASSSGTGASDYT | 0.72014654 |
| NLGAGGTGGNGGNPG | ASSSGTGASDYT | 0.7768914 |
| NTEWPKWETVMAADPVD | ASSSGTGASDYT | 0.8477211 |
| PASAAPAETQP | ASSSGTGASDYT | 0.8370608 |
| QQSSDGKNGVRAAEITGE | ASSSGTGASDYT | 0.7190393 |
| RAPDAVTTSWWKE | ASSSGTGASDYT | 0.8089608 |
| SGFGNVGAHTSGWFNQ | ASSSGTGASDYT | 0.8469478 |
| SGVAGAVGQAASVGGLKV | ASSSGTGASDYT | 0.80957466 |
| SNGGAGTGSAGGNGGT | ASSSGTGASDYT | 0.82780784 |
| TGGAGGEGGAAG | ASSSGTGASDYT | 0.6098815 |
| TGGDGGAAGAGGAPGHG | ASSSGTGASDYT | 0.8637322 |
| TMPPLLAAIANAGRHGV | ASSSGTGASDYT | 0.83679473 |
| TSSTAGNSGGA | ASSSGTGASDYT | 0.8614539 |
| VPATVPVGILC | ASSSGTGASDYT | 0.84440076 |
| VQRPDSGGVVLTGRIS | ASSSGTGASDYT | 0.8012426 |
| WTPNPVPATKGVPAGLVQAT | ASSSGTGASDYT | 0.8216548 |
| YALKRALAKFSDRFDVVII | ASSSGTGASDYT | 0.4696759 |
| YLIGSVIV | ASSSGTGASDYT | 0.70353425 |
| YGQLVAVID | ASSSGTGASDYT | 0.7881117 |
| VTVGGMTLPA | ASSSGTGASDYT | 0.78919125 |
| VLALPAV | ASSSGTGASDYT | 0.8223624 |
| VIPPTLNYET | ASSSGTGASDYT | 0.8514127 |
| VGMGGMHPSSG | ASSSGTGASDYT | 0.6844844 |
| VGLAGGAATA | ASSSGTGASDYT | 0.86084926 |
| VGGAGGAGGSGP | ASSSGTGASDYT | 0.84191024 |
| VFAAFMV | ASSSGTGASDYT | 0.48276055 |
| TLYDKLR | ASSSGTGASDYT | 0.72132844 |
| TLAVFTVL | ASSSGTGASDYT | 0.86036414 |
| TGSVQPARV | ASSSGTGASDYT | 0.7977437 |
| TGGGAGSGGT | ASSSGTGASDYT | 0.7407787 |

| | | |
|---|---|---|
| SHLETSL | ASSSGTGASDYT | 0.81153375 |
| SFTTGMT | ASSSGTGASDYT | 0.8564446 |
| QVEVAKRMGVRQPTVS | ASSSGTGASDYT | 0.7931521 |
| QGSGGPALAM | ASSSGTGASDYT | 0.8702802 |
| MLAPSNP | ASSSGTGASDYT | 0.31756964 |
| LVNNAAI | ASSSGTGASDYT | 0.7959924 |
| LTGEINTLA | ASSSGTGASDYT | 0.75056374 |
| LSIFGPV | ASSSGTGASDYT | 0.4468363 |
| LLALLANL | ASSSGTGASDYT | 0.84992516 |
| LISNASDALDKLR | ASSSGTGASDYT | 0.75686157 |
| LINEAVP | ASSSGTGASDYT | 0.84390867 |
| LGYAFSGL | ASSSGTGASDYT | 0.7260638 |
| LDFVAVAL | ASSSGTGASDYT | 0.849135 |
| LAQPGAE | ASSSGTGASDYT | 0.74385524 |
| ISNLGAVTGF | ASSSGTGASDYT | 0.8089364 |
| HQEDPGRI | ASSSGTGASDYT | 0.7237514 |
| GTPGEPTP | ASSSGTGASDYT | 0.8419164 |
| GFFNSGS | ASSSGTGASDYT | 0.8625681 |
| GAGGGGLTAGAN | ASSSGTGASDYT | 0.8636423 |
| FVDALVNM | ASSSGTGASDYT | 0.4995259 |
| FLTVAPEL | ASSSGTGASDYT | 0.77771807 |
| FETLTAL | ASSSGTGASDYT | 0.7958375 |
| DVINPIEL | ASSSGTGASDYT | 0.36931324 |
| APVSGAPV | ASSSGTGASDYT | 0.44919735 |
| ALVGGLAW | ASSSGTGASDYT | 0.68615943 |
| ALAAAAATAI | ASSSGTGASDYT | 0.64273524 |
| AFGLGLLL | ASSSGTGASDYT | 0.8358667 |
| AFFAAASL | ASSSGTGASDYT | 0.79551566 |
| AAAALPAFSPP | ASSSGTGASDYT | 0.7591833 |
| VVLAPAL | ASSSGTGASDYT | 0.8260733 |
| TIGHVDHGKTTL | ASSSGTGASDYT | 0.7901656 |
| LISNASDALDK | ASSSGTGASDYT | 0.12491536 |
| LISNASDAL | ASSSGTGASDYT | 0.7172539 |
| ISNASDALDK | ASSSGTGASDYT | 0.49893072 |
| NQAETLVYQTE | ASSSGTGASDYT | 0.8734548 |
| TGGTGGTAGAG | ASSSGTGASDYT | 0.6509831 |
| TGSTAGGAGGA | ASSSGTGASDYT | 0.59904826 |
| DPLAVPVAAGPV | ASSSGTGASDYT | 0.7593725 |
| GGTGGTPGNGGN | ASSSGTGASDYT | 0.7463797 |
| PAGLAGVDPATL | ASSSGTGASDYT | 0.7843692 |
| GNNLVVEAPHAGASV | ASSSGTGASDYT | 0.66531336 |
| KVTRIMSASTLEQPA | ASSSGTGASDYT | 0.28894934 |
| LGFALPIALPRRTVV | ASSSGTGASDYT | 0.8445666 |
| TRLGNLTARKLVVLV | ASSSGTGASDYT | 0.5157434 |
| WSQQRRLIYFPSAGP | ASSSGTGASDYT | 0.6710234 |
| GPNGNPDPMAAAVDIR | ASSSGTGASDYT | 0.46818933 |
| TQRCVSVADLRAAASR | ASSSGTGASDYT | 0.6385867 |
| QFVLDVAEHPVEVLTETV | ASSSGTGASDYT | 0.48224753 |
| GGAGGTGGAGGNGGTQAGDG | ASSSGTGASDYT | 0.8612524 |
| PLLATLRAIAVDVDAKPAQV | ASSSGTGASDYT | 0.68558544 |
| APKFRTGGYMIDFWGVGYQVAK | ASSSGTGASDYT | 0.69324106 |

| | | |
|---|---|---|
| GAGGAGGAGGAAGGPGGTGGTG | ASSSGTGASDYT | 0.84823024 |
| IGALGLLATDVAPAGSVAFGVA | ASSSGTGASDYT | 0.74140906 |
| AGGAGGIGGNGGAGGAGGVLMG | ASSSGTGASDYT | 0.72542137 |
| ADELIGLL | ASSSGTGASDYT | 0.709874 |
| ADRALKA | ASSSGTGASDYT | 0.64256364 |
| AGAVGQVGGAGGA | ASSSGTGASDYT | 0.6898488 |
| AGGGGGLGGAP | ASSSGTGASDYT | 0.69886035 |
| AGGLLYGNGGAG | ASSSGTGASDYT | 0.87300587 |
| AGLESIGL | ASSSGTGASDYT | 0.8099062 |
| AGLKTGII | ASSSGTGASDYT | 0.51458466 |
| ASLAALLL | ASSSGTGASDYT | 0.8181937 |
| ASVINISSI | ASSSGTGASDYT | 0.8669552 |
| ATPPGPA | ASSSGTGASDYT | 0.8754563 |
| ATVASALV | ASSSGTGASDYT | 0.6156441 |
| ATVASLIL | ASSSGTGASDYT | 0.79147565 |
| AVNGPVLIRH | ASSSGTGASDYT | 0.77346766 |
| DGAAGAVGI | ASSSGTGASDYT | 0.7338112 |
| DPLLHGI | ASSSGTGASDYT | 0.7473409 |
| ELGAALGAL | ASSSGTGASDYT | 0.6319747 |
| ELGRTQAV | ASSSGTGASDYT | 0.8544214 |
| FVAGGYLS | ASSSGTGASDYT | 0.8840062 |
| GASGGAGGAGGW | ASSSGTGASDYT | 0.77066946 |
| GGAGAGGDGGAGGI | ASSSGTGASDYT | 0.40815338 |
| GTGHNGSQPGMGGNG | ASSSGTGASDYT | 0.7455753 |
| GYAAASASAS | ASSSGTGASDYT | 0.60278034 |
| HQVDVAAVQQG | ASSSGTGASDYT | 0.6856692 |
| IGAAIAHT | ASSSGTGASDYT | 0.7936466 |
| IGMIEVAQL | ASSSGTGASDYT | 0.44112796 |
| IRLAPEHTSAMD | ASSSGTGASDYT | 0.74415386 |
| IYNVSVL | ASSSGTGASDYT | 0.87255925 |
| KQGFLGM | ASSSGTGASDYT | 0.6590858 |
| LESVITQL | ASSSGTGASDYT | 0.79410005 |
| LGLGALVVV | ASSSGTGASDYT | 0.84470326 |
| LIGNGGNGGSGGI | ASSSGTGASDYT | 0.81386167 |
| LNAINAPTL | ASSSGTGASDYT | 0.8618861 |
| LQLLNTGV | ASSSGTGASDYT | 0.8281823 |
| LQTSIEF | ASSSGTGASDYT | 0.82824665 |
| MSFAVSEL | ASSSGTGASDYT | 0.7950597 |
| NTGALGL | ASSSGTGASDYT | 0.7570886 |
| PPGAGGAGGAGTT | ASSSGTGASDYT | 0.77092636 |
| QLVGLGI | ASSSGTGASDYT | 0.75821245 |
| RFLEQVHQL | ASSSGTGASDYT | 0.748382 |
| RGVGYVLR | ASSSGTGASDYT | 0.8327832 |
| SATGVFAGVI | ASSSGTGASDYT | 0.7921479 |
| SLPAITLP | ASSSGTGASDYT | 0.86416906 |
| STALAFTV | ASSSGTGASDYT | 0.5950637 |
| SVLTVGLGK | ASSSGTGASDYT | 0.66160583 |
| TALAGAGSGI | ASSSGTGASDYT | 0.84547174 |
| TGLPFTGI | ASSSGTGASDYT | 0.23626012 |
| VAEHVTV | ASSSGTGASDYT | 0.5457929 |
| VDIVLNSL | ASSSGTGASDYT | 0.7897801 |

| | | |
|---|---|---|
| VLTGGGAVL | ASSSGTGASDYT | 0.86448705 |
| VNVGLAYGI | ASSSGTGASDYT | 0.21433426 |
| VSSAAVGAV | ASSSGTGASDYT | 0.65198326 |
| VTSVYEAGV | ASSSGTGASDYT | 0.59585434 |
| WVALDRGA | ASSSGTGASDYT | 0.84532374 |
| YLVGSVVI | ASSSGTGASDYT | 0.7675281 |
| GPPGTGKTLL | ASSSGTGASDYT | 0.47072554 |
| GVGMTKF | ASSSGTGASDYT | 0.84695643 |
| IGIGGSDL | ASSSGTGASDYT | 0.39070034 |
| IGTLLAEL | ASSSGTGASDYT | 0.60013443 |
| IYTSGTT | ASSSGTGASDYT | 0.8407766 |
| LSFPAGV | ASSSGTGASDYT | 0.8574363 |
| LYTSGTT | ASSSGTGASDYT | 0.5927292 |
| VGAFPFGF | ASSSGTGASDYT | 0.8425941 |
| VGAGTVEF | ASSSGTGASDYT | 0.7060662 |
| VGKSTLLL | ASSSGTGASDYT | 0.88153887 |
| AALAHAGGGIV | ASSSGTGASDYT | 0.7293174 |
| AALVAVSLL | ASSSGTGASDYT | 0.56423825 |
| AAVSRAVAAL | ASSSGTGASDYT | 0.8359203 |
| AGGSSGIGGF | ASSSGTGASDYT | 0.8163545 |
| AGLTVANANG | ASSSGTGASDYT | 0.8384345 |
| AGLVGAVL | ASSSGTGASDYT | 0.82538986 |
| ALSQAFTV | ASSSGTGASDYT | 0.7705537 |
| ALTFSVSL | ASSSGTGASDYT | 0.710194 |
| ALTGAALVAL | ASSSGTGASDYT | 0.8420729 |
| ASFSIGAGVL | ASSSGTGASDYT | 0.82488 |
| DGSVYKI | ASSSGTGASDYT | 0.63125163 |
| EQLAEIV | ASSSGTGASDYT | 0.8589452 |
| FAVSELTVM | ASSSGTGASDYT | 0.7633941 |
| GAAPLAAGL | ASSSGTGASDYT | 0.67342293 |
| GADGGDGGSGGAGG | ASSSGTGASDYT | 0.8711138 |
| GFGLDTGGAGG | ASSSGTGASDYT | 0.8167672 |
| GLASGGLL | ASSSGTGASDYT | 0.8419675 |
| GRIAGAAVLL | ASSSGTGASDYT | 0.8555312 |
| GSAGSLNTSF | ASSSGTGASDYT | 0.8568238 |
| GTGQIGI | ASSSGTGASDYT | 0.56417084 |
| GTLIALIL | ASSSGTGASDYT | 0.7818397 |
| IAATMGFVVG | ASSSGTGASDYT | 0.76426965 |
| IFAQALL | ASSSGTGASDYT | 0.83511144 |
| IGAAVALGF | ASSSGTGASDYT | 0.7733922 |
| IGLTPLAL | ASSSGTGASDYT | 0.5917183 |
| ILGSNTSTL | ASSSGTGASDYT | 0.84317935 |
| IVFSLTVL | ASSSGTGASDYT | 0.85508287 |
| LASKATGF | ASSSGTGASDYT | 0.8015907 |
| LASVALIL | ASSSGTGASDYT | 0.8021934 |
| LFAVGLTL | ASSSGTGASDYT | 0.86690587 |
| LFNGGIGTYI | ASSSGTGASDYT | 0.6972435 |
| LTLGAGII | ASSSGTGASDYT | 0.76976645 |
| LVAAALGDI | ASSSGTGASDYT | 0.8468456 |
| NTAFVQLGI | ASSSGTGASDYT | 0.8509405 |
| QLFSGAEL | ASSSGTGASDYT | 0.79626936 |

| | | |
|---|---|---|
| SAATFITL | ASSSGTGASDYT | 0.86059964 |
| SNYGLANLGV | ASSSGTGASDYT | 0.6409555 |
| SYGGAYAVM | ASSSGTGASDYT | 0.8204019 |
| TTGVGGLTLG | ASSSGTGASDYT | 0.5993876 |
| TTQTIYI | ASSSGTGASDYT | 0.7934707 |
| VAAGAGLGAVI | ASSSGTGASDYT | 0.59640944 |
| VAAVVTLL | ASSSGTGASDYT | 0.80312175 |
| VGFLGGLI | ASSSGTGASDYT | 0.82976687 |
| VGVIPILL | ASSSGTGASDYT | 0.82753205 |
| VIFASSLI | ASSSGTGASDYT | 0.38022795 |
| VKGGDGGAAAG | ASSSGTGASDYT | 0.8110172 |
| VTKTFTNI | ASSSGTGASDYT | 0.83522666 |
| VVAAAILL | ASSSGTGASDYT | 0.87825894 |
| YAALDVEL | ASSSGTGASDYT | 0.7961956 |
| LASVALLL | ASSSGTGASDYT | 0.8412912 |
| LDAVHRV | ASSSGTGASDYT | 0.6991499 |
| QGLSSSIF | ASSSGTGASDYT | 0.7933298 |
| RGTFANIRL | ASSSGTGASDYT | 0.58783144 |
| RQLWWGHRI | ASSSGTGASDYT | 0.7187736 |
| YGPPGTGKTLL | ASSSGTGASDYT | 0.66626775 |
| AALVGAPANVA | ASSSGTGASDYT | 0.620315 |
| AATFRRTE | ASSSGTGASDYT | 0.82884383 |
| AGALGGGI | ASSSGTGASDYT | 0.8462505 |
| AGGAGGAGGIGAG | ASSSGTGASDYT | 0.8263114 |
| AGSGALGL | ASSSGTGASDYT | 0.8472019 |
| AQIADQI | ASSSGTGASDYT | 0.8496847 |
| ASAVAVTYL | ASSSGTGASDYT | 0.8698998 |
| ASTVLGGTGG | ASSSGTGASDYT | 0.70437145 |
| AVGGEPAK | ASSSGTGASDYT | 0.7250833 |
| CEQGTFGGTC | ASSSGTGASDYT | 0.5093304 |
| DEGKAPAG | ASSSGTGASDYT | 0.7274982 |
| DTVVFPAL | ASSSGTGASDYT | 0.7021364 |
| FEGITMAG | ASSSGTGASDYT | 0.78965724 |
| FPLVVQD | ASSSGTGASDYT | 0.19212149 |
| GAFTTPQ | ASSSGTGASDYT | 0.7371629 |
| GFGGFGTAGGVGGA | ASSSGTGASDYT | 0.8696109 |
| GFLAQFL | ASSSGTGASDYT | 0.6910103 |
| GGAGGAGGNAGVA | ASSSGTGASDYT | 0.8327568 |
| GGVGTAGAGGAGGAGG | ASSSGTGASDYT | 0.7602901 |
| GNGGGGDAGNAGSGGN | ASSSGTGASDYT | 0.6979965 |
| GTGGAGGAGNDAGS | ASSSGTGASDYT | 0.82472014 |
| GTTNTGWFNS | ASSSGTGASDYT | 0.6913143 |
| IFMGPLFTL | ASSSGTGASDYT | 0.8543972 |
| ILAHAIEAI | ASSSGTGASDYT | 0.8695713 |
| IRSAIREAL | ASSSGTGASDYT | 0.7812835 |
| ITFAGGG | ASSSGTGASDYT | 0.769325 |
| ITGMSGHQEV | ASSSGTGASDYT | 0.8296829 |
| IVAGLACTL | ASSSGTGASDYT | 0.5038917 |
| IVGGGLAGA | ASSSGTGASDYT | 0.7810619 |
| IVTGVLNI | ASSSGTGASDYT | 0.38319957 |
| LAQQGCI | ASSSGTGASDYT | 0.7434064 |

| | | |
|---|---|---|
| LCLHVEN | ASSSGTGASDYT | 0.69489217 |
| LDAISDGS | ASSSGTGASDYT | 0.79720026 |
| LGGFAAML | ASSSGTGASDYT | 0.7692534 |
| LGGKNPMI | ASSSGTGASDYT | 0.8222969 |
| LGPKGVAAIL | ASSSGTGASDYT | 0.5845866 |
| LMVGVSI | ASSSGTGASDYT | 0.8060931 |
| LVGAAFVL | ASSSGTGASDYT | 0.7495793 |
| LVLAVAPL | ASSSGTGASDYT | 0.57346064 |
| MAAIAVIDAA | ASSSGTGASDYT | 0.8201922 |
| NNGINAELQ | ASSSGTGASDYT | 0.79652196 |
| RGEPLVLVHG | ASSSGTGASDYT | 0.66972524 |
| RGGAGGAAGLI | ASSSGTGASDYT | 0.85447246 |
| RSYRMHPAV | ASSSGTGASDYT | 0.80831003 |
| SAAHLLAGQLA | ASSSGTGASDYT | 0.42685035 |
| SAAVGSY | ASSSGTGASDYT | 0.70188236 |
| SPHAVLHRI | ASSSGTGASDYT | 0.7732674 |
| SQIAKVL | ASSSGTGASDYT | 0.7560097 |
| TALGHLVP | ASSSGTGASDYT | 0.19383787 |
| TAPTTSGAA | ASSSGTGASDYT | 0.80719006 |
| TDGDDGNGGSPGT | ASSSGTGASDYT | 0.67662734 |
| TDGLGEN | ASSSGTGASDYT | 0.49858093 |
| TGGPPLPGQA | ASSSGTGASDYT | 0.70902735 |
| TIPAITI | ASSSGTGASDYT | 0.8285966 |
| TIPGFTI | ASSSGTGASDYT | 0.7385846 |
| TSIASGALA | ASSSGTGASDYT | 0.81298196 |
| TVTLPGAAI | ASSSGTGASDYT | 0.8208855 |
| VASGLSAALP | ASSSGTGASDYT | 0.83302975 |
| VGAGVIGI | ASSSGTGASDYT | 0.8649926 |
| VGPKRFGV | ASSSGTGASDYT | 0.8071546 |
| VNVAFPD | ASSSGTGASDYT | 0.7979411 |
| VSMTNSGVS | ASSSGTGASDYT | 0.84004986 |
| VTDVLGDLSV | ASSSGTGASDYT | 0.78925616 |
| VVVVGAGFAGL | ASSSGTGASDYT | 0.6074781 |
| VYTVGTVV | ASSSGTGASDYT | 0.5310876 |
| WTTFSGP | ASSSGTGASDYT | 0.74740535 |
| YDELVEAF | ASSSGTGASDYT | 0.60033107 |
| YGTGGAGGNGG | ASSSGTGASDYT | 0.79801106 |
| YTNVMAV | ASSSGTGASDYT | 0.874139 |
| FLEQVHQL | ASSSGTGASDYT | 0.7649639 |
| GLSLLAVAV | ASSSGTGASDYT | 0.5824336 |
| LEQVHQL | ASSSGTGASDYT | 0.7198615 |
| LNAAGAI | ASSSGTGASDYT | 0.66294014 |
| PAVHASL | ASSSGTGASDYT | 0.8436537 |
| TELEDDA | ASSSGTGASDYT | 0.84973764 |
| TPPGPPN | ASSSGTGASDYT | 0.86464214 |
| TPPPPPPP | ASSSGTGASDYT | 0.8310769 |
| TQSPALI | ASSRPDWDYAEQF | 0.70175 |
| EFAAAFPQ | ASSRPDWDYAEQF | 0.8340847 |
| LLSGNLDVL | ASSRPDWDYAEQF | 0.75608414 |
| GPANLGNYNI | ASSRPDWDYAEQF | 0.7164955 |
| TLGPTLP | ASSRPDWDYAEQF | 0.80173403 |

| | | |
|---|---|---|
| VYAYSSCCK | ASSRPDWDYAEQF | 0.7793398 |
| APAPAPAPNI | ASSRPDWDYAEQF | 0.7763216 |
| PGGPVPIP | ASSRPDWDYAEQF | 0.7631693 |
| FLTVALEP | ASSRPDWDYAEQF | 0.5022138 |
| SQYFGANL | ASSRPDWDYAEQF | 0.83484167 |
| GGVKPAA | ASSRPDWDYAEQF | 0.8093401 |
| GGTGGAGGAGGW | ASSRPDWDYAEQF | 0.6704105 |
| GVGGVAGLGGAGG | ASSRPDWDYAEQF | 0.7546639 |
| VLLVLATTI | ASSRPDWDYAEQF | 0.8632859 |
| IGIFAAML | ASSRPDWDYAEQF | 0.86268795 |
| AVGDLID | ASSRPDWDYAEQF | 0.66207004 |
| NISPLPL | ASSRPDWDYAEQF | 0.6799468 |
| VTAVFVAI | ASSRPDWDYAEQF | 0.7534768 |
| SGDDNDAAL | ASSRPDWDYAEQF | 0.49933934 |
| AVAALAHS | ASSRPDWDYAEQF | 0.8453007 |
| GGVTVMATP | ASSRPDWDYAEQF | 0.7250342 |
| AGLAANVRD | ASSRPDWDYAEQF | 0.7037494 |
| TNRPDLIDPALLRPG | ASSRPDWDYAEQF | 0.75586474 |
| LANLGVTAL | ASSRPDWDYAEQF | 0.7895604 |
| TGERIEL | ASSRPDWDYAEQF | 0.6578417 |
| NGEELAA | ASSRPDWDYAEQF | 0.8591907 |
| VLSLVWG | ASSRPDWDYAEQF | 0.18381396 |
| IGYALWL | ASSRPDWDYAEQF | 0.81870604 |
| LSLLLAPL | ASSRPDWDYAEQF | 0.6094155 |
| TLDEALK | ASSRPDWDYAEQF | 0.72200286 |
| TDGAGTAEGG | ASSRPDWDYAEQF | 0.47828233 |
| NAPGGAGGNGG | ASSRPDWDYAEQF | 0.75321525 |
| VSVFMGMGGNF | ASSRPDWDYAEQF | 0.8252608 |
| LASYLAAL | ASSRPDWDYAEQF | 0.8483789 |
| LVGGTPAL | ASSRPDWDYAEQF | 0.7036365 |
| VRNADVL | ASSRPDWDYAEQF | 0.8299427 |
| TCSSPFLVSAM | ASSRPDWDYAEQF | 0.8688593 |
| VHITDAE | ASSRPDWDYAEQF | 0.7028905 |
| EYAAPSF | ASSRPDWDYAEQF | 0.45335042 |
| VVDAFHGGL | ASSRPDWDYAEQF | 0.81959814 |
| ASVVDPQ | ASSRPDWDYAEQF | 0.6589582 |
| SYNEFGGLS | ASSRPDWDYAEQF | 0.8275481 |
| DPDGSVY | ASSRPDWDYAEQF | 0.83898306 |
| MVSTIHATE | ASSRPDWDYAEQF | 0.50149894 |
| LAAGVADPV | ASSRPDWDYAEQF | 0.86568356 |
| TKEQIAA | ASSRPDWDYAEQF | 0.830475 |
| SLASPAAL | ASSRPDWDYAEQF | 0.6636104 |
| IGLVFYGV | ASSRPDWDYAEQF | 0.66976774 |
| FLLDASNL | ASSRPDWDYAEQF | 0.68646944 |
| YQVGNFP | ASSRPDWDYAEQF | 0.42585072 |
| ESLLSQST | ASSRPDWDYAEQF | 0.83331424 |
| LFTVGFAAVG | ASSRPDWDYAEQF | 0.6623673 |
| ATPPTPT | ASSRPDWDYAEQF | 0.72645307 |
| TDLATLG | ASSRPDWDYAEQF | 0.78056306 |
| SAGGTGR | ASSRPDWDYAEQF | 0.7458709 |
| GFSNSGS | ASSRPDWDYAEQF | 0.7346214 |

| | | |
|---|---|---|
| GGNGGSGGTSV | ASSRPDWDYAEQF | 0.7328895 |
| GTSVATGGAGN | ASSRPDWDYAEQF | 0.7851828 |
| NGGNGGSGGTS | ASSRPDWDYAEQF | 0.8041781 |
| AVVKFPH | ASSRPDWDYAEQF | 0.7209931 |
| TPPPPPP | ASSRPDWDYAEQF | 0.7713214 |
| AAPAAAAAPA | ASSRPDWDYAEQF | 0.85383123 |
| IALNEDL | ASSRPDWDYAEQF | 0.54311836 |
| IVNVSSM | ASSRPDWDYAEQF | 0.6633053 |
| TGIPPAPR | ASSRPDWDYAEQF | 0.8279773 |
| FDLGGGTF | ASSRPDWDYAEQF | 0.6023074 |
| FDLGGGTFDVS | ASSRPDWDYAEQF | 0.83570176 |
| NEPTAAA | ASSRPDWDYAEQF | 0.7887856 |
| NEPTAAALA | ASSRPDWDYAEQF | 0.80128634 |
| GSIINISSI | ASSRPDWDYAEQF | 0.7520705 |
| VAPGNAGTA | ASSRPDWDYAEQF | 0.86853546 |
| TGGGASLE | ASSRPDWDYAEQF | 0.8368676 |
| VAVYNHYKRI | ASSRPDWDYAEQF | 0.77685165 |
| TDPAPATT | ASSRPDWDYAEQF | 0.7626504 |
| TNRPDILDPALLRPG | ASSRPDWDYAEQF | 0.65332985 |
| LTTTTTS | ASSRPDWDYAEQF | 0.84198356 |
| PTGSGKTL | ASSRPDWDYAEQF | 0.84658945 |
| TGSGKTL | ASSRPDWDYAEQF | 0.53548014 |
| TIGHVDHGK | ASSRPDWDYAEQF | 0.7223387 |
| TIGHVDHGKTT | ASSRPDWDYAEQF | 0.87253064 |
| GHVDHGKTT | ASSRPDWDYAEQF | 0.8465363 |
| AQFEHTLL | ASSRPDWDYAEQF | 0.63769126 |
| IVNQDEL | ASSRPDWDYAEQF | 0.69230115 |
| TNGSQFF | ASSRPDWDYAEQF | 0.5809342 |
| TGKGIVE | ASSRPDWDYAEQF | 0.7761625 |
| LISNASDA | ASSRPDWDYAEQF | 0.57058614 |
| AELRTLGELTKIAKAHGAQ | ASSRPDWDYAEQF | 0.8206341 |
| APNAVAAAEPSW | ASSRPDWDYAEQF | 0.7146115 |
| APQVAGAATAVL | ASSRPDWDYAEQF | 0.8305246 |
| ARRDPAAAAAILPTDA | ASSRPDWDYAEQF | 0.78347385 |
| DAAARDVLAEAGLAEYF | ASSRPDWDYAEQF | 0.8647984 |
| DDQIVLSAKANLAKVEQ | ASSRPDWDYAEQF | 0.715843 |
| DELLPDTGEDSDEDGH | ASSRPDWDYAEQF | 0.7420443 |
| ELPEAPANRATVPQD | ASSRPDWDYAEQF | 0.56958354 |
| FGQSKIERVTGIG | ASSRPDWDYAEQF | 0.669669 |
| GAGGTGSTAGGAGGAGGAGGLY | ASSRPDWDYAEQF | 0.7034078 |
| GHGGTGAAGGNGGTGGAGGSGI | ASSRPDWDYAEQF | 0.74102473 |
| GNGGNGGAGGEGGPGGAGGAGA | ASSRPDWDYAEQF | 0.30813593 |
| GSTAGNGGNGGNATNGTIAGQP | ASSRPDWDYAEQF | 0.85491216 |
| GTGGDGGNGGNGGTGG | ASSRPDWDYAEQF | 0.7925399 |
| GVRRPAGIPADAPV | ASSRPDWDYAEQF | 0.8408542 |
| GYRQVTDAHLLTLARR | ASSRPDWDYAEQF | 0.7269466 |
| IAKMGQDGHDRG | ASSRPDWDYAEQF | 0.7327573 |
| KAEPSIEIQYT | ASSRPDWDYAEQF | 0.6620968 |
| KPVRIRMAADEDTVAVT | ASSRPDWDYAEQF | 0.8278181 |
| MPPRIAGMRLLVIKPEP | ASSRPDWDYAEQF | 0.60365015 |
| MKFYAVALFRSRGVPPTR | ASSRPDWDYAEQF | 0.38828635 |

| | | |
|---|---|---|
| NAGGFGILGAVAHSP | ASSRPDWDYAEQF | 0.7013295 |
| NEGREITLAMAEM | ASSRPDWDYAEQF | 0.43719652 |
| NELVEWSHPLGHRWRWE | ASSRPDWDYAEQF | 0.8674747 |
| NLGAGGTGGNGGNPG | ASSRPDWDYAEQF | 0.84656954 |
| NTEWPKWETVMAADPVD | ASSRPDWDYAEQF | 0.7940458 |
| PASAAPAETQP | ASSRPDWDYAEQF | 0.8697116 |
| QQSSDGKNGVRAAEITGE | ASSRPDWDYAEQF | 0.302721 |
| RAPDAVTTSWWKE | ASSRPDWDYAEQF | 0.67526394 |
| SGFGNVGAHTSGWFNQ | ASSRPDWDYAEQF | 0.8662832 |
| SGVAGAVGQAASVGGLKV | ASSRPDWDYAEQF | 0.8041829 |
| SNGGAGTGSAGGNGGT | ASSRPDWDYAEQF | 0.7474327 |
| TGGAGGEGGAAG | ASSRPDWDYAEQF | 0.862024 |
| TGGDGGAAGAGGAPGHG | ASSRPDWDYAEQF | 0.6591862 |
| TMPPLLAAIANAGRHGV | ASSRPDWDYAEQF | 0.7736669 |
| TSSTAGNSGGA | ASSRPDWDYAEQF | 0.783892 |
| VPATVPVGILC | ASSRPDWDYAEQF | 0.8502646 |
| VQRPDSGGVVLTGRIS | ASSRPDWDYAEQF | 0.81472635 |
| WTPNPVPATKGVPAGLVQAT | ASSRPDWDYAEQF | 0.6182371 |
| YALKRALAKFSDRFDVVII | ASSRPDWDYAEQF | 0.84332854 |
| YLIGSVIV | ASSRPDWDYAEQF | 0.86835736 |
| YGQLVAVID | ASSRPDWDYAEQF | 0.6807945 |
| VTVGGMTLPA | ASSRPDWDYAEQF | 0.8324332 |
| VLALPAV | ASSRPDWDYAEQF | 0.8210965 |
| VIPPTLNYET | ASSRPDWDYAEQF | 0.7773514 |
| VGMGGMHPSSG | ASSRPDWDYAEQF | 0.6284447 |
| VGLAGGAATA | ASSRPDWDYAEQF | 0.63439727 |
| VGGAGGAGGSGP | ASSRPDWDYAEQF | 0.8183266 |
| VFAAFMV | ASSRPDWDYAEQF | 0.5754022 |
| TLYDKLR | ASSRPDWDYAEQF | 0.79013085 |
| TLAVFTVL | ASSRPDWDYAEQF | 0.74543595 |
| TGSVQPARV | ASSRPDWDYAEQF | 0.88742965 |
| TGGGAGSGGT | ASSRPDWDYAEQF | 0.82508427 |
| SHLETSL | ASSRPDWDYAEQF | 0.82048297 |
| SFTTGMT | ASSRPDWDYAEQF | 0.6082467 |
| QVEVAKRMGVRQPTVS | ASSRPDWDYAEQF | 0.8185135 |
| QGSGGPALAM | ASSRPDWDYAEQF | 0.8039248 |
| MLAPSNP | ASSRPDWDYAEQF | 0.51899 |
| LVNNAAI | ASSRPDWDYAEQF | 0.8613087 |
| LTGEINTLA | ASSRPDWDYAEQF | 0.67469245 |
| LSIFGPV | ASSRPDWDYAEQF | 0.7222707 |
| LLALLANL | ASSRPDWDYAEQF | 0.84816223 |
| LISNASDALDKLR | ASSRPDWDYAEQF | 0.85027057 |
| LINEAVP | ASSRPDWDYAEQF | 0.76943946 |
| LGYAFSGL | ASSRPDWDYAEQF | 0.8043522 |
| LDFVAVAL | ASSRPDWDYAEQF | 0.62599075 |
| LAQPGAE | ASSRPDWDYAEQF | 0.8529143 |
| ISNLGAVTGF | ASSRPDWDYAEQF | 0.84753585 |
| HQEDPGRI | ASSRPDWDYAEQF | 0.7519163 |
| GTPGEPTP | ASSRPDWDYAEQF | 0.87065876 |
| GFFNSGS | ASSRPDWDYAEQF | 0.83253753 |
| GAGGGGLTAGAN | ASSRPDWDYAEQF | 0.530598 |

| | | |
|---|---|---|
| FVDALVNM | ASSRPDWDYAEQF | 0.80507314 |
| FLTVAPEL | ASSRPDWDYAEQF | 0.83165956 |
| FETLTAL | ASSRPDWDYAEQF | 0.8431393 |
| DVINPIEL | ASSRPDWDYAEQF | 0.8677368 |
| APVSGAPV | ASSRPDWDYAEQF | 0.8032629 |
| ALVGGLAW | ASSRPDWDYAEQF | 0.6887087 |
| ALAAAAATAI | ASSRPDWDYAEQF | 0.8630358 |
| AFGLGLLL | ASSRPDWDYAEQF | 0.8850571 |
| AFFAAASL | ASSRPDWDYAEQF | 0.8415555 |
| AAAALPAFSPP | ASSRPDWDYAEQF | 0.7479377 |
| VVLAPAL | ASSRPDWDYAEQF | 0.7660767 |
| TIGHVDHGKTTL | ASSRPDWDYAEQF | 0.81928396 |
| LISNASDALDK | ASSRPDWDYAEQF | 0.83397186 |
| LISNASDAL | ASSRPDWDYAEQF | 0.6906811 |
| ISNASDALDK | ASSRPDWDYAEQF | 0.7951764 |
| NQAETLVYQTE | ASSRPDWDYAEQF | 0.8066428 |
| TGGTGGTAGAG | ASSRPDWDYAEQF | 0.43975514 |
| TGSTAGGAGGA | ASSRPDWDYAEQF | 0.7991209 |
| DPLAVPVAAGPV | ASSRPDWDYAEQF | 0.8354487 |
| GGTGGTPGNGGN | ASSRPDWDYAEQF | 0.82157946 |
| PAGLAGVDPATL | ASSRPDWDYAEQF | 0.71102655 |
| GNNLVVEAPHAGASV | ASSRPDWDYAEQF | 0.8467656 |
| KVTRIMSASTLEQPA | ASSRPDWDYAEQF | 0.873294 |
| LGFALPIALPRRTVV | ASSRPDWDYAEQF | 0.87243927 |
| TRLGNLTARKLVVLV | ASSRPDWDYAEQF | 0.4962854 |
| WSQQRRLIYFPSAGP | ASSRPDWDYAEQF | 0.31250268 |
| GPNGNPDPMAAAVDIR | ASSRPDWDYAEQF | 0.735226 |
| TQRCVSVADLRAAASR | ASSRPDWDYAEQF | 0.80173725 |
| QFVLDVAEHPVEVLTETV | ASSRPDWDYAEQF | 0.7444757 |
| GGAGGTGGAGGNGGTQAGDG | ASSRPDWDYAEQF | 0.7108062 |
| PLLATLRAIAVDVDAKPAQV | ASSRPDWDYAEQF | 0.7805519 |
| APKFRTGGYMIDFWGVGYQVAK | ASSRPDWDYAEQF | 0.85113496 |
| GAGGAGGAGGAAGGPGGTGGTG | ASSRPDWDYAEQF | 0.6899829 |
| IGALGLLATDVAPAGSVAFGVA | ASSRPDWDYAEQF | 0.61176264 |
| AGGAGGIGGNGGAGGAGGVLMG | ASSRPDWDYAEQF | 0.798058 |
| ADELIGLL | ASSRPDWDYAEQF | 0.8442638 |
| ADRALKA | ASSRPDWDYAEQF | 0.60270077 |
| AGAVGQVGGAGGA | ASSRPDWDYAEQF | 0.70399404 |
| AGGGGGLGGAP | ASSRPDWDYAEQF | 0.72831506 |
| AGGLLYGNGGAG | ASSRPDWDYAEQF | 0.7333563 |
| AGLESIGL | ASSRPDWDYAEQF | 0.77497977 |
| AGLKTGII | ASSRPDWDYAEQF | 0.62833446 |
| ASLAALLL | ASSRPDWDYAEQF | 0.8166091 |
| ASVINISSI | ASSRPDWDYAEQF | 0.58218706 |
| ATPPGPA | ASSRPDWDYAEQF | 0.84766346 |
| ATVASALV | ASSRPDWDYAEQF | 0.7914325 |
| ATVASLIL | ASSRPDWDYAEQF | 0.8270496 |
| AVNGPVLIRH | ASSRPDWDYAEQF | 0.7202829 |
| DGAAGAVGI | ASSRPDWDYAEQF | 0.65971756 |
| DPLLHGI | ASSRPDWDYAEQF | 0.56410694 |
| ELGAALGAL | ASSRPDWDYAEQF | 0.4811576 |

| | | |
|---|---|---|
| ELGRTQAV | ASSRPDWDYAEQF | 0.8452656 |
| FVAGGYLS | ASSRPDWDYAEQF | 0.5926478 |
| GASGGAGGAGGW | ASSRPDWDYAEQF | 0.8504796 |
| GGAGAGGDGGAGGI | ASSRPDWDYAEQF | 0.834661 |
| GTGHNGSQPGMGGNG | ASSRPDWDYAEQF | 0.6440371 |
| GYAAASASAS | ASSRPDWDYAEQF | 0.82353324 |
| HQVDVAAVQQG | ASSRPDWDYAEQF | 0.6060729 |
| IGAAIAHT | ASSRPDWDYAEQF | 0.7776163 |
| IGMIEVAQL | ASSRPDWDYAEQF | 0.83439386 |
| IRLAPEHTSAMD | ASSRPDWDYAEQF | 0.8443109 |
| IYNVSVL | ASSRPDWDYAEQF | 0.649932 |
| KQGFLGM | ASSRPDWDYAEQF | 0.8403768 |
| LESVITQL | ASSRPDWDYAEQF | 0.79606116 |
| LGLGALVVV | ASSRPDWDYAEQF | 0.69129366 |
| LIGNGGNGGSGGI | ASSRPDWDYAEQF | 0.83039844 |
| LNAINAPTL | ASSRPDWDYAEQF | 0.8451003 |
| LQLLNTGV | ASSRPDWDYAEQF | 0.8452358 |
| LQTSIEF | ASSRPDWDYAEQF | 0.61593765 |
| MSFAVSEL | ASSRPDWDYAEQF | 0.8044735 |
| NTGALGL | ASSRPDWDYAEQF | 0.6222838 |
| PPGAGGAGGAGTT | ASSRPDWDYAEQF | 0.7863102 |
| QLVGLGI | ASSRPDWDYAEQF | 0.78191143 |
| RFLEQVHQL | ASSRPDWDYAEQF | 0.7729061 |
| RGVGYVLR | ASSRPDWDYAEQF | 0.8286591 |
| SATGVFAGVI | ASSRPDWDYAEQF | 0.59116554 |
| SLPAITLP | ASSRPDWDYAEQF | 0.63651574 |
| STALAFTV | ASSRPDWDYAEQF | 0.7734809 |
| SVLTVGLGK | ASSRPDWDYAEQF | 0.79325616 |
| TALAGAGSGI | ASSRPDWDYAEQF | 0.83834517 |
| TGLPFTGI | ASSRPDWDYAEQF | 0.6234306 |
| VAEHVTV | ASSRPDWDYAEQF | 0.79209536 |
| VDIVLNSL | ASSRPDWDYAEQF | 0.75346744 |
| VLTGGGAVL | ASSRPDWDYAEQF | 0.66203946 |
| VNVGLAYGI | ASSRPDWDYAEQF | 0.82791096 |
| VSSAAVGAV | ASSRPDWDYAEQF | 0.7690255 |
| VTSVYEAGV | ASSRPDWDYAEQF | 0.6570611 |
| WVALDRGA | ASSRPDWDYAEQF | 0.8365355 |
| YLVGSVVI | ASSRPDWDYAEQF | 0.6986284 |
| GPPGTGKTLL | ASSRPDWDYAEQF | 0.83644056 |
| GVGMTKF | ASSRPDWDYAEQF | 0.84935045 |
| IGIGGSDL | ASSRPDWDYAEQF | 0.7018364 |
| IGTLLAEL | ASSRPDWDYAEQF | 0.8447837 |
| IYTSGTT | ASSRPDWDYAEQF | 0.7838226 |
| LSFPAGV | ASSRPDWDYAEQF | 0.7911986 |
| LYTSGTT | ASSRPDWDYAEQF | 0.7532327 |
| VGAFPFGF | ASSRPDWDYAEQF | 0.6593534 |
| VGAGTVEF | ASSRPDWDYAEQF | 0.83991 |
| VGKSTLLL | ASSRPDWDYAEQF | 0.82295525 |
| AALAHAGGGIV | ASSRPDWDYAEQF | 0.83932513 |
| AALVAVSLL | ASSRPDWDYAEQF | 0.79403615 |
| AAVSRAVAAL | ASSRPDWDYAEQF | 0.8073684 |

| | | |
|---|---|---|
| AGGSSGIGGF | ASSRPDWDYAEQF | 0.8353589 |
| AGLTVANANG | ASSRPDWDYAEQF | 0.66948205 |
| AGLVGAVL | ASSRPDWDYAEQF | 0.54743433 |
| ALSQAFTV | ASSRPDWDYAEQF | 0.7387721 |
| ALTFSVSL | ASSRPDWDYAEQF | 0.70258045 |
| ALTGAALVAL | ASSRPDWDYAEQF | 0.7205589 |
| ASFSIGAGVL | ASSRPDWDYAEQF | 0.6024229 |
| DGSVYKI | ASSRPDWDYAEQF | 0.84221786 |
| EQLAEIV | ASSRPDWDYAEQF | 0.87691164 |
| FAVSELTVM | ASSRPDWDYAEQF | 0.6634406 |
| GAAPLAAGL | ASSRPDWDYAEQF | 0.8730699 |
| GADGGDGGSGGAGG | ASSRPDWDYAEQF | 0.60743916 |
| GFGLDTGGAGG | ASSRPDWDYAEQF | 0.8401754 |
| GLASGGLL | ASSRPDWDYAEQF | 0.85942185 |
| GRIAGAAVLL | ASSRPDWDYAEQF | 0.56755567 |
| GSAGSLNTSF | ASSRPDWDYAEQF | 0.83690083 |
| GTGQIGI | ASSRPDWDYAEQF | 0.6818584 |
| GTLIALIL | ASSRPDWDYAEQF | 0.72630525 |
| IAATMGFVVG | ASSRPDWDYAEQF | 0.85796404 |
| IFAQALL | ASSRPDWDYAEQF | 0.75791186 |
| IGAAVALGF | ASSRPDWDYAEQF | 0.8310234 |
| IGLTPLAL | ASSRPDWDYAEQF | 0.78288174 |
| ILGSNTSTL | ASSRPDWDYAEQF | 0.6763986 |
| IVFSLTVL | ASSRPDWDYAEQF | 0.8198455 |
| LASKATGF | ASSRPDWDYAEQF | 0.8653157 |
| LASVALIL | ASSRPDWDYAEQF | 0.767024 |
| LFAVGLTL | ASSRPDWDYAEQF | 0.88261473 |
| LFNGGIGTYI | ASSRPDWDYAEQF | 0.6547536 |
| LTLGAGII | ASSRPDWDYAEQF | 0.8491848 |
| LVAAALGDI | ASSRPDWDYAEQF | 0.838098 |
| NTAFVQLGI | ASSRPDWDYAEQF | 0.7515229 |
| QLFSGAEL | ASSRPDWDYAEQF | 0.7636558 |
| SAATFITL | ASSRPDWDYAEQF | 0.8343405 |
| SNYGLANLGV | ASSRPDWDYAEQF | 0.7583162 |
| SYGGAYAVM | ASSRPDWDYAEQF | 0.8221804 |
| TTGVGGLTLG | ASSRPDWDYAEQF | 0.8733739 |
| TTQTIYI | ASSRPDWDYAEQF | 0.3399325 |
| VAAGAGLGAVI | ASSRPDWDYAEQF | 0.6800834 |
| VAAVVTLL | ASSRPDWDYAEQF | 0.5960642 |
| VGFLGGLI | ASSRPDWDYAEQF | 0.42134997 |
| VGVIPILL | ASSRPDWDYAEQF | 0.46539724 |
| VIFASSLI | ASSRPDWDYAEQF | 0.31037986 |
| VKGGDGGAAAG | ASSRPDWDYAEQF | 0.39768678 |
| VTKTFTNI | ASSRPDWDYAEQF | 0.44866872 |
| VVAAAILL | ASSRPDWDYAEQF | 0.6913998 |
| YAALDVEL | ASSRPDWDYAEQF | 0.5388401 |
| LASVALLL | ASSRPDWDYAEQF | 0.4038219 |
| LDAVHRV | ASSRPDWDYAEQF | 0.6608854 |
| QGLSSSIF | ASSRPDWDYAEQF | 0.5187831 |
| RGTFANIRL | ASSRPDWDYAEQF | 0.33995646 |
| RQLWWGHRI | ASSRPDWDYAEQF | 0.4769966 |

| | | |
|---|---|---|
| YGPPGTGKTLL | ASSRPDWDYAEQF | 0.37100345 |
| AALVGAPANVA | ASSRPDWDYAEQF | 0.44070393 |
| AATFRRTE | ASSRPDWDYAEQF | 0.2836887 |
| AGALGGGI | ASSRPDWDYAEQF | 0.5374079 |
| AGGAGGAGGIGAG | ASSRPDWDYAEQF | 0.50331265 |
| AGSGALGL | ASSRPDWDYAEQF | 0.26914746 |
| AQIADQI | ASSRPDWDYAEQF | 0.54626715 |
| ASAVAVTYL | ASSRPDWDYAEQF | 0.477082 |
| ASTVLGGTGG | ASSRPDWDYAEQF | 0.5283818 |
| AVGGEPAK | ASSRPDWDYAEQF | 0.65289664 |
| CEQGTFGGTC | ASSRPDWDYAEQF | 0.3600561 |
| DEGKAPAG | ASSRPDWDYAEQF | 0.364789 |
| DTVVFPAL | ASSRPDWDYAEQF | 0.64744693 |
| FEGITMAG | ASSRPDWDYAEQF | 0.6557757 |
| FPLVVQD | ASSRPDWDYAEQF | 0.34630296 |
| GAFTTPQ | ASSRPDWDYAEQF | 0.5581579 |
| GFGGFGTAGGVGGA | ASSRPDWDYAEQF | 0.6259487 |
| GFLAQFL | ASSRPDWDYAEQF | 0.5534325 |
| GGAGGAGGNAGVA | ASSRPDWDYAEQF | 0.5547449 |
| GGVGTAGAGGAGGAGG | ASSRPDWDYAEQF | 0.57170784 |
| GNGGGGDAGNAGSGGN | ASSRPDWDYAEQF | 0.37294224 |
| GTGGAGGAGNDAGS | ASSRPDWDYAEQF | 0.64845395 |
| GTTNTGWFNS | ASSRPDWDYAEQF | 0.6847573 |
| IFMGPLFTL | ASSRPDWDYAEQF | 0.45348877 |
| ILAHAIEAI | ASSRPDWDYAEQF | 0.46657342 |
| IRSAIREAL | ASSRPDWDYAEQF | 0.33228955 |
| ITFAGGG | ASSRPDWDYAEQF | 0.42785355 |
| ITGMSGHQEV | ASSRPDWDYAEQF | 0.39976057 |
| IVAGLACTL | ASSRPDWDYAEQF | 0.6231667 |
| IVGGGLAGA | ASSRPDWDYAEQF | 0.61305344 |
| IVTGVLNI | ASSRPDWDYAEQF | 0.51327145 |
| LAQQGCI | ASSRPDWDYAEQF | 0.34606668 |
| LCLHVEN | ASSRPDWDYAEQF | 0.32383752 |
| LDAISDGS | ASSRPDWDYAEQF | 0.54201967 |
| LGGFAAML | ASSRPDWDYAEQF | 0.34494823 |
| LGGKNPMI | ASSRPDWDYAEQF | 0.30466932 |
| LGPKGVAAIL | ASSRPDWDYAEQF | 0.47088075 |
| LMVGVSI | ASSRPDWDYAEQF | 0.64754283 |
| LVGAAFVL | ASSRPDWDYAEQF | 0.58431745 |
| LVLAVAPL | ASSRPDWDYAEQF | 0.47355363 |
| MAAIAVIDAA | ASSRPDWDYAEQF | 0.6748386 |
| NNGINAELQ | ASSRPDWDYAEQF | 0.6574806 |
| RGEPLVLVHG | ASSRPDWDYAEQF | 0.5668236 |
| RGGAGGAAGLI | ASSRPDWDYAEQF | 0.38159695 |
| RSYRMHPAV | ASSRPDWDYAEQF | 0.3580523 |
| SAAHLLAGQLA | ASSRPDWDYAEQF | 0.6954449 |
| SAAVGSY | ASSRPDWDYAEQF | 0.57110727 |
| SPHAVLHRI | ASSRPDWDYAEQF | 0.5050386 |
| SQIAKVL | ASSRPDWDYAEQF | 0.6005391 |
| TALGHLVP | ASSRPDWDYAEQF | 0.53029186 |
| TAPTTSGAA | ASSRPDWDYAEQF | 0.4399864 |

| | | |
|---|---|---|
| TDGDDGNGGSPGT | ASSRPDWDYAEQF | 0.61101854 |
| TDGLGEN | ASSRPDWDYAEQF | 0.53173697 |
| TGGPPLPGQA | ASSRPDWDYAEQF | 0.4013658 |
| TIPAITI | ASSRPDWDYAEQF | 0.3936022 |
| TIPGFTI | ASSRPDWDYAEQF | 0.41454354 |
| TSIASGALA | ASSRPDWDYAEQF | 0.5174542 |
| TVTLPGAAI | ASSRPDWDYAEQF | 0.6432452 |
| VASGLSAALP | ASSRPDWDYAEQF | 0.4740978 |
| VGAGVIGI | ASSRPDWDYAEQF | 0.36675742 |
| VGPKRFGV | ASSRPDWDYAEQF | 0.60453224 |
| VNVAFPD | ASSRPDWDYAEQF | 0.49100354 |
| VSMTNSGVS | ASSRPDWDYAEQF | 0.48735517 |
| VTDVLGDLSV | ASSRPDWDYAEQF | 0.42868227 |
| VVVVGAGFAGL | ASSRPDWDYAEQF | 0.505663 |
| VYTVGTVV | ASSRPDWDYAEQF | 0.52907217 |
| WTTFSGP | ASSRPDWDYAEQF | 0.53295285 |
| YDELVEAF | ASSRPDWDYAEQF | 0.46007684 |
| YGTGGAGGNGG | ASSRPDWDYAEQF | 0.5176677 |
| YTNVMAV | ASSRPDWDYAEQF | 0.52547425 |
| FLEQVHQL | ASSRPDWDYAEQF | 0.42328423 |
| GLSLLAVAV | ASSRPDWDYAEQF | 0.33097714 |
| LEQVHQL | ASSRPDWDYAEQF | 0.36392742 |
| LNAAGAI | ASSRPDWDYAEQF | 0.57964164 |
| PAVHASL | ASSRPDWDYAEQF | 0.5786128 |
| TELEDDA | ASSRPDWDYAEQF | 0.6235031 |
| TPPGPPN | ASSRPDWDYAEQF | 0.43528748 |
| TPPPPPPP | ASSRPDWDYAEQF | 0.44418132 |
| TQSPALI | ASSPGQANTEVF | 0.5113239 |
| EFAAAFPQ | ASSPGQANTEVF | 0.61642814 |
| LLSGNLDVL | ASSPGQANTEVF | 0.34900105 |
| GPANLGNYNI | ASSPGQANTEVF | 0.4466036 |
| TLGPTLP | ASSPGQANTEVF | 0.7042997 |
| VYAYSSCCK | ASSPGQANTEVF | 0.62477565 |
| APAPAPAPNI | ASSPGQANTEVF | 0.47312057 |
| PGGPVPIP | ASSPGQANTEVF | 0.36456752 |
| FLTVALEP | ASSPGQANTEVF | 0.70433253 |
| SQYFGANL | ASSPGQANTEVF | 0.40791443 |
| GGVKPAA | ASSPGQANTEVF | 0.69679785 |
| GGTGGAGGAGGW | ASSPGQANTEVF | 0.28506362 |
| GVGGVAGLGGAGG | ASSPGQANTEVF | 0.35893637 |
| VLLVLATTI | ASSPGQANTEVF | 0.3241621 |
| IGIFAAML | ASSPGQANTEVF | 0.28692007 |
| AVGDLID | ASSPGQANTEVF | 0.6205871 |
| NISPLPL | ASSPGQANTEVF | 0.35993266 |
| VTAVFVAI | ASSPGQANTEVF | 0.55247504 |
| SGDDNDAAL | ASSPGQANTEVF | 0.4729744 |
| AVAALAHS | ASSPGQANTEVF | 0.63318413 |
| GGVTVMATP | ASSPGQANTEVF | 0.5621024 |
| AGLAANVRD | ASSPGQANTEVF | 0.43673941 |
| TNRPDLIDPALLRPG | ASSPGQANTEVF | 0.35170883 |
| LANLGVTAL | ASSPGQANTEVF | 0.59559137 |

| | | |
|---|---|---|
| TGERIEL | ASSPGQANTEVF | 0.5823816 |
| NGEELAA | ASSPGQANTEVF | 0.40800193 |
| VLSLVWG | ASSPGQANTEVF | 0.31060332 |
| IGYALWL | ASSPGQANTEVF | 0.5575249 |
| LSLLLAPL | ASSPGQANTEVF | 0.5204614 |
| TLDEALK | ASSPGQANTEVF | 0.3082045 |
| TDGAGTAEGG | ASSPGQANTEVF | 0.28302452 |
| NAPGGAGGNGG | ASSPGQANTEVF | 0.53045815 |
| VSVFMGMGGNF | ASSPGQANTEVF | 0.52204067 |
| LASYLAAL | ASSPGQANTEVF | 0.64277065 |
| LVGGTPAL | ASSPGQANTEVF | 0.65495956 |
| VRNADVL | ASSPGQANTEVF | 0.49780166 |
| TCSSPFLVSAM | ASSPGQANTEVF | 0.5341743 |
| VHITDAE | ASSPGQANTEVF | 0.6023034 |
| EYAAPSF | ASSPGQANTEVF | 0.25619408 |
| VVDAFHGGL | ASSPGQANTEVF | 0.3517634 |
| ASVVDPQ | ASSPGQANTEVF | 0.33356565 |
| SYNEFGGLS | ASSPGQANTEVF | 0.5534203 |
| DPDGSVY | ASSPGQANTEVF | 0.46341556 |
| MVSTIHATE | ASSPGQANTEVF | 0.49889237 |
| LAAGVADPV | ASSPGQANTEVF | 0.54089665 |
| TKEQIAA | ASSPGQANTEVF | 0.66310245 |
| SLASPAAL | ASSPGQANTEVF | 0.48664427 |
| IGLVFYGV | ASSPGQANTEVF | 0.33897942 |
| FLLDASNL | ASSPGQANTEVF | 0.29179865 |
| YQVGNFP | ASSPGQANTEVF | 0.57948565 |
| ESLLSQST | ASSPGQANTEVF | 0.36706594 |
| LFTVGFAAVG | ASSPGQANTEVF | 0.37332034 |
| ATPPTPT | ASSPGQANTEVF | 0.29538116 |
| TDLATLG | ASSPGQANTEVF | 0.58907735 |
| SAGGTGR | ASSPGQANTEVF | 0.2953354 |
| GFSNSGS | ASSPGQANTEVF | 0.48264724 |
| GGNGGSGGTSV | ASSPGQANTEVF | 0.3991797 |
| GTSVATGGAGN | ASSPGQANTEVF | 0.5549711 |
| NGGNGGSGGTS | ASSPGQANTEVF | 0.6944741 |
| AVVKFPH | ASSPGQANTEVF | 0.6649608 |
| TPPPPPP | ASSPGQANTEVF | 0.48103738 |
| AAPAAAAAPA | ASSPGQANTEVF | 0.4138181 |
| IALNEDL | ASSPGQANTEVF | 0.38342184 |
| IVNVSSM | ASSPGQANTEVF | 0.42813495 |
| TGIPPAPR | ASSPGQANTEVF | 0.38571203 |
| FDLGGGTF | ASSPGQANTEVF | 0.6391672 |
| FDLGGGTFDVS | ASSPGQANTEVF | 0.36982685 |
| NEPTAAA | ASSPGQANTEVF | 0.31969512 |
| NEPTAAALA | ASSPGQANTEVF | 0.3777714 |
| GSIINISSI | ASSPGQANTEVF | 0.65979934 |
| VAPGNAGTA | ASSPGQANTEVF | 0.6917869 |
| TGGGASLE | ASSPGQANTEVF | 0.41287205 |
| VAVYNHYKRI | ASSPGQANTEVF | 0.34412864 |
| TDPAPATT | ASSPGQANTEVF | 0.3623547 |
| TNRPDILDPALLRPG | ASSPGQANTEVF | 0.5757339 |

| | | |
|---|---|---|
| LTTTTTS | ASSPGQANTEVF | 0.33413044 |
| PTGSGKTL | ASSPGQANTEVF | 0.35925716 |
| TGSGKTL | ASSPGQANTEVF | 0.49977607 |
| TIGHVDHGK | ASSPGQANTEVF | 0.7048899 |
| TIGHVDHGKTT | ASSPGQANTEVF | 0.42109272 |
| GHVDHGKTT | ASSPGQANTEVF | 0.2910586 |
| AQFEHTLL | ASSPGQANTEVF | 0.5475175 |
| IVNQDEL | ASSPGQANTEVF | 0.36816594 |
| TNGSQFF | ASSPGQANTEVF | 0.47705138 |
| TGKGIVE | ASSPGQANTEVF | 0.37448567 |
| LISNASDA | ASSPGQANTEVF | 0.33187637 |
| AELRTLGELTKIAKAHGAQ | ASSPGQANTEVF | 0.51360345 |
| APNAVAAAEPSW | ASSPGQANTEVF | 0.6759292 |
| APQVAGAATAVL | ASSPGQANTEVF | 0.6058167 |
| ARRDPAAAAAILPTDA | ASSPGQANTEVF | 0.49748707 |
| DAAARDVLAEAGLAEYF | ASSPGQANTEVF | 0.6829728 |
| DDQIVLSAKANLAKVEQ | ASSPGQANTEVF | 0.6165283 |
| DELLPDTGEDSDEDGH | ASSPGQANTEVF | 0.6769891 |
| ELPEAPANRATVPQD | ASSPGQANTEVF | 0.66076887 |
| FGQSKIERVTGIG | ASSPGQANTEVF | 0.44933036 |
| GAGGTGSTAGGAGGAGGAGGLY | ASSPGQANTEVF | 0.42202377 |
| GHGGTGAAGGNGGTGGAGGSGI | ASSPGQANTEVF | 0.6797849 |
| GNGGNGGAGGEGGPGGAGGAGA | ASSPGQANTEVF | 0.5297833 |
| GSTAGNGGNGGNATNGTIAGQP | ASSPGQANTEVF | 0.55167174 |
| GTGGDGGNGGNGGTGG | ASSPGQANTEVF | 0.37659788 |
| GVRRPAGIPADAPV | ASSPGQANTEVF | 0.65145195 |
| GYRQVTDAHLLTLARR | ASSPGQANTEVF | 0.37461773 |
| IAKMGQDGHDRG | ASSPGQANTEVF | 0.6861995 |
| KAEPSIEIQYT | ASSPGQANTEVF | 0.29274085 |
| KPVRIRMAADEDTVAVT | ASSPGQANTEVF | 0.36016843 |
| MPPRIAGMRLLVIKPEP | ASSPGQANTEVF | 0.65289444 |
| MKFYAVALFRSRGVPPTR | ASSPGQANTEVF | 0.2570871 |
| NAGGFGILGAVAHSP | ASSPGQANTEVF | 0.2733225 |
| NEGREITLAMAEM | ASSPGQANTEVF | 0.43797395 |
| NELVEWSHPLGHRWRWE | ASSPGQANTEVF | 0.624724 |
| NLGAGGTGGNGGNPG | ASSPGQANTEVF | 0.26047024 |
| NTEWPKWETVMAADPVD | ASSPGQANTEVF | 0.3684084 |
| PASAAPAETQP | ASSPGQANTEVF | 0.31060815 |
| QQSSDGKNGVRAAEITGE | ASSPGQANTEVF | 0.45147455 |
| RAPDAVTTSWWKE | ASSPGQANTEVF | 0.51870584 |
| SGFGNVGAHTSGWFNQ | ASSPGQANTEVF | 0.3101117 |
| SGVAGAVGQAASVGGLKV | ASSPGQANTEVF | 0.5552478 |
| SNGGAGTGSAGGNGGT | ASSPGQANTEVF | 0.2929028 |
| TGGAGGEGGAAG | ASSPGQANTEVF | 0.5108105 |
| TGGDGGAAGAGGAPGHG | ASSPGQANTEVF | 0.5682057 |
| TMPPLLAAIANAGRHGV | ASSPGQANTEVF | 0.6960358 |
| TSSTAGNSGGA | ASSPGQANTEVF | 0.3566605 |
| VPATVPVGILC | ASSPGQANTEVF | 0.5648997 |
| VQRPDSGGVVLTGRIS | ASSPGQANTEVF | 0.36677444 |
| WTPNPVPATKGVPAGLVQAT | ASSPGQANTEVF | 0.70929456 |
| YALKRALAKFSDRFDVVII | ASSPGQANTEVF | 0.46159375 |

| | | |
|---|---|---|
| YLIGSVIV | ASSPGQANTEVF | 0.30762142 |
| YGQLVAVID | ASSPGQANTEVF | 0.62577003 |
| VTVGGMTLPA | ASSPGQANTEVF | 0.51682526 |
| VLALPAV | ASSPGQANTEVF | 0.44772887 |
| VIPPTLNYET | ASSPGQANTEVF | 0.6765335 |
| VGMGGMHPSSG | ASSPGQANTEVF | 0.37449452 |
| VGLAGGAATA | ASSPGQANTEVF | 0.55287 |
| VGGAGGAGGSGP | ASSPGQANTEVF | 0.6474711 |
| VFAAFMV | ASSPGQANTEVF | 0.63655037 |
| TLYDKLR | ASSPGQANTEVF | 0.38055873 |
| TLAVFTVL | ASSPGQANTEVF | 0.501096 |
| TGSVQPARV | ASSPGQANTEVF | 0.55562395 |
| TGGGAGSGGT | ASSPGQANTEVF | 0.37427998 |
| SHLETSL | ASSPGQANTEVF | 0.6642556 |
| SFTTGMT | ASSPGQANTEVF | 0.40403008 |
| QVEVAKRMGVRQPTVS | ASSPGQANTEVF | 0.5884694 |
| QGSGGPALAM | ASSPGQANTEVF | 0.54895294 |
| MLAPSNP | ASSPGQANTEVF | 0.69251585 |
| LVNNAAI | ASSPGQANTEVF | 0.38724792 |
| LTGEINTLA | ASSPGQANTEVF | 0.48700055 |
| LSIFGPV | ASSPGQANTEVF | 0.52770764 |
| LLALLANL | ASSPGQANTEVF | 0.5718785 |
| LISNASDALDKLR | ASSPGQANTEVF | 0.368817 |
| LINEAVP | ASSPGQANTEVF | 0.55507714 |
| LGYAFSGL | ASSPGQANTEVF | 0.51900977 |
| LDFVAVAL | ASSPGQANTEVF | 0.6498624 |
| LAQPGAE | ASSPGQANTEVF | 0.4480768 |
| ISNLGAVTGF | ASSPGQANTEVF | 0.46639785 |
| HQEDPGRI | ASSPGQANTEVF | 0.6905532 |
| GTPGEPTP | ASSPGQANTEVF | 0.3791882 |
| GFFNSGS | ASSPGQANTEVF | 0.48844504 |
| GAGGGGLTAGAN | ASSPGQANTEVF | 0.56861454 |
| FVDALVNM | ASSPGQANTEVF | 0.61046743 |
| FLTVAPEL | ASSPGQANTEVF | 0.41405112 |
| FETLTAL | ASSPGQANTEVF | 0.6414416 |
| DVINPIEL | ASSPGQANTEVF | 0.3705221 |
| APVSGAPV | ASSPGQANTEVF | 0.6451682 |
| ALVGGLAW | ASSPGQANTEVF | 0.4791245 |
| ALAAAAATAI | ASSPGQANTEVF | 0.47493318 |
| AFGLGLLL | ASSPGQANTEVF | 0.50149375 |
| AFFAAASL | ASSPGQANTEVF | 0.6630542 |
| AAAALPAFSPP | ASSPGQANTEVF | 0.6455525 |
| VVLAPAL | ASSPGQANTEVF | 0.6425234 |
| TIGHVDHGKTTL | ASSPGQANTEVF | 0.272864 |
| LISNASDALDK | ASSPGQANTEVF | 0.6348046 |
| LISNASDAL | ASSPGQANTEVF | 0.6017566 |
| ISNASDALDK | ASSPGQANTEVF | 0.7096646 |
| NQAETLVYQTE | ASSPGQANTEVF | 0.4130758 |
| TGGTGGTAGAG | ASSPGQANTEVF | 0.4669816 |
| TGSTAGGAGGA | ASSPGQANTEVF | 0.46297643 |
| DPLAVPVAAGPV | ASSPGQANTEVF | 0.4381763 |

| | | |
|---|---|---|
| GGTGGTPGNGGN | ASSPGQANTEVF | 0.33225587 |
| PAGLAGVDPATL | ASSPGQANTEVF | 0.40345678 |
| GNNLVVEAPHAGASV | ASSPGQANTEVF | 0.5093372 |
| KVTRIMSASTLEQPA | ASSPGQANTEVF | 0.48140436 |
| LGFALPIALPRRTVV | ASSPGQANTEVF | 0.685781 |
| TRLGNLTARKLVVLV | ASSPGQANTEVF | 0.6444727 |
| WSQQRRLIYFPSAGP | ASSPGQANTEVF | 0.664328 |
| GPNGNPDPMAAAVDIR | ASSPGQANTEVF | 0.6259265 |
| TQRCVSVADLRAAASR | ASSPGQANTEVF | 0.66592574 |
| QFVLDVAEHPVEVLTETV | ASSPGQANTEVF | 0.48255605 |
| GGAGGTGGAGGNGGTQAGDG | ASSPGQANTEVF | 0.45338607 |
| PLLATLRAIAVDVDAKPAQV | ASSPGQANTEVF | 0.4721407 |
| APKFRTGGYMIDFWGVGYQVAK | ASSPGQANTEVF | 0.3791865 |
| GAGGAGGAGGAAGGPGGTGGTG | ASSPGQANTEVF | 0.44763082 |
| IGALGLLATDVAPAGSVAFGVA | ASSPGQANTEVF | 0.4793259 |
| AGGAGGIGGNGGAGGAGGVLMG | ASSPGQANTEVF | 0.5326202 |
| ADELIGLL | ASSPGQANTEVF | 0.2584284 |
| ADRALKA | ASSPGQANTEVF | 0.6592705 |
| AGAVGQVGGAGGA | ASSPGQANTEVF | 0.6752363 |
| AGGGGGLGGAP | ASSPGQANTEVF | 0.5852046 |
| AGGLLYGNGGAG | ASSPGQANTEVF | 0.5246956 |
| AGLESIGL | ASSPGQANTEVF | 0.644177 |
| AGLKTGII | ASSPGQANTEVF | 0.3801829 |
| ASLAALLL | ASSPGQANTEVF | 0.5492553 |
| ASVINISSI | ASSPGQANTEVF | 0.36504763 |
| ATPPGPA | ASSPGQANTEVF | 0.5102112 |
| ATVASALV | ASSPGQANTEVF | 0.6583023 |
| ATVASLIL | ASSPGQANTEVF | 0.55582047 |
| AVNGPVLIRH | ASSPGQANTEVF | 0.3817285 |
| DGAAGAVGI | ASSPGQANTEVF | 0.39116153 |
| DPLLHGI | ASSPGQANTEVF | 0.31870538 |
| ELGAALGAL | ASSPGQANTEVF | 0.47570318 |
| ELGRTQAV | ASSPGQANTEVF | 0.45848128 |
| FVAGGYLS | ASSPGQANTEVF | 0.5317632 |
| GASGGAGGAGGW | ASSPGQANTEVF | 0.38824534 |
| GGAGAGGDGGAGGI | ASSPGQANTEVF | 0.59653485 |
| GTGHNGSQPGMGGNG | ASSPGQANTEVF | 0.6057856 |
| GYAAASASAS | ASSPGQANTEVF | 0.47191626 |
| HQVDVAAVQQG | ASSPGQANTEVF | 0.39033303 |
| IGAAIAHT | ASSPGQANTEVF | 0.58847463 |
| IGMIEVAQL | ASSPGQANTEVF | 0.56367904 |
| IRLAPEHTSAMD | ASSPGQANTEVF | 0.3268746 |
| IYNVSVL | ASSPGQANTEVF | 0.58206 |
| KQGFLGM | ASSPGQANTEVF | 0.37649646 |
| LESVITQL | ASSPGQANTEVF | 0.5406681 |
| LGLGALVVV | ASSPGQANTEVF | 0.6485559 |
| LIGNGGNGGSGGI | ASSPGQANTEVF | 0.54024553 |
| LNAINAPTL | ASSPGQANTEVF | 0.29737547 |
| LQLLNTGV | ASSPGQANTEVF | 0.56326175 |
| LQTSIEF | ASSPGQANTEVF | 0.5366532 |
| MSFAVSEL | ASSPGQANTEVF | 0.50914395 |

| | | |
|---|---|---|
| NTGALGL | ASSPGQANTEVF | 0.26311478 |
| PPGAGGAGGAGTT | ASSPGQANTEVF | 0.5649965 |
| QLVGLGI | ASSPGQANTEVF | 0.37254846 |
| RFLEQVHQL | ASSPGQANTEVF | 0.42423642 |
| RGVGYVLR | ASSPGQANTEVF | 0.4897377 |
| SATGVFAGVI | ASSPGQANTEVF | 0.6532656 |
| SLPAITLP | ASSPGQANTEVF | 0.54195106 |
| STALAFTV | ASSPGQANTEVF | 0.3912285 |
| SVLTVGLGK | ASSPGQANTEVF | 0.61286175 |
| TALAGAGSGI | ASSPGQANTEVF | 0.56630355 |
| TGLPFTGI | ASSPGQANTEVF | 0.66751444 |
| VAEHVTV | ASSPGQANTEVF | 0.571628 |
| VDIVLNSL | ASSPGQANTEVF | 0.37844008 |
| VLTGGGAVL | ASSPGQANTEVF | 0.45075893 |
| VNVGLAYGI | ASSPGQANTEVF | 0.3336473 |
| VSSAAVGAV | ASSPGQANTEVF | 0.37192637 |
| VTSVYEAGV | ASSPGQANTEVF | 0.33493492 |
| WVALDRGA | ASSPGQANTEVF | 0.3792739 |
| YLVGSVVI | ASSPGQANTEVF | 0.31516337 |
| GPPGTGKTLL | ASSPGQANTEVF | 0.5589347 |
| GVGMTKF | ASSPGQANTEVF | 0.47583956 |
| IGIGGSDL | ASSPGQANTEVF | 0.43725857 |
| IGTLLAEL | ASSPGQANTEVF | 0.3106303 |
| IYTSGTT | ASSPGQANTEVF | 0.40208364 |
| LSFPAGV | ASSPGQANTEVF | 0.47471058 |
| LYTSGTT | ASSPGQANTEVF | 0.6262523 |
| VGAFPFGF | ASSPGQANTEVF | 0.50854474 |
| VGAGTVEF | ASSPGQANTEVF | 0.7046871 |
| VGKSTLLL | ASSPGQANTEVF | 0.664839 |
| AALAHAGGGIV | ASSPGQANTEVF | 0.48905027 |
| AALVAVSLL | ASSPGQANTEVF | 0.6374825 |
| AAVSRAVAAL | ASSPGQANTEVF | 0.4005403 |
| AGGSSGIGGF | ASSPGQANTEVF | 0.44778425 |
| AGLTVANANG | ASSPGQANTEVF | 0.42435652 |
| AGLVGAVL | ASSPGQANTEVF | 0.46681917 |
| ALSQAFTV | ASSPGQANTEVF | 0.42595148 |
| ALTFSVSL | ASSPGQANTEVF | 0.4104865 |
| ALTGAALVAL | ASSPGQANTEVF | 0.29293913 |
| ASFSIGAGVL | ASSPGQANTEVF | 0.5312173 |
| DGSVYKI | ASSPGQANTEVF | 0.45332772 |
| EQLAEIV | ASSPGQANTEVF | 0.29889357 |
| FAVSELTVM | ASSPGQANTEVF | 0.37750557 |
| GAAPLAAGL | ASSPGQANTEVF | 0.70120037 |
| GADGGDGGSGGAGG | ASSPGQANTEVF | 0.59873646 |
| GFGLDTGGAGG | ASSPGQANTEVF | 0.44396985 |
| GLASGGLL | ASSPGQANTEVF | 0.37668294 |
| GRIAGAAVLL | ASSPGQANTEVF | 0.66947186 |
| GSAGSLNTSF | ASSPGQANTEVF | 0.40863812 |
| GTGQIGI | ASSPGQANTEVF | 0.4655877 |
| GTLIALIL | ASSPGQANTEVF | 0.596657 |
| IAATMGFVVG | ASSPGQANTEVF | 0.59295875 |

| | | |
|---|---|---|
| IFAQALL | ASSPGQANTEVF | 0.3338024 |
| IGAAVALGF | ASSPGQANTEVF | 0.5740417 |
| IGLTPLAL | ASSPGQANTEVF | 0.57222205 |
| ILGSNTSTL | ASSPGQANTEVF | 0.41651273 |
| IVFSLTVL | ASSPGQANTEVF | 0.2763899 |
| LASKATGF | ASSPGQANTEVF | 0.51470876 |
| LASVALIL | ASSPGQANTEVF | 0.42718977 |
| LFAVGLTL | ASSPGQANTEVF | 0.3370317 |
| LFNGGIGTYI | ASSPGQANTEVF | 0.27796635 |
| LTLGAGII | ASSPGQANTEVF | 0.42192963 |
| LVAAALGDI | ASSPGQANTEVF | 0.64210796 |
| NTAFVQLGI | ASSPGQANTEVF | 0.6612506 |
| QLFSGAEL | ASSPGQANTEVF | 0.44510022 |
| SAATFITL | ASSPGQANTEVF | 0.63266796 |
| SNYGLANLGV | ASSPGQANTEVF | 0.6309979 |
| SYGGAYAVM | ASSPGQANTEVF | 0.4753828 |
| TTGVGGLTLG | ASSPGQANTEVF | 0.30168575 |
| TTQTIYI | ASSPGQANTEVF | 0.4762841 |
| VAAGAGLGAVI | ASSPGQANTEVF | 0.59532315 |
| VAAVVTLL | ASSPGQANTEVF | 0.5561226 |
| VGFLGGLI | ASSPGQANTEVF | 0.6499466 |
| VGVIPILL | ASSPGQANTEVF | 0.38600898 |
| VIFASSLI | ASSPGQANTEVF | 0.5000854 |
| VKGGDGGAAAG | ASSPGQANTEVF | 0.572589 |
| VTKTFTNI | ASSPGQANTEVF | 0.32536507 |
| VVAAAILL | ASSPGQANTEVF | 0.32969558 |
| YAALDVEL | ASSPGQANTEVF | 0.37361532 |
| LASVALLL | ASSPGQANTEVF | 0.35366562 |
| LDAVHRV | ASSPGQANTEVF | 0.56871426 |
| QGLSSSIF | ASSPGQANTEVF | 0.4947182 |
| RGTFANIRL | ASSPGQANTEVF | 0.40805596 |
| RQLWWGHRI | ASSPGQANTEVF | 0.51359975 |
| YGPPGTGKTLL | ASSPGQANTEVF | 0.3988877 |
| AALVGAPANVA | ASSPGQANTEVF | 0.38987398 |
| AATFRRTE | ASSPGQANTEVF | 0.47331062 |
| AGALGGGI | ASSPGQANTEVF | 0.53643847 |
| AGGAGGAGGIGAG | ASSPGQANTEVF | 0.61131465 |
| AGSGALGL | ASSPGQANTEVF | 0.49583727 |
| AQIADQI | ASSPGQANTEVF | 0.38807487 |
| ASAVAVTYL | ASSPGQANTEVF | 0.69393444 |
| ASTVLGGTGG | ASSPGQANTEVF | 0.33560926 |
| AVGGEPAK | ASSPGQANTEVF | 0.40565053 |
| CEQGTFGGTC | ASSPGQANTEVF | 0.5960884 |
| DEGKAPAG | ASSPGQANTEVF | 0.4104458 |
| DTVVFPAL | ASSPGQANTEVF | 0.5862629 |
| FEGITMAG | ASSPGQANTEVF | 0.60154223 |
| FPLVVQD | ASSPGQANTEVF | 0.5952909 |
| GAFTTPQ | ASSPGQANTEVF | 0.5159427 |
| GFGGFGTAGGVGGA | ASSPGQANTEVF | 0.6979444 |
| GFLAQFL | ASSPGQANTEVF | 0.6027977 |
| GGAGGAGGNAGVA | ASSPGQANTEVF | 0.3950294 |

| | | |
|---|---|---|
| GGVGTAGAGGAGGAGG | ASSPGQANTEVF | 0.6937323 |
| GNGGGGDAGNAGSGGN | ASSPGQANTEVF | 0.3230386 |
| GTGGAGGAGNDAGS | ASSPGQANTEVF | 0.6095276 |
| GTTNTGWFNS | ASSPGQANTEVF | 0.52035534 |
| IFMGPLFTL | ASSPGQANTEVF | 0.41705424 |
| ILAHAIEAI | ASSPGQANTEVF | 0.55047464 |
| IRSAIREAL | ASSPGQANTEVF | 0.6052613 |
| ITFAGGG | ASSPGQANTEVF | 0.67255914 |
| ITGMSGHQEV | ASSPGQANTEVF | 0.33790535 |
| IVAGLACTL | ASSPGQANTEVF | 0.4545686 |
| IVGGGLAGA | ASSPGQANTEVF | 0.32576302 |
| IVTGVLNI | ASSPGQANTEVF | 0.3361643 |
| LAQQGCI | ASSPGQANTEVF | 0.33457792 |
| LCLHVEN | ASSPGQANTEVF | 0.55478 |
| LDAISDGS | ASSPGQANTEVF | 0.4656546 |
| LGGFAAML | ASSPGQANTEVF | 0.67237926 |
| LGGKNPMI | ASSPGQANTEVF | 0.5807315 |
| LGPKGVAAIL | ASSPGQANTEVF | 0.64727044 |
| LMVGVSI | ASSPGQANTEVF | 0.38313273 |
| LVGAAFVL | ASSPGQANTEVF | 0.5174135 |
| LVLAVAPL | ASSPGQANTEVF | 0.29832017 |
| MAAIAVIDAA | ASSPGQANTEVF | 0.37914023 |
| NNGINAELQ | ASSPGQANTEVF | 0.32848534 |
| RGEPLVLVHG | ASSPGQANTEVF | 0.49024487 |
| RGGAGGAAGLI | ASSPGQANTEVF | 0.29932797 |
| RSYRMHPAV | ASSPGQANTEVF | 0.6336904 |
| SAAHLLAGQLA | ASSPGQANTEVF | 0.4400298 |
| SAAVGSY | ASSPGQANTEVF | 0.59171486 |
| SPHAVLHRI | ASSPGQANTEVF | 0.393174 |
| SQIAKVL | ASSPGQANTEVF | 0.36880365 |
| TALGHLVP | ASSPGQANTEVF | 0.5328889 |
| TAPTTSGAA | ASSPGQANTEVF | 0.60860544 |
| TDGDDGNGGSPGT | ASSPGQANTEVF | 0.45044276 |
| TDGLGEN | ASSPGQANTEVF | 0.2794904 |
| TGGPPLPGQA | ASSPGQANTEVF | 0.32559404 |
| TIPAITI | ASSPGQANTEVF | 0.32271945 |
| TIPGFTI | ASSPGQANTEVF | 0.6859876 |
| TSIASGALA | ASSPGQANTEVF | 0.6115748 |
| TVTLPGAAI | ASSPGQANTEVF | 0.5415194 |
| VASGLSAALP | ASSPGQANTEVF | 0.68837875 |
| VGAGVIGI | ASSPGQANTEVF | 0.26498032 |
| VGPKRFGV | ASSPGQANTEVF | 0.42596257 |
| VNVAFPD | ASSPGQANTEVF | 0.65895647 |
| VSMTNSGVS | ASSPGQANTEVF | 0.5553897 |
| VTDVLGDLSV | ASSPGQANTEVF | 0.5078434 |
| VVVVGAGFAGL | ASSPGQANTEVF | 0.62931323 |
| VYTVGTVV | ASSPGQANTEVF | 0.46998262 |
| WTTFSGP | ASSPGQANTEVF | 0.48621425 |
| YDELVEAF | ASSPGQANTEVF | 0.5757975 |
| YGTGGAGGNGG | ASSPGQANTEVF | 0.47722393 |
| YTNVMAV | ASSPGQANTEVF | 0.3968837 |

| | | |
|---|---|---|
| FLEQVHQL | ASSPGQANTEVF | 0.31997228 |
| GLSLLAVAV | ASSPGQANTEVF | 0.43804222 |
| LEQVHQL | ASSPGQANTEVF | 0.7151761 |
| LNAAGAI | ASSPGQANTEVF | 0.5399976 |
| PAVHASL | ASSPGQANTEVF | 0.5887366 |
| TELEDDA | ASSPGQANTEVF | 0.49471056 |
| TPPGPPN | ASSPGQANTEVF | 0.6318096 |
| TPPPPPPP | ASSPGQANTEVF | 0.4210077 |
| TQSPALI | ASSQPGQGAETLY | 0.39163584 |
| EFAAAFPQ | ASSQPGQGAETLY | 0.5710338 |
| LLSGNLDVL | ASSQPGQGAETLY | 0.36290413 |
| GPANLGNYNI | ASSQPGQGAETLY | 0.42528448 |
| TLGPTLP | ASSQPGQGAETLY | 0.4084416 |
| VYAYSSCCK | ASSQPGQGAETLY | 0.71441126 |
| APAPAPAPNI | ASSQPGQGAETLY | 0.5493217 |
| PGGPVPIP | ASSQPGQGAETLY | 0.5744151 |
| FLTVALEP | ASSQPGQGAETLY | 0.4760373 |
| SQYFGANL | ASSQPGQGAETLY | 0.42312974 |
| GGVKPAA | ASSQPGQGAETLY | 0.60279745 |
| GGTGGAGGAGGW | ASSQPGQGAETLY | 0.3111732 |
| GVGGVAGLGGAGG | ASSQPGQGAETLY | 0.6101943 |
| VLLVLATTI | ASSQPGQGAETLY | 0.36885977 |
| IGIFAAML | ASSQPGQGAETLY | 0.49063337 |
| AVGDLID | ASSQPGQGAETLY | 0.5293107 |
| NISPLPL | ASSQPGQGAETLY | 0.6995183 |
| VTAVFVAI | ASSQPGQGAETLY | 0.4697268 |
| SGDDNDAAL | ASSQPGQGAETLY | 0.55491734 |
| AVAALAHS | ASSQPGQGAETLY | 0.4578985 |
| GGVTVMATP | ASSQPGQGAETLY | 0.52250385 |
| AGLAANVRD | ASSQPGQGAETLY | 0.6825758 |
| TNRPDLIDPALLRPG | ASSQPGQGAETLY | 0.6014473 |
| LANLGVTAL | ASSQPGQGAETLY | 0.71055734 |
| TGERIEL | ASSQPGQGAETLY | 0.3907631 |
| NGEELAA | ASSQPGQGAETLY | 0.30814093 |
| VLSLVWG | ASSQPGQGAETLY | 0.4125665 |
| IGYALWL | ASSQPGQGAETLY | 0.58746606 |
| LSLLLAPL | ASSQPGQGAETLY | 0.6838558 |
| TLDEALK | ASSQPGQGAETLY | 0.6274725 |
| TDGAGTAEGG | ASSQPGQGAETLY | 0.52923834 |
| NAPGGAGGNGG | ASSQPGQGAETLY | 0.4633574 |
| VSVFMGMGGNF | ASSQPGQGAETLY | 0.6338172 |
| LASYLAAL | ASSQPGQGAETLY | 0.636598 |
| LVGGTPAL | ASSQPGQGAETLY | 0.5001149 |
| VRNADVL | ASSQPGQGAETLY | 0.3319139 |
| TCSSPFLVSAM | ASSQPGQGAETLY | 0.31542662 |
| VHITDAE | ASSQPGQGAETLY | 0.5571187 |
| EYAAPSF | ASSQPGQGAETLY | 0.6737306 |
| VVDAFHGGL | ASSQPGQGAETLY | 0.31721005 |
| ASVVDPQ | ASSQPGQGAETLY | 0.5554377 |
| SYNEFGGLS | ASSQPGQGAETLY | 0.56140006 |
| DPDGSVY | ASSQPGQGAETLY | 0.3076979 |

| | | |
|---|---|---|
| MVSTIHATE | ASSQPGQGAETLY | 0.5121989 |
| LAAGVADPV | ASSQPGQGAETLY | 0.68452275 |
| TKEQIAA | ASSQPGQGAETLY | 0.42719257 |
| SLASPAAL | ASSQPGQGAETLY | 0.32681027 |
| IGLVFYGV | ASSQPGQGAETLY | 0.6468871 |
| FLLDASNL | ASSQPGQGAETLY | 0.7029152 |
| YQVGNFP | ASSQPGQGAETLY | 0.6179986 |
| ESLLSQST | ASSQPGQGAETLY | 0.3000081 |
| LFTVGFAAVG | ASSQPGQGAETLY | 0.2675956 |
| ATPPTPT | ASSQPGQGAETLY | 0.57859695 |
| TDLATLG | ASSQPGQGAETLY | 0.6734165 |
| SAGGTGR | ASSQPGQGAETLY | 0.43289906 |
| GFSNSGS | ASSQPGQGAETLY | 0.42053044 |
| GGNGGSGGTSV | ASSQPGQGAETLY | 0.60648656 |
| GTSVATGGAGN | ASSQPGQGAETLY | 0.61647207 |
| NGGNGGSGGTS | ASSQPGQGAETLY | 0.3612118 |
| AVVKFPH | ASSQPGQGAETLY | 0.31537414 |
| TPPPPPP | ASSQPGQGAETLY | 0.541525 |
| AAPAAAAAPA | ASSQPGQGAETLY | 0.56122506 |
| IALNEDL | ASSQPGQGAETLY | 0.3948169 |
| IVNVSSM | ASSQPGQGAETLY | 0.5463599 |
| TGIPPAPR | ASSQPGQGAETLY | 0.62679064 |
| FDLGGGTF | ASSQPGQGAETLY | 0.42641985 |
| FDLGGGTFDVS | ASSQPGQGAETLY | 0.575101 |
| NEPTAAA | ASSQPGQGAETLY | 0.3595789 |
| NEPTAAALA | ASSQPGQGAETLY | 0.6878141 |
| GSIINISSI | ASSQPGQGAETLY | 0.34853324 |
| VAPGNAGTA | ASSQPGQGAETLY | 0.5515515 |
| TGGGASLE | ASSQPGQGAETLY | 0.57757944 |
| VAVYNHYKRI | ASSQPGQGAETLY | 0.51342213 |
| TDPAPATT | ASSQPGQGAETLY | 0.5095335 |
| TNRPDILDPALLRPG | ASSQPGQGAETLY | 0.3498923 |
| LTTTTTS | ASSQPGQGAETLY | 0.38749444 |
| PTGSGKTL | ASSQPGQGAETLY | 0.33570293 |
| TGSGKTL | ASSQPGQGAETLY | 0.66117483 |
| TIGHVDHGK | ASSQPGQGAETLY | 0.6009698 |
| TIGHVDHGKTT | ASSQPGQGAETLY | 0.6422838 |
| GHVDHGKTT | ASSQPGQGAETLY | 0.5540837 |
| AQFEHTLL | ASSQPGQGAETLY | 0.42090583 |
| IVNQDEL | ASSQPGQGAETLY | 0.51877534 |
| TNGSQFF | ASSQPGQGAETLY | 0.39734417 |
| TGKGIVE | ASSQPGQGAETLY | 0.40755063 |
| LISNASDA | ASSQPGQGAETLY | 0.5235338 |
| AELRTLGELTKIAKAHGAQ | ASSQPGQGAETLY | 0.53902984 |
| APNAVAAAEPSW | ASSQPGQGAETLY | 0.38196653 |
| APQVAGAATAVL | ASSQPGQGAETLY | 0.49662212 |
| ARRDPAAAAAILPTDA | ASSQPGQGAETLY | 0.5412946 |
| DAAARDVLAEAGLAEYF | ASSQPGQGAETLY | 0.31447116 |
| DDQIVLSAKANLAKVEQ | ASSQPGQGAETLY | 0.4229295 |
| DELLPDTGEDSDEDGH | ASSQPGQGAETLY | 0.6505859 |
| ELPEAPANRATVPQD | ASSQPGQGAETLY | 0.4263851 |

| | | |
|---|---|---|
| FGQSKIERVTGIG | ASSQPGQGAETLY | 0.3467881 |
| GAGGTGSTAGGAGGAGGAGGLY | ASSQPGQGAETLY | 0.57636017 |
| GHGGTGAAGGNGGTGGAGGSGI | ASSQPGQGAETLY | 0.42198873 |
| GNGGNGGAGGEGGPGGAGGAGA | ASSQPGQGAETLY | 0.61286944 |
| GSTAGNGGNGGNATNGTIAGQP | ASSQPGQGAETLY | 0.54450977 |
| GTGGDGGNGGNGGTGG | ASSQPGQGAETLY | 0.37364167 |
| GVRRPAGIPADAPV | ASSQPGQGAETLY | 0.57084835 |
| GYRQVTDAHLLTLARR | ASSQPGQGAETLY | 0.29597738 |
| IAKMGQDGHDRG | ASSQPGQGAETLY | 0.38614154 |
| KAEPSIEIQYT | ASSQPGQGAETLY | 0.42730612 |
| KPVRIRMAADEDTVAVT | ASSQPGQGAETLY | 0.5659819 |
| MPPRIAGMRLLVIKPEP | ASSQPGQGAETLY | 0.553976 |
| MKFYAVALFRSRGVPPTR | ASSQPGQGAETLY | 0.37676066 |
| NAGGFGILGAVAHSP | ASSQPGQGAETLY | 0.47912115 |
| NEGREITLAMAEM | ASSQPGQGAETLY | 0.44818726 |
| NELVEWSHPLGHRWRWE | ASSQPGQGAETLY | 0.36702797 |
| NLGAGGTGGNGGNPG | ASSQPGQGAETLY | 0.39117208 |
| NTEWPKWETVMAADPVD | ASSQPGQGAETLY | 0.3471474 |
| PASAAPAETQP | ASSQPGQGAETLY | 0.31402966 |
| QQSSDGKNGVRAAEITGE | ASSQPGQGAETLY | 0.5003019 |
| RAPDAVTTSWWKE | ASSQPGQGAETLY | 0.39396626 |
| SGFGNVGAHTSGWFNQ | ASSQPGQGAETLY | 0.4886598 |
| SGVAGAVGQAASVGGLKV | ASSQPGQGAETLY | 0.6464659 |
| SNGGAGTGSAGGNGGT | ASSQPGQGAETLY | 0.32411745 |
| TGGAGGEGGAAG | ASSQPGQGAETLY | 0.5147053 |
| TGGDGGAAGAGGAPGHG | ASSQPGQGAETLY | 0.52008903 |
| TMPPLLAAIANAGRHGV | ASSQPGQGAETLY | 0.59921014 |
| TSSTAGNSGGA | ASSQPGQGAETLY | 0.5272335 |
| VPATVPVGILC | ASSQPGQGAETLY | 0.3790947 |
| VQRPDSGGVVLTGRIS | ASSQPGQGAETLY | 0.63802123 |
| WTPNPVPATKGVPAGLVQAT | ASSQPGQGAETLY | 0.5772509 |
| YALKRALAKFSDRFDVVII | ASSQPGQGAETLY | 0.44017717 |
| YLIGSVIV | ASSQPGQGAETLY | 0.5425684 |
| YGQLVAVID | ASSQPGQGAETLY | 0.6666032 |
| VTVGGMTLPA | ASSQPGQGAETLY | 0.5937836 |
| VLALPAV | ASSQPGQGAETLY | 0.45845598 |
| VIPPTLNYET | ASSQPGQGAETLY | 0.29545188 |
| VGMGGMHPSSG | ASSQPGQGAETLY | 0.50038123 |
| VGLAGGAATA | ASSQPGQGAETLY | 0.5779745 |
| VGGAGGAGGSGP | ASSQPGQGAETLY | 0.3474645 |
| VFAAFMV | ASSQPGQGAETLY | 0.28699037 |
| TLYDKLR | ASSQPGQGAETLY | 0.65323865 |
| TLAVFTVL | ASSQPGQGAETLY | 0.7056992 |
| TGSVQPARV | ASSQPGQGAETLY | 0.37286344 |
| TGGGAGSGGT | ASSQPGQGAETLY | 0.68711436 |
| SHLETSL | ASSQPGQGAETLY | 0.30531082 |
| SFTTGMT | ASSQPGQGAETLY | 0.5906421 |
| QVEVAKRMGVRQPTVS | ASSQPGQGAETLY | 0.6546147 |
| QGSGGPALAM | ASSQPGQGAETLY | 0.37621176 |
| MLAPSNP | ASSQPGQGAETLY | 0.55143994 |
| LVNNAAI | ASSQPGQGAETLY | 0.38149068 |

| | | |
|---|---|---|
| LTGEINTLA | ASSQPGQGAETLY | 0.43542886 |
| LSIFGPV | ASSQPGQGAETLY | 0.6473963 |
| LLALLANL | ASSQPGQGAETLY | 0.42350486 |
| LISNASDALDKLR | ASSQPGQGAETLY | 0.50604326 |
| LINEAVP | ASSQPGQGAETLY | 0.4651845 |
| LGYAFSGL | ASSQPGQGAETLY | 0.41784152 |
| LDFVAVAL | ASSQPGQGAETLY | 0.49194258 |
| LAQPGAE | ASSQPGQGAETLY | 0.5380974 |
| ISNLGAVTGF | ASSQPGQGAETLY | 0.41884542 |
| HQEDPGRI | ASSQPGQGAETLY | 0.5987247 |
| GTPGEPTP | ASSQPGQGAETLY | 0.3961156 |
| GFFNSGS | ASSQPGQGAETLY | 0.7111086 |
| GAGGGGLTAGAN | ASSQPGQGAETLY | 0.6914885 |
| FVDALVNM | ASSQPGQGAETLY | 0.61996907 |
| FLTVAPEL | ASSQPGQGAETLY | 0.37947032 |
| FETLTAL | ASSQPGQGAETLY | 0.6597456 |
| DVINPIEL | ASSQPGQGAETLY | 0.43092623 |
| APVSGAPV | ASSQPGQGAETLY | 0.4957676 |
| ALVGGLAW | ASSQPGQGAETLY | 0.5224622 |
| ALAAAAATAI | ASSQPGQGAETLY | 0.6249099 |
| AFGLGLLL | ASSQPGQGAETLY | 0.75784826 |
| AFFAAASL | ASSQPGQGAETLY | 0.73261905 |
| AAAALPAFSPP | ASSQPGQGAETLY | 0.73721325 |
| VVLAPAL | ASSQPGQGAETLY | 0.6758143 |
| TIGHVDHGKTTL | ASSQPGQGAETLY | 0.51167786 |
| LISNASDALDK | ASSQPGQGAETLY | 0.6954475 |
| LISNASDAL | ASSQPGQGAETLY | 0.63678545 |
| ISNASDALDK | ASSQPGQGAETLY | 0.7653372 |
| NQAETLVYQTE | ASSQPGQGAETLY | 0.75556767 |
| TGGTGGTAGAG | ASSQPGQGAETLY | 0.6914431 |
| TGSTAGGAGGA | ASSQPGQGAETLY | 0.76072526 |
| DPLAVPVAAGPV | ASSQPGQGAETLY | 0.75167525 |
| GGTGGTPGNGGN | ASSQPGQGAETLY | 0.60137796 |
| PAGLAGVDPATL | ASSQPGQGAETLY | 0.72030807 |
| GNNLVVEAPHAGASV | ASSQPGQGAETLY | 0.701311 |
| KVTRIMSASTLEQPA | ASSQPGQGAETLY | 0.7150358 |
| LGFALPIALPRRTVV | ASSQPGQGAETLY | 0.53046674 |
| TRLGNLTARKLVVLV | ASSQPGQGAETLY | 0.7455046 |
| WSQQRRLIYFPSAGP | ASSQPGQGAETLY | 0.7042983 |
| GPNGNPDPMAAAVDIR | ASSQPGQGAETLY | 0.378348 |
| TQRCVSVADLRAAASR | ASSQPGQGAETLY | 0.74261844 |
| QFVLDVAEHPVEVLTETV | ASSQPGQGAETLY | 0.7223187 |
| GGAGGTGGAGGNGGTQAGDG | ASSQPGQGAETLY | 0.7200216 |
| PLLATLRAIAVDVDAKPAQV | ASSQPGQGAETLY | 0.75613153 |
| APKFRTGGYMIDFWGVGYQVAK | ASSQPGQGAETLY | 0.5746236 |
| GAGGAGGAGGAAGGPGGTGGTG | ASSQPGQGAETLY | 0.6851226 |
| IGALGLLATDVAPAGSVAFGVA | ASSQPGQGAETLY | 0.75253713 |
| AGGAGGIGGNGGAGGAGGVLMG | ASSQPGQGAETLY | 0.7589647 |
| ADELIGLL | ASSQPGQGAETLY | 0.61444575 |
| ADRALKA | ASSQPGQGAETLY | 0.737865 |
| AGAVGQVGGAGGA | ASSQPGQGAETLY | 0.7607922 |

| | | |
|---|---|---|
| AGGGGGLGGAP | ASSQPGQGAETLY | 0.7290839 |
| AGGLLYGNGGAG | ASSQPGQGAETLY | 0.7615898 |
| AGLESIGL | ASSQPGQGAETLY | 0.7620045 |
| AGLKTGII | ASSQPGQGAETLY | 0.63356507 |
| ASLAALLL | ASSQPGQGAETLY | 0.7632501 |
| ASVINISSI | ASSQPGQGAETLY | 0.76805985 |
| ATPPGPA | ASSQPGQGAETLY | 0.70471025 |
| ATVASALV | ASSQPGQGAETLY | 0.7291684 |
| ATVASLIL | ASSQPGQGAETLY | 0.6215528 |
| AVNGPVLIRH | ASSQPGQGAETLY | 0.7193028 |
| DGAAGAVGI | ASSQPGQGAETLY | 0.46439287 |
| DPLLHGI | ASSQPGQGAETLY | 0.7397833 |
| ELGAALGAL | ASSQPGQGAETLY | 0.76266134 |
| ELGRTQAV | ASSQPGQGAETLY | 0.7189208 |
| FVAGGYLS | ASSQPGQGAETLY | 0.6110887 |
| GASGGAGGAGGW | ASSQPGQGAETLY | 0.60175633 |
| GGAGAGGDGGAGGI | ASSQPGQGAETLY | 0.68750715 |
| GTGHNGSQPGMGGNG | ASSQPGQGAETLY | 0.64259994 |
| GYAAASASAS | ASSQPGQGAETLY | 0.53171724 |
| HQVDVAAVQQG | ASSQPGQGAETLY | 0.6062815 |
| IGAAIAHT | ASSQPGQGAETLY | 0.7553704 |
| IGMIEVAQL | ASSQPGQGAETLY | 0.75960714 |
| IRLAPEHTSAMD | ASSQPGQGAETLY | 0.6248504 |
| IYNVSVL | ASSQPGQGAETLY | 0.7642584 |
| KQGFLGM | ASSQPGQGAETLY | 0.75073516 |
| LESVITQL | ASSQPGQGAETLY | 0.75572073 |
| LGLGALVVV | ASSQPGQGAETLY | 0.72264373 |
| LIGNGGNGGSGGI | ASSQPGQGAETLY | 0.61173344 |
| LNAINAPTL | ASSQPGQGAETLY | 0.7617073 |
| LQLLNTGV | ASSQPGQGAETLY | 0.69779515 |
| LQTSIEF | ASSQPGQGAETLY | 0.7319126 |
| MSFAVSEL | ASSQPGQGAETLY | 0.75542295 |
| NTGALGL | ASSQPGQGAETLY | 0.7202898 |
| PPGAGGAGGAGTT | ASSQPGQGAETLY | 0.67474246 |
| QLVGLGI | ASSQPGQGAETLY | 0.7157279 |
| RFLEQVHQL | ASSQPGQGAETLY | 0.7551059 |
| RGVGYVLR | ASSQPGQGAETLY | 0.66067564 |
| SATGVFAGVI | ASSQPGQGAETLY | 0.48417413 |
| SLPAITLP | ASSQPGQGAETLY | 0.6926708 |
| STALAFTV | ASSQPGQGAETLY | 0.710022 |
| SVLTVGLGK | ASSQPGQGAETLY | 0.7614721 |
| TALAGAGSGI | ASSQPGQGAETLY | 0.73884904 |
| TGLPFTGI | ASSQPGQGAETLY | 0.7016669 |
| VAEHVTV | ASSQPGQGAETLY | 0.7484466 |
| VDIVLNSL | ASSQPGQGAETLY | 0.74449146 |
| VLTGGGAVL | ASSQPGQGAETLY | 0.72036654 |
| VNVGLAYGI | ASSQPGQGAETLY | 0.73643243 |
| VSSAAVGAV | ASSQPGQGAETLY | 0.6893492 |
| VTSVYEAGV | ASSQPGQGAETLY | 0.75243247 |
| WVALDRGA | ASSQPGQGAETLY | 0.75244987 |
| YLVGSVVI | ASSQPGQGAETLY | 0.7487043 |

| | | |
|---|---|---|
| GPPGTGKTLL | ASSQPGQGAETLY | 0.75797975 |
| GVGMTKF | ASSQPGQGAETLY | 0.7258589 |
| IGIGGSDL | ASSQPGQGAETLY | 0.72143316 |
| IGTLLAEL | ASSQPGQGAETLY | 0.51922655 |
| IYTSGTT | ASSQPGQGAETLY | 0.6469863 |
| LSFPAGV | ASSQPGQGAETLY | 0.7348725 |
| LYTSGTT | ASSQPGQGAETLY | 0.7410151 |
| VGAFPFGF | ASSQPGQGAETLY | 0.7549732 |
| VGAGTVEF | ASSQPGQGAETLY | 0.7429065 |
| VGKSTLLL | ASSQPGQGAETLY | 0.6794721 |
| AALAHAGGGIV | ASSQPGQGAETLY | 0.7463542 |
| AALVAVSLL | ASSQPGQGAETLY | 0.7625893 |
| AAVSRAVAAL | ASSQPGQGAETLY | 0.60584366 |
| AGGSSGIGGF | ASSQPGQGAETLY | 0.69778866 |
| AGLTVANANG | ASSQPGQGAETLY | 0.7685368 |
| AGLVGAVL | ASSQPGQGAETLY | 0.7421377 |
| ALSQAFTV | ASSQPGQGAETLY | 0.72372127 |
| ALTFSVSL | ASSQPGQGAETLY | 0.7154325 |
| ALTGAALVAL | ASSQPGQGAETLY | 0.7681203 |
| ASFSIGAGVL | ASSQPGQGAETLY | 0.7167357 |
| DGSVYKI | ASSQPGQGAETLY | 0.7668873 |
| EQLAEIV | ASSQPGQGAETLY | 0.43104935 |
| FAVSELTVM | ASSQPGQGAETLY | 0.6901981 |
| GAAPLAAGL | ASSQPGQGAETLY | 0.49928844 |
| GADGGDGGSGGAGG | ASSQPGQGAETLY | 0.31739977 |
| GFGLDTGGAGG | ASSQPGQGAETLY | 0.75565886 |
| GLASGGLL | ASSQPGQGAETLY | 0.63679004 |
| GRIAGAAVLL | ASSQPGQGAETLY | 0.74308264 |
| GSAGSLNTSF | ASSQPGQGAETLY | 0.717479 |
| GTGQIGI | ASSQPGQGAETLY | 0.7558759 |
| GTLIALIL | ASSQPGQGAETLY | 0.73360056 |
| IAATMGFVVG | ASSQPGQGAETLY | 0.7090405 |
| IFAQALL | ASSQPGQGAETLY | 0.64881575 |
| IGAAVALGF | ASSQPGQGAETLY | 0.7482352 |
| IGLTPLAL | ASSQPGQGAETLY | 0.7533522 |
| ILGSNTSTL | ASSQPGQGAETLY | 0.7523282 |
| IVFSLTVL | ASSQPGQGAETLY | 0.47692955 |
| LASKATGF | ASSQPGQGAETLY | 0.7398662 |
| LASVALIL | ASSQPGQGAETLY | 0.7089367 |
| LFAVGLTL | ASSQPGQGAETLY | 0.45373297 |
| LFNGGIGTYI | ASSQPGQGAETLY | 0.4416896 |
| LTLGAGII | ASSQPGQGAETLY | 0.66502726 |
| LVAAALGDI | ASSQPGQGAETLY | 0.6819015 |
| NTAFVQLGI | ASSQPGQGAETLY | 0.7559227 |
| QLFSGAEL | ASSQPGQGAETLY | 0.7562198 |
| SAATFITL | ASSQPGQGAETLY | 0.71543264 |
| SNYGLANLGV | ASSQPGQGAETLY | 0.74841726 |
| SYGGAYAVM | ASSQPGQGAETLY | 0.71592176 |
| TTGVGGLTLG | ASSQPGQGAETLY | 0.21146554 |
| TTQTIYI | ASSQPGQGAETLY | 0.6913072 |
| VAAGAGLGAVI | ASSQPGQGAETLY | 0.6192882 |

| | | |
|---|---|---|
| VAAVVTLL | ASSQPGQGAETLY | 0.7606561 |
| VGFLGGLI | ASSQPGQGAETLY | 0.6545824 |
| VGVIPILL | ASSQPGQGAETLY | 0.7123666 |
| VIFASSLI | ASSQPGQGAETLY | 0.73111904 |
| VKGGDGGAAAG | ASSQPGQGAETLY | 0.7471417 |
| VTKTFTNI | ASSQPGQGAETLY | 0.7276413 |
| VVAAAILL | ASSQPGQGAETLY | 0.65957785 |
| YAALDVEL | ASSQPGQGAETLY | 0.36620724 |
| LASVALLL | ASSQPGQGAETLY | 0.75032055 |
| LDAVHRV | ASSQPGQGAETLY | 0.6697842 |
| QGLSSSIF | ASSQPGQGAETLY | 0.62077165 |
| RGTFANIRL | ASSQPGQGAETLY | 0.5913743 |
| RQLWWGHRI | ASSQPGQGAETLY | 0.7301408 |
| YGPPGTGKTLL | ASSQPGQGAETLY | 0.57070696 |
| AALVGAPANVA | ASSQPGQGAETLY | 0.7594293 |
| AATFRRTE | ASSQPGQGAETLY | 0.65769285 |
| AGALGGGI | ASSQPGQGAETLY | 0.7130282 |
| AGGAGGAGGIGAG | ASSQPGQGAETLY | 0.76788425 |
| AGSGALGL | ASSQPGQGAETLY | 0.7385192 |
| AQIADQI | ASSQPGQGAETLY | 0.7134732 |
| ASAVAVTYL | ASSQPGQGAETLY | 0.69304687 |
| ASTVLGGTGG | ASSQPGQGAETLY | 0.6680602 |
| AVGGEPAK | ASSQPGQGAETLY | 0.67185915 |
| CEQGTFGGTC | ASSQPGQGAETLY | 0.6897438 |
| DEGKAPAG | ASSQPGQGAETLY | 0.7651254 |
| DTVVFPAL | ASSQPGQGAETLY | 0.70699346 |
| FEGITMAG | ASSQPGQGAETLY | 0.6243785 |
| FPLVVQD | ASSQPGQGAETLY | 0.711092 |
| GAFTTPQ | ASSQPGQGAETLY | 0.7666421 |
| GFGGFGTAGGVGGA | ASSQPGQGAETLY | 0.76637065 |
| GFLAQFL | ASSQPGQGAETLY | 0.6549296 |
| GGAGGAGGNAGVA | ASSQPGQGAETLY | 0.6811927 |
| GGVGTAGAGGAGGAGG | ASSQPGQGAETLY | 0.6878046 |
| GNGGGGDAGNAGSGGN | ASSQPGQGAETLY | 0.7178681 |
| GTGGAGGAGNDAGS | ASSQPGQGAETLY | 0.62747705 |
| GTTNTGWFNS | ASSQPGQGAETLY | 0.6413591 |
| IFMGPLFTL | ASSQPGQGAETLY | 0.7509724 |
| ILAHAIEAI | ASSQPGQGAETLY | 0.76976204 |
| IRSAIREAL | ASSQPGQGAETLY | 0.6972964 |
| ITFAGGG | ASSQPGQGAETLY | 0.6378987 |
| ITGMSGHQEV | ASSQPGQGAETLY | 0.72290766 |
| IVAGLACTL | ASSQPGQGAETLY | 0.6730148 |
| IVGGGLAGA | ASSQPGQGAETLY | 0.6833912 |
| IVTGVLNI | ASSQPGQGAETLY | 0.69784063 |
| LAQQGCI | ASSQPGQGAETLY | 0.5892725 |
| LCLHVEN | ASSQPGQGAETLY | 0.72783893 |
| LDAISDGS | ASSQPGQGAETLY | 0.7634915 |
| LGGFAAML | ASSQPGQGAETLY | 0.72278476 |
| LGGKNPMI | ASSQPGQGAETLY | 0.7335448 |
| LGPKGVAAIL | ASSQPGQGAETLY | 0.7660054 |
| LMVGVSI | ASSQPGQGAETLY | 0.75296354 |

| | | |
|---|---|---|
| LVGAAFVL | ASSQPGQGAETLY | 0.76156914 |
| LVLAVAPL | ASSQPGQGAETLY | 0.7580075 |
| MAAIAVIDAA | ASSQPGQGAETLY | 0.71711457 |
| NNGINAELQ | ASSQPGQGAETLY | 0.7008335 |
| RGEPLVLVHG | ASSQPGQGAETLY | 0.7601125 |
| RGGAGGAAGLI | ASSQPGQGAETLY | 0.7276666 |
| RSYRMHPAV | ASSQPGQGAETLY | 0.7398864 |
| SAAHLLAGQLA | ASSQPGQGAETLY | 0.7158605 |
| SAAVGSY | ASSQPGQGAETLY | 0.75712174 |
| SPHAVLHRI | ASSQPGQGAETLY | 0.6952663 |
| SQIAKVL | ASSQPGQGAETLY | 0.76360565 |
| TALGHLVP | ASSQPGQGAETLY | 0.34021562 |
| TAPTTSGAA | ASSQPGQGAETLY | 0.6340107 |
| TDGDDGNGGSPGT | ASSQPGQGAETLY | 0.75600183 |
| TDGLGEN | ASSQPGQGAETLY | 0.24023758 |
| TGGPPLPGQA | ASSQPGQGAETLY | 0.45287284 |
| TIPAITI | ASSQPGQGAETLY | 0.7311448 |
| TIPGFTI | ASSQPGQGAETLY | 0.75218606 |
| TSIASGALA | ASSQPGQGAETLY | 0.24996205 |
| TVTLPGAAI | ASSQPGQGAETLY | 0.6910526 |
| VASGLSAALP | ASSQPGQGAETLY | 0.51712656 |
| VGAGVIGI | ASSQPGQGAETLY | 0.7201514 |
| VGPKRFGV | ASSQPGQGAETLY | 0.7316067 |
| VNVAFPD | ASSQPGQGAETLY | 0.5040136 |
| VSMTNSGVS | ASSQPGQGAETLY | 0.75078356 |
| VTDVLGDLSV | ASSQPGQGAETLY | 0.62482023 |
| VVVVGAGFAGL | ASSQPGQGAETLY | 0.7088406 |
| VYTVGTVV | ASSQPGQGAETLY | 0.75837666 |
| WTTFSGP | ASSQPGQGAETLY | 0.76696765 |
| YDELVEAF | ASSQPGQGAETLY | 0.6140946 |
| YGTGGAGGNGG | ASSQPGQGAETLY | 0.7536036 |
| YTNVMAV | ASSQPGQGAETLY | 0.68290937 |
| FLEQVHQL | ASSQPGQGAETLY | 0.77078915 |
| GLSLLAVAV | ASSQPGQGAETLY | 0.70135546 |
| LEQVHQL | ASSQPGQGAETLY | 0.48862636 |
| LNAAGAI | ASSQPGQGAETLY | 0.7560402 |
| PAVHASL | ASSQPGQGAETLY | 0.7364915 |
| TELEDDA | ASSQPGQGAETLY | 0.70366824 |
| TPPGPPN | ASSQPGQGAETLY | 0.7521411 |
| TPPPPPPP | ASSQPGQGAETLY | 0.56453973 |
| TQSPALI | ASSLLDRGEVF | 0.72580075 |
| EFAAAFPQ | ASSLLDRGEVF | 0.7596723 |
| LLSGNLDVL | ASSLLDRGEVF | 0.75123644 |
| GPANLGNYNI | ASSLLDRGEVF | 0.6100265 |
| TLGPTLP | ASSLLDRGEVF | 0.7593607 |
| VYAYSSCCK | ASSLLDRGEVF | 0.7298374 |
| APAPAPAPNI | ASSLLDRGEVF | 0.6382694 |
| PGGPVPIP | ASSLLDRGEVF | 0.7637185 |
| FLTVALEP | ASSLLDRGEVF | 0.72643906 |
| SQYFGANL | ASSLLDRGEVF | 0.75463545 |
| GGVKPAA | ASSLLDRGEVF | 0.74856144 |

| | | |
|---|---|---|
| GGTGGAGGAGGW | ASSLLDRGEVF | 0.7620925 |
| GVGGVAGLGGAGG | ASSLLDRGEVF | 0.595224 |
| VLLVLATTI | ASSLLDRGEVF | 0.7042247 |
| IGIFAAML | ASSLLDRGEVF | 0.7280897 |
| AVGDLID | ASSLLDRGEVF | 0.75606424 |
| NISPLPL | ASSLLDRGEVF | 0.68353915 |
| VTAVFVAI | ASSLLDRGEVF | 0.71514475 |
| SGDDNDAAL | ASSLLDRGEVF | 0.74355674 |
| AVAALAHS | ASSLLDRGEVF | 0.7640501 |
| GGVTVMATP | ASSLLDRGEVF | 0.71084964 |
| AGLAANVRD | ASSLLDRGEVF | 0.7340563 |
| TNRPDLIDPALLRPG | ASSLLDRGEVF | 0.765864 |
| LANLGVTAL | ASSLLDRGEVF | 0.6530751 |
| TGERIEL | ASSLLDRGEVF | 0.7260128 |
| NGEELAA | ASSLLDRGEVF | 0.7499103 |
| VLSLVWG | ASSLLDRGEVF | 0.75507045 |
| IGYALWL | ASSLLDRGEVF | 0.7124135 |
| LSLLLAPL | ASSLLDRGEVF | 0.761495 |
| TLDEALK | ASSLLDRGEVF | 0.67676055 |
| TDGAGTAEGG | ASSLLDRGEVF | 0.75044733 |
| NAPGGAGGNGG | ASSLLDRGEVF | 0.6699926 |
| VSVFMGMGGNF | ASSLLDRGEVF | 0.73372674 |
| LASYLAAL | ASSLLDRGEVF | 0.63016176 |
| LVGGTPAL | ASSLLDRGEVF | 0.7493242 |
| VRNADVL | ASSLLDRGEVF | 0.758278 |
| TCSSPFLVSAM | ASSLLDRGEVF | 0.7544776 |
| VHITDAE | ASSLLDRGEVF | 0.4578229 |
| EYAAPSF | ASSLLDRGEVF | 0.7501899 |
| VVDAFHGGL | ASSLLDRGEVF | 0.75374246 |
| ASVVDPQ | ASSLLDRGEVF | 0.76869017 |
| SYNEFGGLS | ASSLLDRGEVF | 0.69356835 |
| DPDGSVY | ASSLLDRGEVF | 0.7403965 |
| MVSTIHATE | ASSLLDRGEVF | 0.6925206 |
| LAAGVADPV | ASSLLDRGEVF | 0.7227781 |
| TKEQIAA | ASSLLDRGEVF | 0.60732114 |
| SLASPAAL | ASSLLDRGEVF | 0.61552435 |
| IGLVFYGV | ASSLLDRGEVF | 0.71071875 |
| FLLDASNL | ASSLLDRGEVF | 0.6741864 |
| YQVGNFP | ASSLLDRGEVF | 0.75586927 |
| ESLLSQST | ASSLLDRGEVF | 0.7613653 |
| LFTVGFAAVG | ASSLLDRGEVF | 0.75325286 |
| ATPPTPT | ASSLLDRGEVF | 0.76175636 |
| TDLATLG | ASSLLDRGEVF | 0.7623419 |
| SAGGTGR | ASSLLDRGEVF | 0.7529521 |
| GFSNSGS | ASSLLDRGEVF | 0.68616617 |
| GGNGGSGGTSV | ASSLLDRGEVF | 0.7271658 |
| GTSVATGGAGN | ASSLLDRGEVF | 0.5914186 |
| NGGNGGSGGTS | ASSLLDRGEVF | 0.7199258 |
| AVVKFPH | ASSLLDRGEVF | 0.70091116 |
| TPPPPPP | ASSLLDRGEVF | 0.74164915 |
| AAPAAAAPA | ASSLLDRGEVF | 0.24197084 |

| | | |
|---|---|---|
| IALNEDL | ASSLLDRGEVF | 0.73836064 |
| IVNVSSM | ASSLLDRGEVF | 0.76629275 |
| TGIPPAPR | ASSLLDRGEVF | 0.69132876 |
| FDLGGGTF | ASSLLDRGEVF | 0.74633276 |
| FDLGGGTFDVS | ASSLLDRGEVF | 0.7484692 |
| NEPTAAA | ASSLLDRGEVF | 0.6854949 |
| NEPTAAALA | ASSLLDRGEVF | 0.75138164 |
| GSIINISSI | ASSLLDRGEVF | 0.6799098 |
| VAPGNAGTA | ASSLLDRGEVF | 0.7540921 |
| TGGGASLE | ASSLLDRGEVF | 0.7625724 |
| VAVYNHYKRI | ASSLLDRGEVF | 0.7428949 |
| TDPAPATT | ASSLLDRGEVF | 0.6963723 |
| TNRPDILDPALLRPG | ASSLLDRGEVF | 0.7275528 |
| LTTTTTS | ASSLLDRGEVF | 0.5731077 |
| PTGSGKTL | ASSLLDRGEVF | 0.7129427 |
| TGSGKTL | ASSLLDRGEVF | 0.5686641 |
| TIGHVDHGK | ASSLLDRGEVF | 0.7333503 |
| TIGHVDHGKTT | ASSLLDRGEVF | 0.5860342 |
| GHVDHGKTT | ASSLLDRGEVF | 0.7411684 |
| AQFEHTLL | ASSLLDRGEVF | 0.7281579 |
| IVNQDEL | ASSLLDRGEVF | 0.73200625 |
| TNGSQFF | ASSLLDRGEVF | 0.6930566 |
| TGKGIVE | ASSLLDRGEVF | 0.74664664 |
| LISNASDA | ASSLLDRGEVF | 0.72615075 |
| AELRTLGELTKIAKAHGAQ | ASSLLDRGEVF | 0.58906174 |
| APNAVAAAEPSW | ASSLLDRGEVF | 0.7474106 |
| APQVAGAATAVL | ASSLLDRGEVF | 0.7003157 |
| ARRDPAAAAAILPTDA | ASSLLDRGEVF | 0.69627404 |
| DAAARDVLAEAGLAEYF | ASSLLDRGEVF | 0.7578366 |
| DDQIVLSAKANLAKVEQ | ASSLLDRGEVF | 0.7415177 |
| DELLPDTGEDSDEDGH | ASSLLDRGEVF | 0.5997075 |
| ELPEAPANRATVPQD | ASSLLDRGEVF | 0.70712745 |
| FGQSKIERVTGIG | ASSLLDRGEVF | 0.73828804 |
| GAGGTGSTAGGAGGAGGAGGLY | ASSLLDRGEVF | 0.69184834 |
| GHGGTGAAGGNGGTGGAGGSGI | ASSLLDRGEVF | 0.2777139 |
| GNGGNGGAGGEGGPGGAGGAGA | ASSLLDRGEVF | 0.7347034 |
| GSTAGNGGNGGNATNGTIAGQP | ASSLLDRGEVF | 0.62957346 |
| GTGGDGGNGGNGGTGG | ASSLLDRGEVF | 0.6701481 |
| GVRRPAGIPADAPV | ASSLLDRGEVF | 0.7104584 |
| GYRQVTDAHLLTLARR | ASSLLDRGEVF | 0.75699353 |
| IAKMGQDGHDRG | ASSLLDRGEVF | 0.69426167 |
| KAEPSIEIQYT | ASSLLDRGEVF | 0.65026724 |
| KPVRIRMAADEDTVAVT | ASSLLDRGEVF | 0.75245464 |
| MPPRIAGMRLLVIKPEP | ASSLLDRGEVF | 0.7427075 |
| MKFYAVALFRSRGVPPTR | ASSLLDRGEVF | 0.7566273 |
| NAGGFGILGAVAHSP | ASSLLDRGEVF | 0.7362623 |
| NEGREITLAMAEM | ASSLLDRGEVF | 0.6284501 |
| NELVEWSHPLGHRWRWE | ASSLLDRGEVF | 0.72213286 |
| NLGAGGTGGNGGNPG | ASSLLDRGEVF | 0.6822864 |
| NTEWPKWETVMAADPVD | ASSLLDRGEVF | 0.679803 |
| PASAAPAETQP | ASSLLDRGEVF | 0.61948824 |

| | | |
|---|---|---|
| QQSSDGKNGVRAAEITGE | ASSLLDRGEVF | 0.7151989 |
| RAPDAVTTSWWKE | ASSLLDRGEVF | 0.6633713 |
| SGFGNVGAHTSGWFNQ | ASSLLDRGEVF | 0.7322451 |
| SGVAGAVGQAASVGGLKV | ASSLLDRGEVF | 0.7647747 |
| SNGGAGTGSAGGNGGT | ASSLLDRGEVF | 0.6419455 |
| TGGAGGEGGAAG | ASSLLDRGEVF | 0.4845075 |
| TGGDGGAAGAGGAPGHG | ASSLLDRGEVF | 0.69892883 |
| TMPPLLAAIANAGRHGV | ASSLLDRGEVF | 0.69754386 |
| TSSTAGNSGGA | ASSLLDRGEVF | 0.7526231 |
| VPATVPVGILC | ASSLLDRGEVF | 0.6958028 |
| VQRPDSGGVVLTGRIS | ASSLLDRGEVF | 0.769505 |
| WTPNPVPATKGVPAGLVQAT | ASSLLDRGEVF | 0.7521985 |
| YALKRALAKFSDRFDVVII | ASSLLDRGEVF | 0.6924436 |
| YLIGSVIV | ASSLLDRGEVF | 0.7551637 |
| YGQLVAVID | ASSLLDRGEVF | 0.698118 |
| VTVGGMTLPA | ASSLLDRGEVF | 0.69283164 |
| VLALPAV | ASSLLDRGEVF | 0.7267865 |
| VIPPTLNYET | ASSLLDRGEVF | 0.72429335 |
| VGMGGMHPSSG | ASSLLDRGEVF | 0.702812 |
| VGLAGGAATA | ASSLLDRGEVF | 0.71003556 |
| VGGAGGAGGSGP | ASSLLDRGEVF | 0.5340184 |
| VFAAFMV | ASSLLDRGEVF | 0.75480306 |
| TLYDKLR | ASSLLDRGEVF | 0.6692865 |
| TLAVFTVL | ASSLLDRGEVF | 0.57712847 |
| TGSVQPARV | ASSLLDRGEVF | 0.6881287 |
| TGGGAGSGGT | ASSLLDRGEVF | 0.7647815 |
| SHLETSL | ASSLLDRGEVF | 0.7639508 |
| SFTTGMT | ASSLLDRGEVF | 0.68951964 |
| QVEVAKRMGVRQPTVS | ASSLLDRGEVF | 0.66290486 |
| QGSGGPALAM | ASSLLDRGEVF | 0.745613 |
| MLAPSNP | ASSLLDRGEVF | 0.6139991 |
| LVNNAAI | ASSLLDRGEVF | 0.7385343 |
| LTGEINTLA | ASSLLDRGEVF | 0.7214925 |
| LSIFGPV | ASSLLDRGEVF | 0.71150374 |
| LLALLANL | ASSLLDRGEVF | 0.62631303 |
| LISNASDALDKLR | ASSLLDRGEVF | 0.7485131 |
| LINEAVP | ASSLLDRGEVF | 0.7079635 |
| LGYAFSGL | ASSLLDRGEVF | 0.74191993 |
| LDFVAVAL | ASSLLDRGEVF | 0.39353424 |
| LAQPGAE | ASSLLDRGEVF | 0.7400481 |
| ISNLGAVTGF | ASSLLDRGEVF | 0.6666907 |
| HQEDPGRI | ASSLLDRGEVF | 0.61052513 |
| GTPGEPTP | ASSLLDRGEVF | 0.322627 |
| GFFNSGS | ASSLLDRGEVF | 0.70225394 |
| GAGGGGLTAGAN | ASSLLDRGEVF | 0.7530197 |
| FVDALVNM | ASSLLDRGEVF | 0.7615495 |
| FLTVAPEL | ASSLLDRGEVF | 0.70818585 |
| FETLTAL | ASSLLDRGEVF | 0.75655854 |
| DVINPIEL | ASSLLDRGEVF | 0.7623615 |
| APVSGAPV | ASSLLDRGEVF | 0.7151154 |
| ALVGGLAW | ASSLLDRGEVF | 0.34837237 |

| | | |
|---|---|---|
| ALAAAAATAI | ASSLLDRGEVF | 0.73898506 |
| AFGLGLLL | ASSLLDRGEVF | 0.724348 |
| AFFAAASL | ASSLLDRGEVF | 0.7489835 |
| AAAALPAFSPP | ASSLLDRGEVF | 0.7469071 |
| VVLAPAL | ASSLLDRGEVF | 0.6360903 |
| TIGHVDHGKTTL | ASSLLDRGEVF | 0.7344153 |
| LISNASDALDK | ASSLLDRGEVF | 0.7451454 |
| LISNASDAL | ASSLLDRGEVF | 0.6089864 |
| ISNASDALDK | ASSLLDRGEVF | 0.66634816 |
| NQAETLVYQTE | ASSLLDRGEVF | 0.6495694 |
| TGGTGGTAGAG | ASSLLDRGEVF | 0.45268887 |
| TGSTAGGAGGA | ASSLLDRGEVF | 0.74618006 |
| DPLAVPVAAGPV | ASSLLDRGEVF | 0.70865333 |
| GGTGGTPGNGGN | ASSLLDRGEVF | 0.70982504 |
| PAGLAGVDPATL | ASSLLDRGEVF | 0.7248978 |
| GNNLVVEAPHAGASV | ASSLLDRGEVF | 0.69106406 |
| KVTRIMSASTLEQPA | ASSLLDRGEVF | 0.7023037 |
| LGFALPIALPRRTVV | ASSLLDRGEVF | 0.7076267 |
| TRLGNLTARKLVVLV | ASSLLDRGEVF | 0.74729645 |
| WSQQRRLIYFPSAGP | ASSLLDRGEVF | 0.7509 |
| GPNGNPDPMAAAVDIR | ASSLLDRGEVF | 0.7225695 |
| TQRCVSVADLRAAASR | ASSLLDRGEVF | 0.7231865 |
| QFVLDVAEHPVEVLTETV | ASSLLDRGEVF | 0.76605964 |
| GGAGGTGGAGGNGGTQAGDG | ASSLLDRGEVF | 0.59163463 |
| PLLATLRAIAVDVDAKPAQV | ASSLLDRGEVF | 0.6598358 |
| APKFRTGGYMIDFWGVGYQVAK | ASSLLDRGEVF | 0.74315536 |
| GAGGAGGAGGAAGGPGGTGGTG | ASSLLDRGEVF | 0.6716951 |
| IGALGLLATDVAPAGSVAFGVA | ASSLLDRGEVF | 0.75503767 |
| AGGAGGIGGNGGAGGAGGVLMG | ASSLLDRGEVF | 0.72892547 |
| ADELIGLL | ASSLLDRGEVF | 0.74660504 |
| ADRALKA | ASSLLDRGEVF | 0.7370343 |
| AGAVGQVGGAGGA | ASSLLDRGEVF | 0.7690189 |
| AGGGGGLGGAP | ASSLLDRGEVF | 0.74894273 |
| AGGLLYGNGGAG | ASSLLDRGEVF | 0.6861166 |
| AGLESIGL | ASSLLDRGEVF | 0.75744665 |
| AGLKTGII | ASSLLDRGEVF | 0.49994934 |
| ASLAALLL | ASSLLDRGEVF | 0.75565827 |
| ASVINISSI | ASSLLDRGEVF | 0.7518555 |
| ATPPGPA | ASSLLDRGEVF | 0.66022384 |
| ATVASALV | ASSLLDRGEVF | 0.7257754 |
| ATVASLIL | ASSLLDRGEVF | 0.7587174 |
| AVNGPVLIRH | ASSLLDRGEVF | 0.76393056 |
| DGAAGAVGI | ASSLLDRGEVF | 0.59520096 |
| DPLLHGI | ASSLLDRGEVF | 0.6899335 |
| ELGAALGAL | ASSLLDRGEVF | 0.52868116 |
| ELGRTQAV | ASSLLDRGEVF | 0.5264363 |
| FVAGGYLS | ASSLLDRGEVF | 0.46703255 |
| GASGGAGGAGGW | ASSLLDRGEVF | 0.7478107 |
| GGAGAGGDGGAGGI | ASSLLDRGEVF | 0.7249447 |
| GTGHNGSQPGMGGNG | ASSLLDRGEVF | 0.7540662 |
| GYAAASASAS | ASSLLDRGEVF | 0.7405646 |

| | | |
|---|---|---|
| HQVDVAAVQQG | ASSLLDRGEVF | 0.76163065 |
| IGAAIAHT | ASSLLDRGEVF | 0.6909441 |
| IGMIEVAQL | ASSLLDRGEVF | 0.71380377 |
| IRLAPEHTSAMD | ASSLLDRGEVF | 0.50045 |
| IYNVSVL | ASSLLDRGEVF | 0.5150858 |
| KQGFLGM | ASSLLDRGEVF | 0.58552283 |
| LESVITQL | ASSLLDRGEVF | 0.7122477 |
| LGLGALVVV | ASSLLDRGEVF | 0.45964205 |
| LIGNGGNGGSGGI | ASSLLDRGEVF | 0.7619444 |
| LNAINAPTL | ASSLLDRGEVF | 0.7244183 |
| LQLLNTGV | ASSLLDRGEVF | 0.7539883 |
| LQTSIEF | ASSLLDRGEVF | 0.6956117 |
| MSFAVSEL | ASSLLDRGEVF | 0.64092433 |
| NTGALGL | ASSLLDRGEVF | 0.69470286 |
| PPGAGGAGGAGTT | ASSLLDRGEVF | 0.7450368 |
| QLVGLGI | ASSLLDRGEVF | 0.61299294 |
| RFLEQVHQL | ASSLLDRGEVF | 0.53962874 |
| RGVGYVLR | ASSLLDRGEVF | 0.67051375 |
| SATGVFAGVI | ASSLLDRGEVF | 0.7006175 |
| SLPAITLP | ASSLLDRGEVF | 0.7656922 |
| STALAFTV | ASSLLDRGEVF | 0.7621838 |
| SVLTVGLGK | ASSLLDRGEVF | 0.7429073 |
| TALAGAGSGI | ASSLLDRGEVF | 0.7677593 |
| TGLPFTGI | ASSLLDRGEVF | 0.2534304 |
| VAEHVTV | ASSLLDRGEVF | 0.7163454 |
| VDIVLNSL | ASSLLDRGEVF | 0.7612214 |
| VLTGGGAVL | ASSLLDRGEVF | 0.73830104 |
| VNVGLAYGI | ASSLLDRGEVF | 0.7342448 |
| VSSAAVGAV | ASSLLDRGEVF | 0.76132286 |
| VTSVYEAGV | ASSLLDRGEVF | 0.6703897 |
| WVALDRGA | ASSLLDRGEVF | 0.69565094 |
| YLVGSVVI | ASSLLDRGEVF | 0.7290667 |
| GPPGTGKTLL | ASSLLDRGEVF | 0.7461164 |
| GVGMTKF | ASSLLDRGEVF | 0.68597054 |
| IGIGGSDL | ASSLLDRGEVF | 0.52291906 |
| IGTLLAEL | ASSLLDRGEVF | 0.73797953 |
| IYTSGTT | ASSLLDRGEVF | 0.77270085 |
| LSFPAGV | ASSLLDRGEVF | 0.7160386 |
| LYTSGTT | ASSLLDRGEVF | 0.7555611 |
| VGAFPFGF | ASSLLDRGEVF | 0.7361942 |
| VGAGTVEF | ASSLLDRGEVF | 0.71535945 |
| VGKSTLLL | ASSLLDRGEVF | 0.6983398 |
| AALAHAGGGIV | ASSLLDRGEVF | 0.6743124 |
| AALVAVSLL | ASSLLDRGEVF | 0.74393404 |
| AAVSRAVAAL | ASSLLDRGEVF | 0.6269419 |
| AGGSSGIGGF | ASSLLDRGEVF | 0.7257165 |
| AGLTVANANG | ASSLLDRGEVF | 0.70420474 |
| AGLVGAVL | ASSLLDRGEVF | 0.77144384 |
| ALSQAFTV | ASSLLDRGEVF | 0.74737716 |
| ALTFSVSL | ASSLLDRGEVF | 0.74881506 |
| ALTGAALVAL | ASSLLDRGEVF | 0.6459352 |

| | | |
|---|---|---|
| ASFSIGAGVL | ASSLLDRGEVF | 0.7227752 |
| DGSVYKI | ASSLLDRGEVF | 0.73865575 |
| EQLAEIV | ASSLLDRGEVF | 0.55720246 |
| FAVSELTVM | ASSLLDRGEVF | 0.7597155 |
| GAAPLAAGL | ASSLLDRGEVF | 0.6650418 |
| GADGGDGGSGGAGG | ASSLLDRGEVF | 0.6964992 |
| GFGLDTGGAGG | ASSLLDRGEVF | 0.7454128 |
| GLASGGLL | ASSLLDRGEVF | 0.76157415 |
| GRIAGAAVLL | ASSLLDRGEVF | 0.71224064 |
| GSAGSLNTSF | ASSLLDRGEVF | 0.7410008 |
| GTGQIGI | ASSLLDRGEVF | 0.6689317 |
| GTLIALIL | ASSLLDRGEVF | 0.7526723 |
| IAATMGFVVG | ASSLLDRGEVF | 0.760985 |
| IFAQALL | ASSLLDRGEVF | 0.7342156 |
| IGAAVALGF | ASSLLDRGEVF | 0.7695361 |
| IGLTPLAL | ASSLLDRGEVF | 0.7006439 |
| ILGSNTSTL | ASSLLDRGEVF | 0.395012 |
| IVFSLTVL | ASSLLDRGEVF | 0.70482016 |
| LASKATGF | ASSLLDRGEVF | 0.7578529 |
| LASVALIL | ASSLLDRGEVF | 0.76055634 |
| LFAVGLTL | ASSLLDRGEVF | 0.7629249 |
| LFNGGIGTYI | ASSLLDRGEVF | 0.727818 |
| LTLGAGII | ASSLLDRGEVF | 0.6867819 |
| LVAAALGDI | ASSLLDRGEVF | 0.7647398 |
| NTAFVQLGI | ASSLLDRGEVF | 0.7679 |
| QLFSGAEL | ASSLLDRGEVF | 0.75375926 |
| SAATFITL | ASSLLDRGEVF | 0.67407453 |
| SNYGLANLGV | ASSLLDRGEVF | 0.62596536 |
| SYGGAYAVM | ASSLLDRGEVF | 0.74273056 |
| TTGVGGLTLG | ASSLLDRGEVF | 0.74732786 |
| TTQTIYI | ASSLLDRGEVF | 0.606935 |
| VAAGAGLGAVI | ASSLLDRGEVF | 0.7319832 |
| VAAVVTLL | ASSLLDRGEVF | 0.73099184 |
| VGFLGGLI | ASSLLDRGEVF | 0.44098142 |
| VGVIPILL | ASSLLDRGEVF | 0.74383664 |
| VIFASSLI | ASSLLDRGEVF | 0.7656981 |
| VKGGDGGAAAG | ASSLLDRGEVF | 0.73403084 |
| VTKTFTNI | ASSLLDRGEVF | 0.603758 |
| VVAAAILL | ASSLLDRGEVF | 0.7603136 |
| YAALDVEL | ASSLLDRGEVF | 0.76849616 |
| LASVALLL | ASSLLDRGEVF | 0.76297915 |
| LDAVHRV | ASSLLDRGEVF | 0.54533696 |
| QGLSSSIF | ASSLLDRGEVF | 0.32383615 |
| RGTFANIRL | ASSLLDRGEVF | 0.710808 |
| RQLWWGHRI | ASSLLDRGEVF | 0.7523143 |
| YGPPGTGKTLL | ASSLLDRGEVF | 0.7097511 |
| AALVGAPANVA | ASSLLDRGEVF | 0.6813833 |
| AATFRRTE | ASSLLDRGEVF | 0.7343838 |
| AGALGGGI | ASSLLDRGEVF | 0.7575863 |
| AGGAGGAGGIGAG | ASSLLDRGEVF | 0.62356937 |
| AGSGALGL | ASSLLDRGEVF | 0.49774885 |

| | | |
|---|---|---|
| AQIADQI | ASSLLDRGEVF | 0.738765 |
| ASAVAVTYL | ASSLLDRGEVF | 0.7574376 |
| ASTVLGGTGG | ASSLLDRGEVF | 0.6697082 |
| AVGGEPAK | ASSLLDRGEVF | 0.71397126 |
| CEQGTFGGTC | ASSLLDRGEVF | 0.72339064 |
| DEGKAPAG | ASSLLDRGEVF | 0.6819947 |
| DTVVFPAL | ASSLLDRGEVF | 0.7319702 |
| FEGITMAG | ASSLLDRGEVF | 0.60198414 |
| FPLVVQD | ASSLLDRGEVF | 0.76206493 |
| GAFTTPQ | ASSLLDRGEVF | 0.63845944 |
| GFGGFGTAGGVGGA | ASSLLDRGEVF | 0.7445698 |
| GFLAQFL | ASSLLDRGEVF | 0.73594 |
| GGAGGAGGNAGVA | ASSLLDRGEVF | 0.7407869 |
| GGVGTAGAGGAGGAGG | ASSLLDRGEVF | 0.715056 |
| GNGGGGDAGNAGSGGN | ASSLLDRGEVF | 0.69084454 |
| GTGGAGGAGNDAGS | ASSLLDRGEVF | 0.60438263 |
| GTTNTGWFNS | ASSLLDRGEVF | 0.6016529 |
| IFMGPLFTL | ASSLLDRGEVF | 0.7577344 |
| ILAHAIEAI | ASSLLDRGEVF | 0.7150057 |
| IRSAIREAL | ASSLLDRGEVF | 0.7617883 |
| ITFAGGG | ASSLLDRGEVF | 0.7444334 |
| ITGMSGHQEV | ASSLLDRGEVF | 0.6825962 |
| IVAGLACTL | ASSLLDRGEVF | 0.74159616 |
| IVGGGLAGA | ASSLLDRGEVF | 0.554412 |
| IVTGVLNI | ASSLLDRGEVF | 0.72667444 |
| LAQQGCI | ASSLLDRGEVF | 0.71567106 |
| LCLHVEN | ASSLLDRGEVF | 0.74642956 |
| LDAISDGS | ASSLLDRGEVF | 0.7074838 |
| LGGFAAML | ASSLLDRGEVF | 0.7430794 |
| LGGKNPMI | ASSLLDRGEVF | 0.73975754 |
| LGPKGVAAIL | ASSLLDRGEVF | 0.5813407 |
| LMVGVSI | ASSLLDRGEVF | 0.7214129 |
| LVGAAFVL | ASSLLDRGEVF | 0.7606645 |
| LVLAVAPL | ASSLLDRGEVF | 0.74209577 |
| MAAIAVIDAA | ASSLLDRGEVF | 0.6006839 |
| NNGINAELQ | ASSLLDRGEVF | 0.7438327 |
| RGEPLVLVHG | ASSLLDRGEVF | 0.6800169 |
| RGGAGGAAGLI | ASSLLDRGEVF | 0.7439723 |
| RSYRMHPAV | ASSLLDRGEVF | 0.73146963 |
| SAAHLLAGQLA | ASSLLDRGEVF | 0.68951523 |
| SAAVGSY | ASSLLDRGEVF | 0.74144626 |
| SPHAVLHRI | ASSLLDRGEVF | 0.47506756 |
| SQIAKVL | ASSLLDRGEVF | 0.68420637 |
| TALGHLVP | ASSLLDRGEVF | 0.6952466 |
| TAPTTSGAA | ASSLLDRGEVF | 0.7305188 |
| TDGDDGNGGSPGT | ASSLLDRGEVF | 0.7522431 |
| TDGLGEN | ASSLLDRGEVF | 0.64742196 |
| TGGPPLPGQA | ASSLLDRGEVF | 0.7354513 |
| TIPAITI | ASSLLDRGEVF | 0.6877501 |
| TIPGFTI | ASSLLDRGEVF | 0.6228728 |
| TSIASGALA | ASSLLDRGEVF | 0.7130654 |

| | | |
|---|---|---|
| TVTLPGAAI | ASSLLDRGEVF | 0.68939304 |
| VASGLSAALP | ASSLLDRGEVF | 0.5763838 |
| VGAGVIGI | ASSLLDRGEVF | 0.75195575 |
| VGPKRFGV | ASSLLDRGEVF | 0.62998664 |
| VNVAFPD | ASSLLDRGEVF | 0.731688 |
| VSMTNSGVS | ASSLLDRGEVF | 0.7255977 |
| VTDVLGDLSV | ASSLLDRGEVF | 0.5456768 |
| VVVVGAGFAGL | ASSLLDRGEVF | 0.74087 |
| VYTVGTVV | ASSLLDRGEVF | 0.7271966 |
| WTTFSGP | ASSLLDRGEVF | 0.74221367 |
| YDELVEAF | ASSLLDRGEVF | 0.7229095 |
| YGTGGAGGNGG | ASSLLDRGEVF | 0.6651404 |
| YTNVMAV | ASSLLDRGEVF | 0.7600422 |
| FLEQVHQL | ASSLLDRGEVF | 0.7517761 |
| GLSLLAVAV | ASSLLDRGEVF | 0.7414924 |
| LEQVHQL | ASSLLDRGEVF | 0.7177058 |
| LNAAGAI | ASSLLDRGEVF | 0.7525571 |
| PAVHASL | ASSLLDRGEVF | 0.7542622 |
| TELEDDA | ASSLLDRGEVF | 0.6933079 |
| TPPGPPN | ASSLLDRGEVF | 0.4407534 |
| TPPPPPPP | ASSLLDRGEVF | 0.7217654 |
| TQSPALI | ASSGARDRGDQAPL | 0.70103747 |
| EFAAAFPQ | ASSGARDRGDQAPL | 0.6838329 |
| LLSGNLDVL | ASSGARDRGDQAPL | 0.43544316 |
| GPANLGNYNI | ASSGARDRGDQAPL | 0.7538338 |
| TLGPTLP | ASSGARDRGDQAPL | 0.7715362 |
| VYAYSSCCK | ASSGARDRGDQAPL | 0.7036555 |
| APAPAPAPNI | ASSGARDRGDQAPL | 0.77067447 |
| PGGPVPIP | ASSGARDRGDQAPL | 0.55751127 |
| FLTVALEP | ASSGARDRGDQAPL | 0.74460727 |
| SQYFGANL | ASSGARDRGDQAPL | 0.7566563 |
| GGVKPAA | ASSGARDRGDQAPL | 0.66703594 |
| GGTGGAGGAGGW | ASSGARDRGDQAPL | 0.74615633 |
| GVGGVAGLGGAGG | ASSGARDRGDQAPL | 0.6506413 |
| VLLVLATTI | ASSGARDRGDQAPL | 0.6717509 |
| IGIFAAML | ASSGARDRGDQAPL | 0.7604745 |
| AVGDLID | ASSGARDRGDQAPL | 0.6814873 |
| NISPLPL | ASSGARDRGDQAPL | 0.7420758 |
| VTAVFVAI | ASSGARDRGDQAPL | 0.7189759 |
| SGDDNDAAL | ASSGARDRGDQAPL | 0.6167133 |
| AVAALAHS | ASSGARDRGDQAPL | 0.74097544 |
| GGVTVMATP | ASSGARDRGDQAPL | 0.7575414 |
| AGLAANVRD | ASSGARDRGDQAPL | 0.71929014 |
| TNRPDLIDPALLRPG | ASSGARDRGDQAPL | 0.7679235 |
| LANLGVTAL | ASSGARDRGDQAPL | 0.7044759 |
| TGERIEL | ASSGARDRGDQAPL | 0.7723057 |
| NGEELAA | ASSGARDRGDQAPL | 0.76319396 |
| VLSLVWG | ASSGARDRGDQAPL | 0.73298705 |
| IGYALWL | ASSGARDRGDQAPL | 0.7011665 |
| LSLLLAPL | ASSGARDRGDQAPL | 0.7600943 |
| TLDEALK | ASSGARDRGDQAPL | 0.705986 |

| | | |
|---|---|---|
| TDGAGTAEGG | ASSGARDRGDQAPL | 0.7404136 |
| NAPGGAGGNGG | ASSGARDRGDQAPL | 0.7597004 |
| VSVFMGMGGNF | ASSGARDRGDQAPL | 0.39876473 |
| LASYLAAL | ASSGARDRGDQAPL | 0.76535285 |
| LVGGTPAL | ASSGARDRGDQAPL | 0.7286924 |
| VRNADVL | ASSGARDRGDQAPL | 0.6979226 |
| TCSSPFLVSAM | ASSGARDRGDQAPL | 0.54658186 |
| VHITDAE | ASSGARDRGDQAPL | 0.29429227 |
| EYAAPSF | ASSGARDRGDQAPL | 0.51537466 |
| VVDAFHGGL | ASSGARDRGDQAPL | 0.4862939 |
| ASVVDPQ | ASSGARDRGDQAPL | 0.73560506 |
| SYNEFGGLS | ASSGARDRGDQAPL | 0.73540634 |
| DPDGSVY | ASSGARDRGDQAPL | 0.48734462 |
| MVSTIHATE | ASSGARDRGDQAPL | 0.724548 |
| LAAGVADPV | ASSGARDRGDQAPL | 0.72753155 |
| TKEQIAA | ASSGARDRGDQAPL | 0.32450932 |
| SLASPAAL | ASSGARDRGDQAPL | 0.6752447 |
| IGLVFYGV | ASSGARDRGDQAPL | 0.4031443 |
| FLLDASNL | ASSGARDRGDQAPL | 0.6378825 |
| YQVGNFP | ASSGARDRGDQAPL | 0.2453101 |
| ESLLSQST | ASSGARDRGDQAPL | 0.71966803 |
| LFTVGFAAVG | ASSGARDRGDQAPL | 0.5860512 |
| ATPPTPT | ASSGARDRGDQAPL | 0.20031911 |
| TDLATLG | ASSGARDRGDQAPL | 0.6998693 |
| SAGGTGR | ASSGARDRGDQAPL | 0.5580393 |
| GFSNSGS | ASSGARDRGDQAPL | 0.614153 |
| GGNGGSGGTSV | ASSGARDRGDQAPL | 0.7352576 |
| GTSVATGGAGN | ASSGARDRGDQAPL | 0.48649943 |
| NGGNGGSGGTS | ASSGARDRGDQAPL | 0.43083057 |
| AVVKFPH | ASSGARDRGDQAPL | 0.73094505 |
| TPPPPPP | ASSGARDRGDQAPL | 0.7569529 |
| AAPAAAAAPA | ASSGARDRGDQAPL | 0.39217383 |
| IALNEDL | ASSGARDRGDQAPL | 0.69545114 |
| IVNVSSM | ASSGARDRGDQAPL | 0.7610048 |
| TGIPPAPR | ASSGARDRGDQAPL | 0.65589774 |
| FDLGGGTF | ASSGARDRGDQAPL | 0.74715984 |
| FDLGGGTFDVS | ASSGARDRGDQAPL | 0.76181364 |
| NEPTAAA | ASSGARDRGDQAPL | 0.4811448 |
| NEPTAAALA | ASSGARDRGDQAPL | 0.7561491 |
| GSIINISSI | ASSGARDRGDQAPL | 0.7421545 |
| VAPGNAGTA | ASSGARDRGDQAPL | 0.589172 |
| TGGGASLE | ASSGARDRGDQAPL | 0.6118041 |
| VAVYNHYKRI | ASSGARDRGDQAPL | 0.35975736 |
| TDPAPATT | ASSGARDRGDQAPL | 0.6250807 |
| TNRPDILDPALLRPG | ASSGARDRGDQAPL | 0.3616954 |
| LTTTTTS | ASSGARDRGDQAPL | 0.71677554 |
| PTGSGKTL | ASSGARDRGDQAPL | 0.7534137 |
| TGSGKTL | ASSGARDRGDQAPL | 0.627503 |
| TIGHVDHGK | ASSGARDRGDQAPL | 0.43543756 |
| TIGHVDHGKTT | ASSGARDRGDQAPL | 0.3758338 |
| GHVDHGKTT | ASSGARDRGDQAPL | 0.6977149 |

| | | |
|---|---|---|
| AQFEHTLL | ASSGARDRGDQAPL | 0.46275496 |
| IVNQDEL | ASSGARDRGDQAPL | 0.25818568 |
| TNGSQFF | ASSGARDRGDQAPL | 0.64489967 |
| TGKGIVE | ASSGARDRGDQAPL | 0.7292775 |
| LISNASDA | ASSGARDRGDQAPL | 0.73822117 |
| AELRTLGELTKIAKAHGAQ | ASSGARDRGDQAPL | 0.42022967 |
| APNAVAAAEPSW | ASSGARDRGDQAPL | 0.7697098 |
| APQVAGAATAVL | ASSGARDRGDQAPL | 0.72995245 |
| ARRDPAAAAAILPTDA | ASSGARDRGDQAPL | 0.75348985 |
| DAAARDVLAEAGLAEYF | ASSGARDRGDQAPL | 0.59719 |
| DDQIVLSAKANLAKVEQ | ASSGARDRGDQAPL | 0.45090494 |
| DELLPDTGEDSDEDGH | ASSGARDRGDQAPL | 0.7406437 |
| ELPEAPANRATVPQD | ASSGARDRGDQAPL | 0.65630466 |
| FGQSKIERVTGIG | ASSGARDRGDQAPL | 0.5484177 |
| GAGGTGSTAGGAGGAGGAGGLY | ASSGARDRGDQAPL | 0.7266196 |
| GHGGTGAAGGNGGTGGAGGSGI | ASSGARDRGDQAPL | 0.60833323 |
| GNGGNGGAGGEGGPGGAGGAGA | ASSGARDRGDQAPL | 0.4984997 |
| GSTAGNGGNGGNATNGTIAGQP | ASSGARDRGDQAPL | 0.687205 |
| GTGGDGGNGGNGGTGG | ASSGARDRGDQAPL | 0.7609848 |
| GVRRPAGIPADAPV | ASSGARDRGDQAPL | 0.5251138 |
| GYRQVTDAHLLTLARR | ASSGARDRGDQAPL | 0.44157034 |
| IAKMGQDGHDRG | ASSGARDRGDQAPL | 0.5076794 |
| KAEPSIEIQYT | ASSGARDRGDQAPL | 0.63667107 |
| KPVRIRMAADEDTVAVT | ASSGARDRGDQAPL | 0.75727034 |
| MPPRIAGMRLLVIKPEP | ASSGARDRGDQAPL | 0.6767563 |
| MKFYAVALFRSRGVPPTR | ASSGARDRGDQAPL | 0.62817127 |
| NAGGFGILGAVAHSP | ASSGARDRGDQAPL | 0.7131944 |
| NEGREITLAMAEM | ASSGARDRGDQAPL | 0.7065472 |
| NELVEWSHPLGHRWRWE | ASSGARDRGDQAPL | 0.66141963 |
| NLGAGGTGGNGGNPG | ASSGARDRGDQAPL | 0.7022513 |
| NTEWPKWETVMAADPVD | ASSGARDRGDQAPL | 0.5806587 |
| PASAAPAETQP | ASSGARDRGDQAPL | 0.7561213 |
| QQSSDGKNGVRAAEITGE | ASSGARDRGDQAPL | 0.71968883 |
| RAPDAVTTSWWKE | ASSGARDRGDQAPL | 0.71858954 |
| SGFGNVGAHTSGWFNQ | ASSGARDRGDQAPL | 0.74883497 |
| SGVAGAVGQAASVGGLKV | ASSGARDRGDQAPL | 0.6913911 |
| SNGGAGTGSAGGNGGT | ASSGARDRGDQAPL | 0.6688787 |
| TGGAGGEGGAAG | ASSGARDRGDQAPL | 0.32823282 |
| TGGDGGAAGAGGAPGHG | ASSGARDRGDQAPL | 0.42135435 |
| TMPPLLAAIANAGRHGV | ASSGARDRGDQAPL | 0.69073176 |
| TSSTAGNSGGA | ASSGARDRGDQAPL | 0.72377765 |
| VPATVPVGILC | ASSGARDRGDQAPL | 0.73158646 |
| VQRPDSGGVVLTGRIS | ASSGARDRGDQAPL | 0.6983892 |
| WTPNPVPATKGVPAGLVQAT | ASSGARDRGDQAPL | 0.55266255 |
| YALKRALAKFSDRFDVVII | ASSGARDRGDQAPL | 0.7510228 |
| YLIGSVIV | ASSGARDRGDQAPL | 0.75543064 |
| YGQLVAVID | ASSGARDRGDQAPL | 0.3406921 |
| VTVGGMTLPA | ASSGARDRGDQAPL | 0.6109141 |
| VLALPAV | ASSGARDRGDQAPL | 0.7692547 |
| VIPPTLNYET | ASSGARDRGDQAPL | 0.68415856 |
| VGMGGMHPSSG | ASSGARDRGDQAPL | 0.56784785 |

| | | |
|---|---|---|
| VGLAGGAATA | ASSGARDRGDQAPL | 0.56683147 |
| VGGAGGAGGSGP | ASSGARDRGDQAPL | 0.75749767 |
| VFAAFMV | ASSGARDRGDQAPL | 0.5053071 |
| TLYDKLR | ASSGARDRGDQAPL | 0.7652877 |
| TLAVFTVL | ASSGARDRGDQAPL | 0.24692585 |
| TGSVQPARV | ASSGARDRGDQAPL | 0.57380486 |
| TGGGAGSGGT | ASSGARDRGDQAPL | 0.56421375 |
| SHLETSL | ASSGARDRGDQAPL | 0.2383517 |
| SFTTGMT | ASSGARDRGDQAPL | 0.7288335 |
| QVEVAKRMGVRQPTVS | ASSGARDRGDQAPL | 0.45115447 |
| QGSGGPALAM | ASSGARDRGDQAPL | 0.72241545 |
| MLAPSNP | ASSGARDRGDQAPL | 0.53890747 |
| LVNNAAI | ASSGARDRGDQAPL | 0.7493854 |
| LTGEINTLA | ASSGARDRGDQAPL | 0.67879647 |
| LSIFGPV | ASSGARDRGDQAPL | 0.62836695 |
| LLALLANL | ASSGARDRGDQAPL | 0.43813846 |
| LISNASDALDKLR | ASSGARDRGDQAPL | 0.7039039 |
| LINEAVP | ASSGARDRGDQAPL | 0.757265 |
| LGYAFSGL | ASSGARDRGDQAPL | 0.75097096 |
| LDFVAVAL | ASSGARDRGDQAPL | 0.29081163 |
| LAQPGAE | ASSGARDRGDQAPL | 0.6852435 |
| ISNLGAVTGF | ASSGARDRGDQAPL | 0.6399702 |
| HQEDPGRI | ASSGARDRGDQAPL | 0.27490687 |
| GTPGEPTP | ASSGARDRGDQAPL | 0.22121845 |
| GFFNSGS | ASSGARDRGDQAPL | 0.6845746 |
| GAGGGGLTAGAN | ASSGARDRGDQAPL | 0.64975363 |
| FVDALVNM | ASSGARDRGDQAPL | 0.7285156 |
| FLTVAPEL | ASSGARDRGDQAPL | 0.7363833 |
| FETLTAL | ASSGARDRGDQAPL | 0.5401484 |
| DVINPIEL | ASSGARDRGDQAPL | 0.72388315 |
| APVSGAPV | ASSGARDRGDQAPL | 0.6643915 |
| ALVGGLAW | ASSGARDRGDQAPL | 0.18547316 |
| ALAAAAATAI | ASSGARDRGDQAPL | 0.47704774 |
| AFGLGLLL | ASSGARDRGDQAPL | 0.34110916 |
| AFFAAASL | ASSGARDRGDQAPL | 0.75369227 |
| AAAALPAFSPP | ASSGARDRGDQAPL | 0.4813874 |
| VVLAPAL | ASSGARDRGDQAPL | 0.54531413 |
| TIGHVDHGKTTL | ASSGARDRGDQAPL | 0.6371131 |
| LISNASDALDK | ASSGARDRGDQAPL | 0.71823406 |
| LISNASDAL | ASSGARDRGDQAPL | 0.6652443 |
| ISNASDALDK | ASSGARDRGDQAPL | 0.5272592 |
| NQAETLVYQTE | ASSGARDRGDQAPL | 0.32871857 |
| TGGTGGTAGAG | ASSGARDRGDQAPL | 0.6977604 |
| TGSTAGGAGGA | ASSGARDRGDQAPL | 0.45684677 |
| DPLAVPVAAGPV | ASSGARDRGDQAPL | 0.4615604 |
| GGTGGTPGNGGN | ASSGARDRGDQAPL | 0.32088047 |
| PAGLAGVDPATL | ASSGARDRGDQAPL | 0.66946316 |
| GNNLVVEAPHAGASV | ASSGARDRGDQAPL | 0.2721997 |
| KVTRIMSASTLEQPA | ASSGARDRGDQAPL | 0.7273323 |
| LGFALPIALPRRTVV | ASSGARDRGDQAPL | 0.5572145 |
| TRLGNLTARKLVVLV | ASSGARDRGDQAPL | 0.6296107 |

| | | |
|---|---|---|
| WSQQRRLIYFPSAGP | ASSGARDRGDQAPL | 0.77419126 |
| GPNGNPDPMAAAVDIR | ASSGARDRGDQAPL | 0.6986584 |
| TQRCVSVADLRAAASR | ASSGARDRGDQAPL | 0.6407479 |
| QFVLDVAEHPVEVLTETV | ASSGARDRGDQAPL | 0.49389642 |
| GGAGGTGGAGGNGGTQAGDG | ASSGARDRGDQAPL | 0.42336714 |
| PLLATLRAIAVDVDAKPAQV | ASSGARDRGDQAPL | 0.5158234 |
| APKFRTGGYMIDFWGVGYQVAK | ASSGARDRGDQAPL | 0.50780135 |
| GAGGAGGAGGAAGGPGGTGGTG | ASSGARDRGDQAPL | 0.7600275 |
| IGALGLLATDVAPAGSVAFGVA | ASSGARDRGDQAPL | 0.6454597 |
| AGGAGGIGGNGGAGGAGGVLMG | ASSGARDRGDQAPL | 0.34264576 |
| ADELIGLL | ASSGARDRGDQAPL | 0.67898 |
| ADRALKA | ASSGARDRGDQAPL | 0.75933826 |
| AGAVGQVGGAGGA | ASSGARDRGDQAPL | 0.77267426 |
| AGGGGGLGGAP | ASSGARDRGDQAPL | 0.49353808 |
| AGGLLYGNGGAG | ASSGARDRGDQAPL | 0.512117 |
| AGLESIGL | ASSGARDRGDQAPL | 0.58884025 |
| AGLKTGII | ASSGARDRGDQAPL | 0.64430785 |
| ASLAALLL | ASSGARDRGDQAPL | 0.4788679 |
| ASVINISSI | ASSGARDRGDQAPL | 0.410174 |
| ATPPGPA | ASSGARDRGDQAPL | 0.74783146 |
| ATVASALV | ASSGARDRGDQAPL | 0.77194405 |
| ATVASLIL | ASSGARDRGDQAPL | 0.57518125 |
| AVNGPVLIRH | ASSGARDRGDQAPL | 0.2635971 |
| DGAAGAVGI | ASSGARDRGDQAPL | 0.63719624 |
| DPLLHGI | ASSGARDRGDQAPL | 0.5154246 |
| ELGAALGAL | ASSGARDRGDQAPL | 0.5404478 |
| ELGRTQAV | ASSGARDRGDQAPL | 0.58533716 |
| FVAGGYLS | ASSGARDRGDQAPL | 0.32220918 |
| GASGGAGGAGGW | ASSGARDRGDQAPL | 0.6208009 |
| GGAGAGGDGGAGGI | ASSGARDRGDQAPL | 0.775563 |
| GTGHNGSQPGMGGNG | ASSGARDRGDQAPL | 0.63341224 |
| GYAAASASAS | ASSGARDRGDQAPL | 0.71106935 |
| HQVDVAAVQQG | ASSGARDRGDQAPL | 0.75959265 |
| IGAAIAHT | ASSGARDRGDQAPL | 0.7276873 |
| IGMIEVAQL | ASSGARDRGDQAPL | 0.74841356 |
| IRLAPEHTSAMD | ASSGARDRGDQAPL | 0.7318461 |
| IYNVSVL | ASSGARDRGDQAPL | 0.6096153 |
| KQGFLGM | ASSGARDRGDQAPL | 0.6477263 |
| LESVITQL | ASSGARDRGDQAPL | 0.73427206 |
| LGLGALVVV | ASSGARDRGDQAPL | 0.6226691 |
| LIGNGGNGGSGGI | ASSGARDRGDQAPL | 0.64394134 |
| LNAINAPTL | ASSGARDRGDQAPL | 0.574223 |
| LQLLNTGV | ASSGARDRGDQAPL | 0.74079406 |
| LQTSIEF | ASSGARDRGDQAPL | 0.6161915 |
| MSFAVSEL | ASSGARDRGDQAPL | 0.7525009 |
| NTGALGL | ASSGARDRGDQAPL | 0.251157 |
| PPGAGGAGGAGTT | ASSGARDRGDQAPL | 0.40411735 |
| QLVGLGI | ASSGARDRGDQAPL | 0.74760735 |
| RFLEQVHQL | ASSGARDRGDQAPL | 0.2016255 |
| RGVGYVLR | ASSGARDRGDQAPL | 0.2556504 |
| SATGVFAGVI | ASSGARDRGDQAPL | 0.6661804 |

| | | |
|---|---|---|
| SLPAITLP | ASSGARDRGDQAPL | 0.7569735 |
| STALAFTV | ASSGARDRGDQAPL | 0.21559072 |
| SVLTVGLGK | ASSGARDRGDQAPL | 0.41532475 |
| TALAGAGSGI | ASSGARDRGDQAPL | 0.35513797 |
| TGLPFTGI | ASSGARDRGDQAPL | 0.73097134 |
| VAEHVTV | ASSGARDRGDQAPL | 0.59807634 |
| VDIVLNSL | ASSGARDRGDQAPL | 0.26891905 |
| VLTGGGAVL | ASSGARDRGDQAPL | 0.7241517 |
| VNVGLAYGI | ASSGARDRGDQAPL | 0.28191766 |
| VSSAAVGAV | ASSGARDRGDQAPL | 0.65109885 |
| VTSVYEAGV | ASSGARDRGDQAPL | 0.7412283 |
| WVALDRGA | ASSGARDRGDQAPL | 0.76142704 |
| YLVGSVVI | ASSGARDRGDQAPL | 0.37707716 |
| GPPGTGKTLL | ASSGARDRGDQAPL | 0.71907306 |
| GVGMTKF | ASSGARDRGDQAPL | 0.59579825 |
| IGIGGSDL | ASSGARDRGDQAPL | 0.7781839 |
| IGTLLAEL | ASSGARDRGDQAPL | 0.5831197 |
| IYTSGTT | ASSGARDRGDQAPL | 0.31638616 |
| LSFPAGV | ASSGARDRGDQAPL | 0.7458012 |
| LYTSGTT | ASSGARDRGDQAPL | 0.7242032 |
| VGAFPFGF | ASSGARDRGDQAPL | 0.6161401 |
| VGAGTVEF | ASSGARDRGDQAPL | 0.73433876 |
| VGKSTLLL | ASSGARDRGDQAPL | 0.6477951 |
| AALAHAGGGIV | ASSGARDRGDQAPL | 0.6200273 |
| AALVAVSLL | ASSGARDRGDQAPL | 0.7402071 |
| AAVSRAVAAL | ASSGARDRGDQAPL | 0.7335999 |
| AGGSSGIGGF | ASSGARDRGDQAPL | 0.459865 |
| AGLTVANANG | ASSGARDRGDQAPL | 0.7504647 |
| AGLVGAVL | ASSGARDRGDQAPL | 0.64499307 |
| ALSQAFTV | ASSGARDRGDQAPL | 0.40336913 |
| ALTFSVSL | ASSGARDRGDQAPL | 0.77638304 |
| ALTGAALVAL | ASSGARDRGDQAPL | 0.58220685 |
| ASFSIGAGVL | ASSGARDRGDQAPL | 0.7312728 |
| DGSVYKI | ASSGARDRGDQAPL | 0.7149341 |
| EQLAEIV | ASSGARDRGDQAPL | 0.7570407 |
| FAVSELTVM | ASSGARDRGDQAPL | 0.34818727 |
| GAAPLAAGL | ASSGARDRGDQAPL | 0.59656113 |
| GADGGDGGSGGAGG | ASSGARDRGDQAPL | 0.649179 |
| GFGLDTGGAGG | ASSGARDRGDQAPL | 0.73811054 |
| GLASGGLL | ASSGARDRGDQAPL | 0.60174 |
| GRIAGAAVLL | ASSGARDRGDQAPL | 0.64113116 |
| GSAGSLNTSF | ASSGARDRGDQAPL | 0.7217565 |
| GTGQIGI | ASSGARDRGDQAPL | 0.75431633 |
| GTLIALIL | ASSGARDRGDQAPL | 0.6711724 |
| IAATMGFVVG | ASSGARDRGDQAPL | 0.68423146 |
| IFAQALL | ASSGARDRGDQAPL | 0.7626318 |
| IGAAVALGF | ASSGARDRGDQAPL | 0.38889346 |
| IGLTPLAL | ASSGARDRGDQAPL | 0.62237227 |
| ILGSNTSTL | ASSGARDRGDQAPL | 0.7375475 |
| IVFSLTVL | ASSGARDRGDQAPL | 0.75065327 |
| LASKATGF | ASSGARDRGDQAPL | 0.6478987 |

| | | |
|---|---|---|
| LASVALIL | ASSGARDRGDQAPL | 0.75520825 |
| LFAVGLTL | ASSGARDRGDQAPL | 0.46505105 |
| LFNGGIGTYI | ASSGARDRGDQAPL | 0.7175046 |
| LTLGAGII | ASSGARDRGDQAPL | 0.62701434 |
| LVAAALGDI | ASSGARDRGDQAPL | 0.60239995 |
| NTAFVQLGI | ASSGARDRGDQAPL | 0.5164522 |
| QLFSGAEL | ASSGARDRGDQAPL | 0.6955978 |
| SAATFITL | ASSGARDRGDQAPL | 0.735395 |
| SNYGLANLGV | ASSGARDRGDQAPL | 0.70423985 |
| SYGGAYAVM | ASSGARDRGDQAPL | 0.22036341 |
| TTGVGGLTLG | ASSGARDRGDQAPL | 0.7116002 |
| TTQTIYI | ASSGARDRGDQAPL | 0.72500247 |
| VAAGAGLGAVI | ASSGARDRGDQAPL | 0.76328707 |
| VAAVVTLL | ASSGARDRGDQAPL | 0.6234057 |
| VGFLGGLI | ASSGARDRGDQAPL | 0.7276663 |
| VGVIPILL | ASSGARDRGDQAPL | 0.53578687 |
| VIFASSLI | ASSGARDRGDQAPL | 0.6506034 |
| VKGGDGGAAAG | ASSGARDRGDQAPL | 0.41358674 |
| VTKTFTNI | ASSGARDRGDQAPL | 0.49622762 |
| VVAAAILL | ASSGARDRGDQAPL | 0.57554436 |
| YAALDVEL | ASSGARDRGDQAPL | 0.554762 |
| LASVALLL | ASSGARDRGDQAPL | 0.74053085 |
| LDAVHRV | ASSGARDRGDQAPL | 0.74956274 |
| QGLSSSIF | ASSGARDRGDQAPL | 0.74221706 |
| RGTFANIRL | ASSGARDRGDQAPL | 0.7524617 |
| RQLWWGHRI | ASSGARDRGDQAPL | 0.76215494 |
| YGPPGTGKTLL | ASSGARDRGDQAPL | 0.72635645 |
| AALVGAPANVA | ASSGARDRGDQAPL | 0.5860994 |
| AATFRRTE | ASSGARDRGDQAPL | 0.5344427 |
| AGALGGGI | ASSGARDRGDQAPL | 0.37579876 |
| AGGAGGAGGIGAG | ASSGARDRGDQAPL | 0.55036587 |
| AGSGALGL | ASSGARDRGDQAPL | 0.6001074 |
| AQIADQI | ASSGARDRGDQAPL | 0.64662015 |
| ASAVAVTYL | ASSGARDRGDQAPL | 0.1967136 |
| ASTVLGGTGG | ASSGARDRGDQAPL | 0.7053919 |
| AVGGEPAK | ASSGARDRGDQAPL | 0.76373523 |
| CEQGTFGGTC | ASSGARDRGDQAPL | 0.645769 |
| DEGKAPAG | ASSGARDRGDQAPL | 0.6955013 |
| DTVVFPAL | ASSGARDRGDQAPL | 0.7191631 |
| FEGITMAG | ASSGARDRGDQAPL | 0.63086826 |
| FPLVVQD | ASSGARDRGDQAPL | 0.7151007 |
| GAFTTPQ | ASSGARDRGDQAPL | 0.5321692 |
| GFGGFGTAGGVGGA | ASSGARDRGDQAPL | 0.7222425 |
| GFLAQFL | ASSGARDRGDQAPL | 0.77226555 |
| GGAGGAGGNAGVA | ASSGARDRGDQAPL | 0.6919122 |
| GGVGTAGAGGAGGAGG | ASSGARDRGDQAPL | 0.63805294 |
| GNGGGGDAGNAGSGGN | ASSGARDRGDQAPL | 0.6447244 |
| GTGGAGGAGNDAGS | ASSGARDRGDQAPL | 0.28314877 |
| GTTNTGWFNS | ASSGARDRGDQAPL | 0.67673516 |
| IFMGPLFTL | ASSGARDRGDQAPL | 0.42841685 |
| ILAHAIEAI | ASSGARDRGDQAPL | 0.67146254 |

| | | |
|---|---|---|
| IRSAIREAL | ASSGARDRGDQAPL | 0.47179276 |
| ITFAGGG | ASSGARDRGDQAPL | 0.70214766 |
| ITGMSGHQEV | ASSGARDRGDQAPL | 0.6422906 |
| IVAGLACTL | ASSGARDRGDQAPL | 0.7119773 |
| IVGGGLAGA | ASSGARDRGDQAPL | 0.45801604 |
| IVTGVLNI | ASSGARDRGDQAPL | 0.6495513 |
| LAQQGCI | ASSGARDRGDQAPL | 0.6196847 |
| LCLHVEN | ASSGARDRGDQAPL | 0.37855312 |
| LDAISDGS | ASSGARDRGDQAPL | 0.7353124 |
| LGGFAAML | ASSGARDRGDQAPL | 0.64302826 |
| LGGKNPMI | ASSGARDRGDQAPL | 0.66512215 |
| LGPKGVAAIL | ASSGARDRGDQAPL | 0.74030626 |
| LMVGVSI | ASSGARDRGDQAPL | 0.68121266 |
| LVGAAFVL | ASSGARDRGDQAPL | 0.26554003 |
| LVLAVAPL | ASSGARDRGDQAPL | 0.6253919 |
| MAAIAVIDAA | ASSGARDRGDQAPL | 0.64834845 |
| NNGINAELQ | ASSGARDRGDQAPL | 0.63162833 |
| RGEPLVLVHG | ASSGARDRGDQAPL | 0.20500982 |
| RGGAGGAAGLI | ASSGARDRGDQAPL | 0.7270526 |
| RSYRMHPAV | ASSGARDRGDQAPL | 0.39589992 |
| SAAHLLAGQLA | ASSGARDRGDQAPL | 0.5220655 |
| SAAVGSY | ASSGARDRGDQAPL | 0.5541827 |
| SPHAVLHRI | ASSGARDRGDQAPL | 0.7378346 |
| SQIAKVL | ASSGARDRGDQAPL | 0.68567073 |
| TALGHLVP | ASSGARDRGDQAPL | 0.6067641 |
| TAPTTSGAA | ASSGARDRGDQAPL | 0.71470803 |
| TDGDDGNGGSPGT | ASSGARDRGDQAPL | 0.62859225 |
| TDGLGEN | ASSGARDRGDQAPL | 0.751079 |
| TGGPPLPGQA | ASSGARDRGDQAPL | 0.67938995 |
| TIPAITI | ASSGARDRGDQAPL | 0.6679117 |
| TIPGFTI | ASSGARDRGDQAPL | 0.6584584 |
| TSIASGALA | ASSGARDRGDQAPL | 0.47258723 |
| TVTLPGAAI | ASSGARDRGDQAPL | 0.3746495 |
| VASGLSAALP | ASSGARDRGDQAPL | 0.32905325 |
| VGAGVIGI | ASSGARDRGDQAPL | 0.45223233 |
| VGPKRFGV | ASSGARDRGDQAPL | 0.3173411 |
| VNVAFPD | ASSGARDRGDQAPL | 0.6857469 |
| VSMTNSGVS | ASSGARDRGDQAPL | 0.74947965 |
| VTDVLGDLSV | ASSGARDRGDQAPL | 0.665774 |
| VVVVGAGFAGL | ASSGARDRGDQAPL | 0.30627158 |
| VYTVGTVV | ASSGARDRGDQAPL | 0.6473471 |
| WTTFSGP | ASSGARDRGDQAPL | 0.46467787 |
| YDELVEAF | ASSGARDRGDQAPL | 0.7145674 |
| YGTGGAGGNGG | ASSGARDRGDQAPL | 0.7458533 |
| YTNVMAV | ASSGARDRGDQAPL | 0.75757015 |
| FLEQVHQL | ASSGARDRGDQAPL | 0.73630404 |
| GLSLLAVAV | ASSGARDRGDQAPL | 0.57171714 |
| LEQVHQL | ASSGARDRGDQAPL | 0.7369244 |
| LNAAGAI | ASSGARDRGDQAPL | 0.5931218 |
| PAVHASL | ASSGARDRGDQAPL | 0.61842614 |
| TELEDDA | ASSGARDRGDQAPL | 0.65829456 |

| | | |
|---|---|---|
| TPPGPPN | ASSGARDRGDQAPL | 0.71664166 |
| TPPPPPPP | ASSGARDRGDQAPL | 0.59307116 |
| TQSPALI | ASSFNERLF | 0.6195029 |
| EFAAAFPQ | ASSFNERLF | 0.29236704 |
| LLSGNLDVL | ASSFNERLF | 0.7271929 |
| GPANLGNYNI | ASSFNERLF | 0.4743957 |
| TLGPTLP | ASSFNERLF | 0.29437596 |
| VYAYSSCCK | ASSFNERLF | 0.5968112 |
| APAPAPAPNI | ASSFNERLF | 0.76770866 |
| PGGPVPIP | ASSFNERLF | 0.7547141 |
| FLTVALEP | ASSFNERLF | 0.56575704 |
| SQYFGANL | ASSFNERLF | 0.51344126 |
| GGVKPAA | ASSFNERLF | 0.7217346 |
| GGTGGAGGAGGW | ASSFNERLF | 0.42999852 |
| GVGGVAGLGGAGG | ASSFNERLF | 0.7207236 |
| VLLVLATTI | ASSFNERLF | 0.65567243 |
| IGIFAAML | ASSFNERLF | 0.6574081 |
| AVGDLID | ASSFNERLF | 0.5383965 |
| NISPLPL | ASSFNERLF | 0.7381626 |
| VTAVFVAI | ASSFNERLF | 0.6325215 |
| SGDDNDAAL | ASSFNERLF | 0.6928297 |
| AVAALAHS | ASSFNERLF | 0.21806236 |
| GGVTVMATP | ASSFNERLF | 0.74200934 |
| AGLAANVRD | ASSFNERLF | 0.48251727 |
| TNRPDLIDPALLRPG | ASSFNERLF | 0.44781142 |
| LANLGVTAL | ASSFNERLF | 0.24559975 |
| TGERIEL | ASSFNERLF | 0.55624205 |
| NGEELAA | ASSFNERLF | 0.72231907 |
| VLSLVWG | ASSFNERLF | 0.7637973 |
| IGYALWL | ASSFNERLF | 0.57200295 |
| LSLLLAPL | ASSFNERLF | 0.73704994 |
| TLDEALK | ASSFNERLF | 0.7580999 |
| TDGAGTAEGG | ASSFNERLF | 0.5822426 |
| NAPGGAGGNGG | ASSFNERLF | 0.26916704 |
| VSVFMGMGGNF | ASSFNERLF | 0.6794052 |
| LASYLAAL | ASSFNERLF | 0.5706878 |
| LVGGTPAL | ASSFNERLF | 0.75314593 |
| VRNADVL | ASSFNERLF | 0.7198875 |
| TCSSPFLVSAM | ASSFNERLF | 0.37832144 |
| VHITDAE | ASSFNERLF | 0.7154446 |
| EYAAPSF | ASSFNERLF | 0.7463899 |
| VVDAFHGGL | ASSFNERLF | 0.4395589 |
| ASVVDPQ | ASSFNERLF | 0.38082355 |
| SYNEFGGLS | ASSFNERLF | 0.39667568 |
| DPDGSVY | ASSFNERLF | 0.32940018 |
| MVSTIHATE | ASSFNERLF | 0.66736513 |
| LAAGVADPV | ASSFNERLF | 0.6212082 |
| TKEQIAA | ASSFNERLF | 0.49491167 |
| SLASPAAL | ASSFNERLF | 0.63444865 |
| IGLVFYGV | ASSFNERLF | 0.5000915 |
| FLLDASNL | ASSFNERLF | 0.47730255 |

| | | |
|---|---|---|
| YQVGNFP | ASSFNERLF | 0.5452702 |
| ESLLSQST | ASSFNERLF | 0.6774578 |
| LFTVGFAAVG | ASSFNERLF | 0.7447712 |
| ATPPTPT | ASSFNERLF | 0.6042845 |
| TDLATLG | ASSFNERLF | 0.5853956 |
| SAGGTGR | ASSFNERLF | 0.7624861 |
| GFSNSGS | ASSFNERLF | 0.35395607 |
| GGNGGSGGTSV | ASSFNERLF | 0.41647473 |
| GTSVATGGAGN | ASSFNERLF | 0.71844673 |
| NGGNGGSGGTS | ASSFNERLF | 0.46630058 |
| AVVKFPH | ASSFNERLF | 0.73588216 |
| TPPPPPP | ASSFNERLF | 0.6616334 |
| AAPAAAAAPA | ASSFNERLF | 0.6827444 |
| IALNEDL | ASSFNERLF | 0.60353637 |
| IVNVSSM | ASSFNERLF | 0.77136624 |
| TGIPPAPR | ASSFNERLF | 0.7151396 |
| FDLGGGTF | ASSFNERLF | 0.56245565 |
| FDLGGGTFDVS | ASSFNERLF | 0.71242684 |
| NEPTAAA | ASSFNERLF | 0.3151791 |
| NEPTAAALA | ASSFNERLF | 0.7409611 |
| GSIINISSI | ASSFNERLF | 0.7305158 |
| VAPGNAGTA | ASSFNERLF | 0.48830855 |
| TGGGASLE | ASSFNERLF | 0.6327573 |
| VAVYNHYKRI | ASSFNERLF | 0.7639444 |
| TDPAPATT | ASSFNERLF | 0.75103366 |
| TNRPDILDPALLRPG | ASSFNERLF | 0.37499607 |
| LTTTTTS | ASSFNERLF | 0.5389712 |
| PTGSGKTL | ASSFNERLF | 0.34250325 |
| TGSGKTL | ASSFNERLF | 0.569209 |
| TIGHVDHGK | ASSFNERLF | 0.28501582 |
| TIGHVDHGKTT | ASSFNERLF | 0.71501225 |
| GHVDHGKTT | ASSFNERLF | 0.53901386 |
| AQFEHTLL | ASSFNERLF | 0.7366226 |
| IVNQDEL | ASSFNERLF | 0.695569 |
| TNGSQFF | ASSFNERLF | 0.7497864 |
| TGKGIVE | ASSFNERLF | 0.53315234 |
| LISNASDA | ASSFNERLF | 0.578692 |
| AELRTLGELTKIAKAHGAQ | ASSFNERLF | 0.33759084 |
| APNAVAAAEPSW | ASSFNERLF | 0.54064375 |
| APQVAGAATAVL | ASSFNERLF | 0.500038 |
| ARRDPAAAAAILPTDA | ASSFNERLF | 0.6255381 |
| DAAARDVLAEAGLAEYF | ASSFNERLF | 0.27499267 |
| DDQIVLSAKANLAKVEQ | ASSFNERLF | 0.7663456 |
| DELLPDTGEDSDEDGH | ASSFNERLF | 0.6107667 |
| ELPEAPANRATVPQD | ASSFNERLF | 0.71372205 |
| FGQSKIERVTGIG | ASSFNERLF | 0.4523338 |
| GAGGTGSTAGGAGGAGGAGGLY | ASSFNERLF | 0.44685405 |
| GHGGTGAAGGNGGTGGAGGSGI | ASSFNERLF | 0.5726551 |
| GNGGNGGAGGEGGPGGAGGAGA | ASSFNERLF | 0.72941375 |
| GSTAGNGGNGGNATNGTIAGQP | ASSFNERLF | 0.43140942 |
| GTGGDGGNGGNGGTGG | ASSFNERLF | 0.23665631 |

| | | |
|---|---|---|
| GVRRPAGIPADAPV | ASSFNERLF | 0.4357971 |
| GYRQVTDAHLLTLARR | ASSFNERLF | 0.32813147 |
| IAKMGQDGHDRG | ASSFNERLF | 0.76823974 |
| KAEPSIEIQYT | ASSFNERLF | 0.75193226 |
| KPVRIRMAADEDTVAVT | ASSFNERLF | 0.6944959 |
| MPPRIAGMRLLVIKPEP | ASSFNERLF | 0.7667796 |
| MKFYAVALFRSRGVPPTR | ASSFNERLF | 0.20443334 |
| NAGGFGILGAVAHSP | ASSFNERLF | 0.47350693 |
| NEGREITLAMAEM | ASSFNERLF | 0.7595694 |
| NELVEWSHPLGHRWRWE | ASSFNERLF | 0.71696067 |
| NLGAGGTGGNGGNPG | ASSFNERLF | 0.5974964 |
| NTEWPKWETVMAADPVD | ASSFNERLF | 0.74014294 |
| PASAAPAETQP | ASSFNERLF | 0.50264525 |
| QQSSDGKNGVRAAEITGE | ASSFNERLF | 0.623248 |
| RAPDAVTTSWWKE | ASSFNERLF | 0.6531025 |
| SGFGNVGAHTSGWFNQ | ASSFNERLF | 0.7293179 |
| SGVAGAVGQAASVGGLKV | ASSFNERLF | 0.6699572 |
| SNGGAGTGSAGGNGGT | ASSFNERLF | 0.34541637 |
| TGGAGGEGGAAG | ASSFNERLF | 0.6914964 |
| TGGDGGAAGAGGAPGHG | ASSFNERLF | 0.7749249 |
| TMPPLLAAIANAGRHGV | ASSFNERLF | 0.61572576 |
| TSSTAGNSGGA | ASSFNERLF | 0.7237061 |
| VPATVPVGILC | ASSFNERLF | 0.66353923 |
| VQRPDSGGVVLTGRIS | ASSFNERLF | 0.6846324 |
| WTPNPVPATKGVPAGLVQAT | ASSFNERLF | 0.4921978 |
| YALKRALAKFSDRFDVVII | ASSFNERLF | 0.5121414 |
| YLIGSVIV | ASSFNERLF | 0.6540799 |
| YGQLVAVID | ASSFNERLF | 0.4088404 |
| VTVGGMTLPA | ASSFNERLF | 0.6394665 |
| VLALPAV | ASSFNERLF | 0.54047257 |
| VIPPTLNYET | ASSFNERLF | 0.7850673 |
| VGMGGMHPSSG | ASSFNERLF | 0.7118378 |
| VGLAGGAATA | ASSFNERLF | 0.69410944 |
| VGGAGGAGGSGP | ASSFNERLF | 0.49691153 |
| VFAAFMV | ASSFNERLF | 0.7081435 |
| TLYDKLR | ASSFNERLF | 0.7180858 |
| TLAVFTVL | ASSFNERLF | 0.30256683 |
| TGSVQPARV | ASSFNERLF | 0.74452484 |
| TGGGAGSGGT | ASSFNERLF | 0.418551 |
| SHLETSL | ASSFNERLF | 0.7031639 |
| SFTTGMT | ASSFNERLF | 0.6792036 |
| QVEVAKRMGVRQPTVS | ASSFNERLF | 0.7462792 |
| QGSGGPALAM | ASSFNERLF | 0.60825104 |
| MLAPSNP | ASSFNERLF | 0.61587954 |
| LVNNAAI | ASSFNERLF | 0.44398528 |
| LTGEINTLA | ASSFNERLF | 0.7329913 |
| LSIFGPV | ASSFNERLF | 0.756318 |
| LLALLANL | ASSFNERLF | 0.6867065 |
| LISNASDALDKLR | ASSFNERLF | 0.77037746 |
| LINEAVP | ASSFNERLF | 0.69578576 |
| LGYAFSGL | ASSFNERLF | 0.2702895 |

| | | |
|---|---|---|
| LDFVAVAL | ASSFNERLF | 0.64645284 |
| LAQPGAE | ASSFNERLF | 0.7506008 |
| ISNLGAVTGF | ASSFNERLF | 0.7590442 |
| HQEDPGRI | ASSFNERLF | 0.7387498 |
| GTPGEPTP | ASSFNERLF | 0.698905 |
| GFFNSGS | ASSFNERLF | 0.6377009 |
| GAGGGGLTAGAN | ASSFNERLF | 0.76456696 |
| FVDALVNM | ASSFNERLF | 0.78107655 |
| FLTVAPEL | ASSFNERLF | 0.7213874 |
| FETLTAL | ASSFNERLF | 0.5408097 |
| DVINPIEL | ASSFNERLF | 0.4563809 |
| APVSGAPV | ASSFNERLF | 0.7087498 |
| ALVGGLAW | ASSFNERLF | 0.74906373 |
| ALAAAAATAI | ASSFNERLF | 0.39392686 |
| AFGLGLLL | ASSFNERLF | 0.6523324 |
| AFFAAASL | ASSFNERLF | 0.67249215 |
| AAAALPAFSPP | ASSFNERLF | 0.25402018 |
| VVLAPAL | ASSFNERLF | 0.68546414 |
| TIGHVDHGKTTL | ASSFNERLF | 0.75358176 |
| LISNASDALDK | ASSFNERLF | 0.6782055 |
| LISNASDAL | ASSFNERLF | 0.4145391 |
| ISNASDALDK | ASSFNERLF | 0.74899757 |
| NQAETLVYQTE | ASSFNERLF | 0.77193904 |
| TGGTGGTAGAG | ASSFNERLF | 0.7573239 |
| TGSTAGGAGGA | ASSFNERLF | 0.27800933 |
| DPLAVPVAAGPV | ASSFNERLF | 0.21461658 |
| GGTGGTPGNGGN | ASSFNERLF | 0.6610575 |
| PAGLAGVDPATL | ASSFNERLF | 0.7164327 |
| GNNLVVEAPHAGASV | ASSFNERLF | 0.55299187 |
| KVTRIMSASTLEQPA | ASSFNERLF | 0.55054605 |
| LGFALPIALPRRTVV | ASSFNERLF | 0.67623574 |
| TRLGNLTARKLVVLV | ASSFNERLF | 0.7464454 |
| WSQQRRLIYFPSAGP | ASSFNERLF | 0.37402296 |
| GPNGNPDPMAAAVDIR | ASSFNERLF | 0.40199143 |
| TQRCVSVADLRAAASR | ASSFNERLF | 0.69581413 |
| QFVLDVAEHPVEVLTETV | ASSFNERLF | 0.7397772 |
| GGAGGTGGAGGNGGTQAGDG | ASSFNERLF | 0.5292795 |
| PLLATLRAIAVDVDAKPAQV | ASSFNERLF | 0.6547079 |
| APKFRTGGYMIDFWGVGYQVAK | ASSFNERLF | 0.6428158 |
| GAGGAGGAGGAAGGPGGTGGTG | ASSFNERLF | 0.603618 |
| IGALGLLATDVAPAGSVAFGVA | ASSFNERLF | 0.6971078 |
| AGGAGGIGGNGGAGGAGGVLMG | ASSFNERLF | 0.3865158 |
| ADELIGLL | ASSFNERLF | 0.7534439 |
| ADRALKA | ASSFNERLF | 0.32149926 |
| AGAVGQVGGAGGA | ASSFNERLF | 0.7344891 |
| AGGGGGLGGAP | ASSFNERLF | 0.698671 |
| AGGLLYGNGGAG | ASSFNERLF | 0.70985067 |
| AGLESIGL | ASSFNERLF | 0.5757408 |
| AGLKTGII | ASSFNERLF | 0.4072738 |
| ASLAALLL | ASSFNERLF | 0.40855634 |
| ASVINISSI | ASSFNERLF | 0.36088628 |

| | | |
|---|---|---|
| ATPPGPA | ASSFNERLF | 0.752241 |
| ATVASALV | ASSFNERLF | 0.65804875 |
| ATVASLIL | ASSFNERLF | 0.74763477 |
| AVNGPVLIRH | ASSFNERLF | 0.6556742 |
| DGAAGAVGI | ASSFNERLF | 0.45264235 |
| DPLLHGI | ASSFNERLF | 0.7225593 |
| ELGAALGAL | ASSFNERLF | 0.41587043 |
| ELGRTQAV | ASSFNERLF | 0.5772844 |
| FVAGGYLS | ASSFNERLF | 0.6805262 |
| GASGGAGGAGGW | ASSFNERLF | 0.7068714 |
| GGAGAGGDGGAGGI | ASSFNERLF | 0.40731263 |
| GTGHNGSQPGMGGNG | ASSFNERLF | 0.69814605 |
| GYAAASASAS | ASSFNERLF | 0.70333683 |
| HQVDVAAVQQG | ASSFNERLF | 0.40323132 |
| IGAAIAHT | ASSFNERLF | 0.69064504 |
| IGMIEVAQL | ASSFNERLF | 0.7426044 |
| IRLAPEHTSAMD | ASSFNERLF | 0.67389286 |
| IYNVSVL | ASSFNERLF | 0.3585871 |
| KQGFLGM | ASSFNERLF | 0.6735625 |
| LESVITQL | ASSFNERLF | 0.5549313 |
| LGLGALVVV | ASSFNERLF | 0.70209396 |
| LIGNGGNGGSGGI | ASSFNERLF | 0.6690843 |
| LNAINAPTL | ASSFNERLF | 0.5254171 |
| LQLLNTGV | ASSFNERLF | 0.74821615 |
| LQTSIEF | ASSFNERLF | 0.34801075 |
| MSFAVSEL | ASSFNERLF | 0.56047356 |
| NTGALGL | ASSFNERLF | 0.58670944 |
| PPGAGGAGGAGTT | ASSFNERLF | 0.65023834 |
| QLVGLGI | ASSFNERLF | 0.71413076 |
| RFLEQVHQL | ASSFNERLF | 0.43450212 |
| RGVGYVLR | ASSFNERLF | 0.68021756 |
| SATGVFAGVI | ASSFNERLF | 0.58830976 |
| SLPAITLP | ASSFNERLF | 0.46826184 |
| STALAFTV | ASSFNERLF | 0.64771646 |
| SVLTVGLGK | ASSFNERLF | 0.550398 |
| TALAGAGSGI | ASSFNERLF | 0.3585136 |
| TGLPFTGI | ASSFNERLF | 0.72913516 |
| VAEHVTV | ASSFNERLF | 0.54661644 |
| VDIVLNSL | ASSFNERLF | 0.66117734 |
| VLTGGGAVL | ASSFNERLF | 0.74191594 |
| VNVGLAYGI | ASSFNERLF | 0.40916562 |
| VSSAAVGAV | ASSFNERLF | 0.7246 |
| VTSVYEAGV | ASSFNERLF | 0.6995896 |
| WVALDRGA | ASSFNERLF | 0.7026272 |
| YLVGSVVI | ASSFNERLF | 0.5893012 |
| GPPGTGKTLL | ASSFNERLF | 0.42924216 |
| GVGMTKF | ASSFNERLF | 0.75270224 |
| IGIGGSDL | ASSFNERLF | 0.70497894 |
| IGTLLAEL | ASSFNERLF | 0.6986649 |
| IYTSGTT | ASSFNERLF | 0.71853554 |
| LSFPAGV | ASSFNERLF | 0.6987723 |

| | | |
|---|---|---|
| LYTSGTT | ASSFNERLF | 0.7345289 |
| VGAFPFGF | ASSFNERLF | 0.55912024 |
| VGAGTVEF | ASSFNERLF | 0.4083738 |
| VGKSTLLL | ASSFNERLF | 0.61411905 |
| AALAHAGGGIV | ASSFNERLF | 0.5756634 |
| AALVAVSLL | ASSFNERLF | 0.4364285 |
| AAVSRAVAAL | ASSFNERLF | 0.30784556 |
| AGGSSGIGGF | ASSFNERLF | 0.7406712 |
| AGLTVANANG | ASSFNERLF | 0.78061366 |
| AGLVGAVL | ASSFNERLF | 0.46458966 |
| ALSQAFTV | ASSFNERLF | 0.7731576 |
| ALTFSVSL | ASSFNERLF | 0.36039168 |
| ALTGAALVAL | ASSFNERLF | 0.71940744 |
| ASFSIGAGVL | ASSFNERLF | 0.7386417 |
| DGSVYKI | ASSFNERLF | 0.44788206 |
| EQLAEIV | ASSFNERLF | 0.664752 |
| FAVSELTVM | ASSFNERLF | 0.37957555 |
| GAAPLAAGL | ASSFNERLF | 0.4686354 |
| GADGGDGGSGGAGG | ASSFNERLF | 0.75253916 |
| GFGLDTGGAGG | ASSFNERLF | 0.4877955 |
| GLASGGLL | ASSFNERLF | 0.67617536 |
| GRIAGAAVLL | ASSFNERLF | 0.5824704 |
| GSAGSLNTSF | ASSFNERLF | 0.48468512 |
| GTGQIGI | ASSFNERLF | 0.7106825 |
| GTLIALIL | ASSFNERLF | 0.7318275 |
| IAATMGFVVG | ASSFNERLF | 0.5656816 |
| IFAQALL | ASSFNERLF | 0.77878773 |
| IGAAVALGF | ASSFNERLF | 0.45081034 |
| IGLTPLAL | ASSFNERLF | 0.7632654 |
| ILGSNTSTL | ASSFNERLF | 0.75105184 |
| IVFSLTVL | ASSFNERLF | 0.6925045 |
| LASKATGF | ASSFNERLF | 0.59502155 |
| LASVALIL | ASSFNERLF | 0.7349367 |
| LFAVGLTL | ASSFNERLF | 0.62954664 |
| LFNGGIGTYI | ASSFNERLF | 0.6992341 |
| LTLGAGII | ASSFNERLF | 0.75858927 |
| LVAAALGDI | ASSFNERLF | 0.4485668 |
| NTAFVQLGI | ASSFNERLF | 0.72295165 |
| QLFSGAEL | ASSFNERLF | 0.6676024 |
| SAATFITL | ASSFNERLF | 0.68233305 |
| SNYGLANLGV | ASSFNERLF | 0.4083729 |
| SYGGAYAVM | ASSFNERLF | 0.2846161 |
| TTGVGGLTLG | ASSFNERLF | 0.62176174 |
| TTQTIYI | ASSFNERLF | 0.45253807 |
| VAAGAGLGAVI | ASSFNERLF | 0.73345685 |
| VAAVVTLL | ASSFNERLF | 0.7073653 |
| VGFLGGLI | ASSFNERLF | 0.48557195 |
| VGVIPILL | ASSFNERLF | 0.7205669 |
| VIFASSLI | ASSFNERLF | 0.7198662 |
| VKGGDGGAAAG | ASSFNERLF | 0.36423987 |
| VTKTFTNI | ASSFNERLF | 0.6523345 |

| | | |
|---|---|---|
| VVAAAILL | ASSFNERLF | 0.3665527 |
| YAALDVEL | ASSFNERLF | 0.6408974 |
| LASVALLL | ASSFNERLF | 0.3532718 |
| LDAVHRV | ASSFNERLF | 0.69490147 |
| QGLSSSIF | ASSFNERLF | 0.66062766 |
| RGTFANIRL | ASSFNERLF | 0.31189284 |
| RQLWWGHRI | ASSFNERLF | 0.6789005 |
| YGPPGTGKTLL | ASSFNERLF | 0.61475426 |
| AALVGAPANVA | ASSFNERLF | 0.62549746 |
| AATFRRTE | ASSFNERLF | 0.7149794 |
| AGALGGGI | ASSFNERLF | 0.6506228 |
| AGGAGGAGGIGAG | ASSFNERLF | 0.5347346 |
| AGSGALGL | ASSFNERLF | 0.68581116 |
| AQIADQI | ASSFNERLF | 0.71926075 |
| ASAVAVTYL | ASSFNERLF | 0.47862983 |
| ASTVLGGTGG | ASSFNERLF | 0.6877291 |
| AVGGEPAK | ASSFNERLF | 0.7383244 |
| CEQGTFGGTC | ASSFNERLF | 0.6009802 |
| DEGKAPAG | ASSFNERLF | 0.7314395 |
| DTVVFPAL | ASSFNERLF | 0.7351948 |
| FEGITMAG | ASSFNERLF | 0.5315888 |
| FPLVVQD | ASSFNERLF | 0.7148124 |
| GAFTTPQ | ASSFNERLF | 0.72297955 |
| GFGGFGTAGGVGGA | ASSFNERLF | 0.6002949 |
| GFLAQFL | ASSFNERLF | 0.6548412 |
| GGAGGAGGNAGVA | ASSFNERLF | 0.41279656 |
| GGVGTAGAGGAGGAGG | ASSFNERLF | 0.64583325 |
| GNGGGGDAGNAGSGGN | ASSFNERLF | 0.3134554 |
| GTGGAGGAGNDAGS | ASSFNERLF | 0.69166434 |
| GTTNTGWFNS | ASSFNERLF | 0.7242465 |
| IFMGPLFTL | ASSFNERLF | 0.63820887 |
| ILAHAIEAI | ASSFNERLF | 0.45494765 |
| IRSAIREAL | ASSFNERLF | 0.38069966 |
| ITFAGGG | ASSFNERLF | 0.68342507 |
| ITGMSGHQEV | ASSFNERLF | 0.53170264 |
| IVAGLACTL | ASSFNERLF | 0.4596144 |
| IVGGGLAGA | ASSFNERLF | 0.4326608 |
| IVTGVLNI | ASSFNERLF | 0.70259225 |
| LAQQGCI | ASSFNERLF | 0.713498 |
| LCLHVEN | ASSFNERLF | 0.52303374 |
| LDAISDGS | ASSFNERLF | 0.7249472 |
| LGGFAAML | ASSFNERLF | 0.7323957 |
| LGGKNPMI | ASSFNERLF | 0.68752456 |
| LGPKGVAAIL | ASSFNERLF | 0.6401965 |
| LMVGVSI | ASSFNERLF | 0.4355809 |
| LVGAAFVL | ASSFNERLF | 0.7377011 |
| LVLAVAPL | ASSFNERLF | 0.5536796 |
| MAAIAVIDAA | ASSFNERLF | 0.677982 |
| NNGINAELQ | ASSFNERLF | 0.70123225 |
| RGEPLVLVHG | ASSFNERLF | 0.59741086 |
| RGGAGGAAGLI | ASSFNERLF | 0.51824427 |

| | | |
|---|---|---|
| RSYRMHPAV | ASSFNERLF | 0.70382214 |
| SAAHLLAGQLA | ASSFNERLF | 0.7323068 |
| SAAVGSY | ASSFNERLF | 0.4503916 |
| SPHAVLHRI | ASSFNERLF | 0.3397494 |
| SQIAKVL | ASSFNERLF | 0.66795063 |
| TALGHLVP | ASSFNERLF | 0.6604526 |
| TAPTTSGAA | ASSFNERLF | 0.716962 |
| TDGDDGNGGSPGT | ASSFNERLF | 0.67494595 |
| TDGLGEN | ASSFNERLF | 0.6750104 |
| TGGPPLPGQA | ASSFNERLF | 0.7156958 |
| TIPAITI | ASSFNERLF | 0.69920087 |
| TIPGFTI | ASSFNERLF | 0.6214428 |
| TSIASGALA | ASSFNERLF | 0.7117028 |
| TVTLPGAAI | ASSFNERLF | 0.58154887 |
| VASGLSAALP | ASSFNERLF | 0.7253195 |
| VGAGVIGI | ASSFNERLF | 0.71036744 |
| VGPKRFGV | ASSFNERLF | 0.7052777 |
| VNVAFPD | ASSFNERLF | 0.7280102 |
| VSMTNSGVS | ASSFNERLF | 0.64910316 |
| VTDVLGDLSV | ASSFNERLF | 0.6536586 |
| VVVVGAGFAGL | ASSFNERLF | 0.32458317 |
| VYTVGTVV | ASSFNERLF | 0.55469966 |
| WTTFSGP | ASSFNERLF | 0.68134236 |
| YDELVEAF | ASSFNERLF | 0.6872413 |
| YGTGGAGGNGG | ASSFNERLF | 0.72009766 |
| YTNVMAV | ASSFNERLF | 0.6984345 |
| FLEQVHQL | ASSFNERLF | 0.55627465 |
| GLSLLAVAV | ASSFNERLF | 0.73438275 |
| LEQVHQL | ASSFNERLF | 0.73716533 |
| LNAAGAI | ASSFNERLF | 0.36857748 |
| PAVHASL | ASSFNERLF | 0.60749674 |
| TELEDDA | ASSFNERLF | 0.7323835 |
| TPPGPPN | ASSFNERLF | 0.7150891 |
| TPPPPPPP | ASSFNERLF | 0.5497086 |
| TQSPALI | ASGAPRDRGYNNQAPL | 0.67839324 |
| EFAAAFPQ | ASGAPRDRGYNNQAPL | 0.7416508 |
| LLSGNLDVL | ASGAPRDRGYNNQAPL | 0.6926242 |
| GPANLGNYNI | ASGAPRDRGYNNQAPL | 0.73876274 |
| TLGPTLP | ASGAPRDRGYNNQAPL | 0.3088975 |
| VYAYSSCCK | ASGAPRDRGYNNQAPL | 0.6574288 |
| APAPAPAPNI | ASGAPRDRGYNNQAPL | 0.5267336 |
| PGGPVPIP | ASGAPRDRGYNNQAPL | 0.2813699 |
| FLTVALEP | ASGAPRDRGYNNQAPL | 0.6833086 |
| SQYFGANL | ASGAPRDRGYNNQAPL | 0.6200168 |
| GGVKPAA | ASGAPRDRGYNNQAPL | 0.72205615 |
| GGTGGAGGAGGW | ASGAPRDRGYNNQAPL | 0.5669995 |
| GVGGVAGLGGAGG | ASGAPRDRGYNNQAPL | 0.7043472 |
| VLLVLATTI | ASGAPRDRGYNNQAPL | 0.6642815 |
| IGIFAAML | ASGAPRDRGYNNQAPL | 0.6746547 |
| AVGDLID | ASGAPRDRGYNNQAPL | 0.5147733 |
| NISPLPL | ASGAPRDRGYNNQAPL | 0.63341165 |

| | | |
|---|---|---|
| VTAVFVAI | ASGAPRDRGYNNQAPL | 0.73271036 |
| SGDDNDAAL | ASGAPRDRGYNNQAPL | 0.72292113 |
| AVAALAHS | ASGAPRDRGYNNQAPL | 0.32321972 |
| GGVTVMATP | ASGAPRDRGYNNQAPL | 0.71829903 |
| AGLAANVRD | ASGAPRDRGYNNQAPL | 0.5400108 |
| TNRPDLIDPALLRPG | ASGAPRDRGYNNQAPL | 0.38761222 |
| LANLGVTAL | ASGAPRDRGYNNQAPL | 0.4258079 |
| TGERIEL | ASGAPRDRGYNNQAPL | 0.55971473 |
| NGEELAA | ASGAPRDRGYNNQAPL | 0.6342657 |
| VLSLVWG | ASGAPRDRGYNNQAPL | 0.69876873 |
| IGYALWL | ASGAPRDRGYNNQAPL | 0.7040516 |
| LSLLLAPL | ASGAPRDRGYNNQAPL | 0.5898223 |
| TLDEALK | ASGAPRDRGYNNQAPL | 0.719727 |
| TDGAGTAEGG | ASGAPRDRGYNNQAPL | 0.6533924 |
| NAPGGAGGNGG | ASGAPRDRGYNNQAPL | 0.23171175 |
| VSVFMGMGGNF | ASGAPRDRGYNNQAPL | 0.5625646 |
| LASYLAAL | ASGAPRDRGYNNQAPL | 0.36161554 |
| LVGGTPAL | ASGAPRDRGYNNQAPL | 0.7302855 |
| VRNADVL | ASGAPRDRGYNNQAPL | 0.513394 |
| TCSSPFLVSAM | ASGAPRDRGYNNQAPL | 0.6114596 |
| VHITDAE | ASGAPRDRGYNNQAPL | 0.64426595 |
| EYAAPSF | ASGAPRDRGYNNQAPL | 0.7184874 |
| VVDAFHGGL | ASGAPRDRGYNNQAPL | 0.64422786 |
| ASVVDPQ | ASGAPRDRGYNNQAPL | 0.5462129 |
| SYNEFGGLS | ASGAPRDRGYNNQAPL | 0.31222656 |
| DPDGSVY | ASGAPRDRGYNNQAPL | 0.6605339 |
| MVSTIHATE | ASGAPRDRGYNNQAPL | 0.49354392 |
| LAAGVADPV | ASGAPRDRGYNNQAPL | 0.47604704 |
| TKEQIAA | ASGAPRDRGYNNQAPL | 0.5589633 |
| SLASPAAL | ASGAPRDRGYNNQAPL | 0.6842569 |
| IGLVFYGV | ASGAPRDRGYNNQAPL | 0.36785114 |
| FLLDASNL | ASGAPRDRGYNNQAPL | 0.72716 |
| YQVGNFP | ASGAPRDRGYNNQAPL | 0.52038026 |
| ESLLSQST | ASGAPRDRGYNNQAPL | 0.59998554 |
| LFTVGFAAVG | ASGAPRDRGYNNQAPL | 0.71939576 |
| ATPPTPT | ASGAPRDRGYNNQAPL | 0.71079737 |
| TDLATLG | ASGAPRDRGYNNQAPL | 0.570053 |
| SAGGTGR | ASGAPRDRGYNNQAPL | 0.5908716 |
| GFSNSGS | ASGAPRDRGYNNQAPL | 0.42083 |
| GGNGGSGGTSV | ASGAPRDRGYNNQAPL | 0.51356375 |
| GTSVATGGAGN | ASGAPRDRGYNNQAPL | 0.5180372 |
| NGGNGGSGGTS | ASGAPRDRGYNNQAPL | 0.72883165 |
| AVVKFPH | ASGAPRDRGYNNQAPL | 0.67501825 |
| TPPPPPP | ASGAPRDRGYNNQAPL | 0.40603334 |
| AAPAAAAAPA | ASGAPRDRGYNNQAPL | 0.6927563 |
| IALNEDL | ASGAPRDRGYNNQAPL | 0.7350773 |
| IVNVSSM | ASGAPRDRGYNNQAPL | 0.7263242 |
| TGIPPAPR | ASGAPRDRGYNNQAPL | 0.5466045 |
| FDLGGGTF | ASGAPRDRGYNNQAPL | 0.5626643 |
| FDLGGGTFDVS | ASGAPRDRGYNNQAPL | 0.61871636 |
| NEPTAAA | ASGAPRDRGYNNQAPL | 0.6220181 |

| | | |
|---|---|---|
| NEPTAAALA | ASGAPRDRGYNNQAPL | 0.6053251 |
| GSIINISSI | ASGAPRDRGYNNQAPL | 0.46506816 |
| VAPGNAGTA | ASGAPRDRGYNNQAPL | 0.72753435 |
| TGGGASLE | ASGAPRDRGYNNQAPL | 0.7487885 |
| VAVYNHYKRI | ASGAPRDRGYNNQAPL | 0.6006336 |
| TDPAPATT | ASGAPRDRGYNNQAPL | 0.2831195 |
| TNRPDILDPALLRPG | ASGAPRDRGYNNQAPL | 0.6178449 |
| LTTTTTS | ASGAPRDRGYNNQAPL | 0.48097587 |
| PTGSGKTL | ASGAPRDRGYNNQAPL | 0.5414362 |
| TGSGKTL | ASGAPRDRGYNNQAPL | 0.5971626 |
| TIGHVDHGK | ASGAPRDRGYNNQAPL | 0.39585826 |
| TIGHVDHGKTT | ASGAPRDRGYNNQAPL | 0.6265936 |
| GHVDHGKTT | ASGAPRDRGYNNQAPL | 0.7195053 |
| AQFEHTLL | ASGAPRDRGYNNQAPL | 0.56330144 |
| IVNQDEL | ASGAPRDRGYNNQAPL | 0.68578315 |
| TNGSQFF | ASGAPRDRGYNNQAPL | 0.7264716 |
| TGKGIVE | ASGAPRDRGYNNQAPL | 0.70212924 |
| LISNASDA | ASGAPRDRGYNNQAPL | 0.72648335 |
| AELRTLGELTKIAKAHGAQ | ASGAPRDRGYNNQAPL | 0.73440766 |
| APNAVAAAEPSW | ASGAPRDRGYNNQAPL | 0.64611506 |
| APQVAGAATAVL | ASGAPRDRGYNNQAPL | 0.668041 |
| ARRDPAAAAAILPTDA | ASGAPRDRGYNNQAPL | 0.72190917 |
| DAAARDVLAEAGLAEYF | ASGAPRDRGYNNQAPL | 0.6358959 |
| DDQIVLSAKANLAKVEQ | ASGAPRDRGYNNQAPL | 0.621084 |
| DELLPDTGEDSDEDGH | ASGAPRDRGYNNQAPL | 0.59985924 |
| ELPEAPANRATVPQD | ASGAPRDRGYNNQAPL | 0.71087474 |
| FGQSKIERVTGIG | ASGAPRDRGYNNQAPL | 0.6663165 |
| GAGGTGSTAGGAGGAGGAGGLY | ASGAPRDRGYNNQAPL | 0.7408997 |
| GHGGTGAAGGNGGTGGAGGSGI | ASGAPRDRGYNNQAPL | 0.30400562 |
| GNGGNGGAGGEGGPGGAGGAGA | ASGAPRDRGYNNQAPL | 0.5046162 |
| GSTAGNGGNGGNATNGTIAGQP | ASGAPRDRGYNNQAPL | 0.71942633 |
| GTGGDGGNGGNGGTGG | ASGAPRDRGYNNQAPL | 0.2648235 |
| GVRRPAGIPADAPV | ASGAPRDRGYNNQAPL | 0.5328867 |
| GYRQVTDAHLLTLARR | ASGAPRDRGYNNQAPL | 0.6934474 |
| IAKMGQDGHDRG | ASGAPRDRGYNNQAPL | 0.72203356 |
| KAEPSIEIQYT | ASGAPRDRGYNNQAPL | 0.28102282 |
| KPVRIRMAADEDTVAVT | ASGAPRDRGYNNQAPL | 0.5826926 |
| MPPRIAGMRLLVIKPEP | ASGAPRDRGYNNQAPL | 0.36808363 |
| MKFYAVALFRSRGVPPTR | ASGAPRDRGYNNQAPL | 0.70250964 |
| NAGGFGILGAVAHSP | ASGAPRDRGYNNQAPL | 0.6120926 |
| NEGREITLAMAEM | ASGAPRDRGYNNQAPL | 0.29057056 |
| NELVEWSHPLGHRWRWE | ASGAPRDRGYNNQAPL | 0.7133738 |
| NLGAGGTGGNGGNPG | ASGAPRDRGYNNQAPL | 0.35491982 |
| NTEWPKWETVMAADPVD | ASGAPRDRGYNNQAPL | 0.6880355 |
| PASAAPAETQP | ASGAPRDRGYNNQAPL | 0.7070621 |
| QQSSDGKNGVRAAEITGE | ASGAPRDRGYNNQAPL | 0.7418009 |
| RAPDAVTTSWWKE | ASGAPRDRGYNNQAPL | 0.45528477 |
| SGFGNVGAHTSGWFNQ | ASGAPRDRGYNNQAPL | 0.70366246 |
| SGVAGAVGQAASVGGLKV | ASGAPRDRGYNNQAPL | 0.6340414 |
| SNGGAGTGSAGGNGGT | ASGAPRDRGYNNQAPL | 0.7369535 |
| TGGAGGEGGAAG | ASGAPRDRGYNNQAPL | 0.4223267 |

| | | |
|---|---|---|
| TGGDGGAAGAGGAPGHG | ASGAPRDRGYNNQAPL | 0.400157 |
| TMPPLLAAIANAGRHGV | ASGAPRDRGYNNQAPL | 0.7111044 |
| TSSTAGNSGGA | ASGAPRDRGYNNQAPL | 0.681047 |
| VPATVPVGILC | ASGAPRDRGYNNQAPL | 0.6133734 |
| VQRPDSGGVVLTGRIS | ASGAPRDRGYNNQAPL | 0.7300298 |
| WTPNPVPATKGVPAGLVQAT | ASGAPRDRGYNNQAPL | 0.6211045 |
| YALKRALAKFSDRFDVVII | ASGAPRDRGYNNQAPL | 0.6157518 |
| YLIGSVIV | ASGAPRDRGYNNQAPL | 0.71368766 |
| YGQLVAVID | ASGAPRDRGYNNQAPL | 0.7083421 |
| VTVGGMTLPA | ASGAPRDRGYNNQAPL | 0.5216985 |
| VLALPAV | ASGAPRDRGYNNQAPL | 0.70909667 |
| VIPPTLNYET | ASGAPRDRGYNNQAPL | 0.6616982 |
| VGMGGMHPSSG | ASGAPRDRGYNNQAPL | 0.5017302 |
| VGLAGGAATA | ASGAPRDRGYNNQAPL | 0.71885204 |
| VGGAGGAGGSGP | ASGAPRDRGYNNQAPL | 0.64282554 |
| VFAAFMV | ASGAPRDRGYNNQAPL | 0.7152021 |
| TLYDKLR | ASGAPRDRGYNNQAPL | 0.6411096 |
| TLAVFTVL | ASGAPRDRGYNNQAPL | 0.74437726 |
| TGSVQPARV | ASGAPRDRGYNNQAPL | 0.29466763 |
| TGGGAGSGGT | ASGAPRDRGYNNQAPL | 0.6671232 |
| SHLETSL | ASGAPRDRGYNNQAPL | 0.61435205 |
| SFTTGMT | ASGAPRDRGYNNQAPL | 0.7087444 |
| QVEVAKRMGVRQPTVS | ASGAPRDRGYNNQAPL | 0.6263554 |
| QGSGGPALAM | ASGAPRDRGYNNQAPL | 0.5291656 |
| MLAPSNP | ASGAPRDRGYNNQAPL | 0.68539447 |
| LVNNAAI | ASGAPRDRGYNNQAPL | 0.7238735 |
| LTGEINTLA | ASGAPRDRGYNNQAPL | 0.68440783 |
| LSIFGPV | ASGAPRDRGYNNQAPL | 0.69183296 |
| LLALLANL | ASGAPRDRGYNNQAPL | 0.7330692 |
| LISNASDALDKLR | ASGAPRDRGYNNQAPL | 0.48581064 |
| LINEAVP | ASGAPRDRGYNNQAPL | 0.65997446 |
| LGYAFSGL | ASGAPRDRGYNNQAPL | 0.7102964 |
| LDFVAVAL | ASGAPRDRGYNNQAPL | 0.7271819 |
| LAQPGAE | ASGAPRDRGYNNQAPL | 0.6423288 |
| ISNLGAVTGF | ASGAPRDRGYNNQAPL | 0.73588675 |
| HQEDPGRI | ASGAPRDRGYNNQAPL | 0.51622903 |
| GTPGEPTP | ASGAPRDRGYNNQAPL | 0.7113695 |
| GFFNSGS | ASGAPRDRGYNNQAPL | 0.61280936 |
| GAGGGGLTAGAN | ASGAPRDRGYNNQAPL | 0.67054546 |
| FVDALVNM | ASGAPRDRGYNNQAPL | 0.52455807 |
| FLTVAPEL | ASGAPRDRGYNNQAPL | 0.6738125 |
| FETLTAL | ASGAPRDRGYNNQAPL | 0.70780015 |
| DVINPIEL | ASGAPRDRGYNNQAPL | 0.71294945 |
| APVSGAPV | ASGAPRDRGYNNQAPL | 0.37638748 |
| ALVGGLAW | ASGAPRDRGYNNQAPL | 0.7026448 |
| ALAAAAATAI | ASGAPRDRGYNNQAPL | 0.72282815 |
| AFGLGLLL | ASGAPRDRGYNNQAPL | 0.7434337 |
| AFFAAASL | ASGAPRDRGYNNQAPL | 0.6067748 |
| AAAALPAFSPP | ASGAPRDRGYNNQAPL | 0.7005545 |
| VVLAPAL | ASGAPRDRGYNNQAPL | 0.6118233 |
| TIGHVDHGKTTL | ASGAPRDRGYNNQAPL | 0.6687592 |

| | | |
|---|---|---|
| LISNASDALDK | ASGAPRDRGYNNQAPL | 0.5034044 |
| LISNASDAL | ASGAPRDRGYNNQAPL | 0.5406991 |
| ISNASDALDK | ASGAPRDRGYNNQAPL | 0.65432024 |
| NQAETLVYQTE | ASGAPRDRGYNNQAPL | 0.5431787 |
| TGGTGGTAGAG | ASGAPRDRGYNNQAPL | 0.73268414 |
| TGSTAGGAGGA | ASGAPRDRGYNNQAPL | 0.725425 |
| DPLAVPVAAGPV | ASGAPRDRGYNNQAPL | 0.671093 |
| GGTGGTPGNGGN | ASGAPRDRGYNNQAPL | 0.72708404 |
| PAGLAGVDPATL | ASGAPRDRGYNNQAPL | 0.7305746 |
| GNNLVVEAPHAGASV | ASGAPRDRGYNNQAPL | 0.72490585 |
| KVTRIMSASTLEQPA | ASGAPRDRGYNNQAPL | 0.5205321 |
| LGFALPIALPRRTVV | ASGAPRDRGYNNQAPL | 0.6003042 |
| TRLGNLTARKLVVLV | ASGAPRDRGYNNQAPL | 0.42936456 |
| WSQQRRLIYFPSAGP | ASGAPRDRGYNNQAPL | 0.60770535 |
| GPNGNPDPMAAAVDIR | ASGAPRDRGYNNQAPL | 0.5415522 |
| TQRCVSVADLRAAASR | ASGAPRDRGYNNQAPL | 0.67465544 |
| QFVLDVAEHPVEVLTETV | ASGAPRDRGYNNQAPL | 0.25082445 |
| GGAGGTGGAGGNGGTQAGDG | ASGAPRDRGYNNQAPL | 0.72024924 |
| PLLATLRAIAVDVDAKPAQV | ASGAPRDRGYNNQAPL | 0.7303544 |
| APKFRTGGYMIDFWGVGYQVAK | ASGAPRDRGYNNQAPL | 0.52425987 |
| GAGGAGGAGGAAGGPGGTGGTG | ASGAPRDRGYNNQAPL | 0.6973774 |
| IGALGLLATDVAPAGSVAFGVA | ASGAPRDRGYNNQAPL | 0.68854433 |
| AGGAGGIGGNGGAGGAGGVLMG | ASGAPRDRGYNNQAPL | 0.60069215 |
| ADELIGLL | ASGAPRDRGYNNQAPL | 0.70495594 |
| ADRALKA | ASGAPRDRGYNNQAPL | 0.5762781 |
| AGAVGQVGGAGGA | ASGAPRDRGYNNQAPL | 0.7209337 |
| AGGGGGLGGAP | ASGAPRDRGYNNQAPL | 0.69493854 |
| AGGLLYGNGGAG | ASGAPRDRGYNNQAPL | 0.65623844 |
| AGLESIGL | ASGAPRDRGYNNQAPL | 0.6558644 |
| AGLKTGII | ASGAPRDRGYNNQAPL | 0.6783501 |
| ASLAALLL | ASGAPRDRGYNNQAPL | 0.27993837 |
| ASVINISSI | ASGAPRDRGYNNQAPL | 0.6732284 |
| ATPPGPA | ASGAPRDRGYNNQAPL | 0.42766243 |
| ATVASALV | ASGAPRDRGYNNQAPL | 0.6357884 |
| ATVASLIL | ASGAPRDRGYNNQAPL | 0.37793067 |
| AVNGPVLIRH | ASGAPRDRGYNNQAPL | 0.6855675 |
| DGAAGAVGI | ASGAPRDRGYNNQAPL | 0.6388843 |
| DPLLHGI | ASGAPRDRGYNNQAPL | 0.710243 |
| ELGAALGAL | ASGAPRDRGYNNQAPL | 0.37063625 |
| ELGRTQAV | ASGAPRDRGYNNQAPL | 0.7147232 |
| FVAGGYLS | ASGAPRDRGYNNQAPL | 0.7022728 |
| GASGGAGGAGGW | ASGAPRDRGYNNQAPL | 0.37066585 |
| GGAGAGGDGGAGGI | ASGAPRDRGYNNQAPL | 0.70638454 |
| GTGHNGSQPGMGGNG | ASGAPRDRGYNNQAPL | 0.6587963 |
| GYAAASASAS | ASGAPRDRGYNNQAPL | 0.59152734 |
| HQVDVAAVQQG | ASGAPRDRGYNNQAPL | 0.70371145 |
| IGAAIAHT | ASGAPRDRGYNNQAPL | 0.61862767 |
| IGMIEVAQL | ASGAPRDRGYNNQAPL | 0.30829644 |
| IRLAPEHTSAMD | ASGAPRDRGYNNQAPL | 0.5736915 |
| IYNVSVL | ASGAPRDRGYNNQAPL | 0.66809076 |
| KQGFLGM | ASGAPRDRGYNNQAPL | 0.40421373 |

| | | |
|---|---|---|
| LESVITQL | ASGAPRDRGYNNQAPL | 0.29275185 |
| LGLGALVVV | ASGAPRDRGYNNQAPL | 0.68480074 |
| LIGNGGNGGSGGI | ASGAPRDRGYNNQAPL | 0.5056505 |
| LNAINAPTL | ASGAPRDRGYNNQAPL | 0.454155 |
| LQLLNTGV | ASGAPRDRGYNNQAPL | 0.6065155 |
| LQTSIEF | ASGAPRDRGYNNQAPL | 0.7211003 |
| MSFAVSEL | ASGAPRDRGYNNQAPL | 0.6407523 |
| NTGALGL | ASGAPRDRGYNNQAPL | 0.6914331 |
| PPGAGGAGGAGTT | ASGAPRDRGYNNQAPL | 0.7047445 |
| QLVGLGI | ASGAPRDRGYNNQAPL | 0.6381669 |
| RFLEQVHQL | ASGAPRDRGYNNQAPL | 0.72698903 |
| RGVGYVLR | ASGAPRDRGYNNQAPL | 0.69384676 |
| SATGVFAGVI | ASGAPRDRGYNNQAPL | 0.6484869 |
| SLPAITLP | ASGAPRDRGYNNQAPL | 0.69297355 |
| STALAFTV | ASGAPRDRGYNNQAPL | 0.6501994 |
| SVLTVGLGK | ASGAPRDRGYNNQAPL | 0.64314646 |
| TALAGAGSGI | ASGAPRDRGYNNQAPL | 0.6020361 |
| TGLPFTGI | ASGAPRDRGYNNQAPL | 0.45242667 |
| VAEHVTV | ASGAPRDRGYNNQAPL | 0.59999824 |
| VDIVLNSL | ASGAPRDRGYNNQAPL | 0.594387 |
| VLTGGGAVL | ASGAPRDRGYNNQAPL | 0.7287588 |
| VNVGLAYGI | ASGAPRDRGYNNQAPL | 0.60088086 |
| VSSAAVGAV | ASGAPRDRGYNNQAPL | 0.3866721 |
| VTSVYEAGV | ASGAPRDRGYNNQAPL | 0.63043463 |
| WVALDRGA | ASGAPRDRGYNNQAPL | 0.33769718 |
| YLVGSVVI | ASGAPRDRGYNNQAPL | 0.7075573 |
| GPPGTGKTLL | ASGAPRDRGYNNQAPL | 0.62294054 |
| GVGMTKF | ASGAPRDRGYNNQAPL | 0.74610955 |
| IGIGGSDL | ASGAPRDRGYNNQAPL | 0.71707666 |
| IGTLLAEL | ASGAPRDRGYNNQAPL | 0.58300567 |
| IYTSGTT | ASGAPRDRGYNNQAPL | 0.71910113 |
| LSFPAGV | ASGAPRDRGYNNQAPL | 0.5325037 |
| LYTSGTT | ASGAPRDRGYNNQAPL | 0.6553236 |
| VGAFPFGF | ASGAPRDRGYNNQAPL | 0.6518202 |
| VGAGTVEF | ASGAPRDRGYNNQAPL | 0.67946255 |
| VGKSTLLL | ASGAPRDRGYNNQAPL | 0.6329603 |
| AALAHAGGGIV | ASGAPRDRGYNNQAPL | 0.62631726 |
| AALVAVSLL | ASGAPRDRGYNNQAPL | 0.29162952 |
| AAVSRAVAAL | ASGAPRDRGYNNQAPL | 0.7325099 |
| AGGSSGIGGF | ASGAPRDRGYNNQAPL | 0.3681544 |
| AGLTVANANG | ASGAPRDRGYNNQAPL | 0.5658363 |
| AGLVGAVL | ASGAPRDRGYNNQAPL | 0.566836 |
| ALSQAFTV | ASGAPRDRGYNNQAPL | 0.72670776 |
| ALTFSVSL | ASGAPRDRGYNNQAPL | 0.73483074 |
| ALTGAALVAL | ASGAPRDRGYNNQAPL | 0.529596 |
| ASFSIGAGVL | ASGAPRDRGYNNQAPL | 0.44897467 |
| DGSVYKI | ASGAPRDRGYNNQAPL | 0.72276926 |
| EQLAEIV | ASGAPRDRGYNNQAPL | 0.51157725 |
| FAVSELTVM | ASGAPRDRGYNNQAPL | 0.70137584 |
| GAAPLAAGL | ASGAPRDRGYNNQAPL | 0.6514504 |
| GADGGDGGSGGAGG | ASGAPRDRGYNNQAPL | 0.6128001 |

| | | |
|---|---|---|
| GFGLDTGGAGG | ASGAPRDRGYNNQAPL | 0.57184565 |
| GLASGGLL | ASGAPRDRGYNNQAPL | 0.711378 |
| GRIAGAAVLL | ASGAPRDRGYNNQAPL | 0.5669999 |
| GSAGSLNTSF | ASGAPRDRGYNNQAPL | 0.6958925 |
| GTGQIGI | ASGAPRDRGYNNQAPL | 0.30853412 |
| GTLIALIL | ASGAPRDRGYNNQAPL | 0.68510306 |
| IAATMGFVVG | ASGAPRDRGYNNQAPL | 0.51983297 |
| IFAQALL | ASGAPRDRGYNNQAPL | 0.54467684 |
| IGAAVALGF | ASGAPRDRGYNNQAPL | 0.33709452 |
| IGLTPLAL | ASGAPRDRGYNNQAPL | 0.64950764 |
| ILGSNTSTL | ASGAPRDRGYNNQAPL | 0.6876555 |
| IVFSLTVL | ASGAPRDRGYNNQAPL | 0.7257881 |
| LASKATGF | ASGAPRDRGYNNQAPL | 0.66917574 |
| LASVALIL | ASGAPRDRGYNNQAPL | 0.7074005 |
| LFAVGLTL | ASGAPRDRGYNNQAPL | 0.7160604 |
| LFNGGIGTYI | ASGAPRDRGYNNQAPL | 0.5902705 |
| LTLGAGII | ASGAPRDRGYNNQAPL | 0.27980542 |
| LVAAALGDI | ASGAPRDRGYNNQAPL | 0.640678 |
| NTAFVQLGI | ASGAPRDRGYNNQAPL | 0.6984994 |
| QLFSGAEL | ASGAPRDRGYNNQAPL | 0.71009266 |
| SAATFITL | ASGAPRDRGYNNQAPL | 0.6584488 |
| SNYGLANLGV | ASGAPRDRGYNNQAPL | 0.43097734 |
| SYGGAYAVM | ASGAPRDRGYNNQAPL | 0.67670697 |
| TTGVGGLTLG | ASGAPRDRGYNNQAPL | 0.69703937 |
| TTQTIYI | ASGAPRDRGYNNQAPL | 0.412732 |
| VAAGAGLGAVI | ASGAPRDRGYNNQAPL | 0.56651175 |
| VAAVVTLL | ASGAPRDRGYNNQAPL | 0.5582833 |
| VGFLGGLI | ASGAPRDRGYNNQAPL | 0.38744652 |
| VGVIPILL | ASGAPRDRGYNNQAPL | 0.653365 |
| VIFASSLI | ASGAPRDRGYNNQAPL | 0.5883957 |
| VKGGDGGAAAG | ASGAPRDRGYNNQAPL | 0.6287447 |
| VTKTFTNI | ASGAPRDRGYNNQAPL | 0.6174482 |
| VVAAAILL | ASGAPRDRGYNNQAPL | 0.5563544 |
| YAALDVEL | ASGAPRDRGYNNQAPL | 0.61686647 |
| LASVALLL | ASGAPRDRGYNNQAPL | 0.59473944 |
| LDAVHRV | ASGAPRDRGYNNQAPL | 0.59734344 |
| QGLSSSIF | ASGAPRDRGYNNQAPL | 0.71289134 |
| RGTFANIRL | ASGAPRDRGYNNQAPL | 0.6034919 |
| RQLWWGHRI | ASGAPRDRGYNNQAPL | 0.68849254 |
| YGPPGTGKTLL | ASGAPRDRGYNNQAPL | 0.7291954 |
| AALVGAPANVA | ASGAPRDRGYNNQAPL | 0.42727545 |
| AATFRRTE | ASGAPRDRGYNNQAPL | 0.5723096 |
| AGALGGGI | ASGAPRDRGYNNQAPL | 0.70575905 |
| AGGAGGAGGIGAG | ASGAPRDRGYNNQAPL | 0.57199514 |
| AGSGALGL | ASGAPRDRGYNNQAPL | 0.7268337 |
| AQIADQI | ASGAPRDRGYNNQAPL | 0.6603079 |
| ASAVAVTYL | ASGAPRDRGYNNQAPL | 0.69147164 |
| ASTVLGGTGG | ASGAPRDRGYNNQAPL | 0.70165277 |
| AVGGEPAK | ASGAPRDRGYNNQAPL | 0.732908 |
| CEQGTFGGTC | ASGAPRDRGYNNQAPL | 0.69578516 |
| DEGKAPAG | ASGAPRDRGYNNQAPL | 0.6252557 |

| | | |
|---|---|---|
| DTVVFPAL | ASGAPRDRGYNNQAPL | 0.7250135 |
| FEGITMAG | ASGAPRDRGYNNQAPL | 0.4994967 |
| FPLVVQD | ASGAPRDRGYNNQAPL | 0.71867967 |
| GAFTTPQ | ASGAPRDRGYNNQAPL | 0.71671534 |
| GFGGFGTAGGVGGA | ASGAPRDRGYNNQAPL | 0.39313048 |
| GFLAQFL | ASGAPRDRGYNNQAPL | 0.6232939 |
| GGAGGAGGNAGVA | ASGAPRDRGYNNQAPL | 0.73563075 |
| GGVGTAGAGGAGGAGG | ASGAPRDRGYNNQAPL | 0.72589034 |
| GNGGGGDAGNAGSGGN | ASGAPRDRGYNNQAPL | 0.45219195 |
| GTGGAGGAGNDAGS | ASGAPRDRGYNNQAPL | 0.5919274 |
| GTTNTGWFNS | ASGAPRDRGYNNQAPL | 0.51958954 |
| IFMGPLFTL | ASGAPRDRGYNNQAPL | 0.5937914 |
| ILAHAIEAI | ASGAPRDRGYNNQAPL | 0.3802849 |
| IRSAIREAL | ASGAPRDRGYNNQAPL | 0.70041895 |
| ITFAGGG | ASGAPRDRGYNNQAPL | 0.5805334 |
| ITGMSGHQEV | ASGAPRDRGYNNQAPL | 0.7387433 |
| IVAGLACTL | ASGAPRDRGYNNQAPL | 0.59659314 |
| IVGGGLAGA | ASGAPRDRGYNNQAPL | 0.6994091 |
| IVTGVLNI | ASGAPRDRGYNNQAPL | 0.5638014 |
| LAQQGCI | ASGAPRDRGYNNQAPL | 0.6155055 |
| LCLHVEN | ASGAPRDRGYNNQAPL | 0.3887553 |
| LDAISDGS | ASGAPRDRGYNNQAPL | 0.3411997 |
| LGGFAAML | ASGAPRDRGYNNQAPL | 0.56772935 |
| LGGKNPMI | ASGAPRDRGYNNQAPL | 0.65027934 |
| LGPKGVAAIL | ASGAPRDRGYNNQAPL | 0.314476 |
| LMVGVSI | ASGAPRDRGYNNQAPL | 0.7311126 |
| LVGAAFVL | ASGAPRDRGYNNQAPL | 0.65636146 |
| LVLAVAPL | ASGAPRDRGYNNQAPL | 0.6703086 |
| MAAIAVIDAA | ASGAPRDRGYNNQAPL | 0.5176103 |
| NNGINAELQ | ASGAPRDRGYNNQAPL | 0.55106604 |
| RGEPLVLVHG | ASGAPRDRGYNNQAPL | 0.5645455 |
| RGGAGGAAGLI | ASGAPRDRGYNNQAPL | 0.721184 |
| RSYRMHPAV | ASGAPRDRGYNNQAPL | 0.40169585 |
| SAAHLLAGQLA | ASGAPRDRGYNNQAPL | 0.28789148 |
| SAAVGSY | ASGAPRDRGYNNQAPL | 0.6339203 |
| SPHAVLHRI | ASGAPRDRGYNNQAPL | 0.43210715 |
| SQIAKVL | ASGAPRDRGYNNQAPL | 0.73095715 |
| TALGHLVP | ASGAPRDRGYNNQAPL | 0.7162477 |
| TAPTTSGAA | ASGAPRDRGYNNQAPL | 0.6718694 |
| TDGDDGNGGSPGT | ASGAPRDRGYNNQAPL | 0.7404984 |
| TDGLGEN | ASGAPRDRGYNNQAPL | 0.2556188 |
| TGGPPLPGQA | ASGAPRDRGYNNQAPL | 0.5840708 |
| TIPAITI | ASGAPRDRGYNNQAPL | 0.7342286 |
| TIPGFTI | ASGAPRDRGYNNQAPL | 0.6513747 |
| TSIASGALA | ASGAPRDRGYNNQAPL | 0.58943224 |
| TVTLPGAAI | ASGAPRDRGYNNQAPL | 0.72274935 |
| VASGLSAALP | ASGAPRDRGYNNQAPL | 0.5331464 |
| VGAGVIGI | ASGAPRDRGYNNQAPL | 0.4355383 |
| VGPKRFGV | ASGAPRDRGYNNQAPL | 0.6689886 |
| VNVAFPD | ASGAPRDRGYNNQAPL | 0.68973804 |
| VSMTNSGVS | ASGAPRDRGYNNQAPL | 0.68514776 |

| | | |
|---|---|---|
| VTDVLGDLSV | ASGAPRDRGYNNQAPL | 0.37229425 |
| VVVVGAGFAGL | ASGAPRDRGYNNQAPL | 0.68691206 |
| VYTVGTVV | ASGAPRDRGYNNQAPL | 0.73981404 |
| WTTFSGP | ASGAPRDRGYNNQAPL | 0.6218069 |
| YDELVEAF | ASGAPRDRGYNNQAPL | 0.6347741 |
| YGTGGAGGNGG | ASGAPRDRGYNNQAPL | 0.66599274 |
| YTNVMAV | ASGAPRDRGYNNQAPL | 0.66061723 |
| FLEQVHQL | ASGAPRDRGYNNQAPL | 0.47646385 |
| GLSLLAVAV | ASGAPRDRGYNNQAPL | 0.47973824 |
| LEQVHQL | ASGAPRDRGYNNQAPL | 0.712538 |
| LNAAGAI | ASGAPRDRGYNNQAPL | 0.5313567 |
| PAVHASL | ASGAPRDRGYNNQAPL | 0.67889905 |
| TELEDDA | ASGAPRDRGYNNQAPL | 0.60802597 |
| TPPGPPN | ASGAPRDRGYNNQAPL | 0.7441633 |
| TPPPPPPP | ASGAPRDRGYNNQAPL | 0.7127665 |
| TQSPALI | ASSYRNTEVF | 0.6843228 |
| EFAAAFPQ | ASSYRNTEVF | 0.5555885 |
| LLSGNLDVL | ASSYRNTEVF | 0.69850016 |
| GPANLGNYNI | ASSYRNTEVF | 0.697824 |
| TLGPTLP | ASSYRNTEVF | 0.27865532 |
| VYAYSSCCK | ASSYRNTEVF | 0.7306784 |
| APAPAPAPNI | ASSYRNTEVF | 0.48500663 |
| PGGPVPIP | ASSYRNTEVF | 0.6394136 |
| FLTVALEP | ASSYRNTEVF | 0.71192694 |
| SQYFGANL | ASSYRNTEVF | 0.74287176 |
| GGVKPAA | ASSYRNTEVF | 0.64732254 |
| GGTGGAGGAGGW | ASSYRNTEVF | 0.7130866 |
| GVGGVAGLGGAGG | ASSYRNTEVF | 0.521212 |
| VLLVLATTI | ASSYRNTEVF | 0.7157248 |
| IGIFAAML | ASSYRNTEVF | 0.72114855 |
| AVGDLID | ASSYRNTEVF | 0.68496597 |
| NISPLPL | ASSYRNTEVF | 0.7410628 |
| VTAVFVAI | ASSYRNTEVF | 0.66522366 |
| SGDDNDAAL | ASSYRNTEVF | 0.347962 |
| AVAALAHS | ASSYRNTEVF | 0.61449546 |
| GGVTVMATP | ASSYRNTEVF | 0.7072556 |
| AGLAANVRD | ASSYRNTEVF | 0.72715294 |
| TNRPDLIDPALLRPG | ASSYRNTEVF | 0.66541773 |
| LANLGVTAL | ASSYRNTEVF | 0.71275544 |
| TGERIEL | ASSYRNTEVF | 0.5116075 |
| NGEELAA | ASSYRNTEVF | 0.7287781 |
| VLSLVWG | ASSYRNTEVF | 0.7376816 |
| IGYALWL | ASSYRNTEVF | 0.68833005 |
| LSLLLAPL | ASSYRNTEVF | 0.6077639 |
| TLDEALK | ASSYRNTEVF | 0.5470214 |
| TDGAGTAEGG | ASSYRNTEVF | 0.69578624 |
| NAPGGAGGNGG | ASSYRNTEVF | 0.7069737 |
| VSVFMGMGGNF | ASSYRNTEVF | 0.5099704 |
| LASYLAAL | ASSYRNTEVF | 0.67512333 |
| LVGGTPAL | ASSYRNTEVF | 0.6595446 |
| VRNADVL | ASSYRNTEVF | 0.32953766 |

| | | |
|---|---|---|
| TCSSPFLVSAM | ASSYRNTEVF | 0.67114615 |
| VHITDAE | ASSYRNTEVF | 0.7258885 |
| EYAAPSF | ASSYRNTEVF | 0.6686628 |
| VVDAFHGGL | ASSYRNTEVF | 0.49281877 |
| ASVVDPQ | ASSYRNTEVF | 0.7327614 |
| SYNEFGGLS | ASSYRNTEVF | 0.7325422 |
| DPDGSVY | ASSYRNTEVF | 0.7337703 |
| MVSTIHATE | ASSYRNTEVF | 0.42019123 |
| LAAGVADPV | ASSYRNTEVF | 0.28593114 |
| TKEQIAA | ASSYRNTEVF | 0.6875391 |
| SLASPAAL | ASSYRNTEVF | 0.7297791 |
| IGLVFYGV | ASSYRNTEVF | 0.5885887 |
| FLLDASNL | ASSYRNTEVF | 0.5411848 |
| YQVGNFP | ASSYRNTEVF | 0.6574007 |
| ESLLSQST | ASSYRNTEVF | 0.70918113 |
| LFTVGFAAVG | ASSYRNTEVF | 0.48030096 |
| ATPPTPT | ASSYRNTEVF | 0.4547627 |
| TDLATLG | ASSYRNTEVF | 0.6825663 |
| SAGGTGR | ASSYRNTEVF | 0.7250136 |
| GFSNSGS | ASSYRNTEVF | 0.580118 |
| GGNGGSGGTSV | ASSYRNTEVF | 0.5929735 |
| GTSVATGGAGN | ASSYRNTEVF | 0.65748805 |
| NGGNGGSGGTS | ASSYRNTEVF | 0.6723813 |
| AVVKFPH | ASSYRNTEVF | 0.64740825 |
| TPPPPPP | ASSYRNTEVF | 0.42790276 |
| AAPAAAAAPA | ASSYRNTEVF | 0.70874226 |
| IALNEDL | ASSYRNTEVF | 0.41127056 |
| IVNVSSM | ASSYRNTEVF | 0.7210648 |
| TGIPPAPR | ASSYRNTEVF | 0.68314385 |
| FDLGGGTF | ASSYRNTEVF | 0.642419 |
| FDLGGGTFDVS | ASSYRNTEVF | 0.6092854 |
| NEPTAAA | ASSYRNTEVF | 0.5214226 |
| NEPTAAALA | ASSYRNTEVF | 0.35927045 |
| GSIINISSI | ASSYRNTEVF | 0.5989497 |
| VAPGNAGTA | ASSYRNTEVF | 0.72555757 |
| TGGGASLE | ASSYRNTEVF | 0.58677924 |
| VAVYNHYKRI | ASSYRNTEVF | 0.7119466 |
| TDPAPATT | ASSYRNTEVF | 0.64607537 |
| TNRPDILDPALLRPG | ASSYRNTEVF | 0.4844914 |
| LTTTTTS | ASSYRNTEVF | 0.7075013 |
| PTGSGKTL | ASSYRNTEVF | 0.4514734 |
| TGSGKTL | ASSYRNTEVF | 0.69574153 |
| TIGHVDHGK | ASSYRNTEVF | 0.6387507 |
| TIGHVDHGKTT | ASSYRNTEVF | 0.6988414 |
| GHVDHGKTT | ASSYRNTEVF | 0.50729585 |
| AQFEHTLL | ASSYRNTEVF | 0.7004223 |
| IVNQDEL | ASSYRNTEVF | 0.65906155 |
| TNGSQFF | ASSYRNTEVF | 0.50006187 |
| TGKGIVE | ASSYRNTEVF | 0.67546034 |
| LISNASDA | ASSYRNTEVF | 0.7160008 |
| AELRTLGELTKIAKAHGAQ | ASSYRNTEVF | 0.6853787 |

| | | |
|---|---|---|
| APNAVAAAEPSW | ASSYRNTEVF | 0.38060963 |
| APQVAGAATAVL | ASSYRNTEVF | 0.6309337 |
| ARRDPAAAAAILPTDA | ASSYRNTEVF | 0.56400216 |
| DAAARDVLAEAGLAEYF | ASSYRNTEVF | 0.6640041 |
| DDQIVLSAKANLAKVEQ | ASSYRNTEVF | 0.64845693 |
| DELLPDTGEDSDEDGH | ASSYRNTEVF | 0.619642 |
| ELPEAPANRATVPQD | ASSYRNTEVF | 0.6622175 |
| FGQSKIERVTGIG | ASSYRNTEVF | 0.42071924 |
| GAGGTGSTAGGAGGAGGAGGLY | ASSYRNTEVF | 0.53333294 |
| GHGGTGAAGGNGGTGGAGGSGI | ASSYRNTEVF | 0.60646355 |
| GNGGNGGAGGEGGPGGAGGAGA | ASSYRNTEVF | 0.6668271 |
| GSTAGNGGNGGNATNGTIAGQP | ASSYRNTEVF | 0.6842054 |
| GTGGDGGNGGNGGTGG | ASSYRNTEVF | 0.38863993 |
| GVRRPAGIPADAPV | ASSYRNTEVF | 0.6530577 |
| GYRQVTDAHLLTLARR | ASSYRNTEVF | 0.48929363 |
| IAKMGQDGHDRG | ASSYRNTEVF | 0.35827044 |
| KAEPSIEIQYT | ASSYRNTEVF | 0.6786922 |
| KPVRIRMAADEDTVAVT | ASSYRNTEVF | 0.6547073 |
| MPPRIAGMRLLVIKPEP | ASSYRNTEVF | 0.47247976 |
| MKFYAVALFRSRGVPPTR | ASSYRNTEVF | 0.68994474 |
| NAGGFGILGAVAHSP | ASSYRNTEVF | 0.4790507 |
| NEGREITLAMAEM | ASSYRNTEVF | 0.65635973 |
| NELVEWSHPLGHRWRWE | ASSYRNTEVF | 0.69155025 |
| NLGAGGTGGNGGNPG | ASSYRNTEVF | 0.48365253 |
| NTEWPKWETVMAADPVD | ASSYRNTEVF | 0.71615446 |
| PASAAPAETQP | ASSYRNTEVF | 0.6286328 |
| QQSSDGKNGVRAAEITGE | ASSYRNTEVF | 0.69092035 |
| RAPDAVTTSWWKE | ASSYRNTEVF | 0.62663096 |
| SGFGNVGAHTSGWFNQ | ASSYRNTEVF | 0.46168667 |
| SGVAGAVGQAASVGGLKV | ASSYRNTEVF | 0.7002604 |
| SNGGAGTGSAGGNGGT | ASSYRNTEVF | 0.68227994 |
| TGGAGGEGGAAG | ASSYRNTEVF | 0.7167206 |
| TGGDGGAAGAGGAPGHG | ASSYRNTEVF | 0.6824693 |
| TMPPLLAAIANAGRHGV | ASSYRNTEVF | 0.72997254 |
| TSSTAGNSGGA | ASSYRNTEVF | 0.7157988 |
| VPATVPVGILC | ASSYRNTEVF | 0.5773209 |
| VQRPDSGGVVLTGRIS | ASSYRNTEVF | 0.3577081 |
| WTPNPVPATKGVPAGLVQAT | ASSYRNTEVF | 0.63084567 |
| YALKRALAKFSDRFDVVII | ASSYRNTEVF | 0.66051066 |
| YLIGSVIV | ASSYRNTEVF | 0.53993046 |
| YGQLVAVID | ASSYRNTEVF | 0.44146457 |
| VTVGGMTLPA | ASSYRNTEVF | 0.7188008 |
| VLALPAV | ASSYRNTEVF | 0.7393254 |
| VIPPTLNYET | ASSYRNTEVF | 0.6051073 |
| VGMGGMHPSSG | ASSYRNTEVF | 0.74066496 |
| VGLAGGAATA | ASSYRNTEVF | 0.4537866 |
| VGGAGGAGGSGP | ASSYRNTEVF | 0.6774125 |
| VFAAFMV | ASSYRNTEVF | 0.7343576 |
| TLYDKLR | ASSYRNTEVF | 0.49427497 |
| TLAVFTVL | ASSYRNTEVF | 0.67000985 |
| TGSVQPARV | ASSYRNTEVF | 0.37358826 |

| | | |
|---|---|---|
| TGGGAGSGGT | ASSYRNTEVF | 0.48328596 |
| SHLETSL | ASSYRNTEVF | 0.72429776 |
| SFTTGMT | ASSYRNTEVF | 0.61331224 |
| QVEVAKRMGVRQPTVS | ASSYRNTEVF | 0.700537 |
| QGSGGPALAM | ASSYRNTEVF | 0.64926255 |
| MLAPSNP | ASSYRNTEVF | 0.47592655 |
| LVNNAAI | ASSYRNTEVF | 0.6671082 |
| LTGEINTLA | ASSYRNTEVF | 0.72353065 |
| LSIFGPV | ASSYRNTEVF | 0.691941 |
| LLALLANL | ASSYRNTEVF | 0.73811305 |
| LISNASDALDKLR | ASSYRNTEVF | 0.46947837 |
| LINEAVP | ASSYRNTEVF | 0.72000885 |
| LGYAFSGL | ASSYRNTEVF | 0.73658335 |
| LDFVAVAL | ASSYRNTEVF | 0.6552769 |
| LAQPGAE | ASSYRNTEVF | 0.598354 |
| ISNLGAVTGF | ASSYRNTEVF | 0.7205728 |
| HQEDPGRI | ASSYRNTEVF | 0.6772015 |
| GTPGEPTP | ASSYRNTEVF | 0.7059617 |
| GFFNSGS | ASSYRNTEVF | 0.7278291 |
| GAGGGGLTAGAN | ASSYRNTEVF | 0.36503887 |
| FVDALVNM | ASSYRNTEVF | 0.8159605 |
| FLTVAPEL | ASSYRNTEVF | 0.706936 |
| FETLTAL | ASSYRNTEVF | 0.74690306 |
| DVINPIEL | ASSYRNTEVF | 0.5266179 |
| APVSGAPV | ASSYRNTEVF | 0.26620492 |
| ALVGGLAW | ASSYRNTEVF | 0.6260396 |
| ALAAAAATAI | ASSYRNTEVF | 0.458049 |
| AFGLGLLL | ASSYRNTEVF | 0.7686305 |
| AFFAAASL | ASSYRNTEVF | 0.7918533 |
| AAAALPAFSPP | ASSYRNTEVF | 0.5334992 |
| VVLAPAL | ASSYRNTEVF | 0.7211733 |
| TIGHVDHGKTTL | ASSYRNTEVF | 0.80152655 |
| LISNASDALDK | ASSYRNTEVF | 0.3152762 |
| LISNASDAL | ASSYRNTEVF | 0.6467713 |
| ISNASDALDK | ASSYRNTEVF | 0.34926745 |
| NQAETLVYQTE | ASSYRNTEVF | 0.60214853 |
| TGGTGGTAGAG | ASSYRNTEVF | 0.1920613 |
| TGSTAGGAGGA | ASSYRNTEVF | 0.7444528 |
| DPLAVPVAAGPV | ASSYRNTEVF | 0.5149448 |
| GGTGGTPGNGGN | ASSYRNTEVF | 0.19604093 |
| PAGLAGVDPATL | ASSYRNTEVF | 0.688666 |
| GNNLVVEAPHAGASV | ASSYRNTEVF | 0.54579926 |
| KVTRIMSASTLEQPA | ASSYRNTEVF | 0.55971503 |
| LGFALPIALPRRTVV | ASSYRNTEVF | 0.7540115 |
| TRLGNLTARKLVVLV | ASSYRNTEVF | 0.63939327 |
| WSQQRRLIYFPSAGP | ASSYRNTEVF | 0.4226398 |
| GPNGNPDPMAAAVDIR | ASSYRNTEVF | 0.6496403 |
| TQRCVSVADLRAAASR | ASSYRNTEVF | 0.7875848 |
| QFVLDVAEHPVEVLTETV | ASSYRNTEVF | 0.4420454 |
| GGAGGTGGAGGNGGTQAGDG | ASSYRNTEVF | 0.6483839 |
| PLLATLRAIAVDVDAKPAQV | ASSYRNTEVF | 0.8387672 |

| | | |
|---|---|---|
| APKFRTGGYMIDFWGVGYQVAK | ASSYRNTEVF | 0.67507327 |
| GAGGAGGAGGAAGGPGGTGGTG | ASSYRNTEVF | 0.82483935 |
| IGALGLLATDVAPAGSVAFGVA | ASSYRNTEVF | 0.8478142 |
| AGGAGGIGGNGGAGGAGGVLMG | ASSYRNTEVF | 0.5766659 |
| ADELIGLL | ASSYRNTEVF | 0.78095204 |
| ADRALKA | ASSYRNTEVF | 0.7105729 |
| AGAVGQVGGAGGA | ASSYRNTEVF | 0.66041327 |
| AGGGGGLGGAP | ASSYRNTEVF | 0.5889554 |
| AGGLLYGNGGAG | ASSYRNTEVF | 0.36499646 |
| AGLESIGL | ASSYRNTEVF | 0.6428097 |
| AGLKTGII | ASSYRNTEVF | 0.39682275 |
| ASLAALLL | ASSYRNTEVF | 0.6610279 |
| ASVINISSI | ASSYRNTEVF | 0.82528484 |
| ATPPGPA | ASSYRNTEVF | 0.603297 |
| ATVASALV | ASSYRNTEVF | 0.29026395 |
| ATVASLIL | ASSYRNTEVF | 0.24973471 |
| AVNGPVLIRH | ASSYRNTEVF | 0.7458645 |
| DGAAGAVGI | ASSYRNTEVF | 0.41874978 |
| DPLLHGI | ASSYRNTEVF | 0.42003733 |
| ELGAALGAL | ASSYRNTEVF | 0.52087486 |
| ELGRTQAV | ASSYRNTEVF | 0.7751832 |
| FVAGGYLS | ASSYRNTEVF | 0.79679453 |
| GASGGAGGAGGW | ASSYRNTEVF | 0.46421635 |
| GGAGAGGDGGAGGI | ASSYRNTEVF | 0.8142755 |
| GTGHNGSQPGMGGNG | ASSYRNTEVF | 0.8010686 |
| GYAAASASAS | ASSYRNTEVF | 0.81809723 |
| HQVDVAAVQQG | ASSYRNTEVF | 0.7271711 |
| IGAAIAHT | ASSYRNTEVF | 0.37312382 |
| IGMIEVAQL | ASSYRNTEVF | 0.79608923 |
| IRLAPEHTSAMD | ASSYRNTEVF | 0.50484246 |
| IYNVSVL | ASSYRNTEVF | 0.7106885 |
| KQGFLGM | ASSYRNTEVF | 0.7486621 |
| LESVITQL | ASSYRNTEVF | 0.60030305 |
| LGLGALVVV | ASSYRNTEVF | 0.4601171 |
| LIGNGGNGGSGGI | ASSYRNTEVF | 0.6854687 |
| LNAINAPTL | ASSYRNTEVF | 0.84657407 |
| LQLLNTGV | ASSYRNTEVF | 0.34984386 |
| LQTSIEF | ASSYRNTEVF | 0.4257709 |
| MSFAVSEL | ASSYRNTEVF | 0.62582785 |
| NTGALGL | ASSYRNTEVF | 0.6365901 |
| PPGAGGAGGAGTT | ASSYRNTEVF | 0.80018055 |
| QLVGLGI | ASSYRNTEVF | 0.7710234 |
| RFLEQVHQL | ASSYRNTEVF | 0.65114486 |
| RGVGYVLR | ASSYRNTEVF | 0.6905619 |
| SATGVFAGVI | ASSYRNTEVF | 0.79723287 |
| SLPAITLP | ASSYRNTEVF | 0.6730839 |
| STALAFTV | ASSYRNTEVF | 0.71149254 |
| SVLTVGLGK | ASSYRNTEVF | 0.43485844 |
| TALAGAGSGI | ASSYRNTEVF | 0.8237015 |
| TGLPFTGI | ASSYRNTEVF | 0.7673373 |
| VAEHVTV | ASSYRNTEVF | 0.80563354 |

| | | |
|---|---|---|
| VDIVLNSL | ASSYRNTEVF | 0.81403744 |
| VLTGGGAVL | ASSYRNTEVF | 0.7259116 |
| VNVGLAYGI | ASSYRNTEVF | 0.73181087 |
| VSSAAVGAV | ASSYRNTEVF | 0.3077404 |
| VTSVYEAGV | ASSYRNTEVF | 0.5712882 |
| WVALDRGA | ASSYRNTEVF | 0.59695715 |
| YLVGSVVI | ASSYRNTEVF | 0.6873096 |
| GPPGTGKTLL | ASSYRNTEVF | 0.7177429 |
| GVGMTKF | ASSYRNTEVF | 0.8137667 |
| IGIGGSDL | ASSYRNTEVF | 0.51965797 |
| IGTLLAEL | ASSYRNTEVF | 0.8107356 |
| IYTSGTT | ASSYRNTEVF | 0.81379175 |
| LSFPAGV | ASSYRNTEVF | 0.3763396 |
| LYTSGTT | ASSYRNTEVF | 0.55502754 |
| VGAFPFGF | ASSYRNTEVF | 0.80946815 |
| VGAGTVEF | ASSYRNTEVF | 0.73591703 |
| VGKSTLLL | ASSYRNTEVF | 0.5384857 |
| AALAHAGGGIV | ASSYRNTEVF | 0.6648302 |
| AALVAVSLL | ASSYRNTEVF | 0.8356161 |
| AAVSRAVAAL | ASSYRNTEVF | 0.6707158 |
| AGGSSGIGGF | ASSYRNTEVF | 0.8210553 |
| AGLTVANANG | ASSYRNTEVF | 0.22947904 |
| AGLVGAVL | ASSYRNTEVF | 0.71015704 |
| ALSQAFTV | ASSYRNTEVF | 0.6345663 |
| ALTFSVSL | ASSYRNTEVF | 0.18152216 |
| ALTGAALVAL | ASSYRNTEVF | 0.75932485 |
| ASFSIGAGVL | ASSYRNTEVF | 0.6005588 |
| DGSVYKI | ASSYRNTEVF | 0.756546 |
| EQLAEIV | ASSYRNTEVF | 0.5059521 |
| FAVSELTVM | ASSYRNTEVF | 0.7894842 |
| GAAPLAAGL | ASSYRNTEVF | 0.6234964 |
| GADGGDGGSGGAGG | ASSYRNTEVF | 0.6444875 |
| GFGLDTGGAGG | ASSYRNTEVF | 0.5116662 |
| GLASGGLL | ASSYRNTEVF | 0.7041507 |
| GRIAGAAVLL | ASSYRNTEVF | 0.7965509 |
| GSAGSLNTSF | ASSYRNTEVF | 0.8537767 |
| GTGQIGI | ASSYRNTEVF | 0.24801184 |
| GTLIALIL | ASSYRNTEVF | 0.7164531 |
| IAATMGFVVG | ASSYRNTEVF | 0.6313744 |
| IFAQALL | ASSYRNTEVF | 0.3526409 |
| IGAAVALGF | ASSYRNTEVF | 0.21843015 |
| IGLTPLAL | ASSYRNTEVF | 0.5414738 |
| ILGSNTSTL | ASSYRNTEVF | 0.59140515 |
| IVFSLTVL | ASSYRNTEVF | 0.6805906 |
| LASKATGF | ASSYRNTEVF | 0.68270475 |
| LASVALIL | ASSYRNTEVF | 0.5587938 |
| LFAVGLTL | ASSYRNTEVF | 0.7105324 |
| LFNGGIGTYI | ASSYRNTEVF | 0.6227397 |
| LTLGAGII | ASSYRNTEVF | 0.09356713 |
| LVAAALGDI | ASSYRNTEVF | 0.5029378 |
| NTAFVQLGI | ASSYRNTEVF | 0.28814656 |

| | | |
|---|---|---|
| QLFSGAEL | ASSYRNTEVF | 0.84759784 |
| SAATFITL | ASSYRNTEVF | 0.4054296 |
| SNYGLANLGV | ASSYRNTEVF | 0.5037396 |
| SYGGAYAVM | ASSYRNTEVF | 0.626238 |
| TTGVGGLTLG | ASSYRNTEVF | 0.73507756 |
| TTQTIYI | ASSYRNTEVF | 0.6584839 |
| VAAGAGLGAVI | ASSYRNTEVF | 0.54398483 |
| VAAVVTLL | ASSYRNTEVF | 0.21743502 |
| VGFLGGLI | ASSYRNTEVF | 0.7284071 |
| VGVIPILL | ASSYRNTEVF | 0.5289693 |
| VIFASSLI | ASSYRNTEVF | 0.45107627 |
| VKGGDGGAAAG | ASSYRNTEVF | 0.42403215 |
| VTKTFTNI | ASSYRNTEVF | 0.4929861 |
| VVAAAILL | ASSYRNTEVF | 0.27061462 |
| YAALDVEL | ASSYRNTEVF | 0.8165704 |
| LASVALLL | ASSYRNTEVF | 0.4014327 |
| LDAVHRV | ASSYRNTEVF | 0.5364878 |
| QGLSSSIF | ASSYRNTEVF | 0.8253939 |
| RGTFANIRL | ASSYRNTEVF | 0.6887269 |
| RQLWWGHRI | ASSYRNTEVF | 0.541187 |
| YGPPGTGKTLL | ASSYRNTEVF | 0.56989616 |
| AALVGAPANVA | ASSYRNTEVF | 0.38807157 |
| AATFRRTE | ASSYRNTEVF | 0.4500156 |
| AGALGGGI | ASSYRNTEVF | 0.49753094 |
| AGGAGGAGGIGAG | ASSYRNTEVF | 0.80002016 |
| AGSGALGL | ASSYRNTEVF | 0.7408807 |
| AQIADQI | ASSYRNTEVF | 0.40382898 |
| ASAVAVTYL | ASSYRNTEVF | 0.83452725 |
| ASTVLGGTGG | ASSYRNTEVF | 0.8158877 |
| AVGGEPAK | ASSYRNTEVF | 0.809161 |
| CEQGTFGGTC | ASSYRNTEVF | 0.40005976 |
| DEGKAPAG | ASSYRNTEVF | 0.65851486 |
| DTVVFPAL | ASSYRNTEVF | 0.62080204 |
| FEGITMAG | ASSYRNTEVF | 0.5728417 |
| FPLVVQD | ASSYRNTEVF | 0.60008967 |
| GAFTTPQ | ASSYRNTEVF | 0.45119423 |
| GFGGFGTAGGVGGA | ASSYRNTEVF | 0.818745 |
| GFLAQFL | ASSYRNTEVF | 0.8799559 |
| GGAGGAGGNAGVA | ASSYRNTEVF | 0.6300199 |
| GGVGTAGAGGAGGAGG | ASSYRNTEVF | 0.16239856 |
| GNGGGGDAGNAGSGGN | ASSYRNTEVF | 0.5714815 |
| GTGGAGGAGNDAGS | ASSYRNTEVF | 0.386624 |
| GTTNTGWFNS | ASSYRNTEVF | 0.45662323 |
| IFMGPLFTL | ASSYRNTEVF | 0.61006504 |
| ILAHAIEAI | ASSYRNTEVF | 0.30324933 |
| IRSAIREAL | ASSYRNTEVF | 0.64263165 |
| ITFAGGG | ASSYRNTEVF | 0.83329463 |
| ITGMSGHQEV | ASSYRNTEVF | 0.4920291 |
| IVAGLACTL | ASSYRNTEVF | 0.7274664 |
| IVGGGLAGA | ASSYRNTEVF | 0.7845249 |
| IVTGVLNI | ASSYRNTEVF | 0.70571786 |

| | | |
|---|---|---|
| LAQQGCI | ASSYRNTEVF | 0.7963184 |
| LCLHVEN | ASSYRNTEVF | 0.7946609 |
| LDAISDGS | ASSYRNTEVF | 0.69641864 |
| LGGFAAML | ASSYRNTEVF | 0.70407695 |
| LGGKNPMI | ASSYRNTEVF | 0.7366036 |
| LGPKGVAAIL | ASSYRNTEVF | 0.5580492 |
| LMVGVSI | ASSYRNTEVF | 0.6350885 |
| LVGAAFVL | ASSYRNTEVF | 0.62720954 |
| LVLAVAPL | ASSYRNTEVF | 0.7116219 |
| MAAIAVIDAA | ASSYRNTEVF | 0.69293725 |
| NNGINAELQ | ASSYRNTEVF | 0.85825956 |
| RGEPLVLVHG | ASSYRNTEVF | 0.23124376 |
| RGGAGGAAGLI | ASSYRNTEVF | 0.46011692 |
| RSYRMHPAV | ASSYRNTEVF | 0.79346067 |
| SAAHLLAGQLA | ASSYRNTEVF | 0.16448724 |
| SAAVGSY | ASSYRNTEVF | 0.37589484 |
| SPHAVLHRI | ASSYRNTEVF | 0.71145856 |
| SQIAKVL | ASSYRNTEVF | 0.78939766 |
| TALGHLVP | ASSYRNTEVF | 0.19329968 |
| TAPTTSGAA | ASSYRNTEVF | 0.5556939 |
| TDGDDGNGGSPGT | ASSYRNTEVF | 0.35143024 |
| TDGLGEN | ASSYRNTEVF | 0.8399098 |
| TGGPPLPGQA | ASSYRNTEVF | 0.60596687 |
| TIPAITI | ASSYRNTEVF | 0.30142462 |
| TIPGFTI | ASSYRNTEVF | 0.78310794 |
| TSIASGALA | ASSYRNTEVF | 0.22954352 |
| TVTLPGAAI | ASSYRNTEVF | 0.6494893 |
| VASGLSAALP | ASSYRNTEVF | 0.75576866 |
| VGAGVIGI | ASSYRNTEVF | 0.84521425 |
| VGPKRFGV | ASSYRNTEVF | 0.35715616 |
| VNVAFPD | ASSYRNTEVF | 0.7607318 |
| VSMTNSGVS | ASSYRNTEVF | 0.5950241 |
| VTDVLGDLSV | ASSYRNTEVF | 0.8287157 |
| VVVVGAGFAGL | ASSYRNTEVF | 0.5449374 |
| VYTVGTVV | ASSYRNTEVF | 0.34645063 |
| WTTFSGP | ASSYRNTEVF | 0.78483456 |
| YDELVEAF | ASSYRNTEVF | 0.74308366 |
| YGTGGAGGNGG | ASSYRNTEVF | 0.6753932 |
| YTNVMAV | ASSYRNTEVF | 0.77942824 |
| FLEQVHQL | ASSYRNTEVF | 0.7379956 |
| GLSLLAVAV | ASSYRNTEVF | 0.57513636 |
| LEQVHQL | ASSYRNTEVF | 0.7355132 |
| LNAAGAI | ASSYRNTEVF | 0.6992323 |
| PAVHASL | ASSYRNTEVF | 0.49416804 |
| TELEDDA | ASSYRNTEVF | 0.8210218 |
| TPPGPPN | ASSYRNTEVF | 0.6335056 |
| TPPPPPPP | ASSYRNTEVF | 0.40984187 |
| TQSPALI | ASGDNGLGDPETLY | 0.8446759 |
| EFAAAFPQ | ASGDNGLGDPETLY | 0.7142885 |
| LLSGNLDVL | ASGDNGLGDPETLY | 0.7501446 |
| GPANLGNYNI | ASGDNGLGDPETLY | 0.76905286 |

| | | |
|---|---|---|
| TLGPTLP | ASGDNGLGDPETLY | 0.8817444 |
| VYAYSSCCK | ASGDNGLGDPETLY | 0.34142047 |
| APAPAPAPNI | ASGDNGLGDPETLY | 0.68824536 |
| PGGPVPIP | ASGDNGLGDPETLY | 0.6139082 |
| FLTVALEP | ASGDNGLGDPETLY | 0.78173 |
| SQYFGANL | ASGDNGLGDPETLY | 0.69791114 |
| GGVKPAA | ASGDNGLGDPETLY | 0.49535257 |
| GGTGGAGGAGGW | ASGDNGLGDPETLY | 0.7574959 |
| GVGGVAGLGGAGG | ASGDNGLGDPETLY | 0.8160199 |
| VLLVLATTI | ASGDNGLGDPETLY | 0.69292754 |
| IGIFAAML | ASGDNGLGDPETLY | 0.6781185 |
| AVGDLID | ASGDNGLGDPETLY | 0.8183947 |
| NISPLPL | ASGDNGLGDPETLY | 0.41831815 |
| VTAVFVAI | ASGDNGLGDPETLY | 0.6576674 |
| SGDDNDAAL | ASGDNGLGDPETLY | 0.8164853 |
| AVAALAHS | ASGDNGLGDPETLY | 0.78265786 |
| GGVTVMATP | ASGDNGLGDPETLY | 0.6928757 |
| AGLAANVRD | ASGDNGLGDPETLY | 0.8125011 |
| TNRPDLIDPALLRPG | ASGDNGLGDPETLY | 0.5178944 |
| LANLGVTAL | ASGDNGLGDPETLY | 0.6742975 |
| TGERIEL | ASGDNGLGDPETLY | 0.564048 |
| NGEELAA | ASGDNGLGDPETLY | 0.5184084 |
| VLSLVWG | ASGDNGLGDPETLY | 0.36737767 |
| IGYALWL | ASGDNGLGDPETLY | 0.5197571 |
| LSLLLAPL | ASGDNGLGDPETLY | 0.7239127 |
| TLDEALK | ASGDNGLGDPETLY | 0.6661061 |
| TDGAGTAEGG | ASGDNGLGDPETLY | 0.2545722 |
| NAPGGAGGNGG | ASGDNGLGDPETLY | 0.68677306 |
| VSVFMGMGGNF | ASGDNGLGDPETLY | 0.7163319 |
| LASYLAAL | ASGDNGLGDPETLY | 0.8504976 |
| LVGGTPAL | ASGDNGLGDPETLY | 0.6437523 |
| VRNADVL | ASGDNGLGDPETLY | 0.78749114 |
| TCSSPFLVSAM | ASGDNGLGDPETLY | 0.51406294 |
| VHITDAE | ASGDNGLGDPETLY | 0.70949495 |
| EYAAPSF | ASGDNGLGDPETLY | 0.30800948 |
| VVDAFHGGL | ASGDNGLGDPETLY | 0.45332277 |
| ASVVDPQ | ASGDNGLGDPETLY | 0.550799 |
| SYNEFGGLS | ASGDNGLGDPETLY | 0.40557563 |
| DPDGSVY | ASGDNGLGDPETLY | 0.75693345 |
| MVSTIHATE | ASGDNGLGDPETLY | 0.7474859 |
| LAAGVADPV | ASGDNGLGDPETLY | 0.7642846 |
| TKEQIAA | ASGDNGLGDPETLY | 0.7931644 |
| SLASPAAL | ASGDNGLGDPETLY | 0.8099568 |
| IGLVFYGV | ASGDNGLGDPETLY | 0.83155215 |
| FLLDASNL | ASGDNGLGDPETLY | 0.5137483 |
| YQVGNFP | ASGDNGLGDPETLY | 0.5711167 |
| ESLLSQST | ASGDNGLGDPETLY | 0.26303282 |
| LFTVGFAAVG | ASGDNGLGDPETLY | 0.58270454 |
| ATPPTPT | ASGDNGLGDPETLY | 0.5549458 |
| TDLATLG | ASGDNGLGDPETLY | 0.6341937 |
| SAGGTGR | ASGDNGLGDPETLY | 0.15668887 |

| | | |
|---|---|---|
| GFSNSGS | ASGDNGLGDPETLY | 0.7067644 |
| GGNGGSGGTSV | ASGDNGLGDPETLY | 0.8050316 |
| GTSVATGGAGN | ASGDNGLGDPETLY | 0.42438668 |
| NGGNGGSGGTS | ASGDNGLGDPETLY | 0.74881554 |
| AVVKFPH | ASGDNGLGDPETLY | 0.71283996 |
| TPPPPPP | ASGDNGLGDPETLY | 0.49138874 |
| AAPAAAAAPA | ASGDNGLGDPETLY | 0.72989154 |
| IALNEDL | ASGDNGLGDPETLY | 0.5630747 |
| IVNVSSM | ASGDNGLGDPETLY | 0.8002761 |
| TGIPPAPR | ASGDNGLGDPETLY | 0.8199168 |
| FDLGGGTF | ASGDNGLGDPETLY | 0.69911623 |
| FDLGGGTFDVS | ASGDNGLGDPETLY | 0.5880306 |
| NEPTAAA | ASGDNGLGDPETLY | 0.7251557 |
| NEPTAAALA | ASGDNGLGDPETLY | 0.26120353 |
| GSIINISSI | ASGDNGLGDPETLY | 0.60167974 |
| VAPGNAGTA | ASGDNGLGDPETLY | 0.22748779 |
| TGGGASLE | ASGDNGLGDPETLY | 0.64079916 |
| VAVYNHYKRI | ASGDNGLGDPETLY | 0.45837027 |
| TDPAPATT | ASGDNGLGDPETLY | 0.68064034 |
| TNRPDILDPALLRPG | ASGDNGLGDPETLY | 0.60713977 |
| LTTTTTS | ASGDNGLGDPETLY | 0.74111116 |
| PTGSGKTL | ASGDNGLGDPETLY | 0.29736626 |
| TGSGKTL | ASGDNGLGDPETLY | 0.75936526 |
| TIGHVDHGK | ASGDNGLGDPETLY | 0.702194 |
| TIGHVDHGKTT | ASGDNGLGDPETLY | 0.31025702 |
| GHVDHGKTT | ASGDNGLGDPETLY | 0.7520201 |
| AQFEHTLL | ASGDNGLGDPETLY | 0.7128734 |
| IVNQDEL | ASGDNGLGDPETLY | 0.5724792 |
| TNGSQFF | ASGDNGLGDPETLY | 0.7894471 |
| TGKGIVE | ASGDNGLGDPETLY | 0.66369706 |
| LISNASDA | ASGDNGLGDPETLY | 0.21173406 |
| AELRTLGELTKIAKAHGAQ | ASGDNGLGDPETLY | 0.49406672 |
| APNAVAAAEPSW | ASGDNGLGDPETLY | 0.6689116 |
| APQVAGAATAVL | ASGDNGLGDPETLY | 0.5572739 |
| ARRDPAAAAAILPTDA | ASGDNGLGDPETLY | 0.18359247 |
| DAAARDVLAEAGLAEYF | ASGDNGLGDPETLY | 0.7483092 |
| DDQIVLSAKANLAKVEQ | ASGDNGLGDPETLY | 0.41449818 |
| DELLPDTGEDSDEDGH | ASGDNGLGDPETLY | 0.39932013 |
| ELPEAPANRATVPQD | ASGDNGLGDPETLY | 0.55876213 |
| FGQSKIERVTGIG | ASGDNGLGDPETLY | 0.7613898 |
| GAGGTGSTAGGAGGAGGAGGLY | ASGDNGLGDPETLY | 0.66623247 |
| GHGGTGAAGGNGGTGGAGGSGI | ASGDNGLGDPETLY | 0.688079 |
| GNGGNGGAGGEGGPGGAGGAGA | ASGDNGLGDPETLY | 0.6843872 |
| GSTAGNGGNGGNATNGTIAGQP | ASGDNGLGDPETLY | 0.6890025 |
| GTGGDGGNGGNGGTGG | ASGDNGLGDPETLY | 0.7865842 |
| GVRRPAGIPADAPV | ASGDNGLGDPETLY | 0.7148454 |
| GYRQVTDAHLLTLARR | ASGDNGLGDPETLY | 0.72662795 |
| IAKMGQDGHDRG | ASGDNGLGDPETLY | 0.7422807 |
| KAEPSIEIQYT | ASGDNGLGDPETLY | 0.6502316 |
| KPVRIRMAADEDTVAVT | ASGDNGLGDPETLY | 0.44761646 |
| MPPRIAGMRLLVIKPEP | ASGDNGLGDPETLY | 0.49084437 |

| | | |
|---|---|---|
| MKFYAVALFRSRGVPPTR | ASGDNGLGDPETLY | 0.36757055 |
| NAGGFGILGAVAHSP | ASGDNGLGDPETLY | 0.42109424 |
| NEGREITLAMAEM | ASGDNGLGDPETLY | 0.70557517 |
| NELVEWSHPLGHRWRWE | ASGDNGLGDPETLY | 0.85689735 |
| NLGAGGTGGNGGNPG | ASGDNGLGDPETLY | 0.73283863 |
| NTEWPKWETVMAADPVD | ASGDNGLGDPETLY | 0.3124508 |
| PASAAPAETQP | ASGDNGLGDPETLY | 0.59506685 |
| QQSSDGKNGVRAAEITGE | ASGDNGLGDPETLY | 0.45475554 |
| RAPDAVTTSWWKE | ASGDNGLGDPETLY | 0.7095084 |
| SGFGNVGAHTSGWFNQ | ASGDNGLGDPETLY | 0.8144796 |
| SGVAGAVGQAASVGGLKV | ASGDNGLGDPETLY | 0.85746515 |
| SNGGAGTGSAGGNGGT | ASGDNGLGDPETLY | 0.74192023 |
| TGGAGGEGGAAG | ASGDNGLGDPETLY | 0.50059384 |
| TGGDGGAAGAGGAPGHG | ASGDNGLGDPETLY | 0.7634436 |
| TMPPLLAAIANAGRHGV | ASGDNGLGDPETLY | 0.6101842 |
| TSSTAGNSGGA | ASGDNGLGDPETLY | 0.6392182 |
| VPATVPVGILC | ASGDNGLGDPETLY | 0.68180966 |
| VQRPDSGGVVLTGRIS | ASGDNGLGDPETLY | 0.7472796 |
| WTPNPVPATKGVPAGLVQAT | ASGDNGLGDPETLY | 0.60297966 |
| YALKRALAKFSDRFDVVII | ASGDNGLGDPETLY | 0.5719388 |
| YLIGSVIV | ASGDNGLGDPETLY | 0.25357658 |
| YGQLVAVID | ASGDNGLGDPETLY | 0.8188917 |
| VTVGGMTLPA | ASGDNGLGDPETLY | 0.28820586 |
| VLALPAV | ASGDNGLGDPETLY | 0.5622592 |
| VIPPTLNYET | ASGDNGLGDPETLY | 0.5939071 |
| VGMGGMHPSSG | ASGDNGLGDPETLY | 0.82585657 |
| VGLAGGAATA | ASGDNGLGDPETLY | 0.8251245 |
| VGGAGGAGGSGP | ASGDNGLGDPETLY | 0.5178889 |
| VFAAFMV | ASGDNGLGDPETLY | 0.45320755 |
| TLYDKLR | ASGDNGLGDPETLY | 0.737291 |
| TLAVFTVL | ASGDNGLGDPETLY | 0.49770463 |
| TGSVQPARV | ASGDNGLGDPETLY | 0.80025184 |
| TGGGAGSGGT | ASGDNGLGDPETLY | 0.6023021 |
| SHLETSL | ASGDNGLGDPETLY | 0.54308426 |
| SFTTGMT | ASGDNGLGDPETLY | 0.53834444 |
| QVEVAKRMGVRQPTVS | ASGDNGLGDPETLY | 0.74664474 |
| QGSGGPALAM | ASGDNGLGDPETLY | 0.43642747 |
| MLAPSNP | ASGDNGLGDPETLY | 0.7957195 |
| LVNNAAI | ASGDNGLGDPETLY | 0.15908033 |
| LTGEINTLA | ASGDNGLGDPETLY | 0.7895242 |
| LSIFGPV | ASGDNGLGDPETLY | 0.3886833 |
| LLALLANL | ASGDNGLGDPETLY | 0.5554802 |
| LISNASDALDKLR | ASGDNGLGDPETLY | 0.26445588 |
| LINEAVP | ASGDNGLGDPETLY | 0.63143945 |
| LGYAFSGL | ASGDNGLGDPETLY | 0.6316707 |
| LDFVAVAL | ASGDNGLGDPETLY | 0.80422854 |
| LAQPGAE | ASGDNGLGDPETLY | 0.629305 |
| ISNLGAVTGF | ASGDNGLGDPETLY | 0.68371 |
| HQEDPGRI | ASGDNGLGDPETLY | 0.75152254 |
| GTPGEPTP | ASGDNGLGDPETLY | 0.47143322 |
| GFFNSGS | ASGDNGLGDPETLY | 0.22926834 |

| | | |
|---|---|---|
| GAGGGGLTAGAN | ASGDNGLGDPETLY | 0.7056681 |
| FVDALVNM | ASGDNGLGDPETLY | 0.660656 |
| FLTVAPEL | ASGDNGLGDPETLY | 0.8029853 |
| FETLTAL | ASGDNGLGDPETLY | 0.7032052 |
| DVINPIEL | ASGDNGLGDPETLY | 0.24932653 |
| APVSGAPV | ASGDNGLGDPETLY | 0.7767385 |
| ALVGGLAW | ASGDNGLGDPETLY | 0.74776137 |
| ALAAAAATAI | ASGDNGLGDPETLY | 0.37240613 |
| AFGLGLLL | ASGDNGLGDPETLY | 0.4650834 |
| AFFAAASL | ASGDNGLGDPETLY | 0.52339256 |
| AAAALPAFSPP | ASGDNGLGDPETLY | 0.32322866 |
| VVLAPAL | ASGDNGLGDPETLY | 0.70977026 |
| TIGHVDHGKTTL | ASGDNGLGDPETLY | 0.523605 |
| LISNASDALDK | ASGDNGLGDPETLY | 0.621835 |
| LISNASDAL | ASGDNGLGDPETLY | 0.5932871 |
| ISNASDALDK | ASGDNGLGDPETLY | 0.54394615 |
| NQAETLVYQTE | ASGDNGLGDPETLY | 0.6065059 |
| TGGTGGTAGAG | ASGDNGLGDPETLY | 0.58380836 |
| TGSTAGGAGGA | ASGDNGLGDPETLY | 0.6433417 |
| DPLAVPVAAGPV | ASGDNGLGDPETLY | 0.75757074 |
| GGTGGTPGNGGN | ASGDNGLGDPETLY | 0.589967 |
| PAGLAGVDPATL | ASGDNGLGDPETLY | 0.72301465 |
| GNNLVVEAPHAGASV | ASGDNGLGDPETLY | 0.7670044 |
| KVTRIMSASTLEQPA | ASGDNGLGDPETLY | 0.30867907 |
| LGFALPIALPRRTVV | ASGDNGLGDPETLY | 0.46918604 |
| TRLGNLTARKLVVLV | ASGDNGLGDPETLY | 0.7735466 |
| WSQQRRLIYFPSAGP | ASGDNGLGDPETLY | 0.5843499 |
| GPNGNPDPMAAAVDIR | ASGDNGLGDPETLY | 0.79847574 |
| TQRCVSVADLRAAASR | ASGDNGLGDPETLY | 0.6573754 |
| QFVLDVAEHPVEVLTETV | ASGDNGLGDPETLY | 0.68059635 |
| GGAGGTGGAGGNGGTQAGDG | ASGDNGLGDPETLY | 0.6747433 |
| PLLATLRAIAVDVDAKPAQV | ASGDNGLGDPETLY | 0.81911373 |
| APKFRTGGYMIDFWGVGYQVAK | ASGDNGLGDPETLY | 0.7269243 |
| GAGGAGGAGGAAGGPGGTGGTG | ASGDNGLGDPETLY | 0.6375386 |
| IGALGLLATDVAPAGSVAFGVA | ASGDNGLGDPETLY | 0.7365665 |
| AGGAGGIGGNGGAGGAGGVLMG | ASGDNGLGDPETLY | 0.46524316 |
| ADELIGLL | ASGDNGLGDPETLY | 0.73319066 |
| ADRALKA | ASGDNGLGDPETLY | 0.7896089 |
| AGAVGQVGGAGGA | ASGDNGLGDPETLY | 0.35161787 |
| AGGGGGLGGAP | ASGDNGLGDPETLY | 0.57986724 |
| AGGLLYGNGGAG | ASGDNGLGDPETLY | 0.8359196 |
| AGLESIGL | ASGDNGLGDPETLY | 0.76338494 |
| AGLKTGII | ASGDNGLGDPETLY | 0.5306302 |
| ASLAALLL | ASGDNGLGDPETLY | 0.5129303 |
| ASVINISSI | ASGDNGLGDPETLY | 0.56377155 |
| ATPPGPA | ASGDNGLGDPETLY | 0.6664579 |
| ATVASALV | ASGDNGLGDPETLY | 0.3328039 |
| ATVASLIL | ASGDNGLGDPETLY | 0.73953485 |
| AVNGPVLIRH | ASGDNGLGDPETLY | 0.5229801 |
| DGAAGAVGI | ASGDNGLGDPETLY | 0.8188806 |
| DPLLHGI | ASGDNGLGDPETLY | 0.6967719 |

| | | |
|---|---|---|
| ELGAALGAL | ASGDNGLGDPETLY | 0.78836215 |
| ELGRTQAV | ASGDNGLGDPETLY | 0.55164695 |
| FVAGGYLS | ASGDNGLGDPETLY | 0.5956529 |
| GASGGAGGAGGW | ASGDNGLGDPETLY | 0.3481814 |
| GGAGAGGDGGAGGI | ASGDNGLGDPETLY | 0.33089975 |
| GTGHNGSQPGMGGNG | ASGDNGLGDPETLY | 0.54594225 |
| GYAAASASAS | ASGDNGLGDPETLY | 0.6206613 |
| HQVDVAAVQQG | ASGDNGLGDPETLY | 0.21383908 |
| IGAAIAHT | ASGDNGLGDPETLY | 0.83119863 |
| IGMIEVAQL | ASGDNGLGDPETLY | 0.56110895 |
| IRLAPEHTSAMD | ASGDNGLGDPETLY | 0.7421275 |
| IYNVSVL | ASGDNGLGDPETLY | 0.564088 |
| KQGFLGM | ASGDNGLGDPETLY | 0.42108697 |
| LESVITQL | ASGDNGLGDPETLY | 0.5196886 |
| LGLGALVVV | ASGDNGLGDPETLY | 0.74541736 |
| LIGNGGNGGSGGI | ASGDNGLGDPETLY | 0.33802664 |
| LNAINAPTL | ASGDNGLGDPETLY | 0.17443144 |
| LQLLNTGV | ASGDNGLGDPETLY | 0.5176801 |
| LQTSIEF | ASGDNGLGDPETLY | 0.2615798 |
| MSFAVSEL | ASGDNGLGDPETLY | 0.84558487 |
| NTGALGL | ASGDNGLGDPETLY | 0.791599 |
| PPGAGGAGGAGTT | ASGDNGLGDPETLY | 0.6488197 |
| QLVGLGI | ASGDNGLGDPETLY | 0.86123145 |
| RFLEQVHQL | ASGDNGLGDPETLY | 0.15368725 |
| RGVGYVLR | ASGDNGLGDPETLY | 0.60592896 |
| SATGVFAGVI | ASGDNGLGDPETLY | 0.81423616 |
| SLPAITLP | ASGDNGLGDPETLY | 0.69216204 |
| STALAFTV | ASGDNGLGDPETLY | 0.5658748 |
| SVLTVGLGK | ASGDNGLGDPETLY | 0.77787185 |
| TALAGAGSGI | ASGDNGLGDPETLY | 0.45871088 |
| TGLPFTGI | ASGDNGLGDPETLY | 0.5670109 |
| VAEHVTV | ASGDNGLGDPETLY | 0.6210889 |
| VDIVLNSL | ASGDNGLGDPETLY | 0.7632124 |
| VLTGGGAVL | ASGDNGLGDPETLY | 0.7625452 |
| VNVGLAYGI | ASGDNGLGDPETLY | 0.30120444 |
| VSSAAVGAV | ASGDNGLGDPETLY | 0.7842926 |
| VTSVYEAGV | ASGDNGLGDPETLY | 0.8294816 |
| WVALDRGA | ASGDNGLGDPETLY | 0.4827212 |
| YLVGSVVI | ASGDNGLGDPETLY | 0.68859017 |
| GPPGTGKTLL | ASGDNGLGDPETLY | 0.68620956 |
| GVGMTKF | ASGDNGLGDPETLY | 0.43866777 |
| IGIGGSDL | ASGDNGLGDPETLY | 0.439043 |
| IGTLLAEL | ASGDNGLGDPETLY | 0.3397844 |
| IYTSGTT | ASGDNGLGDPETLY | 0.75612295 |
| LSFPAGV | ASGDNGLGDPETLY | 0.3641678 |
| LYTSGTT | ASGDNGLGDPETLY | 0.7007915 |
| VGAFPFGF | ASGDNGLGDPETLY | 0.6554086 |
| VGAGTVEF | ASGDNGLGDPETLY | 0.85462725 |
| VGKSTLLL | ASGDNGLGDPETLY | 0.7167139 |
| AALAHAGGGIV | ASGDNGLGDPETLY | 0.7107036 |
| AALVAVSLL | ASGDNGLGDPETLY | 0.45859432 |

| | | |
|---|---|---|
| AAVSRAVAAL | ASGDNGLGDPETLY | 0.72540355 |
| AGGSSGIGGF | ASGDNGLGDPETLY | 0.72204936 |
| AGLTVANANG | ASGDNGLGDPETLY | 0.25450778 |
| AGLVGAVL | ASGDNGLGDPETLY | 0.7949575 |
| ALSQAFTV | ASGDNGLGDPETLY | 0.43084288 |
| ALTFSVSL | ASGDNGLGDPETLY | 0.6868614 |
| ALTGAALVAL | ASGDNGLGDPETLY | 0.78254235 |
| ASFSIGAGVL | ASGDNGLGDPETLY | 0.82837296 |
| DGSVYKI | ASGDNGLGDPETLY | 0.65100074 |
| EQLAEIV | ASGDNGLGDPETLY | 0.74950945 |
| FAVSELTVM | ASGDNGLGDPETLY | 0.392501 |
| GAAPLAAGL | ASGDNGLGDPETLY | 0.79339796 |
| GADGGDGGSGGAGG | ASGDNGLGDPETLY | 0.7680232 |
| GFGLDTGGAGG | ASGDNGLGDPETLY | 0.6900749 |
| GLASGGLL | ASGDNGLGDPETLY | 0.8523602 |
| GRIAGAAVLL | ASGDNGLGDPETLY | 0.8299042 |
| GSAGSLNTSF | ASGDNGLGDPETLY | 0.3283686 |
| GTGQIGI | ASGDNGLGDPETLY | 0.6409148 |
| GTLIALIL | ASGDNGLGDPETLY | 0.79854053 |
| IAATMGFVVG | ASGDNGLGDPETLY | 0.8006879 |
| IFAQALL | ASGDNGLGDPETLY | 0.7283801 |
| IGAAVALGF | ASGDNGLGDPETLY | 0.73863685 |
| IGLTPLAL | ASGDNGLGDPETLY | 0.48898423 |
| ILGSNTSTL | ASGDNGLGDPETLY | 0.8124627 |
| IVFSLTVL | ASGDNGLGDPETLY | 0.88810754 |
| LASKATGF | ASGDNGLGDPETLY | 0.7429971 |
| LASVALIL | ASGDNGLGDPETLY | 0.5656705 |
| LFAVGLTL | ASGDNGLGDPETLY | 0.62129533 |
| LFNGGIGTYI | ASGDNGLGDPETLY | 0.75350845 |
| LTLGAGII | ASGDNGLGDPETLY | 0.76819265 |
| LVAAALGDI | ASGDNGLGDPETLY | 0.44689268 |
| NTAFVQLGI | ASGDNGLGDPETLY | 0.6333291 |
| QLFSGAEL | ASGDNGLGDPETLY | 0.6653141 |
| SAATFITL | ASGDNGLGDPETLY | 0.19782393 |
| SNYGLANLGV | ASGDNGLGDPETLY | 0.73260224 |
| SYGGAYAVM | ASGDNGLGDPETLY | 0.7820263 |
| TTGVGGLTLG | ASGDNGLGDPETLY | 0.75144213 |
| TTQTIYI | ASGDNGLGDPETLY | 0.40504795 |
| VAAGAGLGAVI | ASGDNGLGDPETLY | 0.7941092 |
| VAAVVTLL | ASGDNGLGDPETLY | 0.7770572 |
| VGFLGGLI | ASGDNGLGDPETLY | 0.8407464 |
| VGVIPILL | ASGDNGLGDPETLY | 0.32286754 |
| VIFASSLI | ASGDNGLGDPETLY | 0.18801513 |
| VKGGDGGAAAG | ASGDNGLGDPETLY | 0.67703235 |
| VTKTFTNI | ASGDNGLGDPETLY | 0.77386236 |
| VVAAAILL | ASGDNGLGDPETLY | 0.5735713 |
| YAALDVEL | ASGDNGLGDPETLY | 0.46826386 |
| LASVALLL | ASGDNGLGDPETLY | 0.62397087 |
| LDAVHRV | ASGDNGLGDPETLY | 0.787106 |
| QGLSSSIF | ASGDNGLGDPETLY | 0.39581084 |
| RGTFANIRL | ASGDNGLGDPETLY | 0.42275354 |

| | | |
|---|---|---|
| RQLWWGHRI | ASGDNGLGDPETLY | 0.72167534 |
| YGPPGTGKTLL | ASGDNGLGDPETLY | 0.8072505 |
| AALVGAPANVA | ASGDNGLGDPETLY | 0.41350013 |
| AATFRRTE | ASGDNGLGDPETLY | 0.48711255 |
| AGALGGGI | ASGDNGLGDPETLY | 0.70594865 |
| AGGAGGAGGIGAG | ASGDNGLGDPETLY | 0.777779 |
| AGSGALGL | ASGDNGLGDPETLY | 0.6517067 |
| AQIADQI | ASGDNGLGDPETLY | 0.4213767 |
| ASAVAVTYL | ASGDNGLGDPETLY | 0.6997372 |
| ASTVLGGTGG | ASGDNGLGDPETLY | 0.32819897 |
| AVGGEPAK | ASGDNGLGDPETLY | 0.7919196 |
| CEQGTFGGTC | ASGDNGLGDPETLY | 0.6169714 |
| DEGKAPAG | ASGDNGLGDPETLY | 0.72868305 |
| DTVVFPAL | ASGDNGLGDPETLY | 0.56583023 |
| FEGITMAG | ASGDNGLGDPETLY | 0.34610575 |
| FPLVVQD | ASGDNGLGDPETLY | 0.38503385 |
| GAFTTPQ | ASGDNGLGDPETLY | 0.4261555 |
| GFGGFGTAGGVGGA | ASGDNGLGDPETLY | 0.7443694 |
| GFLAQFL | ASGDNGLGDPETLY | 0.26706964 |
| GGAGGAGGNAGVA | ASGDNGLGDPETLY | 0.7483481 |
| GGVGTAGAGGAGGAGG | ASGDNGLGDPETLY | 0.64636624 |
| GNGGGGDAGNAGSGGN | ASGDNGLGDPETLY | 0.42744452 |
| GTGGAGGAGNDAGS | ASGDNGLGDPETLY | 0.7570151 |
| GTTNTGWFNS | ASGDNGLGDPETLY | 0.32876503 |
| IFMGPLFTL | ASGDNGLGDPETLY | 0.7062832 |
| ILAHAIEAI | ASGDNGLGDPETLY | 0.6625631 |
| IRSAIREAL | ASGDNGLGDPETLY | 0.7713089 |
| ITFAGGG | ASGDNGLGDPETLY | 0.44053882 |
| ITGMSGHQEV | ASGDNGLGDPETLY | 0.7839377 |
| IVAGLACTL | ASGDNGLGDPETLY | 0.7175665 |
| IVGGGLAGA | ASGDNGLGDPETLY | 0.43111834 |
| IVTGVLNI | ASGDNGLGDPETLY | 0.7353969 |
| LAQQGCI | ASGDNGLGDPETLY | 0.7692299 |
| LCLHVEN | ASGDNGLGDPETLY | 0.75020283 |
| LDAISDGS | ASGDNGLGDPETLY | 0.24940413 |
| LGGFAAML | ASGDNGLGDPETLY | 0.6886494 |
| LGGKNPMI | ASGDNGLGDPETLY | 0.45933273 |
| LGPKGVAAIL | ASGDNGLGDPETLY | 0.66986895 |
| LMVGVSI | ASGDNGLGDPETLY | 0.63688457 |
| LVGAAFVL | ASGDNGLGDPETLY | 0.611821 |
| LVLAVAPL | ASGDNGLGDPETLY | 0.7608686 |
| MAAIAVIDAA | ASGDNGLGDPETLY | 0.48423028 |
| NNGINAELQ | ASGDNGLGDPETLY | 0.46798754 |
| RGEPLVLVHG | ASGDNGLGDPETLY | 0.624893 |
| RGGAGGAAGLI | ASGDNGLGDPETLY | 0.692647 |
| RSYRMHPAV | ASGDNGLGDPETLY | 0.74731183 |
| SAAHLLAGQLA | ASGDNGLGDPETLY | 0.39676046 |
| SAAVGSY | ASGDNGLGDPETLY | 0.70017964 |
| SPHAVLHRI | ASGDNGLGDPETLY | 0.6051599 |
| SQIAKVL | ASGDNGLGDPETLY | 0.42369366 |
| TALGHLVP | ASGDNGLGDPETLY | 0.7546553 |

| | | |
|---|---|---|
| TAPTTSGAA | ASGDNGLGDPETLY | 0.61398935 |
| TDGDDGNGGSPGT | ASGDNGLGDPETLY | 0.30931166 |
| TDGLGEN | ASGDNGLGDPETLY | 0.7579265 |
| TGGPPLPGQA | ASGDNGLGDPETLY | 0.4603808 |
| TIPAITI | ASGDNGLGDPETLY | 0.7430495 |
| TIPGFTI | ASGDNGLGDPETLY | 0.8335515 |
| TSIASGALA | ASGDNGLGDPETLY | 0.35822254 |
| TVTLPGAAI | ASGDNGLGDPETLY | 0.78292584 |
| VASGLSAALP | ASGDNGLGDPETLY | 0.6332624 |
| VGAGVIGI | ASGDNGLGDPETLY | 0.6317998 |
| VGPKRFGV | ASGDNGLGDPETLY | 0.6315141 |
| VNVAFPD | ASGDNGLGDPETLY | 0.44856644 |
| VSMTNSGVS | ASGDNGLGDPETLY | 0.82021576 |
| VTDVLGDLSV | ASGDNGLGDPETLY | 0.7335331 |
| VVVVGAGFAGL | ASGDNGLGDPETLY | 0.7285124 |
| VYTVGTVV | ASGDNGLGDPETLY | 0.7079097 |
| WTTFSGP | ASGDNGLGDPETLY | 0.7560861 |
| YDELVEAF | ASGDNGLGDPETLY | 0.7741648 |
| YGTGGAGGNGG | ASGDNGLGDPETLY | 0.4626956 |
| YTNVMAV | ASGDNGLGDPETLY | 0.31259894 |
| FLEQVHQL | ASGDNGLGDPETLY | 0.5987519 |
| GLSLLAVAV | ASGDNGLGDPETLY | 0.5681639 |
| LEQVHQL | ASGDNGLGDPETLY | 0.5221 |
| LNAAGAI | ASGDNGLGDPETLY | 0.4349603 |
| PAVHASL | ASGDNGLGDPETLY | 0.7523821 |
| TELEDDA | ASGDNGLGDPETLY | 0.84789705 |
| TPPGPPN | ASGDNGLGDPETLY | 0.5594564 |
| TPPPPPPP | ASGDNGLGDPETLY | 0.859934 |
| TQSPALI | ASSMREVF | 0.3980676 |
| EFAAAFPQ | ASSMREVF | 0.75513077 |
| LLSGNLDVL | ASSMREVF | 0.78826845 |
| GPANLGNYNI | ASSMREVF | 0.4009158 |
| TLGPTLP | ASSMREVF | 0.73100114 |
| VYAYSSCCK | ASSMREVF | 0.3152549 |
| APAPAPAPNI | ASSMREVF | 0.45765218 |
| PGGPVPIP | ASSMREVF | 0.7626147 |
| FLTVALEP | ASSMREVF | 0.61821294 |
| SQYFGANL | ASSMREVF | 0.7355456 |
| GGVKPAA | ASSMREVF | 0.61481583 |
| GGTGGAGGAGGW | ASSMREVF | 0.44832072 |
| GVGGVAGLGGAGG | ASSMREVF | 0.71214724 |
| VLLVLATTI | ASSMREVF | 0.7817558 |
| IGIFAAML | ASSMREVF | 0.7143578 |
| AVGDLID | ASSMREVF | 0.887329 |
| NISPLPL | ASSMREVF | 0.37001437 |
| VTAVFVAI | ASSMREVF | 0.7800056 |
| SGDDNDAAL | ASSMREVF | 0.8218472 |
| AVAALAHS | ASSMREVF | 0.6492518 |
| GGVTVMATP | ASSMREVF | 0.6094394 |
| AGLAANVRD | ASSMREVF | 0.7392792 |
| TNRPDLIDPALLRPG | ASSMREVF | 0.61159366 |

| | | |
|---|---|---|
| LANLGVTAL | ASSMREVF | 0.70309556 |
| TGERIEL | ASSMREVF | 0.83801615 |
| NGEELAA | ASSMREVF | 0.5292405 |
| VLSLVWG | ASSMREVF | 0.83931494 |
| IGYALWL | ASSMREVF | 0.81255877 |
| LSLLLAPL | ASSMREVF | 0.8325723 |
| TLDEALK | ASSMREVF | 0.6876208 |
| TDGAGTAEGG | ASSMREVF | 0.4456582 |
| NAPGGAGGNGG | ASSMREVF | 0.7805652 |
| VSVFMGMGGNF | ASSMREVF | 0.62783146 |
| LASYLAAL | ASSMREVF | 0.8463845 |
| LVGGTPAL | ASSMREVF | 0.8394445 |
| VRNADVL | ASSMREVF | 0.70593476 |
| TCSSPFLVSAM | ASSMREVF | 0.82881725 |
| VHITDAE | ASSMREVF | 0.8428565 |
| EYAAPSF | ASSMREVF | 0.44465074 |
| VVDAFHGGL | ASSMREVF | 0.7867242 |
| ASVVDPQ | ASSMREVF | 0.790772 |
| SYNEFGGLS | ASSMREVF | 0.76587677 |
| DPDGSVY | ASSMREVF | 0.43709964 |
| MVSTIHATE | ASSMREVF | 0.82644665 |
| LAAGVADPV | ASSMREVF | 0.7595655 |
| TKEQIAA | ASSMREVF | 0.4720208 |
| SLASPAAL | ASSMREVF | 0.8163514 |
| IGLVFYGV | ASSMREVF | 0.7349107 |
| FLLDASNL | ASSMREVF | 0.77774894 |
| YQVGNFP | ASSMREVF | 0.8395931 |
| ESLLSQST | ASSMREVF | 0.8126674 |
| LFTVGFAAVG | ASSMREVF | 0.7134107 |
| ATPPTPT | ASSMREVF | 0.8082538 |
| TDLATLG | ASSMREVF | 0.844995 |
| SAGGTGR | ASSMREVF | 0.66982794 |
| GFSNSGS | ASSMREVF | 0.8094704 |
| GGNGGSGGTSV | ASSMREVF | 0.85239017 |
| GTSVATGGAGN | ASSMREVF | 0.78029275 |
| NGGNGGSGGTS | ASSMREVF | 0.85576475 |
| AVVKFPH | ASSMREVF | 0.8578619 |
| TPPPPPP | ASSMREVF | 0.7453907 |
| AAPAAAAAPA | ASSMREVF | 0.841626 |
| IALNEDL | ASSMREVF | 0.83066154 |
| IVNVSSM | ASSMREVF | 0.75995505 |
| TGIPPAPR | ASSMREVF | 0.78790224 |
| FDLGGGTF | ASSMREVF | 0.50510144 |
| FDLGGGTFDVS | ASSMREVF | 0.7985829 |
| NEPTAAA | ASSMREVF | 0.63083065 |
| NEPTAAALA | ASSMREVF | 0.8190011 |
| GSIINISSI | ASSMREVF | 0.8457676 |
| VAPGNAGTA | ASSMREVF | 0.7918545 |
| TGGGASLE | ASSMREVF | 0.5884493 |
| VAVYNHYKRI | ASSMREVF | 0.5509485 |
| TDPAPATT | ASSMREVF | 0.8207288 |

| | | |
|---|---|---|
| TNRPDILDPALLRPG | ASSMREVF | 0.64720196 |
| LTTTTTS | ASSMREVF | 0.6391566 |
| PTGSGKTL | ASSMREVF | 0.708138 |
| TGSGKTL | ASSMREVF | 0.83922595 |
| TIGHVDHGK | ASSMREVF | 0.84344655 |
| TIGHVDHGKTT | ASSMREVF | 0.6999524 |
| GHVDHGKTT | ASSMREVF | 0.8421844 |
| AQFEHTLL | ASSMREVF | 0.8439346 |
| IVNQDEL | ASSMREVF | 0.8470132 |
| TNGSQFF | ASSMREVF | 0.7946982 |
| TGKGIVE | ASSMREVF | 0.5211172 |
| LISNASDA | ASSMREVF | 0.84293544 |
| AELRTLGELTKIAKAHGAQ | ASSMREVF | 0.7587865 |
| APNAVAAAEPSW | ASSMREVF | 0.8505647 |
| APQVAGAATAVL | ASSMREVF | 0.8419179 |
| ARRDPAAAAAILPTDA | ASSMREVF | 0.76562834 |
| DAAARDVLAEAGLAEYF | ASSMREVF | 0.6877104 |
| DDQIVLSAKANLAKVEQ | ASSMREVF | 0.8143357 |
| DELLPDTGEDSDEDGH | ASSMREVF | 0.85512483 |
| ELPEAPANRATVPQD | ASSMREVF | 0.6573843 |
| FGQSKIERVTGIG | ASSMREVF | 0.49491754 |
| GAGGTGSTAGGAGGAGGAGGLY | ASSMREVF | 0.8186619 |
| GHGGTGAAGGNGGTGGAGGSGI | ASSMREVF | 0.79458433 |
| GNGGNGGAGGEGGPGGAGGAGA | ASSMREVF | 0.84893245 |
| GSTAGNGGNGGNATNGTIAGQP | ASSMREVF | 0.8154141 |
| GTGGDGGNGGNGGTGG | ASSMREVF | 0.7969405 |
| GVRRPAGIPADAPV | ASSMREVF | 0.8259244 |
| GYRQVTDAHLLTLARR | ASSMREVF | 0.8376001 |
| IAKMGQDGHDRG | ASSMREVF | 0.80286527 |
| KAEPSIEIQYT | ASSMREVF | 0.8299391 |
| KPVRIRMAADEDTVAVT | ASSMREVF | 0.71594656 |
| MPPRIAGMRLLVIKPEP | ASSMREVF | 0.8513236 |
| MKFYAVALFRSRGVPPTR | ASSMREVF | 0.84375703 |
| NAGGFGILGAVAHSP | ASSMREVF | 0.8466083 |
| NEGREITLAMAEM | ASSMREVF | 0.8498765 |
| NELVEWSHPLGHRWRWE | ASSMREVF | 0.79208624 |
| NLGAGGTGGNGGNPG | ASSMREVF | 0.8138954 |
| NTEWPKWETVMAADPVD | ASSMREVF | 0.6227908 |
| PASAAPAETQP | ASSMREVF | 0.7760783 |
| QQSSDGKNGVRAAEITGE | ASSMREVF | 0.79465014 |
| RAPDAVTTSWWKE | ASSMREVF | 0.82060814 |
| SGFGNVGAHTSGWFNQ | ASSMREVF | 0.83266824 |
| SGVAGAVGQAASVGGLKV | ASSMREVF | 0.8366884 |
| SNGGAGTGSAGGNGGT | ASSMREVF | 0.714339 |
| TGGAGGEGGAAG | ASSMREVF | 0.85410726 |
| TGGDGGAAGAGGAPGHG | ASSMREVF | 0.84520257 |
| TMPPLLAAIANAGRHGV | ASSMREVF | 0.5817467 |
| TSSTAGNSGGA | ASSMREVF | 0.77740914 |
| VPATVPVGILC | ASSMREVF | 0.8409256 |
| VQRPDSGGVVLTGRIS | ASSMREVF | 0.83205897 |
| WTPNPVPATKGVPAGLVQAT | ASSMREVF | 0.76661587 |

| | | |
|---|---|---|
| YALKRALAKFSDRFDVVII | ASSMREVF | 0.8112122 |
| YLIGSVIV | ASSMREVF | 0.8494959 |
| YGQLVAVID | ASSMREVF | 0.84482646 |
| VTVGGMTLPA | ASSMREVF | 0.85043895 |
| VLALPAV | ASSMREVF | 0.44971636 |
| VIPPTLNYET | ASSMREVF | 0.7996351 |
| VGMGGMHPSSG | ASSMREVF | 0.72929966 |
| VGLAGGAATA | ASSMREVF | 0.22726801 |
| VGGAGGAGGSGP | ASSMREVF | 0.84057194 |
| VFAAFMV | ASSMREVF | 0.6830156 |
| TLYDKLR | ASSMREVF | 0.83164406 |
| TLAVFTVL | ASSMREVF | 0.7249752 |
| TGSVQPARV | ASSMREVF | 0.843117 |
| TGGGAGSGGT | ASSMREVF | 0.8067261 |
| SHLETSL | ASSMREVF | 0.7903601 |
| SFTTGMT | ASSMREVF | 0.6794057 |
| QVEVAKRMGVRQPTVS | ASSMREVF | 0.82290006 |
| QGSGGPALAM | ASSMREVF | 0.8527403 |
| MLAPSNP | ASSMREVF | 0.85317254 |
| LVNNAAI | ASSMREVF | 0.43925214 |
| LTGEINTLA | ASSMREVF | 0.8466577 |
| LSIFGPV | ASSMREVF | 0.78638744 |
| LLALLANL | ASSMREVF | 0.48153228 |
| LISNASDALDKLR | ASSMREVF | 0.6725051 |
| LINEAVP | ASSMREVF | 0.7418389 |
| LGYAFSGL | ASSMREVF | 0.77865857 |
| LDFVAVAL | ASSMREVF | 0.8219962 |
| LAQPGAE | ASSMREVF | 0.8314847 |
| ISNLGAVTGF | ASSMREVF | 0.7559424 |
| HQEDPGRI | ASSMREVF | 0.8264278 |
| GTPGEPTP | ASSMREVF | 0.7691504 |
| GFFNSGS | ASSMREVF | 0.13935094 |
| GAGGGGLTAGAN | ASSMREVF | 0.72858244 |
| FVDALVNM | ASSMREVF | 0.46127322 |
| FLTVAPEL | ASSMREVF | 0.8556708 |
| FETLTAL | ASSMREVF | 0.64442515 |
| DVINPIEL | ASSMREVF | 0.76956284 |
| APVSGAPV | ASSMREVF | 0.78708816 |
| ALVGGLAW | ASSMREVF | 0.8233255 |
| ALAAAAATAI | ASSMREVF | 0.81237036 |
| AFGLGLLL | ASSMREVF | 0.71750116 |
| AFFAAASL | ASSMREVF | 0.6334028 |
| AAAALPAFSPP | ASSMREVF | 0.82403654 |
| VVLAPAL | ASSMREVF | 0.807482 |
| TIGHVDHGKTTL | ASSMREVF | 0.6912131 |
| LISNASDALDK | ASSMREVF | 0.7641984 |
| LISNASDAL | ASSMREVF | 0.8423208 |
| ISNASDALDK | ASSMREVF | 0.37328172 |
| NQAETLVYQTE | ASSMREVF | 0.8513006 |
| TGGTGGTAGAG | ASSMREVF | 0.66210794 |
| TGSTAGGAGGA | ASSMREVF | 0.75655526 |

| | | |
|---|---|---|
| DPLAVPVAAGPV | ASSMREVF | 0.8398472 |
| GGTGGTPGNGGN | ASSMREVF | 0.8144028 |
| PAGLAGVDPATL | ASSMREVF | 0.7923853 |
| GNNLVVEAPHAGASV | ASSMREVF | 0.71139044 |
| KVTRIMSASTLEQPA | ASSMREVF | 0.64596164 |
| LGFALPIALPRRTVV | ASSMREVF | 0.68731886 |
| TRLGNLTARKLVVLV | ASSMREVF | 0.69562155 |
| WSQQRRLIYFPSAGP | ASSMREVF | 0.85053587 |
| GPNGNPDPMAAAVDIR | ASSMREVF | 0.8285137 |
| TQRCVSVADLRAAASR | ASSMREVF | 0.4943897 |
| QFVLDVAEHPVEVLTETV | ASSMREVF | 0.8441287 |
| GGAGGTGGAGGNGGTQAGDG | ASSMREVF | 0.85042506 |
| PLLATLRAIAVDVDAKPAQV | ASSMREVF | 0.8487551 |
| APKFRTGGYMIDFWGVGYQVAK | ASSMREVF | 0.625452 |
| GAGGAGGAGGAAGGPGGTGGTG | ASSMREVF | 0.76101744 |
| IGALGLLATDVAPAGSVAFGVA | ASSMREVF | 0.7835887 |
| AGGAGGIGGNGGAGGAGGVLMG | ASSMREVF | 0.783695 |
| ADELIGLL | ASSMREVF | 0.75749135 |
| ADRALKA | ASSMREVF | 0.6481226 |
| AGAVGQVGGAGGA | ASSMREVF | 0.8496064 |
| AGGGGGLGGAP | ASSMREVF | 0.86041355 |
| AGGLLYGNGGAG | ASSMREVF | 0.75207317 |
| AGLESIGL | ASSMREVF | 0.46701205 |
| AGLKTGII | ASSMREVF | 0.7918103 |
| ASLAALLL | ASSMREVF | 0.68018985 |
| ASVINISSI | ASSMREVF | 0.7132391 |
| ATPPGPA | ASSMREVF | 0.7403871 |
| ATVASALV | ASSMREVF | 0.4807821 |
| ATVASLIL | ASSMREVF | 0.78308755 |
| AVNGPVLIRH | ASSMREVF | 0.8467655 |
| DGAAGAVGI | ASSMREVF | 0.75043654 |
| DPLLHGI | ASSMREVF | 0.7833136 |
| ELGAALGAL | ASSMREVF | 0.8408761 |
| ELGRTQAV | ASSMREVF | 0.8326987 |
| FVAGGYLS | ASSMREVF | 0.84620595 |
| GASGGAGGAGGW | ASSMREVF | 0.845376 |
| GGAGAGGDGGAGGI | ASSMREVF | 0.80172527 |
| GTGHNGSQPGMGGNG | ASSMREVF | 0.8090819 |
| GYAAASASAS | ASSMREVF | 0.8259061 |
| HQVDVAAVQQG | ASSMREVF | 0.78527004 |
| IGAAIAHT | ASSMREVF | 0.806936 |
| IGMIEVAQL | ASSMREVF | 0.7989898 |
| IRLAPEHTSAMD | ASSMREVF | 0.8150512 |
| IYNVSVL | ASSMREVF | 0.80967 |
| KQGFLGM | ASSMREVF | 0.8552847 |
| LESVITQL | ASSMREVF | 0.3123111 |
| LGLGALVVV | ASSMREVF | 0.7011016 |
| LIGNGGNGGSGGI | ASSMREVF | 0.8477264 |
| LNAINAPTL | ASSMREVF | 0.2507122 |
| LQLLNTGV | ASSMREVF | 0.71781766 |
| LQTSIEF | ASSMREVF | 0.7971065 |

| | | |
|---|---|---|
| MSFAVSEL | ASSMREVF | 0.8435457 |
| NTGALGL | ASSMREVF | 0.23406261 |
| PPGAGGAGGAGTT | ASSMREVF | 0.776221 |
| QLVGLGI | ASSMREVF | 0.50755 |
| RFLEQVHQL | ASSMREVF | 0.8516842 |
| RGVGYVLR | ASSMREVF | 0.7553058 |
| SATGVFAGVI | ASSMREVF | 0.5016036 |
| SLPAITLP | ASSMREVF | 0.84405386 |
| STALAFTV | ASSMREVF | 0.6638503 |
| SVLTVGLGK | ASSMREVF | 0.8295008 |
| TALAGAGSGI | ASSMREVF | 0.83965683 |
| TGLPFTGI | ASSMREVF | 0.8570695 |
| VAEHVTV | ASSMREVF | 0.59016085 |
| VDIVLNSL | ASSMREVF | 0.83517635 |
| VLTGGGAVL | ASSMREVF | 0.7807653 |
| VNVGLAYGI | ASSMREVF | 0.8556499 |
| VSSAAVGAV | ASSMREVF | 0.72972226 |
| VTSVYEAGV | ASSMREVF | 0.5205604 |
| WVALDRGA | ASSMREVF | 0.8374695 |
| YLVGSVVI | ASSMREVF | 0.8174789 |
| GPPGTGKTLL | ASSMREVF | 0.7732562 |
| GVGMTKF | ASSMREVF | 0.8384557 |
| IGIGGSDL | ASSMREVF | 0.8121543 |
| IGTLLAEL | ASSMREVF | 0.77112263 |
| IYTSGTT | ASSMREVF | 0.82711244 |
| LSFPAGV | ASSMREVF | 0.8261093 |
| LYTSGTT | ASSMREVF | 0.70001465 |
| VGAFPFGF | ASSMREVF | 0.84678906 |
| VGAGTVEF | ASSMREVF | 0.7993025 |
| VGKSTLLL | ASSMREVF | 0.6629364 |
| AALAHAGGGIV | ASSMREVF | 0.8502538 |
| AALVAVSLL | ASSMREVF | 0.8035209 |
| AAVSRAVAAL | ASSMREVF | 0.8348154 |
| AGGSSGIGGF | ASSMREVF | 0.82596624 |
| AGLTVANANG | ASSMREVF | 0.8546557 |
| AGLVGAVL | ASSMREVF | 0.7008294 |
| ALSQAFTV | ASSMREVF | 0.8104746 |
| ALTFSVSL | ASSMREVF | 0.79080045 |
| ALTGAALVAL | ASSMREVF | 0.8426305 |
| ASFSIGAGVL | ASSMREVF | 0.798264 |
| DGSVYKI | ASSMREVF | 0.8189014 |
| EQLAEIV | ASSMREVF | 0.8335805 |
| FAVSELTVM | ASSMREVF | 0.8438083 |
| GAAPLAAGL | ASSMREVF | 0.8032027 |
| GADGGDGGSGGAGG | ASSMREVF | 0.8172631 |
| GFGLDTGGAGG | ASSMREVF | 0.8530232 |
| GLASGGLL | ASSMREVF | 0.5625725 |
| GRIAGAAVLL | ASSMREVF | 0.7926934 |
| GSAGSLNTSF | ASSMREVF | 0.8372529 |
| GTGQIGI | ASSMREVF | 0.8464231 |
| GTLIALIL | ASSMREVF | 0.80160856 |

| | | |
|---|---|---|
| IAATMGFVVG | ASSMREVF | 0.8460212 |
| IFAQALL | ASSMREVF | 0.7678484 |
| IGAAVALGF | ASSMREVF | 0.8123795 |
| IGLTPLAL | ASSMREVF | 0.7557833 |
| ILGSNTSTL | ASSMREVF | 0.7755211 |
| IVFSLTVL | ASSMREVF | 0.6995114 |
| LASKATGF | ASSMREVF | 0.7848917 |
| LASVALIL | ASSMREVF | 0.8305973 |
| LFAVGLTL | ASSMREVF | 0.8158159 |
| LFNGGIGTYI | ASSMREVF | 0.64669514 |
| LTLGAGII | ASSMREVF | 0.8181608 |
| LVAAALGDI | ASSMREVF | 0.83243406 |
| NTAFVQLGI | ASSMREVF | 0.8556493 |
| QLFSGAEL | ASSMREVF | 0.8073244 |
| SAATFITL | ASSMREVF | 0.8374435 |
| SNYGLANLGV | ASSMREVF | 0.76159763 |
| SYGGAYAVM | ASSMREVF | 0.8214538 |
| TTGVGGLTLG | ASSMREVF | 0.6478541 |
| TTQTIYI | ASSMREVF | 0.59626776 |
| VAAGAGLGAVI | ASSMREVF | 0.79223186 |
| VAAVVTLL | ASSMREVF | 0.71152544 |
| VGFLGGLI | ASSMREVF | 0.83426416 |
| VGVIPILL | ASSMREVF | 0.84265435 |
| VIFASSLI | ASSMREVF | 0.82410765 |
| VKGGDGGAAAG | ASSMREVF | 0.84363544 |
| VTKTFTNI | ASSMREVF | 0.8397909 |
| VVAAAILL | ASSMREVF | 0.85247254 |
| YAALDVEL | ASSMREVF | 0.72382617 |
| LASVALLL | ASSMREVF | 0.78783643 |
| LDAVHRV | ASSMREVF | 0.4895653 |
| QGLSSSIF | ASSMREVF | 0.72734696 |
| RGTFANIRL | ASSMREVF | 0.71258926 |
| RQLWWGHRI | ASSMREVF | 0.8026147 |
| YGPPGTGKTLL | ASSMREVF | 0.21186101 |
| AALVGAPANVA | ASSMREVF | 0.8166656 |
| AATFRRTE | ASSMREVF | 0.8494321 |
| AGALGGGI | ASSMREVF | 0.7960434 |
| AGGAGGAGGIGAG | ASSMREVF | 0.8306732 |
| AGSGALGL | ASSMREVF | 0.8142378 |
| AQIADQI | ASSMREVF | 0.7576835 |
| ASAVAVTYL | ASSMREVF | 0.8310896 |
| ASTVLGGTGG | ASSMREVF | 0.7554933 |
| AVGGEPAK | ASSMREVF | 0.8472017 |
| CEQGTFGGTC | ASSMREVF | 0.8418951 |
| DEGKAPAG | ASSMREVF | 0.8100567 |
| DTVVFPAL | ASSMREVF | 0.79099035 |
| FEGITMAG | ASSMREVF | 0.83137554 |
| FPLVVQD | ASSMREVF | 0.7301078 |
| GAFTTPQ | ASSMREVF | 0.8057493 |
| GFGGFGTAGGVGGA | ASSMREVF | 0.78789806 |
| GFLAQFL | ASSMREVF | 0.7990029 |

| | | |
|---|---|---|
| GGAGGAGGNAGVA | ASSMREVF | 0.6392034 |
| GGVGTAGAGGAGGAGG | ASSMREVF | 0.82260275 |
| GNGGGGDAGNAGSGGN | ASSMREVF | 0.7832407 |
| GTGGAGGAGNDAGS | ASSMREVF | 0.8270635 |
| GTTNTGWFNS | ASSMREVF | 0.65945697 |
| IFMGPLFTL | ASSMREVF | 0.85446686 |
| ILAHAIEAI | ASSMREVF | 0.83953536 |
| IRSAIREAL | ASSMREVF | 0.54584706 |
| ITFAGGG | ASSMREVF | 0.83328646 |
| ITGMSGHQEV | ASSMREVF | 0.77607703 |
| IVAGLACTL | ASSMREVF | 0.76811624 |
| IVGGGLAGA | ASSMREVF | 0.8374796 |
| IVTGVLNI | ASSMREVF | 0.7974946 |
| LAQQGCI | ASSMREVF | 0.65385514 |
| LCLHVEN | ASSMREVF | 0.7350226 |
| LDAISDGS | ASSMREVF | 0.7977465 |
| LGGFAAML | ASSMREVF | 0.7458908 |
| LGGKNPMI | ASSMREVF | 0.49298722 |
| LGPKGVAAIL | ASSMREVF | 0.81556815 |
| LMVGVSI | ASSMREVF | 0.66439724 |
| LVGAAFVL | ASSMREVF | 0.71990573 |
| LVLAVAPL | ASSMREVF | 0.74898314 |
| MAAIAVIDAA | ASSMREVF | 0.8419477 |
| NNGINAELQ | ASSMREVF | 0.8342937 |
| RGEPLVLVHG | ASSMREVF | 0.7508482 |
| RGGAGGAAGLI | ASSMREVF | 0.81696135 |
| RSYRMHPAV | ASSMREVF | 0.7900432 |
| SAAHLLAGQLA | ASSMREVF | 0.83959943 |
| SAAVGSY | ASSMREVF | 0.81542516 |
| SPHAVLHRI | ASSMREVF | 0.809073 |
| SQIAKVL | ASSMREVF | 0.8321512 |
| TALGHLVP | ASSMREVF | 0.7985605 |
| TAPTTSGAA | ASSMREVF | 0.7971134 |
| TDGDDGNGGSPGT | ASSMREVF | 0.8042424 |
| TDGLGEN | ASSMREVF | 0.7185673 |
| TGGPPLPGQA | ASSMREVF | 0.776634 |
| TIPAITI | ASSMREVF | 0.80836886 |
| TIPGFTI | ASSMREVF | 0.8563912 |
| TSIASGALA | ASSMREVF | 0.81068385 |
| TVTLPGAAI | ASSMREVF | 0.462384 |
| VASGLSAALP | ASSMREVF | 0.75237626 |
| VGAGVIGI | ASSMREVF | 0.77142334 |
| VGPKRFGV | ASSMREVF | 0.83646876 |
| VNVAFPD | ASSMREVF | 0.8417412 |
| VSMTNSGVS | ASSMREVF | 0.853604 |
| VTDVLGDLSV | ASSMREVF | 0.8307607 |
| VVVVGAGFAGL | ASSMREVF | 0.7266164 |
| VYTVGTVV | ASSMREVF | 0.84149987 |
| WTTFSGP | ASSMREVF | 0.779571 |
| YDELVEAF | ASSMREVF | 0.80236334 |
| YGTGGAGGNGG | ASSMREVF | 0.8056514 |

| | | |
|---|---|---|
| YTNVMAV | ASSMREVF | 0.81760836 |
| FLEQVHQL | ASSMREVF | 0.7537416 |
| GLSLLAVAV | ASSMREVF | 0.77788806 |
| LEQVHQL | ASSMREVF | 0.57040536 |
| LNAAGAI | ASSMREVF | 0.8425647 |
| PAVHASL | ASSMREVF | 0.7297528 |
| TELEDDA | ASSMREVF | 0.81592476 |
| TPPGPPN | ASSMREVF | 0.7833106 |
| TPPPPPPP | ASSMREVF | 0.84463465 |
| TQSPALI | ASSPRTGGPLY | 0.8480833 |
| EFAAAFPQ | ASSPRTGGPLY | 0.74540937 |
| LLSGNLDVL | ASSPRTGGPLY | 0.6718049 |
| GPANLGNYNI | ASSPRTGGPLY | 0.8294653 |
| TLGPTLP | ASSPRTGGPLY | 0.7334357 |
| VYAYSSCCK | ASSPRTGGPLY | 0.84323704 |
| APAPAPAPNI | ASSPRTGGPLY | 0.79524386 |
| PGGPVPIP | ASSPRTGGPLY | 0.7730788 |
| FLTVALEP | ASSPRTGGPLY | 0.7515309 |
| SQYFGANL | ASSPRTGGPLY | 0.8349807 |
| GGVKPAA | ASSPRTGGPLY | 0.7260803 |
| GGTGGAGGAGGW | ASSPRTGGPLY | 0.8400878 |
| GVGGVAGLGGAGG | ASSPRTGGPLY | 0.41151437 |
| VLLVLATTI | ASSPRTGGPLY | 0.8378085 |
| IGIFAAML | ASSPRTGGPLY | 0.67142004 |
| AVGDLID | ASSPRTGGPLY | 0.72448826 |
| NISPLPL | ASSPRTGGPLY | 0.33741784 |
| VTAVFVAI | ASSPRTGGPLY | 0.8039859 |
| SGDDNDAAL | ASSPRTGGPLY | 0.8116542 |
| AVAALAHS | ASSPRTGGPLY | 0.8485197 |
| GGVTVMATP | ASSPRTGGPLY | 0.83652484 |
| AGLAANVRD | ASSPRTGGPLY | 0.8188888 |
| TNRPDLIDPALLRPG | ASSPRTGGPLY | 0.8471776 |
| LANLGVTAL | ASSPRTGGPLY | 0.7540324 |
| TGERIEL | ASSPRTGGPLY | 0.39149016 |
| NGEELAA | ASSPRTGGPLY | 0.80363053 |
| VLSLVWG | ASSPRTGGPLY | 0.8008356 |
| IGYALWL | ASSPRTGGPLY | 0.8483711 |
| LSLLLAPL | ASSPRTGGPLY | 0.81812644 |
| TLDEALK | ASSPRTGGPLY | 0.56313074 |
| TDGAGTAEGG | ASSPRTGGPLY | 0.8354444 |
| NAPGGAGGNGG | ASSPRTGGPLY | 0.8109237 |
| VSVFMGMGGNF | ASSPRTGGPLY | 0.6484548 |
| LASYLAAL | ASSPRTGGPLY | 0.7274283 |
| LVGGTPAL | ASSPRTGGPLY | 0.7221128 |
| VRNADVL | ASSPRTGGPLY | 0.691785 |
| TCSSPFLVSAM | ASSPRTGGPLY | 0.79125273 |
| VHITDAE | ASSPRTGGPLY | 0.75246704 |
| EYAAPSF | ASSPRTGGPLY | 0.7875716 |
| VVDAFHGGL | ASSPRTGGPLY | 0.7823551 |
| ASVVDPQ | ASSPRTGGPLY | 0.7042748 |
| SYNEFGGLS | ASSPRTGGPLY | 0.7756268 |

| | | |
|---|---|---|
| DPDGSVY | ASSPRTGGPLY | 0.76049197 |
| MVSTIHATE | ASSPRTGGPLY | 0.81630147 |
| LAAGVADPV | ASSPRTGGPLY | 0.83945143 |
| TKEQIAA | ASSPRTGGPLY | 0.751723 |
| SLASPAAL | ASSPRTGGPLY | 0.81341976 |
| IGLVFYGV | ASSPRTGGPLY | 0.8406961 |
| FLLDASNL | ASSPRTGGPLY | 0.57403624 |
| YQVGNFP | ASSPRTGGPLY | 0.7257337 |
| ESLLSQST | ASSPRTGGPLY | 0.83228314 |
| LFTVGFAAVG | ASSPRTGGPLY | 0.80305433 |
| ATPPTPT | ASSPRTGGPLY | 0.8429859 |
| TDLATLG | ASSPRTGGPLY | 0.8046265 |
| SAGGTGR | ASSPRTGGPLY | 0.8191574 |
| GFSNSGS | ASSPRTGGPLY | 0.8140447 |
| GGNGGSGGTSV | ASSPRTGGPLY | 0.844056 |
| GTSVATGGAGN | ASSPRTGGPLY | 0.8251321 |
| NGGNGGSGGTS | ASSPRTGGPLY | 0.76828575 |
| AVVKFPH | ASSPRTGGPLY | 0.8336453 |
| TPPPPPP | ASSPRTGGPLY | 0.58172256 |
| AAPAAAAAPA | ASSPRTGGPLY | 0.8330836 |
| IALNEDL | ASSPRTGGPLY | 0.8400545 |
| IVNVSSM | ASSPRTGGPLY | 0.6744498 |
| TGIPPAPR | ASSPRTGGPLY | 0.7781108 |
| FDLGGGTF | ASSPRTGGPLY | 0.85556424 |
| FDLGGGTFDVS | ASSPRTGGPLY | 0.8353815 |
| NEPTAAA | ASSPRTGGPLY | 0.8031953 |
| NEPTAAALA | ASSPRTGGPLY | 0.6985015 |
| GSIINISSI | ASSPRTGGPLY | 0.75717956 |
| VAPGNAGTA | ASSPRTGGPLY | 0.77637285 |
| TGGGASLE | ASSPRTGGPLY | 0.39606166 |
| VAVYNHYKRI | ASSPRTGGPLY | 0.83258843 |
| TDPAPATT | ASSPRTGGPLY | 0.753238 |
| TNRPDILDPALLRPG | ASSPRTGGPLY | 0.8435038 |
| LTTTTTS | ASSPRTGGPLY | 0.8096521 |
| PTGSGKTL | ASSPRTGGPLY | 0.8445475 |
| TGSGKTL | ASSPRTGGPLY | 0.7505827 |
| TIGHVDHGK | ASSPRTGGPLY | 0.7647723 |
| TIGHVDHGKTT | ASSPRTGGPLY | 0.39067048 |
| GHVDHGKTT | ASSPRTGGPLY | 0.7765406 |
| AQFEHTLL | ASSPRTGGPLY | 0.7282672 |
| IVNQDEL | ASSPRTGGPLY | 0.8012564 |
| TNGSQFF | ASSPRTGGPLY | 0.48373267 |
| TGKGIVE | ASSPRTGGPLY | 0.84855926 |
| LISNASDA | ASSPRTGGPLY | 0.77395713 |
| AELRTLGELTKIAKAHGAQ | ASSPRTGGPLY | 0.82818484 |
| APNAVAAAEPSW | ASSPRTGGPLY | 0.6959169 |
| APQVAGAATAVL | ASSPRTGGPLY | 0.64198893 |
| ARRDPAAAAAILPTDA | ASSPRTGGPLY | 0.72309375 |
| DAAARDVLAEAGLAEYF | ASSPRTGGPLY | 0.8350492 |
| DDQIVLSAKANLAKVEQ | ASSPRTGGPLY | 0.5780022 |
| DELLPDTGEDSDEDGH | ASSPRTGGPLY | 0.3987821 |

| | | |
|---|---|---|
| ELPEAPANRATVPQD | ASSPRTGGPLY | 0.7313335 |
| FGQSKIERVTGIG | ASSPRTGGPLY | 0.8022648 |
| GAGGTGSTAGGAGGAGGAGGLY | ASSPRTGGPLY | 0.84856653 |
| GHGGTGAAGGNGGTGGAGGSGI | ASSPRTGGPLY | 0.8397844 |
| GNGGNGGAGGEGGPGGAGGAGA | ASSPRTGGPLY | 0.81234264 |
| GSTAGNGGNGGNATNGTIAGQP | ASSPRTGGPLY | 0.85691476 |
| GTGGDGGNGGNGGTGG | ASSPRTGGPLY | 0.17582496 |
| GVRRPAGIPADAPV | ASSPRTGGPLY | 0.72463375 |
| GYRQVTDAHLLTLARR | ASSPRTGGPLY | 0.84930384 |
| IAKMGQDGHDRG | ASSPRTGGPLY | 0.8074044 |
| KAEPSIEIQYT | ASSPRTGGPLY | 0.7770339 |
| KPVRIRMAADEDTVAVT | ASSPRTGGPLY | 0.844213 |
| MPPRIAGMRLLVIKPEP | ASSPRTGGPLY | 0.68657136 |
| MKFYAVALFRSRGVPPTR | ASSPRTGGPLY | 0.7583082 |
| NAGGFGILGAVAHSP | ASSPRTGGPLY | 0.8051625 |
| NEGREITLAMAEM | ASSPRTGGPLY | 0.83900017 |
| NELVEWSHPLGHRWRWE | ASSPRTGGPLY | 0.8256656 |
| NLGAGGTGGNGGNPG | ASSPRTGGPLY | 0.46425068 |
| NTEWPKWETVMAADPVD | ASSPRTGGPLY | 0.83393335 |
| PASAAPAETQP | ASSPRTGGPLY | 0.8472768 |
| QQSSDGKNGVRAAEITGE | ASSPRTGGPLY | 0.74546343 |
| RAPDAVTTSWWKE | ASSPRTGGPLY | 0.8394189 |
| SGFGNVGAHTSGWFNQ | ASSPRTGGPLY | 0.80487573 |
| SGVAGAVGQAASVGGLKV | ASSPRTGGPLY | 0.8011586 |
| SNGGAGTGSAGGNGGT | ASSPRTGGPLY | 0.6982852 |
| TGGAGGEGGAAG | ASSPRTGGPLY | 0.6702606 |
| TGGDGGAAGAGGAPGHG | ASSPRTGGPLY | 0.8544612 |
| TMPPLLAAIANAGRHGV | ASSPRTGGPLY | 0.69810444 |
| TSSTAGNSGGA | ASSPRTGGPLY | 0.8122866 |
| VPATVPVGILC | ASSPRTGGPLY | 0.7870499 |
| VQRPDSGGVVLTGRIS | ASSPRTGGPLY | 0.8542563 |
| WTPNPVPATKGVPAGLVQAT | ASSPRTGGPLY | 0.82298267 |
| YALKRALAKFSDRFDVVII | ASSPRTGGPLY | 0.8247088 |
| YLIGSVIV | ASSPRTGGPLY | 0.73198843 |
| YGQLVAVID | ASSPRTGGPLY | 0.8255187 |
| VTVGGMTLPA | ASSPRTGGPLY | 0.8266387 |
| VLALPAV | ASSPRTGGPLY | 0.57666135 |
| VIPPTLNYET | ASSPRTGGPLY | 0.8454194 |
| VGMGGMHPSSG | ASSPRTGGPLY | 0.6600359 |
| VGLAGGAATA | ASSPRTGGPLY | 0.80265546 |
| VGGAGGAGGSGP | ASSPRTGGPLY | 0.8299463 |
| VFAAFMV | ASSPRTGGPLY | 0.8453319 |
| TLYDKLR | ASSPRTGGPLY | 0.79459333 |
| TLAVFTVL | ASSPRTGGPLY | 0.853463 |
| TGSVQPARV | ASSPRTGGPLY | 0.6779406 |
| TGGGAGSGGT | ASSPRTGGPLY | 0.8436915 |
| SHLETSL | ASSPRTGGPLY | 0.8421507 |
| SFTTGMT | ASSPRTGGPLY | 0.8277456 |
| QVEVAKRMGVRQPTVS | ASSPRTGGPLY | 0.85600793 |
| QGSGGPALAM | ASSPRTGGPLY | 0.8445595 |
| MLAPSNP | ASSPRTGGPLY | 0.35066098 |

| | | |
|---|---|---|
| LVNNAAI | ASSPRTGGPLY | 0.79938567 |
| LTGEINTLA | ASSPRTGGPLY | 0.8419958 |
| LSIFGPV | ASSPRTGGPLY | 0.8472529 |
| LLALLANL | ASSPRTGGPLY | 0.8385989 |
| LISNASDALDKLR | ASSPRTGGPLY | 0.8241615 |
| LINEAVP | ASSPRTGGPLY | 0.77371013 |
| LGYAFSGL | ASSPRTGGPLY | 0.84859097 |
| LDFVAVAL | ASSPRTGGPLY | 0.8614982 |
| LAQPGAE | ASSPRTGGPLY | 0.82894796 |
| ISNLGAVTGF | ASSPRTGGPLY | 0.75875133 |
| HQEDPGRI | ASSPRTGGPLY | 0.7408366 |
| GTPGEPTP | ASSPRTGGPLY | 0.8279114 |
| GFFNSGS | ASSPRTGGPLY | 0.8332459 |
| GAGGGGLTAGAN | ASSPRTGGPLY | 0.64013493 |
| FVDALVNM | ASSPRTGGPLY | 0.7946566 |
| FLTVAPEL | ASSPRTGGPLY | 0.81332904 |
| FETLTAL | ASSPRTGGPLY | 0.30043888 |
| DVINPIEL | ASSPRTGGPLY | 0.8087604 |
| APVSGAPV | ASSPRTGGPLY | 0.834797 |
| ALVGGLAW | ASSPRTGGPLY | 0.83099586 |
| ALAAAAATAI | ASSPRTGGPLY | 0.67197657 |
| AFGLGLLL | ASSPRTGGPLY | 0.8412169 |
| AFFAAASL | ASSPRTGGPLY | 0.83513814 |
| AAAALPAFSPP | ASSPRTGGPLY | 0.85534143 |
| VVLAPAL | ASSPRTGGPLY | 0.549367 |
| TIGHVDHGKTTL | ASSPRTGGPLY | 0.27555218 |
| LISNASDALDK | ASSPRTGGPLY | 0.82647604 |
| LISNASDAL | ASSPRTGGPLY | 0.8285654 |
| ISNASDALDK | ASSPRTGGPLY | 0.7382275 |
| NQAETLVYQTE | ASSPRTGGPLY | 0.6463011 |
| TGGTGGTAGAG | ASSPRTGGPLY | 0.80169773 |
| TGSTAGGAGGA | ASSPRTGGPLY | 0.838014 |
| DPLAVPVAAGPV | ASSPRTGGPLY | 0.5519778 |
| GGTGGTPGNGGN | ASSPRTGGPLY | 0.60716736 |
| PAGLAGVDPATL | ASSPRTGGPLY | 0.81445956 |
| GNNLVVEAPHAGASV | ASSPRTGGPLY | 0.8464076 |
| KVTRIMSASTLEQPA | ASSPRTGGPLY | 0.67412883 |
| LGFALPIALPRRTVV | ASSPRTGGPLY | 0.74651486 |
| TRLGNLTARKLVVLV | ASSPRTGGPLY | 0.8208026 |
| WSQQRRLIYFPSAGP | ASSPRTGGPLY | 0.84374416 |
| GPNGNPDPMAAAVDIR | ASSPRTGGPLY | 0.8099625 |
| TQRCVSVADLRAAASR | ASSPRTGGPLY | 0.6190036 |
| QFVLDVAEHPVEVLTETV | ASSPRTGGPLY | 0.8142102 |
| GGAGGTGGAGGNGGTQAGDG | ASSPRTGGPLY | 0.62687415 |
| PLLATLRAIAVDVDAKPAQV | ASSPRTGGPLY | 0.8434689 |
| APKFRTGGYMIDFWGVGYQVAK | ASSPRTGGPLY | 0.8091797 |
| GAGGAGGAGGAAGGPGGTGGTG | ASSPRTGGPLY | 0.8087977 |
| IGALGLLATDVAPAGSVAFGVA | ASSPRTGGPLY | 0.7668334 |
| AGGAGGIGGNGGAGGAGGVLMG | ASSPRTGGPLY | 0.65663993 |
| ADELIGLL | ASSPRTGGPLY | 0.55331165 |
| ADRALKA | ASSPRTGGPLY | 0.7791689 |

| | | |
|---|---|---|
| AGAVGQVGGAGGA | ASSPRTGGPLY | 0.84237903 |
| AGGGGGLGGAP | ASSPRTGGPLY | 0.8285034 |
| AGGLLYGNGGAG | ASSPRTGGPLY | 0.8345127 |
| AGLESIGL | ASSPRTGGPLY | 0.8083596 |
| AGLKTGII | ASSPRTGGPLY | 0.6376956 |
| ASLAALLL | ASSPRTGGPLY | 0.83518946 |
| ASVINISSI | ASSPRTGGPLY | 0.53099024 |
| ATPPGPA | ASSPRTGGPLY | 0.8381508 |
| ATVASALV | ASSPRTGGPLY | 0.7813934 |
| ATVASLIL | ASSPRTGGPLY | 0.8374976 |
| AVNGPVLIRH | ASSPRTGGPLY | 0.7263968 |
| DGAAGAVGI | ASSPRTGGPLY | 0.8409301 |
| DPLLHGI | ASSPRTGGPLY | 0.81866735 |
| ELGAALGAL | ASSPRTGGPLY | 0.6676886 |
| ELGRTQAV | ASSPRTGGPLY | 0.8265369 |
| FVAGGYLS | ASSPRTGGPLY | 0.8350765 |
| GASGGAGGAGGW | ASSPRTGGPLY | 0.8373733 |
| GGAGAGGDGGAGGI | ASSPRTGGPLY | 0.4912098 |
| GTGHNGSQPGMGGNG | ASSPRTGGPLY | 0.7844776 |
| GYAAASASAS | ASSPRTGGPLY | 0.74031544 |
| HQVDVAAVQQG | ASSPRTGGPLY | 0.8125305 |
| IGAAIAHT | ASSPRTGGPLY | 0.8016063 |
| IGMIEVAQL | ASSPRTGGPLY | 0.7931683 |
| IRLAPEHTSAMD | ASSPRTGGPLY | 0.80408764 |
| IYNVSVL | ASSPRTGGPLY | 0.5999257 |
| KQGFLGM | ASSPRTGGPLY | 0.72580326 |
| LESVITQL | ASSPRTGGPLY | 0.7646363 |
| LGLGALVVV | ASSPRTGGPLY | 0.8046024 |
| LIGNGGNGGSGGI | ASSPRTGGPLY | 0.8268282 |
| LNAINAPTL | ASSPRTGGPLY | 0.6175191 |
| LQLLNTGV | ASSPRTGGPLY | 0.8095204 |
| LQTSIEF | ASSPRTGGPLY | 0.7671038 |
| MSFAVSEL | ASSPRTGGPLY | 0.6626738 |
| NTGALGL | ASSPRTGGPLY | 0.82596576 |
| PPGAGGAGGAGTT | ASSPRTGGPLY | 0.77792776 |
| QLVGLGI | ASSPRTGGPLY | 0.52178884 |
| RFLEQVHQL | ASSPRTGGPLY | 0.8373946 |
| RGVGYVLR | ASSPRTGGPLY | 0.64176404 |
| SATGVFAGVI | ASSPRTGGPLY | 0.8037017 |
| SLPAITLP | ASSPRTGGPLY | 0.84670746 |
| STALAFTV | ASSPRTGGPLY | 0.5852599 |
| SVLTVGLGK | ASSPRTGGPLY | 0.8395035 |
| TALAGAGSGI | ASSPRTGGPLY | 0.78751683 |
| TGLPFTGI | ASSPRTGGPLY | 0.8059155 |
| VAEHVTV | ASSPRTGGPLY | 0.79329586 |
| VDIVLNSL | ASSPRTGGPLY | 0.7095784 |
| VLTGGGAVL | ASSPRTGGPLY | 0.84107697 |
| VNVGLAYGI | ASSPRTGGPLY | 0.8219018 |
| VSSAAVGAV | ASSPRTGGPLY | 0.827736 |
| VTSVYEAGV | ASSPRTGGPLY | 0.81414664 |
| WVALDRGA | ASSPRTGGPLY | 0.8398007 |

| | | |
|---|---|---|
| YLVGSVVI | ASSPRTGGPLY | 0.8429376 |
| GPPGTGKTLL | ASSPRTGGPLY | 0.7535929 |
| GVGMTKF | ASSPRTGGPLY | 0.49784383 |
| IGIGGSDL | ASSPRTGGPLY | 0.78113705 |
| IGTLLAEL | ASSPRTGGPLY | 0.78050566 |
| IYTSGTT | ASSPRTGGPLY | 0.71923554 |
| LSFPAGV | ASSPRTGGPLY | 0.6004641 |
| LYTSGTT | ASSPRTGGPLY | 0.8312616 |
| VGAFPFGF | ASSPRTGGPLY | 0.8543448 |
| VGAGTVEF | ASSPRTGGPLY | 0.77285784 |
| VGKSTLLL | ASSPRTGGPLY | 0.8573499 |
| AALAHAGGGIV | ASSPRTGGPLY | 0.6344023 |
| AALVAVSLL | ASSPRTGGPLY | 0.8251343 |
| AAVSRAVAAL | ASSPRTGGPLY | 0.84457725 |
| AGGSSGIGGF | ASSPRTGGPLY | 0.66296387 |
| AGLTVANANG | ASSPRTGGPLY | 0.8189409 |
| AGLVGAVL | ASSPRTGGPLY | 0.7215264 |
| ALSQAFTV | ASSPRTGGPLY | 0.6921471 |
| ALTFSVSL | ASSPRTGGPLY | 0.83924794 |
| ALTGAALVAL | ASSPRTGGPLY | 0.7751082 |
| ASFSIGAGVL | ASSPRTGGPLY | 0.83785194 |
| DGSVYKI | ASSPRTGGPLY | 0.77094734 |
| EQLAEIV | ASSPRTGGPLY | 0.6504794 |
| FAVSELTVM | ASSPRTGGPLY | 0.822007 |
| GAAPLAAGL | ASSPRTGGPLY | 0.8429761 |
| GADGGDGGSGGAGG | ASSPRTGGPLY | 0.8421575 |
| GFGLDTGGAGG | ASSPRTGGPLY | 0.8610841 |
| GLASGGLL | ASSPRTGGPLY | 0.7089276 |
| GRIAGAAVLL | ASSPRTGGPLY | 0.84268653 |
| GSAGSLNTSF | ASSPRTGGPLY | 0.84832007 |
| GTGQIGI | ASSPRTGGPLY | 0.8071302 |
| GTLIALIL | ASSPRTGGPLY | 0.7955239 |
| IAATMGFVVG | ASSPRTGGPLY | 0.83266985 |
| IFAQALL | ASSPRTGGPLY | 0.7942517 |
| IGAAVALGF | ASSPRTGGPLY | 0.8259795 |
| IGLTPLAL | ASSPRTGGPLY | 0.8552129 |
| ILGSNTSTL | ASSPRTGGPLY | 0.20411605 |
| IVFSLTVL | ASSPRTGGPLY | 0.7865993 |
| LASKATGF | ASSPRTGGPLY | 0.58956885 |
| LASVALIL | ASSPRTGGPLY | 0.445921 |
| LFAVGLTL | ASSPRTGGPLY | 0.26415992 |
| LFNGGIGTYI | ASSPRTGGPLY | 0.14745824 |
| LTLGAGII | ASSPRTGGPLY | 0.39434344 |
| LVAAALGDI | ASSPRTGGPLY | 0.22602135 |
| NTAFVQLGI | ASSPRTGGPLY | 0.69413227 |
| QLFSGAEL | ASSPRTGGPLY | 0.5465445 |
| SAATFITL | ASSPRTGGPLY | 0.30791092 |
| SNYGLANLGV | ASSPRTGGPLY | 0.7851919 |
| SYGGAYAVM | ASSPRTGGPLY | 0.80312073 |
| TTGVGGLTLG | ASSPRTGGPLY | 0.15506202 |
| TTQTIYI | ASSPRTGGPLY | 0.4369481 |

| | | |
|---|---|---|
| VAAGAGLGAVI | ASSPRTGGPLY | 0.26409557 |
| VAAVVTLL | ASSPRTGGPLY | 0.28645855 |
| VGFLGGLI | ASSPRTGGPLY | 0.13155958 |
| VGVIPILL | ASSPRTGGPLY | 0.5666398 |
| VIFASSLI | ASSPRTGGPLY | 0.29633048 |
| VKGGDGGAAAG | ASSPRTGGPLY | 0.11248512 |
| VTKTFTNI | ASSPRTGGPLY | 0.3695196 |
| VVAAAILL | ASSPRTGGPLY | 0.41360998 |
| YAALDVEL | ASSPRTGGPLY | 0.26504514 |
| LASVALLL | ASSPRTGGPLY | 0.76144385 |
| LDAVHRV | ASSPRTGGPLY | 0.14807665 |
| QGLSSSIF | ASSPRTGGPLY | 0.22454484 |
| RGTFANIRL | ASSPRTGGPLY | 0.70839536 |
| RQLWWGHRI | ASSPRTGGPLY | 0.8119595 |
| YGPPGTGKTLL | ASSPRTGGPLY | 0.187847 |
| AALVGAPANVA | ASSPRTGGPLY | 0.60693574 |
| AATFRRTE | ASSPRTGGPLY | 0.8144506 |
| AGALGGGI | ASSPRTGGPLY | 0.51137316 |
| AGGAGGAGGIGAG | ASSPRTGGPLY | 0.69023407 |
| AGSGALGL | ASSPRTGGPLY | 0.76899755 |
| AQIADQI | ASSPRTGGPLY | 0.25413746 |
| ASAVAVTYL | ASSPRTGGPLY | 0.75251514 |
| ASTVLGGTGG | ASSPRTGGPLY | 0.7894403 |
| AVGGEPAK | ASSPRTGGPLY | 0.38040137 |
| CEQGTFGGTC | ASSPRTGGPLY | 0.4038236 |
| DEGKAPAG | ASSPRTGGPLY | 0.20646095 |
| DTVVFPAL | ASSPRTGGPLY | 0.35112864 |
| FEGITMAG | ASSPRTGGPLY | 0.12746504 |
| FPLVVQD | ASSPRTGGPLY | 0.6994114 |
| GAFTTPQ | ASSPRTGGPLY | 0.63721436 |
| GFGGFGTAGGVGGA | ASSPRTGGPLY | 0.3143729 |
| GFLAQFL | ASSPRTGGPLY | 0.18566833 |
| GGAGGAGGNAGVA | ASSPRTGGPLY | 0.20027521 |
| GGVGTAGAGGAGGAGG | ASSPRTGGPLY | 0.2663167 |
| GNGGGGDAGNAGSGGN | ASSPRTGGPLY | 0.38490766 |
| GTGGAGGAGNDAGS | ASSPRTGGPLY | 0.1342805 |
| GTTNTGWFNS | ASSPRTGGPLY | 0.25832218 |
| IFMGPLFTL | ASSPRTGGPLY | 0.7000343 |
| ILAHAIEAI | ASSPRTGGPLY | 0.7149645 |
| IRSAIREAL | ASSPRTGGPLY | 0.1408285 |
| ITFAGGG | ASSPRTGGPLY | 0.81416583 |
| ITGMSGHQEV | ASSPRTGGPLY | 0.69791293 |
| IVAGLACTL | ASSPRTGGPLY | 0.7555058 |
| IVGGGLAGA | ASSPRTGGPLY | 0.40456378 |
| IVTGVLNI | ASSPRTGGPLY | 0.22538939 |
| LAQQGCI | ASSPRTGGPLY | 0.7766002 |
| LCLHVEN | ASSPRTGGPLY | 0.45084786 |
| LDAISDGS | ASSPRTGGPLY | 0.2104589 |
| LGGFAAML | ASSPRTGGPLY | 0.6162317 |
| LGGKNPMI | ASSPRTGGPLY | 0.3066405 |
| LGPKGVAAIL | ASSPRTGGPLY | 0.29612008 |

| | | |
|---|---|---|
| LMVGVSI | ASSPRTGGPLY | 0.6489781 |
| LVGAAFVL | ASSPRTGGPLY | 0.57320976 |
| LVLAVAPL | ASSPRTGGPLY | 0.21054205 |
| MAAIAVIDAA | ASSPRTGGPLY | 0.18942584 |
| NNGINAELQ | ASSPRTGGPLY | 0.19092576 |
| RGEPLVLVHG | ASSPRTGGPLY | 0.5104064 |
| RGGAGGAAGLI | ASSPRTGGPLY | 0.7692777 |
| RSYRMHPAV | ASSPRTGGPLY | 0.38780853 |
| SAAHLLAGQLA | ASSPRTGGPLY | 0.38395655 |
| SAAVGSY | ASSPRTGGPLY | 0.72284865 |
| SPHAVLHRI | ASSPRTGGPLY | 0.55566794 |
| SQIAKVL | ASSPRTGGPLY | 0.3864502 |
| TALGHLVP | ASSPRTGGPLY | 0.7573443 |
| TAPTTSGAA | ASSPRTGGPLY | 0.37777573 |
| TDGDDGNGGSPGT | ASSPRTGGPLY | 0.5701109 |
| TDGLGEN | ASSPRTGGPLY | 0.52525496 |
| TGGPPLPGQA | ASSPRTGGPLY | 0.61838037 |
| TIPAITI | ASSPRTGGPLY | 0.7102347 |
| TIPGFTI | ASSPRTGGPLY | 0.5983007 |
| TSIASGALA | ASSPRTGGPLY | 0.45157072 |
| TVTLPGAAI | ASSPRTGGPLY | 0.12847348 |
| VASGLSAALP | ASSPRTGGPLY | 0.37776268 |
| VGAGVIGI | ASSPRTGGPLY | 0.6712911 |
| VGPKRFGV | ASSPRTGGPLY | 0.6079713 |
| VNVAFPD | ASSPRTGGPLY | 0.66034997 |
| VSMTNSGVS | ASSPRTGGPLY | 0.51339114 |
| VTDVLGDLSV | ASSPRTGGPLY | 0.29792935 |
| VVVVGAGFAGL | ASSPRTGGPLY | 0.7603961 |
| VYTVGTVV | ASSPRTGGPLY | 0.76012313 |
| WTTFSGP | ASSPRTGGPLY | 0.15333292 |
| YDELVEAF | ASSPRTGGPLY | 0.39987814 |
| YGTGGAGGNGG | ASSPRTGGPLY | 0.80142653 |
| YTNVMAV | ASSPRTGGPLY | 0.7137573 |
| FLEQVHQL | ASSPRTGGPLY | 0.39499304 |
| GLSLLAVAV | ASSPRTGGPLY | 0.2743094 |
| LEQVHQL | ASSPRTGGPLY | 0.81835437 |
| LNAAGAI | ASSPRTGGPLY | 0.2353791 |
| PAVHASL | ASSPRTGGPLY | 0.797603 |
| TELEDDA | ASSPRTGGPLY | 0.120680034 |
| TPPGPPN | ASSPRTGGPLY | 0.29172075 |
| TPPPPPPP | ASSPRTGGPLY | 0.2018223 |
| TQSPALI | ASSFGTEVF | 0.13384548 |
| EFAAAFPQ | ASSFGTEVF | 0.62385386 |
| LLSGNLDVL | ASSFGTEVF | 0.2601951 |
| GPANLGNYNI | ASSFGTEVF | 0.7367486 |
| TLGPTLP | ASSFGTEVF | 0.4345631 |
| VYAYSSCCK | ASSFGTEVF | 0.61851096 |
| APAPAPAPNI | ASSFGTEVF | 0.58018875 |
| PGGPVPIP | ASSFGTEVF | 0.485997 |
| FLTVALEP | ASSFGTEVF | 0.19358516 |
| SQYFGANL | ASSFGTEVF | 0.6001081 |

| | | |
|---|---|---|
| GGVKPAA | ASSFGTEVF | 0.76949596 |
| GGTGGAGGAGGW | ASSFGTEVF | 0.39890677 |
| GVGGVAGLGGAGG | ASSFGTEVF | 0.17042325 |
| VLLVLATTI | ASSFGTEVF | 0.35451293 |
| IGIFAAML | ASSFGTEVF | 0.38382682 |
| AVGDLID | ASSFGTEVF | 0.123416826 |
| NISPLPL | ASSFGTEVF | 0.117101915 |
| VTAVFVAI | ASSFGTEVF | 0.25458172 |
| SGDDNDAAL | ASSFGTEVF | 0.22635242 |
| AVAALAHS | ASSFGTEVF | 0.776512 |
| GGVTVMATP | ASSFGTEVF | 0.72981143 |
| AGLAANVRD | ASSFGTEVF | 0.360349 |
| TNRPDLIDPALLRPG | ASSFGTEVF | 0.7279725 |
| LANLGVTAL | ASSFGTEVF | 0.6598599 |
| TGERIEL | ASSFGTEVF | 0.10932581 |
| NGEELAA | ASSFGTEVF | 0.31701928 |
| VLSLVWG | ASSFGTEVF | 0.15738931 |
| IGYALWL | ASSFGTEVF | 0.72827303 |
| LSLLLAPL | ASSFGTEVF | 0.2055166 |
| TLDEALK | ASSFGTEVF | 0.29623538 |
| TDGAGTAEGG | ASSFGTEVF | 0.43623856 |
| NAPGGAGGNGG | ASSFGTEVF | 0.713847 |
| VSVFMGMGGNF | ASSFGTEVF | 0.5713029 |
| LASYLAAL | ASSFGTEVF | 0.21222374 |
| LVGGTPAL | ASSFGTEVF | 0.11269767 |
| VRNADVL | ASSFGTEVF | 0.58725095 |
| TCSSPFLVSAM | ASSFGTEVF | 0.16205415 |
| VHITDAE | ASSFGTEVF | 0.20336565 |
| EYAAPSF | ASSFGTEVF | 0.13128076 |
| VVDAFHGGL | ASSFGTEVF | 0.12056157 |
| ASVVDPQ | ASSFGTEVF | 0.16364555 |
| SYNEFGGLS | ASSFGTEVF | 0.63222027 |
| DPDGSVY | ASSFGTEVF | 0.48044086 |
| MVSTIHATE | ASSFGTEVF | 0.3880167 |
| LAAGVADPV | ASSFGTEVF | 0.8089588 |
| TKEQIAA | ASSFGTEVF | 0.7564622 |
| SLASPAAL | ASSFGTEVF | 0.3812489 |
| IGLVFYGV | ASSFGTEVF | 0.2988003 |
| FLLDASNL | ASSFGTEVF | 0.20269527 |
| YQVGNFP | ASSFGTEVF | 0.19512719 |
| ESLLSQST | ASSFGTEVF | 0.20334011 |
| LFTVGFAAVG | ASSFGTEVF | 0.82433957 |
| ATPPTPT | ASSFGTEVF | 0.2748994 |
| TDLATLG | ASSFGTEVF | 0.16745894 |
| SAGGTGR | ASSFGTEVF | 0.39660436 |
| GFSNSGS | ASSFGTEVF | 0.7569755 |
| GGNGGSGGTSV | ASSFGTEVF | 0.8275445 |
| GTSVATGGAGN | ASSFGTEVF | 0.19949584 |
| NGGNGGSGGTS | ASSFGTEVF | 0.35642773 |
| AVVKFPH | ASSFGTEVF | 0.3265493 |
| TPPPPPP | ASSFGTEVF | 0.5725915 |

| | | |
|---|---|---|
| AAPAAAAAPA | ASSFGTEVF | 0.28652877 |
| IALNEDL | ASSFGTEVF | 0.17376658 |
| IVNVSSM | ASSFGTEVF | 0.55006766 |
| TGIPPAPR | ASSFGTEVF | 0.83934927 |
| FDLGGGTF | ASSFGTEVF | 0.26913178 |
| FDLGGGTFDVS | ASSFGTEVF | 0.16258681 |
| NEPTAAA | ASSFGTEVF | 0.5212846 |
| NEPTAAALA | ASSFGTEVF | 0.2630582 |
| GSIINISSI | ASSFGTEVF | 0.3181801 |
| VAPGNAGTA | ASSFGTEVF | 0.58238804 |
| TGGGASLE | ASSFGTEVF | 0.16202107 |
| VAVYNHYKRI | ASSFGTEVF | 0.28783822 |
| TDPAPATT | ASSFGTEVF | 0.8348337 |
| TNRPDILDPALLRPG | ASSFGTEVF | 0.49863386 |
| LTTTTTS | ASSFGTEVF | 0.682372 |
| PTGSGKTL | ASSFGTEVF | 0.7597967 |
| TGSGKTL | ASSFGTEVF | 0.63879913 |
| TIGHVDHGK | ASSFGTEVF | 0.80683804 |
| TIGHVDHGKTT | ASSFGTEVF | 0.7744465 |
| GHVDHGKTT | ASSFGTEVF | 0.30630058 |
| AQFEHTLL | ASSFGTEVF | 0.5002302 |
| IVNQDEL | ASSFGTEVF | 0.4397988 |
| TNGSQFF | ASSFGTEVF | 0.5877415 |
| TGKGIVE | ASSFGTEVF | 0.42757258 |
| LISNASDA | ASSFGTEVF | 0.29054224 |
| AELRTLGELTKIAKAHGAQ | ASSFGTEVF | 0.7736799 |
| APNAVAAAEPSW | ASSFGTEVF | 0.5935936 |
| APQVAGAATAVL | ASSFGTEVF | 0.6012512 |
| ARRDPAAAAAILPTDA | ASSFGTEVF | 0.19565827 |
| DAAARDVLAEAGLAEYF | ASSFGTEVF | 0.19586404 |
| DDQIVLSAKANLAKVEQ | ASSFGTEVF | 0.7491803 |
| DELLPDTGEDSDEDGH | ASSFGTEVF | 0.11549501 |
| ELPEAPANRATVPQD | ASSFGTEVF | 0.12889433 |
| FGQSKIERVTGIG | ASSFGTEVF | 0.4263265 |
| GAGGTGSTAGGAGGAGGAGGLY | ASSFGTEVF | 0.81688255 |
| GHGGTGAAGGNGGTGGAGGSGI | ASSFGTEVF | 0.115136124 |
| GNGGNGGAGGEGGPGGAGGAGA | ASSFGTEVF | 0.2117588 |
| GSTAGNGGNGGNATNGTIAGQP | ASSFGTEVF | 0.18807521 |
| GTGGDGGNGGNGGTGG | ASSFGTEVF | 0.51232123 |
| GVRRPAGIPADAPV | ASSFGTEVF | 0.44973063 |
| GYRQVTDAHLLTLARR | ASSFGTEVF | 0.15145004 |
| IAKMGQDGHDRG | ASSFGTEVF | 0.5396112 |
| KAEPSIEIQYT | ASSFGTEVF | 0.13958219 |
| KPVRIRMAADEDTVAVT | ASSFGTEVF | 0.14217913 |
| MPPRIAGMRLLVIKPEP | ASSFGTEVF | 0.6102891 |
| MKFYAVALFRSRGVPPTR | ASSFGTEVF | 0.8049947 |
| NAGGFGILGAVAHSP | ASSFGTEVF | 0.19009142 |
| NEGREITLAMAEM | ASSFGTEVF | 0.7382552 |
| NELVEWSHPLGHRWRWE | ASSFGTEVF | 0.27615827 |
| NLGAGGTGGNGGNPG | ASSFGTEVF | 0.83499783 |
| NTEWPKWETVMAADPVD | ASSFGTEVF | 0.31270573 |

| | | |
|---|---|---|
| PASAAPAETQP | ASSFGTEVF | 0.14481366 |
| QQSSDGKNGVRAAEITGE | ASSFGTEVF | 0.6883272 |
| RAPDAVTTSWWKE | ASSFGTEVF | 0.547861 |
| SGFGNVGAHTSGWFNQ | ASSFGTEVF | 0.39933214 |
| SGVAGAVGQAASVGGLKV | ASSFGTEVF | 0.7211715 |
| SNGGAGTGSAGGNGGT | ASSFGTEVF | 0.25645185 |
| TGGAGGEGGAAG | ASSFGTEVF | 0.41917193 |
| TGGDGGAAGAGGAPGHG | ASSFGTEVF | 0.7704179 |
| TMPPLLAAIANAGRHGV | ASSFGTEVF | 0.74161106 |
| TSSTAGNSGGA | ASSFGTEVF | 0.18067448 |
| VPATVPVGILC | ASSFGTEVF | 0.53415895 |
| VQRPDSGGVVLTGRIS | ASSFGTEVF | 0.49841303 |
| WTPNPVPATKGVPAGLVQAT | ASSFGTEVF | 0.23053971 |
| YALKRALAKFSDRFDVVII | ASSFGTEVF | 0.80564415 |
| YLIGSVIV | ASSFGTEVF | 0.4155836 |
| YGQLVAVID | ASSFGTEVF | 0.6852293 |
| VTVGGMTLPA | ASSFGTEVF | 0.45249188 |
| VLALPAV | ASSFGTEVF | 0.8345742 |
| VIPPTLNYET | ASSFGTEVF | 0.16329941 |
| VGMGGMHPSSG | ASSFGTEVF | 0.6724709 |
| VGLAGGAATA | ASSFGTEVF | 0.51129246 |
| VGGAGGAGGSGP | ASSFGTEVF | 0.6978743 |
| VFAAFMV | ASSFGTEVF | 0.39793783 |
| TLYDKLR | ASSFGTEVF | 0.15633571 |
| TLAVFTVL | ASSFGTEVF | 0.38962466 |
| TGSVQPARV | ASSFGTEVF | 0.7483412 |
| TGGGAGSGGT | ASSFGTEVF | 0.3227878 |
| SHLETSL | ASSFGTEVF | 0.644098 |
| SFTTGMT | ASSFGTEVF | 0.8242946 |
| QVEVAKRMGVRQPTVS | ASSFGTEVF | 0.19975597 |
| QGSGGPALAM | ASSFGTEVF | 0.35010993 |
| MLAPSNP | ASSFGTEVF | 0.58602864 |
| LVNNAAI | ASSFGTEVF | 0.7758392 |
| LTGEINTLA | ASSFGTEVF | 0.32278982 |
| LSIFGPV | ASSFGTEVF | 0.79295665 |
| LLALLANL | ASSFGTEVF | 0.27667886 |
| LISNASDALDKLR | ASSFGTEVF | 0.71969277 |
| LINEAVP | ASSFGTEVF | 0.24847797 |
| LGYAFSGL | ASSFGTEVF | 0.59803116 |
| LDFVAVAL | ASSFGTEVF | 0.21656483 |
| LAQPGAE | ASSFGTEVF | 0.70182765 |
| ISNLGAVTGF | ASSFGTEVF | 0.75843596 |
| HQEDPGRI | ASSFGTEVF | 0.7533345 |
| GTPGEPTP | ASSFGTEVF | 0.11981154 |
| GFFNSGS | ASSFGTEVF | 0.7319113 |
| GAGGGGLTAGAN | ASSFGTEVF | 0.77606267 |
| FVDALVNM | ASSFGTEVF | 0.84236526 |
| FLTVAPEL | ASSFGTEVF | 0.528908 |
| FETLTAL | ASSFGTEVF | 0.6235601 |
| DVINPIEL | ASSFGTEVF | 0.363342 |
| APVSGAPV | ASSFGTEVF | 0.3259942 |

| | | |
|---|---|---|
| ALVGGLAW | ASSFGTEVF | 0.25452635 |
| ALAAAAATAI | ASSFGTEVF | 0.3822051 |
| AFGLGLLL | ASSFGTEVF | 0.49145174 |
| AFFAAASL | ASSFGTEVF | 0.24765447 |
| AAAALPAFSPP | ASSFGTEVF | 0.78602225 |
| VVLAPAL | ASSFGTEVF | 0.7830816 |
| TIGHVDHGKTTL | ASSFGTEVF | 0.76973027 |
| LISNASDALDK | ASSFGTEVF | 0.8012179 |
| LISNASDAL | ASSFGTEVF | 0.7740966 |
| ISNASDALDK | ASSFGTEVF | 0.5783986 |
| NQAETLVYQTE | ASSFGTEVF | 0.28771964 |
| TGGTGGTAGAG | ASSFGTEVF | 0.42921627 |
| TGSTAGGAGGA | ASSFGTEVF | 0.16929036 |
| DPLAVPVAAGPV | ASSFGTEVF | 0.3265653 |
| GGTGGTPGNGGN | ASSFGTEVF | 0.27488238 |
| PAGLAGVDPATL | ASSFGTEVF | 0.5432471 |
| GNNLVVEAPHAGASV | ASSFGTEVF | 0.11148515 |
| KVTRIMSASTLEQPA | ASSFGTEVF | 0.6654947 |
| LGFALPIALPRRTVV | ASSFGTEVF | 0.81336224 |
| TRLGNLTARKLVVLV | ASSFGTEVF | 0.32602882 |
| WSQQRRLIYFPSAGP | ASSFGTEVF | 0.6078875 |
| GPNGNPDPMAAAVDIR | ASSFGTEVF | 0.4681921 |
| TQRCVSVADLRAAASR | ASSFGTEVF | 0.26032725 |
| QFVLDVAEHPVEVLTETV | ASSFGTEVF | 0.76229364 |
| GGAGGTGGAGGNGGTQAGDG | ASSFGTEVF | 0.1987125 |
| PLLATLRAIAVDVDAKPAQV | ASSFGTEVF | 0.54519176 |
| APKFRTGGYMIDFWGVGYQVAK | ASSFGTEVF | 0.79000044 |
| GAGGAGGAGGAAGGPGGTGGTG | ASSFGTEVF | 0.50105375 |
| IGALGLLATDVAPAGSVAFGVA | ASSFGTEVF | 0.60905135 |
| AGGAGGIGGNGGAGGAGGVLMG | ASSFGTEVF | 0.34464076 |
| ADELIGLL | ASSFGTEVF | 0.15356524 |
| ADRALKA | ASSFGTEVF | 0.6679387 |
| AGAVGQVGGAGGA | ASSFGTEVF | 0.111365415 |
| AGGGGGLGGAP | ASSFGTEVF | 0.5983475 |
| AGGLLYGNGGAG | ASSFGTEVF | 0.178624 |
| AGLESIGL | ASSFGTEVF | 0.52090067 |
| AGLKTGII | ASSFGTEVF | 0.4953986 |
| ASLAALLL | ASSFGTEVF | 0.5652969 |
| ASVINISSI | ASSFGTEVF | 0.2591196 |
| ATPPGPA | ASSFGTEVF | 0.3743593 |
| ATVASALV | ASSFGTEVF | 0.24858636 |
| ATVASLIL | ASSFGTEVF | 0.16667967 |
| AVNGPVLIRH | ASSFGTEVF | 0.67017025 |
| DGAAGAVGI | ASSFGTEVF | 0.26058623 |
| DPLLHGI | ASSFGTEVF | 0.39090353 |
| ELGAALGAL | ASSFGTEVF | 0.6911751 |
| ELGRTQAV | ASSFGTEVF | 0.45836475 |
| FVAGGYLS | ASSFGTEVF | 0.16175748 |
| GASGGAGGAGGW | ASSFGTEVF | 0.61380744 |
| GGAGAGGDGGAGGI | ASSFGTEVF | 0.5040964 |
| GTGHNGSQPGMGGNG | ASSFGTEVF | 0.32095072 |

| | | |
|---|---|---|
| GYAAASASAS | ASSFGTEVF | 0.11059529 |
| HQVDVAAVQQG | ASSFGTEVF | 0.49349958 |
| IGAAIAHT | ASSFGTEVF | 0.24032903 |
| IGMIEVAQL | ASSFGTEVF | 0.3497293 |
| IRLAPEHTSAMD | ASSFGTEVF | 0.24731523 |
| IYNVSVL | ASSFGTEVF | 0.7453008 |
| KQGFLGM | ASSFGTEVF | 0.17821854 |
| LESVITQL | ASSFGTEVF | 0.44719133 |
| LGLGALVVV | ASSFGTEVF | 0.72800875 |
| LIGNGGNGGSGGI | ASSFGTEVF | 0.6687502 |
| LNAINAPTL | ASSFGTEVF | 0.812685 |
| LQLLNTGV | ASSFGTEVF | 0.56514347 |
| LQTSIEF | ASSFGTEVF | 0.31424603 |
| MSFAVSEL | ASSFGTEVF | 0.5737565 |
| NTGALGL | ASSFGTEVF | 0.44698983 |
| PPGAGGAGGAGTT | ASSFGTEVF | 0.12669353 |
| QLVGLGI | ASSFGTEVF | 0.12938443 |
| RFLEQVHQL | ASSFGTEVF | 0.56780225 |
| RGVGYVLR | ASSFGTEVF | 0.12685663 |
| SATGVFAGVI | ASSFGTEVF | 0.53845406 |
| SLPAITLP | ASSFGTEVF | 0.6601399 |
| STALAFTV | ASSFGTEVF | 0.47247607 |
| SVLTVGLGK | ASSFGTEVF | 0.16289154 |
| TALAGAGSGI | ASSFGTEVF | 0.38131583 |
| TGLPFTGI | ASSFGTEVF | 0.32890987 |
| VAEHVTV | ASSFGTEVF | 0.6851364 |
| VDIVLNSL | ASSFGTEVF | 0.4492594 |
| VLTGGGAVL | ASSFGTEVF | 0.823465 |
| VNVGLAYGI | ASSFGTEVF | 0.7788899 |
| VSSAAVGAV | ASSFGTEVF | 0.35015225 |
| VTSVYEAGV | ASSFGTEVF | 0.6980424 |
| WVALDRGA | ASSFGTEVF | 0.3054177 |
| YLVGSVVI | ASSFGTEVF | 0.37764955 |
| GPPGTGKTLL | ASSFGTEVF | 0.30473515 |
| GVGMTKF | ASSFGTEVF | 0.41698697 |
| IGIGGSDL | ASSFGTEVF | 0.43356046 |
| IGTLLAEL | ASSFGTEVF | 0.28109 |
| IYTSGTT | ASSFGTEVF | 0.15951584 |
| LSFPAGV | ASSFGTEVF | 0.5371437 |
| LYTSGTT | ASSFGTEVF | 0.5082194 |
| VGAFPFGF | ASSFGTEVF | 0.13885257 |
| VGAGTVEF | ASSFGTEVF | 0.37824333 |
| VGKSTLLL | ASSFGTEVF | 0.837423 |
| AALAHAGGGIV | ASSFGTEVF | 0.83024335 |
| AALVAVSLL | ASSFGTEVF | 0.23475415 |
| AAVSRAVAAL | ASSFGTEVF | 0.2325432 |
| AGGSSGIGGF | ASSFGTEVF | 0.7447885 |
| AGLTVANANG | ASSFGTEVF | 0.14123584 |
| AGLVGAVL | ASSFGTEVF | 0.4524098 |
| ALSQAFTV | ASSFGTEVF | 0.65364605 |
| ALTFSVSL | ASSFGTEVF | 0.4676469 |

| | | |
|---|---|---|
| ALTGAALVAL | ASSFGTEVF | 0.24068287 |
| ASFSIGAGVL | ASSFGTEVF | 0.5291593 |
| DGSVYKI | ASSFGTEVF | 0.36138475 |
| EQLAEIV | ASSFGTEVF | 0.62068576 |
| FAVSELTVM | ASSFGTEVF | 0.11323458 |
| GAAPLAAGL | ASSFGTEVF | 0.45524937 |
| GADGGDGGSGGAGG | ASSFGTEVF | 0.25246212 |
| GFGLDTGGAGG | ASSFGTEVF | 0.2996652 |
| GLASGGLL | ASSFGTEVF | 0.14002812 |
| GRIAGAAVLL | ASSFGTEVF | 0.37498504 |
| GSAGSLNTSF | ASSFGTEVF | 0.7605468 |
| GTGQIGI | ASSFGTEVF | 0.7725297 |
| GTLIALIL | ASSFGTEVF | 0.15524018 |
| IAATMGFVVG | ASSFGTEVF | 0.77250385 |
| IFAQALL | ASSFGTEVF | 0.83304244 |
| IGAAVALGF | ASSFGTEVF | 0.3705092 |
| IGLTPLAL | ASSFGTEVF | 0.14818476 |
| ILGSNTSTL | ASSFGTEVF | 0.5938407 |
| IVFSLTVL | ASSFGTEVF | 0.46178353 |
| LASKATGF | ASSFGTEVF | 0.61285037 |
| LASVALIL | ASSFGTEVF | 0.7868107 |
| LFAVGLTL | ASSFGTEVF | 0.17178488 |
| LFNGGIGTYI | ASSFGTEVF | 0.6769413 |
| LTLGAGII | ASSFGTEVF | 0.75865376 |
| LVAAALGDI | ASSFGTEVF | 0.29619783 |
| NTAFVQLGI | ASSFGTEVF | 0.22814447 |
| QLFSGAEL | ASSFGTEVF | 0.23251048 |
| SAATFITL | ASSFGTEVF | 0.12245626 |
| SNYGLANLGV | ASSFGTEVF | 0.64354694 |
| SYGGAYAVM | ASSFGTEVF | 0.33111763 |
| TTGVGGLTLG | ASSFGTEVF | 0.27555466 |
| TTQTIYI | ASSFGTEVF | 0.39201504 |
| VAAGAGLGAVI | ASSFGTEVF | 0.23757583 |
| VAAVVTLL | ASSFGTEVF | 0.25613976 |
| VGFLGGLI | ASSFGTEVF | 0.27255028 |
| VGVIPILL | ASSFGTEVF | 0.39544687 |
| VIFASSLI | ASSFGTEVF | 0.6239729 |
| VKGGDGGAAAG | ASSFGTEVF | 0.48254803 |
| VTKTFTNI | ASSFGTEVF | 0.5078125 |
| VVAAAILL | ASSFGTEVF | 0.8034952 |
| YAALDVEL | ASSFGTEVF | 0.17364927 |
| LASVALLL | ASSFGTEVF | 0.23187 |
| LDAVHRV | ASSFGTEVF | 0.6453668 |
| QGLSSSIF | ASSFGTEVF | 0.1525538 |
| RGTFANIRL | ASSFGTEVF | 0.7139894 |
| RQLWWGHRI | ASSFGTEVF | 0.60815644 |
| YGPPGTGKTLL | ASSFGTEVF | 0.5805762 |
| AALVGAPANVA | ASSFGTEVF | 0.61968416 |
| AATFRRTE | ASSFGTEVF | 0.83730114 |
| AGALGGGI | ASSFGTEVF | 0.6638129 |
| AGGAGGAGGIGAG | ASSFGTEVF | 0.27156758 |

| | | |
|---|---|---|
| AGSGALGL | ASSFGTEVF | 0.42196876 |
| AQIADQI | ASSFGTEVF | 0.22666268 |
| ASAVAVTYL | ASSFGTEVF | 0.7364733 |
| ASTVLGGTGG | ASSFGTEVF | 0.73294115 |
| AVGGEPAK | ASSFGTEVF | 0.23053107 |
| CEQGTFGGTC | ASSFGTEVF | 0.44375962 |
| DEGKAPAG | ASSFGTEVF | 0.7044995 |
| DTVVFPAL | ASSFGTEVF | 0.7677313 |
| FEGITMAG | ASSFGTEVF | 0.13637924 |
| FPLVVQD | ASSFGTEVF | 0.27416247 |
| GAFTTPQ | ASSFGTEVF | 0.15416683 |
| GFGGFGTAGGVGGA | ASSFGTEVF | 0.4563707 |
| GFLAQFL | ASSFGTEVF | 0.18965472 |
| GGAGGAGGNAGVA | ASSFGTEVF | 0.694976 |
| GGVGTAGAGGAGGAGG | ASSFGTEVF | 0.38207182 |
| GNGGGGDAGNAGSGGN | ASSFGTEVF | 0.7514975 |
| GTGGAGGAGNDAGS | ASSFGTEVF | 0.4864638 |
| GTTNTGWFNS | ASSFGTEVF | 0.69722384 |
| IFMGPLFTL | ASSFGTEVF | 0.34407634 |
| ILAHAIEAI | ASSFGTEVF | 0.41818023 |
| IRSAIREAL | ASSFGTEVF | 0.15626298 |
| ITFAGGG | ASSFGTEVF | 0.14694881 |
| ITGMSGHQEV | ASSFGTEVF | 0.19410849 |
| IVAGLACTL | ASSFGTEVF | 0.35908526 |
| IVGGGLAGA | ASSFGTEVF | 0.120028295 |
| IVTGVLNI | ASSFGTEVF | 0.75297564 |
| LAQQGCI | ASSFGTEVF | 0.5718943 |
| LCLHVEN | ASSFGTEVF | 0.5876634 |
| LDAISDGS | ASSFGTEVF | 0.34766015 |
| LGGFAAML | ASSFGTEVF | 0.3094271 |
| LGGKNPMI | ASSFGTEVF | 0.2784124 |
| LGPKGVAAIL | ASSFGTEVF | 0.7654408 |
| LMVGVSI | ASSFGTEVF | 0.19493662 |
| LVGAAFVL | ASSFGTEVF | 0.13690642 |
| LVLAVAPL | ASSFGTEVF | 0.29294458 |
| MAAIAVIDAA | ASSFGTEVF | 0.17487587 |
| NNGINAELQ | ASSFGTEVF | 0.75972134 |
| RGEPLVLVHG | ASSFGTEVF | 0.69705045 |
| RGGAGGAAGLI | ASSFGTEVF | 0.597817 |
| RSYRMHPAV | ASSFGTEVF | 0.6343185 |
| SAAHLLAGQLA | ASSFGTEVF | 0.12553538 |
| SAAVGSY | ASSFGTEVF | 0.35067856 |
| SPHAVLHRI | ASSFGTEVF | 0.82204044 |
| SQIAKVL | ASSFGTEVF | 0.5147337 |
| TALGHLVP | ASSFGTEVF | 0.41938156 |
| TAPTTSGAA | ASSFGTEVF | 0.7589595 |
| TDGDDGNGGSPGT | ASSFGTEVF | 0.29835284 |
| TDGLGEN | ASSFGTEVF | 0.37005252 |
| TGGPPLPGQA | ASSFGTEVF | 0.36511663 |
| TIPAITI | ASSFGTEVF | 0.4548189 |
| TIPGFTI | ASSFGTEVF | 0.34503084 |

| | | |
|---|---|---|
| TSIASGALA | ASSFGTEVF | 0.20318721 |
| TVTLPGAAI | ASSFGTEVF | 0.517236 |
| VASGLSAALP | ASSFGTEVF | 0.8425745 |
| VGAGVIGI | ASSFGTEVF | 0.38254982 |
| VGPKRFGV | ASSFGTEVF | 0.7517434 |
| VNVAFPD | ASSFGTEVF | 0.6157483 |
| VSMTNSGVS | ASSFGTEVF | 0.6901145 |
| VTDVLGDLSV | ASSFGTEVF | 0.25288123 |
| VVVVGAGFAGL | ASSFGTEVF | 0.2766648 |
| VYTVGTVV | ASSFGTEVF | 0.3841842 |
| WTTFSGP | ASSFGTEVF | 0.2744694 |
| YDELVEAF | ASSFGTEVF | 0.47851276 |
| YGTGGAGGNGG | ASSFGTEVF | 0.25994503 |
| YTNVMAV | ASSFGTEVF | 0.85362285 |
| FLEQVHQL | ASSFGTEVF | 0.67668545 |
| GLSLLAVAV | ASSFGTEVF | 0.5946821 |
| LEQVHQL | ASSFGTEVF | 0.21343955 |
| LNAAGAI | ASSFGTEVF | 0.36080873 |
| PAVHASL | ASSFGTEVF | 0.63233024 |
| TELEDDA | ASSFGTEVF | 0.15866922 |
| TPPGPPN | ASSFGTEVF | 0.80244684 |
| TPPPPPPP | ASSFGTEVF | 0.25348312 |
| TQSPALI | TCSGTGKDERLF | 0.3585062 |
| EFAAAFPQ | TCSGTGKDERLF | 0.65375954 |
| LLSGNLDVL | TCSGTGKDERLF | 0.80729914 |
| GPANLGNYNI | TCSGTGKDERLF | 0.27580535 |
| TLGPTLP | TCSGTGKDERLF | 0.3183398 |
| VYAYSSCCK | TCSGTGKDERLF | 0.19215012 |
| APAPAPAPNI | TCSGTGKDERLF | 0.7079399 |
| PGGPVPIP | TCSGTGKDERLF | 0.77023697 |
| FLTVALEP | TCSGTGKDERLF | 0.42802688 |
| SQYFGANL | TCSGTGKDERLF | 0.80350626 |
| GGVKPAA | TCSGTGKDERLF | 0.42245024 |
| GGTGGAGGAGGW | TCSGTGKDERLF | 0.17068925 |
| GVGGVAGLGGAGG | TCSGTGKDERLF | 0.37948972 |
| VLLVLATTI | TCSGTGKDERLF | 0.6698828 |
| IGIFAAML | TCSGTGKDERLF | 0.79416037 |
| AVGDLID | TCSGTGKDERLF | 0.7809216 |
| NISPLPL | TCSGTGKDERLF | 0.67106277 |
| VTAVFVAI | TCSGTGKDERLF | 0.36742866 |
| SGDDNDAAL | TCSGTGKDERLF | 0.78184056 |
| AVAALAHS | TCSGTGKDERLF | 0.81502396 |
| GGVTVMATP | TCSGTGKDERLF | 0.73983717 |
| AGLAANVRD | TCSGTGKDERLF | 0.382766 |
| TNRPDLIDPALLRPG | TCSGTGKDERLF | 0.32065248 |
| LANLGVTAL | TCSGTGKDERLF | 0.35300815 |
| TGERIEL | TCSGTGKDERLF | 0.72681904 |
| NGEELAA | TCSGTGKDERLF | 0.22186686 |
| VLSLVWG | TCSGTGKDERLF | 0.52452993 |
| IGYALWL | TCSGTGKDERLF | 0.5559977 |
| LSLLLAPL | TCSGTGKDERLF | 0.15359566 |

| | | |
|---|---|---|
| TLDEALK | TCSGTGKDERLF | 0.52985454 |
| TDGAGTAEGG | TCSGTGKDERLF | 0.80878806 |
| NAPGGAGGNGG | TCSGTGKDERLF | 0.27177033 |
| VSVFMGMGGNF | TCSGTGKDERLF | 0.30709666 |
| LASYLAAL | TCSGTGKDERLF | 0.7879852 |
| LVGGTPAL | TCSGTGKDERLF | 0.85015523 |
| VRNADVL | TCSGTGKDERLF | 0.61169213 |
| TCSSPFLVSAM | TCSGTGKDERLF | 0.15964943 |
| VHITDAE | TCSGTGKDERLF | 0.1258338 |
| EYAAPSF | TCSGTGKDERLF | 0.27640706 |
| VVDAFHGGL | TCSGTGKDERLF | 0.7672744 |
| ASVVDPQ | TCSGTGKDERLF | 0.27844182 |
| SYNEFGGLS | TCSGTGKDERLF | 0.23247616 |
| DPDGSVY | TCSGTGKDERLF | 0.61055666 |
| MVSTIHATE | TCSGTGKDERLF | 0.7331038 |
| LAAGVADPV | TCSGTGKDERLF | 0.20521554 |
| TKEQIAA | TCSGTGKDERLF | 0.17512412 |
| SLASPAAL | TCSGTGKDERLF | 0.5227376 |
| IGLVFYGV | TCSGTGKDERLF | 0.71575814 |
| FLLDASNL | TCSGTGKDERLF | 0.21914738 |
| YQVGNFP | TCSGTGKDERLF | 0.3969077 |
| ESLLSQST | TCSGTGKDERLF | 0.23913169 |
| LFTVGFAAVG | TCSGTGKDERLF | 0.2008207 |
| ATPPTPT | TCSGTGKDERLF | 0.670407 |
| TDLATLG | TCSGTGKDERLF | 0.2279867 |
| SAGGTGR | TCSGTGKDERLF | 0.77018535 |
| GFSNSGS | TCSGTGKDERLF | 0.19496492 |
| GGNGGSGGTSV | TCSGTGKDERLF | 0.64039004 |
| GTSVATGGAGN | TCSGTGKDERLF | 0.6406137 |
| NGGNGGSGGTS | TCSGTGKDERLF | 0.5025109 |
| AVVKFPH | TCSGTGKDERLF | 0.38634822 |
| TPPPPPP | TCSGTGKDERLF | 0.2927274 |
| AAPAAAAAPA | TCSGTGKDERLF | 0.18446508 |
| IALNEDL | TCSGTGKDERLF | 0.13364023 |
| IVNVSSM | TCSGTGKDERLF | 0.78618854 |
| TGIPPAPR | TCSGTGKDERLF | 0.11545461 |
| FDLGGGTF | TCSGTGKDERLF | 0.7468276 |
| FDLGGGTFDVS | TCSGTGKDERLF | 0.4996488 |
| NEPTAAA | TCSGTGKDERLF | 0.1908572 |
| NEPTAAALA | TCSGTGKDERLF | 0.4665479 |
| GSIINISSI | TCSGTGKDERLF | 0.20863983 |
| VAPGNAGTA | TCSGTGKDERLF | 0.1716979 |
| TGGGASLE | TCSGTGKDERLF | 0.7537594 |
| VAVYNHYKRI | TCSGTGKDERLF | 0.4947819 |
| TDPAPATT | TCSGTGKDERLF | 0.2814433 |
| TNRPDILDPALLRPG | TCSGTGKDERLF | 0.45023265 |
| LTTTTTS | TCSGTGKDERLF | 0.5562788 |
| PTGSGKTL | TCSGTGKDERLF | 0.3054129 |
| TGSGKTL | TCSGTGKDERLF | 0.27510327 |
| TIGHVDHGK | TCSGTGKDERLF | 0.7676998 |
| TIGHVDHGKTT | TCSGTGKDERLF | 0.37423557 |

| | | |
|---|---|---|
| GHVDHGKTT | TCSGTGKDERLF | 0.21590658 |
| AQFEHTLL | TCSGTGKDERLF | 0.70184255 |
| IVNQDEL | TCSGTGKDERLF | 0.23657702 |
| TNGSQFF | TCSGTGKDERLF | 0.6179119 |
| TGKGIVE | TCSGTGKDERLF | 0.35983926 |
| LISNASDA | TCSGTGKDERLF | 0.4818179 |
| AELRTLGELTKIAKAHGAQ | TCSGTGKDERLF | 0.6321709 |
| APNAVAAAEPSW | TCSGTGKDERLF | 0.19275755 |
| APQVAGAATAVL | TCSGTGKDERLF | 0.28683907 |
| ARRDPAAAAAILPTDA | TCSGTGKDERLF | 0.3391204 |
| DAAARDVLAEAGLAEYF | TCSGTGKDERLF | 0.42505002 |
| DDQIVLSAKANLAKVEQ | TCSGTGKDERLF | 0.71719027 |
| DELLPDTGEDSDEDGH | TCSGTGKDERLF | 0.23353428 |
| ELPEAPANRATVPQD | TCSGTGKDERLF | 0.54402167 |
| FGQSKIERVTGIG | TCSGTGKDERLF | 0.29108232 |
| GAGGTGSTAGGAGGAGGAGGLY | TCSGTGKDERLF | 0.22253358 |
| GHGGTGAAGGNGGTGGAGGSGI | TCSGTGKDERLF | 0.32091397 |
| GNGGNGGAGGEGGPGGAGGAGA | TCSGTGKDERLF | 0.27302724 |
| GSTAGNGGNGGNATNGTIAGQP | TCSGTGKDERLF | 0.27139607 |
| GTGGDGGNGGNGGTGG | TCSGTGKDERLF | 0.5561764 |
| GVRRPAGIPADAPV | TCSGTGKDERLF | 0.20213245 |
| GYRQVTDAHLLTLARR | TCSGTGKDERLF | 0.5391019 |
| IAKMGQDGHDRG | TCSGTGKDERLF | 0.7773811 |
| KAEPSIEIQYT | TCSGTGKDERLF | 0.40033382 |
| KPVRIRMAADEDTVAVT | TCSGTGKDERLF | 0.5472395 |
| MPPRIAGMRLLVIKPEP | TCSGTGKDERLF | 0.6235192 |
| MKFYAVALFRSRGVPPTR | TCSGTGKDERLF | 0.6732816 |
| NAGGFGILGAVAHSP | TCSGTGKDERLF | 0.30331993 |
| NEGREITLAMAEM | TCSGTGKDERLF | 0.29025942 |
| NELVEWSHPLGHRWRWE | TCSGTGKDERLF | 0.7497572 |
| NLGAGGTGGNGGNPG | TCSGTGKDERLF | 0.6738616 |
| NTEWPKWETVMAADPVD | TCSGTGKDERLF | 0.75364196 |
| PASAAPAETQP | TCSGTGKDERLF | 0.5943339 |
| QQSSDGKNGVRAAEITGE | TCSGTGKDERLF | 0.69952726 |
| RAPDAVTTSWWKE | TCSGTGKDERLF | 0.63972056 |
| SGFGNVGAHTSGWFNQ | TCSGTGKDERLF | 0.46899962 |
| SGVAGAVGQAASVGGLKV | TCSGTGKDERLF | 0.1643391 |
| SNGGAGTGSAGGNGGT | TCSGTGKDERLF | 0.30275837 |
| TGGAGGEGGAAG | TCSGTGKDERLF | 0.40894097 |
| TGGDGGAAGAGGAPGHG | TCSGTGKDERLF | 0.24510393 |
| TMPPLLAAIANAGRHGV | TCSGTGKDERLF | 0.16084833 |
| TSSTAGNSGGA | TCSGTGKDERLF | 0.783725 |
| VPATVPVGILC | TCSGTGKDERLF | 0.85145295 |
| VQRPDSGGVVLTGRIS | TCSGTGKDERLF | 0.27437496 |
| WTPNPVPATKGVPAGLVQAT | TCSGTGKDERLF | 0.76329964 |
| YALKRALAKFSDRFDVVII | TCSGTGKDERLF | 0.16001329 |
| YLIGSVIV | TCSGTGKDERLF | 0.664189 |
| YGQLVAVID | TCSGTGKDERLF | 0.78659964 |
| VTVGGMTLPA | TCSGTGKDERLF | 0.24036205 |
| VLALPAV | TCSGTGKDERLF | 0.57817465 |
| VIPPTLNYET | TCSGTGKDERLF | 0.23696168 |

| | | |
|---|---|---|
| VGMGGMHPSSG | TCSGTGKDERLF | 0.2975954 |
| VGLAGGAATA | TCSGTGKDERLF | 0.8089739 |
| VGGAGGAGGSGP | TCSGTGKDERLF | 0.3721761 |
| VFAAFMV | TCSGTGKDERLF | 0.4293937 |
| TLYDKLR | TCSGTGKDERLF | 0.28553462 |
| TLAVFTVL | TCSGTGKDERLF | 0.20704073 |
| TGSVQPARV | TCSGTGKDERLF | 0.34264496 |
| TGGGAGSGGT | TCSGTGKDERLF | 0.8088182 |
| SHLETSL | TCSGTGKDERLF | 0.21497221 |
| SFTTGMT | TCSGTGKDERLF | 0.8057163 |
| QVEVAKRMGVRQPTVS | TCSGTGKDERLF | 0.3155462 |
| QGSGGPALAM | TCSGTGKDERLF | 0.82510567 |
| MLAPSNP | TCSGTGKDERLF | 0.7976874 |
| LVNNAAI | TCSGTGKDERLF | 0.55217195 |
| LTGEINTLA | TCSGTGKDERLF | 0.32426077 |
| LSIFGPV | TCSGTGKDERLF | 0.797126 |
| LLALLANL | TCSGTGKDERLF | 0.4540149 |
| LISNASDALDKLR | TCSGTGKDERLF | 0.73297393 |
| LINEAVP | TCSGTGKDERLF | 0.7000596 |
| LGYAFSGL | TCSGTGKDERLF | 0.35646418 |
| LDFVAVAL | TCSGTGKDERLF | 0.79522145 |
| LAQPGAE | TCSGTGKDERLF | 0.75560474 |
| ISNLGAVTGF | TCSGTGKDERLF | 0.6866178 |
| HQEDPGRI | TCSGTGKDERLF | 0.61461234 |
| GTPGEPTP | TCSGTGKDERLF | 0.28863606 |
| GFFNSGS | TCSGTGKDERLF | 0.52138555 |
| GAGGGGLTAGAN | TCSGTGKDERLF | 0.57715714 |
| FVDALVNM | TCSGTGKDERLF | 0.8352254 |
| FLTVAPEL | TCSGTGKDERLF | 0.7227925 |
| FETLTAL | TCSGTGKDERLF | 0.5107166 |
| DVINPIEL | TCSGTGKDERLF | 0.82874596 |
| APVSGAPV | TCSGTGKDERLF | 0.8022951 |
| ALVGGLAW | TCSGTGKDERLF | 0.3275007 |
| ALAAAAATAI | TCSGTGKDERLF | 0.7120926 |
| AFGLGLLL | TCSGTGKDERLF | 0.32123688 |
| AFFAAASL | TCSGTGKDERLF | 0.50556386 |
| AAAALPAFSPP | TCSGTGKDERLF | 0.16409172 |
| VVLAPAL | TCSGTGKDERLF | 0.7148955 |
| TIGHVDHGKTTL | TCSGTGKDERLF | 0.7636657 |
| LISNASDALDK | TCSGTGKDERLF | 0.24190545 |
| LISNASDAL | TCSGTGKDERLF | 0.6913099 |
| ISNASDALDK | TCSGTGKDERLF | 0.6383623 |
| NQAETLVYQTE | TCSGTGKDERLF | 0.70564073 |
| TGGTGGTAGAG | TCSGTGKDERLF | 0.80265003 |
| TGSTAGGAGGA | TCSGTGKDERLF | 0.26797366 |
| DPLAVPVAAGPV | TCSGTGKDERLF | 0.5987342 |
| GGTGGTPGNGGN | TCSGTGKDERLF | 0.8049007 |
| PAGLAGVDPATL | TCSGTGKDERLF | 0.83484906 |
| GNNLVVEAPHAGASV | TCSGTGKDERLF | 0.48310953 |
| KVTRIMSASTLEQPA | TCSGTGKDERLF | 0.7416825 |
| LGFALPIALPRRTVV | TCSGTGKDERLF | 0.8211436 |

| | | |
|---|---|---|
| TRLGNLTARKLVVLV | TCSGTGKDERLF | 0.7493932 |
| WSQQRRLIYFPSAGP | TCSGTGKDERLF | 0.7181978 |
| GPNGNPDPMAAAVDIR | TCSGTGKDERLF | 0.76913816 |
| TQRCVSVADLRAAASR | TCSGTGKDERLF | 0.402558 |
| QFVLDVAEHPVEVLTETV | TCSGTGKDERLF | 0.82423687 |
| GGAGGTGGAGGNGGTQAGDG | TCSGTGKDERLF | 0.8217529 |
| PLLATLRAIAVDVDAKPAQV | TCSGTGKDERLF | 0.5895518 |
| APKFRTGGYMIDFWGVGYQVAK | TCSGTGKDERLF | 0.6454061 |
| GAGGAGGAGGAAGGPGGTGGTG | TCSGTGKDERLF | 0.34779662 |
| IGALGLLATDVAPAGSVAFGVA | TCSGTGKDERLF | 0.5602043 |
| AGGAGGIGGNGGAGGAGGVLMG | TCSGTGKDERLF | 0.3095839 |
| ADELIGLL | TCSGTGKDERLF | 0.77700436 |
| ADRALKA | TCSGTGKDERLF | 0.79381454 |
| AGAVGQVGGAGGA | TCSGTGKDERLF | 0.715818 |
| AGGGGGLGGAP | TCSGTGKDERLF | 0.321253 |
| AGGLLYGNGGAG | TCSGTGKDERLF | 0.42114323 |
| AGLESIGL | TCSGTGKDERLF | 0.68390155 |
| AGLKTGII | TCSGTGKDERLF | 0.42127296 |
| ASLAALLL | TCSGTGKDERLF | 0.2895168 |
| ASVINISSI | TCSGTGKDERLF | 0.35328826 |
| ATPPGPA | TCSGTGKDERLF | 0.82507175 |
| ATVASALV | TCSGTGKDERLF | 0.8155962 |
| ATVASLIL | TCSGTGKDERLF | 0.6388698 |
| AVNGPVLIRH | TCSGTGKDERLF | 0.80532944 |
| DGAAGAVGI | TCSGTGKDERLF | 0.82464933 |
| DPLLHGI | TCSGTGKDERLF | 0.7364262 |
| ELGAALGAL | TCSGTGKDERLF | 0.46215624 |
| ELGRTQAV | TCSGTGKDERLF | 0.293092 |
| FVAGGYLS | TCSGTGKDERLF | 0.8356405 |
| GASGGAGGAGGW | TCSGTGKDERLF | 0.74358225 |
| GGAGAGGDGGAGGI | TCSGTGKDERLF | 0.72481966 |
| GTGHNGSQPGMGGNG | TCSGTGKDERLF | 0.7592864 |
| GYAAASASAS | TCSGTGKDERLF | 0.7229197 |
| HQVDVAAVQQG | TCSGTGKDERLF | 0.57003754 |
| IGAAIAHT | TCSGTGKDERLF | 0.7788566 |
| IGMIEVAQL | TCSGTGKDERLF | 0.7204834 |
| IRLAPEHTSAMD | TCSGTGKDERLF | 0.5357668 |
| IYNVSVL | TCSGTGKDERLF | 0.35962412 |
| KQGFLGM | TCSGTGKDERLF | 0.67058784 |
| LESVITQL | TCSGTGKDERLF | 0.70329773 |
| LGLGALVVV | TCSGTGKDERLF | 0.82265234 |
| LIGNGGNGGSGGI | TCSGTGKDERLF | 0.55148864 |
| LNAINAPTL | TCSGTGKDERLF | 0.4069531 |
| LQLLNTGV | TCSGTGKDERLF | 0.7993717 |
| LQTSIEF | TCSGTGKDERLF | 0.6285962 |
| MSFAVSEL | TCSGTGKDERLF | 0.6112707 |
| NTGALGL | TCSGTGKDERLF | 0.7345528 |
| PPGAGGAGGAGTT | TCSGTGKDERLF | 0.66184103 |
| QLVGLGI | TCSGTGKDERLF | 0.6932316 |
| RFLEQVHQL | TCSGTGKDERLF | 0.7267065 |
| RGVGYVLR | TCSGTGKDERLF | 0.55296886 |

| | | |
|---|---|---|
| SATGVFAGVI | TCSGTGKDERLF | 0.7149097 |
| SLPAITLP | TCSGTGKDERLF | 0.6393696 |
| STALAFTV | TCSGTGKDERLF | 0.5253464 |
| SVLTVGLGK | TCSGTGKDERLF | 0.52357167 |
| TALAGAGSGI | TCSGTGKDERLF | 0.46612358 |
| TGLPFTGI | TCSGTGKDERLF | 0.77815306 |
| VAEHVTV | TCSGTGKDERLF | 0.76291883 |
| VDIVLNSL | TCSGTGKDERLF | 0.7936687 |
| VLTGGGAVL | TCSGTGKDERLF | 0.653381 |
| VNVGLAYGI | TCSGTGKDERLF | 0.62737924 |
| VSSAAVGAV | TCSGTGKDERLF | 0.76927984 |
| VTSVYEAGV | TCSGTGKDERLF | 0.80312634 |
| WVALDRGA | TCSGTGKDERLF | 0.3108461 |
| YLVGSVVI | TCSGTGKDERLF | 0.6187178 |
| GPPGTGKTLL | TCSGTGKDERLF | 0.8264698 |
| GVGMTKF | TCSGTGKDERLF | 0.8147499 |
| IGIGGSDL | TCSGTGKDERLF | 0.6659532 |
| IGTLLAEL | TCSGTGKDERLF | 0.44741946 |
| IYTSGTT | TCSGTGKDERLF | 0.8486625 |
| LSFPAGV | TCSGTGKDERLF | 0.6994555 |
| LYTSGTT | TCSGTGKDERLF | 0.8361008 |
| VGAFPFGF | TCSGTGKDERLF | 0.29759187 |
| VGAGTVEF | TCSGTGKDERLF | 0.36204436 |
| VGKSTLLL | TCSGTGKDERLF | 0.29340124 |
| AALAHAGGGIV | TCSGTGKDERLF | 0.20868471 |
| AALVAVSLL | TCSGTGKDERLF | 0.80127096 |
| AAVSRAVAAL | TCSGTGKDERLF | 0.45381412 |
| AGGSSGIGGF | TCSGTGKDERLF | 0.7555828 |
| AGLTVANANG | TCSGTGKDERLF | 0.65850884 |
| AGLVGAVL | TCSGTGKDERLF | 0.7751678 |
| ALSQAFTV | TCSGTGKDERLF | 0.6203701 |
| ALTFSVSL | TCSGTGKDERLF | 0.6192785 |
| ALTGAALVAL | TCSGTGKDERLF | 0.2832182 |
| ASFSIGAGVL | TCSGTGKDERLF | 0.80520296 |
| DGSVYKI | TCSGTGKDERLF | 0.8037393 |
| EQLAEIV | TCSGTGKDERLF | 0.60198283 |
| FAVSELTVM | TCSGTGKDERLF | 0.34286496 |
| GAAPLAAGL | TCSGTGKDERLF | 0.7521054 |
| GADGGDGGSGGAGG | TCSGTGKDERLF | 0.73294556 |
| GFGLDTGGAGG | TCSGTGKDERLF | 0.21495453 |
| GLASGGLL | TCSGTGKDERLF | 0.23224325 |
| GRIAGAAVLL | TCSGTGKDERLF | 0.7124251 |
| GSAGSLNTSF | TCSGTGKDERLF | 0.59062576 |
| GTGQIGI | TCSGTGKDERLF | 0.81401134 |
| GTLIALIL | TCSGTGKDERLF | 0.80913675 |
| IAATMGFVVG | TCSGTGKDERLF | 0.63911366 |
| IFAQALL | TCSGTGKDERLF | 0.76847076 |
| IGAAVALGF | TCSGTGKDERLF | 0.77403253 |
| IGLTPLAL | TCSGTGKDERLF | 0.13904418 |
| ILGSNTSTL | TCSGTGKDERLF | 0.35356086 |
| IVFSLTVL | TCSGTGKDERLF | 0.33085567 |

| | | |
|---|---|---|
| LASKATGF | TCSGTGKDERLF | 0.742568 |
| LASVALIL | TCSGTGKDERLF | 0.46907553 |
| LFAVGLTL | TCSGTGKDERLF | 0.63676316 |
| LFNGGIGTYI | TCSGTGKDERLF | 0.7203603 |
| LTLGAGII | TCSGTGKDERLF | 0.81990397 |
| LVAAALGDI | TCSGTGKDERLF | 0.6887186 |
| NTAFVQLGI | TCSGTGKDERLF | 0.2728864 |
| QLFSGAEL | TCSGTGKDERLF | 0.1837036 |
| SAATFITL | TCSGTGKDERLF | 0.79358596 |
| SNYGLANLGV | TCSGTGKDERLF | 0.30907914 |
| SYGGAYAVM | TCSGTGKDERLF | 0.5381552 |
| TTGVGGLTLG | TCSGTGKDERLF | 0.2293098 |
| TTQTIYI | TCSGTGKDERLF | 0.56625986 |
| VAAGAGLGAVI | TCSGTGKDERLF | 0.23637182 |
| VAAVVTLL | TCSGTGKDERLF | 0.6335082 |
| VGFLGGLI | TCSGTGKDERLF | 0.4468099 |
| VGVIPILL | TCSGTGKDERLF | 0.7138365 |
| VIFASSLI | TCSGTGKDERLF | 0.83231854 |
| VKGGDGGAAAG | TCSGTGKDERLF | 0.8109677 |
| VTKTFTNI | TCSGTGKDERLF | 0.62710166 |
| VVAAAILL | TCSGTGKDERLF | 0.5948529 |
| YAALDVEL | TCSGTGKDERLF | 0.40146115 |
| LASVALLL | TCSGTGKDERLF | 0.59537303 |
| LDAVHRV | TCSGTGKDERLF | 0.5638586 |
| QGLSSSIF | TCSGTGKDERLF | 0.824574 |
| RGTFANIRL | TCSGTGKDERLF | 0.36440197 |
| RQLWWGHRI | TCSGTGKDERLF | 0.24661015 |
| YGPPGTGKTLL | TCSGTGKDERLF | 0.3517759 |
| AALVGAPANVA | TCSGTGKDERLF | 0.8129795 |
| AATFRRTE | TCSGTGKDERLF | 0.8324443 |
| AGALGGGI | TCSGTGKDERLF | 0.58342767 |
| AGGAGGAGGIGAG | TCSGTGKDERLF | 0.33994526 |
| AGSGALGL | TCSGTGKDERLF | 0.5096783 |
| AQIADQI | TCSGTGKDERLF | 0.74700767 |
| ASAVAVTYL | TCSGTGKDERLF | 0.27651843 |
| ASTVLGGTGG | TCSGTGKDERLF | 0.37235737 |
| AVGGEPAK | TCSGTGKDERLF | 0.68065387 |
| CEQGTFGGTC | TCSGTGKDERLF | 0.8506361 |
| DEGKAPAG | TCSGTGKDERLF | 0.49029493 |
| DTVVFPAL | TCSGTGKDERLF | 0.18428284 |
| FEGITMAG | TCSGTGKDERLF | 0.7410343 |
| FPLVVQD | TCSGTGKDERLF | 0.5780772 |
| GAFTTPQ | TCSGTGKDERLF | 0.6056917 |
| GFGGFGTAGGVGGA | TCSGTGKDERLF | 0.72019905 |
| GFLAQFL | TCSGTGKDERLF | 0.26129717 |
| GGAGGAGGNAGVA | TCSGTGKDERLF | 0.6682098 |
| GGVGTAGAGGAGGAGG | TCSGTGKDERLF | 0.820873 |
| GNGGGGDAGNAGSGGN | TCSGTGKDERLF | 0.7966504 |
| GTGGAGGAGNDAGS | TCSGTGKDERLF | 0.6118436 |
| GTTNTGWFNS | TCSGTGKDERLF | 0.82469845 |
| IFMGPLFTL | TCSGTGKDERLF | 0.7891262 |

| | | |
|---|---|---|
| ILAHAIEAI | TCSGTGKDERLF | 0.82695544 |
| IRSAIREAL | TCSGTGKDERLF | 0.8314912 |
| ITFAGGG | TCSGTGKDERLF | 0.44057718 |
| ITGMSGHQEV | TCSGTGKDERLF | 0.54323584 |
| IVAGLACTL | TCSGTGKDERLF | 0.7905443 |
| IVGGGLAGA | TCSGTGKDERLF | 0.74788487 |
| IVTGVLNI | TCSGTGKDERLF | 0.73425937 |
| LAQQGCI | TCSGTGKDERLF | 0.41002113 |
| LCLHVEN | TCSGTGKDERLF | 0.82901853 |
| LDAISDGS | TCSGTGKDERLF | 0.4577914 |
| LGGFAAML | TCSGTGKDERLF | 0.8335756 |
| LGGKNPMI | TCSGTGKDERLF | 0.21680236 |
| LGPKGVAAIL | TCSGTGKDERLF | 0.48216045 |
| LMVGVSI | TCSGTGKDERLF | 0.7910277 |
| LVGAAFVL | TCSGTGKDERLF | 0.1574979 |
| LVLAVAPL | TCSGTGKDERLF | 0.25430515 |
| MAAIAVIDAA | TCSGTGKDERLF | 0.6231617 |
| NNGINAELQ | TCSGTGKDERLF | 0.7766092 |
| RGEPLVLVHG | TCSGTGKDERLF | 0.15316978 |
| RGGAGGAAGLI | TCSGTGKDERLF | 0.5890833 |
| RSYRMHPAV | TCSGTGKDERLF | 0.2703095 |
| SAAHLLAGQLA | TCSGTGKDERLF | 0.5683102 |
| SAAVGSY | TCSGTGKDERLF | 0.74642056 |
| SPHAVLHRI | TCSGTGKDERLF | 0.21800101 |
| SQIAKVL | TCSGTGKDERLF | 0.71482956 |
| TALGHLVP | TCSGTGKDERLF | 0.1947698 |
| TAPTTSGAA | TCSGTGKDERLF | 0.46806148 |
| TDGDDGNGGSPGT | TCSGTGKDERLF | 0.7497079 |
| TDGLGEN | TCSGTGKDERLF | 0.8402247 |
| TGGPPLPGQA | TCSGTGKDERLF | 0.36284015 |
| TIPAITI | TCSGTGKDERLF | 0.789477 |
| TIPGFTI | TCSGTGKDERLF | 0.37695968 |
| TSIASGALA | TCSGTGKDERLF | 0.841969 |
| TVTLPGAAI | TCSGTGKDERLF | 0.6512602 |
| VASGLSAALP | TCSGTGKDERLF | 0.20792706 |
| VGAGVIGI | TCSGTGKDERLF | 0.80309117 |
| VGPKRFGV | TCSGTGKDERLF | 0.5092557 |
| VNVAFPD | TCSGTGKDERLF | 0.66361415 |
| VSMTNSGVS | TCSGTGKDERLF | 0.82992595 |
| VTDVLGDLSV | TCSGTGKDERLF | 0.34357285 |
| VVVVGAGFAGL | TCSGTGKDERLF | 0.73143893 |
| VYTVGTVV | TCSGTGKDERLF | 0.81966037 |
| WTTFSGP | TCSGTGKDERLF | 0.7891987 |
| YDELVEAF | TCSGTGKDERLF | 0.3991252 |
| YGTGGAGGNGG | TCSGTGKDERLF | 0.7388848 |
| YTNVMAV | TCSGTGKDERLF | 0.75784695 |
| FLEQVHQL | TCSGTGKDERLF | 0.42392296 |
| GLSLLAVAV | TCSGTGKDERLF | 0.81433904 |
| LEQVHQL | TCSGTGKDERLF | 0.41158128 |
| LNAAGAI | TCSGTGKDERLF | 0.7696338 |
| PAVHASL | TCSGTGKDERLF | 0.7718902 |

| | | |
|---|---|---|
| TELEDDA | TCSGTGKDERLF | 0.8453746 |
| TPPGPPN | TCSGTGKDERLF | 0.37833506 |
| TPPPPPPP | TCSGTGKDERLF | 0.5394424 |
| TQSPALI | ASSDQGPGY | 0.72657394 |
| EFAAAFPQ | ASSDQGPGY | 0.73561674 |
| LLSGNLDVL | ASSDQGPGY | 0.34067956 |
| GPANLGNYNI | ASSDQGPGY | 0.53759015 |
| TLGPTLP | ASSDQGPGY | 0.6206115 |
| VYAYSSCCK | ASSDQGPGY | 0.81973904 |
| APAPAPAPNI | ASSDQGPGY | 0.47166163 |
| PGGPVPIP | ASSDQGPGY | 0.6587122 |
| FLTVALEP | ASSDQGPGY | 0.8366352 |
| SQYFGANL | ASSDQGPGY | 0.4540648 |
| GGVKPAA | ASSDQGPGY | 0.68839085 |
| GGTGGAGGAGGW | ASSDQGPGY | 0.7444383 |
| GVGGVAGLGGAGG | ASSDQGPGY | 0.8159209 |
| VLLVLATTI | ASSDQGPGY | 0.38633233 |
| IGIFAAML | ASSDQGPGY | 0.81252426 |
| AVGDLID | ASSDQGPGY | 0.4964289 |
| NISPLPL | ASSDQGPGY | 0.8053516 |
| VTAVFVAI | ASSDQGPGY | 0.4807504 |
| SGDDNDAAL | ASSDQGPGY | 0.7016129 |
| AVAALAHS | ASSDQGPGY | 0.64927816 |
| GGVTVMATP | ASSDQGPGY | 0.8099369 |
| AGLAANVRD | ASSDQGPGY | 0.807479 |
| TNRPDLIDPALLRPG | ASSDQGPGY | 0.8113382 |
| LANLGVTAL | ASSDQGPGY | 0.22609943 |
| TGERIEL | ASSDQGPGY | 0.80119896 |
| NGEELAA | ASSDQGPGY | 0.8012313 |
| VLSLVWG | ASSDQGPGY | 0.8506906 |
| IGYALWL | ASSDQGPGY | 0.56307757 |
| LSLLLAPL | ASSDQGPGY | 0.7588874 |
| TLDEALK | ASSDQGPGY | 0.72870255 |
| TDGAGTAEGG | ASSDQGPGY | 0.50954163 |
| NAPGGAGGNGG | ASSDQGPGY | 0.44223452 |
| VSVFMGMGGNF | ASSDQGPGY | 0.4507668 |
| LASYLAAL | ASSDQGPGY | 0.6807852 |
| LVGGTPAL | ASSDQGPGY | 0.61566496 |
| VRNADVL | ASSDQGPGY | 0.8339736 |
| TCSSPFLVSAM | ASSDQGPGY | 0.80728567 |
| VHITDAE | ASSDQGPGY | 0.8038688 |
| EYAAPSF | ASSDQGPGY | 0.82558954 |
| VVDAFHGGL | ASSDQGPGY | 0.801311 |
| ASVVDPQ | ASSDQGPGY | 0.58310515 |
| SYNEFGGLS | ASSDQGPGY | 0.64374083 |
| DPDGSVY | ASSDQGPGY | 0.62768793 |
| MVSTIHATE | ASSDQGPGY | 0.3987434 |
| LAAGVADPV | ASSDQGPGY | 0.5292747 |
| TKEQIAA | ASSDQGPGY | 0.6044885 |
| SLASPAAL | ASSDQGPGY | 0.72885233 |
| IGLVFYGV | ASSDQGPGY | 0.14848527 |

| | | |
|---|---|---|
| FLLDASNL | ASSDQGPGY | 0.8115957 |
| YQVGNFP | ASSDQGPGY | 0.8261126 |
| ESLLSQST | ASSDQGPGY | 0.7330958 |
| LFTVGFAAVG | ASSDQGPGY | 0.7287891 |
| ATPPTPT | ASSDQGPGY | 0.7345164 |
| TDLATLG | ASSDQGPGY | 0.4082951 |
| SAGGTGR | ASSDQGPGY | 0.7845083 |
| GFSNSGS | ASSDQGPGY | 0.45455787 |
| GGNGGSGGTSV | ASSDQGPGY | 0.66654795 |
| GTSVATGGAGN | ASSDQGPGY | 0.81664544 |
| NGGNGGSGGTS | ASSDQGPGY | 0.7307092 |
| AVVKFPH | ASSDQGPGY | 0.6989531 |
| TPPPPPP | ASSDQGPGY | 0.53086495 |
| AAPAAAAAPA | ASSDQGPGY | 0.30670387 |
| IALNEDL | ASSDQGPGY | 0.74054396 |
| IVNVSSM | ASSDQGPGY | 0.25787485 |
| TGIPPAPR | ASSDQGPGY | 0.7094052 |
| FDLGGGTF | ASSDQGPGY | 0.42042762 |
| FDLGGGTFDVS | ASSDQGPGY | 0.8002837 |
| NEPTAAA | ASSDQGPGY | 0.76635265 |
| NEPTAAALA | ASSDQGPGY | 0.67671466 |
| GSIINISSI | ASSDQGPGY | 0.5658065 |
| VAPGNAGTA | ASSDQGPGY | 0.77251625 |
| TGGGASLE | ASSDQGPGY | 0.77186126 |
| VAVYNHYKRI | ASSDQGPGY | 0.45847392 |
| TDPAPATT | ASSDQGPGY | 0.6354015 |
| TNRPDILDPALLRPG | ASSDQGPGY | 0.44018152 |
| LTTTTTS | ASSDQGPGY | 0.6215724 |
| PTGSGKTL | ASSDQGPGY | 0.8226776 |
| TGSGKTL | ASSDQGPGY | 0.7496722 |
| TIGHVDHGK | ASSDQGPGY | 0.2022998 |
| TIGHVDHGKTT | ASSDQGPGY | 0.7386074 |
| GHVDHGKTT | ASSDQGPGY | 0.74375933 |
| AQFEHTLL | ASSDQGPGY | 0.6974041 |
| IVNQDEL | ASSDQGPGY | 0.15062793 |
| TNGSQFF | ASSDQGPGY | 0.63388 |
| TGKGIVE | ASSDQGPGY | 0.40936288 |
| LISNASDA | ASSDQGPGY | 0.41149443 |
| AELRTLGELTKIAKAHGAQ | ASSDQGPGY | 0.47955963 |
| APNAVAAAEPSW | ASSDQGPGY | 0.8188206 |
| APQVAGAATAVL | ASSDQGPGY | 0.6808635 |
| ARRDPAAAAAILPTDA | ASSDQGPGY | 0.6694792 |
| DAAARDVLAEAGLAEYF | ASSDQGPGY | 0.7986166 |
| DDQIVLSAKANLAKVEQ | ASSDQGPGY | 0.78798467 |
| DELLPDTGEDSDEDGH | ASSDQGPGY | 0.827131 |
| ELPEAPANRATVPQD | ASSDQGPGY | 0.73275566 |
| FGQSKIERVTGIG | ASSDQGPGY | 0.43914056 |
| GAGGTGSTAGGAGGAGGAGGLY | ASSDQGPGY | 0.68692774 |
| GHGGTGAAGGNGGTGGAGGSGI | ASSDQGPGY | 0.36246693 |
| GNGGNGGAGGEGGPGGAGGAGA | ASSDQGPGY | 0.49481985 |
| GSTAGNGGNGGNATNGTIAGQP | ASSDQGPGY | 0.373687 |

| | | |
|---|---|---|
| GTGGDGGNGGNGGTGG | ASSDQGPGY | 0.6183367 |
| GVRRPAGIPADAPV | ASSDQGPGY | 0.50645036 |
| GYRQVTDAHLLTLARR | ASSDQGPGY | 0.76217407 |
| IAKMGQDGHDRG | ASSDQGPGY | 0.6933495 |
| KAEPSIEIQYT | ASSDQGPGY | 0.5504665 |
| KPVRIRMAADEDTVAVT | ASSDQGPGY | 0.23924965 |
| MPPRIAGMRLLVIKPEP | ASSDQGPGY | 0.6187272 |
| MKFYAVALFRSRGVPPTR | ASSDQGPGY | 0.47660574 |
| NAGGFGILGAVAHSP | ASSDQGPGY | 0.7958467 |
| NEGREITLAMAEM | ASSDQGPGY | 0.7078787 |
| NELVEWSHPLGHRWRWE | ASSDQGPGY | 0.848693 |
| NLGAGGTGGNGGNPG | ASSDQGPGY | 0.82079875 |
| NTEWPKWETVMAADPVD | ASSDQGPGY | 0.6998543 |
| PASAAPAETQP | ASSDQGPGY | 0.7960259 |
| QQSSDGKNGVRAAEITGE | ASSDQGPGY | 0.40140545 |
| RAPDAVTTSWWKE | ASSDQGPGY | 0.4496761 |
| SGFGNVGAHTSGWFNQ | ASSDQGPGY | 0.4430915 |
| SGVAGAVGQAASVGGLKV | ASSDQGPGY | 0.6566606 |
| SNGGAGTGSAGGNGGT | ASSDQGPGY | 0.6476705 |
| TGGAGGEGGAAG | ASSDQGPGY | 0.43726408 |
| TGGDGGAAGAGGAPGHG | ASSDQGPGY | 0.20263982 |
| TMPPLLAAIANAGRHGV | ASSDQGPGY | 0.81203663 |
| TSSTAGNSGGA | ASSDQGPGY | 0.77046704 |
| VPATVPVGILC | ASSDQGPGY | 0.25885418 |
| VQRPDSGGVVLTGRIS | ASSDQGPGY | 0.47383654 |
| WTPNPVPATKGVPAGLVQAT | ASSDQGPGY | 0.83692 |
| YALKRALAKFSDRFDVVII | ASSDQGPGY | 0.82028437 |
| YLIGSVIV | ASSDQGPGY | 0.6202983 |
| YGQLVAVID | ASSDQGPGY | 0.5384733 |
| VTVGGMTLPA | ASSDQGPGY | 0.8146871 |
| VLALPAV | ASSDQGPGY | 0.60375416 |
| VIPPTLNYET | ASSDQGPGY | 0.504982 |
| VGMGGMHPSSG | ASSDQGPGY | 0.73944354 |
| VGLAGGAATA | ASSDQGPGY | 0.65916765 |
| VGGAGGAGGSGP | ASSDQGPGY | 0.35440308 |
| VFAAFMV | ASSDQGPGY | 0.76279896 |
| TLYDKLR | ASSDQGPGY | 0.6681537 |
| TLAVFTVL | ASSDQGPGY | 0.50737023 |
| TGSVQPARV | ASSDQGPGY | 0.17082834 |
| TGGGAGSGGT | ASSDQGPGY | 0.5708014 |
| SHLETSL | ASSDQGPGY | 0.5909837 |
| SFTTGMT | ASSDQGPGY | 0.33476382 |
| QVEVAKRMGVRQPTVS | ASSDQGPGY | 0.18528204 |
| QGSGGPALAM | ASSDQGPGY | 0.5159028 |
| MLAPSNP | ASSDQGPGY | 0.8017726 |
| LVNNAAI | ASSDQGPGY | 0.82014245 |
| LTGEINTLA | ASSDQGPGY | 0.56387603 |
| LSIFGPV | ASSDQGPGY | 0.8049882 |
| LLALLANL | ASSDQGPGY | 0.7986174 |
| LISNASDALDKLR | ASSDQGPGY | 0.6281097 |
| LINEAVP | ASSDQGPGY | 0.2578254 |

| | | |
|---|---|---|
| LGYAFSGL | ASSDQGPGY | 0.75556123 |
| LDFVAVAL | ASSDQGPGY | 0.72852486 |
| LAQPGAE | ASSDQGPGY | 0.6962162 |
| ISNLGAVTGF | ASSDQGPGY | 0.81224227 |
| HQEDPGRI | ASSDQGPGY | 0.46238816 |
| GTPGEPTP | ASSDQGPGY | 0.7858425 |
| GFFNSGS | ASSDQGPGY | 0.7092494 |
| GAGGGGLTAGAN | ASSDQGPGY | 0.44237483 |
| FVDALVNM | ASSDQGPGY | 0.28993922 |
| FLTVAPEL | ASSDQGPGY | 0.30413005 |
| FETLTAL | ASSDQGPGY | 0.2289064 |
| DVINPIEL | ASSDQGPGY | 0.79252434 |
| APVSGAPV | ASSDQGPGY | 0.6735592 |
| ALVGGLAW | ASSDQGPGY | 0.6713394 |
| ALAAAAATAI | ASSDQGPGY | 0.6546179 |
| AFGLGLLL | ASSDQGPGY | 0.61296916 |
| AFFAAASL | ASSDQGPGY | 0.6035522 |
| AAAALPAFSPP | ASSDQGPGY | 0.7124851 |
| VVLAPAL | ASSDQGPGY | 0.6038567 |
| TIGHVDHGKTTL | ASSDQGPGY | 0.76928616 |
| LISNASDALDK | ASSDQGPGY | 0.42540514 |
| LISNASDAL | ASSDQGPGY | 0.39505374 |
| ISNASDALDK | ASSDQGPGY | 0.83613443 |
| NQAETLVYQTE | ASSDQGPGY | 0.38077724 |
| TGGTGGTAGAG | ASSDQGPGY | 0.4457603 |
| TGSTAGGAGGA | ASSDQGPGY | 0.69555134 |
| DPLAVPVAAGPV | ASSDQGPGY | 0.5223936 |
| GGTGGTPGNGGN | ASSDQGPGY | 0.75355387 |
| PAGLAGVDPATL | ASSDQGPGY | 0.7720864 |
| GNNLVVEAPHAGASV | ASSDQGPGY | 0.76304376 |
| KVTRIMSASTLEQPA | ASSDQGPGY | 0.7860438 |
| LGFALPIALPRRTVV | ASSDQGPGY | 0.8361896 |
| TRLGNLTARKLVVLV | ASSDQGPGY | 0.80259645 |
| WSQQRRLIYFPSAGP | ASSDQGPGY | 0.3961708 |
| GPNGNPDPMAAAVDIR | ASSDQGPGY | 0.84087425 |
| TQRCVSVADLRAAASR | ASSDQGPGY | 0.25602493 |
| QFVLDVAEHPVEVLTETV | ASSDQGPGY | 0.7721405 |
| GGAGGTGGAGGNGGTQAGDG | ASSDQGPGY | 0.7430855 |
| PLLATLRAIAVDVDAKPAQV | ASSDQGPGY | 0.5296842 |
| APKFRTGGYMIDFWGVGYQVAK | ASSDQGPGY | 0.73555946 |
| GAGGAGGAGGAAGGPGGTGGTG | ASSDQGPGY | 0.77098864 |
| IGALGLLATDVAPAGSVAFGVA | ASSDQGPGY | 0.8194537 |
| AGGAGGIGGNGGAGGAGGVLMG | ASSDQGPGY | 0.50051796 |
| ADELIGLL | ASSDQGPGY | 0.56542164 |
| ADRALKA | ASSDQGPGY | 0.45629302 |
| AGAVGQVGGAGGA | ASSDQGPGY | 0.30249682 |
| AGGGGGLGGAP | ASSDQGPGY | 0.25194263 |
| AGGLLYGNGGAG | ASSDQGPGY | 0.73902816 |
| AGLESIGL | ASSDQGPGY | 0.6619636 |
| AGLKTGII | ASSDQGPGY | 0.8272699 |
| ASLAALLL | ASSDQGPGY | 0.7749113 |

| | | |
|---|---|---|
| ASVINISSI | ASSDQGPGY | 0.82801175 |
| ATPPGPA | ASSDQGPGY | 0.53530836 |
| ATVASALV | ASSDQGPGY | 0.72100973 |
| ATVASLIL | ASSDQGPGY | 0.20882726 |
| AVNGPVLIRH | ASSDQGPGY | 0.5462121 |
| DGAAGAVGI | ASSDQGPGY | 0.31717873 |
| DPLLHGI | ASSDQGPGY | 0.7065261 |
| ELGAALGAL | ASSDQGPGY | 0.2211701 |
| ELGRTQAV | ASSDQGPGY | 0.7743801 |
| FVAGGYLS | ASSDQGPGY | 0.71409965 |
| GASGGAGGAGGW | ASSDQGPGY | 0.80108523 |
| GGAGAGGDGGAGGI | ASSDQGPGY | 0.5347129 |
| GTGHNGSQPGMGGNG | ASSDQGPGY | 0.39570808 |
| GYAAASASAS | ASSDQGPGY | 0.6470128 |
| HQVDVAAVQQG | ASSDQGPGY | 0.81551117 |
| IGAAIAHT | ASSDQGPGY | 0.46698475 |
| IGMIEVAQL | ASSDQGPGY | 0.17852592 |
| IRLAPEHTSAMD | ASSDQGPGY | 0.3313881 |
| IYNVSVL | ASSDQGPGY | 0.23480688 |
| KQGFLGM | ASSDQGPGY | 0.8263345 |
| LESVITQL | ASSDQGPGY | 0.7942107 |
| LGLGALVVV | ASSDQGPGY | 0.7670059 |
| LIGNGGNGGSGGI | ASSDQGPGY | 0.83902687 |
| LNAINAPTL | ASSDQGPGY | 0.15566128 |
| LQLLNTGV | ASSDQGPGY | 0.5556992 |
| LQTSIEF | ASSDQGPGY | 0.8346654 |
| MSFAVSEL | ASSDQGPGY | 0.7526771 |
| NTGALGL | ASSDQGPGY | 0.6942198 |
| PPGAGGAGGAGTT | ASSDQGPGY | 0.8116467 |
| QLVGLGI | ASSDQGPGY | 0.6891943 |
| RFLEQVHQL | ASSDQGPGY | 0.70880806 |
| RGVGYVLR | ASSDQGPGY | 0.727987 |
| SATGVFAGVI | ASSDQGPGY | 0.5843812 |
| SLPAITLP | ASSDQGPGY | 0.44251543 |
| STALAFTV | ASSDQGPGY | 0.3342908 |
| SVLTVGLGK | ASSDQGPGY | 0.6849443 |
| TALAGAGSGI | ASSDQGPGY | 0.8470197 |
| TGLPFTGI | ASSDQGPGY | 0.7546941 |
| VAEHVTV | ASSDQGPGY | 0.7223572 |
| VDIVLNSL | ASSDQGPGY | 0.73476976 |
| VLTGGGAVL | ASSDQGPGY | 0.819296 |
| VNVGLAYGI | ASSDQGPGY | 0.57645804 |
| VSSAAVGAV | ASSDQGPGY | 0.5314019 |
| VTSVYEAGV | ASSDQGPGY | 0.7816405 |
| WVALDRGA | ASSDQGPGY | 0.44230086 |
| YLVGSVVI | ASSDQGPGY | 0.532487 |
| GPPGTGKTLL | ASSDQGPGY | 0.3691721 |
| GVGMTKF | ASSDQGPGY | 0.85091263 |
| IGIGGSDL | ASSDQGPGY | 0.74275863 |
| IGTLLAEL | ASSDQGPGY | 0.72998405 |
| IYTSGTT | ASSDQGPGY | 0.6849744 |

| | | |
|---|---|---|
| LSFPAGV | ASSDQGPGY | 0.5683313 |
| LYTSGTT | ASSDQGPGY | 0.7519163 |
| VGAFPFGF | ASSDQGPGY | 0.20661697 |
| VGAGTVEF | ASSDQGPGY | 0.8099918 |
| VGKSTLLL | ASSDQGPGY | 0.5163393 |
| AALAHAGGGIV | ASSDQGPGY | 0.583471 |
| AALVAVSLL | ASSDQGPGY | 0.67360556 |
| AAVSRAVAAL | ASSDQGPGY | 0.8415034 |
| AGGSSGIGGF | ASSDQGPGY | 0.48710918 |
| AGLTVANANG | ASSDQGPGY | 0.7558905 |
| AGLVGAVL | ASSDQGPGY | 0.6074484 |
| ALSQAFTV | ASSDQGPGY | 0.7143637 |
| ALTFSVSL | ASSDQGPGY | 0.83334124 |
| ALTGAALVAL | ASSDQGPGY | 0.80838394 |
| ASFSIGAGVL | ASSDQGPGY | 0.8481908 |
| DGSVYKI | ASSDQGPGY | 0.4241953 |
| EQLAEIV | ASSDQGPGY | 0.21606496 |
| FAVSELTVM | ASSDQGPGY | 0.50233364 |
| GAAPLAAGL | ASSDQGPGY | 0.65774405 |
| GADGGDGGSGGAGG | ASSDQGPGY | 0.8049001 |
| GFGLDTGGAGG | ASSDQGPGY | 0.8038268 |
| GLASGGLL | ASSDQGPGY | 0.6871578 |
| GRIAGAAVLL | ASSDQGPGY | 0.6091722 |
| GSAGSLNTSF | ASSDQGPGY | 0.811321 |
| GTGQIGI | ASSDQGPGY | 0.78374976 |
| GTLIALIL | ASSDQGPGY | 0.8023282 |
| IAATMGFVVG | ASSDQGPGY | 0.37008655 |
| IFAQALL | ASSDQGPGY | 0.29789466 |
| IGAAVALGF | ASSDQGPGY | 0.63576156 |
| IGLTPLAL | ASSDQGPGY | 0.79718375 |
| ILGSNTSTL | ASSDQGPGY | 0.28337544 |
| IVFSLTVL | ASSDQGPGY | 0.7187855 |
| LASKATGF | ASSDQGPGY | 0.767555 |
| LASVALIL | ASSDQGPGY | 0.23373537 |
| LFAVGLTL | ASSDQGPGY | 0.40534884 |
| LFNGGIGTYI | ASSDQGPGY | 0.8195194 |
| LTLGAGII | ASSDQGPGY | 0.43760815 |
| LVAAALGDI | ASSDQGPGY | 0.4601851 |
| NTAFVQLGI | ASSDQGPGY | 0.8143367 |
| QLFSGAEL | ASSDQGPGY | 0.8445332 |
| SAATFITL | ASSDQGPGY | 0.766507 |
| SNYGLANLGV | ASSDQGPGY | 0.25104374 |
| SYGGAYAVM | ASSDQGPGY | 0.1503115 |
| TTGVGGLTLG | ASSDQGPGY | 0.7809502 |
| TTQTIYI | ASSDQGPGY | 0.8253058 |
| VAAGAGLGAVI | ASSDQGPGY | 0.6765882 |
| VAAVVTLL | ASSDQGPGY | 0.6214807 |
| VGFLGGLI | ASSDQGPGY | 0.5855048 |
| VGVIPILL | ASSDQGPGY | 0.7808437 |
| VIFASSLI | ASSDQGPGY | 0.43008274 |
| VKGGDGGAAAG | ASSDQGPGY | 0.26451063 |

| | | |
|---|---|---|
| VTKTFTNI | ASSDQGPGY | 0.5243162 |
| VVAAAILL | ASSDQGPGY | 0.7762549 |
| YAALDVEL | ASSDQGPGY | 0.48439392 |
| LASVALLL | ASSDQGPGY | 0.7249572 |
| LDAVHRV | ASSDQGPGY | 0.7951808 |
| QGLSSSIF | ASSDQGPGY | 0.3580625 |
| RGTFANIRL | ASSDQGPGY | 0.75380963 |
| RQLWWGHRI | ASSDQGPGY | 0.44748798 |
| YGPPGTGKTLL | ASSDQGPGY | 0.80578077 |
| AALVGAPANVA | ASSDQGPGY | 0.3580112 |
| AATFRRTE | ASSDQGPGY | 0.70854867 |
| AGALGGGI | ASSDQGPGY | 0.7746491 |
| AGGAGGAGGIGAG | ASSDQGPGY | 0.7060497 |
| AGSGALGL | ASSDQGPGY | 0.6748571 |
| AQIADQI | ASSDQGPGY | 0.477222 |
| ASAVAVTYL | ASSDQGPGY | 0.4758277 |
| ASTVLGGTGG | ASSDQGPGY | 0.4111488 |
| AVGGEPAK | ASSDQGPGY | 0.81478935 |
| CEQGTFGGTC | ASSDQGPGY | 0.61469585 |
| DEGKAPAG | ASSDQGPGY | 0.80021536 |
| DTVVFPAL | ASSDQGPGY | 0.7859947 |
| FEGITMAG | ASSDQGPGY | 0.43801656 |
| FPLVVQD | ASSDQGPGY | 0.6387056 |
| GAFTTPQ | ASSDQGPGY | 0.54656553 |
| GFGGFGTAGGVGGA | ASSDQGPGY | 0.6958927 |
| GFLAQFL | ASSDQGPGY | 0.661821 |
| GGAGGAGGNAGVA | ASSDQGPGY | 0.6532752 |
| GGVGTAGAGGAGGAGG | ASSDQGPGY | 0.5126223 |
| GNGGGGDAGNAGSGGN | ASSDQGPGY | 0.5285214 |
| GTGGAGGAGNDAGS | ASSDQGPGY | 0.7173101 |
| GTTNTGWFNS | ASSDQGPGY | 0.3372599 |
| IFMGPLFTL | ASSDQGPGY | 0.5332993 |
| ILAHAIEAI | ASSDQGPGY | 0.81700873 |
| IRSAIREAL | ASSDQGPGY | 0.5242381 |
| ITFAGGG | ASSDQGPGY | 0.4720732 |
| ITGMSGHQEV | ASSDQGPGY | 0.7591104 |
| IVAGLACTL | ASSDQGPGY | 0.49118906 |
| IVGGGLAGA | ASSDQGPGY | 0.7766937 |
| IVTGVLNI | ASSDQGPGY | 0.5844328 |
| LAQQGCI | ASSDQGPGY | 0.5361874 |
| LCLHVEN | ASSDQGPGY | 0.6521352 |
| LDAISDGS | ASSDQGPGY | 0.1841084 |
| LGGFAAML | ASSDQGPGY | 0.5192748 |
| LGGKNPMI | ASSDQGPGY | 0.5491261 |
| LGPKGVAAIL | ASSDQGPGY | 0.7875649 |
| LMVGVSI | ASSDQGPGY | 0.7902065 |
| LVGAAFVL | ASSDQGPGY | 0.5310135 |
| LVLAVAPL | ASSDQGPGY | 0.6985687 |
| MAAIAVIDAA | ASSDQGPGY | 0.61128855 |
| NNGINAELQ | ASSDQGPGY | 0.5312071 |
| RGEPLVLVHG | ASSDQGPGY | 0.3953179 |

| | | |
|---|---|---|
| RGGAGGAAGLI | ASSDQGPGY | 0.33557793 |
| RSYRMHPAV | ASSDQGPGY | 0.3624556 |
| SAAHLLAGQLA | ASSDQGPGY | 0.6631108 |
| SAAVGSY | ASSDQGPGY | 0.4580541 |
| SPHAVLHRI | ASSDQGPGY | 0.70370734 |
| SQIAKVL | ASSDQGPGY | 0.79973376 |
| TALGHLVP | ASSDQGPGY | 0.5210136 |
| TAPTTSGAA | ASSDQGPGY | 0.6468273 |
| TDGDDGNGGSPGT | ASSDQGPGY | 0.690635 |
| TDGLGEN | ASSDQGPGY | 0.7791897 |
| TGGPPLPGQA | ASSDQGPGY | 0.7425722 |
| TIPAITI | ASSDQGPGY | 0.48832908 |
| TIPGFTI | ASSDQGPGY | 0.7126919 |
| TSIASGALA | ASSDQGPGY | 0.8055657 |
| TVTLPGAAI | ASSDQGPGY | 0.7699722 |
| VASGLSAALP | ASSDQGPGY | 0.6947141 |
| VGAGVIGI | ASSDQGPGY | 0.81874704 |
| VGPKRFGV | ASSDQGPGY | 0.7577285 |
| VNVAFPD | ASSDQGPGY | 0.6658546 |
| VSMTNSGVS | ASSDQGPGY | 0.20637792 |
| VTDVLGDLSV | ASSDQGPGY | 0.7356585 |
| VVVVGAGFAGL | ASSDQGPGY | 0.75214154 |
| VYTVGTVV | ASSDQGPGY | 0.32275823 |
| WTTFSGP | ASSDQGPGY | 0.19381219 |
| YDELVEAF | ASSDQGPGY | 0.8164661 |
| YGTGGAGGNGG | ASSDQGPGY | 0.844031 |
| YTNVMAV | ASSDQGPGY | 0.6428671 |
| FLEQVHQL | ASSDQGPGY | 0.83046126 |
| GLSLLAVAV | ASSDQGPGY | 0.23109722 |
| LEQVHQL | ASSDQGPGY | 0.8024169 |
| LNAAGAI | ASSDQGPGY | 0.8078169 |
| PAVHASL | ASSDQGPGY | 0.5548015 |
| TELEDDA | ASSDQGPGY | 0.7772744 |
| TPPGPPN | ASSDQGPGY | 0.44724956 |
| TPPPPPPP | ASSDQGPGY | 0.53345037 |
| TQSPALI | ASSFSAEVF | 0.83166575 |
| EFAAAFPQ | ASSFSAEVF | 0.7026924 |
| LLSGNLDVL | ASSFSAEVF | 0.5871934 |
| GPANLGNYNI | ASSFSAEVF | 0.6981776 |
| TLGPTLP | ASSFSAEVF | 0.49356887 |
| VYAYSSCCK | ASSFSAEVF | 0.5653231 |
| APAPAPAPNI | ASSFSAEVF | 0.78858036 |
| PGGPVPIP | ASSFSAEVF | 0.6285429 |
| FLTVALEP | ASSFSAEVF | 0.7658749 |
| SQYFGANL | ASSFSAEVF | 0.5118534 |
| GGVKPAA | ASSFSAEVF | 0.8226656 |
| GGTGGAGGAGGW | ASSFSAEVF | 0.83796346 |
| GVGGVAGLGGAGG | ASSFSAEVF | 0.7757355 |
| VLLVLATTI | ASSFSAEVF | 0.42482376 |
| IGIFAAML | ASSFSAEVF | 0.8251284 |
| AVGDLID | ASSFSAEVF | 0.68506473 |

| | | |
|---|---|---|
| NISPLPL | ASSFSAEVF | 0.7364217 |
| VTAVFVAI | ASSFSAEVF | 0.68107885 |
| SGDDNDAAL | ASSFSAEVF | 0.5253011 |
| AVAALAHS | ASSFSAEVF | 0.8202691 |
| GGVTVMATP | ASSFSAEVF | 0.7395005 |
| AGLAANVRD | ASSFSAEVF | 0.6891881 |
| TNRPDLIDPALLRPG | ASSFSAEVF | 0.4676392 |
| LANLGVTAL | ASSFSAEVF | 0.38221353 |
| TGERIEL | ASSFSAEVF | 0.6597899 |
| NGEELAA | ASSFSAEVF | 0.3899417 |
| VLSLVWG | ASSFSAEVF | 0.7684734 |
| IGYALWL | ASSFSAEVF | 0.7847444 |
| LSLLLAPL | ASSFSAEVF | 0.5147744 |
| TLDEALK | ASSFSAEVF | 0.82473755 |
| TDGAGTAEGG | ASSFSAEVF | 0.8026418 |
| NAPGGAGGNGG | ASSFSAEVF | 0.3665918 |
| VSVFMGMGGNF | ASSFSAEVF | 0.62719333 |
| LASYLAAL | ASSFSAEVF | 0.60185504 |
| LVGGTPAL | ASSFSAEVF | 0.6507801 |
| VRNADVL | ASSFSAEVF | 0.3524899 |
| TCSSPFLVSAM | ASSFSAEVF | 0.7760373 |
| VHITDAE | ASSFSAEVF | 0.5859254 |
| EYAAPSF | ASSFSAEVF | 0.16964734 |
| VVDAFHGGL | ASSFSAEVF | 0.7476579 |
| ASVVDPQ | ASSFSAEVF | 0.6808641 |
| SYNEFGGLS | ASSFSAEVF | 0.5411689 |
| DPDGSVY | ASSFSAEVF | 0.8087518 |
| MVSTIHATE | ASSFSAEVF | 0.41655105 |
| LAAGVADPV | ASSFSAEVF | 0.4714377 |
| TKEQIAA | ASSFSAEVF | 0.78591585 |
| SLASPAAL | ASSFSAEVF | 0.80780774 |
| IGLVFYGV | ASSFSAEVF | 0.42831376 |
| FLLDASNL | ASSFSAEVF | 0.7459221 |
| YQVGNFP | ASSFSAEVF | 0.8094139 |
| ESLLSQST | ASSFSAEVF | 0.74572384 |
| LFTVGFAAVG | ASSFSAEVF | 0.79802513 |
| ATPPTPT | ASSFSAEVF | 0.8163048 |
| TDLATLG | ASSFSAEVF | 0.67318463 |
| SAGGTGR | ASSFSAEVF | 0.799057 |
| GFSNSGS | ASSFSAEVF | 0.8364587 |
| GGNGGSGGTSV | ASSFSAEVF | 0.53690857 |
| GTSVATGGAGN | ASSFSAEVF | 0.7223489 |
| NGGNGGSGGTS | ASSFSAEVF | 0.49109983 |
| AVVKFPH | ASSFSAEVF | 0.69801825 |
| TPPPPPP | ASSFSAEVF | 0.2580393 |
| AAPAAAAAPA | ASSFSAEVF | 0.7564149 |
| IALNEDL | ASSFSAEVF | 0.8058576 |
| IVNVSSM | ASSFSAEVF | 0.6465281 |
| TGIPPAPR | ASSFSAEVF | 0.50268275 |
| FDLGGGTF | ASSFSAEVF | 0.3767562 |
| FDLGGGTFDVS | ASSFSAEVF | 0.67601776 |

| | | |
|---|---|---|
| NEPTAAA | ASSFSAEVF | 0.5541589 |
| NEPTAAALA | ASSFSAEVF | 0.26828098 |
| GSIINISSI | ASSFSAEVF | 0.739802 |
| VAPGNAGTA | ASSFSAEVF | 0.79168177 |
| TGGGASLE | ASSFSAEVF | 0.7861173 |
| VAVYNHYKRI | ASSFSAEVF | 0.39174294 |
| TDPAPATT | ASSFSAEVF | 0.83238757 |
| TNRPDILDPALLRPG | ASSFSAEVF | 0.7752297 |
| LTTTTTS | ASSFSAEVF | 0.8188261 |
| PTGSGKTL | ASSFSAEVF | 0.60739744 |
| TGSGKTL | ASSFSAEVF | 0.5098263 |
| TIGHVDHGK | ASSFSAEVF | 0.80641663 |
| TIGHVDHGKTT | ASSFSAEVF | 0.67536664 |
| GHVDHGKTT | ASSFSAEVF | 0.5777489 |
| AQFEHTLL | ASSFSAEVF | 0.80725694 |
| IVNQDEL | ASSFSAEVF | 0.65496886 |
| TNGSQFF | ASSFSAEVF | 0.554158 |
| TGKGIVE | ASSFSAEVF | 0.6589182 |
| LISNASDA | ASSFSAEVF | 0.78703094 |
| AELRTLGELTKIAKAHGAQ | ASSFSAEVF | 0.5255048 |
| APNAVAAAEPSW | ASSFSAEVF | 0.4260525 |
| APQVAGAATAVL | ASSFSAEVF | 0.50672054 |
| ARRDPAAAAAILPTDA | ASSFSAEVF | 0.69947326 |
| DAAARDVLAEAGLAEYF | ASSFSAEVF | 0.79931426 |
| DDQIVLSAKANLAKVEQ | ASSFSAEVF | 0.7265492 |
| DELLPDTGEDSDEDGH | ASSFSAEVF | 0.638589 |
| ELPEAPANRATVPQD | ASSFSAEVF | 0.7611497 |
| FGQSKIERVTGIG | ASSFSAEVF | 0.7022252 |
| GAGGTGSTAGGAGGAGGAGGLY | ASSFSAEVF | 0.71421945 |
| GHGGTGAAGGNGGTGGAGGSGI | ASSFSAEVF | 0.78194547 |
| GNGGNGGAGGEGGPGGAGGAGA | ASSFSAEVF | 0.5760479 |
| GSTAGNGGNGGNATNGTIAGQP | ASSFSAEVF | 0.79000574 |
| GTGGDGGNGGNGGTGG | ASSFSAEVF | 0.7559022 |
| GVRRPAGIPADAPV | ASSFSAEVF | 0.7535652 |
| GYRQVTDAHLLTLARR | ASSFSAEVF | 0.8032383 |
| IAKMGQDGHDRG | ASSFSAEVF | 0.7387556 |
| KAEPSIEIQYT | ASSFSAEVF | 0.6988435 |
| KPVRIRMAADEDTVAVT | ASSFSAEVF | 0.23008166 |
| MPPRIAGMRLLVIKPEP | ASSFSAEVF | 0.7198869 |
| MKFYAVALFRSRGVPPTR | ASSFSAEVF | 0.7676739 |
| NAGGFGILGAVAHSP | ASSFSAEVF | 0.74078274 |
| NEGREITLAMAEM | ASSFSAEVF | 0.7845955 |
| NELVEWSHPLGHRWRWE | ASSFSAEVF | 0.7431484 |
| NLGAGGTGGNGGNPG | ASSFSAEVF | 0.6008518 |
| NTEWPKWETVMAADPVD | ASSFSAEVF | 0.79934806 |
| PASAAPAETQP | ASSFSAEVF | 0.8068075 |
| QQSSDGKNGVRAAEITGE | ASSFSAEVF | 0.33545572 |
| RAPDAVTTSWWKE | ASSFSAEVF | 0.6596248 |
| SGFGNVGAHTSGWFNQ | ASSFSAEVF | 0.8384814 |
| SGVAGAVGQAASVGGLKV | ASSFSAEVF | 0.7589946 |
| SNGGAGTGSAGGNGGT | ASSFSAEVF | 0.6581025 |

| | | |
|---|---|---|
| TGGAGGEGGAAG | ASSFSAEVF | 0.58177507 |
| TGGDGGAAGAGGAPGHG | ASSFSAEVF | 0.8194581 |
| TMPPLLAAIANAGRHGV | ASSFSAEVF | 0.55713916 |
| TSSTAGNSGGA | ASSFSAEVF | 0.8314241 |
| VPATVPVGILC | ASSFSAEVF | 0.20552813 |
| VQRPDSGGVVLTGRIS | ASSFSAEVF | 0.5073225 |
| WTPNPVPATKGVPAGLVQAT | ASSFSAEVF | 0.4061458 |
| YALKRALAKFSDRFDVVII | ASSFSAEVF | 0.24827191 |
| YLIGSVIV | ASSFSAEVF | 0.7797121 |
| YGQLVAVID | ASSFSAEVF | 0.60916615 |
| VTVGGMTLPA | ASSFSAEVF | 0.7864595 |
| VLALPAV | ASSFSAEVF | 0.7437978 |
| VIPPTLNYET | ASSFSAEVF | 0.7976932 |
| VGMGGMHPSSG | ASSFSAEVF | 0.7399576 |
| VGLAGGAATA | ASSFSAEVF | 0.7230673 |
| VGGAGGAGGSGP | ASSFSAEVF | 0.4915743 |
| VFAAFMV | ASSFSAEVF | 0.75629854 |
| TLYDKLR | ASSFSAEVF | 0.81609523 |
| TLAVFTVL | ASSFSAEVF | 0.714727 |
| TGSVQPARV | ASSFSAEVF | 0.35617346 |
| TGGGAGSGGT | ASSFSAEVF | 0.6716598 |
| SHLETSL | ASSFSAEVF | 0.6224021 |
| SFTTGMT | ASSFSAEVF | 0.24184743 |
| QVEVAKRMGVRQPTVS | ASSFSAEVF | 0.21162724 |
| QGSGGPALAM | ASSFSAEVF | 0.5867879 |
| MLAPSNP | ASSFSAEVF | 0.636867 |
| LVNNAAI | ASSFSAEVF | 0.7984108 |
| LTGEINTLA | ASSFSAEVF | 0.78806543 |
| LSIFGPV | ASSFSAEVF | 0.65024567 |
| LLALLANL | ASSFSAEVF | 0.78975916 |
| LISNASDALDKLR | ASSFSAEVF | 0.73593146 |
| LINEAVP | ASSFSAEVF | 0.13973336 |
| LGYAFSGL | ASSFSAEVF | 0.5547909 |
| LDFVAVAL | ASSFSAEVF | 0.3903914 |
| LAQPGAE | ASSFSAEVF | 0.80840755 |
| ISNLGAVTGF | ASSFSAEVF | 0.5879376 |
| HQEDPGRI | ASSFSAEVF | 0.6095207 |
| GTPGEPTP | ASSFSAEVF | 0.69523394 |
| GFFNSGS | ASSFSAEVF | 0.73984635 |
| GAGGGGLTAGAN | ASSFSAEVF | 0.71398795 |
| FVDALVNM | ASSFSAEVF | 0.40740287 |
| FLTVAPEL | ASSFSAEVF | 0.18871117 |
| FETLTAL | ASSFSAEVF | 0.76762027 |
| DVINPIEL | ASSFSAEVF | 0.35850316 |
| APVSGAPV | ASSFSAEVF | 0.4581033 |
| ALVGGLAW | ASSFSAEVF | 0.27107137 |
| ALAAAAATAI | ASSFSAEVF | 0.25160608 |
| AFGLGLLL | ASSFSAEVF | 0.31467962 |
| AFFAAASL | ASSFSAEVF | 0.7925029 |
| AAAALPAFSPP | ASSFSAEVF | 0.6223737 |
| VVLAPAL | ASSFSAEVF | 0.69853806 |

| | | |
|---|---|---|
| TIGHVDHGKTTL | ASSFSAEVF | 0.8400434 |
| LISNASDALDK | ASSFSAEVF | 0.72036016 |
| LISNASDAL | ASSFSAEVF | 0.68407345 |
| ISNASDALDK | ASSFSAEVF | 0.49298346 |
| NQAETLVYQTE | ASSFSAEVF | 0.47252747 |
| TGGTGGTAGAG | ASSFSAEVF | 0.4587906 |
| TGSTAGGAGGA | ASSFSAEVF | 0.49504247 |
| DPLAVPVAAGPV | ASSFSAEVF | 0.82778645 |
| GGTGGTPGNGGN | ASSFSAEVF | 0.51324356 |
| PAGLAGVDPATL | ASSFSAEVF | 0.39529094 |
| GNNLVVEAPHAGASV | ASSFSAEVF | 0.59013546 |
| KVTRIMSASTLEQPA | ASSFSAEVF | 0.8243774 |
| LGFALPIALPRRTVV | ASSFSAEVF | 0.83574224 |
| TRLGNLTARKLVVLV | ASSFSAEVF | 0.5686383 |
| WSQQRRLIYFPSAGP | ASSFSAEVF | 0.5767504 |
| GPNGNPDPMAAAVDIR | ASSFSAEVF | 0.589802 |
| TQRCVSVADLRAAASR | ASSFSAEVF | 0.7054478 |
| QFVLDVAEHPVEVLTETV | ASSFSAEVF | 0.6232359 |
| GGAGGTGGAGGNGGTQAGDG | ASSFSAEVF | 0.36551988 |
| PLLATLRAIAVDVDAKPAQV | ASSFSAEVF | 0.7558967 |
| APKFRTGGYMIDFWGVGYQVAK | ASSFSAEVF | 0.8240645 |
| GAGGAGGAGGAAGGPGGTGGTG | ASSFSAEVF | 0.6339386 |
| IGALGLLATDVAPAGSVAFGVA | ASSFSAEVF | 0.2968419 |
| AGGAGGIGGNGGAGGAGGVLMG | ASSFSAEVF | 0.6768001 |
| ADELIGLL | ASSFSAEVF | 0.48544392 |
| ADRALKA | ASSFSAEVF | 0.593169 |
| AGAVGQVGGAGGA | ASSFSAEVF | 0.6337291 |
| AGGGGGLGGAP | ASSFSAEVF | 0.36964697 |
| AGGLLYGNGGAG | ASSFSAEVF | 0.6815494 |
| AGLESIGL | ASSFSAEVF | 0.8394781 |
| AGLKTGII | ASSFSAEVF | 0.6277907 |
| ASLAALLL | ASSFSAEVF | 0.76209515 |
| ASVINISSI | ASSFSAEVF | 0.8192291 |
| ATPPGPA | ASSFSAEVF | 0.7773339 |
| ATVASALV | ASSFSAEVF | 0.8166712 |
| ATVASLIL | ASSFSAEVF | 0.7855593 |
| AVNGPVLIRH | ASSFSAEVF | 0.71415335 |
| DGAAGAVGI | ASSFSAEVF | 0.6846229 |
| DPLLHGI | ASSFSAEVF | 0.78019035 |
| ELGAALGAL | ASSFSAEVF | 0.69450617 |
| ELGRTQAV | ASSFSAEVF | 0.68756664 |
| FVAGGYLS | ASSFSAEVF | 0.5365424 |
| GASGGAGGAGGW | ASSFSAEVF | 0.8048104 |
| GGAGAGGDGGAGGI | ASSFSAEVF | 0.56943035 |
| GTGHNGSQPGMGGNG | ASSFSAEVF | 0.792168 |
| GYAAASASAS | ASSFSAEVF | 0.34557486 |
| HQVDVAAVQQG | ASSFSAEVF | 0.3773823 |
| IGAAIAHT | ASSFSAEVF | 0.79092914 |
| IGMIEVAQL | ASSFSAEVF | 0.15402976 |
| IRLAPEHTSAMD | ASSFSAEVF | 0.18768138 |
| IYNVSVL | ASSFSAEVF | 0.6715512 |

| | | |
|---|---|---|
| KQGFLGM | ASSFSAEVF | 0.8190104 |
| LESVITQL | ASSFSAEVF | 0.1657733 |
| LGLGALVVV | ASSFSAEVF | 0.47631547 |
| LIGNGGNGGSGGI | ASSFSAEVF | 0.36281833 |
| LNAINAPTL | ASSFSAEVF | 0.7371343 |
| LQLLNTGV | ASSFSAEVF | 0.7071672 |
| LQTSIEF | ASSFSAEVF | 0.35811535 |
| MSFAVSEL | ASSFSAEVF | 0.7561629 |
| NTGALGL | ASSFSAEVF | 0.31027955 |
| PPGAGGAGGAGTT | ASSFSAEVF | 0.4310103 |
| QLVGLGI | ASSFSAEVF | 0.7816061 |
| RFLEQVHQL | ASSFSAEVF | 0.80987334 |
| RGVGYVLR | ASSFSAEVF | 0.5031058 |
| SATGVFAGVI | ASSFSAEVF | 0.76919967 |
| SLPAITLP | ASSFSAEVF | 0.5613607 |
| STALAFTV | ASSFSAEVF | 0.8443781 |
| SVLTVGLGK | ASSFSAEVF | 0.58858883 |
| TALAGAGSGI | ASSFSAEVF | 0.3371329 |
| TGLPFTGI | ASSFSAEVF | 0.7945396 |
| VAEHVTV | ASSFSAEVF | 0.78100747 |
| VDIVLNSL | ASSFSAEVF | 0.69043076 |
| VLTGGGAVL | ASSFSAEVF | 0.7873597 |
| VNVGLAYGI | ASSFSAEVF | 0.46523777 |
| VSSAAVGAV | ASSFSAEVF | 0.65990436 |
| VTSVYEAGV | ASSFSAEVF | 0.8080711 |
| WVALDRGA | ASSFSAEVF | 0.8075372 |
| YLVGSVVI | ASSFSAEVF | 0.4102806 |
| GPPGTGKTLL | ASSFSAEVF | 0.7998128 |
| GVGMTKF | ASSFSAEVF | 0.6881486 |
| IGIGGSDL | ASSFSAEVF | 0.5590457 |
| IGTLLAEL | ASSFSAEVF | 0.8306557 |
| IYTSGTT | ASSFSAEVF | 0.6521348 |
| LSFPAGV | ASSFSAEVF | 0.7864414 |
| LYTSGTT | ASSFSAEVF | 0.7515855 |
| VGAFPFGF | ASSFSAEVF | 0.77180845 |
| VGAGTVEF | ASSFSAEVF | 0.5001288 |
| VGKSTLLL | ASSFSAEVF | 0.77162737 |
| AALAHAGGGIV | ASSFSAEVF | 0.7292331 |
| AALVAVSLL | ASSFSAEVF | 0.7737293 |
| AAVSRAVAAL | ASSFSAEVF | 0.68078536 |
| AGGSSGIGGF | ASSFSAEVF | 0.5075271 |
| AGLTVANANG | ASSFSAEVF | 0.7566329 |
| AGLVGAVL | ASSFSAEVF | 0.82199657 |
| ALSQAFTV | ASSFSAEVF | 0.6964378 |
| ALTFSVSL | ASSFSAEVF | 0.75171316 |
| ALTGAALVAL | ASSFSAEVF | 0.8226663 |
| ASFSIGAGVL | ASSFSAEVF | 0.508163 |
| DGSVYKI | ASSFSAEVF | 0.6551855 |
| EQLAEIV | ASSFSAEVF | 0.75151765 |
| FAVSELTVM | ASSFSAEVF | 0.8105681 |
| GAAPLAAGL | ASSFSAEVF | 0.60590535 |

| | | |
|---|---|---|
| GADGGDGGSGGAGG | ASSFSAEVF | 0.8275494 |
| GFGLDTGGAGG | ASSFSAEVF | 0.65918314 |
| GLASGGLL | ASSFSAEVF | 0.7923159 |
| GRIAGAAVLL | ASSFSAEVF | 0.68114555 |
| GSAGSLNTSF | ASSFSAEVF | 0.73704326 |
| GTGQIGI | ASSFSAEVF | 0.49222508 |
| GTLIALIL | ASSFSAEVF | 0.7892748 |
| IAATMGFVVG | ASSFSAEVF | 0.80037916 |
| IFAQALL | ASSFSAEVF | 0.8054544 |
| IGAAVALGF | ASSFSAEVF | 0.2067959 |
| IGLTPLAL | ASSFSAEVF | 0.7856183 |
| ILGSNTSTL | ASSFSAEVF | 0.77230906 |
| IVFSLTVL | ASSFSAEVF | 0.8389267 |
| LASKATGF | ASSFSAEVF | 0.63817286 |
| LASVALIL | ASSFSAEVF | 0.7442059 |
| LFAVGLTL | ASSFSAEVF | 0.59955347 |
| LFNGGIGTYI | ASSFSAEVF | 0.5701559 |
| LTLGAGII | ASSFSAEVF | 0.41222358 |
| LVAAALGDI | ASSFSAEVF | 0.62261784 |
| NTAFVQLGI | ASSFSAEVF | 0.670353 |
| QLFSGAEL | ASSFSAEVF | 0.53768706 |
| SAATFITL | ASSFSAEVF | 0.79023623 |
| SNYGLANLGV | ASSFSAEVF | 0.8167553 |
| SYGGAYAVM | ASSFSAEVF | 0.8184241 |
| TTGVGGLTLG | ASSFSAEVF | 0.80298054 |
| TTQTIYI | ASSFSAEVF | 0.82513165 |
| VAAGAGLGAVI | ASSFSAEVF | 0.76914555 |
| VAAVVTLL | ASSFSAEVF | 0.49691808 |
| VGFLGGLI | ASSFSAEVF | 0.7108319 |
| VGVIPILL | ASSFSAEVF | 0.3572501 |
| VIFASSLI | ASSFSAEVF | 0.6787126 |
| VKGGDGGAAAG | ASSFSAEVF | 0.5460421 |
| VTKTFTNI | ASSFSAEVF | 0.7402491 |
| VVAAAILL | ASSFSAEVF | 0.14796668 |
| YAALDVEL | ASSFSAEVF | 0.74892735 |
| LASVALLL | ASSFSAEVF | 0.83273846 |
| LDAVHRV | ASSFSAEVF | 0.6052604 |
| QGLSSSIF | ASSFSAEVF | 0.75337267 |
| RGTFANIRL | ASSFSAEVF | 0.77801824 |
| RQLWWGHRI | ASSFSAEVF | 0.5725083 |
| YGPPGTGKTLL | ASSFSAEVF | 0.7718458 |
| AALVGAPANVA | ASSFSAEVF | 0.47762606 |
| AATFRRTE | ASSFSAEVF | 0.7544635 |
| AGALGGGI | ASSFSAEVF | 0.8261453 |
| AGGAGGAGGIGAG | ASSFSAEVF | 0.749061 |
| AGSGALGL | ASSFSAEVF | 0.7386278 |
| AQIADQI | ASSFSAEVF | 0.6658753 |
| ASAVAVTYL | ASSFSAEVF | 0.36816207 |
| ASTVLGGTGG | ASSFSAEVF | 0.74553454 |
| AVGGEPAK | ASSFSAEVF | 0.15659304 |
| CEQGTFGGTC | ASSFSAEVF | 0.7049725 |

| | | |
|---|---|---|
| DEGKAPAG | ASSFSAEVF | 0.41659182 |
| DTVVFPAL | ASSFSAEVF | 0.73079777 |
| FEGITMAG | ASSFSAEVF | 0.73069334 |
| FPLVVQD | ASSFSAEVF | 0.72600305 |
| GAFTTPQ | ASSFSAEVF | 0.50477624 |
| GFGGFGTAGGVGGA | ASSFSAEVF | 0.6939928 |
| GFLAQFL | ASSFSAEVF | 0.64025736 |
| GGAGGAGGNAGVA | ASSFSAEVF | 0.34927282 |
| GGVGTAGAGGAGGAGG | ASSFSAEVF | 0.81779116 |
| GNGGGGDAGNAGSGGN | ASSFSAEVF | 0.56163305 |
| GTGGAGGAGNDAGS | ASSFSAEVF | 0.7171315 |
| GTTNTGWFNS | ASSFSAEVF | 0.8023119 |
| IFMGPLFTL | ASSFSAEVF | 0.7043359 |
| ILAHAIEAI | ASSFSAEVF | 0.28178686 |
| IRSAIREAL | ASSFSAEVF | 0.71339107 |
| ITFAGGG | ASSFSAEVF | 0.70669186 |
| ITGMSGHQEV | ASSFSAEVF | 0.58381706 |
| IVAGLACTL | ASSFSAEVF | 0.15254627 |
| IVGGGLAGA | ASSFSAEVF | 0.7847253 |
| IVTGVLNI | ASSFSAEVF | 0.56639206 |
| LAQQGCI | ASSFSAEVF | 0.5368893 |
| LCLHVEN | ASSFSAEVF | 0.5950962 |
| LDAISDGS | ASSFSAEVF | 0.7926043 |
| LGGFAAML | ASSFSAEVF | 0.5782435 |
| LGGKNPMI | ASSFSAEVF | 0.55644387 |
| LGPKGVAAIL | ASSFSAEVF | 0.7894521 |
| LMVGVSI | ASSFSAEVF | 0.78229755 |
| LVGAAFVL | ASSFSAEVF | 0.8242409 |
| LVLAVAPL | ASSFSAEVF | 0.68066573 |
| MAAIAVIDAA | ASSFSAEVF | 0.5570464 |
| NNGINAELQ | ASSFSAEVF | 0.76380527 |
| RGEPLVLVHG | ASSFSAEVF | 0.61587024 |
| RGGAGGAAGLI | ASSFSAEVF | 0.3292402 |
| RSYRMHPAV | ASSFSAEVF | 0.2479308 |
| SAAHLLAGQLA | ASSFSAEVF | 0.64820313 |
| SAAVGSY | ASSFSAEVF | 0.2923963 |
| SPHAVLHRI | ASSFSAEVF | 0.6826475 |
| SQIAKVL | ASSFSAEVF | 0.78494227 |
| TALGHLVP | ASSFSAEVF | 0.69023263 |
| TAPTTSGAA | ASSFSAEVF | 0.39058477 |
| TDGDDGNGGSPGT | ASSFSAEVF | 0.5965327 |
| TDGLGEN | ASSFSAEVF | 0.70539093 |
| TGGPPLPGQA | ASSFSAEVF | 0.7759837 |
| TIPAITI | ASSFSAEVF | 0.757058 |
| TIPGFTI | ASSFSAEVF | 0.8290908 |
| TSIASGALA | ASSFSAEVF | 0.7931719 |
| TVTLPGAAI | ASSFSAEVF | 0.6004811 |
| VASGLSAALP | ASSFSAEVF | 0.78568465 |
| VGAGVIGI | ASSFSAEVF | 0.6320182 |
| VGPKRFGV | ASSFSAEVF | 0.74627846 |
| VNVAFPD | ASSFSAEVF | 0.7260066 |

| | | |
|---|---|---|
| VSMTNSGVS | ASSFSAEVF | 0.6983691 |
| VTDVLGDLSV | ASSFSAEVF | 0.6793629 |
| VVVVGAGFAGL | ASSFSAEVF | 0.68262154 |
| VYTVGTVV | ASSFSAEVF | 0.26721576 |
| WTTFSGP | ASSFSAEVF | 0.7290211 |
| YDELVEAF | ASSFSAEVF | 0.6970398 |
| YGTGGAGGNGG | ASSFSAEVF | 0.27973285 |
| YTNVMAV | ASSFSAEVF | 0.6112088 |
| FLEQVHQL | ASSFSAEVF | 0.83173305 |
| GLSLLAVAV | ASSFSAEVF | 0.81495905 |
| LEQVHQL | ASSFSAEVF | 0.5198149 |
| LNAAGAI | ASSFSAEVF | 0.43694586 |
| PAVHASL | ASSFSAEVF | 0.74890214 |
| TELEDDA | ASSFSAEVF | 0.30230483 |
| TPPGPPN | ASSFSAEVF | 0.711143 |
| TPPPPPPP | ASSFSAEVF | 0.6910118 |
| TQSPALI | AWSPPGGIYEQY | 0.7416514 |
| EFAAAFPQ | AWSPPGGIYEQY | 0.5286172 |
| LLSGNLDVL | AWSPPGGIYEQY | 0.73891914 |
| GPANLGNYNI | AWSPPGGIYEQY | 0.60517335 |
| TLGPTLP | AWSPPGGIYEQY | 0.75323176 |
| VYAYSSCCK | AWSPPGGIYEQY | 0.1589942 |
| APAPAPAPNI | AWSPPGGIYEQY | 0.7577382 |
| PGGPVPIP | AWSPPGGIYEQY | 0.5543659 |
| FLTVALEP | AWSPPGGIYEQY | 0.45818412 |
| SQYFGANL | AWSPPGGIYEQY | 0.32108903 |
| GGVKPAA | AWSPPGGIYEQY | 0.6960198 |
| GGTGGAGGAGGW | AWSPPGGIYEQY | 0.79065275 |
| GVGGVAGLGGAGG | AWSPPGGIYEQY | 0.7923471 |
| VLLVLATTI | AWSPPGGIYEQY | 0.56636953 |
| IGIFAAML | AWSPPGGIYEQY | 0.79990983 |
| AVGDLID | AWSPPGGIYEQY | 0.8171576 |
| NISPLPL | AWSPPGGIYEQY | 0.6799755 |
| VTAVFVAI | AWSPPGGIYEQY | 0.35585046 |
| SGDDNDAAL | AWSPPGGIYEQY | 0.7076156 |
| AVAALAHS | AWSPPGGIYEQY | 0.6784909 |
| GGVTVMATP | AWSPPGGIYEQY | 0.7735219 |
| AGLAANVRD | AWSPPGGIYEQY | 0.7833481 |
| TNRPDLIDPALLRPG | AWSPPGGIYEQY | 0.43024632 |
| LANLGVTAL | AWSPPGGIYEQY | 0.76767915 |
| TGERIEL | AWSPPGGIYEQY | 0.79739785 |
| NGEELAA | AWSPPGGIYEQY | 0.54920626 |
| VLSLVWG | AWSPPGGIYEQY | 0.5188887 |
| IGYALWL | AWSPPGGIYEQY | 0.5626098 |
| LSLLLAPL | AWSPPGGIYEQY | 0.29974493 |
| TLDEALK | AWSPPGGIYEQY | 0.7948015 |
| TDGAGTAEGG | AWSPPGGIYEQY | 0.644903 |
| NAPGGAGGNGG | AWSPPGGIYEQY | 0.5698812 |
| VSVFMGMGGNF | AWSPPGGIYEQY | 0.69951296 |
| LASYLAAL | AWSPPGGIYEQY | 0.56640685 |
| LVGGTPAL | AWSPPGGIYEQY | 0.5388192 |

| | | |
|---|---|---|
| VRNADVL | AWSPPGGIYEQY | 0.6104597 |
| TCSSPFLVSAM | AWSPPGGIYEQY | 0.72112954 |
| VHITDAE | AWSPPGGIYEQY | 0.7579085 |
| EYAAPSF | AWSPPGGIYEQY | 0.73880756 |
| VVDAFHGGL | AWSPPGGIYEQY | 0.737308 |
| ASVVDPQ | AWSPPGGIYEQY | 0.82787275 |
| SYNEFGGLS | AWSPPGGIYEQY | 0.37346926 |
| DPDGSVY | AWSPPGGIYEQY | 0.5817044 |
| MVSTIHATE | AWSPPGGIYEQY | 0.75718385 |
| LAAGVADPV | AWSPPGGIYEQY | 0.326147 |
| TKEQIAA | AWSPPGGIYEQY | 0.79878575 |
| SLASPAAL | AWSPPGGIYEQY | 0.72965074 |
| IGLVFYGV | AWSPPGGIYEQY | 0.75411236 |
| FLLDASNL | AWSPPGGIYEQY | 0.6996169 |
| YQVGNFP | AWSPPGGIYEQY | 0.8390903 |
| ESLLSQST | AWSPPGGIYEQY | 0.77634776 |
| LFTVGFAAVG | AWSPPGGIYEQY | 0.6976125 |
| ATPPTPT | AWSPPGGIYEQY | 0.79192746 |
| TDLATLG | AWSPPGGIYEQY | 0.39080513 |
| SAGGTGR | AWSPPGGIYEQY | 0.8021076 |
| GFSNSGS | AWSPPGGIYEQY | 0.74097395 |
| GGNGGSGGTSV | AWSPPGGIYEQY | 0.46446735 |
| GTSVATGGAGN | AWSPPGGIYEQY | 0.666865 |
| NGGNGGSGGTS | AWSPPGGIYEQY | 0.8164996 |
| AVVKFPH | AWSPPGGIYEQY | 0.81545484 |
| TPPPPPP | AWSPPGGIYEQY | 0.27401453 |
| AAPAAAAAPA | AWSPPGGIYEQY | 0.6473364 |
| IALNEDL | AWSPPGGIYEQY | 0.3143528 |
| IVNVSSM | AWSPPGGIYEQY | 0.41283232 |
| TGIPPAPR | AWSPPGGIYEQY | 0.42092746 |
| FDLGGGTF | AWSPPGGIYEQY | 0.7717694 |
| FDLGGGTFDVS | AWSPPGGIYEQY | 0.6445631 |
| NEPTAAA | AWSPPGGIYEQY | 0.7525655 |
| NEPTAAALA | AWSPPGGIYEQY | 0.6665213 |
| GSIINISSI | AWSPPGGIYEQY | 0.7986733 |
| VAPGNAGTA | AWSPPGGIYEQY | 0.53038436 |
| TGGGASLE | AWSPPGGIYEQY | 0.62638366 |
| VAVYNHYKRI | AWSPPGGIYEQY | 0.34974656 |
| TDPAPATT | AWSPPGGIYEQY | 0.35075432 |
| TNRPDILDPALLRPG | AWSPPGGIYEQY | 0.539063 |
| LTTTTTS | AWSPPGGIYEQY | 0.6218723 |
| PTGSGKTL | AWSPPGGIYEQY | 0.20238109 |
| TGSGKTL | AWSPPGGIYEQY | 0.8178292 |
| TIGHVDHGK | AWSPPGGIYEQY | 0.6690507 |
| TIGHVDHGKTT | AWSPPGGIYEQY | 0.76151466 |
| GHVDHGKTT | AWSPPGGIYEQY | 0.6058012 |
| AQFEHTLL | AWSPPGGIYEQY | 0.65859854 |
| IVNQDEL | AWSPPGGIYEQY | 0.49144232 |
| TNGSQFF | AWSPPGGIYEQY | 0.76161504 |
| TGKGIVE | AWSPPGGIYEQY | 0.5438055 |
| LISNASDA | AWSPPGGIYEQY | 0.2317781 |

| | | |
|---|---|---|
| AELRTLGELTKIAKAHGAQ | AWSPPGGIYEQY | 0.5934839 |
| APNAVAAAEPSW | AWSPPGGIYEQY | 0.40487945 |
| APQVAGAATAVL | AWSPPGGIYEQY | 0.8276138 |
| ARRDPAAAAAILPTDA | AWSPPGGIYEQY | 0.79780173 |
| DAAARDVLAEAGLAEYF | AWSPPGGIYEQY | 0.71399033 |
| DDQIVLSAKANLAKVEQ | AWSPPGGIYEQY | 0.78790927 |
| DELLPDTGEDSDEDGH | AWSPPGGIYEQY | 0.20645003 |
| ELPEAPANRATVPQD | AWSPPGGIYEQY | 0.6157252 |
| FGQSKIERVTGIG | AWSPPGGIYEQY | 0.8253201 |
| GAGGTGSTAGGAGGAGGAGGLY | AWSPPGGIYEQY | 0.72527254 |
| GHGGTGAAGGNGGTGGAGGSGI | AWSPPGGIYEQY | 0.69485676 |
| GNGGNGGAGGEGGPGGAGGAGA | AWSPPGGIYEQY | 0.8068774 |
| GSTAGNGGNGGNATNGTIAGQP | AWSPPGGIYEQY | 0.564757 |
| GTGGDGGNGGNGGTGG | AWSPPGGIYEQY | 0.58881295 |
| GVRRPAGIPADAPV | AWSPPGGIYEQY | 0.6803255 |
| GYRQVTDAHLLTLARR | AWSPPGGIYEQY | 0.7809731 |
| IAKMGQDGHDRG | AWSPPGGIYEQY | 0.7427772 |
| KAEPSIEIQYT | AWSPPGGIYEQY | 0.5202604 |
| KPVRIRMAADEDTVAVT | AWSPPGGIYEQY | 0.7279216 |
| MPPRIAGMRLLVIKPEP | AWSPPGGIYEQY | 0.8446512 |
| MKFYAVALFRSRGVPPTR | AWSPPGGIYEQY | 0.6130659 |
| NAGGFGILGAVAHSP | AWSPPGGIYEQY | 0.79090333 |
| NEGREITLAMAEM | AWSPPGGIYEQY | 0.7311485 |
| NELVEWSHPLGHRWRWE | AWSPPGGIYEQY | 0.72529376 |
| NLGAGGTGGNGGNPG | AWSPPGGIYEQY | 0.5069845 |
| NTEWPKWETVMAADPVD | AWSPPGGIYEQY | 0.5647404 |
| PASAAPAETQP | AWSPPGGIYEQY | 0.69670546 |
| QQSSDGKNGVRAAEITGE | AWSPPGGIYEQY | 0.54851675 |
| RAPDAVTTSWWKE | AWSPPGGIYEQY | 0.64639366 |
| SGFGNVGAHTSGWFNQ | AWSPPGGIYEQY | 0.61868227 |
| SGVAGAVGQAASVGGLKV | AWSPPGGIYEQY | 0.8461629 |
| SNGGAGTGSAGGNGGT | AWSPPGGIYEQY | 0.76336634 |
| TGGAGGEGGAAG | AWSPPGGIYEQY | 0.75051343 |
| TGGDGGAAGAGGAPGHG | AWSPPGGIYEQY | 0.43983787 |
| TMPPLLAAIANAGRHGV | AWSPPGGIYEQY | 0.7140001 |
| TSSTAGNSGGA | AWSPPGGIYEQY | 0.7599471 |
| VPATVPVGILC | AWSPPGGIYEQY | 0.32856092 |
| VQRPDSGGVVLTGRIS | AWSPPGGIYEQY | 0.82391745 |
| WTPNPVPATKGVPAGLVQAT | AWSPPGGIYEQY | 0.4902125 |
| YALKRALAKFSDRFDVVII | AWSPPGGIYEQY | 0.65951055 |
| YLIGSVIV | AWSPPGGIYEQY | 0.7591927 |
| YGQLVAVID | AWSPPGGIYEQY | 0.8063405 |
| VTVGGMTLPA | AWSPPGGIYEQY | 0.6521828 |
| VLALPAV | AWSPPGGIYEQY | 0.6880493 |
| VIPPTLNYET | AWSPPGGIYEQY | 0.47523177 |
| VGMGGMHPSSG | AWSPPGGIYEQY | 0.77306384 |
| VGLAGGAATA | AWSPPGGIYEQY | 0.7969383 |
| VGGAGGAGGSGP | AWSPPGGIYEQY | 0.6721965 |
| VFAAFMV | AWSPPGGIYEQY | 0.82811695 |
| TLYDKLR | AWSPPGGIYEQY | 0.681679 |
| TLAVFTVL | AWSPPGGIYEQY | 0.3130784 |

| | | |
|---|---|---|
| TGSVQPARV | AWSPPGGIYEQY | 0.65776163 |
| TGGGAGSGGT | AWSPPGGIYEQY | 0.8114743 |
| SHLETSL | AWSPPGGIYEQY | 0.81099856 |
| SFTTGMT | AWSPPGGIYEQY | 0.82206666 |
| QVEVAKRMGVRQPTVS | AWSPPGGIYEQY | 0.7967072 |
| QGSGGPALAM | AWSPPGGIYEQY | 0.6021768 |
| MLAPSNP | AWSPPGGIYEQY | 0.81357896 |
| LVNNAAI | AWSPPGGIYEQY | 0.8300678 |
| LTGEINTLA | AWSPPGGIYEQY | 0.7398089 |
| LSIFGPV | AWSPPGGIYEQY | 0.49210888 |
| LLALLANL | AWSPPGGIYEQY | 0.4070347 |
| LISNASDALDKLR | AWSPPGGIYEQY | 0.75309795 |
| LINEAVP | AWSPPGGIYEQY | 0.789461 |
| LGYAFSGL | AWSPPGGIYEQY | 0.36814782 |
| LDFVAVAL | AWSPPGGIYEQY | 0.74800146 |
| LAQPGAE | AWSPPGGIYEQY | 0.7316232 |
| ISNLGAVTGF | AWSPPGGIYEQY | 0.33481997 |
| HQEDPGRI | AWSPPGGIYEQY | 0.779932 |
| GTPGEPTP | AWSPPGGIYEQY | 0.8297298 |
| GFFNSGS | AWSPPGGIYEQY | 0.6900099 |
| GAGGGGLTAGAN | AWSPPGGIYEQY | 0.5738597 |
| FVDALVNM | AWSPPGGIYEQY | 0.8200203 |
| FLTVAPEL | AWSPPGGIYEQY | 0.8382327 |
| FETLTAL | AWSPPGGIYEQY | 0.81121194 |
| DVINPIEL | AWSPPGGIYEQY | 0.32529137 |
| APVSGAPV | AWSPPGGIYEQY | 0.19860071 |
| ALVGGLAW | AWSPPGGIYEQY | 0.6665058 |
| ALAAAAATAI | AWSPPGGIYEQY | 0.77704144 |
| AFGLGLLL | AWSPPGGIYEQY | 0.61227775 |
| AFFAAASL | AWSPPGGIYEQY | 0.44093686 |
| AAAALPAFSPP | AWSPPGGIYEQY | 0.77209485 |
| VVLAPAL | AWSPPGGIYEQY | 0.79700685 |
| TIGHVDHGKTTL | AWSPPGGIYEQY | 0.4947114 |
| LISNASDALDK | AWSPPGGIYEQY | 0.3077967 |
| LISNASDAL | AWSPPGGIYEQY | 0.79281783 |
| ISNASDALDK | AWSPPGGIYEQY | 0.7358839 |
| NQAETLVYQTE | AWSPPGGIYEQY | 0.5878403 |
| TGGTGGTAGAG | AWSPPGGIYEQY | 0.6451847 |
| TGSTAGGAGGA | AWSPPGGIYEQY | 0.67110443 |
| DPLAVPVAAGPV | AWSPPGGIYEQY | 0.50217164 |
| GGTGGTPGNGGN | AWSPPGGIYEQY | 0.74242043 |
| PAGLAGVDPATL | AWSPPGGIYEQY | 0.34664068 |
| GNNLVVEAPHAGASV | AWSPPGGIYEQY | 0.8257735 |
| KVTRIMSASTLEQPA | AWSPPGGIYEQY | 0.39737213 |
| LGFALPIALPRRTVV | AWSPPGGIYEQY | 0.78737843 |
| TRLGNLTARKLVVLV | AWSPPGGIYEQY | 0.73961127 |
| WSQQRRLIYFPSAGP | AWSPPGGIYEQY | 0.75851595 |
| GPNGNPDPMAAAVDIR | AWSPPGGIYEQY | 0.6530074 |
| TQRCVSVADLRAAASR | AWSPPGGIYEQY | 0.5361634 |
| QFVLDVAEHPVEVLTETV | AWSPPGGIYEQY | 0.48596734 |
| GGAGGTGGAGGNGGTQAGDG | AWSPPGGIYEQY | 0.23438014 |

| | | |
|---|---|---|
| PLLATLRAIAVDVDAKPAQV | AWSPPGGIYEQY | 0.8191255 |
| APKFRTGGYMIDFWGVGYQVAK | AWSPPGGIYEQY | 0.2572489 |
| GAGGAGGAGGAAGGPGGTGGTG | AWSPPGGIYEQY | 0.8078939 |
| IGALGLLATDVAPAGSVAFGVA | AWSPPGGIYEQY | 0.7144965 |
| AGGAGGIGGNGGAGGAGGVLMG | AWSPPGGIYEQY | 0.52672863 |
| ADELIGLL | AWSPPGGIYEQY | 0.7412126 |
| ADRALKA | AWSPPGGIYEQY | 0.39284933 |
| AGAVGQVGGAGGA | AWSPPGGIYEQY | 0.5848373 |
| AGGGGGLGGAP | AWSPPGGIYEQY | 0.7804336 |
| AGGLLYGNGGAG | AWSPPGGIYEQY | 0.75913787 |
| AGLESIGL | AWSPPGGIYEQY | 0.5190132 |
| AGLKTGII | AWSPPGGIYEQY | 0.726161 |
| ASLAALLL | AWSPPGGIYEQY | 0.75474584 |
| ASVINISSI | AWSPPGGIYEQY | 0.54756546 |
| ATPPGPA | AWSPPGGIYEQY | 0.6837709 |
| ATVASALV | AWSPPGGIYEQY | 0.80201536 |
| ATVASLIL | AWSPPGGIYEQY | 0.69814503 |
| AVNGPVLIRH | AWSPPGGIYEQY | 0.44559157 |
| DGAAGAVGI | AWSPPGGIYEQY | 0.78571576 |
| DPLLHGI | AWSPPGGIYEQY | 0.51666737 |
| ELGAALGAL | AWSPPGGIYEQY | 0.7514646 |
| ELGRTQAV | AWSPPGGIYEQY | 0.7253778 |
| FVAGGYLS | AWSPPGGIYEQY | 0.63402426 |
| GASGGAGGAGGW | AWSPPGGIYEQY | 0.804847 |
| GGAGAGGDGGAGGI | AWSPPGGIYEQY | 0.40092602 |
| GTGHNGSQPGMGGNG | AWSPPGGIYEQY | 0.5749577 |
| GYAAASASAS | AWSPPGGIYEQY | 0.64774406 |
| HQVDVAAVQQG | AWSPPGGIYEQY | 0.68014693 |
| IGAAIAHT | AWSPPGGIYEQY | 0.77742314 |
| IGMIEVAQL | AWSPPGGIYEQY | 0.4711735 |
| IRLAPEHTSAMD | AWSPPGGIYEQY | 0.71327037 |
| IYNVSVL | AWSPPGGIYEQY | 0.58293915 |
| KQGFLGM | AWSPPGGIYEQY | 0.40860492 |
| LESVITQL | AWSPPGGIYEQY | 0.59303904 |
| LGLGALVVV | AWSPPGGIYEQY | 0.6569836 |
| LIGNGGNGGSGGI | AWSPPGGIYEQY | 0.47286513 |
| LNAINAPTL | AWSPPGGIYEQY | 0.7782131 |
| LQLLNTGV | AWSPPGGIYEQY | 0.47381058 |
| LQTSIEF | AWSPPGGIYEQY | 0.7368218 |
| MSFAVSEL | AWSPPGGIYEQY | 0.8418356 |
| NTGALGL | AWSPPGGIYEQY | 0.5691744 |
| PPGAGGAGGAGTT | AWSPPGGIYEQY | 0.76183236 |
| QLVGLGI | AWSPPGGIYEQY | 0.72956085 |
| RFLEQVHQL | AWSPPGGIYEQY | 0.78216773 |
| RGVGYVLR | AWSPPGGIYEQY | 0.61073714 |
| SATGVFAGVI | AWSPPGGIYEQY | 0.5832494 |
| SLPAITLP | AWSPPGGIYEQY | 0.8232614 |
| STALAFTV | AWSPPGGIYEQY | 0.776703 |
| SVLTVGLGK | AWSPPGGIYEQY | 0.79308206 |
| TALAGAGSGI | AWSPPGGIYEQY | 0.7775618 |
| TGLPFTGI | AWSPPGGIYEQY | 0.78357893 |

| | | |
|---|---|---|
| VAEHVTV | AWSPPGGIYEQY | 0.7821393 |
| VDIVLNSL | AWSPPGGIYEQY | 0.6034509 |
| VLTGGGAVL | AWSPPGGIYEQY | 0.31439593 |
| VNVGLAYGI | AWSPPGGIYEQY | 0.61557174 |
| VSSAAVGAV | AWSPPGGIYEQY | 0.6695721 |
| VTSVYEAGV | AWSPPGGIYEQY | 0.5649994 |
| WVALDRGA | AWSPPGGIYEQY | 0.24799633 |
| YLVGSVVI | AWSPPGGIYEQY | 0.80800647 |
| GPPGTGKTLL | AWSPPGGIYEQY | 0.839105 |
| GVGMTKF | AWSPPGGIYEQY | 0.5157756 |
| IGIGGSDL | AWSPPGGIYEQY | 0.8348334 |
| IGTLLAEL | AWSPPGGIYEQY | 0.38474432 |
| IYTSGTT | AWSPPGGIYEQY | 0.7828443 |
| LSFPAGV | AWSPPGGIYEQY | 0.8179858 |
| LYTSGTT | AWSPPGGIYEQY | 0.41690797 |
| VGAFPFGF | AWSPPGGIYEQY | 0.7492312 |
| VGAGTVEF | AWSPPGGIYEQY | 0.60480356 |
| VGKSTLLL | AWSPPGGIYEQY | 0.6503626 |
| AALAHAGGGIV | AWSPPGGIYEQY | 0.8212266 |
| AALVAVSLL | AWSPPGGIYEQY | 0.6090827 |
| AAVSRAVAAL | AWSPPGGIYEQY | 0.7847719 |
| AGGSSGIGGF | AWSPPGGIYEQY | 0.5581743 |
| AGLTVANANG | AWSPPGGIYEQY | 0.41955692 |
| AGLVGAVL | AWSPPGGIYEQY | 0.7469685 |
| ALSQAFTV | AWSPPGGIYEQY | 0.7938726 |
| ALTFSVSL | AWSPPGGIYEQY | 0.51640254 |
| ALTGAALVAL | AWSPPGGIYEQY | 0.82483876 |
| ASFSIGAGVL | AWSPPGGIYEQY | 0.5475709 |
| DGSVYKI | AWSPPGGIYEQY | 0.84257525 |
| EQLAEIV | AWSPPGGIYEQY | 0.7983904 |
| FAVSELTVM | AWSPPGGIYEQY | 0.7089484 |
| GAAPLAAGL | AWSPPGGIYEQY | 0.61508036 |
| GADGGDGGSGGAGG | AWSPPGGIYEQY | 0.80238056 |
| GFGLDTGGAGG | AWSPPGGIYEQY | 0.5497476 |
| GLASGGLL | AWSPPGGIYEQY | 0.7928976 |
| GRIAGAAVLL | AWSPPGGIYEQY | 0.80202913 |
| GSAGSLNTSF | AWSPPGGIYEQY | 0.72954196 |
| GTGQIGI | AWSPPGGIYEQY | 0.90572333 |
| GTLIALIL | AWSPPGGIYEQY | 0.8587637 |
| IAATMGFVVG | AWSPPGGIYEQY | 0.8479506 |
| IFAQALL | AWSPPGGIYEQY | 0.6949158 |
| IGAAVALGF | AWSPPGGIYEQY | 0.45992678 |
| IGLTPLAL | AWSPPGGIYEQY | 0.8293993 |
| ILGSNTSTL | AWSPPGGIYEQY | 0.65597403 |
| IVFSLTVL | AWSPPGGIYEQY | 0.85846025 |
| LASKATGF | AWSPPGGIYEQY | 0.8941881 |
| LASVALIL | AWSPPGGIYEQY | 0.74438524 |
| LFAVGLTL | AWSPPGGIYEQY | 0.8989721 |
| LFNGGIGTYI | AWSPPGGIYEQY | 0.8866348 |
| LTLGAGII | AWSPPGGIYEQY | 0.5079814 |
| LVAAALGDI | AWSPPGGIYEQY | 0.8324097 |

| | | |
|---|---|---|
| NTAFVQLGI | AWSPPGGIYEQY | 0.7583561 |
| QLFSGAEL | AWSPPGGIYEQY | 0.83001775 |
| SAATFITL | AWSPPGGIYEQY | 0.46951473 |
| SNYGLANLGV | AWSPPGGIYEQY | 0.8820876 |
| SYGGAYAVM | AWSPPGGIYEQY | 0.772501 |
| TTGVGGLTLG | AWSPPGGIYEQY | 0.16080785 |
| TTQTIYI | AWSPPGGIYEQY | 0.8664175 |
| VAAGAGLGAVI | AWSPPGGIYEQY | 0.8274903 |
| VAAVVTLL | AWSPPGGIYEQY | 0.7853861 |
| VGFLGGLI | AWSPPGGIYEQY | 0.89341044 |
| VGVIPILL | AWSPPGGIYEQY | 0.5914865 |
| VIFASSLI | AWSPPGGIYEQY | 0.7394198 |
| VKGGDGGAAAG | AWSPPGGIYEQY | 0.88773465 |
| VTKTFTNI | AWSPPGGIYEQY | 0.9014744 |
| VVAAAILL | AWSPPGGIYEQY | 0.6512933 |
| YAALDVEL | AWSPPGGIYEQY | 0.8647707 |
| LASVALLL | AWSPPGGIYEQY | 0.89258635 |
| LDAVHRV | AWSPPGGIYEQY | 0.8403092 |
| QGLSSSIF | AWSPPGGIYEQY | 0.89696056 |
| RGTFANIRL | AWSPPGGIYEQY | 0.9039823 |
| RQLWWGHRI | AWSPPGGIYEQY | 0.77362275 |
| YGPPGTGKTLL | AWSPPGGIYEQY | 0.89654726 |
| AALVGAPANVA | AWSPPGGIYEQY | 0.9114591 |
| AATFRRTE | AWSPPGGIYEQY | 0.8098525 |
| AGALGGGI | AWSPPGGIYEQY | 0.83847445 |
| AGGAGGAGGIGAG | AWSPPGGIYEQY | 0.6588281 |
| AGSGALGL | AWSPPGGIYEQY | 0.80990404 |
| AQIADQI | AWSPPGGIYEQY | 0.31745434 |
| ASAVAVTYL | AWSPPGGIYEQY | 0.85705304 |
| ASTVLGGTGG | AWSPPGGIYEQY | 0.8956872 |
| AVGGEPAK | AWSPPGGIYEQY | 0.8062635 |
| CEQGTFGGTC | AWSPPGGIYEQY | 0.7199687 |
| DEGKAPAG | AWSPPGGIYEQY | 0.6936226 |
| DTVVFPAL | AWSPPGGIYEQY | 0.858376 |
| FEGITMAG | AWSPPGGIYEQY | 0.76667213 |
| FPLVVQD | AWSPPGGIYEQY | 0.3765049 |
| GAFTTPQ | AWSPPGGIYEQY | 0.8287916 |
| GFGGFGTAGGVGGA | AWSPPGGIYEQY | 0.88228834 |
| GFLAQFL | AWSPPGGIYEQY | 0.881024 |
| GGAGGAGGNAGVA | AWSPPGGIYEQY | 0.5971879 |
| GGVGTAGAGGAGGAGG | AWSPPGGIYEQY | 0.9081168 |
| GNGGGGDAGNAGSGGN | AWSPPGGIYEQY | 0.873742 |
| GTGGAGGAGNDAGS | AWSPPGGIYEQY | 0.897636 |
| GTTNTGWFNS | AWSPPGGIYEQY | 0.8223267 |
| IFMGPLFTL | AWSPPGGIYEQY | 0.7177097 |
| ILAHAIEAI | AWSPPGGIYEQY | 0.8848561 |
| IRSAIREAL | AWSPPGGIYEQY | 0.8002064 |
| ITFAGGG | AWSPPGGIYEQY | 0.7431521 |
| ITGMSGHQEV | AWSPPGGIYEQY | 0.88468623 |
| IVAGLACTL | AWSPPGGIYEQY | 0.81490433 |
| IVGGGLAGA | AWSPPGGIYEQY | 0.756667 |

| | | |
|---|---|---|
| IVTGVLNI | AWSPPGGIYEQY | 0.7649551 |
| LAQQGCI | AWSPPGGIYEQY | 0.9003464 |
| LCLHVEN | AWSPPGGIYEQY | 0.7177329 |
| LDAISDGS | AWSPPGGIYEQY | 0.58255935 |
| LGGFAAML | AWSPPGGIYEQY | 0.6887847 |
| LGGKNPMI | AWSPPGGIYEQY | 0.81589913 |
| LGPKGVAAIL | AWSPPGGIYEQY | 0.89881355 |
| LMVGVSI | AWSPPGGIYEQY | 0.86804295 |
| LVGAAFVL | AWSPPGGIYEQY | 0.80247104 |
| LVLAVAPL | AWSPPGGIYEQY | 0.86823905 |
| MAAIAVIDAA | AWSPPGGIYEQY | 0.85524344 |
| NNGINAELQ | AWSPPGGIYEQY | 0.8458775 |
| RGEPLVLVHG | AWSPPGGIYEQY | 0.8776059 |
| RGGAGGAAGLI | AWSPPGGIYEQY | 0.75816685 |
| RSYRMHPAV | AWSPPGGIYEQY | 0.90175945 |
| SAAHLLAGQLA | AWSPPGGIYEQY | 0.87124616 |
| SAAVGSY | AWSPPGGIYEQY | 0.8845578 |
| SPHAVLHRI | AWSPPGGIYEQY | 0.9004478 |
| SQIAKVL | AWSPPGGIYEQY | 0.858227 |
| TALGHLVP | AWSPPGGIYEQY | 0.84458673 |
| TAPTTSGAA | AWSPPGGIYEQY | 0.3849191 |
| TDGDDGNGGSPGT | AWSPPGGIYEQY | 0.67500067 |
| TDGLGEN | AWSPPGGIYEQY | 0.8671636 |
| TGGPPLPGQA | AWSPPGGIYEQY | 0.8619654 |
| TIPAITI | AWSPPGGIYEQY | 0.8804793 |
| TIPGFTI | AWSPPGGIYEQY | 0.8769182 |
| TSIASGALA | AWSPPGGIYEQY | 0.77163744 |
| TVTLPGAAI | AWSPPGGIYEQY | 0.8922099 |
| VASGLSAALP | AWSPPGGIYEQY | 0.8922168 |
| VGAGVIGI | AWSPPGGIYEQY | 0.49823126 |
| VGPKRFGV | AWSPPGGIYEQY | 0.8161653 |
| VNVAFPD | AWSPPGGIYEQY | 0.9131714 |
| VSMTNSGVS | AWSPPGGIYEQY | 0.85552704 |
| VTDVLGDLSV | AWSPPGGIYEQY | 0.8079784 |
| VVVVGAGFAGL | AWSPPGGIYEQY | 0.77531314 |
| VYTVGTVV | AWSPPGGIYEQY | 0.8948303 |
| WTTFSGP | AWSPPGGIYEQY | 0.749895 |
| YDELVEAF | AWSPPGGIYEQY | 0.912045 |
| YGTGGAGGNGG | AWSPPGGIYEQY | 0.30455184 |
| YTNVMAV | AWSPPGGIYEQY | 0.78099537 |
| FLEQVHQL | AWSPPGGIYEQY | 0.6496879 |
| GLSLLAVAV | AWSPPGGIYEQY | 0.3032798 |
| LEQVHQL | AWSPPGGIYEQY | 0.87911296 |
| LNAAGAI | AWSPPGGIYEQY | 0.801514 |
| PAVHASL | AWSPPGGIYEQY | 0.87258184 |
| TELEDDA | AWSPPGGIYEQY | 0.83715165 |
| TPPGPPN | AWSPPGGIYEQY | 0.89906085 |
| TPPPPPPP | AWSPPGGIYEQY | 0.85464525 |
| TQSPALI | ASSRTGHNNQAPL | 0.82298416 |
| EFAAAFPQ | ASSRTGHNNQAPL | 0.66055024 |
| LLSGNLDVL | ASSRTGHNNQAPL | 0.87184507 |

| | | |
|---|---|---|
| GPANLGNYNI | ASSRTGHNNQAPL | 0.9011005 |
| TLGPTLP | ASSRTGHNNQAPL | 0.8824533 |
| VYAYSSCCK | ASSRTGHNNQAPL | 0.53529775 |
| APAPAPAPNI | ASSRTGHNNQAPL | 0.8298765 |
| PGGPVPIP | ASSRTGHNNQAPL | 0.81577134 |
| FLTVALEP | ASSRTGHNNQAPL | 0.24243985 |
| SQYFGANL | ASSRTGHNNQAPL | 0.2378772 |
| GGVKPAA | ASSRTGHNNQAPL | 0.8287232 |
| GGTGGAGGAGGW | ASSRTGHNNQAPL | 0.7987213 |
| GVGGVAGLGGAGG | ASSRTGHNNQAPL | 0.8908123 |
| VLLVLATTI | ASSRTGHNNQAPL | 0.8879135 |
| IGIFAAML | ASSRTGHNNQAPL | 0.8111011 |
| AVGDLID | ASSRTGHNNQAPL | 0.88313895 |
| NISPLPL | ASSRTGHNNQAPL | 0.842263 |
| VTAVFVAI | ASSRTGHNNQAPL | 0.08346907 |
| SGDDNDAAL | ASSRTGHNNQAPL | 0.712155 |
| AVAALAHS | ASSRTGHNNQAPL | 0.543854 |
| GGVTVMATP | ASSRTGHNNQAPL | 0.9030212 |
| AGLAANVRD | ASSRTGHNNQAPL | 0.76020116 |
| TNRPDLIDPALLRPG | ASSRTGHNNQAPL | 0.76717067 |
| LANLGVTAL | ASSRTGHNNQAPL | 0.8319452 |
| TGERIEL | ASSRTGHNNQAPL | 0.8397987 |
| NGEELAA | ASSRTGHNNQAPL | 0.83044744 |
| VLSLVWG | ASSRTGHNNQAPL | 0.6793889 |
| IGYALWL | ASSRTGHNNQAPL | 0.16032942 |
| LSLLLAPL | ASSRTGHNNQAPL | 0.87459886 |
| TLDEALK | ASSRTGHNNQAPL | 0.51026565 |
| TDGAGTAEGG | ASSRTGHNNQAPL | 0.713248 |
| NAPGGAGGNGG | ASSRTGHNNQAPL | 0.47787425 |
| VSVFMGMGGNF | ASSRTGHNNQAPL | 0.26968092 |
| LASYLAAL | ASSRTGHNNQAPL | 0.57374537 |
| LVGGTPAL | ASSRTGHNNQAPL | 0.89098287 |
| VRNADVL | ASSRTGHNNQAPL | 0.8019693 |
| TCSSPFLVSAM | ASSRTGHNNQAPL | 0.8198745 |
| VHITDAE | ASSRTGHNNQAPL | 0.9112508 |
| EYAAPSF | ASSRTGHNNQAPL | 0.80995417 |
| VVDAFHGGL | ASSRTGHNNQAPL | 0.8187947 |
| ASVVDPQ | ASSRTGHNNQAPL | 0.7385504 |
| SYNEFGGLS | ASSRTGHNNQAPL | 0.6962403 |
| DPDGSVY | ASSRTGHNNQAPL | 0.72438425 |
| MVSTIHATE | ASSRTGHNNQAPL | 0.7367034 |
| LAAGVADPV | ASSRTGHNNQAPL | 0.90840906 |
| TKEQIAA | ASSRTGHNNQAPL | 0.80038846 |
| SLASPAAL | ASSRTGHNNQAPL | 0.59612894 |
| IGLVFYGV | ASSRTGHNNQAPL | 0.8397892 |
| FLLDASNL | ASSRTGHNNQAPL | 0.9027048 |
| YQVGNFP | ASSRTGHNNQAPL | 0.9118786 |
| ESLLSQST | ASSRTGHNNQAPL | 0.72155696 |
| LFTVGFAAVG | ASSRTGHNNQAPL | 0.7617255 |
| ATPPTPT | ASSRTGHNNQAPL | 0.8072902 |
| TDLATLG | ASSRTGHNNQAPL | 0.8277792 |

| | | |
|---|---|---|
| SAGGTGR | ASSRTGHNNQAPL | 0.75935566 |
| GFSNSGS | ASSRTGHNNQAPL | 0.58865905 |
| GGNGGSGGTSV | ASSRTGHNNQAPL | 0.8888591 |
| GTSVATGGAGN | ASSRTGHNNQAPL | 0.89931464 |
| NGGNGGSGGTS | ASSRTGHNNQAPL | 0.80149865 |
| AVVKFPH | ASSRTGHNNQAPL | 0.54228354 |
| TPPPPPP | ASSRTGHNNQAPL | 0.8181269 |
| AAPAAAAAPA | ASSRTGHNNQAPL | 0.7418287 |
| IALNEDL | ASSRTGHNNQAPL | 0.77235115 |
| IVNVSSM | ASSRTGHNNQAPL | 0.8087137 |
| TGIPPAPR | ASSRTGHNNQAPL | 0.5911422 |
| FDLGGGTF | ASSRTGHNNQAPL | 0.82525307 |
| FDLGGGTFDVS | ASSRTGHNNQAPL | 0.9103707 |
| NEPTAAA | ASSRTGHNNQAPL | 0.7656205 |
| NEPTAAALA | ASSRTGHNNQAPL | 0.87353885 |
| GSIINISSI | ASSRTGHNNQAPL | 0.89993775 |
| VAPGNAGTA | ASSRTGHNNQAPL | 0.87721705 |
| TGGGASLE | ASSRTGHNNQAPL | 0.8995443 |
| VAVYNHYKRI | ASSRTGHNNQAPL | 0.8833787 |
| TDPAPATT | ASSRTGHNNQAPL | 0.8387165 |
| TNRPDILDPALLRPG | ASSRTGHNNQAPL | 0.8464868 |
| LTTTTTS | ASSRTGHNNQAPL | 0.8612435 |
| PTGSGKTL | ASSRTGHNNQAPL | 0.82810926 |
| TGSGKTL | ASSRTGHNNQAPL | 0.82551175 |
| TIGHVDHGK | ASSRTGHNNQAPL | 0.7823937 |
| TIGHVDHGKTT | ASSRTGHNNQAPL | 0.88680387 |
| GHVDHGKTT | ASSRTGHNNQAPL | 0.816151 |
| AQFEHTLL | ASSRTGHNNQAPL | 0.8830339 |
| IVNQDEL | ASSRTGHNNQAPL | 0.4610057 |
| TNGSQFF | ASSRTGHNNQAPL | 0.602285 |
| TGKGIVE | ASSRTGHNNQAPL | 0.8871238 |
| LISNASDA | ASSRTGHNNQAPL | 0.11511542 |
| AELRTLGELTKIAKAHGAQ | ASSRTGHNNQAPL | 0.2617065 |
| APNAVAAAEPSW | ASSRTGHNNQAPL | 0.86652946 |
| APQVAGAATAVL | ASSRTGHNNQAPL | 0.90025336 |
| ARRDPAAAAAILPTDA | ASSRTGHNNQAPL | 0.14091747 |
| DAAARDVLAEAGLAEYF | ASSRTGHNNQAPL | 0.6991818 |
| DDQIVLSAKANLAKVEQ | ASSRTGHNNQAPL | 0.6082789 |
| DELLPDTGEDSDEDGH | ASSRTGHNNQAPL | 0.88227886 |
| ELPEAPANRATVPQD | ASSRTGHNNQAPL | 0.83460367 |
| FGQSKIERVTGIG | ASSRTGHNNQAPL | 0.40158892 |
| GAGGTGSTAGGAGGAGGAGGLY | ASSRTGHNNQAPL | 0.8796463 |
| GHGGTGAAGGNGGTGGAGGSGI | ASSRTGHNNQAPL | 0.52190137 |
| GNGGNGGAGGEGGPGGAGGAGA | ASSRTGHNNQAPL | 0.6266569 |
| GSTAGNGGNGGNATNGTIAGQP | ASSRTGHNNQAPL | 0.8964977 |
| GTGGDGGNGGNGGTGG | ASSRTGHNNQAPL | 0.8966594 |
| GVRRPAGIPADAPV | ASSRTGHNNQAPL | 0.7163369 |
| GYRQVTDAHLLTLARR | ASSRTGHNNQAPL | 0.88413596 |
| IAKMGQDGHDRG | ASSRTGHNNQAPL | 0.7907262 |
| KAEPSIEIQYT | ASSRTGHNNQAPL | 0.91077787 |
| KPVRIRMAADEDTVAVT | ASSRTGHNNQAPL | 0.75597537 |

| | | |
|---|---|---|
| MPPRIAGMRLLVIKPEP | ASSRTGHNNQAPL | 0.50838387 |
| MKFYAVALFRSRGVPPTR | ASSRTGHNNQAPL | 0.8909167 |
| NAGGFGILGAVAHSP | ASSRTGHNNQAPL | 0.8816332 |
| NEGREITLAMAEM | ASSRTGHNNQAPL | 0.84285045 |
| NELVEWSHPLGHRWRWE | ASSRTGHNNQAPL | 0.88037175 |
| NLGAGGTGGNGGNPG | ASSRTGHNNQAPL | 0.752021 |
| NTEWPKWETVMAADPVD | ASSRTGHNNQAPL | 0.810391 |
| PASAAPAETQP | ASSRTGHNNQAPL | 0.89293325 |
| QQSSDGKNGVRAAEITGE | ASSRTGHNNQAPL | 0.89544624 |
| RAPDAVTTSWWKE | ASSRTGHNNQAPL | 0.58217156 |
| SGFGNVGAHTSGWFNQ | ASSRTGHNNQAPL | 0.89906776 |
| SGVAGAVGQAASVGGLKV | ASSRTGHNNQAPL | 0.8181549 |
| SNGGAGTGSAGGNGGT | ASSRTGHNNQAPL | 0.70504045 |
| TGGAGGEGGAAG | ASSRTGHNNQAPL | 0.9078071 |
| TGGDGGAAGAGGAPGHG | ASSRTGHNNQAPL | 0.82635915 |
| TMPPLLAAIANAGRHGV | ASSRTGHNNQAPL | 0.8834552 |
| TSSTAGNSGGA | ASSRTGHNNQAPL | 0.8676724 |
| VPATVPVGILC | ASSRTGHNNQAPL | 0.8496523 |
| VQRPDSGGVVLTGRIS | ASSRTGHNNQAPL | 0.47624746 |
| WTPNPVPATKGVPAGLVQAT | ASSRTGHNNQAPL | 0.8481231 |
| YALKRALAKFSDRFDVVII | ASSRTGHNNQAPL | 0.83474064 |
| YLIGSVIV | ASSRTGHNNQAPL | 0.8906256 |
| YGQLVAVID | ASSRTGHNNQAPL | 0.8291005 |
| VTVGGMTLPA | ASSRTGHNNQAPL | 0.6941147 |
| VLALPAV | ASSRTGHNNQAPL | 0.8648696 |
| VIPPTLNYET | ASSRTGHNNQAPL | 0.9028162 |
| VGMGGMHPSSG | ASSRTGHNNQAPL | 0.86333597 |
| VGLAGGAATA | ASSRTGHNNQAPL | 0.86631876 |
| VGGAGGAGGSGP | ASSRTGHNNQAPL | 0.90367746 |
| VFAAFMV | ASSRTGHNNQAPL | 0.6945131 |
| TLYDKLR | ASSRTGHNNQAPL | 0.8202383 |
| TLAVFTVL | ASSRTGHNNQAPL | 0.8930745 |
| TGSVQPARV | ASSRTGHNNQAPL | 0.89433765 |
| TGGGAGSGGT | ASSRTGHNNQAPL | 0.8155735 |
| SHLETSL | ASSRTGHNNQAPL | 0.90498984 |
| SFTTGMT | ASSRTGHNNQAPL | 0.7438046 |
| QVEVAKRMGVRQPTVS | ASSRTGHNNQAPL | 0.8783425 |
| QGSGGPALAM | ASSRTGHNNQAPL | 0.7874106 |
| MLAPSNP | ASSRTGHNNQAPL | 0.8520699 |
| LVNNAAI | ASSRTGHNNQAPL | 0.71607864 |
| LTGEINTLA | ASSRTGHNNQAPL | 0.88456535 |
| LSIFGPV | ASSRTGHNNQAPL | 0.8900716 |
| LLALLANL | ASSRTGHNNQAPL | 0.8842956 |
| LISNASDALDKLR | ASSRTGHNNQAPL | 0.22753891 |
| LINEAVP | ASSRTGHNNQAPL | 0.8869874 |
| LGYAFSGL | ASSRTGHNNQAPL | 0.86156046 |
| LDFVAVAL | ASSRTGHNNQAPL | 0.91035384 |
| LAQPGAE | ASSRTGHNNQAPL | 0.8061763 |
| ISNLGAVTGF | ASSRTGHNNQAPL | 0.8744277 |
| HQEDPGRI | ASSRTGHNNQAPL | 0.77783763 |
| GTPGEPTP | ASSRTGHNNQAPL | 0.7827562 |

| | | |
|---|---|---|
| GFFNSGS | ASSRTGHNNQAPL | 0.6400646 |
| GAGGGGLTAGAN | ASSRTGHNNQAPL | 0.75040513 |
| FVDALVNM | ASSRTGHNNQAPL | 0.79043317 |
| FLTVAPEL | ASSRTGHNNQAPL | 0.7327776 |
| FETLTAL | ASSRTGHNNQAPL | 0.8779124 |
| DVINPIEL | ASSRTGHNNQAPL | 0.89573777 |
| APVSGAPV | ASSRTGHNNQAPL | 0.8943317 |
| ALVGGLAW | ASSRTGHNNQAPL | 0.8950996 |
| ALAAAAATAI | ASSRTGHNNQAPL | 0.9004469 |
| AFGLGLLL | ASSRTGHNNQAPL | 0.8865757 |
| AFFAAASL | ASSRTGHNNQAPL | 0.7483611 |
| AAAALPAFSPP | ASSRTGHNNQAPL | 0.7978985 |
| VVLAPAL | ASSRTGHNNQAPL | 0.62182987 |
| TIGHVDHGKTTL | ASSRTGHNNQAPL | 0.8270507 |
| LISNASDALDK | ASSRTGHNNQAPL | 0.76276505 |
| LISNASDAL | ASSRTGHNNQAPL | 0.85423267 |
| ISNASDALDK | ASSRTGHNNQAPL | 0.11115354 |
| NQAETLVYQTE | ASSRTGHNNQAPL | 0.84594786 |
| TGGTGGTAGAG | ASSRTGHNNQAPL | 0.9093492 |
| TGSTAGGAGGA | ASSRTGHNNQAPL | 0.7208501 |
| DPLAVPVAAGPV | ASSRTGHNNQAPL | 0.855176 |
| GGTGGTPGNGGN | ASSRTGHNNQAPL | 0.8625978 |
| PAGLAGVDPATL | ASSRTGHNNQAPL | 0.7917925 |
| GNNLVVEAPHAGASV | ASSRTGHNNQAPL | 0.8764705 |
| KVTRIMSASTLEQPA | ASSRTGHNNQAPL | 0.7226145 |
| LGFALPIALPRRTVV | ASSRTGHNNQAPL | 0.87138546 |
| TRLGNLTARKLVVLV | ASSRTGHNNQAPL | 0.9033928 |
| WSQQRRLIYFPSAGP | ASSRTGHNNQAPL | 0.8621991 |
| GPNGNPDPMAAAVDIR | ASSRTGHNNQAPL | 0.8446822 |
| TQRCVSVADLRAAASR | ASSRTGHNNQAPL | 0.83120817 |
| QFVLDVAEHPVEVLTETV | ASSRTGHNNQAPL | 0.34923533 |
| GGAGGTGGAGGNGGTQAGDG | ASSRTGHNNQAPL | 0.8629221 |
| PLLATLRAIAVDVDAKPAQV | ASSRTGHNNQAPL | 0.10415435 |
| APKFRTGGYMIDFWGVGYQVAK | ASSRTGHNNQAPL | 0.8316791 |
| GAGGAGGAGGAAGGPGGTGGTG | ASSRTGHNNQAPL | 0.63240516 |
| IGALGLLATDVAPAGSVAFGVA | ASSRTGHNNQAPL | 0.8555797 |
| AGGAGGIGGNGGAGGAGGVLMG | ASSRTGHNNQAPL | 0.8451188 |
| ADELIGLL | ASSRTGHNNQAPL | 0.8820507 |
| ADRALKA | ASSRTGHNNQAPL | 0.70365226 |
| AGAVGQVGGAGGA | ASSRTGHNNQAPL | 0.8227527 |
| AGGGGGLGGAP | ASSRTGHNNQAPL | 0.7820235 |
| AGGLLYGNGGAG | ASSRTGHNNQAPL | 0.6284827 |
| AGLESIGL | ASSRTGHNNQAPL | 0.89585876 |
| AGLKTGII | ASSRTGHNNQAPL | 0.8278063 |
| ASLAALLL | ASSRTGHNNQAPL | 0.8380183 |
| ASVINISSI | ASSRTGHNNQAPL | 0.8888992 |
| ATPPGPA | ASSRTGHNNQAPL | 0.8510909 |
| ATVASALV | ASSRTGHNNQAPL | 0.42426765 |
| ATVASLIL | ASSRTGHNNQAPL | 0.81195855 |
| AVNGPVLIRH | ASSRTGHNNQAPL | 0.8492789 |
| DGAAGAVGI | ASSRTGHNNQAPL | 0.75070834 |

| | | |
|---|---|---|
| DPLLHGI | ASSRTGHNNQAPL | 0.1205381 |
| ELGAALGAL | ASSRTGHNNQAPL | 0.8820761 |
| ELGRTQAV | ASSRTGHNNQAPL | 0.7026781 |
| FVAGGYLS | ASSRTGHNNQAPL | 0.7190024 |
| GASGGAGGAGGW | ASSRTGHNNQAPL | 0.8073592 |
| GGAGAGGDGGAGGI | ASSRTGHNNQAPL | 0.8845265 |
| GTGHNGSQPGMGGNG | ASSRTGHNNQAPL | 0.69230217 |
| GYAAASASAS | ASSRTGHNNQAPL | 0.71352005 |
| HQVDVAAVQQG | ASSRTGHNNQAPL | 0.88047814 |
| IGAAIAHT | ASSRTGHNNQAPL | 0.8760265 |
| IGMIEVAQL | ASSRTGHNNQAPL | 0.8962829 |
| IRLAPEHTSAMD | ASSRTGHNNQAPL | 0.8553574 |
| IYNVSVL | ASSRTGHNNQAPL | 0.8043046 |
| KQGFLGM | ASSRTGHNNQAPL | 0.85626394 |
| LESVITQL | ASSRTGHNNQAPL | 0.76620924 |
| LGLGALVVV | ASSRTGHNNQAPL | 0.53585374 |
| LIGNGGNGGSGGI | ASSRTGHNNQAPL | 0.40536058 |
| LNAINAPTL | ASSRTGHNNQAPL | 0.8165181 |
| LQLLNTGV | ASSRTGHNNQAPL | 0.47201782 |
| LQTSIEF | ASSRTGHNNQAPL | 0.8251537 |
| MSFAVSEL | ASSRTGHNNQAPL | 0.9020078 |
| NTGALGL | ASSRTGHNNQAPL | 0.8363708 |
| PPGAGGAGGAGTT | ASSRTGHNNQAPL | 0.56505066 |
| QLVGLGI | ASSRTGHNNQAPL | 0.83427835 |
| RFLEQVHQL | ASSRTGHNNQAPL | 0.7394685 |
| RGVGYVLR | ASSRTGHNNQAPL | 0.8658074 |
| SATGVFAGVI | ASSRTGHNNQAPL | 0.87101996 |
| SLPAITLP | ASSRTGHNNQAPL | 0.9031134 |
| STALAFTV | ASSRTGHNNQAPL | 0.88627017 |
| SVLTVGLGK | ASSRTGHNNQAPL | 0.7759378 |
| TALAGAGSGI | ASSRTGHNNQAPL | 0.8866635 |
| TGLPFTGI | ASSRTGHNNQAPL | 0.78366953 |
| VAEHVTV | ASSRTGHNNQAPL | 0.8561923 |
| VDIVLNSL | ASSRTGHNNQAPL | 0.8449299 |
| VLTGGGAVL | ASSRTGHNNQAPL | 0.87122345 |
| VNVGLAYGI | ASSRTGHNNQAPL | 0.82154584 |
| VSSAAVGAV | ASSRTGHNNQAPL | 0.8126564 |
| VTSVYEAGV | ASSRTGHNNQAPL | 0.4916576 |
| WVALDRGA | ASSRTGHNNQAPL | 0.8785159 |
| YLVGSVVI | ASSRTGHNNQAPL | 0.8412371 |
| GPPGTGKTLL | ASSRTGHNNQAPL | 0.41960865 |
| GVGMTKF | ASSRTGHNNQAPL | 0.7791828 |
| IGIGGSDL | ASSRTGHNNQAPL | 0.9091331 |
| IGTLLAEL | ASSRTGHNNQAPL | 0.8988849 |
| IYTSGTT | ASSRTGHNNQAPL | 0.73225445 |
| LSFPAGV | ASSRTGHNNQAPL | 0.7074051 |
| LYTSGTT | ASSRTGHNNQAPL | 0.8522178 |
| VGAFPFGF | ASSRTGHNNQAPL | 0.5337126 |
| VGAGTVEF | ASSRTGHNNQAPL | 0.87089413 |
| VGKSTLLL | ASSRTGHNNQAPL | 0.8248134 |
| AALAHAGGGIV | ASSRTGHNNQAPL | 0.83907163 |

| | | |
|---|---|---|
| AALVAVSLL | ASSRTGHNNQAPL | 0.7822629 |
| AAVSRAVAAL | ASSRTGHNNQAPL | 0.8626405 |
| AGGSSGIGGF | ASSRTGHNNQAPL | 0.80491686 |
| AGLTVANANG | ASSRTGHNNQAPL | 0.874704 |
| AGLVGAVL | ASSRTGHNNQAPL | 0.15605287 |
| ALSQAFTV | ASSRTGHNNQAPL | 0.88156384 |
| ALTFSVSL | ASSRTGHNNQAPL | 0.72442496 |
| ALTGAALVAL | ASSRTGHNNQAPL | 0.66988313 |
| ASFSIGAGVL | ASSRTGHNNQAPL | 0.353316 |
| DGSVYKI | ASSRTGHNNQAPL | 0.845203 |
| EQLAEIV | ASSRTGHNNQAPL | 0.8836188 |
| FAVSELTVM | ASSRTGHNNQAPL | 0.89182335 |
| GAAPLAAGL | ASSRTGHNNQAPL | 0.7060515 |
| GADGGDGGSGGAGG | ASSRTGHNNQAPL | 0.89425707 |
| GFGLDTGGAGG | ASSRTGHNNQAPL | 0.90570444 |
| GLASGGLL | ASSRTGHNNQAPL | 0.8064497 |
| GRIAGAAVLL | ASSRTGHNNQAPL | 0.34932673 |
| GSAGSLNTSF | ASSRTGHNNQAPL | 0.85240793 |
| GTGQIGI | ASSRTGHNNQAPL | 0.8302711 |
| GTLIALIL | ASSRTGHNNQAPL | 0.88692075 |
| IAATMGFVVG | ASSRTGHNNQAPL | 0.8505085 |
| IFAQALL | ASSRTGHNNQAPL | 0.64179873 |
| IGAAVALGF | ASSRTGHNNQAPL | 0.88100016 |
| IGLTPLAL | ASSRTGHNNQAPL | 0.89822257 |
| ILGSNTSTL | ASSRTGHNNQAPL | 0.73634887 |
| IVFSLTVL | ASSRTGHNNQAPL | 0.7653805 |
| LASKATGF | ASSRTGHNNQAPL | 0.7511878 |
| LASVALIL | ASSRTGHNNQAPL | 0.37991 |
| LFAVGLTL | ASSRTGHNNQAPL | 0.8793312 |
| LFNGGIGTYI | ASSRTGHNNQAPL | 0.7932831 |
| LTLGAGII | ASSRTGHNNQAPL | 0.7436706 |
| LVAAALGDI | ASSRTGHNNQAPL | 0.83475274 |
| NTAFVQLGI | ASSRTGHNNQAPL | 0.7889702 |
| QLFSGAEL | ASSRTGHNNQAPL | 0.7474977 |
| SAATFITL | ASSRTGHNNQAPL | 0.7994374 |
| SNYGLANLGV | ASSRTGHNNQAPL | 0.8362642 |
| SYGGAYAVM | ASSRTGHNNQAPL | 0.8730673 |
| TTGVGGLTLG | ASSRTGHNNQAPL | 0.8255714 |
| TTQTIYI | ASSRTGHNNQAPL | 0.84889776 |
| VAAGAGLGAVI | ASSRTGHNNQAPL | 0.9049926 |
| VAAVVTLL | ASSRTGHNNQAPL | 0.58034736 |
| VGFLGGLI | ASSRTGHNNQAPL | 0.72728586 |
| VGVIPILL | ASSRTGHNNQAPL | 0.87446296 |
| VIFASSLI | ASSRTGHNNQAPL | 0.5275277 |
| VKGGDGGAAAG | ASSRTGHNNQAPL | 0.8883221 |
| VTKTFTNI | ASSRTGHNNQAPL | 0.8470454 |
| VVAAAILL | ASSRTGHNNQAPL | 0.8593657 |
| YAALDVEL | ASSRTGHNNQAPL | 0.82463515 |
| LASVALLL | ASSRTGHNNQAPL | 0.9145628 |
| LDAVHRV | ASSRTGHNNQAPL | 0.88163644 |
| QGLSSSIF | ASSRTGHNNQAPL | 0.8495705 |

| | | |
|---|---|---|
| RGTFANIRL | ASSRTGHNNQAPL | 0.8759684 |
| RQLWWGHRI | ASSRTGHNNQAPL | 0.6556901 |
| YGPPGTGKTLL | ASSRTGHNNQAPL | 0.8966119 |
| AALVGAPANVA | ASSRTGHNNQAPL | 0.8725459 |
| AATFRRTE | ASSRTGHNNQAPL | 0.6437162 |
| AGALGGGI | ASSRTGHNNQAPL | 0.81770164 |
| AGGAGGAGGIGAG | ASSRTGHNNQAPL | 0.9042183 |
| AGSGALGL | ASSRTGHNNQAPL | 0.9001868 |
| AQIADQI | ASSRTGHNNQAPL | 0.38011253 |
| ASAVAVTYL | ASSRTGHNNQAPL | 0.7772579 |
| ASTVLGGTGG | ASSRTGHNNQAPL | 0.45667094 |
| AVGGEPAK | ASSRTGHNNQAPL | 0.7656274 |
| CEQGTFGGTC | ASSRTGHNNQAPL | 0.52535546 |
| DEGKAPAG | ASSRTGHNNQAPL | 0.8728002 |
| DTVVFPAL | ASSRTGHNNQAPL | 0.79701364 |
| FEGITMAG | ASSRTGHNNQAPL | 0.8436533 |
| FPLVVQD | ASSRTGHNNQAPL | 0.8290019 |
| GAFTTPQ | ASSRTGHNNQAPL | 0.8922713 |
| GFGGFGTAGGVGGA | ASSRTGHNNQAPL | 0.77946013 |
| GFLAQFL | ASSRTGHNNQAPL | 0.80499256 |
| GGAGGAGGNAGVA | ASSRTGHNNQAPL | 0.5878942 |
| GGVGTAGAGGAGGAGG | ASSRTGHNNQAPL | 0.41045582 |
| GNGGGGDAGNAGSGGN | ASSRTGHNNQAPL | 0.76234233 |
| GTGGAGGAGNDAGS | ASSRTGHNNQAPL | 0.7940894 |
| GTTNTGWFNS | ASSRTGHNNQAPL | 0.22701737 |
| IFMGPLFTL | ASSRTGHNNQAPL | 0.90048784 |
| ILAHAIEAI | ASSRTGHNNQAPL | 0.8448962 |
| IRSAIREAL | ASSRTGHNNQAPL | 0.86739993 |
| ITFAGGG | ASSRTGHNNQAPL | 0.7847223 |
| ITGMSGHQEV | ASSRTGHNNQAPL | 0.77727646 |
| IVAGLACTL | ASSRTGHNNQAPL | 0.73347676 |
| IVGGGLAGA | ASSRTGHNNQAPL | 0.8776953 |
| IVTGVLNI | ASSRTGHNNQAPL | 0.68095744 |
| LAQQGCI | ASSRTGHNNQAPL | 0.4419453 |
| LCLHVEN | ASSRTGHNNQAPL | 0.7152009 |
| LDAISDGS | ASSRTGHNNQAPL | 0.6803704 |
| LGGFAAML | ASSRTGHNNQAPL | 0.90106416 |
| LGGKNPMI | ASSRTGHNNQAPL | 0.89024454 |
| LGPKGVAAIL | ASSRTGHNNQAPL | 0.8469782 |
| LMVGVSI | ASSRTGHNNQAPL | 0.8982256 |
| LVGAAFVL | ASSRTGHNNQAPL | 0.2210003 |
| LVLAVAPL | ASSRTGHNNQAPL | 0.7809199 |
| MAAIAVIDAA | ASSRTGHNNQAPL | 0.8995396 |
| NNGINAELQ | ASSRTGHNNQAPL | 0.85373396 |
| RGEPLVLVHG | ASSRTGHNNQAPL | 0.8252442 |
| RGGAGGAAGLI | ASSRTGHNNQAPL | 0.89657474 |
| RSYRMHPAV | ASSRTGHNNQAPL | 0.7479888 |
| SAAHLLAGQLA | ASSRTGHNNQAPL | 0.7550731 |
| SAAVGSY | ASSRTGHNNQAPL | 0.8323356 |
| SPHAVLHRI | ASSRTGHNNQAPL | 0.88482946 |
| SQIAKVL | ASSRTGHNNQAPL | 0.8539389 |

| | | |
|---|---|---|
| TALGHLVP | ASSRTGHNNQAPL | 0.665166 |
| TAPTTSGAA | ASSRTGHNNQAPL | 0.86282337 |
| TDGDDGNGGSPGT | ASSRTGHNNQAPL | 0.9164253 |
| TDGLGEN | ASSRTGHNNQAPL | 0.7874545 |
| TGGPPLPGQA | ASSRTGHNNQAPL | 0.8858837 |
| TIPAITI | ASSRTGHNNQAPL | 0.86020625 |
| TIPGFTI | ASSRTGHNNQAPL | 0.8121474 |
| TSIASGALA | ASSRTGHNNQAPL | 0.71350026 |
| TVTLPGAAI | ASSRTGHNNQAPL | 0.75994474 |
| VASGLSAALP | ASSRTGHNNQAPL | 0.8281579 |
| VGAGVIGI | ASSRTGHNNQAPL | 0.6784824 |
| VGPKRFGV | ASSRTGHNNQAPL | 0.83417135 |
| VNVAFPD | ASSRTGHNNQAPL | 0.7961184 |
| VSMTNSGVS | ASSRTGHNNQAPL | 0.9178941 |
| VTDVLGDLSV | ASSRTGHNNQAPL | 0.86135674 |
| VVVVGAGFAGL | ASSRTGHNNQAPL | 0.8660331 |
| VYTVGTVV | ASSRTGHNNQAPL | 0.6986823 |
| WTTFSGP | ASSRTGHNNQAPL | 0.8694089 |
| YDELVEAF | ASSRTGHNNQAPL | 0.8714062 |
| YGTGGAGGNGG | ASSRTGHNNQAPL | 0.5606622 |
| YTNVMAV | ASSRTGHNNQAPL | 0.9014332 |
| FLEQVHQL | ASSRTGHNNQAPL | 0.7284714 |
| GLSLLAVAV | ASSRTGHNNQAPL | 0.82628995 |
| LEQVHQL | ASSRTGHNNQAPL | 0.867088 |
| LNAAGAI | ASSRTGHNNQAPL | 0.88688725 |
| PAVHASL | ASSRTGHNNQAPL | 0.8113328 |
| TELEDDA | ASSRTGHNNQAPL | 0.8188629 |
| TPPGPPN | ASSRTGHNNQAPL | 0.6811505 |
| TPPPPPPP | ASSRTGHNNQAPL | 0.8811534 |
| TQSPALI | ASSPDSSAETLY | 0.88942856 |
| EFAAAFPQ | ASSPDSSAETLY | 0.786008 |
| LLSGNLDVL | ASSPDSSAETLY | 0.9032844 |
| GPANLGNYNI | ASSPDSSAETLY | 0.8466102 |
| TLGPTLP | ASSPDSSAETLY | 0.39694422 |
| VYAYSSCCK | ASSPDSSAETLY | 0.8181509 |
| APAPAPAPNI | ASSPDSSAETLY | 0.90013754 |
| PGGPVPIP | ASSPDSSAETLY | 0.89696336 |
| FLTVALEP | ASSPDSSAETLY | 0.9035779 |
| SQYFGANL | ASSPDSSAETLY | 0.8784198 |
| GGVKPAA | ASSPDSSAETLY | 0.8092444 |
| GGTGGAGGAGGW | ASSPDSSAETLY | 0.9011683 |
| GVGGVAGLGGAGG | ASSPDSSAETLY | 0.9123423 |
| VLLVLATTI | ASSPDSSAETLY | 0.8783355 |
| IGIFAAML | ASSPDSSAETLY | 0.7337055 |
| AVGDLID | ASSPDSSAETLY | 0.62163913 |
| NISPLPL | ASSPDSSAETLY | 0.8634232 |
| VTAVFVAI | ASSPDSSAETLY | 0.89089036 |
| SGDDNDAAL | ASSPDSSAETLY | 0.67483187 |
| AVAALAHS | ASSPDSSAETLY | 0.8465321 |
| GGVTVMATP | ASSPDSSAETLY | 0.84571576 |
| AGLAANVRD | ASSPDSSAETLY | 0.41759095 |

| | | |
|---|---|---|
| TNRPDLIDPALLRPG | ASSPDSSAETLY | 0.8692211 |
| LANLGVTAL | ASSPDSSAETLY | 0.907037 |
| TGERIEL | ASSPDSSAETLY | 0.8497648 |
| NGEELAA | ASSPDSSAETLY | 0.76370394 |
| VLSLVWG | ASSPDSSAETLY | 0.90040505 |
| IGYALWL | ASSPDSSAETLY | 0.91457754 |
| LSLLLAPL | ASSPDSSAETLY | 0.8997145 |
| TLDEALK | ASSPDSSAETLY | 0.45152575 |
| TDGAGTAEGG | ASSPDSSAETLY | 0.25911123 |
| NAPGGAGGNGG | ASSPDSSAETLY | 0.8029645 |
| VSVFMGMGGNF | ASSPDSSAETLY | 0.8621098 |
| LASYLAAL | ASSPDSSAETLY | 0.79791385 |
| LVGGTPAL | ASSPDSSAETLY | 0.74338853 |
| VRNADVL | ASSPDSSAETLY | 0.8613181 |
| TCSSPFLVSAM | ASSPDSSAETLY | 0.8917021 |
| VHITDAE | ASSPDSSAETLY | 0.6754776 |
| EYAAPSF | ASSPDSSAETLY | 0.4747069 |
| VVDAFHGGL | ASSPDSSAETLY | 0.88493 |
| ASVVDPQ | ASSPDSSAETLY | 0.8773347 |
| SYNEFGGLS | ASSPDSSAETLY | 0.7200972 |
| DPDGSVY | ASSPDSSAETLY | 0.80816936 |
| MVSTIHATE | ASSPDSSAETLY | 0.8053198 |
| LAAGVADPV | ASSPDSSAETLY | 0.71999776 |
| TKEQIAA | ASSPDSSAETLY | 0.8596644 |
| SLASPAAL | ASSPDSSAETLY | 0.6207456 |
| IGLVFYGV | ASSPDSSAETLY | 0.9085526 |
| FLLDASNL | ASSPDSSAETLY | 0.64397615 |
| YQVGNFP | ASSPDSSAETLY | 0.88753706 |
| ESLLSQST | ASSPDSSAETLY | 0.86254466 |
| LFTVGFAAVG | ASSPDSSAETLY | 0.8689176 |
| ATPPTPT | ASSPDSSAETLY | 0.7902103 |
| TDLATLG | ASSPDSSAETLY | 0.7243418 |
| SAGGTGR | ASSPDSSAETLY | 0.6579081 |
| GFSNSGS | ASSPDSSAETLY | 0.44998658 |
| GGNGGSGGTSV | ASSPDSSAETLY | 0.8971291 |
| GTSVATGGAGN | ASSPDSSAETLY | 0.1747244 |
| NGGNGGSGGTS | ASSPDSSAETLY | 0.8961323 |
| AVVKFPH | ASSPDSSAETLY | 0.84881926 |
| TPPPPPP | ASSPDSSAETLY | 0.7055533 |
| AAPAAAAAPA | ASSPDSSAETLY | 0.8709829 |
| IALNEDL | ASSPDSSAETLY | 0.60426736 |
| IVNVSSM | ASSPDSSAETLY | 0.7396402 |
| TGIPPAPR | ASSPDSSAETLY | 0.86585355 |
| FDLGGGTF | ASSPDSSAETLY | 0.87936854 |
| FDLGGGTFDVS | ASSPDSSAETLY | 0.7310246 |
| NEPTAAA | ASSPDSSAETLY | 0.8672786 |
| NEPTAAALA | ASSPDSSAETLY | 0.8657994 |
| GSIINISSI | ASSPDSSAETLY | 0.70860064 |
| VAPGNAGTA | ASSPDSSAETLY | 0.8582877 |
| TGGGASLE | ASSPDSSAETLY | 0.8937775 |
| VAVYNHYKRI | ASSPDSSAETLY | 0.86568177 |

| | | |
|---|---|---|
| TDPAPATT | ASSPDSSAETLY | 0.5970197 |
| TNRPDILDPALLRPG | ASSPDSSAETLY | 0.878783 |
| LTTTTTS | ASSPDSSAETLY | 0.69255453 |
| PTGSGKTL | ASSPDSSAETLY | 0.86165774 |
| TGSGKTL | ASSPDSSAETLY | 0.8474522 |
| TIGHVDHGK | ASSPDSSAETLY | 0.78872883 |
| TIGHVDHGKTT | ASSPDSSAETLY | 0.8905084 |
| GHVDHGKTT | ASSPDSSAETLY | 0.43210155 |
| AQFEHTLL | ASSPDSSAETLY | 0.72907805 |
| IVNQDEL | ASSPDSSAETLY | 0.78407776 |
| TNGSQFF | ASSPDSSAETLY | 0.8421875 |
| TGKGIVE | ASSPDSSAETLY | 0.88131505 |
| LISNASDA | ASSPDSSAETLY | 0.6406783 |
| AELRTLGELTKIAKAHGAQ | ASSPDSSAETLY | 0.8476573 |
| APNAVAAAEPSW | ASSPDSSAETLY | 0.7393881 |
| APQVAGAATAVL | ASSPDSSAETLY | 0.5719316 |
| ARRDPAAAAAILPTDA | ASSPDSSAETLY | 0.81876075 |
| DAAARDVLAEAGLAEYF | ASSPDSSAETLY | 0.7939358 |
| DDQIVLSAKANLAKVEQ | ASSPDSSAETLY | 0.69034326 |
| DELLPDTGEDSDEDGH | ASSPDSSAETLY | 0.87958515 |
| ELPEAPANRATVPQD | ASSPDSSAETLY | 0.6801089 |
| FGQSKIERVTGIG | ASSPDSSAETLY | 0.8740026 |
| GAGGTGSTAGGAGGAGGAGGLY | ASSPDSSAETLY | 0.9104271 |
| GHGGTGAAGGNGGTGGAGGSGI | ASSPDSSAETLY | 0.67404765 |
| GNGGNGGAGGEGGPGGAGGAGA | ASSPDSSAETLY | 0.8855236 |
| GSTAGNGGNGGNATNGTIAGQP | ASSPDSSAETLY | 0.875548 |
| GTGGDGGNGGNGGTGG | ASSPDSSAETLY | 0.87583876 |
| GVRRPAGIPADAPV | ASSPDSSAETLY | 0.8053225 |
| GYRQVTDAHLLTLARR | ASSPDSSAETLY | 0.7468926 |
| IAKMGQDGHDRG | ASSPDSSAETLY | 0.90336996 |
| KAEPSIEIQYT | ASSPDSSAETLY | 0.88195825 |
| KPVRIRMAADEDTVAVT | ASSPDSSAETLY | 0.8737865 |
| MPPRIAGMRLLVIKPEP | ASSPDSSAETLY | 0.8804277 |
| MKFYAVALFRSRGVPPTR | ASSPDSSAETLY | 0.8768598 |
| NAGGFGILGAVAHSP | ASSPDSSAETLY | 0.8832144 |
| NEGREITLAMAEM | ASSPDSSAETLY | 0.7711138 |
| NELVEWSHPLGHRWRWE | ASSPDSSAETLY | 0.5628699 |
| NLGAGGTGGNGGNPG | ASSPDSSAETLY | 0.79481006 |
| NTEWPKWETVMAADPVD | ASSPDSSAETLY | 0.8298458 |
| PASAAPAETQP | ASSPDSSAETLY | 0.79405653 |
| QQSSDGKNGVRAAEITGE | ASSPDSSAETLY | 0.3817054 |
| RAPDAVTTSWWKE | ASSPDSSAETLY | 0.88767225 |
| SGFGNVGAHTSGWFNQ | ASSPDSSAETLY | 0.9150766 |
| SGVAGAVGQAASVGGLKV | ASSPDSSAETLY | 0.7520826 |
| SNGGAGTGSAGGNGGT | ASSPDSSAETLY | 0.9110955 |
| TGGAGGEGGAAG | ASSPDSSAETLY | 0.6126487 |
| TGGDGGAAGAGGAPGHG | ASSPDSSAETLY | 0.8742903 |
| TMPPLLAAIANAGRHGV | ASSPDSSAETLY | 0.8920529 |
| TSSTAGNSGGA | ASSPDSSAETLY | 0.69111425 |
| VPATVPVGILC | ASSPDSSAETLY | 0.85900116 |
| VQRPDSGGVVLTGRIS | ASSPDSSAETLY | 0.66726863 |

| | | |
|---|---|---|
| WTPNPVPATKGVPAGLVQAT | ASSPDSSAETLY | 0.79784894 |
| YALKRALAKFSDRFDVVII | ASSPDSSAETLY | 0.9007564 |
| YLIGSVIV | ASSPDSSAETLY | 0.77725905 |
| YGQLVAVID | ASSPDSSAETLY | 0.86854833 |
| VTVGGMTLPA | ASSPDSSAETLY | 0.79169774 |
| VLALPAV | ASSPDSSAETLY | 0.64604867 |
| VIPPTLNYET | ASSPDSSAETLY | 0.8625908 |
| VGMGGMHPSSG | ASSPDSSAETLY | 0.89523053 |
| VGLAGGAATA | ASSPDSSAETLY | 0.7263658 |
| VGGAGGAGGSGP | ASSPDSSAETLY | 0.91030264 |
| VFAAFMV | ASSPDSSAETLY | 0.76181453 |
| TLYDKLR | ASSPDSSAETLY | 0.91326237 |
| TLAVFTVL | ASSPDSSAETLY | 0.8957193 |
| TGSVQPARV | ASSPDSSAETLY | 0.8312706 |
| TGGGAGSGGT | ASSPDSSAETLY | 0.7956742 |
| SHLETSL | ASSPDSSAETLY | 0.8853002 |
| SFTTGMT | ASSPDSSAETLY | 0.77164924 |
| QVEVAKRMGVRQPTVS | ASSPDSSAETLY | 0.8779534 |
| QGSGGPALAM | ASSPDSSAETLY | 0.9032966 |
| MLAPSNP | ASSPDSSAETLY | 0.510038 |
| LVNNAAI | ASSPDSSAETLY | 0.7119061 |
| LTGEINTLA | ASSPDSSAETLY | 0.6814888 |
| LSIFGPV | ASSPDSSAETLY | 0.6637105 |
| LLALLANL | ASSPDSSAETLY | 0.60026884 |
| LISNASDALDKLR | ASSPDSSAETLY | 0.38889518 |
| LINEAVP | ASSPDSSAETLY | 0.6171143 |
| LGYAFSGL | ASSPDSSAETLY | 0.53943974 |
| LDFVAVAL | ASSPDSSAETLY | 0.70441854 |
| LAQPGAE | ASSPDSSAETLY | 0.70145035 |
| ISNLGAVTGF | ASSPDSSAETLY | 0.5850339 |
| HQEDPGRI | ASSPDSSAETLY | 0.69397646 |
| GTPGEPTP | ASSPDSSAETLY | 0.60900223 |
| GFFNSGS | ASSPDSSAETLY | 0.47172505 |
| GAGGGGLTAGAN | ASSPDSSAETLY | 0.6642712 |
| FVDALVNM | ASSPDSSAETLY | 0.5972978 |
| FLTVAPEL | ASSPDSSAETLY | 0.66927695 |
| FETLTAL | ASSPDSSAETLY | 0.4367581 |
| DVINPIEL | ASSPDSSAETLY | 0.69623244 |
| APVSGAPV | ASSPDSSAETLY | 0.6118877 |
| ALVGGLAW | ASSPDSSAETLY | 0.29491386 |
| ALAAAAATAI | ASSPDSSAETLY | 0.694292 |
| AFGLGLLL | ASSPDSSAETLY | 0.60976756 |
| AFFAAASL | ASSPDSSAETLY | 0.6144967 |
| AAAALPAFSPP | ASSPDSSAETLY | 0.7023272 |
| VVLAPAL | ASSPDSSAETLY | 0.43848157 |
| TIGHVDHGKTTL | ASSPDSSAETLY | 0.4932375 |
| LISNASDALDK | ASSPDSSAETLY | 0.6975205 |
| LISNASDAL | ASSPDSSAETLY | 0.7017671 |
| ISNASDALDK | ASSPDSSAETLY | 0.43667433 |
| NQAETLVYQTE | ASSPDSSAETLY | 0.6724737 |
| TGGTGGTAGAG | ASSPDSSAETLY | 0.7018378 |

| | | |
|---|---|---|
| TGSTAGGAGGA | ASSPDSSAETLY | 0.6548269 |
| DPLAVPVAAGPV | ASSPDSSAETLY | 0.7111784 |
| GGTGGTPGNGGN | ASSPDSSAETLY | 0.71404743 |
| PAGLAGVDPATL | ASSPDSSAETLY | 0.5999216 |
| GNNLVVEAPHAGASV | ASSPDSSAETLY | 0.71118706 |
| KVTRIMSASTLEQPA | ASSPDSSAETLY | 0.713864 |
| LGFALPIALPRRTVV | ASSPDSSAETLY | 0.6068125 |
| TRLGNLTARKLVVLV | ASSPDSSAETLY | 0.6610527 |
| WSQQRRLIYFPSAGP | ASSPDSSAETLY | 0.46504432 |
| GPNGNPDPMAAAVDIR | ASSPDSSAETLY | 0.61567825 |
| TQRCVSVADLRAAASR | ASSPDSSAETLY | 0.43902472 |
| QFVLDVAEHPVEVLTETV | ASSPDSSAETLY | 0.6841546 |
| GGAGGTGGAGGNGGTQAGDG | ASSPDSSAETLY | 0.71112895 |
| PLLATLRAIAVDVDAKPAQV | ASSPDSSAETLY | 0.65970016 |
| APKFRTGGYMIDFWGVGYQVAK | ASSPDSSAETLY | 0.50178874 |
| GAGGAGGAGGAAGGPGGTGGTG | ASSPDSSAETLY | 0.521899 |
| IGALGLLATDVAPAGSVAFGVA | ASSPDSSAETLY | 0.67223656 |
| AGGAGGIGGNGGAGGAGGVLMG | ASSPDSSAETLY | 0.4917391 |
| ADELIGLL | ASSPDSSAETLY | 0.39081788 |
| ADRALKA | ASSPDSSAETLY | 0.62764233 |
| AGAVGQVGGAGGA | ASSPDSSAETLY | 0.6873264 |
| AGGGGGLGGAP | ASSPDSSAETLY | 0.6930723 |
| AGGLLYGNGGAG | ASSPDSSAETLY | 0.58032703 |
| AGLESIGL | ASSPDSSAETLY | 0.7157267 |
| AGLKTGII | ASSPDSSAETLY | 0.6999347 |
| ASLAALLL | ASSPDSSAETLY | 0.7099883 |
| ASVINISSI | ASSPDSSAETLY | 0.5681216 |
| ATPPGPA | ASSPDSSAETLY | 0.51230127 |
| ATVASALV | ASSPDSSAETLY | 0.70349437 |
| ATVASLIL | ASSPDSSAETLY | 0.61222595 |
| AVNGPVLIRH | ASSPDSSAETLY | 0.657984 |
| DGAAGAVGI | ASSPDSSAETLY | 0.7031981 |
| DPLLHGI | ASSPDSSAETLY | 0.5787732 |
| ELGAALGAL | ASSPDSSAETLY | 0.5597464 |
| ELGRTQAV | ASSPDSSAETLY | 0.6591574 |
| FVAGGYLS | ASSPDSSAETLY | 0.7183567 |
| GASGGAGGAGGW | ASSPDSSAETLY | 0.57722765 |
| GGAGAGGDGGAGGI | ASSPDSSAETLY | 0.45985243 |
| GTGHNGSQPGMGGNG | ASSPDSSAETLY | 0.5913867 |
| GYAAASASAS | ASSPDSSAETLY | 0.59820515 |
| HQVDVAAVQQG | ASSPDSSAETLY | 0.70981884 |
| IGAAIAHT | ASSPDSSAETLY | 0.6821194 |
| IGMIEVAQL | ASSPDSSAETLY | 0.62459403 |
| IRLAPEHTSAMD | ASSPDSSAETLY | 0.68509066 |
| IYNVSVL | ASSPDSSAETLY | 0.6874455 |
| KQGFLGM | ASSPDSSAETLY | 0.65288824 |
| LESVITQL | ASSPDSSAETLY | 0.6156772 |
| LGLGALVVV | ASSPDSSAETLY | 0.61910033 |
| LIGNGGNGGSGGI | ASSPDSSAETLY | 0.7123994 |
| LNAINAPTL | ASSPDSSAETLY | 0.69968295 |
| LQLLNTGV | ASSPDSSAETLY | 0.6995288 |

| | | |
|---|---|---|
| LQTSIEF | ASSPDSSAETLY | 0.708177 |
| MSFAVSEL | ASSPDSSAETLY | 0.6186769 |
| NTGALGL | ASSPDSSAETLY | 0.6763687 |
| PPGAGGAGGAGTT | ASSPDSSAETLY | 0.4753111 |
| QLVGLGI | ASSPDSSAETLY | 0.43953425 |
| RFLEQVHQL | ASSPDSSAETLY | 0.6578443 |
| RGVGYVLR | ASSPDSSAETLY | 0.6854929 |
| SATGVFAGVI | ASSPDSSAETLY | 0.6977482 |
| SLPAITLP | ASSPDSSAETLY | 0.66272056 |
| STALAFTV | ASSPDSSAETLY | 0.5846993 |
| SVLTVGLGK | ASSPDSSAETLY | 0.68694127 |
| TALAGAGSGI | ASSPDSSAETLY | 0.7065171 |
| TGLPFTGI | ASSPDSSAETLY | 0.5344621 |
| VAEHVTV | ASSPDSSAETLY | 0.62450504 |
| VDIVLNSL | ASSPDSSAETLY | 0.72013855 |
| VLTGGGAVL | ASSPDSSAETLY | 0.68281937 |
| VNVGLAYGI | ASSPDSSAETLY | 0.6428165 |
| VSSAAVGAV | ASSPDSSAETLY | 0.6287406 |
| VTSVYEAGV | ASSPDSSAETLY | 0.7102879 |
| WVALDRGA | ASSPDSSAETLY | 0.63994807 |
| YLVGSVVI | ASSPDSSAETLY | 0.7193486 |
| GPPGTGKTLL | ASSPDSSAETLY | 0.379082 |
| GVGMTKF | ASSPDSSAETLY | 0.6545429 |
| IGIGGSDL | ASSPDSSAETLY | 0.55225104 |
| IGTLLAEL | ASSPDSSAETLY | 0.31489265 |
| IYTSGTT | ASSPDSSAETLY | 0.7058742 |
| LSFPAGV | ASSPDSSAETLY | 0.54709804 |
| LYTSGTT | ASSPDSSAETLY | 0.65498054 |
| VGAFPFGF | ASSPDSSAETLY | 0.6008681 |
| VGAGTVEF | ASSPDSSAETLY | 0.7108658 |
| VGKSTLLL | ASSPDSSAETLY | 0.6883181 |
| AALAHAGGGIV | ASSPDSSAETLY | 0.5464754 |
| AALVAVSLL | ASSPDSSAETLY | 0.5491548 |
| AAVSRAVAAL | ASSPDSSAETLY | 0.68869793 |
| AGGSSGIGGF | ASSPDSSAETLY | 0.7004736 |
| AGLTVANANG | ASSPDSSAETLY | 0.70908856 |
| AGLVGAVL | ASSPDSSAETLY | 0.35027206 |
| ALSQAFTV | ASSPDSSAETLY | 0.6890435 |
| ALTFSVSL | ASSPDSSAETLY | 0.6537677 |
| ALTGAALVAL | ASSPDSSAETLY | 0.362812 |
| ASFSIGAGVL | ASSPDSSAETLY | 0.3751741 |
| DGSVYKI | ASSPDSSAETLY | 0.5933404 |
| EQLAEIV | ASSPDSSAETLY | 0.63899624 |
| FAVSELTVM | ASSPDSSAETLY | 0.7001631 |
| GAAPLAAGL | ASSPDSSAETLY | 0.6976156 |
| GADGGDGGSGGAGG | ASSPDSSAETLY | 0.6440861 |
| GFGLDTGGAGG | ASSPDSSAETLY | 0.68007934 |
| GLASGGLL | ASSPDSSAETLY | 0.6269699 |
| GRIAGAAVLL | ASSPDSSAETLY | 0.24989873 |
| GSAGSLNTSF | ASSPDSSAETLY | 0.51986986 |
| GTGQIGI | ASSPDSSAETLY | 0.46848983 |

| | | |
|---|---|---|
| GTLIALIL | ASSPDSSAETLY | 0.7134511 |
| IAATMGFVVG | ASSPDSSAETLY | 0.55206656 |
| IFAQALL | ASSPDSSAETLY | 0.6353007 |
| IGAAVALGF | ASSPDSSAETLY | 0.6681708 |
| IGLTPLAL | ASSPDSSAETLY | 0.68387866 |
| ILGSNTSTL | ASSPDSSAETLY | 0.66100353 |
| IVFSLTVL | ASSPDSSAETLY | 0.52447975 |
| LASKATGF | ASSPDSSAETLY | 0.40037164 |
| LASVALIL | ASSPDSSAETLY | 0.6879202 |
| LFAVGLTL | ASSPDSSAETLY | 0.55037844 |
| LFNGGIGTYI | ASSPDSSAETLY | 0.4668118 |
| LTLGAGII | ASSPDSSAETLY | 0.53656274 |
| LVAAALGDI | ASSPDSSAETLY | 0.6515815 |
| NTAFVQLGI | ASSPDSSAETLY | 0.41367662 |
| QLFSGAEL | ASSPDSSAETLY | 0.70067817 |
| SAATFITL | ASSPDSSAETLY | 0.56774247 |
| SNYGLANLGV | ASSPDSSAETLY | 0.6362588 |
| SYGGAYAVM | ASSPDSSAETLY | 0.71681786 |
| TTGVGGLTLG | ASSPDSSAETLY | 0.67045426 |
| TTQTIYI | ASSPDSSAETLY | 0.6571716 |
| VAAGAGLGAVI | ASSPDSSAETLY | 0.55068594 |
| VAAVVTLL | ASSPDSSAETLY | 0.5906668 |
| VGFLGGLI | ASSPDSSAETLY | 0.5549646 |
| VGVIPILL | ASSPDSSAETLY | 0.5584386 |
| VIFASSLI | ASSPDSSAETLY | 0.6971619 |
| VKGGDGGAAAG | ASSPDSSAETLY | 0.6843716 |
| VTKTFTNI | ASSPDSSAETLY | 0.47808406 |
| VVAAAILL | ASSPDSSAETLY | 0.6859823 |
| YAALDVEL | ASSPDSSAETLY | 0.71714056 |
| LASVALLL | ASSPDSSAETLY | 0.7204022 |
| LDAVHRV | ASSPDSSAETLY | 0.57622665 |
| QGLSSSIF | ASSPDSSAETLY | 0.50188076 |
| RGTFANIRL | ASSPDSSAETLY | 0.5321152 |
| RQLWWGHRI | ASSPDSSAETLY | 0.6558373 |
| YGPPGTGKTLL | ASSPDSSAETLY | 0.49387935 |
| AALVGAPANVA | ASSPDSSAETLY | 0.4843803 |
| AATFRRTE | ASSPDSSAETLY | 0.7070867 |
| AGALGGGI | ASSPDSSAETLY | 0.71873707 |
| AGGAGGAGGIGAG | ASSPDSSAETLY | 0.6032926 |
| AGSGALGL | ASSPDSSAETLY | 0.4772507 |
| AQIADQI | ASSPDSSAETLY | 0.6563834 |
| ASAVAVTYL | ASSPDSSAETLY | 0.5558123 |
| ASTVLGGTGG | ASSPDSSAETLY | 0.602824 |
| AVGGEPAK | ASSPDSSAETLY | 0.49220258 |
| CEQGTFGGTC | ASSPDSSAETLY | 0.48191875 |
| DEGKAPAG | ASSPDSSAETLY | 0.6355208 |
| DTVVFPAL | ASSPDSSAETLY | 0.71360785 |
| FEGITMAG | ASSPDSSAETLY | 0.64607483 |
| FPLVVQD | ASSPDSSAETLY | 0.6710611 |
| GAFTTPQ | ASSPDSSAETLY | 0.7160541 |
| GFGGFGTAGGVGGA | ASSPDSSAETLY | 0.7010913 |

| | | |
|---|---|---|
| GFLAQFL | ASSPDSSAETLY | 0.71146655 |
| GGAGGAGGNAGVA | ASSPDSSAETLY | 0.69979525 |
| GGVGTAGAGGAGGAGG | ASSPDSSAETLY | 0.67602134 |
| GNGGGGDAGNAGSGGN | ASSPDSSAETLY | 0.655713 |
| GTGGAGGAGNDAGS | ASSPDSSAETLY | 0.7083131 |
| GTTNTGWFNS | ASSPDSSAETLY | 0.6506157 |
| IFMGPLFTL | ASSPDSSAETLY | 0.6722051 |
| ILAHAIEAI | ASSPDSSAETLY | 0.6558851 |
| IRSAIREAL | ASSPDSSAETLY | 0.6974791 |
| ITFAGGG | ASSPDSSAETLY | 0.58294725 |
| ITGMSGHQEV | ASSPDSSAETLY | 0.7152145 |
| IVAGLACTL | ASSPDSSAETLY | 0.31712043 |
| IVGGGLAGA | ASSPDSSAETLY | 0.44563404 |
| IVTGVLNI | ASSPDSSAETLY | 0.7092087 |
| LAQQGCI | ASSPDSSAETLY | 0.2629856 |
| LCLHVEN | ASSPDSSAETLY | 0.37485188 |
| LDAISDGS | ASSPDSSAETLY | 0.6711684 |
| LGGFAAML | ASSPDSSAETLY | 0.70633477 |
| LGGKNPMI | ASSPDSSAETLY | 0.27657112 |
| LGPKGVAAIL | ASSPDSSAETLY | 0.5278014 |
| LMVGVSI | ASSPDSSAETLY | 0.45154235 |
| LVGAAFVL | ASSPDSSAETLY | 0.682179 |
| LVLAVAPL | ASSPDSSAETLY | 0.6215178 |
| MAAIAVIDAA | ASSPDSSAETLY | 0.3965494 |
| NNGINAELQ | ASSPDSSAETLY | 0.7033898 |
| RGEPLVLVHG | ASSPDSSAETLY | 0.5487919 |
| RGGAGGAAGLI | ASSPDSSAETLY | 0.6655833 |
| RSYRMHPAV | ASSPDSSAETLY | 0.7086202 |
| SAAHLLAGQLA | ASSPDSSAETLY | 0.71766555 |
| SAAVGSY | ASSPDSSAETLY | 0.50097805 |
| SPHAVLHRI | ASSPDSSAETLY | 0.689519 |
| SQIAKVL | ASSPDSSAETLY | 0.61003935 |
| TALGHLVP | ASSPDSSAETLY | 0.72422826 |
| TAPTTSGAA | ASSPDSSAETLY | 0.6190255 |
| TDGDDGNGGSPGT | ASSPDSSAETLY | 0.46528038 |
| TDGLGEN | ASSPDSSAETLY | 0.7029885 |
| TGGPPLPGQA | ASSPDSSAETLY | 0.7052127 |
| TIPAITI | ASSPDSSAETLY | 0.60060364 |
| TIPGFTI | ASSPDSSAETLY | 0.6988795 |
| TSIASGALA | ASSPDSSAETLY | 0.6256794 |
| TVTLPGAAI | ASSPDSSAETLY | 0.6569078 |
| VASGLSAALP | ASSPDSSAETLY | 0.7010089 |
| VGAGVIGI | ASSPDSSAETLY | 0.6943358 |
| VGPKRFGV | ASSPDSSAETLY | 0.5038991 |
| VNVAFPD | ASSPDSSAETLY | 0.70729893 |
| VSMTNSGVS | ASSPDSSAETLY | 0.6560682 |
| VTDVLGDLSV | ASSPDSSAETLY | 0.5017412 |
| VVVVGAGFAGL | ASSPDSSAETLY | 0.72045374 |
| VYTVGTVV | ASSPDSSAETLY | 0.6130135 |
| WTTFSGP | ASSPDSSAETLY | 0.70204616 |
| YDELVEAF | ASSPDSSAETLY | 0.7003044 |

| | | |
|---|---|---|
| YGTGGAGGNGG | ASSPDSSAETLY | 0.7047954 |
| YTNVMAV | ASSPDSSAETLY | 0.46849936 |
| FLEQVHQL | ASSPDSSAETLY | 0.64067423 |
| GLSLLAVAV | ASSPDSSAETLY | 0.61284524 |
| LEQVHQL | ASSPDSSAETLY | 0.6979375 |
| LNAAGAI | ASSPDSSAETLY | 0.6408231 |
| PAVHASL | ASSPDSSAETLY | 0.6620047 |
| TELEDDA | ASSPDSSAETLY | 0.7021159 |
| TPPGPPN | ASSPDSSAETLY | 0.7101946 |
| TPPPPPPP | ASSPDSSAETLY | 0.6714933 |
| TQSPALI | ASSPRDRSDYT | 0.61380064 |
| EFAAAFPQ | ASSPRDRSDYT | 0.7171838 |
| LLSGNLDVL | ASSPRDRSDYT | 0.56982136 |
| GPANLGNYNI | ASSPRDRSDYT | 0.6563405 |
| TLGPTLP | ASSPRDRSDYT | 0.7003307 |
| VYAYSSCCK | ASSPRDRSDYT | 0.69079393 |
| APAPAPAPNI | ASSPRDRSDYT | 0.6655437 |
| PGGPVPIP | ASSPRDRSDYT | 0.7050773 |
| FLTVALEP | ASSPRDRSDYT | 0.5031725 |
| SQYFGANL | ASSPRDRSDYT | 0.6818149 |
| GGVKPAA | ASSPRDRSDYT | 0.6446465 |
| GGTGGAGGAGGW | ASSPRDRSDYT | 0.58489674 |
| GVGGVAGLGGAGG | ASSPRDRSDYT | 0.54193985 |
| VLLVLATTI | ASSPRDRSDYT | 0.69116455 |
| IGIFAAML | ASSPRDRSDYT | 0.69912875 |
| AVGDLID | ASSPRDRSDYT | 0.6828967 |
| NISPLPL | ASSPRDRSDYT | 0.3475323 |
| VTAVFVAI | ASSPRDRSDYT | 0.68754554 |
| SGDDNDAAL | ASSPRDRSDYT | 0.6706017 |
| AVAALAHS | ASSPRDRSDYT | 0.7191512 |
| GGVTVMATP | ASSPRDRSDYT | 0.62183297 |
| AGLAANVRD | ASSPRDRSDYT | 0.65738976 |
| TNRPDLIDPALLRPG | ASSPRDRSDYT | 0.59409595 |
| LANLGVTAL | ASSPRDRSDYT | 0.6727445 |
| TGERIEL | ASSPRDRSDYT | 0.46722305 |
| NGEELAA | ASSPRDRSDYT | 0.55882794 |
| VLSLVWG | ASSPRDRSDYT | 0.6156683 |
| IGYALWL | ASSPRDRSDYT | 0.6050543 |
| LSLLLAPL | ASSPRDRSDYT | 0.6982892 |
| TLDEALK | ASSPRDRSDYT | 0.70443094 |
| TDGAGTAEGG | ASSPRDRSDYT | 0.6857411 |
| NAPGGAGGNGG | ASSPRDRSDYT | 0.70620453 |
| VSVFMGMGGNF | ASSPRDRSDYT | 0.71624166 |
| LASYLAAL | ASSPRDRSDYT | 0.70553577 |
| LVGGTPAL | ASSPRDRSDYT | 0.6302775 |
| VRNADVL | ASSPRDRSDYT | 0.5635126 |
| TCSSPFLVSAM | ASSPRDRSDYT | 0.49217087 |
| VHITDAE | ASSPRDRSDYT | 0.5833626 |
| EYAAPSF | ASSPRDRSDYT | 0.6249957 |
| VVDAFHGGL | ASSPRDRSDYT | 0.6479198 |
| ASVVDPQ | ASSPRDRSDYT | 0.26387095 |

| | | |
|---|---|---|
| SYNEFGGLS | ASSPRDRSDYT | 0.67809105 |
| DPDGSVY | ASSPRDRSDYT | 0.70888 |
| MVSTIHATE | ASSPRDRSDYT | 0.620159 |
| LAAGVADPV | ASSPRDRSDYT | 0.6739744 |
| TKEQIAA | ASSPRDRSDYT | 0.6910741 |
| SLASPAAL | ASSPRDRSDYT | 0.6274775 |
| IGLVFYGV | ASSPRDRSDYT | 0.6846982 |
| FLLDASNL | ASSPRDRSDYT | 0.60418975 |
| YQVGNFP | ASSPRDRSDYT | 0.70294654 |
| ESLLSQST | ASSPRDRSDYT | 0.7124965 |
| LFTVGFAAVG | ASSPRDRSDYT | 0.687378 |
| ATPPTPT | ASSPRDRSDYT | 0.5781752 |
| TDLATLG | ASSPRDRSDYT | 0.64679164 |
| SAGGTGR | ASSPRDRSDYT | 0.4233573 |
| GFSNSGS | ASSPRDRSDYT | 0.61605185 |
| GGNGGSGGTSV | ASSPRDRSDYT | 0.5386555 |
| GTSVATGGAGN | ASSPRDRSDYT | 0.6679441 |
| NGGNGGSGGTS | ASSPRDRSDYT | 0.5710776 |
| AVVKFPH | ASSPRDRSDYT | 0.6806606 |
| TPPPPPP | ASSPRDRSDYT | 0.6592362 |
| AAPAAAAAPA | ASSPRDRSDYT | 0.6811344 |
| IALNEDL | ASSPRDRSDYT | 0.5854638 |
| IVNVSSM | ASSPRDRSDYT | 0.6903466 |
| TGIPPAPR | ASSPRDRSDYT | 0.66809 |
| FDLGGGTF | ASSPRDRSDYT | 0.4218117 |
| FDLGGGTFDVS | ASSPRDRSDYT | 0.70950174 |
| NEPTAAA | ASSPRDRSDYT | 0.64809257 |
| NEPTAAALA | ASSPRDRSDYT | 0.6509256 |
| GSIINISSI | ASSPRDRSDYT | 0.70038545 |
| VAPGNAGTA | ASSPRDRSDYT | 0.6600374 |
| TGGGASLE | ASSPRDRSDYT | 0.53404665 |
| VAVYNHYKRI | ASSPRDRSDYT | 0.6044606 |
| TDPAPATT | ASSPRDRSDYT | 0.67799497 |
| TNRPDILDPALLRPG | ASSPRDRSDYT | 0.6297488 |
| LTTTTTS | ASSPRDRSDYT | 0.29897264 |
| PTGSGKTL | ASSPRDRSDYT | 0.6949737 |
| TGSGKTL | ASSPRDRSDYT | 0.49706352 |
| TIGHVDHGK | ASSPRDRSDYT | 0.6009922 |
| TIGHVDHGKTT | ASSPRDRSDYT | 0.6545228 |
| GHVDHGKTT | ASSPRDRSDYT | 0.70303833 |
| AQFEHTLL | ASSPRDRSDYT | 0.59504586 |
| IVNQDEL | ASSPRDRSDYT | 0.51603967 |
| TNGSQFF | ASSPRDRSDYT | 0.6780261 |
| TGKGIVE | ASSPRDRSDYT | 0.61302304 |
| LISNASDA | ASSPRDRSDYT | 0.66020995 |
| AELRTLGELTKIAKAHGAQ | ASSPRDRSDYT | 0.6385418 |
| APNAVAAAEPSW | ASSPRDRSDYT | 0.63913465 |
| APQVAGAATAVL | ASSPRDRSDYT | 0.626369 |
| ARRDPAAAAAILPTDA | ASSPRDRSDYT | 0.49420887 |
| DAAARDVLAEAGLAEYF | ASSPRDRSDYT | 0.59303427 |
| DDQIVLSAKANLAKVEQ | ASSPRDRSDYT | 0.5159706 |

| | | |
|---|---|---|
| DELLPDTGEDSDEDGH | ASSPRDRSDYT | 0.6425455 |
| ELPEAPANRATVPQD | ASSPRDRSDYT | 0.54189426 |
| FGQSKIERVTGIG | ASSPRDRSDYT | 0.66683114 |
| GAGGTGSTAGGAGGAGGAGGLY | ASSPRDRSDYT | 0.70945907 |
| GHGGTGAAGGNGGTGGAGGSGI | ASSPRDRSDYT | 0.6244588 |
| GNGGNGGAGGEGGPGGAGGAGA | ASSPRDRSDYT | 0.42341027 |
| GSTAGNGGNGGNATNGTIAGQP | ASSPRDRSDYT | 0.6313933 |
| GTGGDGGNGGNGGTGG | ASSPRDRSDYT | 0.61659956 |
| GVRRPAGIPADAPV | ASSPRDRSDYT | 0.6945964 |
| GYRQVTDAHLLTLARR | ASSPRDRSDYT | 0.69528687 |
| IAKMGQDGHDRG | ASSPRDRSDYT | 0.7134179 |
| KAEPSIEIQYT | ASSPRDRSDYT | 0.69751775 |
| KPVRIRMAADEDTVAVT | ASSPRDRSDYT | 0.62313664 |
| MPPRIAGMRLLVIKPEP | ASSPRDRSDYT | 0.7069566 |
| MKFYAVALFRSRGVPPTR | ASSPRDRSDYT | 0.63468987 |
| NAGGFGILGAVAHSP | ASSPRDRSDYT | 0.66678643 |
| NEGREITLAMAEM | ASSPRDRSDYT | 0.68336105 |
| NELVEWSHPLGHRWRWE | ASSPRDRSDYT | 0.69023716 |
| NLGAGGTGGNGGNPG | ASSPRDRSDYT | 0.51831573 |
| NTEWPKWETVMAADPVD | ASSPRDRSDYT | 0.6636392 |
| PASAAPAETQP | ASSPRDRSDYT | 0.48977965 |
| QQSSDGKNGVRAAEITGE | ASSPRDRSDYT | 0.702703 |
| RAPDAVTTSWWKE | ASSPRDRSDYT | 0.49466497 |
| SGFGNVGAHTSGWFNQ | ASSPRDRSDYT | 0.4629736 |
| SGVAGAVGQAASVGGLKV | ASSPRDRSDYT | 0.648842 |
| SNGGAGTGSAGGNGGT | ASSPRDRSDYT | 0.71545684 |
| TGGAGGEGGAAG | ASSPRDRSDYT | 0.6977613 |
| TGGDGGAAGAGGAPGHG | ASSPRDRSDYT | 0.6159129 |
| TMPPLLAAIANAGRHGV | ASSPRDRSDYT | 0.5831257 |
| TSSTAGNSGGA | ASSPRDRSDYT | 0.68746525 |
| VPATVPVGILC | ASSPRDRSDYT | 0.57603747 |
| VQRPDSGGVVLTGRIS | ASSPRDRSDYT | 0.7029312 |
| WTPNPVPATKGVPAGLVQAT | ASSPRDRSDYT | 0.65255225 |
| YALKRALAKFSDRFDVVII | ASSPRDRSDYT | 0.66057 |
| YLIGSVIV | ASSPRDRSDYT | 0.5057836 |
| YGQLVAVID | ASSPRDRSDYT | 0.6996517 |
| VTVGGMTLPA | ASSPRDRSDYT | 0.6412066 |
| VLALPAV | ASSPRDRSDYT | 0.6736573 |
| VIPPTLNYET | ASSPRDRSDYT | 0.30731335 |
| VGMGGMHPSSG | ASSPRDRSDYT | 0.700105 |
| VGLAGGAATA | ASSPRDRSDYT | 0.5559901 |
| VGGAGGAGGSGP | ASSPRDRSDYT | 0.53081906 |
| VFAAFMV | ASSPRDRSDYT | 0.3334695 |
| TLYDKLR | ASSPRDRSDYT | 0.6416962 |
| TLAVFTVL | ASSPRDRSDYT | 0.69039106 |
| TGSVQPARV | ASSPRDRSDYT | 0.712716 |
| TGGGAGSGGT | ASSPRDRSDYT | 0.6689802 |
| SHLETSL | ASSPRDRSDYT | 0.6578344 |
| SFTTGMT | ASSPRDRSDYT | 0.7073647 |
| QVEVAKRMGVRQPTVS | ASSPRDRSDYT | 0.63603497 |
| QGSGGPALAM | ASSPRDRSDYT | 0.35074174 |

| | | |
|---|---|---|
| MLAPSNP | ASSPRDRSDYT | 0.6356847 |
| LVNNAAI | ASSPRDRSDYT | 0.6751645 |
| LTGEINTLA | ASSPRDRSDYT | 0.7104406 |
| LSIFGPV | ASSPRDRSDYT | 0.683545 |
| LLALLANL | ASSPRDRSDYT | 0.52276516 |
| LISNASDALDKLR | ASSPRDRSDYT | 0.6487031 |
| LINEAVP | ASSPRDRSDYT | 0.67725503 |
| LGYAFSGL | ASSPRDRSDYT | 0.4457457 |
| LDFVAVAL | ASSPRDRSDYT | 0.5906192 |
| LAQPGAE | ASSPRDRSDYT | 0.56102216 |
| ISNLGAVTGF | ASSPRDRSDYT | 0.47787958 |
| HQEDPGRI | ASSPRDRSDYT | 0.6536716 |
| GTPGEPTP | ASSPRDRSDYT | 0.6380969 |
| GFFNSGS | ASSPRDRSDYT | 0.52282584 |
| GAGGGGLTAGAN | ASSPRDRSDYT | 0.6728232 |
| FVDALVNM | ASSPRDRSDYT | 0.5437619 |
| FLTVAPEL | ASSPRDRSDYT | 0.56400096 |
| FETLTAL | ASSPRDRSDYT | 0.5659809 |
| DVINPIEL | ASSPRDRSDYT | 0.67399246 |
| APVSGAPV | ASSPRDRSDYT | 0.70298123 |
| ALVGGLAW | ASSPRDRSDYT | 0.64030457 |
| ALAAAAATAI | ASSPRDRSDYT | 0.64953005 |
| AFGLGLLL | ASSPRDRSDYT | 0.7136165 |
| AFFAAASL | ASSPRDRSDYT | 0.38102543 |
| AAAALPAFSPP | ASSPRDRSDYT | 0.54509866 |
| VVLAPAL | ASSPRDRSDYT | 0.67757493 |
| TIGHVDHGKTTL | ASSPRDRSDYT | 0.59021664 |
| LISNASDALDK | ASSPRDRSDYT | 0.7019921 |
| LISNASDAL | ASSPRDRSDYT | 0.6637411 |
| ISNASDALDK | ASSPRDRSDYT | 0.673633 |
| NQAETLVYQTE | ASSPRDRSDYT | 0.6474857 |
| TGGTGGTAGAG | ASSPRDRSDYT | 0.71461266 |
| TGSTAGGAGGA | ASSPRDRSDYT | 0.6874782 |
| DPLAVPVAAGPV | ASSPRDRSDYT | 0.62868416 |
| GGTGGTPGNGGN | ASSPRDRSDYT | 0.70346457 |
| PAGLAGVDPATL | ASSPRDRSDYT | 0.4122223 |
| GNNLVVEAPHAGASV | ASSPRDRSDYT | 0.678241 |
| KVTRIMSASTLEQPA | ASSPRDRSDYT | 0.696857 |
| LGFALPIALPRRTVV | ASSPRDRSDYT | 0.56026053 |
| TRLGNLTARKLVVLV | ASSPRDRSDYT | 0.6600414 |
| WSQQRRLIYFPSAGP | ASSPRDRSDYT | 0.71740836 |
| GPNGNPDPMAAAVDIR | ASSPRDRSDYT | 0.71043795 |
| TQRCVSVADLRAAASR | ASSPRDRSDYT | 0.5522444 |
| QFVLDVAEHPVEVLTETV | ASSPRDRSDYT | 0.60736525 |
| GGAGGTGGAGGNGGTQAGDG | ASSPRDRSDYT | 0.45691317 |
| PLLATLRAIAVDVDAKPAQV | ASSPRDRSDYT | 0.55237573 |
| APKFRTGGYMIDFWGVGYQVAK | ASSPRDRSDYT | 0.38827062 |
| GAGGAGGAGGAAGGPGGTGGTG | ASSPRDRSDYT | 0.6812053 |
| IGALGLLATDVAPAGSVAFGVA | ASSPRDRSDYT | 0.6278717 |
| AGGAGGIGGNGGAGGAGGVLMG | ASSPRDRSDYT | 0.6974349 |
| ADELIGLL | ASSPRDRSDYT | 0.6764844 |

| | | |
|---|---|---|
| ADRALKA | ASSPRDRSDYT | 0.7109605 |
| AGAVGQVGGAGGA | ASSPRDRSDYT | 0.6326498 |
| AGGGGGLGGAP | ASSPRDRSDYT | 0.6158782 |
| AGGLLYGNGGAG | ASSPRDRSDYT | 0.43267366 |
| AGLESIGL | ASSPRDRSDYT | 0.44441873 |
| AGLKTGII | ASSPRDRSDYT | 0.5777023 |
| ASLAALLL | ASSPRDRSDYT | 0.59529483 |
| ASVINISSI | ASSPRDRSDYT | 0.32690936 |
| ATPPGPA | ASSPRDRSDYT | 0.71727204 |
| ATVASALV | ASSPRDRSDYT | 0.606887 |
| ATVASLIL | ASSPRDRSDYT | 0.6921501 |
| AVNGPVLIRH | ASSPRDRSDYT | 0.5631279 |
| DGAAGAVGI | ASSPRDRSDYT | 0.48363844 |
| DPLLHGI | ASSPRDRSDYT | 0.6154103 |
| ELGAALGAL | ASSPRDRSDYT | 0.67941093 |
| ELGRTQAV | ASSPRDRSDYT | 0.5101448 |
| FVAGGYLS | ASSPRDRSDYT | 0.45674264 |
| GASGGAGGAGGW | ASSPRDRSDYT | 0.46485677 |
| GGAGAGGDGGAGGI | ASSPRDRSDYT | 0.64947516 |
| GTGHNGSQPGMGGNG | ASSPRDRSDYT | 0.71996194 |
| GYAAASASAS | ASSPRDRSDYT | 0.7103182 |
| HQVDVAAVQQG | ASSPRDRSDYT | 0.67973626 |
| IGAAIAHT | ASSPRDRSDYT | 0.72175545 |
| IGMIEVAQL | ASSPRDRSDYT | 0.27568373 |
| IRLAPEHTSAMD | ASSPRDRSDYT | 0.5720872 |
| IYNVSVL | ASSPRDRSDYT | 0.6891166 |
| KQGFLGM | ASSPRDRSDYT | 0.6819329 |
| LESVITQL | ASSPRDRSDYT | 0.6506556 |
| LGLGALVVV | ASSPRDRSDYT | 0.71119654 |
| LIGNGGNGGSGGI | ASSPRDRSDYT | 0.5424243 |
| LNAINAPTL | ASSPRDRSDYT | 0.6223043 |
| LQLLNTGV | ASSPRDRSDYT | 0.6401709 |
| LQTSIEF | ASSPRDRSDYT | 0.7100576 |
| MSFAVSEL | ASSPRDRSDYT | 0.6794058 |
| NTGALGL | ASSPRDRSDYT | 0.46517727 |
| PPGAGGAGGAGTT | ASSPRDRSDYT | 0.6719359 |
| QLVGLGI | ASSPRDRSDYT | 0.72472346 |
| RFLEQVHQL | ASSPRDRSDYT | 0.6295016 |
| RGVGYVLR | ASSPRDRSDYT | 0.7048274 |
| SATGVFAGVI | ASSPRDRSDYT | 0.6528479 |
| SLPAITLP | ASSPRDRSDYT | 0.58616567 |
| STALAFTV | ASSPRDRSDYT | 0.60217905 |
| SVLTVGLGK | ASSPRDRSDYT | 0.569553 |
| TALAGAGSGI | ASSPRDRSDYT | 0.69001675 |
| TGLPFTGI | ASSPRDRSDYT | 0.4735217 |
| VAEHVTV | ASSPRDRSDYT | 0.6567955 |
| VDIVLNSL | ASSPRDRSDYT | 0.62891185 |
| VLTGGGAVL | ASSPRDRSDYT | 0.72366804 |
| VNVGLAYGI | ASSPRDRSDYT | 0.6882931 |
| VSSAAVGAV | ASSPRDRSDYT | 0.6819838 |
| VTSVYEAGV | ASSPRDRSDYT | 0.5838884 |

| | | |
|---|---|---|
| WVALDRGA | ASSPRDRSDYT | 0.68990993 |
| YLVGSVVI | ASSPRDRSDYT | 0.69264305 |
| GPPGTGKTLL | ASSPRDRSDYT | 0.5173699 |
| GVGMTKF | ASSPRDRSDYT | 0.6975915 |
| IGIGGSDL | ASSPRDRSDYT | 0.51598847 |
| IGTLLAEL | ASSPRDRSDYT | 0.67453945 |
| IYTSGTT | ASSPRDRSDYT | 0.61870706 |
| LSFPAGV | ASSPRDRSDYT | 0.70955527 |
| LYTSGTT | ASSPRDRSDYT | 0.6668666 |
| VGAFPFGF | ASSPRDRSDYT | 0.6825866 |
| VGAGTVEF | ASSPRDRSDYT | 0.5742339 |
| VGKSTLLL | ASSPRDRSDYT | 0.6902249 |
| AALAHAGGGIV | ASSPRDRSDYT | 0.71049595 |
| AALVAVSLL | ASSPRDRSDYT | 0.66138864 |
| AAVSRAVAAL | ASSPRDRSDYT | 0.7220601 |
| AGGSSGIGGF | ASSPRDRSDYT | 0.63898337 |
| AGLTVANANG | ASSPRDRSDYT | 0.34888387 |
| AGLVGAVL | ASSPRDRSDYT | 0.6526464 |
| ALSQAFTV | ASSPRDRSDYT | 0.70950186 |
| ALTFSVSL | ASSPRDRSDYT | 0.7138411 |
| ALTGAALVAL | ASSPRDRSDYT | 0.7014744 |
| ASFSIGAGVL | ASSPRDRSDYT | 0.6726483 |
| DGSVYKI | ASSPRDRSDYT | 0.64416397 |
| EQLAEIV | ASSPRDRSDYT | 0.7159837 |
| FAVSELTVM | ASSPRDRSDYT | 0.7223009 |
| GAAPLAAGL | ASSPRDRSDYT | 0.66252553 |
| GADGGDGGSGGAGG | ASSPRDRSDYT | 0.51982516 |
| GFGLDTGGAGG | ASSPRDRSDYT | 0.470426 |
| GLASGGLL | ASSPRDRSDYT | 0.6978146 |
| GRIAGAAVLL | ASSPRDRSDYT | 0.7067195 |
| GSAGSLNTSF | ASSPRDRSDYT | 0.5097412 |
| GTGQIGI | ASSPRDRSDYT | 0.63646066 |
| GTLIALIL | ASSPRDRSDYT | 0.648908 |
| IAATMGFVVG | ASSPRDRSDYT | 0.34863412 |
| IFAQALL | ASSPRDRSDYT | 0.6845838 |
| IGAAVALGF | ASSPRDRSDYT | 0.7049135 |
| IGLTPLAL | ASSPRDRSDYT | 0.6948844 |
| ILGSNTSTL | ASSPRDRSDYT | 0.4368919 |
| IVFSLTVL | ASSPRDRSDYT | 0.70149666 |
| LASKATGF | ASSPRDRSDYT | 0.71393704 |
| LASVALIL | ASSPRDRSDYT | 0.71967554 |
| LFAVGLTL | ASSPRDRSDYT | 0.39322138 |
| LFNGGIGTYI | ASSPRDRSDYT | 0.30583295 |
| LTLGAGII | ASSPRDRSDYT | 0.66064054 |
| LVAAALGDI | ASSPRDRSDYT | 0.6899013 |
| NTAFVQLGI | ASSPRDRSDYT | 0.5851916 |
| QLFSGAEL | ASSPRDRSDYT | 0.52746224 |
| SAATFITL | ASSPRDRSDYT | 0.6657565 |
| SNYGLANLGV | ASSPRDRSDYT | 0.7093203 |
| SYGGAYAVM | ASSPRDRSDYT | 0.5361709 |
| TTGVGGLTLG | ASSPRDRSDYT | 0.4515114 |

| | | |
|---|---|---|
| TTQTIYI | ASSPRDRSDYT | 0.70345914 |
| VAAGAGLGAVI | ASSPRDRSDYT | 0.7047611 |
| VAAVVTLL | ASSPRDRSDYT | 0.5827324 |
| VGFLGGLI | ASSPRDRSDYT | 0.64894736 |
| VGVIPILL | ASSPRDRSDYT | 0.67064923 |
| VIFASSLI | ASSPRDRSDYT | 0.6539182 |
| VKGGDGGAAAG | ASSPRDRSDYT | 0.6755636 |
| VTKTFTNI | ASSPRDRSDYT | 0.49518478 |
| VVAAAILL | ASSPRDRSDYT | 0.71280134 |
| YAALDVEL | ASSPRDRSDYT | 0.55069184 |
| LASVALLL | ASSPRDRSDYT | 0.69169265 |
| LDAVHRV | ASSPRDRSDYT | 0.66258585 |
| QGLSSSIF | ASSPRDRSDYT | 0.68991846 |
| RGTFANIRL | ASSPRDRSDYT | 0.64261127 |
| RQLWWGHRI | ASSPRDRSDYT | 0.48603666 |
| YGPPGTGKTLL | ASSPRDRSDYT | 0.5249002 |
| AALVGAPANVA | ASSPRDRSDYT | 0.4769109 |
| AATFRRTE | ASSPRDRSDYT | 0.7044891 |
| AGALGGGI | ASSPRDRSDYT | 0.60360444 |
| AGGAGGAGGIGAG | ASSPRDRSDYT | 0.707079 |
| AGSGALGL | ASSPRDRSDYT | 0.6718104 |
| AQIADQI | ASSPRDRSDYT | 0.5788078 |
| ASAVAVTYL | ASSPRDRSDYT | 0.6966373 |
| ASTVLGGTGG | ASSPRDRSDYT | 0.39512277 |
| AVGGEPAK | ASSPRDRSDYT | 0.6770663 |
| CEQGTFGGTC | ASSPRDRSDYT | 0.6402277 |
| DEGKAPAG | ASSPRDRSDYT | 0.6798351 |
| DTVVFPAL | ASSPRDRSDYT | 0.58718693 |
| FEGITMAG | ASSPRDRSDYT | 0.69844115 |
| FPLVVQD | ASSPRDRSDYT | 0.6879841 |
| GAFTTPQ | ASSPRDRSDYT | 0.41458613 |
| GFGGFGTAGGVGGA | ASSPRDRSDYT | 0.6819474 |
| GFLAQFL | ASSPRDRSDYT | 0.70697296 |
| GGAGGAGGNAGVA | ASSPRDRSDYT | 0.684612 |
| GGVGTAGAGGAGGAGG | ASSPRDRSDYT | 0.44981417 |
| GNGGGGDAGNAGSGGN | ASSPRDRSDYT | 0.65357554 |
| GTGGAGGAGNDAGS | ASSPRDRSDYT | 0.60161173 |
| GTTNTGWFNS | ASSPRDRSDYT | 0.68540186 |
| IFMGPLFTL | ASSPRDRSDYT | 0.68216723 |
| ILAHAIEAI | ASSPRDRSDYT | 0.5271544 |
| IRSAIREAL | ASSPRDRSDYT | 0.68591917 |
| ITFAGGG | ASSPRDRSDYT | 0.3924256 |
| ITGMSGHQEV | ASSPRDRSDYT | 0.57025045 |
| IVAGLACTL | ASSPRDRSDYT | 0.6299131 |
| IVGGGLAGA | ASSPRDRSDYT | 0.6599488 |
| IVTGVLNI | ASSPRDRSDYT | 0.67945206 |
| LAQQGCI | ASSPRDRSDYT | 0.5064111 |
| LCLHVEN | ASSPRDRSDYT | 0.6683887 |
| LDAISDGS | ASSPRDRSDYT | 0.63001215 |
| LGGFAAML | ASSPRDRSDYT | 0.5379109 |
| LGGKNPMI | ASSPRDRSDYT | 0.6836535 |

| | | |
|---|---|---|
| LGPKGVAAIL | ASSPRDRSDYT | 0.5813142 |
| LMVGVSI | ASSPRDRSDYT | 0.38125834 |
| LVGAAFVL | ASSPRDRSDYT | 0.7055119 |
| LVLAVAPL | ASSPRDRSDYT | 0.5092566 |
| MAAIAVIDAA | ASSPRDRSDYT | 0.64684916 |
| NNGINAELQ | ASSPRDRSDYT | 0.7063881 |
| RGEPLVLVHG | ASSPRDRSDYT | 0.3611472 |
| RGGAGGAAGLI | ASSPRDRSDYT | 0.69274783 |
| RSYRMHPAV | ASSPRDRSDYT | 0.6595485 |
| SAAHLLAGQLA | ASSPRDRSDYT | 0.66582984 |
| SAAVGSY | ASSPRDRSDYT | 0.6173516 |
| SPHAVLHRI | ASSPRDRSDYT | 0.5660861 |
| SQIAKVL | ASSPRDRSDYT | 0.7120221 |
| TALGHLVP | ASSPRDRSDYT | 0.6611296 |
| TAPTTSGAA | ASSPRDRSDYT | 0.4931605 |
| TDGDDGNGGSPGT | ASSPRDRSDYT | 0.67488873 |
| TDGLGEN | ASSPRDRSDYT | 0.6975032 |
| TGGPPLPGQA | ASSPRDRSDYT | 0.702316 |
| TIPAITI | ASSPRDRSDYT | 0.58014417 |
| TIPGFTI | ASSPRDRSDYT | 0.38833642 |
| TSIASGALA | ASSPRDRSDYT | 0.64099056 |
| TVTLPGAAI | ASSPRDRSDYT | 0.62442607 |
| VASGLSAALP | ASSPRDRSDYT | 0.57323295 |
| VGAGVIGI | ASSPRDRSDYT | 0.37369788 |
| VGPKRFGV | ASSPRDRSDYT | 0.6826672 |
| VNVAFPD | ASSPRDRSDYT | 0.7237309 |
| VSMTNSGVS | ASSPRDRSDYT | 0.5032108 |
| VTDVLGDLSV | ASSPRDRSDYT | 0.72255945 |
| VVVVGAGFAGL | ASSPRDRSDYT | 0.49453366 |
| VYTVGTVV | ASSPRDRSDYT | 0.67121965 |
| WTTFSGP | ASSPRDRSDYT | 0.68670756 |
| YDELVEAF | ASSPRDRSDYT | 0.51212525 |
| YGTGGAGGNGG | ASSPRDRSDYT | 0.5899426 |
| YTNVMAV | ASSPRDRSDYT | 0.5908153 |
| FLEQVHQL | ASSPRDRSDYT | 0.5577142 |
| GLSLLAVAV | ASSPRDRSDYT | 0.7021564 |
| LEQVHQL | ASSPRDRSDYT | 0.48088488 |
| LNAAGAI | ASSPRDRSDYT | 0.6834179 |
| PAVHASL | ASSPRDRSDYT | 0.6113551 |
| TELEDDA | ASSPRDRSDYT | 0.5691733 |
| TPPGPPN | ASSPRDRSDYT | 0.7006831 |
| TPPPPPPP | ASSPRDRSDYT | 0.67231137 |
| TQSPALI | ASSDGAGGAYEQY | 0.65364707 |
| EFAAAFPQ | ASSDGAGGAYEQY | 0.72130096 |
| LLSGNLDVL | ASSDGAGGAYEQY | 0.59443873 |
| GPANLGNYNI | ASSDGAGGAYEQY | 0.7232814 |
| TLGPTLP | ASSDGAGGAYEQY | 0.71085525 |
| VYAYSSCCK | ASSDGAGGAYEQY | 0.6770389 |
| APAPAPAPNI | ASSDGAGGAYEQY | 0.63964194 |
| PGGPVPIP | ASSDGAGGAYEQY | 0.6959356 |
| FLTVALEP | ASSDGAGGAYEQY | 0.6076149 |

| | | |
|---|---|---|
| SQYFGANL | ASSDGAGGAYEQY | 0.6355373 |
| GGVKPAA | ASSDGAGGAYEQY | 0.71605265 |
| GGTGGAGGAGGW | ASSDGAGGAYEQY | 0.49205524 |
| GVGGVAGLGGAGG | ASSDGAGGAYEQY | 0.6735749 |
| VLLVLATTI | ASSDGAGGAYEQY | 0.6444932 |
| IGIFAAML | ASSDGAGGAYEQY | 0.62798 |
| AVGDLID | ASSDGAGGAYEQY | 0.55379236 |
| NISPLPL | ASSDGAGGAYEQY | 0.44851804 |
| VTAVFVAI | ASSDGAGGAYEQY | 0.6158432 |
| SGDDNDAAL | ASSDGAGGAYEQY | 0.499513 |
| AVAALAHS | ASSDGAGGAYEQY | 0.68334424 |
| GGVTVMATP | ASSDGAGGAYEQY | 0.6710776 |
| AGLAANVRD | ASSDGAGGAYEQY | 0.5490235 |
| TNRPDLIDPALLRPG | ASSDGAGGAYEQY | 0.6694387 |
| LANLGVTAL | ASSDGAGGAYEQY | 0.66668165 |
| TGERIEL | ASSDGAGGAYEQY | 0.46739638 |
| NGEELAA | ASSDGAGGAYEQY | 0.60651255 |
| VLSLVWG | ASSDGAGGAYEQY | 0.60872936 |
| IGYALWL | ASSDGAGGAYEQY | 0.60990715 |
| LSLLLAPL | ASSDGAGGAYEQY | 0.44117114 |
| TLDEALK | ASSDGAGGAYEQY | 0.6629109 |
| TDGAGTAEGG | ASSDGAGGAYEQY | 0.5944133 |
| NAPGGAGGNGG | ASSDGAGGAYEQY | 0.3632111 |
| VSVFMGMGGNF | ASSDGAGGAYEQY | 0.6359693 |
| LASYLAAL | ASSDGAGGAYEQY | 0.6108804 |
| LVGGTPAL | ASSDGAGGAYEQY | 0.5770156 |
| VRNADVL | ASSDGAGGAYEQY | 0.66952175 |
| TCSSPFLVSAM | ASSDGAGGAYEQY | 0.6389522 |
| VHITDAE | ASSDGAGGAYEQY | 0.52828944 |
| EYAAPSF | ASSDGAGGAYEQY | 0.6602992 |
| VVDAFHGGL | ASSDGAGGAYEQY | 0.672556 |
| ASVVDPQ | ASSDGAGGAYEQY | 0.4668181 |
| SYNEFGGLS | ASSDGAGGAYEQY | 0.6461264 |
| DPDGSVY | ASSDGAGGAYEQY | 0.6756009 |
| MVSTIHATE | ASSDGAGGAYEQY | 0.63258284 |
| LAAGVADPV | ASSDGAGGAYEQY | 0.6802248 |
| TKEQIAA | ASSDGAGGAYEQY | 0.68319887 |
| SLASPAAL | ASSDGAGGAYEQY | 0.5992127 |
| IGLVFYGV | ASSDGAGGAYEQY | 0.6755956 |
| FLLDASNL | ASSDGAGGAYEQY | 0.68095756 |
| YQVGNFP | ASSDGAGGAYEQY | 0.5799465 |
| ESLLSQST | ASSDGAGGAYEQY | 0.6194322 |
| LFTVGFAAVG | ASSDGAGGAYEQY | 0.47831264 |
| ATPPTPT | ASSDGAGGAYEQY | 0.63390005 |
| TDLATLG | ASSDGAGGAYEQY | 0.5453854 |
| SAGGTGR | ASSDGAGGAYEQY | 0.6539427 |
| GFSNSGS | ASSDGAGGAYEQY | 0.67905056 |
| GGNGGSGGTSV | ASSDGAGGAYEQY | 0.61176115 |
| GTSVATGGAGN | ASSDGAGGAYEQY | 0.4801156 |
| NGGNGGSGGTS | ASSDGAGGAYEQY | 0.51472926 |
| AVVKFPH | ASSDGAGGAYEQY | 0.6516546 |

| | | |
|---|---|---|
| TPPPPPP | ASSDGAGGAYEQY | 0.5127753 |
| AAPAAAAAPA | ASSDGAGGAYEQY | 0.51702553 |
| IALNEDL | ASSDGAGGAYEQY | 0.60923976 |
| IVNVSSM | ASSDGAGGAYEQY | 0.6669593 |
| TGIPPAPR | ASSDGAGGAYEQY | 0.66561544 |
| FDLGGGTF | ASSDGAGGAYEQY | 0.5222528 |
| FDLGGGTFDVS | ASSDGAGGAYEQY | 0.68069345 |
| NEPTAAA | ASSDGAGGAYEQY | 0.6778494 |
| NEPTAAALA | ASSDGAGGAYEQY | 0.6789204 |
| GSIINISSI | ASSDGAGGAYEQY | 0.62192345 |
| VAPGNAGTA | ASSDGAGGAYEQY | 0.4569962 |
| TGGGASLE | ASSDGAGGAYEQY | 0.6767496 |
| VAVYNHYKRI | ASSDGAGGAYEQY | 0.58892477 |
| TDPAPATT | ASSDGAGGAYEQY | 0.63442147 |
| TNRPDILDPALLRPG | ASSDGAGGAYEQY | 0.66440463 |
| LTTTTTS | ASSDGAGGAYEQY | 0.6250522 |
| PTGSGKTL | ASSDGAGGAYEQY | 0.5420367 |
| TGSGKTL | ASSDGAGGAYEQY | 0.6358213 |
| TIGHVDHGK | ASSDGAGGAYEQY | 0.6804706 |
| TIGHVDHGKTT | ASSDGAGGAYEQY | 0.53437686 |
| GHVDHGKTT | ASSDGAGGAYEQY | 0.48261857 |
| AQFEHTLL | ASSDGAGGAYEQY | 0.6140998 |
| IVNQDEL | ASSDGAGGAYEQY | 0.61422247 |
| TNGSQFF | ASSDGAGGAYEQY | 0.6725029 |
| TGKGIVE | ASSDGAGGAYEQY | 0.655285 |
| LISNASDA | ASSDGAGGAYEQY | 0.60669667 |
| AELRTLGELTKIAKAHGAQ | ASSDGAGGAYEQY | 0.6518975 |
| APNAVAAAEPSW | ASSDGAGGAYEQY | 0.6577897 |
| APQVAGAATAVL | ASSDGAGGAYEQY | 0.6358528 |
| ARRDPAAAAAILPTDA | ASSDGAGGAYEQY | 0.65515494 |
| DAAARDVLAEAGLAEYF | ASSDGAGGAYEQY | 0.55673033 |
| DDQIVLSAKANLAKVEQ | ASSDGAGGAYEQY | 0.67605704 |
| DELLPDTGEDSDEDGH | ASSDGAGGAYEQY | 0.6629348 |
| ELPEAPANRATVPQD | ASSDGAGGAYEQY | 0.66804385 |
| FGQSKIERVTGIG | ASSDGAGGAYEQY | 0.6815363 |
| GAGGTGSTAGGAGGAGGAGGLY | ASSDGAGGAYEQY | 0.63815266 |
| GHGGTGAAGGNGGTGGAGGSGI | ASSDGAGGAYEQY | 0.6410529 |
| GNGGNGGAGGEGGPGGAGGAGA | ASSDGAGGAYEQY | 0.42416304 |
| GSTAGNGGNGGNATNGTIAGQP | ASSDGAGGAYEQY | 0.6255015 |
| GTGGDGGNGGNGGTGG | ASSDGAGGAYEQY | 0.63636357 |
| GVRRPAGIPADAPV | ASSDGAGGAYEQY | 0.6449945 |
| GYRQVTDAHLLTLARR | ASSDGAGGAYEQY | 0.6678021 |
| IAKMGQDGHDRG | ASSDGAGGAYEQY | 0.65090454 |
| KAEPSIEIQYT | ASSDGAGGAYEQY | 0.5621839 |
| KPVRIRMAADEDTVAVT | ASSDGAGGAYEQY | 0.67704654 |
| MPPRIAGMRLLVIKPEP | ASSDGAGGAYEQY | 0.67559975 |
| MKFYAVALFRSRGVPPTR | ASSDGAGGAYEQY | 0.48140335 |
| NAGGFGILGAVAHSP | ASSDGAGGAYEQY | 0.5901688 |
| NEGREITLAMAEM | ASSDGAGGAYEQY | 0.68086654 |
| NELVEWSHPLGHRWRWE | ASSDGAGGAYEQY | 0.6598363 |
| NLGAGGTGGNGGNPG | ASSDGAGGAYEQY | 0.5976971 |

| | | |
|---|---|---|
| NTEWPKWETVMAADPVD | ASSDGAGGAYEQY | 0.59120655 |
| PASAAPAETQP | ASSDGAGGAYEQY | 0.68842757 |
| QQSSDGKNGVRAAEITGE | ASSDGAGGAYEQY | 0.6393843 |
| RAPDAVTTSWWKE | ASSDGAGGAYEQY | 0.68054694 |
| SGFGNVGAHTSGWFNQ | ASSDGAGGAYEQY | 0.39147562 |
| SGVAGAVGQAASVGGLKV | ASSDGAGGAYEQY | 0.6125364 |
| SNGGAGTGSAGGNGGT | ASSDGAGGAYEQY | 0.556753 |
| TGGAGGEGGAAG | ASSDGAGGAYEQY | 0.3321166 |
| TGGDGGAAGAGGAPGHG | ASSDGAGGAYEQY | 0.6647625 |
| TMPPLLAAIANAGRHGV | ASSDGAGGAYEQY | 0.57542473 |
| TSSTAGNSGGA | ASSDGAGGAYEQY | 0.65431035 |
| VPATVPVGILC | ASSDGAGGAYEQY | 0.6313969 |
| VQRPDSGGVVLTGRIS | ASSDGAGGAYEQY | 0.6730143 |
| WTPNPVPATKGVPAGLVQAT | ASSDGAGGAYEQY | 0.64247763 |
| YALKRALAKFSDRFDVVII | ASSDGAGGAYEQY | 0.6295867 |
| YLIGSVIV | ASSDGAGGAYEQY | 0.5249614 |
| YGQLVAVID | ASSDGAGGAYEQY | 0.6522261 |
| VTVGGMTLPA | ASSDGAGGAYEQY | 0.67878294 |
| VLALPAV | ASSDGAGGAYEQY | 0.6760094 |
| VIPPTLNYET | ASSDGAGGAYEQY | 0.38457552 |
| VGMGGMHPSSG | ASSDGAGGAYEQY | 0.65368277 |
| VGLAGGAATA | ASSDGAGGAYEQY | 0.59639966 |
| VGGAGGAGGSGP | ASSDGAGGAYEQY | 0.5547112 |
| VFAAFMV | ASSDGAGGAYEQY | 0.4018493 |
| TLYDKLR | ASSDGAGGAYEQY | 0.5982918 |
| TLAVFTVL | ASSDGAGGAYEQY | 0.6123614 |
| TGSVQPARV | ASSDGAGGAYEQY | 0.6601685 |
| TGGGAGSGGT | ASSDGAGGAYEQY | 0.656172 |
| SHLETSL | ASSDGAGGAYEQY | 0.59864664 |
| SFTTGMT | ASSDGAGGAYEQY | 0.65987295 |
| QVEVAKRMGVRQPTVS | ASSDGAGGAYEQY | 0.6118007 |
| QGSGGPALAM | ASSDGAGGAYEQY | 0.2872221 |
| MLAPSNP | ASSDGAGGAYEQY | 0.59297824 |
| LVNNAAI | ASSDGAGGAYEQY | 0.46129185 |
| LTGEINTLA | ASSDGAGGAYEQY | 0.68327105 |
| LSIFGPV | ASSDGAGGAYEQY | 0.5350875 |
| LLALLANL | ASSDGAGGAYEQY | 0.58172446 |
| LISNASDALDKLR | ASSDGAGGAYEQY | 0.62016773 |
| LINEAVP | ASSDGAGGAYEQY | 0.6635012 |
| LGYAFSGL | ASSDGAGGAYEQY | 0.6274252 |
| LDFVAVAL | ASSDGAGGAYEQY | 0.5566834 |
| LAQPGAE | ASSDGAGGAYEQY | 0.5307499 |
| ISNLGAVTGF | ASSDGAGGAYEQY | 0.6562189 |
| HQEDPGRI | ASSDGAGGAYEQY | 0.6323543 |
| GTPGEPTP | ASSDGAGGAYEQY | 0.47034252 |
| GFFNSGS | ASSDGAGGAYEQY | 0.5338041 |
| GAGGGGLTAGAN | ASSDGAGGAYEQY | 0.66387033 |
| FVDALVNM | ASSDGAGGAYEQY | 0.43545097 |
| FLTVAPEL | ASSDGAGGAYEQY | 0.6732451 |
| FETLTAL | ASSDGAGGAYEQY | 0.574495 |
| DVINPIEL | ASSDGAGGAYEQY | 0.60017854 |

| | | |
|---|---|---|
| APVSGAPV | ASSDGAGGAYEQY | 0.6841109 |
| ALVGGLAW | ASSDGAGGAYEQY | 0.65395594 |
| ALAAAAATAI | ASSDGAGGAYEQY | 0.61086345 |
| AFGLGLLL | ASSDGAGGAYEQY | 0.5248115 |
| AFFAAASL | ASSDGAGGAYEQY | 0.5376535 |
| AAAALPAFSPP | ASSDGAGGAYEQY | 0.50881135 |
| VVLAPAL | ASSDGAGGAYEQY | 0.54886144 |
| TIGHVDHGKTTL | ASSDGAGGAYEQY | 0.6773389 |
| LISNASDALDK | ASSDGAGGAYEQY | 0.631877 |
| LISNASDAL | ASSDGAGGAYEQY | 0.5192339 |
| ISNASDALDK | ASSDGAGGAYEQY | 0.6537476 |
| NQAETLVYQTE | ASSDGAGGAYEQY | 0.6822628 |
| TGGTGGTAGAG | ASSDGAGGAYEQY | 0.6847534 |
| TGSTAGGAGGA | ASSDGAGGAYEQY | 0.5370747 |
| DPLAVPVAAGPV | ASSDGAGGAYEQY | 0.5986953 |
| GGTGGTPGNGGN | ASSDGAGGAYEQY | 0.55548 |
| PAGLAGVDPATL | ASSDGAGGAYEQY | 0.6110232 |
| GNNLVVEAPHAGASV | ASSDGAGGAYEQY | 0.5480521 |
| KVTRIMSASTLEQPA | ASSDGAGGAYEQY | 0.48045522 |
| LGFALPIALPRRTVV | ASSDGAGGAYEQY | 0.66983616 |
| TRLGNLTARKLVVLV | ASSDGAGGAYEQY | 0.6928767 |
| WSQQRRLIYFPSAGP | ASSDGAGGAYEQY | 0.60405463 |
| GPNGNPDPMAAAVDIR | ASSDGAGGAYEQY | 0.48552215 |
| TQRCVSVADLRAAASR | ASSDGAGGAYEQY | 0.60682285 |
| QFVLDVAEHPVEVLTETV | ASSDGAGGAYEQY | 0.56521183 |
| GGAGGTGGAGGNGGTQAGDG | ASSDGAGGAYEQY | 0.5458737 |
| PLLATLRAIAVDVDAKPAQV | ASSDGAGGAYEQY | 0.6003303 |
| APKFRTGGYMIDFWGVGYQVAK | ASSDGAGGAYEQY | 0.4443441 |
| GAGGAGGAGGAAGGPGGTGGTG | ASSDGAGGAYEQY | 0.6145651 |
| IGALGLLATDVAPAGSVAFGVA | ASSDGAGGAYEQY | 0.68100184 |
| AGGAGGIGGNGGAGGAGGVLMG | ASSDGAGGAYEQY | 0.61505437 |
| ADELIGLL | ASSDGAGGAYEQY | 0.65209043 |
| ADRALKA | ASSDGAGGAYEQY | 0.6800007 |
| AGAVGQVGGAGGA | ASSDGAGGAYEQY | 0.661272 |
| AGGGGGLGGAP | ASSDGAGGAYEQY | 0.6750876 |
| AGGLLYGNGGAG | ASSDGAGGAYEQY | 0.68186605 |
| AGLESIGL | ASSDGAGGAYEQY | 0.62414235 |
| AGLKTGII | ASSDGAGGAYEQY | 0.63732433 |
| ASLAALLL | ASSDGAGGAYEQY | 0.67545164 |
| ASVINISSI | ASSDGAGGAYEQY | 0.6088729 |
| ATPPGPA | ASSDGAGGAYEQY | 0.62660146 |
| ATVASALV | ASSDGAGGAYEQY | 0.58731914 |
| ATVASLIL | ASSDGAGGAYEQY | 0.6658282 |
| AVNGPVLIRH | ASSDGAGGAYEQY | 0.60850066 |
| DGAAGAVGI | ASSDGAGGAYEQY | 0.68639326 |
| DPLLHGI | ASSDGAGGAYEQY | 0.37759078 |
| ELGAALGAL | ASSDGAGGAYEQY | 0.50069034 |
| ELGRTQAV | ASSDGAGGAYEQY | 0.67254514 |
| FVAGGYLS | ASSDGAGGAYEQY | 0.30713412 |
| GASGGAGGAGGW | ASSDGAGGAYEQY | 0.41989505 |
| GGAGAGGDGGAGGI | ASSDGAGGAYEQY | 0.63768697 |

| | | |
|---|---|---|
| GTGHNGSQPGMGGNG | ASSDGAGGAYEQY | 0.67489713 |
| GYAAASASAS | ASSDGAGGAYEQY | 0.37413013 |
| HQVDVAAVQQG | ASSDGAGGAYEQY | 0.56696355 |
| IGAAIAHT | ASSDGAGGAYEQY | 0.4643005 |
| IGMIEVAQL | ASSDGAGGAYEQY | 0.6687157 |
| IRLAPEHTSAMD | ASSDGAGGAYEQY | 0.60938805 |
| IYNVSVL | ASSDGAGGAYEQY | 0.46268472 |
| KQGFLGM | ASSDGAGGAYEQY | 0.6639028 |
| LESVITQL | ASSDGAGGAYEQY | 0.48299313 |
| LGLGALVVV | ASSDGAGGAYEQY | 0.6580756 |
| LIGNGGNGGSGGI | ASSDGAGGAYEQY | 0.67045 |
| LNAINAPTL | ASSDGAGGAYEQY | 0.6870948 |
| LQLLNTGV | ASSDGAGGAYEQY | 0.4859839 |
| LQTSIEF | ASSDGAGGAYEQY | 0.658877 |
| MSFAVSEL | ASSDGAGGAYEQY | 0.60386544 |
| NTGALGL | ASSDGAGGAYEQY | 0.68811256 |
| PPGAGGAGGAGTT | ASSDGAGGAYEQY | 0.58735377 |
| QLVGLGI | ASSDGAGGAYEQY | 0.45891306 |
| RFLEQVHQL | ASSDGAGGAYEQY | 0.6689462 |
| RGVGYVLR | ASSDGAGGAYEQY | 0.65956163 |
| SATGVFAGVI | ASSDGAGGAYEQY | 0.57324463 |
| SLPAITLP | ASSDGAGGAYEQY | 0.67423844 |
| STALAFTV | ASSDGAGGAYEQY | 0.61837566 |
| SVLTVGLGK | ASSDGAGGAYEQY | 0.6130393 |
| TALAGAGSGI | ASSDGAGGAYEQY | 0.6748136 |
| TGLPFTGI | ASSDGAGGAYEQY | 0.6637356 |
| VAEHVTV | ASSDGAGGAYEQY | 0.53133833 |
| VDIVLNSL | ASSDGAGGAYEQY | 0.67499804 |
| VLTGGGAVL | ASSDGAGGAYEQY | 0.61579776 |
| VNVGLAYGI | ASSDGAGGAYEQY | 0.5144353 |
| VSSAAVGAV | ASSDGAGGAYEQY | 0.6845832 |
| VTSVYEAGV | ASSDGAGGAYEQY | 0.6337462 |
| WVALDRGA | ASSDGAGGAYEQY | 0.66615236 |
| YLVGSVVI | ASSDGAGGAYEQY | 0.66013575 |
| GPPGTGKTLL | ASSDGAGGAYEQY | 0.6869626 |
| GVGMTKF | ASSDGAGGAYEQY | 0.5104605 |
| IGIGGSDL | ASSDGAGGAYEQY | 0.64690787 |
| IGTLLAEL | ASSDGAGGAYEQY | 0.61685777 |
| IYTSGTT | ASSDGAGGAYEQY | 0.6693646 |
| LSFPAGV | ASSDGAGGAYEQY | 0.6407479 |
| LYTSGTT | ASSDGAGGAYEQY | 0.6554524 |
| VGAFPFGF | ASSDGAGGAYEQY | 0.6607882 |
| VGAGTVEF | ASSDGAGGAYEQY | 0.67584914 |
| VGKSTLLL | ASSDGAGGAYEQY | 0.6306392 |
| AALAHAGGGIV | ASSDGAGGAYEQY | 0.64838105 |
| AALVAVSLL | ASSDGAGGAYEQY | 0.67917395 |
| AAVSRAVAAL | ASSDGAGGAYEQY | 0.5337211 |
| AGGSSGIGGF | ASSDGAGGAYEQY | 0.61326784 |
| AGLTVANANG | ASSDGAGGAYEQY | 0.66351235 |
| AGLVGAVL | ASSDGAGGAYEQY | 0.6714474 |
| ALSQAFTV | ASSDGAGGAYEQY | 0.61936355 |

| | | |
|---|---|---|
| ALTFSVSL | ASSDGAGGAYEQY | 0.6801208 |
| ALTGAALVAL | ASSDGAGGAYEQY | 0.6003928 |
| ASFSIGAGVL | ASSDGAGGAYEQY | 0.6622711 |
| DGSVYKI | ASSDGAGGAYEQY | 0.6239389 |
| EQLAEIV | ASSDGAGGAYEQY | 0.6254304 |
| FAVSELTVM | ASSDGAGGAYEQY | 0.5206853 |
| GAAPLAAGL | ASSDGAGGAYEQY | 0.64519787 |
| GADGGDGGSGGAGG | ASSDGAGGAYEQY | 0.66748756 |
| GFGLDTGGAGG | ASSDGAGGAYEQY | 0.6578684 |
| GLASGGLL | ASSDGAGGAYEQY | 0.3896746 |
| GRIAGAAVLL | ASSDGAGGAYEQY | 0.6585149 |
| GSAGSLNTSF | ASSDGAGGAYEQY | 0.66138387 |
| GTGQIGI | ASSDGAGGAYEQY | 0.68818176 |
| GTLIALIL | ASSDGAGGAYEQY | 0.6182035 |
| IAATMGFVVG | ASSDGAGGAYEQY | 0.6421503 |
| IFAQALL | ASSDGAGGAYEQY | 0.5619032 |
| IGAAVALGF | ASSDGAGGAYEQY | 0.6257988 |
| IGLTPLAL | ASSDGAGGAYEQY | 0.457452 |
| ILGSNTSTL | ASSDGAGGAYEQY | 0.58107156 |
| IVFSLTVL | ASSDGAGGAYEQY | 0.6086103 |
| LASKATGF | ASSDGAGGAYEQY | 0.5519684 |
| LASVALIL | ASSDGAGGAYEQY | 0.6683401 |
| LFAVGLTL | ASSDGAGGAYEQY | 0.6738819 |
| LFNGGIGTYI | ASSDGAGGAYEQY | 0.6684936 |
| LTLGAGII | ASSDGAGGAYEQY | 0.67512923 |
| LVAAALGDI | ASSDGAGGAYEQY | 0.6792655 |
| NTAFVQLGI | ASSDGAGGAYEQY | 0.6746795 |
| QLFSGAEL | ASSDGAGGAYEQY | 0.5512625 |
| SAATFITL | ASSDGAGGAYEQY | 0.6393354 |
| SNYGLANLGV | ASSDGAGGAYEQY | 0.44043747 |
| SYGGAYAVM | ASSDGAGGAYEQY | 0.61588776 |
| TTGVGGLTLG | ASSDGAGGAYEQY | 0.5759163 |
| TTQTIYI | ASSDGAGGAYEQY | 0.63862014 |
| VAAGAGLGAVI | ASSDGAGGAYEQY | 0.32253042 |
| VAAVVTLL | ASSDGAGGAYEQY | 0.65778047 |
| VGFLGGLI | ASSDGAGGAYEQY | 0.6805804 |
| VGVIPILL | ASSDGAGGAYEQY | 0.60769176 |
| VIFASSLI | ASSDGAGGAYEQY | 0.6472634 |
| VKGGDGGAAAG | ASSDGAGGAYEQY | 0.6644951 |
| VTKTFTNI | ASSDGAGGAYEQY | 0.5782622 |
| VVAAAILL | ASSDGAGGAYEQY | 0.6553731 |
| YAALDVEL | ASSDGAGGAYEQY | 0.58377254 |
| LASVALLL | ASSDGAGGAYEQY | 0.66645694 |
| LDAVHRV | ASSDGAGGAYEQY | 0.6785839 |
| QGLSSSIF | ASSDGAGGAYEQY | 0.64628446 |
| RGTFANIRL | ASSDGAGGAYEQY | 0.62683356 |
| RQLWWGHRI | ASSDGAGGAYEQY | 0.6275612 |
| YGPPGTGKTLL | ASSDGAGGAYEQY | 0.4707923 |
| AALVGAPANVA | ASSDGAGGAYEQY | 0.6341622 |
| AATFRRTE | ASSDGAGGAYEQY | 0.6115774 |
| AGALGGGI | ASSDGAGGAYEQY | 0.63341326 |

| | | |
|---|---|---|
| AGGAGGAGGIGAG | ASSDGAGGAYEQY | 0.548283 |
| AGSGALGL | ASSDGAGGAYEQY | 0.6326419 |
| AQIADQI | ASSDGAGGAYEQY | 0.6152657 |
| ASAVAVTYL | ASSDGAGGAYEQY | 0.6465221 |
| ASTVLGGTGG | ASSDGAGGAYEQY | 0.52536297 |
| AVGGEPAK | ASSDGAGGAYEQY | 0.66379184 |
| CEQGTFGGTC | ASSDGAGGAYEQY | 0.647413 |
| DEGKAPAG | ASSDGAGGAYEQY | 0.44993708 |
| DTVVFPAL | ASSDGAGGAYEQY | 0.67112726 |
| FEGITMAG | ASSDGAGGAYEQY | 0.61049056 |
| FPLVVQD | ASSDGAGGAYEQY | 0.6242492 |
| GAFTTPQ | ASSDGAGGAYEQY | 0.6731381 |
| GFGGFGTAGGVGGA | ASSDGAGGAYEQY | 0.6117921 |
| GFLAQFL | ASSDGAGGAYEQY | 0.45649794 |
| GGAGGAGGNAGVA | ASSDGAGGAYEQY | 0.6067568 |
| GGVGTAGAGGAGGAGG | ASSDGAGGAYEQY | 0.63258827 |
| GNGGGGDAGNAGSGGN | ASSDGAGGAYEQY | 0.5952237 |
| GTGGAGGAGNDAGS | ASSDGAGGAYEQY | 0.43084776 |
| GTTNTGWFNS | ASSDGAGGAYEQY | 0.6609672 |
| IFMGPLFTL | ASSDGAGGAYEQY | 0.5115279 |
| ILAHAIEAI | ASSDGAGGAYEQY | 0.57400686 |
| IRSAIREAL | ASSDGAGGAYEQY | 0.57184744 |
| ITFAGGG | ASSDGAGGAYEQY | 0.6662089 |
| ITGMSGHQEV | ASSDGAGGAYEQY | 0.6464584 |
| IVAGLACTL | ASSDGAGGAYEQY | 0.58451456 |
| IVGGGLAGA | ASSDGAGGAYEQY | 0.6563225 |
| IVTGVLNI | ASSDGAGGAYEQY | 0.63032526 |
| LAQQGCI | ASSDGAGGAYEQY | 0.6739528 |
| LCLHVEN | ASSDGAGGAYEQY | 0.61994237 |
| LDAISDGS | ASSDGAGGAYEQY | 0.62294596 |
| LGGFAAML | ASSDGAGGAYEQY | 0.6439347 |
| LGGKNPMI | ASSDGAGGAYEQY | 0.5927471 |
| LGPKGVAAIL | ASSDGAGGAYEQY | 0.5617736 |
| LMVGVSI | ASSDGAGGAYEQY | 0.5214734 |
| LVGAAFVL | ASSDGAGGAYEQY | 0.6063416 |
| LVLAVAPL | ASSDGAGGAYEQY | 0.50843483 |
| MAAIAVIDAA | ASSDGAGGAYEQY | 0.63066036 |
| NNGINAELQ | ASSDGAGGAYEQY | 0.6820924 |
| RGEPLVLVHG | ASSDGAGGAYEQY | 0.6396896 |
| RGGAGGAAGLI | ASSDGAGGAYEQY | 0.44919506 |
| RSYRMHPAV | ASSDGAGGAYEQY | 0.59259593 |
| SAAHLLAGQLA | ASSDGAGGAYEQY | 0.6396353 |
| SAAVGSY | ASSDGAGGAYEQY | 0.6641102 |
| SPHAVLHRI | ASSDGAGGAYEQY | 0.67420554 |
| SQIAKVL | ASSDGAGGAYEQY | 0.6912025 |
| TALGHLVP | ASSDGAGGAYEQY | 0.6606275 |
| TAPTTSGAA | ASSDGAGGAYEQY | 0.564377 |
| TDGDDGNGGSPGT | ASSDGAGGAYEQY | 0.66107523 |
| TDGLGEN | ASSDGAGGAYEQY | 0.6125157 |
| TGGPPLPGQA | ASSDGAGGAYEQY | 0.6566231 |
| TIPAITI | ASSDGAGGAYEQY | 0.64275455 |

| | | |
|---|---|---|
| TIPGFTI | ASSDGAGGAYEQY | 0.6564585 |
| TSIASGALA | ASSDGAGGAYEQY | 0.5786874 |
| TVTLPGAAI | ASSDGAGGAYEQY | 0.5885308 |
| VASGLSAALP | ASSDGAGGAYEQY | 0.43790242 |
| VGAGVIGI | ASSDGAGGAYEQY | 0.6753083 |
| VGPKRFGV | ASSDGAGGAYEQY | 0.586366 |
| VNVAFPD | ASSDGAGGAYEQY | 0.5786841 |
| VSMTNSGVS | ASSDGAGGAYEQY | 0.5998885 |
| VTDVLGDLSV | ASSDGAGGAYEQY | 0.68181336 |
| VVVVGAGFAGL | ASSDGAGGAYEQY | 0.6771996 |
| VYTVGTVV | ASSDGAGGAYEQY | 0.57206744 |
| WTTFSGP | ASSDGAGGAYEQY | 0.5460124 |
| YDELVEAF | ASSDGAGGAYEQY | 0.6566881 |
| YGTGGAGGNGG | ASSDGAGGAYEQY | 0.5071627 |
| YTNVMAV | ASSDGAGGAYEQY | 0.65851295 |
| FLEQVHQL | ASSDGAGGAYEQY | 0.63511986 |
| GLSLLAVAV | ASSDGAGGAYEQY | 0.6207383 |
| LEQVHQL | ASSDGAGGAYEQY | 0.5456453 |
| LNAAGAI | ASSDGAGGAYEQY | 0.65410876 |
| PAVHASL | ASSDGAGGAYEQY | 0.5793874 |
| TELEDDA | ASSDGAGGAYEQY | 0.6600894 |
| TPPGPPN | ASSDGAGGAYEQY | 0.4774825 |
| TPPPPPPP | ASSDGAGGAYEQY | 0.6678192 |
| TQSPALI | ASSHKTTGGSAETLY | 0.52132994 |
| EFAAAFPQ | ASSHKTTGGSAETLY | 0.5474878 |
| LLSGNLDVL | ASSHKTTGGSAETLY | 0.37703693 |
| GPANLGNYNI | ASSHKTTGGSAETLY | 0.61953384 |
| TLGPTLP | ASSHKTTGGSAETLY | 0.662251 |
| VYAYSSCCK | ASSHKTTGGSAETLY | 0.67109215 |
| APAPAPAPNI | ASSHKTTGGSAETLY | 0.63364184 |
| PGGPVPIP | ASSHKTTGGSAETLY | 0.6633115 |
| FLTVALEP | ASSHKTTGGSAETLY | 0.67637587 |
| SQYFGANL | ASSHKTTGGSAETLY | 0.591509 |
| GGVKPAA | ASSHKTTGGSAETLY | 0.38420832 |
| GGTGGAGGAGGW | ASSHKTTGGSAETLY | 0.6382742 |
| GVGGVAGLGGAGG | ASSHKTTGGSAETLY | 0.64497316 |
| VLLVLATTI | ASSHKTTGGSAETLY | 0.6763104 |
| IGIFAAML | ASSHKTTGGSAETLY | 0.651847 |
| AVGDLID | ASSHKTTGGSAETLY | 0.48261014 |
| NISPLPL | ASSHKTTGGSAETLY | 0.6472119 |
| VTAVFVAI | ASSHKTTGGSAETLY | 0.6651405 |
| SGDDNDAAL | ASSHKTTGGSAETLY | 0.50406945 |
| AVAALAHS | ASSHKTTGGSAETLY | 0.5562809 |
| GGVTVMATP | ASSHKTTGGSAETLY | 0.5402456 |
| AGLAANVRD | ASSHKTTGGSAETLY | 0.5769577 |
| TNRPDLIDPALLRPG | ASSHKTTGGSAETLY | 0.65337574 |
| LANLGVTAL | ASSHKTTGGSAETLY | 0.59480584 |
| TGERIEL | ASSHKTTGGSAETLY | 0.58237237 |
| NGEELAA | ASSHKTTGGSAETLY | 0.61784136 |
| VLSLVWG | ASSHKTTGGSAETLY | 0.542024 |
| IGYALWL | ASSHKTTGGSAETLY | 0.5796865 |

| | | |
|---|---|---|
| LSLLLAPL | ASSHKTTGGSAETLY | 0.59640855 |
| TLDEALK | ASSHKTTGGSAETLY | 0.6508517 |
| TDGAGTAEGG | ASSHKTTGGSAETLY | 0.66355574 |
| NAPGGAGGNGG | ASSHKTTGGSAETLY | 0.6310736 |
| VSVFMGMGGNF | ASSHKTTGGSAETLY | 0.6526457 |
| LASYLAAL | ASSHKTTGGSAETLY | 0.6781906 |
| LVGGTPAL | ASSHKTTGGSAETLY | 0.4710992 |
| VRNADVL | ASSHKTTGGSAETLY | 0.52997386 |
| TCSSPFLVSAM | ASSHKTTGGSAETLY | 0.65461475 |
| VHITDAE | ASSHKTTGGSAETLY | 0.57712334 |
| EYAAPSF | ASSHKTTGGSAETLY | 0.6750356 |
| VVDAFHGGL | ASSHKTTGGSAETLY | 0.6245539 |
| ASVVDPQ | ASSHKTTGGSAETLY | 0.6493673 |
| SYNEFGGLS | ASSHKTTGGSAETLY | 0.6498437 |
| DPDGSVY | ASSHKTTGGSAETLY | 0.6848419 |
| MVSTIHATE | ASSHKTTGGSAETLY | 0.65607727 |
| LAAGVADPV | ASSHKTTGGSAETLY | 0.62273896 |
| TKEQIAA | ASSHKTTGGSAETLY | 0.67218876 |
| SLASPAAL | ASSHKTTGGSAETLY | 0.458563 |
| IGLVFYGV | ASSHKTTGGSAETLY | 0.6699494 |
| FLLDASNL | ASSHKTTGGSAETLY | 0.6576134 |
| YQVGNFP | ASSHKTTGGSAETLY | 0.52878904 |
| ESLLSQST | ASSHKTTGGSAETLY | 0.6175747 |
| LFTVGFAAVG | ASSHKTTGGSAETLY | 0.68079436 |
| ATPPTPT | ASSHKTTGGSAETLY | 0.67398953 |
| TDLATLG | ASSHKTTGGSAETLY | 0.55568916 |
| SAGGTGR | ASSHKTTGGSAETLY | 0.5827799 |
| GFSNSGS | ASSHKTTGGSAETLY | 0.50382906 |
| GGNGGSGGTSV | ASSHKTTGGSAETLY | 0.5414718 |
| GTSVATGGAGN | ASSHKTTGGSAETLY | 0.41029537 |
| NGGNGGSGGTS | ASSHKTTGGSAETLY | 0.6631737 |
| AVVKFPH | ASSHKTTGGSAETLY | 0.58382374 |
| TPPPPPP | ASSHKTTGGSAETLY | 0.6734425 |
| AAPAAAAAPA | ASSHKTTGGSAETLY | 0.6349043 |
| IALNEDL | ASSHKTTGGSAETLY | 0.67099917 |
| IVNVSSM | ASSHKTTGGSAETLY | 0.5752975 |
| TGIPPAPR | ASSHKTTGGSAETLY | 0.5880381 |
| FDLGGGTF | ASSHKTTGGSAETLY | 0.41009623 |
| FDLGGGTFDVS | ASSHKTTGGSAETLY | 0.5943446 |
| NEPTAAA | ASSHKTTGGSAETLY | 0.57952076 |
| NEPTAAALA | ASSHKTTGGSAETLY | 0.5992042 |
| GSIINISSI | ASSHKTTGGSAETLY | 0.5353342 |
| VAPGNAGTA | ASSHKTTGGSAETLY | 0.6795043 |
| TGGGASLE | ASSHKTTGGSAETLY | 0.61297435 |
| VAVYNHYKRI | ASSHKTTGGSAETLY | 0.65891224 |
| TDPAPATT | ASSHKTTGGSAETLY | 0.5772865 |
| TNRPDILDPALLRPG | ASSHKTTGGSAETLY | 0.55456537 |
| LTTTTTS | ASSHKTTGGSAETLY | 0.5482507 |
| PTGSGKTL | ASSHKTTGGSAETLY | 0.6568024 |
| TGSGKTL | ASSHKTTGGSAETLY | 0.5088196 |
| TIGHVDHGK | ASSHKTTGGSAETLY | 0.39770192 |

| | | |
|---|---|---|
| TIGHVDHGKTT | ASSHKTTGGSAETLY | 0.54874635 |
| GHVDHGKTT | ASSHKTTGGSAETLY | 0.5684752 |
| AQFEHTLL | ASSHKTTGGSAETLY | 0.6836175 |
| IVNQDEL | ASSHKTTGGSAETLY | 0.67696464 |
| TNGSQFF | ASSHKTTGGSAETLY | 0.6374047 |
| TGKGIVE | ASSHKTTGGSAETLY | 0.684572 |
| LISNASDA | ASSHKTTGGSAETLY | 0.32451758 |
| AELRTLGELTKIAKAHGAQ | ASSHKTTGGSAETLY | 0.6079674 |
| APNAVAAAEPSW | ASSHKTTGGSAETLY | 0.6774281 |
| APQVAGAATAVL | ASSHKTTGGSAETLY | 0.65312606 |
| ARRDPAAAAAILPTDA | ASSHKTTGGSAETLY | 0.61099905 |
| DAAARDVLAEAGLAEYF | ASSHKTTGGSAETLY | 0.67538416 |
| DDQIVLSAKANLAKVEQ | ASSHKTTGGSAETLY | 0.5165879 |
| DELLPDTGEDSDEDGH | ASSHKTTGGSAETLY | 0.5885166 |
| ELPEAPANRATVPQD | ASSHKTTGGSAETLY | 0.6115756 |
| FGQSKIERVTGIG | ASSHKTTGGSAETLY | 0.66344833 |
| GAGGTGSTAGGAGGAGGAGGLY | ASSHKTTGGSAETLY | 0.66897875 |
| GHGGTGAAGGNGGTGGAGGSGI | ASSHKTTGGSAETLY | 0.4599473 |
| GNGGNGGAGGEGGPGGAGGAGA | ASSHKTTGGSAETLY | 0.6460033 |
| GSTAGNGGNGGNATNGTIAGQP | ASSHKTTGGSAETLY | 0.6870159 |
| GTGGDGGNGGNGGTGG | ASSHKTTGGSAETLY | 0.57221794 |
| GVRRPAGIPADAPV | ASSHKTTGGSAETLY | 0.6710903 |
| GYRQVTDAHLLTLARR | ASSHKTTGGSAETLY | 0.6406448 |
| IAKMGQDGHDRG | ASSHKTTGGSAETLY | 0.641835 |
| KAEPSIEIQYT | ASSHKTTGGSAETLY | 0.55517507 |
| KPVRIRMAADEDTVAVT | ASSHKTTGGSAETLY | 0.5337852 |
| MPPRIAGMRLLVIKPEP | ASSHKTTGGSAETLY | 0.6625552 |
| MKFYAVALFRSRGVPPTR | ASSHKTTGGSAETLY | 0.5468015 |
| NAGGFGILGAVAHSP | ASSHKTTGGSAETLY | 0.6146211 |
| NEGREITLAMAEM | ASSHKTTGGSAETLY | 0.62790966 |
| NELVEWSHPLGHRWRWE | ASSHKTTGGSAETLY | 0.68894565 |
| NLGAGGTGGNGGNPG | ASSHKTTGGSAETLY | 0.6508826 |
| NTEWPKWETVMAADPVD | ASSHKTTGGSAETLY | 0.6497439 |
| PASAAPAETQP | ASSHKTTGGSAETLY | 0.52376413 |
| QQSSDGKNGVRAAEITGE | ASSHKTTGGSAETLY | 0.6532827 |
| RAPDAVTTSWWKE | ASSHKTTGGSAETLY | 0.6627569 |
| SGFGNVGAHTSGWFNQ | ASSHKTTGGSAETLY | 0.46278542 |
| SGVAGAVGQAASVGGLKV | ASSHKTTGGSAETLY | 0.6745471 |
| SNGGAGTGSAGGNGGT | ASSHKTTGGSAETLY | 0.5193002 |
| TGGAGGEGGAAG | ASSHKTTGGSAETLY | 0.636327 |
| TGGDGGAAGAGGAPGHG | ASSHKTTGGSAETLY | 0.65398836 |
| TMPPLLAAIANAGRHGV | ASSHKTTGGSAETLY | 0.684541 |
| TSSTAGNSGGA | ASSHKTTGGSAETLY | 0.6188595 |
| VPATVPVGILC | ASSHKTTGGSAETLY | 0.6597065 |
| VQRPDSGGVVLTGRIS | ASSHKTTGGSAETLY | 0.4889955 |
| WTPNPVPATKGVPAGLVQAT | ASSHKTTGGSAETLY | 0.6690281 |
| YALKRALAKFSDRFDVVII | ASSHKTTGGSAETLY | 0.6670356 |
| YLIGSVIV | ASSHKTTGGSAETLY | 0.6655675 |
| YGQLVAVID | ASSHKTTGGSAETLY | 0.6878753 |
| VTVGGMTLPA | ASSHKTTGGSAETLY | 0.66638166 |
| VLALPAV | ASSHKTTGGSAETLY | 0.3741935 |

| | | |
|---|---|---|
| VIPPTLNYET | ASSHKTTGGSAETLY | 0.6157217 |
| VGMGGMHPSSG | ASSHKTTGGSAETLY | 0.67579705 |
| VGLAGGAATA | ASSHKTTGGSAETLY | 0.6770639 |
| VGGAGGAGGSGP | ASSHKTTGGSAETLY | 0.6781541 |
| VFAAFMV | ASSHKTTGGSAETLY | 0.66392404 |
| TLYDKLR | ASSHKTTGGSAETLY | 0.59536093 |
| TLAVFTVL | ASSHKTTGGSAETLY | 0.6806725 |
| TGSVQPARV | ASSHKTTGGSAETLY | 0.6919167 |
| TGGGAGSGGT | ASSHKTTGGSAETLY | 0.658522 |
| SHLETSL | ASSHKTTGGSAETLY | 0.5529376 |
| SFTTGMT | ASSHKTTGGSAETLY | 0.5371622 |
| QVEVAKRMGVRQPTVS | ASSHKTTGGSAETLY | 0.65293247 |
| QGSGGPALAM | ASSHKTTGGSAETLY | 0.6659813 |
| MLAPSNP | ASSHKTTGGSAETLY | 0.4787467 |
| LVNNAAI | ASSHKTTGGSAETLY | 0.63246137 |
| LTGEINTLA | ASSHKTTGGSAETLY | 0.63687074 |
| LSIFGPV | ASSHKTTGGSAETLY | 0.37309504 |
| LLALLANL | ASSHKTTGGSAETLY | 0.6681622 |
| LISNASDALDKLR | ASSHKTTGGSAETLY | 0.67944056 |
| LINEAVP | ASSHKTTGGSAETLY | 0.65165603 |
| LGYAFSGL | ASSHKTTGGSAETLY | 0.5315096 |
| LDFVAVAL | ASSHKTTGGSAETLY | 0.67511266 |
| LAQPGAE | ASSHKTTGGSAETLY | 0.6847785 |
| ISNLGAVTGF | ASSHKTTGGSAETLY | 0.6848891 |
| HQEDPGRI | ASSHKTTGGSAETLY | 0.40603986 |
| GTPGEPTP | ASSHKTTGGSAETLY | 0.33667156 |
| GFFNSGS | ASSHKTTGGSAETLY | 0.6325698 |
| GAGGGGLTAGAN | ASSHKTTGGSAETLY | 0.66889286 |
| FVDALVNM | ASSHKTTGGSAETLY | 0.58525383 |
| FLTVAPEL | ASSHKTTGGSAETLY | 0.5430962 |
| FETLTAL | ASSHKTTGGSAETLY | 0.6536243 |
| DVINPIEL | ASSHKTTGGSAETLY | 0.6751945 |
| APVSGAPV | ASSHKTTGGSAETLY | 0.5078877 |
| ALVGGLAW | ASSHKTTGGSAETLY | 0.44493145 |
| ALAAAAATAI | ASSHKTTGGSAETLY | 0.671314 |
| AFGLGLLL | ASSHKTTGGSAETLY | 0.6631284 |
| AFFAAASL | ASSHKTTGGSAETLY | 0.5466775 |
| AAAALPAFSPP | ASSHKTTGGSAETLY | 0.5902153 |
| VVLAPAL | ASSHKTTGGSAETLY | 0.63411564 |
| TIGHVDHGKTTL | ASSHKTTGGSAETLY | 0.66956544 |
| LISNASDALDK | ASSHKTTGGSAETLY | 0.64499485 |
| LISNASDAL | ASSHKTTGGSAETLY | 0.47005308 |
| ISNASDALDK | ASSHKTTGGSAETLY | 0.66515183 |
| NQAETLVYQTE | ASSHKTTGGSAETLY | 0.49346542 |
| TGGTGGTAGAG | ASSHKTTGGSAETLY | 0.6640849 |
| TGSTAGGAGGA | ASSHKTTGGSAETLY | 0.6235896 |
| DPLAVPVAAGPV | ASSHKTTGGSAETLY | 0.65142953 |
| GGTGGTPGNGGN | ASSHKTTGGSAETLY | 0.5985074 |
| PAGLAGVDPATL | ASSHKTTGGSAETLY | 0.54822826 |
| GNNLVVEAPHAGASV | ASSHKTTGGSAETLY | 0.5037769 |
| KVTRIMSASTLEQPA | ASSHKTTGGSAETLY | 0.47499737 |

| | | |
|---|---|---|
| LGFALPIALPRRTVV | ASSHKTTGGSAETLY | 0.67318594 |
| TRLGNLTARKLVVLV | ASSHKTTGGSAETLY | 0.65230787 |
| WSQQRRLIYFPSAGP | ASSHKTTGGSAETLY | 0.6700674 |
| GPNGNPDPMAAAVDIR | ASSHKTTGGSAETLY | 0.63682795 |
| TQRCVSVADLRAAASR | ASSHKTTGGSAETLY | 0.49815518 |
| QFVLDVAEHPVEVLTETV | ASSHKTTGGSAETLY | 0.6587012 |
| GGAGGTGGAGGNGGTQAGDG | ASSHKTTGGSAETLY | 0.464055 |
| PLLATLRAIAVDVDAKPAQV | ASSHKTTGGSAETLY | 0.6531954 |
| APKFRTGGYMIDFWGVGYQVAK | ASSHKTTGGSAETLY | 0.63032347 |
| GAGGAGGAGGAAGGPGGTGGTG | ASSHKTTGGSAETLY | 0.6601418 |
| IGALGLLATDVAPAGSVAFGVA | ASSHKTTGGSAETLY | 0.5527333 |
| AGGAGGIGGNGGAGGAGGVLMG | ASSHKTTGGSAETLY | 0.6583338 |
| ADELIGLL | ASSHKTTGGSAETLY | 0.65075785 |
| ADRALKA | ASSHKTTGGSAETLY | 0.551451 |
| AGAVGQVGGAGGA | ASSHKTTGGSAETLY | 0.6415881 |
| AGGGGGLGGAP | ASSHKTTGGSAETLY | 0.67366993 |
| AGGLLYGNGGAG | ASSHKTTGGSAETLY | 0.6442047 |
| AGLESIGL | ASSHKTTGGSAETLY | 0.4624101 |
| AGLKTGII | ASSHKTTGGSAETLY | 0.6480467 |
| ASLAALLL | ASSHKTTGGSAETLY | 0.5545303 |
| ASVINISSI | ASSHKTTGGSAETLY | 0.6485998 |
| ATPPGPA | ASSHKTTGGSAETLY | 0.62381434 |
| ATVASALV | ASSHKTTGGSAETLY | 0.5586891 |
| ATVASLIL | ASSHKTTGGSAETLY | 0.657809 |
| AVNGPVLIRH | ASSHKTTGGSAETLY | 0.48686847 |
| DGAAGAVGI | ASSHKTTGGSAETLY | 0.5595173 |
| DPLLHGI | ASSHKTTGGSAETLY | 0.59474385 |
| ELGAALGAL | ASSHKTTGGSAETLY | 0.61950296 |
| ELGRTQAV | ASSHKTTGGSAETLY | 0.6642289 |
| FVAGGYLS | ASSHKTTGGSAETLY | 0.5092038 |
| GASGGAGGAGGW | ASSHKTTGGSAETLY | 0.6407944 |
| GGAGAGGDGGAGGI | ASSHKTTGGSAETLY | 0.594588 |
| GTGHNGSQPGMGGNG | ASSHKTTGGSAETLY | 0.5080832 |
| GYAAASASAS | ASSHKTTGGSAETLY | 0.6323406 |
| HQVDVAAVQQG | ASSHKTTGGSAETLY | 0.61845505 |
| IGAAIAHT | ASSHKTTGGSAETLY | 0.48745918 |
| IGMIEVAQL | ASSHKTTGGSAETLY | 0.66074264 |
| IRLAPEHTSAMD | ASSHKTTGGSAETLY | 0.5084102 |
| IYNVSVL | ASSHKTTGGSAETLY | 0.6386188 |
| KQGFLGM | ASSHKTTGGSAETLY | 0.68226504 |
| LESVITQL | ASSHKTTGGSAETLY | 0.4921202 |
| LGLGALVVV | ASSHKTTGGSAETLY | 0.6611191 |
| LIGNGGNGGSGGI | ASSHKTTGGSAETLY | 0.6437706 |
| LNAINAPTL | ASSHKTTGGSAETLY | 0.6487777 |
| LQLLNTGV | ASSHKTTGGSAETLY | 0.60296184 |
| LQTSIEF | ASSHKTTGGSAETLY | 0.5765761 |
| MSFAVSEL | ASSHKTTGGSAETLY | 0.6804335 |
| NTGALGL | ASSHKTTGGSAETLY | 0.6515871 |
| PPGAGGAGGAGTT | ASSHKTTGGSAETLY | 0.649633 |
| QLVGLGI | ASSHKTTGGSAETLY | 0.6537873 |
| RFLEQVHQL | ASSHKTTGGSAETLY | 0.6729233 |

| | | |
|---|---|---|
| RGVGYVLR | ASSHKTTGGSAETLY | 0.6650922 |
| SATGVFAGVI | ASSHKTTGGSAETLY | 0.577588 |
| SLPAITLP | ASSHKTTGGSAETLY | 0.4212714 |
| STALAFTV | ASSHKTTGGSAETLY | 0.59461564 |
| SVLTVGLGK | ASSHKTTGGSAETLY | 0.624461 |
| TALAGAGSGI | ASSHKTTGGSAETLY | 0.5873383 |
| TGLPFTGI | ASSHKTTGGSAETLY | 0.41776693 |
| VAEHVTV | ASSHKTTGGSAETLY | 0.6686803 |
| VDIVLNSL | ASSHKTTGGSAETLY | 0.6882769 |
| VLTGGGAVL | ASSHKTTGGSAETLY | 0.56088215 |
| VNVGLAYGI | ASSHKTTGGSAETLY | 0.6901678 |
| VSSAAVGAV | ASSHKTTGGSAETLY | 0.49437162 |
| VTSVYEAGV | ASSHKTTGGSAETLY | 0.65253115 |
| WVALDRGA | ASSHKTTGGSAETLY | 0.6733513 |
| YLVGSVVI | ASSHKTTGGSAETLY | 0.5081662 |
| GPPGTGKTLL | ASSHKTTGGSAETLY | 0.6555888 |
| GVGMTKF | ASSHKTTGGSAETLY | 0.6045972 |
| IGIGGSDL | ASSHKTTGGSAETLY | 0.5757791 |
| IGTLLAEL | ASSHKTTGGSAETLY | 0.67323136 |
| IYTSGTT | ASSHKTTGGSAETLY | 0.5620906 |
| LSFPAGV | ASSHKTTGGSAETLY | 0.6526149 |
| LYTSGTT | ASSHKTTGGSAETLY | 0.5878403 |
| VGAFPFGF | ASSHKTTGGSAETLY | 0.50670886 |
| VGAGTVEF | ASSHKTTGGSAETLY | 0.65151185 |
| VGKSTLLL | ASSHKTTGGSAETLY | 0.66369224 |
| AALAHAGGGIV | ASSHKTTGGSAETLY | 0.64359635 |
| AALVAVSLL | ASSHKTTGGSAETLY | 0.6913077 |
| AAVSRAVAAL | ASSHKTTGGSAETLY | 0.5350736 |
| AGGSSGIGGF | ASSHKTTGGSAETLY | 0.68622375 |
| AGLTVANANG | ASSHKTTGGSAETLY | 0.67190754 |
| AGLVGAVL | ASSHKTTGGSAETLY | 0.63328743 |
| ALSQAFTV | ASSHKTTGGSAETLY | 0.59793264 |
| ALTFSVSL | ASSHKTTGGSAETLY | 0.6698617 |
| ALTGAALVAL | ASSHKTTGGSAETLY | 0.56426007 |
| ASFSIGAGVL | ASSHKTTGGSAETLY | 0.6552253 |
| DGSVYKI | ASSHKTTGGSAETLY | 0.68486416 |
| EQLAEIV | ASSHKTTGGSAETLY | 0.64872146 |
| FAVSELTVM | ASSHKTTGGSAETLY | 0.78049874 |
| GAAPLAAGL | ASSHKTTGGSAETLY | 0.7550124 |
| GADGGDGGSGGAGG | ASSHKTTGGSAETLY | 0.7537369 |
| GFGLDTGGAGG | ASSHKTTGGSAETLY | 0.69318235 |
| GLASGGLL | ASSHKTTGGSAETLY | 0.55975485 |
| GRIAGAAVLL | ASSHKTTGGSAETLY | 0.7440885 |
| GSAGSLNTSF | ASSHKTTGGSAETLY | 0.6372767 |
| GTGQIGI | ASSHKTTGGSAETLY | 0.7868584 |
| GTLIALIL | ASSHKTTGGSAETLY | 0.7798952 |
| IAATMGFVVG | ASSHKTTGGSAETLY | 0.67738307 |
| IFAQALL | ASSHKTTGGSAETLY | 0.7794342 |
| IGAAVALGF | ASSHKTTGGSAETLY | 0.7828816 |
| IGLTPLAL | ASSHKTTGGSAETLY | 0.5678879 |
| ILGSNTSTL | ASSHKTTGGSAETLY | 0.73662484 |

| | | |
|---|---|---|
| IVFSLTVL | ASSHKTTGGSAETLY | 0.73446107 |
| LASKATGF | ASSHKTTGGSAETLY | 0.73274267 |
| LASVALIL | ASSHKTTGGSAETLY | 0.565333 |
| LFAVGLTL | ASSHKTTGGSAETLY | 0.77389497 |
| LFNGGIGTYI | ASSHKTTGGSAETLY | 0.72049123 |
| LTLGAGII | ASSHKTTGGSAETLY | 0.27928725 |
| LVAAALGDI | ASSHKTTGGSAETLY | 0.76081663 |
| NTAFVQLGI | ASSHKTTGGSAETLY | 0.740631 |
| QLFSGAEL | ASSHKTTGGSAETLY | 0.7175326 |
| SAATFITL | ASSHKTTGGSAETLY | 0.7791476 |
| SNYGLANLGV | ASSHKTTGGSAETLY | 0.67197526 |
| SYGGAYAVM | ASSHKTTGGSAETLY | 0.6813507 |
| TTGVGGLTLG | ASSHKTTGGSAETLY | 0.77034396 |
| TTQTIYI | ASSHKTTGGSAETLY | 0.7817303 |
| VAAGAGLGAVI | ASSHKTTGGSAETLY | 0.5827886 |
| VAAVVTLL | ASSHKTTGGSAETLY | 0.76080394 |
| VGFLGGLI | ASSHKTTGGSAETLY | 0.78342354 |
| VGVIPILL | ASSHKTTGGSAETLY | 0.7465496 |
| VIFASSLI | ASSHKTTGGSAETLY | 0.78646487 |
| VKGGDGGAAAG | ASSHKTTGGSAETLY | 0.7883078 |
| VTKTFTNI | ASSHKTTGGSAETLY | 0.71247387 |
| VVAAAILL | ASSHKTTGGSAETLY | 0.7807253 |
| YAALDVEL | ASSHKTTGGSAETLY | 0.78769684 |
| LASVALLL | ASSHKTTGGSAETLY | 0.7131339 |
| LDAVHRV | ASSHKTTGGSAETLY | 0.74184704 |
| QGLSSSIF | ASSHKTTGGSAETLY | 0.64949656 |
| RGTFANIRL | ASSHKTTGGSAETLY | 0.74746734 |
| RQLWWGHRI | ASSHKTTGGSAETLY | 0.5535922 |
| YGPPGTGKTLL | ASSHKTTGGSAETLY | 0.7562181 |
| AALVGAPANVA | ASSHKTTGGSAETLY | 0.7829163 |
| AATFRRTE | ASSHKTTGGSAETLY | 0.718228 |
| AGALGGGI | ASSHKTTGGSAETLY | 0.62791693 |
| AGGAGGAGGIGAG | ASSHKTTGGSAETLY | 0.63512 |
| AGSGALGL | ASSHKTTGGSAETLY | 0.748495 |
| AQIADQI | ASSHKTTGGSAETLY | 0.69969726 |
| ASAVAVTYL | ASSHKTTGGSAETLY | 0.56664467 |
| ASTVLGGTGG | ASSHKTTGGSAETLY | 0.7163398 |
| AVGGEPAK | ASSHKTTGGSAETLY | 0.77600884 |
| CEQGTFGGTC | ASSHKTTGGSAETLY | 0.77296144 |
| DEGKAPAG | ASSHKTTGGSAETLY | 0.5779842 |
| DTVVFPAL | ASSHKTTGGSAETLY | 0.78834563 |
| FEGITMAG | ASSHKTTGGSAETLY | 0.78665453 |
| FPLVVQD | ASSHKTTGGSAETLY | 0.7839309 |
| GAFTTPQ | ASSHKTTGGSAETLY | 0.75096214 |
| GFGGFGTAGGVGGA | ASSHKTTGGSAETLY | 0.6097866 |
| GFLAQFL | ASSHKTTGGSAETLY | 0.78431106 |
| GGAGGAGGNAGVA | ASSHKTTGGSAETLY | 0.7002907 |
| GGVGTAGAGGAGGAGG | ASSHKTTGGSAETLY | 0.7135147 |
| GNGGGGDAGNAGSGGN | ASSHKTTGGSAETLY | 0.7765717 |
| GTGGAGGAGNDAGS | ASSHKTTGGSAETLY | 0.72675526 |
| GTTNTGWFNS | ASSHKTTGGSAETLY | 0.6901939 |

| | | |
|---|---|---|
| IFMGPLFTL | ASSHKTTGGSAETLY | 0.74792624 |
| ILAHAIEAI | ASSHKTTGGSAETLY | 0.78725106 |
| IRSAIREAL | ASSHKTTGGSAETLY | 0.6668418 |
| ITFAGGG | ASSHKTTGGSAETLY | 0.5505392 |
| ITGMSGHQEV | ASSHKTTGGSAETLY | 0.6959541 |
| IVAGLACTL | ASSHKTTGGSAETLY | 0.730335 |
| IVGGGLAGA | ASSHKTTGGSAETLY | 0.77969337 |
| IVTGVLNI | ASSHKTTGGSAETLY | 0.7688805 |
| LAQQGCI | ASSHKTTGGSAETLY | 0.7239363 |
| LCLHVEN | ASSHKTTGGSAETLY | 0.76220095 |
| LDAISDGS | ASSHKTTGGSAETLY | 0.7672067 |
| LGGFAAML | ASSHKTTGGSAETLY | 0.7542279 |
| LGGKNPMI | ASSHKTTGGSAETLY | 0.7800095 |
| LGPKGVAAIL | ASSHKTTGGSAETLY | 0.68037367 |
| LMVGVSI | ASSHKTTGGSAETLY | 0.7866753 |
| LVGAAFVL | ASSHKTTGGSAETLY | 0.7713374 |
| LVLAVAPL | ASSHKTTGGSAETLY | 0.78201807 |
| MAAIAVIDAA | ASSHKTTGGSAETLY | 0.7860104 |
| NNGINAELQ | ASSHKTTGGSAETLY | 0.74711835 |
| RGEPLVLVHG | ASSHKTTGGSAETLY | 0.7544079 |
| RGGAGGAAGLI | ASSHKTTGGSAETLY | 0.46416152 |
| RSYRMHPAV | ASSHKTTGGSAETLY | 0.73214203 |
| SAAHLLAGQLA | ASSHKTTGGSAETLY | 0.7591769 |
| SAAVGSY | ASSHKTTGGSAETLY | 0.7615771 |
| SPHAVLHRI | ASSHKTTGGSAETLY | 0.7759764 |
| SQIAKVL | ASSHKTTGGSAETLY | 0.7749244 |
| TALGHLVP | ASSHKTTGGSAETLY | 0.6977793 |
| TAPTTSGAA | ASSHKTTGGSAETLY | 0.78480685 |
| TDGDDGNGGSPGT | ASSHKTTGGSAETLY | 0.7804694 |
| TDGLGEN | ASSHKTTGGSAETLY | 0.5922102 |
| TGGPPLPGQA | ASSHKTTGGSAETLY | 0.71191734 |
| TIPAITI | ASSHKTTGGSAETLY | 0.78770566 |
| TIPGFTI | ASSHKTTGGSAETLY | 0.7736248 |
| TSIASGALA | ASSHKTTGGSAETLY | 0.71514654 |
| TVTLPGAAI | ASSHKTTGGSAETLY | 0.7288221 |
| VASGLSAALP | ASSHKTTGGSAETLY | 0.79276896 |
| VGAGVIGI | ASSHKTTGGSAETLY | 0.7405058 |
| VGPKRFGV | ASSHKTTGGSAETLY | 0.78686464 |
| VNVAFPD | ASSHKTTGGSAETLY | 0.39001185 |
| VSMTNSGVS | ASSHKTTGGSAETLY | 0.7389155 |
| VTDVLGDLSV | ASSHKTTGGSAETLY | 0.6873016 |
| VVVVGAGFAGL | ASSHKTTGGSAETLY | 0.3270114 |
| VYTVGTVV | ASSHKTTGGSAETLY | 0.77344185 |
| WTTFSGP | ASSHKTTGGSAETLY | 0.7142826 |
| YDELVEAF | ASSHKTTGGSAETLY | 0.7725246 |
| YGTGGAGGNGG | ASSHKTTGGSAETLY | 0.74095345 |
| YTNVMAV | ASSHKTTGGSAETLY | 0.78128195 |
| FLEQVHQL | ASSHKTTGGSAETLY | 0.7502942 |
| GLSLLAVAV | ASSHKTTGGSAETLY | 0.760427 |
| LEQVHQL | ASSHKTTGGSAETLY | 0.6133155 |
| LNAAGAI | ASSHKTTGGSAETLY | 0.7666553 |

| | | |
|---|---|---|
| PAVHASL | ASSHKTTGGSAETLY | 0.7874789 |
| TELEDDA | ASSHKTTGGSAETLY | 0.7842388 |
| TPPGPPN | ASSHKTTGGSAETLY | 0.45010412 |
| TPPPPPPP | ASSHKTTGGSAETLY | 0.74444354 |
| TQSPALI | ASGRDSLSNERLF | 0.731009 |
| EFAAAFPQ | ASGRDSLSNERLF | 0.36637434 |
| LLSGNLDVL | ASGRDSLSNERLF | 0.4062758 |
| GPANLGNYNI | ASGRDSLSNERLF | 0.7293477 |
| TLGPTLP | ASGRDSLSNERLF | 0.7160444 |
| VYAYSSCCK | ASGRDSLSNERLF | 0.7719939 |
| APAPAPAPNI | ASGRDSLSNERLF | 0.76746905 |
| PGGPVPIP | ASGRDSLSNERLF | 0.7171836 |
| FLTVALEP | ASGRDSLSNERLF | 0.76462543 |
| SQYFGANL | ASGRDSLSNERLF | 0.7279832 |
| GGVKPAA | ASGRDSLSNERLF | 0.20387119 |
| GGTGGAGGAGGW | ASGRDSLSNERLF | 0.7087515 |
| GVGGVAGLGGAGG | ASGRDSLSNERLF | 0.5937432 |
| VLLVLATTI | ASGRDSLSNERLF | 0.79081076 |
| IGIFAAML | ASGRDSLSNERLF | 0.65955436 |
| AVGDLID | ASGRDSLSNERLF | 0.70536256 |
| NISPLPL | ASGRDSLSNERLF | 0.72899604 |
| VTAVFVAI | ASGRDSLSNERLF | 0.7715505 |
| SGDDNDAAL | ASGRDSLSNERLF | 0.7411514 |
| AVAALAHS | ASGRDSLSNERLF | 0.68912137 |
| GGVTVMATP | ASGRDSLSNERLF | 0.37652245 |
| AGLAANVRD | ASGRDSLSNERLF | 0.76668775 |
| TNRPDLIDPALLRPG | ASGRDSLSNERLF | 0.6719926 |
| LANLGVTAL | ASGRDSLSNERLF | 0.60267717 |
| TGERIEL | ASGRDSLSNERLF | 0.6101881 |
| NGEELAA | ASGRDSLSNERLF | 0.5666349 |
| VLSLVWG | ASGRDSLSNERLF | 0.5459447 |
| IGYALWL | ASGRDSLSNERLF | 0.78312206 |
| LSLLLAPL | ASGRDSLSNERLF | 0.71251035 |
| TLDEALK | ASGRDSLSNERLF | 0.7232728 |
| TDGAGTAEGG | ASGRDSLSNERLF | 0.7897216 |
| NAPGGAGGNGG | ASGRDSLSNERLF | 0.7574189 |
| VSVFMGMGGNF | ASGRDSLSNERLF | 0.72107744 |
| LASYLAAL | ASGRDSLSNERLF | 0.63636506 |
| LVGGTPAL | ASGRDSLSNERLF | 0.6466695 |
| VRNADVL | ASGRDSLSNERLF | 0.6527977 |
| TCSSPFLVSAM | ASGRDSLSNERLF | 0.6680485 |
| VHITDAE | ASGRDSLSNERLF | 0.78705895 |
| EYAAPSF | ASGRDSLSNERLF | 0.7483323 |
| VVDAFHGGL | ASGRDSLSNERLF | 0.6500704 |
| ASVVDPQ | ASGRDSLSNERLF | 0.76599205 |
| SYNEFGGLS | ASGRDSLSNERLF | 0.7866501 |
| DPDGSVY | ASGRDSLSNERLF | 0.78873223 |
| MVSTIHATE | ASGRDSLSNERLF | 0.6505066 |
| LAAGVADPV | ASGRDSLSNERLF | 0.7381182 |
| TKEQIAA | ASGRDSLSNERLF | 0.72862095 |
| SLASPAAL | ASGRDSLSNERLF | 0.7301921 |

| | | |
|---|---|---|
| IGLVFYGV | ASGRDSLSNERLF | 0.7109171 |
| FLLDASNL | ASGRDSLSNERLF | 0.6211506 |
| YQVGNFP | ASGRDSLSNERLF | 0.78133947 |
| ESLLSQST | ASGRDSLSNERLF | 0.7970744 |
| LFTVGFAAVG | ASGRDSLSNERLF | 0.71514285 |
| ATPPTPT | ASGRDSLSNERLF | 0.61963844 |
| TDLATLG | ASGRDSLSNERLF | 0.7228582 |
| SAGGTGR | ASGRDSLSNERLF | 0.6828364 |
| GFSNSGS | ASGRDSLSNERLF | 0.6777421 |
| GGNGGSGGTSV | ASGRDSLSNERLF | 0.73370767 |
| GTSVATGGAGN | ASGRDSLSNERLF | 0.56399703 |
| NGGNGGSGGTS | ASGRDSLSNERLF | 0.73757106 |
| AVVKFPH | ASGRDSLSNERLF | 0.78567183 |
| TPPPPPP | ASGRDSLSNERLF | 0.72698426 |
| AAPAAAAAPA | ASGRDSLSNERLF | 0.76765186 |
| IALNEDL | ASGRDSLSNERLF | 0.78315675 |
| IVNVSSM | ASGRDSLSNERLF | 0.77103233 |
| TGIPPAPR | ASGRDSLSNERLF | 0.7841667 |
| FDLGGGTF | ASGRDSLSNERLF | 0.7887403 |
| FDLGGGTFDVS | ASGRDSLSNERLF | 0.7457991 |
| NEPTAAA | ASGRDSLSNERLF | 0.754582 |
| NEPTAAALA | ASGRDSLSNERLF | 0.7660456 |
| GSIINISSI | ASGRDSLSNERLF | 0.7268882 |
| VAPGNAGTA | ASGRDSLSNERLF | 0.7296685 |
| TGGGASLE | ASGRDSLSNERLF | 0.72173905 |
| VAVYNHYKRI | ASGRDSLSNERLF | 0.7751807 |
| TDPAPATT | ASGRDSLSNERLF | 0.7430828 |
| TNRPDILDPALLRPG | ASGRDSLSNERLF | 0.78892314 |
| LTTTTTS | ASGRDSLSNERLF | 0.4434726 |
| PTGSGKTL | ASGRDSLSNERLF | 0.612388 |
| TGSGKTL | ASGRDSLSNERLF | 0.77679956 |
| TIGHVDHGK | ASGRDSLSNERLF | 0.24950556 |
| TIGHVDHGKTT | ASGRDSLSNERLF | 0.43023288 |
| GHVDHGKTT | ASGRDSLSNERLF | 0.7579463 |
| AQFEHTLL | ASGRDSLSNERLF | 0.77887917 |
| IVNQDEL | ASGRDSLSNERLF | 0.26078278 |
| TNGSQFF | ASGRDSLSNERLF | 0.67345893 |
| TGKGIVE | ASGRDSLSNERLF | 0.58416986 |
| LISNASDA | ASGRDSLSNERLF | 0.77752566 |
| AELRTLGELTKIAKAHGAQ | ASGRDSLSNERLF | 0.7387289 |
| APNAVAAAEPSW | ASGRDSLSNERLF | 0.5084702 |
| APQVAGAATAVL | ASGRDSLSNERLF | 0.7772056 |
| ARRDPAAAAAILPTDA | ASGRDSLSNERLF | 0.58950627 |
| DAAARDVLAEAGLAEYF | ASGRDSLSNERLF | 0.6987919 |
| DDQIVLSAKANLAKVEQ | ASGRDSLSNERLF | 0.7803769 |
| DELLPDTGEDSDEDGH | ASGRDSLSNERLF | 0.7910973 |
| ELPEAPANRATVPQD | ASGRDSLSNERLF | 0.59754384 |
| FGQSKIERVTGIG | ASGRDSLSNERLF | 0.7721299 |
| GAGGTGSTAGGAGGAGGAGGLY | ASGRDSLSNERLF | 0.71282053 |
| GHGGTGAAGGNGGTGGAGGSGI | ASGRDSLSNERLF | 0.7899549 |
| GNGGNGGAGGEGGPGGAGGAGA | ASGRDSLSNERLF | 0.70940006 |

| | | |
|---|---|---|
| GSTAGNGGNGGNATNGTIAGQP | ASGRDSLSNERLF | 0.5870981 |
| GTGGDGGNGGNGGTGG | ASGRDSLSNERLF | 0.77914584 |
| GVRRPAGIPADAPV | ASGRDSLSNERLF | 0.77398586 |
| GYRQVTDAHLLTLARR | ASGRDSLSNERLF | 0.7279265 |
| IAKMGQDGHDRG | ASGRDSLSNERLF | 0.78437984 |
| KAEPSIEIQYT | ASGRDSLSNERLF | 0.73184234 |
| KPVRIRMAADEDTVAVT | ASGRDSLSNERLF | 0.7261236 |
| MPPRIAGMRLLVIKPEP | ASGRDSLSNERLF | 0.7801486 |
| MKFYAVALFRSRGVPPTR | ASGRDSLSNERLF | 0.7777273 |
| NAGGFGILGAVAHSP | ASGRDSLSNERLF | 0.64665693 |
| NEGREITLAMAEM | ASGRDSLSNERLF | 0.7845075 |
| NELVEWSHPLGHRWRWE | ASGRDSLSNERLF | 0.7278701 |
| NLGAGGTGGNGGNPG | ASGRDSLSNERLF | 0.65883195 |
| NTEWPKWETVMAADPVD | ASGRDSLSNERLF | 0.7884376 |
| PASAAPAETQP | ASGRDSLSNERLF | 0.7609209 |
| QQSSDGKNGVRAAEITGE | ASGRDSLSNERLF | 0.7719146 |
| RAPDAVTTSWWKE | ASGRDSLSNERLF | 0.7707819 |
| SGFGNVGAHTSGWFNQ | ASGRDSLSNERLF | 0.7928207 |
| SGVAGAVGQAASVGGLKV | ASGRDSLSNERLF | 0.59858793 |
| SNGGAGTGSAGGNGGT | ASGRDSLSNERLF | 0.76106954 |
| TGGAGGEGGAAG | ASGRDSLSNERLF | 0.735759 |
| TGGDGGAAGAGGAPGHG | ASGRDSLSNERLF | 0.7805425 |
| TMPPLLAAIANAGRHGV | ASGRDSLSNERLF | 0.75856847 |
| TSSTAGNSGGA | ASGRDSLSNERLF | 0.6999488 |
| VPATVPVGILC | ASGRDSLSNERLF | 0.77005166 |
| VQRPDSGGVVLTGRIS | ASGRDSLSNERLF | 0.78338456 |
| WTPNPVPATKGVPAGLVQAT | ASGRDSLSNERLF | 0.76510525 |
| YALKRALAKFSDRFDVVII | ASGRDSLSNERLF | 0.772257 |
| YLIGSVIV | ASGRDSLSNERLF | 0.7868582 |
| YGQLVAVID | ASGRDSLSNERLF | 0.6604782 |
| VTVGGMTLPA | ASGRDSLSNERLF | 0.72681844 |
| VLALPAV | ASGRDSLSNERLF | 0.77672625 |
| VIPPTLNYET | ASGRDSLSNERLF | 0.7834476 |
| VGMGGMHPSSG | ASGRDSLSNERLF | 0.74750614 |
| VGLAGGAATA | ASGRDSLSNERLF | 0.7856872 |
| VGGAGGAGGSGP | ASGRDSLSNERLF | 0.6937293 |
| VFAAFMV | ASGRDSLSNERLF | 0.76785654 |
| TLYDKLR | ASGRDSLSNERLF | 0.68933684 |
| TLAVFTVL | ASGRDSLSNERLF | 0.75201845 |
| TGSVQPARV | ASGRDSLSNERLF | 0.63865876 |
| TGGGAGSGGT | ASGRDSLSNERLF | 0.76421875 |
| SHLETSL | ASGRDSLSNERLF | 0.77347696 |
| SFTTGMT | ASGRDSLSNERLF | 0.7710047 |
| QVEVAKRMGVRQPTVS | ASGRDSLSNERLF | 0.3834372 |
| QGSGGPALAM | ASGRDSLSNERLF | 0.7671273 |
| MLAPSNP | ASGRDSLSNERLF | 0.77092147 |
| LVNNAAI | ASGRDSLSNERLF | 0.79484904 |
| LTGEINTLA | ASGRDSLSNERLF | 0.73859334 |
| LSIFGPV | ASGRDSLSNERLF | 0.7711315 |
| LLALLANL | ASGRDSLSNERLF | 0.6812522 |
| LISNASDALDKLR | ASGRDSLSNERLF | 0.74987423 |

| | | |
|---|---|---|
| LINEAVP | ASGRDSLSNERLF | 0.60165757 |
| LGYAFSGL | ASGRDSLSNERLF | 0.70930314 |
| LDFVAVAL | ASGRDSLSNERLF | 0.70900697 |
| LAQPGAE | ASGRDSLSNERLF | 0.6731068 |
| ISNLGAVTGF | ASGRDSLSNERLF | 0.7748183 |
| HQEDPGRI | ASGRDSLSNERLF | 0.7836529 |
| GTPGEPTP | ASGRDSLSNERLF | 0.7765534 |
| GFFNSGS | ASGRDSLSNERLF | 0.78093356 |
| GAGGGGLTAGAN | ASGRDSLSNERLF | 0.7827362 |
| FVDALVNM | ASGRDSLSNERLF | 0.7856821 |
| FLTVAPEL | ASGRDSLSNERLF | 0.69427156 |
| FETLTAL | ASGRDSLSNERLF | 0.7426076 |
| DVINPIEL | ASGRDSLSNERLF | 0.5640011 |
| APVSGAPV | ASGRDSLSNERLF | 0.7372506 |
| ALVGGLAW | ASGRDSLSNERLF | 0.70784295 |
| ALAAAAATAI | ASGRDSLSNERLF | 0.75194395 |
| AFGLGLLL | ASGRDSLSNERLF | 0.23533115 |
| AFFAAASL | ASGRDSLSNERLF | 0.7679022 |
| AAAALPAFSPP | ASGRDSLSNERLF | 0.7878039 |
| VVLAPAL | ASGRDSLSNERLF | 0.6744759 |
| TIGHVDHGKTTL | ASGRDSLSNERLF | 0.75905865 |
| LISNASDALDK | ASGRDSLSNERLF | 0.764482 |
| LISNASDAL | ASGRDSLSNERLF | 0.71953416 |
| ISNASDALDK | ASGRDSLSNERLF | 0.7674862 |
| NQAETLVYQTE | ASGRDSLSNERLF | 0.69049716 |
| TGGTGGTAGAG | ASGRDSLSNERLF | 0.77377963 |
| TGSTAGGAGGA | ASGRDSLSNERLF | 0.78701735 |
| DPLAVPVAAGPV | ASGRDSLSNERLF | 0.75747615 |
| GGTGGTPGNGGN | ASGRDSLSNERLF | 0.75494087 |
| PAGLAGVDPATL | ASGRDSLSNERLF | 0.7505982 |
| GNNLVVEAPHAGASV | ASGRDSLSNERLF | 0.56626385 |
| KVTRIMSASTLEQPA | ASGRDSLSNERLF | 0.764923 |
| LGFALPIALPRRTVV | ASGRDSLSNERLF | 0.25997958 |
| TRLGNLTARKLVVLV | ASGRDSLSNERLF | 0.74717677 |
| WSQQRRLIYFPSAGP | ASGRDSLSNERLF | 0.6533935 |
| GPNGNPDPMAAAVDIR | ASGRDSLSNERLF | 0.74765384 |
| TQRCVSVADLRAAASR | ASGRDSLSNERLF | 0.73587227 |
| QFVLDVAEHPVEVLTETV | ASGRDSLSNERLF | 0.7686805 |
| GGAGGTGGAGGNGGTQAGDG | ASGRDSLSNERLF | 0.6568162 |
| PLLATLRAIAVDVDAKPAQV | ASGRDSLSNERLF | 0.7562226 |
| APKFRTGGYMIDFWGVGYQVAK | ASGRDSLSNERLF | 0.7200285 |
| GAGGAGGAGGAAGGPGGTGGTG | ASGRDSLSNERLF | 0.5779834 |
| IGALGLLATDVAPAGSVAFGVA | ASGRDSLSNERLF | 0.77869904 |
| AGGAGGIGGNGGAGGAGGVLMG | ASGRDSLSNERLF | 0.74682415 |
| ADELIGLL | ASGRDSLSNERLF | 0.7324972 |
| ADRALKA | ASGRDSLSNERLF | 0.78020287 |
| AGAVGQVGGAGGA | ASGRDSLSNERLF | 0.7502217 |
| AGGGGGLGGAP | ASGRDSLSNERLF | 0.58877873 |
| AGGLLYGNGGAG | ASGRDSLSNERLF | 0.717679 |
| AGLESIGL | ASGRDSLSNERLF | 0.7434157 |
| AGLKTGII | ASGRDSLSNERLF | 0.7115427 |

| | | |
|---|---|---|
| ASLAALLL | ASGRDSLSNERLF | 0.29656345 |
| ASVINISSI | ASGRDSLSNERLF | 0.76837957 |
| ATPPGPA | ASGRDSLSNERLF | 0.6533599 |
| ATVASALV | ASGRDSLSNERLF | 0.6845481 |
| ATVASLIL | ASGRDSLSNERLF | 0.7184809 |
| AVNGPVLIRH | ASGRDSLSNERLF | 0.7745891 |
| DGAAGAVGI | ASGRDSLSNERLF | 0.70866954 |
| DPLLHGI | ASGRDSLSNERLF | 0.7288171 |
| ELGAALGAL | ASGRDSLSNERLF | 0.77629286 |
| ELGRTQAV | ASGRDSLSNERLF | 0.7579266 |
| FVAGGYLS | ASGRDSLSNERLF | 0.77922976 |
| GASGGAGGAGGW | ASGRDSLSNERLF | 0.74437916 |
| GGAGAGGDGGAGGI | ASGRDSLSNERLF | 0.7516111 |
| GTGHNGSQPGMGGNG | ASGRDSLSNERLF | 0.7703494 |
| GYAAASASAS | ASGRDSLSNERLF | 0.72968894 |
| HQVDVAAVQQG | ASGRDSLSNERLF | 0.63272536 |
| IGAAIAHT | ASGRDSLSNERLF | 0.5838827 |
| IGMIEVAQL | ASGRDSLSNERLF | 0.72343475 |
| IRLAPEHTSAMD | ASGRDSLSNERLF | 0.5164282 |
| IYNVSVL | ASGRDSLSNERLF | 0.7451615 |
| KQGFLGM | ASGRDSLSNERLF | 0.7903018 |
| LESVITQL | ASGRDSLSNERLF | 0.7432258 |
| LGLGALVVV | ASGRDSLSNERLF | 0.554685 |
| LIGNGGNGGSGGI | ASGRDSLSNERLF | 0.7268878 |
| LNAINAPTL | ASGRDSLSNERLF | 0.73625875 |
| LQLLNTGV | ASGRDSLSNERLF | 0.77063084 |
| LQTSIEF | ASGRDSLSNERLF | 0.77561116 |
| MSFAVSEL | ASGRDSLSNERLF | 0.7945322 |
| NTGALGL | ASGRDSLSNERLF | 0.7699439 |
| PPGAGGAGGAGTT | ASGRDSLSNERLF | 0.681821 |
| QLVGLGI | ASGRDSLSNERLF | 0.7708439 |
| RFLEQVHQL | ASGRDSLSNERLF | 0.7432414 |
| RGVGYVLR | ASGRDSLSNERLF | 0.75902563 |
| SATGVFAGVI | ASGRDSLSNERLF | 0.7601123 |
| SLPAITLP | ASGRDSLSNERLF | 0.76435053 |
| STALAFTV | ASGRDSLSNERLF | 0.719285 |
| SVLTVGLGK | ASGRDSLSNERLF | 0.73158085 |
| TALAGAGSGI | ASGRDSLSNERLF | 0.5417258 |
| TGLPFTGI | ASGRDSLSNERLF | 0.7795932 |
| VAEHVTV | ASGRDSLSNERLF | 0.7428629 |
| VDIVLNSL | ASGRDSLSNERLF | 0.67312574 |
| VLTGGGAVL | ASGRDSLSNERLF | 0.71779513 |
| VNVGLAYGI | ASGRDSLSNERLF | 0.788604 |
| VSSAAVGAV | ASGRDSLSNERLF | 0.78336143 |
| VTSVYEAGV | ASGRDSLSNERLF | 0.69471705 |
| WVALDRGA | ASGRDSLSNERLF | 0.683472 |
| YLVGSVVI | ASGRDSLSNERLF | 0.7660879 |
| GPPGTGKTLL | ASGRDSLSNERLF | 0.5643502 |
| GVGMTKF | ASGRDSLSNERLF | 0.7759689 |
| IGIGGSDL | ASGRDSLSNERLF | 0.73809326 |
| IGTLLAEL | ASGRDSLSNERLF | 0.7218591 |

| | | |
|---|---|---|
| IYTSGTT | ASGRDSLSNERLF | 0.711321 |
| LSFPAGV | ASGRDSLSNERLF | 0.7631795 |
| LYTSGTT | ASGRDSLSNERLF | 0.69378555 |
| VGAFPFGF | ASGRDSLSNERLF | 0.7811692 |
| VGAGTVEF | ASGRDSLSNERLF | 0.3137501 |
| VGKSTLLL | ASGRDSLSNERLF | 0.77568614 |
| AALAHAGGGIV | ASGRDSLSNERLF | 0.6586305 |
| AALVAVSLL | ASGRDSLSNERLF | 0.68181086 |
| AAVSRAVAAL | ASGRDSLSNERLF | 0.4335413 |
| AGGSSGIGGF | ASGRDSLSNERLF | 0.75038147 |
| AGLTVANANG | ASGRDSLSNERLF | 0.7682432 |
| AGLVGAVL | ASGRDSLSNERLF | 0.7800224 |
| ALSQAFTV | ASGRDSLSNERLF | 0.7219268 |
| ALTFSVSL | ASGRDSLSNERLF | 0.778304 |
| ALTGAALVAL | ASGRDSLSNERLF | 0.78430945 |
| ASFSIGAGVL | ASGRDSLSNERLF | 0.7198258 |
| DGSVYKI | ASGRDSLSNERLF | 0.40171373 |
| EQLAEIV | ASGRDSLSNERLF | 0.759193 |
| FAVSELTVM | ASGRDSLSNERLF | 0.7573772 |
| GAAPLAAGL | ASGRDSLSNERLF | 0.77982175 |
| GADGGDGGSGGAGG | ASGRDSLSNERLF | 0.75904536 |
| GFGLDTGGAGG | ASGRDSLSNERLF | 0.5943376 |
| GLASGGLL | ASGRDSLSNERLF | 0.7746386 |
| GRIAGAAVLL | ASGRDSLSNERLF | 0.7768587 |
| GSAGSLNTSF | ASGRDSLSNERLF | 0.6517266 |
| GTGQIGI | ASGRDSLSNERLF | 0.7035503 |
| GTLIALIL | ASGRDSLSNERLF | 0.69065726 |
| IAATMGFVVG | ASGRDSLSNERLF | 0.5295522 |
| IFAQALL | ASGRDSLSNERLF | 0.7700604 |
| IGAAVALGF | ASGRDSLSNERLF | 0.71109474 |
| IGLTPLAL | ASGRDSLSNERLF | 0.69321036 |
| ILGSNTSTL | ASGRDSLSNERLF | 0.7440144 |
| IVFSLTVL | ASGRDSLSNERLF | 0.6869912 |
| LASKATGF | ASGRDSLSNERLF | 0.6917447 |
| LASVALIL | ASGRDSLSNERLF | 0.6994424 |
| LFAVGLTL | ASGRDSLSNERLF | 0.7601143 |
| LFNGGIGTYI | ASGRDSLSNERLF | 0.7702123 |
| LTLGAGII | ASGRDSLSNERLF | 0.74987096 |
| LVAAALGDI | ASGRDSLSNERLF | 0.76380193 |
| NTAFVQLGI | ASGRDSLSNERLF | 0.78185356 |
| QLFSGAEL | ASGRDSLSNERLF | 0.57407933 |
| SAATFITL | ASGRDSLSNERLF | 0.6641881 |
| SNYGLANLGV | ASGRDSLSNERLF | 0.7677251 |
| SYGGAYAVM | ASGRDSLSNERLF | 0.6516626 |
| TTGVGGLTLG | ASGRDSLSNERLF | 0.7770858 |
| TTQTIYI | ASGRDSLSNERLF | 0.7376603 |
| VAAGAGLGAVI | ASGRDSLSNERLF | 0.75920045 |
| VAAVVTLL | ASGRDSLSNERLF | 0.756624 |
| VGFLGGLI | ASGRDSLSNERLF | 0.7906373 |
| VGVIPILL | ASGRDSLSNERLF | 0.7723551 |
| VIFASSLI | ASGRDSLSNERLF | 0.75695324 |

| | | |
|---|---|---|
| VKGGDGGAAAG | ASGRDSLSNERLF | 0.7803459 |
| VTKTFTNI | ASGRDSLSNERLF | 0.6243271 |
| VVAAAILL | ASGRDSLSNERLF | 0.7798032 |
| YAALDVEL | ASGRDSLSNERLF | 0.76971424 |
| LASVALLL | ASGRDSLSNERLF | 0.64203244 |
| LDAVHRV | ASGRDSLSNERLF | 0.729298 |
| QGLSSSIF | ASGRDSLSNERLF | 0.7865709 |
| RGTFANIRL | ASGRDSLSNERLF | 0.7839145 |
| RQLWWGHRI | ASGRDSLSNERLF | 0.6017386 |
| YGPPGTGKTLL | ASGRDSLSNERLF | 0.7085165 |
| AALVGAPANVA | ASGRDSLSNERLF | 0.5659247 |
| AATFRRTE | ASGRDSLSNERLF | 0.71161866 |
| AGALGGGI | ASGRDSLSNERLF | 0.49311593 |
| AGGAGGAGGIGAG | ASGRDSLSNERLF | 0.7696891 |
| AGSGALGL | ASGRDSLSNERLF | 0.70049715 |
| AQIADQI | ASGRDSLSNERLF | 0.7831562 |
| ASAVAVTYL | ASGRDSLSNERLF | 0.7480533 |
| ASTVLGGTGG | ASGRDSLSNERLF | 0.7801151 |
| AVGGEPAK | ASGRDSLSNERLF | 0.69959676 |
| CEQGTFGGTC | ASGRDSLSNERLF | 0.70654595 |
| DEGKAPAG | ASGRDSLSNERLF | 0.5111011 |
| DTVVFPAL | ASGRDSLSNERLF | 0.6168096 |
| FEGITMAG | ASGRDSLSNERLF | 0.7002299 |
| FPLVVQD | ASGRDSLSNERLF | 0.7105503 |
| GAFTTPQ | ASGRDSLSNERLF | 0.41286445 |
| GFGGFGTAGGVGGA | ASGRDSLSNERLF | 0.7818417 |
| GFLAQFL | ASGRDSLSNERLF | 0.7443321 |
| GGAGGAGGNAGVA | ASGRDSLSNERLF | 0.76742417 |
| GGVGTAGAGGAGGAGG | ASGRDSLSNERLF | 0.7114827 |
| GNGGGGDAGNAGSGGN | ASGRDSLSNERLF | 0.71191305 |
| GTGGAGGAGNDAGS | ASGRDSLSNERLF | 0.6802492 |
| GTTNTGWFNS | ASGRDSLSNERLF | 0.76496637 |
| IFMGPLFTL | ASGRDSLSNERLF | 0.61614984 |
| ILAHAIEAI | ASGRDSLSNERLF | 0.5518237 |
| IRSAIREAL | ASGRDSLSNERLF | 0.7024213 |
| ITFAGGG | ASGRDSLSNERLF | 0.6927923 |
| ITGMSGHQEV | ASGRDSLSNERLF | 0.7864151 |
| IVAGLACTL | ASGRDSLSNERLF | 0.77874035 |
| IVGGGLAGA | ASGRDSLSNERLF | 0.7564881 |
| IVTGVLNI | ASGRDSLSNERLF | 0.7876428 |
| LAQQGCI | ASGRDSLSNERLF | 0.28542435 |
| LCLHVEN | ASGRDSLSNERLF | 0.7327722 |
| LDAISDGS | ASGRDSLSNERLF | 0.7854721 |
| LGGFAAML | ASGRDSLSNERLF | 0.76483107 |
| LGGKNPMI | ASGRDSLSNERLF | 0.72517836 |
| LGPKGVAAIL | ASGRDSLSNERLF | 0.78196096 |
| LMVGVSI | ASGRDSLSNERLF | 0.65086544 |
| LVGAAFVL | ASGRDSLSNERLF | 0.70209247 |
| LVLAVAPL | ASGRDSLSNERLF | 0.73881555 |
| MAAIAVIDAA | ASGRDSLSNERLF | 0.77820456 |
| NNGINAELQ | ASGRDSLSNERLF | 0.7643565 |

| | | |
|---|---|---|
| RGEPLVLVHG | ASGRDSLSNERLF | 0.6208658 |
| RGGAGGAAGLI | ASGRDSLSNERLF | 0.76596963 |
| RSYRMHPAV | ASGRDSLSNERLF | 0.7920613 |
| SAAHLLAGQLA | ASGRDSLSNERLF | 0.7049813 |
| SAAVGSY | ASGRDSLSNERLF | 0.7768048 |
| SPHAVLHRI | ASGRDSLSNERLF | 0.76209056 |
| SQIAKVL | ASGRDSLSNERLF | 0.74286306 |
| TALGHLVP | ASGRDSLSNERLF | 0.6609888 |
| TAPTTSGAA | ASGRDSLSNERLF | 0.6668631 |
| TDGDDGNGGSPGT | ASGRDSLSNERLF | 0.75236833 |
| TDGLGEN | ASGRDSLSNERLF | 0.63100266 |
| TGGPPLPGQA | ASGRDSLSNERLF | 0.7405013 |
| TIPAITI | ASGRDSLSNERLF | 0.7389606 |
| TIPGFTI | ASGRDSLSNERLF | 0.79198515 |
| TSIASGALA | ASGRDSLSNERLF | 0.7632771 |
| TVTLPGAAI | ASGRDSLSNERLF | 0.76044416 |
| VASGLSAALP | ASGRDSLSNERLF | 0.62434316 |
| VGAGVIGI | ASGRDSLSNERLF | 0.76718295 |
| VGPKRFGV | ASGRDSLSNERLF | 0.7634181 |
| VNVAFPD | ASGRDSLSNERLF | 0.59102654 |
| VSMTNSGVS | ASGRDSLSNERLF | 0.7815635 |
| VTDVLGDLSV | ASGRDSLSNERLF | 0.65752447 |
| VVVVGAGFAGL | ASGRDSLSNERLF | 0.7296011 |
| VYTVGTVV | ASGRDSLSNERLF | 0.76599157 |
| WTTFSGP | ASGRDSLSNERLF | 0.79227436 |
| YDELVEAF | ASGRDSLSNERLF | 0.72744006 |
| YGTGGAGGNGG | ASGRDSLSNERLF | 0.7487063 |
| YTNVMAV | ASGRDSLSNERLF | 0.6022053 |
| FLEQVHQL | ASGRDSLSNERLF | 0.77826893 |
| GLSLLAVAV | ASGRDSLSNERLF | 0.77648425 |
| LEQVHQL | ASGRDSLSNERLF | 0.76535165 |
| LNAAGAI | ASGRDSLSNERLF | 0.79292965 |
| PAVHASL | ASGRDSLSNERLF | 0.7749058 |
| TELEDDA | ASGRDSLSNERLF | 0.41948038 |
| TPPGPPN | ASGRDSLSNERLF | 0.73967314 |
| TPPPPPPP | ASGRDSLSNERLF | 0.7822102 |
| TQSPALI | ASSRSRQGNSQNTLY | 0.78005266 |
| EFAAAFPQ | ASSRSRQGNSQNTLY | 0.78690267 |
| LLSGNLDVL | ASSRSRQGNSQNTLY | 0.77459157 |
| GPANLGNYNI | ASSRSRQGNSQNTLY | 0.71510506 |
| TLGPTLP | ASSRSRQGNSQNTLY | 0.78522795 |
| VYAYSSCCK | ASSRSRQGNSQNTLY | 0.7932288 |
| APAPAPAPNI | ASSRSRQGNSQNTLY | 0.77296644 |
| PGGPVPIP | ASSRSRQGNSQNTLY | 0.7064662 |
| FLTVALEP | ASSRSRQGNSQNTLY | 0.67419386 |
| SQYFGANL | ASSRSRQGNSQNTLY | 0.7593396 |
| GGVKPAA | ASSRSRQGNSQNTLY | 0.7735666 |
| GGTGGAGGAGGW | ASSRSRQGNSQNTLY | 0.6282371 |
| GVGGVAGLGGAGG | ASSRSRQGNSQNTLY | 0.74390316 |
| VLLVLATTI | ASSRSRQGNSQNTLY | 0.74848795 |
| IGIFAAML | ASSRSRQGNSQNTLY | 0.45884198 |

| | | |
|---|---|---|
| AVGDLID | ASSRSRQGNSQNTLY | 0.7736014 |
| NISPLPL | ASSRSRQGNSQNTLY | 0.7874117 |
| VTAVFVAI | ASSRSRQGNSQNTLY | 0.76569414 |
| SGDDNDAAL | ASSRSRQGNSQNTLY | 0.6832715 |
| AVAALAHS | ASSRSRQGNSQNTLY | 0.7791147 |
| GGVTVMATP | ASSRSRQGNSQNTLY | 0.78897274 |
| AGLAANVRD | ASSRSRQGNSQNTLY | 0.7877097 |
| TNRPDLIDPALLRPG | ASSRSRQGNSQNTLY | 0.50584066 |
| LANLGVTAL | ASSRSRQGNSQNTLY | 0.37154648 |
| TGERIEL | ASSRSRQGNSQNTLY | 0.71754646 |
| NGEELAA | ASSRSRQGNSQNTLY | 0.778685 |
| VLSLVWG | ASSRSRQGNSQNTLY | 0.7092154 |
| IGYALWL | ASSRSRQGNSQNTLY | 0.67926705 |
| LSLLLAPL | ASSRSRQGNSQNTLY | 0.75927925 |
| TLDEALK | ASSRSRQGNSQNTLY | 0.78162193 |
| TDGAGTAEGG | ASSRSRQGNSQNTLY | 0.63693935 |
| NAPGGAGGNGG | ASSRSRQGNSQNTLY | 0.55946064 |
| VSVFMGMGGNF | ASSRSRQGNSQNTLY | 0.7751876 |
| LASYLAAL | ASSRSRQGNSQNTLY | 0.7743508 |
| LVGGTPAL | ASSRSRQGNSQNTLY | 0.68129414 |
| VRNADVL | ASSRSRQGNSQNTLY | 0.7237756 |
| TCSSPFLVSAM | ASSRSRQGNSQNTLY | 0.72671294 |
| VHITDAE | ASSRSRQGNSQNTLY | 0.7300243 |
| EYAAPSF | ASSRSRQGNSQNTLY | 0.7562207 |
| VVDAFHGGL | ASSRSRQGNSQNTLY | 0.60705364 |
| ASVVDPQ | ASSRSRQGNSQNTLY | 0.7789015 |
| SYNEFGGLS | ASSRSRQGNSQNTLY | 0.6149492 |
| DPDGSVY | ASSRSRQGNSQNTLY | 0.77702326 |
| MVSTIHATE | ASSRSRQGNSQNTLY | 0.7538943 |
| LAAGVADPV | ASSRSRQGNSQNTLY | 0.77253926 |
| TKEQIAA | ASSRSRQGNSQNTLY | 0.7059288 |
| SLASPAAL | ASSRSRQGNSQNTLY | 0.70349294 |
| IGLVFYGV | ASSRSRQGNSQNTLY | 0.62563074 |
| FLLDASNL | ASSRSRQGNSQNTLY | 0.5727896 |
| YQVGNFP | ASSRSRQGNSQNTLY | 0.7821651 |
| ESLLSQST | ASSRSRQGNSQNTLY | 0.48465037 |
| LFTVGFAAVG | ASSRSRQGNSQNTLY | 0.78557456 |
| ATPPTPT | ASSRSRQGNSQNTLY | 0.75677574 |
| TDLATLG | ASSRSRQGNSQNTLY | 0.63714916 |
| SAGGTGR | ASSRSRQGNSQNTLY | 0.77105474 |
| GFSNSGS | ASSRSRQGNSQNTLY | 0.5837631 |
| GGNGGSGGTSV | ASSRSRQGNSQNTLY | 0.72839 |
| GTSVATGGAGN | ASSRSRQGNSQNTLY | 0.7607252 |
| NGGNGGSGGTS | ASSRSRQGNSQNTLY | 0.777228 |
| AVVKFPH | ASSRSRQGNSQNTLY | 0.6776314 |
| TPPPPPP | ASSRSRQGNSQNTLY | 0.7713607 |
| AAPAAAAAPA | ASSRSRQGNSQNTLY | 0.7631587 |
| IALNEDL | ASSRSRQGNSQNTLY | 0.6601225 |
| IVNVSSM | ASSRSRQGNSQNTLY | 0.7645639 |
| TGIPPAPR | ASSRSRQGNSQNTLY | 0.7770262 |
| FDLGGGTF | ASSRSRQGNSQNTLY | 0.7661306 |

| | | |
|---|---|---|
| FDLGGGTFDVS | ASSRSRQGNSQNTLY | 0.56208754 |
| NEPTAAA | ASSRSRQGNSQNTLY | 0.7630483 |
| NEPTAAALA | ASSRSRQGNSQNTLY | 0.6779238 |
| GSIINISSI | ASSRSRQGNSQNTLY | 0.75788045 |
| VAPGNAGTA | ASSRSRQGNSQNTLY | 0.7437644 |
| TGGGASLE | ASSRSRQGNSQNTLY | 0.7044678 |
| VAVYNHYKRI | ASSRSRQGNSQNTLY | 0.77399385 |
| TDPAPATT | ASSRSRQGNSQNTLY | 0.60208666 |
| TNRPDILDPALLRPG | ASSRSRQGNSQNTLY | 0.679308 |
| LTTTTTS | ASSRSRQGNSQNTLY | 0.7168962 |
| PTGSGKTL | ASSRSRQGNSQNTLY | 0.72578514 |
| TGSGKTL | ASSRSRQGNSQNTLY | 0.77468187 |
| TIGHVDHGK | ASSRSRQGNSQNTLY | 0.6429148 |
| TIGHVDHGKTT | ASSRSRQGNSQNTLY | 0.7540071 |
| GHVDHGKTT | ASSRSRQGNSQNTLY | 0.71089494 |
| AQFEHTLL | ASSRSRQGNSQNTLY | 0.62069315 |
| IVNQDEL | ASSRSRQGNSQNTLY | 0.75923693 |
| TNGSQFF | ASSRSRQGNSQNTLY | 0.7345834 |
| TGKGIVE | ASSRSRQGNSQNTLY | 0.6100154 |
| LISNASDA | ASSRSRQGNSQNTLY | 0.7760409 |
| AELRTLGELTKIAKAHGAQ | ASSRSRQGNSQNTLY | 0.6422895 |
| APNAVAAAEPSW | ASSRSRQGNSQNTLY | 0.762377 |
| APQVAGAATAVL | ASSRSRQGNSQNTLY | 0.7874398 |
| ARRDPAAAAAILPTDA | ASSRSRQGNSQNTLY | 0.65368533 |
| DAAARDVLAEAGLAEYF | ASSRSRQGNSQNTLY | 0.77523124 |
| DDQIVLSAKANLAKVEQ | ASSRSRQGNSQNTLY | 0.7587911 |
| DELLPDTGEDSDEDGH | ASSRSRQGNSQNTLY | 0.76756394 |
| ELPEAPANRATVPQD | ASSRSRQGNSQNTLY | 0.71685636 |
| FGQSKIERVTGIG | ASSRSRQGNSQNTLY | 0.6851427 |
| GAGGTGSTAGGAGGAGGAGGLY | ASSRSRQGNSQNTLY | 0.786134 |
| GHGGTGAAGGNGGTGGAGGSGI | ASSRSRQGNSQNTLY | 0.76493824 |
| GNGGNGGAGGEGGPGGAGGAGA | ASSRSRQGNSQNTLY | 0.77604437 |
| GSTAGNGGNGGNATNGTIAGQP | ASSRSRQGNSQNTLY | 0.7662457 |
| GTGGDGGNGGNGGTGG | ASSRSRQGNSQNTLY | 0.7817977 |
| GVRRPAGIPADAPV | ASSRSRQGNSQNTLY | 0.7742218 |
| GYRQVTDAHLLTLARR | ASSRSRQGNSQNTLY | 0.68504465 |
| IAKMGQDGHDRG | ASSRSRQGNSQNTLY | 0.5302216 |
| KAEPSIEIQYT | ASSRSRQGNSQNTLY | 0.70136 |
| KPVRIRMAADEDTVAVT | ASSRSRQGNSQNTLY | 0.73312247 |
| MPPRIAGMRLLVIKPEP | ASSRSRQGNSQNTLY | 0.7263359 |
| MKFYAVALFRSRGVPPTR | ASSRSRQGNSQNTLY | 0.53718436 |
| NAGGFGILGAVAHSP | ASSRSRQGNSQNTLY | 0.7769096 |
| NEGREITLAMAEM | ASSRSRQGNSQNTLY | 0.79221493 |
| NELVEWSHPLGHRWRWE | ASSRSRQGNSQNTLY | 0.69135505 |
| NLGAGGTGGNGGNPG | ASSRSRQGNSQNTLY | 0.79096913 |
| NTEWPKWETVMAADPVD | ASSRSRQGNSQNTLY | 0.635129 |
| PASAAPAETQP | ASSRSRQGNSQNTLY | 0.76306564 |
| QQSSDGKNGVRAAEITGE | ASSRSRQGNSQNTLY | 0.7772429 |
| RAPDAVTTSWWKE | ASSRSRQGNSQNTLY | 0.6498406 |
| SGFGNVGAHTSGWFNQ | ASSRSRQGNSQNTLY | 0.7693182 |
| SGVAGAVGQAASVGGLKV | ASSRSRQGNSQNTLY | 0.6922072 |

| | | |
|---|---|---|
| SNGGAGTGSAGGNGGT | ASSRSRQGNSQNTLY | 0.6995609 |
| TGGAGGEGGAAG | ASSRSRQGNSQNTLY | 0.78045964 |
| TGGDGGAAGAGGAPGHG | ASSRSRQGNSQNTLY | 0.6830605 |
| TMPPLLAAIANAGRHGV | ASSRSRQGNSQNTLY | 0.7709055 |
| TSSTAGNSGGA | ASSRSRQGNSQNTLY | 0.73115987 |
| VPATVPVGILC | ASSRSRQGNSQNTLY | 0.6372161 |
| VQRPDSGGVVLTGRIS | ASSRSRQGNSQNTLY | 0.7692349 |
| WTPNPVPATKGVPAGLVQAT | ASSRSRQGNSQNTLY | 0.77991104 |
| YALKRALAKFSDRFDVVII | ASSRSRQGNSQNTLY | 0.7340438 |
| YLIGSVIV | ASSRSRQGNSQNTLY | 0.7918669 |
| YGQLVAVID | ASSRSRQGNSQNTLY | 0.66514945 |
| VTVGGMTLPA | ASSRSRQGNSQNTLY | 0.7917656 |
| VLALPAV | ASSRSRQGNSQNTLY | 0.78027153 |
| VIPPTLNYET | ASSRSRQGNSQNTLY | 0.74506694 |
| VGMGGMHPSSG | ASSRSRQGNSQNTLY | 0.71829003 |
| VGLAGGAATA | ASSRSRQGNSQNTLY | 0.7747414 |
| VGGAGGAGGSGP | ASSRSRQGNSQNTLY | 0.705567 |
| VFAAFMV | ASSRSRQGNSQNTLY | 0.777556 |
| TLYDKLR | ASSRSRQGNSQNTLY | 0.7899159 |
| TLAVFTVL | ASSRSRQGNSQNTLY | 0.67667913 |
| TGSVQPARV | ASSRSRQGNSQNTLY | 0.79073334 |
| TGGGAGSGGT | ASSRSRQGNSQNTLY | 0.76771146 |
| SHLETSL | ASSRSRQGNSQNTLY | 0.7036314 |
| SFTTGMT | ASSRSRQGNSQNTLY | 0.69367373 |
| QVEVAKRMGVRQPTVS | ASSRSRQGNSQNTLY | 0.35985854 |
| QGSGGPALAM | ASSRSRQGNSQNTLY | 0.7151083 |
| MLAPSNP | ASSRSRQGNSQNTLY | 0.6360665 |
| LVNNAAI | ASSRSRQGNSQNTLY | 0.7442483 |
| LTGEINTLA | ASSRSRQGNSQNTLY | 0.7670405 |
| LSIFGPV | ASSRSRQGNSQNTLY | 0.6553248 |
| LLALLANL | ASSRSRQGNSQNTLY | 0.7907399 |
| LISNASDALDKLR | ASSRSRQGNSQNTLY | 0.77973014 |
| LINEAVP | ASSRSRQGNSQNTLY | 0.56962264 |
| LGYAFSGL | ASSRSRQGNSQNTLY | 0.73637545 |
| LDFVAVAL | ASSRSRQGNSQNTLY | 0.6805813 |
| LAQPGAE | ASSRSRQGNSQNTLY | 0.72451365 |
| ISNLGAVTGF | ASSRSRQGNSQNTLY | 0.44706848 |
| HQEDPGRI | ASSRSRQGNSQNTLY | 0.77996683 |
| GTPGEPTP | ASSRSRQGNSQNTLY | 0.7107384 |
| GFFNSGS | ASSRSRQGNSQNTLY | 0.20935155 |
| GAGGGGLTAGAN | ASSRSRQGNSQNTLY | 0.7262778 |
| FVDALVNM | ASSRSRQGNSQNTLY | 0.73869514 |
| FLTVAPEL | ASSRSRQGNSQNTLY | 0.74400866 |
| FETLTAL | ASSRSRQGNSQNTLY | 0.7858807 |
| DVINPIEL | ASSRSRQGNSQNTLY | 0.23047605 |
| APVSGAPV | ASSRSRQGNSQNTLY | 0.6128499 |
| ALVGGLAW | ASSRSRQGNSQNTLY | 0.78743213 |
| ALAAAAATAI | ASSRSRQGNSQNTLY | 0.79115117 |
| AFGLGLLL | ASSRSRQGNSQNTLY | 0.49474108 |
| AFFAAASL | ASSRSRQGNSQNTLY | 0.78147274 |
| AAAALPAFSPP | ASSRSRQGNSQNTLY | 0.7746438 |

| | | |
|---|---|---|
| VVLAPAL | ASSRSRQGNSQNTLY | 0.7632581 |
| TIGHVDHGKTTL | ASSRSRQGNSQNTLY | 0.7782892 |
| LISNASDALDK | ASSRSRQGNSQNTLY | 0.78578323 |
| LISNASDAL | ASSRSRQGNSQNTLY | 0.5843266 |
| ISNASDALDK | ASSRSRQGNSQNTLY | 0.7895106 |
| NQAETLVYQTE | ASSRSRQGNSQNTLY | 0.78781974 |
| TGGTGGTAGAG | ASSRSRQGNSQNTLY | 0.63750935 |
| TGSTAGGAGGA | ASSRSRQGNSQNTLY | 0.7621387 |
| DPLAVPVAAGPV | ASSRSRQGNSQNTLY | 0.6210607 |
| GGTGGTPGNGGN | ASSRSRQGNSQNTLY | 0.6108558 |
| PAGLAGVDPATL | ASSRSRQGNSQNTLY | 0.29297522 |
| GNNLVVEAPHAGASV | ASSRSRQGNSQNTLY | 0.77901983 |
| KVTRIMSASTLEQPA | ASSRSRQGNSQNTLY | 0.7732763 |
| LGFALPIALPRRTVV | ASSRSRQGNSQNTLY | 0.7402571 |
| TRLGNLTARKLVVLV | ASSRSRQGNSQNTLY | 0.6607863 |
| WSQQRRLIYFPSAGP | ASSRSRQGNSQNTLY | 0.663657 |
| GPNGNPDPMAAAVDIR | ASSRSRQGNSQNTLY | 0.73108804 |
| TQRCVSVADLRAAASR | ASSRSRQGNSQNTLY | 0.7348726 |
| QFVLDVAEHPVEVLTETV | ASSRSRQGNSQNTLY | 0.3179354 |
| GGAGGTGGAGGNGGTQAGDG | ASSRSRQGNSQNTLY | 0.6640135 |
| PLLATLRAIAVDVDAKPAQV | ASSRSRQGNSQNTLY | 0.7819414 |
| APKFRTGGYMIDFWGVGYQVAK | ASSRSRQGNSQNTLY | 0.77317214 |
| GAGGAGGAGGAAGGPGGTGGTG | ASSRSRQGNSQNTLY | 0.39216566 |
| IGALGLLATDVAPAGSVAFGVA | ASSRSRQGNSQNTLY | 0.7906885 |
| AGGAGGIGGNGGAGGAGGVLMG | ASSRSRQGNSQNTLY | 0.7678046 |
| ADELIGLL | ASSRSRQGNSQNTLY | 0.7845372 |
| ADRALKA | ASSRSRQGNSQNTLY | 0.6840265 |
| AGAVGQVGGAGGA | ASSRSRQGNSQNTLY | 0.6854721 |
| AGGGGGLGGAP | ASSRSRQGNSQNTLY | 0.7703911 |
| AGGLLYGNGGAG | ASSRSRQGNSQNTLY | 0.74499977 |
| AGLESIGL | ASSRSRQGNSQNTLY | 0.40812644 |
| AGLKTGII | ASSRSRQGNSQNTLY | 0.77284193 |
| ASLAALLL | ASSRSRQGNSQNTLY | 0.72558993 |
| ASVINISSI | ASSRSRQGNSQNTLY | 0.721424 |
| ATPPGPA | ASSRSRQGNSQNTLY | 0.6944128 |
| ATVASALV | ASSRSRQGNSQNTLY | 0.7736262 |
| ATVASLIL | ASSRSRQGNSQNTLY | 0.62912285 |
| AVNGPVLIRH | ASSRSRQGNSQNTLY | 0.55627143 |
| DGAAGAVGI | ASSRSRQGNSQNTLY | 0.52161205 |
| DPLLHGI | ASSRSRQGNSQNTLY | 0.76095283 |
| ELGAALGAL | ASSRSRQGNSQNTLY | 0.78771675 |
| ELGRTQAV | ASSRSRQGNSQNTLY | 0.7477633 |
| FVAGGYLS | ASSRSRQGNSQNTLY | 0.6594247 |
| GASGGAGGAGGW | ASSRSRQGNSQNTLY | 0.7781137 |
| GGAGAGGDGGAGGI | ASSRSRQGNSQNTLY | 0.7323544 |
| GTGHNGSQPGMGGNG | ASSRSRQGNSQNTLY | 0.7436996 |
| GYAAASASAS | ASSRSRQGNSQNTLY | 0.75550777 |
| HQVDVAAVQQG | ASSRSRQGNSQNTLY | 0.7133684 |
| IGAAIAHT | ASSRSRQGNSQNTLY | 0.7796874 |
| IGMIEVAQL | ASSRSRQGNSQNTLY | 0.75157154 |
| IRLAPEHTSAMD | ASSRSRQGNSQNTLY | 0.77614313 |

| | | |
|---|---|---|
| IYNVSVL | ASSRSRQGNSQNTLY | 0.7788756 |
| KQGFLGM | ASSRSRQGNSQNTLY | 0.75023925 |
| LESVITQL | ASSRSRQGNSQNTLY | 0.7179 |
| LGLGALVVV | ASSRSRQGNSQNTLY | 0.26557773 |
| LIGNGGNGGSGGI | ASSRSRQGNSQNTLY | 0.62050617 |
| LNAINAPTL | ASSRSRQGNSQNTLY | 0.7711661 |
| LQLLNTGV | ASSRSRQGNSQNTLY | 0.76640075 |
| LQTSIEF | ASSRSRQGNSQNTLY | 0.7746284 |
| MSFAVSEL | ASSRSRQGNSQNTLY | 0.72521573 |
| NTGALGL | ASSRSRQGNSQNTLY | 0.66627693 |
| PPGAGGAGGAGTT | ASSRSRQGNSQNTLY | 0.7639462 |
| QLVGLGI | ASSRSRQGNSQNTLY | 0.7724951 |
| RFLEQVHQL | ASSRSRQGNSQNTLY | 0.51652956 |
| RGVGYVLR | ASSRSRQGNSQNTLY | 0.71707284 |
| SATGVFAGVI | ASSRSRQGNSQNTLY | 0.79340386 |
| SLPAITLP | ASSRSRQGNSQNTLY | 0.75961655 |
| STALAFTV | ASSRSRQGNSQNTLY | 0.73534286 |
| SVLTVGLGK | ASSRSRQGNSQNTLY | 0.56773067 |
| TALAGAGSGI | ASSRSRQGNSQNTLY | 0.77964807 |
| TGLPFTGI | ASSRSRQGNSQNTLY | 0.65905297 |
| VAEHVTV | ASSRSRQGNSQNTLY | 0.7910197 |
| VDIVLNSL | ASSRSRQGNSQNTLY | 0.2260381 |
| VLTGGGAVL | ASSRSRQGNSQNTLY | 0.5961193 |
| VNVGLAYGI | ASSRSRQGNSQNTLY | 0.44863167 |
| VSSAAVGAV | ASSRSRQGNSQNTLY | 0.52551615 |
| VTSVYEAGV | ASSRSRQGNSQNTLY | 0.7830031 |
| WVALDRGA | ASSRSRQGNSQNTLY | 0.63982034 |
| YLVGSVVI | ASSRSRQGNSQNTLY | 0.76451635 |
| GPPGTGKTLL | ASSRSRQGNSQNTLY | 0.7632873 |
| GVGMTKF | ASSRSRQGNSQNTLY | 0.78136414 |
| IGIGGSDL | ASSRSRQGNSQNTLY | 0.76675093 |
| IGTLLAEL | ASSRSRQGNSQNTLY | 0.72832024 |
| IYTSGTT | ASSRSRQGNSQNTLY | 0.5157991 |
| LSFPAGV | ASSRSRQGNSQNTLY | 0.77683306 |
| LYTSGTT | ASSRSRQGNSQNTLY | 0.77868676 |
| VGAFPFGF | ASSRSRQGNSQNTLY | 0.7368665 |
| VGAGTVEF | ASSRSRQGNSQNTLY | 0.4562298 |
| VGKSTLLL | ASSRSRQGNSQNTLY | 0.72245866 |
| AALAHAGGGIV | ASSRSRQGNSQNTLY | 0.7582891 |
| AALVAVSLL | ASSRSRQGNSQNTLY | 0.23082536 |
| AAVSRAVAAL | ASSRSRQGNSQNTLY | 0.24869303 |
| AGGSSGIGGF | ASSRSRQGNSQNTLY | 0.74496245 |
| AGLTVANANG | ASSRSRQGNSQNTLY | 0.66361284 |
| AGLVGAVL | ASSRSRQGNSQNTLY | 0.79101545 |
| ALSQAFTV | ASSRSRQGNSQNTLY | 0.78795767 |
| ALTFSVSL | ASSRSRQGNSQNTLY | 0.74983996 |
| ALTGAALVAL | ASSRSRQGNSQNTLY | 0.76364005 |
| ASFSIGAGVL | ASSRSRQGNSQNTLY | 0.75819004 |
| DGSVYKI | ASSRSRQGNSQNTLY | 0.19389349 |
| EQLAEIV | ASSRSRQGNSQNTLY | 0.6117922 |
| FAVSELTVM | ASSRSRQGNSQNTLY | 0.6109264 |

| | | |
|---|---|---|
| GAAPLAAGL | ASSRSRQGNSQNTLY | 0.78612745 |
| GADGGDGGSGGAGG | ASSRSRQGNSQNTLY | 0.70409214 |
| GFGLDTGGAGG | ASSRSRQGNSQNTLY | 0.69032776 |
| GLASGGLL | ASSRSRQGNSQNTLY | 0.7664337 |
| GRIAGAAVLL | ASSRSRQGNSQNTLY | 0.75304943 |
| GSAGSLNTSF | ASSRSRQGNSQNTLY | 0.7301134 |
| GTGQIGI | ASSRSRQGNSQNTLY | 0.5795025 |
| GTLIALIL | ASSRSRQGNSQNTLY | 0.20073377 |
| IAATMGFVVG | ASSRSRQGNSQNTLY | 0.77249336 |
| IFAQALL | ASSRSRQGNSQNTLY | 0.31635427 |
| IGAAVALGF | ASSRSRQGNSQNTLY | 0.56245 |
| IGLTPLAL | ASSRSRQGNSQNTLY | 0.24543811 |
| ILGSNTSTL | ASSRSRQGNSQNTLY | 0.21932079 |
| IVFSLTVL | ASSRSRQGNSQNTLY | 0.35322642 |
| LASKATGF | ASSRSRQGNSQNTLY | 0.7675314 |
| LASVALIL | ASSRSRQGNSQNTLY | 0.741163 |
| LFAVGLTL | ASSRSRQGNSQNTLY | 0.7233014 |
| LFNGGIGTYI | ASSRSRQGNSQNTLY | 0.7922045 |
| LTLGAGII | ASSRSRQGNSQNTLY | 0.75494605 |
| LVAAALGDI | ASSRSRQGNSQNTLY | 0.73122543 |
| NTAFVQLGI | ASSRSRQGNSQNTLY | 0.57115924 |
| QLFSGAEL | ASSRSRQGNSQNTLY | 0.6326016 |
| SAATFITL | ASSRSRQGNSQNTLY | 0.6423063 |
| SNYGLANLGV | ASSRSRQGNSQNTLY | 0.55727965 |
| SYGGAYAVM | ASSRSRQGNSQNTLY | 0.79184043 |
| TTGVGGLTLG | ASSRSRQGNSQNTLY | 0.63161445 |
| TTQTIYI | ASSRSRQGNSQNTLY | 0.46322048 |
| VAAGAGLGAVI | ASSRSRQGNSQNTLY | 0.6752454 |
| VAAVVTLL | ASSRSRQGNSQNTLY | 0.7878428 |
| VGFLGGLI | ASSRSRQGNSQNTLY | 0.7932186 |
| VGVIPILL | ASSRSRQGNSQNTLY | 0.7079539 |
| VIFASSLI | ASSRSRQGNSQNTLY | 0.55661225 |
| VKGGDGGAAAG | ASSRSRQGNSQNTLY | 0.63804054 |
| VTKTFTNI | ASSRSRQGNSQNTLY | 0.77154136 |
| VVAAAILL | ASSRSRQGNSQNTLY | 0.59997785 |
| YAALDVEL | ASSRSRQGNSQNTLY | 0.44322655 |
| LASVALLL | ASSRSRQGNSQNTLY | 0.76830065 |
| LDAVHRV | ASSRSRQGNSQNTLY | 0.78437996 |
| QGLSSSIF | ASSRSRQGNSQNTLY | 0.6466121 |
| RGTFANIRL | ASSRSRQGNSQNTLY | 0.6051221 |
| RQLWWGHRI | ASSRSRQGNSQNTLY | 0.76091015 |
| YGPPGTGKTLL | ASSRSRQGNSQNTLY | 0.68717444 |
| AALVGAPANVA | ASSRSRQGNSQNTLY | 0.75982165 |
| AATFRRTE | ASSRSRQGNSQNTLY | 0.7451203 |
| AGALGGGI | ASSRSRQGNSQNTLY | 0.5483209 |
| AGGAGGAGGIGAG | ASSRSRQGNSQNTLY | 0.6561548 |
| AGSGALGL | ASSRSRQGNSQNTLY | 0.7908889 |
| AQIADQI | ASSRSRQGNSQNTLY | 0.7290691 |
| ASAVAVTYL | ASSRSRQGNSQNTLY | 0.75359905 |
| ASTVLGGTGG | ASSRSRQGNSQNTLY | 0.7902934 |
| AVGGEPAK | ASSRSRQGNSQNTLY | 0.78027034 |

| | | |
|---|---|---|
| CEQGTFGGTC | ASSRSRQGNSQNTLY | 0.79190624 |
| DEGKAPAG | ASSRSRQGNSQNTLY | 0.77443695 |
| DTVVFPAL | ASSRSRQGNSQNTLY | 0.61854655 |
| FEGITMAG | ASSRSRQGNSQNTLY | 0.6968068 |
| FPLVVQD | ASSRSRQGNSQNTLY | 0.76099 |
| GAFTTPQ | ASSRSRQGNSQNTLY | 0.7734634 |
| GFGGFGTAGGVGGA | ASSRSRQGNSQNTLY | 0.7415124 |
| GFLAQFL | ASSRSRQGNSQNTLY | 0.5979038 |
| GGAGGAGGNAGVA | ASSRSRQGNSQNTLY | 0.7908716 |
| GGVGTAGAGGAGGAGG | ASSRSRQGNSQNTLY | 0.702533 |
| GNGGGGDAGNAGSGGN | ASSRSRQGNSQNTLY | 0.76753 |
| GTGGAGGAGNDAGS | ASSRSRQGNSQNTLY | 0.56486964 |
| GTTNTGWFNS | ASSRSRQGNSQNTLY | 0.5162267 |
| IFMGPLFTL | ASSRSRQGNSQNTLY | 0.7838275 |
| ILAHAIEAI | ASSRSRQGNSQNTLY | 0.19505714 |
| IRSAIREAL | ASSRSRQGNSQNTLY | 0.2350126 |
| ITFAGGG | ASSRSRQGNSQNTLY | 0.7458278 |
| ITGMSGHQEV | ASSRSRQGNSQNTLY | 0.7860482 |
| IVAGLACTL | ASSRSRQGNSQNTLY | 0.20470965 |
| IVGGGLAGA | ASSRSRQGNSQNTLY | 0.51591325 |
| IVTGVLNI | ASSRSRQGNSQNTLY | 0.44105315 |
| LAQQGCI | ASSRSRQGNSQNTLY | 0.7501853 |
| LCLHVEN | ASSRSRQGNSQNTLY | 0.7427349 |
| LDAISDGS | ASSRSRQGNSQNTLY | 0.4605705 |
| LGGFAAML | ASSRSRQGNSQNTLY | 0.7778586 |
| LGGKNPMI | ASSRSRQGNSQNTLY | 0.48376927 |
| LGPKGVAAIL | ASSRSRQGNSQNTLY | 0.3716745 |
| LMVGVSI | ASSRSRQGNSQNTLY | 0.77891743 |
| LVGAAFVL | ASSRSRQGNSQNTLY | 0.7809598 |
| LVLAVAPL | ASSRSRQGNSQNTLY | 0.6310862 |
| MAAIAVIDAA | ASSRSRQGNSQNTLY | 0.7845917 |
| NNGINAELQ | ASSRSRQGNSQNTLY | 0.667747 |
| RGEPLVLVHG | ASSRSRQGNSQNTLY | 0.79092586 |
| RGGAGGAAGLI | ASSRSRQGNSQNTLY | 0.6920055 |
| RSYRMHPAV | ASSRSRQGNSQNTLY | 0.41834533 |
| SAAHLLAGQLA | ASSRSRQGNSQNTLY | 0.7825552 |
| SAAVGSY | ASSRSRQGNSQNTLY | 0.76521236 |
| SPHAVLHRI | ASSRSRQGNSQNTLY | 0.6773231 |
| SQIAKVL | ASSRSRQGNSQNTLY | 0.77682465 |
| TALGHLVP | ASSRSRQGNSQNTLY | 0.6009895 |
| TAPTTSGAA | ASSRSRQGNSQNTLY | 0.76054835 |
| TDGDDGNGGSPGT | ASSRSRQGNSQNTLY | 0.79075027 |
| TDGLGEN | ASSRSRQGNSQNTLY | 0.78676516 |
| TGGPPLPGQA | ASSRSRQGNSQNTLY | 0.5817435 |
| TIPAITI | ASSRSRQGNSQNTLY | 0.7681863 |
| TIPGFTI | ASSRSRQGNSQNTLY | 0.75013125 |
| TSIASGALA | ASSRSRQGNSQNTLY | 0.6333638 |
| TVTLPGAAI | ASSRSRQGNSQNTLY | 0.78930855 |
| VASGLSAALP | ASSRSRQGNSQNTLY | 0.7199158 |
| VGAGVIGI | ASSRSRQGNSQNTLY | 0.7750833 |
| VGPKRFGV | ASSRSRQGNSQNTLY | 0.7594393 |

| | | |
|---|---|---|
| VNVAFPD | ASSRSRQGNSQNTLY | 0.75437945 |
| VSMTNSGVS | ASSRSRQGNSQNTLY | 0.4204886 |
| VTDVLGDLSV | ASSRSRQGNSQNTLY | 0.7298499 |
| VVVVGAGFAGL | ASSRSRQGNSQNTLY | 0.7380043 |
| VYTVGTVV | ASSRSRQGNSQNTLY | 0.7770956 |
| WTTFSGP | ASSRSRQGNSQNTLY | 0.7404246 |
| YDELVEAF | ASSRSRQGNSQNTLY | 0.3790887 |
| YGTGGAGGNGG | ASSRSRQGNSQNTLY | 0.754115 |
| YTNVMAV | ASSRSRQGNSQNTLY | 0.7813306 |
| FLEQVHQL | ASSRSRQGNSQNTLY | 0.724177 |
| GLSLLAVAV | ASSRSRQGNSQNTLY | 0.7215282 |
| LEQVHQL | ASSRSRQGNSQNTLY | 0.79301006 |
| LNAAGAI | ASSRSRQGNSQNTLY | 0.63815045 |
| PAVHASL | ASSRSRQGNSQNTLY | 0.70958865 |
| TELEDDA | ASSRSRQGNSQNTLY | 0.7594393 |
| TPPGPPN | ASSRSRQGNSQNTLY | 0.77689815 |
| TPPPPPPP | ASSRSRQGNSQNTLY | 0.6962677 |
| TQSPALI | ASSQEGGNSYNSPLY | 0.7884044 |
| EFAAAFPQ | ASSQEGGNSYNSPLY | 0.6258099 |
| LLSGNLDVL | ASSQEGGNSYNSPLY | 0.78532815 |
| GPANLGNYNI | ASSQEGGNSYNSPLY | 0.7004821 |
| TLGPTLP | ASSQEGGNSYNSPLY | 0.7768379 |
| VYAYSSCCK | ASSQEGGNSYNSPLY | 0.73268074 |
| APAPAPAPNI | ASSQEGGNSYNSPLY | 0.78760964 |
| PGGPVPIP | ASSQEGGNSYNSPLY | 0.7857036 |
| FLTVALEP | ASSQEGGNSYNSPLY | 0.7853893 |
| SQYFGANL | ASSQEGGNSYNSPLY | 0.22469643 |
| GGVKPAA | ASSQEGGNSYNSPLY | 0.7898823 |
| GGTGGAGGAGGW | ASSQEGGNSYNSPLY | 0.77866024 |
| GVGGVAGLGGAGG | ASSQEGGNSYNSPLY | 0.7929348 |
| VLLVLATTI | ASSQEGGNSYNSPLY | 0.6844063 |
| IGIFAAML | ASSQEGGNSYNSPLY | 0.7500951 |
| AVGDLID | ASSQEGGNSYNSPLY | 0.7165737 |
| NISPLPL | ASSQEGGNSYNSPLY | 0.70717955 |
| VTAVFVAI | ASSQEGGNSYNSPLY | 0.6591925 |
| SGDDNDAAL | ASSQEGGNSYNSPLY | 0.6833577 |
| AVAALAHS | ASSQEGGNSYNSPLY | 0.75446707 |
| GGVTVMATP | ASSQEGGNSYNSPLY | 0.7291355 |
| AGLAANVRD | ASSQEGGNSYNSPLY | 0.7782141 |
| TNRPDLIDPALLRPG | ASSQEGGNSYNSPLY | 0.7793791 |
| LANLGVTAL | ASSQEGGNSYNSPLY | 0.7792225 |
| TGERIEL | ASSQEGGNSYNSPLY | 0.78967065 |
| NGEELAA | ASSQEGGNSYNSPLY | 0.78542495 |
| VLSLVWG | ASSQEGGNSYNSPLY | 0.773331 |
| IGYALWL | ASSQEGGNSYNSPLY | 0.6540603 |
| LSLLLAPL | ASSQEGGNSYNSPLY | 0.72331494 |
| TLDEALK | ASSQEGGNSYNSPLY | 0.69063663 |
| TDGAGTAEGG | ASSQEGGNSYNSPLY | 0.70603627 |
| NAPGGAGGNGG | ASSQEGGNSYNSPLY | 0.68768275 |
| VSVFMGMGGNF | ASSQEGGNSYNSPLY | 0.7669822 |
| LASYLAAL | ASSQEGGNSYNSPLY | 0.1918461 |

| | | |
|---|---|---|
| LVGGTPAL | ASSQEGGNSYNSPLY | 0.74973977 |
| VRNADVL | ASSQEGGNSYNSPLY | 0.7907707 |
| TCSSPFLVSAM | ASSQEGGNSYNSPLY | 0.66980207 |
| VHITDAE | ASSQEGGNSYNSPLY | 0.7344992 |
| EYAAPSF | ASSQEGGNSYNSPLY | 0.7338007 |
| VVDAFHGGL | ASSQEGGNSYNSPLY | 0.6898459 |
| ASVVDPQ | ASSQEGGNSYNSPLY | 0.7845167 |
| SYNEFGGLS | ASSQEGGNSYNSPLY | 0.5505645 |
| DPDGSVY | ASSQEGGNSYNSPLY | 0.76151335 |
| MVSTIHATE | ASSQEGGNSYNSPLY | 0.7878356 |
| LAAGVADPV | ASSQEGGNSYNSPLY | 0.7410898 |
| TKEQIAA | ASSQEGGNSYNSPLY | 0.72070634 |
| SLASPAAL | ASSQEGGNSYNSPLY | 0.6973785 |
| IGLVFYGV | ASSQEGGNSYNSPLY | 0.33583763 |
| FLLDASNL | ASSQEGGNSYNSPLY | 0.7552147 |
| YQVGNFP | ASSQEGGNSYNSPLY | 0.18864103 |
| ESLLSQST | ASSQEGGNSYNSPLY | 0.7561791 |
| LFTVGFAAVG | ASSQEGGNSYNSPLY | 0.5316296 |
| ATPPTPT | ASSQEGGNSYNSPLY | 0.77309763 |
| TDLATLG | ASSQEGGNSYNSPLY | 0.7640383 |
| SAGGTGR | ASSQEGGNSYNSPLY | 0.75365984 |
| GFSNSGS | ASSQEGGNSYNSPLY | 0.68388367 |
| GGNGGSGGTSV | ASSQEGGNSYNSPLY | 0.7371881 |
| GTSVATGGAGN | ASSQEGGNSYNSPLY | 0.54841536 |
| NGGNGGSGGTS | ASSQEGGNSYNSPLY | 0.46075535 |
| AVVKFPH | ASSQEGGNSYNSPLY | 0.77907664 |
| TPPPPPP | ASSQEGGNSYNSPLY | 0.66280556 |
| AAPAAAAAPA | ASSQEGGNSYNSPLY | 0.7133764 |
| IALNEDL | ASSQEGGNSYNSPLY | 0.77910024 |
| IVNVSSM | ASSQEGGNSYNSPLY | 0.7387336 |
| TGIPPAPR | ASSQEGGNSYNSPLY | 0.40366358 |
| FDLGGGTF | ASSQEGGNSYNSPLY | 0.76187503 |
| FDLGGGTFDVS | ASSQEGGNSYNSPLY | 0.73777306 |
| NEPTAAA | ASSQEGGNSYNSPLY | 0.71396804 |
| NEPTAAALA | ASSQEGGNSYNSPLY | 0.18859823 |
| GSIINISSI | ASSQEGGNSYNSPLY | 0.7522716 |
| VAPGNAGTA | ASSQEGGNSYNSPLY | 0.6365912 |
| TGGGASLE | ASSQEGGNSYNSPLY | 0.6935029 |
| VAVYNHYKRI | ASSQEGGNSYNSPLY | 0.6910223 |
| TDPAPATT | ASSQEGGNSYNSPLY | 0.78442687 |
| TNRPDILDPALLRPG | ASSQEGGNSYNSPLY | 0.26028144 |
| LTTTTTS | ASSQEGGNSYNSPLY | 0.74023044 |
| PTGSGKTL | ASSQEGGNSYNSPLY | 0.7792894 |
| TGSGKTL | ASSQEGGNSYNSPLY | 0.7667314 |
| TIGHVDHGK | ASSQEGGNSYNSPLY | 0.78994614 |
| TIGHVDHGKTT | ASSQEGGNSYNSPLY | 0.7411952 |
| GHVDHGKTT | ASSQEGGNSYNSPLY | 0.5889042 |
| AQFEHTLL | ASSQEGGNSYNSPLY | 0.7174978 |
| IVNQDEL | ASSQEGGNSYNSPLY | 0.66526586 |
| TNGSQFF | ASSQEGGNSYNSPLY | 0.3561306 |
| TGKGIVE | ASSQEGGNSYNSPLY | 0.28913587 |

| | | |
|---|---|---|
| LISNASDA | ASSQEGGNSYNSPLY | 0.7668276 |
| AELRTLGELTKIAKAHGAQ | ASSQEGGNSYNSPLY | 0.2454448 |
| APNAVAAAEPSW | ASSQEGGNSYNSPLY | 0.7374065 |
| APQVAGAATAVL | ASSQEGGNSYNSPLY | 0.7779217 |
| ARRDPAAAAAILPTDA | ASSQEGGNSYNSPLY | 0.72468853 |
| DAAARDVLAEAGLAEYF | ASSQEGGNSYNSPLY | 0.44032988 |
| DDQIVLSAKANLAKVEQ | ASSQEGGNSYNSPLY | 0.7224681 |
| DELLPDTGEDSDEDGH | ASSQEGGNSYNSPLY | 0.7350013 |
| ELPEAPANRATVPQD | ASSQEGGNSYNSPLY | 0.7796759 |
| FGQSKIERVTGIG | ASSQEGGNSYNSPLY | 0.73693097 |
| GAGGTGSTAGGAGGAGGAGGLY | ASSQEGGNSYNSPLY | 0.78603494 |
| GHGGTGAAGGNGGTGGAGGSGI | ASSQEGGNSYNSPLY | 0.7851957 |
| GNGGNGGAGGEGGPGGAGGAGA | ASSQEGGNSYNSPLY | 0.7346294 |
| GSTAGNGGNGGNATNGTIAGQP | ASSQEGGNSYNSPLY | 0.77810454 |
| GTGGDGGNGGNGGTGG | ASSQEGGNSYNSPLY | 0.6998204 |
| GVRRPAGIPADAPV | ASSQEGGNSYNSPLY | 0.70598614 |
| GYRQVTDAHLLTLARR | ASSQEGGNSYNSPLY | 0.6915591 |
| IAKMGQDGHDRG | ASSQEGGNSYNSPLY | 0.7505169 |
| KAEPSIEIQYT | ASSQEGGNSYNSPLY | 0.6978783 |
| KPVRIRMAADEDTVAVT | ASSQEGGNSYNSPLY | 0.61431485 |
| MPPRIAGMRLLVIKPEP | ASSQEGGNSYNSPLY | 0.37998426 |
| MKFYAVALFRSRGVPPTR | ASSQEGGNSYNSPLY | 0.75550884 |
| NAGGFGILGAVAHSP | ASSQEGGNSYNSPLY | 0.7331239 |
| NEGREITLAMAEM | ASSQEGGNSYNSPLY | 0.34937876 |
| NELVEWSHPLGHRWRWE | ASSQEGGNSYNSPLY | 0.6637504 |
| NLGAGGTGGNGGNPG | ASSQEGGNSYNSPLY | 0.7941598 |
| NTEWPKWETVMAADPVD | ASSQEGGNSYNSPLY | 0.78833985 |
| PASAAPAETQP | ASSQEGGNSYNSPLY | 0.698665 |
| QQSSDGKNGVRAAEITGE | ASSQEGGNSYNSPLY | 0.5933677 |
| RAPDAVTTSWWKE | ASSQEGGNSYNSPLY | 0.7581076 |
| SGFGNVGAHTSGWFNQ | ASSQEGGNSYNSPLY | 0.29056868 |
| SGVAGAVGQAASVGGLKV | ASSQEGGNSYNSPLY | 0.7496964 |
| SNGGAGTGSAGGNGGT | ASSQEGGNSYNSPLY | 0.76554525 |
| TGGAGGEGGAAG | ASSQEGGNSYNSPLY | 0.7567018 |
| TGGDGGAAGAGGAPGHG | ASSQEGGNSYNSPLY | 0.5335434 |
| TMPPLLAAIANAGRHGV | ASSQEGGNSYNSPLY | 0.75156325 |
| TSSTAGNSGGA | ASSQEGGNSYNSPLY | 0.71259284 |
| VPATVPVGILC | ASSQEGGNSYNSPLY | 0.74872136 |
| VQRPDSGGVVLTGRIS | ASSQEGGNSYNSPLY | 0.21355876 |
| WTPNPVPATKGVPAGLVQAT | ASSQEGGNSYNSPLY | 0.7567433 |
| YALKRALAKFSDRFDVVII | ASSQEGGNSYNSPLY | 0.6751455 |
| YLIGSVIV | ASSQEGGNSYNSPLY | 0.5568179 |
| YGQLVAVID | ASSQEGGNSYNSPLY | 0.35371545 |
| VTVGGMTLPA | ASSQEGGNSYNSPLY | 0.7110218 |
| VLALPAV | ASSQEGGNSYNSPLY | 0.78574544 |
| VIPPTLNYET | ASSQEGGNSYNSPLY | 0.7829454 |
| VGMGGMHPSSG | ASSQEGGNSYNSPLY | 0.50383306 |
| VGLAGGAATA | ASSQEGGNSYNSPLY | 0.78820354 |
| VGGAGGAGGSGP | ASSQEGGNSYNSPLY | 0.7910953 |
| VFAAFMV | ASSQEGGNSYNSPLY | 0.69641185 |
| TLYDKLR | ASSQEGGNSYNSPLY | 0.32668528 |

| | | |
|---|---|---|
| TLAVFTVL | ASSQEGGNSYNSPLY | 0.7683365 |
| TGSVQPARV | ASSQEGGNSYNSPLY | 0.70446694 |
| TGGGAGSGGT | ASSQEGGNSYNSPLY | 0.7767224 |
| SHLETSL | ASSQEGGNSYNSPLY | 0.7441652 |
| SFTTGMT | ASSQEGGNSYNSPLY | 0.6579139 |
| QVEVAKRMGVRQPTVS | ASSQEGGNSYNSPLY | 0.7689029 |
| QGSGGPALAM | ASSQEGGNSYNSPLY | 0.7797494 |
| MLAPSNP | ASSQEGGNSYNSPLY | 0.669726 |
| LVNNAAI | ASSQEGGNSYNSPLY | 0.61371136 |
| LTGEINTLA | ASSQEGGNSYNSPLY | 0.5446766 |
| LSIFGPV | ASSQEGGNSYNSPLY | 0.2915644 |
| LLALLANL | ASSQEGGNSYNSPLY | 0.7543148 |
| LISNASDALDKLR | ASSQEGGNSYNSPLY | 0.6989015 |
| LINEAVP | ASSQEGGNSYNSPLY | 0.6470624 |
| LGYAFSGL | ASSQEGGNSYNSPLY | 0.7729653 |
| LDFVAVAL | ASSQEGGNSYNSPLY | 0.61512434 |
| LAQPGAE | ASSQEGGNSYNSPLY | 0.64477456 |
| ISNLGAVTGF | ASSQEGGNSYNSPLY | 0.66975063 |
| HQEDPGRI | ASSQEGGNSYNSPLY | 0.7544206 |
| GTPGEPTP | ASSQEGGNSYNSPLY | 0.7667345 |
| GFFNSGS | ASSQEGGNSYNSPLY | 0.7565826 |
| GAGGGGLTAGAN | ASSQEGGNSYNSPLY | 0.7176348 |
| FVDALVNM | ASSQEGGNSYNSPLY | 0.7919395 |
| FLTVAPEL | ASSQEGGNSYNSPLY | 0.58659935 |
| FETLTAL | ASSQEGGNSYNSPLY | 0.62159103 |
| DVINPIEL | ASSQEGGNSYNSPLY | 0.77383965 |
| APVSGAPV | ASSQEGGNSYNSPLY | 0.39336252 |
| ALVGGLAW | ASSQEGGNSYNSPLY | 0.7782818 |
| ALAAAAATAI | ASSQEGGNSYNSPLY | 0.75499743 |
| AFGLGLLL | ASSQEGGNSYNSPLY | 0.7627115 |
| AFFAAASL | ASSQEGGNSYNSPLY | 0.7280817 |
| AAAALPAFSPP | ASSQEGGNSYNSPLY | 0.7935396 |
| VVLAPAL | ASSQEGGNSYNSPLY | 0.7862453 |
| TIGHVDHGKTTL | ASSQEGGNSYNSPLY | 0.7165165 |
| LISNASDALDK | ASSQEGGNSYNSPLY | 0.74162364 |
| LISNASDAL | ASSQEGGNSYNSPLY | 0.44702613 |
| ISNASDALDK | ASSQEGGNSYNSPLY | 0.784461 |
| NQAETLVYQTE | ASSQEGGNSYNSPLY | 0.7542855 |
| TGGTGGTAGAG | ASSQEGGNSYNSPLY | 0.67413515 |
| TGSTAGGAGGA | ASSQEGGNSYNSPLY | 0.7630503 |
| DPLAVPVAAGPV | ASSQEGGNSYNSPLY | 0.78100795 |
| GGTGGTPGNGGN | ASSQEGGNSYNSPLY | 0.7875416 |
| PAGLAGVDPATL | ASSQEGGNSYNSPLY | 0.2728649 |
| GNNLVVEAPHAGASV | ASSQEGGNSYNSPLY | 0.75136065 |
| KVTRIMSASTLEQPA | ASSQEGGNSYNSPLY | 0.24391292 |
| LGFALPIALPRRTVV | ASSQEGGNSYNSPLY | 0.4806142 |
| TRLGNLTARKLVVLV | ASSQEGGNSYNSPLY | 0.471613 |
| WSQQRRLIYFPSAGP | ASSQEGGNSYNSPLY | 0.7635939 |
| GPNGNPDPMAAAVDIR | ASSQEGGNSYNSPLY | 0.7193778 |
| TQRCVSVADLRAAASR | ASSQEGGNSYNSPLY | 0.7482155 |
| QFVLDVAEHPVEVLTETV | ASSQEGGNSYNSPLY | 0.76038694 |

| | | |
|---|---|---|
| GGAGGTGGAGGNGGTQAGDG | ASSQEGGNSYNSPLY | 0.7884095 |
| PLLATLRAIAVDVDAKPAQV | ASSQEGGNSYNSPLY | 0.6343894 |
| APKFRTGGYMIDFWGVGYQVAK | ASSQEGGNSYNSPLY | 0.7103343 |
| GAGGAGGAGGAAGGPGGTGGTG | ASSQEGGNSYNSPLY | 0.38077828 |
| IGALGLLATDVAPAGSVAFGVA | ASSQEGGNSYNSPLY | 0.2761985 |
| AGGAGGIGGNGGAGGAGGVLMG | ASSQEGGNSYNSPLY | 0.6385603 |
| ADELIGLL | ASSQEGGNSYNSPLY | 0.7196251 |
| ADRALKA | ASSQEGGNSYNSPLY | 0.23919168 |
| AGAVGQVGGAGGA | ASSQEGGNSYNSPLY | 0.77936375 |
| AGGGGGLGGAP | ASSQEGGNSYNSPLY | 0.77876866 |
| AGGLLYGNGGAG | ASSQEGGNSYNSPLY | 0.77138597 |
| AGLESIGL | ASSQEGGNSYNSPLY | 0.555884 |
| AGLKTGII | ASSQEGGNSYNSPLY | 0.7190982 |
| ASLAALLL | ASSQEGGNSYNSPLY | 0.6991247 |
| ASVINISSI | ASSQEGGNSYNSPLY | 0.7797326 |
| ATPPGPA | ASSQEGGNSYNSPLY | 0.7078061 |
| ATVASALV | ASSQEGGNSYNSPLY | 0.46300483 |
| ATVASLIL | ASSQEGGNSYNSPLY | 0.6049689 |
| AVNGPVLIRH | ASSQEGGNSYNSPLY | 0.6488364 |
| DGAAGAVGI | ASSQEGGNSYNSPLY | 0.7848981 |
| DPLLHGI | ASSQEGGNSYNSPLY | 0.7776069 |
| ELGAALGAL | ASSQEGGNSYNSPLY | 0.77790105 |
| ELGRTQAV | ASSQEGGNSYNSPLY | 0.7753147 |
| FVAGGYLS | ASSQEGGNSYNSPLY | 0.27669567 |
| GASGGAGGAGGW | ASSQEGGNSYNSPLY | 0.6303722 |
| GGAGAGGDGGAGGI | ASSQEGGNSYNSPLY | 0.7896674 |
| GTGHNGSQPGMGGNG | ASSQEGGNSYNSPLY | 0.7536989 |
| GYAAASASAS | ASSQEGGNSYNSPLY | 0.7259666 |
| HQVDVAAVQQG | ASSQEGGNSYNSPLY | 0.7893915 |
| IGAAIAHT | ASSQEGGNSYNSPLY | 0.6884149 |
| IGMIEVAQL | ASSQEGGNSYNSPLY | 0.6958769 |
| IRLAPEHTSAMD | ASSQEGGNSYNSPLY | 0.7523382 |
| IYNVSVL | ASSQEGGNSYNSPLY | 0.74173653 |
| KQGFLGM | ASSQEGGNSYNSPLY | 0.73381364 |
| LESVITQL | ASSQEGGNSYNSPLY | 0.62733525 |
| LGLGALVVV | ASSQEGGNSYNSPLY | 0.7320198 |
| LIGNGGNGGSGGI | ASSQEGGNSYNSPLY | 0.7944006 |
| LNAINAPTL | ASSQEGGNSYNSPLY | 0.764009 |
| LQLLNTGV | ASSQEGGNSYNSPLY | 0.78380287 |
| LQTSIEF | ASSQEGGNSYNSPLY | 0.77009964 |
| MSFAVSEL | ASSQEGGNSYNSPLY | 0.7490091 |
| NTGALGL | ASSQEGGNSYNSPLY | 0.6903571 |
| PPGAGGAGGAGTT | ASSQEGGNSYNSPLY | 0.73746645 |
| QLVGLGI | ASSQEGGNSYNSPLY | 0.73634124 |
| RFLEQVHQL | ASSQEGGNSYNSPLY | 0.71864676 |
| RGVGYVLR | ASSQEGGNSYNSPLY | 0.6798186 |
| SATGVFAGVI | ASSQEGGNSYNSPLY | 0.6744375 |
| SLPAITLP | ASSQEGGNSYNSPLY | 0.7954731 |
| STALAFTV | ASSQEGGNSYNSPLY | 0.7620788 |
| SVLTVGLGK | ASSQEGGNSYNSPLY | 0.77087486 |
| TALAGAGSGI | ASSQEGGNSYNSPLY | 0.6973864 |

| | | |
|---|---|---|
| TGLPFTGI | ASSQEGGNSYNSPLY | 0.7370033 |
| VAEHVTV | ASSQEGGNSYNSPLY | 0.77367854 |
| VDIVLNSL | ASSQEGGNSYNSPLY | 0.441337 |
| VLTGGGAVL | ASSQEGGNSYNSPLY | 0.7870151 |
| VNVGLAYGI | ASSQEGGNSYNSPLY | 0.67942405 |
| VSSAAVGAV | ASSQEGGNSYNSPLY | 0.66209304 |
| VTSVYEAGV | ASSQEGGNSYNSPLY | 0.7636043 |
| WVALDRGA | ASSQEGGNSYNSPLY | 0.77869844 |
| YLVGSVVI | ASSQEGGNSYNSPLY | 0.6625155 |
| GPPGTGKTLL | ASSQEGGNSYNSPLY | 0.6776022 |
| GVGMTKF | ASSQEGGNSYNSPLY | 0.69884384 |
| IGIGGSDL | ASSQEGGNSYNSPLY | 0.769691 |
| IGTLLAEL | ASSQEGGNSYNSPLY | 0.7899549 |
| IYTSGTT | ASSQEGGNSYNSPLY | 0.6601901 |
| LSFPAGV | ASSQEGGNSYNSPLY | 0.78654826 |
| LYTSGTT | ASSQEGGNSYNSPLY | 0.73185486 |
| VGAFPFGF | ASSQEGGNSYNSPLY | 0.34594202 |
| VGAGTVEF | ASSQEGGNSYNSPLY | 0.7166759 |
| VGKSTLLL | ASSQEGGNSYNSPLY | 0.78129756 |
| AALAHAGGGIV | ASSQEGGNSYNSPLY | 0.7804848 |
| AALVAVSLL | ASSQEGGNSYNSPLY | 0.79220176 |
| AAVSRAVAAL | ASSQEGGNSYNSPLY | 0.7552359 |
| AGGSSGIGGF | ASSQEGGNSYNSPLY | 0.706844 |
| AGLTVANANG | ASSQEGGNSYNSPLY | 0.78822577 |
| AGLVGAVL | ASSQEGGNSYNSPLY | 0.78644514 |
| ALSQAFTV | ASSQEGGNSYNSPLY | 0.7828748 |
| ALTFSVSL | ASSQEGGNSYNSPLY | 0.6541772 |
| ALTGAALVAL | ASSQEGGNSYNSPLY | 0.47451335 |
| ASFSIGAGVL | ASSQEGGNSYNSPLY | 0.71881664 |
| DGSVYKI | ASSQEGGNSYNSPLY | 0.78647894 |
| EQLAEIV | ASSQEGGNSYNSPLY | 0.51336944 |
| FAVSELTVM | ASSQEGGNSYNSPLY | 0.7426946 |
| GAAPLAAGL | ASSQEGGNSYNSPLY | 0.7688192 |
| GADGGDGGSGGAGG | ASSQEGGNSYNSPLY | 0.43544328 |
| GFGLDTGGAGG | ASSQEGGNSYNSPLY | 0.76185095 |
| GLASGGLL | ASSQEGGNSYNSPLY | 0.79005253 |
| GRIAGAAVLL | ASSQEGGNSYNSPLY | 0.6816389 |
| GSAGSLNTSF | ASSQEGGNSYNSPLY | 0.73905396 |
| GTGQIGI | ASSQEGGNSYNSPLY | 0.7830659 |
| GTLIALIL | ASSQEGGNSYNSPLY | 0.7963735 |
| IAATMGFVVG | ASSQEGGNSYNSPLY | 0.786922 |
| IFAQALL | ASSQEGGNSYNSPLY | 0.38324997 |
| IGAAVALGF | ASSQEGGNSYNSPLY | 0.3006248 |
| IGLTPLAL | ASSQEGGNSYNSPLY | 0.6574327 |
| ILGSNTSTL | ASSQEGGNSYNSPLY | 0.7598989 |
| IVFSLTVL | ASSQEGGNSYNSPLY | 0.66853213 |
| LASKATGF | ASSQEGGNSYNSPLY | 0.711079 |
| LASVALIL | ASSQEGGNSYNSPLY | 0.77657306 |
| LFAVGLTL | ASSQEGGNSYNSPLY | 0.7847283 |
| LFNGGIGTYI | ASSQEGGNSYNSPLY | 0.58952975 |
| LTLGAGII | ASSQEGGNSYNSPLY | 0.40549234 |

| | | |
|---|---|---|
| LVAAALGDI | ASSQEGGNSYNSPLY | 0.7559367 |
| NTAFVQLGI | ASSQEGGNSYNSPLY | 0.7691163 |
| QLFSGAEL | ASSQEGGNSYNSPLY | 0.7092465 |
| SAATFITL | ASSQEGGNSYNSPLY | 0.7643571 |
| SNYGLANLGV | ASSQEGGNSYNSPLY | 0.51495755 |
| SYGGAYAVM | ASSQEGGNSYNSPLY | 0.25711003 |
| TTGVGGLTLG | ASSQEGGNSYNSPLY | 0.77396697 |
| TTQTIYI | ASSQEGGNSYNSPLY | 0.6404117 |
| VAAGAGLGAVI | ASSQEGGNSYNSPLY | 0.7917569 |
| VAAVVTLL | ASSQEGGNSYNSPLY | 0.70586777 |
| VGFLGGLI | ASSQEGGNSYNSPLY | 0.7710276 |
| VGVIPILL | ASSQEGGNSYNSPLY | 0.77180386 |
| VIFASSLI | ASSQEGGNSYNSPLY | 0.7486117 |
| VKGGDGGAAAG | ASSQEGGNSYNSPLY | 0.7282211 |
| VTKTFTNI | ASSQEGGNSYNSPLY | 0.6591042 |
| VVAAAILL | ASSQEGGNSYNSPLY | 0.57426906 |
| YAALDVEL | ASSQEGGNSYNSPLY | 0.34028226 |
| LASVALLL | ASSQEGGNSYNSPLY | 0.78266776 |
| LDAVHRV | ASSQEGGNSYNSPLY | 0.20128791 |
| QGLSSSIF | ASSQEGGNSYNSPLY | 0.7857131 |
| RGTFANIRL | ASSQEGGNSYNSPLY | 0.7782563 |
| RQLWWGHRI | ASSQEGGNSYNSPLY | 0.64255697 |
| YGPPGTGKTLL | ASSQEGGNSYNSPLY | 0.75748277 |
| AALVGAPANVA | ASSQEGGNSYNSPLY | 0.72876084 |
| AATFRRTE | ASSQEGGNSYNSPLY | 0.53392595 |
| AGALGGGI | ASSQEGGNSYNSPLY | 0.77297294 |
| AGGAGGAGGIGAG | ASSQEGGNSYNSPLY | 0.7700251 |
| AGSGALGL | ASSQEGGNSYNSPLY | 0.6830689 |
| AQIADQI | ASSQEGGNSYNSPLY | 0.7381631 |
| ASAVAVTYL | ASSQEGGNSYNSPLY | 0.7567755 |
| ASTVLGGTGG | ASSQEGGNSYNSPLY | 0.6207422 |
| AVGGEPAK | ASSQEGGNSYNSPLY | 0.6984006 |
| CEQGTFGGTC | ASSQEGGNSYNSPLY | 0.78824437 |
| DEGKAPAG | ASSQEGGNSYNSPLY | 0.71952534 |
| DTVVFPAL | ASSQEGGNSYNSPLY | 0.721501 |
| FEGITMAG | ASSQEGGNSYNSPLY | 0.7661295 |
| FPLVVQD | ASSQEGGNSYNSPLY | 0.5617387 |
| GAFTTPQ | ASSQEGGNSYNSPLY | 0.7824562 |
| GFGGFGTAGGVGGA | ASSQEGGNSYNSPLY | 0.71254 |
| GFLAQFL | ASSQEGGNSYNSPLY | 0.66564614 |
| GGAGGAGGNAGVA | ASSQEGGNSYNSPLY | 0.7662397 |
| GGVGTAGAGGAGGAGG | ASSQEGGNSYNSPLY | 0.4045602 |
| GNGGGGDAGNAGSGGN | ASSQEGGNSYNSPLY | 0.6556636 |
| GTGGAGGAGNDAGS | ASSQEGGNSYNSPLY | 0.68556154 |
| GTTNTGWFNS | ASSQEGGNSYNSPLY | 0.7430637 |
| IFMGPLFTL | ASSQEGGNSYNSPLY | 0.7835845 |
| ILAHAIEAI | ASSQEGGNSYNSPLY | 0.68525195 |
| IRSAIREAL | ASSQEGGNSYNSPLY | 0.75161564 |
| ITFAGGG | ASSQEGGNSYNSPLY | 0.61651444 |
| ITGMSGHQEV | ASSQEGGNSYNSPLY | 0.5569381 |
| IVAGLACTL | ASSQEGGNSYNSPLY | 0.66679376 |

| | | |
|---|---|---|
| IVGGGLAGA | ASSQEGGNSYNSPLY | 0.64617455 |
| IVTGVLNI | ASSQEGGNSYNSPLY | 0.69149107 |
| LAQQGCI | ASSQEGGNSYNSPLY | 0.75733757 |
| LCLHVEN | ASSQEGGNSYNSPLY | 0.6483889 |
| LDAISDGS | ASSQEGGNSYNSPLY | 0.6964831 |
| LGGFAAML | ASSQEGGNSYNSPLY | 0.78445613 |
| LGGKNPMI | ASSQEGGNSYNSPLY | 0.7332495 |
| LGPKGVAAIL | ASSQEGGNSYNSPLY | 0.7676935 |
| LMVGVSI | ASSQEGGNSYNSPLY | 0.77855873 |
| LVGAAFVL | ASSQEGGNSYNSPLY | 0.7723543 |
| LVLAVAPL | ASSQEGGNSYNSPLY | 0.68622756 |
| MAAIAVIDAA | ASSQEGGNSYNSPLY | 0.7196372 |
| NNGINAELQ | ASSQEGGNSYNSPLY | 0.7914969 |
| RGEPLVLVHG | ASSQEGGNSYNSPLY | 0.77306247 |
| RGGAGGAAGLI | ASSQEGGNSYNSPLY | 0.75525343 |
| RSYRMHPAV | ASSQEGGNSYNSPLY | 0.772174 |
| SAAHLLAGQLA | ASSQEGGNSYNSPLY | 0.76523423 |
| SAAVGSY | ASSQEGGNSYNSPLY | 0.7743681 |
| SPHAVLHRI | ASSQEGGNSYNSPLY | 0.74925923 |
| SQIAKVL | ASSQEGGNSYNSPLY | 0.47701582 |
| TALGHLVP | ASSQEGGNSYNSPLY | 0.68456936 |
| TAPTTSGAA | ASSQEGGNSYNSPLY | 0.74614346 |
| TDGDDGNGGSPGT | ASSQEGGNSYNSPLY | 0.67795783 |
| TDGLGEN | ASSQEGGNSYNSPLY | 0.35097915 |
| TGGPPLPGQA | ASSQEGGNSYNSPLY | 0.7868292 |
| TIPAITI | ASSQEGGNSYNSPLY | 0.7953817 |
| TIPGFTI | ASSQEGGNSYNSPLY | 0.55168295 |
| TSIASGALA | ASSQEGGNSYNSPLY | 0.78868914 |
| TVTLPGAAI | ASSQEGGNSYNSPLY | 0.52823275 |
| VASGLSAALP | ASSQEGGNSYNSPLY | 0.7617357 |
| VGAGVIGI | ASSQEGGNSYNSPLY | 0.7848705 |
| VGPKRFGV | ASSQEGGNSYNSPLY | 0.67171013 |
| VNVAFPD | ASSQEGGNSYNSPLY | 0.761302 |
| VSMTNSGVS | ASSQEGGNSYNSPLY | 0.66329944 |
| VTDVLGDLSV | ASSQEGGNSYNSPLY | 0.7433846 |
| VVVVGAGFAGL | ASSQEGGNSYNSPLY | 0.78987336 |
| VYTVGTVV | ASSQEGGNSYNSPLY | 0.6646869 |
| WTTFSGP | ASSQEGGNSYNSPLY | 0.71455884 |
| YDELVEAF | ASSQEGGNSYNSPLY | 0.73076093 |
| YGTGGAGGNGG | ASSQEGGNSYNSPLY | 0.58564276 |
| YTNVMAV | ASSQEGGNSYNSPLY | 0.73752284 |
| FLEQVHQL | ASSQEGGNSYNSPLY | 0.7829777 |
| GLSLLAVAV | ASSQEGGNSYNSPLY | 0.56902987 |
| LEQVHQL | ASSQEGGNSYNSPLY | 0.7851886 |
| LNAAGAI | ASSQEGGNSYNSPLY | 0.6777663 |
| PAVHASL | ASSQEGGNSYNSPLY | 0.7954911 |
| TELEDDA | ASSQEGGNSYNSPLY | 0.7794192 |
| TPPGPPN | ASSQEGGNSYNSPLY | 0.7728672 |
| TPPPPPPP | ASSQEGGNSYNSPLY | 0.66854143 |
| TQSPALI | AWSRTGVNQDTQY | 0.7917584 |
| EFAAAFPQ | AWSRTGVNQDTQY | 0.72751296 |

| | | |
|---|---|---|
| LLSGNLDVL | AWSRTGVNQDTQY | 0.7583077 |
| GPANLGNYNI | AWSRTGVNQDTQY | 0.78570056 |
| TLGPTLP | AWSRTGVNQDTQY | 0.5633472 |
| VYAYSSCCK | AWSRTGVNQDTQY | 0.8771636 |
| APAPAPAPNI | AWSRTGVNQDTQY | 0.7936882 |
| PGGPVPIP | AWSRTGVNQDTQY | 0.78877497 |
| FLTVALEP | AWSRTGVNQDTQY | 0.67266434 |
| SQYFGANL | AWSRTGVNQDTQY | 0.43825096 |
| GGVKPAA | AWSRTGVNQDTQY | 0.7347398 |
| GGTGGAGGAGGW | AWSRTGVNQDTQY | 0.5587428 |
| GVGGVAGLGGAGG | AWSRTGVNQDTQY | 0.79213893 |
| VLLVLATTI | AWSRTGVNQDTQY | 0.8527374 |
| IGIFAAML | AWSRTGVNQDTQY | 0.64983356 |
| AVGDLID | AWSRTGVNQDTQY | 0.862279 |
| NISPLPL | AWSRTGVNQDTQY | 0.85687995 |
| VTAVFVAI | AWSRTGVNQDTQY | 0.38766 |
| SGDDNDAAL | AWSRTGVNQDTQY | 0.7344569 |
| AVAALAHS | AWSRTGVNQDTQY | 0.65908104 |
| GGVTVMATP | AWSRTGVNQDTQY | 0.7248452 |
| AGLAANVRD | AWSRTGVNQDTQY | 0.3011339 |
| TNRPDLIDPALLRPG | AWSRTGVNQDTQY | 0.82800806 |
| LANLGVTAL | AWSRTGVNQDTQY | 0.64918596 |
| TGERIEL | AWSRTGVNQDTQY | 0.14779349 |
| NGEELAA | AWSRTGVNQDTQY | 0.78982365 |
| VLSLVWG | AWSRTGVNQDTQY | 0.7544745 |
| IGYALWL | AWSRTGVNQDTQY | 0.69540167 |
| LSLLLAPL | AWSRTGVNQDTQY | 0.84216106 |
| TLDEALK | AWSRTGVNQDTQY | 0.5827973 |
| TDGAGTAEGG | AWSRTGVNQDTQY | 0.63770664 |
| NAPGGAGGNGG | AWSRTGVNQDTQY | 0.8329162 |
| VSVFMGMGGNF | AWSRTGVNQDTQY | 0.8622478 |
| LASYLAAL | AWSRTGVNQDTQY | 0.5245755 |
| LVGGTPAL | AWSRTGVNQDTQY | 0.7872468 |
| VRNADVL | AWSRTGVNQDTQY | 0.85816646 |
| TCSSPFLVSAM | AWSRTGVNQDTQY | 0.78985596 |
| VHITDAE | AWSRTGVNQDTQY | 0.86653924 |
| EYAAPSF | AWSRTGVNQDTQY | 0.8772284 |
| VVDAFHGGL | AWSRTGVNQDTQY | 0.6719535 |
| ASVVDPQ | AWSRTGVNQDTQY | 0.8573462 |
| SYNEFGGLS | AWSRTGVNQDTQY | 0.87590504 |
| DPDGSVY | AWSRTGVNQDTQY | 0.75414443 |
| MVSTIHATE | AWSRTGVNQDTQY | 0.75569713 |
| LAAGVADPV | AWSRTGVNQDTQY | 0.53619146 |
| TKEQIAA | AWSRTGVNQDTQY | 0.7470686 |
| SLASPAAL | AWSRTGVNQDTQY | 0.38875377 |
| IGLVFYGV | AWSRTGVNQDTQY | 0.78547144 |
| FLLDASNL | AWSRTGVNQDTQY | 0.8598783 |
| YQVGNFP | AWSRTGVNQDTQY | 0.69952196 |
| ESLLSQST | AWSRTGVNQDTQY | 0.5283793 |
| LFTVGFAAVG | AWSRTGVNQDTQY | 0.55395764 |
| ATPPTPT | AWSRTGVNQDTQY | 0.79511714 |

| | | |
|---|---|---|
| TDLATLG | AWSRTGVNQDTQY | 0.6575211 |
| SAGGTGR | AWSRTGVNQDTQY | 0.36979234 |
| GFSNSGS | AWSRTGVNQDTQY | 0.7480892 |
| GGNGGSGGTSV | AWSRTGVNQDTQY | 0.8529791 |
| GTSVATGGAGN | AWSRTGVNQDTQY | 0.8494096 |
| NGGNGGSGGTS | AWSRTGVNQDTQY | 0.44195116 |
| AVVKFPH | AWSRTGVNQDTQY | 0.87825733 |
| TPPPPPP | AWSRTGVNQDTQY | 0.82700145 |
| AAPAAAAAPA | AWSRTGVNQDTQY | 0.87097 |
| IALNEDL | AWSRTGVNQDTQY | 0.78043216 |
| IVNVSSM | AWSRTGVNQDTQY | 0.58691883 |
| TGIPPAPR | AWSRTGVNQDTQY | 0.8419714 |
| FDLGGGTF | AWSRTGVNQDTQY | 0.69769204 |
| FDLGGGTFDVS | AWSRTGVNQDTQY | 0.671666 |
| NEPTAAA | AWSRTGVNQDTQY | 0.8519647 |
| NEPTAAALA | AWSRTGVNQDTQY | 0.7353756 |
| GSIINISSI | AWSRTGVNQDTQY | 0.63799286 |
| VAPGNAGTA | AWSRTGVNQDTQY | 0.65974915 |
| TGGGASLE | AWSRTGVNQDTQY | 0.86982405 |
| VAVYNHYKRI | AWSRTGVNQDTQY | 0.5720872 |
| TDPAPATT | AWSRTGVNQDTQY | 0.5583977 |
| TNRPDILDPALLRPG | AWSRTGVNQDTQY | 0.6299283 |
| LTTTTTS | AWSRTGVNQDTQY | 0.73420566 |
| PTGSGKTL | AWSRTGVNQDTQY | 0.8602236 |
| TGSGKTL | AWSRTGVNQDTQY | 0.826434 |
| TIGHVDHGK | AWSRTGVNQDTQY | 0.7142211 |
| TIGHVDHGKTT | AWSRTGVNQDTQY | 0.79866505 |
| GHVDHGKTT | AWSRTGVNQDTQY | 0.83483 |
| AQFEHTLL | AWSRTGVNQDTQY | 0.77843344 |
| IVNQDEL | AWSRTGVNQDTQY | 0.8347945 |
| TNGSQFF | AWSRTGVNQDTQY | 0.6348948 |
| TGKGIVE | AWSRTGVNQDTQY | 0.8696947 |
| LISNASDA | AWSRTGVNQDTQY | 0.82915413 |
| AELRTLGELTKIAKAHGAQ | AWSRTGVNQDTQY | 0.8551822 |
| APNAVAAAEPSW | AWSRTGVNQDTQY | 0.86305034 |
| APQVAGAATAVL | AWSRTGVNQDTQY | 0.8232135 |
| ARRDPAAAAAILPTDA | AWSRTGVNQDTQY | 0.79562056 |
| DAAARDVLAEAGLAEYF | AWSRTGVNQDTQY | 0.3314487 |
| DDQIVLSAKANLAKVEQ | AWSRTGVNQDTQY | 0.6747016 |
| DELLPDTGEDSDEDGH | AWSRTGVNQDTQY | 0.7986097 |
| ELPEAPANRATVPQD | AWSRTGVNQDTQY | 0.7978746 |
| FGQSKIERVTGIG | AWSRTGVNQDTQY | 0.83272535 |
| GAGGTGSTAGGAGGAGGAGGLY | AWSRTGVNQDTQY | 0.8209079 |
| GHGGTGAAGGNGGTGGAGGSGI | AWSRTGVNQDTQY | 0.6700165 |
| GNGGNGGAGGEGGPGGAGGAGA | AWSRTGVNQDTQY | 0.86952496 |
| GSTAGNGGNGGNATNGTIAGQP | AWSRTGVNQDTQY | 0.85097986 |
| GTGGDGGNGGNGGTGG | AWSRTGVNQDTQY | 0.40423787 |
| GVRRPAGIPADAPV | AWSRTGVNQDTQY | 0.7165934 |
| GYRQVTDAHLLTLARR | AWSRTGVNQDTQY | 0.8872708 |
| IAKMGQDGHDRG | AWSRTGVNQDTQY | 0.77873456 |
| KAEPSIEIQYT | AWSRTGVNQDTQY | 0.71271735 |

| | | |
|---|---|---|
| KPVRIRMAADEDTVAVT | AWSRTGVNQDTQY | 0.7043897 |
| MPPRIAGMRLLVIKPEP | AWSRTGVNQDTQY | 0.8554116 |
| MKFYAVALFRSRGVPPTR | AWSRTGVNQDTQY | 0.6864755 |
| NAGGFGILGAVAHSP | AWSRTGVNQDTQY | 0.8783729 |
| NEGREITLAMAEM | AWSRTGVNQDTQY | 0.23319283 |
| NELVEWSHPLGHRWRWE | AWSRTGVNQDTQY | 0.72291464 |
| NLGAGGTGGNGGNPG | AWSRTGVNQDTQY | 0.6417927 |
| NTEWPKWETVMAADPVD | AWSRTGVNQDTQY | 0.25533667 |
| PASAAPAETQP | AWSRTGVNQDTQY | 0.8464128 |
| QQSSDGKNGVRAAEITGE | AWSRTGVNQDTQY | 0.6521485 |
| RAPDAVTTSWWKE | AWSRTGVNQDTQY | 0.8301164 |
| SGFGNVGAHTSGWFNQ | AWSRTGVNQDTQY | 0.73938864 |
| SGVAGAVGQAASVGGLKV | AWSRTGVNQDTQY | 0.86341846 |
| SNGGAGTGSAGGNGGT | AWSRTGVNQDTQY | 0.76720953 |
| TGGAGGEGGAAG | AWSRTGVNQDTQY | 0.77867657 |
| TGGDGGAAGAGGAPGHG | AWSRTGVNQDTQY | 0.6051038 |
| TMPPLLAAIANAGRHGV | AWSRTGVNQDTQY | 0.8377522 |
| TSSTAGNSGGA | AWSRTGVNQDTQY | 0.87813425 |
| VPATVPVGILC | AWSRTGVNQDTQY | 0.852684 |
| VQRPDSGGVVLTGRIS | AWSRTGVNQDTQY | 0.40006614 |
| WTPNPVPATKGVPAGLVQAT | AWSRTGVNQDTQY | 0.74123275 |
| YALKRALAKFSDRFDVVII | AWSRTGVNQDTQY | 0.7597572 |
| YLIGSVIV | AWSRTGVNQDTQY | 0.2781798 |
| YGQLVAVID | AWSRTGVNQDTQY | 0.22653753 |
| VTVGGMTLPA | AWSRTGVNQDTQY | 0.75301576 |
| VLALPAV | AWSRTGVNQDTQY | 0.6970992 |
| VIPPTLNYET | AWSRTGVNQDTQY | 0.8438891 |
| VGMGGMHPSSG | AWSRTGVNQDTQY | 0.8302864 |
| VGLAGGAATA | AWSRTGVNQDTQY | 0.7036544 |
| VGGAGGAGGSGP | AWSRTGVNQDTQY | 0.8293394 |
| VFAAFMV | AWSRTGVNQDTQY | 0.77755237 |
| TLYDKLR | AWSRTGVNQDTQY | 0.09986039 |
| TLAVFTVL | AWSRTGVNQDTQY | 0.69446105 |
| TGSVQPARV | AWSRTGVNQDTQY | 0.4019643 |
| TGGGAGSGGT | AWSRTGVNQDTQY | 0.8824222 |
| SHLETSL | AWSRTGVNQDTQY | 0.6090276 |
| SFTTGMT | AWSRTGVNQDTQY | 0.6171584 |
| QVEVAKRMGVRQPTVS | AWSRTGVNQDTQY | 0.73743117 |
| QGSGGPALAM | AWSRTGVNQDTQY | 0.7656017 |
| MLAPSNP | AWSRTGVNQDTQY | 0.75343215 |
| LVNNAAI | AWSRTGVNQDTQY | 0.6236839 |
| LTGEINTLA | AWSRTGVNQDTQY | 0.21296862 |
| LSIFGPV | AWSRTGVNQDTQY | 0.8418463 |
| LLALLANL | AWSRTGVNQDTQY | 0.55173755 |
| LISNASDALDKLR | AWSRTGVNQDTQY | 0.56669194 |
| LINEAVP | AWSRTGVNQDTQY | 0.3845673 |
| LGYAFSGL | AWSRTGVNQDTQY | 0.30702382 |
| LDFVAVAL | AWSRTGVNQDTQY | 0.40173262 |
| LAQPGAE | AWSRTGVNQDTQY | 0.84728783 |
| ISNLGAVTGF | AWSRTGVNQDTQY | 0.71564615 |
| HQEDPGRI | AWSRTGVNQDTQY | 0.71217704 |

| | | |
|---|---|---|
| GTPGEPTP | AWSRTGVNQDTQY | 0.8822827 |
| GFFNSGS | AWSRTGVNQDTQY | 0.7341461 |
| GAGGGGLTAGAN | AWSRTGVNQDTQY | 0.7122566 |
| FVDALVNM | AWSRTGVNQDTQY | 0.67128587 |
| FLTVAPEL | AWSRTGVNQDTQY | 0.5486034 |
| FETLTAL | AWSRTGVNQDTQY | 0.6176399 |
| DVINPIEL | AWSRTGVNQDTQY | 0.635409 |
| APVSGAPV | AWSRTGVNQDTQY | 0.8802893 |
| ALVGGLAW | AWSRTGVNQDTQY | 0.7336636 |
| ALAAAAATAI | AWSRTGVNQDTQY | 0.47401989 |
| AFGLGLLL | AWSRTGVNQDTQY | 0.8005562 |
| AFFAAASL | AWSRTGVNQDTQY | 0.8691714 |
| AAAALPAFSPP | AWSRTGVNQDTQY | 0.88719463 |
| VVLAPAL | AWSRTGVNQDTQY | 0.5624029 |
| TIGHVDHGKTTL | AWSRTGVNQDTQY | 0.74166405 |
| LISNASDALDK | AWSRTGVNQDTQY | 0.7119509 |
| LISNASDAL | AWSRTGVNQDTQY | 0.72904605 |
| ISNASDALDK | AWSRTGVNQDTQY | 0.6905221 |
| NQAETLVYQTE | AWSRTGVNQDTQY | 0.5101973 |
| TGGTGGTAGAG | AWSRTGVNQDTQY | 0.85081816 |
| TGSTAGGAGGA | AWSRTGVNQDTQY | 0.8831003 |
| DPLAVPVAAGPV | AWSRTGVNQDTQY | 0.7166702 |
| GGTGGTPGNGGN | AWSRTGVNQDTQY | 0.37917334 |
| PAGLAGVDPATL | AWSRTGVNQDTQY | 0.7339558 |
| GNNLVVEAPHAGASV | AWSRTGVNQDTQY | 0.6204541 |
| KVTRIMSASTLEQPA | AWSRTGVNQDTQY | 0.623866 |
| LGFALPIALPRRTVV | AWSRTGVNQDTQY | 0.73829997 |
| TRLGNLTARKLVVLV | AWSRTGVNQDTQY | 0.38955033 |
| WSQQRRLIYFPSAGP | AWSRTGVNQDTQY | 0.74901843 |
| GPNGNPDPMAAAVDIR | AWSRTGVNQDTQY | 0.88107383 |
| TQRCVSVADLRAAASR | AWSRTGVNQDTQY | 0.68867826 |
| QFVLDVAEHPVEVLTETV | AWSRTGVNQDTQY | 0.8205386 |
| GGAGGTGGAGGNGGTQAGDG | AWSRTGVNQDTQY | 0.8528273 |
| PLLATLRAIAVDVDAKPAQV | AWSRTGVNQDTQY | 0.81672853 |
| APKFRTGGYMIDFWGVGYQVAK | AWSRTGVNQDTQY | 0.85984635 |
| GAGGAGGAGGAAGGPGGTGGTG | AWSRTGVNQDTQY | 0.8398687 |
| IGALGLLATDVAPAGSVAFGVA | AWSRTGVNQDTQY | 0.78508854 |
| AGGAGGIGGNGGAGGAGGVLMG | AWSRTGVNQDTQY | 0.7942414 |
| ADELIGLL | AWSRTGVNQDTQY | 0.80093163 |
| ADRALKA | AWSRTGVNQDTQY | 0.75180876 |
| AGAVGQVGGAGGA | AWSRTGVNQDTQY | 0.7285464 |
| AGGGGGLGGAP | AWSRTGVNQDTQY | 0.7106669 |
| AGGLLYGNGGAG | AWSRTGVNQDTQY | 0.84327084 |
| AGLESIGL | AWSRTGVNQDTQY | 0.7767242 |
| AGLKTGII | AWSRTGVNQDTQY | 0.8484427 |
| ASLAALLL | AWSRTGVNQDTQY | 0.38805145 |
| ASVINISSI | AWSRTGVNQDTQY | 0.5367141 |
| ATPPGPA | AWSRTGVNQDTQY | 0.84319735 |
| ATVASALV | AWSRTGVNQDTQY | 0.1476072 |
| ATVASLIL | AWSRTGVNQDTQY | 0.2748722 |
| AVNGPVLIRH | AWSRTGVNQDTQY | 0.7753078 |

| | | |
|---|---|---|
| DGAAGAVGI | AWSRTGVNQDTQY | 0.86671793 |
| DPLLHGI | AWSRTGVNQDTQY | 0.16946316 |
| ELGAALGAL | AWSRTGVNQDTQY | 0.5923251 |
| ELGRTQAV | AWSRTGVNQDTQY | 0.51595616 |
| FVAGGYLS | AWSRTGVNQDTQY | 0.8482516 |
| GASGGAGGAGGW | AWSRTGVNQDTQY | 0.7563441 |
| GGAGAGGDGGAGGI | AWSRTGVNQDTQY | 0.40099162 |
| GTGHNGSQPGMGGNG | AWSRTGVNQDTQY | 0.837073 |
| GYAAASASAS | AWSRTGVNQDTQY | 0.33729813 |
| HQVDVAAVQQG | AWSRTGVNQDTQY | 0.6217184 |
| IGAAIAHT | AWSRTGVNQDTQY | 0.85540307 |
| IGMIEVAQL | AWSRTGVNQDTQY | 0.8664233 |
| IRLAPEHTSAMD | AWSRTGVNQDTQY | 0.49147314 |
| IYNVSVL | AWSRTGVNQDTQY | 0.8412852 |
| KQGFLGM | AWSRTGVNQDTQY | 0.66477454 |
| LESVITQL | AWSRTGVNQDTQY | 0.8903374 |
| LGLGALVVV | AWSRTGVNQDTQY | 0.7171769 |
| LIGNGGNGGSGGI | AWSRTGVNQDTQY | 0.4333139 |
| LNAINAPTL | AWSRTGVNQDTQY | 0.8402326 |
| LQLLNTGV | AWSRTGVNQDTQY | 0.83179533 |
| LQTSIEF | AWSRTGVNQDTQY | 0.7446735 |
| MSFAVSEL | AWSRTGVNQDTQY | 0.82400286 |
| NTGALGL | AWSRTGVNQDTQY | 0.7103821 |
| PPGAGGAGGAGTT | AWSRTGVNQDTQY | 0.7154251 |
| QLVGLGI | AWSRTGVNQDTQY | 0.85113144 |
| RFLEQVHQL | AWSRTGVNQDTQY | 0.8492586 |
| RGVGYVLR | AWSRTGVNQDTQY | 0.5706593 |
| SATGVFAGVI | AWSRTGVNQDTQY | 0.8558427 |
| SLPAITLP | AWSRTGVNQDTQY | 0.7323329 |
| STALAFTV | AWSRTGVNQDTQY | 0.58170104 |
| SVLTVGLGK | AWSRTGVNQDTQY | 0.8834539 |
| TALAGAGSGI | AWSRTGVNQDTQY | 0.77399814 |
| TGLPFTGI | AWSRTGVNQDTQY | 0.83031034 |
| VAEHVTV | AWSRTGVNQDTQY | 0.8372946 |
| VDIVLNSL | AWSRTGVNQDTQY | 0.8328843 |
| VLTGGGAVL | AWSRTGVNQDTQY | 0.5115081 |
| VNVGLAYGI | AWSRTGVNQDTQY | 0.81559676 |
| VSSAAVGAV | AWSRTGVNQDTQY | 0.7400151 |
| VTSVYEAGV | AWSRTGVNQDTQY | 0.85235894 |
| WVALDRGA | AWSRTGVNQDTQY | 0.7781676 |
| YLVGSVVI | AWSRTGVNQDTQY | 0.6346653 |
| GPPGTGKTLL | AWSRTGVNQDTQY | 0.8194896 |
| GVGMTKF | AWSRTGVNQDTQY | 0.8638433 |
| IGIGGSDL | AWSRTGVNQDTQY | 0.7976315 |
| IGTLLAEL | AWSRTGVNQDTQY | 0.81263125 |
| IYTSGTT | AWSRTGVNQDTQY | 0.871467 |
| LSFPAGV | AWSRTGVNQDTQY | 0.58027637 |
| LYTSGTT | AWSRTGVNQDTQY | 0.7347249 |
| VGAFPFGF | AWSRTGVNQDTQY | 0.84930515 |
| VGAGTVEF | AWSRTGVNQDTQY | 0.86160326 |
| VGKSTLLL | AWSRTGVNQDTQY | 0.759128 |

| | | |
|---|---|---|
| AALAHAGGGIV | AWSRTGVNQDTQY | 0.8729844 |
| AALVAVSLL | AWSRTGVNQDTQY | 0.6573014 |
| AAVSRAVAAL | AWSRTGVNQDTQY | 0.82296914 |
| AGGSSGIGGF | AWSRTGVNQDTQY | 0.6507858 |
| AGLTVANANG | AWSRTGVNQDTQY | 0.7935575 |
| AGLVGAVL | AWSRTGVNQDTQY | 0.6149613 |
| ALSQAFTV | AWSRTGVNQDTQY | 0.82186675 |
| ALTFSVSL | AWSRTGVNQDTQY | 0.8331424 |
| ALTGAALVAL | AWSRTGVNQDTQY | 0.84131193 |
| ASFSIGAGVL | AWSRTGVNQDTQY | 0.23516494 |
| DGSVYKI | AWSRTGVNQDTQY | 0.829564 |
| EQLAEIV | AWSRTGVNQDTQY | 0.83836526 |
| FAVSELTVM | AWSRTGVNQDTQY | 0.883754 |
| GAAPLAAGL | AWSRTGVNQDTQY | 0.76137674 |
| GADGGDGGSGGAGG | AWSRTGVNQDTQY | 0.8239391 |
| GFGLDTGGAGG | AWSRTGVNQDTQY | 0.691449 |
| GLASGGLL | AWSRTGVNQDTQY | 0.7552183 |
| GRIAGAAVLL | AWSRTGVNQDTQY | 0.48783374 |
| GSAGSLNTSF | AWSRTGVNQDTQY | 0.6660084 |
| GTGQIGI | AWSRTGVNQDTQY | 0.66609013 |
| GTLIALIL | AWSRTGVNQDTQY | 0.6053554 |
| IAATMGFVVG | AWSRTGVNQDTQY | 0.81939983 |
| IFAQALL | AWSRTGVNQDTQY | 0.8536019 |
| IGAAVALGF | AWSRTGVNQDTQY | 0.8535516 |
| IGLTPLAL | AWSRTGVNQDTQY | 0.85955596 |
| ILGSNTSTL | AWSRTGVNQDTQY | 0.86101794 |
| IVFSLTVL | AWSRTGVNQDTQY | 0.8605032 |
| LASKATGF | AWSRTGVNQDTQY | 0.6649897 |
| LASVALIL | AWSRTGVNQDTQY | 0.7261905 |
| LFAVGLTL | AWSRTGVNQDTQY | 0.44672245 |
| LFNGGIGTYI | AWSRTGVNQDTQY | 0.74114144 |
| LTLGAGII | AWSRTGVNQDTQY | 0.6824644 |
| LVAAALGDI | AWSRTGVNQDTQY | 0.785154 |
| NTAFVQLGI | AWSRTGVNQDTQY | 0.13211718 |
| QLFSGAEL | AWSRTGVNQDTQY | 0.7565979 |
| SAATFITL | AWSRTGVNQDTQY | 0.879449 |
| SNYGLANLGV | AWSRTGVNQDTQY | 0.6711278 |
| SYGGAYAVM | AWSRTGVNQDTQY | 0.7961692 |
| TTGVGGLTLG | AWSRTGVNQDTQY | 0.8037015 |
| TTQTIYI | AWSRTGVNQDTQY | 0.6797174 |
| VAAGAGLGAVI | AWSRTGVNQDTQY | 0.8267853 |
| VAAVVTLL | AWSRTGVNQDTQY | 0.62857556 |
| VGFLGGLI | AWSRTGVNQDTQY | 0.8241745 |
| VGVIPILL | AWSRTGVNQDTQY | 0.88163805 |
| VIFASSLI | AWSRTGVNQDTQY | 0.7819741 |
| VKGGDGGAAAG | AWSRTGVNQDTQY | 0.78073144 |
| VTKTFTNI | AWSRTGVNQDTQY | 0.76549256 |
| VVAAAILL | AWSRTGVNQDTQY | 0.4160607 |
| YAALDVEL | AWSRTGVNQDTQY | 0.8065526 |
| LASVALLL | AWSRTGVNQDTQY | 0.13316292 |
| LDAVHRV | AWSRTGVNQDTQY | 0.755664 |

| | | |
|---|---|---|
| QGLSSSIF | AWSRTGVNQDTQY | 0.595869 |
| RGTFANIRL | AWSRTGVNQDTQY | 0.79102504 |
| RQLWWGHRI | AWSRTGVNQDTQY | 0.7721101 |
| YGPPGTGKTLL | AWSRTGVNQDTQY | 0.82240045 |
| AALVGAPANVA | AWSRTGVNQDTQY | 0.5630717 |
| AATFRRTE | AWSRTGVNQDTQY | 0.7643293 |
| AGALGGGI | AWSRTGVNQDTQY | 0.6863812 |
| AGGAGGAGGIGAG | AWSRTGVNQDTQY | 0.5062968 |
| AGSGALGL | AWSRTGVNQDTQY | 0.83969045 |
| AQIADQI | AWSRTGVNQDTQY | 0.74279726 |
| ASAVAVTYL | AWSRTGVNQDTQY | 0.7380628 |
| ASTVLGGTGG | AWSRTGVNQDTQY | 0.86221087 |
| AVGGEPAK | AWSRTGVNQDTQY | 0.79324526 |
| CEQGTFGGTC | AWSRTGVNQDTQY | 0.36837506 |
| DEGKAPAG | AWSRTGVNQDTQY | 0.7217004 |
| DTVVFPAL | AWSRTGVNQDTQY | 0.7564523 |
| FEGITMAG | AWSRTGVNQDTQY | 0.72505665 |
| FPLVVQD | AWSRTGVNQDTQY | 0.13623072 |
| GAFTTPQ | AWSRTGVNQDTQY | 0.8410504 |
| GFGGFGTAGGVGGA | AWSRTGVNQDTQY | 0.5883328 |
| GFLAQFL | AWSRTGVNQDTQY | 0.61410093 |
| GGAGGAGGNAGVA | AWSRTGVNQDTQY | 0.66540974 |
| GGVGTAGAGGAGGAGG | AWSRTGVNQDTQY | 0.83339393 |
| GNGGGGDAGNAGSGGN | AWSRTGVNQDTQY | 0.70568955 |
| GTGGAGGAGNDAGS | AWSRTGVNQDTQY | 0.7234454 |
| GTTNTGWFNS | AWSRTGVNQDTQY | 0.8257968 |
| IFMGPLFTL | AWSRTGVNQDTQY | 0.8191171 |
| ILAHAIEAI | AWSRTGVNQDTQY | 0.86121905 |
| IRSAIREAL | AWSRTGVNQDTQY | 0.80714184 |
| ITFAGGG | AWSRTGVNQDTQY | 0.7829333 |
| ITGMSGHQEV | AWSRTGVNQDTQY | 0.8157097 |
| IVAGLACTL | AWSRTGVNQDTQY | 0.739429 |
| IVGGGLAGA | AWSRTGVNQDTQY | 0.4392462 |
| IVTGVLNI | AWSRTGVNQDTQY | 0.39527327 |
| LAQQGCI | AWSRTGVNQDTQY | 0.7521287 |
| LCLHVEN | AWSRTGVNQDTQY | 0.351418 |
| LDAISDGS | AWSRTGVNQDTQY | 0.7879093 |
| LGGFAAML | AWSRTGVNQDTQY | 0.8726641 |
| LGGKNPMI | AWSRTGVNQDTQY | 0.78874063 |
| LGPKGVAAIL | AWSRTGVNQDTQY | 0.4595355 |
| LMVGVSI | AWSRTGVNQDTQY | 0.727794 |
| LVGAAFVL | AWSRTGVNQDTQY | 0.71882945 |
| LVLAVAPL | AWSRTGVNQDTQY | 0.8205655 |
| MAAIAVIDAA | AWSRTGVNQDTQY | 0.8555613 |
| NNGINAELQ | AWSRTGVNQDTQY | 0.86820734 |
| RGEPLVLVHG | AWSRTGVNQDTQY | 0.8424894 |
| RGGAGGAAGLI | AWSRTGVNQDTQY | 0.6626892 |
| RSYRMHPAV | AWSRTGVNQDTQY | 0.8289387 |
| SAAHLLAGQLA | AWSRTGVNQDTQY | 0.73016423 |
| SAAVGSY | AWSRTGVNQDTQY | 0.7750768 |
| SPHAVLHRI | AWSRTGVNQDTQY | 0.775949 |

| | | |
|---|---|---|
| SQIAKVL | AWSRTGVNQDTQY | 0.80285954 |
| TALGHLVP | AWSRTGVNQDTQY | 0.7497162 |
| TAPTTSGAA | AWSRTGVNQDTQY | 0.72837937 |
| TDGDDGNGGSPGT | AWSRTGVNQDTQY | 0.4263842 |
| TDGLGEN | AWSRTGVNQDTQY | 0.8160469 |
| TGGPPLPGQA | AWSRTGVNQDTQY | 0.8242761 |
| TIPAITI | AWSRTGVNQDTQY | 0.4836316 |
| TIPGFTI | AWSRTGVNQDTQY | 0.7176839 |
| TSIASGALA | AWSRTGVNQDTQY | 0.8801311 |
| TVTLPGAAI | AWSRTGVNQDTQY | 0.8665618 |
| VASGLSAALP | AWSRTGVNQDTQY | 0.63370734 |
| VGAGVIGI | AWSRTGVNQDTQY | 0.64189434 |
| VGPKRFGV | AWSRTGVNQDTQY | 0.7811807 |
| VNVAFPD | AWSRTGVNQDTQY | 0.42023242 |
| VSMTNSGVS | AWSRTGVNQDTQY | 0.8252278 |
| VTDVLGDLSV | AWSRTGVNQDTQY | 0.74423194 |
| VVVVGAGFAGL | AWSRTGVNQDTQY | 0.7348267 |
| VYTVGTVV | AWSRTGVNQDTQY | 0.6675439 |
| WTTFSGP | AWSRTGVNQDTQY | 0.7980851 |
| YDELVEAF | AWSRTGVNQDTQY | 0.6993436 |
| YGTGGAGGNGG | AWSRTGVNQDTQY | 0.8357146 |
| YTNVMAV | AWSRTGVNQDTQY | 0.1647364 |
| FLEQVHQL | AWSRTGVNQDTQY | 0.8382572 |
| GLSLLAVAV | AWSRTGVNQDTQY | 0.5719097 |
| LEQVHQL | AWSRTGVNQDTQY | 0.64654255 |
| LNAAGAI | AWSRTGVNQDTQY | 0.32306188 |
| PAVHASL | AWSRTGVNQDTQY | 0.7447108 |
| TELEDDA | AWSRTGVNQDTQY | 0.83481616 |
| TPPGPPN | AWSRTGVNQDTQY | 0.85522014 |
| TPPPPPPP | AWSRTGVNQDTQY | 0.63649386 |
| TQSPALI | ASSRGQLGTEVF | 0.8460803 |
| EFAAAFPQ | ASSRGQLGTEVF | 0.88114095 |
| LLSGNLDVL | ASSRGQLGTEVF | 0.6946913 |
| GPANLGNYNI | ASSRGQLGTEVF | 0.34798864 |
| TLGPTLP | ASSRGQLGTEVF | 0.7943926 |
| VYAYSSCCK | ASSRGQLGTEVF | 0.7261975 |
| APAPAPAPNI | ASSRGQLGTEVF | 0.85128987 |
| PGGPVPIP | ASSRGQLGTEVF | 0.8322972 |
| FLTVALEP | ASSRGQLGTEVF | 0.46828008 |
| SQYFGANL | ASSRGQLGTEVF | 0.8222201 |
| GGVKPAA | ASSRGQLGTEVF | 0.8634938 |
| GGTGGAGGAGGW | ASSRGQLGTEVF | 0.61427915 |
| GVGGVAGLGGAGG | ASSRGQLGTEVF | 0.61656415 |
| VLLVLATTI | ASSRGQLGTEVF | 0.66435444 |
| IGIFAAML | ASSRGQLGTEVF | 0.3697363 |
| AVGDLID | ASSRGQLGTEVF | 0.8288375 |
| NISPLPL | ASSRGQLGTEVF | 0.6765301 |
| VTAVFVAI | ASSRGQLGTEVF | 0.6455098 |
| SGDDNDAAL | ASSRGQLGTEVF | 0.7497157 |
| AVAALAHS | ASSRGQLGTEVF | 0.6669662 |
| GGVTVMATP | ASSRGQLGTEVF | 0.66149217 |

| | | |
|---|---|---|
| AGLAANVRD | ASSRGQLGTEVF | 0.7132616 |
| TNRPDLIDPALLRPG | ASSRGQLGTEVF | 0.79390466 |
| LANLGVTAL | ASSRGQLGTEVF | 0.813809 |
| TGERIEL | ASSRGQLGTEVF | 0.7477758 |
| NGEELAA | ASSRGQLGTEVF | 0.77006614 |
| VLSLVWG | ASSRGQLGTEVF | 0.86549747 |
| IGYALWL | ASSRGQLGTEVF | 0.523049 |
| LSLLLAPL | ASSRGQLGTEVF | 0.6239005 |
| TLDEALK | ASSRGQLGTEVF | 0.8176159 |
| TDGAGTAEGG | ASSRGQLGTEVF | 0.5322365 |
| NAPGGAGGNGG | ASSRGQLGTEVF | 0.8299028 |
| VSVFMGMGGNF | ASSRGQLGTEVF | 0.7852716 |
| LASYLAAL | ASSRGQLGTEVF | 0.7967117 |
| LVGGTPAL | ASSRGQLGTEVF | 0.76064324 |
| VRNADVL | ASSRGQLGTEVF | 0.89069366 |
| TCSSPFLVSAM | ASSRGQLGTEVF | 0.8309562 |
| VHITDAE | ASSRGQLGTEVF | 0.7603494 |
| EYAAPSF | ASSRGQLGTEVF | 0.7802368 |
| VVDAFHGGL | ASSRGQLGTEVF | 0.55868506 |
| ASVVDPQ | ASSRGQLGTEVF | 0.85724777 |
| SYNEFGGLS | ASSRGQLGTEVF | 0.830377 |
| DPDGSVY | ASSRGQLGTEVF | 0.5582793 |
| MVSTIHATE | ASSRGQLGTEVF | 0.72940826 |
| LAAGVADPV | ASSRGQLGTEVF | 0.8715131 |
| TKEQIAA | ASSRGQLGTEVF | 0.85891056 |
| SLASPAAL | ASSRGQLGTEVF | 0.43845877 |
| IGLVFYGV | ASSRGQLGTEVF | 0.6432884 |
| FLLDASNL | ASSRGQLGTEVF | 0.46886027 |
| YQVGNFP | ASSRGQLGTEVF | 0.725454 |
| ESLLSQST | ASSRGQLGTEVF | 0.45569247 |
| LFTVGFAAVG | ASSRGQLGTEVF | 0.81440187 |
| ATPPTPT | ASSRGQLGTEVF | 0.68336594 |
| TDLATLG | ASSRGQLGTEVF | 0.79562294 |
| SAGGTGR | ASSRGQLGTEVF | 0.7807051 |
| GFSNSGS | ASSRGQLGTEVF | 0.8632494 |
| GGNGGSGGTSV | ASSRGQLGTEVF | 0.6639339 |
| GTSVATGGAGN | ASSRGQLGTEVF | 0.73602873 |
| NGGNGGSGGTS | ASSRGQLGTEVF | 0.40303427 |
| AVVKFPH | ASSRGQLGTEVF | 0.6191231 |
| TPPPPPP | ASSRGQLGTEVF | 0.63760054 |
| AAPAAAAAPA | ASSRGQLGTEVF | 0.72764134 |
| IALNEDL | ASSRGQLGTEVF | 0.2857601 |
| IVNVSSM | ASSRGQLGTEVF | 0.8747215 |
| TGIPPAPR | ASSRGQLGTEVF | 0.75650156 |
| FDLGGGTF | ASSRGQLGTEVF | 0.82882625 |
| FDLGGGTFDVS | ASSRGQLGTEVF | 0.6500365 |
| NEPTAAA | ASSRGQLGTEVF | 0.6818019 |
| NEPTAAALA | ASSRGQLGTEVF | 0.63619155 |
| GSIINISSI | ASSRGQLGTEVF | 0.8151978 |
| VAPGNAGTA | ASSRGQLGTEVF | 0.56792605 |
| TGGGASLE | ASSRGQLGTEVF | 0.31514764 |

| | | |
|---|---|---|
| VAVYNHYKRI | ASSRGQLGTEVF | 0.6328723 |
| TDPAPATT | ASSRGQLGTEVF | 0.46737763 |
| TNRPDILDPALLRPG | ASSRGQLGTEVF | 0.8708881 |
| LTTTTTS | ASSRGQLGTEVF | 0.8487037 |
| PTGSGKTL | ASSRGQLGTEVF | 0.7879164 |
| TGSGKTL | ASSRGQLGTEVF | 0.8710207 |
| TIGHVDHGK | ASSRGQLGTEVF | 0.22229248 |
| TIGHVDHGKTT | ASSRGQLGTEVF | 0.6772351 |
| GHVDHGKTT | ASSRGQLGTEVF | 0.8692764 |
| AQFEHTLL | ASSRGQLGTEVF | 0.80629057 |
| IVNQDEL | ASSRGQLGTEVF | 0.73650885 |
| TNGSQFF | ASSRGQLGTEVF | 0.8553412 |
| TGKGIVE | ASSRGQLGTEVF | 0.6119856 |
| LISNASDA | ASSRGQLGTEVF | 0.7254338 |
| AELRTLGELTKIAKAHGAQ | ASSRGQLGTEVF | 0.74222493 |
| APNAVAAAEPSW | ASSRGQLGTEVF | 0.8450922 |
| APQVAGAATAVL | ASSRGQLGTEVF | 0.8004569 |
| ARRDPAAAAAILPTDA | ASSRGQLGTEVF | 0.5127198 |
| DAAARDVLAEAGLAEYF | ASSRGQLGTEVF | 0.8083579 |
| DDQIVLSAKANLAKVEQ | ASSRGQLGTEVF | 0.8920448 |
| DELLPDTGEDSDEDGH | ASSRGQLGTEVF | 0.6828984 |
| ELPEAPANRATVPQD | ASSRGQLGTEVF | 0.8418039 |
| FGQSKIERVTGIG | ASSRGQLGTEVF | 0.801739 |
| GAGGTGSTAGGAGGAGGAGGLY | ASSRGQLGTEVF | 0.7714002 |
| GHGGTGAAGGNGGTGGAGGSGI | ASSRGQLGTEVF | 0.59504396 |
| GNGGNGGAGGEGGPGGAGGAGA | ASSRGQLGTEVF | 0.6009717 |
| GSTAGNGGNGGNATNGTIAGQP | ASSRGQLGTEVF | 0.7639454 |
| GTGGDGGNGGNGGTGG | ASSRGQLGTEVF | 0.5439669 |
| GVRRPAGIPADAPV | ASSRGQLGTEVF | 0.76171887 |
| GYRQVTDAHLLTLARR | ASSRGQLGTEVF | 0.7016532 |
| IAKMGQDGHDRG | ASSRGQLGTEVF | 0.89698625 |
| KAEPSIEIQYT | ASSRGQLGTEVF | 0.8054948 |
| KPVRIRMAADEDTVAVT | ASSRGQLGTEVF | 0.8102001 |
| MPPRIAGMRLLVIKPEP | ASSRGQLGTEVF | 0.5430654 |
| MKFYAVALFRSRGVPPTR | ASSRGQLGTEVF | 0.80948734 |
| NAGGFGILGAVAHSP | ASSRGQLGTEVF | 0.8064153 |
| NEGREITLAMAEM | ASSRGQLGTEVF | 0.44801706 |
| NELVEWSHPLGHRWRWE | ASSRGQLGTEVF | 0.86518407 |
| NLGAGGTGGNGGNPG | ASSRGQLGTEVF | 0.62172425 |
| NTEWPKWETVMAADPVD | ASSRGQLGTEVF | 0.75149167 |
| PASAAPAETQP | ASSRGQLGTEVF | 0.83935606 |
| QQSSDGKNGVRAAEITGE | ASSRGQLGTEVF | 0.8427832 |
| RAPDAVTTSWWKE | ASSRGQLGTEVF | 0.721131 |
| SGFGNVGAHTSGWFNQ | ASSRGQLGTEVF | 0.75003886 |
| SGVAGAVGQAASVGGLKV | ASSRGQLGTEVF | 0.52798593 |
| SNGGAGTGSAGGNGGT | ASSRGQLGTEVF | 0.84270805 |
| TGGAGGEGGAAG | ASSRGQLGTEVF | 0.84670997 |
| TGGDGGAAGAGGAPGHG | ASSRGQLGTEVF | 0.7149987 |
| TMPPLLAAIANAGRHGV | ASSRGQLGTEVF | 0.8784555 |
| TSSTAGNSGGA | ASSRGQLGTEVF | 0.82531166 |
| VPATVPVGILC | ASSRGQLGTEVF | 0.3813102 |

| | | |
|---|---|---|
| VQRPDSGGVVLTGRIS | ASSRGQLGTEVF | 0.75708807 |
| WTPNPVPATKGVPAGLVQAT | ASSRGQLGTEVF | 0.8624952 |
| YALKRALAKFSDRFDVVII | ASSRGQLGTEVF | 0.85286844 |
| YLIGSVIV | ASSRGQLGTEVF | 0.87414587 |
| YGQLVAVID | ASSRGQLGTEVF | 0.8427861 |
| VTVGGMTLPA | ASSRGQLGTEVF | 0.7022098 |
| VLALPAV | ASSRGQLGTEVF | 0.8701428 |
| VIPPTLNYET | ASSRGQLGTEVF | 0.89733946 |
| VGMGGMHPSSG | ASSRGQLGTEVF | 0.8338667 |
| VGLAGGAATA | ASSRGQLGTEVF | 0.66832995 |
| VGGAGGAGGSGP | ASSRGQLGTEVF | 0.6595843 |
| VFAAFMV | ASSRGQLGTEVF | 0.78718746 |
| TLYDKLR | ASSRGQLGTEVF | 0.8354845 |
| TLAVFTVL | ASSRGQLGTEVF | 0.5493231 |
| TGSVQPARV | ASSRGQLGTEVF | 0.7785908 |
| TGGGAGSGGT | ASSRGQLGTEVF | 0.77883816 |
| SHLETSL | ASSRGQLGTEVF | 0.33529603 |
| SFTTGMT | ASSRGQLGTEVF | 0.8430051 |
| QVEVAKRMGVRQPTVS | ASSRGQLGTEVF | 0.8698794 |
| QGSGGPALAM | ASSRGQLGTEVF | 0.78472507 |
| MLAPSNP | ASSRGQLGTEVF | 0.6379087 |
| LVNNAAI | ASSRGQLGTEVF | 0.8560831 |
| LTGEINTLA | ASSRGQLGTEVF | 0.89052904 |
| LSIFGPV | ASSRGQLGTEVF | 0.8693299 |
| LLALLANL | ASSRGQLGTEVF | 0.36983347 |
| LISNASDALDKLR | ASSRGQLGTEVF | 0.21337228 |
| LINEAVP | ASSRGQLGTEVF | 0.69402885 |
| LGYAFSGL | ASSRGQLGTEVF | 0.80866575 |
| LDFVAVAL | ASSRGQLGTEVF | 0.7126205 |
| LAQPGAE | ASSRGQLGTEVF | 0.6286391 |
| ISNLGAVTGF | ASSRGQLGTEVF | 0.788343 |
| HQEDPGRI | ASSRGQLGTEVF | 0.85649174 |
| GTPGEPTP | ASSRGQLGTEVF | 0.5440355 |
| GFFNSGS | ASSRGQLGTEVF | 0.5041313 |
| GAGGGGLTAGAN | ASSRGQLGTEVF | 0.8225619 |
| FVDALVNM | ASSRGQLGTEVF | 0.8379128 |
| FLTVAPEL | ASSRGQLGTEVF | 0.5879899 |
| FETLTAL | ASSRGQLGTEVF | 0.6966975 |
| DVINPIEL | ASSRGQLGTEVF | 0.728184 |
| APVSGAPV | ASSRGQLGTEVF | 0.71171784 |
| ALVGGLAW | ASSRGQLGTEVF | 0.7930578 |
| ALAAAAATAI | ASSRGQLGTEVF | 0.5480056 |
| AFGLGLLL | ASSRGQLGTEVF | 0.86990726 |
| AFFAAASL | ASSRGQLGTEVF | 0.48062462 |
| AAAALPAFSPP | ASSRGQLGTEVF | 0.8403044 |
| VVLAPAL | ASSRGQLGTEVF | 0.7941033 |
| TIGHVDHGKTTL | ASSRGQLGTEVF | 0.81649625 |
| LISNASDALDK | ASSRGQLGTEVF | 0.68653196 |
| LISNASDAL | ASSRGQLGTEVF | 0.6450703 |
| ISNASDALDK | ASSRGQLGTEVF | 0.55384904 |
| NQAETLVYQTE | ASSRGQLGTEVF | 0.3993767 |

| | | |
|---|---|---|
| TGGTGGTAGAG | ASSRGQLGTEVF | 0.86020446 |
| TGSTAGGAGGA | ASSRGQLGTEVF | 0.26448476 |
| DPLAVPVAAGPV | ASSRGQLGTEVF | 0.86292636 |
| GGTGGTPGNGGN | ASSRGQLGTEVF | 0.8185698 |
| PAGLAGVDPATL | ASSRGQLGTEVF | 0.5925118 |
| GNNLVVEAPHAGASV | ASSRGQLGTEVF | 0.8173986 |
| KVTRIMSASTLEQPA | ASSRGQLGTEVF | 0.51082736 |
| LGFALPIALPRRTVV | ASSRGQLGTEVF | 0.6657156 |
| TRLGNLTARKLVVLV | ASSRGQLGTEVF | 0.8311415 |
| WSQQRRLIYFPSAGP | ASSRGQLGTEVF | 0.83147717 |
| GPNGNPDPMAAAVDIR | ASSRGQLGTEVF | 0.5900321 |
| TQRCVSVADLRAAASR | ASSRGQLGTEVF | 0.82025194 |
| QFVLDVAEHPVEVLTETV | ASSRGQLGTEVF | 0.7972087 |
| GGAGGTGGAGGNGGTQAGDG | ASSRGQLGTEVF | 0.62997985 |
| PLLATLRAIAVDVDAKPAQV | ASSRGQLGTEVF | 0.7902762 |
| APKFRTGGYMIDFWGVGYQVAK | ASSRGQLGTEVF | 0.85235226 |
| GAGGAGGAGGAAGGPGGTGGTG | ASSRGQLGTEVF | 0.809788 |
| IGALGLLATDVAPAGSVAFGVA | ASSRGQLGTEVF | 0.47227183 |
| AGGAGGIGGNGGAGGAGGVLMG | ASSRGQLGTEVF | 0.8200754 |
| ADELIGLL | ASSRGQLGTEVF | 0.6005402 |
| ADRALKA | ASSRGQLGTEVF | 0.7911109 |
| AGAVGQVGGAGGA | ASSRGQLGTEVF | 0.7739687 |
| AGGGGGLGGAP | ASSRGQLGTEVF | 0.70556414 |
| AGGLLYGNGGAG | ASSRGQLGTEVF | 0.84412044 |
| AGLESIGL | ASSRGQLGTEVF | 0.58924603 |
| AGLKTGII | ASSRGQLGTEVF | 0.61000884 |
| ASLAALLL | ASSRGQLGTEVF | 0.6947966 |
| ASVINISSI | ASSRGQLGTEVF | 0.77289265 |
| ATPPGPA | ASSRGQLGTEVF | 0.83486915 |
| ATVASALV | ASSRGQLGTEVF | 0.54998934 |
| ATVASLIL | ASSRGQLGTEVF | 0.78594494 |
| AVNGPVLIRH | ASSRGQLGTEVF | 0.71172667 |
| DGAAGAVGI | ASSRGQLGTEVF | 0.58325565 |
| DPLLHGI | ASSRGQLGTEVF | 0.77750325 |
| ELGAALGAL | ASSRGQLGTEVF | 0.7360172 |
| ELGRTQAV | ASSRGQLGTEVF | 0.53516984 |
| FVAGGYLS | ASSRGQLGTEVF | 0.83955044 |
| GASGGAGGAGGW | ASSRGQLGTEVF | 0.62547404 |
| GGAGAGGDGGAGGI | ASSRGQLGTEVF | 0.81783193 |
| GTGHNGSQPGMGGNG | ASSRGQLGTEVF | 0.879478 |
| GYAAASASAS | ASSRGQLGTEVF | 0.5865435 |
| HQVDVAAVQQG | ASSRGQLGTEVF | 0.8361019 |
| IGAAIAHT | ASSRGQLGTEVF | 0.80598634 |
| IGMIEVAQL | ASSRGQLGTEVF | 0.82267076 |
| IRLAPEHTSAMD | ASSRGQLGTEVF | 0.73729086 |
| IYNVSVL | ASSRGQLGTEVF | 0.62237966 |
| KQGFLGM | ASSRGQLGTEVF | 0.86616635 |
| LESVITQL | ASSRGQLGTEVF | 0.82588375 |
| LGLGALVVV | ASSRGQLGTEVF | 0.83249843 |
| LIGNGGNGGSGGI | ASSRGQLGTEVF | 0.82932293 |
| LNAINAPTL | ASSRGQLGTEVF | 0.8110663 |

| | | |
|---|---|---|
| LQLLNTGV | ASSRGQLGTEVF | 0.82631314 |
| LQTSIEF | ASSRGQLGTEVF | 0.63256586 |
| MSFAVSEL | ASSRGQLGTEVF | 0.44118255 |
| NTGALGL | ASSRGQLGTEVF | 0.70798635 |
| PPGAGGAGGAGTT | ASSRGQLGTEVF | 0.77346534 |
| QLVGLGI | ASSRGQLGTEVF | 0.6796738 |
| RFLEQVHQL | ASSRGQLGTEVF | 0.4172123 |
| RGVGYVLR | ASSRGQLGTEVF | 0.8471441 |
| SATGVFAGVI | ASSRGQLGTEVF | 0.8910482 |
| SLPAITLP | ASSRGQLGTEVF | 0.65714765 |
| STALAFTV | ASSRGQLGTEVF | 0.88600606 |
| SVLTVGLGK | ASSRGQLGTEVF | 0.5033749 |
| TALAGAGSGI | ASSRGQLGTEVF | 0.8279846 |
| TGLPFTGI | ASSRGQLGTEVF | 0.8525033 |
| VAEHVTV | ASSRGQLGTEVF | 0.5828981 |
| VDIVLNSL | ASSRGQLGTEVF | 0.82570946 |
| VLTGGGAVL | ASSRGQLGTEVF | 0.59372556 |
| VNVGLAYGI | ASSRGQLGTEVF | 0.6957772 |
| VSSAAVGAV | ASSRGQLGTEVF | 0.8577715 |
| VTSVYEAGV | ASSRGQLGTEVF | 0.68123174 |
| WVALDRGA | ASSRGQLGTEVF | 0.8226032 |
| YLVGSVVI | ASSRGQLGTEVF | 0.6980734 |
| GPPGTGKTLL | ASSRGQLGTEVF | 0.57356554 |
| GVGMTKF | ASSRGQLGTEVF | 0.80833066 |
| IGIGGSDL | ASSRGQLGTEVF | 0.8603065 |
| IGTLLAEL | ASSRGQLGTEVF | 0.7042111 |
| IYTSGTT | ASSRGQLGTEVF | 0.8927244 |
| LSFPAGV | ASSRGQLGTEVF | 0.61916244 |
| LYTSGTT | ASSRGQLGTEVF | 0.88605905 |
| VGAFPFGF | ASSRGQLGTEVF | 0.8493145 |
| VGAGTVEF | ASSRGQLGTEVF | 0.75611347 |
| VGKSTLLL | ASSRGQLGTEVF | 0.70296496 |
| AALAHAGGGIV | ASSRGQLGTEVF | 0.8389641 |
| AALVAVSLL | ASSRGQLGTEVF | 0.68221736 |
| AAVSRAVAAL | ASSRGQLGTEVF | 0.827858 |
| AGGSSGIGGF | ASSRGQLGTEVF | 0.87872005 |
| AGLTVANANG | ASSRGQLGTEVF | 0.7788925 |
| AGLVGAVL | ASSRGQLGTEVF | 0.9078114 |
| ALSQAFTV | ASSRGQLGTEVF | 0.8650037 |
| ALTFSVSL | ASSRGQLGTEVF | 0.83690417 |
| ALTGAALVAL | ASSRGQLGTEVF | 0.8161556 |
| ASFSIGAGVL | ASSRGQLGTEVF | 0.5333842 |
| DGSVYKI | ASSRGQLGTEVF | 0.8198179 |
| EQLAEIV | ASSRGQLGTEVF | 0.76984906 |
| FAVSELTVM | ASSRGQLGTEVF | 0.85962313 |
| GAAPLAAGL | ASSRGQLGTEVF | 0.89936876 |
| GADGGDGGSGGAGG | ASSRGQLGTEVF | 0.8042559 |
| GFGLDTGGAGG | ASSRGQLGTEVF | 0.9085829 |
| GLASGGLL | ASSRGQLGTEVF | 0.8906214 |
| GRIAGAAVLL | ASSRGQLGTEVF | 0.59772706 |
| GSAGSLNTSF | ASSRGQLGTEVF | 0.8224692 |

| | | |
|---|---|---|
| GTGQIGI | ASSRGQLGTEVF | 0.7629729 |
| GTLIALIL | ASSRGQLGTEVF | 0.84162605 |
| IAATMGFVVG | ASSRGQLGTEVF | 0.49269807 |
| IFAQALL | ASSRGQLGTEVF | 0.89786625 |
| IGAAVALGF | ASSRGQLGTEVF | 0.7615944 |
| IGLTPLAL | ASSRGQLGTEVF | 0.2430216 |
| ILGSNTSTL | ASSRGQLGTEVF | 0.86027515 |
| IVFSLTVL | ASSRGQLGTEVF | 0.8628837 |
| LASKATGF | ASSRGQLGTEVF | 0.8577218 |
| LASVALIL | ASSRGQLGTEVF | 0.89211744 |
| LFAVGLTL | ASSRGQLGTEVF | 0.66046953 |
| LFNGGIGTYI | ASSRGQLGTEVF | 0.7757604 |
| LTLGAGII | ASSRGQLGTEVF | 0.8944941 |
| LVAAALGDI | ASSRGQLGTEVF | 0.907109 |
| NTAFVQLGI | ASSRGQLGTEVF | 0.71431154 |
| QLFSGAEL | ASSRGQLGTEVF | 0.8745916 |
| SAATFITL | ASSRGQLGTEVF | 0.89295816 |
| SNYGLANLGV | ASSRGQLGTEVF | 0.8751819 |
| SYGGAYAVM | ASSRGQLGTEVF | 0.89868593 |
| TTGVGGLTLG | ASSRGQLGTEVF | 0.9114028 |
| TTQTIYI | ASSRGQLGTEVF | 0.74948084 |
| VAAGAGLGAVI | ASSRGQLGTEVF | 0.90825284 |
| VAAVVTLL | ASSRGQLGTEVF | 0.9054444 |
| VGFLGGLI | ASSRGQLGTEVF | 0.84000546 |
| VGVIPILL | ASSRGQLGTEVF | 0.85174906 |
| VIFASSLI | ASSRGQLGTEVF | 0.70694494 |
| VKGGDGGAAAG | ASSRGQLGTEVF | 0.83247036 |
| VTKTFTNI | ASSRGQLGTEVF | 0.5846265 |
| VVAAAILL | ASSRGQLGTEVF | 0.8679742 |
| YAALDVEL | ASSRGQLGTEVF | 0.9045594 |
| LASVALLL | ASSRGQLGTEVF | 0.8174277 |
| LDAVHRV | ASSRGQLGTEVF | 0.7413 |
| QGLSSSIF | ASSRGQLGTEVF | 0.7490034 |
| RGTFANIRL | ASSRGQLGTEVF | 0.8419078 |
| RQLWWGHRI | ASSRGQLGTEVF | 0.8340715 |
| YGPPGTGKTLL | ASSRGQLGTEVF | 0.53237426 |
| AALVGAPANVA | ASSRGQLGTEVF | 0.78066695 |
| AATFRRTE | ASSRGQLGTEVF | 0.89796627 |
| AGALGGGI | ASSRGQLGTEVF | 0.89377475 |
| AGGAGGAGGIGAG | ASSRGQLGTEVF | 0.5825145 |
| AGSGALGL | ASSRGQLGTEVF | 0.90518665 |
| AQIADQI | ASSRGQLGTEVF | 0.8841852 |
| ASAVAVTYL | ASSRGQLGTEVF | 0.90049434 |
| ASTVLGGTGG | ASSRGQLGTEVF | 0.836968 |
| AVGGEPAK | ASSRGQLGTEVF | 0.7549648 |
| CEQGTFGGTC | ASSRGQLGTEVF | 0.8741952 |
| DEGKAPAG | ASSRGQLGTEVF | 0.79924023 |
| DTVVFPAL | ASSRGQLGTEVF | 0.78724074 |
| FEGITMAG | ASSRGQLGTEVF | 0.89247584 |
| FPLVVQD | ASSRGQLGTEVF | 0.8360658 |
| GAFTTPQ | ASSRGQLGTEVF | 0.7903752 |

| | | |
|---|---|---|
| GFGGFGTAGGVGGA | ASSRGQLGTEVF | 0.7284812 |
| GFLAQFL | ASSRGQLGTEVF | 0.9090549 |
| GGAGGAGGNAGVA | ASSRGQLGTEVF | 0.71278036 |
| GGVGTAGAGGAGGAGG | ASSRGQLGTEVF | 0.77888274 |
| GNGGGGDAGNAGSGGN | ASSRGQLGTEVF | 0.76570755 |
| GTGGAGGAGNDAGS | ASSRGQLGTEVF | 0.85193 |
| GTTNTGWFNS | ASSRGQLGTEVF | 0.9010726 |
| IFMGPLFTL | ASSRGQLGTEVF | 0.8849847 |
| ILAHAIEAI | ASSRGQLGTEVF | 0.7986704 |
| IRSAIREAL | ASSRGQLGTEVF | 0.8738686 |
| ITFAGGG | ASSRGQLGTEVF | 0.8871365 |
| ITGMSGHQEV | ASSRGQLGTEVF | 0.86509055 |
| IVAGLACTL | ASSRGQLGTEVF | 0.8898922 |
| IVGGGLAGA | ASSRGQLGTEVF | 0.7652807 |
| IVTGVLNI | ASSRGQLGTEVF | 0.90745044 |
| LAQQGCI | ASSRGQLGTEVF | 0.8756057 |
| LCLHVEN | ASSRGQLGTEVF | 0.9038479 |
| LDAISDGS | ASSRGQLGTEVF | 0.900927 |
| LGGFAAML | ASSRGQLGTEVF | 0.88471216 |
| LGGKNPMI | ASSRGQLGTEVF | 0.8681429 |
| LGPKGVAAIL | ASSRGQLGTEVF | 0.4713775 |
| LMVGVSI | ASSRGQLGTEVF | 0.74676895 |
| LVGAAFVL | ASSRGQLGTEVF | 0.8735244 |
| LVLAVAPL | ASSRGQLGTEVF | 0.86122227 |
| MAAIAVIDAA | ASSRGQLGTEVF | 0.8894361 |
| NNGINAELQ | ASSRGQLGTEVF | 0.87523806 |
| RGEPLVLVHG | ASSRGQLGTEVF | 0.7967934 |
| RGGAGGAAGLI | ASSRGQLGTEVF | 0.8940512 |
| RSYRMHPAV | ASSRGQLGTEVF | 0.89038455 |
| SAAHLLAGQLA | ASSRGQLGTEVF | 0.5966081 |
| SAAVGSY | ASSRGQLGTEVF | 0.8262598 |
| SPHAVLHRI | ASSRGQLGTEVF | 0.9163811 |
| SQIAKVL | ASSRGQLGTEVF | 0.84078455 |
| TALGHLVP | ASSRGQLGTEVF | 0.82743543 |
| TAPTTSGAA | ASSRGQLGTEVF | 0.78732145 |
| TDGDDGNGGSPGT | ASSRGQLGTEVF | 0.9035623 |
| TDGLGEN | ASSRGQLGTEVF | 0.82021046 |
| TGGPPLPGQA | ASSRGQLGTEVF | 0.9118199 |
| TIPAITI | ASSRGQLGTEVF | 0.29519838 |
| TIPGFTI | ASSRGQLGTEVF | 0.80862975 |
| TSIASGALA | ASSRGQLGTEVF | 0.73005605 |
| TVTLPGAAI | ASSRGQLGTEVF | 0.5670137 |
| VASGLSAALP | ASSRGQLGTEVF | 0.9045597 |
| VGAGVIGI | ASSRGQLGTEVF | 0.7617605 |
| VGPKRFGV | ASSRGQLGTEVF | 0.8815249 |
| VNVAFPD | ASSRGQLGTEVF | 0.8414134 |
| VSMTNSGVS | ASSRGQLGTEVF | 0.90700805 |
| VTDVLGDLSV | ASSRGQLGTEVF | 0.8373134 |
| VVVVGAGFAGL | ASSRGQLGTEVF | 0.8562759 |
| VYTVGTVV | ASSRGQLGTEVF | 0.7832633 |
| WTTFSGP | ASSRGQLGTEVF | 0.8964777 |

| | | |
|---|---|---|
| YDELVEAF | ASSRGQLGTEVF | 0.9037255 |
| YGTGGAGGNGG | ASSRGQLGTEVF | 0.8880069 |
| YTNVMAV | ASSRGQLGTEVF | 0.65456665 |
| FLEQVHQL | ASSRGQLGTEVF | 0.8242809 |
| GLSLLAVAV | ASSRGQLGTEVF | 0.87274295 |
| LEQVHQL | ASSRGQLGTEVF | 0.31193063 |
| LNAAGAI | ASSRGQLGTEVF | 0.42579204 |
| PAVHASL | ASSRGQLGTEVF | 0.8416711 |
| TELEDDA | ASSRGQLGTEVF | 0.75996995 |
| TPPGPPN | ASSRGQLGTEVF | 0.90521926 |
| TPPPPPPP | ASSRGQLGTEVF | 0.8880156 |
| TQSPALI | ASTPGQQDTQY | 0.8297552 |
| EFAAAFPQ | ASTPGQQDTQY | 0.8873804 |
| LLSGNLDVL | ASTPGQQDTQY | 0.8588732 |
| GPANLGNYNI | ASTPGQQDTQY | 0.10198938 |
| TLGPTLP | ASTPGQQDTQY | 0.7548659 |
| VYAYSSCCK | ASTPGQQDTQY | 0.625757 |
| APAPAPAPNI | ASTPGQQDTQY | 0.91475475 |
| PGGPVPIP | ASTPGQQDTQY | 0.7882924 |
| FLTVALEP | ASTPGQQDTQY | 0.73075557 |
| SQYFGANL | ASTPGQQDTQY | 0.84488225 |
| GGVKPAA | ASTPGQQDTQY | 0.8372561 |
| GGTGGAGGAGGW | ASTPGQQDTQY | 0.8285612 |
| GVGGVAGLGGAGG | ASTPGQQDTQY | 0.7324852 |
| VLLVLATTI | ASTPGQQDTQY | 0.23715276 |
| IGIFAAML | ASTPGQQDTQY | 0.8954917 |
| AVGDLID | ASTPGQQDTQY | 0.57549673 |
| NISPLPL | ASTPGQQDTQY | 0.76267904 |
| VTAVFVAI | ASTPGQQDTQY | 0.3533013 |
| SGDDNDAAL | ASTPGQQDTQY | 0.7534833 |
| AVAALAHS | ASTPGQQDTQY | 0.5417584 |
| GGVTVMATP | ASTPGQQDTQY | 0.89207625 |
| AGLAANVRD | ASTPGQQDTQY | 0.82118046 |
| TNRPDLIDPALLRPG | ASTPGQQDTQY | 0.80238223 |
| LANLGVTAL | ASTPGQQDTQY | 0.9120795 |
| TGERIEL | ASTPGQQDTQY | 0.8167056 |
| NGEELAA | ASTPGQQDTQY | 0.8021066 |
| VLSLVWG | ASTPGQQDTQY | 0.81139004 |
| IGYALWL | ASTPGQQDTQY | 0.7393931 |
| LSLLLAPL | ASTPGQQDTQY | 0.7861022 |
| TLDEALK | ASTPGQQDTQY | 0.80164415 |
| TDGAGTAEGG | ASTPGQQDTQY | 0.91354823 |
| NAPGGAGGNGG | ASTPGQQDTQY | 0.8370714 |
| VSVFMGMGGNF | ASTPGQQDTQY | 0.57235545 |
| LASYLAAL | ASTPGQQDTQY | 0.8475361 |
| LVGGTPAL | ASTPGQQDTQY | 0.913059 |
| VRNADVL | ASTPGQQDTQY | 0.9162177 |
| TCSSPFLVSAM | ASTPGQQDTQY | 0.770419 |
| VHITDAE | ASTPGQQDTQY | 0.7821181 |
| EYAAPSF | ASTPGQQDTQY | 0.7789593 |
| VVDAFHGGL | ASTPGQQDTQY | 0.8393637 |

| | | |
|---|---|---|
| ASVVDPQ | ASTPGQQDTQY | 0.78288656 |
| SYNEFGGLS | ASTPGQQDTQY | 0.70658696 |
| DPDGSVY | ASTPGQQDTQY | 0.8946136 |
| MVSTIHATE | ASTPGQQDTQY | 0.9128496 |
| LAAGVADPV | ASTPGQQDTQY | 0.8321453 |
| TKEQIAA | ASTPGQQDTQY | 0.5266097 |
| SLASPAAL | ASTPGQQDTQY | 0.8466485 |
| IGLVFYGV | ASTPGQQDTQY | 0.7609718 |
| FLLDASNL | ASTPGQQDTQY | 0.8094486 |
| YQVGNFP | ASTPGQQDTQY | 0.8484129 |
| ESLLSQST | ASTPGQQDTQY | 0.5730529 |
| LFTVGFAAVG | ASTPGQQDTQY | 0.83338416 |
| ATPPTPT | ASTPGQQDTQY | 0.91440344 |
| TDLATLG | ASTPGQQDTQY | 0.7930534 |
| SAGGTGR | ASTPGQQDTQY | 0.8495383 |
| GFSNSGS | ASTPGQQDTQY | 0.9038129 |
| GGNGGSGGTSV | ASTPGQQDTQY | 0.8861354 |
| GTSVATGGAGN | ASTPGQQDTQY | 0.90459085 |
| NGGNGGSGGTS | ASTPGQQDTQY | 0.8908042 |
| AVVKFPH | ASTPGQQDTQY | 0.86285174 |
| TPPPPPP | ASTPGQQDTQY | 0.86572367 |
| AAPAAAAAPA | ASTPGQQDTQY | 0.8639207 |
| IALNEDL | ASTPGQQDTQY | 0.8603483 |
| IVNVSSM | ASTPGQQDTQY | 0.8149917 |
| TGIPPAPR | ASTPGQQDTQY | 0.7603166 |
| FDLGGGTF | ASTPGQQDTQY | 0.90290284 |
| FDLGGGTFDVS | ASTPGQQDTQY | 0.8365365 |
| NEPTAAA | ASTPGQQDTQY | 0.8937106 |
| NEPTAAALA | ASTPGQQDTQY | 0.66960657 |
| GSIINISSI | ASTPGQQDTQY | 0.74790084 |
| VAPGNAGTA | ASTPGQQDTQY | 0.89670336 |
| TGGGASLE | ASTPGQQDTQY | 0.13911436 |
| VAVYNHYKRI | ASTPGQQDTQY | 0.40405244 |
| TDPAPATT | ASTPGQQDTQY | 0.85587597 |
| TNRPDILDPALLRPG | ASTPGQQDTQY | 0.90531886 |
| LTTTTTS | ASTPGQQDTQY | 0.17638215 |
| PTGSGKTL | ASTPGQQDTQY | 0.7358849 |
| TGSGKTL | ASTPGQQDTQY | 0.6435509 |
| TIGHVDHGK | ASTPGQQDTQY | 0.8815967 |
| TIGHVDHGKTT | ASTPGQQDTQY | 0.86505103 |
| GHVDHGKTT | ASTPGQQDTQY | 0.56146914 |
| AQFEHTLL | ASTPGQQDTQY | 0.89527965 |
| IVNQDEL | ASTPGQQDTQY | 0.50339866 |
| TNGSQFF | ASTPGQQDTQY | 0.7086201 |
| TGKGIVE | ASTPGQQDTQY | 0.90291435 |
| LISNASDA | ASTPGQQDTQY | 0.90261555 |
| AELRTLGELTKIAKAHGAQ | ASTPGQQDTQY | 0.66668546 |
| APNAVAAAEPSW | ASTPGQQDTQY | 0.89665765 |
| APQVAGAATAVL | ASTPGQQDTQY | 0.7879087 |
| ARRDPAAAAAILPTDA | ASTPGQQDTQY | 0.91452813 |
| DAAARDVLAEAGLAEYF | ASTPGQQDTQY | 0.8166305 |

| | | |
|---|---|---|
| DDQIVLSAKANLAKVEQ | ASTPGQQDTQY | 0.60087186 |
| DELLPDTGEDSDEDGH | ASTPGQQDTQY | 0.8927696 |
| ELPEAPANRATVPQD | ASTPGQQDTQY | 0.87046045 |
| FGQSKIERVTGIG | ASTPGQQDTQY | 0.866047 |
| GAGGTGSTAGGAGGAGGAGGLY | ASTPGQQDTQY | 0.8814087 |
| GHGGTGAAGGNGGTGGAGGSGI | ASTPGQQDTQY | 0.7882293 |
| GNGGNGGAGGEGGPGGAGGAGA | ASTPGQQDTQY | 0.8368453 |
| GSTAGNGGNGGNATNGTIAGQP | ASTPGQQDTQY | 0.9011639 |
| GTGGDGGNGGNGGTGG | ASTPGQQDTQY | 0.8995226 |
| GVRRPAGIPADAPV | ASTPGQQDTQY | 0.74963224 |
| GYRQVTDAHLLTLARR | ASTPGQQDTQY | 0.9012646 |
| IAKMGQDGHDRG | ASTPGQQDTQY | 0.8205662 |
| KAEPSIEIQYT | ASTPGQQDTQY | 0.7504215 |
| KPVRIRMAADEDTVAVT | ASTPGQQDTQY | 0.91202766 |
| MPPRIAGMRLLVIKPEP | ASTPGQQDTQY | 0.8476836 |
| MKFYAVALFRSRGVPPTR | ASTPGQQDTQY | 0.8890958 |
| NAGGFGILGAVAHSP | ASTPGQQDTQY | 0.90322256 |
| NEGREITLAMAEM | ASTPGQQDTQY | 0.8781808 |
| NELVEWSHPLGHRWRWE | ASTPGQQDTQY | 0.6285236 |
| NLGAGGTGGNGGNPG | ASTPGQQDTQY | 0.8486283 |
| NTEWPKWETVMAADPVD | ASTPGQQDTQY | 0.8281837 |
| PASAAPAETQP | ASTPGQQDTQY | 0.89358556 |
| QQSSDGKNGVRAAEITGE | ASTPGQQDTQY | 0.8416647 |
| RAPDAVTTSWWKE | ASTPGQQDTQY | 0.70357263 |
| SGFGNVGAHTSGWFNQ | ASTPGQQDTQY | 0.8862133 |
| SGVAGAVGQAASVGGLKV | ASTPGQQDTQY | 0.9024827 |
| SNGGAGTGSAGGNGGT | ASTPGQQDTQY | 0.86990345 |
| TGGAGGEGGAAG | ASTPGQQDTQY | 0.865421 |
| TGGDGGAAGAGGAPGHG | ASTPGQQDTQY | 0.9095268 |
| TMPPLLAAIANAGRHGV | ASTPGQQDTQY | 0.78204954 |
| TSSTAGNSGGA | ASTPGQQDTQY | 0.83095264 |
| VPATVPVGILC | ASTPGQQDTQY | 0.8943804 |
| VQRPDSGGVVLTGRIS | ASTPGQQDTQY | 0.894859 |
| WTPNPVPATKGVPAGLVQAT | ASTPGQQDTQY | 0.843542 |
| YALKRALAKFSDRFDVVII | ASTPGQQDTQY | 0.90865886 |
| YLIGSVIV | ASTPGQQDTQY | 0.75219953 |
| YGQLVAVID | ASTPGQQDTQY | 0.8892928 |
| VTVGGMTLPA | ASTPGQQDTQY | 0.73482835 |
| VLALPAV | ASTPGQQDTQY | 0.8747821 |
| VIPPTLNYET | ASTPGQQDTQY | 0.7728356 |
| VGMGGMHPSSG | ASTPGQQDTQY | 0.8883401 |
| VGLAGGAATA | ASTPGQQDTQY | 0.88829595 |
| VGGAGGAGGSGP | ASTPGQQDTQY | 0.8926146 |
| VFAAFMV | ASTPGQQDTQY | 0.25449035 |
| TLYDKLR | ASTPGQQDTQY | 0.8983621 |
| TLAVFTVL | ASTPGQQDTQY | 0.87904865 |
| TGSVQPARV | ASTPGQQDTQY | 0.9141536 |
| TGGGAGSGGT | ASTPGQQDTQY | 0.847893 |
| SHLETSL | ASTPGQQDTQY | 0.8561542 |
| SFTTGMT | ASTPGQQDTQY | 0.81912553 |
| QVEVAKRMGVRQPTVS | ASTPGQQDTQY | 0.84376985 |

| | | |
|---|---|---|
| QGSGGPALAM | ASTPGQQDTQY | 0.69683325 |
| MLAPSNP | ASTPGQQDTQY | 0.77502143 |
| LVNNAAI | ASTPGQQDTQY | 0.78573513 |
| LTGEINTLA | ASTPGQQDTQY | 0.76947415 |
| LSIFGPV | ASTPGQQDTQY | 0.88441736 |
| LLALLANL | ASTPGQQDTQY | 0.8974625 |
| LISNASDALDKLR | ASTPGQQDTQY | 0.89379895 |
| LINEAVP | ASTPGQQDTQY | 0.90506566 |
| LGYAFSGL | ASTPGQQDTQY | 0.90112984 |
| LDFVAVAL | ASTPGQQDTQY | 0.90484047 |
| LAQPGAE | ASTPGQQDTQY | 0.81346494 |
| ISNLGAVTGF | ASTPGQQDTQY | 0.81651056 |
| HQEDPGRI | ASTPGQQDTQY | 0.70202994 |
| GTPGEPTP | ASTPGQQDTQY | 0.83908284 |
| GFFNSGS | ASTPGQQDTQY | 0.8290814 |
| GAGGGGLTAGAN | ASTPGQQDTQY | 0.8699604 |
| FVDALVNM | ASTPGQQDTQY | 0.1339121 |
| FLTVAPEL | ASTPGQQDTQY | 0.83972275 |
| FETLTAL | ASTPGQQDTQY | 0.9100158 |
| DVINPIEL | ASTPGQQDTQY | 0.7283612 |
| APVSGAPV | ASTPGQQDTQY | 0.86019707 |
| ALVGGLAW | ASTPGQQDTQY | 0.8405842 |
| ALAAAAATAI | ASTPGQQDTQY | 0.80107594 |
| AFGLGLLL | ASTPGQQDTQY | 0.8897279 |
| AFFAAASL | ASTPGQQDTQY | 0.79197395 |
| AAAALPAFSPP | ASTPGQQDTQY | 0.8781643 |
| VVLAPAL | ASTPGQQDTQY | 0.90853155 |
| TIGHVDHGKTTL | ASTPGQQDTQY | 0.85355294 |
| LISNASDALDK | ASTPGQQDTQY | 0.8531698 |
| LISNASDAL | ASTPGQQDTQY | 0.83996654 |
| ISNASDALDK | ASTPGQQDTQY | 0.50528145 |
| NQAETLVYQTE | ASTPGQQDTQY | 0.87324905 |
| TGGTGGTAGAG | ASTPGQQDTQY | 0.48544836 |
| TGSTAGGAGGA | ASTPGQQDTQY | 0.8354336 |
| DPLAVPVAAGPV | ASTPGQQDTQY | 0.7885387 |
| GGTGGTPGNGGN | ASTPGQQDTQY | 0.87179506 |
| PAGLAGVDPATL | ASTPGQQDTQY | 0.84915125 |
| GNNLVVEAPHAGASV | ASTPGQQDTQY | 0.87788033 |
| KVTRIMSASTLEQPA | ASTPGQQDTQY | 0.7192703 |
| LGFALPIALPRRTVV | ASTPGQQDTQY | 0.8624146 |
| TRLGNLTARKLVVLV | ASTPGQQDTQY | 0.8138663 |
| WSQQRRLIYFPSAGP | ASTPGQQDTQY | 0.7138182 |
| GPNGNPDPMAAAVDIR | ASTPGQQDTQY | 0.8830798 |
| TQRCVSVADLRAAASR | ASTPGQQDTQY | 0.85054225 |
| QFVLDVAEHPVEVLTETV | ASTPGQQDTQY | 0.8153734 |
| GGAGGTGGAGGNGGTQAGDG | ASTPGQQDTQY | 0.9049741 |
| PLLATLRAIAVDVDAKPAQV | ASTPGQQDTQY | 0.8768908 |
| APKFRTGGYMIDFWGVGYQVAK | ASTPGQQDTQY | 0.5417181 |
| GAGGAGGAGGAAGGPGGTGGTG | ASTPGQQDTQY | 0.81288135 |
| IGALGLLATDVAPAGSVAFGVA | ASTPGQQDTQY | 0.8386135 |
| AGGAGGIGGNGGAGGAGGVLMG | ASTPGQQDTQY | 0.8366759 |

| | | |
|---|---|---|
| ADELIGLL | ASTPGQQDTQY | 0.1682985 |
| ADRALKA | ASTPGQQDTQY | 0.8829664 |
| AGAVGQVGGAGGA | ASTPGQQDTQY | 0.75577617 |
| AGGGGGLGGAP | ASTPGQQDTQY | 0.7312335 |
| AGGLLYGNGGAG | ASTPGQQDTQY | 0.8138098 |
| AGLESIGL | ASTPGQQDTQY | 0.88568586 |
| AGLKTGII | ASTPGQQDTQY | 0.7819169 |
| ASLAALLL | ASTPGQQDTQY | 0.84056455 |
| ASVINISSI | ASTPGQQDTQY | 0.8794323 |
| ATPPGPA | ASTPGQQDTQY | 0.8931534 |
| ATVASALV | ASTPGQQDTQY | 0.9067502 |
| ATVASLIL | ASTPGQQDTQY | 0.8849466 |
| AVNGPVLIRH | ASTPGQQDTQY | 0.8328569 |
| DGAAGAVGI | ASTPGQQDTQY | 0.873373 |
| DPLLHGI | ASTPGQQDTQY | 0.81199074 |
| ELGAALGAL | ASTPGQQDTQY | 0.7224108 |
| ELGRTQAV | ASTPGQQDTQY | 0.5979388 |
| FVAGGYLS | ASTPGQQDTQY | 0.85389954 |
| GASGGAGGAGGW | ASTPGQQDTQY | 0.6139229 |
| GGAGAGGDGGAGGI | ASTPGQQDTQY | 0.8603691 |
| GTGHNGSQPGMGGNG | ASTPGQQDTQY | 0.9058963 |
| GYAAASASAS | ASTPGQQDTQY | 0.8230874 |
| HQVDVAAVQQG | ASTPGQQDTQY | 0.6423134 |
| IGAAIAHT | ASTPGQQDTQY | 0.8244084 |
| IGMIEVAQL | ASTPGQQDTQY | 0.786154 |
| IRLAPEHTSAMD | ASTPGQQDTQY | 0.8855237 |
| IYNVSVL | ASTPGQQDTQY | 0.8967486 |
| KQGFLGM | ASTPGQQDTQY | 0.9130835 |
| LESVITQL | ASTPGQQDTQY | 0.90179396 |
| LGLGALVVV | ASTPGQQDTQY | 0.7979309 |
| LIGNGGNGGSGGI | ASTPGQQDTQY | 0.87808853 |
| LNAINAPTL | ASTPGQQDTQY | 0.8222146 |
| LQLLNTGV | ASTPGQQDTQY | 0.7989973 |
| LQTSIEF | ASTPGQQDTQY | 0.8562981 |
| MSFAVSEL | ASTPGQQDTQY | 0.8631026 |
| NTGALGL | ASTPGQQDTQY | 0.8608544 |
| PPGAGGAGGAGTT | ASTPGQQDTQY | 0.81492245 |
| QLVGLGI | ASTPGQQDTQY | 0.5251614 |
| RFLEQVHQL | ASTPGQQDTQY | 0.88487923 |
| RGVGYVLR | ASTPGQQDTQY | 0.88460517 |
| SATGVFAGVI | ASTPGQQDTQY | 0.6387023 |
| SLPAITLP | ASTPGQQDTQY | 0.822295 |
| STALAFTV | ASTPGQQDTQY | 0.9129095 |
| SVLTVGLGK | ASTPGQQDTQY | 0.9025812 |
| TALAGAGSGI | ASTPGQQDTQY | 0.76294416 |
| TGLPFTGI | ASTPGQQDTQY | 0.81160074 |
| VAEHVTV | ASTPGQQDTQY | 0.82810926 |
| VDIVLNSL | ASTPGQQDTQY | 0.53999776 |
| VLTGGGAVL | ASTPGQQDTQY | 0.88405263 |
| VNVGLAYGI | ASTPGQQDTQY | 0.8403595 |
| VSSAAVGAV | ASTPGQQDTQY | 0.8366536 |

| | | |
|---|---|---|
| VTSVYEAGV | ASTPGQQDTQY | 0.75778127 |
| WVALDRGA | ASTPGQQDTQY | 0.85607576 |
| YLVGSVVI | ASTPGQQDTQY | 0.8079732 |
| GPPGTGKTLL | ASTPGQQDTQY | 0.89031154 |
| GVGMTKF | ASTPGQQDTQY | 0.15571377 |
| IGIGGSDL | ASTPGQQDTQY | 0.8923584 |
| IGTLLAEL | ASTPGQQDTQY | 0.7351062 |
| IYTSGTT | ASTPGQQDTQY | 0.7627038 |
| LSFPAGV | ASTPGQQDTQY | 0.45558286 |
| LYTSGTT | ASTPGQQDTQY | 0.818484 |
| VGAFPFGF | ASTPGQQDTQY | 0.88520736 |
| VGAGTVEF | ASTPGQQDTQY | 0.8951448 |
| VGKSTLLL | ASTPGQQDTQY | 0.75480115 |
| AALAHAGGGIV | ASTPGQQDTQY | 0.89714503 |
| AALVAVSLL | ASTPGQQDTQY | 0.91333807 |
| AAVSRAVAAL | ASTPGQQDTQY | 0.8020096 |
| AGGSSGIGGF | ASTPGQQDTQY | 0.4880527 |
| AGLTVANANG | ASTPGQQDTQY | 0.88360465 |
| AGLVGAVL | ASTPGQQDTQY | 0.8022319 |
| ALSQAFTV | ASTPGQQDTQY | 0.89309645 |
| ALTFSVSL | ASTPGQQDTQY | 0.8670987 |
| ALTGAALVAL | ASTPGQQDTQY | 0.68293655 |
| ASFSIGAGVL | ASTPGQQDTQY | 0.8889487 |
| DGSVYKI | ASTPGQQDTQY | 0.8993615 |
| EQLAEIV | ASTPGQQDTQY | 0.760791 |
| FAVSELTVM | ASTPGQQDTQY | 0.73450184 |
| GAAPLAAGL | ASTPGQQDTQY | 0.7458441 |
| GADGGDGGSGGAGG | ASTPGQQDTQY | 0.4068964 |
| GFGLDTGGAGG | ASTPGQQDTQY | 0.8969292 |
| GLASGGLL | ASTPGQQDTQY | 0.77690864 |
| GRIAGAAVLL | ASTPGQQDTQY | 0.7839967 |
| GSAGSLNTSF | ASTPGQQDTQY | 0.8668698 |
| GTGQIGI | ASTPGQQDTQY | 0.8165455 |
| GTLIALIL | ASTPGQQDTQY | 0.80776787 |
| IAATMGFVVG | ASTPGQQDTQY | 0.82471025 |
| IFAQALL | ASTPGQQDTQY | 0.86987203 |
| IGAAVALGF | ASTPGQQDTQY | 0.87008286 |
| IGLTPLAL | ASTPGQQDTQY | 0.8260701 |
| ILGSNTSTL | ASTPGQQDTQY | 0.7971549 |
| IVFSLTVL | ASTPGQQDTQY | 0.90606755 |
| LASKATGF | ASTPGQQDTQY | 0.7014122 |
| LASVALIL | ASTPGQQDTQY | 0.8059006 |
| LFAVGLTL | ASTPGQQDTQY | 0.86940265 |
| LFNGGIGTYI | ASTPGQQDTQY | 0.72165906 |
| LTLGAGII | ASTPGQQDTQY | 0.88461506 |
| LVAAALGDI | ASTPGQQDTQY | 0.8512511 |
| NTAFVQLGI | ASTPGQQDTQY | 0.8639405 |
| QLFSGAEL | ASTPGQQDTQY | 0.8199725 |
| SAATFITL | ASTPGQQDTQY | 0.91325104 |
| SNYGLANLGV | ASTPGQQDTQY | 0.89440715 |
| SYGGAYAVM | ASTPGQQDTQY | 0.834944 |

| | | |
|---|---|---|
| TTGVGGLTLG | ASTPGQQDTQY | 0.86613214 |
| TTQTIYI | ASTPGQQDTQY | 0.626347 |
| VAAGAGLGAVI | ASTPGQQDTQY | 0.9014394 |
| VAAVVTLL | ASTPGQQDTQY | 0.8753457 |
| VGFLGGLI | ASTPGQQDTQY | 0.72812307 |
| VGVIPILL | ASTPGQQDTQY | 0.8477471 |
| VIFASSLI | ASTPGQQDTQY | 0.90927577 |
| VKGGDGGAAAG | ASTPGQQDTQY | 0.8967229 |
| VTKTFTNI | ASTPGQQDTQY | 0.6892437 |
| VVAAAILL | ASTPGQQDTQY | 0.7973056 |
| YAALDVEL | ASTPGQQDTQY | 0.49106437 |
| LASVALLL | ASTPGQQDTQY | 0.7769604 |
| LDAVHRV | ASTPGQQDTQY | 0.65405244 |
| QGLSSSIF | ASTPGQQDTQY | 0.8785542 |
| RGTFANIRL | ASTPGQQDTQY | 0.7968653 |
| RQLWWGHRI | ASTPGQQDTQY | 0.84761024 |
| YGPPGTGKTLL | ASTPGQQDTQY | 0.8615991 |
| AALVGAPANVA | ASTPGQQDTQY | 0.9106389 |
| AATFRRTE | ASTPGQQDTQY | 0.7873807 |
| AGALGGGI | ASTPGQQDTQY | 0.84920084 |
| AGGAGGAGGIGAG | ASTPGQQDTQY | 0.59512955 |
| AGSGALGL | ASTPGQQDTQY | 0.6914773 |
| AQIADQI | ASTPGQQDTQY | 0.77703285 |
| ASAVAVTYL | ASTPGQQDTQY | 0.8487098 |
| ASTVLGGTGG | ASTPGQQDTQY | 0.31150877 |
| AVGGEPAK | ASTPGQQDTQY | 0.90550154 |
| CEQGTFGGTC | ASTPGQQDTQY | 0.86369157 |
| DEGKAPAG | ASTPGQQDTQY | 0.88472813 |
| DTVVFPAL | ASTPGQQDTQY | 0.8184674 |
| FEGITMAG | ASTPGQQDTQY | 0.815011 |
| FPLVVQD | ASTPGQQDTQY | 0.8140253 |
| GAFTTPQ | ASTPGQQDTQY | 0.879374 |
| GFGGFGTAGGVGGA | ASTPGQQDTQY | 0.77179503 |
| GFLAQFL | ASTPGQQDTQY | 0.5086654 |
| GGAGGAGGNAGVA | ASTPGQQDTQY | 0.7206777 |
| GGVGTAGAGGAGGAGG | ASTPGQQDTQY | 0.6253513 |
| GNGGGGDAGNAGSGGN | ASTPGQQDTQY | 0.9075838 |
| GTGGAGGAGNDAGS | ASTPGQQDTQY | 0.9005766 |
| GTTNTGWFNS | ASTPGQQDTQY | 0.8791638 |
| IFMGPLFTL | ASTPGQQDTQY | 0.90296865 |
| ILAHAIEAI | ASTPGQQDTQY | 0.2787677 |
| IRSAIREAL | ASTPGQQDTQY | 0.81287813 |
| ITFAGGG | ASTPGQQDTQY | 0.90627366 |
| ITGMSGHQEV | ASTPGQQDTQY | 0.8788368 |
| IVAGLACTL | ASTPGQQDTQY | 0.81725496 |
| IVGGGLAGA | ASTPGQQDTQY | 0.9053443 |
| IVTGVLNI | ASTPGQQDTQY | 0.7735723 |
| LAQQGCI | ASTPGQQDTQY | 0.83275896 |
| LCLHVEN | ASTPGQQDTQY | 0.85692537 |
| LDAISDGS | ASTPGQQDTQY | 0.894304 |
| LGGFAAML | ASTPGQQDTQY | 0.8578923 |

| | | |
|---|---|---|
| LGGKNPMI | ASTPGQQDTQY | 0.7368836 |
| LGPKGVAAIL | ASTPGQQDTQY | 0.8593357 |
| LMVGVSI | ASTPGQQDTQY | 0.9173372 |
| LVGAAFVL | ASTPGQQDTQY | 0.8156704 |
| LVLAVAPL | ASTPGQQDTQY | 0.8893658 |
| MAAIAVIDAA | ASTPGQQDTQY | 0.87374705 |
| NNGINAELQ | ASTPGQQDTQY | 0.85410327 |
| RGEPLVLVHG | ASTPGQQDTQY | 0.74674964 |
| RGGAGGAAGLI | ASTPGQQDTQY | 0.79286754 |
| RSYRMHPAV | ASTPGQQDTQY | 0.8764204 |
| SAAHLLAGQLA | ASTPGQQDTQY | 0.7249306 |
| SAAVGSY | ASTPGQQDTQY | 0.82617676 |
| SPHAVLHRI | ASTPGQQDTQY | 0.80310553 |
| SQIAKVL | ASTPGQQDTQY | 0.9220876 |
| TALGHLVP | ASTPGQQDTQY | 0.8730329 |
| TAPTTSGAA | ASTPGQQDTQY | 0.8856277 |
| TDGDDGNGGSPGT | ASTPGQQDTQY | 0.7807988 |
| TDGLGEN | ASTPGQQDTQY | 0.8803197 |
| TGGPPLPGQA | ASTPGQQDTQY | 0.88201904 |
| TIPAITI | ASTPGQQDTQY | 0.58185273 |
| TIPGFTI | ASTPGQQDTQY | 0.9077169 |
| TSIASGALA | ASTPGQQDTQY | 0.7786459 |
| TVTLPGAAI | ASTPGQQDTQY | 0.80623615 |
| VASGLSAALP | ASTPGQQDTQY | 0.8938035 |
| VGAGVIGI | ASTPGQQDTQY | 0.90157616 |
| VGPKRFGV | ASTPGQQDTQY | 0.82374924 |
| VNVAFPD | ASTPGQQDTQY | 0.8228518 |
| VSMTNSGVS | ASTPGQQDTQY | 0.7733423 |
| VTDVLGDLSV | ASTPGQQDTQY | 0.89729816 |
| VVVVGAGFAGL | ASTPGQQDTQY | 0.9013965 |
| VYTVGTVV | ASTPGQQDTQY | 0.82037306 |
| WTTFSGP | ASTPGQQDTQY | 0.9118427 |
| YDELVEAF | ASTPGQQDTQY | 0.8741214 |
| YGTGGAGGNGG | ASTPGQQDTQY | 0.5115181 |
| YTNVMAV | ASTPGQQDTQY | 0.85117626 |
| FLEQVHQL | ASTPGQQDTQY | 0.89591897 |
| GLSLLAVAV | ASTPGQQDTQY | 0.89178205 |
| LEQVHQL | ASTPGQQDTQY | 0.9137577 |
| LNAAGAI | ASTPGQQDTQY | 0.8721562 |
| PAVHASL | ASTPGQQDTQY | 0.81359375 |
| TELEDDA | ASTPGQQDTQY | 0.9099227 |
| TPPGPPN | ASTPGQQDTQY | 0.91670597 |
| TPPPPPPP | ASTPGQQDTQY | 0.88881063 |
| TQSPALI | ASSLSYNSPLY | 0.78389144 |
| EFAAAFPQ | ASSLSYNSPLY | 0.76752126 |
| LLSGNLDVL | ASSLSYNSPLY | 0.8625281 |
| GPANLGNYNI | ASSLSYNSPLY | 0.8971062 |
| TLGPTLP | ASSLSYNSPLY | 0.73349917 |
| VYAYSSCCK | ASSLSYNSPLY | 0.8599091 |
| APAPAPAPNI | ASSLSYNSPLY | 0.8645228 |
| PGGPVPIP | ASSLSYNSPLY | 0.5403054 |

| | | |
|---|---|---|
| FLTVALEP | ASSLSYNSPLY | 0.8804331 |
| SQYFGANL | ASSLSYNSPLY | 0.9028524 |
| GGVKPAA | ASSLSYNSPLY | 0.85096866 |
| GGTGGAGGAGGW | ASSLSYNSPLY | 0.8000778 |
| GVGGVAGLGGAGG | ASSLSYNSPLY | 0.90011036 |
| VLLVLATTI | ASSLSYNSPLY | 0.9221199 |
| IGIFAAML | ASSLSYNSPLY | 0.9091983 |
| AVGDLID | ASSLSYNSPLY | 0.57202387 |
| NISPLPL | ASSLSYNSPLY | 0.34633392 |
| VTAVFVAI | ASSLSYNSPLY | 0.7896822 |
| SGDDNDAAL | ASSLSYNSPLY | 0.8707862 |
| AVAALAHS | ASSLSYNSPLY | 0.84316933 |
| GGVTVMATP | ASSLSYNSPLY | 0.83312416 |
| AGLAANVRD | ASSLSYNSPLY | 0.85855395 |
| TNRPDLIDPALLRPG | ASSLSYNSPLY | 0.90166223 |
| LANLGVTAL | ASSLSYNSPLY | 0.75241953 |
| TGERIEL | ASSLSYNSPLY | 0.6320164 |
| NGEELAA | ASSLSYNSPLY | 0.85176075 |
| VLSLVWG | ASSLSYNSPLY | 0.8840697 |
| IGYALWL | ASSLSYNSPLY | 0.7635108 |
| LSLLLAPL | ASSLSYNSPLY | 0.83313346 |
| TLDEALK | ASSLSYNSPLY | 0.775092 |
| TDGAGTAEGG | ASSLSYNSPLY | 0.79100287 |
| NAPGGAGGNGG | ASSLSYNSPLY | 0.8680391 |
| VSVFMGMGGNF | ASSLSYNSPLY | 0.7567414 |
| LASYLAAL | ASSLSYNSPLY | 0.9070194 |
| LVGGTPAL | ASSLSYNSPLY | 0.74742234 |
| VRNADVL | ASSLSYNSPLY | 0.8883526 |
| TCSSPFLVSAM | ASSLSYNSPLY | 0.86789334 |
| VHITDAE | ASSLSYNSPLY | 0.88084865 |
| EYAAPSF | ASSLSYNSPLY | 0.81201863 |
| VVDAFHGGL | ASSLSYNSPLY | 0.78396946 |
| ASVVDPQ | ASSLSYNSPLY | 0.7002922 |
| SYNEFGGLS | ASSLSYNSPLY | 0.5821362 |
| DPDGSVY | ASSLSYNSPLY | 0.8978723 |
| MVSTIHATE | ASSLSYNSPLY | 0.71031827 |
| LAAGVADPV | ASSLSYNSPLY | 0.9033799 |
| TKEQIAA | ASSLSYNSPLY | 0.89518344 |
| SLASPAAL | ASSLSYNSPLY | 0.76803833 |
| IGLVFYGV | ASSLSYNSPLY | 0.8783436 |
| FLLDASNL | ASSLSYNSPLY | 0.7582238 |
| YQVGNFP | ASSLSYNSPLY | 0.8178133 |
| ESLLSQST | ASSLSYNSPLY | 0.88411266 |
| LFTVGFAAVG | ASSLSYNSPLY | 0.88377064 |
| ATPPTPT | ASSLSYNSPLY | 0.7266735 |
| TDLATLG | ASSLSYNSPLY | 0.8729421 |
| SAGGTGR | ASSLSYNSPLY | 0.8629752 |
| GFSNSGS | ASSLSYNSPLY | 0.69564456 |
| GGNGGSGGTSV | ASSLSYNSPLY | 0.8599328 |
| GTSVATGGAGN | ASSLSYNSPLY | 0.89904314 |
| NGGNGGSGGTS | ASSLSYNSPLY | 0.8789682 |

| | | |
|---|---|---|
| AVVKFPH | ASSLSYNSPLY | 0.7390091 |
| TPPPPPP | ASSLSYNSPLY | 0.8828569 |
| AAPAAAAAPA | ASSLSYNSPLY | 0.7256664 |
| IALNEDL | ASSLSYNSPLY | 0.8698524 |
| IVNVSSM | ASSLSYNSPLY | 0.85489964 |
| TGIPPAPR | ASSLSYNSPLY | 0.8061558 |
| FDLGGGTF | ASSLSYNSPLY | 0.8846246 |
| FDLGGGTFDVS | ASSLSYNSPLY | 0.5599826 |
| NEPTAAA | ASSLSYNSPLY | 0.74735296 |
| NEPTAAALA | ASSLSYNSPLY | 0.82059133 |
| GSIINISSI | ASSLSYNSPLY | 0.8652882 |
| VAPGNAGTA | ASSLSYNSPLY | 0.88980854 |
| TGGGASLE | ASSLSYNSPLY | 0.75178945 |
| VAVYNHYKRI | ASSLSYNSPLY | 0.85569507 |
| TDPAPATT | ASSLSYNSPLY | 0.82351804 |
| TNRPDILDPALLRPG | ASSLSYNSPLY | 0.74839133 |
| LTTTTTS | ASSLSYNSPLY | 0.8614503 |
| PTGSGKTL | ASSLSYNSPLY | 0.8311097 |
| TGSGKTL | ASSLSYNSPLY | 0.73306274 |
| TIGHVDHGK | ASSLSYNSPLY | 0.8879603 |
| TIGHVDHGKTT | ASSLSYNSPLY | 0.782047 |
| GHVDHGKTT | ASSLSYNSPLY | 0.88622576 |
| AQFEHTLL | ASSLSYNSPLY | 0.91330767 |
| IVNQDEL | ASSLSYNSPLY | 0.7671073 |
| TNGSQFF | ASSLSYNSPLY | 0.8865968 |
| TGKGIVE | ASSLSYNSPLY | 0.8765087 |
| LISNASDA | ASSLSYNSPLY | 0.87431943 |
| AELRTLGELTKIAKAHGAQ | ASSLSYNSPLY | 0.84025496 |
| APNAVAAAEPSW | ASSLSYNSPLY | 0.78828025 |
| APQVAGAATAVL | ASSLSYNSPLY | 0.9023051 |
| ARRDPAAAAAILPTDA | ASSLSYNSPLY | 0.8886978 |
| DAAARDVLAEAGLAEYF | ASSLSYNSPLY | 0.87290573 |
| DDQIVLSAKANLAKVEQ | ASSLSYNSPLY | 0.8794846 |
| DELLPDTGEDSDEDGH | ASSLSYNSPLY | 0.8602543 |
| ELPEAPANRATVPQD | ASSLSYNSPLY | 0.8780575 |
| FGQSKIERVTGIG | ASSLSYNSPLY | 0.7852812 |
| GAGGTGSTAGGAGGAGGAGGLY | ASSLSYNSPLY | 0.67323667 |
| GHGGTGAAGGNGGTGGAGGSGI | ASSLSYNSPLY | 0.8416679 |
| GNGGNGGAGGEGGPGGAGGAGA | ASSLSYNSPLY | 0.85295707 |
| GSTAGNGGNGGNATNGTIAGQP | ASSLSYNSPLY | 0.80062973 |
| GTGGDGGNGGNGGTGG | ASSLSYNSPLY | 0.54294086 |
| GVRRPAGIPADAPV | ASSLSYNSPLY | 0.89538896 |
| GYRQVTDAHLLTLARR | ASSLSYNSPLY | 0.91945875 |
| IAKMGQDGHDRG | ASSLSYNSPLY | 0.7731923 |
| KAEPSIEIQYT | ASSLSYNSPLY | 0.91665053 |
| KPVRIRMAADEDTVAVT | ASSLSYNSPLY | 0.6895106 |
| MPPRIAGMRLLVIKPEP | ASSLSYNSPLY | 0.89116657 |
| MKFYAVALFRSRGVPPTR | ASSLSYNSPLY | 0.8981171 |
| NAGGFGILGAVAHSP | ASSLSYNSPLY | 0.7408811 |
| NEGREITLAMAEM | ASSLSYNSPLY | 0.8908478 |
| NELVEWSHPLGHRWRWE | ASSLSYNSPLY | 0.74740994 |

| | | |
|---|---|---|
| NLGAGGTGGNGGNPG | ASSLSYNSPLY | 0.82201767 |
| NTEWPKWETVMAADPVD | ASSLSYNSPLY | 0.9050007 |
| PASAAPAETQP | ASSLSYNSPLY | 0.8089264 |
| QQSSDGKNGVRAAEITGE | ASSLSYNSPLY | 0.8729102 |
| RAPDAVTTSWWKE | ASSLSYNSPLY | 0.86108315 |
| SGFGNVGAHTSGWFNQ | ASSLSYNSPLY | 0.8023491 |
| SGVAGAVGQAASVGGLKV | ASSLSYNSPLY | 0.8775718 |
| SNGGAGTGSAGGNGGT | ASSLSYNSPLY | 0.9000899 |
| TGGAGGEGGAAG | ASSLSYNSPLY | 0.81916916 |
| TGGDGGAAGAGGAPGHG | ASSLSYNSPLY | 0.91484857 |
| TMPPLLAAIANAGRHGV | ASSLSYNSPLY | 0.7726933 |
| TSSTAGNSGGA | ASSLSYNSPLY | 0.9147078 |
| VPATVPVGILC | ASSLSYNSPLY | 0.8759023 |
| VQRPDSGGVVLTGRIS | ASSLSYNSPLY | 0.8543699 |
| WTPNPVPATKGVPAGLVQAT | ASSLSYNSPLY | 0.8194952 |
| YALKRALAKFSDRFDVVII | ASSLSYNSPLY | 0.90150243 |
| YLIGSVIV | ASSLSYNSPLY | 0.8097052 |
| YGQLVAVID | ASSLSYNSPLY | 0.8823315 |
| VTVGGMTLPA | ASSLSYNSPLY | 0.91443026 |
| VLALPAV | ASSLSYNSPLY | 0.5732472 |
| VIPPTLNYET | ASSLSYNSPLY | 0.82279885 |
| VGMGGMHPSSG | ASSLSYNSPLY | 0.74985284 |
| VGLAGGAATA | ASSLSYNSPLY | 0.6411073 |
| VGGAGGAGGSGP | ASSLSYNSPLY | 0.5927025 |
| VFAAFMV | ASSLSYNSPLY | 0.34914154 |
| TLYDKLR | ASSLSYNSPLY | 0.65791255 |
| TLAVFTVL | ASSLSYNSPLY | 0.5321849 |
| TGSVQPARV | ASSLSYNSPLY | 0.7473687 |
| TGGGAGSGGT | ASSLSYNSPLY | 0.7776046 |
| SHLETSL | ASSLSYNSPLY | 0.52209485 |
| SFTTGMT | ASSLSYNSPLY | 0.8320271 |
| QVEVAKRMGVRQPTVS | ASSLSYNSPLY | 0.7963781 |
| QGSGGPALAM | ASSLSYNSPLY | 0.40911174 |
| MLAPSNP | ASSLSYNSPLY | 0.6245333 |
| LVNNAAI | ASSLSYNSPLY | 0.534176 |
| LTGEINTLA | ASSLSYNSPLY | 0.65035444 |
| LSIFGPV | ASSLSYNSPLY | 0.26182964 |
| LLALLANL | ASSLSYNSPLY | 0.78942955 |
| LISNASDALDKLR | ASSLSYNSPLY | 0.54450065 |
| LINEAVP | ASSLSYNSPLY | 0.12864839 |
| LGYAFSGL | ASSLSYNSPLY | 0.7174038 |
| LDFVAVAL | ASSLSYNSPLY | 0.63081795 |
| LAQPGAE | ASSLSYNSPLY | 0.69935715 |
| ISNLGAVTGF | ASSLSYNSPLY | 0.8002999 |
| HQEDPGRI | ASSLSYNSPLY | 0.39333892 |
| GTPGEPTP | ASSLSYNSPLY | 0.5076116 |
| GFFNSGS | ASSLSYNSPLY | 0.785802 |
| GAGGGGLTAGAN | ASSLSYNSPLY | 0.8357459 |
| FVDALVNM | ASSLSYNSPLY | 0.44283763 |
| FLTVAPEL | ASSLSYNSPLY | 0.7441289 |
| FETLTAL | ASSLSYNSPLY | 0.8047995 |

| | | |
|---|---|---|
| DVINPIEL | ASSLSYNSPLY | 0.7728939 |
| APVSGAPV | ASSLSYNSPLY | 0.80739623 |
| ALVGGLAW | ASSLSYNSPLY | 0.83805573 |
| ALAAAAATAI | ASSLSYNSPLY | 0.40858674 |
| AFGLGLLL | ASSLSYNSPLY | 0.815729 |
| AFFAAASL | ASSLSYNSPLY | 0.841887 |
| AAAALPAFSPP | ASSLSYNSPLY | 0.6179603 |
| VVLAPAL | ASSLSYNSPLY | 0.6835861 |
| TIGHVDHGKTTL | ASSLSYNSPLY | 0.53454494 |
| LISNASDALDK | ASSLSYNSPLY | 0.5588334 |
| LISNASDAL | ASSLSYNSPLY | 0.2613375 |
| ISNASDALDK | ASSLSYNSPLY | 0.7446047 |
| NQAETLVYQTE | ASSLSYNSPLY | 0.8195522 |
| TGGTGGTAGAG | ASSLSYNSPLY | 0.6090202 |
| TGSTAGGAGGA | ASSLSYNSPLY | 0.45871058 |
| DPLAVPVAAGPV | ASSLSYNSPLY | 0.47121835 |
| GGTGGTPGNGGN | ASSLSYNSPLY | 0.59609324 |
| PAGLAGVDPATL | ASSLSYNSPLY | 0.6118321 |
| GNNLVVEAPHAGASV | ASSLSYNSPLY | 0.26704156 |
| KVTRIMSASTLEQPA | ASSLSYNSPLY | 0.54870856 |
| LGFALPIALPRRTVV | ASSLSYNSPLY | 0.81857884 |
| TRLGNLTARKLVVLV | ASSLSYNSPLY | 0.82783484 |
| WSQQRRLIYFPSAGP | ASSLSYNSPLY | 0.21955493 |
| GPNGNPDPMAAAVDIR | ASSLSYNSPLY | 0.82136524 |
| TQRCVSVADLRAAASR | ASSLSYNSPLY | 0.76340944 |
| QFVLDVAEHPVEVLTETV | ASSLSYNSPLY | 0.83009714 |
| GGAGGTGGAGGNGGTQAGDG | ASSLSYNSPLY | 0.6489123 |
| PLLATLRAIAVDVDAKPAQV | ASSLSYNSPLY | 0.5732831 |
| APKFRTGGYMIDFWGVGYQVAK | ASSLSYNSPLY | 0.7393069 |
| GAGGAGGAGGAAGGPGGTGGTG | ASSLSYNSPLY | 0.5954099 |
| IGALGLLATDVAPAGSVAFGVA | ASSLSYNSPLY | 0.40934002 |
| AGGAGGIGGNGGAGGAGGVLMG | ASSLSYNSPLY | 0.7939888 |
| ADELIGLL | ASSLSYNSPLY | 0.66838294 |
| ADRALKA | ASSLSYNSPLY | 0.54358554 |
| AGAVGQVGGAGGA | ASSLSYNSPLY | 0.52562857 |
| AGGGGGLGGAP | ASSLSYNSPLY | 0.81871927 |
| AGGLLYGNGGAG | ASSLSYNSPLY | 0.3538666 |
| AGLESIGL | ASSLSYNSPLY | 0.5369057 |
| AGLKTGII | ASSLSYNSPLY | 0.3773747 |
| ASLAALLL | ASSLSYNSPLY | 0.67611516 |
| ASVINISSI | ASSLSYNSPLY | 0.81262136 |
| ATPPGPA | ASSLSYNSPLY | 0.77200466 |
| ATVASALV | ASSLSYNSPLY | 0.5478881 |
| ATVASLIL | ASSLSYNSPLY | 0.7519859 |
| AVNGPVLIRH | ASSLSYNSPLY | 0.7415162 |
| DGAAGAVGI | ASSLSYNSPLY | 0.66830826 |
| DPLLHGI | ASSLSYNSPLY | 0.77462006 |
| ELGAALGAL | ASSLSYNSPLY | 0.52292496 |
| ELGRTQAV | ASSLSYNSPLY | 0.8197877 |
| FVAGGYLS | ASSLSYNSPLY | 0.71924406 |
| GASGGAGGAGGW | ASSLSYNSPLY | 0.787676 |

| | | |
|---|---|---|
| GGAGAGGDGGAGGI | ASSLSYNSPLY | 0.80102026 |
| GTGHNGSQPGMGGNG | ASSLSYNSPLY | 0.7422701 |
| GYAAASASAS | ASSLSYNSPLY | 0.65004456 |
| HQVDVAAVQQG | ASSLSYNSPLY | 0.20880827 |
| IGAAIAHT | ASSLSYNSPLY | 0.5220874 |
| IGMIEVAQL | ASSLSYNSPLY | 0.74681604 |
| IRLAPEHTSAMD | ASSLSYNSPLY | 0.71504176 |
| IYNVSVL | ASSLSYNSPLY | 0.7808986 |
| KQGFLGM | ASSLSYNSPLY | 0.76368386 |
| LESVITQL | ASSLSYNSPLY | 0.54807484 |
| LGLGALVVV | ASSLSYNSPLY | 0.7612299 |
| LIGNGGNGGSGGI | ASSLSYNSPLY | 0.78724104 |
| LNAINAPTL | ASSLSYNSPLY | 0.31073263 |
| LQLLNTGV | ASSLSYNSPLY | 0.6132077 |
| LQTSIEF | ASSLSYNSPLY | 0.86766654 |
| MSFAVSEL | ASSLSYNSPLY | 0.6953165 |
| NTGALGL | ASSLSYNSPLY | 0.5886052 |
| PPGAGGAGGAGTT | ASSLSYNSPLY | 0.5035385 |
| QLVGLGI | ASSLSYNSPLY | 0.83566725 |
| RFLEQVHQL | ASSLSYNSPLY | 0.50322723 |
| RGVGYVLR | ASSLSYNSPLY | 0.8344282 |
| SATGVFAGVI | ASSLSYNSPLY | 0.15163074 |
| SLPAITLP | ASSLSYNSPLY | 0.54060143 |
| STALAFTV | ASSLSYNSPLY | 0.46110886 |
| SVLTVGLGK | ASSLSYNSPLY | 0.40550208 |
| TALAGAGSGI | ASSLSYNSPLY | 0.80064774 |
| TGLPFTGI | ASSLSYNSPLY | 0.5643985 |
| VAEHVTV | ASSLSYNSPLY | 0.76323813 |
| VDIVLNSL | ASSLSYNSPLY | 0.6866642 |
| VLTGGGAVL | ASSLSYNSPLY | 0.8097521 |
| VNVGLAYGI | ASSLSYNSPLY | 0.6807929 |
| VSSAAVGAV | ASSLSYNSPLY | 0.69642794 |
| VTSVYEAGV | ASSLSYNSPLY | 0.5979004 |
| WVALDRGA | ASSLSYNSPLY | 0.7783232 |
| YLVGSVVI | ASSLSYNSPLY | 0.79728425 |
| GPPGTGKTLL | ASSLSYNSPLY | 0.76145345 |
| GVGMTKF | ASSLSYNSPLY | 0.36646062 |
| IGIGGSDL | ASSLSYNSPLY | 0.5725639 |
| IGTLLAEL | ASSLSYNSPLY | 0.7259397 |
| IYTSGTT | ASSLSYNSPLY | 0.2503309 |
| LSFPAGV | ASSLSYNSPLY | 0.19750908 |
| LYTSGTT | ASSLSYNSPLY | 0.5937309 |
| VGAFPFGF | ASSLSYNSPLY | 0.47837156 |
| VGAGTVEF | ASSLSYNSPLY | 0.80975956 |
| VGKSTLLL | ASSLSYNSPLY | 0.7907859 |
| AALAHAGGGIV | ASSLSYNSPLY | 0.6443684 |
| AALVAVSLL | ASSLSYNSPLY | 0.74774724 |
| AAVSRAVAAL | ASSLSYNSPLY | 0.6493603 |
| AGGSSGIGGF | ASSLSYNSPLY | 0.11386221 |
| AGLTVANANG | ASSLSYNSPLY | 0.45931885 |
| AGLVGAVL | ASSLSYNSPLY | 0.42102793 |

| | | |
|---|---|---|
| ALSQAFTV | ASSLSYNSPLY | 0.8224998 |
| ALTFSVSL | ASSLSYNSPLY | 0.5967436 |
| ALTGAALVAL | ASSLSYNSPLY | 0.45386928 |
| ASFSIGAGVL | ASSLSYNSPLY | 0.6658689 |
| DGSVYKI | ASSLSYNSPLY | 0.65881896 |
| EQLAEIV | ASSLSYNSPLY | 0.6077355 |
| FAVSELTVM | ASSLSYNSPLY | 0.44803807 |
| GAAPLAAGL | ASSLSYNSPLY | 0.14805518 |
| GADGGDGGSGGAGG | ASSLSYNSPLY | 0.7886618 |
| GFGLDTGGAGG | ASSLSYNSPLY | 0.37892509 |
| GLASGGLL | ASSLSYNSPLY | 0.523097 |
| GRIAGAAVLL | ASSLSYNSPLY | 0.1839001 |
| GSAGSLNTSF | ASSLSYNSPLY | 0.48371887 |
| GTGQIGI | ASSLSYNSPLY | 0.30973673 |
| GTLIALIL | ASSLSYNSPLY | 0.79986995 |
| IAATMGFVVG | ASSLSYNSPLY | 0.5955722 |
| IFAQALL | ASSLSYNSPLY | 0.54002666 |
| IGAAVALGF | ASSLSYNSPLY | 0.8480583 |
| IGLTPLAL | ASSLSYNSPLY | 0.64616454 |
| ILGSNTSTL | ASSLSYNSPLY | 0.5713287 |
| IVFSLTVL | ASSLSYNSPLY | 0.5578551 |
| LASKATGF | ASSLSYNSPLY | 0.47502536 |
| LASVALIL | ASSLSYNSPLY | 0.54394597 |
| LFAVGLTL | ASSLSYNSPLY | 0.5071145 |
| LFNGGIGTYI | ASSLSYNSPLY | 0.84397376 |
| LTLGAGII | ASSLSYNSPLY | 0.52884823 |
| LVAAALGDI | ASSLSYNSPLY | 0.42404014 |
| NTAFVQLGI | ASSLSYNSPLY | 0.60690546 |
| QLFSGAEL | ASSLSYNSPLY | 0.8216611 |
| SAATFITL | ASSLSYNSPLY | 0.86222434 |
| SNYGLANLGV | ASSLSYNSPLY | 0.60122585 |
| SYGGAYAVM | ASSLSYNSPLY | 0.5493201 |
| TTGVGGLTLG | ASSLSYNSPLY | 0.5355343 |
| TTQTIYI | ASSLSYNSPLY | 0.6841 |
| VAAGAGLGAVI | ASSLSYNSPLY | 0.5413356 |
| VAAVVTLL | ASSLSYNSPLY | 0.4523612 |
| VGFLGGLI | ASSLSYNSPLY | 0.7854339 |
| VGVIPILL | ASSLSYNSPLY | 0.8469465 |
| VIFASSLI | ASSLSYNSPLY | 0.58838964 |
| VKGGDGGAAAG | ASSLSYNSPLY | 0.37023574 |
| VTKTFTNI | ASSLSYNSPLY | 0.6235814 |
| VVAAAILL | ASSLSYNSPLY | 0.4873404 |
| YAALDVEL | ASSLSYNSPLY | 0.6174495 |
| LASVALLL | ASSLSYNSPLY | 0.7321174 |
| LDAVHRV | ASSLSYNSPLY | 0.30259323 |
| QGLSSSIF | ASSLSYNSPLY | 0.55850387 |
| RGTFANIRL | ASSLSYNSPLY | 0.861417 |
| RQLWWGHRI | ASSLSYNSPLY | 0.58523405 |
| YGPPGTGKTLL | ASSLSYNSPLY | 0.7113496 |
| AALVGAPANVA | ASSLSYNSPLY | 0.8188964 |
| AATFRRTE | ASSLSYNSPLY | 0.75293684 |

| | | |
|---|---|---|
| AGALGGGI | ASSLSYNSPLY | 0.829191 |
| AGGAGGAGGIGAG | ASSLSYNSPLY | 0.799519 |
| AGSGALGL | ASSLSYNSPLY | 0.6523901 |
| AQIADQI | ASSLSYNSPLY | 0.60918474 |
| ASAVAVTYL | ASSLSYNSPLY | 0.7208251 |
| ASTVLGGTGG | ASSLSYNSPLY | 0.65934116 |
| AVGGEPAK | ASSLSYNSPLY | 0.62522906 |
| CEQGTFGGTC | ASSLSYNSPLY | 0.39948547 |
| DEGKAPAG | ASSLSYNSPLY | 0.8092745 |
| DTVVFPAL | ASSLSYNSPLY | 0.6251291 |
| FEGITMAG | ASSLSYNSPLY | 0.78379965 |
| FPLVVQD | ASSLSYNSPLY | 0.42908475 |
| GAFTTPQ | ASSLSYNSPLY | 0.48606387 |
| GFGGFGTAGGVGGA | ASSLSYNSPLY | 0.7880934 |
| GFLAQFL | ASSLSYNSPLY | 0.12747525 |
| GGAGGAGGNAGVA | ASSLSYNSPLY | 0.1848859 |
| GGVGTAGAGGAGGAGG | ASSLSYNSPLY | 0.71294725 |
| GNGGGGDAGNAGSGGN | ASSLSYNSPLY | 0.81472266 |
| GTGGAGGAGNDAGS | ASSLSYNSPLY | 0.17045853 |
| GTTNTGWFNS | ASSLSYNSPLY | 0.46374175 |
| IFMGPLFTL | ASSLSYNSPLY | 0.34233296 |
| ILAHAIEAI | ASSLSYNSPLY | 0.75349426 |
| IRSAIREAL | ASSLSYNSPLY | 0.73079634 |
| ITFAGGG | ASSLSYNSPLY | 0.31034565 |
| ITGMSGHQEV | ASSLSYNSPLY | 0.78498435 |
| IVAGLACTL | ASSLSYNSPLY | 0.24602064 |
| IVGGGLAGA | ASSLSYNSPLY | 0.38342953 |
| IVTGVLNI | ASSLSYNSPLY | 0.79844815 |
| LAQQGCI | ASSLSYNSPLY | 0.8189799 |
| LCLHVEN | ASSLSYNSPLY | 0.48771805 |
| LDAISDGS | ASSLSYNSPLY | 0.7870264 |
| LGGFAAML | ASSLSYNSPLY | 0.4861402 |
| LGGKNPMI | ASSLSYNSPLY | 0.857394 |
| LGPKGVAAIL | ASSLSYNSPLY | 0.5807922 |
| LMVGVSI | ASSLSYNSPLY | 0.3602247 |
| LVGAAFVL | ASSLSYNSPLY | 0.8002795 |
| LVLAVAPL | ASSLSYNSPLY | 0.7382566 |
| MAAIAVIDAA | ASSLSYNSPLY | 0.72941947 |
| NNGINAELQ | ASSLSYNSPLY | 0.7857857 |
| RGEPLVLVHG | ASSLSYNSPLY | 0.5454507 |
| RGGAGGAAGLI | ASSLSYNSPLY | 0.61512417 |
| RSYRMHPAV | ASSLSYNSPLY | 0.836506 |
| SAAHLLAGQLA | ASSLSYNSPLY | 0.80405897 |
| SAAVGSY | ASSLSYNSPLY | 0.47275847 |
| SPHAVLHRI | ASSLSYNSPLY | 0.8208067 |
| SQIAKVL | ASSLSYNSPLY | 0.64400876 |
| TALGHLVP | ASSLSYNSPLY | 0.5103444 |
| TAPTTSGAA | ASSLSYNSPLY | 0.84375364 |
| TDGDDGNGGSPGT | ASSLSYNSPLY | 0.710727 |
| TDGLGEN | ASSLSYNSPLY | 0.774343 |
| TGGPPLPGQA | ASSLSYNSPLY | 0.7882424 |

| | | |
|---|---|---|
| TIPAITI | ASSLSYNSPLY | 0.7998955 |
| TIPGFTI | ASSLSYNSPLY | 0.4297088 |
| TSIASGALA | ASSLSYNSPLY | 0.6806581 |
| TVTLPGAAI | ASSLSYNSPLY | 0.6540931 |
| VASGLSAALP | ASSLSYNSPLY | 0.7955501 |
| VGAGVIGI | ASSLSYNSPLY | 0.67634124 |
| VGPKRFGV | ASSLSYNSPLY | 0.527185 |
| VNVAFPD | ASSLSYNSPLY | 0.71735686 |
| VSMTNSGVS | ASSLSYNSPLY | 0.8196841 |
| VTDVLGDLSV | ASSLSYNSPLY | 0.66088486 |
| VVVVGAGFAGL | ASSLSYNSPLY | 0.66967565 |
| VYTVGTVV | ASSLSYNSPLY | 0.84847856 |
| WTTFSGP | ASSLSYNSPLY | 0.6087754 |
| YDELVEAF | ASSLSYNSPLY | 0.604368 |
| YGTGGAGGNGG | ASSLSYNSPLY | 0.7730451 |
| YTNVMAV | ASSLSYNSPLY | 0.78767294 |
| FLEQVHQL | ASSLSYNSPLY | 0.62532455 |
| GLSLLAVAV | ASSLSYNSPLY | 0.83969915 |
| LEQVHQL | ASSLSYNSPLY | 0.48268428 |
| LNAAGAI | ASSLSYNSPLY | 0.78675795 |
| PAVHASL | ASSLSYNSPLY | 0.5331519 |
| TELEDDA | ASSLSYNSPLY | 0.7219393 |
| TPPGPPN | ASSLSYNSPLY | 0.49515706 |
| TPPPPPPP | ASSLSYNSPLY | 0.7706137 |
| TQSPALI | ASGWYSGNTLY | 0.7963725 |
| EFAAAFPQ | ASGWYSGNTLY | 0.79264724 |
| LLSGNLDVL | ASGWYSGNTLY | 0.16498792 |
| GPANLGNYNI | ASGWYSGNTLY | 0.8113702 |
| TLGPTLP | ASGWYSGNTLY | 0.78618085 |
| VYAYSSCCK | ASGWYSGNTLY | 0.8567588 |
| APAPAPAPNI | ASGWYSGNTLY | 0.6292801 |
| PGGPVPIP | ASGWYSGNTLY | 0.7147572 |
| FLTVALEP | ASGWYSGNTLY | 0.636446 |
| SQYFGANL | ASGWYSGNTLY | 0.6491958 |
| GGVKPAA | ASGWYSGNTLY | 0.39711168 |
| GGTGGAGGAGGW | ASGWYSGNTLY | 0.62660366 |
| GVGGVAGLGGAGG | ASGWYSGNTLY | 0.5882168 |
| VLLVLATTI | ASGWYSGNTLY | 0.54119194 |
| IGIFAAML | ASGWYSGNTLY | 0.7592237 |
| AVGDLID | ASGWYSGNTLY | 0.8204783 |
| NISPLPL | ASGWYSGNTLY | 0.8079759 |
| VTAVFVAI | ASGWYSGNTLY | 0.81517106 |
| SGDDNDAAL | ASGWYSGNTLY | 0.81547165 |
| AVAALAHS | ASGWYSGNTLY | 0.8205009 |
| GGVTVMATP | ASGWYSGNTLY | 0.560642 |
| AGLAANVRD | ASGWYSGNTLY | 0.6001492 |
| TNRPDLIDPALLRPG | ASGWYSGNTLY | 0.47834265 |
| LANLGVTAL | ASGWYSGNTLY | 0.56835663 |
| TGERIEL | ASGWYSGNTLY | 0.57443374 |
| NGEELAA | ASGWYSGNTLY | 0.71841043 |
| VLSLVWG | ASGWYSGNTLY | 0.14177212 |

| | | |
|---|---|---|
| IGYALWL | ASGWYSGNTLY | 0.65854037 |
| LSLLLAPL | ASGWYSGNTLY | 0.83418435 |
| TLDEALK | ASGWYSGNTLY | 0.46763152 |
| TDGAGTAEGG | ASGWYSGNTLY | 0.7185642 |
| NAPGGAGGNGG | ASGWYSGNTLY | 0.62343454 |
| VSVFMGMGGNF | ASGWYSGNTLY | 0.51782167 |
| LASYLAAL | ASGWYSGNTLY | 0.7739133 |
| LVGGTPAL | ASGWYSGNTLY | 0.44906092 |
| VRNADVL | ASGWYSGNTLY | 0.74101883 |
| TCSSPFLVSAM | ASGWYSGNTLY | 0.8300728 |
| VHITDAE | ASGWYSGNTLY | 0.6861268 |
| EYAAPSF | ASGWYSGNTLY | 0.6548155 |
| VVDAFHGGL | ASGWYSGNTLY | 0.6116624 |
| ASVVDPQ | ASGWYSGNTLY | 0.3565715 |
| SYNEFGGLS | ASGWYSGNTLY | 0.7210463 |
| DPDGSVY | ASGWYSGNTLY | 0.2665627 |
| MVSTIHATE | ASGWYSGNTLY | 0.6544754 |
| LAAGVADPV | ASGWYSGNTLY | 0.48988962 |
| TKEQIAA | ASGWYSGNTLY | 0.7292504 |
| SLASPAAL | ASGWYSGNTLY | 0.65966403 |
| IGLVFYGV | ASGWYSGNTLY | 0.70631474 |
| FLLDASNL | ASGWYSGNTLY | 0.41909125 |
| YQVGNFP | ASGWYSGNTLY | 0.61782104 |
| ESLLSQST | ASGWYSGNTLY | 0.42229348 |
| LFTVGFAAVG | ASGWYSGNTLY | 0.3991204 |
| ATPPTPT | ASGWYSGNTLY | 0.78532934 |
| TDLATLG | ASGWYSGNTLY | 0.65234447 |
| SAGGTGR | ASGWYSGNTLY | 0.587584 |
| GFSNSGS | ASGWYSGNTLY | 0.82421744 |
| GGNGGSGGTSV | ASGWYSGNTLY | 0.7371293 |
| GTSVATGGAGN | ASGWYSGNTLY | 0.35615593 |
| NGGNGGSGGTS | ASGWYSGNTLY | 0.6361904 |
| AVVKFPH | ASGWYSGNTLY | 0.6372778 |
| TPPPPPP | ASGWYSGNTLY | 0.65934736 |
| AAPAAAAAPA | ASGWYSGNTLY | 0.15839103 |
| IALNEDL | ASGWYSGNTLY | 0.74503565 |
| IVNVSSM | ASGWYSGNTLY | 0.50318533 |
| TGIPPAPR | ASGWYSGNTLY | 0.42618376 |
| FDLGGGTF | ASGWYSGNTLY | 0.57186836 |
| FDLGGGTFDVS | ASGWYSGNTLY | 0.78366995 |
| NEPTAAA | ASGWYSGNTLY | 0.5907129 |
| NEPTAAALA | ASGWYSGNTLY | 0.6812606 |
| GSIINISSI | ASGWYSGNTLY | 0.8046851 |
| VAPGNAGTA | ASGWYSGNTLY | 0.78781855 |
| TGGGASLE | ASGWYSGNTLY | 0.8318627 |
| VAVYNHYKRI | ASGWYSGNTLY | 0.75366694 |
| TDPAPATT | ASGWYSGNTLY | 0.5513175 |
| TNRPDILDPALLRPG | ASGWYSGNTLY | 0.6901705 |
| LTTTTTS | ASGWYSGNTLY | 0.6342559 |
| PTGSGKTL | ASGWYSGNTLY | 0.28120768 |
| TGSGKTL | ASGWYSGNTLY | 0.19217384 |

| | | |
|---|---|---|
| TIGHVDHGK | ASGWYSGNTLY | 0.7052512 |
| TIGHVDHGKTT | ASGWYSGNTLY | 0.20719843 |
| GHVDHGKTT | ASGWYSGNTLY | 0.66740507 |
| AQFEHTLL | ASGWYSGNTLY | 0.8304881 |
| IVNQDEL | ASGWYSGNTLY | 0.6594914 |
| TNGSQFF | ASGWYSGNTLY | 0.39187437 |
| TGKGIVE | ASGWYSGNTLY | 0.58964044 |
| LISNASDA | ASGWYSGNTLY | 0.6301665 |
| AELRTLGELTKIAKAHGAQ | ASGWYSGNTLY | 0.79303193 |
| APNAVAAAEPSW | ASGWYSGNTLY | 0.7565353 |
| APQVAGAATAVL | ASGWYSGNTLY | 0.8496965 |
| ARRDPAAAAAILPTDA | ASGWYSGNTLY | 0.81021 |
| DAAARDVLAEAGLAEYF | ASGWYSGNTLY | 0.54821455 |
| DDQIVLSAKANLAKVEQ | ASGWYSGNTLY | 0.7658646 |
| DELLPDTGEDSDEDGH | ASGWYSGNTLY | 0.55768347 |
| ELPEAPANRATVPQD | ASGWYSGNTLY | 0.5475655 |
| FGQSKIERVTGIG | ASGWYSGNTLY | 0.6400772 |
| GAGGTGSTAGGAGGAGGAGGLY | ASGWYSGNTLY | 0.7239197 |
| GHGGTGAAGGNGGTGGAGGSGI | ASGWYSGNTLY | 0.70453215 |
| GNGGNGGAGGEGGPGGAGGAGA | ASGWYSGNTLY | 0.5935625 |
| GSTAGNGGNGGNATNGTIAGQP | ASGWYSGNTLY | 0.29687265 |
| GTGGDGGNGGNGGTGG | ASGWYSGNTLY | 0.7668158 |
| GVRRPAGIPADAPV | ASGWYSGNTLY | 0.7319893 |
| GYRQVTDAHLLTLARR | ASGWYSGNTLY | 0.2981271 |
| IAKMGQDGHDRG | ASGWYSGNTLY | 0.44879335 |
| KAEPSIEIQYT | ASGWYSGNTLY | 0.8584174 |
| KPVRIRMAADEDTVAVT | ASGWYSGNTLY | 0.8380667 |
| MPPRIAGMRLLVIKPEP | ASGWYSGNTLY | 0.5176331 |
| MKFYAVALFRSRGVPPTR | ASGWYSGNTLY | 0.5235832 |
| NAGGFGILGAVAHSP | ASGWYSGNTLY | 0.6513015 |
| NEGREITLAMAEM | ASGWYSGNTLY | 0.23592496 |
| NELVEWSHPLGHRWRWE | ASGWYSGNTLY | 0.72939634 |
| NLGAGGTGGNGGNPG | ASGWYSGNTLY | 0.6540947 |
| NTEWPKWETVMAADPVD | ASGWYSGNTLY | 0.6334159 |
| PASAAPAETQP | ASGWYSGNTLY | 0.4353516 |
| QQSSDGKNGVRAAEITGE | ASGWYSGNTLY | 0.66538984 |
| RAPDAVTTSWWKE | ASGWYSGNTLY | 0.5388767 |
| SGFGNVGAHTSGWFNQ | ASGWYSGNTLY | 0.8008741 |
| SGVAGAVGQAASVGGLKV | ASGWYSGNTLY | 0.14950943 |
| SNGGAGTGSAGGNGGT | ASGWYSGNTLY | 0.7620041 |
| TGGAGGEGGAAG | ASGWYSGNTLY | 0.4177215 |
| TGGDGGAAGAGGAPGHG | ASGWYSGNTLY | 0.5236144 |
| TMPPLLAAIANAGRHGV | ASGWYSGNTLY | 0.33802322 |
| TSSTAGNSGGA | ASGWYSGNTLY | 0.58211684 |
| VPATVPVGILC | ASGWYSGNTLY | 0.7930958 |
| VQRPDSGGVVLTGRIS | ASGWYSGNTLY | 0.7971546 |
| WTPNPVPATKGVPAGLVQAT | ASGWYSGNTLY | 0.38174015 |
| YALKRALAKFSDRFDVVII | ASGWYSGNTLY | 0.80944407 |
| YLIGSVIV | ASGWYSGNTLY | 0.85257685 |
| YGQLVAVID | ASGWYSGNTLY | 0.54913926 |
| VTVGGMTLPA | ASGWYSGNTLY | 0.3879217 |

| | | |
|---|---|---|
| VLALPAV | ASGWYSGNTLY | 0.7715485 |
| VIPPTLNYET | ASGWYSGNTLY | 0.5935958 |
| VGMGGMHPSSG | ASGWYSGNTLY | 0.7781031 |
| VGLAGGAATA | ASGWYSGNTLY | 0.7302675 |
| VGGAGGAGGSGP | ASGWYSGNTLY | 0.40385863 |
| VFAAFMV | ASGWYSGNTLY | 0.7649516 |
| TLYDKLR | ASGWYSGNTLY | 0.8170624 |
| TLAVFTVL | ASGWYSGNTLY | 0.47497624 |
| TGSVQPARV | ASGWYSGNTLY | 0.43899292 |
| TGGGAGSGGT | ASGWYSGNTLY | 0.4667307 |
| SHLETSL | ASGWYSGNTLY | 0.264811 |
| SFTTGMT | ASGWYSGNTLY | 0.771019 |
| QVEVAKRMGVRQPTVS | ASGWYSGNTLY | 0.47870266 |
| QGSGGPALAM | ASGWYSGNTLY | 0.5809072 |
| MLAPSNP | ASGWYSGNTLY | 0.69278264 |
| LVNNAAI | ASGWYSGNTLY | 0.58291674 |
| LTGEINTLA | ASGWYSGNTLY | 0.59976345 |
| LSIFGPV | ASGWYSGNTLY | 0.61251056 |
| LLALLANL | ASGWYSGNTLY | 0.7140668 |
| LISNASDALDKLR | ASGWYSGNTLY | 0.71988815 |
| LINEAVP | ASGWYSGNTLY | 0.6431838 |
| LGYAFSGL | ASGWYSGNTLY | 0.6433359 |
| LDFVAVAL | ASGWYSGNTLY | 0.8228334 |
| LAQPGAE | ASGWYSGNTLY | 0.417369 |
| ISNLGAVTGF | ASGWYSGNTLY | 0.44080365 |
| HQEDPGRI | ASGWYSGNTLY | 0.7655502 |
| GTPGEPTP | ASGWYSGNTLY | 0.31767395 |
| GFFNSGS | ASGWYSGNTLY | 0.77537626 |
| GAGGGGLTAGAN | ASGWYSGNTLY | 0.63295364 |
| FVDALVNM | ASGWYSGNTLY | 0.7157314 |
| FLTVAPEL | ASGWYSGNTLY | 0.66060925 |
| FETLTAL | ASGWYSGNTLY | 0.8569573 |
| DVINPIEL | ASGWYSGNTLY | 0.809076 |
| APVSGAPV | ASGWYSGNTLY | 0.5834173 |
| ALVGGLAW | ASGWYSGNTLY | 0.7904271 |
| ALAAAAATAI | ASGWYSGNTLY | 0.4038164 |
| AFGLGLLL | ASGWYSGNTLY | 0.79254305 |
| AFFAAASL | ASGWYSGNTLY | 0.74079835 |
| AAAALPAFSPP | ASGWYSGNTLY | 0.4087168 |
| VVLAPAL | ASGWYSGNTLY | 0.6230334 |
| TIGHVDHGKTTL | ASGWYSGNTLY | 0.8224664 |
| LISNASDALDK | ASGWYSGNTLY | 0.82572794 |
| LISNASDAL | ASGWYSGNTLY | 0.24474108 |
| ISNASDALDK | ASGWYSGNTLY | 0.63448906 |
| NQAETLVYQTE | ASGWYSGNTLY | 0.22803125 |
| TGGTGGTAGAG | ASGWYSGNTLY | 0.4510569 |
| TGSTAGGAGGA | ASGWYSGNTLY | 0.42277938 |
| DPLAVPVAAGPV | ASGWYSGNTLY | 0.76067024 |
| GGTGGTPGNGGN | ASGWYSGNTLY | 0.56449574 |
| PAGLAGVDPATL | ASGWYSGNTLY | 0.69887984 |
| GNNLVVEAPHAGASV | ASGWYSGNTLY | 0.713384 |

| | | |
|---|---|---|
| KVTRIMSASTLEQPA | ASGWYSGNTLY | 0.8388979 |
| LGFALPIALPRRTVV | ASGWYSGNTLY | 0.48967102 |
| TRLGNLTARKLVVLV | ASGWYSGNTLY | 0.6237966 |
| WSQQRRLIYFPSAGP | ASGWYSGNTLY | 0.3397821 |
| GPNGNPDPMAAAVDIR | ASGWYSGNTLY | 0.4108187 |
| TQRCVSVADLRAAASR | ASGWYSGNTLY | 0.5324863 |
| QFVLDVAEHPVEVLTETV | ASGWYSGNTLY | 0.595371 |
| GGAGGTGGAGGNGGTQAGDG | ASGWYSGNTLY | 0.21986538 |
| PLLATLRAIAVDVDAKPAQV | ASGWYSGNTLY | 0.8244615 |
| APKFRTGGYMIDFWGVGYQVAK | ASGWYSGNTLY | 0.719813 |
| GAGGAGGAGGAAGGPGGTGGTG | ASGWYSGNTLY | 0.8000357 |
| IGALGLLATDVAPAGSVAFGVA | ASGWYSGNTLY | 0.5632956 |
| AGGAGGIGGNGGAGGAGGVLMG | ASGWYSGNTLY | 0.64935017 |
| ADELIGLL | ASGWYSGNTLY | 0.5835749 |
| ADRALKA | ASGWYSGNTLY | 0.7617492 |
| AGAVGQVGGAGGA | ASGWYSGNTLY | 0.57650715 |
| AGGGGGLGGAP | ASGWYSGNTLY | 0.2647549 |
| AGGLLYGNGGAG | ASGWYSGNTLY | 0.5665521 |
| AGLESIGL | ASGWYSGNTLY | 0.3554371 |
| AGLKTGII | ASGWYSGNTLY | 0.8294767 |
| ASLAALLL | ASGWYSGNTLY | 0.8067601 |
| ASVINISSI | ASGWYSGNTLY | 0.7463186 |
| ATPPGPA | ASGWYSGNTLY | 0.7884106 |
| ATVASALV | ASGWYSGNTLY | 0.23689827 |
| ATVASLIL | ASGWYSGNTLY | 0.61357474 |
| AVNGPVLIRH | ASGWYSGNTLY | 0.844988 |
| DGAAGAVGI | ASGWYSGNTLY | 0.76144356 |
| DPLLHGI | ASGWYSGNTLY | 0.58382 |
| ELGAALGAL | ASGWYSGNTLY | 0.8274825 |
| ELGRTQAV | ASGWYSGNTLY | 0.5115715 |
| FVAGGYLS | ASGWYSGNTLY | 0.6371918 |
| GASGGAGGAGGW | ASGWYSGNTLY | 0.7267908 |
| GGAGAGGDGGAGGI | ASGWYSGNTLY | 0.7466103 |
| GTGHNGSQPGMGGNG | ASGWYSGNTLY | 0.738909 |
| GYAAASASAS | ASGWYSGNTLY | 0.54744977 |
| HQVDVAAVQQG | ASGWYSGNTLY | 0.72070104 |
| IGAAIAHT | ASGWYSGNTLY | 0.8648191 |
| IGMIEVAQL | ASGWYSGNTLY | 0.65600765 |
| IRLAPEHTSAMD | ASGWYSGNTLY | 0.80673045 |
| IYNVSVL | ASGWYSGNTLY | 0.7378933 |
| KQGFLGM | ASGWYSGNTLY | 0.7238246 |
| LESVITQL | ASGWYSGNTLY | 0.48759574 |
| LGLGALVVV | ASGWYSGNTLY | 0.60146856 |
| LIGNGGNGGSGGI | ASGWYSGNTLY | 0.5848574 |
| LNAINAPTL | ASGWYSGNTLY | 0.4790131 |
| LQLLNTGV | ASGWYSGNTLY | 0.59747416 |
| LQTSIEF | ASGWYSGNTLY | 0.5426153 |
| MSFAVSEL | ASGWYSGNTLY | 0.8734746 |
| NTGALGL | ASGWYSGNTLY | 0.7169659 |
| PPGAGGAGGAGTT | ASGWYSGNTLY | 0.76065385 |
| QLVGLGI | ASGWYSGNTLY | 0.54530233 |

| | | |
|---|---|---|
| RFLEQVHQL | ASGWYSGNTLY | 0.6926435 |
| RGVGYVLR | ASGWYSGNTLY | 0.751995 |
| SATGVFAGVI | ASGWYSGNTLY | 0.36306265 |
| SLPAITLP | ASGWYSGNTLY | 0.8144077 |
| STALAFTV | ASGWYSGNTLY | 0.50173694 |
| SVLTVGLGK | ASGWYSGNTLY | 0.5693971 |
| TALAGAGSGI | ASGWYSGNTLY | 0.76162136 |
| TGLPFTGI | ASGWYSGNTLY | 0.8076812 |
| VAEHVTV | ASGWYSGNTLY | 0.5591155 |
| VDIVLNSL | ASGWYSGNTLY | 0.4863149 |
| VLTGGGAVL | ASGWYSGNTLY | 0.5350811 |
| VNVGLAYGI | ASGWYSGNTLY | 0.7972043 |
| VSSAAVGAV | ASGWYSGNTLY | 0.8278388 |
| VTSVYEAGV | ASGWYSGNTLY | 0.7118578 |
| WVALDRGA | ASGWYSGNTLY | 0.8556696 |
| YLVGSVVI | ASGWYSGNTLY | 0.731956 |
| GPPGTGKTLL | ASGWYSGNTLY | 0.33677253 |
| GVGMTKF | ASGWYSGNTLY | 0.681846 |
| IGIGGSDL | ASGWYSGNTLY | 0.79695904 |
| IGTLLAEL | ASGWYSGNTLY | 0.7964258 |
| IYTSGTT | ASGWYSGNTLY | 0.85512865 |
| LSFPAGV | ASGWYSGNTLY | 0.7504939 |
| LYTSGTT | ASGWYSGNTLY | 0.5357758 |
| VGAFPFGF | ASGWYSGNTLY | 0.8263832 |
| VGAGTVEF | ASGWYSGNTLY | 0.85422355 |
| VGKSTLLL | ASGWYSGNTLY | 0.8031281 |
| AALAHAGGGIV | ASGWYSGNTLY | 0.5959944 |
| AALVAVSLL | ASGWYSGNTLY | 0.5052301 |
| AAVSRAVAAL | ASGWYSGNTLY | 0.65868914 |
| AGGSSGIGGF | ASGWYSGNTLY | 0.8077525 |
| AGLTVANANG | ASGWYSGNTLY | 0.41733727 |
| AGLVGAVL | ASGWYSGNTLY | 0.64745605 |
| ALSQAFTV | ASGWYSGNTLY | 0.70506394 |
| ALTFSVSL | ASGWYSGNTLY | 0.3225641 |
| ALTGAALVAL | ASGWYSGNTLY | 0.730389 |
| ASFSIGAGVL | ASGWYSGNTLY | 0.8351954 |
| DGSVYKI | ASGWYSGNTLY | 0.5848369 |
| EQLAEIV | ASGWYSGNTLY | 0.56962967 |
| FAVSELTVM | ASGWYSGNTLY | 0.7945233 |
| GAAPLAAGL | ASGWYSGNTLY | 0.87510896 |
| GADGGDGGSGGAGG | ASGWYSGNTLY | 0.8166108 |
| GFGLDTGGAGG | ASGWYSGNTLY | 0.3449507 |
| GLASGGLL | ASGWYSGNTLY | 0.18354742 |
| GRIAGAAVLL | ASGWYSGNTLY | 0.54880434 |
| GSAGSLNTSF | ASGWYSGNTLY | 0.76668763 |
| GTGQIGI | ASGWYSGNTLY | 0.59544 |
| GTLIALIL | ASGWYSGNTLY | 0.6097547 |
| IAATMGFVVG | ASGWYSGNTLY | 0.73072124 |
| IFAQALL | ASGWYSGNTLY | 0.8067967 |
| IGAAVALGF | ASGWYSGNTLY | 0.5600665 |
| IGLTPLAL | ASGWYSGNTLY | 0.42884594 |

| | | |
|---|---|---|
| ILGSNTSTL | ASGWYSGNTLY | 0.7034113 |
| IVFSLTVL | ASGWYSGNTLY | 0.7720213 |
| LASKATGF | ASGWYSGNTLY | 0.523455 |
| LASVALIL | ASGWYSGNTLY | 0.6794795 |
| LFAVGLTL | ASGWYSGNTLY | 0.47135907 |
| LFNGGIGTYI | ASGWYSGNTLY | 0.50747347 |
| LTLGAGII | ASGWYSGNTLY | 0.7119464 |
| LVAAALGDI | ASGWYSGNTLY | 0.44244742 |
| NTAFVQLGI | ASGWYSGNTLY | 0.8322381 |
| QLFSGAEL | ASGWYSGNTLY | 0.4648642 |
| SAATFITL | ASGWYSGNTLY | 0.78599536 |
| SNYGLANLGV | ASGWYSGNTLY | 0.7385508 |
| SYGGAYAVM | ASGWYSGNTLY | 0.7653545 |
| TTGVGGLTLG | ASGWYSGNTLY | 0.5762615 |
| TTQTIYI | ASGWYSGNTLY | 0.4787225 |
| VAAGAGLGAVI | ASGWYSGNTLY | 0.41903937 |
| VAAVVTLL | ASGWYSGNTLY | 0.2067671 |
| VGFLGGLI | ASGWYSGNTLY | 0.80664384 |
| VGVIPILL | ASGWYSGNTLY | 0.4824971 |
| VIFASSLI | ASGWYSGNTLY | 0.8386129 |
| VKGGDGGAAAG | ASGWYSGNTLY | 0.78222185 |
| VTKTFTNI | ASGWYSGNTLY | 0.5287564 |
| VVAAAILL | ASGWYSGNTLY | 0.7404797 |
| YAALDVEL | ASGWYSGNTLY | 0.53938186 |
| LASVALLL | ASGWYSGNTLY | 0.41011834 |
| LDAVHRV | ASGWYSGNTLY | 0.77224046 |
| QGLSSSIF | ASGWYSGNTLY | 0.76511574 |
| RGTFANIRL | ASGWYSGNTLY | 0.45127815 |
| RQLWWGHRI | ASGWYSGNTLY | 0.68516594 |
| YGPPGTGKTLL | ASGWYSGNTLY | 0.7019255 |
| AALVGAPANVA | ASGWYSGNTLY | 0.44338682 |
| AATFRRTE | ASGWYSGNTLY | 0.59114355 |
| AGALGGGI | ASGWYSGNTLY | 0.8170613 |
| AGGAGGAGGIGAG | ASGWYSGNTLY | 0.6577996 |
| AGSGALGL | ASGWYSGNTLY | 0.59123814 |
| AQIADQI | ASGWYSGNTLY | 0.7761121 |
| ASAVAVTYL | ASGWYSGNTLY | 0.4112871 |
| ASTVLGGTGG | ASGWYSGNTLY | 0.7533477 |
| AVGGEPAK | ASGWYSGNTLY | 0.69264215 |
| CEQGTFGGTC | ASGWYSGNTLY | 0.55234957 |
| DEGKAPAG | ASGWYSGNTLY | 0.78816694 |
| DTVVFPAL | ASGWYSGNTLY | 0.43197393 |
| FEGITMAG | ASGWYSGNTLY | 0.5056356 |
| FPLVVQD | ASGWYSGNTLY | 0.5903996 |
| GAFTTPQ | ASGWYSGNTLY | 0.7419611 |
| GFGGFGTAGGVGGA | ASGWYSGNTLY | 0.805053 |
| GFLAQFL | ASGWYSGNTLY | 0.51682407 |
| GGAGGAGGNAGVA | ASGWYSGNTLY | 0.7099498 |
| GGVGTAGAGGAGGAGG | ASGWYSGNTLY | 0.5207716 |
| GNGGGGDAGNAGSGGN | ASGWYSGNTLY | 0.45994735 |
| GTGGAGGAGNDAGS | ASGWYSGNTLY | 0.60236263 |

| | | |
|---|---|---|
| GTTNTGWFNS | ASGWYSGNTLY | 0.60479295 |
| IFMGPLFTL | ASGWYSGNTLY | 0.579359 |
| ILAHAIEAI | ASGWYSGNTLY | 0.73159873 |
| IRSAIREAL | ASGWYSGNTLY | 0.5269237 |
| ITFAGGG | ASGWYSGNTLY | 0.73106384 |
| ITGMSGHQEV | ASGWYSGNTLY | 0.8432961 |
| IVAGLACTL | ASGWYSGNTLY | 0.65117085 |
| IVGGGLAGA | ASGWYSGNTLY | 0.77632576 |
| IVTGVLNI | ASGWYSGNTLY | 0.7490289 |
| LAQQGCI | ASGWYSGNTLY | 0.75556445 |
| LCLHVEN | ASGWYSGNTLY | 0.6148918 |
| LDAISDGS | ASGWYSGNTLY | 0.47530007 |
| LGGFAAML | ASGWYSGNTLY | 0.8303175 |
| LGGKNPMI | ASGWYSGNTLY | 0.79289484 |
| LGPKGVAAIL | ASGWYSGNTLY | 0.75582767 |
| LMVGVSI | ASGWYSGNTLY | 0.7530912 |
| LVGAAFVL | ASGWYSGNTLY | 0.72583175 |
| LVLAVAPL | ASGWYSGNTLY | 0.77604634 |
| MAAIAVIDAA | ASGWYSGNTLY | 0.58916426 |
| NNGINAELQ | ASGWYSGNTLY | 0.32361758 |
| RGEPLVLVHG | ASGWYSGNTLY | 0.6271448 |
| RGGAGGAAGLI | ASGWYSGNTLY | 0.5796995 |
| RSYRMHPAV | ASGWYSGNTLY | 0.5927353 |
| SAAHLLAGQLA | ASGWYSGNTLY | 0.3384296 |
| SAAVGSY | ASGWYSGNTLY | 0.8128339 |
| SPHAVLHRI | ASGWYSGNTLY | 0.8713893 |
| SQIAKVL | ASGWYSGNTLY | 0.5349624 |
| TALGHLVP | ASGWYSGNTLY | 0.85687786 |
| TAPTTSGAA | ASGWYSGNTLY | 0.35154325 |
| TDGDDGNGGSPGT | ASGWYSGNTLY | 0.7878841 |
| TDGLGEN | ASGWYSGNTLY | 0.8068383 |
| TGGPPLPGQA | ASGWYSGNTLY | 0.46776232 |
| TIPAITI | ASGWYSGNTLY | 0.7799872 |
| TIPGFTI | ASGWYSGNTLY | 0.5085843 |
| TSIASGALA | ASGWYSGNTLY | 0.65734094 |
| TVTLPGAAI | ASGWYSGNTLY | 0.84064364 |
| VASGLSAALP | ASGWYSGNTLY | 0.59723777 |
| VGAGVIGI | ASGWYSGNTLY | 0.65520847 |
| VGPKRFGV | ASGWYSGNTLY | 0.72378004 |
| VNVAFPD | ASGWYSGNTLY | 0.53044146 |
| VSMTNSGVS | ASGWYSGNTLY | 0.74272263 |
| VTDVLGDLSV | ASGWYSGNTLY | 0.77478886 |
| VVVVGAGFAGL | ASGWYSGNTLY | 0.45170382 |
| VYTVGTVV | ASGWYSGNTLY | 0.8502766 |
| WTTFSGP | ASGWYSGNTLY | 0.48443997 |
| YDELVEAF | ASGWYSGNTLY | 0.85290366 |
| YGTGGAGGNGG | ASGWYSGNTLY | 0.763926 |
| YTNVMAV | ASGWYSGNTLY | 0.70757383 |
| FLEQVHQL | ASGWYSGNTLY | 0.6053899 |
| GLSLLAVAV | ASGWYSGNTLY | 0.8138113 |
| LEQVHQL | ASGWYSGNTLY | 0.63352185 |

| | | |
|---|---|---|
| LNAAGAI | ASGWYSGNTLY | 0.7518891 |
| PAVHASL | ASGWYSGNTLY | 0.83264875 |
| TELEDDA | ASGWYSGNTLY | 0.6809429 |
| TPPGPPN | ASGWYSGNTLY | 0.72261375 |
| TPPPPPPP | ASGWYSGNTLY | 0.70959115 |
| TQSPALI | ASSELGGYAEQF | 0.7008206 |
| EFAAAFPQ | ASSELGGYAEQF | 0.6876199 |
| LLSGNLDVL | ASSELGGYAEQF | 0.49084592 |
| GPANLGNYNI | ASSELGGYAEQF | 0.69717443 |
| TLGPTLP | ASSELGGYAEQF | 0.663756 |
| VYAYSSCCK | ASSELGGYAEQF | 0.7094543 |
| APAPAPAPNI | ASSELGGYAEQF | 0.71766645 |
| PGGPVPIP | ASSELGGYAEQF | 0.6770966 |
| FLTVALEP | ASSELGGYAEQF | 0.7236626 |
| SQYFGANL | ASSELGGYAEQF | 0.71841687 |
| GGVKPAA | ASSELGGYAEQF | 0.5849328 |
| GGTGGAGGAGGW | ASSELGGYAEQF | 0.6996908 |
| GVGGVAGLGGAGG | ASSELGGYAEQF | 0.6777228 |
| VLLVLATTI | ASSELGGYAEQF | 0.70111585 |
| IGIFAAML | ASSELGGYAEQF | 0.55860084 |
| AVGDLID | ASSELGGYAEQF | 0.71927196 |
| NISPLPL | ASSELGGYAEQF | 0.6856004 |
| VTAVFVAI | ASSELGGYAEQF | 0.31638294 |
| SGDDNDAAL | ASSELGGYAEQF | 0.70919585 |
| AVAALAHS | ASSELGGYAEQF | 0.7075248 |
| GGVTVMATP | ASSELGGYAEQF | 0.69780165 |
| AGLAANVRD | ASSELGGYAEQF | 0.7189381 |
| TNRPDLIDPALLRPG | ASSELGGYAEQF | 0.578591 |
| LANLGVTAL | ASSELGGYAEQF | 0.66304964 |
| TGERIEL | ASSELGGYAEQF | 0.71911454 |
| NGEELAA | ASSELGGYAEQF | 0.7237711 |
| VLSLVWG | ASSELGGYAEQF | 0.6099521 |
| IGYALWL | ASSELGGYAEQF | 0.7142968 |
| LSLLLAPL | ASSELGGYAEQF | 0.7197349 |
| TLDEALK | ASSELGGYAEQF | 0.7135086 |
| TDGAGTAEGG | ASSELGGYAEQF | 0.7200906 |
| NAPGGAGGNGG | ASSELGGYAEQF | 0.72320193 |
| VSVFMGMGGNF | ASSELGGYAEQF | 0.6613102 |
| LASYLAAL | ASSELGGYAEQF | 0.7226691 |
| LVGGTPAL | ASSELGGYAEQF | 0.72157705 |
| VRNADVL | ASSELGGYAEQF | 0.6924777 |
| TCSSPFLVSAM | ASSELGGYAEQF | 0.7077362 |
| VHITDAE | ASSELGGYAEQF | 0.6550976 |
| EYAAPSF | ASSELGGYAEQF | 0.6898942 |
| VVDAFHGGL | ASSELGGYAEQF | 0.49167663 |
| ASVVDPQ | ASSELGGYAEQF | 0.7140809 |
| SYNEFGGLS | ASSELGGYAEQF | 0.720555 |
| DPDGSVY | ASSELGGYAEQF | 0.6959197 |
| MVSTIHATE | ASSELGGYAEQF | 0.65990764 |
| LAAGVADPV | ASSELGGYAEQF | 0.6706565 |
| TKEQIAA | ASSELGGYAEQF | 0.6982716 |

| | | |
|---|---|---|
| SLASPAAL | ASSELGGYAEQF | 0.69561625 |
| IGLVFYGV | ASSELGGYAEQF | 0.49281886 |
| FLLDASNL | ASSELGGYAEQF | 0.68037826 |
| YQVGNFP | ASSELGGYAEQF | 0.72092706 |
| ESLLSQST | ASSELGGYAEQF | 0.72057295 |
| LFTVGFAAVG | ASSELGGYAEQF | 0.5873663 |
| ATPPTPT | ASSELGGYAEQF | 0.7224368 |
| TDLATLG | ASSELGGYAEQF | 0.7185714 |
| SAGGTGR | ASSELGGYAEQF | 0.7206006 |
| GFSNSGS | ASSELGGYAEQF | 0.69563174 |
| GGNGGSGGTSV | ASSELGGYAEQF | 0.67813134 |
| GTSVATGGAGN | ASSELGGYAEQF | 0.71615934 |
| NGGNGGSGGTS | ASSELGGYAEQF | 0.6933242 |
| AVVKFPH | ASSELGGYAEQF | 0.64575857 |
| TPPPPPP | ASSELGGYAEQF | 0.71929306 |
| AAPAAAAAPA | ASSELGGYAEQF | 0.6901876 |
| IALNEDL | ASSELGGYAEQF | 0.6857157 |
| IVNVSSM | ASSELGGYAEQF | 0.67862344 |
| TGIPPAPR | ASSELGGYAEQF | 0.7213615 |
| FDLGGGTF | ASSELGGYAEQF | 0.6537974 |
| FDLGGGTFDVS | ASSELGGYAEQF | 0.6348604 |
| NEPTAAA | ASSELGGYAEQF | 0.6621499 |
| NEPTAAALA | ASSELGGYAEQF | 0.70585954 |
| GSIINISSI | ASSELGGYAEQF | 0.72189355 |
| VAPGNAGTA | ASSELGGYAEQF | 0.71049595 |
| TGGGASLE | ASSELGGYAEQF | 0.6899241 |
| VAVYNHYKRI | ASSELGGYAEQF | 0.7127551 |
| TDPAPATT | ASSELGGYAEQF | 0.7127229 |
| TNRPDILDPALLRPG | ASSELGGYAEQF | 0.709578 |
| LTTTTTS | ASSELGGYAEQF | 0.7155281 |
| PTGSGKTL | ASSELGGYAEQF | 0.6803074 |
| TGSGKTL | ASSELGGYAEQF | 0.7219496 |
| TIGHVDHGK | ASSELGGYAEQF | 0.71350586 |
| TIGHVDHGKTT | ASSELGGYAEQF | 0.71896917 |
| GHVDHGKTT | ASSELGGYAEQF | 0.7205181 |
| AQFEHTLL | ASSELGGYAEQF | 0.71080405 |
| IVNQDEL | ASSELGGYAEQF | 0.7077203 |
| TNGSQFF | ASSELGGYAEQF | 0.47853065 |
| TGKGIVE | ASSELGGYAEQF | 0.66705173 |
| LISNASDA | ASSELGGYAEQF | 0.7140765 |
| AELRTLGELTKIAKAHGAQ | ASSELGGYAEQF | 0.71083105 |
| APNAVAAAEPSW | ASSELGGYAEQF | 0.7174918 |
| APQVAGAATAVL | ASSELGGYAEQF | 0.70594895 |
| ARRDPAAAAAILPTDA | ASSELGGYAEQF | 0.68329316 |
| DAAARDVLAEAGLAEYF | ASSELGGYAEQF | 0.7171274 |
| DDQIVLSAKANLAKVEQ | ASSELGGYAEQF | 0.71974707 |
| DELLPDTGEDSDEDGH | ASSELGGYAEQF | 0.5775374 |
| ELPEAPANRATVPQD | ASSELGGYAEQF | 0.6946881 |
| FGQSKIERVTGIG | ASSELGGYAEQF | 0.7262187 |
| GAGGTGSTAGGAGGAGGAGGLY | ASSELGGYAEQF | 0.7078246 |
| GHGGTGAAGGNGGTGGAGGSGI | ASSELGGYAEQF | 0.6944281 |

| | | |
|---|---|---|
| GNGGNGGAGGEGGPGGAGGAGA | ASSELGGYAEQF | 0.6661361 |
| GSTAGNGGNGGNATNGTIAGQP | ASSELGGYAEQF | 0.72322667 |
| GTGGDGGNGGNGGTGG | ASSELGGYAEQF | 0.68549883 |
| GVRRPAGIPADAPV | ASSELGGYAEQF | 0.7241393 |
| GYRQVTDAHLLTLARR | ASSELGGYAEQF | 0.3765236 |
| IAKMGQDGHDRG | ASSELGGYAEQF | 0.6668974 |
| KAEPSIEIQYT | ASSELGGYAEQF | 0.6138852 |
| KPVRIRMAADEDTVAVT | ASSELGGYAEQF | 0.5423814 |
| MPPRIAGMRLLVIKPEP | ASSELGGYAEQF | 0.7202772 |
| MKFYAVALFRSRGVPPTR | ASSELGGYAEQF | 0.66303045 |
| NAGGFGILGAVAHSP | ASSELGGYAEQF | 0.7150115 |
| NEGREITLAMAEM | ASSELGGYAEQF | 0.7059287 |
| NELVEWSHPLGHRWRWE | ASSELGGYAEQF | 0.722316 |
| NLGAGGTGGNGGNPG | ASSELGGYAEQF | 0.70665765 |
| NTEWPKWETVMAADPVD | ASSELGGYAEQF | 0.7085739 |
| PASAAPAETQP | ASSELGGYAEQF | 0.64625466 |
| QQSSDGKNGVRAAEITGE | ASSELGGYAEQF | 0.7188648 |
| RAPDAVTTSWWKE | ASSELGGYAEQF | 0.7207062 |
| SGFGNVGAHTSGWFNQ | ASSELGGYAEQF | 0.7134919 |
| SGVAGAVGQAASVGGLKV | ASSELGGYAEQF | 0.56677675 |
| SNGGAGTGSAGGNGGT | ASSELGGYAEQF | 0.69894195 |
| TGGAGGEGGAAG | ASSELGGYAEQF | 0.7103885 |
| TGGDGGAAGAGGAPGHG | ASSELGGYAEQF | 0.37503847 |
| TMPPLLAAIANAGRHGV | ASSELGGYAEQF | 0.418531 |
| TSSTAGNSGGA | ASSELGGYAEQF | 0.69739896 |
| VPATVPVGILC | ASSELGGYAEQF | 0.6649051 |
| VQRPDSGGVVLTGRIS | ASSELGGYAEQF | 0.72202766 |
| WTPNPVPATKGVPAGLVQAT | ASSELGGYAEQF | 0.71881104 |
| YALKRALAKFSDRFDVVII | ASSELGGYAEQF | 0.7012273 |
| YLIGSVIV | ASSELGGYAEQF | 0.71666235 |
| YGQLVAVID | ASSELGGYAEQF | 0.70473444 |
| VTVGGMTLPA | ASSELGGYAEQF | 0.2680575 |
| VLALPAV | ASSELGGYAEQF | 0.6719779 |
| VIPPTLNYET | ASSELGGYAEQF | 0.6138115 |
| VGMGGMHPSSG | ASSELGGYAEQF | 0.7232033 |
| VGLAGGAATA | ASSELGGYAEQF | 0.68000394 |
| VGGAGGAGGSGP | ASSELGGYAEQF | 0.6683005 |
| VFAAFMV | ASSELGGYAEQF | 0.7049562 |
| TLYDKLR | ASSELGGYAEQF | 0.70523906 |
| TLAVFTVL | ASSELGGYAEQF | 0.70066595 |
| TGSVQPARV | ASSELGGYAEQF | 0.6627666 |
| TGGGAGSGGT | ASSELGGYAEQF | 0.30772144 |
| SHLETSL | ASSELGGYAEQF | 0.7190964 |
| SFTTGMT | ASSELGGYAEQF | 0.53388005 |
| QVEVAKRMGVRQPTVS | ASSELGGYAEQF | 0.6435579 |
| QGSGGPALAM | ASSELGGYAEQF | 0.4636513 |
| MLAPSNP | ASSELGGYAEQF | 0.38487768 |
| LVNNAAI | ASSELGGYAEQF | 0.48471332 |
| LTGEINTLA | ASSELGGYAEQF | 0.71775436 |
| LSIFGPV | ASSELGGYAEQF | 0.6918986 |
| LLALLANL | ASSELGGYAEQF | 0.6901983 |

| | | |
|---|---|---|
| LISNASDALDKLR | ASSELGGYAEQF | 0.7243314 |
| LINEAVP | ASSELGGYAEQF | 0.702621 |
| LGYAFSGL | ASSELGGYAEQF | 0.6930632 |
| LDFVAVAL | ASSELGGYAEQF | 0.6746309 |
| LAQPGAE | ASSELGGYAEQF | 0.6695335 |
| ISNLGAVTGF | ASSELGGYAEQF | 0.65914184 |
| HQEDPGRI | ASSELGGYAEQF | 0.64766395 |
| GTPGEPTP | ASSELGGYAEQF | 0.7248486 |
| GFFNSGS | ASSELGGYAEQF | 0.67230046 |
| GAGGGGLTAGAN | ASSELGGYAEQF | 0.54177505 |
| FVDALVNM | ASSELGGYAEQF | 0.68080056 |
| FLTVAPEL | ASSELGGYAEQF | 0.7236857 |
| FETLTAL | ASSELGGYAEQF | 0.7256199 |
| DVINPIEL | ASSELGGYAEQF | 0.68131995 |
| APVSGAPV | ASSELGGYAEQF | 0.6536371 |
| ALVGGLAW | ASSELGGYAEQF | 0.6828561 |
| ALAAAAATAI | ASSELGGYAEQF | 0.7062777 |
| AFGLGLLL | ASSELGGYAEQF | 0.6719414 |
| AFFAAASL | ASSELGGYAEQF | 0.57067335 |
| AAAALPAFSPP | ASSELGGYAEQF | 0.7182971 |
| VVLAPAL | ASSELGGYAEQF | 0.72542113 |
| TIGHVDHGKTTL | ASSELGGYAEQF | 0.68693507 |
| LISNASDALDK | ASSELGGYAEQF | 0.5836586 |
| LISNASDAL | ASSELGGYAEQF | 0.70307946 |
| ISNASDALDK | ASSELGGYAEQF | 0.6766211 |
| NQAETLVYQTE | ASSELGGYAEQF | 0.6946705 |
| TGGTGGTAGAG | ASSELGGYAEQF | 0.7044369 |
| TGSTAGGAGGA | ASSELGGYAEQF | 0.5731576 |
| DPLAVPVAAGPV | ASSELGGYAEQF | 0.6875142 |
| GGTGGTPGNGGN | ASSELGGYAEQF | 0.72461236 |
| PAGLAGVDPATL | ASSELGGYAEQF | 0.69400495 |
| GNNLVVEAPHAGASV | ASSELGGYAEQF | 0.70623577 |
| KVTRIMSASTLEQPA | ASSELGGYAEQF | 0.722568 |
| LGFALPIALPRRTVV | ASSELGGYAEQF | 0.715421 |
| TRLGNLTARKLVVLV | ASSELGGYAEQF | 0.72260773 |
| WSQQRRLIYFPSAGP | ASSELGGYAEQF | 0.71999717 |
| GPNGNPDPMAAAVDIR | ASSELGGYAEQF | 0.69406796 |
| TQRCVSVADLRAAASR | ASSELGGYAEQF | 0.7052207 |
| QFVLDVAEHPVEVLTETV | ASSELGGYAEQF | 0.7072526 |
| GGAGGTGGAGGNGGTQAGDG | ASSELGGYAEQF | 0.70672107 |
| PLLATLRAIAVDVDAKPAQV | ASSELGGYAEQF | 0.69752204 |
| APKFRTGGYMIDFWGVGYQVAK | ASSELGGYAEQF | 0.66683435 |
| GAGGAGGAGGAAGGPGGTGGTG | ASSELGGYAEQF | 0.72068626 |
| IGALGLLATDVAPAGSVAFGVA | ASSELGGYAEQF | 0.6979125 |
| AGGAGGIGGNGGAGGAGGVLMG | ASSELGGYAEQF | 0.7202684 |
| ADELIGLL | ASSELGGYAEQF | 0.6192724 |
| ADRALKA | ASSELGGYAEQF | 0.630216 |
| AGAVGQVGGAGGA | ASSELGGYAEQF | 0.71874756 |
| AGGGGGLGGAP | ASSELGGYAEQF | 0.280047 |
| AGGLLYGNGGAG | ASSELGGYAEQF | 0.35611722 |
| AGLESIGL | ASSELGGYAEQF | 0.7063247 |

| | | |
|---|---|---|
| AGLKTGII | ASSELGGYAEQF | 0.7216239 |
| ASLAALLL | ASSELGGYAEQF | 0.2919 |
| ASVINISSI | ASSELGGYAEQF | 0.6436571 |
| ATPPGPA | ASSELGGYAEQF | 0.62514675 |
| ATVASALV | ASSELGGYAEQF | 0.7103944 |
| ATVASLIL | ASSELGGYAEQF | 0.70762825 |
| AVNGPVLIRH | ASSELGGYAEQF | 0.5740803 |
| DGAAGAVGI | ASSELGGYAEQF | 0.71805525 |
| DPLLHGI | ASSELGGYAEQF | 0.5577984 |
| ELGAALGAL | ASSELGGYAEQF | 0.6462964 |
| ELGRTQAV | ASSELGGYAEQF | 0.72027636 |
| FVAGGYLS | ASSELGGYAEQF | 0.72264206 |
| GASGGAGGAGGW | ASSELGGYAEQF | 0.64423704 |
| GGAGAGGDGGAGGI | ASSELGGYAEQF | 0.72017896 |
| GTGHNGSQPGMGGNG | ASSELGGYAEQF | 0.68373454 |
| GYAAASASAS | ASSELGGYAEQF | 0.7248999 |
| HQVDVAAVQQG | ASSELGGYAEQF | 0.6917215 |
| IGAAIAHT | ASSELGGYAEQF | 0.5399842 |
| IGMIEVAQL | ASSELGGYAEQF | 0.7180222 |
| IRLAPEHTSAMD | ASSELGGYAEQF | 0.710379 |
| IYNVSVL | ASSELGGYAEQF | 0.6997191 |
| KQGFLGM | ASSELGGYAEQF | 0.7192565 |
| LESVITQL | ASSELGGYAEQF | 0.6540265 |
| LGLGALVVV | ASSELGGYAEQF | 0.7000416 |
| LIGNGGNGGSGGI | ASSELGGYAEQF | 0.7236579 |
| LNAINAPTL | ASSELGGYAEQF | 0.72165394 |
| LQLLNTGV | ASSELGGYAEQF | 0.61185837 |
| LQTSIEF | ASSELGGYAEQF | 0.7191052 |
| MSFAVSEL | ASSELGGYAEQF | 0.69877124 |
| NTGALGL | ASSELGGYAEQF | 0.6644387 |
| PPGAGGAGGAGTT | ASSELGGYAEQF | 0.7234324 |
| QLVGLGI | ASSELGGYAEQF | 0.70081437 |
| RFLEQVHQL | ASSELGGYAEQF | 0.7182306 |
| RGVGYVLR | ASSELGGYAEQF | 0.71737766 |
| SATGVFAGVI | ASSELGGYAEQF | 0.71467125 |
| SLPAITLP | ASSELGGYAEQF | 0.56262136 |
| STALAFTV | ASSELGGYAEQF | 0.70022464 |
| SVLTVGLGK | ASSELGGYAEQF | 0.70126903 |
| TALAGAGSGI | ASSELGGYAEQF | 0.719936 |
| TGLPFTGI | ASSELGGYAEQF | 0.69721353 |
| VAEHVTV | ASSELGGYAEQF | 0.6036636 |
| VDIVLNSL | ASSELGGYAEQF | 0.7122732 |
| VLTGGGAVL | ASSELGGYAEQF | 0.7214458 |
| VNVGLAYGI | ASSELGGYAEQF | 0.7065363 |
| VSSAAVGAV | ASSELGGYAEQF | 0.7068154 |
| VTSVYEAGV | ASSELGGYAEQF | 0.7241067 |
| WVALDRGA | ASSELGGYAEQF | 0.672642 |
| YLVGSVVI | ASSELGGYAEQF | 0.6952286 |
| GPPGTGKTLL | ASSELGGYAEQF | 0.7160863 |
| GVGMTKF | ASSELGGYAEQF | 0.7193478 |
| IGIGGSDL | ASSELGGYAEQF | 0.69683635 |

| | | |
|---|---|---|
| IGTLLAEL | ASSELGGYAEQF | 0.7236485 |
| IYTSGTT | ASSELGGYAEQF | 0.6693363 |
| LSFPAGV | ASSELGGYAEQF | 0.71958387 |
| LYTSGTT | ASSELGGYAEQF | 0.67035794 |
| VGAFPFGF | ASSELGGYAEQF | 0.7137246 |
| VGAGTVEF | ASSELGGYAEQF | 0.67336047 |
| VGKSTLLL | ASSELGGYAEQF | 0.71884847 |
| AALAHAGGGIV | ASSELGGYAEQF | 0.7190384 |
| AALVAVSLL | ASSELGGYAEQF | 0.7202782 |
| AAVSRAVAAL | ASSELGGYAEQF | 0.34489843 |
| AGGSSGIGGF | ASSELGGYAEQF | 0.7206842 |
| AGLTVANANG | ASSELGGYAEQF | 0.7189423 |
| AGLVGAVL | ASSELGGYAEQF | 0.72578883 |
| ALSQAFTV | ASSELGGYAEQF | 0.7027519 |
| ALTFSVSL | ASSELGGYAEQF | 0.71048146 |
| ALTGAALVAL | ASSELGGYAEQF | 0.69317925 |
| ASFSIGAGVL | ASSELGGYAEQF | 0.6948905 |
| DGSVYKI | ASSELGGYAEQF | 0.65892357 |
| EQLAEIV | ASSELGGYAEQF | 0.68113744 |
| FAVSELTVM | ASSELGGYAEQF | 0.6902386 |
| GAAPLAAGL | ASSELGGYAEQF | 0.68077767 |
| GADGGDGGSGGAGG | ASSELGGYAEQF | 0.71649504 |
| GFGLDTGGAGG | ASSELGGYAEQF | 0.7203697 |
| GLASGGLL | ASSELGGYAEQF | 0.7185396 |
| GRIAGAAVLL | ASSELGGYAEQF | 0.7223314 |
| GSAGSLNTSF | ASSELGGYAEQF | 0.7212956 |
| GTGQIGI | ASSELGGYAEQF | 0.71979237 |
| GTLIALIL | ASSELGGYAEQF | 0.6813866 |
| IAATMGFVVG | ASSELGGYAEQF | 0.6954707 |
| IFAQALL | ASSELGGYAEQF | 0.6499908 |
| IGAAVALGF | ASSELGGYAEQF | 0.69686055 |
| IGLTPLAL | ASSELGGYAEQF | 0.68522096 |
| ILGSNTSTL | ASSELGGYAEQF | 0.71235293 |
| IVFSLTVL | ASSELGGYAEQF | 0.2714878 |
| LASKATGF | ASSELGGYAEQF | 0.7053578 |
| LASVALIL | ASSELGGYAEQF | 0.72378886 |
| LFAVGLTL | ASSELGGYAEQF | 0.66931635 |
| LFNGGIGTYI | ASSELGGYAEQF | 0.70787275 |
| LTLGAGII | ASSELGGYAEQF | 0.69871485 |
| LVAAALGDI | ASSELGGYAEQF | 0.68736327 |
| NTAFVQLGI | ASSELGGYAEQF | 0.7179377 |
| QLFSGAEL | ASSELGGYAEQF | 0.66334105 |
| SAATFITL | ASSELGGYAEQF | 0.71357906 |
| SNYGLANLGV | ASSELGGYAEQF | 0.7224968 |
| SYGGAYAVM | ASSELGGYAEQF | 0.7069398 |
| TTGVGGLTLG | ASSELGGYAEQF | 0.7005296 |
| TTQTIYI | ASSELGGYAEQF | 0.6968396 |
| VAAGAGLGAVI | ASSELGGYAEQF | 0.49859363 |
| VAAVVTLL | ASSELGGYAEQF | 0.7123173 |
| VGFLGGLI | ASSELGGYAEQF | 0.2659474 |
| VGVIPILL | ASSELGGYAEQF | 0.7032117 |

| | | |
|---|---|---|
| VIFASSLI | ASSELGGYAEQF | 0.6493541 |
| VKGGDGGAAAG | ASSELGGYAEQF | 0.7107221 |
| VTKTFTNI | ASSELGGYAEQF | 0.7043041 |
| VVAAAILL | ASSELGGYAEQF | 0.7137342 |
| YAALDVEL | ASSELGGYAEQF | 0.6568742 |
| LASVALLL | ASSELGGYAEQF | 0.70263386 |
| LDAVHRV | ASSELGGYAEQF | 0.6735267 |
| QGLSSSIF | ASSELGGYAEQF | 0.60183215 |
| RGTFANIRL | ASSELGGYAEQF | 0.7155354 |
| RQLWWGHRI | ASSELGGYAEQF | 0.6835022 |
| YGPPGTGKTLL | ASSELGGYAEQF | 0.69093215 |
| AALVGAPANVA | ASSELGGYAEQF | 0.7218857 |
| AATFRRTE | ASSELGGYAEQF | 0.70857257 |
| AGALGGGI | ASSELGGYAEQF | 0.5135257 |
| AGGAGGAGGIGAG | ASSELGGYAEQF | 0.6983641 |
| AGSGALGL | ASSELGGYAEQF | 0.70353365 |
| AQIADQI | ASSELGGYAEQF | 0.69923985 |
| ASAVAVTYL | ASSELGGYAEQF | 0.26627892 |
| ASTVLGGTGG | ASSELGGYAEQF | 0.7122674 |
| AVGGEPAK | ASSELGGYAEQF | 0.66234785 |
| CEQGTFGGTC | ASSELGGYAEQF | 0.6630254 |
| DEGKAPAG | ASSELGGYAEQF | 0.68145937 |
| DTVVFPAL | ASSELGGYAEQF | 0.71754694 |
| FEGITMAG | ASSELGGYAEQF | 0.65010035 |
| FPLVVQD | ASSELGGYAEQF | 0.69327164 |
| GAFTTPQ | ASSELGGYAEQF | 0.7175853 |
| GFGGFGTAGGVGGA | ASSELGGYAEQF | 0.71641076 |
| GFLAQFL | ASSELGGYAEQF | 0.72312534 |
| GGAGGAGGNAGVA | ASSELGGYAEQF | 0.70986587 |
| GGVGTAGAGGAGGAGG | ASSELGGYAEQF | 0.6838106 |
| GNGGGGDAGNAGSGGN | ASSELGGYAEQF | 0.7099217 |
| GTGGAGGAGNDAGS | ASSELGGYAEQF | 0.69149315 |
| GTTNTGWFNS | ASSELGGYAEQF | 0.5899708 |
| IFMGPLFTL | ASSELGGYAEQF | 0.49453074 |
| ILAHAIEAI | ASSELGGYAEQF | 0.70748675 |
| IRSAIREAL | ASSELGGYAEQF | 0.46512607 |
| ITFAGGG | ASSELGGYAEQF | 0.7056585 |
| ITGMSGHQEV | ASSELGGYAEQF | 0.7215252 |
| IVAGLACTL | ASSELGGYAEQF | 0.6923436 |
| IVGGGLAGA | ASSELGGYAEQF | 0.56960154 |
| IVTGVLNI | ASSELGGYAEQF | 0.6963875 |
| LAQQGCI | ASSELGGYAEQF | 0.69600654 |
| LCLHVEN | ASSELGGYAEQF | 0.7171142 |
| LDAISDGS | ASSELGGYAEQF | 0.71339536 |
| LGGFAAML | ASSELGGYAEQF | 0.7250962 |
| LGGKNPMI | ASSELGGYAEQF | 0.72072065 |
| LGPKGVAAIL | ASSELGGYAEQF | 0.686613 |
| LMVGVSI | ASSELGGYAEQF | 0.71613973 |
| LVGAAFVL | ASSELGGYAEQF | 0.6940684 |
| LVLAVAPL | ASSELGGYAEQF | 0.68337 |
| MAAIAVIDAA | ASSELGGYAEQF | 0.6998995 |

| | | |
|---|---|---|
| NNGINAELQ | ASSELGGYAEQF | 0.7084851 |
| RGEPLVLVHG | ASSELGGYAEQF | 0.7005761 |
| RGGAGGAAGLI | ASSELGGYAEQF | 0.6874164 |
| RSYRMHPAV | ASSELGGYAEQF | 0.5417553 |
| SAAHLLAGQLA | ASSELGGYAEQF | 0.71811616 |
| SAAVGSY | ASSELGGYAEQF | 0.70933765 |
| SPHAVLHRI | ASSELGGYAEQF | 0.5925523 |
| SQIAKVL | ASSELGGYAEQF | 0.68624717 |
| TALGHLVP | ASSELGGYAEQF | 0.7251792 |
| TAPTTSGAA | ASSELGGYAEQF | 0.72233784 |
| TDGDDGNGGSPGT | ASSELGGYAEQF | 0.68125093 |
| TDGLGEN | ASSELGGYAEQF | 0.6737758 |
| TGGPPLPGQA | ASSELGGYAEQF | 0.7061746 |
| TIPAITI | ASSELGGYAEQF | 0.54770976 |
| TIPGFTI | ASSELGGYAEQF | 0.7121878 |
| TSIASGALA | ASSELGGYAEQF | 0.7039827 |
| TVTLPGAAI | ASSELGGYAEQF | 0.6987611 |
| VASGLSAALP | ASSELGGYAEQF | 0.64822066 |
| VGAGVIGI | ASSELGGYAEQF | 0.70683473 |
| VGPKRFGV | ASSELGGYAEQF | 0.68713653 |
| VNVAFPD | ASSELGGYAEQF | 0.71552837 |
| VSMTNSGVS | ASSELGGYAEQF | 0.31256577 |
| VTDVLGDLSV | ASSELGGYAEQF | 0.7101073 |
| VVVVGAGFAGL | ASSELGGYAEQF | 0.664803 |
| VYTVGTVV | ASSELGGYAEQF | 0.65863025 |
| WTTFSGP | ASSELGGYAEQF | 0.48232204 |
| YDELVEAF | ASSELGGYAEQF | 0.6968298 |
| YGTGGAGGNGG | ASSELGGYAEQF | 0.7173123 |
| YTNVMAV | ASSELGGYAEQF | 0.71947426 |
| FLEQVHQL | ASSELGGYAEQF | 0.66043526 |
| GLSLLAVAV | ASSELGGYAEQF | 0.72091746 |
| LEQVHQL | ASSELGGYAEQF | 0.7248059 |
| LNAAGAI | ASSELGGYAEQF | 0.6901232 |
| PAVHASL | ASSELGGYAEQF | 0.45582628 |
| TELEDDA | ASSELGGYAEQF | 0.7131762 |
| TPPGPPN | ASSELGGYAEQF | 0.6958176 |
| TPPPPPPP | ASSELGGYAEQF | 0.7166631 |
| TQSPALI | ASSMGLGVQDTQY | 0.7100483 |
| EFAAAFPQ | ASSMGLGVQDTQY | 0.652748 |
| LLSGNLDVL | ASSMGLGVQDTQY | 0.7145366 |
| GPANLGNYNI | ASSMGLGVQDTQY | 0.72047055 |
| TLGPTLP | ASSMGLGVQDTQY | 0.6743219 |
| VYAYSSCCK | ASSMGLGVQDTQY | 0.6481843 |
| APAPAPAPNI | ASSMGLGVQDTQY | 0.6418532 |
| PGGPVPIP | ASSMGLGVQDTQY | 0.44799992 |
| FLTVALEP | ASSMGLGVQDTQY | 0.71535724 |
| SQYFGANL | ASSMGLGVQDTQY | 0.68015146 |
| GGVKPAA | ASSMGLGVQDTQY | 0.66875434 |
| GGTGGAGGAGGW | ASSMGLGVQDTQY | 0.70998573 |
| GVGGVAGLGGAGG | ASSMGLGVQDTQY | 0.6606052 |
| VLLVLATTI | ASSMGLGVQDTQY | 0.67969215 |

| | | |
|---|---|---|
| IGIFAAML | ASSMGLGVQDTQY | 0.68545014 |
| AVGDLID | ASSMGLGVQDTQY | 0.70205635 |
| NISPLPL | ASSMGLGVQDTQY | 0.7120212 |
| VTAVFVAI | ASSMGLGVQDTQY | 0.70097107 |
| SGDDNDAAL | ASSMGLGVQDTQY | 0.6937741 |
| AVAALAHS | ASSMGLGVQDTQY | 0.7229613 |
| GGVTVMATP | ASSMGLGVQDTQY | 0.6119163 |
| AGLAANVRD | ASSMGLGVQDTQY | 0.6674926 |
| TNRPDLIDPALLRPG | ASSMGLGVQDTQY | 0.71364427 |
| LANLGVTAL | ASSMGLGVQDTQY | 0.59024644 |
| TGERIEL | ASSMGLGVQDTQY | 0.7165104 |
| NGEELAA | ASSMGLGVQDTQY | 0.70423734 |
| VLSLVWG | ASSMGLGVQDTQY | 0.71193254 |
| IGYALWL | ASSMGLGVQDTQY | 0.7040895 |
| LSLLLAPL | ASSMGLGVQDTQY | 0.7251907 |
| TLDEALK | ASSMGLGVQDTQY | 0.7190218 |
| TDGAGTAEGG | ASSMGLGVQDTQY | 0.6935922 |
| NAPGGAGGNGG | ASSMGLGVQDTQY | 0.7074257 |
| VSVFMGMGGNF | ASSMGLGVQDTQY | 0.5960539 |
| LASYLAAL | ASSMGLGVQDTQY | 0.7212034 |
| LVGGTPAL | ASSMGLGVQDTQY | 0.71287537 |
| VRNADVL | ASSMGLGVQDTQY | 0.6534244 |
| TCSSPFLVSAM | ASSMGLGVQDTQY | 0.7028582 |
| VHITDAE | ASSMGLGVQDTQY | 0.7227329 |
| EYAAPSF | ASSMGLGVQDTQY | 0.72207034 |
| VVDAFHGGL | ASSMGLGVQDTQY | 0.5254789 |
| ASVVDPQ | ASSMGLGVQDTQY | 0.6956551 |
| SYNEFGGLS | ASSMGLGVQDTQY | 0.49892077 |
| DPDGSVY | ASSMGLGVQDTQY | 0.6478342 |
| MVSTIHATE | ASSMGLGVQDTQY | 0.57020783 |
| LAAGVADPV | ASSMGLGVQDTQY | 0.7154012 |
| TKEQIAA | ASSMGLGVQDTQY | 0.68612206 |
| SLASPAAL | ASSMGLGVQDTQY | 0.71100914 |
| IGLVFYGV | ASSMGLGVQDTQY | 0.708747 |
| FLLDASNL | ASSMGLGVQDTQY | 0.72356796 |
| YQVGNFP | ASSMGLGVQDTQY | 0.6767286 |
| ESLLSQST | ASSMGLGVQDTQY | 0.69721127 |
| LFTVGFAAVG | ASSMGLGVQDTQY | 0.5422394 |
| ATPPTPT | ASSMGLGVQDTQY | 0.50525427 |
| TDLATLG | ASSMGLGVQDTQY | 0.66864806 |
| SAGGTGR | ASSMGLGVQDTQY | 0.6841061 |
| GFSNSGS | ASSMGLGVQDTQY | 0.36632425 |
| GGNGGSGGTSV | ASSMGLGVQDTQY | 0.72167856 |
| GTSVATGGAGN | ASSMGLGVQDTQY | 0.71146524 |
| NGGNGGSGGTS | ASSMGLGVQDTQY | 0.71803296 |
| AVVKFPH | ASSMGLGVQDTQY | 0.67541534 |
| TPPPPPP | ASSMGLGVQDTQY | 0.6942812 |
| AAPAAAAAPA | ASSMGLGVQDTQY | 0.6832041 |
| IALNEDL | ASSMGLGVQDTQY | 0.71459436 |
| IVNVSSM | ASSMGLGVQDTQY | 0.681097 |
| TGIPPAPR | ASSMGLGVQDTQY | 0.52375674 |

| | | |
|---|---|---|
| FDLGGGTF | ASSMGLGVQDTQY | 0.662858 |
| FDLGGGTFDVS | ASSMGLGVQDTQY | 0.6274079 |
| NEPTAAA | ASSMGLGVQDTQY | 0.72157586 |
| NEPTAAALA | ASSMGLGVQDTQY | 0.7200762 |
| GSIINISSI | ASSMGLGVQDTQY | 0.71431494 |
| VAPGNAGTA | ASSMGLGVQDTQY | 0.7214408 |
| TGGGASLE | ASSMGLGVQDTQY | 0.36447167 |
| VAVYNHYKRI | ASSMGLGVQDTQY | 0.6767219 |
| TDPAPATT | ASSMGLGVQDTQY | 0.7237382 |
| TNRPDILDPALLRPG | ASSMGLGVQDTQY | 0.7136344 |
| LTTTTTS | ASSMGLGVQDTQY | 0.6935927 |
| PTGSGKTL | ASSMGLGVQDTQY | 0.7223815 |
| TGSGKTL | ASSMGLGVQDTQY | 0.67069715 |
| TIGHVDHGK | ASSMGLGVQDTQY | 0.69646764 |
| TIGHVDHGKTT | ASSMGLGVQDTQY | 0.7028574 |
| GHVDHGKTT | ASSMGLGVQDTQY | 0.7156774 |
| AQFEHTLL | ASSMGLGVQDTQY | 0.7045896 |
| IVNQDEL | ASSMGLGVQDTQY | 0.6694362 |
| TNGSQFF | ASSMGLGVQDTQY | 0.7098388 |
| TGKGIVE | ASSMGLGVQDTQY | 0.7253798 |
| LISNASDA | ASSMGLGVQDTQY | 0.6956319 |
| AELRTLGELTKIAKAHGAQ | ASSMGLGVQDTQY | 0.7193393 |
| APNAVAAAEPSW | ASSMGLGVQDTQY | 0.71319246 |
| APQVAGAATAVL | ASSMGLGVQDTQY | 0.705786 |
| ARRDPAAAAAILPTDA | ASSMGLGVQDTQY | 0.671687 |
| DAAARDVLAEAGLAEYF | ASSMGLGVQDTQY | 0.6919489 |
| DDQIVLSAKANLAKVEQ | ASSMGLGVQDTQY | 0.70342994 |
| DELLPDTGEDSDEDGH | ASSMGLGVQDTQY | 0.6695997 |
| ELPEAPANRATVPQD | ASSMGLGVQDTQY | 0.69075346 |
| FGQSKIERVTGIG | ASSMGLGVQDTQY | 0.686855 |
| GAGGTGSTAGGAGGAGGAGGLY | ASSMGLGVQDTQY | 0.7265979 |
| GHGGTGAAGGNGGTGGAGGSGI | ASSMGLGVQDTQY | 0.71248114 |
| GNGGNGGAGGEGGPGGAGGAGA | ASSMGLGVQDTQY | 0.71479493 |
| GSTAGNGGNGGNATNGTIAGQP | ASSMGLGVQDTQY | 0.67399776 |
| GTGGDGGNGGNGGTGG | ASSMGLGVQDTQY | 0.71037346 |
| GVRRPAGIPADAPV | ASSMGLGVQDTQY | 0.71500355 |
| GYRQVTDAHLLTLARR | ASSMGLGVQDTQY | 0.5680584 |
| IAKMGQDGHDRG | ASSMGLGVQDTQY | 0.72324276 |
| KAEPSIEIQYT | ASSMGLGVQDTQY | 0.6768034 |
| KPVRIRMAADEDTVAVT | ASSMGLGVQDTQY | 0.68479514 |
| MPPRIAGMRLLVIKPEP | ASSMGLGVQDTQY | 0.7173524 |
| MKFYAVALFRSRGVPPTR | ASSMGLGVQDTQY | 0.7225332 |
| NAGGFGILGAVAHSP | ASSMGLGVQDTQY | 0.6943717 |
| NEGREITLAMAEM | ASSMGLGVQDTQY | 0.6940826 |
| NELVEWSHPLGHRWRWE | ASSMGLGVQDTQY | 0.6718197 |
| NLGAGGTGGNGGNPG | ASSMGLGVQDTQY | 0.71972656 |
| NTEWPKWETVMAADPVD | ASSMGLGVQDTQY | 0.72141385 |
| PASAAPAETQP | ASSMGLGVQDTQY | 0.69382375 |
| QQSSDGKNGVRAAEITGE | ASSMGLGVQDTQY | 0.72492135 |
| RAPDAVTTSWWKE | ASSMGLGVQDTQY | 0.6982264 |
| SGFGNVGAHTSGWFNQ | ASSMGLGVQDTQY | 0.5080619 |

| | | |
|---|---|---|
| SGVAGAVGQAASVGGLKV | ASSMGLGVQDTQY | 0.70318747 |
| SNGGAGTGSAGGNGGT | ASSMGLGVQDTQY | 0.7196658 |
| TGGAGGEGGAAG | ASSMGLGVQDTQY | 0.7185329 |
| TGGDGGAAGAGGAPGHG | ASSMGLGVQDTQY | 0.72373974 |
| TMPPLLAAIANAGRHGV | ASSMGLGVQDTQY | 0.7089925 |
| TSSTAGNSGGA | ASSMGLGVQDTQY | 0.692155 |
| VPATVPVGILC | ASSMGLGVQDTQY | 0.7237609 |
| VQRPDSGGVVLTGRIS | ASSMGLGVQDTQY | 0.7243224 |
| WTPNPVPATKGVPAGLVQAT | ASSMGLGVQDTQY | 0.7190463 |
| YALKRALAKFSDRFDVVII | ASSMGLGVQDTQY | 0.6902007 |
| YLIGSVIV | ASSMGLGVQDTQY | 0.6504289 |
| YGQLVAVID | ASSMGLGVQDTQY | 0.7023792 |
| VTVGGMTLPA | ASSMGLGVQDTQY | 0.7182387 |
| VLALPAV | ASSMGLGVQDTQY | 0.6282932 |
| VIPPTLNYET | ASSMGLGVQDTQY | 0.7071613 |
| VGMGGMHPSSG | ASSMGLGVQDTQY | 0.7101456 |
| VGLAGGAATA | ASSMGLGVQDTQY | 0.5426289 |
| VGGAGGAGGSGP | ASSMGLGVQDTQY | 0.7156164 |
| VFAAFMV | ASSMGLGVQDTQY | 0.72219074 |
| TLYDKLR | ASSMGLGVQDTQY | 0.69365156 |
| TLAVFTVL | ASSMGLGVQDTQY | 0.69169706 |
| TGSVQPARV | ASSMGLGVQDTQY | 0.7211857 |
| TGGGAGSGGT | ASSMGLGVQDTQY | 0.7266671 |
| SHLETSL | ASSMGLGVQDTQY | 0.72259367 |
| SFTTGMT | ASSMGLGVQDTQY | 0.5102823 |
| QVEVAKRMGVRQPTVS | ASSMGLGVQDTQY | 0.43865293 |
| QGSGGPALAM | ASSMGLGVQDTQY | 0.68612075 |
| MLAPSNP | ASSMGLGVQDTQY | 0.7127211 |
| LVNNAAI | ASSMGLGVQDTQY | 0.6812124 |
| LTGEINTLA | ASSMGLGVQDTQY | 0.67687654 |
| LSIFGPV | ASSMGLGVQDTQY | 0.7091861 |
| LLALLANL | ASSMGLGVQDTQY | 0.7219725 |
| LISNASDALDKLR | ASSMGLGVQDTQY | 0.6615507 |
| LINEAVP | ASSMGLGVQDTQY | 0.56946146 |
| LGYAFSGL | ASSMGLGVQDTQY | 0.70507276 |
| LDFVAVAL | ASSMGLGVQDTQY | 0.71621597 |
| LAQPGAE | ASSMGLGVQDTQY | 0.680967 |
| ISNLGAVTGF | ASSMGLGVQDTQY | 0.7021853 |
| HQEDPGRI | ASSMGLGVQDTQY | 0.67747235 |
| GTPGEPTP | ASSMGLGVQDTQY | 0.62308955 |
| GFFNSGS | ASSMGLGVQDTQY | 0.711736 |
| GAGGGGLTAGAN | ASSMGLGVQDTQY | 0.66100067 |
| FVDALVNM | ASSMGLGVQDTQY | 0.723424 |
| FLTVAPEL | ASSMGLGVQDTQY | 0.6615068 |
| FETLTAL | ASSMGLGVQDTQY | 0.7178569 |
| DVINPIEL | ASSMGLGVQDTQY | 0.7124272 |
| APVSGAPV | ASSMGLGVQDTQY | 0.7139248 |
| ALVGGLAW | ASSMGLGVQDTQY | 0.6915834 |
| ALAAAAATAI | ASSMGLGVQDTQY | 0.68156767 |
| AFGLGLLL | ASSMGLGVQDTQY | 0.64637804 |
| AFFAASL | ASSMGLGVQDTQY | 0.5548884 |

| | | |
|---|---|---|
| AAAALPAFSPP | ASSMGLGVQDTQY | 0.7202566 |
| VVLAPAL | ASSMGLGVQDTQY | 0.31684494 |
| TIGHVDHGKTTL | ASSMGLGVQDTQY | 0.72276306 |
| LISNASDALDK | ASSMGLGVQDTQY | 0.71728444 |
| LISNASDAL | ASSMGLGVQDTQY | 0.6724684 |
| ISNASDALDK | ASSMGLGVQDTQY | 0.7146584 |
| NQAETLVYQTE | ASSMGLGVQDTQY | 0.6621733 |
| TGGTGGTAGAG | ASSMGLGVQDTQY | 0.6792595 |
| TGSTAGGAGGA | ASSMGLGVQDTQY | 0.7160686 |
| DPLAVPVAAGPV | ASSMGLGVQDTQY | 0.7157336 |
| GGTGGTPGNGGN | ASSMGLGVQDTQY | 0.66334736 |
| PAGLAGVDPATL | ASSMGLGVQDTQY | 0.70901537 |
| GNNLVVEAPHAGASV | ASSMGLGVQDTQY | 0.71089876 |
| KVTRIMSASTLEQPA | ASSMGLGVQDTQY | 0.6668432 |
| LGFALPIALPRRTVV | ASSMGLGVQDTQY | 0.69306725 |
| TRLGNLTARKLVVLV | ASSMGLGVQDTQY | 0.7217888 |
| WSQQRRLIYFPSAGP | ASSMGLGVQDTQY | 0.70610785 |
| GPNGNPDPMAAAVDIR | ASSMGLGVQDTQY | 0.67894554 |
| TQRCVSVADLRAAASR | ASSMGLGVQDTQY | 0.71327996 |
| QFVLDVAEHPVEVLTETV | ASSMGLGVQDTQY | 0.6655296 |
| GGAGGTGGAGGNGGTQAGDG | ASSMGLGVQDTQY | 0.7139724 |
| PLLATLRAIAVDVDAKPAQV | ASSMGLGVQDTQY | 0.70514226 |
| APKFRTGGYMIDFWGVGYQVAK | ASSMGLGVQDTQY | 0.6851369 |
| GAGGAGGAGGAAGGPGGTGGTG | ASSMGLGVQDTQY | 0.71477693 |
| IGALGLLATDVAPAGSVAFGVA | ASSMGLGVQDTQY | 0.5191886 |
| AGGAGGIGGNGGAGGAGGVLMG | ASSMGLGVQDTQY | 0.66242445 |
| ADELIGLL | ASSMGLGVQDTQY | 0.696257 |
| ADRALKA | ASSMGLGVQDTQY | 0.7060887 |
| AGAVGQVGGAGGA | ASSMGLGVQDTQY | 0.72016513 |
| AGGGGGLGGAP | ASSMGLGVQDTQY | 0.66935384 |
| AGGLLYGNGGAG | ASSMGLGVQDTQY | 0.71026814 |
| AGLESIGL | ASSMGLGVQDTQY | 0.68330646 |
| AGLKTGII | ASSMGLGVQDTQY | 0.6507883 |
| ASLAALLL | ASSMGLGVQDTQY | 0.6905658 |
| ASVINISSI | ASSMGLGVQDTQY | 0.68141615 |
| ATPPGPA | ASSMGLGVQDTQY | 0.6789702 |
| ATVASALV | ASSMGLGVQDTQY | 0.7156949 |
| ATVASLIL | ASSMGLGVQDTQY | 0.63921106 |
| AVNGPVLIRH | ASSMGLGVQDTQY | 0.70865 |
| DGAAGAVGI | ASSMGLGVQDTQY | 0.7228991 |
| DPLLHGI | ASSMGLGVQDTQY | 0.69314593 |
| ELGAALGAL | ASSMGLGVQDTQY | 0.716696 |
| ELGRTQAV | ASSMGLGVQDTQY | 0.71455055 |
| FVAGGYLS | ASSMGLGVQDTQY | 0.7149826 |
| GASGGAGGAGGW | ASSMGLGVQDTQY | 0.68890166 |
| GGAGAGGDGGAGGI | ASSMGLGVQDTQY | 0.6819704 |
| GTGHNGSQPGMGGNG | ASSMGLGVQDTQY | 0.7224139 |
| GYAAASASAS | ASSMGLGVQDTQY | 0.7169327 |
| HQVDVAAVQQG | ASSMGLGVQDTQY | 0.7075304 |
| IGAAIAHT | ASSMGLGVQDTQY | 0.71454036 |
| IGMIEVAQL | ASSMGLGVQDTQY | 0.7150047 |

| | | |
|---|---|---|
| IRLAPEHTSAMD | ASSMGLGVQDTQY | 0.7170603 |
| IYNVSVL | ASSMGLGVQDTQY | 0.6871239 |
| KQGFLGM | ASSMGLGVQDTQY | 0.6037687 |
| LESVITQL | ASSMGLGVQDTQY | 0.69251597 |
| LGLGALVVV | ASSMGLGVQDTQY | 0.7063688 |
| LIGNGGNGGSGGI | ASSMGLGVQDTQY | 0.6770414 |
| LNAINAPTL | ASSMGLGVQDTQY | 0.5032033 |
| LQLLNTGV | ASSMGLGVQDTQY | 0.72135186 |
| LQTSIEF | ASSMGLGVQDTQY | 0.726681 |
| MSFAVSEL | ASSMGLGVQDTQY | 0.6627065 |
| NTGALGL | ASSMGLGVQDTQY | 0.7245287 |
| PPGAGGAGGAGTT | ASSMGLGVQDTQY | 0.60845286 |
| QLVGLGI | ASSMGLGVQDTQY | 0.7153387 |
| RFLEQVHQL | ASSMGLGVQDTQY | 0.72082555 |
| RGVGYVLR | ASSMGLGVQDTQY | 0.66795516 |
| SATGVFAGVI | ASSMGLGVQDTQY | 0.71680295 |
| SLPAITLP | ASSMGLGVQDTQY | 0.6749493 |
| STALAFTV | ASSMGLGVQDTQY | 0.6971946 |
| SVLTVGLGK | ASSMGLGVQDTQY | 0.7233534 |
| TALAGAGSGI | ASSMGLGVQDTQY | 0.6795465 |
| TGLPFTGI | ASSMGLGVQDTQY | 0.7064312 |
| VAEHVTV | ASSMGLGVQDTQY | 0.695371 |
| VDIVLNSL | ASSMGLGVQDTQY | 0.6613991 |
| VLTGGGAVL | ASSMGLGVQDTQY | 0.71155685 |
| VNVGLAYGI | ASSMGLGVQDTQY | 0.7204883 |
| VSSAAVGAV | ASSMGLGVQDTQY | 0.6850678 |
| VTSVYEAGV | ASSMGLGVQDTQY | 0.72344005 |
| WVALDRGA | ASSMGLGVQDTQY | 0.673992 |
| YLVGSVVI | ASSMGLGVQDTQY | 0.7258428 |
| GPPGTGKTLL | ASSMGLGVQDTQY | 0.7167847 |
| GVGMTKF | ASSMGLGVQDTQY | 0.7081558 |
| IGIGGSDL | ASSMGLGVQDTQY | 0.68989843 |
| IGTLLAEL | ASSMGLGVQDTQY | 0.7211527 |
| IYTSGTT | ASSMGLGVQDTQY | 0.68996 |
| LSFPAGV | ASSMGLGVQDTQY | 0.7138783 |
| LYTSGTT | ASSMGLGVQDTQY | 0.7234169 |
| VGAFPFGF | ASSMGLGVQDTQY | 0.13315591 |
| VGAGTVEF | ASSMGLGVQDTQY | 0.4488443 |
| VGKSTLLL | ASSMGLGVQDTQY | 0.23997231 |
| AALAHAGGGIV | ASSMGLGVQDTQY | 0.28623822 |
| AALVAVSLL | ASSMGLGVQDTQY | 0.20484436 |
| AAVSRAVAAL | ASSMGLGVQDTQY | 0.1410948 |
| AGGSSGIGGF | ASSMGLGVQDTQY | 0.2010633 |
| AGLTVANANG | ASSMGLGVQDTQY | 0.16704452 |
| AGLVGAVL | ASSMGLGVQDTQY | 0.81533146 |
| ALSQAFTV | ASSMGLGVQDTQY | 0.31571528 |
| ALTFSVSL | ASSMGLGVQDTQY | 0.20018758 |
| ALTGAALVAL | ASSMGLGVQDTQY | 0.4517886 |
| ASFSIGAGVL | ASSMGLGVQDTQY | 0.42323354 |
| DGSVYKI | ASSMGLGVQDTQY | 0.122929566 |
| EQLAEIV | ASSMGLGVQDTQY | 0.15906394 |

| | | |
|---|---|---|
| FAVSELTVM | ASSMGLGVQDTQY | 0.17542551 |
| GAAPLAAGL | ASSMGLGVQDTQY | 0.23812011 |
| GADGGDGGSGGAGG | ASSMGLGVQDTQY | 0.117763475 |
| GFGLDTGGAGG | ASSMGLGVQDTQY | 0.2839682 |
| GLASGGLL | ASSMGLGVQDTQY | 0.14081676 |
| GRIAGAAVLL | ASSMGLGVQDTQY | 0.10878537 |
| GSAGSLNTSF | ASSMGLGVQDTQY | 0.28562686 |
| GTGQIGI | ASSMGLGVQDTQY | 0.21335793 |
| GTLIALIL | ASSMGLGVQDTQY | 0.17776337 |
| IAATMGFVVG | ASSMGLGVQDTQY | 0.33717814 |
| IFAQALL | ASSMGLGVQDTQY | 0.16673909 |
| IGAAVALGF | ASSMGLGVQDTQY | 0.14468741 |
| IGLTPLAL | ASSMGLGVQDTQY | 0.2753513 |
| ILGSNTSTL | ASSMGLGVQDTQY | 0.35440022 |
| IVFSLTVL | ASSMGLGVQDTQY | 0.15195453 |
| LASKATGF | ASSMGLGVQDTQY | 0.24919745 |
| LASVALIL | ASSMGLGVQDTQY | 0.52445 |
| LFAVGLTL | ASSMGLGVQDTQY | 0.2647127 |
| LFNGGIGTYI | ASSMGLGVQDTQY | 0.37209654 |
| LTLGAGII | ASSMGLGVQDTQY | 0.41735893 |
| LVAAALGDI | ASSMGLGVQDTQY | 0.18138726 |
| NTAFVQLGI | ASSMGLGVQDTQY | 0.45973605 |
| QLFSGAEL | ASSMGLGVQDTQY | 0.607557 |
| SAATFITL | ASSMGLGVQDTQY | 0.20140505 |
| SNYGLANLGV | ASSMGLGVQDTQY | 0.30480713 |
| SYGGAYAVM | ASSMGLGVQDTQY | 0.13990338 |
| TTGVGGLTLG | ASSMGLGVQDTQY | 0.21595179 |
| TTQTIYI | ASSMGLGVQDTQY | 0.13584127 |
| VAAGAGLGAVI | ASSMGLGVQDTQY | 0.3401788 |
| VAAVVTLL | ASSMGLGVQDTQY | 0.37151498 |
| VGFLGGLI | ASSMGLGVQDTQY | 0.20663194 |
| VGVIPILL | ASSMGLGVQDTQY | 0.11870482 |
| VIFASSLI | ASSMGLGVQDTQY | 0.1239521 |
| VKGGDGGAAAG | ASSMGLGVQDTQY | 0.2537085 |
| VTKTFTNI | ASSMGLGVQDTQY | 0.1475603 |
| VVAAAILL | ASSMGLGVQDTQY | 0.13393654 |
| YAALDVEL | ASSMGLGVQDTQY | 0.18574241 |
| LASVALLL | ASSMGLGVQDTQY | 0.34943855 |
| LDAVHRV | ASSMGLGVQDTQY | 0.4289577 |
| QGLSSSIF | ASSMGLGVQDTQY | 0.16916138 |
| RGTFANIRL | ASSMGLGVQDTQY | 0.6150979 |
| RQLWWGHRI | ASSMGLGVQDTQY | 0.6714821 |
| YGPPGTGKTLL | ASSMGLGVQDTQY | 0.5825504 |
| AALVGAPANVA | ASSMGLGVQDTQY | 0.19247289 |
| AATFRRTE | ASSMGLGVQDTQY | 0.13636532 |
| AGALGGGI | ASSMGLGVQDTQY | 0.7788547 |
| AGGAGGAGGIGAG | ASSMGLGVQDTQY | 0.25520682 |
| AGSGALGL | ASSMGLGVQDTQY | 0.4010387 |
| AQIADQI | ASSMGLGVQDTQY | 0.36330932 |
| ASAVAVTYL | ASSMGLGVQDTQY | 0.18909185 |
| ASTVLGGTGG | ASSMGLGVQDTQY | 0.1631047 |

| | | |
|---|---|---|
| AVGGEPAK | ASSMGLGVQDTQY | 0.5721128 |
| CEQGTFGGTC | ASSMGLGVQDTQY | 0.37286693 |
| DEGKAPAG | ASSMGLGVQDTQY | 0.13113724 |
| DTVVFPAL | ASSMGLGVQDTQY | 0.1510675 |
| FEGITMAG | ASSMGLGVQDTQY | 0.1745573 |
| FPLVVQD | ASSMGLGVQDTQY | 0.21923971 |
| GAFTTPQ | ASSMGLGVQDTQY | 0.36105925 |
| GFGGFGTAGGVGGA | ASSMGLGVQDTQY | 0.21669875 |
| GFLAQFL | ASSMGLGVQDTQY | 0.18331374 |
| GGAGGAGGNAGVA | ASSMGLGVQDTQY | 0.308568 |
| GGVGTAGAGGAGGAGG | ASSMGLGVQDTQY | 0.2860378 |
| GNGGGGDAGNAGSGGN | ASSMGLGVQDTQY | 0.29109132 |
| GTGGAGGAGNDAGS | ASSMGLGVQDTQY | 0.2437849 |
| GTTNTGWFNS | ASSMGLGVQDTQY | 0.213225 |
| IFMGPLFTL | ASSMGLGVQDTQY | 0.355788 |
| ILAHAIEAI | ASSMGLGVQDTQY | 0.43085438 |
| IRSAIREAL | ASSMGLGVQDTQY | 0.2834221 |
| ITFAGGG | ASSMGLGVQDTQY | 0.33604228 |
| ITGMSGHQEV | ASSMGLGVQDTQY | 0.21393228 |
| IVAGLACTL | ASSMGLGVQDTQY | 0.24816065 |
| IVGGGLAGA | ASSMGLGVQDTQY | 0.1332132 |
| IVTGVLNI | ASSMGLGVQDTQY | 0.22382948 |
| LAQQGCI | ASSMGLGVQDTQY | 0.2539089 |
| LCLHVEN | ASSMGLGVQDTQY | 0.19955772 |
| LDAISDGS | ASSMGLGVQDTQY | 0.4687613 |
| LGGFAAML | ASSMGLGVQDTQY | 0.23317763 |
| LGGKNPMI | ASSMGLGVQDTQY | 0.16517416 |
| LGPKGVAAIL | ASSMGLGVQDTQY | 0.29967168 |
| LMVGVSI | ASSMGLGVQDTQY | 0.5529542 |
| LVGAAFVL | ASSMGLGVQDTQY | 0.15495859 |
| LVLAVAPL | ASSMGLGVQDTQY | 0.20173317 |
| MAAIAVIDAA | ASSMGLGVQDTQY | 0.6540319 |
| NNGINAELQ | ASSMGLGVQDTQY | 0.5481763 |
| RGEPLVLVHG | ASSMGLGVQDTQY | 0.22379366 |
| RGGAGGAAGLI | ASSMGLGVQDTQY | 0.26864183 |
| RSYRMHPAV | ASSMGLGVQDTQY | 0.82481945 |
| SAAHLLAGQLA | ASSMGLGVQDTQY | 0.28823155 |
| SAAVGSY | ASSMGLGVQDTQY | 0.50981736 |
| SPHAVLHRI | ASSMGLGVQDTQY | 0.12712865 |
| SQIAKVL | ASSMGLGVQDTQY | 0.18129177 |
| TALGHLVP | ASSMGLGVQDTQY | 0.14896493 |
| TAPTTSGAA | ASSMGLGVQDTQY | 0.10941826 |
| TDGDDGNGGSPGT | ASSMGLGVQDTQY | 0.5168849 |
| TDGLGEN | ASSMGLGVQDTQY | 0.18144031 |
| TGGPPLPGQA | ASSMGLGVQDTQY | 0.36517423 |
| TIPAITI | ASSMGLGVQDTQY | 0.19938508 |
| TIPGFTI | ASSMGLGVQDTQY | 0.41466144 |
| TSIASGALA | ASSMGLGVQDTQY | 0.45338276 |
| TVTLPGAAI | ASSMGLGVQDTQY | 0.22524571 |
| VASGLSAALP | ASSMGLGVQDTQY | 0.16205496 |
| VGAGVIGI | ASSMGLGVQDTQY | 0.32100868 |

| | | |
|---|---|---|
| VGPKRFGV | ASSMGLGVQDTQY | 0.3674838 |
| VNVAFPD | ASSMGLGVQDTQY | 0.3042862 |
| VSMTNSGVS | ASSMGLGVQDTQY | 0.12539981 |
| VTDVLGDLSV | ASSMGLGVQDTQY | 0.45233023 |
| VVVVGAGFAGL | ASSMGLGVQDTQY | 0.24196324 |
| VYTVGTVV | ASSMGLGVQDTQY | 0.13376784 |
| WTTFSGP | ASSMGLGVQDTQY | 0.12044782 |
| YDELVEAF | ASSMGLGVQDTQY | 0.12992145 |
| YGTGGAGGNGG | ASSMGLGVQDTQY | 0.19390126 |
| YTNVMAV | ASSMGLGVQDTQY | 0.36135387 |
| FLEQVHQL | ASSMGLGVQDTQY | 0.34854114 |
| GLSLLAVAV | ASSMGLGVQDTQY | 0.21514714 |
| LEQVHQL | ASSMGLGVQDTQY | 0.34410185 |
| LNAAGAI | ASSMGLGVQDTQY | 0.2778897 |
| PAVHASL | ASSMGLGVQDTQY | 0.099834375 |
| TELEDDA | ASSMGLGVQDTQY | 0.16915636 |
| TPPGPPN | ASSMGLGVQDTQY | 0.12798591 |
| TPPPPPPP | ASSMGLGVQDTQY | 0.40077496 |
| TQSPALI | ASSYNNERLF | 0.14396136 |
| EFAAAFPQ | ASSYNNERLF | 0.20620954 |
| LLSGNLDVL | ASSYNNERLF | 0.26482457 |
| GPANLGNYNI | ASSYNNERLF | 0.7127594 |
| TLGPTLP | ASSYNNERLF | 0.30406904 |
| VYAYSSCCK | ASSYNNERLF | 0.14056613 |
| APAPAPAPNI | ASSYNNERLF | 0.12116666 |
| PGGPVPIP | ASSYNNERLF | 0.2995566 |
| FLTVALEP | ASSYNNERLF | 0.20281363 |
| SQYFGANL | ASSYNNERLF | 0.1420277 |
| GGVKPAA | ASSYNNERLF | 0.15847355 |
| GGTGGAGGAGGW | ASSYNNERLF | 0.2854986 |
| GVGGVAGLGGAGG | ASSYNNERLF | 0.1286263 |
| VLLVLATTI | ASSYNNERLF | 0.31350285 |
| IGIFAAML | ASSYNNERLF | 0.15289785 |
| AVGDLID | ASSYNNERLF | 0.19679157 |
| NISPLPL | ASSYNNERLF | 0.53462636 |
| VTAVFVAI | ASSYNNERLF | 0.70842767 |
| SGDDNDAAL | ASSYNNERLF | 0.27772838 |
| AVAALAHS | ASSYNNERLF | 0.16837254 |
| GGVTVMATP | ASSYNNERLF | 0.1497044 |
| AGLAANVRD | ASSYNNERLF | 0.15103082 |
| TNRPDLIDPALLRPG | ASSYNNERLF | 0.16200039 |
| LANLGVTAL | ASSYNNERLF | 0.45758152 |
| TGERIEL | ASSYNNERLF | 0.27953568 |
| NGEELAA | ASSYNNERLF | 0.15009679 |
| VLSLVWG | ASSYNNERLF | 0.33371848 |
| IGYALWL | ASSYNNERLF | 0.60072696 |
| LSLLLAPL | ASSYNNERLF | 0.6447414 |
| TLDEALK | ASSYNNERLF | 0.1348222 |
| TDGAGTAEGG | ASSYNNERLF | 0.17597744 |
| NAPGGAGGNGG | ASSYNNERLF | 0.20565788 |
| VSVFMGMGGNF | ASSYNNERLF | 0.20233554 |

| | | |
|---|---|---|
| LASYLAAL | ASSYNNERLF | 0.16004875 |
| LVGGTPAL | ASSYNNERLF | 0.17100316 |
| VRNADVL | ASSYNNERLF | 0.30478093 |
| TCSSPFLVSAM | ASSYNNERLF | 0.85088676 |
| VHITDAE | ASSYNNERLF | 0.20538732 |
| EYAAPSF | ASSYNNERLF | 0.11624942 |
| VVDAFHGGL | ASSYNNERLF | 0.21497157 |
| ASVVDPQ | ASSYNNERLF | 0.13481483 |
| SYNEFGGLS | ASSYNNERLF | 0.1800785 |
| DPDGSVY | ASSYNNERLF | 0.1761964 |
| MVSTIHATE | ASSYNNERLF | 0.1251044 |
| LAAGVADPV | ASSYNNERLF | 0.21091534 |
| TKEQIAA | ASSYNNERLF | 0.5488486 |
| SLASPAAL | ASSYNNERLF | 0.30911848 |
| IGLVFYGV | ASSYNNERLF | 0.31961063 |
| FLLDASNL | ASSYNNERLF | 0.6135558 |
| YQVGNFP | ASSYNNERLF | 0.29045206 |
| ESLLSQST | ASSYNNERLF | 0.52680284 |
| LFTVGFAAVG | ASSYNNERLF | 0.5745257 |
| ATPPTPT | ASSYNNERLF | 0.2282579 |
| TDLATLG | ASSYNNERLF | 0.19247246 |
| SAGGTGR | ASSYNNERLF | 0.5374645 |
| GFSNSGS | ASSYNNERLF | 0.2636025 |
| GGNGGSGGTSV | ASSYNNERLF | 0.28444615 |
| GTSVATGGAGN | ASSYNNERLF | 0.31748796 |
| NGGNGGSGGTS | ASSYNNERLF | 0.50208336 |
| AVVKFPH | ASSYNNERLF | 0.17997074 |
| TPPPPPP | ASSYNNERLF | 0.79536694 |
| AAPAAAAAPA | ASSYNNERLF | 0.12070175 |
| IALNEDL | ASSYNNERLF | 0.15629895 |
| IVNVSSM | ASSYNNERLF | 0.4399954 |
| TGIPPAPR | ASSYNNERLF | 0.104550235 |
| FDLGGGTF | ASSYNNERLF | 0.13545436 |
| FDLGGGTFDVS | ASSYNNERLF | 0.19366665 |
| NEPTAAA | ASSYNNERLF | 0.5806611 |
| NEPTAAALA | ASSYNNERLF | 0.106038556 |
| GSIINISSI | ASSYNNERLF | 0.17523003 |
| VAPGNAGTA | ASSYNNERLF | 0.1294402 |
| TGGGASLE | ASSYNNERLF | 0.26722085 |
| VAVYNHYKRI | ASSYNNERLF | 0.24618629 |
| TDPAPATT | ASSYNNERLF | 0.1428931 |
| TNRPDILDPALLRPG | ASSYNNERLF | 0.32091096 |
| LTTTTTS | ASSYNNERLF | 0.1287247 |
| PTGSGKTL | ASSYNNERLF | 0.30950868 |
| TGSGKTL | ASSYNNERLF | 0.36573863 |
| TIGHVDHGK | ASSYNNERLF | 0.7542175 |
| TIGHVDHGKTT | ASSYNNERLF | 0.13524085 |
| GHVDHGKTT | ASSYNNERLF | 0.2661707 |
| AQFEHTLL | ASSYNNERLF | 0.1546069 |
| IVNQDEL | ASSYNNERLF | 0.7865038 |
| TNGSQFF | ASSYNNERLF | 0.20916604 |

| | | |
|---|---|---|
| TGKGIVE | ASSYNNERLF | 0.12421402 |
| LISNASDA | ASSYNNERLF | 0.3205648 |
| AELRTLGELTKIAKAHGAQ | ASSYNNERLF | 0.45844883 |
| APNAVAAAEPSW | ASSYNNERLF | 0.19026768 |
| APQVAGAATAVL | ASSYNNERLF | 0.6341524 |
| ARRDPAAAAAILPTDA | ASSYNNERLF | 0.17905605 |
| DAAARDVLAEAGLAEYF | ASSYNNERLF | 0.23758754 |
| DDQIVLSAKANLAKVEQ | ASSYNNERLF | 0.59998465 |
| DELLPDTGEDSDEDGH | ASSYNNERLF | 0.31434304 |
| ELPEAPANRATVPQD | ASSYNNERLF | 0.17603631 |
| FGQSKIERVTGIG | ASSYNNERLF | 0.460031 |
| GAGGTGSTAGGAGGAGGAGGLY | ASSYNNERLF | 0.2646408 |
| GHGGTGAAGGNGGTGGAGGSGI | ASSYNNERLF | 0.13989475 |
| GNGGNGGAGGEGGPGGAGGAGA | ASSYNNERLF | 0.7080229 |
| GSTAGNGGNGGNATNGTIAGQP | ASSYNNERLF | 0.17748708 |
| GTGGDGGNGGNGGTGG | ASSYNNERLF | 0.4515553 |
| GVRRPAGIPADAPV | ASSYNNERLF | 0.31446132 |
| GYRQVTDAHLLTLARR | ASSYNNERLF | 0.8256149 |
| IAKMGQDGHDRG | ASSYNNERLF | 0.16021082 |
| KAEPSIEIQYT | ASSYNNERLF | 0.2771824 |
| KPVRIRMAADEDTVAVT | ASSYNNERLF | 0.2349379 |
| MPPRIAGMRLLVIKPEP | ASSYNNERLF | 0.27698174 |
| MKFYAVALFRSRGVPPTR | ASSYNNERLF | 0.2346093 |
| NAGGFGILGAVAHSP | ASSYNNERLF | 0.22171211 |
| NEGREITLAMAEM | ASSYNNERLF | 0.38683933 |
| NELVEWSHPLGHRWRWE | ASSYNNERLF | 0.43247634 |
| NLGAGGTGGNGGNPG | ASSYNNERLF | 0.18635859 |
| NTEWPKWETVMAADPVD | ASSYNNERLF | 0.19038242 |
| PASAAPAETQP | ASSYNNERLF | 0.6340511 |
| QQSSDGKNGVRAAEITGE | ASSYNNERLF | 0.14501964 |
| RAPDAVTTSWWKE | ASSYNNERLF | 0.28586853 |
| SGFGNVGAHTSGWFNQ | ASSYNNERLF | 0.27557802 |
| SGVAGAVGQAASVGGLKV | ASSYNNERLF | 0.34265393 |
| SNGGAGTGSAGGNGGT | ASSYNNERLF | 0.20006476 |
| TGGAGGEGGAAG | ASSYNNERLF | 0.52174616 |
| TGGDGGAAGAGGAPGHG | ASSYNNERLF | 0.18162952 |
| TMPPLLAAIANAGRHGV | ASSYNNERLF | 0.40914634 |
| TSSTAGNSGGA | ASSYNNERLF | 0.25899655 |
| VPATVPVGILC | ASSYNNERLF | 0.25122362 |
| VQRPDSGGVVLTGRIS | ASSYNNERLF | 0.12688819 |
| WTPNPVPATKGVPAGLVQAT | ASSYNNERLF | 0.25234407 |
| YALKRALAKFSDRFDVVII | ASSYNNERLF | 0.36272675 |
| YLIGSVIV | ASSYNNERLF | 0.41263533 |
| YGQLVAVID | ASSYNNERLF | 0.12971862 |
| VTVGGMTLPA | ASSYNNERLF | 0.37537324 |
| VLALPAV | ASSYNNERLF | 0.40247408 |
| VIPPTLNYET | ASSYNNERLF | 0.8212695 |
| VGMGGMHPSSG | ASSYNNERLF | 0.19519801 |
| VGLAGGAATA | ASSYNNERLF | 0.35551703 |
| VGGAGGAGGSGP | ASSYNNERLF | 0.16598128 |
| VFAAFMV | ASSYNNERLF | 0.19694011 |

| | | |
|---|---|---|
| TLYDKLR | ASSYNNERLF | 0.12025743 |
| TLAVFTVL | ASSYNNERLF | 0.14540908 |
| TGSVQPARV | ASSYNNERLF | 0.25058955 |
| TGGGAGSGGT | ASSYNNERLF | 0.14792125 |
| SHLETSL | ASSYNNERLF | 0.76877093 |
| SFTTGMT | ASSYNNERLF | 0.4306001 |
| QVEVAKRMGVRQPTVS | ASSYNNERLF | 0.34723923 |
| QGSGGPALAM | ASSYNNERLF | 0.44631106 |
| MLAPSNP | ASSYNNERLF | 0.6428766 |
| LVNNAAI | ASSYNNERLF | 0.30987898 |
| LTGEINTLA | ASSYNNERLF | 0.20551097 |
| LSIFGPV | ASSYNNERLF | 0.26392525 |
| LLALLANL | ASSYNNERLF | 0.1189304 |
| LISNASDALDKLR | ASSYNNERLF | 0.18659331 |
| LINEAVP | ASSYNNERLF | 0.18597612 |
| LGYAFSGL | ASSYNNERLF | 0.30146992 |
| LDFVAVAL | ASSYNNERLF | 0.104049735 |
| LAQPGAE | ASSYNNERLF | 0.6419893 |
| ISNLGAVTGF | ASSYNNERLF | 0.5165783 |
| HQEDPGRI | ASSYNNERLF | 0.28693762 |
| GTPGEPTP | ASSYNNERLF | 0.38969144 |
| GFFNSGS | ASSYNNERLF | 0.4758841 |
| GAGGGGLTAGAN | ASSYNNERLF | 0.13665661 |
| FVDALVNM | ASSYNNERLF | 0.30507708 |
| FLTVAPEL | ASSYNNERLF | 0.15670219 |
| FETLTAL | ASSYNNERLF | 0.38602582 |
| DVINPIEL | ASSYNNERLF | 0.48289007 |
| APVSGAPV | ASSYNNERLF | 0.32536933 |
| ALVGGLAW | ASSYNNERLF | 0.16513833 |
| ALAAAAATAI | ASSYNNERLF | 0.28166467 |
| AFGLGLLL | ASSYNNERLF | 0.15989721 |
| AFFAAASL | ASSYNNERLF | 0.17908077 |
| AAAALPAFSPP | ASSYNNERLF | 0.14178498 |
| VVLAPAL | ASSYNNERLF | 0.33915275 |
| TIGHVDHGKTTL | ASSYNNERLF | 0.1571929 |
| LISNASDALDK | ASSYNNERLF | 0.21745446 |
| LISNASDAL | ASSYNNERLF | 0.258151 |
| ISNASDALDK | ASSYNNERLF | 0.199189 |
| NQAETLVYQTE | ASSYNNERLF | 0.16442287 |
| TGGTGGTAGAG | ASSYNNERLF | 0.3748778 |
| TGSTAGGAGGA | ASSYNNERLF | 0.36833346 |
| DPLAVPVAAGPV | ASSYNNERLF | 0.12962395 |
| GGTGGTPGNGGN | ASSYNNERLF | 0.428888 |
| PAGLAGVDPATL | ASSYNNERLF | 0.179222 |
| GNNLVVEAPHAGASV | ASSYNNERLF | 0.28750008 |
| KVTRIMSASTLEQPA | ASSYNNERLF | 0.39796254 |
| LGFALPIALPRRTVV | ASSYNNERLF | 0.21350294 |
| TRLGNLTARKLVVLV | ASSYNNERLF | 0.14487159 |
| WSQQRRLIYFPSAGP | ASSYNNERLF | 0.18699984 |
| GPNGNPDPMAAAVDIR | ASSYNNERLF | 0.22407046 |
| TQRCVSVADLRAAASR | ASSYNNERLF | 0.21204332 |

| | | |
|---|---|---|
| QFVLDVAEHPVEVLTETV | ASSYNNERLF | 0.11706513 |
| GGAGGTGGAGGNGGTQAGDG | ASSYNNERLF | 0.3457775 |
| PLLATLRAIAVDVDAKPAQV | ASSYNNERLF | 0.14001034 |
| APKFRTGGYMIDFWGVGYQVAK | ASSYNNERLF | 0.20897087 |
| GAGGAGGAGGAAGGPGGTGGTG | ASSYNNERLF | 0.1951593 |
| IGALGLLATDVAPAGSVAFGVA | ASSYNNERLF | 0.42339718 |
| AGGAGGIGGNGGAGGAGGVLMG | ASSYNNERLF | 0.27957094 |
| ADELIGLL | ASSYNNERLF | 0.18989278 |
| ADRALKA | ASSYNNERLF | 0.29325637 |
| AGAVGQVGGAGGA | ASSYNNERLF | 0.25903162 |
| AGGGGGLGGAP | ASSYNNERLF | 0.5791739 |
| AGGLLYGNGGAG | ASSYNNERLF | 0.20733872 |
| AGLESIGL | ASSYNNERLF | 0.17939971 |
| AGLKTGII | ASSYNNERLF | 0.29565954 |
| ASLAALLL | ASSYNNERLF | 0.20363794 |
| ASVINISSI | ASSYNNERLF | 0.16129997 |
| ATPPGPA | ASSYNNERLF | 0.14693296 |
| ATVASALV | ASSYNNERLF | 0.15834919 |
| ATVASLIL | ASSYNNERLF | 0.1554399 |
| AVNGPVLIRH | ASSYNNERLF | 0.343345 |
| DGAAGAVGI | ASSYNNERLF | 0.42031378 |
| DPLLHGI | ASSYNNERLF | 0.20871097 |
| ELGAALGAL | ASSYNNERLF | 0.12434069 |
| ELGRTQAV | ASSYNNERLF | 0.13820463 |
| FVAGGYLS | ASSYNNERLF | 0.22712645 |
| GASGGAGGAGGW | ASSYNNERLF | 0.3575496 |
| GGAGAGGDGGAGGI | ASSYNNERLF | 0.28661942 |
| GTGHNGSQPGMGGNG | ASSYNNERLF | 0.8361391 |
| GYAAASASAS | ASSYNNERLF | 0.39593714 |
| HQVDVAAVQQG | ASSYNNERLF | 0.1978015 |
| IGAAIAHT | ASSYNNERLF | 0.35601896 |
| IGMIEVAQL | ASSYNNERLF | 0.22715427 |
| IRLAPEHTSAMD | ASSYNNERLF | 0.21257949 |
| IYNVSVL | ASSYNNERLF | 0.23382849 |
| KQGFLGM | ASSYNNERLF | 0.28091493 |
| LESVITQL | ASSYNNERLF | 0.16028793 |
| LGLGALVVV | ASSYNNERLF | 0.20552105 |
| LIGNGGNGGSGGI | ASSYNNERLF | 0.13785297 |
| LNAINAPTL | ASSYNNERLF | 0.32999128 |
| LQLLNTGV | ASSYNNERLF | 0.14581297 |
| LQTSIEF | ASSYNNERLF | 0.14530225 |
| MSFAVSEL | ASSYNNERLF | 0.21472894 |
| NTGALGL | ASSYNNERLF | 0.6122726 |
| PPGAGGAGGAGTT | ASSYNNERLF | 0.43578577 |
| QLVGLGI | ASSYNNERLF | 0.16421092 |
| RFLEQVHQL | ASSYNNERLF | 0.16418056 |
| RGVGYVLR | ASSYNNERLF | 0.6459925 |
| SATGVFAGVI | ASSYNNERLF | 0.1615099 |
| SLPAITLP | ASSYNNERLF | 0.30363744 |
| STALAFTV | ASSYNNERLF | 0.3205514 |
| SVLTVGLGK | ASSYNNERLF | 0.31037498 |

| | | |
|---|---|---|
| TALAGAGSGI | ASSYNNERLF | 0.18420371 |
| TGLPFTGI | ASSYNNERLF | 0.22932842 |
| VAEHVTV | ASSYNNERLF | 0.20756248 |
| VDIVLNSL | ASSYNNERLF | 0.2277877 |
| VLTGGGAVL | ASSYNNERLF | 0.115168765 |
| VNVGLAYGI | ASSYNNERLF | 0.38580987 |
| VSSAAVGAV | ASSYNNERLF | 0.15082566 |
| VTSVYEAGV | ASSYNNERLF | 0.14804032 |
| WVALDRGA | ASSYNNERLF | 0.11218428 |
| YLVGSVVI | ASSYNNERLF | 0.20622358 |
| GPPGTGKTLL | ASSYNNERLF | 0.3394003 |
| GVGMTKF | ASSYNNERLF | 0.3871274 |
| IGIGGSDL | ASSYNNERLF | 0.33524567 |
| IGTLLAEL | ASSYNNERLF | 0.3484643 |
| IYTSGTT | ASSYNNERLF | 0.4865585 |
| LSFPAGV | ASSYNNERLF | 0.23511383 |
| LYTSGTT | ASSYNNERLF | 0.12262846 |
| VGAFPFGF | ASSYNNERLF | 0.2423392 |
| VGAGTVEF | ASSYNNERLF | 0.43016654 |
| VGKSTLLL | ASSYNNERLF | 0.36822987 |
| AALAHAGGGIV | ASSYNNERLF | 0.45859855 |
| AALVAVSLL | ASSYNNERLF | 0.12164142 |
| AAVSRAVAAL | ASSYNNERLF | 0.3056716 |
| AGGSSGIGGF | ASSYNNERLF | 0.30977857 |
| AGLTVANANG | ASSYNNERLF | 0.13120118 |
| AGLVGAVL | ASSYNNERLF | 0.16628084 |
| ALSQAFTV | ASSYNNERLF | 0.18225566 |
| ALTFSVSL | ASSYNNERLF | 0.12961274 |
| ALTGAALVAL | ASSYNNERLF | 0.25801194 |
| ASFSIGAGVL | ASSYNNERLF | 0.18111135 |
| DGSVYKI | ASSYNNERLF | 0.18874915 |
| EQLAEIV | ASSYNNERLF | 0.19417205 |
| FAVSELTVM | ASSYNNERLF | 0.16634679 |
| GAAPLAAGL | ASSYNNERLF | 0.21184967 |
| GADGGDGGSGGAGG | ASSYNNERLF | 0.19445452 |
| GFGLDTGGAGG | ASSYNNERLF | 0.40269703 |
| GLASGGLL | ASSYNNERLF | 0.25786597 |
| GRIAGAAVLL | ASSYNNERLF | 0.28090453 |
| GSAGSLNTSF | ASSYNNERLF | 0.28369465 |
| GTGQIGI | ASSYNNERLF | 0.61320984 |
| GTLIALIL | ASSYNNERLF | 0.13515878 |
| IAATMGFVVG | ASSYNNERLF | 0.1768495 |
| IFAQALL | ASSYNNERLF | 0.23181446 |
| IGAAVALGF | ASSYNNERLF | 0.17317288 |
| IGLTPLAL | ASSYNNERLF | 0.3732596 |
| ILGSNTSTL | ASSYNNERLF | 0.24312004 |
| IVFSLTVL | ASSYNNERLF | 0.32618177 |
| LASKATGF | ASSYNNERLF | 0.21781334 |
| LASVALIL | ASSYNNERLF | 0.6561022 |
| LFAVGLTL | ASSYNNERLF | 0.2970272 |
| LFNGGIGTYI | ASSYNNERLF | 0.21355921 |

| | | |
|---|---|---|
| LTLGAGII | ASSYNNERLF | 0.51725036 |
| LVAAALGDI | ASSYNNERLF | 0.14931874 |
| NTAFVQLGI | ASSYNNERLF | 0.38759676 |
| QLFSGAEL | ASSYNNERLF | 0.34682828 |
| SAATFITL | ASSYNNERLF | 0.16244689 |
| SNYGLANLGV | ASSYNNERLF | 0.2372919 |
| SYGGAYAVM | ASSYNNERLF | 0.4440337 |
| TTGVGGLTLG | ASSYNNERLF | 0.5618715 |
| TTQTIYI | ASSYNNERLF | 0.16383049 |
| VAAGAGLGAVI | ASSYNNERLF | 0.15010719 |
| VAAVVTLL | ASSYNNERLF | 0.13580738 |
| VGFLGGLI | ASSYNNERLF | 0.14805558 |
| VGVIPILL | ASSYNNERLF | 0.13414611 |
| VIFASSLI | ASSYNNERLF | 0.29851037 |
| VKGGDGGAAAG | ASSYNNERLF | 0.22237182 |
| VTKTFTNI | ASSYNNERLF | 0.76915073 |
| VVAAAILL | ASSYNNERLF | 0.34656858 |
| YAALDVEL | ASSYNNERLF | 0.47860047 |
| LASVALLL | ASSYNNERLF | 0.16651343 |
| LDAVHRV | ASSYNNERLF | 0.20103139 |
| QGLSSSIF | ASSYNNERLF | 0.12481088 |
| RGTFANIRL | ASSYNNERLF | 0.142841 |
| RQLWWGHRI | ASSYNNERLF | 0.14272508 |
| YGPPGTGKTLL | ASSYNNERLF | 0.20257112 |
| AALVGAPANVA | ASSYNNERLF | 0.12025135 |
| AATFRRTE | ASSYNNERLF | 0.68347514 |
| AGALGGGI | ASSYNNERLF | 0.18784979 |
| AGGAGGAGGIGAG | ASSYNNERLF | 0.334578 |
| AGSGALGL | ASSYNNERLF | 0.18464932 |
| AQIADQI | ASSYNNERLF | 0.14983405 |
| ASAVAVTYL | ASSYNNERLF | 0.19198145 |
| ASTVLGGTGG | ASSYNNERLF | 0.30043054 |
| AVGGEPAK | ASSYNNERLF | 0.13055538 |
| CEQGTFGGTC | ASSYNNERLF | 0.123449184 |
| DEGKAPAG | ASSYNNERLF | 0.16494955 |
| DTVVFPAL | ASSYNNERLF | 0.14215294 |
| FEGITMAG | ASSYNNERLF | 0.78391373 |
| FPLVVQD | ASSYNNERLF | 0.44310296 |
| GAFTTPQ | ASSYNNERLF | 0.22997665 |
| GFGGFGTAGGVGGA | ASSYNNERLF | 0.77610266 |
| GFLAQFL | ASSYNNERLF | 0.105119124 |
| GGAGGAGGNAGVA | ASSYNNERLF | 0.19857723 |
| GGVGTAGAGGAGGAGG | ASSYNNERLF | 0.63126314 |
| GNGGGGDAGNAGSGGN | ASSYNNERLF | 0.23606347 |
| GTGGAGGAGNDAGS | ASSYNNERLF | 0.2353228 |
| GTTNTGWFNS | ASSYNNERLF | 0.3970446 |
| IFMGPLFTL | ASSYNNERLF | 0.16256726 |
| ILAHAIEAI | ASSYNNERLF | 0.21271297 |
| IRSAIREAL | ASSYNNERLF | 0.18872803 |
| ITFAGGG | ASSYNNERLF | 0.34923857 |
| ITGMSGHQEV | ASSYNNERLF | 0.26285473 |

| | | |
|---|---|---|
| IVAGLACTL | ASSYNNERLF | 0.13293934 |
| IVGGGLAGA | ASSYNNERLF | 0.3353594 |
| IVTGVLNI | ASSYNNERLF | 0.7870334 |
| LAQQGCI | ASSYNNERLF | 0.16528189 |
| LCLHVEN | ASSYNNERLF | 0.2910264 |
| LDAISDGS | ASSYNNERLF | 0.25026494 |
| LGGFAAML | ASSYNNERLF | 0.16192906 |
| LGGKNPMI | ASSYNNERLF | 0.20217228 |
| LGPKGVAAIL | ASSYNNERLF | 0.13173622 |
| LMVGVSI | ASSYNNERLF | 0.44612888 |
| LVGAAFVL | ASSYNNERLF | 0.15205023 |
| LVLAVAPL | ASSYNNERLF | 0.19961387 |
| MAAIAVIDAA | ASSYNNERLF | 0.294022 |
| NNGINAELQ | ASSYNNERLF | 0.79623926 |
| RGEPLVLVHG | ASSYNNERLF | 0.43638432 |
| RGGAGGAAGLI | ASSYNNERLF | 0.263652 |
| RSYRMHPAV | ASSYNNERLF | 0.14250019 |
| SAAHLLAGQLA | ASSYNNERLF | 0.22118542 |
| SAAVGSY | ASSYNNERLF | 0.32647914 |
| SPHAVLHRI | ASSYNNERLF | 0.14807452 |
| SQIAKVL | ASSYNNERLF | 0.45106903 |
| TALGHLVP | ASSYNNERLF | 0.15806049 |
| TAPTTSGAA | ASSYNNERLF | 0.33954608 |
| TDGDDGNGGSPGT | ASSYNNERLF | 0.3287341 |
| TDGLGEN | ASSYNNERLF | 0.823306 |
| TGGPPLPGQA | ASSYNNERLF | 0.32159448 |
| TIPAITI | ASSYNNERLF | 0.4725731 |
| TIPGFTI | ASSYNNERLF | 0.13836603 |
| TSIASGALA | ASSYNNERLF | 0.32887787 |
| TVTLPGAAI | ASSYNNERLF | 0.48074338 |
| VASGLSAALP | ASSYNNERLF | 0.45866215 |
| VGAGVIGI | ASSYNNERLF | 0.8222932 |
| VGPKRFGV | ASSYNNERLF | 0.28523386 |
| VNVAFPD | ASSYNNERLF | 0.12246637 |
| VSMTNSGVS | ASSYNNERLF | 0.19334392 |
| VTDVLGDLSV | ASSYNNERLF | 0.47696924 |
| VVVVGAGFAGL | ASSYNNERLF | 0.48753673 |
| VYTVGTVV | ASSYNNERLF | 0.42656687 |
| WTTFSGP | ASSYNNERLF | 0.3293536 |
| YDELVEAF | ASSYNNERLF | 0.19223839 |
| YGTGGAGGNGG | ASSYNNERLF | 0.5438967 |
| YTNVMAV | ASSYNNERLF | 0.7695353 |
| FLEQVHQL | ASSYNNERLF | 0.3086655 |
| GLSLLAVAV | ASSYNNERLF | 0.1561929 |
| LEQVHQL | ASSYNNERLF | 0.15217265 |
| LNAAGAI | ASSYNNERLF | 0.3357528 |
| PAVHASL | ASSYNNERLF | 0.46567163 |
| TELEDDA | ASSYNNERLF | 0.13776164 |
| TPPGPPN | ASSYNNERLF | 0.28365946 |
| TPPPPPPP | ASSYNNERLF | 0.22546662 |
| TQSPALI | ASSQEWGGSRDTQY | 0.113533735 |

| | | |
|---|---|---|
| EFAAAFPQ | ASSQEWGGSRDTQY | 0.41040808 |
| LLSGNLDVL | ASSQEWGGSRDTQY | 0.6224301 |
| GPANLGNYNI | ASSQEWGGSRDTQY | 0.33829027 |
| TLGPTLP | ASSQEWGGSRDTQY | 0.13249946 |
| VYAYSSCCK | ASSQEWGGSRDTQY | 0.5044388 |
| APAPAPAPNI | ASSQEWGGSRDTQY | 0.65428305 |
| PGGPVPIP | ASSQEWGGSRDTQY | 0.46517307 |
| FLTVALEP | ASSQEWGGSRDTQY | 0.14139463 |
| SQYFGANL | ASSQEWGGSRDTQY | 0.10683746 |
| GGVKPAA | ASSQEWGGSRDTQY | 0.26256025 |
| GGTGGAGGAGGW | ASSQEWGGSRDTQY | 0.6057893 |
| GVGGVAGLGGAGG | ASSQEWGGSRDTQY | 0.17864563 |
| VLLVLATTI | ASSQEWGGSRDTQY | 0.1407976 |
| IGIFAAML | ASSQEWGGSRDTQY | 0.21460499 |
| AVGDLID | ASSQEWGGSRDTQY | 0.5078056 |
| NISPLPL | ASSQEWGGSRDTQY | 0.13936128 |
| VTAVFVAI | ASSQEWGGSRDTQY | 0.1327224 |
| SGDDNDAAL | ASSQEWGGSRDTQY | 0.2661411 |
| AVAALAHS | ASSQEWGGSRDTQY | 0.34818432 |
| GGVTVMATP | ASSQEWGGSRDTQY | 0.15321276 |
| AGLAANVRD | ASSQEWGGSRDTQY | 0.15564683 |
| TNRPDLIDPALLRPG | ASSQEWGGSRDTQY | 0.2837149 |
| LANLGVTAL | ASSQEWGGSRDTQY | 0.26931366 |
| TGERIEL | ASSQEWGGSRDTQY | 0.26632187 |
| NGEELAA | ASSQEWGGSRDTQY | 0.13982473 |
| VLSLVWG | ASSQEWGGSRDTQY | 0.4422965 |
| IGYALWL | ASSQEWGGSRDTQY | 0.15050843 |
| LSLLLAPL | ASSQEWGGSRDTQY | 0.33746076 |
| TLDEALK | ASSQEWGGSRDTQY | 0.21716934 |
| TDGAGTAEGG | ASSQEWGGSRDTQY | 0.26632053 |
| NAPGGAGGNGG | ASSQEWGGSRDTQY | 0.25057095 |
| VSVFMGMGGNF | ASSQEWGGSRDTQY | 0.15676054 |
| LASYLAAL | ASSQEWGGSRDTQY | 0.1575533 |
| LVGGTPAL | ASSQEWGGSRDTQY | 0.121495515 |
| VRNADVL | ASSQEWGGSRDTQY | 0.43717742 |
| TCSSPFLVSAM | ASSQEWGGSRDTQY | 0.21079703 |
| VHITDAE | ASSQEWGGSRDTQY | 0.39231023 |
| EYAAPSF | ASSQEWGGSRDTQY | 0.33448863 |
| VVDAFHGGL | ASSQEWGGSRDTQY | 0.14836091 |
| ASVVDPQ | ASSQEWGGSRDTQY | 0.26438677 |
| SYNEFGGLS | ASSQEWGGSRDTQY | 0.12175816 |
| DPDGSVY | ASSQEWGGSRDTQY | 0.37038016 |
| MVSTIHATE | ASSQEWGGSRDTQY | 0.20751385 |
| LAAGVADPV | ASSQEWGGSRDTQY | 0.30415228 |
| TKEQIAA | ASSQEWGGSRDTQY | 0.19017102 |
| SLASPAAL | ASSQEWGGSRDTQY | 0.25377277 |
| IGLVFYGV | ASSQEWGGSRDTQY | 0.35730135 |
| FLLDASNL | ASSQEWGGSRDTQY | 0.14240645 |
| YQVGNFP | ASSQEWGGSRDTQY | 0.34518456 |
| ESLLSQST | ASSQEWGGSRDTQY | 0.49999967 |
| LFTVGFAAVG | ASSQEWGGSRDTQY | 0.3819814 |

| | | |
|---|---|---|
| ATPPTPT | ASSQEWGGSRDTQY | 0.123822175 |
| TDLATLG | ASSQEWGGSRDTQY | 0.28767684 |
| SAGGTGR | ASSQEWGGSRDTQY | 0.15872942 |
| GFSNSGS | ASSQEWGGSRDTQY | 0.33080882 |
| GGNGGSGGTSV | ASSQEWGGSRDTQY | 0.23325756 |
| GTSVATGGAGN | ASSQEWGGSRDTQY | 0.18836883 |
| NGGNGGSGGTS | ASSQEWGGSRDTQY | 0.26932245 |
| AVVKFPH | ASSQEWGGSRDTQY | 0.16743265 |
| TPPPPPP | ASSQEWGGSRDTQY | 0.15091294 |
| AAPAAAAAPA | ASSQEWGGSRDTQY | 0.21535432 |
| IALNEDL | ASSQEWGGSRDTQY | 0.26611528 |
| IVNVSSM | ASSQEWGGSRDTQY | 0.32335746 |
| TGIPPAPR | ASSQEWGGSRDTQY | 0.1550133 |
| FDLGGGTF | ASSQEWGGSRDTQY | 0.2713926 |
| FDLGGGTFDVS | ASSQEWGGSRDTQY | 0.17322294 |
| NEPTAAA | ASSQEWGGSRDTQY | 0.17270081 |
| NEPTAAALA | ASSQEWGGSRDTQY | 0.2450152 |
| GSIINISSI | ASSQEWGGSRDTQY | 0.21129203 |
| VAPGNAGTA | ASSQEWGGSRDTQY | 0.12464438 |
| TGGGASLE | ASSQEWGGSRDTQY | 0.3381502 |
| VAVYNHYKRI | ASSQEWGGSRDTQY | 0.15314735 |
| TDPAPATT | ASSQEWGGSRDTQY | 0.20593959 |
| TNRPDILDPALLRPG | ASSQEWGGSRDTQY | 0.58606005 |
| LTTTTTS | ASSQEWGGSRDTQY | 0.12575126 |
| PTGSGKTL | ASSQEWGGSRDTQY | 0.26312757 |
| TGSGKTL | ASSQEWGGSRDTQY | 0.21832442 |
| TIGHVDHGK | ASSQEWGGSRDTQY | 0.29590544 |
| TIGHVDHGKTT | ASSQEWGGSRDTQY | 0.20309639 |
| GHVDHGKTT | ASSQEWGGSRDTQY | 0.17212918 |
| AQFEHTLL | ASSQEWGGSRDTQY | 0.45680875 |
| IVNQDEL | ASSQEWGGSRDTQY | 0.3055897 |
| TNGSQFF | ASSQEWGGSRDTQY | 0.31277937 |
| TGKGIVE | ASSQEWGGSRDTQY | 0.28071612 |
| LISNASDA | ASSQEWGGSRDTQY | 0.6126399 |
| AELRTLGELTKIAKAHGAQ | ASSQEWGGSRDTQY | 0.35907963 |
| APNAVAAAEPSW | ASSQEWGGSRDTQY | 0.17584786 |
| APQVAGAATAVL | ASSQEWGGSRDTQY | 0.12018401 |
| ARRDPAAAAAILPTDA | ASSQEWGGSRDTQY | 0.21433727 |
| DAAARDVLAEAGLAEYF | ASSQEWGGSRDTQY | 0.22536784 |
| DDQIVLSAKANLAKVEQ | ASSQEWGGSRDTQY | 0.1795691 |
| DELLPDTGEDSDEDGH | ASSQEWGGSRDTQY | 0.1296881 |
| ELPEAPANRATVPQD | ASSQEWGGSRDTQY | 0.58251643 |
| FGQSKIERVTGIG | ASSQEWGGSRDTQY | 0.73690367 |
| GAGGTGSTAGGAGGAGGAGGLY | ASSQEWGGSRDTQY | 0.1558626 |
| GHGGTGAAGGNGGTGGAGGSGI | ASSQEWGGSRDTQY | 0.7409717 |
| GNGGNGGAGGEGGPGGAGGAGA | ASSQEWGGSRDTQY | 0.14179493 |
| GSTAGNGGNGGNATNGTIAGQP | ASSQEWGGSRDTQY | 0.29847068 |
| GTGGDGGNGGNGGTGG | ASSQEWGGSRDTQY | 0.60537285 |
| GVRRPAGIPADAPV | ASSQEWGGSRDTQY | 0.14345199 |
| GYRQVTDAHLLTLARR | ASSQEWGGSRDTQY | 0.29561567 |
| IAKMGQDGHDRG | ASSQEWGGSRDTQY | 0.17532931 |

| | | |
|---|---|---|
| KAEPSIEIQYT | ASSQEWGGSRDTQY | 0.15830822 |
| KPVRIRMAADEDTVAVT | ASSQEWGGSRDTQY | 0.527532 |
| MPPRIAGMRLLVIKPEP | ASSQEWGGSRDTQY | 0.17865984 |
| MKFYAVALFRSRGVPPTR | ASSQEWGGSRDTQY | 0.27130964 |
| NAGGFGILGAVAHSP | ASSQEWGGSRDTQY | 0.19297545 |
| NEGREITLAMAEM | ASSQEWGGSRDTQY | 0.14878681 |
| NELVEWSHPLGHRWRWE | ASSQEWGGSRDTQY | 0.29958165 |
| NLGAGGTGGNGGNPG | ASSQEWGGSRDTQY | 0.43296653 |
| NTEWPKWETVMAADPVD | ASSQEWGGSRDTQY | 0.2380042 |
| PASAAPAETQP | ASSQEWGGSRDTQY | 0.6895514 |
| QQSSDGKNGVRAAEITGE | ASSQEWGGSRDTQY | 0.17251262 |
| RAPDAVTTSWWKE | ASSQEWGGSRDTQY | 0.77333164 |
| SGFGNVGAHTSGWFNQ | ASSQEWGGSRDTQY | 0.71773446 |
| SGVAGAVGQAASVGGLKV | ASSQEWGGSRDTQY | 0.24761882 |
| SNGGAGTGSAGGNGGT | ASSQEWGGSRDTQY | 0.17017335 |
| TGGAGGEGGAAG | ASSQEWGGSRDTQY | 0.5966912 |
| TGGDGGAAGAGGAPGHG | ASSQEWGGSRDTQY | 0.17812325 |
| TMPPLLAAIANAGRHGV | ASSQEWGGSRDTQY | 0.22191063 |
| TSSTAGNSGGA | ASSQEWGGSRDTQY | 0.3806321 |
| VPATVPVGILC | ASSQEWGGSRDTQY | 0.63602686 |
| VQRPDSGGVVLTGRIS | ASSQEWGGSRDTQY | 0.7550899 |
| WTPNPVPATKGVPAGLVQAT | ASSQEWGGSRDTQY | 0.73030776 |
| YALKRALAKFSDRFDVVII | ASSQEWGGSRDTQY | 0.669463 |
| YLIGSVIV | ASSQEWGGSRDTQY | 0.5614331 |
| YGQLVAVID | ASSQEWGGSRDTQY | 0.36536396 |
| VTVGGMTLPA | ASSQEWGGSRDTQY | 0.6214069 |
| VLALPAV | ASSQEWGGSRDTQY | 0.47458315 |
| VIPPTLNYET | ASSQEWGGSRDTQY | 0.7374969 |
| VGMGGMHPSSG | ASSQEWGGSRDTQY | 0.80035233 |
| VGLAGGAATA | ASSQEWGGSRDTQY | 0.5330715 |
| VGGAGGAGGSGP | ASSQEWGGSRDTQY | 0.80314976 |
| VFAAFMV | ASSQEWGGSRDTQY | 0.7540887 |
| TLYDKLR | ASSQEWGGSRDTQY | 0.6033262 |
| TLAVFTVL | ASSQEWGGSRDTQY | 0.7142316 |
| TGSVQPARV | ASSQEWGGSRDTQY | 0.5396999 |
| TGGGAGSGGT | ASSQEWGGSRDTQY | 0.71239924 |
| SHLETSL | ASSQEWGGSRDTQY | 0.39060783 |
| SFTTGMT | ASSQEWGGSRDTQY | 0.7676765 |
| QVEVAKRMGVRQPTVS | ASSQEWGGSRDTQY | 0.66829467 |
| QGSGGPALAM | ASSQEWGGSRDTQY | 0.27251807 |
| MLAPSNP | ASSQEWGGSRDTQY | 0.7975915 |
| LVNNAAI | ASSQEWGGSRDTQY | 0.6872943 |
| LTGEINTLA | ASSQEWGGSRDTQY | 0.7508216 |
| LSIFGPV | ASSQEWGGSRDTQY | 0.8028288 |
| LLALLANL | ASSQEWGGSRDTQY | 0.62703544 |
| LISNASDALDKLR | ASSQEWGGSRDTQY | 0.4831993 |
| LINEAVP | ASSQEWGGSRDTQY | 0.7268893 |
| LGYAFSGL | ASSQEWGGSRDTQY | 0.80203307 |
| LDFVAVAL | ASSQEWGGSRDTQY | 0.450912 |
| LAQPGAE | ASSQEWGGSRDTQY | 0.73228955 |
| ISNLGAVTGF | ASSQEWGGSRDTQY | 0.754343 |

| | | |
|---|---|---|
| HQEDPGRI | ASSQEWGGSRDTQY | 0.70522684 |
| GTPGEPTP | ASSQEWGGSRDTQY | 0.80953157 |
| GFFNSGS | ASSQEWGGSRDTQY | 0.8149706 |
| GAGGGGLTAGAN | ASSQEWGGSRDTQY | 0.5957273 |
| FVDALVNM | ASSQEWGGSRDTQY | 0.78836286 |
| FLTVAPEL | ASSQEWGGSRDTQY | 0.8010225 |
| FETLTAL | ASSQEWGGSRDTQY | 0.60698956 |
| DVINPIEL | ASSQEWGGSRDTQY | 0.646309 |
| APVSGAPV | ASSQEWGGSRDTQY | 0.5672478 |
| ALVGGLAW | ASSQEWGGSRDTQY | 0.7413392 |
| ALAAAAATAI | ASSQEWGGSRDTQY | 0.3463837 |
| AFGLGLLL | ASSQEWGGSRDTQY | 0.7385733 |
| AFFAAASL | ASSQEWGGSRDTQY | 0.7894645 |
| AAAALPAFSPP | ASSQEWGGSRDTQY | 0.70926833 |
| VVLAPAL | ASSQEWGGSRDTQY | 0.56877875 |
| TIGHVDHGKTTL | ASSQEWGGSRDTQY | 0.37628454 |
| LISNASDALDK | ASSQEWGGSRDTQY | 0.55689377 |
| LISNASDAL | ASSQEWGGSRDTQY | 0.6306738 |
| ISNASDALDK | ASSQEWGGSRDTQY | 0.2984913 |
| NQAETLVYQTE | ASSQEWGGSRDTQY | 0.7271911 |
| TGGTGGTAGAG | ASSQEWGGSRDTQY | 0.7923559 |
| TGSTAGGAGGA | ASSQEWGGSRDTQY | 0.7624504 |
| DPLAVPVAAGPV | ASSQEWGGSRDTQY | 0.4130848 |
| GGTGGTPGNGGN | ASSQEWGGSRDTQY | 0.80708337 |
| PAGLAGVDPATL | ASSQEWGGSRDTQY | 0.73683053 |
| GNNLVVEAPHAGASV | ASSQEWGGSRDTQY | 0.6702305 |
| KVTRIMSASTLEQPA | ASSQEWGGSRDTQY | 0.56057 |
| LGFALPIALPRRTVV | ASSQEWGGSRDTQY | 0.5790674 |
| TRLGNLTARKLVVLV | ASSQEWGGSRDTQY | 0.7546511 |
| WSQQRRLIYFPSAGP | ASSQEWGGSRDTQY | 0.63753825 |
| GPNGNPDPMAAAVDIR | ASSQEWGGSRDTQY | 0.67789954 |
| TQRCVSVADLRAAASR | ASSQEWGGSRDTQY | 0.81507385 |
| QFVLDVAEHPVEVLTETV | ASSQEWGGSRDTQY | 0.64648056 |
| GGAGGTGGAGGNGGTQAGDG | ASSQEWGGSRDTQY | 0.5130868 |
| PLLATLRAIAVDVDAKPAQV | ASSQEWGGSRDTQY | 0.6070202 |
| APKFRTGGYMIDFWGVGYQVAK | ASSQEWGGSRDTQY | 0.7922143 |
| GAGGAGGAGGAAGGPGGTGGTG | ASSQEWGGSRDTQY | 0.40836686 |
| IGALGLLATDVAPAGSVAFGVA | ASSQEWGGSRDTQY | 0.5718933 |
| AGGAGGIGGNGGAGGAGGVLMG | ASSQEWGGSRDTQY | 0.51601005 |
| ADELIGLL | ASSQEWGGSRDTQY | 0.61423695 |
| ADRALKA | ASSQEWGGSRDTQY | 0.79618406 |
| AGAVGQVGGAGGA | ASSQEWGGSRDTQY | 0.76711446 |
| AGGGGGLGGAP | ASSQEWGGSRDTQY | 0.5475521 |
| AGGLLYGNGGAG | ASSQEWGGSRDTQY | 0.7847583 |
| AGLESIGL | ASSQEWGGSRDTQY | 0.66181374 |
| AGLKTGII | ASSQEWGGSRDTQY | 0.6034869 |
| ASLAALLL | ASSQEWGGSRDTQY | 0.77955616 |
| ASVINISSI | ASSQEWGGSRDTQY | 0.55498874 |
| ATPPGPA | ASSQEWGGSRDTQY | 0.7856698 |
| ATVASALV | ASSQEWGGSRDTQY | 0.72234446 |
| ATVASLIL | ASSQEWGGSRDTQY | 0.785596 |

| | | |
|---|---|---|
| AVNGPVLIRH | ASSQEWGGSRDTQY | 0.7961982 |
| DGAAGAVGI | ASSQEWGGSRDTQY | 0.6402228 |
| DPLLHGI | ASSQEWGGSRDTQY | 0.61118686 |
| ELGAALGAL | ASSQEWGGSRDTQY | 0.32125232 |
| ELGRTQAV | ASSQEWGGSRDTQY | 0.5915441 |
| FVAGGYLS | ASSQEWGGSRDTQY | 0.77598834 |
| GASGGAGGAGGW | ASSQEWGGSRDTQY | 0.75192606 |
| GGAGAGGDGGAGGI | ASSQEWGGSRDTQY | 0.8024702 |
| GTGHNGSQPGMGGNG | ASSQEWGGSRDTQY | 0.74782324 |
| GYAAASASAS | ASSQEWGGSRDTQY | 0.5745882 |
| HQVDVAAVQQG | ASSQEWGGSRDTQY | 0.79349136 |
| IGAAIAHT | ASSQEWGGSRDTQY | 0.7593048 |
| IGMIEVAQL | ASSQEWGGSRDTQY | 0.3222907 |
| IRLAPEHTSAMD | ASSQEWGGSRDTQY | 0.66167074 |
| IYNVSVL | ASSQEWGGSRDTQY | 0.829314 |
| KQGFLGM | ASSQEWGGSRDTQY | 0.7084619 |
| LESVITQL | ASSQEWGGSRDTQY | 0.66269416 |
| LGLGALVVV | ASSQEWGGSRDTQY | 0.6051968 |
| LIGNGGNGGSGGI | ASSQEWGGSRDTQY | 0.7885714 |
| LNAINAPTL | ASSQEWGGSRDTQY | 0.7004901 |
| LQLLNTGV | ASSQEWGGSRDTQY | 0.8135967 |
| LQTSIEF | ASSQEWGGSRDTQY | 0.26720127 |
| MSFAVSEL | ASSQEWGGSRDTQY | 0.59953356 |
| NTGALGL | ASSQEWGGSRDTQY | 0.3357879 |
| PPGAGGAGGAGTT | ASSQEWGGSRDTQY | 0.38299286 |
| QLVGLGI | ASSQEWGGSRDTQY | 0.76160944 |
| RFLEQVHQL | ASSQEWGGSRDTQY | 0.63620317 |
| RGVGYVLR | ASSQEWGGSRDTQY | 0.78210413 |
| SATGVFAGVI | ASSQEWGGSRDTQY | 0.7084953 |
| SLPAITLP | ASSQEWGGSRDTQY | 0.7817894 |
| STALAFTV | ASSQEWGGSRDTQY | 0.58165956 |
| SVLTVGLGK | ASSQEWGGSRDTQY | 0.7644029 |
| TALAGAGSGI | ASSQEWGGSRDTQY | 0.73117435 |
| TGLPFTGI | ASSQEWGGSRDTQY | 0.74626005 |
| VAEHVTV | ASSQEWGGSRDTQY | 0.8138021 |
| VDIVLNSL | ASSQEWGGSRDTQY | 0.7772759 |
| VLTGGGAVL | ASSQEWGGSRDTQY | 0.3653059 |
| VNVGLAYGI | ASSQEWGGSRDTQY | 0.6896603 |
| VSSAAVGAV | ASSQEWGGSRDTQY | 0.6356044 |
| VTSVYEAGV | ASSQEWGGSRDTQY | 0.3352895 |
| WVALDRGA | ASSQEWGGSRDTQY | 0.40698218 |
| YLVGSVVI | ASSQEWGGSRDTQY | 0.59429 |
| GPPGTGKTLL | ASSQEWGGSRDTQY | 0.60206044 |
| GVGMTKF | ASSQEWGGSRDTQY | 0.7972163 |
| IGIGGSDL | ASSQEWGGSRDTQY | 0.74050283 |
| IGTLLAEL | ASSQEWGGSRDTQY | 0.60951906 |
| IYTSGTT | ASSQEWGGSRDTQY | 0.6909109 |
| LSFPAGV | ASSQEWGGSRDTQY | 0.68134844 |
| LYTSGTT | ASSQEWGGSRDTQY | 0.18639454 |
| VGAFPFGF | ASSQEWGGSRDTQY | 0.45331234 |
| VGAGTVEF | ASSQEWGGSRDTQY | 0.63525563 |

| | | |
|---|---|---|
| VGKSTLLL | ASSQEWGGSRDTQY | 0.80577016 |
| AALAHAGGGIV | ASSQEWGGSRDTQY | 0.71243286 |
| AALVAVSLL | ASSQEWGGSRDTQY | 0.58266896 |
| AAVSRAVAAL | ASSQEWGGSRDTQY | 0.64264035 |
| AGGSSGIGGF | ASSQEWGGSRDTQY | 0.66653514 |
| AGLTVANANG | ASSQEWGGSRDTQY | 0.6377394 |
| AGLVGAVL | ASSQEWGGSRDTQY | 0.36733893 |
| ALSQAFTV | ASSQEWGGSRDTQY | 0.26592007 |
| ALTFSVSL | ASSQEWGGSRDTQY | 0.74694777 |
| ALTGAALVAL | ASSQEWGGSRDTQY | 0.4830848 |
| ASFSIGAGVL | ASSQEWGGSRDTQY | 0.6314988 |
| DGSVYKI | ASSQEWGGSRDTQY | 0.4016587 |
| EQLAEIV | ASSQEWGGSRDTQY | 0.698917 |
| FAVSELTVM | ASSQEWGGSRDTQY | 0.2861129 |
| GAAPLAAGL | ASSQEWGGSRDTQY | 0.79960334 |
| GADGGDGGSGGAGG | ASSQEWGGSRDTQY | 0.67123795 |
| GFGLDTGGAGG | ASSQEWGGSRDTQY | 0.6209807 |
| GLASGGLL | ASSQEWGGSRDTQY | 0.78970623 |
| GRIAGAAVLL | ASSQEWGGSRDTQY | 0.6547334 |
| GSAGSLNTSF | ASSQEWGGSRDTQY | 0.65039134 |
| GTGQIGI | ASSQEWGGSRDTQY | 0.56189 |
| GTLIALIL | ASSQEWGGSRDTQY | 0.6310664 |
| IAATMGFVVG | ASSQEWGGSRDTQY | 0.49639976 |
| IFAQALL | ASSQEWGGSRDTQY | 0.5328934 |
| IGAAVALGF | ASSQEWGGSRDTQY | 0.81773376 |
| IGLTPLAL | ASSQEWGGSRDTQY | 0.66289574 |
| ILGSNTSTL | ASSQEWGGSRDTQY | 0.40503955 |
| IVFSLTVL | ASSQEWGGSRDTQY | 0.6077837 |
| LASKATGF | ASSQEWGGSRDTQY | 0.7757298 |
| LASVALIL | ASSQEWGGSRDTQY | 0.789454 |
| LFAVGLTL | ASSQEWGGSRDTQY | 0.72402894 |
| LFNGGIGTYI | ASSQEWGGSRDTQY | 0.4357056 |
| LTLGAGII | ASSQEWGGSRDTQY | 0.5142733 |
| LVAAALGDI | ASSQEWGGSRDTQY | 0.7013962 |
| NTAFVQLGI | ASSQEWGGSRDTQY | 0.582749 |
| QLFSGAEL | ASSQEWGGSRDTQY | 0.43144372 |
| SAATFITL | ASSQEWGGSRDTQY | 0.7734885 |
| SNYGLANLGV | ASSQEWGGSRDTQY | 0.80396175 |
| SYGGAYAVM | ASSQEWGGSRDTQY | 0.54260045 |
| TTGVGGLTLG | ASSQEWGGSRDTQY | 0.44736904 |
| TTQTIYI | ASSQEWGGSRDTQY | 0.6775942 |
| VAAGAGLGAVI | ASSQEWGGSRDTQY | 0.43672204 |
| VAAVVTLL | ASSQEWGGSRDTQY | 0.5511105 |
| VGFLGGLI | ASSQEWGGSRDTQY | 0.64473146 |
| VGVIPILL | ASSQEWGGSRDTQY | 0.35672486 |
| VIFASSLI | ASSQEWGGSRDTQY | 0.5969197 |
| VKGGDGGAAAG | ASSQEWGGSRDTQY | 0.77621764 |
| VTKTFTNI | ASSQEWGGSRDTQY | 0.6149664 |
| VVAAAILL | ASSQEWGGSRDTQY | 0.7330973 |
| YAALDVEL | ASSQEWGGSRDTQY | 0.78760076 |
| LASVALLL | ASSQEWGGSRDTQY | 0.7930087 |

| | | |
|---|---|---|
| LDAVHRV | ASSQEWGGSRDTQY | 0.8112409 |
| QGLSSSIF | ASSQEWGGSRDTQY | 0.75995326 |
| RGTFANIRL | ASSQEWGGSRDTQY | 0.77756214 |
| RQLWWGHRI | ASSQEWGGSRDTQY | 0.5561453 |
| YGPPGTGKTLL | ASSQEWGGSRDTQY | 0.75828993 |
| AALVGAPANVA | ASSQEWGGSRDTQY | 0.69715947 |
| AATFRRTE | ASSQEWGGSRDTQY | 0.69408953 |
| AGALGGGI | ASSQEWGGSRDTQY | 0.5432769 |
| AGGAGGAGGIGAG | ASSQEWGGSRDTQY | 0.7622669 |
| AGSGALGL | ASSQEWGGSRDTQY | 0.5254327 |
| AQIADQI | ASSQEWGGSRDTQY | 0.7467872 |
| ASAVAVTYL | ASSQEWGGSRDTQY | 0.42109728 |
| ASTVLGGTGG | ASSQEWGGSRDTQY | 0.57558656 |
| AVGGEPAK | ASSQEWGGSRDTQY | 0.7757535 |
| CEQGTFGGTC | ASSQEWGGSRDTQY | 0.17569807 |
| DEGKAPAG | ASSQEWGGSRDTQY | 0.40351677 |
| DTVVFPAL | ASSQEWGGSRDTQY | 0.72763294 |
| FEGITMAG | ASSQEWGGSRDTQY | 0.73174375 |
| FPLVVQD | ASSQEWGGSRDTQY | 0.24047562 |
| GAFTTPQ | ASSQEWGGSRDTQY | 0.49396893 |
| GFGGFGTAGGVGGA | ASSQEWGGSRDTQY | 0.49745166 |
| GFLAQFL | ASSQEWGGSRDTQY | 0.6924057 |
| GGAGGAGGNAGVA | ASSQEWGGSRDTQY | 0.66364825 |
| GGVGTAGAGGAGGAGG | ASSQEWGGSRDTQY | 0.30344787 |
| GNGGGGDAGNAGSGGN | ASSQEWGGSRDTQY | 0.8005233 |
| GTGGAGGAGNDAGS | ASSQEWGGSRDTQY | 0.26661634 |
| GTTNTGWFNS | ASSQEWGGSRDTQY | 0.59574646 |
| IFMGPLFTL | ASSQEWGGSRDTQY | 0.7751739 |
| ILAHAIEAI | ASSQEWGGSRDTQY | 0.80069506 |
| IRSAIREAL | ASSQEWGGSRDTQY | 0.68687433 |
| ITFAGGG | ASSQEWGGSRDTQY | 0.7498777 |
| ITGMSGHQEV | ASSQEWGGSRDTQY | 0.5266554 |
| IVAGLACTL | ASSQEWGGSRDTQY | 0.8081031 |
| IVGGGLAGA | ASSQEWGGSRDTQY | 0.6454011 |
| IVTGVLNI | ASSQEWGGSRDTQY | 0.46749854 |
| LAQQGCI | ASSQEWGGSRDTQY | 0.7788045 |
| LCLHVEN | ASSQEWGGSRDTQY | 0.7434503 |
| LDAISDGS | ASSQEWGGSRDTQY | 0.761726 |
| LGGFAAML | ASSQEWGGSRDTQY | 0.75518763 |
| LGGKNPMI | ASSQEWGGSRDTQY | 0.48666093 |
| LGPKGVAAIL | ASSQEWGGSRDTQY | 0.58889604 |
| LMVGVSI | ASSQEWGGSRDTQY | 0.7872325 |
| LVGAAFVL | ASSQEWGGSRDTQY | 0.79376173 |
| LVLAVAPL | ASSQEWGGSRDTQY | 0.45840812 |
| MAAIAVIDAA | ASSQEWGGSRDTQY | 0.79645646 |
| NNGINAELQ | ASSQEWGGSRDTQY | 0.67157847 |
| RGEPLVLVHG | ASSQEWGGSRDTQY | 0.603459 |
| RGGAGGAAGLI | ASSQEWGGSRDTQY | 0.772854 |
| RSYRMHPAV | ASSQEWGGSRDTQY | 0.70724213 |
| SAAHLLAGQLA | ASSQEWGGSRDTQY | 0.73271465 |
| SAAVGSY | ASSQEWGGSRDTQY | 0.69592017 |

| | | |
|---|---|---|
| SPHAVLHRI | ASSQEWGGSRDTQY | 0.7382458 |
| SQIAKVL | ASSQEWGGSRDTQY | 0.5020567 |
| TALGHLVP | ASSQEWGGSRDTQY | 0.66983396 |
| TAPTTSGAA | ASSQEWGGSRDTQY | 0.65466547 |
| TDGDDGNGGSPGT | ASSQEWGGSRDTQY | 0.79616153 |
| TDGLGEN | ASSQEWGGSRDTQY | 0.6241395 |
| TGGPPLPGQA | ASSQEWGGSRDTQY | 0.6725708 |
| TIPAITI | ASSQEWGGSRDTQY | 0.7669686 |
| TIPGFTI | ASSQEWGGSRDTQY | 0.7742214 |
| TSIASGALA | ASSQEWGGSRDTQY | 0.74477136 |
| TVTLPGAAI | ASSQEWGGSRDTQY | 0.7456411 |
| VASGLSAALP | ASSQEWGGSRDTQY | 0.81920326 |
| VGAGVIGI | ASSQEWGGSRDTQY | 0.6740537 |
| VGPKRFGV | ASSQEWGGSRDTQY | 0.6310665 |
| VNVAFPD | ASSQEWGGSRDTQY | 0.78453964 |
| VSMTNSGVS | ASSQEWGGSRDTQY | 0.81352246 |
| VTDVLGDLSV | ASSQEWGGSRDTQY | 0.6753475 |
| VVVVGAGFAGL | ASSQEWGGSRDTQY | 0.7983942 |
| VYTVGTVV | ASSQEWGGSRDTQY | 0.5626235 |
| WTTFSGP | ASSQEWGGSRDTQY | 0.78352135 |
| YDELVEAF | ASSQEWGGSRDTQY | 0.49753374 |
| YGTGGAGGNGG | ASSQEWGGSRDTQY | 0.6916044 |
| YTNVMAV | ASSQEWGGSRDTQY | 0.6597055 |
| FLEQVHQL | ASSQEWGGSRDTQY | 0.7794645 |
| GLSLLAVAV | ASSQEWGGSRDTQY | 0.73025966 |
| LEQVHQL | ASSQEWGGSRDTQY | 0.781108 |
| LNAAGAI | ASSQEWGGSRDTQY | 0.29738697 |
| PAVHASL | ASSQEWGGSRDTQY | 0.7375733 |
| TELEDDA | ASSQEWGGSRDTQY | 0.77745926 |
| TPPGPPN | ASSQEWGGSRDTQY | 0.81855285 |
| TPPPPPPP | ASSQEWGGSRDTQY | 0.5612537 |
| TQSPALI | ASSLDRDNTGQLY | 0.6978231 |
| EFAAAFPQ | ASSLDRDNTGQLY | 0.60744476 |
| LLSGNLDVL | ASSLDRDNTGQLY | 0.7278372 |
| GPANLGNYNI | ASSLDRDNTGQLY | 0.46340975 |
| TLGPTLP | ASSLDRDNTGQLY | 0.6348495 |
| VYAYSSCCK | ASSLDRDNTGQLY | 0.6107626 |
| APAPAPAPNI | ASSLDRDNTGQLY | 0.6721771 |
| PGGPVPIP | ASSLDRDNTGQLY | 0.74893385 |
| FLTVALEP | ASSLDRDNTGQLY | 0.81590706 |
| SQYFGANL | ASSLDRDNTGQLY | 0.74945813 |
| GGVKPAA | ASSLDRDNTGQLY | 0.75764114 |
| GGTGGAGGAGGW | ASSLDRDNTGQLY | 0.7582089 |
| GVGGVAGLGGAGG | ASSLDRDNTGQLY | 0.807518 |
| VLLVLATTI | ASSLDRDNTGQLY | 0.5616492 |
| IGIFAAML | ASSLDRDNTGQLY | 0.7076882 |
| AVGDLID | ASSLDRDNTGQLY | 0.5643807 |
| NISPLPL | ASSLDRDNTGQLY | 0.5879315 |
| VTAVFVAI | ASSLDRDNTGQLY | 0.5510529 |
| SGDDNDAAL | ASSLDRDNTGQLY | 0.6959325 |
| AVAALAHS | ASSLDRDNTGQLY | 0.19766454 |

| | | |
|---|---|---|
| GGVTVMATP | ASSLDRDNTGQLY | 0.64893734 |
| AGLAANVRD | ASSLDRDNTGQLY | 0.8193165 |
| TNRPDLIDPALLRPG | ASSLDRDNTGQLY | 0.58651537 |
| LANLGVTAL | ASSLDRDNTGQLY | 0.6866112 |
| TGERIEL | ASSLDRDNTGQLY | 0.6904718 |
| NGEELAA | ASSLDRDNTGQLY | 0.62624085 |
| VLSLVWG | ASSLDRDNTGQLY | 0.73005426 |
| IGYALWL | ASSLDRDNTGQLY | 0.492029 |
| LSLLLAPL | ASSLDRDNTGQLY | 0.78671885 |
| TLDEALK | ASSLDRDNTGQLY | 0.7732817 |
| TDGAGTAEGG | ASSLDRDNTGQLY | 0.6687585 |
| NAPGGAGGNGG | ASSLDRDNTGQLY | 0.6876756 |
| VSVFMGMGGNF | ASSLDRDNTGQLY | 0.6346379 |
| LASYLAAL | ASSLDRDNTGQLY | 0.43384635 |
| LVGGTPAL | ASSLDRDNTGQLY | 0.7453022 |
| VRNADVL | ASSLDRDNTGQLY | 0.6749172 |
| TCSSPFLVSAM | ASSLDRDNTGQLY | 0.6365817 |
| VHITDAE | ASSLDRDNTGQLY | 0.4776634 |
| EYAAPSF | ASSLDRDNTGQLY | 0.80155903 |
| VVDAFHGGL | ASSLDRDNTGQLY | 0.67222774 |
| ASVVDPQ | ASSLDRDNTGQLY | 0.74271894 |
| SYNEFGGLS | ASSLDRDNTGQLY | 0.5154445 |
| DPDGSVY | ASSLDRDNTGQLY | 0.71024686 |
| MVSTIHATE | ASSLDRDNTGQLY | 0.63974905 |
| LAAGVADPV | ASSLDRDNTGQLY | 0.37267792 |
| TKEQIAA | ASSLDRDNTGQLY | 0.7484926 |
| SLASPAAL | ASSLDRDNTGQLY | 0.68377566 |
| IGLVFYGV | ASSLDRDNTGQLY | 0.53638935 |
| FLLDASNL | ASSLDRDNTGQLY | 0.7830645 |
| YQVGNFP | ASSLDRDNTGQLY | 0.7605366 |
| ESLLSQST | ASSLDRDNTGQLY | 0.3468933 |
| LFTVGFAAVG | ASSLDRDNTGQLY | 0.540727 |
| ATPPTPT | ASSLDRDNTGQLY | 0.6736599 |
| TDLATLG | ASSLDRDNTGQLY | 0.6487612 |
| SAGGTGR | ASSLDRDNTGQLY | 0.23360276 |
| GFSNSGS | ASSLDRDNTGQLY | 0.7744998 |
| GGNGGSGGTSV | ASSLDRDNTGQLY | 0.610556 |
| GTSVATGGAGN | ASSLDRDNTGQLY | 0.40715405 |
| NGGNGGSGGTS | ASSLDRDNTGQLY | 0.5882877 |
| AVVKFPH | ASSLDRDNTGQLY | 0.8078644 |
| TPPPPPP | ASSLDRDNTGQLY | 0.6452067 |
| AAPAAAAAPA | ASSLDRDNTGQLY | 0.6852331 |
| IALNEDL | ASSLDRDNTGQLY | 0.7898524 |
| IVNVSSM | ASSLDRDNTGQLY | 0.7405447 |
| TGIPPAPR | ASSLDRDNTGQLY | 0.79576623 |
| FDLGGGTF | ASSLDRDNTGQLY | 0.7894968 |
| FDLGGGTFDVS | ASSLDRDNTGQLY | 0.46222666 |
| NEPTAAA | ASSLDRDNTGQLY | 0.77720326 |
| NEPTAAALA | ASSLDRDNTGQLY | 0.56918705 |
| GSIINISSI | ASSLDRDNTGQLY | 0.68132913 |
| VAPGNAGTA | ASSLDRDNTGQLY | 0.51521367 |

| | | |
|---|---|---|
| TGGGASLE | ASSLDRDNTGQLY | 0.5629897 |
| VAVYNHYKRI | ASSLDRDNTGQLY | 0.5812173 |
| TDPAPATT | ASSLDRDNTGQLY | 0.7067122 |
| TNRPDILDPALLRPG | ASSLDRDNTGQLY | 0.79346377 |
| LTTTTTS | ASSLDRDNTGQLY | 0.6569483 |
| PTGSGKTL | ASSLDRDNTGQLY | 0.44060153 |
| TGSGKTL | ASSLDRDNTGQLY | 0.6881387 |
| TIGHVDHGK | ASSLDRDNTGQLY | 0.5838777 |
| TIGHVDHGKTT | ASSLDRDNTGQLY | 0.7995703 |
| GHVDHGKTT | ASSLDRDNTGQLY | 0.5333649 |
| AQFEHTLL | ASSLDRDNTGQLY | 0.8015696 |
| IVNQDEL | ASSLDRDNTGQLY | 0.7653278 |
| TNGSQFF | ASSLDRDNTGQLY | 0.5561125 |
| TGKGIVE | ASSLDRDNTGQLY | 0.7889471 |
| LISNASDA | ASSLDRDNTGQLY | 0.49742097 |
| AELRTLGELTKIAKAHGAQ | ASSLDRDNTGQLY | 0.60193384 |
| APNAVAAAEPSW | ASSLDRDNTGQLY | 0.77135515 |
| APQVAGAATAVL | ASSLDRDNTGQLY | 0.7005092 |
| ARRDPAAAAAILPTDA | ASSLDRDNTGQLY | 0.71123815 |
| DAAARDVLAEAGLAEYF | ASSLDRDNTGQLY | 0.7672298 |
| DDQIVLSAKANLAKVEQ | ASSLDRDNTGQLY | 0.3217395 |
| DELLPDTGEDSDEDGH | ASSLDRDNTGQLY | 0.76192635 |
| ELPEAPANRATVPQD | ASSLDRDNTGQLY | 0.48406404 |
| FGQSKIERVTGIG | ASSLDRDNTGQLY | 0.3607247 |
| GAGGTGSTAGGAGGAGGAGGLY | ASSLDRDNTGQLY | 0.671121 |
| GHGGTGAAGGNGGTGGAGGSGI | ASSLDRDNTGQLY | 0.8153146 |
| GNGGNGGAGGEGGPGGAGGAGA | ASSLDRDNTGQLY | 0.77855504 |
| GSTAGNGGNGGNATNGTIAGQP | ASSLDRDNTGQLY | 0.48682868 |
| GTGGDGGNGGNGGTGG | ASSLDRDNTGQLY | 0.47260365 |
| GVRRPAGIPADAPV | ASSLDRDNTGQLY | 0.6699556 |
| GYRQVTDAHLLTLARR | ASSLDRDNTGQLY | 0.52179277 |
| IAKMGQDGHDRG | ASSLDRDNTGQLY | 0.7371658 |
| KAEPSIEIQYT | ASSLDRDNTGQLY | 0.63311744 |
| KPVRIRMAADEDTVAVT | ASSLDRDNTGQLY | 0.57975733 |
| MPPRIAGMRLLVIKPEP | ASSLDRDNTGQLY | 0.44317442 |
| MKFYAVALFRSRGVPPTR | ASSLDRDNTGQLY | 0.75327486 |
| NAGGFGILGAVAHSP | ASSLDRDNTGQLY | 0.534925 |
| NEGREITLAMAEM | ASSLDRDNTGQLY | 0.7546419 |
| NELVEWSHPLGHRWRWE | ASSLDRDNTGQLY | 0.2336011 |
| NLGAGGTGGNGGNPG | ASSLDRDNTGQLY | 0.73572063 |
| NTEWPKWETVMAADPVD | ASSLDRDNTGQLY | 0.523226 |
| PASAAPAETQP | ASSLDRDNTGQLY | 0.41721895 |
| QQSSDGKNGVRAAEITGE | ASSLDRDNTGQLY | 0.33957928 |
| RAPDAVTTSWWKE | ASSLDRDNTGQLY | 0.6073324 |
| SGFGNVGAHTSGWFNQ | ASSLDRDNTGQLY | 0.68948495 |
| SGVAGAVGQAASVGGLKV | ASSLDRDNTGQLY | 0.7987962 |
| SNGGAGTGSAGGNGGT | ASSLDRDNTGQLY | 0.70490384 |
| TGGAGGEGGAAG | ASSLDRDNTGQLY | 0.7932559 |
| TGGDGGAAGAGGAPGHG | ASSLDRDNTGQLY | 0.7444362 |
| TMPPLLAAIANAGRHGV | ASSLDRDNTGQLY | 0.622476 |
| TSSTAGNSGGA | ASSLDRDNTGQLY | 0.620475 |

| | | |
|---|---|---|
| VPATVPVGILC | ASSLDRDNTGQLY | 0.72087616 |
| VQRPDSGGVVLTGRIS | ASSLDRDNTGQLY | 0.6238309 |
| WTPNPVPATKGVPAGLVQAT | ASSLDRDNTGQLY | 0.7769387 |
| YALKRALAKFSDRFDVVII | ASSLDRDNTGQLY | 0.74200976 |
| YLIGSVIV | ASSLDRDNTGQLY | 0.57386804 |
| YGQLVAVID | ASSLDRDNTGQLY | 0.7649007 |
| VTVGGMTLPA | ASSLDRDNTGQLY | 0.7824371 |
| VLALPAV | ASSLDRDNTGQLY | 0.45289347 |
| VIPPTLNYET | ASSLDRDNTGQLY | 0.47175446 |
| VGMGGMHPSSG | ASSLDRDNTGQLY | 0.51184607 |
| VGLAGGAATA | ASSLDRDNTGQLY | 0.41545844 |
| VGGAGGAGGSGP | ASSLDRDNTGQLY | 0.712487 |
| VFAAFMV | ASSLDRDNTGQLY | 0.54754174 |
| TLYDKLR | ASSLDRDNTGQLY | 0.5841632 |
| TLAVFTVL | ASSLDRDNTGQLY | 0.72184014 |
| TGSVQPARV | ASSLDRDNTGQLY | 0.58268136 |
| TGGGAGSGGT | ASSLDRDNTGQLY | 0.60636336 |
| SHLETSL | ASSLDRDNTGQLY | 0.61497957 |
| SFTTGMT | ASSLDRDNTGQLY | 0.6654836 |
| QVEVAKRMGVRQPTVS | ASSLDRDNTGQLY | 0.7664749 |
| QGSGGPALAM | ASSLDRDNTGQLY | 0.51289856 |
| MLAPSNP | ASSLDRDNTGQLY | 0.55783767 |
| LVNNAAI | ASSLDRDNTGQLY | 0.79740953 |
| LTGEINTLA | ASSLDRDNTGQLY | 0.37528852 |
| LSIFGPV | ASSLDRDNTGQLY | 0.44403976 |
| LLALLANL | ASSLDRDNTGQLY | 0.73277676 |
| LISNASDALDKLR | ASSLDRDNTGQLY | 0.46874636 |
| LINEAVP | ASSLDRDNTGQLY | 0.7862309 |
| LGYAFSGL | ASSLDRDNTGQLY | 0.6924908 |
| LDFVAVAL | ASSLDRDNTGQLY | 0.7161987 |
| LAQPGAE | ASSLDRDNTGQLY | 0.6665977 |
| ISNLGAVTGF | ASSLDRDNTGQLY | 0.8195295 |
| HQEDPGRI | ASSLDRDNTGQLY | 0.7950994 |
| GTPGEPTP | ASSLDRDNTGQLY | 0.5269221 |
| GFFNSGS | ASSLDRDNTGQLY | 0.7679868 |
| GAGGGGLTAGAN | ASSLDRDNTGQLY | 0.38162392 |
| FVDALVNM | ASSLDRDNTGQLY | 0.7813475 |
| FLTVAPEL | ASSLDRDNTGQLY | 0.78727055 |
| FETLTAL | ASSLDRDNTGQLY | 0.4169718 |
| DVINPIEL | ASSLDRDNTGQLY | 0.60514694 |
| APVSGAPV | ASSLDRDNTGQLY | 0.80921775 |
| ALVGGLAW | ASSLDRDNTGQLY | 0.8147045 |
| ALAAAAATAI | ASSLDRDNTGQLY | 0.42812586 |
| AFGLGLLL | ASSLDRDNTGQLY | 0.7076721 |
| AFFAAASL | ASSLDRDNTGQLY | 0.36389259 |
| AAAALPAFSPP | ASSLDRDNTGQLY | 0.37662703 |
| VVLAPAL | ASSLDRDNTGQLY | 0.47223467 |
| TIGHVDHGKTTL | ASSLDRDNTGQLY | 0.7555698 |
| LISNASDALDK | ASSLDRDNTGQLY | 0.66203666 |
| LISNASDAL | ASSLDRDNTGQLY | 0.7078049 |
| ISNASDALDK | ASSLDRDNTGQLY | 0.7127288 |

| | | |
|---|---|---|
| NQAETLVYQTE | ASSLDRDNTGQLY | 0.7973286 |
| TGGTGGTAGAG | ASSLDRDNTGQLY | 0.5309621 |
| TGSTAGGAGGA | ASSLDRDNTGQLY | 0.6765447 |
| DPLAVPVAAGPV | ASSLDRDNTGQLY | 0.43667954 |
| GGTGGTPGNGGN | ASSLDRDNTGQLY | 0.5846876 |
| PAGLAGVDPATL | ASSLDRDNTGQLY | 0.64062315 |
| GNNLVVEAPHAGASV | ASSLDRDNTGQLY | 0.6023916 |
| KVTRIMSASTLEQPA | ASSLDRDNTGQLY | 0.31348628 |
| LGFALPIALPRRTVV | ASSLDRDNTGQLY | 0.7484108 |
| TRLGNLTARKLVVLV | ASSLDRDNTGQLY | 0.7354865 |
| WSQQRRLIYFPSAGP | ASSLDRDNTGQLY | 0.74877214 |
| GPNGNPDPMAAAVDIR | ASSLDRDNTGQLY | 0.6402214 |
| TQRCVSVADLRAAASR | ASSLDRDNTGQLY | 0.6885092 |
| QFVLDVAEHPVEVLTETV | ASSLDRDNTGQLY | 0.564547 |
| GGAGGTGGAGGNGGTQAGDG | ASSLDRDNTGQLY | 0.7798986 |
| PLLATLRAIAVDVDAKPAQV | ASSLDRDNTGQLY | 0.71105003 |
| APKFRTGGYMIDFWGVGYQVAK | ASSLDRDNTGQLY | 0.2462146 |
| GAGGAGGAGGAAGGPGGTGGTG | ASSLDRDNTGQLY | 0.49514806 |
| IGALGLLATDVAPAGSVAFGVA | ASSLDRDNTGQLY | 0.36789152 |
| AGGAGGIGGNGGAGGAGGVLMG | ASSLDRDNTGQLY | 0.76335543 |
| ADELIGLL | ASSLDRDNTGQLY | 0.75055194 |
| ADRALKA | ASSLDRDNTGQLY | 0.7764591 |
| AGAVGQVGGAGGA | ASSLDRDNTGQLY | 0.7660886 |
| AGGGGGLGGAP | ASSLDRDNTGQLY | 0.23316605 |
| AGGLLYGNGGAG | ASSLDRDNTGQLY | 0.59327227 |
| AGLESIGL | ASSLDRDNTGQLY | 0.7970357 |
| AGLKTGII | ASSLDRDNTGQLY | 0.65596044 |
| ASLAALLL | ASSLDRDNTGQLY | 0.690462 |
| ASVINISSI | ASSLDRDNTGQLY | 0.79236543 |
| ATPPGPA | ASSLDRDNTGQLY | 0.52850026 |
| ATVASALV | ASSLDRDNTGQLY | 0.65499663 |
| ATVASLIL | ASSLDRDNTGQLY | 0.75447357 |
| AVNGPVLIRH | ASSLDRDNTGQLY | 0.8093446 |
| DGAAGAVGI | ASSLDRDNTGQLY | 0.6988611 |
| DPLLHGI | ASSLDRDNTGQLY | 0.6879761 |
| ELGAALGAL | ASSLDRDNTGQLY | 0.69709444 |
| ELGRTQAV | ASSLDRDNTGQLY | 0.82510227 |
| FVAGGYLS | ASSLDRDNTGQLY | 0.74399126 |
| GASGGAGGAGGW | ASSLDRDNTGQLY | 0.71960115 |
| GGAGAGGDGGAGGI | ASSLDRDNTGQLY | 0.7157116 |
| GTGHNGSQPGMGGNG | ASSLDRDNTGQLY | 0.63454413 |
| GYAAASASAS | ASSLDRDNTGQLY | 0.53771174 |
| HQVDVAAVQQG | ASSLDRDNTGQLY | 0.6858601 |
| IGAAIAHT | ASSLDRDNTGQLY | 0.6955456 |
| IGMIEVAQL | ASSLDRDNTGQLY | 0.47581625 |
| IRLAPEHTSAMD | ASSLDRDNTGQLY | 0.70359147 |
| IYNVSVL | ASSLDRDNTGQLY | 0.50240487 |
| KQGFLGM | ASSLDRDNTGQLY | 0.81658727 |
| LESVITQL | ASSLDRDNTGQLY | 0.71357584 |
| LGLGALVVV | ASSLDRDNTGQLY | 0.7590625 |
| LIGNGGNGGSGGI | ASSLDRDNTGQLY | 0.616809 |

| | | |
|---|---|---|
| LNAINAPTL | ASSLDRDNTGQLY | 0.7293911 |
| LQLLNTGV | ASSLDRDNTGQLY | 0.69709235 |
| LQTSIEF | ASSLDRDNTGQLY | 0.34669363 |
| MSFAVSEL | ASSLDRDNTGQLY | 0.7662884 |
| NTGALGL | ASSLDRDNTGQLY | 0.52972984 |
| PPGAGGAGGAGTT | ASSLDRDNTGQLY | 0.52560586 |
| QLVGLGI | ASSLDRDNTGQLY | 0.71634626 |
| RFLEQVHQL | ASSLDRDNTGQLY | 0.76936364 |
| RGVGYVLR | ASSLDRDNTGQLY | 0.61446315 |
| SATGVFAGVI | ASSLDRDNTGQLY | 0.6644081 |
| SLPAITLP | ASSLDRDNTGQLY | 0.6681963 |
| STALAFTV | ASSLDRDNTGQLY | 0.73869073 |
| SVLTVGLGK | ASSLDRDNTGQLY | 0.81950724 |
| TALAGAGSGI | ASSLDRDNTGQLY | 0.6712702 |
| TGLPFTGI | ASSLDRDNTGQLY | 0.8234935 |
| VAEHVTV | ASSLDRDNTGQLY | 0.6819955 |
| VDIVLNSL | ASSLDRDNTGQLY | 0.36469415 |
| VLTGGGAVL | ASSLDRDNTGQLY | 0.7223995 |
| VNVGLAYGI | ASSLDRDNTGQLY | 0.7730856 |
| VSSAAVGAV | ASSLDRDNTGQLY | 0.80278885 |
| VTSVYEAGV | ASSLDRDNTGQLY | 0.787935 |
| WVALDRGA | ASSLDRDNTGQLY | 0.7704553 |
| YLVGSVVI | ASSLDRDNTGQLY | 0.5200265 |
| GPPGTGKTLL | ASSLDRDNTGQLY | 0.7627158 |
| GVGMTKF | ASSLDRDNTGQLY | 0.7402718 |
| IGIGGSDL | ASSLDRDNTGQLY | 0.7517934 |
| IGTLLAEL | ASSLDRDNTGQLY | 0.5395803 |
| IYTSGTT | ASSLDRDNTGQLY | 0.4589203 |
| LSFPAGV | ASSLDRDNTGQLY | 0.6551772 |
| LYTSGTT | ASSLDRDNTGQLY | 0.7462715 |
| VGAFPFGF | ASSLDRDNTGQLY | 0.50541973 |
| VGAGTVEF | ASSLDRDNTGQLY | 0.7133354 |
| VGKSTLLL | ASSLDRDNTGQLY | 0.674263 |
| AALAHAGGGIV | ASSLDRDNTGQLY | 0.39291972 |
| AALVAVSLL | ASSLDRDNTGQLY | 0.664238 |
| AAVSRAVAAL | ASSLDRDNTGQLY | 0.80151665 |
| AGGSSGIGGF | ASSLDRDNTGQLY | 0.65245515 |
| AGLTVANANG | ASSLDRDNTGQLY | 0.5966848 |
| AGLVGAVL | ASSLDRDNTGQLY | 0.7600744 |
| ALSQAFTV | ASSLDRDNTGQLY | 0.8264111 |
| ALTFSVSL | ASSLDRDNTGQLY | 0.81196916 |
| ALTGAALVAL | ASSLDRDNTGQLY | 0.38722754 |
| ASFSIGAGVL | ASSLDRDNTGQLY | 0.21207467 |
| DGSVYKI | ASSLDRDNTGQLY | 0.71193343 |
| EQLAEIV | ASSLDRDNTGQLY | 0.73512614 |
| FAVSELTVM | ASSLDRDNTGQLY | 0.56778467 |
| GAAPLAAGL | ASSLDRDNTGQLY | 0.6312604 |
| GADGGDGGSGGAGG | ASSLDRDNTGQLY | 0.75903124 |
| GFGLDTGGAGG | ASSLDRDNTGQLY | 0.7235614 |
| GLASGGLL | ASSLDRDNTGQLY | 0.6400529 |
| GRIAGAAVLL | ASSLDRDNTGQLY | 0.34087133 |

| | | |
|---|---|---|
| GSAGSLNTSF | ASSLDRDNTGQLY | 0.70741045 |
| GTGQIGI | ASSLDRDNTGQLY | 0.7850802 |
| GTLIALIL | ASSLDRDNTGQLY | 0.6149292 |
| IAATMGFVVG | ASSLDRDNTGQLY | 0.71908545 |
| IFAQALL | ASSLDRDNTGQLY | 0.67809886 |
| IGAAVALGF | ASSLDRDNTGQLY | 0.59423053 |
| IGLTPLAL | ASSLDRDNTGQLY | 0.6621475 |
| ILGSNTSTL | ASSLDRDNTGQLY | 0.37289062 |
| IVFSLTVL | ASSLDRDNTGQLY | 0.79221684 |
| LASKATGF | ASSLDRDNTGQLY | 0.39098388 |
| LASVALIL | ASSLDRDNTGQLY | 0.7796323 |
| LFAVGLTL | ASSLDRDNTGQLY | 0.77871144 |
| LFNGGIGTYI | ASSLDRDNTGQLY | 0.7874475 |
| LTLGAGII | ASSLDRDNTGQLY | 0.64312583 |
| LVAAALGDI | ASSLDRDNTGQLY | 0.529477 |
| NTAFVQLGI | ASSLDRDNTGQLY | 0.4089555 |
| QLFSGAEL | ASSLDRDNTGQLY | 0.40049326 |
| SAATFITL | ASSLDRDNTGQLY | 0.81423867 |
| SNYGLANLGV | ASSLDRDNTGQLY | 0.63087785 |
| SYGGAYAVM | ASSLDRDNTGQLY | 0.81717503 |
| TTGVGGLTLG | ASSLDRDNTGQLY | 0.72863376 |
| TTQTIYI | ASSLDRDNTGQLY | 0.71648103 |
| VAAGAGLGAVI | ASSLDRDNTGQLY | 0.7635477 |
| VAAVVTLL | ASSLDRDNTGQLY | 0.5724371 |
| VGFLGGLI | ASSLDRDNTGQLY | 0.70273066 |
| VGVIPILL | ASSLDRDNTGQLY | 0.6403274 |
| VIFASSLI | ASSLDRDNTGQLY | 0.7605482 |
| VKGGDGGAAAG | ASSLDRDNTGQLY | 0.5728706 |
| VTKTFTNI | ASSLDRDNTGQLY | 0.75805354 |
| VVAAAILL | ASSLDRDNTGQLY | 0.6838012 |
| YAALDVEL | ASSLDRDNTGQLY | 0.48585272 |
| LASVALLL | ASSLDRDNTGQLY | 0.62885046 |
| LDAVHRV | ASSLDRDNTGQLY | 0.7673115 |
| QGLSSSIF | ASSLDRDNTGQLY | 0.70733094 |
| RGTFANIRL | ASSLDRDNTGQLY | 0.71887857 |
| RQLWWGHRI | ASSLDRDNTGQLY | 0.7141433 |
| YGPPGTGKTLL | ASSLDRDNTGQLY | 0.68618584 |
| AALVGAPANVA | ASSLDRDNTGQLY | 0.71606314 |
| AATFRRTE | ASSLDRDNTGQLY | 0.782952 |
| AGALGGGI | ASSLDRDNTGQLY | 0.5378852 |
| AGGAGGAGGIGAG | ASSLDRDNTGQLY | 0.7834544 |
| AGSGALGL | ASSLDRDNTGQLY | 0.3503506 |
| AQIADQI | ASSLDRDNTGQLY | 0.47408915 |
| ASAVAVTYL | ASSLDRDNTGQLY | 0.69337827 |
| ASTVLGGTGG | ASSLDRDNTGQLY | 0.7135954 |
| AVGGEPAK | ASSLDRDNTGQLY | 0.7618996 |
| CEQGTFGGTC | ASSLDRDNTGQLY | 0.5535526 |
| DEGKAPAG | ASSLDRDNTGQLY | 0.6652635 |
| DTVVFPAL | ASSLDRDNTGQLY | 0.5834553 |
| FEGITMAG | ASSLDRDNTGQLY | 0.5462784 |
| FPLVVQD | ASSLDRDNTGQLY | 0.69102156 |

| | | |
|---|---|---|
| GAFTTPQ | ASSLDRDNTGQLY | 0.5499601 |
| GFGGFGTAGGVGGA | ASSLDRDNTGQLY | 0.55860996 |
| GFLAQFL | ASSLDRDNTGQLY | 0.80377024 |
| GGAGGAGGNAGVA | ASSLDRDNTGQLY | 0.6521585 |
| GGVGTAGAGGAGGAGG | ASSLDRDNTGQLY | 0.6627278 |
| GNGGGGDAGNAGSGGN | ASSLDRDNTGQLY | 0.7613201 |
| GTGGAGGAGNDAGS | ASSLDRDNTGQLY | 0.6201947 |
| GTTNTGWFNS | ASSLDRDNTGQLY | 0.77096164 |
| IFMGPLFTL | ASSLDRDNTGQLY | 0.67606944 |
| ILAHAIEAI | ASSLDRDNTGQLY | 0.78871095 |
| IRSAIREAL | ASSLDRDNTGQLY | 0.6501893 |
| ITFAGGG | ASSLDRDNTGQLY | 0.52427983 |
| ITGMSGHQEV | ASSLDRDNTGQLY | 0.770115 |
| IVAGLACTL | ASSLDRDNTGQLY | 0.73511887 |
| IVGGGLAGA | ASSLDRDNTGQLY | 0.7151382 |
| IVTGVLNI | ASSLDRDNTGQLY | 0.75703794 |
| LAQQGCI | ASSLDRDNTGQLY | 0.7394762 |
| LCLHVEN | ASSLDRDNTGQLY | 0.78751206 |
| LDAISDGS | ASSLDRDNTGQLY | 0.6405762 |
| LGGFAAML | ASSLDRDNTGQLY | 0.40561062 |
| LGGKNPMI | ASSLDRDNTGQLY | 0.64187086 |
| LGPKGVAAIL | ASSLDRDNTGQLY | 0.62121296 |
| LMVGVSI | ASSLDRDNTGQLY | 0.5583832 |
| LVGAAFVL | ASSLDRDNTGQLY | 0.32619572 |
| LVLAVAPL | ASSLDRDNTGQLY | 0.77097255 |
| MAAIAVIDAA | ASSLDRDNTGQLY | 0.8210609 |
| NNGINAELQ | ASSLDRDNTGQLY | 0.55047005 |
| RGEPLVLVHG | ASSLDRDNTGQLY | 0.82417756 |
| RGGAGGAAGLI | ASSLDRDNTGQLY | 0.32231236 |
| RSYRMHPAV | ASSLDRDNTGQLY | 0.7289422 |
| SAAHLLAGQLA | ASSLDRDNTGQLY | 0.7802553 |
| SAAVGSY | ASSLDRDNTGQLY | 0.45683318 |
| SPHAVLHRI | ASSLDRDNTGQLY | 0.7290236 |
| SQIAKVL | ASSLDRDNTGQLY | 0.6218296 |
| TALGHLVP | ASSLDRDNTGQLY | 0.6249669 |
| TAPTTSGAA | ASSLDRDNTGQLY | 0.8071208 |
| TDGDDGNGGSPGT | ASSLDRDNTGQLY | 0.6545124 |
| TDGLGEN | ASSLDRDNTGQLY | 0.7889009 |
| TGGPPLPGQA | ASSLDRDNTGQLY | 0.7482274 |
| TIPAITI | ASSLDRDNTGQLY | 0.66483366 |
| TIPGFTI | ASSLDRDNTGQLY | 0.8069842 |
| TSIASGALA | ASSLDRDNTGQLY | 0.7884555 |
| TVTLPGAAI | ASSLDRDNTGQLY | 0.603608 |
| VASGLSAALP | ASSLDRDNTGQLY | 0.71782583 |
| VGAGVIGI | ASSLDRDNTGQLY | 0.6604092 |
| VGPKRFGV | ASSLDRDNTGQLY | 0.79336965 |
| VNVAFPD | ASSLDRDNTGQLY | 0.7789341 |
| VSMTNSGVS | ASSLDRDNTGQLY | 0.7072809 |
| VTDVLGDLSV | ASSLDRDNTGQLY | 0.6932397 |
| VVVVGAGFAGL | ASSLDRDNTGQLY | 0.7870712 |
| VYTVGTVV | ASSLDRDNTGQLY | 0.74820244 |

| | | |
|---|---|---|
| WTTFSGP | ASSLDRDNTGQLY | 0.7722872 |
| YDELVEAF | ASSLDRDNTGQLY | 0.810228 |
| YGTGGAGGNGG | ASSLDRDNTGQLY | 0.44725463 |
| YTNVMAV | ASSLDRDNTGQLY | 0.7671683 |
| FLEQVHQL | ASSLDRDNTGQLY | 0.68513805 |
| GLSLLAVAV | ASSLDRDNTGQLY | 0.6216662 |
| LEQVHQL | ASSLDRDNTGQLY | 0.56856424 |
| LNAAGAI | ASSLDRDNTGQLY | 0.40432525 |
| PAVHASL | ASSLDRDNTGQLY | 0.57611483 |
| TELEDDA | ASSLDRDNTGQLY | 0.49218524 |
| TPPGPPN | ASSLDRDNTGQLY | 0.7464086 |
| TPPPPPPP | ASSLDRDNTGQLY | 0.74038184 |
| TQSPALI | ASSLPGTGQDTQY | 0.537651 |
| EFAAAFPQ | ASSLPGTGQDTQY | 0.76882803 |
| LLSGNLDVL | ASSLPGTGQDTQY | 0.6894552 |
| GPANLGNYNI | ASSLPGTGQDTQY | 0.39858532 |
| TLGPTLP | ASSLPGTGQDTQY | 0.6010841 |
| VYAYSSCCK | ASSLPGTGQDTQY | 0.5831021 |
| APAPAPAPNI | ASSLPGTGQDTQY | 0.61138165 |
| PGGPVPIP | ASSLPGTGQDTQY | 0.32043722 |
| FLTVALEP | ASSLPGTGQDTQY | 0.7254807 |
| SQYFGANL | ASSLPGTGQDTQY | 0.5632081 |
| GGVKPAA | ASSLPGTGQDTQY | 0.25463519 |
| GGTGGAGGAGGW | ASSLPGTGQDTQY | 0.70660496 |
| GVGGVAGLGGAGG | ASSLPGTGQDTQY | 0.666447 |
| VLLVLATTI | ASSLPGTGQDTQY | 0.62455803 |
| IGIFAAML | ASSLPGTGQDTQY | 0.7617322 |
| AVGDLID | ASSLPGTGQDTQY | 0.54116243 |
| NISPLPL | ASSLPGTGQDTQY | 0.51656294 |
| VTAVFVAI | ASSLPGTGQDTQY | 0.74380684 |
| SGDDNDAAL | ASSLPGTGQDTQY | 0.76760375 |
| AVAALAHS | ASSLPGTGQDTQY | 0.45237887 |
| GGVTVMATP | ASSLPGTGQDTQY | 0.70040965 |
| AGLAANVRD | ASSLPGTGQDTQY | 0.74677026 |
| TNRPDLIDPALLRPG | ASSLPGTGQDTQY | 0.7152275 |
| LANLGVTAL | ASSLPGTGQDTQY | 0.756567 |
| TGERIEL | ASSLPGTGQDTQY | 0.7652651 |
| NGEELAA | ASSLPGTGQDTQY | 0.505759 |
| VLSLVWG | ASSLPGTGQDTQY | 0.7649685 |
| IGYALWL | ASSLPGTGQDTQY | 0.7716671 |
| LSLLLAPL | ASSLPGTGQDTQY | 0.60244995 |
| TLDEALK | ASSLPGTGQDTQY | 0.6664423 |
| TDGAGTAEGG | ASSLPGTGQDTQY | 0.441374 |
| NAPGGAGGNGG | ASSLPGTGQDTQY | 0.6151639 |
| VSVFMGMGGNF | ASSLPGTGQDTQY | 0.39674062 |
| LASYLAAL | ASSLPGTGQDTQY | 0.72261286 |
| LVGGTPAL | ASSLPGTGQDTQY | 0.7709527 |
| VRNADVL | ASSLPGTGQDTQY | 0.6631774 |
| TCSSPFLVSAM | ASSLPGTGQDTQY | 0.43825865 |
| VHITDAE | ASSLPGTGQDTQY | 0.4449324 |
| EYAAPSF | ASSLPGTGQDTQY | 0.6444597 |

| | | |
|---|---|---|
| VVDAFHGGL | ASSLPGTGQDTQY | 0.49503046 |
| ASVVDPQ | ASSLPGTGQDTQY | 0.3435918 |
| SYNEFGGLS | ASSLPGTGQDTQY | 0.6181092 |
| DPDGSVY | ASSLPGTGQDTQY | 0.7556557 |
| MVSTIHATE | ASSLPGTGQDTQY | 0.74902004 |
| LAAGVADPV | ASSLPGTGQDTQY | 0.45863593 |
| TKEQIAA | ASSLPGTGQDTQY | 0.77472603 |
| SLASPAAL | ASSLPGTGQDTQY | 0.7166698 |
| IGLVFYGV | ASSLPGTGQDTQY | 0.7335409 |
| FLLDASNL | ASSLPGTGQDTQY | 0.61153203 |
| YQVGNFP | ASSLPGTGQDTQY | 0.4659498 |
| ESLLSQST | ASSLPGTGQDTQY | 0.749033 |
| LFTVGFAAVG | ASSLPGTGQDTQY | 0.66348755 |
| ATPPTPT | ASSLPGTGQDTQY | 0.6637327 |
| TDLATLG | ASSLPGTGQDTQY | 0.7451049 |
| SAGGTGR | ASSLPGTGQDTQY | 0.66018367 |
| GFSNSGS | ASSLPGTGQDTQY | 0.59567356 |
| GGNGGSGGTSV | ASSLPGTGQDTQY | 0.6452637 |
| GTSVATGGAGN | ASSLPGTGQDTQY | 0.73977804 |
| NGGNGGSGGTS | ASSLPGTGQDTQY | 0.53022516 |
| AVVKFPH | ASSLPGTGQDTQY | 0.43973163 |
| TPPPPPP | ASSLPGTGQDTQY | 0.5878372 |
| AAPAAAAAPA | ASSLPGTGQDTQY | 0.67396927 |
| IALNEDL | ASSLPGTGQDTQY | 0.76438314 |
| IVNVSSM | ASSLPGTGQDTQY | 0.682094 |
| TGIPPAPR | ASSLPGTGQDTQY | 0.5844024 |
| FDLGGGTF | ASSLPGTGQDTQY | 0.7264441 |
| FDLGGGTFDVS | ASSLPGTGQDTQY | 0.7083903 |
| NEPTAAA | ASSLPGTGQDTQY | 0.6672844 |
| NEPTAAALA | ASSLPGTGQDTQY | 0.6576533 |
| GSIINISSI | ASSLPGTGQDTQY | 0.59174263 |
| VAPGNAGTA | ASSLPGTGQDTQY | 0.7309088 |
| TGGGASLE | ASSLPGTGQDTQY | 0.7256316 |
| VAVYNHYKRI | ASSLPGTGQDTQY | 0.7176889 |
| TDPAPATT | ASSLPGTGQDTQY | 0.7476673 |
| TNRPDILDPALLRPG | ASSLPGTGQDTQY | 0.69407237 |
| LTTTTTS | ASSLPGTGQDTQY | 0.66034824 |
| PTGSGKTL | ASSLPGTGQDTQY | 0.3563382 |
| TGSGKTL | ASSLPGTGQDTQY | 0.5548389 |
| TIGHVDHGK | ASSLPGTGQDTQY | 0.70169127 |
| TIGHVDHGKTT | ASSLPGTGQDTQY | 0.70839465 |
| GHVDHGKTT | ASSLPGTGQDTQY | 0.7469505 |
| AQFEHTLL | ASSLPGTGQDTQY | 0.65873086 |
| IVNQDEL | ASSLPGTGQDTQY | 0.5787933 |
| TNGSQFF | ASSLPGTGQDTQY | 0.7256645 |
| TGKGIVE | ASSLPGTGQDTQY | 0.747099 |
| LISNASDA | ASSLPGTGQDTQY | 0.40911463 |
| AELRTLGELTKIAKAHGAQ | ASSLPGTGQDTQY | 0.6199796 |
| APNAVAAAEPSW | ASSLPGTGQDTQY | 0.7869624 |
| APQVAGAATAVL | ASSLPGTGQDTQY | 0.7024783 |
| ARRDPAAAAAILPTDA | ASSLPGTGQDTQY | 0.6496409 |

| | | |
|---|---|---|
| DAAARDVLAEAGLAEYF | ASSLPGTGQDTQY | 0.5373771 |
| DDQIVLSAKANLAKVEQ | ASSLPGTGQDTQY | 0.7654034 |
| DELLPDTGEDSDEDGH | ASSLPGTGQDTQY | 0.6112443 |
| ELPEAPANRATVPQD | ASSLPGTGQDTQY | 0.7755376 |
| FGQSKIERVTGIG | ASSLPGTGQDTQY | 0.28406546 |
| GAGGTGSTAGGAGGAGGAGGLY | ASSLPGTGQDTQY | 0.5112489 |
| GHGGTGAAGGNGGTGGAGGSGI | ASSLPGTGQDTQY | 0.43172145 |
| GNGGNGGAGGEGGPGGAGGAGA | ASSLPGTGQDTQY | 0.29004478 |
| GSTAGNGGNGGNATNGTIAGQP | ASSLPGTGQDTQY | 0.75915396 |
| GTGGDGGNGGNGGTGG | ASSLPGTGQDTQY | 0.47552606 |
| GVRRPAGIPADAPV | ASSLPGTGQDTQY | 0.7046436 |
| GYRQVTDAHLLTLARR | ASSLPGTGQDTQY | 0.663138 |
| IAKMGQDGHDRG | ASSLPGTGQDTQY | 0.757444 |
| KAEPSIEIQYT | ASSLPGTGQDTQY | 0.67640424 |
| KPVRIRMAADEDTVAVT | ASSLPGTGQDTQY | 0.61905307 |
| MPPRIAGMRLLVIKPEP | ASSLPGTGQDTQY | 0.5297686 |
| MKFYAVALFRSRGVPPTR | ASSLPGTGQDTQY | 0.74079525 |
| NAGGFGILGAVAHSP | ASSLPGTGQDTQY | 0.75112915 |
| NEGREITLAMAEM | ASSLPGTGQDTQY | 0.6993358 |
| NELVEWSHPLGHRWRWE | ASSLPGTGQDTQY | 0.36335024 |
| NLGAGGTGGNGGNPG | ASSLPGTGQDTQY | 0.67892206 |
| NTEWPKWETVMAADPVD | ASSLPGTGQDTQY | 0.6639979 |
| PASAAPAETQP | ASSLPGTGQDTQY | 0.36188847 |
| QQSSDGKNGVRAAEITGE | ASSLPGTGQDTQY | 0.29560432 |
| RAPDAVTTSWWKE | ASSLPGTGQDTQY | 0.5235121 |
| SGFGNVGAHTSGWFNQ | ASSLPGTGQDTQY | 0.6302775 |
| SGVAGAVGQAASVGGLKV | ASSLPGTGQDTQY | 0.75531995 |
| SNGGAGTGSAGGNGGT | ASSLPGTGQDTQY | 0.746627 |
| TGGAGGEGGAAG | ASSLPGTGQDTQY | 0.61904144 |
| TGGDGGAAGAGGAPGHG | ASSLPGTGQDTQY | 0.72100705 |
| TMPPLLAAIANAGRHGV | ASSLPGTGQDTQY | 0.6890075 |
| TSSTAGNSGGA | ASSLPGTGQDTQY | 0.19934087 |
| VPATVPVGILC | ASSLPGTGQDTQY | 0.500619 |
| VQRPDSGGVVLTGRIS | ASSLPGTGQDTQY | 0.42332911 |
| WTPNPVPATKGVPAGLVQAT | ASSLPGTGQDTQY | 0.75745296 |
| YALKRALAKFSDRFDVVII | ASSLPGTGQDTQY | 0.5155384 |
| YLIGSVIV | ASSLPGTGQDTQY | 0.6082911 |
| YGQLVAVID | ASSLPGTGQDTQY | 0.67576635 |
| VTVGGMTLPA | ASSLPGTGQDTQY | 0.6968144 |
| VLALPAV | ASSLPGTGQDTQY | 0.67493534 |
| VIPPTLNYET | ASSLPGTGQDTQY | 0.44597882 |
| VGMGGMHPSSG | ASSLPGTGQDTQY | 0.31495836 |
| VGLAGGAATA | ASSLPGTGQDTQY | 0.740702 |
| VGGAGGAGGSGP | ASSLPGTGQDTQY | 0.5386403 |
| VFAAFMV | ASSLPGTGQDTQY | 0.47631136 |
| TLYDKLR | ASSLPGTGQDTQY | 0.3482405 |
| TLAVFTVL | ASSLPGTGQDTQY | 0.71421057 |
| TGSVQPARV | ASSLPGTGQDTQY | 0.32260212 |
| TGGGAGSGGT | ASSLPGTGQDTQY | 0.7273722 |
| SHLETSL | ASSLPGTGQDTQY | 0.5975305 |
| SFTTGMT | ASSLPGTGQDTQY | 0.67501783 |

| | | |
|---|---|---|
| QVEVAKRMGVRQPTVS | ASSLPGTGQDTQY | 0.78364617 |
| QGSGGPALAM | ASSLPGTGQDTQY | 0.68833005 |
| MLAPSNP | ASSLPGTGQDTQY | 0.6529627 |
| LVNNAAI | ASSLPGTGQDTQY | 0.5270206 |
| LTGEINTLA | ASSLPGTGQDTQY | 0.47357857 |
| LSIFGPV | ASSLPGTGQDTQY | 0.53215355 |
| LLALLANL | ASSLPGTGQDTQY | 0.5495565 |
| LISNASDALDKLR | ASSLPGTGQDTQY | 0.77388525 |
| LINEAVP | ASSLPGTGQDTQY | 0.5956546 |
| LGYAFSGL | ASSLPGTGQDTQY | 0.39263165 |
| LDFVAVAL | ASSLPGTGQDTQY | 0.550621 |
| LAQPGAE | ASSLPGTGQDTQY | 0.7735228 |
| ISNLGAVTGF | ASSLPGTGQDTQY | 0.77884555 |
| HQEDPGRI | ASSLPGTGQDTQY | 0.54650456 |
| GTPGEPTP | ASSLPGTGQDTQY | 0.48513883 |
| GFFNSGS | ASSLPGTGQDTQY | 0.49089316 |
| GAGGGGLTAGAN | ASSLPGTGQDTQY | 0.6717224 |
| FVDALVNM | ASSLPGTGQDTQY | 0.46287364 |
| FLTVAPEL | ASSLPGTGQDTQY | 0.46381167 |
| FETLTAL | ASSLPGTGQDTQY | 0.7080457 |
| DVINPIEL | ASSLPGTGQDTQY | 0.7777469 |
| APVSGAPV | ASSLPGTGQDTQY | 0.60511535 |
| ALVGGLAW | ASSLPGTGQDTQY | 0.37150332 |
| ALAAAAATAI | ASSLPGTGQDTQY | 0.66835845 |
| AFGLGLLL | ASSLPGTGQDTQY | 0.53648734 |
| AFFAAASL | ASSLPGTGQDTQY | 0.58229697 |
| AAAALPAFSPP | ASSLPGTGQDTQY | 0.60166097 |
| VVLAPAL | ASSLPGTGQDTQY | 0.37872022 |
| TIGHVDHGKTTL | ASSLPGTGQDTQY | 0.6566225 |
| LISNASDALDK | ASSLPGTGQDTQY | 0.7768202 |
| LISNASDAL | ASSLPGTGQDTQY | 0.63062847 |
| ISNASDALDK | ASSLPGTGQDTQY | 0.68069446 |
| NQAETLVYQTE | ASSLPGTGQDTQY | 0.7744689 |
| TGGTGGTAGAG | ASSLPGTGQDTQY | 0.74057734 |
| TGSTAGGAGGA | ASSLPGTGQDTQY | 0.76898205 |
| DPLAVPVAAGPV | ASSLPGTGQDTQY | 0.73709905 |
| GGTGGTPGNGGN | ASSLPGTGQDTQY | 0.6486267 |
| PAGLAGVDPATL | ASSLPGTGQDTQY | 0.62097573 |
| GNNLVVEAPHAGASV | ASSLPGTGQDTQY | 0.75787306 |
| KVTRIMSASTLEQPA | ASSLPGTGQDTQY | 0.67407286 |
| LGFALPIALPRRTVV | ASSLPGTGQDTQY | 0.6668309 |
| TRLGNLTARKLVVLV | ASSLPGTGQDTQY | 0.5429962 |
| WSQQRRLIYFPSAGP | ASSLPGTGQDTQY | 0.7487261 |
| GPNGNPDPMAAAVDIR | ASSLPGTGQDTQY | 0.57335526 |
| TQRCVSVADLRAAASR | ASSLPGTGQDTQY | 0.75390047 |
| QFVLDVAEHPVEVLTETV | ASSLPGTGQDTQY | 0.32656136 |
| GGAGGTGGAGGNGGTQAGDG | ASSLPGTGQDTQY | 0.48960584 |
| PLLATLRAIAVDVDAKPAQV | ASSLPGTGQDTQY | 0.7528992 |
| APKFRTGGYMIDFWGVGYQVAK | ASSLPGTGQDTQY | 0.21157701 |
| GAGGAGGAGGAAGGPGGTGGTG | ASSLPGTGQDTQY | 0.28834796 |
| IGALGLLATDVAPAGSVAFGVA | ASSLPGTGQDTQY | 0.6125226 |

| | | |
|---|---|---|
| AGGAGGIGGNGGAGGAGGVLMG | ASSLPGTGQDTQY | 0.7532409 |
| ADELIGLL | ASSLPGTGQDTQY | 0.22863679 |
| ADRALKA | ASSLPGTGQDTQY | 0.5087658 |
| AGAVGQVGGAGGA | ASSLPGTGQDTQY | 0.3949788 |
| AGGGGGLGGAP | ASSLPGTGQDTQY | 0.6697166 |
| AGGLLYGNGGAG | ASSLPGTGQDTQY | 0.66559666 |
| AGLESIGL | ASSLPGTGQDTQY | 0.369468 |
| AGLKTGII | ASSLPGTGQDTQY | 0.728534 |
| ASLAALLL | ASSLPGTGQDTQY | 0.3228552 |
| ASVINISSI | ASSLPGTGQDTQY | 0.6665356 |
| ATPPGPA | ASSLPGTGQDTQY | 0.74687105 |
| ATVASALV | ASSLPGTGQDTQY | 0.76860833 |
| ATVASLIL | ASSLPGTGQDTQY | 0.4713925 |
| AVNGPVLIRH | ASSLPGTGQDTQY | 0.7362708 |
| DGAAGAVGI | ASSLPGTGQDTQY | 0.5425043 |
| DPLLHGI | ASSLPGTGQDTQY | 0.787297 |
| ELGAALGAL | ASSLPGTGQDTQY | 0.6312826 |
| ELGRTQAV | ASSLPGTGQDTQY | 0.37130526 |
| FVAGGYLS | ASSLPGTGQDTQY | 0.7454556 |
| GASGGAGGAGGW | ASSLPGTGQDTQY | 0.6997764 |
| GGAGAGGDGGAGGI | ASSLPGTGQDTQY | 0.59243417 |
| GTGHNGSQPGMGGNG | ASSLPGTGQDTQY | 0.72577584 |
| GYAAASASAS | ASSLPGTGQDTQY | 0.49724567 |
| HQVDVAAVQQG | ASSLPGTGQDTQY | 0.64246833 |
| IGAAIAHT | ASSLPGTGQDTQY | 0.75933033 |
| IGMIEVAQL | ASSLPGTGQDTQY | 0.73981273 |
| IRLAPEHTSAMD | ASSLPGTGQDTQY | 0.51160026 |
| IYNVSVL | ASSLPGTGQDTQY | 0.74280745 |
| KQGFLGM | ASSLPGTGQDTQY | 0.67203957 |
| LESVITQL | ASSLPGTGQDTQY | 0.46547723 |
| LGLGALVVV | ASSLPGTGQDTQY | 0.7772447 |
| LIGNGGNGGSGGI | ASSLPGTGQDTQY | 0.5927279 |
| LNAINAPTL | ASSLPGTGQDTQY | 0.7427207 |
| LQLLNTGV | ASSLPGTGQDTQY | 0.7197045 |
| LQTSIEF | ASSLPGTGQDTQY | 0.7470883 |
| MSFAVSEL | ASSLPGTGQDTQY | 0.50220275 |
| NTGALGL | ASSLPGTGQDTQY | 0.65046465 |
| PPGAGGAGGAGTT | ASSLPGTGQDTQY | 0.6574249 |
| QLVGLGI | ASSLPGTGQDTQY | 0.7310096 |
| RFLEQVHQL | ASSLPGTGQDTQY | 0.5881882 |
| RGVGYVLR | ASSLPGTGQDTQY | 0.68204737 |
| SATGVFAGVI | ASSLPGTGQDTQY | 0.7161771 |
| SLPAITLP | ASSLPGTGQDTQY | 0.76893777 |
| STALAFTV | ASSLPGTGQDTQY | 0.6109385 |
| SVLTVGLGK | ASSLPGTGQDTQY | 0.64811563 |
| TALAGAGSGI | ASSLPGTGQDTQY | 0.77801657 |
| TGLPFTGI | ASSLPGTGQDTQY | 0.49944526 |
| VAEHVTV | ASSLPGTGQDTQY | 0.6555085 |
| VDIVLNSL | ASSLPGTGQDTQY | 0.7055554 |
| VLTGGGAVL | ASSLPGTGQDTQY | 0.75308853 |
| VNVGLAYGI | ASSLPGTGQDTQY | 0.59089863 |

| | | |
|---|---|---|
| VSSAAVGAV | ASSLPGTGQDTQY | 0.7574681 |
| VTSVYEAGV | ASSLPGTGQDTQY | 0.58597064 |
| WVALDRGA | ASSLPGTGQDTQY | 0.74425507 |
| YLVGSVVI | ASSLPGTGQDTQY | 0.613099 |
| GPPGTGKTLL | ASSLPGTGQDTQY | 0.66238123 |
| GVGMTKF | ASSLPGTGQDTQY | 0.5476589 |
| IGIGGSDL | ASSLPGTGQDTQY | 0.73563933 |
| IGTLLAEL | ASSLPGTGQDTQY | 0.7488575 |
| IYTSGTT | ASSLPGTGQDTQY | 0.746776 |
| LSFPAGV | ASSLPGTGQDTQY | 0.27278486 |
| LYTSGTT | ASSLPGTGQDTQY | 0.74908245 |
| VGAFPFGF | ASSLPGTGQDTQY | 0.7400254 |
| VGAGTVEF | ASSLPGTGQDTQY | 0.77917063 |
| VGKSTLLL | ASSLPGTGQDTQY | 0.5966928 |
| AALAHAGGGIV | ASSLPGTGQDTQY | 0.66139627 |
| AALVAVSLL | ASSLPGTGQDTQY | 0.59120274 |
| AAVSRAVAAL | ASSLPGTGQDTQY | 0.6362603 |
| AGGSSGIGGF | ASSLPGTGQDTQY | 0.4141835 |
| AGLTVANANG | ASSLPGTGQDTQY | 0.53118527 |
| AGLVGAVL | ASSLPGTGQDTQY | 0.6572888 |
| ALSQAFTV | ASSLPGTGQDTQY | 0.5797899 |
| ALTFSVSL | ASSLPGTGQDTQY | 0.7405716 |
| ALTGAALVAL | ASSLPGTGQDTQY | 0.75895 |
| ASFSIGAGVL | ASSLPGTGQDTQY | 0.75917584 |
| DGSVYKI | ASSLPGTGQDTQY | 0.759259 |
| EQLAEIV | ASSLPGTGQDTQY | 0.7680082 |
| FAVSELTVM | ASSLPGTGQDTQY | 0.7378776 |
| GAAPLAAGL | ASSLPGTGQDTQY | 0.5877454 |
| GADGGDGGSGGAGG | ASSLPGTGQDTQY | 0.66998625 |
| GFGLDTGGAGG | ASSLPGTGQDTQY | 0.42990083 |
| GLASGGLL | ASSLPGTGQDTQY | 0.6430118 |
| GRIAGAAVLL | ASSLPGTGQDTQY | 0.5956813 |
| GSAGSLNTSF | ASSLPGTGQDTQY | 0.71341825 |
| GTGQIGI | ASSLPGTGQDTQY | 0.21574786 |
| GTLIALIL | ASSLPGTGQDTQY | 0.69099224 |
| IAATMGFVVG | ASSLPGTGQDTQY | 0.77812386 |
| IFAQALL | ASSLPGTGQDTQY | 0.63100535 |
| IGAAVALGF | ASSLPGTGQDTQY | 0.69428694 |
| IGLTPLAL | ASSLPGTGQDTQY | 0.7273653 |
| ILGSNTSTL | ASSLPGTGQDTQY | 0.5527322 |
| IVFSLTVL | ASSLPGTGQDTQY | 0.7305771 |
| LASKATGF | ASSLPGTGQDTQY | 0.5555121 |
| LASVALIL | ASSLPGTGQDTQY | 0.7355875 |
| LFAVGLTL | ASSLPGTGQDTQY | 0.7698704 |
| LFNGGIGTYI | ASSLPGTGQDTQY | 0.69315434 |
| LTLGAGII | ASSLPGTGQDTQY | 0.56817317 |
| LVAAALGDI | ASSLPGTGQDTQY | 0.60939705 |
| NTAFVQLGI | ASSLPGTGQDTQY | 0.4164248 |
| QLFSGAEL | ASSLPGTGQDTQY | 0.65841836 |
| SAATFITL | ASSLPGTGQDTQY | 0.5414358 |
| SNYGLANLGV | ASSLPGTGQDTQY | 0.67279553 |

| | | |
|---|---|---|
| SYGGAYAVM | ASSLPGTGQDTQY | 0.5281529 |
| TTGVGGLTLG | ASSLPGTGQDTQY | 0.718549 |
| TTQTIYI | ASSLPGTGQDTQY | 0.7034564 |
| VAAGAGLGAVI | ASSLPGTGQDTQY | 0.6515564 |
| VAAVVTLL | ASSLPGTGQDTQY | 0.50403166 |
| VGFLGGLI | ASSLPGTGQDTQY | 0.71523786 |
| VGVIPILL | ASSLPGTGQDTQY | 0.65948766 |
| VIFASSLI | ASSLPGTGQDTQY | 0.42125547 |
| VKGGDGGAAAG | ASSLPGTGQDTQY | 0.7357497 |
| VTKTFTNI | ASSLPGTGQDTQY | 0.55827427 |
| VVAAAILL | ASSLPGTGQDTQY | 0.6436354 |
| YAALDVEL | ASSLPGTGQDTQY | 0.75647354 |
| LASVALLL | ASSLPGTGQDTQY | 0.6855774 |
| LDAVHRV | ASSLPGTGQDTQY | 0.36198643 |
| QGLSSSIF | ASSLPGTGQDTQY | 0.6280385 |
| RGTFANIRL | ASSLPGTGQDTQY | 0.66919243 |
| RQLWWGHRI | ASSLPGTGQDTQY | 0.64643395 |
| YGPPGTGKTLL | ASSLPGTGQDTQY | 0.24697933 |
| AALVGAPANVA | ASSLPGTGQDTQY | 0.7200844 |
| AATFRRTE | ASSLPGTGQDTQY | 0.46886677 |
| AGALGGGI | ASSLPGTGQDTQY | 0.5083133 |
| AGGAGGAGGIGAG | ASSLPGTGQDTQY | 0.55714995 |
| AGSGALGL | ASSLPGTGQDTQY | 0.7565417 |
| AQIADQI | ASSLPGTGQDTQY | 0.6770085 |
| ASAVAVTYL | ASSLPGTGQDTQY | 0.5550167 |
| ASTVLGGTGG | ASSLPGTGQDTQY | 0.72430336 |
| AVGGEPAK | ASSLPGTGQDTQY | 0.7130009 |
| CEQGTFGGTC | ASSLPGTGQDTQY | 0.76616997 |
| DEGKAPAG | ASSLPGTGQDTQY | 0.68489456 |
| DTVVFPAL | ASSLPGTGQDTQY | 0.5803444 |
| FEGITMAG | ASSLPGTGQDTQY | 0.6543928 |
| FPLVVQD | ASSLPGTGQDTQY | 0.543545 |
| GAFTTPQ | ASSLPGTGQDTQY | 0.49909896 |
| GFGGFGTAGGVGGA | ASSLPGTGQDTQY | 0.3958975 |
| GFLAQFL | ASSLPGTGQDTQY | 0.6042078 |
| GGAGGAGGNAGVA | ASSLPGTGQDTQY | 0.42786127 |
| GGVGTAGAGGAGGAGG | ASSLPGTGQDTQY | 0.6885766 |
| GNGGGGDAGNAGSGGN | ASSLPGTGQDTQY | 0.73765534 |
| GTGGAGGAGNDAGS | ASSLPGTGQDTQY | 0.5948652 |
| GTTNTGWFNS | ASSLPGTGQDTQY | 0.37133306 |
| IFMGPLFTL | ASSLPGTGQDTQY | 0.58755875 |
| ILAHAIEAI | ASSLPGTGQDTQY | 0.62556565 |
| IRSAIREAL | ASSLPGTGQDTQY | 0.75075054 |
| ITFAGGG | ASSLPGTGQDTQY | 0.67443883 |
| ITGMSGHQEV | ASSLPGTGQDTQY | 0.7778449 |
| IVAGLACTL | ASSLPGTGQDTQY | 0.75287884 |
| IVGGGLAGA | ASSLPGTGQDTQY | 0.60659766 |
| IVTGVLNI | ASSLPGTGQDTQY | 0.74052703 |
| LAQQGCI | ASSLPGTGQDTQY | 0.5811367 |
| LCLHVEN | ASSLPGTGQDTQY | 0.6246172 |
| LDAISDGS | ASSLPGTGQDTQY | 0.66570497 |

| | | |
|---|---|---|
| LGGFAAML | ASSLPGTGQDTQY | 0.6523901 |
| LGGKNPMI | ASSLPGTGQDTQY | 0.5786506 |
| LGPKGVAAIL | ASSLPGTGQDTQY | 0.61690503 |
| LMVGVSI | ASSLPGTGQDTQY | 0.35251123 |
| LVGAAFVL | ASSLPGTGQDTQY | 0.70808285 |
| LVLAVAPL | ASSLPGTGQDTQY | 0.47949392 |
| MAAIAVIDAA | ASSLPGTGQDTQY | 0.3662963 |
| NNGINAELQ | ASSLPGTGQDTQY | 0.60421216 |
| RGEPLVLVHG | ASSLPGTGQDTQY | 0.7799728 |
| RGGAGGAAGLI | ASSLPGTGQDTQY | 0.76739025 |
| RSYRMHPAV | ASSLPGTGQDTQY | 0.58156335 |
| SAAHLLAGQLA | ASSLPGTGQDTQY | 0.519008 |
| SAAVGSY | ASSLPGTGQDTQY | 0.7032316 |
| SPHAVLHRI | ASSLPGTGQDTQY | 0.43695903 |
| SQIAKVL | ASSLPGTGQDTQY | 0.68501425 |
| TALGHLVP | ASSLPGTGQDTQY | 0.6847563 |
| TAPTTSGAA | ASSLPGTGQDTQY | 0.66685724 |
| TDGDDGNGGSPGT | ASSLPGTGQDTQY | 0.42255276 |
| TDGLGEN | ASSLPGTGQDTQY | 0.7129697 |
| TGGPPLPGQA | ASSLPGTGQDTQY | 0.6112211 |
| TIPAITI | ASSLPGTGQDTQY | 0.6782183 |
| TIPGFTI | ASSLPGTGQDTQY | 0.3322931 |
| TSIASGALA | ASSLPGTGQDTQY | 0.7295183 |
| TVTLPGAAI | ASSLPGTGQDTQY | 0.5528234 |
| VASGLSAALP | ASSLPGTGQDTQY | 0.46420982 |
| VGAGVIGI | ASSLPGTGQDTQY | 0.28419253 |
| VGPKRFGV | ASSLPGTGQDTQY | 0.591661 |
| VNVAFPD | ASSLPGTGQDTQY | 0.7252283 |
| VSMTNSGVS | ASSLPGTGQDTQY | 0.75858223 |
| VTDVLGDLSV | ASSLPGTGQDTQY | 0.6222131 |
| VVVVGAGFAGL | ASSLPGTGQDTQY | 0.7463083 |
| VYTVGTVV | ASSLPGTGQDTQY | 0.7645572 |
| WTTFSGP | ASSLPGTGQDTQY | 0.66401076 |
| YDELVEAF | ASSLPGTGQDTQY | 0.38467857 |
| YGTGGAGGNGG | ASSLPGTGQDTQY | 0.69293046 |
| YTNVMAV | ASSLPGTGQDTQY | 0.62268376 |
| FLEQVHQL | ASSLPGTGQDTQY | 0.7529385 |
| GLSLLAVAV | ASSLPGTGQDTQY | 0.71654046 |
| LEQVHQL | ASSLPGTGQDTQY | 0.44038588 |
| LNAAGAI | ASSLPGTGQDTQY | 0.66272414 |
| PAVHASL | ASSLPGTGQDTQY | 0.73887527 |
| TELEDDA | ASSLPGTGQDTQY | 0.4513851 |
| TPPGPPN | ASSLPGTGQDTQY | 0.44480264 |
| TPPPPPPP | ASSLPGTGQDTQY | 0.48511225 |
| TQSPALI | ASSSPGQNTGQLY | 0.4002607 |
| EFAAAFPQ | ASSSPGQNTGQLY | 0.7082076 |
| LLSGNLDVL | ASSSPGQNTGQLY | 0.63158053 |
| GPANLGNYNI | ASSSPGQNTGQLY | 0.5955592 |
| TLGPTLP | ASSSPGQNTGQLY | 0.6567987 |
| VYAYSSCCK | ASSSPGQNTGQLY | 0.530869 |
| APAPAPAPNI | ASSSPGQNTGQLY | 0.58371276 |

| | | |
|---|---|---|
| PGGPVPIP | ASSSPGQNTGQLY | 0.61992383 |
| FLTVALEP | ASSSPGQNTGQLY | 0.6871874 |
| SQYFGANL | ASSSPGQNTGQLY | 0.73230064 |
| GGVKPAA | ASSSPGQNTGQLY | 0.6233847 |
| GGTGGAGGAGGW | ASSSPGQNTGQLY | 0.56210124 |
| GVGGVAGLGGAGG | ASSSPGQNTGQLY | 0.77855146 |
| VLLVLATTI | ASSSPGQNTGQLY | 0.4250363 |
| IGIFAAML | ASSSPGQNTGQLY | 0.50590503 |
| AVGDLID | ASSSPGQNTGQLY | 0.7122382 |
| NISPLPL | ASSSPGQNTGQLY | 0.5001598 |
| VTAVFVAI | ASSSPGQNTGQLY | 0.74541056 |
| SGDDNDAAL | ASSSPGQNTGQLY | 0.6968513 |
| AVAALAHS | ASSSPGQNTGQLY | 0.72128373 |
| GGVTVMATP | ASSSPGQNTGQLY | 0.6557671 |
| AGLAANVRD | ASSSPGQNTGQLY | 0.7821063 |
| TNRPDLIDPALLRPG | ASSSPGQNTGQLY | 0.74683565 |
| LANLGVTAL | ASSSPGQNTGQLY | 0.5489111 |
| TGERIEL | ASSSPGQNTGQLY | 0.7415832 |
| NGEELAA | ASSSPGQNTGQLY | 0.3762507 |
| VLSLVWG | ASSSPGQNTGQLY | 0.74559104 |
| IGYALWL | ASSSPGQNTGQLY | 0.7075925 |
| LSLLLAPL | ASSSPGQNTGQLY | 0.5110179 |
| TLDEALK | ASSSPGQNTGQLY | 0.655708 |
| TDGAGTAEGG | ASSSPGQNTGQLY | 0.75552183 |
| NAPGGAGGNGG | ASSSPGQNTGQLY | 0.76626444 |
| VSVFMGMGGNF | ASSSPGQNTGQLY | 0.47339684 |
| LASYLAAL | ASSSPGQNTGQLY | 0.5970279 |
| LVGGTPAL | ASSSPGQNTGQLY | 0.3897041 |
| VRNADVL | ASSSPGQNTGQLY | 0.42312273 |
| TCSSPFLVSAM | ASSSPGQNTGQLY | 0.37671557 |
| VHITDAE | ASSSPGQNTGQLY | 0.7233056 |
| EYAAPSF | ASSSPGQNTGQLY | 0.6295581 |
| VVDAFHGGL | ASSSPGQNTGQLY | 0.72117436 |
| ASVVDPQ | ASSSPGQNTGQLY | 0.70002604 |
| SYNEFGGLS | ASSSPGQNTGQLY | 0.765044 |
| DPDGSVY | ASSSPGQNTGQLY | 0.55160356 |
| MVSTIHATE | ASSSPGQNTGQLY | 0.6436049 |
| LAAGVADPV | ASSSPGQNTGQLY | 0.33000314 |
| TKEQIAA | ASSSPGQNTGQLY | 0.5363506 |
| SLASPAAL | ASSSPGQNTGQLY | 0.47935924 |
| IGLVFYGV | ASSSPGQNTGQLY | 0.62113255 |
| FLLDASNL | ASSSPGQNTGQLY | 0.38561994 |
| YQVGNFP | ASSSPGQNTGQLY | 0.7672481 |
| ESLLSQST | ASSSPGQNTGQLY | 0.6508326 |
| LFTVGFAAVG | ASSSPGQNTGQLY | 0.73625547 |
| ATPPTPT | ASSSPGQNTGQLY | 0.5594287 |
| TDLATLG | ASSSPGQNTGQLY | 0.5145705 |
| SAGGTGR | ASSSPGQNTGQLY | 0.58790815 |
| GFSNSGS | ASSSPGQNTGQLY | 0.7256433 |
| GGNGGSGGTSV | ASSSPGQNTGQLY | 0.4892359 |
| GTSVATGGAGN | ASSSPGQNTGQLY | 0.28364184 |

| | | |
|---|---|---|
| NGGNGGSGGTS | ASSSPGQNTGQLY | 0.46569568 |
| AVVKFPH | ASSSPGQNTGQLY | 0.44931448 |
| TPPPPPP | ASSSPGQNTGQLY | 0.7689125 |
| AAPAAAAAPA | ASSSPGQNTGQLY | 0.7611599 |
| IALNEDL | ASSSPGQNTGQLY | 0.6964747 |
| IVNVSSM | ASSSPGQNTGQLY | 0.7594256 |
| TGIPPAPR | ASSSPGQNTGQLY | 0.23956029 |
| FDLGGGTF | ASSSPGQNTGQLY | 0.591985 |
| FDLGGGTFDVS | ASSSPGQNTGQLY | 0.76395524 |
| NEPTAAA | ASSSPGQNTGQLY | 0.71642476 |
| NEPTAAALA | ASSSPGQNTGQLY | 0.66462946 |
| GSIINISSI | ASSSPGQNTGQLY | 0.7657495 |
| VAPGNAGTA | ASSSPGQNTGQLY | 0.52490634 |
| TGGGASLE | ASSSPGQNTGQLY | 0.63923866 |
| VAVYNHYKRI | ASSSPGQNTGQLY | 0.6955277 |
| TDPAPATT | ASSSPGQNTGQLY | 0.72905314 |
| TNRPDILDPALLRPG | ASSSPGQNTGQLY | 0.6628524 |
| LTTTTTS | ASSSPGQNTGQLY | 0.44043124 |
| PTGSGKTL | ASSSPGQNTGQLY | 0.65563285 |
| TGSGKTL | ASSSPGQNTGQLY | 0.7934382 |
| TIGHVDHGK | ASSSPGQNTGQLY | 0.63654494 |
| TIGHVDHGKTT | ASSSPGQNTGQLY | 0.7472136 |
| GHVDHGKTT | ASSSPGQNTGQLY | 0.6951864 |
| AQFEHTLL | ASSSPGQNTGQLY | 0.68191624 |
| IVNQDEL | ASSSPGQNTGQLY | 0.53292805 |
| TNGSQFF | ASSSPGQNTGQLY | 0.5562674 |
| TGKGIVE | ASSSPGQNTGQLY | 0.7065672 |
| LISNASDA | ASSSPGQNTGQLY | 0.5502906 |
| AELRTLGELTKIAKAHGAQ | ASSSPGQNTGQLY | 0.58112997 |
| APNAVAAAEPSW | ASSSPGQNTGQLY | 0.57970047 |
| APQVAGAATAVL | ASSSPGQNTGQLY | 0.7909975 |
| ARRDPAAAAAILPTDA | ASSSPGQNTGQLY | 0.71413255 |
| DAAARDVLAEAGLAEYF | ASSSPGQNTGQLY | 0.7428944 |
| DDQIVLSAKANLAKVEQ | ASSSPGQNTGQLY | 0.5380772 |
| DELLPDTGEDSDEDGH | ASSSPGQNTGQLY | 0.65483475 |
| ELPEAPANRATVPQD | ASSSPGQNTGQLY | 0.72901267 |
| FGQSKIERVTGIG | ASSSPGQNTGQLY | 0.38748646 |
| GAGGTGSTAGGAGGAGGAGGLY | ASSSPGQNTGQLY | 0.7607553 |
| GHGGTGAAGGNGGTGGAGGSGI | ASSSPGQNTGQLY | 0.54955554 |
| GNGGNGGAGGEGGPGGAGGAGA | ASSSPGQNTGQLY | 0.639428 |
| GSTAGNGGNGGNATNGTIAGQP | ASSSPGQNTGQLY | 0.7022599 |
| GTGGDGGNGGNGGTGG | ASSSPGQNTGQLY | 0.7620013 |
| GVRRPAGIPADAPV | ASSSPGQNTGQLY | 0.63071394 |
| GYRQVTDAHLLTLARR | ASSSPGQNTGQLY | 0.69454485 |
| IAKMGQDGHDRG | ASSSPGQNTGQLY | 0.52981496 |
| KAEPSIEIQYT | ASSSPGQNTGQLY | 0.7182247 |
| KPVRIRMAADEDTVAVT | ASSSPGQNTGQLY | 0.7659769 |
| MPPRIAGMRLLVIKPEP | ASSSPGQNTGQLY | 0.66504145 |
| MKFYAVALFRSRGVPPTR | ASSSPGQNTGQLY | 0.7814441 |
| NAGGFGILGAVAHSP | ASSSPGQNTGQLY | 0.6356019 |
| NEGREITLAMAEM | ASSSPGQNTGQLY | 0.32449168 |

| | | |
|---|---|---|
| NELVEWSHPLGHRWRWE | ASSSPGQNTGQLY | 0.6452 |
| NLGAGGTGGNGGNPG | ASSSPGQNTGQLY | 0.7531401 |
| NTEWPKWETVMAADPVD | ASSSPGQNTGQLY | 0.7686181 |
| PASAAPAETQP | ASSSPGQNTGQLY | 0.75727755 |
| QQSSDGKNGVRAAEITGE | ASSSPGQNTGQLY | 0.69621956 |
| RAPDAVTTSWWKE | ASSSPGQNTGQLY | 0.61127806 |
| SGFGNVGAHTSGWFNQ | ASSSPGQNTGQLY | 0.768772 |
| SGVAGAVGQAASVGGLKV | ASSSPGQNTGQLY | 0.7742244 |
| SNGGAGTGSAGGNGGT | ASSSPGQNTGQLY | 0.7243314 |
| TGGAGGEGGAAG | ASSSPGQNTGQLY | 0.4954243 |
| TGGDGGAAGAGGAPGHG | ASSSPGQNTGQLY | 0.44224718 |
| TMPPLLAAIANAGRHGV | ASSSPGQNTGQLY | 0.6971892 |
| TSSTAGNSGGA | ASSSPGQNTGQLY | 0.7486683 |
| VPATVPVGILC | ASSSPGQNTGQLY | 0.39302674 |
| VQRPDSGGVVLTGRIS | ASSSPGQNTGQLY | 0.71165776 |
| WTPNPVPATKGVPAGLVQAT | ASSSPGQNTGQLY | 0.7027155 |
| YALKRALAKFSDRFDVVII | ASSSPGQNTGQLY | 0.31699434 |
| YLIGSVIV | ASSSPGQNTGQLY | 0.70216596 |
| YGQLVAVID | ASSSPGQNTGQLY | 0.76684225 |
| VTVGGMTLPA | ASSSPGQNTGQLY | 0.65841186 |
| VLALPAV | ASSSPGQNTGQLY | 0.47739446 |
| VIPPTLNYET | ASSSPGQNTGQLY | 0.7606647 |
| VGMGGMHPSSG | ASSSPGQNTGQLY | 0.7871995 |
| VGLAGGAATA | ASSSPGQNTGQLY | 0.7725986 |
| VGGAGGAGGSGP | ASSSPGQNTGQLY | 0.36836597 |
| VFAAFMV | ASSSPGQNTGQLY | 0.24111606 |
| TLYDKLR | ASSSPGQNTGQLY | 0.6655028 |
| TLAVFTVL | ASSSPGQNTGQLY | 0.7266482 |
| TGSVQPARV | ASSSPGQNTGQLY | 0.5940568 |
| TGGGAGSGGT | ASSSPGQNTGQLY | 0.52087635 |
| SHLETSL | ASSSPGQNTGQLY | 0.7144444 |
| SFTTGMT | ASSSPGQNTGQLY | 0.75197697 |
| QVEVAKRMGVRQPTVS | ASSSPGQNTGQLY | 0.45611927 |
| QGSGGPALAM | ASSSPGQNTGQLY | 0.36687806 |
| MLAPSNP | ASSSPGQNTGQLY | 0.7206695 |
| LVNNAAI | ASSSPGQNTGQLY | 0.71238494 |
| LTGEINTLA | ASSSPGQNTGQLY | 0.54197717 |
| LSIFGPV | ASSSPGQNTGQLY | 0.6382321 |
| LLALLANL | ASSSPGQNTGQLY | 0.69807816 |
| LISNASDALDKLR | ASSSPGQNTGQLY | 0.66088104 |
| LINEAVP | ASSSPGQNTGQLY | 0.70938456 |
| LGYAFSGL | ASSSPGQNTGQLY | 0.43072268 |
| LDFVAVAL | ASSSPGQNTGQLY | 0.7690908 |
| LAQPGAE | ASSSPGQNTGQLY | 0.44649088 |
| ISNLGAVTGF | ASSSPGQNTGQLY | 0.70783246 |
| HQEDPGRI | ASSSPGQNTGQLY | 0.70474416 |
| GTPGEPTP | ASSSPGQNTGQLY | 0.6953974 |
| GFFNSGS | ASSSPGQNTGQLY | 0.6365855 |
| GAGGGGLTAGAN | ASSSPGQNTGQLY | 0.545839 |
| FVDALVNM | ASSSPGQNTGQLY | 0.44634286 |
| FLTVAPEL | ASSSPGQNTGQLY | 0.36614048 |

| | | |
|---|---|---|
| FETLTAL | ASSSPGQNTGQLY | 0.76212335 |
| DVINPIEL | ASSSPGQNTGQLY | 0.6962944 |
| APVSGAPV | ASSSPGQNTGQLY | 0.756069 |
| ALVGGLAW | ASSSPGQNTGQLY | 0.72619665 |
| ALAAAAATAI | ASSSPGQNTGQLY | 0.57166624 |
| AFGLGLLL | ASSSPGQNTGQLY | 0.696322 |
| AFFAAASL | ASSSPGQNTGQLY | 0.472186 |
| AAAALPAFSPP | ASSSPGQNTGQLY | 0.65247875 |
| VVLAPAL | ASSSPGQNTGQLY | 0.6230264 |
| TIGHVDHGKTTL | ASSSPGQNTGQLY | 0.6966181 |
| LISNASDALDK | ASSSPGQNTGQLY | 0.5273651 |
| LISNASDAL | ASSSPGQNTGQLY | 0.6930284 |
| ISNASDALDK | ASSSPGQNTGQLY | 0.7122195 |
| NQAETLVYQTE | ASSSPGQNTGQLY | 0.4635225 |
| TGGTGGTAGAG | ASSSPGQNTGQLY | 0.62432545 |
| TGSTAGGAGGA | ASSSPGQNTGQLY | 0.76096797 |
| DPLAVPVAAGPV | ASSSPGQNTGQLY | 0.667976 |
| GGTGGTPGNGGN | ASSSPGQNTGQLY | 0.46876895 |
| PAGLAGVDPATL | ASSSPGQNTGQLY | 0.72917426 |
| GNNLVVEAPHAGASV | ASSSPGQNTGQLY | 0.5416842 |
| KVTRIMSASTLEQPA | ASSSPGQNTGQLY | 0.70685154 |
| LGFALPIALPRRTVV | ASSSPGQNTGQLY | 0.70193744 |
| TRLGNLTARKLVVLV | ASSSPGQNTGQLY | 0.52621275 |
| WSQQRRLIYFPSAGP | ASSSPGQNTGQLY | 0.731865 |
| GPNGNPDPMAAAVDIR | ASSSPGQNTGQLY | 0.3844132 |
| TQRCVSVADLRAAASR | ASSSPGQNTGQLY | 0.52799517 |
| QFVLDVAEHPVEVLTETV | ASSSPGQNTGQLY | 0.6105983 |
| GGAGGTGGAGGNGGTQAGDG | ASSSPGQNTGQLY | 0.6831104 |
| PLLATLRAIAVDVDAKPAQV | ASSSPGQNTGQLY | 0.74400914 |
| APKFRTGGYMIDFWGVGYQVAK | ASSSPGQNTGQLY | 0.51140255 |
| GAGGAGGAGGAAGGPGGTGGTG | ASSSPGQNTGQLY | 0.6919207 |
| IGALGLLATDVAPAGSVAFGVA | ASSSPGQNTGQLY | 0.59687173 |
| AGGAGGIGGNGGAGGAGGVLMG | ASSSPGQNTGQLY | 0.5137083 |
| ADELIGLL | ASSSPGQNTGQLY | 0.59559524 |
| ADRALKA | ASSSPGQNTGQLY | 0.5599947 |
| AGAVGQVGGAGGA | ASSSPGQNTGQLY | 0.4184498 |
| AGGGGGLGGAP | ASSSPGQNTGQLY | 0.72969294 |
| AGGLLYGNGGAG | ASSSPGQNTGQLY | 0.51850134 |
| AGLESIGL | ASSSPGQNTGQLY | 0.67864645 |
| AGLKTGII | ASSSPGQNTGQLY | 0.7603723 |
| ASLAALLL | ASSSPGQNTGQLY | 0.42281723 |
| ASVINISSI | ASSSPGQNTGQLY | 0.6896682 |
| ATPPGPA | ASSSPGQNTGQLY | 0.693774 |
| ATVASALV | ASSSPGQNTGQLY | 0.716803 |
| ATVASLIL | ASSSPGQNTGQLY | 0.64008856 |
| AVNGPVLIRH | ASSSPGQNTGQLY | 0.54990023 |
| DGAAGAVGI | ASSSPGQNTGQLY | 0.7628052 |
| DPLLHGI | ASSSPGQNTGQLY | 0.73706055 |
| ELGAALGAL | ASSSPGQNTGQLY | 0.5988151 |
| ELGRTQAV | ASSSPGQNTGQLY | 0.7132944 |
| FVAGGYLS | ASSSPGQNTGQLY | 0.72013277 |

| | | |
|---|---|---|
| GASGGAGGAGGW | ASSSPGQNTGQLY | 0.7419168 |
| GGAGAGGDGGAGGI | ASSSPGQNTGQLY | 0.61392856 |
| GTGHNGSQPGMGGNG | ASSSPGQNTGQLY | 0.34749967 |
| GYAAASASAS | ASSSPGQNTGQLY | 0.6445978 |
| HQVDVAAVQQG | ASSSPGQNTGQLY | 0.616338 |
| IGAAIAHT | ASSSPGQNTGQLY | 0.52477294 |
| IGMIEVAQL | ASSSPGQNTGQLY | 0.33812946 |
| IRLAPEHTSAMD | ASSSPGQNTGQLY | 0.75061035 |
| IYNVSVL | ASSSPGQNTGQLY | 0.7873169 |
| KQGFLGM | ASSSPGQNTGQLY | 0.5754754 |
| LESVITQL | ASSSPGQNTGQLY | 0.78411424 |
| LGLGALVVV | ASSSPGQNTGQLY | 0.38104582 |
| LIGNGGNGGSGGI | ASSSPGQNTGQLY | 0.74011195 |
| LNAINAPTL | ASSSPGQNTGQLY | 0.75167817 |
| LQLLNTGV | ASSSPGQNTGQLY | 0.53627586 |
| LQTSIEF | ASSSPGQNTGQLY | 0.7121692 |
| MSFAVSEL | ASSSPGQNTGQLY | 0.58505684 |
| NTGALGL | ASSSPGQNTGQLY | 0.586259 |
| PPGAGGAGGAGTT | ASSSPGQNTGQLY | 0.7588512 |
| QLVGLGI | ASSSPGQNTGQLY | 0.5519869 |
| RFLEQVHQL | ASSSPGQNTGQLY | 0.6947814 |
| RGVGYVLR | ASSSPGQNTGQLY | 0.60206014 |
| SATGVFAGVI | ASSSPGQNTGQLY | 0.5172549 |
| SLPAITLP | ASSSPGQNTGQLY | 0.704454 |
| STALAFTV | ASSSPGQNTGQLY | 0.7204194 |
| SVLTVGLGK | ASSSPGQNTGQLY | 0.58592653 |
| TALAGAGSGI | ASSSPGQNTGQLY | 0.7690783 |
| TGLPFTGI | ASSSPGQNTGQLY | 0.554345 |
| VAEHVTV | ASSSPGQNTGQLY | 0.7784753 |
| VDIVLNSL | ASSSPGQNTGQLY | 0.74623567 |
| VLTGGGAVL | ASSSPGQNTGQLY | 0.6939248 |
| VNVGLAYGI | ASSSPGQNTGQLY | 0.5859195 |
| VSSAAVGAV | ASSSPGQNTGQLY | 0.7641274 |
| VTSVYEAGV | ASSSPGQNTGQLY | 0.56672645 |
| WVALDRGA | ASSSPGQNTGQLY | 0.67734283 |
| YLVGSVVI | ASSSPGQNTGQLY | 0.76186705 |
| GPPGTGKTLL | ASSSPGQNTGQLY | 0.33667532 |
| GVGMTKF | ASSSPGQNTGQLY | 0.4677181 |
| IGIGGSDL | ASSSPGQNTGQLY | 0.41772103 |
| IGTLLAEL | ASSSPGQNTGQLY | 0.46856242 |
| IYTSGTT | ASSSPGQNTGQLY | 0.3435094 |
| LSFPAGV | ASSSPGQNTGQLY | 0.28158912 |
| LYTSGTT | ASSSPGQNTGQLY | 0.3947282 |
| VGAFPFGF | ASSSPGQNTGQLY | 0.29616123 |
| VGAGTVEF | ASSSPGQNTGQLY | 0.5643904 |
| VGKSTLLL | ASSSPGQNTGQLY | 0.5235324 |
| AALAHAGGGIV | ASSSPGQNTGQLY | 0.34080768 |
| AALVAVSLL | ASSSPGQNTGQLY | 0.53285444 |
| AAVSRAVAAL | ASSSPGQNTGQLY | 0.49744123 |
| AGGSSGIGGF | ASSSPGQNTGQLY | 0.270041 |
| AGLTVANANG | ASSSPGQNTGQLY | 0.35865372 |

| | | |
|---|---|---|
| AGLVGAVL | ASSSPGQNTGQLY | 0.313676 |
| ALSQAFTV | ASSSPGQNTGQLY | 0.41173536 |
| ALTFSVSL | ASSSPGQNTGQLY | 0.24745853 |
| ALTGAALVAL | ASSSPGQNTGQLY | 0.43823105 |
| ASFSIGAGVL | ASSSPGQNTGQLY | 0.27910027 |
| DGSVYKI | ASSSPGQNTGQLY | 0.21756814 |
| EQLAEIV | ASSSPGQNTGQLY | 0.4600273 |
| FAVSELTVM | ASSSPGQNTGQLY | 0.37570527 |
| GAAPLAAGL | ASSSPGQNTGQLY | 0.42150038 |
| GADGGDGGSGGAGG | ASSSPGQNTGQLY | 0.5213355 |
| GFGLDTGGAGG | ASSSPGQNTGQLY | 0.31756338 |
| GLASGGLL | ASSSPGQNTGQLY | 0.30761248 |
| GRIAGAAVLL | ASSSPGQNTGQLY | 0.41661838 |
| GSAGSLNTSF | ASSSPGQNTGQLY | 0.623493 |
| GTGQIGI | ASSSPGQNTGQLY | 0.3102495 |
| GTLIALIL | ASSSPGQNTGQLY | 0.4203674 |
| IAATMGFVVG | ASSSPGQNTGQLY | 0.5352262 |
| IFAQALL | ASSSPGQNTGQLY | 0.4322415 |
| IGAAVALGF | ASSSPGQNTGQLY | 0.54702747 |
| IGLTPLAL | ASSSPGQNTGQLY | 0.5791867 |
| ILGSNTSTL | ASSSPGQNTGQLY | 0.3176505 |
| IVFSLTVL | ASSSPGQNTGQLY | 0.57581687 |
| LASKATGF | ASSSPGQNTGQLY | 0.5490393 |
| LASVALIL | ASSSPGQNTGQLY | 0.3793161 |
| LFAVGLTL | ASSSPGQNTGQLY | 0.3695617 |
| LFNGGIGTYI | ASSSPGQNTGQLY | 0.32992557 |
| LTLGAGII | ASSSPGQNTGQLY | 0.37672532 |
| LVAAALGDI | ASSSPGQNTGQLY | 0.24807814 |
| NTAFVQLGI | ASSSPGQNTGQLY | 0.4075415 |
| QLFSGAEL | ASSSPGQNTGQLY | 0.5213658 |
| SAATFITL | ASSSPGQNTGQLY | 0.35449946 |
| SNYGLANLGV | ASSSPGQNTGQLY | 0.27381742 |
| SYGGAYAVM | ASSSPGQNTGQLY | 0.27142617 |
| TTGVGGLTLG | ASSSPGQNTGQLY | 0.34830114 |
| TTQTIYI | ASSSPGQNTGQLY | 0.35418996 |
| VAAGAGLGAVI | ASSSPGQNTGQLY | 0.2562123 |
| VAAVVTLL | ASSSPGQNTGQLY | 0.3514728 |
| VGFLGGLI | ASSSPGQNTGQLY | 0.51628965 |
| VGVIPILL | ASSSPGQNTGQLY | 0.5013668 |
| VIFASSLI | ASSSPGQNTGQLY | 0.24837746 |
| VKGGDGGAAAG | ASSSPGQNTGQLY | 0.5958853 |
| VTKTFTNI | ASSSPGQNTGQLY | 0.42289394 |
| VVAAAILL | ASSSPGQNTGQLY | 0.47246853 |
| YAALDVEL | ASSSPGQNTGQLY | 0.38718683 |
| LASVALLL | ASSSPGQNTGQLY | 0.34030327 |
| LDAVHRV | ASSSPGQNTGQLY | 0.5222217 |
| QGLSSSIF | ASSSPGQNTGQLY | 0.32958338 |
| RGTFANIRL | ASSSPGQNTGQLY | 0.37352455 |
| RQLWWGHRI | ASSSPGQNTGQLY | 0.50038296 |
| YGPPGTGKTLL | ASSSPGQNTGQLY | 0.36965528 |
| AALVGAPANVA | ASSSPGQNTGQLY | 0.29306865 |

| | | |
|---|---|---|
| AATFRRTE | ASSSPGQNTGQLY | 0.3329749 |
| AGALGGGI | ASSSPGQNTGQLY | 0.50497186 |
| AGGAGGAGGIGAG | ASSSPGQNTGQLY | 0.24809651 |
| AGSGALGL | ASSSPGQNTGQLY | 0.36534548 |
| AQIADQI | ASSSPGQNTGQLY | 0.29000524 |
| ASAVAVTYL | ASSSPGQNTGQLY | 0.33221245 |
| ASTVLGGTGG | ASSSPGQNTGQLY | 0.5878441 |
| AVGGEPAK | ASSSPGQNTGQLY | 0.4662868 |
| CEQGTFGGTC | ASSSPGQNTGQLY | 0.33865523 |
| DEGKAPAG | ASSSPGQNTGQLY | 0.4989042 |
| DTVVFPAL | ASSSPGQNTGQLY | 0.41052207 |
| FEGITMAG | ASSSPGQNTGQLY | 0.38092342 |
| FPLVVQD | ASSSPGQNTGQLY | 0.46965146 |
| GAFTTPQ | ASSSPGQNTGQLY | 0.28679976 |
| GFGGFGTAGGVGGA | ASSSPGQNTGQLY | 0.48681128 |
| GFLAQFL | ASSSPGQNTGQLY | 0.4340957 |
| GGAGGAGGNAGVA | ASSSPGQNTGQLY | 0.51370984 |
| GGVGTAGAGGAGGAGG | ASSSPGQNTGQLY | 0.52388287 |
| GNGGGGDAGNAGSGGN | ASSSPGQNTGQLY | 0.39753854 |
| GTGGAGGAGNDAGS | ASSSPGQNTGQLY | 0.3674327 |
| GTTNTGWFNS | ASSSPGQNTGQLY | 0.2518845 |
| IFMGPLFTL | ASSSPGQNTGQLY | 0.32467285 |
| ILAHAIEAI | ASSSPGQNTGQLY | 0.45151874 |
| IRSAIREAL | ASSSPGQNTGQLY | 0.38623437 |
| ITFAGGG | ASSSPGQNTGQLY | 0.5202638 |
| ITGMSGHQEV | ASSSPGQNTGQLY | 0.48264173 |
| IVAGLACTL | ASSSPGQNTGQLY | 0.30200937 |
| IVGGGLAGA | ASSSPGQNTGQLY | 0.5059046 |
| IVTGVLNI | ASSSPGQNTGQLY | 0.52543175 |
| LAQQGCI | ASSSPGQNTGQLY | 0.23965567 |
| LCLHVEN | ASSSPGQNTGQLY | 0.3705146 |
| LDAISDGS | ASSSPGQNTGQLY | 0.64027864 |
| LGGFAAML | ASSSPGQNTGQLY | 0.46904725 |
| LGGKNPMI | ASSSPGQNTGQLY | 0.36478907 |
| LGPKGVAAIL | ASSSPGQNTGQLY | 0.42746386 |
| LMVGVSI | ASSSPGQNTGQLY | 0.6436838 |
| LVGAAFVL | ASSSPGQNTGQLY | 0.4069071 |
| LVLAVAPL | ASSSPGQNTGQLY | 0.5619534 |
| MAAIAVIDAA | ASSSPGQNTGQLY | 0.2310418 |
| NNGINAELQ | ASSSPGQNTGQLY | 0.39035356 |
| RGEPLVLVHG | ASSSPGQNTGQLY | 0.29013804 |
| RGGAGGAAGLI | ASSSPGQNTGQLY | 0.2615539 |
| RSYRMHPAV | ASSSPGQNTGQLY | 0.4790822 |
| SAAHLLAGQLA | ASSSPGQNTGQLY | 0.394869 |
| SAAVGSY | ASSSPGQNTGQLY | 0.5001258 |
| SPHAVLHRI | ASSSPGQNTGQLY | 0.3610875 |
| SQIAKVL | ASSSPGQNTGQLY | 0.54756737 |
| TALGHLVP | ASSSPGQNTGQLY | 0.33004016 |
| TAPTTSGAA | ASSSPGQNTGQLY | 0.42975408 |
| TDGDDGNGGSPGT | ASSSPGQNTGQLY | 0.40305623 |
| TDGLGEN | ASSSPGQNTGQLY | 0.46188048 |

| | | |
|---|---|---|
| TGGPPLPGQA | ASSSPGQNTGQLY | 0.5352088 |
| TIPAITI | ASSSPGQNTGQLY | 0.5159303 |
| TIPGFTI | ASSSPGQNTGQLY | 0.27216005 |
| TSIASGALA | ASSSPGQNTGQLY | 0.41236797 |
| TVTLPGAAI | ASSSPGQNTGQLY | 0.4058566 |
| VASGLSAALP | ASSSPGQNTGQLY | 0.2648856 |
| VGAGVIGI | ASSSPGQNTGQLY | 0.2719405 |
| VGPKRFGV | ASSSPGQNTGQLY | 0.31786326 |
| VNVAFPD | ASSSPGQNTGQLY | 0.29604578 |
| VSMTNSGVS | ASSSPGQNTGQLY | 0.5097318 |
| VTDVLGDLSV | ASSSPGQNTGQLY | 0.43613574 |
| VVVVGAGFAGL | ASSSPGQNTGQLY | 0.35682157 |
| VYTVGTVV | ASSSPGQNTGQLY | 0.42122808 |
| WTTFSGP | ASSSPGQNTGQLY | 0.38053986 |
| YDELVEAF | ASSSPGQNTGQLY | 0.20224589 |
| YGTGGAGGNGG | ASSSPGQNTGQLY | 0.30814132 |
| YTNVMAV | ASSSPGQNTGQLY | 0.28016478 |
| FLEQVHQL | ASSSPGQNTGQLY | 0.54326034 |
| GLSLLAVAV | ASSSPGQNTGQLY | 0.34136716 |
| LEQVHQL | ASSSPGQNTGQLY | 0.28063416 |
| LNAAGAI | ASSSPGQNTGQLY | 0.36414447 |
| PAVHASL | ASSSPGQNTGQLY | 0.44060454 |
| TELEDDA | ASSSPGQNTGQLY | 0.35606217 |
| TPPGPPN | ASSSPGQNTGQLY | 0.28090262 |
| TPPPPPPP | ASSSPGQNTGQLY | 0.22750512 |
| TQSPALI | ALMERGGDTDKLV | 0.4832701 |
| EFAAAFPQ | ALMERGGDTDKLV | 0.29765475 |
| LLSGNLDVL | ALMERGGDTDKLV | 0.34573877 |
| GPANLGNYNI | ALMERGGDTDKLV | 0.25916663 |
| TLGPTLP | ALMERGGDTDKLV | 0.33204862 |
| VYAYSSCCK | ALMERGGDTDKLV | 0.24286988 |
| APAPAPAPNI | ALMERGGDTDKLV | 0.5361373 |
| PGGPVPIP | ALMERGGDTDKLV | 0.3393228 |
| FLTVALEP | ALMERGGDTDKLV | 0.31754816 |
| SQYFGANL | ALMERGGDTDKLV | 0.53666496 |
| GGVKPAA | ALMERGGDTDKLV | 0.40217635 |
| GGTGGAGGAGGW | ALMERGGDTDKLV | 0.34374046 |
| GVGGVAGLGGAGG | ALMERGGDTDKLV | 0.34795383 |
| VLLVLATTI | ALMERGGDTDKLV | 0.31415606 |
| IGIFAAML | ALMERGGDTDKLV | 0.30832532 |
| AVGDLID | ALMERGGDTDKLV | 0.33059815 |
| NISPLPL | ALMERGGDTDKLV | 0.5767859 |
| VTAVFVAI | ALMERGGDTDKLV | 0.4321924 |
| SGDDNDAAL | ALMERGGDTDKLV | 0.29533496 |
| AVAALAHS | ALMERGGDTDKLV | 0.45969817 |
| GGVTVMATP | ALMERGGDTDKLV | 0.49763435 |
| AGLAANVRD | ALMERGGDTDKLV | 0.5761807 |
| TNRPDLIDPALLRPG | ALMERGGDTDKLV | 0.34230098 |
| LANLGVTAL | ALMERGGDTDKLV | 0.33042583 |
| TGERIEL | ALMERGGDTDKLV | 0.4057939 |
| NGEELAA | ALMERGGDTDKLV | 0.35799822 |

| | | |
|---|---|---|
| VLSLVWG | ALMERGGDTDKLV | 0.37270492 |
| IGYALWL | ALMERGGDTDKLV | 0.32264692 |
| LSLLLAPL | ALMERGGDTDKLV | 0.4756055 |
| TLDEALK | ALMERGGDTDKLV | 0.67088175 |
| TDGAGTAEGG | ALMERGGDTDKLV | 0.3359997 |
| NAPGGAGGNGG | ALMERGGDTDKLV | 0.24019358 |
| VSVFMGMGGNF | ALMERGGDTDKLV | 0.3664344 |
| LASYLAAL | ALMERGGDTDKLV | 0.2600206 |
| LVGGTPAL | ALMERGGDTDKLV | 0.31018034 |
| VRNADVL | ALMERGGDTDKLV | 0.42475402 |
| TCSSPFLVSAM | ALMERGGDTDKLV | 0.251258 |
| VHITDAE | ALMERGGDTDKLV | 0.3573342 |
| EYAAPSF | ALMERGGDTDKLV | 0.54409045 |
| VVDAFHGGL | ALMERGGDTDKLV | 0.3046993 |
| ASVVDPQ | ALMERGGDTDKLV | 0.46994182 |
| SYNEFGGLS | ALMERGGDTDKLV | 0.4991741 |
| DPDGSVY | ALMERGGDTDKLV | 0.45482296 |
| MVSTIHATE | ALMERGGDTDKLV | 0.5851388 |
| LAAGVADPV | ALMERGGDTDKLV | 0.52074534 |
| TKEQIAA | ALMERGGDTDKLV | 0.4574639 |
| SLASPAAL | ALMERGGDTDKLV | 0.33787978 |
| IGLVFYGV | ALMERGGDTDKLV | 0.4566512 |
| FLLDASNL | ALMERGGDTDKLV | 0.38797942 |
| YQVGNFP | ALMERGGDTDKLV | 0.3814363 |
| ESLLSQST | ALMERGGDTDKLV | 0.35071158 |
| LFTVGFAAVG | ALMERGGDTDKLV | 0.47077957 |
| ATPPTPT | ALMERGGDTDKLV | 0.3271305 |
| TDLATLG | ALMERGGDTDKLV | 0.5286998 |
| SAGGTGR | ALMERGGDTDKLV | 0.2931909 |
| GFSNSGS | ALMERGGDTDKLV | 0.32395333 |
| GGNGGSGGTSV | ALMERGGDTDKLV | 0.50387746 |
| GTSVATGGAGN | ALMERGGDTDKLV | 0.21175092 |
| NGGNGGSGGTS | ALMERGGDTDKLV | 0.29415035 |
| AVVKFPH | ALMERGGDTDKLV | 0.3906726 |
| TPPPPPP | ALMERGGDTDKLV | 0.45255774 |
| AAPAAAAAPA | ALMERGGDTDKLV | 0.21962766 |
| IALNEDL | ALMERGGDTDKLV | 0.30928728 |
| IVNVSSM | ALMERGGDTDKLV | 0.29716793 |
| TGIPPAPR | ALMERGGDTDKLV | 0.50107694 |
| FDLGGGTF | ALMERGGDTDKLV | 0.39452016 |
| FDLGGGTFDVS | ALMERGGDTDKLV | 0.25049412 |
| NEPTAAA | ALMERGGDTDKLV | 0.5136127 |
| NEPTAAALA | ALMERGGDTDKLV | 0.22712617 |
| GSIINISSI | ALMERGGDTDKLV | 0.3401702 |
| VAPGNAGTA | ALMERGGDTDKLV | 0.5254793 |
| TGGGASLE | ALMERGGDTDKLV | 0.6166849 |
| VAVYNHYKRI | ALMERGGDTDKLV | 0.34480032 |
| TDPAPATT | ALMERGGDTDKLV | 0.5147791 |
| TNRPDILDPALLRPG | ALMERGGDTDKLV | 0.28733042 |
| LTTTTTS | ALMERGGDTDKLV | 0.64349395 |
| PTGSGKTL | ALMERGGDTDKLV | 0.37534055 |

| | | |
|---|---|---|
| TGSGKTL | ALMERGGDTDKLV | 0.2894495 |
| TIGHVDHGK | ALMERGGDTDKLV | 0.48703653 |
| TIGHVDHGKTT | ALMERGGDTDKLV | 0.47620884 |
| GHVDHGKTT | ALMERGGDTDKLV | 0.4574668 |
| AQFEHTLL | ALMERGGDTDKLV | 0.45948386 |
| IVNQDEL | ALMERGGDTDKLV | 0.3744291 |
| TNGSQFF | ALMERGGDTDKLV | 0.3346189 |
| TGKGIVE | ALMERGGDTDKLV | 0.54106736 |
| LISNASDA | ALMERGGDTDKLV | 0.49213675 |
| AELRTLGELTKIAKAHGAQ | ALMERGGDTDKLV | 0.34356272 |
| APNAVAAAEPSW | ALMERGGDTDKLV | 0.58124906 |
| APQVAGAATAVL | ALMERGGDTDKLV | 0.3801506 |
| ARRDPAAAAAILPTDA | ALMERGGDTDKLV | 0.34839696 |
| DAAARDVLAEAGLAEYF | ALMERGGDTDKLV | 0.554764 |
| DDQIVLSAKANLAKVEQ | ALMERGGDTDKLV | 0.4375141 |
| DELLPDTGEDSDEDGH | ALMERGGDTDKLV | 0.44333836 |
| ELPEAPANRATVPQD | ALMERGGDTDKLV | 0.450762 |
| FGQSKIERVTGIG | ALMERGGDTDKLV | 0.6170107 |
| GAGGTGSTAGGAGGAGGAGGLY | ALMERGGDTDKLV | 0.2980016 |
| GHGGTGAAGGNGGTGGAGGSGI | ALMERGGDTDKLV | 0.4172105 |
| GNGGNGGAGGEGGPGGAGGAGA | ALMERGGDTDKLV | 0.37452117 |
| GSTAGNGGNGGNATNGTIAGQP | ALMERGGDTDKLV | 0.5344239 |
| GTGGDGGNGGNGGTGG | ALMERGGDTDKLV | 0.41010222 |
| GVRRPAGIPADAPV | ALMERGGDTDKLV | 0.36376554 |
| GYRQVTDAHLLTLARR | ALMERGGDTDKLV | 0.44807258 |
| IAKMGQDGHDRG | ALMERGGDTDKLV | 0.49232206 |
| KAEPSIEIQYT | ALMERGGDTDKLV | 0.40116686 |
| KPVRIRMAADEDTVAVT | ALMERGGDTDKLV | 0.42422107 |
| MPPRIAGMRLLVIKPEP | ALMERGGDTDKLV | 0.620438 |
| MKFYAVALFRSRGVPPTR | ALMERGGDTDKLV | 0.37977433 |
| NAGGFGILGAVAHSP | ALMERGGDTDKLV | 0.36118475 |
| NEGREITLAMAEM | ALMERGGDTDKLV | 0.47195107 |
| NELVEWSHPLGHRWRWE | ALMERGGDTDKLV | 0.5299804 |
| NLGAGGTGGNGGNPG | ALMERGGDTDKLV | 0.41811535 |
| NTEWPKWETVMAADPVD | ALMERGGDTDKLV | 0.57210857 |
| PASAAPAETQP | ALMERGGDTDKLV | 0.2696814 |
| QQSSDGKNGVRAAEITGE | ALMERGGDTDKLV | 0.50973296 |
| RAPDAVTTSWWKE | ALMERGGDTDKLV | 0.27397826 |
| SGFGNVGAHTSGWFNQ | ALMERGGDTDKLV | 0.4246943 |
| SGVAGAVGQAASVGGLKV | ALMERGGDTDKLV | 0.309686 |
| SNGGAGTGSAGGNGGT | ALMERGGDTDKLV | 0.47392058 |
| TGGAGGEGGAAG | ALMERGGDTDKLV | 0.42645982 |
| TGGDGGAAGAGGAPGHG | ALMERGGDTDKLV | 0.486931 |
| TMPPLLAAIANAGRHGV | ALMERGGDTDKLV | 0.24445547 |
| TSSTAGNSGGA | ALMERGGDTDKLV | 0.44475293 |
| VPATVPVGILC | ALMERGGDTDKLV | 0.49622574 |
| VQRPDSGGVVLTGRIS | ALMERGGDTDKLV | 0.6473871 |
| WTPNPVPATKGVPAGLVQAT | ALMERGGDTDKLV | 0.3349097 |
| YALKRALAKFSDRFDVVII | ALMERGGDTDKLV | 0.5032922 |
| YLIGSVIV | ALMERGGDTDKLV | 0.33581918 |
| YGQLVAVID | ALMERGGDTDKLV | 0.3978187 |

| | | |
|---|---|---|
| VTVGGMTLPA | ALMERGGDTDKLV | 0.25861633 |
| VLALPAV | ALMERGGDTDKLV | 0.36034298 |
| VIPPTLNYET | ALMERGGDTDKLV | 0.30463818 |
| VGMGGMHPSSG | ALMERGGDTDKLV | 0.30930102 |
| VGLAGGAATA | ALMERGGDTDKLV | 0.50724185 |
| VGGAGGAGGSGP | ALMERGGDTDKLV | 0.54790837 |
| VFAAFMV | ALMERGGDTDKLV | 0.48170742 |
| TLYDKLR | ALMERGGDTDKLV | 0.46424657 |
| TLAVFTVL | ALMERGGDTDKLV | 0.49322468 |
| TGSVQPARV | ALMERGGDTDKLV | 0.54516685 |
| TGGGAGSGGT | ALMERGGDTDKLV | 0.31977898 |
| SHLETSL | ALMERGGDTDKLV | 0.38399208 |
| SFTTGMT | ALMERGGDTDKLV | 0.26843405 |
| QVEVAKRMGVRQPTVS | ALMERGGDTDKLV | 0.3287836 |
| QGSGGPALAM | ALMERGGDTDKLV | 0.30997872 |
| MLAPSNP | ALMERGGDTDKLV | 0.3979045 |
| LVNNAAI | ALMERGGDTDKLV | 0.22251227 |
| LTGEINTLA | ALMERGGDTDKLV | 0.36610523 |
| LSIFGPV | ALMERGGDTDKLV | 0.5850012 |
| LLALLANL | ALMERGGDTDKLV | 0.32646486 |
| LISNASDALDKLR | ALMERGGDTDKLV | 0.4142687 |
| LINEAVP | ALMERGGDTDKLV | 0.39556155 |
| LGYAFSGL | ALMERGGDTDKLV | 0.30151764 |
| LDFVAVAL | ALMERGGDTDKLV | 0.42717156 |
| LAQPGAE | ALMERGGDTDKLV | 0.29950917 |
| ISNLGAVTGF | ALMERGGDTDKLV | 0.49954775 |
| HQEDPGRI | ALMERGGDTDKLV | 0.54039335 |
| GTPGEPTP | ALMERGGDTDKLV | 0.3997818 |
| GFFNSGS | ALMERGGDTDKLV | 0.38083676 |
| GAGGGGLTAGAN | ALMERGGDTDKLV | 0.45008448 |
| FVDALVNM | ALMERGGDTDKLV | 0.2862642 |
| FLTVAPEL | ALMERGGDTDKLV | 0.43880826 |
| FETLTAL | ALMERGGDTDKLV | 0.3111003 |
| DVINPIEL | ALMERGGDTDKLV | 0.34989855 |
| APVSGAPV | ALMERGGDTDKLV | 0.29376176 |
| ALVGGLAW | ALMERGGDTDKLV | 0.47988254 |
| ALAAAAATAI | ALMERGGDTDKLV | 0.3738149 |
| AFGLGLLL | ALMERGGDTDKLV | 0.40668774 |
| AFFAAASL | ALMERGGDTDKLV | 0.28433916 |
| AAAALPAFSPP | ALMERGGDTDKLV | 0.4317031 |
| VVLAPAL | ALMERGGDTDKLV | 0.32966906 |
| TIGHVDHGKTTL | ALMERGGDTDKLV | 0.28163296 |
| LISNASDALDK | ALMERGGDTDKLV | 0.47332639 |
| LISNASDAL | ALMERGGDTDKLV | 0.37655044 |
| ISNASDALDK | ALMERGGDTDKLV | 0.30700988 |
| NQAETLVYQTE | ALMERGGDTDKLV | 0.49252146 |
| TGGTGGTAGAG | ALMERGGDTDKLV | 0.46036044 |
| TGSTAGGAGGA | ALMERGGDTDKLV | 0.2511761 |
| DPLAVPVAAGPV | ALMERGGDTDKLV | 0.30376932 |
| GGTGGTPGNGGN | ALMERGGDTDKLV | 0.37565437 |
| PAGLAGVDPATL | ALMERGGDTDKLV | 0.370394 |

| | | |
|---|---|---|
| GNNLVVEAPHAGASV | ALMERGGDTDKLV | 0.22198531 |
| KVTRIMSASTLEQPA | ALMERGGDTDKLV | 0.44105017 |
| LGFALPIALPRRTVV | ALMERGGDTDKLV | 0.35178733 |
| TRLGNLTARKLVVLV | ALMERGGDTDKLV | 0.25949925 |
| WSQQRRLIYFPSAGP | ALMERGGDTDKLV | 0.36095864 |
| GPNGNPDPMAAAVDIR | ALMERGGDTDKLV | 0.53656256 |
| TQRCVSVADLRAAASR | ALMERGGDTDKLV | 0.4345875 |
| QFVLDVAEHPVEVLTETV | ALMERGGDTDKLV | 0.44234967 |
| GGAGGTGGAGGNGGTQAGDG | ALMERGGDTDKLV | 0.4691103 |
| PLLATLRAIAVDVDAKPAQV | ALMERGGDTDKLV | 0.45849243 |
| APKFRTGGYMIDFWGVGYQVAK | ALMERGGDTDKLV | 0.56673115 |
| GAGGAGGAGGAAGGPGGTGGTG | ALMERGGDTDKLV | 0.49563292 |
| IGALGLLATDVAPAGSVAFGVA | ALMERGGDTDKLV | 0.34524956 |
| AGGAGGIGGNGGAGGAGGVLMG | ALMERGGDTDKLV | 0.44306853 |
| ADELIGLL | ALMERGGDTDKLV | 0.38248783 |
| ADRALKA | ALMERGGDTDKLV | 0.32796437 |
| AGAVGQVGGAGGA | ALMERGGDTDKLV | 0.26230982 |
| AGGGGGLGGAP | ALMERGGDTDKLV | 0.35924593 |
| AGGLLYGNGGAG | ALMERGGDTDKLV | 0.29258537 |
| AGLESIGL | ALMERGGDTDKLV | 0.41624632 |
| AGLKTGII | ALMERGGDTDKLV | 0.5626617 |
| ASLAALLL | ALMERGGDTDKLV | 0.34830707 |
| ASVINISSI | ALMERGGDTDKLV | 0.28851238 |
| ATPPGPA | ALMERGGDTDKLV | 0.32034898 |
| ATVASALV | ALMERGGDTDKLV | 0.31414333 |
| ATVASLIL | ALMERGGDTDKLV | 0.5097673 |
| AVNGPVLIRH | ALMERGGDTDKLV | 0.40368354 |
| DGAAGAVGI | ALMERGGDTDKLV | 0.63385737 |
| DPLLHGI | ALMERGGDTDKLV | 0.5010497 |
| ELGAALGAL | ALMERGGDTDKLV | 0.3005744 |
| ELGRTQAV | ALMERGGDTDKLV | 0.50453323 |
| FVAGGYLS | ALMERGGDTDKLV | 0.33737585 |
| GASGGAGGAGGW | ALMERGGDTDKLV | 0.37005013 |
| GGAGAGGDGGAGGI | ALMERGGDTDKLV | 0.41479257 |
| GTGHNGSQPGMGGNG | ALMERGGDTDKLV | 0.40141612 |
| GYAAASASAS | ALMERGGDTDKLV | 0.41909954 |
| HQVDVAAVQQG | ALMERGGDTDKLV | 0.42980227 |
| IGAAIAHT | ALMERGGDTDKLV | 0.26257566 |
| IGMIEVAQL | ALMERGGDTDKLV | 0.44663334 |
| IRLAPEHTSAMD | ALMERGGDTDKLV | 0.36116582 |
| IYNVSVL | ALMERGGDTDKLV | 0.28089094 |
| KQGFLGM | ALMERGGDTDKLV | 0.34984103 |
| LESVITQL | ALMERGGDTDKLV | 0.669674 |
| LGLGALVVV | ALMERGGDTDKLV | 0.5100027 |
| LIGNGGNGGSGGI | ALMERGGDTDKLV | 0.27582422 |
| LNAINAPTL | ALMERGGDTDKLV | 0.29972568 |
| LQLLNTGV | ALMERGGDTDKLV | 0.38799855 |
| LQTSIEF | ALMERGGDTDKLV | 0.2763681 |
| MSFAVSEL | ALMERGGDTDKLV | 0.47598475 |
| NTGALGL | ALMERGGDTDKLV | 0.31845495 |
| PPGAGGAGGAGTT | ALMERGGDTDKLV | 0.33421388 |

| | | |
|---|---|---|
| QLVGLGI | ALMERGGDTDKLV | 0.3783572 |
| RFLEQVHQL | ALMERGGDTDKLV | 0.41267917 |
| RGVGYVLR | ALMERGGDTDKLV | 0.3197502 |
| SATGVFAGVI | ALMERGGDTDKLV | 0.5083464 |
| SLPAITLP | ALMERGGDTDKLV | 0.21050845 |
| STALAFTV | ALMERGGDTDKLV | 0.48757595 |
| SVLTVGLGK | ALMERGGDTDKLV | 0.2760665 |
| TALAGAGSGI | ALMERGGDTDKLV | 0.29890656 |
| TGLPFTGI | ALMERGGDTDKLV | 0.26130506 |
| VAEHVTV | ALMERGGDTDKLV | 0.3862392 |
| VDIVLNSL | ALMERGGDTDKLV | 0.42348623 |
| VLTGGGAVL | ALMERGGDTDKLV | 0.51990473 |
| VNVGLAYGI | ALMERGGDTDKLV | 0.35136104 |
| VSSAAVGAV | ALMERGGDTDKLV | 0.5245619 |
| VTSVYEAGV | ALMERGGDTDKLV | 0.5208857 |
| WVALDRGA | ALMERGGDTDKLV | 0.31929353 |
| YLVGSVVI | ALMERGGDTDKLV | 0.3040075 |
| GPPGTGKTLL | ALMERGGDTDKLV | 0.4593909 |
| GVGMTKF | ALMERGGDTDKLV | 0.32180604 |
| IGIGGSDL | ALMERGGDTDKLV | 0.50407887 |
| IGTLLAEL | ALMERGGDTDKLV | 0.4799477 |
| IYTSGTT | ALMERGGDTDKLV | 0.2597976 |
| LSFPAGV | ALMERGGDTDKLV | 0.51692206 |
| LYTSGTT | ALMERGGDTDKLV | 0.5054286 |
| VGAFPFGF | ALMERGGDTDKLV | 0.27654323 |
| VGAGTVEF | ALMERGGDTDKLV | 0.3092499 |
| VGKSTLLL | ALMERGGDTDKLV | 0.3569399 |
| AALAHAGGGIV | ALMERGGDTDKLV | 0.26913625 |
| AALVAVSLL | ALMERGGDTDKLV | 0.4279369 |
| AAVSRAVAAL | ALMERGGDTDKLV | 0.28098994 |
| AGGSSGIGGF | ALMERGGDTDKLV | 0.34221134 |
| AGLTVANANG | ALMERGGDTDKLV | 0.37248176 |
| AGLVGAVL | ALMERGGDTDKLV | 0.3524409 |
| ALSQAFTV | ALMERGGDTDKLV | 0.4003501 |
| ALTFSVSL | ALMERGGDTDKLV | 0.36089635 |
| ALTGAALVAL | ALMERGGDTDKLV | 0.4120533 |
| ASFSIGAGVL | ALMERGGDTDKLV | 0.42526767 |
| DGSVYKI | ALMERGGDTDKLV | 0.29531226 |
| EQLAEIV | ALMERGGDTDKLV | 0.40146282 |
| FAVSELTVM | ALMERGGDTDKLV | 0.5353437 |
| GAAPLAAGL | ALMERGGDTDKLV | 0.29432783 |
| GADGGDGGSGGAGG | ALMERGGDTDKLV | 0.31445873 |
| GFGLDTGGAGG | ALMERGGDTDKLV | 0.44192275 |
| GLASGGLL | ALMERGGDTDKLV | 0.27615327 |
| GRIAGAAVLL | ALMERGGDTDKLV | 0.51268125 |
| GSAGSLNTSF | ALMERGGDTDKLV | 0.37654716 |
| GTGQIGI | ALMERGGDTDKLV | 0.40183377 |
| GTLIALIL | ALMERGGDTDKLV | 0.30753008 |
| IAATMGFVVG | ALMERGGDTDKLV | 0.6283748 |
| IFAQALL | ALMERGGDTDKLV | 0.49143362 |
| IGAAVALGF | ALMERGGDTDKLV | 0.365446 |

| | | |
|---|---|---|
| IGLTPLAL | ALMERGGDTDKLV | 0.5366152 |
| ILGSNTSTL | ALMERGGDTDKLV | 0.3223498 |
| IVFSLTVL | ALMERGGDTDKLV | 0.489878 |
| LASKATGF | ALMERGGDTDKLV | 0.49119252 |
| LASVALIL | ALMERGGDTDKLV | 0.260617 |
| LFAVGLTL | ALMERGGDTDKLV | 0.33616394 |
| LFNGGIGTYI | ALMERGGDTDKLV | 0.5222806 |
| LTLGAGII | ALMERGGDTDKLV | 0.59900594 |
| LVAAALGDI | ALMERGGDTDKLV | 0.28912938 |
| NTAFVQLGI | ALMERGGDTDKLV | 0.32393232 |
| QLFSGAEL | ALMERGGDTDKLV | 0.25723448 |
| SAATFITL | ALMERGGDTDKLV | 0.31438383 |
| SNYGLANLGV | ALMERGGDTDKLV | 0.33438355 |
| SYGGAYAVM | ALMERGGDTDKLV | 0.4219046 |
| TTGVGGLTLG | ALMERGGDTDKLV | 0.36338523 |
| TTQTIYI | ALMERGGDTDKLV | 0.53702503 |
| VAAGAGLGAVI | ALMERGGDTDKLV | 0.40942967 |
| VAAVVTLL | ALMERGGDTDKLV | 0.5217661 |
| VGFLGGLI | ALMERGGDTDKLV | 0.3185973 |
| VGVIPILL | ALMERGGDTDKLV | 0.37615073 |
| VIFASSLI | ALMERGGDTDKLV | 0.2974821 |
| VKGGDGGAAAG | ALMERGGDTDKLV | 0.39096642 |
| VTKTFTNI | ALMERGGDTDKLV | 0.35088608 |
| VVAAAILL | ALMERGGDTDKLV | 0.34804803 |
| YAALDVEL | ALMERGGDTDKLV | 0.2361318 |
| LASVALLL | ALMERGGDTDKLV | 0.5180401 |
| LDAVHRV | ALMERGGDTDKLV | 0.40465644 |
| QGLSSSIF | ALMERGGDTDKLV | 0.4786584 |
| RGTFANIRL | ALMERGGDTDKLV | 0.34316742 |
| RQLWWGHRI | ALMERGGDTDKLV | 0.4063972 |
| YGPPGTGKTLL | ALMERGGDTDKLV | 0.31083792 |
| AALVGAPANVA | ALMERGGDTDKLV | 0.5139691 |
| AATFRRTE | ALMERGGDTDKLV | 0.31970558 |
| AGALGGGI | ALMERGGDTDKLV | 0.23446342 |
| AGGAGGAGGIGAG | ALMERGGDTDKLV | 0.34239385 |
| AGSGALGL | ALMERGGDTDKLV | 0.23702748 |
| AQIADQI | ALMERGGDTDKLV | 0.51950455 |
| ASAVAVTYL | ALMERGGDTDKLV | 0.45888114 |
| ASTVLGGTGG | ALMERGGDTDKLV | 0.43358713 |
| AVGGEPAK | ALMERGGDTDKLV | 0.5637374 |
| CEQGTFGGTC | ALMERGGDTDKLV | 0.23152664 |
| DEGKAPAG | ALMERGGDTDKLV | 0.35423154 |
| DTVVFPAL | ALMERGGDTDKLV | 0.5663899 |
| FEGITMAG | ALMERGGDTDKLV | 0.36154288 |
| FPLVVQD | ALMERGGDTDKLV | 0.38860813 |
| GAFTTPQ | ALMERGGDTDKLV | 0.57016546 |
| GFGGFGTAGGVGGA | ALMERGGDTDKLV | 0.2923218 |
| GFLAQFL | ALMERGGDTDKLV | 0.38527927 |
| GGAGGAGGNAGVA | ALMERGGDTDKLV | 0.42391518 |
| GGVGTAGAGGAGGAGG | ALMERGGDTDKLV | 0.5039931 |
| GNGGGGDAGNAGSGGN | ALMERGGDTDKLV | 0.41957664 |

| | | |
|---|---|---|
| GTGGAGGAGNDAGS | ALMERGGDTDKLV | 0.34606752 |
| GTTNTGWFNS | ALMERGGDTDKLV | 0.498483 |
| IFMGPLFTL | ALMERGGDTDKLV | 0.657918 |
| ILAHAIEAI | ALMERGGDTDKLV | 0.37318078 |
| IRSAIREAL | ALMERGGDTDKLV | 0.46006516 |
| ITFAGGG | ALMERGGDTDKLV | 0.4698677 |
| ITGMSGHQEV | ALMERGGDTDKLV | 0.36283064 |
| IVAGLACTL | ALMERGGDTDKLV | 0.31224945 |
| IVGGGLAGA | ALMERGGDTDKLV | 0.3213718 |
| IVTGVLNI | ALMERGGDTDKLV | 0.42869648 |
| LAQQGCI | ALMERGGDTDKLV | 0.25923628 |
| LCLHVEN | ALMERGGDTDKLV | 0.42122737 |
| LDAISDGS | ALMERGGDTDKLV | 0.33278897 |
| LGGFAAML | ALMERGGDTDKLV | 0.66130996 |
| LGGKNPMI | ALMERGGDTDKLV | 0.43359262 |
| LGPKGVAAIL | ALMERGGDTDKLV | 0.4987032 |
| LMVGVSI | ALMERGGDTDKLV | 0.3107531 |
| LVGAAFVL | ALMERGGDTDKLV | 0.40245804 |
| LVLAVAPL | ALMERGGDTDKLV | 0.36682126 |
| MAAIAVIDAA | ALMERGGDTDKLV | 0.27479422 |
| NNGINAELQ | ALMERGGDTDKLV | 0.48551136 |
| RGEPLVLVHG | ALMERGGDTDKLV | 0.32478428 |
| RGGAGGAAGLI | ALMERGGDTDKLV | 0.31670603 |
| RSYRMHPAV | ALMERGGDTDKLV | 0.45571494 |
| SAAHLLAGQLA | ALMERGGDTDKLV | 0.59600675 |
| SAAVGSY | ALMERGGDTDKLV | 0.366757 |
| SPHAVLHRI | ALMERGGDTDKLV | 0.3529963 |
| SQIAKVL | ALMERGGDTDKLV | 0.32098576 |
| TALGHLVP | ALMERGGDTDKLV | 0.4377859 |
| TAPTTSGAA | ALMERGGDTDKLV | 0.6000872 |
| TDGDDGNGGSPGT | ALMERGGDTDKLV | 0.38619712 |
| TDGLGEN | ALMERGGDTDKLV | 0.65068114 |
| TGGPPLPGQA | ALMERGGDTDKLV | 0.45505098 |
| TIPAITI | ALMERGGDTDKLV | 0.29872024 |
| TIPGFTI | ALMERGGDTDKLV | 0.4360871 |
| TSIASGALA | ALMERGGDTDKLV | 0.52798486 |
| TVTLPGAAI | ALMERGGDTDKLV | 0.49936536 |
| VASGLSAALP | ALMERGGDTDKLV | 0.54791194 |
| VGAGVIGI | ALMERGGDTDKLV | 0.47421435 |
| VGPKRFGV | ALMERGGDTDKLV | 0.30834514 |
| VNVAFPD | ALMERGGDTDKLV | 0.49203512 |
| VSMTNSGVS | ALMERGGDTDKLV | 0.52765834 |
| VTDVLGDLSV | ALMERGGDTDKLV | 0.5177048 |
| VVVVGAGFAGL | ALMERGGDTDKLV | 0.3344008 |
| VYTVGTVV | ALMERGGDTDKLV | 0.3019876 |
| WTTFSGP | ALMERGGDTDKLV | 0.39067638 |
| YDELVEAF | ALMERGGDTDKLV | 0.5331736 |
| YGTGGAGGNGG | ALMERGGDTDKLV | 0.3435128 |
| YTNVMAV | ALMERGGDTDKLV | 0.4000507 |
| FLEQVHQL | ALMERGGDTDKLV | 0.3620035 |
| GLSLLAVAV | ALMERGGDTDKLV | 0.27286112 |

| | | |
|---|---|---|
| LEQVHQL | ALMERGGDTDKLV | 0.39638758 |
| LNAAGAI | ALMERGGDTDKLV | 0.53963524 |
| PAVHASL | ALMERGGDTDKLV | 0.44753307 |
| TELEDDA | ALMERGGDTDKLV | 0.33253267 |
| TPPGPPN | ALMERGGDTDKLV | 0.48286918 |
| TPPPPPPP | ALMERGGDTDKLV | 0.6375482 |
| TQSPALI | AWSAQRTGGTGQLY | 0.5481908 |
| EFAAAFPQ | AWSAQRTGGTGQLY | 0.29055378 |
| LLSGNLDVL | AWSAQRTGGTGQLY | 0.21226946 |
| GPANLGNYNI | AWSAQRTGGTGQLY | 0.40887558 |
| TLGPTLP | AWSAQRTGGTGQLY | 0.48964405 |
| VYAYSSCCK | AWSAQRTGGTGQLY | 0.3428254 |
| APAPAPAPNI | AWSAQRTGGTGQLY | 0.34618643 |
| PGGPVPIP | AWSAQRTGGTGQLY | 0.43850005 |
| FLTVALEP | AWSAQRTGGTGQLY | 0.45281374 |
| SQYFGANL | AWSAQRTGGTGQLY | 0.35965428 |
| GGVKPAA | AWSAQRTGGTGQLY | 0.28287327 |
| GGTGGAGGAGGW | AWSAQRTGGTGQLY | 0.4294027 |
| GVGGVAGLGGAGG | AWSAQRTGGTGQLY | 0.5204062 |
| VLLVLATTI | AWSAQRTGGTGQLY | 0.3207123 |
| IGIFAAML | AWSAQRTGGTGQLY | 0.35498416 |
| AVGDLID | AWSAQRTGGTGQLY | 0.34804493 |
| NISPLPL | AWSAQRTGGTGQLY | 0.34438282 |
| VTAVFVAI | AWSAQRTGGTGQLY | 0.34848443 |
| SGDDNDAAL | AWSAQRTGGTGQLY | 0.2710148 |
| AVAALAHS | AWSAQRTGGTGQLY | 0.5495533 |
| GGVTVMATP | AWSAQRTGGTGQLY | 0.26339072 |
| AGLAANVRD | AWSAQRTGGTGQLY | 0.46811736 |
| TNRPDLIDPALLRPG | AWSAQRTGGTGQLY | 0.4489505 |
| LANLGVTAL | AWSAQRTGGTGQLY | 0.45774975 |
| TGERIEL | AWSAQRTGGTGQLY | 0.3466903 |
| NGEELAA | AWSAQRTGGTGQLY | 0.3185448 |
| VLSLVWG | AWSAQRTGGTGQLY | 0.269226 |
| IGYALWL | AWSAQRTGGTGQLY | 0.24362737 |
| LSLLLAPL | AWSAQRTGGTGQLY | 0.5367434 |
| TLDEALK | AWSAQRTGGTGQLY | 0.32979092 |
| TDGAGTAEGG | AWSAQRTGGTGQLY | 0.5606373 |
| NAPGGAGGNGG | AWSAQRTGGTGQLY | 0.4521908 |
| VSVFMGMGGNF | AWSAQRTGGTGQLY | 0.34105307 |
| LASYLAAL | AWSAQRTGGTGQLY | 0.42830655 |
| LVGGTPAL | AWSAQRTGGTGQLY | 0.31732774 |
| VRNADVL | AWSAQRTGGTGQLY | 0.3783796 |
| TCSSPFLVSAM | AWSAQRTGGTGQLY | 0.40401003 |
| VHITDAE | AWSAQRTGGTGQLY | 0.49150538 |
| EYAAPSF | AWSAQRTGGTGQLY | 0.32189798 |
| VVDAFHGGL | AWSAQRTGGTGQLY | 0.47541636 |
| ASVVDPQ | AWSAQRTGGTGQLY | 0.40326285 |
| SYNEFGGLS | AWSAQRTGGTGQLY | 0.3112603 |
| DPDGSVY | AWSAQRTGGTGQLY | 0.4623177 |
| MVSTIHATE | AWSAQRTGGTGQLY | 0.47362906 |
| LAAGVADPV | AWSAQRTGGTGQLY | 0.47486135 |

| | | |
|---|---|---|
| TKEQIAA | AWSAQRTGGTGQLY | 0.3323834 |
| SLASPAAL | AWSAQRTGGTGQLY | 0.41475463 |
| IGLVFYGV | AWSAQRTGGTGQLY | 0.33107832 |
| FLLDASNL | AWSAQRTGGTGQLY | 0.39799094 |
| YQVGNFP | AWSAQRTGGTGQLY | 0.42871734 |
| ESLLSQST | AWSAQRTGGTGQLY | 0.37399772 |
| LFTVGFAAVG | AWSAQRTGGTGQLY | 0.49539095 |
| ATPPTPT | AWSAQRTGGTGQLY | 0.30296972 |
| TDLATLG | AWSAQRTGGTGQLY | 0.28542185 |
| SAGGTGR | AWSAQRTGGTGQLY | 0.37451786 |
| GFSNSGS | AWSAQRTGGTGQLY | 0.4163208 |
| GGNGGSGGTSV | AWSAQRTGGTGQLY | 0.5078026 |
| GTSVATGGAGN | AWSAQRTGGTGQLY | 0.27468422 |
| NGGNGGSGGTS | AWSAQRTGGTGQLY | 0.391185 |
| AVVKFPH | AWSAQRTGGTGQLY | 0.3524287 |
| TPPPPPP | AWSAQRTGGTGQLY | 0.32373068 |
| AAPAAAAAPA | AWSAQRTGGTGQLY | 0.43655947 |
| IALNEDL | AWSAQRTGGTGQLY | 0.3778537 |
| IVNVSSM | AWSAQRTGGTGQLY | 0.32990986 |
| TGIPPAPR | AWSAQRTGGTGQLY | 0.49418476 |
| FDLGGGTF | AWSAQRTGGTGQLY | 0.36869776 |
| FDLGGGTFDVS | AWSAQRTGGTGQLY | 0.4075367 |
| NEPTAAA | AWSAQRTGGTGQLY | 0.41695684 |
| NEPTAAALA | AWSAQRTGGTGQLY | 0.33609557 |
| GSIINISSI | AWSAQRTGGTGQLY | 0.4373562 |
| VAPGNAGTA | AWSAQRTGGTGQLY | 0.4352134 |
| TGGGASLE | AWSAQRTGGTGQLY | 0.4732482 |
| VAVYNHYKRI | AWSAQRTGGTGQLY | 0.36913237 |
| TDPAPATT | AWSAQRTGGTGQLY | 0.29590705 |
| TNRPDILDPALLRPG | AWSAQRTGGTGQLY | 0.5206999 |
| LTTTTTS | AWSAQRTGGTGQLY | 0.45444363 |
| PTGSGKTL | AWSAQRTGGTGQLY | 0.46884182 |
| TGSGKTL | AWSAQRTGGTGQLY | 0.46812838 |
| TIGHVDHGK | AWSAQRTGGTGQLY | 0.45367318 |
| TIGHVDHGKTT | AWSAQRTGGTGQLY | 0.5028981 |
| GHVDHGKTT | AWSAQRTGGTGQLY | 0.30059502 |
| AQFEHTLL | AWSAQRTGGTGQLY | 0.27127963 |
| IVNQDEL | AWSAQRTGGTGQLY | 0.35644236 |
| TNGSQFF | AWSAQRTGGTGQLY | 0.32300106 |
| TGKGIVE | AWSAQRTGGTGQLY | 0.37973708 |
| LISNASDA | AWSAQRTGGTGQLY | 0.28224263 |
| AELRTLGELTKIAKAHGAQ | AWSAQRTGGTGQLY | 0.51590973 |
| APNAVAAAEPSW | AWSAQRTGGTGQLY | 0.6976873 |
| APQVAGAATAVL | AWSAQRTGGTGQLY | 0.3225598 |
| ARRDPAAAAAILPTDA | AWSAQRTGGTGQLY | 0.5763674 |
| DAAARDVLAEAGLAEYF | AWSAQRTGGTGQLY | 0.2676497 |
| DDQIVLSAKANLAKVEQ | AWSAQRTGGTGQLY | 0.44470662 |
| DELLPDTGEDSDEDGH | AWSAQRTGGTGQLY | 0.52451813 |
| ELPEAPANRATVPQD | AWSAQRTGGTGQLY | 0.27199382 |
| FGQSKIERVTGIG | AWSAQRTGGTGQLY | 0.45369294 |
| GAGGTGSTAGGAGGAGGAGGLY | AWSAQRTGGTGQLY | 0.32505703 |

| | | |
|---|---|---|
| GHGGTGAAGGNGGTGGAGGSGI | AWSAQRTGGTGQLY | 0.3758314 |
| GNGGNGGAGGEGGPGGAGGAGA | AWSAQRTGGTGQLY | 0.5738121 |
| GSTAGNGGNGGNATNGTIAGQP | AWSAQRTGGTGQLY | 0.39041087 |
| GTGGDGGNGGNGGTGG | AWSAQRTGGTGQLY | 0.4699784 |
| GVRRPAGIPADAPV | AWSAQRTGGTGQLY | 0.43552333 |
| GYRQVTDAHLLTLARR | AWSAQRTGGTGQLY | 0.34256384 |
| IAKMGQDGHDRG | AWSAQRTGGTGQLY | 0.48372367 |
| KAEPSIEIQYT | AWSAQRTGGTGQLY | 0.50740063 |
| KPVRIRMAADEDTVAVT | AWSAQRTGGTGQLY | 0.32593256 |
| MPPRIAGMRLLVIKPEP | AWSAQRTGGTGQLY | 0.5207513 |
| MKFYAVALFRSRGVPPTR | AWSAQRTGGTGQLY | 0.33226028 |
| NAGGFGILGAVAHSP | AWSAQRTGGTGQLY | 0.5313602 |
| NEGREITLAMAEM | AWSAQRTGGTGQLY | 0.5938894 |
| NELVEWSHPLGHRWRWE | AWSAQRTGGTGQLY | 0.3647393 |
| NLGAGGTGGNGGNPG | AWSAQRTGGTGQLY | 0.38399816 |
| NTEWPKWETVMAADPVD | AWSAQRTGGTGQLY | 0.51608294 |
| PASAAPAETQP | AWSAQRTGGTGQLY | 0.41177234 |
| QQSSDGKNGVRAAEITGE | AWSAQRTGGTGQLY | 0.46251965 |
| RAPDAVTTSWWKE | AWSAQRTGGTGQLY | 0.5385546 |
| SGFGNVGAHTSGWFNQ | AWSAQRTGGTGQLY | 0.29469714 |
| SGVAGAVGQAASVGGLKV | AWSAQRTGGTGQLY | 0.48846498 |
| SNGGAGTGSAGGNGGT | AWSAQRTGGTGQLY | 0.30866724 |
| TGGAGGEGGAAG | AWSAQRTGGTGQLY | 0.3525597 |
| TGGDGGAAGAGGAPGHG | AWSAQRTGGTGQLY | 0.30380815 |
| TMPPLLAAIANAGRHGV | AWSAQRTGGTGQLY | 0.26606604 |
| TSSTAGNSGGA | AWSAQRTGGTGQLY | 0.33823925 |
| VPATVPVGILC | AWSAQRTGGTGQLY | 0.2908618 |
| VQRPDSGGVVLTGRIS | AWSAQRTGGTGQLY | 0.5481603 |
| WTPNPVPATKGVPAGLVQAT | AWSAQRTGGTGQLY | 0.36880285 |
| YALKRALAKFSDRFDVVII | AWSAQRTGGTGQLY | 0.31584254 |
| YLIGSVIV | AWSAQRTGGTGQLY | 0.5851705 |
| YGQLVAVID | AWSAQRTGGTGQLY | 0.570277 |
| VTVGGMTLPA | AWSAQRTGGTGQLY | 0.2736802 |
| VLALPAV | AWSAQRTGGTGQLY | 0.30721465 |
| VIPPTLNYET | AWSAQRTGGTGQLY | 0.29152295 |
| VGMGGMHPSSG | AWSAQRTGGTGQLY | 0.32223135 |
| VGLAGGAATA | AWSAQRTGGTGQLY | 0.25837395 |
| VGGAGGAGGSGP | AWSAQRTGGTGQLY | 0.37457657 |
| VFAAFMV | AWSAQRTGGTGQLY | 0.2798328 |
| TLYDKLR | AWSAQRTGGTGQLY | 0.25125062 |
| TLAVFTVL | AWSAQRTGGTGQLY | 0.32690203 |
| TGSVQPARV | AWSAQRTGGTGQLY | 0.33921683 |
| TGGGAGSGGT | AWSAQRTGGTGQLY | 0.32713205 |
| SHLETSL | AWSAQRTGGTGQLY | 0.5121865 |
| SFTTGMT | AWSAQRTGGTGQLY | 0.27086195 |
| QVEVAKRMGVRQPTVS | AWSAQRTGGTGQLY | 0.3086611 |
| QGSGGPALAM | AWSAQRTGGTGQLY | 0.43473235 |
| MLAPSNP | AWSAQRTGGTGQLY | 0.48409778 |
| LVNNAAI | AWSAQRTGGTGQLY | 0.29822424 |
| LTGEINTLA | AWSAQRTGGTGQLY | 0.38615426 |
| LSIFGPV | AWSAQRTGGTGQLY | 0.5469488 |

| | | |
|---|---|---|
| LLALLANL | AWSAQRTGGTGQLY | 0.3271094 |
| LISNASDALDKLR | AWSAQRTGGTGQLY | 0.35434327 |
| LINEAVP | AWSAQRTGGTGQLY | 0.37027043 |
| LGYAFSGL | AWSAQRTGGTGQLY | 0.28256515 |
| LDFVAVAL | AWSAQRTGGTGQLY | 0.47769934 |
| LAQPGAE | AWSAQRTGGTGQLY | 0.59608704 |
| ISNLGAVTGF | AWSAQRTGGTGQLY | 0.31357434 |
| HQEDPGRI | AWSAQRTGGTGQLY | 0.35786074 |
| GTPGEPTP | AWSAQRTGGTGQLY | 0.30197906 |
| GFFNSGS | AWSAQRTGGTGQLY | 0.31105396 |
| GAGGGGLTAGAN | AWSAQRTGGTGQLY | 0.25881833 |
| FVDALVNM | AWSAQRTGGTGQLY | 0.40532064 |
| FLTVAPEL | AWSAQRTGGTGQLY | 0.37048185 |
| FETLTAL | AWSAQRTGGTGQLY | 0.32337558 |
| DVINPIEL | AWSAQRTGGTGQLY | 0.27575538 |
| APVSGAPV | AWSAQRTGGTGQLY | 0.27731854 |
| ALVGGLAW | AWSAQRTGGTGQLY | 0.30626273 |
| ALAAAAATAI | AWSAQRTGGTGQLY | 0.30954844 |
| AFGLGLLL | AWSAQRTGGTGQLY | 0.2600583 |
| AFFAAASL | AWSAQRTGGTGQLY | 0.27474427 |
| AAAALPAFSPP | AWSAQRTGGTGQLY | 0.4202505 |
| VVLAPAL | AWSAQRTGGTGQLY | 0.38969335 |
| TIGHVDHGKTTL | AWSAQRTGGTGQLY | 0.28312582 |
| LISNASDALDK | AWSAQRTGGTGQLY | 0.51308596 |
| LISNASDAL | AWSAQRTGGTGQLY | 0.5069883 |
| ISNASDALDK | AWSAQRTGGTGQLY | 0.3325667 |
| NQAETLVYQTE | AWSAQRTGGTGQLY | 0.31741205 |
| TGGTGGTAGAG | AWSAQRTGGTGQLY | 0.29055917 |
| TGSTAGGAGGA | AWSAQRTGGTGQLY | 0.6526551 |
| DPLAVPVAAGPV | AWSAQRTGGTGQLY | 0.30700547 |
| GGTGGTPGNGGN | AWSAQRTGGTGQLY | 0.3018383 |
| PAGLAGVDPATL | AWSAQRTGGTGQLY | 0.37470868 |
| GNNLVVEAPHAGASV | AWSAQRTGGTGQLY | 0.33477834 |
| KVTRIMSASTLEQPA | AWSAQRTGGTGQLY | 0.29633394 |
| LGFALPIALPRRTVV | AWSAQRTGGTGQLY | 0.40866196 |
| TRLGNLTARKLVVLV | AWSAQRTGGTGQLY | 0.36156702 |
| WSQQRRLIYFPSAGP | AWSAQRTGGTGQLY | 0.2734504 |
| GPNGNPDPMAAAVDIR | AWSAQRTGGTGQLY | 0.296509 |
| TQRCVSVADLRAAASR | AWSAQRTGGTGQLY | 0.29031047 |
| QFVLDVAEHPVEVLTETV | AWSAQRTGGTGQLY | 0.38655168 |
| GGAGGTGGAGGNGGTQAGDG | AWSAQRTGGTGQLY | 0.47440028 |
| PLLATLRAIAVDVDAKPAQV | AWSAQRTGGTGQLY | 0.3146619 |
| APKFRTGGYMIDFWGVGYQVAK | AWSAQRTGGTGQLY | 0.28992364 |
| GAGGAGGAGGAAGGPGGTGGTG | AWSAQRTGGTGQLY | 0.48012614 |
| IGALGLLATDVAPAGSVAFGVA | AWSAQRTGGTGQLY | 0.33491835 |
| AGGAGGIGGNGGAGGAGGVLMG | AWSAQRTGGTGQLY | 0.34319246 |
| ADELIGLL | AWSAQRTGGTGQLY | 0.40756872 |
| ADRALKA | AWSAQRTGGTGQLY | 0.31229573 |
| AGAVGQVGGAGGA | AWSAQRTGGTGQLY | 0.36707756 |
| AGGGGGLGGAP | AWSAQRTGGTGQLY | 0.35666794 |
| AGGLLYGNGGAG | AWSAQRTGGTGQLY | 0.34315765 |

| | | |
|---|---|---|
| AGLESIGL | AWSAQRTGGTGQLY | 0.36103785 |
| AGLKTGII | AWSAQRTGGTGQLY | 0.33688772 |
| ASLAALLL | AWSAQRTGGTGQLY | 0.32477036 |
| ASVINISSI | AWSAQRTGGTGQLY | 0.25981024 |
| ATPPGPA | AWSAQRTGGTGQLY | 0.28552595 |
| ATVASALV | AWSAQRTGGTGQLY | 0.43854553 |
| ATVASLIL | AWSAQRTGGTGQLY | 0.33771166 |
| AVNGPVLIRH | AWSAQRTGGTGQLY | 0.50834453 |
| DGAAGAVGI | AWSAQRTGGTGQLY | 0.34543726 |
| DPLLHGI | AWSAQRTGGTGQLY | 0.29822522 |
| ELGAALGAL | AWSAQRTGGTGQLY | 0.43678284 |
| ELGRTQAV | AWSAQRTGGTGQLY | 0.46131104 |
| FVAGGYLS | AWSAQRTGGTGQLY | 0.27061203 |
| GASGGAGGAGGW | AWSAQRTGGTGQLY | 0.30657682 |
| GGAGAGGDGGAGGI | AWSAQRTGGTGQLY | 0.5365795 |
| GTGHNGSQPGMGGNG | AWSAQRTGGTGQLY | 0.5173356 |
| GYAAASASAS | AWSAQRTGGTGQLY | 0.3272694 |
| HQVDVAAVQQG | AWSAQRTGGTGQLY | 0.31889075 |
| IGAAIAHT | AWSAQRTGGTGQLY | 0.64604574 |
| IGMIEVAQL | AWSAQRTGGTGQLY | 0.29553843 |
| IRLAPEHTSAMD | AWSAQRTGGTGQLY | 0.52256113 |
| IYNVSVL | AWSAQRTGGTGQLY | 0.25494382 |
| KQGFLGM | AWSAQRTGGTGQLY | 0.2964776 |
| LESVITQL | AWSAQRTGGTGQLY | 0.28342295 |
| LGLGALVVV | AWSAQRTGGTGQLY | 0.2580571 |
| LIGNGGNGGSGGI | AWSAQRTGGTGQLY | 0.40015015 |
| LNAINAPTL | AWSAQRTGGTGQLY | 0.3173211 |
| LQLLNTGV | AWSAQRTGGTGQLY | 0.4386611 |
| LQTSIEF | AWSAQRTGGTGQLY | 0.30839416 |
| MSFAVSEL | AWSAQRTGGTGQLY | 0.37212086 |
| NTGALGL | AWSAQRTGGTGQLY | 0.36204982 |
| PPGAGGAGGAGTT | AWSAQRTGGTGQLY | 0.32298112 |
| QLVGLGI | AWSAQRTGGTGQLY | 0.28782448 |
| RFLEQVHQL | AWSAQRTGGTGQLY | 0.3763403 |
| RGVGYVLR | AWSAQRTGGTGQLY | 0.50050074 |
| SATGVFAGVI | AWSAQRTGGTGQLY | 0.3307248 |
| SLPAITLP | AWSAQRTGGTGQLY | 0.27845842 |
| STALAFTV | AWSAQRTGGTGQLY | 0.2919063 |
| SVLTVGLGK | AWSAQRTGGTGQLY | 0.3328137 |
| TALAGAGSGI | AWSAQRTGGTGQLY | 0.26469162 |
| TGLPFTGI | AWSAQRTGGTGQLY | 0.25603762 |
| VAEHVTV | AWSAQRTGGTGQLY | 0.27994263 |
| VDIVLNSL | AWSAQRTGGTGQLY | 0.29026407 |
| VLTGGGAVL | AWSAQRTGGTGQLY | 0.51044357 |
| VNVGLAYGI | AWSAQRTGGTGQLY | 0.49811774 |
| VSSAAVGAV | AWSAQRTGGTGQLY | 0.33834317 |
| VTSVYEAGV | AWSAQRTGGTGQLY | 0.40888515 |
| WVALDRGA | AWSAQRTGGTGQLY | 0.43531144 |
| YLVGSVVI | AWSAQRTGGTGQLY | 0.24717878 |
| GPPGTGKTLL | AWSAQRTGGTGQLY | 0.29207236 |
| GVGMTKF | AWSAQRTGGTGQLY | 0.28116074 |

| | | |
|---|---|---|
| IGIGGSDL | AWSAQRTGGTGQLY | 0.38963893 |
| IGTLLAEL | AWSAQRTGGTGQLY | 0.28126603 |
| IYTSGTT | AWSAQRTGGTGQLY | 0.31461877 |
| LSFPAGV | AWSAQRTGGTGQLY | 0.37657335 |
| LYTSGTT | AWSAQRTGGTGQLY | 0.5382817 |
| VGAFPFGF | AWSAQRTGGTGQLY | 0.32282236 |
| VGAGTVEF | AWSAQRTGGTGQLY | 0.27138975 |
| VGKSTLLL | AWSAQRTGGTGQLY | 0.25074512 |
| AALAHAGGGIV | AWSAQRTGGTGQLY | 0.3744927 |
| AALVAVSLL | AWSAQRTGGTGQLY | 0.2707606 |
| AAVSRAVAAL | AWSAQRTGGTGQLY | 0.30584323 |
| AGGSSGIGGF | AWSAQRTGGTGQLY | 0.2597651 |
| AGLTVANANG | AWSAQRTGGTGQLY | 0.2901786 |
| AGLVGAVL | AWSAQRTGGTGQLY | 0.2671182 |
| ALSQAFTV | AWSAQRTGGTGQLY | 0.34856611 |
| ALTFSVSL | AWSAQRTGGTGQLY | 0.30294982 |
| ALTGAALVAL | AWSAQRTGGTGQLY | 0.3375513 |
| ASFSIGAGVL | AWSAQRTGGTGQLY | 0.4806305 |
| DGSVYKI | AWSAQRTGGTGQLY | 0.6229954 |
| EQLAEIV | AWSAQRTGGTGQLY | 0.3093005 |
| FAVSELTVM | AWSAQRTGGTGQLY | 0.30728647 |
| GAAPLAAGL | AWSAQRTGGTGQLY | 0.28752953 |
| GADGGDGGSGGAGG | AWSAQRTGGTGQLY | 0.2987293 |
| GFGLDTGGAGG | AWSAQRTGGTGQLY | 0.30398765 |
| GLASGGLL | AWSAQRTGGTGQLY | 0.57679284 |
| GRIAGAAVLL | AWSAQRTGGTGQLY | 0.30644023 |
| GSAGSLNTSF | AWSAQRTGGTGQLY | 0.28257743 |
| GTGQIGI | AWSAQRTGGTGQLY | 0.32991704 |
| GTLIALIL | AWSAQRTGGTGQLY | 0.48817524 |
| IAATMGFVVG | AWSAQRTGGTGQLY | 0.50065017 |
| IFAQALL | AWSAQRTGGTGQLY | 0.28697708 |
| IGAAVALGF | AWSAQRTGGTGQLY | 0.29168358 |
| IGLTPLAL | AWSAQRTGGTGQLY | 0.32482725 |
| ILGSNTSTL | AWSAQRTGGTGQLY | 0.41377836 |
| IVFSLTVL | AWSAQRTGGTGQLY | 0.29628944 |
| LASKATGF | AWSAQRTGGTGQLY | 0.29020864 |
| LASVALIL | AWSAQRTGGTGQLY | 0.3482807 |
| LFAVGLTL | AWSAQRTGGTGQLY | 0.6855956 |
| LFNGGIGTYI | AWSAQRTGGTGQLY | 0.32095563 |
| LTLGAGII | AWSAQRTGGTGQLY | 0.26282233 |
| LVAAALGDI | AWSAQRTGGTGQLY | 0.37509072 |
| NTAFVQLGI | AWSAQRTGGTGQLY | 0.27540615 |
| QLFSGAEL | AWSAQRTGGTGQLY | 0.32558998 |
| SAATFITL | AWSAQRTGGTGQLY | 0.36224854 |
| SNYGLANLGV | AWSAQRTGGTGQLY | 0.27274716 |
| SYGGAYAVM | AWSAQRTGGTGQLY | 0.32194185 |
| TTGVGGLTLG | AWSAQRTGGTGQLY | 0.4796673 |
| TTQTIYI | AWSAQRTGGTGQLY | 0.29974645 |
| VAAGAGLGAVI | AWSAQRTGGTGQLY | 0.41350222 |
| VAAVVTLL | AWSAQRTGGTGQLY | 0.5100786 |
| VGFLGGLI | AWSAQRTGGTGQLY | 0.42786932 |

| | | |
|---|---|---|
| VGVIPILL | AWSAQRTGGTGQLY | 0.584713 |
| VIFASSLI | AWSAQRTGGTGQLY | 0.4868219 |
| VKGGDGGAAAG | AWSAQRTGGTGQLY | 0.31306285 |
| VTKTFTNI | AWSAQRTGGTGQLY | 0.31234953 |
| VVAAAILL | AWSAQRTGGTGQLY | 0.4654521 |
| YAALDVEL | AWSAQRTGGTGQLY | 0.39729267 |
| LASVALLL | AWSAQRTGGTGQLY | 0.32888925 |
| LDAVHRV | AWSAQRTGGTGQLY | 0.2884648 |
| QGLSSSIF | AWSAQRTGGTGQLY | 0.603908 |
| RGTFANIRL | AWSAQRTGGTGQLY | 0.32015687 |
| RQLWWGHRI | AWSAQRTGGTGQLY | 0.5424031 |
| YGPPGTGKTLL | AWSAQRTGGTGQLY | 0.27694273 |
| AALVGAPANVA | AWSAQRTGGTGQLY | 0.2916862 |
| AATFRRTE | AWSAQRTGGTGQLY | 0.43155754 |
| AGALGGGI | AWSAQRTGGTGQLY | 0.2495768 |
| AGGAGGAGGIGAG | AWSAQRTGGTGQLY | 0.25470597 |
| AGSGALGL | AWSAQRTGGTGQLY | 0.32776582 |
| AQIADQI | AWSAQRTGGTGQLY | 0.43760625 |
| ASAVAVTYL | AWSAQRTGGTGQLY | 0.24984823 |
| ASTVLGGTGG | AWSAQRTGGTGQLY | 0.31315976 |
| AVGGEPAK | AWSAQRTGGTGQLY | 0.27440834 |
| CEQGTFGGTC | AWSAQRTGGTGQLY | 0.3232809 |
| DEGKAPAG | AWSAQRTGGTGQLY | 0.36677572 |
| DTVVFPAL | AWSAQRTGGTGQLY | 0.26425022 |
| FEGITMAG | AWSAQRTGGTGQLY | 0.36698806 |
| FPLVVQD | AWSAQRTGGTGQLY | 0.26613036 |
| GAFTTPQ | AWSAQRTGGTGQLY | 0.28388605 |
| GFGGFGTAGGVGGA | AWSAQRTGGTGQLY | 0.3534053 |
| GFLAQFL | AWSAQRTGGTGQLY | 0.59119403 |
| GGAGGAGGNAGVA | AWSAQRTGGTGQLY | 0.29017964 |
| GGVGTAGAGGAGGAGG | AWSAQRTGGTGQLY | 0.40778324 |
| GNGGGGDAGNAGSGGN | AWSAQRTGGTGQLY | 0.28533977 |
| GTGGAGGAGNDAGS | AWSAQRTGGTGQLY | 0.5706092 |
| GTTNTGWFNS | AWSAQRTGGTGQLY | 0.3190254 |
| IFMGPLFTL | AWSAQRTGGTGQLY | 0.2661699 |
| ILAHAIEAI | AWSAQRTGGTGQLY | 0.48142612 |
| IRSAIREAL | AWSAQRTGGTGQLY | 0.36827505 |
| ITFAGGG | AWSAQRTGGTGQLY | 0.3475365 |
| ITGMSGHQEV | AWSAQRTGGTGQLY | 0.47553736 |
| IVAGLACTL | AWSAQRTGGTGQLY | 0.2932134 |
| IVGGGLAGA | AWSAQRTGGTGQLY | 0.3693883 |
| IVTGVLNI | AWSAQRTGGTGQLY | 0.5121748 |
| LAQQGCI | AWSAQRTGGTGQLY | 0.502526 |
| LCLHVEN | AWSAQRTGGTGQLY | 0.29637846 |
| LDAISDGS | AWSAQRTGGTGQLY | 0.36216548 |
| LGGFAAML | AWSAQRTGGTGQLY | 0.41463926 |
| LGGKNPMI | AWSAQRTGGTGQLY | 0.30783856 |
| LGPKGVAAIL | AWSAQRTGGTGQLY | 0.40746507 |
| LMVGVSI | AWSAQRTGGTGQLY | 0.3150882 |
| LVGAAFVL | AWSAQRTGGTGQLY | 0.4013015 |
| LVLAVAPL | AWSAQRTGGTGQLY | 0.3456917 |

| | | |
|---|---|---|
| MAAIAVIDAA | AWSAQRTGGTGQLY | 0.66106284 |
| NNGINAELQ | AWSAQRTGGTGQLY | 0.2680326 |
| RGEPLVLVHG | AWSAQRTGGTGQLY | 0.35107815 |
| RGGAGGAAGLI | AWSAQRTGGTGQLY | 0.34627065 |
| RSYRMHPAV | AWSAQRTGGTGQLY | 0.3627912 |
| SAAHLLAGQLA | AWSAQRTGGTGQLY | 0.30528003 |
| SAAVGSY | AWSAQRTGGTGQLY | 0.27396297 |
| SPHAVLHRI | AWSAQRTGGTGQLY | 0.3393409 |
| SQIAKVL | AWSAQRTGGTGQLY | 0.48564693 |
| TALGHLVP | AWSAQRTGGTGQLY | 0.3144488 |
| TAPTTSGAA | AWSAQRTGGTGQLY | 0.39108828 |
| TDGDDGNGGSPGT | AWSAQRTGGTGQLY | 0.6489618 |
| TDGLGEN | AWSAQRTGGTGQLY | 0.2964704 |
| TGGPPLPGQA | AWSAQRTGGTGQLY | 0.35284394 |
| TIPAITI | AWSAQRTGGTGQLY | 0.37636846 |
| TIPGFTI | AWSAQRTGGTGQLY | 0.5577935 |
| TSIASGALA | AWSAQRTGGTGQLY | 0.3036421 |
| TVTLPGAAI | AWSAQRTGGTGQLY | 0.46081054 |
| VASGLSAALP | AWSAQRTGGTGQLY | 0.27692583 |
| VGAGVIGI | AWSAQRTGGTGQLY | 0.4740876 |
| VGPKRFGV | AWSAQRTGGTGQLY | 0.28299478 |
| VNVAFPD | AWSAQRTGGTGQLY | 0.35517293 |
| VSMTNSGVS | AWSAQRTGGTGQLY | 0.28312042 |
| VTDVLGDLSV | AWSAQRTGGTGQLY | 0.4242605 |
| VVVVGAGFAGL | AWSAQRTGGTGQLY | 0.47690678 |
| VYTVGTVV | AWSAQRTGGTGQLY | 0.5382078 |
| WTTFSGP | AWSAQRTGGTGQLY | 0.2567626 |
| YDELVEAF | AWSAQRTGGTGQLY | 0.49121767 |
| YGTGGAGGNGG | AWSAQRTGGTGQLY | 0.48415756 |
| YTNVMAV | AWSAQRTGGTGQLY | 0.6923968 |
| FLEQVHQL | AWSAQRTGGTGQLY | 0.29862922 |
| GLSLLAVAV | AWSAQRTGGTGQLY | 0.42632258 |
| LEQVHQL | AWSAQRTGGTGQLY | 0.33433697 |
| LNAAGAI | AWSAQRTGGTGQLY | 0.3000481 |
| PAVHASL | AWSAQRTGGTGQLY | 0.27388656 |
| TELEDDA | AWSAQRTGGTGQLY | 0.3278191 |
| TPPGPPN | AWSAQRTGGTGQLY | 0.34846845 |
| TPPPPPPP | AWSAQRTGGTGQLY | 0.299435 |
| TQSPALI | ASSPGTNTEVF | 0.5730637 |
| EFAAAFPQ | ASSPGTNTEVF | 0.49913812 |
| LLSGNLDVL | ASSPGTNTEVF | 0.44223675 |
| GPANLGNYNI | ASSPGTNTEVF | 0.55881494 |
| TLGPTLP | ASSPGTNTEVF | 0.52347225 |
| VYAYSSCCK | ASSPGTNTEVF | 0.34061676 |
| APAPAPAPNI | ASSPGTNTEVF | 0.3004737 |
| PGGPVPIP | ASSPGTNTEVF | 0.3068556 |
| FLTVALEP | ASSPGTNTEVF | 0.2734898 |
| SQYFGANL | ASSPGTNTEVF | 0.33914095 |
| GGVKPAA | ASSPGTNTEVF | 0.29932848 |
| GGTGGAGGAGGW | ASSPGTNTEVF | 0.3932463 |
| GVGGVAGLGGAGG | ASSPGTNTEVF | 0.24854046 |

| | | |
|---|---|---|
| VLLVLATTI | ASSPGTNTEVF | 0.5027399 |
| IGIFAAML | ASSPGTNTEVF | 0.5811427 |
| AVGDLID | ASSPGTNTEVF | 0.27652672 |
| NISPLPL | ASSPGTNTEVF | 0.38417584 |
| VTAVFVAI | ASSPGTNTEVF | 0.36928967 |
| SGDDNDAAL | ASSPGTNTEVF | 0.27287385 |
| AVAALAHS | ASSPGTNTEVF | 0.47225982 |
| GGVTVMATP | ASSPGTNTEVF | 0.2785871 |
| AGLAANVRD | ASSPGTNTEVF | 0.35174778 |
| TNRPDLIDPALLRPG | ASSPGTNTEVF | 0.4423193 |
| LANLGVTAL | ASSPGTNTEVF | 0.36578786 |
| TGERIEL | ASSPGTNTEVF | 0.3506569 |
| NGEELAA | ASSPGTNTEVF | 0.32982686 |
| VLSLVWG | ASSPGTNTEVF | 0.26523983 |
| IGYALWL | ASSPGTNTEVF | 0.374319 |
| LSLLLAPL | ASSPGTNTEVF | 0.26026422 |
| TLDEALK | ASSPGTNTEVF | 0.33397874 |
| TDGAGTAEGG | ASSPGTNTEVF | 0.28722966 |
| NAPGGAGGNGG | ASSPGTNTEVF | 0.4166737 |
| VSVFMGMGGNF | ASSPGTNTEVF | 0.44762975 |
| LASYLAAL | ASSPGTNTEVF | 0.34167027 |
| LVGGTPAL | ASSPGTNTEVF | 0.28410435 |
| VRNADVL | ASSPGTNTEVF | 0.30436555 |
| TCSSPFLVSAM | ASSPGTNTEVF | 0.29998896 |
| VHITDAE | ASSPGTNTEVF | 0.2799256 |
| EYAAPSF | ASSPGTNTEVF | 0.36487514 |
| VVDAFHGGL | ASSPGTNTEVF | 0.31501853 |
| ASVVDPQ | ASSPGTNTEVF | 0.30457523 |
| SYNEFGGLS | ASSPGTNTEVF | 0.4129265 |
| DPDGSVY | ASSPGTNTEVF | 0.35115942 |
| MVSTIHATE | ASSPGTNTEVF | 0.26858613 |
| LAAGVADPV | ASSPGTNTEVF | 0.38096833 |
| TKEQIAA | ASSPGTNTEVF | 0.34690994 |
| SLASPAAL | ASSPGTNTEVF | 0.30678323 |
| IGLVFYGV | ASSPGTNTEVF | 0.25035083 |
| FLLDASNL | ASSPGTNTEVF | 0.32529923 |
| YQVGNFP | ASSPGTNTEVF | 0.30991355 |
| ESLLSQST | ASSPGTNTEVF | 0.2849366 |
| LFTVGFAAVG | ASSPGTNTEVF | 0.30691037 |
| ATPPTPT | ASSPGTNTEVF | 0.53218 |
| TDLATLG | ASSPGTNTEVF | 0.27699652 |
| SAGGTGR | ASSPGTNTEVF | 0.3505773 |
| GFSNSGS | ASSPGTNTEVF | 0.46505052 |
| GGNGGSGGTSV | ASSPGTNTEVF | 0.43210262 |
| GTSVATGGAGN | ASSPGTNTEVF | 0.5220915 |
| NGGNGGSGGTS | ASSPGTNTEVF | 0.36748627 |
| AVVKFPH | ASSPGTNTEVF | 0.30611962 |
| TPPPPPP | ASSPGTNTEVF | 0.34592256 |
| AAPAAAAAPA | ASSPGTNTEVF | 0.31260458 |
| IALNEDL | ASSPGTNTEVF | 0.26646012 |
| IVNVSSM | ASSPGTNTEVF | 0.257704 |

| | | |
|---|---|---|
| TGIPPAPR | ASSPGTNTEVF | 0.30681857 |
| FDLGGGTF | ASSPGTNTEVF | 0.2612361 |
| FDLGGGTFDVS | ASSPGTNTEVF | 0.34151837 |
| NEPTAAA | ASSPGTNTEVF | 0.386786 |
| NEPTAAALA | ASSPGTNTEVF | 0.29169595 |
| GSIINISSI | ASSPGTNTEVF | 0.2844014 |
| VAPGNAGTA | ASSPGTNTEVF | 0.30818993 |
| TGGGASLE | ASSPGTNTEVF | 0.27770713 |
| VAVYNHYKRI | ASSPGTNTEVF | 0.46114698 |
| TDPAPATT | ASSPGTNTEVF | 0.30650502 |
| TNRPDILDPALLRPG | ASSPGTNTEVF | 0.6890533 |
| LTTTTTS | ASSPGTNTEVF | 0.5479288 |
| PTGSGKTL | ASSPGTNTEVF | 0.35574746 |
| TGSGKTL | ASSPGTNTEVF | 0.4481557 |
| TIGHVDHGK | ASSPGTNTEVF | 0.29422772 |
| TIGHVDHGKTT | ASSPGTNTEVF | 0.3263261 |
| GHVDHGKTT | ASSPGTNTEVF | 0.30437553 |
| AQFEHTLL | ASSPGTNTEVF | 0.31662527 |
| IVNQDEL | ASSPGTNTEVF | 0.31403455 |
| TNGSQFF | ASSPGTNTEVF | 0.2990688 |
| TGKGIVE | ASSPGTNTEVF | 0.26234645 |
| LISNASDA | ASSPGTNTEVF | 0.36419433 |
| AELRTLGELTKIAKAHGAQ | ASSPGTNTEVF | 0.30236197 |
| APNAVAAAEPSW | ASSPGTNTEVF | 0.26032066 |
| APQVAGAATAVL | ASSPGTNTEVF | 0.29072168 |
| ARRDPAAAAAILPTDA | ASSPGTNTEVF | 0.5445985 |
| DAAARDVLAEAGLAEYF | ASSPGTNTEVF | 0.5671943 |
| DDQIVLSAKANLAKVEQ | ASSPGTNTEVF | 0.29255867 |
| DELLPDTGEDSDEDGH | ASSPGTNTEVF | 0.2840606 |
| ELPEAPANRATVPQD | ASSPGTNTEVF | 0.57161146 |
| FGQSKIERVTGIG | ASSPGTNTEVF | 0.26873523 |
| GAGGTGSTAGGAGGAGGAGGLY | ASSPGTNTEVF | 0.3222522 |
| GHGGTGAAGGNGGTGGAGGSGI | ASSPGTNTEVF | 0.3505477 |
| GNGGNGGAGGEGGPGGAGGAGA | ASSPGTNTEVF | 0.38687482 |
| GSTAGNGGNGGNATNGTIAGQP | ASSPGTNTEVF | 0.2974086 |
| GTGGDGGNGGNGGTGG | ASSPGTNTEVF | 0.32895887 |
| GVRRPAGIPADAPV | ASSPGTNTEVF | 0.33439866 |
| GYRQVTDAHLLTLARR | ASSPGTNTEVF | 0.33342427 |
| IAKMGQDGHDRG | ASSPGTNTEVF | 0.25024536 |
| KAEPSIEIQYT | ASSPGTNTEVF | 0.325216 |
| KPVRIRMAADEDTVAVT | ASSPGTNTEVF | 0.30175504 |
| MPPRIAGMRLLVIKPEP | ASSPGTNTEVF | 0.28882188 |
| MKFYAVALFRSRGVPPTR | ASSPGTNTEVF | 0.2605863 |
| NAGGFGILGAVAHSP | ASSPGTNTEVF | 0.3322963 |
| NEGREITLAMAEM | ASSPGTNTEVF | 0.41090068 |
| NELVEWSHPLGHRWRWE | ASSPGTNTEVF | 0.46245497 |
| NLGAGGTGGNGGNPG | ASSPGTNTEVF | 0.26981956 |
| NTEWPKWETVMAADPVD | ASSPGTNTEVF | 0.4456282 |
| PASAAPAETQP | ASSPGTNTEVF | 0.39534023 |
| QQSSDGKNGVRAAEITGE | ASSPGTNTEVF | 0.31067696 |
| RAPDAVTTSWWKE | ASSPGTNTEVF | 0.2633161 |

| | | |
|---|---|---|
| SGFGNVGAHTSGWFNQ | ASSPGTNTEVF | 0.39197254 |
| SGVAGAVGQAASVGGLKV | ASSPGTNTEVF | 0.33115748 |
| SNGGAGTGSAGGNGGT | ASSPGTNTEVF | 0.33685246 |
| TGGAGGEGGAAG | ASSPGTNTEVF | 0.36636293 |
| TGGDGGAAGAGGAPGHG | ASSPGTNTEVF | 0.2775984 |
| TMPPLLAAIANAGRHGV | ASSPGTNTEVF | 0.47457433 |
| TSSTAGNSGGA | ASSPGTNTEVF | 0.39635265 |
| VPATVPVGILC | ASSPGTNTEVF | 0.28122768 |
| VQRPDSGGVVLTGRIS | ASSPGTNTEVF | 0.29703736 |
| WTPNPVPATKGVPAGLVQAT | ASSPGTNTEVF | 0.3022118 |
| YALKRALAKFSDRFDVVII | ASSPGTNTEVF | 0.25246143 |
| YLIGSVIV | ASSPGTNTEVF | 0.4223749 |
| YGQLVAVID | ASSPGTNTEVF | 0.3053253 |
| VTVGGMTLPA | ASSPGTNTEVF | 0.32831722 |
| VLALPAV | ASSPGTNTEVF | 0.3653777 |
| VIPPTLNYET | ASSPGTNTEVF | 0.3167779 |
| VGMGGMHPSSG | ASSPGTNTEVF | 0.3236231 |
| VGLAGGAATA | ASSPGTNTEVF | 0.33967397 |
| VGGAGGAGGSGP | ASSPGTNTEVF | 0.32299733 |
| VFAAFMV | ASSPGTNTEVF | 0.34815544 |
| TLYDKLR | ASSPGTNTEVF | 0.30773214 |
| TLAVFTVL | ASSPGTNTEVF | 0.34719148 |
| TGSVQPARV | ASSPGTNTEVF | 0.6177348 |
| TGGGAGSGGT | ASSPGTNTEVF | 0.28000608 |
| SHLETSL | ASSPGTNTEVF | 0.3133627 |
| SFTTGMT | ASSPGTNTEVF | 0.382882 |
| QVEVAKRMGVRQPTVS | ASSPGTNTEVF | 0.27066028 |
| QGSGGPALAM | ASSPGTNTEVF | 0.43657106 |
| MLAPSNP | ASSPGTNTEVF | 0.44754428 |
| LVNNAAI | ASSPGTNTEVF | 0.39721158 |
| LTGEINTLA | ASSPGTNTEVF | 0.30465254 |
| LSIFGPV | ASSPGTNTEVF | 0.64923733 |
| LLALLANL | ASSPGTNTEVF | 0.46924806 |
| LISNASDALDKLR | ASSPGTNTEVF | 0.32118258 |
| LINEAVP | ASSPGTNTEVF | 0.5562562 |
| LGYAFSGL | ASSPGTNTEVF | 0.2942419 |
| LDFVAVAL | ASSPGTNTEVF | 0.40020043 |
| LAQPGAE | ASSPGTNTEVF | 0.36530206 |
| ISNLGAVTGF | ASSPGTNTEVF | 0.2758914 |
| HQEDPGRI | ASSPGTNTEVF | 0.37481868 |
| GTPGEPTP | ASSPGTNTEVF | 0.37877026 |
| GFFNSGS | ASSPGTNTEVF | 0.51304245 |
| GAGGGGLTAGAN | ASSPGTNTEVF | 0.26690453 |
| FVDALVNM | ASSPGTNTEVF | 0.3164751 |
| FLTVAPEL | ASSPGTNTEVF | 0.2718206 |
| FETLTAL | ASSPGTNTEVF | 0.2961496 |
| DVINPIEL | ASSPGTNTEVF | 0.29560307 |
| APVSGAPV | ASSPGTNTEVF | 0.38123378 |
| ALVGGLAW | ASSPGTNTEVF | 0.32640895 |
| ALAAAAATAI | ASSPGTNTEVF | 0.5971828 |
| AFGLGLLL | ASSPGTNTEVF | 0.34105563 |

| | | |
|---|---|---|
| AFFAAASL | ASSPGTNTEVF | 0.4260341 |
| AAAALPAFSPP | ASSPGTNTEVF | 0.29756317 |
| VVLAPAL | ASSPGTNTEVF | 0.34992418 |
| TIGHVDHGKTTL | ASSPGTNTEVF | 0.26521805 |
| LISNASDALDK | ASSPGTNTEVF | 0.26632816 |
| LISNASDAL | ASSPGTNTEVF | 0.28464156 |
| ISNASDALDK | ASSPGTNTEVF | 0.3369248 |
| NQAETLVYQTE | ASSPGTNTEVF | 0.25467038 |
| TGGTGGTAGAG | ASSPGTNTEVF | 0.39674047 |
| TGSTAGGAGGA | ASSPGTNTEVF | 0.3816851 |
| DPLAVPVAAGPV | ASSPGTNTEVF | 0.36171913 |
| GGTGGTPGNGGN | ASSPGTNTEVF | 0.32660884 |
| PAGLAGVDPATL | ASSPGTNTEVF | 0.31269756 |
| GNNLVVEAPHAGASV | ASSPGTNTEVF | 0.30206558 |
| KVTRIMSASTLEQPA | ASSPGTNTEVF | 0.49580547 |
| LGFALPIALPRRTVV | ASSPGTNTEVF | 0.27317354 |
| TRLGNLTARKLVVLV | ASSPGTNTEVF | 0.27071145 |
| WSQQRRLIYFPSAGP | ASSPGTNTEVF | 0.2904163 |
| GPNGNPDPMAAAVDIR | ASSPGTNTEVF | 0.27219486 |
| TQRCVSVADLRAAASR | ASSPGTNTEVF | 0.53308445 |
| QFVLDVAEHPVEVLTETV | ASSPGTNTEVF | 0.39556822 |
| GGAGGTGGAGGNGGTQAGDG | ASSPGTNTEVF | 0.3990664 |
| PLLATLRAIAVDVDAKPAQV | ASSPGTNTEVF | 0.5266074 |
| APKFRTGGYMIDFWGVGYQVAK | ASSPGTNTEVF | 0.25715828 |
| GAGGAGGAGGAAGGPGGTGGTG | ASSPGTNTEVF | 0.3188285 |
| IGALGLLATDVAPAGSVAFGVA | ASSPGTNTEVF | 0.6295428 |
| AGGAGGIGGNGGAGGAGGVLMG | ASSPGTNTEVF | 0.3136835 |
| ADELIGLL | ASSPGTNTEVF | 0.334821 |
| ADRALKA | ASSPGTNTEVF | 0.41986173 |
| AGAVGQVGGAGGA | ASSPGTNTEVF | 0.34279847 |
| AGGGGGLGGAP | ASSPGTNTEVF | 0.30713487 |
| AGGLLYGNGGAG | ASSPGTNTEVF | 0.35034764 |
| AGLESIGL | ASSPGTNTEVF | 0.32901126 |
| AGLKTGII | ASSPGTNTEVF | 0.31171072 |
| ASLAALLL | ASSPGTNTEVF | 0.27184737 |
| ASVINISSI | ASSPGTNTEVF | 0.37140772 |
| ATPPGPA | ASSPGTNTEVF | 0.69818807 |
| ATVASALV | ASSPGTNTEVF | 0.35623908 |
| ATVASLIL | ASSPGTNTEVF | 0.34381616 |
| AVNGPVLIRH | ASSPGTNTEVF | 0.40081564 |
| DGAAGAVGI | ASSPGTNTEVF | 0.36691827 |
| DPLLHGI | ASSPGTNTEVF | 0.29814 |
| ELGAALGAL | ASSPGTNTEVF | 0.2905349 |
| ELGRTQAV | ASSPGTNTEVF | 0.31646183 |
| FVAGGYLS | ASSPGTNTEVF | 0.28022718 |
| GASGGAGGAGGW | ASSPGTNTEVF | 0.3194632 |
| GGAGAGGDGGAGGI | ASSPGTNTEVF | 0.31267217 |
| GTGHNGSQPGMGGNG | ASSPGTNTEVF | 0.6823618 |
| GYAAASASAS | ASSPGTNTEVF | 0.40097088 |
| HQVDVAAVQQG | ASSPGTNTEVF | 0.3589469 |
| IGAAIAHT | ASSPGTNTEVF | 0.29999775 |

| | | |
|---|---|---|
| IGMIEVAQL | ASSPGTNTEVF | 0.31211686 |
| IRLAPEHTSAMD | ASSPGTNTEVF | 0.33228582 |
| IYNVSVL | ASSPGTNTEVF | 0.26489827 |
| KQGFLGM | ASSPGTNTEVF | 0.5089136 |
| LESVITQL | ASSPGTNTEVF | 0.29463953 |
| LGLGALVVV | ASSPGTNTEVF | 0.29247132 |
| LIGNGGNGGSGGI | ASSPGTNTEVF | 0.36472464 |
| LNAINAPTL | ASSPGTNTEVF | 0.6757293 |
| LQLLNTGV | ASSPGTNTEVF | 0.3045388 |
| LQTSIEF | ASSPGTNTEVF | 0.3134255 |
| MSFAVSEL | ASSPGTNTEVF | 0.30842167 |
| NTGALGL | ASSPGTNTEVF | 0.37578917 |
| PPGAGGAGGAGTT | ASSPGTNTEVF | 0.5893893 |
| QLVGLGI | ASSPGTNTEVF | 0.29542196 |
| RFLEQVHQL | ASSPGTNTEVF | 0.6361214 |
| RGVGYVLR | ASSPGTNTEVF | 0.322597 |
| SATGVFAGVI | ASSPGTNTEVF | 0.2763719 |
| SLPAITLP | ASSPGTNTEVF | 0.33068988 |
| STALAFTV | ASSPGTNTEVF | 0.34160918 |
| SVLTVGLGK | ASSPGTNTEVF | 0.4782235 |
| TALAGAGSGI | ASSPGTNTEVF | 0.4018442 |
| TGLPFTGI | ASSPGTNTEVF | 0.45449537 |
| VAEHVTV | ASSPGTNTEVF | 0.28579226 |
| VDIVLNSL | ASSPGTNTEVF | 0.4473708 |
| VLTGGGAVL | ASSPGTNTEVF | 0.38157517 |
| VNVGLAYGI | ASSPGTNTEVF | 0.40629205 |
| VSSAAVGAV | ASSPGTNTEVF | 0.29735956 |
| VTSVYEAGV | ASSPGTNTEVF | 0.2895925 |
| WVALDRGA | ASSPGTNTEVF | 0.33542812 |
| YLVGSVVI | ASSPGTNTEVF | 0.4407689 |
| GPPGTGKTLL | ASSPGTNTEVF | 0.27956846 |
| GVGMTKF | ASSPGTNTEVF | 0.34806305 |
| IGIGGSDL | ASSPGTNTEVF | 0.40580183 |
| IGTLLAEL | ASSPGTNTEVF | 0.27717212 |
| IYTSGTT | ASSPGTNTEVF | 0.32749206 |
| LSFPAGV | ASSPGTNTEVF | 0.57721186 |
| LYTSGTT | ASSPGTNTEVF | 0.31877655 |
| VGAFPFGF | ASSPGTNTEVF | 0.2976068 |
| VGAGTVEF | ASSPGTNTEVF | 0.52979726 |
| VGKSTLLL | ASSPGTNTEVF | 0.6849668 |
| AALAHAGGGIV | ASSPGTNTEVF | 0.3918604 |
| AALVAVSLL | ASSPGTNTEVF | 0.2814542 |
| AAVSRAVAAL | ASSPGTNTEVF | 0.2563074 |
| AGGSSGIGGF | ASSPGTNTEVF | 0.2968931 |
| AGLTVANANG | ASSPGTNTEVF | 0.5232248 |
| AGLVGAVL | ASSPGTNTEVF | 0.3396606 |
| ALSQAFTV | ASSPGTNTEVF | 0.3003055 |
| ALTFSVSL | ASSPGTNTEVF | 0.38217753 |
| ALTGAALVAL | ASSPGTNTEVF | 0.41258398 |
| ASFSIGAGVL | ASSPGTNTEVF | 0.297312 |
| DGSVYKI | ASSPGTNTEVF | 0.26984498 |

| | | |
|---|---|---|
| EQLAEIV | ASSPGTNTEVF | 0.37194994 |
| FAVSELTVM | ASSPGTNTEVF | 0.39279655 |
| GAAPLAAGL | ASSPGTNTEVF | 0.2872425 |
| GADGGDGGSGGAGG | ASSPGTNTEVF | 0.31954688 |
| GFGLDTGGAGG | ASSPGTNTEVF | 0.30651623 |
| GLASGGLL | ASSPGTNTEVF | 0.28416508 |
| GRIAGAAVLL | ASSPGTNTEVF | 0.34383157 |
| GSAGSLNTSF | ASSPGTNTEVF | 0.29159915 |
| GTGQIGI | ASSPGTNTEVF | 0.4357239 |
| GTLIALIL | ASSPGTNTEVF | 0.29651353 |
| IAATMGFVVG | ASSPGTNTEVF | 0.34945452 |
| IFAQALL | ASSPGTNTEVF | 0.42529848 |
| IGAAVALGF | ASSPGTNTEVF | 0.34162626 |
| IGLTPLAL | ASSPGTNTEVF | 0.31822485 |
| ILGSNTSTL | ASSPGTNTEVF | 0.3011798 |
| IVFSLTVL | ASSPGTNTEVF | 0.27029136 |
| LASKATGF | ASSPGTNTEVF | 0.25645584 |
| LASVALIL | ASSPGTNTEVF | 0.56335753 |
| LFAVGLTL | ASSPGTNTEVF | 0.26262933 |
| LFNGGIGTYI | ASSPGTNTEVF | 0.47820085 |
| LTLGAGII | ASSPGTNTEVF | 0.35986102 |
| LVAAALGDI | ASSPGTNTEVF | 0.28475037 |
| NTAFVQLGI | ASSPGTNTEVF | 0.3276557 |
| QLFSGAEL | ASSPGTNTEVF | 0.28605336 |
| SAATFITL | ASSPGTNTEVF | 0.27927148 |
| SNYGLANLGV | ASSPGTNTEVF | 0.3521618 |
| SYGGAYAVM | ASSPGTNTEVF | 0.35411206 |
| TTGVGGLTLG | ASSPGTNTEVF | 0.30488783 |
| TTQTIYI | ASSPGTNTEVF | 0.3325733 |
| VAAGAGLGAVI | ASSPGTNTEVF | 0.37251332 |
| VAAVVTLL | ASSPGTNTEVF | 0.28938076 |
| VGFLGGLI | ASSPGTNTEVF | 0.3329569 |
| VGVIPILL | ASSPGTNTEVF | 0.49623838 |
| VIFASSLI | ASSPGTNTEVF | 0.339983 |
| VKGGDGGAAAG | ASSPGTNTEVF | 0.27823666 |
| VTKTFTNI | ASSPGTNTEVF | 0.3987528 |
| VVAAAILL | ASSPGTNTEVF | 0.28582415 |
| YAALDVEL | ASSPGTNTEVF | 0.47268945 |
| LASVALLL | ASSPGTNTEVF | 0.31291443 |
| LDAVHRV | ASSPGTNTEVF | 0.32688978 |
| QGLSSSIF | ASSPGTNTEVF | 0.3645921 |
| RGTFANIRL | ASSPGTNTEVF | 0.27126527 |
| RQLWWGHRI | ASSPGTNTEVF | 0.2842492 |
| YGPPGTGKTLL | ASSPGTNTEVF | 0.29873794 |
| AALVGAPANVA | ASSPGTNTEVF | 0.37935877 |
| AATFRRTE | ASSPGTNTEVF | 0.42151892 |
| AGALGGGI | ASSPGTNTEVF | 0.28607225 |
| AGGAGGAGGIGAG | ASSPGTNTEVF | 0.3558287 |
| AGSGALGL | ASSPGTNTEVF | 0.30907017 |
| AQIADQI | ASSPGTNTEVF | 0.29183757 |
| ASAVAVTYL | ASSPGTNTEVF | 0.3197012 |

| | | |
|---|---|---|
| ASTVLGGTGG | ASSPGTNTEVF | 0.2976929 |
| AVGGEPAK | ASSPGTNTEVF | 0.29637754 |
| CEQGTFGGTC | ASSPGTNTEVF | 0.34307542 |
| DEGKAPAG | ASSPGTNTEVF | 0.30915502 |
| DTVVFPAL | ASSPGTNTEVF | 0.34963298 |
| FEGITMAG | ASSPGTNTEVF | 0.35699072 |
| FPLVVQD | ASSPGTNTEVF | 0.30192843 |
| GAFTTPQ | ASSPGTNTEVF | 0.33683446 |
| GFGGFGTAGGVGGA | ASSPGTNTEVF | 0.334846 |
| GFLAQFL | ASSPGTNTEVF | 0.44170955 |
| GGAGGAGGNAGVA | ASSPGTNTEVF | 0.3340298 |
| GGVGTAGAGGAGGAGG | ASSPGTNTEVF | 0.28487542 |
| GNGGGGDAGNAGSGGN | ASSPGTNTEVF | 0.39449167 |
| GTGGAGGAGNDAGS | ASSPGTNTEVF | 0.42006394 |
| GTTNTGWFNS | ASSPGTNTEVF | 0.449616 |
| IFMGPLFTL | ASSPGTNTEVF | 0.36884925 |
| ILAHAIEAI | ASSPGTNTEVF | 0.50692403 |
| IRSAIREAL | ASSPGTNTEVF | 0.40292826 |
| ITFAGGG | ASSPGTNTEVF | 0.33442575 |
| ITGMSGHQEV | ASSPGTNTEVF | 0.26535103 |
| IVAGLACTL | ASSPGTNTEVF | 0.32920673 |
| IVGGGLAGA | ASSPGTNTEVF | 0.32239446 |
| IVTGVLNI | ASSPGTNTEVF | 0.31023294 |
| LAQQGCI | ASSPGTNTEVF | 0.26413298 |
| LCLHVEN | ASSPGTNTEVF | 0.5389014 |
| LDAISDGS | ASSPGTNTEVF | 0.640887 |
| LGGFAAML | ASSPGTNTEVF | 0.31748092 |
| LGGKNPMI | ASSPGTNTEVF | 0.46805665 |
| LGPKGVAAIL | ASSPGTNTEVF | 0.27185145 |
| LMVGVSI | ASSPGTNTEVF | 0.37618548 |
| LVGAAFVL | ASSPGTNTEVF | 0.51388204 |
| LVLAVAPL | ASSPGTNTEVF | 0.2798346 |
| MAAIAVIDAA | ASSPGTNTEVF | 0.37146503 |
| NNGINAELQ | ASSPGTNTEVF | 0.27333525 |
| RGEPLVLVHG | ASSPGTNTEVF | 0.29677108 |
| RGGAGGAAGLI | ASSPGTNTEVF | 0.55193627 |
| RSYRMHPAV | ASSPGTNTEVF | 0.35487837 |
| SAAHLLAGQLA | ASSPGTNTEVF | 0.33921134 |
| SAAVGSY | ASSPGTNTEVF | 0.3294933 |
| SPHAVLHRI | ASSPGTNTEVF | 0.293944 |
| SQIAKVL | ASSPGTNTEVF | 0.31219217 |
| TALGHLVP | ASSPGTNTEVF | 0.49883527 |
| TAPTTSGAA | ASSPGTNTEVF | 0.27783313 |
| TDGDDGNGGSPGT | ASSPGTNTEVF | 0.3813092 |
| TDGLGEN | ASSPGTNTEVF | 0.30307794 |
| TGGPPLPGQA | ASSPGTNTEVF | 0.6096132 |
| TIPAITI | ASSPGTNTEVF | 0.63461053 |
| TIPGFTI | ASSPGTNTEVF | 0.39943984 |
| TSIASGALA | ASSPGTNTEVF | 0.2945923 |
| TVTLPGAAI | ASSPGTNTEVF | 0.62367797 |
| VASGLSAALP | ASSPGTNTEVF | 0.3419734 |

| | | |
|---|---|---|
| VGAGVIGI | ASSPGTNTEVF | 0.41485313 |
| VGPKRFGV | ASSPGTNTEVF | 0.35357675 |
| VNVAFPD | ASSPGTNTEVF | 0.2204306 |
| VSMTNSGVS | ASSPGTNTEVF | 0.81709856 |
| VTDVLGDLSV | ASSPGTNTEVF | 0.5890101 |
| VVVVGAGFAGL | ASSPGTNTEVF | 0.41943765 |
| VYTVGTVV | ASSPGTNTEVF | 0.27564853 |
| WTTFSGP | ASSPGTNTEVF | 0.15886356 |
| YDELVEAF | ASSPGTNTEVF | 0.31378546 |
| YGTGGAGGNGG | ASSPGTNTEVF | 0.24049568 |
| YTNVMAV | ASSPGTNTEVF | 0.6969093 |
| FLEQVHQL | ASSPGTNTEVF | 0.52139056 |
| GLSLLAVAV | ASSPGTNTEVF | 0.27117255 |
| LEQVHQL | ASSPGTNTEVF | 0.6687956 |
| LNAAGAI | ASSPGTNTEVF | 0.72910047 |
| PAVHASL | ASSPGTNTEVF | 0.15839131 |
| TELEDDA | ASSPGTNTEVF | 0.33980036 |
| TPPGPPN | ASSPGTNTEVF | 0.23184331 |
| TPPPPPPP | ASSPGTNTEVF | 0.317691 |
| TQSPALI | GARRDTANSDYT | 0.11247396 |
| EFAAAFPQ | GARRDTANSDYT | 0.55274606 |
| LLSGNLDVL | GARRDTANSDYT | 0.26589206 |
| GPANLGNYNI | GARRDTANSDYT | 0.08848254 |
| TLGPTLP | GARRDTANSDYT | 0.3318608 |
| VYAYSSCCK | GARRDTANSDYT | 0.30145234 |
| APAPAPAPNI | GARRDTANSDYT | 0.31362003 |
| PGGPVPIP | GARRDTANSDYT | 0.6804925 |
| FLTVALEP | GARRDTANSDYT | 0.2312487 |
| SQYFGANL | GARRDTANSDYT | 0.23682913 |
| GGVKPAA | GARRDTANSDYT | 0.60654926 |
| GGTGGAGGAGGW | GARRDTANSDYT | 0.7886273 |
| GVGGVAGLGGAGG | GARRDTANSDYT | 0.21718162 |
| VLLVLATTI | GARRDTANSDYT | 0.5349489 |
| IGIFAAML | GARRDTANSDYT | 0.80056715 |
| AVGDLID | GARRDTANSDYT | 0.4986435 |
| NISPLPL | GARRDTANSDYT | 0.68757594 |
| VTAVFVAI | GARRDTANSDYT | 0.75803983 |
| SGDDNDAAL | GARRDTANSDYT | 0.20297973 |
| AVAALAHS | GARRDTANSDYT | 0.69516426 |
| GGVTVMATP | GARRDTANSDYT | 0.748826 |
| AGLAANVRD | GARRDTANSDYT | 0.36306798 |
| TNRPDLIDPALLRPG | GARRDTANSDYT | 0.3689602 |
| LANLGVTAL | GARRDTANSDYT | 0.226355 |
| TGERIEL | GARRDTANSDYT | 0.26192963 |
| NGEELAA | GARRDTANSDYT | 0.14220613 |
| VLSLVWG | GARRDTANSDYT | 0.589225 |
| IGYALWL | GARRDTANSDYT | 0.6540107 |
| LSLLLAPL | GARRDTANSDYT | 0.3127016 |
| TLDEALK | GARRDTANSDYT | 0.16172063 |
| TDGAGTAEGG | GARRDTANSDYT | 0.16600026 |
| NAPGGAGGNGG | GARRDTANSDYT | 0.47352624 |

| | | |
|---|---|---|
| VSVFMGMGGNF | GARRDTANSDYT | 0.283944 |
| LASYLAAL | GARRDTANSDYT | 0.16318065 |
| LVGGTPAL | GARRDTANSDYT | 0.28671378 |
| VRNADVL | GARRDTANSDYT | 0.6572081 |
| TCSSPFLVSAM | GARRDTANSDYT | 0.687074 |
| VHITDAE | GARRDTANSDYT | 0.15142122 |
| EYAAPSF | GARRDTANSDYT | 0.7883326 |
| VVDAFHGGL | GARRDTANSDYT | 0.72131246 |
| ASVVDPQ | GARRDTANSDYT | 0.8105443 |
| SYNEFGGLS | GARRDTANSDYT | 0.3882004 |
| DPDGSVY | GARRDTANSDYT | 0.23733976 |
| MVSTIHATE | GARRDTANSDYT | 0.733185 |
| LAAGVADPV | GARRDTANSDYT | 0.350238 |
| TKEQIAA | GARRDTANSDYT | 0.2373052 |
| SLASPAAL | GARRDTANSDYT | 0.5157351 |
| IGLVFYGV | GARRDTANSDYT | 0.3421921 |
| FLLDASNL | GARRDTANSDYT | 0.26491043 |
| YQVGNFP | GARRDTANSDYT | 0.50600123 |
| ESLLSQST | GARRDTANSDYT | 0.68617547 |
| LFTVGFAAVG | GARRDTANSDYT | 0.15544845 |
| ATPPTPT | GARRDTANSDYT | 0.2701462 |
| TDLATLG | GARRDTANSDYT | 0.2051447 |
| SAGGTGR | GARRDTANSDYT | 0.41100544 |
| GFSNSGS | GARRDTANSDYT | 0.7295307 |
| GGNGGSGGTSV | GARRDTANSDYT | 0.44601944 |
| GTSVATGGAGN | GARRDTANSDYT | 0.40327322 |
| NGGNGGSGGTS | GARRDTANSDYT | 0.5955045 |
| AVVKFPH | GARRDTANSDYT | 0.5992668 |
| TPPPPPP | GARRDTANSDYT | 0.39483136 |
| AAPAAAAAPA | GARRDTANSDYT | 0.6069324 |
| IALNEDL | GARRDTANSDYT | 0.27557287 |
| IVNVSSM | GARRDTANSDYT | 0.6830299 |
| TGIPPAPR | GARRDTANSDYT | 0.5012258 |
| FDLGGGTF | GARRDTANSDYT | 0.5925335 |
| FDLGGGTFDVS | GARRDTANSDYT | 0.6811725 |
| NEPTAAA | GARRDTANSDYT | 0.6581354 |
| NEPTAAALA | GARRDTANSDYT | 0.40658924 |
| GSIINISSI | GARRDTANSDYT | 0.13411468 |
| VAPGNAGTA | GARRDTANSDYT | 0.24491431 |
| TGGGASLE | GARRDTANSDYT | 0.44767743 |
| VAVYNHYKRI | GARRDTANSDYT | 0.5097854 |
| TDPAPATT | GARRDTANSDYT | 0.5039097 |
| TNRPDILDPALLRPG | GARRDTANSDYT | 0.54101443 |
| LTTTTTS | GARRDTANSDYT | 0.22859848 |
| PTGSGKTL | GARRDTANSDYT | 0.64565665 |
| TGSGKTL | GARRDTANSDYT | 0.73392725 |
| TIGHVDHGK | GARRDTANSDYT | 0.16164732 |
| TIGHVDHGKTT | GARRDTANSDYT | 0.31206313 |
| GHVDHGKTT | GARRDTANSDYT | 0.828511 |
| AQFEHTLL | GARRDTANSDYT | 0.5967938 |
| IVNQDEL | GARRDTANSDYT | 0.26073143 |

| | | |
|---|---|---|
| TNGSQFF | GARRDTANSDYT | 0.31161004 |
| TGKGIVE | GARRDTANSDYT | 0.7981183 |
| LISNASDA | GARRDTANSDYT | 0.31212988 |
| AELRTLGELTKIAKAHGAQ | GARRDTANSDYT | 0.7488718 |
| APNAVAAAEPSW | GARRDTANSDYT | 0.107887805 |
| APQVAGAATAVL | GARRDTANSDYT | 0.3377055 |
| ARRDPAAAAAILPTDA | GARRDTANSDYT | 0.30811605 |
| DAAARDVLAEAGLAEYF | GARRDTANSDYT | 0.14225237 |
| DDQIVLSAKANLAKVEQ | GARRDTANSDYT | 0.5999748 |
| DELLPDTGEDSDEDGH | GARRDTANSDYT | 0.25679782 |
| ELPEAPANRATVPQD | GARRDTANSDYT | 0.60863715 |
| FGQSKIERVTGIG | GARRDTANSDYT | 0.27123493 |
| GAGGTGSTAGGAGGAGGAGGLY | GARRDTANSDYT | 0.6362508 |
| GHGGTGAAGGNGGTGGAGGSGI | GARRDTANSDYT | 0.5649409 |
| GNGGNGGAGGEGGPGGAGGAGA | GARRDTANSDYT | 0.35174742 |
| GSTAGNGGNGGNATNGTIAGQP | GARRDTANSDYT | 0.2594128 |
| GTGGDGGNGGNGGTGG | GARRDTANSDYT | 0.5135355 |
| GVRRPAGIPADAPV | GARRDTANSDYT | 0.6257753 |
| GYRQVTDAHLLTLARR | GARRDTANSDYT | 0.61352587 |
| IAKMGQDGHDRG | GARRDTANSDYT | 0.16548613 |
| KAEPSIEIQYT | GARRDTANSDYT | 0.39898133 |
| KPVRIRMAADEDTVAVT | GARRDTANSDYT | 0.4049457 |
| MPPRIAGMRLLVIKPEP | GARRDTANSDYT | 0.13808775 |
| MKFYAVALFRSRGVPPTR | GARRDTANSDYT | 0.10445813 |
| NAGGFGILGAVAHSP | GARRDTANSDYT | 0.35809425 |
| NEGREITLAMAEM | GARRDTANSDYT | 0.30522758 |
| NELVEWSHPLGHRWRWE | GARRDTANSDYT | 0.6344253 |
| NLGAGGTGGNGGNPG | GARRDTANSDYT | 0.6471621 |
| NTEWPKWETVMAADPVD | GARRDTANSDYT | 0.29618877 |
| PASAAPAETQP | GARRDTANSDYT | 0.6648227 |
| QQSSDGKNGVRAAEITGE | GARRDTANSDYT | 0.48983508 |
| RAPDAVTTSWWKE | GARRDTANSDYT | 0.08034459 |
| SGFGNVGAHTSGWFNQ | GARRDTANSDYT | 0.2815352 |
| SGVAGAVGQAASVGGLKV | GARRDTANSDYT | 0.14965667 |
| SNGGAGTGSAGGNGGT | GARRDTANSDYT | 0.677145 |
| TGGAGGEGGAAG | GARRDTANSDYT | 0.20394373 |
| TGGDGGAAGAGGAPGHG | GARRDTANSDYT | 0.20198503 |
| TMPPLLAAIANAGRHGV | GARRDTANSDYT | 0.3738079 |
| TSSTAGNSGGA | GARRDTANSDYT | 0.6450299 |
| VPATVPVGILC | GARRDTANSDYT | 0.4079795 |
| VQRPDSGGVVLTGRIS | GARRDTANSDYT | 0.25130945 |
| WTPNPVPATKGVPAGLVQAT | GARRDTANSDYT | 0.09507762 |
| YALKRALAKFSDRFDVVII | GARRDTANSDYT | 0.5598985 |
| YLIGSVIV | GARRDTANSDYT | 0.231832 |
| YGQLVAVID | GARRDTANSDYT | 0.24288993 |
| VTVGGMTLPA | GARRDTANSDYT | 0.12726831 |
| VLALPAV | GARRDTANSDYT | 0.11527407 |
| VIPPTLNYET | GARRDTANSDYT | 0.1742384 |
| VGMGGMHPSSG | GARRDTANSDYT | 0.621095 |
| VGLAGGAATA | GARRDTANSDYT | 0.31949234 |
| VGGAGGAGGSGP | GARRDTANSDYT | 0.27397257 |

| | | |
|---|---|---|
| VFAAFMV | GARRDTANSDYT | 0.7858887 |
| TLYDKLR | GARRDTANSDYT | 0.6595869 |
| TLAVFTVL | GARRDTANSDYT | 0.29261357 |
| TGSVQPARV | GARRDTANSDYT | 0.29516333 |
| TGGGAGSGGT | GARRDTANSDYT | 0.20084417 |
| SHLETSL | GARRDTANSDYT | 0.2331061 |
| SFTTGMT | GARRDTANSDYT | 0.23972087 |
| QVEVAKRMGVRQPTVS | GARRDTANSDYT | 0.75450796 |
| QGSGGPALAM | GARRDTANSDYT | 0.34433645 |
| MLAPSNP | GARRDTANSDYT | 0.18747453 |
| LVNNAAI | GARRDTANSDYT | 0.6429656 |
| LTGEINTLA | GARRDTANSDYT | 0.7315233 |
| LSIFGPV | GARRDTANSDYT | 0.8277285 |
| LLALLANL | GARRDTANSDYT | 0.19611083 |
| LISNASDALDKLR | GARRDTANSDYT | 0.3159607 |
| LINEAVP | GARRDTANSDYT | 0.3217218 |
| LGYAFSGL | GARRDTANSDYT | 0.40726867 |
| LDFVAVAL | GARRDTANSDYT | 0.2811705 |
| LAQPGAE | GARRDTANSDYT | 0.23396301 |
| ISNLGAVTGF | GARRDTANSDYT | 0.65914714 |
| HQEDPGRI | GARRDTANSDYT | 0.8450348 |
| GTPGEPTP | GARRDTANSDYT | 0.30327487 |
| GFFNSGS | GARRDTANSDYT | 0.14215201 |
| GAGGGGLTAGAN | GARRDTANSDYT | 0.3029144 |
| FVDALVNM | GARRDTANSDYT | 0.21066166 |
| FLTVAPEL | GARRDTANSDYT | 0.2548193 |
| FETLTAL | GARRDTANSDYT | 0.5136907 |
| DVINPIEL | GARRDTANSDYT | 0.14965744 |
| APVSGAPV | GARRDTANSDYT | 0.2760077 |
| ALVGGLAW | GARRDTANSDYT | 0.8401803 |
| ALAAAAATAI | GARRDTANSDYT | 0.34917742 |
| AFGLGLLL | GARRDTANSDYT | 0.59344697 |
| AFFAAASL | GARRDTANSDYT | 0.73121357 |
| AAAALPAFSPP | GARRDTANSDYT | 0.5502192 |
| VVLAPAL | GARRDTANSDYT | 0.7867876 |
| TIGHVDHGKTTL | GARRDTANSDYT | 0.7492054 |
| LISNASDALDK | GARRDTANSDYT | 0.33977187 |
| LISNASDAL | GARRDTANSDYT | 0.4338548 |
| ISNASDALDK | GARRDTANSDYT | 0.48620927 |
| NQAETLVYQTE | GARRDTANSDYT | 0.33522093 |
| TGGTGGTAGAG | GARRDTANSDYT | 0.30973163 |
| TGSTAGGAGGA | GARRDTANSDYT | 0.28460526 |
| DPLAVPVAAGPV | GARRDTANSDYT | 0.64153904 |
| GGTGGTPGNGGN | GARRDTANSDYT | 0.51931846 |
| PAGLAGVDPATL | GARRDTANSDYT | 0.67280555 |
| GNNLVVEAPHAGASV | GARRDTANSDYT | 0.16195594 |
| KVTRIMSASTLEQPA | GARRDTANSDYT | 0.21528035 |
| LGFALPIALPRRTVV | GARRDTANSDYT | 0.72828764 |
| TRLGNLTARKLVVLV | GARRDTANSDYT | 0.10159969 |
| WSQQRRLIYFPSAGP | GARRDTANSDYT | 0.14224407 |
| GPNGNPDPMAAAVDIR | GARRDTANSDYT | 0.48437786 |

| | | |
|---|---|---|
| TQRCVSVADLRAAASR | GARRDTANSDYT | 0.8074213 |
| QFVLDVAEHPVEVLTETV | GARRDTANSDYT | 0.11347224 |
| GGAGGTGGAGGNGGTQAGDG | GARRDTANSDYT | 0.23559201 |
| PLLATLRAIAVDVDAKPAQV | GARRDTANSDYT | 0.16790047 |
| APKFRTGGYMIDFWGVGYQVAK | GARRDTANSDYT | 0.7142267 |
| GAGGAGGAGGAAGGPGGTGGTG | GARRDTANSDYT | 0.4220959 |
| IGALGLLATDVAPAGSVAFGVA | GARRDTANSDYT | 0.15041049 |
| AGGAGGIGGNGGAGGAGGVLMG | GARRDTANSDYT | 0.55470645 |
| ADELIGLL | GARRDTANSDYT | 0.12354983 |
| ADRALKA | GARRDTANSDYT | 0.24439365 |
| AGAVGQVGGAGGA | GARRDTANSDYT | 0.5969868 |
| AGGGGGLGGAP | GARRDTANSDYT | 0.8020688 |
| AGGLLYGNGGAG | GARRDTANSDYT | 0.18159701 |
| AGLESIGL | GARRDTANSDYT | 0.6947427 |
| AGLKTGII | GARRDTANSDYT | 0.2758809 |
| ASLAALLL | GARRDTANSDYT | 0.8408226 |
| ASVINISSI | GARRDTANSDYT | 0.27084684 |
| ATPPGPA | GARRDTANSDYT | 0.1578172 |
| ATVASALV | GARRDTANSDYT | 0.63911843 |
| ATVASLIL | GARRDTANSDYT | 0.62942684 |
| AVNGPVLIRH | GARRDTANSDYT | 0.40627122 |
| DGAAGAVGI | GARRDTANSDYT | 0.7238747 |
| DPLLHGI | GARRDTANSDYT | 0.43284553 |
| ELGAALGAL | GARRDTANSDYT | 0.31730175 |
| ELGRTQAV | GARRDTANSDYT | 0.7039452 |
| FVAGGYLS | GARRDTANSDYT | 0.6263511 |
| GASGGAGGAGGW | GARRDTANSDYT | 0.24955271 |
| GGAGAGGDGGAGGI | GARRDTANSDYT | 0.5709492 |
| GTGHNGSQPGMGGNG | GARRDTANSDYT | 0.39445436 |
| GYAAASASAS | GARRDTANSDYT | 0.22568552 |
| HQVDVAAVQQG | GARRDTANSDYT | 0.8191838 |
| IGAAIAHT | GARRDTANSDYT | 0.43605542 |
| IGMIEVAQL | GARRDTANSDYT | 0.64227164 |
| IRLAPEHTSAMD | GARRDTANSDYT | 0.5006348 |
| IYNVSVL | GARRDTANSDYT | 0.8194481 |
| KQGFLGM | GARRDTANSDYT | 0.17698842 |
| LESVITQL | GARRDTANSDYT | 0.5365427 |
| LGLGALVVV | GARRDTANSDYT | 0.38337338 |
| LIGNGGNGGSGGI | GARRDTANSDYT | 0.66223335 |
| LNAINAPTL | GARRDTANSDYT | 0.4913419 |
| LQLLNTGV | GARRDTANSDYT | 0.21585992 |
| LQTSIEF | GARRDTANSDYT | 0.39419502 |
| MSFAVSEL | GARRDTANSDYT | 0.69928837 |
| NTGALGL | GARRDTANSDYT | 0.39918005 |
| PPGAGGAGGAGTT | GARRDTANSDYT | 0.4830822 |
| QLVGLGI | GARRDTANSDYT | 0.81351256 |
| RFLEQVHQL | GARRDTANSDYT | 0.21822141 |
| RGVGYVLR | GARRDTANSDYT | 0.35624877 |
| SATGVFAGVI | GARRDTANSDYT | 0.62267727 |
| SLPAITLP | GARRDTANSDYT | 0.65770614 |
| STALAFTV | GARRDTANSDYT | 0.37160006 |

| | | |
|---|---|---|
| SVLTVGLGK | GARRDTANSDYT | 0.72566473 |
| TALAGAGSGI | GARRDTANSDYT | 0.22164859 |
| TGLPFTGI | GARRDTANSDYT | 0.61850667 |
| VAEHVTV | GARRDTANSDYT | 0.26698434 |
| VDIVLNSL | GARRDTANSDYT | 0.44834325 |
| VLTGGGAVL | GARRDTANSDYT | 0.21728024 |
| VNVGLAYGI | GARRDTANSDYT | 0.51529044 |
| VSSAAVGAV | GARRDTANSDYT | 0.68218607 |
| VTSVYEAGV | GARRDTANSDYT | 0.60972714 |
| WVALDRGA | GARRDTANSDYT | 0.103677504 |
| YLVGSVVI | GARRDTANSDYT | 0.6786514 |
| GPPGTGKTLL | GARRDTANSDYT | 0.6626371 |
| GVGMTKF | GARRDTANSDYT | 0.84492856 |
| IGIGGSDL | GARRDTANSDYT | 0.4643969 |
| IGTLLAEL | GARRDTANSDYT | 0.5443363 |
| IYTSGTT | GARRDTANSDYT | 0.30518836 |
| LSFPAGV | GARRDTANSDYT | 0.40152377 |
| LYTSGTT | GARRDTANSDYT | 0.16884004 |
| VGAFPFGF | GARRDTANSDYT | 0.3334407 |
| VGAGTVEF | GARRDTANSDYT | 0.3490379 |
| VGKSTLLL | GARRDTANSDYT | 0.2167002 |
| AALAHAGGGIV | GARRDTANSDYT | 0.74885756 |
| AALVAVSLL | GARRDTANSDYT | 0.6941487 |
| AAVSRAVAAL | GARRDTANSDYT | 0.73421884 |
| AGGSSGIGGF | GARRDTANSDYT | 0.7521657 |
| AGLTVANANG | GARRDTANSDYT | 0.76872724 |
| AGLVGAVL | GARRDTANSDYT | 0.62598807 |
| ALSQAFTV | GARRDTANSDYT | 0.28180742 |
| ALTFSVSL | GARRDTANSDYT | 0.2606158 |
| ALTGAALVAL | GARRDTANSDYT | 0.16905709 |
| ASFSIGAGVL | GARRDTANSDYT | 0.2678341 |
| DGSVYKI | GARRDTANSDYT | 0.30286294 |
| EQLAEIV | GARRDTANSDYT | 0.4284371 |
| FAVSELTVM | GARRDTANSDYT | 0.097853504 |
| GAAPLAAGL | GARRDTANSDYT | 0.62384653 |
| GADGGDGGSGGAGG | GARRDTANSDYT | 0.7489205 |
| GFGLDTGGAGG | GARRDTANSDYT | 0.42207456 |
| GLASGGLL | GARRDTANSDYT | 0.47911787 |
| GRIAGAAVLL | GARRDTANSDYT | 0.46145087 |
| GSAGSLNTSF | GARRDTANSDYT | 0.26849675 |
| GTGQIGI | GARRDTANSDYT | 0.65586543 |
| GTLIALIL | GARRDTANSDYT | 0.22145101 |
| IAATMGFVVG | GARRDTANSDYT | 0.5745362 |
| IFAQALL | GARRDTANSDYT | 0.78473556 |
| IGAAVALGF | GARRDTANSDYT | 0.40100196 |
| IGLTPLAL | GARRDTANSDYT | 0.41100958 |
| ILGSNTSTL | GARRDTANSDYT | 0.39211264 |
| IVFSLTVL | GARRDTANSDYT | 0.15854874 |
| LASKATGF | GARRDTANSDYT | 0.45312202 |
| LASVALIL | GARRDTANSDYT | 0.086030535 |
| LFAVGLTL | GARRDTANSDYT | 0.43180084 |

| | | |
|---|---|---|
| LFNGGIGTYI | GARRDTANSDYT | 0.2095859 |
| LTLGAGII | GARRDTANSDYT | 0.41428715 |
| LVAAALGDI | GARRDTANSDYT | 0.4103945 |
| NTAFVQLGI | GARRDTANSDYT | 0.5603397 |
| QLFSGAEL | GARRDTANSDYT | 0.16294076 |
| SAATFITL | GARRDTANSDYT | 0.48140192 |
| SNYGLANLGV | GARRDTANSDYT | 0.28624022 |
| SYGGAYAVM | GARRDTANSDYT | 0.18751574 |
| TTGVGGLTLG | GARRDTANSDYT | 0.6707802 |
| TTQTIYI | GARRDTANSDYT | 0.3752172 |
| VAAGAGLGAVI | GARRDTANSDYT | 0.47995788 |
| VAAVVTLL | GARRDTANSDYT | 0.6594292 |
| VGFLGGLI | GARRDTANSDYT | 0.4460262 |
| VGVIPILL | GARRDTANSDYT | 0.17726414 |
| VIFASSLI | GARRDTANSDYT | 0.43535858 |
| VKGGDGGAAAG | GARRDTANSDYT | 0.3411854 |
| VTKTFTNI | GARRDTANSDYT | 0.31445047 |
| VVAAAILL | GARRDTANSDYT | 0.08880078 |
| YAALDVEL | GARRDTANSDYT | 0.5061762 |
| LASVALLL | GARRDTANSDYT | 0.22720844 |
| LDAVHRV | GARRDTANSDYT | 0.25639716 |
| QGLSSSIF | GARRDTANSDYT | 0.28291065 |
| RGTFANIRL | GARRDTANSDYT | 0.68426645 |
| RQLWWGHRI | GARRDTANSDYT | 0.4298789 |
| YGPPGTGKTLL | GARRDTANSDYT | 0.4670663 |
| AALVGAPANVA | GARRDTANSDYT | 0.5832876 |
| AATFRRTE | GARRDTANSDYT | 0.4992506 |
| AGALGGGI | GARRDTANSDYT | 0.7776576 |
| AGGAGGAGGIGAG | GARRDTANSDYT | 0.5254911 |
| AGSGALGL | GARRDTANSDYT | 0.4005907 |
| AQIADQI | GARRDTANSDYT | 0.40459403 |
| ASAVAVTYL | GARRDTANSDYT | 0.36673197 |
| ASTVLGGTGG | GARRDTANSDYT | 0.13149287 |
| AVGGEPAK | GARRDTANSDYT | 0.13950863 |
| CEQGTFGGTC | GARRDTANSDYT | 0.40559644 |
| DEGKAPAG | GARRDTANSDYT | 0.12350769 |
| DTVVFPAL | GARRDTANSDYT | 0.421542 |
| FEGITMAG | GARRDTANSDYT | 0.69330764 |
| FPLVVQD | GARRDTANSDYT | 0.5292542 |
| GAFTTPQ | GARRDTANSDYT | 0.17028534 |
| GFGGFGTAGGVGGA | GARRDTANSDYT | 0.30224037 |
| GFLAQFL | GARRDTANSDYT | 0.27000418 |
| GGAGGAGGNAGVA | GARRDTANSDYT | 0.5901375 |
| GGVGTAGAGGAGGAGG | GARRDTANSDYT | 0.6479226 |
| GNGGGGDAGNAGSGGN | GARRDTANSDYT | 0.79892564 |
| GTGGAGGAGNDAGS | GARRDTANSDYT | 0.69601744 |
| GTTNTGWFNS | GARRDTANSDYT | 0.28515327 |
| IFMGPLFTL | GARRDTANSDYT | 0.6084037 |
| ILAHAIEAI | GARRDTANSDYT | 0.3016362 |
| IRSAIREAL | GARRDTANSDYT | 0.3335096 |
| ITFAGGG | GARRDTANSDYT | 0.29389867 |

| | | |
|---|---|---|
| ITGMSGHQEV | GARRDTANSDYT | 0.65459573 |
| IVAGLACTL | GARRDTANSDYT | 0.37075853 |
| IVGGGLAGA | GARRDTANSDYT | 0.25456867 |
| IVTGVLNI | GARRDTANSDYT | 0.17565587 |
| LAQQGCI | GARRDTANSDYT | 0.56557345 |
| LCLHVEN | GARRDTANSDYT | 0.63787806 |
| LDAISDGS | GARRDTANSDYT | 0.18940313 |
| LGGFAAML | GARRDTANSDYT | 0.23699059 |
| LGGKNPMI | GARRDTANSDYT | 0.8400396 |
| LGPKGVAAIL | GARRDTANSDYT | 0.8016579 |
| LMVGVSI | GARRDTANSDYT | 0.22808595 |
| LVGAAFVL | GARRDTANSDYT | 0.24358292 |
| LVLAVAPL | GARRDTANSDYT | 0.6268322 |
| MAAIAVIDAA | GARRDTANSDYT | 0.15840751 |
| NNGINAELQ | GARRDTANSDYT | 0.4900347 |
| RGEPLVLVHG | GARRDTANSDYT | 0.52000916 |
| RGGAGGAAGLI | GARRDTANSDYT | 0.5219225 |
| RSYRMHPAV | GARRDTANSDYT | 0.2475993 |
| SAAHLLAGQLA | GARRDTANSDYT | 0.50003314 |
| SAAVGSY | GARRDTANSDYT | 0.37392908 |
| SPHAVLHRI | GARRDTANSDYT | 0.5817003 |
| SQIAKVL | GARRDTANSDYT | 0.10106999 |
| TALGHLVP | GARRDTANSDYT | 0.58828616 |
| TAPTTSGAA | GARRDTANSDYT | 0.186304 |
| TDGDDGNGGSPGT | GARRDTANSDYT | 0.30786222 |
| TDGLGEN | GARRDTANSDYT | 0.14788963 |
| TGGPPLPGQA | GARRDTANSDYT | 0.3012926 |
| TIPAITI | GARRDTANSDYT | 0.72801447 |
| TIPGFTI | GARRDTANSDYT | 0.74976075 |
| TSIASGALA | GARRDTANSDYT | 0.17832209 |
| TVTLPGAAI | GARRDTANSDYT | 0.6399709 |
| VASGLSAALP | GARRDTANSDYT | 0.83446455 |
| VGAGVIGI | GARRDTANSDYT | 0.2531098 |
| VGPKRFGV | GARRDTANSDYT | 0.14788814 |
| VNVAFPD | GARRDTANSDYT | 0.5668317 |
| VSMTNSGVS | GARRDTANSDYT | 0.5349956 |
| VTDVLGDLSV | GARRDTANSDYT | 0.74319243 |
| VVVVGAGFAGL | GARRDTANSDYT | 0.73457074 |
| VYTVGTVV | GARRDTANSDYT | 0.14568752 |
| WTTFSGP | GARRDTANSDYT | 0.59436315 |
| YDELVEAF | GARRDTANSDYT | 0.7419871 |
| YGTGGAGGNGG | GARRDTANSDYT | 0.21740341 |
| YTNVMAV | GARRDTANSDYT | 0.2273784 |
| FLEQVHQL | GARRDTANSDYT | 0.24936226 |
| GLSLLAVAV | GARRDTANSDYT | 0.13524625 |
| LEQVHQL | GARRDTANSDYT | 0.51013666 |
| LNAAGAI | GARRDTANSDYT | 0.22253159 |
| PAVHASL | GARRDTANSDYT | 0.2942299 |
| TELEDDA | GARRDTANSDYT | 0.36147147 |
| TPPGPPN | GARRDTANSDYT | 0.26907834 |
| TPPPPPPP | GARRDTANSDYT | 0.29630044 |

| | | |
|---|---|---|
| TQSPALI | ASSPLGEQNTLY | 0.30532134 |
| EFAAAFPQ | ASSPLGEQNTLY | 0.4564955 |
| LLSGNLDVL | ASSPLGEQNTLY | 0.58785874 |
| GPANLGNYNI | ASSPLGEQNTLY | 0.47504437 |
| TLGPTLP | ASSPLGEQNTLY | 0.44446597 |
| VYAYSSCCK | ASSPLGEQNTLY | 0.7551999 |
| APAPAPAPNI | ASSPLGEQNTLY | 0.20726316 |
| PGGPVPIP | ASSPLGEQNTLY | 0.24651685 |
| FLTVALEP | ASSPLGEQNTLY | 0.669062 |
| SQYFGANL | ASSPLGEQNTLY | 0.1904338 |
| GGVKPAA | ASSPLGEQNTLY | 0.68897176 |
| GGTGGAGGAGGW | ASSPLGEQNTLY | 0.41746354 |
| GVGGVAGLGGAGG | ASSPLGEQNTLY | 0.47186223 |
| VLLVLATTI | ASSPLGEQNTLY | 0.5047753 |
| IGIFAAML | ASSPLGEQNTLY | 0.83363503 |
| AVGDLID | ASSPLGEQNTLY | 0.58016706 |
| NISPLPL | ASSPLGEQNTLY | 0.29179865 |
| VTAVFVAI | ASSPLGEQNTLY | 0.5386033 |
| SGDDNDAAL | ASSPLGEQNTLY | 0.24589945 |
| AVAALAHS | ASSPLGEQNTLY | 0.6204114 |
| GGVTVMATP | ASSPLGEQNTLY | 0.5228002 |
| AGLAANVRD | ASSPLGEQNTLY | 0.18918553 |
| TNRPDLIDPALLRPG | ASSPLGEQNTLY | 0.324343 |
| LANLGVTAL | ASSPLGEQNTLY | 0.74923885 |
| TGERIEL | ASSPLGEQNTLY | 0.7197143 |
| NGEELAA | ASSPLGEQNTLY | 0.15193619 |
| VLSLVWG | ASSPLGEQNTLY | 0.2662864 |
| IGYALWL | ASSPLGEQNTLY | 0.16754504 |
| LSLLLAPL | ASSPLGEQNTLY | 0.40508926 |
| TLDEALK | ASSPLGEQNTLY | 0.21329342 |
| TDGAGTAEGG | ASSPLGEQNTLY | 0.5793086 |
| NAPGGAGGNGG | ASSPLGEQNTLY | 0.2529917 |
| VSVFMGMGGNF | ASSPLGEQNTLY | 0.71397585 |
| LASYLAAL | ASSPLGEQNTLY | 0.42774364 |
| LVGGTPAL | ASSPLGEQNTLY | 0.6570853 |
| VRNADVL | ASSPLGEQNTLY | 0.24129187 |
| TCSSPFLVSAM | ASSPLGEQNTLY | 0.3144996 |
| VHITDAE | ASSPLGEQNTLY | 0.1702931 |
| EYAAPSF | ASSPLGEQNTLY | 0.20930474 |
| VVDAFHGGL | ASSPLGEQNTLY | 0.24527586 |
| ASVVDPQ | ASSPLGEQNTLY | 0.33282644 |
| SYNEFGGLS | ASSPLGEQNTLY | 0.1307249 |
| DPDGSVY | ASSPLGEQNTLY | 0.8113715 |
| MVSTIHATE | ASSPLGEQNTLY | 0.4606117 |
| LAAGVADPV | ASSPLGEQNTLY | 0.55845135 |
| TKEQIAA | ASSPLGEQNTLY | 0.28366962 |
| SLASPAAL | ASSPLGEQNTLY | 0.25406176 |
| IGLVFYGV | ASSPLGEQNTLY | 0.3030267 |
| FLLDASNL | ASSPLGEQNTLY | 0.6648383 |
| YQVGNFP | ASSPLGEQNTLY | 0.19007622 |
| ESLLSQST | ASSPLGEQNTLY | 0.13198885 |

| | | |
|---|---|---|
| LFTVGFAAVG | ASSPLGEQNTLY | 0.26704067 |
| ATPPTPT | ASSPLGEQNTLY | 0.14850575 |
| TDLATLG | ASSPLGEQNTLY | 0.7858296 |
| SAGGTGR | ASSPLGEQNTLY | 0.661644 |
| GFSNSGS | ASSPLGEQNTLY | 0.47259125 |
| GGNGGSGGTSV | ASSPLGEQNTLY | 0.69848454 |
| GTSVATGGAGN | ASSPLGEQNTLY | 0.11696954 |
| NGGNGGSGGTS | ASSPLGEQNTLY | 0.312535 |
| AVVKFPH | ASSPLGEQNTLY | 0.7964833 |
| TPPPPPP | ASSPLGEQNTLY | 0.5200055 |
| AAPAAAAAPA | ASSPLGEQNTLY | 0.2755782 |
| IALNEDL | ASSPLGEQNTLY | 0.74440974 |
| IVNVSSM | ASSPLGEQNTLY | 0.24857017 |
| TGIPPAPR | ASSPLGEQNTLY | 0.3190196 |
| FDLGGGTF | ASSPLGEQNTLY | 0.405181 |
| FDLGGGTFDVS | ASSPLGEQNTLY | 0.37515956 |
| NEPTAAA | ASSPLGEQNTLY | 0.51342666 |
| NEPTAAALA | ASSPLGEQNTLY | 0.19183005 |
| GSIINISSI | ASSPLGEQNTLY | 0.49262616 |
| VAPGNAGTA | ASSPLGEQNTLY | 0.82992196 |
| TGGGASLE | ASSPLGEQNTLY | 0.30416933 |
| VAVYNHYKRI | ASSPLGEQNTLY | 0.73509896 |
| TDPAPATT | ASSPLGEQNTLY | 0.56109035 |
| TNRPDILDPALLRPG | ASSPLGEQNTLY | 0.6727644 |
| LTTTTTS | ASSPLGEQNTLY | 0.20615593 |
| PTGSGKTL | ASSPLGEQNTLY | 0.21781221 |
| TGSGKTL | ASSPLGEQNTLY | 0.4625644 |
| TIGHVDHGK | ASSPLGEQNTLY | 0.20529753 |
| TIGHVDHGKTT | ASSPLGEQNTLY | 0.38368407 |
| GHVDHGKTT | ASSPLGEQNTLY | 0.30311507 |
| AQFEHTLL | ASSPLGEQNTLY | 0.85862815 |
| IVNQDEL | ASSPLGEQNTLY | 0.56786835 |
| TNGSQFF | ASSPLGEQNTLY | 0.6221349 |
| TGKGIVE | ASSPLGEQNTLY | 0.20691851 |
| LISNASDA | ASSPLGEQNTLY | 0.47228295 |
| AELRTLGELTKIAKAHGAQ | ASSPLGEQNTLY | 0.57204056 |
| APNAVAAAEPSW | ASSPLGEQNTLY | 0.1872598 |
| APQVAGAATAVL | ASSPLGEQNTLY | 0.75350463 |
| ARRDPAAAAAILPTDA | ASSPLGEQNTLY | 0.23211461 |
| DAAARDVLAEAGLAEYF | ASSPLGEQNTLY | 0.50716937 |
| DDQIVLSAKANLAKVEQ | ASSPLGEQNTLY | 0.58803904 |
| DELLPDTGEDSDEDGH | ASSPLGEQNTLY | 0.7647135 |
| ELPEAPANRATVPQD | ASSPLGEQNTLY | 0.28636694 |
| FGQSKIERVTGIG | ASSPLGEQNTLY | 0.37697858 |
| GAGGTGSTAGGAGGAGGAGGLY | ASSPLGEQNTLY | 0.21494392 |
| GHGGTGAAGGNGGTGGAGGSGI | ASSPLGEQNTLY | 0.6648502 |
| GNGGNGGAGGEGGPGGAGGAGA | ASSPLGEQNTLY | 0.7086715 |
| GSTAGNGGNGGNATNGTIAGQP | ASSPLGEQNTLY | 0.54715323 |
| GTGGDGGNGGNGGTGG | ASSPLGEQNTLY | 0.7812726 |
| GVRRPAGIPADAPV | ASSPLGEQNTLY | 0.63906753 |
| GYRQVTDAHLLTLARR | ASSPLGEQNTLY | 0.2061639 |

| | | |
|---|---|---|
| IAKMGQDGHDRG | ASSPLGEQNTLY | 0.34013242 |
| KAEPSIEIQYT | ASSPLGEQNTLY | 0.67715955 |
| KPVRIRMAADEDTVAVT | ASSPLGEQNTLY | 0.77170295 |
| MPPRIAGMRLLVIKPEP | ASSPLGEQNTLY | 0.6963939 |
| MKFYAVALFRSRGVPPTR | ASSPLGEQNTLY | 0.6280959 |
| NAGGFGILGAVAHSP | ASSPLGEQNTLY | 0.27588743 |
| NEGREITLAMAEM | ASSPLGEQNTLY | 0.77332854 |
| NELVEWSHPLGHRWRWE | ASSPLGEQNTLY | 0.8625542 |
| NLGAGGTGGNGGNPG | ASSPLGEQNTLY | 0.69369066 |
| NTEWPKWETVMAADPVD | ASSPLGEQNTLY | 0.30785167 |
| PASAAPAETQP | ASSPLGEQNTLY | 0.32194293 |
| QQSSDGKNGVRAAEITGE | ASSPLGEQNTLY | 0.38825613 |
| RAPDAVTTSWWKE | ASSPLGEQNTLY | 0.7693368 |
| SGFGNVGAHTSGWFNQ | ASSPLGEQNTLY | 0.22338611 |
| SGVAGAVGQAASVGGLKV | ASSPLGEQNTLY | 0.40437335 |
| SNGGAGTGSAGGNGGT | ASSPLGEQNTLY | 0.41972566 |
| TGGAGGEGGAAG | ASSPLGEQNTLY | 0.14071456 |
| TGGDGGAAGAGGAPGHG | ASSPLGEQNTLY | 0.5983649 |
| TMPPLLAAIANAGRHGV | ASSPLGEQNTLY | 0.77097964 |
| TSSTAGNSGGA | ASSPLGEQNTLY | 0.3489878 |
| VPATVPVGILC | ASSPLGEQNTLY | 0.24725333 |
| VQRPDSGGVVLTGRIS | ASSPLGEQNTLY | 0.73977137 |
| WTPNPVPATKGVPAGLVQAT | ASSPLGEQNTLY | 0.836025 |
| YALKRALAKFSDRFDVVII | ASSPLGEQNTLY | 0.670058 |
| YLIGSVIV | ASSPLGEQNTLY | 0.1694008 |
| YGQLVAVID | ASSPLGEQNTLY | 0.10973946 |
| VTVGGMTLPA | ASSPLGEQNTLY | 0.27981946 |
| VLALPAV | ASSPLGEQNTLY | 0.74312925 |
| VIPPTLNYET | ASSPLGEQNTLY | 0.2994819 |
| VGMGGMHPSSG | ASSPLGEQNTLY | 0.25748488 |
| VGLAGGAATA | ASSPLGEQNTLY | 0.60994214 |
| VGGAGGAGGSGP | ASSPLGEQNTLY | 0.7227651 |
| VFAAFMV | ASSPLGEQNTLY | 0.21036151 |
| TLYDKLR | ASSPLGEQNTLY | 0.22666451 |
| TLAVFTVL | ASSPLGEQNTLY | 0.5299374 |
| TGSVQPARV | ASSPLGEQNTLY | 0.6331178 |
| TGGGAGSGGT | ASSPLGEQNTLY | 0.18343705 |
| SHLETSL | ASSPLGEQNTLY | 0.31122303 |
| SFTTGMT | ASSPLGEQNTLY | 0.29093048 |
| QVEVAKRMGVRQPTVS | ASSPLGEQNTLY | 0.52202994 |
| QGSGGPALAM | ASSPLGEQNTLY | 0.5292884 |
| MLAPSNP | ASSPLGEQNTLY | 0.2266707 |
| LVNNAAI | ASSPLGEQNTLY | 0.77440447 |
| LTGEINTLA | ASSPLGEQNTLY | 0.17701831 |
| LSIFGPV | ASSPLGEQNTLY | 0.69265246 |
| LLALLANL | ASSPLGEQNTLY | 0.40897793 |
| LISNASDALDKLR | ASSPLGEQNTLY | 0.3922027 |
| LINEAVP | ASSPLGEQNTLY | 0.27625242 |
| LGYAFSGL | ASSPLGEQNTLY | 0.21291795 |
| LDFVAVAL | ASSPLGEQNTLY | 0.17983112 |
| LAQPGAE | ASSPLGEQNTLY | 0.13160144 |

| | | |
|---|---|---|
| ISNLGAVTGF | ASSPLGEQNTLY | 0.6914779 |
| HQEDPGRI | ASSPLGEQNTLY | 0.10368023 |
| GTPGEPTP | ASSPLGEQNTLY | 0.59991425 |
| GFFNSGS | ASSPLGEQNTLY | 0.48969975 |
| GAGGGGLTAGAN | ASSPLGEQNTLY | 0.20871247 |
| FVDALVNM | ASSPLGEQNTLY | 0.52272093 |
| FLTVAPEL | ASSPLGEQNTLY | 0.22997695 |
| FETLTAL | ASSPLGEQNTLY | 0.23134628 |
| DVINPIEL | ASSPLGEQNTLY | 0.7406733 |
| APVSGAPV | ASSPLGEQNTLY | 0.53153515 |
| ALVGGLAW | ASSPLGEQNTLY | 0.20303257 |
| ALAAAAATAI | ASSPLGEQNTLY | 0.44889438 |
| AFGLGLLL | ASSPLGEQNTLY | 0.4612465 |
| AFFAAASL | ASSPLGEQNTLY | 0.21605773 |
| AAAALPAFSPP | ASSPLGEQNTLY | 0.338457 |
| VVLAPAL | ASSPLGEQNTLY | 0.7177683 |
| TIGHVDHGKTTL | ASSPLGEQNTLY | 0.39636397 |
| LISNASDALDK | ASSPLGEQNTLY | 0.19288206 |
| LISNASDAL | ASSPLGEQNTLY | 0.6566094 |
| ISNASDALDK | ASSPLGEQNTLY | 0.1933572 |
| NQAETLVYQTE | ASSPLGEQNTLY | 0.5202439 |
| TGGTGGTAGAG | ASSPLGEQNTLY | 0.34664115 |
| TGSTAGGAGGA | ASSPLGEQNTLY | 0.3048952 |
| DPLAVPVAAGPV | ASSPLGEQNTLY | 0.6733868 |
| GGTGGTPGNGGN | ASSPLGEQNTLY | 0.2534375 |
| PAGLAGVDPATL | ASSPLGEQNTLY | 0.22055092 |
| GNNLVVEAPHAGASV | ASSPLGEQNTLY | 0.30053645 |
| KVTRIMSASTLEQPA | ASSPLGEQNTLY | 0.39197445 |
| LGFALPIALPRRTVV | ASSPLGEQNTLY | 0.6840767 |
| TRLGNLTARKLVVLV | ASSPLGEQNTLY | 0.20190966 |
| WSQQRRLIYFPSAGP | ASSPLGEQNTLY | 0.46008658 |
| GPNGNPDPMAAAVDIR | ASSPLGEQNTLY | 0.27115947 |
| TQRCVSVADLRAAASR | ASSPLGEQNTLY | 0.20082675 |
| QFVLDVAEHPVEVLTETV | ASSPLGEQNTLY | 0.3804558 |
| GGAGGTGGAGGNGGTQAGDG | ASSPLGEQNTLY | 0.27728426 |
| PLLATLRAIAVDVDAKPAQV | ASSPLGEQNTLY | 0.24540997 |
| APKFRTGGYMIDFWGVGYQVAK | ASSPLGEQNTLY | 0.40547127 |
| GAGGAGGAGGAAGGPGGTGGTG | ASSPLGEQNTLY | 0.26164314 |
| IGALGLLATDVAPAGSVAFGVA | ASSPLGEQNTLY | 0.49914044 |
| AGGAGGIGGNGGAGGAGGVLMG | ASSPLGEQNTLY | 0.8169311 |
| ADELIGLL | ASSPLGEQNTLY | 0.2697329 |
| ADRALKA | ASSPLGEQNTLY | 0.635846 |
| AGAVGQVGGAGGA | ASSPLGEQNTLY | 0.67813766 |
| AGGGGGLGGAP | ASSPLGEQNTLY | 0.50052583 |
| AGGLLYGNGGAG | ASSPLGEQNTLY | 0.31123292 |
| AGLESIGL | ASSPLGEQNTLY | 0.21641955 |
| AGLKTGII | ASSPLGEQNTLY | 0.7719493 |
| ASLAALLL | ASSPLGEQNTLY | 0.6150537 |
| ASVINISSI | ASSPLGEQNTLY | 0.6275359 |
| ATPPGPA | ASSPLGEQNTLY | 0.652512 |
| ATVASALV | ASSPLGEQNTLY | 0.5733936 |

| | | |
|---|---|---|
| ATVASLIL | ASSPLGEQNTLY | 0.61045086 |
| AVNGPVLIRH | ASSPLGEQNTLY | 0.24290967 |
| DGAAGAVGI | ASSPLGEQNTLY | 0.17055137 |
| DPLLHGI | ASSPLGEQNTLY | 0.3201772 |
| ELGAALGAL | ASSPLGEQNTLY | 0.3776762 |
| ELGRTQAV | ASSPLGEQNTLY | 0.25750387 |
| FVAGGYLS | ASSPLGEQNTLY | 0.21240154 |
| GASGGAGGAGGW | ASSPLGEQNTLY | 0.71266186 |
| GGAGAGGDGGAGGI | ASSPLGEQNTLY | 0.8567791 |
| GTGHNGSQPGMGGNG | ASSPLGEQNTLY | 0.28221607 |
| GYAAASASAS | ASSPLGEQNTLY | 0.8164555 |
| HQVDVAAVQQG | ASSPLGEQNTLY | 0.17351511 |
| IGAAIAHT | ASSPLGEQNTLY | 0.6246946 |
| IGMIEVAQL | ASSPLGEQNTLY | 0.73001593 |
| IRLAPEHTSAMD | ASSPLGEQNTLY | 0.24022526 |
| IYNVSVL | ASSPLGEQNTLY | 0.53186363 |
| KQGFLGM | ASSPLGEQNTLY | 0.19187397 |
| LESVITQL | ASSPLGEQNTLY | 0.29461503 |
| LGLGALVVV | ASSPLGEQNTLY | 0.7387567 |
| LIGNGGNGGSGGI | ASSPLGEQNTLY | 0.32054806 |
| LNAINAPTL | ASSPLGEQNTLY | 0.3383762 |
| LQLLNTGV | ASSPLGEQNTLY | 0.33105487 |
| LQTSIEF | ASSPLGEQNTLY | 0.24576068 |
| MSFAVSEL | ASSPLGEQNTLY | 0.35577396 |
| NTGALGL | ASSPLGEQNTLY | 0.6426798 |
| PPGAGGAGGAGTT | ASSPLGEQNTLY | 0.30111793 |
| QLVGLGI | ASSPLGEQNTLY | 0.8573962 |
| RFLEQVHQL | ASSPLGEQNTLY | 0.18308897 |
| RGVGYVLR | ASSPLGEQNTLY | 0.81602925 |
| SATGVFAGVI | ASSPLGEQNTLY | 0.7770504 |
| SLPAITLP | ASSPLGEQNTLY | 0.45061523 |
| STALAFTV | ASSPLGEQNTLY | 0.30211315 |
| SVLTVGLGK | ASSPLGEQNTLY | 0.71061146 |
| TALAGAGSGI | ASSPLGEQNTLY | 0.300097 |
| TGLPFTGI | ASSPLGEQNTLY | 0.56561184 |
| VAEHVTV | ASSPLGEQNTLY | 0.6732818 |
| VDIVLNSL | ASSPLGEQNTLY | 0.23243575 |
| VLTGGGAVL | ASSPLGEQNTLY | 0.7758148 |
| VNVGLAYGI | ASSPLGEQNTLY | 0.54019606 |
| VSSAAVGAV | ASSPLGEQNTLY | 0.5224798 |
| VTSVYEAGV | ASSPLGEQNTLY | 0.31321782 |
| WVALDRGA | ASSPLGEQNTLY | 0.19570011 |
| YLVGSVVI | ASSPLGEQNTLY | 0.32404768 |
| GPPGTGKTLL | ASSPLGEQNTLY | 0.27346995 |
| GVGMTKF | ASSPLGEQNTLY | 0.717457 |
| IGIGGSDL | ASSPLGEQNTLY | 0.5957439 |
| IGTLLAEL | ASSPLGEQNTLY | 0.3355044 |
| IYTSGTT | ASSPLGEQNTLY | 0.7012712 |
| LSFPAGV | ASSPLGEQNTLY | 0.74879974 |
| LYTSGTT | ASSPLGEQNTLY | 0.1807797 |
| VGAFPFGF | ASSPLGEQNTLY | 0.40539363 |

| | | |
|---|---|---|
| VGAGTVEF | ASSPLGEQNTLY | 0.24591866 |
| VGKSTLLL | ASSPLGEQNTLY | 0.34492296 |
| AALAHAGGGIV | ASSPLGEQNTLY | 0.122405276 |
| AALVAVSLL | ASSPLGEQNTLY | 0.56983924 |
| AAVSRAVAAL | ASSPLGEQNTLY | 0.269941 |
| AGGSSGIGGF | ASSPLGEQNTLY | 0.107350625 |
| AGLTVANANG | ASSPLGEQNTLY | 0.4461956 |
| AGLVGAVL | ASSPLGEQNTLY | 0.3895749 |
| ALSQAFTV | ASSPLGEQNTLY | 0.3683208 |
| ALTFSVSL | ASSPLGEQNTLY | 0.6574953 |
| ALTGAALVAL | ASSPLGEQNTLY | 0.28719342 |
| ASFSIGAGVL | ASSPLGEQNTLY | 0.26609343 |
| DGSVYKI | ASSPLGEQNTLY | 0.56966627 |
| EQLAEIV | ASSPLGEQNTLY | 0.76600456 |
| FAVSELTVM | ASSPLGEQNTLY | 0.26046097 |
| GAAPLAAGL | ASSPLGEQNTLY | 0.4763366 |
| GADGGDGGSGGAGG | ASSPLGEQNTLY | 0.79021704 |
| GFGLDTGGAGG | ASSPLGEQNTLY | 0.48235875 |
| GLASGGLL | ASSPLGEQNTLY | 0.67980593 |
| GRIAGAAVLL | ASSPLGEQNTLY | 0.7567342 |
| GSAGSLNTSF | ASSPLGEQNTLY | 0.25785205 |
| GTGQIGI | ASSPLGEQNTLY | 0.68904513 |
| GTLIALIL | ASSPLGEQNTLY | 0.74886346 |
| IAATMGFVVG | ASSPLGEQNTLY | 0.4377116 |
| IFAQALL | ASSPLGEQNTLY | 0.36544004 |
| IGAAVALGF | ASSPLGEQNTLY | 0.23348744 |
| IGLTPLAL | ASSPLGEQNTLY | 0.42995065 |
| ILGSNTSTL | ASSPLGEQNTLY | 0.16760004 |
| IVFSLTVL | ASSPLGEQNTLY | 0.5579231 |
| LASKATGF | ASSPLGEQNTLY | 0.70025945 |
| LASVALIL | ASSPLGEQNTLY | 0.34225553 |
| LFAVGLTL | ASSPLGEQNTLY | 0.16749768 |
| LFNGGIGTYI | ASSPLGEQNTLY | 0.16843303 |
| LTLGAGII | ASSPLGEQNTLY | 0.44717026 |
| LVAAALGDI | ASSPLGEQNTLY | 0.28764713 |
| NTAFVQLGI | ASSPLGEQNTLY | 0.16664608 |
| QLFSGAEL | ASSPLGEQNTLY | 0.36920017 |
| SAATFITL | ASSPLGEQNTLY | 0.6801866 |
| SNYGLANLGV | ASSPLGEQNTLY | 0.69930184 |
| SYGGAYAVM | ASSPLGEQNTLY | 0.2050826 |
| TTGVGGLTLG | ASSPLGEQNTLY | 0.78529406 |
| TTQTIYI | ASSPLGEQNTLY | 0.66329753 |
| VAAGAGLGAVI | ASSPLGEQNTLY | 0.5852169 |
| VAAVVTLL | ASSPLGEQNTLY | 0.37638798 |
| VGFLGGLI | ASSPLGEQNTLY | 0.23520423 |
| VGVIPILL | ASSPLGEQNTLY | 0.75720656 |
| VIFASSLI | ASSPLGEQNTLY | 0.41879624 |
| VKGGDGGAAAG | ASSPLGEQNTLY | 0.36569583 |
| VTKTFTNI | ASSPLGEQNTLY | 0.60203266 |
| VVAAAILL | ASSPLGEQNTLY | 0.44315434 |
| YAALDVEL | ASSPLGEQNTLY | 0.29302317 |

| | | |
|---|---|---|
| LASVALLL | ASSPLGEQNTLY | 0.5687257 |
| LDAVHRV | ASSPLGEQNTLY | 0.60851777 |
| QGLSSSIF | ASSPLGEQNTLY | 0.18951379 |
| RGTFANIRL | ASSPLGEQNTLY | 0.2835112 |
| RQLWWGHRI | ASSPLGEQNTLY | 0.31430623 |
| YGPPGTGKTLL | ASSPLGEQNTLY | 0.46264207 |
| AALVGAPANVA | ASSPLGEQNTLY | 0.7244782 |
| AATFRRTE | ASSPLGEQNTLY | 0.46060404 |
| AGALGGGI | ASSPLGEQNTLY | 0.34779432 |
| AGGAGGAGGIGAG | ASSPLGEQNTLY | 0.6329546 |
| AGSGALGL | ASSPLGEQNTLY | 0.6032315 |
| AQIADQI | ASSPLGEQNTLY | 0.4303838 |
| ASAVAVTYL | ASSPLGEQNTLY | 0.6994199 |
| ASTVLGGTGG | ASSPLGEQNTLY | 0.30102485 |
| AVGGEPAK | ASSPLGEQNTLY | 0.5634439 |
| CEQGTFGGTC | ASSPLGEQNTLY | 0.5544112 |
| DEGKAPAG | ASSPLGEQNTLY | 0.5779246 |
| DTVVFPAL | ASSPLGEQNTLY | 0.6892937 |
| FEGITMAG | ASSPLGEQNTLY | 0.6570512 |
| FPLVVQD | ASSPLGEQNTLY | 0.44591486 |
| GAFTTPQ | ASSPLGEQNTLY | 0.153 |
| GFGGFGTAGGVGGA | ASSPLGEQNTLY | 0.30476868 |
| GFLAQFL | ASSPLGEQNTLY | 0.5806589 |
| GGAGGAGGNAGVA | ASSPLGEQNTLY | 0.5136206 |
| GGVGTAGAGGAGGAGG | ASSPLGEQNTLY | 0.6473714 |
| GNGGGGDAGNAGSGGN | ASSPLGEQNTLY | 0.58324665 |
| GTGGAGGAGNDAGS | ASSPLGEQNTLY | 0.3015271 |
| GTTNTGWFNS | ASSPLGEQNTLY | 0.75802064 |
| IFMGPLFTL | ASSPLGEQNTLY | 0.71613026 |
| ILAHAIEAI | ASSPLGEQNTLY | 0.17363137 |
| IRSAIREAL | ASSPLGEQNTLY | 0.42290848 |
| ITFAGGG | ASSPLGEQNTLY | 0.7946434 |
| ITGMSGHQEV | ASSPLGEQNTLY | 0.6404294 |
| IVAGLACTL | ASSPLGEQNTLY | 0.37108243 |
| IVGGGLAGA | ASSPLGEQNTLY | 0.37604934 |
| IVTGVLNI | ASSPLGEQNTLY | 0.7905066 |
| LAQQGCI | ASSPLGEQNTLY | 0.34579438 |
| LCLHVEN | ASSPLGEQNTLY | 0.76633537 |
| LDAISDGS | ASSPLGEQNTLY | 0.13108419 |
| LGGFAAML | ASSPLGEQNTLY | 0.35631517 |
| LGGKNPMI | ASSPLGEQNTLY | 0.2635448 |
| LGPKGVAAIL | ASSPLGEQNTLY | 0.14168482 |
| LMVGVSI | ASSPLGEQNTLY | 0.61575955 |
| LVGAAFVL | ASSPLGEQNTLY | 0.29688236 |
| LVLAVAPL | ASSPLGEQNTLY | 0.6587567 |
| MAAIAVIDAA | ASSPLGEQNTLY | 0.31794682 |
| NNGINAELQ | ASSPLGEQNTLY | 0.6548139 |
| RGEPLVLVHG | ASSPLGEQNTLY | 0.37549233 |
| RGGAGGAAGLI | ASSPLGEQNTLY | 0.48785886 |
| RSYRMHPAV | ASSPLGEQNTLY | 0.27546227 |
| SAAHLLAGQLA | ASSPLGEQNTLY | 0.60622567 |

| | | |
|---|---|---|
| SAAVGSY | ASSPLGEQNTLY | 0.75393414 |
| SPHAVLHRI | ASSPLGEQNTLY | 0.5136616 |
| SQIAKVL | ASSPLGEQNTLY | 0.18833266 |
| TALGHLVP | ASSPLGEQNTLY | 0.44727677 |
| TAPTTSGAA | ASSPLGEQNTLY | 0.40751612 |
| TDGDDGNGGSPGT | ASSPLGEQNTLY | 0.16169767 |
| TDGLGEN | ASSPLGEQNTLY | 0.12664947 |
| TGGPPLPGQA | ASSPLGEQNTLY | 0.2543255 |
| TIPAITI | ASSPLGEQNTLY | 0.4075992 |
| TIPGFTI | ASSPLGEQNTLY | 0.6470065 |
| TSIASGALA | ASSPLGEQNTLY | 0.61991596 |
| TVTLPGAAI | ASSPLGEQNTLY | 0.30763018 |
| VASGLSAALP | ASSPLGEQNTLY | 0.63035595 |
| VGAGVIGI | ASSPLGEQNTLY | 0.6340983 |
| VGPKRFGV | ASSPLGEQNTLY | 0.094880186 |
| VNVAFPD | ASSPLGEQNTLY | 0.30024332 |
| VSMTNSGVS | ASSPLGEQNTLY | 0.17722647 |
| VTDVLGDLSV | ASSPLGEQNTLY | 0.7174629 |
| VVVVGAGFAGL | ASSPLGEQNTLY | 0.25802404 |
| VYTVGTVV | ASSPLGEQNTLY | 0.26720187 |
| WTTFSGP | ASSPLGEQNTLY | 0.39009106 |
| YDELVEAF | ASSPLGEQNTLY | 0.6753955 |
| YGTGGAGGNGG | ASSPLGEQNTLY | 0.49480093 |
| YTNVMAV | ASSPLGEQNTLY | 0.23329914 |
| FLEQVHQL | ASSPLGEQNTLY | 0.13341364 |
| GLSLLAVAV | ASSPLGEQNTLY | 0.58829737 |
| LEQVHQL | ASSPLGEQNTLY | 0.28225693 |
| LNAAGAI | ASSPLGEQNTLY | 0.2795446 |
| PAVHASL | ASSPLGEQNTLY | 0.15123926 |
| TELEDDA | ASSPLGEQNTLY | 0.46611947 |
| TPPGPPN | ASSPLGEQNTLY | 0.20404556 |
| TPPPPPPP | ASSPLGEQNTLY | 0.62920976 |
| TQSPALI | ASGPGGANTGQLY | 0.28267398 |
| EFAAAFPQ | ASGPGGANTGQLY | 0.36895785 |
| LLSGNLDVL | ASGPGGANTGQLY | 0.773952 |
| GPANLGNYNI | ASGPGGANTGQLY | 0.6711641 |
| TLGPTLP | ASGPGGANTGQLY | 0.3494817 |
| VYAYSSCCK | ASGPGGANTGQLY | 0.3774322 |
| APAPAPAPNI | ASGPGGANTGQLY | 0.22490707 |
| PGGPVPIP | ASGPGGANTGQLY | 0.25670245 |
| FLTVALEP | ASGPGGANTGQLY | 0.302311 |
| SQYFGANL | ASGPGGANTGQLY | 0.7600701 |
| GGVKPAA | ASGPGGANTGQLY | 0.35002807 |
| GGTGGAGGAGGW | ASGPGGANTGQLY | 0.20102198 |
| GVGGVAGLGGAGG | ASGPGGANTGQLY | 0.48116007 |
| VLLVLATTI | ASGPGGANTGQLY | 0.74629945 |
| IGIFAAML | ASGPGGANTGQLY | 0.7932318 |
| AVGDLID | ASGPGGANTGQLY | 0.22614086 |
| NISPLPL | ASGPGGANTGQLY | 0.3375299 |
| VTAVFVAI | ASGPGGANTGQLY | 0.3383216 |
| SGDDNDAAL | ASGPGGANTGQLY | 0.39640445 |

| | | |
|---|---|---|
| AVAALAHS | ASGPGGANTGQLY | 0.3079794 |
| GGVTVMATP | ASGPGGANTGQLY | 0.25938672 |
| AGLAANVRD | ASGPGGANTGQLY | 0.54966587 |
| TNRPDLIDPALLRPG | ASGPGGANTGQLY | 0.82730937 |
| LANLGVTAL | ASGPGGANTGQLY | 0.38772124 |
| TGERIEL | ASGPGGANTGQLY | 0.14031844 |
| NGEELAA | ASGPGGANTGQLY | 0.42088193 |
| VLSLVWG | ASGPGGANTGQLY | 0.23783895 |
| IGYALWL | ASGPGGANTGQLY | 0.25167462 |
| LSLLLAPL | ASGPGGANTGQLY | 0.47780854 |
| TLDEALK | ASGPGGANTGQLY | 0.16674936 |
| TDGAGTAEGG | ASGPGGANTGQLY | 0.38074493 |
| NAPGGAGGNGG | ASGPGGANTGQLY | 0.8031964 |
| VSVFMGMGGNF | ASGPGGANTGQLY | 0.4260434 |
| LASYLAAL | ASGPGGANTGQLY | 0.5959848 |
| LVGGTPAL | ASGPGGANTGQLY | 0.7103612 |
| VRNADVL | ASGPGGANTGQLY | 0.54541075 |
| TCSSPFLVSAM | ASGPGGANTGQLY | 0.74743086 |
| VHITDAE | ASGPGGANTGQLY | 0.7162381 |
| EYAAPSF | ASGPGGANTGQLY | 0.45569116 |
| VVDAFHGGL | ASGPGGANTGQLY | 0.4762581 |
| ASVVDPQ | ASGPGGANTGQLY | 0.683576 |
| SYNEFGGLS | ASGPGGANTGQLY | 0.46149063 |
| DPDGSVY | ASGPGGANTGQLY | 0.4186723 |
| MVSTIHATE | ASGPGGANTGQLY | 0.28642753 |
| LAAGVADPV | ASGPGGANTGQLY | 0.70494473 |
| TKEQIAA | ASGPGGANTGQLY | 0.4639026 |
| SLASPAAL | ASGPGGANTGQLY | 0.73349446 |
| IGLVFYGV | ASGPGGANTGQLY | 0.19975516 |
| FLLDASNL | ASGPGGANTGQLY | 0.2588372 |
| YQVGNFP | ASGPGGANTGQLY | 0.7149422 |
| ESLLSQST | ASGPGGANTGQLY | 0.11762859 |
| LFTVGFAAVG | ASGPGGANTGQLY | 0.15476716 |
| ATPPTPT | ASGPGGANTGQLY | 0.36871237 |
| TDLATLG | ASGPGGANTGQLY | 0.7683425 |
| SAGGTGR | ASGPGGANTGQLY | 0.115643315 |
| GFSNSGS | ASGPGGANTGQLY | 0.2893935 |
| GGNGGSGGTSV | ASGPGGANTGQLY | 0.19682416 |
| GTSVATGGAGN | ASGPGGANTGQLY | 0.6211672 |
| NGGNGGSGGTS | ASGPGGANTGQLY | 0.45260698 |
| AVVKFPH | ASGPGGANTGQLY | 0.18993473 |
| TPPPPPP | ASGPGGANTGQLY | 0.5840831 |
| AAPAAAAAPA | ASGPGGANTGQLY | 0.13793138 |
| IALNEDL | ASGPGGANTGQLY | 0.32427156 |
| IVNVSSM | ASGPGGANTGQLY | 0.59819925 |
| TGIPPAPR | ASGPGGANTGQLY | 0.78051394 |
| FDLGGGTF | ASGPGGANTGQLY | 0.21277961 |
| FDLGGGTFDVS | ASGPGGANTGQLY | 0.6356889 |
| NEPTAAA | ASGPGGANTGQLY | 0.27085805 |
| NEPTAAALA | ASGPGGANTGQLY | 0.8153798 |
| GSIINISSI | ASGPGGANTGQLY | 0.3533492 |

| | | |
|---|---|---|
| VAPGNAGTA | ASGPGGANTGQLY | 0.17628093 |
| TGGGASLE | ASGPGGANTGQLY | 0.6633211 |
| VAVYNHYKRI | ASGPGGANTGQLY | 0.54416674 |
| TDPAPATT | ASGPGGANTGQLY | 0.49760658 |
| TNRPDILDPALLRPG | ASGPGGANTGQLY | 0.66748977 |
| LTTTTTS | ASGPGGANTGQLY | 0.3965159 |
| PTGSGKTL | ASGPGGANTGQLY | 0.3244976 |
| TGSGKTL | ASGPGGANTGQLY | 0.6836754 |
| TIGHVDHGK | ASGPGGANTGQLY | 0.6630399 |
| TIGHVDHGKTT | ASGPGGANTGQLY | 0.28084612 |
| GHVDHGKTT | ASGPGGANTGQLY | 0.65773815 |
| AQFEHTLL | ASGPGGANTGQLY | 0.5117376 |
| IVNQDEL | ASGPGGANTGQLY | 0.26194918 |
| TNGSQFF | ASGPGGANTGQLY | 0.7878438 |
| TGKGIVE | ASGPGGANTGQLY | 0.40415406 |
| LISNASDA | ASGPGGANTGQLY | 0.6198465 |
| AELRTLGELTKIAKAHGAQ | ASGPGGANTGQLY | 0.5388446 |
| APNAVAAAEPSW | ASGPGGANTGQLY | 0.78152776 |
| APQVAGAATAVL | ASGPGGANTGQLY | 0.2285639 |
| ARRDPAAAAAILPTDA | ASGPGGANTGQLY | 0.58821535 |
| DAAARDVLAEAGLAEYF | ASGPGGANTGQLY | 0.45439106 |
| DDQIVLSAKANLAKVEQ | ASGPGGANTGQLY | 0.64146066 |
| DELLPDTGEDSDEDGH | ASGPGGANTGQLY | 0.38079366 |
| ELPEAPANRATVPQD | ASGPGGANTGQLY | 0.37524575 |
| FGQSKIERVTGIG | ASGPGGANTGQLY | 0.5000978 |
| GAGGTGSTAGGAGGAGGAGGLY | ASGPGGANTGQLY | 0.72184765 |
| GHGGTGAAGGNGGTGGAGGSGI | ASGPGGANTGQLY | 0.3644035 |
| GNGGNGGAGGEGGPGGAGGAGA | ASGPGGANTGQLY | 0.62282777 |
| GSTAGNGGNGGNATNGTIAGQP | ASGPGGANTGQLY | 0.793556 |
| GTGGDGGNGGNGGTGG | ASGPGGANTGQLY | 0.2449612 |
| GVRRPAGIPADAPV | ASGPGGANTGQLY | 0.4395696 |
| GYRQVTDAHLLTLARR | ASGPGGANTGQLY | 0.57560307 |
| IAKMGQDGHDRG | ASGPGGANTGQLY | 0.73830277 |
| KAEPSIEIQYT | ASGPGGANTGQLY | 0.374914 |
| KPVRIRMAADEDTVAVT | ASGPGGANTGQLY | 0.74357307 |
| MPPRIAGMRLLVIKPEP | ASGPGGANTGQLY | 0.35566515 |
| MKFYAVALFRSRGVPPTR | ASGPGGANTGQLY | 0.59648895 |
| NAGGFGILGAVAHSP | ASGPGGANTGQLY | 0.27760056 |
| NEGREITLAMAEM | ASGPGGANTGQLY | 0.43062097 |
| NELVEWSHPLGHRWRWE | ASGPGGANTGQLY | 0.25739792 |
| NLGAGGTGGNGGNPG | ASGPGGANTGQLY | 0.52618265 |
| NTEWPKWETVMAADPVD | ASGPGGANTGQLY | 0.640206 |
| PASAAPAETQP | ASGPGGANTGQLY | 0.64870286 |
| QQSSDGKNGVRAAEITGE | ASGPGGANTGQLY | 0.12367408 |
| RAPDAVTTSWWKE | ASGPGGANTGQLY | 0.6199771 |
| SGFGNVGAHTSGWFNQ | ASGPGGANTGQLY | 0.6704805 |
| SGVAGAVGQAASVGGLKV | ASGPGGANTGQLY | 0.8240459 |
| SNGGAGTGSAGGNGGT | ASGPGGANTGQLY | 0.46335757 |
| TGGAGGEGGAAG | ASGPGGANTGQLY | 0.6058763 |
| TGGDGGAAGAGGAPGHG | ASGPGGANTGQLY | 0.3598484 |
| TMPPLLAAIANAGRHGV | ASGPGGANTGQLY | 0.38668418 |

| | | |
|---|---|---|
| TSSTAGNSGGA | ASGPGGANTGQLY | 0.18923645 |
| VPATVPVGILC | ASGPGGANTGQLY | 0.332359 |
| VQRPDSGGVVLTGRIS | ASGPGGANTGQLY | 0.34760803 |
| WTPNPVPATKGVPAGLVQAT | ASGPGGANTGQLY | 0.2323204 |
| YALKRALAKFSDRFDVVII | ASGPGGANTGQLY | 0.748564 |
| YLIGSVIV | ASGPGGANTGQLY | 0.7248864 |
| YGQLVAVID | ASGPGGANTGQLY | 0.7237326 |
| VTVGGMTLPA | ASGPGGANTGQLY | 0.7425312 |
| VLALPAV | ASGPGGANTGQLY | 0.74278593 |
| VIPPTLNYET | ASGPGGANTGQLY | 0.6115227 |
| VGMGGMHPSSG | ASGPGGANTGQLY | 0.33817446 |
| VGLAGGAATA | ASGPGGANTGQLY | 0.3356973 |
| VGGAGGAGGSGP | ASGPGGANTGQLY | 0.18137585 |
| VFAAFMV | ASGPGGANTGQLY | 0.32889915 |
| TLYDKLR | ASGPGGANTGQLY | 0.34183908 |
| TLAVFTVL | ASGPGGANTGQLY | 0.44537663 |
| TGSVQPARV | ASGPGGANTGQLY | 0.11561008 |
| TGGGAGSGGT | ASGPGGANTGQLY | 0.62030596 |
| SHLETSL | ASGPGGANTGQLY | 0.76578414 |
| SFTTGMT | ASGPGGANTGQLY | 0.4437182 |
| QVEVAKRMGVRQPTVS | ASGPGGANTGQLY | 0.6032245 |
| QGSGGPALAM | ASGPGGANTGQLY | 0.5788994 |
| MLAPSNP | ASGPGGANTGQLY | 0.2714444 |
| LVNNAAI | ASGPGGANTGQLY | 0.67090464 |
| LTGEINTLA | ASGPGGANTGQLY | 0.27871543 |
| LSIFGPV | ASGPGGANTGQLY | 0.5989558 |
| LLALLANL | ASGPGGANTGQLY | 0.74646276 |
| LISNASDALDKLR | ASGPGGANTGQLY | 0.46344435 |
| LINEAVP | ASGPGGANTGQLY | 0.53298837 |
| LGYAFSGL | ASGPGGANTGQLY | 0.44573155 |
| LDFVAVAL | ASGPGGANTGQLY | 0.19508734 |
| LAQPGAE | ASGPGGANTGQLY | 0.54882175 |
| ISNLGAVTGF | ASGPGGANTGQLY | 0.19473183 |
| HQEDPGRI | ASGPGGANTGQLY | 0.43991274 |
| GTPGEPTP | ASGPGGANTGQLY | 0.24024393 |
| GFFNSGS | ASGPGGANTGQLY | 0.5292618 |
| GAGGGGLTAGAN | ASGPGGANTGQLY | 0.48753357 |
| FVDALVNM | ASGPGGANTGQLY | 0.46804786 |
| FLTVAPEL | ASGPGGANTGQLY | 0.22376412 |
| FETLTAL | ASGPGGANTGQLY | 0.47529775 |
| DVINPIEL | ASGPGGANTGQLY | 0.39191815 |
| APVSGAPV | ASGPGGANTGQLY | 0.20686418 |
| ALVGGLAW | ASGPGGANTGQLY | 0.53447914 |
| ALAAAAATAI | ASGPGGANTGQLY | 0.342688 |
| AFGLGLLL | ASGPGGANTGQLY | 0.42874622 |
| AFFAAASL | ASGPGGANTGQLY | 0.68575 |
| AAAALPAFSPP | ASGPGGANTGQLY | 0.48720214 |
| VVLAPAL | ASGPGGANTGQLY | 0.16958459 |
| TIGHVDHGKTTL | ASGPGGANTGQLY | 0.3710668 |
| LISNASDALDK | ASGPGGANTGQLY | 0.45813382 |
| LISNASDAL | ASGPGGANTGQLY | 0.37989852 |

| | | |
|---|---|---|
| ISNASDALDK | ASGPGGANTGQLY | 0.11227201 |
| NQAETLVYQTE | ASGPGGANTGQLY | 0.6042503 |
| TGGTGGTAGAG | ASGPGGANTGQLY | 0.2696242 |
| TGSTAGGAGGA | ASGPGGANTGQLY | 0.22238678 |
| DPLAVPVAAGPV | ASGPGGANTGQLY | 0.315074 |
| GGTGGTPGNGGN | ASGPGGANTGQLY | 0.66549814 |
| PAGLAGVDPATL | ASGPGGANTGQLY | 0.4385779 |
| GNNLVVEAPHAGASV | ASGPGGANTGQLY | 0.4734149 |
| KVTRIMSASTLEQPA | ASGPGGANTGQLY | 0.6411965 |
| LGFALPIALPRRTVV | ASGPGGANTGQLY | 0.6300465 |
| TRLGNLTARKLVVLV | ASGPGGANTGQLY | 0.7624327 |
| WSQQRRLIYFPSAGP | ASGPGGANTGQLY | 0.611887 |
| GPNGNPDPMAAAVDIR | ASGPGGANTGQLY | 0.3978601 |
| TQRCVSVADLRAAASR | ASGPGGANTGQLY | 0.5862056 |
| QFVLDVAEHPVEVLTETV | ASGPGGANTGQLY | 0.3771123 |
| GGAGGTGGAGGNGGTQAGDG | ASGPGGANTGQLY | 0.1617702 |
| PLLATLRAIAVDVDAKPAQV | ASGPGGANTGQLY | 0.15967378 |
| APKFRTGGYMIDFWGVGYQVAK | ASGPGGANTGQLY | 0.28818417 |
| GAGGAGGAGGAAGGPGGTGGTG | ASGPGGANTGQLY | 0.1578822 |
| IGALGLLATDVAPAGSVAFGVA | ASGPGGANTGQLY | 0.5504464 |
| AGGAGGIGGNGGAGGAGGVLMG | ASGPGGANTGQLY | 0.68315065 |
| ADELIGLL | ASGPGGANTGQLY | 0.48582426 |
| ADRALKA | ASGPGGANTGQLY | 0.1864236 |
| AGAVGQVGGAGGA | ASGPGGANTGQLY | 0.33749276 |
| AGGGGGLGGAP | ASGPGGANTGQLY | 0.32040942 |
| AGGLLYGNGGAG | ASGPGGANTGQLY | 0.62490916 |
| AGLESIGL | ASGPGGANTGQLY | 0.49849772 |
| AGLKTGII | ASGPGGANTGQLY | 0.8051795 |
| ASLAALLL | ASGPGGANTGQLY | 0.718983 |
| ASVINISSI | ASGPGGANTGQLY | 0.34520203 |
| ATPPGPA | ASGPGGANTGQLY | 0.6300324 |
| ATVASALV | ASGPGGANTGQLY | 0.297147 |
| ATVASLIL | ASGPGGANTGQLY | 0.3752141 |
| AVNGPVLIRH | ASGPGGANTGQLY | 0.37235743 |
| DGAAGAVGI | ASGPGGANTGQLY | 0.4651811 |
| DPLLHGI | ASGPGGANTGQLY | 0.5104113 |
| ELGAALGAL | ASGPGGANTGQLY | 0.33222073 |
| ELGRTQAV | ASGPGGANTGQLY | 0.18061435 |
| FVAGGYLS | ASGPGGANTGQLY | 0.57257414 |
| GASGGAGGAGGW | ASGPGGANTGQLY | 0.29980105 |
| GGAGAGGDGGAGGI | ASGPGGANTGQLY | 0.17720288 |
| GTGHNGSQPGMGGNG | ASGPGGANTGQLY | 0.38809967 |
| GYAAASASAS | ASGPGGANTGQLY | 0.8149158 |
| HQVDVAAVQQG | ASGPGGANTGQLY | 0.79293466 |
| IGAAIAHT | ASGPGGANTGQLY | 0.25268823 |
| IGMIEVAQL | ASGPGGANTGQLY | 0.28616422 |
| IRLAPEHTSAMD | ASGPGGANTGQLY | 0.68240285 |
| IYNVSVL | ASGPGGANTGQLY | 0.2059838 |
| KQGFLGM | ASGPGGANTGQLY | 0.49866268 |
| LESVITQL | ASGPGGANTGQLY | 0.514543 |
| LGLGALVVV | ASGPGGANTGQLY | 0.4320296 |

| | | |
|---|---|---|
| LIGNGGNGGSGGI | ASGPGGANTGQLY | 0.29393613 |
| LNAINAPTL | ASGPGGANTGQLY | 0.52361673 |
| LQLLNTGV | ASGPGGANTGQLY | 0.3739317 |
| LQTSIEF | ASGPGGANTGQLY | 0.57476926 |
| MSFAVSEL | ASGPGGANTGQLY | 0.1180259 |
| NTGALGL | ASGPGGANTGQLY | 0.60687834 |
| PPGAGGAGGAGTT | ASGPGGANTGQLY | 0.23863593 |
| QLVGLGI | ASGPGGANTGQLY | 0.30442783 |
| RFLEQVHQL | ASGPGGANTGQLY | 0.1471302 |
| RGVGYVLR | ASGPGGANTGQLY | 0.31715468 |
| SATGVFAGVI | ASGPGGANTGQLY | 0.5997695 |
| SLPAITLP | ASGPGGANTGQLY | 0.7202091 |
| STALAFTV | ASGPGGANTGQLY | 0.2598821 |
| SVLTVGLGK | ASGPGGANTGQLY | 0.66253495 |
| TALAGAGSGI | ASGPGGANTGQLY | 0.69022286 |
| TGLPFTGI | ASGPGGANTGQLY | 0.31070027 |
| VAEHVTV | ASGPGGANTGQLY | 0.17443623 |
| VDIVLNSL | ASGPGGANTGQLY | 0.59633553 |
| VLTGGGAVL | ASGPGGANTGQLY | 0.44582912 |
| VNVGLAYGI | ASGPGGANTGQLY | 0.69613147 |
| VSSAAVGAV | ASGPGGANTGQLY | 0.7239293 |
| VTSVYEAGV | ASGPGGANTGQLY | 0.16615462 |
| WVALDRGA | ASGPGGANTGQLY | 0.68128145 |
| YLVGSVVI | ASGPGGANTGQLY | 0.6010237 |
| GPPGTGKTLL | ASGPGGANTGQLY | 0.23812467 |
| GVGMTKF | ASGPGGANTGQLY | 0.22359261 |
| IGIGGSDL | ASGPGGANTGQLY | 0.27978987 |
| IGTLLAEL | ASGPGGANTGQLY | 0.14844838 |
| IYTSGTT | ASGPGGANTGQLY | 0.62271726 |
| LSFPAGV | ASGPGGANTGQLY | 0.3111009 |
| LYTSGTT | ASGPGGANTGQLY | 0.3699123 |
| VGAFPFGF | ASGPGGANTGQLY | 0.35842803 |
| VGAGTVEF | ASGPGGANTGQLY | 0.352694 |
| VGKSTLLL | ASGPGGANTGQLY | 0.3416554 |
| AALAHAGGGIV | ASGPGGANTGQLY | 0.36117426 |
| AALVAVSLL | ASGPGGANTGQLY | 0.45823306 |
| AAVSRAVAAL | ASGPGGANTGQLY | 0.580428 |
| AGGSSGIGGF | ASGPGGANTGQLY | 0.35156712 |
| AGLTVANANG | ASGPGGANTGQLY | 0.43615875 |
| AGLVGAVL | ASGPGGANTGQLY | 0.73146385 |
| ALSQAFTV | ASGPGGANTGQLY | 0.2207962 |
| ALTFSVSL | ASGPGGANTGQLY | 0.28958565 |
| ALTGAALVAL | ASGPGGANTGQLY | 0.63554657 |
| ASFSIGAGVL | ASGPGGANTGQLY | 0.23291004 |
| DGSVYKI | ASGPGGANTGQLY | 0.6755457 |
| EQLAEIV | ASGPGGANTGQLY | 0.58737886 |
| FAVSELTVM | ASGPGGANTGQLY | 0.5145768 |
| GAAPLAAGL | ASGPGGANTGQLY | 0.49875742 |
| GADGGDGGSGGAGG | ASGPGGANTGQLY | 0.80553764 |
| GFGLDTGGAGG | ASGPGGANTGQLY | 0.5766734 |
| GLASGGLL | ASGPGGANTGQLY | 0.296779 |

| | | |
|---|---|---|
| GRIAGAAVLL | ASGPGGANTGQLY | 0.6773056 |
| GSAGSLNTSF | ASGPGGANTGQLY | 0.2647295 |
| GTGQIGI | ASGPGGANTGQLY | 0.6460236 |
| GTLIALIL | ASGPGGANTGQLY | 0.6620976 |
| IAATMGFVVG | ASGPGGANTGQLY | 0.21369061 |
| IFAQALL | ASGPGGANTGQLY | 0.3507377 |
| IGAAVALGF | ASGPGGANTGQLY | 0.6803994 |
| IGLTPLAL | ASGPGGANTGQLY | 0.68708634 |
| ILGSNTSTL | ASGPGGANTGQLY | 0.18108995 |
| IVFSLTVL | ASGPGGANTGQLY | 0.25967547 |
| LASKATGF | ASGPGGANTGQLY | 0.2247879 |
| LASVALIL | ASGPGGANTGQLY | 0.3881505 |
| LFAVGLTL | ASGPGGANTGQLY | 0.2402361 |
| LFNGGIGTYI | ASGPGGANTGQLY | 0.59858525 |
| LTLGAGII | ASGPGGANTGQLY | 0.33586216 |
| LVAAALGDI | ASGPGGANTGQLY | 0.72280085 |
| NTAFVQLGI | ASGPGGANTGQLY | 0.5286187 |
| QLFSGAEL | ASGPGGANTGQLY | 0.7037715 |
| SAATFITL | ASGPGGANTGQLY | 0.31619716 |
| SNYGLANLGV | ASGPGGANTGQLY | 0.47263414 |
| SYGGAYAVM | ASGPGGANTGQLY | 0.17586522 |
| TTGVGGLTLG | ASGPGGANTGQLY | 0.23482513 |
| TTQTIYI | ASGPGGANTGQLY | 0.22366822 |
| VAAGAGLGAVI | ASGPGGANTGQLY | 0.43836474 |
| VAAVVTLL | ASGPGGANTGQLY | 0.14284423 |
| VGFLGGLI | ASGPGGANTGQLY | 0.7734479 |
| VGVIPILL | ASGPGGANTGQLY | 0.3931315 |
| VIFASSLI | ASGPGGANTGQLY | 0.5902394 |
| VKGGDGGAAAG | ASGPGGANTGQLY | 0.37151563 |
| VTKTFTNI | ASGPGGANTGQLY | 0.2982561 |
| VVAAAILL | ASGPGGANTGQLY | 0.32702604 |
| YAALDVEL | ASGPGGANTGQLY | 0.7060671 |
| LASVALLL | ASGPGGANTGQLY | 0.21836853 |
| LDAVHRV | ASGPGGANTGQLY | 0.12688342 |
| QGLSSSIF | ASGPGGANTGQLY | 0.25744748 |
| RGTFANIRL | ASGPGGANTGQLY | 0.16162401 |
| RQLWWGHRI | ASGPGGANTGQLY | 0.76750624 |
| YGPPGTGKTLL | ASGPGGANTGQLY | 0.676615 |
| AALVGAPANVA | ASGPGGANTGQLY | 0.43544877 |
| AATFRRTE | ASGPGGANTGQLY | 0.7413881 |
| AGALGGGI | ASGPGGANTGQLY | 0.119874306 |
| AGGAGGAGGIGAG | ASGPGGANTGQLY | 0.3670496 |
| AGSGALGL | ASGPGGANTGQLY | 0.7895641 |
| AQIADQI | ASGPGGANTGQLY | 0.48405612 |
| ASAVAVTYL | ASGPGGANTGQLY | 0.40021843 |
| ASTVLGGTGG | ASGPGGANTGQLY | 0.72802913 |
| AVGGEPAK | ASGPGGANTGQLY | 0.32831994 |
| CEQGTFGGTC | ASGPGGANTGQLY | 0.41427642 |
| DEGKAPAG | ASGPGGANTGQLY | 0.44914973 |
| DTVVFPAL | ASGPGGANTGQLY | 0.5275717 |
| FEGITMAG | ASGPGGANTGQLY | 0.38546744 |

| | | |
|---|---|---|
| FPLVVQD | ASGPGGANTGQLY | 0.2122865 |
| GAFTTPQ | ASGPGGANTGQLY | 0.554078 |
| GFGGFGTAGGVGGA | ASGPGGANTGQLY | 0.81242085 |
| GFLAQFL | ASGPGGANTGQLY | 0.31696332 |
| GGAGGAGGNAGVA | ASGPGGANTGQLY | 0.6211512 |
| GGVGTAGAGGAGGAGG | ASGPGGANTGQLY | 0.52393216 |
| GNGGGGDAGNAGSGGN | ASGPGGANTGQLY | 0.567503 |
| GTGGAGGAGNDAGS | ASGPGGANTGQLY | 0.2508416 |
| GTTNTGWFNS | ASGPGGANTGQLY | 0.23207659 |
| IFMGPLFTL | ASGPGGANTGQLY | 0.4868018 |
| ILAHAIEAI | ASGPGGANTGQLY | 0.20635322 |
| IRSAIREAL | ASGPGGANTGQLY | 0.41793352 |
| ITFAGGG | ASGPGGANTGQLY | 0.31243595 |
| ITGMSGHQEV | ASGPGGANTGQLY | 0.8395542 |
| IVAGLACTL | ASGPGGANTGQLY | 0.60842013 |
| IVGGGLAGA | ASGPGGANTGQLY | 0.5928815 |
| IVTGVLNI | ASGPGGANTGQLY | 0.23944521 |
| LAQQGCI | ASGPGGANTGQLY | 0.36851922 |
| LCLHVEN | ASGPGGANTGQLY | 0.6222372 |
| LDAISDGS | ASGPGGANTGQLY | 0.19186632 |
| LGGFAAML | ASGPGGANTGQLY | 0.75110996 |
| LGGKNPMI | ASGPGGANTGQLY | 0.2718073 |
| LGPKGVAAIL | ASGPGGANTGQLY | 0.48393124 |
| LMVGVSI | ASGPGGANTGQLY | 0.6595748 |
| LVGAAFVL | ASGPGGANTGQLY | 0.7585536 |
| LVLAVAPL | ASGPGGANTGQLY | 0.32021064 |
| MAAIAVIDAA | ASGPGGANTGQLY | 0.4099214 |
| NNGINAELQ | ASGPGGANTGQLY | 0.24125797 |
| RGEPLVLVHG | ASGPGGANTGQLY | 0.63494456 |
| RGGAGGAAGLI | ASGPGGANTGQLY | 0.7216941 |
| RSYRMHPAV | ASGPGGANTGQLY | 0.5565692 |
| SAAHLLAGQLA | ASGPGGANTGQLY | 0.8007782 |
| SAAVGSY | ASGPGGANTGQLY | 0.56672764 |
| SPHAVLHRI | ASGPGGANTGQLY | 0.21377675 |
| SQIAKVL | ASGPGGANTGQLY | 0.4020299 |
| TALGHLVP | ASGPGGANTGQLY | 0.66699183 |
| TAPTTSGAA | ASGPGGANTGQLY | 0.72008836 |
| TDGDDGNGGSPGT | ASGPGGANTGQLY | 0.71433306 |
| TDGLGEN | ASGPGGANTGQLY | 0.6821039 |
| TGGPPLPGQA | ASGPGGANTGQLY | 0.28242743 |
| TIPAITI | ASGPGGANTGQLY | 0.7214396 |
| TIPGFTI | ASGPGGANTGQLY | 0.791543 |
| TSIASGALA | ASGPGGANTGQLY | 0.6400957 |
| TVTLPGAAI | ASGPGGANTGQLY | 0.3048101 |
| VASGLSAALP | ASGPGGANTGQLY | 0.3818124 |
| VGAGVIGI | ASGPGGANTGQLY | 0.47458348 |
| VGPKRFGV | ASGPGGANTGQLY | 0.7346858 |
| VNVAFPD | ASGPGGANTGQLY | 0.22594208 |
| VSMTNSGVS | ASGPGGANTGQLY | 0.49574852 |
| VTDVLGDLSV | ASGPGGANTGQLY | 0.5279333 |
| VVVVGAGFAGL | ASGPGGANTGQLY | 0.15851898 |

| | | |
|---|---|---|
| VYTVGTVV | ASGPGGANTGQLY | 0.45767298 |
| WTTFSGP | ASGPGGANTGQLY | 0.76620126 |
| YDELVEAF | ASGPGGANTGQLY | 0.40871522 |
| YGTGGAGGNGG | ASGPGGANTGQLY | 0.25446075 |
| YTNVMAV | ASGPGGANTGQLY | 0.73322284 |
| FLEQVHQL | ASGPGGANTGQLY | 0.81036025 |
| GLSLLAVAV | ASGPGGANTGQLY | 0.6903909 |
| LEQVHQL | ASGPGGANTGQLY | 0.18746178 |
| LNAAGAI | ASGPGGANTGQLY | 0.116111636 |
| PAVHASL | ASGPGGANTGQLY | 0.42022917 |
| TELEDDA | ASGPGGANTGQLY | 0.7330381 |
| TPPGPPN | ASGPGGANTGQLY | 0.40709305 |
| TPPPPPPP | ASGPGGANTGQLY | 0.29050004 |
| TQSPALI | ASSQKTGVGAETLY | 0.5466093 |
| EFAAAFPQ | ASSQKTGVGAETLY | 0.65265864 |
| LLSGNLDVL | ASSQKTGVGAETLY | 0.245765 |
| GPANLGNYNI | ASSQKTGVGAETLY | 0.22127074 |
| TLGPTLP | ASSQKTGVGAETLY | 0.5201527 |
| VYAYSSCCK | ASSQKTGVGAETLY | 0.6599337 |
| APAPAPAPNI | ASSQKTGVGAETLY | 0.20505291 |
| PGGPVPIP | ASSQKTGVGAETLY | 0.29254693 |
| FLTVALEP | ASSQKTGVGAETLY | 0.4194277 |
| SQYFGANL | ASSQKTGVGAETLY | 0.3731233 |
| GGVKPAA | ASSQKTGVGAETLY | 0.561219 |
| GGTGGAGGAGGW | ASSQKTGVGAETLY | 0.24964201 |
| GVGGVAGLGGAGG | ASSQKTGVGAETLY | 0.7099207 |
| VLLVLATTI | ASSQKTGVGAETLY | 0.18351525 |
| IGIFAAML | ASSQKTGVGAETLY | 0.58676577 |
| AVGDLID | ASSQKTGVGAETLY | 0.5394938 |
| NISPLPL | ASSQKTGVGAETLY | 0.49174672 |
| VTAVFVAI | ASSQKTGVGAETLY | 0.36496186 |
| SGDDNDAAL | ASSQKTGVGAETLY | 0.23236993 |
| AVAALAHS | ASSQKTGVGAETLY | 0.21491188 |
| GGVTVMATP | ASSQKTGVGAETLY | 0.16202025 |
| AGLAANVRD | ASSQKTGVGAETLY | 0.72371906 |
| TNRPDLIDPALLRPG | ASSQKTGVGAETLY | 0.36525145 |
| LANLGVTAL | ASSQKTGVGAETLY | 0.69367886 |
| TGERIEL | ASSQKTGVGAETLY | 0.4814381 |
| NGEELAA | ASSQKTGVGAETLY | 0.24231389 |
| VLSLVWG | ASSQKTGVGAETLY | 0.47837538 |
| IGYALWL | ASSQKTGVGAETLY | 0.24860084 |
| LSLLLAPL | ASSQKTGVGAETLY | 0.29077804 |
| TLDEALK | ASSQKTGVGAETLY | 0.5789305 |
| TDGAGTAEGG | ASSQKTGVGAETLY | 0.58638394 |
| NAPGGAGGNGG | ASSQKTGVGAETLY | 0.250391 |
| VSVFMGMGGNF | ASSQKTGVGAETLY | 0.48318654 |
| LASYLAAL | ASSQKTGVGAETLY | 0.49682713 |
| LVGGTPAL | ASSQKTGVGAETLY | 0.24669628 |
| VRNADVL | ASSQKTGVGAETLY | 0.42764255 |
| TCSSPFLVSAM | ASSQKTGVGAETLY | 0.71222055 |
| VHITDAE | ASSQKTGVGAETLY | 0.47040683 |

| | | |
|---|---|---|
| EYAAPSF | ASSQKTGVGAETLY | 0.189748 |
| VVDAFHGGL | ASSQKTGVGAETLY | 0.59037983 |
| ASVVDPQ | ASSQKTGVGAETLY | 0.24560197 |
| SYNEFGGLS | ASSQKTGVGAETLY | 0.51552385 |
| DPDGSVY | ASSQKTGVGAETLY | 0.44432396 |
| MVSTIHATE | ASSQKTGVGAETLY | 0.44388872 |
| LAAGVADPV | ASSQKTGVGAETLY | 0.6199604 |
| TKEQIAA | ASSQKTGVGAETLY | 0.23875932 |
| SLASPAAL | ASSQKTGVGAETLY | 0.2500209 |
| IGLVFYGV | ASSQKTGVGAETLY | 0.36516553 |
| FLLDASNL | ASSQKTGVGAETLY | 0.48488796 |
| YQVGNFP | ASSQKTGVGAETLY | 0.63759243 |
| ESLLSQST | ASSQKTGVGAETLY | 0.22545329 |
| LFTVGFAAVG | ASSQKTGVGAETLY | 0.46786582 |
| ATPPTPT | ASSQKTGVGAETLY | 0.34978282 |
| TDLATLG | ASSQKTGVGAETLY | 0.26332307 |
| SAGGTGR | ASSQKTGVGAETLY | 0.40046483 |
| GFSNSGS | ASSQKTGVGAETLY | 0.293437 |
| GGNGGSGGTSV | ASSQKTGVGAETLY | 0.22791056 |
| GTSVATGGAGN | ASSQKTGVGAETLY | 0.5794753 |
| NGGNGGSGGTS | ASSQKTGVGAETLY | 0.3103642 |
| AVVKFPH | ASSQKTGVGAETLY | 0.5811979 |
| TPPPPPP | ASSQKTGVGAETLY | 0.6988039 |
| AAPAAAAAPA | ASSQKTGVGAETLY | 0.295953 |
| IALNEDL | ASSQKTGVGAETLY | 0.49347752 |
| IVNVSSM | ASSQKTGVGAETLY | 0.5545038 |
| TGIPPAPR | ASSQKTGVGAETLY | 0.5950216 |
| FDLGGGTF | ASSQKTGVGAETLY | 0.40729713 |
| FDLGGGTFDVS | ASSQKTGVGAETLY | 0.24527049 |
| NEPTAAA | ASSQKTGVGAETLY | 0.7030687 |
| NEPTAAALA | ASSQKTGVGAETLY | 0.6209656 |
| GSIINISSI | ASSQKTGVGAETLY | 0.6720872 |
| VAPGNAGTA | ASSQKTGVGAETLY | 0.65809697 |
| TGGGASLE | ASSQKTGVGAETLY | 0.6254641 |
| VAVYNHYKRI | ASSQKTGVGAETLY | 0.5845356 |
| TDPAPATT | ASSQKTGVGAETLY | 0.35076612 |
| TNRPDILDPALLRPG | ASSQKTGVGAETLY | 0.18015341 |
| LTTTTTS | ASSQKTGVGAETLY | 0.4100901 |
| PTGSGKTL | ASSQKTGVGAETLY | 0.37589645 |
| TGSGKTL | ASSQKTGVGAETLY | 0.25418323 |
| TIGHVDHGK | ASSQKTGVGAETLY | 0.21321973 |
| TIGHVDHGKTT | ASSQKTGVGAETLY | 0.70376456 |
| GHVDHGKTT | ASSQKTGVGAETLY | 0.83479476 |
| AQFEHTLL | ASSQKTGVGAETLY | 0.33372065 |
| IVNQDEL | ASSQKTGVGAETLY | 0.77696323 |
| TNGSQFF | ASSQKTGVGAETLY | 0.18073578 |
| TGKGIVE | ASSQKTGVGAETLY | 0.67804205 |
| LISNASDA | ASSQKTGVGAETLY | 0.7292891 |
| AELRTLGELTKIAKAHGAQ | ASSQKTGVGAETLY | 0.24486107 |
| APNAVAAAEPSW | ASSQKTGVGAETLY | 0.5923307 |
| APQVAGAATAVL | ASSQKTGVGAETLY | 0.21819712 |

| | | |
|---|---|---|
| ARRDPAAAAAILPTDA | ASSQKTGVGAETLY | 0.3200197 |
| DAAARDVLAEAGLAEYF | ASSQKTGVGAETLY | 0.7485578 |
| DDQIVLSAKANLAKVEQ | ASSQKTGVGAETLY | 0.4604752 |
| DELLPDTGEDSDEDGH | ASSQKTGVGAETLY | 0.5290104 |
| ELPEAPANRATVPQD | ASSQKTGVGAETLY | 0.40431604 |
| FGQSKIERVTGIG | ASSQKTGVGAETLY | 0.28767827 |
| GAGGTGSTAGGAGGAGGAGGLY | ASSQKTGVGAETLY | 0.45303333 |
| GHGGTGAAGGNGGTGGAGGSGI | ASSQKTGVGAETLY | 0.7544111 |
| GNGGNGGAGGEGGPGGAGGAGA | ASSQKTGVGAETLY | 0.32412505 |
| GSTAGNGGNGGNATNGTIAGQP | ASSQKTGVGAETLY | 0.75167483 |
| GTGGDGGNGGNGGTGG | ASSQKTGVGAETLY | 0.2505629 |
| GVRRPAGIPADAPV | ASSQKTGVGAETLY | 0.7657418 |
| GYRQVTDAHLLTLARR | ASSQKTGVGAETLY | 0.76393604 |
| IAKMGQDGHDRG | ASSQKTGVGAETLY | 0.46463656 |
| KAEPSIEIQYT | ASSQKTGVGAETLY | 0.35139707 |
| KPVRIRMAADEDTVAVT | ASSQKTGVGAETLY | 0.72430015 |
| MPPRIAGMRLLVIKPEP | ASSQKTGVGAETLY | 0.4333086 |
| MKFYAVALFRSRGVPPTR | ASSQKTGVGAETLY | 0.65834916 |
| NAGGFGILGAVAHSP | ASSQKTGVGAETLY | 0.6931838 |
| NEGREITLAMAEM | ASSQKTGVGAETLY | 0.4183538 |
| NELVEWSHPLGHRWRWE | ASSQKTGVGAETLY | 0.6436066 |
| NLGAGGTGGNGGNPG | ASSQKTGVGAETLY | 0.5627527 |
| NTEWPKWETVMAADPVD | ASSQKTGVGAETLY | 0.507551 |
| PASAAPAETQP | ASSQKTGVGAETLY | 0.5391313 |
| QQSSDGKNGVRAAEITGE | ASSQKTGVGAETLY | 0.35002387 |
| RAPDAVTTSWWKE | ASSQKTGVGAETLY | 0.4928794 |
| SGFGNVGAHTSGWFNQ | ASSQKTGVGAETLY | 0.48918322 |
| SGVAGAVGQAASVGGLKV | ASSQKTGVGAETLY | 0.6161789 |
| SNGGAGTGSAGGNGGT | ASSQKTGVGAETLY | 0.54272825 |
| TGGAGGEGGAAG | ASSQKTGVGAETLY | 0.46252802 |
| TGGDGGAAGAGGAPGHG | ASSQKTGVGAETLY | 0.6630273 |
| TMPPLLAAIANAGRHGV | ASSQKTGVGAETLY | 0.62456065 |
| TSSTAGNSGGA | ASSQKTGVGAETLY | 0.36906254 |
| VPATVPVGILC | ASSQKTGVGAETLY | 0.5162915 |
| VQRPDSGGVVLTGRIS | ASSQKTGVGAETLY | 0.41042733 |
| WTPNPVPATKGVPAGLVQAT | ASSQKTGVGAETLY | 0.52957845 |
| YALKRALAKFSDRFDVVII | ASSQKTGVGAETLY | 0.3047535 |
| YLIGSVIV | ASSQKTGVGAETLY | 0.61145085 |
| YGQLVAVID | ASSQKTGVGAETLY | 0.38702902 |
| VTVGGMTLPA | ASSQKTGVGAETLY | 0.28753036 |
| VLALPAV | ASSQKTGVGAETLY | 0.53916836 |
| VIPPTLNYET | ASSQKTGVGAETLY | 0.55157554 |
| VGMGGMHPSSG | ASSQKTGVGAETLY | 0.53568506 |
| VGLAGGAATA | ASSQKTGVGAETLY | 0.63453233 |
| VGGAGGAGGSGP | ASSQKTGVGAETLY | 0.3139682 |
| VFAAFMV | ASSQKTGVGAETLY | 0.4207259 |
| TLYDKLR | ASSQKTGVGAETLY | 0.6343913 |
| TLAVFTVL | ASSQKTGVGAETLY | 0.6598419 |
| TGSVQPARV | ASSQKTGVGAETLY | 0.3855243 |
| TGGGAGSGGT | ASSQKTGVGAETLY | 0.5952512 |
| SHLETSL | ASSQKTGVGAETLY | 0.63487625 |

| | | |
|---|---|---|
| SFTTGMT | ASSQKTGVGAETLY | 0.5961734 |
| QVEVAKRMGVRQPTVS | ASSQKTGVGAETLY | 0.61042714 |
| QGSGGPALAM | ASSQKTGVGAETLY | 0.6367647 |
| MLAPSNP | ASSQKTGVGAETLY | 0.34563914 |
| LVNNAAI | ASSQKTGVGAETLY | 0.6540257 |
| LTGEINTLA | ASSQKTGVGAETLY | 0.658222 |
| LSIFGPV | ASSQKTGVGAETLY | 0.46441478 |
| LLALLANL | ASSQKTGVGAETLY | 0.5769784 |
| LISNASDALDKLR | ASSQKTGVGAETLY | 0.3914117 |
| LINEAVP | ASSQKTGVGAETLY | 0.40740797 |
| LGYAFSGL | ASSQKTGVGAETLY | 0.3060592 |
| LDFVAVAL | ASSQKTGVGAETLY | 0.6164477 |
| LAQPGAE | ASSQKTGVGAETLY | 0.6174101 |
| ISNLGAVTGF | ASSQKTGVGAETLY | 0.56501025 |
| HQEDPGRI | ASSQKTGVGAETLY | 0.35854214 |
| GTPGEPTP | ASSQKTGVGAETLY | 0.44871512 |
| GFFNSGS | ASSQKTGVGAETLY | 0.37523538 |
| GAGGGGLTAGAN | ASSQKTGVGAETLY | 0.4982389 |
| FVDALVNM | ASSQKTGVGAETLY | 0.31034023 |
| FLTVAPEL | ASSQKTGVGAETLY | 0.3720831 |
| FETLTAL | ASSQKTGVGAETLY | 0.657135 |
| DVINPIEL | ASSQKTGVGAETLY | 0.65017366 |
| APVSGAPV | ASSQKTGVGAETLY | 0.34121498 |
| ALVGGLAW | ASSQKTGVGAETLY | 0.6564391 |
| ALAAAAATAI | ASSQKTGVGAETLY | 0.6171229 |
| AFGLGLLL | ASSQKTGVGAETLY | 0.6469544 |
| AFFAAASL | ASSQKTGVGAETLY | 0.46954256 |
| AAAALPAFSPP | ASSQKTGVGAETLY | 0.41680816 |
| VVLAPAL | ASSQKTGVGAETLY | 0.6230516 |
| TIGHVDHGKTTL | ASSQKTGVGAETLY | 0.55783343 |
| LISNASDALDK | ASSQKTGVGAETLY | 0.3647683 |
| LISNASDAL | ASSQKTGVGAETLY | 0.6291054 |
| ISNASDALDK | ASSQKTGVGAETLY | 0.46869653 |
| NQAETLVYQTE | ASSQKTGVGAETLY | 0.5028661 |
| TGGTGGTAGAG | ASSQKTGVGAETLY | 0.5440242 |
| TGSTAGGAGGA | ASSQKTGVGAETLY | 0.5553551 |
| DPLAVPVAAGPV | ASSQKTGVGAETLY | 0.39876103 |
| GGTGGTPGNGGN | ASSQKTGVGAETLY | 0.3810307 |
| PAGLAGVDPATL | ASSQKTGVGAETLY | 0.3598622 |
| GNNLVVEAPHAGASV | ASSQKTGVGAETLY | 0.58956695 |
| KVTRIMSASTLEQPA | ASSQKTGVGAETLY | 0.6523679 |
| LGFALPIALPRRTVV | ASSQKTGVGAETLY | 0.54314226 |
| TRLGNLTARKLVVLV | ASSQKTGVGAETLY | 0.38381538 |
| WSQQRRLIYFPSAGP | ASSQKTGVGAETLY | 0.6294912 |
| GPNGNPDPMAAAVDIR | ASSQKTGVGAETLY | 0.5660195 |
| TQRCVSVADLRAAASR | ASSQKTGVGAETLY | 0.52821857 |
| QFVLDVAEHPVEVLTETV | ASSQKTGVGAETLY | 0.49184006 |
| GGAGGTGGAGGNGGTQAGDG | ASSQKTGVGAETLY | 0.4997005 |
| PLLATLRAIAVDVDAKPAQV | ASSQKTGVGAETLY | 0.5846523 |
| APKFRTGGYMIDFWGVGYQVAK | ASSQKTGVGAETLY | 0.580856 |
| GAGGAGGAGGAAGGPGGTGGTG | ASSQKTGVGAETLY | 0.5905096 |

| | | |
|---|---|---|
| IGALGLLATDVAPAGSVAFGVA | ASSQKTGVGAETLY | 0.5920093 |
| AGGAGGIGGNGGAGGAGGVLMG | ASSQKTGVGAETLY | 0.47306567 |
| ADELIGLL | ASSQKTGVGAETLY | 0.49200895 |
| ADRALKA | ASSQKTGVGAETLY | 0.33096647 |
| AGAVGQVGGAGGA | ASSQKTGVGAETLY | 0.3623901 |
| AGGGGGLGGAP | ASSQKTGVGAETLY | 0.5757914 |
| AGGLLYGNGGAG | ASSQKTGVGAETLY | 0.57017636 |
| AGLESIGL | ASSQKTGVGAETLY | 0.6219994 |
| AGLKTGII | ASSQKTGVGAETLY | 0.5019202 |
| ASLAALLL | ASSQKTGVGAETLY | 0.39886063 |
| ASVINISSI | ASSQKTGVGAETLY | 0.57880986 |
| ATPPGPA | ASSQKTGVGAETLY | 0.628498 |
| ATVASALV | ASSQKTGVGAETLY | 0.38264373 |
| ATVASLIL | ASSQKTGVGAETLY | 0.44547462 |
| AVNGPVLIRH | ASSQKTGVGAETLY | 0.66931355 |
| DGAAGAVGI | ASSQKTGVGAETLY | 0.5864905 |
| DPLLHGI | ASSQKTGVGAETLY | 0.5378089 |
| ELGAALGAL | ASSQKTGVGAETLY | 0.43888053 |
| ELGRTQAV | ASSQKTGVGAETLY | 0.66547215 |
| FVAGGYLS | ASSQKTGVGAETLY | 0.38248834 |
| GASGGAGGAGGW | ASSQKTGVGAETLY | 0.6573526 |
| GGAGAGGDGGAGGI | ASSQKTGVGAETLY | 0.3079254 |
| GTGHNGSQPGMGGNG | ASSQKTGVGAETLY | 0.42660305 |
| GYAAASASAS | ASSQKTGVGAETLY | 0.38103995 |
| HQVDVAAVQQG | ASSQKTGVGAETLY | 0.3390963 |
| IGAAIAHT | ASSQKTGVGAETLY | 0.6508428 |
| IGMIEVAQL | ASSQKTGVGAETLY | 0.43997657 |
| IRLAPEHTSAMD | ASSQKTGVGAETLY | 0.6257268 |
| IYNVSVL | ASSQKTGVGAETLY | 0.4896092 |
| KQGFLGM | ASSQKTGVGAETLY | 0.63752836 |
| LESVITQL | ASSQKTGVGAETLY | 0.59155285 |
| LGLGALVVV | ASSQKTGVGAETLY | 0.4859893 |
| LIGNGGNGGSGGI | ASSQKTGVGAETLY | 0.4013207 |
| LNAINAPTL | ASSQKTGVGAETLY | 0.6432463 |
| LQLLNTGV | ASSQKTGVGAETLY | 0.60740024 |
| LQTSIEF | ASSQKTGVGAETLY | 0.5162421 |
| MSFAVSEL | ASSQKTGVGAETLY | 0.36183923 |
| NTGALGL | ASSQKTGVGAETLY | 0.43307033 |
| PPGAGGAGGAGTT | ASSQKTGVGAETLY | 0.6271097 |
| QLVGLGI | ASSQKTGVGAETLY | 0.29955786 |
| RFLEQVHQL | ASSQKTGVGAETLY | 0.299273 |
| RGVGYVLR | ASSQKTGVGAETLY | 0.39276105 |
| SATGVFAGVI | ASSQKTGVGAETLY | 0.35591328 |
| SLPAITLP | ASSQKTGVGAETLY | 0.66213924 |
| STALAFTV | ASSQKTGVGAETLY | 0.6411313 |
| SVLTVGLGK | ASSQKTGVGAETLY | 0.58051616 |
| TALAGAGSGI | ASSQKTGVGAETLY | 0.5186088 |
| TGLPFTGI | ASSQKTGVGAETLY | 0.5714438 |
| VAEHVTV | ASSQKTGVGAETLY | 0.28091088 |
| VDIVLNSL | ASSQKTGVGAETLY | 0.38084543 |
| VLTGGGAVL | ASSQKTGVGAETLY | 0.3915547 |

| | | |
|---|---|---|
| VNVGLAYGI | ASSQKTGVGAETLY | 0.62833047 |
| VSSAAVGAV | ASSQKTGVGAETLY | 0.40623075 |
| VTSVYEAGV | ASSQKTGVGAETLY | 0.4264657 |
| WVALDRGA | ASSQKTGVGAETLY | 0.6106794 |
| YLVGSVVI | ASSQKTGVGAETLY | 0.60311 |
| GPPGTGKTLL | ASSQKTGVGAETLY | 0.5374741 |
| GVGMTKF | ASSQKTGVGAETLY | 0.394549 |
| IGIGGSDL | ASSQKTGVGAETLY | 0.28860277 |
| IGTLLAEL | ASSQKTGVGAETLY | 0.64467776 |
| IYTSGTT | ASSQKTGVGAETLY | 0.32163808 |
| LSFPAGV | ASSQKTGVGAETLY | 0.4031439 |
| LYTSGTT | ASSQKTGVGAETLY | 0.30099428 |
| VGAFPFGF | ASSQKTGVGAETLY | 0.3216777 |
| VGAGTVEF | ASSQKTGVGAETLY | 0.34814948 |
| VGKSTLLL | ASSQKTGVGAETLY | 0.566257 |
| AALAHAGGGIV | ASSQKTGVGAETLY | 0.53786683 |
| AALVAVSLL | ASSQKTGVGAETLY | 0.5378164 |
| AAVSRAVAAL | ASSQKTGVGAETLY | 0.6654076 |
| AGGSSGIGGF | ASSQKTGVGAETLY | 0.6268715 |
| AGLTVANANG | ASSQKTGVGAETLY | 0.51087075 |
| AGLVGAVL | ASSQKTGVGAETLY | 0.45134956 |
| ALSQAFTV | ASSQKTGVGAETLY | 0.41731042 |
| ALTFSVSL | ASSQKTGVGAETLY | 0.4010414 |
| ALTGAALVAL | ASSQKTGVGAETLY | 0.41958258 |
| ASFSIGAGVL | ASSQKTGVGAETLY | 0.6644579 |
| DGSVYKI | ASSQKTGVGAETLY | 0.43283328 |
| EQLAEIV | ASSQKTGVGAETLY | 0.36582345 |
| FAVSELTVM | ASSQKTGVGAETLY | 0.43852776 |
| GAAPLAAGL | ASSQKTGVGAETLY | 0.64881057 |
| GADGGDGGSGGAGG | ASSQKTGVGAETLY | 0.6683749 |
| GFGLDTGGAGG | ASSQKTGVGAETLY | 0.4041385 |
| GLASGGLL | ASSQKTGVGAETLY | 0.425918 |
| GRIAGAAVLL | ASSQKTGVGAETLY | 0.43168676 |
| GSAGSLNTSF | ASSQKTGVGAETLY | 0.6163713 |
| GTGQIGI | ASSQKTGVGAETLY | 0.3859234 |
| GTLIALIL | ASSQKTGVGAETLY | 0.39320344 |
| IAATMGFVVG | ASSQKTGVGAETLY | 0.541832 |
| IFAQALL | ASSQKTGVGAETLY | 0.6762488 |
| IGAAVALGF | ASSQKTGVGAETLY | 0.41363826 |
| IGLTPLAL | ASSQKTGVGAETLY | 0.3406684 |
| ILGSNTSTL | ASSQKTGVGAETLY | 0.60277224 |
| IVFSLTVL | ASSQKTGVGAETLY | 0.43821874 |
| LASKATGF | ASSQKTGVGAETLY | 0.5584908 |
| LASVALIL | ASSQKTGVGAETLY | 0.47089285 |
| LFAVGLTL | ASSQKTGVGAETLY | 0.36249837 |
| LFNGGIGTYI | ASSQKTGVGAETLY | 0.46162784 |
| LTLGAGII | ASSQKTGVGAETLY | 0.6643345 |
| LVAAALGDI | ASSQKTGVGAETLY | 0.57338727 |
| NTAFVQLGI | ASSQKTGVGAETLY | 0.56526124 |
| QLFSGAEL | ASSQKTGVGAETLY | 0.6562703 |
| SAATFITL | ASSQKTGVGAETLY | 0.6177768 |

| | | |
|---|---|---|
| SNYGLANLGV | ASSQKTGVGAETLY | 0.65706867 |
| SYGGAYAVM | ASSQKTGVGAETLY | 0.6188945 |
| TTGVGGLTLG | ASSQKTGVGAETLY | 0.48760027 |
| TTQTIYI | ASSQKTGVGAETLY | 0.48869085 |
| VAAGAGLGAVI | ASSQKTGVGAETLY | 0.5882199 |
| VAAVVTLL | ASSQKTGVGAETLY | 0.62171894 |
| VGFLGGLI | ASSQKTGVGAETLY | 0.5559046 |
| VGVIPILL | ASSQKTGVGAETLY | 0.406714 |
| VIFASSLI | ASSQKTGVGAETLY | 0.660874 |
| VKGGDGGAAAG | ASSQKTGVGAETLY | 0.4620821 |
| VTKTFTNI | ASSQKTGVGAETLY | 0.6130604 |
| VVAAAILL | ASSQKTGVGAETLY | 0.36504418 |
| YAALDVEL | ASSQKTGVGAETLY | 0.3860985 |
| LASVALLL | ASSQKTGVGAETLY | 0.6268306 |
| LDAVHRV | ASSQKTGVGAETLY | 0.28661698 |
| QGLSSSIF | ASSQKTGVGAETLY | 0.30492648 |
| RGTFANIRL | ASSQKTGVGAETLY | 0.49302143 |
| RQLWWGHRI | ASSQKTGVGAETLY | 0.6372894 |
| YGPPGTGKTLL | ASSQKTGVGAETLY | 0.28546083 |
| AALVGAPANVA | ASSQKTGVGAETLY | 0.39202943 |
| AATFRRTE | ASSQKTGVGAETLY | 0.349128 |
| AGALGGGI | ASSQKTGVGAETLY | 0.49154526 |
| AGGAGGAGGIGAG | ASSQKTGVGAETLY | 0.5548713 |
| AGSGALGL | ASSQKTGVGAETLY | 0.34699398 |
| AQIADQI | ASSQKTGVGAETLY | 0.614138 |
| ASAVAVTYL | ASSQKTGVGAETLY | 0.3205344 |
| ASTVLGGTGG | ASSQKTGVGAETLY | 0.32739234 |
| AVGGEPAK | ASSQKTGVGAETLY | 0.6098112 |
| CEQGTFGGTC | ASSQKTGVGAETLY | 0.6508755 |
| DEGKAPAG | ASSQKTGVGAETLY | 0.36348236 |
| DTVVFPAL | ASSQKTGVGAETLY | 0.6241367 |
| FEGITMAG | ASSQKTGVGAETLY | 0.39099002 |
| FPLVVQD | ASSQKTGVGAETLY | 0.66450405 |
| GAFTTPQ | ASSQKTGVGAETLY | 0.537116 |
| GFGGFGTAGGVGGA | ASSQKTGVGAETLY | 0.34346625 |
| GFLAQFL | ASSQKTGVGAETLY | 0.6282059 |
| GGAGGAGGNAGVA | ASSQKTGVGAETLY | 0.57764643 |
| GGVGTAGAGGAGGAGG | ASSQKTGVGAETLY | 0.48470104 |
| GNGGGGDAGNAGSGGN | ASSQKTGVGAETLY | 0.6249214 |
| GTGGAGGAGNDAGS | ASSQKTGVGAETLY | 0.38328308 |
| GTTNTGWFNS | ASSQKTGVGAETLY | 0.594381 |
| IFMGPLFTL | ASSQKTGVGAETLY | 0.6645943 |
| ILAHAIEAI | ASSQKTGVGAETLY | 0.62644297 |
| IRSAIREAL | ASSQKTGVGAETLY | 0.3833807 |
| ITFAGGG | ASSQKTGVGAETLY | 0.57082 |
| ITGMSGHQEV | ASSQKTGVGAETLY | 0.58328557 |
| IVAGLACTL | ASSQKTGVGAETLY | 0.44320142 |
| IVGGGLAGA | ASSQKTGVGAETLY | 0.6552563 |
| IVTGVLNI | ASSQKTGVGAETLY | 0.4479016 |
| LAQQGCI | ASSQKTGVGAETLY | 0.6117982 |
| LCLHVEN | ASSQKTGVGAETLY | 0.62616825 |

| | | |
|---|---|---|
| LDAISDGS | ASSQKTGVGAETLY | 0.66245866 |
| LGGFAAML | ASSQKTGVGAETLY | 0.35526982 |
| LGGKNPMI | ASSQKTGVGAETLY | 0.4828041 |
| LGPKGVAAIL | ASSQKTGVGAETLY | 0.52082187 |
| LMVGVSI | ASSQKTGVGAETLY | 0.61832416 |
| LVGAAFVL | ASSQKTGVGAETLY | 0.46558517 |
| LVLAVAPL | ASSQKTGVGAETLY | 0.33490568 |
| MAAIAVIDAA | ASSQKTGVGAETLY | 0.5339107 |
| NNGINAELQ | ASSQKTGVGAETLY | 0.6306238 |
| RGEPLVLVHG | ASSQKTGVGAETLY | 0.46922338 |
| RGGAGGAAGLI | ASSQKTGVGAETLY | 0.47068104 |
| RSYRMHPAV | ASSQKTGVGAETLY | 0.6671324 |
| SAAHLLAGQLA | ASSQKTGVGAETLY | 0.43488815 |
| SAAVGSY | ASSQKTGVGAETLY | 0.5390178 |
| SPHAVLHRI | ASSQKTGVGAETLY | 0.5834185 |
| SQIAKVL | ASSQKTGVGAETLY | 0.62481344 |
| TALGHLVP | ASSQKTGVGAETLY | 0.47689283 |
| TAPTTSGAA | ASSQKTGVGAETLY | 0.6226285 |
| TDGDDGNGGSPGT | ASSQKTGVGAETLY | 0.34606314 |
| TDGLGEN | ASSQKTGVGAETLY | 0.6310625 |
| TGGPPLPGQA | ASSQKTGVGAETLY | 0.36383313 |
| TIPAITI | ASSQKTGVGAETLY | 0.6035012 |
| TIPGFTI | ASSQKTGVGAETLY | 0.40830207 |
| TSIASGALA | ASSQKTGVGAETLY | 0.6436513 |
| TVTLPGAAI | ASSQKTGVGAETLY | 0.6257122 |
| VASGLSAALP | ASSQKTGVGAETLY | 0.65233374 |
| VGAGVIGI | ASSQKTGVGAETLY | 0.29677722 |
| VGPKRFGV | ASSQKTGVGAETLY | 0.64197123 |
| VNVAFPD | ASSQKTGVGAETLY | 0.65380585 |
| VSMTNSGVS | ASSQKTGVGAETLY | 0.67065454 |
| VTDVLGDLSV | ASSQKTGVGAETLY | 0.495911 |
| VVVVGAGFAGL | ASSQKTGVGAETLY | 0.5591222 |
| VYTVGTVV | ASSQKTGVGAETLY | 0.4594864 |
| WTTFSGP | ASSQKTGVGAETLY | 0.4702855 |
| YDELVEAF | ASSQKTGVGAETLY | 0.39345068 |
| YGTGGAGGNGG | ASSQKTGVGAETLY | 0.45849147 |
| YTNVMAV | ASSQKTGVGAETLY | 0.5143672 |
| FLEQVHQL | ASSQKTGVGAETLY | 0.5363405 |
| GLSLLAVAV | ASSQKTGVGAETLY | 0.63613594 |
| LEQVHQL | ASSQKTGVGAETLY | 0.6332605 |
| LNAAGAI | ASSQKTGVGAETLY | 0.62548405 |
| PAVHASL | ASSQKTGVGAETLY | 0.65305525 |
| TELEDDA | ASSQKTGVGAETLY | 0.64279664 |
| TPPGPPN | ASSQKTGVGAETLY | 0.5730217 |
| TPPPPPPP | ASSQKTGVGAETLY | 0.49449176 |
| TQSPALI | ASSFSYNSPLY | 0.4216374 |
| EFAAAFPQ | ASSFSYNSPLY | 0.41416386 |
| LLSGNLDVL | ASSFSYNSPLY | 0.45370555 |
| GPANLGNYNI | ASSFSYNSPLY | 0.46731323 |
| TLGPTLP | ASSFSYNSPLY | 0.6056886 |
| VYAYSSCCK | ASSFSYNSPLY | 0.28712153 |

| | | |
|---|---|---|
| APAPAPAPNI | ASSFSYNSPLY | 0.5428499 |
| PGGPVPIP | ASSFSYNSPLY | 0.66153026 |
| FLTVALEP | ASSFSYNSPLY | 0.39811525 |
| SQYFGANL | ASSFSYNSPLY | 0.56831217 |
| GGVKPAA | ASSFSYNSPLY | 0.5247145 |
| GGTGGAGGAGGW | ASSFSYNSPLY | 0.41187987 |
| GVGGVAGLGGAGG | ASSFSYNSPLY | 0.63619363 |
| VLLVLATTI | ASSFSYNSPLY | 0.39478013 |
| IGIFAAML | ASSFSYNSPLY | 0.5754678 |
| AVGDLID | ASSFSYNSPLY | 0.6622879 |
| NISPLPL | ASSFSYNSPLY | 0.5605012 |
| VTAVFVAI | ASSFSYNSPLY | 0.43819845 |
| SGDDNDAAL | ASSFSYNSPLY | 0.48674095 |
| AVAALAHS | ASSFSYNSPLY | 0.35006395 |
| GGVTVMATP | ASSFSYNSPLY | 0.5376916 |
| AGLAANVRD | ASSFSYNSPLY | 0.29176557 |
| TNRPDLIDPALLRPG | ASSFSYNSPLY | 0.57459825 |
| LANLGVTAL | ASSFSYNSPLY | 0.3939277 |
| TGERIEL | ASSFSYNSPLY | 0.618312 |
| NGEELAA | ASSFSYNSPLY | 0.5946427 |
| VLSLVWG | ASSFSYNSPLY | 0.52206576 |
| IGYALWL | ASSFSYNSPLY | 0.4710837 |
| LSLLLAPL | ASSFSYNSPLY | 0.46210974 |
| TLDEALK | ASSFSYNSPLY | 0.36617735 |
| TDGAGTAEGG | ASSFSYNSPLY | 0.37475386 |
| NAPGGAGGNGG | ASSFSYNSPLY | 0.6017181 |
| VSVFMGMGGNF | ASSFSYNSPLY | 0.43309194 |
| LASYLAAL | ASSFSYNSPLY | 0.4298765 |
| LVGGTPAL | ASSFSYNSPLY | 0.6512984 |
| VRNADVL | ASSFSYNSPLY | 0.5348535 |
| TCSSPFLVSAM | ASSFSYNSPLY | 0.3412699 |
| VHITDAE | ASSFSYNSPLY | 0.5536941 |
| EYAAPSF | ASSFSYNSPLY | 0.54165256 |
| VVDAFHGGL | ASSFSYNSPLY | 0.5610811 |
| ASVVDPQ | ASSFSYNSPLY | 0.28464362 |
| SYNEFGGLS | ASSFSYNSPLY | 0.47492594 |
| DPDGSVY | ASSFSYNSPLY | 0.4395771 |
| MVSTIHATE | ASSFSYNSPLY | 0.422401 |
| LAAGVADPV | ASSFSYNSPLY | 0.45632187 |
| TKEQIAA | ASSFSYNSPLY | 0.64910674 |
| SLASPAAL | ASSFSYNSPLY | 0.3695908 |
| IGLVFYGV | ASSFSYNSPLY | 0.54711473 |
| FLLDASNL | ASSFSYNSPLY | 0.6008382 |
| YQVGNFP | ASSFSYNSPLY | 0.591779 |
| ESLLSQST | ASSFSYNSPLY | 0.66449285 |
| LFTVGFAAVG | ASSFSYNSPLY | 0.5273421 |
| ATPPTPT | ASSFSYNSPLY | 0.43818787 |
| TDLATLG | ASSFSYNSPLY | 0.5253699 |
| SAGGTGR | ASSFSYNSPLY | 0.42618424 |
| GFSNSGS | ASSFSYNSPLY | 0.3140642 |
| GGNGGSGGTSV | ASSFSYNSPLY | 0.3005481 |

| | | |
|---|---|---|
| GTSVATGGAGN | ASSFSYNSPLY | 0.59714866 |
| NGGNGGSGGTS | ASSFSYNSPLY | 0.30962762 |
| AVVKFPH | ASSFSYNSPLY | 0.59425133 |
| TPPPPPP | ASSFSYNSPLY | 0.5866742 |
| AAPAAAAAPA | ASSFSYNSPLY | 0.3841479 |
| IALNEDL | ASSFSYNSPLY | 0.34880754 |
| IVNVSSM | ASSFSYNSPLY | 0.44187808 |
| TGIPPAPR | ASSFSYNSPLY | 0.44088015 |
| FDLGGGTF | ASSFSYNSPLY | 0.62414944 |
| FDLGGGTFDVS | ASSFSYNSPLY | 0.5244014 |
| NEPTAAA | ASSFSYNSPLY | 0.66287744 |
| NEPTAAALA | ASSFSYNSPLY | 0.6539964 |
| GSIINISSI | ASSFSYNSPLY | 0.5488894 |
| VAPGNAGTA | ASSFSYNSPLY | 0.622397 |
| TGGGASLE | ASSFSYNSPLY | 0.4721239 |
| VAVYNHYKRI | ASSFSYNSPLY | 0.41669324 |
| TDPAPATT | ASSFSYNSPLY | 0.4577843 |
| TNRPDILDPALLRPG | ASSFSYNSPLY | 0.4837833 |
| LTTTTTS | ASSFSYNSPLY | 0.46181354 |
| PTGSGKTL | ASSFSYNSPLY | 0.4520104 |
| TGSGKTL | ASSFSYNSPLY | 0.3571425 |
| TIGHVDHGK | ASSFSYNSPLY | 0.5214469 |
| TIGHVDHGKTT | ASSFSYNSPLY | 0.5279232 |
| GHVDHGKTT | ASSFSYNSPLY | 0.3158753 |
| AQFEHTLL | ASSFSYNSPLY | 0.46494523 |
| IVNQDEL | ASSFSYNSPLY | 0.6722752 |
| TNGSQFF | ASSFSYNSPLY | 0.6494086 |
| TGKGIVE | ASSFSYNSPLY | 0.5065665 |
| LISNASDA | ASSFSYNSPLY | 0.44418696 |
| AELRTLGELTKIAKAHGAQ | ASSFSYNSPLY | 0.5593197 |
| APNAVAAAEPSW | ASSFSYNSPLY | 0.33392286 |
| APQVAGAATAVL | ASSFSYNSPLY | 0.4991936 |
| ARRDPAAAAAILPTDA | ASSFSYNSPLY | 0.59888864 |
| DAAARDVLAEAGLAEYF | ASSFSYNSPLY | 0.52181435 |
| DDQIVLSAKANLAKVEQ | ASSFSYNSPLY | 0.39050424 |
| DELLPDTGEDSDEDGH | ASSFSYNSPLY | 0.55316997 |
| ELPEAPANRATVPQD | ASSFSYNSPLY | 0.48982653 |
| FGQSKIERVTGIG | ASSFSYNSPLY | 0.5224487 |
| GAGGTGSTAGGAGGAGGAGGLY | ASSFSYNSPLY | 0.28444338 |
| GHGGTGAAGGNGGTGGAGGSGI | ASSFSYNSPLY | 0.5208654 |
| GNGGNGGAGGEGGPGGAGGAGA | ASSFSYNSPLY | 0.41447055 |
| GSTAGNGGNGGNATNGTIAGQP | ASSFSYNSPLY | 0.39782184 |
| GTGGDGGNGGNGGTGG | ASSFSYNSPLY | 0.31861472 |
| GVRRPAGIPADAPV | ASSFSYNSPLY | 0.47368932 |
| GYRQVTDAHLLTLARR | ASSFSYNSPLY | 0.6326963 |
| IAKMGQDGHDRG | ASSFSYNSPLY | 0.6328137 |
| KAEPSIEIQYT | ASSFSYNSPLY | 0.32845268 |
| KPVRIRMAADEDTVAVT | ASSFSYNSPLY | 0.64745766 |
| MPPRIAGMRLLVIKPEP | ASSFSYNSPLY | 0.6592107 |
| MKFYAVALFRSRGVPPTR | ASSFSYNSPLY | 0.4713773 |
| NAGGFGILGAVAHSP | ASSFSYNSPLY | 0.33839038 |

| | | |
|---|---|---|
| NEGREITLAMAEM | ASSFSYNSPLY | 0.59021205 |
| NELVEWSHPLGHRWRWE | ASSFSYNSPLY | 0.45722282 |
| NLGAGGTGGNGGNPG | ASSFSYNSPLY | 0.58821744 |
| NTEWPKWETVMAADPVD | ASSFSYNSPLY | 0.5629661 |
| PASAAPAETQP | ASSFSYNSPLY | 0.42296958 |
| QQSSDGKNGVRAAEITGE | ASSFSYNSPLY | 0.57198304 |
| RAPDAVTTSWWKE | ASSFSYNSPLY | 0.6219281 |
| SGFGNVGAHTSGWFNQ | ASSFSYNSPLY | 0.38147384 |
| SGVAGAVGQAASVGGLKV | ASSFSYNSPLY | 0.3683449 |
| SNGGAGTGSAGGNGGT | ASSFSYNSPLY | 0.3871086 |
| TGGAGGEGGAAG | ASSFSYNSPLY | 0.29930004 |
| TGGDGGAAGAGGAPGHG | ASSFSYNSPLY | 0.5650112 |
| TMPPLLAAIANAGRHGV | ASSFSYNSPLY | 0.47679698 |
| TSSTAGNSGGA | ASSFSYNSPLY | 0.44780537 |
| VPATVPVGILC | ASSFSYNSPLY | 0.5955663 |
| VQRPDSGGVVLTGRIS | ASSFSYNSPLY | 0.43085903 |
| WTPNPVPATKGVPAGLVQAT | ASSFSYNSPLY | 0.49387276 |
| YALKRALAKFSDRFDVVII | ASSFSYNSPLY | 0.48396653 |
| YLIGSVIV | ASSFSYNSPLY | 0.5349326 |
| YGQLVAVID | ASSFSYNSPLY | 0.5593778 |
| VTVGGMTLPA | ASSFSYNSPLY | 0.48394355 |
| VLALPAV | ASSFSYNSPLY | 0.500936 |
| VIPPTLNYET | ASSFSYNSPLY | 0.66329175 |
| VGMGGMHPSSG | ASSFSYNSPLY | 0.3606003 |
| VGLAGGAATA | ASSFSYNSPLY | 0.43680012 |
| VGGAGGAGGSGP | ASSFSYNSPLY | 0.5730255 |
| VFAAFMV | ASSFSYNSPLY | 0.34159464 |
| TLYDKLR | ASSFSYNSPLY | 0.57791936 |
| TLAVFTVL | ASSFSYNSPLY | 0.5749291 |
| TGSVQPARV | ASSFSYNSPLY | 0.60280186 |
| TGGGAGSGGT | ASSFSYNSPLY | 0.41214725 |
| SHLETSL | ASSFSYNSPLY | 0.6661558 |
| SFTTGMT | ASSFSYNSPLY | 0.63900065 |
| QVEVAKRMGVRQPTVS | ASSFSYNSPLY | 0.46065086 |
| QGSGGPALAM | ASSFSYNSPLY | 0.49363843 |
| MLAPSNP | ASSFSYNSPLY | 0.36978105 |
| LVNNAAI | ASSFSYNSPLY | 0.6299369 |
| LTGEINTLA | ASSFSYNSPLY | 0.5884981 |
| LSIFGPV | ASSFSYNSPLY | 0.448981 |
| LLALLANL | ASSFSYNSPLY | 0.6020485 |
| LISNASDALDKLR | ASSFSYNSPLY | 0.6238534 |
| LINEAVP | ASSFSYNSPLY | 0.6570811 |
| LGYAFSGL | ASSFSYNSPLY | 0.32626995 |
| LDFVAVAL | ASSFSYNSPLY | 0.5348109 |
| LAQPGAE | ASSFSYNSPLY | 0.31204382 |
| ISNLGAVTGF | ASSFSYNSPLY | 0.38492194 |
| HQEDPGRI | ASSFSYNSPLY | 0.3845875 |
| GTPGEPTP | ASSFSYNSPLY | 0.5846552 |
| GFFNSGS | ASSFSYNSPLY | 0.52116525 |
| GAGGGGLTAGAN | ASSFSYNSPLY | 0.6175845 |
| FVDALVNM | ASSFSYNSPLY | 0.60712475 |

| | | |
|---|---|---|
| FLTVAPEL | ASSFSYNSPLY | 0.6666194 |
| FETLTAL | ASSFSYNSPLY | 0.44584942 |
| DVINPIEL | ASSFSYNSPLY | 0.5278852 |
| APVSGAPV | ASSFSYNSPLY | 0.33854878 |
| ALVGGLAW | ASSFSYNSPLY | 0.3340508 |
| ALAAAAATAI | ASSFSYNSPLY | 0.37353677 |
| AFGLGLLL | ASSFSYNSPLY | 0.50137043 |
| AFFAAASL | ASSFSYNSPLY | 0.29324707 |
| AAAALPAFSPP | ASSFSYNSPLY | 0.6266589 |
| VVLAPAL | ASSFSYNSPLY | 0.59264636 |
| TIGHVDHGKTTL | ASSFSYNSPLY | 0.6449858 |
| LISNASDALDK | ASSFSYNSPLY | 0.4059645 |
| LISNASDAL | ASSFSYNSPLY | 0.4505853 |
| ISNASDALDK | ASSFSYNSPLY | 0.47987768 |
| NQAETLVYQTE | ASSFSYNSPLY | 0.6313604 |
| TGGTGGTAGAG | ASSFSYNSPLY | 0.39927626 |
| TGSTAGGAGGA | ASSFSYNSPLY | 0.34108233 |
| DPLAVPVAAGPV | ASSFSYNSPLY | 0.3999805 |
| GGTGGTPGNGGN | ASSFSYNSPLY | 0.3524733 |
| PAGLAGVDPATL | ASSFSYNSPLY | 0.6413239 |
| GNNLVVEAPHAGASV | ASSFSYNSPLY | 0.63815343 |
| KVTRIMSASTLEQPA | ASSFSYNSPLY | 0.61997294 |
| LGFALPIALPRRTVV | ASSFSYNSPLY | 0.60050017 |
| TRLGNLTARKLVVLV | ASSFSYNSPLY | 0.3009946 |
| WSQQRRLIYFPSAGP | ASSFSYNSPLY | 0.40825194 |
| GPNGNPDPMAAAVDIR | ASSFSYNSPLY | 0.6681612 |
| TQRCVSVADLRAAASR | ASSFSYNSPLY | 0.5682893 |
| QFVLDVAEHPVEVLTETV | ASSFSYNSPLY | 0.5424974 |
| GGAGGTGGAGGNGGTQAGDG | ASSFSYNSPLY | 0.64767987 |
| PLLATLRAIAVDVDAKPAQV | ASSFSYNSPLY | 0.4997175 |
| APKFRTGGYMIDFWGVGYQVAK | ASSFSYNSPLY | 0.55727094 |
| GAGGAGGAGGAAGGPGGTGGTG | ASSFSYNSPLY | 0.5287149 |
| IGALGLLATDVAPAGSVAFGVA | ASSFSYNSPLY | 0.53254974 |
| AGGAGGIGGNGGAGGAGGVLMG | ASSFSYNSPLY | 0.470983 |
| ADELIGLL | ASSFSYNSPLY | 0.39791158 |
| ADRALKA | ASSFSYNSPLY | 0.5335804 |
| AGAVGQVGGAGGA | ASSFSYNSPLY | 0.67260265 |
| AGGGGGLGGAP | ASSFSYNSPLY | 0.5911678 |
| AGGLLYGNGGAG | ASSFSYNSPLY | 0.62586594 |
| AGLESIGL | ASSFSYNSPLY | 0.59307194 |
| AGLKTGII | ASSFSYNSPLY | 0.62239033 |
| ASLAALLL | ASSFSYNSPLY | 0.5036843 |
| ASVINISSI | ASSFSYNSPLY | 0.4777387 |
| ATPPGPA | ASSFSYNSPLY | 0.47088176 |
| ATVASALV | ASSFSYNSPLY | 0.39167106 |
| ATVASLIL | ASSFSYNSPLY | 0.47548804 |
| AVNGPVLIRH | ASSFSYNSPLY | 0.455927 |
| DGAAGAVGI | ASSFSYNSPLY | 0.6790585 |
| DPLLHGI | ASSFSYNSPLY | 0.5941582 |
| ELGAALGAL | ASSFSYNSPLY | 0.55118763 |
| ELGRTQAV | ASSFSYNSPLY | 0.4927929 |

| | | |
|---|---|---|
| FVAGGYLS | ASSFSYNSPLY | 0.49973777 |
| GASGGAGGAGGW | ASSFSYNSPLY | 0.5810022 |
| GGAGAGGDGGAGGI | ASSFSYNSPLY | 0.37238762 |
| GTGHNGSQPGMGGNG | ASSFSYNSPLY | 0.63003504 |
| GYAAASASAS | ASSFSYNSPLY | 0.46651018 |
| HQVDVAAVQQG | ASSFSYNSPLY | 0.42726496 |
| IGAAIAHT | ASSFSYNSPLY | 0.6136608 |
| IGMIEVAQL | ASSFSYNSPLY | 0.6662584 |
| IRLAPEHTSAMD | ASSFSYNSPLY | 0.45932916 |
| IYNVSVL | ASSFSYNSPLY | 0.41411427 |
| KQGFLGM | ASSFSYNSPLY | 0.473646 |
| LESVITQL | ASSFSYNSPLY | 0.56923246 |
| LGLGALVVV | ASSFSYNSPLY | 0.66329944 |
| LIGNGGNGGSGGI | ASSFSYNSPLY | 0.44737896 |
| LNAINAPTL | ASSFSYNSPLY | 0.6578145 |
| LQLLNTGV | ASSFSYNSPLY | 0.46166426 |
| LQTSIEF | ASSFSYNSPLY | 0.34787297 |
| MSFAVSEL | ASSFSYNSPLY | 0.5017771 |
| NTGALGL | ASSFSYNSPLY | 0.6173151 |
| PPGAGGAGGAGTT | ASSFSYNSPLY | 0.6255654 |
| QLVGLGI | ASSFSYNSPLY | 0.6692009 |
| RFLEQVHQL | ASSFSYNSPLY | 0.49682456 |
| RGVGYVLR | ASSFSYNSPLY | 0.48075494 |
| SATGVFAGVI | ASSFSYNSPLY | 0.65677035 |
| SLPAITLP | ASSFSYNSPLY | 0.6448473 |
| STALAFTV | ASSFSYNSPLY | 0.63415956 |
| SVLTVGLGK | ASSFSYNSPLY | 0.45531452 |
| TALAGAGSGI | ASSFSYNSPLY | 0.38703358 |
| TGLPFTGI | ASSFSYNSPLY | 0.51472783 |
| VAEHVTV | ASSFSYNSPLY | 0.63041514 |
| VDIVLNSL | ASSFSYNSPLY | 0.38206023 |
| VLTGGGAVL | ASSFSYNSPLY | 0.5631693 |
| VNVGLAYGI | ASSFSYNSPLY | 0.6117042 |
| VSSAAVGAV | ASSFSYNSPLY | 0.34640822 |
| VTSVYEAGV | ASSFSYNSPLY | 0.49939838 |
| WVALDRGA | ASSFSYNSPLY | 0.6452462 |
| YLVGSVVI | ASSFSYNSPLY | 0.46239403 |
| GPPGTGKTLL | ASSFSYNSPLY | 0.42723405 |
| GVGMTKF | ASSFSYNSPLY | 0.6234896 |
| IGIGGSDL | ASSFSYNSPLY | 0.67988175 |
| IGTLLAEL | ASSFSYNSPLY | 0.63278943 |
| IYTSGTT | ASSFSYNSPLY | 0.3603626 |
| LSFPAGV | ASSFSYNSPLY | 0.30601254 |
| LYTSGTT | ASSFSYNSPLY | 0.45978034 |
| VGAFPFGF | ASSFSYNSPLY | 0.62268966 |
| VGAGTVEF | ASSFSYNSPLY | 0.47731227 |
| VGKSTLLL | ASSFSYNSPLY | 0.45970434 |
| AALAHAGGGIV | ASSFSYNSPLY | 0.58311987 |
| AALVAVSLL | ASSFSYNSPLY | 0.65178204 |
| AAVSRAVAAL | ASSFSYNSPLY | 0.41867784 |
| AGGSSGIGGF | ASSFSYNSPLY | 0.35748544 |

| | | |
|---|---|---|
| AGLTVANANG | ASSFSYNSPLY | 0.54090506 |
| AGLVGAVL | ASSFSYNSPLY | 0.6076712 |
| ALSQAFTV | ASSFSYNSPLY | 0.47798592 |
| ALTFSVSL | ASSFSYNSPLY | 0.5856974 |
| ALTGAALVAL | ASSFSYNSPLY | 0.42521155 |
| ASFSIGAGVL | ASSFSYNSPLY | 0.35295442 |
| DGSVYKI | ASSFSYNSPLY | 0.59858465 |
| EQLAEIV | ASSFSYNSPLY | 0.4525866 |
| FAVSELTVM | ASSFSYNSPLY | 0.6445441 |
| GAAPLAAGL | ASSFSYNSPLY | 0.46638927 |
| GADGGDGGSGGAGG | ASSFSYNSPLY | 0.51204824 |
| GFGLDTGGAGG | ASSFSYNSPLY | 0.62021554 |
| GLASGGLL | ASSFSYNSPLY | 0.56394875 |
| GRIAGAAVLL | ASSFSYNSPLY | 0.52920824 |
| GSAGSLNTSF | ASSFSYNSPLY | 0.4546401 |
| GTGQIGI | ASSFSYNSPLY | 0.3797738 |
| GTLIALIL | ASSFSYNSPLY | 0.30153677 |
| IAATMGFVVG | ASSFSYNSPLY | 0.6392597 |
| IFAQALL | ASSFSYNSPLY | 0.30735683 |
| IGAAVALGF | ASSFSYNSPLY | 0.64918476 |
| IGLTPLAL | ASSFSYNSPLY | 0.63828194 |
| ILGSNTSTL | ASSFSYNSPLY | 0.43121207 |
| IVFSLTVL | ASSFSYNSPLY | 0.55805784 |
| LASKATGF | ASSFSYNSPLY | 0.38161463 |
| LASVALIL | ASSFSYNSPLY | 0.36226046 |
| LFAVGLTL | ASSFSYNSPLY | 0.59193176 |
| LFNGGIGTYI | ASSFSYNSPLY | 0.59078443 |
| LTLGAGII | ASSFSYNSPLY | 0.43101823 |
| LVAAALGDI | ASSFSYNSPLY | 0.50062954 |
| NTAFVQLGI | ASSFSYNSPLY | 0.5798424 |
| QLFSGAEL | ASSFSYNSPLY | 0.37866834 |
| SAATFITL | ASSFSYNSPLY | 0.47802797 |
| SNYGLANLGV | ASSFSYNSPLY | 0.6552216 |
| SYGGAYAVM | ASSFSYNSPLY | 0.50578034 |
| TTGVGGLTLG | ASSFSYNSPLY | 0.44714588 |
| TTQTIYI | ASSFSYNSPLY | 0.5821606 |
| VAAGAGLGAVI | ASSFSYNSPLY | 0.40021658 |
| VAAVVTLL | ASSFSYNSPLY | 0.62873644 |
| VGFLGGLI | ASSFSYNSPLY | 0.47484303 |
| VGVIPILL | ASSFSYNSPLY | 0.47292095 |
| VIFASSLI | ASSFSYNSPLY | 0.58304244 |
| VKGGDGGAAAG | ASSFSYNSPLY | 0.34861505 |
| VTKTFTNI | ASSFSYNSPLY | 0.39653292 |
| VVAAAILL | ASSFSYNSPLY | 0.47196496 |
| YAALDVEL | ASSFSYNSPLY | 0.5820855 |
| LASVALLL | ASSFSYNSPLY | 0.625425 |
| LDAVHRV | ASSFSYNSPLY | 0.4893815 |
| QGLSSSIF | ASSFSYNSPLY | 0.5530287 |
| RGTFANIRL | ASSFSYNSPLY | 0.4683941 |
| RQLWWGHRI | ASSFSYNSPLY | 0.4585188 |
| YGPPGTGKTLL | ASSFSYNSPLY | 0.47057575 |

| | | |
|---|---|---|
| AALVGAPANVA | ASSFSYNSPLY | 0.42119315 |
| AATFRRTE | ASSFSYNSPLY | 0.3676722 |
| AGALGGGI | ASSFSYNSPLY | 0.5557599 |
| AGGAGGAGGIGAG | ASSFSYNSPLY | 0.39603794 |
| AGSGALGL | ASSFSYNSPLY | 0.49551573 |
| AQIADQI | ASSFSYNSPLY | 0.5852008 |
| ASAVAVTYL | ASSFSYNSPLY | 0.43424454 |
| ASTVLGGTGG | ASSFSYNSPLY | 0.53754675 |
| AVGGEPAK | ASSFSYNSPLY | 0.58288485 |
| CEQGTFGGTC | ASSFSYNSPLY | 0.5909345 |
| DEGKAPAG | ASSFSYNSPLY | 0.5089386 |
| DTVVFPAL | ASSFSYNSPLY | 0.4360274 |
| FEGITMAG | ASSFSYNSPLY | 0.6479306 |
| FPLVVQD | ASSFSYNSPLY | 0.6126987 |
| GAFTTPQ | ASSFSYNSPLY | 0.56051075 |
| GFGGFGTAGGVGGA | ASSFSYNSPLY | 0.56331193 |
| GFLAQFL | ASSFSYNSPLY | 0.6037939 |
| GGAGGAGGNAGVA | ASSFSYNSPLY | 0.6161773 |
| GGVGTAGAGGAGGAGG | ASSFSYNSPLY | 0.54211754 |
| GNGGGGDAGNAGSGGN | ASSFSYNSPLY | 0.34239304 |
| GTGGAGGAGNDAGS | ASSFSYNSPLY | 0.51283664 |
| GTTNTGWFNS | ASSFSYNSPLY | 0.52106434 |
| IFMGPLFTL | ASSFSYNSPLY | 0.42853868 |
| ILAHAIEAI | ASSFSYNSPLY | 0.3395818 |
| IRSAIREAL | ASSFSYNSPLY | 0.6593535 |
| ITFAGGG | ASSFSYNSPLY | 0.6791706 |
| ITGMSGHQEV | ASSFSYNSPLY | 0.39557606 |
| IVAGLACTL | ASSFSYNSPLY | 0.65692335 |
| IVGGGLAGA | ASSFSYNSPLY | 0.34780127 |
| IVTGVLNI | ASSFSYNSPLY | 0.596079 |
| LAQQGCI | ASSFSYNSPLY | 0.6526243 |
| LCLHVEN | ASSFSYNSPLY | 0.45790598 |
| LDAISDGS | ASSFSYNSPLY | 0.6249363 |
| LGGFAAML | ASSFSYNSPLY | 0.3960774 |
| LGGKNPMI | ASSFSYNSPLY | 0.45474547 |
| LGPKGVAAIL | ASSFSYNSPLY | 0.6574547 |
| LMVGVSI | ASSFSYNSPLY | 0.46739635 |
| LVGAAFVL | ASSFSYNSPLY | 0.50699186 |
| LVLAVAPL | ASSFSYNSPLY | 0.54792595 |
| MAAIAVIDAA | ASSFSYNSPLY | 0.4450289 |
| NNGINAELQ | ASSFSYNSPLY | 0.53174233 |
| RGEPLVLVHG | ASSFSYNSPLY | 0.63411754 |
| RGGAGGAAGLI | ASSFSYNSPLY | 0.35554937 |
| RSYRMHPAV | ASSFSYNSPLY | 0.6408931 |
| SAAHLLAGQLA | ASSFSYNSPLY | 0.40234542 |
| SAAVGSY | ASSFSYNSPLY | 0.67368245 |
| SPHAVLHRI | ASSFSYNSPLY | 0.5998664 |
| SQIAKVL | ASSFSYNSPLY | 0.61413974 |
| TALGHLVP | ASSFSYNSPLY | 0.43562582 |
| TAPTTSGAA | ASSFSYNSPLY | 0.6610392 |
| TDGDDGNGGSPGT | ASSFSYNSPLY | 0.5756347 |

| | | |
|---|---|---|
| TDGLGEN | ASSFSYNSPLY | 0.52202594 |
| TGGPPLPGQA | ASSFSYNSPLY | 0.62160337 |
| TIPAITI | ASSFSYNSPLY | 0.12253698 |
| TIPGFTI | ASSFSYNSPLY | 0.51302254 |
| TSIASGALA | ASSFSYNSPLY | 0.3773335 |
| TVTLPGAAI | ASSFSYNSPLY | 0.28737092 |
| VASGLSAALP | ASSFSYNSPLY | 0.16126953 |
| VGAGVIGI | ASSFSYNSPLY | 0.13118011 |
| VGPKRFGV | ASSFSYNSPLY | 0.1695633 |
| VNVAFPD | ASSFSYNSPLY | 0.13546486 |
| VSMTNSGVS | ASSFSYNSPLY | 0.74474335 |
| VTDVLGDLSV | ASSFSYNSPLY | 0.29387784 |
| VVVVGAGFAGL | ASSFSYNSPLY | 0.16491103 |
| VYTVGTVV | ASSFSYNSPLY | 0.27061778 |
| WTTFSGP | ASSFSYNSPLY | 0.36274314 |
| YDELVEAF | ASSFSYNSPLY | 0.1198569 |
| YGTGGAGGNGG | ASSFSYNSPLY | 0.25370985 |
| YTNVMAV | ASSFSYNSPLY | 0.18953252 |
| FLEQVHQL | ASSFSYNSPLY | 0.17787476 |
| GLSLLAVAV | ASSFSYNSPLY | 0.107636176 |
| LEQVHQL | ASSFSYNSPLY | 0.3572241 |
| LNAAGAI | ASSFSYNSPLY | 0.1825085 |
| PAVHASL | ASSFSYNSPLY | 0.10397892 |
| TELEDDA | ASSFSYNSPLY | 0.2540617 |
| TPPGPPN | ASSFSYNSPLY | 0.15716843 |
| TPPPPPPP | ASSFSYNSPLY | 0.16056152 |
| TQSPALI | ASSATGGASQNTLY | 0.27396396 |
| EFAAAFPQ | ASSATGGASQNTLY | 0.27980858 |
| LLSGNLDVL | ASSATGGASQNTLY | 0.16078416 |
| GPANLGNYNI | ASSATGGASQNTLY | 0.3695811 |
| TLGPTLP | ASSATGGASQNTLY | 0.42024928 |
| VYAYSSCCK | ASSATGGASQNTLY | 0.14374457 |
| APAPAPAPNI | ASSATGGASQNTLY | 0.22706904 |
| PGGPVPIP | ASSATGGASQNTLY | 0.66731066 |
| FLTVALEP | ASSATGGASQNTLY | 0.2082885 |
| SQYFGANL | ASSATGGASQNTLY | 0.53885365 |
| GGVKPAA | ASSATGGASQNTLY | 0.5722527 |
| GGTGGAGGAGGW | ASSATGGASQNTLY | 0.19244899 |
| GVGGVAGLGGAGG | ASSATGGASQNTLY | 0.40958112 |
| VLLVLATTI | ASSATGGASQNTLY | 0.4320331 |
| IGIFAAML | ASSATGGASQNTLY | 0.22986218 |
| AVGDLID | ASSATGGASQNTLY | 0.2054656 |
| NISPLPL | ASSATGGASQNTLY | 0.12616551 |
| VTAVFVAI | ASSATGGASQNTLY | 0.24447823 |
| SGDDNDAAL | ASSATGGASQNTLY | 0.14534703 |
| AVAALAHS | ASSATGGASQNTLY | 0.35842282 |
| GGVTVMATP | ASSATGGASQNTLY | 0.5825019 |
| AGLAANVRD | ASSATGGASQNTLY | 0.19287816 |
| TNRPDLIDPALLRPG | ASSATGGASQNTLY | 0.1144369 |
| LANLGVTAL | ASSATGGASQNTLY | 0.12226441 |
| TGERIEL | ASSATGGASQNTLY | 0.32684922 |

| | | |
|---|---|---|
| NGEELAA | ASSATGGASQNTLY | 0.12487801 |
| VLSLVWG | ASSATGGASQNTLY | 0.13994369 |
| IGYALWL | ASSATGGASQNTLY | 0.14906284 |
| LSLLLAPL | ASSATGGASQNTLY | 0.37033075 |
| TLDEALK | ASSATGGASQNTLY | 0.40976125 |
| TDGAGTAEGG | ASSATGGASQNTLY | 0.14431219 |
| NAPGGAGGNGG | ASSATGGASQNTLY | 0.5634254 |
| VSVFMGMGGNF | ASSATGGASQNTLY | 0.7532216 |
| LASYLAAL | ASSATGGASQNTLY | 0.7578147 |
| LVGGTPAL | ASSATGGASQNTLY | 0.28189027 |
| VRNADVL | ASSATGGASQNTLY | 0.124376245 |
| TCSSPFLVSAM | ASSATGGASQNTLY | 0.6703124 |
| VHITDAE | ASSATGGASQNTLY | 0.1583733 |
| EYAAPSF | ASSATGGASQNTLY | 0.36831516 |
| VVDAFHGGL | ASSATGGASQNTLY | 0.32217082 |
| ASVVDPQ | ASSATGGASQNTLY | 0.19317095 |
| SYNEFGGLS | ASSATGGASQNTLY | 0.14184083 |
| DPDGSVY | ASSATGGASQNTLY | 0.6523339 |
| MVSTIHATE | ASSATGGASQNTLY | 0.5810022 |
| LAAGVADPV | ASSATGGASQNTLY | 0.13310385 |
| TKEQIAA | ASSATGGASQNTLY | 0.1427693 |
| SLASPAAL | ASSATGGASQNTLY | 0.22569561 |
| IGLVFYGV | ASSATGGASQNTLY | 0.19727275 |
| FLLDASNL | ASSATGGASQNTLY | 0.37676382 |
| YQVGNFP | ASSATGGASQNTLY | 0.30472815 |
| ESLLSQST | ASSATGGASQNTLY | 0.35517052 |
| LFTVGFAAVG | ASSATGGASQNTLY | 0.2849189 |
| ATPPTPT | ASSATGGASQNTLY | 0.3898207 |
| TDLATLG | ASSATGGASQNTLY | 0.23757926 |
| SAGGTGR | ASSATGGASQNTLY | 0.26237446 |
| GFSNSGS | ASSATGGASQNTLY | 0.13889107 |
| GGNGGSGGTSV | ASSATGGASQNTLY | 0.5372514 |
| GTSVATGGAGN | ASSATGGASQNTLY | 0.45271584 |
| NGGNGGSGGTS | ASSATGGASQNTLY | 0.36601835 |
| AVVKFPH | ASSATGGASQNTLY | 0.57449675 |
| TPPPPPP | ASSATGGASQNTLY | 0.30597305 |
| AAPAAAAAPA | ASSATGGASQNTLY | 0.33945736 |
| IALNEDL | ASSATGGASQNTLY | 0.12772292 |
| IVNVSSM | ASSATGGASQNTLY | 0.19999026 |
| TGIPPAPR | ASSATGGASQNTLY | 0.1523692 |
| FDLGGGTF | ASSATGGASQNTLY | 0.3499211 |
| FDLGGGTFDVS | ASSATGGASQNTLY | 0.27579862 |
| NEPTAAA | ASSATGGASQNTLY | 0.44521576 |
| NEPTAAALA | ASSATGGASQNTLY | 0.17477548 |
| GSIINISSI | ASSATGGASQNTLY | 0.43455324 |
| VAPGNAGTA | ASSATGGASQNTLY | 0.6273333 |
| TGGGASLE | ASSATGGASQNTLY | 0.13409351 |
| VAVYNHYKRI | ASSATGGASQNTLY | 0.16311349 |
| TDPAPATT | ASSATGGASQNTLY | 0.47824913 |
| TNRPDILDPALLRPG | ASSATGGASQNTLY | 0.3501901 |
| LTTTTTS | ASSATGGASQNTLY | 0.1697011 |

| | | |
|---|---|---|
| PTGSGKTL | ASSATGGASQNTLY | 0.20254004 |
| TGSGKTL | ASSATGGASQNTLY | 0.7933758 |
| TIGHVDHGK | ASSATGGASQNTLY | 0.21922919 |
| TIGHVDHGKTT | ASSATGGASQNTLY | 0.6274669 |
| GHVDHGKTT | ASSATGGASQNTLY | 0.117941864 |
| AQFEHTLL | ASSATGGASQNTLY | 0.24484332 |
| IVNQDEL | ASSATGGASQNTLY | 0.23260263 |
| TNGSQFF | ASSATGGASQNTLY | 0.10143521 |
| TGKGIVE | ASSATGGASQNTLY | 0.36297193 |
| LISNASDA | ASSATGGASQNTLY | 0.15152186 |
| AELRTLGELTKIAKAHGAQ | ASSATGGASQNTLY | 0.2753948 |
| APNAVAAAEPSW | ASSATGGASQNTLY | 0.16275325 |
| APQVAGAATAVL | ASSATGGASQNTLY | 0.38169092 |
| ARRDPAAAAAILPTDA | ASSATGGASQNTLY | 0.39392632 |
| DAAARDVLAEAGLAEYF | ASSATGGASQNTLY | 0.19220486 |
| DDQIVLSAKANLAKVEQ | ASSATGGASQNTLY | 0.15531974 |
| DELLPDTGEDSDEDGH | ASSATGGASQNTLY | 0.22287387 |
| ELPEAPANRATVPQD | ASSATGGASQNTLY | 0.39443287 |
| FGQSKIERVTGIG | ASSATGGASQNTLY | 0.5380929 |
| GAGGTGSTAGGAGGAGGAGGLY | ASSATGGASQNTLY | 0.1218595 |
| GHGGTGAAGGNGGTGGAGGSGI | ASSATGGASQNTLY | 0.30358493 |
| GNGGNGGAGGEGGPGGAGGAGA | ASSATGGASQNTLY | 0.18750755 |
| GSTAGNGGNGGNATNGTIAGQP | ASSATGGASQNTLY | 0.11948622 |
| GTGGDGGNGGNGGTGG | ASSATGGASQNTLY | 0.10279157 |
| GVRRPAGIPADAPV | ASSATGGASQNTLY | 0.22026029 |
| GYRQVTDAHLLTLARR | ASSATGGASQNTLY | 0.1805973 |
| IAKMGQDGHDRG | ASSATGGASQNTLY | 0.2243338 |
| KAEPSIEIQYT | ASSATGGASQNTLY | 0.42603436 |
| KPVRIRMAADEDTVAVT | ASSATGGASQNTLY | 0.1482814 |
| MPPRIAGMRLLVIKPEP | ASSATGGASQNTLY | 0.46006706 |
| MKFYAVALFRSRGVPPTR | ASSATGGASQNTLY | 0.22180805 |
| NAGGFGILGAVAHSP | ASSATGGASQNTLY | 0.092489816 |
| NEGREITLAMAEM | ASSATGGASQNTLY | 0.18040983 |
| NELVEWSHPLGHRWRWE | ASSATGGASQNTLY | 0.11234609 |
| NLGAGGTGGNGGNPG | ASSATGGASQNTLY | 0.5470294 |
| NTEWPKWETVMAADPVD | ASSATGGASQNTLY | 0.12441352 |
| PASAAPAETQP | ASSATGGASQNTLY | 0.17080069 |
| QQSSDGKNGVRAAEITGE | ASSATGGASQNTLY | 0.1752219 |
| RAPDAVTTSWWKE | ASSATGGASQNTLY | 0.7292682 |
| SGFGNVGAHTSGWFNQ | ASSATGGASQNTLY | 0.29671818 |
| SGVAGAVGQAASVGGLKV | ASSATGGASQNTLY | 0.19258186 |
| SNGGAGTGSAGGNGGT | ASSATGGASQNTLY | 0.11615661 |
| TGGAGGEGGAAG | ASSATGGASQNTLY | 0.24791437 |
| TGGDGGAAGAGGAPGHG | ASSATGGASQNTLY | 0.3693666 |
| TMPPLLAAIANAGRHGV | ASSATGGASQNTLY | 0.15586501 |
| TSSTAGNSGGA | ASSATGGASQNTLY | 0.13088298 |
| VPATVPVGILC | ASSATGGASQNTLY | 0.23965274 |
| VQRPDSGGVVLTGRIS | ASSATGGASQNTLY | 0.13035251 |
| WTPNPVPATKGVPAGLVQAT | ASSATGGASQNTLY | 0.44933867 |
| YALKRALAKFSDRFDVVII | ASSATGGASQNTLY | 0.1356466 |
| YLIGSVIV | ASSATGGASQNTLY | 0.1522856 |

| | | |
|---|---|---|
| YGQLVAVID | ASSATGGASQNTLY | 0.65642655 |
| VTVGGMTLPA | ASSATGGASQNTLY | 0.6059317 |
| VLALPAV | ASSATGGASQNTLY | 0.18061364 |
| VIPPTLNYET | ASSATGGASQNTLY | 0.17602693 |
| VGMGGMHPSSG | ASSATGGASQNTLY | 0.13745712 |
| VGLAGGAATA | ASSATGGASQNTLY | 0.1449925 |
| VGGAGGAGGSGP | ASSATGGASQNTLY | 0.1670702 |
| VFAAFMV | ASSATGGASQNTLY | 0.30307215 |
| TLYDKLR | ASSATGGASQNTLY | 0.26100844 |
| TLAVFTVL | ASSATGGASQNTLY | 0.1304993 |
| TGSVQPARV | ASSATGGASQNTLY | 0.6568538 |
| TGGGAGSGGT | ASSATGGASQNTLY | 0.62324595 |
| SHLETSL | ASSATGGASQNTLY | 0.70129144 |
| SFTTGMT | ASSATGGASQNTLY | 0.1277191 |
| QVEVAKRMGVRQPTVS | ASSATGGASQNTLY | 0.2523979 |
| QGSGGPALAM | ASSATGGASQNTLY | 0.19468506 |
| MLAPSNP | ASSATGGASQNTLY | 0.18501481 |
| LVNNAAI | ASSATGGASQNTLY | 0.15973277 |
| LTGEINTLA | ASSATGGASQNTLY | 0.1520585 |
| LSIFGPV | ASSATGGASQNTLY | 0.57303154 |
| LLALLANL | ASSATGGASQNTLY | 0.84170556 |
| LISNASDALDKLR | ASSATGGASQNTLY | 0.20414156 |
| LINEAVP | ASSATGGASQNTLY | 0.110544994 |
| LGYAFSGL | ASSATGGASQNTLY | 0.17103207 |
| LDFVAVAL | ASSATGGASQNTLY | 0.15225616 |
| LAQPGAE | ASSATGGASQNTLY | 0.1336461 |
| ISNLGAVTGF | ASSATGGASQNTLY | 0.19795221 |
| HQEDPGRI | ASSATGGASQNTLY | 0.11301273 |
| GTPGEPTP | ASSATGGASQNTLY | 0.20379214 |
| GFFNSGS | ASSATGGASQNTLY | 0.6789501 |
| GAGGGGLTAGAN | ASSATGGASQNTLY | 0.23620437 |
| FVDALVNM | ASSATGGASQNTLY | 0.26941168 |
| FLTVAPEL | ASSATGGASQNTLY | 0.56791836 |
| FETLTAL | ASSATGGASQNTLY | 0.3291477 |
| DVINPIEL | ASSATGGASQNTLY | 0.4600861 |
| APVSGAPV | ASSATGGASQNTLY | 0.7232801 |
| ALVGGLAW | ASSATGGASQNTLY | 0.22330754 |
| ALAAAAATAI | ASSATGGASQNTLY | 0.34116846 |
| AFGLGLLL | ASSATGGASQNTLY | 0.40321958 |
| AFFAAASL | ASSATGGASQNTLY | 0.16684628 |
| AAAALPAFSPP | ASSATGGASQNTLY | 0.23112741 |
| VVLAPAL | ASSATGGASQNTLY | 0.28385508 |
| TIGHVDHGKTTL | ASSATGGASQNTLY | 0.29434383 |
| LISNASDALDK | ASSATGGASQNTLY | 0.29371288 |
| LISNASDAL | ASSATGGASQNTLY | 0.72689587 |
| ISNASDALDK | ASSATGGASQNTLY | 0.10676523 |
| NQAETLVYQTE | ASSATGGASQNTLY | 0.14187293 |
| TGGTGGTAGAG | ASSATGGASQNTLY | 0.5612248 |
| TGSTAGGAGGA | ASSATGGASQNTLY | 0.10738382 |
| DPLAVPVAAGPV | ASSATGGASQNTLY | 0.13168237 |
| GGTGGTPGNGGN | ASSATGGASQNTLY | 0.30799657 |

| | | |
|---|---|---|
| PAGLAGVDPATL | ASSATGGASQNTLY | 0.71456647 |
| GNNLVVEAPHAGASV | ASSATGGASQNTLY | 0.10959159 |
| KVTRIMSASTLEQPA | ASSATGGASQNTLY | 0.21381097 |
| LGFALPIALPRRTVV | ASSATGGASQNTLY | 0.12825912 |
| TRLGNLTARKLVVLV | ASSATGGASQNTLY | 0.5768528 |
| WSQQRRLIYFPSAGP | ASSATGGASQNTLY | 0.19703043 |
| GPNGNPDPMAAAVDIR | ASSATGGASQNTLY | 0.12205449 |
| TQRCVSVADLRAAASR | ASSATGGASQNTLY | 0.33502623 |
| QFVLDVAEHPVEVLTETV | ASSATGGASQNTLY | 0.12732205 |
| GGAGGTGGAGGNGGTQAGDG | ASSATGGASQNTLY | 0.33395863 |
| PLLATLRAIAVDVDAKPAQV | ASSATGGASQNTLY | 0.35711145 |
| APKFRTGGYMIDFWGVGYQVAK | ASSATGGASQNTLY | 0.78973955 |
| GAGGAGGAGGAAGGPGGTGGTG | ASSATGGASQNTLY | 0.11894694 |
| IGALGLLATDVAPAGSVAFGVA | ASSATGGASQNTLY | 0.3081977 |
| AGGAGGIGGNGGAGGAGGVLMG | ASSATGGASQNTLY | 0.20028493 |
| ADELIGLL | ASSATGGASQNTLY | 0.80332464 |
| ADRALKA | ASSATGGASQNTLY | 0.16656128 |
| AGAVGQVGGAGGA | ASSATGGASQNTLY | 0.12323647 |
| AGGGGGLGGAP | ASSATGGASQNTLY | 0.4105079 |
| AGGLLYGNGGAG | ASSATGGASQNTLY | 0.34317133 |
| AGLESIGL | ASSATGGASQNTLY | 0.22050461 |
| AGLKTGII | ASSATGGASQNTLY | 0.70709044 |
| ASLAALLL | ASSATGGASQNTLY | 0.32341576 |
| ASVINISSI | ASSATGGASQNTLY | 0.18695086 |
| ATPPGPA | ASSATGGASQNTLY | 0.32591614 |
| ATVASALV | ASSATGGASQNTLY | 0.23519263 |
| ATVASLIL | ASSATGGASQNTLY | 0.15840736 |
| AVNGPVLIRH | ASSATGGASQNTLY | 0.39987648 |
| DGAAGAVGI | ASSATGGASQNTLY | 0.20993784 |
| DPLLHGI | ASSATGGASQNTLY | 0.130572 |
| ELGAALGAL | ASSATGGASQNTLY | 0.7577219 |
| ELGRTQAV | ASSATGGASQNTLY | 0.24506207 |
| FVAGGYLS | ASSATGGASQNTLY | 0.40832275 |
| GASGGAGGAGGW | ASSATGGASQNTLY | 0.30670306 |
| GGAGAGGDGGAGGI | ASSATGGASQNTLY | 0.83361226 |
| GTGHNGSQPGMGGNG | ASSATGGASQNTLY | 0.14655246 |
| GYAAASASAS | ASSATGGASQNTLY | 0.25323987 |
| HQVDVAAVQQG | ASSATGGASQNTLY | 0.25437772 |
| IGAAIAHT | ASSATGGASQNTLY | 0.44713777 |
| IGMIEVAQL | ASSATGGASQNTLY | 0.2728772 |
| IRLAPEHTSAMD | ASSATGGASQNTLY | 0.38725623 |
| IYNVSVL | ASSATGGASQNTLY | 0.37320697 |
| KQGFLGM | ASSATGGASQNTLY | 0.46179825 |
| LESVITQL | ASSATGGASQNTLY | 0.2576114 |
| LGLGALVVV | ASSATGGASQNTLY | 0.21634376 |
| LIGNGGNGGSGGI | ASSATGGASQNTLY | 0.5933919 |
| LNAINAPTL | ASSATGGASQNTLY | 0.12953107 |
| LQLLNTGV | ASSATGGASQNTLY | 0.2184639 |
| LQTSIEF | ASSATGGASQNTLY | 0.52777123 |
| MSFAVSEL | ASSATGGASQNTLY | 0.2967074 |
| NTGALGL | ASSATGGASQNTLY | 0.2406778 |

| | | |
|---|---|---|
| PPGAGGAGGAGTT | ASSATGGASQNTLY | 0.44618696 |
| QLVGLGI | ASSATGGASQNTLY | 0.19078124 |
| RFLEQVHQL | ASSATGGASQNTLY | 0.25712833 |
| RGVGYVLR | ASSATGGASQNTLY | 0.30166936 |
| SATGVFAGVI | ASSATGGASQNTLY | 0.1621768 |
| SLPAITLP | ASSATGGASQNTLY | 0.12206145 |
| STALAFTV | ASSATGGASQNTLY | 0.1711275 |
| SVLTVGLGK | ASSATGGASQNTLY | 0.45400748 |
| TALAGAGSGI | ASSATGGASQNTLY | 0.24307853 |
| TGLPFTGI | ASSATGGASQNTLY | 0.10900947 |
| VAEHVTV | ASSATGGASQNTLY | 0.27725253 |
| VDIVLNSL | ASSATGGASQNTLY | 0.33766437 |
| VLTGGGAVL | ASSATGGASQNTLY | 0.7631161 |
| VNVGLAYGI | ASSATGGASQNTLY | 0.31899434 |
| VSSAAVGAV | ASSATGGASQNTLY | 0.3912333 |
| VTSVYEAGV | ASSATGGASQNTLY | 0.18711956 |
| WVALDRGA | ASSATGGASQNTLY | 0.30024463 |
| YLVGSVVI | ASSATGGASQNTLY | 0.119667634 |
| GPPGTGKTLL | ASSATGGASQNTLY | 0.1454916 |
| GVGMTKF | ASSATGGASQNTLY | 0.21054111 |
| IGIGGSDL | ASSATGGASQNTLY | 0.12869278 |
| IGTLLAEL | ASSATGGASQNTLY | 0.6673846 |
| IYTSGTT | ASSATGGASQNTLY | 0.35889515 |
| LSFPAGV | ASSATGGASQNTLY | 0.4069545 |
| LYTSGTT | ASSATGGASQNTLY | 0.6370125 |
| VGAFPFGF | ASSATGGASQNTLY | 0.65004635 |
| VGAGTVEF | ASSATGGASQNTLY | 0.34792072 |
| VGKSTLLL | ASSATGGASQNTLY | 0.15500715 |
| AALAHAGGGIV | ASSATGGASQNTLY | 0.19127804 |
| AALVAVSLL | ASSATGGASQNTLY | 0.10992428 |
| AAVSRAVAAL | ASSATGGASQNTLY | 0.18143456 |
| AGGSSGIGGF | ASSATGGASQNTLY | 0.17479238 |
| AGLTVANANG | ASSATGGASQNTLY | 0.19598556 |
| AGLVGAVL | ASSATGGASQNTLY | 0.10328136 |
| ALSQAFTV | ASSATGGASQNTLY | 0.6505474 |
| ALTFSVSL | ASSATGGASQNTLY | 0.2945658 |
| ALTGAALVAL | ASSATGGASQNTLY | 0.21601072 |
| ASFSIGAGVL | ASSATGGASQNTLY | 0.38526335 |
| DGSVYKI | ASSATGGASQNTLY | 0.3587948 |
| EQLAEIV | ASSATGGASQNTLY | 0.17085503 |
| FAVSELTVM | ASSATGGASQNTLY | 0.424538 |
| GAAPLAAGL | ASSATGGASQNTLY | 0.19639187 |
| GADGGDGGSGGAGG | ASSATGGASQNTLY | 0.4483352 |
| GFGLDTGGAGG | ASSATGGASQNTLY | 0.4666046 |
| GLASGGLL | ASSATGGASQNTLY | 0.34789962 |
| GRIAGAAVLL | ASSATGGASQNTLY | 0.1769105 |
| GSAGSLNTSF | ASSATGGASQNTLY | 0.2591535 |
| GTGQIGI | ASSATGGASQNTLY | 0.15172482 |
| GTLIALIL | ASSATGGASQNTLY | 0.18280149 |
| IAATMGFVVG | ASSATGGASQNTLY | 0.11457605 |
| IFAQALL | ASSATGGASQNTLY | 0.35777724 |

| | | |
|---|---|---|
| IGAAVALGF | ASSATGGASQNTLY | 0.15439612 |
| IGLTPLAL | ASSATGGASQNTLY | 0.24780689 |
| ILGSNTSTL | ASSATGGASQNTLY | 0.19062455 |
| IVFSLTVL | ASSATGGASQNTLY | 0.39548784 |
| LASKATGF | ASSATGGASQNTLY | 0.13256082 |
| LASVALIL | ASSATGGASQNTLY | 0.33004278 |
| LFAVGLTL | ASSATGGASQNTLY | 0.46252802 |
| LFNGGIGTYI | ASSATGGASQNTLY | 0.13471602 |
| LTLGAGII | ASSATGGASQNTLY | 0.3255024 |
| LVAAALGDI | ASSATGGASQNTLY | 0.2700986 |
| NTAFVQLGI | ASSATGGASQNTLY | 0.19276506 |
| QLFSGAEL | ASSATGGASQNTLY | 0.46922252 |
| SAATFITL | ASSATGGASQNTLY | 0.21684149 |
| SNYGLANLGV | ASSATGGASQNTLY | 0.13350286 |
| SYGGAYAVM | ASSATGGASQNTLY | 0.17789297 |
| TTGVGGLTLG | ASSATGGASQNTLY | 0.2837154 |
| TTQTIYI | ASSATGGASQNTLY | 0.18988308 |
| VAAGAGLGAVI | ASSATGGASQNTLY | 0.0999047 |
| VAAVVTLL | ASSATGGASQNTLY | 0.29216194 |
| VGFLGGLI | ASSATGGASQNTLY | 0.13198549 |
| VGVIPILL | ASSATGGASQNTLY | 0.14595617 |
| VIFASSLI | ASSATGGASQNTLY | 0.15022351 |
| VKGGDGGAAAG | ASSATGGASQNTLY | 0.37070048 |
| VTKTFTNI | ASSATGGASQNTLY | 0.46955857 |
| VVAAAILL | ASSATGGASQNTLY | 0.2385584 |
| YAALDVEL | ASSATGGASQNTLY | 0.22972208 |
| LASVALLL | ASSATGGASQNTLY | 0.25763723 |
| LDAVHRV | ASSATGGASQNTLY | 0.36178252 |
| QGLSSSIF | ASSATGGASQNTLY | 0.25841057 |
| RGTFANIRL | ASSATGGASQNTLY | 0.32858342 |
| RQLWWGHRI | ASSATGGASQNTLY | 0.33024612 |
| YGPPGTGKTLL | ASSATGGASQNTLY | 0.20685034 |
| AALVGAPANVA | ASSATGGASQNTLY | 0.13056426 |
| AATFRRTE | ASSATGGASQNTLY | 0.1848359 |
| AGALGGGI | ASSATGGASQNTLY | 0.15784648 |
| AGGAGGAGGIGAG | ASSATGGASQNTLY | 0.14252424 |
| AGSGALGL | ASSATGGASQNTLY | 0.3004567 |
| AQIADQI | ASSATGGASQNTLY | 0.56442904 |
| ASAVAVTYL | ASSATGGASQNTLY | 0.37512335 |
| ASTVLGGTGG | ASSATGGASQNTLY | 0.118470356 |
| AVGGEPAK | ASSATGGASQNTLY | 0.19855615 |
| CEQGTFGGTC | ASSATGGASQNTLY | 0.20376623 |
| DEGKAPAG | ASSATGGASQNTLY | 0.25297529 |
| DTVVFPAL | ASSATGGASQNTLY | 0.5588125 |
| FEGITMAG | ASSATGGASQNTLY | 0.8203634 |
| FPLVVQD | ASSATGGASQNTLY | 0.388798 |
| GAFTTPQ | ASSATGGASQNTLY | 0.14980271 |
| GFGGFGTAGGVGGA | ASSATGGASQNTLY | 0.30839825 |
| GFLAQFL | ASSATGGASQNTLY | 0.20964083 |
| GGAGGAGGNAGVA | ASSATGGASQNTLY | 0.16939719 |
| GGVGTAGAGGAGGAGG | ASSATGGASQNTLY | 0.24508324 |

| | | |
|---|---|---|
| GNGGGGDAGNAGSGGN | ASSATGGASQNTLY | 0.48240453 |
| GTGGAGGAGNDAGS | ASSATGGASQNTLY | 0.17471385 |
| GTTNTGWFNS | ASSATGGASQNTLY | 0.16808577 |
| IFMGPLFTL | ASSATGGASQNTLY | 0.1379631 |
| ILAHAIEAI | ASSATGGASQNTLY | 0.46973613 |
| IRSAIREAL | ASSATGGASQNTLY | 0.1608591 |
| ITFAGGG | ASSATGGASQNTLY | 0.183629 |
| ITGMSGHQEV | ASSATGGASQNTLY | 0.20868063 |
| IVAGLACTL | ASSATGGASQNTLY | 0.65269125 |
| IVGGGLAGA | ASSATGGASQNTLY | 0.70780313 |
| IVTGVLNI | ASSATGGASQNTLY | 0.18366669 |
| LAQQGCI | ASSATGGASQNTLY | 0.15925154 |
| LCLHVEN | ASSATGGASQNTLY | 0.66434884 |
| LDAISDGS | ASSATGGASQNTLY | 0.16109824 |
| LGGFAAML | ASSATGGASQNTLY | 0.36780262 |
| LGGKNPMI | ASSATGGASQNTLY | 0.27797037 |
| LGPKGVAAIL | ASSATGGASQNTLY | 0.1708138 |
| LMVGVSI | ASSATGGASQNTLY | 0.17383792 |
| LVGAAFVL | ASSATGGASQNTLY | 0.4491409 |
| LVLAVAPL | ASSATGGASQNTLY | 0.15563084 |
| MAAIAVIDAA | ASSATGGASQNTLY | 0.38073716 |
| NNGINAELQ | ASSATGGASQNTLY | 0.117607825 |
| RGEPLVLVHG | ASSATGGASQNTLY | 0.3501982 |
| RGGAGGAAGLI | ASSATGGASQNTLY | 0.12363499 |
| RSYRMHPAV | ASSATGGASQNTLY | 0.17844611 |
| SAAHLLAGQLA | ASSATGGASQNTLY | 0.10869008 |
| SAAVGSY | ASSATGGASQNTLY | 0.168353 |
| SPHAVLHRI | ASSATGGASQNTLY | 0.40447402 |
| SQIAKVL | ASSATGGASQNTLY | 0.6598962 |
| TALGHLVP | ASSATGGASQNTLY | 0.2492395 |
| TAPTTSGAA | ASSATGGASQNTLY | 0.19061954 |
| TDGDDGNGGSPGT | ASSATGGASQNTLY | 0.7311077 |
| TDGLGEN | ASSATGGASQNTLY | 0.14138004 |
| TGGPPLPGQA | ASSATGGASQNTLY | 0.11038949 |
| TIPAITI | ASSATGGASQNTLY | 0.23860618 |
| TIPGFTI | ASSATGGASQNTLY | 0.45298874 |
| TSIASGALA | ASSATGGASQNTLY | 0.59850883 |
| TVTLPGAAI | ASSATGGASQNTLY | 0.5998646 |
| VASGLSAALP | ASSATGGASQNTLY | 0.109439015 |
| VGAGVIGI | ASSATGGASQNTLY | 0.36882862 |
| VGPKRFGV | ASSATGGASQNTLY | 0.3986195 |
| VNVAFPD | ASSATGGASQNTLY | 0.13847741 |
| VSMTNSGVS | ASSATGGASQNTLY | 0.13719937 |
| VTDVLGDLSV | ASSATGGASQNTLY | 0.15277535 |
| VVVVGAGFAGL | ASSATGGASQNTLY | 0.113061786 |
| VYTVGTVV | ASSATGGASQNTLY | 0.30598396 |
| WTTFSGP | ASSATGGASQNTLY | 0.17049588 |
| YDELVEAF | ASSATGGASQNTLY | 0.24108313 |
| YGTGGAGGNGG | ASSATGGASQNTLY | 0.18438348 |
| YTNVMAV | ASSATGGASQNTLY | 0.17945649 |
| FLEQVHQL | ASSATGGASQNTLY | 0.19148582 |

| | | |
|---|---|---|
| GLSLLAVAV | ASSATGGASQNTLY | 0.18150535 |
| LEQVHQL | ASSATGGASQNTLY | 0.27314466 |
| LNAAGAI | ASSATGGASQNTLY | 0.51410604 |
| PAVHASL | ASSATGGASQNTLY | 0.1972039 |
| TELEDDA | ASSATGGASQNTLY | 0.252582 |
| TPPGPPN | ASSATGGASQNTLY | 0.41993046 |
| TPPPPPPP | ASSATGGASQNTLY | 0.13265324 |
| TQSPALI | ASSGTGGALEQY | 0.14861968 |
| EFAAAFPQ | ASSGTGGALEQY | 0.3673057 |
| LLSGNLDVL | ASSGTGGALEQY | 0.20467852 |
| GPANLGNYNI | ASSGTGGALEQY | 0.5925603 |
| TLGPTLP | ASSGTGGALEQY | 0.22840273 |
| VYAYSSCCK | ASSGTGGALEQY | 0.2335938 |
| APAPAPAPNI | ASSGTGGALEQY | 0.45485625 |
| PGGPVPIP | ASSGTGGALEQY | 0.5400221 |
| FLTVALEP | ASSGTGGALEQY | 0.23779196 |
| SQYFGANL | ASSGTGGALEQY | 0.16751575 |
| GGVKPAA | ASSGTGGALEQY | 0.5481523 |
| GGTGGAGGAGGW | ASSGTGGALEQY | 0.14247805 |
| GVGGVAGLGGAGG | ASSGTGGALEQY | 0.2870642 |
| VLLVLATTI | ASSGTGGALEQY | 0.4344083 |
| IGIFAAML | ASSGTGGALEQY | 0.12509914 |
| AVGDLID | ASSGTGGALEQY | 0.19097124 |
| NISPLPL | ASSGTGGALEQY | 0.6379783 |
| VTAVFVAI | ASSGTGGALEQY | 0.31086028 |
| SGDDNDAAL | ASSGTGGALEQY | 0.18034607 |
| AVAALAHS | ASSGTGGALEQY | 0.1444904 |
| GGVTVMATP | ASSGTGGALEQY | 0.18922825 |
| AGLAANVRD | ASSGTGGALEQY | 0.31481653 |
| TNRPDLIDPALLRPG | ASSGTGGALEQY | 0.123254284 |
| LANLGVTAL | ASSGTGGALEQY | 0.41656616 |
| TGERIEL | ASSGTGGALEQY | 0.17365234 |
| NGEELAA | ASSGTGGALEQY | 0.7515617 |
| VLSLVWG | ASSGTGGALEQY | 0.27289632 |
| IGYALWL | ASSGTGGALEQY | 0.366544 |
| LSLLLAPL | ASSGTGGALEQY | 0.21123949 |
| TLDEALK | ASSGTGGALEQY | 0.18304002 |
| TDGAGTAEGG | ASSGTGGALEQY | 0.12906215 |
| NAPGGAGGNGG | ASSGTGGALEQY | 0.14903766 |
| VSVFMGMGGNF | ASSGTGGALEQY | 0.153141 |
| LASYLAAL | ASSGTGGALEQY | 0.23091352 |
| LVGGTPAL | ASSGTGGALEQY | 0.1325253 |
| VRNADVL | ASSGTGGALEQY | 0.6714453 |
| TCSSPFLVSAM | ASSGTGGALEQY | 0.1537248 |
| VHITDAE | ASSGTGGALEQY | 0.2795626 |
| EYAAPSF | ASSGTGGALEQY | 0.21197465 |
| VVDAFHGGL | ASSGTGGALEQY | 0.12097918 |
| ASVVDPQ | ASSGTGGALEQY | 0.15132546 |
| SYNEFGGLS | ASSGTGGALEQY | 0.41222566 |
| DPDGSVY | ASSGTGGALEQY | 0.12024749 |
| MVSTIHATE | ASSGTGGALEQY | 0.11682173 |

| | | |
|---|---|---|
| LAAGVADPV | ASSGTGGALEQY | 0.16467188 |
| TKEQIAA | ASSGTGGALEQY | 0.22088039 |
| SLASPAAL | ASSGTGGALEQY | 0.7538774 |
| IGLVFYGV | ASSGTGGALEQY | 0.5587361 |
| FLLDASNL | ASSGTGGALEQY | 0.2507527 |
| YQVGNFP | ASSGTGGALEQY | 0.74937856 |
| ESLLSQST | ASSGTGGALEQY | 0.10248514 |
| LFTVGFAAVG | ASSGTGGALEQY | 0.21280035 |
| ATPPTPT | ASSGTGGALEQY | 0.57228506 |
| TDLATLG | ASSGTGGALEQY | 0.33922789 |
| SAGGTGR | ASSGTGGALEQY | 0.1791727 |
| GFSNSGS | ASSGTGGALEQY | 0.3829383 |
| GGNGGSGGTSV | ASSGTGGALEQY | 0.14754859 |
| GTSVATGGAGN | ASSGTGGALEQY | 0.18802375 |
| NGGNGGSGGTS | ASSGTGGALEQY | 0.18998429 |
| AVVKFPH | ASSGTGGALEQY | 0.3293216 |
| TPPPPPP | ASSGTGGALEQY | 0.31931162 |
| AAPAAAAAPA | ASSGTGGALEQY | 0.117470205 |
| IALNEDL | ASSGTGGALEQY | 0.33766308 |
| IVNVSSM | ASSGTGGALEQY | 0.76840997 |
| TGIPPAPR | ASSGTGGALEQY | 0.14676689 |
| FDLGGGTF | ASSGTGGALEQY | 0.4885543 |
| FDLGGGTFDVS | ASSGTGGALEQY | 0.2044536 |
| NEPTAAA | ASSGTGGALEQY | 0.45490986 |
| NEPTAAALA | ASSGTGGALEQY | 0.15123077 |
| GSIINISSI | ASSGTGGALEQY | 0.11956435 |
| VAPGNAGTA | ASSGTGGALEQY | 0.35966116 |
| TGGGASLE | ASSGTGGALEQY | 0.13160884 |
| VAVYNHYKRI | ASSGTGGALEQY | 0.29040253 |
| TDPAPATT | ASSGTGGALEQY | 0.25653508 |
| TNRPDILDPALLRPG | ASSGTGGALEQY | 0.80695724 |
| LTTTTTS | ASSGTGGALEQY | 0.4018292 |
| PTGSGKTL | ASSGTGGALEQY | 0.23890042 |
| TGSGKTL | ASSGTGGALEQY | 0.1354289 |
| TIGHVDHGK | ASSGTGGALEQY | 0.3132714 |
| TIGHVDHGKTT | ASSGTGGALEQY | 0.4640195 |
| GHVDHGKTT | ASSGTGGALEQY | 0.14102904 |
| AQFEHTLL | ASSGTGGALEQY | 0.5547477 |
| IVNQDEL | ASSGTGGALEQY | 0.13109498 |
| TNGSQFF | ASSGTGGALEQY | 0.3386126 |
| TGKGIVE | ASSGTGGALEQY | 0.32964385 |
| LISNASDA | ASSGTGGALEQY | 0.79759264 |
| AELRTLGELTKIAKAHGAQ | ASSGTGGALEQY | 0.2123163 |
| APNAVAAAEPSW | ASSGTGGALEQY | 0.4285796 |
| APQVAGAATAVL | ASSGTGGALEQY | 0.12784387 |
| ARRDPAAAAAILPTDA | ASSGTGGALEQY | 0.5610368 |
| DAAARDVLAEAGLAEYF | ASSGTGGALEQY | 0.44573438 |
| DDQIVLSAKANLAKVEQ | ASSGTGGALEQY | 0.4843871 |
| DELLPDTGEDSDEDGH | ASSGTGGALEQY | 0.785643 |
| ELPEAPANRATVPQD | ASSGTGGALEQY | 0.47024915 |
| FGQSKIERVTGIG | ASSGTGGALEQY | 0.12829384 |

| | | |
|---|---|---|
| GAGGTGSTAGGAGGAGGAGGLY | ASSGTGGALEQY | 0.19537568 |
| GHGGTGAAGGNGGTGGAGGSGI | ASSGTGGALEQY | 0.44322437 |
| GNGGNGGAGGEGGPGGAGGAGA | ASSGTGGALEQY | 0.72897613 |
| GSTAGNGGNGGNATNGTIAGQP | ASSGTGGALEQY | 0.38007194 |
| GTGGDGGNGGNGGTGG | ASSGTGGALEQY | 0.22735378 |
| GVRRPAGIPADAPV | ASSGTGGALEQY | 0.23342228 |
| GYRQVTDAHLLTLARR | ASSGTGGALEQY | 0.6535203 |
| IAKMGQDGHDRG | ASSGTGGALEQY | 0.85188025 |
| KAEPSIEIQYT | ASSGTGGALEQY | 0.30786553 |
| KPVRIRMAADEDTVAVT | ASSGTGGALEQY | 0.17678618 |
| MPPRIAGMRLLVIKPEP | ASSGTGGALEQY | 0.2165876 |
| MKFYAVALFRSRGVPPTR | ASSGTGGALEQY | 0.3646332 |
| NAGGFGILGAVAHSP | ASSGTGGALEQY | 0.38747662 |
| NEGREITLAMAEM | ASSGTGGALEQY | 0.14692302 |
| NELVEWSHPLGHRWRWE | ASSGTGGALEQY | 0.19497412 |
| NLGAGGTGGNGGNPG | ASSGTGGALEQY | 0.21330056 |
| NTEWPKWETVMAADPVD | ASSGTGGALEQY | 0.1056957 |
| PASAAPAETQP | ASSGTGGALEQY | 0.5068369 |
| QQSSDGKNGVRAAEITGE | ASSGTGGALEQY | 0.5127755 |
| RAPDAVTTSWWKE | ASSGTGGALEQY | 0.25863606 |
| SGFGNVGAHTSGWFNQ | ASSGTGGALEQY | 0.12477993 |
| SGVAGAVGQAASVGGLKV | ASSGTGGALEQY | 0.5703995 |
| SNGGAGTGSAGGNGGT | ASSGTGGALEQY | 0.48359472 |
| TGGAGGEGGAAG | ASSGTGGALEQY | 0.51099646 |
| TGGDGGAAGAGGAPGHG | ASSGTGGALEQY | 0.1288076 |
| TMPPLLAAIANAGRHGV | ASSGTGGALEQY | 0.10320672 |
| TSSTAGNSGGA | ASSGTGGALEQY | 0.20412558 |
| VPATVPVGILC | ASSGTGGALEQY | 0.7120187 |
| VQRPDSGGVVLTGRIS | ASSGTGGALEQY | 0.20555404 |
| WTPNPVPATKGVPAGLVQAT | ASSGTGGALEQY | 0.15334111 |
| YALKRALAKFSDRFDVVII | ASSGTGGALEQY | 0.19143042 |
| YLIGSVIV | ASSGTGGALEQY | 0.45342267 |
| YGQLVAVID | ASSGTGGALEQY | 0.12963396 |
| VTVGGMTLPA | ASSGTGGALEQY | 0.1465222 |
| VLALPAV | ASSGTGGALEQY | 0.19721489 |
| VIPPTLNYET | ASSGTGGALEQY | 0.5288735 |
| VGMGGMHPSSG | ASSGTGGALEQY | 0.12453246 |
| VGLAGGAATA | ASSGTGGALEQY | 0.16163121 |
| VGGAGGAGGSGP | ASSGTGGALEQY | 0.272317 |
| VFAAFMV | ASSGTGGALEQY | 0.63726497 |
| TLYDKLR | ASSGTGGALEQY | 0.3084611 |
| TLAVFTVL | ASSGTGGALEQY | 0.1339351 |
| TGSVQPARV | ASSGTGGALEQY | 0.3048985 |
| TGGGAGSGGT | ASSGTGGALEQY | 0.12507251 |
| SHLETSL | ASSGTGGALEQY | 0.55092734 |
| SFTTGMT | ASSGTGGALEQY | 0.16518547 |
| QVEVAKRMGVRQPTVS | ASSGTGGALEQY | 0.29281378 |
| QGSGGPALAM | ASSGTGGALEQY | 0.16944575 |
| MLAPSNP | ASSGTGGALEQY | 0.13479038 |
| LVNNAAI | ASSGTGGALEQY | 0.13667648 |
| LTGEINTLA | ASSGTGGALEQY | 0.15378764 |

| | | |
|---|---|---|
| LSIFGPV | ASSGTGGALEQY | 0.3343169 |
| LLALLANL | ASSGTGGALEQY | 0.11686107 |
| LISNASDALDKLR | ASSGTGGALEQY | 0.24228671 |
| LINEAVP | ASSGTGGALEQY | 0.18899076 |
| LGYAFSGL | ASSGTGGALEQY | 0.13498494 |
| LDFVAVAL | ASSGTGGALEQY | 0.3886282 |
| LAQPGAE | ASSGTGGALEQY | 0.11896395 |
| ISNLGAVTGF | ASSGTGGALEQY | 0.3058291 |
| HQEDPGRI | ASSGTGGALEQY | 0.4423025 |
| GTPGEPTP | ASSGTGGALEQY | 0.2820248 |
| GFFNSGS | ASSGTGGALEQY | 0.17141773 |
| GAGGGGLTAGAN | ASSGTGGALEQY | 0.30809444 |
| FVDALVNM | ASSGTGGALEQY | 0.39001057 |
| FLTVAPEL | ASSGTGGALEQY | 0.15567943 |
| FETLTAL | ASSGTGGALEQY | 0.24348399 |
| DVINPIEL | ASSGTGGALEQY | 0.46903133 |
| APVSGAPV | ASSGTGGALEQY | 0.2565404 |
| ALVGGLAW | ASSGTGGALEQY | 0.11071285 |
| ALAAAAATAI | ASSGTGGALEQY | 0.26906994 |
| AFGLGLLL | ASSGTGGALEQY | 0.14687851 |
| AFFAAASL | ASSGTGGALEQY | 0.28437364 |
| AAAALPAFSPP | ASSGTGGALEQY | 0.20201983 |
| VVLAPAL | ASSGTGGALEQY | 0.23360628 |
| TIGHVDHGKTTL | ASSGTGGALEQY | 0.34384227 |
| LISNASDALDK | ASSGTGGALEQY | 0.14366983 |
| LISNASDAL | ASSGTGGALEQY | 0.17496139 |
| ISNASDALDK | ASSGTGGALEQY | 0.2024596 |
| NQAETLVYQTE | ASSGTGGALEQY | 0.24434638 |
| TGGTGGTAGAG | ASSGTGGALEQY | 0.29849708 |
| TGSTAGGAGGA | ASSGTGGALEQY | 0.12962328 |
| DPLAVPVAAGPV | ASSGTGGALEQY | 0.3746035 |
| GGTGGTPGNGGN | ASSGTGGALEQY | 0.21414384 |
| PAGLAGVDPATL | ASSGTGGALEQY | 0.1380465 |
| GNNLVVEAPHAGASV | ASSGTGGALEQY | 0.26082444 |
| KVTRIMSASTLEQPA | ASSGTGGALEQY | 0.17905268 |
| LGFALPIALPRRTVV | ASSGTGGALEQY | 0.1270587 |
| TRLGNLTARKLVVLV | ASSGTGGALEQY | 0.3994181 |
| WSQQRRLIYFPSAGP | ASSGTGGALEQY | 0.15674332 |
| GPNGNPDPMAAAVDIR | ASSGTGGALEQY | 0.35627472 |
| TQRCVSVADLRAAASR | ASSGTGGALEQY | 0.74388486 |
| QFVLDVAEHPVEVLTETV | ASSGTGGALEQY | 0.11949133 |
| GGAGGTGGAGGNGGTQAGDG | ASSGTGGALEQY | 0.461975 |
| PLLATLRAIAVDVDAKPAQV | ASSGTGGALEQY | 0.19504131 |
| APKFRTGGYMIDFWGVGYQVAK | ASSGTGGALEQY | 0.17365605 |
| GAGGAGGAGGAAGGPGGTGGTG | ASSGTGGALEQY | 0.1993494 |
| IGALGLLATDVAPAGSVAFGVA | ASSGTGGALEQY | 0.15576388 |
| AGGAGGIGGNGGAGGAGGVLMG | ASSGTGGALEQY | 0.43093893 |
| ADELIGLL | ASSGTGGALEQY | 0.29568395 |
| ADRALKA | ASSGTGGALEQY | 0.21616504 |
| AGAVGQVGGAGGA | ASSGTGGALEQY | 0.24625917 |
| AGGGGGLGGAP | ASSGTGGALEQY | 0.32926226 |

| | | |
|---|---|---|
| AGGLLYGNGGAG | ASSGTGGALEQY | 0.36390674 |
| AGLESIGL | ASSGTGGALEQY | 0.13957942 |
| AGLKTGII | ASSGTGGALEQY | 0.12134596 |
| ASLAALLL | ASSGTGGALEQY | 0.18840742 |
| ASVINISSI | ASSGTGGALEQY | 0.18848446 |
| ATPPGPA | ASSGTGGALEQY | 0.14075473 |
| ATVASALV | ASSGTGGALEQY | 0.15046845 |
| ATVASLIL | ASSGTGGALEQY | 0.3737184 |
| AVNGPVLIRH | ASSGTGGALEQY | 0.75458896 |
| DGAAGAVGI | ASSGTGGALEQY | 0.20041922 |
| DPLLHGI | ASSGTGGALEQY | 0.7970382 |
| ELGAALGAL | ASSGTGGALEQY | 0.13085991 |
| ELGRTQAV | ASSGTGGALEQY | 0.36462334 |
| FVAGGYLS | ASSGTGGALEQY | 0.44089884 |
| GASGGAGGAGGW | ASSGTGGALEQY | 0.13068673 |
| GGAGAGGDGGAGGI | ASSGTGGALEQY | 0.25967887 |
| GTGHNGSQPGMGGNG | ASSGTGGALEQY | 0.13768065 |
| GYAAASASAS | ASSGTGGALEQY | 0.14040843 |
| HQVDVAAVQQG | ASSGTGGALEQY | 0.28513443 |
| IGAAIAHT | ASSGTGGALEQY | 0.22031948 |
| IGMIEVAQL | ASSGTGGALEQY | 0.34616607 |
| IRLAPEHTSAMD | ASSGTGGALEQY | 0.174355 |
| IYNVSVL | ASSGTGGALEQY | 0.13642891 |
| KQGFLGM | ASSGTGGALEQY | 0.24456176 |
| LESVITQL | ASSGTGGALEQY | 0.28939515 |
| LGLGALVVV | ASSGTGGALEQY | 0.35343602 |
| LIGNGGNGGSGGI | ASSGTGGALEQY | 0.8491416 |
| LNAINAPTL | ASSGTGGALEQY | 0.13296673 |
| LQLLNTGV | ASSGTGGALEQY | 0.69957936 |
| LQTSIEF | ASSGTGGALEQY | 0.73365104 |
| MSFAVSEL | ASSGTGGALEQY | 0.2699021 |
| NTGALGL | ASSGTGGALEQY | 0.2231212 |
| PPGAGGAGGAGTT | ASSGTGGALEQY | 0.5002708 |
| QLVGLGI | ASSGTGGALEQY | 0.19115464 |
| RFLEQVHQL | ASSGTGGALEQY | 0.23032117 |
| RGVGYVLR | ASSGTGGALEQY | 0.50202626 |
| SATGVFAGVI | ASSGTGGALEQY | 0.51143557 |
| SLPAITLP | ASSGTGGALEQY | 0.8228514 |
| STALAFTV | ASSGTGGALEQY | 0.7610291 |
| SVLTVGLGK | ASSGTGGALEQY | 0.69670576 |
| TALAGAGSGI | ASSGTGGALEQY | 0.65405536 |
| TGLPFTGI | ASSGTGGALEQY | 0.29560748 |
| VAEHVTV | ASSGTGGALEQY | 0.6653888 |
| VDIVLNSL | ASSGTGGALEQY | 0.56736624 |
| VLTGGGAVL | ASSGTGGALEQY | 0.78483 |
| VNVGLAYGI | ASSGTGGALEQY | 0.77175754 |
| VSSAAVGAV | ASSGTGGALEQY | 0.64146924 |
| VTSVYEAGV | ASSGTGGALEQY | 0.84675574 |
| WVALDRGA | ASSGTGGALEQY | 0.8452234 |
| YLVGSVVI | ASSGTGGALEQY | 0.3977646 |
| GPPGTGKTLL | ASSGTGGALEQY | 0.6717566 |

| | | |
|---|---|---|
| GVGMTKF | ASSGTGGALEQY | 0.5802673 |
| IGIGGSDL | ASSGTGGALEQY | 0.71095103 |
| IGTLLAEL | ASSGTGGALEQY | 0.22175452 |
| IYTSGTT | ASSGTGGALEQY | 0.8103274 |
| LSFPAGV | ASSGTGGALEQY | 0.56987524 |
| LYTSGTT | ASSGTGGALEQY | 0.11752848 |
| VGAFPFGF | ASSGTGGALEQY | 0.7237557 |
| VGAGTVEF | ASSGTGGALEQY | 0.7168614 |
| VGKSTLLL | ASSGTGGALEQY | 0.719746 |
| AALAHAGGGIV | ASSGTGGALEQY | 0.8309438 |
| AALVAVSLL | ASSGTGGALEQY | 0.2386358 |
| AAVSRAVAAL | ASSGTGGALEQY | 0.5299225 |
| AGGSSGIGGF | ASSGTGGALEQY | 0.8069128 |
| AGLTVANANG | ASSGTGGALEQY | 0.8432469 |
| AGLVGAVL | ASSGTGGALEQY | 0.44315362 |
| ALSQAFTV | ASSGTGGALEQY | 0.77918303 |
| ALTFSVSL | ASSGTGGALEQY | 0.84241366 |
| ALTGAALVAL | ASSGTGGALEQY | 0.7860222 |
| ASFSIGAGVL | ASSGTGGALEQY | 0.8056625 |
| DGSVYKI | ASSGTGGALEQY | 0.83959556 |
| EQLAEIV | ASSGTGGALEQY | 0.4118507 |
| FAVSELTVM | ASSGTGGALEQY | 0.8376094 |
| GAAPLAAGL | ASSGTGGALEQY | 0.84652966 |
| GADGGDGGSGGAGG | ASSGTGGALEQY | 0.62894917 |
| GFGLDTGGAGG | ASSGTGGALEQY | 0.7301359 |
| GLASGGLL | ASSGTGGALEQY | 0.53156775 |
| GRIAGAAVLL | ASSGTGGALEQY | 0.5763366 |
| GSAGSLNTSF | ASSGTGGALEQY | 0.21673238 |
| GTGQIGI | ASSGTGGALEQY | 0.7750536 |
| GTLIALIL | ASSGTGGALEQY | 0.82809967 |
| IAATMGFVVG | ASSGTGGALEQY | 0.69758475 |
| IFAQALL | ASSGTGGALEQY | 0.44287467 |
| IGAAVALGF | ASSGTGGALEQY | 0.49588236 |
| IGLTPLAL | ASSGTGGALEQY | 0.61956924 |
| ILGSNTSTL | ASSGTGGALEQY | 0.7103408 |
| IVFSLTVL | ASSGTGGALEQY | 0.18711796 |
| LASKATGF | ASSGTGGALEQY | 0.558383 |
| LASVALIL | ASSGTGGALEQY | 0.83342254 |
| LFAVGLTL | ASSGTGGALEQY | 0.8280879 |
| LFNGGIGTYI | ASSGTGGALEQY | 0.37281913 |
| LTLGAGII | ASSGTGGALEQY | 0.84303105 |
| LVAAALGDI | ASSGTGGALEQY | 0.78177965 |
| NTAFVQLGI | ASSGTGGALEQY | 0.8300423 |
| QLFSGAEL | ASSGTGGALEQY | 0.6201759 |
| SAATFITL | ASSGTGGALEQY | 0.53232086 |
| SNYGLANLGV | ASSGTGGALEQY | 0.791098 |
| SYGGAYAVM | ASSGTGGALEQY | 0.69657975 |
| TTGVGGLTLG | ASSGTGGALEQY | 0.43459278 |
| TTQTIYI | ASSGTGGALEQY | 0.82260275 |
| VAAGAGLGAVI | ASSGTGGALEQY | 0.7028128 |
| VAAVVTLL | ASSGTGGALEQY | 0.6273144 |

| | | |
|---|---|---|
| VGFLGGLI | ASSGTGGALEQY | 0.64396006 |
| VGVIPILL | ASSGTGGALEQY | 0.8000243 |
| VIFASSLI | ASSGTGGALEQY | 0.50370204 |
| VKGGDGGAAAG | ASSGTGGALEQY | 0.5519485 |
| VTKTFTNI | ASSGTGGALEQY | 0.4416128 |
| VVAAAILL | ASSGTGGALEQY | 0.7563815 |
| YAALDVEL | ASSGTGGALEQY | 0.8335665 |
| LASVALLL | ASSGTGGALEQY | 0.7117636 |
| LDAVHRV | ASSGTGGALEQY | 0.5595925 |
| QGLSSSIF | ASSGTGGALEQY | 0.79665786 |
| RGTFANIRL | ASSGTGGALEQY | 0.75982857 |
| RQLWWGHRI | ASSGTGGALEQY | 0.7291137 |
| YGPPGTGKTLL | ASSGTGGALEQY | 0.8174089 |
| AALVGAPANVA | ASSGTGGALEQY | 0.6006425 |
| AATFRRTE | ASSGTGGALEQY | 0.788149 |
| AGALGGGI | ASSGTGGALEQY | 0.7733411 |
| AGGAGGAGGIGAG | ASSGTGGALEQY | 0.80371064 |
| AGSGALGL | ASSGTGGALEQY | 0.80063546 |
| AQIADQI | ASSGTGGALEQY | 0.7291726 |
| ASAVAVTYL | ASSGTGGALEQY | 0.67224914 |
| ASTVLGGTGG | ASSGTGGALEQY | 0.24393348 |
| AVGGEPAK | ASSGTGGALEQY | 0.57111835 |
| CEQGTFGGTC | ASSGTGGALEQY | 0.79893804 |
| DEGKAPAG | ASSGTGGALEQY | 0.759489 |
| DTVVFPAL | ASSGTGGALEQY | 0.8121377 |
| FEGITMAG | ASSGTGGALEQY | 0.711915 |
| FPLVVQD | ASSGTGGALEQY | 0.5241512 |
| GAFTTPQ | ASSGTGGALEQY | 0.827532 |
| GFGGFGTAGGVGGA | ASSGTGGALEQY | 0.81373954 |
| GFLAQFL | ASSGTGGALEQY | 0.3807707 |
| GGAGGAGGNAGVA | ASSGTGGALEQY | 0.6152458 |
| GGVGTAGAGGAGGAGG | ASSGTGGALEQY | 0.8535862 |
| GNGGGGDAGNAGSGGN | ASSGTGGALEQY | 0.7530389 |
| GTGGAGGAGNDAGS | ASSGTGGALEQY | 0.70308137 |
| GTTNTGWFNS | ASSGTGGALEQY | 0.60509753 |
| IFMGPLFTL | ASSGTGGALEQY | 0.8479096 |
| ILAHAIEAI | ASSGTGGALEQY | 0.49974716 |
| IRSAIREAL | ASSGTGGALEQY | 0.84364897 |
| ITFAGGG | ASSGTGGALEQY | 0.189582 |
| ITGMSGHQEV | ASSGTGGALEQY | 0.5383737 |
| IVAGLACTL | ASSGTGGALEQY | 0.39915153 |
| IVGGGLAGA | ASSGTGGALEQY | 0.32225847 |
| IVTGVLNI | ASSGTGGALEQY | 0.8235123 |
| LAQQGCI | ASSGTGGALEQY | 0.5637186 |
| LCLHVEN | ASSGTGGALEQY | 0.8254295 |
| LDAISDGS | ASSGTGGALEQY | 0.68859303 |
| LGGFAAML | ASSGTGGALEQY | 0.8131689 |
| LGGKNPMI | ASSGTGGALEQY | 0.7510174 |
| LGPKGVAAIL | ASSGTGGALEQY | 0.7669748 |
| LMVGVSI | ASSGTGGALEQY | 0.46802956 |
| LVGAAFVL | ASSGTGGALEQY | 0.8128763 |

| | | |
|---|---|---|
| LVLAVAPL | ASSGTGGALEQY | 0.83384836 |
| MAAIAVIDAA | ASSGTGGALEQY | 0.7087151 |
| NNGINAELQ | ASSGTGGALEQY | 0.3897935 |
| RGEPLVLVHG | ASSGTGGALEQY | 0.55789936 |
| RGGAGGAAGLI | ASSGTGGALEQY | 0.7604773 |
| RSYRMHPAV | ASSGTGGALEQY | 0.18252362 |
| SAAHLLAGQLA | ASSGTGGALEQY | 0.15365106 |
| SAAVGSY | ASSGTGGALEQY | 0.60528076 |
| SPHAVLHRI | ASSGTGGALEQY | 0.49782795 |
| SQIAKVL | ASSGTGGALEQY | 0.8421358 |
| TALGHLVP | ASSGTGGALEQY | 0.8076109 |
| TAPTTSGAA | ASSGTGGALEQY | 0.7418101 |
| TDGDDGNGGSPGT | ASSGTGGALEQY | 0.79223263 |
| TDGLGEN | ASSGTGGALEQY | 0.7415444 |
| TGGPPLPGQA | ASSGTGGALEQY | 0.103687435 |
| TIPAITI | ASSGTGGALEQY | 0.53573 |
| TIPGFTI | ASSGTGGALEQY | 0.4349223 |
| TSIASGALA | ASSGTGGALEQY | 0.83834326 |
| TVTLPGAAI | ASSGTGGALEQY | 0.53646755 |
| VASGLSAALP | ASSGTGGALEQY | 0.5511061 |
| VGAGVIGI | ASSGTGGALEQY | 0.7628225 |
| VGPKRFGV | ASSGTGGALEQY | 0.739685 |
| VNVAFPD | ASSGTGGALEQY | 0.69034624 |
| VSMTNSGVS | ASSGTGGALEQY | 0.46633002 |
| VTDVLGDLSV | ASSGTGGALEQY | 0.13010344 |
| VVVVGAGFAGL | ASSGTGGALEQY | 0.8265599 |
| VYTVGTVV | ASSGTGGALEQY | 0.2734375 |
| WTTFSGP | ASSGTGGALEQY | 0.48747095 |
| YDELVEAF | ASSGTGGALEQY | 0.18236129 |
| YGTGGAGGNGG | ASSGTGGALEQY | 0.24915199 |
| YTNVMAV | ASSGTGGALEQY | 0.2930251 |
| FLEQVHQL | ASSGTGGALEQY | 0.7963221 |
| GLSLLAVAV | ASSGTGGALEQY | 0.71959156 |
| LEQVHQL | ASSGTGGALEQY | 0.65103173 |
| LNAAGAI | ASSGTGGALEQY | 0.8420025 |
| PAVHASL | ASSGTGGALEQY | 0.7693573 |
| TELEDDA | ASSGTGGALEQY | 0.67017615 |
| TPPGPPN | ASSGTGGALEQY | 0.6143108 |
| TPPPPPPP | ASSGTGGALEQY | 0.5148947 |
| TQSPALI | ASTGGAYEQY | 0.52162325 |
| EFAAAFPQ | ASTGGAYEQY | 0.51625276 |
| LLSGNLDVL | ASTGGAYEQY | 0.86007845 |
| GPANLGNYNI | ASTGGAYEQY | 0.5755032 |
| TLGPTLP | ASTGGAYEQY | 0.37177548 |
| VYAYSSCCK | ASTGGAYEQY | 0.5715247 |
| APAPAPAPNI | ASTGGAYEQY | 0.84794796 |
| PGGPVPIP | ASTGGAYEQY | 0.86801326 |
| FLTVALEP | ASTGGAYEQY | 0.5795289 |
| SQYFGANL | ASTGGAYEQY | 0.570777 |
| GGVKPAA | ASTGGAYEQY | 0.58088064 |
| GGTGGAGGAGGW | ASTGGAYEQY | 0.731879 |

| | | |
|---|---|---|
| GVGGVAGLGGAGG | ASTGGAYEQY | 0.5577508 |
| VLLVLATTI | ASTGGAYEQY | 0.41838822 |
| IGIFAAML | ASTGGAYEQY | 0.75712836 |
| AVGDLID | ASTGGAYEQY | 0.8677021 |
| NISPLPL | ASTGGAYEQY | 0.5801561 |
| VTAVFVAI | ASTGGAYEQY | 0.292616 |
| SGDDNDAAL | ASTGGAYEQY | 0.7295112 |
| AVAALAHS | ASTGGAYEQY | 0.5671577 |
| GGVTVMATP | ASTGGAYEQY | 0.6838279 |
| AGLAANVRD | ASTGGAYEQY | 0.73428446 |
| TNRPDLIDPALLRPG | ASTGGAYEQY | 0.37295145 |
| LANLGVTAL | ASTGGAYEQY | 0.6062517 |
| TGERIEL | ASTGGAYEQY | 0.8524089 |
| NGEELAA | ASTGGAYEQY | 0.6886927 |
| VLSLVWG | ASTGGAYEQY | 0.7598046 |
| IGYALWL | ASTGGAYEQY | 0.8340144 |
| LSLLLAPL | ASTGGAYEQY | 0.79885423 |
| TLDEALK | ASTGGAYEQY | 0.8443576 |
| TDGAGTAEGG | ASTGGAYEQY | 0.7974756 |
| NAPGGAGGNGG | ASTGGAYEQY | 0.66362935 |
| VSVFMGMGGNF | ASTGGAYEQY | 0.6467807 |
| LASYLAAL | ASTGGAYEQY | 0.76069736 |
| LVGGTPAL | ASTGGAYEQY | 0.7561004 |
| VRNADVL | ASTGGAYEQY | 0.7118639 |
| TCSSPFLVSAM | ASTGGAYEQY | 0.465859 |
| VHITDAE | ASTGGAYEQY | 0.8396257 |
| EYAAPSF | ASTGGAYEQY | 0.63437206 |
| VVDAFHGGL | ASTGGAYEQY | 0.81959426 |
| ASVVDPQ | ASTGGAYEQY | 0.45504093 |
| SYNEFGGLS | ASTGGAYEQY | 0.48659393 |
| DPDGSVY | ASTGGAYEQY | 0.80934465 |
| MVSTIHATE | ASTGGAYEQY | 0.11919693 |
| LAAGVADPV | ASTGGAYEQY | 0.17432638 |
| TKEQIAA | ASTGGAYEQY | 0.6838855 |
| SLASPAAL | ASTGGAYEQY | 0.8453771 |
| IGLVFYGV | ASTGGAYEQY | 0.12059772 |
| FLLDASNL | ASTGGAYEQY | 0.45435387 |
| YQVGNFP | ASTGGAYEQY | 0.41558993 |
| ESLLSQST | ASTGGAYEQY | 0.72837275 |
| LFTVGFAAVG | ASTGGAYEQY | 0.7474899 |
| ATPPTPT | ASTGGAYEQY | 0.35307404 |
| TDLATLG | ASTGGAYEQY | 0.80878794 |
| SAGGTGR | ASTGGAYEQY | 0.28465074 |
| GFSNSGS | ASTGGAYEQY | 0.33310127 |
| GGNGGSGGTSV | ASTGGAYEQY | 0.7999499 |
| GTSVATGGAGN | ASTGGAYEQY | 0.82213086 |
| NGGNGGSGGTS | ASTGGAYEQY | 0.40126213 |
| AVVKFPH | ASTGGAYEQY | 0.8173224 |
| TPPPPPP | ASTGGAYEQY | 0.50508964 |
| AAPAAAAAPA | ASTGGAYEQY | 0.86727786 |
| IALNEDL | ASTGGAYEQY | 0.6542375 |

| | | |
|---|---|---|
| IVNVSSM | ASTGGAYEQY | 0.2890709 |
| TGIPPAPR | ASTGGAYEQY | 0.798035 |
| FDLGGGTF | ASTGGAYEQY | 0.791325 |
| FDLGGGTFDVS | ASTGGAYEQY | 0.7357513 |
| NEPTAAA | ASTGGAYEQY | 0.78997016 |
| NEPTAAALA | ASTGGAYEQY | 0.4815522 |
| GSIINISSI | ASTGGAYEQY | 0.7216028 |
| VAPGNAGTA | ASTGGAYEQY | 0.8421377 |
| TGGGASLE | ASTGGAYEQY | 0.8257125 |
| VAVYNHYKRI | ASTGGAYEQY | 0.4867829 |
| TDPAPATT | ASTGGAYEQY | 0.7934489 |
| TNRPDILDPALLRPG | ASTGGAYEQY | 0.7313953 |
| LTTTTTS | ASTGGAYEQY | 0.5702582 |
| PTGSGKTL | ASTGGAYEQY | 0.8386446 |
| TGSGKTL | ASTGGAYEQY | 0.6809894 |
| TIGHVDHGK | ASTGGAYEQY | 0.81460166 |
| TIGHVDHGKTT | ASTGGAYEQY | 0.809116 |
| GHVDHGKTT | ASTGGAYEQY | 0.8652543 |
| AQFEHTLL | ASTGGAYEQY | 0.37276074 |
| IVNQDEL | ASTGGAYEQY | 0.7668244 |
| TNGSQFF | ASTGGAYEQY | 0.71245354 |
| TGKGIVE | ASTGGAYEQY | 0.7998198 |
| LISNASDA | ASTGGAYEQY | 0.6391469 |
| AELRTLGELTKIAKAHGAQ | ASTGGAYEQY | 0.33275682 |
| APNAVAAAEPSW | ASTGGAYEQY | 0.75244033 |
| APQVAGAATAVL | ASTGGAYEQY | 0.8310635 |
| ARRDPAAAAAILPTDA | ASTGGAYEQY | 0.69731534 |
| DAAARDVLAEAGLAEYF | ASTGGAYEQY | 0.7547189 |
| DDQIVLSAKANLAKVEQ | ASTGGAYEQY | 0.85562485 |
| DELLPDTGEDSDEDGH | ASTGGAYEQY | 0.6013772 |
| ELPEAPANRATVPQD | ASTGGAYEQY | 0.7204422 |
| FGQSKIERVTGIG | ASTGGAYEQY | 0.76648825 |
| GAGGTGSTAGGAGGAGGAGGLY | ASTGGAYEQY | 0.8271587 |
| GHGGTGAAGGNGGTGGAGGSGI | ASTGGAYEQY | 0.63766503 |
| GNGGNGGAGGEGGPGGAGGAGA | ASTGGAYEQY | 0.8391963 |
| GSTAGNGGNGGNATNGTIAGQP | ASTGGAYEQY | 0.4388837 |
| GTGGDGGNGGNGGTGG | ASTGGAYEQY | 0.81816757 |
| GVRRPAGIPADAPV | ASTGGAYEQY | 0.46167636 |
| GYRQVTDAHLLTLARR | ASTGGAYEQY | 0.7871575 |
| IAKMGQDGHDRG | ASTGGAYEQY | 0.5945511 |
| KAEPSIEIQYT | ASTGGAYEQY | 0.8059411 |
| KPVRIRMAADEDTVAVT | ASTGGAYEQY | 0.8116454 |
| MPPRIAGMRLLVIKPEP | ASTGGAYEQY | 0.82705235 |
| MKFYAVALFRSRGVPPTR | ASTGGAYEQY | 0.1479374 |
| NAGGFGILGAVAHSP | ASTGGAYEQY | 0.8180175 |
| NEGREITLAMAEM | ASTGGAYEQY | 0.82069004 |
| NELVEWSHPLGHRWRWE | ASTGGAYEQY | 0.86054116 |
| NLGAGGTGGNGGNPG | ASTGGAYEQY | 0.6621825 |
| NTEWPKWETVMAADPVD | ASTGGAYEQY | 0.75500685 |
| PASAAPAETQP | ASTGGAYEQY | 0.66985106 |
| QQSSDGKNGVRAAEITGE | ASTGGAYEQY | 0.6334486 |

| | | |
|---|---|---|
| RAPDAVTTSWWKE | ASTGGAYEQY | 0.53474796 |
| SGFGNVGAHTSGWFNQ | ASTGGAYEQY | 0.6940898 |
| SGVAGAVGQAASVGGLKV | ASTGGAYEQY | 0.6595666 |
| SNGGAGTGSAGGNGGT | ASTGGAYEQY | 0.64698386 |
| TGGAGGEGGAAG | ASTGGAYEQY | 0.799794 |
| TGGDGGAAGAGGAPGHG | ASTGGAYEQY | 0.8306373 |
| TMPPLLAAIANAGRHGV | ASTGGAYEQY | 0.80300426 |
| TSSTAGNSGGA | ASTGGAYEQY | 0.82840335 |
| VPATVPVGILC | ASTGGAYEQY | 0.84561205 |
| VQRPDSGGVVLTGRIS | ASTGGAYEQY | 0.8135367 |
| WTPNPVPATKGVPAGLVQAT | ASTGGAYEQY | 0.6455461 |
| YALKRALAKFSDRFDVVII | ASTGGAYEQY | 0.6686342 |
| YLIGSVIV | ASTGGAYEQY | 0.4911151 |
| YGQLVAVID | ASTGGAYEQY | 0.5914944 |
| VTVGGMTLPA | ASTGGAYEQY | 0.64768505 |
| VLALPAV | ASTGGAYEQY | 0.76695174 |
| VIPPTLNYET | ASTGGAYEQY | 0.12575172 |
| VGMGGMHPSSG | ASTGGAYEQY | 0.6863778 |
| VGLAGGAATA | ASTGGAYEQY | 0.8566711 |
| VGGAGGAGGSGP | ASTGGAYEQY | 0.6031353 |
| VFAAFMV | ASTGGAYEQY | 0.7726142 |
| TLYDKLR | ASTGGAYEQY | 0.7125468 |
| TLAVFTVL | ASTGGAYEQY | 0.5597793 |
| TGSVQPARV | ASTGGAYEQY | 0.80931306 |
| TGGGAGSGGT | ASTGGAYEQY | 0.5031953 |
| SHLETSL | ASTGGAYEQY | 0.78061974 |
| SFTTGMT | ASTGGAYEQY | 0.84564877 |
| QVEVAKRMGVRQPTVS | ASTGGAYEQY | 0.7253414 |
| QGSGGPALAM | ASTGGAYEQY | 0.7318289 |
| MLAPSNP | ASTGGAYEQY | 0.67389107 |
| LVNNAAI | ASTGGAYEQY | 0.3236022 |
| LTGEINTLA | ASTGGAYEQY | 0.7664362 |
| LSIFGPV | ASTGGAYEQY | 0.19192341 |
| LLALLANL | ASTGGAYEQY | 0.7116767 |
| LISNASDALDKLR | ASTGGAYEQY | 0.46703944 |
| LINEAVP | ASTGGAYEQY | 0.7808496 |
| LGYAFSGL | ASTGGAYEQY | 0.74874234 |
| LDFVAVAL | ASTGGAYEQY | 0.7436431 |
| LAQPGAE | ASTGGAYEQY | 0.5685748 |
| ISNLGAVTGF | ASTGGAYEQY | 0.6525152 |
| HQEDPGRI | ASTGGAYEQY | 0.51513064 |
| GTPGEPTP | ASTGGAYEQY | 0.38436615 |
| GFFNSGS | ASTGGAYEQY | 0.80011284 |
| GAGGGGLTAGAN | ASTGGAYEQY | 0.57061064 |
| FVDALVNM | ASTGGAYEQY | 0.6143221 |
| FLTVAPEL | ASTGGAYEQY | 0.8300836 |
| FETLTAL | ASTGGAYEQY | 0.7502285 |
| DVINPIEL | ASTGGAYEQY | 0.2975695 |
| APVSGAPV | ASTGGAYEQY | 0.723497 |
| ALVGGLAW | ASTGGAYEQY | 0.71457195 |
| ALAAAAATAI | ASTGGAYEQY | 0.71945536 |

| | | |
|---|---|---|
| AFGLGLLL | ASTGGAYEQY | 0.11496531 |
| AFFAAASL | ASTGGAYEQY | 0.72202325 |
| AAAALPAFSPP | ASTGGAYEQY | 0.57402897 |
| VVLAPAL | ASTGGAYEQY | 0.5216489 |
| TIGHVDHGKTTL | ASTGGAYEQY | 0.5912571 |
| LISNASDALDK | ASTGGAYEQY | 0.82750934 |
| LISNASDAL | ASTGGAYEQY | 0.52930886 |
| ISNASDALDK | ASTGGAYEQY | 0.7703877 |
| NQAETLVYQTE | ASTGGAYEQY | 0.8089924 |
| TGGTGGTAGAG | ASTGGAYEQY | 0.79345405 |
| TGSTAGGAGGA | ASTGGAYEQY | 0.853807 |
| DPLAVPVAAGPV | ASTGGAYEQY | 0.794389 |
| GGTGGTPGNGGN | ASTGGAYEQY | 0.56512463 |
| PAGLAGVDPATL | ASTGGAYEQY | 0.73360336 |
| GNNLVVEAPHAGASV | ASTGGAYEQY | 0.6111492 |
| KVTRIMSASTLEQPA | ASTGGAYEQY | 0.19624016 |
| LGFALPIALPRRTVV | ASTGGAYEQY | 0.18804966 |
| TRLGNLTARKLVVLV | ASTGGAYEQY | 0.76174176 |
| WSQQRRLIYFPSAGP | ASTGGAYEQY | 0.21102332 |
| GPNGNPDPMAAAVDIR | ASTGGAYEQY | 0.7470498 |
| TQRCVSVADLRAAASR | ASTGGAYEQY | 0.81325257 |
| QFVLDVAEHPVEVLTETV | ASTGGAYEQY | 0.5766009 |
| GGAGGTGGAGGNGGTQAGDG | ASTGGAYEQY | 0.35628298 |
| PLLATLRAIAVDVDAKPAQV | ASTGGAYEQY | 0.5771029 |
| APKFRTGGYMIDFWGVGYQVAK | ASTGGAYEQY | 0.6434915 |
| GAGGAGGAGGAAGGPGGTGGTG | ASTGGAYEQY | 0.82465994 |
| IGALGLLATDVAPAGSVAFGVA | ASTGGAYEQY | 0.74726117 |
| AGGAGGIGGNGGAGGAGGVLMG | ASTGGAYEQY | 0.8416456 |
| ADELIGLL | ASTGGAYEQY | 0.8192922 |
| ADRALKA | ASTGGAYEQY | 0.6819416 |
| AGAVGQVGGAGGA | ASTGGAYEQY | 0.8094133 |
| AGGGGGLGGAP | ASTGGAYEQY | 0.6037092 |
| AGGLLYGNGGAG | ASTGGAYEQY | 0.6429002 |
| AGLESIGL | ASTGGAYEQY | 0.6217882 |
| AGLKTGII | ASTGGAYEQY | 0.70561767 |
| ASLAALLL | ASTGGAYEQY | 0.7200897 |
| ASVINISSI | ASTGGAYEQY | 0.60141516 |
| ATPPGPA | ASTGGAYEQY | 0.3069535 |
| ATVASALV | ASTGGAYEQY | 0.7707003 |
| ATVASLIL | ASTGGAYEQY | 0.7913183 |
| AVNGPVLIRH | ASTGGAYEQY | 0.20466885 |
| DGAAGAVGI | ASTGGAYEQY | 0.5525947 |
| DPLLHGI | ASTGGAYEQY | 0.8644974 |
| ELGAALGAL | ASTGGAYEQY | 0.8402777 |
| ELGRTQAV | ASTGGAYEQY | 0.54702216 |
| FVAGGYLS | ASTGGAYEQY | 0.5732184 |
| GASGGAGGAGGW | ASTGGAYEQY | 0.6768104 |
| GGAGAGGDGGAGGI | ASTGGAYEQY | 0.2966673 |
| GTGHNGSQPGMGGNG | ASTGGAYEQY | 0.7181392 |
| GYAAASASAS | ASTGGAYEQY | 0.71130633 |
| HQVDVAAVQQG | ASTGGAYEQY | 0.7152591 |

| | | |
|---|---|---|
| IGAAIAHT | ASTGGAYEQY | 0.47810006 |
| IGMIEVAQL | ASTGGAYEQY | 0.71636206 |
| IRLAPEHTSAMD | ASTGGAYEQY | 0.6408521 |
| IYNVSVL | ASTGGAYEQY | 0.7765598 |
| KQGFLGM | ASTGGAYEQY | 0.12170998 |
| LESVITQL | ASTGGAYEQY | 0.72748566 |
| LGLGALVVV | ASTGGAYEQY | 0.5745357 |
| LIGNGGNGGSGGI | ASTGGAYEQY | 0.47114316 |
| LNAINAPTL | ASTGGAYEQY | 0.2755338 |
| LQLLNTGV | ASTGGAYEQY | 0.671317 |
| LQTSIEF | ASTGGAYEQY | 0.8175584 |
| MSFAVSEL | ASTGGAYEQY | 0.8101531 |
| NTGALGL | ASTGGAYEQY | 0.31830305 |
| PPGAGGAGGAGTT | ASTGGAYEQY | 0.8264215 |
| QLVGLGI | ASTGGAYEQY | 0.8604188 |
| RFLEQVHQL | ASTGGAYEQY | 0.6220341 |
| RGVGYVLR | ASTGGAYEQY | 0.28876615 |
| SATGVFAGVI | ASTGGAYEQY | 0.79822737 |
| SLPAITLP | ASTGGAYEQY | 0.5998678 |
| STALAFTV | ASTGGAYEQY | 0.76311743 |
| SVLTVGLGK | ASTGGAYEQY | 0.7986377 |
| TALAGAGSGI | ASTGGAYEQY | 0.5081665 |
| TGLPFTGI | ASTGGAYEQY | 0.81248677 |
| VAEHVTV | ASTGGAYEQY | 0.81018984 |
| VDIVLNSL | ASTGGAYEQY | 0.533722 |
| VLTGGGAVL | ASTGGAYEQY | 0.47060025 |
| VNVGLAYGI | ASTGGAYEQY | 0.47499645 |
| VSSAAVGAV | ASTGGAYEQY | 0.18544321 |
| VTSVYEAGV | ASTGGAYEQY | 0.79153717 |
| WVALDRGA | ASTGGAYEQY | 0.62641186 |
| YLVGSVVI | ASTGGAYEQY | 0.5931098 |
| GPPGTGKTLL | ASTGGAYEQY | 0.77647924 |
| GVGMTKF | ASTGGAYEQY | 0.5339999 |
| IGIGGSDL | ASTGGAYEQY | 0.6922147 |
| IGTLLAEL | ASTGGAYEQY | 0.64605904 |
| IYTSGTT | ASTGGAYEQY | 0.74160016 |
| LSFPAGV | ASTGGAYEQY | 0.7474809 |
| LYTSGTT | ASTGGAYEQY | 0.6983143 |
| VGAFPFGF | ASTGGAYEQY | 0.6949849 |
| VGAGTVEF | ASTGGAYEQY | 0.8489957 |
| VGKSTLLL | ASTGGAYEQY | 0.42947507 |
| AALAHAGGGIV | ASTGGAYEQY | 0.57958853 |
| AALVAVSLL | ASTGGAYEQY | 0.76901364 |
| AAVSRAVAAL | ASTGGAYEQY | 0.32512677 |
| AGGSSGIGGF | ASTGGAYEQY | 0.7913984 |
| AGLTVANANG | ASTGGAYEQY | 0.7317374 |
| AGLVGAVL | ASTGGAYEQY | 0.78302705 |
| ALSQAFTV | ASTGGAYEQY | 0.5849184 |
| ALTFSVSL | ASTGGAYEQY | 0.8592557 |
| ALTGAALVAL | ASTGGAYEQY | 0.8186577 |
| ASFSIGAGVL | ASTGGAYEQY | 0.6409606 |

| | | |
|---|---|---|
| DGSVYKI | ASTGGAYEQY | 0.7759737 |
| EQLAEIV | ASTGGAYEQY | 0.42395994 |
| FAVSELTVM | ASTGGAYEQY | 0.8231801 |
| GAAPLAAGL | ASTGGAYEQY | 0.783142 |
| GADGGDGGSGGAGG | ASTGGAYEQY | 0.5323031 |
| GFGLDTGGAGG | ASTGGAYEQY | 0.7264675 |
| GLASGGLL | ASTGGAYEQY | 0.8217669 |
| GRIAGAAVLL | ASTGGAYEQY | 0.83934045 |
| GSAGSLNTSF | ASTGGAYEQY | 0.25144428 |
| GTGQIGI | ASTGGAYEQY | 0.6899841 |
| GTLIALIL | ASTGGAYEQY | 0.23872189 |
| IAATMGFVVG | ASTGGAYEQY | 0.45678043 |
| IFAQALL | ASTGGAYEQY | 0.48529857 |
| IGAAVALGF | ASTGGAYEQY | 0.7825178 |
| IGLTPLAL | ASTGGAYEQY | 0.67007965 |
| ILGSNTSTL | ASTGGAYEQY | 0.78352314 |
| IVFSLTVL | ASTGGAYEQY | 0.7548628 |
| LASKATGF | ASTGGAYEQY | 0.8450106 |
| LASVALIL | ASTGGAYEQY | 0.57563233 |
| LFAVGLTL | ASTGGAYEQY | 0.7190047 |
| LFNGGIGTYI | ASTGGAYEQY | 0.3431364 |
| LTLGAGII | ASTGGAYEQY | 0.34214705 |
| LVAAALGDI | ASTGGAYEQY | 0.45216066 |
| NTAFVQLGI | ASTGGAYEQY | 0.673743 |
| QLFSGAEL | ASTGGAYEQY | 0.17493348 |
| SAATFITL | ASTGGAYEQY | 0.8303636 |
| SNYGLANLGV | ASTGGAYEQY | 0.76339686 |
| SYGGAYAVM | ASTGGAYEQY | 0.8217943 |
| TTGVGGLTLG | ASTGGAYEQY | 0.5490834 |
| TTQTIYI | ASTGGAYEQY | 0.67355406 |
| VAAGAGLGAVI | ASTGGAYEQY | 0.627964 |
| VAAVVTLL | ASTGGAYEQY | 0.79797846 |
| VGFLGGLI | ASTGGAYEQY | 0.539618 |
| VGVIPILL | ASTGGAYEQY | 0.27987403 |
| VIFASSLI | ASTGGAYEQY | 0.4998502 |
| VKGGDGGAAAG | ASTGGAYEQY | 0.33902222 |
| VTKTFTNI | ASTGGAYEQY | 0.8331599 |
| VVAAAILL | ASTGGAYEQY | 0.8267875 |
| YAALDVEL | ASTGGAYEQY | 0.7653637 |
| LASVALLL | ASTGGAYEQY | 0.81069136 |
| LDAVHRV | ASTGGAYEQY | 0.18717456 |
| QGLSSSIF | ASTGGAYEQY | 0.5907658 |
| RGTFANIRL | ASTGGAYEQY | 0.8568864 |
| RQLWWGHRI | ASTGGAYEQY | 0.7505584 |
| YGPPGTGKTLL | ASTGGAYEQY | 0.71031284 |
| AALVGAPANVA | ASTGGAYEQY | 0.8438735 |
| AATFRRTE | ASTGGAYEQY | 0.62678444 |
| AGALGGGI | ASTGGAYEQY | 0.7175738 |
| AGGAGGAGGIGAG | ASTGGAYEQY | 0.72696894 |
| AGSGALGL | ASTGGAYEQY | 0.7210498 |
| AQIADQI | ASTGGAYEQY | 0.6598319 |

| | | |
|---|---|---|
| ASAVAVTYL | ASTGGAYEQY | 0.47192988 |
| ASTVLGGTGG | ASTGGAYEQY | 0.73753875 |
| AVGGEPAK | ASTGGAYEQY | 0.85860807 |
| CEQGTFGGTC | ASTGGAYEQY | 0.71747464 |
| DEGKAPAG | ASTGGAYEQY | 0.8308948 |
| DTVVFPAL | ASTGGAYEQY | 0.79077697 |
| FEGITMAG | ASTGGAYEQY | 0.81266284 |
| FPLVVQD | ASTGGAYEQY | 0.5938914 |
| GAFTTPQ | ASTGGAYEQY | 0.6468834 |
| GFGGFGTAGGVGGA | ASTGGAYEQY | 0.64274335 |
| GFLAQFL | ASTGGAYEQY | 0.57384 |
| GGAGGAGGNAGVA | ASTGGAYEQY | 0.6609731 |
| GGVGTAGAGGAGGAGG | ASTGGAYEQY | 0.6015924 |
| GNGGGGDAGNAGSGGN | ASTGGAYEQY | 0.87834775 |
| GTGGAGGAGNDAGS | ASTGGAYEQY | 0.796512 |
| GTTNTGWFNS | ASTGGAYEQY | 0.7859405 |
| IFMGPLFTL | ASTGGAYEQY | 0.57408726 |
| ILAHAIEAI | ASTGGAYEQY | 0.69109225 |
| IRSAIREAL | ASTGGAYEQY | 0.74912447 |
| ITFAGGG | ASTGGAYEQY | 0.34203503 |
| ITGMSGHQEV | ASTGGAYEQY | 0.84415704 |
| IVAGLACTL | ASTGGAYEQY | 0.63217545 |
| IVGGGLAGA | ASTGGAYEQY | 0.5515379 |
| IVTGVLNI | ASTGGAYEQY | 0.80873245 |
| LAQQGCI | ASTGGAYEQY | 0.84141487 |
| LCLHVEN | ASTGGAYEQY | 0.63392603 |
| LDAISDGS | ASTGGAYEQY | 0.60114753 |
| LGGFAAML | ASTGGAYEQY | 0.6303873 |
| LGGKNPMI | ASTGGAYEQY | 0.79597634 |
| LGPKGVAAIL | ASTGGAYEQY | 0.8444458 |
| LMVGVSI | ASTGGAYEQY | 0.69133204 |
| LVGAAFVL | ASTGGAYEQY | 0.84958655 |
| LVLAVAPL | ASTGGAYEQY | 0.66436964 |
| MAAIAVIDAA | ASTGGAYEQY | 0.35239542 |
| NNGINAELQ | ASTGGAYEQY | 0.71999705 |
| RGEPLVLVHG | ASTGGAYEQY | 0.80524665 |
| RGGAGGAAGLI | ASTGGAYEQY | 0.8152636 |
| RSYRMHPAV | ASTGGAYEQY | 0.86339253 |
| SAAHLLAGQLA | ASTGGAYEQY | 0.78789496 |
| SAAVGSY | ASTGGAYEQY | 0.559225 |
| SPHAVLHRI | ASTGGAYEQY | 0.84101075 |
| SQIAKVL | ASTGGAYEQY | 0.84768116 |
| TALGHLVP | ASTGGAYEQY | 0.8167418 |
| TAPTTSGAA | ASTGGAYEQY | 0.63997984 |
| TDGDDGNGGSPGT | ASTGGAYEQY | 0.47869688 |
| TDGLGEN | ASTGGAYEQY | 0.6932835 |
| TGGPPLPGQA | ASTGGAYEQY | 0.80714405 |
| TIPAITI | ASTGGAYEQY | 0.47219834 |
| TIPGFTI | ASTGGAYEQY | 0.75676394 |
| TSIASGALA | ASTGGAYEQY | 0.78303105 |
| TVTLPGAAI | ASTGGAYEQY | 0.38529333 |

| | | |
|---|---|---|
| VASGLSAALP | ASTGGAYEQY | 0.7035645 |
| VGAGVIGI | ASTGGAYEQY | 0.8389197 |
| VGPKRFGV | ASTGGAYEQY | 0.63621366 |
| VNVAFPD | ASTGGAYEQY | 0.58737653 |
| VSMTNSGVS | ASTGGAYEQY | 0.8241478 |
| VTDVLGDLSV | ASTGGAYEQY | 0.87587994 |
| VVVVGAGFAGL | ASTGGAYEQY | 0.8313855 |
| VYTVGTVV | ASTGGAYEQY | 0.35876167 |
| WTTFSGP | ASTGGAYEQY | 0.19899036 |
| YDELVEAF | ASTGGAYEQY | 0.61331606 |
| YGTGGAGGNGG | ASTGGAYEQY | 0.78395104 |
| YTNVMAV | ASTGGAYEQY | 0.63356173 |
| FLEQVHQL | ASTGGAYEQY | 0.6323964 |
| GLSLLAVAV | ASTGGAYEQY | 0.7822061 |
| LEQVHQL | ASTGGAYEQY | 0.83460546 |
| LNAAGAI | ASTGGAYEQY | 0.5407127 |
| PAVHASL | ASTGGAYEQY | 0.36557645 |
| TELEDDA | ASTGGAYEQY | 0.73363143 |
| TPPGPPN | ASTGGAYEQY | 0.79463613 |
| TPPPPPPP | ASTGGAYEQY | 0.5860907 |
| TQSPALI | ASSLGGSQNTLY | 0.7262815 |
| EFAAAFPQ | ASSLGGSQNTLY | 0.55900323 |
| LLSGNLDVL | ASSLGGSQNTLY | 0.30131355 |
| GPANLGNYNI | ASSLGGSQNTLY | 0.7477307 |
| TLGPTLP | ASSLGGSQNTLY | 0.51560485 |
| VYAYSSCCK | ASSLGGSQNTLY | 0.81182057 |
| APAPAPAPNI | ASSLGGSQNTLY | 0.5388903 |
| PGGPVPIP | ASSLGGSQNTLY | 0.77802366 |
| FLTVALEP | ASSLGGSQNTLY | 0.80330336 |
| SQYFGANL | ASSLGGSQNTLY | 0.76233906 |
| GGVKPAA | ASSLGGSQNTLY | 0.689727 |
| GGTGGAGGAGGW | ASSLGGSQNTLY | 0.62104654 |
| GVGGVAGLGGAGG | ASSLGGSQNTLY | 0.43164638 |
| VLLVLATTI | ASSLGGSQNTLY | 0.21625136 |
| IGIFAAML | ASSLGGSQNTLY | 0.8316721 |
| AVGDLID | ASSLGGSQNTLY | 0.32568568 |
| NISPLPL | ASSLGGSQNTLY | 0.8424109 |
| VTAVFVAI | ASSLGGSQNTLY | 0.8120803 |
| SGDDNDAAL | ASSLGGSQNTLY | 0.5711196 |
| AVAALAHS | ASSLGGSQNTLY | 0.7581091 |
| GGVTVMATP | ASSLGGSQNTLY | 0.5682163 |
| AGLAANVRD | ASSLGGSQNTLY | 0.37121546 |
| TNRPDLIDPALLRPG | ASSLGGSQNTLY | 0.80713093 |
| LANLGVTAL | ASSLGGSQNTLY | 0.7975745 |
| TGERIEL | ASSLGGSQNTLY | 0.53799117 |
| NGEELAA | ASSLGGSQNTLY | 0.7486238 |
| VLSLVWG | ASSLGGSQNTLY | 0.7288471 |
| IGYALWL | ASSLGGSQNTLY | 0.45981878 |
| LSLLLAPL | ASSLGGSQNTLY | 0.64798546 |
| TLDEALK | ASSLGGSQNTLY | 0.8140434 |
| TDGAGTAEGG | ASSLGGSQNTLY | 0.7160094 |

| | | |
|---|---|---|
| NAPGGAGGNGG | ASSLGGSQNTLY | 0.5641772 |
| VSVFMGMGGNF | ASSLGGSQNTLY | 0.75175273 |
| LASYLAAL | ASSLGGSQNTLY | 0.5373075 |
| LVGGTPAL | ASSLGGSQNTLY | 0.78420126 |
| VRNADVL | ASSLGGSQNTLY | 0.6987986 |
| TCSSPFLVSAM | ASSLGGSQNTLY | 0.67422783 |
| VHITDAE | ASSLGGSQNTLY | 0.78043437 |
| EYAAPSF | ASSLGGSQNTLY | 0.34539634 |
| VVDAFHGGL | ASSLGGSQNTLY | 0.44791186 |
| ASVVDPQ | ASSLGGSQNTLY | 0.6332991 |
| SYNEFGGLS | ASSLGGSQNTLY | 0.74609727 |
| DPDGSVY | ASSLGGSQNTLY | 0.8159133 |
| MVSTIHATE | ASSLGGSQNTLY | 0.55622536 |
| LAAGVADPV | ASSLGGSQNTLY | 0.7087952 |
| TKEQIAA | ASSLGGSQNTLY | 0.65164196 |
| SLASPAAL | ASSLGGSQNTLY | 0.5295896 |
| IGLVFYGV | ASSLGGSQNTLY | 0.6089593 |
| FLLDASNL | ASSLGGSQNTLY | 0.550225 |
| YQVGNFP | ASSLGGSQNTLY | 0.5481682 |
| ESLLSQST | ASSLGGSQNTLY | 0.746889 |
| LFTVGFAAVG | ASSLGGSQNTLY | 0.55510664 |
| ATPPTPT | ASSLGGSQNTLY | 0.6905855 |
| TDLATLG | ASSLGGSQNTLY | 0.7440635 |
| SAGGTGR | ASSLGGSQNTLY | 0.6656157 |
| GFSNSGS | ASSLGGSQNTLY | 0.7556433 |
| GGNGGSGGTSV | ASSLGGSQNTLY | 0.75328904 |
| GTSVATGGAGN | ASSLGGSQNTLY | 0.7983464 |
| NGGNGGSGGTS | ASSLGGSQNTLY | 0.6737362 |
| AVVKFPH | ASSLGGSQNTLY | 0.5933769 |
| TPPPPPP | ASSLGGSQNTLY | 0.82903624 |
| AAPAAAAAPA | ASSLGGSQNTLY | 0.7990775 |
| IALNEDL | ASSLGGSQNTLY | 0.8204633 |
| IVNVSSM | ASSLGGSQNTLY | 0.76984024 |
| TGIPPAPR | ASSLGGSQNTLY | 0.74878156 |
| FDLGGGTF | ASSLGGSQNTLY | 0.80480766 |
| FDLGGGTFDVS | ASSLGGSQNTLY | 0.6623064 |
| NEPTAAA | ASSLGGSQNTLY | 0.3857755 |
| NEPTAAALA | ASSLGGSQNTLY | 0.66435015 |
| GSIINISSI | ASSLGGSQNTLY | 0.7235192 |
| VAPGNAGTA | ASSLGGSQNTLY | 0.5770175 |
| TGGGASLE | ASSLGGSQNTLY | 0.27688426 |
| VAVYNHYKRI | ASSLGGSQNTLY | 0.8381853 |
| TDPAPATT | ASSLGGSQNTLY | 0.87525666 |
| TNRPDILDPALLRPG | ASSLGGSQNTLY | 0.50964487 |
| LTTTTTS | ASSLGGSQNTLY | 0.8627981 |
| PTGSGKTL | ASSLGGSQNTLY | 0.31409085 |
| TGSGKTL | ASSLGGSQNTLY | 0.7857885 |
| TIGHVDHGK | ASSLGGSQNTLY | 0.8388653 |
| TIGHVDHGKTT | ASSLGGSQNTLY | 0.56270975 |
| GHVDHGKTT | ASSLGGSQNTLY | 0.8104449 |
| AQFEHTLL | ASSLGGSQNTLY | 0.52419305 |

| | | |
|---|---|---|
| IVNQDEL | ASSLGGSQNTLY | 0.6441766 |
| TNGSQFF | ASSLGGSQNTLY | 0.84019613 |
| TGKGIVE | ASSLGGSQNTLY | 0.65903354 |
| LISNASDA | ASSLGGSQNTLY | 0.7188219 |
| AELRTLGELTKIAKAHGAQ | ASSLGGSQNTLY | 0.743294 |
| APNAVAAAEPSW | ASSLGGSQNTLY | 0.5960772 |
| APQVAGAATAVL | ASSLGGSQNTLY | 0.72180057 |
| ARRDPAAAAAILPTDA | ASSLGGSQNTLY | 0.8451994 |
| DAAARDVLAEAGLAEYF | ASSLGGSQNTLY | 0.4116646 |
| DDQIVLSAKANLAKVEQ | ASSLGGSQNTLY | 0.839701 |
| DELLPDTGEDSDEDGH | ASSLGGSQNTLY | 0.47932112 |
| ELPEAPANRATVPQD | ASSLGGSQNTLY | 0.86680615 |
| FGQSKIERVTGIG | ASSLGGSQNTLY | 0.76331174 |
| GAGGTGSTAGGAGGAGGAGGLY | ASSLGGSQNTLY | 0.7493672 |
| GHGGTGAAGGNGGTGGAGGSGI | ASSLGGSQNTLY | 0.6043512 |
| GNGGNGGAGGEGGPGGAGGAGA | ASSLGGSQNTLY | 0.8410932 |
| GSTAGNGGNGGNATNGTIAGQP | ASSLGGSQNTLY | 0.7364942 |
| GTGGDGGNGGNGGTGG | ASSLGGSQNTLY | 0.79500955 |
| GVRRPAGIPADAPV | ASSLGGSQNTLY | 0.8235311 |
| GYRQVTDAHLLTLARR | ASSLGGSQNTLY | 0.12848319 |
| IAKMGQDGHDRG | ASSLGGSQNTLY | 0.3488812 |
| KAEPSIEIQYT | ASSLGGSQNTLY | 0.15249327 |
| KPVRIRMAADEDTVAVT | ASSLGGSQNTLY | 0.17410721 |
| MPPRIAGMRLLVIKPEP | ASSLGGSQNTLY | 0.14429787 |
| MKFYAVALFRSRGVPPTR | ASSLGGSQNTLY | 0.123976484 |
| NAGGFGILGAVAHSP | ASSLGGSQNTLY | 0.13540766 |
| NEGREITLAMAEM | ASSLGGSQNTLY | 0.1372863 |
| NELVEWSHPLGHRWRWE | ASSLGGSQNTLY | 0.20708743 |
| NLGAGGTGGNGGNPG | ASSLGGSQNTLY | 0.15739681 |
| NTEWPKWETVMAADPVD | ASSLGGSQNTLY | 0.14702922 |
| PASAAPAETQP | ASSLGGSQNTLY | 0.36176994 |
| QQSSDGKNGVRAAEITGE | ASSLGGSQNTLY | 0.45285222 |
| RAPDAVTTSWWKE | ASSLGGSQNTLY | 0.12321333 |
| SGFGNVGAHTSGWFNQ | ASSLGGSQNTLY | 0.1442275 |
| SGVAGAVGQAASVGGLKV | ASSLGGSQNTLY | 0.13055252 |
| SNGGAGTGSAGGNGGT | ASSLGGSQNTLY | 0.14792445 |
| TGGAGGEGGAAG | ASSLGGSQNTLY | 0.11791953 |
| TGGDGGAAGAGGAPGHG | ASSLGGSQNTLY | 0.17306437 |
| TMPPLLAAIANAGRHGV | ASSLGGSQNTLY | 0.12389421 |
| TSSTAGNSGGA | ASSLGGSQNTLY | 0.11702884 |
| VPATVPVGILC | ASSLGGSQNTLY | 0.1525121 |
| VQRPDSGGVVLTGRIS | ASSLGGSQNTLY | 0.14949052 |
| WTPNPVPATKGVPAGLVQAT | ASSLGGSQNTLY | 0.1429102 |
| YALKRALAKFSDRFDVVII | ASSLGGSQNTLY | 0.28529632 |
| YLIGSVIV | ASSLGGSQNTLY | 0.120327644 |
| YGQLVAVID | ASSLGGSQNTLY | 0.12934364 |
| VTVGGMTLPA | ASSLGGSQNTLY | 0.20585757 |
| VLALPAV | ASSLGGSQNTLY | 0.3177197 |
| VIPPTLNYET | ASSLGGSQNTLY | 0.1315871 |
| VGMGGMHPSSG | ASSLGGSQNTLY | 0.1682787 |
| VGLAGGAATA | ASSLGGSQNTLY | 0.28637716 |

| | | |
|---|---|---|
| VGGAGGAGGSGP | ASSLGGSQNTLY | 0.1439709 |
| VFAAFMV | ASSLGGSQNTLY | 0.16801012 |
| TLYDKLR | ASSLGGSQNTLY | 0.20955066 |
| TLAVFTVL | ASSLGGSQNTLY | 0.12891282 |
| TGSVQPARV | ASSLGGSQNTLY | 0.2607904 |
| TGGGAGSGGT | ASSLGGSQNTLY | 0.3784426 |
| SHLETSL | ASSLGGSQNTLY | 0.14466704 |
| SFTTGMT | ASSLGGSQNTLY | 0.1603716 |
| QVEVAKRMGVRQPTVS | ASSLGGSQNTLY | 0.12574191 |
| QGSGGPALAM | ASSLGGSQNTLY | 0.14330283 |
| MLAPSNP | ASSLGGSQNTLY | 0.1200442 |
| LVNNAAI | ASSLGGSQNTLY | 0.22161898 |
| LTGEINTLA | ASSLGGSQNTLY | 0.18211818 |
| LSIFGPV | ASSLGGSQNTLY | 0.14770067 |
| LLALLANL | ASSLGGSQNTLY | 0.12156318 |
| LISNASDALDKLR | ASSLGGSQNTLY | 0.1276523 |
| LINEAVP | ASSLGGSQNTLY | 0.12803566 |
| LGYAFSGL | ASSLGGSQNTLY | 0.14059833 |
| LDFVAVAL | ASSLGGSQNTLY | 0.11930557 |
| LAQPGAE | ASSLGGSQNTLY | 0.14702114 |
| ISNLGAVTGF | ASSLGGSQNTLY | 0.21030454 |
| HQEDPGRI | ASSLGGSQNTLY | 0.19329676 |
| GTPGEPTP | ASSLGGSQNTLY | 0.12695533 |
| GFFNSGS | ASSLGGSQNTLY | 0.41100496 |
| GAGGGGLTAGAN | ASSLGGSQNTLY | 0.16708466 |
| FVDALVNM | ASSLGGSQNTLY | 0.15985231 |
| FLTVAPEL | ASSLGGSQNTLY | 0.14109322 |
| FETLTAL | ASSLGGSQNTLY | 0.12770776 |
| DVINPIEL | ASSLGGSQNTLY | 0.43663073 |
| APVSGAPV | ASSLGGSQNTLY | 0.1447787 |
| ALVGGLAW | ASSLGGSQNTLY | 0.13009053 |
| ALAAAAATAI | ASSLGGSQNTLY | 0.21447986 |
| AFGLGLLL | ASSLGGSQNTLY | 0.1469302 |
| AFFAAASL | ASSLGGSQNTLY | 0.13532151 |
| AAAALPAFSPP | ASSLGGSQNTLY | 0.18149698 |
| VVLAPAL | ASSLGGSQNTLY | 0.1594896 |
| TIGHVDHGKTTL | ASSLGGSQNTLY | 0.12617229 |
| LISNASDALDK | ASSLGGSQNTLY | 0.13733263 |
| LISNASDAL | ASSLGGSQNTLY | 0.12796463 |
| ISNASDALDK | ASSLGGSQNTLY | 0.15304022 |
| NQAETLVYQTE | ASSLGGSQNTLY | 0.27962887 |
| TGGTGGTAGAG | ASSLGGSQNTLY | 0.14776528 |
| TGSTAGGAGGA | ASSLGGSQNTLY | 0.13196622 |
| DPLAVPVAAGPV | ASSLGGSQNTLY | 0.25144333 |
| GGTGGTPGNGGN | ASSLGGSQNTLY | 0.15574804 |
| PAGLAGVDPATL | ASSLGGSQNTLY | 0.15849179 |
| GNNLVVEAPHAGASV | ASSLGGSQNTLY | 0.29659522 |
| KVTRIMSASTLEQPA | ASSLGGSQNTLY | 0.13566196 |
| LGFALPIALPRRTVV | ASSLGGSQNTLY | 0.16034935 |
| TRLGNLTARKLVVLV | ASSLGGSQNTLY | 0.19295785 |
| WSQQRRLIYFPSAGP | ASSLGGSQNTLY | 0.17910042 |

| | | |
|---|---|---|
| GPNGNPDPMAAAVDIR | ASSLGGSQNTLY | 0.1782712 |
| TQRCVSVADLRAAASR | ASSLGGSQNTLY | 0.14917925 |
| QFVLDVAEHPVEVLTETV | ASSLGGSQNTLY | 0.15021028 |
| GGAGGTGGAGGNGGTQAGDG | ASSLGGSQNTLY | 0.12188868 |
| PLLATLRAIAVDVDAKPAQV | ASSLGGSQNTLY | 0.12782462 |
| APKFRTGGYMIDFWGVGYQVAK | ASSLGGSQNTLY | 0.25785136 |
| GAGGAGGAGGAAGGPGGTGGTG | ASSLGGSQNTLY | 0.1729191 |
| IGALGLLATDVAPAGSVAFGVA | ASSLGGSQNTLY | 0.24743186 |
| AGGAGGIGGNGGAGGAGGVLMG | ASSLGGSQNTLY | 0.16724595 |
| ADELIGLL | ASSLGGSQNTLY | 0.12601562 |
| ADRALKA | ASSLGGSQNTLY | 0.27465403 |
| AGAVGQVGGAGGA | ASSLGGSQNTLY | 0.26176375 |
| AGGGGGLGGAP | ASSLGGSQNTLY | 0.12496837 |
| AGGLLYGNGGAG | ASSLGGSQNTLY | 0.14830941 |
| AGLESIGL | ASSLGGSQNTLY | 0.48161632 |
| AGLKTGII | ASSLGGSQNTLY | 0.1791313 |
| ASLAALLL | ASSLGGSQNTLY | 0.16172753 |
| ASVINISSI | ASSLGGSQNTLY | 0.14124082 |
| ATPPGPA | ASSLGGSQNTLY | 0.2677474 |
| ATVASALV | ASSLGGSQNTLY | 0.12302346 |
| ATVASLIL | ASSLGGSQNTLY | 0.32033992 |
| AVNGPVLIRH | ASSLGGSQNTLY | 0.12068397 |
| DGAAGAVGI | ASSLGGSQNTLY | 0.1347672 |
| DPLLHGI | ASSLGGSQNTLY | 0.124204464 |
| ELGAALGAL | ASSLGGSQNTLY | 0.11897688 |
| ELGRTQAV | ASSLGGSQNTLY | 0.1815005 |
| FVAGGYLS | ASSLGGSQNTLY | 0.13448752 |
| GASGGAGGAGGW | ASSLGGSQNTLY | 0.28188387 |
| GGAGAGGDGGAGGI | ASSLGGSQNTLY | 0.13715519 |
| GTGHNGSQPGMGGNG | ASSLGGSQNTLY | 0.1978004 |
| GYAAASASAS | ASSLGGSQNTLY | 0.17224035 |
| HQVDVAAVQQG | ASSLGGSQNTLY | 0.17573588 |
| IGAAIAHT | ASSLGGSQNTLY | 0.13006581 |
| IGMIEVAQL | ASSLGGSQNTLY | 0.19252853 |
| IRLAPEHTSAMD | ASSLGGSQNTLY | 0.2920783 |
| IYNVSVL | ASSLGGSQNTLY | 0.15289575 |
| KQGFLGM | ASSLGGSQNTLY | 0.12736863 |
| LESVITQL | ASSLGGSQNTLY | 0.12588818 |
| LGLGALVVV | ASSLGGSQNTLY | 0.16562274 |
| LIGNGGNGGSGGI | ASSLGGSQNTLY | 0.11885078 |
| LNAINAPTL | ASSLGGSQNTLY | 0.117675945 |
| LQLLNTGV | ASSLGGSQNTLY | 0.12377866 |
| LQTSIEF | ASSLGGSQNTLY | 0.1292079 |
| MSFAVSEL | ASSLGGSQNTLY | 0.37378755 |
| NTGALGL | ASSLGGSQNTLY | 0.22057626 |
| PPGAGGAGGAGTT | ASSLGGSQNTLY | 0.15445277 |
| QLVGLGI | ASSLGGSQNTLY | 0.18124762 |
| RFLEQVHQL | ASSLGGSQNTLY | 0.19572227 |
| RGVGYVLR | ASSLGGSQNTLY | 0.11617648 |
| SATGVFAGVI | ASSLGGSQNTLY | 0.13009632 |
| SLPAITLP | ASSLGGSQNTLY | 0.12582009 |

| | | |
|---|---|---|
| STALAFTV | ASSLGGSQNTLY | 0.26285076 |
| SVLTVGLGK | ASSLGGSQNTLY | 0.1293824 |
| TALAGAGSGI | ASSLGGSQNTLY | 0.13154083 |
| TGLPFTGI | ASSLGGSQNTLY | 0.17344818 |
| VAEHVTV | ASSLGGSQNTLY | 0.16063744 |
| VDIVLNSL | ASSLGGSQNTLY | 0.15169235 |
| VLTGGGAVL | ASSLGGSQNTLY | 0.12430717 |
| VNVGLAYGI | ASSLGGSQNTLY | 0.117487 |
| VSSAAVGAV | ASSLGGSQNTLY | 0.18048865 |
| VTSVYEAGV | ASSLGGSQNTLY | 0.12091434 |
| WVALDRGA | ASSLGGSQNTLY | 0.139405 |
| YLVGSVVI | ASSLGGSQNTLY | 0.11898912 |
| GPPGTGKTLL | ASSLGGSQNTLY | 0.12196295 |
| GVGMTKF | ASSLGGSQNTLY | 0.1262142 |
| IGIGGSDL | ASSLGGSQNTLY | 0.17475967 |
| IGTLLAEL | ASSLGGSQNTLY | 0.16259208 |
| IYTSGTT | ASSLGGSQNTLY | 0.15112124 |
| LSFPAGV | ASSLGGSQNTLY | 0.36807764 |
| LYTSGTT | ASSLGGSQNTLY | 0.32712927 |
| VGAFPFGF | ASSLGGSQNTLY | 0.14956716 |
| VGAGTVEF | ASSLGGSQNTLY | 0.14513633 |
| VGKSTLLL | ASSLGGSQNTLY | 0.13101462 |
| AALAHAGGGIV | ASSLGGSQNTLY | 0.12891819 |
| AALVAVSLL | ASSLGGSQNTLY | 0.13573587 |
| AAVSRAVAAL | ASSLGGSQNTLY | 0.48376995 |
| AGGSSGIGGF | ASSLGGSQNTLY | 0.13733925 |
| AGLTVANANG | ASSLGGSQNTLY | 0.12424951 |
| AGLVGAVL | ASSLGGSQNTLY | 0.1482351 |
| ALSQAFTV | ASSLGGSQNTLY | 0.32298976 |
| ALTFSVSL | ASSLGGSQNTLY | 0.43390256 |
| ALTGAALVAL | ASSLGGSQNTLY | 0.12697649 |
| ASFSIGAGVL | ASSLGGSQNTLY | 0.13717015 |
| DGSVYKI | ASSLGGSQNTLY | 0.14496753 |
| EQLAEIV | ASSLGGSQNTLY | 0.19388789 |
| FAVSELTVM | ASSLGGSQNTLY | 0.12615429 |
| GAAPLAAGL | ASSLGGSQNTLY | 0.13422999 |
| GADGGDGGSGGAGG | ASSLGGSQNTLY | 0.15503368 |
| GFGLDTGGAGG | ASSLGGSQNTLY | 0.5702205 |
| GLASGGLL | ASSLGGSQNTLY | 0.1373995 |
| GRIAGAAVLL | ASSLGGSQNTLY | 0.11953822 |
| GSAGSLNTSF | ASSLGGSQNTLY | 0.19844376 |
| GTGQIGI | ASSLGGSQNTLY | 0.12665318 |
| GTLIALIL | ASSLGGSQNTLY | 0.13978776 |
| IAATMGFVVG | ASSLGGSQNTLY | 0.14025173 |
| IFAQALL | ASSLGGSQNTLY | 0.1233943 |
| IGAAVALGF | ASSLGGSQNTLY | 0.14018379 |
| IGLTPLAL | ASSLGGSQNTLY | 0.3015079 |
| ILGSNTSTL | ASSLGGSQNTLY | 0.14220937 |
| IVFSLTVL | ASSLGGSQNTLY | 0.2563997 |
| LASKATGF | ASSLGGSQNTLY | 0.2612639 |
| LASVALIL | ASSLGGSQNTLY | 0.23478669 |

| | | |
|---|---|---|
| LFAVGLTL | ASSLGGSQNTLY | 0.35576296 |
| LFNGGIGTYI | ASSLGGSQNTLY | 0.19418532 |
| LTLGAGII | ASSLGGSQNTLY | 0.15153003 |
| LVAAALGDI | ASSLGGSQNTLY | 0.13806897 |
| NTAFVQLGI | ASSLGGSQNTLY | 0.19062214 |
| QLFSGAEL | ASSLGGSQNTLY | 0.22991112 |
| SAATFITL | ASSLGGSQNTLY | 0.17543626 |
| SNYGLANLGV | ASSLGGSQNTLY | 0.13108423 |
| SYGGAYAVM | ASSLGGSQNTLY | 0.376261 |
| TTGVGGLTLG | ASSLGGSQNTLY | 0.1340211 |
| TTQTIYI | ASSLGGSQNTLY | 0.20738395 |
| VAAGAGLGAVI | ASSLGGSQNTLY | 0.12576585 |
| VAAVVTLL | ASSLGGSQNTLY | 0.13198261 |
| VGFLGGLI | ASSLGGSQNTLY | 0.29726285 |
| VGVIPILL | ASSLGGSQNTLY | 0.11725307 |
| VIFASSLI | ASSLGGSQNTLY | 0.11925948 |
| VKGGDGGAAAG | ASSLGGSQNTLY | 0.13833635 |
| VTKTFTNI | ASSLGGSQNTLY | 0.35572833 |
| VVAAAILL | ASSLGGSQNTLY | 0.116448 |
| YAALDVEL | ASSLGGSQNTLY | 0.13310486 |
| LASVALLL | ASSLGGSQNTLY | 0.12367022 |
| LDAVHRV | ASSLGGSQNTLY | 0.14833798 |
| QGLSSSIF | ASSLGGSQNTLY | 0.1509434 |
| RGTFANIRL | ASSLGGSQNTLY | 0.12299151 |
| RQLWWGHRI | ASSLGGSQNTLY | 0.19281842 |
| YGPPGTGKTLL | ASSLGGSQNTLY | 0.11999775 |
| AALVGAPANVA | ASSLGGSQNTLY | 0.12473207 |
| AATFRRTE | ASSLGGSQNTLY | 0.16808435 |
| AGALGGGI | ASSLGGSQNTLY | 0.27921104 |
| AGGAGGAGGIGAG | ASSLGGSQNTLY | 0.12301518 |
| AGSGALGL | ASSLGGSQNTLY | 0.17295238 |
| AQIADQI | ASSLGGSQNTLY | 0.12895742 |
| ASAVAVTYL | ASSLGGSQNTLY | 0.4772269 |
| ASTVLGGTGG | ASSLGGSQNTLY | 0.15051368 |
| AVGGEPAK | ASSLGGSQNTLY | 0.12030005 |
| CEQGTFGGTC | ASSLGGSQNTLY | 0.19719346 |
| DEGKAPAG | ASSLGGSQNTLY | 0.21113718 |
| DTVVFPAL | ASSLGGSQNTLY | 0.16613829 |
| FEGITMAG | ASSLGGSQNTLY | 0.16326919 |
| FPLVVQD | ASSLGGSQNTLY | 0.12971412 |
| GAFTTPQ | ASSLGGSQNTLY | 0.16303208 |
| GFGGFGTAGGVGGA | ASSLGGSQNTLY | 0.28058797 |
| GFLAQFL | ASSLGGSQNTLY | 0.3348797 |
| GGAGGAGGNAGVA | ASSLGGSQNTLY | 0.13333115 |
| GGVGTAGAGGAGGAGG | ASSLGGSQNTLY | 0.16483493 |
| GNGGGGDAGNAGSGGN | ASSLGGSQNTLY | 0.17461817 |
| GTGGAGGAGNDAGS | ASSLGGSQNTLY | 0.13780646 |
| GTTNTGWFNS | ASSLGGSQNTLY | 0.2532094 |
| IFMGPLFTL | ASSLGGSQNTLY | 0.14029773 |
| ILAHAIEAI | ASSLGGSQNTLY | 0.20849302 |
| IRSAIREAL | ASSLGGSQNTLY | 0.16661116 |

| | | |
|---|---|---|
| ITFAGGG | ASSLGGSQNTLY | 0.548136 |
| ITGMSGHQEV | ASSLGGSQNTLY | 0.12512349 |
| IVAGLACTL | ASSLGGSQNTLY | 0.16323338 |
| IVGGGLAGA | ASSLGGSQNTLY | 0.15999381 |
| IVTGVLNI | ASSLGGSQNTLY | 0.20961565 |
| LAQQGCI | ASSLGGSQNTLY | 0.12919733 |
| LCLHVEN | ASSLGGSQNTLY | 0.12287497 |
| LDAISDGS | ASSLGGSQNTLY | 0.1565968 |
| LGGFAAML | ASSLGGSQNTLY | 0.23240578 |
| LGGKNPMI | ASSLGGSQNTLY | 0.13651606 |
| LGPKGVAAIL | ASSLGGSQNTLY | 0.20157501 |
| LMVGVSI | ASSLGGSQNTLY | 0.4637887 |
| LVGAAFVL | ASSLGGSQNTLY | 0.12842803 |
| LVLAVAPL | ASSLGGSQNTLY | 0.16013144 |
| MAAIAVIDAA | ASSLGGSQNTLY | 0.16914797 |
| NNGINAELQ | ASSLGGSQNTLY | 0.34514543 |
| RGEPLVLVHG | ASSLGGSQNTLY | 0.14400041 |
| RGGAGGAAGLI | ASSLGGSQNTLY | 0.30385357 |
| RSYRMHPAV | ASSLGGSQNTLY | 0.12382921 |
| SAAHLLAGQLA | ASSLGGSQNTLY | 0.2560869 |
| SAAVGSY | ASSLGGSQNTLY | 0.12479888 |
| SPHAVLHRI | ASSLGGSQNTLY | 0.17617892 |
| SQIAKVL | ASSLGGSQNTLY | 0.12557107 |
| TALGHLVP | ASSLGGSQNTLY | 0.25620085 |
| TAPTTSGAA | ASSLGGSQNTLY | 0.2587386 |
| TDGDDGNGGSPGT | ASSLGGSQNTLY | 0.31597367 |
| TDGLGEN | ASSLGGSQNTLY | 0.117584474 |
| TGGPPLPGQA | ASSLGGSQNTLY | 0.18818781 |
| TIPAITI | ASSLGGSQNTLY | 0.26947445 |
| TIPGFTI | ASSLGGSQNTLY | 0.628976 |
| TSIASGALA | ASSLGGSQNTLY | 0.13410453 |
| TVTLPGAAI | ASSLGGSQNTLY | 0.2275118 |
| VASGLSAALP | ASSLGGSQNTLY | 0.13466634 |
| VGAGVIGI | ASSLGGSQNTLY | 0.13622418 |
| VGPKRFGV | ASSLGGSQNTLY | 0.124906644 |
| VNVAFPD | ASSLGGSQNTLY | 0.13494736 |
| VSMTNSGVS | ASSLGGSQNTLY | 0.13367268 |
| VTDVLGDLSV | ASSLGGSQNTLY | 0.133007 |
| VVVVGAGFAGL | ASSLGGSQNTLY | 0.3053791 |
| VYTVGTVV | ASSLGGSQNTLY | 0.35555992 |
| WTTFSGP | ASSLGGSQNTLY | 0.20758659 |
| YDELVEAF | ASSLGGSQNTLY | 0.36103344 |
| YGTGGAGGNGG | ASSLGGSQNTLY | 0.41227084 |
| YTNVMAV | ASSLGGSQNTLY | 0.16050895 |
| FLEQVHQL | ASSLGGSQNTLY | 0.13825677 |
| GLSLLAVAV | ASSLGGSQNTLY | 0.14093938 |
| LEQVHQL | ASSLGGSQNTLY | 0.12456913 |
| LNAAGAI | ASSLGGSQNTLY | 0.13546059 |
| PAVHASL | ASSLGGSQNTLY | 0.14135332 |
| TELEDDA | ASSLGGSQNTLY | 0.16521639 |
| TPPGPPN | ASSLGGSQNTLY | 0.11640864 |

| | | |
|---|---|---|
| TPPPPPPP | ASSLGGSQNTLY | 0.20151597 |
| TQSPALI | ASSLRDKPTEVF | 0.48436102 |
| EFAAAFPQ | ASSLRDKPTEVF | 0.12712532 |
| LLSGNLDVL | ASSLRDKPTEVF | 0.17694442 |
| GPANLGNYNI | ASSLRDKPTEVF | 0.17475411 |
| TLGPTLP | ASSLRDKPTEVF | 0.12924433 |
| VYAYSSCCK | ASSLRDKPTEVF | 0.28541142 |
| APAPAPAPNI | ASSLRDKPTEVF | 0.12962969 |
| PGGPVPIP | ASSLRDKPTEVF | 0.17572097 |
| FLTVALEP | ASSLRDKPTEVF | 0.30187988 |
| SQYFGANL | ASSLRDKPTEVF | 0.16635582 |
| GGVKPAA | ASSLRDKPTEVF | 0.19625677 |
| GGTGGAGGAGGW | ASSLRDKPTEVF | 0.144555 |
| GVGGVAGLGGAGG | ASSLRDKPTEVF | 0.12528951 |
| VLLVLATTI | ASSLRDKPTEVF | 0.20052686 |
| IGIFAAML | ASSLRDKPTEVF | 0.11827184 |
| AVGDLID | ASSLRDKPTEVF | 0.15539283 |
| NISPLPL | ASSLRDKPTEVF | 0.12784968 |
| VTAVFVAI | ASSLRDKPTEVF | 0.25333288 |
| SGDDNDAAL | ASSLRDKPTEVF | 0.1684066 |
| AVAALAHS | ASSLRDKPTEVF | 0.1461595 |
| GGVTVMATP | ASSLRDKPTEVF | 0.14156899 |
| AGLAANVRD | ASSLRDKPTEVF | 0.13350627 |
| TNRPDLIDPALLRPG | ASSLRDKPTEVF | 0.12692274 |
| LANLGVTAL | ASSLRDKPTEVF | 0.12822579 |
| TGERIEL | ASSLRDKPTEVF | 0.16837715 |
| NGEELAA | ASSLRDKPTEVF | 0.13609706 |
| VLSLVWG | ASSLRDKPTEVF | 0.15547788 |
| IGYALWL | ASSLRDKPTEVF | 0.20317078 |
| LSLLLAPL | ASSLRDKPTEVF | 0.15598938 |
| TLDEALK | ASSLRDKPTEVF | 0.12314204 |
| TDGAGTAEGG | ASSLRDKPTEVF | 0.16133521 |
| NAPGGAGGNGG | ASSLRDKPTEVF | 0.19243896 |
| VSVFMGMGGNF | ASSLRDKPTEVF | 0.14130421 |
| LASYLAAL | ASSLRDKPTEVF | 0.11631089 |
| LVGGTPAL | ASSLRDKPTEVF | 0.18901013 |
| VRNADVL | ASSLRDKPTEVF | 0.14103618 |
| TCSSPFLVSAM | ASSLRDKPTEVF | 0.13223785 |
| VHITDAE | ASSLRDKPTEVF | 0.1399068 |
| EYAAPSF | ASSLRDKPTEVF | 0.31513616 |
| VVDAFHGGL | ASSLRDKPTEVF | 0.1254774 |
| ASVVDPQ | ASSLRDKPTEVF | 0.14152299 |
| SYNEFGGLS | ASSLRDKPTEVF | 0.23197997 |
| DPDGSVY | ASSLRDKPTEVF | 0.19751059 |
| MVSTIHATE | ASSLRDKPTEVF | 0.40169343 |
| LAAGVADPV | ASSLRDKPTEVF | 0.18121122 |
| TKEQIAA | ASSLRDKPTEVF | 0.12989043 |
| SLASPAAL | ASSLRDKPTEVF | 0.17956305 |
| IGLVFYGV | ASSLRDKPTEVF | 0.13438699 |
| FLLDASNL | ASSLRDKPTEVF | 0.12122828 |
| YQVGNFP | ASSLRDKPTEVF | 0.11789928 |

| | | |
|---|---|---|
| ESLLSQST | ASSLRDKPTEVF | 0.14514011 |
| LFTVGFAAVG | ASSLRDKPTEVF | 0.12370058 |
| ATPPTPT | ASSLRDKPTEVF | 0.17986497 |
| TDLATLG | ASSLRDKPTEVF | 0.17901248 |
| SAGGTGR | ASSLRDKPTEVF | 0.13179891 |
| GFSNSGS | ASSLRDKPTEVF | 0.12159599 |
| GGNGGSGGTSV | ASSLRDKPTEVF | 0.13508244 |
| GTSVATGGAGN | ASSLRDKPTEVF | 0.13196805 |
| NGGNGGSGGTS | ASSLRDKPTEVF | 0.20587608 |
| AVVKFPH | ASSLRDKPTEVF | 0.13704608 |
| TPPPPPP | ASSLRDKPTEVF | 0.67048824 |
| AAPAAAAAPA | ASSLRDKPTEVF | 0.23409833 |
| IALNEDL | ASSLRDKPTEVF | 0.13908333 |
| IVNVSSM | ASSLRDKPTEVF | 0.2319772 |
| TGIPPAPR | ASSLRDKPTEVF | 0.13323796 |
| FDLGGGTF | ASSLRDKPTEVF | 0.1337597 |
| FDLGGGTFDVS | ASSLRDKPTEVF | 0.13944425 |
| NEPTAAA | ASSLRDKPTEVF | 0.14129107 |
| NEPTAAALA | ASSLRDKPTEVF | 0.14592147 |
| GSIINISSI | ASSLRDKPTEVF | 0.13850552 |
| VAPGNAGTA | ASSLRDKPTEVF | 0.12366523 |
| TGGGASLE | ASSLRDKPTEVF | 0.15218623 |
| VAVYNHYKRI | ASSLRDKPTEVF | 0.13204516 |
| TDPAPATT | ASSLRDKPTEVF | 0.119897895 |
| TNRPDILDPALLRPG | ASSLRDKPTEVF | 0.14330727 |
| LTTTTTS | ASSLRDKPTEVF | 0.40223795 |
| PTGSGKTL | ASSLRDKPTEVF | 0.32289732 |
| TGSGKTL | ASSLRDKPTEVF | 0.13509497 |
| TIGHVDHGK | ASSLRDKPTEVF | 0.14036462 |
| TIGHVDHGKTT | ASSLRDKPTEVF | 0.26962775 |
| GHVDHGKTT | ASSLRDKPTEVF | 0.12563053 |
| AQFEHTLL | ASSLRDKPTEVF | 0.14854968 |
| IVNQDEL | ASSLRDKPTEVF | 0.17622295 |
| TNGSQFF | ASSLRDKPTEVF | 0.18955135 |
| TGKGIVE | ASSLRDKPTEVF | 0.1371092 |
| LISNASDA | ASSLRDKPTEVF | 0.15366545 |
| AELRTLGELTKIAKAHGAQ | ASSLRDKPTEVF | 0.14299554 |
| APNAVAAAEPSW | ASSLRDKPTEVF | 0.15842316 |
| APQVAGAATAVL | ASSLRDKPTEVF | 0.11629916 |
| ARRDPAAAAAILPTDA | ASSLRDKPTEVF | 0.15139623 |
| DAAARDVLAEAGLAEYF | ASSLRDKPTEVF | 0.1319678 |
| DDQIVLSAKANLAKVEQ | ASSLRDKPTEVF | 0.12558293 |
| DELLPDTGEDSDEDGH | ASSLRDKPTEVF | 0.11915054 |
| ELPEAPANRATVPQD | ASSLRDKPTEVF | 0.13679865 |
| FGQSKIERVTGIG | ASSLRDKPTEVF | 0.22042364 |
| GAGGTGSTAGGAGGAGGAGGLY | ASSLRDKPTEVF | 0.2721025 |
| GHGGTGAAGGNGGTGGAGGSGI | ASSLRDKPTEVF | 0.12255346 |
| GNGGNGGAGGEGGPGGAGGAGA | ASSLRDKPTEVF | 0.34589714 |
| GSTAGNGGNGGNATNGTIAGQP | ASSLRDKPTEVF | 0.22972399 |
| GTGGDGGNGGNGGTGG | ASSLRDKPTEVF | 0.1451948 |
| GVRRPAGIPADAPV | ASSLRDKPTEVF | 0.12235422 |

| | | |
|---|---|---|
| GYRQVTDAHLLTLARR | ASSLRDKPTEVF | 0.17302148 |
| IAKMGQDGHDRG | ASSLRDKPTEVF | 0.13675022 |
| KAEPSIEIQYT | ASSLRDKPTEVF | 0.2007465 |
| KPVRIRMAADEDTVAVT | ASSLRDKPTEVF | 0.31423387 |
| MPPRIAGMRLLVIKPEP | ASSLRDKPTEVF | 0.122961044 |
| MKFYAVALFRSRGVPPTR | ASSLRDKPTEVF | 0.23670253 |
| NAGGFGILGAVAHSP | ASSLRDKPTEVF | 0.21642709 |
| NEGREITLAMAEM | ASSLRDKPTEVF | 0.1270021 |
| NELVEWSHPLGHRWRWE | ASSLRDKPTEVF | 0.12365231 |
| NLGAGGTGGNGGNPG | ASSLRDKPTEVF | 0.13092345 |
| NTEWPKWETVMAADPVD | ASSLRDKPTEVF | 0.118002445 |
| PASAAPAETQP | ASSLRDKPTEVF | 0.19100514 |
| QQSSDGKNGVRAAEITGE | ASSLRDKPTEVF | 0.14217375 |
| RAPDAVTTSWWKE | ASSLRDKPTEVF | 0.13565046 |
| SGFGNVGAHTSGWFNQ | ASSLRDKPTEVF | 0.17269093 |
| SGVAGAVGQAASVGGLKV | ASSLRDKPTEVF | 0.14006405 |
| SNGGAGTGSAGGNGGT | ASSLRDKPTEVF | 0.14840521 |
| TGGAGGEGGAAG | ASSLRDKPTEVF | 0.14843428 |
| TGGDGGAAGAGGAPGHG | ASSLRDKPTEVF | 0.1417782 |
| TMPPLLAAIANAGRHGV | ASSLRDKPTEVF | 0.1657953 |
| TSSTAGNSGGA | ASSLRDKPTEVF | 0.13950051 |
| VPATVPVGILC | ASSLRDKPTEVF | 0.13735025 |
| VQRPDSGGVVLTGRIS | ASSLRDKPTEVF | 0.3839325 |
| WTPNPVPATKGVPAGLVQAT | ASSLRDKPTEVF | 0.12569767 |
| YALKRALAKFSDRFDVVII | ASSLRDKPTEVF | 0.13799733 |
| YLIGSVIV | ASSLRDKPTEVF | 0.17272827 |
| YGQLVAVID | ASSLRDKPTEVF | 0.12454333 |
| VTVGGMTLPA | ASSLRDKPTEVF | 0.18984446 |
| VLALPAV | ASSLRDKPTEVF | 0.17973812 |
| VIPPTLNYET | ASSLRDKPTEVF | 0.18239851 |
| VGMGGMHPSSG | ASSLRDKPTEVF | 0.12444109 |
| VGLAGGAATA | ASSLRDKPTEVF | 0.5140816 |
| VGGAGGAGGSGP | ASSLRDKPTEVF | 0.19694819 |
| VFAAFMV | ASSLRDKPTEVF | 0.13317525 |
| TLYDKLR | ASSLRDKPTEVF | 0.17531015 |
| TLAVFTVL | ASSLRDKPTEVF | 0.12804446 |
| TGSVQPARV | ASSLRDKPTEVF | 0.2727248 |
| TGGGAGSGGT | ASSLRDKPTEVF | 0.24568757 |
| SHLETSL | ASSLRDKPTEVF | 0.12847514 |
| SFTTGMT | ASSLRDKPTEVF | 0.17393512 |
| QVEVAKRMGVRQPTVS | ASSLRDKPTEVF | 0.1844685 |
| QGSGGPALAM | ASSLRDKPTEVF | 0.26367983 |
| MLAPSNP | ASSLRDKPTEVF | 0.1237048 |
| LVNNAAI | ASSLRDKPTEVF | 0.1358049 |
| LTGEINTLA | ASSLRDKPTEVF | 0.122954026 |
| LSIFGPV | ASSLRDKPTEVF | 0.12912017 |
| LLALLANL | ASSLRDKPTEVF | 0.13404474 |
| LISNASDALDKLR | ASSLRDKPTEVF | 0.18320477 |
| LINEAVP | ASSLRDKPTEVF | 0.15181327 |
| LGYAFSGL | ASSLRDKPTEVF | 0.2711965 |
| LDFVAVAL | ASSLRDKPTEVF | 0.1773054 |

| | | |
|---|---|---|
| LAQPGAE | ASSLRDKPTEVF | 0.21524854 |
| ISNLGAVTGF | ASSLRDKPTEVF | 0.136397 |
| HQEDPGRI | ASSLRDKPTEVF | 0.15884125 |
| GTPGEPTP | ASSLRDKPTEVF | 0.12199582 |
| GFFNSGS | ASSLRDKPTEVF | 0.12062306 |
| GAGGGGLTAGAN | ASSLRDKPTEVF | 0.12321895 |
| FVDALVNM | ASSLRDKPTEVF | 0.15626225 |
| FLTVAPEL | ASSLRDKPTEVF | 0.11770486 |
| FETLTAL | ASSLRDKPTEVF | 0.25608638 |
| DVINPIEL | ASSLRDKPTEVF | 0.16048272 |
| APVSGAPV | ASSLRDKPTEVF | 0.17767955 |
| ALVGGLAW | ASSLRDKPTEVF | 0.13718925 |
| ALAAAAATAI | ASSLRDKPTEVF | 0.13422988 |
| AFGLGLLL | ASSLRDKPTEVF | 0.1397985 |
| AFFAAASL | ASSLRDKPTEVF | 0.28219396 |
| AAAALPAFSPP | ASSLRDKPTEVF | 0.12723178 |
| VVLAPAL | ASSLRDKPTEVF | 0.12018698 |
| TIGHVDHGKTTL | ASSLRDKPTEVF | 0.12376402 |
| LISNASDALDK | ASSLRDKPTEVF | 0.12109025 |
| LISNASDAL | ASSLRDKPTEVF | 0.3441152 |
| ISNASDALDK | ASSLRDKPTEVF | 0.23428285 |
| NQAETLVYQTE | ASSLRDKPTEVF | 0.194713 |
| TGGTGGTAGAG | ASSLRDKPTEVF | 0.21644425 |
| TGSTAGGAGGA | ASSLRDKPTEVF | 0.117951356 |
| DPLAVPVAAGPV | ASSLRDKPTEVF | 0.1329479 |
| GGTGGTPGNGGN | ASSLRDKPTEVF | 0.47836244 |
| PAGLAGVDPATL | ASSLRDKPTEVF | 0.1424606 |
| GNNLVVEAPHAGASV | ASSLRDKPTEVF | 0.17246714 |
| KVTRIMSASTLEQPA | ASSLRDKPTEVF | 0.22003788 |
| LGFALPIALPRRTVV | ASSLRDKPTEVF | 0.14087364 |
| TRLGNLTARKLVVLV | ASSLRDKPTEVF | 0.14524077 |
| WSQQRRLIYFPSAGP | ASSLRDKPTEVF | 0.13904645 |
| GPNGNPDPMAAAVDIR | ASSLRDKPTEVF | 0.1633779 |
| TQRCVSVADLRAAASR | ASSLRDKPTEVF | 0.13079953 |
| QFVLDVAEHPVEVLTETV | ASSLRDKPTEVF | 0.12624452 |
| GGAGGTGGAGGNGGTQAGDG | ASSLRDKPTEVF | 0.17414188 |
| PLLATLRAIAVDVDAKPAQV | ASSLRDKPTEVF | 0.61197156 |
| APKFRTGGYMIDFWGVGYQVAK | ASSLRDKPTEVF | 0.17153518 |
| GAGGAGGAGGAAGGPGGTGGTG | ASSLRDKPTEVF | 0.18014014 |
| IGALGLLATDVAPAGSVAFGVA | ASSLRDKPTEVF | 0.15089288 |
| AGGAGGIGGNGGAGGAGGVLMG | ASSLRDKPTEVF | 0.21636811 |
| ADELIGLL | ASSLRDKPTEVF | 0.14230356 |
| ADRALKA | ASSLRDKPTEVF | 0.13426116 |
| AGAVGQVGGAGGA | ASSLRDKPTEVF | 0.13724218 |
| AGGGGGLGGAP | ASSLRDKPTEVF | 0.12340259 |
| AGGLLYGNGGAG | ASSLRDKPTEVF | 0.15585423 |
| AGLESIGL | ASSLRDKPTEVF | 0.1392712 |
| AGLKTGII | ASSLRDKPTEVF | 0.6879462 |
| ASLAALLL | ASSLRDKPTEVF | 0.18898046 |
| ASVINISSI | ASSLRDKPTEVF | 0.15536144 |
| ATPPGPA | ASSLRDKPTEVF | 0.12961106 |

| | | |
|---|---|---|
| ATVASALV | ASSLRDKPTEVF | 0.13477063 |
| ATVASLIL | ASSLRDKPTEVF | 0.17497297 |
| AVNGPVLIRH | ASSLRDKPTEVF | 0.12551963 |
| DGAAGAVGI | ASSLRDKPTEVF | 0.2719782 |
| DPLLHGI | ASSLRDKPTEVF | 0.13396868 |
| ELGAALGAL | ASSLRDKPTEVF | 0.14041781 |
| ELGRTQAV | ASSLRDKPTEVF | 0.19278082 |
| FVAGGYLS | ASSLRDKPTEVF | 0.637133 |
| GASGGAGGAGGW | ASSLRDKPTEVF | 0.13790716 |
| GGAGAGGDGGAGGI | ASSLRDKPTEVF | 0.1318709 |
| GTGHNGSQPGMGGNG | ASSLRDKPTEVF | 0.13414115 |
| GYAAASASAS | ASSLRDKPTEVF | 0.18201935 |
| HQVDVAAVQQG | ASSLRDKPTEVF | 0.40820658 |
| IGAAIAHT | ASSLRDKPTEVF | 0.13709918 |
| IGMIEVAQL | ASSLRDKPTEVF | 0.45848882 |
| IRLAPEHTSAMD | ASSLRDKPTEVF | 0.14272308 |
| IYNVSVL | ASSLRDKPTEVF | 0.124980085 |
| KQGFLGM | ASSLRDKPTEVF | 0.15049812 |
| LESVITQL | ASSLRDKPTEVF | 0.16834085 |
| LGLGALVVV | ASSLRDKPTEVF | 0.41148597 |
| LIGNGGNGGSGGI | ASSLRDKPTEVF | 0.24338952 |
| LNAINAPTL | ASSLRDKPTEVF | 0.151094 |
| LQLLNTGV | ASSLRDKPTEVF | 0.13194793 |
| LQTSIEF | ASSLRDKPTEVF | 0.2537257 |
| MSFAVSEL | ASSLRDKPTEVF | 0.20502497 |
| NTGALGL | ASSLRDKPTEVF | 0.18507439 |
| PPGAGGAGGAGTT | ASSLRDKPTEVF | 0.13300908 |
| QLVGLGI | ASSLRDKPTEVF | 0.13287717 |
| RFLEQVHQL | ASSLRDKPTEVF | 0.15127707 |
| RGVGYVLR | ASSLRDKPTEVF | 0.3285485 |
| SATGVFAGVI | ASSLRDKPTEVF | 0.12787922 |
| SLPAITLP | ASSLRDKPTEVF | 0.16974998 |
| STALAFTV | ASSLRDKPTEVF | 0.18213502 |
| SVLTVGLGK | ASSLRDKPTEVF | 0.12355841 |
| TALAGAGSGI | ASSLRDKPTEVF | 0.13884285 |
| TGLPFTGI | ASSLRDKPTEVF | 0.39671424 |
| VAEHVTV | ASSLRDKPTEVF | 0.14324468 |
| VDIVLNSL | ASSLRDKPTEVF | 0.12738428 |
| VLTGGGAVL | ASSLRDKPTEVF | 0.27515844 |
| VNVGLAYGI | ASSLRDKPTEVF | 0.7094983 |
| VSSAAVGAV | ASSLRDKPTEVF | 0.21115252 |
| VTSVYEAGV | ASSLRDKPTEVF | 0.12555347 |
| WVALDRGA | ASSLRDKPTEVF | 0.117662445 |
| YLVGSVVI | ASSLRDKPTEVF | 0.13800879 |
| GPPGTGKTLL | ASSLRDKPTEVF | 0.17380713 |
| GVGMTKF | ASSLRDKPTEVF | 0.14422363 |
| IGIGGSDL | ASSLRDKPTEVF | 0.13383457 |
| IGTLLAEL | ASSLRDKPTEVF | 0.19637266 |
| IYTSGTT | ASSLRDKPTEVF | 0.22559197 |
| LSFPAGV | ASSLRDKPTEVF | 0.13020317 |
| LYTSGTT | ASSLRDKPTEVF | 0.1229524 |

| | | |
|---|---|---|
| VGAFPFGF | ASSLRDKPTEVF | 0.14193402 |
| VGAGTVEF | ASSLRDKPTEVF | 0.24046408 |
| VGKSTLLL | ASSLRDKPTEVF | 0.1275404 |
| AALAHAGGGIV | ASSLRDKPTEVF | 0.1479834 |
| AALVAVSLL | ASSLRDKPTEVF | 0.13903086 |
| AAVSRAVAAL | ASSLRDKPTEVF | 0.12735389 |
| AGGSSGIGGF | ASSLRDKPTEVF | 0.18109457 |
| AGLTVANANG | ASSLRDKPTEVF | 0.13253894 |
| AGLVGAVL | ASSLRDKPTEVF | 0.40330535 |
| ALSQAFTV | ASSLRDKPTEVF | 0.1291517 |
| ALTFSVSL | ASSLRDKPTEVF | 0.16661884 |
| ALTGAALVAL | ASSLRDKPTEVF | 0.2231116 |
| ASFSIGAGVL | ASSLRDKPTEVF | 0.17397842 |
| DGSVYKI | ASSLRDKPTEVF | 0.14685819 |
| EQLAEIV | ASSLRDKPTEVF | 0.13069275 |
| FAVSELTVM | ASSLRDKPTEVF | 0.12710187 |
| GAAPLAAGL | ASSLRDKPTEVF | 0.1177328 |
| GADGGDGGSGGAGG | ASSLRDKPTEVF | 0.3706197 |
| GFGLDTGGAGG | ASSLRDKPTEVF | 0.12231166 |
| GLASGGLL | ASSLRDKPTEVF | 0.3263758 |
| GRIAGAAVLL | ASSLRDKPTEVF | 0.16224602 |
| GSAGSLNTSF | ASSLRDKPTEVF | 0.13179992 |
| GTGQIGI | ASSLRDKPTEVF | 0.15337317 |
| GTLIALIL | ASSLRDKPTEVF | 0.123171225 |
| IAATMGFVVG | ASSLRDKPTEVF | 0.12790883 |
| IFAQALL | ASSLRDKPTEVF | 0.19881502 |
| IGAAVALGF | ASSLRDKPTEVF | 0.18608178 |
| IGLTPLAL | ASSLRDKPTEVF | 0.1323462 |
| ILGSNTSTL | ASSLRDKPTEVF | 0.1602438 |
| IVFSLTVL | ASSLRDKPTEVF | 0.17418815 |
| LASKATGF | ASSLRDKPTEVF | 0.1287487 |
| LASVALIL | ASSLRDKPTEVF | 0.14900614 |
| LFAVGLTL | ASSLRDKPTEVF | 0.22998583 |
| LFNGGIGTYI | ASSLRDKPTEVF | 0.15702438 |
| LTLGAGII | ASSLRDKPTEVF | 0.12566365 |
| LVAAALGDI | ASSLRDKPTEVF | 0.16190813 |
| NTAFVQLGI | ASSLRDKPTEVF | 0.13304266 |
| QLFSGAEL | ASSLRDKPTEVF | 0.21122693 |
| SAATFITL | ASSLRDKPTEVF | 0.15132706 |
| SNYGLANLGV | ASSLRDKPTEVF | 0.15300186 |
| SYGGAYAVM | ASSLRDKPTEVF | 0.17089255 |
| TTGVGGLTLG | ASSLRDKPTEVF | 0.121951915 |
| TTQTIYI | ASSLRDKPTEVF | 0.12543677 |
| VAAGAGLGAVI | ASSLRDKPTEVF | 0.14218557 |
| VAAVVTLL | ASSLRDKPTEVF | 0.16504343 |
| VGFLGGLI | ASSLRDKPTEVF | 0.17642266 |
| VGVIPILL | ASSLRDKPTEVF | 0.13095969 |
| VIFASSLI | ASSLRDKPTEVF | 0.16051343 |
| VKGGDGGAAAG | ASSLRDKPTEVF | 0.14603797 |
| VTKTFTNI | ASSLRDKPTEVF | 0.13381442 |
| VVAAAILL | ASSLRDKPTEVF | 0.1496452 |

| | | |
|---|---|---|
| YAALDVEL | ASSLRDKPTEVF | 0.12760524 |
| LASVALLL | ASSLRDKPTEVF | 0.12683587 |
| LDAVHRV | ASSLRDKPTEVF | 0.19460768 |
| QGLSSSIF | ASSLRDKPTEVF | 0.13549684 |
| RGTFANIRL | ASSLRDKPTEVF | 0.155818 |
| RQLWWGHRI | ASSLRDKPTEVF | 0.33559418 |
| YGPPGTGKTLL | ASSLRDKPTEVF | 0.13251278 |
| AALVGAPANVA | ASSLRDKPTEVF | 0.14508298 |
| AATFRRTE | ASSLRDKPTEVF | 0.1725863 |
| AGALGGGI | ASSLRDKPTEVF | 0.24335846 |
| AGGAGGAGGIGAG | ASSLRDKPTEVF | 0.15061864 |
| AGSGALGL | ASSLRDKPTEVF | 0.13280687 |
| AQIADQI | ASSLRDKPTEVF | 0.21768206 |
| ASAVAVTYL | ASSLRDKPTEVF | 0.17458354 |
| ASTVLGGTGG | ASSLRDKPTEVF | 0.31016344 |
| AVGGEPAK | ASSLRDKPTEVF | 0.25477138 |
| CEQGTFGGTC | ASSLRDKPTEVF | 0.18089113 |
| DEGKAPAG | ASSLRDKPTEVF | 0.18965441 |
| DTVVFPAL | ASSLRDKPTEVF | 0.14557895 |
| FEGITMAG | ASSLRDKPTEVF | 0.121158585 |
| FPLVVQD | ASSLRDKPTEVF | 0.14031962 |
| GAFTTPQ | ASSLRDKPTEVF | 0.13653871 |
| GFGGFGTAGGVGGA | ASSLRDKPTEVF | 0.1288794 |
| GFLAQFL | ASSLRDKPTEVF | 0.12146362 |
| GGAGGAGGNAGVA | ASSLRDKPTEVF | 0.33044755 |
| GGVGTAGAGGAGGAGG | ASSLRDKPTEVF | 0.6132263 |
| GNGGGGDAGNAGSGGN | ASSLRDKPTEVF | 0.12944272 |
| GTGGAGGAGNDAGS | ASSLRDKPTEVF | 0.30684447 |
| GTTNTGWFNS | ASSLRDKPTEVF | 0.12107257 |
| IFMGPLFTL | ASSLRDKPTEVF | 0.15434705 |
| ILAHAIEAI | ASSLRDKPTEVF | 0.28876185 |
| IRSAIREAL | ASSLRDKPTEVF | 0.12825738 |
| ITFAGGG | ASSLRDKPTEVF | 0.17639932 |
| ITGMSGHQEV | ASSLRDKPTEVF | 0.12678726 |
| IVAGLACTL | ASSLRDKPTEVF | 0.13702615 |
| IVGGGLAGA | ASSLRDKPTEVF | 0.42226171 |
| IVTGVLNI | ASSLRDKPTEVF | 0.14845589 |
| LAQQGCI | ASSLRDKPTEVF | 0.16528344 |
| LCLHVEN | ASSLRDKPTEVF | 0.14538191 |
| LDAISDGS | ASSLRDKPTEVF | 0.13539104 |
| LGGFAAML | ASSLRDKPTEVF | 0.15352982 |
| LGGKNPMI | ASSLRDKPTEVF | 0.37673193 |
| LGPKGVAAIL | ASSLRDKPTEVF | 0.12556326 |
| LMVGVSI | ASSLRDKPTEVF | 0.18735558 |
| LVGAAFVL | ASSLRDKPTEVF | 0.13139431 |
| LVLAVAPL | ASSLRDKPTEVF | 0.55250204 |
| MAAIAVIDAA | ASSLRDKPTEVF | 0.23207077 |
| NNGINAELQ | ASSLRDKPTEVF | 0.19234107 |
| RGEPLVLVHG | ASSLRDKPTEVF | 0.1369186 |
| RGGAGGAAGLI | ASSLRDKPTEVF | 0.30187482 |
| RSYRMHPAV | ASSLRDKPTEVF | 0.18609136 |

| | | |
|---|---|---|
| SAAHLLAGQLA | ASSLRDKPTEVF | 0.22543815 |
| SAAVGSY | ASSLRDKPTEVF | 0.18515657 |
| SPHAVLHRI | ASSLRDKPTEVF | 0.08738741 |
| SQIAKVL | ASSLRDKPTEVF | 0.13202733 |
| TALGHLVP | ASSLRDKPTEVF | 0.07830211 |
| TAPTTSGAA | ASSLRDKPTEVF | 0.088182874 |
| TDGDDGNGGSPGT | ASSLRDKPTEVF | 0.11678923 |
| TDGLGEN | ASSLRDKPTEVF | 0.07893671 |
| TGGPPLPGQA | ASSLRDKPTEVF | 0.1055716 |
| TIPAITI | ASSLRDKPTEVF | 0.1049426 |
| TIPGFTI | ASSLRDKPTEVF | 0.0730805 |
| TSIASGALA | ASSLRDKPTEVF | 0.0914054 |
| TVTLPGAAI | ASSLRDKPTEVF | 0.11535096 |
| VASGLSAALP | ASSLRDKPTEVF | 0.22220235 |
| VGAGVIGI | ASSLRDKPTEVF | 0.118641324 |
| VGPKRFGV | ASSLRDKPTEVF | 0.09065094 |
| VNVAFPD | ASSLRDKPTEVF | 0.08969024 |
| VSMTNSGVS | ASSLRDKPTEVF | 0.105231054 |
| VTDVLGDLSV | ASSLRDKPTEVF | 0.089851245 |
| VVVVGAGFAGL | ASSLRDKPTEVF | 0.069031976 |
| VYTVGTVV | ASSLRDKPTEVF | 0.09723326 |
| WTTFSGP | ASSLRDKPTEVF | 0.071913406 |
| YDELVEAF | ASSLRDKPTEVF | 0.08226422 |
| YGTGGAGGNGG | ASSLRDKPTEVF | 0.12454893 |
| YTNVMAV | ASSLRDKPTEVF | 0.11095439 |
| FLEQVHQL | ASSLRDKPTEVF | 0.11257763 |
| GLSLLAVAV | ASSLRDKPTEVF | 0.16487655 |
| LEQVHQL | ASSLRDKPTEVF | 0.16684937 |
| LNAAGAI | ASSLRDKPTEVF | 0.10301056 |
| PAVHASL | ASSLRDKPTEVF | 0.1429326 |
| TELEDDA | ASSLRDKPTEVF | 0.15934725 |
| TPPGPPN | ASSLRDKPTEVF | 0.09638847 |
| TPPPPPPP | ASSLRDKPTEVF | 0.083394036 |
| TQSPALI | ASSGQGAAETLY | 0.13562213 |
| EFAAAFPQ | ASSGQGAAETLY | 0.10683093 |
| LLSGNLDVL | ASSGQGAAETLY | 0.08790237 |
| GPANLGNYNI | ASSGQGAAETLY | 0.0898487 |
| TLGPTLP | ASSGQGAAETLY | 0.073789306 |
| VYAYSSCCK | ASSGQGAAETLY | 0.1351759 |
| APAPAPAPNI | ASSGQGAAETLY | 0.11794012 |
| PGGPVPIP | ASSGQGAAETLY | 0.12848914 |
| FLTVALEP | ASSGQGAAETLY | 0.109626696 |
| SQYFGANL | ASSGQGAAETLY | 0.08666982 |
| GGVKPAA | ASSGQGAAETLY | 0.09193416 |
| GGTGGAGGAGGW | ASSGQGAAETLY | 0.12126251 |
| GVGGVAGLGGAGG | ASSGQGAAETLY | 0.105357505 |
| VLLVLATTI | ASSGQGAAETLY | 0.12073361 |
| IGIFAAML | ASSGQGAAETLY | 0.11290114 |
| AVGDLID | ASSGQGAAETLY | 0.075934574 |
| NISPLPL | ASSGQGAAETLY | 0.09177072 |
| VTAVFVAI | ASSGQGAAETLY | 0.06842562 |

| | | |
|---|---|---|
| SGDDNDAAL | ASSGQGAAETLY | 0.07912164 |
| AVAALAHS | ASSGQGAAETLY | 0.117372565 |
| GGVTVMATP | ASSGQGAAETLY | 0.0857299 |
| AGLAANVRD | ASSGQGAAETLY | 0.18999647 |
| TNRPDLIDPALLRPG | ASSGQGAAETLY | 0.15430191 |
| LANLGVTAL | ASSGQGAAETLY | 0.1371691 |
| TGERIEL | ASSGQGAAETLY | 0.12617774 |
| NGEELAA | ASSGQGAAETLY | 0.0750621 |
| VLSLVWG | ASSGQGAAETLY | 0.078023605 |
| IGYALWL | ASSGQGAAETLY | 0.093931474 |
| LSLLLAPL | ASSGQGAAETLY | 0.09724579 |
| TLDEALK | ASSGQGAAETLY | 0.12859936 |
| TDGAGTAEGG | ASSGQGAAETLY | 0.0944426 |
| NAPGGAGGNGG | ASSGQGAAETLY | 0.18330158 |
| VSVFMGMGGNF | ASSGQGAAETLY | 0.17790753 |
| LASYLAAL | ASSGQGAAETLY | 0.11785231 |
| LVGGTPAL | ASSGQGAAETLY | 0.1035325 |
| VRNADVL | ASSGQGAAETLY | 0.076376066 |
| TCSSPFLVSAM | ASSGQGAAETLY | 0.08181367 |
| VHITDAE | ASSGQGAAETLY | 0.085666426 |
| EYAAPSF | ASSGQGAAETLY | 0.096015394 |
| VVDAFHGGL | ASSGQGAAETLY | 0.16291796 |
| ASVVDPQ | ASSGQGAAETLY | 0.11297186 |
| SYNEFGGLS | ASSGQGAAETLY | 0.2017568 |
| DPDGSVY | ASSGQGAAETLY | 0.09199191 |
| MVSTIHATE | ASSGQGAAETLY | 0.07470777 |
| LAAGVADPV | ASSGQGAAETLY | 0.12806907 |
| TKEQIAA | ASSGQGAAETLY | 0.09422881 |
| SLASPAAL | ASSGQGAAETLY | 0.107146695 |
| IGLVFYGV | ASSGQGAAETLY | 0.116461344 |
| FLLDASNL | ASSGQGAAETLY | 0.09412508 |
| YQVGNFP | ASSGQGAAETLY | 0.093339175 |
| ESLLSQST | ASSGQGAAETLY | 0.12939742 |
| LFTVGFAAVG | ASSGQGAAETLY | 0.0953687 |
| ATPPTPT | ASSGQGAAETLY | 0.08845264 |
| TDLATLG | ASSGQGAAETLY | 0.09598567 |
| SAGGTGR | ASSGQGAAETLY | 0.08692251 |
| GFSNSGS | ASSGQGAAETLY | 0.14036477 |
| GGNGGSGGTSV | ASSGQGAAETLY | 0.07641707 |
| GTSVATGGAGN | ASSGQGAAETLY | 0.10901715 |
| NGGNGGSGGTS | ASSGQGAAETLY | 0.10085776 |
| AVVKFPH | ASSGQGAAETLY | 0.0914703 |
| TPPPPPP | ASSGQGAAETLY | 0.083343804 |
| AAPAAAAAPA | ASSGQGAAETLY | 0.091391794 |
| IALNEDL | ASSGQGAAETLY | 0.09214303 |
| IVNVSSM | ASSGQGAAETLY | 0.10533155 |
| TGIPPAPR | ASSGQGAAETLY | 0.079718925 |
| FDLGGGTF | ASSGQGAAETLY | 0.1021565 |
| FDLGGGTFDVS | ASSGQGAAETLY | 0.12163208 |
| NEPTAAA | ASSGQGAAETLY | 0.102804124 |
| NEPTAAALA | ASSGQGAAETLY | 0.14378159 |

| | | |
|---|---|---|
| GSIINISSI | ASSGQGAAETLY | 0.07058683 |
| VAPGNAGTA | ASSGQGAAETLY | 0.08914217 |
| TGGGASLE | ASSGQGAAETLY | 0.15862633 |
| VAVYNHYKRI | ASSGQGAAETLY | 0.13330366 |
| TDPAPATT | ASSGQGAAETLY | 0.11419432 |
| TNRPDILDPALLRPG | ASSGQGAAETLY | 0.070295736 |
| LTTTTTS | ASSGQGAAETLY | 0.07020561 |
| PTGSGKTL | ASSGQGAAETLY | 0.07169137 |
| TGSGKTL | ASSGQGAAETLY | 0.15020977 |
| TIGHVDHGK | ASSGQGAAETLY | 0.07588807 |
| TIGHVDHGKTT | ASSGQGAAETLY | 0.10614305 |
| GHVDHGKTT | ASSGQGAAETLY | 0.09513405 |
| AQFEHTLL | ASSGQGAAETLY | 0.112644345 |
| IVNQDEL | ASSGQGAAETLY | 0.08770477 |
| TNGSQFF | ASSGQGAAETLY | 0.09941876 |
| TGKGIVE | ASSGQGAAETLY | 0.08962989 |
| LISNASDA | ASSGQGAAETLY | 0.14472663 |
| AELRTLGELTKIAKAHGAQ | ASSGQGAAETLY | 0.10924979 |
| APNAVAAAEPSW | ASSGQGAAETLY | 0.0771934 |
| APQVAGAATAVL | ASSGQGAAETLY | 0.092160985 |
| ARRDPAAAAAILPTDA | ASSGQGAAETLY | 0.19342288 |
| DAAARDVLAEAGLAEYF | ASSGQGAAETLY | 0.12634411 |
| DDQIVLSAKANLAKVEQ | ASSGQGAAETLY | 0.1422852 |
| DELLPDTGEDSDEDGH | ASSGQGAAETLY | 0.11276827 |
| ELPEAPANRATVPQD | ASSGQGAAETLY | 0.06837968 |
| FGQSKIERVTGIG | ASSGQGAAETLY | 0.07274974 |
| GAGGTGSTAGGAGGAGGAGGLY | ASSGQGAAETLY | 0.22557023 |
| GHGGTGAAGGNGGTGGAGGSGI | ASSGQGAAETLY | 0.18838793 |
| GNGGNGGAGGEGGPGGAGGAGA | ASSGQGAAETLY | 0.11505467 |
| GSTAGNGGNGGNATNGTIAGQP | ASSGQGAAETLY | 0.10180744 |
| GTGGDGGNGGNGGTGG | ASSGQGAAETLY | 0.13151379 |
| GVRRPAGIPADAPV | ASSGQGAAETLY | 0.070242144 |
| GYRQVTDAHLLTLARR | ASSGQGAAETLY | 0.08193877 |
| IAKMGQDGHDRG | ASSGQGAAETLY | 0.09869793 |
| KAEPSIEIQYT | ASSGQGAAETLY | 0.0964087 |
| KPVRIRMAADEDTVAVT | ASSGQGAAETLY | 0.08189804 |
| MPPRIAGMRLLVIKPEP | ASSGQGAAETLY | 0.09970135 |
| MKFYAVALFRSRGVPPTR | ASSGQGAAETLY | 0.15524499 |
| NAGGFGILGAVAHSP | ASSGQGAAETLY | 0.07454615 |
| NEGREITLAMAEM | ASSGQGAAETLY | 0.09196431 |
| NELVEWSHPLGHRWRWE | ASSGQGAAETLY | 0.06873709 |
| NLGAGGTGGNGGNPG | ASSGQGAAETLY | 0.11192946 |
| NTEWPKWETVMAADPVD | ASSGQGAAETLY | 0.17294532 |
| PASAAPAETQP | ASSGQGAAETLY | 0.13119934 |
| QQSSDGKNGVRAAEITGE | ASSGQGAAETLY | 0.10762407 |
| RAPDAVTTSWWKE | ASSGQGAAETLY | 0.1392246 |
| SGFGNVGAHTSGWFNQ | ASSGQGAAETLY | 0.27370763 |
| SGVAGAVGQAASVGGLKV | ASSGQGAAETLY | 0.077193126 |
| SNGGAGTGSAGGNGGT | ASSGQGAAETLY | 0.087188505 |
| TGGAGGEGGAAG | ASSGQGAAETLY | 0.10267203 |
| TGGDGGAAGAGGAPGHG | ASSGQGAAETLY | 0.13731764 |

| | | |
|---|---|---|
| TMPPLLAAIANAGRHGV | ASSGQGAAETLY | 0.15317227 |
| TSSTAGNSGGA | ASSGQGAAETLY | 0.11742963 |
| VPATVPVGILC | ASSGQGAAETLY | 0.12952185 |
| VQRPDSGGVVLTGRIS | ASSGQGAAETLY | 0.11835709 |
| WTPNPVPATKGVPAGLVQAT | ASSGQGAAETLY | 0.098536834 |
| YALKRALAKFSDRFDVVII | ASSGQGAAETLY | 0.09822283 |
| YLIGSVIV | ASSGQGAAETLY | 0.10125448 |
| YGQLVAVID | ASSGQGAAETLY | 0.16349891 |
| VTVGGMTLPA | ASSGQGAAETLY | 0.06856205 |
| VLALPAV | ASSGQGAAETLY | 0.08297324 |
| VIPPTLNYET | ASSGQGAAETLY | 0.069522105 |
| VGMGGMHPSSG | ASSGQGAAETLY | 0.10943529 |
| VGLAGGAATA | ASSGQGAAETLY | 0.10955103 |
| VGGAGGAGGSGP | ASSGQGAAETLY | 0.1016628 |
| VFAAFMV | ASSGQGAAETLY | 0.083212085 |
| TLYDKLR | ASSGQGAAETLY | 0.07961266 |
| TLAVFTVL | ASSGQGAAETLY | 0.11092612 |
| TGSVQPARV | ASSGQGAAETLY | 0.073799 |
| TGGGAGSGGT | ASSGQGAAETLY | 0.093217455 |
| SHLETSL | ASSGQGAAETLY | 0.084640495 |
| SFTTGMT | ASSGQGAAETLY | 0.12114598 |
| QVEVAKRMGVRQPTVS | ASSGQGAAETLY | 0.08992098 |
| QGSGGPALAM | ASSGQGAAETLY | 0.07852098 |
| MLAPSNP | ASSGQGAAETLY | 0.1494755 |
| LVNNAAI | ASSGQGAAETLY | 0.086287715 |
| LTGEINTLA | ASSGQGAAETLY | 0.118907146 |
| LSIFGPV | ASSGQGAAETLY | 0.10189769 |
| LLALLANL | ASSGQGAAETLY | 0.07703611 |
| LISNASDALDKLR | ASSGQGAAETLY | 0.09062273 |
| LINEAVP | ASSGQGAAETLY | 0.12751682 |
| LGYAFSGL | ASSGQGAAETLY | 0.099894986 |
| LDFVAVAL | ASSGQGAAETLY | 0.10217443 |
| LAQPGAE | ASSGQGAAETLY | 0.1519293 |
| ISNLGAVTGF | ASSGQGAAETLY | 0.12667872 |
| HQEDPGRI | ASSGQGAAETLY | 0.13973278 |
| GTPGEPTP | ASSGQGAAETLY | 0.07353964 |
| GFFNSGS | ASSGQGAAETLY | 0.09868555 |
| GAGGGGLTAGAN | ASSGQGAAETLY | 0.08456423 |
| FVDALVNM | ASSGQGAAETLY | 0.09539463 |
| FLTVAPEL | ASSGQGAAETLY | 0.15861417 |
| FETLTAL | ASSGQGAAETLY | 0.14648192 |
| DVINPIEL | ASSGQGAAETLY | 0.072477505 |
| APVSGAPV | ASSGQGAAETLY | 0.23409906 |
| ALVGGLAW | ASSGQGAAETLY | 0.09427841 |
| ALAAAAATAI | ASSGQGAAETLY | 0.07874435 |
| AFGLGLLL | ASSGQGAAETLY | 0.07862621 |
| AFFAAASL | ASSGQGAAETLY | 0.11070229 |
| AAAALPAFSPP | ASSGQGAAETLY | 0.09183051 |
| VVLAPAL | ASSGQGAAETLY | 0.08285574 |
| TIGHVDHGKTTL | ASSGQGAAETLY | 0.11537066 |
| LISNASDALDK | ASSGQGAAETLY | 0.086558454 |

| | | |
|---|---|---|
| LISNASDAL | ASSGQGAAETLY | 0.08855331 |
| ISNASDALDK | ASSGQGAAETLY | 0.07833562 |
| NQAETLVYQTE | ASSGQGAAETLY | 0.115797974 |
| TGGTGGTAGAG | ASSGQGAAETLY | 0.08022429 |
| TGSTAGGAGGA | ASSGQGAAETLY | 0.06962125 |
| DPLAVPVAAGPV | ASSGQGAAETLY | 0.10565521 |
| GGTGGTPGNGGN | ASSGQGAAETLY | 0.0844599 |
| PAGLAGVDPATL | ASSGQGAAETLY | 0.09882908 |
| GNNLVVEAPHAGASV | ASSGQGAAETLY | 0.078515396 |
| KVTRIMSASTLEQPA | ASSGQGAAETLY | 0.20105422 |
| LGFALPIALPRRTVV | ASSGQGAAETLY | 0.10463324 |
| TRLGNLTARKLVVLV | ASSGQGAAETLY | 0.08930951 |
| WSQQRRLIYFPSAGP | ASSGQGAAETLY | 0.08636994 |
| GPNGNPDPMAAAVDIR | ASSGQGAAETLY | 0.11158682 |
| TQRCVSVADLRAAASR | ASSGQGAAETLY | 0.06931542 |
| QFVLDVAEHPVEVLTETV | ASSGQGAAETLY | 0.10296329 |
| GGAGGTGGAGGNGGTQAGDG | ASSGQGAAETLY | 0.14317083 |
| PLLATLRAIAVDVDAKPAQV | ASSGQGAAETLY | 0.06837848 |
| APKFRTGGYMIDFWGVGYQVAK | ASSGQGAAETLY | 0.11599399 |
| GAGGAGGAGGAAGGPGGTGGTG | ASSGQGAAETLY | 0.08255068 |
| IGALGLLATDVAPAGSVAFGVA | ASSGQGAAETLY | 0.12760474 |
| AGGAGGIGGNGGAGGAGGVLMG | ASSGQGAAETLY | 0.082281426 |
| ADELIGLL | ASSGQGAAETLY | 0.06834055 |
| ADRALKA | ASSGQGAAETLY | 0.12503389 |
| AGAVGQVGGAGGA | ASSGQGAAETLY | 0.14102425 |
| AGGGGGLGGAP | ASSGQGAAETLY | 0.1460495 |
| AGGLLYGNGGAG | ASSGQGAAETLY | 0.08730116 |
| AGLESIGL | ASSGQGAAETLY | 0.09231717 |
| AGLKTGII | ASSGQGAAETLY | 0.121962056 |
| ASLAALLL | ASSGQGAAETLY | 0.12424028 |
| ASVINISSI | ASSGQGAAETLY | 0.09364586 |
| ATPPGPA | ASSGQGAAETLY | 0.08138044 |
| ATVASALV | ASSGQGAAETLY | 0.118165426 |
| ATVASLIL | ASSGQGAAETLY | 0.105115466 |
| AVNGPVLIRH | ASSGQGAAETLY | 0.07368911 |
| DGAAGAVGI | ASSGQGAAETLY | 0.089503616 |
| DPLLHGI | ASSGQGAAETLY | 0.08206674 |
| ELGAALGAL | ASSGQGAAETLY | 0.1019965 |
| ELGRTQAV | ASSGQGAAETLY | 0.113643 |
| FVAGGYLS | ASSGQGAAETLY | 0.06944843 |
| GASGGAGGAGGW | ASSGQGAAETLY | 0.19313791 |
| GGAGAGGDGGAGGI | ASSGQGAAETLY | 0.0895459 |
| GTGHNGSQPGMGGNG | ASSGQGAAETLY | 0.1636163 |
| GYAAASASAS | ASSGQGAAETLY | 0.08432138 |
| HQVDVAAVQQG | ASSGQGAAETLY | 0.10663868 |
| IGAAIAHT | ASSGQGAAETLY | 0.21063945 |
| IGMIEVAQL | ASSGQGAAETLY | 0.085902 |
| IRLAPEHTSAMD | ASSGQGAAETLY | 0.12345433 |
| IYNVSVL | ASSGQGAAETLY | 0.115277074 |
| KQGFLGM | ASSGQGAAETLY | 0.14625818 |
| LESVITQL | ASSGQGAAETLY | 0.08139497 |

| | | |
|---|---|---|
| LGLGALVVV | ASSGQGAAETLY | 0.09652547 |
| LIGNGGNGGSGGI | ASSGQGAAETLY | 0.083973244 |
| LNAINAPTL | ASSGQGAAETLY | 0.1935609 |
| LQLLNTGV | ASSGQGAAETLY | 0.06756707 |
| LQTSIEF | ASSGQGAAETLY | 0.09258938 |
| MSFAVSEL | ASSGQGAAETLY | 0.083523475 |
| NTGALGL | ASSGQGAAETLY | 0.20335911 |
| PPGAGGAGGAGTT | ASSGQGAAETLY | 0.1357171 |
| QLVGLGI | ASSGQGAAETLY | 0.164514 |
| RFLEQVHQL | ASSGQGAAETLY | 0.101666585 |
| RGVGYVLR | ASSGQGAAETLY | 0.16946024 |
| SATGVFAGVI | ASSGQGAAETLY | 0.20130935 |
| SLPAITLP | ASSGQGAAETLY | 0.17435059 |
| STALAFTV | ASSGQGAAETLY | 0.08580757 |
| SVLTVGLGK | ASSGQGAAETLY | 0.107979536 |
| TALAGAGSGI | ASSGQGAAETLY | 0.11813023 |
| TGLPFTGI | ASSGQGAAETLY | 0.082893826 |
| VAEHVTV | ASSGQGAAETLY | 0.074864514 |
| VDIVLNSL | ASSGQGAAETLY | 0.091445364 |
| VLTGGGAVL | ASSGQGAAETLY | 0.09235797 |
| VNVGLAYGI | ASSGQGAAETLY | 0.10172511 |
| VSSAAVGAV | ASSGQGAAETLY | 0.1048873 |
| VTSVYEAGV | ASSGQGAAETLY | 0.1308302 |
| WVALDRGA | ASSGQGAAETLY | 0.13764848 |
| YLVGSVVI | ASSGQGAAETLY | 0.16339955 |
| GPPGTGKTLL | ASSGQGAAETLY | 0.1249236 |
| GVGMTKF | ASSGQGAAETLY | 0.07967657 |
| IGIGGSDL | ASSGQGAAETLY | 0.10896321 |
| IGTLLAEL | ASSGQGAAETLY | 0.089629374 |
| IYTSGTT | ASSGQGAAETLY | 0.08529657 |
| LSFPAGV | ASSGQGAAETLY | 0.09190863 |
| LYTSGTT | ASSGQGAAETLY | 0.10486832 |
| VGAFPFGF | ASSGQGAAETLY | 0.15297917 |
| VGAGTVEF | ASSGQGAAETLY | 0.065338 |
| VGKSTLLL | ASSGQGAAETLY | 0.07693491 |
| AALAHAGGGIV | ASSGQGAAETLY | 0.15068662 |
| AALVAVSLL | ASSGQGAAETLY | 0.11218796 |
| AAVSRAVAAL | ASSGQGAAETLY | 0.09912891 |
| AGGSSGIGGF | ASSGQGAAETLY | 0.095560916 |
| AGLTVANANG | ASSGQGAAETLY | 0.06992321 |
| AGLVGAVL | ASSGQGAAETLY | 0.16117692 |
| ALSQAFTV | ASSGQGAAETLY | 0.079462126 |
| ALTFSVSL | ASSGQGAAETLY | 0.08787197 |
| ALTGAALVAL | ASSGQGAAETLY | 0.11319169 |
| ASFSIGAGVL | ASSGQGAAETLY | 0.113905095 |
| DGSVYKI | ASSGQGAAETLY | 0.088000596 |
| EQLAEIV | ASSGQGAAETLY | 0.07348956 |
| FAVSELTVM | ASSGQGAAETLY | 0.09376083 |
| GAAPLAAGL | ASSGQGAAETLY | 0.101384506 |
| GADGGDGGSGGAGG | ASSGQGAAETLY | 0.15678205 |
| GFGLDTGGAGG | ASSGQGAAETLY | 0.11190847 |

| | | |
|---|---|---|
| GLASGGLL | ASSGQGAAETLY | 0.08353436 |
| GRIAGAAVLL | ASSGQGAAETLY | 0.21489684 |
| GSAGSLNTSF | ASSGQGAAETLY | 0.14648284 |
| GTGQIGI | ASSGQGAAETLY | 0.09879401 |
| GTLIALIL | ASSGQGAAETLY | 0.10721375 |
| IAATMGFVVG | ASSGQGAAETLY | 0.20890905 |
| IFAQALL | ASSGQGAAETLY | 0.17370583 |
| IGAAVALGF | ASSGQGAAETLY | 0.0927857 |
| IGLTPLAL | ASSGQGAAETLY | 0.081957884 |
| ILGSNTSTL | ASSGQGAAETLY | 0.08051775 |
| IVFSLTVL | ASSGQGAAETLY | 0.09156826 |
| LASKATGF | ASSGQGAAETLY | 0.14966857 |
| LASVALIL | ASSGQGAAETLY | 0.12101618 |
| LFAVGLTL | ASSGQGAAETLY | 0.084589124 |
| LFNGGIGTYI | ASSGQGAAETLY | 0.090170406 |
| LTLGAGII | ASSGQGAAETLY | 0.119948715 |
| LVAAALGDI | ASSGQGAAETLY | 0.11734572 |
| NTAFVQLGI | ASSGQGAAETLY | 0.0927603 |
| QLFSGAEL | ASSGQGAAETLY | 0.09249755 |
| SAATFITL | ASSGQGAAETLY | 0.1287224 |
| SNYGLANLGV | ASSGQGAAETLY | 0.09019912 |
| SYGGAYAVM | ASSGQGAAETLY | 0.08369362 |
| TTGVGGLTLG | ASSGQGAAETLY | 0.1901083 |
| TTQTIYI | ASSGQGAAETLY | 0.12802121 |
| VAAGAGLGAVI | ASSGQGAAETLY | 0.081356294 |
| VAAVVTLL | ASSGQGAAETLY | 0.12013169 |
| VGFLGGLI | ASSGQGAAETLY | 0.19113593 |
| VGVIPILL | ASSGQGAAETLY | 0.118025914 |
| VIFASSLI | ASSGQGAAETLY | 0.14627162 |
| VKGGDGGAAAG | ASSGQGAAETLY | 0.08043748 |
| VTKTFTNI | ASSGQGAAETLY | 0.121705584 |
| VVAAAILL | ASSGQGAAETLY | 0.09539378 |
| YAALDVEL | ASSGQGAAETLY | 0.17030655 |
| LASVALLL | ASSGQGAAETLY | 0.1215707 |
| LDAVHRV | ASSGQGAAETLY | 0.09539582 |
| QGLSSSIF | ASSGQGAAETLY | 0.15649569 |
| RGTFANIRL | ASSGQGAAETLY | 0.1576088 |
| RQLWWGHRI | ASSGQGAAETLY | 0.07764888 |
| YGPPGTGKTLL | ASSGQGAAETLY | 0.071181916 |
| AALVGAPANVA | ASSGQGAAETLY | 0.07501829 |
| AATFRRTE | ASSGQGAAETLY | 0.08961776 |
| AGALGGGI | ASSGQGAAETLY | 0.10411472 |
| AGGAGGAGGIGAG | ASSGQGAAETLY | 0.1383088 |
| AGSGALGL | ASSGQGAAETLY | 0.07224467 |
| AQIADQI | ASSGQGAAETLY | 0.220131 |
| ASAVAVTYL | ASSGQGAAETLY | 0.22958475 |
| ASTVLGGTGG | ASSGQGAAETLY | 0.10555752 |
| AVGGEPAK | ASSGQGAAETLY | 0.15733725 |
| CEQGTFGGTC | ASSGQGAAETLY | 0.08124354 |
| DEGKAPAG | ASSGQGAAETLY | 0.07529078 |
| DTVVFPAL | ASSGQGAAETLY | 0.08529949 |

| | | |
|---|---|---|
| FEGITMAG | ASSGQGAAETLY | 0.0743553 |
| FPLVVQD | ASSGQGAAETLY | 0.0904504 |
| GAFTTPQ | ASSGQGAAETLY | 0.08275066 |
| GFGGFGTAGGVGGA | ASSGQGAAETLY | 0.07901871 |
| GFLAQFL | ASSGQGAAETLY | 0.068865344 |
| GGAGGAGGNAGVA | ASSGQGAAETLY | 0.07704397 |
| GGVGTAGAGGAGGAGG | ASSGQGAAETLY | 0.12858886 |
| GNGGGGDAGNAGSGGN | ASSGQGAAETLY | 0.085703194 |
| GTGGAGGAGNDAGS | ASSGQGAAETLY | 0.1343021 |
| GTTNTGWFNS | ASSGQGAAETLY | 0.2038332 |
| IFMGPLFTL | ASSGQGAAETLY | 0.08434412 |
| ILAHAIEAI | ASSGQGAAETLY | 0.10674745 |
| IRSAIREAL | ASSGQGAAETLY | 0.090421125 |
| ITFAGGG | ASSGQGAAETLY | 0.14723712 |
| ITGMSGHQEV | ASSGQGAAETLY | 0.07343778 |
| IVAGLACTL | ASSGQGAAETLY | 0.107430995 |
| IVGGGLAGA | ASSGQGAAETLY | 0.1044457 |
| IVTGVLNI | ASSGQGAAETLY | 0.0701623 |
| LAQQGCI | ASSGQGAAETLY | 0.078528985 |
| LCLHVEN | ASSGQGAAETLY | 0.099492006 |
| LDAISDGS | ASSGQGAAETLY | 0.08637082 |
| LGGFAAML | ASSGQGAAETLY | 0.07470251 |
| LGGKNPMI | ASSGQGAAETLY | 0.08656228 |
| LGPKGVAAIL | ASSGQGAAETLY | 0.09829526 |
| LMVGVSI | ASSGQGAAETLY | 0.08237501 |
| LVGAAFVL | ASSGQGAAETLY | 0.06833139 |
| LVLAVAPL | ASSGQGAAETLY | 0.109803766 |
| MAAIAVIDAA | ASSGQGAAETLY | 0.09715152 |
| NNGINAELQ | ASSGQGAAETLY | 0.12518598 |
| RGEPLVLVHG | ASSGQGAAETLY | 0.21818708 |
| RGGAGGAAGLI | ASSGQGAAETLY | 0.097934954 |
| RSYRMHPAV | ASSGQGAAETLY | 0.08632426 |
| SAAHLLAGQLA | ASSGQGAAETLY | 0.11735633 |
| SAAVGSY | ASSGQGAAETLY | 0.09944234 |
| SPHAVLHRI | ASSGQGAAETLY | 0.12496612 |
| SQIAKVL | ASSGQGAAETLY | 0.07238214 |
| TALGHLVP | ASSGQGAAETLY | 0.08393393 |
| TAPTTSGAA | ASSGQGAAETLY | 0.10360486 |
| TDGDDGNGGSPGT | ASSGQGAAETLY | 0.083974265 |
| TDGLGEN | ASSGQGAAETLY | 0.091666356 |
| TGGPPLPGQA | ASSGQGAAETLY | 0.18229859 |
| TIPAITI | ASSGQGAAETLY | 0.08037514 |
| TIPGFTI | ASSGQGAAETLY | 0.07925887 |
| TSIASGALA | ASSGQGAAETLY | 0.09164075 |
| TVTLPGAAI | ASSGQGAAETLY | 0.11930039 |
| VASGLSAALP | ASSGQGAAETLY | 0.15655597 |
| VGAGVIGI | ASSGQGAAETLY | 0.10223948 |
| VGPKRFGV | ASSGQGAAETLY | 0.1053275 |
| VNVAFPD | ASSGQGAAETLY | 0.110759795 |
| VSMTNSGVS | ASSGQGAAETLY | 0.10046706 |
| VTDVLGDLSV | ASSGQGAAETLY | 0.12458384 |

| | | |
|---|---|---|
| VVVVGAGFAGL | ASSGQGAAETLY | 0.1328984 |
| VYTVGTVV | ASSGQGAAETLY | 0.08567087 |
| WTTFSGP | ASSGQGAAETLY | 0.08598205 |
| YDELVEAF | ASSGQGAAETLY | 0.07567561 |
| YGTGGAGGNGG | ASSGQGAAETLY | 0.075520605 |
| YTNVMAV | ASSGQGAAETLY | 0.22052981 |
| FLEQVHQL | ASSGQGAAETLY | 0.08059386 |
| GLSLLAVAV | ASSGQGAAETLY | 0.08686876 |
| LEQVHQL | ASSGQGAAETLY | 0.09144948 |
| LNAAGAI | ASSGQGAAETLY | 0.13055441 |
| PAVHASL | ASSGQGAAETLY | 0.0933378 |
| TELEDDA | ASSGQGAAETLY | 0.1608917 |
| TPPGPPN | ASSGQGAAETLY | 0.14863725 |
| TPPPPPPP | ASSGQGAAETLY | 0.073067166 |
| TQSPALI | ASSGTGGKDTQY | 0.09491478 |
| EFAAAFPQ | ASSGTGGKDTQY | 0.16238946 |
| LLSGNLDVL | ASSGTGGKDTQY | 0.07737943 |
| GPANLGNYNI | ASSGTGGKDTQY | 0.0779439 |
| TLGPTLP | ASSGTGGKDTQY | 0.08064649 |
| VYAYSSCCK | ASSGTGGKDTQY | 0.08536228 |
| APAPAPAPNI | ASSGTGGKDTQY | 0.09587159 |
| PGGPVPIP | ASSGTGGKDTQY | 0.09428317 |
| FLTVALEP | ASSGTGGKDTQY | 0.124814294 |
| SQYFGANL | ASSGTGGKDTQY | 0.094709806 |
| GGVKPAA | ASSGTGGKDTQY | 0.12594439 |
| GGTGGAGGAGGW | ASSGTGGKDTQY | 0.15722895 |
| GVGGVAGLGGAGG | ASSGTGGKDTQY | 0.115459375 |
| VLLVLATTI | ASSGTGGKDTQY | 0.16598386 |
| IGIFAAML | ASSGTGGKDTQY | 0.07730852 |
| AVGDLID | ASSGTGGKDTQY | 0.10514158 |
| NISPLPL | ASSGTGGKDTQY | 0.11527437 |
| VTAVFVAI | ASSGTGGKDTQY | 0.13259542 |
| SGDDNDAAL | ASSGTGGKDTQY | 0.07500028 |
| AVAALAHS | ASSGTGGKDTQY | 0.16075075 |
| GGVTVMATP | ASSGTGGKDTQY | 0.19401765 |
| AGLAANVRD | ASSGTGGKDTQY | 0.087710805 |
| TNRPDLIDPALLRPG | ASSGTGGKDTQY | 0.16227905 |
| LANLGVTAL | ASSGTGGKDTQY | 0.068736084 |
| TGERIEL | ASSGTGGKDTQY | 0.091483876 |
| NGEELAA | ASSGTGGKDTQY | 0.071552336 |
| VLSLVWG | ASSGTGGKDTQY | 0.12899962 |
| IGYALWL | ASSGTGGKDTQY | 0.092443526 |
| LSLLLAPL | ASSGTGGKDTQY | 0.08680963 |
| TLDEALK | ASSGTGGKDTQY | 0.101371065 |
| TDGAGTAEGG | ASSGTGGKDTQY | 0.15459785 |
| NAPGGAGGNGG | ASSGTGGKDTQY | 0.111398034 |
| VSVFMGMGGNF | ASSGTGGKDTQY | 0.089043394 |
| LASYLAAL | ASSGTGGKDTQY | 0.107566975 |
| LVGGTPAL | ASSGTGGKDTQY | 0.1452885 |
| VRNADVL | ASSGTGGKDTQY | 0.09382711 |
| TCSSPFLVSAM | ASSGTGGKDTQY | 0.07498838 |

| | | |
|---|---|---|
| VHITDAE | ASSGTGGKDTQY | 0.07796944 |
| EYAAPSF | ASSGTGGKDTQY | 0.08773326 |
| VVDAFHGGL | ASSGTGGKDTQY | 0.09977075 |
| ASVVDPQ | ASSGTGGKDTQY | 0.11748957 |
| SYNEFGGLS | ASSGTGGKDTQY | 0.12417324 |
| DPDGSVY | ASSGTGGKDTQY | 0.07662642 |
| MVSTIHATE | ASSGTGGKDTQY | 0.0660908 |
| LAAGVADPV | ASSGTGGKDTQY | 0.10456203 |
| TKEQIAA | ASSGTGGKDTQY | 0.2055108 |
| SLASPAAL | ASSGTGGKDTQY | 0.096488565 |
| IGLVFYGV | ASSGTGGKDTQY | 0.15718822 |
| FLLDASNL | ASSGTGGKDTQY | 0.12516008 |
| YQVGNFP | ASSGTGGKDTQY | 0.107997276 |
| ESLLSQST | ASSGTGGKDTQY | 0.11757079 |
| LFTVGFAAVG | ASSGTGGKDTQY | 0.1199175 |
| ATPPTPT | ASSGTGGKDTQY | 0.09569753 |
| TDLATLG | ASSGTGGKDTQY | 0.07189797 |
| SAGGTGR | ASSGTGGKDTQY | 0.08788883 |
| GFSNSGS | ASSGTGGKDTQY | 0.09352367 |
| GGNGGSGGTSV | ASSGTGGKDTQY | 0.19379985 |
| GTSVATGGAGN | ASSGTGGKDTQY | 0.12496491 |
| NGGNGGSGGTS | ASSGTGGKDTQY | 0.120688975 |
| AVVKFPH | ASSGTGGKDTQY | 0.098569535 |
| TPPPPPP | ASSGTGGKDTQY | 0.111326784 |
| AAPAAAAAPA | ASSGTGGKDTQY | 0.10515561 |
| IALNEDL | ASSGTGGKDTQY | 0.09417793 |
| IVNVSSM | ASSGTGGKDTQY | 0.20675917 |
| TGIPPAPR | ASSGTGGKDTQY | 0.07893004 |
| FDLGGGTF | ASSGTGGKDTQY | 0.086534195 |
| FDLGGGTFDVS | ASSGTGGKDTQY | 0.09015246 |
| NEPTAAA | ASSGTGGKDTQY | 0.14346889 |
| NEPTAAALA | ASSGTGGKDTQY | 0.095104106 |
| GSIINISSI | ASSGTGGKDTQY | 0.13210201 |
| VAPGNAGTA | ASSGTGGKDTQY | 0.10195594 |
| TGGGASLE | ASSGTGGKDTQY | 0.075231925 |
| VAVYNHYKRI | ASSGTGGKDTQY | 0.087662235 |
| TDPAPATT | ASSGTGGKDTQY | 0.08302423 |
| TNRPDILDPALLRPG | ASSGTGGKDTQY | 0.17507027 |
| LTTTTTS | ASSGTGGKDTQY | 0.10057166 |
| PTGSGKTL | ASSGTGGKDTQY | 0.07320246 |
| TGSGKTL | ASSGTGGKDTQY | 0.09347167 |
| TIGHVDHGK | ASSGTGGKDTQY | 0.20421736 |
| TIGHVDHGKTT | ASSGTGGKDTQY | 0.09012631 |
| GHVDHGKTT | ASSGTGGKDTQY | 0.22592004 |
| AQFEHTLL | ASSGTGGKDTQY | 0.107751 |
| IVNQDEL | ASSGTGGKDTQY | 0.09564293 |
| TNGSQFF | ASSGTGGKDTQY | 0.23027825 |
| TGKGIVE | ASSGTGGKDTQY | 0.0686625 |
| LISNASDA | ASSGTGGKDTQY | 0.1294089 |
| AELRTLGELTKIAKAHGAQ | ASSGTGGKDTQY | 0.073189504 |
| APNAVAAAEPSW | ASSGTGGKDTQY | 0.080191396 |

| | | |
|---|---|---|
| APQVAGAATAVL | ASSGTGGKDTQY | 0.109635584 |
| ARRDPAAAAAILPTDA | ASSGTGGKDTQY | 0.09342355 |
| DAAARDVLAEAGLAEYF | ASSGTGGKDTQY | 0.105181634 |
| DDQIVLSAKANLAKVEQ | ASSGTGGKDTQY | 0.10467447 |
| DELLPDTGEDSDEDGH | ASSGTGGKDTQY | 0.0713991 |
| ELPEAPANRATVPQD | ASSGTGGKDTQY | 0.08585931 |
| FGQSKIERVTGIG | ASSGTGGKDTQY | 0.12610085 |
| GAGGTGSTAGGAGGAGGAGGLY | ASSGTGGKDTQY | 0.07968222 |
| GHGGTGAAGGNGGTGGAGGSGI | ASSGTGGKDTQY | 0.128744 |
| GNGGNGGAGGEGGPGGAGGAGA | ASSGTGGKDTQY | 0.07903684 |
| GSTAGNGGNGGNATNGTIAGQP | ASSGTGGKDTQY | 0.077535555 |
| GTGGDGGNGGNGGTGG | ASSGTGGKDTQY | 0.08261178 |
| GVRRPAGIPADAPV | ASSGTGGKDTQY | 0.09727507 |
| GYRQVTDAHLLTLARR | ASSGTGGKDTQY | 0.07255848 |
| IAKMGQDGHDRG | ASSGTGGKDTQY | 0.10280905 |
| KAEPSIEIQYT | ASSGTGGKDTQY | 0.1425328 |
| KPVRIRMAADEDTVAVT | ASSGTGGKDTQY | 0.086050294 |
| MPPRIAGMRLLVIKPEP | ASSGTGGKDTQY | 0.08271681 |
| MKFYAVALFRSRGVPPTR | ASSGTGGKDTQY | 0.09522984 |
| NAGGFGILGAVAHSP | ASSGTGGKDTQY | 0.07481298 |
| NEGREITLAMAEM | ASSGTGGKDTQY | 0.07412592 |
| NELVEWSHPLGHRWRWE | ASSGTGGKDTQY | 0.14292625 |
| NLGAGGTGGNGGNPG | ASSGTGGKDTQY | 0.2742745 |
| NTEWPKWETVMAADPVD | ASSGTGGKDTQY | 0.09186422 |
| PASAAPAETQP | ASSGTGGKDTQY | 0.08433499 |
| QQSSDGKNGVRAAEITGE | ASSGTGGKDTQY | 0.06717796 |
| RAPDAVTTSWWKE | ASSGTGGKDTQY | 0.23323421 |
| SGFGNVGAHTSGWFNQ | ASSGTGGKDTQY | 0.0938981 |
| SGVAGAVGQAASVGGLKV | ASSGTGGKDTQY | 0.118872814 |
| SNGGAGTGSAGGNGGT | ASSGTGGKDTQY | 0.08552215 |
| TGGAGGEGGAAG | ASSGTGGKDTQY | 0.08843854 |
| TGGDGGAAGAGGAPGHG | ASSGTGGKDTQY | 0.1286901 |
| TMPPLLAAIANAGRHGV | ASSGTGGKDTQY | 0.08517947 |
| TSSTAGNSGGA | ASSGTGGKDTQY | 0.0725105 |
| VPATVPVGILC | ASSGTGGKDTQY | 0.077069856 |
| VQRPDSGGVVLTGRIS | ASSGTGGKDTQY | 0.11673022 |
| WTPNPVPATKGVPAGLVQAT | ASSGTGGKDTQY | 0.077569745 |
| YALKRALAKFSDRFDVVII | ASSGTGGKDTQY | 0.11271656 |
| YLIGSVIV | ASSGTGGKDTQY | 0.2594571 |
| YGQLVAVID | ASSGTGGKDTQY | 0.21088535 |
| VTVGGMTLPA | ASSGTGGKDTQY | 0.10281529 |
| VLALPAV | ASSGTGGKDTQY | 0.09466086 |
| VIPPTLNYET | ASSGTGGKDTQY | 0.132507 |
| VGMGGMHPSSG | ASSGTGGKDTQY | 0.10283206 |
| VGLAGGAATA | ASSGTGGKDTQY | 0.08669415 |
| VGGAGGAGGSGP | ASSGTGGKDTQY | 0.14672361 |
| VFAAFMV | ASSGTGGKDTQY | 0.1330816 |
| TLYDKLR | ASSGTGGKDTQY | 0.11174652 |
| TLAVFTVL | ASSGTGGKDTQY | 0.0837583 |
| TGSVQPARV | ASSGTGGKDTQY | 0.091363624 |
| TGGGAGSGGT | ASSGTGGKDTQY | 0.09652462 |

| | | |
|---|---|---|
| SHLETSL | ASSGTGGKDTQY | 0.14034864 |
| SFTTGMT | ASSGTGGKDTQY | 0.16605566 |
| QVEVAKRMGVRQPTVS | ASSGTGGKDTQY | 0.13915144 |
| QGSGGPALAM | ASSGTGGKDTQY | 0.1314307 |
| MLAPSNP | ASSGTGGKDTQY | 0.1127926 |
| LVNNAAI | ASSGTGGKDTQY | 0.095779285 |
| LTGEINTLA | ASSGTGGKDTQY | 0.079048775 |
| LSIFGPV | ASSGTGGKDTQY | 0.21616739 |
| LLALLANL | ASSGTGGKDTQY | 0.07777607 |
| LISNASDALDKLR | ASSGTGGKDTQY | 0.083988406 |
| LINEAVP | ASSGTGGKDTQY | 0.10565388 |
| LGYAFSGL | ASSGTGGKDTQY | 0.08947621 |
| LDFVAVAL | ASSGTGGKDTQY | 0.12942375 |
| LAQPGAE | ASSGTGGKDTQY | 0.0876125 |
| ISNLGAVTGF | ASSGTGGKDTQY | 0.07022352 |
| HQEDPGRI | ASSGTGGKDTQY | 0.17115167 |
| GTPGEPTP | ASSGTGGKDTQY | 0.08372402 |
| GFFNSGS | ASSGTGGKDTQY | 0.09446893 |
| GAGGGGLTAGAN | ASSGTGGKDTQY | 0.12978347 |
| FVDALVNM | ASSGTGGKDTQY | 0.08118815 |
| FLTVAPEL | ASSGTGGKDTQY | 0.13333333 |
| FETLTAL | ASSGTGGKDTQY | 0.12837313 |
| DVINPIEL | ASSGTGGKDTQY | 0.10537401 |
| APVSGAPV | ASSGTGGKDTQY | 0.1199247 |
| ALVGGLAW | ASSGTGGKDTQY | 0.06822764 |
| ALAAAAATAI | ASSGTGGKDTQY | 0.08329993 |
| AFGLGLLL | ASSGTGGKDTQY | 0.08273463 |
| AFFAAASL | ASSGTGGKDTQY | 0.1712129 |
| AAAALPAFSPP | ASSGTGGKDTQY | 0.13143013 |
| VVLAPAL | ASSGTGGKDTQY | 0.09078972 |
| TIGHVDHGKTTL | ASSGTGGKDTQY | 0.1099789 |
| LISNASDALDK | ASSGTGGKDTQY | 0.10340156 |
| LISNASDAL | ASSGTGGKDTQY | 0.10589739 |
| ISNASDALDK | ASSGTGGKDTQY | 0.07773991 |
| NQAETLVYQTE | ASSGTGGKDTQY | 0.07457846 |
| TGGTGGTAGAG | ASSGTGGKDTQY | 0.08360832 |
| TGSTAGGAGGA | ASSGTGGKDTQY | 0.10433825 |
| DPLAVPVAAGPV | ASSGTGGKDTQY | 0.09875556 |
| GGTGGTPGNGGN | ASSGTGGKDTQY | 0.115089506 |
| PAGLAGVDPATL | ASSGTGGKDTQY | 0.08369771 |
| GNNLVVEAPHAGASV | ASSGTGGKDTQY | 0.089220695 |
| KVTRIMSASTLEQPA | ASSGTGGKDTQY | 0.08872073 |
| LGFALPIALPRRTVV | ASSGTGGKDTQY | 0.094106875 |
| TRLGNLTARKLVVLV | ASSGTGGKDTQY | 0.11630714 |
| WSQQRRLIYFPSAGP | ASSGTGGKDTQY | 0.12849352 |
| GPNGNPDPMAAAVDIR | ASSGTGGKDTQY | 0.09358958 |
| TQRCVSVADLRAAASR | ASSGTGGKDTQY | 0.10710613 |
| QFVLDVAEHPVEVLTETV | ASSGTGGKDTQY | 0.16346723 |
| GGAGGTGGAGGNGGTQAGDG | ASSGTGGKDTQY | 0.081256315 |
| PLLATLRAIAVDVDAKPAQV | ASSGTGGKDTQY | 0.081686184 |
| APKFRTGGYMIDFWGVGYQVAK | ASSGTGGKDTQY | 0.08645677 |

| | | |
|---|---|---|
| GAGGAGGAGGAAGGPGGTGGTG | ASSGTGGKDTQY | 0.12990162 |
| IGALGLLATDVAPAGSVAFGVA | ASSGTGGKDTQY | 0.11077625 |
| AGGAGGIGGNGGAGGAGGVLMG | ASSGTGGKDTQY | 0.070445664 |
| ADELIGLL | ASSGTGGKDTQY | 0.12949328 |
| ADRALKA | ASSGTGGKDTQY | 0.13039766 |
| AGAVGQVGGAGGA | ASSGTGGKDTQY | 0.08940054 |
| AGGGGGLGGAP | ASSGTGGKDTQY | 0.06608299 |
| AGGLLYGNGGAG | ASSGTGGKDTQY | 0.1294187 |
| AGLESIGL | ASSGTGGKDTQY | 0.15065295 |
| AGLKTGII | ASSGTGGKDTQY | 0.10307852 |
| ASLAALLL | ASSGTGGKDTQY | 0.08171998 |
| ASVINISSI | ASSGTGGKDTQY | 0.068342134 |
| ATPPGPA | ASSGTGGKDTQY | 0.10667099 |
| ATVASALV | ASSGTGGKDTQY | 0.1052125 |
| ATVASLIL | ASSGTGGKDTQY | 0.086415105 |
| AVNGPVLIRH | ASSGTGGKDTQY | 0.12632509 |
| DGAAGAVGI | ASSGTGGKDTQY | 0.08563902 |
| DPLLHGI | ASSGTGGKDTQY | 0.089996286 |
| ELGAALGAL | ASSGTGGKDTQY | 0.1286175 |
| ELGRTQAV | ASSGTGGKDTQY | 0.129573 |
| FVAGGYLS | ASSGTGGKDTQY | 0.086843066 |
| GASGGAGGAGGW | ASSGTGGKDTQY | 0.100244366 |
| GGAGAGGDGGAGGI | ASSGTGGKDTQY | 0.114582375 |
| GTGHNGSQPGMGGNG | ASSGTGGKDTQY | 0.1034728 |
| GYAAASASAS | ASSGTGGKDTQY | 0.10588904 |
| HQVDVAAVQQG | ASSGTGGKDTQY | 0.17947237 |
| IGAAIAHT | ASSGTGGKDTQY | 0.081131436 |
| IGMIEVAQL | ASSGTGGKDTQY | 0.11380243 |
| IRLAPEHTSAMD | ASSGTGGKDTQY | 0.18395343 |
| IYNVSVL | ASSGTGGKDTQY | 0.0774085 |
| KQGFLGM | ASSGTGGKDTQY | 0.12214786 |
| LESVITQL | ASSGTGGKDTQY | 0.086754575 |
| LGLGALVVV | ASSGTGGKDTQY | 0.20583895 |
| LIGNGGNGGSGGI | ASSGTGGKDTQY | 0.12568079 |
| LNAINAPTL | ASSGTGGKDTQY | 0.111917004 |
| LQLLNTGV | ASSGTGGKDTQY | 0.087866545 |
| LQTSIEF | ASSGTGGKDTQY | 0.35011196 |
| MSFAVSEL | ASSGTGGKDTQY | 0.71908027 |
| NTGALGL | ASSGTGGKDTQY | 0.5161454 |
| PPGAGGAGGAGTT | ASSGTGGKDTQY | 0.6297279 |
| QLVGLGI | ASSGTGGKDTQY | 0.33730674 |
| RFLEQVHQL | ASSGTGGKDTQY | 0.10162102 |
| RGVGYVLR | ASSGTGGKDTQY | 0.4790328 |
| SATGVFAGVI | ASSGTGGKDTQY | 0.22081578 |
| SLPAITLP | ASSGTGGKDTQY | 0.16649926 |
| STALAFTV | ASSGTGGKDTQY | 0.56940913 |
| SVLTVGLGK | ASSGTGGKDTQY | 0.3501616 |
| TALAGAGSGI | ASSGTGGKDTQY | 0.80556554 |
| TGLPFTGI | ASSGTGGKDTQY | 0.7514742 |
| VAEHVTV | ASSGTGGKDTQY | 0.1901629 |
| VDIVLNSL | ASSGTGGKDTQY | 0.6036516 |

| | | |
|---|---|---|
| VLTGGGAVL | ASSGTGGKDTQY | 0.22033103 |
| VNVGLAYGI | ASSGTGGKDTQY | 0.5624706 |
| VSSAAVGAV | ASSGTGGKDTQY | 0.08974631 |
| VTSVYEAGV | ASSGTGGKDTQY | 0.66236365 |
| WVALDRGA | ASSGTGGKDTQY | 0.29937148 |
| YLVGSVVI | ASSGTGGKDTQY | 0.07127275 |
| GPPGTGKTLL | ASSGTGGKDTQY | 0.65338844 |
| GVGMTKF | ASSGTGGKDTQY | 0.43889347 |
| IGIGGSDL | ASSGTGGKDTQY | 0.4323957 |
| IGTLLAEL | ASSGTGGKDTQY | 0.8069042 |
| IYTSGTT | ASSGTGGKDTQY | 0.08318804 |
| LSFPAGV | ASSGTGGKDTQY | 0.16810161 |
| LYTSGTT | ASSGTGGKDTQY | 0.71406394 |
| VGAFPFGF | ASSGTGGKDTQY | 0.83058214 |
| VGAGTVEF | ASSGTGGKDTQY | 0.15108287 |
| VGKSTLLL | ASSGTGGKDTQY | 0.71742034 |
| AALAHAGGGIV | ASSGTGGKDTQY | 0.8322508 |
| AALVAVSLL | ASSGTGGKDTQY | 0.48661086 |
| AAVSRAVAAL | ASSGTGGKDTQY | 0.71391904 |
| AGGSSGIGGF | ASSGTGGKDTQY | 0.7759179 |
| AGLTVANANG | ASSGTGGKDTQY | 0.14399105 |
| AGLVGAVL | ASSGTGGKDTQY | 0.77786726 |
| ALSQAFTV | ASSGTGGKDTQY | 0.6577743 |
| ALTFSVSL | ASSGTGGKDTQY | 0.4037559 |
| ALTGAALVAL | ASSGTGGKDTQY | 0.60253465 |
| ASFSIGAGVL | ASSGTGGKDTQY | 0.2766728 |
| DGSVYKI | ASSGTGGKDTQY | 0.41362715 |
| EQLAEIV | ASSGTGGKDTQY | 0.0797077 |
| FAVSELTVM | ASSGTGGKDTQY | 0.7182817 |
| GAAPLAAGL | ASSGTGGKDTQY | 0.5925957 |
| GADGGDGGSGGAGG | ASSGTGGKDTQY | 0.5921564 |
| GFGLDTGGAGG | ASSGTGGKDTQY | 0.25798813 |
| GLASGGLL | ASSGTGGKDTQY | 0.3262807 |
| GRIAGAAVLL | ASSGTGGKDTQY | 0.29347965 |
| GSAGSLNTSF | ASSGTGGKDTQY | 0.51933897 |
| GTGQIGI | ASSGTGGKDTQY | 0.09516031 |
| GTLIALIL | ASSGTGGKDTQY | 0.21667129 |
| IAATMGFVVG | ASSGTGGKDTQY | 0.74441063 |
| IFAQALL | ASSGTGGKDTQY | 0.74152625 |
| IGAAVALGF | ASSGTGGKDTQY | 0.13596483 |
| IGLTPLAL | ASSGTGGKDTQY | 0.76988435 |
| ILGSNTSTL | ASSGTGGKDTQY | 0.6653576 |
| IVFSLTVL | ASSGTGGKDTQY | 0.58805835 |
| LASKATGF | ASSGTGGKDTQY | 0.38568366 |
| LASVALIL | ASSGTGGKDTQY | 0.4175732 |
| LFAVGLTL | ASSGTGGKDTQY | 0.7069933 |
| LFNGGIGTYI | ASSGTGGKDTQY | 0.5459888 |
| LTLGAGII | ASSGTGGKDTQY | 0.13246828 |
| LVAAALGDI | ASSGTGGKDTQY | 0.7989938 |
| NTAFVQLGI | ASSGTGGKDTQY | 0.22830111 |
| QLFSGAEL | ASSGTGGKDTQY | 0.3392579 |

| | | |
|---|---|---|
| SAATFITL | ASSGTGGKDTQY | 0.57718945 |
| SNYGLANLGV | ASSGTGGKDTQY | 0.7743919 |
| SYGGAYAVM | ASSGTGGKDTQY | 0.24655947 |
| TTGVGGLTLG | ASSGTGGKDTQY | 0.16316712 |
| TTQTIYI | ASSGTGGKDTQY | 0.16158146 |
| VAAGAGLGAVI | ASSGTGGKDTQY | 0.58590436 |
| VAAVVTLL | ASSGTGGKDTQY | 0.82750463 |
| VGFLGGLI | ASSGTGGKDTQY | 0.48150808 |
| VGVIPILL | ASSGTGGKDTQY | 0.31881526 |
| VIFASSLI | ASSGTGGKDTQY | 0.76918536 |
| VKGGDGGAAAG | ASSGTGGKDTQY | 0.53024495 |
| VTKTFTNI | ASSGTGGKDTQY | 0.5134489 |
| VVAAAILL | ASSGTGGKDTQY | 0.6656284 |
| YAALDVEL | ASSGTGGKDTQY | 0.42650557 |
| LASVALLL | ASSGTGGKDTQY | 0.7836026 |
| LDAVHRV | ASSGTGGKDTQY | 0.70665294 |
| QGLSSSIF | ASSGTGGKDTQY | 0.7132415 |
| RGTFANIRL | ASSGTGGKDTQY | 0.69306636 |
| RQLWWGHRI | ASSGTGGKDTQY | 0.27701515 |
| YGPPGTGKTLL | ASSGTGGKDTQY | 0.6298511 |
| AALVGAPANVA | ASSGTGGKDTQY | 0.11230221 |
| AATFRRTE | ASSGTGGKDTQY | 0.17433922 |
| AGALGGGI | ASSGTGGKDTQY | 0.67132413 |
| AGGAGGAGGIGAG | ASSGTGGKDTQY | 0.7250916 |
| AGSGALGL | ASSGTGGKDTQY | 0.7121421 |
| AQIADQI | ASSGTGGKDTQY | 0.56599677 |
| ASAVAVTYL | ASSGTGGKDTQY | 0.28986874 |
| ASTVLGGTGG | ASSGTGGKDTQY | 0.71041316 |
| AVGGEPAK | ASSGTGGKDTQY | 0.7964387 |
| CEQGTFGGTC | ASSGTGGKDTQY | 0.16851678 |
| DEGKAPAG | ASSGTGGKDTQY | 0.33082157 |
| DTVVFPAL | ASSGTGGKDTQY | 0.80442667 |
| FEGITMAG | ASSGTGGKDTQY | 0.647695 |
| FPLVVQD | ASSGTGGKDTQY | 0.5172372 |
| GAFTTPQ | ASSGTGGKDTQY | 0.37442768 |
| GFGGFGTAGGVGGA | ASSGTGGKDTQY | 0.55132806 |
| GFLAQFL | ASSGTGGKDTQY | 0.17786539 |
| GGAGGAGGNAGVA | ASSGTGGKDTQY | 0.81530386 |
| GGVGTAGAGGAGGAGG | ASSGTGGKDTQY | 0.0906651 |
| GNGGGGDAGNAGSGGN | ASSGTGGKDTQY | 0.48882332 |
| GTGGAGGAGNDAGS | ASSGTGGKDTQY | 0.18089889 |
| GTTNTGWFNS | ASSGTGGKDTQY | 0.14341009 |
| IFMGPLFTL | ASSGTGGKDTQY | 0.62629074 |
| ILAHAIEAI | ASSGTGGKDTQY | 0.33432645 |
| IRSAIREAL | ASSGTGGKDTQY | 0.78290045 |
| ITFAGGG | ASSGTGGKDTQY | 0.34470823 |
| ITGMSGHQEV | ASSGTGGKDTQY | 0.68132937 |
| IVAGLACTL | ASSGTGGKDTQY | 0.63273555 |
| IVGGGLAGA | ASSGTGGKDTQY | 0.5893598 |
| IVTGVLNI | ASSGTGGKDTQY | 0.3145891 |
| LAQQGCI | ASSGTGGKDTQY | 0.60322744 |

| | | |
|---|---|---|
| LCLHVEN | ASSGTGGKDTQY | 0.7680937 |
| LDAISDGS | ASSGTGGKDTQY | 0.7269776 |
| LGGFAAML | ASSGTGGKDTQY | 0.115775615 |
| LGGKNPMI | ASSGTGGKDTQY | 0.42787883 |
| LGPKGVAAIL | ASSGTGGKDTQY | 0.5187872 |
| LMVGVSI | ASSGTGGKDTQY | 0.07672198 |
| LVGAAFVL | ASSGTGGKDTQY | 0.0847041 |
| LVLAVAPL | ASSGTGGKDTQY | 0.24612868 |
| MAAIAVIDAA | ASSGTGGKDTQY | 0.26819167 |
| NNGINAELQ | ASSGTGGKDTQY | 0.801077 |
| RGEPLVLVHG | ASSGTGGKDTQY | 0.75549686 |
| RGGAGGAAGLI | ASSGTGGKDTQY | 0.57427865 |
| RSYRMHPAV | ASSGTGGKDTQY | 0.6603915 |
| SAAHLLAGQLA | ASSGTGGKDTQY | 0.4929166 |
| SAAVGSY | ASSGTGGKDTQY | 0.06826978 |
| SPHAVLHRI | ASSGTGGKDTQY | 0.19680724 |
| SQIAKVL | ASSGTGGKDTQY | 0.23502368 |
| TALGHLVP | ASSGTGGKDTQY | 0.7460918 |
| TAPTTSGAA | ASSGTGGKDTQY | 0.25020766 |
| TDGDDGNGGSPGT | ASSGTGGKDTQY | 0.35206103 |
| TDGLGEN | ASSGTGGKDTQY | 0.6554863 |
| TGGPPLPGQA | ASSGTGGKDTQY | 0.50996226 |
| TIPAITI | ASSGTGGKDTQY | 0.5800195 |
| TIPGFTI | ASSGTGGKDTQY | 0.19728191 |
| TSIASGALA | ASSGTGGKDTQY | 0.07290746 |
| TVTLPGAAI | ASSGTGGKDTQY | 0.6602274 |
| VASGLSAALP | ASSGTGGKDTQY | 0.08114286 |
| VGAGVIGI | ASSGTGGKDTQY | 0.26173833 |
| VGPKRFGV | ASSGTGGKDTQY | 0.087839186 |
| VNVAFPD | ASSGTGGKDTQY | 0.072728686 |
| VSMTNSGVS | ASSGTGGKDTQY | 0.09705605 |
| VTDVLGDLSV | ASSGTGGKDTQY | 0.6824726 |
| VVVVGAGFAGL | ASSGTGGKDTQY | 0.65164995 |
| VYTVGTVV | ASSGTGGKDTQY | 0.5355034 |
| WTTFSGP | ASSGTGGKDTQY | 0.7446442 |
| YDELVEAF | ASSGTGGKDTQY | 0.64328074 |
| YGTGGAGGNGG | ASSGTGGKDTQY | 0.60197204 |
| YTNVMAV | ASSGTGGKDTQY | 0.24809453 |
| FLEQVHQL | ASSGTGGKDTQY | 0.31388652 |
| GLSLLAVAV | ASSGTGGKDTQY | 0.20501669 |
| LEQVHQL | ASSGTGGKDTQY | 0.16626994 |
| LNAAGAI | ASSGTGGKDTQY | 0.8405809 |
| PAVHASL | ASSGTGGKDTQY | 0.40122342 |
| TELEDDA | ASSGTGGKDTQY | 0.12743072 |
| TPPGPPN | ASSGTGGKDTQY | 0.5212921 |
| TPPPPPPP | ASSGTGGKDTQY | 0.7534143 |
| TQSPALI | ASSLRDSNTEVF | 0.7790394 |
| EFAAAFPQ | ASSLRDSNTEVF | 0.43678683 |
| LLSGNLDVL | ASSLRDSNTEVF | 0.2481535 |
| GPANLGNYNI | ASSLRDSNTEVF | 0.5385984 |
| TLGPTLP | ASSLRDSNTEVF | 0.66185886 |

| | | |
|---|---|---|
| VYAYSSCCK | ASSLRDSNTEVF | 0.25658095 |
| APAPAPAPNI | ASSLRDSNTEVF | 0.15988451 |
| PGGPVPIP | ASSLRDSNTEVF | 0.7747834 |
| FLTVALEP | ASSLRDSNTEVF | 0.7822017 |
| SQYFGANL | ASSLRDSNTEVF | 0.31432486 |
| GGVKPAA | ASSLRDSNTEVF | 0.11142885 |
| GGTGGAGGAGGW | ASSLRDSNTEVF | 0.64024174 |
| GVGGVAGLGGAGG | ASSLRDSNTEVF | 0.289303 |
| VLLVLATTI | ASSLRDSNTEVF | 0.592489 |
| IGIFAAML | ASSLRDSNTEVF | 0.37829703 |
| AVGDLID | ASSLRDSNTEVF | 0.159268 |
| NISPLPL | ASSLRDSNTEVF | 0.3215425 |
| VTAVFVAI | ASSLRDSNTEVF | 0.7563643 |
| SGDDNDAAL | ASSLRDSNTEVF | 0.46169752 |
| AVAALAHS | ASSLRDSNTEVF | 0.7002518 |
| GGVTVMATP | ASSLRDSNTEVF | 0.7991327 |
| AGLAANVRD | ASSLRDSNTEVF | 0.7747665 |
| TNRPDLIDPALLRPG | ASSLRDSNTEVF | 0.8267298 |
| LANLGVTAL | ASSLRDSNTEVF | 0.64481163 |
| TGERIEL | ASSLRDSNTEVF | 0.40284207 |
| NGEELAA | ASSLRDSNTEVF | 0.50163186 |
| VLSLVWG | ASSLRDSNTEVF | 0.62075615 |
| IGYALWL | ASSLRDSNTEVF | 0.65923285 |
| LSLLLAPL | ASSLRDSNTEVF | 0.63273275 |
| TLDEALK | ASSLRDSNTEVF | 0.3006308 |
| TDGAGTAEGG | ASSLRDSNTEVF | 0.7725433 |
| NAPGGAGGNGG | ASSLRDSNTEVF | 0.464511 |
| VSVFMGMGGNF | ASSLRDSNTEVF | 0.66305155 |
| LASYLAAL | ASSLRDSNTEVF | 0.17001583 |
| LVGGTPAL | ASSLRDSNTEVF | 0.1763353 |
| VRNADVL | ASSLRDSNTEVF | 0.7818182 |
| TCSSPFLVSAM | ASSLRDSNTEVF | 0.08254368 |
| VHITDAE | ASSLRDSNTEVF | 0.10704262 |
| EYAAPSF | ASSLRDSNTEVF | 0.6975247 |
| VVDAFHGGL | ASSLRDSNTEVF | 0.6959062 |
| ASVVDPQ | ASSLRDSNTEVF | 0.07051927 |
| SYNEFGGLS | ASSLRDSNTEVF | 0.19677843 |
| DPDGSVY | ASSLRDSNTEVF | 0.321822 |
| MVSTIHATE | ASSLRDSNTEVF | 0.46184987 |
| LAAGVADPV | ASSLRDSNTEVF | 0.37401438 |
| TKEQIAA | ASSLRDSNTEVF | 0.111155696 |
| SLASPAAL | ASSLRDSNTEVF | 0.74575496 |
| IGLVFYGV | ASSLRDSNTEVF | 0.115373224 |
| FLLDASNL | ASSLRDSNTEVF | 0.112085596 |
| YQVGNFP | ASSLRDSNTEVF | 0.6521972 |
| ESLLSQST | ASSLRDSNTEVF | 0.7610164 |
| LFTVGFAAVG | ASSLRDSNTEVF | 0.22168887 |
| ATPPTPT | ASSLRDSNTEVF | 0.73384124 |
| TDLATLG | ASSLRDSNTEVF | 0.31759405 |
| SAGGTGR | ASSLRDSNTEVF | 0.7459966 |
| GFSNSGS | ASSLRDSNTEVF | 0.4312268 |

| | | |
|---|---|---|
| GGNGGSGGTSV | ASSLRDSNTEVF | 0.12771758 |
| GTSVATGGAGN | ASSLRDSNTEVF | 0.74504656 |
| NGGNGGSGGTS | ASSLRDSNTEVF | 0.6848176 |
| AVVKFPH | ASSLRDSNTEVF | 0.57075524 |
| TPPPPPP | ASSLRDSNTEVF | 0.72371143 |
| AAPAAAAAPA | ASSLRDSNTEVF | 0.33092952 |
| IALNEDL | ASSLRDSNTEVF | 0.6346263 |
| IVNVSSM | ASSLRDSNTEVF | 0.8009527 |
| TGIPPAPR | ASSLRDSNTEVF | 0.734391 |
| FDLGGGTF | ASSLRDSNTEVF | 0.14994736 |
| FDLGGGTFDVS | ASSLRDSNTEVF | 0.6855601 |
| NEPTAAA | ASSLRDSNTEVF | 0.5317325 |
| NEPTAAALA | ASSLRDSNTEVF | 0.4696939 |
| GSIINISSI | ASSLRDSNTEVF | 0.6236701 |
| VAPGNAGTA | ASSLRDSNTEVF | 0.4887479 |
| TGGGASLE | ASSLRDSNTEVF | 0.740134 |
| VAVYNHYKRI | ASSLRDSNTEVF | 0.5154464 |
| TDPAPATT | ASSLRDSNTEVF | 0.68877566 |
| TNRPDILDPALLRPG | ASSLRDSNTEVF | 0.117946036 |
| LTTTTTS | ASSLRDSNTEVF | 0.3034674 |
| PTGSGKTL | ASSLRDSNTEVF | 0.5866783 |
| TGSGKTL | ASSLRDSNTEVF | 0.7619533 |
| TIGHVDHGK | ASSLRDSNTEVF | 0.24962172 |
| TIGHVDHGKTT | ASSLRDSNTEVF | 0.083619 |
| GHVDHGKTT | ASSLRDSNTEVF | 0.6071203 |
| AQFEHTLL | ASSLRDSNTEVF | 0.74513316 |
| IVNQDEL | ASSLRDSNTEVF | 0.5025451 |
| TNGSQFF | ASSLRDSNTEVF | 0.4215968 |
| TGKGIVE | ASSLRDSNTEVF | 0.8628611 |
| LISNASDA | ASSLRDSNTEVF | 0.42722142 |
| AELRTLGELTKIAKAHGAQ | ASSLRDSNTEVF | 0.423782 |
| APNAVAAAEPSW | ASSLRDSNTEVF | 0.6627381 |
| APQVAGAATAVL | ASSLRDSNTEVF | 0.792331 |
| ARRDPAAAAAILPTDA | ASSLRDSNTEVF | 0.5097559 |
| DAAARDVLAEAGLAEYF | ASSLRDSNTEVF | 0.7753676 |
| DDQIVLSAKANLAKVEQ | ASSLRDSNTEVF | 0.16562192 |
| DELLPDTGEDSDEDGH | ASSLRDSNTEVF | 0.72974324 |
| ELPEAPANRATVPQD | ASSLRDSNTEVF | 0.1561437 |
| FGQSKIERVTGIG | ASSLRDSNTEVF | 0.6226859 |
| GAGGTGSTAGGAGGAGGAGGLY | ASSLRDSNTEVF | 0.23422036 |
| GHGGTGAAGGNGGTGGAGGSGI | ASSLRDSNTEVF | 0.73081064 |
| GNGGNGGAGGEGGPGGAGGAGA | ASSLRDSNTEVF | 0.7520217 |
| GSTAGNGGNGGNATNGTIAGQP | ASSLRDSNTEVF | 0.76100564 |
| GTGGDGGNGGNGGTGG | ASSLRDSNTEVF | 0.07581581 |
| GVRRPAGIPADAPV | ASSLRDSNTEVF | 0.76355433 |
| GYRQVTDAHLLTLARR | ASSLRDSNTEVF | 0.78417754 |
| IAKMGQDGHDRG | ASSLRDSNTEVF | 0.8371428 |
| KAEPSIEIQYT | ASSLRDSNTEVF | 0.48740065 |
| KPVRIRMAADEDTVAVT | ASSLRDSNTEVF | 0.7871548 |
| MPPRIAGMRLLVIKPEP | ASSLRDSNTEVF | 0.3139939 |
| MKFYAVALFRSRGVPPTR | ASSLRDSNTEVF | 0.51654446 |

| | | |
|---|---|---|
| NAGGFGILGAVAHSP | ASSLRDSNTEVF | 0.32117617 |
| NEGREITLAMAEM | ASSLRDSNTEVF | 0.49036747 |
| NELVEWSHPLGHRWRWE | ASSLRDSNTEVF | 0.49569446 |
| NLGAGGTGGNGGNPG | ASSLRDSNTEVF | 0.5387487 |
| NTEWPKWETVMAADPVD | ASSLRDSNTEVF | 0.5976571 |
| PASAAPAETQP | ASSLRDSNTEVF | 0.7749994 |
| QQSSDGKNGVRAAEITGE | ASSLRDSNTEVF | 0.63438725 |
| RAPDAVTTSWWKE | ASSLRDSNTEVF | 0.81318134 |
| SGFGNVGAHTSGWFNQ | ASSLRDSNTEVF | 0.772983 |
| SGVAGAVGQAASVGGLKV | ASSLRDSNTEVF | 0.74912226 |
| SNGGAGTGSAGGNGGT | ASSLRDSNTEVF | 0.2704232 |
| TGGAGGEGGAAG | ASSLRDSNTEVF | 0.2490573 |
| TGGDGGAAGAGGAPGHG | ASSLRDSNTEVF | 0.26532406 |
| TMPPLLAAIANAGRHGV | ASSLRDSNTEVF | 0.31476462 |
| TSSTAGNSGGA | ASSLRDSNTEVF | 0.24649042 |
| VPATVPVGILC | ASSLRDSNTEVF | 0.692534 |
| VQRPDSGGVVLTGRIS | ASSLRDSNTEVF | 0.07001904 |
| WTPNPVPATKGVPAGLVQAT | ASSLRDSNTEVF | 0.63241833 |
| YALKRALAKFSDRFDVVII | ASSLRDSNTEVF | 0.81807804 |
| YLIGSVIV | ASSLRDSNTEVF | 0.117857076 |
| YGQLVAVID | ASSLRDSNTEVF | 0.5795965 |
| VTVGGMTLPA | ASSLRDSNTEVF | 0.46978623 |
| VLALPAV | ASSLRDSNTEVF | 0.37223268 |
| VIPPTLNYET | ASSLRDSNTEVF | 0.78373456 |
| VGMGGMHPSSG | ASSLRDSNTEVF | 0.22302385 |
| VGLAGGAATA | ASSLRDSNTEVF | 0.73857474 |
| VGGAGGAGGSGP | ASSLRDSNTEVF | 0.6600542 |
| VFAAFMV | ASSLRDSNTEVF | 0.6495115 |
| TLYDKLR | ASSLRDSNTEVF | 0.5313417 |
| TLAVFTVL | ASSLRDSNTEVF | 0.5106652 |
| TGSVQPARV | ASSLRDSNTEVF | 0.11191498 |
| TGGGAGSGGT | ASSLRDSNTEVF | 0.6949005 |
| SHLETSL | ASSLRDSNTEVF | 0.06950696 |
| SFTTGMT | ASSLRDSNTEVF | 0.6749801 |
| QVEVAKRMGVRQPTVS | ASSLRDSNTEVF | 0.17767034 |
| QGSGGPALAM | ASSLRDSNTEVF | 0.78945595 |
| MLAPSNP | ASSLRDSNTEVF | 0.5335701 |
| LVNNAAI | ASSLRDSNTEVF | 0.6916995 |
| LTGEINTLA | ASSLRDSNTEVF | 0.316589 |
| LSIFGPV | ASSLRDSNTEVF | 0.31060678 |
| LLALLANL | ASSLRDSNTEVF | 0.1286101 |
| LISNASDALDKLR | ASSLRDSNTEVF | 0.13944344 |
| LINEAVP | ASSLRDSNTEVF | 0.5866524 |
| LGYAFSGL | ASSLRDSNTEVF | 0.3865256 |
| LDFVAVAL | ASSLRDSNTEVF | 0.40249625 |
| LAQPGAE | ASSLRDSNTEVF | 0.68859935 |
| ISNLGAVTGF | ASSLRDSNTEVF | 0.58413196 |
| HQEDPGRI | ASSLRDSNTEVF | 0.121503554 |
| GTPGEPTP | ASSLRDSNTEVF | 0.5949547 |
| GFFNSGS | ASSLRDSNTEVF | 0.6377022 |
| GAGGGGLTAGAN | ASSLRDSNTEVF | 0.4482546 |

| | | |
|---|---|---|
| FVDALVNM | ASSLRDSNTEVF | 0.0681402 |
| FLTVAPEL | ASSLRDSNTEVF | 0.531458 |
| FETLTAL | ASSLRDSNTEVF | 0.46493632 |
| DVINPIEL | ASSLRDSNTEVF | 0.35672128 |
| APVSGAPV | ASSLRDSNTEVF | 0.2809583 |
| ALVGGLAW | ASSLRDSNTEVF | 0.8350107 |
| ALAAAAATAI | ASSLRDSNTEVF | 0.08685279 |
| AFGLGLLL | ASSLRDSNTEVF | 0.36838335 |
| AFFAAASL | ASSLRDSNTEVF | 0.7707088 |
| AAAALPAFSPP | ASSLRDSNTEVF | 0.5682193 |
| VVLAPAL | ASSLRDSNTEVF | 0.8001948 |
| TIGHVDHGKTTL | ASSLRDSNTEVF | 0.4168648 |
| LISNASDALDK | ASSLRDSNTEVF | 0.27859473 |
| LISNASDAL | ASSLRDSNTEVF | 0.6570863 |
| ISNASDALDK | ASSLRDSNTEVF | 0.30488458 |
| NQAETLVYQTE | ASSLRDSNTEVF | 0.12252736 |
| TGGTGGTAGAG | ASSLRDSNTEVF | 0.0873038 |
| TGSTAGGAGGA | ASSLRDSNTEVF | 0.5000814 |
| DPLAVPVAAGPV | ASSLRDSNTEVF | 0.10782387 |
| GGTGGTPGNGGN | ASSLRDSNTEVF | 0.5373683 |
| PAGLAGVDPATL | ASSLRDSNTEVF | 0.7300515 |
| GNNLVVEAPHAGASV | ASSLRDSNTEVF | 0.26880938 |
| KVTRIMSASTLEQPA | ASSLRDSNTEVF | 0.1540317 |
| LGFALPIALPRRTVV | ASSLRDSNTEVF | 0.5258828 |
| TRLGNLTARKLVVLV | ASSLRDSNTEVF | 0.22989339 |
| WSQQRRLIYFPSAGP | ASSLRDSNTEVF | 0.78145427 |
| GPNGNPDPMAAAVDIR | ASSLRDSNTEVF | 0.20272608 |
| TQRCVSVADLRAAASR | ASSLRDSNTEVF | 0.7517611 |
| QFVLDVAEHPVEVLTETV | ASSLRDSNTEVF | 0.7631821 |
| GGAGGTGGAGGNGGTQAGDG | ASSLRDSNTEVF | 0.3693688 |
| PLLATLRAIAVDVDAKPAQV | ASSLRDSNTEVF | 0.80470407 |
| APKFRTGGYMIDFWGVGYQVAK | ASSLRDSNTEVF | 0.42159933 |
| GAGGAGGAGGAAGGPGGTGGTG | ASSLRDSNTEVF | 0.22390583 |
| IGALGLLATDVAPAGSVAFGVA | ASSLRDSNTEVF | 0.48482478 |
| AGGAGGIGGNGGAGGAGGVLMG | ASSLRDSNTEVF | 0.5658088 |
| ADELIGLL | ASSLRDSNTEVF | 0.31444836 |
| ADRALKA | ASSLRDSNTEVF | 0.50682926 |
| AGAVGQVGGAGGA | ASSLRDSNTEVF | 0.12005598 |
| AGGGGGLGGAP | ASSLRDSNTEVF | 0.630594 |
| AGGLLYGNGGAG | ASSLRDSNTEVF | 0.10961503 |
| AGLESIGL | ASSLRDSNTEVF | 0.114724666 |
| AGLKTGII | ASSLRDSNTEVF | 0.3589667 |
| ASLAALLL | ASSLRDSNTEVF | 0.7909121 |
| ASVINISSI | ASSLRDSNTEVF | 0.833828 |
| ATPPGPA | ASSLRDSNTEVF | 0.47716606 |
| ATVASALV | ASSLRDSNTEVF | 0.23976384 |
| ATVASLIL | ASSLRDSNTEVF | 0.66277224 |
| AVNGPVLIRH | ASSLRDSNTEVF | 0.1320512 |
| DGAAGAVGI | ASSLRDSNTEVF | 0.6247157 |
| DPLLHGI | ASSLRDSNTEVF | 0.5913831 |
| ELGAALGAL | ASSLRDSNTEVF | 0.44530195 |

| | | |
|---|---|---|
| ELGRTQAV | ASSLRDSNTEVF | 0.3563773 |
| FVAGGYLS | ASSLRDSNTEVF | 0.66880524 |
| GASGGAGGAGGW | ASSLRDSNTEVF | 0.30701 |
| GGAGAGGDGGAGGI | ASSLRDSNTEVF | 0.68463624 |
| GTGHNGSQPGMGGNG | ASSLRDSNTEVF | 0.06934669 |
| GYAAASASAS | ASSLRDSNTEVF | 0.2762673 |
| HQVDVAAVQQG | ASSLRDSNTEVF | 0.28805104 |
| IGAAIAHT | ASSLRDSNTEVF | 0.2468265 |
| IGMIEVAQL | ASSLRDSNTEVF | 0.10385352 |
| IRLAPEHTSAMD | ASSLRDSNTEVF | 0.40532064 |
| IYNVSVL | ASSLRDSNTEVF | 0.6912123 |
| KQGFLGM | ASSLRDSNTEVF | 0.81675005 |
| LESVITQL | ASSLRDSNTEVF | 0.2342848 |
| LGLGALVVV | ASSLRDSNTEVF | 0.7540174 |
| LIGNGGNGGSGGI | ASSLRDSNTEVF | 0.7624432 |
| LNAINAPTL | ASSLRDSNTEVF | 0.50965977 |
| LQLLNTGV | ASSLRDSNTEVF | 0.12297036 |
| LQTSIEF | ASSLRDSNTEVF | 0.49399543 |
| MSFAVSEL | ASSLRDSNTEVF | 0.16591124 |
| NTGALGL | ASSLRDSNTEVF | 0.6849558 |
| PPGAGGAGGAGTT | ASSLRDSNTEVF | 0.61215794 |
| QLVGLGI | ASSLRDSNTEVF | 0.29858518 |
| RFLEQVHQL | ASSLRDSNTEVF | 0.6665729 |
| RGVGYVLR | ASSLRDSNTEVF | 0.7762328 |
| SATGVFAGVI | ASSLRDSNTEVF | 0.29853106 |
| SLPAITLP | ASSLRDSNTEVF | 0.18575417 |
| STALAFTV | ASSLRDSNTEVF | 0.17226905 |
| SVLTVGLGK | ASSLRDSNTEVF | 0.08074175 |
| TALAGAGSGI | ASSLRDSNTEVF | 0.47902143 |
| TGLPFTGI | ASSLRDSNTEVF | 0.4213472 |
| VAEHVTV | ASSLRDSNTEVF | 0.19690248 |
| VDIVLNSL | ASSLRDSNTEVF | 0.6616249 |
| VLTGGGAVL | ASSLRDSNTEVF | 0.19168541 |
| VNVGLAYGI | ASSLRDSNTEVF | 0.3025844 |
| VSSAAVGAV | ASSLRDSNTEVF | 0.2747857 |
| VTSVYEAGV | ASSLRDSNTEVF | 0.15345821 |
| WVALDRGA | ASSLRDSNTEVF | 0.73158085 |
| YLVGSVVI | ASSLRDSNTEVF | 0.19301835 |
| GPPGTGKTLL | ASSLRDSNTEVF | 0.40742877 |
| GVGMTKF | ASSLRDSNTEVF | 0.8147129 |
| IGIGGSDL | ASSLRDSNTEVF | 0.11109053 |
| IGTLLAEL | ASSLRDSNTEVF | 0.19613671 |
| IYTSGTT | ASSLRDSNTEVF | 0.72036004 |
| LSFPAGV | ASSLRDSNTEVF | 0.09848559 |
| LYTSGTT | ASSLRDSNTEVF | 0.720014 |
| VGAFPFGF | ASSLRDSNTEVF | 0.5013425 |
| VGAGTVEF | ASSLRDSNTEVF | 0.70373 |
| VGKSTLLL | ASSLRDSNTEVF | 0.26495433 |
| AALAHAGGGIV | ASSLRDSNTEVF | 0.8039232 |
| AALVAVSLL | ASSLRDSNTEVF | 0.7865616 |
| AAVSRAVAAL | ASSLRDSNTEVF | 0.2520755 |

| | | |
|---|---|---|
| AGGSSGIGGF | ASSLRDSNTEVF | 0.1154248 |
| AGLTVANANG | ASSLRDSNTEVF | 0.17138727 |
| AGLVGAVL | ASSLRDSNTEVF | 0.7146554 |
| ALSQAFTV | ASSLRDSNTEVF | 0.77327687 |
| ALTFSVSL | ASSLRDSNTEVF | 0.39049384 |
| ALTGAALVAL | ASSLRDSNTEVF | 0.62957287 |
| ASFSIGAGVL | ASSLRDSNTEVF | 0.7754056 |
| DGSVYKI | ASSLRDSNTEVF | 0.8348835 |
| EQLAEIV | ASSLRDSNTEVF | 0.104338996 |
| FAVSELTVM | ASSLRDSNTEVF | 0.5472735 |
| GAAPLAAGL | ASSLRDSNTEVF | 0.1474255 |
| GADGGDGGSGGAGG | ASSLRDSNTEVF | 0.39691657 |
| GFGLDTGGAGG | ASSLRDSNTEVF | 0.21684088 |
| GLASGGLL | ASSLRDSNTEVF | 0.7336234 |
| GRIAGAAVLL | ASSLRDSNTEVF | 0.4778444 |
| GSAGSLNTSF | ASSLRDSNTEVF | 0.66136885 |
| GTGQIGI | ASSLRDSNTEVF | 0.628126 |
| GTLIALIL | ASSLRDSNTEVF | 0.77735615 |
| IAATMGFVVG | ASSLRDSNTEVF | 0.4443389 |
| IFAQALL | ASSLRDSNTEVF | 0.3296479 |
| IGAAVALGF | ASSLRDSNTEVF | 0.15456301 |
| IGLTPLAL | ASSLRDSNTEVF | 0.07748608 |
| ILGSNTSTL | ASSLRDSNTEVF | 0.22564152 |
| IVFSLTVL | ASSLRDSNTEVF | 0.23401004 |
| LASKATGF | ASSLRDSNTEVF | 0.07114494 |
| LASVALIL | ASSLRDSNTEVF | 0.66022307 |
| LFAVGLTL | ASSLRDSNTEVF | 0.71041226 |
| LFNGGIGTYI | ASSLRDSNTEVF | 0.694813 |
| LTLGAGII | ASSLRDSNTEVF | 0.20588678 |
| LVAAALGDI | ASSLRDSNTEVF | 0.2732393 |
| NTAFVQLGI | ASSLRDSNTEVF | 0.2803352 |
| QLFSGAEL | ASSLRDSNTEVF | 0.80688554 |
| SAATFITL | ASSLRDSNTEVF | 0.27976322 |
| SNYGLANLGV | ASSLRDSNTEVF | 0.121281266 |
| SYGGAYAVM | ASSLRDSNTEVF | 0.19280516 |
| TTGVGGLTLG | ASSLRDSNTEVF | 0.17050342 |
| TTQTIYI | ASSLRDSNTEVF | 0.768508 |
| VAAGAGLGAVI | ASSLRDSNTEVF | 0.7076083 |
| VAAVVTLL | ASSLRDSNTEVF | 0.7701219 |
| VGFLGGLI | ASSLRDSNTEVF | 0.67572224 |
| VGVIPILL | ASSLRDSNTEVF | 0.09749211 |
| VIFASSLI | ASSLRDSNTEVF | 0.19523856 |
| VKGGDGGAAAG | ASSLRDSNTEVF | 0.854735 |
| VTKTFTNI | ASSLRDSNTEVF | 0.49781328 |
| VVAAAILL | ASSLRDSNTEVF | 0.555936 |
| YAALDVEL | ASSLRDSNTEVF | 0.7560619 |
| LASVALLL | ASSLRDSNTEVF | 0.25900316 |
| LDAVHRV | ASSLRDSNTEVF | 0.3855897 |
| QGLSSSIF | ASSLRDSNTEVF | 0.4910125 |
| RGTFANIRL | ASSLRDSNTEVF | 0.69345737 |
| RQLWWGHRI | ASSLRDSNTEVF | 0.33295602 |

| | | |
|---|---|---|
| YGPPGTGKTLL | ASSLRDSNTEVF | 0.21334758 |
| AALVGAPANVA | ASSLRDSNTEVF | 0.7241714 |
| AATFRRTE | ASSLRDSNTEVF | 0.80551505 |
| AGALGGGI | ASSLRDSNTEVF | 0.71365696 |
| AGGAGGAGGIGAG | ASSLRDSNTEVF | 0.6384319 |
| AGSGALGL | ASSLRDSNTEVF | 0.5886637 |
| AQIADQI | ASSLRDSNTEVF | 0.28254163 |
| ASAVAVTYL | ASSLRDSNTEVF | 0.4095348 |
| ASTVLGGTGG | ASSLRDSNTEVF | 0.55204624 |
| AVGGEPAK | ASSLRDSNTEVF | 0.26598188 |
| CEQGTFGGTC | ASSLRDSNTEVF | 0.26074097 |
| DEGKAPAG | ASSLRDSNTEVF | 0.560818 |
| DTVVFPAL | ASSLRDSNTEVF | 0.28668988 |
| FEGITMAG | ASSLRDSNTEVF | 0.82043904 |
| FPLVVQD | ASSLRDSNTEVF | 0.6601811 |
| GAFTTPQ | ASSLRDSNTEVF | 0.63292634 |
| GFGGFGTAGGVGGA | ASSLRDSNTEVF | 0.2806049 |
| GFLAQFL | ASSLRDSNTEVF | 0.557434 |
| GGAGGAGGNAGVA | ASSLRDSNTEVF | 0.6047494 |
| GGVGTAGAGGAGGAGG | ASSLRDSNTEVF | 0.12394563 |
| GNGGGGDAGNAGSGGN | ASSLRDSNTEVF | 0.7893636 |
| GTGGAGGAGNDAGS | ASSLRDSNTEVF | 0.28393215 |
| GTTNTGWFNS | ASSLRDSNTEVF | 0.34892797 |
| IFMGPLFTL | ASSLRDSNTEVF | 0.59107816 |
| ILAHAIEAI | ASSLRDSNTEVF | 0.731485 |
| IRSAIREAL | ASSLRDSNTEVF | 0.3843244 |
| ITFAGGG | ASSLRDSNTEVF | 0.28270662 |
| ITGMSGHQEV | ASSLRDSNTEVF | 0.2621302 |
| IVAGLACTL | ASSLRDSNTEVF | 0.73687166 |
| IVGGGLAGA | ASSLRDSNTEVF | 0.85293996 |
| IVTGVLNI | ASSLRDSNTEVF | 0.11595368 |
| LAQQGCI | ASSLRDSNTEVF | 0.81118405 |
| LCLHVEN | ASSLRDSNTEVF | 0.25370806 |
| LDAISDGS | ASSLRDSNTEVF | 0.17849018 |
| LGGFAAML | ASSLRDSNTEVF | 0.7066064 |
| LGGKNPMI | ASSLRDSNTEVF | 0.57007223 |
| LGPKGVAAIL | ASSLRDSNTEVF | 0.7839073 |
| LMVGVSI | ASSLRDSNTEVF | 0.4554059 |
| LVGAAFVL | ASSLRDSNTEVF | 0.45587915 |
| LVLAVAPL | ASSLRDSNTEVF | 0.55694896 |
| MAAIAVIDAA | ASSLRDSNTEVF | 0.8033386 |
| NNGINAELQ | ASSLRDSNTEVF | 0.69092214 |
| RGEPLVLVHG | ASSLRDSNTEVF | 0.7510762 |
| RGGAGGAAGLI | ASSLRDSNTEVF | 0.49187854 |
| RSYRMHPAV | ASSLRDSNTEVF | 0.26447794 |
| SAAHLLAGQLA | ASSLRDSNTEVF | 0.42884678 |
| SAAVGSY | ASSLRDSNTEVF | 0.69587046 |
| SPHAVLHRI | ASSLRDSNTEVF | 0.309326 |
| SQIAKVL | ASSLRDSNTEVF | 0.5443872 |
| TALGHLVP | ASSLRDSNTEVF | 0.645889 |
| TAPTTSGAA | ASSLRDSNTEVF | 0.1702117 |

| | | |
|---|---|---|
| TDGDDGNGGSPGT | ASSLRDSNTEVF | 0.28677315 |
| TDGLGEN | ASSLRDSNTEVF | 0.7460782 |
| TGGPPLPGQA | ASSLRDSNTEVF | 0.47933853 |
| TIPAITI | ASSLRDSNTEVF | 0.31739414 |
| TIPGFTI | ASSLRDSNTEVF | 0.76003885 |
| TSIASGALA | ASSLRDSNTEVF | 0.8474454 |
| TVTLPGAAI | ASSLRDSNTEVF | 0.7897924 |
| VASGLSAALP | ASSLRDSNTEVF | 0.11675085 |
| VGAGVIGI | ASSLRDSNTEVF | 0.09453245 |
| VGPKRFGV | ASSLRDSNTEVF | 0.47219568 |
| VNVAFPD | ASSLRDSNTEVF | 0.32343435 |
| VSMTNSGVS | ASSLRDSNTEVF | 0.20949529 |
| VTDVLGDLSV | ASSLRDSNTEVF | 0.2674342 |
| VVVVGAGFAGL | ASSLRDSNTEVF | 0.61242884 |
| VYTVGTVV | ASSLRDSNTEVF | 0.73345983 |
| WTTFSGP | ASSLRDSNTEVF | 0.37163305 |
| YDELVEAF | ASSLRDSNTEVF | 0.13656566 |
| YGTGGAGGNGG | ASSLRDSNTEVF | 0.3718492 |
| YTNVMAV | ASSLRDSNTEVF | 0.8146039 |
| FLEQVHQL | ASSLRDSNTEVF | 0.40684885 |
| GLSLLAVAV | ASSLRDSNTEVF | 0.7037492 |
| LEQVHQL | ASSLRDSNTEVF | 0.29284805 |
| LNAAGAI | ASSLRDSNTEVF | 0.20519985 |
| PAVHASL | ASSLRDSNTEVF | 0.63614964 |
| TELEDDA | ASSLRDSNTEVF | 0.33134648 |
| TPPGPPN | ASSLRDSNTEVF | 0.72298694 |
| TPPPPPPP | ASSLRDSNTEVF | 0.3057107 |
| TQSPALI | GANPGTGENTEVF | 0.7197466 |
| EFAAAFPQ | GANPGTGENTEVF | 0.7151955 |
| LLSGNLDVL | GANPGTGENTEVF | 0.69403607 |
| GPANLGNYNI | GANPGTGENTEVF | 0.49166334 |
| TLGPTLP | GANPGTGENTEVF | 0.3332073 |
| VYAYSSCCK | GANPGTGENTEVF | 0.16999657 |
| APAPAPAPNI | GANPGTGENTEVF | 0.08862559 |
| PGGPVPIP | GANPGTGENTEVF | 0.7779973 |
| FLTVALEP | GANPGTGENTEVF | 0.06897713 |
| SQYFGANL | GANPGTGENTEVF | 0.8276658 |
| GGVKPAA | GANPGTGENTEVF | 0.56177354 |
| GGTGGAGGAGGW | GANPGTGENTEVF | 0.34513605 |
| GVGGVAGLGGAGG | GANPGTGENTEVF | 0.6567498 |
| VLLVLATTI | GANPGTGENTEVF | 0.19599853 |
| IGIFAAML | GANPGTGENTEVF | 0.27928454 |
| AVGDLID | GANPGTGENTEVF | 0.5997863 |
| NISPLPL | GANPGTGENTEVF | 0.67946994 |
| VTAVFVAI | GANPGTGENTEVF | 0.31586385 |
| SGDDNDAAL | GANPGTGENTEVF | 0.6393922 |
| AVAALAHS | GANPGTGENTEVF | 0.73491657 |
| GGVTVMATP | GANPGTGENTEVF | 0.27178794 |
| AGLAANVRD | GANPGTGENTEVF | 0.42718434 |
| TNRPDLIDPALLRPG | GANPGTGENTEVF | 0.77983105 |
| LANLGVTAL | GANPGTGENTEVF | 0.5740615 |

| | | |
|---|---|---|
| TGERIEL | GANPGTGENTEVF | 0.32275122 |
| NGEELAA | GANPGTGENTEVF | 0.43427342 |
| VLSLVWG | GANPGTGENTEVF | 0.43130663 |
| IGYALWL | GANPGTGENTEVF | 0.7280039 |
| LSLLLAPL | GANPGTGENTEVF | 0.5993692 |
| TLDEALK | GANPGTGENTEVF | 0.4721269 |
| TDGAGTAEGG | GANPGTGENTEVF | 0.66592455 |
| NAPGGAGGNGG | GANPGTGENTEVF | 0.095905215 |
| VSVFMGMGGNF | GANPGTGENTEVF | 0.34949625 |
| LASYLAAL | GANPGTGENTEVF | 0.41410828 |
| LVGGTPAL | GANPGTGENTEVF | 0.5247637 |
| VRNADVL | GANPGTGENTEVF | 0.70237815 |
| TCSSPFLVSAM | GANPGTGENTEVF | 0.22775663 |
| VHITDAE | GANPGTGENTEVF | 0.73015404 |
| EYAAPSF | GANPGTGENTEVF | 0.2860428 |
| VVDAFHGGL | GANPGTGENTEVF | 0.22675008 |
| ASVVDPQ | GANPGTGENTEVF | 0.54703915 |
| SYNEFGGLS | GANPGTGENTEVF | 0.2016732 |
| DPDGSVY | GANPGTGENTEVF | 0.39725888 |
| MVSTIHATE | GANPGTGENTEVF | 0.6958392 |
| LAAGVADPV | GANPGTGENTEVF | 0.20138942 |
| TKEQIAA | GANPGTGENTEVF | 0.43296665 |
| SLASPAAL | GANPGTGENTEVF | 0.3270209 |
| IGLVFYGV | GANPGTGENTEVF | 0.35553834 |
| FLLDASNL | GANPGTGENTEVF | 0.6925934 |
| YQVGNFP | GANPGTGENTEVF | 0.5714835 |
| ESLLSQST | GANPGTGENTEVF | 0.7052206 |
| LFTVGFAAVG | GANPGTGENTEVF | 0.2564132 |
| ATPPTPT | GANPGTGENTEVF | 0.18588832 |
| TDLATLG | GANPGTGENTEVF | 0.56440544 |
| SAGGTGR | GANPGTGENTEVF | 0.6615724 |
| GFSNSGS | GANPGTGENTEVF | 0.6586325 |
| GGNGGSGGTSV | GANPGTGENTEVF | 0.5744819 |
| GTSVATGGAGN | GANPGTGENTEVF | 0.6091033 |
| NGGNGGSGGTS | GANPGTGENTEVF | 0.79831535 |
| AVVKFPH | GANPGTGENTEVF | 0.5375403 |
| TPPPPPP | GANPGTGENTEVF | 0.19278958 |
| AAPAAAAAPA | GANPGTGENTEVF | 0.25951934 |
| IALNEDL | GANPGTGENTEVF | 0.23030034 |
| IVNVSSM | GANPGTGENTEVF | 0.25633773 |
| TGIPPAPR | GANPGTGENTEVF | 0.14294909 |
| FDLGGGTF | GANPGTGENTEVF | 0.77635765 |
| FDLGGGTFDVS | GANPGTGENTEVF | 0.83810186 |
| NEPTAAA | GANPGTGENTEVF | 0.1841832 |
| NEPTAAALA | GANPGTGENTEVF | 0.79814893 |
| GSIINISSI | GANPGTGENTEVF | 0.12103385 |
| VAPGNAGTA | GANPGTGENTEVF | 0.4936182 |
| TGGGASLE | GANPGTGENTEVF | 0.75475395 |
| VAVYNHYKRI | GANPGTGENTEVF | 0.2824508 |
| TDPAPATT | GANPGTGENTEVF | 0.44876066 |
| TNRPDILDPALLRPG | GANPGTGENTEVF | 0.20210081 |

| | | |
|---|---|---|
| LTTTTTS | GANPGTGENTEVF | 0.23807362 |
| PTGSGKTL | GANPGTGENTEVF | 0.8046255 |
| TGSGKTL | GANPGTGENTEVF | 0.23599008 |
| TIGHVDHGK | GANPGTGENTEVF | 0.5355443 |
| TIGHVDHGKTT | GANPGTGENTEVF | 0.4323392 |
| GHVDHGKTT | GANPGTGENTEVF | 0.2096934 |
| AQFEHTLL | GANPGTGENTEVF | 0.6456968 |
| IVNQDEL | GANPGTGENTEVF | 0.750677 |
| TNGSQFF | GANPGTGENTEVF | 0.21516794 |
| TGKGIVE | GANPGTGENTEVF | 0.70842445 |
| LISNASDA | GANPGTGENTEVF | 0.26969713 |
| AELRTLGELTKIAKAHGAQ | GANPGTGENTEVF | 0.73912406 |
| APNAVAAAEPSW | GANPGTGENTEVF | 0.745342 |
| APQVAGAATAVL | GANPGTGENTEVF | 0.712302 |
| ARRDPAAAAAILPTDA | GANPGTGENTEVF | 0.374303 |
| DAAARDVLAEAGLAEYF | GANPGTGENTEVF | 0.7814146 |
| DDQIVLSAKANLAKVEQ | GANPGTGENTEVF | 0.7102899 |
| DELLPDTGEDSDEDGH | GANPGTGENTEVF | 0.6034275 |
| ELPEAPANRATVPQD | GANPGTGENTEVF | 0.7467414 |
| FGQSKIERVTGIG | GANPGTGENTEVF | 0.15630406 |
| GAGGTGSTAGGAGGAGGAGGLY | GANPGTGENTEVF | 0.18293324 |
| GHGGTGAAGGNGGTGGAGGSGI | GANPGTGENTEVF | 0.15861489 |
| GNGGNGGAGGEGGPGGAGGAGA | GANPGTGENTEVF | 0.17629062 |
| GSTAGNGGNGGNATNGTIAGQP | GANPGTGENTEVF | 0.16737586 |
| GTGGDGGNGGNGGTGG | GANPGTGENTEVF | 0.14334351 |
| GVRRPAGIPADAPV | GANPGTGENTEVF | 0.18062098 |
| GYRQVTDAHLLTLARR | GANPGTGENTEVF | 0.15426262 |
| IAKMGQDGHDRG | GANPGTGENTEVF | 0.14524366 |
| KAEPSIEIQYT | GANPGTGENTEVF | 0.18058941 |
| KPVRIRMAADEDTVAVT | GANPGTGENTEVF | 0.16776608 |
| MPPRIAGMRLLVIKPEP | GANPGTGENTEVF | 0.23259825 |
| MKFYAVALFRSRGVPPTR | GANPGTGENTEVF | 0.2165847 |
| NAGGFGILGAVAHSP | GANPGTGENTEVF | 0.14686653 |
| NEGREITLAMAEM | GANPGTGENTEVF | 0.16290003 |
| NELVEWSHPLGHRWRWE | GANPGTGENTEVF | 0.15709136 |
| NLGAGGTGGNGGNPG | GANPGTGENTEVF | 0.16290338 |
| NTEWPKWETVMAADPVD | GANPGTGENTEVF | 0.1404376 |
| PASAAPAETQP | GANPGTGENTEVF | 0.16613637 |
| QQSSDGKNGVRAAEITGE | GANPGTGENTEVF | 0.14119294 |
| RAPDAVTTSWWKE | GANPGTGENTEVF | 0.1369963 |
| SGFGNVGAHTSGWFNQ | GANPGTGENTEVF | 0.17273584 |
| SGVAGAVGQAASVGGLKV | GANPGTGENTEVF | 0.15963271 |
| SNGGAGTGSAGGNGGT | GANPGTGENTEVF | 0.156828 |
| TGGAGGEGGAAG | GANPGTGENTEVF | 0.19687094 |
| TGGDGGAAGAGGAPGHG | GANPGTGENTEVF | 0.13909669 |
| TMPPLLAAIANAGRHGV | GANPGTGENTEVF | 0.14890279 |
| TSSTAGNSGGA | GANPGTGENTEVF | 0.17400289 |
| VPATVPVGILC | GANPGTGENTEVF | 0.31673908 |
| VQRPDSGGVVLTGRIS | GANPGTGENTEVF | 0.14827748 |
| WTPNPVPATKGVPAGLVQAT | GANPGTGENTEVF | 0.16059431 |
| YALKRALAKFSDRFDVVII | GANPGTGENTEVF | 0.22809295 |

| | | |
|---|---|---|
| YLIGSVIV | GANPGTGENTEVF | 0.17068505 |
| YGQLVAVID | GANPGTGENTEVF | 0.18628567 |
| VTVGGMTLPA | GANPGTGENTEVF | 0.19171505 |
| VLALPAV | GANPGTGENTEVF | 0.149692 |
| VIPPTLNYET | GANPGTGENTEVF | 0.2538637 |
| VGMGGMHPSSG | GANPGTGENTEVF | 0.2418729 |
| VGLAGGAATA | GANPGTGENTEVF | 0.17731008 |
| VGGAGGAGGSGP | GANPGTGENTEVF | 0.1668051 |
| VFAAFMV | GANPGTGENTEVF | 0.15486561 |
| TLYDKLR | GANPGTGENTEVF | 0.1587742 |
| TLAVFTVL | GANPGTGENTEVF | 0.13829032 |
| TGSVQPARV | GANPGTGENTEVF | 0.16807786 |
| TGGGAGSGGT | GANPGTGENTEVF | 0.18984373 |
| SHLETSL | GANPGTGENTEVF | 0.16103446 |
| SFTTGMT | GANPGTGENTEVF | 0.14450115 |
| QVEVAKRMGVRQPTVS | GANPGTGENTEVF | 0.15110649 |
| QGSGGPALAM | GANPGTGENTEVF | 0.14800546 |
| MLAPSNP | GANPGTGENTEVF | 0.1524099 |
| LVNNAAI | GANPGTGENTEVF | 0.13834663 |
| LTGEINTLA | GANPGTGENTEVF | 0.15853852 |
| LSIFGPV | GANPGTGENTEVF | 0.2070056 |
| LLALLANL | GANPGTGENTEVF | 0.20061764 |
| LISNASDALDKLR | GANPGTGENTEVF | 0.14938073 |
| LINEAVP | GANPGTGENTEVF | 0.30896336 |
| LGYAFSGL | GANPGTGENTEVF | 0.14727974 |
| LDFVAVAL | GANPGTGENTEVF | 0.18116713 |
| LAQPGAE | GANPGTGENTEVF | 0.16523072 |
| ISNLGAVTGF | GANPGTGENTEVF | 0.15679373 |
| HQEDPGRI | GANPGTGENTEVF | 0.18335724 |
| GTPGEPTP | GANPGTGENTEVF | 0.15658662 |
| GFFNSGS | GANPGTGENTEVF | 0.14529109 |
| GAGGGGLTAGAN | GANPGTGENTEVF | 0.18842708 |
| FVDALVNM | GANPGTGENTEVF | 0.15563975 |
| FLTVAPEL | GANPGTGENTEVF | 0.15163091 |
| FETLTAL | GANPGTGENTEVF | 0.14335133 |
| DVINPIEL | GANPGTGENTEVF | 0.18427663 |
| APVSGAPV | GANPGTGENTEVF | 0.14241728 |
| ALVGGLAW | GANPGTGENTEVF | 0.15437578 |
| ALAAAAATAI | GANPGTGENTEVF | 0.13978496 |
| AFGLGLLL | GANPGTGENTEVF | 0.15648675 |
| AFFAAASL | GANPGTGENTEVF | 0.31136692 |
| AAAALPAFSPP | GANPGTGENTEVF | 0.17766716 |
| VVLAPAL | GANPGTGENTEVF | 0.14995551 |
| TIGHVDHGKTTL | GANPGTGENTEVF | 0.19800718 |
| LISNASDALDK | GANPGTGENTEVF | 0.16325301 |
| LISNASDAL | GANPGTGENTEVF | 0.16034682 |
| ISNASDALDK | GANPGTGENTEVF | 0.1851 |
| NQAETLVYQTE | GANPGTGENTEVF | 0.1500747 |
| TGGTGGTAGAG | GANPGTGENTEVF | 0.18866114 |
| TGSTAGGAGGA | GANPGTGENTEVF | 0.1755828 |
| DPLAVPVAAGPV | GANPGTGENTEVF | 0.19099237 |

| | | |
|---|---|---|
| GGTGGTPGNGGN | GANPGTGENTEVF | 0.18877977 |
| PAGLAGVDPATL | GANPGTGENTEVF | 0.16497515 |
| GNNLVVEAPHAGASV | GANPGTGENTEVF | 0.16040085 |
| KVTRIMSASTLEQPA | GANPGTGENTEVF | 0.14901245 |
| LGFALPIALPRRTVV | GANPGTGENTEVF | 0.15541533 |
| TRLGNLTARKLVVLV | GANPGTGENTEVF | 0.18917131 |
| WSQQRRLIYFPSAGP | GANPGTGENTEVF | 0.16444129 |
| GPNGNPDPMAAAVDIR | GANPGTGENTEVF | 0.19009283 |
| TQRCVSVADLRAAASR | GANPGTGENTEVF | 0.1880159 |
| QFVLDVAEHPVEVLTETV | GANPGTGENTEVF | 0.14932647 |
| GGAGGTGGAGGNGGTQAGDG | GANPGTGENTEVF | 0.19076833 |
| PLLATLRAIAVDVDAKPAQV | GANPGTGENTEVF | 0.25631812 |
| APKFRTGGYMIDFWGVGYQVAK | GANPGTGENTEVF | 0.14426845 |
| GAGGAGGAGGAAGGPGGTGGTG | GANPGTGENTEVF | 0.16286281 |
| IGALGLLATDVAPAGSVAFGVA | GANPGTGENTEVF | 0.2420207 |
| AGGAGGIGGNGGAGGAGGVLMG | GANPGTGENTEVF | 0.1616257 |
| ADELIGLL | GANPGTGENTEVF | 0.1704599 |
| ADRALKA | GANPGTGENTEVF | 0.15771733 |
| AGAVGQVGGAGGA | GANPGTGENTEVF | 0.16296631 |
| AGGGGGLGGAP | GANPGTGENTEVF | 0.14164567 |
| AGGLLYGNGGAG | GANPGTGENTEVF | 0.21019845 |
| AGLESIGL | GANPGTGENTEVF | 0.14461412 |
| AGLKTGII | GANPGTGENTEVF | 0.16540077 |
| ASLAALLL | GANPGTGENTEVF | 0.14908764 |
| ASVINISSI | GANPGTGENTEVF | 0.14102896 |
| ATPPGPA | GANPGTGENTEVF | 0.18794467 |
| ATVASALV | GANPGTGENTEVF | 0.16825326 |
| ATVASLIL | GANPGTGENTEVF | 0.1921099 |
| AVNGPVLIRH | GANPGTGENTEVF | 0.1615051 |
| DGAAGAVGI | GANPGTGENTEVF | 0.20023517 |
| DPLLHGI | GANPGTGENTEVF | 0.15885668 |
| ELGAALGAL | GANPGTGENTEVF | 0.16356336 |
| ELGRTQAV | GANPGTGENTEVF | 0.15701719 |
| FVAGGYLS | GANPGTGENTEVF | 0.1818305 |
| GASGGAGGAGGW | GANPGTGENTEVF | 0.22473593 |
| GGAGAGGDGGAGGI | GANPGTGENTEVF | 0.1848145 |
| GTGHNGSQPGMGGNG | GANPGTGENTEVF | 0.14246029 |
| GYAAASASAS | GANPGTGENTEVF | 0.14109613 |
| HQVDVAAVQQG | GANPGTGENTEVF | 0.17661168 |
| IGAAIAHT | GANPGTGENTEVF | 0.13852279 |
| IGMIEVAQL | GANPGTGENTEVF | 0.13705137 |
| IRLAPEHTSAMD | GANPGTGENTEVF | 0.14360529 |
| IYNVSVL | GANPGTGENTEVF | 0.14860597 |
| KQGFLGM | GANPGTGENTEVF | 0.22616819 |
| LESVITQL | GANPGTGENTEVF | 0.19123957 |
| LGLGALVVV | GANPGTGENTEVF | 0.16056918 |
| LIGNGGNGGSGGI | GANPGTGENTEVF | 0.18154731 |
| LNAINAPTL | GANPGTGENTEVF | 0.16157594 |
| LQLLNTGV | GANPGTGENTEVF | 0.13618122 |
| LQTSIEF | GANPGTGENTEVF | 0.155415 |
| MSFAVSEL | GANPGTGENTEVF | 0.14922476 |

| | | |
|---|---|---|
| NTGALGL | GANPGTGENTEVF | 0.19865137 |
| PPGAGGAGGAGTT | GANPGTGENTEVF | 0.15234794 |
| QLVGLGI | GANPGTGENTEVF | 0.15051877 |
| RFLEQVHQL | GANPGTGENTEVF | 0.16582546 |
| RGVGYVLR | GANPGTGENTEVF | 0.14679138 |
| SATGVFAGVI | GANPGTGENTEVF | 0.15547907 |
| SLPAITLP | GANPGTGENTEVF | 0.1436638 |
| STALAFTV | GANPGTGENTEVF | 0.13672496 |
| SVLTVGLGK | GANPGTGENTEVF | 0.1951525 |
| TALAGAGSGI | GANPGTGENTEVF | 0.14031766 |
| TGLPFTGI | GANPGTGENTEVF | 0.15530463 |
| VAEHVTV | GANPGTGENTEVF | 0.14131969 |
| VDIVLNSL | GANPGTGENTEVF | 0.1373925 |
| VLTGGGAVL | GANPGTGENTEVF | 0.14257117 |
| VNVGLAYGI | GANPGTGENTEVF | 0.18382102 |
| VSSAAVGAV | GANPGTGENTEVF | 0.16651101 |
| VTSVYEAGV | GANPGTGENTEVF | 0.16313758 |
| WVALDRGA | GANPGTGENTEVF | 0.21968451 |
| YLVGSVVI | GANPGTGENTEVF | 0.15663779 |
| GPPGTGKTLL | GANPGTGENTEVF | 0.16462427 |
| GVGMTKF | GANPGTGENTEVF | 0.15689057 |
| IGIGGSDL | GANPGTGENTEVF | 0.15721792 |
| IGTLLAEL | GANPGTGENTEVF | 0.14787662 |
| IYTSGTT | GANPGTGENTEVF | 0.14963722 |
| LSFPAGV | GANPGTGENTEVF | 0.25966612 |
| LYTSGTT | GANPGTGENTEVF | 0.15895565 |
| VGAFPFGF | GANPGTGENTEVF | 0.1505946 |
| VGAGTVEF | GANPGTGENTEVF | 0.18595408 |
| VGKSTLLL | GANPGTGENTEVF | 0.19104241 |
| AALAHAGGGIV | GANPGTGENTEVF | 0.22523007 |
| AALVAVSLL | GANPGTGENTEVF | 0.1538624 |
| AAVSRAVAAL | GANPGTGENTEVF | 0.15893836 |
| AGGSSGIGGF | GANPGTGENTEVF | 0.17237893 |
| AGLTVANANG | GANPGTGENTEVF | 0.16264716 |
| AGLVGAVL | GANPGTGENTEVF | 0.15197192 |
| ALSQAFTV | GANPGTGENTEVF | 0.14890215 |
| ALTFSVSL | GANPGTGENTEVF | 0.18042962 |
| ALTGAALVAL | GANPGTGENTEVF | 0.21278976 |
| ASFSIGAGVL | GANPGTGENTEVF | 0.15271178 |
| DGSVYKI | GANPGTGENTEVF | 0.14675574 |
| EQLAEIV | GANPGTGENTEVF | 0.1678142 |
| FAVSELTVM | GANPGTGENTEVF | 0.1452415 |
| GAAPLAAGL | GANPGTGENTEVF | 0.15666114 |
| GADGGDGGSGGAGG | GANPGTGENTEVF | 0.17722413 |
| GFGLDTGGAGG | GANPGTGENTEVF | 0.14521451 |
| GLASGGLL | GANPGTGENTEVF | 0.15547016 |
| GRIAGAAVLL | GANPGTGENTEVF | 0.22950208 |
| GSAGSLNTSF | GANPGTGENTEVF | 0.15315808 |
| GTGQIGI | GANPGTGENTEVF | 0.26420122 |
| GTLIALIL | GANPGTGENTEVF | 0.21256007 |
| IAATMGFVVG | GANPGTGENTEVF | 0.1810639 |

| | | |
|---|---|---|
| IFAQALL | GANPGTGENTEVF | 0.22118646 |
| IGAAVALGF | GANPGTGENTEVF | 0.15558155 |
| IGLTPLAL | GANPGTGENTEVF | 0.16145767 |
| ILGSNTSTL | GANPGTGENTEVF | 0.15564974 |
| IVFSLTVL | GANPGTGENTEVF | 0.16819748 |
| LASKATGF | GANPGTGENTEVF | 0.18156764 |
| LASVALIL | GANPGTGENTEVF | 0.17694454 |
| LFAVGLTL | GANPGTGENTEVF | 0.16141766 |
| LFNGGIGTYI | GANPGTGENTEVF | 0.22537927 |
| LTLGAGII | GANPGTGENTEVF | 0.15387744 |
| LVAAALGDI | GANPGTGENTEVF | 0.15199444 |
| NTAFVQLGI | GANPGTGENTEVF | 0.14549384 |
| QLFSGAEL | GANPGTGENTEVF | 0.14889657 |
| SAATFITL | GANPGTGENTEVF | 0.18114135 |
| SNYGLANLGV | GANPGTGENTEVF | 0.13904326 |
| SYGGAYAVM | GANPGTGENTEVF | 0.13798808 |
| TTGVGGLTLG | GANPGTGENTEVF | 0.17941038 |
| TTQTIYI | GANPGTGENTEVF | 0.17483473 |
| VAAGAGLGAVI | GANPGTGENTEVF | 0.1365564 |
| VAAVVTLL | GANPGTGENTEVF | 0.15338822 |
| VGFLGGLI | GANPGTGENTEVF | 0.14543581 |
| VGVIPILL | GANPGTGENTEVF | 0.16384085 |
| VIFASSLI | GANPGTGENTEVF | 0.16784048 |
| VKGGDGGAAAG | GANPGTGENTEVF | 0.14263257 |
| VTKTFTNI | GANPGTGENTEVF | 0.18100995 |
| VVAAAILL | GANPGTGENTEVF | 0.14271729 |
| YAALDVEL | GANPGTGENTEVF | 0.14076008 |
| LASVALLL | GANPGTGENTEVF | 0.1933883 |
| LDAVHRV | GANPGTGENTEVF | 0.18057932 |
| QGLSSSIF | GANPGTGENTEVF | 0.15267602 |
| RGTFANIRL | GANPGTGENTEVF | 0.20892763 |
| RQLWWGHRI | GANPGTGENTEVF | 0.14620818 |
| YGPPGTGKTLL | GANPGTGENTEVF | 0.25105608 |
| AALVGAPANVA | GANPGTGENTEVF | 0.18255474 |
| AATFRRTE | GANPGTGENTEVF | 0.14294371 |
| AGALGGGI | GANPGTGENTEVF | 0.1838516 |
| AGGAGGAGGIGAG | GANPGTGENTEVF | 0.17894931 |
| AGSGALGL | GANPGTGENTEVF | 0.184814 |
| AQIADQI | GANPGTGENTEVF | 0.15638725 |
| ASAVAVTYL | GANPGTGENTEVF | 0.15460719 |
| ASTVLGGTGG | GANPGTGENTEVF | 0.1650687 |
| AVGGEPAK | GANPGTGENTEVF | 0.2517945 |
| CEQGTFGGTC | GANPGTGENTEVF | 0.19548197 |
| DEGKAPAG | GANPGTGENTEVF | 0.14717561 |
| DTVVFPAL | GANPGTGENTEVF | 0.19909537 |
| FEGITMAG | GANPGTGENTEVF | 0.16071405 |
| FPLVVQD | GANPGTGENTEVF | 0.161366 |
| GAFTTPQ | GANPGTGENTEVF | 0.20189232 |
| GFGGFGTAGGVGGA | GANPGTGENTEVF | 0.16837746 |
| GFLAQFL | GANPGTGENTEVF | 0.19034518 |
| GGAGGAGGNAGVA | GANPGTGENTEVF | 0.16948348 |

| | | |
|---|---|---|
| GGVGTAGAGGAGGAGG | GANPGTGENTEVF | 0.15126504 |
| GNGGGGDAGNAGSGGN | GANPGTGENTEVF | 0.14501329 |
| GTGGAGGAGNDAGS | GANPGTGENTEVF | 0.17778978 |
| GTTNTGWFNS | GANPGTGENTEVF | 0.17109834 |
| IFMGPLFTL | GANPGTGENTEVF | 0.22157425 |
| ILAHAIEAI | GANPGTGENTEVF | 0.15444084 |
| IRSAIREAL | GANPGTGENTEVF | 0.14239068 |
| ITFAGGG | GANPGTGENTEVF | 0.16622488 |
| ITGMSGHQEV | GANPGTGENTEVF | 0.20761064 |
| IVAGLACTL | GANPGTGENTEVF | 0.16126339 |
| IVGGGLAGA | GANPGTGENTEVF | 0.16811676 |
| IVTGVLNI | GANPGTGENTEVF | 0.25205234 |
| LAQQGCI | GANPGTGENTEVF | 0.15905266 |
| LCLHVEN | GANPGTGENTEVF | 0.16601934 |
| LDAISDGS | GANPGTGENTEVF | 0.19564784 |
| LGGFAAML | GANPGTGENTEVF | 0.2474044 |
| LGGKNPMI | GANPGTGENTEVF | 0.16723853 |
| LGPKGVAAIL | GANPGTGENTEVF | 0.2125391 |
| LMVGVSI | GANPGTGENTEVF | 0.14671971 |
| LVGAAFVL | GANPGTGENTEVF | 0.21893124 |
| LVLAVAPL | GANPGTGENTEVF | 0.14341767 |
| MAAIAVIDAA | GANPGTGENTEVF | 0.1747278 |
| NNGINAELQ | GANPGTGENTEVF | 0.14195402 |
| RGEPLVLVHG | GANPGTGENTEVF | 0.18953426 |
| RGGAGGAAGLI | GANPGTGENTEVF | 0.18818593 |
| RSYRMHPAV | GANPGTGENTEVF | 0.21268167 |
| SAAHLLAGQLA | GANPGTGENTEVF | 0.13834715 |
| SAAVGSY | GANPGTGENTEVF | 0.194269 |
| SPHAVLHRI | GANPGTGENTEVF | 0.20669866 |
| SQIAKVL | GANPGTGENTEVF | 0.2305663 |
| TALGHLVP | GANPGTGENTEVF | 0.15496604 |
| TAPTTSGAA | GANPGTGENTEVF | 0.23964694 |
| TDGDDGNGGSPGT | GANPGTGENTEVF | 0.16306143 |
| TDGLGEN | GANPGTGENTEVF | 0.16122767 |
| TGGPPLPGQA | GANPGTGENTEVF | 0.14228156 |
| TIPAITI | GANPGTGENTEVF | 0.16078815 |
| TIPGFTI | GANPGTGENTEVF | 0.15189737 |
| TSIASGALA | GANPGTGENTEVF | 0.15612824 |
| TVTLPGAAI | GANPGTGENTEVF | 0.16416156 |
| VASGLSAALP | GANPGTGENTEVF | 0.21060771 |
| VGAGVIGI | GANPGTGENTEVF | 0.20908004 |
| VGPKRFGV | GANPGTGENTEVF | 0.22885242 |
| VNVAFPD | GANPGTGENTEVF | 0.2556204 |
| VSMTNSGVS | GANPGTGENTEVF | 0.18771392 |
| VTDVLGDLSV | GANPGTGENTEVF | 0.15152279 |
| VVVVGAGFAGL | GANPGTGENTEVF | 0.16612767 |
| VYTVGTVV | GANPGTGENTEVF | 0.14628525 |
| WTTFSGP | GANPGTGENTEVF | 0.15631276 |
| YDELVEAF | GANPGTGENTEVF | 0.14924762 |
| YGTGGAGGNGG | GANPGTGENTEVF | 0.1837584 |
| YTNVMAV | GANPGTGENTEVF | 0.13606165 |

| | | |
|---|---|---|
| FLEQVHQL | GANPGTGENTEVF | 0.14766951 |
| GLSLLAVAV | GANPGTGENTEVF | 0.2804341 |
| LEQVHQL | GANPGTGENTEVF | 0.13862921 |
| LNAAGAI | GANPGTGENTEVF | 0.16778108 |
| PAVHASL | GANPGTGENTEVF | 0.16951783 |
| TELEDDA | GANPGTGENTEVF | 0.14418103 |
| TPPGPPN | GANPGTGENTEVF | 0.20593119 |
| TPPPPPPP | GANPGTGENTEVF | 0.1470866 |
| TQSPALI | ASSLDWGSNTGQLY | 0.17391376 |
| EFAAAFPQ | ASSLDWGSNTGQLY | 0.2130613 |
| LLSGNLDVL | ASSLDWGSNTGQLY | 0.17675497 |
| GPANLGNYNI | ASSLDWGSNTGQLY | 0.16864635 |
| TLGPTLP | ASSLDWGSNTGQLY | 0.16050288 |
| VYAYSSCCK | ASSLDWGSNTGQLY | 0.14620917 |
| APAPAPAPNI | ASSLDWGSNTGQLY | 0.18851419 |
| PGGPVPIP | ASSLDWGSNTGQLY | 0.13656847 |
| FLTVALEP | ASSLDWGSNTGQLY | 0.17101626 |
| SQYFGANL | ASSLDWGSNTGQLY | 0.14974155 |
| GGVKPAA | ASSLDWGSNTGQLY | 0.19753098 |
| GGTGGAGGAGGW | ASSLDWGSNTGQLY | 0.1740775 |
| GVGGVAGLGGAGG | ASSLDWGSNTGQLY | 0.16674332 |
| VLLVLATTI | ASSLDWGSNTGQLY | 0.15703486 |
| IGIFAAML | ASSLDWGSNTGQLY | 0.14441179 |
| AVGDLID | ASSLDWGSNTGQLY | 0.14162901 |
| NISPLPL | ASSLDWGSNTGQLY | 0.14620887 |
| VTAVFVAI | ASSLDWGSNTGQLY | 0.1872585 |
| SGDDNDAAL | ASSLDWGSNTGQLY | 0.16496988 |
| AVAALAHS | ASSLDWGSNTGQLY | 0.15640618 |
| GGVTVMATP | ASSLDWGSNTGQLY | 0.19564004 |
| AGLAANVRD | ASSLDWGSNTGQLY | 0.18268748 |
| TNRPDLIDPALLRPG | ASSLDWGSNTGQLY | 0.14840403 |
| LANLGVTAL | ASSLDWGSNTGQLY | 0.15267858 |
| TGERIEL | ASSLDWGSNTGQLY | 0.18075047 |
| NGEELAA | ASSLDWGSNTGQLY | 0.16353804 |
| VLSLVWG | ASSLDWGSNTGQLY | 0.13613307 |
| IGYALWL | ASSLDWGSNTGQLY | 0.17627461 |
| LSLLLAPL | ASSLDWGSNTGQLY | 0.16045275 |
| TLDEALK | ASSLDWGSNTGQLY | 0.14806524 |
| TDGAGTAEGG | ASSLDWGSNTGQLY | 0.1565609 |
| NAPGGAGGNGG | ASSLDWGSNTGQLY | 0.2483308 |
| VSVFMGMGGNF | ASSLDWGSNTGQLY | 0.13811553 |
| LASYLAAL | ASSLDWGSNTGQLY | 0.1588233 |
| LVGGTPAL | ASSLDWGSNTGQLY | 0.20231311 |
| VRNADVL | ASSLDWGSNTGQLY | 0.1942782 |
| TCSSPFLVSAM | ASSLDWGSNTGQLY | 0.21920505 |
| VHITDAE | ASSLDWGSNTGQLY | 0.19371492 |
| EYAAPSF | ASSLDWGSNTGQLY | 0.15851371 |
| VVDAFHGGL | ASSLDWGSNTGQLY | 0.18022431 |
| ASVVDPQ | ASSLDWGSNTGQLY | 0.16815332 |
| SYNEFGGLS | ASSLDWGSNTGQLY | 0.14078844 |
| DPDGSVY | ASSLDWGSNTGQLY | 0.13802576 |

| | | |
|---|---|---|
| MVSTIHATE | ASSLDWGSNTGQLY | 0.16004181 |
| LAAGVADPV | ASSLDWGSNTGQLY | 0.13875125 |
| TKEQIAA | ASSLDWGSNTGQLY | 0.19283842 |
| SLASPAAL | ASSLDWGSNTGQLY | 0.20398597 |
| IGLVFYGV | ASSLDWGSNTGQLY | 0.15506604 |
| FLLDASNL | ASSLDWGSNTGQLY | 0.14713095 |
| YQVGNFP | ASSLDWGSNTGQLY | 0.16065371 |
| ESLLSQST | ASSLDWGSNTGQLY | 0.16256827 |
| LFTVGFAAVG | ASSLDWGSNTGQLY | 0.18853608 |
| ATPPTPT | ASSLDWGSNTGQLY | 0.16185717 |
| TDLATLG | ASSLDWGSNTGQLY | 0.3366412 |
| SAGGTGR | ASSLDWGSNTGQLY | 0.19423448 |
| GFSNSGS | ASSLDWGSNTGQLY | 0.15305775 |
| GGNGGSGGTSV | ASSLDWGSNTGQLY | 0.19788818 |
| GTSVATGGAGN | ASSLDWGSNTGQLY | 0.1555271 |
| NGGNGGSGGTS | ASSLDWGSNTGQLY | 0.15989509 |
| AVVKFPH | ASSLDWGSNTGQLY | 0.16164526 |
| TPPPPPP | ASSLDWGSNTGQLY | 0.15846163 |
| AAPAAAAAPA | ASSLDWGSNTGQLY | 0.15680301 |
| IALNEDL | ASSLDWGSNTGQLY | 0.15844108 |
| IVNVSSM | ASSLDWGSNTGQLY | 0.14592768 |
| TGIPPAPR | ASSLDWGSNTGQLY | 0.14655681 |
| FDLGGGTF | ASSLDWGSNTGQLY | 0.14858112 |
| FDLGGGTFDVS | ASSLDWGSNTGQLY | 0.13819653 |
| NEPTAAA | ASSLDWGSNTGQLY | 0.15267286 |
| NEPTAAALA | ASSLDWGSNTGQLY | 0.3609053 |
| GSIINISSI | ASSLDWGSNTGQLY | 0.25806555 |
| VAPGNAGTA | ASSLDWGSNTGQLY | 0.14960523 |
| TGGGASLE | ASSLDWGSNTGQLY | 0.15261735 |
| VAVYNHYKRI | ASSLDWGSNTGQLY | 0.15970701 |
| TDPAPATT | ASSLDWGSNTGQLY | 0.15147054 |
| TNRPDILDPALLRPG | ASSLDWGSNTGQLY | 0.17480855 |
| LTTTTTS | ASSLDWGSNTGQLY | 0.15825711 |
| PTGSGKTL | ASSLDWGSNTGQLY | 0.15498653 |
| TGSGKTL | ASSLDWGSNTGQLY | 0.15703143 |
| TIGHVDHGK | ASSLDWGSNTGQLY | 0.15884313 |
| TIGHVDHGKTT | ASSLDWGSNTGQLY | 0.15347494 |
| GHVDHGKTT | ASSLDWGSNTGQLY | 0.17912783 |
| AQFEHTLL | ASSLDWGSNTGQLY | 0.13647918 |
| IVNQDEL | ASSLDWGSNTGQLY | 0.16854 |
| TNGSQFF | ASSLDWGSNTGQLY | 0.14638416 |
| TGKGIVE | ASSLDWGSNTGQLY | 0.14797036 |
| LISNASDA | ASSLDWGSNTGQLY | 0.14153075 |
| AELRTLGELTKIAKAHGAQ | ASSLDWGSNTGQLY | 0.16907676 |
| APNAVAAAEPSW | ASSLDWGSNTGQLY | 0.19346264 |
| APQVAGAATAVL | ASSLDWGSNTGQLY | 0.19900985 |
| ARRDPAAAAAILPTDA | ASSLDWGSNTGQLY | 0.14007406 |
| DAAARDVLAEAGLAEYF | ASSLDWGSNTGQLY | 0.2345585 |
| DDQIVLSAKANLAKVEQ | ASSLDWGSNTGQLY | 0.19031328 |
| DELLPDTGEDSDEDGH | ASSLDWGSNTGQLY | 0.15757342 |
| ELPEAPANRATVPQD | ASSLDWGSNTGQLY | 0.14536975 |

| | | |
|---|---|---|
| FGQSKIERVTGIG | ASSLDWGSNTGQLY | 0.18309061 |
| GAGGTGSTAGGAGGAGGAGGLY | ASSLDWGSNTGQLY | 0.1437894 |
| GHGGTGAAGGNGGTGGAGGSGI | ASSLDWGSNTGQLY | 0.17298348 |
| GNGGNGGAGGEGGPGGAGGAGA | ASSLDWGSNTGQLY | 0.20358688 |
| GSTAGNGGNGGNATNGTIAGQP | ASSLDWGSNTGQLY | 0.14514978 |
| GTGGDGGNGGNGGTGG | ASSLDWGSNTGQLY | 0.1961159 |
| GVRRPAGIPADAPV | ASSLDWGSNTGQLY | 0.25772846 |
| GYRQVTDAHLLTLARR | ASSLDWGSNTGQLY | 0.14489403 |
| IAKMGQDGHDRG | ASSLDWGSNTGQLY | 0.15724358 |
| KAEPSIEIQYT | ASSLDWGSNTGQLY | 0.16128917 |
| KPVRIRMAADEDTVAVT | ASSLDWGSNTGQLY | 0.13973598 |
| MPPRIAGMRLLVIKPEP | ASSLDWGSNTGQLY | 0.17995052 |
| MKFYAVALFRSRGVPPTR | ASSLDWGSNTGQLY | 0.14769498 |
| NAGGFGILGAVAHSP | ASSLDWGSNTGQLY | 0.1507339 |
| NEGREITLAMAEM | ASSLDWGSNTGQLY | 0.1582185 |
| NELVEWSHPLGHRWRWE | ASSLDWGSNTGQLY | 0.15051703 |
| NLGAGGTGGNGGNPG | ASSLDWGSNTGQLY | 0.16051365 |
| NTEWPKWETVMAADPVD | ASSLDWGSNTGQLY | 0.16175109 |
| PASAAPAETQP | ASSLDWGSNTGQLY | 0.15777619 |
| QQSSDGKNGVRAAEITGE | ASSLDWGSNTGQLY | 0.16483459 |
| RAPDAVTTSWWKE | ASSLDWGSNTGQLY | 0.14823078 |
| SGFGNVGAHTSGWFNQ | ASSLDWGSNTGQLY | 0.16037455 |
| SGVAGAVGQAASVGGLKV | ASSLDWGSNTGQLY | 0.24541503 |
| SNGGAGTGSAGGNGGT | ASSLDWGSNTGQLY | 0.14259286 |
| TGGAGGEGGAAG | ASSLDWGSNTGQLY | 0.1466936 |
| TGGDGGAAGAGGAPGHG | ASSLDWGSNTGQLY | 0.16571493 |
| TMPPLLAAIANAGRHGV | ASSLDWGSNTGQLY | 0.13933334 |
| TSSTAGNSGGA | ASSLDWGSNTGQLY | 0.17255022 |
| VPATVPVGILC | ASSLDWGSNTGQLY | 0.16295226 |
| VQRPDSGGVVLTGRIS | ASSLDWGSNTGQLY | 0.17426275 |
| WTPNPVPATKGVPAGLVQAT | ASSLDWGSNTGQLY | 0.1417815 |
| YALKRALAKFSDRFDVVII | ASSLDWGSNTGQLY | 0.3613196 |
| YLIGSVIV | ASSLDWGSNTGQLY | 0.19913055 |
| YGQLVAVID | ASSLDWGSNTGQLY | 0.15648173 |
| VTVGGMTLPA | ASSLDWGSNTGQLY | 0.14326458 |
| VLALPAV | ASSLDWGSNTGQLY | 0.15520316 |
| VIPPTLNYET | ASSLDWGSNTGQLY | 0.18995808 |
| VGMGGMHPSSG | ASSLDWGSNTGQLY | 0.22122307 |
| VGLAGGAATA | ASSLDWGSNTGQLY | 0.14922127 |
| VGGAGGAGGSGP | ASSLDWGSNTGQLY | 0.16311847 |
| VFAAFMV | ASSLDWGSNTGQLY | 0.18278877 |
| TLYDKLR | ASSLDWGSNTGQLY | 0.25913078 |
| TLAVFTVL | ASSLDWGSNTGQLY | 0.13963236 |
| TGSVQPARV | ASSLDWGSNTGQLY | 0.15108724 |
| TGGGAGSGGT | ASSLDWGSNTGQLY | 0.14772935 |
| SHLETSL | ASSLDWGSNTGQLY | 0.15515757 |
| SFTTGMT | ASSLDWGSNTGQLY | 0.15228789 |
| QVEVAKRMGVRQPTVS | ASSLDWGSNTGQLY | 0.17808738 |
| QGSGGPALAM | ASSLDWGSNTGQLY | 0.17167191 |
| MLAPSNP | ASSLDWGSNTGQLY | 0.15346865 |
| LVNNAAI | ASSLDWGSNTGQLY | 0.19154301 |

| | | |
|---|---|---|
| LTGEINTLA | ASSLDWGSNTGQLY | 0.20283063 |
| LSIFGPV | ASSLDWGSNTGQLY | 0.15659532 |
| LLALLANL | ASSLDWGSNTGQLY | 0.17392305 |
| LISNASDALDKLR | ASSLDWGSNTGQLY | 0.14610896 |
| LINEAVP | ASSLDWGSNTGQLY | 0.1396726 |
| LGYAFSGL | ASSLDWGSNTGQLY | 0.14723293 |
| LDFVAVAL | ASSLDWGSNTGQLY | 0.14913519 |
| LAQPGAE | ASSLDWGSNTGQLY | 0.1370104 |
| ISNLGAVTGF | ASSLDWGSNTGQLY | 0.18208759 |
| HQEDPGRI | ASSLDWGSNTGQLY | 0.16559353 |
| GTPGEPTP | ASSLDWGSNTGQLY | 0.18867093 |
| GFFNSGS | ASSLDWGSNTGQLY | 0.15685862 |
| GAGGGGLTAGAN | ASSLDWGSNTGQLY | 0.16249411 |
| FVDALVNM | ASSLDWGSNTGQLY | 0.15134023 |
| FLTVAPEL | ASSLDWGSNTGQLY | 0.26384646 |
| FETLTAL | ASSLDWGSNTGQLY | 0.14995444 |
| DVINPIEL | ASSLDWGSNTGQLY | 0.14310452 |
| APVSGAPV | ASSLDWGSNTGQLY | 0.15183213 |
| ALVGGLAW | ASSLDWGSNTGQLY | 0.14555994 |
| ALAAAAATAI | ASSLDWGSNTGQLY | 0.19763465 |
| AFGLGLLL | ASSLDWGSNTGQLY | 0.17919685 |
| AFFAAASL | ASSLDWGSNTGQLY | 0.17874436 |
| AAAALPAFSPP | ASSLDWGSNTGQLY | 0.15856855 |
| VVLAPAL | ASSLDWGSNTGQLY | 0.13998581 |
| TIGHVDHGKTTL | ASSLDWGSNTGQLY | 0.15482014 |
| LISNASDALDK | ASSLDWGSNTGQLY | 0.2837978 |
| LISNASDAL | ASSLDWGSNTGQLY | 0.162483 |
| ISNASDALDK | ASSLDWGSNTGQLY | 0.17706417 |
| NQAETLVYQTE | ASSLDWGSNTGQLY | 0.21229503 |
| TGGTGGTAGAG | ASSLDWGSNTGQLY | 0.1527486 |
| TGSTAGGAGGA | ASSLDWGSNTGQLY | 0.16420005 |
| DPLAVPVAAGPV | ASSLDWGSNTGQLY | 0.15943803 |
| GGTGGTPGNGGN | ASSLDWGSNTGQLY | 0.17647734 |
| PAGLAGVDPATL | ASSLDWGSNTGQLY | 0.15581784 |
| GNNLVVEAPHAGASV | ASSLDWGSNTGQLY | 0.15037984 |
| KVTRIMSASTLEQPA | ASSLDWGSNTGQLY | 0.19804671 |
| LGFALPIALPRRTVV | ASSLDWGSNTGQLY | 0.2986993 |
| TRLGNLTARKLVVLV | ASSLDWGSNTGQLY | 0.17397104 |
| WSQQRRLIYFPSAGP | ASSLDWGSNTGQLY | 0.17185088 |
| GPNGNPDPMAAAVDIR | ASSLDWGSNTGQLY | 0.1912187 |
| TQRCVSVADLRAAASR | ASSLDWGSNTGQLY | 0.14344068 |
| QFVLDVAEHPVEVLTETV | ASSLDWGSNTGQLY | 0.16600893 |
| GGAGGTGGAGGNGGTQAGDG | ASSLDWGSNTGQLY | 0.1487712 |
| PLLATLRAIAVDVDAKPAQV | ASSLDWGSNTGQLY | 0.1440796 |
| APKFRTGGYMIDFWGVGYQVAK | ASSLDWGSNTGQLY | 0.1438475 |
| GAGGAGGAGGAAGGPGGTGGTG | ASSLDWGSNTGQLY | 0.18080077 |
| IGALGLLATDVAPAGSVAFGVA | ASSLDWGSNTGQLY | 0.15693897 |
| AGGAGGIGGNGGAGGAGGVLMG | ASSLDWGSNTGQLY | 0.37524742 |
| ADELIGLL | ASSLDWGSNTGQLY | 0.17775422 |
| ADRALKA | ASSLDWGSNTGQLY | 0.18825376 |
| AGAVGQVGGAGGA | ASSLDWGSNTGQLY | 0.1461383 |

| | | |
|---|---|---|
| AGGGGGLGGAP | ASSLDWGSNTGQLY | 0.1576063 |
| AGGLLYGNGGAG | ASSLDWGSNTGQLY | 0.15389955 |
| AGLESIGL | ASSLDWGSNTGQLY | 0.14966592 |
| AGLKTGII | ASSLDWGSNTGQLY | 0.21116734 |
| ASLAALLL | ASSLDWGSNTGQLY | 0.153809 |
| ASVINISSI | ASSLDWGSNTGQLY | 0.1527221 |
| ATPPGPA | ASSLDWGSNTGQLY | 0.1665114 |
| ATVASALV | ASSLDWGSNTGQLY | 0.22173348 |
| ATVASLIL | ASSLDWGSNTGQLY | 0.16081832 |
| AVNGPVLIRH | ASSLDWGSNTGQLY | 0.14249615 |
| DGAAGAVGI | ASSLDWGSNTGQLY | 0.14590497 |
| DPLLHGI | ASSLDWGSNTGQLY | 0.178267 |
| ELGAALGAL | ASSLDWGSNTGQLY | 0.2348195 |
| ELGRTQAV | ASSLDWGSNTGQLY | 0.14011373 |
| FVAGGYLS | ASSLDWGSNTGQLY | 0.22198284 |
| GASGGAGGAGGW | ASSLDWGSNTGQLY | 0.16697326 |
| GGAGAGGDGGAGGI | ASSLDWGSNTGQLY | 0.14767113 |
| GTGHNGSQPGMGGNG | ASSLDWGSNTGQLY | 0.16526006 |
| GYAAASASAS | ASSLDWGSNTGQLY | 0.18959895 |
| HQVDVAAVQQG | ASSLDWGSNTGQLY | 0.20982733 |
| IGAAIAHT | ASSLDWGSNTGQLY | 0.18395135 |
| IGMIEVAQL | ASSLDWGSNTGQLY | 0.17998819 |
| IRLAPEHTSAMD | ASSLDWGSNTGQLY | 0.15427417 |
| IYNVSVL | ASSLDWGSNTGQLY | 0.19355902 |
| KQGFLGM | ASSLDWGSNTGQLY | 0.2014519 |
| LESVITQL | ASSLDWGSNTGQLY | 0.21355417 |
| LGLGALVVV | ASSLDWGSNTGQLY | 0.15479411 |
| LIGNGGNGGSGGI | ASSLDWGSNTGQLY | 0.15122606 |
| LNAINAPTL | ASSLDWGSNTGQLY | 0.1582541 |
| LQLLNTGV | ASSLDWGSNTGQLY | 0.19147363 |
| LQTSIEF | ASSLDWGSNTGQLY | 0.15469228 |
| MSFAVSEL | ASSLDWGSNTGQLY | 0.16641364 |
| NTGALGL | ASSLDWGSNTGQLY | 0.15957049 |
| PPGAGGAGGAGTT | ASSLDWGSNTGQLY | 0.14560343 |
| QLVGLGI | ASSLDWGSNTGQLY | 0.1695232 |
| RFLEQVHQL | ASSLDWGSNTGQLY | 0.30428267 |
| RGVGYVLR | ASSLDWGSNTGQLY | 0.16181429 |
| SATGVFAGVI | ASSLDWGSNTGQLY | 0.14520372 |
| SLPAITLP | ASSLDWGSNTGQLY | 0.22199228 |
| STALAFTV | ASSLDWGSNTGQLY | 0.5116056 |
| SVLTVGLGK | ASSLDWGSNTGQLY | 0.18905726 |
| TALAGAGSGI | ASSLDWGSNTGQLY | 0.14442909 |
| TGLPFTGI | ASSLDWGSNTGQLY | 0.13787352 |
| VAEHVTV | ASSLDWGSNTGQLY | 0.17573343 |
| VDIVLNSL | ASSLDWGSNTGQLY | 0.15510447 |
| VLTGGGAVL | ASSLDWGSNTGQLY | 0.15319891 |
| VNVGLAYGI | ASSLDWGSNTGQLY | 0.14773747 |
| VSSAAVGAV | ASSLDWGSNTGQLY | 0.1606318 |
| VTSVYEAGV | ASSLDWGSNTGQLY | 0.18558781 |
| WVALDRGA | ASSLDWGSNTGQLY | 0.15944287 |
| YLVGSVVI | ASSLDWGSNTGQLY | 0.14551033 |

| | | |
|---|---|---|
| GPPGTGKTLL | ASSLDWGSNTGQLY | 0.16463903 |
| GVGMTKF | ASSLDWGSNTGQLY | 0.22601146 |
| IGIGGSDL | ASSLDWGSNTGQLY | 0.14602049 |
| IGTLLAEL | ASSLDWGSNTGQLY | 0.1609692 |
| IYTSGTT | ASSLDWGSNTGQLY | 0.17707393 |
| LSFPAGV | ASSLDWGSNTGQLY | 0.1394029 |
| LYTSGTT | ASSLDWGSNTGQLY | 0.15454708 |
| VGAFPFGF | ASSLDWGSNTGQLY | 0.15130413 |
| VGAGTVEF | ASSLDWGSNTGQLY | 0.21035837 |
| VGKSTLLL | ASSLDWGSNTGQLY | 0.15044296 |
| AALAHAGGGIV | ASSLDWGSNTGQLY | 0.17488389 |
| AALVAVSLL | ASSLDWGSNTGQLY | 0.18481565 |
| AAVSRAVAAL | ASSLDWGSNTGQLY | 0.18514018 |
| AGGSSGIGGF | ASSLDWGSNTGQLY | 0.16352934 |
| AGLTVANANG | ASSLDWGSNTGQLY | 0.15365659 |
| AGLVGAVL | ASSLDWGSNTGQLY | 0.14960913 |
| ALSQAFTV | ASSLDWGSNTGQLY | 0.1370262 |
| ALTFSVSL | ASSLDWGSNTGQLY | 0.24179175 |
| ALTGAALVAL | ASSLDWGSNTGQLY | 0.13712241 |
| ASFSIGAGVL | ASSLDWGSNTGQLY | 0.24102873 |
| DGSVYKI | ASSLDWGSNTGQLY | 0.18248329 |
| EQLAEIV | ASSLDWGSNTGQLY | 0.15149951 |
| FAVSELTVM | ASSLDWGSNTGQLY | 0.17153189 |
| GAAPLAAGL | ASSLDWGSNTGQLY | 0.14223577 |
| GADGGDGGSGGAGG | ASSLDWGSNTGQLY | 0.14184356 |
| GFGLDTGGAGG | ASSLDWGSNTGQLY | 0.16564621 |
| GLASGGLL | ASSLDWGSNTGQLY | 0.17614377 |
| GRIAGAAVLL | ASSLDWGSNTGQLY | 0.1621236 |
| GSAGSLNTSF | ASSLDWGSNTGQLY | 0.17746307 |
| GTGQIGI | ASSLDWGSNTGQLY | 0.18701828 |
| GTLIALIL | ASSLDWGSNTGQLY | 0.14971846 |
| IAATMGFVVG | ASSLDWGSNTGQLY | 0.16025138 |
| IFAQALL | ASSLDWGSNTGQLY | 0.2197885 |
| IGAAVALGF | ASSLDWGSNTGQLY | 0.17291597 |
| IGLTPLAL | ASSLDWGSNTGQLY | 0.15228468 |
| ILGSNTSTL | ASSLDWGSNTGQLY | 0.18701456 |
| IVFSLTVL | ASSLDWGSNTGQLY | 0.15367989 |
| LASKATGF | ASSLDWGSNTGQLY | 0.17400658 |
| LASVALIL | ASSLDWGSNTGQLY | 0.16624826 |
| LFAVGLTL | ASSLDWGSNTGQLY | 0.17419845 |
| LFNGGIGTYI | ASSLDWGSNTGQLY | 0.20622295 |
| LTLGAGII | ASSLDWGSNTGQLY | 0.14733846 |
| LVAAALGDI | ASSLDWGSNTGQLY | 0.14787802 |
| NTAFVQLGI | ASSLDWGSNTGQLY | 0.15612352 |
| QLFSGAEL | ASSLDWGSNTGQLY | 0.20110507 |
| SAATFITL | ASSLDWGSNTGQLY | 0.21168429 |
| SNYGLANLGV | ASSLDWGSNTGQLY | 0.14712153 |
| SYGGAYAVM | ASSLDWGSNTGQLY | 0.17824002 |
| TTGVGGLTLG | ASSLDWGSNTGQLY | 0.15831247 |
| TTQTIYI | ASSLDWGSNTGQLY | 0.16032077 |
| VAAGAGLGAVI | ASSLDWGSNTGQLY | 0.17527854 |

| | | |
|---|---|---|
| VAAVVTLL | ASSLDWGSNTGQLY | 0.15434718 |
| VGFLGGLI | ASSLDWGSNTGQLY | 0.15295848 |
| VGVIPILL | ASSLDWGSNTGQLY | 0.1731792 |
| VIFASSLI | ASSLDWGSNTGQLY | 0.15478171 |
| VKGGDGGAAAG | ASSLDWGSNTGQLY | 0.17501044 |
| VTKTFTNI | ASSLDWGSNTGQLY | 0.18241297 |
| VVAAAILL | ASSLDWGSNTGQLY | 0.15248808 |
| YAALDVEL | ASSLDWGSNTGQLY | 0.17383863 |
| LASVALLL | ASSLDWGSNTGQLY | 0.18176357 |
| LDAVHRV | ASSLDWGSNTGQLY | 0.19965482 |
| QGLSSSIF | ASSLDWGSNTGQLY | 0.1548999 |
| RGTFANIRL | ASSLDWGSNTGQLY | 0.14795554 |
| RQLWWGHRI | ASSLDWGSNTGQLY | 0.21149352 |
| YGPPGTGKTLL | ASSLDWGSNTGQLY | 0.20370485 |
| AALVGAPANVA | ASSLDWGSNTGQLY | 0.19210988 |
| AATFRRTE | ASSLDWGSNTGQLY | 0.17947008 |
| AGALGGGI | ASSLDWGSNTGQLY | 0.16809256 |
| AGGAGGAGGIGAG | ASSLDWGSNTGQLY | 0.1865044 |
| AGSGALGL | ASSLDWGSNTGQLY | 0.1532181 |
| AQIADQI | ASSLDWGSNTGQLY | 0.14275521 |
| ASAVAVTYL | ASSLDWGSNTGQLY | 0.15951937 |
| ASTVLGGTGG | ASSLDWGSNTGQLY | 0.15478957 |
| AVGGEPAK | ASSLDWGSNTGQLY | 0.16379653 |
| CEQGTFGGTC | ASSLDWGSNTGQLY | 0.1417342 |
| DEGKAPAG | ASSLDWGSNTGQLY | 0.23094791 |
| DTVVFPAL | ASSLDWGSNTGQLY | 0.43445852 |
| FEGITMAG | ASSLDWGSNTGQLY | 0.15062739 |
| FPLVVQD | ASSLDWGSNTGQLY | 0.19829564 |
| GAFTTPQ | ASSLDWGSNTGQLY | 0.14280327 |
| GFGGFGTAGGVGGA | ASSLDWGSNTGQLY | 0.17103252 |
| GFLAQFL | ASSLDWGSNTGQLY | 0.21068688 |
| GGAGGAGGNAGVA | ASSLDWGSNTGQLY | 0.14832537 |
| GGVGTAGAGGAGGAGG | ASSLDWGSNTGQLY | 0.1754049 |
| GNGGGGDAGNAGSGGN | ASSLDWGSNTGQLY | 0.15217587 |
| GTGGAGGAGNDAGS | ASSLDWGSNTGQLY | 0.16036226 |
| GTTNTGWFNS | ASSLDWGSNTGQLY | 0.28551933 |
| IFMGPLFTL | ASSLDWGSNTGQLY | 0.15880029 |
| ILAHAIEAI | ASSLDWGSNTGQLY | 0.15916413 |
| IRSAIREAL | ASSLDWGSNTGQLY | 0.15930869 |
| ITFAGGG | ASSLDWGSNTGQLY | 0.14894995 |
| ITGMSGHQEV | ASSLDWGSNTGQLY | 0.1708085 |
| IVAGLACTL | ASSLDWGSNTGQLY | 0.23131683 |
| IVGGGLAGA | ASSLDWGSNTGQLY | 0.14034505 |
| IVTGVLNI | ASSLDWGSNTGQLY | 0.20030843 |
| LAQQGCI | ASSLDWGSNTGQLY | 0.15776823 |
| LCLHVEN | ASSLDWGSNTGQLY | 0.33929414 |
| LDAISDGS | ASSLDWGSNTGQLY | 0.1931381 |
| LGGFAAML | ASSLDWGSNTGQLY | 0.16640171 |
| LGGKNPMI | ASSLDWGSNTGQLY | 0.15546332 |
| LGPKGVAAIL | ASSLDWGSNTGQLY | 0.21417181 |
| LMVGVSI | ASSLDWGSNTGQLY | 0.17872275 |

| | | |
|---|---|---|
| LVGAAFVL | ASSLDWGSNTGQLY | 0.17936824 |
| LVLAVAPL | ASSLDWGSNTGQLY | 0.19401403 |
| MAAIAVIDAA | ASSLDWGSNTGQLY | 0.20361447 |
| NNGINAELQ | ASSLDWGSNTGQLY | 0.23621945 |
| RGEPLVLVHG | ASSLDWGSNTGQLY | 0.2218714 |
| RGGAGGAAGLI | ASSLDWGSNTGQLY | 0.24386244 |
| RSYRMHPAV | ASSLDWGSNTGQLY | 0.20672622 |
| SAAHLLAGQLA | ASSLDWGSNTGQLY | 0.20263183 |
| SAAVGSY | ASSLDWGSNTGQLY | 0.23148179 |
| SPHAVLHRI | ASSLDWGSNTGQLY | 0.1981474 |
| SQIAKVL | ASSLDWGSNTGQLY | 0.26620823 |
| TALGHLVP | ASSLDWGSNTGQLY | 0.23649718 |
| TAPTTSGAA | ASSLDWGSNTGQLY | 0.20761174 |
| TDGDDGNGGSPGT | ASSLDWGSNTGQLY | 0.27774265 |
| TDGLGEN | ASSLDWGSNTGQLY | 0.27042392 |
| TGGPPLPGQA | ASSLDWGSNTGQLY | 0.20350078 |
| TIPAITI | ASSLDWGSNTGQLY | 0.21193907 |
| TIPGFTI | ASSLDWGSNTGQLY | 0.22666468 |
| TSIASGALA | ASSLDWGSNTGQLY | 0.21416152 |
| TVTLPGAAI | ASSLDWGSNTGQLY | 0.19556893 |
| VASGLSAALP | ASSLDWGSNTGQLY | 0.2224459 |
| VGAGVIGI | ASSLDWGSNTGQLY | 0.20294854 |
| VGPKRFGV | ASSLDWGSNTGQLY | 0.19370528 |
| VNVAFPD | ASSLDWGSNTGQLY | 0.2717178 |
| VSMTNSGVS | ASSLDWGSNTGQLY | 0.21028726 |
| VTDVLGDLSV | ASSLDWGSNTGQLY | 0.20691322 |
| VVVVGAGFAGL | ASSLDWGSNTGQLY | 0.25457355 |
| VYTVGTVV | ASSLDWGSNTGQLY | 0.19943392 |
| WTTFSGP | ASSLDWGSNTGQLY | 0.20495787 |
| YDELVEAF | ASSLDWGSNTGQLY | 0.25101557 |
| YGTGGAGGNGG | ASSLDWGSNTGQLY | 0.2612001 |
| YTNVMAV | ASSLDWGSNTGQLY | 0.20314236 |
| FLEQVHQL | ASSLDWGSNTGQLY | 0.21240611 |
| GLSLLAVAV | ASSLDWGSNTGQLY | 0.3133635 |
| LEQVHQL | ASSLDWGSNTGQLY | 0.2261922 |
| LNAAGAI | ASSLDWGSNTGQLY | 0.26264927 |
| PAVHASL | ASSLDWGSNTGQLY | 0.26760736 |
| TELEDDA | ASSLDWGSNTGQLY | 0.20885018 |
| TPPGPPN | ASSLDWGSNTGQLY | 0.25464094 |
| TPPPPPPP | ASSLDWGSNTGQLY | 0.30009073 |
| TQSPALI | ASSLARGGQNTLY | 0.21783395 |
| EFAAAFPQ | ASSLARGGQNTLY | 0.21707354 |
| LLSGNLDVL | ASSLARGGQNTLY | 0.2067221 |
| GPANLGNYNI | ASSLARGGQNTLY | 0.2395542 |
| TLGPTLP | ASSLARGGQNTLY | 0.19496928 |
| VYAYSSCCK | ASSLARGGQNTLY | 0.23422758 |
| APAPAPAPNI | ASSLARGGQNTLY | 0.26646972 |
| PGGPVPIP | ASSLARGGQNTLY | 0.2210311 |
| FLTVALEP | ASSLARGGQNTLY | 0.19735241 |
| SQYFGANL | ASSLARGGQNTLY | 0.1986042 |
| GGVKPAA | ASSLARGGQNTLY | 0.1997157 |

| | | |
|---|---|---|
| GGTGGAGGAGGW | ASSLARGGQNTLY | 0.20236467 |
| GVGGVAGLGGAGG | ASSLARGGQNTLY | 0.19995372 |
| VLLVLATTI | ASSLARGGQNTLY | 0.20531955 |
| IGIFAAML | ASSLARGGQNTLY | 0.2993018 |
| AVGDLID | ASSLARGGQNTLY | 0.30469245 |
| NISPLPL | ASSLARGGQNTLY | 0.19826373 |
| VTAVFVAI | ASSLARGGQNTLY | 0.24135748 |
| SGDDNDAAL | ASSLARGGQNTLY | 0.29525894 |
| AVAALAHS | ASSLARGGQNTLY | 0.25940144 |
| GGVTVMATP | ASSLARGGQNTLY | 0.22429483 |
| AGLAANVRD | ASSLARGGQNTLY | 0.20009647 |
| TNRPDLIDPALLRPG | ASSLARGGQNTLY | 0.35973787 |
| LANLGVTAL | ASSLARGGQNTLY | 0.21209458 |
| TGERIEL | ASSLARGGQNTLY | 0.24342783 |
| NGEELAA | ASSLARGGQNTLY | 0.29378513 |
| VLSLVWG | ASSLARGGQNTLY | 0.21001364 |
| IGYALWL | ASSLARGGQNTLY | 0.20073225 |
| LSLLLAPL | ASSLARGGQNTLY | 0.27754927 |
| TLDEALK | ASSLARGGQNTLY | 0.23723106 |
| TDGAGTAEGG | ASSLARGGQNTLY | 0.19937932 |
| NAPGGAGGNGG | ASSLARGGQNTLY | 0.20364352 |
| VSVFMGMGGNF | ASSLARGGQNTLY | 0.21583995 |
| LASYLAAL | ASSLARGGQNTLY | 0.23009428 |
| LVGGTPAL | ASSLARGGQNTLY | 0.26321924 |
| VRNADVL | ASSLARGGQNTLY | 0.2291168 |
| TCSSPFLVSAM | ASSLARGGQNTLY | 0.20547307 |
| VHITDAE | ASSLARGGQNTLY | 0.25521216 |
| EYAAPSF | ASSLARGGQNTLY | 0.22343454 |
| VVDAFHGGL | ASSLARGGQNTLY | 0.22489698 |
| ASVVDPQ | ASSLARGGQNTLY | 0.24327362 |
| SYNEFGGLS | ASSLARGGQNTLY | 0.19649549 |
| DPDGSVY | ASSLARGGQNTLY | 0.2409187 |
| MVSTIHATE | ASSLARGGQNTLY | 0.24481031 |
| LAAGVADPV | ASSLARGGQNTLY | 0.23894018 |
| TKEQIAA | ASSLARGGQNTLY | 0.25580555 |
| SLASPAAL | ASSLARGGQNTLY | 0.21239978 |
| IGLVFYGV | ASSLARGGQNTLY | 0.22117501 |
| FLLDASNL | ASSLARGGQNTLY | 0.20122236 |
| YQVGNFP | ASSLARGGQNTLY | 0.22477162 |
| ESLLSQST | ASSLARGGQNTLY | 0.23486476 |
| LFTVGFAAVG | ASSLARGGQNTLY | 0.22450246 |
| ATPPTPT | ASSLARGGQNTLY | 0.27801386 |
| TDLATLG | ASSLARGGQNTLY | 0.24230304 |
| SAGGTGR | ASSLARGGQNTLY | 0.21290089 |
| GFSNSGS | ASSLARGGQNTLY | 0.276971 |
| GGNGGSGGTSV | ASSLARGGQNTLY | 0.26626176 |
| GTSVATGGAGN | ASSLARGGQNTLY | 0.19769925 |
| NGGNGGSGGTS | ASSLARGGQNTLY | 0.20701963 |
| AVVKFPH | ASSLARGGQNTLY | 0.23868755 |
| TPPPPPP | ASSLARGGQNTLY | 0.27110434 |
| AAPAAAAPA | ASSLARGGQNTLY | 0.22516921 |

| | | |
|---|---|---|
| IALNEDL | ASSLARGGQNTLY | 0.22514671 |
| IVNVSSM | ASSLARGGQNTLY | 0.34614438 |
| TGIPPAPR | ASSLARGGQNTLY | 0.21105583 |
| FDLGGGTF | ASSLARGGQNTLY | 0.28453994 |
| FDLGGGTFDVS | ASSLARGGQNTLY | 0.19569494 |
| NEPTAAA | ASSLARGGQNTLY | 0.20804623 |
| NEPTAAALA | ASSLARGGQNTLY | 0.20357686 |
| GSIINISSI | ASSLARGGQNTLY | 0.19296235 |
| VAPGNAGTA | ASSLARGGQNTLY | 0.26433086 |
| TGGGASLE | ASSLARGGQNTLY | 0.2141211 |
| VAVYNHYKRI | ASSLARGGQNTLY | 0.2461367 |
| TDPAPATT | ASSLARGGQNTLY | 0.22411403 |
| TNRPDILDPALLRPG | ASSLARGGQNTLY | 0.23200902 |
| LTTTTTS | ASSLARGGQNTLY | 0.21450461 |
| PTGSGKTL | ASSLARGGQNTLY | 0.2292891 |
| TGSGKTL | ASSLARGGQNTLY | 0.2096953 |
| TIGHVDHGK | ASSLARGGQNTLY | 0.24322733 |
| TIGHVDHGKTT | ASSLARGGQNTLY | 0.29570413 |
| GHVDHGKTT | ASSLARGGQNTLY | 0.23599182 |
| AQFEHTLL | ASSLARGGQNTLY | 0.1983247 |
| IVNQDEL | ASSLARGGQNTLY | 0.21619146 |
| TNGSQFF | ASSLARGGQNTLY | 0.21641472 |
| TGKGIVE | ASSLARGGQNTLY | 0.19534957 |
| LISNASDA | ASSLARGGQNTLY | 0.1963811 |
| AELRTLGELTKIAKAHGAQ | ASSLARGGQNTLY | 0.19679715 |
| APNAVAAAEPSW | ASSLARGGQNTLY | 0.19979526 |
| APQVAGAATAVL | ASSLARGGQNTLY | 0.26270017 |
| ARRDPAAAAAILPTDA | ASSLARGGQNTLY | 0.2891066 |
| DAAARDVLAEAGLAEYF | ASSLARGGQNTLY | 0.21003209 |
| DDQIVLSAKANLAKVEQ | ASSLARGGQNTLY | 0.2611505 |
| DELLPDTGEDSDEDGH | ASSLARGGQNTLY | 0.20976427 |
| ELPEAPANRATVPQD | ASSLARGGQNTLY | 0.18987344 |
| FGQSKIERVTGIG | ASSLARGGQNTLY | 0.21052894 |
| GAGGTGSTAGGAGGAGGAGGLY | ASSLARGGQNTLY | 0.20599699 |
| GHGGTGAAGGNGGTGGAGGSGI | ASSLARGGQNTLY | 0.25469524 |
| GNGGNGGAGGEGGPGGAGGAGA | ASSLARGGQNTLY | 0.19909878 |
| GSTAGNGGNGGNATNGTIAGQP | ASSLARGGQNTLY | 0.2188714 |
| GTGGDGGNGGNGGTGG | ASSLARGGQNTLY | 0.21724851 |
| GVRRPAGIPADAPV | ASSLARGGQNTLY | 0.24834043 |
| GYRQVTDAHLLTLARR | ASSLARGGQNTLY | 0.24641447 |
| IAKMGQDGHDRG | ASSLARGGQNTLY | 0.1995004 |
| KAEPSIEIQYT | ASSLARGGQNTLY | 0.19355202 |
| KPVRIRMAADEDTVAVT | ASSLARGGQNTLY | 0.2444329 |
| MPPRIAGMRLLVIKPEP | ASSLARGGQNTLY | 0.20358396 |
| MKFYAVALFRSRGVPPTR | ASSLARGGQNTLY | 0.20835786 |
| NAGGFGILGAVAHSP | ASSLARGGQNTLY | 0.19905502 |
| NEGREITLAMAEM | ASSLARGGQNTLY | 0.21043944 |
| NELVEWSHPLGHRWRWE | ASSLARGGQNTLY | 0.19715856 |
| NLGAGGTGGNGGNPG | ASSLARGGQNTLY | 0.2539997 |
| NTEWPKWETVMAADPVD | ASSLARGGQNTLY | 0.20645143 |
| PASAAPAETQP | ASSLARGGQNTLY | 0.21937251 |

| | | |
|---|---|---|
| QQSSDGKNGVRAAEITGE | ASSLARGGQNTLY | 0.29267016 |
| RAPDAVTTSWWKE | ASSLARGGQNTLY | 0.26812991 |
| SGFGNVGAHTSGWFNQ | ASSLARGGQNTLY | 0.21243276 |
| SGVAGAVGQAASVGGLKV | ASSLARGGQNTLY | 0.20654552 |
| SNGGAGTGSAGGNGGT | ASSLARGGQNTLY | 0.20821199 |
| TGGAGGEGGAAG | ASSLARGGQNTLY | 0.20362619 |
| TGGDGGAAGAGGAPGHG | ASSLARGGQNTLY | 0.20571902 |
| TMPPLLAAIANAGRHGV | ASSLARGGQNTLY | 0.2861696 |
| TSSTAGNSGGA | ASSLARGGQNTLY | 0.21325392 |
| VPATVPVGILC | ASSLARGGQNTLY | 0.20265095 |
| VQRPDSGGVVLTGRIS | ASSLARGGQNTLY | 0.21263535 |
| WTPNPVPATKGVPAGLVQAT | ASSLARGGQNTLY | 0.27458164 |
| YALKRALAKFSDRFDVVII | ASSLARGGQNTLY | 0.30925834 |
| YLIGSVIV | ASSLARGGQNTLY | 0.21016 |
| YGQLVAVID | ASSLARGGQNTLY | 0.2172642 |
| VTVGGMTLPA | ASSLARGGQNTLY | 0.22255129 |
| VLALPAV | ASSLARGGQNTLY | 0.22049093 |
| VIPPTLNYET | ASSLARGGQNTLY | 0.21080862 |
| VGMGGMHPSSG | ASSLARGGQNTLY | 0.20022348 |
| VGLAGGAATA | ASSLARGGQNTLY | 0.23449212 |
| VGGAGGAGGSGP | ASSLARGGQNTLY | 0.58093095 |
| VFAAFMV | ASSLARGGQNTLY | 0.21055317 |
| TLYDKLR | ASSLARGGQNTLY | 0.19998069 |
| TLAVFTVL | ASSLARGGQNTLY | 0.22179379 |
| TGSVQPARV | ASSLARGGQNTLY | 0.20020021 |
| TGGGAGSGGT | ASSLARGGQNTLY | 0.20533393 |
| SHLETSL | ASSLARGGQNTLY | 0.22150558 |
| SFTTGMT | ASSLARGGQNTLY | 0.1947728 |
| QVEVAKRMGVRQPTVS | ASSLARGGQNTLY | 0.2116458 |
| QGSGGPALAM | ASSLARGGQNTLY | 0.2902655 |
| MLAPSNP | ASSLARGGQNTLY | 0.21766095 |
| LVNNAAI | ASSLARGGQNTLY | 0.2376965 |
| LTGEINTLA | ASSLARGGQNTLY | 0.2859099 |
| LSIFGPV | ASSLARGGQNTLY | 0.23295775 |
| LLALLANL | ASSLARGGQNTLY | 0.2821182 |
| LISNASDALDKLR | ASSLARGGQNTLY | 0.30149633 |
| LINEAVP | ASSLARGGQNTLY | 0.2241817 |
| LGYAFSGL | ASSLARGGQNTLY | 0.2139145 |
| LDFVAVAL | ASSLARGGQNTLY | 0.25714242 |
| LAQPGAE | ASSLARGGQNTLY | 0.2205657 |
| ISNLGAVTGF | ASSLARGGQNTLY | 0.23221658 |
| HQEDPGRI | ASSLARGGQNTLY | 0.20978811 |
| GTPGEPTP | ASSLARGGQNTLY | 0.28082666 |
| GFFNSGS | ASSLARGGQNTLY | 0.21650165 |
| GAGGGGLTAGAN | ASSLARGGQNTLY | 0.26652813 |
| FVDALVNM | ASSLARGGQNTLY | 0.19839765 |
| FLTVAPEL | ASSLARGGQNTLY | 0.20400746 |
| FETLTAL | ASSLARGGQNTLY | 0.25513577 |
| DVINPIEL | ASSLARGGQNTLY | 0.19329214 |
| APVSGAPV | ASSLARGGQNTLY | 0.19788268 |
| ALVGGLAW | ASSLARGGQNTLY | 0.21960427 |

| | | |
|---|---|---|
| ALAAAAATAI | ASSLARGGQNTLY | 0.24243903 |
| AFGLGLLL | ASSLARGGQNTLY | 0.19096461 |
| AFFAAASL | ASSLARGGQNTLY | 0.21090253 |
| AAAALPAFSPP | ASSLARGGQNTLY | 0.20289488 |
| VVLAPAL | ASSLARGGQNTLY | 0.21571286 |
| TIGHVDHGKTTL | ASSLARGGQNTLY | 0.21252915 |
| LISNASDALDK | ASSLARGGQNTLY | 0.19787773 |
| LISNASDAL | ASSLARGGQNTLY | 0.2328676 |
| ISNASDALDK | ASSLARGGQNTLY | 0.20079212 |
| NQAETLVYQTE | ASSLARGGQNTLY | 0.2030806 |
| TGGTGGTAGAG | ASSLARGGQNTLY | 0.24025154 |
| TGSTAGGAGGA | ASSLARGGQNTLY | 0.3822092 |
| DPLAVPVAAGPV | ASSLARGGQNTLY | 0.20320837 |
| GGTGGTPGNGGN | ASSLARGGQNTLY | 0.23385274 |
| PAGLAGVDPATL | ASSLARGGQNTLY | 0.20008826 |
| GNNLVVEAPHAGASV | ASSLARGGQNTLY | 0.46738577 |
| KVTRIMSASTLEQPA | ASSLARGGQNTLY | 0.2200932 |
| LGFALPIALPRRTVV | ASSLARGGQNTLY | 0.19652376 |
| TRLGNLTARKLVVLV | ASSLARGGQNTLY | 0.2571681 |
| WSQQRRLIYFPSAGP | ASSLARGGQNTLY | 0.22792475 |
| GPNGNPDPMAAAVDIR | ASSLARGGQNTLY | 0.23959298 |
| TQRCVSVADLRAAASR | ASSLARGGQNTLY | 0.27189502 |
| QFVLDVAEHPVEVLTETV | ASSLARGGQNTLY | 0.20206957 |
| GGAGGTGGAGGNGGTQAGDG | ASSLARGGQNTLY | 0.21162814 |
| PLLATLRAIAVDVDAKPAQV | ASSLARGGQNTLY | 0.25021008 |
| APKFRTGGYMIDFWGVGYQVAK | ASSLARGGQNTLY | 0.23684725 |
| GAGGAGGAGGAAGGPGGTGGTG | ASSLARGGQNTLY | 0.19822055 |
| IGALGLLATDVAPAGSVAFGVA | ASSLARGGQNTLY | 0.2532966 |
| AGGAGGIGGNGGAGGAGGVLMG | ASSLARGGQNTLY | 0.21539843 |
| ADELIGLL | ASSLARGGQNTLY | 0.21214789 |
| ADRALKA | ASSLARGGQNTLY | 0.2857628 |
| AGAVGQVGGAGGA | ASSLARGGQNTLY | 0.22186534 |
| AGGGGGLGGAP | ASSLARGGQNTLY | 0.25326714 |
| AGGLLYGNGGAG | ASSLARGGQNTLY | 0.23261657 |
| AGLESIGL | ASSLARGGQNTLY | 0.37593502 |
| AGLKTGII | ASSLARGGQNTLY | 0.20673713 |
| ASLAALLL | ASSLARGGQNTLY | 0.22486693 |
| ASVINISSI | ASSLARGGQNTLY | 0.25078684 |
| ATPPGPA | ASSLARGGQNTLY | 0.24334593 |
| ATVASALV | ASSLARGGQNTLY | 0.20632412 |
| ATVASLIL | ASSLARGGQNTLY | 0.21830957 |
| AVNGPVLIRH | ASSLARGGQNTLY | 0.23293985 |
| DGAAGAVGI | ASSLARGGQNTLY | 0.29449165 |
| DPLLHGI | ASSLARGGQNTLY | 0.21529125 |
| ELGAALGAL | ASSLARGGQNTLY | 0.22831309 |
| ELGRTQAV | ASSLARGGQNTLY | 0.34589237 |
| FVAGGYLS | ASSLARGGQNTLY | 0.20782769 |
| GASGGAGGAGGW | ASSLARGGQNTLY | 0.2117692 |
| GGAGAGGDGGAGGI | ASSLARGGQNTLY | 0.25030133 |
| GTGHNGSQPGMGGNG | ASSLARGGQNTLY | 0.3094734 |
| GYAAASASAS | ASSLARGGQNTLY | 0.21159977 |

| | | |
|---|---|---|
| HQVDVAAVQQG | ASSLARGGQNTLY | 0.26299322 |
| IGAAIAHT | ASSLARGGQNTLY | 0.21220396 |
| IGMIEVAQL | ASSLARGGQNTLY | 0.26939434 |
| IRLAPEHTSAMD | ASSLARGGQNTLY | 0.20145854 |
| IYNVSVL | ASSLARGGQNTLY | 0.21761245 |
| KQGFLGM | ASSLARGGQNTLY | 0.19625024 |
| LESVITQL | ASSLARGGQNTLY | 0.2393168 |
| LGLGALVVV | ASSLARGGQNTLY | 0.2759824 |
| LIGNGGNGGSGGI | ASSLARGGQNTLY | 0.24748188 |
| LNAINAPTL | ASSLARGGQNTLY | 0.19960424 |
| LQLLNTGV | ASSLARGGQNTLY | 0.2295655 |
| LQTSIEF | ASSLARGGQNTLY | 0.2607603 |
| MSFAVSEL | ASSLARGGQNTLY | 0.45242348 |
| NTGALGL | ASSLARGGQNTLY | 0.2199865 |
| PPGAGGAGGAGTT | ASSLARGGQNTLY | 0.28167674 |
| QLVGLGI | ASSLARGGQNTLY | 0.2175729 |
| RFLEQVHQL | ASSLARGGQNTLY | 0.21498968 |
| RGVGYVLR | ASSLARGGQNTLY | 0.19672984 |
| SATGVFAGVI | ASSLARGGQNTLY | 0.21224739 |
| SLPAITLP | ASSLARGGQNTLY | 0.21828109 |
| STALAFTV | ASSLARGGQNTLY | 0.20532629 |
| SVLTVGLGK | ASSLARGGQNTLY | 0.2549392 |
| TALAGAGSGI | ASSLARGGQNTLY | 0.3124056 |
| TGLPFTGI | ASSLARGGQNTLY | 0.24692747 |
| VAEHVTV | ASSLARGGQNTLY | 0.30235445 |
| VDIVLNSL | ASSLARGGQNTLY | 0.25065485 |
| VLTGGGAVL | ASSLARGGQNTLY | 0.23635426 |
| VNVGLAYGI | ASSLARGGQNTLY | 0.20110807 |
| VSSAAVGAV | ASSLARGGQNTLY | 0.23456044 |
| VTSVYEAGV | ASSLARGGQNTLY | 0.19795747 |
| WVALDRGA | ASSLARGGQNTLY | 0.20673397 |
| YLVGSVVI | ASSLARGGQNTLY | 0.20177637 |
| GPPGTGKTLL | ASSLARGGQNTLY | 0.22528063 |
| GVGMTKF | ASSLARGGQNTLY | 0.19074191 |
| IGIGGSDL | ASSLARGGQNTLY | 0.29502183 |
| IGTLLAEL | ASSLARGGQNTLY | 0.2938412 |
| IYTSGTT | ASSLARGGQNTLY | 0.19792546 |
| LSFPAGV | ASSLARGGQNTLY | 0.26268208 |
| LYTSGTT | ASSLARGGQNTLY | 0.22482155 |
| VGAFPFGF | ASSLARGGQNTLY | 0.198006 |
| VGAGTVEF | ASSLARGGQNTLY | 0.2717602 |
| VGKSTLLL | ASSLARGGQNTLY | 0.20338084 |
| AALAHAGGGIV | ASSLARGGQNTLY | 0.24673012 |
| AALVAVSLL | ASSLARGGQNTLY | 0.26609266 |
| AAVSRAVAAL | ASSLARGGQNTLY | 0.23953493 |
| AGGSSGIGGF | ASSLARGGQNTLY | 0.22243212 |
| AGLTVANANG | ASSLARGGQNTLY | 0.23003978 |
| AGLVGAVL | ASSLARGGQNTLY | 0.20609947 |
| ALSQAFTV | ASSLARGGQNTLY | 0.22912976 |
| ALTFSVSL | ASSLARGGQNTLY | 0.20127384 |
| ALTGAALVAL | ASSLARGGQNTLY | 0.23667392 |

| | | |
|---|---|---|
| ASFSIGAGVL | ASSLARGGQNTLY | 0.20189385 |
| DGSVYKI | ASSLARGGQNTLY | 0.28633535 |
| EQLAEIV | ASSLARGGQNTLY | 0.2156504 |
| FAVSELTVM | ASSLARGGQNTLY | 0.2279907 |
| GAAPLAAGL | ASSLARGGQNTLY | 0.21001612 |
| GADGGDGGSGGAGG | ASSLARGGQNTLY | 0.22121629 |
| GFGLDTGGAGG | ASSLARGGQNTLY | 0.23122197 |
| GLASGGLL | ASSLARGGQNTLY | 0.19920039 |
| GRIAGAAVLL | ASSLARGGQNTLY | 0.23036541 |
| GSAGSLNTSF | ASSLARGGQNTLY | 0.21592265 |
| GTGQIGI | ASSLARGGQNTLY | 0.20188165 |
| GTLIALIL | ASSLARGGQNTLY | 0.28939 |
| IAATMGFVVG | ASSLARGGQNTLY | 0.22518963 |
| IFAQALL | ASSLARGGQNTLY | 0.20135614 |
| IGAAVALGF | ASSLARGGQNTLY | 0.21464315 |
| IGLTPLAL | ASSLARGGQNTLY | 0.2378699 |
| ILGSNTSTL | ASSLARGGQNTLY | 0.21417838 |
| IVFSLTVL | ASSLARGGQNTLY | 0.19170707 |
| LASKATGF | ASSLARGGQNTLY | 0.21999174 |
| LASVALIL | ASSLARGGQNTLY | 0.21259089 |
| LFAVGLTL | ASSLARGGQNTLY | 0.19682665 |
| LFNGGIGTYI | ASSLARGGQNTLY | 0.19974642 |
| LTLGAGII | ASSLARGGQNTLY | 0.27157444 |
| LVAAALGDI | ASSLARGGQNTLY | 0.20405139 |
| NTAFVQLGI | ASSLARGGQNTLY | 0.21491955 |
| QLFSGAEL | ASSLARGGQNTLY | 0.27220035 |
| SAATFITL | ASSLARGGQNTLY | 0.23500565 |
| SNYGLANLGV | ASSLARGGQNTLY | 0.26198238 |
| SYGGAYAVM | ASSLARGGQNTLY | 0.21792968 |
| TTGVGGLTLG | ASSLARGGQNTLY | 0.21339911 |
| TTQTIYI | ASSLARGGQNTLY | 0.2695304 |
| VAAGAGLGAVI | ASSLARGGQNTLY | 0.22495107 |
| VAAVVTLL | ASSLARGGQNTLY | 0.20447917 |
| VGFLGGLI | ASSLARGGQNTLY | 0.19797045 |
| VGVIPILL | ASSLARGGQNTLY | 0.20905091 |
| VIFASSLI | ASSLARGGQNTLY | 0.20653008 |
| VKGGDGGAAAG | ASSLARGGQNTLY | 0.2378522 |
| VTKTFTNI | ASSLARGGQNTLY | 0.251966 |
| VVAAAILL | ASSLARGGQNTLY | 0.20191455 |
| YAALDVEL | ASSLARGGQNTLY | 0.20091237 |
| LASVALLL | ASSLARGGQNTLY | 0.20789267 |
| LDAVHRV | ASSLARGGQNTLY | 0.21342885 |
| QGLSSSIF | ASSLARGGQNTLY | 0.2652232 |
| RGTFANIRL | ASSLARGGQNTLY | 0.2195985 |
| RQLWWGHRI | ASSLARGGQNTLY | 0.5912136 |
| YGPPGTGKTLL | ASSLARGGQNTLY | 0.25303793 |
| AALVGAPANVA | ASSLARGGQNTLY | 0.20375468 |
| AATFRRTE | ASSLARGGQNTLY | 0.24612272 |
| AGALGGGI | ASSLARGGQNTLY | 0.20661883 |
| AGGAGGAGGIGAG | ASSLARGGQNTLY | 0.21775049 |
| AGSGALGL | ASSLARGGQNTLY | 0.22568375 |

| | | |
|---|---|---|
| AQIADQI | ASSLARGGQNTLY | 0.20757692 |
| ASAVAVTYL | ASSLARGGQNTLY | 0.20695819 |
| ASTVLGGTGG | ASSLARGGQNTLY | 0.22775218 |
| AVGGEPAK | ASSLARGGQNTLY | 0.20062631 |
| CEQGTFGGTC | ASSLARGGQNTLY | 0.22295898 |
| DEGKAPAG | ASSLARGGQNTLY | 0.2005543 |
| DTVVFPAL | ASSLARGGQNTLY | 0.20120277 |
| FEGITMAG | ASSLARGGQNTLY | 0.21433297 |
| FPLVVQD | ASSLARGGQNTLY | 0.27291974 |
| GAFTTPQ | ASSLARGGQNTLY | 0.4211256 |
| GFGGFGTAGGVGGA | ASSLARGGQNTLY | 0.20606044 |
| GFLAQFL | ASSLARGGQNTLY | 0.20215149 |
| GGAGGAGGNAGVA | ASSLARGGQNTLY | 0.32612938 |
| GGVGTAGAGGAGGAGG | ASSLARGGQNTLY | 0.20026033 |
| GNGGGGDAGNAGSGGN | ASSLARGGQNTLY | 0.2186705 |
| GTGGAGGAGNDAGS | ASSLARGGQNTLY | 0.21055382 |
| GTTNTGWFNS | ASSLARGGQNTLY | 0.20180558 |
| IFMGPLFTL | ASSLARGGQNTLY | 0.20711045 |
| ILAHAIEAI | ASSLARGGQNTLY | 0.22377548 |
| IRSAIREAL | ASSLARGGQNTLY | 0.19842194 |
| ITFAGGG | ASSLARGGQNTLY | 0.2372944 |
| ITGMSGHQEV | ASSLARGGQNTLY | 0.1902093 |
| IVAGLACTL | ASSLARGGQNTLY | 0.21457952 |
| IVGGGLAGA | ASSLARGGQNTLY | 0.20487486 |
| IVTGVLNI | ASSLARGGQNTLY | 0.20289403 |
| LAQQGCI | ASSLARGGQNTLY | 0.19606161 |
| LCLHVEN | ASSLARGGQNTLY | 0.22798109 |
| LDAISDGS | ASSLARGGQNTLY | 0.25209615 |
| LGGFAAML | ASSLARGGQNTLY | 0.29748327 |
| LGGKNPMI | ASSLARGGQNTLY | 0.22168133 |
| LGPKGVAAIL | ASSLARGGQNTLY | 0.23974808 |
| LMVGVSI | ASSLARGGQNTLY | 0.27498412 |
| LVGAAFVL | ASSLARGGQNTLY | 0.21075854 |
| LVLAVAPL | ASSLARGGQNTLY | 0.20347726 |
| MAAIAVIDAA | ASSLARGGQNTLY | 0.22175859 |
| NNGINAELQ | ASSLARGGQNTLY | 0.2023909 |
| RGEPLVLVHG | ASSLARGGQNTLY | 0.22180328 |
| RGGAGGAAGLI | ASSLARGGQNTLY | 0.28782004 |
| RSYRMHPAV | ASSLARGGQNTLY | 0.19780606 |
| SAAHLLAGQLA | ASSLARGGQNTLY | 0.25088432 |
| SAAVGSY | ASSLARGGQNTLY | 0.25961152 |
| SPHAVLHRI | ASSLARGGQNTLY | 0.20021467 |
| SQIAKVL | ASSLARGGQNTLY | 0.21063508 |
| TALGHLVP | ASSLARGGQNTLY | 0.20851249 |
| TAPTTSGAA | ASSLARGGQNTLY | 0.19326735 |
| TDGDDGNGGSPGT | ASSLARGGQNTLY | 0.23596917 |
| TDGLGEN | ASSLARGGQNTLY | 0.2085722 |
| TGGPPLPGQA | ASSLARGGQNTLY | 0.2136375 |
| TIPAITI | ASSLARGGQNTLY | 0.21154615 |
| TIPGFTI | ASSLARGGQNTLY | 0.20992953 |
| TSIASGALA | ASSLARGGQNTLY | 0.21042933 |

| | | |
|---|---|---|
| TVTLPGAAI | ASSLARGGQNTLY | 0.21201284 |
| VASGLSAALP | ASSLARGGQNTLY | 0.2193802 |
| VGAGVIGI | ASSLARGGQNTLY | 0.22870496 |
| VGPKRFGV | ASSLARGGQNTLY | 0.20442118 |
| VNVAFPD | ASSLARGGQNTLY | 0.22232324 |
| VSMTNSGVS | ASSLARGGQNTLY | 0.3200167 |
| VTDVLGDLSV | ASSLARGGQNTLY | 0.1952884 |
| VVVVGAGFAGL | ASSLARGGQNTLY | 0.19856982 |
| VYTVGTVV | ASSLARGGQNTLY | 0.21900027 |
| WTTFSGP | ASSLARGGQNTLY | 0.20037657 |
| YDELVEAF | ASSLARGGQNTLY | 0.26569304 |
| YGTGGAGGNGG | ASSLARGGQNTLY | 0.21947707 |
| YTNVMAV | ASSLARGGQNTLY | 0.22729354 |
| FLEQVHQL | ASSLARGGQNTLY | 0.21934514 |
| GLSLLAVAV | ASSLARGGQNTLY | 0.30609277 |
| LEQVHQL | ASSLARGGQNTLY | 0.24559344 |
| LNAAGAI | ASSLARGGQNTLY | 0.21166483 |
| PAVHASL | ASSLARGGQNTLY | 0.22062507 |
| TELEDDA | ASSLARGGQNTLY | 0.20223898 |
| TPPGPPN | ASSLARGGQNTLY | 0.23164922 |
| TPPPPPPP | ASSLARGGQNTLY | 0.25024393 |
| TQSPALI | ASSPSTASSYEQY | 0.1966794 |
| EFAAAFPQ | ASSPSTASSYEQY | 0.21226871 |
| LLSGNLDVL | ASSPSTASSYEQY | 0.2654595 |
| GPANLGNYNI | ASSPSTASSYEQY | 0.26833868 |
| TLGPTLP | ASSPSTASSYEQY | 0.2042433 |
| VYAYSSCCK | ASSPSTASSYEQY | 0.20821379 |
| APAPAPAPNI | ASSPSTASSYEQY | 0.20861025 |
| PGGPVPIP | ASSPSTASSYEQY | 0.20573454 |
| FLTVALEP | ASSPSTASSYEQY | 0.20078026 |
| SQYFGANL | ASSPSTASSYEQY | 0.24772991 |
| GGVKPAA | ASSPSTASSYEQY | 0.2284774 |
| GGTGGAGGAGGW | ASSPSTASSYEQY | 0.31917807 |
| GVGGVAGLGGAGG | ASSPSTASSYEQY | 0.23174511 |
| VLLVLATTI | ASSPSTASSYEQY | 0.28402925 |
| IGIFAAML | ASSPSTASSYEQY | 0.21016467 |
| AVGDLID | ASSPSTASSYEQY | 0.21030821 |
| NISPLPL | ASSPSTASSYEQY | 0.20063962 |
| VTAVFVAI | ASSPSTASSYEQY | 0.20659284 |
| SGDDNDAAL | ASSPSTASSYEQY | 0.2061262 |
| AVAALAHS | ASSPSTASSYEQY | 0.20640056 |
| GGVTVMATP | ASSPSTASSYEQY | 0.19257377 |
| AGLAANVRD | ASSPSTASSYEQY | 0.22990662 |
| TNRPDLIDPALLRPG | ASSPSTASSYEQY | 0.21246065 |
| LANLGVTAL | ASSPSTASSYEQY | 0.2480678 |
| TGERIEL | ASSPSTASSYEQY | 0.21690752 |
| NGEELAA | ASSPSTASSYEQY | 0.21049966 |
| VLSLVWG | ASSPSTASSYEQY | 0.19888884 |
| IGYALWL | ASSPSTASSYEQY | 0.2832334 |
| LSLLLAPL | ASSPSTASSYEQY | 0.19913952 |
| TLDEALK | ASSPSTASSYEQY | 0.19860086 |

| | | |
|---|---|---|
| TDGAGTAEGG | ASSPSTASSYEQY | 0.21173915 |
| NAPGGAGGNGG | ASSPSTASSYEQY | 0.21265288 |
| VSVFMGMGGNF | ASSPSTASSYEQY | 0.30580544 |
| LASYLAAL | ASSPSTASSYEQY | 0.27425697 |
| LVGGTPAL | ASSPSTASSYEQY | 0.22245115 |
| VRNADVL | ASSPSTASSYEQY | 0.26064292 |
| TCSSPFLVSAM | ASSPSTASSYEQY | 0.19328631 |
| VHITDAE | ASSPSTASSYEQY | 0.21069725 |
| EYAAPSF | ASSPSTASSYEQY | 0.35343426 |
| VVDAFHGGL | ASSPSTASSYEQY | 0.22061147 |
| ASVVDPQ | ASSPSTASSYEQY | 0.23347807 |
| SYNEFGGLS | ASSPSTASSYEQY | 0.248256 |
| DPDGSVY | ASSPSTASSYEQY | 0.20278183 |
| MVSTIHATE | ASSPSTASSYEQY | 0.21823224 |
| LAAGVADPV | ASSPSTASSYEQY | 0.21660127 |
| TKEQIAA | ASSPSTASSYEQY | 0.2475228 |
| SLASPAAL | ASSPSTASSYEQY | 0.21444868 |
| IGLVFYGV | ASSPSTASSYEQY | 0.20521241 |
| FLLDASNL | ASSPSTASSYEQY | 0.25755617 |
| YQVGNFP | ASSPSTASSYEQY | 0.4202314 |
| ESLLSQST | ASSPSTASSYEQY | 0.22766262 |
| LFTVGFAAVG | ASSPSTASSYEQY | 0.24454704 |
| ATPPTPT | ASSPSTASSYEQY | 0.22144945 |
| TDLATLG | ASSPSTASSYEQY | 0.2167868 |
| SAGGTGR | ASSPSTASSYEQY | 0.20773242 |
| GFSNSGS | ASSPSTASSYEQY | 0.20248273 |
| GGNGGSGGTSV | ASSPSTASSYEQY | 0.22508238 |
| GTSVATGGAGN | ASSPSTASSYEQY | 0.1992938 |
| NGGNGGSGGTS | ASSPSTASSYEQY | 0.22514509 |
| AVVKFPH | ASSPSTASSYEQY | 0.21168733 |
| TPPPPPP | ASSPSTASSYEQY | 0.45415914 |
| AAPAAAAAPA | ASSPSTASSYEQY | 0.23870794 |
| IALNEDL | ASSPSTASSYEQY | 0.21524297 |
| IVNVSSM | ASSPSTASSYEQY | 0.19941908 |
| TGIPPAPR | ASSPSTASSYEQY | 0.21478765 |
| FDLGGGTF | ASSPSTASSYEQY | 0.22763936 |
| FDLGGGTFDVS | ASSPSTASSYEQY | 0.2026658 |
| NEPTAAA | ASSPSTASSYEQY | 0.29137868 |
| NEPTAAALA | ASSPSTASSYEQY | 0.20261554 |
| GSIINISSI | ASSPSTASSYEQY | 0.20344844 |
| VAPGNAGTA | ASSPSTASSYEQY | 0.21924388 |
| TGGGASLE | ASSPSTASSYEQY | 0.53069556 |
| VAVYNHYKRI | ASSPSTASSYEQY | 0.21613812 |
| TDPAPATT | ASSPSTASSYEQY | 0.23385699 |
| TNRPDILDPALLRPG | ASSPSTASSYEQY | 0.2006748 |
| LTTTTTS | ASSPSTASSYEQY | 0.24085712 |
| PTGSGKTL | ASSPSTASSYEQY | 0.38922915 |
| TGSGKTL | ASSPSTASSYEQY | 0.21303281 |
| TIGHVDHGK | ASSPSTASSYEQY | 0.5158727 |
| TIGHVDHGKTT | ASSPSTASSYEQY | 0.21795136 |
| GHVDHGKTT | ASSPSTASSYEQY | 0.19794306 |

| | | |
|---|---|---|
| AQFEHTLL | ASSPSTASSYEQY | 0.23266783 |
| IVNQDEL | ASSPSTASSYEQY | 0.22623229 |
| TNGSQFF | ASSPSTASSYEQY | 0.2671996 |
| TGKGIVE | ASSPSTASSYEQY | 0.26176912 |
| LISNASDA | ASSPSTASSYEQY | 0.23512287 |
| AELRTLGELTKIAKAHGAQ | ASSPSTASSYEQY | 0.21026693 |
| APNAVAAAEPSW | ASSPSTASSYEQY | 0.27550173 |
| APQVAGAATAVL | ASSPSTASSYEQY | 0.3373638 |
| ARRDPAAAAAILPTDA | ASSPSTASSYEQY | 0.23440279 |
| DAAARDVLAEAGLAEYF | ASSPSTASSYEQY | 0.20703492 |
| DDQIVLSAKANLAKVEQ | ASSPSTASSYEQY | 0.20673211 |
| DELLPDTGEDSDEDGH | ASSPSTASSYEQY | 0.2316467 |
| ELPEAPANRATVPQD | ASSPSTASSYEQY | 0.21471009 |
| FGQSKIERVTGIG | ASSPSTASSYEQY | 0.2024305 |
| GAGGTGSTAGGAGGAGGAGGLY | ASSPSTASSYEQY | 0.21634132 |
| GHGGTGAAGGNGGTGGAGGSGI | ASSPSTASSYEQY | 0.24258837 |
| GNGGNGGAGGEGGPGGAGGAGA | ASSPSTASSYEQY | 0.19839656 |
| GSTAGNGGNGGNATNGTIAGQP | ASSPSTASSYEQY | 0.22354373 |
| GTGGDGGNGGNGGTGG | ASSPSTASSYEQY | 0.2859204 |
| GVRRPAGIPADAPV | ASSPSTASSYEQY | 0.21785289 |
| GYRQVTDAHLLTLARR | ASSPSTASSYEQY | 0.20086804 |
| IAKMGQDGHDRG | ASSPSTASSYEQY | 0.3171296 |
| KAEPSIEIQYT | ASSPSTASSYEQY | 0.42926162 |
| KPVRIRMAADEDTVAVT | ASSPSTASSYEQY | 0.2470039 |
| MPPRIAGMRLLVIKPEP | ASSPSTASSYEQY | 0.20010641 |
| MKFYAVALFRSRGVPPTR | ASSPSTASSYEQY | 0.19276227 |
| NAGGFGILGAVAHSP | ASSPSTASSYEQY | 0.23048005 |
| NEGREITLAMAEM | ASSPSTASSYEQY | 0.23955771 |
| NELVEWSHPLGHRWRWE | ASSPSTASSYEQY | 0.21013267 |
| NLGAGGTGGNGGNPG | ASSPSTASSYEQY | 0.200409 |
| NTEWPKWETVMAADPVD | ASSPSTASSYEQY | 0.21270038 |
| PASAAPAETQP | ASSPSTASSYEQY | 0.23832175 |
| QQSSDGKNGVRAAEITGE | ASSPSTASSYEQY | 0.20892802 |
| RAPDAVTTSWWKE | ASSPSTASSYEQY | 0.19713254 |
| SGFGNVGAHTSGWFNQ | ASSPSTASSYEQY | 0.22242442 |
| SGVAGAVGQAASVGGLKV | ASSPSTASSYEQY | 0.25509453 |
| SNGGAGTGSAGGNGGT | ASSPSTASSYEQY | 0.19984709 |
| TGGAGGEGGAAG | ASSPSTASSYEQY | 0.21020302 |
| TGGDGGAAGAGGAPGHG | ASSPSTASSYEQY | 0.2643767 |
| TMPPLLAAIANAGRHGV | ASSPSTASSYEQY | 0.20619822 |
| TSSTAGNSGGA | ASSPSTASSYEQY | 0.21637036 |
| VPATVPVGILC | ASSPSTASSYEQY | 0.19940981 |
| VQRPDSGGVVLTGRIS | ASSPSTASSYEQY | 0.2188327 |
| WTPNPVPATKGVPAGLVQAT | ASSPSTASSYEQY | 0.20259616 |
| YALKRALAKFSDRFDVVII | ASSPSTASSYEQY | 0.24811305 |
| YLIGSVIV | ASSPSTASSYEQY | 0.24417177 |
| YGQLVAVID | ASSPSTASSYEQY | 0.26012212 |
| VTVGGMTLPA | ASSPSTASSYEQY | 0.21914709 |
| VLALPAV | ASSPSTASSYEQY | 0.20353803 |
| VIPPTLNYET | ASSPSTASSYEQY | 0.20216356 |
| VGMGGMHPSSG | ASSPSTASSYEQY | 0.19387692 |

| | | |
|---|---|---|
| VGLAGGAATA | ASSPSTASSYEQY | 0.3055133 |
| VGGAGGAGGSGP | ASSPSTASSYEQY | 0.20410511 |
| VFAAFMV | ASSPSTASSYEQY | 0.26279908 |
| TLYDKLR | ASSPSTASSYEQY | 0.23450992 |
| TLAVFTVL | ASSPSTASSYEQY | 0.20692815 |
| TGSVQPARV | ASSPSTASSYEQY | 0.22516078 |
| TGGGAGSGGT | ASSPSTASSYEQY | 0.19540069 |
| SHLETSL | ASSPSTASSYEQY | 0.2271948 |
| SFTTGMT | ASSPSTASSYEQY | 0.21043009 |
| QVEVAKRMGVRQPTVS | ASSPSTASSYEQY | 0.21990761 |
| QGSGGPALAM | ASSPSTASSYEQY | 0.22036777 |
| MLAPSNP | ASSPSTASSYEQY | 0.22835457 |
| LVNNAAI | ASSPSTASSYEQY | 0.24347873 |
| LTGEINTLA | ASSPSTASSYEQY | 0.2114977 |
| LSIFGPV | ASSPSTASSYEQY | 0.21890475 |
| LLALLANL | ASSPSTASSYEQY | 0.26704463 |
| LISNASDALDKLR | ASSPSTASSYEQY | 0.23738377 |
| LINEAVP | ASSPSTASSYEQY | 0.20213193 |
| LGYAFSGL | ASSPSTASSYEQY | 0.22712648 |
| LDFVAVAL | ASSPSTASSYEQY | 0.20649649 |
| LAQPGAE | ASSPSTASSYEQY | 0.23109002 |
| ISNLGAVTGF | ASSPSTASSYEQY | 0.22326979 |
| HQEDPGRI | ASSPSTASSYEQY | 0.22633505 |
| GTPGEPTP | ASSPSTASSYEQY | 0.23677056 |
| GFFNSGS | ASSPSTASSYEQY | 0.19986692 |
| GAGGGGLTAGAN | ASSPSTASSYEQY | 0.20819685 |
| FVDALVNM | ASSPSTASSYEQY | 0.2192819 |
| FLTVAPEL | ASSPSTASSYEQY | 0.22894146 |
| FETLTAL | ASSPSTASSYEQY | 0.23145442 |
| DVINPIEL | ASSPSTASSYEQY | 0.19966139 |
| APVSGAPV | ASSPSTASSYEQY | 0.2402312 |
| ALVGGLAW | ASSPSTASSYEQY | 0.21323654 |
| ALAAAAATAI | ASSPSTASSYEQY | 0.20640399 |
| AFGLGLLL | ASSPSTASSYEQY | 0.21514769 |
| AFFAAASL | ASSPSTASSYEQY | 0.21126892 |
| AAAALPAFSPP | ASSPSTASSYEQY | 0.20505057 |
| VVLAPAL | ASSPSTASSYEQY | 0.24463014 |
| TIGHVDHGKTTL | ASSPSTASSYEQY | 0.20713116 |
| LISNASDALDK | ASSPSTASSYEQY | 0.22952813 |
| LISNASDAL | ASSPSTASSYEQY | 0.22234562 |
| ISNASDALDK | ASSPSTASSYEQY | 0.20499443 |
| NQAETLVYQTE | ASSPSTASSYEQY | 0.23623653 |
| TGGTGGTAGAG | ASSPSTASSYEQY | 0.20438245 |
| TGSTAGGAGGA | ASSPSTASSYEQY | 0.24357094 |
| DPLAVPVAAGPV | ASSPSTASSYEQY | 0.21331108 |
| GGTGGTPGNGGN | ASSPSTASSYEQY | 0.20855801 |
| PAGLAGVDPATL | ASSPSTASSYEQY | 0.22558066 |
| GNNLVVEAPHAGASV | ASSPSTASSYEQY | 0.244288 |
| KVTRIMSASTLEQPA | ASSPSTASSYEQY | 0.24429336 |
| LGFALPIALPRRTVV | ASSPSTASSYEQY | 0.21331015 |
| TRLGNLTARKLVVLV | ASSPSTASSYEQY | 0.26978314 |

| | | |
|---|---|---|
| WSQQRRLIYFPSAGP | ASSPSTASSYEQY | 0.24973543 |
| GPNGNPDPMAAAVDIR | ASSPSTASSYEQY | 0.2062322 |
| TQRCVSVADLRAAASR | ASSPSTASSYEQY | 0.19497423 |
| QFVLDVAEHPVEVLTETV | ASSPSTASSYEQY | 0.20987746 |
| GGAGGTGGAGGNGGTQAGDG | ASSPSTASSYEQY | 0.20112056 |
| PLLATLRAIAVDVDAKPAQV | ASSPSTASSYEQY | 0.21452028 |
| APKFRTGGYMIDFWGVGYQVAK | ASSPSTASSYEQY | 0.19605465 |
| GAGGAGGAGGAAGGPGGTGGTG | ASSPSTASSYEQY | 0.25460112 |
| IGALGLLATDVAPAGSVAFGVA | ASSPSTASSYEQY | 0.29773414 |
| AGGAGGIGGNGGAGGAGGVLMG | ASSPSTASSYEQY | 0.20741728 |
| ADELIGLL | ASSPSTASSYEQY | 0.26803026 |
| ADRALKA | ASSPSTASSYEQY | 0.19775936 |
| AGAVGQVGGAGGA | ASSPSTASSYEQY | 0.24532998 |
| AGGGGGLGGAP | ASSPSTASSYEQY | 0.28233218 |
| AGGLLYGNGGAG | ASSPSTASSYEQY | 0.19767593 |
| AGLESIGL | ASSPSTASSYEQY | 0.23207316 |
| AGLKTGII | ASSPSTASSYEQY | 0.22174525 |
| ASLAALLL | ASSPSTASSYEQY | 0.20513308 |
| ASVINISSI | ASSPSTASSYEQY | 0.29674426 |
| ATPPGPA | ASSPSTASSYEQY | 0.21764538 |
| ATVASALV | ASSPSTASSYEQY | 0.24483918 |
| ATVASLIL | ASSPSTASSYEQY | 0.20900413 |
| AVNGPVLIRH | ASSPSTASSYEQY | 0.20266022 |
| DGAAGAVGI | ASSPSTASSYEQY | 0.2352802 |
| DPLLHGI | ASSPSTASSYEQY | 0.25459015 |
| ELGAALGAL | ASSPSTASSYEQY | 0.21380238 |
| ELGRTQAV | ASSPSTASSYEQY | 0.29666921 |
| FVAGGYLS | ASSPSTASSYEQY | 0.21505213 |
| GASGGAGGAGGW | ASSPSTASSYEQY | 0.34743536 |
| GGAGAGGDGGAGGI | ASSPSTASSYEQY | 0.3286512 |
| GTGHNGSQPGMGGNG | ASSPSTASSYEQY | 0.23219353 |
| GYAAASASAS | ASSPSTASSYEQY | 0.21348567 |
| HQVDVAAVQQG | ASSPSTASSYEQY | 0.35469925 |
| IGAAIAHT | ASSPSTASSYEQY | 0.23280911 |
| IGMIEVAQL | ASSPSTASSYEQY | 0.25105643 |
| IRLAPEHTSAMD | ASSPSTASSYEQY | 0.24385154 |
| IYNVSVL | ASSPSTASSYEQY | 0.20450704 |
| KQGFLGM | ASSPSTASSYEQY | 0.570664 |
| LESVITQL | ASSPSTASSYEQY | 0.24511766 |
| LGLGALVVV | ASSPSTASSYEQY | 0.23221357 |
| LIGNGGNGGSGGI | ASSPSTASSYEQY | 0.23421049 |
| LNAINAPTL | ASSPSTASSYEQY | 0.19767937 |
| LQLLNTGV | ASSPSTASSYEQY | 0.21069539 |
| LQTSIEF | ASSPSTASSYEQY | 0.21609397 |
| MSFAVSEL | ASSPSTASSYEQY | 0.5007288 |
| NTGALGL | ASSPSTASSYEQY | 0.26840812 |
| PPGAGGAGGAGTT | ASSPSTASSYEQY | 0.23886652 |
| QLVGLGI | ASSPSTASSYEQY | 0.5290576 |
| RFLEQVHQL | ASSPSTASSYEQY | 0.45194882 |
| RGVGYVLR | ASSPSTASSYEQY | 0.199492 |
| SATGVFAGVI | ASSPSTASSYEQY | 0.2405353 |

| | | |
|---|---|---|
| SLPAITLP | ASSPSTASSYEQY | 0.20668577 |
| STALAFTV | ASSPSTASSYEQY | 0.24211235 |
| SVLTVGLGK | ASSPSTASSYEQY | 0.19258475 |
| TALAGAGSGI | ASSPSTASSYEQY | 0.3467666 |
| TGLPFTGI | ASSPSTASSYEQY | 0.21187603 |
| VAEHVTV | ASSPSTASSYEQY | 0.19232461 |
| VDIVLNSL | ASSPSTASSYEQY | 0.22750998 |
| VLTGGGAVL | ASSPSTASSYEQY | 0.2535172 |
| VNVGLAYGI | ASSPSTASSYEQY | 0.23671538 |
| VSSAAVGAV | ASSPSTASSYEQY | 0.55887556 |
| VTSVYEAGV | ASSPSTASSYEQY | 0.20874214 |
| WVALDRGA | ASSPSTASSYEQY | 0.20513743 |
| YLVGSVVI | ASSPSTASSYEQY | 0.39503676 |
| GPPGTGKTLL | ASSPSTASSYEQY | 0.45354512 |
| GVGMTKF | ASSPSTASSYEQY | 0.20398684 |
| IGIGGSDL | ASSPSTASSYEQY | 0.38635397 |
| IGTLLAEL | ASSPSTASSYEQY | 0.40023795 |
| IYTSGTT | ASSPSTASSYEQY | 0.2540801 |
| LSFPAGV | ASSPSTASSYEQY | 0.26799428 |
| LYTSGTT | ASSPSTASSYEQY | 0.3141923 |
| VGAFPFGF | ASSPSTASSYEQY | 0.20425488 |
| VGAGTVEF | ASSPSTASSYEQY | 0.4909442 |
| VGKSTLLL | ASSPSTASSYEQY | 0.5968788 |
| AALAHAGGGIV | ASSPSTASSYEQY | 0.22712517 |
| AALVAVSLL | ASSPSTASSYEQY | 0.25028387 |
| AAVSRAVAAL | ASSPSTASSYEQY | 0.20282379 |
| AGGSSGIGGF | ASSPSTASSYEQY | 0.23376217 |
| AGLTVANANG | ASSPSTASSYEQY | 0.20134751 |
| AGLVGAVL | ASSPSTASSYEQY | 0.44137624 |
| ALSQAFTV | ASSPSTASSYEQY | 0.45807624 |
| ALTFSVSL | ASSPSTASSYEQY | 0.26458994 |
| ALTGAALVAL | ASSPSTASSYEQY | 0.19931585 |
| ASFSIGAGVL | ASSPSTASSYEQY | 0.2138472 |
| DGSVYKI | ASSPSTASSYEQY | 0.22347777 |
| EQLAEIV | ASSPSTASSYEQY | 0.23776962 |
| FAVSELTVM | ASSPSTASSYEQY | 0.19898516 |
| GAAPLAAGL | ASSPSTASSYEQY | 0.21953689 |
| GADGGDGGSGGAGG | ASSPSTASSYEQY | 0.44516098 |
| GFGLDTGGAGG | ASSPSTASSYEQY | 0.3754113 |
| GLASGGLL | ASSPSTASSYEQY | 0.23209316 |
| GRIAGAAVLL | ASSPSTASSYEQY | 0.6282022 |
| GSAGSLNTSF | ASSPSTASSYEQY | 0.32713413 |
| GTGQIGI | ASSPSTASSYEQY | 0.35638765 |
| GTLIALIL | ASSPSTASSYEQY | 0.22864494 |
| IAATMGFVVG | ASSPSTASSYEQY | 0.20750716 |
| IFAQALL | ASSPSTASSYEQY | 0.55901104 |
| IGAAVALGF | ASSPSTASSYEQY | 0.25740603 |
| IGLTPLAL | ASSPSTASSYEQY | 0.21730998 |
| ILGSNTSTL | ASSPSTASSYEQY | 0.3180935 |
| IVFSLTVL | ASSPSTASSYEQY | 0.23713645 |
| LASKATGF | ASSPSTASSYEQY | 0.22938138 |

| | | |
|---|---|---|
| LASVALIL | ASSPSTASSYEQY | 0.4168088 |
| LFAVGLTL | ASSPSTASSYEQY | 0.30353197 |
| LFNGGIGTYI | ASSPSTASSYEQY | 0.20560022 |
| LTLGAGII | ASSPSTASSYEQY | 0.22516191 |
| LVAAALGDI | ASSPSTASSYEQY | 0.22151726 |
| NTAFVQLGI | ASSPSTASSYEQY | 0.27306938 |
| QLFSGAEL | ASSPSTASSYEQY | 0.45524567 |
| SAATFITL | ASSPSTASSYEQY | 0.24968106 |
| SNYGLANLGV | ASSPSTASSYEQY | 0.22952288 |
| SYGGAYAVM | ASSPSTASSYEQY | 0.40250668 |
| TTGVGGLTLG | ASSPSTASSYEQY | 0.29215834 |
| TTQTIYI | ASSPSTASSYEQY | 0.26063284 |
| VAAGAGLGAVI | ASSPSTASSYEQY | 0.31533086 |
| VAAVVTLL | ASSPSTASSYEQY | 0.22982931 |
| VGFLGGLI | ASSPSTASSYEQY | 0.29177248 |
| VGVIPILL | ASSPSTASSYEQY | 0.2964344 |
| VIFASSLI | ASSPSTASSYEQY | 0.30641732 |
| VKGGDGGAAAG | ASSPSTASSYEQY | 0.3128731 |
| VTKTFTNI | ASSPSTASSYEQY | 0.2626104 |
| VVAAAILL | ASSPSTASSYEQY | 0.2642969 |
| YAALDVEL | ASSPSTASSYEQY | 0.21659987 |
| LASVALLL | ASSPSTASSYEQY | 0.20154485 |
| LDAVHRV | ASSPSTASSYEQY | 0.37699127 |
| QGLSSSIF | ASSPSTASSYEQY | 0.36055017 |
| RGTFANIRL | ASSPSTASSYEQY | 0.39175868 |
| RQLWWGHRI | ASSPSTASSYEQY | 0.24178982 |
| YGPPGTGKTLL | ASSPSTASSYEQY | 0.21549983 |
| AALVGAPANVA | ASSPSTASSYEQY | 0.3398708 |
| AATFRRTE | ASSPSTASSYEQY | 0.4254238 |
| AGALGGGI | ASSPSTASSYEQY | 0.20250323 |
| AGGAGGAGGIGAG | ASSPSTASSYEQY | 0.21333104 |
| AGSGALGL | ASSPSTASSYEQY | 0.62475085 |
| AQIADQI | ASSPSTASSYEQY | 0.32468823 |
| ASAVAVTYL | ASSPSTASSYEQY | 0.27449256 |
| ASTVLGGTGG | ASSPSTASSYEQY | 0.21781945 |
| AVGGEPAK | ASSPSTASSYEQY | 0.64190614 |
| CEQGTFGGTC | ASSPSTASSYEQY | 0.21858227 |
| DEGKAPAG | ASSPSTASSYEQY | 0.53327656 |
| DTVVFPAL | ASSPSTASSYEQY | 0.2039828 |
| FEGITMAG | ASSPSTASSYEQY | 0.2273949 |
| FPLVVQD | ASSPSTASSYEQY | 0.2041447 |
| GAFTTPQ | ASSPSTASSYEQY | 0.19553174 |
| GFGGFGTAGGVGGA | ASSPSTASSYEQY | 0.3542347 |
| GFLAQFL | ASSPSTASSYEQY | 0.20388356 |
| GGAGGAGGNAGVA | ASSPSTASSYEQY | 0.44539088 |
| GGVGTAGAGGAGGAGG | ASSPSTASSYEQY | 0.23948926 |
| GNGGGGDAGNAGSGGN | ASSPSTASSYEQY | 0.3220527 |
| GTGGAGGAGNDAGS | ASSPSTASSYEQY | 0.33556822 |
| GTTNTGWFNS | ASSPSTASSYEQY | 0.26632202 |
| IFMGPLFTL | ASSPSTASSYEQY | 0.21148162 |
| ILAHAIEAI | ASSPSTASSYEQY | 0.364923 |

| | | |
|---|---|---|
| IRSAIREAL | ASSPSTASSYEQY | 0.3709102 |
| ITFAGGG | ASSPSTASSYEQY | 0.2483164 |
| ITGMSGHQEV | ASSPSTASSYEQY | 0.19964792 |
| IVAGLACTL | ASSPSTASSYEQY | 0.22922462 |
| IVGGGLAGA | ASSPSTASSYEQY | 0.27113014 |
| IVTGVLNI | ASSPSTASSYEQY | 0.20020343 |
| LAQQGCI | ASSPSTASSYEQY | 0.19636406 |
| LCLHVEN | ASSPSTASSYEQY | 0.21153899 |
| LDAISDGS | ASSPSTASSYEQY | 0.21677251 |
| LGGFAAML | ASSPSTASSYEQY | 0.5400095 |
| LGGKNPMI | ASSPSTASSYEQY | 0.39073664 |
| LGPKGVAAIL | ASSPSTASSYEQY | 0.2798928 |
| LMVGVSI | ASSPSTASSYEQY | 0.3605522 |
| LVGAAFVL | ASSPSTASSYEQY | 0.28942588 |
| LVLAVAPL | ASSPSTASSYEQY | 0.19162686 |
| MAAIAVIDAA | ASSPSTASSYEQY | 0.21080074 |
| NNGINAELQ | ASSPSTASSYEQY | 0.20126085 |
| RGEPLVLVHG | ASSPSTASSYEQY | 0.34710097 |
| RGGAGGAAGLI | ASSPSTASSYEQY | 0.20118162 |
| RSYRMHPAV | ASSPSTASSYEQY | 0.2124269 |
| SAAHLLAGQLA | ASSPSTASSYEQY | 0.3298811 |
| SAAVGSY | ASSPSTASSYEQY | 0.38855007 |
| SPHAVLHRI | ASSPSTASSYEQY | 0.26687688 |
| SQIAKVL | ASSPSTASSYEQY | 0.20495112 |
| TALGHLVP | ASSPSTASSYEQY | 0.1997183 |
| TAPTTSGAA | ASSPSTASSYEQY | 0.35506016 |
| TDGDDGNGGSPGT | ASSPSTASSYEQY | 0.20468959 |
| TDGLGEN | ASSPSTASSYEQY | 0.20622894 |
| TGGPPLPGQA | ASSPSTASSYEQY | 0.19890235 |
| TIPAITI | ASSPSTASSYEQY | 0.22271983 |
| TIPGFTI | ASSPSTASSYEQY | 0.19731419 |
| TSIASGALA | ASSPSTASSYEQY | 0.26863116 |
| TVTLPGAAI | ASSPSTASSYEQY | 0.34791785 |
| VASGLSAALP | ASSPSTASSYEQY | 0.25782257 |
| VGAGVIGI | ASSPSTASSYEQY | 0.6457094 |
| VGPKRFGV | ASSPSTASSYEQY | 0.5664901 |
| VNVAFPD | ASSPSTASSYEQY | 0.27022436 |
| VSMTNSGVS | ASSPSTASSYEQY | 0.2185866 |
| VTDVLGDLSV | ASSPSTASSYEQY | 0.21439014 |
| VVVVGAGFAGL | ASSPSTASSYEQY | 0.20743598 |
| VYTVGTVV | ASSPSTASSYEQY | 0.21130286 |
| WTTFSGP | ASSPSTASSYEQY | 0.5482911 |
| YDELVEAF | ASSPSTASSYEQY | 0.2224417 |
| YGTGGAGGNGG | ASSPSTASSYEQY | 0.1973964 |
| YTNVMAV | ASSPSTASSYEQY | 0.24124745 |
| FLEQVHQL | ASSPSTASSYEQY | 0.5025537 |
| GLSLLAVAV | ASSPSTASSYEQY | 0.62720364 |
| LEQVHQL | ASSPSTASSYEQY | 0.20775507 |
| LNAAGAI | ASSPSTASSYEQY | 0.21794881 |
| PAVHASL | ASSPSTASSYEQY | 0.21222338 |
| TELEDDA | ASSPSTASSYEQY | 0.32304975 |

| | | |
|---|---|---|
| TPPGPPN | ASSPSTASSYEQY | 0.20384085 |
| TPPPPPPP | ASSPSTASSYEQY | 0.2062294 |
| TQSPALI | ASSPGQTNSDYT | 0.28937033 |
| EFAAAFPQ | ASSPGQTNSDYT | 0.6696265 |
| LLSGNLDVL | ASSPGQTNSDYT | 0.22506487 |
| GPANLGNYNI | ASSPGQTNSDYT | 0.19446336 |
| TLGPTLP | ASSPGQTNSDYT | 0.35164607 |
| VYAYSSCCK | ASSPGQTNSDYT | 0.21721908 |
| APAPAPAPNI | ASSPGQTNSDYT | 0.25783086 |
| PGGPVPIP | ASSPGQTNSDYT | 0.24381343 |
| FLTVALEP | ASSPGQTNSDYT | 0.19957438 |
| SQYFGANL | ASSPGQTNSDYT | 0.22428271 |
| GGVKPAA | ASSPGQTNSDYT | 0.5177338 |
| GGTGGAGGAGGW | ASSPGQTNSDYT | 0.27302298 |
| GVGGVAGLGGAGG | ASSPGQTNSDYT | 0.4167995 |
| VLLVLATTI | ASSPGQTNSDYT | 0.59535104 |
| IGIFAAML | ASSPGQTNSDYT | 0.506779 |
| AVGDLID | ASSPGQTNSDYT | 0.5783045 |
| NISPLPL | ASSPGQTNSDYT | 0.36937517 |
| VTAVFVAI | ASSPGQTNSDYT | 0.22802033 |
| SGDDNDAAL | ASSPGQTNSDYT | 0.2492037 |
| AVAALAHS | ASSPGQTNSDYT | 0.46673453 |
| GGVTVMATP | ASSPGQTNSDYT | 0.3612072 |
| AGLAANVRD | ASSPGQTNSDYT | 0.30900997 |
| TNRPDLIDPALLRPG | ASSPGQTNSDYT | 0.20790088 |
| LANLGVTAL | ASSPGQTNSDYT | 0.51467186 |
| TGERIEL | ASSPGQTNSDYT | 0.22336526 |
| NGEELAA | ASSPGQTNSDYT | 0.381267 |
| VLSLVWG | ASSPGQTNSDYT | 0.20411137 |
| IGYALWL | ASSPGQTNSDYT | 0.21099697 |
| LSLLLAPL | ASSPGQTNSDYT | 0.5506412 |
| TLDEALK | ASSPGQTNSDYT | 0.19712713 |
| TDGAGTAEGG | ASSPGQTNSDYT | 0.20080414 |
| NAPGGAGGNGG | ASSPGQTNSDYT | 0.24850087 |
| VSVFMGMGGNF | ASSPGQTNSDYT | 0.5581932 |
| LASYLAAL | ASSPGQTNSDYT | 0.19286294 |
| LVGGTPAL | ASSPGQTNSDYT | 0.20354559 |
| VRNADVL | ASSPGQTNSDYT | 0.21229136 |
| TCSSPFLVSAM | ASSPGQTNSDYT | 0.23692629 |
| VHITDAE | ASSPGQTNSDYT | 0.23848882 |
| EYAAPSF | ASSPGQTNSDYT | 0.19857949 |
| VVDAFHGGL | ASSPGQTNSDYT | 0.3425569 |
| ASVVDPQ | ASSPGQTNSDYT | 0.1959992 |
| SYNEFGGLS | ASSPGQTNSDYT | 0.2384469 |
| DPDGSVY | ASSPGQTNSDYT | 0.27138206 |
| MVSTIHATE | ASSPGQTNSDYT | 0.5295104 |
| LAAGVADPV | ASSPGQTNSDYT | 0.1975877 |
| TKEQIAA | ASSPGQTNSDYT | 0.3699053 |
| SLASPAAL | ASSPGQTNSDYT | 0.21093665 |
| IGLVFYGV | ASSPGQTNSDYT | 0.62604415 |
| FLLDASNL | ASSPGQTNSDYT | 0.24380468 |

| | | |
|---|---|---|
| YQVGNFP | ASSPGQTNSDYT | 0.19663319 |
| ESLLSQST | ASSPGQTNSDYT | 0.40011072 |
| LFTVGFAAVG | ASSPGQTNSDYT | 0.40219295 |
| ATPPTPT | ASSPGQTNSDYT | 0.23985308 |
| TDLATLG | ASSPGQTNSDYT | 0.37192452 |
| SAGGTGR | ASSPGQTNSDYT | 0.2249451 |
| GFSNSGS | ASSPGQTNSDYT | 0.33285788 |
| GGNGGSGGTSV | ASSPGQTNSDYT | 0.49677432 |
| GTSVATGGAGN | ASSPGQTNSDYT | 0.40612906 |
| NGGNGGSGGTS | ASSPGQTNSDYT | 0.21033522 |
| AVVKFPH | ASSPGQTNSDYT | 0.2949045 |
| TPPPPPP | ASSPGQTNSDYT | 0.31287932 |
| AAPAAAAAPA | ASSPGQTNSDYT | 0.20820612 |
| IALNEDL | ASSPGQTNSDYT | 0.43913358 |
| IVNVSSM | ASSPGQTNSDYT | 0.2408298 |
| TGIPPAPR | ASSPGQTNSDYT | 0.4594772 |
| FDLGGGTF | ASSPGQTNSDYT | 0.31445128 |
| FDLGGGTFDVS | ASSPGQTNSDYT | 0.67102253 |
| NEPTAAA | ASSPGQTNSDYT | 0.19808461 |
| NEPTAAALA | ASSPGQTNSDYT | 0.22018321 |
| GSIINISSI | ASSPGQTNSDYT | 0.23102675 |
| VAPGNAGTA | ASSPGQTNSDYT | 0.3292178 |
| TGGGASLE | ASSPGQTNSDYT | 0.21478179 |
| VAVYNHYKRI | ASSPGQTNSDYT | 0.20905715 |
| TDPAPATT | ASSPGQTNSDYT | 0.28197894 |
| TNRPDILDPALLRPG | ASSPGQTNSDYT | 0.4048647 |
| LTTTTTS | ASSPGQTNSDYT | 0.22354689 |
| PTGSGKTL | ASSPGQTNSDYT | 0.24928704 |
| TGSGKTL | ASSPGQTNSDYT | 0.639393 |
| TIGHVDHGK | ASSPGQTNSDYT | 0.2111706 |
| TIGHVDHGKTT | ASSPGQTNSDYT | 0.25583076 |
| GHVDHGKTT | ASSPGQTNSDYT | 0.25012636 |
| AQFEHTLL | ASSPGQTNSDYT | 0.43846714 |
| IVNQDEL | ASSPGQTNSDYT | 0.2389437 |
| TNGSQFF | ASSPGQTNSDYT | 0.5092443 |
| TGKGIVE | ASSPGQTNSDYT | 0.20456287 |
| LISNASDA | ASSPGQTNSDYT | 0.4953165 |
| AELRTLGELTKIAKAHGAQ | ASSPGQTNSDYT | 0.24836922 |
| APNAVAAAEPSW | ASSPGQTNSDYT | 0.2668711 |
| APQVAGAATAVL | ASSPGQTNSDYT | 0.20967872 |
| ARRDPAAAAAILPTDA | ASSPGQTNSDYT | 0.38965753 |
| DAAARDVLAEAGLAEYF | ASSPGQTNSDYT | 0.49679828 |
| DDQIVLSAKANLAKVEQ | ASSPGQTNSDYT | 0.5025833 |
| DELLPDTGEDSDEDGH | ASSPGQTNSDYT | 0.19432414 |
| ELPEAPANRATVPQD | ASSPGQTNSDYT | 0.45592928 |
| FGQSKIERVTGIG | ASSPGQTNSDYT | 0.44152802 |
| GAGGTGSTAGGAGGAGGAGGLY | ASSPGQTNSDYT | 0.70789844 |
| GHGGTGAAGGNGGTGGAGGSGI | ASSPGQTNSDYT | 0.22934829 |
| GNGGNGGAGGEGGPGGAGGAGA | ASSPGQTNSDYT | 0.28169203 |
| GSTAGNGGNGGNATNGTIAGQP | ASSPGQTNSDYT | 0.22576249 |
| GTGGDGGNGGNGGTGG | ASSPGQTNSDYT | 0.23771058 |

| | | |
|---|---|---|
| GVRRPAGIPADAPV | ASSPGQTNSDYT | 0.20950955 |
| GYRQVTDAHLLTLARR | ASSPGQTNSDYT | 0.22374147 |
| IAKMGQDGHDRG | ASSPGQTNSDYT | 0.2267029 |
| KAEPSIEIQYT | ASSPGQTNSDYT | 0.23337053 |
| KPVRIRMAADEDTVAVT | ASSPGQTNSDYT | 0.45688933 |
| MPPRIAGMRLLVIKPEP | ASSPGQTNSDYT | 0.4186486 |
| MKFYAVALFRSRGVPPTR | ASSPGQTNSDYT | 0.3947584 |
| NAGGFGILGAVAHSP | ASSPGQTNSDYT | 0.58578694 |
| NEGREITLAMAEM | ASSPGQTNSDYT | 0.6665878 |
| NELVEWSHPLGHRWRWE | ASSPGQTNSDYT | 0.2938454 |
| NLGAGGTGGNGGNPG | ASSPGQTNSDYT | 0.22976825 |
| NTEWPKWETVMAADPVD | ASSPGQTNSDYT | 0.2232412 |
| PASAAPAETQP | ASSPGQTNSDYT | 0.2008085 |
| QQSSDGKNGVRAAEITGE | ASSPGQTNSDYT | 0.21855213 |
| RAPDAVTTSWWKE | ASSPGQTNSDYT | 0.23476824 |
| SGFGNVGAHTSGWFNQ | ASSPGQTNSDYT | 0.2797399 |
| SGVAGAVGQAASVGGLKV | ASSPGQTNSDYT | 0.19270435 |
| SNGGAGTGSAGGNGGT | ASSPGQTNSDYT | 0.37354136 |
| TGGAGGEGGAAG | ASSPGQTNSDYT | 0.58742815 |
| TGGDGGAAGAGGAPGHG | ASSPGQTNSDYT | 0.22179201 |
| TMPPLLAAIANAGRHGV | ASSPGQTNSDYT | 0.28325897 |
| TSSTAGNSGGA | ASSPGQTNSDYT | 0.371907 |
| VPATVPVGILC | ASSPGQTNSDYT | 0.21451236 |
| VQRPDSGGVVLTGRIS | ASSPGQTNSDYT | 0.48819354 |
| WTPNPVPATKGVPAGLVQAT | ASSPGQTNSDYT | 0.21722321 |
| YALKRALAKFSDRFDVVII | ASSPGQTNSDYT | 0.35852858 |
| YLIGSVIV | ASSPGQTNSDYT | 0.51971865 |
| YGQLVAVID | ASSPGQTNSDYT | 0.2734943 |
| VTVGGMTLPA | ASSPGQTNSDYT | 0.25791067 |
| VLALPAV | ASSPGQTNSDYT | 0.22839528 |
| VIPPTLNYET | ASSPGQTNSDYT | 0.19915281 |
| VGMGGMHPSSG | ASSPGQTNSDYT | 0.2739761 |
| VGLAGGAATA | ASSPGQTNSDYT | 0.20822099 |
| VGGAGGAGGSGP | ASSPGQTNSDYT | 0.30417567 |
| VFAAFMV | ASSPGQTNSDYT | 0.21301295 |
| TLYDKLR | ASSPGQTNSDYT | 0.3697518 |
| TLAVFTVL | ASSPGQTNSDYT | 0.29493445 |
| TGSVQPARV | ASSPGQTNSDYT | 0.24567379 |
| TGGGAGSGGT | ASSPGQTNSDYT | 0.22194009 |
| SHLETSL | ASSPGQTNSDYT | 0.2514804 |
| SFTTGMT | ASSPGQTNSDYT | 0.22765906 |
| QVEVAKRMGVRQPTVS | ASSPGQTNSDYT | 0.20079628 |
| QGSGGPALAM | ASSPGQTNSDYT | 0.28928098 |
| MLAPSNP | ASSPGQTNSDYT | 0.21913083 |
| LVNNAAI | ASSPGQTNSDYT | 0.27102846 |
| LTGEINTLA | ASSPGQTNSDYT | 0.4513775 |
| LSIFGPV | ASSPGQTNSDYT | 0.24805394 |
| LLALLANL | ASSPGQTNSDYT | 0.19849952 |
| LISNASDALDKLR | ASSPGQTNSDYT | 0.33425522 |
| LINEAVP | ASSPGQTNSDYT | 0.32191122 |
| LGYAFSGL | ASSPGQTNSDYT | 0.23935387 |

| | | |
|---|---|---|
| LDFVAVAL | ASSPGQTNSDYT | 0.19268237 |
| LAQPGAE | ASSPGQTNSDYT | 0.3063642 |
| ISNLGAVTGF | ASSPGQTNSDYT | 0.21007952 |
| HQEDPGRI | ASSPGQTNSDYT | 0.22148287 |
| GTPGEPTP | ASSPGQTNSDYT | 0.21626014 |
| GFFNSGS | ASSPGQTNSDYT | 0.5611637 |
| GAGGGGLTAGAN | ASSPGQTNSDYT | 0.23792093 |
| FVDALVNM | ASSPGQTNSDYT | 0.23946498 |
| FLTVAPEL | ASSPGQTNSDYT | 0.3840307 |
| FETLTAL | ASSPGQTNSDYT | 0.2863248 |
| DVINPIEL | ASSPGQTNSDYT | 0.49844557 |
| APVSGAPV | ASSPGQTNSDYT | 0.29986987 |
| ALVGGLAW | ASSPGQTNSDYT | 0.22014067 |
| ALAAAAATAI | ASSPGQTNSDYT | 0.25562224 |
| AFGLGLLL | ASSPGQTNSDYT | 0.20958129 |
| AFFAAASL | ASSPGQTNSDYT | 0.20076935 |
| AAAALPAFSPP | ASSPGQTNSDYT | 0.19725189 |
| VVLAPAL | ASSPGQTNSDYT | 0.28405538 |
| TIGHVDHGKTTL | ASSPGQTNSDYT | 0.20953703 |
| LISNASDALDK | ASSPGQTNSDYT | 0.29650217 |
| LISNASDAL | ASSPGQTNSDYT | 0.2815687 |
| ISNASDALDK | ASSPGQTNSDYT | 0.22682032 |
| NQAETLVYQTE | ASSPGQTNSDYT | 0.19851464 |
| TGGTGGTAGAG | ASSPGQTNSDYT | 0.2189331 |
| TGSTAGGAGGA | ASSPGQTNSDYT | 0.21682385 |
| DPLAVPVAAGPV | ASSPGQTNSDYT | 0.5001917 |
| GGTGGTPGNGGN | ASSPGQTNSDYT | 0.24724293 |
| PAGLAGVDPATL | ASSPGQTNSDYT | 0.69653386 |
| GNNLVVEAPHAGASV | ASSPGQTNSDYT | 0.45408756 |
| KVTRIMSASTLEQPA | ASSPGQTNSDYT | 0.24662027 |
| LGFALPIALPRRTVV | ASSPGQTNSDYT | 0.48485872 |
| TRLGNLTARKLVVLV | ASSPGQTNSDYT | 0.23063628 |
| WSQQRRLIYFPSAGP | ASSPGQTNSDYT | 0.21858989 |
| GPNGNPDPMAAAVDIR | ASSPGQTNSDYT | 0.2290671 |
| TQRCVSVADLRAAASR | ASSPGQTNSDYT | 0.254302 |
| QFVLDVAEHPVEVLTETV | ASSPGQTNSDYT | 0.22821964 |
| GGAGGTGGAGGNGGTQAGDG | ASSPGQTNSDYT | 0.21631691 |
| PLLATLRAIAVDVDAKPAQV | ASSPGQTNSDYT | 0.19914299 |
| APKFRTGGYMIDFWGVGYQVAK | ASSPGQTNSDYT | 0.2589997 |
| GAGGAGGAGGAAGGPGGTGGTG | ASSPGQTNSDYT | 0.23138335 |
| IGALGLLATDVAPAGSVAFGVA | ASSPGQTNSDYT | 0.20545255 |
| AGGAGGIGGNGGAGGAGGVLMG | ASSPGQTNSDYT | 0.22370237 |
| ADELIGLL | ASSPGQTNSDYT | 0.5575998 |
| ADRALKA | ASSPGQTNSDYT | 0.59587955 |
| AGAVGQVGGAGGA | ASSPGQTNSDYT | 0.22501773 |
| AGGGGGLGGAP | ASSPGQTNSDYT | 0.22860004 |
| AGGLLYGNGGAG | ASSPGQTNSDYT | 0.40178508 |
| AGLESIGL | ASSPGQTNSDYT | 0.22145785 |
| AGLKTGII | ASSPGQTNSDYT | 0.26770285 |
| ASLAALLL | ASSPGQTNSDYT | 0.35006225 |
| ASVINISSI | ASSPGQTNSDYT | 0.33671087 |

| | | |
|---|---|---|
| ATPPGPA | ASSPGQTNSDYT | 0.20657054 |
| ATVASALV | ASSPGQTNSDYT | 0.27709112 |
| ATVASLIL | ASSPGQTNSDYT | 0.2510783 |
| AVNGPVLIRH | ASSPGQTNSDYT | 0.29192817 |
| DGAAGAVGI | ASSPGQTNSDYT | 0.19231927 |
| DPLLHGI | ASSPGQTNSDYT | 0.22113368 |
| ELGAALGAL | ASSPGQTNSDYT | 0.21417892 |
| ELGRTQAV | ASSPGQTNSDYT | 0.21218929 |
| FVAGGYLS | ASSPGQTNSDYT | 0.1951017 |
| GASGGAGGAGGW | ASSPGQTNSDYT | 0.20804803 |
| GGAGAGGDGGAGGI | ASSPGQTNSDYT | 0.42072034 |
| GTGHNGSQPGMGGNG | ASSPGQTNSDYT | 0.47797927 |
| GYAAASASAS | ASSPGQTNSDYT | 0.21079206 |
| HQVDVAAVQQG | ASSPGQTNSDYT | 0.4167297 |
| IGAAIAHT | ASSPGQTNSDYT | 0.61984754 |
| IGMIEVAQL | ASSPGQTNSDYT | 0.23555145 |
| IRLAPEHTSAMD | ASSPGQTNSDYT | 0.19691691 |
| IYNVSVL | ASSPGQTNSDYT | 0.29754966 |
| KQGFLGM | ASSPGQTNSDYT | 0.2204256 |
| LESVITQL | ASSPGQTNSDYT | 0.31293002 |
| LGLGALVVV | ASSPGQTNSDYT | 0.40470594 |
| LIGNGGNGGSGGI | ASSPGQTNSDYT | 0.20419827 |
| LNAINAPTL | ASSPGQTNSDYT | 0.334844 |
| LQLLNTGV | ASSPGQTNSDYT | 0.42205346 |
| LQTSIEF | ASSPGQTNSDYT | 0.21250091 |
| MSFAVSEL | ASSPGQTNSDYT | 0.19822511 |
| NTGALGL | ASSPGQTNSDYT | 0.20169377 |
| PPGAGGAGGAGTT | ASSPGQTNSDYT | 0.20568755 |
| QLVGLGI | ASSPGQTNSDYT | 0.28574085 |
| RFLEQVHQL | ASSPGQTNSDYT | 0.24652252 |
| RGVGYVLR | ASSPGQTNSDYT | 0.21057703 |
| SATGVFAGVI | ASSPGQTNSDYT | 0.32354456 |
| SLPAITLP | ASSPGQTNSDYT | 0.21304584 |
| STALAFTV | ASSPGQTNSDYT | 0.22262128 |
| SVLTVGLGK | ASSPGQTNSDYT | 0.2214489 |
| TALAGAGSGI | ASSPGQTNSDYT | 0.21919957 |
| TGLPFTGI | ASSPGQTNSDYT | 0.3639493 |
| VAEHVTV | ASSPGQTNSDYT | 0.21920466 |
| VDIVLNSL | ASSPGQTNSDYT | 0.21898702 |
| VLTGGGAVL | ASSPGQTNSDYT | 0.61351925 |
| VNVGLAYGI | ASSPGQTNSDYT | 0.19999397 |
| VSSAAVGAV | ASSPGQTNSDYT | 0.21094465 |
| VTSVYEAGV | ASSPGQTNSDYT | 0.4039011 |
| WVALDRGA | ASSPGQTNSDYT | 0.20644805 |
| YLVGSVVI | ASSPGQTNSDYT | 0.46101612 |
| GPPGTGKTLL | ASSPGQTNSDYT | 0.31812608 |
| GVGMTKF | ASSPGQTNSDYT | 0.34531224 |
| IGIGGSDL | ASSPGQTNSDYT | 0.21340941 |
| IGTLLAEL | ASSPGQTNSDYT | 0.6325033 |
| IYTSGTT | ASSPGQTNSDYT | 0.39182493 |
| LSFPAGV | ASSPGQTNSDYT | 0.20921943 |

| | | |
|---|---|---|
| LYTSGTT | ASSPGQTNSDYT | 0.46666783 |
| VGAFPFGF | ASSPGQTNSDYT | 0.19941987 |
| VGAGTVEF | ASSPGQTNSDYT | 0.39914435 |
| VGKSTLLL | ASSPGQTNSDYT | 0.42582947 |
| AALAHAGGGIV | ASSPGQTNSDYT | 0.22125691 |
| AALVAVSLL | ASSPGQTNSDYT | 0.34794253 |
| AAVSRAVAAL | ASSPGQTNSDYT | 0.3010832 |
| AGGSSGIGGF | ASSPGQTNSDYT | 0.5627074 |
| AGLTVANANG | ASSPGQTNSDYT | 0.20483442 |
| AGLVGAVL | ASSPGQTNSDYT | 0.23686351 |
| ALSQAFTV | ASSPGQTNSDYT | 0.22206667 |
| ALTFSVSL | ASSPGQTNSDYT | 0.2133253 |
| ALTGAALVAL | ASSPGQTNSDYT | 0.20354809 |
| ASFSIGAGVL | ASSPGQTNSDYT | 0.37578112 |
| DGSVYKI | ASSPGQTNSDYT | 0.24631545 |
| EQLAEIV | ASSPGQTNSDYT | 0.46555477 |
| FAVSELTVM | ASSPGQTNSDYT | 0.302682 |
| GAAPLAAGL | ASSPGQTNSDYT | 0.4321149 |
| GADGGDGGSGGAGG | ASSPGQTNSDYT | 0.23503354 |
| GFGLDTGGAGG | ASSPGQTNSDYT | 0.24772143 |
| GLASGGLL | ASSPGQTNSDYT | 0.19898763 |
| GRIAGAAVLL | ASSPGQTNSDYT | 0.20257804 |
| GSAGSLNTSF | ASSPGQTNSDYT | 0.20342033 |
| GTGQIGI | ASSPGQTNSDYT | 0.23123059 |
| GTLIALIL | ASSPGQTNSDYT | 0.19616553 |
| IAATMGFVVG | ASSPGQTNSDYT | 0.48659468 |
| IFAQALL | ASSPGQTNSDYT | 0.30018938 |
| IGAAVALGF | ASSPGQTNSDYT | 0.3762503 |
| IGLTPLAL | ASSPGQTNSDYT | 0.21221551 |
| ILGSNTSTL | ASSPGQTNSDYT | 0.20682484 |
| IVFSLTVL | ASSPGQTNSDYT | 0.23043984 |
| LASKATGF | ASSPGQTNSDYT | 0.44704837 |
| LASVALIL | ASSPGQTNSDYT | 0.20161653 |
| LFAVGLTL | ASSPGQTNSDYT | 0.19515276 |
| LFNGGIGTYI | ASSPGQTNSDYT | 0.19912289 |
| LTLGAGII | ASSPGQTNSDYT | 0.19582166 |
| LVAAALGDI | ASSPGQTNSDYT | 0.6522449 |
| NTAFVQLGI | ASSPGQTNSDYT | 0.48964942 |
| QLFSGAEL | ASSPGQTNSDYT | 0.33529449 |
| SAATFITL | ASSPGQTNSDYT | 0.34661186 |
| SNYGLANLGV | ASSPGQTNSDYT | 0.19336006 |
| SYGGAYAVM | ASSPGQTNSDYT | 0.20975514 |
| TTGVGGLTLG | ASSPGQTNSDYT | 0.58293307 |
| TTQTIYI | ASSPGQTNSDYT | 0.27715328 |
| VAAGAGLGAVI | ASSPGQTNSDYT | 0.27900884 |
| VAAVVTLL | ASSPGQTNSDYT | 0.43140674 |
| VGFLGGLI | ASSPGQTNSDYT | 0.22180575 |
| VGVIPILL | ASSPGQTNSDYT | 0.23439367 |
| VIFASSLI | ASSPGQTNSDYT | 0.23350231 |
| VKGGDGGAAAG | ASSPGQTNSDYT | 0.2570446 |
| VTKTFTNI | ASSPGQTNSDYT | 0.2342193 |

| | | |
|---|---|---|
| VVAAAILL | ASSPGQTNSDYT | 0.20312029 |
| YAALDVEL | ASSPGQTNSDYT | 0.26056683 |
| LASVALLL | ASSPGQTNSDYT | 0.7072617 |
| LDAVHRV | ASSPGQTNSDYT | 0.27790785 |
| QGLSSSIF | ASSPGQTNSDYT | 0.36689937 |
| RGTFANIRL | ASSPGQTNSDYT | 0.304188 |
| RQLWWGHRI | ASSPGQTNSDYT | 0.57875985 |
| YGPPGTGKTLL | ASSPGQTNSDYT | 0.23670526 |
| AALVGAPANVA | ASSPGQTNSDYT | 0.23243427 |
| AATFRRTE | ASSPGQTNSDYT | 0.2538904 |
| AGALGGGI | ASSPGQTNSDYT | 0.20423818 |
| AGGAGGAGGIGAG | ASSPGQTNSDYT | 0.22869673 |
| AGSGALGL | ASSPGQTNSDYT | 0.22027323 |
| AQIADQI | ASSPGQTNSDYT | 0.7213711 |
| ASAVAVTYL | ASSPGQTNSDYT | 0.36632693 |
| ASTVLGGTGG | ASSPGQTNSDYT | 0.30549723 |
| AVGGEPAK | ASSPGQTNSDYT | 0.21332417 |
| CEQGTFGGTC | ASSPGQTNSDYT | 0.22373089 |
| DEGKAPAG | ASSPGQTNSDYT | 0.3605851 |
| DTVVFPAL | ASSPGQTNSDYT | 0.20038919 |
| FEGITMAG | ASSPGQTNSDYT | 0.46628347 |
| FPLVVQD | ASSPGQTNSDYT | 0.22029407 |
| GAFTTPQ | ASSPGQTNSDYT | 0.24299477 |
| GFGGFGTAGGVGGA | ASSPGQTNSDYT | 0.32216913 |
| GFLAQFL | ASSPGQTNSDYT | 0.67365766 |
| GGAGGAGGNAGVA | ASSPGQTNSDYT | 0.22204475 |
| GGVGTAGAGGAGGAGG | ASSPGQTNSDYT | 0.23608837 |
| GNGGGGDAGNAGSGGN | ASSPGQTNSDYT | 0.21213706 |
| GTGGAGGAGNDAGS | ASSPGQTNSDYT | 0.35257503 |
| GTTNTGWFNS | ASSPGQTNSDYT | 0.58309126 |
| IFMGPLFTL | ASSPGQTNSDYT | 0.3004542 |
| ILAHAIEAI | ASSPGQTNSDYT | 0.62883663 |
| IRSAIREAL | ASSPGQTNSDYT | 0.22565545 |
| ITFAGGG | ASSPGQTNSDYT | 0.1978439 |
| ITGMSGHQEV | ASSPGQTNSDYT | 0.26981542 |
| IVAGLACTL | ASSPGQTNSDYT | 0.25983903 |
| IVGGGLAGA | ASSPGQTNSDYT | 0.6344496 |
| IVTGVLNI | ASSPGQTNSDYT | 0.42041665 |
| LAQQGCI | ASSPGQTNSDYT | 0.24905655 |
| LCLHVEN | ASSPGQTNSDYT | 0.22558707 |
| LDAISDGS | ASSPGQTNSDYT | 0.52725244 |
| LGGFAAML | ASSPGQTNSDYT | 0.4675483 |
| LGGKNPMI | ASSPGQTNSDYT | 0.33108273 |
| LGPKGVAAIL | ASSPGQTNSDYT | 0.21484888 |
| LMVGVSI | ASSPGQTNSDYT | 0.21380809 |
| LVGAAFVL | ASSPGQTNSDYT | 0.24881011 |
| LVLAVAPL | ASSPGQTNSDYT | 0.49547637 |
| MAAIAVIDAA | ASSPGQTNSDYT | 0.20636782 |
| NNGINAELQ | ASSPGQTNSDYT | 0.26582175 |
| RGEPLVLVHG | ASSPGQTNSDYT | 0.36634976 |
| RGGAGGAAGLI | ASSPGQTNSDYT | 0.19647548 |

| | | |
|---|---|---|
| RSYRMHPAV | ASSPGQTNSDYT | 0.2383533 |
| SAAHLLAGQLA | ASSPGQTNSDYT | 0.57342243 |
| SAAVGSY | ASSPGQTNSDYT | 0.23353976 |
| SPHAVLHRI | ASSPGQTNSDYT | 0.2092014 |
| SQIAKVL | ASSPGQTNSDYT | 0.5106806 |
| TALGHLVP | ASSPGQTNSDYT | 0.72897124 |
| TAPTTSGAA | ASSPGQTNSDYT | 0.41877067 |
| TDGDDGNGGSPGT | ASSPGQTNSDYT | 0.19788955 |
| TDGLGEN | ASSPGQTNSDYT | 0.19311734 |
| TGGPPLPGQA | ASSPGQTNSDYT | 0.43835008 |
| TIPAITI | ASSPGQTNSDYT | 0.43282264 |
| TIPGFTI | ASSPGQTNSDYT | 0.22317015 |
| TSIASGALA | ASSPGQTNSDYT | 0.21714926 |
| TVTLPGAAI | ASSPGQTNSDYT | 0.40936017 |
| VASGLSAALP | ASSPGQTNSDYT | 0.44937444 |
| VGAGVIGI | ASSPGQTNSDYT | 0.20632376 |
| VGPKRFGV | ASSPGQTNSDYT | 0.20049566 |
| VNVAFPD | ASSPGQTNSDYT | 0.22345586 |
| VSMTNSGVS | ASSPGQTNSDYT | 0.37610415 |
| VTDVLGDLSV | ASSPGQTNSDYT | 0.21687649 |
| VVVVGAGFAGL | ASSPGQTNSDYT | 0.29095128 |
| VYTVGTVV | ASSPGQTNSDYT | 0.34356222 |
| WTTFSGP | ASSPGQTNSDYT | 0.24107867 |
| YDELVEAF | ASSPGQTNSDYT | 0.3544125 |
| YGTGGAGGNGG | ASSPGQTNSDYT | 0.21786302 |
| YTNVMAV | ASSPGQTNSDYT | 0.5313683 |
| FLEQVHQL | ASSPGQTNSDYT | 0.21420239 |
| GLSLLAVAV | ASSPGQTNSDYT | 0.26815075 |
| LEQVHQL | ASSPGQTNSDYT | 0.34183028 |
| LNAAGAI | ASSPGQTNSDYT | 0.28435653 |
| PAVHASL | ASSPGQTNSDYT | 0.2401653 |
| TELEDDA | ASSPGQTNSDYT | 0.21208501 |
| TPPGPPN | ASSPGQTNSDYT | 0.20203732 |
| TPPPPPPP | ASSPGQTNSDYT | 0.19661315 |
| TQSPALI | ASSSRLVNYAEQF | 0.37833896 |
| EFAAAFPQ | ASSSRLVNYAEQF | 0.21374114 |
| LLSGNLDVL | ASSSRLVNYAEQF | 0.4858029 |
| GPANLGNYNI | ASSSRLVNYAEQF | 0.29909828 |
| TLGPTLP | ASSSRLVNYAEQF | 0.20802112 |
| VYAYSSCCK | ASSSRLVNYAEQF | 0.29377398 |
| APAPAPAPNI | ASSSRLVNYAEQF | 0.2061258 |
| PGGPVPIP | ASSSRLVNYAEQF | 0.21747121 |
| FLTVALEP | ASSSRLVNYAEQF | 0.42635787 |
| SQYFGANL | ASSSRLVNYAEQF | 0.28708518 |
| GGVKPAA | ASSSRLVNYAEQF | 0.21068181 |
| GGTGGAGGAGGW | ASSSRLVNYAEQF | 0.30056253 |
| GVGGVAGLGGAGG | ASSSRLVNYAEQF | 0.3317635 |
| VLLVLATTI | ASSSRLVNYAEQF | 0.20282139 |
| IGIFAAML | ASSSRLVNYAEQF | 0.22277425 |
| AVGDLID | ASSSRLVNYAEQF | 0.47393692 |
| NISPLPL | ASSSRLVNYAEQF | 0.2544046 |

| | | |
|---|---|---|
| VTAVFVAI | ASSSRLVNYAEQF | 0.20482872 |
| SGDDNDAAL | ASSSRLVNYAEQF | 0.3152124 |
| AVAALAHS | ASSSRLVNYAEQF | 0.21769714 |
| GGVTVMATP | ASSSRLVNYAEQF | 0.3940907 |
| AGLAANVRD | ASSSRLVNYAEQF | 0.25759715 |
| TNRPDLIDPALLRPG | ASSSRLVNYAEQF | 0.2254413 |
| LANLGVTAL | ASSSRLVNYAEQF | 0.32871282 |
| TGERIEL | ASSSRLVNYAEQF | 0.19828916 |
| NGEELAA | ASSSRLVNYAEQF | 0.20949753 |
| VLSLVWG | ASSSRLVNYAEQF | 0.23018053 |
| IGYALWL | ASSSRLVNYAEQF | 0.2576684 |
| LSLLLAPL | ASSSRLVNYAEQF | 0.36166298 |
| TLDEALK | ASSSRLVNYAEQF | 0.21486701 |
| TDGAGTAEGG | ASSSRLVNYAEQF | 0.3087994 |
| NAPGGAGGNGG | ASSSRLVNYAEQF | 0.23598224 |
| VSVFMGMGGNF | ASSSRLVNYAEQF | 0.21595575 |
| LASYLAAL | ASSSRLVNYAEQF | 0.24388412 |
| LVGGTPAL | ASSSRLVNYAEQF | 0.20682207 |
| VRNADVL | ASSSRLVNYAEQF | 0.21083862 |
| TCSSPFLVSAM | ASSSRLVNYAEQF | 0.2886026 |
| VHITDAE | ASSSRLVNYAEQF | 0.21229589 |
| EYAAPSF | ASSSRLVNYAEQF | 0.2406771 |
| VVDAFHGGL | ASSSRLVNYAEQF | 0.34464756 |
| ASVVDPQ | ASSSRLVNYAEQF | 0.20694432 |
| SYNEFGGLS | ASSSRLVNYAEQF | 0.24141696 |
| DPDGSVY | ASSSRLVNYAEQF | 0.29883847 |
| MVSTIHATE | ASSSRLVNYAEQF | 0.41583228 |
| LAAGVADPV | ASSSRLVNYAEQF | 0.227912 |
| TKEQIAA | ASSSRLVNYAEQF | 0.21112972 |
| SLASPAAL | ASSSRLVNYAEQF | 0.30119044 |
| IGLVFYGV | ASSSRLVNYAEQF | 0.28261706 |
| FLLDASNL | ASSSRLVNYAEQF | 0.29277378 |
| YQVGNFP | ASSSRLVNYAEQF | 0.31879455 |
| ESLLSQST | ASSSRLVNYAEQF | 0.24182372 |
| LFTVGFAAVG | ASSSRLVNYAEQF | 0.36028403 |
| ATPPTPT | ASSSRLVNYAEQF | 0.30625764 |
| TDLATLG | ASSSRLVNYAEQF | 0.20302764 |
| SAGGTGR | ASSSRLVNYAEQF | 0.2270656 |
| GFSNSGS | ASSSRLVNYAEQF | 0.23781835 |
| GGNGGSGGTSV | ASSSRLVNYAEQF | 0.20755933 |
| GTSVATGGAGN | ASSSRLVNYAEQF | 0.19751525 |
| NGGNGGSGGTS | ASSSRLVNYAEQF | 0.496664 |
| AVVKFPH | ASSSRLVNYAEQF | 0.6954942 |
| TPPPPPP | ASSSRLVNYAEQF | 0.20740274 |
| AAPAAAAAPA | ASSSRLVNYAEQF | 0.68950504 |
| IALNEDL | ASSSRLVNYAEQF | 0.19694395 |
| IVNVSSM | ASSSRLVNYAEQF | 0.26305947 |
| TGIPPAPR | ASSSRLVNYAEQF | 0.39777863 |
| FDLGGGTF | ASSSRLVNYAEQF | 0.21637967 |
| FDLGGGTFDVS | ASSSRLVNYAEQF | 0.32027692 |
| NEPTAAA | ASSSRLVNYAEQF | 0.20154078 |

| | | |
|---|---|---|
| NEPTAAALA | ASSSRLVNYAEQF | 0.21263634 |
| GSIINISSI | ASSSRLVNYAEQF | 0.47605163 |
| VAPGNAGTA | ASSSRLVNYAEQF | 0.22263448 |
| TGGGASLE | ASSSRLVNYAEQF | 0.24308006 |
| VAVYNHYKRI | ASSSRLVNYAEQF | 0.2319367 |
| TDPAPATT | ASSSRLVNYAEQF | 0.22107661 |
| TNRPDILDPALLRPG | ASSSRLVNYAEQF | 0.25075713 |
| LTTTTTS | ASSSRLVNYAEQF | 0.4539424 |
| PTGSGKTL | ASSSRLVNYAEQF | 0.21923922 |
| TGSGKTL | ASSSRLVNYAEQF | 0.5162784 |
| TIGHVDHGK | ASSSRLVNYAEQF | 0.20765927 |
| TIGHVDHGKTT | ASSSRLVNYAEQF | 0.58336145 |
| GHVDHGKTT | ASSSRLVNYAEQF | 0.40068948 |
| AQFEHTLL | ASSSRLVNYAEQF | 0.39266568 |
| IVNQDEL | ASSSRLVNYAEQF | 0.22950356 |
| TNGSQFF | ASSSRLVNYAEQF | 0.5395032 |
| TGKGIVE | ASSSRLVNYAEQF | 0.27779847 |
| LISNASDA | ASSSRLVNYAEQF | 0.25274608 |
| AELRTLGELTKIAKAHGAQ | ASSSRLVNYAEQF | 0.32250854 |
| APNAVAAAEPSW | ASSSRLVNYAEQF | 0.15661827 |
| APQVAGAATAVL | ASSSRLVNYAEQF | 0.17650023 |
| ARRDPAAAAAILPTDA | ASSSRLVNYAEQF | 0.14789455 |
| DAAARDVLAEAGLAEYF | ASSSRLVNYAEQF | 0.16410178 |
| DDQIVLSAKANLAKVEQ | ASSSRLVNYAEQF | 0.1644545 |
| DELLPDTGEDSDEDGH | ASSSRLVNYAEQF | 0.13982208 |
| ELPEAPANRATVPQD | ASSSRLVNYAEQF | 0.16768089 |
| FGQSKIERVTGIG | ASSSRLVNYAEQF | 0.15377384 |
| GAGGTGSTAGGAGGAGGAGGLY | ASSSRLVNYAEQF | 0.17049992 |
| GHGGTGAAGGNGGTGGAGGSGI | ASSSRLVNYAEQF | 0.17565139 |
| GNGGNGGAGGEGGPGGAGGAGA | ASSSRLVNYAEQF | 0.16170934 |
| GSTAGNGGNGGNATNGTIAGQP | ASSSRLVNYAEQF | 0.20293845 |
| GTGGDGGNGGNGGTGG | ASSSRLVNYAEQF | 0.1928407 |
| GVRRPAGIPADAPV | ASSSRLVNYAEQF | 0.15062983 |
| GYRQVTDAHLLTLARR | ASSSRLVNYAEQF | 0.15841207 |
| IAKMGQDGHDRG | ASSSRLVNYAEQF | 0.15378603 |
| KAEPSIEIQYT | ASSSRLVNYAEQF | 0.16535768 |
| KPVRIRMAADEDTVAVT | ASSSRLVNYAEQF | 0.13197957 |
| MPPRIAGMRLLVIKPEP | ASSSRLVNYAEQF | 0.1734193 |
| MKFYAVALFRSRGVPPTR | ASSSRLVNYAEQF | 0.14139271 |
| NAGGFGILGAVAHSP | ASSSRLVNYAEQF | 0.12920345 |
| NEGREITLAMAEM | ASSSRLVNYAEQF | 0.19220954 |
| NELVEWSHPLGHRWRWE | ASSSRLVNYAEQF | 0.16656463 |
| NLGAGGTGGNGGNPG | ASSSRLVNYAEQF | 0.18737167 |
| NTEWPKWETVMAADPVD | ASSSRLVNYAEQF | 0.21690741 |
| PASAAPAETQP | ASSSRLVNYAEQF | 0.13499354 |
| QQSSDGKNGVRAAEITGE | ASSSRLVNYAEQF | 0.14553975 |
| RAPDAVTTSWWKE | ASSSRLVNYAEQF | 0.1648761 |
| SGFGNVGAHTSGWFNQ | ASSSRLVNYAEQF | 0.22746822 |
| SGVAGAVGQAASVGGLKV | ASSSRLVNYAEQF | 0.14655733 |
| SNGGAGTGSAGGNGGT | ASSSRLVNYAEQF | 0.16270655 |
| TGGAGGEGGAAG | ASSSRLVNYAEQF | 0.19530267 |

| | | |
|---|---|---|
| TGGDGGAAGAGGAPGHG | ASSSRLVNYAEQF | 0.16461323 |
| TMPPLLAAIANAGRHGV | ASSSRLVNYAEQF | 0.18142004 |
| TSSTAGNSGGA | ASSSRLVNYAEQF | 0.19291435 |
| VPATVPVGILC | ASSSRLVNYAEQF | 0.13471977 |
| VQRPDSGGVVLTGRIS | ASSSRLVNYAEQF | 0.20195217 |
| WTPNPVPATKGVPAGLVQAT | ASSSRLVNYAEQF | 0.17501667 |
| YALKRALAKFSDRFDVVII | ASSSRLVNYAEQF | 0.16185704 |
| YLIGSVIV | ASSSRLVNYAEQF | 0.1612525 |
| YGQLVAVID | ASSSRLVNYAEQF | 0.15189904 |
| VTVGGMTLPA | ASSSRLVNYAEQF | 0.15528212 |
| VLALPAV | ASSSRLVNYAEQF | 0.13161038 |
| VIPPTLNYET | ASSSRLVNYAEQF | 0.15900353 |
| VGMGGMHPSSG | ASSSRLVNYAEQF | 0.17933412 |
| VGLAGGAATA | ASSSRLVNYAEQF | 0.16458656 |
| VGGAGGAGGSGP | ASSSRLVNYAEQF | 0.1429928 |
| VFAAFMV | ASSSRLVNYAEQF | 0.1436063 |
| TLYDKLR | ASSSRLVNYAEQF | 0.13594942 |
| TLAVFTVL | ASSSRLVNYAEQF | 0.14672312 |
| TGSVQPARV | ASSSRLVNYAEQF | 0.13024107 |
| TGGGAGSGGT | ASSSRLVNYAEQF | 0.15786034 |
| SHLETSL | ASSSRLVNYAEQF | 0.22466454 |
| SFTTGMT | ASSSRLVNYAEQF | 0.2048349 |
| QVEVAKRMGVRQPTVS | ASSSRLVNYAEQF | 0.14610866 |
| QGSGGPALAM | ASSSRLVNYAEQF | 0.211366 |
| MLAPSNP | ASSSRLVNYAEQF | 0.1592944 |
| LVNNAAI | ASSSRLVNYAEQF | 0.15329938 |
| LTGEINTLA | ASSSRLVNYAEQF | 0.15356405 |
| LSIFGPV | ASSSRLVNYAEQF | 0.16394375 |
| LLALLANL | ASSSRLVNYAEQF | 0.25080746 |
| LISNASDALDKLR | ASSSRLVNYAEQF | 0.1512618 |
| LINEAVP | ASSSRLVNYAEQF | 0.1529562 |
| LGYAFSGL | ASSSRLVNYAEQF | 0.24105108 |
| LDFVAVAL | ASSSRLVNYAEQF | 0.15732783 |
| LAQPGAE | ASSSRLVNYAEQF | 0.15077658 |
| ISNLGAVTGF | ASSSRLVNYAEQF | 0.14308655 |
| HQEDPGRI | ASSSRLVNYAEQF | 0.17764418 |
| GTPGEPTP | ASSSRLVNYAEQF | 0.13584527 |
| GFFNSGS | ASSSRLVNYAEQF | 0.15774183 |
| GAGGGGLTAGAN | ASSSRLVNYAEQF | 0.13874635 |
| FVDALVNM | ASSSRLVNYAEQF | 0.1585778 |
| FLTVAPEL | ASSSRLVNYAEQF | 0.2683297 |
| FETLTAL | ASSSRLVNYAEQF | 0.16784434 |
| DVINPIEL | ASSSRLVNYAEQF | 0.13811688 |
| APVSGAPV | ASSSRLVNYAEQF | 0.21083522 |
| ALVGGLAW | ASSSRLVNYAEQF | 0.15708171 |
| ALAAAAATAI | ASSSRLVNYAEQF | 0.15831684 |
| AFGLGLLL | ASSSRLVNYAEQF | 0.18076785 |
| AFFAAASL | ASSSRLVNYAEQF | 0.14475453 |
| AAAALPAFSPP | ASSSRLVNYAEQF | 0.18391131 |
| VVLAPAL | ASSSRLVNYAEQF | 0.17200273 |
| TIGHVDHGKTTL | ASSSRLVNYAEQF | 0.19120589 |

| | | |
|---|---|---|
| LISNASDALDK | ASSSRLVNYAEQF | 0.1814857 |
| LISNASDAL | ASSSRLVNYAEQF | 0.15307122 |
| ISNASDALDK | ASSSRLVNYAEQF | 0.1500297 |
| NQAETLVYQTE | ASSSRLVNYAEQF | 0.13984458 |
| TGGTGGTAGAG | ASSSRLVNYAEQF | 0.13935249 |
| TGSTAGGAGGA | ASSSRLVNYAEQF | 0.17728455 |
| DPLAVPVAAGPV | ASSSRLVNYAEQF | 0.16351525 |
| GGTGGTPGNGGN | ASSSRLVNYAEQF | 0.19406174 |
| PAGLAGVDPATL | ASSSRLVNYAEQF | 0.17859907 |
| GNNLVVEAPHAGASV | ASSSRLVNYAEQF | 0.14186552 |
| KVTRIMSASTLEQPA | ASSSRLVNYAEQF | 0.1757019 |
| LGFALPIALPRRTVV | ASSSRLVNYAEQF | 0.18766901 |
| TRLGNLTARKLVVLV | ASSSRLVNYAEQF | 0.13783503 |
| WSQQRRLIYFPSAGP | ASSSRLVNYAEQF | 0.15757039 |
| GPNGNPDPMAAAVDIR | ASSSRLVNYAEQF | 0.25742993 |
| TQRCVSVADLRAAASR | ASSSRLVNYAEQF | 0.17557894 |
| QFVLDVAEHPVEVLTETV | ASSSRLVNYAEQF | 0.17005733 |
| GGAGGTGGAGGNGGTQAGDG | ASSSRLVNYAEQF | 0.14551185 |
| PLLATLRAIAVDVDAKPAQV | ASSSRLVNYAEQF | 0.2918858 |
| APKFRTGGYMIDFWGVGYQVAK | ASSSRLVNYAEQF | 0.14128451 |
| GAGGAGGAGGAAGGPGGTGGTG | ASSSRLVNYAEQF | 0.18219133 |
| IGALGLLATDVAPAGSVAFGVA | ASSSRLVNYAEQF | 0.1350931 |
| AGGAGGIGGNGGAGGAGGVLMG | ASSSRLVNYAEQF | 0.15182091 |
| ADELIGLL | ASSSRLVNYAEQF | 0.13417184 |
| ADRALKA | ASSSRLVNYAEQF | 0.13648611 |
| AGAVGQVGGAGGA | ASSSRLVNYAEQF | 0.188143 |
| AGGGGGLGGAP | ASSSRLVNYAEQF | 0.1606629 |
| AGGLLYGNGGAG | ASSSRLVNYAEQF | 0.18132618 |
| AGLESIGL | ASSSRLVNYAEQF | 0.15207933 |
| AGLKTGII | ASSSRLVNYAEQF | 0.1890266 |
| ASLAALLL | ASSSRLVNYAEQF | 0.14742872 |
| ASVINISSI | ASSSRLVNYAEQF | 0.16958109 |
| ATPPGPA | ASSSRLVNYAEQF | 0.16895668 |
| ATVASALV | ASSSRLVNYAEQF | 0.17726445 |
| ATVASLIL | ASSSRLVNYAEQF | 0.19173169 |
| AVNGPVLIRH | ASSSRLVNYAEQF | 0.17957951 |
| DGAAGAVGI | ASSSRLVNYAEQF | 0.13856319 |
| DPLLHGI | ASSSRLVNYAEQF | 0.13888212 |
| ELGAALGAL | ASSSRLVNYAEQF | 0.1779835 |
| ELGRTQAV | ASSSRLVNYAEQF | 0.13331023 |
| FVAGGYLS | ASSSRLVNYAEQF | 0.13222143 |
| GASGGAGGAGGW | ASSSRLVNYAEQF | 0.1387212 |
| GGAGAGGDGGAGGI | ASSSRLVNYAEQF | 0.13997735 |
| GTGHNGSQPGMGGNG | ASSSRLVNYAEQF | 0.19787541 |
| GYAAASASAS | ASSSRLVNYAEQF | 0.1929747 |
| HQVDVAAVQQG | ASSSRLVNYAEQF | 0.15961589 |
| IGAAIAHT | ASSSRLVNYAEQF | 0.15144776 |
| IGMIEVAQL | ASSSRLVNYAEQF | 0.15460561 |
| IRLAPEHTSAMD | ASSSRLVNYAEQF | 0.12735267 |
| IYNVSVL | ASSSRLVNYAEQF | 0.1400652 |
| KQGFLGM | ASSSRLVNYAEQF | 0.15719211 |

| | | |
|---|---|---|
| LESVITQL | ASSSRLVNYAEQF | 0.19757366 |
| LGLGALVVV | ASSSRLVNYAEQF | 0.15222906 |
| LIGNGGNGGSGGI | ASSSRLVNYAEQF | 0.14658219 |
| LNAINAPTL | ASSSRLVNYAEQF | 0.17656903 |
| LQLLNTGV | ASSSRLVNYAEQF | 0.15905108 |
| LQTSIEF | ASSSRLVNYAEQF | 0.1492062 |
| MSFAVSEL | ASSSRLVNYAEQF | 0.13335507 |
| NTGALGL | ASSSRLVNYAEQF | 0.12920195 |
| PPGAGGAGGAGTT | ASSSRLVNYAEQF | 0.19878766 |
| QLVGLGI | ASSSRLVNYAEQF | 0.13594408 |
| RFLEQVHQL | ASSSRLVNYAEQF | 0.16493288 |
| RGVGYVLR | ASSSRLVNYAEQF | 0.13112937 |
| SATGVFAGVI | ASSSRLVNYAEQF | 0.13710935 |
| SLPAITLP | ASSSRLVNYAEQF | 0.13443169 |
| STALAFTV | ASSSRLVNYAEQF | 0.17535298 |
| SVLTVGLGK | ASSSRLVNYAEQF | 0.17267677 |
| TALAGAGSGI | ASSSRLVNYAEQF | 0.16389619 |
| TGLPFTGI | ASSSRLVNYAEQF | 0.19578624 |
| VAEHVTV | ASSSRLVNYAEQF | 0.22923288 |
| VDIVLNSL | ASSSRLVNYAEQF | 0.15701996 |
| VLTGGGAVL | ASSSRLVNYAEQF | 0.1574541 |
| VNVGLAYGI | ASSSRLVNYAEQF | 0.1581652 |
| VSSAAVGAV | ASSSRLVNYAEQF | 0.14602336 |
| VTSVYEAGV | ASSSRLVNYAEQF | 0.14616881 |
| WVALDRGA | ASSSRLVNYAEQF | 0.22338857 |
| YLVGSVVI | ASSSRLVNYAEQF | 0.14881936 |
| GPPGTGKTLL | ASSSRLVNYAEQF | 0.14189957 |
| GVGMTKF | ASSSRLVNYAEQF | 0.15504502 |
| IGIGGSDL | ASSSRLVNYAEQF | 0.18036343 |
| IGTLLAEL | ASSSRLVNYAEQF | 0.17964044 |
| IYTSGTT | ASSSRLVNYAEQF | 0.16103914 |
| LSFPAGV | ASSSRLVNYAEQF | 0.14508013 |
| LYTSGTT | ASSSRLVNYAEQF | 0.15328173 |
| VGAFPFGF | ASSSRLVNYAEQF | 0.16297053 |
| VGAGTVEF | ASSSRLVNYAEQF | 0.14685768 |
| VGKSTLLL | ASSSRLVNYAEQF | 0.1471991 |
| AALAHAGGGIV | ASSSRLVNYAEQF | 0.17383865 |
| AALVAVSLL | ASSSRLVNYAEQF | 0.23990868 |
| AAVSRAVAAL | ASSSRLVNYAEQF | 0.14568108 |
| AGGSSGIGGF | ASSSRLVNYAEQF | 0.13842537 |
| AGLTVANANG | ASSSRLVNYAEQF | 0.17111917 |
| AGLVGAVL | ASSSRLVNYAEQF | 0.1379805 |
| ALSQAFTV | ASSSRLVNYAEQF | 0.16081727 |
| ALTFSVSL | ASSSRLVNYAEQF | 0.15775426 |
| ALTGAALVAL | ASSSRLVNYAEQF | 0.13752031 |
| ASFSIGAGVL | ASSSRLVNYAEQF | 0.1504762 |
| DGSVYKI | ASSSRLVNYAEQF | 0.18965432 |
| EQLAEIV | ASSSRLVNYAEQF | 0.1600299 |
| FAVSELTVM | ASSSRLVNYAEQF | 0.19419079 |
| GAAPLAAGL | ASSSRLVNYAEQF | 0.19173056 |
| GADGGDGGSGGAGG | ASSSRLVNYAEQF | 0.19112132 |

| | | |
|---|---|---|
| GFGLDTGGAGG | ASSSRLVNYAEQF | 0.24083714 |
| GLASGGLL | ASSSRLVNYAEQF | 0.16940343 |
| GRIAGAAVLL | ASSSRLVNYAEQF | 0.18979438 |
| GSAGSLNTSF | ASSSRLVNYAEQF | 0.1449457 |
| GTGQIGI | ASSSRLVNYAEQF | 0.17135301 |
| GTLIALIL | ASSSRLVNYAEQF | 0.17157386 |
| IAATMGFVVG | ASSSRLVNYAEQF | 0.18239927 |
| IFAQALL | ASSSRLVNYAEQF | 0.14088081 |
| IGAAVALGF | ASSSRLVNYAEQF | 0.19383085 |
| IGLTPLAL | ASSSRLVNYAEQF | 0.1486834 |
| ILGSNTSTL | ASSSRLVNYAEQF | 0.17175002 |
| IVFSLTVL | ASSSRLVNYAEQF | 0.14206868 |
| LASKATGF | ASSSRLVNYAEQF | 0.14940657 |
| LASVALIL | ASSSRLVNYAEQF | 0.17819265 |
| LFAVGLTL | ASSSRLVNYAEQF | 0.1291035 |
| LFNGGIGTYI | ASSSRLVNYAEQF | 0.13099189 |
| LTLGAGII | ASSSRLVNYAEQF | 0.16316098 |
| LVAAALGDI | ASSSRLVNYAEQF | 0.1553471 |
| NTAFVQLGI | ASSSRLVNYAEQF | 0.12808728 |
| QLFSGAEL | ASSSRLVNYAEQF | 0.15115795 |
| SAATFITL | ASSSRLVNYAEQF | 0.14705278 |
| SNYGLANLGV | ASSSRLVNYAEQF | 0.15926234 |
| SYGGAYAVM | ASSSRLVNYAEQF | 0.16755165 |
| TTGVGGLTLG | ASSSRLVNYAEQF | 0.13995326 |
| TTQTIYI | ASSSRLVNYAEQF | 0.19862303 |
| VAAGAGLGAVI | ASSSRLVNYAEQF | 0.13383411 |
| VAAVVTLL | ASSSRLVNYAEQF | 0.13746765 |
| VGFLGGLI | ASSSRLVNYAEQF | 0.17397025 |
| VGVIPILL | ASSSRLVNYAEQF | 0.26021478 |
| VIFASSLI | ASSSRLVNYAEQF | 0.15214437 |
| VKGGDGGAAAG | ASSSRLVNYAEQF | 0.17916009 |
| VTKTFTNI | ASSSRLVNYAEQF | 0.13550778 |
| VVAAAILL | ASSSRLVNYAEQF | 0.18979059 |
| YAALDVEL | ASSSRLVNYAEQF | 0.17694831 |
| LASVALLL | ASSSRLVNYAEQF | 0.13760716 |
| LDAVHRV | ASSSRLVNYAEQF | 0.1910169 |
| QGLSSSIF | ASSSRLVNYAEQF | 0.1821597 |
| RGTFANIRL | ASSSRLVNYAEQF | 0.18530208 |
| RQLWWGHRI | ASSSRLVNYAEQF | 0.1846571 |
| YGPPGTGKTLL | ASSSRLVNYAEQF | 0.1440079 |
| AALVGAPANVA | ASSSRLVNYAEQF | 0.16459711 |
| AATFRRTE | ASSSRLVNYAEQF | 0.20446607 |
| AGALGGGI | ASSSRLVNYAEQF | 0.20067358 |
| AGGAGGAGGIGAG | ASSSRLVNYAEQF | 0.14722586 |
| AGSGALGL | ASSSRLVNYAEQF | 0.18171361 |
| AQIADQI | ASSSRLVNYAEQF | 0.16496499 |
| ASAVAVTYL | ASSSRLVNYAEQF | 0.17846979 |
| ASTVLGGTGG | ASSSRLVNYAEQF | 0.16861801 |
| AVGGEPAK | ASSSRLVNYAEQF | 0.16210558 |
| CEQGTFGGTC | ASSSRLVNYAEQF | 0.16316852 |
| DEGKAPAG | ASSSRLVNYAEQF | 0.16477141 |

| | | |
|---|---|---|
| DTVVFPAL | ASSSRLVNYAEQF | 0.16861984 |
| FEGITMAG | ASSSRLVNYAEQF | 0.14320536 |
| FPLVVQD | ASSSRLVNYAEQF | 0.14935084 |
| GAFTTPQ | ASSSRLVNYAEQF | 0.15680262 |
| GFGGFGTAGGVGGA | ASSSRLVNYAEQF | 0.21510936 |
| GFLAQFL | ASSSRLVNYAEQF | 0.1452884 |
| GGAGGAGGNAGVA | ASSSRLVNYAEQF | 0.14026344 |
| GGVGTAGAGGAGGAGG | ASSSRLVNYAEQF | 0.16828857 |
| GNGGGGDAGNAGSGGN | ASSSRLVNYAEQF | 0.18547115 |
| GTGGAGGAGNDAGS | ASSSRLVNYAEQF | 0.15710387 |
| GTTNTGWFNS | ASSSRLVNYAEQF | 0.16725086 |
| IFMGPLFTL | ASSSRLVNYAEQF | 0.33306277 |
| ILAHAIEAI | ASSSRLVNYAEQF | 0.16536301 |
| IRSAIREAL | ASSSRLVNYAEQF | 0.16248216 |
| ITFAGGG | ASSSRLVNYAEQF | 0.17976393 |
| ITGMSGHQEV | ASSSRLVNYAEQF | 0.22536299 |
| IVAGLACTL | ASSSRLVNYAEQF | 0.1598345 |
| IVGGGLAGA | ASSSRLVNYAEQF | 0.17338724 |
| IVTGVLNI | ASSSRLVNYAEQF | 0.13645706 |
| LAQQGCI | ASSSRLVNYAEQF | 0.17195575 |
| LCLHVEN | ASSSRLVNYAEQF | 0.13036627 |
| LDAISDGS | ASSSRLVNYAEQF | 0.15743013 |
| LGGFAAML | ASSSRLVNYAEQF | 0.1519372 |
| LGGKNPMI | ASSSRLVNYAEQF | 0.17814341 |
| LGPKGVAAIL | ASSSRLVNYAEQF | 0.16670614 |
| LMVGVSI | ASSSRLVNYAEQF | 0.18382628 |
| LVGAAFVL | ASSSRLVNYAEQF | 0.13088606 |
| LVLAVAPL | ASSSRLVNYAEQF | 0.19846857 |
| MAAIAVIDAA | ASSSRLVNYAEQF | 0.19227493 |
| NNGINAELQ | ASSSRLVNYAEQF | 0.4711083 |
| RGEPLVLVHG | ASSSRLVNYAEQF | 0.14428023 |
| RGGAGGAAGLI | ASSSRLVNYAEQF | 0.18693757 |
| RSYRMHPAV | ASSSRLVNYAEQF | 0.15266171 |
| SAAHLLAGQLA | ASSSRLVNYAEQF | 0.1597856 |
| SAAVGSY | ASSSRLVNYAEQF | 0.13612765 |
| SPHAVLHRI | ASSSRLVNYAEQF | 0.1603341 |
| SQIAKVL | ASSSRLVNYAEQF | 0.14621437 |
| TALGHLVP | ASSSRLVNYAEQF | 0.16752677 |
| TAPTTSGAA | ASSSRLVNYAEQF | 0.20734054 |
| TDGDDGNGGSPGT | ASSSRLVNYAEQF | 0.21721105 |
| TDGLGEN | ASSSRLVNYAEQF | 0.1900456 |
| TGGPPLPGQA | ASSSRLVNYAEQF | 0.19735672 |
| TIPAITI | ASSSRLVNYAEQF | 0.18116763 |
| TIPGFTI | ASSSRLVNYAEQF | 0.188408 |
| TSIASGALA | ASSSRLVNYAEQF | 0.14990054 |
| TVTLPGAAI | ASSSRLVNYAEQF | 0.14921653 |
| VASGLSAALP | ASSSRLVNYAEQF | 0.14980206 |
| VGAGVIGI | ASSSRLVNYAEQF | 0.1508442 |
| VGPKRFGV | ASSSRLVNYAEQF | 0.14707892 |
| VNVAFPD | ASSSRLVNYAEQF | 0.1876996 |
| VSMTNSGVS | ASSSRLVNYAEQF | 0.12865801 |

| | | |
|---|---|---|
| VTDVLGDLSV | ASSSRLVNYAEQF | 0.1496019 |
| VVVVGAGFAGL | ASSSRLVNYAEQF | 0.21887745 |
| VYTVGTVV | ASSSRLVNYAEQF | 0.13504621 |
| WTTFSGP | ASSSRLVNYAEQF | 0.15801409 |
| YDELVEAF | ASSSRLVNYAEQF | 0.16274649 |
| YGTGGAGGNGG | ASSSRLVNYAEQF | 0.13855988 |
| YTNVMAV | ASSSRLVNYAEQF | 0.18978661 |
| FLEQVHQL | ASSSRLVNYAEQF | 0.1380306 |
| GLSLLAVAV | ASSSRLVNYAEQF | 0.17004839 |
| LEQVHQL | ASSSRLVNYAEQF | 0.18549025 |
| LNAAGAI | ASSSRLVNYAEQF | 0.16758482 |
| PAVHASL | ASSSRLVNYAEQF | 0.15195166 |
| TELEDDA | ASSSRLVNYAEQF | 0.15084209 |
| TPPGPPN | ASSSRLVNYAEQF | 0.14362149 |
| TPPPPPPP | ASSSRLVNYAEQF | 0.17475139 |
| TQSPALI | ASSHRDISGNTLY | 0.1368105 |
| EFAAAFPQ | ASSHRDISGNTLY | 0.16290681 |
| LLSGNLDVL | ASSHRDISGNTLY | 0.1411518 |
| GPANLGNYNI | ASSHRDISGNTLY | 0.2702428 |
| TLGPTLP | ASSHRDISGNTLY | 0.17811082 |
| VYAYSSCCK | ASSHRDISGNTLY | 0.16086453 |
| APAPAPAPNI | ASSHRDISGNTLY | 0.1532674 |
| PGGPVPIP | ASSHRDISGNTLY | 0.15200286 |
| FLTVALEP | ASSHRDISGNTLY | 0.13830242 |
| SQYFGANL | ASSHRDISGNTLY | 0.13989845 |
| GGVKPAA | ASSHRDISGNTLY | 0.16023667 |
| GGTGGAGGAGGW | ASSHRDISGNTLY | 0.15181367 |
| GVGGVAGLGGAGG | ASSHRDISGNTLY | 0.14457065 |
| VLLVLATTI | ASSHRDISGNTLY | 0.18336637 |
| IGIFAAML | ASSHRDISGNTLY | 0.19726345 |
| AVGDLID | ASSHRDISGNTLY | 0.13978194 |
| NISPLPL | ASSHRDISGNTLY | 0.14587794 |
| VTAVFVAI | ASSHRDISGNTLY | 0.16141625 |
| SGDDNDAAL | ASSHRDISGNTLY | 0.16102616 |
| AVAALAHS | ASSHRDISGNTLY | 0.12789631 |
| GGVTVMATP | ASSHRDISGNTLY | 0.17098302 |
| AGLAANVRD | ASSHRDISGNTLY | 0.17643122 |
| TNRPDLIDPALLRPG | ASSHRDISGNTLY | 0.1420425 |
| LANLGVTAL | ASSHRDISGNTLY | 0.14806983 |
| TGERIEL | ASSHRDISGNTLY | 0.29059112 |
| NGEELAA | ASSHRDISGNTLY | 0.13438468 |
| VLSLVWG | ASSHRDISGNTLY | 0.15824829 |
| IGYALWL | ASSHRDISGNTLY | 0.20069182 |
| LSLLLAPL | ASSHRDISGNTLY | 0.19468372 |
| TLDEALK | ASSHRDISGNTLY | 0.18592502 |
| TDGAGTAEGG | ASSHRDISGNTLY | 0.20407143 |
| NAPGGAGGNGG | ASSHRDISGNTLY | 0.1405253 |
| VSVFMGMGGNF | ASSHRDISGNTLY | 0.18837044 |
| LASYLAAL | ASSHRDISGNTLY | 0.15233953 |
| LVGGTPAL | ASSHRDISGNTLY | 0.14655744 |
| VRNADVL | ASSHRDISGNTLY | 0.1324504 |

| | | |
|---|---|---|
| TCSSPFLVSAM | ASSHRDISGNTLY | 0.14885548 |
| VHITDAE | ASSHRDISGNTLY | 0.13579768 |
| EYAAPSF | ASSHRDISGNTLY | 0.17401698 |
| VVDAFHGGL | ASSHRDISGNTLY | 0.19302097 |
| ASVVDPQ | ASSHRDISGNTLY | 0.1438447 |
| SYNEFGGLS | ASSHRDISGNTLY | 0.13883652 |
| DPDGSVY | ASSHRDISGNTLY | 0.14792113 |
| MVSTIHATE | ASSHRDISGNTLY | 0.15872954 |
| LAAGVADPV | ASSHRDISGNTLY | 0.19922194 |
| TKEQIAA | ASSHRDISGNTLY | 0.13958865 |
| SLASPAAL | ASSHRDISGNTLY | 0.31152764 |
| IGLVFYGV | ASSHRDISGNTLY | 0.20262249 |
| FLLDASNL | ASSHRDISGNTLY | 0.14942315 |
| YQVGNFP | ASSHRDISGNTLY | 0.21598934 |
| ESLLSQST | ASSHRDISGNTLY | 0.14475366 |
| LFTVGFAAVG | ASSHRDISGNTLY | 0.14550538 |
| ATPPTPT | ASSHRDISGNTLY | 0.16283874 |
| TDLATLG | ASSHRDISGNTLY | 0.15423569 |
| SAGGTGR | ASSHRDISGNTLY | 0.16157365 |
| GFSNSGS | ASSHRDISGNTLY | 0.16978343 |
| GGNGGSGGTSV | ASSHRDISGNTLY | 0.13911843 |
| GTSVATGGAGN | ASSHRDISGNTLY | 0.16255993 |
| NGGNGGSGGTS | ASSHRDISGNTLY | 0.13935918 |
| AVVKFPH | ASSHRDISGNTLY | 0.13142888 |
| TPPPPPP | ASSHRDISGNTLY | 0.14329258 |
| AAPAAAAAPA | ASSHRDISGNTLY | 0.22469227 |
| IALNEDL | ASSHRDISGNTLY | 0.21421848 |
| IVNVSSM | ASSHRDISGNTLY | 0.14165726 |
| TGIPPAPR | ASSHRDISGNTLY | 0.14640659 |
| FDLGGGTF | ASSHRDISGNTLY | 0.17744896 |
| FDLGGGTFDVS | ASSHRDISGNTLY | 0.14788057 |
| NEPTAAA | ASSHRDISGNTLY | 0.16480018 |
| NEPTAAALA | ASSHRDISGNTLY | 0.15644164 |
| GSIINISSI | ASSHRDISGNTLY | 0.15274228 |
| VAPGNAGTA | ASSHRDISGNTLY | 0.14298752 |
| TGGGASLE | ASSHRDISGNTLY | 0.15594485 |
| VAVYNHYKRI | ASSHRDISGNTLY | 0.1479366 |
| TDPAPATT | ASSHRDISGNTLY | 0.17902593 |
| TNRPDILDPALLRPG | ASSHRDISGNTLY | 0.1275424 |
| LTTTTTS | ASSHRDISGNTLY | 0.16325168 |
| PTGSGKTL | ASSHRDISGNTLY | 0.1446741 |
| TGSGKTL | ASSHRDISGNTLY | 0.1405852 |
| TIGHVDHGK | ASSHRDISGNTLY | 0.13287151 |
| TIGHVDHGKTT | ASSHRDISGNTLY | 0.16269268 |
| GHVDHGKTT | ASSHRDISGNTLY | 0.16072784 |
| AQFEHTLL | ASSHRDISGNTLY | 0.20054643 |
| IVNQDEL | ASSHRDISGNTLY | 0.14415063 |
| TNGSQFF | ASSHRDISGNTLY | 0.1777423 |
| TGKGIVE | ASSHRDISGNTLY | 0.15889113 |
| LISNASDA | ASSHRDISGNTLY | 0.1531546 |
| AELRTLGELTKIAKAHGAQ | ASSHRDISGNTLY | 0.14971521 |

| | | |
|---|---|---|
| APNAVAAAEPSW | ASSHRDISGNTLY | 0.18043496 |
| APQVAGAATAVL | ASSHRDISGNTLY | 0.14056858 |
| ARRDPAAAAAILPTDA | ASSHRDISGNTLY | 0.18286395 |
| DAAARDVLAEAGLAEYF | ASSHRDISGNTLY | 0.15962087 |
| DDQIVLSAKANLAKVEQ | ASSHRDISGNTLY | 0.14840217 |
| DELLPDTGEDSDEDGH | ASSHRDISGNTLY | 0.1939846 |
| ELPEAPANRATVPQD | ASSHRDISGNTLY | 0.21051554 |
| FGQSKIERVTGIG | ASSHRDISGNTLY | 0.13769375 |
| GAGGTGSTAGGAGGAGGAGGLY | ASSHRDISGNTLY | 0.14024885 |
| GHGGTGAAGGNGGTGGAGGSGI | ASSHRDISGNTLY | 0.15121913 |
| GNGGNGGAGGEGGPGGAGGAGA | ASSHRDISGNTLY | 0.1299122 |
| GSTAGNGGNGGNATNGTIAGQP | ASSHRDISGNTLY | 0.16715755 |
| GTGGDGGNGGNGGTGG | ASSHRDISGNTLY | 0.14287123 |
| GVRRPAGIPADAPV | ASSHRDISGNTLY | 0.15232149 |
| GYRQVTDAHLLTLARR | ASSHRDISGNTLY | 0.17114769 |
| IAKMGQDGHDRG | ASSHRDISGNTLY | 0.15265639 |
| KAEPSIEIQYT | ASSHRDISGNTLY | 0.16466266 |
| KPVRIRMAADEDTVAVT | ASSHRDISGNTLY | 0.16927305 |
| MPPRIAGMRLLVIKPEP | ASSHRDISGNTLY | 0.15478131 |
| MKFYAVALFRSRGVPPTR | ASSHRDISGNTLY | 0.16247575 |
| NAGGFGILGAVAHSP | ASSHRDISGNTLY | 0.14056768 |
| NEGREITLAMAEM | ASSHRDISGNTLY | 0.15134358 |
| NELVEWSHPLGHRWRWE | ASSHRDISGNTLY | 0.23350316 |
| NLGAGGTGGNGGNPG | ASSHRDISGNTLY | 0.14137891 |
| NTEWPKWETVMAADPVD | ASSHRDISGNTLY | 0.14349912 |
| PASAAPAETQP | ASSHRDISGNTLY | 0.17289706 |
| QQSSDGKNGVRAAEITGE | ASSHRDISGNTLY | 0.13689156 |
| RAPDAVTTSWWKE | ASSHRDISGNTLY | 0.17697164 |
| SGFGNVGAHTSGWFNQ | ASSHRDISGNTLY | 0.15958954 |
| SGVAGAVGQAASVGGLKV | ASSHRDISGNTLY | 0.18139465 |
| SNGGAGTGSAGGNGGT | ASSHRDISGNTLY | 0.14338319 |
| TGGAGGEGGAAG | ASSHRDISGNTLY | 0.20118515 |
| TGGDGGAAGAGGAPGHG | ASSHRDISGNTLY | 0.21939422 |
| TMPPLLAAIANAGRHGV | ASSHRDISGNTLY | 0.14816728 |
| TSSTAGNSGGA | ASSHRDISGNTLY | 0.19054444 |
| VPATVPVGILC | ASSHRDISGNTLY | 0.14354993 |
| VQRPDSGGVVLTGRIS | ASSHRDISGNTLY | 0.16299519 |
| WTPNPVPATKGVPAGLVQAT | ASSHRDISGNTLY | 0.18529943 |
| YALKRALAKFSDRFDVVII | ASSHRDISGNTLY | 0.14598402 |
| YLIGSVIV | ASSHRDISGNTLY | 0.16362715 |
| YGQLVAVID | ASSHRDISGNTLY | 0.17801984 |
| VTVGGMTLPA | ASSHRDISGNTLY | 0.23699595 |
| VLALPAV | ASSHRDISGNTLY | 0.13668272 |
| VIPPTLNYET | ASSHRDISGNTLY | 0.16452153 |
| VGMGGMHPSSG | ASSHRDISGNTLY | 0.13971557 |
| VGLAGGAATA | ASSHRDISGNTLY | 0.13926798 |
| VGGAGGAGGSGP | ASSHRDISGNTLY | 0.15563026 |
| VFAAFMV | ASSHRDISGNTLY | 0.17068475 |
| TLYDKLR | ASSHRDISGNTLY | 0.16637388 |
| TLAVFTVL | ASSHRDISGNTLY | 0.1650245 |
| TGSVQPARV | ASSHRDISGNTLY | 0.19583575 |

| | | |
|---|---|---|
| TGGGAGSGGT | ASSHRDISGNTLY | 0.22743532 |
| SHLETSL | ASSHRDISGNTLY | 0.14864029 |
| SFTTGMT | ASSHRDISGNTLY | 0.16288605 |
| QVEVAKRMGVRQPTVS | ASSHRDISGNTLY | 0.13495865 |
| QGSGGPALAM | ASSHRDISGNTLY | 0.14036717 |
| MLAPSNP | ASSHRDISGNTLY | 0.14466399 |
| LVNNAAI | ASSHRDISGNTLY | 0.15144944 |
| LTGEINTLA | ASSHRDISGNTLY | 0.12937066 |
| LSIFGPV | ASSHRDISGNTLY | 0.17238392 |
| LLALLANL | ASSHRDISGNTLY | 0.17077005 |
| LISNASDALDKLR | ASSHRDISGNTLY | 0.19554864 |
| LINEAVP | ASSHRDISGNTLY | 0.15052937 |
| LGYAFSGL | ASSHRDISGNTLY | 0.16523562 |
| LDFVAVAL | ASSHRDISGNTLY | 0.15436201 |
| LAQPGAE | ASSHRDISGNTLY | 0.20279261 |
| ISNLGAVTGF | ASSHRDISGNTLY | 0.15925002 |
| HQEDPGRI | ASSHRDISGNTLY | 0.13501863 |
| GTPGEPTP | ASSHRDISGNTLY | 0.14503752 |
| GFFNSGS | ASSHRDISGNTLY | 0.1396945 |
| GAGGGGLTAGAN | ASSHRDISGNTLY | 0.16921352 |
| FVDALVNM | ASSHRDISGNTLY | 0.16682306 |
| FLTVAPEL | ASSHRDISGNTLY | 0.20651734 |
| FETLTAL | ASSHRDISGNTLY | 0.1828734 |
| DVINPIEL | ASSHRDISGNTLY | 0.13055368 |
| APVSGAPV | ASSHRDISGNTLY | 0.14897193 |
| ALVGGLAW | ASSHRDISGNTLY | 0.24643274 |
| ALAAAAATAI | ASSHRDISGNTLY | 0.15011649 |
| AFGLGLLL | ASSHRDISGNTLY | 0.17723335 |
| AFFAAASL | ASSHRDISGNTLY | 0.19116071 |
| AAAALPAFSPP | ASSHRDISGNTLY | 0.14601192 |
| VVLAPAL | ASSHRDISGNTLY | 0.15890603 |
| TIGHVDHGKTTL | ASSHRDISGNTLY | 0.17797548 |
| LISNASDALDK | ASSHRDISGNTLY | 0.19187956 |
| LISNASDAL | ASSHRDISGNTLY | 0.14601775 |
| ISNASDALDK | ASSHRDISGNTLY | 0.15732554 |
| NQAETLVYQTE | ASSHRDISGNTLY | 0.17658544 |
| TGGTGGTAGAG | ASSHRDISGNTLY | 0.22502975 |
| TGSTAGGAGGA | ASSHRDISGNTLY | 0.21732563 |
| DPLAVPVAAGPV | ASSHRDISGNTLY | 0.17209496 |
| GGTGGTPGNGGN | ASSHRDISGNTLY | 0.17343499 |
| PAGLAGVDPATL | ASSHRDISGNTLY | 0.14424835 |
| GNNLVVEAPHAGASV | ASSHRDISGNTLY | 0.15857174 |
| KVTRIMSASTLEQPA | ASSHRDISGNTLY | 0.1540384 |
| LGFALPIALPRRTVV | ASSHRDISGNTLY | 0.15110923 |
| TRLGNLTARKLVVLV | ASSHRDISGNTLY | 0.13613431 |
| WSQQRRLIYFPSAGP | ASSHRDISGNTLY | 0.16078857 |
| GPNGNPDPMAAAVDIR | ASSHRDISGNTLY | 0.14801474 |
| TQRCVSVADLRAAASR | ASSHRDISGNTLY | 0.20483719 |
| QFVLDVAEHPVEVLTETV | ASSHRDISGNTLY | 0.15430161 |
| GGAGGTGGAGGNGGTQAGDG | ASSHRDISGNTLY | 0.1866546 |
| PLLATLRAIAVDVDAKPAQV | ASSHRDISGNTLY | 0.15245254 |

| | | |
|---|---|---|
| APKFRTGGYMIDFWGVGYQVAK | ASSHRDISGNTLY | 0.15283926 |
| GAGGAGGAGGAAGGPGGTGGTG | ASSHRDISGNTLY | 0.14661056 |
| IGALGLLATDVAPAGSVAFGVA | ASSHRDISGNTLY | 0.14328025 |
| AGGAGGIGGNGGAGGAGGVLMG | ASSHRDISGNTLY | 0.1780324 |
| ADELIGLL | ASSHRDISGNTLY | 0.1497099 |
| ADRALKA | ASSHRDISGNTLY | 0.13911691 |
| AGAVGQVGGAGGA | ASSHRDISGNTLY | 0.16607991 |
| AGGGGGLGGAP | ASSHRDISGNTLY | 0.25157326 |
| AGGLLYGNGGAG | ASSHRDISGNTLY | 0.15106767 |
| AGLESIGL | ASSHRDISGNTLY | 0.15021828 |
| AGLKTGII | ASSHRDISGNTLY | 0.1654915 |
| ASLAALLL | ASSHRDISGNTLY | 0.16595909 |
| ASVINISSI | ASSHRDISGNTLY | 0.36275825 |
| ATPPGPA | ASSHRDISGNTLY | 0.14194247 |
| ATVASALV | ASSHRDISGNTLY | 0.39598942 |
| ATVASLIL | ASSHRDISGNTLY | 0.15980205 |
| AVNGPVLIRH | ASSHRDISGNTLY | 0.14304723 |
| DGAAGAVGI | ASSHRDISGNTLY | 0.17519979 |
| DPLLHGI | ASSHRDISGNTLY | 0.17535354 |
| ELGAALGAL | ASSHRDISGNTLY | 0.1749137 |
| ELGRTQAV | ASSHRDISGNTLY | 0.17242613 |
| FVAGGYLS | ASSHRDISGNTLY | 0.15684235 |
| GASGGAGGAGGW | ASSHRDISGNTLY | 0.1411964 |
| GGAGAGGDGGAGGI | ASSHRDISGNTLY | 0.182074 |
| GTGHNGSQPGMGGNG | ASSHRDISGNTLY | 0.15077639 |
| GYAAASASAS | ASSHRDISGNTLY | 0.1961962 |
| HQVDVAAVQQG | ASSHRDISGNTLY | 0.14258108 |
| IGAAIAHT | ASSHRDISGNTLY | 0.1410931 |
| IGMIEVAQL | ASSHRDISGNTLY | 0.14836514 |
| IRLAPEHTSAMD | ASSHRDISGNTLY | 0.16679929 |
| IYNVSVL | ASSHRDISGNTLY | 0.14293793 |
| KQGFLGM | ASSHRDISGNTLY | 0.16285841 |
| LESVITQL | ASSHRDISGNTLY | 0.16854905 |
| LGLGALVVV | ASSHRDISGNTLY | 0.14466608 |
| LIGNGGNGGSGGI | ASSHRDISGNTLY | 0.14641409 |
| LNAINAPTL | ASSHRDISGNTLY | 0.18352489 |
| LQLLNTGV | ASSHRDISGNTLY | 0.15454 |
| LQTSIEF | ASSHRDISGNTLY | 0.14532216 |
| MSFAVSEL | ASSHRDISGNTLY | 0.17960371 |
| NTGALGL | ASSHRDISGNTLY | 0.24083066 |
| PPGAGGAGGAGTT | ASSHRDISGNTLY | 0.18846776 |
| QLVGLGI | ASSHRDISGNTLY | 0.13952263 |
| RFLEQVHQL | ASSHRDISGNTLY | 0.12959969 |
| RGVGYVLR | ASSHRDISGNTLY | 0.15543018 |
| SATGVFAGVI | ASSHRDISGNTLY | 0.17777982 |
| SLPAITLP | ASSHRDISGNTLY | 0.15248759 |
| STALAFTV | ASSHRDISGNTLY | 0.1523006 |
| SVLTVGLGK | ASSHRDISGNTLY | 0.16797276 |
| TALAGAGSGI | ASSHRDISGNTLY | 0.17760867 |
| TGLPFTGI | ASSHRDISGNTLY | 0.16326945 |
| VAEHVTV | ASSHRDISGNTLY | 0.13499486 |

| | | |
|---|---|---|
| VDIVLNSL | ASSHRDISGNTLY | 0.15175451 |
| VLTGGGAVL | ASSHRDISGNTLY | 0.21095525 |
| VNVGLAYGI | ASSHRDISGNTLY | 0.14186536 |
| VSSAAVGAV | ASSHRDISGNTLY | 0.16878688 |
| VTSVYEAGV | ASSHRDISGNTLY | 0.1802915 |
| WVALDRGA | ASSHRDISGNTLY | 0.13616674 |
| YLVGSVVI | ASSHRDISGNTLY | 0.15115382 |
| GPPGTGKTLL | ASSHRDISGNTLY | 0.14672038 |
| GVGMTKF | ASSHRDISGNTLY | 0.17906767 |
| IGIGGSDL | ASSHRDISGNTLY | 0.14832295 |
| IGTLLAEL | ASSHRDISGNTLY | 0.15537368 |
| IYTSGTT | ASSHRDISGNTLY | 0.20293494 |
| LSFPAGV | ASSHRDISGNTLY | 0.19380924 |
| LYTSGTT | ASSHRDISGNTLY | 0.16161227 |
| VGAFPFGF | ASSHRDISGNTLY | 0.14777388 |
| VGAGTVEF | ASSHRDISGNTLY | 0.14456044 |
| VGKSTLLL | ASSHRDISGNTLY | 0.13052368 |
| AALAHAGGGIV | ASSHRDISGNTLY | 0.21716258 |
| AALVAVSLL | ASSHRDISGNTLY | 0.13272218 |
| AAVSRAVAAL | ASSHRDISGNTLY | 0.25000367 |
| AGGSSGIGGF | ASSHRDISGNTLY | 0.18014792 |
| AGLTVANANG | ASSHRDISGNTLY | 0.16904455 |
| AGLVGAVL | ASSHRDISGNTLY | 0.169559 |
| ALSQAFTV | ASSHRDISGNTLY | 0.1450416 |
| ALTFSVSL | ASSHRDISGNTLY | 0.14308314 |
| ALTGAALVAL | ASSHRDISGNTLY | 0.14634353 |
| ASFSIGAGVL | ASSHRDISGNTLY | 0.16847879 |
| DGSVYKI | ASSHRDISGNTLY | 0.15212709 |
| EQLAEIV | ASSHRDISGNTLY | 0.17141502 |
| FAVSELTVM | ASSHRDISGNTLY | 0.17077868 |
| GAAPLAAGL | ASSHRDISGNTLY | 0.14352898 |
| GADGGDGGSGGAGG | ASSHRDISGNTLY | 0.15455791 |
| GFGLDTGGAGG | ASSHRDISGNTLY | 0.17965795 |
| GLASGGLL | ASSHRDISGNTLY | 0.16065015 |
| GRIAGAAVLL | ASSHRDISGNTLY | 0.16634637 |
| GSAGSLNTSF | ASSHRDISGNTLY | 0.16764723 |
| GTGQIGI | ASSHRDISGNTLY | 0.15004767 |
| GTLIALIL | ASSHRDISGNTLY | 0.17573217 |
| IAATMGFVVG | ASSHRDISGNTLY | 0.19221342 |
| IFAQALL | ASSHRDISGNTLY | 0.15212922 |
| IGAAVALGF | ASSHRDISGNTLY | 0.20758015 |
| IGLTPLAL | ASSHRDISGNTLY | 0.13527028 |
| ILGSNTSTL | ASSHRDISGNTLY | 0.13823518 |
| IVFSLTVL | ASSHRDISGNTLY | 0.14477149 |
| LASKATGF | ASSHRDISGNTLY | 0.2032932 |
| LASVALIL | ASSHRDISGNTLY | 0.1970339 |
| LFAVGLTL | ASSHRDISGNTLY | 0.14413984 |
| LFNGGIGTYI | ASSHRDISGNTLY | 0.16047008 |
| LTLGAGII | ASSHRDISGNTLY | 0.1474527 |
| LVAAALGDI | ASSHRDISGNTLY | 0.150166 |
| NTAFVQLGI | ASSHRDISGNTLY | 0.16165726 |

| | | |
|---|---|---|
| QLFSGAEL | ASSHRDISGNTLY | 0.14296573 |
| SAATFITL | ASSHRDISGNTLY | 0.14691988 |
| SNYGLANLGV | ASSHRDISGNTLY | 0.1877504 |
| SYGGAYAVM | ASSHRDISGNTLY | 0.1612554 |
| TTGVGGLTLG | ASSHRDISGNTLY | 0.15739363 |
| TTQTIYI | ASSHRDISGNTLY | 0.14477855 |
| VAAGAGLGAVI | ASSHRDISGNTLY | 0.15284745 |
| VAAVVTLL | ASSHRDISGNTLY | 0.16114879 |
| VGFLGGLI | ASSHRDISGNTLY | 0.16278075 |
| VGVIPILL | ASSHRDISGNTLY | 0.1924553 |
| VIFASSLI | ASSHRDISGNTLY | 0.16348062 |
| VKGGDGGAAAG | ASSHRDISGNTLY | 0.14354578 |
| VTKTFTNI | ASSHRDISGNTLY | 0.19282055 |
| VVAAAILL | ASSHRDISGNTLY | 0.21495119 |
| YAALDVEL | ASSHRDISGNTLY | 0.15947953 |
| LASVALLL | ASSHRDISGNTLY | 0.17318699 |
| LDAVHRV | ASSHRDISGNTLY | 0.1721903 |
| QGLSSSIF | ASSHRDISGNTLY | 0.20686544 |
| RGTFANIRL | ASSHRDISGNTLY | 0.15135635 |
| RQLWWGHRI | ASSHRDISGNTLY | 0.13650897 |
| YGPPGTGKTLL | ASSHRDISGNTLY | 0.15988483 |
| AALVGAPANVA | ASSHRDISGNTLY | 0.14829719 |
| AATFRRTE | ASSHRDISGNTLY | 0.15401195 |
| AGALGGGI | ASSHRDISGNTLY | 0.13409339 |
| AGGAGGAGGIGAG | ASSHRDISGNTLY | 0.19492586 |
| AGSGALGL | ASSHRDISGNTLY | 0.23890194 |
| AQIADQI | ASSHRDISGNTLY | 0.14604324 |
| ASAVAVTYL | ASSHRDISGNTLY | 0.1809548 |
| ASTVLGGTGG | ASSHRDISGNTLY | 0.13310449 |
| AVGGEPAK | ASSHRDISGNTLY | 0.16644637 |
| CEQGTFGGTC | ASSHRDISGNTLY | 0.17759317 |
| DEGKAPAG | ASSHRDISGNTLY | 0.14390656 |
| DTVVFPAL | ASSHRDISGNTLY | 0.17441411 |
| FEGITMAG | ASSHRDISGNTLY | 0.14508645 |
| FPLVVQD | ASSHRDISGNTLY | 0.15833235 |
| GAFTTPQ | ASSHRDISGNTLY | 0.2002045 |
| GFGGFGTAGGVGGA | ASSHRDISGNTLY | 0.16006678 |
| GFLAQFL | ASSHRDISGNTLY | 0.17451194 |
| GGAGGAGGNAGVA | ASSHRDISGNTLY | 0.17394406 |
| GGVGTAGAGGAGGAGG | ASSHRDISGNTLY | 0.16438472 |
| GNGGGGDAGNAGSGGN | ASSHRDISGNTLY | 0.19683339 |
| GTGGAGGAGNDAGS | ASSHRDISGNTLY | 0.17565879 |
| GTTNTGWFNS | ASSHRDISGNTLY | 0.13788854 |
| IFMGPLFTL | ASSHRDISGNTLY | 0.15156561 |
| ILAHAIEAI | ASSHRDISGNTLY | 0.14735049 |
| IRSAIREAL | ASSHRDISGNTLY | 0.20705786 |
| ITFAGGG | ASSHRDISGNTLY | 0.22239637 |
| ITGMSGHQEV | ASSHRDISGNTLY | 0.16809864 |
| IVAGLACTL | ASSHRDISGNTLY | 0.1530868 |
| IVGGGLAGA | ASSHRDISGNTLY | 0.20437947 |
| IVTGVLNI | ASSHRDISGNTLY | 0.18590671 |

| | | |
|---|---|---|
| LAQQGCI | ASSHRDISGNTLY | 0.17225626 |
| LCLHVEN | ASSHRDISGNTLY | 0.19738756 |
| LDAISDGS | ASSHRDISGNTLY | 0.084847406 |
| LGGFAAML | ASSHRDISGNTLY | 0.14141132 |
| LGGKNPMI | ASSHRDISGNTLY | 0.08935472 |
| LGPKGVAAIL | ASSHRDISGNTLY | 0.097524114 |
| LMVGVSI | ASSHRDISGNTLY | 0.09419056 |
| LVGAAFVL | ASSHRDISGNTLY | 0.081864506 |
| LVLAVAPL | ASSHRDISGNTLY | 0.08905338 |
| MAAIAVIDAA | ASSHRDISGNTLY | 0.08646086 |
| NNGINAELQ | ASSHRDISGNTLY | 0.083403714 |
| RGEPLVLVHG | ASSHRDISGNTLY | 0.094302274 |
| RGGAGGAAGLI | ASSHRDISGNTLY | 0.09519569 |
| RSYRMHPAV | ASSHRDISGNTLY | 0.34032905 |
| SAAHLLAGQLA | ASSHRDISGNTLY | 0.1110751 |
| SAAVGSY | ASSHRDISGNTLY | 0.084257156 |
| SPHAVLHRI | ASSHRDISGNTLY | 0.093142286 |
| SQIAKVL | ASSHRDISGNTLY | 0.089654915 |
| TALGHLVP | ASSHRDISGNTLY | 0.09720722 |
| TAPTTSGAA | ASSHRDISGNTLY | 0.07981682 |
| TDGDDGNGGSPGT | ASSHRDISGNTLY | 0.102972955 |
| TDGLGEN | ASSHRDISGNTLY | 0.081246324 |
| TGGPPLPGQA | ASSHRDISGNTLY | 0.0787419 |
| TIPAITI | ASSHRDISGNTLY | 0.11998662 |
| TIPGFTI | ASSHRDISGNTLY | 0.0959381 |
| TSIASGALA | ASSHRDISGNTLY | 0.086921 |
| TVTLPGAAI | ASSHRDISGNTLY | 0.14003712 |
| VASGLSAALP | ASSHRDISGNTLY | 0.08300007 |
| VGAGVIGI | ASSHRDISGNTLY | 0.08542325 |
| VGPKRFGV | ASSHRDISGNTLY | 0.13381475 |
| VNVAFPD | ASSHRDISGNTLY | 0.12493319 |
| VSMTNSGVS | ASSHRDISGNTLY | 0.08601194 |
| VTDVLGDLSV | ASSHRDISGNTLY | 0.09527925 |
| VVVVGAGFAGL | ASSHRDISGNTLY | 0.17892018 |
| VYTVGTVV | ASSHRDISGNTLY | 0.0991314 |
| WTTFSGP | ASSHRDISGNTLY | 0.09599276 |
| YDELVEAF | ASSHRDISGNTLY | 0.100322604 |
| YGTGGAGGNGG | ASSHRDISGNTLY | 0.083459646 |
| YTNVMAV | ASSHRDISGNTLY | 0.12611087 |
| FLEQVHQL | ASSHRDISGNTLY | 0.16105705 |
| GLSLLAVAV | ASSHRDISGNTLY | 0.10178214 |
| LEQVHQL | ASSHRDISGNTLY | 0.112530604 |
| LNAAGAI | ASSHRDISGNTLY | 0.08468996 |
| PAVHASL | ASSHRDISGNTLY | 0.09115863 |
| TELEDDA | ASSHRDISGNTLY | 0.080262244 |
| TPPGPPN | ASSHRDISGNTLY | 0.11782122 |
| TPPPPPPP | ASSHRDISGNTLY | 0.112927966 |
| TQSPALI | TCSADGGYEQY | 0.104072675 |
| EFAAAFPQ | TCSADGGYEQY | 0.08232804 |
| LLSGNLDVL | TCSADGGYEQY | 0.08562932 |
| GPANLGNYNI | TCSADGGYEQY | 0.08228014 |

| | | |
|---|---|---|
| TLGPTLP | TCSADGGYEQY | 0.083991915 |
| VYAYSSCCK | TCSADGGYEQY | 0.08234883 |
| APAPAPAPNI | TCSADGGYEQY | 0.08670068 |
| PGGPVPIP | TCSADGGYEQY | 0.13491969 |
| FLTVALEP | TCSADGGYEQY | 0.12405986 |
| SQYFGANL | TCSADGGYEQY | 0.104069844 |
| GGVKPAA | TCSADGGYEQY | 0.16184166 |
| GGTGGAGGAGGW | TCSADGGYEQY | 0.08682481 |
| GVGGVAGLGGAGG | TCSADGGYEQY | 0.09548739 |
| VLLVLATTI | TCSADGGYEQY | 0.092515334 |
| IGIFAAML | TCSADGGYEQY | 0.08709915 |
| AVGDLID | TCSADGGYEQY | 0.17770708 |
| NISPLPL | TCSADGGYEQY | 0.09561344 |
| VTAVFVAI | TCSADGGYEQY | 0.089522466 |
| SGDDNDAAL | TCSADGGYEQY | 0.15573713 |
| AVAALAHS | TCSADGGYEQY | 0.09063568 |
| GGVTVMATP | TCSADGGYEQY | 0.088868044 |
| AGLAANVRD | TCSADGGYEQY | 0.09348411 |
| TNRPDLIDPALLRPG | TCSADGGYEQY | 0.101885155 |
| LANLGVTAL | TCSADGGYEQY | 0.08633896 |
| TGERIEL | TCSADGGYEQY | 0.085169725 |
| NGEELAA | TCSADGGYEQY | 0.088140965 |
| VLSLVWG | TCSADGGYEQY | 0.09877184 |
| IGYALWL | TCSADGGYEQY | 0.13331297 |
| LSLLLAPL | TCSADGGYEQY | 0.097050615 |
| TLDEALK | TCSADGGYEQY | 0.08465561 |
| TDGAGTAEGG | TCSADGGYEQY | 0.124838874 |
| NAPGGAGGNGG | TCSADGGYEQY | 0.09607226 |
| VSVFMGMGGNF | TCSADGGYEQY | 0.099747695 |
| LASYLAAL | TCSADGGYEQY | 0.094201975 |
| LVGGTPAL | TCSADGGYEQY | 0.08639761 |
| VRNADVL | TCSADGGYEQY | 0.107288584 |
| TCSSPFLVSAM | TCSADGGYEQY | 0.123097785 |
| VHITDAE | TCSADGGYEQY | 0.09852855 |
| EYAAPSF | TCSADGGYEQY | 0.095864125 |
| VVDAFHGGL | TCSADGGYEQY | 0.09206292 |
| ASVVDPQ | TCSADGGYEQY | 0.09871318 |
| SYNEFGGLS | TCSADGGYEQY | 0.0954317 |
| DPDGSVY | TCSADGGYEQY | 0.08641497 |
| MVSTIHATE | TCSADGGYEQY | 0.1079307 |
| LAAGVADPV | TCSADGGYEQY | 0.115345985 |
| TKEQIAA | TCSADGGYEQY | 0.12298341 |
| SLASPAAL | TCSADGGYEQY | 0.098626174 |
| IGLVFYGV | TCSADGGYEQY | 0.08697469 |
| FLLDASNL | TCSADGGYEQY | 0.09979395 |
| YQVGNFP | TCSADGGYEQY | 0.22062322 |
| ESLLSQST | TCSADGGYEQY | 0.0833758 |
| LFTVGFAAVG | TCSADGGYEQY | 0.086878404 |
| ATPPTPT | TCSADGGYEQY | 0.28736517 |
| TDLATLG | TCSADGGYEQY | 0.09262992 |
| SAGGTGR | TCSADGGYEQY | 0.115112 |

| | | |
|---|---|---|
| GFSNSGS | TCSADGGYEQY | 0.08634046 |
| GGNGGSGGTSV | TCSADGGYEQY | 0.107636854 |
| GTSVATGGAGN | TCSADGGYEQY | 0.08513249 |
| NGGNGGSGGTS | TCSADGGYEQY | 0.37302634 |
| AVVKFPH | TCSADGGYEQY | 0.08846375 |
| TPPPPPP | TCSADGGYEQY | 0.08357504 |
| AAPAAAAAPA | TCSADGGYEQY | 0.084956706 |
| IALNEDL | TCSADGGYEQY | 0.07895854 |
| IVNVSSM | TCSADGGYEQY | 0.123761676 |
| TGIPPAPR | TCSADGGYEQY | 0.08714846 |
| FDLGGGTF | TCSADGGYEQY | 0.121594675 |
| FDLGGGTFDVS | TCSADGGYEQY | 0.0959981 |
| NEPTAAA | TCSADGGYEQY | 0.10774521 |
| NEPTAAALA | TCSADGGYEQY | 0.109819025 |
| GSIINISSI | TCSADGGYEQY | 0.09198526 |
| VAPGNAGTA | TCSADGGYEQY | 0.08776233 |
| TGGGASLE | TCSADGGYEQY | 0.12102715 |
| VAVYNHYKRI | TCSADGGYEQY | 0.1132132 |
| TDPAPATT | TCSADGGYEQY | 0.093326755 |
| TNRPDILDPALLRPG | TCSADGGYEQY | 0.083130166 |
| LTTTTTS | TCSADGGYEQY | 0.10293718 |
| PTGSGKTL | TCSADGGYEQY | 0.104911804 |
| TGSGKTL | TCSADGGYEQY | 0.08042557 |
| TIGHVDHGK | TCSADGGYEQY | 0.08079813 |
| TIGHVDHGKTT | TCSADGGYEQY | 0.08102216 |
| GHVDHGKTT | TCSADGGYEQY | 0.08432905 |
| AQFEHTLL | TCSADGGYEQY | 0.24109107 |
| IVNQDEL | TCSADGGYEQY | 0.1491722 |
| TNGSQFF | TCSADGGYEQY | 0.1034766 |
| TGKGIVE | TCSADGGYEQY | 0.102701336 |
| LISNASDA | TCSADGGYEQY | 0.096442424 |
| AELRTLGELTKIAKAHGAQ | TCSADGGYEQY | 0.07742881 |
| APNAVAAAEPSW | TCSADGGYEQY | 0.08534759 |
| APQVAGAATAVL | TCSADGGYEQY | 0.08710798 |
| ARRDPAAAAAILPTDA | TCSADGGYEQY | 0.10634423 |
| DAAARDVLAEAGLAEYF | TCSADGGYEQY | 0.08204754 |
| DDQIVLSAKANLAKVEQ | TCSADGGYEQY | 0.09351801 |
| DELLPDTGEDSDEDGH | TCSADGGYEQY | 0.11223417 |
| ELPEAPANRATVPQD | TCSADGGYEQY | 0.08837166 |
| FGQSKIERVTGIG | TCSADGGYEQY | 0.10014718 |
| GAGGTGSTAGGAGGAGGAGGLY | TCSADGGYEQY | 0.08114139 |
| GHGGTGAAGGNGGTGGAGGSGI | TCSADGGYEQY | 0.08154332 |
| GNGGNGGAGGEGGPGGAGGAGA | TCSADGGYEQY | 0.12258622 |
| GSTAGNGGNGGNATNGTIAGQP | TCSADGGYEQY | 0.08004843 |
| GTGGDGGNGGNGGTGG | TCSADGGYEQY | 0.08752705 |
| GVRRPAGIPADAPV | TCSADGGYEQY | 0.08652368 |
| GYRQVTDAHLLTLARR | TCSADGGYEQY | 0.09165038 |
| IAKMGQDGHDRG | TCSADGGYEQY | 0.08091008 |
| KAEPSIEIQYT | TCSADGGYEQY | 0.09862711 |
| KPVRIRMAADEDTVAVT | TCSADGGYEQY | 0.10804379 |
| MPPRIAGMRLLVIKPEP | TCSADGGYEQY | 0.114823416 |

| | | |
|---|---|---|
| MKFYAVALFRSRGVPPTR | TCSADGGYEQY | 0.19441989 |
| NAGGFGILGAVAHSP | TCSADGGYEQY | 0.13040869 |
| NEGREITLAMAEM | TCSADGGYEQY | 0.10743651 |
| NELVEWSHPLGHRWRWE | TCSADGGYEQY | 0.09447087 |
| NLGAGGTGGNGGNPG | TCSADGGYEQY | 0.08733469 |
| NTEWPKWETVMAADPVD | TCSADGGYEQY | 0.08555588 |
| PASAAPAETQP | TCSADGGYEQY | 0.0866298 |
| QQSSDGKNGVRAAEITGE | TCSADGGYEQY | 0.15764883 |
| RAPDAVTTSWWKE | TCSADGGYEQY | 0.0833655 |
| SGFGNVGAHTSGWFNQ | TCSADGGYEQY | 0.084248036 |
| SGVAGAVGQAASVGGLKV | TCSADGGYEQY | 0.08257573 |
| SNGGAGTGSAGGNGGT | TCSADGGYEQY | 0.21503663 |
| TGGAGGEGGAAG | TCSADGGYEQY | 0.1537267 |
| TGGDGGAAGAGGAPGHG | TCSADGGYEQY | 0.08704466 |
| TMPPLLAAIANAGRHGV | TCSADGGYEQY | 0.09234687 |
| TSSTAGNSGGA | TCSADGGYEQY | 0.08835341 |
| VPATVPVGILC | TCSADGGYEQY | 0.111910395 |
| VQRPDSGGVVLTGRIS | TCSADGGYEQY | 0.08237561 |
| WTPNPVPATKGVPAGLVQAT | TCSADGGYEQY | 0.08547556 |
| YALKRALAKFSDRFDVVII | TCSADGGYEQY | 0.09825085 |
| YLIGSVIV | TCSADGGYEQY | 0.18705265 |
| YGQLVAVID | TCSADGGYEQY | 0.08870392 |
| VTVGGMTLPA | TCSADGGYEQY | 0.08109186 |
| VLALPAV | TCSADGGYEQY | 0.11768944 |
| VIPPTLNYET | TCSADGGYEQY | 0.08417435 |
| VGMGGMHPSSG | TCSADGGYEQY | 0.0956817 |
| VGLAGGAATA | TCSADGGYEQY | 0.08898127 |
| VGGAGGAGGSGP | TCSADGGYEQY | 0.0819614 |
| VFAAFMV | TCSADGGYEQY | 0.095371485 |
| TLYDKLR | TCSADGGYEQY | 0.15182912 |
| TLAVFTVL | TCSADGGYEQY | 0.08740049 |
| TGSVQPARV | TCSADGGYEQY | 0.14105925 |
| TGGGAGSGGT | TCSADGGYEQY | 0.33721054 |
| SHLETSL | TCSADGGYEQY | 0.1379313 |
| SFTTGMT | TCSADGGYEQY | 0.14184664 |
| QVEVAKRMGVRQPTVS | TCSADGGYEQY | 0.08461728 |
| QGSGGPALAM | TCSADGGYEQY | 0.0992098 |
| MLAPSNP | TCSADGGYEQY | 0.094014645 |
| LVNNAAI | TCSADGGYEQY | 0.1787194 |
| LTGEINTLA | TCSADGGYEQY | 0.11679314 |
| LSIFGPV | TCSADGGYEQY | 0.12581076 |
| LLALLANL | TCSADGGYEQY | 0.085881576 |
| LISNASDALDKLR | TCSADGGYEQY | 0.23094475 |
| LINEAVP | TCSADGGYEQY | 0.090321764 |
| LGYAFSGL | TCSADGGYEQY | 0.100265466 |
| LDFVAVAL | TCSADGGYEQY | 0.08133369 |
| LAQPGAE | TCSADGGYEQY | 0.08484515 |
| ISNLGAVTGF | TCSADGGYEQY | 0.14034508 |
| HQEDPGRI | TCSADGGYEQY | 0.08159874 |
| GTPGEPTP | TCSADGGYEQY | 0.0813095 |
| GFFNSGS | TCSADGGYEQY | 0.09438615 |

| | | |
|---|---|---|
| GAGGGGLTAGAN | TCSADGGYEQY | 0.115015954 |
| FVDALVNM | TCSADGGYEQY | 0.07772668 |
| FLTVAPEL | TCSADGGYEQY | 0.08754603 |
| FETLTAL | TCSADGGYEQY | 0.08136328 |
| DVINPIEL | TCSADGGYEQY | 0.08404398 |
| APVSGAPV | TCSADGGYEQY | 0.101545915 |
| ALVGGLAW | TCSADGGYEQY | 0.081154935 |
| ALAAAAATAI | TCSADGGYEQY | 0.10127972 |
| AFGLGLLL | TCSADGGYEQY | 0.081892826 |
| AFFAAASL | TCSADGGYEQY | 0.08995461 |
| AAAALPAFSPP | TCSADGGYEQY | 0.10118541 |
| VVLAPAL | TCSADGGYEQY | 0.10296699 |
| TIGHVDHGKTTL | TCSADGGYEQY | 0.08246046 |
| LISNASDALDK | TCSADGGYEQY | 0.101168394 |
| LISNASDAL | TCSADGGYEQY | 0.083495244 |
| ISNASDALDK | TCSADGGYEQY | 0.20390737 |
| NQAETLVYQTE | TCSADGGYEQY | 0.10269725 |
| TGGTGGTAGAG | TCSADGGYEQY | 0.080301754 |
| TGSTAGGAGGA | TCSADGGYEQY | 0.11600035 |
| DPLAVPVAAGPV | TCSADGGYEQY | 0.11419082 |
| GGTGGTPGNGGN | TCSADGGYEQY | 0.099105954 |
| PAGLAGVDPATL | TCSADGGYEQY | 0.09397188 |
| GNNLVVEAPHAGASV | TCSADGGYEQY | 0.08033926 |
| KVTRIMSASTLEQPA | TCSADGGYEQY | 0.111289956 |
| LGFALPIALPRRTVV | TCSADGGYEQY | 0.22656135 |
| TRLGNLTARKLVVLV | TCSADGGYEQY | 0.12317066 |
| WSQQRRLIYFPSAGP | TCSADGGYEQY | 0.08456332 |
| GPNGNPDPMAAAVDIR | TCSADGGYEQY | 0.10133034 |
| TQRCVSVADLRAAASR | TCSADGGYEQY | 0.103458956 |
| QFVLDVAEHPVEVLTETV | TCSADGGYEQY | 0.094149604 |
| GGAGGTGGAGGNGGTQAGDG | TCSADGGYEQY | 0.10753818 |
| PLLATLRAIAVDVDAKPAQV | TCSADGGYEQY | 0.085520394 |
| APKFRTGGYMIDFWGVGYQVAK | TCSADGGYEQY | 0.1407963 |
| GAGGAGGAGGAAGGPGGTGGTG | TCSADGGYEQY | 0.10574468 |
| IGALGLLATDVAPAGSVAFGVA | TCSADGGYEQY | 0.091938935 |
| AGGAGGIGGNGGAGGAGGVLMG | TCSADGGYEQY | 0.082835525 |
| ADELIGLL | TCSADGGYEQY | 0.08258852 |
| ADRALKA | TCSADGGYEQY | 0.10679734 |
| AGAVGQVGGAGGA | TCSADGGYEQY | 0.10006619 |
| AGGGGGLGGAP | TCSADGGYEQY | 0.08049993 |
| AGGLLYGNGGAG | TCSADGGYEQY | 0.08957528 |
| AGLESIGL | TCSADGGYEQY | 0.10910602 |
| AGLKTGII | TCSADGGYEQY | 0.18033653 |
| ASLAALLL | TCSADGGYEQY | 0.08929833 |
| ASVINISSI | TCSADGGYEQY | 0.09255083 |
| ATPPGPA | TCSADGGYEQY | 0.19179761 |
| ATVASALV | TCSADGGYEQY | 0.0859951 |
| ATVASLIL | TCSADGGYEQY | 0.10223477 |
| AVNGPVLIRH | TCSADGGYEQY | 0.1086543 |
| DGAAGAVGI | TCSADGGYEQY | 0.12862423 |
| DPLLHGI | TCSADGGYEQY | 0.08652565 |

| | | |
|---|---|---|
| ELGAALGAL | TCSADGGYEQY | 0.1131584 |
| ELGRTQAV | TCSADGGYEQY | 0.08387062 |
| FVAGGYLS | TCSADGGYEQY | 0.22282484 |
| GASGGAGGAGGW | TCSADGGYEQY | 0.08574742 |
| GGAGAGGDGGAGGI | TCSADGGYEQY | 0.1103016 |
| GTGHNGSQPGMGGNG | TCSADGGYEQY | 0.08143099 |
| GYAAASASAS | TCSADGGYEQY | 0.2971734 |
| HQVDVAAVQQG | TCSADGGYEQY | 0.14582536 |
| IGAAIAHT | TCSADGGYEQY | 0.29927132 |
| IGMIEVAQL | TCSADGGYEQY | 0.081107855 |
| IRLAPEHTSAMD | TCSADGGYEQY | 0.11508796 |
| IYNVSVL | TCSADGGYEQY | 0.25724554 |
| KQGFLGM | TCSADGGYEQY | 0.23882763 |
| LESVITQL | TCSADGGYEQY | 0.089616805 |
| LGLGALVVV | TCSADGGYEQY | 0.11123505 |
| LIGNGGNGGSGGI | TCSADGGYEQY | 0.0929572 |
| LNAINAPTL | TCSADGGYEQY | 0.08573787 |
| LQLLNTGV | TCSADGGYEQY | 0.08202838 |
| LQTSIEF | TCSADGGYEQY | 0.08659498 |
| MSFAVSEL | TCSADGGYEQY | 0.08785634 |
| NTGALGL | TCSADGGYEQY | 0.09131839 |
| PPGAGGAGGAGTT | TCSADGGYEQY | 0.105261594 |
| QLVGLGI | TCSADGGYEQY | 0.1452909 |
| RFLEQVHQL | TCSADGGYEQY | 0.15029381 |
| RGVGYVLR | TCSADGGYEQY | 0.21937943 |
| SATGVFAGVI | TCSADGGYEQY | 0.3297365 |
| SLPAITLP | TCSADGGYEQY | 0.08916311 |
| STALAFTV | TCSADGGYEQY | 0.08610076 |
| SVLTVGLGK | TCSADGGYEQY | 0.09175755 |
| TALAGAGSGI | TCSADGGYEQY | 0.08466736 |
| TGLPFTGI | TCSADGGYEQY | 0.0911051 |
| VAEHVTV | TCSADGGYEQY | 0.08750412 |
| VDIVLNSL | TCSADGGYEQY | 0.11330044 |
| VLTGGGAVL | TCSADGGYEQY | 0.07716551 |
| VNVGLAYGI | TCSADGGYEQY | 0.10191815 |
| VSSAAVGAV | TCSADGGYEQY | 0.19554837 |
| VTSVYEAGV | TCSADGGYEQY | 0.07982521 |
| WVALDRGA | TCSADGGYEQY | 0.11153081 |
| YLVGSVVI | TCSADGGYEQY | 0.16066703 |
| GPPGTGKTLL | TCSADGGYEQY | 0.08347563 |
| GVGMTKF | TCSADGGYEQY | 0.15423627 |
| IGIGGSDL | TCSADGGYEQY | 0.086054385 |
| IGTLLAEL | TCSADGGYEQY | 0.09792151 |
| IYTSGTT | TCSADGGYEQY | 0.1452336 |
| LSFPAGV | TCSADGGYEQY | 0.13443358 |
| LYTSGTT | TCSADGGYEQY | 0.08708315 |
| VGAFPFGF | TCSADGGYEQY | 0.08769911 |
| VGAGTVEF | TCSADGGYEQY | 0.08333152 |
| VGKSTLLL | TCSADGGYEQY | 0.09615063 |
| AALAHAGGGIV | TCSADGGYEQY | 0.084879555 |
| AALVAVSLL | TCSADGGYEQY | 0.12117792 |

| | | |
|---|---|---|
| AAVSRAVAAL | TCSADGGYEQY | 0.08756187 |
| AGGSSGIGGF | TCSADGGYEQY | 0.13788989 |
| AGLTVANANG | TCSADGGYEQY | 0.10784329 |
| AGLVGAVL | TCSADGGYEQY | 0.09270708 |
| ALSQAFTV | TCSADGGYEQY | 0.09459165 |
| ALTFSVSL | TCSADGGYEQY | 0.091662966 |
| ALTGAALVAL | TCSADGGYEQY | 0.08568906 |
| ASFSIGAGVL | TCSADGGYEQY | 0.084018186 |
| DGSVYKI | TCSADGGYEQY | 0.09651385 |
| EQLAEIV | TCSADGGYEQY | 0.089478776 |
| FAVSELTVM | TCSADGGYEQY | 0.09236065 |
| GAAPLAAGL | TCSADGGYEQY | 0.1284351 |
| GADGGDGGSGGAGG | TCSADGGYEQY | 0.100913994 |
| GFGLDTGGAGG | TCSADGGYEQY | 0.08243245 |
| GLASGGLL | TCSADGGYEQY | 0.103683814 |
| GRIAGAAVLL | TCSADGGYEQY | 0.1204686 |
| GSAGSLNTSF | TCSADGGYEQY | 0.094032 |
| GTGQIGI | TCSADGGYEQY | 0.07782581 |
| GTLIALIL | TCSADGGYEQY | 0.09837399 |
| IAATMGFVVG | TCSADGGYEQY | 0.09454238 |
| IFAQALL | TCSADGGYEQY | 0.08549585 |
| IGAAVALGF | TCSADGGYEQY | 0.08488453 |
| IGLTPLAL | TCSADGGYEQY | 0.16797605 |
| ILGSNTSTL | TCSADGGYEQY | 0.082445025 |
| IVFSLTVL | TCSADGGYEQY | 0.08245399 |
| LASKATGF | TCSADGGYEQY | 0.11142637 |
| LASVALIL | TCSADGGYEQY | 0.1311479 |
| LFAVGLTL | TCSADGGYEQY | 0.1740933 |
| LFNGGIGTYI | TCSADGGYEQY | 0.10077142 |
| LTLGAGII | TCSADGGYEQY | 0.09082167 |
| LVAAALGDI | TCSADGGYEQY | 0.10530129 |
| NTAFVQLGI | TCSADGGYEQY | 0.09149168 |
| QLFSGAEL | TCSADGGYEQY | 0.08557694 |
| SAATFITL | TCSADGGYEQY | 0.07918653 |
| SNYGLANLGV | TCSADGGYEQY | 0.08867845 |
| SYGGAYAVM | TCSADGGYEQY | 0.08824654 |
| TTGVGGLTLG | TCSADGGYEQY | 0.10939164 |
| TTQTIYI | TCSADGGYEQY | 0.09717364 |
| VAAGAGLGAVI | TCSADGGYEQY | 0.08248317 |
| VAAVVTLL | TCSADGGYEQY | 0.08126888 |
| VGFLGGLI | TCSADGGYEQY | 0.09044217 |
| VGVIPILL | TCSADGGYEQY | 0.09131458 |
| VIFASSLI | TCSADGGYEQY | 0.11065891 |
| VKGGDGGAAAG | TCSADGGYEQY | 0.08862799 |
| VTKTFTNI | TCSADGGYEQY | 0.545663 |
| VVAAAILL | TCSADGGYEQY | 0.14095409 |
| YAALDVEL | TCSADGGYEQY | 0.09130855 |
| LASVALLL | TCSADGGYEQY | 0.14086103 |
| LDAVHRV | TCSADGGYEQY | 0.09048093 |
| QGLSSSIF | TCSADGGYEQY | 0.08472494 |
| RGTFANIRL | TCSADGGYEQY | 0.09869008 |

| | | |
|---|---|---|
| RQLWWGHRI | TCSADGGYEQY | 0.08541143 |
| YGPPGTGKTLL | TCSADGGYEQY | 0.08892608 |
| AALVGAPANVA | TCSADGGYEQY | 0.09067921 |
| AATFRRTE | TCSADGGYEQY | 0.08129169 |
| AGALGGGI | TCSADGGYEQY | 0.084000066 |
| AGGAGGAGGIGAG | TCSADGGYEQY | 0.08336862 |
| AGSGALGL | TCSADGGYEQY | 0.081817746 |
| AQIADQI | TCSADGGYEQY | 0.09176973 |
| ASAVAVTYL | TCSADGGYEQY | 0.12595022 |
| ASTVLGGTGG | TCSADGGYEQY | 0.16124254 |
| AVGGEPAK | TCSADGGYEQY | 0.088305764 |
| CEQGTFGGTC | TCSADGGYEQY | 0.08597078 |
| DEGKAPAG | TCSADGGYEQY | 0.15581836 |
| DTVVFPAL | TCSADGGYEQY | 0.09616811 |
| FEGITMAG | TCSADGGYEQY | 0.088775516 |
| FPLVVQD | TCSADGGYEQY | 0.09637896 |
| GAFTTPQ | TCSADGGYEQY | 0.096964985 |
| GFGGFGTAGGVGGA | TCSADGGYEQY | 0.082969196 |
| GFLAQFL | TCSADGGYEQY | 0.107400104 |
| GGAGGAGGNAGVA | TCSADGGYEQY | 0.08895008 |
| GGVGTAGAGGAGGAGG | TCSADGGYEQY | 0.08882633 |
| GNGGGGDAGNAGSGGN | TCSADGGYEQY | 0.077557646 |
| GTGGAGGAGNDAGS | TCSADGGYEQY | 0.088264555 |
| GTTNTGWFNS | TCSADGGYEQY | 0.09178598 |
| IFMGPLFTL | TCSADGGYEQY | 0.084377214 |
| ILAHAIEAI | TCSADGGYEQY | 0.07887811 |
| IRSAIREAL | TCSADGGYEQY | 0.09133205 |
| ITFAGGG | TCSADGGYEQY | 0.117818594 |
| ITGMSGHQEV | TCSADGGYEQY | 0.17934671 |
| IVAGLACTL | TCSADGGYEQY | 0.090329625 |
| IVGGGLAGA | TCSADGGYEQY | 0.13291675 |
| IVTGVLNI | TCSADGGYEQY | 0.11337954 |
| LAQQGCI | TCSADGGYEQY | 0.10841875 |
| LCLHVEN | TCSADGGYEQY | 0.0826061 |
| LDAISDGS | TCSADGGYEQY | 0.09681157 |
| LGGFAAML | TCSADGGYEQY | 0.07920697 |
| LGGKNPMI | TCSADGGYEQY | 0.08964756 |
| LGPKGVAAIL | TCSADGGYEQY | 0.096154034 |
| LMVGVSI | TCSADGGYEQY | 0.08297019 |
| LVGAAFVL | TCSADGGYEQY | 0.1022891 |
| LVLAVAPL | TCSADGGYEQY | 0.18478045 |
| MAAIAVIDAA | TCSADGGYEQY | 0.0841055 |
| NNGINAELQ | TCSADGGYEQY | 0.08393513 |
| RGEPLVLVHG | TCSADGGYEQY | 0.086342774 |
| RGGAGGAAGLI | TCSADGGYEQY | 0.08484501 |
| RSYRMHPAV | TCSADGGYEQY | 0.12049337 |
| SAAHLLAGQLA | TCSADGGYEQY | 0.102446415 |
| SAAVGSY | TCSADGGYEQY | 0.087359555 |
| SPHAVLHRI | TCSADGGYEQY | 0.10508836 |
| SQIAKVL | TCSADGGYEQY | 0.08954825 |
| TALGHLVP | TCSADGGYEQY | 0.092151135 |

| | | |
|---|---|---|
| TAPTTSGAA | TCSADGGYEQY | 0.096947566 |
| TDGDDGNGGSPGT | TCSADGGYEQY | 0.08742694 |
| TDGLGEN | TCSADGGYEQY | 0.11829184 |
| TGGPPLPGQA | TCSADGGYEQY | 0.08472765 |
| TIPAITI | TCSADGGYEQY | 0.08622729 |
| TIPGFTI | TCSADGGYEQY | 0.37768596 |
| TSIASGALA | TCSADGGYEQY | 0.08133524 |
| TVTLPGAAI | TCSADGGYEQY | 0.08730426 |
| VASGLSAALP | TCSADGGYEQY | 0.08984193 |
| VGAGVIGI | TCSADGGYEQY | 0.08149313 |
| VGPKRFGV | TCSADGGYEQY | 0.0970658 |
| VNVAFPD | TCSADGGYEQY | 0.10362175 |
| VSMTNSGVS | TCSADGGYEQY | 0.118610494 |
| VTDVLGDLSV | TCSADGGYEQY | 0.08209153 |
| VVVVGAGFAGL | TCSADGGYEQY | 0.23655269 |
| VYTVGTVV | TCSADGGYEQY | 0.11647673 |
| WTTFSGP | TCSADGGYEQY | 0.08680324 |
| YDELVEAF | TCSADGGYEQY | 0.0824443 |
| YGTGGAGGNGG | TCSADGGYEQY | 0.08296643 |
| YTNVMAV | TCSADGGYEQY | 0.11083454 |
| FLEQVHQL | TCSADGGYEQY | 0.105577074 |
| GLSLLAVAV | TCSADGGYEQY | 0.08676836 |
| LEQVHQL | TCSADGGYEQY | 0.11597805 |
| LNAAGAI | TCSADGGYEQY | 0.10926031 |
| PAVHASL | TCSADGGYEQY | 0.2427206 |
| TELEDDA | TCSADGGYEQY | 0.08365823 |
| TPPGPPN | TCSADGGYEQY | 0.09183143 |
| TPPPPPPP | TCSADGGYEQY | 0.09926274 |
| TQSPALI | TCSAGDRPDYT | 0.0865246 |
| EFAAAFPQ | TCSAGDRPDYT | 0.08850678 |
| LLSGNLDVL | TCSAGDRPDYT | 0.11640396 |
| GPANLGNYNI | TCSAGDRPDYT | 0.110579856 |
| TLGPTLP | TCSAGDRPDYT | 0.10370236 |
| VYAYSSCCK | TCSAGDRPDYT | 0.10976163 |
| APAPAPAPNI | TCSAGDRPDYT | 0.13486552 |
| PGGPVPIP | TCSAGDRPDYT | 0.0924376 |
| FLTVALEP | TCSAGDRPDYT | 0.09980465 |
| SQYFGANL | TCSAGDRPDYT | 0.080275424 |
| GGVKPAA | TCSAGDRPDYT | 0.07891962 |
| GGTGGAGGAGGW | TCSAGDRPDYT | 0.08244056 |
| GVGGVAGLGGAGG | TCSAGDRPDYT | 0.09038282 |
| VLLVLATTI | TCSAGDRPDYT | 0.07831509 |
| IGIFAAML | TCSAGDRPDYT | 0.106666535 |
| AVGDLID | TCSAGDRPDYT | 0.092008695 |
| NISPLPL | TCSAGDRPDYT | 0.120701656 |
| VTAVFVAI | TCSAGDRPDYT | 0.09150616 |
| SGDDNDAAL | TCSAGDRPDYT | 0.08718722 |
| AVAALAHS | TCSAGDRPDYT | 0.08759836 |
| GGVTVMATP | TCSAGDRPDYT | 0.12135724 |
| AGLAANVRD | TCSAGDRPDYT | 0.08375745 |
| TNRPDLIDPALLRPG | TCSAGDRPDYT | 0.081375726 |

| | | |
|---|---|---|
| LANLGVTAL | TCSAGDRPDYT | 0.08566739 |
| TGERIEL | TCSAGDRPDYT | 0.084520616 |
| NGEELAA | TCSAGDRPDYT | 0.26116663 |
| VLSLVWG | TCSAGDRPDYT | 0.14095657 |
| IGYALWL | TCSAGDRPDYT | 0.11646105 |
| LSLLLAPL | TCSAGDRPDYT | 0.101814605 |
| TLDEALK | TCSAGDRPDYT | 0.07848949 |
| TDGAGTAEGG | TCSAGDRPDYT | 0.09271529 |
| NAPGGAGGNGG | TCSAGDRPDYT | 0.23792565 |
| VSVFMGMGGNF | TCSAGDRPDYT | 0.09422212 |
| LASYLAAL | TCSAGDRPDYT | 0.12488152 |
| LVGGTPAL | TCSAGDRPDYT | 0.11219313 |
| VRNADVL | TCSAGDRPDYT | 0.0943526 |
| TCSSPFLVSAM | TCSAGDRPDYT | 0.09422978 |
| VHITDAE | TCSAGDRPDYT | 0.08762258 |
| EYAAPSF | TCSAGDRPDYT | 0.11575753 |
| VVDAFHGGL | TCSAGDRPDYT | 0.084554665 |
| ASVVDPQ | TCSAGDRPDYT | 0.08562878 |
| SYNEFGGLS | TCSAGDRPDYT | 0.09952742 |
| DPDGSVY | TCSAGDRPDYT | 0.6538621 |
| MVSTIHATE | TCSAGDRPDYT | 0.10974461 |
| LAAGVADPV | TCSAGDRPDYT | 0.09826584 |
| TKEQIAA | TCSAGDRPDYT | 0.09400713 |
| SLASPAAL | TCSAGDRPDYT | 0.08593024 |
| IGLVFYGV | TCSAGDRPDYT | 0.09545943 |
| FLLDASNL | TCSAGDRPDYT | 0.08743476 |
| YQVGNFP | TCSAGDRPDYT | 0.091906525 |
| ESLLSQST | TCSAGDRPDYT | 0.082569666 |
| LFTVGFAAVG | TCSAGDRPDYT | 0.089237325 |
| ATPPTPT | TCSAGDRPDYT | 0.09104862 |
| TDLATLG | TCSAGDRPDYT | 0.6118901 |
| SAGGTGR | TCSAGDRPDYT | 0.11560713 |
| GFSNSGS | TCSAGDRPDYT | 0.095774606 |
| GGNGGSGGTSV | TCSAGDRPDYT | 0.08442786 |
| GTSVATGGAGN | TCSAGDRPDYT | 0.08864509 |
| NGGNGGSGGTS | TCSAGDRPDYT | 0.1019935 |
| AVVKFPH | TCSAGDRPDYT | 0.08356324 |
| TPPPPPP | TCSAGDRPDYT | 0.14201026 |
| AAPAAAAAPA | TCSAGDRPDYT | 0.090315744 |
| IALNEDL | TCSAGDRPDYT | 0.08599256 |
| IVNVSSM | TCSAGDRPDYT | 0.10285983 |
| TGIPPAPR | TCSAGDRPDYT | 0.35522485 |
| FDLGGGTF | TCSAGDRPDYT | 0.097558565 |
| FDLGGGTFDVS | TCSAGDRPDYT | 0.08800747 |
| NEPTAAA | TCSAGDRPDYT | 0.086272426 |
| NEPTAAALA | TCSAGDRPDYT | 0.11036922 |
| GSIINISSI | TCSAGDRPDYT | 0.21700633 |
| VAPGNAGTA | TCSAGDRPDYT | 0.07997651 |
| TGGGASLE | TCSAGDRPDYT | 0.23122881 |
| VAVYNHYKRI | TCSAGDRPDYT | 0.083934695 |
| TDPAPATT | TCSAGDRPDYT | 0.08238222 |

| | | |
|---|---|---|
| TNRPDILDPALLRPG | TCSAGDRPDYT | 0.09149759 |
| LTTTTTS | TCSAGDRPDYT | 0.095089205 |
| PTGSGKTL | TCSAGDRPDYT | 0.2753883 |
| TGSGKTL | TCSAGDRPDYT | 0.10368123 |
| TIGHVDHGK | TCSAGDRPDYT | 0.08142559 |
| TIGHVDHGKTT | TCSAGDRPDYT | 0.09264949 |
| GHVDHGKTT | TCSAGDRPDYT | 0.17214604 |
| AQFEHTLL | TCSAGDRPDYT | 0.101982065 |
| IVNQDEL | TCSAGDRPDYT | 0.10228038 |
| TNGSQFF | TCSAGDRPDYT | 0.0861011 |
| TGKGIVE | TCSAGDRPDYT | 0.084207855 |
| LISNASDA | TCSAGDRPDYT | 0.114176996 |
| AELRTLGELTKIAKAHGAQ | TCSAGDRPDYT | 0.09297394 |
| APNAVAAAEPSW | TCSAGDRPDYT | 0.08172549 |
| APQVAGAATAVL | TCSAGDRPDYT | 0.10181895 |
| ARRDPAAAAAILPTDA | TCSAGDRPDYT | 0.11756607 |
| DAAARDVLAEAGLAEYF | TCSAGDRPDYT | 0.082869776 |
| DDQIVLSAKANLAKVEQ | TCSAGDRPDYT | 0.09609159 |
| DELLPDTGEDSDEDGH | TCSAGDRPDYT | 0.18003345 |
| ELPEAPANRATVPQD | TCSAGDRPDYT | 0.09321853 |
| FGQSKIERVTGIG | TCSAGDRPDYT | 0.081815384 |
| GAGGTGSTAGGAGGAGGAGGLY | TCSAGDRPDYT | 0.16199894 |
| GHGGTGAAGGNGGTGGAGGSGI | TCSAGDRPDYT | 0.688131 |
| GNGGNGGAGGEGGPGGAGGAGA | TCSAGDRPDYT | 0.117059566 |
| GSTAGNGGNGGNATNGTIAGQP | TCSAGDRPDYT | 0.08198743 |
| GTGGDGGNGGNGGTGG | TCSAGDRPDYT | 0.07849711 |
| GVRRPAGIPADAPV | TCSAGDRPDYT | 0.18251245 |
| GYRQVTDAHLLTLARR | TCSAGDRPDYT | 0.08981578 |
| IAKMGQDGHDRG | TCSAGDRPDYT | 0.09025787 |
| KAEPSIEIQYT | TCSAGDRPDYT | 0.08292537 |
| KPVRIRMAADEDTVAVT | TCSAGDRPDYT | 0.09305034 |
| MPPRIAGMRLLVIKPEP | TCSAGDRPDYT | 0.12942517 |
| MKFYAVALFRSRGVPPTR | TCSAGDRPDYT | 0.08717802 |
| NAGGFGILGAVAHSP | TCSAGDRPDYT | 0.08148285 |
| NEGREITLAMAEM | TCSAGDRPDYT | 0.08787147 |
| NELVEWSHPLGHRWRWE | TCSAGDRPDYT | 0.110036954 |
| NLGAGGTGGNGGNPG | TCSAGDRPDYT | 0.08856077 |
| NTEWPKWETVMAADPVD | TCSAGDRPDYT | 0.09811795 |
| PASAAPAETQP | TCSAGDRPDYT | 0.15941362 |
| QQSSDGKNGVRAAEITGE | TCSAGDRPDYT | 0.090739965 |
| RAPDAVTTSWWKE | TCSAGDRPDYT | 0.10330837 |
| SGFGNVGAHTSGWFNQ | TCSAGDRPDYT | 0.08586734 |
| SGVAGAVGQAASVGGLKV | TCSAGDRPDYT | 0.19927739 |
| SNGGAGTGSAGGNGGT | TCSAGDRPDYT | 0.089915015 |
| TGGAGGEGGAAG | TCSAGDRPDYT | 0.0956223 |
| TGGDGGAAGAGGAPGHG | TCSAGDRPDYT | 0.10928325 |
| TMPPLLAAIANAGRHGV | TCSAGDRPDYT | 0.11777629 |
| TSSTAGNSGGA | TCSAGDRPDYT | 0.106126234 |
| VPATVPVGILC | TCSAGDRPDYT | 0.08596551 |
| VQRPDSGGVVLTGRIS | TCSAGDRPDYT | 0.08472536 |
| WTPNPVPATKGVPAGLVQAT | TCSAGDRPDYT | 0.07859009 |

| | | |
|---|---|---|
| YALKRALAKFSDRFDVVII | TCSAGDRPDYT | 0.13767512 |
| YLIGSVIV | TCSAGDRPDYT | 0.08166494 |
| YGQLVAVID | TCSAGDRPDYT | 0.17679594 |
| VTVGGMTLPA | TCSAGDRPDYT | 0.114797205 |
| VLALPAV | TCSAGDRPDYT | 0.092192575 |
| VIPPTLNYET | TCSAGDRPDYT | 0.09740612 |
| VGMGGMHPSSG | TCSAGDRPDYT | 0.07974681 |
| VGLAGGAATA | TCSAGDRPDYT | 0.09250512 |
| VGGAGGAGGSGP | TCSAGDRPDYT | 0.09304349 |
| VFAAFMV | TCSAGDRPDYT | 0.092124775 |
| TLYDKLR | TCSAGDRPDYT | 0.0920881 |
| TLAVFTVL | TCSAGDRPDYT | 0.09313973 |
| TGSVQPARV | TCSAGDRPDYT | 0.13287996 |
| TGGGAGSGGT | TCSAGDRPDYT | 0.085192434 |
| SHLETSL | TCSAGDRPDYT | 0.09210048 |
| SFTTGMT | TCSAGDRPDYT | 0.20605546 |
| QVEVAKRMGVRQPTVS | TCSAGDRPDYT | 0.097738914 |
| QGSGGPALAM | TCSAGDRPDYT | 0.083930574 |
| MLAPSNP | TCSAGDRPDYT | 0.11228676 |
| LVNNAAI | TCSAGDRPDYT | 0.08568216 |
| LTGEINTLA | TCSAGDRPDYT | 0.12205969 |
| LSIFGPV | TCSAGDRPDYT | 0.100189306 |
| LLALLANL | TCSAGDRPDYT | 0.09919499 |
| LISNASDALDKLR | TCSAGDRPDYT | 0.11257279 |
| LINEAVP | TCSAGDRPDYT | 0.080318175 |
| LGYAFSGL | TCSAGDRPDYT | 0.086705774 |
| LDFVAVAL | TCSAGDRPDYT | 0.09323957 |
| LAQPGAE | TCSAGDRPDYT | 0.122363016 |
| ISNLGAVTGF | TCSAGDRPDYT | 0.1032074 |
| HQEDPGRI | TCSAGDRPDYT | 0.0853246 |
| GTPGEPTP | TCSAGDRPDYT | 0.109712005 |
| GFFNSGS | TCSAGDRPDYT | 0.09615771 |
| GAGGGGLTAGAN | TCSAGDRPDYT | 0.090553194 |
| FVDALVNM | TCSAGDRPDYT | 0.0902118 |
| FLTVAPEL | TCSAGDRPDYT | 0.085911155 |
| FETLTAL | TCSAGDRPDYT | 0.08220678 |
| DVINPIEL | TCSAGDRPDYT | 0.12108862 |
| APVSGAPV | TCSAGDRPDYT | 0.08521974 |
| ALVGGLAW | TCSAGDRPDYT | 0.1062854 |
| ALAAAAATAI | TCSAGDRPDYT | 0.098142244 |
| AFGLGLLL | TCSAGDRPDYT | 0.08416163 |
| AFFAAASL | TCSAGDRPDYT | 0.09471619 |
| AAAALPAFSPP | TCSAGDRPDYT | 0.08678519 |
| VVLAPAL | TCSAGDRPDYT | 0.11615106 |
| TIGHVDHGKTTL | TCSAGDRPDYT | 0.09532114 |
| LISNASDALDK | TCSAGDRPDYT | 0.086737216 |
| LISNASDAL | TCSAGDRPDYT | 0.10258152 |
| ISNASDALDK | TCSAGDRPDYT | 0.12833169 |
| NQAETLVYQTE | TCSAGDRPDYT | 0.09429415 |
| TGGTGGTAGAG | TCSAGDRPDYT | 0.090904385 |
| TGSTAGGAGGA | TCSAGDRPDYT | 0.0935126 |

| | | |
|---|---|---|
| DPLAVPVAAGPV | TCSAGDRPDYT | 0.11485065 |
| GGTGGTPGNGGN | TCSAGDRPDYT | 0.090905264 |
| PAGLAGVDPATL | TCSAGDRPDYT | 0.07980212 |
| GNNLVVEAPHAGASV | TCSAGDRPDYT | 0.09088052 |
| KVTRIMSASTLEQPA | TCSAGDRPDYT | 0.09141383 |
| LGFALPIALPRRTVV | TCSAGDRPDYT | 0.08633627 |
| TRLGNLTARKLVVLV | TCSAGDRPDYT | 0.078982994 |
| WSQQRRLIYFPSAGP | TCSAGDRPDYT | 0.19562194 |
| GPNGNPDPMAAAVDIR | TCSAGDRPDYT | 0.20517293 |
| TQRCVSVADLRAAASR | TCSAGDRPDYT | 0.08586469 |
| QFVLDVAEHPVEVLTETV | TCSAGDRPDYT | 0.12704344 |
| GGAGGTGGAGGNGGTQAGDG | TCSAGDRPDYT | 0.08105715 |
| PLLATLRAIAVDVDAKPAQV | TCSAGDRPDYT | 0.10333602 |
| APKFRTGGYMIDFWGVGYQVAK | TCSAGDRPDYT | 0.17186928 |
| GAGGAGGAGGAAGGPGGTGGTG | TCSAGDRPDYT | 0.08541167 |
| IGALGLLATDVAPAGSVAFGVA | TCSAGDRPDYT | 0.10763908 |
| AGGAGGIGGNGGAGGAGGVLMG | TCSAGDRPDYT | 0.08686596 |
| ADELIGLL | TCSAGDRPDYT | 0.086831465 |
| ADRALKA | TCSAGDRPDYT | 0.14442042 |
| AGAVGQVGGAGGA | TCSAGDRPDYT | 0.093110174 |
| AGGGGGLGGAP | TCSAGDRPDYT | 0.11014855 |
| AGGLLYGNGGAG | TCSAGDRPDYT | 0.086464085 |
| AGLESIGL | TCSAGDRPDYT | 0.08703189 |
| AGLKTGII | TCSAGDRPDYT | 0.106113836 |
| ASLAALLL | TCSAGDRPDYT | 0.109028146 |
| ASVINISSI | TCSAGDRPDYT | 0.08952651 |
| ATPPGPA | TCSAGDRPDYT | 0.10094396 |
| ATVASALV | TCSAGDRPDYT | 0.09493334 |
| ATVASLIL | TCSAGDRPDYT | 0.43453678 |
| AVNGPVLIRH | TCSAGDRPDYT | 0.09960292 |
| DGAAGAVGI | TCSAGDRPDYT | 0.12914638 |
| DPLLHGI | TCSAGDRPDYT | 0.08918793 |
| ELGAALGAL | TCSAGDRPDYT | 0.19337633 |
| ELGRTQAV | TCSAGDRPDYT | 0.0934832 |
| FVAGGYLS | TCSAGDRPDYT | 0.096214294 |
| GASGGAGGAGGW | TCSAGDRPDYT | 0.10275508 |
| GGAGAGGDGGAGGI | TCSAGDRPDYT | 0.08781756 |
| GTGHNGSQPGMGGNG | TCSAGDRPDYT | 0.2739144 |
| GYAAASASAS | TCSAGDRPDYT | 0.11813023 |
| HQVDVAAVQQG | TCSAGDRPDYT | 0.11928801 |
| IGAAIAHT | TCSAGDRPDYT | 0.11315766 |
| IGMIEVAQL | TCSAGDRPDYT | 0.083024345 |
| IRLAPEHTSAMD | TCSAGDRPDYT | 0.10529319 |
| IYNVSVL | TCSAGDRPDYT | 0.09709338 |
| KQGFLGM | TCSAGDRPDYT | 0.08448999 |
| LESVITQL | TCSAGDRPDYT | 0.105295636 |
| LGLGALVVV | TCSAGDRPDYT | 0.11570536 |
| LIGNGGNGGSGGI | TCSAGDRPDYT | 0.3983176 |
| LNAINAPTL | TCSAGDRPDYT | 0.16740242 |
| LQLLNTGV | TCSAGDRPDYT | 0.09142836 |
| LQTSIEF | TCSAGDRPDYT | 0.115891464 |

| | | |
|---|---|---|
| MSFAVSEL | TCSAGDRPDYT | 0.10099685 |
| NTGALGL | TCSAGDRPDYT | 0.101592265 |
| PPGAGGAGGAGTT | TCSAGDRPDYT | 0.0794979 |
| QLVGLGI | TCSAGDRPDYT | 0.13848248 |
| RFLEQVHQL | TCSAGDRPDYT | 0.09051754 |
| RGVGYVLR | TCSAGDRPDYT | 0.07791701 |
| SATGVFAGVI | TCSAGDRPDYT | 0.11544206 |
| SLPAITLP | TCSAGDRPDYT | 0.102825396 |
| STALAFTV | TCSAGDRPDYT | 0.09149786 |
| SVLTVGLGK | TCSAGDRPDYT | 0.25915915 |
| TALAGAGSGI | TCSAGDRPDYT | 0.07694405 |
| TGLPFTGI | TCSAGDRPDYT | 0.10214542 |
| VAEHVTV | TCSAGDRPDYT | 0.28027093 |
| VDIVLNSL | TCSAGDRPDYT | 0.2587378 |
| VLTGGGAVL | TCSAGDRPDYT | 0.095161214 |
| VNVGLAYGI | TCSAGDRPDYT | 0.13398868 |
| VSSAAVGAV | TCSAGDRPDYT | 0.49290714 |
| VTSVYEAGV | TCSAGDRPDYT | 0.13830128 |
| WVALDRGA | TCSAGDRPDYT | 0.13282529 |
| YLVGSVVI | TCSAGDRPDYT | 0.15314373 |
| GPPGTGKTLL | TCSAGDRPDYT | 0.087799996 |
| GVGMTKF | TCSAGDRPDYT | 0.1495437 |
| IGIGGSDL | TCSAGDRPDYT | 0.16342317 |
| IGTLLAEL | TCSAGDRPDYT | 0.13112003 |
| IYTSGTT | TCSAGDRPDYT | 0.14235458 |
| LSFPAGV | TCSAGDRPDYT | 0.09069034 |
| LYTSGTT | TCSAGDRPDYT | 0.09241733 |
| VGAFPFGF | TCSAGDRPDYT | 0.08160341 |
| VGAGTVEF | TCSAGDRPDYT | 0.19906306 |
| VGKSTLLL | TCSAGDRPDYT | 0.1613869 |
| AALAHAGGGIV | TCSAGDRPDYT | 0.14333744 |
| AALVAVSLL | TCSAGDRPDYT | 0.08344924 |
| AAVSRAVAAL | TCSAGDRPDYT | 0.09743878 |
| AGGSSGIGGF | TCSAGDRPDYT | 0.08364845 |
| AGLTVANANG | TCSAGDRPDYT | 0.092150815 |
| AGLVGAVL | TCSAGDRPDYT | 0.09223824 |
| ALSQAFTV | TCSAGDRPDYT | 0.08089699 |
| ALTFSVSL | TCSAGDRPDYT | 0.2749107 |
| ALTGAALVAL | TCSAGDRPDYT | 0.2850096 |
| ASFSIGAGVL | TCSAGDRPDYT | 0.12181169 |
| DGSVYKI | TCSAGDRPDYT | 0.13880453 |
| EQLAEIV | TCSAGDRPDYT | 0.16436915 |
| FAVSELTVM | TCSAGDRPDYT | 0.24193725 |
| GAAPLAAGL | TCSAGDRPDYT | 0.13489835 |
| GADGGDGGSGGAGG | TCSAGDRPDYT | 0.0869008 |
| GFGLDTGGAGG | TCSAGDRPDYT | 0.1782955 |
| GLASGGLL | TCSAGDRPDYT | 0.1430632 |
| GRIAGAAVLL | TCSAGDRPDYT | 0.09742783 |
| GSAGSLNTSF | TCSAGDRPDYT | 0.1799765 |
| GTGQIGI | TCSAGDRPDYT | 0.10150409 |
| GTLIALIL | TCSAGDRPDYT | 0.10640465 |

| | | |
|---|---|---|
| IAATMGFVVG | TCSAGDRPDYT | 0.186784 |
| IFAQALL | TCSAGDRPDYT | 0.15611303 |
| IGAAVALGF | TCSAGDRPDYT | 0.09158067 |
| IGLTPLAL | TCSAGDRPDYT | 0.09095745 |
| ILGSNTSTL | TCSAGDRPDYT | 0.10770332 |
| IVFSLTVL | TCSAGDRPDYT | 0.19358775 |
| LASKATGF | TCSAGDRPDYT | 0.23371607 |
| LASVALIL | TCSAGDRPDYT | 0.13042139 |
| LFAVGLTL | TCSAGDRPDYT | 0.10808976 |
| LFNGGIGTYI | TCSAGDRPDYT | 0.2289546 |
| LTLGAGII | TCSAGDRPDYT | 0.15640806 |
| LVAAALGDI | TCSAGDRPDYT | 0.12481855 |
| NTAFVQLGI | TCSAGDRPDYT | 0.11736638 |
| QLFSGAEL | TCSAGDRPDYT | 0.09313615 |
| SAATFITL | TCSAGDRPDYT | 0.16331196 |
| SNYGLANLGV | TCSAGDRPDYT | 0.19247532 |
| SYGGAYAVM | TCSAGDRPDYT | 0.121017374 |
| TTGVGGLTLG | TCSAGDRPDYT | 0.12803823 |
| TTQTIYI | TCSAGDRPDYT | 0.1231565 |
| VAAGAGLGAVI | TCSAGDRPDYT | 0.124899365 |
| VAAVVTLL | TCSAGDRPDYT | 0.118882954 |
| VGFLGGLI | TCSAGDRPDYT | 0.10119011 |
| VGVIPILL | TCSAGDRPDYT | 0.105490014 |
| VIFASSLI | TCSAGDRPDYT | 0.17401437 |
| VKGGDGGAAAG | TCSAGDRPDYT | 0.09885436 |
| VTKTFTNI | TCSAGDRPDYT | 0.1315963 |
| VVAAAILL | TCSAGDRPDYT | 0.10208884 |
| YAALDVEL | TCSAGDRPDYT | 0.1340635 |
| LASVALLL | TCSAGDRPDYT | 0.23586687 |
| LDAVHRV | TCSAGDRPDYT | 0.09201811 |
| QGLSSSIF | TCSAGDRPDYT | 0.09844301 |
| RGTFANIRL | TCSAGDRPDYT | 0.16487035 |
| RQLWWGHRI | TCSAGDRPDYT | 0.13934231 |
| YGPPGTGKTLL | TCSAGDRPDYT | 0.14848298 |
| AALVGAPANVA | TCSAGDRPDYT | 0.09041036 |
| AATFRRTE | TCSAGDRPDYT | 0.19599295 |
| AGALGGGI | TCSAGDRPDYT | 0.089880615 |
| AGGAGGAGGIGAG | TCSAGDRPDYT | 0.3580696 |
| AGSGALGL | TCSAGDRPDYT | 0.09977469 |
| AQIADQI | TCSAGDRPDYT | 0.096388794 |
| ASAVAVTYL | TCSAGDRPDYT | 0.10018981 |
| ASTVLGGTGG | TCSAGDRPDYT | 0.07861816 |
| AVGGEPAK | TCSAGDRPDYT | 0.20126998 |
| CEQGTFGGTC | TCSAGDRPDYT | 0.08590315 |
| DEGKAPAG | TCSAGDRPDYT | 0.14794302 |
| DTVVFPAL | TCSAGDRPDYT | 0.10607856 |
| FEGITMAG | TCSAGDRPDYT | 0.11730806 |
| FPLVVQD | TCSAGDRPDYT | 0.13819903 |
| GAFTTPQ | TCSAGDRPDYT | 0.0963287 |
| GFGGFGTAGGVGGA | TCSAGDRPDYT | 0.08946167 |
| GFLAQFL | TCSAGDRPDYT | 0.17607054 |

| | | |
|---|---|---|
| GGAGGAGGNAGVA | TCSAGDRPDYT | 0.12439629 |
| GGVGTAGAGGAGGAGG | TCSAGDRPDYT | 0.12380133 |
| GNGGGGDAGNAGSGGN | TCSAGDRPDYT | 0.090243824 |
| GTGGAGGAGNDAGS | TCSAGDRPDYT | 0.10013062 |
| GTTNTGWFNS | TCSAGDRPDYT | 0.13787445 |
| IFMGPLFTL | TCSAGDRPDYT | 0.07885201 |
| ILAHAIEAI | TCSAGDRPDYT | 0.07967293 |
| IRSAIREAL | TCSAGDRPDYT | 0.07952824 |
| ITFAGGG | TCSAGDRPDYT | 0.08149631 |
| ITGMSGHQEV | TCSAGDRPDYT | 0.36125785 |
| IVAGLACTL | TCSAGDRPDYT | 0.4430596 |
| IVGGGLAGA | TCSAGDRPDYT | 0.121176735 |
| IVTGVLNI | TCSAGDRPDYT | 0.2565481 |
| LAQQGCI | TCSAGDRPDYT | 0.15119459 |
| LCLHVEN | TCSAGDRPDYT | 0.07594318 |
| LDAISDGS | TCSAGDRPDYT | 0.10315552 |
| LGGFAAML | TCSAGDRPDYT | 0.08911093 |
| LGGKNPMI | TCSAGDRPDYT | 0.14615242 |
| LGPKGVAAIL | TCSAGDRPDYT | 0.08215219 |
| LMVGVSI | TCSAGDRPDYT | 0.10973274 |
| LVGAAFVL | TCSAGDRPDYT | 0.16937949 |
| LVLAVAPL | TCSAGDRPDYT | 0.12448685 |
| MAAIAVIDAA | TCSAGDRPDYT | 0.172879 |
| NNGINAELQ | TCSAGDRPDYT | 0.094669536 |
| RGEPLVLVHG | TCSAGDRPDYT | 0.0786873 |
| RGGAGGAAGLI | TCSAGDRPDYT | 0.19495958 |
| RSYRMHPAV | TCSAGDRPDYT | 0.08313223 |
| SAAHLLAGQLA | TCSAGDRPDYT | 0.09813793 |
| SAAVGSY | TCSAGDRPDYT | 0.09159227 |
| SPHAVLHRI | TCSAGDRPDYT | 0.07620937 |
| SQIAKVL | TCSAGDRPDYT | 0.08737715 |
| TALGHLVP | TCSAGDRPDYT | 0.12880565 |
| TAPTTSGAA | TCSAGDRPDYT | 0.091853395 |
| TDGDDGNGGSPGT | TCSAGDRPDYT | 0.13568434 |
| TDGLGEN | TCSAGDRPDYT | 0.22990605 |
| TGGPPLPGQA | TCSAGDRPDYT | 0.23439297 |
| TIPAITI | TCSAGDRPDYT | 0.12656656 |
| TIPGFTI | TCSAGDRPDYT | 0.1095618 |
| TSIASGALA | TCSAGDRPDYT | 0.09329976 |
| TVTLPGAAI | TCSAGDRPDYT | 0.09487759 |
| VASGLSAALP | TCSAGDRPDYT | 0.09819067 |
| VGAGVIGI | TCSAGDRPDYT | 0.24819142 |
| VGPKRFGV | TCSAGDRPDYT | 0.08573126 |
| VNVAFPD | TCSAGDRPDYT | 0.08563995 |
| VSMTNSGVS | TCSAGDRPDYT | 0.09474871 |
| VTDVLGDLSV | TCSAGDRPDYT | 0.25068817 |
| VVVVGAGFAGL | TCSAGDRPDYT | 0.28348726 |
| VYTVGTVV | TCSAGDRPDYT | 0.09269099 |
| WTTFSGP | TCSAGDRPDYT | 0.12052186 |
| YDELVEAF | TCSAGDRPDYT | 0.097383596 |
| YGTGGAGGNGG | TCSAGDRPDYT | 0.16041678 |

| | | |
|---|---|---|
| YTNVMAV | TCSAGDRPDYT | 0.08656596 |
| FLEQVHQL | TCSAGDRPDYT | 0.09626874 |
| GLSLLAVAV | TCSAGDRPDYT | 0.17580748 |
| LEQVHQL | TCSAGDRPDYT | 0.51099885 |
| LNAAGAI | TCSAGDRPDYT | 0.10434483 |
| PAVHASL | TCSAGDRPDYT | 0.08782661 |
| TELEDDA | TCSAGDRPDYT | 0.16186602 |
| TPPGPPN | TCSAGDRPDYT | 0.09943385 |
| TPPPPPPP | TCSAGDRPDYT | 0.12874009 |
| TQSPALI | ASSLRSNQDTQY | 0.1352105 |
| EFAAAFPQ | ASSLRSNQDTQY | 0.08322601 |
| LLSGNLDVL | ASSLRSNQDTQY | 0.09451891 |
| GPANLGNYNI | ASSLRSNQDTQY | 0.3499289 |
| TLGPTLP | ASSLRSNQDTQY | 0.11999528 |
| VYAYSSCCK | ASSLRSNQDTQY | 0.09471763 |
| APAPAPAPNI | ASSLRSNQDTQY | 0.30166426 |
| PGGPVPIP | ASSLRSNQDTQY | 0.17407984 |
| FLTVALEP | ASSLRSNQDTQY | 0.33155358 |
| SQYFGANL | ASSLRSNQDTQY | 0.12326194 |
| GGVKPAA | ASSLRSNQDTQY | 0.101191156 |
| GGTGGAGGAGGW | ASSLRSNQDTQY | 0.1169279 |
| GVGGVAGLGGAGG | ASSLRSNQDTQY | 0.087457635 |
| VLLVLATTI | ASSLRSNQDTQY | 0.16373551 |
| IGIFAAML | ASSLRSNQDTQY | 0.17940542 |
| AVGDLID | ASSLRSNQDTQY | 0.095144354 |
| NISPLPL | ASSLRSNQDTQY | 0.40249377 |
| VTAVFVAI | ASSLRSNQDTQY | 0.1649013 |
| SGDDNDAAL | ASSLRSNQDTQY | 0.10213996 |
| AVAALAHS | ASSLRSNQDTQY | 0.08269726 |
| GGVTVMATP | ASSLRSNQDTQY | 0.09430787 |
| AGLAANVRD | ASSLRSNQDTQY | 0.19225293 |
| TNRPDLIDPALLRPG | ASSLRSNQDTQY | 0.10169224 |
| LANLGVTAL | ASSLRSNQDTQY | 0.087646574 |
| TGERIEL | ASSLRSNQDTQY | 0.13897252 |
| NGEELAA | ASSLRSNQDTQY | 0.2905513 |
| VLSLVWG | ASSLRSNQDTQY | 0.07801151 |
| IGYALWL | ASSLRSNQDTQY | 0.105191365 |
| LSLLLAPL | ASSLRSNQDTQY | 0.08324974 |
| TLDEALK | ASSLRSNQDTQY | 0.08950388 |
| TDGAGTAEGG | ASSLRSNQDTQY | 0.12868607 |
| NAPGGAGGNGG | ASSLRSNQDTQY | 0.08583937 |
| VSVFMGMGGNF | ASSLRSNQDTQY | 0.11804778 |
| LASYLAAL | ASSLRSNQDTQY | 0.0859447 |
| LVGGTPAL | ASSLRSNQDTQY | 0.08147014 |
| VRNADVL | ASSLRSNQDTQY | 0.12619984 |
| TCSSPFLVSAM | ASSLRSNQDTQY | 0.14682882 |
| VHITDAE | ASSLRSNQDTQY | 0.08525287 |
| EYAAPSF | ASSLRSNQDTQY | 0.1773033 |
| VVDAFHGGL | ASSLRSNQDTQY | 0.09737455 |
| ASVVDPQ | ASSLRSNQDTQY | 0.28639752 |
| SYNEFGGLS | ASSLRSNQDTQY | 0.11934008 |

| | | |
|---|---|---|
| DPDGSVY | ASSLRSNQDTQY | 0.082629494 |
| MVSTIHATE | ASSLRSNQDTQY | 0.1582761 |
| LAAGVADPV | ASSLRSNQDTQY | 0.08858297 |
| TKEQIAA | ASSLRSNQDTQY | 0.12680106 |
| SLASPAAL | ASSLRSNQDTQY | 0.16835353 |
| IGLVFYGV | ASSLRSNQDTQY | 0.083745666 |
| FLLDASNL | ASSLRSNQDTQY | 0.12765662 |
| YQVGNFP | ASSLRSNQDTQY | 0.24288929 |
| ESLLSQST | ASSLRSNQDTQY | 0.11879535 |
| LFTVGFAAVG | ASSLRSNQDTQY | 0.08753495 |
| ATPPTPT | ASSLRSNQDTQY | 0.10377003 |
| TDLATLG | ASSLRSNQDTQY | 0.14350289 |
| SAGGTGR | ASSLRSNQDTQY | 0.09935417 |
| GFSNSGS | ASSLRSNQDTQY | 0.2083561 |
| GGNGGSGGTSV | ASSLRSNQDTQY | 0.105406396 |
| GTSVATGGAGN | ASSLRSNQDTQY | 0.2014398 |
| NGGNGGSGGTS | ASSLRSNQDTQY | 0.13698821 |
| AVVKFPH | ASSLRSNQDTQY | 0.081783935 |
| TPPPPPP | ASSLRSNQDTQY | 0.08592621 |
| AAPAAAAAPA | ASSLRSNQDTQY | 0.08532003 |
| IALNEDL | ASSLRSNQDTQY | 0.14679188 |
| IVNVSSM | ASSLRSNQDTQY | 0.13357385 |
| TGIPPAPR | ASSLRSNQDTQY | 0.08207773 |
| FDLGGGTF | ASSLRSNQDTQY | 0.08695496 |
| FDLGGGTFDVS | ASSLRSNQDTQY | 0.10996856 |
| NEPTAAA | ASSLRSNQDTQY | 0.26048487 |
| NEPTAAALA | ASSLRSNQDTQY | 0.10439707 |
| GSIINISSI | ASSLRSNQDTQY | 0.104520805 |
| VAPGNAGTA | ASSLRSNQDTQY | 0.5214319 |
| TGGGASLE | ASSLRSNQDTQY | 0.086288765 |
| VAVYNHYKRI | ASSLRSNQDTQY | 0.12455704 |
| TDPAPATT | ASSLRSNQDTQY | 0.1637684 |
| TNRPDILDPALLRPG | ASSLRSNQDTQY | 0.153548 |
| LTTTTTS | ASSLRSNQDTQY | 0.10384959 |
| PTGSGKTL | ASSLRSNQDTQY | 0.16118863 |
| TGSGKTL | ASSLRSNQDTQY | 0.08797317 |
| TIGHVDHGK | ASSLRSNQDTQY | 0.23495284 |
| TIGHVDHGKTT | ASSLRSNQDTQY | 0.08170122 |
| GHVDHGKTT | ASSLRSNQDTQY | 0.17178774 |
| AQFEHTLL | ASSLRSNQDTQY | 0.08071724 |
| IVNQDEL | ASSLRSNQDTQY | 0.2739929 |
| TNGSQFF | ASSLRSNQDTQY | 0.3875192 |
| TGKGIVE | ASSLRSNQDTQY | 0.25651014 |
| LISNASDA | ASSLRSNQDTQY | 0.07999661 |
| AELRTLGELTKIAKAHGAQ | ASSLRSNQDTQY | 0.23144628 |
| APNAVAAAEPSW | ASSLRSNQDTQY | 0.37428367 |
| APQVAGAATAVL | ASSLRSNQDTQY | 0.4837704 |
| ARRDPAAAAAILPTDA | ASSLRSNQDTQY | 0.13143727 |
| DAAARDVLAEAGLAEYF | ASSLRSNQDTQY | 0.142016 |
| DDQIVLSAKANLAKVEQ | ASSLRSNQDTQY | 0.1220808 |
| DELLPDTGEDSDEDGH | ASSLRSNQDTQY | 0.09972093 |

| | | |
|---|---|---|
| ELPEAPANRATVPQD | ASSLRSNQDTQY | 0.096020564 |
| FGQSKIERVTGIG | ASSLRSNQDTQY | 0.092032656 |
| GAGGTGSTAGGAGGAGGAGGLY | ASSLRSNQDTQY | 0.15158212 |
| GHGGTGAAGGNGGTGGAGGSGI | ASSLRSNQDTQY | 0.10026974 |
| GNGGNGGAGGEGGPGGAGGAGA | ASSLRSNQDTQY | 0.24742831 |
| GSTAGNGGNGGNATNGTIAGQP | ASSLRSNQDTQY | 0.18040876 |
| GTGGDGGNGGNGGTGG | ASSLRSNQDTQY | 0.18661425 |
| GVRRPAGIPADAPV | ASSLRSNQDTQY | 0.52659523 |
| GYRQVTDAHLLTLARR | ASSLRSNQDTQY | 0.1541566 |
| IAKMGQDGHDRG | ASSLRSNQDTQY | 0.09856293 |
| KAEPSIEIQYT | ASSLRSNQDTQY | 0.098434635 |
| KPVRIRMAADEDTVAVT | ASSLRSNQDTQY | 0.1041554 |
| MPPRIAGMRLLVIKPEP | ASSLRSNQDTQY | 0.0908613 |
| MKFYAVALFRSRGVPPTR | ASSLRSNQDTQY | 0.10144899 |
| NAGGFGILGAVAHSP | ASSLRSNQDTQY | 0.09739303 |
| NEGREITLAMAEM | ASSLRSNQDTQY | 0.1654588 |
| NELVEWSHPLGHRWRWE | ASSLRSNQDTQY | 0.076119944 |
| NLGAGGTGGNGGNPG | ASSLRSNQDTQY | 0.11969818 |
| NTEWPKWETVMAADPVD | ASSLRSNQDTQY | 0.20809284 |
| PASAAPAETQP | ASSLRSNQDTQY | 0.08432729 |
| QQSSDGKNGVRAAEITGE | ASSLRSNQDTQY | 0.19585982 |
| RAPDAVTTSWWKE | ASSLRSNQDTQY | 0.091406815 |
| SGFGNVGAHTSGWFNQ | ASSLRSNQDTQY | 0.08601648 |
| SGVAGAVGQAASVGGLKV | ASSLRSNQDTQY | 0.45759124 |
| SNGGAGTGSAGGNGGT | ASSLRSNQDTQY | 0.09138243 |
| TGGAGGEGGAAG | ASSLRSNQDTQY | 0.1317055 |
| TGGDGGAAGAGGAPGHG | ASSLRSNQDTQY | 0.2570816 |
| TMPPLLAAIANAGRHGV | ASSLRSNQDTQY | 0.13522379 |
| TSSTAGNSGGA | ASSLRSNQDTQY | 0.09594917 |
| VPATVPVGILC | ASSLRSNQDTQY | 0.09992281 |
| VQRPDSGGVVLTGRIS | ASSLRSNQDTQY | 0.086699516 |
| WTPNPVPATKGVPAGLVQAT | ASSLRSNQDTQY | 0.10256067 |
| YALKRALAKFSDRFDVVII | ASSLRSNQDTQY | 0.075958215 |
| YLIGSVIV | ASSLRSNQDTQY | 0.18626261 |
| YGQLVAVID | ASSLRSNQDTQY | 0.09141001 |
| VTVGGMTLPA | ASSLRSNQDTQY | 0.24113697 |
| VLALPAV | ASSLRSNQDTQY | 0.13627139 |
| VIPPTLNYET | ASSLRSNQDTQY | 0.16459441 |
| VGMGGMHPSSG | ASSLRSNQDTQY | 0.10744274 |
| VGLAGGAATA | ASSLRSNQDTQY | 0.0983884 |
| VGGAGGAGGSGP | ASSLRSNQDTQY | 0.084846616 |
| VFAAFMV | ASSLRSNQDTQY | 0.09084378 |
| TLYDKLR | ASSLRSNQDTQY | 0.087817274 |
| TLAVFTVL | ASSLRSNQDTQY | 0.107316576 |
| TGSVQPARV | ASSLRSNQDTQY | 0.08977683 |
| TGGGAGSGGT | ASSLRSNQDTQY | 0.19816294 |
| SHLETSL | ASSLRSNQDTQY | 0.11202095 |
| SFTTGMT | ASSLRSNQDTQY | 0.08784986 |
| QVEVAKRMGVRQPTVS | ASSLRSNQDTQY | 0.19945621 |
| QGSGGPALAM | ASSLRSNQDTQY | 0.14290737 |
| MLAPSNP | ASSLRSNQDTQY | 0.11531395 |

| | | |
|---|---|---|
| LVNNAAI | ASSLRSNQDTQY | 0.07588802 |
| LTGEINTLA | ASSLRSNQDTQY | 0.09199717 |
| LSIFGPV | ASSLRSNQDTQY | 0.09986388 |
| LLALLANL | ASSLRSNQDTQY | 0.08754253 |
| LISNASDALDKLR | ASSLRSNQDTQY | 0.08801529 |
| LINEAVP | ASSLRSNQDTQY | 0.3434132 |
| LGYAFSGL | ASSLRSNQDTQY | 0.077268004 |
| LDFVAVAL | ASSLRSNQDTQY | 0.09350716 |
| LAQPGAE | ASSLRSNQDTQY | 0.15223952 |
| ISNLGAVTGF | ASSLRSNQDTQY | 0.21131885 |
| HQEDPGRI | ASSLRSNQDTQY | 0.3770268 |
| GTPGEPTP | ASSLRSNQDTQY | 0.11129293 |
| GFFNSGS | ASSLRSNQDTQY | 0.10944823 |
| GAGGGGLTAGAN | ASSLRSNQDTQY | 0.11496077 |
| FVDALVNM | ASSLRSNQDTQY | 0.13414454 |
| FLTVAPEL | ASSLRSNQDTQY | 0.0831881 |
| FETLTAL | ASSLRSNQDTQY | 0.08053109 |
| DVINPIEL | ASSLRSNQDTQY | 0.13158178 |
| APVSGAPV | ASSLRSNQDTQY | 0.0883633 |
| ALVGGLAW | ASSLRSNQDTQY | 0.16431384 |
| ALAAAAATAI | ASSLRSNQDTQY | 0.12595242 |
| AFGLGLLL | ASSLRSNQDTQY | 0.09633729 |
| AFFAAASL | ASSLRSNQDTQY | 0.08309947 |
| AAAALPAFSPP | ASSLRSNQDTQY | 0.11362529 |
| VVLAPAL | ASSLRSNQDTQY | 0.103404835 |
| TIGHVDHGKTTL | ASSLRSNQDTQY | 0.1805647 |
| LISNASDALDK | ASSLRSNQDTQY | 0.102614 |
| LISNASDAL | ASSLRSNQDTQY | 0.52056015 |
| ISNASDALDK | ASSLRSNQDTQY | 0.34466127 |
| NQAETLVYQTE | ASSLRSNQDTQY | 0.10832439 |
| TGGTGGTAGAG | ASSLRSNQDTQY | 0.2169932 |
| TGSTAGGAGGA | ASSLRSNQDTQY | 0.097693406 |
| DPLAVPVAAGPV | ASSLRSNQDTQY | 0.07907844 |
| GGTGGTPGNGGN | ASSLRSNQDTQY | 0.09425194 |
| PAGLAGVDPATL | ASSLRSNQDTQY | 0.09439528 |
| GNNLVVEAPHAGASV | ASSLRSNQDTQY | 0.09377939 |
| KVTRIMSASTLEQPA | ASSLRSNQDTQY | 0.086071484 |
| LGFALPIALPRRTVV | ASSLRSNQDTQY | 0.09041111 |
| TRLGNLTARKLVVLV | ASSLRSNQDTQY | 0.10128417 |
| WSQQRRLIYFPSAGP | ASSLRSNQDTQY | 0.08950615 |
| GPNGNPDPMAAAVDIR | ASSLRSNQDTQY | 0.097128525 |
| TQRCVSVADLRAAASR | ASSLRSNQDTQY | 0.09865636 |
| QFVLDVAEHPVEVLTETV | ASSLRSNQDTQY | 0.20969787 |
| GGAGGTGGAGGNGGTQAGDG | ASSLRSNQDTQY | 0.676889 |
| PLLATLRAIAVDVDAKPAQV | ASSLRSNQDTQY | 0.1062822 |
| APKFRTGGYMIDFWGVGYQVAK | ASSLRSNQDTQY | 0.097603254 |
| GAGGAGGAGGAAGGPGGTGGTG | ASSLRSNQDTQY | 0.1782876 |
| IGALGLLATDVAPAGSVAFGVA | ASSLRSNQDTQY | 0.12145979 |
| AGGAGGIGGNGGAGGAGGVLMG | ASSLRSNQDTQY | 0.09973686 |
| ADELIGLL | ASSLRSNQDTQY | 0.1548902 |
| ADRALKA | ASSLRSNQDTQY | 0.09808736 |

| | | |
|---|---|---|
| AGAVGQVGGAGGA | ASSLRSNQDTQY | 0.085373946 |
| AGGGGGLGGAP | ASSLRSNQDTQY | 0.12918527 |
| AGGLLYGNGGAG | ASSLRSNQDTQY | 0.090634085 |
| AGLESIGL | ASSLRSNQDTQY | 0.124628946 |
| AGLKTGII | ASSLRSNQDTQY | 0.07710211 |
| ASLAALLL | ASSLRSNQDTQY | 0.087873854 |
| ASVINISSI | ASSLRSNQDTQY | 0.10153588 |
| ATPPGPA | ASSLRSNQDTQY | 0.10432835 |
| ATVASALV | ASSLRSNQDTQY | 0.08170517 |
| ATVASLIL | ASSLRSNQDTQY | 0.09981324 |
| AVNGPVLIRH | ASSLRSNQDTQY | 0.27237096 |
| DGAAGAVGI | ASSLRSNQDTQY | 0.36842978 |
| DPLLHGI | ASSLRSNQDTQY | 0.08679849 |
| ELGAALGAL | ASSLRSNQDTQY | 0.16813049 |
| ELGRTQAV | ASSLRSNQDTQY | 0.46256572 |
| FVAGGYLS | ASSLRSNQDTQY | 0.108437836 |
| GASGGAGGAGGW | ASSLRSNQDTQY | 0.08221858 |
| GGAGAGGDGGAGGI | ASSLRSNQDTQY | 0.1269867 |
| GTGHNGSQPGMGGNG | ASSLRSNQDTQY | 0.07840732 |
| GYAAASASAS | ASSLRSNQDTQY | 0.091044776 |
| HQVDVAAVQQG | ASSLRSNQDTQY | 0.16174467 |
| IGAAIAHT | ASSLRSNQDTQY | 0.08514985 |
| IGMIEVAQL | ASSLRSNQDTQY | 0.13623932 |
| IRLAPEHTSAMD | ASSLRSNQDTQY | 0.2105385 |
| IYNVSVL | ASSLRSNQDTQY | 0.09713724 |
| KQGFLGM | ASSLRSNQDTQY | 0.08812396 |
| LESVITQL | ASSLRSNQDTQY | 0.08634955 |
| LGLGALVVV | ASSLRSNQDTQY | 0.08002779 |
| LIGNGGNGGSGGI | ASSLRSNQDTQY | 0.15648249 |
| LNAINAPTL | ASSLRSNQDTQY | 0.10643834 |
| LQLLNTGV | ASSLRSNQDTQY | 0.111614004 |
| LQTSIEF | ASSLRSNQDTQY | 0.11786131 |
| MSFAVSEL | ASSLRSNQDTQY | 0.10339498 |
| NTGALGL | ASSLRSNQDTQY | 0.1102489 |
| PPGAGGAGGAGTT | ASSLRSNQDTQY | 0.113317974 |
| QLVGLGI | ASSLRSNQDTQY | 0.0968802 |
| RFLEQVHQL | ASSLRSNQDTQY | 0.1633243 |
| RGVGYVLR | ASSLRSNQDTQY | 0.10013155 |
| SATGVFAGVI | ASSLRSNQDTQY | 0.09156859 |
| SLPAITLP | ASSLRSNQDTQY | 0.5014895 |
| STALAFTV | ASSLRSNQDTQY | 0.08571373 |
| SVLTVGLGK | ASSLRSNQDTQY | 0.09826085 |
| TALAGAGSGI | ASSLRSNQDTQY | 0.13767664 |
| TGLPFTGI | ASSLRSNQDTQY | 0.08361456 |
| VAEHVTV | ASSLRSNQDTQY | 0.17300622 |
| VDIVLNSL | ASSLRSNQDTQY | 0.14062935 |
| VLTGGGAVL | ASSLRSNQDTQY | 0.16933694 |
| VNVGLAYGI | ASSLRSNQDTQY | 0.11870507 |
| VSSAAVGAV | ASSLRSNQDTQY | 0.16615444 |
| VTSVYEAGV | ASSLRSNQDTQY | 0.19272292 |
| WVALDRGA | ASSLRSNQDTQY | 0.08850862 |

| | | |
|---|---|---|
| YLVGSVVI | ASSLRSNQDTQY | 0.07804428 |
| GPPGTGKTLL | ASSLRSNQDTQY | 0.08672292 |
| GVGMTKF | ASSLRSNQDTQY | 0.12395205 |
| IGIGGSDL | ASSLRSNQDTQY | 0.17473696 |
| IGTLLAEL | ASSLRSNQDTQY | 0.09222251 |
| IYTSGTT | ASSLRSNQDTQY | 0.13115127 |
| LSFPAGV | ASSLRSNQDTQY | 0.19845504 |
| LYTSGTT | ASSLRSNQDTQY | 0.1843989 |
| VGAFPFGF | ASSLRSNQDTQY | 0.08610839 |
| VGAGTVEF | ASSLRSNQDTQY | 0.10701962 |
| VGKSTLLL | ASSLRSNQDTQY | 0.1556778 |
| AALAHAGGGIV | ASSLRSNQDTQY | 0.12597159 |
| AALVAVSLL | ASSLRSNQDTQY | 0.088358104 |
| AAVSRAVAAL | ASSLRSNQDTQY | 0.25442135 |
| AGGSSGIGGF | ASSLRSNQDTQY | 0.14671299 |
| AGLTVANANG | ASSLRSNQDTQY | 0.14629592 |
| AGLVGAVL | ASSLRSNQDTQY | 0.1709675 |
| ALSQAFTV | ASSLRSNQDTQY | 0.27189788 |
| ALTFSVSL | ASSLRSNQDTQY | 0.11815978 |
| ALTGAALVAL | ASSLRSNQDTQY | 0.11789708 |
| ASFSIGAGVL | ASSLRSNQDTQY | 0.083371505 |
| DGSVYKI | ASSLRSNQDTQY | 0.08099083 |
| EQLAEIV | ASSLRSNQDTQY | 0.08640384 |
| FAVSELTVM | ASSLRSNQDTQY | 0.10910188 |
| GAAPLAAGL | ASSLRSNQDTQY | 0.07919985 |
| GADGGDGGSGGAGG | ASSLRSNQDTQY | 0.15443166 |
| GFGLDTGGAGG | ASSLRSNQDTQY | 0.14377525 |
| GLASGGLL | ASSLRSNQDTQY | 0.20530167 |
| GRIAGAAVLL | ASSLRSNQDTQY | 0.11554984 |
| GSAGSLNTSF | ASSLRSNQDTQY | 0.08440884 |
| GTGQIGI | ASSLRSNQDTQY | 0.0978039 |
| GTLIALIL | ASSLRSNQDTQY | 0.30067268 |
| IAATMGFVVG | ASSLRSNQDTQY | 0.084321305 |
| IFAQALL | ASSLRSNQDTQY | 0.08503299 |
| IGAAVALGF | ASSLRSNQDTQY | 0.10995486 |
| IGLTPLAL | ASSLRSNQDTQY | 0.09289095 |
| ILGSNTSTL | ASSLRSNQDTQY | 0.16196616 |
| IVFSLTVL | ASSLRSNQDTQY | 0.24493894 |
| LASKATGF | ASSLRSNQDTQY | 0.16250134 |
| LASVALIL | ASSLRSNQDTQY | 0.103015274 |
| LFAVGLTL | ASSLRSNQDTQY | 0.07952603 |
| LFNGGIGTYI | ASSLRSNQDTQY | 0.10779917 |
| LTLGAGII | ASSLRSNQDTQY | 0.55549055 |
| LVAAALGDI | ASSLRSNQDTQY | 0.13068256 |
| NTAFVQLGI | ASSLRSNQDTQY | 0.14685412 |
| QLFSGAEL | ASSLRSNQDTQY | 0.1846512 |
| SAATFITL | ASSLRSNQDTQY | 0.1022651 |
| SNYGLANLGV | ASSLRSNQDTQY | 0.117014855 |
| SYGGAYAVM | ASSLRSNQDTQY | 0.098792665 |
| TTGVGGLTLG | ASSLRSNQDTQY | 0.102089316 |
| TTQTIYI | ASSLRSNQDTQY | 0.09021354 |

| | | |
|---|---|---|
| VAAGAGLGAVI | ASSLRSNQDTQY | 0.08709636 |
| VAAVVTLL | ASSLRSNQDTQY | 0.13321742 |
| VGFLGGLI | ASSLRSNQDTQY | 0.57914966 |
| VGVIPILL | ASSLRSNQDTQY | 0.18215956 |
| VIFASSLI | ASSLRSNQDTQY | 0.17367399 |
| VKGGDGGAAAG | ASSLRSNQDTQY | 0.11991078 |
| VTKTFTNI | ASSLRSNQDTQY | 0.18748353 |
| VVAAAILL | ASSLRSNQDTQY | 0.108602166 |
| YAALDVEL | ASSLRSNQDTQY | 0.095413655 |
| LASVALLL | ASSLRSNQDTQY | 0.09744061 |
| LDAVHRV | ASSLRSNQDTQY | 0.10020654 |
| QGLSSSIF | ASSLRSNQDTQY | 0.11884492 |
| RGTFANIRL | ASSLRSNQDTQY | 0.11214233 |
| RQLWWGHRI | ASSLRSNQDTQY | 0.5974773 |
| YGPPGTGKTLL | ASSLRSNQDTQY | 0.19398148 |
| AALVGAPANVA | ASSLRSNQDTQY | 0.107209735 |
| AATFRRTE | ASSLRSNQDTQY | 0.09082389 |
| AGALGGGI | ASSLRSNQDTQY | 0.10575523 |
| AGGAGGAGGIGAG | ASSLRSNQDTQY | 0.13013859 |
| AGSGALGL | ASSLRSNQDTQY | 0.08979717 |
| AQIADQI | ASSLRSNQDTQY | 0.4492433 |
| ASAVAVTYL | ASSLRSNQDTQY | 0.101323545 |
| ASTVLGGTGG | ASSLRSNQDTQY | 0.09589924 |
| AVGGEPAK | ASSLRSNQDTQY | 0.14093608 |
| CEQGTFGGTC | ASSLRSNQDTQY | 0.38754958 |
| DEGKAPAG | ASSLRSNQDTQY | 0.10622892 |
| DTVVFPAL | ASSLRSNQDTQY | 0.10216981 |
| FEGITMAG | ASSLRSNQDTQY | 0.091311835 |
| FPLVVQD | ASSLRSNQDTQY | 0.2205129 |
| GAFTTPQ | ASSLRSNQDTQY | 0.46601516 |
| GFGGFGTAGGVGGA | ASSLRSNQDTQY | 0.077650756 |
| GFLAQFL | ASSLRSNQDTQY | 0.17225587 |
| GGAGGAGGNAGVA | ASSLRSNQDTQY | 0.0876498 |
| GGVGTAGAGGAGGAGG | ASSLRSNQDTQY | 0.08836152 |
| GNGGGGDAGNAGSGGN | ASSLRSNQDTQY | 0.107896514 |
| GTGGAGGAGNDAGS | ASSLRSNQDTQY | 0.09949255 |
| GTTNTGWFNS | ASSLRSNQDTQY | 0.44699225 |
| IFMGPLFTL | ASSLRSNQDTQY | 0.15907872 |
| ILAHAIEAI | ASSLRSNQDTQY | 0.07861548 |
| IRSAIREAL | ASSLRSNQDTQY | 0.10481746 |
| ITFAGGG | ASSLRSNQDTQY | 0.37666148 |
| ITGMSGHQEV | ASSLRSNQDTQY | 0.21222998 |
| IVAGLACTL | ASSLRSNQDTQY | 0.19714585 |
| IVGGGLAGA | ASSLRSNQDTQY | 0.123100504 |
| IVTGVLNI | ASSLRSNQDTQY | 0.11406657 |
| LAQQGCI | ASSLRSNQDTQY | 0.1100466 |
| LCLHVEN | ASSLRSNQDTQY | 0.1014822 |
| LDAISDGS | ASSLRSNQDTQY | 0.0897087 |
| LGGFAAML | ASSLRSNQDTQY | 0.1180088 |
| LGGKNPMI | ASSLRSNQDTQY | 0.24437863 |
| LGPKGVAAIL | ASSLRSNQDTQY | 0.08330866 |

| | | |
|---|---|---|
| LMVGVSI | ASSLRSNQDTQY | 0.11420826 |
| LVGAAFVL | ASSLRSNQDTQY | 0.117971465 |
| LVLAVAPL | ASSLRSNQDTQY | 0.08739072 |
| MAAIAVIDAA | ASSLRSNQDTQY | 0.088293515 |
| NNGINAELQ | ASSLRSNQDTQY | 0.35950568 |
| RGEPLVLVHG | ASSLRSNQDTQY | 0.6510361 |
| RGGAGGAAGLI | ASSLRSNQDTQY | 0.12963666 |
| RSYRMHPAV | ASSLRSNQDTQY | 0.08681547 |
| SAAHLLAGQLA | ASSLRSNQDTQY | 0.08089576 |
| SAAVGSY | ASSLRSNQDTQY | 0.22676331 |
| SPHAVLHRI | ASSLRSNQDTQY | 0.15649472 |
| SQIAKVL | ASSLRSNQDTQY | 0.100283004 |
| TALGHLVP | ASSLRSNQDTQY | 0.087066025 |
| TAPTTSGAA | ASSLRSNQDTQY | 0.08942874 |
| TDGDDGNGGSPGT | ASSLRSNQDTQY | 0.25267667 |
| TDGLGEN | ASSLRSNQDTQY | 0.087767854 |
| TGGPPLPGQA | ASSLRSNQDTQY | 0.08935412 |
| TIPAITI | ASSLRSNQDTQY | 0.08186793 |
| TIPGFTI | ASSLRSNQDTQY | 0.23709238 |
| TSIASGALA | ASSLRSNQDTQY | 0.08824684 |
| TVTLPGAAI | ASSLRSNQDTQY | 0.13472681 |
| VASGLSAALP | ASSLRSNQDTQY | 0.24315962 |
| VGAGVIGI | ASSLRSNQDTQY | 0.07719709 |
| VGPKRFGV | ASSLRSNQDTQY | 0.14210573 |
| VNVAFPD | ASSLRSNQDTQY | 0.10562968 |
| VSMTNSGVS | ASSLRSNQDTQY | 0.17284632 |
| VTDVLGDLSV | ASSLRSNQDTQY | 0.100912124 |
| VVVVGAGFAGL | ASSLRSNQDTQY | 0.17495345 |
| VYTVGTVV | ASSLRSNQDTQY | 0.12308731 |
| WTTFSGP | ASSLRSNQDTQY | 0.12479931 |
| YDELVEAF | ASSLRSNQDTQY | 0.13418832 |
| YGTGGAGGNGG | ASSLRSNQDTQY | 0.09738676 |
| YTNVMAV | ASSLRSNQDTQY | 0.08919423 |
| FLEQVHQL | ASSLRSNQDTQY | 0.08112666 |
| GLSLLAVAV | ASSLRSNQDTQY | 0.16533697 |
| LEQVHQL | ASSLRSNQDTQY | 0.07604265 |
| LNAAGAI | ASSLRSNQDTQY | 0.1227621 |
| PAVHASL | ASSLRSNQDTQY | 0.16214234 |
| TELEDDA | ASSLRSNQDTQY | 0.090983115 |
| TPPGPPN | ASSLRSNQDTQY | 0.120963715 |
| TPPPPPPP | ASSLRSNQDTQY | 0.08009161 |
| TQSPALI | ASGRQGVGERLF | 0.096599765 |
| EFAAAFPQ | ASGRQGVGERLF | 0.2303299 |
| LLSGNLDVL | ASGRQGVGERLF | 0.102272645 |
| GPANLGNYNI | ASGRQGVGERLF | 0.112052746 |
| TLGPTLP | ASGRQGVGERLF | 0.10530296 |
| VYAYSSCCK | ASGRQGVGERLF | 0.14495741 |
| APAPAPAPNI | ASGRQGVGERLF | 0.099879615 |
| PGGPVPIP | ASGRQGVGERLF | 0.08877248 |
| FLTVALEP | ASGRQGVGERLF | 0.329612 |
| SQYFGANL | ASGRQGVGERLF | 0.09878598 |

| | | |
|---|---|---|
| GGVKPAA | ASGRQGVGERLF | 0.09009383 |
| GGTGGAGGAGGW | ASGRQGVGERLF | 0.15483218 |
| GVGGVAGLGGAGG | ASGRQGVGERLF | 0.0860032 |
| VLLVLATTI | ASGRQGVGERLF | 0.18857953 |
| IGIFAAML | ASGRQGVGERLF | 0.09740897 |
| AVGDLID | ASGRQGVGERLF | 0.14949465 |
| NISPLPL | ASGRQGVGERLF | 0.115657285 |
| VTAVFVAI | ASGRQGVGERLF | 0.08048115 |
| SGDDNDAAL | ASGRQGVGERLF | 0.101666786 |
| AVAALAHS | ASGRQGVGERLF | 0.10669653 |
| GGVTVMATP | ASGRQGVGERLF | 0.15204124 |
| AGLAANVRD | ASGRQGVGERLF | 0.1536511 |
| TNRPDLIDPALLRPG | ASGRQGVGERLF | 0.10070898 |
| LANLGVTAL | ASGRQGVGERLF | 0.16473994 |
| TGERIEL | ASGRQGVGERLF | 0.10325681 |
| NGEELAA | ASGRQGVGERLF | 0.101482704 |
| VLSLVWG | ASGRQGVGERLF | 0.09844242 |
| IGYALWL | ASGRQGVGERLF | 0.09524917 |
| LSLLLAPL | ASGRQGVGERLF | 0.100540444 |
| TLDEALK | ASGRQGVGERLF | 0.109949805 |
| TDGAGTAEGG | ASGRQGVGERLF | 0.08765229 |
| NAPGGAGGNGG | ASGRQGVGERLF | 0.16142808 |
| VSVFMGMGGNF | ASGRQGVGERLF | 0.084781155 |
| LASYLAAL | ASGRQGVGERLF | 0.09880363 |
| LVGGTPAL | ASGRQGVGERLF | 0.15532634 |
| VRNADVL | ASGRQGVGERLF | 0.09377349 |
| TCSSPFLVSAM | ASGRQGVGERLF | 0.10361564 |
| VHITDAE | ASGRQGVGERLF | 0.11676344 |
| EYAAPSF | ASGRQGVGERLF | 0.09524154 |
| VVDAFHGGL | ASGRQGVGERLF | 0.116384834 |
| ASVVDPQ | ASGRQGVGERLF | 0.23235747 |
| SYNEFGGLS | ASGRQGVGERLF | 0.12901133 |
| DPDGSVY | ASGRQGVGERLF | 0.08386303 |
| MVSTIHATE | ASGRQGVGERLF | 0.09492768 |
| LAAGVADPV | ASGRQGVGERLF | 0.16814488 |
| TKEQIAA | ASGRQGVGERLF | 0.1284153 |
| SLASPAAL | ASGRQGVGERLF | 0.08087146 |
| IGLVFYGV | ASGRQGVGERLF | 0.10389368 |
| FLLDASNL | ASGRQGVGERLF | 0.08665244 |
| YQVGNFP | ASGRQGVGERLF | 0.08999266 |
| ESLLSQST | ASGRQGVGERLF | 0.088569775 |
| LFTVGFAAVG | ASGRQGVGERLF | 0.27364877 |
| ATPPTPT | ASGRQGVGERLF | 0.42718095 |
| TDLATLG | ASGRQGVGERLF | 0.111715555 |
| SAGGTGR | ASGRQGVGERLF | 0.10623625 |
| GFSNSGS | ASGRQGVGERLF | 0.08317027 |
| GGNGGSGGTSV | ASGRQGVGERLF | 0.1573257 |
| GTSVATGGAGN | ASGRQGVGERLF | 0.29361778 |
| NGGNGGSGGTS | ASGRQGVGERLF | 0.1016815 |
| AVVKFPH | ASGRQGVGERLF | 0.13477944 |
| TPPPPPP | ASGRQGVGERLF | 0.09037796 |

| | | |
|---|---|---|
| AAPAAAAAPA | ASGRQGVGERLF | 0.09361823 |
| IALNEDL | ASGRQGVGERLF | 0.19496323 |
| IVNVSSM | ASGRQGVGERLF | 0.12004987 |
| TGIPPAPR | ASGRQGVGERLF | 0.09562342 |
| FDLGGGTF | ASGRQGVGERLF | 0.092808016 |
| FDLGGGTFDVS | ASGRQGVGERLF | 0.0933632 |
| NEPTAAA | ASGRQGVGERLF | 0.09895779 |
| NEPTAAALA | ASGRQGVGERLF | 0.15606907 |
| GSIINISSI | ASGRQGVGERLF | 0.107132345 |
| VAPGNAGTA | ASGRQGVGERLF | 0.22554836 |
| TGGGASLE | ASGRQGVGERLF | 0.10351101 |
| VAVYNHYKRI | ASGRQGVGERLF | 0.29961228 |
| TDPAPATT | ASGRQGVGERLF | 0.16280517 |
| TNRPDILDPALLRPG | ASGRQGVGERLF | 0.19645776 |
| LTTTTTS | ASGRQGVGERLF | 0.09942732 |
| PTGSGKTL | ASGRQGVGERLF | 0.5690564 |
| TGSGKTL | ASGRQGVGERLF | 0.14037529 |
| TIGHVDHGK | ASGRQGVGERLF | 0.119787075 |
| TIGHVDHGKTT | ASGRQGVGERLF | 0.12261409 |
| GHVDHGKTT | ASGRQGVGERLF | 0.285489 |
| AQFEHTLL | ASGRQGVGERLF | 0.2892299 |
| IVNQDEL | ASGRQGVGERLF | 0.28607884 |
| TNGSQFF | ASGRQGVGERLF | 0.28552753 |
| TGKGIVE | ASGRQGVGERLF | 0.28837726 |
| LISNASDA | ASGRQGVGERLF | 0.284691 |
| AELRTLGELTKIAKAHGAQ | ASGRQGVGERLF | 0.28680462 |
| APNAVAAAEPSW | ASGRQGVGERLF | 0.28967983 |
| APQVAGAATAVL | ASGRQGVGERLF | 0.28492093 |
| ARRDPAAAAAILPTDA | ASGRQGVGERLF | 0.286561 |
| DAAARDVLAEAGLAEYF | ASGRQGVGERLF | 0.2880762 |
| DDQIVLSAKANLAKVEQ | ASGRQGVGERLF | 0.29710543 |
| DELLPDTGEDSDEDGH | ASGRQGVGERLF | 0.2915196 |
| ELPEAPANRATVPQD | ASGRQGVGERLF | 0.2857306 |
| FGQSKIERVTGIG | ASGRQGVGERLF | 0.2847772 |
| GAGGTGSTAGGAGGAGGAGGLY | ASGRQGVGERLF | 0.28698415 |
| GHGGTGAAGGNGGTGGAGGSGI | ASGRQGVGERLF | 0.28633532 |
| GNGGNGGAGGEGGPGGAGGAGA | ASGRQGVGERLF | 0.28391543 |
| GSTAGNGGNGGNATNGTIAGQP | ASGRQGVGERLF | 0.2877868 |
| GTGGDGGNGGNGGTGG | ASGRQGVGERLF | 0.28387374 |
| GVRRPAGIPADAPV | ASGRQGVGERLF | 0.28633493 |
| GYRQVTDAHLLTLARR | ASGRQGVGERLF | 0.28778708 |
| IAKMGQDGHDRG | ASGRQGVGERLF | 0.28651336 |
| KAEPSIEIQYT | ASGRQGVGERLF | 0.2859488 |
| KPVRIRMAADEDTVAVT | ASGRQGVGERLF | 0.2911539 |
| MPPRIAGMRLLVIKPEP | ASGRQGVGERLF | 0.2869246 |
| MKFYAVALFRSRGVPPTR | ASGRQGVGERLF | 0.28638688 |
| NAGGFGILGAVAHSP | ASGRQGVGERLF | 0.29124624 |
| NEGREITLAMAEM | ASGRQGVGERLF | 0.29540414 |
| NELVEWSHPLGHRWRWE | ASGRQGVGERLF | 0.28548485 |
| NLGAGGTGGNGGNPG | ASGRQGVGERLF | 0.28618425 |
| NTEWPKWETVMAADPVD | ASGRQGVGERLF | 0.2920959 |

| | | |
|---|---|---|
| PASAAPAETQP | ASGRQGVGERLF | 0.29130697 |
| QQSSDGKNGVRAAEITGE | ASGRQGVGERLF | 0.28709736 |
| RAPDAVTTSWWKE | ASGRQGVGERLF | 0.28759724 |
| SGFGNVGAHTSGWFNQ | ASGRQGVGERLF | 0.2840657 |
| SGVAGAVGQAASVGGLKV | ASGRQGVGERLF | 0.29057673 |
| SNGGAGTGSAGGNGGT | ASGRQGVGERLF | 0.2910821 |
| TGGAGGEGGAAG | ASGRQGVGERLF | 0.2875815 |
| TGGDGGAAGAGGAPGHG | ASGRQGVGERLF | 0.29064623 |
| TMPPLLAAIANAGRHGV | ASGRQGVGERLF | 0.28583598 |
| TSSTAGNSGGA | ASGRQGVGERLF | 0.2846093 |
| VPATVPVGILC | ASGRQGVGERLF | 0.29291412 |
| VQRPDSGGVVLTGRIS | ASGRQGVGERLF | 0.28822017 |
| WTPNPVPATKGVPAGLVQAT | ASGRQGVGERLF | 0.28968257 |
| YALKRALAKFSDRFDVVII | ASGRQGVGERLF | 0.28826648 |
| YLIGSVIV | ASGRQGVGERLF | 0.2850238 |
| YGQLVAVID | ASGRQGVGERLF | 0.2863414 |
| VTVGGMTLPA | ASGRQGVGERLF | 0.283522 |
| VLALPAV | ASGRQGVGERLF | 0.28540742 |
| VIPPTLNYET | ASGRQGVGERLF | 0.29289922 |
| VGMGGMHPSSG | ASGRQGVGERLF | 0.28482196 |
| VGLAGGAATA | ASGRQGVGERLF | 0.29881814 |
| VGGAGGAGGSGP | ASGRQGVGERLF | 0.29332182 |
| VFAAFMV | ASGRQGVGERLF | 0.3092754 |
| TLYDKLR | ASGRQGVGERLF | 0.289614 |
| TLAVFTVL | ASGRQGVGERLF | 0.28420958 |
| TGSVQPARV | ASGRQGVGERLF | 0.2903804 |
| TGGGAGSGGT | ASGRQGVGERLF | 0.28722158 |
| SHLETSL | ASGRQGVGERLF | 0.28573328 |
| SFTTGMT | ASGRQGVGERLF | 0.28629047 |
| QVEVAKRMGVRQPTVS | ASGRQGVGERLF | 0.28785592 |
| QGSGGPALAM | ASGRQGVGERLF | 0.31454065 |
| MLAPSNP | ASGRQGVGERLF | 0.2902448 |
| LVNNAAI | ASGRQGVGERLF | 0.28712118 |
| LTGEINTLA | ASGRQGVGERLF | 0.28751427 |
| LSIFGPV | ASGRQGVGERLF | 0.28341067 |
| LLALLANL | ASGRQGVGERLF | 0.28534937 |
| LISNASDALDKLR | ASGRQGVGERLF | 0.28500083 |
| LINEAVP | ASGRQGVGERLF | 0.2855344 |
| LGYAFSGL | ASGRQGVGERLF | 0.30150464 |
| LDFVAVAL | ASGRQGVGERLF | 0.28925362 |
| LAQPGAE | ASGRQGVGERLF | 0.2950452 |
| ISNLGAVTGF | ASGRQGVGERLF | 0.2873085 |
| HQEDPGRI | ASGRQGVGERLF | 0.28451177 |
| GTPGEPTP | ASGRQGVGERLF | 0.28829288 |
| GFFNSGS | ASGRQGVGERLF | 0.28730345 |
| GAGGGGLTAGAN | ASGRQGVGERLF | 0.28809044 |
| FVDALVNM | ASGRQGVGERLF | 0.28531596 |
| FLTVAPEL | ASGRQGVGERLF | 0.2865791 |
| FETLTAL | ASGRQGVGERLF | 0.2865832 |
| DVINPIEL | ASGRQGVGERLF | 0.28857636 |
| APVSGAPV | ASGRQGVGERLF | 0.28699827 |

| | | |
|---|---|---|
| ALVGGLAW | ASGRQGVGERLF | 0.28620458 |
| ALAAAAATAI | ASGRQGVGERLF | 0.2869346 |
| AFGLGLLL | ASGRQGVGERLF | 0.28528145 |
| AFFAAASL | ASGRQGVGERLF | 0.3066955 |
| AAAALPAFSPP | ASGRQGVGERLF | 0.28590083 |
| VVLAPAL | ASGRQGVGERLF | 0.28561085 |
| TIGHVDHGKTTL | ASGRQGVGERLF | 0.28593543 |
| LISNASDALDK | ASGRQGVGERLF | 0.28548852 |
| LISNASDAL | ASGRQGVGERLF | 0.28621918 |
| ISNASDALDK | ASGRQGVGERLF | 0.2850159 |
| NQAETLVYQTE | ASGRQGVGERLF | 0.28483984 |
| TGGTGGTAGAG | ASGRQGVGERLF | 0.28903416 |
| TGSTAGGAGGA | ASGRQGVGERLF | 0.28617296 |
| DPLAVPVAAGPV | ASGRQGVGERLF | 0.2865073 |
| GGTGGTPGNGGN | ASGRQGVGERLF | 0.28945848 |
| PAGLAGVDPATL | ASGRQGVGERLF | 0.28519684 |
| GNNLVVEAPHAGASV | ASGRQGVGERLF | 0.28920224 |
| KVTRIMSASTLEQPA | ASGRQGVGERLF | 0.28467247 |
| LGFALPIALPRRTVV | ASGRQGVGERLF | 0.28669426 |
| TRLGNLTARKLVVLV | ASGRQGVGERLF | 0.3032204 |
| WSQQRRLIYFPSAGP | ASGRQGVGERLF | 0.2868783 |
| GPNGNPDPMAAAVDIR | ASGRQGVGERLF | 0.29849166 |
| TQRCVSVADLRAAASR | ASGRQGVGERLF | 0.28425905 |
| QFVLDVAEHPVEVLTETV | ASGRQGVGERLF | 0.2846602 |
| GGAGGTGGAGGNGGTQAGDG | ASGRQGVGERLF | 0.2845809 |
| PLLATLRAIAVDVDAKPAQV | ASGRQGVGERLF | 0.29375938 |
| APKFRTGGYMIDFWGVGYQVAK | ASGRQGVGERLF | 0.2848168 |
| GAGGAGGAGGAAGGPGGTGGTG | ASGRQGVGERLF | 0.2874073 |
| IGALGLLATDVAPAGSVAFGVA | ASGRQGVGERLF | 0.28662243 |
| AGGAGGIGGNGGAGGAGGVLMG | ASGRQGVGERLF | 0.28953245 |
| ADELIGLL | ASGRQGVGERLF | 0.2890191 |
| ADRALKA | ASGRQGVGERLF | 0.28559357 |
| AGAVGQVGGAGGA | ASGRQGVGERLF | 0.28514668 |
| AGGGGGLGGAP | ASGRQGVGERLF | 0.29131743 |
| AGGLLYGNGGAG | ASGRQGVGERLF | 0.28544828 |
| AGLESIGL | ASGRQGVGERLF | 0.28533792 |
| AGLKTGII | ASGRQGVGERLF | 0.2854191 |
| ASLAALLL | ASGRQGVGERLF | 0.2991839 |
| ASVINISSI | ASGRQGVGERLF | 0.29151583 |
| ATPPGPA | ASGRQGVGERLF | 0.29035437 |
| ATVASALV | ASGRQGVGERLF | 0.2893388 |
| ATVASLIL | ASGRQGVGERLF | 0.2837371 |
| AVNGPVLIRH | ASGRQGVGERLF | 0.28413686 |
| DGAAGAVGI | ASGRQGVGERLF | 0.28988543 |
| DPLLHGI | ASGRQGVGERLF | 0.29576424 |
| ELGAALGAL | ASGRQGVGERLF | 0.28839538 |
| ELGRTQAV | ASGRQGVGERLF | 0.28848886 |
| FVAGGYLS | ASGRQGVGERLF | 0.288598 |
| GASGGAGGAGGW | ASGRQGVGERLF | 0.28402174 |
| GGAGAGGDGGAGGI | ASGRQGVGERLF | 0.28652027 |
| GTGHNGSQPGMGGNG | ASGRQGVGERLF | 0.2858211 |

| | | |
|---|---|---|
| GYAAASASAS | ASGRQGVGERLF | 0.28726315 |
| HQVDVAAVQQG | ASGRQGVGERLF | 0.28497326 |
| IGAAIAHT | ASGRQGVGERLF | 0.28571284 |
| IGMIEVAQL | ASGRQGVGERLF | 0.29040766 |
| IRLAPEHTSAMD | ASGRQGVGERLF | 0.2842756 |
| IYNVSVL | ASGRQGVGERLF | 0.28531814 |
| KQGFLGM | ASGRQGVGERLF | 0.28406498 |
| LESVITQL | ASGRQGVGERLF | 0.28678367 |
| LGLGALVVV | ASGRQGVGERLF | 0.29573658 |
| LIGNGGNGGSGGI | ASGRQGVGERLF | 0.29646477 |
| LNAINAPTL | ASGRQGVGERLF | 0.28647342 |
| LQLLNTGV | ASGRQGVGERLF | 0.2913232 |
| LQTSIEF | ASGRQGVGERLF | 0.28712493 |
| MSFAVSEL | ASGRQGVGERLF | 0.28475103 |
| NTGALGL | ASGRQGVGERLF | 0.28734076 |
| PPGAGGAGGAGTT | ASGRQGVGERLF | 0.28592795 |
| QLVGLGI | ASGRQGVGERLF | 0.28772098 |
| RFLEQVHQL | ASGRQGVGERLF | 0.29671216 |
| RGVGYVLR | ASGRQGVGERLF | 0.28617185 |
| SATGVFAGVI | ASGRQGVGERLF | 0.28682464 |
| SLPAITLP | ASGRQGVGERLF | 0.28684992 |
| STALAFTV | ASGRQGVGERLF | 0.2857803 |
| SVLTVGLGK | ASGRQGVGERLF | 0.28595605 |
| TALAGAGSGI | ASGRQGVGERLF | 0.28575614 |
| TGLPFTGI | ASGRQGVGERLF | 0.29343456 |
| VAEHVTV | ASGRQGVGERLF | 0.28437406 |
| VDIVLNSL | ASGRQGVGERLF | 0.28508735 |
| VLTGGGAVL | ASGRQGVGERLF | 0.28459978 |
| VNVGLAYGI | ASGRQGVGERLF | 0.2894835 |
| VSSAAVGAV | ASGRQGVGERLF | 0.29591596 |
| VTSVYEAGV | ASGRQGVGERLF | 0.28524768 |
| WVALDRGA | ASGRQGVGERLF | 0.2866298 |
| YLVGSVVI | ASGRQGVGERLF | 0.28518218 |
| GPPGTGKTLL | ASGRQGVGERLF | 0.28943032 |
| GVGMTKF | ASGRQGVGERLF | 0.28524557 |
| IGIGGSDL | ASGRQGVGERLF | 0.2861756 |
| IGTLLAEL | ASGRQGVGERLF | 0.28533444 |
| IYTSGTT | ASGRQGVGERLF | 0.28703976 |
| LSFPAGV | ASGRQGVGERLF | 0.28551215 |
| LYTSGTT | ASGRQGVGERLF | 0.2850936 |
| VGAFPFGF | ASGRQGVGERLF | 0.2880501 |
| VGAGTVEF | ASGRQGVGERLF | 0.2853861 |
| VGKSTLLL | ASGRQGVGERLF | 0.28928503 |
| AALAHAGGGIV | ASGRQGVGERLF | 0.288683 |
| AALVAVSLL | ASGRQGVGERLF | 0.28494313 |
| AAVSRAVAAL | ASGRQGVGERLF | 0.28533328 |
| AGGSSGIGGF | ASGRQGVGERLF | 0.29633966 |
| AGLTVANANG | ASGRQGVGERLF | 0.28563145 |
| AGLVGAVL | ASGRQGVGERLF | 0.28574866 |
| ALSQAFTV | ASGRQGVGERLF | 0.2912183 |
| ALTFSVSL | ASGRQGVGERLF | 0.28695482 |

| | | |
|---|---|---|
| ALTGAALVAL | ASGRQGVGERLF | 0.29201618 |
| ASFSIGAGVL | ASGRQGVGERLF | 0.2842819 |
| DGSVYKI | ASGRQGVGERLF | 0.28551662 |
| EQLAEIV | ASGRQGVGERLF | 0.28587928 |
| FAVSELTVM | ASGRQGVGERLF | 0.28623852 |
| GAAPLAAGL | ASGRQGVGERLF | 0.28905937 |
| GADGGDGGSGGAGG | ASGRQGVGERLF | 0.2886784 |
| GFGLDTGGAGG | ASGRQGVGERLF | 0.2845728 |
| GLASGGLL | ASGRQGVGERLF | 0.29140115 |
| GRIAGAAVLL | ASGRQGVGERLF | 0.28665447 |
| GSAGSLNTSF | ASGRQGVGERLF | 0.284482 |
| GTGQIGI | ASGRQGVGERLF | 0.2847482 |
| GTLIALIL | ASGRQGVGERLF | 0.28578222 |
| IAATMGFVVG | ASGRQGVGERLF | 0.28653267 |
| IFAQALL | ASGRQGVGERLF | 0.2902129 |
| IGAAVALGF | ASGRQGVGERLF | 0.28775826 |
| IGLTPLAL | ASGRQGVGERLF | 0.28559285 |
| ILGSNTSTL | ASGRQGVGERLF | 0.28963277 |
| IVFSLTVL | ASGRQGVGERLF | 0.2854925 |
| LASKATGF | ASGRQGVGERLF | 0.28702334 |
| LASVALIL | ASGRQGVGERLF | 0.28424656 |
| LFAVGLTL | ASGRQGVGERLF | 0.28437108 |
| LFNGGIGTYI | ASGRQGVGERLF | 0.28951403 |
| LTLGAGII | ASGRQGVGERLF | 0.28517503 |
| LVAAALGDI | ASGRQGVGERLF | 0.2864982 |
| NTAFVQLGI | ASGRQGVGERLF | 0.2850233 |
| QLFSGAEL | ASGRQGVGERLF | 0.2921133 |
| SAATFITL | ASGRQGVGERLF | 0.2894051 |
| SNYGLANLGV | ASGRQGVGERLF | 0.28523827 |
| SYGGAYAVM | ASGRQGVGERLF | 0.28501394 |
| TTGVGGLTLG | ASGRQGVGERLF | 0.2897294 |
| TTQTIYI | ASGRQGVGERLF | 0.2841132 |
| VAAGAGLGAVI | ASGRQGVGERLF | 0.29598188 |
| VAAVVTLL | ASGRQGVGERLF | 0.28888768 |
| VGFLGGLI | ASGRQGVGERLF | 0.28432214 |
| VGVIPILL | ASGRQGVGERLF | 0.28702077 |
| VIFASSLI | ASGRQGVGERLF | 0.28583923 |
| VKGGDGGAAAG | ASGRQGVGERLF | 0.28734842 |
| VTKTFTNI | ASGRQGVGERLF | 0.2850976 |
| VVAAAILL | ASGRQGVGERLF | 0.28405556 |
| YAALDVEL | ASGRQGVGERLF | 0.28744796 |
| LASVALLL | ASGRQGVGERLF | 0.29400334 |
| LDAVHRV | ASGRQGVGERLF | 0.28768632 |
| QGLSSSIF | ASGRQGVGERLF | 0.2856241 |
| RGTFANIRL | ASGRQGVGERLF | 0.2874622 |
| RQLWWGHRI | ASGRQGVGERLF | 0.28925753 |
| YGPPGTGKTLL | ASGRQGVGERLF | 0.28640407 |
| AALVGAPANVA | ASGRQGVGERLF | 0.2906029 |
| AATFRRTE | ASGRQGVGERLF | 0.285889 |
| AGALGGGI | ASGRQGVGERLF | 0.28888482 |
| AGGAGGAGGIGAG | ASGRQGVGERLF | 0.28863218 |

| | | |
|---|---|---|
| AGSGALGL | ASGRQGVGERLF | 0.28573132 |
| AQIADQI | ASGRQGVGERLF | 0.2856761 |
| ASAVAVTYL | ASGRQGVGERLF | 0.2865442 |
| ASTVLGGTGG | ASGRQGVGERLF | 0.28800687 |
| AVGGEPAK | ASGRQGVGERLF | 0.28742036 |
| CEQGTFGGTC | ASGRQGVGERLF | 0.2844711 |
| DEGKAPAG | ASGRQGVGERLF | 0.30532023 |
| DTVVFPAL | ASGRQGVGERLF | 0.28555235 |
| FEGITMAG | ASGRQGVGERLF | 0.29242682 |
| FPLVVQD | ASGRQGVGERLF | 0.28539252 |
| GAFTTPQ | ASGRQGVGERLF | 0.285258 |
| GFGGFGTAGGVGGA | ASGRQGVGERLF | 0.29473856 |
| GFLAQFL | ASGRQGVGERLF | 0.28515646 |
| GGAGGAGGNAGVA | ASGRQGVGERLF | 0.28905714 |
| GGVGTAGAGGAGGAGG | ASGRQGVGERLF | 0.28734022 |
| GNGGGGDAGNAGSGGN | ASGRQGVGERLF | 0.28722352 |
| GTGGAGGAGNDAGS | ASGRQGVGERLF | 0.2856485 |
| GTTNTGWFNS | ASGRQGVGERLF | 0.2869167 |
| IFMGPLFTL | ASGRQGVGERLF | 0.28481796 |
| ILAHAIEAI | ASGRQGVGERLF | 0.2897066 |
| IRSAIREAL | ASGRQGVGERLF | 0.28444856 |
| ITFAGGG | ASGRQGVGERLF | 0.28930718 |
| ITGMSGHQEV | ASGRQGVGERLF | 0.28455132 |
| IVAGLACTL | ASGRQGVGERLF | 0.28864977 |
| IVGGGLAGA | ASGRQGVGERLF | 0.29164425 |
| IVTGVLNI | ASGRQGVGERLF | 0.29047257 |
| LAQQGCI | ASGRQGVGERLF | 0.2907806 |
| LCLHVEN | ASGRQGVGERLF | 0.29511932 |
| LDAISDGS | ASGRQGVGERLF | 0.28979418 |
| LGGFAAML | ASGRQGVGERLF | 0.289876 |
| LGGKNPMI | ASGRQGVGERLF | 0.28679296 |
| LGPKGVAAIL | ASGRQGVGERLF | 0.2868986 |
| LMVGVSI | ASGRQGVGERLF | 0.2902837 |
| LVGAAFVL | ASGRQGVGERLF | 0.2852418 |
| LVLAVAPL | ASGRQGVGERLF | 0.2846653 |
| MAAIAVIDAA | ASGRQGVGERLF | 0.28643835 |
| NNGINAELQ | ASGRQGVGERLF | 0.2879818 |
| RGEPLVLVHG | ASGRQGVGERLF | 0.28699693 |
| RGGAGGAAGLI | ASGRQGVGERLF | 0.2868748 |
| RSYRMHPAV | ASGRQGVGERLF | 0.28797877 |
| SAAHLLAGQLA | ASGRQGVGERLF | 0.2917353 |
| SAAVGSY | ASGRQGVGERLF | 0.29076824 |
| SPHAVLHRI | ASGRQGVGERLF | 0.29008174 |
| SQIAKVL | ASGRQGVGERLF | 0.2855258 |
| TALGHLVP | ASGRQGVGERLF | 0.28605428 |
| TAPTTSGAA | ASGRQGVGERLF | 0.28663358 |
| TDGDDGNGGSPGT | ASGRQGVGERLF | 0.28585434 |
| TDGLGEN | ASGRQGVGERLF | 0.28638068 |
| TGGPPLPGQA | ASGRQGVGERLF | 0.28677684 |
| TIPAITI | ASGRQGVGERLF | 0.29104078 |
| TIPGFTI | ASGRQGVGERLF | 0.28325397 |

| | | |
|---|---|---|
| TSIASGALA | ASGRQGVGERLF | 0.28436068 |
| TVTLPGAAI | ASGRQGVGERLF | 0.2917331 |
| VASGLSAALP | ASGRQGVGERLF | 0.3014489 |
| VGAGVIGI | ASGRQGVGERLF | 0.28634632 |
| VGPKRFGV | ASGRQGVGERLF | 0.28636932 |
| VNVAFPD | ASGRQGVGERLF | 0.2842998 |
| VSMTNSGVS | ASGRQGVGERLF | 0.29046524 |
| VTDVLGDLSV | ASGRQGVGERLF | 0.28511423 |
| VVVVGAGFAGL | ASGRQGVGERLF | 0.28647268 |
| VYTVGTVV | ASGRQGVGERLF | 0.29137918 |
| WTTFSGP | ASGRQGVGERLF | 0.28679666 |
| YDELVEAF | ASGRQGVGERLF | 0.2849378 |
| YGTGGAGGNGG | ASGRQGVGERLF | 0.2851223 |
| YTNVMAV | ASGRQGVGERLF | 0.28583628 |
| FLEQVHQL | ASGRQGVGERLF | 0.2855706 |
| GLSLLAVAV | ASGRQGVGERLF | 0.28449583 |
| LEQVHQL | ASGRQGVGERLF | 0.2869588 |
| LNAAGAI | ASGRQGVGERLF | 0.2850004 |
| PAVHASL | ASGRQGVGERLF | 0.29456434 |
| TELEDDA | ASGRQGVGERLF | 0.29073328 |
| TPPGPPN | ASGRQGVGERLF | 0.28580067 |
| TPPPPPPP | ASGRQGVGERLF | 0.28649923 |
| TQSPALI | ASSLYRGDYAEQF | 0.312532 |
| EFAAAFPQ | ASSLYRGDYAEQF | 0.30412602 |
| LLSGNLDVL | ASSLYRGDYAEQF | 0.28558853 |
| GPANLGNYNI | ASSLYRGDYAEQF | 0.28725293 |
| TLGPTLP | ASSLYRGDYAEQF | 0.28522703 |
| VYAYSSCCK | ASSLYRGDYAEQF | 0.28581864 |
| APAPAPAPNI | ASSLYRGDYAEQF | 0.29380846 |
| PGGPVPIP | ASSLYRGDYAEQF | 0.28682858 |
| FLTVALEP | ASSLYRGDYAEQF | 0.2855698 |
| SQYFGANL | ASSLYRGDYAEQF | 0.28706565 |
| GGVKPAA | ASSLYRGDYAEQF | 0.28663832 |
| GGTGGAGGAGGW | ASSLYRGDYAEQF | 0.2881496 |
| GVGGVAGLGGAGG | ASSLYRGDYAEQF | 0.28369197 |
| VLLVLATTI | ASSLYRGDYAEQF | 0.28500494 |
| IGIFAAML | ASSLYRGDYAEQF | 0.28650928 |
| AVGDLID | ASSLYRGDYAEQF | 0.28563118 |
| NISPLPL | ASSLYRGDYAEQF | 0.28555235 |
| VTAVFVAI | ASSLYRGDYAEQF | 0.2914452 |
| SGDDNDAAL | ASSLYRGDYAEQF | 0.28864548 |
| AVAALAHS | ASSLYRGDYAEQF | 0.28553492 |
| GGVTVMATP | ASSLYRGDYAEQF | 0.2869615 |
| AGLAANVRD | ASSLYRGDYAEQF | 0.29620722 |
| TNRPDLIDPALLRPG | ASSLYRGDYAEQF | 0.29626176 |
| LANLGVTAL | ASSLYRGDYAEQF | 0.28761494 |
| TGERIEL | ASSLYRGDYAEQF | 0.2852863 |
| NGEELAA | ASSLYRGDYAEQF | 0.28647137 |
| VLSLVWG | ASSLYRGDYAEQF | 0.2876736 |
| IGYALWL | ASSLYRGDYAEQF | 0.29173622 |
| LSLLLAPL | ASSLYRGDYAEQF | 0.2879798 |

| | | |
|---|---|---|
| TLDEALK | ASSLYRGDYAEQF | 0.2890216 |
| TDGAGTAEGG | ASSLYRGDYAEQF | 0.29168323 |
| NAPGGAGGNGG | ASSLYRGDYAEQF | 0.28910995 |
| VSVFMGMGGNF | ASSLYRGDYAEQF | 0.28678313 |
| LASYLAAL | ASSLYRGDYAEQF | 0.2842877 |
| LVGGTPAL | ASSLYRGDYAEQF | 0.2848384 |
| VRNADVL | ASSLYRGDYAEQF | 0.28486356 |
| TCSSPFLVSAM | ASSLYRGDYAEQF | 0.28949022 |
| VHITDAE | ASSLYRGDYAEQF | 0.29035658 |
| EYAAPSF | ASSLYRGDYAEQF | 0.285867 |
| VVDAFHGGL | ASSLYRGDYAEQF | 0.2876216 |
| ASVVDPQ | ASSLYRGDYAEQF | 0.29381263 |
| SYNEFGGLS | ASSLYRGDYAEQF | 0.28661075 |
| DPDGSVY | ASSLYRGDYAEQF | 0.28812292 |
| MVSTIHATE | ASSLYRGDYAEQF | 0.2857793 |
| LAAGVADPV | ASSLYRGDYAEQF | 0.28496623 |
| TKEQIAA | ASSLYRGDYAEQF | 0.28546774 |
| SLASPAAL | ASSLYRGDYAEQF | 0.28487256 |
| IGLVFYGV | ASSLYRGDYAEQF | 0.28477752 |
| FLLDASNL | ASSLYRGDYAEQF | 0.2845745 |
| YQVGNFP | ASSLYRGDYAEQF | 0.28502387 |
| ESLLSQST | ASSLYRGDYAEQF | 0.28378415 |
| LFTVGFAAVG | ASSLYRGDYAEQF | 0.2857039 |
| ATPPTPT | ASSLYRGDYAEQF | 0.29089576 |
| TDLATLG | ASSLYRGDYAEQF | 0.28435165 |
| SAGGTGR | ASSLYRGDYAEQF | 0.29678258 |
| GFSNSGS | ASSLYRGDYAEQF | 0.29717058 |
| GGNGGSGGTSV | ASSLYRGDYAEQF | 0.28550017 |
| GTSVATGGAGN | ASSLYRGDYAEQF | 0.28657442 |
| NGGNGGSGGTS | ASSLYRGDYAEQF | 0.28425536 |
| AVVKFPH | ASSLYRGDYAEQF | 0.3050593 |
| TPPPPPP | ASSLYRGDYAEQF | 0.28495258 |
| AAPAAAAAPA | ASSLYRGDYAEQF | 0.2883315 |
| IALNEDL | ASSLYRGDYAEQF | 0.28865132 |
| IVNVSSM | ASSLYRGDYAEQF | 0.28417143 |
| TGIPPAPR | ASSLYRGDYAEQF | 0.28434137 |
| FDLGGGTF | ASSLYRGDYAEQF | 0.2858935 |
| FDLGGGTFDVS | ASSLYRGDYAEQF | 0.28591236 |
| NEPTAAA | ASSLYRGDYAEQF | 0.2849576 |
| NEPTAAALA | ASSLYRGDYAEQF | 0.2864755 |
| GSIINISSI | ASSLYRGDYAEQF | 0.28579074 |
| VAPGNAGTA | ASSLYRGDYAEQF | 0.2855297 |
| TGGGASLE | ASSLYRGDYAEQF | 0.2844796 |
| VAVYNHYKRI | ASSLYRGDYAEQF | 0.28649074 |
| TDPAPATT | ASSLYRGDYAEQF | 0.29189757 |
| TNRPDILDPALLRPG | ASSLYRGDYAEQF | 0.28719383 |
| LTTTTTS | ASSLYRGDYAEQF | 0.29709265 |
| PTGSGKTL | ASSLYRGDYAEQF | 0.28816876 |
| TGSGKTL | ASSLYRGDYAEQF | 0.29629138 |
| TIGHVDHGK | ASSLYRGDYAEQF | 0.28715748 |
| TIGHVDHGKTT | ASSLYRGDYAEQF | 0.2850201 |

| | | |
|---|---|---|
| GHVDHGKTT | ASSLYRGDYAEQF | 0.29021123 |
| AQFEHTLL | ASSLYRGDYAEQF | 0.28416353 |
| IVNQDEL | ASSLYRGDYAEQF | 0.28549814 |
| TNGSQFF | ASSLYRGDYAEQF | 0.28831813 |
| TGKGIVE | ASSLYRGDYAEQF | 0.28534782 |
| LISNASDA | ASSLYRGDYAEQF | 0.28736186 |
| AELRTLGELTKIAKAHGAQ | ASSLYRGDYAEQF | 0.29035994 |
| APNAVAAAEPSW | ASSLYRGDYAEQF | 0.28474027 |
| APQVAGAATAVL | ASSLYRGDYAEQF | 0.28526753 |
| ARRDPAAAAAILPTDA | ASSLYRGDYAEQF | 0.2854986 |
| DAAARDVLAEAGLAEYF | ASSLYRGDYAEQF | 0.29077905 |
| DDQIVLSAKANLAKVEQ | ASSLYRGDYAEQF | 0.29067346 |
| DELLPDTGEDSDEDGH | ASSLYRGDYAEQF | 0.28540722 |
| ELPEAPANRATVPQD | ASSLYRGDYAEQF | 0.2866242 |
| FGQSKIERVTGIG | ASSLYRGDYAEQF | 0.2877184 |
| GAGGTGSTAGGAGGAGGAGGLY | ASSLYRGDYAEQF | 0.28533897 |
| GHGGTGAAGGNGGTGGAGGSGI | ASSLYRGDYAEQF | 0.28774244 |
| GNGGNGGAGGEGGPGGAGGAGA | ASSLYRGDYAEQF | 0.28869897 |
| GSTAGNGGNGGNATNGTIAGQP | ASSLYRGDYAEQF | 0.28667426 |
| GTGGDGGNGGNGGTGG | ASSLYRGDYAEQF | 0.28469104 |
| GVRRPAGIPADAPV | ASSLYRGDYAEQF | 0.2873321 |
| GYRQVTDAHLLTLARR | ASSLYRGDYAEQF | 0.28523222 |
| IAKMGQDGHDRG | ASSLYRGDYAEQF | 0.298403 |
| KAEPSIEIQYT | ASSLYRGDYAEQF | 0.28493825 |
| KPVRIRMAADEDTVAVT | ASSLYRGDYAEQF | 0.28596056 |
| MPPRIAGMRLLVIKPEP | ASSLYRGDYAEQF | 0.28541696 |
| MKFYAVALFRSRGVPPTR | ASSLYRGDYAEQF | 0.29799545 |
| NAGGFGILGAVAHSP | ASSLYRGDYAEQF | 0.2865054 |
| NEGREITLAMAEM | ASSLYRGDYAEQF | 0.2873295 |
| NELVEWSHPLGHRWRWE | ASSLYRGDYAEQF | 0.28926864 |
| NLGAGGTGGNGGNPG | ASSLYRGDYAEQF | 0.28437868 |
| NTEWPKWETVMAADPVD | ASSLYRGDYAEQF | 0.29087356 |
| PASAAPAETQP | ASSLYRGDYAEQF | 0.29078066 |
| QQSSDGKNGVRAAEITGE | ASSLYRGDYAEQF | 0.28579962 |
| RAPDAVTTSWWKE | ASSLYRGDYAEQF | 0.2851297 |
| SGFGNVGAHTSGWFNQ | ASSLYRGDYAEQF | 0.2846448 |
| SGVAGAVGQAASVGGLKV | ASSLYRGDYAEQF | 0.28697976 |
| SNGGAGTGSAGGNGGT | ASSLYRGDYAEQF | 0.2856781 |
| TGGAGGEGGAAG | ASSLYRGDYAEQF | 0.28660843 |
| TGGDGGAAGAGGAPGHG | ASSLYRGDYAEQF | 0.28767326 |
| TMPPLLAAIANAGRHGV | ASSLYRGDYAEQF | 0.28674853 |
| TSSTAGNSGGA | ASSLYRGDYAEQF | 0.2888903 |
| VPATVPVGILC | ASSLYRGDYAEQF | 0.2961593 |
| VQRPDSGGVVLTGRIS | ASSLYRGDYAEQF | 0.28783306 |
| WTPNPVPATKGVPAGLVQAT | ASSLYRGDYAEQF | 0.30344784 |
| YALKRALAKFSDRFDVVII | ASSLYRGDYAEQF | 0.2847024 |
| YLIGSVIV | ASSLYRGDYAEQF | 0.28608325 |
| YGQLVAVID | ASSLYRGDYAEQF | 0.28676173 |
| VTVGGMTLPA | ASSLYRGDYAEQF | 0.287925 |
| VLALPAV | ASSLYRGDYAEQF | 0.28464592 |
| VIPPTLNYET | ASSLYRGDYAEQF | 0.29198167 |

| | | |
|---|---|---|
| VGMGGMHPSSG | ASSLYRGDYAEQF | 0.29493514 |
| VGLAGGAATA | ASSLYRGDYAEQF | 0.28510737 |
| VGGAGGAGGSGP | ASSLYRGDYAEQF | 0.29072738 |
| VFAAFMV | ASSLYRGDYAEQF | 0.28407165 |
| TLYDKLR | ASSLYRGDYAEQF | 0.28520527 |
| TLAVFTVL | ASSLYRGDYAEQF | 0.28431618 |
| TGSVQPARV | ASSLYRGDYAEQF | 0.28655088 |
| TGGGAGSGGT | ASSLYRGDYAEQF | 0.28951243 |
| SHLETSL | ASSLYRGDYAEQF | 0.28788805 |
| SFTTGMT | ASSLYRGDYAEQF | 0.28723338 |
| QVEVAKRMGVRQPTVS | ASSLYRGDYAEQF | 0.29366007 |
| QGSGGPALAM | ASSLYRGDYAEQF | 0.2870692 |
| MLAPSNP | ASSLYRGDYAEQF | 0.28544345 |
| LVNNAAI | ASSLYRGDYAEQF | 0.287632 |
| LTGEINTLA | ASSLYRGDYAEQF | 0.28787935 |
| LSIFGPV | ASSLYRGDYAEQF | 0.28560874 |
| LLALLANL | ASSLYRGDYAEQF | 0.28484726 |
| LISNASDALDKLR | ASSLYRGDYAEQF | 0.28600436 |
| LINEAVP | ASSLYRGDYAEQF | 0.28582317 |
| LGYAFSGL | ASSLYRGDYAEQF | 0.2865599 |
| LDFVAVAL | ASSLYRGDYAEQF | 0.29041305 |
| LAQPGAE | ASSLYRGDYAEQF | 0.28712016 |
| ISNLGAVTGF | ASSLYRGDYAEQF | 0.28442174 |
| HQEDPGRI | ASSLYRGDYAEQF | 0.2840324 |
| GTPGEPTP | ASSLYRGDYAEQF | 0.28780544 |
| GFFNSGS | ASSLYRGDYAEQF | 0.30113548 |
| GAGGGGLTAGAN | ASSLYRGDYAEQF | 0.28844166 |
| FVDALVNM | ASSLYRGDYAEQF | 0.2877925 |
| FLTVAPEL | ASSLYRGDYAEQF | 0.29284343 |
| FETLTAL | ASSLYRGDYAEQF | 0.28709465 |
| DVINPIEL | ASSLYRGDYAEQF | 0.28747112 |
| APVSGAPV | ASSLYRGDYAEQF | 0.28722596 |
| ALVGGLAW | ASSLYRGDYAEQF | 0.28536445 |
| ALAAAAATAI | ASSLYRGDYAEQF | 0.2853943 |
| AFGLGLLL | ASSLYRGDYAEQF | 0.28498715 |
| AFFAAASL | ASSLYRGDYAEQF | 0.28639963 |
| AAAALPAFSPP | ASSLYRGDYAEQF | 0.2925847 |
| VVLAPAL | ASSLYRGDYAEQF | 0.28972036 |
| TIGHVDHGKTTL | ASSLYRGDYAEQF | 0.29349017 |
| LISNASDALDK | ASSLYRGDYAEQF | 0.28818876 |
| LISNASDAL | ASSLYRGDYAEQF | 0.28784397 |
| ISNASDALDK | ASSLYRGDYAEQF | 0.28709677 |
| NQAETLVYQTE | ASSLYRGDYAEQF | 0.28596893 |
| TGGTGGTAGAG | ASSLYRGDYAEQF | 0.31194738 |
| TGSTAGGAGGA | ASSLYRGDYAEQF | 0.285864 |
| DPLAVPVAAGPV | ASSLYRGDYAEQF | 0.2855848 |
| GGTGGTPGNGGN | ASSLYRGDYAEQF | 0.28703764 |
| PAGLAGVDPATL | ASSLYRGDYAEQF | 0.29993325 |
| GNNLVVEAPHAGASV | ASSLYRGDYAEQF | 0.28681478 |
| KVTRIMSASTLEQPA | ASSLYRGDYAEQF | 0.2893687 |
| LGFALPIALPRRTVV | ASSLYRGDYAEQF | 0.2874178 |

| | | |
|---|---|---|
| TRLGNLTARKLVVLV | ASSLYRGDYAEQF | 0.28500554 |
| WSQQRRLIYFPSAGP | ASSLYRGDYAEQF | 0.28697214 |
| GPNGNPDPMAAAVDIR | ASSLYRGDYAEQF | 0.28596514 |
| TQRCVSVADLRAAASR | ASSLYRGDYAEQF | 0.29537785 |
| QFVLDVAEHPVEVLTETV | ASSLYRGDYAEQF | 0.28700653 |
| GGAGGTGGAGGNGGTQAGDG | ASSLYRGDYAEQF | 0.28436399 |
| PLLATLRAIAVDVDAKPAQV | ASSLYRGDYAEQF | 0.28735015 |
| APKFRTGGYMIDFWGVGYQVAK | ASSLYRGDYAEQF | 0.28646165 |
| GAGGAGGAGGAAGGPGGTGGTG | ASSLYRGDYAEQF | 0.28702807 |
| IGALGLLATDVAPAGSVAFGVA | ASSLYRGDYAEQF | 0.32010025 |
| AGGAGGIGGNGGAGGAGGVLMG | ASSLYRGDYAEQF | 0.28800008 |
| ADELIGLL | ASSLYRGDYAEQF | 0.2865372 |
| ADRALKA | ASSLYRGDYAEQF | 0.29518497 |
| AGAVGQVGGAGGA | ASSLYRGDYAEQF | 0.2840225 |
| AGGGGGLGGAP | ASSLYRGDYAEQF | 0.28614512 |
| AGGLLYGNGGAG | ASSLYRGDYAEQF | 0.28497872 |
| AGLESIGL | ASSLYRGDYAEQF | 0.28468883 |
| AGLKTGII | ASSLYRGDYAEQF | 0.28617686 |
| ASLAALLL | ASSLYRGDYAEQF | 0.28828713 |
| ASVINISSI | ASSLYRGDYAEQF | 0.28632128 |
| ATPPGPA | ASSLYRGDYAEQF | 0.29310343 |
| ATVASALV | ASSLYRGDYAEQF | 0.28432003 |
| ATVASLIL | ASSLYRGDYAEQF | 0.28545645 |
| AVNGPVLIRH | ASSLYRGDYAEQF | 0.2922971 |
| DGAAGAVGI | ASSLYRGDYAEQF | 0.2886573 |
| DPLLHGI | ASSLYRGDYAEQF | 0.2909468 |
| ELGAALGAL | ASSLYRGDYAEQF | 0.28512475 |
| ELGRTQAV | ASSLYRGDYAEQF | 0.2845762 |
| FVAGGYLS | ASSLYRGDYAEQF | 0.28528142 |
| GASGGAGGAGGW | ASSLYRGDYAEQF | 0.28863242 |
| GGAGAGGDGGAGGI | ASSLYRGDYAEQF | 0.2846637 |
| GTGHNGSQPGMGGNG | ASSLYRGDYAEQF | 0.28803352 |
| GYAAASASAS | ASSLYRGDYAEQF | 0.28926757 |
| HQVDVAAVQQG | ASSLYRGDYAEQF | 0.28508636 |
| IGAAIAHT | ASSLYRGDYAEQF | 0.28839925 |
| IGMIEVAQL | ASSLYRGDYAEQF | 0.28902927 |
| IRLAPEHTSAMD | ASSLYRGDYAEQF | 0.28487074 |
| IYNVSVL | ASSLYRGDYAEQF | 0.2849533 |
| KQGFLGM | ASSLYRGDYAEQF | 0.29235196 |
| LESVITQL | ASSLYRGDYAEQF | 0.29978654 |
| LGLGALVVV | ASSLYRGDYAEQF | 0.28701124 |
| LIGNGGNGGSGGI | ASSLYRGDYAEQF | 0.28513992 |
| LNAINAPTL | ASSLYRGDYAEQF | 0.28401777 |
| LQLLNTGV | ASSLYRGDYAEQF | 0.3314358 |
| LQTSIEF | ASSLYRGDYAEQF | 0.2859486 |
| MSFAVSEL | ASSLYRGDYAEQF | 0.28641945 |
| NTGALGL | ASSLYRGDYAEQF | 0.2853355 |
| PPGAGGAGGAGTT | ASSLYRGDYAEQF | 0.28693125 |
| QLVGLGI | ASSLYRGDYAEQF | 0.29432124 |
| RFLEQVHQL | ASSLYRGDYAEQF | 0.28509176 |
| RGVGYVLR | ASSLYRGDYAEQF | 0.28458574 |

| | | |
|---|---|---|
| SATGVFAGVI | ASSLYRGDYAEQF | 0.28567293 |
| SLPAITLP | ASSLYRGDYAEQF | 0.28714675 |
| STALAFTV | ASSLYRGDYAEQF | 0.28487548 |
| SVLTVGLGK | ASSLYRGDYAEQF | 0.2871845 |
| TALAGAGSGI | ASSLYRGDYAEQF | 0.34559295 |
| TGLPFTGI | ASSLYRGDYAEQF | 0.28676888 |
| VAEHVTV | ASSLYRGDYAEQF | 0.2876477 |
| VDIVLNSL | ASSLYRGDYAEQF | 0.28588197 |
| VLTGGGAVL | ASSLYRGDYAEQF | 0.28909463 |
| VNVGLAYGI | ASSLYRGDYAEQF | 0.28767094 |
| VSSAAVGAV | ASSLYRGDYAEQF | 0.2859565 |
| VTSVYEAGV | ASSLYRGDYAEQF | 0.28651768 |
| WVALDRGA | ASSLYRGDYAEQF | 0.28735846 |
| YLVGSVVI | ASSLYRGDYAEQF | 0.28808662 |
| GPPGTGKTLL | ASSLYRGDYAEQF | 0.28607738 |
| GVGMTKF | ASSLYRGDYAEQF | 0.28586686 |
| IGIGGSDL | ASSLYRGDYAEQF | 0.2879189 |
| IGTLLAEL | ASSLYRGDYAEQF | 0.28855374 |
| IYTSGTT | ASSLYRGDYAEQF | 0.28602174 |
| LSFPAGV | ASSLYRGDYAEQF | 0.28715113 |
| LYTSGTT | ASSLYRGDYAEQF | 0.29154432 |
| VGAFPFGF | ASSLYRGDYAEQF | 0.28582257 |
| VGAGTVEF | ASSLYRGDYAEQF | 0.28602877 |
| VGKSTLLL | ASSLYRGDYAEQF | 0.28458273 |
| AALAHAGGGIV | ASSLYRGDYAEQF | 0.29880753 |
| AALVAVSLL | ASSLYRGDYAEQF | 0.2879694 |
| AAVSRAVAAL | ASSLYRGDYAEQF | 0.28622445 |
| AGGSSGIGGF | ASSLYRGDYAEQF | 0.28605035 |
| AGLTVANANG | ASSLYRGDYAEQF | 0.28602773 |
| AGLVGAVL | ASSLYRGDYAEQF | 0.28703418 |
| ALSQAFTV | ASSLYRGDYAEQF | 0.2849219 |
| ALTFSVSL | ASSLYRGDYAEQF | 0.28463182 |
| ALTGAALVAL | ASSLYRGDYAEQF | 0.29176646 |
| ASFSIGAGVL | ASSLYRGDYAEQF | 0.28564498 |
| DGSVYKI | ASSLYRGDYAEQF | 0.28614122 |
| EQLAEIV | ASSLYRGDYAEQF | 0.29119205 |
| FAVSELTVM | ASSLYRGDYAEQF | 0.28474063 |
| GAAPLAAGL | ASSLYRGDYAEQF | 0.29004744 |
| GADGGDGGSGGAGG | ASSLYRGDYAEQF | 0.28630784 |
| GFGLDTGGAGG | ASSLYRGDYAEQF | 0.28563464 |
| GLASGGLL | ASSLYRGDYAEQF | 0.28757885 |
| GRIAGAAVLL | ASSLYRGDYAEQF | 0.28502452 |
| GSAGSLNTSF | ASSLYRGDYAEQF | 0.28538945 |
| GTGQIGI | ASSLYRGDYAEQF | 0.28635404 |
| GTLIALIL | ASSLYRGDYAEQF | 0.30094528 |
| IAATMGFVVG | ASSLYRGDYAEQF | 0.29207188 |
| IFAQALL | ASSLYRGDYAEQF | 0.2865765 |
| IGAAVALGF | ASSLYRGDYAEQF | 0.28670207 |
| IGLTPLAL | ASSLYRGDYAEQF | 0.2859507 |
| ILGSNTSTL | ASSLYRGDYAEQF | 0.2860192 |
| IVFSLTVL | ASSLYRGDYAEQF | 0.284978 |

| | | |
|---|---|---|
| LASKATGF | ASSLYRGDYAEQF | 0.28604704 |
| LASVALIL | ASSLYRGDYAEQF | 0.28548425 |
| LFAVGLTL | ASSLYRGDYAEQF | 0.28538343 |
| LFNGGIGTYI | ASSLYRGDYAEQF | 0.28518036 |
| LTLGAGII | ASSLYRGDYAEQF | 0.2875436 |
| LVAAALGDI | ASSLYRGDYAEQF | 0.2869996 |
| NTAFVQLGI | ASSLYRGDYAEQF | 0.28532144 |
| QLFSGAEL | ASSLYRGDYAEQF | 0.28691998 |
| SAATFITL | ASSLYRGDYAEQF | 0.28715268 |
| SNYGLANLGV | ASSLYRGDYAEQF | 0.28607008 |
| SYGGAYAVM | ASSLYRGDYAEQF | 0.28771585 |
| TTGVGGLTLG | ASSLYRGDYAEQF | 0.28582814 |
| TTQTIYI | ASSLYRGDYAEQF | 0.28998068 |
| VAAGAGLGAVI | ASSLYRGDYAEQF | 0.2952589 |
| VAAVVTLL | ASSLYRGDYAEQF | 0.28794017 |
| VGFLGGLI | ASSLYRGDYAEQF | 0.2854698 |
| VGVIPILL | ASSLYRGDYAEQF | 0.28534904 |
| VIFASSLI | ASSLYRGDYAEQF | 0.28841236 |
| VKGGDGGAAAG | ASSLYRGDYAEQF | 0.28704047 |
| VTKTFTNI | ASSLYRGDYAEQF | 0.28412896 |
| VVAAAILL | ASSLYRGDYAEQF | 0.28814226 |
| YAALDVEL | ASSLYRGDYAEQF | 0.2872657 |
| LASVALLL | ASSLYRGDYAEQF | 0.2862336 |
| LDAVHRV | ASSLYRGDYAEQF | 0.28343162 |
| QGLSSSIF | ASSLYRGDYAEQF | 0.29375616 |
| RGTFANIRL | ASSLYRGDYAEQF | 0.2989475 |
| RQLWWGHRI | ASSLYRGDYAEQF | 0.28622064 |
| YGPPGTGKTLL | ASSLYRGDYAEQF | 0.28689322 |
| AALVGAPANVA | ASSLYRGDYAEQF | 0.28435785 |
| AATFRRTE | ASSLYRGDYAEQF | 0.28909603 |
| AGALGGGI | ASSLYRGDYAEQF | 0.29216033 |
| AGGAGGAGGIGAG | ASSLYRGDYAEQF | 0.286044 |
| AGSGALGL | ASSLYRGDYAEQF | 0.2919846 |
| AQIADQI | ASSLYRGDYAEQF | 0.28606907 |
| ASAVAVTYL | ASSLYRGDYAEQF | 0.28725854 |
| ASTVLGGTGG | ASSLYRGDYAEQF | 0.29103923 |
| AVGGEPAK | ASSLYRGDYAEQF | 0.28739247 |
| CEQGTFGGTC | ASSLYRGDYAEQF | 0.2849134 |
| DEGKAPAG | ASSLYRGDYAEQF | 0.28579333 |
| DTVVFPAL | ASSLYRGDYAEQF | 0.28655463 |
| FEGITMAG | ASSLYRGDYAEQF | 0.285917 |
| FPLVVQD | ASSLYRGDYAEQF | 0.28677213 |
| GAFTTPQ | ASSLYRGDYAEQF | 0.29812926 |
| GFGGFGTAGGVGGA | ASSLYRGDYAEQF | 0.28823838 |
| GFLAQFL | ASSLYRGDYAEQF | 0.28563964 |
| GGAGGAGGNAGVA | ASSLYRGDYAEQF | 0.29750752 |
| GGVGTAGAGGAGGAGG | ASSLYRGDYAEQF | 0.28459364 |
| GNGGGGDAGNAGSGGN | ASSLYRGDYAEQF | 0.28831968 |
| GTGGAGGAGNDAGS | ASSLYRGDYAEQF | 0.28518412 |
| GTTNTGWFNS | ASSLYRGDYAEQF | 0.29508692 |
| IFMGPLFTL | ASSLYRGDYAEQF | 0.28552446 |

| | | |
|---|---|---|
| ILAHAIEAI | ASSLYRGDYAEQF | 0.28630996 |
| IRSAIREAL | ASSLYRGDYAEQF | 0.28638887 |
| ITFAGGG | ASSLYRGDYAEQF | 0.20752643 |
| ITGMSGHQEV | ASSLYRGDYAEQF | 0.22454771 |
| IVAGLACTL | ASSLYRGDYAEQF | 0.21647394 |
| IVGGGLAGA | ASSLYRGDYAEQF | 0.2527923 |
| IVTGVLNI | ASSLYRGDYAEQF | 0.21821775 |
| LAQQGCI | ASSLYRGDYAEQF | 0.20765312 |
| LCLHVEN | ASSLYRGDYAEQF | 0.22957976 |
| LDAISDGS | ASSLYRGDYAEQF | 0.2049289 |
| LGGFAAML | ASSLYRGDYAEQF | 0.20766425 |
| LGGKNPMI | ASSLYRGDYAEQF | 0.23056343 |
| LGPKGVAAIL | ASSLYRGDYAEQF | 0.21535467 |
| LMVGVSI | ASSLYRGDYAEQF | 0.2702588 |
| LVGAAFVL | ASSLYRGDYAEQF | 0.24485756 |
| LVLAVAPL | ASSLYRGDYAEQF | 0.20509173 |
| MAAIAVIDAA | ASSLYRGDYAEQF | 0.21032827 |
| NNGINAELQ | ASSLYRGDYAEQF | 0.22929627 |
| RGEPLVLVHG | ASSLYRGDYAEQF | 0.22175865 |
| RGGAGGAAGLI | ASSLYRGDYAEQF | 0.19887705 |
| RSYRMHPAV | ASSLYRGDYAEQF | 0.22903952 |
| SAAHLLAGQLA | ASSLYRGDYAEQF | 0.20220251 |
| SAAVGSY | ASSLYRGDYAEQF | 0.19743551 |
| SPHAVLHRI | ASSLYRGDYAEQF | 0.24740382 |
| SQIAKVL | ASSLYRGDYAEQF | 0.21894522 |
| TALGHLVP | ASSLYRGDYAEQF | 0.21103586 |
| TAPTTSGAA | ASSLYRGDYAEQF | 0.24545291 |
| TDGDDGNGGSPGT | ASSLYRGDYAEQF | 0.20099722 |
| TDGLGEN | ASSLYRGDYAEQF | 0.2097438 |
| TGGPPLPGQA | ASSLYRGDYAEQF | 0.23721053 |
| TIPAITI | ASSLYRGDYAEQF | 0.24659988 |
| TIPGFTI | ASSLYRGDYAEQF | 0.21054702 |
| TSIASGALA | ASSLYRGDYAEQF | 0.21996854 |
| TVTLPGAAI | ASSLYRGDYAEQF | 0.30451182 |
| VASGLSAALP | ASSLYRGDYAEQF | 0.23068252 |
| VGAGVIGI | ASSLYRGDYAEQF | 0.24233767 |
| VGPKRFGV | ASSLYRGDYAEQF | 0.25181663 |
| VNVAFPD | ASSLYRGDYAEQF | 0.21207525 |
| VSMTNSGVS | ASSLYRGDYAEQF | 0.2338207 |
| VTDVLGDLSV | ASSLYRGDYAEQF | 0.28363627 |
| VVVVGAGFAGL | ASSLYRGDYAEQF | 0.22060327 |
| VYTVGTVV | ASSLYRGDYAEQF | 0.23213783 |
| WTTFSGP | ASSLYRGDYAEQF | 0.21422629 |
| YDELVEAF | ASSLYRGDYAEQF | 0.22711925 |
| YGTGGAGGNGG | ASSLYRGDYAEQF | 0.198508 |
| YTNVMAV | ASSLYRGDYAEQF | 0.21635954 |
| FLEQVHQL | ASSLYRGDYAEQF | 0.25879163 |
| GLSLLAVAV | ASSLYRGDYAEQF | 0.22676538 |
| LEQVHQL | ASSLYRGDYAEQF | 0.20161085 |
| LNAAGAI | ASSLYRGDYAEQF | 0.20912734 |
| PAVHASL | ASSLYRGDYAEQF | 0.20312153 |

| | | |
|---|---|---|
| TELEDDA | ASSLYRGDYAEQF | 0.21246864 |
| TPPGPPN | ASSLYRGDYAEQF | 0.20154053 |
| TPPPPPPP | ASSLYRGDYAEQF | 0.2015452 |
| TQSPALI | ASGDPTGGNTGQLY | 0.2613624 |
| EFAAAFPQ | ASGDPTGGNTGQLY | 0.26237234 |
| LLSGNLDVL | ASGDPTGGNTGQLY | 0.21153486 |
| GPANLGNYNI | ASGDPTGGNTGQLY | 0.22921866 |
| TLGPTLP | ASGDPTGGNTGQLY | 0.24147537 |
| VYAYSSCCK | ASGDPTGGNTGQLY | 0.2865284 |
| APAPAPAPNI | ASGDPTGGNTGQLY | 0.23575388 |
| PGGPVPIP | ASGDPTGGNTGQLY | 0.20392677 |
| FLTVALEP | ASGDPTGGNTGQLY | 0.28702703 |
| SQYFGANL | ASGDPTGGNTGQLY | 0.21743277 |
| GGVKPAA | ASGDPTGGNTGQLY | 0.21911788 |
| GGTGGAGGAGGW | ASGDPTGGNTGQLY | 0.26399857 |
| GVGGVAGLGGAGG | ASGDPTGGNTGQLY | 0.21172932 |
| VLLVLATTI | ASGDPTGGNTGQLY | 0.20686716 |
| IGIFAAML | ASGDPTGGNTGQLY | 0.24075389 |
| AVGDLID | ASGDPTGGNTGQLY | 0.24888478 |
| NISPLPL | ASGDPTGGNTGQLY | 0.20657629 |
| VTAVFVAI | ASGDPTGGNTGQLY | 0.2043005 |
| SGDDNDAAL | ASGDPTGGNTGQLY | 0.21008015 |
| AVAALAHS | ASGDPTGGNTGQLY | 0.23228544 |
| GGVTVMATP | ASGDPTGGNTGQLY | 0.25810048 |
| AGLAANVRD | ASGDPTGGNTGQLY | 0.22941619 |
| TNRPDLIDPALLRPG | ASGDPTGGNTGQLY | 0.20567912 |
| LANLGVTAL | ASGDPTGGNTGQLY | 0.24119125 |
| TGERIEL | ASGDPTGGNTGQLY | 0.22467138 |
| NGEELAA | ASGDPTGGNTGQLY | 0.22992627 |
| VLSLVWG | ASGDPTGGNTGQLY | 0.23125766 |
| IGYALWL | ASGDPTGGNTGQLY | 0.20035668 |
| LSLLLAPL | ASGDPTGGNTGQLY | 0.24355108 |
| TLDEALK | ASGDPTGGNTGQLY | 0.26116097 |
| TDGAGTAEGG | ASGDPTGGNTGQLY | 0.23630258 |
| NAPGGAGGNGG | ASGDPTGGNTGQLY | 0.23611037 |
| VSVFMGMGGNF | ASGDPTGGNTGQLY | 0.21172951 |
| LASYLAAL | ASGDPTGGNTGQLY | 0.23713487 |
| LVGGTPAL | ASGDPTGGNTGQLY | 0.21360555 |
| VRNADVL | ASGDPTGGNTGQLY | 0.2191899 |
| TCSSPFLVSAM | ASGDPTGGNTGQLY | 0.22989365 |
| VHITDAE | ASGDPTGGNTGQLY | 0.21738133 |
| EYAAPSF | ASGDPTGGNTGQLY | 0.24407196 |
| VVDAFHGGL | ASGDPTGGNTGQLY | 0.23469394 |
| ASVVDPQ | ASGDPTGGNTGQLY | 0.21632467 |
| SYNEFGGLS | ASGDPTGGNTGQLY | 0.27629226 |
| DPDGSVY | ASGDPTGGNTGQLY | 0.27999493 |
| MVSTIHATE | ASGDPTGGNTGQLY | 0.20745125 |
| LAAGVADPV | ASGDPTGGNTGQLY | 0.20891793 |
| TKEQIAA | ASGDPTGGNTGQLY | 0.22448292 |
| SLASPAAL | ASGDPTGGNTGQLY | 0.24472328 |
| IGLVFYGV | ASGDPTGGNTGQLY | 0.23468879 |

| | | |
|---|---|---|
| FLLDASNL | ASGDPTGGNTGQLY | 0.23683989 |
| YQVGNFP | ASGDPTGGNTGQLY | 0.23980418 |
| ESLLSQST | ASGDPTGGNTGQLY | 0.20311552 |
| LFTVGFAAVG | ASGDPTGGNTGQLY | 0.28607202 |
| ATPPTPT | ASGDPTGGNTGQLY | 0.21008828 |
| TDLATLG | ASGDPTGGNTGQLY | 0.21825747 |
| SAGGTGR | ASGDPTGGNTGQLY | 0.21128863 |
| GFSNSGS | ASGDPTGGNTGQLY | 0.19755965 |
| GGNGGSGGTSV | ASGDPTGGNTGQLY | 0.26207826 |
| GTSVATGGAGN | ASGDPTGGNTGQLY | 0.21891478 |
| NGGNGGSGGTS | ASGDPTGGNTGQLY | 0.25160903 |
| AVVKFPH | ASGDPTGGNTGQLY | 0.22014649 |
| TPPPPPP | ASGDPTGGNTGQLY | 0.23238665 |
| AAPAAAAAPA | ASGDPTGGNTGQLY | 0.22967605 |
| IALNEDL | ASGDPTGGNTGQLY | 0.22648637 |
| IVNVSSM | ASGDPTGGNTGQLY | 0.20955056 |
| TGIPPAPR | ASGDPTGGNTGQLY | 0.24482146 |
| FDLGGGTF | ASGDPTGGNTGQLY | 0.2708193 |
| FDLGGGTFDVS | ASGDPTGGNTGQLY | 0.24037907 |
| NEPTAAA | ASGDPTGGNTGQLY | 0.20552367 |
| NEPTAAALA | ASGDPTGGNTGQLY | 0.20694996 |
| GSIINISSI | ASGDPTGGNTGQLY | 0.22692615 |
| VAPGNAGTA | ASGDPTGGNTGQLY | 0.19694811 |
| TGGGASLE | ASGDPTGGNTGQLY | 0.19753711 |
| VAVYNHYKRI | ASGDPTGGNTGQLY | 0.19878866 |
| TDPAPATT | ASGDPTGGNTGQLY | 0.20278616 |
| TNRPDILDPALLRPG | ASGDPTGGNTGQLY | 0.25202018 |
| LTTTTTS | ASGDPTGGNTGQLY | 0.24183686 |
| PTGSGKTL | ASGDPTGGNTGQLY | 0.22760642 |
| TGSGKTL | ASGDPTGGNTGQLY | 0.25506622 |
| TIGHVDHGK | ASGDPTGGNTGQLY | 0.21698391 |
| TIGHVDHGKTT | ASGDPTGGNTGQLY | 0.19526385 |
| GHVDHGKTT | ASGDPTGGNTGQLY | 0.21397182 |
| AQFEHTLL | ASGDPTGGNTGQLY | 0.20900337 |
| IVNQDEL | ASGDPTGGNTGQLY | 0.24674559 |
| TNGSQFF | ASGDPTGGNTGQLY | 0.20099835 |
| TGKGIVE | ASGDPTGGNTGQLY | 0.21823688 |
| LISNASDA | ASGDPTGGNTGQLY | 0.23617917 |
| AELRTLGELTKIAKAHGAQ | ASGDPTGGNTGQLY | 0.21520577 |
| APNAVAAAEPSW | ASGDPTGGNTGQLY | 0.24326697 |
| APQVAGAATAVL | ASGDPTGGNTGQLY | 0.20508923 |
| ARRDPAAAAAILPTDA | ASGDPTGGNTGQLY | 0.1996777 |
| DAAARDVLAEAGLAEYF | ASGDPTGGNTGQLY | 0.24989098 |
| DDQIVLSAKANLAKVEQ | ASGDPTGGNTGQLY | 0.19921425 |
| DELLPDTGEDSDEDGH | ASGDPTGGNTGQLY | 0.21260783 |
| ELPEAPANRATVPQD | ASGDPTGGNTGQLY | 0.20212273 |
| FGQSKIERVTGIG | ASGDPTGGNTGQLY | 0.19770718 |
| GAGGTGSTAGGAGGAGGAGGLY | ASGDPTGGNTGQLY | 0.20156957 |
| GHGGTGAAGGNGGTGGAGGSGI | ASGDPTGGNTGQLY | 0.23823005 |
| GNGGNGGAGGEGGPGGAGGAGA | ASGDPTGGNTGQLY | 0.21455884 |
| GSTAGNGGNGGNATNGTIAGQP | ASGDPTGGNTGQLY | 0.2235364 |

| | | |
|---|---|---|
| GTGGDGGNGGNGGTGG | ASGDPTGGNTGQLY | 0.2635969 |
| GVRRPAGIPADAPV | ASGDPTGGNTGQLY | 0.21623456 |
| GYRQVTDAHLLTLARR | ASGDPTGGNTGQLY | 0.22220786 |
| IAKMGQDGHDRG | ASGDPTGGNTGQLY | 0.20870122 |
| KAEPSIEIQYT | ASGDPTGGNTGQLY | 0.21197088 |
| KPVRIRMAADEDTVAVT | ASGDPTGGNTGQLY | 0.20987085 |
| MPPRIAGMRLLVIKPEP | ASGDPTGGNTGQLY | 0.21333148 |
| MKFYAVALFRSRGVPPTR | ASGDPTGGNTGQLY | 0.27497405 |
| NAGGFGILGAVAHSP | ASGDPTGGNTGQLY | 0.22207947 |
| NEGREITLAMAEM | ASGDPTGGNTGQLY | 0.20630184 |
| NELVEWSHPLGHRWRWE | ASGDPTGGNTGQLY | 0.21323775 |
| NLGAGGTGGNGGNPG | ASGDPTGGNTGQLY | 0.29179695 |
| NTEWPKWETVMAADPVD | ASGDPTGGNTGQLY | 0.29689145 |
| PASAAPAETQP | ASGDPTGGNTGQLY | 0.20765851 |
| QQSSDGKNGVRAAEITGE | ASGDPTGGNTGQLY | 0.23724523 |
| RAPDAVTTSWWKE | ASGDPTGGNTGQLY | 0.23207542 |
| SGFGNVGAHTSGWFNQ | ASGDPTGGNTGQLY | 0.21613348 |
| SGVAGAVGQAASVGGLKV | ASGDPTGGNTGQLY | 0.21240512 |
| SNGGAGTGSAGGNGGT | ASGDPTGGNTGQLY | 0.2091848 |
| TGGAGGEGGAAG | ASGDPTGGNTGQLY | 0.23856525 |
| TGGDGGAAGAGGAPGHG | ASGDPTGGNTGQLY | 0.5081389 |
| TMPPLLAAIANAGRHGV | ASGDPTGGNTGQLY | 0.21618803 |
| TSSTAGNSGGA | ASGDPTGGNTGQLY | 0.20357175 |
| VPATVPVGILC | ASGDPTGGNTGQLY | 0.22779763 |
| VQRPDSGGVVLTGRIS | ASGDPTGGNTGQLY | 0.20532662 |
| WTPNPVPATKGVPAGLVQAT | ASGDPTGGNTGQLY | 0.21758097 |
| YALKRALAKFSDRFDVVII | ASGDPTGGNTGQLY | 0.21846509 |
| YLIGSVIV | ASGDPTGGNTGQLY | 0.20288181 |
| YGQLVAVID | ASGDPTGGNTGQLY | 0.23172584 |
| VTVGGMTLPA | ASGDPTGGNTGQLY | 0.25589326 |
| VLALPAV | ASGDPTGGNTGQLY | 0.21777327 |
| VIPPTLNYET | ASGDPTGGNTGQLY | 0.24983929 |
| VGMGGMHPSSG | ASGDPTGGNTGQLY | 0.2831825 |
| VGLAGGAATA | ASGDPTGGNTGQLY | 0.23679912 |
| VGGAGGAGGSGP | ASGDPTGGNTGQLY | 0.27260503 |
| VFAAFMV | ASGDPTGGNTGQLY | 0.21899788 |
| TLYDKLR | ASGDPTGGNTGQLY | 0.22234155 |
| TLAVFTVL | ASGDPTGGNTGQLY | 0.21920328 |
| TGSVQPARV | ASGDPTGGNTGQLY | 0.25832438 |
| TGGGAGSGGT | ASGDPTGGNTGQLY | 0.23106663 |
| SHLETSL | ASGDPTGGNTGQLY | 0.23750766 |
| SFTTGMT | ASGDPTGGNTGQLY | 0.22552325 |
| QVEVAKRMGVRQPTVS | ASGDPTGGNTGQLY | 0.2789288 |
| QGSGGPALAM | ASGDPTGGNTGQLY | 0.21957022 |
| MLAPSNP | ASGDPTGGNTGQLY | 0.2954054 |
| LVNNAAI | ASGDPTGGNTGQLY | 0.20290367 |
| LTGEINTLA | ASGDPTGGNTGQLY | 0.20949204 |
| LSIFGPV | ASGDPTGGNTGQLY | 0.23709206 |
| LLALLANL | ASGDPTGGNTGQLY | 0.20201029 |
| LISNASDALDKLR | ASGDPTGGNTGQLY | 0.19981892 |
| LINEAVP | ASGDPTGGNTGQLY | 0.21980241 |

| | | |
|---|---|---|
| LGYAFSGL | ASGDPTGGNTGQLY | 0.2370392 |
| LDFVAVAL | ASGDPTGGNTGQLY | 0.19557573 |
| LAQPGAE | ASGDPTGGNTGQLY | 0.21989661 |
| ISNLGAVTGF | ASGDPTGGNTGQLY | 0.2114964 |
| HQEDPGRI | ASGDPTGGNTGQLY | 0.21092413 |
| GTPGEPTP | ASGDPTGGNTGQLY | 0.21807209 |
| GFFNSGS | ASGDPTGGNTGQLY | 0.20408164 |
| GAGGGGLTAGAN | ASGDPTGGNTGQLY | 0.24395968 |
| FVDALVNM | ASGDPTGGNTGQLY | 0.20503591 |
| FLTVAPEL | ASGDPTGGNTGQLY | 0.20058006 |
| FETLTAL | ASGDPTGGNTGQLY | 0.23450688 |
| DVINPIEL | ASGDPTGGNTGQLY | 0.26417476 |
| APVSGAPV | ASGDPTGGNTGQLY | 0.20790628 |
| ALVGGLAW | ASGDPTGGNTGQLY | 0.23370527 |
| ALAAAAATAI | ASGDPTGGNTGQLY | 0.20527418 |
| AFGLGLLL | ASGDPTGGNTGQLY | 0.4230261 |
| AFFAAASL | ASGDPTGGNTGQLY | 0.22910273 |
| AAAALPAFSPP | ASGDPTGGNTGQLY | 0.20504002 |
| VVLAPAL | ASGDPTGGNTGQLY | 0.24417295 |
| TIGHVDHGKTTL | ASGDPTGGNTGQLY | 0.23748538 |
| LISNASDALDK | ASGDPTGGNTGQLY | 0.24657655 |
| LISNASDAL | ASGDPTGGNTGQLY | 0.23943 |
| ISNASDALDK | ASGDPTGGNTGQLY | 0.20415053 |
| NQAETLVYQTE | ASGDPTGGNTGQLY | 0.22475551 |
| TGGTGGTAGAG | ASGDPTGGNTGQLY | 0.24186115 |
| TGSTAGGAGGA | ASGDPTGGNTGQLY | 0.22739924 |
| DPLAVPVAAGPV | ASGDPTGGNTGQLY | 0.20297669 |
| GGTGGTPGNGGN | ASGDPTGGNTGQLY | 0.24190038 |
| PAGLAGVDPATL | ASGDPTGGNTGQLY | 0.21672776 |
| GNNLVVEAPHAGASV | ASGDPTGGNTGQLY | 0.21865879 |
| KVTRIMSASTLEQPA | ASGDPTGGNTGQLY | 0.25384486 |
| LGFALPIALPRRTVV | ASGDPTGGNTGQLY | 0.21937962 |
| TRLGNLTARKLVVLV | ASGDPTGGNTGQLY | 0.2642755 |
| WSQQRRLIYFPSAGP | ASGDPTGGNTGQLY | 0.24956818 |
| GPNGNPDPMAAAVDIR | ASGDPTGGNTGQLY | 0.21753038 |
| TQRCVSVADLRAAASR | ASGDPTGGNTGQLY | 0.20769688 |
| QFVLDVAEHPVEVLTETV | ASGDPTGGNTGQLY | 0.22829454 |
| GGAGGTGGAGGNGGTQAGDG | ASGDPTGGNTGQLY | 0.24493906 |
| PLLATLRAIAVDVDAKPAQV | ASGDPTGGNTGQLY | 0.22829536 |
| APKFRTGGYMIDFWGVGYQVAK | ASGDPTGGNTGQLY | 0.2038043 |
| GAGGAGGAGGAAGGPGGTGGTG | ASGDPTGGNTGQLY | 0.20428047 |
| IGALGLLATDVAPAGSVAFGVA | ASGDPTGGNTGQLY | 0.24847715 |
| AGGAGGIGGNGGAGGAGGVLMG | ASGDPTGGNTGQLY | 0.2874248 |
| ADELIGLL | ASGDPTGGNTGQLY | 0.21522206 |
| ADRALKA | ASGDPTGGNTGQLY | 0.22312735 |
| AGAVGQVGGAGGA | ASGDPTGGNTGQLY | 0.30987728 |
| AGGGGGLGGAP | ASGDPTGGNTGQLY | 0.21348476 |
| AGGLLYGNGGAG | ASGDPTGGNTGQLY | 0.22013839 |
| AGLESIGL | ASGDPTGGNTGQLY | 0.25742447 |
| AGLKTGII | ASGDPTGGNTGQLY | 0.28193814 |
| ASLAALLL | ASGDPTGGNTGQLY | 0.21728256 |

| | | |
|---|---|---|
| ASVINISSI | ASGDPTGGNTGQLY | 0.26758882 |
| ATPPGPA | ASGDPTGGNTGQLY | 0.20733383 |
| ATVASALV | ASGDPTGGNTGQLY | 0.24550137 |
| ATVASLIL | ASGDPTGGNTGQLY | 0.20288812 |
| AVNGPVLIRH | ASGDPTGGNTGQLY | 0.2258203 |
| DGAAGAVGI | ASGDPTGGNTGQLY | 0.19830309 |
| DPLLHGI | ASGDPTGGNTGQLY | 0.24054958 |
| ELGAALGAL | ASGDPTGGNTGQLY | 0.2563841 |
| ELGRTQAV | ASGDPTGGNTGQLY | 0.24863414 |
| FVAGGYLS | ASGDPTGGNTGQLY | 0.1991867 |
| GASGGAGGAGGW | ASGDPTGGNTGQLY | 0.22211379 |
| GGAGAGGDGGAGGI | ASGDPTGGNTGQLY | 0.25261024 |
| GTGHNGSQPGMGGNG | ASGDPTGGNTGQLY | 0.36971423 |
| GYAAASASAS | ASGDPTGGNTGQLY | 0.22223255 |
| HQVDVAAVQQG | ASGDPTGGNTGQLY | 0.2794003 |
| IGAAIAHT | ASGDPTGGNTGQLY | 0.2143519 |
| IGMIEVAQL | ASGDPTGGNTGQLY | 0.21484788 |
| IRLAPEHTSAMD | ASGDPTGGNTGQLY | 0.20387477 |
| IYNVSVL | ASGDPTGGNTGQLY | 0.21634836 |
| KQGFLGM | ASGDPTGGNTGQLY | 0.21755092 |
| LESVITQL | ASGDPTGGNTGQLY | 0.2076013 |
| LGLGALVVV | ASGDPTGGNTGQLY | 0.21911877 |
| LIGNGGNGGSGGI | ASGDPTGGNTGQLY | 0.28163624 |
| LNAINAPTL | ASGDPTGGNTGQLY | 0.24453145 |
| LQLLNTGV | ASGDPTGGNTGQLY | 0.27834207 |
| LQTSIEF | ASGDPTGGNTGQLY | 0.30363482 |
| MSFAVSEL | ASGDPTGGNTGQLY | 0.23705994 |
| NTGALGL | ASGDPTGGNTGQLY | 0.20774697 |
| PPGAGGAGGAGTT | ASGDPTGGNTGQLY | 0.22219373 |
| QLVGLGI | ASGDPTGGNTGQLY | 0.20185937 |
| RFLEQVHQL | ASGDPTGGNTGQLY | 0.22181411 |
| RGVGYVLR | ASGDPTGGNTGQLY | 0.20669445 |
| SATGVFAGVI | ASGDPTGGNTGQLY | 0.24134542 |
| SLPAITLP | ASGDPTGGNTGQLY | 0.19535837 |
| STALAFTV | ASGDPTGGNTGQLY | 0.2502621 |
| SVLTVGLGK | ASGDPTGGNTGQLY | 0.26930577 |
| TALAGAGSGI | ASGDPTGGNTGQLY | 0.19740279 |
| TGLPFTGI | ASGDPTGGNTGQLY | 0.27075732 |
| VAEHVTV | ASGDPTGGNTGQLY | 0.25850013 |
| VDIVLNSL | ASGDPTGGNTGQLY | 0.20128384 |
| VLTGGGAVL | ASGDPTGGNTGQLY | 0.25479135 |
| VNVGLAYGI | ASGDPTGGNTGQLY | 0.20956737 |
| VSSAAVGAV | ASGDPTGGNTGQLY | 0.25394416 |
| VTSVYEAGV | ASGDPTGGNTGQLY | 0.26622456 |
| WVALDRGA | ASGDPTGGNTGQLY | 0.25160944 |
| YLVGSVVI | ASGDPTGGNTGQLY | 0.21978946 |
| GPPGTGKTLL | ASGDPTGGNTGQLY | 0.24693227 |
| GVGMTKF | ASGDPTGGNTGQLY | 0.20960946 |
| IGIGGSDL | ASGDPTGGNTGQLY | 0.22218868 |
| IGTLLAEL | ASGDPTGGNTGQLY | 0.1965828 |
| IYTSGTT | ASGDPTGGNTGQLY | 0.23588794 |

| | | |
|---|---|---|
| LSFPAGV | ASGDPTGGNTGQLY | 0.21549858 |
| LYTSGTT | ASGDPTGGNTGQLY | 0.25337565 |
| VGAFPFGF | ASGDPTGGNTGQLY | 0.21791536 |
| VGAGTVEF | ASGDPTGGNTGQLY | 0.23064801 |
| VGKSTLLL | ASGDPTGGNTGQLY | 0.22054222 |
| AALAHAGGGIV | ASGDPTGGNTGQLY | 0.2074769 |
| AALVAVSLL | ASGDPTGGNTGQLY | 0.20541571 |
| AAVSRAVAAL | ASGDPTGGNTGQLY | 0.20607597 |
| AGGSSGIGGF | ASGDPTGGNTGQLY | 0.23189726 |
| AGLTVANANG | ASGDPTGGNTGQLY | 0.2243686 |
| AGLVGAVL | ASGDPTGGNTGQLY | 0.20355777 |
| ALSQAFTV | ASGDPTGGNTGQLY | 0.25631037 |
| ALTFSVSL | ASGDPTGGNTGQLY | 0.23139417 |
| ALTGAALVAL | ASGDPTGGNTGQLY | 0.20440716 |
| ASFSIGAGVL | ASGDPTGGNTGQLY | 0.22047901 |
| DGSVYKI | ASGDPTGGNTGQLY | 0.23745012 |
| EQLAEIV | ASGDPTGGNTGQLY | 0.22613195 |
| FAVSELTVM | ASGDPTGGNTGQLY | 0.19563098 |
| GAAPLAAGL | ASGDPTGGNTGQLY | 0.2107789 |
| GADGGDGGSGGAGG | ASGDPTGGNTGQLY | 0.2192143 |
| GFGLDTGGAGG | ASGDPTGGNTGQLY | 0.20085055 |
| GLASGGLL | ASGDPTGGNTGQLY | 0.20608966 |
| GRIAGAAVLL | ASGDPTGGNTGQLY | 0.26395816 |
| GSAGSLNTSF | ASGDPTGGNTGQLY | 0.19869778 |
| GTGQIGI | ASGDPTGGNTGQLY | 0.21736488 |
| GTLIALIL | ASGDPTGGNTGQLY | 0.24391854 |
| IAATMGFVVG | ASGDPTGGNTGQLY | 0.2503283 |
| IFAQALL | ASGDPTGGNTGQLY | 0.2765513 |
| IGAAVALGF | ASGDPTGGNTGQLY | 0.21312569 |
| IGLTPLAL | ASGDPTGGNTGQLY | 0.21970513 |
| ILGSNTSTL | ASGDPTGGNTGQLY | 0.24299651 |
| IVFSLTVL | ASGDPTGGNTGQLY | 0.22619297 |
| LASKATGF | ASGDPTGGNTGQLY | 0.20212531 |
| LASVALIL | ASGDPTGGNTGQLY | 0.19985262 |
| LFAVGLTL | ASGDPTGGNTGQLY | 0.21824756 |
| LFNGGIGTYI | ASGDPTGGNTGQLY | 0.20506755 |
| LTLGAGII | ASGDPTGGNTGQLY | 0.24906279 |
| LVAAALGDI | ASGDPTGGNTGQLY | 0.24542688 |
| NTAFVQLGI | ASGDPTGGNTGQLY | 0.2045915 |
| QLFSGAEL | ASGDPTGGNTGQLY | 0.20603728 |
| SAATFITL | ASGDPTGGNTGQLY | 0.21121176 |
| SNYGLANLGV | ASGDPTGGNTGQLY | 0.21096918 |
| SYGGAYAVM | ASGDPTGGNTGQLY | 0.24329215 |
| TTGVGGLTLG | ASGDPTGGNTGQLY | 0.21294422 |
| TTQTIYI | ASGDPTGGNTGQLY | 0.5115372 |
| VAAGAGLGAVI | ASGDPTGGNTGQLY | 0.23228648 |
| VAAVVTLL | ASGDPTGGNTGQLY | 0.20809042 |
| VGFLGGLI | ASGDPTGGNTGQLY | 0.25936264 |
| VGVIPILL | ASGDPTGGNTGQLY | 0.22462578 |
| VIFASSLI | ASGDPTGGNTGQLY | 0.215756 |
| VKGGDGGAAAG | ASGDPTGGNTGQLY | 0.22767505 |

| | | |
|---|---|---|
| VTKTFTNI | ASGDPTGGNTGQLY | 0.20647566 |
| VVAAAILL | ASGDPTGGNTGQLY | 0.20644851 |
| YAALDVEL | ASGDPTGGNTGQLY | 0.22839715 |
| LASVALLL | ASGDPTGGNTGQLY | 0.20757964 |
| LDAVHRV | ASGDPTGGNTGQLY | 0.21605173 |
| QGLSSSIF | ASGDPTGGNTGQLY | 0.20049463 |
| RGTFANIRL | ASGDPTGGNTGQLY | 0.20127344 |
| RQLWWGHRI | ASGDPTGGNTGQLY | 0.22248238 |
| YGPPGTGKTLL | ASGDPTGGNTGQLY | 0.23196915 |
| AALVGAPANVA | ASGDPTGGNTGQLY | 0.31430572 |
| AATFRRTE | ASGDPTGGNTGQLY | 0.21182738 |
| AGALGGGI | ASGDPTGGNTGQLY | 0.20822084 |
| AGGAGGAGGIGAG | ASGDPTGGNTGQLY | 0.25078404 |
| AGSGALGL | ASGDPTGGNTGQLY | 0.21515852 |
| AQIADQI | ASGDPTGGNTGQLY | 0.227573 |
| ASAVAVTYL | ASGDPTGGNTGQLY | 0.2088808 |
| ASTVLGGTGG | ASGDPTGGNTGQLY | 0.20430546 |
| AVGGEPAK | ASGDPTGGNTGQLY | 0.21878882 |
| CEQGTFGGTC | ASGDPTGGNTGQLY | 0.22030216 |
| DEGKAPAG | ASGDPTGGNTGQLY | 0.2019497 |
| DTVVFPAL | ASGDPTGGNTGQLY | 0.23573546 |
| FEGITMAG | ASGDPTGGNTGQLY | 0.19546115 |
| FPLVVQD | ASGDPTGGNTGQLY | 0.21075232 |
| GAFTTPQ | ASGDPTGGNTGQLY | 0.21086401 |
| GFGGFGTAGGVGGA | ASGDPTGGNTGQLY | 0.2066383 |
| GFLAQFL | ASGDPTGGNTGQLY | 0.19881229 |
| GGAGGAGGNAGVA | ASGDPTGGNTGQLY | 0.23089015 |
| GGVGTAGAGGAGGAGG | ASGDPTGGNTGQLY | 0.25030956 |
| GNGGGGDAGNAGSGGN | ASGDPTGGNTGQLY | 0.24827726 |
| GTGGAGGAGNDAGS | ASGDPTGGNTGQLY | 0.20718783 |
| GTTNTGWFNS | ASGDPTGGNTGQLY | 0.24408513 |
| IFMGPLFTL | ASGDPTGGNTGQLY | 0.29273796 |
| ILAHAIEAI | ASGDPTGGNTGQLY | 0.22211398 |
| IRSAIREAL | ASGDPTGGNTGQLY | 0.20269957 |
| ITFAGGG | ASGDPTGGNTGQLY | 0.22146752 |
| ITGMSGHQEV | ASGDPTGGNTGQLY | 0.1989698 |
| IVAGLACTL | ASGDPTGGNTGQLY | 0.21043941 |
| IVGGGLAGA | ASGDPTGGNTGQLY | 0.33303958 |
| IVTGVLNI | ASGDPTGGNTGQLY | 0.2019498 |
| LAQQGCI | ASGDPTGGNTGQLY | 0.23989947 |
| LCLHVEN | ASGDPTGGNTGQLY | 0.25639263 |
| LDAISDGS | ASGDPTGGNTGQLY | 0.20902155 |
| LGGFAAML | ASGDPTGGNTGQLY | 0.21454147 |
| LGGKNPMI | ASGDPTGGNTGQLY | 0.20903102 |
| LGPKGVAAIL | ASGDPTGGNTGQLY | 0.19895364 |
| LMVGVSI | ASGDPTGGNTGQLY | 0.24121504 |
| LVGAAFVL | ASGDPTGGNTGQLY | 0.21426535 |
| LVLAVAPL | ASGDPTGGNTGQLY | 0.21963848 |
| MAAIAVIDAA | ASGDPTGGNTGQLY | 0.22270402 |
| NNGINAELQ | ASGDPTGGNTGQLY | 0.21576224 |
| RGEPLVLVHG | ASGDPTGGNTGQLY | 0.2242401 |

| | | |
|---|---|---|
| RGGAGGAAGLI | ASGDPTGGNTGQLY | 0.21924978 |
| RSYRMHPAV | ASGDPTGGNTGQLY | 0.21409345 |
| SAAHLLAGQLA | ASGDPTGGNTGQLY | 0.2245205 |
| SAAVGSY | ASGDPTGGNTGQLY | 0.20399769 |
| SPHAVLHRI | ASGDPTGGNTGQLY | 0.22368713 |
| SQIAKVL | ASGDPTGGNTGQLY | 0.2968104 |
| TALGHLVP | ASGDPTGGNTGQLY | 0.19950353 |
| TAPTTSGAA | ASGDPTGGNTGQLY | 0.20690225 |
| TDGDDGNGGSPGT | ASGDPTGGNTGQLY | 0.21536106 |
| TDGLGEN | ASGDPTGGNTGQLY | 0.20193873 |
| TGGPPLPGQA | ASGDPTGGNTGQLY | 0.2534609 |
| TIPAITI | ASGDPTGGNTGQLY | 0.21969411 |
| TIPGFTI | ASGDPTGGNTGQLY | 0.24241447 |
| TSIASGALA | ASGDPTGGNTGQLY | 0.20561375 |
| TVTLPGAAI | ASGDPTGGNTGQLY | 0.31425905 |
| VASGLSAALP | ASGDPTGGNTGQLY | 0.24922433 |
| VGAGVIGI | ASGDPTGGNTGQLY | 0.22054553 |
| VGPKRFGV | ASGDPTGGNTGQLY | 0.20429814 |
| VNVAFPD | ASGDPTGGNTGQLY | 0.20735747 |
| VSMTNSGVS | ASGDPTGGNTGQLY | 0.23692682 |
| VTDVLGDLSV | ASGDPTGGNTGQLY | 0.24601495 |
| VVVVGAGFAGL | ASGDPTGGNTGQLY | 0.2026153 |
| VYTVGTVV | ASGDPTGGNTGQLY | 0.2242246 |
| WTTFSGP | ASGDPTGGNTGQLY | 0.26739958 |
| YDELVEAF | ASGDPTGGNTGQLY | 0.26402035 |
| YGTGGAGGNGG | ASGDPTGGNTGQLY | 0.20613785 |
| YTNVMAV | ASGDPTGGNTGQLY | 0.21054803 |
| FLEQVHQL | ASGDPTGGNTGQLY | 0.22118734 |
| GLSLLAVAV | ASGDPTGGNTGQLY | 0.212335 |
| LEQVHQL | ASGDPTGGNTGQLY | 0.20510963 |
| LNAAGAI | ASGDPTGGNTGQLY | 0.24816518 |
| PAVHASL | ASGDPTGGNTGQLY | 0.23562479 |
| TELEDDA | ASGDPTGGNTGQLY | 0.28540865 |
| TPPGPPN | ASGDPTGGNTGQLY | 0.24222429 |
| TPPPPPPP | ASGDPTGGNTGQLY | 0.26851735 |
| TQSPALI | ASSLGEDTQY | 0.22679529 |
| EFAAAFPQ | ASSLGEDTQY | 0.21349661 |
| LLSGNLDVL | ASSLGEDTQY | 0.20474139 |
| GPANLGNYNI | ASSLGEDTQY | 0.20083569 |
| TLGPTLP | ASSLGEDTQY | 0.20614433 |
| VYAYSSCCK | ASSLGEDTQY | 0.21123123 |
| APAPAPAPNI | ASSLGEDTQY | 0.19614573 |
| PGGPVPIP | ASSLGEDTQY | 0.223182 |
| FLTVALEP | ASSLGEDTQY | 0.2238073 |
| SQYFGANL | ASSLGEDTQY | 0.24903667 |
| GGVKPAA | ASSLGEDTQY | 0.22583719 |
| GGTGGAGGAGGW | ASSLGEDTQY | 0.2107787 |
| GVGGVAGLGGAGG | ASSLGEDTQY | 0.20456958 |
| VLLVLATTI | ASSLGEDTQY | 0.26179162 |
| IGIFAAML | ASSLGEDTQY | 0.2001001 |
| AVGDLID | ASSLGEDTQY | 0.20889284 |

| | | |
|---|---|---|
| NISPLPL | ASSLGEDTQY | 0.21402739 |
| VTAVFVAI | ASSLGEDTQY | 0.20895849 |
| SGDDNDAAL | ASSLGEDTQY | 0.45351338 |
| AVAALAHS | ASSLGEDTQY | 0.27126545 |
| GGVTVMATP | ASSLGEDTQY | 0.22964008 |
| AGLAANVRD | ASSLGEDTQY | 0.29145613 |
| TNRPDLIDPALLRPG | ASSLGEDTQY | 0.19830564 |
| LANLGVTAL | ASSLGEDTQY | 0.21309556 |
| TGERIEL | ASSLGEDTQY | 0.3239939 |
| NGEELAA | ASSLGEDTQY | 0.22550963 |
| VLSLVWG | ASSLGEDTQY | 0.23817524 |
| IGYALWL | ASSLGEDTQY | 0.2384141 |
| LSLLLAPL | ASSLGEDTQY | 0.20775259 |
| TLDEALK | ASSLGEDTQY | 0.22805807 |
| TDGAGTAEGG | ASSLGEDTQY | 0.21337989 |
| NAPGGAGGNGG | ASSLGEDTQY | 0.24543208 |
| VSVFMGMGGNF | ASSLGEDTQY | 0.2077202 |
| LASYLAAL | ASSLGEDTQY | 0.20617265 |
| LVGGTPAL | ASSLGEDTQY | 0.26220056 |
| VRNADVL | ASSLGEDTQY | 0.5283977 |
| TCSSPFLVSAM | ASSLGEDTQY | 0.22196004 |
| VHITDAE | ASSLGEDTQY | 0.22242215 |
| EYAAPSF | ASSLGEDTQY | 0.22502357 |
| VVDAFHGGL | ASSLGEDTQY | 0.20801573 |
| ASVVDPQ | ASSLGEDTQY | 0.21413694 |
| SYNEFGGLS | ASSLGEDTQY | 0.20504539 |
| DPDGSVY | ASSLGEDTQY | 0.20905082 |
| MVSTIHATE | ASSLGEDTQY | 0.20590642 |
| LAAGVADPV | ASSLGEDTQY | 0.2292728 |
| TKEQIAA | ASSLGEDTQY | 0.22428843 |
| SLASPAAL | ASSLGEDTQY | 0.48719072 |
| IGLVFYGV | ASSLGEDTQY | 0.26100874 |
| FLLDASNL | ASSLGEDTQY | 0.21504872 |
| YQVGNFP | ASSLGEDTQY | 0.20363379 |
| ESLLSQST | ASSLGEDTQY | 0.21889512 |
| LFTVGFAAVG | ASSLGEDTQY | 0.21456437 |
| ATPPTPT | ASSLGEDTQY | 0.20780888 |
| TDLATLG | ASSLGEDTQY | 0.2862585 |
| SAGGTGR | ASSLGEDTQY | 0.2114929 |
| GFSNSGS | ASSLGEDTQY | 0.20655517 |
| GGNGGSGGTSV | ASSLGEDTQY | 0.22403927 |
| GTSVATGGAGN | ASSLGEDTQY | 0.42593712 |
| NGGNGGSGGTS | ASSLGEDTQY | 0.2348354 |
| AVVKFPH | ASSLGEDTQY | 0.21271822 |
| TPPPPPP | ASSLGEDTQY | 0.20413159 |
| AAPAAAAAPA | ASSLGEDTQY | 0.24446413 |
| IALNEDL | ASSLGEDTQY | 0.30146292 |
| IVNVSSM | ASSLGEDTQY | 0.20197558 |
| TGIPPAPR | ASSLGEDTQY | 0.34826553 |
| FDLGGGTF | ASSLGEDTQY | 0.20798264 |
| FDLGGGTFDVS | ASSLGEDTQY | 0.2034424 |

| | | |
|---|---|---|
| NEPTAAA | ASSLGEDTQY | 0.23602624 |
| NEPTAAALA | ASSLGEDTQY | 0.22203732 |
| GSIINISSI | ASSLGEDTQY | 0.25845358 |
| VAPGNAGTA | ASSLGEDTQY | 0.2474288 |
| TGGGASLE | ASSLGEDTQY | 0.24836425 |
| VAVYNHYKRI | ASSLGEDTQY | 0.21414602 |
| TDPAPATT | ASSLGEDTQY | 0.2783907 |
| TNRPDILDPALLRPG | ASSLGEDTQY | 0.26782855 |
| LTTTTTS | ASSLGEDTQY | 0.24064249 |
| PTGSGKTL | ASSLGEDTQY | 0.2157091 |
| TGSGKTL | ASSLGEDTQY | 0.2110591 |
| TIGHVDHGK | ASSLGEDTQY | 0.25654498 |
| TIGHVDHGKTT | ASSLGEDTQY | 0.21108565 |
| GHVDHGKTT | ASSLGEDTQY | 0.21055497 |
| AQFEHTLL | ASSLGEDTQY | 0.23101072 |
| IVNQDEL | ASSLGEDTQY | 0.23714487 |
| TNGSQFF | ASSLGEDTQY | 0.20449796 |
| TGKGIVE | ASSLGEDTQY | 0.2276862 |
| LISNASDA | ASSLGEDTQY | 0.2983886 |
| AELRTLGELTKIAKAHGAQ | ASSLGEDTQY | 0.24048808 |
| APNAVAAAEPSW | ASSLGEDTQY | 0.20455481 |
| APQVAGAATAVL | ASSLGEDTQY | 0.30822897 |
| ARRDPAAAAAILPTDA | ASSLGEDTQY | 0.36381233 |
| DAAARDVLAEAGLAEYF | ASSLGEDTQY | 0.2578348 |
| DDQIVLSAKANLAKVEQ | ASSLGEDTQY | 0.20761544 |
| DELLPDTGEDSDEDGH | ASSLGEDTQY | 0.19946896 |
| ELPEAPANRATVPQD | ASSLGEDTQY | 0.22585689 |
| FGQSKIERVTGIG | ASSLGEDTQY | 0.22361273 |
| GAGGTGSTAGGAGGAGGAGGLY | ASSLGEDTQY | 0.21703061 |
| GHGGTGAAGGNGGTGGAGGSGI | ASSLGEDTQY | 0.20573701 |
| GNGGNGGAGGEGGPGGAGGAGA | ASSLGEDTQY | 0.21121925 |
| GSTAGNGGNGGNATNGTIAGQP | ASSLGEDTQY | 0.24091502 |
| GTGGDGGNGGNGGTGG | ASSLGEDTQY | 0.21330808 |
| GVRRPAGIPADAPV | ASSLGEDTQY | 0.201481 |
| GYRQVTDAHLLTLARR | ASSLGEDTQY | 0.21527547 |
| IAKMGQDGHDRG | ASSLGEDTQY | 0.26132116 |
| KAEPSIEIQYT | ASSLGEDTQY | 0.20402724 |
| KPVRIRMAADEDTVAVT | ASSLGEDTQY | 0.2144434 |
| MPPRIAGMRLLVIKPEP | ASSLGEDTQY | 0.24004269 |
| MKFYAVALFRSRGVPPTR | ASSLGEDTQY | 0.2042607 |
| NAGGFGILGAVAHSP | ASSLGEDTQY | 0.21249634 |
| NEGREITLAMAEM | ASSLGEDTQY | 0.20724285 |
| NELVEWSHPLGHRWRWE | ASSLGEDTQY | 0.21118386 |
| NLGAGGTGGNGGNPG | ASSLGEDTQY | 0.21419336 |
| NTEWPKWETVMAADPVD | ASSLGEDTQY | 0.24025263 |
| PASAAPAETQP | ASSLGEDTQY | 0.22975639 |
| QQSSDGKNGVRAAEITGE | ASSLGEDTQY | 0.24163747 |
| RAPDAVTTSWWKE | ASSLGEDTQY | 0.22761285 |
| SGFGNVGAHTSGWFNQ | ASSLGEDTQY | 0.21561351 |
| SGVAGAVGQAASVGGLKV | ASSLGEDTQY | 0.2069552 |
| SNGGAGTGSAGGNGGT | ASSLGEDTQY | 0.19795066 |

| | | |
|---|---|---|
| TGGAGGEGGAAG | ASSLGEDTQY | 0.27834997 |
| TGGDGGAAGAGGAPGHG | ASSLGEDTQY | 0.19686462 |
| TMPPLLAAIANAGRHGV | ASSLGEDTQY | 0.25424218 |
| TSSTAGNSGGA | ASSLGEDTQY | 0.24329725 |
| VPATVPVGILC | ASSLGEDTQY | 0.20889616 |
| VQRPDSGGVVLTGRIS | ASSLGEDTQY | 0.2244555 |
| WTPNPVPATKGVPAGLVQAT | ASSLGEDTQY | 0.1981031 |
| YALKRALAKFSDRFDVVII | ASSLGEDTQY | 0.21484901 |
| YLIGSVIV | ASSLGEDTQY | 0.2300517 |
| YGQLVAVID | ASSLGEDTQY | 0.23252107 |
| VTVGGMTLPA | ASSLGEDTQY | 0.23097607 |
| VLALPAV | ASSLGEDTQY | 0.23471838 |
| VIPPTLNYET | ASSLGEDTQY | 0.2538858 |
| VGMGGMHPSSG | ASSLGEDTQY | 0.21536227 |
| VGLAGGAATA | ASSLGEDTQY | 0.23951195 |
| VGGAGGAGGSGP | ASSLGEDTQY | 0.24693958 |
| VFAAFMV | ASSLGEDTQY | 0.25472736 |
| TLYDKLR | ASSLGEDTQY | 0.20322682 |
| TLAVFTVL | ASSLGEDTQY | 0.23062165 |
| TGSVQPARV | ASSLGEDTQY | 0.2071046 |
| TGGGAGSGGT | ASSLGEDTQY | 0.23207135 |
| SHLETSL | ASSLGEDTQY | 0.21358888 |
| SFTTGMT | ASSLGEDTQY | 0.2483544 |
| QVEVAKRMGVRQPTVS | ASSLGEDTQY | 0.23028111 |
| QGSGGPALAM | ASSLGEDTQY | 0.20014505 |
| MLAPSNP | ASSLGEDTQY | 0.21092355 |
| LVNNAAI | ASSLGEDTQY | 0.22566023 |
| LTGEINTLA | ASSLGEDTQY | 0.23386742 |
| LSIFGPV | ASSLGEDTQY | 0.22637455 |
| LLALLANL | ASSLGEDTQY | 0.20644104 |
| LISNASDALDKLR | ASSLGEDTQY | 0.23935026 |
| LINEAVP | ASSLGEDTQY | 0.22389054 |
| LGYAFSGL | ASSLGEDTQY | 0.21727282 |
| LDFVAVAL | ASSLGEDTQY | 0.23119423 |
| LAQPGAE | ASSLGEDTQY | 0.21589194 |
| ISNLGAVTGF | ASSLGEDTQY | 0.20643516 |
| HQEDPGRI | ASSLGEDTQY | 0.24514507 |
| GTPGEPTP | ASSLGEDTQY | 0.20775814 |
| GFFNSGS | ASSLGEDTQY | 0.23994803 |
| GAGGGGLTAGAN | ASSLGEDTQY | 0.20571633 |
| FVDALVNM | ASSLGEDTQY | 0.20636223 |
| FLTVAPEL | ASSLGEDTQY | 0.23400387 |
| FETLTAL | ASSLGEDTQY | 0.20637439 |
| DVINPIEL | ASSLGEDTQY | 0.23144993 |
| APVSGAPV | ASSLGEDTQY | 0.21884942 |
| ALVGGLAW | ASSLGEDTQY | 0.21908624 |
| ALAAAAATAI | ASSLGEDTQY | 0.22020066 |
| AFGLGLLL | ASSLGEDTQY | 0.24623246 |
| AFFAAASL | ASSLGEDTQY | 0.2444083 |
| AAAALPAFSPP | ASSLGEDTQY | 0.20713663 |
| VVLAPAL | ASSLGEDTQY | 0.23185337 |

| | | |
|---|---|---|
| TIGHVDHGKTTL | ASSLGEDTQY | 0.25717497 |
| LISNASDALDK | ASSLGEDTQY | 0.20985854 |
| LISNASDAL | ASSLGEDTQY | 0.20079589 |
| ISNASDALDK | ASSLGEDTQY | 0.20937672 |
| NQAETLVYQTE | ASSLGEDTQY | 0.20025612 |
| TGGTGGTAGAG | ASSLGEDTQY | 0.22056046 |
| TGSTAGGAGGA | ASSLGEDTQY | 0.20218378 |
| DPLAVPVAAGPV | ASSLGEDTQY | 0.24749097 |
| GGTGGTPGNGGN | ASSLGEDTQY | 0.2512053 |
| PAGLAGVDPATL | ASSLGEDTQY | 0.2112813 |
| GNNLVVEAPHAGASV | ASSLGEDTQY | 0.24130696 |
| KVTRIMSASTLEQPA | ASSLGEDTQY | 0.20665997 |
| LGFALPIALPRRTVV | ASSLGEDTQY | 0.24271539 |
| TRLGNLTARKLVVLV | ASSLGEDTQY | 0.27667397 |
| WSQQRRLIYFPSAGP | ASSLGEDTQY | 0.20388806 |
| GPNGNPDPMAAAVDIR | ASSLGEDTQY | 0.22791688 |
| TQRCVSVADLRAAASR | ASSLGEDTQY | 0.21602988 |
| QFVLDVAEHPVEVLTETV | ASSLGEDTQY | 0.20764829 |
| GGAGGTGGAGGNGGTQAGDG | ASSLGEDTQY | 0.2616817 |
| PLLATLRAIAVDVDAKPAQV | ASSLGEDTQY | 0.22282945 |
| APKFRTGGYMIDFWGVGYQVAK | ASSLGEDTQY | 0.2414706 |
| GAGGAGGAGGAAGGPGGTGGTG | ASSLGEDTQY | 0.21490134 |
| IGALGLLATDVAPAGSVAFGVA | ASSLGEDTQY | 0.20540512 |
| AGGAGGIGGNGGAGGAGGVLMG | ASSLGEDTQY | 0.23384133 |
| ADELIGLL | ASSLGEDTQY | 0.2533152 |
| ADRALKA | ASSLGEDTQY | 0.20927724 |
| AGAVGQVGGAGGA | ASSLGEDTQY | 0.260504 |
| AGGGGGLGGAP | ASSLGEDTQY | 0.21636689 |
| AGGLLYGNGGAG | ASSLGEDTQY | 0.2886702 |
| AGLESIGL | ASSLGEDTQY | 0.26086235 |
| AGLKTGII | ASSLGEDTQY | 0.22148411 |
| ASLAALLL | ASSLGEDTQY | 0.21569075 |
| ASVINISSI | ASSLGEDTQY | 0.3030275 |
| ATPPGPA | ASSLGEDTQY | 0.23772116 |
| ATVASALV | ASSLGEDTQY | 0.24244668 |
| ATVASLIL | ASSLGEDTQY | 0.23633628 |
| AVNGPVLIRH | ASSLGEDTQY | 0.1681954 |
| DGAAGAVGI | ASSLGEDTQY | 0.27950346 |
| DPLLHGI | ASSLGEDTQY | 0.4109211 |
| ELGAALGAL | ASSLGEDTQY | 0.44159624 |
| ELGRTQAV | ASSLGEDTQY | 0.1656428 |
| FVAGGYLS | ASSLGEDTQY | 0.11098584 |
| GASGGAGGAGGW | ASSLGEDTQY | 0.3198494 |
| GGAGAGGDGGAGGI | ASSLGEDTQY | 0.12433513 |
| GTGHNGSQPGMGGNG | ASSLGEDTQY | 0.13557169 |
| GYAAASASAS | ASSLGEDTQY | 0.36982626 |
| HQVDVAAVQQG | ASSLGEDTQY | 0.18743074 |
| IGAAIAHT | ASSLGEDTQY | 0.71933323 |
| IGMIEVAQL | ASSLGEDTQY | 0.6329113 |
| IRLAPEHTSAMD | ASSLGEDTQY | 0.13692392 |
| IYNVSVL | ASSLGEDTQY | 0.24976775 |

| | | |
|---|---|---|
| KQGFLGM | ASSLGEDTQY | 0.22474235 |
| LESVITQL | ASSLGEDTQY | 0.26231965 |
| LGLGALVVV | ASSLGEDTQY | 0.102231584 |
| LIGNGGNGGSGGI | ASSLGEDTQY | 0.41879606 |
| LNAINAPTL | ASSLGEDTQY | 0.20269638 |
| LQLLNTGV | ASSLGEDTQY | 0.089601144 |
| LQTSIEF | ASSLGEDTQY | 0.32920712 |
| MSFAVSEL | ASSLGEDTQY | 0.26585793 |
| NTGALGL | ASSLGEDTQY | 0.16526079 |
| PPGAGGAGGAGTT | ASSLGEDTQY | 0.71811074 |
| QLVGLGI | ASSLGEDTQY | 0.08931034 |
| RFLEQVHQL | ASSLGEDTQY | 0.1379482 |
| RGVGYVLR | ASSLGEDTQY | 0.5425591 |
| SATGVFAGVI | ASSLGEDTQY | 0.66874194 |
| SLPAITLP | ASSLGEDTQY | 0.13131338 |
| STALAFTV | ASSLGEDTQY | 0.52921414 |
| SVLTVGLGK | ASSLGEDTQY | 0.7574481 |
| TALAGAGSGI | ASSLGEDTQY | 0.3399334 |
| TGLPFTGI | ASSLGEDTQY | 0.59571815 |
| VAEHVTV | ASSLGEDTQY | 0.68665695 |
| VDIVLNSL | ASSLGEDTQY | 0.19147816 |
| VLTGGGAVL | ASSLGEDTQY | 0.5443722 |
| VNVGLAYGI | ASSLGEDTQY | 0.6915809 |
| VSSAAVGAV | ASSLGEDTQY | 0.20973772 |
| VTSVYEAGV | ASSLGEDTQY | 0.29551983 |
| WVALDRGA | ASSLGEDTQY | 0.22003143 |
| YLVGSVVI | ASSLGEDTQY | 0.32884103 |
| GPPGTGKTLL | ASSLGEDTQY | 0.09043001 |
| GVGMTKF | ASSLGEDTQY | 0.48783726 |
| IGIGGSDL | ASSLGEDTQY | 0.4733305 |
| IGTLLAEL | ASSLGEDTQY | 0.2749597 |
| IYTSGTT | ASSLGEDTQY | 0.14702602 |
| LSFPAGV | ASSLGEDTQY | 0.15216205 |
| LYTSGTT | ASSLGEDTQY | 0.10904916 |
| VGAFPFGF | ASSLGEDTQY | 0.3964308 |
| VGAGTVEF | ASSLGEDTQY | 0.105617076 |
| VGKSTLLL | ASSLGEDTQY | 0.108404435 |
| AALAHAGGGIV | ASSLGEDTQY | 0.4634958 |
| AALVAVSLL | ASSLGEDTQY | 0.47843307 |
| AAVSRAVAAL | ASSLGEDTQY | 0.10088796 |
| AGGSSGIGGF | ASSLGEDTQY | 0.54670537 |
| AGLTVANANG | ASSLGEDTQY | 0.5535588 |
| AGLVGAVL | ASSLGEDTQY | 0.64675355 |
| ALSQAFTV | ASSLGEDTQY | 0.30818176 |
| ALTFSVSL | ASSLGEDTQY | 0.13002081 |
| ALTGAALVAL | ASSLGEDTQY | 0.7292332 |
| ASFSIGAGVL | ASSLGEDTQY | 0.3380263 |
| DGSVYKI | ASSLGEDTQY | 0.12115754 |
| EQLAEIV | ASSLGEDTQY | 0.5482177 |
| FAVSELTVM | ASSLGEDTQY | 0.1472048 |
| GAAPLAAGL | ASSLGEDTQY | 0.15749969 |

| | | |
|---|---|---|
| GADGGDGGSGGAGG | ASSLGEDTQY | 0.56427956 |
| GFGLDTGGAGG | ASSLGEDTQY | 0.49102655 |
| GLASGGLL | ASSLGEDTQY | 0.16669917 |
| GRIAGAAVLL | ASSLGEDTQY | 0.121019065 |
| GSAGSLNTSF | ASSLGEDTQY | 0.14321837 |
| GTGQIGI | ASSLGEDTQY | 0.4251107 |
| GTLIALIL | ASSLGEDTQY | 0.5806705 |
| IAATMGFVVG | ASSLGEDTQY | 0.27896488 |
| IFAQALL | ASSLGEDTQY | 0.21807745 |
| IGAAVALGF | ASSLGEDTQY | 0.6512575 |
| IGLTPLAL | ASSLGEDTQY | 0.3276692 |
| ILGSNTSTL | ASSLGEDTQY | 0.2926873 |
| IVFSLTVL | ASSLGEDTQY | 0.61734843 |
| LASKATGF | ASSLGEDTQY | 0.13167809 |
| LASVALIL | ASSLGEDTQY | 0.44596773 |
| LFAVGLTL | ASSLGEDTQY | 0.46993136 |
| LFNGGIGTYI | ASSLGEDTQY | 0.5550682 |
| LTLGAGII | ASSLGEDTQY | 0.57126415 |
| LVAAALGDI | ASSLGEDTQY | 0.1538328 |
| NTAFVQLGI | ASSLGEDTQY | 0.4291179 |
| QLFSGAEL | ASSLGEDTQY | 0.09931879 |
| SAATFITL | ASSLGEDTQY | 0.3049326 |
| SNYGLANLGV | ASSLGEDTQY | 0.62194324 |
| SYGGAYAVM | ASSLGEDTQY | 0.47419173 |
| TTGVGGLTLG | ASSLGEDTQY | 0.597135 |
| TTQTIYI | ASSLGEDTQY | 0.4364536 |
| VAAGAGLGAVI | ASSLGEDTQY | 0.22674815 |
| VAAVVTLL | ASSLGEDTQY | 0.70684195 |
| VGFLGGLI | ASSLGEDTQY | 0.6827384 |
| VGVIPILL | ASSLGEDTQY | 0.13487038 |
| VIFASSLI | ASSLGEDTQY | 0.18366452 |
| VKGGDGGAAAG | ASSLGEDTQY | 0.28854853 |
| VTKTFTNI | ASSLGEDTQY | 0.6169593 |
| VVAAAILL | ASSLGEDTQY | 0.36192316 |
| YAALDVEL | ASSLGEDTQY | 0.36367106 |
| LASVALLL | ASSLGEDTQY | 0.3234061 |
| LDAVHRV | ASSLGEDTQY | 0.12359601 |
| QGLSSSIF | ASSLGEDTQY | 0.7379945 |
| RGTFANIRL | ASSLGEDTQY | 0.09819078 |
| RQLWWGHRI | ASSLGEDTQY | 0.31912854 |
| YGPPGTGKTLL | ASSLGEDTQY | 0.14891675 |
| AALVGAPANVA | ASSLGEDTQY | 0.10198939 |
| AATFRRTE | ASSLGEDTQY | 0.40398163 |
| AGALGGGI | ASSLGEDTQY | 0.22593698 |
| AGGAGGAGGIGAG | ASSLGEDTQY | 0.6909987 |
| AGSGALGL | ASSLGEDTQY | 0.3080146 |
| AQIADQI | ASSLGEDTQY | 0.35893005 |
| ASAVAVTYL | ASSLGEDTQY | 0.39550132 |
| ASTVLGGTGG | ASSLGEDTQY | 0.35122463 |
| AVGGEPAK | ASSLGEDTQY | 0.15176126 |
| CEQGTFGGTC | ASSLGEDTQY | 0.41144365 |

| | | |
|---|---|---|
| DEGKAPAG | ASSLGEDTQY | 0.703386 |
| DTVVFPAL | ASSLGEDTQY | 0.37900478 |
| FEGITMAG | ASSLGEDTQY | 0.11672615 |
| FPLVVQD | ASSLGEDTQY | 0.16112287 |
| GAFTTPQ | ASSLGEDTQY | 0.23379114 |
| GFGGFGTAGGVGGA | ASSLGEDTQY | 0.086942695 |
| GFLAQFL | ASSLGEDTQY | 0.09442183 |
| GGAGGAGGNAGVA | ASSLGEDTQY | 0.100445256 |
| GGVGTAGAGGAGGAGG | ASSLGEDTQY | 0.10368651 |
| GNGGGGDAGNAGSGGN | ASSLGEDTQY | 0.6695803 |
| GTGGAGGAGNDAGS | ASSLGEDTQY | 0.5244576 |
| GTTNTGWFNS | ASSLGEDTQY | 0.31310415 |
| IFMGPLFTL | ASSLGEDTQY | 0.636971 |
| ILAHAIEAI | ASSLGEDTQY | 0.33210838 |
| IRSAIREAL | ASSLGEDTQY | 0.085346594 |
| ITFAGGG | ASSLGEDTQY | 0.21327595 |
| ITGMSGHQEV | ASSLGEDTQY | 0.16227992 |
| IVAGLACTL | ASSLGEDTQY | 0.65964544 |
| IVGGGLAGA | ASSLGEDTQY | 0.121679865 |
| IVTGVLNI | ASSLGEDTQY | 0.22501038 |
| LAQQGCI | ASSLGEDTQY | 0.31932026 |
| LCLHVEN | ASSLGEDTQY | 0.25075793 |
| LDAISDGS | ASSLGEDTQY | 0.51112235 |
| LGGFAAML | ASSLGEDTQY | 0.14151533 |
| LGGKNPMI | ASSLGEDTQY | 0.08974084 |
| LGPKGVAAIL | ASSLGEDTQY | 0.38046443 |
| LMVGVSI | ASSLGEDTQY | 0.09265043 |
| LVGAAFVL | ASSLGEDTQY | 0.15438068 |
| LVLAVAPL | ASSLGEDTQY | 0.09849175 |
| MAAIAVIDAA | ASSLGEDTQY | 0.085054316 |
| NNGINAELQ | ASSLGEDTQY | 0.112878844 |
| RGEPLVLVHG | ASSLGEDTQY | 0.60936856 |
| RGGAGGAAGLI | ASSLGEDTQY | 0.46095034 |
| RSYRMHPAV | ASSLGEDTQY | 0.30947545 |
| SAAHLLAGQLA | ASSLGEDTQY | 0.5086287 |
| SAAVGSY | ASSLGEDTQY | 0.49613583 |
| SPHAVLHRI | ASSLGEDTQY | 0.30451143 |
| SQIAKVL | ASSLGEDTQY | 0.15129934 |
| TALGHLVP | ASSLGEDTQY | 0.16683188 |
| TAPTTSGAA | ASSLGEDTQY | 0.14290386 |
| TDGDDGNGGSPGT | ASSLGEDTQY | 0.14384715 |
| TDGLGEN | ASSLGEDTQY | 0.7559246 |
| TGGPPLPGQA | ASSLGEDTQY | 0.30245122 |
| TIPAITI | ASSLGEDTQY | 0.12940106 |
| TIPGFTI | ASSLGEDTQY | 0.15730244 |
| TSIASGALA | ASSLGEDTQY | 0.70558465 |
| TVTLPGAAI | ASSLGEDTQY | 0.751518 |
| VASGLSAALP | ASSLGEDTQY | 0.17128567 |
| VGAGVIGI | ASSLGEDTQY | 0.27878994 |
| VGPKRFGV | ASSLGEDTQY | 0.32694846 |
| VNVAFPD | ASSLGEDTQY | 0.3114317 |

| | | |
|---|---|---|
| VSMTNSGVS | ASSLGEDTQY | 0.28155845 |
| VTDVLGDLSV | ASSLGEDTQY | 0.13575086 |
| VVVVGAGFAGL | ASSLGEDTQY | 0.47531003 |
| VYTVGTVV | ASSLGEDTQY | 0.80270267 |
| WTTFSGP | ASSLGEDTQY | 0.17180026 |
| YDELVEAF | ASSLGEDTQY | 0.13106441 |
| YGTGGAGGNGG | ASSLGEDTQY | 0.37309647 |
| YTNVMAV | ASSLGEDTQY | 0.15437838 |
| FLEQVHQL | ASSLGEDTQY | 0.24523927 |
| GLSLLAVAV | ASSLGEDTQY | 0.29074082 |
| LEQVHQL | ASSLGEDTQY | 0.12746459 |
| LNAAGAI | ASSLGEDTQY | 0.23619893 |
| PAVHASL | ASSLGEDTQY | 0.6017338 |
| TELEDDA | ASSLGEDTQY | 0.22345233 |
| TPPGPPN | ASSLGEDTQY | 0.6395966 |
| TPPPPPPP | ASSLGEDTQY | 0.6953548 |
| TQSPALI | ASSEGGWDTQY | 0.5887104 |
| EFAAAFPQ | ASSEGGWDTQY | 0.75121504 |
| LLSGNLDVL | ASSEGGWDTQY | 0.6124648 |
| GPANLGNYNI | ASSEGGWDTQY | 0.21521178 |
| TLGPTLP | ASSEGGWDTQY | 0.30591673 |
| VYAYSSCCK | ASSEGGWDTQY | 0.47166735 |
| APAPAPAPNI | ASSEGGWDTQY | 0.39744306 |
| PGGPVPIP | ASSEGGWDTQY | 0.35692638 |
| FLTVALEP | ASSEGGWDTQY | 0.23485875 |
| SQYFGANL | ASSEGGWDTQY | 0.68255067 |
| GGVKPAA | ASSEGGWDTQY | 0.35898682 |
| GGTGGAGGAGGW | ASSEGGWDTQY | 0.60960484 |
| GVGGVAGLGGAGG | ASSEGGWDTQY | 0.19375357 |
| VLLVLATTI | ASSEGGWDTQY | 0.14457907 |
| IGIFAAML | ASSEGGWDTQY | 0.6550048 |
| AVGDLID | ASSEGGWDTQY | 0.09040137 |
| NISPLPL | ASSEGGWDTQY | 0.10497127 |
| VTAVFVAI | ASSEGGWDTQY | 0.31830218 |
| SGDDNDAAL | ASSEGGWDTQY | 0.57608944 |
| AVAALAHS | ASSEGGWDTQY | 0.08591365 |
| GGVTVMATP | ASSEGGWDTQY | 0.14589995 |
| AGLAANVRD | ASSEGGWDTQY | 0.1950085 |
| TNRPDLIDPALLRPG | ASSEGGWDTQY | 0.16672404 |
| LANLGVTAL | ASSEGGWDTQY | 0.23517752 |
| TGERIEL | ASSEGGWDTQY | 0.12587196 |
| NGEELAA | ASSEGGWDTQY | 0.4690831 |
| VLSLVWG | ASSEGGWDTQY | 0.11901621 |
| IGYALWL | ASSEGGWDTQY | 0.091544025 |
| LSLLLAPL | ASSEGGWDTQY | 0.41209447 |
| TLDEALK | ASSEGGWDTQY | 0.6852554 |
| TDGAGTAEGG | ASSEGGWDTQY | 0.17788208 |
| NAPGGAGGNGG | ASSEGGWDTQY | 0.5261784 |
| VSVFMGMGGNF | ASSEGGWDTQY | 0.21714506 |
| LASYLAAL | ASSEGGWDTQY | 0.78081024 |
| LVGGTPAL | ASSEGGWDTQY | 0.20548841 |

| | | |
|---|---|---|
| VRNADVL | ASSEGGWDTQY | 0.1481396 |
| TCSSPFLVSAM | ASSEGGWDTQY | 0.5605736 |
| VHITDAE | ASSEGGWDTQY | 0.6219216 |
| EYAAPSF | ASSEGGWDTQY | 0.29851544 |
| VVDAFHGGL | ASSEGGWDTQY | 0.59366393 |
| ASVVDPQ | ASSEGGWDTQY | 0.13265814 |
| SYNEFGGLS | ASSEGGWDTQY | 0.30027273 |
| DPDGSVY | ASSEGGWDTQY | 0.60130274 |
| MVSTIHATE | ASSEGGWDTQY | 0.564661 |
| LAAGVADPV | ASSEGGWDTQY | 0.10950776 |
| TKEQIAA | ASSEGGWDTQY | 0.4087538 |
| SLASPAAL | ASSEGGWDTQY | 0.24652442 |
| IGLVFYGV | ASSEGGWDTQY | 0.22038981 |
| FLLDASNL | ASSEGGWDTQY | 0.49336892 |
| YQVGNFP | ASSEGGWDTQY | 0.39334455 |
| ESLLSQST | ASSEGGWDTQY | 0.64108926 |
| LFTVGFAAVG | ASSEGGWDTQY | 0.34000316 |
| ATPPTPT | ASSEGGWDTQY | 0.30280587 |
| TDLATLG | ASSEGGWDTQY | 0.12596218 |
| SAGGTGR | ASSEGGWDTQY | 0.5658723 |
| GFSNSGS | ASSEGGWDTQY | 0.36345008 |
| GGNGGSGGTSV | ASSEGGWDTQY | 0.46929932 |
| GTSVATGGAGN | ASSEGGWDTQY | 0.39108253 |
| NGGNGGSGGTS | ASSEGGWDTQY | 0.09743481 |
| AVVKFPH | ASSEGGWDTQY | 0.43263045 |
| TPPPPPP | ASSEGGWDTQY | 0.6671816 |
| AAPAAAAAPA | ASSEGGWDTQY | 0.22478734 |
| IALNEDL | ASSEGGWDTQY | 0.47288778 |
| IVNVSSM | ASSEGGWDTQY | 0.78164434 |
| TGIPPAPR | ASSEGGWDTQY | 0.21191065 |
| FDLGGGTF | ASSEGGWDTQY | 0.21171455 |
| FDLGGGTFDVS | ASSEGGWDTQY | 0.41239518 |
| NEPTAAA | ASSEGGWDTQY | 0.7261454 |
| NEPTAAALA | ASSEGGWDTQY | 0.27224785 |
| GSIINISSI | ASSEGGWDTQY | 0.70364976 |
| VAPGNAGTA | ASSEGGWDTQY | 0.12941407 |
| TGGGASLE | ASSEGGWDTQY | 0.5991652 |
| VAVYNHYKRI | ASSEGGWDTQY | 0.20530461 |
| TDPAPATT | ASSEGGWDTQY | 0.5149431 |
| TNRPDILDPALLRPG | ASSEGGWDTQY | 0.09789609 |
| LTTTTTS | ASSEGGWDTQY | 0.5776477 |
| PTGSGKTL | ASSEGGWDTQY | 0.6603066 |
| TGSGKTL | ASSEGGWDTQY | 0.6439024 |
| TIGHVDHGK | ASSEGGWDTQY | 0.0925526 |
| TIGHVDHGKTT | ASSEGGWDTQY | 0.45078862 |
| GHVDHGKTT | ASSEGGWDTQY | 0.6655282 |
| AQFEHTLL | ASSEGGWDTQY | 0.818264 |
| IVNQDEL | ASSEGGWDTQY | 0.35481286 |
| TNGSQFF | ASSEGGWDTQY | 0.6013614 |
| TGKGIVE | ASSEGGWDTQY | 0.1707427 |
| LISNASDA | ASSEGGWDTQY | 0.19243862 |

| | | |
|---|---|---|
| AELRTLGELTKIAKAHGAQ | ASSEGGWDTQY | 0.19389272 |
| APNAVAAAEPSW | ASSEGGWDTQY | 0.39073348 |
| APQVAGAATAVL | ASSEGGWDTQY | 0.3004442 |
| ARRDPAAAAAILPTDA | ASSEGGWDTQY | 0.18320033 |
| DAAARDVLAEAGLAEYF | ASSEGGWDTQY | 0.44888288 |
| DDQIVLSAKANLAKVEQ | ASSEGGWDTQY | 0.72484225 |
| DELLPDTGEDSDEDGH | ASSEGGWDTQY | 0.43364695 |
| ELPEAPANRATVPQD | ASSEGGWDTQY | 0.7113855 |
| FGQSKIERVTGIG | ASSEGGWDTQY | 0.748149 |
| GAGGTGSTAGGAGGAGGAGGLY | ASSEGGWDTQY | 0.5450166 |
| GHGGTGAAGGNGGTGGAGGSGI | ASSEGGWDTQY | 0.1403337 |
| GNGGNGGAGGEGGPGGAGGAGA | ASSEGGWDTQY | 0.3327147 |
| GSTAGNGGNGGNATNGTIAGQP | ASSEGGWDTQY | 0.12931283 |
| GTGGDGGNGGNGGTGG | ASSEGGWDTQY | 0.26932293 |
| GVRRPAGIPADAPV | ASSEGGWDTQY | 0.14281636 |
| GYRQVTDAHLLTLARR | ASSEGGWDTQY | 0.3900183 |
| IAKMGQDGHDRG | ASSEGGWDTQY | 0.08567502 |
| KAEPSIEIQYT | ASSEGGWDTQY | 0.5191416 |
| KPVRIRMAADEDTVAVT | ASSEGGWDTQY | 0.7580446 |
| MPPRIAGMRLLVIKPEP | ASSEGGWDTQY | 0.10264421 |
| MKFYAVALFRSRGVPPTR | ASSEGGWDTQY | 0.5607488 |
| NAGGFGILGAVAHSP | ASSEGGWDTQY | 0.59618735 |
| NEGREITLAMAEM | ASSEGGWDTQY | 0.1535878 |
| NELVEWSHPLGHRWRWE | ASSEGGWDTQY | 0.6647328 |
| NLGAGGTGGNGGNPG | ASSEGGWDTQY | 0.1581663 |
| NTEWPKWETVMAADPVD | ASSEGGWDTQY | 0.5771342 |
| PASAAPAETQP | ASSEGGWDTQY | 0.5230256 |
| QQSSDGKNGVRAAEITGE | ASSEGGWDTQY | 0.39873773 |
| RAPDAVTTSWWKE | ASSEGGWDTQY | 0.5127803 |
| SGFGNVGAHTSGWFNQ | ASSEGGWDTQY | 0.40756622 |
| SGVAGAVGQAASVGGLKV | ASSEGGWDTQY | 0.12066549 |
| SNGGAGTGSAGGNGGT | ASSEGGWDTQY | 0.5401027 |
| TGGAGGEGGAAG | ASSEGGWDTQY | 0.0847934 |
| TGGDGGAAGAGGAPGHG | ASSEGGWDTQY | 0.45132148 |
| TMPPLLAAIANAGRHGV | ASSEGGWDTQY | 0.14819327 |
| TSSTAGNSGGA | ASSEGGWDTQY | 0.43608558 |
| VPATVPVGILC | ASSEGGWDTQY | 0.23169616 |
| VQRPDSGGVVLTGRIS | ASSEGGWDTQY | 0.3875675 |
| WTPNPVPATKGVPAGLVQAT | ASSEGGWDTQY | 0.23956549 |
| YALKRALAKFSDRFDVVII | ASSEGGWDTQY | 0.15284753 |
| YLIGSVIV | ASSEGGWDTQY | 0.10659669 |
| YGQLVAVID | ASSEGGWDTQY | 0.11943116 |
| VTVGGMTLPA | ASSEGGWDTQY | 0.60562736 |
| VLALPAV | ASSEGGWDTQY | 0.2251999 |
| VIPPTLNYET | ASSEGGWDTQY | 0.120867655 |
| VGMGGMHPSSG | ASSEGGWDTQY | 0.49983943 |
| VGLAGGAATA | ASSEGGWDTQY | 0.28892663 |
| VGGAGGAGGSGP | ASSEGGWDTQY | 0.11624908 |
| VFAAFMV | ASSEGGWDTQY | 0.5219089 |
| TLYDKLR | ASSEGGWDTQY | 0.41063708 |
| TLAVFTVL | ASSEGGWDTQY | 0.21659228 |

| | | |
|---|---|---|
| TGSVQPARV | ASSEGGWDTQY | 0.0850717 |
| TGGGAGSGGT | ASSEGGWDTQY | 0.13784832 |
| SHLETSL | ASSEGGWDTQY | 0.22720222 |
| SFTTGMT | ASSEGGWDTQY | 0.107030004 |
| QVEVAKRMGVRQPTVS | ASSEGGWDTQY | 0.14749989 |
| QGSGGPALAM | ASSEGGWDTQY | 0.68804604 |
| MLAPSNP | ASSEGGWDTQY | 0.089649774 |
| LVNNAAI | ASSEGGWDTQY | 0.17030141 |
| LTGEINTLA | ASSEGGWDTQY | 0.6773242 |
| LSIFGPV | ASSEGGWDTQY | 0.39206573 |
| LLALLANL | ASSEGGWDTQY | 0.7627078 |
| LISNASDALDKLR | ASSEGGWDTQY | 0.16109377 |
| LINEAVP | ASSEGGWDTQY | 0.18656747 |
| LGYAFSGL | ASSEGGWDTQY | 0.37739483 |
| LDFVAVAL | ASSEGGWDTQY | 0.36272103 |
| LAQPGAE | ASSEGGWDTQY | 0.10578786 |
| ISNLGAVTGF | ASSEGGWDTQY | 0.09230466 |
| HQEDPGRI | ASSEGGWDTQY | 0.35691708 |
| GTPGEPTP | ASSEGGWDTQY | 0.10557811 |
| GFFNSGS | ASSEGGWDTQY | 0.30402508 |
| GAGGGGLTAGAN | ASSEGGWDTQY | 0.60492975 |
| FVDALVNM | ASSEGGWDTQY | 0.12522514 |
| FLTVAPEL | ASSEGGWDTQY | 0.16058154 |
| FETLTAL | ASSEGGWDTQY | 0.25764585 |
| DVINPIEL | ASSEGGWDTQY | 0.21128619 |
| APVSGAPV | ASSEGGWDTQY | 0.64998746 |
| ALVGGLAW | ASSEGGWDTQY | 0.18662778 |
| ALAAAAATAI | ASSEGGWDTQY | 0.77279246 |
| AFGLGLLL | ASSEGGWDTQY | 0.46844086 |
| AFFAAASL | ASSEGGWDTQY | 0.17453751 |
| AAAALPAFSPP | ASSEGGWDTQY | 0.6439636 |
| VVLAPAL | ASSEGGWDTQY | 0.3095777 |
| TIGHVDHGKTTL | ASSEGGWDTQY | 0.38633534 |
| LISNASDALDK | ASSEGGWDTQY | 0.3222579 |
| LISNASDAL | ASSEGGWDTQY | 0.121880986 |
| ISNASDALDK | ASSEGGWDTQY | 0.20456198 |
| NQAETLVYQTE | ASSEGGWDTQY | 0.29029942 |
| TGGTGGTAGAG | ASSEGGWDTQY | 0.1306875 |
| TGSTAGGAGGA | ASSEGGWDTQY | 0.3515625 |
| DPLAVPVAAGPV | ASSEGGWDTQY | 0.12589441 |
| GGTGGTPGNGGN | ASSEGGWDTQY | 0.11271438 |
| PAGLAGVDPATL | ASSEGGWDTQY | 0.35878396 |
| GNNLVVEAPHAGASV | ASSEGGWDTQY | 0.6396961 |
| KVTRIMSASTLEQPA | ASSEGGWDTQY | 0.775085 |
| LGFALPIALPRRTVV | ASSEGGWDTQY | 0.19057396 |
| TRLGNLTARKLVVLV | ASSEGGWDTQY | 0.13500717 |
| WSQQRRLIYFPSAGP | ASSEGGWDTQY | 0.6203139 |
| GPNGNPDPMAAAVDIR | ASSEGGWDTQY | 0.10282402 |
| TQRCVSVADLRAAASR | ASSEGGWDTQY | 0.4355021 |
| QFVLDVAEHPVEVLTETV | ASSEGGWDTQY | 0.3157572 |
| GGAGGTGGAGGNGGTQAGDG | ASSEGGWDTQY | 0.15159287 |

| | | |
|---|---|---|
| PLLATLRAIAVDVDAKPAQV | ASSEGGWDTQY | 0.24904124 |
| APKFRTGGYMIDFWGVGYQVAK | ASSEGGWDTQY | 0.38437915 |
| GAGGAGGAGGAAGGPGGTGGTG | ASSEGGWDTQY | 0.11150244 |
| IGALGLLATDVAPAGSVAFGVA | ASSEGGWDTQY | 0.5609797 |
| AGGAGGIGGNGGAGGAGGVLMG | ASSEGGWDTQY | 0.085512355 |
| ADELIGLL | ASSEGGWDTQY | 0.13759847 |
| ADRALKA | ASSEGGWDTQY | 0.19799867 |
| AGAVGQVGGAGGA | ASSEGGWDTQY | 0.18179283 |
| AGGGGGLGGAP | ASSEGGWDTQY | 0.1340051 |
| AGGLLYGNGGAG | ASSEGGWDTQY | 0.25621474 |
| AGLESIGL | ASSEGGWDTQY | 0.6542577 |
| AGLKTGII | ASSEGGWDTQY | 0.66259867 |
| ASLAALLL | ASSEGGWDTQY | 0.11269895 |
| ASVINISSI | ASSEGGWDTQY | 0.677721 |
| ATPPGPA | ASSEGGWDTQY | 0.7709626 |
| ATVASALV | ASSEGGWDTQY | 0.33041784 |
| ATVASLIL | ASSEGGWDTQY | 0.11426589 |
| AVNGPVLIRH | ASSEGGWDTQY | 0.30143887 |
| DGAAGAVGI | ASSEGGWDTQY | 0.092893355 |
| DPLLHGI | ASSEGGWDTQY | 0.15529746 |
| ELGAALGAL | ASSEGGWDTQY | 0.71825516 |
| ELGRTQAV | ASSEGGWDTQY | 0.12468298 |
| FVAGGYLS | ASSEGGWDTQY | 0.5528302 |
| GASGGAGGAGGW | ASSEGGWDTQY | 0.6506219 |
| GGAGAGGDGGAGGI | ASSEGGWDTQY | 0.20768118 |
| GTGHNGSQPGMGGNG | ASSEGGWDTQY | 0.19086829 |
| GYAAASASAS | ASSEGGWDTQY | 0.13871084 |
| HQVDVAAVQQG | ASSEGGWDTQY | 0.09293239 |
| IGAAIAHT | ASSEGGWDTQY | 0.37006897 |
| IGMIEVAQL | ASSEGGWDTQY | 0.2693474 |
| IRLAPEHTSAMD | ASSEGGWDTQY | 0.15661286 |
| IYNVSVL | ASSEGGWDTQY | 0.31155655 |
| KQGFLGM | ASSEGGWDTQY | 0.16076484 |
| LESVITQL | ASSEGGWDTQY | 0.18841247 |
| LGLGALVVV | ASSEGGWDTQY | 0.1769994 |
| LIGNGGNGGSGGI | ASSEGGWDTQY | 0.12980299 |
| LNAINAPTL | ASSEGGWDTQY | 0.4815891 |
| LQLLNTGV | ASSEGGWDTQY | 0.12562023 |
| LQTSIEF | ASSEGGWDTQY | 0.4205104 |
| MSFAVSEL | ASSEGGWDTQY | 0.75531626 |
| NTGALGL | ASSEGGWDTQY | 0.10012586 |
| PPGAGGAGGAGTT | ASSEGGWDTQY | 0.15571813 |
| QLVGLGI | ASSEGGWDTQY | 0.3364108 |
| RFLEQVHQL | ASSEGGWDTQY | 0.10005785 |
| RGVGYVLR | ASSEGGWDTQY | 0.64379406 |
| SATGVFAGVI | ASSEGGWDTQY | 0.27916414 |
| SLPAITLP | ASSEGGWDTQY | 0.46371496 |
| STALAFTV | ASSEGGWDTQY | 0.27743277 |
| SVLTVGLGK | ASSEGGWDTQY | 0.7487416 |
| TALAGAGSGI | ASSEGGWDTQY | 0.60090196 |
| TGLPFTGI | ASSEGGWDTQY | 0.2301188 |

| | | |
|---|---|---|
| VAEHVTV | ASSEGGWDTQY | 0.1021393 |
| VDIVLNSL | ASSEGGWDTQY | 0.13731652 |
| VLTGGGAVL | ASSEGGWDTQY | 0.6068139 |
| VNVGLAYGI | ASSEGGWDTQY | 0.69086593 |
| VSSAAVGAV | ASSEGGWDTQY | 0.13416208 |
| VTSVYEAGV | ASSEGGWDTQY | 0.31442946 |
| WVALDRGA | ASSEGGWDTQY | 0.5958471 |
| YLVGSVVI | ASSEGGWDTQY | 0.7007223 |
| GPPGTGKTLL | ASSEGGWDTQY | 0.10168698 |
| GVGMTKF | ASSEGGWDTQY | 0.19941892 |
| IGIGGSDL | ASSEGGWDTQY | 0.11416348 |
| IGTLLAEL | ASSEGGWDTQY | 0.27368408 |
| IYTSGTT | ASSEGGWDTQY | 0.13869615 |
| LSFPAGV | ASSEGGWDTQY | 0.64706826 |
| LYTSGTT | ASSEGGWDTQY | 0.32020193 |
| VGAFPFGF | ASSEGGWDTQY | 0.6747117 |
| VGAGTVEF | ASSEGGWDTQY | 0.27682558 |
| VGKSTLLL | ASSEGGWDTQY | 0.4939816 |
| AALAHAGGGIV | ASSEGGWDTQY | 0.29482216 |
| AALVAVSLL | ASSEGGWDTQY | 0.17260246 |
| AAVSRAVAAL | ASSEGGWDTQY | 0.15716147 |
| AGGSSGIGGF | ASSEGGWDTQY | 0.09561042 |
| AGLTVANANG | ASSEGGWDTQY | 0.16455507 |
| AGLVGAVL | ASSEGGWDTQY | 0.1552917 |
| ALSQAFTV | ASSEGGWDTQY | 0.0869504 |
| ALTFSVSL | ASSEGGWDTQY | 0.22939256 |
| ALTGAALVAL | ASSEGGWDTQY | 0.5164709 |
| ASFSIGAGVL | ASSEGGWDTQY | 0.39822116 |
| DGSVYKI | ASSEGGWDTQY | 0.19782905 |
| EQLAEIV | ASSEGGWDTQY | 0.33813357 |
| FAVSELTVM | ASSEGGWDTQY | 0.13062376 |
| GAAPLAAGL | ASSEGGWDTQY | 0.6889854 |
| GADGGDGGSGGAGG | ASSEGGWDTQY | 0.11669506 |
| GFGLDTGGAGG | ASSEGGWDTQY | 0.123950064 |
| GLASGGLL | ASSEGGWDTQY | 0.23095891 |
| GRIAGAAVLL | ASSEGGWDTQY | 0.13214166 |
| GSAGSLNTSF | ASSEGGWDTQY | 0.7246653 |
| GTGQIGI | ASSEGGWDTQY | 0.60979813 |
| GTLIALIL | ASSEGGWDTQY | 0.47500664 |
| IAATMGFVVG | ASSEGGWDTQY | 0.604024 |
| IFAQALL | ASSEGGWDTQY | 0.112034224 |
| IGAAVALGF | ASSEGGWDTQY | 0.16982083 |
| IGLTPLAL | ASSEGGWDTQY | 0.76659036 |
| ILGSNTSTL | ASSEGGWDTQY | 0.29301745 |
| IVFSLTVL | ASSEGGWDTQY | 0.39283523 |
| LASKATGF | ASSEGGWDTQY | 0.6215008 |
| LASVALIL | ASSEGGWDTQY | 0.14363807 |
| LFAVGLTL | ASSEGGWDTQY | 0.22582656 |
| LFNGGIGTYI | ASSEGGWDTQY | 0.18844385 |
| LTLGAGII | ASSEGGWDTQY | 0.44473743 |
| LVAAALGDI | ASSEGGWDTQY | 0.32236812 |

| | | |
|---|---|---|
| NTAFVQLGI | ASSEGGWDTQY | 0.14834191 |
| QLFSGAEL | ASSEGGWDTQY | 0.5922695 |
| SAATFITL | ASSEGGWDTQY | 0.7883545 |
| SNYGLANLGV | ASSEGGWDTQY | 0.30178124 |
| SYGGAYAVM | ASSEGGWDTQY | 0.43750432 |
| TTGVGGLTLG | ASSEGGWDTQY | 0.43982613 |
| TTQTIYI | ASSEGGWDTQY | 0.48585063 |
| VAAGAGLGAVI | ASSEGGWDTQY | 0.18286403 |
| VAAVVTLL | ASSEGGWDTQY | 0.16921116 |
| VGFLGGLI | ASSEGGWDTQY | 0.15040594 |
| VGVIPILL | ASSEGGWDTQY | 0.1906288 |
| VIFASSLI | ASSEGGWDTQY | 0.45893514 |
| VKGGDGGAAAG | ASSEGGWDTQY | 0.21972868 |
| VTKTFTNI | ASSEGGWDTQY | 0.82173085 |
| VVAAAILL | ASSEGGWDTQY | 0.63948923 |
| YAALDVEL | ASSEGGWDTQY | 0.27579814 |
| LASVALLL | ASSEGGWDTQY | 0.120104074 |
| LDAVHRV | ASSEGGWDTQY | 0.26703733 |
| QGLSSSIF | ASSEGGWDTQY | 0.38070574 |
| RGTFANIRL | ASSEGGWDTQY | 0.13155153 |
| RQLWWGHRI | ASSEGGWDTQY | 0.7213492 |
| YGPPGTGKTLL | ASSEGGWDTQY | 0.186448 |
| AALVGAPANVA | ASSEGGWDTQY | 0.11883651 |
| AATFRRTE | ASSEGGWDTQY | 0.37099797 |
| AGALGGGI | ASSEGGWDTQY | 0.7482298 |
| AGGAGGAGGIGAG | ASSEGGWDTQY | 0.30199337 |
| AGSGALGL | ASSEGGWDTQY | 0.14303611 |
| AQIADQI | ASSEGGWDTQY | 0.1273684 |
| ASAVAVTYL | ASSEGGWDTQY | 0.6433189 |
| ASTVLGGTGG | ASSEGGWDTQY | 0.7044578 |
| AVGGEPAK | ASSEGGWDTQY | 0.09161125 |
| CEQGTFGGTC | ASSEGGWDTQY | 0.74027723 |
| DEGKAPAG | ASSEGGWDTQY | 0.14790873 |
| DTVVFPAL | ASSEGGWDTQY | 0.13390994 |
| FEGITMAG | ASSEGGWDTQY | 0.41500217 |
| FPLVVQD | ASSEGGWDTQY | 0.33669904 |
| GAFTTPQ | ASSEGGWDTQY | 0.705644 |
| GFGGFGTAGGVGGA | ASSEGGWDTQY | 0.42979616 |
| GFLAQFL | ASSEGGWDTQY | 0.6525347 |
| GGAGGAGGNAGVA | ASSEGGWDTQY | 0.24478894 |
| GGVGTAGAGGAGGAGG | ASSEGGWDTQY | 0.71592915 |
| GNGGGGDAGNAGSGGN | ASSEGGWDTQY | 0.68035537 |
| GTGGAGGAGNDAGS | ASSEGGWDTQY | 0.4923515 |
| GTTNTGWFNS | ASSEGGWDTQY | 0.34717435 |
| IFMGPLFTL | ASSEGGWDTQY | 0.2039774 |
| ILAHAIEAI | ASSEGGWDTQY | 0.31320116 |
| IRSAIREAL | ASSEGGWDTQY | 0.18058577 |
| ITFAGGG | ASSEGGWDTQY | 0.22984992 |
| ITGMSGHQEV | ASSEGGWDTQY | 0.51113045 |
| IVAGLACTL | ASSEGGWDTQY | 0.49801958 |
| IVGGGLAGA | ASSEGGWDTQY | 0.15237625 |

| | | |
|---|---|---|
| IVTGVLNI | ASSEGGWDTQY | 0.44174808 |
| LAQQGCI | ASSEGGWDTQY | 0.70593256 |
| LCLHVEN | ASSEGGWDTQY | 0.332327 |
| LDAISDGS | ASSEGGWDTQY | 0.24644652 |
| LGGFAAML | ASSEGGWDTQY | 0.7191372 |
| LGGKNPMI | ASSEGGWDTQY | 0.8095021 |
| LGPKGVAAIL | ASSEGGWDTQY | 0.63113666 |
| LMVGVSI | ASSEGGWDTQY | 0.12386572 |
| LVGAAFVL | ASSEGGWDTQY | 0.114611164 |
| LVLAVAPL | ASSEGGWDTQY | 0.17999536 |
| MAAIAVIDAA | ASSEGGWDTQY | 0.20773597 |
| NNGINAELQ | ASSEGGWDTQY | 0.15426569 |
| RGEPLVLVHG | ASSEGGWDTQY | 0.14408147 |
| RGGAGGAAGLI | ASSEGGWDTQY | 0.6071898 |
| RSYRMHPAV | ASSEGGWDTQY | 0.58611953 |
| SAAHLLAGQLA | ASSEGGWDTQY | 0.20720977 |
| SAAVGSY | ASSEGGWDTQY | 0.11709667 |
| SPHAVLHRI | ASSEGGWDTQY | 0.50938255 |
| SQIAKVL | ASSEGGWDTQY | 0.64009786 |
| TALGHLVP | ASSEGGWDTQY | 0.18004693 |
| TAPTTSGAA | ASSEGGWDTQY | 0.31543565 |
| TDGDDGNGGSPGT | ASSEGGWDTQY | 0.1620465 |
| TDGLGEN | ASSEGGWDTQY | 0.090399265 |
| TGGPPLPGQA | ASSEGGWDTQY | 0.3925426 |
| TIPAITI | ASSEGGWDTQY | 0.15429848 |
| TIPGFTI | ASSEGGWDTQY | 0.21672171 |
| TSIASGALA | ASSEGGWDTQY | 0.18341821 |
| TVTLPGAAI | ASSEGGWDTQY | 0.5403286 |
| VASGLSAALP | ASSEGGWDTQY | 0.4671771 |
| VGAGVIGI | ASSEGGWDTQY | 0.44909108 |
| VGPKRFGV | ASSEGGWDTQY | 0.3526281 |
| VNVAFPD | ASSEGGWDTQY | 0.27561724 |
| VSMTNSGVS | ASSEGGWDTQY | 0.13116023 |
| VTDVLGDLSV | ASSEGGWDTQY | 0.09321364 |
| VVVVGAGFAGL | ASSEGGWDTQY | 0.7210645 |
| VYTVGTVV | ASSEGGWDTQY | 0.0850025 |
| WTTFSGP | ASSEGGWDTQY | 0.6948359 |
| YDELVEAF | ASSEGGWDTQY | 0.3456602 |
| YGTGGAGGNGG | ASSEGGWDTQY | 0.1441847 |
| YTNVMAV | ASSEGGWDTQY | 0.29015982 |
| FLEQVHQL | ASSEGGWDTQY | 0.09810524 |
| GLSLLAVAV | ASSEGGWDTQY | 0.13639009 |
| LEQVHQL | ASSEGGWDTQY | 0.70956314 |
| LNAAGAI | ASSEGGWDTQY | 0.4403609 |
| PAVHASL | ASSEGGWDTQY | 0.23275119 |
| TELEDDA | ASSEGGWDTQY | 0.50536066 |
| TPPGPPN | ASSEGGWDTQY | 0.48304912 |
| TPPPPPPP | ASSEGGWDTQY | 0.21632636 |
| TQSPALI | ASSFHWGGRDTQY | 0.3391103 |
| EFAAAFPQ | ASSFHWGGRDTQY | 0.59730256 |
| LLSGNLDVL | ASSFHWGGRDTQY | 0.45479035 |

| | | |
|---|---|---|
| GPANLGNYNI | ASSFHWGGRDTQY | 0.15126649 |
| TLGPTLP | ASSFHWGGRDTQY | 0.2963063 |
| VYAYSSCCK | ASSFHWGGRDTQY | 0.1805723 |
| APAPAPAPNI | ASSFHWGGRDTQY | 0.43561643 |
| PGGPVPIP | ASSFHWGGRDTQY | 0.21583238 |
| FLTVALEP | ASSFHWGGRDTQY | 0.3811776 |
| SQYFGANL | ASSFHWGGRDTQY | 0.30260885 |
| GGVKPAA | ASSFHWGGRDTQY | 0.10211012 |
| GGTGGAGGAGGW | ASSFHWGGRDTQY | 0.18813209 |
| GVGGVAGLGGAGG | ASSFHWGGRDTQY | 0.36391407 |
| VLLVLATTI | ASSFHWGGRDTQY | 0.23505242 |
| IGIFAAML | ASSFHWGGRDTQY | 0.4565431 |
| AVGDLID | ASSFHWGGRDTQY | 0.1461824 |
| NISPLPL | ASSFHWGGRDTQY | 0.45525265 |
| VTAVFVAI | ASSFHWGGRDTQY | 0.1925072 |
| SGDDNDAAL | ASSFHWGGRDTQY | 0.15825509 |
| AVAALAHS | ASSFHWGGRDTQY | 0.3719159 |
| GGVTVMATP | ASSFHWGGRDTQY | 0.25034773 |
| AGLAANVRD | ASSFHWGGRDTQY | 0.23547757 |
| TNRPDLIDPALLRPG | ASSFHWGGRDTQY | 0.5402958 |
| LANLGVTAL | ASSFHWGGRDTQY | 0.13183819 |
| TGERIEL | ASSFHWGGRDTQY | 0.33420038 |
| NGEELAA | ASSFHWGGRDTQY | 0.1904675 |
| VLSLVWG | ASSFHWGGRDTQY | 0.29536325 |
| IGYALWL | ASSFHWGGRDTQY | 0.44366622 |
| LSLLLAPL | ASSFHWGGRDTQY | 0.13639362 |
| TLDEALK | ASSFHWGGRDTQY | 0.60961473 |
| TDGAGTAEGG | ASSFHWGGRDTQY | 0.17998573 |
| NAPGGAGGNGG | ASSFHWGGRDTQY | 0.18856975 |
| VSVFMGMGGNF | ASSFHWGGRDTQY | 0.24904698 |
| LASYLAAL | ASSFHWGGRDTQY | 0.3859945 |
| LVGGTPAL | ASSFHWGGRDTQY | 0.6817234 |
| VRNADVL | ASSFHWGGRDTQY | 0.14750463 |
| TCSSPFLVSAM | ASSFHWGGRDTQY | 0.5201654 |
| VHITDAE | ASSFHWGGRDTQY | 0.597592 |
| EYAAPSF | ASSFHWGGRDTQY | 0.2953072 |
| VVDAFHGGL | ASSFHWGGRDTQY | 0.1249441 |
| ASVVDPQ | ASSFHWGGRDTQY | 0.15784216 |
| SYNEFGGLS | ASSFHWGGRDTQY | 0.115933895 |
| DPDGSVY | ASSFHWGGRDTQY | 0.23920795 |
| MVSTIHATE | ASSFHWGGRDTQY | 0.1259867 |
| LAAGVADPV | ASSFHWGGRDTQY | 0.6136601 |
| TKEQIAA | ASSFHWGGRDTQY | 0.6940414 |
| SLASPAAL | ASSFHWGGRDTQY | 0.14984946 |
| IGLVFYGV | ASSFHWGGRDTQY | 0.72042084 |
| FLLDASNL | ASSFHWGGRDTQY | 0.14606485 |
| YQVGNFP | ASSFHWGGRDTQY | 0.42205262 |
| ESLLSQST | ASSFHWGGRDTQY | 0.73404324 |
| LFTVGFAAVG | ASSFHWGGRDTQY | 0.13956699 |
| ATPPTPT | ASSFHWGGRDTQY | 0.28245905 |
| TDLATLG | ASSFHWGGRDTQY | 0.21164797 |

| | | |
|---|---|---|
| SAGGTGR | ASSFHWGGRDTQY | 0.19305345 |
| GFSNSGS | ASSFHWGGRDTQY | 0.7325529 |
| GGNGGSGGTSV | ASSFHWGGRDTQY | 0.183913 |
| GTSVATGGAGN | ASSFHWGGRDTQY | 0.41684645 |
| NGGNGGSGGTS | ASSFHWGGRDTQY | 0.18349175 |
| AVVKFPH | ASSFHWGGRDTQY | 0.13060081 |
| TPPPPPP | ASSFHWGGRDTQY | 0.30891278 |
| AAPAAAAAPA | ASSFHWGGRDTQY | 0.7287832 |
| IALNEDL | ASSFHWGGRDTQY | 0.14385751 |
| IVNVSSM | ASSFHWGGRDTQY | 0.68388575 |
| TGIPPAPR | ASSFHWGGRDTQY | 0.21416146 |
| FDLGGGTF | ASSFHWGGRDTQY | 0.58068806 |
| FDLGGGTFDVS | ASSFHWGGRDTQY | 0.69926333 |
| NEPTAAA | ASSFHWGGRDTQY | 0.41044283 |
| NEPTAAALA | ASSFHWGGRDTQY | 0.22110254 |
| GSIINISSI | ASSFHWGGRDTQY | 0.70149636 |
| VAPGNAGTA | ASSFHWGGRDTQY | 0.41226345 |
| TGGGASLE | ASSFHWGGRDTQY | 0.65789413 |
| VAVYNHYKRI | ASSFHWGGRDTQY | 0.59907264 |
| TDPAPATT | ASSFHWGGRDTQY | 0.08967576 |
| TNRPDILDPALLRPG | ASSFHWGGRDTQY | 0.093281925 |
| LTTTTTS | ASSFHWGGRDTQY | 0.0880991 |
| PTGSGKTL | ASSFHWGGRDTQY | 0.09770134 |
| TGSGKTL | ASSFHWGGRDTQY | 0.09666215 |
| TIGHVDHGK | ASSFHWGGRDTQY | 0.08715111 |
| TIGHVDHGKTT | ASSFHWGGRDTQY | 0.09838038 |
| GHVDHGKTT | ASSFHWGGRDTQY | 0.09198273 |
| AQFEHTLL | ASSFHWGGRDTQY | 0.08812641 |
| IVNQDEL | ASSFHWGGRDTQY | 0.09171063 |
| TNGSQFF | ASSFHWGGRDTQY | 0.09591951 |
| TGKGIVE | ASSFHWGGRDTQY | 0.12337107 |
| LISNASDA | ASSFHWGGRDTQY | 0.11000244 |
| AELRTLGELTKIAKAHGAQ | ASSFHWGGRDTQY | 0.088384144 |
| APNAVAAAEPSW | ASSFHWGGRDTQY | 0.0921397 |
| APQVAGAATAVL | ASSFHWGGRDTQY | 0.091755666 |
| ARRDPAAAAAILPTDA | ASSFHWGGRDTQY | 0.09060167 |
| DAAARDVLAEAGLAEYF | ASSFHWGGRDTQY | 0.084450245 |
| DDQIVLSAKANLAKVEQ | ASSFHWGGRDTQY | 0.095231876 |
| DELLPDTGEDSDEDGH | ASSFHWGGRDTQY | 0.086192496 |
| ELPEAPANRATVPQD | ASSFHWGGRDTQY | 0.08382284 |
| FGQSKIERVTGIG | ASSFHWGGRDTQY | 0.100082725 |
| GAGGTGSTAGGAGGAGGAGGLY | ASSFHWGGRDTQY | 0.09358762 |
| GHGGTGAAGGNGGTGGAGGSGI | ASSFHWGGRDTQY | 0.0900209 |
| GNGGNGGAGGEGGPGGAGGAGA | ASSFHWGGRDTQY | 0.10166819 |
| GSTAGNGGNGGNATNGTIAGQP | ASSFHWGGRDTQY | 0.084514536 |
| GTGGDGGNGGNGGTGG | ASSFHWGGRDTQY | 0.09098357 |
| GVRRPAGIPADAPV | ASSFHWGGRDTQY | 0.099246256 |
| GYRQVTDAHLLTLARR | ASSFHWGGRDTQY | 0.12576094 |
| IAKMGQDGHDRG | ASSFHWGGRDTQY | 0.08954712 |
| KAEPSIEIQYT | ASSFHWGGRDTQY | 0.089558296 |
| KPVRIRMAADEDTVAVT | ASSFHWGGRDTQY | 0.122733906 |

| | | |
|---|---|---|
| MPPRIAGMRLLVIKPEP | ASSFHWGGRDTQY | 0.097034425 |
| MKFYAVALFRSRGVPPTR | ASSFHWGGRDTQY | 0.09736272 |
| NAGGFGILGAVAHSP | ASSFHWGGRDTQY | 0.100308634 |
| NEGREITLAMAEM | ASSFHWGGRDTQY | 0.087136455 |
| NELVEWSHPLGHRWRWE | ASSFHWGGRDTQY | 0.10376321 |
| NLGAGGTGGNGGNPG | ASSFHWGGRDTQY | 0.10008878 |
| NTEWPKWETVMAADPVD | ASSFHWGGRDTQY | 0.09471191 |
| PASAAPAETQP | ASSFHWGGRDTQY | 0.102402054 |
| QQSSDGKNGVRAAEITGE | ASSFHWGGRDTQY | 0.08953568 |
| RAPDAVTTSWWKE | ASSFHWGGRDTQY | 0.08907771 |
| SGFGNVGAHTSGWFNQ | ASSFHWGGRDTQY | 0.08424272 |
| SGVAGAVGQAASVGGLKV | ASSFHWGGRDTQY | 0.09248952 |
| SNGGAGTGSAGGNGGT | ASSFHWGGRDTQY | 0.09517374 |
| TGGAGGEGGAAG | ASSFHWGGRDTQY | 0.09663769 |
| TGGDGGAAGAGGAPGHG | ASSFHWGGRDTQY | 0.086166546 |
| TMPPLLAAIANAGRHGV | ASSFHWGGRDTQY | 0.091237925 |
| TSSTAGNSGGA | ASSFHWGGRDTQY | 0.08438327 |
| VPATVPVGILC | ASSFHWGGRDTQY | 0.08812582 |
| VQRPDSGGVVLTGRIS | ASSFHWGGRDTQY | 0.08463774 |
| WTPNPVPATKGVPAGLVQAT | ASSFHWGGRDTQY | 0.08521274 |
| YALKRALAKFSDRFDVVII | ASSFHWGGRDTQY | 0.12525818 |
| YLIGSVIV | ASSFHWGGRDTQY | 0.121820346 |
| YGQLVAVID | ASSFHWGGRDTQY | 0.10130867 |
| VTVGGMTLPA | ASSFHWGGRDTQY | 0.094067074 |
| VLALPAV | ASSFHWGGRDTQY | 0.08966668 |
| VIPPTLNYET | ASSFHWGGRDTQY | 0.09749201 |
| VGMGGMHPSSG | ASSFHWGGRDTQY | 0.09610842 |
| VGLAGGAATA | ASSFHWGGRDTQY | 0.08764636 |
| VGGAGGAGGSGP | ASSFHWGGRDTQY | 0.10152119 |
| VFAAFMV | ASSFHWGGRDTQY | 0.092379816 |
| TLYDKLR | ASSFHWGGRDTQY | 0.087691315 |
| TLAVFTVL | ASSFHWGGRDTQY | 0.1174341 |
| TGSVQPARV | ASSFHWGGRDTQY | 0.090202704 |
| TGGGAGSGGT | ASSFHWGGRDTQY | 0.09031947 |
| SHLETSL | ASSFHWGGRDTQY | 0.08574093 |
| SFTTGMT | ASSFHWGGRDTQY | 0.09353808 |
| QVEVAKRMGVRQPTVS | ASSFHWGGRDTQY | 0.086338446 |
| QGSGGPALAM | ASSFHWGGRDTQY | 0.086415395 |
| MLAPSNP | ASSFHWGGRDTQY | 0.086954296 |
| LVNNAAI | ASSFHWGGRDTQY | 0.09799612 |
| LTGEINTLA | ASSFHWGGRDTQY | 0.13022523 |
| LSIFGPV | ASSFHWGGRDTQY | 0.09546052 |
| LLALLANL | ASSFHWGGRDTQY | 0.08684842 |
| LISNASDALDKLR | ASSFHWGGRDTQY | 0.10231824 |
| LINEAVP | ASSFHWGGRDTQY | 0.09171415 |
| LGYAFSGL | ASSFHWGGRDTQY | 0.09461328 |
| LDFVAVAL | ASSFHWGGRDTQY | 0.09321024 |
| LAQPGAE | ASSFHWGGRDTQY | 0.08767613 |
| ISNLGAVTGF | ASSFHWGGRDTQY | 0.0963783 |
| HQEDPGRI | ASSFHWGGRDTQY | 0.10241814 |
| GTPGEPTP | ASSFHWGGRDTQY | 0.096300066 |

| | | |
|---|---|---|
| GFFNSGS | ASSFHWGGRDTQY | 0.093877904 |
| GAGGGGLTAGAN | ASSFHWGGRDTQY | 0.08966323 |
| FVDALVNM | ASSFHWGGRDTQY | 0.092061095 |
| FLTVAPEL | ASSFHWGGRDTQY | 0.09512664 |
| FETLTAL | ASSFHWGGRDTQY | 0.09044656 |
| DVINPIEL | ASSFHWGGRDTQY | 0.0912782 |
| APVSGAPV | ASSFHWGGRDTQY | 0.09130732 |
| ALVGGLAW | ASSFHWGGRDTQY | 0.09091991 |
| ALAAAAATAI | ASSFHWGGRDTQY | 0.097288094 |
| AFGLGLLL | ASSFHWGGRDTQY | 0.08738927 |
| AFFAAASL | ASSFHWGGRDTQY | 0.09527512 |
| AAAALPAFSPP | ASSFHWGGRDTQY | 0.104533486 |
| VVLAPAL | ASSFHWGGRDTQY | 0.08814343 |
| TIGHVDHGKTTL | ASSFHWGGRDTQY | 0.093202814 |
| LISNASDALDK | ASSFHWGGRDTQY | 0.09485239 |
| LISNASDAL | ASSFHWGGRDTQY | 0.09060304 |
| ISNASDALDK | ASSFHWGGRDTQY | 0.10049811 |
| NQAETLVYQTE | ASSFHWGGRDTQY | 0.08903836 |
| TGGTGGTAGAG | ASSFHWGGRDTQY | 0.0984558 |
| TGSTAGGAGGA | ASSFHWGGRDTQY | 0.0853208 |
| DPLAVPVAAGPV | ASSFHWGGRDTQY | 0.10396883 |
| GGTGGTPGNGGN | ASSFHWGGRDTQY | 0.09084145 |
| PAGLAGVDPATL | ASSFHWGGRDTQY | 0.08792286 |
| GNNLVVEAPHAGASV | ASSFHWGGRDTQY | 0.08728333 |
| KVTRIMSASTLEQPA | ASSFHWGGRDTQY | 0.08453549 |
| LGFALPIALPRRTVV | ASSFHWGGRDTQY | 0.10533782 |
| TRLGNLTARKLVVLV | ASSFHWGGRDTQY | 0.09159719 |
| WSQQRRLIYFPSAGP | ASSFHWGGRDTQY | 0.0962946 |
| GPNGNPDPMAAAVDIR | ASSFHWGGRDTQY | 0.09212916 |
| TQRCVSVADLRAAASR | ASSFHWGGRDTQY | 0.10080576 |
| QFVLDVAEHPVEVLTETV | ASSFHWGGRDTQY | 0.09065617 |
| GGAGGTGGAGGNGGTQAGDG | ASSFHWGGRDTQY | 0.09240976 |
| PLLATLRAIAVDVDAKPAQV | ASSFHWGGRDTQY | 0.08790912 |
| APKFRTGGYMIDFWGVGYQVAK | ASSFHWGGRDTQY | 0.10432522 |
| GAGGAGGAGGAAGGPGGTGGTG | ASSFHWGGRDTQY | 0.09868834 |
| IGALGLLATDVAPAGSVAFGVA | ASSFHWGGRDTQY | 0.09534554 |
| AGGAGGIGGNGGAGGAGGVLMG | ASSFHWGGRDTQY | 0.08876023 |
| ADELIGLL | ASSFHWGGRDTQY | 0.08479721 |
| ADRALKA | ASSFHWGGRDTQY | 0.10339231 |
| AGAVGQVGGAGGA | ASSFHWGGRDTQY | 0.08453365 |
| AGGGGGLGGAP | ASSFHWGGRDTQY | 0.083885185 |
| AGGLLYGNGGAG | ASSFHWGGRDTQY | 0.08462439 |
| AGLESIGL | ASSFHWGGRDTQY | 0.08432843 |
| AGLKTGII | ASSFHWGGRDTQY | 0.10786137 |
| ASLAALLL | ASSFHWGGRDTQY | 0.11601785 |
| ASVINISSI | ASSFHWGGRDTQY | 0.09482613 |
| ATPPGPA | ASSFHWGGRDTQY | 0.09632098 |
| ATVASALV | ASSFHWGGRDTQY | 0.09795438 |
| ATVASLIL | ASSFHWGGRDTQY | 0.08357911 |
| AVNGPVLIRH | ASSFHWGGRDTQY | 0.08999598 |
| DGAAGAVGI | ASSFHWGGRDTQY | 0.08954976 |

| | | |
|---|---|---|
| DPLLHGI | ASSFHWGGRDTQY | 0.10240398 |
| ELGAALGAL | ASSFHWGGRDTQY | 0.0868277 |
| ELGRTQAV | ASSFHWGGRDTQY | 0.08871108 |
| FVAGGYLS | ASSFHWGGRDTQY | 0.10251103 |
| GASGGAGGAGGW | ASSFHWGGRDTQY | 0.090099335 |
| GGAGAGGDGGAGGI | ASSFHWGGRDTQY | 0.09185668 |
| GTGHNGSQPGMGGNG | ASSFHWGGRDTQY | 0.08579688 |
| GYAAASASAS | ASSFHWGGRDTQY | 0.08369912 |
| HQVDVAAVQQG | ASSFHWGGRDTQY | 0.114717536 |
| IGAAIAHT | ASSFHWGGRDTQY | 0.08531541 |
| IGMIEVAQL | ASSFHWGGRDTQY | 0.09181941 |
| IRLAPEHTSAMD | ASSFHWGGRDTQY | 0.08412521 |
| IYNVSVL | ASSFHWGGRDTQY | 0.08467392 |
| KQGFLGM | ASSFHWGGRDTQY | 0.08705963 |
| LESVITQL | ASSFHWGGRDTQY | 0.09603296 |
| LGLGALVVV | ASSFHWGGRDTQY | 0.08819259 |
| LIGNGGNGGSGGI | ASSFHWGGRDTQY | 0.094925776 |
| LNAINAPTL | ASSFHWGGRDTQY | 0.10081395 |
| LQLLNTGV | ASSFHWGGRDTQY | 0.11573902 |
| LQTSIEF | ASSFHWGGRDTQY | 0.09171279 |
| MSFAVSEL | ASSFHWGGRDTQY | 0.09274946 |
| NTGALGL | ASSFHWGGRDTQY | 0.08802774 |
| PPGAGGAGGAGTT | ASSFHWGGRDTQY | 0.08785209 |
| QLVGLGI | ASSFHWGGRDTQY | 0.08936039 |
| RFLEQVHQL | ASSFHWGGRDTQY | 0.119190946 |
| RGVGYVLR | ASSFHWGGRDTQY | 0.08695222 |
| SATGVFAGVI | ASSFHWGGRDTQY | 0.08710372 |
| SLPAITLP | ASSFHWGGRDTQY | 0.08826861 |
| STALAFTV | ASSFHWGGRDTQY | 0.10399569 |
| SVLTVGLGK | ASSFHWGGRDTQY | 0.10567194 |
| TALAGAGSGI | ASSFHWGGRDTQY | 0.08864694 |
| TGLPFTGI | ASSFHWGGRDTQY | 0.09383015 |
| VAEHVTV | ASSFHWGGRDTQY | 0.0923928 |
| VDIVLNSL | ASSFHWGGRDTQY | 0.09478401 |
| VLTGGGAVL | ASSFHWGGRDTQY | 0.08764573 |
| VNVGLAYGI | ASSFHWGGRDTQY | 0.09035982 |
| VSSAAVGAV | ASSFHWGGRDTQY | 0.09335908 |
| VTSVYEAGV | ASSFHWGGRDTQY | 0.11026431 |
| WVALDRGA | ASSFHWGGRDTQY | 0.08880105 |
| YLVGSVVI | ASSFHWGGRDTQY | 0.08649716 |
| GPPGTGKTLL | ASSFHWGGRDTQY | 0.10292306 |
| GVGMTKF | ASSFHWGGRDTQY | 0.0872327 |
| IGIGGSDL | ASSFHWGGRDTQY | 0.094308294 |
| IGTLLAEL | ASSFHWGGRDTQY | 0.095858544 |
| IYTSGTT | ASSFHWGGRDTQY | 0.086072974 |
| LSFPAGV | ASSFHWGGRDTQY | 0.0893831 |
| LYTSGTT | ASSFHWGGRDTQY | 0.104765646 |
| VGAFPFGF | ASSFHWGGRDTQY | 0.09204692 |
| VGAGTVEF | ASSFHWGGRDTQY | 0.09268369 |
| VGKSTLLL | ASSFHWGGRDTQY | 0.10365886 |
| AALAHAGGGIV | ASSFHWGGRDTQY | 0.093266614 |

| | | |
|---|---|---|
| AALVAVSLL | ASSFHWGGRDTQY | 0.103684254 |
| AAVSRAVAAL | ASSFHWGGRDTQY | 0.090045564 |
| AGGSSGIGGF | ASSFHWGGRDTQY | 0.093313344 |
| AGLTVANANG | ASSFHWGGRDTQY | 0.09121296 |
| AGLVGAVL | ASSFHWGGRDTQY | 0.089999646 |
| ALSQAFTV | ASSFHWGGRDTQY | 0.10781887 |
| ALTFSVSL | ASSFHWGGRDTQY | 0.10105246 |
| ALTGAALVAL | ASSFHWGGRDTQY | 0.08798416 |
| ASFSIGAGVL | ASSFHWGGRDTQY | 0.11645957 |
| DGSVYKI | ASSFHWGGRDTQY | 0.09281137 |
| EQLAEIV | ASSFHWGGRDTQY | 0.090673245 |
| FAVSELTVM | ASSFHWGGRDTQY | 0.08749233 |
| GAAPLAAGL | ASSFHWGGRDTQY | 0.087770075 |
| GADGGDGGSGGAGG | ASSFHWGGRDTQY | 0.09097687 |
| GFGLDTGGAGG | ASSFHWGGRDTQY | 0.08757929 |
| GLASGGLL | ASSFHWGGRDTQY | 0.08382643 |
| GRIAGAAVLL | ASSFHWGGRDTQY | 0.09280744 |
| GSAGSLNTSF | ASSFHWGGRDTQY | 0.09381606 |
| GTGQIGI | ASSFHWGGRDTQY | 0.083733924 |
| GTLIALIL | ASSFHWGGRDTQY | 0.094060466 |
| IAATMGFVVG | ASSFHWGGRDTQY | 0.085693285 |
| IFAQALL | ASSFHWGGRDTQY | 0.086730145 |
| IGAAVALGF | ASSFHWGGRDTQY | 0.09746963 |
| IGLTPLAL | ASSFHWGGRDTQY | 0.087049745 |
| ILGSNTSTL | ASSFHWGGRDTQY | 0.09393444 |
| IVFSLTVL | ASSFHWGGRDTQY | 0.08583918 |
| LASKATGF | ASSFHWGGRDTQY | 0.08514298 |
| LASVALIL | ASSFHWGGRDTQY | 0.09886278 |
| LFAVGLTL | ASSFHWGGRDTQY | 0.09383555 |
| LFNGGIGTYI | ASSFHWGGRDTQY | 0.089026965 |
| LTLGAGII | ASSFHWGGRDTQY | 0.09961126 |
| LVAAALGDI | ASSFHWGGRDTQY | 0.08587047 |
| NTAFVQLGI | ASSFHWGGRDTQY | 0.108382456 |
| QLFSGAEL | ASSFHWGGRDTQY | 0.10350062 |
| SAATFITL | ASSFHWGGRDTQY | 0.08544145 |
| SNYGLANLGV | ASSFHWGGRDTQY | 0.09470465 |
| SYGGAYAVM | ASSFHWGGRDTQY | 0.08847507 |
| TTGVGGLTLG | ASSFHWGGRDTQY | 0.09805205 |
| TTQTIYI | ASSFHWGGRDTQY | 0.09109803 |
| VAAGAGLGAVI | ASSFHWGGRDTQY | 0.084708005 |
| VAAVVTLL | ASSFHWGGRDTQY | 0.09713813 |
| VGFLGGLI | ASSFHWGGRDTQY | 0.11250939 |
| VGVIPILL | ASSFHWGGRDTQY | 0.09645709 |
| VIFASSLI | ASSFHWGGRDTQY | 0.08833506 |
| VKGGDGGAAAG | ASSFHWGGRDTQY | 0.09420983 |
| VTKTFTNI | ASSFHWGGRDTQY | 0.09727281 |
| VVAAAILL | ASSFHWGGRDTQY | 0.09807856 |
| YAALDVEL | ASSFHWGGRDTQY | 0.095938236 |
| LASVALLL | ASSFHWGGRDTQY | 0.09022803 |
| LDAVHRV | ASSFHWGGRDTQY | 0.09674767 |
| QGLSSSIF | ASSFHWGGRDTQY | 0.09753863 |

| | | |
|---|---|---|
| RGTFANIRL | ASSFHWGGRDTQY | 0.08827976 |
| RQLWWGHRI | ASSFHWGGRDTQY | 0.08802886 |
| YGPPGTGKTLL | ASSFHWGGRDTQY | 0.09122485 |
| AALVGAPANVA | ASSFHWGGRDTQY | 0.09939284 |
| AATFRRTE | ASSFHWGGRDTQY | 0.10037392 |
| AGALGGGI | ASSFHWGGRDTQY | 0.08592765 |
| AGGAGGAGGIGAG | ASSFHWGGRDTQY | 0.0858712 |
| AGSGALGL | ASSFHWGGRDTQY | 0.09264971 |
| AQIADQI | ASSFHWGGRDTQY | 0.10518241 |
| ASAVAVTYL | ASSFHWGGRDTQY | 0.08950309 |
| ASTVLGGTGG | ASSFHWGGRDTQY | 0.09038654 |
| AVGGEPAK | ASSFHWGGRDTQY | 0.11942032 |
| CEQGTFGGTC | ASSFHWGGRDTQY | 0.089989595 |
| DEGKAPAG | ASSFHWGGRDTQY | 0.098523244 |
| DTVVFPAL | ASSFHWGGRDTQY | 0.100398645 |
| FEGITMAG | ASSFHWGGRDTQY | 0.102635734 |
| FPLVVQD | ASSFHWGGRDTQY | 0.08955976 |
| GAFTTPQ | ASSFHWGGRDTQY | 0.09688287 |
| GFGGFGTAGGVGGA | ASSFHWGGRDTQY | 0.086959615 |
| GFLAQFL | ASSFHWGGRDTQY | 0.09880055 |
| GGAGGAGGNAGVA | ASSFHWGGRDTQY | 0.085063666 |
| GGVGTAGAGGAGGAGG | ASSFHWGGRDTQY | 0.096077524 |
| GNGGGGDAGNAGSGGN | ASSFHWGGRDTQY | 0.08541784 |
| GTGGAGGAGNDAGS | ASSFHWGGRDTQY | 0.10031784 |
| GTTNTGWFNS | ASSFHWGGRDTQY | 0.09810974 |
| IFMGPLFTL | ASSFHWGGRDTQY | 0.10679109 |
| ILAHAIEAI | ASSFHWGGRDTQY | 0.084039085 |
| IRSAIREAL | ASSFHWGGRDTQY | 0.1026959 |
| ITFAGGG | ASSFHWGGRDTQY | 0.11087316 |
| ITGMSGHQEV | ASSFHWGGRDTQY | 0.12585165 |
| IVAGLACTL | ASSFHWGGRDTQY | 0.09244647 |
| IVGGGLAGA | ASSFHWGGRDTQY | 0.107846685 |
| IVTGVLNI | ASSFHWGGRDTQY | 0.094564766 |
| LAQQGCI | ASSFHWGGRDTQY | 0.087147035 |
| LCLHVEN | ASSFHWGGRDTQY | 0.08610978 |
| LDAISDGS | ASSFHWGGRDTQY | 0.09098403 |
| LGGFAAML | ASSFHWGGRDTQY | 0.092754565 |
| LGGKNPMI | ASSFHWGGRDTQY | 0.090040416 |
| LGPKGVAAIL | ASSFHWGGRDTQY | 0.10545246 |
| LMVGVSI | ASSFHWGGRDTQY | 0.09954299 |
| LVGAAFVL | ASSFHWGGRDTQY | 0.09881962 |
| LVLAVAPL | ASSFHWGGRDTQY | 0.117559664 |
| MAAIAVIDAA | ASSFHWGGRDTQY | 0.095426306 |
| NNGINAELQ | ASSFHWGGRDTQY | 0.09198977 |
| RGEPLVLVHG | ASSFHWGGRDTQY | 0.08926796 |
| RGGAGGAAGLI | ASSFHWGGRDTQY | 0.090942085 |
| RSYRMHPAV | ASSFHWGGRDTQY | 0.08736594 |
| SAAHLLAGQLA | ASSFHWGGRDTQY | 0.091010354 |
| SAAVGSY | ASSFHWGGRDTQY | 0.089148425 |
| SPHAVLHRI | ASSFHWGGRDTQY | 0.107988976 |
| SQIAKVL | ASSFHWGGRDTQY | 0.08337287 |

| | | |
|---|---|---|
| TALGHLVP | ASSFHWGGRDTQY | 0.0877764 |
| TAPTTSGAA | ASSFHWGGRDTQY | 0.10944985 |
| TDGDDGNGGSPGT | ASSFHWGGRDTQY | 0.08506528 |
| TDGLGEN | ASSFHWGGRDTQY | 0.09843293 |
| TGGPPLPGQA | ASSFHWGGRDTQY | 0.08941753 |
| TIPAITI | ASSFHWGGRDTQY | 0.085588396 |
| TIPGFTI | ASSFHWGGRDTQY | 0.115053385 |
| TSIASGALA | ASSFHWGGRDTQY | 0.08736618 |
| TVTLPGAAI | ASSFHWGGRDTQY | 0.09313361 |
| VASGLSAALP | ASSFHWGGRDTQY | 0.0971958 |
| VGAGVIGI | ASSFHWGGRDTQY | 0.099471375 |
| VGPKRFGV | ASSFHWGGRDTQY | 0.09018 |
| VNVAFPD | ASSFHWGGRDTQY | 0.09185019 |
| VSMTNSGVS | ASSFHWGGRDTQY | 0.08831183 |
| VTDVLGDLSV | ASSFHWGGRDTQY | 0.093143605 |
| VVVVGAGFAGL | ASSFHWGGRDTQY | 0.08368914 |
| VYTVGTVV | ASSFHWGGRDTQY | 0.09437044 |
| WTTFSGP | ASSFHWGGRDTQY | 0.08648263 |
| YDELVEAF | ASSFHWGGRDTQY | 0.12547123 |
| YGTGGAGGNGG | ASSFHWGGRDTQY | 0.099897705 |
| YTNVMAV | ASSFHWGGRDTQY | 0.09567599 |
| FLEQVHQL | ASSFHWGGRDTQY | 0.093189865 |
| GLSLLAVAV | ASSFHWGGRDTQY | 0.08633217 |
| LEQVHQL | ASSFHWGGRDTQY | 0.08590392 |
| LNAAGAI | ASSFHWGGRDTQY | 0.08826495 |
| PAVHASL | ASSFHWGGRDTQY | 0.09029634 |
| TELEDDA | ASSFHWGGRDTQY | 0.08954274 |
| TPPGPPN | ASSFHWGGRDTQY | 0.085694164 |
| TPPPPPPP | ASSFHWGGRDTQY | 0.106499776 |
| TQSPALI | ASNPRGRNSPLY | 0.09466195 |
| EFAAAFPQ | ASNPRGRNSPLY | 0.08642339 |
| LLSGNLDVL | ASNPRGRNSPLY | 0.09014571 |
| GPANLGNYNI | ASNPRGRNSPLY | 0.09758632 |
| TLGPTLP | ASNPRGRNSPLY | 0.09524753 |
| VYAYSSCCK | ASNPRGRNSPLY | 0.083468445 |
| APAPAPAPNI | ASNPRGRNSPLY | 0.0871279 |
| PGGPVPIP | ASNPRGRNSPLY | 0.09840774 |
| FLTVALEP | ASNPRGRNSPLY | 0.08653452 |
| SQYFGANL | ASNPRGRNSPLY | 0.087594025 |
| GGVKPAA | ASNPRGRNSPLY | 0.11654468 |
| GGTGGAGGAGGW | ASNPRGRNSPLY | 0.0855949 |
| GVGGVAGLGGAGG | ASNPRGRNSPLY | 0.08928727 |
| VLLVLATTI | ASNPRGRNSPLY | 0.09566694 |
| IGIFAAML | ASNPRGRNSPLY | 0.124894924 |
| AVGDLID | ASNPRGRNSPLY | 0.113991655 |
| NISPLPL | ASNPRGRNSPLY | 0.09226127 |
| VTAVFVAI | ASNPRGRNSPLY | 0.08863955 |
| SGDDNDAAL | ASNPRGRNSPLY | 0.102152176 |
| AVAALAHS | ASNPRGRNSPLY | 0.099812426 |
| GGVTVMATP | ASNPRGRNSPLY | 0.08448112 |
| AGLAANVRD | ASNPRGRNSPLY | 0.083708115 |

| | | |
|---|---|---|
| TNRPDLIDPALLRPG | ASNPRGRNSPLY | 0.09540419 |
| LANLGVTAL | ASNPRGRNSPLY | 0.08463157 |
| TGERIEL | ASNPRGRNSPLY | 0.11061063 |
| NGEELAA | ASNPRGRNSPLY | 0.09733311 |
| VLSLVWG | ASNPRGRNSPLY | 0.08603128 |
| IGYALWL | ASNPRGRNSPLY | 0.08731156 |
| LSLLLAPL | ASNPRGRNSPLY | 0.09032595 |
| TLDEALK | ASNPRGRNSPLY | 0.093331516 |
| TDGAGTAEGG | ASNPRGRNSPLY | 0.09965093 |
| NAPGGAGGNGG | ASNPRGRNSPLY | 0.08801229 |
| VSVFMGMGGNF | ASNPRGRNSPLY | 0.12572831 |
| LASYLAAL | ASNPRGRNSPLY | 0.11070912 |
| LVGGTPAL | ASNPRGRNSPLY | 0.09120687 |
| VRNADVL | ASNPRGRNSPLY | 0.09853267 |
| TCSSPFLVSAM | ASNPRGRNSPLY | 0.088395685 |
| VHITDAE | ASNPRGRNSPLY | 0.08690784 |
| EYAAPSF | ASNPRGRNSPLY | 0.09080015 |
| VVDAFHGGL | ASNPRGRNSPLY | 0.08665668 |
| ASVVDPQ | ASNPRGRNSPLY | 0.088334315 |
| SYNEFGGLS | ASNPRGRNSPLY | 0.089156434 |
| DPDGSVY | ASNPRGRNSPLY | 0.08744063 |
| MVSTIHATE | ASNPRGRNSPLY | 0.08829526 |
| LAAGVADPV | ASNPRGRNSPLY | 0.09011584 |
| TKEQIAA | ASNPRGRNSPLY | 0.08424527 |
| SLASPAAL | ASNPRGRNSPLY | 0.08934049 |
| IGLVFYGV | ASNPRGRNSPLY | 0.1153608 |
| FLLDASNL | ASNPRGRNSPLY | 0.13637015 |
| YQVGNFP | ASNPRGRNSPLY | 0.090626135 |
| ESLLSQST | ASNPRGRNSPLY | 0.09009484 |
| LFTVGFAAVG | ASNPRGRNSPLY | 0.08908956 |
| ATPPTPT | ASNPRGRNSPLY | 0.09747603 |
| TDLATLG | ASNPRGRNSPLY | 0.08813189 |
| SAGGTGR | ASNPRGRNSPLY | 0.09290277 |
| GFSNSGS | ASNPRGRNSPLY | 0.08852186 |
| GGNGGSGGTSV | ASNPRGRNSPLY | 0.08829349 |
| GTSVATGGAGN | ASNPRGRNSPLY | 0.08887697 |
| NGGNGGSGGTS | ASNPRGRNSPLY | 0.08684253 |
| AVVKFPH | ASNPRGRNSPLY | 0.09266548 |
| TPPPPPP | ASNPRGRNSPLY | 0.08376189 |
| AAPAAAAAPA | ASNPRGRNSPLY | 0.08967344 |
| IALNEDL | ASNPRGRNSPLY | 0.08893815 |
| IVNVSSM | ASNPRGRNSPLY | 0.08812139 |
| TGIPPAPR | ASNPRGRNSPLY | 0.0852924 |
| FDLGGGTF | ASNPRGRNSPLY | 0.094441324 |
| FDLGGGTFDVS | ASNPRGRNSPLY | 0.09822552 |
| NEPTAAA | ASNPRGRNSPLY | 0.09877709 |
| NEPTAAALA | ASNPRGRNSPLY | 0.08452704 |
| GSIINISSI | ASNPRGRNSPLY | 0.09713204 |
| VAPGNAGTA | ASNPRGRNSPLY | 0.09942997 |
| TGGGASLE | ASNPRGRNSPLY | 0.09107924 |
| VAVYNHYKRI | ASNPRGRNSPLY | 0.087093204 |

| | | |
|---|---|---|
| TDPAPATT | ASNPRGRNSPLY | 0.09878143 |
| TNRPDILDPALLRPG | ASNPRGRNSPLY | 0.08535748 |
| LTTTTTS | ASNPRGRNSPLY | 0.088065475 |
| PTGSGKTL | ASNPRGRNSPLY | 0.09736701 |
| TGSGKTL | ASNPRGRNSPLY | 0.08646322 |
| TIGHVDHGK | ASNPRGRNSPLY | 0.102916524 |
| TIGHVDHGKTT | ASNPRGRNSPLY | 0.10344191 |
| GHVDHGKTT | ASNPRGRNSPLY | 0.08730062 |
| AQFEHTLL | ASNPRGRNSPLY | 0.08796777 |
| IVNQDEL | ASNPRGRNSPLY | 0.088810734 |
| TNGSQFF | ASNPRGRNSPLY | 0.08399414 |
| TGKGIVE | ASNPRGRNSPLY | 0.10835371 |
| LISNASDA | ASNPRGRNSPLY | 0.0878242 |
| AELRTLGELTKIAKAHGAQ | ASNPRGRNSPLY | 0.09302269 |
| APNAVAAAEPSW | ASNPRGRNSPLY | 0.092872374 |
| APQVAGAATAVL | ASNPRGRNSPLY | 0.09169866 |
| ARRDPAAAAAILPTDA | ASNPRGRNSPLY | 0.09530216 |
| DAAARDVLAEAGLAEYF | ASNPRGRNSPLY | 0.097325124 |
| DDQIVLSAKANLAKVEQ | ASNPRGRNSPLY | 0.08830417 |
| DELLPDTGEDSDEDGH | ASNPRGRNSPLY | 0.088617966 |
| ELPEAPANRATVPQD | ASNPRGRNSPLY | 0.087844975 |
| FGQSKIERVTGIG | ASNPRGRNSPLY | 0.090064965 |
| GAGGTGSTAGGAGGAGGAGGLY | ASNPRGRNSPLY | 0.12401252 |
| GHGGTGAAGGNGGTGGAGGSGI | ASNPRGRNSPLY | 0.086503655 |
| GNGGNGGAGGEGGPGGAGGAGA | ASNPRGRNSPLY | 0.090560496 |
| GSTAGNGGNGGNATNGTIAGQP | ASNPRGRNSPLY | 0.0882676 |
| GTGGDGGNGGNGGTGG | ASNPRGRNSPLY | 0.08521788 |
| GVRRPAGIPADAPV | ASNPRGRNSPLY | 0.091692775 |
| GYRQVTDAHLLTLARR | ASNPRGRNSPLY | 0.09749652 |
| IAKMGQDGHDRG | ASNPRGRNSPLY | 0.10082204 |
| KAEPSIEIQYT | ASNPRGRNSPLY | 0.09011455 |
| KPVRIRMAADEDTVAVT | ASNPRGRNSPLY | 0.09860312 |
| MPPRIAGMRLLVIKPEP | ASNPRGRNSPLY | 0.102990896 |
| MKFYAVALFRSRGVPPTR | ASNPRGRNSPLY | 0.08987135 |
| NAGGFGILGAVAHSP | ASNPRGRNSPLY | 0.0867703 |
| NEGREITLAMAEM | ASNPRGRNSPLY | 0.08771607 |
| NELVEWSHPLGHRWRWE | ASNPRGRNSPLY | 0.09373041 |
| NLGAGGTGGNGGNPG | ASNPRGRNSPLY | 0.0967988 |
| NTEWPKWETVMAADPVD | ASNPRGRNSPLY | 0.08858026 |
| PASAAPAETQP | ASNPRGRNSPLY | 0.09743502 |
| QQSSDGKNGVRAAEITGE | ASNPRGRNSPLY | 0.09953034 |
| RAPDAVTTSWWKE | ASNPRGRNSPLY | 0.09870693 |
| SGFGNVGAHTSGWFNQ | ASNPRGRNSPLY | 0.08491046 |
| SGVAGAVGQAASVGGLKV | ASNPRGRNSPLY | 0.091220975 |
| SNGGAGTGSAGGNGGT | ASNPRGRNSPLY | 0.10038277 |
| TGGAGGEGGAAG | ASNPRGRNSPLY | 0.09077841 |
| TGGDGGAAGAGGAPGHG | ASNPRGRNSPLY | 0.09197266 |
| TMPPLLAAIANAGRHGV | ASNPRGRNSPLY | 0.09543421 |
| TSSTAGNSGGA | ASNPRGRNSPLY | 0.09919964 |
| VPATVPVGILC | ASNPRGRNSPLY | 0.09001264 |
| VQRPDSGGVVLTGRIS | ASNPRGRNSPLY | 0.11212772 |

| | | |
|---|---|---|
| WTPNPVPATKGVPAGLVQAT | ASNPRGRNSPLY | 0.12161253 |
| YALKRALAKFSDRFDVVII | ASNPRGRNSPLY | 0.08975634 |
| YLIGSVIV | ASNPRGRNSPLY | 0.096820645 |
| YGQLVAVID | ASNPRGRNSPLY | 0.08493849 |
| VTVGGMTLPA | ASNPRGRNSPLY | 0.08493904 |
| VLALPAV | ASNPRGRNSPLY | 0.08680428 |
| VIPPTLNYET | ASNPRGRNSPLY | 0.09235544 |
| VGMGGMHPSSG | ASNPRGRNSPLY | 0.08399573 |
| VGLAGGAATA | ASNPRGRNSPLY | 0.09081964 |
| VGGAGGAGGSGP | ASNPRGRNSPLY | 0.094622515 |
| VFAAFMV | ASNPRGRNSPLY | 0.108999506 |
| TLYDKLR | ASNPRGRNSPLY | 0.09283226 |
| TLAVFTVL | ASNPRGRNSPLY | 0.08998984 |
| TGSVQPARV | ASNPRGRNSPLY | 0.09000234 |
| TGGGAGSGGT | ASNPRGRNSPLY | 0.10690247 |
| SHLETSL | ASNPRGRNSPLY | 0.087609686 |
| SFTTGMT | ASNPRGRNSPLY | 0.08599485 |
| QVEVAKRMGVRQPTVS | ASNPRGRNSPLY | 0.09221993 |
| QGSGGPALAM | ASNPRGRNSPLY | 0.087768964 |
| MLAPSNP | ASNPRGRNSPLY | 0.09481773 |
| LVNNAAI | ASNPRGRNSPLY | 0.098109215 |
| LTGEINTLA | ASNPRGRNSPLY | 0.09892322 |
| LSIFGPV | ASNPRGRNSPLY | 0.09205773 |
| LLALLANL | ASNPRGRNSPLY | 0.08435963 |
| LISNASDALDKLR | ASNPRGRNSPLY | 0.093975134 |
| LINEAVP | ASNPRGRNSPLY | 0.15668239 |
| LGYAFSGL | ASNPRGRNSPLY | 0.091191724 |
| LDFVAVAL | ASNPRGRNSPLY | 0.10057776 |
| LAQPGAE | ASNPRGRNSPLY | 0.10114167 |
| ISNLGAVTGF | ASNPRGRNSPLY | 0.09273174 |
| HQEDPGRI | ASNPRGRNSPLY | 0.09828652 |
| GTPGEPTP | ASNPRGRNSPLY | 0.09060138 |
| GFFNSGS | ASNPRGRNSPLY | 0.09439837 |
| GAGGGGLTAGAN | ASNPRGRNSPLY | 0.08741468 |
| FVDALVNM | ASNPRGRNSPLY | 0.08808425 |
| FLTVAPEL | ASNPRGRNSPLY | 0.10331161 |
| FETLTAL | ASNPRGRNSPLY | 0.10453443 |
| DVINPIEL | ASNPRGRNSPLY | 0.10266057 |
| APVSGAPV | ASNPRGRNSPLY | 0.09744274 |
| ALVGGLAW | ASNPRGRNSPLY | 0.09468247 |
| ALAAAAATAI | ASNPRGRNSPLY | 0.09094773 |
| AFGLGLLL | ASNPRGRNSPLY | 0.094158836 |
| AFFAAASL | ASNPRGRNSPLY | 0.087957226 |
| AAAALPAFSPP | ASNPRGRNSPLY | 0.08628744 |
| VVLAPAL | ASNPRGRNSPLY | 0.08739682 |
| TIGHVDHGKTTL | ASNPRGRNSPLY | 0.09361472 |
| LISNASDALDK | ASNPRGRNSPLY | 0.091437876 |
| LISNASDAL | ASNPRGRNSPLY | 0.11365318 |
| ISNASDALDK | ASNPRGRNSPLY | 0.093459725 |
| NQAETLVYQTE | ASNPRGRNSPLY | 0.09341958 |
| TGGTGGTAGAG | ASNPRGRNSPLY | 0.08921842 |

| | | |
|---|---|---|
| TGSTAGGAGGA | ASNPRGRNSPLY | 0.090223365 |
| DPLAVPVAAGPV | ASNPRGRNSPLY | 0.09093876 |
| GGTGGTPGNGGN | ASNPRGRNSPLY | 0.08767681 |
| PAGLAGVDPATL | ASNPRGRNSPLY | 0.122320965 |
| GNNLVVEAPHAGASV | ASNPRGRNSPLY | 0.09073267 |
| KVTRIMSASTLEQPA | ASNPRGRNSPLY | 0.08524979 |
| LGFALPIALPRRTVV | ASNPRGRNSPLY | 0.09385682 |
| TRLGNLTARKLVVLV | ASNPRGRNSPLY | 0.11283451 |
| WSQQRRLIYFPSAGP | ASNPRGRNSPLY | 0.091351725 |
| GPNGNPDPMAAAVDIR | ASNPRGRNSPLY | 0.08725461 |
| TQRCVSVADLRAAASR | ASNPRGRNSPLY | 0.08996041 |
| QFVLDVAEHPVEVLTETV | ASNPRGRNSPLY | 0.09562586 |
| GGAGGTGGAGGNGGTQAGDG | ASNPRGRNSPLY | 0.12809893 |
| PLLATLRAIAVDVDAKPAQV | ASNPRGRNSPLY | 0.08503107 |
| APKFRTGGYMIDFWGVGYQVAK | ASNPRGRNSPLY | 0.09879515 |
| GAGGAGGAGGAAGGPGGTGGTG | ASNPRGRNSPLY | 0.08734208 |
| IGALGLLATDVAPAGSVAFGVA | ASNPRGRNSPLY | 0.08772832 |
| AGGAGGIGGNGGAGGAGGVLMG | ASNPRGRNSPLY | 0.09395098 |
| ADELIGLL | ASNPRGRNSPLY | 0.09525084 |
| ADRALKA | ASNPRGRNSPLY | 0.09074716 |
| AGAVGQVGGAGGA | ASNPRGRNSPLY | 0.098123394 |
| AGGGGGLGGAP | ASNPRGRNSPLY | 0.08617544 |
| AGGLLYGNGGAG | ASNPRGRNSPLY | 0.08924151 |
| AGLESIGL | ASNPRGRNSPLY | 0.1067255 |
| AGLKTGII | ASNPRGRNSPLY | 0.092760265 |
| ASLAALLL | ASNPRGRNSPLY | 0.10592149 |
| ASVINISSI | ASNPRGRNSPLY | 0.09033638 |
| ATPPGPA | ASNPRGRNSPLY | 0.088365644 |
| ATVASALV | ASNPRGRNSPLY | 0.092714325 |
| ATVASLIL | ASNPRGRNSPLY | 0.089052975 |
| AVNGPVLIRH | ASNPRGRNSPLY | 0.08709006 |
| DGAAGAVGI | ASNPRGRNSPLY | 0.094787054 |
| DPLLHGI | ASNPRGRNSPLY | 0.102299474 |
| ELGAALGAL | ASNPRGRNSPLY | 0.088313416 |
| ELGRTQAV | ASNPRGRNSPLY | 0.09285649 |
| FVAGGYLS | ASNPRGRNSPLY | 0.09422151 |
| GASGGAGGAGGW | ASNPRGRNSPLY | 0.08881053 |
| GGAGAGGDGGAGGI | ASNPRGRNSPLY | 0.08860741 |
| GTGHNGSQPGMGGNG | ASNPRGRNSPLY | 0.10845386 |
| GYAAASASAS | ASNPRGRNSPLY | 0.1160659 |
| HQVDVAAVQQG | ASNPRGRNSPLY | 0.09737744 |
| IGAAIAHT | ASNPRGRNSPLY | 0.08717684 |
| IGMIEVAQL | ASNPRGRNSPLY | 0.08439581 |
| IRLAPEHTSAMD | ASNPRGRNSPLY | 0.10734316 |
| IYNVSVL | ASNPRGRNSPLY | 0.09229066 |
| KQGFLGM | ASNPRGRNSPLY | 0.09208798 |
| LESVITQL | ASNPRGRNSPLY | 0.08781385 |
| LGLGALVVV | ASNPRGRNSPLY | 0.08841151 |
| LIGNGGNGGSGGI | ASNPRGRNSPLY | 0.10355034 |
| LNAINAPTL | ASNPRGRNSPLY | 0.09142834 |
| LQLLNTGV | ASNPRGRNSPLY | 0.085709736 |

| | | |
|---|---|---|
| LQTSIEF | ASNPRGRNSPLY | 0.08822708 |
| MSFAVSEL | ASNPRGRNSPLY | 0.10499397 |
| NTGALGL | ASNPRGRNSPLY | 0.087532766 |
| PPGAGGAGGAGTT | ASNPRGRNSPLY | 0.09091742 |
| QLVGLGI | ASNPRGRNSPLY | 0.12763931 |
| RFLEQVHQL | ASNPRGRNSPLY | 0.08473283 |
| RGVGYVLR | ASNPRGRNSPLY | 0.092275396 |
| SATGVFAGVI | ASNPRGRNSPLY | 0.09040942 |
| SLPAITLP | ASNPRGRNSPLY | 0.09116832 |
| STALAFTV | ASNPRGRNSPLY | 0.09130904 |
| SVLTVGLGK | ASNPRGRNSPLY | 0.092345774 |
| TALAGAGSGI | ASNPRGRNSPLY | 0.097471 |
| TGLPFTGI | ASNPRGRNSPLY | 0.10042694 |
| VAEHVTV | ASNPRGRNSPLY | 0.094561644 |
| VDIVLNSL | ASNPRGRNSPLY | 0.089367084 |
| VLTGGGAVL | ASNPRGRNSPLY | 0.08863036 |
| VNVGLAYGI | ASNPRGRNSPLY | 0.08363139 |
| VSSAAVGAV | ASNPRGRNSPLY | 0.10061093 |
| VTSVYEAGV | ASNPRGRNSPLY | 0.0847059 |
| WVALDRGA | ASNPRGRNSPLY | 0.10072478 |
| YLVGSVVI | ASNPRGRNSPLY | 0.1022849 |
| GPPGTGKTLL | ASNPRGRNSPLY | 0.08919283 |
| GVGMTKF | ASNPRGRNSPLY | 0.09106237 |
| IGIGGSDL | ASNPRGRNSPLY | 0.08533839 |
| IGTLLAEL | ASNPRGRNSPLY | 0.085240334 |
| IYTSGTT | ASNPRGRNSPLY | 0.095081225 |
| LSFPAGV | ASNPRGRNSPLY | 0.09154423 |
| LYTSGTT | ASNPRGRNSPLY | 0.092665456 |
| VGAFPFGF | ASNPRGRNSPLY | 0.09039205 |
| VGAGTVEF | ASNPRGRNSPLY | 0.09692248 |
| VGKSTLLL | ASNPRGRNSPLY | 0.09163203 |
| AALAHAGGGIV | ASNPRGRNSPLY | 0.09011072 |
| AALVAVSLL | ASNPRGRNSPLY | 0.10848734 |
| AAVSRAVAAL | ASNPRGRNSPLY | 0.092798345 |
| AGGSSGIGGF | ASNPRGRNSPLY | 0.08990583 |
| AGLTVANANG | ASNPRGRNSPLY | 0.096732326 |
| AGLVGAVL | ASNPRGRNSPLY | 0.08678451 |
| ALSQAFTV | ASNPRGRNSPLY | 0.09961257 |
| ALTFSVSL | ASNPRGRNSPLY | 0.08970384 |
| ALTGAALVAL | ASNPRGRNSPLY | 0.09693788 |
| ASFSIGAGVL | ASNPRGRNSPLY | 0.09215326 |
| DGSVYKI | ASNPRGRNSPLY | 0.08571975 |
| EQLAEIV | ASNPRGRNSPLY | 0.08806043 |
| FAVSELTVM | ASNPRGRNSPLY | 0.09109698 |
| GAAPLAAGL | ASNPRGRNSPLY | 0.11733785 |
| GADGGDGGSGGAGG | ASNPRGRNSPLY | 0.10315332 |
| GFGLDTGGAGG | ASNPRGRNSPLY | 0.089932516 |
| GLASGGLL | ASNPRGRNSPLY | 0.09661741 |
| GRIAGAAVLL | ASNPRGRNSPLY | 0.09393743 |
| GSAGSLNTSF | ASNPRGRNSPLY | 0.09373762 |
| GTGQIGI | ASNPRGRNSPLY | 0.09035803 |

| | | |
|---|---|---|
| GTLIALIL | ASNPRGRNSPLY | 0.0895637 |
| IAATMGFVVG | ASNPRGRNSPLY | 0.088355154 |
| IFAQALL | ASNPRGRNSPLY | 0.09503788 |
| IGAAVALGF | ASNPRGRNSPLY | 0.087217815 |
| IGLTPLAL | ASNPRGRNSPLY | 0.097884096 |
| ILGSNTSTL | ASNPRGRNSPLY | 0.091918215 |
| IVFSLTVL | ASNPRGRNSPLY | 0.09032556 |
| LASKATGF | ASNPRGRNSPLY | 0.09337097 |
| LASVALIL | ASNPRGRNSPLY | 0.08869356 |
| LFAVGLTL | ASNPRGRNSPLY | 0.09246629 |
| LFNGGIGTYI | ASNPRGRNSPLY | 0.09618825 |
| LTLGAGII | ASNPRGRNSPLY | 0.089664035 |
| LVAAALGDI | ASNPRGRNSPLY | 0.09606266 |
| NTAFVQLGI | ASNPRGRNSPLY | 0.11855139 |
| QLFSGAEL | ASNPRGRNSPLY | 0.09799185 |
| SAATFITL | ASNPRGRNSPLY | 0.08687905 |
| SNYGLANLGV | ASNPRGRNSPLY | 0.09238627 |
| SYGGAYAVM | ASNPRGRNSPLY | 0.10120013 |
| TTGVGGLTLG | ASNPRGRNSPLY | 0.090775594 |
| TTQTIYI | ASNPRGRNSPLY | 0.08479274 |
| VAAGAGLGAVI | ASNPRGRNSPLY | 0.093946025 |
| VAAVVTLL | ASNPRGRNSPLY | 0.091037765 |
| VGFLGGLI | ASNPRGRNSPLY | 0.090674594 |
| VGVIPILL | ASNPRGRNSPLY | 0.08445177 |
| VIFASSLI | ASNPRGRNSPLY | 0.109957516 |
| VKGGDGGAAAG | ASNPRGRNSPLY | 0.11610911 |
| VTKTFTNI | ASNPRGRNSPLY | 0.09090639 |
| VVAAAILL | ASNPRGRNSPLY | 0.08931717 |
| YAALDVEL | ASNPRGRNSPLY | 0.085572466 |
| LASVALLL | ASNPRGRNSPLY | 0.09941676 |
| LDAVHRV | ASNPRGRNSPLY | 0.10718927 |
| QGLSSSIF | ASNPRGRNSPLY | 0.0878843 |
| RGTFANIRL | ASNPRGRNSPLY | 0.102979206 |
| RQLWWGHRI | ASNPRGRNSPLY | 0.08982848 |
| YGPPGTGKTLL | ASNPRGRNSPLY | 0.09128944 |
| AALVGAPANVA | ASNPRGRNSPLY | 0.112455904 |
| AATFRRTE | ASNPRGRNSPLY | 0.0968292 |
| AGALGGGI | ASNPRGRNSPLY | 0.09344568 |
| AGGAGGAGGIGAG | ASNPRGRNSPLY | 0.09045508 |
| AGSGALGL | ASNPRGRNSPLY | 0.089169525 |
| AQIADQI | ASNPRGRNSPLY | 0.09287797 |
| ASAVAVTYL | ASNPRGRNSPLY | 0.098950915 |
| ASTVLGGTGG | ASNPRGRNSPLY | 0.08544295 |
| AVGGEPAK | ASNPRGRNSPLY | 0.09318156 |
| CEQGTFGGTC | ASNPRGRNSPLY | 0.09120711 |
| DEGKAPAG | ASNPRGRNSPLY | 0.107679814 |
| DTVVFPAL | ASNPRGRNSPLY | 0.09359745 |
| FEGITMAG | ASNPRGRNSPLY | 0.09497888 |
| FPLVVQD | ASNPRGRNSPLY | 0.08801444 |
| GAFTTPQ | ASNPRGRNSPLY | 0.119799815 |
| GFGGFGTAGGVGGA | ASNPRGRNSPLY | 0.0961785 |

| | | |
|---|---|---|
| GFLAQFL | ASNPRGRNSPLY | 0.09421084 |
| GGAGGAGGNAGVA | ASNPRGRNSPLY | 0.09681992 |
| GGVGTAGAGGAGGAGG | ASNPRGRNSPLY | 0.086886376 |
| GNGGGGDAGNAGSGGN | ASNPRGRNSPLY | 0.15337783 |
| GTGGAGGAGNDAGS | ASNPRGRNSPLY | 0.12667455 |
| GTTNTGWFNS | ASNPRGRNSPLY | 0.14532268 |
| IFMGPLFTL | ASNPRGRNSPLY | 0.105675444 |
| ILAHAIEAI | ASNPRGRNSPLY | 0.081137374 |
| IRSAIREAL | ASNPRGRNSPLY | 0.105666414 |
| ITFAGGG | ASNPRGRNSPLY | 0.08730608 |
| ITGMSGHQEV | ASNPRGRNSPLY | 0.0846728 |
| IVAGLACTL | ASNPRGRNSPLY | 0.11251675 |
| IVGGGLAGA | ASNPRGRNSPLY | 0.09925671 |
| IVTGVLNI | ASNPRGRNSPLY | 0.28939778 |
| LAQQGCI | ASNPRGRNSPLY | 0.20125078 |
| LCLHVEN | ASNPRGRNSPLY | 0.08543596 |
| LDAISDGS | ASNPRGRNSPLY | 0.11075752 |
| LGGFAAML | ASNPRGRNSPLY | 0.09787749 |
| LGGKNPMI | ASNPRGRNSPLY | 0.10427864 |
| LGPKGVAAIL | ASNPRGRNSPLY | 0.07564469 |
| LMVGVSI | ASNPRGRNSPLY | 0.1294164 |
| LVGAAFVL | ASNPRGRNSPLY | 0.084450856 |
| LVLAVAPL | ASNPRGRNSPLY | 0.07459631 |
| MAAIAVIDAA | ASNPRGRNSPLY | 0.12348509 |
| NNGINAELQ | ASNPRGRNSPLY | 0.102047406 |
| RGEPLVLVHG | ASNPRGRNSPLY | 0.09144975 |
| RGGAGGAAGLI | ASNPRGRNSPLY | 0.22407728 |
| RSYRMHPAV | ASNPRGRNSPLY | 0.07515008 |
| SAAHLLAGQLA | ASNPRGRNSPLY | 0.0943063 |
| SAAVGSY | ASNPRGRNSPLY | 0.19368583 |
| SPHAVLHRI | ASNPRGRNSPLY | 0.28387135 |
| SQIAKVL | ASNPRGRNSPLY | 0.093075484 |
| TALGHLVP | ASNPRGRNSPLY | 0.121797204 |
| TAPTTSGAA | ASNPRGRNSPLY | 0.44941345 |
| TDGDDGNGGSPGT | ASNPRGRNSPLY | 0.12919627 |
| TDGLGEN | ASNPRGRNSPLY | 0.15032603 |
| TGGPPLPGQA | ASNPRGRNSPLY | 0.18832833 |
| TIPAITI | ASNPRGRNSPLY | 0.08306492 |
| TIPGFTI | ASNPRGRNSPLY | 0.15308891 |
| TSIASGALA | ASNPRGRNSPLY | 0.23388432 |
| TVTLPGAAI | ASNPRGRNSPLY | 0.11596376 |
| VASGLSAALP | ASNPRGRNSPLY | 0.124931924 |
| VGAGVIGI | ASNPRGRNSPLY | 0.08989186 |
| VGPKRFGV | ASNPRGRNSPLY | 0.09798904 |
| VNVAFPD | ASNPRGRNSPLY | 0.07747337 |
| VSMTNSGVS | ASNPRGRNSPLY | 0.15228063 |
| VTDVLGDLSV | ASNPRGRNSPLY | 0.14257203 |
| VVVVGAGFAGL | ASNPRGRNSPLY | 0.12918334 |
| VYTVGTVV | ASNPRGRNSPLY | 0.08164777 |
| WTTFSGP | ASNPRGRNSPLY | 0.08960357 |
| YDELVEAF | ASNPRGRNSPLY | 0.08230769 |

| | | |
|---|---|---|
| YGTGGAGGNGG | ASNPRGRNSPLY | 0.09869961 |
| YTNVMAV | ASNPRGRNSPLY | 0.08121024 |
| FLEQVHQL | ASNPRGRNSPLY | 0.07767686 |
| GLSLLAVAV | ASNPRGRNSPLY | 0.21974437 |
| LEQVHQL | ASNPRGRNSPLY | 0.22717974 |
| LNAAGAI | ASNPRGRNSPLY | 0.10203704 |
| PAVHASL | ASNPRGRNSPLY | 0.24789424 |
| TELEDDA | ASNPRGRNSPLY | 0.16195646 |
| TPPGPPN | ASNPRGRNSPLY | 0.18479137 |
| TPPPPPPP | ASNPRGRNSPLY | 0.120038725 |
| TQSPALI | ASSADTSSGNTLY | 0.082464285 |
| EFAAAFPQ | ASSADTSSGNTLY | 0.3064837 |
| LLSGNLDVL | ASSADTSSGNTLY | 0.14678468 |
| GPANLGNYNI | ASSADTSSGNTLY | 0.08658847 |
| TLGPTLP | ASSADTSSGNTLY | 0.20351893 |
| VYAYSSCCK | ASSADTSSGNTLY | 0.09689066 |
| APAPAPAPNI | ASSADTSSGNTLY | 0.095439345 |
| PGGPVPIP | ASSADTSSGNTLY | 0.16555974 |
| FLTVALEP | ASSADTSSGNTLY | 0.14730906 |
| SQYFGANL | ASSADTSSGNTLY | 0.08608175 |
| GGVKPAA | ASSADTSSGNTLY | 0.08422659 |
| GGTGGAGGAGGW | ASSADTSSGNTLY | 0.087086596 |
| GVGGVAGLGGAGG | ASSADTSSGNTLY | 0.15038691 |
| VLLVLATTI | ASSADTSSGNTLY | 0.22430617 |
| IGIFAAML | ASSADTSSGNTLY | 0.12713408 |
| AVGDLID | ASSADTSSGNTLY | 0.106034204 |
| NISPLPL | ASSADTSSGNTLY | 0.21931626 |
| VTAVFVAI | ASSADTSSGNTLY | 0.13011912 |
| SGDDNDAAL | ASSADTSSGNTLY | 0.117762536 |
| AVAALAHS | ASSADTSSGNTLY | 0.14735046 |
| GGVTVMATP | ASSADTSSGNTLY | 0.08355081 |
| AGLAANVRD | ASSADTSSGNTLY | 0.15162103 |
| TNRPDLIDPALLRPG | ASSADTSSGNTLY | 0.20270771 |
| LANLGVTAL | ASSADTSSGNTLY | 0.1358927 |
| TGERIEL | ASSADTSSGNTLY | 0.14288922 |
| NGEELAA | ASSADTSSGNTLY | 0.09983488 |
| VLSLVWG | ASSADTSSGNTLY | 0.12737635 |
| IGYALWL | ASSADTSSGNTLY | 0.09453006 |
| LSLLLAPL | ASSADTSSGNTLY | 0.09163864 |
| TLDEALK | ASSADTSSGNTLY | 0.1308856 |
| TDGAGTAEGG | ASSADTSSGNTLY | 0.13694218 |
| NAPGGAGGNGG | ASSADTSSGNTLY | 0.12507764 |
| VSVFMGMGGNF | ASSADTSSGNTLY | 0.13142106 |
| LASYLAAL | ASSADTSSGNTLY | 0.09551484 |
| LVGGTPAL | ASSADTSSGNTLY | 0.1646712 |
| VRNADVL | ASSADTSSGNTLY | 0.28884047 |
| TCSSPFLVSAM | ASSADTSSGNTLY | 0.08565825 |
| VHITDAE | ASSADTSSGNTLY | 0.08963235 |
| EYAAPSF | ASSADTSSGNTLY | 0.12849158 |
| VVDAFHGGL | ASSADTSSGNTLY | 0.15196693 |
| ASVVDPQ | ASSADTSSGNTLY | 0.1368207 |

| | | |
|---|---|---|
| SYNEFGGLS | ASSADTSSGNTLY | 0.10856299 |
| DPDGSVY | ASSADTSSGNTLY | 0.1447564 |
| MVSTIHATE | ASSADTSSGNTLY | 0.079272635 |
| LAAGVADPV | ASSADTSSGNTLY | 0.2584409 |
| TKEQIAA | ASSADTSSGNTLY | 0.08654693 |
| SLASPAAL | ASSADTSSGNTLY | 0.100623205 |
| IGLVFYGV | ASSADTSSGNTLY | 0.094926044 |
| FLLDASNL | ASSADTSSGNTLY | 0.076106295 |
| YQVGNFP | ASSADTSSGNTLY | 0.15735267 |
| ESLLSQST | ASSADTSSGNTLY | 0.08957031 |
| LFTVGFAAVG | ASSADTSSGNTLY | 0.18498188 |
| ATPPTPT | ASSADTSSGNTLY | 0.09328729 |
| TDLATLG | ASSADTSSGNTLY | 0.13275665 |
| SAGGTGR | ASSADTSSGNTLY | 0.140912 |
| GFSNSGS | ASSADTSSGNTLY | 0.10976822 |
| GGNGGSGGTSV | ASSADTSSGNTLY | 0.086533934 |
| GTSVATGGAGN | ASSADTSSGNTLY | 0.1579673 |
| NGGNGGSGGTS | ASSADTSSGNTLY | 0.16105732 |
| AVVKFPH | ASSADTSSGNTLY | 0.13026722 |
| TPPPPPP | ASSADTSSGNTLY | 0.08589617 |
| AAPAAAAAPA | ASSADTSSGNTLY | 0.084253386 |
| IALNEDL | ASSADTSSGNTLY | 0.12417803 |
| IVNVSSM | ASSADTSSGNTLY | 0.07486318 |
| TGIPPAPR | ASSADTSSGNTLY | 0.07541402 |
| FDLGGGTF | ASSADTSSGNTLY | 0.07860023 |
| FDLGGGTFDVS | ASSADTSSGNTLY | 0.07858052 |
| NEPTAAA | ASSADTSSGNTLY | 0.24596119 |
| NEPTAAALA | ASSADTSSGNTLY | 0.28645915 |
| GSIINISSI | ASSADTSSGNTLY | 0.122447215 |
| VAPGNAGTA | ASSADTSSGNTLY | 0.22135375 |
| TGGGASLE | ASSADTSSGNTLY | 0.121070415 |
| VAVYNHYKRI | ASSADTSSGNTLY | 0.07368716 |
| TDPAPATT | ASSADTSSGNTLY | 0.09145538 |
| TNRPDILDPALLRPG | ASSADTSSGNTLY | 0.08593179 |
| LTTTTTS | ASSADTSSGNTLY | 0.14357677 |
| PTGSGKTL | ASSADTSSGNTLY | 0.07881445 |
| TGSGKTL | ASSADTSSGNTLY | 0.09838886 |
| TIGHVDHGK | ASSADTSSGNTLY | 0.14288308 |
| TIGHVDHGKTT | ASSADTSSGNTLY | 0.1021827 |
| GHVDHGKTT | ASSADTSSGNTLY | 0.14887448 |
| AQFEHTLL | ASSADTSSGNTLY | 0.08906431 |
| IVNQDEL | ASSADTSSGNTLY | 0.0765841 |
| TNGSQFF | ASSADTSSGNTLY | 0.17776342 |
| TGKGIVE | ASSADTSSGNTLY | 0.07711938 |
| LISNASDA | ASSADTSSGNTLY | 0.0976502 |
| AELRTLGELTKIAKAHGAQ | ASSADTSSGNTLY | 0.07964548 |
| APNAVAAAEPSW | ASSADTSSGNTLY | 0.073788695 |
| APQVAGAATAVL | ASSADTSSGNTLY | 0.08129558 |
| ARRDPAAAAAILPTDA | ASSADTSSGNTLY | 0.14562126 |
| DAAARDVLAEAGLAEYF | ASSADTSSGNTLY | 0.096734196 |
| DDQIVLSAKANLAKVEQ | ASSADTSSGNTLY | 0.12954329 |

| | | |
|---|---|---|
| DELLPDTGEDSDEDGH | ASSADTSSGNTLY | 0.209508 |
| ELPEAPANRATVPQD | ASSADTSSGNTLY | 0.16667281 |
| FGQSKIERVTGIG | ASSADTSSGNTLY | 0.1356171 |
| GAGGTGSTAGGAGGAGGAGGLY | ASSADTSSGNTLY | 0.098167926 |
| GHGGTGAAGGNGGTGGAGGSGI | ASSADTSSGNTLY | 0.090187676 |
| GNGGNGGAGGEGGPGGAGGAGA | ASSADTSSGNTLY | 0.091269836 |
| GSTAGNGGNGGNATNGTIAGQP | ASSADTSSGNTLY | 0.09427266 |
| GTGGDGGNGGNGGTGG | ASSADTSSGNTLY | 0.3027004 |
| GVRRPAGIPADAPV | ASSADTSSGNTLY | 0.0952559 |
| GYRQVTDAHLLTLARR | ASSADTSSGNTLY | 0.081930906 |
| IAKMGQDGHDRG | ASSADTSSGNTLY | 0.09751226 |
| KAEPSIEIQYT | ASSADTSSGNTLY | 0.24526452 |
| KPVRIRMAADEDTVAVT | ASSADTSSGNTLY | 0.39987212 |
| MPPRIAGMRLLVIKPEP | ASSADTSSGNTLY | 0.08902266 |
| MKFYAVALFRSRGVPPTR | ASSADTSSGNTLY | 0.10865359 |
| NAGGFGILGAVAHSP | ASSADTSSGNTLY | 0.11101124 |
| NEGREITLAMAEM | ASSADTSSGNTLY | 0.12481378 |
| NELVEWSHPLGHRWRWE | ASSADTSSGNTLY | 0.08943923 |
| NLGAGGTGGNGGNPG | ASSADTSSGNTLY | 0.09033463 |
| NTEWPKWETVMAADPVD | ASSADTSSGNTLY | 0.15719996 |
| PASAAPAETQP | ASSADTSSGNTLY | 0.5609516 |
| QQSSDGKNGVRAAEITGE | ASSADTSSGNTLY | 0.092717424 |
| RAPDAVTTSWWKE | ASSADTSSGNTLY | 0.07984778 |
| SGFGNVGAHTSGWFNQ | ASSADTSSGNTLY | 0.14866325 |
| SGVAGAVGQAASVGGLKV | ASSADTSSGNTLY | 0.08976225 |
| SNGGAGTGSAGGNGGT | ASSADTSSGNTLY | 0.11319257 |
| TGGAGGEGGAAG | ASSADTSSGNTLY | 0.113096654 |
| TGGDGGAAGAGGAPGHG | ASSADTSSGNTLY | 0.08058874 |
| TMPPLLAAIANAGRHGV | ASSADTSSGNTLY | 0.09626076 |
| TSSTAGNSGGA | ASSADTSSGNTLY | 0.4123945 |
| VPATVPVGILC | ASSADTSSGNTLY | 0.104009755 |
| VQRPDSGGVVLTGRIS | ASSADTSSGNTLY | 0.14900859 |
| WTPNPVPATKGVPAGLVQAT | ASSADTSSGNTLY | 0.26567996 |
| YALKRALAKFSDRFDVVII | ASSADTSSGNTLY | 0.15433209 |
| YLIGSVIV | ASSADTSSGNTLY | 0.3324036 |
| YGQLVAVID | ASSADTSSGNTLY | 0.16916692 |
| VTVGGMTLPA | ASSADTSSGNTLY | 0.10374865 |
| VLALPAV | ASSADTSSGNTLY | 0.110953204 |
| VIPPTLNYET | ASSADTSSGNTLY | 0.088553004 |
| VGMGGMHPSSG | ASSADTSSGNTLY | 0.1428855 |
| VGLAGGAATA | ASSADTSSGNTLY | 0.17253663 |
| VGGAGGAGGSGP | ASSADTSSGNTLY | 0.09982774 |
| VFAAFMV | ASSADTSSGNTLY | 0.25693604 |
| TLYDKLR | ASSADTSSGNTLY | 0.12732859 |
| TLAVFTVL | ASSADTSSGNTLY | 0.14068022 |
| TGSVQPARV | ASSADTSSGNTLY | 0.08369061 |
| TGGGAGSGGT | ASSADTSSGNTLY | 0.09258891 |
| SHLETSL | ASSADTSSGNTLY | 0.20317864 |
| SFTTGMT | ASSADTSSGNTLY | 0.08192557 |
| QVEVAKRMGVRQPTVS | ASSADTSSGNTLY | 0.080240265 |
| QGSGGPALAM | ASSADTSSGNTLY | 0.11623471 |

| | | |
|---|---|---|
| MLAPSNP | ASSADTSSGNTLY | 0.41283435 |
| LVNNAAI | ASSADTSSGNTLY | 0.07452871 |
| LTGEINTLA | ASSADTSSGNTLY | 0.09461068 |
| LSIFGPV | ASSADTSSGNTLY | 0.08433834 |
| LLALLANL | ASSADTSSGNTLY | 0.0991925 |
| LISNASDALDKLR | ASSADTSSGNTLY | 0.11261902 |
| LINEAVP | ASSADTSSGNTLY | 0.080086805 |
| LGYAFSGL | ASSADTSSGNTLY | 0.13431261 |
| LDFVAVAL | ASSADTSSGNTLY | 0.07980415 |
| LAQPGAE | ASSADTSSGNTLY | 0.07742051 |
| ISNLGAVTGF | ASSADTSSGNTLY | 0.12986234 |
| HQEDPGRI | ASSADTSSGNTLY | 0.24204254 |
| GTPGEPTP | ASSADTSSGNTLY | 0.08413118 |
| GFFNSGS | ASSADTSSGNTLY | 0.15563864 |
| GAGGGGLTAGAN | ASSADTSSGNTLY | 0.08868052 |
| FVDALVNM | ASSADTSSGNTLY | 0.5304758 |
| FLTVAPEL | ASSADTSSGNTLY | 0.11978582 |
| FETLTAL | ASSADTSSGNTLY | 0.079950325 |
| DVINPIEL | ASSADTSSGNTLY | 0.1490734 |
| APVSGAPV | ASSADTSSGNTLY | 0.11448225 |
| ALVGGLAW | ASSADTSSGNTLY | 0.13400842 |
| ALAAAAATAI | ASSADTSSGNTLY | 0.19258142 |
| AFGLGLLL | ASSADTSSGNTLY | 0.08563284 |
| AFFAAASL | ASSADTSSGNTLY | 0.117621444 |
| AAAALPAFSPP | ASSADTSSGNTLY | 0.22098638 |
| VVLAPAL | ASSADTSSGNTLY | 0.13705407 |
| TIGHVDHGKTTL | ASSADTSSGNTLY | 0.08132232 |
| LISNASDALDK | ASSADTSSGNTLY | 0.12365085 |
| LISNASDAL | ASSADTSSGNTLY | 0.11436783 |
| ISNASDALDK | ASSADTSSGNTLY | 0.098629944 |
| NQAETLVYQTE | ASSADTSSGNTLY | 0.18595257 |
| TGGTGGTAGAG | ASSADTSSGNTLY | 0.10955434 |
| TGSTAGGAGGA | ASSADTSSGNTLY | 0.21907783 |
| DPLAVPVAAGPV | ASSADTSSGNTLY | 0.131081 |
| GGTGGTPGNGGN | ASSADTSSGNTLY | 0.07441386 |
| PAGLAGVDPATL | ASSADTSSGNTLY | 0.08322738 |
| GNNLVVEAPHAGASV | ASSADTSSGNTLY | 0.09975015 |
| KVTRIMSASTLEQPA | ASSADTSSGNTLY | 0.15262595 |
| LGFALPIALPRRTVV | ASSADTSSGNTLY | 0.15120688 |
| TRLGNLTARKLVVLV | ASSADTSSGNTLY | 0.07979833 |
| WSQQRRLIYFPSAGP | ASSADTSSGNTLY | 0.08402145 |
| GPNGNPDPMAAAVDIR | ASSADTSSGNTLY | 0.117717445 |
| TQRCVSVADLRAAASR | ASSADTSSGNTLY | 0.23405907 |
| QFVLDVAEHPVEVLTETV | ASSADTSSGNTLY | 0.10181412 |
| GGAGGTGGAGGNGGTQAGDG | ASSADTSSGNTLY | 0.11240274 |
| PLLATLRAIAVDVDAKPAQV | ASSADTSSGNTLY | 0.45620406 |
| APKFRTGGYMIDFWGVGYQVAK | ASSADTSSGNTLY | 0.08771893 |
| GAGGAGGAGGAAGGPGGTGGTG | ASSADTSSGNTLY | 0.10931651 |
| IGALGLLATDVAPAGSVAFGVA | ASSADTSSGNTLY | 0.15845065 |
| AGGAGGIGGNGGAGGAGGVLMG | ASSADTSSGNTLY | 0.2226935 |
| ADELIGLL | ASSADTSSGNTLY | 0.107749715 |

| | | |
|---|---|---|
| ADRALKA | ASSADTSSGNTLY | 0.20987645 |
| AGAVGQVGGAGGA | ASSADTSSGNTLY | 0.08075625 |
| AGGGGGLGGAP | ASSADTSSGNTLY | 0.21530202 |
| AGGLLYGNGGAG | ASSADTSSGNTLY | 0.076862454 |
| AGLESIGL | ASSADTSSGNTLY | 0.13693856 |
| AGLKTGII | ASSADTSSGNTLY | 0.07710555 |
| ASLAALLL | ASSADTSSGNTLY | 0.17553967 |
| ASVINISSI | ASSADTSSGNTLY | 0.2635737 |
| ATPPGPA | ASSADTSSGNTLY | 0.23357292 |
| ATVASALV | ASSADTSSGNTLY | 0.07532859 |
| ATVASLIL | ASSADTSSGNTLY | 0.14023462 |
| AVNGPVLIRH | ASSADTSSGNTLY | 0.27890104 |
| DGAAGAVGI | ASSADTSSGNTLY | 0.5570862 |
| DPLLHGI | ASSADTSSGNTLY | 0.12663504 |
| ELGAALGAL | ASSADTSSGNTLY | 0.18190615 |
| ELGRTQAV | ASSADTSSGNTLY | 0.102119565 |
| FVAGGYLS | ASSADTSSGNTLY | 0.09742069 |
| GASGGAGGAGGW | ASSADTSSGNTLY | 0.08864363 |
| GGAGAGGDGGAGGI | ASSADTSSGNTLY | 0.088224955 |
| GTGHNGSQPGMGGNG | ASSADTSSGNTLY | 0.11957401 |
| GYAAASASAS | ASSADTSSGNTLY | 0.096582256 |
| HQVDVAAVQQG | ASSADTSSGNTLY | 0.19239563 |
| IGAAIAHT | ASSADTSSGNTLY | 0.24537922 |
| IGMIEVAQL | ASSADTSSGNTLY | 0.19463629 |
| IRLAPEHTSAMD | ASSADTSSGNTLY | 0.38736174 |
| IYNVSVL | ASSADTSSGNTLY | 0.31044406 |
| KQGFLGM | ASSADTSSGNTLY | 0.12649298 |
| LESVITQL | ASSADTSSGNTLY | 0.09137218 |
| LGLGALVVV | ASSADTSSGNTLY | 0.09550782 |
| LIGNGGNGGSGGI | ASSADTSSGNTLY | 0.083478525 |
| LNAINAPTL | ASSADTSSGNTLY | 0.09529769 |
| LQLLNTGV | ASSADTSSGNTLY | 0.0896728 |
| LQTSIEF | ASSADTSSGNTLY | 0.15695503 |
| MSFAVSEL | ASSADTSSGNTLY | 0.074007906 |
| NTGALGL | ASSADTSSGNTLY | 0.17650631 |
| PPGAGGAGGAGTT | ASSADTSSGNTLY | 0.28791702 |
| QLVGLGI | ASSADTSSGNTLY | 0.07822892 |
| RFLEQVHQL | ASSADTSSGNTLY | 0.18060982 |
| RGVGYVLR | ASSADTSSGNTLY | 0.10908676 |
| SATGVFAGVI | ASSADTSSGNTLY | 0.08643161 |
| SLPAITLP | ASSADTSSGNTLY | 0.33351034 |
| STALAFTV | ASSADTSSGNTLY | 0.09074868 |
| SVLTVGLGK | ASSADTSSGNTLY | 0.15007502 |
| TALAGAGSGI | ASSADTSSGNTLY | 0.1857143 |
| TGLPFTGI | ASSADTSSGNTLY | 0.13831417 |
| VAEHVTV | ASSADTSSGNTLY | 0.10922047 |
| VDIVLNSL | ASSADTSSGNTLY | 0.117344946 |
| VLTGGGAVL | ASSADTSSGNTLY | 0.08383912 |
| VNVGLAYGI | ASSADTSSGNTLY | 0.12153924 |
| VSSAAVGAV | ASSADTSSGNTLY | 0.07363717 |
| VTSVYEAGV | ASSADTSSGNTLY | 0.1625184 |

| | | |
|---|---|---|
| WVALDRGA | ASSADTSSGNTLY | 0.090003006 |
| YLVGSVVI | ASSADTSSGNTLY | 0.19511455 |
| GPPGTGKTLL | ASSADTSSGNTLY | 0.110771626 |
| GVGMTKF | ASSADTSSGNTLY | 0.14353642 |
| IGIGGSDL | ASSADTSSGNTLY | 0.09925006 |
| IGTLLAEL | ASSADTSSGNTLY | 0.0829698 |
| IYTSGTT | ASSADTSSGNTLY | 0.08058024 |
| LSFPAGV | ASSADTSSGNTLY | 0.0861664 |
| LYTSGTT | ASSADTSSGNTLY | 0.11464562 |
| VGAFPFGF | ASSADTSSGNTLY | 0.10251904 |
| VGAGTVEF | ASSADTSSGNTLY | 0.08197188 |
| VGKSTLLL | ASSADTSSGNTLY | 0.16492668 |
| AALAHAGGGIV | ASSADTSSGNTLY | 0.115278065 |
| AALVAVSLL | ASSADTSSGNTLY | 0.080399044 |
| AAVSRAVAAL | ASSADTSSGNTLY | 0.15295342 |
| AGGSSGIGGF | ASSADTSSGNTLY | 0.12905298 |
| AGLTVANANG | ASSADTSSGNTLY | 0.120547056 |
| AGLVGAVL | ASSADTSSGNTLY | 0.07360194 |
| ALSQAFTV | ASSADTSSGNTLY | 0.0878786 |
| ALTFSVSL | ASSADTSSGNTLY | 0.099653915 |
| ALTGAALVAL | ASSADTSSGNTLY | 0.07844563 |
| ASFSIGAGVL | ASSADTSSGNTLY | 0.08456559 |
| DGSVYKI | ASSADTSSGNTLY | 0.31192276 |
| EQLAEIV | ASSADTSSGNTLY | 0.07472036 |
| FAVSELTVM | ASSADTSSGNTLY | 0.09472079 |
| GAAPLAAGL | ASSADTSSGNTLY | 0.17094778 |
| GADGGDGGSGGAGG | ASSADTSSGNTLY | 0.16974862 |
| GFGLDTGGAGG | ASSADTSSGNTLY | 0.4202143 |
| GLASGGLL | ASSADTSSGNTLY | 0.09691579 |
| GRIAGAAVLL | ASSADTSSGNTLY | 0.102311775 |
| GSAGSLNTSF | ASSADTSSGNTLY | 0.12783554 |
| GTGQIGI | ASSADTSSGNTLY | 0.1126665 |
| GTLIALIL | ASSADTSSGNTLY | 0.07780751 |
| IAATMGFVVG | ASSADTSSGNTLY | 0.07641513 |
| IFAQALL | ASSADTSSGNTLY | 0.11544731 |
| IGAAVALGF | ASSADTSSGNTLY | 0.08210943 |
| IGLTPLAL | ASSADTSSGNTLY | 0.15016748 |
| ILGSNTSTL | ASSADTSSGNTLY | 0.1465849 |
| IVFSLTVL | ASSADTSSGNTLY | 0.08429851 |
| LASKATGF | ASSADTSSGNTLY | 0.080650784 |
| LASVALIL | ASSADTSSGNTLY | 0.10046839 |
| LFAVGLTL | ASSADTSSGNTLY | 0.08993006 |
| LFNGGIGTYI | ASSADTSSGNTLY | 0.17442554 |
| LTLGAGII | ASSADTSSGNTLY | 0.10010079 |
| LVAAALGDI | ASSADTSSGNTLY | 0.59708 |
| NTAFVQLGI | ASSADTSSGNTLY | 0.18038328 |
| QLFSGAEL | ASSADTSSGNTLY | 0.093602166 |
| SAATFITL | ASSADTSSGNTLY | 0.21188891 |
| SNYGLANLGV | ASSADTSSGNTLY | 0.10235362 |
| SYGGAYAVM | ASSADTSSGNTLY | 0.079763465 |
| TTGVGGLTLG | ASSADTSSGNTLY | 0.100388415 |

| | | |
|---|---|---|
| TTQTIYI | ASSADTSSGNTLY | 0.08654441 |
| VAAGAGLGAVI | ASSADTSSGNTLY | 0.09406166 |
| VAAVVTLL | ASSADTSSGNTLY | 0.09858301 |
| VGFLGGLI | ASSADTSSGNTLY | 0.08796442 |
| VGVIPILL | ASSADTSSGNTLY | 0.1023512 |
| VIFASSLI | ASSADTSSGNTLY | 0.09707485 |
| VKGGDGGAAAG | ASSADTSSGNTLY | 0.080785714 |
| VTKTFTNI | ASSADTSSGNTLY | 0.10825614 |
| VVAAAILL | ASSADTSSGNTLY | 0.3323364 |
| YAALDVEL | ASSADTSSGNTLY | 0.578084 |
| LASVALLL | ASSADTSSGNTLY | 0.10252546 |
| LDAVHRV | ASSADTSSGNTLY | 0.08981418 |
| QGLSSSIF | ASSADTSSGNTLY | 0.2001445 |
| RGTFANIRL | ASSADTSSGNTLY | 0.105343275 |
| RQLWWGHRI | ASSADTSSGNTLY | 0.11012813 |
| YGPPGTGKTLL | ASSADTSSGNTLY | 0.1185958 |
| AALVGAPANVA | ASSADTSSGNTLY | 0.08518419 |
| AATFRRTE | ASSADTSSGNTLY | 0.092234775 |
| AGALGGGI | ASSADTSSGNTLY | 0.116318196 |
| AGGAGGAGGIGAG | ASSADTSSGNTLY | 0.080996156 |
| AGSGALGL | ASSADTSSGNTLY | 0.13760649 |
| AQIADQI | ASSADTSSGNTLY | 0.0739636 |
| ASAVAVTYL | ASSADTSSGNTLY | 0.086185776 |
| ASTVLGGTGG | ASSADTSSGNTLY | 0.099194884 |
| AVGGEPAK | ASSADTSSGNTLY | 0.0933118 |
| CEQGTFGGTC | ASSADTSSGNTLY | 0.0787262 |
| DEGKAPAG | ASSADTSSGNTLY | 0.099009134 |
| DTVVFPAL | ASSADTSSGNTLY | 0.283625 |
| FEGITMAG | ASSADTSSGNTLY | 0.28206128 |
| FPLVVQD | ASSADTSSGNTLY | 0.07983371 |
| GAFTTPQ | ASSADTSSGNTLY | 0.18429886 |
| GFGGFGTAGGVGGA | ASSADTSSGNTLY | 0.54408485 |
| GFLAQFL | ASSADTSSGNTLY | 0.111233585 |
| GGAGGAGGNAGVA | ASSADTSSGNTLY | 0.08081988 |
| GGVGTAGAGGAGGAGG | ASSADTSSGNTLY | 0.12085067 |
| GNGGGGDAGNAGSGGN | ASSADTSSGNTLY | 0.07534616 |
| GTGGAGGAGNDAGS | ASSADTSSGNTLY | 0.09445676 |
| GTTNTGWFNS | ASSADTSSGNTLY | 0.39409763 |
| IFMGPLFTL | ASSADTSSGNTLY | 0.08133617 |
| ILAHAIEAI | ASSADTSSGNTLY | 0.15082316 |
| IRSAIREAL | ASSADTSSGNTLY | 0.25057185 |
| ITFAGGG | ASSADTSSGNTLY | 0.09797727 |
| ITGMSGHQEV | ASSADTSSGNTLY | 0.08533345 |
| IVAGLACTL | ASSADTSSGNTLY | 0.083462924 |
| IVGGGLAGA | ASSADTSSGNTLY | 0.07742786 |
| IVTGVLNI | ASSADTSSGNTLY | 0.13610691 |
| LAQQGCI | ASSADTSSGNTLY | 0.09865551 |
| LCLHVEN | ASSADTSSGNTLY | 0.10240037 |
| LDAISDGS | ASSADTSSGNTLY | 0.11144473 |
| LGGFAAML | ASSADTSSGNTLY | 0.09808664 |
| LGGKNPMI | ASSADTSSGNTLY | 0.10791209 |

| | | |
|---|---|---|
| LGPKGVAAIL | ASSADTSSGNTLY | 0.10831013 |
| LMVGVSI | ASSADTSSGNTLY | 0.087048 |
| LVGAAFVL | ASSADTSSGNTLY | 0.14156568 |
| LVLAVAPL | ASSADTSSGNTLY | 0.08550497 |
| MAAIAVIDAA | ASSADTSSGNTLY | 0.09920708 |
| NNGINAELQ | ASSADTSSGNTLY | 0.41196877 |
| RGEPLVLVHG | ASSADTSSGNTLY | 0.07974285 |
| RGGAGGAAGLI | ASSADTSSGNTLY | 0.089713484 |
| RSYRMHPAV | ASSADTSSGNTLY | 0.11469889 |
| SAAHLLAGQLA | ASSADTSSGNTLY | 0.0788029 |
| SAAVGSY | ASSADTSSGNTLY | 0.18514606 |
| SPHAVLHRI | ASSADTSSGNTLY | 0.117969446 |
| SQIAKVL | ASSADTSSGNTLY | 0.17792028 |
| TALGHLVP | ASSADTSSGNTLY | 0.09361015 |
| TAPTTSGAA | ASSADTSSGNTLY | 0.37396994 |
| TDGDDGNGGSPGT | ASSADTSSGNTLY | 0.16223776 |
| TDGLGEN | ASSADTSSGNTLY | 0.08851845 |
| TGGPPLPGQA | ASSADTSSGNTLY | 0.07434193 |
| TIPAITI | ASSADTSSGNTLY | 0.084113196 |
| TIPGFTI | ASSADTSSGNTLY | 0.1561198 |
| TSIASGALA | ASSADTSSGNTLY | 0.16488685 |
| TVTLPGAAI | ASSADTSSGNTLY | 0.08406314 |
| VASGLSAALP | ASSADTSSGNTLY | 0.11999832 |
| VGAGVIGI | ASSADTSSGNTLY | 0.20972627 |
| VGPKRFGV | ASSADTSSGNTLY | 0.24357733 |
| VNVAFPD | ASSADTSSGNTLY | 0.08104777 |
| VSMTNSGVS | ASSADTSSGNTLY | 0.09445095 |
| VTDVLGDLSV | ASSADTSSGNTLY | 0.11653179 |
| VVVVGAGFAGL | ASSADTSSGNTLY | 0.11691632 |
| VYTVGTVV | ASSADTSSGNTLY | 0.08462675 |
| WTTFSGP | ASSADTSSGNTLY | 0.23457098 |
| YDELVEAF | ASSADTSSGNTLY | 0.13226439 |
| YGTGGAGGNGG | ASSADTSSGNTLY | 0.22458914 |
| YTNVMAV | ASSADTSSGNTLY | 0.15867707 |
| FLEQVHQL | ASSADTSSGNTLY | 0.22225307 |
| GLSLLAVAV | ASSADTSSGNTLY | 0.10751257 |
| LEQVHQL | ASSADTSSGNTLY | 0.10766678 |
| LNAAGAI | ASSADTSSGNTLY | 0.08347165 |
| PAVHASL | ASSADTSSGNTLY | 0.082285136 |
| TELEDDA | ASSADTSSGNTLY | 0.08250213 |
| TPPGPPN | ASSADTSSGNTLY | 0.09715878 |
| TPPPPPPP | ASSADTSSGNTLY | 0.07473294 |
| TQSPALI | ASSRLGGSQNTLY | 0.12990972 |
| EFAAAFPQ | ASSRLGGSQNTLY | 0.12371217 |
| LLSGNLDVL | ASSRLGGSQNTLY | 0.18674469 |
| GPANLGNYNI | ASSRLGGSQNTLY | 0.10351791 |
| TLGPTLP | ASSRLGGSQNTLY | 0.092027865 |
| VYAYSSCCK | ASSRLGGSQNTLY | 0.08694076 |
| APAPAPAPNI | ASSRLGGSQNTLY | 0.31311303 |
| PGGPVPIP | ASSRLGGSQNTLY | 0.078557536 |
| FLTVALEP | ASSRLGGSQNTLY | 0.08084967 |

| | | |
|---|---|---|
| SQYFGANL | ASSRLGGSQNTLY | 0.09282726 |
| GGVKPAA | ASSRLGGSQNTLY | 0.08087843 |
| GGTGGAGGAGGW | ASSRLGGSQNTLY | 0.2849177 |
| GVGGVAGLGGAGG | ASSRLGGSQNTLY | 0.2028338 |
| VLLVLATTI | ASSRLGGSQNTLY | 0.13856633 |
| IGIFAAML | ASSRLGGSQNTLY | 0.14829643 |
| AVGDLID | ASSRLGGSQNTLY | 0.07659666 |
| NISPLPL | ASSRLGGSQNTLY | 0.093567386 |
| VTAVFVAI | ASSRLGGSQNTLY | 0.5250524 |
| SGDDNDAAL | ASSRLGGSQNTLY | 0.111005105 |
| AVAALAHS | ASSRLGGSQNTLY | 0.14402857 |
| GGVTVMATP | ASSRLGGSQNTLY | 0.18223196 |
| AGLAANVRD | ASSRLGGSQNTLY | 0.093924426 |
| TNRPDLIDPALLRPG | ASSRLGGSQNTLY | 0.122553974 |
| LANLGVTAL | ASSRLGGSQNTLY | 0.09560951 |
| TGERIEL | ASSRLGGSQNTLY | 0.11903149 |
| NGEELAA | ASSRLGGSQNTLY | 0.082071275 |
| VLSLVWG | ASSRLGGSQNTLY | 0.08315425 |
| IGYALWL | ASSRLGGSQNTLY | 0.16293381 |
| LSLLLAPL | ASSRLGGSQNTLY | 0.62078285 |
| TLDEALK | ASSRLGGSQNTLY | 0.1335994 |
| TDGAGTAEGG | ASSRLGGSQNTLY | 0.14122218 |
| NAPGGAGGNGG | ASSRLGGSQNTLY | 0.11978851 |
| VSVFMGMGGNF | ASSRLGGSQNTLY | 0.17302854 |
| LASYLAAL | ASSRLGGSQNTLY | 0.1008921 |
| LVGGTPAL | ASSRLGGSQNTLY | 0.089722306 |
| VRNADVL | ASSRLGGSQNTLY | 0.08322223 |
| TCSSPFLVSAM | ASSRLGGSQNTLY | 0.088035084 |
| VHITDAE | ASSRLGGSQNTLY | 0.11826542 |
| EYAAPSF | ASSRLGGSQNTLY | 0.10330769 |
| VVDAFHGGL | ASSRLGGSQNTLY | 0.6733572 |
| ASVVDPQ | ASSRLGGSQNTLY | 0.18864141 |
| SYNEFGGLS | ASSRLGGSQNTLY | 0.09977012 |
| DPDGSVY | ASSRLGGSQNTLY | 0.083934225 |
| MVSTIHATE | ASSRLGGSQNTLY | 0.10186914 |
| LAAGVADPV | ASSRLGGSQNTLY | 0.12696101 |
| TKEQIAA | ASSRLGGSQNTLY | 0.086090155 |
| SLASPAAL | ASSRLGGSQNTLY | 0.36983418 |
| IGLVFYGV | ASSRLGGSQNTLY | 0.09934929 |
| FLLDASNL | ASSRLGGSQNTLY | 0.08836483 |
| YQVGNFP | ASSRLGGSQNTLY | 0.13300627 |
| ESLLSQST | ASSRLGGSQNTLY | 0.43490928 |
| LFTVGFAAVG | ASSRLGGSQNTLY | 0.11332684 |
| ATPPTPT | ASSRLGGSQNTLY | 0.08621547 |
| TDLATLG | ASSRLGGSQNTLY | 0.085141376 |
| SAGGTGR | ASSRLGGSQNTLY | 0.21647832 |
| GFSNSGS | ASSRLGGSQNTLY | 0.36492014 |
| GGNGGSGGTSV | ASSRLGGSQNTLY | 0.07386811 |
| GTSVATGGAGN | ASSRLGGSQNTLY | 0.30024278 |
| NGGNGGSGGTS | ASSRLGGSQNTLY | 0.08991678 |
| AVVKFPH | ASSRLGGSQNTLY | 0.0846869 |

| | | |
|---|---|---|
| TPPPPPP | ASSRLGGSQNTLY | 0.121964544 |
| AAPAAAAAPA | ASSRLGGSQNTLY | 0.11224429 |
| IALNEDL | ASSRLGGSQNTLY | 0.4356056 |
| IVNVSSM | ASSRLGGSQNTLY | 0.13698715 |
| TGIPPAPR | ASSRLGGSQNTLY | 0.094944656 |
| FDLGGGTF | ASSRLGGSQNTLY | 0.1025159 |
| FDLGGGTFDVS | ASSRLGGSQNTLY | 0.29114527 |
| NEPTAAA | ASSRLGGSQNTLY | 0.14741032 |
| NEPTAAALA | ASSRLGGSQNTLY | 0.16130236 |
| GSIINISSI | ASSRLGGSQNTLY | 0.120375514 |
| VAPGNAGTA | ASSRLGGSQNTLY | 0.101963654 |
| TGGGASLE | ASSRLGGSQNTLY | 0.11143511 |
| VAVYNHYKRI | ASSRLGGSQNTLY | 0.094741575 |
| TDPAPATT | ASSRLGGSQNTLY | 0.089725256 |
| TNRPDILDPALLRPG | ASSRLGGSQNTLY | 0.14522687 |
| LTTTTTS | ASSRLGGSQNTLY | 0.17636436 |
| PTGSGKTL | ASSRLGGSQNTLY | 0.082375385 |
| TGSGKTL | ASSRLGGSQNTLY | 0.10387985 |
| TIGHVDHGK | ASSRLGGSQNTLY | 0.2584391 |
| TIGHVDHGKTT | ASSRLGGSQNTLY | 0.100424446 |
| GHVDHGKTT | ASSRLGGSQNTLY | 0.08590324 |
| AQFEHTLL | ASSRLGGSQNTLY | 0.35832414 |
| IVNQDEL | ASSRLGGSQNTLY | 0.6810864 |
| TNGSQFF | ASSRLGGSQNTLY | 0.14858848 |
| TGKGIVE | ASSRLGGSQNTLY | 0.0842802 |
| LISNASDA | ASSRLGGSQNTLY | 0.07765589 |
| AELRTLGELTKIAKAHGAQ | ASSRLGGSQNTLY | 0.13078433 |
| APNAVAAAEPSW | ASSRLGGSQNTLY | 0.14469749 |
| APQVAGAATAVL | ASSRLGGSQNTLY | 0.09706292 |
| ARRDPAAAAAILPTDA | ASSRLGGSQNTLY | 0.08580552 |
| DAAARDVLAEAGLAEYF | ASSRLGGSQNTLY | 0.12788293 |
| DDQIVLSAKANLAKVEQ | ASSRLGGSQNTLY | 0.19274217 |
| DELLPDTGEDSDEDGH | ASSRLGGSQNTLY | 0.08917627 |
| ELPEAPANRATVPQD | ASSRLGGSQNTLY | 0.08338165 |
| FGQSKIERVTGIG | ASSRLGGSQNTLY | 0.08626996 |
| GAGGTGSTAGGAGGAGGAGGLY | ASSRLGGSQNTLY | 0.2197262 |
| GHGGTGAAGGNGGTGGAGGSGI | ASSRLGGSQNTLY | 0.08571574 |
| GNGGNGGAGGEGGPGGAGGAGA | ASSRLGGSQNTLY | 0.1118483 |
| GSTAGNGGNGGNATNGTIAGQP | ASSRLGGSQNTLY | 0.15974078 |
| GTGGDGGNGGNGGTGG | ASSRLGGSQNTLY | 0.07623524 |
| GVRRPAGIPADAPV | ASSRLGGSQNTLY | 0.12675433 |
| GYRQVTDAHLLTLARR | ASSRLGGSQNTLY | 0.09335528 |
| IAKMGQDGHDRG | ASSRLGGSQNTLY | 0.110250294 |
| KAEPSIEIQYT | ASSRLGGSQNTLY | 0.09253344 |
| KPVRIRMAADEDTVAVT | ASSRLGGSQNTLY | 0.18721098 |
| MPPRIAGMRLLVIKPEP | ASSRLGGSQNTLY | 0.13556449 |
| MKFYAVALFRSRGVPPTR | ASSRLGGSQNTLY | 0.13565704 |
| NAGGFGILGAVAHSP | ASSRLGGSQNTLY | 0.1326632 |
| NEGREITLAMAEM | ASSRLGGSQNTLY | 0.09500295 |
| NELVEWSHPLGHRWRWE | ASSRLGGSQNTLY | 0.082990594 |
| NLGAGGTGGNGGNPG | ASSRLGGSQNTLY | 0.07617115 |

| | | |
|---|---|---|
| NTEWPKWETVMAADPVD | ASSRLGGSQNTLY | 0.23066315 |
| PASAAPAETQP | ASSRLGGSQNTLY | 0.073722936 |
| QQSSDGKNGVRAAEITGE | ASSRLGGSQNTLY | 0.17945918 |
| RAPDAVTTSWWKE | ASSRLGGSQNTLY | 0.1402449 |
| SGFGNVGAHTSGWFNQ | ASSRLGGSQNTLY | 0.087283075 |
| SGVAGAVGQAASVGGLKV | ASSRLGGSQNTLY | 0.12196103 |
| SNGGAGTGSAGGNGGT | ASSRLGGSQNTLY | 0.076388 |
| TGGAGGEGGAAG | ASSRLGGSQNTLY | 0.086816974 |
| TGGDGGAAGAGGAPGHG | ASSRLGGSQNTLY | 0.4277917 |
| TMPPLLAAIANAGRHGV | ASSRLGGSQNTLY | 0.12432215 |
| TSSTAGNSGGA | ASSRLGGSQNTLY | 0.10066534 |
| VPATVPVGILC | ASSRLGGSQNTLY | 0.12061591 |
| VQRPDSGGVVLTGRIS | ASSRLGGSQNTLY | 0.16281146 |
| WTPNPVPATKGVPAGLVQAT | ASSRLGGSQNTLY | 0.09265368 |
| YALKRALAKFSDRFDVVII | ASSRLGGSQNTLY | 0.10174222 |
| YLIGSVIV | ASSRLGGSQNTLY | 0.24595916 |
| YGQLVAVID | ASSRLGGSQNTLY | 0.1230091 |
| VTVGGMTLPA | ASSRLGGSQNTLY | 0.084424414 |
| VLALPAV | ASSRLGGSQNTLY | 0.13439761 |
| VIPPTLNYET | ASSRLGGSQNTLY | 0.0888788 |
| VGMGGMHPSSG | ASSRLGGSQNTLY | 0.15892836 |
| VGLAGGAATA | ASSRLGGSQNTLY | 0.09525054 |
| VGGAGGAGGSGP | ASSRLGGSQNTLY | 0.13769706 |
| VFAAFMV | ASSRLGGSQNTLY | 0.12550189 |
| TLYDKLR | ASSRLGGSQNTLY | 0.0776938 |
| TLAVFTVL | ASSRLGGSQNTLY | 0.09383012 |
| TGSVQPARV | ASSRLGGSQNTLY | 0.11790751 |
| TGGGAGSGGT | ASSRLGGSQNTLY | 0.13461365 |
| SHLETSL | ASSRLGGSQNTLY | 0.13750802 |
| SFTTGMT | ASSRLGGSQNTLY | 0.09185437 |
| QVEVAKRMGVRQPTVS | ASSRLGGSQNTLY | 0.159974 |
| QGSGGPALAM | ASSRLGGSQNTLY | 0.103272095 |
| MLAPSNP | ASSRLGGSQNTLY | 0.100989975 |
| LVNNAAI | ASSRLGGSQNTLY | 0.106094174 |
| LTGEINTLA | ASSRLGGSQNTLY | 0.08965384 |
| LSIFGPV | ASSRLGGSQNTLY | 0.09408387 |
| LLALLANL | ASSRLGGSQNTLY | 0.13078187 |
| LISNASDALDKLR | ASSRLGGSQNTLY | 0.086740725 |
| LINEAVP | ASSRLGGSQNTLY | 0.13977 |
| LGYAFSGL | ASSRLGGSQNTLY | 0.07481373 |
| LDFVAVAL | ASSRLGGSQNTLY | 0.09269628 |
| LAQPGAE | ASSRLGGSQNTLY | 0.15079357 |
| ISNLGAVTGF | ASSRLGGSQNTLY | 0.085095674 |
| HQEDPGRI | ASSRLGGSQNTLY | 0.1449147 |
| GTPGEPTP | ASSRLGGSQNTLY | 0.11023487 |
| GFFNSGS | ASSRLGGSQNTLY | 0.087265834 |
| GAGGGGLTAGAN | ASSRLGGSQNTLY | 0.10588397 |
| FVDALVNM | ASSRLGGSQNTLY | 0.18361397 |
| FLTVAPEL | ASSRLGGSQNTLY | 0.17525569 |
| FETLTAL | ASSRLGGSQNTLY | 0.08678467 |
| DVINPIEL | ASSRLGGSQNTLY | 0.11087966 |

| | | |
|---|---|---|
| APVSGAPV | ASSRLGGSQNTLY | 0.1816666 |
| ALVGGLAW | ASSRLGGSQNTLY | 0.11444715 |
| ALAAAAATAI | ASSRLGGSQNTLY | 0.08069539 |
| AFGLGLLL | ASSRLGGSQNTLY | 0.09848421 |
| AFFAAASL | ASSRLGGSQNTLY | 0.08271999 |
| AAAALPAFSPP | ASSRLGGSQNTLY | 0.09187343 |
| VVLAPAL | ASSRLGGSQNTLY | 0.08401023 |
| TIGHVDHGKTTL | ASSRLGGSQNTLY | 0.24523365 |
| LISNASDALDK | ASSRLGGSQNTLY | 0.56450605 |
| LISNASDAL | ASSRLGGSQNTLY | 0.1007327 |
| ISNASDALDK | ASSRLGGSQNTLY | 0.103418946 |
| NQAETLVYQTE | ASSRLGGSQNTLY | 0.08056794 |
| TGGTGGTAGAG | ASSRLGGSQNTLY | 0.16279662 |
| TGSTAGGAGGA | ASSRLGGSQNTLY | 0.30528224 |
| DPLAVPVAAGPV | ASSRLGGSQNTLY | 0.09245415 |
| GGTGGTPGNGGN | ASSRLGGSQNTLY | 0.13023125 |
| PAGLAGVDPATL | ASSRLGGSQNTLY | 0.08929007 |
| GNNLVVEAPHAGASV | ASSRLGGSQNTLY | 0.08702308 |
| KVTRIMSASTLEQPA | ASSRLGGSQNTLY | 0.29998368 |
| LGFALPIALPRRTVV | ASSRLGGSQNTLY | 0.114343256 |
| TRLGNLTARKLVVLV | ASSRLGGSQNTLY | 0.10865658 |
| WSQQRRLIYFPSAGP | ASSRLGGSQNTLY | 0.0950505 |
| GPNGNPDPMAAAVDIR | ASSRLGGSQNTLY | 0.08684169 |
| TQRCVSVADLRAAASR | ASSRLGGSQNTLY | 0.11116347 |
| QFVLDVAEHPVEVLTETV | ASSRLGGSQNTLY | 0.22326706 |
| GGAGGTGGAGGNGGTQAGDG | ASSRLGGSQNTLY | 0.08634343 |
| PLLATLRAIAVDVDAKPAQV | ASSRLGGSQNTLY | 0.13825564 |
| APKFRTGGYMIDFWGVGYQVAK | ASSRLGGSQNTLY | 0.095402054 |
| GAGGAGGAGGAAGGPGGTGGTG | ASSRLGGSQNTLY | 0.29485694 |
| IGALGLLATDVAPAGSVAFGVA | ASSRLGGSQNTLY | 0.23898643 |
| AGGAGGIGGNGGAGGAGGVLMG | ASSRLGGSQNTLY | 0.15574439 |
| ADELIGLL | ASSRLGGSQNTLY | 0.101072885 |
| ADRALKA | ASSRLGGSQNTLY | 0.36836946 |
| AGAVGQVGGAGGA | ASSRLGGSQNTLY | 0.1310787 |
| AGGGGGLGGAP | ASSRLGGSQNTLY | 0.14673606 |
| AGGLLYGNGGAG | ASSRLGGSQNTLY | 0.14140075 |
| AGLESIGL | ASSRLGGSQNTLY | 0.5828046 |
| AGLKTGII | ASSRLGGSQNTLY | 0.82004404 |
| ASLAALLL | ASSRLGGSQNTLY | 0.7723352 |
| ASVINISSI | ASSRLGGSQNTLY | 0.681916 |
| ATPPGPA | ASSRLGGSQNTLY | 0.5489229 |
| ATVASALV | ASSRLGGSQNTLY | 0.26742715 |
| ATVASLIL | ASSRLGGSQNTLY | 0.70997596 |
| AVNGPVLIRH | ASSRLGGSQNTLY | 0.6030314 |
| DGAAGAVGI | ASSRLGGSQNTLY | 0.8129093 |
| DPLLHGI | ASSRLGGSQNTLY | 0.77589756 |
| ELGAALGAL | ASSRLGGSQNTLY | 0.603404 |
| ELGRTQAV | ASSRLGGSQNTLY | 0.8215067 |
| FVAGGYLS | ASSRLGGSQNTLY | 0.8135915 |
| GASGGAGGAGGW | ASSRLGGSQNTLY | 0.4394151 |
| GGAGAGGDGGAGGI | ASSRLGGSQNTLY | 0.7426467 |

| | | |
|---|---|---|
| GTGHNGSQPGMGGNG | ASSRLGGSQNTLY | 0.5325816 |
| GYAAASASAS | ASSRLGGSQNTLY | 0.7207222 |
| HQVDVAAVQQG | ASSRLGGSQNTLY | 0.42645895 |
| IGAAIAHT | ASSRLGGSQNTLY | 0.7761096 |
| IGMIEVAQL | ASSRLGGSQNTLY | 0.72663736 |
| IRLAPEHTSAMD | ASSRLGGSQNTLY | 0.23267028 |
| IYNVSVL | ASSRLGGSQNTLY | 0.7715392 |
| KQGFLGM | ASSRLGGSQNTLY | 0.74105966 |
| LESVITQL | ASSRLGGSQNTLY | 0.7582861 |
| LGLGALVVV | ASSRLGGSQNTLY | 0.8111634 |
| LIGNGGNGGSGGI | ASSRLGGSQNTLY | 0.5534388 |
| LNAINAPTL | ASSRLGGSQNTLY | 0.6480016 |
| LQLLNTGV | ASSRLGGSQNTLY | 0.8079898 |
| LQTSIEF | ASSRLGGSQNTLY | 0.8071936 |
| MSFAVSEL | ASSRLGGSQNTLY | 0.5412947 |
| NTGALGL | ASSRLGGSQNTLY | 0.7822038 |
| PPGAGGAGGAGTT | ASSRLGGSQNTLY | 0.80922407 |
| QLVGLGI | ASSRLGGSQNTLY | 0.703526 |
| RFLEQVHQL | ASSRLGGSQNTLY | 0.78556913 |
| RGVGYVLR | ASSRLGGSQNTLY | 0.7983672 |
| SATGVFAGVI | ASSRLGGSQNTLY | 0.52199996 |
| SLPAITLP | ASSRLGGSQNTLY | 0.8112785 |
| STALAFTV | ASSRLGGSQNTLY | 0.82354844 |
| SVLTVGLGK | ASSRLGGSQNTLY | 0.68014157 |
| TALAGAGSGI | ASSRLGGSQNTLY | 0.71104944 |
| TGLPFTGI | ASSRLGGSQNTLY | 0.48903847 |
| VAEHVTV | ASSRLGGSQNTLY | 0.71901643 |
| VDIVLNSL | ASSRLGGSQNTLY | 0.2848022 |
| VLTGGGAVL | ASSRLGGSQNTLY | 0.7921635 |
| VNVGLAYGI | ASSRLGGSQNTLY | 0.8065858 |
| VSSAAVGAV | ASSRLGGSQNTLY | 0.7302058 |
| VTSVYEAGV | ASSRLGGSQNTLY | 0.5929551 |
| WVALDRGA | ASSRLGGSQNTLY | 0.5136665 |
| YLVGSVVI | ASSRLGGSQNTLY | 0.762076 |
| GPPGTGKTLL | ASSRLGGSQNTLY | 0.55044854 |
| GVGMTKF | ASSRLGGSQNTLY | 0.3483165 |
| IGIGGSDL | ASSRLGGSQNTLY | 0.54203403 |
| IGTLLAEL | ASSRLGGSQNTLY | 0.7958115 |
| IYTSGTT | ASSRLGGSQNTLY | 0.78336334 |
| LSFPAGV | ASSRLGGSQNTLY | 0.65561455 |
| LYTSGTT | ASSRLGGSQNTLY | 0.8163724 |
| VGAFPFGF | ASSRLGGSQNTLY | 0.81077796 |
| VGAGTVEF | ASSRLGGSQNTLY | 0.79405344 |
| VGKSTLLL | ASSRLGGSQNTLY | 0.71675307 |
| AALAHAGGGIV | ASSRLGGSQNTLY | 0.6073576 |
| AALVAVSLL | ASSRLGGSQNTLY | 0.81808233 |
| AAVSRAVAAL | ASSRLGGSQNTLY | 0.7246404 |
| AGGSSGIGGF | ASSRLGGSQNTLY | 0.660201 |
| AGLTVANANG | ASSRLGGSQNTLY | 0.7728306 |
| AGLVGAVL | ASSRLGGSQNTLY | 0.7508514 |
| ALSQAFTV | ASSRLGGSQNTLY | 0.6630758 |

| | | |
|---|---|---|
| ALTFSVSL | ASSRLGGSQNTLY | 0.77448595 |
| ALTGAALVAL | ASSRLGGSQNTLY | 0.7862499 |
| ASFSIGAGVL | ASSRLGGSQNTLY | 0.6084549 |
| DGSVYKI | ASSRLGGSQNTLY | 0.3773135 |
| EQLAEIV | ASSRLGGSQNTLY | 0.6872032 |
| FAVSELTVM | ASSRLGGSQNTLY | 0.7675096 |
| GAAPLAAGL | ASSRLGGSQNTLY | 0.8059883 |
| GADGGDGGSGGAGG | ASSRLGGSQNTLY | 0.7512597 |
| GFGLDTGGAGG | ASSRLGGSQNTLY | 0.6653158 |
| GLASGGLL | ASSRLGGSQNTLY | 0.79830897 |
| GRIAGAAVLL | ASSRLGGSQNTLY | 0.74975574 |
| GSAGSLNTSF | ASSRLGGSQNTLY | 0.7342047 |
| GTGQIGI | ASSRLGGSQNTLY | 0.8148809 |
| GTLIALIL | ASSRLGGSQNTLY | 0.72895813 |
| IAATMGFVVG | ASSRLGGSQNTLY | 0.785789 |
| IFAQALL | ASSRLGGSQNTLY | 0.7619685 |
| IGAAVALGF | ASSRLGGSQNTLY | 0.74238336 |
| IGLTPLAL | ASSRLGGSQNTLY | 0.78529453 |
| ILGSNTSTL | ASSRLGGSQNTLY | 0.72377115 |
| IVFSLTVL | ASSRLGGSQNTLY | 0.6879045 |
| LASKATGF | ASSRLGGSQNTLY | 0.31571046 |
| LASVALIL | ASSRLGGSQNTLY | 0.6525612 |
| LFAVGLTL | ASSRLGGSQNTLY | 0.7988996 |
| LFNGGIGTYI | ASSRLGGSQNTLY | 0.7858522 |
| LTLGAGII | ASSRLGGSQNTLY | 0.8040136 |
| LVAAALGDI | ASSRLGGSQNTLY | 0.7124748 |
| NTAFVQLGI | ASSRLGGSQNTLY | 0.6599164 |
| QLFSGAEL | ASSRLGGSQNTLY | 0.8090921 |
| SAATFITL | ASSRLGGSQNTLY | 0.8090155 |
| SNYGLANLGV | ASSRLGGSQNTLY | 0.3671763 |
| SYGGAYAVM | ASSRLGGSQNTLY | 0.6770379 |
| TTGVGGLTLG | ASSRLGGSQNTLY | 0.82973266 |
| TTQTIYI | ASSRLGGSQNTLY | 0.79587245 |
| VAAGAGLGAVI | ASSRLGGSQNTLY | 0.67025375 |
| VAAVVTLL | ASSRLGGSQNTLY | 0.567634 |
| VGFLGGLI | ASSRLGGSQNTLY | 0.82753134 |
| VGVIPILL | ASSRLGGSQNTLY | 0.66325414 |
| VIFASSLI | ASSRLGGSQNTLY | 0.82739896 |
| VKGGDGGAAAG | ASSRLGGSQNTLY | 0.34850425 |
| VTKTFTNI | ASSRLGGSQNTLY | 0.58995855 |
| VVAAAILL | ASSRLGGSQNTLY | 0.45075986 |
| YAALDVEL | ASSRLGGSQNTLY | 0.29838747 |
| LASVALLL | ASSRLGGSQNTLY | 0.7640629 |
| LDAVHRV | ASSRLGGSQNTLY | 0.6627466 |
| QGLSSSIF | ASSRLGGSQNTLY | 0.79388 |
| RGTFANIRL | ASSRLGGSQNTLY | 0.71848774 |
| RQLWWGHRI | ASSRLGGSQNTLY | 0.7983042 |
| YGPPGTGKTLL | ASSRLGGSQNTLY | 0.77525955 |
| AALVGAPANVA | ASSRLGGSQNTLY | 0.7657821 |
| AATFRRTE | ASSRLGGSQNTLY | 0.50223416 |
| AGALGGGI | ASSRLGGSQNTLY | 0.75968534 |

| | | |
|---|---|---|
| AGGAGGAGGIGAG | ASSRLGGSQNTLY | 0.8105668 |
| AGSGALGL | ASSRLGGSQNTLY | 0.6918262 |
| AQIADQI | ASSRLGGSQNTLY | 0.36120266 |
| ASAVAVTYL | ASSRLGGSQNTLY | 0.7775847 |
| ASTVLGGTGG | ASSRLGGSQNTLY | 0.6517087 |
| AVGGEPAK | ASSRLGGSQNTLY | 0.2706977 |
| CEQGTFGGTC | ASSRLGGSQNTLY | 0.23956946 |
| DEGKAPAG | ASSRLGGSQNTLY | 0.69512933 |
| DTVVFPAL | ASSRLGGSQNTLY | 0.68844604 |
| FEGITMAG | ASSRLGGSQNTLY | 0.8141514 |
| FPLVVQD | ASSRLGGSQNTLY | 0.81031466 |
| GAFTTPQ | ASSRLGGSQNTLY | 0.7235323 |
| GFGGFGTAGGVGGA | ASSRLGGSQNTLY | 0.8024882 |
| GFLAQFL | ASSRLGGSQNTLY | 0.7739713 |
| GGAGGAGGNAGVA | ASSRLGGSQNTLY | 0.16456036 |
| GGVGTAGAGGAGGAGG | ASSRLGGSQNTLY | 0.557814 |
| GNGGGGDAGNAGSGGN | ASSRLGGSQNTLY | 0.47427225 |
| GTGGAGGAGNDAGS | ASSRLGGSQNTLY | 0.7885538 |
| GTTNTGWFNS | ASSRLGGSQNTLY | 0.66330284 |
| IFMGPLFTL | ASSRLGGSQNTLY | 0.71310204 |
| ILAHAIEAI | ASSRLGGSQNTLY | 0.7612729 |
| IRSAIREAL | ASSRLGGSQNTLY | 0.8007238 |
| ITFAGGG | ASSRLGGSQNTLY | 0.706897 |
| ITGMSGHQEV | ASSRLGGSQNTLY | 0.5380813 |
| IVAGLACTL | ASSRLGGSQNTLY | 0.23861352 |
| IVGGGLAGA | ASSRLGGSQNTLY | 0.74435914 |
| IVTGVLNI | ASSRLGGSQNTLY | 0.48594657 |
| LAQQGCI | ASSRLGGSQNTLY | 0.5616154 |
| LCLHVEN | ASSRLGGSQNTLY | 0.44276446 |
| LDAISDGS | ASSRLGGSQNTLY | 0.70503306 |
| LGGFAAML | ASSRLGGSQNTLY | 0.3901543 |
| LGGKNPMI | ASSRLGGSQNTLY | 0.775118 |
| LGPKGVAAIL | ASSRLGGSQNTLY | 0.58015084 |
| LMVGVSI | ASSRLGGSQNTLY | 0.7492664 |
| LVGAAFVL | ASSRLGGSQNTLY | 0.8170127 |
| LVLAVAPL | ASSRLGGSQNTLY | 0.7986084 |
| MAAIAVIDAA | ASSRLGGSQNTLY | 0.68525624 |
| NNGINAELQ | ASSRLGGSQNTLY | 0.67868775 |
| RGEPLVLVHG | ASSRLGGSQNTLY | 0.5484941 |
| RGGAGGAAGLI | ASSRLGGSQNTLY | 0.67121935 |
| RSYRMHPAV | ASSRLGGSQNTLY | 0.6485591 |
| SAAHLLAGQLA | ASSRLGGSQNTLY | 0.82387465 |
| SAAVGSY | ASSRLGGSQNTLY | 0.5886967 |
| SPHAVLHRI | ASSRLGGSQNTLY | 0.5381872 |
| SQIAKVL | ASSRLGGSQNTLY | 0.6309712 |
| TALGHLVP | ASSRLGGSQNTLY | 0.8200504 |
| TAPTTSGAA | ASSRLGGSQNTLY | 0.82126814 |
| TDGDDGNGGSPGT | ASSRLGGSQNTLY | 0.6378672 |
| TDGLGEN | ASSRLGGSQNTLY | 0.7204753 |
| TGGPPLPGQA | ASSRLGGSQNTLY | 0.60877633 |
| TIPAITI | ASSRLGGSQNTLY | 0.766392 |

| | | |
|---|---|---|
| TIPGFTI | ASSRLGGSQNTLY | 0.6101166 |
| TSIASGALA | ASSRLGGSQNTLY | 0.5379471 |
| TVTLPGAAI | ASSRLGGSQNTLY | 0.76185703 |
| VASGLSAALP | ASSRLGGSQNTLY | 0.83112043 |
| VGAGVIGI | ASSRLGGSQNTLY | 0.6746731 |
| VGPKRFGV | ASSRLGGSQNTLY | 0.38598573 |
| VNVAFPD | ASSRLGGSQNTLY | 0.7634355 |
| VSMTNSGVS | ASSRLGGSQNTLY | 0.62164295 |
| VTDVLGDLSV | ASSRLGGSQNTLY | 0.6854031 |
| VVVVGAGFAGL | ASSRLGGSQNTLY | 0.6761131 |
| VYTVGTVV | ASSRLGGSQNTLY | 0.45109355 |
| WTTFSGP | ASSRLGGSQNTLY | 0.7262267 |
| YDELVEAF | ASSRLGGSQNTLY | 0.81888413 |
| YGTGGAGGNGG | ASSRLGGSQNTLY | 0.7627374 |
| YTNVMAV | ASSRLGGSQNTLY | 0.7685139 |
| FLEQVHQL | ASSRLGGSQNTLY | 0.8214014 |
| GLSLLAVAV | ASSRLGGSQNTLY | 0.79878104 |
| LEQVHQL | ASSRLGGSQNTLY | 0.8170903 |
| LNAAGAI | ASSRLGGSQNTLY | 0.8187702 |
| PAVHASL | ASSRLGGSQNTLY | 0.6854458 |
| TELEDDA | ASSRLGGSQNTLY | 0.67428297 |
| TPPGPPN | ASSRLGGSQNTLY | 0.80255544 |
| TPPPPPPP | ASSRLGGSQNTLY | 0.75869423 |
| TQSPALI | GARVQGANERLF | 0.73616815 |
| EFAAAFPQ | GARVQGANERLF | 0.5476703 |
| LLSGNLDVL | GARVQGANERLF | 0.8204423 |
| GPANLGNYNI | GARVQGANERLF | 0.66748524 |
| TLGPTLP | GARVQGANERLF | 0.8176404 |
| VYAYSSCCK | GARVQGANERLF | 0.2900533 |
| APAPAPAPNI | GARVQGANERLF | 0.5952863 |
| PGGPVPIP | GARVQGANERLF | 0.8086612 |
| FLTVALEP | GARVQGANERLF | 0.17522827 |
| SQYFGANL | GARVQGANERLF | 0.24025413 |
| GGVKPAA | GARVQGANERLF | 0.7493378 |
| GGTGGAGGAGGW | GARVQGANERLF | 0.8101977 |
| GVGGVAGLGGAGG | GARVQGANERLF | 0.18388508 |
| VLLVLATTI | GARVQGANERLF | 0.6114347 |
| IGIFAAML | GARVQGANERLF | 0.3435983 |
| AVGDLID | GARVQGANERLF | 0.72186655 |
| NISPLPL | GARVQGANERLF | 0.7385386 |
| VTAVFVAI | GARVQGANERLF | 0.28574598 |
| SGDDNDAAL | GARVQGANERLF | 0.776991 |
| AVAALAHS | GARVQGANERLF | 0.39108384 |
| GGVTVMATP | GARVQGANERLF | 0.6417798 |
| AGLAANVRD | GARVQGANERLF | 0.7838592 |
| TNRPDLIDPALLRPG | GARVQGANERLF | 0.82559764 |
| LANLGVTAL | GARVQGANERLF | 0.49608776 |
| TGERIEL | GARVQGANERLF | 0.77830344 |
| NGEELAA | GARVQGANERLF | 0.61067885 |
| VLSLVWG | GARVQGANERLF | 0.8258583 |
| IGYALWL | GARVQGANERLF | 0.5591328 |

| | | |
|---|---|---|
| LSLLLAPL | GARVQGANERLF | 0.33929932 |
| TLDEALK | GARVQGANERLF | 0.80899894 |
| TDGAGTAEGG | GARVQGANERLF | 0.7517079 |
| NAPGGAGGNGG | GARVQGANERLF | 0.68880224 |
| VSVFMGMGGNF | GARVQGANERLF | 0.81604564 |
| LASYLAAL | GARVQGANERLF | 0.52055764 |
| LVGGTPAL | GARVQGANERLF | 0.7486838 |
| VRNADVL | GARVQGANERLF | 0.8174061 |
| TCSSPFLVSAM | GARVQGANERLF | 0.8141659 |
| VHITDAE | GARVQGANERLF | 0.4748145 |
| EYAAPSF | GARVQGANERLF | 0.7737174 |
| VVDAFHGGL | GARVQGANERLF | 0.7780001 |
| ASVVDPQ | GARVQGANERLF | 0.5595333 |
| SYNEFGGLS | GARVQGANERLF | 0.7997005 |
| DPDGSVY | GARVQGANERLF | 0.63268095 |
| MVSTIHATE | GARVQGANERLF | 0.80446506 |
| LAAGVADPV | GARVQGANERLF | 0.7354686 |
| TKEQIAA | GARVQGANERLF | 0.8225395 |
| SLASPAAL | GARVQGANERLF | 0.31510577 |
| IGLVFYGV | GARVQGANERLF | 0.7716769 |
| FLLDASNL | GARVQGANERLF | 0.7349112 |
| YQVGNFP | GARVQGANERLF | 0.7934644 |
| ESLLSQST | GARVQGANERLF | 0.6525576 |
| LFTVGFAAVG | GARVQGANERLF | 0.66114914 |
| ATPPTPT | GARVQGANERLF | 0.73330116 |
| TDLATLG | GARVQGANERLF | 0.81994486 |
| SAGGTGR | GARVQGANERLF | 0.70905095 |
| GFSNSGS | GARVQGANERLF | 0.784008 |
| GGNGGSGGTSV | GARVQGANERLF | 0.8235359 |
| GTSVATGGAGN | GARVQGANERLF | 0.5841912 |
| NGGNGGSGGTS | GARVQGANERLF | 0.746169 |
| AVVKFPH | GARVQGANERLF | 0.78292716 |
| TPPPPPP | GARVQGANERLF | 0.80947566 |
| AAPAAAAAPA | GARVQGANERLF | 0.66062224 |
| IALNEDL | GARVQGANERLF | 0.82813174 |
| IVNVSSM | GARVQGANERLF | 0.4474091 |
| TGIPPAPR | GARVQGANERLF | 0.81579757 |
| FDLGGGTF | GARVQGANERLF | 0.6223926 |
| FDLGGGTFDVS | GARVQGANERLF | 0.7461795 |
| NEPTAAA | GARVQGANERLF | 0.67919636 |
| NEPTAAALA | GARVQGANERLF | 0.8150784 |
| GSIINISSI | GARVQGANERLF | 0.81285536 |
| VAPGNAGTA | GARVQGANERLF | 0.81402516 |
| TGGGASLE | GARVQGANERLF | 0.27293235 |
| VAVYNHYKRI | GARVQGANERLF | 0.8092177 |
| TDPAPATT | GARVQGANERLF | 0.81382793 |
| TNRPDILDPALLRPG | GARVQGANERLF | 0.8295239 |
| LTTTTTS | GARVQGANERLF | 0.6309247 |
| PTGSGKTL | GARVQGANERLF | 0.75006413 |
| TGSGKTL | GARVQGANERLF | 0.7507386 |
| TIGHVDHGK | GARVQGANERLF | 0.6677687 |

| | | |
|---|---|---|
| TIGHVDHGKTT | GARVQGANERLF | 0.60407305 |
| GHVDHGKTT | GARVQGANERLF | 0.66259575 |
| AQFEHTLL | GARVQGANERLF | 0.71267617 |
| IVNQDEL | GARVQGANERLF | 0.70491827 |
| TNGSQFF | GARVQGANERLF | 0.81540155 |
| TGKGIVE | GARVQGANERLF | 0.81832397 |
| LISNASDA | GARVQGANERLF | 0.7966007 |
| AELRTLGELTKIAKAHGAQ | GARVQGANERLF | 0.8188404 |
| APNAVAAAEPSW | GARVQGANERLF | 0.8190818 |
| APQVAGAATAVL | GARVQGANERLF | 0.7542057 |
| ARRDPAAAAAILPTDA | GARVQGANERLF | 0.7052733 |
| DAAARDVLAEAGLAEYF | GARVQGANERLF | 0.70259273 |
| DDQIVLSAKANLAKVEQ | GARVQGANERLF | 0.6200437 |
| DELLPDTGEDSDEDGH | GARVQGANERLF | 0.71089387 |
| ELPEAPANRATVPQD | GARVQGANERLF | 0.6805371 |
| FGQSKIERVTGIG | GARVQGANERLF | 0.7656195 |
| GAGGTGSTAGGAGGAGGAGGLY | GARVQGANERLF | 0.16052008 |
| GHGGTGAAGGNGGTGGAGGSGI | GARVQGANERLF | 0.79623777 |
| GNGGNGGAGGEGGPGGAGGAGA | GARVQGANERLF | 0.8257185 |
| GSTAGNGGNGGNATNGTIAGQP | GARVQGANERLF | 0.72309035 |
| GTGGDGGNGGNGGTGG | GARVQGANERLF | 0.7612013 |
| GVRRPAGIPADAPV | GARVQGANERLF | 0.78300995 |
| GYRQVTDAHLLTLARR | GARVQGANERLF | 0.6261877 |
| IAKMGQDGHDRG | GARVQGANERLF | 0.8005992 |
| KAEPSIEIQYT | GARVQGANERLF | 0.5584086 |
| KPVRIRMAADEDTVAVT | GARVQGANERLF | 0.7514689 |
| MPPRIAGMRLLVIKPEP | GARVQGANERLF | 0.79625237 |
| MKFYAVALFRSRGVPPTR | GARVQGANERLF | 0.7430823 |
| NAGGFGILGAVAHSP | GARVQGANERLF | 0.7610185 |
| NEGREITLAMAEM | GARVQGANERLF | 0.6082734 |
| NELVEWSHPLGHRWRWE | GARVQGANERLF | 0.23894122 |
| NLGAGGTGGNGGNPG | GARVQGANERLF | 0.77244985 |
| NTEWPKWETVMAADPVD | GARVQGANERLF | 0.56530005 |
| PASAAPAETQP | GARVQGANERLF | 0.7343527 |
| QQSSDGKNGVRAAEITGE | GARVQGANERLF | 0.4250552 |
| RAPDAVTTSWWKE | GARVQGANERLF | 0.78434 |
| SGFGNVGAHTSGWFNQ | GARVQGANERLF | 0.77985954 |
| SGVAGAVGQAASVGGLKV | GARVQGANERLF | 0.75698125 |
| SNGGAGTGSAGGNGGT | GARVQGANERLF | 0.5922836 |
| TGGAGGEGGAAG | GARVQGANERLF | 0.75714886 |
| TGGDGGAAGAGGAPGHG | GARVQGANERLF | 0.75393707 |
| TMPPLLAAIANAGRHGV | GARVQGANERLF | 0.49523932 |
| TSSTAGNSGGA | GARVQGANERLF | 0.8048355 |
| VPATVPVGILC | GARVQGANERLF | 0.6407545 |
| VQRPDSGGVVLTGRIS | GARVQGANERLF | 0.6775272 |
| WTPNPVPATKGVPAGLVQAT | GARVQGANERLF | 0.8009306 |
| YALKRALAKFSDRFDVVII | GARVQGANERLF | 0.7409644 |
| YLIGSVIV | GARVQGANERLF | 0.30351964 |
| YGQLVAVID | GARVQGANERLF | 0.7516111 |
| VTVGGMTLPA | GARVQGANERLF | 0.74856025 |
| VLALPAV | GARVQGANERLF | 0.53959465 |

| | | |
|---|---|---|
| VIPPTLNYET | GARVQGANERLF | 0.25392124 |
| VGMGGMHPSSG | GARVQGANERLF | 0.7596077 |
| VGLAGGAATA | GARVQGANERLF | 0.54193103 |
| VGGAGGAGGSGP | GARVQGANERLF | 0.5571849 |
| VFAAFMV | GARVQGANERLF | 0.7167785 |
| TLYDKLR | GARVQGANERLF | 0.81378424 |
| TLAVFTVL | GARVQGANERLF | 0.5766303 |
| TGSVQPARV | GARVQGANERLF | 0.61646396 |
| TGGGAGSGGT | GARVQGANERLF | 0.80593383 |
| SHLETSL | GARVQGANERLF | 0.7571603 |
| SFTTGMT | GARVQGANERLF | 0.81372106 |
| QVEVAKRMGVRQPTVS | GARVQGANERLF | 0.80019313 |
| QGSGGPALAM | GARVQGANERLF | 0.5915884 |
| MLAPSNP | GARVQGANERLF | 0.7550321 |
| LVNNAAI | GARVQGANERLF | 0.67338485 |
| LTGEINTLA | GARVQGANERLF | 0.47100624 |
| LSIFGPV | GARVQGANERLF | 0.423541 |
| LLALLANL | GARVQGANERLF | 0.64068997 |
| LISNASDALDKLR | GARVQGANERLF | 0.37929666 |
| LINEAVP | GARVQGANERLF | 0.6906457 |
| LGYAFSGL | GARVQGANERLF | 0.7818502 |
| LDFVAVAL | GARVQGANERLF | 0.6110103 |
| LAQPGAE | GARVQGANERLF | 0.39790514 |
| ISNLGAVTGF | GARVQGANERLF | 0.71274847 |
| HQEDPGRI | GARVQGANERLF | 0.40618387 |
| GTPGEPTP | GARVQGANERLF | 0.78375643 |
| GFFNSGS | GARVQGANERLF | 0.6969806 |
| GAGGGGLTAGAN | GARVQGANERLF | 0.8305936 |
| FVDALVNM | GARVQGANERLF | 0.81808436 |
| FLTVAPEL | GARVQGANERLF | 0.7492926 |
| FETLTAL | GARVQGANERLF | 0.8078632 |
| DVINPIEL | GARVQGANERLF | 0.54376113 |
| APVSGAPV | GARVQGANERLF | 0.7331232 |
| ALVGGLAW | GARVQGANERLF | 0.68526965 |
| ALAAAAATAI | GARVQGANERLF | 0.7570107 |
| AFGLGLLL | GARVQGANERLF | 0.7338003 |
| AFFAAASL | GARVQGANERLF | 0.6604537 |
| AAAALPAFSPP | GARVQGANERLF | 0.2709878 |
| VVLAPAL | GARVQGANERLF | 0.80389535 |
| TIGHVDHGKTTL | GARVQGANERLF | 0.6816349 |
| LISNASDALDK | GARVQGANERLF | 0.31347498 |
| LISNASDAL | GARVQGANERLF | 0.5479104 |
| ISNASDALDK | GARVQGANERLF | 0.82568014 |
| NQAETLVYQTE | GARVQGANERLF | 0.8212852 |
| TGGTGGTAGAG | GARVQGANERLF | 0.6844832 |
| TGSTAGGAGGA | GARVQGANERLF | 0.58065355 |
| DPLAVPVAAGPV | GARVQGANERLF | 0.80285406 |
| GGTGGTPGNGGN | GARVQGANERLF | 0.50995195 |
| PAGLAGVDPATL | GARVQGANERLF | 0.7195588 |
| GNNLVVEAPHAGASV | GARVQGANERLF | 0.7616179 |
| KVTRIMSASTLEQPA | GARVQGANERLF | 0.76029754 |

| | | |
|---|---|---|
| LGFALPIALPRRTVV | GARVQGANERLF | 0.476677 |
| TRLGNLTARKLVVLV | GARVQGANERLF | 0.77792215 |
| WSQQRRLIYFPSAGP | GARVQGANERLF | 0.7586935 |
| GPNGNPDPMAAAVDIR | GARVQGANERLF | 0.71784496 |
| TQRCVSVADLRAAASR | GARVQGANERLF | 0.21955512 |
| QFVLDVAEHPVEVLTETV | GARVQGANERLF | 0.715089 |
| GGAGGTGGAGGNGGTQAGDG | GARVQGANERLF | 0.6722397 |
| PLLATLRAIAVDVDAKPAQV | GARVQGANERLF | 0.45698127 |
| APKFRTGGYMIDFWGVGYQVAK | GARVQGANERLF | 0.31078896 |
| GAGGAGGAGGAAGGPGGTGGTG | GARVQGANERLF | 0.7196307 |
| IGALGLLATDVAPAGSVAFGVA | GARVQGANERLF | 0.8104258 |
| AGGAGGIGGNGGAGGAGGVLMG | GARVQGANERLF | 0.8140874 |
| ADELIGLL | GARVQGANERLF | 0.59655184 |
| ADRALKA | GARVQGANERLF | 0.82050836 |
| AGAVGQVGGAGGA | GARVQGANERLF | 0.7852322 |
| AGGGGGLGGAP | GARVQGANERLF | 0.6768666 |
| AGGLLYGNGGAG | GARVQGANERLF | 0.2619794 |
| AGLESIGL | GARVQGANERLF | 0.75272787 |
| AGLKTGII | GARVQGANERLF | 0.7946999 |
| ASLAALLL | GARVQGANERLF | 0.768376 |
| ASVINISSI | GARVQGANERLF | 0.7972692 |
| ATPPGPA | GARVQGANERLF | 0.6063491 |
| ATVASALV | GARVQGANERLF | 0.7503979 |
| ATVASLIL | GARVQGANERLF | 0.77467835 |
| AVNGPVLIRH | GARVQGANERLF | 0.5286586 |
| DGAAGAVGI | GARVQGANERLF | 0.63958436 |
| DPLLHGI | GARVQGANERLF | 0.6393838 |
| ELGAALGAL | GARVQGANERLF | 0.4631948 |
| ELGRTQAV | GARVQGANERLF | 0.7674263 |
| FVAGGYLS | GARVQGANERLF | 0.72394997 |
| GASGGAGGAGGW | GARVQGANERLF | 0.667567 |
| GGAGAGGDGGAGGI | GARVQGANERLF | 0.7562543 |
| GTGHNGSQPGMGGNG | GARVQGANERLF | 0.6768794 |
| GYAAASASAS | GARVQGANERLF | 0.6507623 |
| HQVDVAAVQQG | GARVQGANERLF | 0.7243167 |
| IGAAIAHT | GARVQGANERLF | 0.723226 |
| IGMIEVAQL | GARVQGANERLF | 0.79517823 |
| IRLAPEHTSAMD | GARVQGANERLF | 0.6664344 |
| IYNVSVL | GARVQGANERLF | 0.7226698 |
| KQGFLGM | GARVQGANERLF | 0.82528675 |
| LESVITQL | GARVQGANERLF | 0.4768685 |
| LGLGALVVV | GARVQGANERLF | 0.5558824 |
| LIGNGGNGGSGGI | GARVQGANERLF | 0.78672564 |
| LNAINAPTL | GARVQGANERLF | 0.518686 |
| LQLLNTGV | GARVQGANERLF | 0.79841405 |
| LQTSIEF | GARVQGANERLF | 0.78301066 |
| MSFAVSEL | GARVQGANERLF | 0.7885376 |
| NTGALGL | GARVQGANERLF | 0.7611333 |
| PPGAGGAGGAGTT | GARVQGANERLF | 0.82820064 |
| QLVGLGI | GARVQGANERLF | 0.7946266 |
| RFLEQVHQL | GARVQGANERLF | 0.68645954 |

| | | |
|---|---|---|
| RGVGYVLR | GARVQGANERLF | 0.81831163 |
| SATGVFAGVI | GARVQGANERLF | 0.53143334 |
| SLPAITLP | GARVQGANERLF | 0.81535226 |
| STALAFTV | GARVQGANERLF | 0.7983803 |
| SVLTVGLGK | GARVQGANERLF | 0.48399752 |
| TALAGAGSGI | GARVQGANERLF | 0.75509965 |
| TGLPFTGI | GARVQGANERLF | 0.8058256 |
| VAEHVTV | GARVQGANERLF | 0.8222512 |
| VDIVLNSL | GARVQGANERLF | 0.32532918 |
| VLTGGGAVL | GARVQGANERLF | 0.7076366 |
| VNVGLAYGI | GARVQGANERLF | 0.41909948 |
| VSSAAVGAV | GARVQGANERLF | 0.5226649 |
| VTSVYEAGV | GARVQGANERLF | 0.38747436 |
| WVALDRGA | GARVQGANERLF | 0.7683426 |
| YLVGSVVI | GARVQGANERLF | 0.6749109 |
| GPPGTGKTLL | GARVQGANERLF | 0.80996466 |
| GVGMTKF | GARVQGANERLF | 0.7096666 |
| IGIGGSDL | GARVQGANERLF | 0.7923728 |
| IGTLLAEL | GARVQGANERLF | 0.64298004 |
| IYTSGTT | GARVQGANERLF | 0.746392 |
| LSFPAGV | GARVQGANERLF | 0.36385876 |
| LYTSGTT | GARVQGANERLF | 0.35974988 |
| VGAFPFGF | GARVQGANERLF | 0.48371226 |
| VGAGTVEF | GARVQGANERLF | 0.72666097 |
| VGKSTLLL | GARVQGANERLF | 0.25290725 |
| AALAHAGGGIV | GARVQGANERLF | 0.81169075 |
| AALVAVSLL | GARVQGANERLF | 0.7349272 |
| AAVSRAVAAL | GARVQGANERLF | 0.76310915 |
| AGGSSGIGGF | GARVQGANERLF | 0.56991017 |
| AGLTVANANG | GARVQGANERLF | 0.7180939 |
| AGLVGAVL | GARVQGANERLF | 0.6929673 |
| ALSQAFTV | GARVQGANERLF | 0.80025625 |
| ALTFSVSL | GARVQGANERLF | 0.5631741 |
| ALTGAALVAL | GARVQGANERLF | 0.30268687 |
| ASFSIGAGVL | GARVQGANERLF | 0.6375085 |
| DGSVYKI | GARVQGANERLF | 0.5361555 |
| EQLAEIV | GARVQGANERLF | 0.8152529 |
| FAVSELTVM | GARVQGANERLF | 0.8084483 |
| GAAPLAAGL | GARVQGANERLF | 0.7706574 |
| GADGGDGGSGGAGG | GARVQGANERLF | 0.82112956 |
| GFGLDTGGAGG | GARVQGANERLF | 0.1982787 |
| GLASGGLL | GARVQGANERLF | 0.65295017 |
| GRIAGAAVLL | GARVQGANERLF | 0.819666 |
| GSAGSLNTSF | GARVQGANERLF | 0.74049187 |
| GTGQIGI | GARVQGANERLF | 0.7004999 |
| GTLIALIL | GARVQGANERLF | 0.8070479 |
| IAATMGFVVG | GARVQGANERLF | 0.69653213 |
| IFAQALL | GARVQGANERLF | 0.5473674 |
| IGAAVALGF | GARVQGANERLF | 0.7713666 |
| IGLTPLAL | GARVQGANERLF | 0.75136673 |
| ILGSNTSTL | GARVQGANERLF | 0.7006865 |

| | | |
|---|---|---|
| IVFSLTVL | GARVQGANERLF | 0.4774758 |
| LASKATGF | GARVQGANERLF | 0.70114326 |
| LASVALIL | GARVQGANERLF | 0.8328348 |
| LFAVGLTL | GARVQGANERLF | 0.7597785 |
| LFNGGIGTYI | GARVQGANERLF | 0.7728403 |
| LTLGAGII | GARVQGANERLF | 0.74929774 |
| LVAAALGDI | GARVQGANERLF | 0.76345885 |
| NTAFVQLGI | GARVQGANERLF | 0.6074195 |
| QLFSGAEL | GARVQGANERLF | 0.5752261 |
| SAATFITL | GARVQGANERLF | 0.7538533 |
| SNYGLANLGV | GARVQGANERLF | 0.5378809 |
| SYGGAYAVM | GARVQGANERLF | 0.6971495 |
| TTGVGGLTLG | GARVQGANERLF | 0.6329264 |
| TTQTIYI | GARVQGANERLF | 0.8336248 |
| VAAGAGLGAVI | GARVQGANERLF | 0.80095226 |
| VAAVVTLL | GARVQGANERLF | 0.76474166 |
| VGFLGGLI | GARVQGANERLF | 0.6992743 |
| VGVIPILL | GARVQGANERLF | 0.7030381 |
| VIFASSLI | GARVQGANERLF | 0.7756399 |
| VKGGDGGAAAG | GARVQGANERLF | 0.27646455 |
| VTKTFTNI | GARVQGANERLF | 0.81055784 |
| VVAAAILL | GARVQGANERLF | 0.5891204 |
| YAALDVEL | GARVQGANERLF | 0.6464355 |
| LASVALLL | GARVQGANERLF | 0.78052866 |
| LDAVHRV | GARVQGANERLF | 0.8244658 |
| QGLSSSIF | GARVQGANERLF | 0.65380096 |
| RGTFANIRL | GARVQGANERLF | 0.7357691 |
| RQLWWGHRI | GARVQGANERLF | 0.6916369 |
| YGPPGTGKTLL | GARVQGANERLF | 0.76241106 |
| AALVGAPANVA | GARVQGANERLF | 0.8193164 |
| AATFRRTE | GARVQGANERLF | 0.7685776 |
| AGALGGGI | GARVQGANERLF | 0.8280977 |
| AGGAGGAGGIGAG | GARVQGANERLF | 0.6593429 |
| AGSGALGL | GARVQGANERLF | 0.31500787 |
| AQIADQI | GARVQGANERLF | 0.6530724 |
| ASAVAVTYL | GARVQGANERLF | 0.7683882 |
| ASTVLGGTGG | GARVQGANERLF | 0.817008 |
| AVGGEPAK | GARVQGANERLF | 0.8053802 |
| CEQGTFGGTC | GARVQGANERLF | 0.7758697 |
| DEGKAPAG | GARVQGANERLF | 0.60777926 |
| DTVVFPAL | GARVQGANERLF | 0.8141656 |
| FEGITMAG | GARVQGANERLF | 0.8229662 |
| FPLVVQD | GARVQGANERLF | 0.7697186 |
| GAFTTPQ | GARVQGANERLF | 0.5389961 |
| GFGGFGTAGGVGGA | GARVQGANERLF | 0.474177 |
| GFLAQFL | GARVQGANERLF | 0.7364118 |
| GGAGGAGGNAGVA | GARVQGANERLF | 0.80256414 |
| GGVGTAGAGGAGGAGG | GARVQGANERLF | 0.46903658 |
| GNGGGGDAGNAGSGGN | GARVQGANERLF | 0.73890316 |
| GTGGAGGAGNDAGS | GARVQGANERLF | 0.7869588 |
| GTTNTGWFNS | GARVQGANERLF | 0.34937054 |

| | | |
|---|---|---|
| IFMGPLFTL | GARVQGANERLF | 0.67869496 |
| ILAHAIEAI | GARVQGANERLF | 0.8246002 |
| IRSAIREAL | GARVQGANERLF | 0.64881414 |
| ITFAGGG | GARVQGANERLF | 0.522552 |
| ITGMSGHQEV | GARVQGANERLF | 0.8178745 |
| IVAGLACTL | GARVQGANERLF | 0.8310777 |
| IVGGGLAGA | GARVQGANERLF | 0.7968571 |
| IVTGVLNI | GARVQGANERLF | 0.34769702 |
| LAQQGCI | GARVQGANERLF | 0.25021005 |
| LCLHVEN | GARVQGANERLF | 0.74847025 |
| LDAISDGS | GARVQGANERLF | 0.8118814 |
| LGGFAAML | GARVQGANERLF | 0.7222225 |
| LGGKNPMI | GARVQGANERLF | 0.6858374 |
| LGPKGVAAIL | GARVQGANERLF | 0.777532 |
| LMVGVSI | GARVQGANERLF | 0.80714667 |
| LVGAAFVL | GARVQGANERLF | 0.5555838 |
| LVLAVAPL | GARVQGANERLF | 0.40307027 |
| MAAIAVIDAA | GARVQGANERLF | 0.7789298 |
| NNGINAELQ | GARVQGANERLF | 0.7985707 |
| RGEPLVLVHG | GARVQGANERLF | 0.62650096 |
| RGGAGGAAGLI | GARVQGANERLF | 0.75513405 |
| RSYRMHPAV | GARVQGANERLF | 0.7260541 |
| SAAHLLAGQLA | GARVQGANERLF | 0.6357075 |
| SAAVGSY | GARVQGANERLF | 0.76462674 |
| SPHAVLHRI | GARVQGANERLF | 0.53472555 |
| SQIAKVL | GARVQGANERLF | 0.8210695 |
| TALGHLVP | GARVQGANERLF | 0.53530437 |
| TAPTTSGAA | GARVQGANERLF | 0.7774043 |
| TDGDDGNGGSPGT | GARVQGANERLF | 0.79259914 |
| TDGLGEN | GARVQGANERLF | 0.7455903 |
| TGGPPLPGQA | GARVQGANERLF | 0.69459456 |
| TIPAITI | GARVQGANERLF | 0.5579432 |
| TIPGFTI | GARVQGANERLF | 0.45570984 |
| TSIASGALA | GARVQGANERLF | 0.50961125 |
| TVTLPGAAI | GARVQGANERLF | 0.8134862 |
| VASGLSAALP | GARVQGANERLF | 0.6877752 |
| VGAGVIGI | GARVQGANERLF | 0.81606174 |
| VGPKRFGV | GARVQGANERLF | 0.72331905 |
| VNVAFPD | GARVQGANERLF | 0.623558 |
| VSMTNSGVS | GARVQGANERLF | 0.7713397 |
| VTDVLGDLSV | GARVQGANERLF | 0.5526203 |
| VVVVGAGFAGL | GARVQGANERLF | 0.5303919 |
| VYTVGTVV | GARVQGANERLF | 0.70941603 |
| WTTFSGP | GARVQGANERLF | 0.7696917 |
| YDELVEAF | GARVQGANERLF | 0.573364 |
| YGTGGAGGNGG | GARVQGANERLF | 0.74436104 |
| YTNVMAV | GARVQGANERLF | 0.74826217 |
| FLEQVHQL | GARVQGANERLF | 0.5023619 |
| GLSLLAVAV | GARVQGANERLF | 0.66085947 |
| LEQVHQL | GARVQGANERLF | 0.8152031 |
| LNAAGAI | GARVQGANERLF | 0.70476025 |

| | | |
|---|---|---|
| PAVHASL | GARVQGANERLF | 0.45817214 |
| TELEDDA | GARVQGANERLF | 0.76536363 |
| TPPGPPN | GARVQGANERLF | 0.5804452 |
| TPPPPPPP | GARVQGANERLF | 0.7905634 |
| TQSPALI | ASGDARQGNTEVF | 0.75786394 |
| EFAAAFPQ | ASGDARQGNTEVF | 0.660381 |
| LLSGNLDVL | ASGDARQGNTEVF | 0.75985694 |
| GPANLGNYNI | ASGDARQGNTEVF | 0.3886994 |
| TLGPTLP | ASGDARQGNTEVF | 0.5669043 |
| VYAYSSCCK | ASGDARQGNTEVF | 0.6222047 |
| APAPAPAPNI | ASGDARQGNTEVF | 0.77159965 |
| PGGPVPIP | ASGDARQGNTEVF | 0.7845212 |
| FLTVALEP | ASGDARQGNTEVF | 0.5497297 |
| SQYFGANL | ASGDARQGNTEVF | 0.72493017 |
| GGVKPAA | ASGDARQGNTEVF | 0.51937425 |
| GGTGGAGGAGGW | ASGDARQGNTEVF | 0.40641212 |
| GVGGVAGLGGAGG | ASGDARQGNTEVF | 0.64455795 |
| VLLVLATTI | ASGDARQGNTEVF | 0.6438297 |
| IGIFAAML | ASGDARQGNTEVF | 0.4157436 |
| AVGDLID | ASGDARQGNTEVF | 0.7714989 |
| NISPLPL | ASGDARQGNTEVF | 0.56517303 |
| VTAVFVAI | ASGDARQGNTEVF | 0.72446823 |
| SGDDNDAAL | ASGDARQGNTEVF | 0.8022419 |
| AVAALAHS | ASGDARQGNTEVF | 0.58484817 |
| GGVTVMATP | ASGDARQGNTEVF | 0.750307 |
| AGLAANVRD | ASGDARQGNTEVF | 0.7494366 |
| TNRPDLIDPALLRPG | ASGDARQGNTEVF | 0.7962063 |
| LANLGVTAL | ASGDARQGNTEVF | 0.7437174 |
| TGERIEL | ASGDARQGNTEVF | 0.55734175 |
| NGEELAA | ASGDARQGNTEVF | 0.79949677 |
| VLSLVWG | ASGDARQGNTEVF | 0.7786145 |
| IGYALWL | ASGDARQGNTEVF | 0.8070626 |
| LSLLLAPL | ASGDARQGNTEVF | 0.76846904 |
| TLDEALK | ASGDARQGNTEVF | 0.8148124 |
| TDGAGTAEGG | ASGDARQGNTEVF | 0.7868292 |
| NAPGGAGGNGG | ASGDARQGNTEVF | 0.72526896 |
| VSVFMGMGGNF | ASGDARQGNTEVF | 0.3125232 |
| LASYLAAL | ASGDARQGNTEVF | 0.74706626 |
| LVGGTPAL | ASGDARQGNTEVF | 0.7607137 |
| VRNADVL | ASGDARQGNTEVF | 0.6094963 |
| TCSSPFLVSAM | ASGDARQGNTEVF | 0.29670903 |
| VHITDAE | ASGDARQGNTEVF | 0.81059945 |
| EYAAPSF | ASGDARQGNTEVF | 0.83171844 |
| VVDAFHGGL | ASGDARQGNTEVF | 0.5971491 |
| ASVVDPQ | ASGDARQGNTEVF | 0.81973064 |
| SYNEFGGLS | ASGDARQGNTEVF | 0.39424485 |
| DPDGSVY | ASGDARQGNTEVF | 0.77223027 |
| MVSTIHATE | ASGDARQGNTEVF | 0.81453717 |
| LAAGVADPV | ASGDARQGNTEVF | 0.5403751 |
| TKEQIAA | ASGDARQGNTEVF | 0.7927033 |
| SLASPAAL | ASGDARQGNTEVF | 0.39801443 |

| | | |
|---|---|---|
| IGLVFYGV | ASGDARQGNTEVF | 0.5771728 |
| FLLDASNL | ASGDARQGNTEVF | 0.82059324 |
| YQVGNFP | ASGDARQGNTEVF | 0.6959246 |
| ESLLSQST | ASGDARQGNTEVF | 0.7356721 |
| LFTVGFAAVG | ASGDARQGNTEVF | 0.72078884 |
| ATPPTPT | ASGDARQGNTEVF | 0.60713696 |
| TDLATLG | ASGDARQGNTEVF | 0.7399132 |
| SAGGTGR | ASGDARQGNTEVF | 0.82036185 |
| GFSNSGS | ASGDARQGNTEVF | 0.59860533 |
| GGNGGSGGTSV | ASGDARQGNTEVF | 0.8199868 |
| GTSVATGGAGN | ASGDARQGNTEVF | 0.57875144 |
| NGGNGGSGGTS | ASGDARQGNTEVF | 0.8283947 |
| AVVKFPH | ASGDARQGNTEVF | 0.82223356 |
| TPPPPPP | ASGDARQGNTEVF | 0.78275573 |
| AAPAAAAAPA | ASGDARQGNTEVF | 0.63505167 |
| IALNEDL | ASGDARQGNTEVF | 0.81452185 |
| IVNVSSM | ASGDARQGNTEVF | 0.7137428 |
| TGIPPAPR | ASGDARQGNTEVF | 0.79614925 |
| FDLGGGTF | ASGDARQGNTEVF | 0.783025 |
| FDLGGGTFDVS | ASGDARQGNTEVF | 0.6389041 |
| NEPTAAA | ASGDARQGNTEVF | 0.8053274 |
| NEPTAAALA | ASGDARQGNTEVF | 0.783283 |
| GSIINISSI | ASGDARQGNTEVF | 0.74725986 |
| VAPGNAGTA | ASGDARQGNTEVF | 0.7410134 |
| TGGGASLE | ASGDARQGNTEVF | 0.4410723 |
| VAVYNHYKRI | ASGDARQGNTEVF | 0.71147096 |
| TDPAPATT | ASGDARQGNTEVF | 0.677823 |
| TNRPDILDPALLRPG | ASGDARQGNTEVF | 0.79622936 |
| LTTTTTS | ASGDARQGNTEVF | 0.7959369 |
| PTGSGKTL | ASGDARQGNTEVF | 0.72453773 |
| TGSGKTL | ASGDARQGNTEVF | 0.80354774 |
| TIGHVDHGK | ASGDARQGNTEVF | 0.7868495 |
| TIGHVDHGKTT | ASGDARQGNTEVF | 0.62688375 |
| GHVDHGKTT | ASGDARQGNTEVF | 0.75515175 |
| AQFEHTLL | ASGDARQGNTEVF | 0.5893613 |
| IVNQDEL | ASGDARQGNTEVF | 0.7699324 |
| TNGSQFF | ASGDARQGNTEVF | 0.3965481 |
| TGKGIVE | ASGDARQGNTEVF | 0.7886549 |
| LISNASDA | ASGDARQGNTEVF | 0.71616757 |
| AELRTLGELTKIAKAHGAQ | ASGDARQGNTEVF | 0.2355746 |
| APNAVAAAEPSW | ASGDARQGNTEVF | 0.7762326 |
| APQVAGAATAVL | ASGDARQGNTEVF | 0.76290524 |
| ARRDPAAAAAILPTDA | ASGDARQGNTEVF | 0.7663281 |
| DAAARDVLAEAGLAEYF | ASGDARQGNTEVF | 0.79703677 |
| DDQIVLSAKANLAKVEQ | ASGDARQGNTEVF | 0.6885133 |
| DELLPDTGEDSDEDGH | ASGDARQGNTEVF | 0.6846458 |
| ELPEAPANRATVPQD | ASGDARQGNTEVF | 0.794551 |
| FGQSKIERVTGIG | ASGDARQGNTEVF | 0.81195116 |
| GAGGTGSTAGGAGGAGGAGGLY | ASGDARQGNTEVF | 0.62969136 |
| GHGGTGAAGGNGGTGGAGGSGI | ASGDARQGNTEVF | 0.78605783 |
| GNGGNGGAGGEGGPGGAGGAGA | ASGDARQGNTEVF | 0.7978418 |

| | | |
|---|---|---|
| GSTAGNGGNGGNATNGTIAGQP | ASGDARQGNTEVF | 0.7715824 |
| GTGGDGGNGGNGGTGG | ASGDARQGNTEVF | 0.7965447 |
| GVRRPAGIPADAPV | ASGDARQGNTEVF | 0.80615675 |
| GYRQVTDAHLLTLARR | ASGDARQGNTEVF | 0.61788535 |
| IAKMGQDGHDRG | ASGDARQGNTEVF | 0.8113971 |
| KAEPSIEIQYT | ASGDARQGNTEVF | 0.8092693 |
| KPVRIRMAADEDTVAVT | ASGDARQGNTEVF | 0.7685425 |
| MPPRIAGMRLLVIKPEP | ASGDARQGNTEVF | 0.72842926 |
| MKFYAVALFRSRGVPPTR | ASGDARQGNTEVF | 0.6003896 |
| NAGGFGILGAVAHSP | ASGDARQGNTEVF | 0.74257433 |
| NEGREITLAMAEM | ASGDARQGNTEVF | 0.6391977 |
| NELVEWSHPLGHRWRWE | ASGDARQGNTEVF | 0.77380174 |
| NLGAGGTGGNGGNPG | ASGDARQGNTEVF | 0.8052991 |
| NTEWPKWETVMAADPVD | ASGDARQGNTEVF | 0.7359866 |
| PASAAPAETQP | ASGDARQGNTEVF | 0.612504 |
| QQSSDGKNGVRAAEITGE | ASGDARQGNTEVF | 0.62623346 |
| RAPDAVTTSWWKE | ASGDARQGNTEVF | 0.7537457 |
| SGFGNVGAHTSGWFNQ | ASGDARQGNTEVF | 0.7142266 |
| SGVAGAVGQAASVGGLKV | ASGDARQGNTEVF | 0.50650525 |
| SNGGAGTGSAGGNGGT | ASGDARQGNTEVF | 0.69858944 |
| TGGAGGEGGAAG | ASGDARQGNTEVF | 0.79641414 |
| TGGDGGAAGAGGAPGHG | ASGDARQGNTEVF | 0.8001605 |
| TMPPLLAAIANAGRHGV | ASGDARQGNTEVF | 0.5682938 |
| TSSTAGNSGGA | ASGDARQGNTEVF | 0.8107848 |
| VPATVPVGILC | ASGDARQGNTEVF | 0.7881005 |
| VQRPDSGGVVLTGRIS | ASGDARQGNTEVF | 0.8060324 |
| WTPNPVPATKGVPAGLVQAT | ASGDARQGNTEVF | 0.75331855 |
| YALKRALAKFSDRFDVVII | ASGDARQGNTEVF | 0.64778906 |
| YLIGSVIV | ASGDARQGNTEVF | 0.7961819 |
| YGQLVAVID | ASGDARQGNTEVF | 0.74698746 |
| VTVGGMTLPA | ASGDARQGNTEVF | 0.68165267 |
| VLALPAV | ASGDARQGNTEVF | 0.79317915 |
| VIPPTLNYET | ASGDARQGNTEVF | 0.74190533 |
| VGMGGMHPSSG | ASGDARQGNTEVF | 0.6914884 |
| VGLAGGAATA | ASGDARQGNTEVF | 0.733655 |
| VGGAGGAGGSGP | ASGDARQGNTEVF | 0.80099225 |
| VFAAFMV | ASGDARQGNTEVF | 0.62378407 |
| TLYDKLR | ASGDARQGNTEVF | 0.6963657 |
| TLAVFTVL | ASGDARQGNTEVF | 0.6786745 |
| TGSVQPARV | ASGDARQGNTEVF | 0.764025 |
| TGGGAGSGGT | ASGDARQGNTEVF | 0.8073139 |
| SHLETSL | ASGDARQGNTEVF | 0.78416693 |
| SFTTGMT | ASGDARQGNTEVF | 0.7286417 |
| QVEVAKRMGVRQPTVS | ASGDARQGNTEVF | 0.7969157 |
| QGSGGPALAM | ASGDARQGNTEVF | 0.78168726 |
| MLAPSNP | ASGDARQGNTEVF | 0.7682862 |
| LVNNAAI | ASGDARQGNTEVF | 0.76746815 |
| LTGEINTLA | ASGDARQGNTEVF | 0.69218177 |
| LSIFGPV | ASGDARQGNTEVF | 0.80007625 |
| LLALLANL | ASGDARQGNTEVF | 0.76956975 |
| LISNASDALDKLR | ASGDARQGNTEVF | 0.78865004 |

| | | |
|---|---|---|
| LINEAVP | ASGDARQGNTEVF | 0.7955947 |
| LGYAFSGL | ASGDARQGNTEVF | 0.789419 |
| LDFVAVAL | ASGDARQGNTEVF | 0.77670753 |
| LAQPGAE | ASGDARQGNTEVF | 0.45650196 |
| ISNLGAVTGF | ASGDARQGNTEVF | 0.63265777 |
| HQEDPGRI | ASGDARQGNTEVF | 0.79102784 |
| GTPGEPTP | ASGDARQGNTEVF | 0.77561724 |
| GFFNSGS | ASGDARQGNTEVF | 0.8016596 |
| GAGGGGLTAGAN | ASGDARQGNTEVF | 0.7558069 |
| FVDALVNM | ASGDARQGNTEVF | 0.72148067 |
| FLTVAPEL | ASGDARQGNTEVF | 0.7769905 |
| FETLTAL | ASGDARQGNTEVF | 0.8035077 |
| DVINPIEL | ASGDARQGNTEVF | 0.4885221 |
| APVSGAPV | ASGDARQGNTEVF | 0.73245233 |
| ALVGGLAW | ASGDARQGNTEVF | 0.8167755 |
| ALAAAAATAI | ASGDARQGNTEVF | 0.7716416 |
| AFGLGLLL | ASGDARQGNTEVF | 0.75000226 |
| AFFAAASL | ASGDARQGNTEVF | 0.69315827 |
| AAAALPAFSPP | ASGDARQGNTEVF | 0.81302476 |
| VVLAPAL | ASGDARQGNTEVF | 0.7225789 |
| TIGHVDHGKTTL | ASGDARQGNTEVF | 0.809515 |
| LISNASDALDK | ASGDARQGNTEVF | 0.35137904 |
| LISNASDAL | ASGDARQGNTEVF | 0.7211497 |
| ISNASDALDK | ASGDARQGNTEVF | 0.5392254 |
| NQAETLVYQTE | ASGDARQGNTEVF | 0.48166043 |
| TGGTGGTAGAG | ASGDARQGNTEVF | 0.78961575 |
| TGSTAGGAGGA | ASGDARQGNTEVF | 0.70535266 |
| DPLAVPVAAGPV | ASGDARQGNTEVF | 0.7955102 |
| GGTGGTPGNGGN | ASGDARQGNTEVF | 0.7484923 |
| PAGLAGVDPATL | ASGDARQGNTEVF | 0.7998915 |
| GNNLVVEAPHAGASV | ASGDARQGNTEVF | 0.7510594 |
| KVTRIMSASTLEQPA | ASGDARQGNTEVF | 0.77293336 |
| LGFALPIALPRRTVV | ASGDARQGNTEVF | 0.6955004 |
| TRLGNLTARKLVVLV | ASGDARQGNTEVF | 0.78080577 |
| WSQQRRLIYFPSAGP | ASGDARQGNTEVF | 0.7976501 |
| GPNGNPDPMAAAVDIR | ASGDARQGNTEVF | 0.76889884 |
| TQRCVSVADLRAAASR | ASGDARQGNTEVF | 0.4869768 |
| QFVLDVAEHPVEVLTETV | ASGDARQGNTEVF | 0.7397312 |
| GGAGGTGGAGGNGGTQAGDG | ASGDARQGNTEVF | 0.75792146 |
| PLLATLRAIAVDVDAKPAQV | ASGDARQGNTEVF | 0.36694318 |
| APKFRTGGYMIDFWGVGYQVAK | ASGDARQGNTEVF | 0.33584943 |
| GAGGAGGAGGAAGGPGGTGGTG | ASGDARQGNTEVF | 0.7398145 |
| IGALGLLATDVAPAGSVAFGVA | ASGDARQGNTEVF | 0.69358903 |
| AGGAGGIGGNGGAGGAGGVLMG | ASGDARQGNTEVF | 0.8056777 |
| ADELIGLL | ASGDARQGNTEVF | 0.7943176 |
| ADRALKA | ASGDARQGNTEVF | 0.73672616 |
| AGAVGQVGGAGGA | ASGDARQGNTEVF | 0.7959142 |
| AGGGGGLGGAP | ASGDARQGNTEVF | 0.7664275 |
| AGGLLYGNGGAG | ASGDARQGNTEVF | 0.19755818 |
| AGLESIGL | ASGDARQGNTEVF | 0.6739939 |
| AGLKTGII | ASGDARQGNTEVF | 0.63551354 |

| | | |
|---|---|---|
| ASLAALLL | ASGDARQGNTEVF | 0.8004428 |
| ASVINISSI | ASGDARQGNTEVF | 0.7083291 |
| ATPPGPA | ASGDARQGNTEVF | 0.6759795 |
| ATVASALV | ASGDARQGNTEVF | 0.7650759 |
| ATVASLIL | ASGDARQGNTEVF | 0.7724681 |
| AVNGPVLIRH | ASGDARQGNTEVF | 0.7481798 |
| DGAAGAVGI | ASGDARQGNTEVF | 0.6618355 |
| DPLLHGI | ASGDARQGNTEVF | 0.38902545 |
| ELGAALGAL | ASGDARQGNTEVF | 0.7908565 |
| ELGRTQAV | ASGDARQGNTEVF | 0.6043952 |
| FVAGGYLS | ASGDARQGNTEVF | 0.6909834 |
| GASGGAGGAGGW | ASGDARQGNTEVF | 0.41726822 |
| GGAGAGGDGGAGGI | ASGDARQGNTEVF | 0.66807103 |
| GTGHNGSQPGMGGNG | ASGDARQGNTEVF | 0.38865903 |
| GYAAASASAS | ASGDARQGNTEVF | 0.7873194 |
| HQVDVAAVQQG | ASGDARQGNTEVF | 0.7530986 |
| IGAAIAHT | ASGDARQGNTEVF | 0.72900796 |
| IGMIEVAQL | ASGDARQGNTEVF | 0.81009024 |
| IRLAPEHTSAMD | ASGDARQGNTEVF | 0.76931703 |
| IYNVSVL | ASGDARQGNTEVF | 0.74578357 |
| KQGFLGM | ASGDARQGNTEVF | 0.7295948 |
| LESVITQL | ASGDARQGNTEVF | 0.6883131 |
| LGLGALVVV | ASGDARQGNTEVF | 0.71518797 |
| LIGNGGNGGSGGI | ASGDARQGNTEVF | 0.71259356 |
| LNAINAPTL | ASGDARQGNTEVF | 0.80852115 |
| LQLLNTGV | ASGDARQGNTEVF | 0.70826924 |
| LQTSIEF | ASGDARQGNTEVF | 0.53722125 |
| MSFAVSEL | ASGDARQGNTEVF | 0.7430992 |
| NTGALGL | ASGDARQGNTEVF | 0.80816793 |
| PPGAGGAGGAGTT | ASGDARQGNTEVF | 0.8158226 |
| QLVGLGI | ASGDARQGNTEVF | 0.73051476 |
| RFLEQVHQL | ASGDARQGNTEVF | 0.6053494 |
| RGVGYVLR | ASGDARQGNTEVF | 0.7064394 |
| SATGVFAGVI | ASGDARQGNTEVF | 0.73861635 |
| SLPAITLP | ASGDARQGNTEVF | 0.64111793 |
| STALAFTV | ASGDARQGNTEVF | 0.5996431 |
| SVLTVGLGK | ASGDARQGNTEVF | 0.7948911 |
| TALAGAGSGI | ASGDARQGNTEVF | 0.81430316 |
| TGLPFTGI | ASGDARQGNTEVF | 0.7458988 |
| VAEHVTV | ASGDARQGNTEVF | 0.32136494 |
| VDIVLNSL | ASGDARQGNTEVF | 0.7542864 |
| VLTGGGAVL | ASGDARQGNTEVF | 0.6867055 |
| VNVGLAYGI | ASGDARQGNTEVF | 0.7342708 |
| VSSAAVGAV | ASGDARQGNTEVF | 0.7718172 |
| VTSVYEAGV | ASGDARQGNTEVF | 0.48147792 |
| WVALDRGA | ASGDARQGNTEVF | 0.7378033 |
| YLVGSVVI | ASGDARQGNTEVF | 0.8105403 |
| GPPGTGKTLL | ASGDARQGNTEVF | 0.753152 |
| GVGMTKF | ASGDARQGNTEVF | 0.7933841 |
| IGIGGSDL | ASGDARQGNTEVF | 0.8051846 |
| IGTLLAEL | ASGDARQGNTEVF | 0.78839386 |

| | | |
|---|---|---|
| IYTSGTT | ASGDARQGNTEVF | 0.80668724 |
| LSFPAGV | ASGDARQGNTEVF | 0.8019291 |
| LYTSGTT | ASGDARQGNTEVF | 0.73622525 |
| VGAFPFGF | ASGDARQGNTEVF | 0.7223136 |
| VGAGTVEF | ASGDARQGNTEVF | 0.7924874 |
| VGKSTLLL | ASGDARQGNTEVF | 0.76363313 |
| AALAHAGGGIV | ASGDARQGNTEVF | 0.760035 |
| AALVAVSLL | ASGDARQGNTEVF | 0.6717199 |
| AAVSRAVAAL | ASGDARQGNTEVF | 0.8041786 |
| AGGSSGIGGF | ASGDARQGNTEVF | 0.72043097 |
| AGLTVANANG | ASGDARQGNTEVF | 0.8075989 |
| AGLVGAVL | ASGDARQGNTEVF | 0.47543773 |
| ALSQAFTV | ASGDARQGNTEVF | 0.6945843 |
| ALTFSVSL | ASGDARQGNTEVF | 0.79689336 |
| ALTGAALVAL | ASGDARQGNTEVF | 0.21200082 |
| ASFSIGAGVL | ASGDARQGNTEVF | 0.3501051 |
| DGSVYKI | ASGDARQGNTEVF | 0.77950156 |
| EQLAEIV | ASGDARQGNTEVF | 0.80132604 |
| FAVSELTVM | ASGDARQGNTEVF | 0.24648578 |
| GAAPLAAGL | ASGDARQGNTEVF | 0.64899683 |
| GADGGDGGSGGAGG | ASGDARQGNTEVF | 0.548502 |
| GFGLDTGGAGG | ASGDARQGNTEVF | 0.78153944 |
| GLASGGLL | ASGDARQGNTEVF | 0.752959 |
| GRIAGAAVLL | ASGDARQGNTEVF | 0.46405885 |
| GSAGSLNTSF | ASGDARQGNTEVF | 0.7940105 |
| GTGQIGI | ASGDARQGNTEVF | 0.4077462 |
| GTLIALIL | ASGDARQGNTEVF | 0.70588905 |
| IAATMGFVVG | ASGDARQGNTEVF | 0.79626274 |
| IFAQALL | ASGDARQGNTEVF | 0.8101977 |
| IGAAVALGF | ASGDARQGNTEVF | 0.6191994 |
| IGLTPLAL | ASGDARQGNTEVF | 0.7966684 |
| ILGSNTSTL | ASGDARQGNTEVF | 0.66580856 |
| IVFSLTVL | ASGDARQGNTEVF | 0.8175622 |
| LASKATGF | ASGDARQGNTEVF | 0.75141203 |
| LASVALIL | ASGDARQGNTEVF | 0.4696123 |
| LFAVGLTL | ASGDARQGNTEVF | 0.7974786 |
| LFNGGIGTYI | ASGDARQGNTEVF | 0.7959111 |
| LTLGAGII | ASGDARQGNTEVF | 0.7509022 |
| LVAAALGDI | ASGDARQGNTEVF | 0.7969393 |
| NTAFVQLGI | ASGDARQGNTEVF | 0.7020463 |
| QLFSGAEL | ASGDARQGNTEVF | 0.7639898 |
| SAATFITL | ASGDARQGNTEVF | 0.8080231 |
| SNYGLANLGV | ASGDARQGNTEVF | 0.7980603 |
| SYGGAYAVM | ASGDARQGNTEVF | 0.5892498 |
| TTGVGGLTLG | ASGDARQGNTEVF | 0.79071707 |
| TTQTIYI | ASGDARQGNTEVF | 0.7476108 |
| VAAGAGLGAVI | ASGDARQGNTEVF | 0.6311467 |
| VAAVVTLL | ASGDARQGNTEVF | 0.80465525 |
| VGFLGGLI | ASGDARQGNTEVF | 0.7499795 |
| VGVIPILL | ASGDARQGNTEVF | 0.80001056 |
| VIFASSLI | ASGDARQGNTEVF | 0.78843695 |

| | | |
|---|---|---|
| VKGGDGGAAAG | ASGDARQGNTEVF | 0.804803 |
| VTKTFTNI | ASGDARQGNTEVF | 0.52138096 |
| VVAAAILL | ASGDARQGNTEVF | 0.75299466 |
| YAALDVEL | ASGDARQGNTEVF | 0.765767 |
| LASVALLL | ASGDARQGNTEVF | 0.79913986 |
| LDAVHRV | ASGDARQGNTEVF | 0.7331941 |
| QGLSSSIF | ASGDARQGNTEVF | 0.69083524 |
| RGTFANIRL | ASGDARQGNTEVF | 0.7909956 |
| RQLWWGHRI | ASGDARQGNTEVF | 0.8030968 |
| YGPPGTGKTLL | ASGDARQGNTEVF | 0.74738514 |
| AALVGAPANVA | ASGDARQGNTEVF | 0.7513343 |
| AATFRRTE | ASGDARQGNTEVF | 0.81024224 |
| AGALGGGI | ASGDARQGNTEVF | 0.7208879 |
| AGGAGGAGGIGAG | ASGDARQGNTEVF | 0.74666214 |
| AGSGALGL | ASGDARQGNTEVF | 0.78886455 |
| AQIADQI | ASGDARQGNTEVF | 0.80322415 |
| ASAVAVTYL | ASGDARQGNTEVF | 0.73035514 |
| ASTVLGGTGG | ASGDARQGNTEVF | 0.8080951 |
| AVGGEPAK | ASGDARQGNTEVF | 0.6541996 |
| CEQGTFGGTC | ASGDARQGNTEVF | 0.8080693 |
| DEGKAPAG | ASGDARQGNTEVF | 0.69631827 |
| DTVVFPAL | ASGDARQGNTEVF | 0.7770647 |
| FEGITMAG | ASGDARQGNTEVF | 0.68974483 |
| FPLVVQD | ASGDARQGNTEVF | 0.7970013 |
| GAFTTPQ | ASGDARQGNTEVF | 0.79677236 |
| GFGGFGTAGGVGGA | ASGDARQGNTEVF | 0.799294 |
| GFLAQFL | ASGDARQGNTEVF | 0.29913688 |
| GGAGGAGGNAGVA | ASGDARQGNTEVF | 0.79402024 |
| GGVGTAGAGGAGGAGG | ASGDARQGNTEVF | 0.8029174 |
| GNGGGGDAGNAGSGGN | ASGDARQGNTEVF | 0.81291926 |
| GTGGAGGAGNDAGS | ASGDARQGNTEVF | 0.7356031 |
| GTTNTGWFNS | ASGDARQGNTEVF | 0.7794607 |
| IFMGPLFTL | ASGDARQGNTEVF | 0.7511374 |
| ILAHAIEAI | ASGDARQGNTEVF | 0.7479675 |
| IRSAIREAL | ASGDARQGNTEVF | 0.63710976 |
| ITFAGGG | ASGDARQGNTEVF | 0.7378416 |
| ITGMSGHQEV | ASGDARQGNTEVF | 0.7233168 |
| IVAGLACTL | ASGDARQGNTEVF | 0.732019 |
| IVGGGLAGA | ASGDARQGNTEVF | 0.7918275 |
| IVTGVLNI | ASGDARQGNTEVF | 0.8000418 |
| LAQQGCI | ASGDARQGNTEVF | 0.79604566 |
| LCLHVEN | ASGDARQGNTEVF | 0.8082832 |
| LDAISDGS | ASGDARQGNTEVF | 0.8109647 |
| LGGFAAML | ASGDARQGNTEVF | 0.7989736 |
| LGGKNPMI | ASGDARQGNTEVF | 0.71483696 |
| LGPKGVAAIL | ASGDARQGNTEVF | 0.7524004 |
| LMVGVSI | ASGDARQGNTEVF | 0.6072103 |
| LVGAAFVL | ASGDARQGNTEVF | 0.7207761 |
| LVLAVAPL | ASGDARQGNTEVF | 0.7256141 |
| MAAIAVIDAA | ASGDARQGNTEVF | 0.7714184 |
| NNGINAELQ | ASGDARQGNTEVF | 0.22105184 |

| | | |
|---|---|---|
| RGEPLVLVHG | ASGDARQGNTEVF | 0.7471658 |
| RGGAGGAAGLI | ASGDARQGNTEVF | 0.81117827 |
| RSYRMHPAV | ASGDARQGNTEVF | 0.70657516 |
| SAAHLLAGQLA | ASGDARQGNTEVF | 0.7838825 |
| SAAVGSY | ASGDARQGNTEVF | 0.7697057 |
| SPHAVLHRI | ASGDARQGNTEVF | 0.67053795 |
| SQIAKVL | ASGDARQGNTEVF | 0.7972381 |
| TALGHLVP | ASGDARQGNTEVF | 0.6771959 |
| TAPTTSGAA | ASGDARQGNTEVF | 0.7857334 |
| TDGDDGNGGSPGT | ASGDARQGNTEVF | 0.80156124 |
| TDGLGEN | ASGDARQGNTEVF | 0.7818953 |
| TGGPPLPGQA | ASGDARQGNTEVF | 0.7112714 |
| TIPAITI | ASGDARQGNTEVF | 0.7413907 |
| TIPGFTI | ASGDARQGNTEVF | 0.45528245 |
| TSIASGALA | ASGDARQGNTEVF | 0.7642437 |
| TVTLPGAAI | ASGDARQGNTEVF | 0.6600413 |
| VASGLSAALP | ASGDARQGNTEVF | 0.77019477 |
| VGAGVIGI | ASGDARQGNTEVF | 0.7006564 |
| VGPKRFGV | ASGDARQGNTEVF | 0.7880589 |
| VNVAFPD | ASGDARQGNTEVF | 0.7646663 |
| VSMTNSGVS | ASGDARQGNTEVF | 0.7690642 |
| VTDVLGDLSV | ASGDARQGNTEVF | 0.6798481 |
| VVVVGAGFAGL | ASGDARQGNTEVF | 0.7598809 |
| VYTVGTVV | ASGDARQGNTEVF | 0.71103764 |
| WTTFSGP | ASGDARQGNTEVF | 0.60720223 |
| YDELVEAF | ASGDARQGNTEVF | 0.79004073 |
| YGTGGAGGNGG | ASGDARQGNTEVF | 0.7442338 |
| YTNVMAV | ASGDARQGNTEVF | 0.7148992 |
| FLEQVHQL | ASGDARQGNTEVF | 0.7986725 |
| GLSLLAVAV | ASGDARQGNTEVF | 0.7837938 |
| LEQVHQL | ASGDARQGNTEVF | 0.39018297 |
| LNAAGAI | ASGDARQGNTEVF | 0.76159877 |
| PAVHASL | ASGDARQGNTEVF | 0.76921594 |
| TELEDDA | ASGDARQGNTEVF | 0.7708779 |
| TPPGPPN | ASGDARQGNTEVF | 0.2684374 |
| TPPPPPPP | ASGDARQGNTEVF | 0.7916359 |
| TQSPALI | ASSSGQISNERLF | 0.6194051 |
| EFAAAFPQ | ASSSGQISNERLF | 0.6725492 |
| LLSGNLDVL | ASSSGQISNERLF | 0.69284254 |
| GPANLGNYNI | ASSSGQISNERLF | 0.8024397 |
| TLGPTLP | ASSSGQISNERLF | 0.71855795 |
| VYAYSSCCK | ASSSGQISNERLF | 0.7476253 |
| APAPAPAPNI | ASSSGQISNERLF | 0.79481864 |
| PGGPVPIP | ASSSGQISNERLF | 0.76911384 |
| FLTVALEP | ASSSGQISNERLF | 0.8075021 |
| SQYFGANL | ASSSGQISNERLF | 0.80067647 |
| GGVKPAA | ASSSGQISNERLF | 0.66364443 |
| GGTGGAGGAGGW | ASSSGQISNERLF | 0.7448052 |
| GVGGVAGLGGAGG | ASSSGQISNERLF | 0.64182234 |
| VLLVLATTI | ASSSGQISNERLF | 0.5902143 |
| IGIFAAML | ASSSGQISNERLF | 0.5697231 |

| | | |
|---|---|---|
| AVGDLID | ASSSGQISNERLF | 0.7491729 |
| NISPLPL | ASSSGQISNERLF | 0.4608161 |
| VTAVFVAI | ASSSGQISNERLF | 0.7792882 |
| SGDDNDAAL | ASSSGQISNERLF | 0.8027067 |
| AVAALAHS | ASSSGQISNERLF | 0.7474439 |
| GGVTVMATP | ASSSGQISNERLF | 0.46567684 |
| AGLAANVRD | ASSSGQISNERLF | 0.72081196 |
| TNRPDLIDPALLRPG | ASSSGQISNERLF | 0.6687844 |
| LANLGVTAL | ASSSGQISNERLF | 0.796682 |
| TGERIEL | ASSSGQISNERLF | 0.78479254 |
| NGEELAA | ASSSGQISNERLF | 0.8127425 |
| VLSLVWG | ASSSGQISNERLF | 0.7985201 |
| IGYALWL | ASSSGQISNERLF | 0.72312284 |
| LSLLLAPL | ASSSGQISNERLF | 0.7940303 |
| TLDEALK | ASSSGQISNERLF | 0.70801187 |
| TDGAGTAEGG | ASSSGQISNERLF | 0.69861186 |
| NAPGGAGGNGG | ASSSGQISNERLF | 0.7333254 |
| VSVFMGMGGNF | ASSSGQISNERLF | 0.7702263 |
| LASYLAAL | ASSSGQISNERLF | 0.77841324 |
| LVGGTPAL | ASSSGQISNERLF | 0.71812165 |
| VRNADVL | ASSSGQISNERLF | 0.41598076 |
| TCSSPFLVSAM | ASSSGQISNERLF | 0.7940335 |
| VHITDAE | ASSSGQISNERLF | 0.76824534 |
| EYAAPSF | ASSSGQISNERLF | 0.48321098 |
| VVDAFHGGL | ASSSGQISNERLF | 0.718741 |
| ASVVDPQ | ASSSGQISNERLF | 0.816653 |
| SYNEFGGLS | ASSSGQISNERLF | 0.80926317 |
| DPDGSVY | ASSSGQISNERLF | 0.70207024 |
| MVSTIHATE | ASSSGQISNERLF | 0.6902376 |
| LAAGVADPV | ASSSGQISNERLF | 0.76305866 |
| TKEQIAA | ASSSGQISNERLF | 0.5763744 |
| SLASPAAL | ASSSGQISNERLF | 0.77843404 |
| IGLVFYGV | ASSSGQISNERLF | 0.7433169 |
| FLLDASNL | ASSSGQISNERLF | 0.7460027 |
| YQVGNFP | ASSSGQISNERLF | 0.6010841 |
| ESLLSQST | ASSSGQISNERLF | 0.7711828 |
| LFTVGFAAVG | ASSSGQISNERLF | 0.7394019 |
| ATPPTPT | ASSSGQISNERLF | 0.7836299 |
| TDLATLG | ASSSGQISNERLF | 0.2471607 |
| SAGGTGR | ASSSGQISNERLF | 0.77929056 |
| GFSNSGS | ASSSGQISNERLF | 0.6884086 |
| GGNGGSGGTSV | ASSSGQISNERLF | 0.6604371 |
| GTSVATGGAGN | ASSSGQISNERLF | 0.38054112 |
| NGGNGGSGGTS | ASSSGQISNERLF | 0.6845633 |
| AVVKFPH | ASSSGQISNERLF | 0.8017112 |
| TPPPPPP | ASSSGQISNERLF | 0.8010932 |
| AAPAAAAAPA | ASSSGQISNERLF | 0.6920798 |
| IALNEDL | ASSSGQISNERLF | 0.80294794 |
| IVNVSSM | ASSSGQISNERLF | 0.8092098 |
| TGIPPAPR | ASSSGQISNERLF | 0.7057812 |
| FDLGGGTF | ASSSGQISNERLF | 0.4184174 |

| | | |
|---|---|---|
| FDLGGGTFDVS | ASSSGQISNERLF | 0.7834205 |
| NEPTAAA | ASSSGQISNERLF | 0.7332882 |
| NEPTAAALA | ASSSGQISNERLF | 0.78962815 |
| GSIINISSI | ASSSGQISNERLF | 0.8002889 |
| VAPGNAGTA | ASSSGQISNERLF | 0.6618973 |
| TGGGASLE | ASSSGQISNERLF | 0.7669066 |
| VAVYNHYKRI | ASSSGQISNERLF | 0.81253815 |
| TDPAPATT | ASSSGQISNERLF | 0.68265414 |
| TNRPDILDPALLRPG | ASSSGQISNERLF | 0.5215344 |
| LTTTTTS | ASSSGQISNERLF | 0.5875524 |
| PTGSGKTL | ASSSGQISNERLF | 0.5294053 |
| TGSGKTL | ASSSGQISNERLF | 0.776312 |
| TIGHVDHGK | ASSSGQISNERLF | 0.7121996 |
| TIGHVDHGKTT | ASSSGQISNERLF | 0.7141708 |
| GHVDHGKTT | ASSSGQISNERLF | 0.75927496 |
| AQFEHTLL | ASSSGQISNERLF | 0.72222626 |
| IVNQDEL | ASSSGQISNERLF | 0.7290202 |
| TNGSQFF | ASSSGQISNERLF | 0.7290071 |
| TGKGIVE | ASSSGQISNERLF | 0.7655678 |
| LISNASDA | ASSSGQISNERLF | 0.7841557 |
| AELRTLGELTKIAKAHGAQ | ASSSGQISNERLF | 0.704171 |
| APNAVAAAEPSW | ASSSGQISNERLF | 0.7109221 |
| APQVAGAATAVL | ASSSGQISNERLF | 0.8074616 |
| ARRDPAAAAAILPTDA | ASSSGQISNERLF | 0.6004145 |
| DAAARDVLAEAGLAEYF | ASSSGQISNERLF | 0.6442455 |
| DDQIVLSAKANLAKVEQ | ASSSGQISNERLF | 0.79135287 |
| DELLPDTGEDSDEDGH | ASSSGQISNERLF | 0.6086726 |
| ELPEAPANRATVPQD | ASSSGQISNERLF | 0.7981031 |
| FGQSKIERVTGIG | ASSSGQISNERLF | 0.76067436 |
| GAGGTGSTAGGAGGAGGAGGLY | ASSSGQISNERLF | 0.7765642 |
| GHGGTGAAGGNGGTGGAGGSGI | ASSSGQISNERLF | 0.74186456 |
| GNGGNGGAGGEGGPGGAGGAGA | ASSSGQISNERLF | 0.8138819 |
| GSTAGNGGNGGNATNGTIAGQP | ASSSGQISNERLF | 0.7950858 |
| GTGGDGGNGGNGGTGG | ASSSGQISNERLF | 0.66998523 |
| GVRRPAGIPADAPV | ASSSGQISNERLF | 0.7906941 |
| GYRQVTDAHLLTLARR | ASSSGQISNERLF | 0.5042912 |
| IAKMGQDGHDRG | ASSSGQISNERLF | 0.7995784 |
| KAEPSIEIQYT | ASSSGQISNERLF | 0.7987545 |
| KPVRIRMAADEDTVAVT | ASSSGQISNERLF | 0.65624857 |
| MPPRIAGMRLLVIKPEP | ASSSGQISNERLF | 0.76878655 |
| MKFYAVALFRSRGVPPTR | ASSSGQISNERLF | 0.80244315 |
| NAGGFGILGAVAHSP | ASSSGQISNERLF | 0.8105408 |
| NEGREITLAMAEM | ASSSGQISNERLF | 0.5600726 |
| NELVEWSHPLGHRWRWE | ASSSGQISNERLF | 0.7192652 |
| NLGAGGTGGNGGNPG | ASSSGQISNERLF | 0.45437205 |
| NTEWPKWETVMAADPVD | ASSSGQISNERLF | 0.5741408 |
| PASAAPAETQP | ASSSGQISNERLF | 0.4744283 |
| QQSSDGKNGVRAAEITGE | ASSSGQISNERLF | 0.78886205 |
| RAPDAVTTSWWKE | ASSSGQISNERLF | 0.74217916 |
| SGFGNVGAHTSGWFNQ | ASSSGQISNERLF | 0.7926576 |
| SGVAGAVGQAASVGGLKV | ASSSGQISNERLF | 0.7837843 |

| | | |
|---|---|---|
| SNGGAGTGSAGGNGGT | ASSSGQISNERLF | 0.8048156 |
| TGGAGGEGGAAG | ASSSGQISNERLF | 0.7354646 |
| TGGDGGAAGAGGAPGHG | ASSSGQISNERLF | 0.7623595 |
| TMPPLLAAIANAGRHGV | ASSSGQISNERLF | 0.57144356 |
| TSSTAGNSGGA | ASSSGQISNERLF | 0.60397756 |
| VPATVPVGILC | ASSSGQISNERLF | 0.5738992 |
| VQRPDSGGVVLTGRIS | ASSSGQISNERLF | 0.7346827 |
| WTPNPVPATKGVPAGLVQAT | ASSSGQISNERLF | 0.40904808 |
| YALKRALAKFSDRFDVVII | ASSSGQISNERLF | 0.80582666 |
| YLIGSVIV | ASSSGQISNERLF | 0.77634823 |
| YGQLVAVID | ASSSGQISNERLF | 0.7878482 |
| VTVGGMTLPA | ASSSGQISNERLF | 0.7045979 |
| VLALPAV | ASSSGQISNERLF | 0.694069 |
| VIPPTLNYET | ASSSGQISNERLF | 0.72422445 |
| VGMGGMHPSSG | ASSSGQISNERLF | 0.8004741 |
| VGLAGGAATA | ASSSGQISNERLF | 0.70462704 |
| VGGAGGAGGSGP | ASSSGQISNERLF | 0.35590878 |
| VFAAFMV | ASSSGQISNERLF | 0.6160083 |
| TLYDKLR | ASSSGQISNERLF | 0.40118423 |
| TLAVFTVL | ASSSGQISNERLF | 0.8102404 |
| TGSVQPARV | ASSSGQISNERLF | 0.8018904 |
| TGGGAGSGGT | ASSSGQISNERLF | 0.7890926 |
| SHLETSL | ASSSGQISNERLF | 0.8120382 |
| SFTTGMT | ASSSGQISNERLF | 0.26010332 |
| QVEVAKRMGVRQPTVS | ASSSGQISNERLF | 0.6860689 |
| QGSGGPALAM | ASSSGQISNERLF | 0.80792546 |
| MLAPSNP | ASSSGQISNERLF | 0.78058434 |
| LVNNAAI | ASSSGQISNERLF | 0.76388705 |
| LTGEINTLA | ASSSGQISNERLF | 0.80528224 |
| LSIFGPV | ASSSGQISNERLF | 0.6976832 |
| LLALLANL | ASSSGQISNERLF | 0.77151895 |
| LISNASDALDKLR | ASSSGQISNERLF | 0.7676007 |
| LINEAVP | ASSSGQISNERLF | 0.7830089 |
| LGYAFSGL | ASSSGQISNERLF | 0.74803305 |
| LDFVAVAL | ASSSGQISNERLF | 0.60942507 |
| LAQPGAE | ASSSGQISNERLF | 0.77896243 |
| ISNLGAVTGF | ASSSGQISNERLF | 0.8127644 |
| HQEDPGRI | ASSSGQISNERLF | 0.7674061 |
| GTPGEPTP | ASSSGQISNERLF | 0.79513896 |
| GFFNSGS | ASSSGQISNERLF | 0.7843019 |
| GAGGGGLTAGAN | ASSSGQISNERLF | 0.7724364 |
| FVDALVNM | ASSSGQISNERLF | 0.71235096 |
| FLTVAPEL | ASSSGQISNERLF | 0.68598485 |
| FETLTAL | ASSSGQISNERLF | 0.75456476 |
| DVINPIEL | ASSSGQISNERLF | 0.65617657 |
| APVSGAPV | ASSSGQISNERLF | 0.7443284 |
| ALVGGLAW | ASSSGQISNERLF | 0.6878709 |
| ALAAAAATAI | ASSSGQISNERLF | 0.8192199 |
| AFGLGLLL | ASSSGQISNERLF | 0.7913489 |
| AFFAAASL | ASSSGQISNERLF | 0.7866869 |
| AAAALPAFSPP | ASSSGQISNERLF | 0.6873739 |

| | | |
|---|---|---|
| VVLAPAL | ASSSGQISNERLF | 0.761038 |
| TIGHVDHGKTTL | ASSSGQISNERLF | 0.7691991 |
| LISNASDALDK | ASSSGQISNERLF | 0.4263776 |
| LISNASDAL | ASSSGQISNERLF | 0.80432796 |
| ISNASDALDK | ASSSGQISNERLF | 0.6795243 |
| NQAETLVYQTE | ASSSGQISNERLF | 0.7328323 |
| TGGTGGTAGAG | ASSSGQISNERLF | 0.79393977 |
| TGSTAGGAGGA | ASSSGQISNERLF | 0.80499053 |
| DPLAVPVAAGPV | ASSSGQISNERLF | 0.7405255 |
| GGTGGTPGNGGN | ASSSGQISNERLF | 0.70957756 |
| PAGLAGVDPATL | ASSSGQISNERLF | 0.69272137 |
| GNNLVVEAPHAGASV | ASSSGQISNERLF | 0.79920065 |
| KVTRIMSASTLEQPA | ASSSGQISNERLF | 0.8083365 |
| LGFALPIALPRRTVV | ASSSGQISNERLF | 0.77390736 |
| TRLGNLTARKLVVLV | ASSSGQISNERLF | 0.8168095 |
| WSQQRRLIYFPSAGP | ASSSGQISNERLF | 0.7476413 |
| GPNGNPDPMAAAVDIR | ASSSGQISNERLF | 0.42654356 |
| TQRCVSVADLRAAASR | ASSSGQISNERLF | 0.78151894 |
| QFVLDVAEHPVEVLTETV | ASSSGQISNERLF | 0.7965131 |
| GGAGGTGGAGGNGGTQAGDG | ASSSGQISNERLF | 0.7988409 |
| PLLATLRAIAVDVDAKPAQV | ASSSGQISNERLF | 0.8089313 |
| APKFRTGGYMIDFWGVGYQVAK | ASSSGQISNERLF | 0.7865665 |
| GAGGAGGAGGAAGGPGGTGGTG | ASSSGQISNERLF | 0.72417384 |
| IGALGLLATDVAPAGSVAFGVA | ASSSGQISNERLF | 0.8060293 |
| AGGAGGIGGNGGAGGAGGVLMG | ASSSGQISNERLF | 0.8147826 |
| ADELIGLL | ASSSGQISNERLF | 0.79760206 |
| ADRALKA | ASSSGQISNERLF | 0.7413583 |
| AGAVGQVGGAGGA | ASSSGQISNERLF | 0.6956596 |
| AGGGGGLGGAP | ASSSGQISNERLF | 0.7720082 |
| AGGLLYGNGGAG | ASSSGQISNERLF | 0.7947377 |
| AGLESIGL | ASSSGQISNERLF | 0.60455716 |
| AGLKTGII | ASSSGQISNERLF | 0.7766919 |
| ASLAALLL | ASSSGQISNERLF | 0.7707974 |
| ASVINISSI | ASSSGQISNERLF | 0.4392443 |
| ATPPGPA | ASSSGQISNERLF | 0.7827134 |
| ATVASALV | ASSSGQISNERLF | 0.806333 |
| ATVASLIL | ASSSGQISNERLF | 0.72776055 |
| AVNGPVLIRH | ASSSGQISNERLF | 0.66516024 |
| DGAAGAVGI | ASSSGQISNERLF | 0.80457866 |
| DPLLHGI | ASSSGQISNERLF | 0.81817555 |
| ELGAALGAL | ASSSGQISNERLF | 0.8039359 |
| ELGRTQAV | ASSSGQISNERLF | 0.5041261 |
| FVAGGYLS | ASSSGQISNERLF | 0.31581372 |
| GASGGAGGAGGW | ASSSGQISNERLF | 0.74663234 |
| GGAGAGGDGGAGGI | ASSSGQISNERLF | 0.7967158 |
| GTGHNGSQPGMGGNG | ASSSGQISNERLF | 0.7432047 |
| GYAAASASAS | ASSSGQISNERLF | 0.74313295 |
| HQVDVAAVQQG | ASSSGQISNERLF | 0.77361465 |
| IGAAIAHT | ASSSGQISNERLF | 0.79581785 |
| IGMIEVAQL | ASSSGQISNERLF | 0.6801592 |
| IRLAPEHTSAMD | ASSSGQISNERLF | 0.55480945 |

| | | |
|---|---|---|
| IYNVSVL | ASSSGQISNERLF | 0.7267211 |
| KQGFLGM | ASSSGQISNERLF | 0.80174774 |
| LESVITQL | ASSSGQISNERLF | 0.7027836 |
| LGLGALVVV | ASSSGQISNERLF | 0.7636864 |
| LIGNGGNGGSGGI | ASSSGQISNERLF | 0.7164174 |
| LNAINAPTL | ASSSGQISNERLF | 0.7205696 |
| LQLLNTGV | ASSSGQISNERLF | 0.75444126 |
| LQTSIEF | ASSSGQISNERLF | 0.6894133 |
| MSFAVSEL | ASSSGQISNERLF | 0.80594647 |
| NTGALGL | ASSSGQISNERLF | 0.6143084 |
| PPGAGGAGGAGTT | ASSSGQISNERLF | 0.7890655 |
| QLVGLGI | ASSSGQISNERLF | 0.79059374 |
| RFLEQVHQL | ASSSGQISNERLF | 0.7910068 |
| RGVGYVLR | ASSSGQISNERLF | 0.75491774 |
| SATGVFAGVI | ASSSGQISNERLF | 0.71285903 |
| SLPAITLP | ASSSGQISNERLF | 0.5734818 |
| STALAFTV | ASSSGQISNERLF | 0.58442956 |
| SVLTVGLGK | ASSSGQISNERLF | 0.80464786 |
| TALAGAGSGI | ASSSGQISNERLF | 0.6937454 |
| TGLPFTGI | ASSSGQISNERLF | 0.8100488 |
| VAEHVTV | ASSSGQISNERLF | 0.78091013 |
| VDIVLNSL | ASSSGQISNERLF | 0.7128074 |
| VLTGGGAVL | ASSSGQISNERLF | 0.7847018 |
| VNVGLAYGI | ASSSGQISNERLF | 0.67474675 |
| VSSAAVGAV | ASSSGQISNERLF | 0.70970875 |
| VTSVYEAGV | ASSSGQISNERLF | 0.7911434 |
| WVALDRGA | ASSSGQISNERLF | 0.77704465 |
| YLVGSVVI | ASSSGQISNERLF | 0.6970438 |
| GPPGTGKTLL | ASSSGQISNERLF | 0.76649207 |
| GVGMTKF | ASSSGQISNERLF | 0.78283024 |
| IGIGGSDL | ASSSGQISNERLF | 0.55935717 |
| IGTLLAEL | ASSSGQISNERLF | 0.74496424 |
| IYTSGTT | ASSSGQISNERLF | 0.8003291 |
| LSFPAGV | ASSSGQISNERLF | 0.7640797 |
| LYTSGTT | ASSSGQISNERLF | 0.66524637 |
| VGAFPFGF | ASSSGQISNERLF | 0.77728885 |
| VGAGTVEF | ASSSGQISNERLF | 0.6906141 |
| VGKSTLLL | ASSSGQISNERLF | 0.78371507 |
| AALAHAGGGIV | ASSSGQISNERLF | 0.7705386 |
| AALVAVSLL | ASSSGQISNERLF | 0.7330125 |
| AAVSRAVAAL | ASSSGQISNERLF | 0.79222083 |
| AGGSSGIGGF | ASSSGQISNERLF | 0.42888802 |
| AGLTVANANG | ASSSGQISNERLF | 0.6809825 |
| AGLVGAVL | ASSSGQISNERLF | 0.7430696 |
| ALSQAFTV | ASSSGQISNERLF | 0.7725996 |
| ALTFSVSL | ASSSGQISNERLF | 0.7973519 |
| ALTGAALVAL | ASSSGQISNERLF | 0.66275156 |
| ASFSIGAGVL | ASSSGQISNERLF | 0.781054 |
| DGSVYKI | ASSSGQISNERLF | 0.73699147 |
| EQLAEIV | ASSSGQISNERLF | 0.69561464 |
| FAVSELTVM | ASSSGQISNERLF | 0.72756743 |

| | | |
|---|---|---|
| GAAPLAAGL | ASSSGQISNERLF | 0.6624766 |
| GADGGDGGSGGAGG | ASSSGQISNERLF | 0.6533443 |
| GFGLDTGGAGG | ASSSGQISNERLF | 0.78465134 |
| GLASGGLL | ASSSGQISNERLF | 0.6913099 |
| GRIAGAAVLL | ASSSGQISNERLF | 0.76723194 |
| GSAGSLNTSF | ASSSGQISNERLF | 0.7786522 |
| GTGQIGI | ASSSGQISNERLF | 0.67066205 |
| GTLIALIL | ASSSGQISNERLF | 0.7832875 |
| IAATMGFVVG | ASSSGQISNERLF | 0.78175044 |
| IFAQALL | ASSSGQISNERLF | 0.79672617 |
| IGAAVALGF | ASSSGQISNERLF | 0.7266676 |
| IGLTPLAL | ASSSGQISNERLF | 0.65502334 |
| ILGSNTSTL | ASSSGQISNERLF | 0.8015413 |
| IVFSLTVL | ASSSGQISNERLF | 0.7877872 |
| LASKATGF | ASSSGQISNERLF | 0.74880844 |
| LASVALIL | ASSSGQISNERLF | 0.7917687 |
| LFAVGLTL | ASSSGQISNERLF | 0.78378296 |
| LFNGGIGTYI | ASSSGQISNERLF | 0.7909248 |
| LTLGAGII | ASSSGQISNERLF | 0.7429558 |
| LVAAALGDI | ASSSGQISNERLF | 0.53019905 |
| NTAFVQLGI | ASSSGQISNERLF | 0.74894 |
| QLFSGAEL | ASSSGQISNERLF | 0.75405824 |
| SAATFITL | ASSSGQISNERLF | 0.6980626 |
| SNYGLANLGV | ASSSGQISNERLF | 0.485919 |
| SYGGAYAVM | ASSSGQISNERLF | 0.8030506 |
| TTGVGGLTLG | ASSSGQISNERLF | 0.8187589 |
| TTQTIYI | ASSSGQISNERLF | 0.69614625 |
| VAAGAGLGAVI | ASSSGQISNERLF | 0.81500286 |
| VAAVVTLL | ASSSGQISNERLF | 0.4779508 |
| VGFLGGLI | ASSSGQISNERLF | 0.789573 |
| VGVIPILL | ASSSGQISNERLF | 0.80418044 |
| VIFASSLI | ASSSGQISNERLF | 0.6699431 |
| VKGGDGGAAAG | ASSSGQISNERLF | 0.78492963 |
| VTKTFTNI | ASSSGQISNERLF | 0.61855596 |
| VVAAAILL | ASSSGQISNERLF | 0.70830566 |
| YAALDVEL | ASSSGQISNERLF | 0.8103066 |
| LASVALLL | ASSSGQISNERLF | 0.7069018 |
| LDAVHRV | ASSSGQISNERLF | 0.74690783 |
| QGLSSSIF | ASSSGQISNERLF | 0.76902384 |
| RGTFANIRL | ASSSGQISNERLF | 0.69850683 |
| RQLWWGHRI | ASSSGQISNERLF | 0.77651316 |
| YGPPGTGKTLL | ASSSGQISNERLF | 0.7984736 |
| AALVGAPANVA | ASSSGQISNERLF | 0.7000426 |
| AATFRRTE | ASSSGQISNERLF | 0.81429464 |
| AGALGGGI | ASSSGQISNERLF | 0.6835993 |
| AGGAGGAGGIGAG | ASSSGQISNERLF | 0.81659895 |
| AGSGALGL | ASSSGQISNERLF | 0.80082047 |
| AQIADQI | ASSSGQISNERLF | 0.7718538 |
| ASAVAVTYL | ASSSGQISNERLF | 0.72860754 |
| ASTVLGGTGG | ASSSGQISNERLF | 0.8032806 |
| AVGGEPAK | ASSSGQISNERLF | 0.7659099 |

| | | |
|---|---|---|
| CEQGTFGGTC | ASSSGQISNERLF | 0.7627472 |
| DEGKAPAG | ASSSGQISNERLF | 0.80493367 |
| DTVVFPAL | ASSSGQISNERLF | 0.1648246 |
| FEGITMAG | ASSSGQISNERLF | 0.46829966 |
| FPLVVQD | ASSSGQISNERLF | 0.23362741 |
| GAFTTPQ | ASSSGQISNERLF | 0.18830198 |
| GFGGFGTAGGVGGA | ASSSGQISNERLF | 0.18628016 |
| GFLAQFL | ASSSGQISNERLF | 0.14902298 |
| GGAGGAGGNAGVA | ASSSGQISNERLF | 0.18904692 |
| GGVGTAGAGGAGGAGG | ASSSGQISNERLF | 0.17688383 |
| GNGGGGDAGNAGSGGN | ASSSGQISNERLF | 0.3509729 |
| GTGGAGGAGNDAGS | ASSSGQISNERLF | 0.20272161 |
| GTTNTGWFNS | ASSSGQISNERLF | 0.1728045 |
| IFMGPLFTL | ASSSGQISNERLF | 0.57243305 |
| ILAHAIEAI | ASSSGQISNERLF | 0.49171686 |
| IRSAIREAL | ASSSGQISNERLF | 0.154914 |
| ITFAGGG | ASSSGQISNERLF | 0.19868805 |
| ITGMSGHQEV | ASSSGQISNERLF | 0.16654626 |
| IVAGLACTL | ASSSGQISNERLF | 0.18156865 |
| IVGGGLAGA | ASSSGQISNERLF | 0.14457983 |
| IVTGVLNI | ASSSGQISNERLF | 0.24942546 |
| LAQQGCI | ASSSGQISNERLF | 0.16451594 |
| LCLHVEN | ASSSGQISNERLF | 0.1420798 |
| LDAISDGS | ASSSGQISNERLF | 0.18795967 |
| LGGFAAML | ASSSGQISNERLF | 0.17579882 |
| LGGKNPMI | ASSSGQISNERLF | 0.18704611 |
| LGPKGVAAIL | ASSSGQISNERLF | 0.49765342 |
| LMVGVSI | ASSSGQISNERLF | 0.1467214 |
| LVGAAFVL | ASSSGQISNERLF | 0.16435805 |
| LVLAVAPL | ASSSGQISNERLF | 0.4028498 |
| MAAIAVIDAA | ASSSGQISNERLF | 0.65707964 |
| NNGINAELQ | ASSSGQISNERLF | 0.15779169 |
| RGEPLVLVHG | ASSSGQISNERLF | 0.30190745 |
| RGGAGGAAGLI | ASSSGQISNERLF | 0.5665647 |
| RSYRMHPAV | ASSSGQISNERLF | 0.25102037 |
| SAAHLLAGQLA | ASSSGQISNERLF | 0.27270138 |
| SAAVGSY | ASSSGQISNERLF | 0.31606132 |
| SPHAVLHRI | ASSSGQISNERLF | 0.1528095 |
| SQIAKVL | ASSSGQISNERLF | 0.4271288 |
| TALGHLVP | ASSSGQISNERLF | 0.5056939 |
| TAPTTSGAA | ASSSGQISNERLF | 0.19305712 |
| TDGDDGNGGSPGT | ASSSGQISNERLF | 0.22386591 |
| TDGLGEN | ASSSGQISNERLF | 0.16129383 |
| TGGPPLPGQA | ASSSGQISNERLF | 0.1593045 |
| TIPAITI | ASSSGQISNERLF | 0.14494735 |
| TIPGFTI | ASSSGQISNERLF | 0.40758455 |
| TSIASGALA | ASSSGQISNERLF | 0.27988383 |
| TVTLPGAAI | ASSSGQISNERLF | 0.22072527 |
| VASGLSAALP | ASSSGQISNERLF | 0.15482922 |
| VGAGVIGI | ASSSGQISNERLF | 0.16471545 |
| VGPKRFGV | ASSSGQISNERLF | 0.16327488 |

| | | |
|---|---|---|
| VNVAFPD | ASSSGQISNERLF | 0.18170524 |
| VSMTNSGVS | ASSSGQISNERLF | 0.14667235 |
| VTDVLGDLSV | ASSSGQISNERLF | 0.15217067 |
| VVVVGAGFAGL | ASSSGQISNERLF | 0.47193182 |
| VYTVGTVV | ASSSGQISNERLF | 0.46363342 |
| WTTFSGP | ASSSGQISNERLF | 0.15108587 |
| YDELVEAF | ASSSGQISNERLF | 0.5495498 |
| YGTGGAGGNGG | ASSSGQISNERLF | 0.28162655 |
| YTNVMAV | ASSSGQISNERLF | 0.38891244 |
| FLEQVHQL | ASSSGQISNERLF | 0.19701356 |
| GLSLLAVAV | ASSSGQISNERLF | 0.16433105 |
| LEQVHQL | ASSSGQISNERLF | 0.35014683 |
| LNAAGAI | ASSSGQISNERLF | 0.23807383 |
| PAVHASL | ASSSGQISNERLF | 0.15913238 |
| TELEDDA | ASSSGQISNERLF | 0.27205914 |
| TPPGPPN | ASSSGQISNERLF | 0.18914366 |
| TPPPPPPP | ASSSGQISNERLF | 0.19904338 |
| TQSPALI | ASGDDRVSYNSPLY | 0.25812656 |
| EFAAAFPQ | ASGDDRVSYNSPLY | 0.24402642 |
| LLSGNLDVL | ASGDDRVSYNSPLY | 0.15646043 |
| GPANLGNYNI | ASGDDRVSYNSPLY | 0.15908901 |
| TLGPTLP | ASGDDRVSYNSPLY | 0.15461564 |
| VYAYSSCCK | ASGDDRVSYNSPLY | 0.31253803 |
| APAPAPAPNI | ASGDDRVSYNSPLY | 0.5775478 |
| PGGPVPIP | ASGDDRVSYNSPLY | 0.20456813 |
| FLTVALEP | ASGDDRVSYNSPLY | 0.17499997 |
| SQYFGANL | ASGDDRVSYNSPLY | 0.47666568 |
| GGVKPAA | ASGDDRVSYNSPLY | 0.22718877 |
| GGTGGAGGAGGW | ASGDDRVSYNSPLY | 0.19569962 |
| GVGGVAGLGGAGG | ASGDDRVSYNSPLY | 0.3079077 |
| VLLVLATTI | ASGDDRVSYNSPLY | 0.1942507 |
| IGIFAAML | ASGDDRVSYNSPLY | 0.24992111 |
| AVGDLID | ASGDDRVSYNSPLY | 0.23386271 |
| NISPLPL | ASGDDRVSYNSPLY | 0.23251721 |
| VTAVFVAI | ASGDDRVSYNSPLY | 0.26281136 |
| SGDDNDAAL | ASGDDRVSYNSPLY | 0.2143518 |
| AVAALAHS | ASGDDRVSYNSPLY | 0.19309153 |
| GGVTVMATP | ASGDDRVSYNSPLY | 0.1481102 |
| AGLAANVRD | ASGDDRVSYNSPLY | 0.17124331 |
| TNRPDLIDPALLRPG | ASGDDRVSYNSPLY | 0.24313226 |
| LANLGVTAL | ASGDDRVSYNSPLY | 0.2886294 |
| TGERIEL | ASGDDRVSYNSPLY | 0.22771731 |
| NGEELAA | ASGDDRVSYNSPLY | 0.21647406 |
| VLSLVWG | ASGDDRVSYNSPLY | 0.1645454 |
| IGYALWL | ASGDDRVSYNSPLY | 0.2581588 |
| LSLLLAPL | ASGDDRVSYNSPLY | 0.46775743 |
| TLDEALK | ASGDDRVSYNSPLY | 0.15445983 |
| TDGAGTAEGG | ASGDDRVSYNSPLY | 0.17469585 |
| NAPGGAGGNGG | ASGDDRVSYNSPLY | 0.50488555 |
| VSVFMGMGGNF | ASGDDRVSYNSPLY | 0.30709535 |
| LASYLAAL | ASGDDRVSYNSPLY | 0.19785859 |

| | | |
|---|---|---|
| LVGGTPAL | ASGDDRVSYNSPLY | 0.16740121 |
| VRNADVL | ASGDDRVSYNSPLY | 0.6511434 |
| TCSSPFLVSAM | ASGDDRVSYNSPLY | 0.156943 |
| VHITDAE | ASGDDRVSYNSPLY | 0.45369598 |
| EYAAPSF | ASGDDRVSYNSPLY | 0.14460589 |
| VVDAFHGGL | ASGDDRVSYNSPLY | 0.17122939 |
| ASVVDPQ | ASGDDRVSYNSPLY | 0.16805468 |
| SYNEFGGLS | ASGDDRVSYNSPLY | 0.14711943 |
| DPDGSVY | ASGDDRVSYNSPLY | 0.34830022 |
| MVSTIHATE | ASGDDRVSYNSPLY | 0.16598493 |
| LAAGVADPV | ASGDDRVSYNSPLY | 0.35976 |
| TKEQIAA | ASGDDRVSYNSPLY | 0.18932778 |
| SLASPAAL | ASGDDRVSYNSPLY | 0.26201713 |
| IGLVFYGV | ASGDDRVSYNSPLY | 0.23883164 |
| FLLDASNL | ASGDDRVSYNSPLY | 0.18791284 |
| YQVGNFP | ASGDDRVSYNSPLY | 0.1625279 |
| ESLLSQST | ASGDDRVSYNSPLY | 0.31710976 |
| LFTVGFAAVG | ASGDDRVSYNSPLY | 0.26112893 |
| ATPPTPT | ASGDDRVSYNSPLY | 0.21072228 |
| TDLATLG | ASGDDRVSYNSPLY | 0.15217458 |
| SAGGTGR | ASGDDRVSYNSPLY | 0.16456257 |
| GFSNSGS | ASGDDRVSYNSPLY | 0.28973752 |
| GGNGGSGGTSV | ASGDDRVSYNSPLY | 0.14491034 |
| GTSVATGGAGN | ASGDDRVSYNSPLY | 0.14297874 |
| NGGNGGSGGTS | ASGDDRVSYNSPLY | 0.17572047 |
| AVVKFPH | ASGDDRVSYNSPLY | 0.15109435 |
| TPPPPPP | ASGDDRVSYNSPLY | 0.5384521 |
| AAPAAAAAPA | ASGDDRVSYNSPLY | 0.53946114 |
| IALNEDL | ASGDDRVSYNSPLY | 0.20995879 |
| IVNVSSM | ASGDDRVSYNSPLY | 0.33906353 |
| TGIPPAPR | ASGDDRVSYNSPLY | 0.24834822 |
| FDLGGGTF | ASGDDRVSYNSPLY | 0.14115746 |
| FDLGGGTFDVS | ASGDDRVSYNSPLY | 0.17421032 |
| NEPTAAA | ASGDDRVSYNSPLY | 0.15590015 |
| NEPTAAALA | ASGDDRVSYNSPLY | 0.25879914 |
| GSIINISSI | ASGDDRVSYNSPLY | 0.1557614 |
| VAPGNAGTA | ASGDDRVSYNSPLY | 0.17507139 |
| TGGGASLE | ASGDDRVSYNSPLY | 0.26608863 |
| VAVYNHYKRI | ASGDDRVSYNSPLY | 0.36415493 |
| TDPAPATT | ASGDDRVSYNSPLY | 0.20611693 |
| TNRPDILDPALLRPG | ASGDDRVSYNSPLY | 0.15958686 |
| LTTTTTS | ASGDDRVSYNSPLY | 0.14167711 |
| PTGSGKTL | ASGDDRVSYNSPLY | 0.34579492 |
| TGSGKTL | ASGDDRVSYNSPLY | 0.1486808 |
| TIGHVDHGK | ASGDDRVSYNSPLY | 0.16584118 |
| TIGHVDHGKTT | ASGDDRVSYNSPLY | 0.143824 |
| GHVDHGKTT | ASGDDRVSYNSPLY | 0.14345929 |
| AQFEHTLL | ASGDDRVSYNSPLY | 0.15017816 |
| IVNQDEL | ASGDDRVSYNSPLY | 0.23040223 |
| TNGSQFF | ASGDDRVSYNSPLY | 0.17734563 |
| TGKGIVE | ASGDDRVSYNSPLY | 0.21021283 |

| | | |
|---|---|---|
| LISNASDA | ASGDDRVSYNSPLY | 0.6212645 |
| AELRTLGELTKIAKAHGAQ | ASGDDRVSYNSPLY | 0.59787154 |
| APNAVAAAEPSW | ASGDDRVSYNSPLY | 0.19780649 |
| APQVAGAATAVL | ASGDDRVSYNSPLY | 0.1753129 |
| ARRDPAAAAAILPTDA | ASGDDRVSYNSPLY | 0.1582674 |
| DAAARDVLAEAGLAEYF | ASGDDRVSYNSPLY | 0.16217193 |
| DDQIVLSAKANLAKVEQ | ASGDDRVSYNSPLY | 0.1612701 |
| DELLPDTGEDSDEDGH | ASGDDRVSYNSPLY | 0.5958922 |
| ELPEAPANRATVPQD | ASGDDRVSYNSPLY | 0.16589025 |
| FGQSKIERVTGIG | ASGDDRVSYNSPLY | 0.15380792 |
| GAGGTGSTAGGAGGAGGAGGLY | ASGDDRVSYNSPLY | 0.18238735 |
| GHGGTGAAGGNGGTGGAGGSGI | ASGDDRVSYNSPLY | 0.36047292 |
| GNGGNGGAGGEGGPGGAGGAGA | ASGDDRVSYNSPLY | 0.61327744 |
| GSTAGNGGNGGNATNGTIAGQP | ASGDDRVSYNSPLY | 0.16722709 |
| GTGGDGGNGGNGGTGG | ASGDDRVSYNSPLY | 0.18046266 |
| GVRRPAGIPADAPV | ASGDDRVSYNSPLY | 0.17521289 |
| GYRQVTDAHLLTLARR | ASGDDRVSYNSPLY | 0.3356314 |
| IAKMGQDGHDRG | ASGDDRVSYNSPLY | 0.16610819 |
| KAEPSIEIQYT | ASGDDRVSYNSPLY | 0.16107193 |
| KPVRIRMAADEDTVAVT | ASGDDRVSYNSPLY | 0.24406408 |
| MPPRIAGMRLLVIKPEP | ASGDDRVSYNSPLY | 0.6887771 |
| MKFYAVALFRSRGVPPTR | ASGDDRVSYNSPLY | 0.16808224 |
| NAGGFGILGAVAHSP | ASGDDRVSYNSPLY | 0.1505297 |
| NEGREITLAMAEM | ASGDDRVSYNSPLY | 0.25428936 |
| NELVEWSHPLGHRWRWE | ASGDDRVSYNSPLY | 0.16927911 |
| NLGAGGTGGNGGNPG | ASGDDRVSYNSPLY | 0.22141445 |
| NTEWPKWETVMAADPVD | ASGDDRVSYNSPLY | 0.2505567 |
| PASAAPAETQP | ASGDDRVSYNSPLY | 0.15100838 |
| QQSSDGKNGVRAAEITGE | ASGDDRVSYNSPLY | 0.16170438 |
| RAPDAVTTSWWKE | ASGDDRVSYNSPLY | 0.6261836 |
| SGFGNVGAHTSGWFNQ | ASGDDRVSYNSPLY | 0.22145224 |
| SGVAGAVGQAASVGGLKV | ASGDDRVSYNSPLY | 0.19980916 |
| SNGGAGTGSAGGNGGT | ASGDDRVSYNSPLY | 0.5066928 |
| TGGAGGEGGAAG | ASGDDRVSYNSPLY | 0.3169444 |
| TGGDGGAAGAGGAPGHG | ASGDDRVSYNSPLY | 0.6218183 |
| TMPPLLAAIANAGRHGV | ASGDDRVSYNSPLY | 0.36076957 |
| TSSTAGNSGGA | ASGDDRVSYNSPLY | 0.17217496 |
| VPATVPVGILC | ASGDDRVSYNSPLY | 0.19585794 |
| VQRPDSGGVVLTGRIS | ASGDDRVSYNSPLY | 0.18093285 |
| WTPNPVPATKGVPAGLVQAT | ASGDDRVSYNSPLY | 0.27358437 |
| YALKRALAKFSDRFDVVII | ASGDDRVSYNSPLY | 0.22027147 |
| YLIGSVIV | ASGDDRVSYNSPLY | 0.15974933 |
| YGQLVAVID | ASGDDRVSYNSPLY | 0.5517255 |
| VTVGGMTLPA | ASGDDRVSYNSPLY | 0.22051258 |
| VLALPAV | ASGDDRVSYNSPLY | 0.22630033 |
| VIPPTLNYET | ASGDDRVSYNSPLY | 0.15353301 |
| VGMGGMHPSSG | ASGDDRVSYNSPLY | 0.15978111 |
| VGLAGGAATA | ASGDDRVSYNSPLY | 0.37041256 |
| VGGAGGAGGSGP | ASGDDRVSYNSPLY | 0.14227636 |
| VFAAFMV | ASGDDRVSYNSPLY | 0.14388117 |
| TLYDKLR | ASGDDRVSYNSPLY | 0.20605788 |

| | | |
|---|---|---|
| TLAVFTVL | ASGDDRVSYNSPLY | 0.509793 |
| TGSVQPARV | ASGDDRVSYNSPLY | 0.1422025 |
| TGGGAGSGGT | ASGDDRVSYNSPLY | 0.16683654 |
| SHLETSL | ASGDDRVSYNSPLY | 0.1493063 |
| SFTTGMT | ASGDDRVSYNSPLY | 0.19645384 |
| QVEVAKRMGVRQPTVS | ASGDDRVSYNSPLY | 0.21485847 |
| QGSGGPALAM | ASGDDRVSYNSPLY | 0.14800675 |
| MLAPSNP | ASGDDRVSYNSPLY | 0.24859416 |
| LVNNAAI | ASGDDRVSYNSPLY | 0.14780132 |
| LTGEINTLA | ASGDDRVSYNSPLY | 0.14965881 |
| LSIFGPV | ASGDDRVSYNSPLY | 0.23836815 |
| LLALLANL | ASGDDRVSYNSPLY | 0.45221552 |
| LISNASDALDKLR | ASGDDRVSYNSPLY | 0.15386961 |
| LINEAVP | ASGDDRVSYNSPLY | 0.3727705 |
| LGYAFSGL | ASGDDRVSYNSPLY | 0.16778408 |
| LDFVAVAL | ASGDDRVSYNSPLY | 0.61892 |
| LAQPGAE | ASGDDRVSYNSPLY | 0.20102628 |
| ISNLGAVTGF | ASGDDRVSYNSPLY | 0.14918181 |
| HQEDPGRI | ASGDDRVSYNSPLY | 0.3165415 |
| GTPGEPTP | ASGDDRVSYNSPLY | 0.18564813 |
| GFFNSGS | ASGDDRVSYNSPLY | 0.20433177 |
| GAGGGGLTAGAN | ASGDDRVSYNSPLY | 0.36155334 |
| FVDALVNM | ASGDDRVSYNSPLY | 0.16698375 |
| FLTVAPEL | ASGDDRVSYNSPLY | 0.24609244 |
| FETLTAL | ASGDDRVSYNSPLY | 0.5326131 |
| DVINPIEL | ASGDDRVSYNSPLY | 0.3125816 |
| APVSGAPV | ASGDDRVSYNSPLY | 0.15904336 |
| ALVGGLAW | ASGDDRVSYNSPLY | 0.20725697 |
| ALAAAAATAI | ASGDDRVSYNSPLY | 0.26003093 |
| AFGLGLLL | ASGDDRVSYNSPLY | 0.1733918 |
| AFFAAASL | ASGDDRVSYNSPLY | 0.44754243 |
| AAAALPAFSPP | ASGDDRVSYNSPLY | 0.1957471 |
| VVLAPAL | ASGDDRVSYNSPLY | 0.29464918 |
| TIGHVDHGKTTL | ASGDDRVSYNSPLY | 0.23554835 |
| LISNASDALDK | ASGDDRVSYNSPLY | 0.64850837 |
| LISNASDAL | ASGDDRVSYNSPLY | 0.15302563 |
| ISNASDALDK | ASGDDRVSYNSPLY | 0.19603954 |
| NQAETLVYQTE | ASGDDRVSYNSPLY | 0.23225181 |
| TGGTGGTAGAG | ASGDDRVSYNSPLY | 0.33262837 |
| TGSTAGGAGGA | ASGDDRVSYNSPLY | 0.16592866 |
| DPLAVPVAAGPV | ASGDDRVSYNSPLY | 0.15057224 |
| GGTGGTPGNGGN | ASGDDRVSYNSPLY | 0.18516535 |
| PAGLAGVDPATL | ASGDDRVSYNSPLY | 0.37897894 |
| GNNLVVEAPHAGASV | ASGDDRVSYNSPLY | 0.18162973 |
| KVTRIMSASTLEQPA | ASGDDRVSYNSPLY | 0.21353777 |
| LGFALPIALPRRTVV | ASGDDRVSYNSPLY | 0.70278347 |
| TRLGNLTARKLVVLV | ASGDDRVSYNSPLY | 0.1631942 |
| WSQQRRLIYFPSAGP | ASGDDRVSYNSPLY | 0.19649664 |
| GPNGNPDPMAAAVDIR | ASGDDRVSYNSPLY | 0.2314077 |
| TQRCVSVADLRAAASR | ASGDDRVSYNSPLY | 0.34011737 |
| QFVLDVAEHPVEVLTETV | ASGDDRVSYNSPLY | 0.18389179 |

| | | |
|---|---|---|
| GGAGGTGGAGGNGGTQAGDG | ASGDDRVSYNSPLY | 0.3938102 |
| PLLATLRAIAVDVDAKPAQV | ASGDDRVSYNSPLY | 0.15680607 |
| APKFRTGGYMIDFWGVGYQVAK | ASGDDRVSYNSPLY | 0.4563404 |
| GAGGAGGAGGAAGGPGGTGGTG | ASGDDRVSYNSPLY | 0.15435657 |
| IGALGLLATDVAPAGSVAFGVA | ASGDDRVSYNSPLY | 0.2929504 |
| AGGAGGIGGNGGAGGAGGVLMG | ASGDDRVSYNSPLY | 0.16226226 |
| ADELIGLL | ASGDDRVSYNSPLY | 0.4366937 |
| ADRALKA | ASGDDRVSYNSPLY | 0.5031276 |
| AGAVGQVGGAGGA | ASGDDRVSYNSPLY | 0.48840117 |
| AGGGGGLGGAP | ASGDDRVSYNSPLY | 0.1432536 |
| AGGLLYGNGGAG | ASGDDRVSYNSPLY | 0.4489444 |
| AGLESIGL | ASGDDRVSYNSPLY | 0.5418418 |
| AGLKTGII | ASGDDRVSYNSPLY | 0.7391721 |
| ASLAALLL | ASGDDRVSYNSPLY | 0.204039 |
| ASVINISSI | ASGDDRVSYNSPLY | 0.24786705 |
| ATPPGPA | ASGDDRVSYNSPLY | 0.22347622 |
| ATVASALV | ASGDDRVSYNSPLY | 0.17678739 |
| ATVASLIL | ASGDDRVSYNSPLY | 0.16541739 |
| AVNGPVLIRH | ASGDDRVSYNSPLY | 0.18209167 |
| DGAAGAVGI | ASGDDRVSYNSPLY | 0.26408252 |
| DPLLHGI | ASGDDRVSYNSPLY | 0.2020829 |
| ELGAALGAL | ASGDDRVSYNSPLY | 0.4920707 |
| ELGRTQAV | ASGDDRVSYNSPLY | 0.42289215 |
| FVAGGYLS | ASGDDRVSYNSPLY | 0.3977827 |
| GASGGAGGAGGW | ASGDDRVSYNSPLY | 0.6374454 |
| GGAGAGGDGGAGGI | ASGDDRVSYNSPLY | 0.38119775 |
| GTGHNGSQPGMGGNG | ASGDDRVSYNSPLY | 0.2271385 |
| GYAAASASAS | ASGDDRVSYNSPLY | 0.17155619 |
| HQVDVAAVQQG | ASGDDRVSYNSPLY | 0.18471023 |
| IGAAIAHT | ASGDDRVSYNSPLY | 0.16263351 |
| IGMIEVAQL | ASGDDRVSYNSPLY | 0.1664543 |
| IRLAPEHTSAMD | ASGDDRVSYNSPLY | 0.1775424 |
| IYNVSVL | ASGDDRVSYNSPLY | 0.24998271 |
| KQGFLGM | ASGDDRVSYNSPLY | 0.14162962 |
| LESVITQL | ASGDDRVSYNSPLY | 0.2684914 |
| LGLGALVVV | ASGDDRVSYNSPLY | 0.54608244 |
| LIGNGGNGGSGGI | ASGDDRVSYNSPLY | 0.15933265 |
| LNAINAPTL | ASGDDRVSYNSPLY | 0.22530346 |
| LQLLNTGV | ASGDDRVSYNSPLY | 0.16445231 |
| LQTSIEF | ASGDDRVSYNSPLY | 0.16045012 |
| MSFAVSEL | ASGDDRVSYNSPLY | 0.56741893 |
| NTGALGL | ASGDDRVSYNSPLY | 0.15723296 |
| PPGAGGAGGAGTT | ASGDDRVSYNSPLY | 0.2341741 |
| QLVGLGI | ASGDDRVSYNSPLY | 0.4655481 |
| RFLEQVHQL | ASGDDRVSYNSPLY | 0.2112568 |
| RGVGYVLR | ASGDDRVSYNSPLY | 0.19731212 |
| SATGVFAGVI | ASGDDRVSYNSPLY | 0.177596 |
| SLPAITLP | ASGDDRVSYNSPLY | 0.15212567 |
| STALAFTV | ASGDDRVSYNSPLY | 0.24126121 |
| SVLTVGLGK | ASGDDRVSYNSPLY | 0.14130673 |
| TALAGAGSGI | ASGDDRVSYNSPLY | 0.2592587 |

| | | |
|---|---|---|
| TGLPFTGI | ASGDDRVSYNSPLY | 0.1570231 |
| VAEHVTV | ASGDDRVSYNSPLY | 0.3887644 |
| VDIVLNSL | ASGDDRVSYNSPLY | 0.27343544 |
| VLTGGGAVL | ASGDDRVSYNSPLY | 0.21473925 |
| VNVGLAYGI | ASGDDRVSYNSPLY | 0.16668366 |
| VSSAAVGAV | ASGDDRVSYNSPLY | 0.17961104 |
| VTSVYEAGV | ASGDDRVSYNSPLY | 0.16527274 |
| WVALDRGA | ASGDDRVSYNSPLY | 0.15587214 |
| YLVGSVVI | ASGDDRVSYNSPLY | 0.22784014 |
| GPPGTGKTLL | ASGDDRVSYNSPLY | 0.18146044 |
| GVGMTKF | ASGDDRVSYNSPLY | 0.16812807 |
| IGIGGSDL | ASGDDRVSYNSPLY | 0.3575377 |
| IGTLLAEL | ASGDDRVSYNSPLY | 0.201865 |
| IYTSGTT | ASGDDRVSYNSPLY | 0.15134096 |
| LSFPAGV | ASGDDRVSYNSPLY | 0.29693192 |
| LYTSGTT | ASGDDRVSYNSPLY | 0.21458206 |
| VGAFPFGF | ASGDDRVSYNSPLY | 0.19711062 |
| VGAGTVEF | ASGDDRVSYNSPLY | 0.14106686 |
| VGKSTLLL | ASGDDRVSYNSPLY | 0.17884947 |
| AALAHAGGGIV | ASGDDRVSYNSPLY | 0.17261693 |
| AALVAVSLL | ASGDDRVSYNSPLY | 0.16724752 |
| AAVSRAVAAL | ASGDDRVSYNSPLY | 0.16379829 |
| AGGSSGIGGF | ASGDDRVSYNSPLY | 0.60469556 |
| AGLTVANANG | ASGDDRVSYNSPLY | 0.15131976 |
| AGLVGAVL | ASGDDRVSYNSPLY | 0.24762005 |
| ALSQAFTV | ASGDDRVSYNSPLY | 0.30900264 |
| ALTFSVSL | ASGDDRVSYNSPLY | 0.28537264 |
| ALTGAALVAL | ASGDDRVSYNSPLY | 0.6190331 |
| ASFSIGAGVL | ASGDDRVSYNSPLY | 0.23191535 |
| DGSVYKI | ASGDDRVSYNSPLY | 0.18531454 |
| EQLAEIV | ASGDDRVSYNSPLY | 0.19452491 |
| FAVSELTVM | ASGDDRVSYNSPLY | 0.19870819 |
| GAAPLAAGL | ASGDDRVSYNSPLY | 0.14488433 |
| GADGGDGGSGGAGG | ASGDDRVSYNSPLY | 0.14453225 |
| GFGLDTGGAGG | ASGDDRVSYNSPLY | 0.22683765 |
| GLASGGLL | ASGDDRVSYNSPLY | 0.14405432 |
| GRIAGAAVLL | ASGDDRVSYNSPLY | 0.25067157 |
| GSAGSLNTSF | ASGDDRVSYNSPLY | 0.24345186 |
| GTGQIGI | ASGDDRVSYNSPLY | 0.17833282 |
| GTLIALIL | ASGDDRVSYNSPLY | 0.15380938 |
| IAATMGFVVG | ASGDDRVSYNSPLY | 0.17884135 |
| IFAQALL | ASGDDRVSYNSPLY | 0.18337983 |
| IGAAVALGF | ASGDDRVSYNSPLY | 0.3679865 |
| IGLTPLAL | ASGDDRVSYNSPLY | 0.21077655 |
| ILGSNTSTL | ASGDDRVSYNSPLY | 0.72725123 |
| IVFSLTVL | ASGDDRVSYNSPLY | 0.51767814 |
| LASKATGF | ASGDDRVSYNSPLY | 0.20001537 |
| LASVALIL | ASGDDRVSYNSPLY | 0.39029783 |
| LFAVGLTL | ASGDDRVSYNSPLY | 0.17011578 |
| LFNGGIGTYI | ASGDDRVSYNSPLY | 0.15727006 |
| LTLGAGII | ASGDDRVSYNSPLY | 0.16823424 |

| | | |
|---|---|---|
| LVAAALGDI | ASGDDRVSYNSPLY | 0.1837835 |
| NTAFVQLGI | ASGDDRVSYNSPLY | 0.1783095 |
| QLFSGAEL | ASGDDRVSYNSPLY | 0.15929352 |
| SAATFITL | ASGDDRVSYNSPLY | 0.15098175 |
| SNYGLANLGV | ASGDDRVSYNSPLY | 0.18705893 |
| SYGGAYAVM | ASGDDRVSYNSPLY | 0.24817555 |
| TTGVGGLTLG | ASGDDRVSYNSPLY | 0.14726955 |
| TTQTIYI | ASGDDRVSYNSPLY | 0.15805183 |
| VAAGAGLGAVI | ASGDDRVSYNSPLY | 0.7383097 |
| VAAVVTLL | ASGDDRVSYNSPLY | 0.69973737 |
| VGFLGGLI | ASGDDRVSYNSPLY | 0.18567881 |
| VGVIPILL | ASGDDRVSYNSPLY | 0.16876382 |
| VIFASSLI | ASGDDRVSYNSPLY | 0.31198603 |
| VKGGDGGAAAG | ASGDDRVSYNSPLY | 0.14973433 |
| VTKTFTNI | ASGDDRVSYNSPLY | 0.1880483 |
| VVAAAILL | ASGDDRVSYNSPLY | 0.33684933 |
| YAALDVEL | ASGDDRVSYNSPLY | 0.19689137 |
| LASVALLL | ASGDDRVSYNSPLY | 0.15863186 |
| LDAVHRV | ASGDDRVSYNSPLY | 0.20741436 |
| QGLSSSIF | ASGDDRVSYNSPLY | 0.17880483 |
| RGTFANIRL | ASGDDRVSYNSPLY | 0.2334277 |
| RQLWWGHRI | ASGDDRVSYNSPLY | 0.14176998 |
| YGPPGTGKTLL | ASGDDRVSYNSPLY | 0.18306041 |
| AALVGAPANVA | ASGDDRVSYNSPLY | 0.17006613 |
| AATFRRTE | ASGDDRVSYNSPLY | 0.17144895 |
| AGALGGGI | ASGDDRVSYNSPLY | 0.1507126 |
| AGGAGGAGGIGAG | ASGDDRVSYNSPLY | 0.18087146 |
| AGSGALGL | ASGDDRVSYNSPLY | 0.51817924 |
| AQIADQI | ASGDDRVSYNSPLY | 0.45689997 |
| ASAVAVTYL | ASGDDRVSYNSPLY | 0.14697148 |
| ASTVLGGTGG | ASGDDRVSYNSPLY | 0.4421903 |
| AVGGEPAK | ASGDDRVSYNSPLY | 0.66146165 |
| CEQGTFGGTC | ASGDDRVSYNSPLY | 0.17902076 |
| DEGKAPAG | ASGDDRVSYNSPLY | 0.1495924 |
| DTVVFPAL | ASGDDRVSYNSPLY | 0.2726393 |
| FEGITMAG | ASGDDRVSYNSPLY | 0.1800065 |
| FPLVVQD | ASGDDRVSYNSPLY | 0.24130484 |
| GAFTTPQ | ASGDDRVSYNSPLY | 0.2641703 |
| GFGGFGTAGGVGGA | ASGDDRVSYNSPLY | 0.15572432 |
| GFLAQFL | ASGDDRVSYNSPLY | 0.30898213 |
| GGAGGAGGNAGVA | ASGDDRVSYNSPLY | 0.34636813 |
| GGVGTAGAGGAGGAGG | ASGDDRVSYNSPLY | 0.15994778 |
| GNGGGGDAGNAGSGGN | ASGDDRVSYNSPLY | 0.15901734 |
| GTGGAGGAGNDAGS | ASGDDRVSYNSPLY | 0.15840496 |
| GTTNTGWFNS | ASGDDRVSYNSPLY | 0.14325568 |
| IFMGPLFTL | ASGDDRVSYNSPLY | 0.21903573 |
| ILAHAIEAI | ASGDDRVSYNSPLY | 0.17453577 |
| IRSAIREAL | ASGDDRVSYNSPLY | 0.17386757 |
| ITFAGGG | ASGDDRVSYNSPLY | 0.21456122 |
| ITGMSGHQEV | ASGDDRVSYNSPLY | 0.16573422 |
| IVAGLACTL | ASGDDRVSYNSPLY | 0.17714657 |

| | | |
|---|---|---|
| IVGGGLAGA | ASGDDRVSYNSPLY | 0.18041188 |
| IVTGVLNI | ASGDDRVSYNSPLY | 0.18132317 |
| LAQQGCI | ASGDDRVSYNSPLY | 0.25806963 |
| LCLHVEN | ASGDDRVSYNSPLY | 0.20061538 |
| LDAISDGS | ASGDDRVSYNSPLY | 0.18014775 |
| LGGFAAML | ASGDDRVSYNSPLY | 0.6384409 |
| LGGKNPMI | ASGDDRVSYNSPLY | 0.15334107 |
| LGPKGVAAIL | ASGDDRVSYNSPLY | 0.1607991 |
| LMVGVSI | ASGDDRVSYNSPLY | 0.26900685 |
| LVGAAFVL | ASGDDRVSYNSPLY | 0.15072118 |
| LVLAVAPL | ASGDDRVSYNSPLY | 0.34007123 |
| MAAIAVIDAA | ASGDDRVSYNSPLY | 0.24639757 |
| NNGINAELQ | ASGDDRVSYNSPLY | 0.25215983 |
| RGEPLVLVHG | ASGDDRVSYNSPLY | 0.20829158 |
| RGGAGGAAGLI | ASGDDRVSYNSPLY | 0.61374927 |
| RSYRMHPAV | ASGDDRVSYNSPLY | 0.37566817 |
| SAAHLLAGQLA | ASGDDRVSYNSPLY | 0.16834016 |
| SAAVGSY | ASGDDRVSYNSPLY | 0.19244367 |
| SPHAVLHRI | ASGDDRVSYNSPLY | 0.15794355 |
| SQIAKVL | ASGDDRVSYNSPLY | 0.30842388 |
| TALGHLVP | ASGDDRVSYNSPLY | 0.2390911 |
| TAPTTSGAA | ASGDDRVSYNSPLY | 0.16547735 |
| TDGDDGNGGSPGT | ASGDDRVSYNSPLY | 0.24795076 |
| TDGLGEN | ASGDDRVSYNSPLY | 0.28691092 |
| TGGPPLPGQA | ASGDDRVSYNSPLY | 0.47303587 |
| TIPAITI | ASGDDRVSYNSPLY | 0.14750671 |
| TIPGFTI | ASGDDRVSYNSPLY | 0.20236936 |
| TSIASGALA | ASGDDRVSYNSPLY | 0.1497786 |
| TVTLPGAAI | ASGDDRVSYNSPLY | 0.18792513 |
| VASGLSAALP | ASGDDRVSYNSPLY | 0.15722777 |
| VGAGVIGI | ASGDDRVSYNSPLY | 0.31379092 |
| VGPKRFGV | ASGDDRVSYNSPLY | 0.19313729 |
| VNVAFPD | ASGDDRVSYNSPLY | 0.34954053 |
| VSMTNSGVS | ASGDDRVSYNSPLY | 0.2688055 |
| VTDVLGDLSV | ASGDDRVSYNSPLY | 0.44250333 |
| VVVVGAGFAGL | ASGDDRVSYNSPLY | 0.16714658 |
| VYTVGTVV | ASGDDRVSYNSPLY | 0.20471719 |
| WTTFSGP | ASGDDRVSYNSPLY | 0.14969772 |
| YDELVEAF | ASGDDRVSYNSPLY | 0.1494248 |
| YGTGGAGGNGG | ASGDDRVSYNSPLY | 0.15794857 |
| YTNVMAV | ASGDDRVSYNSPLY | 0.17917702 |
| FLEQVHQL | ASGDDRVSYNSPLY | 0.14346954 |
| GLSLLAVAV | ASGDDRVSYNSPLY | 0.35715798 |
| LEQVHQL | ASGDDRVSYNSPLY | 0.28008685 |
| LNAAGAI | ASGDDRVSYNSPLY | 0.36686435 |
| PAVHASL | ASGDDRVSYNSPLY | 0.16912118 |
| TELEDDA | ASGDDRVSYNSPLY | 0.16949838 |
| TPPGPPN | ASGDDRVSYNSPLY | 0.17868595 |
| TPPPPPPP | ASGDDRVSYNSPLY | 0.55182457 |
| TQSPALI | ASSQDRGEEDTQY | 0.15636085 |
| EFAAAFPQ | ASSQDRGEEDTQY | 0.14785616 |

| | | |
|---|---|---|
| LLSGNLDVL | ASSQDRGEEDTQY | 0.16826014 |
| GPANLGNYNI | ASSQDRGEEDTQY | 0.15604347 |
| TLGPTLP | ASSQDRGEEDTQY | 0.3031113 |
| VYAYSSCCK | ASSQDRGEEDTQY | 0.3361809 |
| APAPAPAPNI | ASSQDRGEEDTQY | 0.29987955 |
| PGGPVPIP | ASSQDRGEEDTQY | 0.21807346 |
| FLTVALEP | ASSQDRGEEDTQY | 0.14527702 |
| SQYFGANL | ASSQDRGEEDTQY | 0.17716056 |
| GGVKPAA | ASSQDRGEEDTQY | 0.70770836 |
| GGTGGAGGAGGW | ASSQDRGEEDTQY | 0.2396904 |
| GVGGVAGLGGAGG | ASSQDRGEEDTQY | 0.2015704 |
| VLLVLATTI | ASSQDRGEEDTQY | 0.40910572 |
| IGIFAAML | ASSQDRGEEDTQY | 0.18110502 |
| AVGDLID | ASSQDRGEEDTQY | 0.1956006 |
| NISPLPL | ASSQDRGEEDTQY | 0.21261573 |
| VTAVFVAI | ASSQDRGEEDTQY | 0.18068622 |
| SGDDNDAAL | ASSQDRGEEDTQY | 0.17354068 |
| AVAALAHS | ASSQDRGEEDTQY | 0.15243408 |
| GGVTVMATP | ASSQDRGEEDTQY | 0.21481784 |
| AGLAANVRD | ASSQDRGEEDTQY | 0.71348685 |
| TNRPDLIDPALLRPG | ASSQDRGEEDTQY | 0.32363522 |
| LANLGVTAL | ASSQDRGEEDTQY | 0.37733126 |
| TGERIEL | ASSQDRGEEDTQY | 0.26255327 |
| NGEELAA | ASSQDRGEEDTQY | 0.55009854 |
| VLSLVWG | ASSQDRGEEDTQY | 0.18101214 |
| IGYALWL | ASSQDRGEEDTQY | 0.17950079 |
| LSLLLAPL | ASSQDRGEEDTQY | 0.17635415 |
| TLDEALK | ASSQDRGEEDTQY | 0.17762457 |
| TDGAGTAEGG | ASSQDRGEEDTQY | 0.18871452 |
| NAPGGAGGNGG | ASSQDRGEEDTQY | 0.17283545 |
| VSVFMGMGGNF | ASSQDRGEEDTQY | 0.7545544 |
| LASYLAAL | ASSQDRGEEDTQY | 0.25238928 |
| LVGGTPAL | ASSQDRGEEDTQY | 0.23993993 |
| VRNADVL | ASSQDRGEEDTQY | 0.17262697 |
| TCSSPFLVSAM | ASSQDRGEEDTQY | 0.1859362 |
| VHITDAE | ASSQDRGEEDTQY | 0.23390298 |
| EYAAPSF | ASSQDRGEEDTQY | 0.15227593 |
| VVDAFHGGL | ASSQDRGEEDTQY | 0.5354179 |
| ASVVDPQ | ASSQDRGEEDTQY | 0.17571425 |
| SYNEFGGLS | ASSQDRGEEDTQY | 0.16816851 |
| DPDGSVY | ASSQDRGEEDTQY | 0.26601166 |
| MVSTIHATE | ASSQDRGEEDTQY | 0.66735196 |
| LAAGVADPV | ASSQDRGEEDTQY | 0.16877571 |
| TKEQIAA | ASSQDRGEEDTQY | 0.1827425 |
| SLASPAAL | ASSQDRGEEDTQY | 0.1676022 |
| IGLVFYGV | ASSQDRGEEDTQY | 0.29421228 |
| FLLDASNL | ASSQDRGEEDTQY | 0.62641174 |
| YQVGNFP | ASSQDRGEEDTQY | 0.18119785 |
| ESLLSQST | ASSQDRGEEDTQY | 0.4387414 |
| LFTVGFAAVG | ASSQDRGEEDTQY | 0.18524368 |
| ATPPTPT | ASSQDRGEEDTQY | 0.1545143 |

| | | |
|---|---|---|
| TDLATLG | ASSQDRGEEDTQY | 0.19024788 |
| SAGGTGR | ASSQDRGEEDTQY | 0.24386877 |
| GFSNSGS | ASSQDRGEEDTQY | 0.55814433 |
| GGNGGSGGTSV | ASSQDRGEEDTQY | 0.39056695 |
| GTSVATGGAGN | ASSQDRGEEDTQY | 0.17716515 |
| NGGNGGSGGTS | ASSQDRGEEDTQY | 0.17028229 |
| AVVKFPH | ASSQDRGEEDTQY | 0.5188854 |
| TPPPPPP | ASSQDRGEEDTQY | 0.45142376 |
| AAPAAAAAPA | ASSQDRGEEDTQY | 0.38902426 |
| IALNEDL | ASSQDRGEEDTQY | 0.18341182 |
| IVNVSSM | ASSQDRGEEDTQY | 0.17154376 |
| TGIPPAPR | ASSQDRGEEDTQY | 0.17864053 |
| FDLGGGTF | ASSQDRGEEDTQY | 0.35382938 |
| FDLGGGTFDVS | ASSQDRGEEDTQY | 0.15732798 |
| NEPTAAA | ASSQDRGEEDTQY | 0.18992046 |
| NEPTAAALA | ASSQDRGEEDTQY | 0.33018914 |
| GSIINISSI | ASSQDRGEEDTQY | 0.15064064 |
| VAPGNAGTA | ASSQDRGEEDTQY | 0.22395515 |
| TGGGASLE | ASSQDRGEEDTQY | 0.47289568 |
| VAVYNHYKRI | ASSQDRGEEDTQY | 0.17101102 |
| TDPAPATT | ASSQDRGEEDTQY | 0.17686914 |
| TNRPDILDPALLRPG | ASSQDRGEEDTQY | 0.49719137 |
| LTTTTTS | ASSQDRGEEDTQY | 0.7615223 |
| PTGSGKTL | ASSQDRGEEDTQY | 0.28761676 |
| TGSGKTL | ASSQDRGEEDTQY | 0.15236719 |
| TIGHVDHGK | ASSQDRGEEDTQY | 0.14446367 |
| TIGHVDHGKTT | ASSQDRGEEDTQY | 0.17113668 |
| GHVDHGKTT | ASSQDRGEEDTQY | 0.38892174 |
| AQFEHTLL | ASSQDRGEEDTQY | 0.16934042 |
| IVNQDEL | ASSQDRGEEDTQY | 0.17006284 |
| TNGSQFF | ASSQDRGEEDTQY | 0.28097674 |
| TGKGIVE | ASSQDRGEEDTQY | 0.4178378 |
| LISNASDA | ASSQDRGEEDTQY | 0.16307463 |
| AELRTLGELTKIAKAHGAQ | ASSQDRGEEDTQY | 0.1575196 |
| APNAVAAAEPSW | ASSQDRGEEDTQY | 0.1775332 |
| APQVAGAATAVL | ASSQDRGEEDTQY | 0.30853096 |
| ARRDPAAAAAILPTDA | ASSQDRGEEDTQY | 0.16458336 |
| DAAARDVLAEAGLAEYF | ASSQDRGEEDTQY | 0.26988563 |
| DDQIVLSAKANLAKVEQ | ASSQDRGEEDTQY | 0.18510476 |
| DELLPDTGEDSDEDGH | ASSQDRGEEDTQY | 0.1507365 |
| ELPEAPANRATVPQD | ASSQDRGEEDTQY | 0.27511266 |
| FGQSKIERVTGIG | ASSQDRGEEDTQY | 0.16900302 |
| GAGGTGSTAGGAGGAGGAGGLY | ASSQDRGEEDTQY | 0.46233425 |
| GHGGTGAAGGNGGTGGAGGSGI | ASSQDRGEEDTQY | 0.1673198 |
| GNGGNGGAGGEGGPGGAGGAGA | ASSQDRGEEDTQY | 0.26094002 |
| GSTAGNGGNGGNATNGTIAGQP | ASSQDRGEEDTQY | 0.3179322 |
| GTGGDGGNGGNGGTGG | ASSQDRGEEDTQY | 0.20099275 |
| GVRRPAGIPADAPV | ASSQDRGEEDTQY | 0.19520384 |
| GYRQVTDAHLLTLARR | ASSQDRGEEDTQY | 0.16096026 |
| IAKMGQDGHDRG | ASSQDRGEEDTQY | 0.15501632 |
| KAEPSIEIQYT | ASSQDRGEEDTQY | 0.14418839 |

| | | |
|---|---|---|
| KPVRIRMAADEDTVAVT | ASSQDRGEEDTQY | 0.409729 |
| MPPRIAGMRLLVIKPEP | ASSQDRGEEDTQY | 0.14210606 |
| MKFYAVALFRSRGVPPTR | ASSQDRGEEDTQY | 0.30255306 |
| NAGGFGILGAVAHSP | ASSQDRGEEDTQY | 0.30083996 |
| NEGREITLAMAEM | ASSQDRGEEDTQY | 0.15949172 |
| NELVEWSHPLGHRWRWE | ASSQDRGEEDTQY | 0.20930293 |
| NLGAGGTGGNGGNPG | ASSQDRGEEDTQY | 0.16186464 |
| NTEWPKWETVMAADPVD | ASSQDRGEEDTQY | 0.15401115 |
| PASAAPAETQP | ASSQDRGEEDTQY | 0.32692474 |
| QQSSDGKNGVRAAEITGE | ASSQDRGEEDTQY | 0.22951935 |
| RAPDAVTTSWWKE | ASSQDRGEEDTQY | 0.16574804 |
| SGFGNVGAHTSGWFNQ | ASSQDRGEEDTQY | 0.19544013 |
| SGVAGAVGQAASVGGLKV | ASSQDRGEEDTQY | 0.255766 |
| SNGGAGTGSAGGNGGT | ASSQDRGEEDTQY | 0.16602531 |
| TGGAGGEGGAAG | ASSQDRGEEDTQY | 0.16952582 |
| TGGDGGAAGAGGAPGHG | ASSQDRGEEDTQY | 0.47075826 |
| TMPPLLAAIANAGRHGV | ASSQDRGEEDTQY | 0.18353805 |
| TSSTAGNSGGA | ASSQDRGEEDTQY | 0.16773118 |
| VPATVPVGILC | ASSQDRGEEDTQY | 0.30673474 |
| VQRPDSGGVVLTGRIS | ASSQDRGEEDTQY | 0.15923089 |
| WTPNPVPATKGVPAGLVQAT | ASSQDRGEEDTQY | 0.34970182 |
| YALKRALAKFSDRFDVVII | ASSQDRGEEDTQY | 0.1749591 |
| YLIGSVIV | ASSQDRGEEDTQY | 0.18723297 |
| YGQLVAVID | ASSQDRGEEDTQY | 0.2591165 |
| VTVGGMTLPA | ASSQDRGEEDTQY | 0.15535803 |
| VLALPAV | ASSQDRGEEDTQY | 0.16346166 |
| VIPPTLNYET | ASSQDRGEEDTQY | 0.17037912 |
| VGMGGMHPSSG | ASSQDRGEEDTQY | 0.26773384 |
| VGLAGGAATA | ASSQDRGEEDTQY | 0.38749316 |
| VGGAGGAGGSGP | ASSQDRGEEDTQY | 0.17362821 |
| VFAAFMV | ASSQDRGEEDTQY | 0.239415 |
| TLYDKLR | ASSQDRGEEDTQY | 0.17080663 |
| TLAVFTVL | ASSQDRGEEDTQY | 0.16482395 |
| TGSVQPARV | ASSQDRGEEDTQY | 0.1814572 |
| TGGGAGSGGT | ASSQDRGEEDTQY | 0.16950971 |
| SHLETSL | ASSQDRGEEDTQY | 0.17555864 |
| SFTTGMT | ASSQDRGEEDTQY | 0.18341988 |
| QVEVAKRMGVRQPTVS | ASSQDRGEEDTQY | 0.16480699 |
| QGSGGPALAM | ASSQDRGEEDTQY | 0.20812833 |
| MLAPSNP | ASSQDRGEEDTQY | 0.42805666 |
| LVNNAAI | ASSQDRGEEDTQY | 0.18737176 |
| LTGEINTLA | ASSQDRGEEDTQY | 0.23125266 |
| LSIFGPV | ASSQDRGEEDTQY | 0.28004354 |
| LLALLANL | ASSQDRGEEDTQY | 0.27177054 |
| LISNASDALDKLR | ASSQDRGEEDTQY | 0.18132268 |
| LINEAVP | ASSQDRGEEDTQY | 0.15857695 |
| LGYAFSGL | ASSQDRGEEDTQY | 0.3623168 |
| LDFVAVAL | ASSQDRGEEDTQY | 0.3486411 |
| LAQPGAE | ASSQDRGEEDTQY | 0.37520885 |
| ISNLGAVTGF | ASSQDRGEEDTQY | 0.21673591 |
| HQEDPGRI | ASSQDRGEEDTQY | 0.25425133 |

| | | |
|---|---|---|
| GTPGEPTP | ASSQDRGEEDTQY | 0.3287539 |
| GFFNSGS | ASSQDRGEEDTQY | 0.22587413 |
| GAGGGGLTAGAN | ASSQDRGEEDTQY | 0.1488191 |
| FVDALVNM | ASSQDRGEEDTQY | 0.18331628 |
| FLTVAPEL | ASSQDRGEEDTQY | 0.2276353 |
| FETLTAL | ASSQDRGEEDTQY | 0.16697882 |
| DVINPIEL | ASSQDRGEEDTQY | 0.14982015 |
| APVSGAPV | ASSQDRGEEDTQY | 0.592985 |
| ALVGGLAW | ASSQDRGEEDTQY | 0.764225 |
| ALAAAAATAI | ASSQDRGEEDTQY | 0.17113411 |
| AFGLGLLL | ASSQDRGEEDTQY | 0.3559807 |
| AFFAAASL | ASSQDRGEEDTQY | 0.14987385 |
| AAAALPAFSPP | ASSQDRGEEDTQY | 0.2581738 |
| VVLAPAL | ASSQDRGEEDTQY | 0.49055067 |
| TIGHVDHGKTTL | ASSQDRGEEDTQY | 0.18075669 |
| LISNASDALDK | ASSQDRGEEDTQY | 0.2844255 |
| LISNASDAL | ASSQDRGEEDTQY | 0.15828572 |
| ISNASDALDK | ASSQDRGEEDTQY | 0.17499906 |
| NQAETLVYQTE | ASSQDRGEEDTQY | 0.51882803 |
| TGGTGGTAGAG | ASSQDRGEEDTQY | 0.18001987 |
| TGSTAGGAGGA | ASSQDRGEEDTQY | 0.16941774 |
| DPLAVPVAAGPV | ASSQDRGEEDTQY | 0.18723914 |
| GGTGGTPGNGGN | ASSQDRGEEDTQY | 0.16195473 |
| PAGLAGVDPATL | ASSQDRGEEDTQY | 0.17764089 |
| GNNLVVEAPHAGASV | ASSQDRGEEDTQY | 0.3483811 |
| KVTRIMSASTLEQPA | ASSQDRGEEDTQY | 0.15318981 |
| LGFALPIALPRRTVV | ASSQDRGEEDTQY | 0.41221878 |
| TRLGNLTARKLVVLV | ASSQDRGEEDTQY | 0.1583383 |
| WSQQRRLIYFPSAGP | ASSQDRGEEDTQY | 0.69343203 |
| GPNGNPDPMAAAVDIR | ASSQDRGEEDTQY | 0.39582193 |
| TQRCVSVADLRAAASR | ASSQDRGEEDTQY | 0.36583418 |
| QFVLDVAEHPVEVLTETV | ASSQDRGEEDTQY | 0.16919716 |
| GGAGGTGGAGGNGGTQAGDG | ASSQDRGEEDTQY | 0.64117104 |
| PLLATLRAIAVDVDAKPAQV | ASSQDRGEEDTQY | 0.2397624 |
| APKFRTGGYMIDFWGVGYQVAK | ASSQDRGEEDTQY | 0.2614153 |
| GAGGAGGAGGAAGGPGGTGGTG | ASSQDRGEEDTQY | 0.23854837 |
| IGALGLLATDVAPAGSVAFGVA | ASSQDRGEEDTQY | 0.21141489 |
| AGGAGGIGGNGGAGGAGGVLMG | ASSQDRGEEDTQY | 0.85085285 |
| ADELIGLL | ASSQDRGEEDTQY | 0.72218865 |
| ADRALKA | ASSQDRGEEDTQY | 0.55924225 |
| AGAVGQVGGAGGA | ASSQDRGEEDTQY | 0.30815637 |
| AGGGGGLGGAP | ASSQDRGEEDTQY | 0.16133004 |
| AGGLLYGNGGAG | ASSQDRGEEDTQY | 0.3546396 |
| AGLESIGL | ASSQDRGEEDTQY | 0.25342217 |
| AGLKTGII | ASSQDRGEEDTQY | 0.78934085 |
| ASLAALLL | ASSQDRGEEDTQY | 0.6823641 |
| ASVINISSI | ASSQDRGEEDTQY | 0.30161548 |
| ATPPGPA | ASSQDRGEEDTQY | 0.78126776 |
| ATVASALV | ASSQDRGEEDTQY | 0.79976094 |
| ATVASLIL | ASSQDRGEEDTQY | 0.15635198 |
| AVNGPVLIRH | ASSQDRGEEDTQY | 0.46543264 |

| | | |
|---|---|---|
| DGAAGAVGI | ASSQDRGEEDTQY | 0.2565696 |
| DPLLHGI | ASSQDRGEEDTQY | 0.3960635 |
| ELGAALGAL | ASSQDRGEEDTQY | 0.10831059 |
| ELGRTQAV | ASSQDRGEEDTQY | 0.720294 |
| FVAGGYLS | ASSQDRGEEDTQY | 0.3842815 |
| GASGGAGGAGGW | ASSQDRGEEDTQY | 0.100752406 |
| GGAGAGGDGGAGGI | ASSQDRGEEDTQY | 0.37071663 |
| GTGHNGSQPGMGGNG | ASSQDRGEEDTQY | 0.37520587 |
| GYAAASASAS | ASSQDRGEEDTQY | 0.36045882 |
| HQVDVAAVQQG | ASSQDRGEEDTQY | 0.77406895 |
| IGAAIAHT | ASSQDRGEEDTQY | 0.34670198 |
| IGMIEVAQL | ASSQDRGEEDTQY | 0.25742966 |
| IRLAPEHTSAMD | ASSQDRGEEDTQY | 0.7656537 |
| IYNVSVL | ASSQDRGEEDTQY | 0.8275894 |
| KQGFLGM | ASSQDRGEEDTQY | 0.22924475 |
| LESVITQL | ASSQDRGEEDTQY | 0.7029284 |
| LGLGALVVV | ASSQDRGEEDTQY | 0.84510505 |
| LIGNGGNGGSGGI | ASSQDRGEEDTQY | 0.5883064 |
| LNAINAPTL | ASSQDRGEEDTQY | 0.78519297 |
| LQLLNTGV | ASSQDRGEEDTQY | 0.8237712 |
| LQTSIEF | ASSQDRGEEDTQY | 0.24850395 |
| MSFAVSEL | ASSQDRGEEDTQY | 0.7943495 |
| NTGALGL | ASSQDRGEEDTQY | 0.8118654 |
| PPGAGGAGGAGTT | ASSQDRGEEDTQY | 0.46842426 |
| QLVGLGI | ASSQDRGEEDTQY | 0.49012738 |
| RFLEQVHQL | ASSQDRGEEDTQY | 0.20496805 |
| RGVGYVLR | ASSQDRGEEDTQY | 0.38667196 |
| SATGVFAGVI | ASSQDRGEEDTQY | 0.19249727 |
| SLPAITLP | ASSQDRGEEDTQY | 0.72672665 |
| STALAFTV | ASSQDRGEEDTQY | 0.79488933 |
| SVLTVGLGK | ASSQDRGEEDTQY | 0.45383933 |
| TALAGAGSGI | ASSQDRGEEDTQY | 0.21549994 |
| TGLPFTGI | ASSQDRGEEDTQY | 0.24527878 |
| VAEHVTV | ASSQDRGEEDTQY | 0.5406948 |
| VDIVLNSL | ASSQDRGEEDTQY | 0.37036547 |
| VLTGGGAVL | ASSQDRGEEDTQY | 0.18340889 |
| VNVGLAYGI | ASSQDRGEEDTQY | 0.26824814 |
| VSSAAVGAV | ASSQDRGEEDTQY | 0.7718803 |
| VTSVYEAGV | ASSQDRGEEDTQY | 0.7880548 |
| WVALDRGA | ASSQDRGEEDTQY | 0.17425045 |
| YLVGSVVI | ASSQDRGEEDTQY | 0.82526803 |
| GPPGTGKTLL | ASSQDRGEEDTQY | 0.787166 |
| GVGMTKF | ASSQDRGEEDTQY | 0.85137254 |
| IGIGGSDL | ASSQDRGEEDTQY | 0.5229804 |
| IGTLLAEL | ASSQDRGEEDTQY | 0.22727203 |
| IYTSGTT | ASSQDRGEEDTQY | 0.8043711 |
| LSFPAGV | ASSQDRGEEDTQY | 0.4978547 |
| LYTSGTT | ASSQDRGEEDTQY | 0.39867648 |
| VGAFPFGF | ASSQDRGEEDTQY | 0.6540199 |
| VGAGTVEF | ASSQDRGEEDTQY | 0.45148498 |
| VGKSTLLL | ASSQDRGEEDTQY | 0.32662517 |

| | | |
|---|---|---|
| AALAHAGGGIV | ASSQDRGEEDTQY | 0.691603 |
| AALVAVSLL | ASSQDRGEEDTQY | 0.75941 |
| AAVSRAVAAL | ASSQDRGEEDTQY | 0.22264826 |
| AGGSSGIGGF | ASSQDRGEEDTQY | 0.26228848 |
| AGLTVANANG | ASSQDRGEEDTQY | 0.34270734 |
| AGLVGAVL | ASSQDRGEEDTQY | 0.57744265 |
| ALSQAFTV | ASSQDRGEEDTQY | 0.79892045 |
| ALTFSVSL | ASSQDRGEEDTQY | 0.5682217 |
| ALTGAALVAL | ASSQDRGEEDTQY | 0.5988786 |
| ASFSIGAGVL | ASSQDRGEEDTQY | 0.7436982 |
| DGSVYKI | ASSQDRGEEDTQY | 0.76648206 |
| EQLAEIV | ASSQDRGEEDTQY | 0.50243694 |
| FAVSELTVM | ASSQDRGEEDTQY | 0.6875501 |
| GAAPLAAGL | ASSQDRGEEDTQY | 0.38492313 |
| GADGGDGGSGGAGG | ASSQDRGEEDTQY | 0.7638908 |
| GFGLDTGGAGG | ASSQDRGEEDTQY | 0.66256344 |
| GLASGGLL | ASSQDRGEEDTQY | 0.72250307 |
| GRIAGAAVLL | ASSQDRGEEDTQY | 0.78693634 |
| GSAGSLNTSF | ASSQDRGEEDTQY | 0.7778644 |
| GTGQIGI | ASSQDRGEEDTQY | 0.5959439 |
| GTLIALIL | ASSQDRGEEDTQY | 0.15302096 |
| IAATMGFVVG | ASSQDRGEEDTQY | 0.3419731 |
| IFAQALL | ASSQDRGEEDTQY | 0.5983069 |
| IGAAVALGF | ASSQDRGEEDTQY | 0.6727582 |
| IGLTPLAL | ASSQDRGEEDTQY | 0.6967336 |
| ILGSNTSTL | ASSQDRGEEDTQY | 0.71325994 |
| IVFSLTVL | ASSQDRGEEDTQY | 0.35640007 |
| LASKATGF | ASSQDRGEEDTQY | 0.76276255 |
| LASVALIL | ASSQDRGEEDTQY | 0.82140243 |
| LFAVGLTL | ASSQDRGEEDTQY | 0.17078716 |
| LFNGGIGTYI | ASSQDRGEEDTQY | 0.3334247 |
| LTLGAGII | ASSQDRGEEDTQY | 0.84028566 |
| LVAAALGDI | ASSQDRGEEDTQY | 0.7509439 |
| NTAFVQLGI | ASSQDRGEEDTQY | 0.37596518 |
| QLFSGAEL | ASSQDRGEEDTQY | 0.43483785 |
| SAATFITL | ASSQDRGEEDTQY | 0.8462351 |
| SNYGLANLGV | ASSQDRGEEDTQY | 0.44194126 |
| SYGGAYAVM | ASSQDRGEEDTQY | 0.8303914 |
| TTGVGGLTLG | ASSQDRGEEDTQY | 0.1204781 |
| TTQTIYI | ASSQDRGEEDTQY | 0.47439367 |
| VAAGAGLGAVI | ASSQDRGEEDTQY | 0.33766004 |
| VAAVVTLL | ASSQDRGEEDTQY | 0.12139787 |
| VGFLGGLI | ASSQDRGEEDTQY | 0.71198976 |
| VGVIPILL | ASSQDRGEEDTQY | 0.29307228 |
| VIFASSLI | ASSQDRGEEDTQY | 0.7526687 |
| VKGGDGGAAAG | ASSQDRGEEDTQY | 0.4332811 |
| VTKTFTNI | ASSQDRGEEDTQY | 0.72838724 |
| VVAAAILL | ASSQDRGEEDTQY | 0.6908107 |
| YAALDVEL | ASSQDRGEEDTQY | 0.43816438 |
| LASVALLL | ASSQDRGEEDTQY | 0.24241847 |
| LDAVHRV | ASSQDRGEEDTQY | 0.6397302 |

| | | |
|---|---|---|
| QGLSSSIF | ASSQDRGEEDTQY | 0.75135607 |
| RGTFANIRL | ASSQDRGEEDTQY | 0.70093036 |
| RQLWWGHRI | ASSQDRGEEDTQY | 0.16766739 |
| YGPPGTGKTLL | ASSQDRGEEDTQY | 0.5257729 |
| AALVGAPANVA | ASSQDRGEEDTQY | 0.5121429 |
| AATFRRTE | ASSQDRGEEDTQY | 0.12655102 |
| AGALGGGI | ASSQDRGEEDTQY | 0.111362115 |
| AGGAGGAGGIGAG | ASSQDRGEEDTQY | 0.50058466 |
| AGSGALGL | ASSQDRGEEDTQY | 0.35496324 |
| AQIADQI | ASSQDRGEEDTQY | 0.7517622 |
| ASAVAVTYL | ASSQDRGEEDTQY | 0.77286685 |
| ASTVLGGTGG | ASSQDRGEEDTQY | 0.388976 |
| AVGGEPAK | ASSQDRGEEDTQY | 0.7733426 |
| CEQGTFGGTC | ASSQDRGEEDTQY | 0.54843074 |
| DEGKAPAG | ASSQDRGEEDTQY | 0.08568986 |
| DTVVFPAL | ASSQDRGEEDTQY | 0.41416344 |
| FEGITMAG | ASSQDRGEEDTQY | 0.14736527 |
| FPLVVQD | ASSQDRGEEDTQY | 0.78429294 |
| GAFTTPQ | ASSQDRGEEDTQY | 0.1906467 |
| GFGGFGTAGGVGGA | ASSQDRGEEDTQY | 0.27026647 |
| GFLAQFL | ASSQDRGEEDTQY | 0.508721 |
| GGAGGAGGNAGVA | ASSQDRGEEDTQY | 0.7639539 |
| GGVGTAGAGGAGGAGG | ASSQDRGEEDTQY | 0.6200258 |
| GNGGGGDAGNAGSGGN | ASSQDRGEEDTQY | 0.36845738 |
| GTGGAGGAGNDAGS | ASSQDRGEEDTQY | 0.101863205 |
| GTTNTGWFNS | ASSQDRGEEDTQY | 0.6921487 |
| IFMGPLFTL | ASSQDRGEEDTQY | 0.40894824 |
| ILAHAIEAI | ASSQDRGEEDTQY | 0.2329481 |
| IRSAIREAL | ASSQDRGEEDTQY | 0.12733012 |
| ITFAGGG | ASSQDRGEEDTQY | 0.14809102 |
| ITGMSGHQEV | ASSQDRGEEDTQY | 0.18579867 |
| IVAGLACTL | ASSQDRGEEDTQY | 0.7332691 |
| IVGGGLAGA | ASSQDRGEEDTQY | 0.46878898 |
| IVTGVLNI | ASSQDRGEEDTQY | 0.40158945 |
| LAQQGCI | ASSQDRGEEDTQY | 0.8417136 |
| LCLHVEN | ASSQDRGEEDTQY | 0.7561975 |
| LDAISDGS | ASSQDRGEEDTQY | 0.443469 |
| LGGFAAML | ASSQDRGEEDTQY | 0.3461718 |
| LGGKNPMI | ASSQDRGEEDTQY | 0.197909 |
| LGPKGVAAIL | ASSQDRGEEDTQY | 0.24058156 |
| LMVGVSI | ASSQDRGEEDTQY | 0.28544462 |
| LVGAAFVL | ASSQDRGEEDTQY | 0.8062334 |
| LVLAVAPL | ASSQDRGEEDTQY | 0.44678104 |
| MAAIAVIDAA | ASSQDRGEEDTQY | 0.170926 |
| NNGINAELQ | ASSQDRGEEDTQY | 0.7032021 |
| RGEPLVLVHG | ASSQDRGEEDTQY | 0.8247076 |
| RGGAGGAAGLI | ASSQDRGEEDTQY | 0.8491725 |
| RSYRMHPAV | ASSQDRGEEDTQY | 0.19978356 |
| SAAHLLAGQLA | ASSQDRGEEDTQY | 0.49328634 |
| SAAVGSY | ASSQDRGEEDTQY | 0.40892333 |
| SPHAVLHRI | ASSQDRGEEDTQY | 0.5623497 |

| | | |
|---|---|---|
| SQIAKVL | ASSQDRGEEDTQY | 0.38987833 |
| TALGHLVP | ASSQDRGEEDTQY | 0.21780545 |
| TAPTTSGAA | ASSQDRGEEDTQY | 0.75856996 |
| TDGDDGNGGSPGT | ASSQDRGEEDTQY | 0.8695896 |
| TDGLGEN | ASSQDRGEEDTQY | 0.34134078 |
| TGGPPLPGQA | ASSQDRGEEDTQY | 0.13426442 |
| TIPAITI | ASSQDRGEEDTQY | 0.43654668 |
| TIPGFTI | ASSQDRGEEDTQY | 0.3120514 |
| TSIASGALA | ASSQDRGEEDTQY | 0.35224342 |
| TVTLPGAAI | ASSQDRGEEDTQY | 0.6420533 |
| VASGLSAALP | ASSQDRGEEDTQY | 0.16193858 |
| VGAGVIGI | ASSQDRGEEDTQY | 0.42553878 |
| VGPKRFGV | ASSQDRGEEDTQY | 0.87117445 |
| VNVAFPD | ASSQDRGEEDTQY | 0.5245762 |
| VSMTNSGVS | ASSQDRGEEDTQY | 0.6930124 |
| VTDVLGDLSV | ASSQDRGEEDTQY | 0.8106335 |
| VVVVGAGFAGL | ASSQDRGEEDTQY | 0.71001375 |
| VYTVGTVV | ASSQDRGEEDTQY | 0.8265878 |
| WTTFSGP | ASSQDRGEEDTQY | 0.8102939 |
| YDELVEAF | ASSQDRGEEDTQY | 0.35515255 |
| YGTGGAGGNGG | ASSQDRGEEDTQY | 0.65342927 |
| YTNVMAV | ASSQDRGEEDTQY | 0.58050156 |
| FLEQVHQL | ASSQDRGEEDTQY | 0.44471925 |
| GLSLLAVAV | ASSQDRGEEDTQY | 0.48081112 |
| LEQVHQL | ASSQDRGEEDTQY | 0.37603414 |
| LNAAGAI | ASSQDRGEEDTQY | 0.78534544 |
| PAVHASL | ASSQDRGEEDTQY | 0.64561856 |
| TELEDDA | ASSQDRGEEDTQY | 0.7915456 |
| TPPGPPN | ASSQDRGEEDTQY | 0.16753726 |
| TPPPPPPP | ASSQDRGEEDTQY | 0.21429743 |
| TQSPALI | ASSSGGGSYEQY | 0.8101044 |
| EFAAAFPQ | ASSSGGGSYEQY | 0.1086936 |
| LLSGNLDVL | ASSSGGGSYEQY | 0.14740847 |
| GPANLGNYNI | ASSSGGGSYEQY | 0.6207755 |
| TLGPTLP | ASSSGGGSYEQY | 0.84422654 |
| VYAYSSCCK | ASSSGGGSYEQY | 0.11425904 |
| APAPAPAPNI | ASSSGGGSYEQY | 0.24673411 |
| PGGPVPIP | ASSSGGGSYEQY | 0.21042031 |
| FLTVALEP | ASSSGGGSYEQY | 0.78416026 |
| SQYFGANL | ASSSGGGSYEQY | 0.5868303 |
| GGVKPAA | ASSSGGGSYEQY | 0.14104797 |
| GGTGGAGGAGGW | ASSSGGGSYEQY | 0.69484735 |
| GVGGVAGLGGAGG | ASSSGGGSYEQY | 0.13181527 |
| VLLVLATTI | ASSSGGGSYEQY | 0.27487382 |
| IGIFAAML | ASSSGGGSYEQY | 0.7011603 |
| AVGDLID | ASSSGGGSYEQY | 0.8345868 |
| NISPLPL | ASSSGGGSYEQY | 0.17291316 |
| VTAVFVAI | ASSSGGGSYEQY | 0.7796161 |
| SGDDNDAAL | ASSSGGGSYEQY | 0.46252996 |
| AVAALAHS | ASSSGGGSYEQY | 0.8624952 |
| GGVTVMATP | ASSSGGGSYEQY | 0.33060497 |

| | | |
|---|---|---|
| AGLAANVRD | ASSSGGGSYEQY | 0.16636336 |
| TNRPDLIDPALLRPG | ASSSGGGSYEQY | 0.76883745 |
| LANLGVTAL | ASSSGGGSYEQY | 0.68938273 |
| TGERIEL | ASSSGGGSYEQY | 0.46423414 |
| NGEELAA | ASSSGGGSYEQY | 0.7873574 |
| VLSLVWG | ASSSGGGSYEQY | 0.5081309 |
| IGYALWL | ASSSGGGSYEQY | 0.5225848 |
| LSLLLAPL | ASSSGGGSYEQY | 0.78519994 |
| TLDEALK | ASSSGGGSYEQY | 0.68890834 |
| TDGAGTAEGG | ASSSGGGSYEQY | 0.25876534 |
| NAPGGAGGNGG | ASSSGGGSYEQY | 0.6635259 |
| VSVFMGMGGNF | ASSSGGGSYEQY | 0.49802953 |
| LASYLAAL | ASSSGGGSYEQY | 0.24478 |
| LVGGTPAL | ASSSGGGSYEQY | 0.8549131 |
| VRNADVL | ASSSGGGSYEQY | 0.56897426 |
| TCSSPFLVSAM | ASSSGGGSYEQY | 0.77166355 |
| VHITDAE | ASSSGGGSYEQY | 0.6820822 |
| EYAAPSF | ASSSGGGSYEQY | 0.86573136 |
| VVDAFHGGL | ASSSGGGSYEQY | 0.16574623 |
| ASVVDPQ | ASSSGGGSYEQY | 0.5749675 |
| SYNEFGGLS | ASSSGGGSYEQY | 0.55541515 |
| DPDGSVY | ASSSGGGSYEQY | 0.76536006 |
| MVSTIHATE | ASSSGGGSYEQY | 0.5688962 |
| LAAGVADPV | ASSSGGGSYEQY | 0.3798513 |
| TKEQIAA | ASSSGGGSYEQY | 0.6038928 |
| SLASPAAL | ASSSGGGSYEQY | 0.8114465 |
| IGLVFYGV | ASSSGGGSYEQY | 0.47212243 |
| FLLDASNL | ASSSGGGSYEQY | 0.60743576 |
| YQVGNFP | ASSSGGGSYEQY | 0.8432604 |
| ESLLSQST | ASSSGGGSYEQY | 0.2276722 |
| LFTVGFAAVG | ASSSGGGSYEQY | 0.4259748 |
| ATPPTPT | ASSSGGGSYEQY | 0.7328199 |
| TDLATLG | ASSSGGGSYEQY | 0.7535114 |
| SAGGTGR | ASSSGGGSYEQY | 0.48721796 |
| GFSNSGS | ASSSGGGSYEQY | 0.804226 |
| GGNGGSGGTSV | ASSSGGGSYEQY | 0.2413359 |
| GTSVATGGAGN | ASSSGGGSYEQY | 0.73303896 |
| NGGNGGSGGTS | ASSSGGGSYEQY | 0.4073431 |
| AVVKFPH | ASSSGGGSYEQY | 0.6408273 |
| TPPPPPP | ASSSGGGSYEQY | 0.28772515 |
| AAPAAAAAPA | ASSSGGGSYEQY | 0.6805643 |
| IALNEDL | ASSSGGGSYEQY | 0.7816118 |
| IVNVSSM | ASSSGGGSYEQY | 0.75048596 |
| TGIPPAPR | ASSSGGGSYEQY | 0.11368054 |
| FDLGGGTF | ASSSGGGSYEQY | 0.7452661 |
| FDLGGGTFDVS | ASSSGGGSYEQY | 0.77514195 |
| NEPTAAA | ASSSGGGSYEQY | 0.86315125 |
| NEPTAAALA | ASSSGGGSYEQY | 0.6308265 |
| GSIINISSI | ASSSGGGSYEQY | 0.66205716 |
| VAPGNAGTA | ASSSGGGSYEQY | 0.35468617 |
| TGGGASLE | ASSSGGGSYEQY | 0.5216596 |

| | | |
|---|---|---|
| VAVYNHYKRI | ASSSGGGSYEQY | 0.26228845 |
| TDPAPATT | ASSSGGGSYEQY | 0.40328485 |
| TNRPDILDPALLRPG | ASSSGGGSYEQY | 0.5276029 |
| LTTTTTS | ASSSGGGSYEQY | 0.290202 |
| PTGSGKTL | ASSSGGGSYEQY | 0.8064927 |
| TGSGKTL | ASSSGGGSYEQY | 0.7651814 |
| TIGHVDHGK | ASSSGGGSYEQY | 0.81809133 |
| TIGHVDHGKTT | ASSSGGGSYEQY | 0.8347539 |
| GHVDHGKTT | ASSSGGGSYEQY | 0.8106169 |
| AQFEHTLL | ASSSGGGSYEQY | 0.7179391 |
| IVNQDEL | ASSSGGGSYEQY | 0.3083958 |
| TNGSQFF | ASSSGGGSYEQY | 0.41094723 |
| TGKGIVE | ASSSGGGSYEQY | 0.16528368 |
| LISNASDA | ASSSGGGSYEQY | 0.39071742 |
| AELRTLGELTKIAKAHGAQ | ASSSGGGSYEQY | 0.39790398 |
| APNAVAAAEPSW | ASSSGGGSYEQY | 0.59612167 |
| APQVAGAATAVL | ASSSGGGSYEQY | 0.10663808 |
| ARRDPAAAAAILPTDA | ASSSGGGSYEQY | 0.73037946 |
| DAAARDVLAEAGLAEYF | ASSSGGGSYEQY | 0.8048285 |
| DDQIVLSAKANLAKVEQ | ASSSGGGSYEQY | 0.47511625 |
| DELLPDTGEDSDEDGH | ASSSGGGSYEQY | 0.6943233 |
| ELPEAPANRATVPQD | ASSSGGGSYEQY | 0.5801667 |
| FGQSKIERVTGIG | ASSSGGGSYEQY | 0.37499413 |
| GAGGTGSTAGGAGGAGGAGGLY | ASSSGGGSYEQY | 0.78764373 |
| GHGGTGAAGGNGGTGGAGGSGI | ASSSGGGSYEQY | 0.2927249 |
| GNGGNGGAGGEGGPGGAGGAGA | ASSSGGGSYEQY | 0.74128133 |
| GSTAGNGGNGGNATNGTIAGQP | ASSSGGGSYEQY | 0.8414873 |
| GTGGDGGNGGNGGTGG | ASSSGGGSYEQY | 0.6251459 |
| GVRRPAGIPADAPV | ASSSGGGSYEQY | 0.569839 |
| GYRQVTDAHLLTLARR | ASSSGGGSYEQY | 0.5407079 |
| IAKMGQDGHDRG | ASSSGGGSYEQY | 0.15086536 |
| KAEPSIEIQYT | ASSSGGGSYEQY | 0.6029072 |
| KPVRIRMAADEDTVAVT | ASSSGGGSYEQY | 0.09288512 |
| MPPRIAGMRLLVIKPEP | ASSSGGGSYEQY | 0.66426444 |
| MKFYAVALFRSRGVPPTR | ASSSGGGSYEQY | 0.22724941 |
| NAGGFGILGAVAHSP | ASSSGGGSYEQY | 0.5847275 |
| NEGREITLAMAEM | ASSSGGGSYEQY | 0.48375812 |
| NELVEWSHPLGHRWRWE | ASSSGGGSYEQY | 0.6651797 |
| NLGAGGTGGNGGNPG | ASSSGGGSYEQY | 0.2187853 |
| NTEWPKWETVMAADPVD | ASSSGGGSYEQY | 0.63967174 |
| PASAAPAETQP | ASSSGGGSYEQY | 0.51910317 |
| QQSSDGKNGVRAAEITGE | ASSSGGGSYEQY | 0.19990355 |
| RAPDAVTTSWWKE | ASSSGGGSYEQY | 0.7027243 |
| SGFGNVGAHTSGWFNQ | ASSSGGGSYEQY | 0.44872525 |
| SGVAGAVGQAASVGGLKV | ASSSGGGSYEQY | 0.5763423 |
| SNGGAGTGSAGGNGGT | ASSSGGGSYEQY | 0.7888652 |
| TGGAGGEGGAAG | ASSSGGGSYEQY | 0.5845215 |
| TGGDGGAAGAGGAPGHG | ASSSGGGSYEQY | 0.14378507 |
| TMPPLLAAIANAGRHGV | ASSSGGGSYEQY | 0.6428547 |
| TSSTAGNSGGA | ASSSGGGSYEQY | 0.52251816 |
| VPATVPVGILC | ASSSGGGSYEQY | 0.4107306 |

| | | |
|---|---|---|
| VQRPDSGGVVLTGRIS | ASSSGGGSYEQY | 0.09840381 |
| WTPNPVPATKGVPAGLVQAT | ASSSGGGSYEQY | 0.58846194 |
| YALKRALAKFSDRFDVVII | ASSSGGGSYEQY | 0.2427871 |
| YLIGSVIV | ASSSGGGSYEQY | 0.3589028 |
| YGQLVAVID | ASSSGGGSYEQY | 0.27397555 |
| VTVGGMTLPA | ASSSGGGSYEQY | 0.7926037 |
| VLALPAV | ASSSGGGSYEQY | 0.54340935 |
| VIPPTLNYET | ASSSGGGSYEQY | 0.62318146 |
| VGMGGMHPSSG | ASSSGGGSYEQY | 0.6753686 |
| VGLAGGAATA | ASSSGGGSYEQY | 0.6080922 |
| VGGAGGAGGSGP | ASSSGGGSYEQY | 0.8288043 |
| VFAAFMV | ASSSGGGSYEQY | 0.6212776 |
| TLYDKLR | ASSSGGGSYEQY | 0.533533 |
| TLAVFTVL | ASSSGGGSYEQY | 0.5125676 |
| TGSVQPARV | ASSSGGGSYEQY | 0.48926893 |
| TGGGAGSGGT | ASSSGGGSYEQY | 0.14546001 |
| SHLETSL | ASSSGGGSYEQY | 0.17938188 |
| SFTTGMT | ASSSGGGSYEQY | 0.60326827 |
| QVEVAKRMGVRQPTVS | ASSSGGGSYEQY | 0.14179075 |
| QGSGGPALAM | ASSSGGGSYEQY | 0.5553209 |
| MLAPSNP | ASSSGGGSYEQY | 0.8014431 |
| LVNNAAI | ASSSGGGSYEQY | 0.63270646 |
| LTGEINTLA | ASSSGGGSYEQY | 0.17378473 |
| LSIFGPV | ASSSGGGSYEQY | 0.39795187 |
| LLALLANL | ASSSGGGSYEQY | 0.32851738 |
| LISNASDALDKLR | ASSSGGGSYEQY | 0.7255238 |
| LINEAVP | ASSSGGGSYEQY | 0.7512977 |
| LGYAFSGL | ASSSGGGSYEQY | 0.8471949 |
| LDFVAVAL | ASSSGGGSYEQY | 0.78857464 |
| LAQPGAE | ASSSGGGSYEQY | 0.3522741 |
| ISNLGAVTGF | ASSSGGGSYEQY | 0.73079216 |
| HQEDPGRI | ASSSGGGSYEQY | 0.4588192 |
| GTPGEPTP | ASSSGGGSYEQY | 0.35854524 |
| GFFNSGS | ASSSGGGSYEQY | 0.40190554 |
| GAGGGGLTAGAN | ASSSGGGSYEQY | 0.65494287 |
| FVDALVNM | ASSSGGGSYEQY | 0.51141274 |
| FLTVAPEL | ASSSGGGSYEQY | 0.3036716 |
| FETLTAL | ASSSGGGSYEQY | 0.1651868 |
| DVINPIEL | ASSSGGGSYEQY | 0.5951179 |
| APVSGAPV | ASSSGGGSYEQY | 0.7902094 |
| ALVGGLAW | ASSSGGGSYEQY | 0.22842449 |
| ALAAAAATAI | ASSSGGGSYEQY | 0.33143923 |
| AFGLGLLL | ASSSGGGSYEQY | 0.85317165 |
| AFFAAASL | ASSSGGGSYEQY | 0.84934527 |
| AAAALPAFSPP | ASSSGGGSYEQY | 0.3645511 |
| VVLAPAL | ASSSGGGSYEQY | 0.3001392 |
| TIGHVDHGKTTL | ASSSGGGSYEQY | 0.7507729 |
| LISNASDALDK | ASSSGGGSYEQY | 0.17786726 |
| LISNASDAL | ASSSGGGSYEQY | 0.64009 |
| ISNASDALDK | ASSSGGGSYEQY | 0.65751255 |
| NQAETLVYQTE | ASSSGGGSYEQY | 0.56429046 |

| | | |
|---|---|---|
| TGGTGGTAGAG | ASSSGGGSYEQY | 0.31935704 |
| TGSTAGGAGGA | ASSSGGGSYEQY | 0.6880018 |
| DPLAVPVAAGPV | ASSSGGGSYEQY | 0.41158256 |
| GGTGGTPGNGGN | ASSSGGGSYEQY | 0.7253778 |
| PAGLAGVDPATL | ASSSGGGSYEQY | 0.103188805 |
| GNNLVVEAPHAGASV | ASSSGGGSYEQY | 0.6199888 |
| KVTRIMSASTLEQPA | ASSSGGGSYEQY | 0.2458436 |
| LGFALPIALPRRTVV | ASSSGGGSYEQY | 0.42282036 |
| TRLGNLTARKLVVLV | ASSSGGGSYEQY | 0.15107399 |
| WSQQRRLIYFPSAGP | ASSSGGGSYEQY | 0.3448537 |
| GPNGNPDPMAAAVDIR | ASSSGGGSYEQY | 0.79118174 |
| TQRCVSVADLRAAASR | ASSSGGGSYEQY | 0.8092852 |
| QFVLDVAEHPVEVLTETV | ASSSGGGSYEQY | 0.23828122 |
| GGAGGTGGAGGNGGTQAGDG | ASSSGGGSYEQY | 0.7312399 |
| PLLATLRAIAVDVDAKPAQV | ASSSGGGSYEQY | 0.85909534 |
| APKFRTGGYMIDFWGVGYQVAK | ASSSGGGSYEQY | 0.38489115 |
| GAGGAGGAGGAAGGPGGTGGTG | ASSSGGGSYEQY | 0.13523553 |
| IGALGLLATDVAPAGSVAFGVA | ASSSGGGSYEQY | 0.6863107 |
| AGGAGGIGGNGGAGGAGGVLMG | ASSSGGGSYEQY | 0.49043038 |
| ADELIGLL | ASSSGGGSYEQY | 0.77971196 |
| ADRALKA | ASSSGGGSYEQY | 0.7953065 |
| AGAVGQVGGAGGA | ASSSGGGSYEQY | 0.16891259 |
| AGGGGGLGGAP | ASSSGGGSYEQY | 0.7657174 |
| AGGLLYGNGGAG | ASSSGGGSYEQY | 0.814863 |
| AGLESIGL | ASSSGGGSYEQY | 0.28652674 |
| AGLKTGII | ASSSGGGSYEQY | 0.20846388 |
| ASLAALLL | ASSSGGGSYEQY | 0.22079809 |
| ASVINISSI | ASSSGGGSYEQY | 0.12798603 |
| ATPPGPA | ASSSGGGSYEQY | 0.63228303 |
| ATVASALV | ASSSGGGSYEQY | 0.37158388 |
| ATVASLIL | ASSSGGGSYEQY | 0.3603704 |
| AVNGPVLIRH | ASSSGGGSYEQY | 0.49640217 |
| DGAAGAVGI | ASSSGGGSYEQY | 0.3032464 |
| DPLLHGI | ASSSGGGSYEQY | 0.32935846 |
| ELGAALGAL | ASSSGGGSYEQY | 0.35431612 |
| ELGRTQAV | ASSSGGGSYEQY | 0.5212668 |
| FVAGGYLS | ASSSGGGSYEQY | 0.73060656 |
| GASGGAGGAGGW | ASSSGGGSYEQY | 0.61195576 |
| GGAGAGGDGGAGGI | ASSSGGGSYEQY | 0.60295737 |
| GTGHNGSQPGMGGNG | ASSSGGGSYEQY | 0.81747526 |
| GYAAASASAS | ASSSGGGSYEQY | 0.24758735 |
| HQVDVAAVQQG | ASSSGGGSYEQY | 0.26480657 |
| IGAAIAHT | ASSSGGGSYEQY | 0.7668868 |
| IGMIEVAQL | ASSSGGGSYEQY | 0.2746316 |
| IRLAPEHTSAMD | ASSSGGGSYEQY | 0.79689294 |
| IYNVSVL | ASSSGGGSYEQY | 0.49708873 |
| KQGFLGM | ASSSGGGSYEQY | 0.6252291 |
| LESVITQL | ASSSGGGSYEQY | 0.6961458 |
| LGLGALVVV | ASSSGGGSYEQY | 0.8324805 |
| LIGNGGNGGSGGI | ASSSGGGSYEQY | 0.7314717 |
| LNAINAPTL | ASSSGGGSYEQY | 0.34206343 |

| | | |
|---|---|---|
| LQLLNTGV | ASSSGGGSYEQY | 0.6926131 |
| LQTSIEF | ASSSGGGSYEQY | 0.21913728 |
| MSFAVSEL | ASSSGGGSYEQY | 0.7343675 |
| NTGALGL | ASSSGGGSYEQY | 0.69571364 |
| PPGAGGAGGAGTT | ASSSGGGSYEQY | 0.2732889 |
| QLVGLGI | ASSSGGGSYEQY | 0.5294045 |
| RFLEQVHQL | ASSSGGGSYEQY | 0.8072728 |
| RGVGYVLR | ASSSGGGSYEQY | 0.7800535 |
| SATGVFAGVI | ASSSGGGSYEQY | 0.18075162 |
| SLPAITLP | ASSSGGGSYEQY | 0.31828177 |
| STALAFTV | ASSSGGGSYEQY | 0.219696 |
| SVLTVGLGK | ASSSGGGSYEQY | 0.574122 |
| TALAGAGSGI | ASSSGGGSYEQY | 0.1751133 |
| TGLPFTGI | ASSSGGGSYEQY | 0.73461777 |
| VAEHVTV | ASSSGGGSYEQY | 0.3718716 |
| VDIVLNSL | ASSSGGGSYEQY | 0.80004364 |
| VLTGGGAVL | ASSSGGGSYEQY | 0.5691536 |
| VNVGLAYGI | ASSSGGGSYEQY | 0.75892615 |
| VSSAAVGAV | ASSSGGGSYEQY | 0.34979197 |
| VTSVYEAGV | ASSSGGGSYEQY | 0.37918445 |
| WVALDRGA | ASSSGGGSYEQY | 0.17598437 |
| YLVGSVVI | ASSSGGGSYEQY | 0.3220424 |
| GPPGTGKTLL | ASSSGGGSYEQY | 0.2771755 |
| GVGMTKF | ASSSGGGSYEQY | 0.48560652 |
| IGIGGSDL | ASSSGGGSYEQY | 0.14906384 |
| IGTLLAEL | ASSSGGGSYEQY | 0.8416452 |
| IYTSGTT | ASSSGGGSYEQY | 0.61621785 |
| LSFPAGV | ASSSGGGSYEQY | 0.6942543 |
| LYTSGTT | ASSSGGGSYEQY | 0.32978857 |
| VGAFPFGF | ASSSGGGSYEQY | 0.26781493 |
| VGAGTVEF | ASSSGGGSYEQY | 0.3291918 |
| VGKSTLLL | ASSSGGGSYEQY | 0.787225 |
| AALAHAGGGIV | ASSSGGGSYEQY | 0.1746472 |
| AALVAVSLL | ASSSGGGSYEQY | 0.12480899 |
| AAVSRAVAAL | ASSSGGGSYEQY | 0.33210623 |
| AGGSSGIGGF | ASSSGGGSYEQY | 0.15104029 |
| AGLTVANANG | ASSSGGGSYEQY | 0.8262818 |
| AGLVGAVL | ASSSGGGSYEQY | 0.80485857 |
| ALSQAFTV | ASSSGGGSYEQY | 0.6721054 |
| ALTFSVSL | ASSSGGGSYEQY | 0.7972057 |
| ALTGAALVAL | ASSSGGGSYEQY | 0.11707838 |
| ASFSIGAGVL | ASSSGGGSYEQY | 0.42248282 |
| DGSVYKI | ASSSGGGSYEQY | 0.83795285 |
| EQLAEIV | ASSSGGGSYEQY | 0.7053253 |
| FAVSELTVM | ASSSGGGSYEQY | 0.4074589 |
| GAAPLAAGL | ASSSGGGSYEQY | 0.7951445 |
| GADGGDGGSGGAGG | ASSSGGGSYEQY | 0.3051272 |
| GFGLDTGGAGG | ASSSGGGSYEQY | 0.42775893 |
| GLASGGLL | ASSSGGGSYEQY | 0.47186667 |
| GRIAGAAVLL | ASSSGGGSYEQY | 0.5231686 |
| GSAGSLNTSF | ASSSGGGSYEQY | 0.5290902 |

| | | |
|---|---|---|
| GTGQIGI | ASSSGGGSYEQY | 0.17129847 |
| GTLIALIL | ASSSGGGSYEQY | 0.65897226 |
| IAATMGFVVG | ASSSGGGSYEQY | 0.84484303 |
| IFAQALL | ASSSGGGSYEQY | 0.45616966 |
| IGAAVALGF | ASSSGGGSYEQY | 0.7774779 |
| IGLTPLAL | ASSSGGGSYEQY | 0.67435676 |
| ILGSNTSTL | ASSSGGGSYEQY | 0.80919844 |
| IVFSLTVL | ASSSGGGSYEQY | 0.28577676 |
| LASKATGF | ASSSGGGSYEQY | 0.23127995 |
| LASVALIL | ASSSGGGSYEQY | 0.6382818 |
| LFAVGLTL | ASSSGGGSYEQY | 0.29119068 |
| LFNGGIGTYI | ASSSGGGSYEQY | 0.5589751 |
| LTLGAGII | ASSSGGGSYEQY | 0.34935737 |
| LVAAALGDI | ASSSGGGSYEQY | 0.8716032 |
| NTAFVQLGI | ASSSGGGSYEQY | 0.74378407 |
| QLFSGAEL | ASSSGGGSYEQY | 0.6649125 |
| SAATFITL | ASSSGGGSYEQY | 0.23484787 |
| SNYGLANLGV | ASSSGGGSYEQY | 0.583495 |
| SYGGAYAVM | ASSSGGGSYEQY | 0.7256932 |
| TTGVGGLTLG | ASSSGGGSYEQY | 0.16360278 |
| TTQTIYI | ASSSGGGSYEQY | 0.82206243 |
| VAAGAGLGAVI | ASSSGGGSYEQY | 0.27348712 |
| VAAVVTLL | ASSSGGGSYEQY | 0.66758496 |
| VGFLGGLI | ASSSGGGSYEQY | 0.75795805 |
| VGVIPILL | ASSSGGGSYEQY | 0.8320974 |
| VIFASSLI | ASSSGGGSYEQY | 0.37829912 |
| VKGGDGGAAAG | ASSSGGGSYEQY | 0.5697051 |
| VTKTFTNI | ASSSGGGSYEQY | 0.2204626 |
| VVAAAILL | ASSSGGGSYEQY | 0.79478794 |
| YAALDVEL | ASSSGGGSYEQY | 0.79372793 |
| LASVALLL | ASSSGGGSYEQY | 0.72177076 |
| LDAVHRV | ASSSGGGSYEQY | 0.83584154 |
| QGLSSSIF | ASSSGGGSYEQY | 0.65785456 |
| RGTFANIRL | ASSSGGGSYEQY | 0.18294269 |
| RQLWWGHRI | ASSSGGGSYEQY | 0.45215273 |
| YGPPGTGKTLL | ASSSGGGSYEQY | 0.75976235 |
| AALVGAPANVA | ASSSGGGSYEQY | 0.81434107 |
| AATFRRTE | ASSSGGGSYEQY | 0.7705302 |
| AGALGGGI | ASSSGGGSYEQY | 0.6340141 |
| AGGAGGAGGIGAG | ASSSGGGSYEQY | 0.4318651 |
| AGSGALGL | ASSSGGGSYEQY | 0.8350947 |
| AQIADQI | ASSSGGGSYEQY | 0.87529874 |
| ASAVAVTYL | ASSSGGGSYEQY | 0.76665735 |
| ASTVLGGTGG | ASSSGGGSYEQY | 0.50420517 |
| AVGGEPAK | ASSSGGGSYEQY | 0.48379984 |
| CEQGTFGGTC | ASSSGGGSYEQY | 0.6043626 |
| DEGKAPAG | ASSSGGGSYEQY | 0.787639 |
| DTVVFPAL | ASSSGGGSYEQY | 0.26259637 |
| FEGITMAG | ASSSGGGSYEQY | 0.57632625 |
| FPLVVQD | ASSSGGGSYEQY | 0.5917388 |
| GAFTTPQ | ASSSGGGSYEQY | 0.13617006 |

| | | |
|---|---|---|
| GFGGFGTAGGVGGA | ASSSGGGSYEQY | 0.70502526 |
| GFLAQFL | ASSSGGGSYEQY | 0.7782293 |
| GGAGGAGGNAGVA | ASSSGGGSYEQY | 0.4931513 |
| GGVGTAGAGGAGGAGG | ASSSGGGSYEQY | 0.37232867 |
| GNGGGGDAGNAGSGGN | ASSSGGGSYEQY | 0.81322116 |
| GTGGAGGAGNDAGS | ASSSGGGSYEQY | 0.8565396 |
| GTTNTGWFNS | ASSSGGGSYEQY | 0.80148876 |
| IFMGPLFTL | ASSSGGGSYEQY | 0.16131644 |
| ILAHAIEAI | ASSSGGGSYEQY | 0.11416646 |
| IRSAIREAL | ASSSGGGSYEQY | 0.42199638 |
| ITFAGGG | ASSSGGGSYEQY | 0.83590883 |
| ITGMSGHQEV | ASSSGGGSYEQY | 0.40424082 |
| IVAGLACTL | ASSSGGGSYEQY | 0.34718806 |
| IVGGGLAGA | ASSSGGGSYEQY | 0.7003562 |
| IVTGVLNI | ASSSGGGSYEQY | 0.80883646 |
| LAQQGCI | ASSSGGGSYEQY | 0.22254474 |
| LCLHVEN | ASSSGGGSYEQY | 0.24512371 |
| LDAISDGS | ASSSGGGSYEQY | 0.58891475 |
| LGGFAAML | ASSSGGGSYEQY | 0.7772684 |
| LGGKNPMI | ASSSGGGSYEQY | 0.26735348 |
| LGPKGVAAIL | ASSSGGGSYEQY | 0.48025644 |
| LMVGVSI | ASSSGGGSYEQY | 0.45667112 |
| LVGAAFVL | ASSSGGGSYEQY | 0.6656774 |
| LVLAVAPL | ASSSGGGSYEQY | 0.68274575 |
| MAAIAVIDAA | ASSSGGGSYEQY | 0.2523684 |
| NNGINAELQ | ASSSGGGSYEQY | 0.80164003 |
| RGEPLVLVHG | ASSSGGGSYEQY | 0.1945479 |
| RGGAGGAAGLI | ASSSGGGSYEQY | 0.7607255 |
| RSYRMHPAV | ASSSGGGSYEQY | 0.5294692 |
| SAAHLLAGQLA | ASSSGGGSYEQY | 0.5411925 |
| SAAVGSY | ASSSGGGSYEQY | 0.4261256 |
| SPHAVLHRI | ASSSGGGSYEQY | 0.31121162 |
| SQIAKVL | ASSSGGGSYEQY | 0.18398201 |
| TALGHLVP | ASSSGGGSYEQY | 0.19409278 |
| TAPTTSGAA | ASSSGGGSYEQY | 0.7646437 |
| TDGDDGNGGSPGT | ASSSGGGSYEQY | 0.10681541 |
| TDGLGEN | ASSSGGGSYEQY | 0.70172834 |
| TGGPPLPGQA | ASSSGGGSYEQY | 0.62004554 |
| TIPAITI | ASSSGGGSYEQY | 0.19202903 |
| TIPGFTI | ASSSGGGSYEQY | 0.6454122 |
| TSIASGALA | ASSSGGGSYEQY | 0.29557866 |
| TVTLPGAAI | ASSSGGGSYEQY | 0.32482937 |
| VASGLSAALP | ASSSGGGSYEQY | 0.80839235 |
| VGAGVIGI | ASSSGGGSYEQY | 0.6774633 |
| VGPKRFGV | ASSSGGGSYEQY | 0.22901689 |
| VNVAFPD | ASSSGGGSYEQY | 0.6134942 |
| VSMTNSGVS | ASSSGGGSYEQY | 0.66363966 |
| VTDVLGDLSV | ASSSGGGSYEQY | 0.2821568 |
| VVVVGAGFAGL | ASSSGGGSYEQY | 0.45036584 |
| VYTVGTVV | ASSSGGGSYEQY | 0.8034412 |
| WTTFSGP | ASSSGGGSYEQY | 0.5347065 |

| | | |
|---|---|---|
| YDELVEAF | ASSSGGGSYEQY | 0.19707024 |
| YGTGGAGGNGG | ASSSGGGSYEQY | 0.7792969 |
| YTNVMAV | ASSSGGGSYEQY | 0.2797733 |
| FLEQVHQL | ASSSGGGSYEQY | 0.6835002 |
| GLSLLAVAV | ASSSGGGSYEQY | 0.36485302 |
| LEQVHQL | ASSSGGGSYEQY | 0.45350847 |
| LNAAGAI | ASSSGGGSYEQY | 0.7888158 |
| PAVHASL | ASSSGGGSYEQY | 0.23225011 |
| TELEDDA | ASSSGGGSYEQY | 0.33990008 |
| TPPGPPN | ASSSGGGSYEQY | 0.41476062 |
| TPPPPPPP | ASSSGGGSYEQY | 0.46646786 |
| TQSPALI | GARRHSETLY | 0.77300155 |
| EFAAAFPQ | GARRHSETLY | 0.25464484 |
| LLSGNLDVL | GARRHSETLY | 0.65946317 |
| GPANLGNYNI | GARRHSETLY | 0.39411083 |
| TLGPTLP | GARRHSETLY | 0.2357131 |
| VYAYSSCCK | GARRHSETLY | 0.47922358 |
| APAPAPAPNI | GARRHSETLY | 0.41378483 |
| PGGPVPIP | GARRHSETLY | 0.28918368 |
| FLTVALEP | GARRHSETLY | 0.56275773 |
| SQYFGANL | GARRHSETLY | 0.27204043 |
| GGVKPAA | GARRHSETLY | 0.7032381 |
| GGTGGAGGAGGW | GARRHSETLY | 0.8141042 |
| GVGGVAGLGGAGG | GARRHSETLY | 0.30320376 |
| VLLVLATTI | GARRHSETLY | 0.727153 |
| IGIFAAML | GARRHSETLY | 0.71812034 |
| AVGDLID | GARRHSETLY | 0.6458944 |
| NISPLPL | GARRHSETLY | 0.36793944 |
| VTAVFVAI | GARRHSETLY | 0.25329366 |
| SGDDNDAAL | GARRHSETLY | 0.7989747 |
| AVAALAHS | GARRHSETLY | 0.74948275 |
| GGVTVMATP | GARRHSETLY | 0.71488607 |
| AGLAANVRD | GARRHSETLY | 0.704543 |
| TNRPDLIDPALLRPG | GARRHSETLY | 0.68477446 |
| LANLGVTAL | GARRHSETLY | 0.7459365 |
| TGERIEL | GARRHSETLY | 0.35611498 |
| NGEELAA | GARRHSETLY | 0.19386017 |
| VLSLVWG | GARRHSETLY | 0.39338237 |
| IGYALWL | GARRHSETLY | 0.5650702 |
| LSLLLAPL | GARRHSETLY | 0.25807214 |
| TLDEALK | GARRHSETLY | 0.21807148 |
| TDGAGTAEGG | GARRHSETLY | 0.798751 |
| NAPGGAGGNGG | GARRHSETLY | 0.8674216 |
| VSVFMGMGGNF | GARRHSETLY | 0.3240291 |
| LASYLAAL | GARRHSETLY | 0.82865363 |
| LVGGTPAL | GARRHSETLY | 0.20651777 |
| VRNADVL | GARRHSETLY | 0.7700227 |
| TCSSPFLVSAM | GARRHSETLY | 0.80812275 |
| VHITDAE | GARRHSETLY | 0.28733867 |
| EYAAPSF | GARRHSETLY | 0.6514678 |
| VVDAFHGGL | GARRHSETLY | 0.2164469 |

| | | |
|---|---|---|
| ASVVDPQ | GARRHSETLY | 0.3058319 |
| SYNEFGGLS | GARRHSETLY | 0.80469155 |
| DPDGSVY | GARRHSETLY | 0.38555247 |
| MVSTIHATE | GARRHSETLY | 0.51060987 |
| LAAGVADPV | GARRHSETLY | 0.39783147 |
| TKEQIAA | GARRHSETLY | 0.23043782 |
| SLASPAAL | GARRHSETLY | 0.42317972 |
| IGLVFYGV | GARRHSETLY | 0.755067 |
| FLLDASNL | GARRHSETLY | 0.45799157 |
| YQVGNFP | GARRHSETLY | 0.8725131 |
| ESLLSQST | GARRHSETLY | 0.20234455 |
| LFTVGFAAVG | GARRHSETLY | 0.83646643 |
| ATPPTPT | GARRHSETLY | 0.8135947 |
| TDLATLG | GARRHSETLY | 0.63585114 |
| SAGGTGR | GARRHSETLY | 0.4400707 |
| GFSNSGS | GARRHSETLY | 0.8080121 |
| GGNGGSGGTSV | GARRHSETLY | 0.43962803 |
| GTSVATGGAGN | GARRHSETLY | 0.65345275 |
| NGGNGGSGGTS | GARRHSETLY | 0.77642655 |
| AVVKFPH | GARRHSETLY | 0.15272884 |
| TPPPPPP | GARRHSETLY | 0.75411 |
| AAPAAAAAPA | GARRHSETLY | 0.5700253 |
| IALNEDL | GARRHSETLY | 0.48543853 |
| IVNVSSM | GARRHSETLY | 0.22438842 |
| TGIPPAPR | GARRHSETLY | 0.12721393 |
| FDLGGGTF | GARRHSETLY | 0.27316898 |
| FDLGGGTFDVS | GARRHSETLY | 0.1775909 |
| NEPTAAA | GARRHSETLY | 0.50592184 |
| NEPTAAALA | GARRHSETLY | 0.4973713 |
| GSIINISSI | GARRHSETLY | 0.22067606 |
| VAPGNAGTA | GARRHSETLY | 0.6924475 |
| TGGGASLE | GARRHSETLY | 0.72121906 |
| VAVYNHYKRI | GARRHSETLY | 0.12759984 |
| TDPAPATT | GARRHSETLY | 0.3435325 |
| TNRPDILDPALLRPG | GARRHSETLY | 0.16938677 |
| LTTTTTS | GARRHSETLY | 0.23880556 |
| PTGSGKTL | GARRHSETLY | 0.10508249 |
| TGSGKTL | GARRHSETLY | 0.51102644 |
| TIGHVDHGK | GARRHSETLY | 0.21485865 |
| TIGHVDHGKTT | GARRHSETLY | 0.10038562 |
| GHVDHGKTT | GARRHSETLY | 0.4051459 |
| AQFEHTLL | GARRHSETLY | 0.29825956 |
| IVNQDEL | GARRHSETLY | 0.23170882 |
| TNGSQFF | GARRHSETLY | 0.6624799 |
| TGKGIVE | GARRHSETLY | 0.11443534 |
| LISNASDA | GARRHSETLY | 0.18119481 |
| AELRTLGELTKIAKAHGAQ | GARRHSETLY | 0.5652046 |
| APNAVAAAEPSW | GARRHSETLY | 0.74875724 |
| APQVAGAATAVL | GARRHSETLY | 0.17107822 |
| ARRDPAAAAAILPTDA | GARRHSETLY | 0.47447383 |
| DAAARDVLAEAGLAEYF | GARRHSETLY | 0.78072643 |

| | | |
|---|---|---|
| DDQIVLSAKANLAKVEQ | GARRHSETLY | 0.38650227 |
| DELLPDTGEDSDEDGH | GARRHSETLY | 0.63149345 |
| ELPEAPANRATVPQD | GARRHSETLY | 0.7379953 |
| FGQSKIERVTGIG | GARRHSETLY | 0.19906501 |
| GAGGTGSTAGGAGGAGGAGGLY | GARRHSETLY | 0.63181615 |
| GHGGTGAAGGNGGTGGAGGSGI | GARRHSETLY | 0.7474251 |
| GNGGNGGAGGEGGPGGAGGAGA | GARRHSETLY | 0.29543442 |
| GSTAGNGGNGGNATNGTIAGQP | GARRHSETLY | 0.30267018 |
| GTGGDGGNGGNGGTGG | GARRHSETLY | 0.16499515 |
| GVRRPAGIPADAPV | GARRHSETLY | 0.32690883 |
| GYRQVTDAHLLTLARR | GARRHSETLY | 0.12866297 |
| IAKMGQDGHDRG | GARRHSETLY | 0.5235635 |
| KAEPSIEIQYT | GARRHSETLY | 0.63371515 |
| KPVRIRMAADEDTVAVT | GARRHSETLY | 0.28511864 |
| MPPRIAGMRLLVIKPEP | GARRHSETLY | 0.1381132 |
| MKFYAVALFRSRGVPPTR | GARRHSETLY | 0.1304803 |
| NAGGFGILGAVAHSP | GARRHSETLY | 0.30933002 |
| NEGREITLAMAEM | GARRHSETLY | 0.3345507 |
| NELVEWSHPLGHRWRWE | GARRHSETLY | 0.11816285 |
| NLGAGGTGGNGGNPG | GARRHSETLY | 0.27637532 |
| NTEWPKWETVMAADPVD | GARRHSETLY | 0.6120468 |
| PASAAPAETQP | GARRHSETLY | 0.6240928 |
| QQSSDGKNGVRAAEITGE | GARRHSETLY | 0.1341075 |
| RAPDAVTTSWWKE | GARRHSETLY | 0.7738045 |
| SGFGNVGAHTSGWFNQ | GARRHSETLY | 0.64890015 |
| SGVAGAVGQAASVGGLKV | GARRHSETLY | 0.7141458 |
| SNGGAGTGSAGGNGGT | GARRHSETLY | 0.27023786 |
| TGGAGGEGGAAG | GARRHSETLY | 0.15835744 |
| TGGDGGAAGAGGAPGHG | GARRHSETLY | 0.74505806 |
| TMPPLLAAIANAGRHGV | GARRHSETLY | 0.40477097 |
| TSSTAGNSGGA | GARRHSETLY | 0.19109304 |
| VPATVPVGILC | GARRHSETLY | 0.5379851 |
| VQRPDSGGVVLTGRIS | GARRHSETLY | 0.39163482 |
| WTPNPVPATKGVPAGLVQAT | GARRHSETLY | 0.22006708 |
| YALKRALAKFSDRFDVVII | GARRHSETLY | 0.6266061 |
| YLIGSVIV | GARRHSETLY | 0.528665 |
| YGQLVAVID | GARRHSETLY | 0.17858425 |
| VTVGGMTLPA | GARRHSETLY | 0.19034402 |
| VLALPAV | GARRHSETLY | 0.20383307 |
| VIPPTLNYET | GARRHSETLY | 0.44723934 |
| VGMGGMHPSSG | GARRHSETLY | 0.6738987 |
| VGLAGGAATA | GARRHSETLY | 0.31245667 |
| VGGAGGAGGSGP | GARRHSETLY | 0.30508143 |
| VFAAFMV | GARRHSETLY | 0.663816 |
| TLYDKLR | GARRHSETLY | 0.5212305 |
| TLAVFTVL | GARRHSETLY | 0.32669997 |
| TGSVQPARV | GARRHSETLY | 0.7058555 |
| TGGGAGSGGT | GARRHSETLY | 0.19676901 |
| SHLETSL | GARRHSETLY | 0.4848122 |
| SFTTGMT | GARRHSETLY | 0.51065826 |
| QVEVAKRMGVRQPTVS | GARRHSETLY | 0.5316316 |

| | | |
|---|---|---|
| QGSGGPALAM | GARRHSETLY | 0.6360252 |
| MLAPSNP | GARRHSETLY | 0.5510381 |
| LVNNAAI | GARRHSETLY | 0.42704076 |
| LTGEINTLA | GARRHSETLY | 0.1200581 |
| LSIFGPV | GARRHSETLY | 0.2428576 |
| LLALLANL | GARRHSETLY | 0.6090307 |
| LISNASDALDKLR | GARRHSETLY | 0.53227687 |
| LINEAVP | GARRHSETLY | 0.64817715 |
| LGYAFSGL | GARRHSETLY | 0.47562283 |
| LDFVAVAL | GARRHSETLY | 0.2354756 |
| LAQPGAE | GARRHSETLY | 0.76317644 |
| ISNLGAVTGF | GARRHSETLY | 0.7107389 |
| HQEDPGRI | GARRHSETLY | 0.13214685 |
| GTPGEPTP | GARRHSETLY | 0.27439746 |
| GFFNSGS | GARRHSETLY | 0.7460917 |
| GAGGGGLTAGAN | GARRHSETLY | 0.60208386 |
| FVDALVNM | GARRHSETLY | 0.31006753 |
| FLTVAPEL | GARRHSETLY | 0.34797826 |
| FETLTAL | GARRHSETLY | 0.7802963 |
| DVINPIEL | GARRHSETLY | 0.2223295 |
| APVSGAPV | GARRHSETLY | 0.7657534 |
| ALVGGLAW | GARRHSETLY | 0.11281365 |
| ALAAAAATAI | GARRHSETLY | 0.28644443 |
| AFGLGLLL | GARRHSETLY | 0.1763716 |
| AFFAAASL | GARRHSETLY | 0.114316426 |
| AAAALPAFSPP | GARRHSETLY | 0.5083302 |
| VVLAPAL | GARRHSETLY | 0.22530138 |
| TIGHVDHGKTTL | GARRHSETLY | 0.68379366 |
| LISNASDALDK | GARRHSETLY | 0.24388799 |
| LISNASDAL | GARRHSETLY | 0.62183195 |
| ISNASDALDK | GARRHSETLY | 0.4009773 |
| NQAETLVYQTE | GARRHSETLY | 0.4310236 |
| TGGTGGTAGAG | GARRHSETLY | 0.16467854 |
| TGSTAGGAGGA | GARRHSETLY | 0.5700599 |
| DPLAVPVAAGPV | GARRHSETLY | 0.75959325 |
| GGTGGTPGNGGN | GARRHSETLY | 0.41293883 |
| PAGLAGVDPATL | GARRHSETLY | 0.14924546 |
| GNNLVVEAPHAGASV | GARRHSETLY | 0.3613981 |
| KVTRIMSASTLEQPA | GARRHSETLY | 0.33044547 |
| LGFALPIALPRRTVV | GARRHSETLY | 0.10634814 |
| TRLGNLTARKLVVLV | GARRHSETLY | 0.104198195 |
| WSQQRRLIYFPSAGP | GARRHSETLY | 0.3446743 |
| GPNGNPDPMAAAVDIR | GARRHSETLY | 0.21676457 |
| TQRCVSVADLRAAASR | GARRHSETLY | 0.6543494 |
| QFVLDVAEHPVEVLTETV | GARRHSETLY | 0.5428517 |
| GGAGGTGGAGGNGGTQAGDG | GARRHSETLY | 0.25259534 |
| PLLATLRAIAVDVDAKPAQV | GARRHSETLY | 0.61497456 |
| APKFRTGGYMIDFWGVGYQVAK | GARRHSETLY | 0.58955085 |
| GAGGAGGAGGAAGGPGGTGGTG | GARRHSETLY | 0.09734856 |
| IGALGLLATDVAPAGSVAFGVA | GARRHSETLY | 0.19117203 |
| AGGAGGIGGNGGAGGAGGVLMG | GARRHSETLY | 0.13217825 |

| | | |
|---|---|---|
| ADELIGLL | GARRHSETLY | 0.67615634 |
| ADRALKA | GARRHSETLY | 0.1537291 |
| AGAVGQVGGAGGA | GARRHSETLY | 0.21024969 |
| AGGGGGLGGAP | GARRHSETLY | 0.3058747 |
| AGGLLYGNGGAG | GARRHSETLY | 0.68614185 |
| AGLESIGL | GARRHSETLY | 0.49473897 |
| AGLKTGII | GARRHSETLY | 0.16895448 |
| ASLAALLL | GARRHSETLY | 0.10099504 |
| ASVINISSI | GARRHSETLY | 0.5034288 |
| ATPPGPA | GARRHSETLY | 0.13410804 |
| ATVASALV | GARRHSETLY | 0.18886586 |
| ATVASLIL | GARRHSETLY | 0.11786731 |
| AVNGPVLIRH | GARRHSETLY | 0.10185458 |
| DGAAGAVGI | GARRHSETLY | 0.13802454 |
| DPLLHGI | GARRHSETLY | 0.6112995 |
| ELGAALGAL | GARRHSETLY | 0.25388092 |
| ELGRTQAV | GARRHSETLY | 0.31798023 |
| FVAGGYLS | GARRHSETLY | 0.7680555 |
| GASGGAGGAGGW | GARRHSETLY | 0.68096036 |
| GGAGAGGDGGAGGI | GARRHSETLY | 0.2912745 |
| GTGHNGSQPGMGGNG | GARRHSETLY | 0.25553915 |
| GYAAASASAS | GARRHSETLY | 0.14920789 |
| HQVDVAAVQQG | GARRHSETLY | 0.17197526 |
| IGAAIAHT | GARRHSETLY | 0.20409265 |
| IGMIEVAQL | GARRHSETLY | 0.76155686 |
| IRLAPEHTSAMD | GARRHSETLY | 0.25505114 |
| IYNVSVL | GARRHSETLY | 0.13366622 |
| KQGFLGM | GARRHSETLY | 0.3685933 |
| LESVITQL | GARRHSETLY | 0.7047807 |
| LGLGALVVV | GARRHSETLY | 0.7773481 |
| LIGNGGNGGSGGI | GARRHSETLY | 0.16124715 |
| LNAINAPTL | GARRHSETLY | 0.3037545 |
| LQLLNTGV | GARRHSETLY | 0.32221022 |
| LQTSIEF | GARRHSETLY | 0.355188 |
| MSFAVSEL | GARRHSETLY | 0.2625742 |
| NTGALGL | GARRHSETLY | 0.15788461 |
| PPGAGGAGGAGTT | GARRHSETLY | 0.49507502 |
| QLVGLGI | GARRHSETLY | 0.8075376 |
| RFLEQVHQL | GARRHSETLY | 0.23987463 |
| RGVGYVLR | GARRHSETLY | 0.11503358 |
| SATGVFAGVI | GARRHSETLY | 0.37909755 |
| SLPAITLP | GARRHSETLY | 0.20732921 |
| STALAFTV | GARRHSETLY | 0.19778316 |
| SVLTVGLGK | GARRHSETLY | 0.37132734 |
| TALAGAGSGI | GARRHSETLY | 0.12668239 |
| TGLPFTGI | GARRHSETLY | 0.26087093 |
| VAEHVTV | GARRHSETLY | 0.8164023 |
| VDIVLNSL | GARRHSETLY | 0.41304404 |
| VLTGGGAVL | GARRHSETLY | 0.63829505 |
| VNVGLAYGI | GARRHSETLY | 0.6727829 |
| VSSAAVGAV | GARRHSETLY | 0.5516639 |

| | | |
|---|---|---|
| VTSVYEAGV | GARRHSETLY | 0.7339251 |
| WVALDRGA | GARRHSETLY | 0.7060349 |
| YLVGSVVI | GARRHSETLY | 0.31170207 |
| GPPGTGKTLL | GARRHSETLY | 0.45426983 |
| GVGMTKF | GARRHSETLY | 0.33197802 |
| IGIGGSDL | GARRHSETLY | 0.43090615 |
| IGTLLAEL | GARRHSETLY | 0.3380515 |
| IYTSGTT | GARRHSETLY | 0.22882658 |
| LSFPAGV | GARRHSETLY | 0.6919126 |
| LYTSGTT | GARRHSETLY | 0.4120092 |
| VGAFPFGF | GARRHSETLY | 0.6271354 |
| VGAGTVEF | GARRHSETLY | 0.15436542 |
| VGKSTLLL | GARRHSETLY | 0.17322682 |
| AALAHAGGGIV | GARRHSETLY | 0.69202304 |
| AALVAVSLL | GARRHSETLY | 0.103372745 |
| AAVSRAVAAL | GARRHSETLY | 0.117250435 |
| AGGSSGIGGF | GARRHSETLY | 0.2913227 |
| AGLTVANANG | GARRHSETLY | 0.7871352 |
| AGLVGAVL | GARRHSETLY | 0.10218653 |
| ALSQAFTV | GARRHSETLY | 0.1754374 |
| ALTFSVSL | GARRHSETLY | 0.15030912 |
| ALTGAALVAL | GARRHSETLY | 0.5819818 |
| ASFSIGAGVL | GARRHSETLY | 0.33624083 |
| DGSVYKI | GARRHSETLY | 0.12644233 |
| EQLAEIV | GARRHSETLY | 0.52337915 |
| FAVSELTVM | GARRHSETLY | 0.110961586 |
| GAAPLAAGL | GARRHSETLY | 0.11769503 |
| GADGGDGGSGGAGG | GARRHSETLY | 0.58687156 |
| GFGLDTGGAGG | GARRHSETLY | 0.75077516 |
| GLASGGLL | GARRHSETLY | 0.13633907 |
| GRIAGAAVLL | GARRHSETLY | 0.61269265 |
| GSAGSLNTSF | GARRHSETLY | 0.2133005 |
| GTGQIGI | GARRHSETLY | 0.7870513 |
| GTLIALIL | GARRHSETLY | 0.2747053 |
| IAATMGFVVG | GARRHSETLY | 0.123899475 |
| IFAQALL | GARRHSETLY | 0.592088 |
| IGAAVALGF | GARRHSETLY | 0.5204179 |
| IGLTPLAL | GARRHSETLY | 0.38598216 |
| ILGSNTSTL | GARRHSETLY | 0.60463834 |
| IVFSLTVL | GARRHSETLY | 0.28626356 |
| LASKATGF | GARRHSETLY | 0.27176487 |
| LASVALIL | GARRHSETLY | 0.62705714 |
| LFAVGLTL | GARRHSETLY | 0.6517689 |
| LFNGGIGTYI | GARRHSETLY | 0.19233479 |
| LTLGAGII | GARRHSETLY | 0.615639 |
| LVAAALGDI | GARRHSETLY | 0.42408282 |
| NTAFVQLGI | GARRHSETLY | 0.18380688 |
| QLFSGAEL | GARRHSETLY | 0.77823997 |
| SAATFITL | GARRHSETLY | 0.31137767 |
| SNYGLANLGV | GARRHSETLY | 0.617408 |
| SYGGAYAVM | GARRHSETLY | 0.49144775 |

| | | |
|---|---|---|
| TTGVGGLTLG | GARRHSETLY | 0.6611617 |
| TTQTIYI | GARRHSETLY | 0.1387483 |
| VAAGAGLGAVI | GARRHSETLY | 0.64335203 |
| VAAVVTLL | GARRHSETLY | 0.4385043 |
| VGFLGGLI | GARRHSETLY | 0.594962 |
| VGVIPILL | GARRHSETLY | 0.3123916 |
| VIFASSLI | GARRHSETLY | 0.15862277 |
| VKGGDGGAAAG | GARRHSETLY | 0.4006313 |
| VTKTFTNI | GARRHSETLY | 0.67832947 |
| VVAAAILL | GARRHSETLY | 0.27288148 |
| YAALDVEL | GARRHSETLY | 0.57580835 |
| LASVALLL | GARRHSETLY | 0.7628504 |
| LDAVHRV | GARRHSETLY | 0.16274792 |
| QGLSSSIF | GARRHSETLY | 0.3282531 |
| RGTFANIRL | GARRHSETLY | 0.50441086 |
| RQLWWGHRI | GARRHSETLY | 0.7468532 |
| YGPPGTGKTLL | GARRHSETLY | 0.27408493 |
| AALVGAPANVA | GARRHSETLY | 0.7311791 |
| AATFRRTE | GARRHSETLY | 0.21080866 |
| AGALGGGI | GARRHSETLY | 0.59041256 |
| AGGAGGAGGIGAG | GARRHSETLY | 0.15733495 |
| AGSGALGL | GARRHSETLY | 0.4329276 |
| AQIADQI | GARRHSETLY | 0.22890657 |
| ASAVAVTYL | GARRHSETLY | 0.5752572 |
| ASTVLGGTGG | GARRHSETLY | 0.63166296 |
| AVGGEPAK | GARRHSETLY | 0.63632506 |
| CEQGTFGGTC | GARRHSETLY | 0.10611467 |
| DEGKAPAG | GARRHSETLY | 0.5467068 |
| DTVVFPAL | GARRHSETLY | 0.6618567 |
| FEGITMAG | GARRHSETLY | 0.8054733 |
| FPLVVQD | GARRHSETLY | 0.46508121 |
| GAFTTPQ | GARRHSETLY | 0.6455498 |
| GFGGFGTAGGVGGA | GARRHSETLY | 0.25763583 |
| GFLAQFL | GARRHSETLY | 0.28196573 |
| GGAGGAGGNAGVA | GARRHSETLY | 0.17548831 |
| GGVGTAGAGGAGGAGG | GARRHSETLY | 0.31527862 |
| GNGGGGDAGNAGSGGN | GARRHSETLY | 0.28425014 |
| GTGGAGGAGNDAGS | GARRHSETLY | 0.17782435 |
| GTTNTGWFNS | GARRHSETLY | 0.75664103 |
| IFMGPLFTL | GARRHSETLY | 0.7478123 |
| ILAHAIEAI | GARRHSETLY | 0.7368101 |
| IRSAIREAL | GARRHSETLY | 0.72357774 |
| ITFAGGG | GARRHSETLY | 0.6878846 |
| ITGMSGHQEV | GARRHSETLY | 0.5316393 |
| IVAGLACTL | GARRHSETLY | 0.24133977 |
| IVGGGLAGA | GARRHSETLY | 0.24095255 |
| IVTGVLNI | GARRHSETLY | 0.13351561 |
| LAQQGCI | GARRHSETLY | 0.2167027 |
| LCLHVEN | GARRHSETLY | 0.2565122 |
| LDAISDGS | GARRHSETLY | 0.353095 |
| LGGFAAML | GARRHSETLY | 0.10235025 |

| | | |
|---|---|---|
| LGGKNPMI | GARRHSETLY | 0.5892806 |
| LGPKGVAAIL | GARRHSETLY | 0.76005536 |
| LMVGVSI | GARRHSETLY | 0.29913557 |
| LVGAAFVL | GARRHSETLY | 0.5653999 |
| LVLAVAPL | GARRHSETLY | 0.46940637 |
| MAAIAVIDAA | GARRHSETLY | 0.21627483 |
| NNGINAELQ | GARRHSETLY | 0.6719856 |
| RGEPLVLVHG | GARRHSETLY | 0.21887526 |
| RGGAGGAAGLI | GARRHSETLY | 0.57577205 |
| RSYRMHPAV | GARRHSETLY | 0.77021533 |
| SAAHLLAGQLA | GARRHSETLY | 0.39055464 |
| SAAVGSY | GARRHSETLY | 0.57744515 |
| SPHAVLHRI | GARRHSETLY | 0.44028753 |
| SQIAKVL | GARRHSETLY | 0.13045952 |
| TALGHLVP | GARRHSETLY | 0.60788476 |
| TAPTTSGAA | GARRHSETLY | 0.09810809 |
| TDGDDGNGGSPGT | GARRHSETLY | 0.43887013 |
| TDGLGEN | GARRHSETLY | 0.1768746 |
| TGGPPLPGQA | GARRHSETLY | 0.5232327 |
| TIPAITI | GARRHSETLY | 0.3754183 |
| TIPGFTI | GARRHSETLY | 0.4240341 |
| TSIASGALA | GARRHSETLY | 0.20855156 |
| TVTLPGAAI | GARRHSETLY | 0.31633174 |
| VASGLSAALP | GARRHSETLY | 0.26795146 |
| VGAGVIGI | GARRHSETLY | 0.149808 |
| VGPKRFGV | GARRHSETLY | 0.57887936 |
| VNVAFPD | GARRHSETLY | 0.21545258 |
| VSMTNSGVS | GARRHSETLY | 0.39443165 |
| VTDVLGDLSV | GARRHSETLY | 0.6612215 |
| VVVVGAGFAGL | GARRHSETLY | 0.3829475 |
| VYTVGTVV | GARRHSETLY | 0.12643804 |
| WTTFSGP | GARRHSETLY | 0.4958204 |
| YDELVEAF | GARRHSETLY | 0.41215897 |
| YGTGGAGGNGG | GARRHSETLY | 0.30693656 |
| YTNVMAV | GARRHSETLY | 0.098311946 |
| FLEQVHQL | GARRHSETLY | 0.4907892 |
| GLSLLAVAV | GARRHSETLY | 0.19607788 |
| LEQVHQL | GARRHSETLY | 0.19062094 |
| LNAAGAI | GARRHSETLY | 0.1993584 |
| PAVHASL | GARRHSETLY | 0.66642845 |
| TELEDDA | GARRHSETLY | 0.14550634 |
| TPPGPPN | GARRHSETLY | 0.51604843 |
| TPPPPPPP | GARRHSETLY | 0.6543672 |
| TQSPALI | ASGGGAEEDEQY | 0.56769013 |
| EFAAAFPQ | ASGGGAEEDEQY | 0.7699289 |
| LLSGNLDVL | ASGGGAEEDEQY | 0.5800806 |
| GPANLGNYNI | ASGGGAEEDEQY | 0.27026775 |
| TLGPTLP | ASGGGAEEDEQY | 0.5248373 |
| VYAYSSCCK | ASGGGAEEDEQY | 0.2933362 |
| APAPAPAPNI | ASGGGAEEDEQY | 0.10983171 |
| PGGPVPIP | ASGGGAEEDEQY | 0.12343323 |

| | | |
|---|---|---|
| FLTVALEP | ASGGGAEEDEQY | 0.26996636 |
| SQYFGANL | ASGGGAEEDEQY | 0.114895195 |
| GGVKPAA | ASGGGAEEDEQY | 0.50001645 |
| GGTGGAGGAGGW | ASGGGAEEDEQY | 0.6065308 |
| GVGGVAGLGGAGG | ASGGGAEEDEQY | 0.45562148 |
| VLLVLATTI | ASGGGAEEDEQY | 0.13318963 |
| IGIFAAML | ASGGGAEEDEQY | 0.3033117 |
| AVGDLID | ASGGGAEEDEQY | 0.1809471 |
| NISPLPL | ASGGGAEEDEQY | 0.5473692 |
| VTAVFVAI | ASGGGAEEDEQY | 0.53618467 |
| SGDDNDAAL | ASGGGAEEDEQY | 0.77849174 |
| AVAALAHS | ASGGGAEEDEQY | 0.63872766 |
| GGVTVMATP | ASGGGAEEDEQY | 0.25814763 |
| AGLAANVRD | ASGGGAEEDEQY | 0.5981821 |
| TNRPDLIDPALLRPG | ASGGGAEEDEQY | 0.25632465 |
| LANLGVTAL | ASGGGAEEDEQY | 0.3285987 |
| TGERIEL | ASGGGAEEDEQY | 0.3163794 |
| NGEELAA | ASGGGAEEDEQY | 0.37429613 |
| VLSLVWG | ASGGGAEEDEQY | 0.3829529 |
| IGYALWL | ASGGGAEEDEQY | 0.2924024 |
| LSLLLAPL | ASGGGAEEDEQY | 0.12856695 |
| TLDEALK | ASGGGAEEDEQY | 0.48209673 |
| TDGAGTAEGG | ASGGGAEEDEQY | 0.56652415 |
| NAPGGAGGNGG | ASGGGAEEDEQY | 0.12429826 |
| VSVFMGMGGNF | ASGGGAEEDEQY | 0.41359678 |
| LASYLAAL | ASGGGAEEDEQY | 0.7906883 |
| LVGGTPAL | ASGGGAEEDEQY | 0.7825663 |
| VRNADVL | ASGGGAEEDEQY | 0.22078979 |
| TCSSPFLVSAM | ASGGGAEEDEQY | 0.20452715 |
| VHITDAE | ASGGGAEEDEQY | 0.66493225 |
| EYAAPSF | ASGGGAEEDEQY | 0.14177836 |
| VVDAFHGGL | ASGGGAEEDEQY | 0.42379266 |
| ASVVDPQ | ASGGGAEEDEQY | 0.48225766 |
| SYNEFGGLS | ASGGGAEEDEQY | 0.30484194 |
| DPDGSVY | ASGGGAEEDEQY | 0.28511322 |
| MVSTIHATE | ASGGGAEEDEQY | 0.5085533 |
| LAAGVADPV | ASGGGAEEDEQY | 0.3166535 |
| TKEQIAA | ASGGGAEEDEQY | 0.55276763 |
| SLASPAAL | ASGGGAEEDEQY | 0.10078364 |
| IGLVFYGV | ASGGGAEEDEQY | 0.45596015 |
| FLLDASNL | ASGGGAEEDEQY | 0.19440198 |
| YQVGNFP | ASGGGAEEDEQY | 0.19777055 |
| ESLLSQST | ASGGGAEEDEQY | 0.120774165 |
| LFTVGFAAVG | ASGGGAEEDEQY | 0.25152633 |
| ATPPTPT | ASGGGAEEDEQY | 0.62839746 |
| TDLATLG | ASGGGAEEDEQY | 0.7192933 |
| SAGGTGR | ASGGGAEEDEQY | 0.15277871 |
| GFSNSGS | ASGGGAEEDEQY | 0.69583696 |
| GGNGGSGGTSV | ASGGGAEEDEQY | 0.792914 |
| GTSVATGGAGN | ASGGGAEEDEQY | 0.25797036 |
| NGGNGGSGGTS | ASGGGAEEDEQY | 0.1276394 |

| | | |
|---|---|---|
| AVVKFPH | ASGGGAEEDEQY | 0.44957262 |
| TPPPPPP | ASGGGAEEDEQY | 0.27577132 |
| AAPAAAAAPA | ASGGGAEEDEQY | 0.6046717 |
| IALNEDL | ASGGGAEEDEQY | 0.765111 |
| IVNVSSM | ASGGGAEEDEQY | 0.12235219 |
| TGIPPAPR | ASGGGAEEDEQY | 0.67403597 |
| FDLGGGTF | ASGGGAEEDEQY | 0.73027444 |
| FDLGGGTFDVS | ASGGGAEEDEQY | 0.21436529 |
| NEPTAAA | ASGGGAEEDEQY | 0.16377816 |
| NEPTAAALA | ASGGGAEEDEQY | 0.17053446 |
| GSIINISSI | ASGGGAEEDEQY | 0.107747965 |
| VAPGNAGTA | ASGGGAEEDEQY | 0.59042954 |
| TGGGASLE | ASGGGAEEDEQY | 0.2859135 |
| VAVYNHYKRI | ASGGGAEEDEQY | 0.22995843 |
| TDPAPATT | ASGGGAEEDEQY | 0.26862198 |
| TNRPDILDPALLRPG | ASGGGAEEDEQY | 0.24094917 |
| LTTTTTS | ASGGGAEEDEQY | 0.22233807 |
| PTGSGKTL | ASGGGAEEDEQY | 0.2385104 |
| TGSGKTL | ASGGGAEEDEQY | 0.3140818 |
| TIGHVDHGK | ASGGGAEEDEQY | 0.5756607 |
| TIGHVDHGKTT | ASGGGAEEDEQY | 0.334461 |
| GHVDHGKTT | ASGGGAEEDEQY | 0.37714994 |
| AQFEHTLL | ASGGGAEEDEQY | 0.7350447 |
| IVNQDEL | ASGGGAEEDEQY | 0.17183742 |
| TNGSQFF | ASGGGAEEDEQY | 0.20077977 |
| TGKGIVE | ASGGGAEEDEQY | 0.60549986 |
| LISNASDA | ASGGGAEEDEQY | 0.15276276 |
| AELRTLGELTKIAKAHGAQ | ASGGGAEEDEQY | 0.6473328 |
| APNAVAAAEPSW | ASGGGAEEDEQY | 0.5142256 |
| APQVAGAATAVL | ASGGGAEEDEQY | 0.44046536 |
| ARRDPAAAAAILPTDA | ASGGGAEEDEQY | 0.41928205 |
| DAAARDVLAEAGLAEYF | ASGGGAEEDEQY | 0.7850838 |
| DDQIVLSAKANLAKVEQ | ASGGGAEEDEQY | 0.51510787 |
| DELLPDTGEDSDEDGH | ASGGGAEEDEQY | 0.22043978 |
| ELPEAPANRATVPQD | ASGGGAEEDEQY | 0.3349597 |
| FGQSKIERVTGIG | ASGGGAEEDEQY | 0.17900854 |
| GAGGTGSTAGGAGGAGGAGGLY | ASGGGAEEDEQY | 0.678534 |
| GHGGTGAAGGNGGTGGAGGSGI | ASGGGAEEDEQY | 0.6845747 |
| GNGGNGGAGGEGGPGGAGGAGA | ASGGGAEEDEQY | 0.15561025 |
| GSTAGNGGNGGNATNGTIAGQP | ASGGGAEEDEQY | 0.28654575 |
| GTGGDGGNGGNGGTGG | ASGGGAEEDEQY | 0.64833677 |
| GVRRPAGIPADAPV | ASGGGAEEDEQY | 0.6787869 |
| GYRQVTDAHLLTLARR | ASGGGAEEDEQY | 0.13093695 |
| IAKMGQDGHDRG | ASGGGAEEDEQY | 0.18903664 |
| KAEPSIEIQYT | ASGGGAEEDEQY | 0.14649664 |
| KPVRIRMAADEDTVAVT | ASGGGAEEDEQY | 0.4077621 |
| MPPRIAGMRLLVIKPEP | ASGGGAEEDEQY | 0.15141614 |
| MKFYAVALFRSRGVPPTR | ASGGGAEEDEQY | 0.5985921 |
| NAGGFGILGAVAHSP | ASGGGAEEDEQY | 0.27581272 |
| NEGREITLAMAEM | ASGGGAEEDEQY | 0.7272179 |
| NELVEWSHPLGHRWRWE | ASGGGAEEDEQY | 0.40347743 |

| | | |
|---|---|---|
| NLGAGGTGGNGGNPG | ASGGGAEEDEQY | 0.6439456 |
| NTEWPKWETVMAADPVD | ASGGGAEEDEQY | 0.26866198 |
| PASAAPAETQP | ASGGGAEEDEQY | 0.31767428 |
| QQSSDGKNGVRAAEITGE | ASGGGAEEDEQY | 0.13854901 |
| RAPDAVTTSWWKE | ASGGGAEEDEQY | 0.15993193 |
| SGFGNVGAHTSGWFNQ | ASGGGAEEDEQY | 0.14708397 |
| SGVAGAVGQAASVGGLKV | ASGGGAEEDEQY | 0.3097453 |
| SNGGAGTGSAGGNGGT | ASGGGAEEDEQY | 0.10766104 |
| TGGAGGEGGAAG | ASGGGAEEDEQY | 0.7366153 |
| TGGDGGAAGAGGAPGHG | ASGGGAEEDEQY | 0.39941454 |
| TMPPLLAAIANAGRHGV | ASGGGAEEDEQY | 0.49153045 |
| TSSTAGNSGGA | ASGGGAEEDEQY | 0.27787638 |
| VPATVPVGILC | ASGGGAEEDEQY | 0.24237344 |
| VQRPDSGGVVLTGRIS | ASGGGAEEDEQY | 0.2115038 |
| WTPNPVPATKGVPAGLVQAT | ASGGGAEEDEQY | 0.7265234 |
| YALKRALAKFSDRFDVVII | ASGGGAEEDEQY | 0.1407374 |
| YLIGSVIV | ASGGGAEEDEQY | 0.111883976 |
| YGQLVAVID | ASGGGAEEDEQY | 0.18785019 |
| VTVGGMTLPA | ASGGGAEEDEQY | 0.11829133 |
| VLALPAV | ASGGGAEEDEQY | 0.70774484 |
| VIPPTLNYET | ASGGGAEEDEQY | 0.6528348 |
| VGMGGMHPSSG | ASGGGAEEDEQY | 0.43264347 |
| VGLAGGAATA | ASGGGAEEDEQY | 0.66903806 |
| VGGAGGAGGSGP | ASGGGAEEDEQY | 0.10688959 |
| VFAAFMV | ASGGGAEEDEQY | 0.21535742 |
| TLYDKLR | ASGGGAEEDEQY | 0.7849891 |
| TLAVFTVL | ASGGGAEEDEQY | 0.4204679 |
| TGSVQPARV | ASGGGAEEDEQY | 0.3525628 |
| TGGGAGSGGT | ASGGGAEEDEQY | 0.67997813 |
| SHLETSL | ASGGGAEEDEQY | 0.2245591 |
| SFTTGMT | ASGGGAEEDEQY | 0.36621368 |
| QVEVAKRMGVRQPTVS | ASGGGAEEDEQY | 0.28260782 |
| QGSGGPALAM | ASGGGAEEDEQY | 0.52390003 |
| MLAPSNP | ASGGGAEEDEQY | 0.23840475 |
| LVNNAAI | ASGGGAEEDEQY | 0.14681199 |
| LTGEINTLA | ASGGGAEEDEQY | 0.54727626 |
| LSIFGPV | ASGGGAEEDEQY | 0.7840573 |
| LLALLANL | ASGGGAEEDEQY | 0.3116982 |
| LISNASDALDKLR | ASGGGAEEDEQY | 0.57801497 |
| LINEAVP | ASGGGAEEDEQY | 0.46742788 |
| LGYAFSGL | ASGGGAEEDEQY | 0.7292901 |
| LDFVAVAL | ASGGGAEEDEQY | 0.18555954 |
| LAQPGAE | ASGGGAEEDEQY | 0.17244065 |
| ISNLGAVTGF | ASGGGAEEDEQY | 0.33923715 |
| HQEDPGRI | ASGGGAEEDEQY | 0.16678116 |
| GTPGEPTP | ASGGGAEEDEQY | 0.404827 |
| GFFNSGS | ASGGGAEEDEQY | 0.21448463 |
| GAGGGGLTAGAN | ASGGGAEEDEQY | 0.8224537 |
| FVDALVNM | ASGGGAEEDEQY | 0.6346868 |
| FLTVAPEL | ASGGGAEEDEQY | 0.42273074 |
| FETLTAL | ASGGGAEEDEQY | 0.15584575 |

| | | |
|---|---|---|
| DVINPIEL | ASGGGAEEDEQY | 0.29168764 |
| APVSGAPV | ASGGGAEEDEQY | 0.58915854 |
| ALVGGLAW | ASGGGAEEDEQY | 0.12809773 |
| ALAAAAATAI | ASGGGAEEDEQY | 0.71553576 |
| AFGLGLLL | ASGGGAEEDEQY | 0.2074173 |
| AFFAAASL | ASGGGAEEDEQY | 0.35475284 |
| AAAALPAFSPP | ASGGGAEEDEQY | 0.5766554 |
| VVLAPAL | ASGGGAEEDEQY | 0.7578681 |
| TIGHVDHGKTTL | ASGGGAEEDEQY | 0.26656616 |
| LISNASDALDK | ASGGGAEEDEQY | 0.29835415 |
| LISNASDAL | ASGGGAEEDEQY | 0.15202466 |
| ISNASDALDK | ASGGGAEEDEQY | 0.6395777 |
| NQAETLVYQTE | ASGGGAEEDEQY | 0.71584487 |
| TGGTGGTAGAG | ASGGGAEEDEQY | 0.18987308 |
| TGSTAGGAGGA | ASGGGAEEDEQY | 0.7564373 |
| DPLAVPVAAGPV | ASGGGAEEDEQY | 0.35087854 |
| GGTGGTPGNGGN | ASGGGAEEDEQY | 0.14299631 |
| PAGLAGVDPATL | ASGGGAEEDEQY | 0.33831358 |
| GNNLVVEAPHAGASV | ASGGGAEEDEQY | 0.6208067 |
| KVTRIMSASTLEQPA | ASGGGAEEDEQY | 0.73693013 |
| LGFALPIALPRRTVV | ASGGGAEEDEQY | 0.7511667 |
| TRLGNLTARKLVVLV | ASGGGAEEDEQY | 0.6400591 |
| WSQQRRLIYFPSAGP | ASGGGAEEDEQY | 0.26170376 |
| GPNGNPDPMAAAVDIR | ASGGGAEEDEQY | 0.7233716 |
| TQRCVSVADLRAAASR | ASGGGAEEDEQY | 0.7769662 |
| QFVLDVAEHPVEVLTETV | ASGGGAEEDEQY | 0.584303 |
| GGAGGTGGAGGNGGTQAGDG | ASGGGAEEDEQY | 0.27916417 |
| PLLATLRAIAVDVDAKPAQV | ASGGGAEEDEQY | 0.2750598 |
| APKFRTGGYMIDFWGVGYQVAK | ASGGGAEEDEQY | 0.3475277 |
| GAGGAGGAGGAAGGPGGTGGTG | ASGGGAEEDEQY | 0.69338167 |
| IGALGLLATDVAPAGSVAFGVA | ASGGGAEEDEQY | 0.1892885 |
| AGGAGGIGGNGGAGGAGGVLMG | ASGGGAEEDEQY | 0.5002169 |
| ADELIGLL | ASGGGAEEDEQY | 0.5202514 |
| ADRALKA | ASGGGAEEDEQY | 0.12881409 |
| AGAVGQVGGAGGA | ASGGGAEEDEQY | 0.4049989 |
| AGGGGGLGGAP | ASGGGAEEDEQY | 0.7429476 |
| AGGLLYGNGGAG | ASGGGAEEDEQY | 0.31104457 |
| AGLESIGL | ASGGGAEEDEQY | 0.18736967 |
| AGLKTGII | ASGGGAEEDEQY | 0.7119749 |
| ASLAALLL | ASGGGAEEDEQY | 0.80632865 |
| ASVINISSI | ASGGGAEEDEQY | 0.623284 |
| ATPPGPA | ASGGGAEEDEQY | 0.14110391 |
| ATVASALV | ASGGGAEEDEQY | 0.10860989 |
| ATVASLIL | ASGGGAEEDEQY | 0.30555093 |
| AVNGPVLIRH | ASGGGAEEDEQY | 0.7437061 |
| DGAAGAVGI | ASGGGAEEDEQY | 0.27674583 |
| DPLLHGI | ASGGGAEEDEQY | 0.21084063 |
| ELGAALGAL | ASGGGAEEDEQY | 0.5760032 |
| ELGRTQAV | ASGGGAEEDEQY | 0.6175814 |
| FVAGGYLS | ASGGGAEEDEQY | 0.16719042 |
| GASGGAGGAGGW | ASGGGAEEDEQY | 0.14208889 |

| | | |
|---|---|---|
| GGAGAGGDGGAGGI | ASGGGAEEDEQY | 0.43389362 |
| GTGHNGSQPGMGGNG | ASGGGAEEDEQY | 0.6244975 |
| GYAAASASAS | ASGGGAEEDEQY | 0.15974687 |
| HQVDVAAVQQG | ASGGGAEEDEQY | 0.31843308 |
| IGAAIAHT | ASGGGAEEDEQY | 0.26268834 |
| IGMIEVAQL | ASGGGAEEDEQY | 0.14391491 |
| IRLAPEHTSAMD | ASGGGAEEDEQY | 0.5405443 |
| IYNVSVL | ASGGGAEEDEQY | 0.1739584 |
| KQGFLGM | ASGGGAEEDEQY | 0.6929628 |
| LESVITQL | ASGGGAEEDEQY | 0.13361847 |
| LGLGALVVV | ASGGGAEEDEQY | 0.5786405 |
| LIGNGGNGGSGGI | ASGGGAEEDEQY | 0.559664 |
| LNAINAPTL | ASGGGAEEDEQY | 0.45731086 |
| LQLLNTGV | ASGGGAEEDEQY | 0.3164553 |
| LQTSIEF | ASGGGAEEDEQY | 0.20939113 |
| MSFAVSEL | ASGGGAEEDEQY | 0.14644384 |
| NTGALGL | ASGGGAEEDEQY | 0.12751184 |
| PPGAGGAGGAGTT | ASGGGAEEDEQY | 0.7460995 |
| QLVGLGI | ASGGGAEEDEQY | 0.10101105 |
| RFLEQVHQL | ASGGGAEEDEQY | 0.70774084 |
| RGVGYVLR | ASGGGAEEDEQY | 0.4126508 |
| SATGVFAGVI | ASGGGAEEDEQY | 0.15399764 |
| SLPAITLP | ASGGGAEEDEQY | 0.3744895 |
| STALAFTV | ASGGGAEEDEQY | 0.19184002 |
| SVLTVGLGK | ASGGGAEEDEQY | 0.17237476 |
| TALAGAGSGI | ASGGGAEEDEQY | 0.7330376 |
| TGLPFTGI | ASGGGAEEDEQY | 0.53854215 |
| VAEHVTV | ASGGGAEEDEQY | 0.19733869 |
| VDIVLNSL | ASGGGAEEDEQY | 0.4386624 |
| VLTGGGAVL | ASGGGAEEDEQY | 0.45485416 |
| VNVGLAYGI | ASGGGAEEDEQY | 0.19135416 |
| VSSAAVGAV | ASGGGAEEDEQY | 0.32180586 |
| VTSVYEAGV | ASGGGAEEDEQY | 0.6439918 |
| WVALDRGA | ASGGGAEEDEQY | 0.4453152 |
| YLVGSVVI | ASGGGAEEDEQY | 0.13937645 |
| GPPGTGKTLL | ASGGGAEEDEQY | 0.553736 |
| GVGMTKF | ASGGGAEEDEQY | 0.19491647 |
| IGIGGSDL | ASGGGAEEDEQY | 0.43402156 |
| IGTLLAEL | ASGGGAEEDEQY | 0.3029726 |
| IYTSGTT | ASGGGAEEDEQY | 0.41160268 |
| LSFPAGV | ASGGGAEEDEQY | 0.6078068 |
| LYTSGTT | ASGGGAEEDEQY | 0.1473353 |
| VGAFPFGF | ASGGGAEEDEQY | 0.19039664 |
| VGAGTVEF | ASGGGAEEDEQY | 0.35971725 |
| VGKSTLLL | ASGGGAEEDEQY | 0.35683885 |
| AALAHAGGGIV | ASGGGAEEDEQY | 0.5471867 |
| AALVAVSLL | ASGGGAEEDEQY | 0.16931652 |
| AAVSRAVAAL | ASGGGAEEDEQY | 0.4108866 |
| AGGSSGIGGF | ASGGGAEEDEQY | 0.2953574 |
| AGLTVANANG | ASGGGAEEDEQY | 0.21113671 |
| AGLVGAVL | ASGGGAEEDEQY | 0.30630964 |

| | | |
|---|---|---|
| ALSQAFTV | ASGGGAEEDEQY | 0.22327273 |
| ALTFSVSL | ASGGGAEEDEQY | 0.15957592 |
| ALTGAALVAL | ASGGGAEEDEQY | 0.59228057 |
| ASFSIGAGVL | ASGGGAEEDEQY | 0.2141775 |
| DGSVYKI | ASGGGAEEDEQY | 0.5379152 |
| EQLAEIV | ASGGGAEEDEQY | 0.5777023 |
| FAVSELTVM | ASGGGAEEDEQY | 0.24012578 |
| GAAPLAAGL | ASGGGAEEDEQY | 0.44159955 |
| GADGGDGGSGGAGG | ASGGGAEEDEQY | 0.5216068 |
| GFGLDTGGAGG | ASGGGAEEDEQY | 0.5968707 |
| GLASGGLL | ASGGGAEEDEQY | 0.27729735 |
| GRIAGAAVLL | ASGGGAEEDEQY | 0.16971906 |
| GSAGSLNTSF | ASGGGAEEDEQY | 0.6252922 |
| GTGQIGI | ASGGGAEEDEQY | 0.524293 |
| GTLIALIL | ASGGGAEEDEQY | 0.7036634 |
| IAATMGFVVG | ASGGGAEEDEQY | 0.60165787 |
| IFAQALL | ASGGGAEEDEQY | 0.60662866 |
| IGAAVALGF | ASGGGAEEDEQY | 0.5682664 |
| IGLTPLAL | ASGGGAEEDEQY | 0.28579575 |
| ILGSNTSTL | ASGGGAEEDEQY | 0.16920975 |
| IVFSLTVL | ASGGGAEEDEQY | 0.23643678 |
| LASKATGF | ASGGGAEEDEQY | 0.44490522 |
| LASVALIL | ASGGGAEEDEQY | 0.18585347 |
| LFAVGLTL | ASGGGAEEDEQY | 0.14371468 |
| LFNGGIGTYI | ASGGGAEEDEQY | 0.6706631 |
| LTLGAGII | ASGGGAEEDEQY | 0.82043225 |
| LVAAALGDI | ASGGGAEEDEQY | 0.20909649 |
| NTAFVQLGI | ASGGGAEEDEQY | 0.68820024 |
| QLFSGAEL | ASGGGAEEDEQY | 0.13489687 |
| SAATFITL | ASGGGAEEDEQY | 0.6448663 |
| SNYGLANLGV | ASGGGAEEDEQY | 0.7274148 |
| SYGGAYAVM | ASGGGAEEDEQY | 0.19881892 |
| TTGVGGLTLG | ASGGGAEEDEQY | 0.43358308 |
| TTQTIYI | ASGGGAEEDEQY | 0.16619061 |
| VAAGAGLGAVI | ASGGGAEEDEQY | 0.20710886 |
| VAAVVTLL | ASGGGAEEDEQY | 0.7518083 |
| VGFLGGLI | ASGGGAEEDEQY | 0.34518933 |
| VGVIPILL | ASGGGAEEDEQY | 0.4714861 |
| VIFASSLI | ASGGGAEEDEQY | 0.26060563 |
| VKGGDGGAAAG | ASGGGAEEDEQY | 0.19672725 |
| VTKTFTNI | ASGGGAEEDEQY | 0.35867524 |
| VVAAAILL | ASGGGAEEDEQY | 0.75380707 |
| YAALDVEL | ASGGGAEEDEQY | 0.22342777 |
| LASVALLL | ASGGGAEEDEQY | 0.75173414 |
| LDAVHRV | ASGGGAEEDEQY | 0.20108357 |
| QGLSSSIF | ASGGGAEEDEQY | 0.74248135 |
| RGTFANIRL | ASGGGAEEDEQY | 0.7351188 |
| RQLWWGHRI | ASGGGAEEDEQY | 0.44279093 |
| YGPPGTGKTLL | ASGGGAEEDEQY | 0.24868351 |
| AALVGAPANVA | ASGGGAEEDEQY | 0.6996678 |
| AATFRRTE | ASGGGAEEDEQY | 0.48343205 |

| | | |
|---|---|---|
| AGALGGGI | ASGGGAEEDEQY | 0.690658 |
| AGGAGGAGGIGAG | ASGGGAEEDEQY | 0.6485027 |
| AGSGALGL | ASGGGAEEDEQY | 0.55925524 |
| AQIADQI | ASGGGAEEDEQY | 0.8204759 |
| ASAVAVTYL | ASGGGAEEDEQY | 0.69412863 |
| ASTVLGGTGG | ASGGGAEEDEQY | 0.7573069 |
| AVGGEPAK | ASGGGAEEDEQY | 0.4780457 |
| CEQGTFGGTC | ASGGGAEEDEQY | 0.3061859 |
| DEGKAPAG | ASGGGAEEDEQY | 0.7126961 |
| DTVVFPAL | ASGGGAEEDEQY | 0.4491324 |
| FEGITMAG | ASGGGAEEDEQY | 0.74030185 |
| FPLVVQD | ASGGGAEEDEQY | 0.7725855 |
| GAFTTPQ | ASGGGAEEDEQY | 0.51435196 |
| GFGGFGTAGGVGGA | ASGGGAEEDEQY | 0.82833993 |
| GFLAQFL | ASGGGAEEDEQY | 0.8225272 |
| GGAGGAGGNAGVA | ASGGGAEEDEQY | 0.356434 |
| GGVGTAGAGGAGGAGG | ASGGGAEEDEQY | 0.6004645 |
| GNGGGGDAGNAGSGGN | ASGGGAEEDEQY | 0.5446045 |
| GTGGAGGAGNDAGS | ASGGGAEEDEQY | 0.6860379 |
| GTTNTGWFNS | ASGGGAEEDEQY | 0.29481846 |
| IFMGPLFTL | ASGGGAEEDEQY | 0.7904754 |
| ILAHAIEAI | ASGGGAEEDEQY | 0.53627414 |
| IRSAIREAL | ASGGGAEEDEQY | 0.1691726 |
| ITFAGGG | ASGGGAEEDEQY | 0.7456248 |
| ITGMSGHQEV | ASGGGAEEDEQY | 0.64791614 |
| IVAGLACTL | ASGGGAEEDEQY | 0.63313484 |
| IVGGGLAGA | ASGGGAEEDEQY | 0.8090087 |
| IVTGVLNI | ASGGGAEEDEQY | 0.5516432 |
| LAQQGCI | ASGGGAEEDEQY | 0.5035806 |
| LCLHVEN | ASGGGAEEDEQY | 0.74417865 |
| LDAISDGS | ASGGGAEEDEQY | 0.80949426 |
| LGGFAAML | ASGGGAEEDEQY | 0.49152425 |
| LGGKNPMI | ASGGGAEEDEQY | 0.70534396 |
| LGPKGVAAIL | ASGGGAEEDEQY | 0.8394568 |
| LMVGVSI | ASGGGAEEDEQY | 0.724746 |
| LVGAAFVL | ASGGGAEEDEQY | 0.81333303 |
| LVLAVAPL | ASGGGAEEDEQY | 0.82436275 |
| MAAIAVIDAA | ASGGGAEEDEQY | 0.53634405 |
| NNGINAELQ | ASGGGAEEDEQY | 0.78028226 |
| RGEPLVLVHG | ASGGGAEEDEQY | 0.827049 |
| RGGAGGAAGLI | ASGGGAEEDEQY | 0.635545 |
| RSYRMHPAV | ASGGGAEEDEQY | 0.7297718 |
| SAAHLLAGQLA | ASGGGAEEDEQY | 0.49413285 |
| SAAVGSY | ASGGGAEEDEQY | 0.6500093 |
| SPHAVLHRI | ASGGGAEEDEQY | 0.22670239 |
| SQIAKVL | ASGGGAEEDEQY | 0.715394 |
| TALGHLVP | ASGGGAEEDEQY | 0.8176676 |
| TAPTTSGAA | ASGGGAEEDEQY | 0.64819336 |
| TDGDDGNGGSPGT | ASGGGAEEDEQY | 0.47964394 |
| TDGLGEN | ASGGGAEEDEQY | 0.3443035 |
| TGGPPLPGQA | ASGGGAEEDEQY | 0.7241127 |

| | | |
|---|---|---|
| TIPAITI | ASGGGAEEDEQY | 0.58794874 |
| TIPGFTI | ASGGGAEEDEQY | 0.34298277 |
| TSIASGALA | ASGGGAEEDEQY | 0.5859581 |
| TVTLPGAAI | ASGGGAEEDEQY | 0.80973303 |
| VASGLSAALP | ASGGGAEEDEQY | 0.81348765 |
| VGAGVIGI | ASGGGAEEDEQY | 0.42315996 |
| VGPKRFGV | ASGGGAEEDEQY | 0.8210036 |
| VNVAFPD | ASGGGAEEDEQY | 0.7629349 |
| VSMTNSGVS | ASGGGAEEDEQY | 0.78305924 |
| VTDVLGDLSV | ASGGGAEEDEQY | 0.7143139 |
| VVVVGAGFAGL | ASGGGAEEDEQY | 0.51382047 |
| VYTVGTVV | ASGGGAEEDEQY | 0.799644 |
| WTTFSGP | ASGGGAEEDEQY | 0.6011306 |
| YDELVEAF | ASGGGAEEDEQY | 0.62477463 |
| YGTGGAGGNGG | ASGGGAEEDEQY | 0.79301953 |
| YTNVMAV | ASGGGAEEDEQY | 0.68047917 |
| FLEQVHQL | ASGGGAEEDEQY | 0.5049838 |
| GLSLLAVAV | ASGGGAEEDEQY | 0.6105928 |
| LEQVHQL | ASGGGAEEDEQY | 0.81636345 |
| LNAAGAI | ASGGGAEEDEQY | 0.41186574 |
| PAVHASL | ASGGGAEEDEQY | 0.45317715 |
| TELEDDA | ASGGGAEEDEQY | 0.5643477 |
| TPPGPPN | ASGGGAEEDEQY | 0.72500306 |
| TPPPPPPP | ASGGGAEEDEQY | 0.79765546 |
| TQSPALI | ASSPDRGGTEVF | 0.75971836 |
| EFAAAFPQ | ASSPDRGGTEVF | 0.64421594 |
| LLSGNLDVL | ASSPDRGGTEVF | 0.7682231 |
| GPANLGNYNI | ASSPDRGGTEVF | 0.7809396 |
| TLGPTLP | ASSPDRGGTEVF | 0.73101574 |
| VYAYSSCCK | ASSPDRGGTEVF | 0.7699653 |
| APAPAPAPNI | ASSPDRGGTEVF | 0.54913384 |
| PGGPVPIP | ASSPDRGGTEVF | 0.8054508 |
| FLTVALEP | ASSPDRGGTEVF | 0.7854708 |
| SQYFGANL | ASSPDRGGTEVF | 0.78179586 |
| GGVKPAA | ASSPDRGGTEVF | 0.7842939 |
| GGTGGAGGAGGW | ASSPDRGGTEVF | 0.737208 |
| GVGGVAGLGGAGG | ASSPDRGGTEVF | 0.7078429 |
| VLLVLATTI | ASSPDRGGTEVF | 0.2286777 |
| IGIFAAML | ASSPDRGGTEVF | 0.6072887 |
| AVGDLID | ASSPDRGGTEVF | 0.71998656 |
| NISPLPL | ASSPDRGGTEVF | 0.70185065 |
| VTAVFVAI | ASSPDRGGTEVF | 0.7500397 |
| SGDDNDAAL | ASSPDRGGTEVF | 0.7884733 |
| AVAALAHS | ASSPDRGGTEVF | 0.5371065 |
| GGVTVMATP | ASSPDRGGTEVF | 0.80897856 |
| AGLAANVRD | ASSPDRGGTEVF | 0.8126643 |
| TNRPDLIDPALLRPG | ASSPDRGGTEVF | 0.32200372 |
| LANLGVTAL | ASSPDRGGTEVF | 0.5568353 |
| TGERIEL | ASSPDRGGTEVF | 0.83979535 |
| NGEELAA | ASSPDRGGTEVF | 0.73441696 |
| VLSLVWG | ASSPDRGGTEVF | 0.61805904 |

| | | |
|---|---|---|
| IGYALWL | ASSPDRGGTEVF | 0.7003722 |
| LSLLLAPL | ASSPDRGGTEVF | 0.83633393 |
| TLDEALK | ASSPDRGGTEVF | 0.629757 |
| TDGAGTAEGG | ASSPDRGGTEVF | 0.8330934 |
| NAPGGAGGNGG | ASSPDRGGTEVF | 0.19945815 |
| VSVFMGMGGNF | ASSPDRGGTEVF | 0.6446769 |
| LASYLAAL | ASSPDRGGTEVF | 0.56852084 |
| LVGGTPAL | ASSPDRGGTEVF | 0.25001094 |
| VRNADVL | ASSPDRGGTEVF | 0.7773773 |
| TCSSPFLVSAM | ASSPDRGGTEVF | 0.69209933 |
| VHITDAE | ASSPDRGGTEVF | 0.8026303 |
| EYAAPSF | ASSPDRGGTEVF | 0.66307336 |
| VVDAFHGGL | ASSPDRGGTEVF | 0.78284585 |
| ASVVDPQ | ASSPDRGGTEVF | 0.688596 |
| SYNEFGGLS | ASSPDRGGTEVF | 0.7148116 |
| DPDGSVY | ASSPDRGGTEVF | 0.56319296 |
| MVSTIHATE | ASSPDRGGTEVF | 0.7594295 |
| LAAGVADPV | ASSPDRGGTEVF | 0.8105213 |
| TKEQIAA | ASSPDRGGTEVF | 0.7927557 |
| SLASPAAL | ASSPDRGGTEVF | 0.31818402 |
| IGLVFYGV | ASSPDRGGTEVF | 0.65043736 |
| FLLDASNL | ASSPDRGGTEVF | 0.6435787 |
| YQVGNFP | ASSPDRGGTEVF | 0.2798518 |
| ESLLSQST | ASSPDRGGTEVF | 0.23730358 |
| LFTVGFAAVG | ASSPDRGGTEVF | 0.541515 |
| ATPPTPT | ASSPDRGGTEVF | 0.6175387 |
| TDLATLG | ASSPDRGGTEVF | 0.79200226 |
| SAGGTGR | ASSPDRGGTEVF | 0.7647631 |
| GFSNSGS | ASSPDRGGTEVF | 0.6544191 |
| GGNGGSGGTSV | ASSPDRGGTEVF | 0.78831226 |
| GTSVATGGAGN | ASSPDRGGTEVF | 0.71578 |
| NGGNGGSGGTS | ASSPDRGGTEVF | 0.11223871 |
| AVVKFPH | ASSPDRGGTEVF | 0.60505295 |
| TPPPPPP | ASSPDRGGTEVF | 0.37877092 |
| AAPAAAAAPA | ASSPDRGGTEVF | 0.82002914 |
| IALNEDL | ASSPDRGGTEVF | 0.4763114 |
| IVNVSSM | ASSPDRGGTEVF | 0.56766987 |
| TGIPPAPR | ASSPDRGGTEVF | 0.68557185 |
| FDLGGGTF | ASSPDRGGTEVF | 0.6923101 |
| FDLGGGTFDVS | ASSPDRGGTEVF | 0.6881903 |
| NEPTAAA | ASSPDRGGTEVF | 0.48937774 |
| NEPTAAALA | ASSPDRGGTEVF | 0.17704052 |
| GSIINISSI | ASSPDRGGTEVF | 0.7928668 |
| VAPGNAGTA | ASSPDRGGTEVF | 0.411303 |
| TGGGASLE | ASSPDRGGTEVF | 0.50409865 |
| VAVYNHYKRI | ASSPDRGGTEVF | 0.38084584 |
| TDPAPATT | ASSPDRGGTEVF | 0.62532586 |
| TNRPDILDPALLRPG | ASSPDRGGTEVF | 0.2975027 |
| LTTTTTS | ASSPDRGGTEVF | 0.8186554 |
| PTGSGKTL | ASSPDRGGTEVF | 0.5521939 |
| TGSGKTL | ASSPDRGGTEVF | 0.6349401 |

| | | |
|---|---|---|
| TIGHVDHGK | ASSPDRGGTEVF | 0.8252517 |
| TIGHVDHGKTT | ASSPDRGGTEVF | 0.69908285 |
| GHVDHGKTT | ASSPDRGGTEVF | 0.6221096 |
| AQFEHTLL | ASSPDRGGTEVF | 0.6140543 |
| IVNQDEL | ASSPDRGGTEVF | 0.43933016 |
| TNGSQFF | ASSPDRGGTEVF | 0.5275012 |
| TGKGIVE | ASSPDRGGTEVF | 0.5798993 |
| LISNASDA | ASSPDRGGTEVF | 0.84227043 |
| AELRTLGELTKIAKAHGAQ | ASSPDRGGTEVF | 0.6607923 |
| APNAVAAAEPSW | ASSPDRGGTEVF | 0.4939724 |
| APQVAGAATAVL | ASSPDRGGTEVF | 0.74857473 |
| ARRDPAAAAAILPTDA | ASSPDRGGTEVF | 0.8343911 |
| DAAARDVLAEAGLAEYF | ASSPDRGGTEVF | 0.832639 |
| DDQIVLSAKANLAKVEQ | ASSPDRGGTEVF | 0.5039935 |
| DELLPDTGEDSDEDGH | ASSPDRGGTEVF | 0.67370486 |
| ELPEAPANRATVPQD | ASSPDRGGTEVF | 0.68299204 |
| FGQSKIERVTGIG | ASSPDRGGTEVF | 0.6535976 |
| GAGGTGSTAGGAGGAGGAGGLY | ASSPDRGGTEVF | 0.6499192 |
| GHGGTGAAGGNGGTGGAGGSGI | ASSPDRGGTEVF | 0.47318137 |
| GNGGNGGAGGEGGPGGAGGAGA | ASSPDRGGTEVF | 0.7883154 |
| GSTAGNGGNGGNATNGTIAGQP | ASSPDRGGTEVF | 0.85587585 |
| GTGGDGGNGGNGGTGG | ASSPDRGGTEVF | 0.68490744 |
| GVRRPAGIPADAPV | ASSPDRGGTEVF | 0.28586778 |
| GYRQVTDAHLLTLARR | ASSPDRGGTEVF | 0.6355523 |
| IAKMGQDGHDRG | ASSPDRGGTEVF | 0.39054972 |
| KAEPSIEIQYT | ASSPDRGGTEVF | 0.5697483 |
| KPVRIRMAADEDTVAVT | ASSPDRGGTEVF | 0.68782526 |
| MPPRIAGMRLLVIKPEP | ASSPDRGGTEVF | 0.36078376 |
| MKFYAVALFRSRGVPPTR | ASSPDRGGTEVF | 0.6759213 |
| NAGGFGILGAVAHSP | ASSPDRGGTEVF | 0.8386963 |
| NEGREITLAMAEM | ASSPDRGGTEVF | 0.47708026 |
| NELVEWSHPLGHRWRWE | ASSPDRGGTEVF | 0.7576777 |
| NLGAGGTGGNGGNPG | ASSPDRGGTEVF | 0.80240816 |
| NTEWPKWETVMAADPVD | ASSPDRGGTEVF | 0.7597499 |
| PASAAPAETQP | ASSPDRGGTEVF | 0.819332 |
| QQSSDGKNGVRAAEITGE | ASSPDRGGTEVF | 0.7620835 |
| RAPDAVTTSWWKE | ASSPDRGGTEVF | 0.7256348 |
| SGFGNVGAHTSGWFNQ | ASSPDRGGTEVF | 0.7196336 |
| SGVAGAVGQAASVGGLKV | ASSPDRGGTEVF | 0.7099918 |
| SNGGAGTGSAGGNGGT | ASSPDRGGTEVF | 0.67425185 |
| TGGAGGEGGAAG | ASSPDRGGTEVF | 0.64672965 |
| TGGDGGAAGAGGAPGHG | ASSPDRGGTEVF | 0.5926411 |
| TMPPLLAAIANAGRHGV | ASSPDRGGTEVF | 0.79696476 |
| TSSTAGNSGGA | ASSPDRGGTEVF | 0.69506323 |
| VPATVPVGILC | ASSPDRGGTEVF | 0.7990304 |
| VQRPDSGGVVLTGRIS | ASSPDRGGTEVF | 0.38952142 |
| WTPNPVPATKGVPAGLVQAT | ASSPDRGGTEVF | 0.5034438 |
| YALKRALAKFSDRFDVVII | ASSPDRGGTEVF | 0.7836282 |
| YLIGSVIV | ASSPDRGGTEVF | 0.1516011 |
| YGQLVAVID | ASSPDRGGTEVF | 0.257256 |
| VTVGGMTLPA | ASSPDRGGTEVF | 0.7086025 |

| | | |
|---|---|---|
| VLALPAV | ASSPDRGGTEVF | 0.8130528 |
| VIPPTLNYET | ASSPDRGGTEVF | 0.16726674 |
| VGMGGMHPSSG | ASSPDRGGTEVF | 0.53107214 |
| VGLAGGAATA | ASSPDRGGTEVF | 0.35736188 |
| VGGAGGAGGSGP | ASSPDRGGTEVF | 0.79928535 |
| VFAAFMV | ASSPDRGGTEVF | 0.70040023 |
| TLYDKLR | ASSPDRGGTEVF | 0.23713997 |
| TLAVFTVL | ASSPDRGGTEVF | 0.7719238 |
| TGSVQPARV | ASSPDRGGTEVF | 0.34797028 |
| TGGGAGSGGT | ASSPDRGGTEVF | 0.5338551 |
| SHLETSL | ASSPDRGGTEVF | 0.77076983 |
| SFTTGMT | ASSPDRGGTEVF | 0.81077296 |
| QVEVAKRMGVRQPTVS | ASSPDRGGTEVF | 0.45419922 |
| QGSGGPALAM | ASSPDRGGTEVF | 0.7635666 |
| MLAPSNP | ASSPDRGGTEVF | 0.5518501 |
| LVNNAAI | ASSPDRGGTEVF | 0.84018975 |
| LTGEINTLA | ASSPDRGGTEVF | 0.5539423 |
| LSIFGPV | ASSPDRGGTEVF | 0.36833355 |
| LLALLANL | ASSPDRGGTEVF | 0.79241014 |
| LISNASDALDKLR | ASSPDRGGTEVF | 0.739557 |
| LINEAVP | ASSPDRGGTEVF | 0.73357415 |
| LGYAFSGL | ASSPDRGGTEVF | 0.7655102 |
| LDFVAVAL | ASSPDRGGTEVF | 0.63165385 |
| LAQPGAE | ASSPDRGGTEVF | 0.6155734 |
| ISNLGAVTGF | ASSPDRGGTEVF | 0.80409974 |
| HQEDPGRI | ASSPDRGGTEVF | 0.77147067 |
| GTPGEPTP | ASSPDRGGTEVF | 0.5242888 |
| GFFNSGS | ASSPDRGGTEVF | 0.79731166 |
| GAGGGGLTAGAN | ASSPDRGGTEVF | 0.6990912 |
| FVDALVNM | ASSPDRGGTEVF | 0.5607721 |
| FLTVAPEL | ASSPDRGGTEVF | 0.82419574 |
| FETLTAL | ASSPDRGGTEVF | 0.699284 |
| DVINPIEL | ASSPDRGGTEVF | 0.80224526 |
| APVSGAPV | ASSPDRGGTEVF | 0.7560651 |
| ALVGGLAW | ASSPDRGGTEVF | 0.8190926 |
| ALAAAAATAI | ASSPDRGGTEVF | 0.34829912 |
| AFGLGLLL | ASSPDRGGTEVF | 0.7621789 |
| AFFAAASL | ASSPDRGGTEVF | 0.71406084 |
| AAAALPAFSPP | ASSPDRGGTEVF | 0.76014704 |
| VVLAPAL | ASSPDRGGTEVF | 0.6678599 |
| TIGHVDHGKTTL | ASSPDRGGTEVF | 0.4517316 |
| LISNASDALDK | ASSPDRGGTEVF | 0.73461825 |
| LISNASDAL | ASSPDRGGTEVF | 0.8328546 |
| ISNASDALDK | ASSPDRGGTEVF | 0.726699 |
| NQAETLVYQTE | ASSPDRGGTEVF | 0.7392029 |
| TGGTGGTAGAG | ASSPDRGGTEVF | 0.8283583 |
| TGSTAGGAGGA | ASSPDRGGTEVF | 0.50016034 |
| DPLAVPVAAGPV | ASSPDRGGTEVF | 0.69857246 |
| GGTGGTPGNGGN | ASSPDRGGTEVF | 0.80349267 |
| PAGLAGVDPATL | ASSPDRGGTEVF | 0.8084514 |
| GNNLVVEAPHAGASV | ASSPDRGGTEVF | 0.6741806 |

| | | |
|---|---|---|
| KVTRIMSASTLEQPA | ASSPDRGGTEVF | 0.8163042 |
| LGFALPIALPRRTVV | ASSPDRGGTEVF | 0.4363945 |
| TRLGNLTARKLVVLV | ASSPDRGGTEVF | 0.77478766 |
| WSQQRRLIYFPSAGP | ASSPDRGGTEVF | 0.535955 |
| GPNGNPDPMAAAVDIR | ASSPDRGGTEVF | 0.73737764 |
| TQRCVSVADLRAAASR | ASSPDRGGTEVF | 0.47922042 |
| QFVLDVAEHPVEVLTETV | ASSPDRGGTEVF | 0.7503034 |
| GGAGGTGGAGGNGGTQAGDG | ASSPDRGGTEVF | 0.78964746 |
| PLLATLRAIAVDVDAKPAQV | ASSPDRGGTEVF | 0.79794955 |
| APKFRTGGYMIDFWGVGYQVAK | ASSPDRGGTEVF | 0.28658915 |
| GAGGAGGAGGAAGGPGGTGGTG | ASSPDRGGTEVF | 0.77814245 |
| IGALGLLATDVAPAGSVAFGVA | ASSPDRGGTEVF | 0.8003438 |
| AGGAGGIGGNGGAGGAGGVLMG | ASSPDRGGTEVF | 0.8536613 |
| ADELIGLL | ASSPDRGGTEVF | 0.6755252 |
| ADRALKA | ASSPDRGGTEVF | 0.79119796 |
| AGAVGQVGGAGGA | ASSPDRGGTEVF | 0.61591434 |
| AGGGGGLGGAP | ASSPDRGGTEVF | 0.67821956 |
| AGGLLYGNGGAG | ASSPDRGGTEVF | 0.45465785 |
| AGLESIGL | ASSPDRGGTEVF | 0.5969601 |
| AGLKTGII | ASSPDRGGTEVF | 0.63135225 |
| ASLAALLL | ASSPDRGGTEVF | 0.5114136 |
| ASVINISSI | ASSPDRGGTEVF | 0.7799041 |
| ATPPGPA | ASSPDRGGTEVF | 0.799558 |
| ATVASALV | ASSPDRGGTEVF | 0.8010278 |
| ATVASLIL | ASSPDRGGTEVF | 0.8064048 |
| AVNGPVLIRH | ASSPDRGGTEVF | 0.82068837 |
| DGAAGAVGI | ASSPDRGGTEVF | 0.79160476 |
| DPLLHGI | ASSPDRGGTEVF | 0.55760133 |
| ELGAALGAL | ASSPDRGGTEVF | 0.63114655 |
| ELGRTQAV | ASSPDRGGTEVF | 0.3926605 |
| FVAGGYLS | ASSPDRGGTEVF | 0.64461637 |
| GASGGAGGAGGW | ASSPDRGGTEVF | 0.5579618 |
| GGAGAGGDGGAGGI | ASSPDRGGTEVF | 0.7333411 |
| GTGHNGSQPGMGGNG | ASSPDRGGTEVF | 0.13534331 |
| GYAAASASAS | ASSPDRGGTEVF | 0.693867 |
| HQVDVAAVQQG | ASSPDRGGTEVF | 0.8421342 |
| IGAAIAHT | ASSPDRGGTEVF | 0.4731093 |
| IGMIEVAQL | ASSPDRGGTEVF | 0.7789217 |
| IRLAPEHTSAMD | ASSPDRGGTEVF | 0.7132119 |
| IYNVSVL | ASSPDRGGTEVF | 0.55941725 |
| KQGFLGM | ASSPDRGGTEVF | 0.78313506 |
| LESVITQL | ASSPDRGGTEVF | 0.494311 |
| LGLGALVVV | ASSPDRGGTEVF | 0.7908883 |
| LIGNGGNGGSGGI | ASSPDRGGTEVF | 0.8227627 |
| LNAINAPTL | ASSPDRGGTEVF | 0.73496974 |
| LQLLNTGV | ASSPDRGGTEVF | 0.67521113 |
| LQTSIEF | ASSPDRGGTEVF | 0.7330981 |
| MSFAVSEL | ASSPDRGGTEVF | 0.2901935 |
| NTGALGL | ASSPDRGGTEVF | 0.6927657 |
| PPGAGGAGGAGTT | ASSPDRGGTEVF | 0.27037898 |
| QLVGLGI | ASSPDRGGTEVF | 0.6418785 |

| | | |
|---|---|---|
| RFLEQVHQL | ASSPDRGGTEVF | 0.45170805 |
| RGVGYVLR | ASSPDRGGTEVF | 0.728988 |
| SATGVFAGVI | ASSPDRGGTEVF | 0.7224349 |
| SLPAITLP | ASSPDRGGTEVF | 0.74847937 |
| STALAFTV | ASSPDRGGTEVF | 0.36883765 |
| SVLTVGLGK | ASSPDRGGTEVF | 0.6605347 |
| TALAGAGSGI | ASSPDRGGTEVF | 0.6587262 |
| TGLPFTGI | ASSPDRGGTEVF | 0.39247996 |
| VAEHVTV | ASSPDRGGTEVF | 0.7720268 |
| VDIVLNSL | ASSPDRGGTEVF | 0.7111837 |
| VLTGGGAVL | ASSPDRGGTEVF | 0.6247254 |
| VNVGLAYGI | ASSPDRGGTEVF | 0.8085846 |
| VSSAAVGAV | ASSPDRGGTEVF | 0.71467984 |
| VTSVYEAGV | ASSPDRGGTEVF | 0.27459586 |
| WVALDRGA | ASSPDRGGTEVF | 0.61845803 |
| YLVGSVVI | ASSPDRGGTEVF | 0.6599901 |
| GPPGTGKTLL | ASSPDRGGTEVF | 0.53083724 |
| GVGMTKF | ASSPDRGGTEVF | 0.16860661 |
| IGIGGSDL | ASSPDRGGTEVF | 0.7523009 |
| IGTLLAEL | ASSPDRGGTEVF | 0.5726789 |
| IYTSGTT | ASSPDRGGTEVF | 0.4169714 |
| LSFPAGV | ASSPDRGGTEVF | 0.5860197 |
| LYTSGTT | ASSPDRGGTEVF | 0.7899608 |
| VGAFPFGF | ASSPDRGGTEVF | 0.5738617 |
| VGAGTVEF | ASSPDRGGTEVF | 0.728111 |
| VGKSTLLL | ASSPDRGGTEVF | 0.749321 |
| AALAHAGGGIV | ASSPDRGGTEVF | 0.78177136 |
| AALVAVSLL | ASSPDRGGTEVF | 0.8335411 |
| AAVSRAVAAL | ASSPDRGGTEVF | 0.7613114 |
| AGGSSGIGGF | ASSPDRGGTEVF | 0.67608416 |
| AGLTVANANG | ASSPDRGGTEVF | 0.7743627 |
| AGLVGAVL | ASSPDRGGTEVF | 0.70935386 |
| ALSQAFTV | ASSPDRGGTEVF | 0.38193005 |
| ALTFSVSL | ASSPDRGGTEVF | 0.29579312 |
| ALTGAALVAL | ASSPDRGGTEVF | 0.57068706 |
| ASFSIGAGVL | ASSPDRGGTEVF | 0.3154747 |
| DGSVYKI | ASSPDRGGTEVF | 0.6707809 |
| EQLAEIV | ASSPDRGGTEVF | 0.8204708 |
| FAVSELTVM | ASSPDRGGTEVF | 0.6193884 |
| GAAPLAAGL | ASSPDRGGTEVF | 0.4111524 |
| GADGGDGGSGGAGG | ASSPDRGGTEVF | 0.60124606 |
| GFGLDTGGAGG | ASSPDRGGTEVF | 0.41973948 |
| GLASGGLL | ASSPDRGGTEVF | 0.7754748 |
| GRIAGAAVLL | ASSPDRGGTEVF | 0.7217827 |
| GSAGSLNTSF | ASSPDRGGTEVF | 0.83745396 |
| GTGQIGI | ASSPDRGGTEVF | 0.79252964 |
| GTLIALIL | ASSPDRGGTEVF | 0.6211237 |
| IAATMGFVVG | ASSPDRGGTEVF | 0.7756041 |
| IFAQALL | ASSPDRGGTEVF | 0.5692049 |
| IGAAVALGF | ASSPDRGGTEVF | 0.7250029 |
| IGLTPLAL | ASSPDRGGTEVF | 0.67959803 |

| | | |
|---|---|---|
| ILGSNTSTL | ASSPDRGGTEVF | 0.7255068 |
| IVFSLTVL | ASSPDRGGTEVF | 0.6465203 |
| LASKATGF | ASSPDRGGTEVF | 0.6428354 |
| LASVALIL | ASSPDRGGTEVF | 0.2549878 |
| LFAVGLTL | ASSPDRGGTEVF | 0.7677467 |
| LFNGGIGTYI | ASSPDRGGTEVF | 0.6128792 |
| LTLGAGII | ASSPDRGGTEVF | 0.38311198 |
| LVAAALGDI | ASSPDRGGTEVF | 0.5719781 |
| NTAFVQLGI | ASSPDRGGTEVF | 0.8264866 |
| QLFSGAEL | ASSPDRGGTEVF | 0.83300763 |
| SAATFITL | ASSPDRGGTEVF | 0.54122114 |
| SNYGLANLGV | ASSPDRGGTEVF | 0.46611077 |
| SYGGAYAVM | ASSPDRGGTEVF | 0.70694566 |
| TTGVGGLTLG | ASSPDRGGTEVF | 0.31391287 |
| TTQTIYI | ASSPDRGGTEVF | 0.7397715 |
| VAAGAGLGAVI | ASSPDRGGTEVF | 0.65038145 |
| VAAVVTLL | ASSPDRGGTEVF | 0.67272854 |
| VGFLGGLI | ASSPDRGGTEVF | 0.60025316 |
| VGVIPILL | ASSPDRGGTEVF | 0.6999632 |
| VIFASSLI | ASSPDRGGTEVF | 0.57154524 |
| VKGGDGGAAAG | ASSPDRGGTEVF | 0.77774084 |
| VTKTFTNI | ASSPDRGGTEVF | 0.14592084 |
| VVAAAILL | ASSPDRGGTEVF | 0.7400628 |
| YAALDVEL | ASSPDRGGTEVF | 0.523732 |
| LASVALLL | ASSPDRGGTEVF | 0.558769 |
| LDAVHRV | ASSPDRGGTEVF | 0.33456302 |
| QGLSSSIF | ASSPDRGGTEVF | 0.73379207 |
| RGTFANIRL | ASSPDRGGTEVF | 0.7468282 |
| RQLWWGHRI | ASSPDRGGTEVF | 0.78109455 |
| YGPPGTGKTLL | ASSPDRGGTEVF | 0.47243333 |
| AALVGAPANVA | ASSPDRGGTEVF | 0.7760898 |
| AATFRRTE | ASSPDRGGTEVF | 0.79070306 |
| AGALGGGI | ASSPDRGGTEVF | 0.6394603 |
| AGGAGGAGGIGAG | ASSPDRGGTEVF | 0.29622906 |
| AGSGALGL | ASSPDRGGTEVF | 0.7586342 |
| AQIADQI | ASSPDRGGTEVF | 0.6608015 |
| ASAVAVTYL | ASSPDRGGTEVF | 0.7696517 |
| ASTVLGGTGG | ASSPDRGGTEVF | 0.73606634 |
| AVGGEPAK | ASSPDRGGTEVF | 0.398195 |
| CEQGTFGGTC | ASSPDRGGTEVF | 0.812291 |
| DEGKAPAG | ASSPDRGGTEVF | 0.79120815 |
| DTVVFPAL | ASSPDRGGTEVF | 0.4631472 |
| FEGITMAG | ASSPDRGGTEVF | 0.6095257 |
| FPLVVQD | ASSPDRGGTEVF | 0.6602874 |
| GAFTTPQ | ASSPDRGGTEVF | 0.32947302 |
| GFGGFGTAGGVGGA | ASSPDRGGTEVF | 0.7709085 |
| GFLAQFL | ASSPDRGGTEVF | 0.57774234 |
| GGAGGAGGNAGVA | ASSPDRGGTEVF | 0.60621697 |
| GGVGTAGAGGAGGAGG | ASSPDRGGTEVF | 0.71604943 |
| GNGGGGDAGNAGSGGN | ASSPDRGGTEVF | 0.6088941 |
| GTGGAGGAGNDAGS | ASSPDRGGTEVF | 0.6106919 |

| | | |
|---|---|---|
| GTTNTGWFNS | ASSPDRGGTEVF | 0.6557586 |
| IFMGPLFTL | ASSPDRGGTEVF | 0.6722811 |
| ILAHAIEAI | ASSPDRGGTEVF | 0.7184415 |
| IRSAIREAL | ASSPDRGGTEVF | 0.6727901 |
| ITFAGGG | ASSPDRGGTEVF | 0.74578357 |
| ITGMSGHQEV | ASSPDRGGTEVF | 0.83130634 |
| IVAGLACTL | ASSPDRGGTEVF | 0.39323896 |
| IVGGGLAGA | ASSPDRGGTEVF | 0.55006194 |
| IVTGVLNI | ASSPDRGGTEVF | 0.75673157 |
| LAQQGCI | ASSPDRGGTEVF | 0.39948365 |
| LCLHVEN | ASSPDRGGTEVF | 0.7850915 |
| LDAISDGS | ASSPDRGGTEVF | 0.72057176 |
| LGGFAAML | ASSPDRGGTEVF | 0.7426827 |
| LGGKNPMI | ASSPDRGGTEVF | 0.6495559 |
| LGPKGVAAIL | ASSPDRGGTEVF | 0.83997035 |
| LMVGVSI | ASSPDRGGTEVF | 0.79490095 |
| LVGAAFVL | ASSPDRGGTEVF | 0.72666276 |
| LVLAVAPL | ASSPDRGGTEVF | 0.7643769 |
| MAAIAVIDAA | ASSPDRGGTEVF | 0.5903667 |
| NNGINAELQ | ASSPDRGGTEVF | 0.78112125 |
| RGEPLVLVHG | ASSPDRGGTEVF | 0.77009916 |
| RGGAGGAAGLI | ASSPDRGGTEVF | 0.36461815 |
| RSYRMHPAV | ASSPDRGGTEVF | 0.6193402 |
| SAAHLLAGQLA | ASSPDRGGTEVF | 0.8287749 |
| SAAVGSY | ASSPDRGGTEVF | 0.81170243 |
| SPHAVLHRI | ASSPDRGGTEVF | 0.31552237 |
| SQIAKVL | ASSPDRGGTEVF | 0.625395 |
| TALGHLVP | ASSPDRGGTEVF | 0.41603374 |
| TAPTTSGAA | ASSPDRGGTEVF | 0.6263039 |
| TDGDDGNGGSPGT | ASSPDRGGTEVF | 0.46360368 |
| TDGLGEN | ASSPDRGGTEVF | 0.7543231 |
| TGGPPLPGQA | ASSPDRGGTEVF | 0.60212886 |
| TIPAITI | ASSPDRGGTEVF | 0.7794558 |
| TIPGFTI | ASSPDRGGTEVF | 0.65509796 |
| TSIASGALA | ASSPDRGGTEVF | 0.80724263 |
| TVTLPGAAI | ASSPDRGGTEVF | 0.54050016 |
| VASGLSAALP | ASSPDRGGTEVF | 0.68500674 |
| VGAGVIGI | ASSPDRGGTEVF | 0.36099634 |
| VGPKRFGV | ASSPDRGGTEVF | 0.35502398 |
| VNVAFPD | ASSPDRGGTEVF | 0.5321314 |
| VSMTNSGVS | ASSPDRGGTEVF | 0.6357994 |
| VTDVLGDLSV | ASSPDRGGTEVF | 0.22509749 |
| VVVVGAGFAGL | ASSPDRGGTEVF | 0.8324138 |
| VYTVGTVV | ASSPDRGGTEVF | 0.70552284 |
| WTTFSGP | ASSPDRGGTEVF | 0.7752384 |
| YDELVEAF | ASSPDRGGTEVF | 0.61362606 |
| YGTGGAGGNGG | ASSPDRGGTEVF | 0.6564925 |
| YTNVMAV | ASSPDRGGTEVF | 0.52457166 |
| FLEQVHQL | ASSPDRGGTEVF | 0.7786585 |
| GLSLLAVAV | ASSPDRGGTEVF | 0.3841325 |
| LEQVHQL | ASSPDRGGTEVF | 0.28477466 |

| | | |
|---|---|---|
| LNAAGAI | ASSPDRGGTEVF | 0.5805929 |
| PAVHASL | ASSPDRGGTEVF | 0.40557796 |
| TELEDDA | ASSPDRGGTEVF | 0.8120129 |
| TPPGPPN | ASSPDRGGTEVF | 0.8002771 |
| TPPPPPPP | ASSPDRGGTEVF | 0.7135514 |
| TQSPALI | ASSPSIQDTQY | 0.7971497 |
| EFAAAFPQ | ASSPSIQDTQY | 0.20455614 |
| LLSGNLDVL | ASSPSIQDTQY | 0.62511456 |
| GPANLGNYNI | ASSPSIQDTQY | 0.8421601 |
| TLGPTLP | ASSPSIQDTQY | 0.7207016 |
| VYAYSSCCK | ASSPSIQDTQY | 0.6467985 |
| APAPAPAPNI | ASSPSIQDTQY | 0.81437206 |
| PGGPVPIP | ASSPSIQDTQY | 0.58390254 |
| FLTVALEP | ASSPSIQDTQY | 0.54980004 |
| SQYFGANL | ASSPSIQDTQY | 0.7062175 |
| GGVKPAA | ASSPSIQDTQY | 0.7553284 |
| GGTGGAGGAGGW | ASSPSIQDTQY | 0.7398559 |
| GVGGVAGLGGAGG | ASSPSIQDTQY | 0.4125641 |
| VLLVLATTI | ASSPSIQDTQY | 0.7741904 |
| IGIFAAML | ASSPSIQDTQY | 0.8475537 |
| AVGDLID | ASSPSIQDTQY | 0.6120596 |
| NISPLPL | ASSPSIQDTQY | 0.7610973 |
| VTAVFVAI | ASSPSIQDTQY | 0.7578703 |
| SGDDNDAAL | ASSPSIQDTQY | 0.6370306 |
| AVAALAHS | ASSPSIQDTQY | 0.458651 |
| GGVTVMATP | ASSPSIQDTQY | 0.5111418 |
| AGLAANVRD | ASSPSIQDTQY | 0.6900816 |
| TNRPDLIDPALLRPG | ASSPSIQDTQY | 0.4772366 |
| LANLGVTAL | ASSPSIQDTQY | 0.7128941 |
| TGERIEL | ASSPSIQDTQY | 0.6696682 |
| NGEELAA | ASSPSIQDTQY | 0.85055774 |
| VLSLVWG | ASSPSIQDTQY | 0.7785822 |
| IGYALWL | ASSPSIQDTQY | 0.74617493 |
| LSLLLAPL | ASSPSIQDTQY | 0.45787847 |
| TLDEALK | ASSPSIQDTQY | 0.7270332 |
| TDGAGTAEGG | ASSPSIQDTQY | 0.7287266 |
| NAPGGAGGNGG | ASSPSIQDTQY | 0.27916 |
| VSVFMGMGGNF | ASSPSIQDTQY | 0.8353671 |
| LASYLAAL | ASSPSIQDTQY | 0.5102728 |
| LVGGTPAL | ASSPSIQDTQY | 0.63980937 |
| VRNADVL | ASSPSIQDTQY | 0.79537356 |
| TCSSPFLVSAM | ASSPSIQDTQY | 0.816599 |
| VHITDAE | ASSPSIQDTQY | 0.66348815 |
| EYAAPSF | ASSPSIQDTQY | 0.7093115 |
| VVDAFHGGL | ASSPSIQDTQY | 0.5175625 |
| ASVVDPQ | ASSPSIQDTQY | 0.7736687 |
| SYNEFGGLS | ASSPSIQDTQY | 0.79890186 |
| DPDGSVY | ASSPSIQDTQY | 0.59656805 |
| MVSTIHATE | ASSPSIQDTQY | 0.81138647 |
| LAAGVADPV | ASSPSIQDTQY | 0.7559165 |
| TKEQIAA | ASSPSIQDTQY | 0.37257507 |

| | | |
|---|---|---|
| SLASPAAL | ASSPSIQDTQY | 0.66112906 |
| IGLVFYGV | ASSPSIQDTQY | 0.7682425 |
| FLLDASNL | ASSPSIQDTQY | 0.78648174 |
| YQVGNFP | ASSPSIQDTQY | 0.8022602 |
| ESLLSQST | ASSPSIQDTQY | 0.7573023 |
| LFTVGFAAVG | ASSPSIQDTQY | 0.4748945 |
| ATPPTPT | ASSPSIQDTQY | 0.8174355 |
| TDLATLG | ASSPSIQDTQY | 0.84516835 |
| SAGGTGR | ASSPSIQDTQY | 0.7690399 |
| GFSNSGS | ASSPSIQDTQY | 0.5729877 |
| GGNGGSGGTSV | ASSPSIQDTQY | 0.5770824 |
| GTSVATGGAGN | ASSPSIQDTQY | 0.7532614 |
| NGGNGGSGGTS | ASSPSIQDTQY | 0.7762341 |
| AVVKFPH | ASSPSIQDTQY | 0.4891202 |
| TPPPPPP | ASSPSIQDTQY | 0.71958673 |
| AAPAAAAAPA | ASSPSIQDTQY | 0.75086254 |
| IALNEDL | ASSPSIQDTQY | 0.32769352 |
| IVNVSSM | ASSPSIQDTQY | 0.74838376 |
| TGIPPAPR | ASSPSIQDTQY | 0.81296027 |
| FDLGGGTF | ASSPSIQDTQY | 0.7337922 |
| FDLGGGTFDVS | ASSPSIQDTQY | 0.53467494 |
| NEPTAAA | ASSPSIQDTQY | 0.8263776 |
| NEPTAAALA | ASSPSIQDTQY | 0.82959235 |
| GSIINISSI | ASSPSIQDTQY | 0.8223748 |
| VAPGNAGTA | ASSPSIQDTQY | 0.36581674 |
| TGGGASLE | ASSPSIQDTQY | 0.23833184 |
| VAVYNHYKRI | ASSPSIQDTQY | 0.644959 |
| TDPAPATT | ASSPSIQDTQY | 0.7866379 |
| TNRPDILDPALLRPG | ASSPSIQDTQY | 0.6525271 |
| LTTTTTS | ASSPSIQDTQY | 0.5303818 |
| PTGSGKTL | ASSPSIQDTQY | 0.7416152 |
| TGSGKTL | ASSPSIQDTQY | 0.8078417 |
| TIGHVDHGK | ASSPSIQDTQY | 0.52480686 |
| TIGHVDHGKTT | ASSPSIQDTQY | 0.37145638 |
| GHVDHGKTT | ASSPSIQDTQY | 0.7834381 |
| AQFEHTLL | ASSPSIQDTQY | 0.77279854 |
| IVNQDEL | ASSPSIQDTQY | 0.5256954 |
| TNGSQFF | ASSPSIQDTQY | 0.6004461 |
| TGKGIVE | ASSPSIQDTQY | 0.6298852 |
| LISNASDA | ASSPSIQDTQY | 0.6565815 |
| AELRTLGELTKIAKAHGAQ | ASSPSIQDTQY | 0.68580884 |
| APNAVAAAEPSW | ASSPSIQDTQY | 0.39563644 |
| APQVAGAATAVL | ASSPSIQDTQY | 0.79175997 |
| ARRDPAAAAAILPTDA | ASSPSIQDTQY | 0.46006107 |
| DAAARDVLAEAGLAEYF | ASSPSIQDTQY | 0.7916859 |
| DDQIVLSAKANLAKVEQ | ASSPSIQDTQY | 0.7359733 |
| DELLPDTGEDSDEDGH | ASSPSIQDTQY | 0.7243794 |
| ELPEAPANRATVPQD | ASSPSIQDTQY | 0.62139994 |
| FGQSKIERVTGIG | ASSPSIQDTQY | 0.50113565 |
| GAGGTGSTAGGAGGAGGAGGLY | ASSPSIQDTQY | 0.33726352 |
| GHGGTGAAGGNGGTGGAGGSGI | ASSPSIQDTQY | 0.30092105 |

| | | |
|---|---|---|
| GNGGNGGAGGEGGPGGAGGAGA | ASSPSIQDTQY | 0.80856025 |
| GSTAGNGGNGGNATNGTIAGQP | ASSPSIQDTQY | 0.47888857 |
| GTGGDGGNGGNGGTGG | ASSPSIQDTQY | 0.7901201 |
| GVRRPAGIPADAPV | ASSPSIQDTQY | 0.745554 |
| GYRQVTDAHLLTLARR | ASSPSIQDTQY | 0.4998238 |
| IAKMGQDGHDRG | ASSPSIQDTQY | 0.7594398 |
| KAEPSIEIQYT | ASSPSIQDTQY | 0.4254731 |
| KPVRIRMAADEDTVAVT | ASSPSIQDTQY | 0.5699167 |
| MPPRIAGMRLLVIKPEP | ASSPSIQDTQY | 0.71072763 |
| MKFYAVALFRSRGVPPTR | ASSPSIQDTQY | 0.78221625 |
| NAGGFGILGAVAHSP | ASSPSIQDTQY | 0.49883455 |
| NEGREITLAMAEM | ASSPSIQDTQY | 0.76659036 |
| NELVEWSHPLGHRWRWE | ASSPSIQDTQY | 0.7287992 |
| NLGAGGTGGNGGNPG | ASSPSIQDTQY | 0.5301827 |
| NTEWPKWETVMAADPVD | ASSPSIQDTQY | 0.7385136 |
| PASAAPAETQP | ASSPSIQDTQY | 0.7956909 |
| QQSSDGKNGVRAAEITGE | ASSPSIQDTQY | 0.76607794 |
| RAPDAVTTSWWKE | ASSPSIQDTQY | 0.39441666 |
| SGFGNVGAHTSGWFNQ | ASSPSIQDTQY | 0.7723732 |
| SGVAGAVGQAASVGGLKV | ASSPSIQDTQY | 0.5203088 |
| SNGGAGTGSAGGNGGT | ASSPSIQDTQY | 0.7335248 |
| TGGAGGEGGAAG | ASSPSIQDTQY | 0.7130662 |
| TGGDGGAAGAGGAPGHG | ASSPSIQDTQY | 0.708984 |
| TMPPLLAAIANAGRHGV | ASSPSIQDTQY | 0.7852833 |
| TSSTAGNSGGA | ASSPSIQDTQY | 0.4514467 |
| VPATVPVGILC | ASSPSIQDTQY | 0.49008036 |
| VQRPDSGGVVLTGRIS | ASSPSIQDTQY | 0.63389367 |
| WTPNPVPATKGVPAGLVQAT | ASSPSIQDTQY | 0.7314782 |
| YALKRALAKFSDRFDVVII | ASSPSIQDTQY | 0.7838803 |
| YLIGSVIV | ASSPSIQDTQY | 0.4911996 |
| YGQLVAVID | ASSPSIQDTQY | 0.6951809 |
| VTVGGMTLPA | ASSPSIQDTQY | 0.56365037 |
| VLALPAV | ASSPSIQDTQY | 0.4010179 |
| VIPPTLNYET | ASSPSIQDTQY | 0.7049265 |
| VGMGGMHPSSG | ASSPSIQDTQY | 0.6561301 |
| VGLAGGAATA | ASSPSIQDTQY | 0.46866688 |
| VGGAGGAGGSGP | ASSPSIQDTQY | 0.75264317 |
| VFAAFMV | ASSPSIQDTQY | 0.5821866 |
| TLYDKLR | ASSPSIQDTQY | 0.7618235 |
| TLAVFTVL | ASSPSIQDTQY | 0.85196763 |
| TGSVQPARV | ASSPSIQDTQY | 0.54115814 |
| TGGGAGSGGT | ASSPSIQDTQY | 0.78097415 |
| SHLETSL | ASSPSIQDTQY | 0.7018298 |
| SFTTGMT | ASSPSIQDTQY | 0.73960143 |
| QVEVAKRMGVRQPTVS | ASSPSIQDTQY | 0.6479137 |
| QGSGGPALAM | ASSPSIQDTQY | 0.46461433 |
| MLAPSNP | ASSPSIQDTQY | 0.79715836 |
| LVNNAAI | ASSPSIQDTQY | 0.78242016 |
| LTGEINTLA | ASSPSIQDTQY | 0.80134726 |
| LSIFGPV | ASSPSIQDTQY | 0.7557144 |
| LLALLANL | ASSPSIQDTQY | 0.79537433 |

| | | |
|---|---|---|
| LISNASDALDKLR | ASSPSIQDTQY | 0.78141004 |
| LINEAVP | ASSPSIQDTQY | 0.55735 |
| LGYAFSGL | ASSPSIQDTQY | 0.30210778 |
| LDFVAVAL | ASSPSIQDTQY | 0.6623075 |
| LAQPGAE | ASSPSIQDTQY | 0.6804832 |
| ISNLGAVTGF | ASSPSIQDTQY | 0.6627103 |
| HQEDPGRI | ASSPSIQDTQY | 0.36515507 |
| GTPGEPTP | ASSPSIQDTQY | 0.80256826 |
| GFFNSGS | ASSPSIQDTQY | 0.8427434 |
| GAGGGGLTAGAN | ASSPSIQDTQY | 0.5304608 |
| FVDALVNM | ASSPSIQDTQY | 0.82017803 |
| FLTVAPEL | ASSPSIQDTQY | 0.41587874 |
| FETLTAL | ASSPSIQDTQY | 0.78208154 |
| DVINPIEL | ASSPSIQDTQY | 0.8333548 |
| APVSGAPV | ASSPSIQDTQY | 0.38894826 |
| ALVGGLAW | ASSPSIQDTQY | 0.7722945 |
| ALAAAAATAI | ASSPSIQDTQY | 0.510176 |
| AFGLGLLL | ASSPSIQDTQY | 0.56184375 |
| AFFAAASL | ASSPSIQDTQY | 0.82563 |
| AAAALPAFSPP | ASSPSIQDTQY | 0.6488644 |
| VVLAPAL | ASSPSIQDTQY | 0.75870615 |
| TIGHVDHGKTTL | ASSPSIQDTQY | 0.6509527 |
| LISNASDALDK | ASSPSIQDTQY | 0.4804567 |
| LISNASDAL | ASSPSIQDTQY | 0.71805876 |
| ISNASDALDK | ASSPSIQDTQY | 0.8155358 |
| NQAETLVYQTE | ASSPSIQDTQY | 0.52169144 |
| TGGTGGTAGAG | ASSPSIQDTQY | 0.8464235 |
| TGSTAGGAGGA | ASSPSIQDTQY | 0.5451604 |
| DPLAVPVAAGPV | ASSPSIQDTQY | 0.82669425 |
| GGTGGTPGNGGN | ASSPSIQDTQY | 0.7927228 |
| PAGLAGVDPATL | ASSPSIQDTQY | 0.658181 |
| GNNLVVEAPHAGASV | ASSPSIQDTQY | 0.6556373 |
| KVTRIMSASTLEQPA | ASSPSIQDTQY | 0.8057375 |
| LGFALPIALPRRTVV | ASSPSIQDTQY | 0.63667834 |
| TRLGNLTARKLVVLV | ASSPSIQDTQY | 0.77449536 |
| WSQQRRLIYFPSAGP | ASSPSIQDTQY | 0.80341876 |
| GPNGNPDPMAAAVDIR | ASSPSIQDTQY | 0.78218865 |
| TQRCVSVADLRAAASR | ASSPSIQDTQY | 0.90327215 |
| QFVLDVAEHPVEVLTETV | ASSPSIQDTQY | 0.86613095 |
| GGAGGTGGAGGNGGTQAGDG | ASSPSIQDTQY | 0.8777454 |
| PLLATLRAIAVDVDAKPAQV | ASSPSIQDTQY | 0.8108117 |
| APKFRTGGYMIDFWGVGYQVAK | ASSPSIQDTQY | 0.49913263 |
| GAGGAGGAGGAAGGPGGTGGTG | ASSPSIQDTQY | 0.86689925 |
| IGALGLLATDVAPAGSVAFGVA | ASSPSIQDTQY | 0.7423948 |
| AGGAGGIGGNGGAGGAGGVLMG | ASSPSIQDTQY | 0.88139015 |
| ADELIGLL | ASSPSIQDTQY | 0.895836 |
| ADRALKA | ASSPSIQDTQY | 0.8106779 |
| AGAVGQVGGAGGA | ASSPSIQDTQY | 0.89348125 |
| AGGGGGLGGAP | ASSPSIQDTQY | 0.8997912 |
| AGGLLYGNGGAG | ASSPSIQDTQY | 0.713572 |
| AGLESIGL | ASSPSIQDTQY | 0.8483814 |

| | | |
|---|---|---|
| AGLKTGII | ASSPSIQDTQY | 0.7931495 |
| ASLAALLL | ASSPSIQDTQY | 0.86175394 |
| ASVINISSI | ASSPSIQDTQY | 0.5809297 |
| ATPPGPA | ASSPSIQDTQY | 0.8919368 |
| ATVASALV | ASSPSIQDTQY | 0.811502 |
| ATVASLIL | ASSPSIQDTQY | 0.18168764 |
| AVNGPVLIRH | ASSPSIQDTQY | 0.8846252 |
| DGAAGAVGI | ASSPSIQDTQY | 0.8469996 |
| DPLLHGI | ASSPSIQDTQY | 0.8235227 |
| ELGAALGAL | ASSPSIQDTQY | 0.8947871 |
| ELGRTQAV | ASSPSIQDTQY | 0.67235494 |
| FVAGGYLS | ASSPSIQDTQY | 0.7404628 |
| GASGGAGGAGGW | ASSPSIQDTQY | 0.88433 |
| GGAGAGGDGGAGGI | ASSPSIQDTQY | 0.90015316 |
| GTGHNGSQPGMGGNG | ASSPSIQDTQY | 0.6405214 |
| GYAAASASAS | ASSPSIQDTQY | 0.87203074 |
| HQVDVAAVQQG | ASSPSIQDTQY | 0.9048165 |
| IGAAIAHT | ASSPSIQDTQY | 0.85857487 |
| IGMIEVAQL | ASSPSIQDTQY | 0.9039103 |
| IRLAPEHTSAMD | ASSPSIQDTQY | 0.90727335 |
| IYNVSVL | ASSPSIQDTQY | 0.82400584 |
| KQGFLGM | ASSPSIQDTQY | 0.89689946 |
| LESVITQL | ASSPSIQDTQY | 0.9043633 |
| LGLGALVVV | ASSPSIQDTQY | 0.835451 |
| LIGNGGNGGSGGI | ASSPSIQDTQY | 0.86022663 |
| LNAINAPTL | ASSPSIQDTQY | 0.80074227 |
| LQLLNTGV | ASSPSIQDTQY | 0.8630669 |
| LQTSIEF | ASSPSIQDTQY | 0.3500688 |
| MSFAVSEL | ASSPSIQDTQY | 0.8661896 |
| NTGALGL | ASSPSIQDTQY | 0.90201694 |
| PPGAGGAGGAGTT | ASSPSIQDTQY | 0.8462658 |
| QLVGLGI | ASSPSIQDTQY | 0.7483095 |
| RFLEQVHQL | ASSPSIQDTQY | 0.739145 |
| RGVGYVLR | ASSPSIQDTQY | 0.8786871 |
| SATGVFAGVI | ASSPSIQDTQY | 0.7915906 |
| SLPAITLP | ASSPSIQDTQY | 0.4503654 |
| STALAFTV | ASSPSIQDTQY | 0.82978404 |
| SVLTVGLGK | ASSPSIQDTQY | 0.90004456 |
| TALAGAGSGI | ASSPSIQDTQY | 0.90036297 |
| TGLPFTGI | ASSPSIQDTQY | 0.6330918 |
| VAEHVTV | ASSPSIQDTQY | 0.9055995 |
| VDIVLNSL | ASSPSIQDTQY | 0.88886803 |
| VLTGGGAVL | ASSPSIQDTQY | 0.90333354 |
| VNVGLAYGI | ASSPSIQDTQY | 0.86617696 |
| VSSAAVGAV | ASSPSIQDTQY | 0.7875484 |
| VTSVYEAGV | ASSPSIQDTQY | 0.89272904 |
| WVALDRGA | ASSPSIQDTQY | 0.8171854 |
| YLVGSVVI | ASSPSIQDTQY | 0.7984388 |
| GPPGTGKTLL | ASSPSIQDTQY | 0.902377 |
| GVGMTKF | ASSPSIQDTQY | 0.8173868 |
| IGIGGSDL | ASSPSIQDTQY | 0.76678705 |

| | | |
|---|---|---|
| IGTLLAEL | ASSPSIQDTQY | 0.82192785 |
| IYTSGTT | ASSPSIQDTQY | 0.9089915 |
| LSFPAGV | ASSPSIQDTQY | 0.7470778 |
| LYTSGTT | ASSPSIQDTQY | 0.719761 |
| VGAFPFGF | ASSPSIQDTQY | 0.74266696 |
| VGAGTVEF | ASSPSIQDTQY | 0.8489846 |
| VGKSTLLL | ASSPSIQDTQY | 0.8994035 |
| AALAHAGGGIV | ASSPSIQDTQY | 0.88886297 |
| AALVAVSLL | ASSPSIQDTQY | 0.83311564 |
| AAVSRAVAAL | ASSPSIQDTQY | 0.88223517 |
| AGGSSGIGGF | ASSPSIQDTQY | 0.8851148 |
| AGLTVANANG | ASSPSIQDTQY | 0.87110525 |
| AGLVGAVL | ASSPSIQDTQY | 0.8960397 |
| ALSQAFTV | ASSPSIQDTQY | 0.77824765 |
| ALTFSVSL | ASSPSIQDTQY | 0.9068862 |
| ALTGAALVAL | ASSPSIQDTQY | 0.888489 |
| ASFSIGAGVL | ASSPSIQDTQY | 0.8971573 |
| DGSVYKI | ASSPSIQDTQY | 0.90029603 |
| EQLAEIV | ASSPSIQDTQY | 0.86850405 |
| FAVSELTVM | ASSPSIQDTQY | 0.8746135 |
| GAAPLAAGL | ASSPSIQDTQY | 0.5243466 |
| GADGGDGGSGGAGG | ASSPSIQDTQY | 0.80446506 |
| GFGLDTGGAGG | ASSPSIQDTQY | 0.8755 |
| GLASGGLL | ASSPSIQDTQY | 0.870159 |
| GRIAGAAVLL | ASSPSIQDTQY | 0.8893695 |
| GSAGSLNTSF | ASSPSIQDTQY | 0.8891901 |
| GTGQIGI | ASSPSIQDTQY | 0.81789553 |
| GTLIALIL | ASSPSIQDTQY | 0.904358 |
| IAATMGFVVG | ASSPSIQDTQY | 0.89761424 |
| IFAQALL | ASSPSIQDTQY | 0.67923677 |
| IGAAVALGF | ASSPSIQDTQY | 0.84317696 |
| IGLTPLAL | ASSPSIQDTQY | 0.90970504 |
| ILGSNTSTL | ASSPSIQDTQY | 0.8653757 |
| IVFSLTVL | ASSPSIQDTQY | 0.84613955 |
| LASKATGF | ASSPSIQDTQY | 0.8493873 |
| LASVALIL | ASSPSIQDTQY | 0.8992027 |
| LFAVGLTL | ASSPSIQDTQY | 0.8134468 |
| LFNGGIGTYI | ASSPSIQDTQY | 0.9070566 |
| LTLGAGII | ASSPSIQDTQY | 0.30280828 |
| LVAAALGDI | ASSPSIQDTQY | 0.85276365 |
| NTAFVQLGI | ASSPSIQDTQY | 0.8074827 |
| QLFSGAEL | ASSPSIQDTQY | 0.31741375 |
| SAATFITL | ASSPSIQDTQY | 0.89204425 |
| SNYGLANLGV | ASSPSIQDTQY | 0.8399553 |
| SYGGAYAVM | ASSPSIQDTQY | 0.89081496 |
| TTGVGGLTLG | ASSPSIQDTQY | 0.8538774 |
| TTQTIYI | ASSPSIQDTQY | 0.89491385 |
| VAAGAGLGAVI | ASSPSIQDTQY | 0.8638411 |
| VAAVVTLL | ASSPSIQDTQY | 0.87692976 |
| VGFLGGLI | ASSPSIQDTQY | 0.81298697 |
| VGVIPILL | ASSPSIQDTQY | 0.88163966 |

| | | |
|---|---|---|
| VIFASSLI | ASSPSIQDTQY | 0.90560347 |
| VKGGDGGAAAG | ASSPSIQDTQY | 0.90305954 |
| VTKTFTNI | ASSPSIQDTQY | 0.4757024 |
| VVAAAILL | ASSPSIQDTQY | 0.8437179 |
| YAALDVEL | ASSPSIQDTQY | 0.84831357 |
| LASVALLL | ASSPSIQDTQY | 0.2935547 |
| LDAVHRV | ASSPSIQDTQY | 0.37355593 |
| QGLSSSIF | ASSPSIQDTQY | 0.82817715 |
| RGTFANIRL | ASSPSIQDTQY | 0.8449729 |
| RQLWWGHRI | ASSPSIQDTQY | 0.8937981 |
| YGPPGTGKTLL | ASSPSIQDTQY | 0.89363885 |
| AALVGAPANVA | ASSPSIQDTQY | 0.83505404 |
| AATFRRTE | ASSPSIQDTQY | 0.89174473 |
| AGALGGGI | ASSPSIQDTQY | 0.84961504 |
| AGGAGGAGGIGAG | ASSPSIQDTQY | 0.08317769 |
| AGSGALGL | ASSPSIQDTQY | 0.8304499 |
| AQIADQI | ASSPSIQDTQY | 0.72818375 |
| ASAVAVTYL | ASSPSIQDTQY | 0.9039917 |
| ASTVLGGTGG | ASSPSIQDTQY | 0.7891962 |
| AVGGEPAK | ASSPSIQDTQY | 0.8040893 |
| CEQGTFGGTC | ASSPSIQDTQY | 0.8614372 |
| DEGKAPAG | ASSPSIQDTQY | 0.85802543 |
| DTVVFPAL | ASSPSIQDTQY | 0.8573918 |
| FEGITMAG | ASSPSIQDTQY | 0.78170425 |
| FPLVVQD | ASSPSIQDTQY | 0.2070436 |
| GAFTTPQ | ASSPSIQDTQY | 0.88968956 |
| GFGGFGTAGGVGGA | ASSPSIQDTQY | 0.63314116 |
| GFLAQFL | ASSPSIQDTQY | 0.72959024 |
| GGAGGAGGNAGVA | ASSPSIQDTQY | 0.6264336 |
| GGVGTAGAGGAGGAGG | ASSPSIQDTQY | 0.526001 |
| GNGGGGDAGNAGSGGN | ASSPSIQDTQY | 0.5590941 |
| GTGGAGGAGNDAGS | ASSPSIQDTQY | 0.90051466 |
| GTTNTGWFNS | ASSPSIQDTQY | 0.830961 |
| IFMGPLFTL | ASSPSIQDTQY | 0.83217776 |
| ILAHAIEAI | ASSPSIQDTQY | 0.90437543 |
| IRSAIREAL | ASSPSIQDTQY | 0.8385999 |
| ITFAGGG | ASSPSIQDTQY | 0.8404989 |
| ITGMSGHQEV | ASSPSIQDTQY | 0.7865404 |
| IVAGLACTL | ASSPSIQDTQY | 0.8017409 |
| IVGGGLAGA | ASSPSIQDTQY | 0.77572143 |
| IVTGVLNI | ASSPSIQDTQY | 0.75285745 |
| LAQQGCI | ASSPSIQDTQY | 0.90589917 |
| LCLHVEN | ASSPSIQDTQY | 0.86122894 |
| LDAISDGS | ASSPSIQDTQY | 0.7012471 |
| LGGFAAML | ASSPSIQDTQY | 0.89141893 |
| LGGKNPMI | ASSPSIQDTQY | 0.90237796 |
| LGPKGVAAIL | ASSPSIQDTQY | 0.91073585 |
| LMVGVSI | ASSPSIQDTQY | 0.80995554 |
| LVGAAFVL | ASSPSIQDTQY | 0.8357701 |
| LVLAVAPL | ASSPSIQDTQY | 0.85215867 |
| MAAIAVIDAA | ASSPSIQDTQY | 0.840495 |

| | | |
|---|---|---|
| NNGINAELQ | ASSPSIQDTQY | 0.8188246 |
| RGEPLVLVHG | ASSPSIQDTQY | 0.6305704 |
| RGGAGGAAGLI | ASSPSIQDTQY | 0.9031217 |
| RSYRMHPAV | ASSPSIQDTQY | 0.90945065 |
| SAAHLLAGQLA | ASSPSIQDTQY | 0.8496167 |
| SAAVGSY | ASSPSIQDTQY | 0.7271018 |
| SPHAVLHRI | ASSPSIQDTQY | 0.84739864 |
| SQIAKVL | ASSPSIQDTQY | 0.74966747 |
| TALGHLVP | ASSPSIQDTQY | 0.82252365 |
| TAPTTSGAA | ASSPSIQDTQY | 0.84984374 |
| TDGDDGNGGSPGT | ASSPSIQDTQY | 0.67920655 |
| TDGLGEN | ASSPSIQDTQY | 0.84258074 |
| TGGPPLPGQA | ASSPSIQDTQY | 0.909848 |
| TIPAITI | ASSPSIQDTQY | 0.8169831 |
| TIPGFTI | ASSPSIQDTQY | 0.880892 |
| TSIASGALA | ASSPSIQDTQY | 0.8986712 |
| TVTLPGAAI | ASSPSIQDTQY | 0.890414 |
| VASGLSAALP | ASSPSIQDTQY | 0.90404004 |
| VGAGVIGI | ASSPSIQDTQY | 0.8906872 |
| VGPKRFGV | ASSPSIQDTQY | 0.87938523 |
| VNVAFPD | ASSPSIQDTQY | 0.86374176 |
| VSMTNSGVS | ASSPSIQDTQY | 0.8747051 |
| VTDVLGDLSV | ASSPSIQDTQY | 0.86070085 |
| VVVVGAGFAGL | ASSPSIQDTQY | 0.8618995 |
| VYTVGTVV | ASSPSIQDTQY | 0.84190917 |
| WTTFSGP | ASSPSIQDTQY | 0.8941412 |
| YDELVEAF | ASSPSIQDTQY | 0.86772245 |
| YGTGGAGGNGG | ASSPSIQDTQY | 0.8975569 |
| YTNVMAV | ASSPSIQDTQY | 0.4519903 |
| FLEQVHQL | ASSPSIQDTQY | 0.6961485 |
| GLSLLAVAV | ASSPSIQDTQY | 0.88991684 |
| LEQVHQL | ASSPSIQDTQY | 0.18248886 |
| LNAAGAI | ASSPSIQDTQY | 0.4392357 |
| PAVHASL | ASSPSIQDTQY | 0.8820913 |
| TELEDDA | ASSPSIQDTQY | 0.9015754 |
| TPPGPPN | ASSPSIQDTQY | 0.1824046 |
| TPPPPPPP | ASSPSIQDTQY | 0.7183845 |
| TQSPALI | ASSRQNTEVF | 0.70746815 |
| EFAAAFPQ | ASSRQNTEVF | 0.8990756 |
| LLSGNLDVL | ASSRQNTEVF | 0.8392849 |
| GPANLGNYNI | ASSRQNTEVF | 0.46512222 |
| TLGPTLP | ASSRQNTEVF | 0.89236104 |
| VYAYSSCCK | ASSRQNTEVF | 0.6809796 |
| APAPAPAPNI | ASSRQNTEVF | 0.7739012 |
| PGGPVPIP | ASSRQNTEVF | 0.89780045 |
| FLTVALEP | ASSRQNTEVF | 0.90189195 |
| SQYFGANL | ASSRQNTEVF | 0.7749945 |
| GGVKPAA | ASSRQNTEVF | 0.88924146 |
| GGTGGAGGAGGW | ASSRQNTEVF | 0.8212384 |
| GVGGVAGLGGAGG | ASSRQNTEVF | 0.90871674 |
| VLLVLATTI | ASSRQNTEVF | 0.83452857 |

| | | |
|---|---|---|
| IGIFAAML | ASSRQNTEVF | 0.6133276 |
| AVGDLID | ASSRQNTEVF | 0.8947768 |
| NISPLPL | ASSRQNTEVF | 0.89044845 |
| VTAVFVAI | ASSRQNTEVF | 0.8731574 |
| SGDDNDAAL | ASSRQNTEVF | 0.89017713 |
| AVAALAHS | ASSRQNTEVF | 0.8576683 |
| GGVTVMATP | ASSRQNTEVF | 0.83515346 |
| AGLAANVRD | ASSRQNTEVF | 0.8910133 |
| TNRPDLIDPALLRPG | ASSRQNTEVF | 0.8941964 |
| LANLGVTAL | ASSRQNTEVF | 0.6299057 |
| TGERIEL | ASSRQNTEVF | 0.9009387 |
| NGEELAA | ASSRQNTEVF | 0.84275854 |
| VLSLVWG | ASSRQNTEVF | 0.81808454 |
| IGYALWL | ASSRQNTEVF | 0.9086792 |
| LSLLLAPL | ASSRQNTEVF | 0.87461627 |
| TLDEALK | ASSRQNTEVF | 0.8933425 |
| TDGAGTAEGG | ASSRQNTEVF | 0.8888601 |
| NAPGGAGGNGG | ASSRQNTEVF | 0.9031684 |
| VSVFMGMGGNF | ASSRQNTEVF | 0.4702365 |
| LASYLAAL | ASSRQNTEVF | 0.86605406 |
| LVGGTPAL | ASSRQNTEVF | 0.8679338 |
| VRNADVL | ASSRQNTEVF | 0.89403343 |
| TCSSPFLVSAM | ASSRQNTEVF | 0.85216814 |
| VHITDAE | ASSRQNTEVF | 0.6814463 |
| EYAAPSF | ASSRQNTEVF | 0.89094234 |
| VVDAFHGGL | ASSRQNTEVF | 0.9013077 |
| ASVVDPQ | ASSRQNTEVF | 0.88503075 |
| SYNEFGGLS | ASSRQNTEVF | 0.88645744 |
| DPDGSVY | ASSRQNTEVF | 0.908731 |
| MVSTIHATE | ASSRQNTEVF | 0.7990689 |
| LAAGVADPV | ASSRQNTEVF | 0.84703153 |
| TKEQIAA | ASSRQNTEVF | 0.9027648 |
| SLASPAAL | ASSRQNTEVF | 0.90335625 |
| IGLVFYGV | ASSRQNTEVF | 0.8605158 |
| FLLDASNL | ASSRQNTEVF | 0.9047873 |
| YQVGNFP | ASSRQNTEVF | 0.7944094 |
| ESLLSQST | ASSRQNTEVF | 0.88429433 |
| LFTVGFAAVG | ASSRQNTEVF | 0.76607144 |
| ATPPTPT | ASSRQNTEVF | 0.8691071 |
| TDLATLG | ASSRQNTEVF | 0.7364732 |
| SAGGTGR | ASSRQNTEVF | 0.88077843 |
| GFSNSGS | ASSRQNTEVF | 0.8906852 |
| GGNGGSGGTSV | ASSRQNTEVF | 0.8827044 |
| GTSVATGGAGN | ASSRQNTEVF | 0.30384496 |
| NGGNGGSGGTS | ASSRQNTEVF | 0.8884809 |
| AVVKFPH | ASSRQNTEVF | 0.88896215 |
| TPPPPPP | ASSRQNTEVF | 0.9097906 |
| AAPAAAAAPA | ASSRQNTEVF | 0.85603595 |
| IALNEDL | ASSRQNTEVF | 0.8904526 |
| IVNVSSM | ASSRQNTEVF | 0.8196049 |
| TGIPPAPR | ASSRQNTEVF | 0.8759797 |

| | | |
|---|---|---|
| FDLGGGTF | ASSRQNTEVF | 0.7482792 |
| FDLGGGTFDVS | ASSRQNTEVF | 0.8305348 |
| NEPTAAA | ASSRQNTEVF | 0.8132977 |
| NEPTAAALA | ASSRQNTEVF | 0.7991478 |
| GSIINISSI | ASSRQNTEVF | 0.88381743 |
| VAPGNAGTA | ASSRQNTEVF | 0.90037906 |
| TGGGASLE | ASSRQNTEVF | 0.89300406 |
| VAVYNHYKRI | ASSRQNTEVF | 0.8934417 |
| TDPAPATT | ASSRQNTEVF | 0.9030938 |
| TNRPDILDPALLRPG | ASSRQNTEVF | 0.90124285 |
| LTTTTTS | ASSRQNTEVF | 0.8150327 |
| PTGSGKTL | ASSRQNTEVF | 0.84621036 |
| TGSGKTL | ASSRQNTEVF | 0.7070708 |
| TIGHVDHGK | ASSRQNTEVF | 0.83195233 |
| TIGHVDHGKTT | ASSRQNTEVF | 0.8081235 |
| GHVDHGKTT | ASSRQNTEVF | 0.8744417 |
| AQFEHTLL | ASSRQNTEVF | 0.13554761 |
| IVNQDEL | ASSRQNTEVF | 0.85194254 |
| TNGSQFF | ASSRQNTEVF | 0.9057283 |
| TGKGIVE | ASSRQNTEVF | 0.75102794 |
| LISNASDA | ASSRQNTEVF | 0.8867835 |
| AELRTLGELTKIAKAHGAQ | ASSRQNTEVF | 0.8721334 |
| APNAVAAAEPSW | ASSRQNTEVF | 0.81924766 |
| APQVAGAATAVL | ASSRQNTEVF | 0.8869921 |
| ARRDPAAAAAILPTDA | ASSRQNTEVF | 0.75345755 |
| DAAARDVLAEAGLAEYF | ASSRQNTEVF | 0.8921852 |
| DDQIVLSAKANLAKVEQ | ASSRQNTEVF | 0.90756404 |
| DELLPDTGEDSDEDGH | ASSRQNTEVF | 0.8737292 |
| ELPEAPANRATVPQD | ASSRQNTEVF | 0.86893374 |
| FGQSKIERVTGIG | ASSRQNTEVF | 0.86566633 |
| GAGGTGSTAGGAGGAGGAGGLY | ASSRQNTEVF | 0.4474783 |
| GHGGTGAAGGNGGTGGAGGSGI | ASSRQNTEVF | 0.8749926 |
| GNGGNGGAGGEGGPGGAGGAGA | ASSRQNTEVF | 0.21598345 |
| GSTAGNGGNGGNATNGTIAGQP | ASSRQNTEVF | 0.85337996 |
| GTGGDGGNGGNGGTGG | ASSRQNTEVF | 0.76138437 |
| GVRRPAGIPADAPV | ASSRQNTEVF | 0.88204414 |
| GYRQVTDAHLLTLARR | ASSRQNTEVF | 0.8546028 |
| IAKMGQDGHDRG | ASSRQNTEVF | 0.8961766 |
| KAEPSIEIQYT | ASSRQNTEVF | 0.78471303 |
| KPVRIRMAADEDTVAVT | ASSRQNTEVF | 0.8446702 |
| MPPRIAGMRLLVIKPEP | ASSRQNTEVF | 0.81233585 |
| MKFYAVALFRSRGVPPTR | ASSRQNTEVF | 0.61269486 |
| NAGGFGILGAVAHSP | ASSRQNTEVF | 0.8916297 |
| NEGREITLAMAEM | ASSRQNTEVF | 0.8773328 |
| NELVEWSHPLGHRWRWE | ASSRQNTEVF | 0.8456422 |
| NLGAGGTGGNGGNPG | ASSRQNTEVF | 0.8991468 |
| NTEWPKWETVMAADPVD | ASSRQNTEVF | 0.87646914 |
| PASAAPAETQP | ASSRQNTEVF | 0.6045858 |
| QQSSDGKNGVRAAEITGE | ASSRQNTEVF | 0.8256242 |
| RAPDAVTTSWWKE | ASSRQNTEVF | 0.8543341 |
| SGFGNVGAHTSGWFNQ | ASSRQNTEVF | 0.8367514 |

| | | |
|---|---|---|
| SGVAGAVGQAASVGGLKV | ASSRQNTEVF | 0.14536694 |
| SNGGAGTGSAGGNGGT | ASSRQNTEVF | 0.89806235 |
| TGGAGGEGGAAG | ASSRQNTEVF | 0.815729 |
| TGGDGGAAGAGGAPGHG | ASSRQNTEVF | 0.77470005 |
| TMPPLLAAIANAGRHGV | ASSRQNTEVF | 0.8227408 |
| TSSTAGNSGGA | ASSRQNTEVF | 0.8964107 |
| VPATVPVGILC | ASSRQNTEVF | 0.73984116 |
| VQRPDSGGVVLTGRIS | ASSRQNTEVF | 0.8402132 |
| WTPNPVPATKGVPAGLVQAT | ASSRQNTEVF | 0.894473 |
| YALKRALAKFSDRFDVVII | ASSRQNTEVF | 0.87669086 |
| YLIGSVIV | ASSRQNTEVF | 0.89732486 |
| YGQLVAVID | ASSRQNTEVF | 0.88369423 |
| VTVGGMTLPA | ASSRQNTEVF | 0.8627057 |
| VLALPAV | ASSRQNTEVF | 0.8915312 |
| VIPPTLNYET | ASSRQNTEVF | 0.85721576 |
| VGMGGMHPSSG | ASSRQNTEVF | 0.69834423 |
| VGLAGGAATA | ASSRQNTEVF | 0.6628693 |
| VGGAGGAGGSGP | ASSRQNTEVF | 0.85446465 |
| VFAAFMV | ASSRQNTEVF | 0.62453634 |
| TLYDKLR | ASSRQNTEVF | 0.8724359 |
| TLAVFTVL | ASSRQNTEVF | 0.9054432 |
| TGSVQPARV | ASSRQNTEVF | 0.8747668 |
| TGGGAGSGGT | ASSRQNTEVF | 0.72903347 |
| SHLETSL | ASSRQNTEVF | 0.8456434 |
| SFTTGMT | ASSRQNTEVF | 0.749329 |
| QVEVAKRMGVRQPTVS | ASSRQNTEVF | 0.8907549 |
| QGSGGPALAM | ASSRQNTEVF | 0.884327 |
| MLAPSNP | ASSRQNTEVF | 0.90384245 |
| LVNNAAI | ASSRQNTEVF | 0.89328563 |
| LTGEINTLA | ASSRQNTEVF | 0.8058834 |
| LSIFGPV | ASSRQNTEVF | 0.8963188 |
| LLALLANL | ASSRQNTEVF | 0.8350504 |
| LISNASDALDKLR | ASSRQNTEVF | 0.8650578 |
| LINEAVP | ASSRQNTEVF | 0.87816197 |
| LGYAFSGL | ASSRQNTEVF | 0.8897725 |
| LDFVAVAL | ASSRQNTEVF | 0.85184 |
| LAQPGAE | ASSRQNTEVF | 0.8595661 |
| ISNLGAVTGF | ASSRQNTEVF | 0.69651866 |
| HQEDPGRI | ASSRQNTEVF | 0.8930818 |
| GTPGEPTP | ASSRQNTEVF | 0.80453265 |
| GFFNSGS | ASSRQNTEVF | 0.6425172 |
| GAGGGGLTAGAN | ASSRQNTEVF | 0.83411753 |
| FVDALVNM | ASSRQNTEVF | 0.9058538 |
| FLTVAPEL | ASSRQNTEVF | 0.90243226 |
| FETLTAL | ASSRQNTEVF | 0.76796675 |
| DVINPIEL | ASSRQNTEVF | 0.76896095 |
| APVSGAPV | ASSRQNTEVF | 0.86136675 |
| ALVGGLAW | ASSRQNTEVF | 0.6545348 |
| ALAAAAATAI | ASSRQNTEVF | 0.8883845 |
| AFGLGLLL | ASSRQNTEVF | 0.835371 |
| AFFAASL | ASSRQNTEVF | 0.8386711 |

| | | |
|---|---|---|
| AAAALPAFSPP | ASSRQNTEVF | 0.8202008 |
| VVLAPAL | ASSRQNTEVF | 0.8775346 |
| TIGHVDHGKTTL | ASSRQNTEVF | 0.82258904 |
| LISNASDALDK | ASSRQNTEVF | 0.89498174 |
| LISNASDAL | ASSRQNTEVF | 0.19709188 |
| ISNASDALDK | ASSRQNTEVF | 0.88435614 |
| NQAETLVYQTE | ASSRQNTEVF | 0.7648661 |
| TGGTGGTAGAG | ASSRQNTEVF | 0.8023657 |
| TGSTAGGAGGA | ASSRQNTEVF | 0.437761 |
| DPLAVPVAAGPV | ASSRQNTEVF | 0.87619644 |
| GGTGGTPGNGGN | ASSRQNTEVF | 0.88199 |
| PAGLAGVDPATL | ASSRQNTEVF | 0.9002309 |
| GNNLVVEAPHAGASV | ASSRQNTEVF | 0.82217443 |
| KVTRIMSASTLEQPA | ASSRQNTEVF | 0.8894982 |
| LGFALPIALPRRTVV | ASSRQNTEVF | 0.90194196 |
| TRLGNLTARKLVVLV | ASSRQNTEVF | 0.82259625 |
| WSQQRRLIYFPSAGP | ASSRQNTEVF | 0.42872623 |
| GPNGNPDPMAAAVDIR | ASSRQNTEVF | 0.8643905 |
| TQRCVSVADLRAAASR | ASSRQNTEVF | 0.85285574 |
| QFVLDVAEHPVEVLTETV | ASSRQNTEVF | 0.89742714 |
| GGAGGTGGAGGNGGTQAGDG | ASSRQNTEVF | 0.88056767 |
| PLLATLRAIAVDVDAKPAQV | ASSRQNTEVF | 0.74585277 |
| APKFRTGGYMIDFWGVGYQVAK | ASSRQNTEVF | 0.8857833 |
| GAGGAGGAGGAAGGPGGTGGTG | ASSRQNTEVF | 0.8957239 |
| IGALGLLATDVAPAGSVAFGVA | ASSRQNTEVF | 0.777181 |
| AGGAGGIGGNGGAGGAGGVLMG | ASSRQNTEVF | 0.8448528 |
| ADELIGLL | ASSRQNTEVF | 0.8224971 |
| ADRALKA | ASSRQNTEVF | 0.350951 |
| AGAVGQVGGAGGA | ASSRQNTEVF | 0.879757 |
| AGGGGGLGGAP | ASSRQNTEVF | 0.80063665 |
| AGGLLYGNGGAG | ASSRQNTEVF | 0.77052593 |
| AGLESIGL | ASSRQNTEVF | 0.8667325 |
| AGLKTGII | ASSRQNTEVF | 0.78065634 |
| ASLAALLL | ASSRQNTEVF | 0.7616962 |
| ASVINISSI | ASSRQNTEVF | 0.7911926 |
| ATPPGPA | ASSRQNTEVF | 0.84499264 |
| ATVASALV | ASSRQNTEVF | 0.884937 |
| ATVASLIL | ASSRQNTEVF | 0.8485426 |
| AVNGPVLIRH | ASSRQNTEVF | 0.8787708 |
| DGAAGAVGI | ASSRQNTEVF | 0.90241206 |
| DPLLHGI | ASSRQNTEVF | 0.60494334 |
| ELGAALGAL | ASSRQNTEVF | 0.7541522 |
| ELGRTQAV | ASSRQNTEVF | 0.8884416 |
| FVAGGYLS | ASSRQNTEVF | 0.67144984 |
| GASGGAGGAGGW | ASSRQNTEVF | 0.8939997 |
| GGAGAGGDGGAGGI | ASSRQNTEVF | 0.8564122 |
| GTGHNGSQPGMGGNG | ASSRQNTEVF | 0.87872213 |
| GYAAASASAS | ASSRQNTEVF | 0.8523585 |
| HQVDVAAVQQG | ASSRQNTEVF | 0.9097365 |
| IGAAIAHT | ASSRQNTEVF | 0.89228046 |
| IGMIEVAQL | ASSRQNTEVF | 0.85947406 |

| | | |
|---|---|---|
| IRLAPEHTSAMD | ASSRQNTEVF | 0.85599613 |
| IYNVSVL | ASSRQNTEVF | 0.74620396 |
| KQGFLGM | ASSRQNTEVF | 0.89162993 |
| LESVITQL | ASSRQNTEVF | 0.89233255 |
| LGLGALVVV | ASSRQNTEVF | 0.74495673 |
| LIGNGGNGGSGGI | ASSRQNTEVF | 0.8375368 |
| LNAINAPTL | ASSRQNTEVF | 0.9095963 |
| LQLLNTGV | ASSRQNTEVF | 0.90195787 |
| LQTSIEF | ASSRQNTEVF | 0.5734292 |
| MSFAVSEL | ASSRQNTEVF | 0.836547 |
| NTGALGL | ASSRQNTEVF | 0.6179072 |
| PPGAGGAGGAGTT | ASSRQNTEVF | 0.8456625 |
| QLVGLGI | ASSRQNTEVF | 0.6246406 |
| RFLEQVHQL | ASSRQNTEVF | 0.8878578 |
| RGVGYVLR | ASSRQNTEVF | 0.8394201 |
| SATGVFAGVI | ASSRQNTEVF | 0.8796425 |
| SLPAITLP | ASSRQNTEVF | 0.87468 |
| STALAFTV | ASSRQNTEVF | 0.9047264 |
| SVLTVGLGK | ASSRQNTEVF | 0.8241935 |
| TALAGAGSGI | ASSRQNTEVF | 0.8259878 |
| TGLPFTGI | ASSRQNTEVF | 0.67011327 |
| VAEHVTV | ASSRQNTEVF | 0.5485741 |
| VDIVLNSL | ASSRQNTEVF | 0.82135236 |
| VLTGGGAVL | ASSRQNTEVF | 0.80287874 |
| VNVGLAYGI | ASSRQNTEVF | 0.28382695 |
| VSSAAVGAV | ASSRQNTEVF | 0.9056017 |
| VTSVYEAGV | ASSRQNTEVF | 0.86512387 |
| WVALDRGA | ASSRQNTEVF | 0.8879032 |
| YLVGSVVI | ASSRQNTEVF | 0.81422037 |
| GPPGTGKTLL | ASSRQNTEVF | 0.7727469 |
| GVGMTKF | ASSRQNTEVF | 0.8004055 |
| IGIGGSDL | ASSRQNTEVF | 0.88927627 |
| IGTLLAEL | ASSRQNTEVF | 0.7535343 |
| IYTSGTT | ASSRQNTEVF | 0.6565385 |
| LSFPAGV | ASSRQNTEVF | 0.82122374 |
| LYTSGTT | ASSRQNTEVF | 0.7834933 |
| VGAFPFGF | ASSRQNTEVF | 0.90689325 |
| VGAGTVEF | ASSRQNTEVF | 0.8938485 |
| VGKSTLLL | ASSRQNTEVF | 0.8768519 |
| AALAHAGGGIV | ASSRQNTEVF | 0.90445244 |
| AALVAVSLL | ASSRQNTEVF | 0.2592514 |
| AAVSRAVAAL | ASSRQNTEVF | 0.7964107 |
| AGGSSGIGGF | ASSRQNTEVF | 0.9041667 |
| AGLTVANANG | ASSRQNTEVF | 0.8789903 |
| AGLVGAVL | ASSRQNTEVF | 0.8552528 |
| ALSQAFTV | ASSRQNTEVF | 0.898691 |
| ALTFSVSL | ASSRQNTEVF | 0.7652527 |
| ALTGAALVAL | ASSRQNTEVF | 0.8419796 |
| ASFSIGAGVL | ASSRQNTEVF | 0.8506153 |
| DGSVYKI | ASSRQNTEVF | 0.896838 |
| EQLAEIV | ASSRQNTEVF | 0.8789593 |

| | | |
|---|---|---|
| FAVSELTVM | ASSRQNTEVF | 0.7541003 |
| GAAPLAAGL | ASSRQNTEVF | 0.88221896 |
| GADGGDGGSGGAGG | ASSRQNTEVF | 0.90912795 |
| GFGLDTGGAGG | ASSRQNTEVF | 0.8472794 |
| GLASGGLL | ASSRQNTEVF | 0.8895741 |
| GRIAGAAVLL | ASSRQNTEVF | 0.8752156 |
| GSAGSLNTSF | ASSRQNTEVF | 0.827276 |
| GTGQIGI | ASSRQNTEVF | 0.7909643 |
| GTLIALIL | ASSRQNTEVF | 0.8069704 |
| IAATMGFVVG | ASSRQNTEVF | 0.8498818 |
| IFAQALL | ASSRQNTEVF | 0.70596135 |
| IGAAVALGF | ASSRQNTEVF | 0.86721915 |
| IGLTPLAL | ASSRQNTEVF | 0.8384036 |
| ILGSNTSTL | ASSRQNTEVF | 0.9140502 |
| IVFSLTVL | ASSRQNTEVF | 0.88643026 |
| LASKATGF | ASSRQNTEVF | 0.88151264 |
| LASVALIL | ASSRQNTEVF | 0.74152565 |
| LFAVGLTL | ASSRQNTEVF | 0.8870164 |
| LFNGGIGTYI | ASSRQNTEVF | 0.86383903 |
| LTLGAGII | ASSRQNTEVF | 0.6975087 |
| LVAAALGDI | ASSRQNTEVF | 0.9027691 |
| NTAFVQLGI | ASSRQNTEVF | 0.7305715 |
| QLFSGAEL | ASSRQNTEVF | 0.85156596 |
| SAATFITL | ASSRQNTEVF | 0.8895008 |
| SNYGLANLGV | ASSRQNTEVF | 0.89056385 |
| SYGGAYAVM | ASSRQNTEVF | 0.83924353 |
| TTGVGGLTLG | ASSRQNTEVF | 0.84727925 |
| TTQTIYI | ASSRQNTEVF | 0.7702118 |
| VAAGAGLGAVI | ASSRQNTEVF | 0.896242 |
| VAAVVTLL | ASSRQNTEVF | 0.9031556 |
| VGFLGGLI | ASSRQNTEVF | 0.8422195 |
| VGVIPILL | ASSRQNTEVF | 0.90916085 |
| VIFASSLI | ASSRQNTEVF | 0.8792079 |
| VKGGDGGAAAG | ASSRQNTEVF | 0.5652826 |
| VTKTFTNI | ASSRQNTEVF | 0.86914515 |
| VVAAAILL | ASSRQNTEVF | 0.898425 |
| YAALDVEL | ASSRQNTEVF | 0.8974378 |
| LASVALLL | ASSRQNTEVF | 0.8996563 |
| LDAVHRV | ASSRQNTEVF | 0.8847413 |
| QGLSSSIF | ASSRQNTEVF | 0.8188371 |
| RGTFANIRL | ASSRQNTEVF | 0.89960206 |
| RQLWWGHRI | ASSRQNTEVF | 0.9124602 |
| YGPPGTGKTLL | ASSRQNTEVF | 0.8910667 |
| AALVGAPANVA | ASSRQNTEVF | 0.83690524 |
| AATFRRTE | ASSRQNTEVF | 0.80584514 |
| AGALGGGI | ASSRQNTEVF | 0.87895066 |
| AGGAGGAGGIGAG | ASSRQNTEVF | 0.89352375 |
| AGSGALGL | ASSRQNTEVF | 0.76570517 |
| AQIADQI | ASSRQNTEVF | 0.85902905 |
| ASAVAVTYL | ASSRQNTEVF | 0.86942255 |
| ASTVLGGTGG | ASSRQNTEVF | 0.61049277 |

| | | |
|---|---|---|
| AVGGEPAK | ASSRQNTEVF | 0.8760588 |
| CEQGTFGGTC | ASSRQNTEVF | 0.90270483 |
| DEGKAPAG | ASSRQNTEVF | 0.87090296 |
| DTVVFPAL | ASSRQNTEVF | 0.7693952 |
| FEGITMAG | ASSRQNTEVF | 0.8992622 |
| FPLVVQD | ASSRQNTEVF | 0.90958023 |
| GAFTTPQ | ASSRQNTEVF | 0.90333134 |
| GFGGFGTAGGVGGA | ASSRQNTEVF | 0.4977945 |
| GFLAQFL | ASSRQNTEVF | 0.3814935 |
| GGAGGAGGNAGVA | ASSRQNTEVF | 0.8125165 |
| GGVGTAGAGGAGGAGG | ASSRQNTEVF | 0.87789404 |
| GNGGGGDAGNAGSGGN | ASSRQNTEVF | 0.805678 |
| GTGGAGGAGNDAGS | ASSRQNTEVF | 0.7643002 |
| GTTNTGWFNS | ASSRQNTEVF | 0.8529029 |
| IFMGPLFTL | ASSRQNTEVF | 0.89452803 |
| ILAHAIEAI | ASSRQNTEVF | 0.7876229 |
| IRSAIREAL | ASSRQNTEVF | 0.7236755 |
| ITFAGGG | ASSRQNTEVF | 0.88316184 |
| ITGMSGHQEV | ASSRQNTEVF | 0.89761424 |
| IVAGLACTL | ASSRQNTEVF | 0.7674718 |
| IVGGGLAGA | ASSRQNTEVF | 0.85225606 |
| IVTGVLNI | ASSRQNTEVF | 0.820762 |
| LAQQGCI | ASSRQNTEVF | 0.7989069 |
| LCLHVEN | ASSRQNTEVF | 0.8524146 |
| LDAISDGS | ASSRQNTEVF | 0.71198523 |
| LGGFAAML | ASSRQNTEVF | 0.9012308 |
| LGGKNPMI | ASSRQNTEVF | 0.7388003 |
| LGPKGVAAIL | ASSRQNTEVF | 0.8996807 |
| LMVGVSI | ASSRQNTEVF | 0.8796188 |
| LVGAAFVL | ASSRQNTEVF | 0.8897033 |
| LVLAVAPL | ASSRQNTEVF | 0.82485807 |
| MAAIAVIDAA | ASSRQNTEVF | 0.80397713 |
| NNGINAELQ | ASSRQNTEVF | 0.70481527 |
| RGEPLVLVHG | ASSRQNTEVF | 0.62739015 |
| RGGAGGAAGLI | ASSRQNTEVF | 0.90205956 |
| RSYRMHPAV | ASSRQNTEVF | 0.35323972 |
| SAAHLLAGQLA | ASSRQNTEVF | 0.8990525 |
| SAAVGSY | ASSRQNTEVF | 0.8803115 |
| SPHAVLHRI | ASSRQNTEVF | 0.79198074 |
| SQIAKVL | ASSRQNTEVF | 0.89287287 |
| TALGHLVP | ASSRQNTEVF | 0.627947 |
| TAPTTSGAA | ASSRQNTEVF | 0.8487459 |
| TDGDDGNGGSPGT | ASSRQNTEVF | 0.878233 |
| TDGLGEN | ASSRQNTEVF | 0.88735235 |
| TGGPPLPGQA | ASSRQNTEVF | 0.81828964 |
| TIPAITI | ASSRQNTEVF | 0.88760674 |
| TIPGFTI | ASSRQNTEVF | 0.8772703 |
| TSIASGALA | ASSRQNTEVF | 0.80854106 |
| TVTLPGAAI | ASSRQNTEVF | 0.8673129 |
| VASGLSAALP | ASSRQNTEVF | 0.8913286 |
| VGAGVIGI | ASSRQNTEVF | 0.88145554 |

| | | |
|---|---|---|
| VGPKRFGV | ASSRQNTEVF | 0.75975096 |
| VNVAFPD | ASSRQNTEVF | 0.8777578 |
| VSMTNSGVS | ASSRQNTEVF | 0.7972196 |
| VTDVLGDLSV | ASSRQNTEVF | 0.87423277 |
| VVVVGAGFAGL | ASSRQNTEVF | 0.8776922 |
| VYTVGTVV | ASSRQNTEVF | 0.86050063 |
| WTTFSGP | ASSRQNTEVF | 0.8972886 |
| YDELVEAF | ASSRQNTEVF | 0.5829048 |
| YGTGGAGGNGG | ASSRQNTEVF | 0.795251 |
| YTNVMAV | ASSRQNTEVF | 0.82815194 |
| FLEQVHQL | ASSRQNTEVF | 0.8550922 |
| GLSLLAVAV | ASSRQNTEVF | 0.8871813 |
| LEQVHQL | ASSRQNTEVF | 0.75077903 |
| LNAAGAI | ASSRQNTEVF | 0.8688479 |
| PAVHASL | ASSRQNTEVF | 0.8203169 |
| TELEDDA | ASSRQNTEVF | 0.75188375 |
| TPPGPPN | ASSRQNTEVF | 0.87118435 |
| TPPPPPPP | ASSRQNTEVF | 0.8262548 |
| TQSPALI | ASGDEENSDYT | 0.7523095 |
| EFAAAFPQ | ASGDEENSDYT | 0.89254254 |
| LLSGNLDVL | ASGDEENSDYT | 0.7684392 |
| GPANLGNYNI | ASGDEENSDYT | 0.8740603 |
| TLGPTLP | ASGDEENSDYT | 0.8994216 |
| VYAYSSCCK | ASGDEENSDYT | 0.77839494 |
| APAPAPAPNI | ASGDEENSDYT | 0.8990374 |
| PGGPVPIP | ASGDEENSDYT | 0.8784583 |
| FLTVALEP | ASGDEENSDYT | 0.88133556 |
| SQYFGANL | ASGDEENSDYT | 0.8029241 |
| GGVKPAA | ASGDEENSDYT | 0.80754393 |
| GGTGGAGGAGGW | ASGDEENSDYT | 0.9012649 |
| GVGGVAGLGGAGG | ASGDEENSDYT | 0.8851652 |
| VLLVLATTI | ASGDEENSDYT | 0.8950673 |
| IGIFAAML | ASGDEENSDYT | 0.8910184 |
| AVGDLID | ASGDEENSDYT | 0.8836549 |
| NISPLPL | ASGDEENSDYT | 0.8939968 |
| VTAVFVAI | ASGDEENSDYT | 0.80167085 |
| SGDDNDAAL | ASGDEENSDYT | 0.63775015 |
| AVAALAHS | ASGDEENSDYT | 0.81886685 |
| GGVTVMATP | ASGDEENSDYT | 0.80203915 |
| AGLAANVRD | ASGDEENSDYT | 0.83960855 |
| TNRPDLIDPALLRPG | ASGDEENSDYT | 0.6075759 |
| LANLGVTAL | ASGDEENSDYT | 0.88879436 |
| TGERIEL | ASGDEENSDYT | 0.9126551 |
| NGEELAA | ASGDEENSDYT | 0.7611289 |
| VLSLVWG | ASGDEENSDYT | 0.9088119 |
| IGYALWL | ASGDEENSDYT | 0.6649189 |
| LSLLLAPL | ASGDEENSDYT | 0.8842217 |
| TLDEALK | ASGDEENSDYT | 0.89833856 |
| TDGAGTAEGG | ASGDEENSDYT | 0.73252404 |
| NAPGGAGGNGG | ASGDEENSDYT | 0.87347484 |
| VSVFMGMGGNF | ASGDEENSDYT | 0.7683643 |

| | | |
|---|---|---|
| LASYLAAL | ASGDEENSDYT | 0.8119377 |
| LVGGTPAL | ASGDEENSDYT | 0.8995593 |
| VRNADVL | ASGDEENSDYT | 0.78072643 |
| TCSSPFLVSAM | ASGDEENSDYT | 0.88785547 |
| VHITDAE | ASGDEENSDYT | 0.82919234 |
| EYAAPSF | ASGDEENSDYT | 0.7881769 |
| VVDAFHGGL | ASGDEENSDYT | 0.8874759 |
| ASVVDPQ | ASGDEENSDYT | 0.90121067 |
| SYNEFGGLS | ASGDEENSDYT | 0.82457006 |
| DPDGSVY | ASGDEENSDYT | 0.9116914 |
| MVSTIHATE | ASGDEENSDYT | 0.81686294 |
| LAAGVADPV | ASGDEENSDYT | 0.9058916 |
| TKEQIAA | ASGDEENSDYT | 0.8986962 |
| SLASPAAL | ASGDEENSDYT | 0.85897243 |
| IGLVFYGV | ASGDEENSDYT | 0.84933484 |
| FLLDASNL | ASGDEENSDYT | 0.89490414 |
| YQVGNFP | ASGDEENSDYT | 0.8582946 |
| ESLLSQST | ASGDEENSDYT | 0.8930297 |
| LFTVGFAAVG | ASGDEENSDYT | 0.9041907 |
| ATPPTPT | ASGDEENSDYT | 0.6053541 |
| TDLATLG | ASGDEENSDYT | 0.66716635 |
| SAGGTGR | ASGDEENSDYT | 0.6476735 |
| GFSNSGS | ASGDEENSDYT | 0.62552315 |
| GGNGGSGGTSV | ASGDEENSDYT | 0.61355597 |
| GTSVATGGAGN | ASGDEENSDYT | 0.50645673 |
| NGGNGGSGGTS | ASGDEENSDYT | 0.6186945 |
| AVVKFPH | ASGDEENSDYT | 0.6018396 |
| TPPPPPP | ASGDEENSDYT | 0.65189296 |
| AAPAAAAAPA | ASGDEENSDYT | 0.6584865 |
| IALNEDL | ASGDEENSDYT | 0.6039463 |
| IVNVSSM | ASGDEENSDYT | 0.66763294 |
| TGIPPAPR | ASGDEENSDYT | 0.6602317 |
| FDLGGGTF | ASGDEENSDYT | 0.5409047 |
| FDLGGGTFDVS | ASGDEENSDYT | 0.6246095 |
| NEPTAAA | ASGDEENSDYT | 0.59811145 |
| NEPTAAALA | ASGDEENSDYT | 0.6304495 |
| GSIINISSI | ASGDEENSDYT | 0.4972306 |
| VAPGNAGTA | ASGDEENSDYT | 0.6583101 |
| TGGGASLE | ASGDEENSDYT | 0.59525806 |
| VAVYNHYKRI | ASGDEENSDYT | 0.32785007 |
| TDPAPATT | ASGDEENSDYT | 0.639585 |
| TNRPDILDPALLRPG | ASGDEENSDYT | 0.6405259 |
| LTTTTTS | ASGDEENSDYT | 0.6467153 |
| PTGSGKTL | ASGDEENSDYT | 0.65856475 |
| TGSGKTL | ASGDEENSDYT | 0.5198308 |
| TIGHVDHGK | ASGDEENSDYT | 0.6153872 |
| TIGHVDHGKTT | ASGDEENSDYT | 0.65897125 |
| GHVDHGKTT | ASGDEENSDYT | 0.66480434 |
| AQFEHTLL | ASGDEENSDYT | 0.58457613 |
| IVNQDEL | ASGDEENSDYT | 0.647191 |
| TNGSQFF | ASGDEENSDYT | 0.65850943 |

| | | |
|---|---|---|
| TGKGIVE | ASGDEENSDYT | 0.65134853 |
| LISNASDA | ASGDEENSDYT | 0.6576692 |
| AELRTLGELTKIAKAHGAQ | ASGDEENSDYT | 0.6642051 |
| APNAVAAAEPSW | ASGDEENSDYT | 0.57857597 |
| APQVAGAATAVL | ASGDEENSDYT | 0.6645318 |
| ARRDPAAAAAILPTDA | ASGDEENSDYT | 0.6671425 |
| DAAARDVLAEAGLAEYF | ASGDEENSDYT | 0.6378961 |
| DDQIVLSAKANLAKVEQ | ASGDEENSDYT | 0.63530725 |
| DELLPDTGEDSDEDGH | ASGDEENSDYT | 0.5633099 |
| ELPEAPANRATVPQD | ASGDEENSDYT | 0.6212131 |
| FGQSKIERVTGIG | ASGDEENSDYT | 0.4817402 |
| GAGGTGSTAGGAGGAGGAGGLY | ASGDEENSDYT | 0.6487851 |
| GHGGTGAAGGNGGTGGAGGSGI | ASGDEENSDYT | 0.66320074 |
| GNGGNGGAGGEGGPGGAGGAGA | ASGDEENSDYT | 0.61955625 |
| GSTAGNGGNGGNATNGTIAGQP | ASGDEENSDYT | 0.5855986 |
| GTGGDGGNGGNGGTGG | ASGDEENSDYT | 0.5856115 |
| GVRRPAGIPADAPV | ASGDEENSDYT | 0.6303659 |
| GYRQVTDAHLLTLARR | ASGDEENSDYT | 0.61912906 |
| IAKMGQDGHDRG | ASGDEENSDYT | 0.40436298 |
| KAEPSIEIQYT | ASGDEENSDYT | 0.6253184 |
| KPVRIRMAADEDTVAVT | ASGDEENSDYT | 0.6627626 |
| MPPRIAGMRLLVIKPEP | ASGDEENSDYT | 0.6599922 |
| MKFYAVALFRSRGVPPTR | ASGDEENSDYT | 0.48740697 |
| NAGGFGILGAVAHSP | ASGDEENSDYT | 0.6657672 |
| NEGREITLAMAEM | ASGDEENSDYT | 0.65120995 |
| NELVEWSHPLGHRWRWE | ASGDEENSDYT | 0.6623254 |
| NLGAGGTGGNGGNPG | ASGDEENSDYT | 0.6328683 |
| NTEWPKWETVMAADPVD | ASGDEENSDYT | 0.60589826 |
| PASAAPAETQP | ASGDEENSDYT | 0.6543596 |
| QQSSDGKNGVRAAEITGE | ASGDEENSDYT | 0.6175777 |
| RAPDAVTTSWWKE | ASGDEENSDYT | 0.54086065 |
| SGFGNVGAHTSGWFNQ | ASGDEENSDYT | 0.65698916 |
| SGVAGAVGQAASVGGLKV | ASGDEENSDYT | 0.64351785 |
| SNGGAGTGSAGGNGGT | ASGDEENSDYT | 0.6089885 |
| TGGAGGEGGAAG | ASGDEENSDYT | 0.5881071 |
| TGGDGGAAGAGGAPGHG | ASGDEENSDYT | 0.65882427 |
| TMPPLLAAIANAGRHGV | ASGDEENSDYT | 0.570462 |
| TSSTAGNSGGA | ASGDEENSDYT | 0.6213609 |
| VPATVPVGILC | ASGDEENSDYT | 0.5523711 |
| VQRPDSGGVVLTGRIS | ASGDEENSDYT | 0.6360937 |
| WTPNPVPATKGVPAGLVQAT | ASGDEENSDYT | 0.66235054 |
| YALKRALAKFSDRFDVVII | ASGDEENSDYT | 0.6519056 |
| YLIGSVIV | ASGDEENSDYT | 0.61030126 |
| YGQLVAVID | ASGDEENSDYT | 0.64624023 |
| VTVGGMTLPA | ASGDEENSDYT | 0.65259427 |
| VLALPAV | ASGDEENSDYT | 0.64211845 |
| VIPPTLNYET | ASGDEENSDYT | 0.65256697 |
| VGMGGMHPSSG | ASGDEENSDYT | 0.60881144 |
| VGLAGGAATA | ASGDEENSDYT | 0.6597258 |
| VGGAGGAGGSGP | ASGDEENSDYT | 0.6439582 |
| VFAAFMV | ASGDEENSDYT | 0.6582901 |

| | | |
|---|---|---|
| TLYDKLR | ASGDEENSDYT | 0.6591925 |
| TLAVFTVL | ASGDEENSDYT | 0.66128886 |
| TGSVQPARV | ASGDEENSDYT | 0.63546014 |
| TGGGAGSGGT | ASGDEENSDYT | 0.41056064 |
| SHLETSL | ASGDEENSDYT | 0.5978335 |
| SFTTGMT | ASGDEENSDYT | 0.6494586 |
| QVEVAKRMGVRQPTVS | ASGDEENSDYT | 0.6429884 |
| QGSGGPALAM | ASGDEENSDYT | 0.6572574 |
| MLAPSNP | ASGDEENSDYT | 0.6487122 |
| LVNNAAI | ASGDEENSDYT | 0.6054058 |
| LTGEINTLA | ASGDEENSDYT | 0.65408635 |
| LSIFGPV | ASGDEENSDYT | 0.6555772 |
| LLALLANL | ASGDEENSDYT | 0.4967095 |
| LISNASDALDKLR | ASGDEENSDYT | 0.6276494 |
| LINEAVP | ASGDEENSDYT | 0.67247915 |
| LGYAFSGL | ASGDEENSDYT | 0.63206 |
| LDFVAVAL | ASGDEENSDYT | 0.6180587 |
| LAQPGAE | ASGDEENSDYT | 0.59216356 |
| ISNLGAVTGF | ASGDEENSDYT | 0.66626704 |
| HQEDPGRI | ASGDEENSDYT | 0.5716493 |
| GTPGEPTP | ASGDEENSDYT | 0.6664262 |
| GFFNSGS | ASGDEENSDYT | 0.3576908 |
| GAGGGGLTAGAN | ASGDEENSDYT | 0.57860965 |
| FVDALVNM | ASGDEENSDYT | 0.5570853 |
| FLTVAPEL | ASGDEENSDYT | 0.5220756 |
| FETLTAL | ASGDEENSDYT | 0.6604662 |
| DVINPIEL | ASGDEENSDYT | 0.58811617 |
| APVSGAPV | ASGDEENSDYT | 0.6540158 |
| ALVGGLAW | ASGDEENSDYT | 0.63598764 |
| ALAAAAATAI | ASGDEENSDYT | 0.6644552 |
| AFGLGLLL | ASGDEENSDYT | 0.6344029 |
| AFFAAASL | ASGDEENSDYT | 0.6445527 |
| AAAALPAFSPP | ASGDEENSDYT | 0.6044157 |
| VVLAPAL | ASGDEENSDYT | 0.6574686 |
| TIGHVDHGKTTL | ASGDEENSDYT | 0.66058195 |
| LISNASDALDK | ASGDEENSDYT | 0.6409799 |
| LISNASDAL | ASGDEENSDYT | 0.5596054 |
| ISNASDALDK | ASGDEENSDYT | 0.5978512 |
| NQAETLVYQTE | ASGDEENSDYT | 0.6472764 |
| TGGTGGTAGAG | ASGDEENSDYT | 0.40317714 |
| TGSTAGGAGGA | ASGDEENSDYT | 0.36171678 |
| DPLAVPVAAGPV | ASGDEENSDYT | 0.6349046 |
| GGTGGTPGNGGN | ASGDEENSDYT | 0.6082447 |
| PAGLAGVDPATL | ASGDEENSDYT | 0.66294205 |
| GNNLVVEAPHAGASV | ASGDEENSDYT | 0.6590649 |
| KVTRIMSASTLEQPA | ASGDEENSDYT | 0.6255331 |
| LGFALPIALPRRTVV | ASGDEENSDYT | 0.6547265 |
| TRLGNLTARKLVVLV | ASGDEENSDYT | 0.6433948 |
| WSQQRRLIYFPSAGP | ASGDEENSDYT | 0.31897312 |
| GPNGNPDPMAAAVDIR | ASGDEENSDYT | 0.61120695 |
| TQRCVSVADLRAAASR | ASGDEENSDYT | 0.54875404 |

| | | |
|---|---|---|
| QFVLDVAEHPVEVLTETV | ASGDEENSDYT | 0.66603434 |
| GGAGGTGGAGGNGGTQAGDG | ASGDEENSDYT | 0.6228241 |
| PLLATLRAIAVDVDAKPAQV | ASGDEENSDYT | 0.5872814 |
| APKFRTGGYMIDFWGVGYQVAK | ASGDEENSDYT | 0.6339702 |
| GAGGAGGAGGAAGGPGGTGGTG | ASGDEENSDYT | 0.6295098 |
| IGALGLLATDVAPAGSVAFGVA | ASGDEENSDYT | 0.6244823 |
| AGGAGGIGGNGGAGGAGGVLMG | ASGDEENSDYT | 0.58525 |
| ADELIGLL | ASGDEENSDYT | 0.35746387 |
| ADRALKA | ASGDEENSDYT | 0.6594273 |
| AGAVGQVGGAGGA | ASGDEENSDYT | 0.5271823 |
| AGGGGGLGGAP | ASGDEENSDYT | 0.61015415 |
| AGGLLYGNGGAG | ASGDEENSDYT | 0.37317622 |
| AGLESIGL | ASGDEENSDYT | 0.5884378 |
| AGLKTGII | ASGDEENSDYT | 0.4900843 |
| ASLAALLL | ASGDEENSDYT | 0.6534837 |
| ASVINISSI | ASGDEENSDYT | 0.6226124 |
| ATPPGPA | ASGDEENSDYT | 0.61565864 |
| ATVASALV | ASGDEENSDYT | 0.66864574 |
| ATVASLIL | ASGDEENSDYT | 0.63106847 |
| AVNGPVLIRH | ASGDEENSDYT | 0.6160288 |
| DGAAGAVGI | ASGDEENSDYT | 0.62046564 |
| DPLLHGI | ASGDEENSDYT | 0.5798139 |
| ELGAALGAL | ASGDEENSDYT | 0.61341953 |
| ELGRTQAV | ASGDEENSDYT | 0.6134758 |
| FVAGGYLS | ASGDEENSDYT | 0.6682985 |
| GASGGAGGAGGW | ASGDEENSDYT | 0.59427136 |
| GGAGAGGDGGAGGI | ASGDEENSDYT | 0.5308261 |
| GTGHNGSQPGMGGNG | ASGDEENSDYT | 0.6130052 |
| GYAAASASAS | ASGDEENSDYT | 0.6665086 |
| HQVDVAAVQQG | ASGDEENSDYT | 0.66891956 |
| IGAAIAHT | ASGDEENSDYT | 0.6064776 |
| IGMIEVAQL | ASGDEENSDYT | 0.60003614 |
| IRLAPEHTSAMD | ASGDEENSDYT | 0.59358263 |
| IYNVSVL | ASGDEENSDYT | 0.6328679 |
| KQGFLGM | ASGDEENSDYT | 0.59778684 |
| LESVITQL | ASGDEENSDYT | 0.56348836 |
| LGLGALVVV | ASGDEENSDYT | 0.65043896 |
| LIGNGGNGGSGGI | ASGDEENSDYT | 0.66866696 |
| LNAINAPTL | ASGDEENSDYT | 0.6235766 |
| LQLLNTGV | ASGDEENSDYT | 0.49951723 |
| LQTSIEF | ASGDEENSDYT | 0.6286268 |
| MSFAVSEL | ASGDEENSDYT | 0.5932628 |
| NTGALGL | ASGDEENSDYT | 0.6174954 |
| PPGAGGAGGAGTT | ASGDEENSDYT | 0.64416885 |
| QLVGLGI | ASGDEENSDYT | 0.49771488 |
| RFLEQVHQL | ASGDEENSDYT | 0.62627506 |
| RGVGYVLR | ASGDEENSDYT | 0.6689301 |
| SATGVFAGVI | ASGDEENSDYT | 0.60984 |
| SLPAITLP | ASGDEENSDYT | 0.64177406 |
| STALAFTV | ASGDEENSDYT | 0.66309834 |
| SVLTVGLGK | ASGDEENSDYT | 0.6508382 |

| | | |
|---|---|---|
| TALAGAGSGI | ASGDEENSDYT | 0.663 |
| TGLPFTGI | ASGDEENSDYT | 0.6577285 |
| VAEHVTV | ASGDEENSDYT | 0.6394551 |
| VDIVLNSL | ASGDEENSDYT | 0.6321999 |
| VLTGGGAVL | ASGDEENSDYT | 0.6473215 |
| VNVGLAYGI | ASGDEENSDYT | 0.63507843 |
| VSSAAVGAV | ASGDEENSDYT | 0.61697555 |
| VTSVYEAGV | ASGDEENSDYT | 0.5702739 |
| WVALDRGA | ASGDEENSDYT | 0.661513 |
| YLVGSVVI | ASGDEENSDYT | 0.6232736 |
| GPPGTGKTLL | ASGDEENSDYT | 0.65711933 |
| GVGMTKF | ASGDEENSDYT | 0.53873664 |
| IGIGGSDL | ASGDEENSDYT | 0.58413035 |
| IGTLLAEL | ASGDEENSDYT | 0.65676403 |
| IYTSGTT | ASGDEENSDYT | 0.3191642 |
| LSFPAGV | ASGDEENSDYT | 0.351696 |
| LYTSGTT | ASGDEENSDYT | 0.63089293 |
| VGAFPFGF | ASGDEENSDYT | 0.66284275 |
| VGAGTVEF | ASGDEENSDYT | 0.33455953 |
| VGKSTLLL | ASGDEENSDYT | 0.59581465 |
| AALAHAGGGIV | ASGDEENSDYT | 0.53696966 |
| AALVAVSLL | ASGDEENSDYT | 0.65546465 |
| AAVSRAVAAL | ASGDEENSDYT | 0.6494554 |
| AGGSSGIGGF | ASGDEENSDYT | 0.5147799 |
| AGLTVANANG | ASGDEENSDYT | 0.65532297 |
| AGLVGAVL | ASGDEENSDYT | 0.47460115 |
| ALSQAFTV | ASGDEENSDYT | 0.5711851 |
| ALTFSVSL | ASGDEENSDYT | 0.6611685 |
| ALTGAALVAL | ASGDEENSDYT | 0.66260874 |
| ASFSIGAGVL | ASGDEENSDYT | 0.56864995 |
| DGSVYKI | ASGDEENSDYT | 0.65718114 |
| EQLAEIV | ASGDEENSDYT | 0.59425974 |
| FAVSELTVM | ASGDEENSDYT | 0.6699733 |
| GAAPLAAGL | ASGDEENSDYT | 0.6187762 |
| GADGGDGGSGGAGG | ASGDEENSDYT | 0.4962871 |
| GFGLDTGGAGG | ASGDEENSDYT | 0.6569755 |
| GLASGGLL | ASGDEENSDYT | 0.6489555 |
| GRIAGAAVLL | ASGDEENSDYT | 0.6426135 |
| GSAGSLNTSF | ASGDEENSDYT | 0.65440774 |
| GTGQIGI | ASGDEENSDYT | 0.5984945 |
| GTLIALIL | ASGDEENSDYT | 0.6241036 |
| IAATMGFVVG | ASGDEENSDYT | 0.66454536 |
| IFAQALL | ASGDEENSDYT | 0.6608084 |
| IGAAVALGF | ASGDEENSDYT | 0.5776126 |
| IGLTPLAL | ASGDEENSDYT | 0.65768385 |
| ILGSNTSTL | ASGDEENSDYT | 0.6286082 |
| IVFSLTVL | ASGDEENSDYT | 0.5813031 |
| LASKATGF | ASGDEENSDYT | 0.66718644 |
| LASVALIL | ASGDEENSDYT | 0.633427 |
| LFAVGLTL | ASGDEENSDYT | 0.65516114 |
| LFNGGIGTYI | ASGDEENSDYT | 0.65812135 |

| | | |
|---|---|---|
| LTLGAGII | ASGDEENSDYT | 0.6568353 |
| LVAAALGDI | ASGDEENSDYT | 0.5548088 |
| NTAFVQLGI | ASGDEENSDYT | 0.6339107 |
| QLFSGAEL | ASGDEENSDYT | 0.62694216 |
| SAATFITL | ASGDEENSDYT | 0.65828586 |
| SNYGLANLGV | ASGDEENSDYT | 0.63185275 |
| SYGGAYAVM | ASGDEENSDYT | 0.58734924 |
| TTGVGGLTLG | ASGDEENSDYT | 0.6490892 |
| TTQTIYI | ASGDEENSDYT | 0.663097 |
| VAAGAGLGAVI | ASGDEENSDYT | 0.6390483 |
| VAAVVTLL | ASGDEENSDYT | 0.64702106 |
| VGFLGGLI | ASGDEENSDYT | 0.66642404 |
| VGVIPILL | ASGDEENSDYT | 0.5933541 |
| VIFASSLI | ASGDEENSDYT | 0.62289536 |
| VKGGDGGAAAG | ASGDEENSDYT | 0.6543524 |
| VTKTFTNI | ASGDEENSDYT | 0.65928566 |
| VVAAAILL | ASGDEENSDYT | 0.6254028 |
| YAALDVEL | ASGDEENSDYT | 0.66324645 |
| LASVALLL | ASGDEENSDYT | 0.5962816 |
| LDAVHRV | ASGDEENSDYT | 0.661255 |
| QGLSSSIF | ASGDEENSDYT | 0.5948955 |
| RGTFANIRL | ASGDEENSDYT | 0.6483874 |
| RQLWWGHRI | ASGDEENSDYT | 0.6168632 |
| YGPPGTGKTLL | ASGDEENSDYT | 0.65885884 |
| AALVGAPANVA | ASGDEENSDYT | 0.6576612 |
| AATFRRTE | ASGDEENSDYT | 0.6616999 |
| AGALGGGI | ASGDEENSDYT | 0.3430684 |
| AGGAGGAGGIGAG | ASGDEENSDYT | 0.6622565 |
| AGSGALGL | ASGDEENSDYT | 0.6617042 |
| AQIADQI | ASGDEENSDYT | 0.66845393 |
| ASAVAVTYL | ASGDEENSDYT | 0.6272927 |
| ASTVLGGTGG | ASGDEENSDYT | 0.63712335 |
| AVGGEPAK | ASGDEENSDYT | 0.6317757 |
| CEQGTFGGTC | ASGDEENSDYT | 0.62464577 |
| DEGKAPAG | ASGDEENSDYT | 0.5661669 |
| DTVVFPAL | ASGDEENSDYT | 0.6107678 |
| FEGITMAG | ASGDEENSDYT | 0.6100672 |
| FPLVVQD | ASGDEENSDYT | 0.6046397 |
| GAFTTPQ | ASGDEENSDYT | 0.65327287 |
| GFGGFGTAGGVGGA | ASGDEENSDYT | 0.6589838 |
| GFLAQFL | ASGDEENSDYT | 0.6614065 |
| GGAGGAGGNAGVA | ASGDEENSDYT | 0.66387284 |
| GGVGTAGAGGAGGAGG | ASGDEENSDYT | 0.66363436 |
| GNGGGGDAGNAGSGGN | ASGDEENSDYT | 0.6595283 |
| GTGGAGGAGNDAGS | ASGDEENSDYT | 0.61941385 |
| GTTNTGWFNS | ASGDEENSDYT | 0.6275949 |
| IFMGPLFTL | ASGDEENSDYT | 0.5682245 |
| ILAHAIEAI | ASGDEENSDYT | 0.6277913 |
| IRSAIREAL | ASGDEENSDYT | 0.6317363 |
| ITFAGGG | ASGDEENSDYT | 0.6466633 |
| ITGMSGHQEV | ASGDEENSDYT | 0.32350978 |

| | | |
|---|---|---|
| IVAGLACTL | ASGDEENSDYT | 0.63099617 |
| IVGGGLAGA | ASGDEENSDYT | 0.6668956 |
| IVTGVLNI | ASGDEENSDYT | 0.58140725 |
| LAQQGCI | ASGDEENSDYT | 0.64057684 |
| LCLHVEN | ASGDEENSDYT | 0.6396735 |
| LDAISDGS | ASGDEENSDYT | 0.6099695 |
| LGGFAAML | ASGDEENSDYT | 0.65546775 |
| LGGKNPMI | ASGDEENSDYT | 0.5857388 |
| LGPKGVAAIL | ASGDEENSDYT | 0.6445011 |
| LMVGVSI | ASGDEENSDYT | 0.66561437 |
| LVGAAFVL | ASGDEENSDYT | 0.63688976 |
| LVLAVAPL | ASGDEENSDYT | 0.62710387 |
| MAAIAVIDAA | ASGDEENSDYT | 0.62360466 |
| NNGINAELQ | ASGDEENSDYT | 0.48437825 |
| RGEPLVLVHG | ASGDEENSDYT | 0.6442955 |
| RGGAGGAAGLI | ASGDEENSDYT | 0.45841914 |
| RSYRMHPAV | ASGDEENSDYT | 0.6286657 |
| SAAHLLAGQLA | ASGDEENSDYT | 0.5945605 |
| SAAVGSY | ASGDEENSDYT | 0.6475146 |
| SPHAVLHRI | ASGDEENSDYT | 0.6415225 |
| SQIAKVL | ASGDEENSDYT | 0.6445671 |
| TALGHLVP | ASGDEENSDYT | 0.5702286 |
| TAPTTSGAA | ASGDEENSDYT | 0.6207899 |
| TDGDDGNGGSPGT | ASGDEENSDYT | 0.5698035 |
| TDGLGEN | ASGDEENSDYT | 0.58258915 |
| TGGPPLPGQA | ASGDEENSDYT | 0.65371764 |
| TIPAITI | ASGDEENSDYT | 0.61885124 |
| TIPGFTI | ASGDEENSDYT | 0.6308909 |
| TSIASGALA | ASGDEENSDYT | 0.6636741 |
| TVTLPGAAI | ASGDEENSDYT | 0.6480543 |
| VASGLSAALP | ASGDEENSDYT | 0.4779303 |
| VGAGVIGI | ASGDEENSDYT | 0.6276716 |
| VGPKRFGV | ASGDEENSDYT | 0.62531185 |
| VNVAFPD | ASGDEENSDYT | 0.6277975 |
| VSMTNSGVS | ASGDEENSDYT | 0.33237058 |
| VTDVLGDLSV | ASGDEENSDYT | 0.6547523 |
| VVVVGAGFAGL | ASGDEENSDYT | 0.5822327 |
| VYTVGTVV | ASGDEENSDYT | 0.5886129 |
| WTTFSGP | ASGDEENSDYT | 0.6145857 |
| YDELVEAF | ASGDEENSDYT | 0.65589654 |
| YGTGGAGGNGG | ASGDEENSDYT | 0.59323156 |
| YTNVMAV | ASGDEENSDYT | 0.64496803 |
| FLEQVHQL | ASGDEENSDYT | 0.65317595 |
| GLSLLAVAV | ASGDEENSDYT | 0.65903676 |
| LEQVHQL | ASGDEENSDYT | 0.66490734 |
| LNAAGAI | ASGDEENSDYT | 0.6614814 |
| PAVHASL | ASGDEENSDYT | 0.6137036 |
| TELEDDA | ASGDEENSDYT | 0.6460358 |
| TPPGPPN | ASGDEENSDYT | 0.6157782 |
| TPPPPPPP | ASGDEENSDYT | 0.46434683 |
| TQSPALI | ASSLGQNNQAPL | 0.38230386 |

| | | |
|---|---|---|
| EFAAAFPQ | ASSLGQNNQAPL | 0.63793296 |
| LLSGNLDVL | ASSLGQNNQAPL | 0.39553905 |
| GPANLGNYNI | ASSLGQNNQAPL | 0.6377618 |
| TLGPTLP | ASSLGQNNQAPL | 0.6613178 |
| VYAYSSCCK | ASSLGQNNQAPL | 0.63071656 |
| APAPAPAPNI | ASSLGQNNQAPL | 0.51278174 |
| PGGPVPIP | ASSLGQNNQAPL | 0.61910033 |
| FLTVALEP | ASSLGQNNQAPL | 0.63212526 |
| SQYFGANL | ASSLGQNNQAPL | 0.6548705 |
| GGVKPAA | ASSLGQNNQAPL | 0.6574939 |
| GGTGGAGGAGGW | ASSLGQNNQAPL | 0.66862214 |
| GVGGVAGLGGAGG | ASSLGQNNQAPL | 0.6641146 |
| VLLVLATTI | ASSLGQNNQAPL | 0.6147742 |
| IGIFAAML | ASSLGQNNQAPL | 0.6498092 |
| AVGDLID | ASSLGQNNQAPL | 0.6169562 |
| NISPLPL | ASSLGQNNQAPL | 0.6197709 |
| VTAVFVAI | ASSLGQNNQAPL | 0.62774754 |
| SGDDNDAAL | ASSLGQNNQAPL | 0.64163417 |
| AVAALAHS | ASSLGQNNQAPL | 0.6465242 |
| GGVTVMATP | ASSLGQNNQAPL | 0.61694336 |
| AGLAANVRD | ASSLGQNNQAPL | 0.45869386 |
| TNRPDLIDPALLRPG | ASSLGQNNQAPL | 0.644268 |
| LANLGVTAL | ASSLGQNNQAPL | 0.65007085 |
| TGERIEL | ASSLGQNNQAPL | 0.40341645 |
| NGEELAA | ASSLGQNNQAPL | 0.6137099 |
| VLSLVWG | ASSLGQNNQAPL | 0.6685624 |
| IGYALWL | ASSLGQNNQAPL | 0.6647694 |
| LSLLLAPL | ASSLGQNNQAPL | 0.59427583 |
| TLDEALK | ASSLGQNNQAPL | 0.60989916 |
| TDGAGTAEGG | ASSLGQNNQAPL | 0.63270974 |
| NAPGGAGGNGG | ASSLGQNNQAPL | 0.44080192 |
| VSVFMGMGGNF | ASSLGQNNQAPL | 0.6426902 |
| LASYLAAL | ASSLGQNNQAPL | 0.6355394 |
| LVGGTPAL | ASSLGQNNQAPL | 0.6395672 |
| VRNADVL | ASSLGQNNQAPL | 0.55763316 |
| TCSSPFLVSAM | ASSLGQNNQAPL | 0.63714665 |
| VHITDAE | ASSLGQNNQAPL | 0.63277507 |
| EYAAPSF | ASSLGQNNQAPL | 0.65057117 |
| VVDAFHGGL | ASSLGQNNQAPL | 0.3440592 |
| ASVVDPQ | ASSLGQNNQAPL | 0.6607443 |
| SYNEFGGLS | ASSLGQNNQAPL | 0.5846729 |
| DPDGSVY | ASSLGQNNQAPL | 0.58409417 |
| MVSTIHATE | ASSLGQNNQAPL | 0.44835612 |
| LAAGVADPV | ASSLGQNNQAPL | 0.6244534 |
| TKEQIAA | ASSLGQNNQAPL | 0.6575344 |
| SLASPAAL | ASSLGQNNQAPL | 0.65665424 |
| IGLVFYGV | ASSLGQNNQAPL | 0.50959414 |
| FLLDASNL | ASSLGQNNQAPL | 0.66159254 |
| YQVGNFP | ASSLGQNNQAPL | 0.6672526 |
| ESLLSQST | ASSLGQNNQAPL | 0.61415803 |
| LFTVGFAAVG | ASSLGQNNQAPL | 0.46804845 |

| | | |
|---|---|---|
| ATPPTPT | ASSLGQNNQAPL | 0.65587676 |
| TDLATLG | ASSLGQNNQAPL | 0.60656464 |
| SAGGTGR | ASSLGQNNQAPL | 0.65690595 |
| GFSNSGS | ASSLGQNNQAPL | 0.64520675 |
| GGNGGSGGTSV | ASSLGQNNQAPL | 0.56855875 |
| GTSVATGGAGN | ASSLGQNNQAPL | 0.65294135 |
| NGGNGGSGGTS | ASSLGQNNQAPL | 0.66379464 |
| AVVKFPH | ASSLGQNNQAPL | 0.58808947 |
| TPPPPPP | ASSLGQNNQAPL | 0.5634173 |
| AAPAAAAAPA | ASSLGQNNQAPL | 0.59144986 |
| IALNEDL | ASSLGQNNQAPL | 0.4379345 |
| IVNVSSM | ASSLGQNNQAPL | 0.66017383 |
| TGIPPAPR | ASSLGQNNQAPL | 0.6019606 |
| FDLGGGTF | ASSLGQNNQAPL | 0.6180348 |
| FDLGGGTFDVS | ASSLGQNNQAPL | 0.6426728 |
| NEPTAAA | ASSLGQNNQAPL | 0.622573 |
| NEPTAAALA | ASSLGQNNQAPL | 0.614445 |
| GSIINISSI | ASSLGQNNQAPL | 0.63235 |
| VAPGNAGTA | ASSLGQNNQAPL | 0.6562673 |
| TGGGASLE | ASSLGQNNQAPL | 0.6439541 |
| VAVYNHYKRI | ASSLGQNNQAPL | 0.6330524 |
| TDPAPATT | ASSLGQNNQAPL | 0.615322 |
| TNRPDILDPALLRPG | ASSLGQNNQAPL | 0.66497535 |
| LTTTTTS | ASSLGQNNQAPL | 0.5873068 |
| PTGSGKTL | ASSLGQNNQAPL | 0.6129152 |
| TGSGKTL | ASSLGQNNQAPL | 0.6475239 |
| TIGHVDHGK | ASSLGQNNQAPL | 0.4780239 |
| TIGHVDHGKTT | ASSLGQNNQAPL | 0.65226454 |
| GHVDHGKTT | ASSLGQNNQAPL | 0.6404462 |
| AQFEHTLL | ASSLGQNNQAPL | 0.6451583 |
| IVNQDEL | ASSLGQNNQAPL | 0.6177033 |
| TNGSQFF | ASSLGQNNQAPL | 0.66907436 |
| TGKGIVE | ASSLGQNNQAPL | 0.65892446 |
| LISNASDA | ASSLGQNNQAPL | 0.62449604 |
| AELRTLGELTKIAKAHGAQ | ASSLGQNNQAPL | 0.6547256 |
| APNAVAAAEPSW | ASSLGQNNQAPL | 0.5505998 |
| APQVAGAATAVL | ASSLGQNNQAPL | 0.6610329 |
| ARRDPAAAAAILPTDA | ASSLGQNNQAPL | 0.64299935 |
| DAAARDVLAEAGLAEYF | ASSLGQNNQAPL | 0.57601017 |
| DDQIVLSAKANLAKVEQ | ASSLGQNNQAPL | 0.627704 |
| DELLPDTGEDSDEDGH | ASSLGQNNQAPL | 0.6606339 |
| ELPEAPANRATVPQD | ASSLGQNNQAPL | 0.6608747 |
| FGQSKIERVTGIG | ASSLGQNNQAPL | 0.4252131 |
| GAGGTGSTAGGAGGAGGAGGLY | ASSLGQNNQAPL | 0.62130374 |
| GHGGTGAAGGNGGTGGAGGSGI | ASSLGQNNQAPL | 0.41746876 |
| GNGGNGGAGGEGGPGGAGGAGA | ASSLGQNNQAPL | 0.5671622 |
| GSTAGNGGNGGNATNGTIAGQP | ASSLGQNNQAPL | 0.5426382 |
| GTGGDGGNGGNGGTGG | ASSLGQNNQAPL | 0.6491748 |
| GVRRPAGIPADAPV | ASSLGQNNQAPL | 0.6033707 |
| GYRQVTDAHLLTLARR | ASSLGQNNQAPL | 0.6404004 |
| IAKMGQDGHDRG | ASSLGQNNQAPL | 0.6385817 |

| | | |
|---|---|---|
| KAEPSIEIQYT | ASSLGQNNQAPL | 0.6649934 |
| KPVRIRMAADEDTVAVT | ASSLGQNNQAPL | 0.58576185 |
| MPPRIAGMRLLVIKPEP | ASSLGQNNQAPL | 0.6440225 |
| MKFYAVALFRSRGVPPTR | ASSLGQNNQAPL | 0.50221014 |
| NAGGFGILGAVAHSP | ASSLGQNNQAPL | 0.561319 |
| NEGREITLAMAEM | ASSLGQNNQAPL | 0.5804892 |
| NELVEWSHPLGHRWRWE | ASSLGQNNQAPL | 0.6406684 |
| NLGAGGTGGNGGNPG | ASSLGQNNQAPL | 0.40644002 |
| NTEWPKWETVMAADPVD | ASSLGQNNQAPL | 0.66643023 |
| PASAAPAETQP | ASSLGQNNQAPL | 0.6388092 |
| QQSSDGKNGVRAAEITGE | ASSLGQNNQAPL | 0.6547948 |
| RAPDAVTTSWWKE | ASSLGQNNQAPL | 0.62630755 |
| SGFGNVGAHTSGWFNQ | ASSLGQNNQAPL | 0.62454665 |
| SGVAGAVGQAASVGGLKV | ASSLGQNNQAPL | 0.62540185 |
| SNGGAGTGSAGGNGGT | ASSLGQNNQAPL | 0.65076584 |
| TGGAGGEGGAAG | ASSLGQNNQAPL | 0.6179935 |
| TGGDGGAAGAGGAPGHG | ASSLGQNNQAPL | 0.4490512 |
| TMPPLLAAIANAGRHGV | ASSLGQNNQAPL | 0.5861574 |
| TSSTAGNSGGA | ASSLGQNNQAPL | 0.53182995 |
| VPATVPVGILC | ASSLGQNNQAPL | 0.66409016 |
| VQRPDSGGVVLTGRIS | ASSLGQNNQAPL | 0.6607456 |
| WTPNPVPATKGVPAGLVQAT | ASSLGQNNQAPL | 0.6474844 |
| YALKRALAKFSDRFDVVII | ASSLGQNNQAPL | 0.6570969 |
| YLIGSVIV | ASSLGQNNQAPL | 0.36766222 |
| YGQLVAVID | ASSLGQNNQAPL | 0.63229376 |
| VTVGGMTLPA | ASSLGQNNQAPL | 0.66709113 |
| VLALPAV | ASSLGQNNQAPL | 0.6494795 |
| VIPPTLNYET | ASSLGQNNQAPL | 0.6169455 |
| VGMGGMHPSSG | ASSLGQNNQAPL | 0.6630577 |
| VGLAGGAATA | ASSLGQNNQAPL | 0.6086079 |
| VGGAGGAGGSGP | ASSLGQNNQAPL | 0.6260472 |
| VFAAFMV | ASSLGQNNQAPL | 0.6480031 |
| TLYDKLR | ASSLGQNNQAPL | 0.65129125 |
| TLAVFTVL | ASSLGQNNQAPL | 0.63354903 |
| TGSVQPARV | ASSLGQNNQAPL | 0.5871969 |
| TGGGAGSGGT | ASSLGQNNQAPL | 0.6348995 |
| SHLETSL | ASSLGQNNQAPL | 0.67147523 |
| SFTTGMT | ASSLGQNNQAPL | 0.6239828 |
| QVEVAKRMGVRQPTVS | ASSLGQNNQAPL | 0.6596558 |
| QGSGGPALAM | ASSLGQNNQAPL | 0.6497923 |
| MLAPSNP | ASSLGQNNQAPL | 0.64563704 |
| LVNNAAI | ASSLGQNNQAPL | 0.5820093 |
| LTGEINTLA | ASSLGQNNQAPL | 0.61190933 |
| LSIFGPV | ASSLGQNNQAPL | 0.6200998 |
| LLALLANL | ASSLGQNNQAPL | 0.5827894 |
| LISNASDALDKLR | ASSLGQNNQAPL | 0.61256987 |
| LINEAVP | ASSLGQNNQAPL | 0.61513764 |
| LGYAFSGL | ASSLGQNNQAPL | 0.6728064 |
| LDFVAVAL | ASSLGQNNQAPL | 0.6470573 |
| LAQPGAE | ASSLGQNNQAPL | 0.65706635 |
| ISNLGAVTGF | ASSLGQNNQAPL | 0.6077714 |

| | | |
|---|---|---|
| HQEDPGRI | ASSLGQNNQAPL | 0.6399565 |
| GTPGEPTP | ASSLGQNNQAPL | 0.6495206 |
| GFFNSGS | ASSLGQNNQAPL | 0.49670506 |
| GAGGGGLTAGAN | ASSLGQNNQAPL | 0.66449 |
| FVDALVNM | ASSLGQNNQAPL | 0.5981851 |
| FLTVAPEL | ASSLGQNNQAPL | 0.62077785 |
| FETLTAL | ASSLGQNNQAPL | 0.6608534 |
| DVINPIEL | ASSLGQNNQAPL | 0.6648777 |
| APVSGAPV | ASSLGQNNQAPL | 0.6132368 |
| ALVGGLAW | ASSLGQNNQAPL | 0.5937811 |
| ALAAAAATAI | ASSLGQNNQAPL | 0.6107311 |
| AFGLGLLL | ASSLGQNNQAPL | 0.65555024 |
| AFFAAASL | ASSLGQNNQAPL | 0.66190195 |
| AAAALPAFSPP | ASSLGQNNQAPL | 0.6326316 |
| VVLAPAL | ASSLGQNNQAPL | 0.66831136 |
| TIGHVDHGKTTL | ASSLGQNNQAPL | 0.6510823 |
| LISNASDALDK | ASSLGQNNQAPL | 0.47835872 |
| LISNASDAL | ASSLGQNNQAPL | 0.63069284 |
| ISNASDALDK | ASSLGQNNQAPL | 0.6604403 |
| NQAETLVYQTE | ASSLGQNNQAPL | 0.6584789 |
| TGGTGGTAGAG | ASSLGQNNQAPL | 0.6704972 |
| TGSTAGGAGGA | ASSLGQNNQAPL | 0.65111655 |
| DPLAVPVAAGPV | ASSLGQNNQAPL | 0.6069778 |
| GGTGGTPGNGGN | ASSLGQNNQAPL | 0.6651092 |
| PAGLAGVDPATL | ASSLGQNNQAPL | 0.67004144 |
| GNNLVVEAPHAGASV | ASSLGQNNQAPL | 0.6552611 |
| KVTRIMSASTLEQPA | ASSLGQNNQAPL | 0.6051549 |
| LGFALPIALPRRTVV | ASSLGQNNQAPL | 0.5880353 |
| TRLGNLTARKLVVLV | ASSLGQNNQAPL | 0.6368246 |
| WSQQRRLIYFPSAGP | ASSLGQNNQAPL | 0.6613084 |
| GPNGNPDPMAAAVDIR | ASSLGQNNQAPL | 0.5292886 |
| TQRCVSVADLRAAASR | ASSLGQNNQAPL | 0.6393223 |
| QFVLDVAEHPVEVLTETV | ASSLGQNNQAPL | 0.64372385 |
| GGAGGTGGAGGNGGTQAGDG | ASSLGQNNQAPL | 0.48807144 |
| PLLATLRAIAVDVDAKPAQV | ASSLGQNNQAPL | 0.65904665 |
| APKFRTGGYMIDFWGVGYQVAK | ASSLGQNNQAPL | 0.66386616 |
| GAGGAGGAGGAAGGPGGTGGTG | ASSLGQNNQAPL | 0.62357396 |
| IGALGLLATDVAPAGSVAFGVA | ASSLGQNNQAPL | 0.6126878 |
| AGGAGGIGGNGGAGGAGGVLMG | ASSLGQNNQAPL | 0.660647 |
| ADELIGLL | ASSLGQNNQAPL | 0.6726011 |
| ADRALKA | ASSLGQNNQAPL | 0.66131246 |
| AGAVGQVGGAGGA | ASSLGQNNQAPL | 0.5181264 |
| AGGGGGLGGAP | ASSLGQNNQAPL | 0.3881764 |
| AGGLLYGNGGAG | ASSLGQNNQAPL | 0.6012564 |
| AGLESIGL | ASSLGQNNQAPL | 0.6524818 |
| AGLKTGII | ASSLGQNNQAPL | 0.6365427 |
| ASLAALLL | ASSLGQNNQAPL | 0.63174415 |
| ASVINISSI | ASSLGQNNQAPL | 0.64517766 |
| ATPPGPA | ASSLGQNNQAPL | 0.6629964 |
| ATVASALV | ASSLGQNNQAPL | 0.58318895 |
| ATVASLIL | ASSLGQNNQAPL | 0.53463423 |

| | | |
|---|---|---|
| AVNGPVLIRH | ASSLGQNNQAPL | 0.6425116 |
| DGAAGAVGI | ASSLGQNNQAPL | 0.6465975 |
| DPLLHGI | ASSLGQNNQAPL | 0.6004029 |
| ELGAALGAL | ASSLGQNNQAPL | 0.6292378 |
| ELGRTQAV | ASSLGQNNQAPL | 0.5935282 |
| FVAGGYLS | ASSLGQNNQAPL | 0.57054245 |
| GASGGAGGAGGW | ASSLGQNNQAPL | 0.6455378 |
| GGAGAGGDGGAGGI | ASSLGQNNQAPL | 0.57936776 |
| GTGHNGSQPGMGGNG | ASSLGQNNQAPL | 0.6677304 |
| GYAAASASAS | ASSLGQNNQAPL | 0.5739558 |
| HQVDVAAVQQG | ASSLGQNNQAPL | 0.64930254 |
| IGAAIAHT | ASSLGQNNQAPL | 0.6485456 |
| IGMIEVAQL | ASSLGQNNQAPL | 0.64921236 |
| IRLAPEHTSAMD | ASSLGQNNQAPL | 0.61377966 |
| IYNVSVL | ASSLGQNNQAPL | 0.602201 |
| KQGFLGM | ASSLGQNNQAPL | 0.55033183 |
| LESVITQL | ASSLGQNNQAPL | 0.40606648 |
| LGLGALVVV | ASSLGQNNQAPL | 0.6605877 |
| LIGNGGNGGSGGI | ASSLGQNNQAPL | 0.5952897 |
| LNAINAPTL | ASSLGQNNQAPL | 0.66384125 |
| LQLLNTGV | ASSLGQNNQAPL | 0.655057 |
| LQTSIEF | ASSLGQNNQAPL | 0.60526353 |
| MSFAVSEL | ASSLGQNNQAPL | 0.64684033 |
| NTGALGL | ASSLGQNNQAPL | 0.6109869 |
| PPGAGGAGGAGTT | ASSLGQNNQAPL | 0.5401905 |
| QLVGLGI | ASSLGQNNQAPL | 0.6506537 |
| RFLEQVHQL | ASSLGQNNQAPL | 0.65214705 |
| RGVGYVLR | ASSLGQNNQAPL | 0.56972134 |
| SATGVFAGVI | ASSLGQNNQAPL | 0.6391705 |
| SLPAITLP | ASSLGQNNQAPL | 0.6383465 |
| STALAFTV | ASSLGQNNQAPL | 0.5663094 |
| SVLTVGLGK | ASSLGQNNQAPL | 0.63014615 |
| TALAGAGSGI | ASSLGQNNQAPL | 0.6612062 |
| TGLPFTGI | ASSLGQNNQAPL | 0.6370779 |
| VAEHVTV | ASSLGQNNQAPL | 0.6015399 |
| VDIVLNSL | ASSLGQNNQAPL | 0.6553432 |
| VLTGGGAVL | ASSLGQNNQAPL | 0.5781199 |
| VNVGLAYGI | ASSLGQNNQAPL | 0.64833516 |
| VSSAAVGAV | ASSLGQNNQAPL | 0.63879603 |
| VTSVYEAGV | ASSLGQNNQAPL | 0.6104025 |
| WVALDRGA | ASSLGQNNQAPL | 0.65787697 |
| YLVGSVVI | ASSLGQNNQAPL | 0.5510398 |
| GPPGTGKTLL | ASSLGQNNQAPL | 0.57868403 |
| GVGMTKF | ASSLGQNNQAPL | 0.61481345 |
| IGIGGSDL | ASSLGQNNQAPL | 0.64719045 |
| IGTLLAEL | ASSLGQNNQAPL | 0.6601389 |
| IYTSGTT | ASSLGQNNQAPL | 0.59584796 |
| LSFPAGV | ASSLGQNNQAPL | 0.6350781 |
| LYTSGTT | ASSLGQNNQAPL | 0.6111835 |
| VGAFPFGF | ASSLGQNNQAPL | 0.57612765 |
| VGAGTVEF | ASSLGQNNQAPL | 0.612848 |

| | | |
|---|---|---|
| VGKSTLLL | ASSLGQNNQAPL | 0.61459386 |
| AALAHAGGGIV | ASSLGQNNQAPL | 0.59437096 |
| AALVAVSLL | ASSLGQNNQAPL | 0.64985925 |
| AAVSRAVAAL | ASSLGQNNQAPL | 0.6217254 |
| AGGSSGIGGF | ASSLGQNNQAPL | 0.65270674 |
| AGLTVANANG | ASSLGQNNQAPL | 0.6648036 |
| AGLVGAVL | ASSLGQNNQAPL | 0.6080357 |
| ALSQAFTV | ASSLGQNNQAPL | 0.64797115 |
| ALTFSVSL | ASSLGQNNQAPL | 0.6520699 |
| ALTGAALVAL | ASSLGQNNQAPL | 0.65125084 |
| ASFSIGAGVL | ASSLGQNNQAPL | 0.63339406 |
| DGSVYKI | ASSLGQNNQAPL | 0.6031884 |
| EQLAEIV | ASSLGQNNQAPL | 0.66524863 |
| FAVSELTVM | ASSLGQNNQAPL | 0.657218 |
| GAAPLAAGL | ASSLGQNNQAPL | 0.6425935 |
| GADGGDGGSGGAGG | ASSLGQNNQAPL | 0.6571946 |
| GFGLDTGGAGG | ASSLGQNNQAPL | 0.6417376 |
| GLASGGLL | ASSLGQNNQAPL | 0.6525742 |
| GRIAGAAVLL | ASSLGQNNQAPL | 0.6102632 |
| GSAGSLNTSF | ASSLGQNNQAPL | 0.5466208 |
| GTGQIGI | ASSLGQNNQAPL | 0.6385324 |
| GTLIALIL | ASSLGQNNQAPL | 0.63690674 |
| IAATMGFVVG | ASSLGQNNQAPL | 0.604754 |
| IFAQALL | ASSLGQNNQAPL | 0.46327516 |
| IGAAVALGF | ASSLGQNNQAPL | 0.6633143 |
| IGLTPLAL | ASSLGQNNQAPL | 0.6711205 |
| ILGSNTSTL | ASSLGQNNQAPL | 0.61029285 |
| IVFSLTVL | ASSLGQNNQAPL | 0.6675483 |
| LASKATGF | ASSLGQNNQAPL | 0.5365226 |
| LASVALIL | ASSLGQNNQAPL | 0.6569767 |
| LFAVGLTL | ASSLGQNNQAPL | 0.6616219 |
| LFNGGIGTYI | ASSLGQNNQAPL | 0.58301616 |
| LTLGAGII | ASSLGQNNQAPL | 0.66122794 |
| LVAAALGDI | ASSLGQNNQAPL | 0.5900414 |
| NTAFVQLGI | ASSLGQNNQAPL | 0.6324359 |
| QLFSGAEL | ASSLGQNNQAPL | 0.6645182 |
| SAATFITL | ASSLGQNNQAPL | 0.6288368 |
| SNYGLANLGV | ASSLGQNNQAPL | 0.6433378 |
| SYGGAYAVM | ASSLGQNNQAPL | 0.64327645 |
| TTGVGGLTLG | ASSLGQNNQAPL | 0.609884 |
| TTQTIYI | ASSLGQNNQAPL | 0.647682 |
| VAAGAGLGAVI | ASSLGQNNQAPL | 0.6601834 |
| VAAVVTLL | ASSLGQNNQAPL | 0.5392424 |
| VGFLGGLI | ASSLGQNNQAPL | 0.66870093 |
| VGVIPILL | ASSLGQNNQAPL | 0.5921048 |
| VIFASSLI | ASSLGQNNQAPL | 0.67230505 |
| VKGGDGGAAAG | ASSLGQNNQAPL | 0.6512531 |
| VTKTFTNI | ASSLGQNNQAPL | 0.6395077 |
| VVAAAILL | ASSLGQNNQAPL | 0.6199186 |
| YAALDVEL | ASSLGQNNQAPL | 0.6618029 |
| LASVALLL | ASSLGQNNQAPL | 0.62165797 |

| | | |
|---|---|---|
| LDAVHRV | ASSLGQNNQAPL | 0.6496712 |
| QGLSSSIF | ASSLGQNNQAPL | 0.6653973 |
| RGTFANIRL | ASSLGQNNQAPL | 0.3566472 |
| RQLWWGHRI | ASSLGQNNQAPL | 0.61770505 |
| YGPPGTGKTLL | ASSLGQNNQAPL | 0.53883725 |
| AALVGAPANVA | ASSLGQNNQAPL | 0.4366845 |
| AATFRRTE | ASSLGQNNQAPL | 0.43996406 |
| AGALGGGI | ASSLGQNNQAPL | 0.35980016 |
| AGGAGGAGGIGAG | ASSLGQNNQAPL | 0.40619195 |
| AGSGALGL | ASSLGQNNQAPL | 0.42176455 |
| AQIADQI | ASSLGQNNQAPL | 0.6046989 |
| ASAVAVTYL | ASSLGQNNQAPL | 0.5358536 |
| ASTVLGGTGG | ASSLGQNNQAPL | 0.41911387 |
| AVGGEPAK | ASSLGQNNQAPL | 0.56235576 |
| CEQGTFGGTC | ASSLGQNNQAPL | 0.43136 |
| DEGKAPAG | ASSLGQNNQAPL | 0.37738276 |
| DTVVFPAL | ASSLGQNNQAPL | 0.46159315 |
| FEGITMAG | ASSLGQNNQAPL | 0.38988099 |
| FPLVVQD | ASSLGQNNQAPL | 0.4550207 |
| GAFTTPQ | ASSLGQNNQAPL | 0.3617906 |
| GFGGFGTAGGVGGA | ASSLGQNNQAPL | 0.5362179 |
| GFLAQFL | ASSLGQNNQAPL | 0.36352822 |
| GGAGGAGGNAGVA | ASSLGQNNQAPL | 0.3116876 |
| GGVGTAGAGGAGGAGG | ASSLGQNNQAPL | 0.52166325 |
| GNGGGGDAGNAGSGGN | ASSLGQNNQAPL | 0.40944418 |
| GTGGAGGAGNDAGS | ASSLGQNNQAPL | 0.45277113 |
| GTTNTGWFNS | ASSLGQNNQAPL | 0.53144467 |
| IFMGPLFTL | ASSLGQNNQAPL | 0.40362003 |
| ILAHAIEAI | ASSLGQNNQAPL | 0.3920832 |
| IRSAIREAL | ASSLGQNNQAPL | 0.577166 |
| ITFAGGG | ASSLGQNNQAPL | 0.5413332 |
| ITGMSGHQEV | ASSLGQNNQAPL | 0.3839112 |
| IVAGLACTL | ASSLGQNNQAPL | 0.46674877 |
| IVGGGLAGA | ASSLGQNNQAPL | 0.5735539 |
| IVTGVLNI | ASSLGQNNQAPL | 0.49769726 |
| LAQQGCI | ASSLGQNNQAPL | 0.5039824 |
| LCLHVEN | ASSLGQNNQAPL | 0.5140424 |
| LDAISDGS | ASSLGQNNQAPL | 0.41370097 |
| LGGFAAML | ASSLGQNNQAPL | 0.5765347 |
| LGGKNPMI | ASSLGQNNQAPL | 0.5589563 |
| LGPKGVAAIL | ASSLGQNNQAPL | 0.45751965 |
| LMVGVSI | ASSLGQNNQAPL | 0.4841605 |
| LVGAAFVL | ASSLGQNNQAPL | 0.34659836 |
| LVLAVAPL | ASSLGQNNQAPL | 0.4106453 |
| MAAIAVIDAA | ASSLGQNNQAPL | 0.39055645 |
| NNGINAELQ | ASSLGQNNQAPL | 0.53412753 |
| RGEPLVLVHG | ASSLGQNNQAPL | 0.581074 |
| RGGAGGAAGLI | ASSLGQNNQAPL | 0.47970226 |
| RSYRMHPAV | ASSLGQNNQAPL | 0.36138478 |
| SAAHLLAGQLA | ASSLGQNNQAPL | 0.3602173 |
| SAAVGSY | ASSLGQNNQAPL | 0.54191995 |

| | | |
|---|---|---|
| SPHAVLHRI | ASSLGQNNQAPL | 0.33971268 |
| SQIAKVL | ASSLGQNNQAPL | 0.3356563 |
| TALGHLVP | ASSLGQNNQAPL | 0.47192627 |
| TAPTTSGAA | ASSLGQNNQAPL | 0.5797641 |
| TDGDDGNGGSPGT | ASSLGQNNQAPL | 0.55837 |
| TDGLGEN | ASSLGQNNQAPL | 0.45365375 |
| TGGPPLPGQA | ASSLGQNNQAPL | 0.57486975 |
| TIPAITI | ASSLGQNNQAPL | 0.57820207 |
| TIPGFTI | ASSLGQNNQAPL | 0.49275637 |
| TSIASGALA | ASSLGQNNQAPL | 0.38077056 |
| TVTLPGAAI | ASSLGQNNQAPL | 0.38772112 |
| VASGLSAALP | ASSLGQNNQAPL | 0.5856245 |
| VGAGVIGI | ASSLGQNNQAPL | 0.4755438 |
| VGPKRFGV | ASSLGQNNQAPL | 0.56116605 |
| VNVAFPD | ASSLGQNNQAPL | 0.53788877 |
| VSMTNSGVS | ASSLGQNNQAPL | 0.43246788 |
| VTDVLGDLSV | ASSLGQNNQAPL | 0.39601874 |
| VVVVGAGFAGL | ASSLGQNNQAPL | 0.5401556 |
| VYTVGTVV | ASSLGQNNQAPL | 0.57279944 |
| WTTFSGP | ASSLGQNNQAPL | 0.42715675 |
| YDELVEAF | ASSLGQNNQAPL | 0.39399266 |
| YGTGGAGGNGG | ASSLGQNNQAPL | 0.42097682 |
| YTNVMAV | ASSLGQNNQAPL | 0.44535047 |
| FLEQVHQL | ASSLGQNNQAPL | 0.5784185 |
| GLSLLAVAV | ASSLGQNNQAPL | 0.4564368 |
| LEQVHQL | ASSLGQNNQAPL | 0.42763492 |
| LNAAGAI | ASSLGQNNQAPL | 0.4807511 |
| PAVHASL | ASSLGQNNQAPL | 0.52206993 |
| TELEDDA | ASSLGQNNQAPL | 0.47059795 |
| TPPGPPN | ASSLGQNNQAPL | 0.46077913 |
| TPPPPPPP | ASSLGQNNQAPL | 0.45734602 |
| TQSPALI | ASSQVQDANTEVF | 0.5569738 |
| EFAAAFPQ | ASSQVQDANTEVF | 0.5275061 |
| LLSGNLDVL | ASSQVQDANTEVF | 0.4648593 |
| GPANLGNYNI | ASSQVQDANTEVF | 0.5189109 |
| TLGPTLP | ASSQVQDANTEVF | 0.4765616 |
| VYAYSSCCK | ASSQVQDANTEVF | 0.44085774 |
| APAPAPAPNI | ASSQVQDANTEVF | 0.3695353 |
| PGGPVPIP | ASSQVQDANTEVF | 0.365115 |
| FLTVALEP | ASSQVQDANTEVF | 0.46032304 |
| SQYFGANL | ASSQVQDANTEVF | 0.51998717 |
| GGVKPAA | ASSQVQDANTEVF | 0.531197 |
| GGTGGAGGAGGW | ASSQVQDANTEVF | 0.4509635 |
| GVGGVAGLGGAGG | ASSQVQDANTEVF | 0.44917765 |
| VLLVLATTI | ASSQVQDANTEVF | 0.48641878 |
| IGIFAAML | ASSQVQDANTEVF | 0.5403682 |
| AVGDLID | ASSQVQDANTEVF | 0.37410146 |
| NISPLPL | ASSQVQDANTEVF | 0.48853827 |
| VTAVFVAI | ASSQVQDANTEVF | 0.6276063 |
| SGDDNDAAL | ASSQVQDANTEVF | 0.49641204 |
| AVAALAHS | ASSQVQDANTEVF | 0.44894886 |

| | | |
|---|---|---|
| GGVTVMATP | ASSQVQDANTEVF | 0.38871697 |
| AGLAANVRD | ASSQVQDANTEVF | 0.613515 |
| TNRPDLIDPALLRPG | ASSQVQDANTEVF | 0.46455947 |
| LANLGVTAL | ASSQVQDANTEVF | 0.62161696 |
| TGERIEL | ASSQVQDANTEVF | 0.32884654 |
| NGEELAA | ASSQVQDANTEVF | 0.37896803 |
| VLSLVWG | ASSQVQDANTEVF | 0.39167887 |
| IGYALWL | ASSQVQDANTEVF | 0.31776512 |
| LSLLLAPL | ASSQVQDANTEVF | 0.5910896 |
| TLDEALK | ASSQVQDANTEVF | 0.35555327 |
| TDGAGTAEGG | ASSQVQDANTEVF | 0.43485725 |
| NAPGGAGGNGG | ASSQVQDANTEVF | 0.4085428 |
| VSVFMGMGGNF | ASSQVQDANTEVF | 0.58241546 |
| LASYLAAL | ASSQVQDANTEVF | 0.5272589 |
| LVGGTPAL | ASSQVQDANTEVF | 0.41112444 |
| VRNADVL | ASSQVQDANTEVF | 0.40547055 |
| TCSSPFLVSAM | ASSQVQDANTEVF | 0.5522694 |
| VHITDAE | ASSQVQDANTEVF | 0.5182034 |
| EYAAPSF | ASSQVQDANTEVF | 0.5059836 |
| VVDAFHGGL | ASSQVQDANTEVF | 0.35125345 |
| ASVVDPQ | ASSQVQDANTEVF | 0.52663326 |
| SYNEFGGLS | ASSQVQDANTEVF | 0.5058582 |
| DPDGSVY | ASSQVQDANTEVF | 0.38918936 |
| MVSTIHATE | ASSQVQDANTEVF | 0.3240553 |
| LAAGVADPV | ASSQVQDANTEVF | 0.4793139 |
| TKEQIAA | ASSQVQDANTEVF | 0.47108322 |
| SLASPAAL | ASSQVQDANTEVF | 0.5699517 |
| IGLVFYGV | ASSQVQDANTEVF | 0.59488523 |
| FLLDASNL | ASSQVQDANTEVF | 0.42263195 |
| YQVGNFP | ASSQVQDANTEVF | 0.5003948 |
| ESLLSQST | ASSQVQDANTEVF | 0.53150916 |
| LFTVGFAAVG | ASSQVQDANTEVF | 0.29276285 |
| ATPPTPT | ASSQVQDANTEVF | 0.37719947 |
| TDLATLG | ASSQVQDANTEVF | 0.36924154 |
| SAGGTGR | ASSQVQDANTEVF | 0.5458787 |
| GFSNSGS | ASSQVQDANTEVF | 0.42770326 |
| GGNGGSGGTSV | ASSQVQDANTEVF | 0.4886173 |
| GTSVATGGAGN | ASSQVQDANTEVF | 0.50496006 |
| NGGNGGSGGTS | ASSQVQDANTEVF | 0.5518285 |
| AVVKFPH | ASSQVQDANTEVF | 0.47161514 |
| TPPPPPP | ASSQVQDANTEVF | 0.37566176 |
| AAPAAAAAPA | ASSQVQDANTEVF | 0.38844952 |
| IALNEDL | ASSQVQDANTEVF | 0.5637202 |
| IVNVSSM | ASSQVQDANTEVF | 0.4618853 |
| TGIPPAPR | ASSQVQDANTEVF | 0.39158407 |
| FDLGGGTF | ASSQVQDANTEVF | 0.39757198 |
| FDLGGGTFDVS | ASSQVQDANTEVF | 0.5876827 |
| NEPTAAA | ASSQVQDANTEVF | 0.33345592 |
| NEPTAAALA | ASSQVQDANTEVF | 0.4563772 |
| GSIINISSI | ASSQVQDANTEVF | 0.36218834 |
| VAPGNAGTA | ASSQVQDANTEVF | 0.5022592 |

| | | |
|---|---|---|
| TGGGASLE | ASSQVQDANTEVF | 0.62484443 |
| VAVYNHYKRI | ASSQVQDANTEVF | 0.5772366 |
| TDPAPATT | ASSQVQDANTEVF | 0.48939323 |
| TNRPDILDPALLRPG | ASSQVQDANTEVF | 0.42845634 |
| LTTTTTS | ASSQVQDANTEVF | 0.38559556 |
| PTGSGKTL | ASSQVQDANTEVF | 0.4213061 |
| TGSGKTL | ASSQVQDANTEVF | 0.41289535 |
| TIGHVDHGK | ASSQVQDANTEVF | 0.5143076 |
| TIGHVDHGKTT | ASSQVQDANTEVF | 0.448376 |
| GHVDHGKTT | ASSQVQDANTEVF | 0.37370217 |
| AQFEHTLL | ASSQVQDANTEVF | 0.47017434 |
| IVNQDEL | ASSQVQDANTEVF | 0.5953058 |
| TNGSQFF | ASSQVQDANTEVF | 0.60057014 |
| TGKGIVE | ASSQVQDANTEVF | 0.4448337 |
| LISNASDA | ASSQVQDANTEVF | 0.37304994 |
| AELRTLGELTKIAKAHGAQ | ASSQVQDANTEVF | 0.35716563 |
| APNAVAAAEPSW | ASSQVQDANTEVF | 0.5046662 |
| APQVAGAATAVL | ASSQVQDANTEVF | 0.35126796 |
| ARRDPAAAAAILPTDA | ASSQVQDANTEVF | 0.38376674 |
| DAAARDVLAEAGLAEYF | ASSQVQDANTEVF | 0.5345089 |
| DDQIVLSAKANLAKVEQ | ASSQVQDANTEVF | 0.6409102 |
| DELLPDTGEDSDEDGH | ASSQVQDANTEVF | 0.43234372 |
| ELPEAPANRATVPQD | ASSQVQDANTEVF | 0.34106475 |
| FGQSKIERVTGIG | ASSQVQDANTEVF | 0.5023632 |
| GAGGTGSTAGGAGGAGGAGGLY | ASSQVQDANTEVF | 0.37971526 |
| GHGGTGAAGGNGGTGGAGGSGI | ASSQVQDANTEVF | 0.41984266 |
| GNGGNGGAGGEGGPGGAGGAGA | ASSQVQDANTEVF | 0.3896598 |
| GSTAGNGGNGGNATNGTIAGQP | ASSQVQDANTEVF | 0.34416163 |
| GTGGDGGNGGNGGTGG | ASSQVQDANTEVF | 0.44062096 |
| GVRRPAGIPADAPV | ASSQVQDANTEVF | 0.6086026 |
| GYRQVTDAHLLTLARR | ASSQVQDANTEVF | 0.48400044 |
| IAKMGQDGHDRG | ASSQVQDANTEVF | 0.40127912 |
| KAEPSIEIQYT | ASSQVQDANTEVF | 0.60547554 |
| KPVRIRMAADEDTVAVT | ASSQVQDANTEVF | 0.5511837 |
| MPPRIAGMRLLVIKPEP | ASSQVQDANTEVF | 0.5678873 |
| MKFYAVALFRSRGVPPTR | ASSQVQDANTEVF | 0.5700859 |
| NAGGFGILGAVAHSP | ASSQVQDANTEVF | 0.49361295 |
| NEGREITLAMAEM | ASSQVQDANTEVF | 0.45256454 |
| NELVEWSHPLGHRWRWE | ASSQVQDANTEVF | 0.5996046 |
| NLGAGGTGGNGGNPG | ASSQVQDANTEVF | 0.49129432 |
| NTEWPKWETVMAADPVD | ASSQVQDANTEVF | 0.5033461 |
| PASAAPAETQP | ASSQVQDANTEVF | 0.3956748 |
| QQSSDGKNGVRAAEITGE | ASSQVQDANTEVF | 0.5837873 |
| RAPDAVTTSWWKE | ASSQVQDANTEVF | 0.4095534 |
| SGFGNVGAHTSGWFNQ | ASSQVQDANTEVF | 0.6024377 |
| SGVAGAVGQAASVGGLKV | ASSQVQDANTEVF | 0.31383747 |
| SNGGAGTGSAGGNGGT | ASSQVQDANTEVF | 0.37644333 |
| TGGAGGEGGAAG | ASSQVQDANTEVF | 0.5842932 |
| TGGDGGAAGAGGAPGHG | ASSQVQDANTEVF | 0.3055343 |
| TMPPLLAAIANAGRHGV | ASSQVQDANTEVF | 0.32406747 |
| TSSTAGNSGGA | ASSQVQDANTEVF | 0.4269687 |

| | | |
|---|---|---|
| VPATVPVGILC | ASSQVQDANTEVF | 0.5696466 |
| VQRPDSGGVVLTGRIS | ASSQVQDANTEVF | 0.31344402 |
| WTPNPVPATKGVPAGLVQAT | ASSQVQDANTEVF | 0.412413 |
| YALKRALAKFSDRFDVVII | ASSQVQDANTEVF | 0.3387837 |
| YLIGSVIV | ASSQVQDANTEVF | 0.507565 |
| YGQLVAVID | ASSQVQDANTEVF | 0.46942097 |
| VTVGGMTLPA | ASSQVQDANTEVF | 0.34736377 |
| VLALPAV | ASSQVQDANTEVF | 0.5340618 |
| VIPPTLNYET | ASSQVQDANTEVF | 0.3580994 |
| VGMGGMHPSSG | ASSQVQDANTEVF | 0.5485085 |
| VGLAGGAATA | ASSQVQDANTEVF | 0.5580789 |
| VGGAGGAGGSGP | ASSQVQDANTEVF | 0.6099293 |
| VFAAFMV | ASSQVQDANTEVF | 0.3737753 |
| TLYDKLR | ASSQVQDANTEVF | 0.48300785 |
| TLAVFTVL | ASSQVQDANTEVF | 0.3956887 |
| TGSVQPARV | ASSQVQDANTEVF | 0.63872504 |
| TGGGAGSGGT | ASSQVQDANTEVF | 0.456012 |
| SHLETSL | ASSQVQDANTEVF | 0.33042818 |
| SFTTGMT | ASSQVQDANTEVF | 0.55392396 |
| QVEVAKRMGVRQPTVS | ASSQVQDANTEVF | 0.47805008 |
| QGSGGPALAM | ASSQVQDANTEVF | 0.4325012 |
| MLAPSNP | ASSQVQDANTEVF | 0.56835514 |
| LVNNAAI | ASSQVQDANTEVF | 0.4143646 |
| LTGEINTLA | ASSQVQDANTEVF | 0.47127926 |
| LSIFGPV | ASSQVQDANTEVF | 0.5302216 |
| LLALLANL | ASSQVQDANTEVF | 0.55826867 |
| LISNASDALDKLR | ASSQVQDANTEVF | 0.419133 |
| LINEAVP | ASSQVQDANTEVF | 0.50290716 |
| LGYAFSGL | ASSQVQDANTEVF | 0.52628374 |
| LDFVAVAL | ASSQVQDANTEVF | 0.37795386 |
| LAQPGAE | ASSQVQDANTEVF | 0.62614846 |
| ISNLGAVTGF | ASSQVQDANTEVF | 0.37839085 |
| HQEDPGRI | ASSQVQDANTEVF | 0.52711976 |
| GTPGEPTP | ASSQVQDANTEVF | 0.5482654 |
| GFFNSGS | ASSQVQDANTEVF | 0.6038778 |
| GAGGGGLTAGAN | ASSQVQDANTEVF | 0.38618585 |
| FVDALVNM | ASSQVQDANTEVF | 0.4438573 |
| FLTVAPEL | ASSQVQDANTEVF | 0.42597982 |
| FETLTAL | ASSQVQDANTEVF | 0.49416244 |
| DVINPIEL | ASSQVQDANTEVF | 0.3909613 |
| APVSGAPV | ASSQVQDANTEVF | 0.5698878 |
| ALVGGLAW | ASSQVQDANTEVF | 0.5435164 |
| ALAAAAATAI | ASSQVQDANTEVF | 0.571648 |
| AFGLGLLL | ASSQVQDANTEVF | 0.46813726 |
| AFFAAASL | ASSQVQDANTEVF | 0.4539579 |
| AAAALPAFSPP | ASSQVQDANTEVF | 0.6078594 |
| VVLAPAL | ASSQVQDANTEVF | 0.36253664 |
| TIGHVDHGKTTL | ASSQVQDANTEVF | 0.50163645 |
| LISNASDALDK | ASSQVQDANTEVF | 0.53371656 |
| LISNASDAL | ASSQVQDANTEVF | 0.4961497 |
| ISNASDALDK | ASSQVQDANTEVF | 0.42540848 |

| | | |
|---|---|---|
| NQAETLVYQTE | ASSQVQDANTEVF | 0.5682631 |
| TGGTGGTAGAG | ASSQVQDANTEVF | 0.41337386 |
| TGSTAGGAGGA | ASSQVQDANTEVF | 0.5638845 |
| DPLAVPVAAGPV | ASSQVQDANTEVF | 0.47511655 |
| GGTGGTPGNGGN | ASSQVQDANTEVF | 0.38092145 |
| PAGLAGVDPATL | ASSQVQDANTEVF | 0.44232732 |
| GNNLVVEAPHAGASV | ASSQVQDANTEVF | 0.5644859 |
| KVTRIMSASTLEQPA | ASSQVQDANTEVF | 0.5399152 |
| LGFALPIALPRRTVV | ASSQVQDANTEVF | 0.5344493 |
| TRLGNLTARKLVVLV | ASSQVQDANTEVF | 0.3305259 |
| WSQQRRLIYFPSAGP | ASSQVQDANTEVF | 0.5580156 |
| GPNGNPDPMAAAVDIR | ASSQVQDANTEVF | 0.54692143 |
| TQRCVSVADLRAAASR | ASSQVQDANTEVF | 0.62069005 |
| QFVLDVAEHPVEVLTETV | ASSQVQDANTEVF | 0.4199985 |
| GGAGGTGGAGGNGGTQAGDG | ASSQVQDANTEVF | 0.41791236 |
| PLLATLRAIAVDVDAKPAQV | ASSQVQDANTEVF | 0.48046818 |
| APKFRTGGYMIDFWGVGYQVAK | ASSQVQDANTEVF | 0.47961527 |
| GAGGAGGAGGAAGGPGGTGGTG | ASSQVQDANTEVF | 0.33883035 |
| IGALGLLATDVAPAGSVAFGVA | ASSQVQDANTEVF | 0.37686968 |
| AGGAGGIGGNGGAGGAGGVLMG | ASSQVQDANTEVF | 0.44147733 |
| ADELIGLL | ASSQVQDANTEVF | 0.4427142 |
| ADRALKA | ASSQVQDANTEVF | 0.59508026 |
| AGAVGQVGGAGGA | ASSQVQDANTEVF | 0.5572027 |
| AGGGGGLGGAP | ASSQVQDANTEVF | 0.5604375 |
| AGGLLYGNGGAG | ASSQVQDANTEVF | 0.56506443 |
| AGLESIGL | ASSQVQDANTEVF | 0.5819811 |
| AGLKTGII | ASSQVQDANTEVF | 0.45721543 |
| ASLAALLL | ASSQVQDANTEVF | 0.46105704 |
| ASVINISSI | ASSQVQDANTEVF | 0.4049316 |
| ATPPGPA | ASSQVQDANTEVF | 0.36496294 |
| ATVASALV | ASSQVQDANTEVF | 0.39566052 |
| ATVASLIL | ASSQVQDANTEVF | 0.46443406 |
| AVNGPVLIRH | ASSQVQDANTEVF | 0.45589545 |
| DGAAGAVGI | ASSQVQDANTEVF | 0.3032578 |
| DPLLHGI | ASSQVQDANTEVF | 0.5654903 |
| ELGAALGAL | ASSQVQDANTEVF | 0.532793 |
| ELGRTQAV | ASSQVQDANTEVF | 0.525594 |
| FVAGGYLS | ASSQVQDANTEVF | 0.47082198 |
| GASGGAGGAGGW | ASSQVQDANTEVF | 0.47338048 |
| GGAGAGGDGGAGGI | ASSQVQDANTEVF | 0.4019494 |
| GTGHNGSQPGMGGNG | ASSQVQDANTEVF | 0.49540284 |
| GYAAASASAS | ASSQVQDANTEVF | 0.40468508 |
| HQVDVAAVQQG | ASSQVQDANTEVF | 0.49270153 |
| IGAAIAHT | ASSQVQDANTEVF | 0.5758213 |
| IGMIEVAQL | ASSQVQDANTEVF | 0.50719965 |
| IRLAPEHTSAMD | ASSQVQDANTEVF | 0.37124756 |
| IYNVSVL | ASSQVQDANTEVF | 0.4036713 |
| KQGFLGM | ASSQVQDANTEVF | 0.36521867 |
| LESVITQL | ASSQVQDANTEVF | 0.3997202 |
| LGLGALVVV | ASSQVQDANTEVF | 0.44041142 |
| LIGNGGNGGSGGI | ASSQVQDANTEVF | 0.49410066 |

| | | |
|---|---|---|
| LNAINAPTL | ASSQVQDANTEVF | 0.42042914 |
| LQLLNTGV | ASSQVQDANTEVF | 0.55671614 |
| LQTSIEF | ASSQVQDANTEVF | 0.5495946 |
| MSFAVSEL | ASSQVQDANTEVF | 0.49268907 |
| NTGALGL | ASSQVQDANTEVF | 0.39706993 |
| PPGAGGAGGAGTT | ASSQVQDANTEVF | 0.5614371 |
| QLVGLGI | ASSQVQDANTEVF | 0.58502686 |
| RFLEQVHQL | ASSQVQDANTEVF | 0.37149355 |
| RGVGYVLR | ASSQVQDANTEVF | 0.503821 |
| SATGVFAGVI | ASSQVQDANTEVF | 0.42112732 |
| SLPAITLP | ASSQVQDANTEVF | 0.5155266 |
| STALAFTV | ASSQVQDANTEVF | 0.5838693 |
| SVLTVGLGK | ASSQVQDANTEVF | 0.46117735 |
| TALAGAGSGI | ASSQVQDANTEVF | 0.36964843 |
| TGLPFTGI | ASSQVQDANTEVF | 0.41128197 |
| VAEHVTV | ASSQVQDANTEVF | 0.4740032 |
| VDIVLNSL | ASSQVQDANTEVF | 0.456781 |
| VLTGGGAVL | ASSQVQDANTEVF | 0.32330436 |
| VNVGLAYGI | ASSQVQDANTEVF | 0.57306176 |
| VSSAAVGAV | ASSQVQDANTEVF | 0.37843654 |
| VTSVYEAGV | ASSQVQDANTEVF | 0.45507115 |
| WVALDRGA | ASSQVQDANTEVF | 0.46281037 |
| YLVGSVVI | ASSQVQDANTEVF | 0.546344 |
| GPPGTGKTLL | ASSQVQDANTEVF | 0.50379986 |
| GVGMTKF | ASSQVQDANTEVF | 0.42086583 |
| IGIGGSDL | ASSQVQDANTEVF | 0.4708287 |
| IGTLLAEL | ASSQVQDANTEVF | 0.5144605 |
| IYTSGTT | ASSQVQDANTEVF | 0.47749174 |
| LSFPAGV | ASSQVQDANTEVF | 0.5366571 |
| LYTSGTT | ASSQVQDANTEVF | 0.46344948 |
| VGAFPFGF | ASSQVQDANTEVF | 0.44127393 |
| VGAGTVEF | ASSQVQDANTEVF | 0.37946275 |
| VGKSTLLL | ASSQVQDANTEVF | 0.40460354 |
| AALAHAGGGIV | ASSQVQDANTEVF | 0.4126225 |
| AALVAVSLL | ASSQVQDANTEVF | 0.38325125 |
| AAVSRAVAAL | ASSQVQDANTEVF | 0.37509844 |
| AGGSSGIGGF | ASSQVQDANTEVF | 0.5250222 |
| AGLTVANANG | ASSQVQDANTEVF | 0.49240172 |
| AGLVGAVL | ASSQVQDANTEVF | 0.42453274 |
| ALSQAFTV | ASSQVQDANTEVF | 0.34147063 |
| ALTFSVSL | ASSQVQDANTEVF | 0.44038713 |
| ALTGAALVAL | ASSQVQDANTEVF | 0.42967603 |
| ASFSIGAGVL | ASSQVQDANTEVF | 0.52210313 |
| DGSVYKI | ASSQVQDANTEVF | 0.57488257 |
| EQLAEIV | ASSQVQDANTEVF | 0.6360433 |
| FAVSELTVM | ASSQVQDANTEVF | 0.61749375 |
| GAAPLAAGL | ASSQVQDANTEVF | 0.49299064 |
| GADGGDGGSGGAGG | ASSQVQDANTEVF | 0.566133 |
| GFGLDTGGAGG | ASSQVQDANTEVF | 0.3986618 |
| GLASGGLL | ASSQVQDANTEVF | 0.43094158 |
| GRIAGAAVLL | ASSQVQDANTEVF | 0.47529688 |

| | | |
|---|---|---|
| GSAGSLNTSF | ASSQVQDANTEVF | 0.5174607 |
| GTGQIGI | ASSQVQDANTEVF | 0.3882379 |
| GTLIALIL | ASSQVQDANTEVF | 0.41995788 |
| IAATMGFVVG | ASSQVQDANTEVF | 0.33975387 |
| IFAQALL | ASSQVQDANTEVF | 0.52141047 |
| IGAAVALGF | ASSQVQDANTEVF | 0.36466604 |
| IGLTPLAL | ASSQVQDANTEVF | 0.33181548 |
| ILGSNTSTL | ASSQVQDANTEVF | 0.45732585 |
| IVFSLTVL | ASSQVQDANTEVF | 0.5951718 |
| LASKATGF | ASSQVQDANTEVF | 0.5319817 |
| LASVALIL | ASSQVQDANTEVF | 0.45433873 |
| LFAVGLTL | ASSQVQDANTEVF | 0.4245249 |
| LFNGGIGTYI | ASSQVQDANTEVF | 0.57119364 |
| LTLGAGII | ASSQVQDANTEVF | 0.4119298 |
| LVAAALGDI | ASSQVQDANTEVF | 0.5020002 |
| NTAFVQLGI | ASSQVQDANTEVF | 0.49052304 |
| QLFSGAEL | ASSQVQDANTEVF | 0.5504408 |
| SAATFITL | ASSQVQDANTEVF | 0.34470853 |
| SNYGLANLGV | ASSQVQDANTEVF | 0.5494771 |
| SYGGAYAVM | ASSQVQDANTEVF | 0.53858197 |
| TTGVGGLTLG | ASSQVQDANTEVF | 0.4177183 |
| TTQTIYI | ASSQVQDANTEVF | 0.33536074 |
| VAAGAGLGAVI | ASSQVQDANTEVF | 0.5928731 |
| VAAVVTLL | ASSQVQDANTEVF | 0.42782485 |
| VGFLGGLI | ASSQVQDANTEVF | 0.38097325 |
| VGVIPILL | ASSQVQDANTEVF | 0.31141064 |
| VIFASSLI | ASSQVQDANTEVF | 0.45793742 |
| VKGGDGGAAAG | ASSQVQDANTEVF | 0.47802797 |
| VTKTFTNI | ASSQVQDANTEVF | 0.59298337 |
| VVAAAILL | ASSQVQDANTEVF | 0.49361622 |
| YAALDVEL | ASSQVQDANTEVF | 0.44421092 |
| LASVALLL | ASSQVQDANTEVF | 0.57081586 |
| LDAVHRV | ASSQVQDANTEVF | 0.44751373 |
| QGLSSSIF | ASSQVQDANTEVF | 0.35457522 |
| RGTFANIRL | ASSQVQDANTEVF | 0.43406412 |
| RQLWWGHRI | ASSQVQDANTEVF | 0.48518324 |
| YGPPGTGKTLL | ASSQVQDANTEVF | 0.5818095 |
| AALVGAPANVA | ASSQVQDANTEVF | 0.55840325 |
| AATFRRTE | ASSQVQDANTEVF | 0.37361988 |
| AGALGGGI | ASSQVQDANTEVF | 0.4460193 |
| AGGAGGAGGIGAG | ASSQVQDANTEVF | 0.5229701 |
| AGSGALGL | ASSQVQDANTEVF | 0.33743587 |
| AQIADQI | ASSQVQDANTEVF | 0.39643195 |
| ASAVAVTYL | ASSQVQDANTEVF | 0.4191596 |
| ASTVLGGTGG | ASSQVQDANTEVF | 0.37248904 |
| AVGGEPAK | ASSQVQDANTEVF | 0.5353865 |
| CEQGTFGGTC | ASSQVQDANTEVF | 0.48136255 |
| DEGKAPAG | ASSQVQDANTEVF | 0.40968218 |
| DTVVFPAL | ASSQVQDANTEVF | 0.49457932 |
| FEGITMAG | ASSQVQDANTEVF | 0.40751413 |
| FPLVVQD | ASSQVQDANTEVF | 0.40529987 |

| | | |
|---|---|---|
| GAFTTPQ | ASSQVQDANTEVF | 0.43342677 |
| GFGGFGTAGGVGGA | ASSQVQDANTEVF | 0.5502203 |
| GFLAQFL | ASSQVQDANTEVF | 0.5650079 |
| GGAGGAGGNAGVA | ASSQVQDANTEVF | 0.459746 |
| GGVGTAGAGGAGGAGG | ASSQVQDANTEVF | 0.3878216 |
| GNGGGGDAGNAGSGGN | ASSQVQDANTEVF | 0.6123489 |
| GTGGAGGAGNDAGS | ASSQVQDANTEVF | 0.37038577 |
| GTTNTGWFNS | ASSQVQDANTEVF | 0.42465076 |
| IFMGPLFTL | ASSQVQDANTEVF | 0.49635822 |
| ILAHAIEAI | ASSQVQDANTEVF | 0.47501832 |
| IRSAIREAL | ASSQVQDANTEVF | 0.5451275 |
| ITFAGGG | ASSQVQDANTEVF | 0.56257874 |
| ITGMSGHQEV | ASSQVQDANTEVF | 0.49188212 |
| IVAGLACTL | ASSQVQDANTEVF | 0.43303555 |
| IVGGGLAGA | ASSQVQDANTEVF | 0.5210651 |
| IVTGVLNI | ASSQVQDANTEVF | 0.50772554 |
| LAQQGCI | ASSQVQDANTEVF | 0.4201222 |
| LCLHVEN | ASSQVQDANTEVF | 0.54420084 |
| LDAISDGS | ASSQVQDANTEVF | 0.35998422 |
| LGGFAAML | ASSQVQDANTEVF | 0.46326077 |
| LGGKNPMI | ASSQVQDANTEVF | 0.47799444 |
| LGPKGVAAIL | ASSQVQDANTEVF | 0.40887994 |
| LMVGVSI | ASSQVQDANTEVF | 0.48609677 |
| LVGAAFVL | ASSQVQDANTEVF | 0.5786987 |
| LVLAVAPL | ASSQVQDANTEVF | 0.54490906 |
| MAAIAVIDAA | ASSQVQDANTEVF | 0.4422615 |
| NNGINAELQ | ASSQVQDANTEVF | 0.4569642 |
| RGEPLVLVHG | ASSQVQDANTEVF | 0.37601694 |
| RGGAGGAAGLI | ASSQVQDANTEVF | 0.39273506 |
| RSYRMHPAV | ASSQVQDANTEVF | 0.3651976 |
| SAAHLLAGQLA | ASSQVQDANTEVF | 0.46780306 |
| SAAVGSY | ASSQVQDANTEVF | 0.42555064 |
| SPHAVLHRI | ASSQVQDANTEVF | 0.58003503 |
| SQIAKVL | ASSQVQDANTEVF | 0.54275537 |
| TALGHLVP | ASSQVQDANTEVF | 0.59991515 |
| TAPTTSGAA | ASSQVQDANTEVF | 0.3998047 |
| TDGDDGNGGSPGT | ASSQVQDANTEVF | 0.5230626 |
| TDGLGEN | ASSQVQDANTEVF | 0.32616827 |
| TGGPPLPGQA | ASSQVQDANTEVF | 0.47191733 |
| TIPAITI | ASSQVQDANTEVF | 0.36866778 |
| TIPGFTI | ASSQVQDANTEVF | 0.47211573 |
| TSIASGALA | ASSQVQDANTEVF | 0.35144204 |
| TVTLPGAAI | ASSQVQDANTEVF | 0.612819 |
| VASGLSAALP | ASSQVQDANTEVF | 0.38235205 |
| VGAGVIGI | ASSQVQDANTEVF | 0.54925394 |
| VGPKRFGV | ASSQVQDANTEVF | 0.42672738 |
| VNVAFPD | ASSQVQDANTEVF | 0.35808197 |
| VSMTNSGVS | ASSQVQDANTEVF | 0.4638356 |
| VTDVLGDLSV | ASSQVQDANTEVF | 0.48445648 |
| VVVVGAGFAGL | ASSQVQDANTEVF | 0.40392005 |
| VYTVGTVV | ASSQVQDANTEVF | 0.32536682 |

| | | |
|---|---|---|
| WTTFSGP | ASSQVQDANTEVF | 0.3659663 |
| YDELVEAF | ASSQVQDANTEVF | 0.429906 |
| YGTGGAGGNGG | ASSQVQDANTEVF | 0.6146387 |
| YTNVMAV | ASSQVQDANTEVF | 0.5916447 |
| FLEQVHQL | ASSQVQDANTEVF | 0.49853504 |
| GLSLLAVAV | ASSQVQDANTEVF | 0.5965445 |
| LEQVHQL | ASSQVQDANTEVF | 0.3008049 |
| LNAAGAI | ASSQVQDANTEVF | 0.4515902 |
| PAVHASL | ASSQVQDANTEVF | 0.5582295 |
| TELEDDA | ASSQVQDANTEVF | 0.52881664 |
| TPPGPPN | ASSQVQDANTEVF | 0.45310494 |
| TPPPPPPP | ASSQVQDANTEVF | 0.53783154 |
| TQSPALI | ASSSPPNERLF | 0.45752257 |
| EFAAAFPQ | ASSSPPNERLF | 0.44505513 |
| LLSGNLDVL | ASSSPPNERLF | 0.46931088 |
| GPANLGNYNI | ASSSPPNERLF | 0.53233945 |
| TLGPTLP | ASSSPPNERLF | 0.4361356 |
| VYAYSSCCK | ASSSPPNERLF | 0.3550155 |
| APAPAPAPNI | ASSSPPNERLF | 0.41384143 |
| PGGPVPIP | ASSSPPNERLF | 0.64386135 |
| FLTVALEP | ASSSPPNERLF | 0.456657 |
| SQYFGANL | ASSSPPNERLF | 0.5315155 |
| GGVKPAA | ASSSPPNERLF | 0.42656386 |
| GGTGGAGGAGGW | ASSSPPNERLF | 0.5186636 |
| GVGGVAGLGGAGG | ASSSPPNERLF | 0.40494004 |
| VLLVLATTI | ASSSPPNERLF | 0.37116417 |
| IGIFAAML | ASSSPPNERLF | 0.5711298 |
| AVGDLID | ASSSPPNERLF | 0.3523493 |
| NISPLPL | ASSSPPNERLF | 0.40109402 |
| VTAVFVAI | ASSSPPNERLF | 0.42911923 |
| SGDDNDAAL | ASSSPPNERLF | 0.6429633 |
| AVAALAHS | ASSSPPNERLF | 0.49764067 |
| GGVTVMATP | ASSSPPNERLF | 0.48602074 |
| AGLAANVRD | ASSSPPNERLF | 0.44717196 |
| TNRPDLIDPALLRPG | ASSSPPNERLF | 0.46341872 |
| LANLGVTAL | ASSSPPNERLF | 0.5507206 |
| TGERIEL | ASSSPPNERLF | 0.35398415 |
| NGEELAA | ASSSPPNERLF | 0.55996925 |
| VLSLVWG | ASSSPPNERLF | 0.36554185 |
| IGYALWL | ASSSPPNERLF | 0.5220907 |
| LSLLLAPL | ASSSPPNERLF | 0.437903 |
| TLDEALK | ASSSPPNERLF | 0.6276111 |
| TDGAGTAEGG | ASSSPPNERLF | 0.44339553 |
| NAPGGAGGNGG | ASSSPPNERLF | 0.554983 |
| VSVFMGMGGNF | ASSSPPNERLF | 0.4292098 |
| LASYLAAL | ASSSPPNERLF | 0.47912148 |
| LVGGTPAL | ASSSPPNERLF | 0.60481906 |
| VRNADVL | ASSSPPNERLF | 0.41871852 |
| TCSSPFLVSAM | ASSSPPNERLF | 0.63963914 |
| VHITDAE | ASSSPPNERLF | 0.45676118 |
| EYAAPSF | ASSSPPNERLF | 0.3366898 |

| | | |
|---|---|---|
| VVDAFHGGL | ASSSPPNERLF | 0.42127272 |
| ASVVDPQ | ASSSPPNERLF | 0.56754863 |
| SYNEFGGLS | ASSSPPNERLF | 0.60706156 |
| DPDGSVY | ASSSPPNERLF | 0.52162087 |
| MVSTIHATE | ASSSPPNERLF | 0.4255231 |
| LAAGVADPV | ASSSPPNERLF | 0.44769508 |
| TKEQIAA | ASSSPPNERLF | 0.60542846 |
| SLASPAAL | ASSSPPNERLF | 0.6125165 |
| IGLVFYGV | ASSSPPNERLF | 0.43596828 |
| FLLDASNL | ASSSPPNERLF | 0.3438767 |
| YQVGNFP | ASSSPPNERLF | 0.34960726 |
| ESLLSQST | ASSSPPNERLF | 0.53846455 |
| LFTVGFAAVG | ASSSPPNERLF | 0.60671055 |
| ATPPTPT | ASSSPPNERLF | 0.33180898 |
| TDLATLG | ASSSPPNERLF | 0.41920927 |
| SAGGTGR | ASSSPPNERLF | 0.4918033 |
| GFSNSGS | ASSSPPNERLF | 0.32913625 |
| GGNGGSGGTSV | ASSSPPNERLF | 0.52130866 |
| GTSVATGGAGN | ASSSPPNERLF | 0.5320447 |
| NGGNGGSGGTS | ASSSPPNERLF | 0.45619115 |
| AVVKFPH | ASSSPPNERLF | 0.3379322 |
| TPPPPPP | ASSSPPNERLF | 0.5590465 |
| AAPAAAAAPA | ASSSPPNERLF | 0.6005527 |
| IALNEDL | ASSSPPNERLF | 0.5818579 |
| IVNVSSM | ASSSPPNERLF | 0.3486625 |
| TGIPPAPR | ASSSPPNERLF | 0.3036237 |
| FDLGGGTF | ASSSPPNERLF | 0.5066248 |
| FDLGGGTFDVS | ASSSPPNERLF | 0.5263839 |
| NEPTAAA | ASSSPPNERLF | 0.4469247 |
| NEPTAAALA | ASSSPPNERLF | 0.3896817 |
| GSIINISSI | ASSSPPNERLF | 0.4311388 |
| VAPGNAGTA | ASSSPPNERLF | 0.5877441 |
| TGGGASLE | ASSSPPNERLF | 0.35743794 |
| VAVYNHYKRI | ASSSPPNERLF | 0.35491934 |
| TDPAPATT | ASSSPPNERLF | 0.4857127 |
| TNRPDILDPALLRPG | ASSSPPNERLF | 0.5187243 |
| LTTTTTS | ASSSPPNERLF | 0.41249657 |
| PTGSGKTL | ASSSPPNERLF | 0.472508 |
| TGSGKTL | ASSSPPNERLF | 0.5702461 |
| TIGHVDHGK | ASSSPPNERLF | 0.54741967 |
| TIGHVDHGKTT | ASSSPPNERLF | 0.5159131 |
| GHVDHGKTT | ASSSPPNERLF | 0.35810074 |
| AQFEHTLL | ASSSPPNERLF | 0.61080277 |
| IVNQDEL | ASSSPPNERLF | 0.35816696 |
| TNGSQFF | ASSSPPNERLF | 0.5242982 |
| TGKGIVE | ASSSPPNERLF | 0.47328803 |
| LISNASDA | ASSSPPNERLF | 0.51842606 |
| AELRTLGELTKIAKAHGAQ | ASSSPPNERLF | 0.4454338 |
| APNAVAAAEPSW | ASSSPPNERLF | 0.33836243 |
| APQVAGAATAVL | ASSSPPNERLF | 0.38090658 |
| ARRDPAAAAAILPTDA | ASSSPPNERLF | 0.35325328 |

| | | |
|---|---|---|
| DAAARDVLAEAGLAEYF | ASSSPPNERLF | 0.54952806 |
| DDQIVLSAKANLAKVEQ | ASSSPPNERLF | 0.56462634 |
| DELLPDTGEDSDEDGH | ASSSPPNERLF | 0.51414967 |
| ELPEAPANRATVPQD | ASSSPPNERLF | 0.5020221 |
| FGQSKIERVTGIG | ASSSPPNERLF | 0.43554136 |
| GAGGTGSTAGGAGGAGGAGGLY | ASSSPPNERLF | 0.5188508 |
| GHGGTGAAGGNGGTGGAGGSGI | ASSSPPNERLF | 0.37491974 |
| GNGGNGGAGGEGGPGGAGGAGA | ASSSPPNERLF | 0.47178277 |
| GSTAGNGGNGGNATNGTIAGQP | ASSSPPNERLF | 0.36554697 |
| GTGGDGGNGGNGGTGG | ASSSPPNERLF | 0.44095573 |
| GVRRPAGIPADAPV | ASSSPPNERLF | 0.38932097 |
| GYRQVTDAHLLTLARR | ASSSPPNERLF | 0.46928144 |
| IAKMGQDGHDRG | ASSSPPNERLF | 0.50164866 |
| KAEPSIEIQYT | ASSSPPNERLF | 0.34599972 |
| KPVRIRMAADEDTVAVT | ASSSPPNERLF | 0.44168603 |
| MPPRIAGMRLLVIKPEP | ASSSPPNERLF | 0.60512376 |
| MKFYAVALFRSRGVPPTR | ASSSPPNERLF | 0.43673253 |
| NAGGFGILGAVAHSP | ASSSPPNERLF | 0.34028345 |
| NEGREITLAMAEM | ASSSPPNERLF | 0.49837723 |
| NELVEWSHPLGHRWRWE | ASSSPPNERLF | 0.41241035 |
| NLGAGGTGGNGGNPG | ASSSPPNERLF | 0.54853284 |
| NTEWPKWETVMAADPVD | ASSSPPNERLF | 0.5324703 |
| PASAAPAETQP | ASSSPPNERLF | 0.36897364 |
| QQSSDGKNGVRAAEITGE | ASSSPPNERLF | 0.5189056 |
| RAPDAVTTSWWKE | ASSSPPNERLF | 0.36281592 |
| SGFGNVGAHTSGWFNQ | ASSSPPNERLF | 0.38673428 |
| SGVAGAVGQAASVGGLKV | ASSSPPNERLF | 0.40183142 |
| SNGGAGTGSAGGNGGT | ASSSPPNERLF | 0.53978324 |
| TGGAGGEGGAAG | ASSSPPNERLF | 0.48154598 |
| TGGDGGAAGAGGAPGHG | ASSSPPNERLF | 0.36584243 |
| TMPPLLAAIANAGRHGV | ASSSPPNERLF | 0.44611126 |
| TSSTAGNSGGA | ASSSPPNERLF | 0.45959526 |
| VPATVPVGILC | ASSSPPNERLF | 0.4083874 |
| VQRPDSGGVVLTGRIS | ASSSPPNERLF | 0.42721146 |
| WTPNPVPATKGVPAGLVQAT | ASSSPPNERLF | 0.38113812 |
| YALKRALAKFSDRFDVVII | ASSSPPNERLF | 0.32569808 |
| YLIGSVIV | ASSSPPNERLF | 0.5429646 |
| YGQLVAVID | ASSSPPNERLF | 0.40204033 |
| VTVGGMTLPA | ASSSPPNERLF | 0.4891985 |
| VLALPAV | ASSSPPNERLF | 0.48247778 |
| VIPPTLNYET | ASSSPPNERLF | 0.33119953 |
| VGMGGMHPSSG | ASSSPPNERLF | 0.4822655 |
| VGLAGGAATA | ASSSPPNERLF | 0.4856044 |
| VGGAGGAGGSGP | ASSSPPNERLF | 0.46995348 |
| VFAAFMV | ASSSPPNERLF | 0.46839327 |
| TLYDKLR | ASSSPPNERLF | 0.4248759 |
| TLAVFTVL | ASSSPPNERLF | 0.56717086 |
| TGSVQPARV | ASSSPPNERLF | 0.5216584 |
| TGGGAGSGGT | ASSSPPNERLF | 0.36854967 |
| SHLETSL | ASSSPPNERLF | 0.5165973 |
| SFTTGMT | ASSSPPNERLF | 0.5497768 |

| | | |
|---|---|---|
| QVEVAKRMGVRQPTVS | ASSSPPNERLF | 0.5387349 |
| QGSGGPALAM | ASSSPPNERLF | 0.41609743 |
| MLAPSNP | ASSSPPNERLF | 0.32661822 |
| LVNNAAI | ASSSPPNERLF | 0.52717996 |
| LTGEINTLA | ASSSPPNERLF | 0.51521105 |
| LSIFGPV | ASSSPPNERLF | 0.3779158 |
| LLALLANL | ASSSPPNERLF | 0.32407707 |
| LISNASDALDKLR | ASSSPPNERLF | 0.52739614 |
| LINEAVP | ASSSPPNERLF | 0.6299875 |
| LGYAFSGL | ASSSPPNERLF | 0.38599065 |
| LDFVAVAL | ASSSPPNERLF | 0.6277253 |
| LAQPGAE | ASSSPPNERLF | 0.33898103 |
| ISNLGAVTGF | ASSSPPNERLF | 0.5123544 |
| HQEDPGRI | ASSSPPNERLF | 0.50619113 |
| GTPGEPTP | ASSSPPNERLF | 0.35564536 |
| GFFNSGS | ASSSPPNERLF | 0.4756082 |
| GAGGGGLTAGAN | ASSSPPNERLF | 0.37603882 |
| FVDALVNM | ASSSPPNERLF | 0.38660985 |
| FLTVAPEL | ASSSPPNERLF | 0.5065613 |
| FETLTAL | ASSSPPNERLF | 0.40908158 |
| DVINPIEL | ASSSPPNERLF | 0.46235293 |
| APVSGAPV | ASSSPPNERLF | 0.4240153 |
| ALVGGLAW | ASSSPPNERLF | 0.42450932 |
| ALAAAAATAI | ASSSPPNERLF | 0.5300836 |
| AFGLGLLL | ASSSPPNERLF | 0.44177115 |
| AFFAAASL | ASSSPPNERLF | 0.40207487 |
| AAAALPAFSPP | ASSSPPNERLF | 0.57045805 |
| VVLAPAL | ASSSPPNERLF | 0.42364222 |
| TIGHVDHGKTTL | ASSSPPNERLF | 0.63818467 |
| LISNASDALDK | ASSSPPNERLF | 0.52262175 |
| LISNASDAL | ASSSPPNERLF | 0.53980803 |
| ISNASDALDK | ASSSPPNERLF | 0.42656046 |
| NQAETLVYQTE | ASSSPPNERLF | 0.58423156 |
| TGGTGGTAGAG | ASSSPPNERLF | 0.42148596 |
| TGSTAGGAGGA | ASSSPPNERLF | 0.44709432 |
| DPLAVPVAAGPV | ASSSPPNERLF | 0.5369175 |
| GGTGGTPGNGGN | ASSSPPNERLF | 0.39167523 |
| PAGLAGVDPATL | ASSSPPNERLF | 0.8152783 |
| GNNLVVEAPHAGASV | ASSSPPNERLF | 0.76690745 |
| KVTRIMSASTLEQPA | ASSSPPNERLF | 0.74527776 |
| LGFALPIALPRRTVV | ASSSPPNERLF | 0.58139676 |
| TRLGNLTARKLVVLV | ASSSPPNERLF | 0.3376869 |
| WSQQRRLIYFPSAGP | ASSSPPNERLF | 0.64197063 |
| GPNGNPDPMAAAVDIR | ASSSPPNERLF | 0.44969004 |
| TQRCVSVADLRAAASR | ASSSPPNERLF | 0.81806207 |
| QFVLDVAEHPVEVLTETV | ASSSPPNERLF | 0.795751 |
| GGAGGTGGAGGNGGTQAGDG | ASSSPPNERLF | 0.55966043 |
| PLLATLRAIAVDVDAKPAQV | ASSSPPNERLF | 0.77885365 |
| APKFRTGGYMIDFWGVGYQVAK | ASSSPPNERLF | 0.7859664 |
| GAGGAGGAGGAAGGPGGTGGTG | ASSSPPNERLF | 0.3729598 |
| IGALGLLATDVAPAGSVAFGVA | ASSSPPNERLF | 0.6962949 |

| | | |
|---|---|---|
| AGGAGGIGGNGGAGGAGGVLMG | ASSSPPNERLF | 0.70016074 |
| ADELIGLL | ASSSPPNERLF | 0.6400317 |
| ADRALKA | ASSSPPNERLF | 0.37627435 |
| AGAVGQVGGAGGA | ASSSPPNERLF | 0.773716 |
| AGGGGGLGGAP | ASSSPPNERLF | 0.6602148 |
| AGGLLYGNGGAG | ASSSPPNERLF | 0.21117792 |
| AGLESIGL | ASSSPPNERLF | 0.74391645 |
| AGLKTGII | ASSSPPNERLF | 0.66461813 |
| ASLAALLL | ASSSPPNERLF | 0.5482038 |
| ASVINISSI | ASSSPPNERLF | 0.7883285 |
| ATPPGPA | ASSSPPNERLF | 0.72845304 |
| ATVASALV | ASSSPPNERLF | 0.5456334 |
| ATVASLIL | ASSSPPNERLF | 0.7803514 |
| AVNGPVLIRH | ASSSPPNERLF | 0.81203175 |
| DGAAGAVGI | ASSSPPNERLF | 0.38841486 |
| DPLLHGI | ASSSPPNERLF | 0.74329245 |
| ELGAALGAL | ASSSPPNERLF | 0.8166371 |
| ELGRTQAV | ASSSPPNERLF | 0.7035483 |
| FVAGGYLS | ASSSPPNERLF | 0.8211783 |
| GASGGAGGAGGW | ASSSPPNERLF | 0.8253045 |
| GGAGAGGDGGAGGI | ASSSPPNERLF | 0.63145995 |
| GTGHNGSQPGMGGNG | ASSSPPNERLF | 0.81190574 |
| GYAAASASAS | ASSSPPNERLF | 0.8063718 |
| HQVDVAAVQQG | ASSSPPNERLF | 0.66505194 |
| IGAAIAHT | ASSSPPNERLF | 0.7040665 |
| IGMIEVAQL | ASSSPPNERLF | 0.36059687 |
| IRLAPEHTSAMD | ASSSPPNERLF | 0.6921935 |
| IYNVSVL | ASSSPPNERLF | 0.49824795 |
| KQGFLGM | ASSSPPNERLF | 0.76670027 |
| LESVITQL | ASSSPPNERLF | 0.8159757 |
| LGLGALVVV | ASSSPPNERLF | 0.67645216 |
| LIGNGGNGGSGGI | ASSSPPNERLF | 0.4059141 |
| LNAINAPTL | ASSSPPNERLF | 0.503174 |
| LQLLNTGV | ASSSPPNERLF | 0.7642666 |
| LQTSIEF | ASSSPPNERLF | 0.43518004 |
| MSFAVSEL | ASSSPPNERLF | 0.41199982 |
| NTGALGL | ASSSPPNERLF | 0.6610633 |
| PPGAGGAGGAGTT | ASSSPPNERLF | 0.7856152 |
| QLVGLGI | ASSSPPNERLF | 0.7970562 |
| RFLEQVHQL | ASSSPPNERLF | 0.5081528 |
| RGVGYVLR | ASSSPPNERLF | 0.8209095 |
| SATGVFAGVI | ASSSPPNERLF | 0.81054187 |
| SLPAITLP | ASSSPPNERLF | 0.82542634 |
| STALAFTV | ASSSPPNERLF | 0.71739185 |
| SVLTVGLGK | ASSSPPNERLF | 0.41247386 |
| TALAGAGSGI | ASSSPPNERLF | 0.8129881 |
| TGLPFTGI | ASSSPPNERLF | 0.63143414 |
| VAEHVTV | ASSSPPNERLF | 0.7445112 |
| VDIVLNSL | ASSSPPNERLF | 0.78814065 |
| VLTGGGAVL | ASSSPPNERLF | 0.6024692 |
| VNVGLAYGI | ASSSPPNERLF | 0.5853508 |

| | | |
|---|---|---|
| VSSAAVGAV | ASSSPPNERLF | 0.7539927 |
| VTSVYEAGV | ASSSPPNERLF | 0.81894946 |
| WVALDRGA | ASSSPPNERLF | 0.5367328 |
| YLVGSVVI | ASSSPPNERLF | 0.40891305 |
| GPPGTGKTLL | ASSSPPNERLF | 0.66258943 |
| GVGMTKF | ASSSPPNERLF | 0.67655426 |
| IGIGGSDL | ASSSPPNERLF | 0.8125068 |
| IGTLLAEL | ASSSPPNERLF | 0.75735784 |
| IYTSGTT | ASSSPPNERLF | 0.73964024 |
| LSFPAGV | ASSSPPNERLF | 0.7635474 |
| LYTSGTT | ASSSPPNERLF | 0.78502285 |
| VGAFPFGF | ASSSPPNERLF | 0.71609366 |
| VGAGTVEF | ASSSPPNERLF | 0.769101 |
| VGKSTLLL | ASSSPPNERLF | 0.59756845 |
| AALAHAGGGIV | ASSSPPNERLF | 0.81718326 |
| AALVAVSLL | ASSSPPNERLF | 0.7953763 |
| AAVSRAVAAL | ASSSPPNERLF | 0.7956667 |
| AGGSSGIGGF | ASSSPPNERLF | 0.820345 |
| AGLTVANANG | ASSSPPNERLF | 0.77655333 |
| AGLVGAVL | ASSSPPNERLF | 0.73780596 |
| ALSQAFTV | ASSSPPNERLF | 0.4020153 |
| ALTFSVSL | ASSSPPNERLF | 0.672189 |
| ALTGAALVAL | ASSSPPNERLF | 0.72492063 |
| ASFSIGAGVL | ASSSPPNERLF | 0.7627154 |
| DGSVYKI | ASSSPPNERLF | 0.7841541 |
| EQLAEIV | ASSSPPNERLF | 0.77409804 |
| FAVSELTVM | ASSSPPNERLF | 0.6019157 |
| GAAPLAAGL | ASSSPPNERLF | 0.79828477 |
| GADGGDGGSGGAGG | ASSSPPNERLF | 0.8122412 |
| GFGLDTGGAGG | ASSSPPNERLF | 0.46726224 |
| GLASGGLL | ASSSPPNERLF | 0.65672255 |
| GRIAGAAVLL | ASSSPPNERLF | 0.8240307 |
| GSAGSLNTSF | ASSSPPNERLF | 0.7802249 |
| GTGQIGI | ASSSPPNERLF | 0.6232472 |
| GTLIALIL | ASSSPPNERLF | 0.63379776 |
| IAATMGFVVG | ASSSPPNERLF | 0.82963365 |
| IFAQALL | ASSSPPNERLF | 0.719305 |
| IGAAVALGF | ASSSPPNERLF | 0.8163814 |
| IGLTPLAL | ASSSPPNERLF | 0.2778937 |
| ILGSNTSTL | ASSSPPNERLF | 0.670013 |
| IVFSLTVL | ASSSPPNERLF | 0.55008155 |
| LASKATGF | ASSSPPNERLF | 0.207118 |
| LASVALIL | ASSSPPNERLF | 0.7831036 |
| LFAVGLTL | ASSSPPNERLF | 0.52913827 |
| LFNGGIGTYI | ASSSPPNERLF | 0.76063275 |
| LTLGAGII | ASSSPPNERLF | 0.6563594 |
| LVAAALGDI | ASSSPPNERLF | 0.8069788 |
| NTAFVQLGI | ASSSPPNERLF | 0.7749461 |
| QLFSGAEL | ASSSPPNERLF | 0.6908959 |
| SAATFITL | ASSSPPNERLF | 0.5176209 |
| SNYGLANLGV | ASSSPPNERLF | 0.7278751 |

| | | |
|---|---|---|
| SYGGAYAVM | ASSSPPNERLF | 0.80374354 |
| TTGVGGLTLG | ASSSPPNERLF | 0.8162694 |
| TTQTIYI | ASSSPPNERLF | 0.29359767 |
| VAAGAGLGAVI | ASSSPPNERLF | 0.772401 |
| VAAVVTLL | ASSSPPNERLF | 0.64293694 |
| VGFLGGLI | ASSSPPNERLF | 0.33525217 |
| VGVIPILL | ASSSPPNERLF | 0.23514758 |
| VIFASSLI | ASSSPPNERLF | 0.64729977 |
| VKGGDGGAAAG | ASSSPPNERLF | 0.7023945 |
| VTKTFTNI | ASSSPPNERLF | 0.7611104 |
| VVAAAILL | ASSSPPNERLF | 0.7994461 |
| YAALDVEL | ASSSPPNERLF | 0.62311 |
| LASVALLL | ASSSPPNERLF | 0.79124343 |
| LDAVHRV | ASSSPPNERLF | 0.69326377 |
| QGLSSSIF | ASSSPPNERLF | 0.14933474 |
| RGTFANIRL | ASSSPPNERLF | 0.64213943 |
| RQLWWGHRI | ASSSPPNERLF | 0.34611243 |
| YGPPGTGKTLL | ASSSPPNERLF | 0.8237301 |
| AALVGAPANVA | ASSSPPNERLF | 0.48963422 |
| AATFRRTE | ASSSPPNERLF | 0.6513029 |
| AGALGGGI | ASSSPPNERLF | 0.7064748 |
| AGGAGGAGGIGAG | ASSSPPNERLF | 0.79362154 |
| AGSGALGL | ASSSPPNERLF | 0.7259747 |
| AQIADQI | ASSSPPNERLF | 0.5847175 |
| ASAVAVTYL | ASSSPPNERLF | 0.27923855 |
| ASTVLGGTGG | ASSSPPNERLF | 0.7579439 |
| AVGGEPAK | ASSSPPNERLF | 0.76002 |
| CEQGTFGGTC | ASSSPPNERLF | 0.41079247 |
| DEGKAPAG | ASSSPPNERLF | 0.48452917 |
| DTVVFPAL | ASSSPPNERLF | 0.6013363 |
| FEGITMAG | ASSSPPNERLF | 0.3261039 |
| FPLVVQD | ASSSPPNERLF | 0.8112011 |
| GAFTTPQ | ASSSPPNERLF | 0.5092979 |
| GFGGFGTAGGVGGA | ASSSPPNERLF | 0.6248647 |
| GFLAQFL | ASSSPPNERLF | 0.8248056 |
| GGAGGAGGNAGVA | ASSSPPNERLF | 0.7721051 |
| GGVGTAGAGGAGGAGG | ASSSPPNERLF | 0.6888491 |
| GNGGGGDAGNAGSGGN | ASSSPPNERLF | 0.5274751 |
| GTGGAGGAGNDAGS | ASSSPPNERLF | 0.5077692 |
| GTTNTGWFNS | ASSSPPNERLF | 0.44595915 |
| IFMGPLFTL | ASSSPPNERLF | 0.5054789 |
| ILAHAIEAI | ASSSPPNERLF | 0.8092796 |
| IRSAIREAL | ASSSPPNERLF | 0.7028707 |
| ITFAGGG | ASSSPPNERLF | 0.44119376 |
| ITGMSGHQEV | ASSSPPNERLF | 0.78963125 |
| IVAGLACTL | ASSSPPNERLF | 0.8168877 |
| IVGGGLAGA | ASSSPPNERLF | 0.83082074 |
| IVTGVLNI | ASSSPPNERLF | 0.5093446 |
| LAQQGCI | ASSSPPNERLF | 0.67097455 |
| LCLHVEN | ASSSPPNERLF | 0.6318153 |
| LDAISDGS | ASSSPPNERLF | 0.6874703 |

| | | |
|---|---|---|
| LGGFAAML | ASSSPPNERLF | 0.5774669 |
| LGGKNPMI | ASSSPPNERLF | 0.43243417 |
| LGPKGVAAIL | ASSSPPNERLF | 0.80573475 |
| LMVGVSI | ASSSPPNERLF | 0.83707446 |
| LVGAAFVL | ASSSPPNERLF | 0.67009187 |
| LVLAVAPL | ASSSPPNERLF | 0.38649824 |
| MAAIAVIDAA | ASSSPPNERLF | 0.6786663 |
| NNGINAELQ | ASSSPPNERLF | 0.606732 |
| RGEPLVLVHG | ASSSPPNERLF | 0.557846 |
| RGGAGGAAGLI | ASSSPPNERLF | 0.6912497 |
| RSYRMHPAV | ASSSPPNERLF | 0.37372744 |
| SAAHLLAGQLA | ASSSPPNERLF | 0.6720197 |
| SAAVGSY | ASSSPPNERLF | 0.82660866 |
| SPHAVLHRI | ASSSPPNERLF | 0.71527183 |
| SQIAKVL | ASSSPPNERLF | 0.71605325 |
| TALGHLVP | ASSSPPNERLF | 0.81053317 |
| TAPTTSGAA | ASSSPPNERLF | 0.7729868 |
| TDGDDGNGGSPGT | ASSSPPNERLF | 0.8113065 |
| TDGLGEN | ASSSPPNERLF | 0.81865644 |
| TGGPPLPGQA | ASSSPPNERLF | 0.70588124 |
| TIPAITI | ASSSPPNERLF | 0.7218064 |
| TIPGFTI | ASSSPPNERLF | 0.7926225 |
| TSIASGALA | ASSSPPNERLF | 0.6704225 |
| TVTLPGAAI | ASSSPPNERLF | 0.7275567 |
| VASGLSAALP | ASSSPPNERLF | 0.68352693 |
| VGAGVIGI | ASSSPPNERLF | 0.7635908 |
| VGPKRFGV | ASSSPPNERLF | 0.7064137 |
| VNVAFPD | ASSSPPNERLF | 0.8278893 |
| VSMTNSGVS | ASSSPPNERLF | 0.24274042 |
| VTDVLGDLSV | ASSSPPNERLF | 0.42120016 |
| VVVVGAGFAGL | ASSSPPNERLF | 0.8096659 |
| VYTVGTVV | ASSSPPNERLF | 0.20350887 |
| WTTFSGP | ASSSPPNERLF | 0.2906956 |
| YDELVEAF | ASSSPPNERLF | 0.7412615 |
| YGTGGAGGNGG | ASSSPPNERLF | 0.8149157 |
| YTNVMAV | ASSSPPNERLF | 0.24015813 |
| FLEQVHQL | ASSSPPNERLF | 0.6413131 |
| GLSLLAVAV | ASSSPPNERLF | 0.3647136 |
| LEQVHQL | ASSSPPNERLF | 0.8108118 |
| LNAAGAI | ASSSPPNERLF | 0.69628155 |
| PAVHASL | ASSSPPNERLF | 0.4107716 |
| TELEDDA | ASSSPPNERLF | 0.7818477 |
| TPPGPPN | ASSSPPNERLF | 0.513348 |
| TPPPPPPP | ASSSPPNERLF | 0.75284827 |
| TQSPALI | TCSAEPDWGGEQY | 0.80587256 |
| EFAAAFPQ | TCSAEPDWGGEQY | 0.8296823 |
| LLSGNLDVL | TCSAEPDWGGEQY | 0.43698877 |
| GPANLGNYNI | TCSAEPDWGGEQY | 0.7895696 |
| TLGPTLP | TCSAEPDWGGEQY | 0.66042286 |
| VYAYSSCCK | TCSAEPDWGGEQY | 0.8358273 |
| APAPAPAPNI | TCSAEPDWGGEQY | 0.5883572 |

| | | |
|---|---|---|
| PGGPVPIP | TCSAEPDWGGEQY | 0.35628688 |
| FLTVALEP | TCSAEPDWGGEQY | 0.7933613 |
| SQYFGANL | TCSAEPDWGGEQY | 0.7836671 |
| GGVKPAA | TCSAEPDWGGEQY | 0.617607 |
| GGTGGAGGAGGW | TCSAEPDWGGEQY | 0.8062618 |
| GVGGVAGLGGAGG | TCSAEPDWGGEQY | 0.7002896 |
| VLLVLATTI | TCSAEPDWGGEQY | 0.686234 |
| IGIFAAML | TCSAEPDWGGEQY | 0.7958029 |
| AVGDLID | TCSAEPDWGGEQY | 0.78992 |
| NISPLPL | TCSAEPDWGGEQY | 0.45712596 |
| VTAVFVAI | TCSAEPDWGGEQY | 0.80439746 |
| SGDDNDAAL | TCSAEPDWGGEQY | 0.7054925 |
| AVAALAHS | TCSAEPDWGGEQY | 0.45584267 |
| GGVTVMATP | TCSAEPDWGGEQY | 0.828879 |
| AGLAANVRD | TCSAEPDWGGEQY | 0.7387073 |
| TNRPDLIDPALLRPG | TCSAEPDWGGEQY | 0.7881573 |
| LANLGVTAL | TCSAEPDWGGEQY | 0.73941934 |
| TGERIEL | TCSAEPDWGGEQY | 0.8323081 |
| NGEELAA | TCSAEPDWGGEQY | 0.49802452 |
| VLSLVWG | TCSAEPDWGGEQY | 0.7279253 |
| IGYALWL | TCSAEPDWGGEQY | 0.71010035 |
| LSLLLAPL | TCSAEPDWGGEQY | 0.8070327 |
| TLDEALK | TCSAEPDWGGEQY | 0.7444618 |
| TDGAGTAEGG | TCSAEPDWGGEQY | 0.7221451 |
| NAPGGAGGNGG | TCSAEPDWGGEQY | 0.76533496 |
| VSVFMGMGGNF | TCSAEPDWGGEQY | 0.8074847 |
| LASYLAAL | TCSAEPDWGGEQY | 0.7279328 |
| LVGGTPAL | TCSAEPDWGGEQY | 0.74652195 |
| VRNADVL | TCSAEPDWGGEQY | 0.8209605 |
| TCSSPFLVSAM | TCSAEPDWGGEQY | 0.43642277 |
| VHITDAE | TCSAEPDWGGEQY | 0.7018415 |
| EYAAPSF | TCSAEPDWGGEQY | 0.79292095 |
| VVDAFHGGL | TCSAEPDWGGEQY | 0.786834 |
| ASVVDPQ | TCSAEPDWGGEQY | 0.6984382 |
| SYNEFGGLS | TCSAEPDWGGEQY | 0.81582344 |
| DPDGSVY | TCSAEPDWGGEQY | 0.6584034 |
| MVSTIHATE | TCSAEPDWGGEQY | 0.77276444 |
| LAAGVADPV | TCSAEPDWGGEQY | 0.7235982 |
| TKEQIAA | TCSAEPDWGGEQY | 0.66601574 |
| SLASPAAL | TCSAEPDWGGEQY | 0.40495276 |
| IGLVFYGV | TCSAEPDWGGEQY | 0.7347457 |
| FLLDASNL | TCSAEPDWGGEQY | 0.7982886 |
| YQVGNFP | TCSAEPDWGGEQY | 0.7592935 |
| ESLLSQST | TCSAEPDWGGEQY | 0.25162563 |
| LFTVGFAAVG | TCSAEPDWGGEQY | 0.769346 |
| ATPPTPT | TCSAEPDWGGEQY | 0.77831244 |
| TDLATLG | TCSAEPDWGGEQY | 0.8298459 |
| SAGGTGR | TCSAEPDWGGEQY | 0.70986295 |
| GFSNSGS | TCSAEPDWGGEQY | 0.7677747 |
| GGNGGSGGTSV | TCSAEPDWGGEQY | 0.6179682 |
| GTSVATGGAGN | TCSAEPDWGGEQY | 0.73342824 |

| | | |
|---|---|---|
| NGGNGGSGGTS | TCSAEPDWGGEQY | 0.42602587 |
| AVVKFPH | TCSAEPDWGGEQY | 0.47267178 |
| TPPPPPP | TCSAEPDWGGEQY | 0.7105459 |
| AAPAAAAAPA | TCSAEPDWGGEQY | 0.51755273 |
| IALNEDL | TCSAEPDWGGEQY | 0.79518455 |
| IVNVSSM | TCSAEPDWGGEQY | 0.80387044 |
| TGIPPAPR | TCSAEPDWGGEQY | 0.8011232 |
| FDLGGGTF | TCSAEPDWGGEQY | 0.8111956 |
| FDLGGGTFDVS | TCSAEPDWGGEQY | 0.81471735 |
| NEPTAAA | TCSAEPDWGGEQY | 0.8113928 |
| NEPTAAALA | TCSAEPDWGGEQY | 0.5400665 |
| GSIINISSI | TCSAEPDWGGEQY | 0.7165327 |
| VAPGNAGTA | TCSAEPDWGGEQY | 0.341783 |
| TGGGASLE | TCSAEPDWGGEQY | 0.66414034 |
| VAVYNHYKRI | TCSAEPDWGGEQY | 0.58531535 |
| TDPAPATT | TCSAEPDWGGEQY | 0.73040485 |
| TNRPDILDPALLRPG | TCSAEPDWGGEQY | 0.18291546 |
| LTTTTTS | TCSAEPDWGGEQY | 0.77703667 |
| PTGSGKTL | TCSAEPDWGGEQY | 0.81011635 |
| TGSGKTL | TCSAEPDWGGEQY | 0.7106571 |
| TIGHVDHGK | TCSAEPDWGGEQY | 0.7513102 |
| TIGHVDHGKTT | TCSAEPDWGGEQY | 0.77232885 |
| GHVDHGKTT | TCSAEPDWGGEQY | 0.63692105 |
| AQFEHTLL | TCSAEPDWGGEQY | 0.77996194 |
| IVNQDEL | TCSAEPDWGGEQY | 0.6259134 |
| TNGSQFF | TCSAEPDWGGEQY | 0.7920202 |
| TGKGIVE | TCSAEPDWGGEQY | 0.79601866 |
| LISNASDA | TCSAEPDWGGEQY | 0.75046563 |
| AELRTLGELTKIAKAHGAQ | TCSAEPDWGGEQY | 0.6765255 |
| APNAVAAAEPSW | TCSAEPDWGGEQY | 0.712232 |
| APQVAGAATAVL | TCSAEPDWGGEQY | 0.4690472 |
| ARRDPAAAAAILPTDA | TCSAEPDWGGEQY | 0.7184272 |
| DAAARDVLAEAGLAEYF | TCSAEPDWGGEQY | 0.2821071 |
| DDQIVLSAKANLAKVEQ | TCSAEPDWGGEQY | 0.74434507 |
| DELLPDTGEDSDEDGH | TCSAEPDWGGEQY | 0.48043203 |
| ELPEAPANRATVPQD | TCSAEPDWGGEQY | 0.72170997 |
| FGQSKIERVTGIG | TCSAEPDWGGEQY | 0.68337166 |
| GAGGTGSTAGGAGGAGGAGGLY | TCSAEPDWGGEQY | 0.77473336 |
| GHGGTGAAGGNGGTGGAGGSGI | TCSAEPDWGGEQY | 0.5264714 |
| GNGGNGGAGGEGGPGGAGGAGA | TCSAEPDWGGEQY | 0.7871796 |
| GSTAGNGGNGGNATNGTIAGQP | TCSAEPDWGGEQY | 0.74449927 |
| GTGGDGGNGGNGGTGG | TCSAEPDWGGEQY | 0.36279252 |
| GVRRPAGIPADAPV | TCSAEPDWGGEQY | 0.80572045 |
| GYRQVTDAHLLTLARR | TCSAEPDWGGEQY | 0.68309426 |
| IAKMGQDGHDRG | TCSAEPDWGGEQY | 0.70930743 |
| KAEPSIEIQYT | TCSAEPDWGGEQY | 0.7874289 |
| KPVRIRMAADEDTVAVT | TCSAEPDWGGEQY | 0.71444285 |
| MPPRIAGMRLLVIKPEP | TCSAEPDWGGEQY | 0.5104108 |
| MKFYAVALFRSRGVPPTR | TCSAEPDWGGEQY | 0.67007273 |
| NAGGFGILGAVAHSP | TCSAEPDWGGEQY | 0.73232025 |
| NEGREITLAMAEM | TCSAEPDWGGEQY | 0.6418562 |

| | | |
|---|---|---|
| NELVEWSHPLGHRWRWE | TCSAEPDWGGEQY | 0.28028992 |
| NLGAGGTGGNGGNPG | TCSAEPDWGGEQY | 0.78848803 |
| NTEWPKWETVMAADPVD | TCSAEPDWGGEQY | 0.45720345 |
| PASAAPAETQP | TCSAEPDWGGEQY | 0.6447673 |
| QQSSDGKNGVRAAEITGE | TCSAEPDWGGEQY | 0.600298 |
| RAPDAVTTSWWKE | TCSAEPDWGGEQY | 0.7962496 |
| SGFGNVGAHTSGWFNQ | TCSAEPDWGGEQY | 0.7903753 |
| SGVAGAVGQAASVGGLKV | TCSAEPDWGGEQY | 0.51366526 |
| SNGGAGTGSAGGNGGT | TCSAEPDWGGEQY | 0.7654653 |
| TGGAGGEGGAAG | TCSAEPDWGGEQY | 0.73642397 |
| TGGDGGAAGAGGAPGHG | TCSAEPDWGGEQY | 0.7966662 |
| TMPPLLAAIANAGRHGV | TCSAEPDWGGEQY | 0.67116237 |
| TSSTAGNSGGA | TCSAEPDWGGEQY | 0.68880194 |
| VPATVPVGILC | TCSAEPDWGGEQY | 0.7372421 |
| VQRPDSGGVVLTGRIS | TCSAEPDWGGEQY | 0.68767595 |
| WTPNPVPATKGVPAGLVQAT | TCSAEPDWGGEQY | 0.463916 |
| YALKRALAKFSDRFDVVII | TCSAEPDWGGEQY | 0.44965017 |
| YLIGSVIV | TCSAEPDWGGEQY | 0.6915678 |
| YGQLVAVID | TCSAEPDWGGEQY | 0.4377805 |
| VTVGGMTLPA | TCSAEPDWGGEQY | 0.7228652 |
| VLALPAV | TCSAEPDWGGEQY | 0.82590824 |
| VIPPTLNYET | TCSAEPDWGGEQY | 0.7692512 |
| VGMGGMHPSSG | TCSAEPDWGGEQY | 0.3188886 |
| VGLAGGAATA | TCSAEPDWGGEQY | 0.64369124 |
| VGGAGGAGGSGP | TCSAEPDWGGEQY | 0.7445803 |
| VFAAFMV | TCSAEPDWGGEQY | 0.7745597 |
| TLYDKLR | TCSAEPDWGGEQY | 0.8074157 |
| TLAVFTVL | TCSAEPDWGGEQY | 0.83359945 |
| TGSVQPARV | TCSAEPDWGGEQY | 0.7854297 |
| TGGGAGSGGT | TCSAEPDWGGEQY | 0.5789188 |
| SHLETSL | TCSAEPDWGGEQY | 0.78736466 |
| SFTTGMT | TCSAEPDWGGEQY | 0.6742884 |
| QVEVAKRMGVRQPTVS | TCSAEPDWGGEQY | 0.682207 |
| QGSGGPALAM | TCSAEPDWGGEQY | 0.7247299 |
| MLAPSNP | TCSAEPDWGGEQY | 0.7866276 |
| LVNNAAI | TCSAEPDWGGEQY | 0.6616002 |
| LTGEINTLA | TCSAEPDWGGEQY | 0.63073635 |
| LSIFGPV | TCSAEPDWGGEQY | 0.41074246 |
| LLALLANL | TCSAEPDWGGEQY | 0.80849385 |
| LISNASDALDKLR | TCSAEPDWGGEQY | 0.6420882 |
| LINEAVP | TCSAEPDWGGEQY | 0.5994613 |
| LGYAFSGL | TCSAEPDWGGEQY | 0.6566485 |
| LDFVAVAL | TCSAEPDWGGEQY | 0.8241765 |
| LAQPGAE | TCSAEPDWGGEQY | 0.8183962 |
| ISNLGAVTGF | TCSAEPDWGGEQY | 0.62142044 |
| HQEDPGRI | TCSAEPDWGGEQY | 0.52951884 |
| GTPGEPTP | TCSAEPDWGGEQY | 0.7873593 |
| GFFNSGS | TCSAEPDWGGEQY | 0.5325425 |
| GAGGGGLTAGAN | TCSAEPDWGGEQY | 0.788798 |
| FVDALVNM | TCSAEPDWGGEQY | 0.73803365 |
| FLTVAPEL | TCSAEPDWGGEQY | 0.7254092 |

| | | |
|---|---|---|
| FETLTAL | TCSAEPDWGGEQY | 0.5685848 |
| DVINPIEL | TCSAEPDWGGEQY | 0.7823764 |
| APVSGAPV | TCSAEPDWGGEQY | 0.6634066 |
| ALVGGLAW | TCSAEPDWGGEQY | 0.79061866 |
| ALAAAAATAI | TCSAEPDWGGEQY | 0.3405791 |
| AFGLGLLL | TCSAEPDWGGEQY | 0.8047997 |
| AFFAAASL | TCSAEPDWGGEQY | 0.4925847 |
| AAAALPAFSPP | TCSAEPDWGGEQY | 0.5957198 |
| VVLAPAL | TCSAEPDWGGEQY | 0.2743764 |
| TIGHVDHGKTTL | TCSAEPDWGGEQY | 0.6796465 |
| LISNASDALDK | TCSAEPDWGGEQY | 0.7941222 |
| LISNASDAL | TCSAEPDWGGEQY | 0.8141661 |
| ISNASDALDK | TCSAEPDWGGEQY | 0.71977246 |
| NQAETLVYQTE | TCSAEPDWGGEQY | 0.77802104 |
| TGGTGGTAGAG | TCSAEPDWGGEQY | 0.82507026 |
| TGSTAGGAGGA | TCSAEPDWGGEQY | 0.5971508 |
| DPLAVPVAAGPV | TCSAEPDWGGEQY | 0.24723841 |
| GGTGGTPGNGGN | TCSAEPDWGGEQY | 0.7334787 |
| PAGLAGVDPATL | TCSAEPDWGGEQY | 0.76964104 |
| GNNLVVEAPHAGASV | TCSAEPDWGGEQY | 0.81569356 |
| KVTRIMSASTLEQPA | TCSAEPDWGGEQY | 0.7808783 |
| LGFALPIALPRRTVV | TCSAEPDWGGEQY | 0.40065527 |
| TRLGNLTARKLVVLV | TCSAEPDWGGEQY | 0.74774194 |
| WSQQRRLIYFPSAGP | TCSAEPDWGGEQY | 0.7736323 |
| GPNGNPDPMAAAVDIR | TCSAEPDWGGEQY | 0.49900788 |
| TQRCVSVADLRAAASR | TCSAEPDWGGEQY | 0.53218186 |
| QFVLDVAEHPVEVLTETV | TCSAEPDWGGEQY | 0.52706665 |
| GGAGGTGGAGGNGGTQAGDG | TCSAEPDWGGEQY | 0.44992256 |
| PLLATLRAIAVDVDAKPAQV | TCSAEPDWGGEQY | 0.7657481 |
| APKFRTGGYMIDFWGVGYQVAK | TCSAEPDWGGEQY | 0.64989704 |
| GAGGAGGAGGAAGGPGGTGGTG | TCSAEPDWGGEQY | 0.63228834 |
| IGALGLLATDVAPAGSVAFGVA | TCSAEPDWGGEQY | 0.686847 |
| AGGAGGIGGNGGAGGAGGVLMG | TCSAEPDWGGEQY | 0.5154183 |
| ADELIGLL | TCSAEPDWGGEQY | 0.58759093 |
| ADRALKA | TCSAEPDWGGEQY | 0.5823505 |
| AGAVGQVGGAGGA | TCSAEPDWGGEQY | 0.77425224 |
| AGGGGGLGGAP | TCSAEPDWGGEQY | 0.7958933 |
| AGGLLYGNGGAG | TCSAEPDWGGEQY | 0.74232066 |
| AGLESIGL | TCSAEPDWGGEQY | 0.7608996 |
| AGLKTGII | TCSAEPDWGGEQY | 0.8036264 |
| ASLAALLL | TCSAEPDWGGEQY | 0.41095233 |
| ASVINISSI | TCSAEPDWGGEQY | 0.56226355 |
| ATPPGPA | TCSAEPDWGGEQY | 0.7784943 |
| ATVASALV | TCSAEPDWGGEQY | 0.5695014 |
| ATVASLIL | TCSAEPDWGGEQY | 0.80918616 |
| AVNGPVLIRH | TCSAEPDWGGEQY | 0.71818537 |
| DGAAGAVGI | TCSAEPDWGGEQY | 0.7604196 |
| DPLLHGI | TCSAEPDWGGEQY | 0.76972955 |
| ELGAALGAL | TCSAEPDWGGEQY | 0.82604533 |
| ELGRTQAV | TCSAEPDWGGEQY | 0.7477972 |
| FVAGGYLS | TCSAEPDWGGEQY | 0.6376332 |

| | | |
|---|---|---|
| GASGGAGGAGGW | TCSAEPDWGGEQY | 0.787182 |
| GGAGAGGDGGAGGI | TCSAEPDWGGEQY | 0.41229373 |
| GTGHNGSQPGMGGNG | TCSAEPDWGGEQY | 0.7937157 |
| GYAAASASAS | TCSAEPDWGGEQY | 0.7811746 |
| HQVDVAAVQQG | TCSAEPDWGGEQY | 0.5169555 |
| IGAAIAHT | TCSAEPDWGGEQY | 0.6942653 |
| IGMIEVAQL | TCSAEPDWGGEQY | 0.82147723 |
| IRLAPEHTSAMD | TCSAEPDWGGEQY | 0.8014692 |
| IYNVSVL | TCSAEPDWGGEQY | 0.50248265 |
| KQGFLGM | TCSAEPDWGGEQY | 0.59160626 |
| LESVITQL | TCSAEPDWGGEQY | 0.49842095 |
| LGLGALVVV | TCSAEPDWGGEQY | 0.596874 |
| LIGNGGNGGSGGI | TCSAEPDWGGEQY | 0.3157418 |
| LNAINAPTL | TCSAEPDWGGEQY | 0.7760523 |
| LQLLNTGV | TCSAEPDWGGEQY | 0.60823864 |
| LQTSIEF | TCSAEPDWGGEQY | 0.8074055 |
| MSFAVSEL | TCSAEPDWGGEQY | 0.74548095 |
| NTGALGL | TCSAEPDWGGEQY | 0.79618514 |
| PPGAGGAGGAGTT | TCSAEPDWGGEQY | 0.62333566 |
| QLVGLGI | TCSAEPDWGGEQY | 0.6074327 |
| RFLEQVHQL | TCSAEPDWGGEQY | 0.32937932 |
| RGVGYVLR | TCSAEPDWGGEQY | 0.42916602 |
| SATGVFAGVI | TCSAEPDWGGEQY | 0.6040552 |
| SLPAITLP | TCSAEPDWGGEQY | 0.66569704 |
| STALAFTV | TCSAEPDWGGEQY | 0.4112451 |
| SVLTVGLGK | TCSAEPDWGGEQY | 0.825541 |
| TALAGAGSGI | TCSAEPDWGGEQY | 0.6460973 |
| TGLPFTGI | TCSAEPDWGGEQY | 0.77011114 |
| VAEHVTV | TCSAEPDWGGEQY | 0.6320408 |
| VDIVLNSL | TCSAEPDWGGEQY | 0.4674498 |
| VLTGGGAVL | TCSAEPDWGGEQY | 0.56676316 |
| VNVGLAYGI | TCSAEPDWGGEQY | 0.77074957 |
| VSSAAVGAV | TCSAEPDWGGEQY | 0.4449254 |
| VTSVYEAGV | TCSAEPDWGGEQY | 0.34005678 |
| WVALDRGA | TCSAEPDWGGEQY | 0.6192642 |
| YLVGSVVI | TCSAEPDWGGEQY | 0.7163471 |
| GPPGTGKTLL | TCSAEPDWGGEQY | 0.82367605 |
| GVGMTKF | TCSAEPDWGGEQY | 0.8091157 |
| IGIGGSDL | TCSAEPDWGGEQY | 0.73610115 |
| IGTLLAEL | TCSAEPDWGGEQY | 0.8302414 |
| IYTSGTT | TCSAEPDWGGEQY | 0.19224179 |
| LSFPAGV | TCSAEPDWGGEQY | 0.7082195 |
| LYTSGTT | TCSAEPDWGGEQY | 0.81498975 |
| VGAFPFGF | TCSAEPDWGGEQY | 0.762067 |
| VGAGTVEF | TCSAEPDWGGEQY | 0.64056075 |
| VGKSTLLL | TCSAEPDWGGEQY | 0.809932 |
| AALAHAGGGIV | TCSAEPDWGGEQY | 0.49940357 |
| AALVAVSLL | TCSAEPDWGGEQY | 0.63082665 |
| AAVSRAVAAL | TCSAEPDWGGEQY | 0.65246 |
| AGGSSGIGGF | TCSAEPDWGGEQY | 0.78250057 |
| AGLTVANANG | TCSAEPDWGGEQY | 0.77618074 |

| | | |
|---|---|---|
| AGLVGAVL | TCSAEPDWGGEQY | 0.38540804 |
| ALSQAFTV | TCSAEPDWGGEQY | 0.7570148 |
| ALTFSVSL | TCSAEPDWGGEQY | 0.8277682 |
| ALTGAALVAL | TCSAEPDWGGEQY | 0.60212684 |
| ASFSIGAGVL | TCSAEPDWGGEQY | 0.81843466 |
| DGSVYKI | TCSAEPDWGGEQY | 0.73015165 |
| EQLAEIV | TCSAEPDWGGEQY | 0.71752906 |
| FAVSELTVM | TCSAEPDWGGEQY | 0.57320416 |
| GAAPLAAGL | TCSAEPDWGGEQY | 0.47702253 |
| GADGGDGGSGGAGG | TCSAEPDWGGEQY | 0.7854043 |
| GFGLDTGGAGG | TCSAEPDWGGEQY | 0.5767767 |
| GLASGGLL | TCSAEPDWGGEQY | 0.7032347 |
| GRIAGAAVLL | TCSAEPDWGGEQY | 0.7099484 |
| GSAGSLNTSF | TCSAEPDWGGEQY | 0.83174205 |
| GTGQIGI | TCSAEPDWGGEQY | 0.76251775 |
| GTLIALIL | TCSAEPDWGGEQY | 0.7685174 |
| IAATMGFVVG | TCSAEPDWGGEQY | 0.5137227 |
| IFAQALL | TCSAEPDWGGEQY | 0.7698065 |
| IGAAVALGF | TCSAEPDWGGEQY | 0.7886977 |
| IGLTPLAL | TCSAEPDWGGEQY | 0.47364938 |
| ILGSNTSTL | TCSAEPDWGGEQY | 0.8095506 |
| IVFSLTVL | TCSAEPDWGGEQY | 0.49025387 |
| LASKATGF | TCSAEPDWGGEQY | 0.7545335 |
| LASVALIL | TCSAEPDWGGEQY | 0.76972014 |
| LFAVGLTL | TCSAEPDWGGEQY | 0.82024497 |
| LFNGGIGTYI | TCSAEPDWGGEQY | 0.6746454 |
| LTLGAGII | TCSAEPDWGGEQY | 0.7716561 |
| LVAAALGDI | TCSAEPDWGGEQY | 0.44648647 |
| NTAFVQLGI | TCSAEPDWGGEQY | 0.8008946 |
| QLFSGAEL | TCSAEPDWGGEQY | 0.7972314 |
| SAATFITL | TCSAEPDWGGEQY | 0.7761766 |
| SNYGLANLGV | TCSAEPDWGGEQY | 0.83430904 |
| SYGGAYAVM | TCSAEPDWGGEQY | 0.79500586 |
| TTGVGGLTLG | TCSAEPDWGGEQY | 0.27891976 |
| TTQTIYI | TCSAEPDWGGEQY | 0.67790633 |
| VAAGAGLGAVI | TCSAEPDWGGEQY | 0.8181279 |
| VAAVVTLL | TCSAEPDWGGEQY | 0.8150418 |
| VGFLGGLI | TCSAEPDWGGEQY | 0.80241895 |
| VGVIPILL | TCSAEPDWGGEQY | 0.75290775 |
| VIFASSLI | TCSAEPDWGGEQY | 0.67198324 |
| VKGGDGGAAAG | TCSAEPDWGGEQY | 0.81905407 |
| VTKTFTNI | TCSAEPDWGGEQY | 0.8400823 |
| VVAAAILL | TCSAEPDWGGEQY | 0.78643 |
| YAALDVEL | TCSAEPDWGGEQY | 0.62024975 |
| LASVALLL | TCSAEPDWGGEQY | 0.5344193 |
| LDAVHRV | TCSAEPDWGGEQY | 0.7610342 |
| QGLSSSIF | TCSAEPDWGGEQY | 0.7941649 |
| RGTFANIRL | TCSAEPDWGGEQY | 0.42632386 |
| RQLWWGHRI | TCSAEPDWGGEQY | 0.7091229 |
| YGPPGTGKTLL | TCSAEPDWGGEQY | 0.7072723 |
| AALVGAPANVA | TCSAEPDWGGEQY | 0.23802352 |

| | | |
|---|---|---|
| AATFRRTE | TCSAEPDWGGEQY | 0.79929173 |
| AGALGGGI | TCSAEPDWGGEQY | 0.8142053 |
| AGGAGGAGGIGAG | TCSAEPDWGGEQY | 0.74117553 |
| AGSGALGL | TCSAEPDWGGEQY | 0.48089278 |
| AQIADQI | TCSAEPDWGGEQY | 0.80706036 |
| ASAVAVTYL | TCSAEPDWGGEQY | 0.8131492 |
| ASTVLGGTGG | TCSAEPDWGGEQY | 0.82030517 |
| AVGGEPAK | TCSAEPDWGGEQY | 0.33452624 |
| CEQGTFGGTC | TCSAEPDWGGEQY | 0.20841733 |
| DEGKAPAG | TCSAEPDWGGEQY | 0.7198914 |
| DTVVFPAL | TCSAEPDWGGEQY | 0.80366147 |
| FEGITMAG | TCSAEPDWGGEQY | 0.5854755 |
| FPLVVQD | TCSAEPDWGGEQY | 0.46183193 |
| GAFTTPQ | TCSAEPDWGGEQY | 0.7338403 |
| GFGGFGTAGGVGGA | TCSAEPDWGGEQY | 0.80654263 |
| GFLAQFL | TCSAEPDWGGEQY | 0.3752822 |
| GGAGGAGGNAGVA | TCSAEPDWGGEQY | 0.44403547 |
| GGVGTAGAGGAGGAGG | TCSAEPDWGGEQY | 0.7683058 |
| GNGGGGDAGNAGSGGN | TCSAEPDWGGEQY | 0.80122703 |
| GTGGAGGAGNDAGS | TCSAEPDWGGEQY | 0.5766351 |
| GTTNTGWFNS | TCSAEPDWGGEQY | 0.6369364 |
| IFMGPLFTL | TCSAEPDWGGEQY | 0.73076427 |
| ILAHAIEAI | TCSAEPDWGGEQY | 0.79867417 |
| IRSAIREAL | TCSAEPDWGGEQY | 0.76639587 |
| ITFAGGG | TCSAEPDWGGEQY | 0.44739583 |
| ITGMSGHQEV | TCSAEPDWGGEQY | 0.8063195 |
| IVAGLACTL | TCSAEPDWGGEQY | 0.45070893 |
| IVGGGLAGA | TCSAEPDWGGEQY | 0.8019662 |
| IVTGVLNI | TCSAEPDWGGEQY | 0.68785703 |
| LAQQGCI | TCSAEPDWGGEQY | 0.7461648 |
| LCLHVEN | TCSAEPDWGGEQY | 0.6277673 |
| LDAISDGS | TCSAEPDWGGEQY | 0.54846287 |
| LGGFAAML | TCSAEPDWGGEQY | 0.4600402 |
| LGGKNPMI | TCSAEPDWGGEQY | 0.4674594 |
| LGPKGVAAIL | TCSAEPDWGGEQY | 0.80037785 |
| LMVGVSI | TCSAEPDWGGEQY | 0.5115943 |
| LVGAAFVL | TCSAEPDWGGEQY | 0.7943907 |
| LVLAVAPL | TCSAEPDWGGEQY | 0.6906755 |
| MAAIAVIDAA | TCSAEPDWGGEQY | 0.50862384 |
| NNGINAELQ | TCSAEPDWGGEQY | 0.78145623 |
| RGEPLVLVHG | TCSAEPDWGGEQY | 0.32700104 |
| RGGAGGAAGLI | TCSAEPDWGGEQY | 0.7327389 |
| RSYRMHPAV | TCSAEPDWGGEQY | 0.72732097 |
| SAAHLLAGQLA | TCSAEPDWGGEQY | 0.7684846 |
| SAAVGSY | TCSAEPDWGGEQY | 0.577752 |
| SPHAVLHRI | TCSAEPDWGGEQY | 0.76521647 |
| SQIAKVL | TCSAEPDWGGEQY | 0.76352257 |
| TALGHLVP | TCSAEPDWGGEQY | 0.51863766 |
| TAPTTSGAA | TCSAEPDWGGEQY | 0.7189918 |
| TDGDDGNGGSPGT | TCSAEPDWGGEQY | 0.8017595 |
| TDGLGEN | TCSAEPDWGGEQY | 0.7311574 |

| | | |
|---|---|---|
| TGGPPLPGQA | TCSAEPDWGGEQY | 0.3504768 |
| TIPAITI | TCSAEPDWGGEQY | 0.75998354 |
| TIPGFTI | TCSAEPDWGGEQY | 0.5764116 |
| TSIASGALA | TCSAEPDWGGEQY | 0.7607801 |
| TVTLPGAAI | TCSAEPDWGGEQY | 0.7210325 |
| VASGLSAALP | TCSAEPDWGGEQY | 0.5754417 |
| VGAGVIGI | TCSAEPDWGGEQY | 0.7725359 |
| VGPKRFGV | TCSAEPDWGGEQY | 0.530972 |
| VNVAFPD | TCSAEPDWGGEQY | 0.5960326 |
| VSMTNSGVS | TCSAEPDWGGEQY | 0.6384388 |
| VTDVLGDLSV | TCSAEPDWGGEQY | 0.675233 |
| VVVVGAGFAGL | TCSAEPDWGGEQY | 0.7871944 |
| VYTVGTVV | TCSAEPDWGGEQY | 0.45265198 |
| WTTFSGP | TCSAEPDWGGEQY | 0.756327 |
| YDELVEAF | TCSAEPDWGGEQY | 0.626315 |
| YGTGGAGGNGG | TCSAEPDWGGEQY | 0.4516495 |
| YTNVMAV | TCSAEPDWGGEQY | 0.70512336 |
| FLEQVHQL | TCSAEPDWGGEQY | 0.67353857 |
| GLSLLAVAV | TCSAEPDWGGEQY | 0.40836853 |
| LEQVHQL | TCSAEPDWGGEQY | 0.78929585 |
| LNAAGAI | TCSAEPDWGGEQY | 0.37395927 |
| PAVHASL | TCSAEPDWGGEQY | 0.75200295 |
| TELEDDA | TCSAEPDWGGEQY | 0.82539964 |
| TPPGPPN | TCSAEPDWGGEQY | 0.43751702 |
| TPPPPPPP | TCSAEPDWGGEQY | 0.79514384 |
| TQSPALI | TCSAEAGGDTGQLY | 0.7470944 |
| EFAAAFPQ | TCSAEAGGDTGQLY | 0.74454355 |
| LLSGNLDVL | TCSAEAGGDTGQLY | 0.63271165 |
| GPANLGNYNI | TCSAEAGGDTGQLY | 0.5988448 |
| TLGPTLP | TCSAEAGGDTGQLY | 0.82177 |
| VYAYSSCCK | TCSAEAGGDTGQLY | 0.7847643 |
| APAPAPAPNI | TCSAEAGGDTGQLY | 0.7353552 |
| PGGPVPIP | TCSAEAGGDTGQLY | 0.7565991 |
| FLTVALEP | TCSAEAGGDTGQLY | 0.79766965 |
| SQYFGANL | TCSAEAGGDTGQLY | 0.7975905 |
| GGVKPAA | TCSAEAGGDTGQLY | 0.6271876 |
| GGTGGAGGAGGW | TCSAEAGGDTGQLY | 0.33783305 |
| GVGGVAGLGGAGG | TCSAEAGGDTGQLY | 0.60802543 |
| VLLVLATTI | TCSAEAGGDTGQLY | 0.6342418 |
| IGIFAAML | TCSAEAGGDTGQLY | 0.5872159 |
| AVGDLID | TCSAEAGGDTGQLY | 0.39387453 |
| NISPLPL | TCSAEAGGDTGQLY | 0.793777 |
| VTAVFVAI | TCSAEAGGDTGQLY | 0.83481574 |
| SGDDNDAAL | TCSAEAGGDTGQLY | 0.61454475 |
| AVAALAHS | TCSAEAGGDTGQLY | 0.83496654 |
| GGVTVMATP | TCSAEAGGDTGQLY | 0.42776445 |
| AGLAANVRD | TCSAEAGGDTGQLY | 0.78103465 |
| TNRPDLIDPALLRPG | TCSAEAGGDTGQLY | 0.7962922 |
| LANLGVTAL | TCSAEAGGDTGQLY | 0.5246388 |
| TGERIEL | TCSAEAGGDTGQLY | 0.748968 |
| NGEELAA | TCSAEAGGDTGQLY | 0.5705502 |

| | | |
|---|---|---|
| VLSLVWG | TCSAEAGGDTGQLY | 0.59690654 |
| IGYALWL | TCSAEAGGDTGQLY | 0.7962061 |
| LSLLLAPL | TCSAEAGGDTGQLY | 0.5375377 |
| TLDEALK | TCSAEAGGDTGQLY | 0.755955 |
| TDGAGTAEGG | TCSAEAGGDTGQLY | 0.5552963 |
| NAPGGAGGNGG | TCSAEAGGDTGQLY | 0.46783137 |
| VSVFMGMGGNF | TCSAEAGGDTGQLY | 0.7571429 |
| LASYLAAL | TCSAEAGGDTGQLY | 0.7817626 |
| LVGGTPAL | TCSAEAGGDTGQLY | 0.74740136 |
| VRNADVL | TCSAEAGGDTGQLY | 0.83864903 |
| TCSSPFLVSAM | TCSAEAGGDTGQLY | 0.54005873 |
| VHITDAE | TCSAEAGGDTGQLY | 0.83214027 |
| EYAAPSF | TCSAEAGGDTGQLY | 0.81868774 |
| VVDAFHGGL | TCSAEAGGDTGQLY | 0.7456825 |
| ASVVDPQ | TCSAEAGGDTGQLY | 0.6686895 |
| SYNEFGGLS | TCSAEAGGDTGQLY | 0.79331267 |
| DPDGSVY | TCSAEAGGDTGQLY | 0.5853698 |
| MVSTIHATE | TCSAEAGGDTGQLY | 0.7581918 |
| LAAGVADPV | TCSAEAGGDTGQLY | 0.82498574 |
| TKEQIAA | TCSAEAGGDTGQLY | 0.7333739 |
| SLASPAAL | TCSAEAGGDTGQLY | 0.90537393 |
| IGLVFYGV | TCSAEAGGDTGQLY | 0.8582618 |
| FLLDASNL | TCSAEAGGDTGQLY | 0.815005 |
| YQVGNFP | TCSAEAGGDTGQLY | 0.75161654 |
| ESLLSQST | TCSAEAGGDTGQLY | 0.42027617 |
| LFTVGFAAVG | TCSAEAGGDTGQLY | 0.781881 |
| ATPPTPT | TCSAEAGGDTGQLY | 0.70208025 |
| TDLATLG | TCSAEAGGDTGQLY | 0.8620033 |
| SAGGTGR | TCSAEAGGDTGQLY | 0.8850529 |
| GFSNSGS | TCSAEAGGDTGQLY | 0.7409971 |
| GGNGGSGGTSV | TCSAEAGGDTGQLY | 0.8951198 |
| GTSVATGGAGN | TCSAEAGGDTGQLY | 0.87541544 |
| NGGNGGSGGTS | TCSAEAGGDTGQLY | 0.5017803 |
| AVVKFPH | TCSAEAGGDTGQLY | 0.8055968 |
| TPPPPPP | TCSAEAGGDTGQLY | 0.7588139 |
| AAPAAAAAPA | TCSAEAGGDTGQLY | 0.8170098 |
| IALNEDL | TCSAEAGGDTGQLY | 0.46162072 |
| IVNVSSM | TCSAEAGGDTGQLY | 0.8903614 |
| TGIPPAPR | TCSAEAGGDTGQLY | 0.77578896 |
| FDLGGGTF | TCSAEAGGDTGQLY | 0.19543104 |
| FDLGGGTFDVS | TCSAEAGGDTGQLY | 0.84341717 |
| NEPTAAA | TCSAEAGGDTGQLY | 0.8480513 |
| NEPTAAALA | TCSAEAGGDTGQLY | 0.8099717 |
| GSIINISSI | TCSAEAGGDTGQLY | 0.8900669 |
| VAPGNAGTA | TCSAEAGGDTGQLY | 0.7265143 |
| TGGGASLE | TCSAEAGGDTGQLY | 0.64380014 |
| VAVYNHYKRI | TCSAEAGGDTGQLY | 0.88210773 |
| TDPAPATT | TCSAEAGGDTGQLY | 0.90083957 |
| TNRPDILDPALLRPG | TCSAEAGGDTGQLY | 0.54397917 |
| LTTTTTS | TCSAEAGGDTGQLY | 0.868314 |
| PTGSGKTL | TCSAEAGGDTGQLY | 0.87821364 |

| | | |
|---|---|---|
| TGSGKTL | TCSAEAGGDTGQLY | 0.859524 |
| TIGHVDHGK | TCSAEAGGDTGQLY | 0.8975396 |
| TIGHVDHGKTT | TCSAEAGGDTGQLY | 0.9079838 |
| GHVDHGKTT | TCSAEAGGDTGQLY | 0.70777667 |
| AQFEHTLL | TCSAEAGGDTGQLY | 0.8976383 |
| IVNQDEL | TCSAEAGGDTGQLY | 0.8983159 |
| TNGSQFF | TCSAEAGGDTGQLY | 0.7737992 |
| TGKGIVE | TCSAEAGGDTGQLY | 0.8378929 |
| LISNASDA | TCSAEAGGDTGQLY | 0.6503963 |
| AELRTLGELTKIAKAHGAQ | TCSAEAGGDTGQLY | 0.8061557 |
| APNAVAAAEPSW | TCSAEAGGDTGQLY | 0.48846298 |
| APQVAGAATAVL | TCSAEAGGDTGQLY | 0.8687244 |
| ARRDPAAAAAILPTDA | TCSAEAGGDTGQLY | 0.89590347 |
| DAAARDVLAEAGLAEYF | TCSAEAGGDTGQLY | 0.80339104 |
| DDQIVLSAKANLAKVEQ | TCSAEAGGDTGQLY | 0.72094846 |
| DELLPDTGEDSDEDGH | TCSAEAGGDTGQLY | 0.6850589 |
| ELPEAPANRATVPQD | TCSAEAGGDTGQLY | 0.8463094 |
| FGQSKIERVTGIG | TCSAEAGGDTGQLY | 0.78685033 |
| GAGGTGSTAGGAGGAGGAGGLY | TCSAEAGGDTGQLY | 0.5405152 |
| GHGGTGAAGGNGGTGGAGGSGI | TCSAEAGGDTGQLY | 0.7922929 |
| GNGGNGGAGGEGGPGGAGGAGA | TCSAEAGGDTGQLY | 0.8805186 |
| GSTAGNGGNGGNATNGTIAGQP | TCSAEAGGDTGQLY | 0.8784864 |
| GTGGDGGNGGNGGTGG | TCSAEAGGDTGQLY | 0.5495314 |
| GVRRPAGIPADAPV | TCSAEAGGDTGQLY | 0.9032059 |
| GYRQVTDAHLLTLARR | TCSAEAGGDTGQLY | 0.87632936 |
| IAKMGQDGHDRG | TCSAEAGGDTGQLY | 0.8968537 |
| KAEPSIEIQYT | TCSAEAGGDTGQLY | 0.8160651 |
| KPVRIRMAADEDTVAVT | TCSAEAGGDTGQLY | 0.72832507 |
| MPPRIAGMRLLVIKPEP | TCSAEAGGDTGQLY | 0.8820354 |
| MKFYAVALFRSRGVPPTR | TCSAEAGGDTGQLY | 0.8071053 |
| NAGGFGILGAVAHSP | TCSAEAGGDTGQLY | 0.7112244 |
| NEGREITLAMAEM | TCSAEAGGDTGQLY | 0.8861362 |
| NELVEWSHPLGHRWRWE | TCSAEAGGDTGQLY | 0.79223907 |
| NLGAGGTGGNGGNPG | TCSAEAGGDTGQLY | 0.7634441 |
| NTEWPKWETVMAADPVD | TCSAEAGGDTGQLY | 0.73570395 |
| PASAAPAETQP | TCSAEAGGDTGQLY | 0.9068531 |
| QQSSDGKNGVRAAEITGE | TCSAEAGGDTGQLY | 0.67370105 |
| RAPDAVTTSWWKE | TCSAEAGGDTGQLY | 0.7158698 |
| SGFGNVGAHTSGWFNQ | TCSAEAGGDTGQLY | 0.7314818 |
| SGVAGAVGQAASVGGLKV | TCSAEAGGDTGQLY | 0.8368667 |
| SNGGAGTGSAGGNGGT | TCSAEAGGDTGQLY | 0.8928654 |
| TGGAGGEGGAAG | TCSAEAGGDTGQLY | 0.8700259 |
| TGGDGGAAGAGGAPGHG | TCSAEAGGDTGQLY | 0.8051972 |
| TMPPLLAAIANAGRHGV | TCSAEAGGDTGQLY | 0.8557659 |
| TSSTAGNSGGA | TCSAEAGGDTGQLY | 0.8752781 |
| VPATVPVGILC | TCSAEAGGDTGQLY | 0.8586204 |
| VQRPDSGGVVLTGRIS | TCSAEAGGDTGQLY | 0.86889076 |
| WTPNPVPATKGVPAGLVQAT | TCSAEAGGDTGQLY | 0.76240456 |
| YALKRALAKFSDRFDVVII | TCSAEAGGDTGQLY | 0.9061726 |
| YLIGSVIV | TCSAEAGGDTGQLY | 0.864987 |
| YGQLVAVID | TCSAEAGGDTGQLY | 0.894284 |

| | | |
|---|---|---|
| VTVGGMTLPA | TCSAEAGGDTGQLY | 0.8982304 |
| VLALPAV | TCSAEAGGDTGQLY | 0.8673521 |
| VIPPTLNYET | TCSAEAGGDTGQLY | 0.85498434 |
| VGMGGMHPSSG | TCSAEAGGDTGQLY | 0.31317592 |
| VGLAGGAATA | TCSAEAGGDTGQLY | 0.7088003 |
| VGGAGGAGGSGP | TCSAEAGGDTGQLY | 0.864539 |
| VFAAFMV | TCSAEAGGDTGQLY | 0.86249995 |
| TLYDKLR | TCSAEAGGDTGQLY | 0.8828355 |
| TLAVFTVL | TCSAEAGGDTGQLY | 0.8526911 |
| TGSVQPARV | TCSAEAGGDTGQLY | 0.76181936 |
| TGGGAGSGGT | TCSAEAGGDTGQLY | 0.88903606 |
| SHLETSL | TCSAEAGGDTGQLY | 0.8831861 |
| SFTTGMT | TCSAEAGGDTGQLY | 0.508038 |
| QVEVAKRMGVRQPTVS | TCSAEAGGDTGQLY | 0.7847306 |
| QGSGGPALAM | TCSAEAGGDTGQLY | 0.91359746 |
| MLAPSNP | TCSAEAGGDTGQLY | 0.8421711 |
| LVNNAAI | TCSAEAGGDTGQLY | 0.80207574 |
| LTGEINTLA | TCSAEAGGDTGQLY | 0.7360128 |
| LSIFGPV | TCSAEAGGDTGQLY | 0.9006438 |
| LLALLANL | TCSAEAGGDTGQLY | 0.78887427 |
| LISNASDALDKLR | TCSAEAGGDTGQLY | 0.9054676 |
| LINEAVP | TCSAEAGGDTGQLY | 0.2121713 |
| LGYAFSGL | TCSAEAGGDTGQLY | 0.7633532 |
| LDFVAVAL | TCSAEAGGDTGQLY | 0.6426407 |
| LAQPGAE | TCSAEAGGDTGQLY | 0.47486624 |
| ISNLGAVTGF | TCSAEAGGDTGQLY | 0.8855045 |
| HQEDPGRI | TCSAEAGGDTGQLY | 0.7134006 |
| GTPGEPTP | TCSAEAGGDTGQLY | 0.85124135 |
| GFFNSGS | TCSAEAGGDTGQLY | 0.83834994 |
| GAGGGGLTAGAN | TCSAEAGGDTGQLY | 0.8968235 |
| FVDALVNM | TCSAEAGGDTGQLY | 0.84026074 |
| FLTVAPEL | TCSAEAGGDTGQLY | 0.83842325 |
| FETLTAL | TCSAEAGGDTGQLY | 0.7292795 |
| DVINPIEL | TCSAEAGGDTGQLY | 0.8719569 |
| APVSGAPV | TCSAEAGGDTGQLY | 0.90157413 |
| ALVGGLAW | TCSAEAGGDTGQLY | 0.8848249 |
| ALAAAAATAI | TCSAEAGGDTGQLY | 0.39005286 |
| AFGLGLLL | TCSAEAGGDTGQLY | 0.818743 |
| AFFAAASL | TCSAEAGGDTGQLY | 0.82887065 |
| AAAALPAFSPP | TCSAEAGGDTGQLY | 0.3729691 |
| VVLAPAL | TCSAEAGGDTGQLY | 0.34698308 |
| TIGHVDHGKTTL | TCSAEAGGDTGQLY | 0.8301871 |
| LISNASDALDK | TCSAEAGGDTGQLY | 0.7771088 |
| LISNASDAL | TCSAEAGGDTGQLY | 0.8909138 |
| ISNASDALDK | TCSAEAGGDTGQLY | 0.87724805 |
| NQAETLVYQTE | TCSAEAGGDTGQLY | 0.8142915 |
| TGGTGGTAGAG | TCSAEAGGDTGQLY | 0.8802762 |
| TGSTAGGAGGA | TCSAEAGGDTGQLY | 0.82961595 |
| DPLAVPVAAGPV | TCSAEAGGDTGQLY | 0.106508926 |
| GGTGGTPGNGGN | TCSAEAGGDTGQLY | 0.7604341 |
| PAGLAGVDPATL | TCSAEAGGDTGQLY | 0.5346716 |

| | | |
|---|---|---|
| GNNLVVEAPHAGASV | TCSAEAGGDTGQLY | 0.9045857 |
| KVTRIMSASTLEQPA | TCSAEAGGDTGQLY | 0.7305869 |
| LGFALPIALPRRTVV | TCSAEAGGDTGQLY | 0.71787417 |
| TRLGNLTARKLVVLV | TCSAEAGGDTGQLY | 0.82955337 |
| WSQQRRLIYFPSAGP | TCSAEAGGDTGQLY | 0.8364758 |
| GPNGNPDPMAAAVDIR | TCSAEAGGDTGQLY | 0.8240831 |
| TQRCVSVADLRAAASR | TCSAEAGGDTGQLY | 0.7185528 |
| QFVLDVAEHPVEVLTETV | TCSAEAGGDTGQLY | 0.25027487 |
| GGAGGTGGAGGNGGTQAGDG | TCSAEAGGDTGQLY | 0.87770265 |
| PLLATLRAIAVDVDAKPAQV | TCSAEAGGDTGQLY | 0.6547281 |
| APKFRTGGYMIDFWGVGYQVAK | TCSAEAGGDTGQLY | 0.5843645 |
| GAGGAGGAGGAAGGPGGTGGTG | TCSAEAGGDTGQLY | 0.38223636 |
| IGALGLLATDVAPAGSVAFGVA | TCSAEAGGDTGQLY | 0.8226724 |
| AGGAGGIGGNGGAGGAGGVLMG | TCSAEAGGDTGQLY | 0.38759667 |
| ADELIGLL | TCSAEAGGDTGQLY | 0.89045763 |
| ADRALKA | TCSAEAGGDTGQLY | 0.7980856 |
| AGAVGQVGGAGGA | TCSAEAGGDTGQLY | 0.7975513 |
| AGGGGGLGGAP | TCSAEAGGDTGQLY | 0.9100474 |
| AGGLLYGNGGAG | TCSAEAGGDTGQLY | 0.80491394 |
| AGLESIGL | TCSAEAGGDTGQLY | 0.80869 |
| AGLKTGII | TCSAEAGGDTGQLY | 0.69307333 |
| ASLAALLL | TCSAEAGGDTGQLY | 0.71631205 |
| ASVINISSI | TCSAEAGGDTGQLY | 0.7035103 |
| ATPPGPA | TCSAEAGGDTGQLY | 0.6607512 |
| ATVASALV | TCSAEAGGDTGQLY | 0.908265 |
| ATVASLIL | TCSAEAGGDTGQLY | 0.78538835 |
| AVNGPVLIRH | TCSAEAGGDTGQLY | 0.4867163 |
| DGAAGAVGI | TCSAEAGGDTGQLY | 0.8267513 |
| DPLLHGI | TCSAEAGGDTGQLY | 0.9080306 |
| ELGAALGAL | TCSAEAGGDTGQLY | 0.91418874 |
| ELGRTQAV | TCSAEAGGDTGQLY | 0.7406217 |
| FVAGGYLS | TCSAEAGGDTGQLY | 0.7061384 |
| GASGGAGGAGGW | TCSAEAGGDTGQLY | 0.7342459 |
| GGAGAGGDGGAGGI | TCSAEAGGDTGQLY | 0.82664084 |
| GTGHNGSQPGMGGNG | TCSAEAGGDTGQLY | 0.7517108 |
| GYAAASASAS | TCSAEAGGDTGQLY | 0.56678784 |
| HQVDVAAVQQG | TCSAEAGGDTGQLY | 0.89559925 |
| IGAAIAHT | TCSAEAGGDTGQLY | 0.9052513 |
| IGMIEVAQL | TCSAEAGGDTGQLY | 0.8036018 |
| IRLAPEHTSAMD | TCSAEAGGDTGQLY | 0.48336512 |
| IYNVSVL | TCSAEAGGDTGQLY | 0.81793016 |
| KQGFLGM | TCSAEAGGDTGQLY | 0.7294246 |
| LESVITQL | TCSAEAGGDTGQLY | 0.796305 |
| LGLGALVVV | TCSAEAGGDTGQLY | 0.82535106 |
| LIGNGGNGGSGGI | TCSAEAGGDTGQLY | 0.5140108 |
| LNAINAPTL | TCSAEAGGDTGQLY | 0.7994864 |
| LQLLNTGV | TCSAEAGGDTGQLY | 0.90999186 |
| LQTSIEF | TCSAEAGGDTGQLY | 0.7715607 |
| MSFAVSEL | TCSAEAGGDTGQLY | 0.848396 |
| NTGALGL | TCSAEAGGDTGQLY | 0.9014766 |
| PPGAGGAGGAGTT | TCSAEAGGDTGQLY | 0.88063717 |

| | | |
|---|---|---|
| QLVGLGI | TCSAEAGGDTGQLY | 0.9031505 |
| RFLEQVHQL | TCSAEAGGDTGQLY | 0.8807776 |
| RGVGYVLR | TCSAEAGGDTGQLY | 0.8288038 |
| SATGVFAGVI | TCSAEAGGDTGQLY | 0.8542572 |
| SLPAITLP | TCSAEAGGDTGQLY | 0.86337405 |
| STALAFTV | TCSAEAGGDTGQLY | 0.83397114 |
| SVLTVGLGK | TCSAEAGGDTGQLY | 0.81404185 |
| TALAGAGSGI | TCSAEAGGDTGQLY | 0.7503495 |
| TGLPFTGI | TCSAEAGGDTGQLY | 0.88841105 |
| VAEHVTV | TCSAEAGGDTGQLY | 0.82508504 |
| VDIVLNSL | TCSAEAGGDTGQLY | 0.89192903 |
| VLTGGGAVL | TCSAEAGGDTGQLY | 0.5891191 |
| VNVGLAYGI | TCSAEAGGDTGQLY | 0.6333449 |
| VSSAAVGAV | TCSAEAGGDTGQLY | 0.8931756 |
| VTSVYEAGV | TCSAEAGGDTGQLY | 0.13685367 |
| WVALDRGA | TCSAEAGGDTGQLY | 0.28418583 |
| YLVGSVVI | TCSAEAGGDTGQLY | 0.8449013 |
| GPPGTGKTLL | TCSAEAGGDTGQLY | 0.8994678 |
| GVGMTKF | TCSAEAGGDTGQLY | 0.18575485 |
| IGIGGSDL | TCSAEAGGDTGQLY | 0.6230624 |
| IGTLLAEL | TCSAEAGGDTGQLY | 0.59601164 |
| IYTSGTT | TCSAEAGGDTGQLY | 0.88056684 |
| LSFPAGV | TCSAEAGGDTGQLY | 0.816998 |
| LYTSGTT | TCSAEAGGDTGQLY | 0.5011667 |
| VGAFPFGF | TCSAEAGGDTGQLY | 0.8828052 |
| VGAGTVEF | TCSAEAGGDTGQLY | 0.4851715 |
| VGKSTLLL | TCSAEAGGDTGQLY | 0.7612265 |
| AALAHAGGGIV | TCSAEAGGDTGQLY | 0.88935137 |
| AALVAVSLL | TCSAEAGGDTGQLY | 0.8987736 |
| AAVSRAVAAL | TCSAEAGGDTGQLY | 0.6396302 |
| AGGSSGIGGF | TCSAEAGGDTGQLY | 0.88290966 |
| AGLTVANANG | TCSAEAGGDTGQLY | 0.78265554 |
| AGLVGAVL | TCSAEAGGDTGQLY | 0.91502285 |
| ALSQAFTV | TCSAEAGGDTGQLY | 0.768098 |
| ALTFSVSL | TCSAEAGGDTGQLY | 0.5356358 |
| ALTGAALVAL | TCSAEAGGDTGQLY | 0.88563395 |
| ASFSIGAGVL | TCSAEAGGDTGQLY | 0.8660958 |
| DGSVYKI | TCSAEAGGDTGQLY | 0.7721803 |
| EQLAEIV | TCSAEAGGDTGQLY | 0.8842542 |
| FAVSELTVM | TCSAEAGGDTGQLY | 0.80071414 |
| GAAPLAAGL | TCSAEAGGDTGQLY | 0.8212209 |
| GADGGDGGSGGAGG | TCSAEAGGDTGQLY | 0.897915 |
| GFGLDTGGAGG | TCSAEAGGDTGQLY | 0.89644265 |
| GLASGGLL | TCSAEAGGDTGQLY | 0.66736025 |
| GRIAGAAVLL | TCSAEAGGDTGQLY | 0.89102626 |
| GSAGSLNTSF | TCSAEAGGDTGQLY | 0.79890454 |
| GTGQIGI | TCSAEAGGDTGQLY | 0.6586443 |
| GTLIALIL | TCSAEAGGDTGQLY | 0.9077978 |
| IAATMGFVVG | TCSAEAGGDTGQLY | 0.8428987 |
| IFAQALL | TCSAEAGGDTGQLY | 0.8868887 |
| IGAAVALGF | TCSAEAGGDTGQLY | 0.8745306 |

| | | |
|---|---|---|
| IGLTPLAL | TCSAEAGGDTGQLY | 0.87014097 |
| ILGSNTSTL | TCSAEAGGDTGQLY | 0.5225866 |
| IVFSLTVL | TCSAEAGGDTGQLY | 0.79517555 |
| LASKATGF | TCSAEAGGDTGQLY | 0.81160206 |
| LASVALIL | TCSAEAGGDTGQLY | 0.88717234 |
| LFAVGLTL | TCSAEAGGDTGQLY | 0.817153 |
| LFNGGIGTYI | TCSAEAGGDTGQLY | 0.72129464 |
| LTLGAGII | TCSAEAGGDTGQLY | 0.88017774 |
| LVAAALGDI | TCSAEAGGDTGQLY | 0.8961667 |
| NTAFVQLGI | TCSAEAGGDTGQLY | 0.8552904 |
| QLFSGAEL | TCSAEAGGDTGQLY | 0.8510534 |
| SAATFITL | TCSAEAGGDTGQLY | 0.9068285 |
| SNYGLANLGV | TCSAEAGGDTGQLY | 0.70918006 |
| SYGGAYAVM | TCSAEAGGDTGQLY | 0.78838515 |
| TTGVGGLTLG | TCSAEAGGDTGQLY | 0.8772346 |
| TTQTIYI | TCSAEAGGDTGQLY | 0.8919765 |
| VAAGAGLGAVI | TCSAEAGGDTGQLY | 0.83121 |
| VAAVVTLL | TCSAEAGGDTGQLY | 0.9030893 |
| VGFLGGLI | TCSAEAGGDTGQLY | 0.7305777 |
| VGVIPILL | TCSAEAGGDTGQLY | 0.88063407 |
| VIFASSLI | TCSAEAGGDTGQLY | 0.7354176 |
| VKGGDGGAAAG | TCSAEAGGDTGQLY | 0.84895325 |
| VTKTFTNI | TCSAEAGGDTGQLY | 0.7552677 |
| VVAAAILL | TCSAEAGGDTGQLY | 0.8695831 |
| YAALDVEL | TCSAEAGGDTGQLY | 0.8879652 |
| LASVALLL | TCSAEAGGDTGQLY | 0.8776081 |
| LDAVHRV | TCSAEAGGDTGQLY | 0.24449304 |
| QGLSSSIF | TCSAEAGGDTGQLY | 0.8947257 |
| RGTFANIRL | TCSAEAGGDTGQLY | 0.8633426 |
| RQLWWGHRI | TCSAEAGGDTGQLY | 0.9117234 |
| YGPPGTGKTLL | TCSAEAGGDTGQLY | 0.84139156 |
| AALVGAPANVA | TCSAEAGGDTGQLY | 0.8451131 |
| AATFRRTE | TCSAEAGGDTGQLY | 0.7592213 |
| AGALGGGI | TCSAEAGGDTGQLY | 0.83126295 |
| AGGAGGAGGIGAG | TCSAEAGGDTGQLY | 0.6723739 |
| AGSGALGL | TCSAEAGGDTGQLY | 0.71831346 |
| AQIADQI | TCSAEAGGDTGQLY | 0.78657377 |
| ASAVAVTYL | TCSAEAGGDTGQLY | 0.75882185 |
| ASTVLGGTGG | TCSAEAGGDTGQLY | 0.879424 |
| AVGGEPAK | TCSAEAGGDTGQLY | 0.89165914 |
| CEQGTFGGTC | TCSAEAGGDTGQLY | 0.8813263 |
| DEGKAPAG | TCSAEAGGDTGQLY | 0.8994207 |
| DTVVFPAL | TCSAEAGGDTGQLY | 0.9018322 |
| FEGITMAG | TCSAEAGGDTGQLY | 0.89760613 |
| FPLVVQD | TCSAEAGGDTGQLY | 0.7378306 |
| GAFTTPQ | TCSAEAGGDTGQLY | 0.8015158 |
| GFGGFGTAGGVGGA | TCSAEAGGDTGQLY | 0.65540564 |
| GFLAQFL | TCSAEAGGDTGQLY | 0.8217213 |
| GGAGGAGGNAGVA | TCSAEAGGDTGQLY | 0.7821902 |
| GGVGTAGAGGAGGAGG | TCSAEAGGDTGQLY | 0.86537045 |
| GNGGGGDAGNAGSGGN | TCSAEAGGDTGQLY | 0.14268056 |

| | | |
|---|---|---|
| GTGGAGGAGNDAGS | TCSAEAGGDTGQLY | 0.83656925 |
| GTTNTGWFNS | TCSAEAGGDTGQLY | 0.9072814 |
| IFMGPLFTL | TCSAEAGGDTGQLY | 0.74844146 |
| ILAHAIEAI | TCSAEAGGDTGQLY | 0.83932495 |
| IRSAIREAL | TCSAEAGGDTGQLY | 0.8536409 |
| ITFAGGG | TCSAEAGGDTGQLY | 0.79421556 |
| ITGMSGHQEV | TCSAEAGGDTGQLY | 0.8788929 |
| IVAGLACTL | TCSAEAGGDTGQLY | 0.7345603 |
| IVGGGLAGA | TCSAEAGGDTGQLY | 0.8715426 |
| IVTGVLNI | TCSAEAGGDTGQLY | 0.9025411 |
| LAQQGCI | TCSAEAGGDTGQLY | 0.84290195 |
| LCLHVEN | TCSAEAGGDTGQLY | 0.8364028 |
| LDAISDGS | TCSAEAGGDTGQLY | 0.8188504 |
| LGGFAAML | TCSAEAGGDTGQLY | 0.41107136 |
| LGGKNPMI | TCSAEAGGDTGQLY | 0.8572614 |
| LGPKGVAAIL | TCSAEAGGDTGQLY | 0.62708306 |
| LMVGVSI | TCSAEAGGDTGQLY | 0.8295666 |
| LVGAAFVL | TCSAEAGGDTGQLY | 0.70229053 |
| LVLAVAPL | TCSAEAGGDTGQLY | 0.8406248 |
| MAAIAVIDAA | TCSAEAGGDTGQLY | 0.8303031 |
| NNGINAELQ | TCSAEAGGDTGQLY | 0.8665106 |
| RGEPLVLVHG | TCSAEAGGDTGQLY | 0.64861095 |
| RGGAGGAAGLI | TCSAEAGGDTGQLY | 0.8379157 |
| RSYRMHPAV | TCSAEAGGDTGQLY | 0.78746563 |
| SAAHLLAGQLA | TCSAEAGGDTGQLY | 0.51030844 |
| SAAVGSY | TCSAEAGGDTGQLY | 0.8733599 |
| SPHAVLHRI | TCSAEAGGDTGQLY | 0.79655063 |
| SQIAKVL | TCSAEAGGDTGQLY | 0.8220346 |
| TALGHLVP | TCSAEAGGDTGQLY | 0.88841033 |
| TAPTTSGAA | TCSAEAGGDTGQLY | 0.83761334 |
| TDGDDGNGGSPGT | TCSAEAGGDTGQLY | 0.43953705 |
| TDGLGEN | TCSAEAGGDTGQLY | 0.812495 |
| TGGPPLPGQA | TCSAEAGGDTGQLY | 0.8214979 |
| TIPAITI | TCSAEAGGDTGQLY | 0.7977914 |
| TIPGFTI | TCSAEAGGDTGQLY | 0.20559926 |
| TSIASGALA | TCSAEAGGDTGQLY | 0.8791344 |
| TVTLPGAAI | TCSAEAGGDTGQLY | 0.6548388 |
| VASGLSAALP | TCSAEAGGDTGQLY | 0.70344853 |
| VGAGVIGI | TCSAEAGGDTGQLY | 0.76473486 |
| VGPKRFGV | TCSAEAGGDTGQLY | 0.87928414 |
| VNVAFPD | TCSAEAGGDTGQLY | 0.79586405 |
| VSMTNSGVS | TCSAEAGGDTGQLY | 0.744235 |
| VTDVLGDLSV | TCSAEAGGDTGQLY | 0.876114 |
| VVVVGAGFAGL | TCSAEAGGDTGQLY | 0.8577323 |
| VYTVGTVV | TCSAEAGGDTGQLY | 0.8975682 |
| WTTFSGP | TCSAEAGGDTGQLY | 0.84654844 |
| YDELVEAF | TCSAEAGGDTGQLY | 0.8024533 |
| YGTGGAGGNGG | TCSAEAGGDTGQLY | 0.8445016 |
| YTNVMAV | TCSAEAGGDTGQLY | 0.7963561 |
| FLEQVHQL | TCSAEAGGDTGQLY | 0.6234458 |
| GLSLLAVAV | TCSAEAGGDTGQLY | 0.48566625 |

| | | |
|---|---|---|
| LEQVHQL | TCSAEAGGDTGQLY | 0.83605516 |
| LNAAGAI | TCSAEAGGDTGQLY | 0.45367873 |
| PAVHASL | TCSAEAGGDTGQLY | 0.8181181 |
| TELEDDA | TCSAEAGGDTGQLY | 0.8992865 |
| TPPGPPN | TCSAEAGGDTGQLY | 0.81039375 |
| TPPPPPPP | TCSAEAGGDTGQLY | 0.5284867 |
| TQSPALI | ASRTGTDHNSPLY | 0.80376935 |
| EFAAAFPQ | ASRTGTDHNSPLY | 0.79929376 |
| LLSGNLDVL | ASRTGTDHNSPLY | 0.8846226 |
| GPANLGNYNI | ASRTGTDHNSPLY | 0.88756967 |
| TLGPTLP | ASRTGTDHNSPLY | 0.90850943 |
| VYAYSSCCK | ASRTGTDHNSPLY | 0.8818165 |
| APAPAPAPNI | ASRTGTDHNSPLY | 0.7560774 |
| PGGPVPIP | ASRTGTDHNSPLY | 0.87976384 |
| FLTVALEP | ASRTGTDHNSPLY | 0.81557083 |
| SQYFGANL | ASRTGTDHNSPLY | 0.7826345 |
| GGVKPAA | ASRTGTDHNSPLY | 0.84275144 |
| GGTGGAGGAGGW | ASRTGTDHNSPLY | 0.85720766 |
| GVGGVAGLGGAGG | ASRTGTDHNSPLY | 0.8347097 |
| VLLVLATTI | ASRTGTDHNSPLY | 0.7699609 |
| IGIFAAML | ASRTGTDHNSPLY | 0.46435326 |
| AVGDLID | ASRTGTDHNSPLY | 0.878539 |
| NISPLPL | ASRTGTDHNSPLY | 0.8820998 |
| VTAVFVAI | ASRTGTDHNSPLY | 0.657774 |
| SGDDNDAAL | ASRTGTDHNSPLY | 0.77600735 |
| AVAALAHS | ASRTGTDHNSPLY | 0.9083729 |
| GGVTVMATP | ASRTGTDHNSPLY | 0.8976458 |
| AGLAANVRD | ASRTGTDHNSPLY | 0.7431015 |
| TNRPDLIDPALLRPG | ASRTGTDHNSPLY | 0.74264073 |
| LANLGVTAL | ASRTGTDHNSPLY | 0.84640217 |
| TGERIEL | ASRTGTDHNSPLY | 0.48902798 |
| NGEELAA | ASRTGTDHNSPLY | 0.8778779 |
| VLSLVWG | ASRTGTDHNSPLY | 0.8380413 |
| IGYALWL | ASRTGTDHNSPLY | 0.8266375 |
| LSLLLAPL | ASRTGTDHNSPLY | 0.6882552 |
| TLDEALK | ASRTGTDHNSPLY | 0.85065186 |
| TDGAGTAEGG | ASRTGTDHNSPLY | 0.78281397 |
| NAPGGAGGNGG | ASRTGTDHNSPLY | 0.88220346 |
| VSVFMGMGGNF | ASRTGTDHNSPLY | 0.18658903 |
| LASYLAAL | ASRTGTDHNSPLY | 0.8738939 |
| LVGGTPAL | ASRTGTDHNSPLY | 0.7072525 |
| VRNADVL | ASRTGTDHNSPLY | 0.74294126 |
| TCSSPFLVSAM | ASRTGTDHNSPLY | 0.43020248 |
| VHITDAE | ASRTGTDHNSPLY | 0.79237586 |
| EYAAPSF | ASRTGTDHNSPLY | 0.8831456 |
| VVDAFHGGL | ASRTGTDHNSPLY | 0.8904268 |
| ASVVDPQ | ASRTGTDHNSPLY | 0.75641936 |
| SYNEFGGLS | ASRTGTDHNSPLY | 0.88958114 |
| DPDGSVY | ASRTGTDHNSPLY | 0.9084853 |
| MVSTIHATE | ASRTGTDHNSPLY | 0.8063832 |
| LAAGVADPV | ASRTGTDHNSPLY | 0.3735406 |

| | | |
|---|---|---|
| TKEQIAA | ASRTGTDHNSPLY | 0.8593482 |
| SLASPAAL | ASRTGTDHNSPLY | 0.7816845 |
| IGLVFYGV | ASRTGTDHNSPLY | 0.88788605 |
| FLLDASNL | ASRTGTDHNSPLY | 0.8493235 |
| YQVGNFP | ASRTGTDHNSPLY | 0.68197787 |
| ESLLSQST | ASRTGTDHNSPLY | 0.86924016 |
| LFTVGFAAVG | ASRTGTDHNSPLY | 0.8922812 |
| ATPPTPT | ASRTGTDHNSPLY | 0.7353176 |
| TDLATLG | ASRTGTDHNSPLY | 0.6691606 |
| SAGGTGR | ASRTGTDHNSPLY | 0.6784284 |
| GFSNSGS | ASRTGTDHNSPLY | 0.47088337 |
| GGNGGSGGTSV | ASRTGTDHNSPLY | 0.8578011 |
| GTSVATGGAGN | ASRTGTDHNSPLY | 0.772988 |
| NGGNGGSGGTS | ASRTGTDHNSPLY | 0.6602255 |
| AVVKFPH | ASRTGTDHNSPLY | 0.84656155 |
| TPPPPPP | ASRTGTDHNSPLY | 0.69156927 |
| AAPAAAAAPA | ASRTGTDHNSPLY | 0.70809484 |
| IALNEDL | ASRTGTDHNSPLY | 0.7571454 |
| IVNVSSM | ASRTGTDHNSPLY | 0.8216029 |
| TGIPPAPR | ASRTGTDHNSPLY | 0.8741128 |
| FDLGGGTF | ASRTGTDHNSPLY | 0.81736064 |
| FDLGGGTFDVS | ASRTGTDHNSPLY | 0.79828966 |
| NEPTAAA | ASRTGTDHNSPLY | 0.9030875 |
| NEPTAAALA | ASRTGTDHNSPLY | 0.5802207 |
| GSIINISSI | ASRTGTDHNSPLY | 0.6953975 |
| VAPGNAGTA | ASRTGTDHNSPLY | 0.86954194 |
| TGGGASLE | ASRTGTDHNSPLY | 0.63576555 |
| VAVYNHYKRI | ASRTGTDHNSPLY | 0.8833009 |
| TDPAPATT | ASRTGTDHNSPLY | 0.824394 |
| TNRPDILDPALLRPG | ASRTGTDHNSPLY | 0.8622446 |
| LTTTTTS | ASRTGTDHNSPLY | 0.82720786 |
| PTGSGKTL | ASRTGTDHNSPLY | 0.91227984 |
| TGSGKTL | ASRTGTDHNSPLY | 0.8849155 |
| TIGHVDHGK | ASRTGTDHNSPLY | 0.81223655 |
| TIGHVDHGKTT | ASRTGTDHNSPLY | 0.87602854 |
| GHVDHGKTT | ASRTGTDHNSPLY | 0.55571616 |
| AQFEHTLL | ASRTGTDHNSPLY | 0.8958827 |
| IVNQDEL | ASRTGTDHNSPLY | 0.86527336 |
| TNGSQFF | ASRTGTDHNSPLY | 0.6952626 |
| TGKGIVE | ASRTGTDHNSPLY | 0.8292223 |
| LISNASDA | ASRTGTDHNSPLY | 0.9077269 |
| AELRTLGELTKIAKAHGAQ | ASRTGTDHNSPLY | 0.89496946 |
| APNAVAAAEPSW | ASRTGTDHNSPLY | 0.49302 |
| APQVAGAATAVL | ASRTGTDHNSPLY | 0.7948437 |
| ARRDPAAAAAILPTDA | ASRTGTDHNSPLY | 0.45572662 |
| DAAARDVLAEAGLAEYF | ASRTGTDHNSPLY | 0.7241333 |
| DDQIVLSAKANLAKVEQ | ASRTGTDHNSPLY | 0.4648844 |
| DELLPDTGEDSDEDGH | ASRTGTDHNSPLY | 0.8782619 |
| ELPEAPANRATVPQD | ASRTGTDHNSPLY | 0.7694836 |
| FGQSKIERVTGIG | ASRTGTDHNSPLY | 0.8377329 |
| GAGGTGSTAGGAGGAGGAGGLY | ASRTGTDHNSPLY | 0.84046793 |

| | | |
|---|---|---|
| GHGGTGAAGGNGGTGGAGGSGI | ASRTGTDHNSPLY | 0.8983916 |
| GNGGNGGAGGEGGPGGAGGAGA | ASRTGTDHNSPLY | 0.73157954 |
| GSTAGNGGNGGNATNGTIAGQP | ASRTGTDHNSPLY | 0.77818125 |
| GTGGDGGNGGNGGTGG | ASRTGTDHNSPLY | 0.48378003 |
| GVRRPAGIPADAPV | ASRTGTDHNSPLY | 0.6482934 |
| GYRQVTDAHLLTLARR | ASRTGTDHNSPLY | 0.77125347 |
| IAKMGQDGHDRG | ASRTGTDHNSPLY | 0.76177764 |
| KAEPSIEIQYT | ASRTGTDHNSPLY | 0.3868357 |
| KPVRIRMAADEDTVAVT | ASRTGTDHNSPLY | 0.9013686 |
| MPPRIAGMRLLVIKPEP | ASRTGTDHNSPLY | 0.85542655 |
| MKFYAVALFRSRGVPPTR | ASRTGTDHNSPLY | 0.87158436 |
| NAGGFGILGAVAHSP | ASRTGTDHNSPLY | 0.69880337 |
| NEGREITLAMAEM | ASRTGTDHNSPLY | 0.798643 |
| NELVEWSHPLGHRWRWE | ASRTGTDHNSPLY | 0.77909124 |
| NLGAGGTGGNGGNPG | ASRTGTDHNSPLY | 0.8616693 |
| NTEWPKWETVMAADPVD | ASRTGTDHNSPLY | 0.72395205 |
| PASAAPAETQP | ASRTGTDHNSPLY | 0.37441638 |
| QQSSDGKNGVRAAEITGE | ASRTGTDHNSPLY | 0.6804669 |
| RAPDAVTTSWWKE | ASRTGTDHNSPLY | 0.62099224 |
| SGFGNVGAHTSGWFNQ | ASRTGTDHNSPLY | 0.90411216 |
| SGVAGAVGQAASVGGLKV | ASRTGTDHNSPLY | 0.89179057 |
| SNGGAGTGSAGGNGGT | ASRTGTDHNSPLY | 0.86425996 |
| TGGAGGEGGAAG | ASRTGTDHNSPLY | 0.90186906 |
| TGGDGGAAGAGGAPGHG | ASRTGTDHNSPLY | 0.2596673 |
| TMPPLLAAIANAGRHGV | ASRTGTDHNSPLY | 0.7221185 |
| TSSTAGNSGGA | ASRTGTDHNSPLY | 0.89219236 |
| VPATVPVGILC | ASRTGTDHNSPLY | 0.867782 |
| VQRPDSGGVVLTGRIS | ASRTGTDHNSPLY | 0.79861087 |
| WTPNPVPATKGVPAGLVQAT | ASRTGTDHNSPLY | 0.9019544 |
| YALKRALAKFSDRFDVVII | ASRTGTDHNSPLY | 0.65137696 |
| YLIGSVIV | ASRTGTDHNSPLY | 0.7730442 |
| YGQLVAVID | ASRTGTDHNSPLY | 0.83445895 |
| VTVGGMTLPA | ASRTGTDHNSPLY | 0.88645864 |
| VLALPAV | ASRTGTDHNSPLY | 0.85239303 |
| VIPPTLNYET | ASRTGTDHNSPLY | 0.7032672 |
| VGMGGMHPSSG | ASRTGTDHNSPLY | 0.84539807 |
| VGLAGGAATA | ASRTGTDHNSPLY | 0.91726303 |
| VGGAGGAGGSGP | ASRTGTDHNSPLY | 0.77655107 |
| VFAAFMV | ASRTGTDHNSPLY | 0.88848215 |
| TLYDKLR | ASRTGTDHNSPLY | 0.863539 |
| TLAVFTVL | ASRTGTDHNSPLY | 0.84809184 |
| TGSVQPARV | ASRTGTDHNSPLY | 0.7125832 |
| TGGGAGSGGT | ASRTGTDHNSPLY | 0.78087294 |
| SHLETSL | ASRTGTDHNSPLY | 0.83792984 |
| SFTTGMT | ASRTGTDHNSPLY | 0.72288585 |
| QVEVAKRMGVRQPTVS | ASRTGTDHNSPLY | 0.78353214 |
| QGSGGPALAM | ASRTGTDHNSPLY | 0.7887838 |
| MLAPSNP | ASRTGTDHNSPLY | 0.9200803 |
| LVNNAAI | ASRTGTDHNSPLY | 0.8605292 |
| LTGEINTLA | ASRTGTDHNSPLY | 0.8806547 |
| LSIFGPV | ASRTGTDHNSPLY | 0.7127842 |

| | | |
|---|---|---|
| LLALLANL | ASRTGTDHNSPLY | 0.8708587 |
| LISNASDALDKLR | ASRTGTDHNSPLY | 0.87623584 |
| LINEAVP | ASRTGTDHNSPLY | 0.513689 |
| LGYAFSGL | ASRTGTDHNSPLY | 0.9043777 |
| LDFVAVAL | ASRTGTDHNSPLY | 0.7299789 |
| LAQPGAE | ASRTGTDHNSPLY | 0.81107044 |
| ISNLGAVTGF | ASRTGTDHNSPLY | 0.87159324 |
| HQEDPGRI | ASRTGTDHNSPLY | 0.8851408 |
| GTPGEPTP | ASRTGTDHNSPLY | 0.81808007 |
| GFFNSGS | ASRTGTDHNSPLY | 0.80835253 |
| GAGGGGLTAGAN | ASRTGTDHNSPLY | 0.70181286 |
| FVDALVNM | ASRTGTDHNSPLY | 0.8953996 |
| FLTVAPEL | ASRTGTDHNSPLY | 0.88981175 |
| FETLTAL | ASRTGTDHNSPLY | 0.8232755 |
| DVINPIEL | ASRTGTDHNSPLY | 0.91134065 |
| APVSGAPV | ASRTGTDHNSPLY | 0.85929537 |
| ALVGGLAW | ASRTGTDHNSPLY | 0.3765089 |
| ALAAAAATAI | ASRTGTDHNSPLY | 0.82515144 |
| AFGLGLLL | ASRTGTDHNSPLY | 0.89092946 |
| AFFAAASL | ASRTGTDHNSPLY | 0.89378107 |
| AAAALPAFSPP | ASRTGTDHNSPLY | 0.9006994 |
| VVLAPAL | ASRTGTDHNSPLY | 0.8612362 |
| TIGHVDHGKTTL | ASRTGTDHNSPLY | 0.78845793 |
| LISNASDALDK | ASRTGTDHNSPLY | 0.9065699 |
| LISNASDAL | ASRTGTDHNSPLY | 0.91229016 |
| ISNASDALDK | ASRTGTDHNSPLY | 0.8777876 |
| NQAETLVYQTE | ASRTGTDHNSPLY | 0.77311647 |
| TGGTGGTAGAG | ASRTGTDHNSPLY | 0.7303194 |
| TGSTAGGAGGA | ASRTGTDHNSPLY | 0.8518158 |
| DPLAVPVAAGPV | ASRTGTDHNSPLY | 0.88992727 |
| GGTGGTPGNGGN | ASRTGTDHNSPLY | 0.6387955 |
| PAGLAGVDPATL | ASRTGTDHNSPLY | 0.85825455 |
| GNNLVVEAPHAGASV | ASRTGTDHNSPLY | 0.8529917 |
| KVTRIMSASTLEQPA | ASRTGTDHNSPLY | 0.36989054 |
| LGFALPIALPRRTVV | ASRTGTDHNSPLY | 0.874712 |
| TRLGNLTARKLVVLV | ASRTGTDHNSPLY | 0.89679396 |
| WSQQRRLIYFPSAGP | ASRTGTDHNSPLY | 0.8367206 |
| GPNGNPDPMAAAVDIR | ASRTGTDHNSPLY | 0.78408045 |
| TQRCVSVADLRAAASR | ASRTGTDHNSPLY | 0.898584 |
| QFVLDVAEHPVEVLTETV | ASRTGTDHNSPLY | 0.91834974 |
| GGAGGTGGAGGNGGTQAGDG | ASRTGTDHNSPLY | 0.9063684 |
| PLLATLRAIAVDVDAKPAQV | ASRTGTDHNSPLY | 0.35097912 |
| APKFRTGGYMIDFWGVGYQVAK | ASRTGTDHNSPLY | 0.28513166 |
| GAGGAGGAGGAAGGPGGTGGTG | ASRTGTDHNSPLY | 0.7891668 |
| IGALGLLATDVAPAGSVAFGVA | ASRTGTDHNSPLY | 0.85840374 |
| AGGAGGIGGNGGAGGAGGVLMG | ASRTGTDHNSPLY | 0.74467206 |
| ADELIGLL | ASRTGTDHNSPLY | 0.73317575 |
| ADRALKA | ASRTGTDHNSPLY | 0.859026 |
| AGAVGQVGGAGGA | ASRTGTDHNSPLY | 0.8979338 |
| AGGGGGLGGAP | ASRTGTDHNSPLY | 0.64866567 |
| AGGLLYGNGGAG | ASRTGTDHNSPLY | 0.5524675 |

| | | |
|---|---|---|
| AGLESIGL | ASRTGTDHNSPLY | 0.84967536 |
| AGLKTGII | ASRTGTDHNSPLY | 0.8715372 |
| ASLAALLL | ASRTGTDHNSPLY | 0.72861516 |
| ASVINISSI | ASRTGTDHNSPLY | 0.81174725 |
| ATPPGPA | ASRTGTDHNSPLY | 0.7489249 |
| ATVASALV | ASRTGTDHNSPLY | 0.8232264 |
| ATVASLIL | ASRTGTDHNSPLY | 0.8614651 |
| AVNGPVLIRH | ASRTGTDHNSPLY | 0.6768018 |
| DGAAGAVGI | ASRTGTDHNSPLY | 0.9022312 |
| DPLLHGI | ASRTGTDHNSPLY | 0.68512094 |
| ELGAALGAL | ASRTGTDHNSPLY | 0.8875351 |
| ELGRTQAV | ASRTGTDHNSPLY | 0.85320187 |
| FVAGGYLS | ASRTGTDHNSPLY | 0.86887217 |
| GASGGAGGAGGW | ASRTGTDHNSPLY | 0.8008921 |
| GGAGAGGDGGAGGI | ASRTGTDHNSPLY | 0.75656056 |
| GTGHNGSQPGMGGNG | ASRTGTDHNSPLY | 0.6352699 |
| GYAAASASAS | ASRTGTDHNSPLY | 0.5242517 |
| HQVDVAAVQQG | ASRTGTDHNSPLY | 0.8961211 |
| IGAAIAHT | ASRTGTDHNSPLY | 0.7767025 |
| IGMIEVAQL | ASRTGTDHNSPLY | 0.9007628 |
| IRLAPEHTSAMD | ASRTGTDHNSPLY | 0.8686718 |
| IYNVSVL | ASRTGTDHNSPLY | 0.7078855 |
| KQGFLGM | ASRTGTDHNSPLY | 0.87796086 |
| LESVITQL | ASRTGTDHNSPLY | 0.7093682 |
| LGLGALVVV | ASRTGTDHNSPLY | 0.7934575 |
| LIGNGGNGGSGGI | ASRTGTDHNSPLY | 0.8752856 |
| LNAINAPTL | ASRTGTDHNSPLY | 0.86995614 |
| LQLLNTGV | ASRTGTDHNSPLY | 0.6862779 |
| LQTSIEF | ASRTGTDHNSPLY | 0.8702087 |
| MSFAVSEL | ASRTGTDHNSPLY | 0.86420596 |
| NTGALGL | ASRTGTDHNSPLY | 0.689957 |
| PPGAGGAGGAGTT | ASRTGTDHNSPLY | 0.8288998 |
| QLVGLGI | ASRTGTDHNSPLY | 0.8953395 |
| RFLEQVHQL | ASRTGTDHNSPLY | 0.8598849 |
| RGVGYVLR | ASRTGTDHNSPLY | 0.7086178 |
| SATGVFAGVI | ASRTGTDHNSPLY | 0.8668809 |
| SLPAITLP | ASRTGTDHNSPLY | 0.7076068 |
| STALAFTV | ASRTGTDHNSPLY | 0.8669323 |
| SVLTVGLGK | ASRTGTDHNSPLY | 0.85336983 |
| TALAGAGSGI | ASRTGTDHNSPLY | 0.7435911 |
| TGLPFTGI | ASRTGTDHNSPLY | 0.8777993 |
| VAEHVTV | ASRTGTDHNSPLY | 0.53824204 |
| VDIVLNSL | ASRTGTDHNSPLY | 0.7188835 |
| VLTGGGAVL | ASRTGTDHNSPLY | 0.8056083 |
| VNVGLAYGI | ASRTGTDHNSPLY | 0.8200692 |
| VSSAAVGAV | ASRTGTDHNSPLY | 0.8887512 |
| VTSVYEAGV | ASRTGTDHNSPLY | 0.6842902 |
| WVALDRGA | ASRTGTDHNSPLY | 0.85306376 |
| YLVGSVVI | ASRTGTDHNSPLY | 0.7365444 |
| GPPGTGKTLL | ASRTGTDHNSPLY | 0.6031165 |
| GVGMTKF | ASRTGTDHNSPLY | 0.83476985 |

| | | |
|---|---|---|
| IGIGGSDL | ASRTGTDHNSPLY | 0.80069596 |
| IGTLLAEL | ASRTGTDHNSPLY | 0.7272334 |
| IYTSGTT | ASRTGTDHNSPLY | 0.8800603 |
| LSFPAGV | ASRTGTDHNSPLY | 0.65979624 |
| LYTSGTT | ASRTGTDHNSPLY | 0.8396379 |
| VGAFPFGF | ASRTGTDHNSPLY | 0.9133327 |
| VGAGTVEF | ASRTGTDHNSPLY | 0.7715515 |
| VGKSTLLL | ASRTGTDHNSPLY | 0.8882523 |
| AALAHAGGGIV | ASRTGTDHNSPLY | 0.87247884 |
| AALVAVSLL | ASRTGTDHNSPLY | 0.87574196 |
| AAVSRAVAAL | ASRTGTDHNSPLY | 0.761271 |
| AGGSSGIGGF | ASRTGTDHNSPLY | 0.74682534 |
| AGLTVANANG | ASRTGTDHNSPLY | 0.90169495 |
| AGLVGAVL | ASRTGTDHNSPLY | 0.8698269 |
| ALSQAFTV | ASRTGTDHNSPLY | 0.8402933 |
| ALTFSVSL | ASRTGTDHNSPLY | 0.8776002 |
| ALTGAALVAL | ASRTGTDHNSPLY | 0.8627931 |
| ASFSIGAGVL | ASRTGTDHNSPLY | 0.87470347 |
| DGSVYKI | ASRTGTDHNSPLY | 0.7741712 |
| EQLAEIV | ASRTGTDHNSPLY | 0.58879876 |
| FAVSELTVM | ASRTGTDHNSPLY | 0.78923607 |
| GAAPLAAGL | ASRTGTDHNSPLY | 0.85307735 |
| GADGGDGGSGGAGG | ASRTGTDHNSPLY | 0.7674333 |
| GFGLDTGGAGG | ASRTGTDHNSPLY | 0.5097809 |
| GLASGGLL | ASRTGTDHNSPLY | 0.8835782 |
| GRIAGAAVLL | ASRTGTDHNSPLY | 0.91572547 |
| GSAGSLNTSF | ASRTGTDHNSPLY | 0.65148354 |
| GTGQIGI | ASRTGTDHNSPLY | 0.91350496 |
| GTLIALIL | ASRTGTDHNSPLY | 0.62087935 |
| IAATMGFVVG | ASRTGTDHNSPLY | 0.86794007 |
| IFAQALL | ASRTGTDHNSPLY | 0.88908815 |
| IGAAVALGF | ASRTGTDHNSPLY | 0.6914562 |
| IGLTPLAL | ASRTGTDHNSPLY | 0.8595611 |
| ILGSNTSTL | ASRTGTDHNSPLY | 0.74578714 |
| IVFSLTVL | ASRTGTDHNSPLY | 0.8073742 |
| LASKATGF | ASRTGTDHNSPLY | 0.9023287 |
| LASVALIL | ASRTGTDHNSPLY | 0.6810209 |
| LFAVGLTL | ASRTGTDHNSPLY | 0.8548924 |
| LFNGGIGTYI | ASRTGTDHNSPLY | 0.7757691 |
| LTLGAGII | ASRTGTDHNSPLY | 0.7196551 |
| LVAAALGDI | ASRTGTDHNSPLY | 0.8697976 |
| NTAFVQLGI | ASRTGTDHNSPLY | 0.8883817 |
| QLFSGAEL | ASRTGTDHNSPLY | 0.8085662 |
| SAATFITL | ASRTGTDHNSPLY | 0.9092057 |
| SNYGLANLGV | ASRTGTDHNSPLY | 0.7176498 |
| SYGGAYAVM | ASRTGTDHNSPLY | 0.91119385 |
| TTGVGGLTLG | ASRTGTDHNSPLY | 0.8821787 |
| TTQTIYI | ASRTGTDHNSPLY | 0.843644 |
| VAAGAGLGAVI | ASRTGTDHNSPLY | 0.8051217 |
| VAAVVTLL | ASRTGTDHNSPLY | 0.8887677 |
| VGFLGGLI | ASRTGTDHNSPLY | 0.71624756 |

| | | |
|---|---|---|
| VGVIPILL | ASRTGTDHNSPLY | 0.8706038 |
| VIFASSLI | ASRTGTDHNSPLY | 0.91017735 |
| VKGGDGGAAAG | ASRTGTDHNSPLY | 0.7051875 |
| VTKTFTNI | ASRTGTDHNSPLY | 0.86938196 |
| VVAAAILL | ASRTGTDHNSPLY | 0.8086798 |
| YAALDVEL | ASRTGTDHNSPLY | 0.7819424 |
| LASVALLL | ASRTGTDHNSPLY | 0.7204435 |
| LDAVHRV | ASRTGTDHNSPLY | 0.5107239 |
| QGLSSSIF | ASRTGTDHNSPLY | 0.75966275 |
| RGTFANIRL | ASRTGTDHNSPLY | 0.651397 |
| RQLWWGHRI | ASRTGTDHNSPLY | 0.78689164 |
| YGPPGTGKTLL | ASRTGTDHNSPLY | 0.84931475 |
| AALVGAPANVA | ASRTGTDHNSPLY | 0.7158053 |
| AATFRRTE | ASRTGTDHNSPLY | 0.8620753 |
| AGALGGGI | ASRTGTDHNSPLY | 0.8336713 |
| AGGAGGAGGIGAG | ASRTGTDHNSPLY | 0.5171122 |
| AGSGALGL | ASRTGTDHNSPLY | 0.74857664 |
| AQIADQI | ASRTGTDHNSPLY | 0.728531 |
| ASAVAVTYL | ASRTGTDHNSPLY | 0.78134024 |
| ASTVLGGTGG | ASRTGTDHNSPLY | 0.40493917 |
| AVGGEPAK | ASRTGTDHNSPLY | 0.84798586 |
| CEQGTFGGTC | ASRTGTDHNSPLY | 0.6842959 |
| DEGKAPAG | ASRTGTDHNSPLY | 0.19607744 |
| DTVVFPAL | ASRTGTDHNSPLY | 0.81255144 |
| FEGITMAG | ASRTGTDHNSPLY | 0.7983861 |
| FPLVVQD | ASRTGTDHNSPLY | 0.7568285 |
| GAFTTPQ | ASRTGTDHNSPLY | 0.847293 |
| GFGGFGTAGGVGGA | ASRTGTDHNSPLY | 0.60103804 |
| GFLAQFL | ASRTGTDHNSPLY | 0.6480038 |
| GGAGGAGGNAGVA | ASRTGTDHNSPLY | 0.83300793 |
| GGVGTAGAGGAGGAGG | ASRTGTDHNSPLY | 0.8639473 |
| GNGGGGDAGNAGSGGN | ASRTGTDHNSPLY | 0.5851801 |
| GTGGAGGAGNDAGS | ASRTGTDHNSPLY | 0.8104528 |
| GTTNTGWFNS | ASRTGTDHNSPLY | 0.8448197 |
| IFMGPLFTL | ASRTGTDHNSPLY | 0.82190084 |
| ILAHAIEAI | ASRTGTDHNSPLY | 0.86433864 |
| IRSAIREAL | ASRTGTDHNSPLY | 0.8768941 |
| ITFAGGG | ASRTGTDHNSPLY | 0.654542 |
| ITGMSGHQEV | ASRTGTDHNSPLY | 0.85649633 |
| IVAGLACTL | ASRTGTDHNSPLY | 0.863647 |
| IVGGGLAGA | ASRTGTDHNSPLY | 0.7645271 |
| IVTGVLNI | ASRTGTDHNSPLY | 0.7892083 |
| LAQQGCI | ASRTGTDHNSPLY | 0.6867855 |
| LCLHVEN | ASRTGTDHNSPLY | 0.7579261 |
| LDAISDGS | ASRTGTDHNSPLY | 0.42942488 |
| LGGFAAML | ASRTGTDHNSPLY | 0.80849487 |
| LGGKNPMI | ASRTGTDHNSPLY | 0.86468536 |
| LGPKGVAAIL | ASRTGTDHNSPLY | 0.7380341 |
| LMVGVSI | ASRTGTDHNSPLY | 0.64072967 |
| LVGAAFVL | ASRTGTDHNSPLY | 0.63202983 |
| LVLAVAPL | ASRTGTDHNSPLY | 0.80201066 |

| | | |
|---|---|---|
| MAAIAVIDAA | ASRTGTDHNSPLY | 0.7439324 |
| NNGINAELQ | ASRTGTDHNSPLY | 0.5090937 |
| RGEPLVLVHG | ASRTGTDHNSPLY | 0.71495545 |
| RGGAGGAAGLI | ASRTGTDHNSPLY | 0.84380835 |
| RSYRMHPAV | ASRTGTDHNSPLY | 0.84135544 |
| SAAHLLAGQLA | ASRTGTDHNSPLY | 0.47038174 |
| SAAVGSY | ASRTGTDHNSPLY | 0.86792827 |
| SPHAVLHRI | ASRTGTDHNSPLY | 0.8194643 |
| SQIAKVL | ASRTGTDHNSPLY | 0.8599664 |
| TALGHLVP | ASRTGTDHNSPLY | 0.7908528 |
| TAPTTSGAA | ASRTGTDHNSPLY | 0.69600224 |
| TDGDDGNGGSPGT | ASRTGTDHNSPLY | 0.83488536 |
| TDGLGEN | ASRTGTDHNSPLY | 0.7358538 |
| TGGPPLPGQA | ASRTGTDHNSPLY | 0.6793337 |
| TIPAITI | ASRTGTDHNSPLY | 0.85335624 |
| TIPGFTI | ASRTGTDHNSPLY | 0.7589066 |
| TSIASGALA | ASRTGTDHNSPLY | 0.68403447 |
| TVTLPGAAI | ASRTGTDHNSPLY | 0.6340106 |
| VASGLSAALP | ASRTGTDHNSPLY | 0.8778619 |
| VGAGVIGI | ASRTGTDHNSPLY | 0.59554595 |
| VGPKRFGV | ASRTGTDHNSPLY | 0.71460867 |
| VNVAFPD | ASRTGTDHNSPLY | 0.6344788 |
| VSMTNSGVS | ASRTGTDHNSPLY | 0.7800069 |
| VTDVLGDLSV | ASRTGTDHNSPLY | 0.8551111 |
| VVVVGAGFAGL | ASRTGTDHNSPLY | 0.83803266 |
| VYTVGTVV | ASRTGTDHNSPLY | 0.7341253 |
| WTTFSGP | ASRTGTDHNSPLY | 0.8040824 |
| YDELVEAF | ASRTGTDHNSPLY | 0.84262073 |
| YGTGGAGGNGG | ASRTGTDHNSPLY | 0.8144354 |
| YTNVMAV | ASRTGTDHNSPLY | 0.841927 |
| FLEQVHQL | ASRTGTDHNSPLY | 0.6616105 |
| GLSLLAVAV | ASRTGTDHNSPLY | 0.8727762 |
| LEQVHQL | ASRTGTDHNSPLY | 0.825884 |
| LNAAGAI | ASRTGTDHNSPLY | 0.85908127 |
| PAVHASL | ASRTGTDHNSPLY | 0.86014295 |
| TELEDDA | ASRTGTDHNSPLY | 0.8263676 |
| TPPGPPN | ASRTGTDHNSPLY | 0.8178575 |
| TPPPPPPP | ASRTGTDHNSPLY | 0.37552387 |
| TQSPALI | TCSAETISGNTLY | 0.6282578 |
| EFAAAFPQ | TCSAETISGNTLY | 0.8176071 |
| LLSGNLDVL | TCSAETISGNTLY | 0.8045936 |
| GPANLGNYNI | TCSAETISGNTLY | 0.8407767 |
| TLGPTLP | TCSAETISGNTLY | 0.82005596 |
| VYAYSSCCK | TCSAETISGNTLY | 0.7053649 |
| APAPAPAPNI | TCSAETISGNTLY | 0.84926003 |
| PGGPVPIP | TCSAETISGNTLY | 0.84147114 |
| FLTVALEP | TCSAETISGNTLY | 0.49586493 |
| SQYFGANL | TCSAETISGNTLY | 0.7259237 |
| GGVKPAA | TCSAETISGNTLY | 0.8782184 |
| GGTGGAGGAGGW | TCSAETISGNTLY | 0.78117704 |
| GVGGVAGLGGAGG | TCSAETISGNTLY | 0.76501286 |

| | | |
|---|---|---|
| VLLVLATTI | TCSAETISGNTLY | 0.7316093 |
| IGIFAAML | TCSAETISGNTLY | 0.85481644 |
| AVGDLID | TCSAETISGNTLY | 0.7015121 |
| NISPLPL | TCSAETISGNTLY | 0.87368035 |
| VTAVFVAI | TCSAETISGNTLY | 0.24358161 |
| SGDDNDAAL | TCSAETISGNTLY | 0.7545029 |
| AVAALAHS | TCSAETISGNTLY | 0.725611 |
| GGVTVMATP | TCSAETISGNTLY | 0.4937145 |
| AGLAANVRD | TCSAETISGNTLY | 0.8554741 |
| TNRPDLIDPALLRPG | TCSAETISGNTLY | 0.68834215 |
| LANLGVTAL | TCSAETISGNTLY | 0.81747097 |
| TGERIEL | TCSAETISGNTLY | 0.7716973 |
| NGEELAA | TCSAETISGNTLY | 0.862249 |
| VLSLVWG | TCSAETISGNTLY | 0.7712865 |
| IGYALWL | TCSAETISGNTLY | 0.7815447 |
| LSLLLAPL | TCSAETISGNTLY | 0.74664116 |
| TLDEALK | TCSAETISGNTLY | 0.84098595 |
| TDGAGTAEGG | TCSAETISGNTLY | 0.86334527 |
| NAPGGAGGNGG | TCSAETISGNTLY | 0.85532993 |
| VSVFMGMGGNF | TCSAETISGNTLY | 0.4545392 |
| LASYLAAL | TCSAETISGNTLY | 0.74839115 |
| LVGGTPAL | TCSAETISGNTLY | 0.7905621 |
| VRNADVL | TCSAETISGNTLY | 0.38771677 |
| TCSSPFLVSAM | TCSAETISGNTLY | 0.35452285 |
| VHITDAE | TCSAETISGNTLY | 0.76389366 |
| EYAAPSF | TCSAETISGNTLY | 0.7027652 |
| VVDAFHGGL | TCSAETISGNTLY | 0.8500153 |
| ASVVDPQ | TCSAETISGNTLY | 0.82779855 |
| SYNEFGGLS | TCSAETISGNTLY | 0.7557778 |
| DPDGSVY | TCSAETISGNTLY | 0.8401694 |
| MVSTIHATE | TCSAETISGNTLY | 0.7784442 |
| LAAGVADPV | TCSAETISGNTLY | 0.11813303 |
| TKEQIAA | TCSAETISGNTLY | 0.72122854 |
| SLASPAAL | TCSAETISGNTLY | 0.5202745 |
| IGLVFYGV | TCSAETISGNTLY | 0.8648077 |
| FLLDASNL | TCSAETISGNTLY | 0.67467284 |
| YQVGNFP | TCSAETISGNTLY | 0.642902 |
| ESLLSQST | TCSAETISGNTLY | 0.7694234 |
| LFTVGFAAVG | TCSAETISGNTLY | 0.7607932 |
| ATPPTPT | TCSAETISGNTLY | 0.77373064 |
| TDLATLG | TCSAETISGNTLY | 0.66241276 |
| SAGGTGR | TCSAETISGNTLY | 0.28172514 |
| GFSNSGS | TCSAETISGNTLY | 0.84567475 |
| GGNGGSGGTSV | TCSAETISGNTLY | 0.5496272 |
| GTSVATGGAGN | TCSAETISGNTLY | 0.6072405 |
| NGGNGGSGGTS | TCSAETISGNTLY | 0.41955316 |
| AVVKFPH | TCSAETISGNTLY | 0.72839576 |
| TPPPPPP | TCSAETISGNTLY | 0.43923572 |
| AAPAAAAAPA | TCSAETISGNTLY | 0.8575889 |
| IALNEDL | TCSAETISGNTLY | 0.72236353 |
| IVNVSSM | TCSAETISGNTLY | 0.73376024 |

| | | |
|---|---|---|
| TGIPPAPR | TCSAETISGNTLY | 0.87272376 |
| FDLGGGTF | TCSAETISGNTLY | 0.7120475 |
| FDLGGGTFDVS | TCSAETISGNTLY | 0.74601066 |
| NEPTAAA | TCSAETISGNTLY | 0.6957059 |
| NEPTAAALA | TCSAETISGNTLY | 0.6809508 |
| GSIINISSI | TCSAETISGNTLY | 0.7155419 |
| VAPGNAGTA | TCSAETISGNTLY | 0.6718482 |
| TGGGASLE | TCSAETISGNTLY | 0.8746871 |
| VAVYNHYKRI | TCSAETISGNTLY | 0.76752645 |
| TDPAPATT | TCSAETISGNTLY | 0.5616625 |
| TNRPDILDPALLRPG | TCSAETISGNTLY | 0.8002879 |
| LTTTTTS | TCSAETISGNTLY | 0.8719009 |
| PTGSGKTL | TCSAETISGNTLY | 0.8807739 |
| TGSGKTL | TCSAETISGNTLY | 0.6751047 |
| TIGHVDHGK | TCSAETISGNTLY | 0.6923139 |
| TIGHVDHGKTT | TCSAETISGNTLY | 0.7103857 |
| GHVDHGKTT | TCSAETISGNTLY | 0.7500417 |
| AQFEHTLL | TCSAETISGNTLY | 0.7323323 |
| IVNQDEL | TCSAETISGNTLY | 0.61618406 |
| TNGSQFF | TCSAETISGNTLY | 0.8600393 |
| TGKGIVE | TCSAETISGNTLY | 0.8641651 |
| LISNASDA | TCSAETISGNTLY | 0.7827461 |
| AELRTLGELTKIAKAHGAQ | TCSAETISGNTLY | 0.46474844 |
| APNAVAAAEPSW | TCSAETISGNTLY | 0.758761 |
| APQVAGAATAVL | TCSAETISGNTLY | 0.6488848 |
| ARRDPAAAAAILPTDA | TCSAETISGNTLY | 0.7225113 |
| DAAARDVLAEAGLAEYF | TCSAETISGNTLY | 0.7883214 |
| DDQIVLSAKANLAKVEQ | TCSAETISGNTLY | 0.48017842 |
| DELLPDTGEDSDEDGH | TCSAETISGNTLY | 0.76438564 |
| ELPEAPANRATVPQD | TCSAETISGNTLY | 0.8798271 |
| FGQSKIERVTGIG | TCSAETISGNTLY | 0.6641426 |
| GAGGTGSTAGGAGGAGGAGGLY | TCSAETISGNTLY | 0.8010954 |
| GHGGTGAAGGNGGTGGAGGSGI | TCSAETISGNTLY | 0.8629426 |
| GNGGNGGAGGEGGPGGAGGAGA | TCSAETISGNTLY | 0.8359498 |
| GSTAGNGGNGGNATNGTIAGQP | TCSAETISGNTLY | 0.861898 |
| GTGGDGGNGGNGGTGG | TCSAETISGNTLY | 0.822895 |
| GVRRPAGIPADAPV | TCSAETISGNTLY | 0.820813 |
| GYRQVTDAHLLTLARR | TCSAETISGNTLY | 0.8156928 |
| IAKMGQDGHDRG | TCSAETISGNTLY | 0.8008947 |
| KAEPSIEIQYT | TCSAETISGNTLY | 0.77665 |
| KPVRIRMAADEDTVAVT | TCSAETISGNTLY | 0.7613313 |
| MPPRIAGMRLLVIKPEP | TCSAETISGNTLY | 0.72659844 |
| MKFYAVALFRSRGVPPTR | TCSAETISGNTLY | 0.83913875 |
| NAGGFGILGAVAHSP | TCSAETISGNTLY | 0.8081555 |
| NEGREITLAMAEM | TCSAETISGNTLY | 0.84530556 |
| NELVEWSHPLGHRWRWE | TCSAETISGNTLY | 0.59814626 |
| NLGAGGTGGNGGNPG | TCSAETISGNTLY | 0.64701587 |
| NTEWPKWETVMAADPVD | TCSAETISGNTLY | 0.84445226 |
| PASAAPAETQP | TCSAETISGNTLY | 0.19249876 |
| QQSSDGKNGVRAAEITGE | TCSAETISGNTLY | 0.3334309 |
| RAPDAVTTSWWKE | TCSAETISGNTLY | 0.8075565 |

| | | |
|---|---|---|
| SGFGNVGAHTSGWFNQ | TCSAETISGNTLY | 0.86472195 |
| SGVAGAVGQAASVGGLKV | TCSAETISGNTLY | 0.21789944 |
| SNGGAGTGSAGGNGGT | TCSAETISGNTLY | 0.6249067 |
| TGGAGGEGGAAG | TCSAETISGNTLY | 0.56598353 |
| TGGDGGAAGAGGAPGHG | TCSAETISGNTLY | 0.8428864 |
| TMPPLLAAIANAGRHGV | TCSAETISGNTLY | 0.7716212 |
| TSSTAGNSGGA | TCSAETISGNTLY | 0.4767798 |
| VPATVPVGILC | TCSAETISGNTLY | 0.8400283 |
| VQRPDSGGVVLTGRIS | TCSAETISGNTLY | 0.4313103 |
| WTPNPVPATKGVPAGLVQAT | TCSAETISGNTLY | 0.6604513 |
| YALKRALAKFSDRFDVVII | TCSAETISGNTLY | 0.8586935 |
| YLIGSVIV | TCSAETISGNTLY | 0.8557505 |
| YGQLVAVID | TCSAETISGNTLY | 0.629218 |
| VTVGGMTLPA | TCSAETISGNTLY | 0.8444352 |
| VLALPAV | TCSAETISGNTLY | 0.69215846 |
| VIPPTLNYET | TCSAETISGNTLY | 0.8822217 |
| VGMGGMHPSSG | TCSAETISGNTLY | 0.7412192 |
| VGLAGGAATA | TCSAETISGNTLY | 0.5712197 |
| VGGAGGAGGSGP | TCSAETISGNTLY | 0.8409154 |
| VFAAFMV | TCSAETISGNTLY | 0.81639385 |
| TLYDKLR | TCSAETISGNTLY | 0.77645373 |
| TLAVFTVL | TCSAETISGNTLY | 0.82945305 |
| TGSVQPARV | TCSAETISGNTLY | 0.7616087 |
| TGGGAGSGGT | TCSAETISGNTLY | 0.7488916 |
| SHLETSL | TCSAETISGNTLY | 0.8522245 |
| SFTTGMT | TCSAETISGNTLY | 0.8552655 |
| QVEVAKRMGVRQPTVS | TCSAETISGNTLY | 0.6757574 |
| QGSGGPALAM | TCSAETISGNTLY | 0.8638041 |
| MLAPSNP | TCSAETISGNTLY | 0.73577 |
| LVNNAAI | TCSAETISGNTLY | 0.6572306 |
| LTGEINTLA | TCSAETISGNTLY | 0.8769459 |
| LSIFGPV | TCSAETISGNTLY | 0.7999574 |
| LLALLANL | TCSAETISGNTLY | 0.849955 |
| LISNASDALDKLR | TCSAETISGNTLY | 0.8474361 |
| LINEAVP | TCSAETISGNTLY | 0.81066567 |
| LGYAFSGL | TCSAETISGNTLY | 0.5809327 |
| LDFVAVAL | TCSAETISGNTLY | 0.75550973 |
| LAQPGAE | TCSAETISGNTLY | 0.76820695 |
| ISNLGAVTGF | TCSAETISGNTLY | 0.84193635 |
| HQEDPGRI | TCSAETISGNTLY | 0.76795983 |
| GTPGEPTP | TCSAETISGNTLY | 0.6382676 |
| GFFNSGS | TCSAETISGNTLY | 0.83996505 |
| GAGGGGLTAGAN | TCSAETISGNTLY | 0.8641787 |
| FVDALVNM | TCSAETISGNTLY | 0.8329769 |
| FLTVAPEL | TCSAETISGNTLY | 0.8190595 |
| FETLTAL | TCSAETISGNTLY | 0.86791736 |
| DVINPIEL | TCSAETISGNTLY | 0.7195946 |
| APVSGAPV | TCSAETISGNTLY | 0.74910444 |
| ALVGGLAW | TCSAETISGNTLY | 0.84985214 |
| ALAAAAATAI | TCSAETISGNTLY | 0.8561787 |
| AFGLGLLL | TCSAETISGNTLY | 0.7879331 |

| | | |
|---|---|---|
| AFFAAASL | TCSAETISGNTLY | 0.86624134 |
| AAAALPAFSPP | TCSAETISGNTLY | 0.6648004 |
| VVLAPAL | TCSAETISGNTLY | 0.81919277 |
| TIGHVDHGKTTL | TCSAETISGNTLY | 0.61425674 |
| LISNASDALDK | TCSAETISGNTLY | 0.8049599 |
| LISNASDAL | TCSAETISGNTLY | 0.66888785 |
| ISNASDALDK | TCSAETISGNTLY | 0.820505 |
| NQAETLVYQTE | TCSAETISGNTLY | 0.84292674 |
| TGGTGGTAGAG | TCSAETISGNTLY | 0.8348892 |
| TGSTAGGAGGA | TCSAETISGNTLY | 0.2950872 |
| DPLAVPVAAGPV | TCSAETISGNTLY | 0.8513246 |
| GGTGGTPGNGGN | TCSAETISGNTLY | 0.7980521 |
| PAGLAGVDPATL | TCSAETISGNTLY | 0.8768535 |
| GNNLVVEAPHAGASV | TCSAETISGNTLY | 0.80561596 |
| KVTRIMSASTLEQPA | TCSAETISGNTLY | 0.80773866 |
| LGFALPIALPRRTVV | TCSAETISGNTLY | 0.7184179 |
| TRLGNLTARKLVVLV | TCSAETISGNTLY | 0.7989446 |
| WSQQRRLIYFPSAGP | TCSAETISGNTLY | 0.5947406 |
| GPNGNPDPMAAAVDIR | TCSAETISGNTLY | 0.68831486 |
| TQRCVSVADLRAAASR | TCSAETISGNTLY | 0.6944919 |
| QFVLDVAEHPVEVLTETV | TCSAETISGNTLY | 0.66700536 |
| GGAGGTGGAGGNGGTQAGDG | TCSAETISGNTLY | 0.8299719 |
| PLLATLRAIAVDVDAKPAQV | TCSAETISGNTLY | 0.859551 |
| APKFRTGGYMIDFWGVGYQVAK | TCSAETISGNTLY | 0.8439405 |
| GAGGAGGAGGAAGGPGGTGGTG | TCSAETISGNTLY | 0.8557563 |
| IGALGLLATDVAPAGSVAFGVA | TCSAETISGNTLY | 0.86461395 |
| AGGAGGIGGNGGAGGAGGVLMG | TCSAETISGNTLY | 0.86333144 |
| ADELIGLL | TCSAETISGNTLY | 0.7113483 |
| ADRALKA | TCSAETISGNTLY | 0.7232976 |
| AGAVGQVGGAGGA | TCSAETISGNTLY | 0.58967537 |
| AGGGGGLGGAP | TCSAETISGNTLY | 0.7630403 |
| AGGLLYGNGGAG | TCSAETISGNTLY | 0.7301596 |
| AGLESIGL | TCSAETISGNTLY | 0.8146374 |
| AGLKTGII | TCSAETISGNTLY | 0.17246613 |
| ASLAALLL | TCSAETISGNTLY | 0.75783277 |
| ASVINISSI | TCSAETISGNTLY | 0.8749181 |
| ATPPGPA | TCSAETISGNTLY | 0.63730586 |
| ATVASALV | TCSAETISGNTLY | 0.80261683 |
| ATVASLIL | TCSAETISGNTLY | 0.79992664 |
| AVNGPVLIRH | TCSAETISGNTLY | 0.7394221 |
| DGAAGAVGI | TCSAETISGNTLY | 0.83275425 |
| DPLLHGI | TCSAETISGNTLY | 0.6964263 |
| ELGAALGAL | TCSAETISGNTLY | 0.83395904 |
| ELGRTQAV | TCSAETISGNTLY | 0.8730909 |
| FVAGGYLS | TCSAETISGNTLY | 0.79851305 |
| GASGGAGGAGGW | TCSAETISGNTLY | 0.7853682 |
| GGAGAGGDGGAGGI | TCSAETISGNTLY | 0.79639584 |
| GTGHNGSQPGMGGNG | TCSAETISGNTLY | 0.50470364 |
| GYAAASASAS | TCSAETISGNTLY | 0.8039713 |
| HQVDVAAVQQG | TCSAETISGNTLY | 0.5072286 |
| IGAAIAHT | TCSAETISGNTLY | 0.7611704 |

| | | |
|---|---|---|
| IGMIEVAQL | TCSAETISGNTLY | 0.72509885 |
| IRLAPEHTSAMD | TCSAETISGNTLY | 0.7965073 |
| IYNVSVL | TCSAETISGNTLY | 0.77737314 |
| KQGFLGM | TCSAETISGNTLY | 0.835992 |
| LESVITQL | TCSAETISGNTLY | 0.5872772 |
| LGLGALVVV | TCSAETISGNTLY | 0.7611301 |
| LIGNGGNGGSGGI | TCSAETISGNTLY | 0.6908974 |
| LNAINAPTL | TCSAETISGNTLY | 0.56747895 |
| LQLLNTGV | TCSAETISGNTLY | 0.83021355 |
| LQTSIEF | TCSAETISGNTLY | 0.80606127 |
| MSFAVSEL | TCSAETISGNTLY | 0.7545493 |
| NTGALGL | TCSAETISGNTLY | 0.851074 |
| PPGAGGAGGAGTT | TCSAETISGNTLY | 0.8126844 |
| QLVGLGI | TCSAETISGNTLY | 0.50962424 |
| RFLEQVHQL | TCSAETISGNTLY | 0.7250414 |
| RGVGYVLR | TCSAETISGNTLY | 0.76802856 |
| SATGVFAGVI | TCSAETISGNTLY | 0.768953 |
| SLPAITLP | TCSAETISGNTLY | 0.20104007 |
| STALAFTV | TCSAETISGNTLY | 0.83854586 |
| SVLTVGLGK | TCSAETISGNTLY | 0.65853244 |
| TALAGAGSGI | TCSAETISGNTLY | 0.61336833 |
| TGLPFTGI | TCSAETISGNTLY | 0.7135508 |
| VAEHVTV | TCSAETISGNTLY | 0.831918 |
| VDIVLNSL | TCSAETISGNTLY | 0.71654564 |
| VLTGGGAVL | TCSAETISGNTLY | 0.72733724 |
| VNVGLAYGI | TCSAETISGNTLY | 0.83373123 |
| VSSAAVGAV | TCSAETISGNTLY | 0.8249683 |
| VTSVYEAGV | TCSAETISGNTLY | 0.863981 |
| WVALDRGA | TCSAETISGNTLY | 0.81524944 |
| YLVGSVVI | TCSAETISGNTLY | 0.80495954 |
| GPPGTGKTLL | TCSAETISGNTLY | 0.8041318 |
| GVGMTKF | TCSAETISGNTLY | 0.77601886 |
| IGIGGSDL | TCSAETISGNTLY | 0.5449109 |
| IGTLLAEL | TCSAETISGNTLY | 0.44460818 |
| IYTSGTT | TCSAETISGNTLY | 0.77113056 |
| LSFPAGV | TCSAETISGNTLY | 0.47109202 |
| LYTSGTT | TCSAETISGNTLY | 0.7815932 |
| VGAFPFGF | TCSAETISGNTLY | 0.8613601 |
| VGAGTVEF | TCSAETISGNTLY | 0.7643489 |
| VGKSTLLL | TCSAETISGNTLY | 0.59489167 |
| AALAHAGGGIV | TCSAETISGNTLY | 0.75347006 |
| AALVAVSLL | TCSAETISGNTLY | 0.71523243 |
| AAVSRAVAAL | TCSAETISGNTLY | 0.84293485 |
| AGGSSGIGGF | TCSAETISGNTLY | 0.8569519 |
| AGLTVANANG | TCSAETISGNTLY | 0.8653995 |
| AGLVGAVL | TCSAETISGNTLY | 0.83968776 |
| ALSQAFTV | TCSAETISGNTLY | 0.69630337 |
| ALTFSVSL | TCSAETISGNTLY | 0.8379708 |
| ALTGAALVAL | TCSAETISGNTLY | 0.7605221 |
| ASFSIGAGVL | TCSAETISGNTLY | 0.7358418 |
| DGSVYKI | TCSAETISGNTLY | 0.81036055 |

| | | |
|---|---|---|
| EQLAEIV | TCSAETISGNTLY | 0.81039786 |
| FAVSELTVM | TCSAETISGNTLY | 0.7984263 |
| GAAPLAAGL | TCSAETISGNTLY | 0.7497097 |
| GADGGDGGSGGAGG | TCSAETISGNTLY | 0.51123536 |
| GFGLDTGGAGG | TCSAETISGNTLY | 0.8268949 |
| GLASGGLL | TCSAETISGNTLY | 0.8324542 |
| GRIAGAAVLL | TCSAETISGNTLY | 0.5726924 |
| GSAGSLNTSF | TCSAETISGNTLY | 0.7360103 |
| GTGQIGI | TCSAETISGNTLY | 0.87048054 |
| GTLIALIL | TCSAETISGNTLY | 0.86103016 |
| IAATMGFVVG | TCSAETISGNTLY | 0.64142245 |
| IFAQALL | TCSAETISGNTLY | 0.69687736 |
| IGAAVALGF | TCSAETISGNTLY | 0.77954286 |
| IGLTPLAL | TCSAETISGNTLY | 0.45352316 |
| ILGSNTSTL | TCSAETISGNTLY | 0.8385678 |
| IVFSLTVL | TCSAETISGNTLY | 0.7596325 |
| LASKATGF | TCSAETISGNTLY | 0.75002766 |
| LASVALIL | TCSAETISGNTLY | 0.6825909 |
| LFAVGLTL | TCSAETISGNTLY | 0.8025191 |
| LFNGGIGTYI | TCSAETISGNTLY | 0.7118326 |
| LTLGAGII | TCSAETISGNTLY | 0.8474659 |
| LVAAALGDI | TCSAETISGNTLY | 0.186293 |
| NTAFVQLGI | TCSAETISGNTLY | 0.839744 |
| QLFSGAEL | TCSAETISGNTLY | 0.6250479 |
| SAATFITL | TCSAETISGNTLY | 0.74921334 |
| SNYGLANLGV | TCSAETISGNTLY | 0.4701512 |
| SYGGAYAVM | TCSAETISGNTLY | 0.7681303 |
| TTGVGGLTLG | TCSAETISGNTLY | 0.8347742 |
| TTQTIYI | TCSAETISGNTLY | 0.85063446 |
| VAAGAGLGAVI | TCSAETISGNTLY | 0.6737904 |
| VAAVVTLL | TCSAETISGNTLY | 0.8431915 |
| VGFLGGLI | TCSAETISGNTLY | 0.87526983 |
| VGVIPILL | TCSAETISGNTLY | 0.74585164 |
| VIFASSLI | TCSAETISGNTLY | 0.49377167 |
| VKGGDGGAAAG | TCSAETISGNTLY | 0.81523 |
| VTKTFTNI | TCSAETISGNTLY | 0.69343185 |
| VVAAAILL | TCSAETISGNTLY | 0.8454019 |
| YAALDVEL | TCSAETISGNTLY | 0.7957377 |
| LASVALLL | TCSAETISGNTLY | 0.5749686 |
| LDAVHRV | TCSAETISGNTLY | 0.81987154 |
| QGLSSSIF | TCSAETISGNTLY | 0.848944 |
| RGTFANIRL | TCSAETISGNTLY | 0.68061674 |
| RQLWWGHRI | TCSAETISGNTLY | 0.7156294 |
| YGPPGTGKTLL | TCSAETISGNTLY | 0.7125342 |
| AALVGAPANVA | TCSAETISGNTLY | 0.41261524 |
| AATFRRTE | TCSAETISGNTLY | 0.8181923 |
| AGALGGGI | TCSAETISGNTLY | 0.69440585 |
| AGGAGGAGGIGAG | TCSAETISGNTLY | 0.6529803 |
| AGSGALGL | TCSAETISGNTLY | 0.7858853 |
| AQIADQI | TCSAETISGNTLY | 0.69950473 |
| ASAVAVTYL | TCSAETISGNTLY | 0.7021782 |

| | | |
|---|---|---|
| ASTVLGGTGG | TCSAETISGNTLY | 0.74198526 |
| AVGGEPAK | TCSAETISGNTLY | 0.7736696 |
| CEQGTFGGTC | TCSAETISGNTLY | 0.82658815 |
| DEGKAPAG | TCSAETISGNTLY | 0.7470835 |
| DTVVFPAL | TCSAETISGNTLY | 0.7430961 |
| FEGITMAG | TCSAETISGNTLY | 0.86847585 |
| FPLVVQD | TCSAETISGNTLY | 0.56345236 |
| GAFTTPQ | TCSAETISGNTLY | 0.65343034 |
| GFGGFGTAGGVGGA | TCSAETISGNTLY | 0.8214497 |
| GFLAQFL | TCSAETISGNTLY | 0.5575237 |
| GGAGGAGGNAGVA | TCSAETISGNTLY | 0.8315141 |
| GGVGTAGAGGAGGAGG | TCSAETISGNTLY | 0.77615064 |
| GNGGGGDAGNAGSGGN | TCSAETISGNTLY | 0.8125717 |
| GTGGAGGAGNDAGS | TCSAETISGNTLY | 0.7514342 |
| GTTNTGWFNS | TCSAETISGNTLY | 0.87880826 |
| IFMGPLFTL | TCSAETISGNTLY | 0.8439269 |
| ILAHAIEAI | TCSAETISGNTLY | 0.7817444 |
| IRSAIREAL | TCSAETISGNTLY | 0.8126373 |
| ITFAGGG | TCSAETISGNTLY | 0.6369081 |
| ITGMSGHQEV | TCSAETISGNTLY | 0.8536552 |
| IVAGLACTL | TCSAETISGNTLY | 0.8213445 |
| IVGGGLAGA | TCSAETISGNTLY | 0.6235871 |
| IVTGVLNI | TCSAETISGNTLY | 0.75567997 |
| LAQQGCI | TCSAETISGNTLY | 0.8797431 |
| LCLHVEN | TCSAETISGNTLY | 0.8569058 |
| LDAISDGS | TCSAETISGNTLY | 0.50367916 |
| LGGFAAML | TCSAETISGNTLY | 0.7202853 |
| LGGKNPMI | TCSAETISGNTLY | 0.52375287 |
| LGPKGVAAIL | TCSAETISGNTLY | 0.76832503 |
| LMVGVSI | TCSAETISGNTLY | 0.5725234 |
| LVGAAFVL | TCSAETISGNTLY | 0.8367971 |
| LVLAVAPL | TCSAETISGNTLY | 0.7300956 |
| MAAIAVIDAA | TCSAETISGNTLY | 0.7812459 |
| NNGINAELQ | TCSAETISGNTLY | 0.7948669 |
| RGEPLVLVHG | TCSAETISGNTLY | 0.8663135 |
| RGGAGGAAGLI | TCSAETISGNTLY | 0.7082479 |
| RSYRMHPAV | TCSAETISGNTLY | 0.74624985 |
| SAAHLLAGQLA | TCSAETISGNTLY | 0.5385798 |
| SAAVGSY | TCSAETISGNTLY | 0.6136844 |
| SPHAVLHRI | TCSAETISGNTLY | 0.74309814 |
| SQIAKVL | TCSAETISGNTLY | 0.71788865 |
| TALGHLVP | TCSAETISGNTLY | 0.34293944 |
| TAPTTSGAA | TCSAETISGNTLY | 0.86457837 |
| TDGDDGNGGSPGT | TCSAETISGNTLY | 0.79833585 |
| TDGLGEN | TCSAETISGNTLY | 0.83509433 |
| TGGPPLPGQA | TCSAETISGNTLY | 0.6835018 |
| TIPAITI | TCSAETISGNTLY | 0.7650623 |
| TIPGFTI | TCSAETISGNTLY | 0.71059406 |
| TSIASGALA | TCSAETISGNTLY | 0.81699634 |
| TVTLPGAAI | TCSAETISGNTLY | 0.64449865 |
| VASGLSAALP | TCSAETISGNTLY | 0.43390527 |

| | | |
|---|---|---|
| VGAGVIGI | TCSAETISGNTLY | 0.6821984 |
| VGPKRFGV | TCSAETISGNTLY | 0.546395 |
| VNVAFPD | TCSAETISGNTLY | 0.86820924 |
| VSMTNSGVS | TCSAETISGNTLY | 0.8500953 |
| VTDVLGDLSV | TCSAETISGNTLY | 0.80554247 |
| VVVVGAGFAGL | TCSAETISGNTLY | 0.8652248 |
| VYTVGTVV | TCSAETISGNTLY | 0.31236982 |
| WTTFSGP | TCSAETISGNTLY | 0.67944366 |
| YDELVEAF | TCSAETISGNTLY | 0.85345805 |
| YGTGGAGGNGG | TCSAETISGNTLY | 0.8278571 |
| YTNVMAV | TCSAETISGNTLY | 0.7566718 |
| FLEQVHQL | TCSAETISGNTLY | 0.8665188 |
| GLSLLAVAV | TCSAETISGNTLY | 0.6119933 |
| LEQVHQL | TCSAETISGNTLY | 0.75446755 |
| LNAAGAI | TCSAETISGNTLY | 0.7845597 |
| PAVHASL | TCSAETISGNTLY | 0.8552793 |
| TELEDDA | TCSAETISGNTLY | 0.8158859 |
| TPPGPPN | TCSAETISGNTLY | 0.6869629 |
| TPPPPPPP | TCSAETISGNTLY | 0.80705386 |
| TQSPALI | ASSPDWASQNTLY | 0.8873112 |
| EFAAAFPQ | ASSPDWASQNTLY | 0.7097976 |
| LLSGNLDVL | ASSPDWASQNTLY | 0.84349114 |
| GPANLGNYNI | ASSPDWASQNTLY | 0.8214375 |
| TLGPTLP | ASSPDWASQNTLY | 0.7547011 |
| VYAYSSCCK | ASSPDWASQNTLY | 0.6571977 |
| APAPAPAPNI | ASSPDWASQNTLY | 0.70510006 |
| PGGPVPIP | ASSPDWASQNTLY | 0.76149046 |
| FLTVALEP | ASSPDWASQNTLY | 0.60066575 |
| SQYFGANL | ASSPDWASQNTLY | 0.75230014 |
| GGVKPAA | ASSPDWASQNTLY | 0.75275636 |
| GGTGGAGGAGGW | ASSPDWASQNTLY | 0.8919132 |
| GVGGVAGLGGAGG | ASSPDWASQNTLY | 0.8265315 |
| VLLVLATTI | ASSPDWASQNTLY | 0.8370117 |
| IGIFAAML | ASSPDWASQNTLY | 0.6268673 |
| AVGDLID | ASSPDWASQNTLY | 0.8387918 |
| NISPLPL | ASSPDWASQNTLY | 0.82361317 |
| VTAVFVAI | ASSPDWASQNTLY | 0.5611615 |
| SGDDNDAAL | ASSPDWASQNTLY | 0.8700754 |
| AVAALAHS | ASSPDWASQNTLY | 0.6904989 |
| GGVTVMATP | ASSPDWASQNTLY | 0.7487842 |
| AGLAANVRD | ASSPDWASQNTLY | 0.8310248 |
| TNRPDLIDPALLRPG | ASSPDWASQNTLY | 0.8370392 |
| LANLGVTAL | ASSPDWASQNTLY | 0.7781072 |
| TGERIEL | ASSPDWASQNTLY | 0.7457017 |
| NGEELAA | ASSPDWASQNTLY | 0.63782907 |
| VLSLVWG | ASSPDWASQNTLY | 0.8531855 |
| IGYALWL | ASSPDWASQNTLY | 0.8485505 |
| LSLLLAPL | ASSPDWASQNTLY | 0.73699355 |
| TLDEALK | ASSPDWASQNTLY | 0.8748276 |
| TDGAGTAEGG | ASSPDWASQNTLY | 0.8198366 |
| NAPGGAGGNGG | ASSPDWASQNTLY | 0.48962826 |

| | | |
|---|---|---|
| VSVFMGMGGNF | ASSPDWASQNTLY | 0.791484 |
| LASYLAAL | ASSPDWASQNTLY | 0.85314465 |
| LVGGTPAL | ASSPDWASQNTLY | 0.8547857 |
| VRNADVL | ASSPDWASQNTLY | 0.86318725 |
| TCSSPFLVSAM | ASSPDWASQNTLY | 0.81247604 |
| VHITDAE | ASSPDWASQNTLY | 0.73435587 |
| EYAAPSF | ASSPDWASQNTLY | 0.8668088 |
| VVDAFHGGL | ASSPDWASQNTLY | 0.8818681 |
| ASVVDPQ | ASSPDWASQNTLY | 0.83516705 |
| SYNEFGGLS | ASSPDWASQNTLY | 0.7263821 |
| DPDGSVY | ASSPDWASQNTLY | 0.71757776 |
| MVSTIHATE | ASSPDWASQNTLY | 0.8212944 |
| LAAGVADPV | ASSPDWASQNTLY | 0.8453494 |
| TKEQIAA | ASSPDWASQNTLY | 0.6496345 |
| SLASPAAL | ASSPDWASQNTLY | 0.8129624 |
| IGLVFYGV | ASSPDWASQNTLY | 0.80439913 |
| FLLDASNL | ASSPDWASQNTLY | 0.44314778 |
| YQVGNFP | ASSPDWASQNTLY | 0.8067695 |
| ESLLSQST | ASSPDWASQNTLY | 0.85969824 |
| LFTVGFAAVG | ASSPDWASQNTLY | 0.8088809 |
| ATPPTPT | ASSPDWASQNTLY | 0.6945999 |
| TDLATLG | ASSPDWASQNTLY | 0.86316323 |
| SAGGTGR | ASSPDWASQNTLY | 0.8871229 |
| GFSNSGS | ASSPDWASQNTLY | 0.87096715 |
| GGNGGSGGTSV | ASSPDWASQNTLY | 0.4486081 |
| GTSVATGGAGN | ASSPDWASQNTLY | 0.3144592 |
| NGGNGGSGGTS | ASSPDWASQNTLY | 0.7202004 |
| AVVKFPH | ASSPDWASQNTLY | 0.7982828 |
| TPPPPPP | ASSPDWASQNTLY | 0.7189646 |
| AAPAAAAAPA | ASSPDWASQNTLY | 0.69789773 |
| IALNEDL | ASSPDWASQNTLY | 0.7944894 |
| IVNVSSM | ASSPDWASQNTLY | 0.85589075 |
| TGIPPAPR | ASSPDWASQNTLY | 0.65379244 |
| FDLGGGTF | ASSPDWASQNTLY | 0.63304424 |
| FDLGGGTFDVS | ASSPDWASQNTLY | 0.8037957 |
| NEPTAAA | ASSPDWASQNTLY | 0.8291909 |
| NEPTAAALA | ASSPDWASQNTLY | 0.64409304 |
| GSIINISSI | ASSPDWASQNTLY | 0.7386072 |
| VAPGNAGTA | ASSPDWASQNTLY | 0.69766474 |
| TGGGASLE | ASSPDWASQNTLY | 0.73821694 |
| VAVYNHYKRI | ASSPDWASQNTLY | 0.80050385 |
| TDPAPATT | ASSPDWASQNTLY | 0.64393497 |
| TNRPDILDPALLRPG | ASSPDWASQNTLY | 0.86692786 |
| LTTTTTS | ASSPDWASQNTLY | 0.6483228 |
| PTGSGKTL | ASSPDWASQNTLY | 0.84570944 |
| TGSGKTL | ASSPDWASQNTLY | 0.81079096 |
| TIGHVDHGK | ASSPDWASQNTLY | 0.8297677 |
| TIGHVDHGKTT | ASSPDWASQNTLY | 0.7461999 |
| GHVDHGKTT | ASSPDWASQNTLY | 0.6975489 |
| AQFEHTLL | ASSPDWASQNTLY | 0.59661275 |
| IVNQDEL | ASSPDWASQNTLY | 0.45141184 |

| | | |
|---|---|---|
| TNGSQFF | ASSPDWASQNTLY | 0.86053073 |
| TGKGIVE | ASSPDWASQNTLY | 0.66738653 |
| LISNASDA | ASSPDWASQNTLY | 0.86263824 |
| AELRTLGELTKIAKAHGAQ | ASSPDWASQNTLY | 0.8406509 |
| APNAVAAAEPSW | ASSPDWASQNTLY | 0.694489 |
| APQVAGAATAVL | ASSPDWASQNTLY | 0.84080017 |
| ARRDPAAAAAILPTDA | ASSPDWASQNTLY | 0.62772155 |
| DAAARDVLAEAGLAEYF | ASSPDWASQNTLY | 0.72133243 |
| DDQIVLSAKANLAKVEQ | ASSPDWASQNTLY | 0.83057964 |
| DELLPDTGEDSDEDGH | ASSPDWASQNTLY | 0.8246451 |
| ELPEAPANRATVPQD | ASSPDWASQNTLY | 0.66460013 |
| FGQSKIERVTGIG | ASSPDWASQNTLY | 0.8347082 |
| GAGGTGSTAGGAGGAGGAGGLY | ASSPDWASQNTLY | 0.81593245 |
| GHGGTGAAGGNGGTGGAGGSGI | ASSPDWASQNTLY | 0.6515186 |
| GNGGNGGAGGEGGPGGAGGAGA | ASSPDWASQNTLY | 0.805733 |
| GSTAGNGGNGGNATNGTIAGQP | ASSPDWASQNTLY | 0.8502714 |
| GTGGDGGNGGNGGTGG | ASSPDWASQNTLY | 0.8380839 |
| GVRRPAGIPADAPV | ASSPDWASQNTLY | 0.64578915 |
| GYRQVTDAHLLTLARR | ASSPDWASQNTLY | 0.83026075 |
| IAKMGQDGHDRG | ASSPDWASQNTLY | 0.6264627 |
| KAEPSIEIQYT | ASSPDWASQNTLY | 0.8013381 |
| KPVRIRMAADEDTVAVT | ASSPDWASQNTLY | 0.8146459 |
| MPPRIAGMRLLVIKPEP | ASSPDWASQNTLY | 0.72833836 |
| MKFYAVALFRSRGVPPTR | ASSPDWASQNTLY | 0.8441274 |
| NAGGFGILGAVAHSP | ASSPDWASQNTLY | 0.54694796 |
| NEGREITLAMAEM | ASSPDWASQNTLY | 0.67410547 |
| NELVEWSHPLGHRWRWE | ASSPDWASQNTLY | 0.751587 |
| NLGAGGTGGNGGNPG | ASSPDWASQNTLY | 0.79579276 |
| NTEWPKWETVMAADPVD | ASSPDWASQNTLY | 0.8443414 |
| PASAAPAETQP | ASSPDWASQNTLY | 0.6166978 |
| QQSSDGKNGVRAAEITGE | ASSPDWASQNTLY | 0.7961588 |
| RAPDAVTTSWWKE | ASSPDWASQNTLY | 0.72716045 |
| SGFGNVGAHTSGWFNQ | ASSPDWASQNTLY | 0.60566723 |
| SGVAGAVGQAASVGGLKV | ASSPDWASQNTLY | 0.80657 |
| SNGGAGTGSAGGNGGT | ASSPDWASQNTLY | 0.7833406 |
| TGGAGGEGGAAG | ASSPDWASQNTLY | 0.6443474 |
| TGGDGGAAGAGGAPGHG | ASSPDWASQNTLY | 0.8407223 |
| TMPPLLAAIANAGRHGV | ASSPDWASQNTLY | 0.69392884 |
| TSSTAGNSGGA | ASSPDWASQNTLY | 0.82221276 |
| VPATVPVGILC | ASSPDWASQNTLY | 0.87630385 |
| VQRPDSGGVVLTGRIS | ASSPDWASQNTLY | 0.6986363 |
| WTPNPVPATKGVPAGLVQAT | ASSPDWASQNTLY | 0.8529128 |
| YALKRALAKFSDRFDVVII | ASSPDWASQNTLY | 0.8184365 |
| YLIGSVIV | ASSPDWASQNTLY | 0.82935673 |
| YGQLVAVID | ASSPDWASQNTLY | 0.7397555 |
| VTVGGMTLPA | ASSPDWASQNTLY | 0.67627954 |
| VLALPAV | ASSPDWASQNTLY | 0.8571604 |
| VIPPTLNYET | ASSPDWASQNTLY | 0.8312772 |
| VGMGGMHPSSG | ASSPDWASQNTLY | 0.8180288 |
| VGLAGGAATA | ASSPDWASQNTLY | 0.8302891 |
| VGGAGGAGGSGP | ASSPDWASQNTLY | 0.804703 |

| | | |
|---|---|---|
| VFAAFMV | ASSPDWASQNTLY | 0.8264653 |
| TLYDKLR | ASSPDWASQNTLY | 0.6788751 |
| TLAVFTVL | ASSPDWASQNTLY | 0.58037347 |
| TGSVQPARV | ASSPDWASQNTLY | 0.7444341 |
| TGGGAGSGGT | ASSPDWASQNTLY | 0.784987 |
| SHLETSL | ASSPDWASQNTLY | 0.7594146 |
| SFTTGMT | ASSPDWASQNTLY | 0.54763395 |
| QVEVAKRMGVRQPTVS | ASSPDWASQNTLY | 0.83964765 |
| QGSGGPALAM | ASSPDWASQNTLY | 0.88401103 |
| MLAPSNP | ASSPDWASQNTLY | 0.65997905 |
| LVNNAAI | ASSPDWASQNTLY | 0.87727726 |
| LTGEINTLA | ASSPDWASQNTLY | 0.59744203 |
| LSIFGPV | ASSPDWASQNTLY | 0.8376648 |
| LLALLANL | ASSPDWASQNTLY | 0.85234785 |
| LISNASDALDKLR | ASSPDWASQNTLY | 0.6180608 |
| LINEAVP | ASSPDWASQNTLY | 0.8063591 |
| LGYAFSGL | ASSPDWASQNTLY | 0.7045486 |
| LDFVAVAL | ASSPDWASQNTLY | 0.74840724 |
| LAQPGAE | ASSPDWASQNTLY | 0.8653946 |
| ISNLGAVTGF | ASSPDWASQNTLY | 0.6635961 |
| HQEDPGRI | ASSPDWASQNTLY | 0.81898284 |
| GTPGEPTP | ASSPDWASQNTLY | 0.75077814 |
| GFFNSGS | ASSPDWASQNTLY | 0.6973343 |
| GAGGGGLTAGAN | ASSPDWASQNTLY | 0.8365907 |
| FVDALVNM | ASSPDWASQNTLY | 0.8518105 |
| FLTVAPEL | ASSPDWASQNTLY | 0.72638416 |
| FETLTAL | ASSPDWASQNTLY | 0.8777658 |
| DVINPIEL | ASSPDWASQNTLY | 0.6945077 |
| APVSGAPV | ASSPDWASQNTLY | 0.8702355 |
| ALVGGLAW | ASSPDWASQNTLY | 0.83803266 |
| ALAAAAATAI | ASSPDWASQNTLY | 0.76582813 |
| AFGLGLLL | ASSPDWASQNTLY | 0.77114356 |
| AFFAAASL | ASSPDWASQNTLY | 0.8456806 |
| AAAALPAFSPP | ASSPDWASQNTLY | 0.6769302 |
| VVLAPAL | ASSPDWASQNTLY | 0.84227073 |
| TIGHVDHGKTTL | ASSPDWASQNTLY | 0.87648165 |
| LISNASDALDK | ASSPDWASQNTLY | 0.20932214 |
| LISNASDAL | ASSPDWASQNTLY | 0.53222543 |
| ISNASDALDK | ASSPDWASQNTLY | 0.33808243 |
| NQAETLVYQTE | ASSPDWASQNTLY | 0.34427726 |
| TGGTGGTAGAG | ASSPDWASQNTLY | 0.31201664 |
| TGSTAGGAGGA | ASSPDWASQNTLY | 0.19244479 |
| DPLAVPVAAGPV | ASSPDWASQNTLY | 0.31401792 |
| GGTGGTPGNGGN | ASSPDWASQNTLY | 0.25948006 |
| PAGLAGVDPATL | ASSPDWASQNTLY | 0.72956455 |
| GNNLVVEAPHAGASV | ASSPDWASQNTLY | 0.46082956 |
| KVTRIMSASTLEQPA | ASSPDWASQNTLY | 0.29946834 |
| LGFALPIALPRRTVV | ASSPDWASQNTLY | 0.5790744 |
| TRLGNLTARKLVVLV | ASSPDWASQNTLY | 0.40844125 |
| WSQQRRLIYFPSAGP | ASSPDWASQNTLY | 0.18960571 |
| GPNGNPDPMAAAVDIR | ASSPDWASQNTLY | 0.2708902 |

| | | |
|---|---|---|
| TQRCVSVADLRAAASR | ASSPDWASQNTLY | 0.29103282 |
| QFVLDVAEHPVEVLTETV | ASSPDWASQNTLY | 0.31501856 |
| GGAGGTGGAGGNGGTQAGDG | ASSPDWASQNTLY | 0.18984324 |
| PLLATLRAIAVDVDAKPAQV | ASSPDWASQNTLY | 0.3650254 |
| APKFRTGGYMIDFWGVGYQVAK | ASSPDWASQNTLY | 0.22425449 |
| GAGGAGGAGGAAGGPGGTGGTG | ASSPDWASQNTLY | 0.1631741 |
| IGALGLLATDVAPAGSVAFGVA | ASSPDWASQNTLY | 0.34977883 |
| AGGAGGIGGNGGAGGAGGVLMG | ASSPDWASQNTLY | 0.28553805 |
| ADELIGLL | ASSPDWASQNTLY | 0.23611343 |
| ADRALKA | ASSPDWASQNTLY | 0.47288543 |
| AGAVGQVGGAGGA | ASSPDWASQNTLY | 0.2242575 |
| AGGGGGLGGAP | ASSPDWASQNTLY | 0.23570503 |
| AGGLLYGNGGAG | ASSPDWASQNTLY | 0.4353059 |
| AGLESIGL | ASSPDWASQNTLY | 0.59888256 |
| AGLKTGII | ASSPDWASQNTLY | 0.21082942 |
| ASLAALLL | ASSPDWASQNTLY | 0.39151126 |
| ASVINISSI | ASSPDWASQNTLY | 0.5848851 |
| ATPPGPA | ASSPDWASQNTLY | 0.32280427 |
| ATVASALV | ASSPDWASQNTLY | 0.61983085 |
| ATVASLIL | ASSPDWASQNTLY | 0.6223699 |
| AVNGPVLIRH | ASSPDWASQNTLY | 0.23890972 |
| DGAAGAVGI | ASSPDWASQNTLY | 0.6704116 |
| DPLLHGI | ASSPDWASQNTLY | 0.67013586 |
| ELGAALGAL | ASSPDWASQNTLY | 0.28064653 |
| ELGRTQAV | ASSPDWASQNTLY | 0.46159014 |
| FVAGGYLS | ASSPDWASQNTLY | 0.20707265 |
| GASGGAGGAGGW | ASSPDWASQNTLY | 0.26621327 |
| GGAGAGGDGGAGGI | ASSPDWASQNTLY | 0.18793291 |
| GTGHNGSQPGMGGNG | ASSPDWASQNTLY | 0.47701487 |
| GYAAASASAS | ASSPDWASQNTLY | 0.44711864 |
| HQVDVAAVQQG | ASSPDWASQNTLY | 0.3520728 |
| IGAAIAHT | ASSPDWASQNTLY | 0.18841489 |
| IGMIEVAQL | ASSPDWASQNTLY | 0.19883135 |
| IRLAPEHTSAMD | ASSPDWASQNTLY | 0.29811767 |
| IYNVSVL | ASSPDWASQNTLY | 0.20949946 |
| KQGFLGM | ASSPDWASQNTLY | 0.18859528 |
| LESVITQL | ASSPDWASQNTLY | 0.23222205 |
| LGLGALVVV | ASSPDWASQNTLY | 0.44367626 |
| LIGNGGNGGSGGI | ASSPDWASQNTLY | 0.44405916 |
| LNAINAPTL | ASSPDWASQNTLY | 0.27411914 |
| LQLLNTGV | ASSPDWASQNTLY | 0.7228887 |
| LQTSIEF | ASSPDWASQNTLY | 0.67028046 |
| MSFAVSEL | ASSPDWASQNTLY | 0.6146462 |
| NTGALGL | ASSPDWASQNTLY | 0.29020935 |
| PPGAGGAGGAGTT | ASSPDWASQNTLY | 0.21717855 |
| QLVGLGI | ASSPDWASQNTLY | 0.75321895 |
| RFLEQVHQL | ASSPDWASQNTLY | 0.3537879 |
| RGVGYVLR | ASSPDWASQNTLY | 0.3879329 |
| SATGVFAGVI | ASSPDWASQNTLY | 0.37495807 |
| SLPAITLP | ASSPDWASQNTLY | 0.2438572 |
| STALAFTV | ASSPDWASQNTLY | 0.29034448 |

| | | |
|---|---|---|
| SVLTVGLGK | ASSPDWASQNTLY | 0.5068772 |
| TALAGAGSGI | ASSPDWASQNTLY | 0.5853303 |
| TGLPFTGI | ASSPDWASQNTLY | 0.20785624 |
| VAEHVTV | ASSPDWASQNTLY | 0.20812184 |
| VDIVLNSL | ASSPDWASQNTLY | 0.21110168 |
| VLTGGGAVL | ASSPDWASQNTLY | 0.25612396 |
| VNVGLAYGI | ASSPDWASQNTLY | 0.5954982 |
| VSSAAVGAV | ASSPDWASQNTLY | 0.34597978 |
| VTSVYEAGV | ASSPDWASQNTLY | 0.310406 |
| WVALDRGA | ASSPDWASQNTLY | 0.42478758 |
| YLVGSVVI | ASSPDWASQNTLY | 0.36327285 |
| GPPGTGKTLL | ASSPDWASQNTLY | 0.33571687 |
| GVGMTKF | ASSPDWASQNTLY | 0.29021755 |
| IGIGGSDL | ASSPDWASQNTLY | 0.32736427 |
| IGTLLAEL | ASSPDWASQNTLY | 0.58774805 |
| IYTSGTT | ASSPDWASQNTLY | 0.4960909 |
| LSFPAGV | ASSPDWASQNTLY | 0.4916609 |
| LYTSGTT | ASSPDWASQNTLY | 0.6318794 |
| VGAFPFGF | ASSPDWASQNTLY | 0.29089594 |
| VGAGTVEF | ASSPDWASQNTLY | 0.33575943 |
| VGKSTLLL | ASSPDWASQNTLY | 0.22062708 |
| AALAHAGGGIV | ASSPDWASQNTLY | 0.24075474 |
| AALVAVSLL | ASSPDWASQNTLY | 0.32271305 |
| AAVSRAVAAL | ASSPDWASQNTLY | 0.39187902 |
| AGGSSGIGGF | ASSPDWASQNTLY | 0.50198627 |
| AGLTVANANG | ASSPDWASQNTLY | 0.3860765 |
| AGLVGAVL | ASSPDWASQNTLY | 0.21373522 |
| ALSQAFTV | ASSPDWASQNTLY | 0.42649126 |
| ALTFSVSL | ASSPDWASQNTLY | 0.66130793 |
| ALTGAALVAL | ASSPDWASQNTLY | 0.24889986 |
| ASFSIGAGVL | ASSPDWASQNTLY | 0.28684533 |
| DGSVYKI | ASSPDWASQNTLY | 0.68863153 |
| EQLAEIV | ASSPDWASQNTLY | 0.5516226 |
| FAVSELTVM | ASSPDWASQNTLY | 0.34775615 |
| GAAPLAAGL | ASSPDWASQNTLY | 0.28912845 |
| GADGGDGGSGGAGG | ASSPDWASQNTLY | 0.7705241 |
| GFGLDTGGAGG | ASSPDWASQNTLY | 0.30830622 |
| GLASGGLL | ASSPDWASQNTLY | 0.5890236 |
| GRIAGAAVLL | ASSPDWASQNTLY | 0.20633374 |
| GSAGSLNTSF | ASSPDWASQNTLY | 0.30852854 |
| GTGQIGI | ASSPDWASQNTLY | 0.21184947 |
| GTLIALIL | ASSPDWASQNTLY | 0.17105734 |
| IAATMGFVVG | ASSPDWASQNTLY | 0.572188 |
| IFAQALL | ASSPDWASQNTLY | 0.25618118 |
| IGAAVALGF | ASSPDWASQNTLY | 0.36063436 |
| IGLTPLAL | ASSPDWASQNTLY | 0.24036117 |
| ILGSNTSTL | ASSPDWASQNTLY | 0.59263694 |
| IVFSLTVL | ASSPDWASQNTLY | 0.6662695 |
| LASKATGF | ASSPDWASQNTLY | 0.23758969 |
| LASVALIL | ASSPDWASQNTLY | 0.24261653 |
| LFAVGLTL | ASSPDWASQNTLY | 0.34889236 |

| | | |
|---|---|---|
| LFNGGIGTYI | ASSPDWASQNTLY | 0.47996137 |
| LTLGAGII | ASSPDWASQNTLY | 0.525589 |
| LVAAALGDI | ASSPDWASQNTLY | 0.19168277 |
| NTAFVQLGI | ASSPDWASQNTLY | 0.55260897 |
| QLFSGAEL | ASSPDWASQNTLY | 0.35975012 |
| SAATFITL | ASSPDWASQNTLY | 0.18600309 |
| SNYGLANLGV | ASSPDWASQNTLY | 0.17657284 |
| SYGGAYAVM | ASSPDWASQNTLY | 0.2126644 |
| TTGVGGLTLG | ASSPDWASQNTLY | 0.27487826 |
| TTQTIYI | ASSPDWASQNTLY | 0.5298232 |
| VAAGAGLGAVI | ASSPDWASQNTLY | 0.6172727 |
| VAAVVTLL | ASSPDWASQNTLY | 0.27690256 |
| VGFLGGLI | ASSPDWASQNTLY | 0.4533131 |
| VGVIPILL | ASSPDWASQNTLY | 0.34246206 |
| VIFASSLI | ASSPDWASQNTLY | 0.15831678 |
| VKGGDGGAAAG | ASSPDWASQNTLY | 0.24062654 |
| VTKTFTNI | ASSPDWASQNTLY | 0.19422396 |
| VVAAAILL | ASSPDWASQNTLY | 0.63161993 |
| YAALDVEL | ASSPDWASQNTLY | 0.20933853 |
| LASVALLL | ASSPDWASQNTLY | 0.38185665 |
| LDAVHRV | ASSPDWASQNTLY | 0.385334 |
| QGLSSSIF | ASSPDWASQNTLY | 0.68687505 |
| RGTFANIRL | ASSPDWASQNTLY | 0.3663897 |
| RQLWWGHRI | ASSPDWASQNTLY | 0.21074846 |
| YGPPGTGKTLL | ASSPDWASQNTLY | 0.17221439 |
| AALVGAPANVA | ASSPDWASQNTLY | 0.40161353 |
| AATFRRTE | ASSPDWASQNTLY | 0.26453155 |
| AGALGGGI | ASSPDWASQNTLY | 0.20394355 |
| AGGAGGAGGIGAG | ASSPDWASQNTLY | 0.24252787 |
| AGSGALGL | ASSPDWASQNTLY | 0.36060858 |
| AQIADQI | ASSPDWASQNTLY | 0.18743601 |
| ASAVAVTYL | ASSPDWASQNTLY | 0.52248955 |
| ASTVLGGTGG | ASSPDWASQNTLY | 0.23069334 |
| AVGGEPAK | ASSPDWASQNTLY | 0.33438152 |
| CEQGTFGGTC | ASSPDWASQNTLY | 0.6053655 |
| DEGKAPAG | ASSPDWASQNTLY | 0.69600946 |
| DTVVFPAL | ASSPDWASQNTLY | 0.43141717 |
| FEGITMAG | ASSPDWASQNTLY | 0.23486713 |
| FPLVVQD | ASSPDWASQNTLY | 0.22700737 |
| GAFTTPQ | ASSPDWASQNTLY | 0.21470729 |
| GFGGFGTAGGVGGA | ASSPDWASQNTLY | 0.20920198 |
| GFLAQFL | ASSPDWASQNTLY | 0.52623606 |
| GGAGGAGGNAGVA | ASSPDWASQNTLY | 0.36181122 |
| GGVGTAGAGGAGGAGG | ASSPDWASQNTLY | 0.22566108 |
| GNGGGGDAGNAGSGGN | ASSPDWASQNTLY | 0.50536877 |
| GTGGAGGAGNDAGS | ASSPDWASQNTLY | 0.64615107 |
| GTTNTGWFNS | ASSPDWASQNTLY | 0.6699803 |
| IFMGPLFTL | ASSPDWASQNTLY | 0.20828798 |
| ILAHAIEAI | ASSPDWASQNTLY | 0.24995574 |
| IRSAIREAL | ASSPDWASQNTLY | 0.31935 |
| ITFAGGG | ASSPDWASQNTLY | 0.40453 |

| | | |
|---|---|---|
| ITGMSGHQEV | ASSPDWASQNTLY | 0.2181583 |
| IVAGLACTL | ASSPDWASQNTLY | 0.24205405 |
| IVGGGLAGA | ASSPDWASQNTLY | 0.54527277 |
| IVTGVLNI | ASSPDWASQNTLY | 0.7890883 |
| LAQQGCI | ASSPDWASQNTLY | 0.25666383 |
| LCLHVEN | ASSPDWASQNTLY | 0.19666 |
| LDAISDGS | ASSPDWASQNTLY | 0.30307895 |
| LGGFAAML | ASSPDWASQNTLY | 0.20971088 |
| LGGKNPMI | ASSPDWASQNTLY | 0.26942772 |
| LGPKGVAAIL | ASSPDWASQNTLY | 0.2656052 |
| LMVGVSI | ASSPDWASQNTLY | 0.20013808 |
| LVGAAFVL | ASSPDWASQNTLY | 0.25915685 |
| LVLAVAPL | ASSPDWASQNTLY | 0.65129673 |
| MAAIAVIDAA | ASSPDWASQNTLY | 0.52575994 |
| NNGINAELQ | ASSPDWASQNTLY | 0.40568477 |
| RGEPLVLVHG | ASSPDWASQNTLY | 0.62193257 |
| RGGAGGAAGLI | ASSPDWASQNTLY | 0.43616337 |
| RSYRMHPAV | ASSPDWASQNTLY | 0.63819027 |
| SAAHLLAGQLA | ASSPDWASQNTLY | 0.7122809 |
| SAAVGSY | ASSPDWASQNTLY | 0.27398884 |
| SPHAVLHRI | ASSPDWASQNTLY | 0.27330118 |
| SQIAKVL | ASSPDWASQNTLY | 0.5931039 |
| TALGHLVP | ASSPDWASQNTLY | 0.38583267 |
| TAPTTSGAA | ASSPDWASQNTLY | 0.3724109 |
| TDGDDGNGGSPGT | ASSPDWASQNTLY | 0.4127344 |
| TDGLGEN | ASSPDWASQNTLY | 0.5313369 |
| TGGPPLPGQA | ASSPDWASQNTLY | 0.22888696 |
| TIPAITI | ASSPDWASQNTLY | 0.7420405 |
| TIPGFTI | ASSPDWASQNTLY | 0.1770176 |
| TSIASGALA | ASSPDWASQNTLY | 0.21285765 |
| TVTLPGAAI | ASSPDWASQNTLY | 0.6807079 |
| VASGLSAALP | ASSPDWASQNTLY | 0.16220646 |
| VGAGVIGI | ASSPDWASQNTLY | 0.19345525 |
| VGPKRFGV | ASSPDWASQNTLY | 0.37022334 |
| VNVAFPD | ASSPDWASQNTLY | 0.6164307 |
| VSMTNSGVS | ASSPDWASQNTLY | 0.16454376 |
| VTDVLGDLSV | ASSPDWASQNTLY | 0.27029017 |
| VVVVGAGFAGL | ASSPDWASQNTLY | 0.19660497 |
| VYTVGTVV | ASSPDWASQNTLY | 0.46421784 |
| WTTFSGP | ASSPDWASQNTLY | 0.35723233 |
| YDELVEAF | ASSPDWASQNTLY | 0.20238377 |
| YGTGGAGGNGG | ASSPDWASQNTLY | 0.41276115 |
| YTNVMAV | ASSPDWASQNTLY | 0.2529059 |
| FLEQVHQL | ASSPDWASQNTLY | 0.46410337 |
| GLSLLAVAV | ASSPDWASQNTLY | 0.57763165 |
| LEQVHQL | ASSPDWASQNTLY | 0.7789348 |
| LNAAGAI | ASSPDWASQNTLY | 0.19935814 |
| PAVHASL | ASSPDWASQNTLY | 0.45531058 |
| TELEDDA | ASSPDWASQNTLY | 0.24679515 |
| TPPGPPN | ASSPDWASQNTLY | 0.7621361 |
| TPPPPPPP | ASSPDWASQNTLY | 0.32903135 |

| | | |
|---|---|---|
| TQSPALI | ASSLDWGEAEQF | 0.18856233 |
| EFAAAFPQ | ASSLDWGEAEQF | 0.42392093 |
| LLSGNLDVL | ASSLDWGEAEQF | 0.5028631 |
| GPANLGNYNI | ASSLDWGEAEQF | 0.25214168 |
| TLGPTLP | ASSLDWGEAEQF | 0.7000425 |
| VYAYSSCCK | ASSLDWGEAEQF | 0.3033921 |
| APAPAPAPNI | ASSLDWGEAEQF | 0.3026966 |
| PGGPVPIP | ASSLDWGEAEQF | 0.6595909 |
| FLTVALEP | ASSLDWGEAEQF | 0.4762271 |
| SQYFGANL | ASSLDWGEAEQF | 0.24048434 |
| GGVKPAA | ASSLDWGEAEQF | 0.5291797 |
| GGTGGAGGAGGW | ASSLDWGEAEQF | 0.32878268 |
| GVGGVAGLGGAGG | ASSLDWGEAEQF | 0.2172642 |
| VLLVLATTI | ASSLDWGEAEQF | 0.7275015 |
| IGIFAAML | ASSLDWGEAEQF | 0.33021426 |
| AVGDLID | ASSLDWGEAEQF | 0.5009689 |
| NISPLPL | ASSLDWGEAEQF | 0.32174024 |
| VTAVFVAI | ASSLDWGEAEQF | 0.76580846 |
| SGDDNDAAL | ASSLDWGEAEQF | 0.21092542 |
| AVAALAHS | ASSLDWGEAEQF | 0.3452953 |
| GGVTVMATP | ASSLDWGEAEQF | 0.3176343 |
| AGLAANVRD | ASSLDWGEAEQF | 0.5964765 |
| TNRPDLIDPALLRPG | ASSLDWGEAEQF | 0.3259848 |
| LANLGVTAL | ASSLDWGEAEQF | 0.3165783 |
| TGERIEL | ASSLDWGEAEQF | 0.4564878 |
| NGEELAA | ASSLDWGEAEQF | 0.49688375 |
| VLSLVWG | ASSLDWGEAEQF | 0.32588723 |
| IGYALWL | ASSLDWGEAEQF | 0.28879112 |
| LSLLLAPL | ASSLDWGEAEQF | 0.694168 |
| TLDEALK | ASSLDWGEAEQF | 0.22014841 |
| TDGAGTAEGG | ASSLDWGEAEQF | 0.3900951 |
| NAPGGAGGNGG | ASSLDWGEAEQF | 0.41976708 |
| VSVFMGMGGNF | ASSLDWGEAEQF | 0.47101462 |
| LASYLAAL | ASSLDWGEAEQF | 0.33493137 |
| LVGGTPAL | ASSLDWGEAEQF | 0.5838468 |
| VRNADVL | ASSLDWGEAEQF | 0.21405588 |
| TCSSPFLVSAM | ASSLDWGEAEQF | 0.5578997 |
| VHITDAE | ASSLDWGEAEQF | 0.29392195 |
| EYAAPSF | ASSLDWGEAEQF | 0.3581251 |
| VVDAFHGGL | ASSLDWGEAEQF | 0.18885037 |
| ASVVDPQ | ASSLDWGEAEQF | 0.4496196 |
| SYNEFGGLS | ASSLDWGEAEQF | 0.5372673 |
| DPDGSVY | ASSLDWGEAEQF | 0.526983 |
| MVSTIHATE | ASSLDWGEAEQF | 0.17907813 |
| LAAGVADPV | ASSLDWGEAEQF | 0.5482744 |
| TKEQIAA | ASSLDWGEAEQF | 0.55201924 |
| SLASPAAL | ASSLDWGEAEQF | 0.7806516 |
| IGLVFYGV | ASSLDWGEAEQF | 0.24393411 |
| FLLDASNL | ASSLDWGEAEQF | 0.5173323 |
| YQVGNFP | ASSLDWGEAEQF | 0.28621116 |
| ESLLSQST | ASSLDWGEAEQF | 0.36138853 |

| | | |
|---|---|---|
| LFTVGFAAVG | ASSLDWGEAEQF | 0.1824426 |
| ATPPTPT | ASSLDWGEAEQF | 0.22493884 |
| TDLATLG | ASSLDWGEAEQF | 0.40481028 |
| SAGGTGR | ASSLDWGEAEQF | 0.22627689 |
| GFSNSGS | ASSLDWGEAEQF | 0.7165525 |
| GGNGGSGGTSV | ASSLDWGEAEQF | 0.5110792 |
| GTSVATGGAGN | ASSLDWGEAEQF | 0.55114305 |
| NGGNGGSGGTS | ASSLDWGEAEQF | 0.60248286 |
| AVVKFPH | ASSLDWGEAEQF | 0.7592232 |
| TPPPPPP | ASSLDWGEAEQF | 0.5040829 |
| AAPAAAAAPA | ASSLDWGEAEQF | 0.293818 |
| IALNEDL | ASSLDWGEAEQF | 0.28504124 |
| IVNVSSM | ASSLDWGEAEQF | 0.19021058 |
| TGIPPAPR | ASSLDWGEAEQF | 0.2592837 |
| FDLGGGTF | ASSLDWGEAEQF | 0.2442613 |
| FDLGGGTFDVS | ASSLDWGEAEQF | 0.44374847 |
| NEPTAAA | ASSLDWGEAEQF | 0.16346774 |
| NEPTAAALA | ASSLDWGEAEQF | 0.6671485 |
| GSIINISSI | ASSLDWGEAEQF | 0.56798947 |
| VAPGNAGTA | ASSLDWGEAEQF | 0.30206782 |
| TGGGASLE | ASSLDWGEAEQF | 0.32794818 |
| VAVYNHYKRI | ASSLDWGEAEQF | 0.6330341 |
| TDPAPATT | ASSLDWGEAEQF | 0.23602636 |
| TNRPDILDPALLRPG | ASSLDWGEAEQF | 0.4138276 |
| LTTTTTS | ASSLDWGEAEQF | 0.21832004 |
| PTGSGKTL | ASSLDWGEAEQF | 0.49019837 |
| TGSGKTL | ASSLDWGEAEQF | 0.56867236 |
| TIGHVDHGK | ASSLDWGEAEQF | 0.3914231 |
| TIGHVDHGKTT | ASSLDWGEAEQF | 0.24999927 |
| GHVDHGKTT | ASSLDWGEAEQF | 0.35322064 |
| AQFEHTLL | ASSLDWGEAEQF | 0.21306288 |
| IVNQDEL | ASSLDWGEAEQF | 0.3183612 |
| TNGSQFF | ASSLDWGEAEQF | 0.20153338 |
| TGKGIVE | ASSLDWGEAEQF | 0.39980018 |
| LISNASDA | ASSLDWGEAEQF | 0.21364482 |
| AELRTLGELTKIAKAHGAQ | ASSLDWGEAEQF | 0.3116656 |
| APNAVAAAEPSW | ASSLDWGEAEQF | 0.41299576 |
| APQVAGAATAVL | ASSLDWGEAEQF | 0.37090206 |
| ARRDPAAAAAILPTDA | ASSLDWGEAEQF | 0.3092655 |
| DAAARDVLAEAGLAEYF | ASSLDWGEAEQF | 0.500331 |
| DDQIVLSAKANLAKVEQ | ASSLDWGEAEQF | 0.28044772 |
| DELLPDTGEDSDEDGH | ASSLDWGEAEQF | 0.19281836 |
| ELPEAPANRATVPQD | ASSLDWGEAEQF | 0.63084006 |
| FGQSKIERVTGIG | ASSLDWGEAEQF | 0.28711322 |
| GAGGTGSTAGGAGGAGGAGGLY | ASSLDWGEAEQF | 0.47542557 |
| GHGGTGAAGGNGGTGGAGGSGI | ASSLDWGEAEQF | 0.37823325 |
| GNGGNGGAGGEGGPGGAGGAGA | ASSLDWGEAEQF | 0.28865016 |
| GSTAGNGGNGGNATNGTIAGQP | ASSLDWGEAEQF | 0.23618664 |
| GTGGDGGNGGNGGTGG | ASSLDWGEAEQF | 0.24641317 |
| GVRRPAGIPADAPV | ASSLDWGEAEQF | 0.33186537 |
| GYRQVTDAHLLTLARR | ASSLDWGEAEQF | 0.28667578 |

| | | |
|---|---|---|
| IAKMGQDGHDRG | ASSLDWGEAEQF | 0.1706413 |
| KAEPSIEIQYT | ASSLDWGEAEQF | 0.44913727 |
| KPVRIRMAADEDTVAVT | ASSLDWGEAEQF | 0.21763453 |
| MPPRIAGMRLLVIKPEP | ASSLDWGEAEQF | 0.3505147 |
| MKFYAVALFRSRGVPPTR | ASSLDWGEAEQF | 0.3003839 |
| NAGGFGILGAVAHSP | ASSLDWGEAEQF | 0.5296848 |
| NEGREITLAMAEM | ASSLDWGEAEQF | 0.37703258 |
| NELVEWSHPLGHRWRWE | ASSLDWGEAEQF | 0.23603518 |
| NLGAGGTGGNGGNPG | ASSLDWGEAEQF | 0.43479383 |
| NTEWPKWETVMAADPVD | ASSLDWGEAEQF | 0.32922423 |
| PASAAPAETQP | ASSLDWGEAEQF | 0.51457274 |
| QQSSDGKNGVRAAEITGE | ASSLDWGEAEQF | 0.29074356 |
| RAPDAVTTSWWKE | ASSLDWGEAEQF | 0.25726658 |
| SGFGNVGAHTSGWFNQ | ASSLDWGEAEQF | 0.2833228 |
| SGVAGAVGQAASVGGLKV | ASSLDWGEAEQF | 0.2408955 |
| SNGGAGTGSAGGNGGT | ASSLDWGEAEQF | 0.20577298 |
| TGGAGGEGGAAG | ASSLDWGEAEQF | 0.1908461 |
| TGGDGGAAGAGGAPGHG | ASSLDWGEAEQF | 0.29826546 |
| TMPPLLAAIANAGRHGV | ASSLDWGEAEQF | 0.19498816 |
| TSSTAGNSGGA | ASSLDWGEAEQF | 0.4901477 |
| VPATVPVGILC | ASSLDWGEAEQF | 0.6133553 |
| VQRPDSGGVVLTGRIS | ASSLDWGEAEQF | 0.2538797 |
| WTPNPVPATKGVPAGLVQAT | ASSLDWGEAEQF | 0.1962179 |
| YALKRALAKFSDRFDVVII | ASSLDWGEAEQF | 0.24466604 |
| YLIGSVIV | ASSLDWGEAEQF | 0.36009848 |
| YGQLVAVID | ASSLDWGEAEQF | 0.42099935 |
| VTVGGMTLPA | ASSLDWGEAEQF | 0.33002415 |
| VLALPAV | ASSLDWGEAEQF | 0.779002 |
| VIPPTLNYET | ASSLDWGEAEQF | 0.54388386 |
| VGMGGMHPSSG | ASSLDWGEAEQF | 0.29211855 |
| VGLAGGAATA | ASSLDWGEAEQF | 0.4244998 |
| VGGAGGAGGSGP | ASSLDWGEAEQF | 0.2952757 |
| VFAAFMV | ASSLDWGEAEQF | 0.33433282 |
| TLYDKLR | ASSLDWGEAEQF | 0.34083152 |
| TLAVFTVL | ASSLDWGEAEQF | 0.4008417 |
| TGSVQPARV | ASSLDWGEAEQF | 0.23538223 |
| TGGGAGSGGT | ASSLDWGEAEQF | 0.28276917 |
| SHLETSL | ASSLDWGEAEQF | 0.21769162 |
| SFTTGMT | ASSLDWGEAEQF | 0.4975237 |
| QVEVAKRMGVRQPTVS | ASSLDWGEAEQF | 0.21831124 |
| QGSGGPALAM | ASSLDWGEAEQF | 0.20895396 |
| MLAPSNP | ASSLDWGEAEQF | 0.28267887 |
| LVNNAAI | ASSLDWGEAEQF | 0.74438953 |
| LTGEINTLA | ASSLDWGEAEQF | 0.55716425 |
| LSIFGPV | ASSLDWGEAEQF | 0.2635121 |
| LLALLANL | ASSLDWGEAEQF | 0.23497663 |
| LISNASDALDKLR | ASSLDWGEAEQF | 0.6517573 |
| LINEAVP | ASSLDWGEAEQF | 0.2818774 |
| LGYAFSGL | ASSLDWGEAEQF | 0.49733424 |
| LDFVAVAL | ASSLDWGEAEQF | 0.5018101 |
| LAQPGAE | ASSLDWGEAEQF | 0.47134557 |

| | | |
|---|---|---|
| ISNLGAVTGF | ASSLDWGEAEQF | 0.27347577 |
| HQEDPGRI | ASSLDWGEAEQF | 0.46006915 |
| GTPGEPTP | ASSLDWGEAEQF | 0.39003214 |
| GFFNSGS | ASSLDWGEAEQF | 0.41868997 |
| GAGGGGLTAGAN | ASSLDWGEAEQF | 0.17726167 |
| FVDALVNM | ASSLDWGEAEQF | 0.48864755 |
| FLTVAPEL | ASSLDWGEAEQF | 0.23310517 |
| FETLTAL | ASSLDWGEAEQF | 0.19956067 |
| DVINPIEL | ASSLDWGEAEQF | 0.17286469 |
| APVSGAPV | ASSLDWGEAEQF | 0.3092141 |
| ALVGGLAW | ASSLDWGEAEQF | 0.46060213 |
| ALAAAAATAI | ASSLDWGEAEQF | 0.6119824 |
| AFGLGLLL | ASSLDWGEAEQF | 0.40257058 |
| AFFAAASL | ASSLDWGEAEQF | 0.43928215 |
| AAAALPAFSPP | ASSLDWGEAEQF | 0.66597456 |
| VVLAPAL | ASSLDWGEAEQF | 0.36138672 |
| TIGHVDHGKTTL | ASSLDWGEAEQF | 0.1766581 |
| LISNASDALDK | ASSLDWGEAEQF | 0.35688707 |
| LISNASDAL | ASSLDWGEAEQF | 0.5342612 |
| ISNASDALDK | ASSLDWGEAEQF | 0.65903056 |
| NQAETLVYQTE | ASSLDWGEAEQF | 0.47478193 |
| TGGTGGTAGAG | ASSLDWGEAEQF | 0.19505729 |
| TGSTAGGAGGA | ASSLDWGEAEQF | 0.3928097 |
| DPLAVPVAAGPV | ASSLDWGEAEQF | 0.4451498 |
| GGTGGTPGNGGN | ASSLDWGEAEQF | 0.1884613 |
| PAGLAGVDPATL | ASSLDWGEAEQF | 0.24764886 |
| GNNLVVEAPHAGASV | ASSLDWGEAEQF | 0.26745397 |
| KVTRIMSASTLEQPA | ASSLDWGEAEQF | 0.20444004 |
| LGFALPIALPRRTVV | ASSLDWGEAEQF | 0.30457556 |
| TRLGNLTARKLVVLV | ASSLDWGEAEQF | 0.2525988 |
| WSQQRRLIYFPSAGP | ASSLDWGEAEQF | 0.25495347 |
| GPNGNPDPMAAAVDIR | ASSLDWGEAEQF | 0.2796104 |
| TQRCVSVADLRAAASR | ASSLDWGEAEQF | 0.21035272 |
| QFVLDVAEHPVEVLTETV | ASSLDWGEAEQF | 0.27581412 |
| GGAGGTGGAGGNGGTQAGDG | ASSLDWGEAEQF | 0.2707153 |
| PLLATLRAIAVDVDAKPAQV | ASSLDWGEAEQF | 0.3988961 |
| APKFRTGGYMIDFWGVGYQVAK | ASSLDWGEAEQF | 0.50620145 |
| GAGGAGGAGGAAGGPGGTGGTG | ASSLDWGEAEQF | 0.3024736 |
| IGALGLLATDVAPAGSVAFGVA | ASSLDWGEAEQF | 0.35901254 |
| AGGAGGIGGNGGAGGAGGVLMG | ASSLDWGEAEQF | 0.7131962 |
| ADELIGLL | ASSLDWGEAEQF | 0.20346336 |
| ADRALKA | ASSLDWGEAEQF | 0.26191956 |
| AGAVGQVGGAGGA | ASSLDWGEAEQF | 0.37151736 |
| AGGGGGLGGAP | ASSLDWGEAEQF | 0.20241249 |
| AGGLLYGNGGAG | ASSLDWGEAEQF | 0.5870997 |
| AGLESIGL | ASSLDWGEAEQF | 0.34390253 |
| AGLKTGII | ASSLDWGEAEQF | 0.42829978 |
| ASLAALLL | ASSLDWGEAEQF | 0.2762412 |
| ASVINISSI | ASSLDWGEAEQF | 0.7273506 |
| ATPPGPA | ASSLDWGEAEQF | 0.3695249 |
| ATVASALV | ASSLDWGEAEQF | 0.29827714 |

| | | |
|---|---|---|
| ATVASLIL | ASSLDWGEAEQF | 0.43616638 |
| AVNGPVLIRH | ASSLDWGEAEQF | 0.22652365 |
| DGAAGAVGI | ASSLDWGEAEQF | 0.48235667 |
| DPLLHGI | ASSLDWGEAEQF | 0.5128455 |
| ELGAALGAL | ASSLDWGEAEQF | 0.2610852 |
| ELGRTQAV | ASSLDWGEAEQF | 0.31165272 |
| FVAGGYLS | ASSLDWGEAEQF | 0.62367404 |
| GASGGAGGAGGW | ASSLDWGEAEQF | 0.59723556 |
| GGAGAGGDGGAGGI | ASSLDWGEAEQF | 0.1977604 |
| GTGHNGSQPGMGGNG | ASSLDWGEAEQF | 0.23172179 |
| GYAAASASAS | ASSLDWGEAEQF | 0.19062218 |
| HQVDVAAVQQG | ASSLDWGEAEQF | 0.21827853 |
| IGAAIAHT | ASSLDWGEAEQF | 0.2073377 |
| IGMIEVAQL | ASSLDWGEAEQF | 0.37134427 |
| IRLAPEHTSAMD | ASSLDWGEAEQF | 0.35499787 |
| IYNVSVL | ASSLDWGEAEQF | 0.6951964 |
| KQGFLGM | ASSLDWGEAEQF | 0.47970062 |
| LESVITQL | ASSLDWGEAEQF | 0.43265855 |
| LGLGALVVV | ASSLDWGEAEQF | 0.24241762 |
| LIGNGGNGGSGGI | ASSLDWGEAEQF | 0.32086667 |
| LNAINAPTL | ASSLDWGEAEQF | 0.20092413 |
| LQLLNTGV | ASSLDWGEAEQF | 0.19700778 |
| LQTSIEF | ASSLDWGEAEQF | 0.21782158 |
| MSFAVSEL | ASSLDWGEAEQF | 0.28237623 |
| NTGALGL | ASSLDWGEAEQF | 0.17776284 |
| PPGAGGAGGAGTT | ASSLDWGEAEQF | 0.6806015 |
| QLVGLGI | ASSLDWGEAEQF | 0.27973932 |
| RFLEQVHQL | ASSLDWGEAEQF | 0.36239427 |
| RGVGYVLR | ASSLDWGEAEQF | 0.27495927 |
| SATGVFAGVI | ASSLDWGEAEQF | 0.21013823 |
| SLPAITLP | ASSLDWGEAEQF | 0.30293915 |
| STALAFTV | ASSLDWGEAEQF | 0.42878726 |
| SVLTVGLGK | ASSLDWGEAEQF | 0.1959355 |
| TALAGAGSGI | ASSLDWGEAEQF | 0.21535343 |
| TGLPFTGI | ASSLDWGEAEQF | 0.21994825 |
| VAEHVTV | ASSLDWGEAEQF | 0.38315403 |
| VDIVLNSL | ASSLDWGEAEQF | 0.74920666 |
| VLTGGGAVL | ASSLDWGEAEQF | 0.539663 |
| VNVGLAYGI | ASSLDWGEAEQF | 0.36014324 |
| VSSAAVGAV | ASSLDWGEAEQF | 0.7453463 |
| VTSVYEAGV | ASSLDWGEAEQF | 0.16940391 |
| WVALDRGA | ASSLDWGEAEQF | 0.3216496 |
| YLVGSVVI | ASSLDWGEAEQF | 0.6665459 |
| GPPGTGKTLL | ASSLDWGEAEQF | 0.3690015 |
| GVGMTKF | ASSLDWGEAEQF | 0.36852276 |
| IGIGGSDL | ASSLDWGEAEQF | 0.5789302 |
| IGTLLAEL | ASSLDWGEAEQF | 0.24959312 |
| IYTSGTT | ASSLDWGEAEQF | 0.29069784 |
| LSFPAGV | ASSLDWGEAEQF | 0.27364263 |
| LYTSGTT | ASSLDWGEAEQF | 0.43927634 |
| VGAFPFGF | ASSLDWGEAEQF | 0.3478744 |

| | | |
|---|---|---|
| VGAGTVEF | ASSLDWGEAEQF | 0.19406888 |
| VGKSTLLL | ASSLDWGEAEQF | 0.46756792 |
| AALAHAGGGIV | ASSLDWGEAEQF | 0.7847004 |
| AALVAVSLL | ASSLDWGEAEQF | 0.23444709 |
| AAVSRAVAAL | ASSLDWGEAEQF | 0.636803 |
| AGGSSGIGGF | ASSLDWGEAEQF | 0.33883092 |
| AGLTVANANG | ASSLDWGEAEQF | 0.24429967 |
| AGLVGAVL | ASSLDWGEAEQF | 0.33586943 |
| ALSQAFTV | ASSLDWGEAEQF | 0.21309893 |
| ALTFSVSL | ASSLDWGEAEQF | 0.47657657 |
| ALTGAALVAL | ASSLDWGEAEQF | 0.25499153 |
| ASFSIGAGVL | ASSLDWGEAEQF | 0.35296834 |
| DGSVYKI | ASSLDWGEAEQF | 0.4102555 |
| EQLAEIV | ASSLDWGEAEQF | 0.780902 |
| FAVSELTVM | ASSLDWGEAEQF | 0.4147801 |
| GAAPLAAGL | ASSLDWGEAEQF | 0.44336736 |
| GADGGDGGSGGAGG | ASSLDWGEAEQF | 0.23949194 |
| GFGLDTGGAGG | ASSLDWGEAEQF | 0.40603793 |
| GLASGGLL | ASSLDWGEAEQF | 0.41405493 |
| GRIAGAAVLL | ASSLDWGEAEQF | 0.24042057 |
| GSAGSLNTSF | ASSLDWGEAEQF | 0.53918463 |
| GTGQIGI | ASSLDWGEAEQF | 0.2441136 |
| GTLIALIL | ASSLDWGEAEQF | 0.45192957 |
| IAATMGFVVG | ASSLDWGEAEQF | 0.35671633 |
| IFAQALL | ASSLDWGEAEQF | 0.7547312 |
| IGAAVALGF | ASSLDWGEAEQF | 0.41861504 |
| IGLTPLAL | ASSLDWGEAEQF | 0.36957422 |
| ILGSNTSTL | ASSLDWGEAEQF | 0.20878589 |
| IVFSLTVL | ASSLDWGEAEQF | 0.44445926 |
| LASKATGF | ASSLDWGEAEQF | 0.5156496 |
| LASVALIL | ASSLDWGEAEQF | 0.34115267 |
| LFAVGLTL | ASSLDWGEAEQF | 0.7827527 |
| LFNGGIGTYI | ASSLDWGEAEQF | 0.36943877 |
| LTLGAGII | ASSLDWGEAEQF | 0.19410434 |
| LVAAALGDI | ASSLDWGEAEQF | 0.28239667 |
| NTAFVQLGI | ASSLDWGEAEQF | 0.608551 |
| QLFSGAEL | ASSLDWGEAEQF | 0.7189371 |
| SAATFITL | ASSLDWGEAEQF | 0.36973897 |
| SNYGLANLGV | ASSLDWGEAEQF | 0.34157026 |
| SYGGAYAVM | ASSLDWGEAEQF | 0.30125445 |
| TTGVGGLTLG | ASSLDWGEAEQF | 0.6435977 |
| TTQTIYI | ASSLDWGEAEQF | 0.75133574 |
| VAAGAGLGAVI | ASSLDWGEAEQF | 0.42653817 |
| VAAVVTLL | ASSLDWGEAEQF | 0.22614874 |
| VGFLGGLI | ASSLDWGEAEQF | 0.20850451 |
| VGVIPILL | ASSLDWGEAEQF | 0.35718632 |
| VIFASSLI | ASSLDWGEAEQF | 0.6242782 |
| VKGGDGGAAAG | ASSLDWGEAEQF | 0.22408386 |
| VTKTFTNI | ASSLDWGEAEQF | 0.37598503 |
| VVAAAILL | ASSLDWGEAEQF | 0.2497279 |
| YAALDVEL | ASSLDWGEAEQF | 0.18267325 |

| | | |
|---|---|---|
| LASVALLL | ASSLDWGEAEQF | 0.4641978 |
| LDAVHRV | ASSLDWGEAEQF | 0.70128864 |
| QGLSSSIF | ASSLDWGEAEQF | 0.43833107 |
| RGTFANIRL | ASSLDWGEAEQF | 0.1968322 |
| RQLWWGHRI | ASSLDWGEAEQF | 0.587116 |
| YGPPGTGKTLL | ASSLDWGEAEQF | 0.734833 |
| AALVGAPANVA | ASSLDWGEAEQF | 0.61484003 |
| AATFRRTE | ASSLDWGEAEQF | 0.21088536 |
| AGALGGGI | ASSLDWGEAEQF | 0.16695382 |
| AGGAGGAGGIGAG | ASSLDWGEAEQF | 0.34196883 |
| AGSGALGL | ASSLDWGEAEQF | 0.6136464 |
| AQIADQI | ASSLDWGEAEQF | 0.22594377 |
| ASAVAVTYL | ASSLDWGEAEQF | 0.2106593 |
| ASTVLGGTGG | ASSLDWGEAEQF | 0.32409084 |
| AVGGEPAK | ASSLDWGEAEQF | 0.5992171 |
| CEQGTFGGTC | ASSLDWGEAEQF | 0.21349305 |
| DEGKAPAG | ASSLDWGEAEQF | 0.19268015 |
| DTVVFPAL | ASSLDWGEAEQF | 0.39150348 |
| FEGITMAG | ASSLDWGEAEQF | 0.5280409 |
| FPLVVQD | ASSLDWGEAEQF | 0.22952363 |
| GAFTTPQ | ASSLDWGEAEQF | 0.26891539 |
| GFGGFGTAGGVGGA | ASSLDWGEAEQF | 0.42605716 |
| GFLAQFL | ASSLDWGEAEQF | 0.4748379 |
| GGAGGAGGNAGVA | ASSLDWGEAEQF | 0.3624504 |
| GGVGTAGAGGAGGAGG | ASSLDWGEAEQF | 0.20559064 |
| GNGGGGDAGNAGSGGN | ASSLDWGEAEQF | 0.6848922 |
| GTGGAGGAGNDAGS | ASSLDWGEAEQF | 0.22986764 |
| GTTNTGWFNS | ASSLDWGEAEQF | 0.51045 |
| IFMGPLFTL | ASSLDWGEAEQF | 0.28937006 |
| ILAHAIEAI | ASSLDWGEAEQF | 0.3928687 |
| IRSAIREAL | ASSLDWGEAEQF | 0.34828836 |
| ITFAGGG | ASSLDWGEAEQF | 0.27223554 |
| ITGMSGHQEV | ASSLDWGEAEQF | 0.21297303 |
| IVAGLACTL | ASSLDWGEAEQF | 0.17739028 |
| IVGGGLAGA | ASSLDWGEAEQF | 0.553699 |
| IVTGVLNI | ASSLDWGEAEQF | 0.23816873 |
| LAQQGCI | ASSLDWGEAEQF | 0.5319097 |
| LCLHVEN | ASSLDWGEAEQF | 0.37713256 |
| LDAISDGS | ASSLDWGEAEQF | 0.22192737 |
| LGGFAAML | ASSLDWGEAEQF | 0.44612843 |
| LGGKNPMI | ASSLDWGEAEQF | 0.18830135 |
| LGPKGVAAIL | ASSLDWGEAEQF | 0.3687584 |
| LMVGVSI | ASSLDWGEAEQF | 0.2600969 |
| LVGAAFVL | ASSLDWGEAEQF | 0.42378166 |
| LVLAVAPL | ASSLDWGEAEQF | 0.367561 |
| MAAIAVIDAA | ASSLDWGEAEQF | 0.43008277 |
| NNGINAELQ | ASSLDWGEAEQF | 0.45484877 |
| RGEPLVLVHG | ASSLDWGEAEQF | 0.20187517 |
| RGGAGGAAGLI | ASSLDWGEAEQF | 0.4262463 |
| RSYRMHPAV | ASSLDWGEAEQF | 0.6472665 |
| SAAHLLAGQLA | ASSLDWGEAEQF | 0.3847468 |

| | | |
|---|---|---|
| SAAVGSY | ASSLDWGEAEQF | 0.18541795 |
| SPHAVLHRI | ASSLDWGEAEQF | 0.4526881 |
| SQIAKVL | ASSLDWGEAEQF | 0.26274022 |
| TALGHLVP | ASSLDWGEAEQF | 0.42044157 |
| TAPTTSGAA | ASSLDWGEAEQF | 0.33326733 |
| TDGDDGNGGSPGT | ASSLDWGEAEQF | 0.24773693 |
| TDGLGEN | ASSLDWGEAEQF | 0.36511683 |
| TGGPPLPGQA | ASSLDWGEAEQF | 0.20599522 |
| TIPAITI | ASSLDWGEAEQF | 0.21655826 |
| TIPGFTI | ASSLDWGEAEQF | 0.35279882 |
| TSIASGALA | ASSLDWGEAEQF | 0.3615147 |
| TVTLPGAAI | ASSLDWGEAEQF | 0.49017483 |
| VASGLSAALP | ASSLDWGEAEQF | 0.2235383 |
| VGAGVIGI | ASSLDWGEAEQF | 0.39502054 |
| VGPKRFGV | ASSLDWGEAEQF | 0.24083783 |
| VNVAFPD | ASSLDWGEAEQF | 0.2574375 |
| VSMTNSGVS | ASSLDWGEAEQF | 0.36176673 |
| VTDVLGDLSV | ASSLDWGEAEQF | 0.24158606 |
| VVVVGAGFAGL | ASSLDWGEAEQF | 0.19668533 |
| VYTVGTVV | ASSLDWGEAEQF | 0.48541817 |
| WTTFSGP | ASSLDWGEAEQF | 0.21085921 |
| YDELVEAF | ASSLDWGEAEQF | 0.28627202 |
| YGTGGAGGNGG | ASSLDWGEAEQF | 0.6971499 |
| YTNVMAV | ASSLDWGEAEQF | 0.18655433 |
| FLEQVHQL | ASSLDWGEAEQF | 0.46876347 |
| GLSLLAVAV | ASSLDWGEAEQF | 0.42049506 |
| LEQVHQL | ASSLDWGEAEQF | 0.36348355 |
| LNAAGAI | ASSLDWGEAEQF | 0.25903738 |
| PAVHASL | ASSLDWGEAEQF | 0.21933223 |
| TELEDDA | ASSLDWGEAEQF | 0.61059964 |
| TPPGPPN | ASSLDWGEAEQF | 0.46815917 |
| TPPPPPPP | ASSLDWGEAEQF | 0.27998874 |
| TQSPALI | ASGDAADRGLEQY | 0.39415476 |
| EFAAAFPQ | ASGDAADRGLEQY | 0.67542994 |
| LLSGNLDVL | ASGDAADRGLEQY | 0.48472577 |
| GPANLGNYNI | ASGDAADRGLEQY | 0.24552508 |
| TLGPTLP | ASGDAADRGLEQY | 0.18615073 |
| VYAYSSCCK | ASGDAADRGLEQY | 0.30316567 |
| APAPAPAPNI | ASGDAADRGLEQY | 0.31590965 |
| PGGPVPIP | ASGDAADRGLEQY | 0.2665281 |
| FLTVALEP | ASGDAADRGLEQY | 0.19644451 |
| SQYFGANL | ASGDAADRGLEQY | 0.6468469 |
| GGVKPAA | ASGDAADRGLEQY | 0.7852105 |
| GGTGGAGGAGGW | ASGDAADRGLEQY | 0.24144825 |
| GVGGVAGLGGAGG | ASGDAADRGLEQY | 0.773399 |
| VLLVLATTI | ASGDAADRGLEQY | 0.18949631 |
| IGIFAAML | ASGDAADRGLEQY | 0.384417 |
| AVGDLID | ASGDAADRGLEQY | 0.5662428 |
| NISPLPL | ASGDAADRGLEQY | 0.24819262 |
| VTAVFVAI | ASGDAADRGLEQY | 0.3421558 |
| SGDDNDAAL | ASGDAADRGLEQY | 0.21371321 |

| | | |
|---|---|---|
| AVAALAHS | ASGDAADRGLEQY | 0.24732457 |
| GGVTVMATP | ASGDAADRGLEQY | 0.5446742 |
| AGLAANVRD | ASGDAADRGLEQY | 0.23567456 |
| TNRPDLIDPALLRPG | ASGDAADRGLEQY | 0.32516629 |
| LANLGVTAL | ASGDAADRGLEQY | 0.23945972 |
| TGERIEL | ASGDAADRGLEQY | 0.22409764 |
| NGEELAA | ASGDAADRGLEQY | 0.37158793 |
| VLSLVWG | ASGDAADRGLEQY | 0.38201562 |
| IGYALWL | ASGDAADRGLEQY | 0.28290594 |
| LSLLLAPL | ASGDAADRGLEQY | 0.74591136 |
| TLDEALK | ASGDAADRGLEQY | 0.24966773 |
| TDGAGTAEGG | ASGDAADRGLEQY | 0.7841016 |
| NAPGGAGGNGG | ASGDAADRGLEQY | 0.7504196 |
| VSVFMGMGGNF | ASGDAADRGLEQY | 0.34522212 |
| LASYLAAL | ASGDAADRGLEQY | 0.30331403 |
| LVGGTPAL | ASGDAADRGLEQY | 0.6701999 |
| VRNADVL | ASGDAADRGLEQY | 0.24600297 |
| TCSSPFLVSAM | ASGDAADRGLEQY | 0.29229814 |
| VHITDAE | ASGDAADRGLEQY | 0.6235346 |
| EYAAPSF | ASGDAADRGLEQY | 0.652177 |
| VVDAFHGGL | ASGDAADRGLEQY | 0.75739336 |
| ASVVDPQ | ASGDAADRGLEQY | 0.7422788 |
| SYNEFGGLS | ASGDAADRGLEQY | 0.7206274 |
| DPDGSVY | ASGDAADRGLEQY | 0.7078614 |
| MVSTIHATE | ASGDAADRGLEQY | 0.43732494 |
| LAAGVADPV | ASGDAADRGLEQY | 0.71010315 |
| TKEQIAA | ASGDAADRGLEQY | 0.68474966 |
| SLASPAAL | ASGDAADRGLEQY | 0.73422253 |
| IGLVFYGV | ASGDAADRGLEQY | 0.7459326 |
| FLLDASNL | ASGDAADRGLEQY | 0.689448 |
| YQVGNFP | ASGDAADRGLEQY | 0.75623214 |
| ESLLSQST | ASGDAADRGLEQY | 0.75674677 |
| LFTVGFAAVG | ASGDAADRGLEQY | 0.5980856 |
| ATPPTPT | ASGDAADRGLEQY | 0.7233918 |
| TDLATLG | ASGDAADRGLEQY | 0.65607446 |
| SAGGTGR | ASGDAADRGLEQY | 0.7204573 |
| GFSNSGS | ASGDAADRGLEQY | 0.4026901 |
| GGNGGSGGTSV | ASGDAADRGLEQY | 0.75049114 |
| GTSVATGGAGN | ASGDAADRGLEQY | 0.68265885 |
| NGGNGGSGGTS | ASGDAADRGLEQY | 0.23947784 |
| AVVKFPH | ASGDAADRGLEQY | 0.7270251 |
| TPPPPPP | ASGDAADRGLEQY | 0.7209339 |
| AAPAAAAAPA | ASGDAADRGLEQY | 0.7248208 |
| IALNEDL | ASGDAADRGLEQY | 0.7487081 |
| IVNVSSM | ASGDAADRGLEQY | 0.5390412 |
| TGIPPAPR | ASGDAADRGLEQY | 0.6806805 |
| FDLGGGTF | ASGDAADRGLEQY | 0.749178 |
| FDLGGGTFDVS | ASGDAADRGLEQY | 0.7576122 |
| NEPTAAA | ASGDAADRGLEQY | 0.6202774 |
| NEPTAAALA | ASGDAADRGLEQY | 0.74128765 |
| GSIINISSI | ASGDAADRGLEQY | 0.757733 |

| | | |
|---|---|---|
| VAPGNAGTA | ASGDAADRGLEQY | 0.7442242 |
| TGGGASLE | ASGDAADRGLEQY | 0.7498303 |
| VAVYNHYKRI | ASGDAADRGLEQY | 0.75618565 |
| TDPAPATT | ASGDAADRGLEQY | 0.6510956 |
| TNRPDILDPALLRPG | ASGDAADRGLEQY | 0.7541375 |
| LTTTTTS | ASGDAADRGLEQY | 0.75636923 |
| PTGSGKTL | ASGDAADRGLEQY | 0.71503884 |
| TGSGKTL | ASGDAADRGLEQY | 0.72914004 |
| TIGHVDHGK | ASGDAADRGLEQY | 0.63223696 |
| TIGHVDHGKTT | ASGDAADRGLEQY | 0.7048594 |
| GHVDHGKTT | ASGDAADRGLEQY | 0.39748114 |
| AQFEHTLL | ASGDAADRGLEQY | 0.73699427 |
| IVNQDEL | ASGDAADRGLEQY | 0.7543287 |
| TNGSQFF | ASGDAADRGLEQY | 0.71494097 |
| TGKGIVE | ASGDAADRGLEQY | 0.61401653 |
| LISNASDA | ASGDAADRGLEQY | 0.67869234 |
| AELRTLGELTKIAKAHGAQ | ASGDAADRGLEQY | 0.7065532 |
| APNAVAAAEPSW | ASGDAADRGLEQY | 0.71976185 |
| APQVAGAATAVL | ASGDAADRGLEQY | 0.3534587 |
| ARRDPAAAAAILPTDA | ASGDAADRGLEQY | 0.6396523 |
| DAAARDVLAEAGLAEYF | ASGDAADRGLEQY | 0.75510705 |
| DDQIVLSAKANLAKVEQ | ASGDAADRGLEQY | 0.75753206 |
| DELLPDTGEDSDEDGH | ASGDAADRGLEQY | 0.5173028 |
| ELPEAPANRATVPQD | ASGDAADRGLEQY | 0.7530261 |
| FGQSKIERVTGIG | ASGDAADRGLEQY | 0.7416779 |
| GAGGTGSTAGGAGGAGGAGGLY | ASGDAADRGLEQY | 0.7595241 |
| GHGGTGAAGGNGGTGGAGGSGI | ASGDAADRGLEQY | 0.7278905 |
| GNGGNGGAGGEGGPGGAGGAGA | ASGDAADRGLEQY | 0.6514719 |
| GSTAGNGGNGGNATNGTIAGQP | ASGDAADRGLEQY | 0.7378992 |
| GTGGDGGNGGNGGTGG | ASGDAADRGLEQY | 0.7130009 |
| GVRRPAGIPADAPV | ASGDAADRGLEQY | 0.5752243 |
| GYRQVTDAHLLTLARR | ASGDAADRGLEQY | 0.74848 |
| IAKMGQDGHDRG | ASGDAADRGLEQY | 0.71938276 |
| KAEPSIEIQYT | ASGDAADRGLEQY | 0.69531834 |
| KPVRIRMAADEDTVAVT | ASGDAADRGLEQY | 0.6973331 |
| MPPRIAGMRLLVIKPEP | ASGDAADRGLEQY | 0.7521676 |
| MKFYAVALFRSRGVPPTR | ASGDAADRGLEQY | 0.6506078 |
| NAGGFGILGAVAHSP | ASGDAADRGLEQY | 0.6183183 |
| NEGREITLAMAEM | ASGDAADRGLEQY | 0.6583236 |
| NELVEWSHPLGHRWRWE | ASGDAADRGLEQY | 0.73620236 |
| NLGAGGTGGNGGNPG | ASGDAADRGLEQY | 0.7536831 |
| NTEWPKWETVMAADPVD | ASGDAADRGLEQY | 0.74020785 |
| PASAAPAETQP | ASGDAADRGLEQY | 0.69530743 |
| QQSSDGKNGVRAAEITGE | ASGDAADRGLEQY | 0.74538136 |
| RAPDAVTTSWWKE | ASGDAADRGLEQY | 0.746099 |
| SGFGNVGAHTSGWFNQ | ASGDAADRGLEQY | 0.73055696 |
| SGVAGAVGQAASVGGLKV | ASGDAADRGLEQY | 0.7413568 |
| SNGGAGTGSAGGNGGT | ASGDAADRGLEQY | 0.68517303 |
| TGGAGGEGGAAG | ASGDAADRGLEQY | 0.7520741 |
| TGGDGGAAGAGGAPGHG | ASGDAADRGLEQY | 0.7407476 |
| TMPPLLAAIANAGRHGV | ASGDAADRGLEQY | 0.7486833 |

| | | |
|---|---|---|
| TSSTAGNSGGA | ASGDAADRGLEQY | 0.7464707 |
| VPATVPVGILC | ASGDAADRGLEQY | 0.7403061 |
| VQRPDSGGVVLTGRIS | ASGDAADRGLEQY | 0.73410106 |
| WTPNPVPATKGVPAGLVQAT | ASGDAADRGLEQY | 0.4410126 |
| YALKRALAKFSDRFDVVII | ASGDAADRGLEQY | 0.6966214 |
| YLIGSVIV | ASGDAADRGLEQY | 0.7343207 |
| YGQLVAVID | ASGDAADRGLEQY | 0.7374231 |
| VTVGGMTLPA | ASGDAADRGLEQY | 0.7421958 |
| VLALPAV | ASGDAADRGLEQY | 0.7333304 |
| VIPPTLNYET | ASGDAADRGLEQY | 0.6946459 |
| VGMGGMHPSSG | ASGDAADRGLEQY | 0.748508 |
| VGLAGGAATA | ASGDAADRGLEQY | 0.7517861 |
| VGGAGGAGGSGP | ASGDAADRGLEQY | 0.5319563 |
| VFAAFMV | ASGDAADRGLEQY | 0.6961982 |
| TLYDKLR | ASGDAADRGLEQY | 0.7594898 |
| TLAVFTVL | ASGDAADRGLEQY | 0.72658455 |
| TGSVQPARV | ASGDAADRGLEQY | 0.719089 |
| TGGGAGSGGT | ASGDAADRGLEQY | 0.69088054 |
| SHLETSL | ASGDAADRGLEQY | 0.7556034 |
| SFTTGMT | ASGDAADRGLEQY | 0.6441753 |
| QVEVAKRMGVRQPTVS | ASGDAADRGLEQY | 0.75983065 |
| QGSGGPALAM | ASGDAADRGLEQY | 0.31917888 |
| MLAPSNP | ASGDAADRGLEQY | 0.6975137 |
| LVNNAAI | ASGDAADRGLEQY | 0.6570273 |
| LTGEINTLA | ASGDAADRGLEQY | 0.4729115 |
| LSIFGPV | ASGDAADRGLEQY | 0.7543439 |
| LLALLANL | ASGDAADRGLEQY | 0.70062447 |
| LISNASDALDKLR | ASGDAADRGLEQY | 0.75120807 |
| LINEAVP | ASGDAADRGLEQY | 0.7226249 |
| LGYAFSGL | ASGDAADRGLEQY | 0.74971545 |
| LDFVAVAL | ASGDAADRGLEQY | 0.7300836 |
| LAQPGAE | ASGDAADRGLEQY | 0.73518896 |
| ISNLGAVTGF | ASGDAADRGLEQY | 0.6030445 |
| HQEDPGRI | ASGDAADRGLEQY | 0.75191957 |
| GTPGEPTP | ASGDAADRGLEQY | 0.75055104 |
| GFFNSGS | ASGDAADRGLEQY | 0.7375053 |
| GAGGGGLTAGAN | ASGDAADRGLEQY | 0.58090127 |
| FVDALVNM | ASGDAADRGLEQY | 0.673922 |
| FLTVAPEL | ASGDAADRGLEQY | 0.7427488 |
| FETLTAL | ASGDAADRGLEQY | 0.2593301 |
| DVINPIEL | ASGDAADRGLEQY | 0.27017683 |
| APVSGAPV | ASGDAADRGLEQY | 0.7032459 |
| ALVGGLAW | ASGDAADRGLEQY | 0.62058043 |
| ALAAAAATAI | ASGDAADRGLEQY | 0.7547438 |
| AFGLGLLL | ASGDAADRGLEQY | 0.74913156 |
| AFFAAASL | ASGDAADRGLEQY | 0.7253534 |
| AAAALPAFSPP | ASGDAADRGLEQY | 0.7474091 |
| VVLAPAL | ASGDAADRGLEQY | 0.7370899 |
| TIGHVDHGKTTL | ASGDAADRGLEQY | 0.2187519 |
| LISNASDALDK | ASGDAADRGLEQY | 0.6988709 |
| LISNASDAL | ASGDAADRGLEQY | 0.6195291 |

| | | |
|---|---|---|
| ISNASDALDK | ASGDAADRGLEQY | 0.7549801 |
| NQAETLVYQTE | ASGDAADRGLEQY | 0.6510347 |
| TGGTGGTAGAG | ASGDAADRGLEQY | 0.6654775 |
| TGSTAGGAGGA | ASGDAADRGLEQY | 0.7303459 |
| DPLAVPVAAGPV | ASGDAADRGLEQY | 0.7259668 |
| GGTGGTPGNGGN | ASGDAADRGLEQY | 0.7224674 |
| PAGLAGVDPATL | ASGDAADRGLEQY | 0.6445404 |
| GNNLVVEAPHAGASV | ASGDAADRGLEQY | 0.25252983 |
| KVTRIMSASTLEQPA | ASGDAADRGLEQY | 0.7523415 |
| LGFALPIALPRRTVV | ASGDAADRGLEQY | 0.4071675 |
| TRLGNLTARKLVVLV | ASGDAADRGLEQY | 0.6628089 |
| WSQQRRLIYFPSAGP | ASGDAADRGLEQY | 0.3151846 |
| GPNGNPDPMAAAVDIR | ASGDAADRGLEQY | 0.55931807 |
| TQRCVSVADLRAAASR | ASGDAADRGLEQY | 0.4757185 |
| QFVLDVAEHPVEVLTETV | ASGDAADRGLEQY | 0.7458651 |
| GGAGGTGGAGGNGGTQAGDG | ASGDAADRGLEQY | 0.7107297 |
| PLLATLRAIAVDVDAKPAQV | ASGDAADRGLEQY | 0.6987341 |
| APKFRTGGYMIDFWGVGYQVAK | ASGDAADRGLEQY | 0.7566651 |
| GAGGAGGAGGAAGGPGGTGGTG | ASGDAADRGLEQY | 0.72963834 |
| IGALGLLATDVAPAGSVAFGVA | ASGDAADRGLEQY | 0.704128 |
| AGGAGGIGGNGGAGGAGGVLMG | ASGDAADRGLEQY | 0.7059221 |
| ADELIGLL | ASGDAADRGLEQY | 0.6522083 |
| ADRALKA | ASGDAADRGLEQY | 0.666496 |
| AGAVGQVGGAGGA | ASGDAADRGLEQY | 0.6668699 |
| AGGGGGLGGAP | ASGDAADRGLEQY | 0.7588799 |
| AGGLLYGNGGAG | ASGDAADRGLEQY | 0.6872381 |
| AGLESIGL | ASGDAADRGLEQY | 0.5224371 |
| AGLKTGII | ASGDAADRGLEQY | 0.70974755 |
| ASLAALLL | ASGDAADRGLEQY | 0.756813 |
| ASVINISSI | ASGDAADRGLEQY | 0.761761 |
| ATPPGPA | ASGDAADRGLEQY | 0.6748906 |
| ATVASALV | ASGDAADRGLEQY | 0.7170055 |
| ATVASLIL | ASGDAADRGLEQY | 0.70095026 |
| AVNGPVLIRH | ASGDAADRGLEQY | 0.729088 |
| DGAAGAVGI | ASGDAADRGLEQY | 0.69058645 |
| DPLLHGI | ASGDAADRGLEQY | 0.5251434 |
| ELGAALGAL | ASGDAADRGLEQY | 0.7479185 |
| ELGRTQAV | ASGDAADRGLEQY | 0.7608106 |
| FVAGGYLS | ASGDAADRGLEQY | 0.68989575 |
| GASGGAGGAGGW | ASGDAADRGLEQY | 0.507351 |
| GGAGAGGDGGAGGI | ASGDAADRGLEQY | 0.7272401 |
| GTGHNGSQPGMGGNG | ASGDAADRGLEQY | 0.68425727 |
| GYAAASASAS | ASGDAADRGLEQY | 0.7131883 |
| HQVDVAAVQQG | ASGDAADRGLEQY | 0.7387186 |
| IGAAIAHT | ASGDAADRGLEQY | 0.51206267 |
| IGMIEVAQL | ASGDAADRGLEQY | 0.68621194 |
| IRLAPEHTSAMD | ASGDAADRGLEQY | 0.76084405 |
| IYNVSVL | ASGDAADRGLEQY | 0.7127865 |
| KQGFLGM | ASGDAADRGLEQY | 0.73542506 |
| LESVITQL | ASGDAADRGLEQY | 0.75279295 |
| LGLGALVVV | ASGDAADRGLEQY | 0.7410028 |

| | | |
|---|---|---|
| LIGNGGNGGSGGI | ASGDAADRGLEQY | 0.7552784 |
| LNAINAPTL | ASGDAADRGLEQY | 0.7439103 |
| LQLLNTGV | ASGDAADRGLEQY | 0.69405997 |
| LQTSIEF | ASGDAADRGLEQY | 0.72797275 |
| MSFAVSEL | ASGDAADRGLEQY | 0.7241022 |
| NTGALGL | ASGDAADRGLEQY | 0.7342228 |
| PPGAGGAGGAGTT | ASGDAADRGLEQY | 0.7208615 |
| QLVGLGI | ASGDAADRGLEQY | 0.6598346 |
| RFLEQVHQL | ASGDAADRGLEQY | 0.7561041 |
| RGVGYVLR | ASGDAADRGLEQY | 0.7292793 |
| SATGVFAGVI | ASGDAADRGLEQY | 0.7479796 |
| SLPAITLP | ASGDAADRGLEQY | 0.6277184 |
| STALAFTV | ASGDAADRGLEQY | 0.6433871 |
| SVLTVGLGK | ASGDAADRGLEQY | 0.7503625 |
| TALAGAGSGI | ASGDAADRGLEQY | 0.24994911 |
| TGLPFTGI | ASGDAADRGLEQY | 0.2967847 |
| VAEHVTV | ASGDAADRGLEQY | 0.7316964 |
| VDIVLNSL | ASGDAADRGLEQY | 0.756798 |
| VLTGGGAVL | ASGDAADRGLEQY | 0.23981935 |
| VNVGLAYGI | ASGDAADRGLEQY | 0.6530107 |
| VSSAAVGAV | ASGDAADRGLEQY | 0.60353106 |
| VTSVYEAGV | ASGDAADRGLEQY | 0.7341318 |
| WVALDRGA | ASGDAADRGLEQY | 0.7277092 |
| YLVGSVVI | ASGDAADRGLEQY | 0.47068697 |
| GPPGTGKTLL | ASGDAADRGLEQY | 0.7476102 |
| GVGMTKF | ASGDAADRGLEQY | 0.44738907 |
| IGIGGSDL | ASGDAADRGLEQY | 0.42862302 |
| IGTLLAEL | ASGDAADRGLEQY | 0.74702746 |
| IYTSGTT | ASGDAADRGLEQY | 0.7508069 |
| LSFPAGV | ASGDAADRGLEQY | 0.57992005 |
| LYTSGTT | ASGDAADRGLEQY | 0.7505355 |
| VGAFPFGF | ASGDAADRGLEQY | 0.68847144 |
| VGAGTVEF | ASGDAADRGLEQY | 0.7611399 |
| VGKSTLLL | ASGDAADRGLEQY | 0.71479315 |
| AALAHAGGGIV | ASGDAADRGLEQY | 0.4623972 |
| AALVAVSLL | ASGDAADRGLEQY | 0.7495891 |
| AAVSRAVAAL | ASGDAADRGLEQY | 0.7381626 |
| AGGSSGIGGF | ASGDAADRGLEQY | 0.7335819 |
| AGLTVANANG | ASGDAADRGLEQY | 0.7420298 |
| AGLVGAVL | ASGDAADRGLEQY | 0.6618879 |
| ALSQAFTV | ASGDAADRGLEQY | 0.71987665 |
| ALTFSVSL | ASGDAADRGLEQY | 0.7559843 |
| ALTGAALVAL | ASGDAADRGLEQY | 0.7453556 |
| ASFSIGAGVL | ASGDAADRGLEQY | 0.5512606 |
| DGSVYKI | ASGDAADRGLEQY | 0.7449518 |
| EQLAEIV | ASGDAADRGLEQY | 0.72960365 |
| FAVSELTVM | ASGDAADRGLEQY | 0.6729439 |
| GAAPLAAGL | ASGDAADRGLEQY | 0.75929767 |
| GADGGDGGSGGAGG | ASGDAADRGLEQY | 0.7190201 |
| GFGLDTGGAGG | ASGDAADRGLEQY | 0.74906176 |
| GLASGGLL | ASGDAADRGLEQY | 0.75315243 |

| | | |
|---|---|---|
| GRIAGAAVLL | ASGDAADRGLEQY | 0.7558396 |
| GSAGSLNTSF | ASGDAADRGLEQY | 0.51065403 |
| GTGQIGI | ASGDAADRGLEQY | 0.7391701 |
| GTLIALIL | ASGDAADRGLEQY | 0.73109823 |
| IAATMGFVVG | ASGDAADRGLEQY | 0.7450844 |
| IFAQALL | ASGDAADRGLEQY | 0.7169136 |
| IGAAVALGF | ASGDAADRGLEQY | 0.5525724 |
| IGLTPLAL | ASGDAADRGLEQY | 0.738907 |
| ILGSNTSTL | ASGDAADRGLEQY | 0.75374746 |
| IVFSLTVL | ASGDAADRGLEQY | 0.7198022 |
| LASKATGF | ASGDAADRGLEQY | 0.72910255 |
| LASVALIL | ASGDAADRGLEQY | 0.75813186 |
| LFAVGLTL | ASGDAADRGLEQY | 0.67589957 |
| LFNGGIGTYI | ASGDAADRGLEQY | 0.7189996 |
| LTLGAGII | ASGDAADRGLEQY | 0.7487958 |
| LVAAALGDI | ASGDAADRGLEQY | 0.74685323 |
| NTAFVQLGI | ASGDAADRGLEQY | 0.7076466 |
| QLFSGAEL | ASGDAADRGLEQY | 0.756619 |
| SAATFITL | ASGDAADRGLEQY | 0.65562224 |
| SNYGLANLGV | ASGDAADRGLEQY | 0.7512504 |
| SYGGAYAVM | ASGDAADRGLEQY | 0.67903066 |
| TTGVGGLTLG | ASGDAADRGLEQY | 0.74287665 |
| TTQTIYI | ASGDAADRGLEQY | 0.6710472 |
| VAAGAGLGAVI | ASGDAADRGLEQY | 0.74819785 |
| VAAVVTLL | ASGDAADRGLEQY | 0.7493205 |
| VGFLGGLI | ASGDAADRGLEQY | 0.7489834 |
| VGVIPILL | ASGDAADRGLEQY | 0.28602698 |
| VIFASSLI | ASGDAADRGLEQY | 0.7501056 |
| VKGGDGGAAAG | ASGDAADRGLEQY | 0.7547992 |
| VTKTFTNI | ASGDAADRGLEQY | 0.7575443 |
| VVAAAILL | ASGDAADRGLEQY | 0.72937596 |
| YAALDVEL | ASGDAADRGLEQY | 0.74262834 |
| LASVALLL | ASGDAADRGLEQY | 0.72424066 |
| LDAVHRV | ASGDAADRGLEQY | 0.7102169 |
| QGLSSSIF | ASGDAADRGLEQY | 0.6736642 |
| RGTFANIRL | ASGDAADRGLEQY | 0.7225102 |
| RQLWWGHRI | ASGDAADRGLEQY | 0.70233786 |
| YGPPGTGKTLL | ASGDAADRGLEQY | 0.6924578 |
| AALVGAPANVA | ASGDAADRGLEQY | 0.74645126 |
| AATFRRTE | ASGDAADRGLEQY | 0.7480804 |
| AGALGGGI | ASGDAADRGLEQY | 0.75022197 |
| AGGAGGAGGIGAG | ASGDAADRGLEQY | 0.7563587 |
| AGSGALGL | ASGDAADRGLEQY | 0.75345266 |
| AQIADQI | ASGDAADRGLEQY | 0.7503137 |
| ASAVAVTYL | ASGDAADRGLEQY | 0.7032901 |
| ASTVLGGTGG | ASGDAADRGLEQY | 0.71983635 |
| AVGGEPAK | ASGDAADRGLEQY | 0.6321715 |
| CEQGTFGGTC | ASGDAADRGLEQY | 0.68750215 |
| DEGKAPAG | ASGDAADRGLEQY | 0.70178825 |
| DTVVFPAL | ASGDAADRGLEQY | 0.73267597 |
| FEGITMAG | ASGDAADRGLEQY | 0.2338139 |

| | | |
|---|---|---|
| FPLVVQD | ASGDAADRGLEQY | 0.71433073 |
| GAFTTPQ | ASGDAADRGLEQY | 0.7566397 |
| GFGGFGTAGGVGGA | ASGDAADRGLEQY | 0.67187166 |
| GFLAQFL | ASGDAADRGLEQY | 0.7451112 |
| GGAGGAGGNAGVA | ASGDAADRGLEQY | 0.71341825 |
| GGVGTAGAGGAGGAGG | ASGDAADRGLEQY | 0.684087 |
| GNGGGGDAGNAGSGGN | ASGDAADRGLEQY | 0.75099 |
| GTGGAGGAGNDAGS | ASGDAADRGLEQY | 0.6523448 |
| GTTNTGWFNS | ASGDAADRGLEQY | 0.7451121 |
| IFMGPLFTL | ASGDAADRGLEQY | 0.75861967 |
| ILAHAIEAI | ASGDAADRGLEQY | 0.7324323 |
| IRSAIREAL | ASGDAADRGLEQY | 0.71997344 |
| ITFAGGG | ASGDAADRGLEQY | 0.71627355 |
| ITGMSGHQEV | ASGDAADRGLEQY | 0.45582992 |
| IVAGLACTL | ASGDAADRGLEQY | 0.72962755 |
| IVGGGLAGA | ASGDAADRGLEQY | 0.28324246 |
| IVTGVLNI | ASGDAADRGLEQY | 0.7269441 |
| LAQQGCI | ASGDAADRGLEQY | 0.6631837 |
| LCLHVEN | ASGDAADRGLEQY | 0.7415266 |
| LDAISDGS | ASGDAADRGLEQY | 0.73106 |
| LGGFAAML | ASGDAADRGLEQY | 0.7399675 |
| LGGKNPMI | ASGDAADRGLEQY | 0.67430854 |
| LGPKGVAAIL | ASGDAADRGLEQY | 0.69368565 |
| LMVGVSI | ASGDAADRGLEQY | 0.6807388 |
| LVGAAFVL | ASGDAADRGLEQY | 0.606879 |
| LVLAVAPL | ASGDAADRGLEQY | 0.7445246 |
| MAAIAVIDAA | ASGDAADRGLEQY | 0.69975203 |
| NNGINAELQ | ASGDAADRGLEQY | 0.6739023 |
| RGEPLVLVHG | ASGDAADRGLEQY | 0.75608504 |
| RGGAGGAAGLI | ASGDAADRGLEQY | 0.7260293 |
| RSYRMHPAV | ASGDAADRGLEQY | 0.43734542 |
| SAAHLLAGQLA | ASGDAADRGLEQY | 0.7198925 |
| SAAVGSY | ASGDAADRGLEQY | 0.73176134 |
| SPHAVLHRI | ASGDAADRGLEQY | 0.727153 |
| SQIAKVL | ASGDAADRGLEQY | 0.23183031 |
| TALGHLVP | ASGDAADRGLEQY | 0.732854 |
| TAPTTSGAA | ASGDAADRGLEQY | 0.6765697 |
| TDGDDGNGGSPGT | ASGDAADRGLEQY | 0.66261774 |
| TDGLGEN | ASGDAADRGLEQY | 0.66936326 |
| TGGPPLPGQA | ASGDAADRGLEQY | 0.74807644 |
| TIPAITI | ASGDAADRGLEQY | 0.6885489 |
| TIPGFTI | ASGDAADRGLEQY | 0.742273 |
| TSIASGALA | ASGDAADRGLEQY | 0.74292827 |
| TVTLPGAAI | ASGDAADRGLEQY | 0.74782515 |
| VASGLSAALP | ASGDAADRGLEQY | 0.7585137 |
| VGAGVIGI | ASGDAADRGLEQY | 0.74497604 |
| VGPKRFGV | ASGDAADRGLEQY | 0.721057 |
| VNVAFPD | ASGDAADRGLEQY | 0.74136055 |
| VSMTNSGVS | ASGDAADRGLEQY | 0.7202852 |
| VTDVLGDLSV | ASGDAADRGLEQY | 0.34954607 |
| VVVVGAGFAGL | ASGDAADRGLEQY | 0.40474042 |

| | | |
|---|---|---|
| VYTVGTVV | ASGDAADRGLEQY | 0.7393599 |
| WTTFSGP | ASGDAADRGLEQY | 0.3193298 |
| YDELVEAF | ASGDAADRGLEQY | 0.7397609 |
| YGTGGAGGNGG | ASGDAADRGLEQY | 0.75264704 |
| YTNVMAV | ASGDAADRGLEQY | 0.71792215 |
| FLEQVHQL | ASGDAADRGLEQY | 0.5329006 |
| GLSLLAVAV | ASGDAADRGLEQY | 0.71302783 |
| LEQVHQL | ASGDAADRGLEQY | 0.66938555 |
| LNAAGAI | ASGDAADRGLEQY | 0.74910784 |
| PAVHASL | ASGDAADRGLEQY | 0.7511514 |
| TELEDDA | ASGDAADRGLEQY | 0.7564205 |
| TPPGPPN | ASGDAADRGLEQY | 0.75520444 |
| TPPPPPPP | ASGDAADRGLEQY | 0.71386933 |
| TQSPALI | ASSLGRDNNERLF | 0.74567354 |
| EFAAAFPQ | ASSLGRDNNERLF | 0.6982341 |
| LLSGNLDVL | ASSLGRDNNERLF | 0.7049077 |
| GPANLGNYNI | ASSLGRDNNERLF | 0.7046212 |
| TLGPTLP | ASSLGRDNNERLF | 0.72766614 |
| VYAYSSCCK | ASSLGRDNNERLF | 0.7175421 |
| APAPAPAPNI | ASSLGRDNNERLF | 0.67177266 |
| PGGPVPIP | ASSLGRDNNERLF | 0.5017384 |
| FLTVALEP | ASSLGRDNNERLF | 0.73600954 |
| SQYFGANL | ASSLGRDNNERLF | 0.7371801 |
| GGVKPAA | ASSLGRDNNERLF | 0.40002394 |
| GGTGGAGGAGGW | ASSLGRDNNERLF | 0.6888628 |
| GVGGVAGLGGAGG | ASSLGRDNNERLF | 0.75999653 |
| VLLVLATTI | ASSLGRDNNERLF | 0.75839686 |
| IGIFAAML | ASSLGRDNNERLF | 0.6874846 |
| AVGDLID | ASSLGRDNNERLF | 0.6839553 |
| NISPLPL | ASSLGRDNNERLF | 0.717023 |
| VTAVFVAI | ASSLGRDNNERLF | 0.48294234 |
| SGDDNDAAL | ASSLGRDNNERLF | 0.73659515 |
| AVAALAHS | ASSLGRDNNERLF | 0.721373 |
| GGVTVMATP | ASSLGRDNNERLF | 0.71024287 |
| AGLAANVRD | ASSLGRDNNERLF | 0.6551442 |
| TNRPDLIDPALLRPG | ASSLGRDNNERLF | 0.73102087 |
| LANLGVTAL | ASSLGRDNNERLF | 0.707767 |
| TGERIEL | ASSLGRDNNERLF | 0.74552715 |
| NGEELAA | ASSLGRDNNERLF | 0.23604478 |
| VLSLVWG | ASSLGRDNNERLF | 0.72659427 |
| IGYALWL | ASSLGRDNNERLF | 0.6670331 |
| LSLLLAPL | ASSLGRDNNERLF | 0.6725484 |
| TLDEALK | ASSLGRDNNERLF | 0.42102057 |
| TDGAGTAEGG | ASSLGRDNNERLF | 0.704301 |
| NAPGGAGGNGG | ASSLGRDNNERLF | 0.7530396 |
| VSVFMGMGGNF | ASSLGRDNNERLF | 0.7515126 |
| LASYLAAL | ASSLGRDNNERLF | 0.51082426 |
| LVGGTPAL | ASSLGRDNNERLF | 0.7503214 |
| VRNADVL | ASSLGRDNNERLF | 0.76143706 |
| TCSSPFLVSAM | ASSLGRDNNERLF | 0.6901739 |
| VHITDAE | ASSLGRDNNERLF | 0.39002672 |

| | | |
|---|---|---|
| EYAAPSF | ASSLGRDNNERLF | 0.7457225 |
| VVDAFHGGL | ASSLGRDNNERLF | 0.6943902 |
| ASVVDPQ | ASSLGRDNNERLF | 0.7394028 |
| SYNEFGGLS | ASSLGRDNNERLF | 0.7486057 |
| DPDGSVY | ASSLGRDNNERLF | 0.6331477 |
| MVSTIHATE | ASSLGRDNNERLF | 0.7552891 |
| LAAGVADPV | ASSLGRDNNERLF | 0.7559058 |
| TKEQIAA | ASSLGRDNNERLF | 0.68051285 |
| SLASPAAL | ASSLGRDNNERLF | 0.6270045 |
| IGLVFYGV | ASSLGRDNNERLF | 0.5961149 |
| FLLDASNL | ASSLGRDNNERLF | 0.3476196 |
| YQVGNFP | ASSLGRDNNERLF | 0.7448021 |
| ESLLSQST | ASSLGRDNNERLF | 0.6864687 |
| LFTVGFAAVG | ASSLGRDNNERLF | 0.69878983 |
| ATPPTPT | ASSLGRDNNERLF | 0.73479605 |
| TDLATLG | ASSLGRDNNERLF | 0.67606425 |
| SAGGTGR | ASSLGRDNNERLF | 0.6993649 |
| GFSNSGS | ASSLGRDNNERLF | 0.7041019 |
| GGNGGSGGTSV | ASSLGRDNNERLF | 0.73421973 |
| GTSVATGGAGN | ASSLGRDNNERLF | 0.7381469 |
| NGGNGGSGGTS | ASSLGRDNNERLF | 0.71523607 |
| AVVKFPH | ASSLGRDNNERLF | 0.72202575 |
| TPPPPPP | ASSLGRDNNERLF | 0.7547243 |
| AAPAAAAAPA | ASSLGRDNNERLF | 0.5749635 |
| IALNEDL | ASSLGRDNNERLF | 0.6565863 |
| IVNVSSM | ASSLGRDNNERLF | 0.74529135 |
| TGIPPAPR | ASSLGRDNNERLF | 0.5404729 |
| FDLGGGTF | ASSLGRDNNERLF | 0.7461212 |
| FDLGGGTFDVS | ASSLGRDNNERLF | 0.7350521 |
| NEPTAAA | ASSLGRDNNERLF | 0.7353245 |
| NEPTAAALA | ASSLGRDNNERLF | 0.71080106 |
| GSIINISSI | ASSLGRDNNERLF | 0.75850666 |
| VAPGNAGTA | ASSLGRDNNERLF | 0.7504615 |
| TGGGASLE | ASSLGRDNNERLF | 0.70338696 |
| VAVYNHYKRI | ASSLGRDNNERLF | 0.7186507 |
| TDPAPATT | ASSLGRDNNERLF | 0.5876324 |
| TNRPDILDPALLRPG | ASSLGRDNNERLF | 0.75186974 |
| LTTTTTS | ASSLGRDNNERLF | 0.7472439 |
| PTGSGKTL | ASSLGRDNNERLF | 0.63991153 |
| TGSGKTL | ASSLGRDNNERLF | 0.7229608 |
| TIGHVDHGK | ASSLGRDNNERLF | 0.75477815 |
| TIGHVDHGKTT | ASSLGRDNNERLF | 0.7534308 |
| GHVDHGKTT | ASSLGRDNNERLF | 0.4275506 |
| AQFEHTLL | ASSLGRDNNERLF | 0.71250916 |
| IVNQDEL | ASSLGRDNNERLF | 0.44291705 |
| TNGSQFF | ASSLGRDNNERLF | 0.7066083 |
| TGKGIVE | ASSLGRDNNERLF | 0.57944787 |
| LISNASDA | ASSLGRDNNERLF | 0.74252903 |
| AELRTLGELTKIAKAHGAQ | ASSLGRDNNERLF | 0.7103514 |
| APNAVAAAEPSW | ASSLGRDNNERLF | 0.73845816 |
| APQVAGAATAVL | ASSLGRDNNERLF | 0.73789036 |

| | | |
|---|---|---|
| ARRDPAAAAAILPTDA | ASSLGRDNNERLF | 0.7586627 |
| DAAARDVLAEAGLAEYF | ASSLGRDNNERLF | 0.7039082 |
| DDQIVLSAKANLAKVEQ | ASSLGRDNNERLF | 0.72903764 |
| DELLPDTGEDSDEDGH | ASSLGRDNNERLF | 0.504472 |
| ELPEAPANRATVPQD | ASSLGRDNNERLF | 0.5129489 |
| FGQSKIERVTGIG | ASSLGRDNNERLF | 0.6402869 |
| GAGGTGSTAGGAGGAGGAGGLY | ASSLGRDNNERLF | 0.71630627 |
| GHGGTGAAGGNGGTGGAGGSGI | ASSLGRDNNERLF | 0.27996242 |
| GNGGNGGAGGEGGPGGAGGAGA | ASSLGRDNNERLF | 0.7556584 |
| GSTAGNGGNGGNATNGTIAGQP | ASSLGRDNNERLF | 0.7376671 |
| GTGGDGGNGGNGGTGG | ASSLGRDNNERLF | 0.7523075 |
| GVRRPAGIPADAPV | ASSLGRDNNERLF | 0.6921426 |
| GYRQVTDAHLLTLARR | ASSLGRDNNERLF | 0.6871016 |
| IAKMGQDGHDRG | ASSLGRDNNERLF | 0.69928914 |
| KAEPSIEIQYT | ASSLGRDNNERLF | 0.75032187 |
| KPVRIRMAADEDTVAVT | ASSLGRDNNERLF | 0.64375937 |
| MPPRIAGMRLLVIKPEP | ASSLGRDNNERLF | 0.48335928 |
| MKFYAVALFRSRGVPPTR | ASSLGRDNNERLF | 0.680684 |
| NAGGFGILGAVAHSP | ASSLGRDNNERLF | 0.5475277 |
| NEGREITLAMAEM | ASSLGRDNNERLF | 0.7558351 |
| NELVEWSHPLGHRWRWE | ASSLGRDNNERLF | 0.75399554 |
| NLGAGGTGGNGGNPG | ASSLGRDNNERLF | 0.7410226 |
| NTEWPKWETVMAADPVD | ASSLGRDNNERLF | 0.7507707 |
| PASAAPAETQP | ASSLGRDNNERLF | 0.31123072 |
| QQSSDGKNGVRAAEITGE | ASSLGRDNNERLF | 0.6776228 |
| RAPDAVTTSWWKE | ASSLGRDNNERLF | 0.7600335 |
| SGFGNVGAHTSGWFNQ | ASSLGRDNNERLF | 0.7438956 |
| SGVAGAVGQAASVGGLKV | ASSLGRDNNERLF | 0.717497 |
| SNGGAGTGSAGGNGGT | ASSLGRDNNERLF | 0.75547063 |
| TGGAGGEGGAAG | ASSLGRDNNERLF | 0.7027054 |
| TGGDGGAAGAGGAPGHG | ASSLGRDNNERLF | 0.7306849 |
| TMPPLLAAIANAGRHGV | ASSLGRDNNERLF | 0.72917426 |
| TSSTAGNSGGA | ASSLGRDNNERLF | 0.7362192 |
| VPATVPVGILC | ASSLGRDNNERLF | 0.7333578 |
| VQRPDSGGVVLTGRIS | ASSLGRDNNERLF | 0.6295705 |
| WTPNPVPATKGVPAGLVQAT | ASSLGRDNNERLF | 0.7346518 |
| YALKRALAKFSDRFDVVII | ASSLGRDNNERLF | 0.75926816 |
| YLIGSVIV | ASSLGRDNNERLF | 0.72163695 |
| YGQLVAVID | ASSLGRDNNERLF | 0.7494402 |
| VTVGGMTLPA | ASSLGRDNNERLF | 0.74034196 |
| VLALPAV | ASSLGRDNNERLF | 0.7452244 |
| VIPPTLNYET | ASSLGRDNNERLF | 0.68361956 |
| VGMGGMHPSSG | ASSLGRDNNERLF | 0.6972485 |
| VGLAGGAATA | ASSLGRDNNERLF | 0.70161986 |
| VGGAGGAGGSGP | ASSLGRDNNERLF | 0.6636895 |
| VFAAFMV | ASSLGRDNNERLF | 0.71997404 |
| TLYDKLR | ASSLGRDNNERLF | 0.6821259 |
| TLAVFTVL | ASSLGRDNNERLF | 0.763794 |
| TGSVQPARV | ASSLGRDNNERLF | 0.7439287 |
| TGGGAGSGGT | ASSLGRDNNERLF | 0.7361238 |
| SHLETSL | ASSLGRDNNERLF | 0.69045144 |

| | | |
|---|---|---|
| SFTTGMT | ASSLGRDNNERLF | 0.7302091 |
| QVEVAKRMGVRQPTVS | ASSLGRDNNERLF | 0.739376 |
| QGSGGPALAM | ASSLGRDNNERLF | 0.47744948 |
| MLAPSNP | ASSLGRDNNERLF | 0.7564868 |
| LVNNAAI | ASSLGRDNNERLF | 0.6845997 |
| LTGEINTLA | ASSLGRDNNERLF | 0.6965481 |
| LSIFGPV | ASSLGRDNNERLF | 0.75352025 |
| LLALLANL | ASSLGRDNNERLF | 0.7528802 |
| LISNASDALDKLR | ASSLGRDNNERLF | 0.69545656 |
| LINEAVP | ASSLGRDNNERLF | 0.6593762 |
| LGYAFSGL | ASSLGRDNNERLF | 0.6789534 |
| LDFVAVAL | ASSLGRDNNERLF | 0.75251883 |
| LAQPGAE | ASSLGRDNNERLF | 0.7548572 |
| ISNLGAVTGF | ASSLGRDNNERLF | 0.7211977 |
| HQEDPGRI | ASSLGRDNNERLF | 0.75601244 |
| GTPGEPTP | ASSLGRDNNERLF | 0.7182921 |
| GFFNSGS | ASSLGRDNNERLF | 0.49064186 |
| GAGGGGLTAGAN | ASSLGRDNNERLF | 0.72994095 |
| FVDALVNM | ASSLGRDNNERLF | 0.74719465 |
| FLTVAPEL | ASSLGRDNNERLF | 0.74956536 |
| FETLTAL | ASSLGRDNNERLF | 0.7620542 |
| DVINPIEL | ASSLGRDNNERLF | 0.73420346 |
| APVSGAPV | ASSLGRDNNERLF | 0.7077571 |
| ALVGGLAW | ASSLGRDNNERLF | 0.75723696 |
| ALAAAAATAI | ASSLGRDNNERLF | 0.7621472 |
| AFGLGLLL | ASSLGRDNNERLF | 0.7520063 |
| AFFAAASL | ASSLGRDNNERLF | 0.7199522 |
| AAAALPAFSPP | ASSLGRDNNERLF | 0.6935004 |
| VVLAPAL | ASSLGRDNNERLF | 0.7272602 |
| TIGHVDHGKTTL | ASSLGRDNNERLF | 0.74845135 |
| LISNASDALDK | ASSLGRDNNERLF | 0.63179183 |
| LISNASDAL | ASSLGRDNNERLF | 0.72566533 |
| ISNASDALDK | ASSLGRDNNERLF | 0.74189377 |
| NQAETLVYQTE | ASSLGRDNNERLF | 0.5547875 |
| TGGTGGTAGAG | ASSLGRDNNERLF | 0.74676025 |
| TGSTAGGAGGA | ASSLGRDNNERLF | 0.7519979 |
| DPLAVPVAAGPV | ASSLGRDNNERLF | 0.69681805 |
| GGTGGTPGNGGN | ASSLGRDNNERLF | 0.7041576 |
| PAGLAGVDPATL | ASSLGRDNNERLF | 0.7525271 |
| GNNLVVEAPHAGASV | ASSLGRDNNERLF | 0.763126 |
| KVTRIMSASTLEQPA | ASSLGRDNNERLF | 0.7546979 |
| LGFALPIALPRRTVV | ASSLGRDNNERLF | 0.51057065 |
| TRLGNLTARKLVVLV | ASSLGRDNNERLF | 0.37209076 |
| WSQQRRLIYFPSAGP | ASSLGRDNNERLF | 0.678231 |
| GPNGNPDPMAAAVDIR | ASSLGRDNNERLF | 0.7417803 |
| TQRCVSVADLRAAASR | ASSLGRDNNERLF | 0.709331 |
| QFVLDVAEHPVEVLTETV | ASSLGRDNNERLF | 0.7086226 |
| GGAGGTGGAGGNGGTQAGDG | ASSLGRDNNERLF | 0.72813153 |
| PLLATLRAIAVDVDAKPAQV | ASSLGRDNNERLF | 0.7549822 |
| APKFRTGGYMIDFWGVGYQVAK | ASSLGRDNNERLF | 0.66899693 |
| GAGGAGGAGGAAGGPGGTGGTG | ASSLGRDNNERLF | 0.58999527 |

| | | |
|---|---|---|
| IGALGLLATDVAPAGSVAFGVA | ASSLGRDNNERLF | 0.7302505 |
| AGGAGGIGGNGGAGGAGGVLMG | ASSLGRDNNERLF | 0.74871445 |
| ADELIGLL | ASSLGRDNNERLF | 0.68520033 |
| ADRALKA | ASSLGRDNNERLF | 0.72440046 |
| AGAVGQVGGAGGA | ASSLGRDNNERLF | 0.6796809 |
| AGGGGGLGGAP | ASSLGRDNNERLF | 0.63584524 |
| AGGLLYGNGGAG | ASSLGRDNNERLF | 0.7352034 |
| AGLESIGL | ASSLGRDNNERLF | 0.6734352 |
| AGLKTGII | ASSLGRDNNERLF | 0.7537092 |
| ASLAALLL | ASSLGRDNNERLF | 0.66399837 |
| ASVINISSI | ASSLGRDNNERLF | 0.7458956 |
| ATPPGPA | ASSLGRDNNERLF | 0.7384306 |
| ATVASALV | ASSLGRDNNERLF | 0.7419975 |
| ATVASLIL | ASSLGRDNNERLF | 0.71401554 |
| AVNGPVLIRH | ASSLGRDNNERLF | 0.68124855 |
| DGAAGAVGI | ASSLGRDNNERLF | 0.6135144 |
| DPLLHGI | ASSLGRDNNERLF | 0.49856758 |
| ELGAALGAL | ASSLGRDNNERLF | 0.74716747 |
| ELGRTQAV | ASSLGRDNNERLF | 0.49013153 |
| FVAGGYLS | ASSLGRDNNERLF | 0.75232595 |
| GASGGAGGAGGW | ASSLGRDNNERLF | 0.74979836 |
| GGAGAGGDGGAGGI | ASSLGRDNNERLF | 0.65155756 |
| GTGHNGSQPGMGGNG | ASSLGRDNNERLF | 0.73831475 |
| GYAAASASAS | ASSLGRDNNERLF | 0.6626958 |
| HQVDVAAVQQG | ASSLGRDNNERLF | 0.58814 |
| IGAAIAHT | ASSLGRDNNERLF | 0.75356203 |
| IGMIEVAQL | ASSLGRDNNERLF | 0.74304616 |
| IRLAPEHTSAMD | ASSLGRDNNERLF | 0.6766496 |
| IYNVSVL | ASSLGRDNNERLF | 0.73227966 |
| KQGFLGM | ASSLGRDNNERLF | 0.73547804 |
| LESVITQL | ASSLGRDNNERLF | 0.6892246 |
| LGLGALVVV | ASSLGRDNNERLF | 0.6992185 |
| LIGNGGNGGSGGI | ASSLGRDNNERLF | 0.7555461 |
| LNAINAPTL | ASSLGRDNNERLF | 0.7254968 |
| LQLLNTGV | ASSLGRDNNERLF | 0.6733573 |
| LQTSIEF | ASSLGRDNNERLF | 0.74312186 |
| MSFAVSEL | ASSLGRDNNERLF | 0.6506345 |
| NTGALGL | ASSLGRDNNERLF | 0.742084 |
| PPGAGGAGGAGTT | ASSLGRDNNERLF | 0.7045794 |
| QLVGLGI | ASSLGRDNNERLF | 0.7266123 |
| RFLEQVHQL | ASSLGRDNNERLF | 0.74006987 |
| RGVGYVLR | ASSLGRDNNERLF | 0.5627795 |
| SATGVFAGVI | ASSLGRDNNERLF | 0.6776968 |
| SLPAITLP | ASSLGRDNNERLF | 0.710703 |
| STALAFTV | ASSLGRDNNERLF | 0.7406658 |
| SVLTVGLGK | ASSLGRDNNERLF | 0.7494515 |
| TALAGAGSGI | ASSLGRDNNERLF | 0.68389994 |
| TGLPFTGI | ASSLGRDNNERLF | 0.7378075 |
| VAEHVTV | ASSLGRDNNERLF | 0.7119698 |
| VDIVLNSL | ASSLGRDNNERLF | 0.6853664 |
| VLTGGGAVL | ASSLGRDNNERLF | 0.71073604 |

| | | |
|---|---|---|
| VNVGLAYGI | ASSLGRDNNERLF | 0.6969229 |
| VSSAAVGAV | ASSLGRDNNERLF | 0.68391246 |
| VTSVYEAGV | ASSLGRDNNERLF | 0.7383578 |
| WVALDRGA | ASSLGRDNNERLF | 0.6521032 |
| YLVGSVVI | ASSLGRDNNERLF | 0.738137 |
| GPPGTGKTLL | ASSLGRDNNERLF | 0.7529644 |
| GVGMTKF | ASSLGRDNNERLF | 0.6964766 |
| IGIGGSDL | ASSLGRDNNERLF | 0.743027 |
| IGTLLAEL | ASSLGRDNNERLF | 0.7372923 |
| IYTSGTT | ASSLGRDNNERLF | 0.7349075 |
| LSFPAGV | ASSLGRDNNERLF | 0.7142416 |
| LYTSGTT | ASSLGRDNNERLF | 0.691318 |
| VGAFPFGF | ASSLGRDNNERLF | 0.7511186 |
| VGAGTVEF | ASSLGRDNNERLF | 0.74560213 |
| VGKSTLLL | ASSLGRDNNERLF | 0.74900174 |
| AALAHAGGGIV | ASSLGRDNNERLF | 0.7370175 |
| AALVAVSLL | ASSLGRDNNERLF | 0.73337406 |
| AAVSRAVAAL | ASSLGRDNNERLF | 0.7466099 |
| AGGSSGIGGF | ASSLGRDNNERLF | 0.7070849 |
| AGLTVANANG | ASSLGRDNNERLF | 0.56086737 |
| AGLVGAVL | ASSLGRDNNERLF | 0.7174358 |
| ALSQAFTV | ASSLGRDNNERLF | 0.7113496 |
| ALTFSVSL | ASSLGRDNNERLF | 0.67074037 |
| ALTGAALVAL | ASSLGRDNNERLF | 0.4641763 |
| ASFSIGAGVL | ASSLGRDNNERLF | 0.7551838 |
| DGSVYKI | ASSLGRDNNERLF | 0.76235443 |
| EQLAEIV | ASSLGRDNNERLF | 0.68374133 |
| FAVSELTVM | ASSLGRDNNERLF | 0.75849706 |
| GAAPLAAGL | ASSLGRDNNERLF | 0.54109275 |
| GADGGDGGSGGAGG | ASSLGRDNNERLF | 0.7464218 |
| GFGLDTGGAGG | ASSLGRDNNERLF | 0.75621617 |
| GLASGGLL | ASSLGRDNNERLF | 0.6733289 |
| GRIAGAAVLL | ASSLGRDNNERLF | 0.7526243 |
| GSAGSLNTSF | ASSLGRDNNERLF | 0.65512043 |
| GTGQIGI | ASSLGRDNNERLF | 0.668129 |
| GTLIALIL | ASSLGRDNNERLF | 0.7558817 |
| IAATMGFVVG | ASSLGRDNNERLF | 0.7180077 |
| IFAQALL | ASSLGRDNNERLF | 0.72356963 |
| IGAAVALGF | ASSLGRDNNERLF | 0.7301717 |
| IGLTPLAL | ASSLGRDNNERLF | 0.67052317 |
| ILGSNTSTL | ASSLGRDNNERLF | 0.7237915 |
| IVFSLTVL | ASSLGRDNNERLF | 0.7556942 |
| LASKATGF | ASSLGRDNNERLF | 0.6290454 |
| LASVALIL | ASSLGRDNNERLF | 0.76165247 |
| LFAVGLTL | ASSLGRDNNERLF | 0.6664977 |
| LFNGGIGTYI | ASSLGRDNNERLF | 0.7609839 |
| LTLGAGII | ASSLGRDNNERLF | 0.74194187 |
| LVAAALGDI | ASSLGRDNNERLF | 0.7337573 |
| NTAFVQLGI | ASSLGRDNNERLF | 0.6927825 |
| QLFSGAEL | ASSLGRDNNERLF | 0.7552254 |
| SAATFITL | ASSLGRDNNERLF | 0.72944367 |

| | | |
|---|---|---|
| SNYGLANLGV | ASSLGRDNNERLF | 0.7358089 |
| SYGGAYAVM | ASSLGRDNNERLF | 0.7542943 |
| TTGVGGLTLG | ASSLGRDNNERLF | 0.5969399 |
| TTQTIYI | ASSLGRDNNERLF | 0.6811336 |
| VAAGAGLGAVI | ASSLGRDNNERLF | 0.6632025 |
| VAAVVTLL | ASSLGRDNNERLF | 0.6624374 |
| VGFLGGLI | ASSLGRDNNERLF | 0.6302942 |
| VGVIPILL | ASSLGRDNNERLF | 0.53663397 |
| VIFASSLI | ASSLGRDNNERLF | 0.65512896 |
| VKGGDGGAAAG | ASSLGRDNNERLF | 0.60809195 |
| VTKTFTNI | ASSLGRDNNERLF | 0.67598253 |
| VVAAAILL | ASSLGRDNNERLF | 0.6754337 |
| YAALDVEL | ASSLGRDNNERLF | 0.63011044 |
| LASVALLL | ASSLGRDNNERLF | 0.679572 |
| LDAVHRV | ASSLGRDNNERLF | 0.6813477 |
| QGLSSSIF | ASSLGRDNNERLF | 0.5460817 |
| RGTFANIRL | ASSLGRDNNERLF | 0.6430604 |
| RQLWWGHRI | ASSLGRDNNERLF | 0.6374661 |
| YGPPGTGKTLL | ASSLGRDNNERLF | 0.6505331 |
| AALVGAPANVA | ASSLGRDNNERLF | 0.48137242 |
| AATFRRTE | ASSLGRDNNERLF | 0.6726711 |
| AGALGGGI | ASSLGRDNNERLF | 0.6235374 |
| AGGAGGAGGIGAG | ASSLGRDNNERLF | 0.3289932 |
| AGSGALGL | ASSLGRDNNERLF | 0.66258144 |
| AQIADQI | ASSLGRDNNERLF | 0.6492244 |
| ASAVAVTYL | ASSLGRDNNERLF | 0.63872945 |
| ASTVLGGTGG | ASSLGRDNNERLF | 0.6728555 |
| AVGGEPAK | ASSLGRDNNERLF | 0.56785417 |
| CEQGTFGGTC | ASSLGRDNNERLF | 0.6233106 |
| DEGKAPAG | ASSLGRDNNERLF | 0.66803086 |
| DTVVFPAL | ASSLGRDNNERLF | 0.68048203 |
| FEGITMAG | ASSLGRDNNERLF | 0.5851121 |
| FPLVVQD | ASSLGRDNNERLF | 0.6568608 |
| GAFTTPQ | ASSLGRDNNERLF | 0.6837557 |
| GFGGFGTAGGVGGA | ASSLGRDNNERLF | 0.6652783 |
| GFLAQFL | ASSLGRDNNERLF | 0.68076515 |
| GGAGGAGGNAGVA | ASSLGRDNNERLF | 0.6845229 |
| GGVGTAGAGGAGGAGG | ASSLGRDNNERLF | 0.6250975 |
| GNGGGGDAGNAGSGGN | ASSLGRDNNERLF | 0.6775507 |
| GTGGAGGAGNDAGS | ASSLGRDNNERLF | 0.6810591 |
| GTTNTGWFNS | ASSLGRDNNERLF | 0.6482478 |
| IFMGPLFTL | ASSLGRDNNERLF | 0.65559125 |
| ILAHAIEAI | ASSLGRDNNERLF | 0.6052638 |
| IRSAIREAL | ASSLGRDNNERLF | 0.65246034 |
| ITFAGGG | ASSLGRDNNERLF | 0.4715373 |
| ITGMSGHQEV | ASSLGRDNNERLF | 0.65761775 |
| IVAGLACTL | ASSLGRDNNERLF | 0.6813388 |
| IVGGGLAGA | ASSLGRDNNERLF | 0.64084506 |
| IVTGVLNI | ASSLGRDNNERLF | 0.5649339 |
| LAQQGCI | ASSLGRDNNERLF | 0.6037104 |
| LCLHVEN | ASSLGRDNNERLF | 0.6532522 |

| | | |
|---|---|---|
| LDAISDGS | ASSLGRDNNERLF | 0.63113356 |
| LGGFAAML | ASSLGRDNNERLF | 0.4491964 |
| LGGKNPMI | ASSLGRDNNERLF | 0.6371669 |
| LGPKGVAAIL | ASSLGRDNNERLF | 0.67870355 |
| LMVGVSI | ASSLGRDNNERLF | 0.67907286 |
| LVGAAFVL | ASSLGRDNNERLF | 0.5685073 |
| LVLAVAPL | ASSLGRDNNERLF | 0.67959213 |
| MAAIAVIDAA | ASSLGRDNNERLF | 0.67337215 |
| NNGINAELQ | ASSLGRDNNERLF | 0.68632287 |
| RGEPLVLVHG | ASSLGRDNNERLF | 0.66322243 |
| RGGAGGAAGLI | ASSLGRDNNERLF | 0.60818493 |
| RSYRMHPAV | ASSLGRDNNERLF | 0.67183316 |
| SAAHLLAGQLA | ASSLGRDNNERLF | 0.63229966 |
| SAAVGSY | ASSLGRDNNERLF | 0.6233286 |
| SPHAVLHRI | ASSLGRDNNERLF | 0.67645895 |
| SQIAKVL | ASSLGRDNNERLF | 0.65133494 |
| TALGHLVP | ASSLGRDNNERLF | 0.61887383 |
| TAPTTSGAA | ASSLGRDNNERLF | 0.6333735 |
| TDGDDGNGGSPGT | ASSLGRDNNERLF | 0.68329155 |
| TDGLGEN | ASSLGRDNNERLF | 0.59403825 |
| TGGPPLPGQA | ASSLGRDNNERLF | 0.61744887 |
| TIPAITI | ASSLGRDNNERLF | 0.6136345 |
| TIPGFTI | ASSLGRDNNERLF | 0.6547357 |
| TSIASGALA | ASSLGRDNNERLF | 0.6790487 |
| TVTLPGAAI | ASSLGRDNNERLF | 0.67378163 |
| VASGLSAALP | ASSLGRDNNERLF | 0.6371093 |
| VGAGVIGI | ASSLGRDNNERLF | 0.66450465 |
| VGPKRFGV | ASSLGRDNNERLF | 0.6757928 |
| VNVAFPD | ASSLGRDNNERLF | 0.66233385 |
| VSMTNSGVS | ASSLGRDNNERLF | 0.6790684 |
| VTDVLGDLSV | ASSLGRDNNERLF | 0.6045346 |
| VVVVGAGFAGL | ASSLGRDNNERLF | 0.6794151 |
| VYTVGTVV | ASSLGRDNNERLF | 0.67235607 |
| WTTFSGP | ASSLGRDNNERLF | 0.6777016 |
| YDELVEAF | ASSLGRDNNERLF | 0.6772124 |
| YGTGGAGGNGG | ASSLGRDNNERLF | 0.66791815 |
| YTNVMAV | ASSLGRDNNERLF | 0.6632042 |
| FLEQVHQL | ASSLGRDNNERLF | 0.5470861 |
| GLSLLAVAV | ASSLGRDNNERLF | 0.645252 |
| LEQVHQL | ASSLGRDNNERLF | 0.6626193 |
| LNAAGAI | ASSLGRDNNERLF | 0.65711313 |
| PAVHASL | ASSLGRDNNERLF | 0.67273426 |
| TELEDDA | ASSLGRDNNERLF | 0.66997266 |
| TPPGPPN | ASSLGRDNNERLF | 0.6182613 |
| TPPPPPPP | ASSLGRDNNERLF | 0.68293995 |
| TQSPALI | ASSHLRDPSAETLY | 0.67770386 |
| EFAAAFPQ | ASSHLRDPSAETLY | 0.5586044 |
| LLSGNLDVL | ASSHLRDPSAETLY | 0.6444568 |
| GPANLGNYNI | ASSHLRDPSAETLY | 0.68392915 |
| TLGPTLP | ASSHLRDPSAETLY | 0.65768707 |
| VYAYSSCCK | ASSHLRDPSAETLY | 0.6443916 |

| | | |
|---|---|---|
| APAPAPAPNI | ASSHLRDPSAETLY | 0.64763653 |
| PGGPVPIP | ASSHLRDPSAETLY | 0.68404627 |
| FLTVALEP | ASSHLRDPSAETLY | 0.6117972 |
| SQYFGANL | ASSHLRDPSAETLY | 0.6844926 |
| GGVKPAA | ASSHLRDPSAETLY | 0.4449576 |
| GGTGGAGGAGGW | ASSHLRDPSAETLY | 0.64292073 |
| GVGGVAGLGGAGG | ASSHLRDPSAETLY | 0.627916 |
| VLLVLATTI | ASSHLRDPSAETLY | 0.43739402 |
| IGIFAAML | ASSHLRDPSAETLY | 0.67633855 |
| AVGDLID | ASSHLRDPSAETLY | 0.6354136 |
| NISPLPL | ASSHLRDPSAETLY | 0.67400134 |
| VTAVFVAI | ASSHLRDPSAETLY | 0.64853203 |
| SGDDNDAAL | ASSHLRDPSAETLY | 0.6814096 |
| AVAALAHS | ASSHLRDPSAETLY | 0.65850806 |
| GGVTVMATP | ASSHLRDPSAETLY | 0.66785336 |
| AGLAANVRD | ASSHLRDPSAETLY | 0.61014336 |
| TNRPDLIDPALLRPG | ASSHLRDPSAETLY | 0.6745444 |
| LANLGVTAL | ASSHLRDPSAETLY | 0.68301475 |
| TGERIEL | ASSHLRDPSAETLY | 0.6781261 |
| NGEELAA | ASSHLRDPSAETLY | 0.548891 |
| VLSLVWG | ASSHLRDPSAETLY | 0.6415013 |
| IGYALWL | ASSHLRDPSAETLY | 0.6609821 |
| LSLLLAPL | ASSHLRDPSAETLY | 0.40848327 |
| TLDEALK | ASSHLRDPSAETLY | 0.37116852 |
| TDGAGTAEGG | ASSHLRDPSAETLY | 0.625407 |
| NAPGGAGGNGG | ASSHLRDPSAETLY | 0.6248063 |
| VSVFMGMGGNF | ASSHLRDPSAETLY | 0.674297 |
| LASYLAAL | ASSHLRDPSAETLY | 0.66875863 |
| LVGGTPAL | ASSHLRDPSAETLY | 0.6467302 |
| VRNADVL | ASSHLRDPSAETLY | 0.6715231 |
| TCSSPFLVSAM | ASSHLRDPSAETLY | 0.6622469 |
| VHITDAE | ASSHLRDPSAETLY | 0.2897155 |
| EYAAPSF | ASSHLRDPSAETLY | 0.63589066 |
| VVDAFHGGL | ASSHLRDPSAETLY | 0.546934 |
| ASVVDPQ | ASSHLRDPSAETLY | 0.68398666 |
| SYNEFGGLS | ASSHLRDPSAETLY | 0.6062614 |
| DPDGSVY | ASSHLRDPSAETLY | 0.6071725 |
| MVSTIHATE | ASSHLRDPSAETLY | 0.65639806 |
| LAAGVADPV | ASSHLRDPSAETLY | 0.6506719 |
| TKEQIAA | ASSHLRDPSAETLY | 0.6444746 |
| SLASPAAL | ASSHLRDPSAETLY | 0.6211477 |
| IGLVFYGV | ASSHLRDPSAETLY | 0.39446375 |
| FLLDASNL | ASSHLRDPSAETLY | 0.6772979 |
| YQVGNFP | ASSHLRDPSAETLY | 0.5454349 |
| ESLLSQST | ASSHLRDPSAETLY | 0.60191864 |
| LFTVGFAAVG | ASSHLRDPSAETLY | 0.5161679 |
| ATPPTPT | ASSHLRDPSAETLY | 0.60393023 |
| TDLATLG | ASSHLRDPSAETLY | 0.528691 |
| SAGGTGR | ASSHLRDPSAETLY | 0.67830133 |
| GFSNSGS | ASSHLRDPSAETLY | 0.6285192 |
| GGNGGSGGTSV | ASSHLRDPSAETLY | 0.63355595 |

| | | |
|---|---|---|
| GTSVATGGAGN | ASSHLRDPSAETLY | 0.68016785 |
| NGGNGGSGGTS | ASSHLRDPSAETLY | 0.65597105 |
| AVVKFPH | ASSHLRDPSAETLY | 0.63502014 |
| TPPPPPP | ASSHLRDPSAETLY | 0.6367213 |
| AAPAAAAAPA | ASSHLRDPSAETLY | 0.59948784 |
| IALNEDL | ASSHLRDPSAETLY | 0.6244873 |
| IVNVSSM | ASSHLRDPSAETLY | 0.6262884 |
| TGIPPAPR | ASSHLRDPSAETLY | 0.68467337 |
| FDLGGGTF | ASSHLRDPSAETLY | 0.64301455 |
| FDLGGGTFDVS | ASSHLRDPSAETLY | 0.5823436 |
| NEPTAAA | ASSHLRDPSAETLY | 0.6561008 |
| NEPTAAALA | ASSHLRDPSAETLY | 0.6825458 |
| GSIINISSI | ASSHLRDPSAETLY | 0.6876811 |
| VAPGNAGTA | ASSHLRDPSAETLY | 0.6156387 |
| TGGGASLE | ASSHLRDPSAETLY | 0.65837 |
| VAVYNHYKRI | ASSHLRDPSAETLY | 0.6451634 |
| TDPAPATT | ASSHLRDPSAETLY | 0.6434164 |
| TNRPDILDPALLRPG | ASSHLRDPSAETLY | 0.6426159 |
| LTTTTTS | ASSHLRDPSAETLY | 0.580984 |
| PTGSGKTL | ASSHLRDPSAETLY | 0.6763374 |
| TGSGKTL | ASSHLRDPSAETLY | 0.6888355 |
| TIGHVDHGK | ASSHLRDPSAETLY | 0.6495446 |
| TIGHVDHGKTT | ASSHLRDPSAETLY | 0.54072744 |
| GHVDHGKTT | ASSHLRDPSAETLY | 0.6501112 |
| AQFEHTLL | ASSHLRDPSAETLY | 0.6114693 |
| IVNQDEL | ASSHLRDPSAETLY | 0.62668127 |
| TNGSQFF | ASSHLRDPSAETLY | 0.65484524 |
| TGKGIVE | ASSHLRDPSAETLY | 0.5281614 |
| LISNASDA | ASSHLRDPSAETLY | 0.6446244 |
| AELRTLGELTKIAKAHGAQ | ASSHLRDPSAETLY | 0.6867756 |
| APNAVAAAEPSW | ASSHLRDPSAETLY | 0.6260814 |
| APQVAGAATAVL | ASSHLRDPSAETLY | 0.66181743 |
| ARRDPAAAAAILPTDA | ASSHLRDPSAETLY | 0.6775256 |
| DAAARDVLAEAGLAEYF | ASSHLRDPSAETLY | 0.66512954 |
| DDQIVLSAKANLAKVEQ | ASSHLRDPSAETLY | 0.67709064 |
| DELLPDTGEDSDEDGH | ASSHLRDPSAETLY | 0.67600954 |
| ELPEAPANRATVPQD | ASSHLRDPSAETLY | 0.66745937 |
| FGQSKIERVTGIG | ASSHLRDPSAETLY | 0.6598704 |
| GAGGTGSTAGGAGGAGGAGGLY | ASSHLRDPSAETLY | 0.6608108 |
| GHGGTGAAGGNGGTGGAGGSGI | ASSHLRDPSAETLY | 0.6560515 |
| GNGGNGGAGGEGGPGGAGGAGA | ASSHLRDPSAETLY | 0.64963126 |
| GSTAGNGGNGGNATNGTIAGQP | ASSHLRDPSAETLY | 0.62704116 |
| GTGGDGGNGGNGGTGG | ASSHLRDPSAETLY | 0.67279327 |
| GVRRPAGIPADAPV | ASSHLRDPSAETLY | 0.6572777 |
| GYRQVTDAHLLTLARR | ASSHLRDPSAETLY | 0.68117166 |
| IAKMGQDGHDRG | ASSHLRDPSAETLY | 0.51293087 |
| KAEPSIEIQYT | ASSHLRDPSAETLY | 0.59753317 |
| KPVRIRMAADEDTVAVT | ASSHLRDPSAETLY | 0.6720443 |
| MPPRIAGMRLLVIKPEP | ASSHLRDPSAETLY | 0.34842715 |
| MKFYAVALFRSRGVPPTR | ASSHLRDPSAETLY | 0.38536295 |
| NAGGFGILGAVAHSP | ASSHLRDPSAETLY | 0.657583 |

| | | |
|---|---|---|
| NEGREITLAMAEM | ASSHLRDPSAETLY | 0.68352354 |
| NELVEWSHPLGHRWRWE | ASSHLRDPSAETLY | 0.33112296 |
| NLGAGGTGGNGGNPG | ASSHLRDPSAETLY | 0.6053525 |
| NTEWPKWETVMAADPVD | ASSHLRDPSAETLY | 0.5745604 |
| PASAAPAETQP | ASSHLRDPSAETLY | 0.67412674 |
| QQSSDGKNGVRAAEITGE | ASSHLRDPSAETLY | 0.6517955 |
| RAPDAVTTSWWKE | ASSHLRDPSAETLY | 0.53574175 |
| SGFGNVGAHTSGWFNQ | ASSHLRDPSAETLY | 0.6742934 |
| SGVAGAVGQAASVGGLKV | ASSHLRDPSAETLY | 0.51118207 |
| SNGGAGTGSAGGNGGT | ASSHLRDPSAETLY | 0.6097329 |
| TGGAGGEGGAAG | ASSHLRDPSAETLY | 0.6787683 |
| TGGDGGAAGAGGAPGHG | ASSHLRDPSAETLY | 0.6831673 |
| TMPPLLAAIANAGRHGV | ASSHLRDPSAETLY | 0.5745163 |
| TSSTAGNSGGA | ASSHLRDPSAETLY | 0.6750073 |
| VPATVPVGILC | ASSHLRDPSAETLY | 0.62563306 |
| VQRPDSGGVVLTGRIS | ASSHLRDPSAETLY | 0.6883048 |
| WTPNPVPATKGVPAGLVQAT | ASSHLRDPSAETLY | 0.6395975 |
| YALKRALAKFSDRFDVVII | ASSHLRDPSAETLY | 0.5289892 |
| YLIGSVIV | ASSHLRDPSAETLY | 0.6724323 |
| YGQLVAVID | ASSHLRDPSAETLY | 0.6680826 |
| VTVGGMTLPA | ASSHLRDPSAETLY | 0.66250646 |
| VLALPAV | ASSHLRDPSAETLY | 0.67046154 |
| VIPPTLNYET | ASSHLRDPSAETLY | 0.6304232 |
| VGMGGMHPSSG | ASSHLRDPSAETLY | 0.6386454 |
| VGLAGGAATA | ASSHLRDPSAETLY | 0.6757643 |
| VGGAGGAGGSGP | ASSHLRDPSAETLY | 0.67512506 |
| VFAAFMV | ASSHLRDPSAETLY | 0.578182 |
| TLYDKLR | ASSHLRDPSAETLY | 0.6808291 |
| TLAVFTVL | ASSHLRDPSAETLY | 0.6535919 |
| TGSVQPARV | ASSHLRDPSAETLY | 0.6245661 |
| TGGGAGSGGT | ASSHLRDPSAETLY | 0.6853994 |
| SHLETSL | ASSHLRDPSAETLY | 0.6627784 |
| SFTTGMT | ASSHLRDPSAETLY | 0.6743909 |
| QVEVAKRMGVRQPTVS | ASSHLRDPSAETLY | 0.67734766 |
| QGSGGPALAM | ASSHLRDPSAETLY | 0.68883395 |
| MLAPSNP | ASSHLRDPSAETLY | 0.5787061 |
| LVNNAAI | ASSHLRDPSAETLY | 0.66986376 |
| LTGEINTLA | ASSHLRDPSAETLY | 0.6571293 |
| LSIFGPV | ASSHLRDPSAETLY | 0.67649746 |
| LLALLANL | ASSHLRDPSAETLY | 0.6555597 |
| LISNASDALDKLR | ASSHLRDPSAETLY | 0.60916626 |
| LINEAVP | ASSHLRDPSAETLY | 0.66848326 |
| LGYAFSGL | ASSHLRDPSAETLY | 0.6801667 |
| LDFVAVAL | ASSHLRDPSAETLY | 0.66341126 |
| LAQPGAE | ASSHLRDPSAETLY | 0.6712267 |
| ISNLGAVTGF | ASSHLRDPSAETLY | 0.68155324 |
| HQEDPGRI | ASSHLRDPSAETLY | 0.6186609 |
| GTPGEPTP | ASSHLRDPSAETLY | 0.6558999 |
| GFFNSGS | ASSHLRDPSAETLY | 0.67671263 |
| GAGGGGLTAGAN | ASSHLRDPSAETLY | 0.6791278 |
| FVDALVNM | ASSHLRDPSAETLY | 0.6477481 |

| | | |
|---|---|---|
| FLTVAPEL | ASSHLRDPSAETLY | 0.683457 |
| FETLTAL | ASSHLRDPSAETLY | 0.6223869 |
| DVINPIEL | ASSHLRDPSAETLY | 0.6701761 |
| APVSGAPV | ASSHLRDPSAETLY | 0.6132879 |
| ALVGGLAW | ASSHLRDPSAETLY | 0.6614003 |
| ALAAAAATAI | ASSHLRDPSAETLY | 0.58996826 |
| AFGLGLLL | ASSHLRDPSAETLY | 0.66556513 |
| AFFAAASL | ASSHLRDPSAETLY | 0.672345 |
| AAAALPAFSPP | ASSHLRDPSAETLY | 0.67246985 |
| VVLAPAL | ASSHLRDPSAETLY | 0.39244282 |
| TIGHVDHGKTTL | ASSHLRDPSAETLY | 0.6706265 |
| LISNASDALDK | ASSHLRDPSAETLY | 0.6727562 |
| LISNASDAL | ASSHLRDPSAETLY | 0.6840931 |
| ISNASDALDK | ASSHLRDPSAETLY | 0.6559789 |
| NQAETLVYQTE | ASSHLRDPSAETLY | 0.6713499 |
| TGGTGGTAGAG | ASSHLRDPSAETLY | 0.6398411 |
| TGSTAGGAGGA | ASSHLRDPSAETLY | 0.65474576 |
| DPLAVPVAAGPV | ASSHLRDPSAETLY | 0.5882603 |
| GGTGGTPGNGGN | ASSHLRDPSAETLY | 0.6448412 |
| PAGLAGVDPATL | ASSHLRDPSAETLY | 0.6276101 |
| GNNLVVEAPHAGASV | ASSHLRDPSAETLY | 0.6036901 |
| KVTRIMSASTLEQPA | ASSHLRDPSAETLY | 0.67055684 |
| LGFALPIALPRRTVV | ASSHLRDPSAETLY | 0.68040186 |
| TRLGNLTARKLVVLV | ASSHLRDPSAETLY | 0.67467415 |
| WSQQRRLIYFPSAGP | ASSHLRDPSAETLY | 0.67797357 |
| GPNGNPDPMAAAVDIR | ASSHLRDPSAETLY | 0.68000364 |
| TQRCVSVADLRAAASR | ASSHLRDPSAETLY | 0.6805484 |
| QFVLDVAEHPVEVLTETV | ASSHLRDPSAETLY | 0.63382226 |
| GGAGGTGGAGGNGGTQAGDG | ASSHLRDPSAETLY | 0.6541878 |
| PLLATLRAIAVDVDAKPAQV | ASSHLRDPSAETLY | 0.5701693 |
| APKFRTGGYMIDFWGVGYQVAK | ASSHLRDPSAETLY | 0.63602656 |
| GAGGAGGAGGAAGGPGGTGGTG | ASSHLRDPSAETLY | 0.62880427 |
| IGALGLLATDVAPAGSVAFGVA | ASSHLRDPSAETLY | 0.6579411 |
| AGGAGGIGGNGGAGGAGGVLMG | ASSHLRDPSAETLY | 0.3153028 |
| ADELIGLL | ASSHLRDPSAETLY | 0.64929795 |
| ADRALKA | ASSHLRDPSAETLY | 0.68354225 |
| AGAVGQVGGAGGA | ASSHLRDPSAETLY | 0.6347666 |
| AGGGGGLGGAP | ASSHLRDPSAETLY | 0.6677157 |
| AGGLLYGNGGAG | ASSHLRDPSAETLY | 0.6583576 |
| AGLESIGL | ASSHLRDPSAETLY | 0.6252371 |
| AGLKTGII | ASSHLRDPSAETLY | 0.6716223 |
| ASLAALLL | ASSHLRDPSAETLY | 0.6200738 |
| ASVINISSI | ASSHLRDPSAETLY | 0.6731802 |
| ATPPGPA | ASSHLRDPSAETLY | 0.6838616 |
| ATVASALV | ASSHLRDPSAETLY | 0.65813446 |
| ATVASLIL | ASSHLRDPSAETLY | 0.6613152 |
| AVNGPVLIRH | ASSHLRDPSAETLY | 0.6607247 |
| DGAAGAVGI | ASSHLRDPSAETLY | 0.5166063 |
| DPLLHGI | ASSHLRDPSAETLY | 0.66428024 |
| ELGAALGAL | ASSHLRDPSAETLY | 0.50090826 |
| ELGRTQAV | ASSHLRDPSAETLY | 0.6395267 |

| | | |
|---|---|---|
| FVAGGYLS | ASSHLRDPSAETLY | 0.6165365 |
| GASGGAGGAGGW | ASSHLRDPSAETLY | 0.6631447 |
| GGAGAGGDGGAGGI | ASSHLRDPSAETLY | 0.6554079 |
| GTGHNGSQPGMGGNG | ASSHLRDPSAETLY | 0.67042387 |
| GYAAASASAS | ASSHLRDPSAETLY | 0.6075787 |
| HQVDVAAVQQG | ASSHLRDPSAETLY | 0.6491536 |
| IGAAIAHT | ASSHLRDPSAETLY | 0.63364375 |
| IGMIEVAQL | ASSHLRDPSAETLY | 0.5868605 |
| IRLAPEHTSAMD | ASSHLRDPSAETLY | 0.6771133 |
| IYNVSVL | ASSHLRDPSAETLY | 0.65371513 |
| KQGFLGM | ASSHLRDPSAETLY | 0.6342701 |
| LESVITQL | ASSHLRDPSAETLY | 0.6790866 |
| LGLGALVVV | ASSHLRDPSAETLY | 0.66026103 |
| LIGNGGNGGSGGI | ASSHLRDPSAETLY | 0.5319443 |
| LNAINAPTL | ASSHLRDPSAETLY | 0.6408274 |
| LQLLNTGV | ASSHLRDPSAETLY | 0.6531129 |
| LQTSIEF | ASSHLRDPSAETLY | 0.6473075 |
| MSFAVSEL | ASSHLRDPSAETLY | 0.3454231 |
| NTGALGL | ASSHLRDPSAETLY | 0.67101115 |
| PPGAGGAGGAGTT | ASSHLRDPSAETLY | 0.6268824 |
| QLVGLGI | ASSHLRDPSAETLY | 0.59992385 |
| RFLEQVHQL | ASSHLRDPSAETLY | 0.63052917 |
| RGVGYVLR | ASSHLRDPSAETLY | 0.6722901 |
| SATGVFAGVI | ASSHLRDPSAETLY | 0.6518806 |
| SLPAITLP | ASSHLRDPSAETLY | 0.6571453 |
| STALAFTV | ASSHLRDPSAETLY | 0.670891 |
| SVLTVGLGK | ASSHLRDPSAETLY | 0.6743716 |
| TALAGAGSGI | ASSHLRDPSAETLY | 0.68196964 |
| TGLPFTGI | ASSHLRDPSAETLY | 0.6687685 |
| VAEHVTV | ASSHLRDPSAETLY | 0.6584413 |
| VDIVLNSL | ASSHLRDPSAETLY | 0.6730218 |
| VLTGGGAVL | ASSHLRDPSAETLY | 0.6620275 |
| VNVGLAYGI | ASSHLRDPSAETLY | 0.4870571 |
| VSSAAVGAV | ASSHLRDPSAETLY | 0.48607057 |
| VTSVYEAGV | ASSHLRDPSAETLY | 0.65828776 |
| WVALDRGA | ASSHLRDPSAETLY | 0.49036595 |
| YLVGSVVI | ASSHLRDPSAETLY | 0.6585058 |
| GPPGTGKTLL | ASSHLRDPSAETLY | 0.68164164 |
| GVGMTKF | ASSHLRDPSAETLY | 0.6572776 |
| IGIGGSDL | ASSHLRDPSAETLY | 0.56466794 |
| IGTLLAEL | ASSHLRDPSAETLY | 0.63728344 |
| IYTSGTT | ASSHLRDPSAETLY | 0.6383772 |
| LSFPAGV | ASSHLRDPSAETLY | 0.6706636 |
| LYTSGTT | ASSHLRDPSAETLY | 0.6794238 |
| VGAFPFGF | ASSHLRDPSAETLY | 0.68474054 |
| VGAGTVEF | ASSHLRDPSAETLY | 0.6760401 |
| VGKSTLLL | ASSHLRDPSAETLY | 0.6324185 |
| AALAHAGGGIV | ASSHLRDPSAETLY | 0.6705997 |
| AALVAVSLL | ASSHLRDPSAETLY | 0.63239527 |
| AAVSRAVAAL | ASSHLRDPSAETLY | 0.6545464 |
| AGGSSGIGGF | ASSHLRDPSAETLY | 0.65891147 |

| | | |
|---|---|---|
| AGLTVANANG | ASSHLRDPSAETLY | 0.66655695 |
| AGLVGAVL | ASSHLRDPSAETLY | 0.6559276 |
| ALSQAFTV | ASSHLRDPSAETLY | 0.64436615 |
| ALTFSVSL | ASSHLRDPSAETLY | 0.53896666 |
| ALTGAALVAL | ASSHLRDPSAETLY | 0.6737555 |
| ASFSIGAGVL | ASSHLRDPSAETLY | 0.6558127 |
| DGSVYKI | ASSHLRDPSAETLY | 0.49116814 |
| EQLAEIV | ASSHLRDPSAETLY | 0.6427522 |
| FAVSELTVM | ASSHLRDPSAETLY | 0.6835286 |
| GAAPLAAGL | ASSHLRDPSAETLY | 0.6820755 |
| GADGGDGGSGGAGG | ASSHLRDPSAETLY | 0.60945624 |
| GFGLDTGGAGG | ASSHLRDPSAETLY | 0.6176182 |
| GLASGGLL | ASSHLRDPSAETLY | 0.65001905 |
| GRIAGAAVLL | ASSHLRDPSAETLY | 0.5578817 |
| GSAGSLNTSF | ASSHLRDPSAETLY | 0.665497 |
| GTGQIGI | ASSHLRDPSAETLY | 0.64548075 |
| GTLIALIL | ASSHLRDPSAETLY | 0.6458483 |
| IAATMGFVVG | ASSHLRDPSAETLY | 0.6370478 |
| IFAQALL | ASSHLRDPSAETLY | 0.65808845 |
| IGAAVALGF | ASSHLRDPSAETLY | 0.63944006 |
| IGLTPLAL | ASSHLRDPSAETLY | 0.67700505 |
| ILGSNTSTL | ASSHLRDPSAETLY | 0.34680352 |
| IVFSLTVL | ASSHLRDPSAETLY | 0.6751247 |
| LASKATGF | ASSHLRDPSAETLY | 0.6008324 |
| LASVALIL | ASSHLRDPSAETLY | 0.636868 |
| LFAVGLTL | ASSHLRDPSAETLY | 0.48677248 |
| LFNGGIGTYI | ASSHLRDPSAETLY | 0.65818995 |
| LTLGAGII | ASSHLRDPSAETLY | 0.67426836 |
| LVAAALGDI | ASSHLRDPSAETLY | 0.6749182 |
| NTAFVQLGI | ASSHLRDPSAETLY | 0.59160304 |
| QLFSGAEL | ASSHLRDPSAETLY | 0.67383033 |
| SAATFITL | ASSHLRDPSAETLY | 0.68661094 |
| SNYGLANLGV | ASSHLRDPSAETLY | 0.62438744 |
| SYGGAYAVM | ASSHLRDPSAETLY | 0.48045605 |
| TTGVGGLTLG | ASSHLRDPSAETLY | 0.6650512 |
| TTQTIYI | ASSHLRDPSAETLY | 0.6576392 |
| VAAGAGLGAVI | ASSHLRDPSAETLY | 0.67697954 |
| VAAVVTLL | ASSHLRDPSAETLY | 0.67686164 |
| VGFLGGLI | ASSHLRDPSAETLY | 0.5578778 |
| VGVIPILL | ASSHLRDPSAETLY | 0.67216194 |
| VIFASSLI | ASSHLRDPSAETLY | 0.67471635 |
| VKGGDGGAAAG | ASSHLRDPSAETLY | 0.5996868 |
| VTKTFTNI | ASSHLRDPSAETLY | 0.63315797 |
| VVAAAILL | ASSHLRDPSAETLY | 0.63255835 |
| YAALDVEL | ASSHLRDPSAETLY | 0.5014959 |
| LASVALLL | ASSHLRDPSAETLY | 0.6763332 |
| LDAVHRV | ASSHLRDPSAETLY | 0.6217423 |
| QGLSSSIF | ASSHLRDPSAETLY | 0.631531 |
| RGTFANIRL | ASSHLRDPSAETLY | 0.6549349 |
| RQLWWGHRI | ASSHLRDPSAETLY | 0.63247687 |
| YGPPGTGKTLL | ASSHLRDPSAETLY | 0.6443083 |

| | | |
|---|---|---|
| AALVGAPANVA | ASSHLRDPSAETLY | 0.64165837 |
| AATFRRTE | ASSHLRDPSAETLY | 0.6644075 |
| AGALGGGI | ASSHLRDPSAETLY | 0.6634257 |
| AGGAGGAGGIGAG | ASSHLRDPSAETLY | 0.65107346 |
| AGSGALGL | ASSHLRDPSAETLY | 0.6647128 |
| AQIADQI | ASSHLRDPSAETLY | 0.6787993 |
| ASAVAVTYL | ASSHLRDPSAETLY | 0.5766886 |
| ASTVLGGTGG | ASSHLRDPSAETLY | 0.6288583 |
| AVGGEPAK | ASSHLRDPSAETLY | 0.6695739 |
| CEQGTFGGTC | ASSHLRDPSAETLY | 0.56355983 |
| DEGKAPAG | ASSHLRDPSAETLY | 0.6710821 |
| DTVVFPAL | ASSHLRDPSAETLY | 0.66003674 |
| FEGITMAG | ASSHLRDPSAETLY | 0.66326296 |
| FPLVVQD | ASSHLRDPSAETLY | 0.65479827 |
| GAFTTPQ | ASSHLRDPSAETLY | 0.68426526 |
| GFGGFGTAGGVGGA | ASSHLRDPSAETLY | 0.66999996 |
| GFLAQFL | ASSHLRDPSAETLY | 0.6531625 |
| GGAGGAGGNAGVA | ASSHLRDPSAETLY | 0.66508484 |
| GGVGTAGAGGAGGAGG | ASSHLRDPSAETLY | 0.6108391 |
| GNGGGGDAGNAGSGGN | ASSHLRDPSAETLY | 0.67735565 |
| GTGGAGGAGNDAGS | ASSHLRDPSAETLY | 0.6722921 |
| GTTNTGWFNS | ASSHLRDPSAETLY | 0.59112924 |
| IFMGPLFTL | ASSHLRDPSAETLY | 0.6431179 |
| ILAHAIEAI | ASSHLRDPSAETLY | 0.6838229 |
| IRSAIREAL | ASSHLRDPSAETLY | 0.6767732 |
| ITFAGGG | ASSHLRDPSAETLY | 0.54432666 |
| ITGMSGHQEV | ASSHLRDPSAETLY | 0.63127387 |
| IVAGLACTL | ASSHLRDPSAETLY | 0.56932 |
| IVGGGLAGA | ASSHLRDPSAETLY | 0.6485905 |
| IVTGVLNI | ASSHLRDPSAETLY | 0.5781036 |
| LAQQGCI | ASSHLRDPSAETLY | 0.66762835 |
| LCLHVEN | ASSHLRDPSAETLY | 0.6358392 |
| LDAISDGS | ASSHLRDPSAETLY | 0.6740536 |
| LGGFAAML | ASSHLRDPSAETLY | 0.6575952 |
| LGGKNPMI | ASSHLRDPSAETLY | 0.6820599 |
| LGPKGVAAIL | ASSHLRDPSAETLY | 0.6319953 |
| LMVGVSI | ASSHLRDPSAETLY | 0.64919156 |
| LVGAAFVL | ASSHLRDPSAETLY | 0.5355885 |
| LVLAVAPL | ASSHLRDPSAETLY | 0.61754787 |
| MAAIAVIDAA | ASSHLRDPSAETLY | 0.6265357 |
| NNGINAELQ | ASSHLRDPSAETLY | 0.6455422 |
| RGEPLVLVHG | ASSHLRDPSAETLY | 0.4057129 |
| RGGAGGAAGLI | ASSHLRDPSAETLY | 0.68315077 |
| RSYRMHPAV | ASSHLRDPSAETLY | 0.6497942 |
| SAAHLLAGQLA | ASSHLRDPSAETLY | 0.67546517 |
| SAAVGSY | ASSHLRDPSAETLY | 0.63219595 |
| SPHAVLHRI | ASSHLRDPSAETLY | 0.6394121 |
| SQIAKVL | ASSHLRDPSAETLY | 0.62482405 |
| TALGHLVP | ASSHLRDPSAETLY | 0.6709448 |
| TAPTTSGAA | ASSHLRDPSAETLY | 0.5987655 |
| TDGDDGNGGSPGT | ASSHLRDPSAETLY | 0.5038211 |

| | | |
|---|---|---|
| TDGLGEN | ASSHLRDPSAETLY | 0.64044625 |
| TGGPPLPGQA | ASSHLRDPSAETLY | 0.5651251 |
| TIPAITI | ASSHLRDPSAETLY | 0.6825774 |
| TIPGFTI | ASSHLRDPSAETLY | 0.67881763 |
| TSIASGALA | ASSHLRDPSAETLY | 0.65495276 |
| TVTLPGAAI | ASSHLRDPSAETLY | 0.68046796 |
| VASGLSAALP | ASSHLRDPSAETLY | 0.354584 |
| VGAGVIGI | ASSHLRDPSAETLY | 0.63135004 |
| VGPKRFGV | ASSHLRDPSAETLY | 0.68347573 |
| VNVAFPD | ASSHLRDPSAETLY | 0.6661489 |
| VSMTNSGVS | ASSHLRDPSAETLY | 0.64624125 |
| VTDVLGDLSV | ASSHLRDPSAETLY | 0.6796565 |
| VVVVGAGFAGL | ASSHLRDPSAETLY | 0.6217802 |
| VYTVGTVV | ASSHLRDPSAETLY | 0.64977336 |
| WTTFSGP | ASSHLRDPSAETLY | 0.6495851 |
| YDELVEAF | ASSHLRDPSAETLY | 0.6741384 |
| YGTGGAGGNGG | ASSHLRDPSAETLY | 0.67112106 |
| YTNVMAV | ASSHLRDPSAETLY | 0.5927773 |
| FLEQVHQL | ASSHLRDPSAETLY | 0.66768765 |
| GLSLLAVAV | ASSHLRDPSAETLY | 0.6839899 |
| LEQVHQL | ASSHLRDPSAETLY | 0.6308849 |
| LNAAGAI | ASSHLRDPSAETLY | 0.67902076 |
| PAVHASL | ASSHLRDPSAETLY | 0.6644115 |
| TELEDDA | ASSHLRDPSAETLY | 0.66302705 |
| TPPGPPN | ASSHLRDPSAETLY | 0.6105717 |
| TPPPPPPP | ASSHLRDPSAETLY | 0.6148543 |
| TQSPALI | ASSQPGGRGQY | 0.6524453 |
| EFAAAFPQ | ASSQPGGRGQY | 0.5968504 |
| LLSGNLDVL | ASSQPGGRGQY | 0.65338945 |
| GPANLGNYNI | ASSQPGGRGQY | 0.6432235 |
| TLGPTLP | ASSQPGGRGQY | 0.6903877 |
| VYAYSSCCK | ASSQPGGRGQY | 0.6700299 |
| APAPAPAPNI | ASSQPGGRGQY | 0.6666951 |
| PGGPVPIP | ASSQPGGRGQY | 0.60113025 |
| FLTVALEP | ASSQPGGRGQY | 0.6647846 |
| SQYFGANL | ASSQPGGRGQY | 0.66328967 |
| GGVKPAA | ASSQPGGRGQY | 0.5459441 |
| GGTGGAGGAGGW | ASSQPGGRGQY | 0.68096197 |
| GVGGVAGLGGAGG | ASSQPGGRGQY | 0.61657625 |
| VLLVLATTI | ASSQPGGRGQY | 0.64031005 |
| IGIFAAML | ASSQPGGRGQY | 0.67673963 |
| AVGDLID | ASSQPGGRGQY | 0.6816611 |
| NISPLPL | ASSQPGGRGQY | 0.64402485 |
| VTAVFVAI | ASSQPGGRGQY | 0.6502707 |
| SGDDNDAAL | ASSQPGGRGQY | 0.5995904 |
| AVAALAHS | ASSQPGGRGQY | 0.67436117 |
| GGVTVMATP | ASSQPGGRGQY | 0.67617035 |
| AGLAANVRD | ASSQPGGRGQY | 0.66048396 |
| TNRPDLIDPALLRPG | ASSQPGGRGQY | 0.68509865 |
| LANLGVTAL | ASSQPGGRGQY | 0.66680634 |
| TGERIEL | ASSQPGGRGQY | 0.508062 |

| | | |
|---|---|---|
| NGEELAA | ASSQPGGRGQY | 0.6615741 |
| VLSLVWG | ASSQPGGRGQY | 0.6795695 |
| IGYALWL | ASSQPGGRGQY | 0.6763947 |
| LSLLLAPL | ASSQPGGRGQY | 0.6864371 |
| TLDEALK | ASSQPGGRGQY | 0.66936684 |
| TDGAGTAEGG | ASSQPGGRGQY | 0.6324338 |
| NAPGGAGGNGG | ASSQPGGRGQY | 0.68031466 |
| VSVFMGMGGNF | ASSQPGGRGQY | 0.6906322 |
| LASYLAAL | ASSQPGGRGQY | 0.67317724 |
| LVGGTPAL | ASSQPGGRGQY | 0.6376299 |
| VRNADVL | ASSQPGGRGQY | 0.62150097 |
| TCSSPFLVSAM | ASSQPGGRGQY | 0.664374 |
| VHITDAE | ASSQPGGRGQY | 0.6740369 |
| EYAAPSF | ASSQPGGRGQY | 0.5985042 |
| VVDAFHGGL | ASSQPGGRGQY | 0.65638256 |
| ASVVDPQ | ASSQPGGRGQY | 0.6634374 |
| SYNEFGGLS | ASSQPGGRGQY | 0.5410521 |
| DPDGSVY | ASSQPGGRGQY | 0.6709137 |
| MVSTIHATE | ASSQPGGRGQY | 0.67875904 |
| LAAGVADPV | ASSQPGGRGQY | 0.6577671 |
| TKEQIAA | ASSQPGGRGQY | 0.61283123 |
| SLASPAAL | ASSQPGGRGQY | 0.6767627 |
| IGLVFYGV | ASSQPGGRGQY | 0.6864519 |
| FLLDASNL | ASSQPGGRGQY | 0.68191063 |
| YQVGNFP | ASSQPGGRGQY | 0.5200883 |
| ESLLSQST | ASSQPGGRGQY | 0.41528606 |
| LFTVGFAAVG | ASSQPGGRGQY | 0.6388458 |
| ATPPTPT | ASSQPGGRGQY | 0.6711012 |
| TDLATLG | ASSQPGGRGQY | 0.6464484 |
| SAGGTGR | ASSQPGGRGQY | 0.6253693 |
| GFSNSGS | ASSQPGGRGQY | 0.654732 |
| GGNGGSGGTSV | ASSQPGGRGQY | 0.67840624 |
| GTSVATGGAGN | ASSQPGGRGQY | 0.60644233 |
| NGGNGGSGGTS | ASSQPGGRGQY | 0.59760445 |
| AVVKFPH | ASSQPGGRGQY | 0.66735363 |
| TPPPPPP | ASSQPGGRGQY | 0.6726477 |
| AAPAAAAAPA | ASSQPGGRGQY | 0.6023628 |
| IALNEDL | ASSQPGGRGQY | 0.6357471 |
| IVNVSSM | ASSQPGGRGQY | 0.6457758 |
| TGIPPAPR | ASSQPGGRGQY | 0.63595384 |
| FDLGGGTF | ASSQPGGRGQY | 0.6559484 |
| FDLGGGTFDVS | ASSQPGGRGQY | 0.59531474 |
| NEPTAAA | ASSQPGGRGQY | 0.6786078 |
| NEPTAAALA | ASSQPGGRGQY | 0.5885813 |
| GSIINISSI | ASSQPGGRGQY | 0.6749681 |
| VAPGNAGTA | ASSQPGGRGQY | 0.66271776 |
| TGGGASLE | ASSQPGGRGQY | 0.667084 |
| VAVYNHYKRI | ASSQPGGRGQY | 0.63677084 |
| TDPAPATT | ASSQPGGRGQY | 0.62398386 |
| TNRPDILDPALLRPG | ASSQPGGRGQY | 0.59054196 |
| LTTTTTS | ASSQPGGRGQY | 0.50935197 |

| | | |
|---|---|---|
| PTGSGKTL | ASSQPGGRGQY | 0.67912424 |
| TGSGKTL | ASSQPGGRGQY | 0.6229712 |
| TIGHVDHGK | ASSQPGGRGQY | 0.6778623 |
| TIGHVDHGKTT | ASSQPGGRGQY | 0.6701108 |
| GHVDHGKTT | ASSQPGGRGQY | 0.6140424 |
| AQFEHTLL | ASSQPGGRGQY | 0.6698599 |
| IVNQDEL | ASSQPGGRGQY | 0.5663731 |
| TNGSQFF | ASSQPGGRGQY | 0.62813896 |
| TGKGIVE | ASSQPGGRGQY | 0.6721938 |
| LISNASDA | ASSQPGGRGQY | 0.6734818 |
| AELRTLGELTKIAKAHGAQ | ASSQPGGRGQY | 0.61133033 |
| APNAVAAAEPSW | ASSQPGGRGQY | 0.6679734 |
| APQVAGAATAVL | ASSQPGGRGQY | 0.6595619 |
| ARRDPAAAAAILPTDA | ASSQPGGRGQY | 0.63320225 |
| DAAARDVLAEAGLAEYF | ASSQPGGRGQY | 0.65906227 |
| DDQIVLSAKANLAKVEQ | ASSQPGGRGQY | 0.6779611 |
| DELLPDTGEDSDEDGH | ASSQPGGRGQY | 0.6676474 |
| ELPEAPANRATVPQD | ASSQPGGRGQY | 0.59240854 |
| FGQSKIERVTGIG | ASSQPGGRGQY | 0.6668525 |
| GAGGTGSTAGGAGGAGGAGGLY | ASSQPGGRGQY | 0.60330975 |
| GHGGTGAAGGNGGTGGAGGSGI | ASSQPGGRGQY | 0.65971386 |
| GNGGNGGAGGEGGPGGAGGAGA | ASSQPGGRGQY | 0.6577561 |
| GSTAGNGGNGGNATNGTIAGQP | ASSQPGGRGQY | 0.6478886 |
| GTGGDGGNGGNGGTGG | ASSQPGGRGQY | 0.6724759 |
| GVRRPAGIPADAPV | ASSQPGGRGQY | 0.6109258 |
| GYRQVTDAHLLTLARR | ASSQPGGRGQY | 0.6092429 |
| IAKMGQDGHDRG | ASSQPGGRGQY | 0.64029753 |
| KAEPSIEIQYT | ASSQPGGRGQY | 0.65826464 |
| KPVRIRMAADEDTVAVT | ASSQPGGRGQY | 0.67109656 |
| MPPRIAGMRLLVIKPEP | ASSQPGGRGQY | 0.5982709 |
| MKFYAVALFRSRGVPPTR | ASSQPGGRGQY | 0.6568082 |
| NAGGFGILGAVAHSP | ASSQPGGRGQY | 0.6412925 |
| NEGREITLAMAEM | ASSQPGGRGQY | 0.60383034 |
| NELVEWSHPLGHRWRWE | ASSQPGGRGQY | 0.6512087 |
| NLGAGGTGGNGGNPG | ASSQPGGRGQY | 0.65433073 |
| NTEWPKWETVMAADPVD | ASSQPGGRGQY | 0.5804159 |
| PASAAPAETQP | ASSQPGGRGQY | 0.6726545 |
| QQSSDGKNGVRAAEITGE | ASSQPGGRGQY | 0.6140245 |
| RAPDAVTTSWWKE | ASSQPGGRGQY | 0.67177415 |
| SGFGNVGAHTSGWFNQ | ASSQPGGRGQY | 0.6810287 |
| SGVAGAVGQAASVGGLKV | ASSQPGGRGQY | 0.6261723 |
| SNGGAGTGSAGGNGGT | ASSQPGGRGQY | 0.67388284 |
| TGGAGGEGGAAG | ASSQPGGRGQY | 0.6636802 |
| TGGDGGAAGAGGAPGHG | ASSQPGGRGQY | 0.66544056 |
| TMPPLLAAIANAGRHGV | ASSQPGGRGQY | 0.64845586 |
| TSSTAGNSGGA | ASSQPGGRGQY | 0.6296498 |
| VPATVPVGILC | ASSQPGGRGQY | 0.6782499 |
| VQRPDSGGVVLTGRIS | ASSQPGGRGQY | 0.6715017 |
| WTPNPVPATKGVPAGLVQAT | ASSQPGGRGQY | 0.67630947 |
| YALKRALAKFSDRFDVVII | ASSQPGGRGQY | 0.6701782 |
| YLIGSVIV | ASSQPGGRGQY | 0.6707891 |

| | | |
|---|---|---|
| YGQLVAVID | ASSQPGGRGQY | 0.6709415 |
| VTVGGMTLPA | ASSQPGGRGQY | 0.62542975 |
| VLALPAV | ASSQPGGRGQY | 0.55923986 |
| VIPPTLNYET | ASSQPGGRGQY | 0.64104813 |
| VGMGGMHPSSG | ASSQPGGRGQY | 0.62710834 |
| VGLAGGAATA | ASSQPGGRGQY | 0.64768475 |
| VGGAGGAGGSGP | ASSQPGGRGQY | 0.50968516 |
| VFAAFMV | ASSQPGGRGQY | 0.6759275 |
| TLYDKLR | ASSQPGGRGQY | 0.68858075 |
| TLAVFTVL | ASSQPGGRGQY | 0.6223323 |
| TGSVQPARV | ASSQPGGRGQY | 0.68504655 |
| TGGGAGSGGT | ASSQPGGRGQY | 0.5602042 |
| SHLETSL | ASSQPGGRGQY | 0.6750039 |
| SFTTGMT | ASSQPGGRGQY | 0.6790987 |
| QVEVAKRMGVRQPTVS | ASSQPGGRGQY | 0.604838 |
| QGSGGPALAM | ASSQPGGRGQY | 0.6730785 |
| MLAPSNP | ASSQPGGRGQY | 0.629236 |
| LVNNAAI | ASSQPGGRGQY | 0.6449734 |
| LTGEINTLA | ASSQPGGRGQY | 0.6789162 |
| LSIFGPV | ASSQPGGRGQY | 0.64053416 |
| LLALLANL | ASSQPGGRGQY | 0.6703057 |
| LISNASDALDKLR | ASSQPGGRGQY | 0.6492348 |
| LINEAVP | ASSQPGGRGQY | 0.6189605 |
| LGYAFSGL | ASSQPGGRGQY | 0.6676721 |
| LDFVAVAL | ASSQPGGRGQY | 0.68146706 |
| LAQPGAE | ASSQPGGRGQY | 0.6241925 |
| ISNLGAVTGF | ASSQPGGRGQY | 0.68986046 |
| HQEDPGRI | ASSQPGGRGQY | 0.6123895 |
| GTPGEPTP | ASSQPGGRGQY | 0.68426144 |
| GFFNSGS | ASSQPGGRGQY | 0.6681907 |
| GAGGGGLTAGAN | ASSQPGGRGQY | 0.64718425 |
| FVDALVNM | ASSQPGGRGQY | 0.64100766 |
| FLTVAPEL | ASSQPGGRGQY | 0.6739675 |
| FETLTAL | ASSQPGGRGQY | 0.6463523 |
| DVINPIEL | ASSQPGGRGQY | 0.6735782 |
| APVSGAPV | ASSQPGGRGQY | 0.6826384 |
| ALVGGLAW | ASSQPGGRGQY | 0.5977822 |
| ALAAAAATAI | ASSQPGGRGQY | 0.83835906 |
| AFGLGLLL | ASSQPGGRGQY | 0.7388257 |
| AFFAAASL | ASSQPGGRGQY | 0.67942727 |
| AAAALPAFSPP | ASSQPGGRGQY | 0.5396136 |
| VVLAPAL | ASSQPGGRGQY | 0.39158726 |
| TIGHVDHGKTTL | ASSQPGGRGQY | 0.6988573 |
| LISNASDALDK | ASSQPGGRGQY | 0.4971808 |
| LISNASDAL | ASSQPGGRGQY | 0.7284367 |
| ISNASDALDK | ASSQPGGRGQY | 0.7613485 |
| NQAETLVYQTE | ASSQPGGRGQY | 0.60318583 |
| TGGTGGTAGAG | ASSQPGGRGQY | 0.83595586 |
| TGSTAGGAGGA | ASSQPGGRGQY | 0.84203994 |
| DPLAVPVAAGPV | ASSQPGGRGQY | 0.42937005 |
| GGTGGTPGNGGN | ASSQPGGRGQY | 0.6042103 |

| | | |
|---|---|---|
| PAGLAGVDPATL | ASSQPGGRGQY | 0.59315556 |
| GNNLVVEAPHAGASV | ASSQPGGRGQY | 0.66159475 |
| KVTRIMSASTLEQPA | ASSQPGGRGQY | 0.33051318 |
| LGFALPIALPRRTVV | ASSQPGGRGQY | 0.7927468 |
| TRLGNLTARKLVVLV | ASSQPGGRGQY | 0.54651153 |
| WSQQRRLIYFPSAGP | ASSQPGGRGQY | 0.15734798 |
| GPNGNPDPMAAAVDIR | ASSQPGGRGQY | 0.694298 |
| TQRCVSVADLRAAASR | ASSQPGGRGQY | 0.697968 |
| QFVLDVAEHPVEVLTETV | ASSQPGGRGQY | 0.63309073 |
| GGAGGTGGAGGNGGTQAGDG | ASSQPGGRGQY | 0.82512176 |
| PLLATLRAIAVDVDAKPAQV | ASSQPGGRGQY | 0.3870891 |
| APKFRTGGYMIDFWGVGYQVAK | ASSQPGGRGQY | 0.5351763 |
| GAGGAGGAGGAAGGPGGTGGTG | ASSQPGGRGQY | 0.78989583 |
| IGALGLLATDVAPAGSVAFGVA | ASSQPGGRGQY | 0.83511496 |
| AGGAGGIGGNGGAGGAGGVLMG | ASSQPGGRGQY | 0.459303 |
| ADELIGLL | ASSQPGGRGQY | 0.76026183 |
| ADRALKA | ASSQPGGRGQY | 0.8267658 |
| AGAVGQVGGAGGA | ASSQPGGRGQY | 0.7660102 |
| AGGGGGLGGAP | ASSQPGGRGQY | 0.8012832 |
| AGGLLYGNGGAG | ASSQPGGRGQY | 0.8306954 |
| AGLESIGL | ASSQPGGRGQY | 0.60203844 |
| AGLKTGII | ASSQPGGRGQY | 0.8113351 |
| ASLAALLL | ASSQPGGRGQY | 0.8364521 |
| ASVINISSI | ASSQPGGRGQY | 0.6635296 |
| ATPPGPA | ASSQPGGRGQY | 0.7230977 |
| ATVASALV | ASSQPGGRGQY | 0.57051873 |
| ATVASLIL | ASSQPGGRGQY | 0.6224907 |
| AVNGPVLIRH | ASSQPGGRGQY | 0.2587348 |
| DGAAGAVGI | ASSQPGGRGQY | 0.7718796 |
| DPLLHGI | ASSQPGGRGQY | 0.8071456 |
| ELGAALGAL | ASSQPGGRGQY | 0.6207657 |
| ELGRTQAV | ASSQPGGRGQY | 0.50532037 |
| FVAGGYLS | ASSQPGGRGQY | 0.4880412 |
| GASGGAGGAGGW | ASSQPGGRGQY | 0.6714432 |
| GGAGAGGDGGAGGI | ASSQPGGRGQY | 0.6425427 |
| GTGHNGSQPGMGGNG | ASSQPGGRGQY | 0.28819054 |
| GYAAASASAS | ASSQPGGRGQY | 0.66768956 |
| HQVDVAAVQQG | ASSQPGGRGQY | 0.8175087 |
| IGAAIAHT | ASSQPGGRGQY | 0.8255879 |
| IGMIEVAQL | ASSQPGGRGQY | 0.28741387 |
| IRLAPEHTSAMD | ASSQPGGRGQY | 0.83790517 |
| IYNVSVL | ASSQPGGRGQY | 0.7538748 |
| KQGFLGM | ASSQPGGRGQY | 0.8402668 |
| LESVITQL | ASSQPGGRGQY | 0.67729545 |
| LGLGALVVV | ASSQPGGRGQY | 0.58295476 |
| LIGNGGNGGSGGI | ASSQPGGRGQY | 0.7856346 |
| LNAINAPTL | ASSQPGGRGQY | 0.66421616 |
| LQLLNTGV | ASSQPGGRGQY | 0.42100307 |
| LQTSIEF | ASSQPGGRGQY | 0.81362003 |
| MSFAVSEL | ASSQPGGRGQY | 0.6851066 |
| NTGALGL | ASSQPGGRGQY | 0.5713729 |

| | | |
|---|---|---|
| PPGAGGAGGAGTT | ASSQPGGRGQY | 0.53537726 |
| QLVGLGI | ASSQPGGRGQY | 0.7763023 |
| RFLEQVHQL | ASSQPGGRGQY | 0.43139586 |
| RGVGYVLR | ASSQPGGRGQY | 0.56370294 |
| SATGVFAGVI | ASSQPGGRGQY | 0.44767502 |
| SLPAITLP | ASSQPGGRGQY | 0.73191303 |
| STALAFTV | ASSQPGGRGQY | 0.8160545 |
| SVLTVGLGK | ASSQPGGRGQY | 0.75011075 |
| TALAGAGSGI | ASSQPGGRGQY | 0.6245338 |
| TGLPFTGI | ASSQPGGRGQY | 0.7682708 |
| VAEHVTV | ASSQPGGRGQY | 0.7565994 |
| VDIVLNSL | ASSQPGGRGQY | 0.7371901 |
| VLTGGGAVL | ASSQPGGRGQY | 0.81370413 |
| VNVGLAYGI | ASSQPGGRGQY | 0.58686996 |
| VSSAAVGAV | ASSQPGGRGQY | 0.7857106 |
| VTSVYEAGV | ASSQPGGRGQY | 0.7399451 |
| WVALDRGA | ASSQPGGRGQY | 0.7873392 |
| YLVGSVVI | ASSQPGGRGQY | 0.7899685 |
| GPPGTGKTLL | ASSQPGGRGQY | 0.7774956 |
| GVGMTKF | ASSQPGGRGQY | 0.6843442 |
| IGIGGSDL | ASSQPGGRGQY | 0.21579807 |
| IGTLLAEL | ASSQPGGRGQY | 0.66888463 |
| IYTSGTT | ASSQPGGRGQY | 0.7787035 |
| LSFPAGV | ASSQPGGRGQY | 0.72827893 |
| LYTSGTT | ASSQPGGRGQY | 0.78310937 |
| VGAFPFGF | ASSQPGGRGQY | 0.7631134 |
| VGAGTVEF | ASSQPGGRGQY | 0.579837 |
| VGKSTLLL | ASSQPGGRGQY | 0.81907725 |
| AALAHAGGGIV | ASSQPGGRGQY | 0.7947361 |
| AALVAVSLL | ASSQPGGRGQY | 0.37127247 |
| AAVSRAVAAL | ASSQPGGRGQY | 0.6674055 |
| AGGSSGIGGF | ASSQPGGRGQY | 0.8585024 |
| AGLTVANANG | ASSQPGGRGQY | 0.7290932 |
| AGLVGAVL | ASSQPGGRGQY | 0.6540396 |
| ALSQAFTV | ASSQPGGRGQY | 0.56463623 |
| ALTFSVSL | ASSQPGGRGQY | 0.8228137 |
| ALTGAALVAL | ASSQPGGRGQY | 0.5549685 |
| ASFSIGAGVL | ASSQPGGRGQY | 0.83587277 |
| DGSVYKI | ASSQPGGRGQY | 0.1866997 |
| EQLAEIV | ASSQPGGRGQY | 0.54489464 |
| FAVSELTVM | ASSQPGGRGQY | 0.5250441 |
| GAAPLAAGL | ASSQPGGRGQY | 0.38493937 |
| GADGGDGGSGGAGG | ASSQPGGRGQY | 0.7892526 |
| GFGLDTGGAGG | ASSQPGGRGQY | 0.6371563 |
| GLASGGLL | ASSQPGGRGQY | 0.8021677 |
| GRIAGAAVLL | ASSQPGGRGQY | 0.7360652 |
| GSAGSLNTSF | ASSQPGGRGQY | 0.77957445 |
| GTGQIGI | ASSQPGGRGQY | 0.7315437 |
| GTLIALIL | ASSQPGGRGQY | 0.77364266 |
| IAATMGFVVG | ASSQPGGRGQY | 0.5842675 |
| IFAQALL | ASSQPGGRGQY | 0.7732315 |

| | | |
|---|---|---|
| IGAAVALGF | ASSQPGGRGQY | 0.82703483 |
| IGLTPLAL | ASSQPGGRGQY | 0.71624106 |
| ILGSNTSTL | ASSQPGGRGQY | 0.40246487 |
| IVFSLTVL | ASSQPGGRGQY | 0.5422765 |
| LASKATGF | ASSQPGGRGQY | 0.7042502 |
| LASVALIL | ASSQPGGRGQY | 0.27357203 |
| LFAVGLTL | ASSQPGGRGQY | 0.22404815 |
| LFNGGIGTYI | ASSQPGGRGQY | 0.62286353 |
| LTLGAGII | ASSQPGGRGQY | 0.54202783 |
| LVAAALGDI | ASSQPGGRGQY | 0.8202243 |
| NTAFVQLGI | ASSQPGGRGQY | 0.79906934 |
| QLFSGAEL | ASSQPGGRGQY | 0.6841431 |
| SAATFITL | ASSQPGGRGQY | 0.803846 |
| SNYGLANLGV | ASSQPGGRGQY | 0.74132216 |
| SYGGAYAVM | ASSQPGGRGQY | 0.13670135 |
| TTGVGGLTLG | ASSQPGGRGQY | 0.672703 |
| TTQTIYI | ASSQPGGRGQY | 0.44014943 |
| VAAGAGLGAVI | ASSQPGGRGQY | 0.8201935 |
| VAAVVTLL | ASSQPGGRGQY | 0.5820708 |
| VGFLGGLI | ASSQPGGRGQY | 0.5194591 |
| VGVIPILL | ASSQPGGRGQY | 0.7155557 |
| VIFASSLI | ASSQPGGRGQY | 0.6773517 |
| VKGGDGGAAAG | ASSQPGGRGQY | 0.7056734 |
| VTKTFTNI | ASSQPGGRGQY | 0.48601797 |
| VVAAAILL | ASSQPGGRGQY | 0.16405565 |
| YAALDVEL | ASSQPGGRGQY | 0.7965849 |
| LASVALLL | ASSQPGGRGQY | 0.3643014 |
| LDAVHRV | ASSQPGGRGQY | 0.51326454 |
| QGLSSSIF | ASSQPGGRGQY | 0.24475853 |
| RGTFANIRL | ASSQPGGRGQY | 0.17918535 |
| RQLWWGHRI | ASSQPGGRGQY | 0.4032754 |
| YGPPGTGKTLL | ASSQPGGRGQY | 0.7960712 |
| AALVGAPANVA | ASSQPGGRGQY | 0.65381855 |
| AATFRRTE | ASSQPGGRGQY | 0.6172961 |
| AGALGGGI | ASSQPGGRGQY | 0.8437869 |
| AGGAGGAGGIGAG | ASSQPGGRGQY | 0.70399857 |
| AGSGALGL | ASSQPGGRGQY | 0.6600067 |
| AQIADQI | ASSQPGGRGQY | 0.579012 |
| ASAVAVTYL | ASSQPGGRGQY | 0.5221879 |
| ASTVLGGTGG | ASSQPGGRGQY | 0.5302415 |
| AVGGEPAK | ASSQPGGRGQY | 0.49981213 |
| CEQGTFGGTC | ASSQPGGRGQY | 0.8485752 |
| DEGKAPAG | ASSQPGGRGQY | 0.54896003 |
| DTVVFPAL | ASSQPGGRGQY | 0.481371 |
| FEGITMAG | ASSQPGGRGQY | 0.6549331 |
| FPLVVQD | ASSQPGGRGQY | 0.83542854 |
| GAFTTPQ | ASSQPGGRGQY | 0.86074877 |
| GFGGFGTAGGVGGA | ASSQPGGRGQY | 0.5877435 |
| GFLAQFL | ASSQPGGRGQY | 0.6699326 |
| GGAGGAGGNAGVA | ASSQPGGRGQY | 0.6031117 |
| GGVGTAGAGGAGGAGG | ASSQPGGRGQY | 0.71753913 |

| | | |
|---|---|---|
| GNGGGGDAGNAGSGGN | ASSQPGGRGQY | 0.628654 |
| GTGGAGGAGNDAGS | ASSQPGGRGQY | 0.4730211 |
| GTTNTGWFNS | ASSQPGGRGQY | 0.7525793 |
| IFMGPLFTL | ASSQPGGRGQY | 0.84489053 |
| ILAHAIEAI | ASSQPGGRGQY | 0.65322125 |
| IRSAIREAL | ASSQPGGRGQY | 0.3854576 |
| ITFAGGG | ASSQPGGRGQY | 0.6864361 |
| ITGMSGHQEV | ASSQPGGRGQY | 0.4915896 |
| IVAGLACTL | ASSQPGGRGQY | 0.6471282 |
| IVGGGLAGA | ASSQPGGRGQY | 0.7687262 |
| IVTGVLNI | ASSQPGGRGQY | 0.37783718 |
| LAQQGCI | ASSQPGGRGQY | 0.57348096 |
| LCLHVEN | ASSQPGGRGQY | 0.8566596 |
| LDAISDGS | ASSQPGGRGQY | 0.592988 |
| LGGFAAML | ASSQPGGRGQY | 0.7289307 |
| LGGKNPMI | ASSQPGGRGQY | 0.82024693 |
| LGPKGVAAIL | ASSQPGGRGQY | 0.7630681 |
| LMVGVSI | ASSQPGGRGQY | 0.84058636 |
| LVGAAFVL | ASSQPGGRGQY | 0.77989113 |
| LVLAVAPL | ASSQPGGRGQY | 0.6942433 |
| MAAIAVIDAA | ASSQPGGRGQY | 0.70061404 |
| NNGINAELQ | ASSQPGGRGQY | 0.7270348 |
| RGEPLVLVHG | ASSQPGGRGQY | 0.7310663 |
| RGGAGGAAGLI | ASSQPGGRGQY | 0.6563134 |
| RSYRMHPAV | ASSQPGGRGQY | 0.5037403 |
| SAAHLLAGQLA | ASSQPGGRGQY | 0.82327235 |
| SAAVGSY | ASSQPGGRGQY | 0.70943415 |
| SPHAVLHRI | ASSQPGGRGQY | 0.78053296 |
| SQIAKVL | ASSQPGGRGQY | 0.54375523 |
| TALGHLVP | ASSQPGGRGQY | 0.4799495 |
| TAPTTSGAA | ASSQPGGRGQY | 0.79976505 |
| TDGDDGNGGSPGT | ASSQPGGRGQY | 0.17064384 |
| TDGLGEN | ASSQPGGRGQY | 0.21722786 |
| TGGPPLPGQA | ASSQPGGRGQY | 0.6728207 |
| TIPAITI | ASSQPGGRGQY | 0.83941543 |
| TIPGFTI | ASSQPGGRGQY | 0.17930356 |
| TSIASGALA | ASSQPGGRGQY | 0.5177658 |
| TVTLPGAAI | ASSQPGGRGQY | 0.39733502 |
| VASGLSAALP | ASSQPGGRGQY | 0.7850961 |
| VGAGVIGI | ASSQPGGRGQY | 0.7180928 |
| VGPKRFGV | ASSQPGGRGQY | 0.42985708 |
| VNVAFPD | ASSQPGGRGQY | 0.7685491 |
| VSMTNSGVS | ASSQPGGRGQY | 0.3186105 |
| VTDVLGDLSV | ASSQPGGRGQY | 0.37875435 |
| VVVVGAGFAGL | ASSQPGGRGQY | 0.7710128 |
| VYTVGTVV | ASSQPGGRGQY | 0.8095775 |
| WTTFSGP | ASSQPGGRGQY | 0.50429237 |
| YDELVEAF | ASSQPGGRGQY | 0.79786265 |
| YGTGGAGGNGG | ASSQPGGRGQY | 0.5698764 |
| YTNVMAV | ASSQPGGRGQY | 0.85777235 |
| FLEQVHQL | ASSQPGGRGQY | 0.59457713 |

| | | |
|---|---|---|
| GLSLLAVAV | ASSQPGGRGQY | 0.40223235 |
| LEQVHQL | ASSQPGGRGQY | 0.7948235 |
| LNAAGAI | ASSQPGGRGQY | 0.77035373 |
| PAVHASL | ASSQPGGRGQY | 0.73592025 |
| TELEDDA | ASSQPGGRGQY | 0.7916434 |
| TPPGPPN | ASSQPGGRGQY | 0.57784706 |
| TPPPPPPP | ASSQPGGRGQY | 0.6676722 |
| TQSPALI | ASSGTGGGNEQY | 0.8317703 |
| EFAAAFPQ | ASSGTGGGNEQY | 0.8157362 |
| LLSGNLDVL | ASSGTGGGNEQY | 0.54103637 |
| GPANLGNYNI | ASSGTGGGNEQY | 0.7806517 |
| TLGPTLP | ASSGTGGGNEQY | 0.69801295 |
| VYAYSSCCK | ASSGTGGGNEQY | 0.5764338 |
| APAPAPAPNI | ASSGTGGGNEQY | 0.8329165 |
| PGGPVPIP | ASSGTGGGNEQY | 0.7192877 |
| FLTVALEP | ASSGTGGGNEQY | 0.80141217 |
| SQYFGANL | ASSGTGGGNEQY | 0.74520123 |
| GGVKPAA | ASSGTGGGNEQY | 0.7911787 |
| GGTGGAGGAGGW | ASSGTGGGNEQY | 0.48629212 |
| GVGGVAGLGGAGG | ASSGTGGGNEQY | 0.77992344 |
| VLLVLATTI | ASSGTGGGNEQY | 0.71725595 |
| IGIFAAML | ASSGTGGGNEQY | 0.77822596 |
| AVGDLID | ASSGTGGGNEQY | 0.70684046 |
| NISPLPL | ASSGTGGGNEQY | 0.40387312 |
| VTAVFVAI | ASSGTGGGNEQY | 0.72601485 |
| SGDDNDAAL | ASSGTGGGNEQY | 0.8256103 |
| AVAALAHS | ASSGTGGGNEQY | 0.6630106 |
| GGVTVMATP | ASSGTGGGNEQY | 0.75753814 |
| AGLAANVRD | ASSGTGGGNEQY | 0.84607583 |
| TNRPDLIDPALLRPG | ASSGTGGGNEQY | 0.57773525 |
| LANLGVTAL | ASSGTGGGNEQY | 0.65612406 |
| TGERIEL | ASSGTGGGNEQY | 0.7391759 |
| NGEELAA | ASSGTGGGNEQY | 0.81515723 |
| VLSLVWG | ASSGTGGGNEQY | 0.62609935 |
| IGYALWL | ASSGTGGGNEQY | 0.8427962 |
| LSLLLAPL | ASSGTGGGNEQY | 0.6034966 |
| TLDEALK | ASSGTGGGNEQY | 0.80351543 |
| TDGAGTAEGG | ASSGTGGGNEQY | 0.57228035 |
| NAPGGAGGNGG | ASSGTGGGNEQY | 0.77174056 |
| VSVFMGMGGNF | ASSGTGGGNEQY | 0.5736126 |
| LASYLAAL | ASSGTGGGNEQY | 0.7907359 |
| LVGGTPAL | ASSGTGGGNEQY | 0.8133431 |
| VRNADVL | ASSGTGGGNEQY | 0.80539954 |
| TCSSPFLVSAM | ASSGTGGGNEQY | 0.2163445 |
| VHITDAE | ASSGTGGGNEQY | 0.82328147 |
| EYAAPSF | ASSGTGGGNEQY | 0.8097482 |
| VVDAFHGGL | ASSGTGGGNEQY | 0.8589816 |
| ASVVDPQ | ASSGTGGGNEQY | 0.7118834 |
| SYNEFGGLS | ASSGTGGGNEQY | 0.7583057 |
| DPDGSVY | ASSGTGGGNEQY | 0.68411404 |
| MVSTIHATE | ASSGTGGGNEQY | 0.61663383 |

| | | |
|---|---|---|
| LAAGVADPV | ASSGTGGGNEQY | 0.50478184 |
| TKEQIAA | ASSGTGGGNEQY | 0.6837672 |
| SLASPAAL | ASSGTGGGNEQY | 0.66461957 |
| IGLVFYGV | ASSGTGGGNEQY | 0.5752183 |
| FLLDASNL | ASSGTGGGNEQY | 0.78960466 |
| YQVGNFP | ASSGTGGGNEQY | 0.8103018 |
| ESLLSQST | ASSGTGGGNEQY | 0.8151666 |
| LFTVGFAAVG | ASSGTGGGNEQY | 0.8177564 |
| ATPPTPT | ASSGTGGGNEQY | 0.8279826 |
| TDLATLG | ASSGTGGGNEQY | 0.7903505 |
| SAGGTGR | ASSGTGGGNEQY | 0.51195765 |
| GFSNSGS | ASSGTGGGNEQY | 0.7072998 |
| GGNGGSGGTSV | ASSGTGGGNEQY | 0.43584415 |
| GTSVATGGAGN | ASSGTGGGNEQY | 0.6370174 |
| NGGNGGSGGTS | ASSGTGGGNEQY | 0.5649803 |
| AVVKFPH | ASSGTGGGNEQY | 0.763284 |
| TPPPPPP | ASSGTGGGNEQY | 0.14487338 |
| AAPAAAAAPA | ASSGTGGGNEQY | 0.7057306 |
| IALNEDL | ASSGTGGGNEQY | 0.84372264 |
| IVNVSSM | ASSGTGGGNEQY | 0.5269808 |
| TGIPPAPR | ASSGTGGGNEQY | 0.77321213 |
| FDLGGGTF | ASSGTGGGNEQY | 0.6963648 |
| FDLGGGTFDVS | ASSGTGGGNEQY | 0.54070604 |
| NEPTAAA | ASSGTGGGNEQY | 0.7944276 |
| NEPTAAALA | ASSGTGGGNEQY | 0.46711957 |
| GSIINISSI | ASSGTGGGNEQY | 0.74626106 |
| VAPGNAGTA | ASSGTGGGNEQY | 0.8234458 |
| TGGGASLE | ASSGTGGGNEQY | 0.70545554 |
| VAVYNHYKRI | ASSGTGGGNEQY | 0.74194485 |
| TDPAPATT | ASSGTGGGNEQY | 0.64763415 |
| TNRPDILDPALLRPG | ASSGTGGGNEQY | 0.43056148 |
| LTTTTTS | ASSGTGGGNEQY | 0.7548911 |
| PTGSGKTL | ASSGTGGGNEQY | 0.1317879 |
| TGSGKTL | ASSGTGGGNEQY | 0.697716 |
| TIGHVDHGK | ASSGTGGGNEQY | 0.4823712 |
| TIGHVDHGKTT | ASSGTGGGNEQY | 0.71920663 |
| GHVDHGKTT | ASSGTGGGNEQY | 0.7317021 |
| AQFEHTLL | ASSGTGGGNEQY | 0.7422172 |
| IVNQDEL | ASSGTGGGNEQY | 0.47984797 |
| TNGSQFF | ASSGTGGGNEQY | 0.62491655 |
| TGKGIVE | ASSGTGGGNEQY | 0.48466545 |
| LISNASDA | ASSGTGGGNEQY | 0.4307963 |
| AELRTLGELTKIAKAHGAQ | ASSGTGGGNEQY | 0.80298 |
| APNAVAAAEPSW | ASSGTGGGNEQY | 0.6088588 |
| APQVAGAATAVL | ASSGTGGGNEQY | 0.6696106 |
| ARRDPAAAAAILPTDA | ASSGTGGGNEQY | 0.8132713 |
| DAAARDVLAEAGLAEYF | ASSGTGGGNEQY | 0.6905939 |
| DDQIVLSAKANLAKVEQ | ASSGTGGGNEQY | 0.39540625 |
| DELLPDTGEDSDEDGH | ASSGTGGGNEQY | 0.7141311 |
| ELPEAPANRATVPQD | ASSGTGGGNEQY | 0.6759505 |
| FGQSKIERVTGIG | ASSGTGGGNEQY | 0.6062863 |

| | | |
|---|---|---|
| GAGGTGSTAGGAGGAGGAGGLY | ASSGTGGGNEQY | 0.13588482 |
| GHGGTGAAGGNGGTGGAGGSGI | ASSGTGGGNEQY | 0.7713653 |
| GNGGNGGAGGEGGPGGAGGAGA | ASSGTGGGNEQY | 0.5822636 |
| GSTAGNGGNGGNATNGTIAGQP | ASSGTGGGNEQY | 0.5363803 |
| GTGGDGGNGGNGGTGG | ASSGTGGGNEQY | 0.55050856 |
| GVRRPAGIPADAPV | ASSGTGGGNEQY | 0.8002438 |
| GYRQVTDAHLLTLARR | ASSGTGGGNEQY | 0.4533511 |
| IAKMGQDGHDRG | ASSGTGGGNEQY | 0.7421875 |
| KAEPSIEIQYT | ASSGTGGGNEQY | 0.8064504 |
| KPVRIRMAADEDTVAVT | ASSGTGGGNEQY | 0.8013034 |
| MPPRIAGMRLLVIKPEP | ASSGTGGGNEQY | 0.8420688 |
| MKFYAVALFRSRGVPPTR | ASSGTGGGNEQY | 0.77145404 |
| NAGGFGILGAVAHSP | ASSGTGGGNEQY | 0.6138459 |
| NEGREITLAMAEM | ASSGTGGGNEQY | 0.7788357 |
| NELVEWSHPLGHRWRWE | ASSGTGGGNEQY | 0.73989993 |
| NLGAGGTGGNGGNPG | ASSGTGGGNEQY | 0.2601911 |
| NTEWPKWETVMAADPVD | ASSGTGGGNEQY | 0.22037457 |
| PASAAPAETQP | ASSGTGGGNEQY | 0.7398789 |
| QQSSDGKNGVRAAEITGE | ASSGTGGGNEQY | 0.24742694 |
| RAPDAVTTSWWKE | ASSGTGGGNEQY | 0.6818005 |
| SGFGNVGAHTSGWFNQ | ASSGTGGGNEQY | 0.8101192 |
| SGVAGAVGQAASVGGLKV | ASSGTGGGNEQY | 0.7215153 |
| SNGGAGTGSAGGNGGT | ASSGTGGGNEQY | 0.48530447 |
| TGGAGGEGGAAG | ASSGTGGGNEQY | 0.5745882 |
| TGGDGGAAGAGGAPGHG | ASSGTGGGNEQY | 0.7199981 |
| TMPPLLAAIANAGRHGV | ASSGTGGGNEQY | 0.8105973 |
| TSSTAGNSGGA | ASSGTGGGNEQY | 0.7810737 |
| VPATVPVGILC | ASSGTGGGNEQY | 0.83415353 |
| VQRPDSGGVVLTGRIS | ASSGTGGGNEQY | 0.819036 |
| WTPNPVPATKGVPAGLVQAT | ASSGTGGGNEQY | 0.62630445 |
| YALKRALAKFSDRFDVVII | ASSGTGGGNEQY | 0.7864941 |
| YLIGSVIV | ASSGTGGGNEQY | 0.62712467 |
| YGQLVAVID | ASSGTGGGNEQY | 0.6732596 |
| VTVGGMTLPA | ASSGTGGGNEQY | 0.67358655 |
| VLALPAV | ASSGTGGGNEQY | 0.7095725 |
| VIPPTLNYET | ASSGTGGGNEQY | 0.72913325 |
| VGMGGMHPSSG | ASSGTGGGNEQY | 0.5923129 |
| VGLAGGAATA | ASSGTGGGNEQY | 0.3782749 |
| VGGAGGAGGSGP | ASSGTGGGNEQY | 0.7089283 |
| VFAAFMV | ASSGTGGGNEQY | 0.7598062 |
| TLYDKLR | ASSGTGGGNEQY | 0.30126557 |
| TLAVFTVL | ASSGTGGGNEQY | 0.568344 |
| TGSVQPARV | ASSGTGGGNEQY | 0.8566877 |
| TGGGAGSGGT | ASSGTGGGNEQY | 0.83596754 |
| SHLETSL | ASSGTGGGNEQY | 0.49953556 |
| SFTTGMT | ASSGTGGGNEQY | 0.5024931 |
| QVEVAKRMGVRQPTVS | ASSGTGGGNEQY | 0.6875031 |
| QGSGGPALAM | ASSGTGGGNEQY | 0.25304833 |
| MLAPSNP | ASSGTGGGNEQY | 0.7034507 |
| LVNNAAI | ASSGTGGGNEQY | 0.70964056 |
| LTGEINTLA | ASSGTGGGNEQY | 0.7181356 |

| | | |
|---|---|---|
| LSIFGPV | ASSGTGGGNEQY | 0.5194906 |
| LLALLANL | ASSGTGGGNEQY | 0.65859354 |
| LISNASDALDKLR | ASSGTGGGNEQY | 0.60907865 |
| LINEAVP | ASSGTGGGNEQY | 0.78276443 |
| LGYAFSGL | ASSGTGGGNEQY | 0.16868982 |
| LDFVAVAL | ASSGTGGGNEQY | 0.73404664 |
| LAQPGAE | ASSGTGGGNEQY | 0.50998676 |
| ISNLGAVTGF | ASSGTGGGNEQY | 0.61683285 |
| HQEDPGRI | ASSGTGGGNEQY | 0.44052953 |
| GTPGEPTP | ASSGTGGGNEQY | 0.69366324 |
| GFFNSGS | ASSGTGGGNEQY | 0.8123934 |
| GAGGGGLTAGAN | ASSGTGGGNEQY | 0.7912767 |
| FVDALVNM | ASSGTGGGNEQY | 0.35715967 |
| FLTVAPEL | ASSGTGGGNEQY | 0.8136423 |
| FETLTAL | ASSGTGGGNEQY | 0.8514615 |
| DVINPIEL | ASSGTGGGNEQY | 0.6167371 |
| APVSGAPV | ASSGTGGGNEQY | 0.4314146 |
| ALVGGLAW | ASSGTGGGNEQY | 0.77578294 |
| ALAAAAATAI | ASSGTGGGNEQY | 0.65540445 |
| AFGLGLLL | ASSGTGGGNEQY | 0.78300154 |
| AFFAAASL | ASSGTGGGNEQY | 0.8022573 |
| AAAALPAFSPP | ASSGTGGGNEQY | 0.47221535 |
| VVLAPAL | ASSGTGGGNEQY | 0.8187399 |
| TIGHVDHGKTTL | ASSGTGGGNEQY | 0.8215115 |
| LISNASDALDK | ASSGTGGGNEQY | 0.55175924 |
| LISNASDAL | ASSGTGGGNEQY | 0.5652771 |
| ISNASDALDK | ASSGTGGGNEQY | 0.5981656 |
| NQAETLVYQTE | ASSGTGGGNEQY | 0.27205318 |
| TGGTGGTAGAG | ASSGTGGGNEQY | 0.7832072 |
| TGSTAGGAGGA | ASSGTGGGNEQY | 0.5520738 |
| DPLAVPVAAGPV | ASSGTGGGNEQY | 0.574199 |
| GGTGGTPGNGGN | ASSGTGGGNEQY | 0.71732765 |
| PAGLAGVDPATL | ASSGTGGGNEQY | 0.568869 |
| GNNLVVEAPHAGASV | ASSGTGGGNEQY | 0.58919054 |
| KVTRIMSASTLEQPA | ASSGTGGGNEQY | 0.61180055 |
| LGFALPIALPRRTVV | ASSGTGGGNEQY | 0.7308321 |
| TRLGNLTARKLVVLV | ASSGTGGGNEQY | 0.7154013 |
| WSQQRRLIYFPSAGP | ASSGTGGGNEQY | 0.7485622 |
| GPNGNPDPMAAAVDIR | ASSGTGGGNEQY | 0.73594844 |
| TQRCVSVADLRAAASR | ASSGTGGGNEQY | 0.8359314 |
| QFVLDVAEHPVEVLTETV | ASSGTGGGNEQY | 0.47192377 |
| GGAGGTGGAGGNGGTQAGDG | ASSGTGGGNEQY | 0.55519074 |
| PLLATLRAIAVDVDAKPAQV | ASSGTGGGNEQY | 0.7796254 |
| APKFRTGGYMIDFWGVGYQVAK | ASSGTGGGNEQY | 0.31534195 |
| GAGGAGGAGGAAGGPGGTGGTG | ASSGTGGGNEQY | 0.79710054 |
| IGALGLLATDVAPAGSVAFGVA | ASSGTGGGNEQY | 0.72248083 |
| AGGAGGIGGNGGAGGAGGVLMG | ASSGTGGGNEQY | 0.7621066 |
| ADELIGLL | ASSGTGGGNEQY | 0.6998645 |
| ADRALKA | ASSGTGGGNEQY | 0.8579652 |
| AGAVGQVGGAGGA | ASSGTGGGNEQY | 0.81201553 |
| AGGGGGLGGAP | ASSGTGGGNEQY | 0.68290377 |

| | | |
|---|---|---|
| AGGLLYGNGGAG | ASSGTGGGNEQY | 0.7501738 |
| AGLESIGL | ASSGTGGGNEQY | 0.5631984 |
| AGLKTGII | ASSGTGGGNEQY | 0.811966 |
| ASLAALLL | ASSGTGGGNEQY | 0.74988294 |
| ASVINISSI | ASSGTGGGNEQY | 0.48305416 |
| ATPPGPA | ASSGTGGGNEQY | 0.67388403 |
| ATVASALV | ASSGTGGGNEQY | 0.81564367 |
| ATVASLIL | ASSGTGGGNEQY | 0.8179397 |
| AVNGPVLIRH | ASSGTGGGNEQY | 0.25946712 |
| DGAAGAVGI | ASSGTGGGNEQY | 0.6778727 |
| DPLLHGI | ASSGTGGGNEQY | 0.31021845 |
| ELGAALGAL | ASSGTGGGNEQY | 0.617803 |
| ELGRTQAV | ASSGTGGGNEQY | 0.5327303 |
| FVAGGYLS | ASSGTGGGNEQY | 0.7758143 |
| GASGGAGGAGGW | ASSGTGGGNEQY | 0.6134776 |
| GGAGAGGDGGAGGI | ASSGTGGGNEQY | 0.73393255 |
| GTGHNGSQPGMGGNG | ASSGTGGGNEQY | 0.68221134 |
| GYAAASASAS | ASSGTGGGNEQY | 0.8204946 |
| HQVDVAAVQQG | ASSGTGGGNEQY | 0.52407527 |
| IGAAIAHT | ASSGTGGGNEQY | 0.6647291 |
| IGMIEVAQL | ASSGTGGGNEQY | 0.40249103 |
| IRLAPEHTSAMD | ASSGTGGGNEQY | 0.34276745 |
| IYNVSVL | ASSGTGGGNEQY | 0.5426388 |
| KQGFLGM | ASSGTGGGNEQY | 0.62324834 |
| LESVITQL | ASSGTGGGNEQY | 0.24653244 |
| LGLGALVVV | ASSGTGGGNEQY | 0.83654684 |
| LIGNGGNGGSGGI | ASSGTGGGNEQY | 0.7509437 |
| LNAINAPTL | ASSGTGGGNEQY | 0.78619397 |
| LQLLNTGV | ASSGTGGGNEQY | 0.6056136 |
| LQTSIEF | ASSGTGGGNEQY | 0.6828487 |
| MSFAVSEL | ASSGTGGGNEQY | 0.54321027 |
| NTGALGL | ASSGTGGGNEQY | 0.7743455 |
| PPGAGGAGGAGTT | ASSGTGGGNEQY | 0.5813716 |
| QLVGLGI | ASSGTGGGNEQY | 0.32410687 |
| RFLEQVHQL | ASSGTGGGNEQY | 0.6527548 |
| RGVGYVLR | ASSGTGGGNEQY | 0.42390892 |
| SATGVFAGVI | ASSGTGGGNEQY | 0.82723916 |
| SLPAITLP | ASSGTGGGNEQY | 0.7920154 |
| STALAFTV | ASSGTGGGNEQY | 0.74647784 |
| SVLTVGLGK | ASSGTGGGNEQY | 0.7736386 |
| TALAGAGSGI | ASSGTGGGNEQY | 0.3077884 |
| TGLPFTGI | ASSGTGGGNEQY | 0.6719153 |
| VAEHVTV | ASSGTGGGNEQY | 0.84560585 |
| VDIVLNSL | ASSGTGGGNEQY | 0.7524655 |
| VLTGGGAVL | ASSGTGGGNEQY | 0.65213954 |
| VNVGLAYGI | ASSGTGGGNEQY | 0.83241624 |
| VSSAAVGAV | ASSGTGGGNEQY | 0.53968793 |
| VTSVYEAGV | ASSGTGGGNEQY | 0.6270232 |
| WVALDRGA | ASSGTGGGNEQY | 0.7095127 |
| YLVGSVVI | ASSGTGGGNEQY | 0.74953425 |
| GPPGTGKTLL | ASSGTGGGNEQY | 0.7447363 |

| | | |
|---|---|---|
| GVGMTKF | ASSGTGGGNEQY | 0.5839294 |
| IGIGGSDL | ASSGTGGGNEQY | 0.7297149 |
| IGTLLAEL | ASSGTGGGNEQY | 0.84924906 |
| IYTSGTT | ASSGTGGGNEQY | 0.634754 |
| LSFPAGV | ASSGTGGGNEQY | 0.82388705 |
| LYTSGTT | ASSGTGGGNEQY | 0.77772796 |
| VGAFPFGF | ASSGTGGGNEQY | 0.7856453 |
| VGAGTVEF | ASSGTGGGNEQY | 0.5240898 |
| VGKSTLLL | ASSGTGGGNEQY | 0.61407954 |
| AALAHAGGGIV | ASSGTGGGNEQY | 0.6166346 |
| AALVAVSLL | ASSGTGGGNEQY | 0.5637526 |
| AAVSRAVAAL | ASSGTGGGNEQY | 0.62704223 |
| AGGSSGIGGF | ASSGTGGGNEQY | 0.62345254 |
| AGLTVANANG | ASSGTGGGNEQY | 0.8647082 |
| AGLVGAVL | ASSGTGGGNEQY | 0.7707541 |
| ALSQAFTV | ASSGTGGGNEQY | 0.78331995 |
| ALTFSVSL | ASSGTGGGNEQY | 0.5120766 |
| ALTGAALVAL | ASSGTGGGNEQY | 0.68110204 |
| ASFSIGAGVL | ASSGTGGGNEQY | 0.7618163 |
| DGSVYKI | ASSGTGGGNEQY | 0.44102624 |
| EQLAEIV | ASSGTGGGNEQY | 0.8430939 |
| FAVSELTVM | ASSGTGGGNEQY | 0.54701513 |
| GAAPLAAGL | ASSGTGGGNEQY | 0.61978734 |
| GADGGDGGSGGAGG | ASSGTGGGNEQY | 0.80173826 |
| GFGLDTGGAGG | ASSGTGGGNEQY | 0.8104379 |
| GLASGGLL | ASSGTGGGNEQY | 0.597852 |
| GRIAGAAVLL | ASSGTGGGNEQY | 0.57967496 |
| GSAGSLNTSF | ASSGTGGGNEQY | 0.53756255 |
| GTGQIGI | ASSGTGGGNEQY | 0.79982865 |
| GTLIALIL | ASSGTGGGNEQY | 0.81653273 |
| IAATMGFVVG | ASSGTGGGNEQY | 0.64894813 |
| IFAQALL | ASSGTGGGNEQY | 0.8310339 |
| IGAAVALGF | ASSGTGGGNEQY | 0.75859445 |
| IGLTPLAL | ASSGTGGGNEQY | 0.4447034 |
| ILGSNTSTL | ASSGTGGGNEQY | 0.6958114 |
| IVFSLTVL | ASSGTGGGNEQY | 0.79067206 |
| LASKATGF | ASSGTGGGNEQY | 0.8041753 |
| LASVALIL | ASSGTGGGNEQY | 0.8492625 |
| LFAVGLTL | ASSGTGGGNEQY | 0.78743905 |
| LFNGGIGTYI | ASSGTGGGNEQY | 0.58674556 |
| LTLGAGII | ASSGTGGGNEQY | 0.8319845 |
| LVAAALGDI | ASSGTGGGNEQY | 0.8553096 |
| NTAFVQLGI | ASSGTGGGNEQY | 0.8087442 |
| QLFSGAEL | ASSGTGGGNEQY | 0.6314297 |
| SAATFITL | ASSGTGGGNEQY | 0.59865856 |
| SNYGLANLGV | ASSGTGGGNEQY | 0.67546993 |
| SYGGAYAVM | ASSGTGGGNEQY | 0.8086293 |
| TTGVGGLTLG | ASSGTGGGNEQY | 0.45539698 |
| TTQTIYI | ASSGTGGGNEQY | 0.72754145 |
| VAAGAGLGAVI | ASSGTGGGNEQY | 0.75689626 |
| VAAVVTLL | ASSGTGGGNEQY | 0.3909724 |

| | | |
|---|---|---|
| VGFLGGLI | ASSGTGGGNEQY | 0.7881228 |
| VGVIPILL | ASSGTGGGNEQY | 0.83677953 |
| VIFASSLI | ASSGTGGGNEQY | 0.60985094 |
| VKGGDGGAAAG | ASSGTGGGNEQY | 0.6611162 |
| VTKTFTNI | ASSGTGGGNEQY | 0.82694745 |
| VVAAAILL | ASSGTGGGNEQY | 0.8643812 |
| YAALDVEL | ASSGTGGGNEQY | 0.8123447 |
| LASVALLL | ASSGTGGGNEQY | 0.41035202 |
| LDAVHRV | ASSGTGGGNEQY | 0.2612694 |
| QGLSSSIF | ASSGTGGGNEQY | 0.50351834 |
| RGTFANIRL | ASSGTGGGNEQY | 0.7727027 |
| RQLWWGHRI | ASSGTGGGNEQY | 0.5842229 |
| YGPPGTGKTLL | ASSGTGGGNEQY | 0.5641645 |
| AALVGAPANVA | ASSGTGGGNEQY | 0.77655643 |
| AATFRRTE | ASSGTGGGNEQY | 0.8218231 |
| AGALGGGI | ASSGTGGGNEQY | 0.57821536 |
| AGGAGGAGGIGAG | ASSGTGGGNEQY | 0.46985015 |
| AGSGALGL | ASSGTGGGNEQY | 0.76617384 |
| AQIADQI | ASSGTGGGNEQY | 0.7617999 |
| ASAVAVTYL | ASSGTGGGNEQY | 0.54435825 |
| ASTVLGGTGG | ASSGTGGGNEQY | 0.65194255 |
| AVGGEPAK | ASSGTGGGNEQY | 0.46459565 |
| CEQGTFGGTC | ASSGTGGGNEQY | 0.3651638 |
| DEGKAPAG | ASSGTGGGNEQY | 0.7480286 |
| DTVVFPAL | ASSGTGGGNEQY | 0.5277784 |
| FEGITMAG | ASSGTGGGNEQY | 0.83769625 |
| FPLVVQD | ASSGTGGGNEQY | 0.5147344 |
| GAFTTPQ | ASSGTGGGNEQY | 0.7907001 |
| GFGGFGTAGGVGGA | ASSGTGGGNEQY | 0.7602227 |
| GFLAQFL | ASSGTGGGNEQY | 0.76242757 |
| GGAGGAGGNAGVA | ASSGTGGGNEQY | 0.63077855 |
| GGVGTAGAGGAGGAGG | ASSGTGGGNEQY | 0.57977927 |
| GNGGGGDAGNAGSGGN | ASSGTGGGNEQY | 0.45577636 |
| GTGGAGGAGNDAGS | ASSGTGGGNEQY | 0.23230352 |
| GTTNTGWFNS | ASSGTGGGNEQY | 0.8188343 |
| IFMGPLFTL | ASSGTGGGNEQY | 0.17458369 |
| ILAHAIEAI | ASSGTGGGNEQY | 0.820826 |
| IRSAIREAL | ASSGTGGGNEQY | 0.76522744 |
| ITFAGGG | ASSGTGGGNEQY | 0.52798784 |
| ITGMSGHQEV | ASSGTGGGNEQY | 0.7279315 |
| IVAGLACTL | ASSGTGGGNEQY | 0.5905807 |
| IVGGGLAGA | ASSGTGGGNEQY | 0.40901694 |
| IVTGVLNI | ASSGTGGGNEQY | 0.8167322 |
| LAQQGCI | ASSGTGGGNEQY | 0.7600076 |
| LCLHVEN | ASSGTGGGNEQY | 0.5162403 |
| LDAISDGS | ASSGTGGGNEQY | 0.704903 |
| LGGFAAML | ASSGTGGGNEQY | 0.72801024 |
| LGGKNPMI | ASSGTGGGNEQY | 0.6098401 |
| LGPKGVAAIL | ASSGTGGGNEQY | 0.61930156 |
| LMVGVSI | ASSGTGGGNEQY | 0.81978357 |
| LVGAAFVL | ASSGTGGGNEQY | 0.67605245 |

| | | |
|---|---|---|
| LVLAVAPL | ASSGTGGGNEQY | 0.5821451 |
| MAAIAVIDAA | ASSGTGGGNEQY | 0.79198366 |
| NNGINAELQ | ASSGTGGGNEQY | 0.47828948 |
| RGEPLVLVHG | ASSGTGGGNEQY | 0.7754227 |
| RGGAGGAAGLI | ASSGTGGGNEQY | 0.7044575 |
| RSYRMHPAV | ASSGTGGGNEQY | 0.6771024 |
| SAAHLLAGQLA | ASSGTGGGNEQY | 0.79520845 |
| SAAVGSY | ASSGTGGGNEQY | 0.5710956 |
| SPHAVLHRI | ASSGTGGGNEQY | 0.5492102 |
| SQIAKVL | ASSGTGGGNEQY | 0.64923185 |
| TALGHLVP | ASSGTGGGNEQY | 0.7441999 |
| TAPTTSGAA | ASSGTGGGNEQY | 0.8194103 |
| TDGDDGNGGSPGT | ASSGTGGGNEQY | 0.5029451 |
| TDGLGEN | ASSGTGGGNEQY | 0.7316143 |
| TGGPPLPGQA | ASSGTGGGNEQY | 0.5378111 |
| TIPAITI | ASSGTGGGNEQY | 0.4555371 |
| TIPGFTI | ASSGTGGGNEQY | 0.61787283 |
| TSIASGALA | ASSGTGGGNEQY | 0.66117233 |
| TVTLPGAAI | ASSGTGGGNEQY | 0.6337186 |
| VASGLSAALP | ASSGTGGGNEQY | 0.75339276 |
| VGAGVIGI | ASSGTGGGNEQY | 0.5791541 |
| VGPKRFGV | ASSGTGGGNEQY | 0.7380846 |
| VNVAFPD | ASSGTGGGNEQY | 0.8449513 |
| VSMTNSGVS | ASSGTGGGNEQY | 0.7200768 |
| VTDVLGDLSV | ASSGTGGGNEQY | 0.7649948 |
| VVVVGAGFAGL | ASSGTGGGNEQY | 0.7805978 |
| VYTVGTVV | ASSGTGGGNEQY | 0.7777099 |
| WTTFSGP | ASSGTGGGNEQY | 0.5933267 |
| YDELVEAF | ASSGTGGGNEQY | 0.5793948 |
| YGTGGAGGNGG | ASSGTGGGNEQY | 0.8391317 |
| YTNVMAV | ASSGTGGGNEQY | 0.7924788 |
| FLEQVHQL | ASSGTGGGNEQY | 0.81495404 |
| GLSLLAVAV | ASSGTGGGNEQY | 0.8003534 |
| LEQVHQL | ASSGTGGGNEQY | 0.7696277 |
| LNAAGAI | ASSGTGGGNEQY | 0.78963053 |
| PAVHASL | ASSGTGGGNEQY | 0.64283186 |
| TELEDDA | ASSGTGGGNEQY | 0.38613516 |
| TPPGPPN | ASSGTGGGNEQY | 0.63650304 |
| TPPPPPPP | ASSGTGGGNEQY | 0.74519914 |
| TQSPALI | ASRATGNNQAPL | 0.62719476 |
| EFAAAFPQ | ASRATGNNQAPL | 0.39767686 |
| LLSGNLDVL | ASRATGNNQAPL | 0.8255233 |
| GPANLGNYNI | ASRATGNNQAPL | 0.86368996 |
| TLGPTLP | ASRATGNNQAPL | 0.54249996 |
| VYAYSSCCK | ASRATGNNQAPL | 0.8439772 |
| APAPAPAPNI | ASRATGNNQAPL | 0.4113614 |
| PGGPVPIP | ASRATGNNQAPL | 0.78364503 |
| FLTVALEP | ASRATGNNQAPL | 0.8334865 |
| SQYFGANL | ASRATGNNQAPL | 0.46128252 |
| GGVKPAA | ASRATGNNQAPL | 0.8042655 |
| GGTGGAGGAGGW | ASRATGNNQAPL | 0.5923177 |

| | | |
|---|---|---|
| GVGGVAGLGGAGG | ASRATGNNQAPL | 0.6792914 |
| VLLVLATTI | ASRATGNNQAPL | 0.8369709 |
| IGIFAAML | ASRATGNNQAPL | 0.6271556 |
| AVGDLID | ASRATGNNQAPL | 0.7289421 |
| NISPLPL | ASRATGNNQAPL | 0.64311934 |
| VTAVFVAI | ASRATGNNQAPL | 0.5132008 |
| SGDDNDAAL | ASRATGNNQAPL | 0.7238737 |
| AVAALAHS | ASRATGNNQAPL | 0.8291743 |
| GGVTVMATP | ASRATGNNQAPL | 0.42988458 |
| AGLAANVRD | ASRATGNNQAPL | 0.84894633 |
| TNRPDLIDPALLRPG | ASRATGNNQAPL | 0.51480657 |
| LANLGVTAL | ASRATGNNQAPL | 0.8599377 |
| TGERIEL | ASRATGNNQAPL | 0.7895052 |
| NGEELAA | ASRATGNNQAPL | 0.71327144 |
| VLSLVWG | ASRATGNNQAPL | 0.62017494 |
| IGYALWL | ASRATGNNQAPL | 0.82318634 |
| LSLLLAPL | ASRATGNNQAPL | 0.60888773 |
| TLDEALK | ASRATGNNQAPL | 0.79475445 |
| TDGAGTAEGG | ASRATGNNQAPL | 0.81378335 |
| NAPGGAGGNGG | ASRATGNNQAPL | 0.5452554 |
| VSVFMGMGGNF | ASRATGNNQAPL | 0.8605636 |
| LASYLAAL | ASRATGNNQAPL | 0.7343471 |
| LVGGTPAL | ASRATGNNQAPL | 0.7130515 |
| VRNADVL | ASRATGNNQAPL | 0.51562357 |
| TCSSPFLVSAM | ASRATGNNQAPL | 0.42581278 |
| VHITDAE | ASRATGNNQAPL | 0.7346822 |
| EYAAPSF | ASRATGNNQAPL | 0.4157545 |
| VVDAFHGGL | ASRATGNNQAPL | 0.80399615 |
| ASVVDPQ | ASRATGNNQAPL | 0.8118303 |
| SYNEFGGLS | ASRATGNNQAPL | 0.53585446 |
| DPDGSVY | ASRATGNNQAPL | 0.8096143 |
| MVSTIHATE | ASRATGNNQAPL | 0.7947335 |
| LAAGVADPV | ASRATGNNQAPL | 0.4631873 |
| TKEQIAA | ASRATGNNQAPL | 0.6361854 |
| SLASPAAL | ASRATGNNQAPL | 0.6616703 |
| IGLVFYGV | ASRATGNNQAPL | 0.6912506 |
| FLLDASNL | ASRATGNNQAPL | 0.41437793 |
| YQVGNFP | ASRATGNNQAPL | 0.7768892 |
| ESLLSQST | ASRATGNNQAPL | 0.5586637 |
| LFTVGFAAVG | ASRATGNNQAPL | 0.16786994 |
| ATPPTPT | ASRATGNNQAPL | 0.7798873 |
| TDLATLG | ASRATGNNQAPL | 0.6625462 |
| SAGGTGR | ASRATGNNQAPL | 0.6024058 |
| GFSNSGS | ASRATGNNQAPL | 0.83187604 |
| GGNGGSGGTSV | ASRATGNNQAPL | 0.6251177 |
| GTSVATGGAGN | ASRATGNNQAPL | 0.48398262 |
| NGGNGGSGGTS | ASRATGNNQAPL | 0.78653246 |
| AVVKFPH | ASRATGNNQAPL | 0.8300558 |
| TPPPPPP | ASRATGNNQAPL | 0.40427703 |
| AAPAAAAAPA | ASRATGNNQAPL | 0.7326409 |
| IALNEDL | ASRATGNNQAPL | 0.817195 |

| | | |
|---|---|---|
| IVNVSSM | ASRATGNNQAPL | 0.69809014 |
| TGIPPAPR | ASRATGNNQAPL | 0.8559103 |
| FDLGGGTF | ASRATGNNQAPL | 0.8605487 |
| FDLGGGTFDVS | ASRATGNNQAPL | 0.6705593 |
| NEPTAAA | ASRATGNNQAPL | 0.82443273 |
| NEPTAAALA | ASRATGNNQAPL | 0.8624552 |
| GSIINISSI | ASRATGNNQAPL | 0.66814315 |
| VAPGNAGTA | ASRATGNNQAPL | 0.68500286 |
| TGGGASLE | ASRATGNNQAPL | 0.5283253 |
| VAVYNHYKRI | ASRATGNNQAPL | 0.720517 |
| TDPAPATT | ASRATGNNQAPL | 0.3457536 |
| TNRPDILDPALLRPG | ASRATGNNQAPL | 0.7278574 |
| LTTTTTS | ASRATGNNQAPL | 0.83618355 |
| PTGSGKTL | ASRATGNNQAPL | 0.6026156 |
| TGSGKTL | ASRATGNNQAPL | 0.46770406 |
| TIGHVDHGK | ASRATGNNQAPL | 0.42232567 |
| TIGHVDHGKTT | ASRATGNNQAPL | 0.78109825 |
| GHVDHGKTT | ASRATGNNQAPL | 0.5080018 |
| AQFEHTLL | ASRATGNNQAPL | 0.40660375 |
| IVNQDEL | ASRATGNNQAPL | 0.7609049 |
| TNGSQFF | ASRATGNNQAPL | 0.78118885 |
| TGKGIVE | ASRATGNNQAPL | 0.80419564 |
| LISNASDA | ASRATGNNQAPL | 0.35583863 |
| AELRTLGELTKIAKAHGAQ | ASRATGNNQAPL | 0.8706329 |
| APNAVAAAEPSW | ASRATGNNQAPL | 0.7752266 |
| APQVAGAATAVL | ASRATGNNQAPL | 0.8599207 |
| ARRDPAAAAAILPTDA | ASRATGNNQAPL | 0.7014352 |
| DAAARDVLAEAGLAEYF | ASRATGNNQAPL | 0.597973 |
| DDQIVLSAKANLAKVEQ | ASRATGNNQAPL | 0.8150172 |
| DELLPDTGEDSDEDGH | ASRATGNNQAPL | 0.5738479 |
| ELPEAPANRATVPQD | ASRATGNNQAPL | 0.61323416 |
| FGQSKIERVTGIG | ASRATGNNQAPL | 0.8320682 |
| GAGGTGSTAGGAGGAGGAGGLY | ASRATGNNQAPL | 0.57363796 |
| GHGGTGAAGGNGGTGGAGGSGI | ASRATGNNQAPL | 0.4753016 |
| GNGGNGGAGGEGGPGGAGGAGA | ASRATGNNQAPL | 0.59229076 |
| GSTAGNGGNGGNATNGTIAGQP | ASRATGNNQAPL | 0.85649264 |
| GTGGDGGNGGNGGTGG | ASRATGNNQAPL | 0.4676451 |
| GVRRPAGIPADAPV | ASRATGNNQAPL | 0.53278035 |
| GYRQVTDAHLLTLARR | ASRATGNNQAPL | 0.5047983 |
| IAKMGQDGHDRG | ASRATGNNQAPL | 0.64633906 |
| KAEPSIEIQYT | ASRATGNNQAPL | 0.8370348 |
| KPVRIRMAADEDTVAVT | ASRATGNNQAPL | 0.7910085 |
| MPPRIAGMRLLVIKPEP | ASRATGNNQAPL | 0.71103525 |
| MKFYAVALFRSRGVPPTR | ASRATGNNQAPL | 0.77814615 |
| NAGGFGILGAVAHSP | ASRATGNNQAPL | 0.77083504 |
| NEGREITLAMAEM | ASRATGNNQAPL | 0.73034996 |
| NELVEWSHPLGHRWRWE | ASRATGNNQAPL | 0.74329066 |
| NLGAGGTGGNGGNPG | ASRATGNNQAPL | 0.51182145 |
| NTEWPKWETVMAADPVD | ASRATGNNQAPL | 0.85651094 |
| PASAAPAETQP | ASRATGNNQAPL | 0.7547153 |
| QQSSDGKNGVRAAEITGE | ASRATGNNQAPL | 0.8286371 |

| | | |
|---|---|---|
| RAPDAVTTSWWKE | ASRATGNNQAPL | 0.85055566 |
| SGFGNVGAHTSGWFNQ | ASRATGNNQAPL | 0.7724726 |
| SGVAGAVGQAASVGGLKV | ASRATGNNQAPL | 0.7547133 |
| SNGGAGTGSAGGNGGT | ASRATGNNQAPL | 0.28788808 |
| TGGAGGEGGAAG | ASRATGNNQAPL | 0.6466846 |
| TGGDGGAAGAGGAPGHG | ASRATGNNQAPL | 0.7741794 |
| TMPPLLAAIANAGRHGV | ASRATGNNQAPL | 0.7256448 |
| TSSTAGNSGGA | ASRATGNNQAPL | 0.8060768 |
| VPATVPVGILC | ASRATGNNQAPL | 0.8147596 |
| VQRPDSGGVVLTGRIS | ASRATGNNQAPL | 0.61487234 |
| WTPNPVPATKGVPAGLVQAT | ASRATGNNQAPL | 0.8207789 |
| YALKRALAKFSDRFDVVII | ASRATGNNQAPL | 0.8300115 |
| YLIGSVIV | ASRATGNNQAPL | 0.36018506 |
| YGQLVAVID | ASRATGNNQAPL | 0.68943346 |
| VTVGGMTLPA | ASRATGNNQAPL | 0.88401353 |
| VLALPAV | ASRATGNNQAPL | 0.75664955 |
| VIPPTLNYET | ASRATGNNQAPL | 0.617717 |
| VGMGGMHPSSG | ASRATGNNQAPL | 0.6203966 |
| VGLAGGAATA | ASRATGNNQAPL | 0.83867514 |
| VGGAGGAGGSGP | ASRATGNNQAPL | 0.6342035 |
| VFAAFMV | ASRATGNNQAPL | 0.85161734 |
| TLYDKLR | ASRATGNNQAPL | 0.2251236 |
| TLAVFTVL | ASRATGNNQAPL | 0.65066284 |
| TGSVQPARV | ASRATGNNQAPL | 0.58182806 |
| TGGGAGSGGT | ASRATGNNQAPL | 0.27342921 |
| SHLETSL | ASRATGNNQAPL | 0.80175644 |
| SFTTGMT | ASRATGNNQAPL | 0.64601094 |
| QVEVAKRMGVRQPTVS | ASRATGNNQAPL | 0.7979256 |
| QGSGGPALAM | ASRATGNNQAPL | 0.74149215 |
| MLAPSNP | ASRATGNNQAPL | 0.8233568 |
| LVNNAAI | ASRATGNNQAPL | 0.75096405 |
| LTGEINTLA | ASRATGNNQAPL | 0.72187513 |
| LSIFGPV | ASRATGNNQAPL | 0.6523202 |
| LLALLANL | ASRATGNNQAPL | 0.7179598 |
| LISNASDALDKLR | ASRATGNNQAPL | 0.8460322 |
| LINEAVP | ASRATGNNQAPL | 0.84813946 |
| LGYAFSGL | ASRATGNNQAPL | 0.30585843 |
| LDFVAVAL | ASRATGNNQAPL | 0.70581996 |
| LAQPGAE | ASRATGNNQAPL | 0.62660575 |
| ISNLGAVTGF | ASRATGNNQAPL | 0.35739094 |
| HQEDPGRI | ASRATGNNQAPL | 0.2580074 |
| GTPGEPTP | ASRATGNNQAPL | 0.6364404 |
| GFFNSGS | ASRATGNNQAPL | 0.7035902 |
| GAGGGGLTAGAN | ASRATGNNQAPL | 0.7865968 |
| FVDALVNM | ASRATGNNQAPL | 0.80442035 |
| FLTVAPEL | ASRATGNNQAPL | 0.5704747 |
| FETLTAL | ASRATGNNQAPL | 0.80498517 |
| DVINPIEL | ASRATGNNQAPL | 0.61622596 |
| APVSGAPV | ASRATGNNQAPL | 0.09908585 |
| ALVGGLAW | ASRATGNNQAPL | 0.6449996 |
| ALAAAAATAI | ASRATGNNQAPL | 0.47821474 |

| | | |
|---|---|---|
| AFGLGLLL | ASRATGNNQAPL | 0.84810126 |
| AFFAAASL | ASRATGNNQAPL | 0.5612415 |
| AAAALPAFSPP | ASRATGNNQAPL | 0.6061964 |
| VVLAPAL | ASRATGNNQAPL | 0.6392162 |
| TIGHVDHGKTTL | ASRATGNNQAPL | 0.7075834 |
| LISNASDALDK | ASRATGNNQAPL | 0.7016356 |
| LISNASDAL | ASRATGNNQAPL | 0.50752413 |
| ISNASDALDK | ASRATGNNQAPL | 0.25773498 |
| NQAETLVYQTE | ASRATGNNQAPL | 0.75998455 |
| TGGTGGTAGAG | ASRATGNNQAPL | 0.5772522 |
| TGSTAGGAGGA | ASRATGNNQAPL | 0.5053096 |
| DPLAVPVAAGPV | ASRATGNNQAPL | 0.45112512 |
| GGTGGTPGNGGN | ASRATGNNQAPL | 0.6866815 |
| PAGLAGVDPATL | ASRATGNNQAPL | 0.35768753 |
| GNNLVVEAPHAGASV | ASRATGNNQAPL | 0.8498024 |
| KVTRIMSASTLEQPA | ASRATGNNQAPL | 0.6434774 |
| LGFALPIALPRRTVV | ASRATGNNQAPL | 0.6236479 |
| TRLGNLTARKLVVLV | ASRATGNNQAPL | 0.8701957 |
| WSQQRRLIYFPSAGP | ASRATGNNQAPL | 0.6394359 |
| GPNGNPDPMAAAVDIR | ASRATGNNQAPL | 0.65494215 |
| TQRCVSVADLRAAASR | ASRATGNNQAPL | 0.56335145 |
| QFVLDVAEHPVEVLTETV | ASRATGNNQAPL | 0.55512524 |
| GGAGGTGGAGGNGGTQAGDG | ASRATGNNQAPL | 0.50768834 |
| PLLATLRAIAVDVDAKPAQV | ASRATGNNQAPL | 0.50224257 |
| APKFRTGGYMIDFWGVGYQVAK | ASRATGNNQAPL | 0.85072047 |
| GAGGAGGAGGAAGGPGGTGGTG | ASRATGNNQAPL | 0.69172114 |
| IGALGLLATDVAPAGSVAFGVA | ASRATGNNQAPL | 0.52950746 |
| AGGAGGIGGNGGAGGAGGVLMG | ASRATGNNQAPL | 0.8011179 |
| ADELIGLL | ASRATGNNQAPL | 0.84814876 |
| ADRALKA | ASRATGNNQAPL | 0.8751385 |
| AGAVGQVGGAGGA | ASRATGNNQAPL | 0.5952108 |
| AGGGGGLGGAP | ASRATGNNQAPL | 0.6279444 |
| AGGLLYGNGGAG | ASRATGNNQAPL | 0.64530146 |
| AGLESIGL | ASRATGNNQAPL | 0.63077444 |
| AGLKTGII | ASRATGNNQAPL | 0.636798 |
| ASLAALLL | ASRATGNNQAPL | 0.44268063 |
| ASVINISSI | ASRATGNNQAPL | 0.8349026 |
| ATPPGPA | ASRATGNNQAPL | 0.8353123 |
| ATVASALV | ASRATGNNQAPL | 0.68665105 |
| ATVASLIL | ASRATGNNQAPL | 0.46078336 |
| AVNGPVLIRH | ASRATGNNQAPL | 0.58288646 |
| DGAAGAVGI | ASRATGNNQAPL | 0.44171542 |
| DPLLHGI | ASRATGNNQAPL | 0.53069675 |
| ELGAALGAL | ASRATGNNQAPL | 0.69995207 |
| ELGRTQAV | ASRATGNNQAPL | 0.3935802 |
| FVAGGYLS | ASRATGNNQAPL | 0.66132087 |
| GASGGAGGAGGW | ASRATGNNQAPL | 0.8575437 |
| GGAGAGGDGGAGGI | ASRATGNNQAPL | 0.5886804 |
| GTGHNGSQPGMGGNG | ASRATGNNQAPL | 0.79401815 |
| GYAAASASAS | ASRATGNNQAPL | 0.8438932 |
| HQVDVAAVQQG | ASRATGNNQAPL | 0.7791109 |

| | | |
|---|---|---|
| IGAAIAHT | ASRATGNNQAPL | 0.84540206 |
| IGMIEVAQL | ASRATGNNQAPL | 0.79649323 |
| IRLAPEHTSAMD | ASRATGNNQAPL | 0.7718653 |
| IYNVSVL | ASRATGNNQAPL | 0.7464156 |
| KQGFLGM | ASRATGNNQAPL | 0.8273979 |
| LESVITQL | ASRATGNNQAPL | 0.61323965 |
| LGLGALVVV | ASRATGNNQAPL | 0.63020617 |
| LIGNGGNGGSGGI | ASRATGNNQAPL | 0.6555982 |
| LNAINAPTL | ASRATGNNQAPL | 0.78255343 |
| LQLLNTGV | ASRATGNNQAPL | 0.69269115 |
| LQTSIEF | ASRATGNNQAPL | 0.82067716 |
| MSFAVSEL | ASRATGNNQAPL | 0.3812429 |
| NTGALGL | ASRATGNNQAPL | 0.41415223 |
| PPGAGGAGGAGTT | ASRATGNNQAPL | 0.8249279 |
| QLVGLGI | ASRATGNNQAPL | 0.15233049 |
| RFLEQVHQL | ASRATGNNQAPL | 0.2648061 |
| RGVGYVLR | ASRATGNNQAPL | 0.7410896 |
| SATGVFAGVI | ASRATGNNQAPL | 0.82059896 |
| SLPAITLP | ASRATGNNQAPL | 0.1919487 |
| STALAFTV | ASRATGNNQAPL | 0.518533 |
| SVLTVGLGK | ASRATGNNQAPL | 0.46644706 |
| TALAGAGSGI | ASRATGNNQAPL | 0.85301805 |
| TGLPFTGI | ASRATGNNQAPL | 0.64206576 |
| VAEHVTV | ASRATGNNQAPL | 0.37246054 |
| VDIVLNSL | ASRATGNNQAPL | 0.79432654 |
| VLTGGGAVL | ASRATGNNQAPL | 0.42722973 |
| VNVGLAYGI | ASRATGNNQAPL | 0.70544493 |
| VSSAAVGAV | ASRATGNNQAPL | 0.82813776 |
| VTSVYEAGV | ASRATGNNQAPL | 0.8503028 |
| WVALDRGA | ASRATGNNQAPL | 0.5368557 |
| YLVGSVVI | ASRATGNNQAPL | 0.791777 |
| GPPGTGKTLL | ASRATGNNQAPL | 0.5981519 |
| GVGMTKF | ASRATGNNQAPL | 0.8788302 |
| IGIGGSDL | ASRATGNNQAPL | 0.59118515 |
| IGTLLAEL | ASRATGNNQAPL | 0.4165759 |
| IYTSGTT | ASRATGNNQAPL | 0.8179586 |
| LSFPAGV | ASRATGNNQAPL | 0.8306887 |
| LYTSGTT | ASRATGNNQAPL | 0.691589 |
| VGAFPFGF | ASRATGNNQAPL | 0.79864264 |
| VGAGTVEF | ASRATGNNQAPL | 0.6879637 |
| VGKSTLLL | ASRATGNNQAPL | 0.62572896 |
| AALAHAGGGIV | ASRATGNNQAPL | 0.8199395 |
| AALVAVSLL | ASRATGNNQAPL | 0.8211272 |
| AAVSRAVAAL | ASRATGNNQAPL | 0.48114073 |
| AGGSSGIGGF | ASRATGNNQAPL | 0.8402847 |
| AGLTVANANG | ASRATGNNQAPL | 0.59202445 |
| AGLVGAVL | ASRATGNNQAPL | 0.565025 |
| ALSQAFTV | ASRATGNNQAPL | 0.8554627 |
| ALTFSVSL | ASRATGNNQAPL | 0.7549832 |
| ALTGAALVAL | ASRATGNNQAPL | 0.7971146 |
| ASFSIGAGVL | ASRATGNNQAPL | 0.7372644 |

| | | |
|---|---|---|
| DGSVYKI | ASRATGNNQAPL | 0.81120086 |
| EQLAEIV | ASRATGNNQAPL | 0.49551266 |
| FAVSELTVM | ASRATGNNQAPL | 0.7395537 |
| GAAPLAAGL | ASRATGNNQAPL | 0.68789315 |
| GADGGDGGSGGAGG | ASRATGNNQAPL | 0.8235324 |
| GFGLDTGGAGG | ASRATGNNQAPL | 0.7462727 |
| GLASGGLL | ASRATGNNQAPL | 0.6437092 |
| GRIAGAAVLL | ASRATGNNQAPL | 0.8074331 |
| GSAGSLNTSF | ASRATGNNQAPL | 0.83872086 |
| GTGQIGI | ASRATGNNQAPL | 0.7516584 |
| GTLIALIL | ASRATGNNQAPL | 0.74984986 |
| IAATMGFVVG | ASRATGNNQAPL | 0.85980755 |
| IFAQALL | ASRATGNNQAPL | 0.5639955 |
| IGAAVALGF | ASRATGNNQAPL | 0.63925135 |
| IGLTPLAL | ASRATGNNQAPL | 0.80888414 |
| ILGSNTSTL | ASRATGNNQAPL | 0.85004115 |
| IVFSLTVL | ASRATGNNQAPL | 0.72008234 |
| LASKATGF | ASRATGNNQAPL | 0.8517946 |
| LASVALIL | ASRATGNNQAPL | 0.68465215 |
| LFAVGLTL | ASRATGNNQAPL | 0.8032843 |
| LFNGGIGTYI | ASRATGNNQAPL | 0.6614826 |
| LTLGAGII | ASRATGNNQAPL | 0.6986848 |
| LVAAALGDI | ASRATGNNQAPL | 0.49272665 |
| NTAFVQLGI | ASRATGNNQAPL | 0.77911586 |
| QLFSGAEL | ASRATGNNQAPL | 0.8019644 |
| SAATFITL | ASRATGNNQAPL | 0.7728333 |
| SNYGLANLGV | ASRATGNNQAPL | 0.28125402 |
| SYGGAYAVM | ASRATGNNQAPL | 0.7922583 |
| TTGVGGLTLG | ASRATGNNQAPL | 0.79000163 |
| TTQTIYI | ASRATGNNQAPL | 0.87395227 |
| VAAGAGLGAVI | ASRATGNNQAPL | 0.71054757 |
| VAAVVTLL | ASRATGNNQAPL | 0.77859783 |
| VGFLGGLI | ASRATGNNQAPL | 0.6298138 |
| VGVIPILL | ASRATGNNQAPL | 0.712224 |
| VIFASSLI | ASRATGNNQAPL | 0.3842566 |
| VKGGDGGAAAG | ASRATGNNQAPL | 0.62175703 |
| VTKTFTNI | ASRATGNNQAPL | 0.62143826 |
| VVAAAILL | ASRATGNNQAPL | 0.55455154 |
| YAALDVEL | ASRATGNNQAPL | 0.76257634 |
| LASVALLL | ASRATGNNQAPL | 0.82725966 |
| LDAVHRV | ASRATGNNQAPL | 0.8203562 |
| QGLSSSIF | ASRATGNNQAPL | 0.8171928 |
| RGTFANIRL | ASRATGNNQAPL | 0.8596437 |
| RQLWWGHRI | ASRATGNNQAPL | 0.84037507 |
| YGPPGTGKTLL | ASRATGNNQAPL | 0.47728717 |
| AALVGAPANVA | ASRATGNNQAPL | 0.7177632 |
| AATFRRTE | ASRATGNNQAPL | 0.37878335 |
| AGALGGGI | ASRATGNNQAPL | 0.66774 |
| AGGAGGAGGIGAG | ASRATGNNQAPL | 0.53848004 |
| AGSGALGL | ASRATGNNQAPL | 0.7300931 |
| AQIADQI | ASRATGNNQAPL | 0.18374036 |

| | | |
|---|---|---|
| ASAVAVTYL | ASRATGNNQAPL | 0.7245259 |
| ASTVLGGTGG | ASRATGNNQAPL | 0.8588533 |
| AVGGEPAK | ASRATGNNQAPL | 0.5036385 |
| CEQGTFGGTC | ASRATGNNQAPL | 0.760505 |
| DEGKAPAG | ASRATGNNQAPL | 0.7884519 |
| DTVVFPAL | ASRATGNNQAPL | 0.55812114 |
| FEGITMAG | ASRATGNNQAPL | 0.75279886 |
| FPLVVQD | ASRATGNNQAPL | 0.49774963 |
| GAFTTPQ | ASRATGNNQAPL | 0.80679893 |
| GFGGFGTAGGVGGA | ASRATGNNQAPL | 0.8489523 |
| GFLAQFL | ASRATGNNQAPL | 0.7577629 |
| GGAGGAGGNAGVA | ASRATGNNQAPL | 0.65893304 |
| GGVGTAGAGGAGGAGG | ASRATGNNQAPL | 0.7021181 |
| GNGGGGDAGNAGSGGN | ASRATGNNQAPL | 0.42794502 |
| GTGGAGGAGNDAGS | ASRATGNNQAPL | 0.7176899 |
| GTTNTGWFNS | ASRATGNNQAPL | 0.46407205 |
| IFMGPLFTL | ASRATGNNQAPL | 0.70980525 |
| ILAHAIEAI | ASRATGNNQAPL | 0.5118098 |
| IRSAIREAL | ASRATGNNQAPL | 0.7343312 |
| ITFAGGG | ASRATGNNQAPL | 0.6692331 |
| ITGMSGHQEV | ASRATGNNQAPL | 0.7702879 |
| IVAGLACTL | ASRATGNNQAPL | 0.47212982 |
| IVGGGLAGA | ASRATGNNQAPL | 0.72200716 |
| IVTGVLNI | ASRATGNNQAPL | 0.6774825 |
| LAQQGCI | ASRATGNNQAPL | 0.38244998 |
| LCLHVEN | ASRATGNNQAPL | 0.8436128 |
| LDAISDGS | ASRATGNNQAPL | 0.706784 |
| LGGFAAML | ASRATGNNQAPL | 0.70175713 |
| LGGKNPMI | ASRATGNNQAPL | 0.80338705 |
| LGPKGVAAIL | ASRATGNNQAPL | 0.73818797 |
| LMVGVSI | ASRATGNNQAPL | 0.4560062 |
| LVGAAFVL | ASRATGNNQAPL | 0.6507957 |
| LVLAVAPL | ASRATGNNQAPL | 0.6999915 |
| MAAIAVIDAA | ASRATGNNQAPL | 0.56471264 |
| NNGINAELQ | ASRATGNNQAPL | 0.20974381 |
| RGEPLVLVHG | ASRATGNNQAPL | 0.85255766 |
| RGGAGGAAGLI | ASRATGNNQAPL | 0.57018805 |
| RSYRMHPAV | ASRATGNNQAPL | 0.54333854 |
| SAAHLLAGQLA | ASRATGNNQAPL | 0.61225414 |
| SAAVGSY | ASRATGNNQAPL | 0.81670034 |
| SPHAVLHRI | ASRATGNNQAPL | 0.5903302 |
| SQIAKVL | ASRATGNNQAPL | 0.6353714 |
| TALGHLVP | ASRATGNNQAPL | 0.8023312 |
| TAPTTSGAA | ASRATGNNQAPL | 0.6976773 |
| TDGDDGNGGSPGT | ASRATGNNQAPL | 0.8021364 |
| TDGLGEN | ASRATGNNQAPL | 0.73952687 |
| TGGPPLPGQA | ASRATGNNQAPL | 0.71621764 |
| TIPAITI | ASRATGNNQAPL | 0.7828838 |
| TIPGFTI | ASRATGNNQAPL | 0.64981496 |
| TSIASGALA | ASRATGNNQAPL | 0.42267978 |
| TVTLPGAAI | ASRATGNNQAPL | 0.39145812 |

| | | |
|---|---|---|
| VASGLSAALP | ASRATGNNQAPL | 0.6842667 |
| VGAGVIGI | ASRATGNNQAPL | 0.35126865 |
| VGPKRFGV | ASRATGNNQAPL | 0.6968804 |
| VNVAFPD | ASRATGNNQAPL | 0.86609226 |
| VSMTNSGVS | ASRATGNNQAPL | 0.7670378 |
| VTDVLGDLSV | ASRATGNNQAPL | 0.47052804 |
| VVVVGAGFAGL | ASRATGNNQAPL | 0.62176484 |
| VYTVGTVV | ASRATGNNQAPL | 0.6966418 |
| WTTFSGP | ASRATGNNQAPL | 0.8060378 |
| YDELVEAF | ASRATGNNQAPL | 0.83673626 |
| YGTGGAGGNGG | ASRATGNNQAPL | 0.8484508 |
| YTNVMAV | ASRATGNNQAPL | 0.78796715 |
| FLEQVHQL | ASRATGNNQAPL | 0.52600837 |
| GLSLLAVAV | ASRATGNNQAPL | 0.8149203 |
| LEQVHQL | ASRATGNNQAPL | 0.6607608 |
| LNAAGAI | ASRATGNNQAPL | 0.7895356 |
| PAVHASL | ASRATGNNQAPL | 0.77834946 |
| TELEDDA | ASRATGNNQAPL | 0.7857459 |
| TPPGPPN | ASRATGNNQAPL | 0.6880258 |
| TPPPPPPP | ASRATGNNQAPL | 0.7233931 |
| TQSPALI | AWSLGERLF | 0.41309577 |
| EFAAAFPQ | AWSLGERLF | 0.78302693 |
| LLSGNLDVL | AWSLGERLF | 0.575461 |
| GPANLGNYNI | AWSLGERLF | 0.42807582 |
| TLGPTLP | AWSLGERLF | 0.6158151 |
| VYAYSSCCK | AWSLGERLF | 0.8565691 |
| APAPAPAPNI | AWSLGERLF | 0.8348918 |
| PGGPVPIP | AWSLGERLF | 0.4931875 |
| FLTVALEP | AWSLGERLF | 0.4928298 |
| SQYFGANL | AWSLGERLF | 0.7578976 |
| GGVKPAA | AWSLGERLF | 0.33833483 |
| GGTGGAGGAGGW | AWSLGERLF | 0.8001714 |
| GVGGVAGLGGAGG | AWSLGERLF | 0.6359028 |
| VLLVLATTI | AWSLGERLF | 0.6845227 |
| IGIFAAML | AWSLGERLF | 0.5470045 |
| AVGDLID | AWSLGERLF | 0.75421417 |
| NISPLPL | AWSLGERLF | 0.6274893 |
| VTAVFVAI | AWSLGERLF | 0.81910145 |
| SGDDNDAAL | AWSLGERLF | 0.2048623 |
| AVAALAHS | AWSLGERLF | 0.81176114 |
| GGVTVMATP | AWSLGERLF | 0.46491593 |
| AGLAANVRD | AWSLGERLF | 0.59299874 |
| TNRPDLIDPALLRPG | AWSLGERLF | 0.3409294 |
| LANLGVTAL | AWSLGERLF | 0.7333847 |
| TGERIEL | AWSLGERLF | 0.8002588 |
| NGEELAA | AWSLGERLF | 0.820424 |
| VLSLVWG | AWSLGERLF | 0.6710975 |
| IGYALWL | AWSLGERLF | 0.80329317 |
| LSLLLAPL | AWSLGERLF | 0.8409478 |
| TLDEALK | AWSLGERLF | 0.6382756 |
| TDGAGTAEGG | AWSLGERLF | 0.41948962 |

| | | |
|---|---|---|
| NAPGGAGGNGG | AWSLGERLF | 0.7335359 |
| VSVFMGMGGNF | AWSLGERLF | 0.718548 |
| LASYLAAL | AWSLGERLF | 0.84484565 |
| LVGGTPAL | AWSLGERLF | 0.762467 |
| VRNADVL | AWSLGERLF | 0.43312383 |
| TCSSPFLVSAM | AWSLGERLF | 0.75403655 |
| VHITDAE | AWSLGERLF | 0.8434148 |
| EYAAPSF | AWSLGERLF | 0.5217669 |
| VVDAFHGGL | AWSLGERLF | 0.61793655 |
| ASVVDPQ | AWSLGERLF | 0.6367306 |
| SYNEFGGLS | AWSLGERLF | 0.39169672 |
| DPDGSVY | AWSLGERLF | 0.7374963 |
| MVSTIHATE | AWSLGERLF | 0.58910775 |
| LAAGVADPV | AWSLGERLF | 0.53465533 |
| TKEQIAA | AWSLGERLF | 0.66900724 |
| SLASPAAL | AWSLGERLF | 0.55924267 |
| IGLVFYGV | AWSLGERLF | 0.54079914 |
| FLLDASNL | AWSLGERLF | 0.56043196 |
| YQVGNFP | AWSLGERLF | 0.7725902 |
| ESLLSQST | AWSLGERLF | 0.78640914 |
| LFTVGFAAVG | AWSLGERLF | 0.69628245 |
| ATPPTPT | AWSLGERLF | 0.74656403 |
| TDLATLG | AWSLGERLF | 0.84229875 |
| SAGGTGR | AWSLGERLF | 0.39017287 |
| GFSNSGS | AWSLGERLF | 0.46323016 |
| GGNGGSGGTSV | AWSLGERLF | 0.79119265 |
| GTSVATGGAGN | AWSLGERLF | 0.41129643 |
| NGGNGGSGGTS | AWSLGERLF | 0.8406633 |
| AVVKFPH | AWSLGERLF | 0.6599287 |
| TPPPPPP | AWSLGERLF | 0.7264086 |
| AAPAAAAAPA | AWSLGERLF | 0.7367648 |
| IALNEDL | AWSLGERLF | 0.8787236 |
| IVNVSSM | AWSLGERLF | 0.809716 |
| TGIPPAPR | AWSLGERLF | 0.7246475 |
| FDLGGGTF | AWSLGERLF | 0.8185277 |
| FDLGGGTFDVS | AWSLGERLF | 0.4935846 |
| NEPTAAA | AWSLGERLF | 0.80939305 |
| NEPTAAALA | AWSLGERLF | 0.7843278 |
| GSIINISSI | AWSLGERLF | 0.41323304 |
| VAPGNAGTA | AWSLGERLF | 0.62707984 |
| TGGGASLE | AWSLGERLF | 0.8582697 |
| VAVYNHYKRI | AWSLGERLF | 0.8548986 |
| TDPAPATT | AWSLGERLF | 0.36749154 |
| TNRPDILDPALLRPG | AWSLGERLF | 0.6441456 |
| LTTTTTS | AWSLGERLF | 0.44757557 |
| PTGSGKTL | AWSLGERLF | 0.63920355 |
| TGSGKTL | AWSLGERLF | 0.47661513 |
| TIGHVDHGK | AWSLGERLF | 0.7977122 |
| TIGHVDHGKTT | AWSLGERLF | 0.60927653 |
| GHVDHGKTT | AWSLGERLF | 0.7344328 |
| AQFEHTLL | AWSLGERLF | 0.6856145 |

| | | |
|---|---|---|
| IVNQDEL | AWSLGERLF | 0.8063197 |
| TNGSQFF | AWSLGERLF | 0.5719594 |
| TGKGIVE | AWSLGERLF | 0.58104336 |
| LISNASDA | AWSLGERLF | 0.43820083 |
| AELRTLGELTKIAKAHGAQ | AWSLGERLF | 0.42411706 |
| APNAVAAAEPSW | AWSLGERLF | 0.6504802 |
| APQVAGAATAVL | AWSLGERLF | 0.54343307 |
| ARRDPAAAAAILPTDA | AWSLGERLF | 0.2826087 |
| DAAARDVLAEAGLAEYF | AWSLGERLF | 0.85915905 |
| DDQIVLSAKANLAKVEQ | AWSLGERLF | 0.6928728 |
| DELLPDTGEDSDEDGH | AWSLGERLF | 0.7387875 |
| ELPEAPANRATVPQD | AWSLGERLF | 0.6271907 |
| FGQSKIERVTGIG | AWSLGERLF | 0.59006727 |
| GAGGTGSTAGGAGGAGGAGGLY | AWSLGERLF | 0.48158422 |
| GHGGTGAAGGNGGTGGAGGSGI | AWSLGERLF | 0.7736515 |
| GNGGNGGAGGEGGPGGAGGAGA | AWSLGERLF | 0.52967024 |
| GSTAGNGGNGGNATNGTIAGQP | AWSLGERLF | 0.40231794 |
| GTGGDGGNGGNGGTGG | AWSLGERLF | 0.59811693 |
| GVRRPAGIPADAPV | AWSLGERLF | 0.63524514 |
| GYRQVTDAHLLTLARR | AWSLGERLF | 0.87001145 |
| IAKMGQDGHDRG | AWSLGERLF | 0.8157283 |
| KAEPSIEIQYT | AWSLGERLF | 0.7290243 |
| KPVRIRMAADEDTVAVT | AWSLGERLF | 0.84769 |
| MPPRIAGMRLLVIKPEP | AWSLGERLF | 0.25683743 |
| MKFYAVALFRSRGVPPTR | AWSLGERLF | 0.66456586 |
| NAGGFGILGAVAHSP | AWSLGERLF | 0.8319042 |
| NEGREITLAMAEM | AWSLGERLF | 0.75750226 |
| NELVEWSHPLGHRWRWE | AWSLGERLF | 0.6589308 |
| NLGAGGTGGNGGNPG | AWSLGERLF | 0.8393235 |
| NTEWPKWETVMAADPVD | AWSLGERLF | 0.44833064 |
| PASAAPAETQP | AWSLGERLF | 0.5812523 |
| QQSSDGKNGVRAAEITGE | AWSLGERLF | 0.7135843 |
| RAPDAVTTSWWKE | AWSLGERLF | 0.8379525 |
| SGFGNVGAHTSGWFNQ | AWSLGERLF | 0.8017346 |
| SGVAGAVGQAASVGGLKV | AWSLGERLF | 0.5500974 |
| SNGGAGTGSAGGNGGT | AWSLGERLF | 0.76487756 |
| TGGAGGEGGAAG | AWSLGERLF | 0.8798693 |
| TGGDGGAAGAGGAPGHG | AWSLGERLF | 0.61587644 |
| TMPPLLAAIANAGRHGV | AWSLGERLF | 0.81235635 |
| TSSTAGNSGGA | AWSLGERLF | 0.75412023 |
| VPATVPVGILC | AWSLGERLF | 0.5555 |
| VQRPDSGGVVLTGRIS | AWSLGERLF | 0.5010664 |
| WTPNPVPATKGVPAGLVQAT | AWSLGERLF | 0.5137171 |
| YALKRALAKFSDRFDVVII | AWSLGERLF | 0.7081524 |
| YLIGSVIV | AWSLGERLF | 0.46214178 |
| YGQLVAVID | AWSLGERLF | 0.7133939 |
| VTVGGMTLPA | AWSLGERLF | 0.68217367 |
| VLALPAV | AWSLGERLF | 0.8816015 |
| VIPPTLNYET | AWSLGERLF | 0.7650908 |
| VGMGGMHPSSG | AWSLGERLF | 0.7961905 |
| VGLAGGAATA | AWSLGERLF | 0.40428236 |

| | | |
|---|---|---|
| VGGAGGAGGSGP | AWSLGERLF | 0.7837582 |
| VFAAFMV | AWSLGERLF | 0.75026935 |
| TLYDKLR | AWSLGERLF | 0.46077392 |
| TLAVFTVL | AWSLGERLF | 0.8362746 |
| TGSVQPARV | AWSLGERLF | 0.44725826 |
| TGGGAGSGGT | AWSLGERLF | 0.7521899 |
| SHLETSL | AWSLGERLF | 0.7694115 |
| SFTTGMT | AWSLGERLF | 0.7848835 |
| QVEVAKRMGVRQPTVS | AWSLGERLF | 0.66954035 |
| QGSGGPALAM | AWSLGERLF | 0.69643617 |
| MLAPSNP | AWSLGERLF | 0.46382982 |
| LVNNAAI | AWSLGERLF | 0.81632197 |
| LTGEINTLA | AWSLGERLF | 0.8293332 |
| LSIFGPV | AWSLGERLF | 0.73550206 |
| LLALLANL | AWSLGERLF | 0.8695775 |
| LISNASDALDKLR | AWSLGERLF | 0.80920756 |
| LINEAVP | AWSLGERLF | 0.38586167 |
| LGYAFSGL | AWSLGERLF | 0.7054739 |
| LDFVAVAL | AWSLGERLF | 0.830109 |
| LAQPGAE | AWSLGERLF | 0.83999807 |
| ISNLGAVTGF | AWSLGERLF | 0.8529971 |
| HQEDPGRI | AWSLGERLF | 0.832113 |
| GTPGEPTP | AWSLGERLF | 0.6279467 |
| GFFNSGS | AWSLGERLF | 0.83883107 |
| GAGGGGLTAGAN | AWSLGERLF | 0.88802147 |
| FVDALVNM | AWSLGERLF | 0.79025936 |
| FLTVAPEL | AWSLGERLF | 0.578925 |
| FETLTAL | AWSLGERLF | 0.5466207 |
| DVINPIEL | AWSLGERLF | 0.78443813 |
| APVSGAPV | AWSLGERLF | 0.8227607 |
| ALVGGLAW | AWSLGERLF | 0.4618398 |
| ALAAAAATAI | AWSLGERLF | 0.6860574 |
| AFGLGLLL | AWSLGERLF | 0.6829741 |
| AFFAAASL | AWSLGERLF | 0.33808026 |
| AAAALPAFSPP | AWSLGERLF | 0.83696187 |
| VVLAPAL | AWSLGERLF | 0.85959387 |
| TIGHVDHGKTTL | AWSLGERLF | 0.75647944 |
| LISNASDALDK | AWSLGERLF | 0.54086393 |
| LISNASDAL | AWSLGERLF | 0.83762157 |
| ISNASDALDK | AWSLGERLF | 0.8659835 |
| NQAETLVYQTE | AWSLGERLF | 0.8477561 |
| TGGTGGTAGAG | AWSLGERLF | 0.36254662 |
| TGSTAGGAGGA | AWSLGERLF | 0.2091749 |
| DPLAVPVAAGPV | AWSLGERLF | 0.64355266 |
| GGTGGTPGNGGN | AWSLGERLF | 0.77205527 |
| PAGLAGVDPATL | AWSLGERLF | 0.5665978 |
| GNNLVVEAPHAGASV | AWSLGERLF | 0.49356142 |
| KVTRIMSASTLEQPA | AWSLGERLF | 0.7718227 |
| LGFALPIALPRRTVV | AWSLGERLF | 0.8109897 |
| TRLGNLTARKLVVLV | AWSLGERLF | 0.5384592 |
| WSQQRRLIYFPSAGP | AWSLGERLF | 0.45457736 |

| | | |
|---|---|---|
| GPNGNPDPMAAAVDIR | AWSLGERLF | 0.82605135 |
| TQRCVSVADLRAAASR | AWSLGERLF | 0.8053427 |
| QFVLDVAEHPVEVLTETV | AWSLGERLF | 0.5300535 |
| GGAGGTGGAGGNGGTQAGDG | AWSLGERLF | 0.59534705 |
| PLLATLRAIAVDVDAKPAQV | AWSLGERLF | 0.635405 |
| APKFRTGGYMIDFWGVGYQVAK | AWSLGERLF | 0.72646594 |
| GAGGAGGAGGAAGGPGGTGGTG | AWSLGERLF | 0.70256126 |
| IGALGLLATDVAPAGSVAFGVA | AWSLGERLF | 0.40278414 |
| AGGAGGIGGNGGAGGAGGVLMG | AWSLGERLF | 0.85404646 |
| ADELIGLL | AWSLGERLF | 0.4337397 |
| ADRALKA | AWSLGERLF | 0.817252 |
| AGAVGQVGGAGGA | AWSLGERLF | 0.7569536 |
| AGGGGGLGGAP | AWSLGERLF | 0.80564356 |
| AGGLLYGNGGAG | AWSLGERLF | 0.5953539 |
| AGLESIGL | AWSLGERLF | 0.53052974 |
| AGLKTGII | AWSLGERLF | 0.47963816 |
| ASLAALLL | AWSLGERLF | 0.30028862 |
| ASVINISSI | AWSLGERLF | 0.84757304 |
| ATPPGPA | AWSLGERLF | 0.48101214 |
| ATVASALV | AWSLGERLF | 0.8433405 |
| ATVASLIL | AWSLGERLF | 0.73011994 |
| AVNGPVLIRH | AWSLGERLF | 0.59163916 |
| DGAAGAVGI | AWSLGERLF | 0.7990215 |
| DPLLHGI | AWSLGERLF | 0.42802963 |
| ELGAALGAL | AWSLGERLF | 0.68525124 |
| ELGRTQAV | AWSLGERLF | 0.74848855 |
| FVAGGYLS | AWSLGERLF | 0.78869194 |
| GASGGAGGAGGW | AWSLGERLF | 0.5843195 |
| GGAGAGGDGGAGGI | AWSLGERLF | 0.7944188 |
| GTGHNGSQPGMGGNG | AWSLGERLF | 0.7820523 |
| GYAAASASAS | AWSLGERLF | 0.5985472 |
| HQVDVAAVQQG | AWSLGERLF | 0.74212265 |
| IGAAIAHT | AWSLGERLF | 0.81868684 |
| IGMIEVAQL | AWSLGERLF | 0.76368594 |
| IRLAPEHTSAMD | AWSLGERLF | 0.4597977 |
| IYNVSVL | AWSLGERLF | 0.76623905 |
| KQGFLGM | AWSLGERLF | 0.5340184 |
| LESVITQL | AWSLGERLF | 0.742023 |
| LGLGALVVV | AWSLGERLF | 0.7814041 |
| LIGNGGNGGSGGI | AWSLGERLF | 0.6261057 |
| LNAINAPTL | AWSLGERLF | 0.8327961 |
| LQLLNTGV | AWSLGERLF | 0.4923599 |
| LQTSIEF | AWSLGERLF | 0.5929891 |
| MSFAVSEL | AWSLGERLF | 0.6393048 |
| NTGALGL | AWSLGERLF | 0.692512 |
| PPGAGGAGGAGTT | AWSLGERLF | 0.8111105 |
| QLVGLGI | AWSLGERLF | 0.4014875 |
| RFLEQVHQL | AWSLGERLF | 0.7496455 |
| RGVGYVLR | AWSLGERLF | 0.5735165 |
| SATGVFAGVI | AWSLGERLF | 0.4250673 |
| SLPAITLP | AWSLGERLF | 0.7308426 |

| | | |
|---|---|---|
| STALAFTV | AWSLGERLF | 0.6744239 |
| SVLTVGLGK | AWSLGERLF | 0.5335227 |
| TALAGAGSGI | AWSLGERLF | 0.81740856 |
| TGLPFTGI | AWSLGERLF | 0.5764199 |
| VAEHVTV | AWSLGERLF | 0.749213 |
| VDIVLNSL | AWSLGERLF | 0.8906615 |
| VLTGGGAVL | AWSLGERLF | 0.4910417 |
| VNVGLAYGI | AWSLGERLF | 0.8155264 |
| VSSAAVGAV | AWSLGERLF | 0.7745882 |
| VTSVYEAGV | AWSLGERLF | 0.7999697 |
| WVALDRGA | AWSLGERLF | 0.53605026 |
| YLVGSVVI | AWSLGERLF | 0.59512293 |
| GPPGTGKTLL | AWSLGERLF | 0.8576182 |
| GVGMTKF | AWSLGERLF | 0.7631504 |
| IGIGGSDL | AWSLGERLF | 0.70772666 |
| IGTLLAEL | AWSLGERLF | 0.84546757 |
| IYTSGTT | AWSLGERLF | 0.7960917 |
| LSFPAGV | AWSLGERLF | 0.8163448 |
| LYTSGTT | AWSLGERLF | 0.519568 |
| VGAFPFGF | AWSLGERLF | 0.37763026 |
| VGAGTVEF | AWSLGERLF | 0.58656496 |
| VGKSTLLL | AWSLGERLF | 0.588586 |
| AALAHAGGGIV | AWSLGERLF | 0.67201054 |
| AALVAVSLL | AWSLGERLF | 0.3844754 |
| AAVSRAVAAL | AWSLGERLF | 0.7977332 |
| AGGSSGIGGF | AWSLGERLF | 0.8820257 |
| AGLTVANANG | AWSLGERLF | 0.4895453 |
| AGLVGAVL | AWSLGERLF | 0.882008 |
| ALSQAFTV | AWSLGERLF | 0.4167128 |
| ALTFSVSL | AWSLGERLF | 0.7537763 |
| ALTGAALVAL | AWSLGERLF | 0.8183814 |
| ASFSIGAGVL | AWSLGERLF | 0.3801404 |
| DGSVYKI | AWSLGERLF | 0.73337734 |
| EQLAEIV | AWSLGERLF | 0.6479427 |
| FAVSELTVM | AWSLGERLF | 0.6368637 |
| GAAPLAAGL | AWSLGERLF | 0.84287894 |
| GADGGDGGSGGAGG | AWSLGERLF | 0.51256704 |
| GFGLDTGGAGG | AWSLGERLF | 0.7958808 |
| GLASGGLL | AWSLGERLF | 0.60451984 |
| GRIAGAAVLL | AWSLGERLF | 0.48891187 |
| GSAGSLNTSF | AWSLGERLF | 0.81257105 |
| GTGQIGI | AWSLGERLF | 0.80486166 |
| GTLIALIL | AWSLGERLF | 0.63507116 |
| IAATMGFVVG | AWSLGERLF | 0.8854031 |
| IFAQALL | AWSLGERLF | 0.58715475 |
| IGAAVALGF | AWSLGERLF | 0.87465894 |
| IGLTPLAL | AWSLGERLF | 0.83877903 |
| ILGSNTSTL | AWSLGERLF | 0.6677999 |
| IVFSLTVL | AWSLGERLF | 0.6739186 |
| LASKATGF | AWSLGERLF | 0.79526246 |
| LASVALIL | AWSLGERLF | 0.5821907 |

| | | |
|---|---|---|
| LFAVGLTL | AWSLGERLF | 0.76390916 |
| LFNGGIGTYI | AWSLGERLF | 0.8576586 |
| LTLGAGII | AWSLGERLF | 0.63587296 |
| LVAAALGDI | AWSLGERLF | 0.874691 |
| NTAFVQLGI | AWSLGERLF | 0.8588046 |
| QLFSGAEL | AWSLGERLF | 0.8282596 |
| SAATFITL | AWSLGERLF | 0.7280197 |
| SNYGLANLGV | AWSLGERLF | 0.43900743 |
| SYGGAYAVM | AWSLGERLF | 0.7748468 |
| TTGVGGLTLG | AWSLGERLF | 0.7076358 |
| TTQTIYI | AWSLGERLF | 0.8606318 |
| VAAGAGLGAVI | AWSLGERLF | 0.8642776 |
| VAAVVTLL | AWSLGERLF | 0.6986916 |
| VGFLGGLI | AWSLGERLF | 0.86810136 |
| VGVIPILL | AWSLGERLF | 0.8699672 |
| VIFASSLI | AWSLGERLF | 0.3877843 |
| VKGGDGGAAAG | AWSLGERLF | 0.8077979 |
| VTKTFTNI | AWSLGERLF | 0.7320342 |
| VVAAAILL | AWSLGERLF | 0.77800864 |
| YAALDVEL | AWSLGERLF | 0.4056462 |
| LASVALLL | AWSLGERLF | 0.85634255 |
| LDAVHRV | AWSLGERLF | 0.7922789 |
| QGLSSSIF | AWSLGERLF | 0.1530141 |
| RGTFANIRL | AWSLGERLF | 0.82102835 |
| RQLWWGHRI | AWSLGERLF | 0.7770148 |
| YGPPGTGKTLL | AWSLGERLF | 0.7962158 |
| AALVGAPANVA | AWSLGERLF | 0.8639969 |
| AATFRRTE | AWSLGERLF | 0.63846314 |
| AGALGGGI | AWSLGERLF | 0.75602156 |
| AGGAGGAGGIGAG | AWSLGERLF | 0.8592172 |
| AGSGALGL | AWSLGERLF | 0.87216413 |
| AQIADQI | AWSLGERLF | 0.6603223 |
| ASAVAVTYL | AWSLGERLF | 0.8419899 |
| ASTVLGGTGG | AWSLGERLF | 0.87781155 |
| AVGGEPAK | AWSLGERLF | 0.83901834 |
| CEQGTFGGTC | AWSLGERLF | 0.8759187 |
| DEGKAPAG | AWSLGERLF | 0.87818867 |
| DTVVFPAL | AWSLGERLF | 0.734352 |
| FEGITMAG | AWSLGERLF | 0.8679229 |
| FPLVVQD | AWSLGERLF | 0.8783103 |
| GAFTTPQ | AWSLGERLF | 0.7730633 |
| GFGGFGTAGGVGGA | AWSLGERLF | 0.8148439 |
| GFLAQFL | AWSLGERLF | 0.6868732 |
| GGAGGAGGNAGVA | AWSLGERLF | 0.8073095 |
| GGVGTAGAGGAGGAGG | AWSLGERLF | 0.48106498 |
| GNGGGDAGNAGSGGN | AWSLGERLF | 0.84089744 |
| GTGGAGGAGNDAGS | AWSLGERLF | 0.87319595 |
| GTTNTGWFNS | AWSLGERLF | 0.7855774 |
| IFMGPLFTL | AWSLGERLF | 0.6800547 |
| ILAHAIEAI | AWSLGERLF | 0.6654384 |
| IRSAIREAL | AWSLGERLF | 0.8297701 |

| | | |
|---|---|---|
| ITFAGGG | AWSLGERLF | 0.75409764 |
| ITGMSGHQEV | AWSLGERLF | 0.35790175 |
| IVAGLACTL | AWSLGERLF | 0.7585908 |
| IVGGGLAGA | AWSLGERLF | 0.8644402 |
| IVTGVLNI | AWSLGERLF | 0.86709905 |
| LAQQGCI | AWSLGERLF | 0.502874 |
| LCLHVEN | AWSLGERLF | 0.8751703 |
| LDAISDGS | AWSLGERLF | 0.86062443 |
| LGGFAAML | AWSLGERLF | 0.8820118 |
| LGGKNPMI | AWSLGERLF | 0.8341295 |
| LGPKGVAAIL | AWSLGERLF | 0.6495886 |
| LMVGVSI | AWSLGERLF | 0.85746104 |
| LVGAAFVL | AWSLGERLF | 0.78764176 |
| LVLAVAPL | AWSLGERLF | 0.7894206 |
| MAAIAVIDAA | AWSLGERLF | 0.85780025 |
| NNGINAELQ | AWSLGERLF | 0.8259863 |
| RGEPLVLVHG | AWSLGERLF | 0.73768955 |
| RGGAGGAAGLI | AWSLGERLF | 0.80700684 |
| RSYRMHPAV | AWSLGERLF | 0.8789795 |
| SAAHLLAGQLA | AWSLGERLF | 0.66935205 |
| SAAVGSY | AWSLGERLF | 0.6786469 |
| SPHAVLHRI | AWSLGERLF | 0.71449196 |
| SQIAKVL | AWSLGERLF | 0.82298756 |
| TALGHLVP | AWSLGERLF | 0.8700032 |
| TAPTTSGAA | AWSLGERLF | 0.84896535 |
| TDGDDGNGGSPGT | AWSLGERLF | 0.79871386 |
| TDGLGEN | AWSLGERLF | 0.85339975 |
| TGGPPLPGQA | AWSLGERLF | 0.8597064 |
| TIPAITI | AWSLGERLF | 0.8278059 |
| TIPGFTI | AWSLGERLF | 0.8687434 |
| TSIASGALA | AWSLGERLF | 0.729529 |
| TVTLPGAAI | AWSLGERLF | 0.87666714 |
| VASGLSAALP | AWSLGERLF | 0.8508345 |
| VGAGVIGI | AWSLGERLF | 0.86526 |
| VGPKRFGV | AWSLGERLF | 0.86993295 |
| VNVAFPD | AWSLGERLF | 0.8591616 |
| VSMTNSGVS | AWSLGERLF | 0.83709025 |
| VTDVLGDLSV | AWSLGERLF | 0.44380975 |
| VVVVGAGFAGL | AWSLGERLF | 0.7610773 |
| VYTVGTVV | AWSLGERLF | 0.8470297 |
| WTTFSGP | AWSLGERLF | 0.8452399 |
| YDELVEAF | AWSLGERLF | 0.8618159 |
| YGTGGAGGNGG | AWSLGERLF | 0.8680844 |
| YTNVMAV | AWSLGERLF | 0.7449013 |
| FLEQVHQL | AWSLGERLF | 0.87634474 |
| GLSLLAVAV | AWSLGERLF | 0.8685219 |
| LEQVHQL | AWSLGERLF | 0.51096773 |
| LNAAGAI | AWSLGERLF | 0.7894303 |
| PAVHASL | AWSLGERLF | 0.87545794 |
| TELEDDA | AWSLGERLF | 0.84028757 |
| TPPGPPN | AWSLGERLF | 0.76252747 |

| | | |
|---|---|---|
| TPPPPPPP | AWSLGERLF | 0.7888266 |
| TQSPALI | ASSDSLEQY | 0.8743873 |
| EFAAAFPQ | ASSDSLEQY | 0.73400617 |
| LLSGNLDVL | ASSDSLEQY | 0.8782313 |
| GPANLGNYNI | ASSDSLEQY | 0.2345998 |
| TLGPTLP | ASSDSLEQY | 0.7756539 |
| VYAYSSCCK | ASSDSLEQY | 0.7839181 |
| APAPAPAPNI | ASSDSLEQY | 0.32630694 |
| PGGPVPIP | ASSDSLEQY | 0.8652203 |
| FLTVALEP | ASSDSLEQY | 0.6720444 |
| SQYFGANL | ASSDSLEQY | 0.858815 |
| GGVKPAA | ASSDSLEQY | 0.8200735 |
| GGTGGAGGAGGW | ASSDSLEQY | 0.87090826 |
| GVGGVAGLGGAGG | ASSDSLEQY | 0.8380518 |
| VLLVLATTI | ASSDSLEQY | 0.8054811 |
| IGIFAAML | ASSDSLEQY | 0.5753838 |
| AVGDLID | ASSDSLEQY | 0.85599583 |
| NISPLPL | ASSDSLEQY | 0.87791777 |
| VTAVFVAI | ASSDSLEQY | 0.86999345 |
| SGDDNDAAL | ASSDSLEQY | 0.5214539 |
| AVAALAHS | ASSDSLEQY | 0.8186481 |
| GGVTVMATP | ASSDSLEQY | 0.8271624 |
| AGLAANVRD | ASSDSLEQY | 0.39440244 |
| TNRPDLIDPALLRPG | ASSDSLEQY | 0.21106878 |
| LANLGVTAL | ASSDSLEQY | 0.8036252 |
| TGERIEL | ASSDSLEQY | 0.78791296 |
| NGEELAA | ASSDSLEQY | 0.86067927 |
| VLSLVWG | ASSDSLEQY | 0.8629089 |
| IGYALWL | ASSDSLEQY | 0.7779727 |
| LSLLLAPL | ASSDSLEQY | 0.8603345 |
| TLDEALK | ASSDSLEQY | 0.8353593 |
| TDGAGTAEGG | ASSDSLEQY | 0.1169412 |
| NAPGGAGGNGG | ASSDSLEQY | 0.7596813 |
| VSVFMGMGGNF | ASSDSLEQY | 0.44096276 |
| LASYLAAL | ASSDSLEQY | 0.8759881 |
| LVGGTPAL | ASSDSLEQY | 0.69914687 |
| VRNADVL | ASSDSLEQY | 0.71498024 |
| TCSSPFLVSAM | ASSDSLEQY | 0.8050646 |
| VHITDAE | ASSDSLEQY | 0.83219874 |
| EYAAPSF | ASSDSLEQY | 0.82042664 |
| VVDAFHGGL | ASSDSLEQY | 0.7503263 |
| ASVVDPQ | ASSDSLEQY | 0.27746204 |
| SYNEFGGLS | ASSDSLEQY | 0.8567565 |
| DPDGSVY | ASSDSLEQY | 0.65640724 |
| MVSTIHATE | ASSDSLEQY | 0.66839623 |
| LAAGVADPV | ASSDSLEQY | 0.32263058 |
| TKEQIAA | ASSDSLEQY | 0.78901684 |
| SLASPAAL | ASSDSLEQY | 0.5021544 |
| IGLVFYGV | ASSDSLEQY | 0.8727119 |
| FLLDASNL | ASSDSLEQY | 0.7567182 |
| YQVGNFP | ASSDSLEQY | 0.78060377 |

| | | |
|---|---|---|
| ESLLSQST | ASSDSLEQY | 0.87800145 |
| LFTVGFAAVG | ASSDSLEQY | 0.82452387 |
| ATPPTPT | ASSDSLEQY | 0.77527493 |
| TDLATLG | ASSDSLEQY | 0.7339334 |
| SAGGTGR | ASSDSLEQY | 0.60161114 |
| GFSNSGS | ASSDSLEQY | 0.73651135 |
| GGNGGSGGTSV | ASSDSLEQY | 0.7464359 |
| GTSVATGGAGN | ASSDSLEQY | 0.87812686 |
| NGGNGGSGGTS | ASSDSLEQY | 0.7827853 |
| AVVKFPH | ASSDSLEQY | 0.4547999 |
| TPPPPPP | ASSDSLEQY | 0.83794314 |
| AAPAAAAAPA | ASSDSLEQY | 0.8752715 |
| IALNEDL | ASSDSLEQY | 0.88377064 |
| IVNVSSM | ASSDSLEQY | 0.6343953 |
| TGIPPAPR | ASSDSLEQY | 0.81270933 |
| FDLGGGTF | ASSDSLEQY | 0.7923633 |
| FDLGGGTFDVS | ASSDSLEQY | 0.816486 |
| NEPTAAA | ASSDSLEQY | 0.78682363 |
| NEPTAAALA | ASSDSLEQY | 0.60041416 |
| GSIINISSI | ASSDSLEQY | 0.87084496 |
| VAPGNAGTA | ASSDSLEQY | 0.8795496 |
| TGGGASLE | ASSDSLEQY | 0.782081 |
| VAVYNHYKRI | ASSDSLEQY | 0.6267562 |
| TDPAPATT | ASSDSLEQY | 0.794986 |
| TNRPDILDPALLRPG | ASSDSLEQY | 0.7307102 |
| LTTTTTS | ASSDSLEQY | 0.7577331 |
| PTGSGKTL | ASSDSLEQY | 0.8312621 |
| TGSGKTL | ASSDSLEQY | 0.44975215 |
| TIGHVDHGK | ASSDSLEQY | 0.807057 |
| TIGHVDHGKTT | ASSDSLEQY | 0.8846656 |
| GHVDHGKTT | ASSDSLEQY | 0.7761687 |
| AQFEHTLL | ASSDSLEQY | 0.8371946 |
| IVNQDEL | ASSDSLEQY | 0.8708875 |
| TNGSQFF | ASSDSLEQY | 0.8517132 |
| TGKGIVE | ASSDSLEQY | 0.8705348 |
| LISNASDA | ASSDSLEQY | 0.8634843 |
| AELRTLGELTKIAKAHGAQ | ASSDSLEQY | 0.77295655 |
| APNAVAAAEPSW | ASSDSLEQY | 0.83800256 |
| APQVAGAATAVL | ASSDSLEQY | 0.8400956 |
| ARRDPAAAAAILPTDA | ASSDSLEQY | 0.80374295 |
| DAAARDVLAEAGLAEYF | ASSDSLEQY | 0.7920489 |
| DDQIVLSAKANLAKVEQ | ASSDSLEQY | 0.76141274 |
| DELLPDTGEDSDEDGH | ASSDSLEQY | 0.861629 |
| ELPEAPANRATVPQD | ASSDSLEQY | 0.83235556 |
| FGQSKIERVTGIG | ASSDSLEQY | 0.86552685 |
| GAGGTGSTAGGAGGAGGAGGLY | ASSDSLEQY | 0.3988371 |
| GHGGTGAAGGNGGTGGAGGSGI | ASSDSLEQY | 0.63285285 |
| GNGGNGGAGGEGGPGGAGGAGA | ASSDSLEQY | 0.86182904 |
| GSTAGNGGNGGNATNGTIAGQP | ASSDSLEQY | 0.22023687 |
| GTGGDGGNGGNGGTGG | ASSDSLEQY | 0.24869251 |
| GVRRPAGIPADAPV | ASSDSLEQY | 0.8434537 |

| | | |
|---|---|---|
| GYRQVTDAHLLTLARR | ASSDSLEQY | 0.87624246 |
| IAKMGQDGHDRG | ASSDSLEQY | 0.21527459 |
| KAEPSIEIQYT | ASSDSLEQY | 0.68626493 |
| KPVRIRMAADEDTVAVT | ASSDSLEQY | 0.4811858 |
| MPPRIAGMRLLVIKPEP | ASSDSLEQY | 0.87030846 |
| MKFYAVALFRSRGVPPTR | ASSDSLEQY | 0.83776397 |
| NAGGFGILGAVAHSP | ASSDSLEQY | 0.3327294 |
| NEGREITLAMAEM | ASSDSLEQY | 0.85730165 |
| NELVEWSHPLGHRWRWE | ASSDSLEQY | 0.50663143 |
| NLGAGGTGGNGGNPG | ASSDSLEQY | 0.7603308 |
| NTEWPKWETVMAADPVD | ASSDSLEQY | 0.8709583 |
| PASAAPAETQP | ASSDSLEQY | 0.87483597 |
| QQSSDGKNGVRAAEITGE | ASSDSLEQY | 0.5492984 |
| RAPDAVTTSWWKE | ASSDSLEQY | 0.8646618 |
| SGFGNVGAHTSGWFNQ | ASSDSLEQY | 0.7858119 |
| SGVAGAVGQAASVGGLKV | ASSDSLEQY | 0.88589174 |
| SNGGAGTGSAGGNGGT | ASSDSLEQY | 0.7558106 |
| TGGAGGEGGAAG | ASSDSLEQY | 0.49956685 |
| TGGDGGAAGAGGAPGHG | ASSDSLEQY | 0.86781687 |
| TMPPLLAAIANAGRHGV | ASSDSLEQY | 0.84797555 |
| TSSTAGNSGGA | ASSDSLEQY | 0.83576095 |
| VPATVPVGILC | ASSDSLEQY | 0.86156356 |
| VQRPDSGGVVLTGRIS | ASSDSLEQY | 0.79194283 |
| WTPNPVPATKGVPAGLVQAT | ASSDSLEQY | 0.78981465 |
| YALKRALAKFSDRFDVVII | ASSDSLEQY | 0.8598956 |
| YLIGSVIV | ASSDSLEQY | 0.8608942 |
| YGQLVAVID | ASSDSLEQY | 0.61614347 |
| VTVGGMTLPA | ASSDSLEQY | 0.8769687 |
| VLALPAV | ASSDSLEQY | 0.81145334 |
| VIPPTLNYET | ASSDSLEQY | 0.6679412 |
| VGMGGMHPSSG | ASSDSLEQY | 0.8813335 |
| VGLAGGAATA | ASSDSLEQY | 0.84241897 |
| VGGAGGAGGSGP | ASSDSLEQY | 0.86361325 |
| VFAAFMV | ASSDSLEQY | 0.85996497 |
| TLYDKLR | ASSDSLEQY | 0.8757021 |
| TLAVFTVL | ASSDSLEQY | 0.45492107 |
| TGSVQPARV | ASSDSLEQY | 0.8425416 |
| TGGGAGSGGT | ASSDSLEQY | 0.8277043 |
| SHLETSL | ASSDSLEQY | 0.8670311 |
| SFTTGMT | ASSDSLEQY | 0.83611864 |
| QVEVAKRMGVRQPTVS | ASSDSLEQY | 0.79229414 |
| QGSGGPALAM | ASSDSLEQY | 0.8446169 |
| MLAPSNP | ASSDSLEQY | 0.87504184 |
| LVNNAAI | ASSDSLEQY | 0.8388068 |
| LTGEINTLA | ASSDSLEQY | 0.8501705 |
| LSIFGPV | ASSDSLEQY | 0.8756391 |
| LLALLANL | ASSDSLEQY | 0.63241124 |
| LISNASDALDKLR | ASSDSLEQY | 0.80296266 |
| LINEAVP | ASSDSLEQY | 0.86847746 |
| LGYAFSGL | ASSDSLEQY | 0.87476766 |
| LDFVAVAL | ASSDSLEQY | 0.80164707 |

| | | |
|---|---|---|
| LAQPGAE | ASSDSLEQY | 0.8734733 |
| ISNLGAVTGF | ASSDSLEQY | 0.7372378 |
| HQEDPGRI | ASSDSLEQY | 0.85323155 |
| GTPGEPTP | ASSDSLEQY | 0.79985315 |
| GFFNSGS | ASSDSLEQY | 0.8354625 |
| GAGGGGLTAGAN | ASSDSLEQY | 0.7137184 |
| FVDALVNM | ASSDSLEQY | 0.8422682 |
| FLTVAPEL | ASSDSLEQY | 0.8659028 |
| FETLTAL | ASSDSLEQY | 0.8537167 |
| DVINPIEL | ASSDSLEQY | 0.19070867 |
| APVSGAPV | ASSDSLEQY | 0.8553242 |
| ALVGGLAW | ASSDSLEQY | 0.85038203 |
| ALAAAAATAI | ASSDSLEQY | 0.87916446 |
| AFGLGLLL | ASSDSLEQY | 0.8268833 |
| AFFAAASL | ASSDSLEQY | 0.8541666 |
| AAAALPAFSPP | ASSDSLEQY | 0.7892715 |
| VVLAPAL | ASSDSLEQY | 0.82549083 |
| TIGHVDHGKTTL | ASSDSLEQY | 0.6556748 |
| LISNASDALDK | ASSDSLEQY | 0.8010788 |
| LISNASDAL | ASSDSLEQY | 0.7835945 |
| ISNASDALDK | ASSDSLEQY | 0.70818216 |
| NQAETLVYQTE | ASSDSLEQY | 0.8589622 |
| TGGTGGTAGAG | ASSDSLEQY | 0.8753682 |
| TGSTAGGAGGA | ASSDSLEQY | 0.8722446 |
| DPLAVPVAAGPV | ASSDSLEQY | 0.8714416 |
| GGTGGTPGNGGN | ASSDSLEQY | 0.87181777 |
| PAGLAGVDPATL | ASSDSLEQY | 0.87258905 |
| GNNLVVEAPHAGASV | ASSDSLEQY | 0.7670011 |
| KVTRIMSASTLEQPA | ASSDSLEQY | 0.79408705 |
| LGFALPIALPRRTVV | ASSDSLEQY | 0.6314584 |
| TRLGNLTARKLVVLV | ASSDSLEQY | 0.777635 |
| WSQQRRLIYFPSAGP | ASSDSLEQY | 0.77967036 |
| GPNGNPDPMAAAVDIR | ASSDSLEQY | 0.8254262 |
| TQRCVSVADLRAAASR | ASSDSLEQY | 0.13628851 |
| QFVLDVAEHPVEVLTETV | ASSDSLEQY | 0.83588547 |
| GGAGGTGGAGGNGGTQAGDG | ASSDSLEQY | 0.8781527 |
| PLLATLRAIAVDVDAKPAQV | ASSDSLEQY | 0.7881924 |
| APKFRTGGYMIDFWGVGYQVAK | ASSDSLEQY | 0.85156465 |
| GAGGAGGAGGAAGGPGGTGGTG | ASSDSLEQY | 0.8297929 |
| IGALGLLATDVAPAGSVAFGVA | ASSDSLEQY | 0.78981507 |
| AGGAGGIGGNGGAGGAGGVLMG | ASSDSLEQY | 0.86081797 |
| ADELIGLL | ASSDSLEQY | 0.7164721 |
| ADRALKA | ASSDSLEQY | 0.86040914 |
| AGAVGQVGGAGGA | ASSDSLEQY | 0.8810392 |
| AGGGGGLGGAP | ASSDSLEQY | 0.8412902 |
| AGGLLYGNGGAG | ASSDSLEQY | 0.84403384 |
| AGLESIGL | ASSDSLEQY | 0.8315095 |
| AGLKTGII | ASSDSLEQY | 0.36704183 |
| ASLAALLL | ASSDSLEQY | 0.85049415 |
| ASVINISSI | ASSDSLEQY | 0.41142535 |
| ATPPGPA | ASSDSLEQY | 0.8112942 |

| | | |
|---|---|---|
| ATVASALV | ASSDSLEQY | 0.6753346 |
| ATVASLIL | ASSDSLEQY | 0.83755606 |
| AVNGPVLIRH | ASSDSLEQY | 0.82106066 |
| DGAAGAVGI | ASSDSLEQY | 0.85678554 |
| DPLLHGI | ASSDSLEQY | 0.63880295 |
| ELGAALGAL | ASSDSLEQY | 0.8240639 |
| ELGRTQAV | ASSDSLEQY | 0.7936609 |
| FVAGGYLS | ASSDSLEQY | 0.64190865 |
| GASGGAGGAGGW | ASSDSLEQY | 0.86452305 |
| GGAGAGGDGGAGGI | ASSDSLEQY | 0.8322186 |
| GTGHNGSQPGMGGNG | ASSDSLEQY | 0.8175662 |
| GYAAASASAS | ASSDSLEQY | 0.8684394 |
| HQVDVAAVQQG | ASSDSLEQY | 0.8466855 |
| IGAAIAHT | ASSDSLEQY | 0.52495146 |
| IGMIEVAQL | ASSDSLEQY | 0.82167137 |
| IRLAPEHTSAMD | ASSDSLEQY | 0.81507444 |
| IYNVSVL | ASSDSLEQY | 0.79013443 |
| KQGFLGM | ASSDSLEQY | 0.19920048 |
| LESVITQL | ASSDSLEQY | 0.8477716 |
| LGLGALVVV | ASSDSLEQY | 0.66230655 |
| LIGNGGNGGSGGI | ASSDSLEQY | 0.74270695 |
| LNAINAPTL | ASSDSLEQY | 0.78510314 |
| LQLLNTGV | ASSDSLEQY | 0.86308014 |
| LQTSIEF | ASSDSLEQY | 0.83425367 |
| MSFAVSEL | ASSDSLEQY | 0.8355635 |
| NTGALGL | ASSDSLEQY | 0.8648259 |
| PPGAGGAGGAGTT | ASSDSLEQY | 0.8587736 |
| QLVGLGI | ASSDSLEQY | 0.8726996 |
| RFLEQVHQL | ASSDSLEQY | 0.84810007 |
| RGVGYVLR | ASSDSLEQY | 0.8409544 |
| SATGVFAGVI | ASSDSLEQY | 0.8320517 |
| SLPAITLP | ASSDSLEQY | 0.83075786 |
| STALAFTV | ASSDSLEQY | 0.4208705 |
| SVLTVGLGK | ASSDSLEQY | 0.41317642 |
| TALAGAGSGI | ASSDSLEQY | 0.8281579 |
| TGLPFTGI | ASSDSLEQY | 0.38040543 |
| VAEHVTV | ASSDSLEQY | 0.8187058 |
| VDIVLNSL | ASSDSLEQY | 0.8748729 |
| VLTGGGAVL | ASSDSLEQY | 0.8459161 |
| VNVGLAYGI | ASSDSLEQY | 0.5369945 |
| VSSAAVGAV | ASSDSLEQY | 0.8033005 |
| VTSVYEAGV | ASSDSLEQY | 0.7048424 |
| WVALDRGA | ASSDSLEQY | 0.8588811 |
| YLVGSVVI | ASSDSLEQY | 0.8744153 |
| GPPGTGKTLL | ASSDSLEQY | 0.8756117 |
| GVGMTKF | ASSDSLEQY | 0.86322916 |
| IGIGGSDL | ASSDSLEQY | 0.76856625 |
| IGTLLAEL | ASSDSLEQY | 0.8546201 |
| IYTSGTT | ASSDSLEQY | 0.7781857 |
| LSFPAGV | ASSDSLEQY | 0.80815303 |
| LYTSGTT | ASSDSLEQY | 0.8094535 |

| | | |
|---|---|---|
| VGAFPFGF | ASSDSLEQY | 0.8520342 |
| VGAGTVEF | ASSDSLEQY | 0.8214785 |
| VGKSTLLL | ASSDSLEQY | 0.7646283 |
| AALAHAGGGIV | ASSDSLEQY | 0.45721307 |
| AALVAVSLL | ASSDSLEQY | 0.85760283 |
| AAVSRAVAAL | ASSDSLEQY | 0.82315934 |
| AGGSSGIGGF | ASSDSLEQY | 0.47111446 |
| AGLTVANANG | ASSDSLEQY | 0.7371669 |
| AGLVGAVL | ASSDSLEQY | 0.8765762 |
| ALSQAFTV | ASSDSLEQY | 0.8776523 |
| ALTFSVSL | ASSDSLEQY | 0.7362397 |
| ALTGAALVAL | ASSDSLEQY | 0.7098696 |
| ASFSIGAGVL | ASSDSLEQY | 0.83569956 |
| DGSVYKI | ASSDSLEQY | 0.47641277 |
| EQLAEIV | ASSDSLEQY | 0.8475615 |
| FAVSELTVM | ASSDSLEQY | 0.8147819 |
| GAAPLAAGL | ASSDSLEQY | 0.82160956 |
| GADGGDGGSGGAGG | ASSDSLEQY | 0.74970603 |
| GFGLDTGGAGG | ASSDSLEQY | 0.85204935 |
| GLASGGLL | ASSDSLEQY | 0.7945585 |
| GRIAGAAVLL | ASSDSLEQY | 0.86977565 |
| GSAGSLNTSF | ASSDSLEQY | 0.21821809 |
| GTGQIGI | ASSDSLEQY | 0.8653686 |
| GTLIALIL | ASSDSLEQY | 0.6543636 |
| IAATMGFVVG | ASSDSLEQY | 0.7319366 |
| IFAQALL | ASSDSLEQY | 0.47103804 |
| IGAAVALGF | ASSDSLEQY | 0.79301035 |
| IGLTPLAL | ASSDSLEQY | 0.8704762 |
| ILGSNTSTL | ASSDSLEQY | 0.87123525 |
| IVFSLTVL | ASSDSLEQY | 0.6500784 |
| LASKATGF | ASSDSLEQY | 0.86302805 |
| LASVALIL | ASSDSLEQY | 0.8846418 |
| LFAVGLTL | ASSDSLEQY | 0.7729507 |
| LFNGGIGTYI | ASSDSLEQY | 0.27029568 |
| LTLGAGII | ASSDSLEQY | 0.84921545 |
| LVAAALGDI | ASSDSLEQY | 0.8081059 |
| NTAFVQLGI | ASSDSLEQY | 0.86939836 |
| QLFSGAEL | ASSDSLEQY | 0.86411 |
| SAATFITL | ASSDSLEQY | 0.576833 |
| SNYGLANLGV | ASSDSLEQY | 0.8645058 |
| SYGGAYAVM | ASSDSLEQY | 0.8623884 |
| TTGVGGLTLG | ASSDSLEQY | 0.7071372 |
| TTQTIYI | ASSDSLEQY | 0.74365175 |
| VAAGAGLGAVI | ASSDSLEQY | 0.6313449 |
| VAAVVTLL | ASSDSLEQY | 0.4863441 |
| VGFLGGLI | ASSDSLEQY | 0.86716455 |
| VGVIPILL | ASSDSLEQY | 0.75105953 |
| VIFASSLI | ASSDSLEQY | 0.79716724 |
| VKGGDGGAAAG | ASSDSLEQY | 0.80284286 |
| VTKTFTNI | ASSDSLEQY | 0.7740537 |
| VVAAAILL | ASSDSLEQY | 0.73929954 |

| | | |
|---|---|---|
| YAALDVEL | ASSDSLEQY | 0.77559125 |
| LASVALLL | ASSDSLEQY | 0.8482718 |
| LDAVHRV | ASSDSLEQY | 0.85686344 |
| QGLSSSIF | ASSDSLEQY | 0.82660556 |
| RGTFANIRL | ASSDSLEQY | 0.8304614 |
| RQLWWGHRI | ASSDSLEQY | 0.8731973 |
| YGPPGTGKTLL | ASSDSLEQY | 0.69348603 |
| AALVGAPANVA | ASSDSLEQY | 0.7143319 |
| AATFRRTE | ASSDSLEQY | 0.8564526 |
| AGALGGGI | ASSDSLEQY | 0.59355915 |
| AGGAGGAGGIGAG | ASSDSLEQY | 0.8604865 |
| AGSGALGL | ASSDSLEQY | 0.8212611 |
| AQIADQI | ASSDSLEQY | 0.8331907 |
| ASAVAVTYL | ASSDSLEQY | 0.8353936 |
| ASTVLGGTGG | ASSDSLEQY | 0.8813194 |
| AVGGEPAK | ASSDSLEQY | 0.8609756 |
| CEQGTFGGTC | ASSDSLEQY | 0.7934424 |
| DEGKAPAG | ASSDSLEQY | 0.8593343 |
| DTVVFPAL | ASSDSLEQY | 0.57874364 |
| FEGITMAG | ASSDSLEQY | 0.86587656 |
| FPLVVQD | ASSDSLEQY | 0.84869224 |
| GAFTTPQ | ASSDSLEQY | 0.69075674 |
| GFGGFGTAGGVGGA | ASSDSLEQY | 0.7948272 |
| GFLAQFL | ASSDSLEQY | 0.8790864 |
| GGAGGAGGNAGVA | ASSDSLEQY | 0.87036586 |
| GGVGTAGAGGAGGAGG | ASSDSLEQY | 0.50787103 |
| GNGGGGDAGNAGSGGN | ASSDSLEQY | 0.7235484 |
| GTGGAGGAGNDAGS | ASSDSLEQY | 0.49010873 |
| GTTNTGWFNS | ASSDSLEQY | 0.80553705 |
| IFMGPLFTL | ASSDSLEQY | 0.44568175 |
| ILAHAIEAI | ASSDSLEQY | 0.85185134 |
| IRSAIREAL | ASSDSLEQY | 0.7503727 |
| ITFAGGG | ASSDSLEQY | 0.85532707 |
| ITGMSGHQEV | ASSDSLEQY | 0.8221392 |
| IVAGLACTL | ASSDSLEQY | 0.87103045 |
| IVGGGLAGA | ASSDSLEQY | 0.7426059 |
| IVTGVLNI | ASSDSLEQY | 0.77664477 |
| LAQQGCI | ASSDSLEQY | 0.44493064 |
| LCLHVEN | ASSDSLEQY | 0.7239718 |
| LDAISDGS | ASSDSLEQY | 0.7881951 |
| LGGFAAML | ASSDSLEQY | 0.82189405 |
| LGGKNPMI | ASSDSLEQY | 0.37977025 |
| LGPKGVAAIL | ASSDSLEQY | 0.8744473 |
| LMVGVSI | ASSDSLEQY | 0.82171655 |
| LVGAAFVL | ASSDSLEQY | 0.85548604 |
| LVLAVAPL | ASSDSLEQY | 0.76539594 |
| MAAIAVIDAA | ASSDSLEQY | 0.77581096 |
| NNGINAELQ | ASSDSLEQY | 0.7650915 |
| RGEPLVLVHG | ASSDSLEQY | 0.8564395 |
| RGGAGGAAGLI | ASSDSLEQY | 0.62514555 |
| RSYRMHPAV | ASSDSLEQY | 0.41604227 |

| | | |
|---|---|---|
| SAAHLLAGQLA | ASSDSLEQY | 0.8129978 |
| SAAVGSY | ASSDSLEQY | 0.62319076 |
| SPHAVLHRI | ASSDSLEQY | 0.87900186 |
| SQIAKVL | ASSDSLEQY | 0.8707076 |
| TALGHLVP | ASSDSLEQY | 0.83316875 |
| TAPTTSGAA | ASSDSLEQY | 0.8675721 |
| TDGDDGNGGSPGT | ASSDSLEQY | 0.18363896 |
| TDGLGEN | ASSDSLEQY | 0.7838877 |
| TGGPPLPGQA | ASSDSLEQY | 0.87515926 |
| TIPAITI | ASSDSLEQY | 0.8539654 |
| TIPGFTI | ASSDSLEQY | 0.768221 |
| TSIASGALA | ASSDSLEQY | 0.8714522 |
| TVTLPGAAI | ASSDSLEQY | 0.7542614 |
| VASGLSAALP | ASSDSLEQY | 0.7854165 |
| VGAGVIGI | ASSDSLEQY | 0.83724403 |
| VGPKRFGV | ASSDSLEQY | 0.8487155 |
| VNVAFPD | ASSDSLEQY | 0.860873 |
| VSMTNSGVS | ASSDSLEQY | 0.6022808 |
| VTDVLGDLSV | ASSDSLEQY | 0.838262 |
| VVVVGAGFAGL | ASSDSLEQY | 0.883749 |
| VYTVGTVV | ASSDSLEQY | 0.7992252 |
| WTTFSGP | ASSDSLEQY | 0.8684756 |
| YDELVEAF | ASSDSLEQY | 0.84663725 |
| YGTGGAGGNGG | ASSDSLEQY | 0.8523302 |
| YTNVMAV | ASSDSLEQY | 0.6803721 |
| FLEQVHQL | ASSDSLEQY | 0.7424089 |
| GLSLLAVAV | ASSDSLEQY | 0.823263 |
| LEQVHQL | ASSDSLEQY | 0.68935627 |
| LNAAGAI | ASSDSLEQY | 0.8066807 |
| PAVHASL | ASSDSLEQY | 0.7593393 |
| TELEDDA | ASSDSLEQY | 0.8865425 |
| TPPGPPN | ASSDSLEQY | 0.85382515 |
| TPPPPPPP | ASSDSLEQY | 0.849458 |
| TQSPALI | ASSMGGANSDYT | 0.7059367 |
| EFAAAFPQ | ASSMGGANSDYT | 0.8530842 |
| LLSGNLDVL | ASSMGGANSDYT | 0.8478899 |
| GPANLGNYNI | ASSMGGANSDYT | 0.48035094 |
| TLGPTLP | ASSMGGANSDYT | 0.87109756 |
| VYAYSSCCK | ASSMGGANSDYT | 0.6756133 |
| APAPAPAPNI | ASSMGGANSDYT | 0.82663554 |
| PGGPVPIP | ASSMGGANSDYT | 0.8546475 |
| FLTVALEP | ASSMGGANSDYT | 0.8667718 |
| SQYFGANL | ASSMGGANSDYT | 0.7797142 |
| GGVKPAA | ASSMGGANSDYT | 0.8064315 |
| GGTGGAGGAGGW | ASSMGGANSDYT | 0.64373815 |
| GVGGVAGLGGAGG | ASSMGGANSDYT | 0.8618513 |
| VLLVLATTI | ASSMGGANSDYT | 0.86506677 |
| IGIFAAML | ASSMGGANSDYT | 0.8463047 |
| AVGDLID | ASSMGGANSDYT | 0.8770172 |
| NISPLPL | ASSMGGANSDYT | 0.8620313 |
| VTAVFVAI | ASSMGGANSDYT | 0.4029041 |

| | | |
|---|---|---|
| SGDDNDAAL | ASSMGGANSDYT | 0.80145746 |
| AVAALAHS | ASSMGGANSDYT | 0.87042314 |
| GGVTVMATP | ASSMGGANSDYT | 0.8735995 |
| AGLAANVRD | ASSMGGANSDYT | 0.8782809 |
| TNRPDLIDPALLRPG | ASSMGGANSDYT | 0.86182725 |
| LANLGVTAL | ASSMGGANSDYT | 0.7898561 |
| TGERIEL | ASSMGGANSDYT | 0.8738665 |
| NGEELAA | ASSMGGANSDYT | 0.88741326 |
| VLSLVWG | ASSMGGANSDYT | 0.8574333 |
| IGYALWL | ASSMGGANSDYT | 0.76234305 |
| LSLLLAPL | ASSMGGANSDYT | 0.70794535 |
| TLDEALK | ASSMGGANSDYT | 0.85279274 |
| TDGAGTAEGG | ASSMGGANSDYT | 0.86829734 |
| NAPGGAGGNGG | ASSMGGANSDYT | 0.5971286 |
| VSVFMGMGGNF | ASSMGGANSDYT | 0.82175046 |
| LASYLAAL | ASSMGGANSDYT | 0.8324505 |
| LVGGTPAL | ASSMGGANSDYT | 0.4009527 |
| VRNADVL | ASSMGGANSDYT | 0.8505474 |
| TCSSPFLVSAM | ASSMGGANSDYT | 0.8735364 |
| VHITDAE | ASSMGGANSDYT | 0.82677245 |
| EYAAPSF | ASSMGGANSDYT | 0.7213076 |
| VVDAFHGGL | ASSMGGANSDYT | 0.8679465 |
| ASVVDPQ | ASSMGGANSDYT | 0.8816955 |
| SYNEFGGLS | ASSMGGANSDYT | 0.8743154 |
| DPDGSVY | ASSMGGANSDYT | 0.4585354 |
| MVSTIHATE | ASSMGGANSDYT | 0.2859351 |
| LAAGVADPV | ASSMGGANSDYT | 0.77744937 |
| TKEQIAA | ASSMGGANSDYT | 0.8644475 |
| SLASPAAL | ASSMGGANSDYT | 0.802899 |
| IGLVFYGV | ASSMGGANSDYT | 0.7587866 |
| FLLDASNL | ASSMGGANSDYT | 0.8476581 |
| YQVGNFP | ASSMGGANSDYT | 0.86755073 |
| ESLLSQST | ASSMGGANSDYT | 0.5888647 |
| LFTVGFAAVG | ASSMGGANSDYT | 0.6734072 |
| ATPPTPT | ASSMGGANSDYT | 0.8575514 |
| TDLATLG | ASSMGGANSDYT | 0.85673547 |
| SAGGTGR | ASSMGGANSDYT | 0.7038461 |
| GFSNSGS | ASSMGGANSDYT | 0.8063605 |
| GGNGGSGGTSV | ASSMGGANSDYT | 0.78337896 |
| GTSVATGGAGN | ASSMGGANSDYT | 0.7829746 |
| NGGNGGSGGTS | ASSMGGANSDYT | 0.8352888 |
| AVVKFPH | ASSMGGANSDYT | 0.626409 |
| TPPPPPP | ASSMGGANSDYT | 0.8644631 |
| AAPAAAAAPA | ASSMGGANSDYT | 0.5862365 |
| IALNEDL | ASSMGGANSDYT | 0.86683273 |
| IVNVSSM | ASSMGGANSDYT | 0.84148705 |
| TGIPPAPR | ASSMGGANSDYT | 0.84265035 |
| FDLGGGTF | ASSMGGANSDYT | 0.76823425 |
| FDLGGGTFDVS | ASSMGGANSDYT | 0.7278412 |
| NEPTAAA | ASSMGGANSDYT | 0.54895437 |
| NEPTAAALA | ASSMGGANSDYT | 0.4660218 |

| | | |
|---|---|---|
| GSIINISSI | ASSMGGANSDYT | 0.874757 |
| VAPGNAGTA | ASSMGGANSDYT | 0.8138329 |
| TGGGASLE | ASSMGGANSDYT | 0.8675941 |
| VAVYNHYKRI | ASSMGGANSDYT | 0.85158587 |
| TDPAPATT | ASSMGGANSDYT | 0.5945372 |
| TNRPDILDPALLRPG | ASSMGGANSDYT | 0.85788167 |
| LTTTTTS | ASSMGGANSDYT | 0.68489456 |
| PTGSGKTL | ASSMGGANSDYT | 0.6661663 |
| TGSGKTL | ASSMGGANSDYT | 0.859205 |
| TIGHVDHGK | ASSMGGANSDYT | 0.8594631 |
| TIGHVDHGKTT | ASSMGGANSDYT | 0.6473967 |
| GHVDHGKTT | ASSMGGANSDYT | 0.84106964 |
| AQFEHTLL | ASSMGGANSDYT | 0.8400365 |
| IVNQDEL | ASSMGGANSDYT | 0.7348182 |
| TNGSQFF | ASSMGGANSDYT | 0.8253981 |
| TGKGIVE | ASSMGGANSDYT | 0.86289144 |
| LISNASDA | ASSMGGANSDYT | 0.8353044 |
| AELRTLGELTKIAKAHGAQ | ASSMGGANSDYT | 0.6448496 |
| APNAVAAAEPSW | ASSMGGANSDYT | 0.8652396 |
| APQVAGAATAVL | ASSMGGANSDYT | 0.6779804 |
| ARRDPAAAAAILPTDA | ASSMGGANSDYT | 0.8326456 |
| DAAARDVLAEAGLAEYF | ASSMGGANSDYT | 0.7904432 |
| DDQIVLSAKANLAKVEQ | ASSMGGANSDYT | 0.7933526 |
| DELLPDTGEDSDEDGH | ASSMGGANSDYT | 0.8686332 |
| ELPEAPANRATVPQD | ASSMGGANSDYT | 0.79770744 |
| FGQSKIERVTGIG | ASSMGGANSDYT | 0.76937366 |
| GAGGTGSTAGGAGGAGGAGGLY | ASSMGGANSDYT | 0.76290584 |
| GHGGTGAAGGNGGTGGAGGSGI | ASSMGGANSDYT | 0.82005554 |
| GNGGNGGAGGEGGPGGAGGAGA | ASSMGGANSDYT | 0.85748154 |
| GSTAGNGGNGGNATNGTIAGQP | ASSMGGANSDYT | 0.6749969 |
| GTGGDGGNGGNGGTGG | ASSMGGANSDYT | 0.83648413 |
| GVRRPAGIPADAPV | ASSMGGANSDYT | 0.78360355 |
| GYRQVTDAHLLTLARR | ASSMGGANSDYT | 0.6539948 |
| IAKMGQDGHDRG | ASSMGGANSDYT | 0.80582356 |
| KAEPSIEIQYT | ASSMGGANSDYT | 0.8288181 |
| KPVRIRMAADEDTVAVT | ASSMGGANSDYT | 0.6850852 |
| MPPRIAGMRLLVIKPEP | ASSMGGANSDYT | 0.8458701 |
| MKFYAVALFRSRGVPPTR | ASSMGGANSDYT | 0.7389452 |
| NAGGFGILGAVAHSP | ASSMGGANSDYT | 0.8622273 |
| NEGREITLAMAEM | ASSMGGANSDYT | 0.8767576 |
| NELVEWSHPLGHRWRWE | ASSMGGANSDYT | 0.7775445 |
| NLGAGGTGGNGGNPG | ASSMGGANSDYT | 0.86590075 |
| NTEWPKWETVMAADPVD | ASSMGGANSDYT | 0.8471641 |
| PASAAPAETQP | ASSMGGANSDYT | 0.8503164 |
| QQSSDGKNGVRAAEITGE | ASSMGGANSDYT | 0.78906286 |
| RAPDAVTTSWWKE | ASSMGGANSDYT | 0.6930126 |
| SGFGNVGAHTSGWFNQ | ASSMGGANSDYT | 0.86914253 |
| SGVAGAVGQAASVGGLKV | ASSMGGANSDYT | 0.8655541 |
| SNGGAGTGSAGGNGGT | ASSMGGANSDYT | 0.8595965 |
| TGGAGGEGGAAG | ASSMGGANSDYT | 0.855369 |
| TGGDGGAAGAGGAPGHG | ASSMGGANSDYT | 0.84670645 |

| | | |
|---|---|---|
| TMPPLLAAIANAGRHGV | ASSMGGANSDYT | 0.85884583 |
| TSSTAGNSGGA | ASSMGGANSDYT | 0.7513447 |
| VPATVPVGILC | ASSMGGANSDYT | 0.5259321 |
| VQRPDSGGVVLTGRIS | ASSMGGANSDYT | 0.7882179 |
| WTPNPVPATKGVPAGLVQAT | ASSMGGANSDYT | 0.7041651 |
| YALKRALAKFSDRFDVVII | ASSMGGANSDYT | 0.7333184 |
| YLIGSVIV | ASSMGGANSDYT | 0.45989615 |
| YGQLVAVID | ASSMGGANSDYT | 0.8619466 |
| VTVGGMTLPA | ASSMGGANSDYT | 0.88366723 |
| VLALPAV | ASSMGGANSDYT | 0.7930366 |
| VIPPTLNYET | ASSMGGANSDYT | 0.8809006 |
| VGMGGMHPSSG | ASSMGGANSDYT | 0.6406597 |
| VGLAGGAATA | ASSMGGANSDYT | 0.8670583 |
| VGGAGGAGGSGP | ASSMGGANSDYT | 0.8685138 |
| VFAAFMV | ASSMGGANSDYT | 0.7315257 |
| TLYDKLR | ASSMGGANSDYT | 0.8516778 |
| TLAVFTVL | ASSMGGANSDYT | 0.672077 |
| TGSVQPARV | ASSMGGANSDYT | 0.78774303 |
| TGGGAGSGGT | ASSMGGANSDYT | 0.8729354 |
| SHLETSL | ASSMGGANSDYT | 0.7588289 |
| SFTTGMT | ASSMGGANSDYT | 0.84144676 |
| QVEVAKRMGVRQPTVS | ASSMGGANSDYT | 0.7995802 |
| QGSGGPALAM | ASSMGGANSDYT | 0.68489677 |
| MLAPSNP | ASSMGGANSDYT | 0.8173853 |
| LVNNAAI | ASSMGGANSDYT | 0.8706221 |
| LTGEINTLA | ASSMGGANSDYT | 0.7349372 |
| LSIFGPV | ASSMGGANSDYT | 0.88697267 |
| LLALLANL | ASSMGGANSDYT | 0.70486045 |
| LISNASDALDKLR | ASSMGGANSDYT | 0.87657106 |
| LINEAVP | ASSMGGANSDYT | 0.85402966 |
| LGYAFSGL | ASSMGGANSDYT | 0.8154913 |
| LDFVAVAL | ASSMGGANSDYT | 0.7717342 |
| LAQPGAE | ASSMGGANSDYT | 0.8664694 |
| ISNLGAVTGF | ASSMGGANSDYT | 0.7765143 |
| HQEDPGRI | ASSMGGANSDYT | 0.8653908 |
| GTPGEPTP | ASSMGGANSDYT | 0.8747282 |
| GFFNSGS | ASSMGGANSDYT | 0.22006996 |
| GAGGGGLTAGAN | ASSMGGANSDYT | 0.79531467 |
| FVDALVNM | ASSMGGANSDYT | 0.6025281 |
| FLTVAPEL | ASSMGGANSDYT | 0.42012668 |
| FETLTAL | ASSMGGANSDYT | 0.28895587 |
| DVINPIEL | ASSMGGANSDYT | 0.18375738 |
| APVSGAPV | ASSMGGANSDYT | 0.43044278 |
| ALVGGLAW | ASSMGGANSDYT | 0.27108347 |
| ALAAAAATAI | ASSMGGANSDYT | 0.8159423 |
| AFGLGLLL | ASSMGGANSDYT | 0.62212974 |
| AFFAAASL | ASSMGGANSDYT | 0.29338476 |
| AAAALPAFSPP | ASSMGGANSDYT | 0.7533247 |
| VVLAPAL | ASSMGGANSDYT | 0.73990387 |
| TIGHVDHGKTTL | ASSMGGANSDYT | 0.17570224 |
| LISNASDALDK | ASSMGGANSDYT | 0.36494648 |

| | | |
|---|---|---|
| LISNASDAL | ASSMGGANSDYT | 0.31155095 |
| ISNASDALDK | ASSMGGANSDYT | 0.30327302 |
| NQAETLVYQTE | ASSMGGANSDYT | 0.15044823 |
| TGGTGGTAGAG | ASSMGGANSDYT | 0.60681945 |
| TGSTAGGAGGA | ASSMGGANSDYT | 0.44000688 |
| DPLAVPVAAGPV | ASSMGGANSDYT | 0.1217359 |
| GGTGGTPGNGGN | ASSMGGANSDYT | 0.42974728 |
| PAGLAGVDPATL | ASSMGGANSDYT | 0.37134016 |
| GNNLVVEAPHAGASV | ASSMGGANSDYT | 0.36533535 |
| KVTRIMSASTLEQPA | ASSMGGANSDYT | 0.72933584 |
| LGFALPIALPRRTVV | ASSMGGANSDYT | 0.38745767 |
| TRLGNLTARKLVVLV | ASSMGGANSDYT | 0.29738364 |
| WSQQRRLIYFPSAGP | ASSMGGANSDYT | 0.7010964 |
| GPNGNPDPMAAAVDIR | ASSMGGANSDYT | 0.7853015 |
| TQRCVSVADLRAAASR | ASSMGGANSDYT | 0.24745864 |
| QFVLDVAEHPVEVLTETV | ASSMGGANSDYT | 0.58774984 |
| GGAGGTGGAGGNGGTQAGDG | ASSMGGANSDYT | 0.81847346 |
| PLLATLRAIAVDVDAKPAQV | ASSMGGANSDYT | 0.5538757 |
| APKFRTGGYMIDFWGVGYQVAK | ASSMGGANSDYT | 0.7463261 |
| GAGGAGGAGGAAGGPGGTGGTG | ASSMGGANSDYT | 0.76612604 |
| IGALGLLATDVAPAGSVAFGVA | ASSMGGANSDYT | 0.2532695 |
| AGGAGGIGGNGGAGGAGGVLMG | ASSMGGANSDYT | 0.7392857 |
| ADELIGLL | ASSMGGANSDYT | 0.83167744 |
| ADRALKA | ASSMGGANSDYT | 0.35518724 |
| AGAVGQVGGAGGA | ASSMGGANSDYT | 0.3967448 |
| AGGGGGLGGAP | ASSMGGANSDYT | 0.2681863 |
| AGGLLYGNGGAG | ASSMGGANSDYT | 0.3865175 |
| AGLESIGL | ASSMGGANSDYT | 0.17481995 |
| AGLKTGII | ASSMGGANSDYT | 0.69627434 |
| ASLAALLL | ASSMGGANSDYT | 0.7583132 |
| ASVINISSI | ASSMGGANSDYT | 0.42002013 |
| ATPPGPA | ASSMGGANSDYT | 0.20064166 |
| ATVASALV | ASSMGGANSDYT | 0.19580755 |
| ATVASLIL | ASSMGGANSDYT | 0.504259 |
| AVNGPVLIRH | ASSMGGANSDYT | 0.28400522 |
| DGAAGAVGI | ASSMGGANSDYT | 0.1665333 |
| DPLLHGI | ASSMGGANSDYT | 0.27960715 |
| ELGAALGAL | ASSMGGANSDYT | 0.740945 |
| ELGRTQAV | ASSMGGANSDYT | 0.74774563 |
| FVAGGYLS | ASSMGGANSDYT | 0.18171634 |
| GASGGAGGAGGW | ASSMGGANSDYT | 0.80928975 |
| GGAGAGGDGGAGGI | ASSMGGANSDYT | 0.8040552 |
| GTGHNGSQPGMGGNG | ASSMGGANSDYT | 0.7525169 |
| GYAAASASAS | ASSMGGANSDYT | 0.45199373 |
| HQVDVAAVQQG | ASSMGGANSDYT | 0.2383508 |
| IGAAIAHT | ASSMGGANSDYT | 0.81726205 |
| IGMIEVAQL | ASSMGGANSDYT | 0.45109698 |
| IRLAPEHTSAMD | ASSMGGANSDYT | 0.4321431 |
| IYNVSVL | ASSMGGANSDYT | 0.5834768 |
| KQGFLGM | ASSMGGANSDYT | 0.5377176 |
| LESVITQL | ASSMGGANSDYT | 0.33592504 |

| | | |
|---|---|---|
| LGLGALVVV | ASSMGGANSDYT | 0.7241141 |
| LIGNGGNGGSGGI | ASSMGGANSDYT | 0.65388644 |
| LNAINAPTL | ASSMGGANSDYT | 0.24644414 |
| LQLLNTGV | ASSMGGANSDYT | 0.23434258 |
| LQTSIEF | ASSMGGANSDYT | 0.27148598 |
| MSFAVSEL | ASSMGGANSDYT | 0.50129557 |
| NTGALGL | ASSMGGANSDYT | 0.7590953 |
| PPGAGGAGGAGTT | ASSMGGANSDYT | 0.47439528 |
| QLVGLGI | ASSMGGANSDYT | 0.481579 |
| RFLEQVHQL | ASSMGGANSDYT | 0.6841411 |
| RGVGYVLR | ASSMGGANSDYT | 0.66427416 |
| SATGVFAGVI | ASSMGGANSDYT | 0.4266361 |
| SLPAITLP | ASSMGGANSDYT | 0.70347846 |
| STALAFTV | ASSMGGANSDYT | 0.38758314 |
| SVLTVGLGK | ASSMGGANSDYT | 0.68289244 |
| TALAGAGSGI | ASSMGGANSDYT | 0.5760163 |
| TGLPFTGI | ASSMGGANSDYT | 0.61645883 |
| VAEHVTV | ASSMGGANSDYT | 0.73309356 |
| VDIVLNSL | ASSMGGANSDYT | 0.67248774 |
| VLTGGGAVL | ASSMGGANSDYT | 0.41282326 |
| VNVGLAYGI | ASSMGGANSDYT | 0.1603534 |
| VSSAAVGAV | ASSMGGANSDYT | 0.37045312 |
| VTSVYEAGV | ASSMGGANSDYT | 0.6040617 |
| WVALDRGA | ASSMGGANSDYT | 0.5915915 |
| YLVGSVVI | ASSMGGANSDYT | 0.67401916 |
| GPPGTGKTLL | ASSMGGANSDYT | 0.6464102 |
| GVGMTKF | ASSMGGANSDYT | 0.32551 |
| IGIGGSDL | ASSMGGANSDYT | 0.7483527 |
| IGTLLAEL | ASSMGGANSDYT | 0.78314614 |
| IYTSGTT | ASSMGGANSDYT | 0.17920995 |
| LSFPAGV | ASSMGGANSDYT | 0.36172155 |
| LYTSGTT | ASSMGGANSDYT | 0.7969318 |
| VGAFPFGF | ASSMGGANSDYT | 0.7599462 |
| VGAGTVEF | ASSMGGANSDYT | 0.3752942 |
| VGKSTLLL | ASSMGGANSDYT | 0.3638785 |
| AALAHAGGGIV | ASSMGGANSDYT | 0.84514374 |
| AALVAVSLL | ASSMGGANSDYT | 0.3373043 |
| AAVSRAVAAL | ASSMGGANSDYT | 0.7907669 |
| AGGSSGIGGF | ASSMGGANSDYT | 0.1344535 |
| AGLTVANANG | ASSMGGANSDYT | 0.32759282 |
| AGLVGAVL | ASSMGGANSDYT | 0.2861218 |
| ALSQAFTV | ASSMGGANSDYT | 0.13617197 |
| ALTFSVSL | ASSMGGANSDYT | 0.71433574 |
| ALTGAALVAL | ASSMGGANSDYT | 0.27387318 |
| ASFSIGAGVL | ASSMGGANSDYT | 0.6713316 |
| DGSVYKI | ASSMGGANSDYT | 0.47686037 |
| EQLAEIV | ASSMGGANSDYT | 0.67188096 |
| FAVSELTVM | ASSMGGANSDYT | 0.5587349 |
| GAAPLAAGL | ASSMGGANSDYT | 0.3878637 |
| GADGGDGGSGGAGG | ASSMGGANSDYT | 0.22683908 |
| GFGLDTGGAGG | ASSMGGANSDYT | 0.65104014 |

| | | |
|---|---|---|
| GLASGGLL | ASSMGGANSDYT | 0.7405391 |
| GRIAGAAVLL | ASSMGGANSDYT | 0.5682559 |
| GSAGSLNTSF | ASSMGGANSDYT | 0.1897253 |
| GTGQIGI | ASSMGGANSDYT | 0.44074583 |
| GTLIALIL | ASSMGGANSDYT | 0.46913734 |
| IAATMGFVVG | ASSMGGANSDYT | 0.18615918 |
| IFAQALL | ASSMGGANSDYT | 0.13223767 |
| IGAAVALGF | ASSMGGANSDYT | 0.32601827 |
| IGLTPLAL | ASSMGGANSDYT | 0.482503 |
| ILGSNTSTL | ASSMGGANSDYT | 0.71394336 |
| IVFSLTVL | ASSMGGANSDYT | 0.7396077 |
| LASKATGF | ASSMGGANSDYT | 0.32141578 |
| LASVALIL | ASSMGGANSDYT | 0.7480586 |
| LFAVGLTL | ASSMGGANSDYT | 0.60502553 |
| LFNGGIGTYI | ASSMGGANSDYT | 0.10892276 |
| LTLGAGII | ASSMGGANSDYT | 0.3881082 |
| LVAAALGDI | ASSMGGANSDYT | 0.17415674 |
| NTAFVQLGI | ASSMGGANSDYT | 0.7080369 |
| QLFSGAEL | ASSMGGANSDYT | 0.21362318 |
| SAATFITL | ASSMGGANSDYT | 0.3776303 |
| SNYGLANLGV | ASSMGGANSDYT | 0.41531825 |
| SYGGAYAVM | ASSMGGANSDYT | 0.78317326 |
| TTGVGGLTLG | ASSMGGANSDYT | 0.61492074 |
| TTQTIYI | ASSMGGANSDYT | 0.27794388 |
| VAAGAGLGAVI | ASSMGGANSDYT | 0.14094277 |
| VAAVVTLL | ASSMGGANSDYT | 0.640271 |
| VGFLGGLI | ASSMGGANSDYT | 0.41866034 |
| VGVIPILL | ASSMGGANSDYT | 0.22814327 |
| VIFASSLI | ASSMGGANSDYT | 0.16889822 |
| VKGGDGGAAAG | ASSMGGANSDYT | 0.6147011 |
| VTKTFTNI | ASSMGGANSDYT | 0.23314306 |
| VVAAAILL | ASSMGGANSDYT | 0.7148955 |
| YAALDVEL | ASSMGGANSDYT | 0.3173852 |
| LASVALLL | ASSMGGANSDYT | 0.44066668 |
| LDAVHRV | ASSMGGANSDYT | 0.82185566 |
| QGLSSSIF | ASSMGGANSDYT | 0.77645445 |
| RGTFANIRL | ASSMGGANSDYT | 0.3654604 |
| RQLWWGHRI | ASSMGGANSDYT | 0.30929106 |
| YGPPGTGKTLL | ASSMGGANSDYT | 0.20790648 |
| AALVGAPANVA | ASSMGGANSDYT | 0.2897462 |
| AATFRRTE | ASSMGGANSDYT | 0.29146826 |
| AGALGGGI | ASSMGGANSDYT | 0.76218 |
| AGGAGGAGGIGAG | ASSMGGANSDYT | 0.3204909 |
| AGSGALGL | ASSMGGANSDYT | 0.20541553 |
| AQIADQI | ASSMGGANSDYT | 0.563076 |
| ASAVAVTYL | ASSMGGANSDYT | 0.7853626 |
| ASTVLGGTGG | ASSMGGANSDYT | 0.8420869 |
| AVGGEPAK | ASSMGGANSDYT | 0.21682975 |
| CEQGTFGGTC | ASSMGGANSDYT | 0.4051713 |
| DEGKAPAG | ASSMGGANSDYT | 0.31693858 |
| DTVVFPAL | ASSMGGANSDYT | 0.5212413 |

| | | |
|---|---|---|
| FEGITMAG | ASSMGGANSDYT | 0.3120173 |
| FPLVVQD | ASSMGGANSDYT | 0.22558603 |
| GAFTTPQ | ASSMGGANSDYT | 0.6607072 |
| GFGGFGTAGGVGGA | ASSMGGANSDYT | 0.86043507 |
| GFLAQFL | ASSMGGANSDYT | 0.3023836 |
| GGAGGAGGNAGVA | ASSMGGANSDYT | 0.25384519 |
| GGVGTAGAGGAGGAGG | ASSMGGANSDYT | 0.411389 |
| GNGGGGDAGNAGSGGN | ASSMGGANSDYT | 0.3055447 |
| GTGGAGGAGNDAGS | ASSMGGANSDYT | 0.3099653 |
| GTTNTGWFNS | ASSMGGANSDYT | 0.5933243 |
| IFMGPLFTL | ASSMGGANSDYT | 0.1629098 |
| ILAHAIEAI | ASSMGGANSDYT | 0.4045887 |
| IRSAIREAL | ASSMGGANSDYT | 0.83734745 |
| ITFAGGG | ASSMGGANSDYT | 0.5331204 |
| ITGMSGHQEV | ASSMGGANSDYT | 0.56006503 |
| IVAGLACTL | ASSMGGANSDYT | 0.77244836 |
| IVGGGLAGA | ASSMGGANSDYT | 0.63441306 |
| IVTGVLNI | ASSMGGANSDYT | 0.7835431 |
| LAQQGCI | ASSMGGANSDYT | 0.8144978 |
| LCLHVEN | ASSMGGANSDYT | 0.31568754 |
| LDAISDGS | ASSMGGANSDYT | 0.54557014 |
| LGGFAAML | ASSMGGANSDYT | 0.75215113 |
| LGGKNPMI | ASSMGGANSDYT | 0.43727145 |
| LGPKGVAAIL | ASSMGGANSDYT | 0.41569197 |
| LMVGVSI | ASSMGGANSDYT | 0.359353 |
| LVGAAFVL | ASSMGGANSDYT | 0.7316164 |
| LVLAVAPL | ASSMGGANSDYT | 0.63515747 |
| MAAIAVIDAA | ASSMGGANSDYT | 0.7588441 |
| NNGINAELQ | ASSMGGANSDYT | 0.16760388 |
| RGEPLVLVHG | ASSMGGANSDYT | 0.24737027 |
| RGGAGGAAGLI | ASSMGGANSDYT | 0.7652611 |
| RSYRMHPAV | ASSMGGANSDYT | 0.12951942 |
| SAAHLLAGQLA | ASSMGGANSDYT | 0.14963293 |
| SAAVGSY | ASSMGGANSDYT | 0.49629584 |
| SPHAVLHRI | ASSMGGANSDYT | 0.81259423 |
| SQIAKVL | ASSMGGANSDYT | 0.13361433 |
| TALGHLVP | ASSMGGANSDYT | 0.27326623 |
| TAPTTSGAA | ASSMGGANSDYT | 0.18288809 |
| TDGDDGNGGSPGT | ASSMGGANSDYT | 0.65336525 |
| TDGLGEN | ASSMGGANSDYT | 0.5428973 |
| TGGPPLPGQA | ASSMGGANSDYT | 0.15649146 |
| TIPAITI | ASSMGGANSDYT | 0.5591991 |
| TIPGFTI | ASSMGGANSDYT | 0.18070453 |
| TSIASGALA | ASSMGGANSDYT | 0.41019136 |
| TVTLPGAAI | ASSMGGANSDYT | 0.7011353 |
| VASGLSAALP | ASSMGGANSDYT | 0.83222675 |
| VGAGVIGI | ASSMGGANSDYT | 0.18921515 |
| VGPKRFGV | ASSMGGANSDYT | 0.72775656 |
| VNVAFPD | ASSMGGANSDYT | 0.37424585 |
| VSMTNSGVS | ASSMGGANSDYT | 0.8474985 |
| VTDVLGDLSV | ASSMGGANSDYT | 0.32258713 |

| | | |
|---|---|---|
| VVVVGAGFAGL | ASSMGGANSDYT | 0.17127079 |
| VYTVGTVV | ASSMGGANSDYT | 0.69341284 |
| WTTFSGP | ASSMGGANSDYT | 0.5100814 |
| YDELVEAF | ASSMGGANSDYT | 0.4112869 |
| YGTGGAGGNGG | ASSMGGANSDYT | 0.78112125 |
| YTNVMAV | ASSMGGANSDYT | 0.3781554 |
| FLEQVHQL | ASSMGGANSDYT | 0.3962348 |
| GLSLLAVAV | ASSMGGANSDYT | 0.7498015 |
| LEQVHQL | ASSMGGANSDYT | 0.66321456 |
| LNAAGAI | ASSMGGANSDYT | 0.25972426 |
| PAVHASL | ASSMGGANSDYT | 0.66216594 |
| TELEDDA | ASSMGGANSDYT | 0.531142 |
| TPPGPPN | ASSMGGANSDYT | 0.2718384 |
| TPPPPPPP | ASSMGGANSDYT | 0.82257533 |
| TQSPALI | ASSSGLGGGETLY | 0.45560688 |
| EFAAAFPQ | ASSSGLGGGETLY | 0.7139112 |
| LLSGNLDVL | ASSSGLGGGETLY | 0.58637226 |
| GPANLGNYNI | ASSSGLGGGETLY | 0.84870225 |
| TLGPTLP | ASSSGLGGGETLY | 0.19424407 |
| VYAYSSCCK | ASSSGLGGGETLY | 0.6797436 |
| APAPAPAPNI | ASSSGLGGGETLY | 0.46703762 |
| PGGPVPIP | ASSSGLGGGETLY | 0.711312 |
| FLTVALEP | ASSSGLGGGETLY | 0.5098167 |
| SQYFGANL | ASSSGLGGGETLY | 0.50892234 |
| GGVKPAA | ASSSGLGGGETLY | 0.46822894 |
| GGTGGAGGAGGW | ASSSGLGGGETLY | 0.7725374 |
| GVGGVAGLGGAGG | ASSSGLGGGETLY | 0.38607994 |
| VLLVLATTI | ASSSGLGGGETLY | 0.62607235 |
| IGIFAAML | ASSSGLGGGETLY | 0.8124606 |
| AVGDLID | ASSSGLGGGETLY | 0.23407586 |
| NISPLPL | ASSSGLGGGETLY | 0.44604313 |
| VTAVFVAI | ASSSGLGGGETLY | 0.6511858 |
| SGDDNDAAL | ASSSGLGGGETLY | 0.76260656 |
| AVAALAHS | ASSSGLGGGETLY | 0.37834644 |
| GGVTVMATP | ASSSGLGGGETLY | 0.74296355 |
| AGLAANVRD | ASSSGLGGGETLY | 0.38459945 |
| TNRPDLIDPALLRPG | ASSSGLGGGETLY | 0.6482301 |
| LANLGVTAL | ASSSGLGGGETLY | 0.44332346 |
| TGERIEL | ASSSGLGGGETLY | 0.5982262 |
| NGEELAA | ASSSGLGGGETLY | 0.28191262 |
| VLSLVWG | ASSSGLGGGETLY | 0.679019 |
| IGYALWL | ASSSGLGGGETLY | 0.7662612 |
| LSLLLAPL | ASSSGLGGGETLY | 0.70598924 |
| TLDEALK | ASSSGLGGGETLY | 0.13974002 |
| TDGAGTAEGG | ASSSGLGGGETLY | 0.7197868 |
| NAPGGAGGNGG | ASSSGLGGGETLY | 0.71555746 |
| VSVFMGMGGNF | ASSSGLGGGETLY | 0.84786975 |
| LASYLAAL | ASSSGLGGGETLY | 0.55071 |
| LVGGTPAL | ASSSGLGGGETLY | 0.56510043 |
| VRNADVL | ASSSGLGGGETLY | 0.42995375 |
| TCSSPFLVSAM | ASSSGLGGGETLY | 0.48086888 |

| | | |
|---|---|---|
| VHITDAE | ASSSGLGGGETLY | 0.22715029 |
| EYAAPSF | ASSSGLGGGETLY | 0.47394463 |
| VVDAFHGGL | ASSSGLGGGETLY | 0.47525388 |
| ASVVDPQ | ASSSGLGGGETLY | 0.2692625 |
| SYNEFGGLS | ASSSGLGGGETLY | 0.8134519 |
| DPDGSVY | ASSSGLGGGETLY | 0.7800982 |
| MVSTIHATE | ASSSGLGGGETLY | 0.77670884 |
| LAAGVADPV | ASSSGLGGGETLY | 0.78454345 |
| TKEQIAA | ASSSGLGGGETLY | 0.8013559 |
| SLASPAAL | ASSSGLGGGETLY | 0.5879689 |
| IGLVFYGV | ASSSGLGGGETLY | 0.37492898 |
| FLLDASNL | ASSSGLGGGETLY | 0.44943726 |
| YQVGNFP | ASSSGLGGGETLY | 0.1999866 |
| ESLLSQST | ASSSGLGGGETLY | 0.3824997 |
| LFTVGFAAVG | ASSSGLGGGETLY | 0.41658342 |
| ATPPTPT | ASSSGLGGGETLY | 0.50971997 |
| TDLATLG | ASSSGLGGGETLY | 0.1239138 |
| SAGGTGR | ASSSGLGGGETLY | 0.75802517 |
| GFSNSGS | ASSSGLGGGETLY | 0.77435875 |
| GGNGGSGGTSV | ASSSGLGGGETLY | 0.48957416 |
| GTSVATGGAGN | ASSSGLGGGETLY | 0.61487365 |
| NGGNGGSGGTS | ASSSGLGGGETLY | 0.6464275 |
| AVVKFPH | ASSSGLGGGETLY | 0.3038932 |
| TPPPPPP | ASSSGLGGGETLY | 0.72550327 |
| AAPAAAAAPA | ASSSGLGGGETLY | 0.25627878 |
| IALNEDL | ASSSGLGGGETLY | 0.6528408 |
| IVNVSSM | ASSSGLGGGETLY | 0.8025174 |
| TGIPPAPR | ASSSGLGGGETLY | 0.4840867 |
| FDLGGGTF | ASSSGLGGGETLY | 0.5467398 |
| FDLGGGTFDVS | ASSSGLGGGETLY | 0.42349848 |
| NEPTAAA | ASSSGLGGGETLY | 0.18450391 |
| NEPTAAALA | ASSSGLGGGETLY | 0.56985897 |
| GSIINISSI | ASSSGLGGGETLY | 0.17080808 |
| VAPGNAGTA | ASSSGLGGGETLY | 0.56652987 |
| TGGGASLE | ASSSGLGGGETLY | 0.21783389 |
| VAVYNHYKRI | ASSSGLGGGETLY | 0.5505025 |
| TDPAPATT | ASSSGLGGGETLY | 0.5304401 |
| TNRPDILDPALLRPG | ASSSGLGGGETLY | 0.54436815 |
| LTTTTTS | ASSSGLGGGETLY | 0.2512617 |
| PTGSGKTL | ASSSGLGGGETLY | 0.5325544 |
| TGSGKTL | ASSSGLGGGETLY | 0.41705036 |
| TIGHVDHGK | ASSSGLGGGETLY | 0.2135411 |
| TIGHVDHGKTT | ASSSGLGGGETLY | 0.69774693 |
| GHVDHGKTT | ASSSGLGGGETLY | 0.36126843 |
| AQFEHTLL | ASSSGLGGGETLY | 0.5459466 |
| IVNQDEL | ASSSGLGGGETLY | 0.75817025 |
| TNGSQFF | ASSSGLGGGETLY | 0.5211705 |
| TGKGIVE | ASSSGLGGGETLY | 0.19618864 |
| LISNASDA | ASSSGLGGGETLY | 0.53299534 |
| AELRTLGELTKIAKAHGAQ | ASSSGLGGGETLY | 0.42504483 |
| APNAVAAAEPSW | ASSSGLGGGETLY | 0.34293503 |

| | | |
|---|---|---|
| APQVAGAATAVL | ASSSGLGGGETLY | 0.12963264 |
| ARRDPAAAAAILPTDA | ASSSGLGGGETLY | 0.48015898 |
| DAAARDVLAEAGLAEYF | ASSSGLGGGETLY | 0.27675343 |
| DDQIVLSAKANLAKVEQ | ASSSGLGGGETLY | 0.30982736 |
| DELLPDTGEDSDEDGH | ASSSGLGGGETLY | 0.28983027 |
| ELPEAPANRATVPQD | ASSSGLGGGETLY | 0.71894646 |
| FGQSKIERVTGIG | ASSSGLGGGETLY | 0.6554987 |
| GAGGTGSTAGGAGGAGGAGGLY | ASSSGLGGGETLY | 0.56018734 |
| GHGGTGAAGGNGGTGGAGGSGI | ASSSGLGGGETLY | 0.7256137 |
| GNGGNGGAGGEGGPGGAGGAGA | ASSSGLGGGETLY | 0.6165987 |
| GSTAGNGGNGGNATNGTIAGQP | ASSSGLGGGETLY | 0.788225 |
| GTGGDGGNGGNGGTGG | ASSSGLGGGETLY | 0.61260706 |
| GVRRPAGIPADAPV | ASSSGLGGGETLY | 0.5019134 |
| GYRQVTDAHLLTLARR | ASSSGLGGGETLY | 0.47152567 |
| IAKMGQDGHDRG | ASSSGLGGGETLY | 0.51412135 |
| KAEPSIEIQYT | ASSSGLGGGETLY | 0.15461983 |
| KPVRIRMAADEDTVAVT | ASSSGLGGGETLY | 0.17105769 |
| MPPRIAGMRLLVIKPEP | ASSSGLGGGETLY | 0.45503774 |
| MKFYAVALFRSRGVPPTR | ASSSGLGGGETLY | 0.14622702 |
| NAGGFGILGAVAHSP | ASSSGLGGGETLY | 0.5166038 |
| NEGREITLAMAEM | ASSSGLGGGETLY | 0.7200442 |
| NELVEWSHPLGHRWRWE | ASSSGLGGGETLY | 0.51058227 |
| NLGAGGTGGNGGNPG | ASSSGLGGGETLY | 0.19919822 |
| NTEWPKWETVMAADPVD | ASSSGLGGGETLY | 0.33862486 |
| PASAAPAETQP | ASSSGLGGGETLY | 0.3215562 |
| QQSSDGKNGVRAAEITGE | ASSSGLGGGETLY | 0.7160362 |
| RAPDAVTTSWWKE | ASSSGLGGGETLY | 0.64482 |
| SGFGNVGAHTSGWFNQ | ASSSGLGGGETLY | 0.8441334 |
| SGVAGAVGQAASVGGLKV | ASSSGLGGGETLY | 0.76823604 |
| SNGGAGTGSAGGNGGT | ASSSGLGGGETLY | 0.37942326 |
| TGGAGGEGGAAG | ASSSGLGGGETLY | 0.65813994 |
| TGGDGGAAGAGGAPGHG | ASSSGLGGGETLY | 0.35182434 |
| TMPPLLAAIANAGRHGV | ASSSGLGGGETLY | 0.4942515 |
| TSSTAGNSGGA | ASSSGLGGGETLY | 0.35149857 |
| VPATVPVGILC | ASSSGLGGGETLY | 0.5876899 |
| VQRPDSGGVVLTGRIS | ASSSGLGGGETLY | 0.40641534 |
| WTPNPVPATKGVPAGLVQAT | ASSSGLGGGETLY | 0.26439166 |
| YALKRALAKFSDRFDVVII | ASSSGLGGGETLY | 0.19068629 |
| YLIGSVIV | ASSSGLGGGETLY | 0.5526907 |
| YGQLVAVID | ASSSGLGGGETLY | 0.33932608 |
| VTVGGMTLPA | ASSSGLGGGETLY | 0.19998801 |
| VLALPAV | ASSSGLGGGETLY | 0.30143526 |
| VIPPTLNYET | ASSSGLGGGETLY | 0.83265764 |
| VGMGGMHPSSG | ASSSGLGGGETLY | 0.827705 |
| VGLAGGAATA | ASSSGLGGGETLY | 0.29636204 |
| VGGAGGAGGSGP | ASSSGLGGGETLY | 0.29958674 |
| VFAAFMV | ASSSGLGGGETLY | 0.7792827 |
| TLYDKLR | ASSSGLGGGETLY | 0.18742207 |
| TLAVFTVL | ASSSGLGGGETLY | 0.4984799 |
| TGSVQPARV | ASSSGLGGGETLY | 0.65304846 |
| TGGGAGSGGT | ASSSGLGGGETLY | 0.45499977 |

| | | |
|---|---|---|
| SHLETSL | ASSSGLGGGETLY | 0.23260668 |
| SFTTGMT | ASSSGLGGGETLY | 0.59245837 |
| QVEVAKRMGVRQPTVS | ASSSGLGGGETLY | 0.4404076 |
| QGSGGPALAM | ASSSGLGGGETLY | 0.6498759 |
| MLAPSNP | ASSSGLGGGETLY | 0.20032124 |
| LVNNAAI | ASSSGLGGGETLY | 0.64802647 |
| LTGEINTLA | ASSSGLGGGETLY | 0.26084056 |
| LSIFGPV | ASSSGLGGGETLY | 0.3839004 |
| LLALLANL | ASSSGLGGGETLY | 0.1538843 |
| LISNASDALDKLR | ASSSGLGGGETLY | 0.4032852 |
| LINEAVP | ASSSGLGGGETLY | 0.782935 |
| LGYAFSGL | ASSSGLGGGETLY | 0.7745266 |
| LDFVAVAL | ASSSGLGGGETLY | 0.23610301 |
| LAQPGAE | ASSSGLGGGETLY | 0.714295 |
| ISNLGAVTGF | ASSSGLGGGETLY | 0.8330946 |
| HQEDPGRI | ASSSGLGGGETLY | 0.42689407 |
| GTPGEPTP | ASSSGLGGGETLY | 0.15899415 |
| GFFNSGS | ASSSGLGGGETLY | 0.6354195 |
| GAGGGGLTAGAN | ASSSGLGGGETLY | 0.6450895 |
| FVDALVNM | ASSSGLGGGETLY | 0.74118304 |
| FLTVAPEL | ASSSGLGGGETLY | 0.78188443 |
| FETLTAL | ASSSGLGGGETLY | 0.18140154 |
| DVINPIEL | ASSSGLGGGETLY | 0.69025373 |
| APVSGAPV | ASSSGLGGGETLY | 0.65908694 |
| ALVGGLAW | ASSSGLGGGETLY | 0.21700364 |
| ALAAAAATAI | ASSSGLGGGETLY | 0.2927312 |
| AFGLGLLL | ASSSGLGGGETLY | 0.271441 |
| AFFAAASL | ASSSGLGGGETLY | 0.1713742 |
| AAAALPAFSPP | ASSSGLGGGETLY | 0.61729777 |
| VVLAPAL | ASSSGLGGGETLY | 0.37584287 |
| TIGHVDHGKTTL | ASSSGLGGGETLY | 0.41490072 |
| LISNASDALDK | ASSSGLGGGETLY | 0.35558674 |
| LISNASDAL | ASSSGLGGGETLY | 0.31806564 |
| ISNASDALDK | ASSSGLGGGETLY | 0.32166398 |
| NQAETLVYQTE | ASSSGLGGGETLY | 0.3496763 |
| TGGTGGTAGAG | ASSSGLGGGETLY | 0.57423913 |
| TGSTAGGAGGA | ASSSGLGGGETLY | 0.6618358 |
| DPLAVPVAAGPV | ASSSGLGGGETLY | 0.5882292 |
| GGTGGTPGNGGN | ASSSGLGGGETLY | 0.59390086 |
| PAGLAGVDPATL | ASSSGLGGGETLY | 0.79893816 |
| GNNLVVEAPHAGASV | ASSSGLGGGETLY | 0.24351646 |
| KVTRIMSASTLEQPA | ASSSGLGGGETLY | 0.22933576 |
| LGFALPIALPRRTVV | ASSSGLGGGETLY | 0.683615 |
| TRLGNLTARKLVVLV | ASSSGLGGGETLY | 0.22877079 |
| WSQQRRLIYFPSAGP | ASSSGLGGGETLY | 0.7439482 |
| GPNGNPDPMAAAVDIR | ASSSGLGGGETLY | 0.5378461 |
| TQRCVSVADLRAAASR | ASSSGLGGGETLY | 0.5754074 |
| QFVLDVAEHPVEVLTETV | ASSSGLGGGETLY | 0.7236101 |
| GGAGGTGGAGGNGGTQAGDG | ASSSGLGGGETLY | 0.82451606 |
| PLLATLRAIAVDVDAKPAQV | ASSSGLGGGETLY | 0.65232027 |
| APKFRTGGYMIDFWGVGYQVAK | ASSSGLGGGETLY | 0.33521754 |

| | | |
|---|---|---|
| GAGGAGGAGGAAGGPGGTGGTG | ASSSGLGGGETLY | 0.7655933 |
| IGALGLLATDVAPAGSVAFGVA | ASSSGLGGGETLY | 0.23012766 |
| AGGAGGIGGNGGAGGAGGVLMG | ASSSGLGGGETLY | 0.6686479 |
| ADELIGLL | ASSSGLGGGETLY | 0.6629985 |
| ADRALKA | ASSSGLGGGETLY | 0.24764368 |
| AGAVGQVGGAGGA | ASSSGLGGGETLY | 0.41045853 |
| AGGGGGLGGAP | ASSSGLGGGETLY | 0.72908896 |
| AGGLLYGNGGAG | ASSSGLGGGETLY | 0.7389493 |
| AGLESIGL | ASSSGLGGGETLY | 0.16873725 |
| AGLKTGII | ASSSGLGGGETLY | 0.29297447 |
| ASLAALLL | ASSSGLGGGETLY | 0.21364474 |
| ASVINISSI | ASSSGLGGGETLY | 0.42649424 |
| ATPPGPA | ASSSGLGGGETLY | 0.19641134 |
| ATVASALV | ASSSGLGGGETLY | 0.6813195 |
| ATVASLIL | ASSSGLGGGETLY | 0.35774726 |
| AVNGPVLIRH | ASSSGLGGGETLY | 0.8074659 |
| DGAAGAVGI | ASSSGLGGGETLY | 0.520419 |
| DPLLHGI | ASSSGLGGGETLY | 0.7429166 |
| ELGAALGAL | ASSSGLGGGETLY | 0.28872097 |
| ELGRTQAV | ASSSGLGGGETLY | 0.42043936 |
| FVAGGYLS | ASSSGLGGGETLY | 0.16836712 |
| GASGGAGGAGGW | ASSSGLGGGETLY | 0.2656211 |
| GGAGAGGDGGAGGI | ASSSGLGGGETLY | 0.24862817 |
| GTGHNGSQPGMGGNG | ASSSGLGGGETLY | 0.45646983 |
| GYAAASASAS | ASSSGLGGGETLY | 0.23214391 |
| HQVDVAAVQQG | ASSSGLGGGETLY | 0.7913933 |
| IGAAIAHT | ASSSGLGGGETLY | 0.47730517 |
| IGMIEVAQL | ASSSGLGGGETLY | 0.6647544 |
| IRLAPEHTSAMD | ASSSGLGGGETLY | 0.3754846 |
| IYNVSVL | ASSSGLGGGETLY | 0.30416656 |
| KQGFLGM | ASSSGLGGGETLY | 0.3738382 |
| LESVITQL | ASSSGLGGGETLY | 0.72496 |
| LGLGALVVV | ASSSGLGGGETLY | 0.18996744 |
| LIGNGGNGGSGGI | ASSSGLGGGETLY | 0.15859018 |
| LNAINAPTL | ASSSGLGGGETLY | 0.40676332 |
| LQLLNTGV | ASSSGLGGGETLY | 0.2618195 |
| LQTSIEF | ASSSGLGGGETLY | 0.8025911 |
| MSFAVSEL | ASSSGLGGGETLY | 0.7458065 |
| NTGALGL | ASSSGLGGGETLY | 0.5228897 |
| PPGAGGAGGAGTT | ASSSGLGGGETLY | 0.7475339 |
| QLVGLGI | ASSSGLGGGETLY | 0.13401915 |
| RFLEQVHQL | ASSSGLGGGETLY | 0.42384234 |
| RGVGYVLR | ASSSGLGGGETLY | 0.8151529 |
| SATGVFAGVI | ASSSGLGGGETLY | 0.64653325 |
| SLPAITLP | ASSSGLGGGETLY | 0.39785686 |
| STALAFTV | ASSSGLGGGETLY | 0.76456994 |
| SVLTVGLGK | ASSSGLGGGETLY | 0.30974573 |
| TALAGAGSGI | ASSSGLGGGETLY | 0.35331813 |
| TGLPFTGI | ASSSGLGGGETLY | 0.5454288 |
| VAEHVTV | ASSSGLGGGETLY | 0.42070568 |
| VDIVLNSL | ASSSGLGGGETLY | 0.58977497 |

| | | |
|---|---|---|
| VLTGGGAVL | ASSSGLGGGETLY | 0.19421306 |
| VNVGLAYGI | ASSSGLGGGETLY | 0.5235982 |
| VSSAAVGAV | ASSSGLGGGETLY | 0.84370387 |
| VTSVYEAGV | ASSSGLGGGETLY | 0.42921484 |
| WVALDRGA | ASSSGLGGGETLY | 0.7663451 |
| YLVGSVVI | ASSSGLGGGETLY | 0.61314553 |
| GPPGTGKTLL | ASSSGLGGGETLY | 0.79800755 |
| GVGMTKF | ASSSGLGGGETLY | 0.26377565 |
| IGIGGSDL | ASSSGLGGGETLY | 0.24002022 |
| IGTLLAEL | ASSSGLGGGETLY | 0.5289414 |
| IYTSGTT | ASSSGLGGGETLY | 0.24381836 |
| LSFPAGV | ASSSGLGGGETLY | 0.47590828 |
| LYTSGTT | ASSSGLGGGETLY | 0.3061695 |
| VGAFPFGF | ASSSGLGGGETLY | 0.8544219 |
| VGAGTVEF | ASSSGLGGGETLY | 0.6983942 |
| VGKSTLLL | ASSSGLGGGETLY | 0.6460395 |
| AALAHAGGGIV | ASSSGLGGGETLY | 0.26468801 |
| AALVAVSLL | ASSSGLGGGETLY | 0.43471557 |
| AAVSRAVAAL | ASSSGLGGGETLY | 0.7011554 |
| AGGSSGIGGF | ASSSGLGGGETLY | 0.1912639 |
| AGLTVANANG | ASSSGLGGGETLY | 0.77929103 |
| AGLVGAVL | ASSSGLGGGETLY | 0.27646303 |
| ALSQAFTV | ASSSGLGGGETLY | 0.47406307 |
| ALTFSVSL | ASSSGLGGGETLY | 0.6385492 |
| ALTGAALVAL | ASSSGLGGGETLY | 0.8354677 |
| ASFSIGAGVL | ASSSGLGGGETLY | 0.323041 |
| DGSVYKI | ASSSGLGGGETLY | 0.48750138 |
| EQLAEIV | ASSSGLGGGETLY | 0.22348037 |
| FAVSELTVM | ASSSGLGGGETLY | 0.71495485 |
| GAAPLAAGL | ASSSGLGGGETLY | 0.7698896 |
| GADGGDGGSGGAGG | ASSSGLGGGETLY | 0.7391318 |
| GFGLDTGGAGG | ASSSGLGGGETLY | 0.8259698 |
| GLASGGLL | ASSSGLGGGETLY | 0.62780595 |
| GRIAGAAVLL | ASSSGLGGGETLY | 0.18577409 |
| GSAGSLNTSF | ASSSGLGGGETLY | 0.33759314 |
| GTGQIGI | ASSSGLGGGETLY | 0.7150688 |
| GTLIALIL | ASSSGLGGGETLY | 0.79720616 |
| IAATMGFVVG | ASSSGLGGGETLY | 0.78065556 |
| IFAQALL | ASSSGLGGGETLY | 0.6863011 |
| IGAAVALGF | ASSSGLGGGETLY | 0.3625143 |
| IGLTPLAL | ASSSGLGGGETLY | 0.7881695 |
| ILGSNTSTL | ASSSGLGGGETLY | 0.84086114 |
| IVFSLTVL | ASSSGLGGGETLY | 0.72094977 |
| LASKATGF | ASSSGLGGGETLY | 0.35878736 |
| LASVALIL | ASSSGLGGGETLY | 0.33837065 |
| LFAVGLTL | ASSSGLGGGETLY | 0.45402795 |
| LFNGGIGTYI | ASSSGLGGGETLY | 0.75433564 |
| LTLGAGII | ASSSGLGGGETLY | 0.22927001 |
| LVAAALGDI | ASSSGLGGGETLY | 0.5335348 |
| NTAFVQLGI | ASSSGLGGGETLY | 0.5634589 |
| QLFSGAEL | ASSSGLGGGETLY | 0.17136857 |

| | | |
|---|---|---|
| SAATFITL | ASSSGLGGGETLY | 0.6363951 |
| SNYGLANLGV | ASSSGLGGGETLY | 0.8073418 |
| SYGGAYAVM | ASSSGLGGGETLY | 0.35998648 |
| TTGVGGLTLG | ASSSGLGGGETLY | 0.2594455 |
| TTQTIYI | ASSSGLGGGETLY | 0.7905307 |
| VAAGAGLGAVI | ASSSGLGGGETLY | 0.84467506 |
| VAAVVTLL | ASSSGLGGGETLY | 0.66831714 |
| VGFLGGLI | ASSSGLGGGETLY | 0.19110438 |
| VGVIPILL | ASSSGLGGGETLY | 0.13764854 |
| VIFASSLI | ASSSGLGGGETLY | 0.32669193 |
| VKGGDGGAAAG | ASSSGLGGGETLY | 0.80780554 |
| VTKTFTNI | ASSSGLGGGETLY | 0.42143303 |
| VVAAAILL | ASSSGLGGGETLY | 0.33265164 |
| YAALDVEL | ASSSGLGGGETLY | 0.6732932 |
| LASVALLL | ASSSGLGGGETLY | 0.75218034 |
| LDAVHRV | ASSSGLGGGETLY | 0.22757602 |
| QGLSSSIF | ASSSGLGGGETLY | 0.23297992 |
| RGTFANIRL | ASSSGLGGGETLY | 0.68628854 |
| RQLWWGHRI | ASSSGLGGGETLY | 0.69250333 |
| YGPPGTGKTLL | ASSSGLGGGETLY | 0.23846196 |
| AALVGAPANVA | ASSSGLGGGETLY | 0.39951718 |
| AATFRRTE | ASSSGLGGGETLY | 0.4278474 |
| AGALGGGI | ASSSGLGGGETLY | 0.5469916 |
| AGGAGGAGGIGAG | ASSSGLGGGETLY | 0.6623799 |
| AGSGALGL | ASSSGLGGGETLY | 0.22048256 |
| AQIADQI | ASSSGLGGGETLY | 0.78490394 |
| ASAVAVTYL | ASSSGLGGGETLY | 0.19756365 |
| ASTVLGGTGG | ASSSGLGGGETLY | 0.6843826 |
| AVGGEPAK | ASSSGLGGGETLY | 0.5045296 |
| CEQGTFGGTC | ASSSGLGGGETLY | 0.46791244 |
| DEGKAPAG | ASSSGLGGGETLY | 0.3593869 |
| DTVVFPAL | ASSSGLGGGETLY | 0.33667958 |
| FEGITMAG | ASSSGLGGGETLY | 0.20783775 |
| FPLVVQD | ASSSGLGGGETLY | 0.17701817 |
| GAFTTPQ | ASSSGLGGGETLY | 0.7742346 |
| GFGGFGTAGGVGGA | ASSSGLGGGETLY | 0.5653391 |
| GFLAQFL | ASSSGLGGGETLY | 0.690907 |
| GGAGGAGGNAGVA | ASSSGLGGGETLY | 0.614076 |
| GGVGTAGAGGAGGAGG | ASSSGLGGGETLY | 0.20971298 |
| GNGGGGDAGNAGSGGN | ASSSGLGGGETLY | 0.5712358 |
| GTGGAGGAGNDAGS | ASSSGLGGGETLY | 0.28055814 |
| GTTNTGWFNS | ASSSGLGGGETLY | 0.24677718 |
| IFMGPLFTL | ASSSGLGGGETLY | 0.72167563 |
| ILAHAIEAI | ASSSGLGGGETLY | 0.5771067 |
| IRSAIREAL | ASSSGLGGGETLY | 0.26672637 |
| ITFAGGG | ASSSGLGGGETLY | 0.5159839 |
| ITGMSGHQEV | ASSSGLGGGETLY | 0.5309453 |
| IVAGLACTL | ASSSGLGGGETLY | 0.2611604 |
| IVGGGLAGA | ASSSGLGGGETLY | 0.3149311 |
| IVTGVLNI | ASSSGLGGGETLY | 0.773239 |
| LAQQGCI | ASSSGLGGGETLY | 0.4399871 |

| | | |
|---|---|---|
| LCLHVEN | ASSSGLGGGETLY | 0.1864139 |
| LDAISDGS | ASSSGLGGGETLY | 0.7243373 |
| LGGFAAML | ASSSGLGGGETLY | 0.26196712 |
| LGGKNPMI | ASSSGLGGGETLY | 0.595278 |
| LGPKGVAAIL | ASSSGLGGGETLY | 0.40340692 |
| LMVGVSI | ASSSGLGGGETLY | 0.3719133 |
| LVGAAFVL | ASSSGLGGGETLY | 0.6629696 |
| LVLAVAPL | ASSSGLGGGETLY | 0.33374894 |
| MAAIAVIDAA | ASSSGLGGGETLY | 0.32889426 |
| NNGINAELQ | ASSSGLGGGETLY | 0.33696973 |
| RGEPLVLVHG | ASSSGLGGGETLY | 0.49314833 |
| RGGAGGAAGLI | ASSSGLGGGETLY | 0.71223044 |
| RSYRMHPAV | ASSSGLGGGETLY | 0.25135103 |
| SAAHLLAGQLA | ASSSGLGGGETLY | 0.50337696 |
| SAAVGSY | ASSSGLGGGETLY | 0.30853015 |
| SPHAVLHRI | ASSSGLGGGETLY | 0.23719841 |
| SQIAKVL | ASSSGLGGGETLY | 0.36426893 |
| TALGHLVP | ASSSGLGGGETLY | 0.39854527 |
| TAPTTSGAA | ASSSGLGGGETLY | 0.24121282 |
| TDGDDGNGGSPGT | ASSSGLGGGETLY | 0.46517232 |
| TDGLGEN | ASSSGLGGGETLY | 0.2916239 |
| TGGPPLPGQA | ASSSGLGGGETLY | 0.6009241 |
| TIPAITI | ASSSGLGGGETLY | 0.74466515 |
| TIPGFTI | ASSSGLGGGETLY | 0.2969849 |
| TSIASGALA | ASSSGLGGGETLY | 0.631583 |
| TVTLPGAAI | ASSSGLGGGETLY | 0.65796274 |
| VASGLSAALP | ASSSGLGGGETLY | 0.63001287 |
| VGAGVIGI | ASSSGLGGGETLY | 0.38902104 |
| VGPKRFGV | ASSSGLGGGETLY | 0.25022945 |
| VNVAFPD | ASSSGLGGGETLY | 0.75733054 |
| VSMTNSGVS | ASSSGLGGGETLY | 0.68950737 |
| VTDVLGDLSV | ASSSGLGGGETLY | 0.68947935 |
| VVVVGAGFAGL | ASSSGLGGGETLY | 0.5993943 |
| VYTVGTVV | ASSSGLGGGETLY | 0.7632017 |
| WTTFSGP | ASSSGLGGGETLY | 0.63275194 |
| YDELVEAF | ASSSGLGGGETLY | 0.4068989 |
| YGTGGAGGNGG | ASSSGLGGGETLY | 0.16953477 |
| YTNVMAV | ASSSGLGGGETLY | 0.43827972 |
| FLEQVHQL | ASSSGLGGGETLY | 0.35366708 |
| GLSLLAVAV | ASSSGLGGGETLY | 0.30472094 |
| LEQVHQL | ASSSGLGGGETLY | 0.18659258 |
| LNAAGAI | ASSSGLGGGETLY | 0.7698456 |
| PAVHASL | ASSSGLGGGETLY | 0.8485109 |
| TELEDDA | ASSSGLGGGETLY | 0.3874362 |
| TPPGPPN | ASSSGLGGGETLY | 0.8302666 |
| TPPPPPPP | ASSSGLGGGETLY | 0.19403829 |
| TQSPALI | ASSLDGGENTGQLY | 0.72042245 |
| EFAAAFPQ | ASSLDGGENTGQLY | 0.7733123 |
| LLSGNLDVL | ASSLDGGENTGQLY | 0.29030395 |
| GPANLGNYNI | ASSLDGGENTGQLY | 0.62682307 |
| TLGPTLP | ASSLDGGENTGQLY | 0.2271649 |

| | | |
|---|---|---|
| VYAYSSCCK | ASSLDGGENTGQLY | 0.31895807 |
| APAPAPAPNI | ASSLDGGENTGQLY | 0.73726344 |
| PGGPVPIP | ASSLDGGENTGQLY | 0.41212058 |
| FLTVALEP | ASSLDGGENTGQLY | 0.42998683 |
| SQYFGANL | ASSLDGGENTGQLY | 0.39556348 |
| GGVKPAA | ASSLDGGENTGQLY | 0.2546131 |
| GGTGGAGGAGGW | ASSLDGGENTGQLY | 0.37519276 |
| GVGGVAGLGGAGG | ASSLDGGENTGQLY | 0.6985321 |
| VLLVLATTI | ASSLDGGENTGQLY | 0.37968183 |
| IGIFAAML | ASSLDGGENTGQLY | 0.8347114 |
| AVGDLID | ASSLDGGENTGQLY | 0.25790283 |
| NISPLPL | ASSLDGGENTGQLY | 0.8245354 |
| VTAVFVAI | ASSLDGGENTGQLY | 0.8048773 |
| SGDDNDAAL | ASSLDGGENTGQLY | 0.57368827 |
| AVAALAHS | ASSLDGGENTGQLY | 0.35948765 |
| GGVTVMATP | ASSLDGGENTGQLY | 0.79475534 |
| AGLAANVRD | ASSLDGGENTGQLY | 0.41544652 |
| TNRPDLIDPALLRPG | ASSLDGGENTGQLY | 0.6915729 |
| LANLGVTAL | ASSLDGGENTGQLY | 0.68645394 |
| TGERIEL | ASSLDGGENTGQLY | 0.16183971 |
| NGEELAA | ASSLDGGENTGQLY | 0.417988 |
| VLSLVWG | ASSLDGGENTGQLY | 0.21067518 |
| IGYALWL | ASSLDGGENTGQLY | 0.29552495 |
| LSLLLAPL | ASSLDGGENTGQLY | 0.18477331 |
| TLDEALK | ASSLDGGENTGQLY | 0.12803282 |
| TDGAGTAEGG | ASSLDGGENTGQLY | 0.2223175 |
| NAPGGAGGNGG | ASSLDGGENTGQLY | 0.14932953 |
| VSVFMGMGGNF | ASSLDGGENTGQLY | 0.16435179 |
| LASYLAAL | ASSLDGGENTGQLY | 0.26729664 |
| LVGGTPAL | ASSLDGGENTGQLY | 0.21322429 |
| VRNADVL | ASSLDGGENTGQLY | 0.61679304 |
| TCSSPFLVSAM | ASSLDGGENTGQLY | 0.4770139 |
| VHITDAE | ASSLDGGENTGQLY | 0.13772754 |
| EYAAPSF | ASSLDGGENTGQLY | 0.24779232 |
| VVDAFHGGL | ASSLDGGENTGQLY | 0.1567648 |
| ASVVDPQ | ASSLDGGENTGQLY | 0.21921478 |
| SYNEFGGLS | ASSLDGGENTGQLY | 0.11541279 |
| DPDGSVY | ASSLDGGENTGQLY | 0.32950547 |
| MVSTIHATE | ASSLDGGENTGQLY | 0.16478291 |
| LAAGVADPV | ASSLDGGENTGQLY | 0.11307197 |
| TKEQIAA | ASSLDGGENTGQLY | 0.2340273 |
| SLASPAAL | ASSLDGGENTGQLY | 0.20101358 |
| IGLVFYGV | ASSLDGGENTGQLY | 0.19739142 |
| FLLDASNL | ASSLDGGENTGQLY | 0.60893416 |
| YQVGNFP | ASSLDGGENTGQLY | 0.11634115 |
| ESLLSQST | ASSLDGGENTGQLY | 0.16285485 |
| LFTVGFAAVG | ASSLDGGENTGQLY | 0.4547633 |
| ATPPTPT | ASSLDGGENTGQLY | 0.5966752 |
| TDLATLG | ASSLDGGENTGQLY | 0.15477328 |
| SAGGTGR | ASSLDGGENTGQLY | 0.41532597 |
| GFSNSGS | ASSLDGGENTGQLY | 0.6915924 |

| | | |
|---|---|---|
| GGNGGSGGTSV | ASSLDGGENTGQLY | 0.27959326 |
| GTSVATGGAGN | ASSLDGGENTGQLY | 0.34942663 |
| NGGNGGSGGTS | ASSLDGGENTGQLY | 0.44565272 |
| AVVKFPH | ASSLDGGENTGQLY | 0.14941192 |
| TPPPPPP | ASSLDGGENTGQLY | 0.4936325 |
| AAPAAAAAPA | ASSLDGGENTGQLY | 0.4547312 |
| IALNEDL | ASSLDGGENTGQLY | 0.2400702 |
| IVNVSSM | ASSLDGGENTGQLY | 0.23851398 |
| TGIPPAPR | ASSLDGGENTGQLY | 0.172865 |
| FDLGGGTF | ASSLDGGENTGQLY | 0.21723855 |
| FDLGGGTFDVS | ASSLDGGENTGQLY | 0.115771726 |
| NEPTAAA | ASSLDGGENTGQLY | 0.4118225 |
| NEPTAAALA | ASSLDGGENTGQLY | 0.45983028 |
| GSIINISSI | ASSLDGGENTGQLY | 0.25272962 |
| VAPGNAGTA | ASSLDGGENTGQLY | 0.13645302 |
| TGGGASLE | ASSLDGGENTGQLY | 0.17250054 |
| VAVYNHYKRI | ASSLDGGENTGQLY | 0.1511474 |
| TDPAPATT | ASSLDGGENTGQLY | 0.2390158 |
| TNRPDILDPALLRPG | ASSLDGGENTGQLY | 0.12829123 |
| LTTTTTS | ASSLDGGENTGQLY | 0.12372841 |
| PTGSGKTL | ASSLDGGENTGQLY | 0.47948876 |
| TGSGKTL | ASSLDGGENTGQLY | 0.517029 |
| TIGHVDHGK | ASSLDGGENTGQLY | 0.13387056 |
| TIGHVDHGKTT | ASSLDGGENTGQLY | 0.459537 |
| GHVDHGKTT | ASSLDGGENTGQLY | 0.52917457 |
| AQFEHTLL | ASSLDGGENTGQLY | 0.3194011 |
| IVNQDEL | ASSLDGGENTGQLY | 0.2600902 |
| TNGSQFF | ASSLDGGENTGQLY | 0.15130544 |
| TGKGIVE | ASSLDGGENTGQLY | 0.58901316 |
| LISNASDA | ASSLDGGENTGQLY | 0.26583633 |
| AELRTLGELTKIAKAHGAQ | ASSLDGGENTGQLY | 0.15403345 |
| APNAVAAAEPSW | ASSLDGGENTGQLY | 0.37343025 |
| APQVAGAATAVL | ASSLDGGENTGQLY | 0.19002715 |
| ARRDPAAAAAILPTDA | ASSLDGGENTGQLY | 0.16032115 |
| DAAARDVLAEAGLAEYF | ASSLDGGENTGQLY | 0.49283454 |
| DDQIVLSAKANLAKVEQ | ASSLDGGENTGQLY | 0.4496511 |
| DELLPDTGEDSDEDGH | ASSLDGGENTGQLY | 0.14311782 |
| ELPEAPANRATVPQD | ASSLDGGENTGQLY | 0.13932241 |
| FGQSKIERVTGIG | ASSLDGGENTGQLY | 0.17075133 |
| GAGGTGSTAGGAGGAGGAGGLY | ASSLDGGENTGQLY | 0.38439244 |
| GHGGTGAAGGNGGTGGAGGSGI | ASSLDGGENTGQLY | 0.54604626 |
| GNGGNGGAGGEGGPGGAGGAGA | ASSLDGGENTGQLY | 0.2531282 |
| GSTAGNGGNGGNATNGTIAGQP | ASSLDGGENTGQLY | 0.19349432 |
| GTGGDGGNGGNGGTGG | ASSLDGGENTGQLY | 0.52683425 |
| GVRRPAGIPADAPV | ASSLDGGENTGQLY | 0.3339605 |
| GYRQVTDAHLLTLARR | ASSLDGGENTGQLY | 0.23623459 |
| IAKMGQDGHDRG | ASSLDGGENTGQLY | 0.4534787 |
| KAEPSIEIQYT | ASSLDGGENTGQLY | 0.16079001 |
| KPVRIRMAADEDTVAVT | ASSLDGGENTGQLY | 0.4050814 |
| MPPRIAGMRLLVIKPEP | ASSLDGGENTGQLY | 0.33134142 |
| MKFYAVALFRSRGVPPTR | ASSLDGGENTGQLY | 0.39063907 |

| | | |
|---|---|---|
| NAGGFGILGAVAHSP | ASSLDGGENTGQLY | 0.38681176 |
| NEGREITLAMAEM | ASSLDGGENTGQLY | 0.20622051 |
| NELVEWSHPLGHRWRWE | ASSLDGGENTGQLY | 0.34070212 |
| NLGAGGTGGNGGNPG | ASSLDGGENTGQLY | 0.1274258 |
| NTEWPKWETVMAADPVD | ASSLDGGENTGQLY | 0.15442389 |
| PASAAPAETQP | ASSLDGGENTGQLY | 0.43686634 |
| QQSSDGKNGVRAAEITGE | ASSLDGGENTGQLY | 0.39861614 |
| RAPDAVTTSWWKE | ASSLDGGENTGQLY | 0.4269855 |
| SGFGNVGAHTSGWFNQ | ASSLDGGENTGQLY | 0.31848353 |
| SGVAGAVGQAASVGGLKV | ASSLDGGENTGQLY | 0.18468328 |
| SNGGAGTGSAGGNGGT | ASSLDGGENTGQLY | 0.5729956 |
| TGGAGGEGGAAG | ASSLDGGENTGQLY | 0.6134478 |
| TGGDGGAAGAGGAPGHG | ASSLDGGENTGQLY | 0.1381458 |
| TMPPLLAAIANAGRHGV | ASSLDGGENTGQLY | 0.16722608 |
| TSSTAGNSGGA | ASSLDGGENTGQLY | 0.45415598 |
| VPATVPVGILC | ASSLDGGENTGQLY | 0.50618386 |
| VQRPDSGGVVLTGRIS | ASSLDGGENTGQLY | 0.30524105 |
| WTPNPVPATKGVPAGLVQAT | ASSLDGGENTGQLY | 0.2417865 |
| YALKRALAKFSDRFDVVII | ASSLDGGENTGQLY | 0.52039754 |
| YLIGSVIV | ASSLDGGENTGQLY | 0.14743079 |
| YGQLVAVID | ASSLDGGENTGQLY | 0.66702825 |
| VTVGGMTLPA | ASSLDGGENTGQLY | 0.12145107 |
| VLALPAV | ASSLDGGENTGQLY | 0.2714331 |
| VIPPTLNYET | ASSLDGGENTGQLY | 0.1665574 |
| VGMGGMHPSSG | ASSLDGGENTGQLY | 0.11943161 |
| VGLAGGAATA | ASSLDGGENTGQLY | 0.41397095 |
| VGGAGGAGGSGP | ASSLDGGENTGQLY | 0.17934375 |
| VFAAFMV | ASSLDGGENTGQLY | 0.54405713 |
| TLYDKLR | ASSLDGGENTGQLY | 0.18603298 |
| TLAVFTVL | ASSLDGGENTGQLY | 0.26828033 |
| TGSVQPARV | ASSLDGGENTGQLY | 0.27247778 |
| TGGGAGSGGT | ASSLDGGENTGQLY | 0.24201208 |
| SHLETSL | ASSLDGGENTGQLY | 0.16040829 |
| SFTTGMT | ASSLDGGENTGQLY | 0.42529643 |
| QVEVAKRMGVRQPTVS | ASSLDGGENTGQLY | 0.55872536 |
| QGSGGPALAM | ASSLDGGENTGQLY | 0.33962148 |
| MLAPSNP | ASSLDGGENTGQLY | 0.13787274 |
| LVNNAAI | ASSLDGGENTGQLY | 0.18866602 |
| LTGEINTLA | ASSLDGGENTGQLY | 0.25239843 |
| LSIFGPV | ASSLDGGENTGQLY | 0.11455758 |
| LLALLANL | ASSLDGGENTGQLY | 0.117729686 |
| LISNASDALDKLR | ASSLDGGENTGQLY | 0.13971765 |
| LINEAVP | ASSLDGGENTGQLY | 0.1320632 |
| LGYAFSGL | ASSLDGGENTGQLY | 0.60659444 |
| LDFVAVAL | ASSLDGGENTGQLY | 0.46565634 |
| LAQPGAE | ASSLDGGENTGQLY | 0.2638973 |
| ISNLGAVTGF | ASSLDGGENTGQLY | 0.49970654 |
| HQEDPGRI | ASSLDGGENTGQLY | 0.31164414 |
| GTPGEPTP | ASSLDGGENTGQLY | 0.11063983 |
| GFFNSGS | ASSLDGGENTGQLY | 0.1690384 |
| GAGGGGLTAGAN | ASSLDGGENTGQLY | 0.14612214 |

| | | |
|---|---|---|
| FVDALVNM | ASSLDGGENTGQLY | 0.45890233 |
| FLTVAPEL | ASSLDGGENTGQLY | 0.1309801 |
| FETLTAL | ASSLDGGENTGQLY | 0.17469649 |
| DVINPIEL | ASSLDGGENTGQLY | 0.2865303 |
| APVSGAPV | ASSLDGGENTGQLY | 0.32471234 |
| ALVGGLAW | ASSLDGGENTGQLY | 0.35881308 |
| ALAAAAATAI | ASSLDGGENTGQLY | 0.15470462 |
| AFGLGLLL | ASSLDGGENTGQLY | 0.11241581 |
| AFFAAASL | ASSLDGGENTGQLY | 0.37716454 |
| AAAALPAFSPP | ASSLDGGENTGQLY | 0.11870752 |
| VVLAPAL | ASSLDGGENTGQLY | 0.17567076 |
| TIGHVDHGKTTL | ASSLDGGENTGQLY | 0.12033656 |
| LISNASDALDK | ASSLDGGENTGQLY | 0.11660248 |
| LISNASDAL | ASSLDGGENTGQLY | 0.12749133 |
| ISNASDALDK | ASSLDGGENTGQLY | 0.3417116 |
| NQAETLVYQTE | ASSLDGGENTGQLY | 0.36012018 |
| TGGTGGTAGAG | ASSLDGGENTGQLY | 0.23467624 |
| TGSTAGGAGGA | ASSLDGGENTGQLY | 0.5213613 |
| DPLAVPVAAGPV | ASSLDGGENTGQLY | 0.51613 |
| GGTGGTPGNGGN | ASSLDGGENTGQLY | 0.2289335 |
| PAGLAGVDPATL | ASSLDGGENTGQLY | 0.19969493 |
| GNNLVVEAPHAGASV | ASSLDGGENTGQLY | 0.15695663 |
| KVTRIMSASTLEQPA | ASSLDGGENTGQLY | 0.17015445 |
| LGFALPIALPRRTVV | ASSLDGGENTGQLY | 0.16675806 |
| TRLGNLTARKLVVLV | ASSLDGGENTGQLY | 0.64949036 |
| WSQQRRLIYFPSAGP | ASSLDGGENTGQLY | 0.22930086 |
| GPNGNPDPMAAAVDIR | ASSLDGGENTGQLY | 0.1323326 |
| TQRCVSVADLRAAASR | ASSLDGGENTGQLY | 0.22905727 |
| QFVLDVAEHPVEVLTETV | ASSLDGGENTGQLY | 0.60376626 |
| GGAGGTGGAGGNGGTQAGDG | ASSLDGGENTGQLY | 0.58646727 |
| PLLATLRAIAVDVDAKPAQV | ASSLDGGENTGQLY | 0.15661371 |
| APKFRTGGYMIDFWGVGYQVAK | ASSLDGGENTGQLY | 0.22822714 |
| GAGGAGGAGGAAGGPGGTGGTG | ASSLDGGENTGQLY | 0.22599967 |
| IGALGLLATDVAPAGSVAFGVA | ASSLDGGENTGQLY | 0.2771333 |
| AGGAGGIGGNGGAGGAGGVLMG | ASSLDGGENTGQLY | 0.16286439 |
| ADELIGLL | ASSLDGGENTGQLY | 0.15339965 |
| ADRALKA | ASSLDGGENTGQLY | 0.4365734 |
| AGAVGQVGGAGGA | ASSLDGGENTGQLY | 0.7510266 |
| AGGGGGLGGAP | ASSLDGGENTGQLY | 0.17665477 |
| AGGLLYGNGGAG | ASSLDGGENTGQLY | 0.12256801 |
| AGLESIGL | ASSLDGGENTGQLY | 0.35014102 |
| AGLKTGII | ASSLDGGENTGQLY | 0.16408128 |
| ASLAALLL | ASSLDGGENTGQLY | 0.24656186 |
| ASVINISSI | ASSLDGGENTGQLY | 0.28406757 |
| ATPPGPA | ASSLDGGENTGQLY | 0.13097374 |
| ATVASALV | ASSLDGGENTGQLY | 0.18185264 |
| ATVASLIL | ASSLDGGENTGQLY | 0.5542293 |
| AVNGPVLIRH | ASSLDGGENTGQLY | 0.2889486 |
| DGAAGAVGI | ASSLDGGENTGQLY | 0.5009101 |
| DPLLHGI | ASSLDGGENTGQLY | 0.6078937 |
| ELGAALGAL | ASSLDGGENTGQLY | 0.4791011 |

| | | |
|---|---|---|
| ELGRTQAV | ASSLDGGENTGQLY | 0.6388139 |
| FVAGGYLS | ASSLDGGENTGQLY | 0.5003138 |
| GASGGAGGAGGW | ASSLDGGENTGQLY | 0.20300859 |
| GGAGAGGDGGAGGI | ASSLDGGENTGQLY | 0.29835594 |
| GTGHNGSQPGMGGNG | ASSLDGGENTGQLY | 0.33634034 |
| GYAAASASAS | ASSLDGGENTGQLY | 0.3603394 |
| HQVDVAAVQQG | ASSLDGGENTGQLY | 0.31048334 |
| IGAAIAHT | ASSLDGGENTGQLY | 0.18123394 |
| IGMIEVAQL | ASSLDGGENTGQLY | 0.62446976 |
| IRLAPEHTSAMD | ASSLDGGENTGQLY | 0.31269413 |
| IYNVSVL | ASSLDGGENTGQLY | 0.50983447 |
| KQGFLGM | ASSLDGGENTGQLY | 0.14540121 |
| LESVITQL | ASSLDGGENTGQLY | 0.1682899 |
| LGLGALVVV | ASSLDGGENTGQLY | 0.5619991 |
| LIGNGGNGGSGGI | ASSLDGGENTGQLY | 0.11506417 |
| LNAINAPTL | ASSLDGGENTGQLY | 0.1280563 |
| LQLLNTGV | ASSLDGGENTGQLY | 0.32506293 |
| LQTSIEF | ASSLDGGENTGQLY | 0.44759494 |
| MSFAVSEL | ASSLDGGENTGQLY | 0.11159668 |
| NTGALGL | ASSLDGGENTGQLY | 0.16195326 |
| PPGAGGAGGAGTT | ASSLDGGENTGQLY | 0.16433397 |
| QLVGLGI | ASSLDGGENTGQLY | 0.21830364 |
| RFLEQVHQL | ASSLDGGENTGQLY | 0.2316538 |
| RGVGYVLR | ASSLDGGENTGQLY | 0.12832472 |
| SATGVFAGVI | ASSLDGGENTGQLY | 0.3272868 |
| SLPAITLP | ASSLDGGENTGQLY | 0.12519273 |
| STALAFTV | ASSLDGGENTGQLY | 0.1230333 |
| SVLTVGLGK | ASSLDGGENTGQLY | 0.28449717 |
| TALAGAGSGI | ASSLDGGENTGQLY | 0.62193966 |
| TGLPFTGI | ASSLDGGENTGQLY | 0.14414339 |
| VAEHVTV | ASSLDGGENTGQLY | 0.49372327 |
| VDIVLNSL | ASSLDGGENTGQLY | 0.1803594 |
| VLTGGGAVL | ASSLDGGENTGQLY | 0.5131114 |
| VNVGLAYGI | ASSLDGGENTGQLY | 0.22631778 |
| VSSAAVGAV | ASSLDGGENTGQLY | 0.13443421 |
| VTSVYEAGV | ASSLDGGENTGQLY | 0.46974587 |
| WVALDRGA | ASSLDGGENTGQLY | 0.35023284 |
| YLVGSVVI | ASSLDGGENTGQLY | 0.27491385 |
| GPPGTGKTLL | ASSLDGGENTGQLY | 0.5552118 |
| GVGMTKF | ASSLDGGENTGQLY | 0.18123968 |
| IGIGGSDL | ASSLDGGENTGQLY | 0.2918096 |
| IGTLLAEL | ASSLDGGENTGQLY | 0.53152114 |
| IYTSGTT | ASSLDGGENTGQLY | 0.3957979 |
| LSFPAGV | ASSLDGGENTGQLY | 0.1418237 |
| LYTSGTT | ASSLDGGENTGQLY | 0.29700285 |
| VGAFPFGF | ASSLDGGENTGQLY | 0.27282935 |
| VGAGTVEF | ASSLDGGENTGQLY | 0.1943893 |
| VGKSTLLL | ASSLDGGENTGQLY | 0.37991756 |
| AALAHAGGGIV | ASSLDGGENTGQLY | 0.2817685 |
| AALVAVSLL | ASSLDGGENTGQLY | 0.53261733 |
| AAVSRAVAAL | ASSLDGGENTGQLY | 0.3351995 |

| | | |
|---|---|---|
| AGGSSGIGGF | ASSLDGGENTGQLY | 0.6200361 |
| AGLTVANANG | ASSLDGGENTGQLY | 0.13040197 |
| AGLVGAVL | ASSLDGGENTGQLY | 0.18036564 |
| ALSQAFTV | ASSLDGGENTGQLY | 0.28734428 |
| ALTFSVSL | ASSLDGGENTGQLY | 0.37247336 |
| ALTGAALVAL | ASSLDGGENTGQLY | 0.14609772 |
| ASFSIGAGVL | ASSLDGGENTGQLY | 0.11888075 |
| DGSVYKI | ASSLDGGENTGQLY | 0.29858652 |
| EQLAEIV | ASSLDGGENTGQLY | 0.60150886 |
| FAVSELTVM | ASSLDGGENTGQLY | 0.21059288 |
| GAAPLAAGL | ASSLDGGENTGQLY | 0.3015214 |
| GADGGDGGSGGAGG | ASSLDGGENTGQLY | 0.716422 |
| GFGLDTGGAGG | ASSLDGGENTGQLY | 0.19705528 |
| GLASGGLL | ASSLDGGENTGQLY | 0.2150723 |
| GRIAGAAVLL | ASSLDGGENTGQLY | 0.33250773 |
| GSAGSLNTSF | ASSLDGGENTGQLY | 0.6019012 |
| GTGQIGI | ASSLDGGENTGQLY | 0.22349784 |
| GTLIALIL | ASSLDGGENTGQLY | 0.57984686 |
| IAATMGFVVG | ASSLDGGENTGQLY | 0.13639635 |
| IFAQALL | ASSLDGGENTGQLY | 0.48491257 |
| IGAAVALGF | ASSLDGGENTGQLY | 0.13863239 |
| IGLTPLAL | ASSLDGGENTGQLY | 0.35106754 |
| ILGSNTSTL | ASSLDGGENTGQLY | 0.12942414 |
| IVFSLTVL | ASSLDGGENTGQLY | 0.49091282 |
| LASKATGF | ASSLDGGENTGQLY | 0.5815623 |
| LASVALIL | ASSLDGGENTGQLY | 0.5388683 |
| LFAVGLTL | ASSLDGGENTGQLY | 0.1155484 |
| LFNGGIGTYI | ASSLDGGENTGQLY | 0.4994505 |
| LTLGAGII | ASSLDGGENTGQLY | 0.5901066 |
| LVAAALGDI | ASSLDGGENTGQLY | 0.7511428 |
| NTAFVQLGI | ASSLDGGENTGQLY | 0.30607063 |
| QLFSGAEL | ASSLDGGENTGQLY | 0.42846668 |
| SAATFITL | ASSLDGGENTGQLY | 0.23070788 |
| SNYGLANLGV | ASSLDGGENTGQLY | 0.2049642 |
| SYGGAYAVM | ASSLDGGENTGQLY | 0.1687893 |
| TTGVGGLTLG | ASSLDGGENTGQLY | 0.30086446 |
| TTQTIYI | ASSLDGGENTGQLY | 0.22617638 |
| VAAGAGLGAVI | ASSLDGGENTGQLY | 0.19419667 |
| VAAVVTLL | ASSLDGGENTGQLY | 0.43842196 |
| VGFLGGLI | ASSLDGGENTGQLY | 0.5780647 |
| VGVIPILL | ASSLDGGENTGQLY | 0.42544618 |
| VIFASSLI | ASSLDGGENTGQLY | 0.6967107 |
| VKGGDGGAAAG | ASSLDGGENTGQLY | 0.646263 |
| VTKTFTNI | ASSLDGGENTGQLY | 0.40343395 |
| VVAAAILL | ASSLDGGENTGQLY | 0.17039151 |
| YAALDVEL | ASSLDGGENTGQLY | 0.19682476 |
| LASVALLL | ASSLDGGENTGQLY | 0.13716726 |
| LDAVHRV | ASSLDGGENTGQLY | 0.188643 |
| QGLSSSIF | ASSLDGGENTGQLY | 0.17204484 |
| RGTFANIRL | ASSLDGGENTGQLY | 0.3002836 |
| RQLWWGHRI | ASSLDGGENTGQLY | 0.11211878 |

| | | |
|---|---|---|
| YGPPGTGKTLL | ASSLDGGENTGQLY | 0.40476638 |
| AALVGAPANVA | ASSLDGGENTGQLY | 0.6373457 |
| AATFRRTE | ASSLDGGENTGQLY | 0.12239616 |
| AGALGGGI | ASSLDGGENTGQLY | 0.4477446 |
| AGGAGGAGGIGAG | ASSLDGGENTGQLY | 0.267901 |
| AGSGALGL | ASSLDGGENTGQLY | 0.15184638 |
| AQIADQI | ASSLDGGENTGQLY | 0.6454755 |
| ASAVAVTYL | ASSLDGGENTGQLY | 0.1588082 |
| ASTVLGGTGG | ASSLDGGENTGQLY | 0.39979696 |
| AVGGEPAK | ASSLDGGENTGQLY | 0.5386342 |
| CEQGTFGGTC | ASSLDGGENTGQLY | 0.33339727 |
| DEGKAPAG | ASSLDGGENTGQLY | 0.32493907 |
| DTVVFPAL | ASSLDGGENTGQLY | 0.2615585 |
| FEGITMAG | ASSLDGGENTGQLY | 0.13237965 |
| FPLVVQD | ASSLDGGENTGQLY | 0.4116844 |
| GAFTTPQ | ASSLDGGENTGQLY | 0.113755874 |
| GFGGFGTAGGVGGA | ASSLDGGENTGQLY | 0.37067944 |
| GFLAQFL | ASSLDGGENTGQLY | 0.15219858 |
| GGAGGAGGNAGVA | ASSLDGGENTGQLY | 0.44609672 |
| GGVGTAGAGGAGGAGG | ASSLDGGENTGQLY | 0.2664112 |
| GNGGGGDAGNAGSGGN | ASSLDGGENTGQLY | 0.29935002 |
| GTGGAGGAGNDAGS | ASSLDGGENTGQLY | 0.1935138 |
| GTTNTGWFNS | ASSLDGGENTGQLY | 0.18452325 |
| IFMGPLFTL | ASSLDGGENTGQLY | 0.14894845 |
| ILAHAIEAI | ASSLDGGENTGQLY | 0.14123866 |
| IRSAIREAL | ASSLDGGENTGQLY | 0.2235115 |
| ITFAGGG | ASSLDGGENTGQLY | 0.22439688 |
| ITGMSGHQEV | ASSLDGGENTGQLY | 0.1515381 |
| IVAGLACTL | ASSLDGGENTGQLY | 0.45976043 |
| IVGGGLAGA | ASSLDGGENTGQLY | 0.2666755 |
| IVTGVLNI | ASSLDGGENTGQLY | 0.13059378 |
| LAQQGCI | ASSLDGGENTGQLY | 0.42148805 |
| LCLHVEN | ASSLDGGENTGQLY | 0.37264812 |
| LDAISDGS | ASSLDGGENTGQLY | 0.23197696 |
| LGGFAAML | ASSLDGGENTGQLY | 0.11078574 |
| LGGKNPMI | ASSLDGGENTGQLY | 0.17424 |
| LGPKGVAAIL | ASSLDGGENTGQLY | 0.19774099 |
| LMVGVSI | ASSLDGGENTGQLY | 0.14354561 |
| LVGAAFVL | ASSLDGGENTGQLY | 0.15778561 |
| LVLAVAPL | ASSLDGGENTGQLY | 0.62567675 |
| MAAIAVIDAA | ASSLDGGENTGQLY | 0.124768384 |
| NNGINAELQ | ASSLDGGENTGQLY | 0.26942042 |
| RGEPLVLVHG | ASSLDGGENTGQLY | 0.53982484 |
| RGGAGGAAGLI | ASSLDGGENTGQLY | 0.37454945 |
| RSYRMHPAV | ASSLDGGENTGQLY | 0.6253898 |
| SAAHLLAGQLA | ASSLDGGENTGQLY | 0.2496111 |
| SAAVGSY | ASSLDGGENTGQLY | 0.20859852 |
| SPHAVLHRI | ASSLDGGENTGQLY | 0.25466266 |
| SQIAKVL | ASSLDGGENTGQLY | 0.20974123 |
| TALGHLVP | ASSLDGGENTGQLY | 0.12758955 |
| TAPTTSGAA | ASSLDGGENTGQLY | 0.12223594 |

| | | |
|---|---|---|
| TDGDDGNGGSPGT | ASSLDGGENTGQLY | 0.2389524 |
| TDGLGEN | ASSLDGGENTGQLY | 0.12881601 |
| TGGPPLPGQA | ASSLDGGENTGQLY | 0.33272588 |
| TIPAITI | ASSLDGGENTGQLY | 0.42674053 |
| TIPGFTI | ASSLDGGENTGQLY | 0.16003278 |
| TSIASGALA | ASSLDGGENTGQLY | 0.14713039 |
| TVTLPGAAI | ASSLDGGENTGQLY | 0.22649276 |
| VASGLSAALP | ASSLDGGENTGQLY | 0.15421735 |
| VGAGVIGI | ASSLDGGENTGQLY | 0.55662584 |
| VGPKRFGV | ASSLDGGENTGQLY | 0.17398643 |
| VNVAFPD | ASSLDGGENTGQLY | 0.69887733 |
| VSMTNSGVS | ASSLDGGENTGQLY | 0.534207 |
| VTDVLGDLSV | ASSLDGGENTGQLY | 0.2086688 |
| VVVVGAGFAGL | ASSLDGGENTGQLY | 0.52882576 |
| VYTVGTVV | ASSLDGGENTGQLY | 0.21077168 |
| WTTFSGP | ASSLDGGENTGQLY | 0.15590203 |
| YDELVEAF | ASSLDGGENTGQLY | 0.21283928 |
| YGTGGAGGNGG | ASSLDGGENTGQLY | 0.17862675 |
| YTNVMAV | ASSLDGGENTGQLY | 0.22438656 |
| FLEQVHQL | ASSLDGGENTGQLY | 0.20284133 |
| GLSLLAVAV | ASSLDGGENTGQLY | 0.13590008 |
| LEQVHQL | ASSLDGGENTGQLY | 0.3706658 |
| LNAAGAI | ASSLDGGENTGQLY | 0.14213903 |
| PAVHASL | ASSLDGGENTGQLY | 0.13695525 |
| TELEDDA | ASSLDGGENTGQLY | 0.21306315 |
| TPPGPPN | ASSLDGGENTGQLY | 0.52546173 |
| TPPPPPPP | ASSLDGGENTGQLY | 0.74018335 |
| TQSPALI | ASGPGTANERLF | 0.19911033 |
| EFAAAFPQ | ASGPGTANERLF | 0.1696347 |
| LLSGNLDVL | ASGPGTANERLF | 0.5233063 |
| GPANLGNYNI | ASGPGTANERLF | 0.1288238 |
| TLGPTLP | ASGPGTANERLF | 0.3083221 |
| VYAYSSCCK | ASGPGTANERLF | 0.27852657 |
| APAPAPAPNI | ASGPGTANERLF | 0.17489116 |
| PGGPVPIP | ASGPGTANERLF | 0.18965027 |
| FLTVALEP | ASGPGTANERLF | 0.30016476 |
| SQYFGANL | ASGPGTANERLF | 0.16303381 |
| GGVKPAA | ASGPGTANERLF | 0.42138642 |
| GGTGGAGGAGGW | ASGPGTANERLF | 0.11140271 |
| GVGGVAGLGGAGG | ASGPGTANERLF | 0.15199542 |
| VLLVLATTI | ASGPGTANERLF | 0.16903205 |
| IGIFAAML | ASGPGTANERLF | 0.18293501 |
| AVGDLID | ASGPGTANERLF | 0.12556636 |
| NISPLPL | ASGPGTANERLF | 0.18726069 |
| VTAVFVAI | ASGPGTANERLF | 0.49377328 |
| SGDDNDAAL | ASGPGTANERLF | 0.583746 |
| AVAALAHS | ASGPGTANERLF | 0.13435827 |
| GGVTVMATP | ASGPGTANERLF | 0.5068495 |
| AGLAANVRD | ASGPGTANERLF | 0.63852024 |
| TNRPDLIDPALLRPG | ASGPGTANERLF | 0.20357276 |
| LANLGVTAL | ASGPGTANERLF | 0.1286479 |

| | | |
|---|---|---|
| TGERIEL | ASGPGTANERLF | 0.28651324 |
| NGEELAA | ASGPGTANERLF | 0.12550849 |
| VLSLVWG | ASGPGTANERLF | 0.22151367 |
| IGYALWL | ASGPGTANERLF | 0.49351442 |
| LSLLLAPL | ASGPGTANERLF | 0.13385363 |
| TLDEALK | ASGPGTANERLF | 0.43212092 |
| TDGAGTAEGG | ASGPGTANERLF | 0.54175735 |
| NAPGGAGGNGG | ASGPGTANERLF | 0.17837392 |
| VSVFMGMGGNF | ASGPGTANERLF | 0.14692546 |
| LASYLAAL | ASGPGTANERLF | 0.1422546 |
| LVGGTPAL | ASGPGTANERLF | 0.11241431 |
| VRNADVL | ASGPGTANERLF | 0.33020478 |
| TCSSPFLVSAM | ASGPGTANERLF | 0.19756322 |
| VHITDAE | ASGPGTANERLF | 0.19090271 |
| EYAAPSF | ASGPGTANERLF | 0.26935607 |
| VVDAFHGGL | ASGPGTANERLF | 0.18280461 |
| ASVVDPQ | ASGPGTANERLF | 0.19805582 |
| SYNEFGGLS | ASGPGTANERLF | 0.20852461 |
| DPDGSVY | ASGPGTANERLF | 0.14177825 |
| MVSTIHATE | ASGPGTANERLF | 0.39624023 |
| LAAGVADPV | ASGPGTANERLF | 0.13355264 |
| TKEQIAA | ASGPGTANERLF | 0.20903942 |
| SLASPAAL | ASGPGTANERLF | 0.67924666 |
| IGLVFYGV | ASGPGTANERLF | 0.12694691 |
| FLLDASNL | ASGPGTANERLF | 0.16165727 |
| YQVGNFP | ASGPGTANERLF | 0.37306342 |
| ESLLSQST | ASGPGTANERLF | 0.13120721 |
| LFTVGFAAVG | ASGPGTANERLF | 0.5064504 |
| ATPPTPT | ASGPGTANERLF | 0.31323293 |
| TDLATLG | ASGPGTANERLF | 0.35329115 |
| SAGGTGR | ASGPGTANERLF | 0.21096054 |
| GFSNSGS | ASGPGTANERLF | 0.6364821 |
| GGNGGSGGTSV | ASGPGTANERLF | 0.5182421 |
| GTSVATGGAGN | ASGPGTANERLF | 0.17674197 |
| NGGNGGSGGTS | ASGPGTANERLF | 0.14237788 |
| AVVKFPH | ASGPGTANERLF | 0.15707088 |
| TPPPPPP | ASGPGTANERLF | 0.44304812 |
| AAPAAAAAPA | ASGPGTANERLF | 0.57153296 |
| IALNEDL | ASGPGTANERLF | 0.16136107 |
| IVNVSSM | ASGPGTANERLF | 0.30464283 |
| TGIPPAPR | ASGPGTANERLF | 0.439727 |
| FDLGGGTF | ASGPGTANERLF | 0.6014039 |
| FDLGGGTFDVS | ASGPGTANERLF | 0.13225071 |
| NEPTAAA | ASGPGTANERLF | 0.176554 |
| NEPTAAALA | ASGPGTANERLF | 0.14898705 |
| GSIINISSI | ASGPGTANERLF | 0.2997377 |
| VAPGNAGTA | ASGPGTANERLF | 0.15831263 |
| TGGGASLE | ASGPGTANERLF | 0.5633775 |
| VAVYNHYKRI | ASGPGTANERLF | 0.27455014 |
| TDPAPATT | ASGPGTANERLF | 0.6189065 |
| TNRPDILDPALLRPG | ASGPGTANERLF | 0.29572695 |

| | | |
|---|---|---|
| LTTTTTS | ASGPGTANERLF | 0.49353755 |
| PTGSGKTL | ASGPGTANERLF | 0.27395952 |
| TGSGKTL | ASGPGTANERLF | 0.22861966 |
| TIGHVDHGK | ASGPGTANERLF | 0.14257662 |
| TIGHVDHGKTT | ASGPGTANERLF | 0.12038337 |
| GHVDHGKTT | ASGPGTANERLF | 0.15292408 |
| AQFEHTLL | ASGPGTANERLF | 0.18909231 |
| IVNQDEL | ASGPGTANERLF | 0.11266139 |
| TNGSQFF | ASGPGTANERLF | 0.4082783 |
| TGKGIVE | ASGPGTANERLF | 0.3553387 |
| LISNASDA | ASGPGTANERLF | 0.42344493 |
| AELRTLGELTKIAKAHGAQ | ASGPGTANERLF | 0.1886302 |
| APNAVAAAEPSW | ASGPGTANERLF | 0.16667053 |
| APQVAGAATAVL | ASGPGTANERLF | 0.16141836 |
| ARRDPAAAAAILPTDA | ASGPGTANERLF | 0.63288987 |
| DAAARDVLAEAGLAEYF | ASGPGTANERLF | 0.1283574 |
| DDQIVLSAKANLAKVEQ | ASGPGTANERLF | 0.12752375 |
| DELLPDTGEDSDEDGH | ASGPGTANERLF | 0.15989345 |
| ELPEAPANRATVPQD | ASGPGTANERLF | 0.12857018 |
| FGQSKIERVTGIG | ASGPGTANERLF | 0.59968734 |
| GAGGTGSTAGGAGGAGGAGGLY | ASGPGTANERLF | 0.53196573 |
| GHGGTGAAGGNGGTGGAGGSGI | ASGPGTANERLF | 0.389247 |
| GNGGNGGAGGEGGPGGAGGAGA | ASGPGTANERLF | 0.51354563 |
| GSTAGNGGNGGNATNGTIAGQP | ASGPGTANERLF | 0.12230243 |
| GTGGDGGNGGNGGTGG | ASGPGTANERLF | 0.16840363 |
| GVRRPAGIPADAPV | ASGPGTANERLF | 0.69833606 |
| GYRQVTDAHLLTLARR | ASGPGTANERLF | 0.30969018 |
| IAKMGQDGHDRG | ASGPGTANERLF | 0.32003158 |
| KAEPSIEIQYT | ASGPGTANERLF | 0.44303104 |
| KPVRIRMAADEDTVAVT | ASGPGTANERLF | 0.17460734 |
| MPPRIAGMRLLVIKPEP | ASGPGTANERLF | 0.2388548 |
| MKFYAVALFRSRGVPPTR | ASGPGTANERLF | 0.22641814 |
| NAGGFGILGAVAHSP | ASGPGTANERLF | 0.27549553 |
| NEGREITLAMAEM | ASGPGTANERLF | 0.17099586 |
| NELVEWSHPLGHRWRWE | ASGPGTANERLF | 0.13741785 |
| NLGAGGTGGNGGNPG | ASGPGTANERLF | 0.39001176 |
| NTEWPKWETVMAADPVD | ASGPGTANERLF | 0.7035519 |
| PASAAPAETQP | ASGPGTANERLF | 0.3431828 |
| QQSSDGKNGVRAAEITGE | ASGPGTANERLF | 0.25140217 |
| RAPDAVTTSWWKE | ASGPGTANERLF | 0.3357103 |
| SGFGNVGAHTSGWFNQ | ASGPGTANERLF | 0.29901212 |
| SGVAGAVGQAASVGGLKV | ASGPGTANERLF | 0.19724707 |
| SNGGAGTGSAGGNGGT | ASGPGTANERLF | 0.17110139 |
| TGGAGGEGGAAG | ASGPGTANERLF | 0.17639843 |
| TGGDGGAAGAGGAPGHG | ASGPGTANERLF | 0.14641422 |
| TMPPLLAAIANAGRHGV | ASGPGTANERLF | 0.27778617 |
| TSSTAGNSGGA | ASGPGTANERLF | 0.178372 |
| VPATVPVGILC | ASGPGTANERLF | 0.726341 |
| VQRPDSGGVVLTGRIS | ASGPGTANERLF | 0.51260495 |
| WTPNPVPATKGVPAGLVQAT | ASGPGTANERLF | 0.27712327 |
| YALKRALAKFSDRFDVVII | ASGPGTANERLF | 0.15082452 |

| | | |
|---|---|---|
| YLIGSVIV | ASGPGTANERLF | 0.23179142 |
| YGQLVAVID | ASGPGTANERLF | 0.29633588 |
| VTVGGMTLPA | ASGPGTANERLF | 0.13120738 |
| VLALPAV | ASGPGTANERLF | 0.6215022 |
| VIPPTLNYET | ASGPGTANERLF | 0.17686087 |
| VGMGGMHPSSG | ASGPGTANERLF | 0.16112953 |
| VGLAGGAATA | ASGPGTANERLF | 0.35355914 |
| VGGAGGAGGSGP | ASGPGTANERLF | 0.6856744 |
| VFAAFMV | ASGPGTANERLF | 0.19121021 |
| TLYDKLR | ASGPGTANERLF | 0.17903188 |
| TLAVFTVL | ASGPGTANERLF | 0.15146385 |
| TGSVQPARV | ASGPGTANERLF | 0.5703305 |
| TGGGAGSGGT | ASGPGTANERLF | 0.6765311 |
| SHLETSL | ASGPGTANERLF | 0.14693469 |
| SFTTGMT | ASGPGTANERLF | 0.6797005 |
| QVEVAKRMGVRQPTVS | ASGPGTANERLF | 0.16330013 |
| QGSGGPALAM | ASGPGTANERLF | 0.15557781 |
| MLAPSNP | ASGPGTANERLF | 0.3172406 |
| LVNNAAI | ASGPGTANERLF | 0.1972666 |
| LTGEINTLA | ASGPGTANERLF | 0.62815464 |
| LSIFGPV | ASGPGTANERLF | 0.3161059 |
| LLALLANL | ASGPGTANERLF | 0.18718818 |
| LISNASDALDKLR | ASGPGTANERLF | 0.21366796 |
| LINEAVP | ASGPGTANERLF | 0.64507794 |
| LGYAFSGL | ASGPGTANERLF | 0.43095315 |
| LDFVAVAL | ASGPGTANERLF | 0.39591828 |
| LAQPGAE | ASGPGTANERLF | 0.2627741 |
| ISNLGAVTGF | ASGPGTANERLF | 0.22786096 |
| HQEDPGRI | ASGPGTANERLF | 0.25152928 |
| GTPGEPTP | ASGPGTANERLF | 0.3043272 |
| GFFNSGS | ASGPGTANERLF | 0.18426931 |
| GAGGGGLTAGAN | ASGPGTANERLF | 0.25441465 |
| FVDALVNM | ASGPGTANERLF | 0.48605543 |
| FLTVAPEL | ASGPGTANERLF | 0.14324707 |
| FETLTAL | ASGPGTANERLF | 0.17885402 |
| DVINPIEL | ASGPGTANERLF | 0.5673538 |
| APVSGAPV | ASGPGTANERLF | 0.23445258 |
| ALVGGLAW | ASGPGTANERLF | 0.17461313 |
| ALAAAAATAI | ASGPGTANERLF | 0.64051604 |
| AFGLGLLL | ASGPGTANERLF | 0.7494483 |
| AFFAAASL | ASGPGTANERLF | 0.46027192 |
| AAAALPAFSPP | ASGPGTANERLF | 0.13929296 |
| VVLAPAL | ASGPGTANERLF | 0.1228117 |
| TIGHVDHGKTTL | ASGPGTANERLF | 0.20259188 |
| LISNASDALDK | ASGPGTANERLF | 0.3510906 |
| LISNASDAL | ASGPGTANERLF | 0.18324357 |
| ISNASDALDK | ASGPGTANERLF | 0.17822017 |
| NQAETLVYQTE | ASGPGTANERLF | 0.23259062 |
| TGGTGGTAGAG | ASGPGTANERLF | 0.45013344 |
| TGSTAGGAGGA | ASGPGTANERLF | 0.19202366 |
| DPLAVPVAAGPV | ASGPGTANERLF | 0.14221172 |

| | | |
|---|---|---|
| GGTGGTPGNGGN | ASGPGTANERLF | 0.15241867 |
| PAGLAGVDPATL | ASGPGTANERLF | 0.5171157 |
| GNNLVVEAPHAGASV | ASGPGTANERLF | 0.1611155 |
| KVTRIMSASTLEQPA | ASGPGTANERLF | 0.30854988 |
| LGFALPIALPRRTVV | ASGPGTANERLF | 0.21378247 |
| TRLGNLTARKLVVLV | ASGPGTANERLF | 0.13062808 |
| WSQQRRLIYFPSAGP | ASGPGTANERLF | 0.32661813 |
| GPNGNPDPMAAAVDIR | ASGPGTANERLF | 0.18588534 |
| TQRCVSVADLRAAASR | ASGPGTANERLF | 0.3522876 |
| QFVLDVAEHPVEVLTETV | ASGPGTANERLF | 0.16992049 |
| GGAGGTGGAGGNGGTQAGDG | ASGPGTANERLF | 0.34034157 |
| PLLATLRAIAVDVDAKPAQV | ASGPGTANERLF | 0.4326504 |
| APKFRTGGYMIDFWGVGYQVAK | ASGPGTANERLF | 0.32370666 |
| GAGGAGGAGGAAGGPGGTGGTG | ASGPGTANERLF | 0.25717226 |
| IGALGLLATDVAPAGSVAFGVA | ASGPGTANERLF | 0.1776291 |
| AGGAGGIGGNGGAGGAGGVLMG | ASGPGTANERLF | 0.13766861 |
| ADELIGLL | ASGPGTANERLF | 0.12512657 |
| ADRALKA | ASGPGTANERLF | 0.53332007 |
| AGAVGQVGGAGGA | ASGPGTANERLF | 0.11407326 |
| AGGGGGLGGAP | ASGPGTANERLF | 0.58572567 |
| AGGLLYGNGGAG | ASGPGTANERLF | 0.37008983 |
| AGLESIGL | ASGPGTANERLF | 0.15059027 |
| AGLKTGII | ASGPGTANERLF | 0.27218702 |
| ASLAALLL | ASGPGTANERLF | 0.13114756 |
| ASVINISSI | ASGPGTANERLF | 0.16255327 |
| ATPPGPA | ASGPGTANERLF | 0.4612822 |
| ATVASALV | ASGPGTANERLF | 0.28981125 |
| ATVASLIL | ASGPGTANERLF | 0.19221 |
| AVNGPVLIRH | ASGPGTANERLF | 0.3279189 |
| DGAAGAVGI | ASGPGTANERLF | 0.3886968 |
| DPLLHGI | ASGPGTANERLF | 0.17629774 |
| ELGAALGAL | ASGPGTANERLF | 0.24362193 |
| ELGRTQAV | ASGPGTANERLF | 0.537411 |
| FVAGGYLS | ASGPGTANERLF | 0.2947464 |
| GASGGAGGAGGW | ASGPGTANERLF | 0.14951067 |
| GGAGAGGDGGAGGI | ASGPGTANERLF | 0.31844702 |
| GTGHNGSQPGMGGNG | ASGPGTANERLF | 0.16215447 |
| GYAAASASAS | ASGPGTANERLF | 0.38552088 |
| HQVDVAAVQQG | ASGPGTANERLF | 0.19787331 |
| IGAAIAHT | ASGPGTANERLF | 0.32530192 |
| IGMIEVAQL | ASGPGTANERLF | 0.28732765 |
| IRLAPEHTSAMD | ASGPGTANERLF | 0.120678775 |
| IYNVSVL | ASGPGTANERLF | 0.18039957 |
| KQGFLGM | ASGPGTANERLF | 0.21710119 |
| LESVITQL | ASGPGTANERLF | 0.31034172 |
| LGLGALVVV | ASGPGTANERLF | 0.3593529 |
| LIGNGGNGGSGGI | ASGPGTANERLF | 0.15123947 |
| LNAINAPTL | ASGPGTANERLF | 0.4039333 |
| LQLLNTGV | ASGPGTANERLF | 0.2086897 |
| LQTSIEF | ASGPGTANERLF | 0.1768076 |
| MSFAVSEL | ASGPGTANERLF | 0.27088884 |

| | | |
|---|---|---|
| NTGALGL | ASGPGTANERLF | 0.16444969 |
| PPGAGGAGGAGTT | ASGPGTANERLF | 0.18440726 |
| QLVGLGI | ASGPGTANERLF | 0.32780957 |
| RFLEQVHQL | ASGPGTANERLF | 0.15369587 |
| RGVGYVLR | ASGPGTANERLF | 0.30662954 |
| SATGVFAGVI | ASGPGTANERLF | 0.141728 |
| SLPAITLP | ASGPGTANERLF | 0.18343863 |
| STALAFTV | ASGPGTANERLF | 0.30956274 |
| SVLTVGLGK | ASGPGTANERLF | 0.19480263 |
| TALAGAGSGI | ASGPGTANERLF | 0.41585487 |
| TGLPFTGI | ASGPGTANERLF | 0.21272355 |
| VAEHVTV | ASGPGTANERLF | 0.15440524 |
| VDIVLNSL | ASGPGTANERLF | 0.19194318 |
| VLTGGGAVL | ASGPGTANERLF | 0.38333452 |
| VNVGLAYGI | ASGPGTANERLF | 0.39906076 |
| VSSAAVGAV | ASGPGTANERLF | 0.18712418 |
| VTSVYEAGV | ASGPGTANERLF | 0.26964486 |
| WVALDRGA | ASGPGTANERLF | 0.43446907 |
| YLVGSVVI | ASGPGTANERLF | 0.25928235 |
| GPPGTGKTLL | ASGPGTANERLF | 0.13270603 |
| GVGMTKF | ASGPGTANERLF | 0.19467387 |
| IGIGGSDL | ASGPGTANERLF | 0.14990214 |
| IGTLLAEL | ASGPGTANERLF | 0.1788427 |
| IYTSGTT | ASGPGTANERLF | 0.14761613 |
| LSFPAGV | ASGPGTANERLF | 0.5370636 |
| LYTSGTT | ASGPGTANERLF | 0.7021786 |
| VGAFPFGF | ASGPGTANERLF | 0.19188648 |
| VGAGTVEF | ASGPGTANERLF | 0.5745381 |
| VGKSTLLL | ASGPGTANERLF | 0.13363555 |
| AALAHAGGGIV | ASGPGTANERLF | 0.34546268 |
| AALVAVSLL | ASGPGTANERLF | 0.55854064 |
| AAVSRAVAAL | ASGPGTANERLF | 0.1574186 |
| AGGSSGIGGF | ASGPGTANERLF | 0.28211614 |
| AGLTVANANG | ASGPGTANERLF | 0.14060031 |
| AGLVGAVL | ASGPGTANERLF | 0.15629967 |
| ALSQAFTV | ASGPGTANERLF | 0.5711863 |
| ALTFSVSL | ASGPGTANERLF | 0.21803881 |
| ALTGAALVAL | ASGPGTANERLF | 0.2390202 |
| ASFSIGAGVL | ASGPGTANERLF | 0.20665874 |
| DGSVYKI | ASGPGTANERLF | 0.15222162 |
| EQLAEIV | ASGPGTANERLF | 0.222912 |
| FAVSELTVM | ASGPGTANERLF | 0.6221281 |
| GAAPLAAGL | ASGPGTANERLF | 0.17476802 |
| GADGGDGGSGGAGG | ASGPGTANERLF | 0.5229687 |
| GFGLDTGGAGG | ASGPGTANERLF | 0.1824156 |
| GLASGGLL | ASGPGTANERLF | 0.36461666 |
| GRIAGAAVLL | ASGPGTANERLF | 0.6075314 |
| GSAGSLNTSF | ASGPGTANERLF | 0.38597804 |
| GTGQIGI | ASGPGTANERLF | 0.1969285 |
| GTLIALIL | ASGPGTANERLF | 0.65387625 |
| IAATMGFVVG | ASGPGTANERLF | 0.4219575 |

| | | |
|---|---|---|
| IFAQALL | ASGPGTANERLF | 0.40165848 |
| IGAAVALGF | ASGPGTANERLF | 0.42128062 |
| IGLTPLAL | ASGPGTANERLF | 0.098762095 |
| ILGSNTSTL | ASGPGTANERLF | 0.12931211 |
| IVFSLTVL | ASGPGTANERLF | 0.122909784 |
| LASKATGF | ASGPGTANERLF | 0.1879669 |
| LASVALIL | ASGPGTANERLF | 0.1035324 |
| LFAVGLTL | ASGPGTANERLF | 0.09213409 |
| LFNGGIGTYI | ASGPGTANERLF | 0.14375846 |
| LTLGAGII | ASGPGTANERLF | 0.09209757 |
| LVAAALGDI | ASGPGTANERLF | 0.10871619 |
| NTAFVQLGI | ASGPGTANERLF | 0.1274961 |
| QLFSGAEL | ASGPGTANERLF | 0.108967915 |
| SAATFITL | ASGPGTANERLF | 0.19310184 |
| SNYGLANLGV | ASGPGTANERLF | 0.1776436 |
| SYGGAYAVM | ASGPGTANERLF | 0.09602413 |
| TTGVGGLTLG | ASGPGTANERLF | 0.1158407 |
| TTQTIYI | ASGPGTANERLF | 0.14133586 |
| VAAGAGLGAVI | ASGPGTANERLF | 0.11646019 |
| VAAVVTLL | ASGPGTANERLF | 0.08911217 |
| VGFLGGLI | ASGPGTANERLF | 0.124527164 |
| VGVIPILL | ASGPGTANERLF | 0.09258249 |
| VIFASSLI | ASGPGTANERLF | 0.08431972 |
| VKGGDGGAAAG | ASGPGTANERLF | 0.16411293 |
| VTKTFTNI | ASGPGTANERLF | 0.11147321 |
| VVAAAILL | ASGPGTANERLF | 0.097876444 |
| YAALDVEL | ASGPGTANERLF | 0.18477188 |
| LASVALLL | ASGPGTANERLF | 0.10903248 |
| LDAVHRV | ASGPGTANERLF | 0.09831002 |
| QGLSSSIF | ASGPGTANERLF | 0.13686576 |
| RGTFANIRL | ASGPGTANERLF | 0.14799985 |
| RQLWWGHRI | ASGPGTANERLF | 0.099300146 |
| YGPPGTGKTLL | ASGPGTANERLF | 0.12305079 |
| AALVGAPANVA | ASGPGTANERLF | 0.30333996 |
| AATFRRTE | ASGPGTANERLF | 0.11110795 |
| AGALGGGI | ASGPGTANERLF | 0.28506157 |
| AGGAGGAGGIGAG | ASGPGTANERLF | 0.3329235 |
| AGSGALGL | ASGPGTANERLF | 0.105834216 |
| AQIADQI | ASGPGTANERLF | 0.13048251 |
| ASAVAVTYL | ASGPGTANERLF | 0.15237841 |
| ASTVLGGTGG | ASGPGTANERLF | 0.12304609 |
| AVGGEPAK | ASGPGTANERLF | 0.12239507 |
| CEQGTFGGTC | ASGPGTANERLF | 0.10571158 |
| DEGKAPAG | ASGPGTANERLF | 0.14709781 |
| DTVVFPAL | ASGPGTANERLF | 0.086464025 |
| FEGITMAG | ASGPGTANERLF | 0.11929642 |
| FPLVVQD | ASGPGTANERLF | 0.15588771 |
| GAFTTPQ | ASGPGTANERLF | 0.12655653 |
| GFGGFGTAGGVGGA | ASGPGTANERLF | 0.09084479 |
| GFLAQFL | ASGPGTANERLF | 0.09466359 |
| GGAGGAGGNAGVA | ASGPGTANERLF | 0.09223276 |

| | | |
|---|---|---|
| GGVGTAGAGGAGGAGG | ASGPGTANERLF | 0.110215396 |
| GNGGGGDAGNAGSGGN | ASGPGTANERLF | 0.09431776 |
| GTGGAGGAGNDAGS | ASGPGTANERLF | 0.0955658 |
| GTTNTGWFNS | ASGPGTANERLF | 0.15969911 |
| IFMGPLFTL | ASGPGTANERLF | 0.15730637 |
| ILAHAIEAI | ASGPGTANERLF | 0.10926457 |
| IRSAIREAL | ASGPGTANERLF | 0.1388472 |
| ITFAGGG | ASGPGTANERLF | 0.2663287 |
| ITGMSGHQEV | ASGPGTANERLF | 0.2661087 |
| IVAGLACTL | ASGPGTANERLF | 0.12732352 |
| IVGGGLAGA | ASGPGTANERLF | 0.09260823 |
| IVTGVLNI | ASGPGTANERLF | 0.23540735 |
| LAQQGCI | ASGPGTANERLF | 0.114167064 |
| LCLHVEN | ASGPGTANERLF | 0.12152032 |
| LDAISDGS | ASGPGTANERLF | 0.21223944 |
| LGGFAAML | ASGPGTANERLF | 0.10194995 |
| LGGKNPMI | ASGPGTANERLF | 0.09572125 |
| LGPKGVAAIL | ASGPGTANERLF | 0.25205296 |
| LMVGVSI | ASGPGTANERLF | 0.20659669 |
| LVGAAFVL | ASGPGTANERLF | 0.09756065 |
| LVLAVAPL | ASGPGTANERLF | 0.09672648 |
| MAAIAVIDAA | ASGPGTANERLF | 0.107385725 |
| NNGINAELQ | ASGPGTANERLF | 0.13394454 |
| RGEPLVLVHG | ASGPGTANERLF | 0.1601662 |
| RGGAGGAAGLI | ASGPGTANERLF | 0.13092501 |
| RSYRMHPAV | ASGPGTANERLF | 0.10441495 |
| SAAHLLAGQLA | ASGPGTANERLF | 0.1796844 |
| SAAVGSY | ASGPGTANERLF | 0.12789872 |
| SPHAVLHRI | ASGPGTANERLF | 0.12374454 |
| SQIAKVL | ASGPGTANERLF | 0.20492846 |
| TALGHLVP | ASGPGTANERLF | 0.089885175 |
| TAPTTSGAA | ASGPGTANERLF | 0.17637666 |
| TDGDDGNGGSPGT | ASGPGTANERLF | 0.2091756 |
| TDGLGEN | ASGPGTANERLF | 0.21535687 |
| TGGPPLPGQA | ASGPGTANERLF | 0.32130522 |
| TIPAITI | ASGPGTANERLF | 0.106908135 |
| TIPGFTI | ASGPGTANERLF | 0.16538554 |
| TSIASGALA | ASGPGTANERLF | 0.09899263 |
| TVTLPGAAI | ASGPGTANERLF | 0.11885375 |
| VASGLSAALP | ASGPGTANERLF | 0.18515807 |
| VGAGVIGI | ASGPGTANERLF | 0.13924451 |
| VGPKRFGV | ASGPGTANERLF | 0.28514722 |
| VNVAFPD | ASGPGTANERLF | 0.19513443 |
| VSMTNSGVS | ASGPGTANERLF | 0.11089742 |
| VTDVLGDLSV | ASGPGTANERLF | 0.5150027 |
| VVVVGAGFAGL | ASGPGTANERLF | 0.2572037 |
| VYTVGTVV | ASGPGTANERLF | 0.09469885 |
| WTTFSGP | ASGPGTANERLF | 0.09926499 |
| YDELVEAF | ASGPGTANERLF | 0.12615141 |
| YGTGGAGGNGG | ASGPGTANERLF | 0.1682119 |
| YTNVMAV | ASGPGTANERLF | 0.15084004 |

| | | |
|---|---|---|
| FLEQVHQL | ASGPGTANERLF | 0.15993758 |
| GLSLLAVAV | ASGPGTANERLF | 0.12599325 |
| LEQVHQL | ASGPGTANERLF | 0.0949862 |
| LNAAGAI | ASGPGTANERLF | 0.2873658 |
| PAVHASL | ASGPGTANERLF | 0.09446708 |
| TELEDDA | ASGPGTANERLF | 0.13146493 |
| TPPGPPN | ASGPGTANERLF | 0.09697978 |
| TPPPPPPP | ASGPGTANERLF | 0.08416229 |
| TQSPALI | ASSLGRGANTEVF | 0.13868387 |
| EFAAAFPQ | ASSLGRGANTEVF | 0.12427412 |
| LLSGNLDVL | ASSLGRGANTEVF | 0.19725586 |
| GPANLGNYNI | ASSLGRGANTEVF | 0.11345083 |
| TLGPTLP | ASSLGRGANTEVF | 0.12998861 |
| VYAYSSCCK | ASSLGRGANTEVF | 0.14710887 |
| APAPAPAPNI | ASSLGRGANTEVF | 0.12877274 |
| PGGPVPIP | ASSLGRGANTEVF | 0.10106409 |
| FLTVALEP | ASSLGRGANTEVF | 0.12533385 |
| SQYFGANL | ASSLGRGANTEVF | 0.45719874 |
| GGVKPAA | ASSLGRGANTEVF | 0.18586193 |
| GGTGGAGGAGGW | ASSLGRGANTEVF | 0.0940356 |
| GVGGVAGLGGAGG | ASSLGRGANTEVF | 0.12189533 |
| VLLVLATTI | ASSLGRGANTEVF | 0.10614702 |
| IGIFAAML | ASSLGRGANTEVF | 0.08488947 |
| AVGDLID | ASSLGRGANTEVF | 0.08927815 |
| NISPLPL | ASSLGRGANTEVF | 0.086218126 |
| VTAVFVAI | ASSLGRGANTEVF | 0.09266478 |
| SGDDNDAAL | ASSLGRGANTEVF | 0.17002244 |
| AVAALAHS | ASSLGRGANTEVF | 0.17523853 |
| GGVTVMATP | ASSLGRGANTEVF | 0.12612456 |
| AGLAANVRD | ASSLGRGANTEVF | 0.18573487 |
| TNRPDLIDPALLRPG | ASSLGRGANTEVF | 0.12336469 |
| LANLGVTAL | ASSLGRGANTEVF | 0.08177381 |
| TGERIEL | ASSLGRGANTEVF | 0.10110153 |
| NGEELAA | ASSLGRGANTEVF | 0.10275052 |
| VLSLVWG | ASSLGRGANTEVF | 0.3644831 |
| IGYALWL | ASSLGRGANTEVF | 0.088879466 |
| LSLLLAPL | ASSLGRGANTEVF | 0.121076815 |
| TLDEALK | ASSLGRGANTEVF | 0.13631658 |
| TDGAGTAEGG | ASSLGRGANTEVF | 0.09977667 |
| NAPGGAGGNGG | ASSLGRGANTEVF | 0.17424709 |
| VSVFMGMGGNF | ASSLGRGANTEVF | 0.09603644 |
| LASYLAAL | ASSLGRGANTEVF | 0.100708514 |
| LVGGTPAL | ASSLGRGANTEVF | 0.134325 |
| VRNADVL | ASSLGRGANTEVF | 0.092351414 |
| TCSSPFLVSAM | ASSLGRGANTEVF | 0.102817774 |
| VHITDAE | ASSLGRGANTEVF | 0.11544674 |
| EYAAPSF | ASSLGRGANTEVF | 0.12104711 |
| VVDAFHGGL | ASSLGRGANTEVF | 0.09137128 |
| ASVVDPQ | ASSLGRGANTEVF | 0.2741429 |
| SYNEFGGLS | ASSLGRGANTEVF | 0.12636885 |
| DPDGSVY | ASSLGRGANTEVF | 0.122949444 |

| | | |
|---|---|---|
| MVSTIHATE | ASSLGRGANTEVF | 0.13307925 |
| LAAGVADPV | ASSLGRGANTEVF | 0.106015585 |
| TKEQIAA | ASSLGRGANTEVF | 0.1354824 |
| SLASPAAL | ASSLGRGANTEVF | 0.1050574 |
| IGLVFYGV | ASSLGRGANTEVF | 0.10514709 |
| FLLDASNL | ASSLGRGANTEVF | 0.096419394 |
| YQVGNFP | ASSLGRGANTEVF | 0.10023474 |
| ESLLSQST | ASSLGRGANTEVF | 0.28783825 |
| LFTVGFAAVG | ASSLGRGANTEVF | 0.13114768 |
| ATPPTPT | ASSLGRGANTEVF | 0.09768968 |
| TDLATLG | ASSLGRGANTEVF | 0.10771542 |
| SAGGTGR | ASSLGRGANTEVF | 0.3013908 |
| GFSNSGS | ASSLGRGANTEVF | 0.34374863 |
| GGNGGSGGTSV | ASSLGRGANTEVF | 0.10436835 |
| GTSVATGGAGN | ASSLGRGANTEVF | 0.12360092 |
| NGGNGGSGGTS | ASSLGRGANTEVF | 0.14353299 |
| AVVKFPH | ASSLGRGANTEVF | 0.108047776 |
| TPPPPPP | ASSLGRGANTEVF | 0.114271946 |
| AAPAAAAAPA | ASSLGRGANTEVF | 0.09646963 |
| IALNEDL | ASSLGRGANTEVF | 0.20620054 |
| IVNVSSM | ASSLGRGANTEVF | 0.64673626 |
| TGIPPAPR | ASSLGRGANTEVF | 0.1039496 |
| FDLGGGTF | ASSLGRGANTEVF | 0.09352403 |
| FDLGGGTFDVS | ASSLGRGANTEVF | 0.13952455 |
| NEPTAAA | ASSLGRGANTEVF | 0.09309883 |
| NEPTAAALA | ASSLGRGANTEVF | 0.11464435 |
| GSIINISSI | ASSLGRGANTEVF | 0.10558544 |
| VAPGNAGTA | ASSLGRGANTEVF | 0.08855109 |
| TGGGASLE | ASSLGRGANTEVF | 0.11526081 |
| VAVYNHYKRI | ASSLGRGANTEVF | 0.16107395 |
| TDPAPATT | ASSLGRGANTEVF | 0.099514894 |
| TNRPDILDPALLRPG | ASSLGRGANTEVF | 0.23168144 |
| LTTTTTS | ASSLGRGANTEVF | 0.22026561 |
| PTGSGKTL | ASSLGRGANTEVF | 0.15667418 |
| TGSGKTL | ASSLGRGANTEVF | 0.20547149 |
| TIGHVDHGK | ASSLGRGANTEVF | 0.37867424 |
| TIGHVDHGKTT | ASSLGRGANTEVF | 0.12462823 |
| GHVDHGKTT | ASSLGRGANTEVF | 0.11901831 |
| AQFEHTLL | ASSLGRGANTEVF | 0.16129039 |
| IVNQDEL | ASSLGRGANTEVF | 0.15462385 |
| TNGSQFF | ASSLGRGANTEVF | 0.16406575 |
| TGKGIVE | ASSLGRGANTEVF | 0.12790197 |
| LISNASDA | ASSLGRGANTEVF | 0.16996914 |
| AELRTLGELTKIAKAHGAQ | ASSLGRGANTEVF | 0.11377151 |
| APNAVAAAEPSW | ASSLGRGANTEVF | 0.27808577 |
| APQVAGAATAVL | ASSLGRGANTEVF | 0.09732874 |
| ARRDPAAAAAILPTDA | ASSLGRGANTEVF | 0.10023222 |
| DAAARDVLAEAGLAEYF | ASSLGRGANTEVF | 0.20985627 |
| DDQIVLSAKANLAKVEQ | ASSLGRGANTEVF | 0.09329171 |
| DELLPDTGEDSDEDGH | ASSLGRGANTEVF | 0.10122889 |
| ELPEAPANRATVPQD | ASSLGRGANTEVF | 0.1295321 |

| | | |
|---|---|---|
| FGQSKIERVTGIG | ASSLGRGANTEVF | 0.18047258 |
| GAGGTGSTAGGAGGAGGAGGLY | ASSLGRGANTEVF | 0.08226506 |
| GHGGTGAAGGNGGTGGAGGSGI | ASSLGRGANTEVF | 0.11092068 |
| GNGGNGGAGGEGGPGGAGGAGA | ASSLGRGANTEVF | 0.11351117 |
| GSTAGNGGNGGNATNGTIAGQP | ASSLGRGANTEVF | 0.12828977 |
| GTGGDGGNGGNGGTGG | ASSLGRGANTEVF | 0.10603194 |
| GVRRPAGIPADAPV | ASSLGRGANTEVF | 0.094661824 |
| GYRQVTDAHLLTLARR | ASSLGRGANTEVF | 0.14546601 |
| IAKMGQDGHDRG | ASSLGRGANTEVF | 0.101224 |
| KAEPSIEIQYT | ASSLGRGANTEVF | 0.092346705 |
| KPVRIRMAADEDTVAVT | ASSLGRGANTEVF | 0.13726845 |
| MPPRIAGMRLLVIKPEP | ASSLGRGANTEVF | 0.5093087 |
| MKFYAVALFRSRGVPPTR | ASSLGRGANTEVF | 0.101325974 |
| NAGGFGILGAVAHSP | ASSLGRGANTEVF | 0.13192046 |
| NEGREITLAMAEM | ASSLGRGANTEVF | 0.10170887 |
| NELVEWSHPLGHRWRWE | ASSLGRGANTEVF | 0.62801373 |
| NLGAGGTGGNGGNPG | ASSLGRGANTEVF | 0.1156922 |
| NTEWPKWETVMAADPVD | ASSLGRGANTEVF | 0.09595262 |
| PASAAPAETQP | ASSLGRGANTEVF | 0.16135195 |
| QQSSDGKNGVRAAEITGE | ASSLGRGANTEVF | 0.20273307 |
| RAPDAVTTSWWKE | ASSLGRGANTEVF | 0.14022726 |
| SGFGNVGAHTSGWFNQ | ASSLGRGANTEVF | 0.20008662 |
| SGVAGAVGQAASVGGLKV | ASSLGRGANTEVF | 0.10142278 |
| SNGGAGTGSAGGNGGT | ASSLGRGANTEVF | 0.11324985 |
| TGGAGGEGGAAG | ASSLGRGANTEVF | 0.13655423 |
| TGGDGGAAGAGGAPGHG | ASSLGRGANTEVF | 0.14794797 |
| TMPPLLAAIANAGRHGV | ASSLGRGANTEVF | 0.09176508 |
| TSSTAGNSGGA | ASSLGRGANTEVF | 0.15474027 |
| VPATVPVGILC | ASSLGRGANTEVF | 0.11360889 |
| VQRPDSGGVVLTGRIS | ASSLGRGANTEVF | 0.12006892 |
| WTPNPVPATKGVPAGLVQAT | ASSLGRGANTEVF | 0.1320313 |
| YALKRALAKFSDRFDVVII | ASSLGRGANTEVF | 0.14650908 |
| YLIGSVIV | ASSLGRGANTEVF | 0.1849978 |
| YGQLVAVID | ASSLGRGANTEVF | 0.12284144 |
| VTVGGMTLPA | ASSLGRGANTEVF | 0.5214956 |
| VLALPAV | ASSLGRGANTEVF | 0.09428925 |
| VIPPTLNYET | ASSLGRGANTEVF | 0.30676144 |
| VGMGGMHPSSG | ASSLGRGANTEVF | 0.13928168 |
| VGLAGGAATA | ASSLGRGANTEVF | 0.17771317 |
| VGGAGGAGGSGP | ASSLGRGANTEVF | 0.1209307 |
| VFAAFMV | ASSLGRGANTEVF | 0.10701629 |
| TLYDKLR | ASSLGRGANTEVF | 0.17927651 |
| TLAVFTVL | ASSLGRGANTEVF | 0.1956291 |
| TGSVQPARV | ASSLGRGANTEVF | 0.11158396 |
| TGGGAGSGGT | ASSLGRGANTEVF | 0.16187239 |
| SHLETSL | ASSLGRGANTEVF | 0.28073752 |
| SFTTGMT | ASSLGRGANTEVF | 0.1098238 |
| QVEVAKRMGVRQPTVS | ASSLGRGANTEVF | 0.11443583 |
| QGSGGPALAM | ASSLGRGANTEVF | 0.17514765 |
| MLAPSNP | ASSLGRGANTEVF | 0.41611418 |
| LVNNAAI | ASSLGRGANTEVF | 0.11763437 |

| | | |
|---|---|---|
| LTGEINTLA | ASSLGRGANTEVF | 0.27960742 |
| LSIFGPV | ASSLGRGANTEVF | 0.09315523 |
| LLALLANL | ASSLGRGANTEVF | 0.17672333 |
| LISNASDALDKLR | ASSLGRGANTEVF | 0.09682781 |
| LINEAVP | ASSLGRGANTEVF | 0.13106506 |
| LGYAFSGL | ASSLGRGANTEVF | 0.0873006 |
| LDFVAVAL | ASSLGRGANTEVF | 0.14213249 |
| LAQPGAE | ASSLGRGANTEVF | 0.19006954 |
| ISNLGAVTGF | ASSLGRGANTEVF | 0.17720649 |
| HQEDPGRI | ASSLGRGANTEVF | 0.09087262 |
| GTPGEPTP | ASSLGRGANTEVF | 0.11596032 |
| GFFNSGS | ASSLGRGANTEVF | 0.20970869 |
| GAGGGGLTAGAN | ASSLGRGANTEVF | 0.38555226 |
| FVDALVNM | ASSLGRGANTEVF | 0.12106368 |
| FLTVAPEL | ASSLGRGANTEVF | 0.22586095 |
| FETLTAL | ASSLGRGANTEVF | 0.1088446 |
| DVINPIEL | ASSLGRGANTEVF | 0.10691714 |
| APVSGAPV | ASSLGRGANTEVF | 0.097744346 |
| ALVGGLAW | ASSLGRGANTEVF | 0.121159 |
| ALAAAAATAI | ASSLGRGANTEVF | 0.11979952 |
| AFGLGLLL | ASSLGRGANTEVF | 0.0986859 |
| AFFAAASL | ASSLGRGANTEVF | 0.105670266 |
| AAAALPAFSPP | ASSLGRGANTEVF | 0.43106207 |
| VVLAPAL | ASSLGRGANTEVF | 0.13644657 |
| TIGHVDHGKTTL | ASSLGRGANTEVF | 0.31506357 |
| LISNASDALDK | ASSLGRGANTEVF | 0.39357987 |
| LISNASDAL | ASSLGRGANTEVF | 0.20721427 |
| ISNASDALDK | ASSLGRGANTEVF | 0.09632151 |
| NQAETLVYQTE | ASSLGRGANTEVF | 0.12944975 |
| TGGTGGTAGAG | ASSLGRGANTEVF | 0.08988641 |
| TGSTAGGAGGA | ASSLGRGANTEVF | 0.10705509 |
| DPLAVPVAAGPV | ASSLGRGANTEVF | 0.092816696 |
| GGTGGTPGNGGN | ASSLGRGANTEVF | 0.13685156 |
| PAGLAGVDPATL | ASSLGRGANTEVF | 0.08299738 |
| GNNLVVEAPHAGASV | ASSLGRGANTEVF | 0.1377557 |
| KVTRIMSASTLEQPA | ASSLGRGANTEVF | 0.2678738 |
| LGFALPIALPRRTVV | ASSLGRGANTEVF | 0.088843435 |
| TRLGNLTARKLVVLV | ASSLGRGANTEVF | 0.20939943 |
| WSQQRRLIYFPSAGP | ASSLGRGANTEVF | 0.21876207 |
| GPNGNPDPMAAAVDIR | ASSLGRGANTEVF | 0.09104056 |
| TQRCVSVADLRAAASR | ASSLGRGANTEVF | 0.22766194 |
| QFVLDVAEHPVEVLTETV | ASSLGRGANTEVF | 0.098325066 |
| GGAGGTGGAGGNGGTQAGDG | ASSLGRGANTEVF | 0.2244852 |
| PLLATLRAIAVDVDAKPAQV | ASSLGRGANTEVF | 0.14195068 |
| APKFRTGGYMIDFWGVGYQVAK | ASSLGRGANTEVF | 0.15960252 |
| GAGGAGGAGGAAGGPGGTGGTG | ASSLGRGANTEVF | 0.17781276 |
| IGALGLLATDVAPAGSVAFGVA | ASSLGRGANTEVF | 0.1822978 |
| AGGAGGIGGNGGAGGAGGVLMG | ASSLGRGANTEVF | 0.09768246 |
| ADELIGLL | ASSLGRGANTEVF | 0.19297405 |
| ADRALKA | ASSLGRGANTEVF | 0.10616094 |
| AGAVGQVGGAGGA | ASSLGRGANTEVF | 0.14624879 |

| | | |
|---|---|---|
| AGGGGGLGGAP | ASSLGRGANTEVF | 0.0988131 |
| AGGLLYGNGGAG | ASSLGRGANTEVF | 0.18025936 |
| AGLESIGL | ASSLGRGANTEVF | 0.11186065 |
| AGLKTGII | ASSLGRGANTEVF | 0.15622458 |
| ASLAALLL | ASSLGRGANTEVF | 0.12061457 |
| ASVINISSI | ASSLGRGANTEVF | 0.10145159 |
| ATPPGPA | ASSLGRGANTEVF | 0.09650253 |
| ATVASALV | ASSLGRGANTEVF | 0.09084251 |
| ATVASLIL | ASSLGRGANTEVF | 0.22409278 |
| AVNGPVLIRH | ASSLGRGANTEVF | 0.11738272 |
| DGAAGAVGI | ASSLGRGANTEVF | 0.097180754 |
| DPLLHGI | ASSLGRGANTEVF | 0.1481952 |
| ELGAALGAL | ASSLGRGANTEVF | 0.117455766 |
| ELGRTQAV | ASSLGRGANTEVF | 0.09671921 |
| FVAGGYLS | ASSLGRGANTEVF | 0.12383676 |
| GASGGAGGAGGW | ASSLGRGANTEVF | 0.14288802 |
| GGAGAGGDGGAGGI | ASSLGRGANTEVF | 0.11206576 |
| GTGHNGSQPGMGGNG | ASSLGRGANTEVF | 0.0860955 |
| GYAAASASAS | ASSLGRGANTEVF | 0.10775117 |
| HQVDVAAVQQG | ASSLGRGANTEVF | 0.12405988 |
| IGAAIAHT | ASSLGRGANTEVF | 0.09195262 |
| IGMIEVAQL | ASSLGRGANTEVF | 0.09306157 |
| IRLAPEHTSAMD | ASSLGRGANTEVF | 0.20314072 |
| IYNVSVL | ASSLGRGANTEVF | 0.15567093 |
| KQGFLGM | ASSLGRGANTEVF | 0.11040195 |
| LESVITQL | ASSLGRGANTEVF | 0.22840878 |
| LGLGALVVV | ASSLGRGANTEVF | 0.18989775 |
| LIGNGGNGGSGGI | ASSLGRGANTEVF | 0.1930239 |
| LNAINAPTL | ASSLGRGANTEVF | 0.11434626 |
| LQLLNTGV | ASSLGRGANTEVF | 0.10803862 |
| LQTSIEF | ASSLGRGANTEVF | 0.16078515 |
| MSFAVSEL | ASSLGRGANTEVF | 0.13562474 |
| NTGALGL | ASSLGRGANTEVF | 0.09482696 |
| PPGAGGAGGAGTT | ASSLGRGANTEVF | 0.08823969 |
| QLVGLGI | ASSLGRGANTEVF | 0.11477336 |
| RFLEQVHQL | ASSLGRGANTEVF | 0.1087669 |
| RGVGYVLR | ASSLGRGANTEVF | 0.1385563 |
| SATGVFAGVI | ASSLGRGANTEVF | 0.30261958 |
| SLPAITLP | ASSLGRGANTEVF | 0.09772772 |
| STALAFTV | ASSLGRGANTEVF | 0.09596281 |
| SVLTVGLGK | ASSLGRGANTEVF | 0.110917136 |
| TALAGAGSGI | ASSLGRGANTEVF | 0.10183133 |
| TGLPFTGI | ASSLGRGANTEVF | 0.18623698 |
| VAEHVTV | ASSLGRGANTEVF | 0.105237156 |
| VDIVLNSL | ASSLGRGANTEVF | 0.5822352 |
| VLTGGGAVL | ASSLGRGANTEVF | 0.13864672 |
| VNVGLAYGI | ASSLGRGANTEVF | 0.09870557 |
| VSSAAVGAV | ASSLGRGANTEVF | 0.2019105 |
| VTSVYEAGV | ASSLGRGANTEVF | 0.114060186 |
| WVALDRGA | ASSLGRGANTEVF | 0.17515904 |
| YLVGSVVI | ASSLGRGANTEVF | 0.1449495 |

| | | |
|---|---|---|
| GPPGTGKTLL | ASSLGRGANTEVF | 0.09349415 |
| GVGMTKF | ASSLGRGANTEVF | 0.11297473 |
| IGIGGSDL | ASSLGRGANTEVF | 0.15761521 |
| IGTLLAEL | ASSLGRGANTEVF | 0.10120998 |
| IYTSGTT | ASSLGRGANTEVF | 0.17597947 |
| LSFPAGV | ASSLGRGANTEVF | 0.09000621 |
| LYTSGTT | ASSLGRGANTEVF | 0.099965155 |
| VGAFPFGF | ASSLGRGANTEVF | 0.20121895 |
| VGAGTVEF | ASSLGRGANTEVF | 0.13697873 |
| VGKSTLLL | ASSLGRGANTEVF | 0.43331483 |
| AALAHAGGGIV | ASSLGRGANTEVF | 0.10519041 |
| AALVAVSLL | ASSLGRGANTEVF | 0.09431097 |
| AAVSRAVAAL | ASSLGRGANTEVF | 0.24607465 |
| AGGSSGIGGF | ASSLGRGANTEVF | 0.111850604 |
| AGLTVANANG | ASSLGRGANTEVF | 0.1497638 |
| AGLVGAVL | ASSLGRGANTEVF | 0.09856233 |
| ALSQAFTV | ASSLGRGANTEVF | 0.1003421 |
| ALTFSVSL | ASSLGRGANTEVF | 0.117293246 |
| ALTGAALVAL | ASSLGRGANTEVF | 0.14555499 |
| ASFSIGAGVL | ASSLGRGANTEVF | 0.09253027 |
| DGSVYKI | ASSLGRGANTEVF | 0.23716849 |
| EQLAEIV | ASSLGRGANTEVF | 0.081979066 |
| FAVSELTVM | ASSLGRGANTEVF | 0.10706143 |
| GAAPLAAGL | ASSLGRGANTEVF | 0.11070982 |
| GADGGDGGSGGAGG | ASSLGRGANTEVF | 0.0985677 |
| GFGLDTGGAGG | ASSLGRGANTEVF | 0.086978324 |
| GLASGGLL | ASSLGRGANTEVF | 0.141636 |
| GRIAGAAVLL | ASSLGRGANTEVF | 0.21635146 |
| GSAGSLNTSF | ASSLGRGANTEVF | 0.23791622 |
| GTGQIGI | ASSLGRGANTEVF | 0.11383721 |
| GTLIALIL | ASSLGRGANTEVF | 0.17773308 |
| IAATMGFVVG | ASSLGRGANTEVF | 0.4017992 |
| IFAQALL | ASSLGRGANTEVF | 0.12207317 |
| IGAAVALGF | ASSLGRGANTEVF | 0.090747036 |
| IGLTPLAL | ASSLGRGANTEVF | 0.1121076 |
| ILGSNTSTL | ASSLGRGANTEVF | 0.088355295 |
| IVFSLTVL | ASSLGRGANTEVF | 0.11041799 |
| LASKATGF | ASSLGRGANTEVF | 0.656648 |
| LASVALIL | ASSLGRGANTEVF | 0.08971162 |
| LFAVGLTL | ASSLGRGANTEVF | 0.14757705 |
| LFNGGIGTYI | ASSLGRGANTEVF | 0.19326553 |
| LTLGAGII | ASSLGRGANTEVF | 0.11197299 |
| LVAAALGDI | ASSLGRGANTEVF | 0.11207567 |
| NTAFVQLGI | ASSLGRGANTEVF | 0.103377044 |
| QLFSGAEL | ASSLGRGANTEVF | 0.097751476 |
| SAATFITL | ASSLGRGANTEVF | 0.16515224 |
| SNYGLANLGV | ASSLGRGANTEVF | 0.113069326 |
| SYGGAYAVM | ASSLGRGANTEVF | 0.11034742 |
| TTGVGGLTLG | ASSLGRGANTEVF | 0.11543725 |
| TTQTIYI | ASSLGRGANTEVF | 0.10656484 |
| VAAGAGLGAVI | ASSLGRGANTEVF | 0.10993991 |

| | | |
|---|---|---|
| VAAVVTLL | ASSLGRGANTEVF | 0.10480399 |
| VGFLGGLI | ASSLGRGANTEVF | 0.106983244 |
| VGVIPILL | ASSLGRGANTEVF | 0.16596392 |
| VIFASSLI | ASSLGRGANTEVF | 0.09088692 |
| VKGGDGGAAAG | ASSLGRGANTEVF | 0.18624279 |
| VTKTFTNI | ASSLGRGANTEVF | 0.221113 |
| VVAAAILL | ASSLGRGANTEVF | 0.0888514 |
| YAALDVEL | ASSLGRGANTEVF | 0.1004282 |
| LASVALLL | ASSLGRGANTEVF | 0.11741204 |
| LDAVHRV | ASSLGRGANTEVF | 0.08976478 |
| QGLSSSIF | ASSLGRGANTEVF | 0.23709793 |
| RGTFANIRL | ASSLGRGANTEVF | 0.1249372 |
| RQLWWGHRI | ASSLGRGANTEVF | 0.1514144 |
| YGPPGTGKTLL | ASSLGRGANTEVF | 0.09794718 |
| AALVGAPANVA | ASSLGRGANTEVF | 0.44098637 |
| AATFRRTE | ASSLGRGANTEVF | 0.14634623 |
| AGALGGGI | ASSLGRGANTEVF | 0.11776763 |
| AGGAGGAGGIGAG | ASSLGRGANTEVF | 0.09418483 |
| AGSGALGL | ASSLGRGANTEVF | 0.10275002 |
| AQIADQI | ASSLGRGANTEVF | 0.1792289 |
| ASAVAVTYL | ASSLGRGANTEVF | 0.2994541 |
| ASTVLGGTGG | ASSLGRGANTEVF | 0.091364056 |
| AVGGEPAK | ASSLGRGANTEVF | 0.11509639 |
| CEQGTFGGTC | ASSLGRGANTEVF | 0.24945359 |
| DEGKAPAG | ASSLGRGANTEVF | 0.19782005 |
| DTVVFPAL | ASSLGRGANTEVF | 0.095595144 |
| FEGITMAG | ASSLGRGANTEVF | 0.103033684 |
| FPLVVQD | ASSLGRGANTEVF | 0.14152701 |
| GAFTTPQ | ASSLGRGANTEVF | 0.119426675 |
| GFGGFGTAGGVGGA | ASSLGRGANTEVF | 0.09599392 |
| GFLAQFL | ASSLGRGANTEVF | 0.22239661 |
| GGAGGAGGNAGVA | ASSLGRGANTEVF | 0.15160096 |
| GGVGTAGAGGAGGAGG | ASSLGRGANTEVF | 0.32266802 |
| GNGGGGDAGNAGSGGN | ASSLGRGANTEVF | 0.1327617 |
| GTGGAGGAGNDAGS | ASSLGRGANTEVF | 0.15810692 |
| GTTNTGWFNS | ASSLGRGANTEVF | 0.13811097 |
| IFMGPLFTL | ASSLGRGANTEVF | 0.10851953 |
| ILAHAIEAI | ASSLGRGANTEVF | 0.10069709 |
| IRSAIREAL | ASSLGRGANTEVF | 0.090328634 |
| ITFAGGG | ASSLGRGANTEVF | 0.09729123 |
| ITGMSGHQEV | ASSLGRGANTEVF | 0.10798856 |
| IVAGLACTL | ASSLGRGANTEVF | 0.083287135 |
| IVGGGLAGA | ASSLGRGANTEVF | 0.118827626 |
| IVTGVLNI | ASSLGRGANTEVF | 0.11727154 |
| LAQQGCI | ASSLGRGANTEVF | 0.13694775 |
| LCLHVEN | ASSLGRGANTEVF | 0.12564397 |
| LDAISDGS | ASSLGRGANTEVF | 0.11744885 |
| LGGFAAML | ASSLGRGANTEVF | 0.091655694 |
| LGGKNPMI | ASSLGRGANTEVF | 0.24581897 |
| LGPKGVAAIL | ASSLGRGANTEVF | 0.08852011 |
| LMVGVSI | ASSLGRGANTEVF | 0.10245657 |

| | | |
|---|---|---|
| LVGAAFVL | ASSLGRGANTEVF | 0.11656616 |
| LVLAVAPL | ASSLGRGANTEVF | 0.1217555 |
| MAAIAVIDAA | ASSLGRGANTEVF | 0.57221156 |
| NNGINAELQ | ASSLGRGANTEVF | 0.20897877 |
| RGEPLVLVHG | ASSLGRGANTEVF | 0.13158108 |
| RGGAGGAAGLI | ASSLGRGANTEVF | 0.31741917 |
| RSYRMHPAV | ASSLGRGANTEVF | 0.08719896 |
| SAAHLLAGQLA | ASSLGRGANTEVF | 0.10496471 |
| SAAVGSY | ASSLGRGANTEVF | 0.27344695 |
| SPHAVLHRI | ASSLGRGANTEVF | 0.10428735 |
| SQIAKVL | ASSLGRGANTEVF | 0.16115303 |
| TALGHLVP | ASSLGRGANTEVF | 0.13390361 |
| TAPTTSGAA | ASSLGRGANTEVF | 0.10126042 |
| TDGDDGNGGSPGT | ASSLGRGANTEVF | 0.11401427 |
| TDGLGEN | ASSLGRGANTEVF | 0.09919879 |
| TGGPPLPGQA | ASSLGRGANTEVF | 0.1886213 |
| TIPAITI | ASSLGRGANTEVF | 0.13071221 |
| TIPGFTI | ASSLGRGANTEVF | 0.095879056 |
| TSIASGALA | ASSLGRGANTEVF | 0.2253943 |
| TVTLPGAAI | ASSLGRGANTEVF | 0.62856454 |
| VASGLSAALP | ASSLGRGANTEVF | 0.115283936 |
| VGAGVIGI | ASSLGRGANTEVF | 0.14353308 |
| VGPKRFGV | ASSLGRGANTEVF | 0.11370184 |
| VNVAFPD | ASSLGRGANTEVF | 0.1255694 |
| VSMTNSGVS | ASSLGRGANTEVF | 0.10679177 |
| VTDVLGDLSV | ASSLGRGANTEVF | 0.09778356 |
| VVVVGAGFAGL | ASSLGRGANTEVF | 0.10188238 |
| VYTVGTVV | ASSLGRGANTEVF | 0.099927284 |
| WTTFSGP | ASSLGRGANTEVF | 0.14977232 |
| YDELVEAF | ASSLGRGANTEVF | 0.116809495 |
| YGTGGAGGNGG | ASSLGRGANTEVF | 0.6536509 |
| YTNVMAV | ASSLGRGANTEVF | 0.17644495 |
| FLEQVHQL | ASSLGRGANTEVF | 0.104740076 |
| GLSLLAVAV | ASSLGRGANTEVF | 0.0914615 |
| LEQVHQL | ASSLGRGANTEVF | 0.1208306 |
| LNAAGAI | ASSLGRGANTEVF | 0.11019774 |
| PAVHASL | ASSLGRGANTEVF | 0.101426244 |
| TELEDDA | ASSLGRGANTEVF | 0.22776188 |
| TPPGPPN | ASSLGRGANTEVF | 0.101157695 |
| TPPPPPPP | ASSLGRGANTEVF | 0.10160768 |
| TQSPALI | ASGDRDWGIAEQF | 0.13071978 |
| EFAAAFPQ | ASGDRDWGIAEQF | 0.43622482 |
| LLSGNLDVL | ASGDRDWGIAEQF | 0.13465983 |
| GPANLGNYNI | ASGDRDWGIAEQF | 0.11308525 |
| TLGPTLP | ASGDRDWGIAEQF | 0.09179733 |
| VYAYSSCCK | ASGDRDWGIAEQF | 0.22127351 |
| APAPAPAPNI | ASGDRDWGIAEQF | 0.23463395 |
| PGGPVPIP | ASGDRDWGIAEQF | 0.15072449 |
| FLTVALEP | ASGDRDWGIAEQF | 0.45895928 |
| SQYFGANL | ASGDRDWGIAEQF | 0.11065272 |
| GGVKPAA | ASGDRDWGIAEQF | 0.095868595 |

| | | |
|---|---|---|
| GGTGGAGGAGGW | ASGDRDWGIAEQF | 0.16196749 |
| GVGGVAGLGGAGG | ASGDRDWGIAEQF | 0.13353491 |
| VLLVLATTI | ASGDRDWGIAEQF | 0.37936398 |
| IGIFAAML | ASGDRDWGIAEQF | 0.1282183 |
| AVGDLID | ASGDRDWGIAEQF | 0.46611047 |
| NISPLPL | ASGDRDWGIAEQF | 0.1129599 |
| VTAVFVAI | ASGDRDWGIAEQF | 0.24598135 |
| SGDDNDAAL | ASGDRDWGIAEQF | 0.30387983 |
| AVAALAHS | ASGDRDWGIAEQF | 0.12729773 |
| GGVTVMATP | ASGDRDWGIAEQF | 0.12157514 |
| AGLAANVRD | ASGDRDWGIAEQF | 0.11143682 |
| TNRPDLIDPALLRPG | ASGDRDWGIAEQF | 0.14860499 |
| LANLGVTAL | ASGDRDWGIAEQF | 0.10282978 |
| TGERIEL | ASGDRDWGIAEQF | 0.11143365 |
| NGEELAA | ASGDRDWGIAEQF | 0.13194765 |
| VLSLVWG | ASGDRDWGIAEQF | 0.15795466 |
| IGYALWL | ASGDRDWGIAEQF | 0.097542405 |
| LSLLLAPL | ASGDRDWGIAEQF | 0.183718 |
| TLDEALK | ASGDRDWGIAEQF | 0.38339457 |
| TDGAGTAEGG | ASGDRDWGIAEQF | 0.15017201 |
| NAPGGAGGNGG | ASGDRDWGIAEQF | 0.09571147 |
| VSVFMGMGGNF | ASGDRDWGIAEQF | 0.25872245 |
| LASYLAAL | ASGDRDWGIAEQF | 0.40141004 |
| LVGGTPAL | ASGDRDWGIAEQF | 0.17772648 |
| VRNADVL | ASGDRDWGIAEQF | 0.095268905 |
| TCSSPFLVSAM | ASGDRDWGIAEQF | 0.08909218 |
| VHITDAE | ASGDRDWGIAEQF | 0.17023112 |
| EYAAPSF | ASGDRDWGIAEQF | 0.118946776 |
| VVDAFHGGL | ASGDRDWGIAEQF | 0.1058555 |
| ASVVDPQ | ASGDRDWGIAEQF | 0.09042663 |
| SYNEFGGLS | ASGDRDWGIAEQF | 0.1267737 |
| DPDGSVY | ASGDRDWGIAEQF | 0.13941553 |
| MVSTIHATE | ASGDRDWGIAEQF | 0.1098695 |
| LAAGVADPV | ASGDRDWGIAEQF | 0.090087004 |
| TKEQIAA | ASGDRDWGIAEQF | 0.15613127 |
| SLASPAAL | ASGDRDWGIAEQF | 0.22013289 |
| IGLVFYGV | ASGDRDWGIAEQF | 0.096595645 |
| FLLDASNL | ASGDRDWGIAEQF | 0.10690508 |
| YQVGNFP | ASGDRDWGIAEQF | 0.14575921 |
| ESLLSQST | ASGDRDWGIAEQF | 0.13390183 |
| LFTVGFAAVG | ASGDRDWGIAEQF | 0.10686668 |
| ATPPTPT | ASGDRDWGIAEQF | 0.09896359 |
| TDLATLG | ASGDRDWGIAEQF | 0.10866126 |
| SAGGTGR | ASGDRDWGIAEQF | 0.10783933 |
| GFSNSGS | ASGDRDWGIAEQF | 0.18981123 |
| GGNGGSGGTSV | ASGDRDWGIAEQF | 0.18100742 |
| GTSVATGGAGN | ASGDRDWGIAEQF | 0.16329059 |
| NGGNGGSGGTS | ASGDRDWGIAEQF | 0.13241258 |
| AVVKFPH | ASGDRDWGIAEQF | 0.10955043 |
| TPPPPPP | ASGDRDWGIAEQF | 0.09321083 |
| AAPAAAAAPA | ASGDRDWGIAEQF | 0.086560585 |

| | | |
|---|---|---|
| IALNEDL | ASGDRDWGIAEQF | 0.38129464 |
| IVNVSSM | ASGDRDWGIAEQF | 0.10832972 |
| TGIPPAPR | ASGDRDWGIAEQF | 0.23649824 |
| FDLGGGTF | ASGDRDWGIAEQF | 0.114337675 |
| FDLGGGTFDVS | ASGDRDWGIAEQF | 0.101332456 |
| NEPTAAA | ASGDRDWGIAEQF | 0.12384532 |
| NEPTAAALA | ASGDRDWGIAEQF | 0.085198715 |
| GSIINISSI | ASGDRDWGIAEQF | 0.12053607 |
| VAPGNAGTA | ASGDRDWGIAEQF | 0.21432808 |
| TGGGASLE | ASGDRDWGIAEQF | 0.1307668 |
| VAVYNHYKRI | ASGDRDWGIAEQF | 0.14236699 |
| TDPAPATT | ASGDRDWGIAEQF | 0.16641825 |
| TNRPDILDPALLRPG | ASGDRDWGIAEQF | 0.17933944 |
| LTTTTTS | ASGDRDWGIAEQF | 0.11269531 |
| PTGSGKTL | ASGDRDWGIAEQF | 0.14710619 |
| TGSGKTL | ASGDRDWGIAEQF | 0.16366935 |
| TIGHVDHGK | ASGDRDWGIAEQF | 0.19553429 |
| TIGHVDHGKTT | ASGDRDWGIAEQF | 0.09129901 |
| GHVDHGKTT | ASGDRDWGIAEQF | 0.1166725 |
| AQFEHTLL | ASGDRDWGIAEQF | 0.100485526 |
| IVNQDEL | ASGDRDWGIAEQF | 0.13379821 |
| TNGSQFF | ASGDRDWGIAEQF | 0.11555134 |
| TGKGIVE | ASGDRDWGIAEQF | 0.17425963 |
| LISNASDA | ASGDRDWGIAEQF | 0.12258812 |
| AELRTLGELTKIAKAHGAQ | ASGDRDWGIAEQF | 0.09190094 |
| APNAVAAAEPSW | ASGDRDWGIAEQF | 0.10293416 |
| APQVAGAATAVL | ASGDRDWGIAEQF | 0.1389066 |
| ARRDPAAAAAILPTDA | ASGDRDWGIAEQF | 0.1299289 |
| DAAARDVLAEAGLAEYF | ASGDRDWGIAEQF | 0.11414464 |
| DDQIVLSAKANLAKVEQ | ASGDRDWGIAEQF | 0.095708616 |
| DELLPDTGEDSDEDGH | ASGDRDWGIAEQF | 0.17049146 |
| ELPEAPANRATVPQD | ASGDRDWGIAEQF | 0.10799116 |
| FGQSKIERVTGIG | ASGDRDWGIAEQF | 0.10589576 |
| GAGGTGSTAGGAGGAGGAGGLY | ASGDRDWGIAEQF | 0.14384979 |
| GHGGTGAAGGNGGTGGAGGSGI | ASGDRDWGIAEQF | 0.11159672 |
| GNGGNGGAGGEGGPGGAGGAGA | ASGDRDWGIAEQF | 0.10515733 |
| GSTAGNGGNGGNATNGTIAGQP | ASGDRDWGIAEQF | 0.3516701 |
| GTGGDGGNGGNGGTGG | ASGDRDWGIAEQF | 0.09782062 |
| GVRRPAGIPADAPV | ASGDRDWGIAEQF | 0.13753553 |
| GYRQVTDAHLLTLARR | ASGDRDWGIAEQF | 0.13826191 |
| IAKMGQDGHDRG | ASGDRDWGIAEQF | 0.107900135 |
| KAEPSIEIQYT | ASGDRDWGIAEQF | 0.15641363 |
| KPVRIRMAADEDTVAVT | ASGDRDWGIAEQF | 0.09388303 |
| MPPRIAGMRLLVIKPEP | ASGDRDWGIAEQF | 0.2008166 |
| MKFYAVALFRSRGVPPTR | ASGDRDWGIAEQF | 0.10838703 |
| NAGGFGILGAVAHSP | ASGDRDWGIAEQF | 0.10692551 |
| NEGREITLAMAEM | ASGDRDWGIAEQF | 0.10936011 |
| NELVEWSHPLGHRWRWE | ASGDRDWGIAEQF | 0.137311 |
| NLGAGGTGGNGGNPG | ASGDRDWGIAEQF | 0.17717497 |
| NTEWPKWETVMAADPVD | ASGDRDWGIAEQF | 0.108518295 |
| PASAAPAETQP | ASGDRDWGIAEQF | 0.1594993 |

| | | |
|---|---|---|
| QQSSDGKNGVRAAEITGE | ASGDRDWGIAEQF | 0.18643475 |
| RAPDAVTTSWWKE | ASGDRDWGIAEQF | 0.13190736 |
| SGFGNVGAHTSGWFNQ | ASGDRDWGIAEQF | 0.09030158 |
| SGVAGAVGQAASVGGLKV | ASGDRDWGIAEQF | 0.101412036 |
| SNGGAGTGSAGGNGGT | ASGDRDWGIAEQF | 0.096402116 |
| TGGAGGEGGAAG | ASGDRDWGIAEQF | 0.12040653 |
| TGGDGGAAGAGGAPGHG | ASGDRDWGIAEQF | 0.0959996 |
| TMPPLLAAIANAGRHGV | ASGDRDWGIAEQF | 0.14785615 |
| TSSTAGNSGGA | ASGDRDWGIAEQF | 0.1827355 |
| VPATVPVGILC | ASGDRDWGIAEQF | 0.10539416 |
| VQRPDSGGVVLTGRIS | ASGDRDWGIAEQF | 0.40087277 |
| WTPNPVPATKGVPAGLVQAT | ASGDRDWGIAEQF | 0.09138594 |
| YALKRALAKFSDRFDVVII | ASGDRDWGIAEQF | 0.12934743 |
| YLIGSVIV | ASGDRDWGIAEQF | 0.22920434 |
| YGQLVAVID | ASGDRDWGIAEQF | 0.0924273 |
| VTVGGMTLPA | ASGDRDWGIAEQF | 0.119826004 |
| VLALPAV | ASGDRDWGIAEQF | 0.1116846 |
| VIPPTLNYET | ASGDRDWGIAEQF | 0.09462438 |
| VGMGGMHPSSG | ASGDRDWGIAEQF | 0.20649025 |
| VGLAGGAATA | ASGDRDWGIAEQF | 0.11772521 |
| VGGAGGAGGSGP | ASGDRDWGIAEQF | 0.20265707 |
| VFAAFMV | ASGDRDWGIAEQF | 0.10122581 |
| TLYDKLR | ASGDRDWGIAEQF | 0.09343695 |
| TLAVFTVL | ASGDRDWGIAEQF | 0.14192715 |
| TGSVQPARV | ASGDRDWGIAEQF | 0.4734707 |
| TGGGAGSGGT | ASGDRDWGIAEQF | 0.11156316 |
| SHLETSL | ASGDRDWGIAEQF | 0.3450055 |
| SFTTGMT | ASGDRDWGIAEQF | 0.14542118 |
| QVEVAKRMGVRQPTVS | ASGDRDWGIAEQF | 0.18312304 |
| QGSGGPALAM | ASGDRDWGIAEQF | 0.38131577 |
| MLAPSNP | ASGDRDWGIAEQF | 0.12429322 |
| LVNNAAI | ASGDRDWGIAEQF | 0.1148947 |
| LTGEINTLA | ASGDRDWGIAEQF | 0.25071537 |
| LSIFGPV | ASGDRDWGIAEQF | 0.15348244 |
| LLALLANL | ASGDRDWGIAEQF | 0.2523813 |
| LISNASDALDKLR | ASGDRDWGIAEQF | 0.26850018 |
| LINEAVP | ASGDRDWGIAEQF | 0.4271407 |
| LGYAFSGL | ASGDRDWGIAEQF | 0.7728678 |
| LDFVAVAL | ASGDRDWGIAEQF | 0.646101 |
| LAQPGAE | ASGDRDWGIAEQF | 0.55691546 |
| ISNLGAVTGF | ASGDRDWGIAEQF | 0.41881555 |
| HQEDPGRI | ASGDRDWGIAEQF | 0.33218604 |
| GTPGEPTP | ASGDRDWGIAEQF | 0.5628005 |
| GFFNSGS | ASGDRDWGIAEQF | 0.44455293 |
| GAGGGGLTAGAN | ASGDRDWGIAEQF | 0.69085634 |
| FVDALVNM | ASGDRDWGIAEQF | 0.70632774 |
| FLTVAPEL | ASGDRDWGIAEQF | 0.4482243 |
| FETLTAL | ASGDRDWGIAEQF | 0.79794246 |
| DVINPIEL | ASGDRDWGIAEQF | 0.7917893 |
| APVSGAPV | ASGDRDWGIAEQF | 0.29982737 |
| ALVGGLAW | ASGDRDWGIAEQF | 0.44696978 |

| | | |
|---|---|---|
| ALAAAAATAI | ASGDRDWGIAEQF | 0.4501739 |
| AFGLGLLL | ASGDRDWGIAEQF | 0.5066176 |
| AFFAAASL | ASGDRDWGIAEQF | 0.19208795 |
| AAAALPAFSPP | ASGDRDWGIAEQF | 0.74102056 |
| VVLAPAL | ASGDRDWGIAEQF | 0.40016925 |
| TIGHVDHGKTTL | ASGDRDWGIAEQF | 0.15369532 |
| LISNASDALDK | ASGDRDWGIAEQF | 0.5470737 |
| LISNASDAL | ASGDRDWGIAEQF | 0.52684206 |
| ISNASDALDK | ASGDRDWGIAEQF | 0.5782639 |
| NQAETLVYQTE | ASGDRDWGIAEQF | 0.7640538 |
| TGGTGGTAGAG | ASGDRDWGIAEQF | 0.3552821 |
| TGSTAGGAGGA | ASGDRDWGIAEQF | 0.44114533 |
| DPLAVPVAAGPV | ASGDRDWGIAEQF | 0.7273388 |
| GGTGGTPGNGGN | ASGDRDWGIAEQF | 0.8122403 |
| PAGLAGVDPATL | ASGDRDWGIAEQF | 0.40019745 |
| GNNLVVEAPHAGASV | ASGDRDWGIAEQF | 0.66310734 |
| KVTRIMSASTLEQPA | ASGDRDWGIAEQF | 0.7908315 |
| LGFALPIALPRRTVV | ASGDRDWGIAEQF | 0.705799 |
| TRLGNLTARKLVVLV | ASGDRDWGIAEQF | 0.72837895 |
| WSQQRRLIYFPSAGP | ASGDRDWGIAEQF | 0.78306484 |
| GPNGNPDPMAAAVDIR | ASGDRDWGIAEQF | 0.38478687 |
| TQRCVSVADLRAAASR | ASGDRDWGIAEQF | 0.7716861 |
| QFVLDVAEHPVEVLTETV | ASGDRDWGIAEQF | 0.80167496 |
| GGAGGTGGAGGNGGTQAGDG | ASGDRDWGIAEQF | 0.49762753 |
| PLLATLRAIAVDVDAKPAQV | ASGDRDWGIAEQF | 0.64737093 |
| APKFRTGGYMIDFWGVGYQVAK | ASGDRDWGIAEQF | 0.43373764 |
| GAGGAGGAGGAAGGPGGTGGTG | ASGDRDWGIAEQF | 0.4725971 |
| IGALGLLATDVAPAGSVAFGVA | ASGDRDWGIAEQF | 0.2026875 |
| AGGAGGIGGNGGAGGAGGVLMG | ASGDRDWGIAEQF | 0.68880296 |
| ADELIGLL | ASGDRDWGIAEQF | 0.7577637 |
| ADRALKA | ASGDRDWGIAEQF | 0.55203843 |
| AGAVGQVGGAGGA | ASGDRDWGIAEQF | 0.36552718 |
| AGGGGGLGGAP | ASGDRDWGIAEQF | 0.37417185 |
| AGGLLYGNGGAG | ASGDRDWGIAEQF | 0.44383416 |
| AGLESIGL | ASGDRDWGIAEQF | 0.5382969 |
| AGLKTGII | ASGDRDWGIAEQF | 0.279781 |
| ASLAALLL | ASGDRDWGIAEQF | 0.35941014 |
| ASVINISSI | ASGDRDWGIAEQF | 0.78113985 |
| ATPPGPA | ASGDRDWGIAEQF | 0.79529655 |
| ATVASALV | ASGDRDWGIAEQF | 0.2022209 |
| ATVASLIL | ASGDRDWGIAEQF | 0.7692733 |
| AVNGPVLIRH | ASGDRDWGIAEQF | 0.6586832 |
| DGAAGAVGI | ASGDRDWGIAEQF | 0.7869747 |
| DPLLHGI | ASGDRDWGIAEQF | 0.58906686 |
| ELGAALGAL | ASGDRDWGIAEQF | 0.37027112 |
| ELGRTQAV | ASGDRDWGIAEQF | 0.6960765 |
| FVAGGYLS | ASGDRDWGIAEQF | 0.52461696 |
| GASGGAGGAGGW | ASGDRDWGIAEQF | 0.4648298 |
| GGAGAGGDGGAGGI | ASGDRDWGIAEQF | 0.71030015 |
| GTGHNGSQPGMGGNG | ASGDRDWGIAEQF | 0.57689524 |
| GYAAASASAS | ASGDRDWGIAEQF | 0.43447143 |

| | | |
|---|---|---|
| HQVDVAAVQQG | ASGDRDWGIAEQF | 0.41111314 |
| IGAAIAHT | ASGDRDWGIAEQF | 0.69115484 |
| IGMIEVAQL | ASGDRDWGIAEQF | 0.28254995 |
| IRLAPEHTSAMD | ASGDRDWGIAEQF | 0.45852694 |
| IYNVSVL | ASGDRDWGIAEQF | 0.39815262 |
| KQGFLGM | ASGDRDWGIAEQF | 0.6630626 |
| LESVITQL | ASGDRDWGIAEQF | 0.7788545 |
| LGLGALVVV | ASGDRDWGIAEQF | 0.6486982 |
| LIGNGGNGGSGGI | ASGDRDWGIAEQF | 0.39801106 |
| LNAINAPTL | ASGDRDWGIAEQF | 0.7177747 |
| LQLLNTGV | ASGDRDWGIAEQF | 0.6629603 |
| LQTSIEF | ASGDRDWGIAEQF | 0.58762705 |
| MSFAVSEL | ASGDRDWGIAEQF | 0.64876735 |
| NTGALGL | ASGDRDWGIAEQF | 0.44416383 |
| PPGAGGAGGAGTT | ASGDRDWGIAEQF | 0.6795767 |
| QLVGLGI | ASGDRDWGIAEQF | 0.63824743 |
| RFLEQVHQL | ASGDRDWGIAEQF | 0.71140414 |
| RGVGYVLR | ASGDRDWGIAEQF | 0.6774368 |
| SATGVFAGVI | ASGDRDWGIAEQF | 0.6642002 |
| SLPAITLP | ASGDRDWGIAEQF | 0.5409507 |
| STALAFTV | ASGDRDWGIAEQF | 0.18658924 |
| SVLTVGLGK | ASGDRDWGIAEQF | 0.4657827 |
| TALAGAGSGI | ASGDRDWGIAEQF | 0.62529075 |
| TGLPFTGI | ASGDRDWGIAEQF | 0.61246073 |
| VAEHVTV | ASGDRDWGIAEQF | 0.6726002 |
| VDIVLNSL | ASGDRDWGIAEQF | 0.6878457 |
| VLTGGGAVL | ASGDRDWGIAEQF | 0.42591125 |
| VNVGLAYGI | ASGDRDWGIAEQF | 0.72276795 |
| VSSAAVGAV | ASGDRDWGIAEQF | 0.728135 |
| VTSVYEAGV | ASGDRDWGIAEQF | 0.28248376 |
| WVALDRGA | ASGDRDWGIAEQF | 0.5038428 |
| YLVGSVVI | ASGDRDWGIAEQF | 0.7942234 |
| GPPGTGKTLL | ASGDRDWGIAEQF | 0.6230638 |
| GVGMTKF | ASGDRDWGIAEQF | 0.52132255 |
| IGIGGSDL | ASGDRDWGIAEQF | 0.4593612 |
| IGTLLAEL | ASGDRDWGIAEQF | 0.7923949 |
| IYTSGTT | ASGDRDWGIAEQF | 0.47294438 |
| LSFPAGV | ASGDRDWGIAEQF | 0.7937098 |
| LYTSGTT | ASGDRDWGIAEQF | 0.14045934 |
| VGAFPFGF | ASGDRDWGIAEQF | 0.4237836 |
| VGAGTVEF | ASGDRDWGIAEQF | 0.3928146 |
| VGKSTLLL | ASGDRDWGIAEQF | 0.28133717 |
| AALAHAGGGIV | ASGDRDWGIAEQF | 0.7761611 |
| AALVAVSLL | ASGDRDWGIAEQF | 0.4572098 |
| AAVSRAVAAL | ASGDRDWGIAEQF | 0.7153315 |
| AGGSSGIGGF | ASGDRDWGIAEQF | 0.58121264 |
| AGLTVANANG | ASGDRDWGIAEQF | 0.7099354 |
| AGLVGAVL | ASGDRDWGIAEQF | 0.61939603 |
| ALSQAFTV | ASGDRDWGIAEQF | 0.51726747 |
| ALTFSVSL | ASGDRDWGIAEQF | 0.44530442 |
| ALTGAALVAL | ASGDRDWGIAEQF | 0.7398207 |

| | | |
|---|---|---|
| ASFSIGAGVL | ASGDRDWGIAEQF | 0.73579955 |
| DGSVYKI | ASGDRDWGIAEQF | 0.6211964 |
| EQLAEIV | ASGDRDWGIAEQF | 0.3575182 |
| FAVSELTVM | ASGDRDWGIAEQF | 0.4130934 |
| GAAPLAAGL | ASGDRDWGIAEQF | 0.65771806 |
| GADGGDGGSGGAGG | ASGDRDWGIAEQF | 0.23846613 |
| GFGLDTGGAGG | ASGDRDWGIAEQF | 0.21058352 |
| GLASGGLL | ASGDRDWGIAEQF | 0.42020985 |
| GRIAGAAVLL | ASGDRDWGIAEQF | 0.34690702 |
| GSAGSLNTSF | ASGDRDWGIAEQF | 0.7764069 |
| GTGQIGI | ASGDRDWGIAEQF | 0.7507962 |
| GTLIALIL | ASGDRDWGIAEQF | 0.5622573 |
| IAATMGFVVG | ASGDRDWGIAEQF | 0.7179262 |
| IFAQALL | ASGDRDWGIAEQF | 0.6698358 |
| IGAAVALGF | ASGDRDWGIAEQF | 0.121700525 |
| IGLTPLAL | ASGDRDWGIAEQF | 0.46649885 |
| ILGSNTSTL | ASGDRDWGIAEQF | 0.32114807 |
| IVFSLTVL | ASGDRDWGIAEQF | 0.7701132 |
| LASKATGF | ASGDRDWGIAEQF | 0.40103954 |
| LASVALIL | ASGDRDWGIAEQF | 0.35179356 |
| LFAVGLTL | ASGDRDWGIAEQF | 0.6051705 |
| LFNGGIGTYI | ASGDRDWGIAEQF | 0.5713529 |
| LTLGAGII | ASGDRDWGIAEQF | 0.5667385 |
| LVAAALGDI | ASGDRDWGIAEQF | 0.3459873 |
| NTAFVQLGI | ASGDRDWGIAEQF | 0.14802937 |
| QLFSGAEL | ASGDRDWGIAEQF | 0.7519235 |
| SAATFITL | ASGDRDWGIAEQF | 0.30659628 |
| SNYGLANLGV | ASGDRDWGIAEQF | 0.4458533 |
| SYGGAYAVM | ASGDRDWGIAEQF | 0.1698286 |
| TTGVGGLTLG | ASGDRDWGIAEQF | 0.4729698 |
| TTQTIYI | ASGDRDWGIAEQF | 0.34123275 |
| VAAGAGLGAVI | ASGDRDWGIAEQF | 0.729679 |
| VAAVVTLL | ASGDRDWGIAEQF | 0.47937852 |
| VGFLGGLI | ASGDRDWGIAEQF | 0.4537034 |
| VGVIPILL | ASGDRDWGIAEQF | 0.7952025 |
| VIFASSLI | ASGDRDWGIAEQF | 0.62872666 |
| VKGGDGGAAAG | ASGDRDWGIAEQF | 0.45919663 |
| VTKTFTNI | ASGDRDWGIAEQF | 0.46887055 |
| VVAAAILL | ASGDRDWGIAEQF | 0.36384726 |
| YAALDVEL | ASGDRDWGIAEQF | 0.45058843 |
| LASVALLL | ASGDRDWGIAEQF | 0.4447145 |
| LDAVHRV | ASGDRDWGIAEQF | 0.8124206 |
| QGLSSSIF | ASGDRDWGIAEQF | 0.4426499 |
| RGTFANIRL | ASGDRDWGIAEQF | 0.34444663 |
| RQLWWGHRI | ASGDRDWGIAEQF | 0.494402 |
| YGPPGTGKTLL | ASGDRDWGIAEQF | 0.79786235 |
| AALVGAPANVA | ASGDRDWGIAEQF | 0.8433588 |
| AATFRRTE | ASGDRDWGIAEQF | 0.37135857 |
| AGALGGGI | ASGDRDWGIAEQF | 0.55353713 |
| AGGAGGAGGIGAG | ASGDRDWGIAEQF | 0.47027537 |
| AGSGALGL | ASGDRDWGIAEQF | 0.6308972 |

| | | |
|---|---|---|
| AQIADQI | ASGDRDWGIAEQF | 0.45382828 |
| ASAVAVTYL | ASGDRDWGIAEQF | 0.4379788 |
| ASTVLGGTGG | ASGDRDWGIAEQF | 0.64026123 |
| AVGGEPAK | ASGDRDWGIAEQF | 0.836588 |
| CEQGTFGGTC | ASGDRDWGIAEQF | 0.48966503 |
| DEGKAPAG | ASGDRDWGIAEQF | 0.32803932 |
| DTVVFPAL | ASGDRDWGIAEQF | 0.5772265 |
| FEGITMAG | ASGDRDWGIAEQF | 0.37326363 |
| FPLVVQD | ASGDRDWGIAEQF | 0.5281015 |
| GAFTTPQ | ASGDRDWGIAEQF | 0.7040474 |
| GFGGFGTAGGVGGA | ASGDRDWGIAEQF | 0.24713738 |
| GFLAQFL | ASGDRDWGIAEQF | 0.45267534 |
| GGAGGAGGNAGVA | ASGDRDWGIAEQF | 0.84287375 |
| GGVGTAGAGGAGGAGG | ASGDRDWGIAEQF | 0.45196784 |
| GNGGGGDAGNAGSGGN | ASGDRDWGIAEQF | 0.5556883 |
| GTGGAGGAGNDAGS | ASGDRDWGIAEQF | 0.77707624 |
| GTTNTGWFNS | ASGDRDWGIAEQF | 0.6856398 |
| IFMGPLFTL | ASGDRDWGIAEQF | 0.8005061 |
| ILAHAIEAI | ASGDRDWGIAEQF | 0.70121264 |
| IRSAIREAL | ASGDRDWGIAEQF | 0.5259396 |
| ITFAGGG | ASGDRDWGIAEQF | 0.52947813 |
| ITGMSGHQEV | ASGDRDWGIAEQF | 0.6411408 |
| IVAGLACTL | ASGDRDWGIAEQF | 0.6392346 |
| IVGGGLAGA | ASGDRDWGIAEQF | 0.51420474 |
| IVTGVLNI | ASGDRDWGIAEQF | 0.3588918 |
| LAQQGCI | ASGDRDWGIAEQF | 0.7882174 |
| LCLHVEN | ASGDRDWGIAEQF | 0.5692755 |
| LDAISDGS | ASGDRDWGIAEQF | 0.70490414 |
| LGGFAAML | ASGDRDWGIAEQF | 0.3954279 |
| LGGKNPMI | ASGDRDWGIAEQF | 0.43240753 |
| LGPKGVAAIL | ASGDRDWGIAEQF | 0.7391769 |
| LMVGVSI | ASGDRDWGIAEQF | 0.1355916 |
| LVGAAFVL | ASGDRDWGIAEQF | 0.1973297 |
| LVLAVAPL | ASGDRDWGIAEQF | 0.5264835 |
| MAAIAVIDAA | ASGDRDWGIAEQF | 0.7949351 |
| NNGINAELQ | ASGDRDWGIAEQF | 0.14789878 |
| RGEPLVLVHG | ASGDRDWGIAEQF | 0.43606633 |
| RGGAGGAAGLI | ASGDRDWGIAEQF | 0.28263542 |
| RSYRMHPAV | ASGDRDWGIAEQF | 0.6095177 |
| SAAHLLAGQLA | ASGDRDWGIAEQF | 0.65178645 |
| SAAVGSY | ASGDRDWGIAEQF | 0.31847942 |
| SPHAVLHRI | ASGDRDWGIAEQF | 0.69256365 |
| SQIAKVL | ASGDRDWGIAEQF | 0.24033162 |
| TALGHLVP | ASGDRDWGIAEQF | 0.34036383 |
| TAPTTSGAA | ASGDRDWGIAEQF | 0.7097597 |
| TDGDDGNGGSPGT | ASGDRDWGIAEQF | 0.754193 |
| TDGLGEN | ASGDRDWGIAEQF | 0.35818735 |
| TGGPPLPGQA | ASGDRDWGIAEQF | 0.750569 |
| TIPAITI | ASGDRDWGIAEQF | 0.41825053 |
| TIPGFTI | ASGDRDWGIAEQF | 0.83881986 |
| TSIASGALA | ASGDRDWGIAEQF | 0.4607142 |

| | | |
|---|---|---|
| TVTLPGAAI | ASGDRDWGIAEQF | 0.3027463 |
| VASGLSAALP | ASGDRDWGIAEQF | 0.73825204 |
| VGAGVIGI | ASGDRDWGIAEQF | 0.57459086 |
| VGPKRFGV | ASGDRDWGIAEQF | 0.67229444 |
| VNVAFPD | ASGDRDWGIAEQF | 0.69389653 |
| VSMTNSGVS | ASGDRDWGIAEQF | 0.42770365 |
| VTDVLGDLSV | ASGDRDWGIAEQF | 0.5503078 |
| VVVVGAGFAGL | ASGDRDWGIAEQF | 0.77285373 |
| VYTVGTVV | ASGDRDWGIAEQF | 0.6868288 |
| WTTFSGP | ASGDRDWGIAEQF | 0.44261256 |
| YDELVEAF | ASGDRDWGIAEQF | 0.7018122 |
| YGTGGAGGNGG | ASGDRDWGIAEQF | 0.60134274 |
| YTNVMAV | ASGDRDWGIAEQF | 0.4156189 |
| FLEQVHQL | ASGDRDWGIAEQF | 0.80082715 |
| GLSLLAVAV | ASGDRDWGIAEQF | 0.58751124 |
| LEQVHQL | ASGDRDWGIAEQF | 0.73887306 |
| LNAAGAI | ASGDRDWGIAEQF | 0.72829884 |
| PAVHASL | ASGDRDWGIAEQF | 0.8017435 |
| TELEDDA | ASGDRDWGIAEQF | 0.41057065 |
| TPPGPPN | ASGDRDWGIAEQF | 0.6300746 |
| TPPPPPPP | ASGDRDWGIAEQF | 0.5888903 |
| TQSPALI | ASSDAGRDTEVF | 0.66486996 |
| EFAAAFPQ | ASSDAGRDTEVF | 0.50648546 |
| LLSGNLDVL | ASSDAGRDTEVF | 0.44736296 |
| GPANLGNYNI | ASSDAGRDTEVF | 0.6103783 |
| TLGPTLP | ASSDAGRDTEVF | 0.78349084 |
| VYAYSSCCK | ASSDAGRDTEVF | 0.50203496 |
| APAPAPAPNI | ASSDAGRDTEVF | 0.5892472 |
| PGGPVPIP | ASSDAGRDTEVF | 0.81899154 |
| FLTVALEP | ASSDAGRDTEVF | 0.4356178 |
| SQYFGANL | ASSDAGRDTEVF | 0.5792604 |
| GGVKPAA | ASSDAGRDTEVF | 0.6542079 |
| GGTGGAGGAGGW | ASSDAGRDTEVF | 0.7312027 |
| GVGGVAGLGGAGG | ASSDAGRDTEVF | 0.50317746 |
| VLLVLATTI | ASSDAGRDTEVF | 0.7638808 |
| IGIFAAML | ASSDAGRDTEVF | 0.4009768 |
| AVGDLID | ASSDAGRDTEVF | 0.7208099 |
| NISPLPL | ASSDAGRDTEVF | 0.4262913 |
| VTAVFVAI | ASSDAGRDTEVF | 0.68558556 |
| SGDDNDAAL | ASSDAGRDTEVF | 0.41771954 |
| AVAALAHS | ASSDAGRDTEVF | 0.70660734 |
| GGVTVMATP | ASSDAGRDTEVF | 0.7652912 |
| AGLAANVRD | ASSDAGRDTEVF | 0.71552825 |
| TNRPDLIDPALLRPG | ASSDAGRDTEVF | 0.18534909 |
| LANLGVTAL | ASSDAGRDTEVF | 0.7742609 |
| TGERIEL | ASSDAGRDTEVF | 0.72701854 |
| NGEELAA | ASSDAGRDTEVF | 0.83775324 |
| VLSLVWG | ASSDAGRDTEVF | 0.55608594 |
| IGYALWL | ASSDAGRDTEVF | 0.65690273 |
| LSLLLAPL | ASSDAGRDTEVF | 0.5485751 |
| TLDEALK | ASSDAGRDTEVF | 0.5210287 |

| | | |
|---|---|---|
| TDGAGTAEGG | ASSDAGRDTEVF | 0.338826 |
| NAPGGAGGNGG | ASSDAGRDTEVF | 0.54238045 |
| VSVFMGMGGNF | ASSDAGRDTEVF | 0.5663444 |
| LASYLAAL | ASSDAGRDTEVF | 0.42983755 |
| LVGGTPAL | ASSDAGRDTEVF | 0.7290285 |
| VRNADVL | ASSDAGRDTEVF | 0.7401958 |
| TCSSPFLVSAM | ASSDAGRDTEVF | 0.76834667 |
| VHITDAE | ASSDAGRDTEVF | 0.78587997 |
| EYAAPSF | ASSDAGRDTEVF | 0.78336406 |
| VVDAFHGGL | ASSDAGRDTEVF | 0.70589936 |
| ASVVDPQ | ASSDAGRDTEVF | 0.48159897 |
| SYNEFGGLS | ASSDAGRDTEVF | 0.5525764 |
| DPDGSVY | ASSDAGRDTEVF | 0.3347451 |
| MVSTIHATE | ASSDAGRDTEVF | 0.48040372 |
| LAAGVADPV | ASSDAGRDTEVF | 0.51012933 |
| TKEQIAA | ASSDAGRDTEVF | 0.65584296 |
| SLASPAAL | ASSDAGRDTEVF | 0.14681542 |
| IGLVFYGV | ASSDAGRDTEVF | 0.6000394 |
| FLLDASNL | ASSDAGRDTEVF | 0.79703796 |
| YQVGNFP | ASSDAGRDTEVF | 0.37707505 |
| ESLLSQST | ASSDAGRDTEVF | 0.6689239 |
| LFTVGFAAVG | ASSDAGRDTEVF | 0.54269576 |
| ATPPTPT | ASSDAGRDTEVF | 0.37995136 |
| TDLATLG | ASSDAGRDTEVF | 0.75133306 |
| SAGGTGR | ASSDAGRDTEVF | 0.3955181 |
| GFSNSGS | ASSDAGRDTEVF | 0.6956974 |
| GGNGGSGGTSV | ASSDAGRDTEVF | 0.7953063 |
| GTSVATGGAGN | ASSDAGRDTEVF | 0.5739209 |
| NGGNGGSGGTS | ASSDAGRDTEVF | 0.57097614 |
| AVVKFPH | ASSDAGRDTEVF | 0.5457626 |
| TPPPPPP | ASSDAGRDTEVF | 0.36623257 |
| AAPAAAAAPA | ASSDAGRDTEVF | 0.61210525 |
| IALNEDL | ASSDAGRDTEVF | 0.1922934 |
| IVNVSSM | ASSDAGRDTEVF | 0.53619343 |
| TGIPPAPR | ASSDAGRDTEVF | 0.35988584 |
| FDLGGGTF | ASSDAGRDTEVF | 0.666846 |
| FDLGGGTFDVS | ASSDAGRDTEVF | 0.6370597 |
| NEPTAAA | ASSDAGRDTEVF | 0.58741415 |
| NEPTAAALA | ASSDAGRDTEVF | 0.33981368 |
| GSIINISSI | ASSDAGRDTEVF | 0.58112526 |
| VAPGNAGTA | ASSDAGRDTEVF | 0.43005306 |
| TGGGASLE | ASSDAGRDTEVF | 0.36568975 |
| VAVYNHYKRI | ASSDAGRDTEVF | 0.6860433 |
| TDPAPATT | ASSDAGRDTEVF | 0.51654404 |
| TNRPDILDPALLRPG | ASSDAGRDTEVF | 0.4801002 |
| LTTTTTS | ASSDAGRDTEVF | 0.7764557 |
| PTGSGKTL | ASSDAGRDTEVF | 0.64152247 |
| TGSGKTL | ASSDAGRDTEVF | 0.3288029 |
| TIGHVDHGK | ASSDAGRDTEVF | 0.58642274 |
| TIGHVDHGKTT | ASSDAGRDTEVF | 0.5393318 |
| GHVDHGKTT | ASSDAGRDTEVF | 0.51619023 |

| | | |
|---|---|---|
| AQFEHTLL | ASSDAGRDTEVF | 0.15105784 |
| IVNQDEL | ASSDAGRDTEVF | 0.54389507 |
| TNGSQFF | ASSDAGRDTEVF | 0.4204217 |
| TGKGIVE | ASSDAGRDTEVF | 0.34022436 |
| LISNASDA | ASSDAGRDTEVF | 0.43766555 |
| AELRTLGELTKIAKAHGAQ | ASSDAGRDTEVF | 0.7428464 |
| APNAVAAAEPSW | ASSDAGRDTEVF | 0.520639 |
| APQVAGAATAVL | ASSDAGRDTEVF | 0.6729625 |
| ARRDPAAAAAILPTDA | ASSDAGRDTEVF | 0.69981503 |
| DAAARDVLAEAGLAEYF | ASSDAGRDTEVF | 0.7602099 |
| DDQIVLSAKANLAKVEQ | ASSDAGRDTEVF | 0.8074233 |
| DELLPDTGEDSDEDGH | ASSDAGRDTEVF | 0.6347648 |
| ELPEAPANRATVPQD | ASSDAGRDTEVF | 0.46172878 |
| FGQSKIERVTGIG | ASSDAGRDTEVF | 0.61857486 |
| GAGGTGSTAGGAGGAGGAGGLY | ASSDAGRDTEVF | 0.60317856 |
| GHGGTGAAGGNGGTGGAGGSGI | ASSDAGRDTEVF | 0.27214038 |
| GNGGNGGAGGEGGPGGAGGAGA | ASSDAGRDTEVF | 0.19475712 |
| GSTAGNGGNGGNATNGTIAGQP | ASSDAGRDTEVF | 0.6766572 |
| GTGGDGGNGGNGGTGG | ASSDAGRDTEVF | 0.22510725 |
| GVRRPAGIPADAPV | ASSDAGRDTEVF | 0.5663747 |
| GYRQVTDAHLLTLARR | ASSDAGRDTEVF | 0.73655975 |
| IAKMGQDGHDRG | ASSDAGRDTEVF | 0.50986755 |
| KAEPSIEIQYT | ASSDAGRDTEVF | 0.35209465 |
| KPVRIRMAADEDTVAVT | ASSDAGRDTEVF | 0.44788447 |
| MPPRIAGMRLLVIKPEP | ASSDAGRDTEVF | 0.5599651 |
| MKFYAVALFRSRGVPPTR | ASSDAGRDTEVF | 0.7594468 |
| NAGGFGILGAVAHSP | ASSDAGRDTEVF | 0.6506453 |
| NEGREITLAMAEM | ASSDAGRDTEVF | 0.81241906 |
| NELVEWSHPLGHRWRWE | ASSDAGRDTEVF | 0.76801634 |
| NLGAGGTGGNGGNPG | ASSDAGRDTEVF | 0.52597225 |
| NTEWPKWETVMAADPVD | ASSDAGRDTEVF | 0.683356 |
| PASAAPAETQP | ASSDAGRDTEVF | 0.4604247 |
| QQSSDGKNGVRAAEITGE | ASSDAGRDTEVF | 0.4755901 |
| RAPDAVTTSWWKE | ASSDAGRDTEVF | 0.5054128 |
| SGFGNVGAHTSGWFNQ | ASSDAGRDTEVF | 0.51431656 |
| SGVAGAVGQAASVGGLKV | ASSDAGRDTEVF | 0.51339555 |
| SNGGAGTGSAGGNGGT | ASSDAGRDTEVF | 0.42065248 |
| TGGAGGEGGAAG | ASSDAGRDTEVF | 0.26537043 |
| TGGDGGAAGAGGAPGHG | ASSDAGRDTEVF | 0.58094156 |
| TMPPLLAAIANAGRHGV | ASSDAGRDTEVF | 0.65455914 |
| TSSTAGNSGGA | ASSDAGRDTEVF | 0.30514726 |
| VPATVPVGILC | ASSDAGRDTEVF | 0.40138152 |
| VQRPDSGGVVLTGRIS | ASSDAGRDTEVF | 0.8337001 |
| WTPNPVPATKGVPAGLVQAT | ASSDAGRDTEVF | 0.8219447 |
| YALKRALAKFSDRFDVVII | ASSDAGRDTEVF | 0.4229828 |
| YLIGSVIV | ASSDAGRDTEVF | 0.45101327 |
| YGQLVAVID | ASSDAGRDTEVF | 0.57700825 |
| VTVGGMTLPA | ASSDAGRDTEVF | 0.18502559 |
| VLALPAV | ASSDAGRDTEVF | 0.5657362 |
| VIPPTLNYET | ASSDAGRDTEVF | 0.61777186 |
| VGMGGMHPSSG | ASSDAGRDTEVF | 0.59052503 |

| | | |
|---|---|---|
| VGLAGGAATA | ASSDAGRDTEVF | 0.401785 |
| VGGAGGAGGSGP | ASSDAGRDTEVF | 0.5358672 |
| VFAAFMV | ASSDAGRDTEVF | 0.47504926 |
| TLYDKLR | ASSDAGRDTEVF | 0.6828011 |
| TLAVFTVL | ASSDAGRDTEVF | 0.14055294 |
| TGSVQPARV | ASSDAGRDTEVF | 0.6533389 |
| TGGGAGSGGT | ASSDAGRDTEVF | 0.34132773 |
| SHLETSL | ASSDAGRDTEVF | 0.47709253 |
| SFTTGMT | ASSDAGRDTEVF | 0.2839811 |
| QVEVAKRMGVRQPTVS | ASSDAGRDTEVF | 0.50116944 |
| QGSGGPALAM | ASSDAGRDTEVF | 0.7722856 |
| MLAPSNP | ASSDAGRDTEVF | 0.72200984 |
| LVNNAAI | ASSDAGRDTEVF | 0.29666713 |
| LTGEINTLA | ASSDAGRDTEVF | 0.73623776 |
| LSIFGPV | ASSDAGRDTEVF | 0.8306894 |
| LLALLANL | ASSDAGRDTEVF | 0.48354888 |
| LISNASDALDKLR | ASSDAGRDTEVF | 0.34418783 |
| LINEAVP | ASSDAGRDTEVF | 0.7407014 |
| LGYAFSGL | ASSDAGRDTEVF | 0.4439209 |
| LDFVAVAL | ASSDAGRDTEVF | 0.67167825 |
| LAQPGAE | ASSDAGRDTEVF | 0.68106407 |
| ISNLGAVTGF | ASSDAGRDTEVF | 0.30186218 |
| HQEDPGRI | ASSDAGRDTEVF | 0.7938849 |
| GTPGEPTP | ASSDAGRDTEVF | 0.71874535 |
| GFFNSGS | ASSDAGRDTEVF | 0.34707507 |
| GAGGGGLTAGAN | ASSDAGRDTEVF | 0.39881697 |
| FVDALVNM | ASSDAGRDTEVF | 0.40954277 |
| FLTVAPEL | ASSDAGRDTEVF | 0.21038932 |
| FETLTAL | ASSDAGRDTEVF | 0.73707926 |
| DVINPIEL | ASSDAGRDTEVF | 0.38387355 |
| APVSGAPV | ASSDAGRDTEVF | 0.5140527 |
| ALVGGLAW | ASSDAGRDTEVF | 0.6204531 |
| ALAAAAATAI | ASSDAGRDTEVF | 0.50774556 |
| AFGLGLLL | ASSDAGRDTEVF | 0.5161758 |
| AFFAAASL | ASSDAGRDTEVF | 0.52643055 |
| AAAALPAFSPP | ASSDAGRDTEVF | 0.6422349 |
| VVLAPAL | ASSDAGRDTEVF | 0.5831674 |
| TIGHVDHGKTTL | ASSDAGRDTEVF | 0.58058715 |
| LISNASDALDK | ASSDAGRDTEVF | 0.54527724 |
| LISNASDAL | ASSDAGRDTEVF | 0.80868703 |
| ISNASDALDK | ASSDAGRDTEVF | 0.35390455 |
| NQAETLVYQTE | ASSDAGRDTEVF | 0.4549838 |
| TGGTGGTAGAG | ASSDAGRDTEVF | 0.66210604 |
| TGSTAGGAGGA | ASSDAGRDTEVF | 0.31506112 |
| DPLAVPVAAGPV | ASSDAGRDTEVF | 0.72227305 |
| GGTGGTPGNGGN | ASSDAGRDTEVF | 0.61710656 |
| PAGLAGVDPATL | ASSDAGRDTEVF | 0.63126254 |
| GNNLVVEAPHAGASV | ASSDAGRDTEVF | 0.5694732 |
| KVTRIMSASTLEQPA | ASSDAGRDTEVF | 0.79656446 |
| LGFALPIALPRRTVV | ASSDAGRDTEVF | 0.7695489 |
| TRLGNLTARKLVVLV | ASSDAGRDTEVF | 0.52071625 |

| | | |
|---|---|---|
| WSQQRRLIYFPSAGP | ASSDAGRDTEVF | 0.7407688 |
| GPNGNPDPMAAAVDIR | ASSDAGRDTEVF | 0.35492137 |
| TQRCVSVADLRAAASR | ASSDAGRDTEVF | 0.71735317 |
| QFVLDVAEHPVEVLTETV | ASSDAGRDTEVF | 0.6453423 |
| GGAGGTGGAGGNGGTQAGDG | ASSDAGRDTEVF | 0.33793527 |
| PLLATLRAIAVDVDAKPAQV | ASSDAGRDTEVF | 0.5571935 |
| APKFRTGGYMIDFWGVGYQVAK | ASSDAGRDTEVF | 0.74484026 |
| GAGGAGGAGGAAGGPGGTGGTG | ASSDAGRDTEVF | 0.72632164 |
| IGALGLLATDVAPAGSVAFGVA | ASSDAGRDTEVF | 0.24917595 |
| AGGAGGIGGNGGAGGAGGVLMG | ASSDAGRDTEVF | 0.52228695 |
| ADELIGLL | ASSDAGRDTEVF | 0.22748265 |
| ADRALKA | ASSDAGRDTEVF | 0.44246083 |
| AGAVGQVGGAGGA | ASSDAGRDTEVF | 0.37398466 |
| AGGGGGLGGAP | ASSDAGRDTEVF | 0.69001186 |
| AGGLLYGNGGAG | ASSDAGRDTEVF | 0.49188262 |
| AGLESIGL | ASSDAGRDTEVF | 0.6752396 |
| AGLKTGII | ASSDAGRDTEVF | 0.58489835 |
| ASLAALLL | ASSDAGRDTEVF | 0.80516577 |
| ASVINISSI | ASSDAGRDTEVF | 0.37209415 |
| ATPPGPA | ASSDAGRDTEVF | 0.5701722 |
| ATVASALV | ASSDAGRDTEVF | 0.26616254 |
| ATVASLIL | ASSDAGRDTEVF | 0.40136135 |
| AVNGPVLIRH | ASSDAGRDTEVF | 0.3910675 |
| DGAAGAVGI | ASSDAGRDTEVF | 0.55922496 |
| DPLLHGI | ASSDAGRDTEVF | 0.20508216 |
| ELGAALGAL | ASSDAGRDTEVF | 0.7593639 |
| ELGRTQAV | ASSDAGRDTEVF | 0.66698575 |
| FVAGGYLS | ASSDAGRDTEVF | 0.7446894 |
| GASGGAGGAGGW | ASSDAGRDTEVF | 0.5453982 |
| GGAGAGGDGGAGGI | ASSDAGRDTEVF | 0.500981 |
| GTGHNGSQPGMGGNG | ASSDAGRDTEVF | 0.5002398 |
| GYAAASASAS | ASSDAGRDTEVF | 0.7300266 |
| HQVDVAAVQQG | ASSDAGRDTEVF | 0.40249664 |
| IGAAIAHT | ASSDAGRDTEVF | 0.24012293 |
| IGMIEVAQL | ASSDAGRDTEVF | 0.47742608 |
| IRLAPEHTSAMD | ASSDAGRDTEVF | 0.3318737 |
| IYNVSVL | ASSDAGRDTEVF | 0.764257 |
| KQGFLGM | ASSDAGRDTEVF | 0.7500841 |
| LESVITQL | ASSDAGRDTEVF | 0.6852809 |
| LGLGALVVV | ASSDAGRDTEVF | 0.6643765 |
| LIGNGGNGGSGGI | ASSDAGRDTEVF | 0.20537737 |
| LNAINAPTL | ASSDAGRDTEVF | 0.5378632 |
| LQLLNTGV | ASSDAGRDTEVF | 0.817665 |
| LQTSIEF | ASSDAGRDTEVF | 0.6794039 |
| MSFAVSEL | ASSDAGRDTEVF | 0.50285494 |
| NTGALGL | ASSDAGRDTEVF | 0.79557407 |
| PPGAGGAGGAGTT | ASSDAGRDTEVF | 0.49154806 |
| QLVGLGI | ASSDAGRDTEVF | 0.5240912 |
| RFLEQVHQL | ASSDAGRDTEVF | 0.6511921 |
| RGVGYVLR | ASSDAGRDTEVF | 0.59010196 |
| SATGVFAGVI | ASSDAGRDTEVF | 0.6002973 |

| | | |
|---|---|---|
| SLPAITLP | ASSDAGRDTEVF | 0.3416222 |
| STALAFTV | ASSDAGRDTEVF | 0.6453088 |
| SVLTVGLGK | ASSDAGRDTEVF | 0.8257577 |
| TALAGAGSGI | ASSDAGRDTEVF | 0.5574174 |
| TGLPFTGI | ASSDAGRDTEVF | 0.77779526 |
| VAEHVTV | ASSDAGRDTEVF | 0.69157064 |
| VDIVLNSL | ASSDAGRDTEVF | 0.75247204 |
| VLTGGGAVL | ASSDAGRDTEVF | 0.37412438 |
| VNVGLAYGI | ASSDAGRDTEVF | 0.47123268 |
| VSSAAVGAV | ASSDAGRDTEVF | 0.5920991 |
| VTSVYEAGV | ASSDAGRDTEVF | 0.4235992 |
| WVALDRGA | ASSDAGRDTEVF | 0.5128181 |
| YLVGSVVI | ASSDAGRDTEVF | 0.4762051 |
| GPPGTGKTLL | ASSDAGRDTEVF | 0.8568076 |
| GVGMTKF | ASSDAGRDTEVF | 0.6722872 |
| IGIGGSDL | ASSDAGRDTEVF | 0.71107817 |
| IGTLLAEL | ASSDAGRDTEVF | 0.4560886 |
| IYTSGTT | ASSDAGRDTEVF | 0.5474397 |
| LSFPAGV | ASSDAGRDTEVF | 0.70229554 |
| LYTSGTT | ASSDAGRDTEVF | 0.3191083 |
| VGAFPFGF | ASSDAGRDTEVF | 0.8025407 |
| VGAGTVEF | ASSDAGRDTEVF | 0.4337508 |
| VGKSTLLL | ASSDAGRDTEVF | 0.41809928 |
| AALAHAGGGIV | ASSDAGRDTEVF | 0.6708697 |
| AALVAVSLL | ASSDAGRDTEVF | 0.78484726 |
| AAVSRAVAAL | ASSDAGRDTEVF | 0.4739288 |
| AGGSSGIGGF | ASSDAGRDTEVF | 0.5091505 |
| AGLTVANANG | ASSDAGRDTEVF | 0.4647906 |
| AGLVGAVL | ASSDAGRDTEVF | 0.72023684 |
| ALSQAFTV | ASSDAGRDTEVF | 0.7883464 |
| ALTFSVSL | ASSDAGRDTEVF | 0.59781367 |
| ALTGAALVAL | ASSDAGRDTEVF | 0.7806697 |
| ASFSIGAGVL | ASSDAGRDTEVF | 0.643659 |
| DGSVYKI | ASSDAGRDTEVF | 0.29048058 |
| EQLAEIV | ASSDAGRDTEVF | 0.5712104 |
| FAVSELTVM | ASSDAGRDTEVF | 0.71682835 |
| GAAPLAAGL | ASSDAGRDTEVF | 0.73888814 |
| GADGGDGGSGGAGG | ASSDAGRDTEVF | 0.82561916 |
| GFGLDTGGAGG | ASSDAGRDTEVF | 0.5553226 |
| GLASGGLL | ASSDAGRDTEVF | 0.4582988 |
| GRIAGAAVLL | ASSDAGRDTEVF | 0.7966331 |
| GSAGSLNTSF | ASSDAGRDTEVF | 0.82851446 |
| GTGQIGI | ASSDAGRDTEVF | 0.7606688 |
| GTLIALIL | ASSDAGRDTEVF | 0.47534624 |
| IAATMGFVVG | ASSDAGRDTEVF | 0.44751158 |
| IFAQALL | ASSDAGRDTEVF | 0.55844104 |
| IGAAVALGF | ASSDAGRDTEVF | 0.7574904 |
| IGLTPLAL | ASSDAGRDTEVF | 0.3582953 |
| ILGSNTSTL | ASSDAGRDTEVF | 0.5994466 |
| IVFSLTVL | ASSDAGRDTEVF | 0.68638366 |
| LASKATGF | ASSDAGRDTEVF | 0.28152722 |

| | | |
|---|---|---|
| LASVALIL | ASSDAGRDTEVF | 0.6145752 |
| LFAVGLTL | ASSDAGRDTEVF | 0.7629069 |
| LFNGGIGTYI | ASSDAGRDTEVF | 0.49480462 |
| LTLGAGII | ASSDAGRDTEVF | 0.5108449 |
| LVAAALGDI | ASSDAGRDTEVF | 0.7719066 |
| NTAFVQLGI | ASSDAGRDTEVF | 0.85211855 |
| QLFSGAEL | ASSDAGRDTEVF | 0.7750368 |
| SAATFITL | ASSDAGRDTEVF | 0.35579726 |
| SNYGLANLGV | ASSDAGRDTEVF | 0.1873754 |
| SYGGAYAVM | ASSDAGRDTEVF | 0.36739883 |
| TTGVGGLTLG | ASSDAGRDTEVF | 0.7157426 |
| TTQTIYI | ASSDAGRDTEVF | 0.5083731 |
| VAAGAGLGAVI | ASSDAGRDTEVF | 0.5177379 |
| VAAVVTLL | ASSDAGRDTEVF | 0.6820028 |
| VGFLGGLI | ASSDAGRDTEVF | 0.76941514 |
| VGVIPILL | ASSDAGRDTEVF | 0.4261674 |
| VIFASSLI | ASSDAGRDTEVF | 0.3661423 |
| VKGGDGGAAAG | ASSDAGRDTEVF | 0.60254616 |
| VTKTFTNI | ASSDAGRDTEVF | 0.66851175 |
| VVAAAILL | ASSDAGRDTEVF | 0.3697866 |
| YAALDVEL | ASSDAGRDTEVF | 0.55320245 |
| LASVALLL | ASSDAGRDTEVF | 0.3620471 |
| LDAVHRV | ASSDAGRDTEVF | 0.452007 |
| QGLSSSIF | ASSDAGRDTEVF | 0.65781915 |
| RGTFANIRL | ASSDAGRDTEVF | 0.3836409 |
| RQLWWGHRI | ASSDAGRDTEVF | 0.7414382 |
| YGPPGTGKTLL | ASSDAGRDTEVF | 0.4024254 |
| AALVGAPANVA | ASSDAGRDTEVF | 0.67471105 |
| AATFRRTE | ASSDAGRDTEVF | 0.63819575 |
| AGALGGGI | ASSDAGRDTEVF | 0.63150746 |
| AGGAGGAGGIGAG | ASSDAGRDTEVF | 0.4959546 |
| AGSGALGL | ASSDAGRDTEVF | 0.4011703 |
| AQIADQI | ASSDAGRDTEVF | 0.32260737 |
| ASAVAVTYL | ASSDAGRDTEVF | 0.18631865 |
| ASTVLGGTGG | ASSDAGRDTEVF | 0.73940957 |
| AVGGEPAK | ASSDAGRDTEVF | 0.42980486 |
| CEQGTFGGTC | ASSDAGRDTEVF | 0.73592246 |
| DEGKAPAG | ASSDAGRDTEVF | 0.7483309 |
| DTVVFPAL | ASSDAGRDTEVF | 0.3571993 |
| FEGITMAG | ASSDAGRDTEVF | 0.58622676 |
| FPLVVQD | ASSDAGRDTEVF | 0.41588807 |
| GAFTTPQ | ASSDAGRDTEVF | 0.38229692 |
| GFGGFGTAGGVGGA | ASSDAGRDTEVF | 0.722139 |
| GFLAQFL | ASSDAGRDTEVF | 0.6541999 |
| GGAGGAGGNAGVA | ASSDAGRDTEVF | 0.37649277 |
| GGVGTAGAGGAGGAGG | ASSDAGRDTEVF | 0.5838425 |
| GNGGGGDAGNAGSGGN | ASSDAGRDTEVF | 0.6017914 |
| GTGGAGGAGNDAGS | ASSDAGRDTEVF | 0.4181719 |
| GTTNTGWFNS | ASSDAGRDTEVF | 0.5152808 |
| IFMGPLFTL | ASSDAGRDTEVF | 0.77116007 |
| ILAHAIEAI | ASSDAGRDTEVF | 0.5837172 |

| | | |
|---|---|---|
| IRSAIREAL | ASSDAGRDTEVF | 0.4544728 |
| ITFAGGG | ASSDAGRDTEVF | 0.75341463 |
| ITGMSGHQEV | ASSDAGRDTEVF | 0.309572 |
| IVAGLACTL | ASSDAGRDTEVF | 0.6887905 |
| IVGGGLAGA | ASSDAGRDTEVF | 0.5036756 |
| IVTGVLNI | ASSDAGRDTEVF | 0.5355712 |
| LAQQGCI | ASSDAGRDTEVF | 0.7155316 |
| LCLHVEN | ASSDAGRDTEVF | 0.37470415 |
| LDAISDGS | ASSDAGRDTEVF | 0.41338924 |
| LGGFAAML | ASSDAGRDTEVF | 0.5057612 |
| LGGKNPMI | ASSDAGRDTEVF | 0.66437846 |
| LGPKGVAAIL | ASSDAGRDTEVF | 0.7582809 |
| LMVGVSI | ASSDAGRDTEVF | 0.42971945 |
| LVGAAFVL | ASSDAGRDTEVF | 0.60862964 |
| LVLAVAPL | ASSDAGRDTEVF | 0.43703768 |
| MAAIAVIDAA | ASSDAGRDTEVF | 0.3615805 |
| NNGINAELQ | ASSDAGRDTEVF | 0.53414756 |
| RGEPLVLVHG | ASSDAGRDTEVF | 0.52600867 |
| RGGAGGAAGLI | ASSDAGRDTEVF | 0.48076475 |
| RSYRMHPAV | ASSDAGRDTEVF | 0.5865447 |
| SAAHLLAGQLA | ASSDAGRDTEVF | 0.45547044 |
| SAAVGSY | ASSDAGRDTEVF | 0.6777245 |
| SPHAVLHRI | ASSDAGRDTEVF | 0.81448567 |
| SQIAKVL | ASSDAGRDTEVF | 0.50552076 |
| TALGHLVP | ASSDAGRDTEVF | 0.6571023 |
| TAPTTSGAA | ASSDAGRDTEVF | 0.68691456 |
| TDGDDGNGGSPGT | ASSDAGRDTEVF | 0.62361616 |
| TDGLGEN | ASSDAGRDTEVF | 0.5333726 |
| TGGPPLPGQA | ASSDAGRDTEVF | 0.43443504 |
| TIPAITI | ASSDAGRDTEVF | 0.77495533 |
| TIPGFTI | ASSDAGRDTEVF | 0.7655679 |
| TSIASGALA | ASSDAGRDTEVF | 0.73544526 |
| TVTLPGAAI | ASSDAGRDTEVF | 0.63748163 |
| VASGLSAALP | ASSDAGRDTEVF | 0.6549635 |
| VGAGVIGI | ASSDAGRDTEVF | 0.71485406 |
| VGPKRFGV | ASSDAGRDTEVF | 0.5295875 |
| VNVAFPD | ASSDAGRDTEVF | 0.26482192 |
| VSMTNSGVS | ASSDAGRDTEVF | 0.5578078 |
| VTDVLGDLSV | ASSDAGRDTEVF | 0.44799548 |
| VVVVGAGFAGL | ASSDAGRDTEVF | 0.5041392 |
| VYTVGTVV | ASSDAGRDTEVF | 0.30907544 |
| WTTFSGP | ASSDAGRDTEVF | 0.75650823 |
| YDELVEAF | ASSDAGRDTEVF | 0.8496379 |
| YGTGGAGGNGG | ASSDAGRDTEVF | 0.48903936 |
| YTNVMAV | ASSDAGRDTEVF | 0.8059703 |
| FLEQVHQL | ASSDAGRDTEVF | 0.29564896 |
| GLSLLAVAV | ASSDAGRDTEVF | 0.76160145 |
| LEQVHQL | ASSDAGRDTEVF | 0.7703688 |
| LNAAGAI | ASSDAGRDTEVF | 0.35062224 |
| PAVHASL | ASSDAGRDTEVF | 0.7297435 |
| TELEDDA | ASSDAGRDTEVF | 0.4503888 |

| | | |
|---|---|---|
| TPPGPPN | ASSDAGRDTEVF | 0.5474892 |
| TPPPPPPP | ASSDAGRDTEVF | 0.7938609 |
| TQSPALI | ASSETGDANTEVF | 0.55213356 |
| EFAAAFPQ | ASSETGDANTEVF | 0.5726924 |
| LLSGNLDVL | ASSETGDANTEVF | 0.6012954 |
| GPANLGNYNI | ASSETGDANTEVF | 0.45015886 |
| TLGPTLP | ASSETGDANTEVF | 0.53739387 |
| VYAYSSCCK | ASSETGDANTEVF | 0.74602926 |
| APAPAPAPNI | ASSETGDANTEVF | 0.40123105 |
| PGGPVPIP | ASSETGDANTEVF | 0.820082 |
| FLTVALEP | ASSETGDANTEVF | 0.38170266 |
| SQYFGANL | ASSETGDANTEVF | 0.8105786 |
| GGVKPAA | ASSETGDANTEVF | 0.68028855 |
| GGTGGAGGAGGW | ASSETGDANTEVF | 0.6314791 |
| GVGGVAGLGGAGG | ASSETGDANTEVF | 0.46439627 |
| VLLVLATTI | ASSETGDANTEVF | 0.7960292 |
| IGIFAAML | ASSETGDANTEVF | 0.51000077 |
| AVGDLID | ASSETGDANTEVF | 0.6742154 |
| NISPLPL | ASSETGDANTEVF | 0.7444345 |
| VTAVFVAI | ASSETGDANTEVF | 0.16060765 |
| SGDDNDAAL | ASSETGDANTEVF | 0.45042157 |
| AVAALAHS | ASSETGDANTEVF | 0.22485562 |
| GGVTVMATP | ASSETGDANTEVF | 0.21978493 |
| AGLAANVRD | ASSETGDANTEVF | 0.18631813 |
| TNRPDLIDPALLRPG | ASSETGDANTEVF | 0.16118574 |
| LANLGVTAL | ASSETGDANTEVF | 0.19960013 |
| TGERIEL | ASSETGDANTEVF | 0.1718202 |
| NGEELAA | ASSETGDANTEVF | 0.7798295 |
| VLSLVWG | ASSETGDANTEVF | 0.265952 |
| IGYALWL | ASSETGDANTEVF | 0.17783113 |
| LSLLLAPL | ASSETGDANTEVF | 0.47941393 |
| TLDEALK | ASSETGDANTEVF | 0.25620115 |
| TDGAGTAEGG | ASSETGDANTEVF | 0.1557015 |
| NAPGGAGGNGG | ASSETGDANTEVF | 0.19863491 |
| VSVFMGMGGNF | ASSETGDANTEVF | 0.18362924 |
| LASYLAAL | ASSETGDANTEVF | 0.20857467 |
| LVGGTPAL | ASSETGDANTEVF | 0.14157747 |
| VRNADVL | ASSETGDANTEVF | 0.26895416 |
| TCSSPFLVSAM | ASSETGDANTEVF | 0.24364752 |
| VHITDAE | ASSETGDANTEVF | 0.14042562 |
| EYAAPSF | ASSETGDANTEVF | 0.23385406 |
| VVDAFHGGL | ASSETGDANTEVF | 0.23248735 |
| ASVVDPQ | ASSETGDANTEVF | 0.22165182 |
| SYNEFGGLS | ASSETGDANTEVF | 0.30437747 |
| DPDGSVY | ASSETGDANTEVF | 0.16725814 |
| MVSTIHATE | ASSETGDANTEVF | 0.17707823 |
| LAAGVADPV | ASSETGDANTEVF | 0.36994052 |
| TKEQIAA | ASSETGDANTEVF | 0.31763586 |
| SLASPAAL | ASSETGDANTEVF | 0.16072638 |
| IGLVFYGV | ASSETGDANTEVF | 0.23177253 |
| FLLDASNL | ASSETGDANTEVF | 0.3531236 |

| | | |
|---|---|---|
| YQVGNFP | ASSETGDANTEVF | 0.22466074 |
| ESLLSQST | ASSETGDANTEVF | 0.3016671 |
| LFTVGFAAVG | ASSETGDANTEVF | 0.3377661 |
| ATPPTPT | ASSETGDANTEVF | 0.16212144 |
| TDLATLG | ASSETGDANTEVF | 0.39239588 |
| SAGGTGR | ASSETGDANTEVF | 0.5352008 |
| GFSNSGS | ASSETGDANTEVF | 0.21285133 |
| GGNGGSGGTSV | ASSETGDANTEVF | 0.21812437 |
| GTSVATGGAGN | ASSETGDANTEVF | 0.16078462 |
| NGGNGGSGGTS | ASSETGDANTEVF | 0.19687004 |
| AVVKFPH | ASSETGDANTEVF | 0.16525641 |
| TPPPPPP | ASSETGDANTEVF | 0.28559914 |
| AAPAAAAAPA | ASSETGDANTEVF | 0.29902264 |
| IALNEDL | ASSETGDANTEVF | 0.20061214 |
| IVNVSSM | ASSETGDANTEVF | 0.15052089 |
| TGIPPAPR | ASSETGDANTEVF | 0.16246407 |
| FDLGGGTF | ASSETGDANTEVF | 0.18078196 |
| FDLGGGTFDVS | ASSETGDANTEVF | 0.15902308 |
| NEPTAAA | ASSETGDANTEVF | 0.15369122 |
| NEPTAAALA | ASSETGDANTEVF | 0.17724238 |
| GSIINISSI | ASSETGDANTEVF | 0.32867938 |
| VAPGNAGTA | ASSETGDANTEVF | 0.3156352 |
| TGGGASLE | ASSETGDANTEVF | 0.17668839 |
| VAVYNHYKRI | ASSETGDANTEVF | 0.37614417 |
| TDPAPATT | ASSETGDANTEVF | 0.72560644 |
| TNRPDILDPALLRPG | ASSETGDANTEVF | 0.25372526 |
| LTTTTTS | ASSETGDANTEVF | 0.19465213 |
| PTGSGKTL | ASSETGDANTEVF | 0.15850396 |
| TGSGKTL | ASSETGDANTEVF | 0.78283644 |
| TIGHVDHGK | ASSETGDANTEVF | 0.2203221 |
| TIGHVDHGKTT | ASSETGDANTEVF | 0.24807397 |
| GHVDHGKTT | ASSETGDANTEVF | 0.2972831 |
| AQFEHTLL | ASSETGDANTEVF | 0.24005371 |
| IVNQDEL | ASSETGDANTEVF | 0.16391908 |
| TNGSQFF | ASSETGDANTEVF | 0.5463928 |
| TGKGIVE | ASSETGDANTEVF | 0.235339 |
| LISNASDA | ASSETGDANTEVF | 0.16817543 |
| AELRTLGELTKIAKAHGAQ | ASSETGDANTEVF | 0.1610721 |
| APNAVAAAEPSW | ASSETGDANTEVF | 0.23095894 |
| APQVAGAATAVL | ASSETGDANTEVF | 0.22090922 |
| ARRDPAAAAAILPTDA | ASSETGDANTEVF | 0.31393245 |
| DAAARDVLAEAGLAEYF | ASSETGDANTEVF | 0.21021381 |
| DDQIVLSAKANLAKVEQ | ASSETGDANTEVF | 0.15740792 |
| DELLPDTGEDSDEDGH | ASSETGDANTEVF | 0.36144102 |
| ELPEAPANRATVPQD | ASSETGDANTEVF | 0.26458842 |
| FGQSKIERVTGIG | ASSETGDANTEVF | 0.21145071 |
| GAGGTGSTAGGAGGAGGAGGLY | ASSETGDANTEVF | 0.20119376 |
| GHGGTGAAGGNGGTGGAGGSGI | ASSETGDANTEVF | 0.20622954 |
| GNGGNGGAGGEGGPGGAGGAGA | ASSETGDANTEVF | 0.2531066 |
| GSTAGNGGNGGNATNGTIAGQP | ASSETGDANTEVF | 0.29736114 |
| GTGGDGGNGGNGGTGG | ASSETGDANTEVF | 0.22840403 |

| | | |
|---|---|---|
| GVRRPAGIPADAPV | ASSETGDANTEVF | 0.28233472 |
| GYRQVTDAHLLTLARR | ASSETGDANTEVF | 0.20166224 |
| IAKMGQDGHDRG | ASSETGDANTEVF | 0.20186187 |
| KAEPSIEIQYT | ASSETGDANTEVF | 0.15327641 |
| KPVRIRMAADEDTVAVT | ASSETGDANTEVF | 0.17579037 |
| MPPRIAGMRLLVIKPEP | ASSETGDANTEVF | 0.24653152 |
| MKFYAVALFRSRGVPPTR | ASSETGDANTEVF | 0.22280353 |
| NAGGFGILGAVAHSP | ASSETGDANTEVF | 0.41165367 |
| NEGREITLAMAEM | ASSETGDANTEVF | 0.20297536 |
| NELVEWSHPLGHRWRWE | ASSETGDANTEVF | 0.17401141 |
| NLGAGGTGGNGGNPG | ASSETGDANTEVF | 0.28669783 |
| NTEWPKWETVMAADPVD | ASSETGDANTEVF | 0.34509283 |
| PASAAPAETQP | ASSETGDANTEVF | 0.15738404 |
| QQSSDGKNGVRAAEITGE | ASSETGDANTEVF | 0.18217118 |
| RAPDAVTTSWWKE | ASSETGDANTEVF | 0.5839044 |
| SGFGNVGAHTSGWFNQ | ASSETGDANTEVF | 0.57373875 |
| SGVAGAVGQAASVGGLKV | ASSETGDANTEVF | 0.20386967 |
| SNGGAGTGSAGGNGGT | ASSETGDANTEVF | 0.2161692 |
| TGGAGGEGGAAG | ASSETGDANTEVF | 0.7968409 |
| TGGDGGAAGAGGAPGHG | ASSETGDANTEVF | 0.22350466 |
| TMPPLLAAIANAGRHGV | ASSETGDANTEVF | 0.5396338 |
| TSSTAGNSGGA | ASSETGDANTEVF | 0.14565785 |
| VPATVPVGILC | ASSETGDANTEVF | 0.18271983 |
| VQRPDSGGVVLTGRIS | ASSETGDANTEVF | 0.16366625 |
| WTPNPVPATKGVPAGLVQAT | ASSETGDANTEVF | 0.13849874 |
| YALKRALAKFSDRFDVVII | ASSETGDANTEVF | 0.32572275 |
| YLIGSVIV | ASSETGDANTEVF | 0.18473084 |
| YGQLVAVID | ASSETGDANTEVF | 0.29463232 |
| VTVGGMTLPA | ASSETGDANTEVF | 0.22807242 |
| VLALPAV | ASSETGDANTEVF | 0.31235033 |
| VIPPTLNYET | ASSETGDANTEVF | 0.21526016 |
| VGMGGMHPSSG | ASSETGDANTEVF | 0.19123141 |
| VGLAGGAATA | ASSETGDANTEVF | 0.17399743 |
| VGGAGGAGGSGP | ASSETGDANTEVF | 0.27839452 |
| VFAAFMV | ASSETGDANTEVF | 0.31400612 |
| TLYDKLR | ASSETGDANTEVF | 0.21242052 |
| TLAVFTVL | ASSETGDANTEVF | 0.14930454 |
| TGSVQPARV | ASSETGDANTEVF | 0.31409338 |
| TGGGAGSGGT | ASSETGDANTEVF | 0.20595661 |
| SHLETSL | ASSETGDANTEVF | 0.15711918 |
| SFTTGMT | ASSETGDANTEVF | 0.15078577 |
| QVEVAKRMGVRQPTVS | ASSETGDANTEVF | 0.17529513 |
| QGSGGPALAM | ASSETGDANTEVF | 0.16866542 |
| MLAPSNP | ASSETGDANTEVF | 0.3404838 |
| LVNNAAI | ASSETGDANTEVF | 0.379062 |
| LTGEINTLA | ASSETGDANTEVF | 0.19735044 |
| LSIFGPV | ASSETGDANTEVF | 0.2380656 |
| LLALLANL | ASSETGDANTEVF | 0.22197443 |
| LISNASDALDKLR | ASSETGDANTEVF | 0.13369617 |
| LINEAVP | ASSETGDANTEVF | 0.18039331 |
| LGYAFSGL | ASSETGDANTEVF | 0.16176936 |

| | | |
|---|---|---|
| LDFVAVAL | ASSETGDANTEVF | 0.32790267 |
| LAQPGAE | ASSETGDANTEVF | 0.16360746 |
| ISNLGAVTGF | ASSETGDANTEVF | 0.21255732 |
| HQEDPGRI | ASSETGDANTEVF | 0.22293903 |
| GTPGEPTP | ASSETGDANTEVF | 0.69035375 |
| GFFNSGS | ASSETGDANTEVF | 0.268149 |
| GAGGGGLTAGAN | ASSETGDANTEVF | 0.15994664 |
| FVDALVNM | ASSETGDANTEVF | 0.14995131 |
| FLTVAPEL | ASSETGDANTEVF | 0.26901525 |
| FETLTAL | ASSETGDANTEVF | 0.18807624 |
| DVINPIEL | ASSETGDANTEVF | 0.16577004 |
| APVSGAPV | ASSETGDANTEVF | 0.1601568 |
| ALVGGLAW | ASSETGDANTEVF | 0.3370401 |
| ALAAAAATAI | ASSETGDANTEVF | 0.14972256 |
| AFGLGLLL | ASSETGDANTEVF | 0.23387384 |
| AFFAAASL | ASSETGDANTEVF | 0.19350152 |
| AAAALPAFSPP | ASSETGDANTEVF | 0.21795905 |
| VVLAPAL | ASSETGDANTEVF | 0.60187316 |
| TIGHVDHGKTTL | ASSETGDANTEVF | 0.7274054 |
| LISNASDALDK | ASSETGDANTEVF | 0.18653467 |
| LISNASDAL | ASSETGDANTEVF | 0.18881774 |
| ISNASDALDK | ASSETGDANTEVF | 0.1685219 |
| NQAETLVYQTE | ASSETGDANTEVF | 0.17211325 |
| TGGTGGTAGAG | ASSETGDANTEVF | 0.1846858 |
| TGSTAGGAGGA | ASSETGDANTEVF | 0.35957655 |
| DPLAVPVAAGPV | ASSETGDANTEVF | 0.19116464 |
| GGTGGTPGNGGN | ASSETGDANTEVF | 0.16075276 |
| PAGLAGVDPATL | ASSETGDANTEVF | 0.19664223 |
| GNNLVVEAPHAGASV | ASSETGDANTEVF | 0.48647016 |
| KVTRIMSASTLEQPA | ASSETGDANTEVF | 0.39717114 |
| LGFALPIALPRRTVV | ASSETGDANTEVF | 0.16331 |
| TRLGNLTARKLVVLV | ASSETGDANTEVF | 0.18318686 |
| WSQQRRLIYFPSAGP | ASSETGDANTEVF | 0.18222487 |
| GPNGNPDPMAAAVDIR | ASSETGDANTEVF | 0.21634531 |
| TQRCVSVADLRAAASR | ASSETGDANTEVF | 0.1625782 |
| QFVLDVAEHPVEVLTETV | ASSETGDANTEVF | 0.1701378 |
| GGAGGTGGAGGNGGTQAGDG | ASSETGDANTEVF | 0.2222959 |
| PLLATLRAIAVDVDAKPAQV | ASSETGDANTEVF | 0.81735665 |
| APKFRTGGYMIDFWGVGYQVAK | ASSETGDANTEVF | 0.18063533 |
| GAGGAGGAGGAAGGPGGTGGTG | ASSETGDANTEVF | 0.14905927 |
| IGALGLLATDVAPAGSVAFGVA | ASSETGDANTEVF | 0.24546468 |
| AGGAGGIGGNGGAGGAGGVLMG | ASSETGDANTEVF | 0.16000444 |
| ADELIGLL | ASSETGDANTEVF | 0.16609457 |
| ADRALKA | ASSETGDANTEVF | 0.17394072 |
| AGAVGQVGGAGGA | ASSETGDANTEVF | 0.14692241 |
| AGGGGGLGGAP | ASSETGDANTEVF | 0.22976339 |
| AGGLLYGNGGAG | ASSETGDANTEVF | 0.41899997 |
| AGLESIGL | ASSETGDANTEVF | 0.44388887 |
| AGLKTGII | ASSETGDANTEVF | 0.24493583 |
| ASLAALLL | ASSETGDANTEVF | 0.47207484 |
| ASVINISSI | ASSETGDANTEVF | 0.2752491 |

| | | |
|---|---|---|
| ATPPGPA | ASSETGDANTEVF | 0.461029 |
| ATVASALV | ASSETGDANTEVF | 0.67070657 |
| ATVASLIL | ASSETGDANTEVF | 0.21683535 |
| AVNGPVLIRH | ASSETGDANTEVF | 0.1991113 |
| DGAAGAVGI | ASSETGDANTEVF | 0.5681716 |
| DPLLHGI | ASSETGDANTEVF | 0.23494324 |
| ELGAALGAL | ASSETGDANTEVF | 0.22706774 |
| ELGRTQAV | ASSETGDANTEVF | 0.19194415 |
| FVAGGYLS | ASSETGDANTEVF | 0.55161625 |
| GASGGAGGAGGW | ASSETGDANTEVF | 0.1863469 |
| GGAGAGGDGGAGGI | ASSETGDANTEVF | 0.737813 |
| GTGHNGSQPGMGGNG | ASSETGDANTEVF | 0.14622237 |
| GYAAASASAS | ASSETGDANTEVF | 0.16028196 |
| HQVDVAAVQQG | ASSETGDANTEVF | 0.35669273 |
| IGAAIAHT | ASSETGDANTEVF | 0.13559873 |
| IGMIEVAQL | ASSETGDANTEVF | 0.1515403 |
| IRLAPEHTSAMD | ASSETGDANTEVF | 0.19368331 |
| IYNVSVL | ASSETGDANTEVF | 0.29937452 |
| KQGFLGM | ASSETGDANTEVF | 0.13622378 |
| LESVITQL | ASSETGDANTEVF | 0.18237773 |
| LGLGALVVV | ASSETGDANTEVF | 0.15815547 |
| LIGNGGNGGSGGI | ASSETGDANTEVF | 0.20089254 |
| LNAINAPTL | ASSETGDANTEVF | 0.21125335 |
| LQLLNTGV | ASSETGDANTEVF | 0.15098539 |
| LQTSIEF | ASSETGDANTEVF | 0.28088513 |
| MSFAVSEL | ASSETGDANTEVF | 0.14898656 |
| NTGALGL | ASSETGDANTEVF | 0.2721027 |
| PPGAGGAGGAGTT | ASSETGDANTEVF | 0.29288262 |
| QLVGLGI | ASSETGDANTEVF | 0.74577594 |
| RFLEQVHQL | ASSETGDANTEVF | 0.15949453 |
| RGVGYVLR | ASSETGDANTEVF | 0.24711257 |
| SATGVFAGVI | ASSETGDANTEVF | 0.15795742 |
| SLPAITLP | ASSETGDANTEVF | 0.6642554 |
| STALAFTV | ASSETGDANTEVF | 0.20343834 |
| SVLTVGLGK | ASSETGDANTEVF | 0.14972521 |
| TALAGAGSGI | ASSETGDANTEVF | 0.33516812 |
| TGLPFTGI | ASSETGDANTEVF | 0.23347864 |
| VAEHVTV | ASSETGDANTEVF | 0.1849582 |
| VDIVLNSL | ASSETGDANTEVF | 0.68536186 |
| VLTGGGAVL | ASSETGDANTEVF | 0.15842207 |
| VNVGLAYGI | ASSETGDANTEVF | 0.26393744 |
| VSSAAVGAV | ASSETGDANTEVF | 0.3905312 |
| VTSVYEAGV | ASSETGDANTEVF | 0.2917956 |
| WVALDRGA | ASSETGDANTEVF | 0.17777964 |
| YLVGSVVI | ASSETGDANTEVF | 0.22785589 |
| GPPGTGKTLL | ASSETGDANTEVF | 0.22030173 |
| GVGMTKF | ASSETGDANTEVF | 0.17477003 |
| IGIGGSDL | ASSETGDANTEVF | 0.4062456 |
| IGTLLAEL | ASSETGDANTEVF | 0.17295377 |
| IYTSGTT | ASSETGDANTEVF | 0.27435815 |
| LSFPAGV | ASSETGDANTEVF | 0.261067 |

| | | |
|---|---|---|
| LYTSGTT | ASSETGDANTEVF | 0.78141236 |
| VGAFPFGF | ASSETGDANTEVF | 0.16741866 |
| VGAGTVEF | ASSETGDANTEVF | 0.19530925 |
| VGKSTLLL | ASSETGDANTEVF | 0.2186298 |
| AALAHAGGGIV | ASSETGDANTEVF | 0.27558947 |
| AALVAVSLL | ASSETGDANTEVF | 0.17567684 |
| AAVSRAVAAL | ASSETGDANTEVF | 0.27557185 |
| AGGSSGIGGF | ASSETGDANTEVF | 0.2711622 |
| AGLTVANANG | ASSETGDANTEVF | 0.33820772 |
| AGLVGAVL | ASSETGDANTEVF | 0.19479878 |
| ALSQAFTV | ASSETGDANTEVF | 0.22351441 |
| ALTFSVSL | ASSETGDANTEVF | 0.5546284 |
| ALTGAALVAL | ASSETGDANTEVF | 0.17476366 |
| ASFSIGAGVL | ASSETGDANTEVF | 0.200026 |
| DGSVYKI | ASSETGDANTEVF | 0.25244248 |
| EQLAEIV | ASSETGDANTEVF | 0.33322987 |
| FAVSELTVM | ASSETGDANTEVF | 0.18491776 |
| GAAPLAAGL | ASSETGDANTEVF | 0.29882818 |
| GADGGDGGSGGAGG | ASSETGDANTEVF | 0.17096303 |
| GFGLDTGGAGG | ASSETGDANTEVF | 0.29602757 |
| GLASGGLL | ASSETGDANTEVF | 0.18506783 |
| GRIAGAAVLL | ASSETGDANTEVF | 0.21362631 |
| GSAGSLNTSF | ASSETGDANTEVF | 0.17580357 |
| GTGQIGI | ASSETGDANTEVF | 0.335752 |
| GTLIALIL | ASSETGDANTEVF | 0.2783775 |
| IAATMGFVVG | ASSETGDANTEVF | 0.39951155 |
| IFAQALL | ASSETGDANTEVF | 0.15141891 |
| IGAAVALGF | ASSETGDANTEVF | 0.34526455 |
| IGLTPLAL | ASSETGDANTEVF | 0.43366402 |
| ILGSNTSTL | ASSETGDANTEVF | 0.8010349 |
| IVFSLTVL | ASSETGDANTEVF | 0.18897164 |
| LASKATGF | ASSETGDANTEVF | 0.23986459 |
| LASVALIL | ASSETGDANTEVF | 0.18171598 |
| LFAVGLTL | ASSETGDANTEVF | 0.1868067 |
| LFNGGIGTYI | ASSETGDANTEVF | 0.14834051 |
| LTLGAGII | ASSETGDANTEVF | 0.18244624 |
| LVAAALGDI | ASSETGDANTEVF | 0.22588092 |
| NTAFVQLGI | ASSETGDANTEVF | 0.19019842 |
| QLFSGAEL | ASSETGDANTEVF | 0.7465807 |
| SAATFITL | ASSETGDANTEVF | 0.34181112 |
| SNYGLANLGV | ASSETGDANTEVF | 0.3063801 |
| SYGGAYAVM | ASSETGDANTEVF | 0.41402477 |
| TTGVGGLTLG | ASSETGDANTEVF | 0.35016733 |
| TTQTIYI | ASSETGDANTEVF | 0.22720142 |
| VAAGAGLGAVI | ASSETGDANTEVF | 0.1816421 |
| VAAVVTLL | ASSETGDANTEVF | 0.2265214 |
| VGFLGGLI | ASSETGDANTEVF | 0.14984587 |
| VGVIPILL | ASSETGDANTEVF | 0.22248714 |
| VIFASSLI | ASSETGDANTEVF | 0.19320768 |
| VKGGDGGAAAG | ASSETGDANTEVF | 0.26496875 |
| VTKTFTNI | ASSETGDANTEVF | 0.137076 |

| | | |
|---|---|---|
| VVAAAILL | ASSETGDANTEVF | 0.6811705 |
| YAALDVEL | ASSETGDANTEVF | 0.4349916 |
| LASVALLL | ASSETGDANTEVF | 0.24057041 |
| LDAVHRV | ASSETGDANTEVF | 0.303688 |
| QGLSSSIF | ASSETGDANTEVF | 0.37107933 |
| RGTFANIRL | ASSETGDANTEVF | 0.15899454 |
| RQLWWGHRI | ASSETGDANTEVF | 0.2660987 |
| YGPPGTGKTLL | ASSETGDANTEVF | 0.16765071 |
| AALVGAPANVA | ASSETGDANTEVF | 0.32664055 |
| AATFRRTE | ASSETGDANTEVF | 0.3629769 |
| AGALGGGI | ASSETGDANTEVF | 0.27769184 |
| AGGAGGAGGIGAG | ASSETGDANTEVF | 0.17296821 |
| AGSGALGL | ASSETGDANTEVF | 0.20996726 |
| AQIADQI | ASSETGDANTEVF | 0.1621148 |
| ASAVAVTYL | ASSETGDANTEVF | 0.19995897 |
| ASTVLGGTGG | ASSETGDANTEVF | 0.18949753 |
| AVGGEPAK | ASSETGDANTEVF | 0.27706978 |
| CEQGTFGGTC | ASSETGDANTEVF | 0.16565107 |
| DEGKAPAG | ASSETGDANTEVF | 0.3036785 |
| DTVVFPAL | ASSETGDANTEVF | 0.24935725 |
| FEGITMAG | ASSETGDANTEVF | 0.2119785 |
| FPLVVQD | ASSETGDANTEVF | 0.18086763 |
| GAFTTPQ | ASSETGDANTEVF | 0.31819344 |
| GFGGFGTAGGVGGA | ASSETGDANTEVF | 0.3188312 |
| GFLAQFL | ASSETGDANTEVF | 0.15840746 |
| GGAGGAGGNAGVA | ASSETGDANTEVF | 0.26316756 |
| GGVGTAGAGGAGGAGG | ASSETGDANTEVF | 0.17480974 |
| GNGGGGDAGNAGSGGN | ASSETGDANTEVF | 0.19830397 |
| GTGGAGGAGNDAGS | ASSETGDANTEVF | 0.36609674 |
| GTTNTGWFNS | ASSETGDANTEVF | 0.20300226 |
| IFMGPLFTL | ASSETGDANTEVF | 0.15021272 |
| ILAHAIEAI | ASSETGDANTEVF | 0.21818912 |
| IRSAIREAL | ASSETGDANTEVF | 0.26448575 |
| ITFAGGG | ASSETGDANTEVF | 0.19364895 |
| ITGMSGHQEV | ASSETGDANTEVF | 0.14055274 |
| IVAGLACTL | ASSETGDANTEVF | 0.23064592 |
| IVGGGLAGA | ASSETGDANTEVF | 0.1728213 |
| IVTGVLNI | ASSETGDANTEVF | 0.19503266 |
| LAQQGCI | ASSETGDANTEVF | 0.18223573 |
| LCLHVEN | ASSETGDANTEVF | 0.4037845 |
| LDAISDGS | ASSETGDANTEVF | 0.2045117 |
| LGGFAAML | ASSETGDANTEVF | 0.21872984 |
| LGGKNPMI | ASSETGDANTEVF | 0.26901668 |
| LGPKGVAAIL | ASSETGDANTEVF | 0.21794435 |
| LMVGVSI | ASSETGDANTEVF | 0.393214 |
| LVGAAFVL | ASSETGDANTEVF | 0.2554714 |
| LVLAVAPL | ASSETGDANTEVF | 0.18721567 |
| MAAIAVIDAA | ASSETGDANTEVF | 0.2119121 |
| NNGINAELQ | ASSETGDANTEVF | 0.17807515 |
| RGEPLVLVHG | ASSETGDANTEVF | 0.19502085 |
| RGGAGGAAGLI | ASSETGDANTEVF | 0.16461352 |

| | | |
|---|---|---|
| RSYRMHPAV | ASSETGDANTEVF | 0.1779771 |
| SAAHLLAGQLA | ASSETGDANTEVF | 0.18157409 |
| SAAVGSY | ASSETGDANTEVF | 0.279849 |
| SPHAVLHRI | ASSETGDANTEVF | 0.24393731 |
| SQIAKVL | ASSETGDANTEVF | 0.1619296 |
| TALGHLVP | ASSETGDANTEVF | 0.15328999 |
| TAPTTSGAA | ASSETGDANTEVF | 0.17745939 |
| TDGDDGNGGSPGT | ASSETGDANTEVF | 0.19313748 |
| TDGLGEN | ASSETGDANTEVF | 0.33882475 |
| TGGPPLPGQA | ASSETGDANTEVF | 0.23952731 |
| TIPAITI | ASSETGDANTEVF | 0.8038628 |
| TIPGFTI | ASSETGDANTEVF | 0.4949031 |
| TSIASGALA | ASSETGDANTEVF | 0.19472799 |
| TVTLPGAAI | ASSETGDANTEVF | 0.28342646 |
| VASGLSAALP | ASSETGDANTEVF | 0.1776104 |
| VGAGVIGI | ASSETGDANTEVF | 0.20977473 |
| VGPKRFGV | ASSETGDANTEVF | 0.20104156 |
| VNVAFPD | ASSETGDANTEVF | 0.17241412 |
| VSMTNSGVS | ASSETGDANTEVF | 0.18532445 |
| VTDVLGDLSV | ASSETGDANTEVF | 0.20154607 |
| VVVVGAGFAGL | ASSETGDANTEVF | 0.15029839 |
| VYTVGTVV | ASSETGDANTEVF | 0.35613483 |
| WTTFSGP | ASSETGDANTEVF | 0.18874587 |
| YDELVEAF | ASSETGDANTEVF | 0.15289195 |
| YGTGGAGGNGG | ASSETGDANTEVF | 0.18355274 |
| YTNVMAV | ASSETGDANTEVF | 0.43836868 |
| FLEQVHQL | ASSETGDANTEVF | 0.35619688 |
| GLSLLAVAV | ASSETGDANTEVF | 0.19004878 |
| LEQVHQL | ASSETGDANTEVF | 0.16799122 |
| LNAAGAI | ASSETGDANTEVF | 0.721727 |
| PAVHASL | ASSETGDANTEVF | 0.16586718 |
| TELEDDA | ASSETGDANTEVF | 0.19458531 |
| TPPGPPN | ASSETGDANTEVF | 0.25630733 |
| TPPPPPPP | ASSETGDANTEVF | 0.19926229 |
| TQSPALI | ASSQSAWGKNTLY | 0.16907963 |
| EFAAAFPQ | ASSQSAWGKNTLY | 0.27988204 |
| LLSGNLDVL | ASSQSAWGKNTLY | 0.21681538 |
| GPANLGNYNI | ASSQSAWGKNTLY | 0.18718015 |
| TLGPTLP | ASSQSAWGKNTLY | 0.13930905 |
| VYAYSSCCK | ASSQSAWGKNTLY | 0.23403001 |
| APAPAPAPNI | ASSQSAWGKNTLY | 0.16555554 |
| PGGPVPIP | ASSQSAWGKNTLY | 0.15887722 |
| FLTVALEP | ASSQSAWGKNTLY | 0.13880718 |
| SQYFGANL | ASSQSAWGKNTLY | 0.19832239 |
| GGVKPAA | ASSQSAWGKNTLY | 0.31692928 |
| GGTGGAGGAGGW | ASSQSAWGKNTLY | 0.4106997 |
| GVGGVAGLGGAGG | ASSQSAWGKNTLY | 0.20381607 |
| VLLVLATTI | ASSQSAWGKNTLY | 0.31357166 |
| IGIFAAML | ASSQSAWGKNTLY | 0.30071583 |
| AVGDLID | ASSQSAWGKNTLY | 0.20065027 |
| NISPLPL | ASSQSAWGKNTLY | 0.15412906 |

| | | |
|---|---|---|
| VTAVFVAI | ASSQSAWGKNTLY | 0.20148093 |
| SGDDNDAAL | ASSQSAWGKNTLY | 0.3772425 |
| AVAALAHS | ASSQSAWGKNTLY | 0.2540841 |
| GGVTVMATP | ASSQSAWGKNTLY | 0.2648926 |
| AGLAANVRD | ASSQSAWGKNTLY | 0.16425526 |
| TNRPDLIDPALLRPG | ASSQSAWGKNTLY | 0.214011 |
| LANLGVTAL | ASSQSAWGKNTLY | 0.2939062 |
| TGERIEL | ASSQSAWGKNTLY | 0.15101749 |
| NGEELAA | ASSQSAWGKNTLY | 0.16672322 |
| VLSLVWG | ASSQSAWGKNTLY | 0.18227187 |
| IGYALWL | ASSQSAWGKNTLY | 0.15198515 |
| LSLLLAPL | ASSQSAWGKNTLY | 0.23131976 |
| TLDEALK | ASSQSAWGKNTLY | 0.20715418 |
| TDGAGTAEGG | ASSQSAWGKNTLY | 0.19246846 |
| NAPGGAGGNGG | ASSQSAWGKNTLY | 0.19715571 |
| VSVFMGMGGNF | ASSQSAWGKNTLY | 0.18254505 |
| LASYLAAL | ASSQSAWGKNTLY | 0.18682353 |
| LVGGTPAL | ASSQSAWGKNTLY | 0.20701785 |
| VRNADVL | ASSQSAWGKNTLY | 0.30040592 |
| TCSSPFLVSAM | ASSQSAWGKNTLY | 0.25767726 |
| VHITDAE | ASSQSAWGKNTLY | 0.21306159 |
| EYAAPSF | ASSQSAWGKNTLY | 0.19306223 |
| VVDAFHGGL | ASSQSAWGKNTLY | 0.60899264 |
| ASVVDPQ | ASSQSAWGKNTLY | 0.1598429 |
| SYNEFGGLS | ASSQSAWGKNTLY | 0.17233628 |
| DPDGSVY | ASSQSAWGKNTLY | 0.22359933 |
| MVSTIHATE | ASSQSAWGKNTLY | 0.19447064 |
| LAAGVADPV | ASSQSAWGKNTLY | 0.26093575 |
| TKEQIAA | ASSQSAWGKNTLY | 0.25784206 |
| SLASPAAL | ASSQSAWGKNTLY | 0.28744924 |
| IGLVFYGV | ASSQSAWGKNTLY | 0.4234465 |
| FLLDASNL | ASSQSAWGKNTLY | 0.53093475 |
| YQVGNFP | ASSQSAWGKNTLY | 0.30690977 |
| ESLLSQST | ASSQSAWGKNTLY | 0.19420129 |
| LFTVGFAAVG | ASSQSAWGKNTLY | 0.610235 |
| ATPPTPT | ASSQSAWGKNTLY | 0.16132557 |
| TDLATLG | ASSQSAWGKNTLY | 0.3036674 |
| SAGGTGR | ASSQSAWGKNTLY | 0.26566312 |
| GFSNSGS | ASSQSAWGKNTLY | 0.17059726 |
| GGNGGSGGTSV | ASSQSAWGKNTLY | 0.26754963 |
| GTSVATGGAGN | ASSQSAWGKNTLY | 0.3641234 |
| NGGNGGSGGTS | ASSQSAWGKNTLY | 0.4062944 |
| AVVKFPH | ASSQSAWGKNTLY | 0.17808293 |
| TPPPPPP | ASSQSAWGKNTLY | 0.1764086 |
| AAPAAAAAPA | ASSQSAWGKNTLY | 0.1620694 |
| IALNEDL | ASSQSAWGKNTLY | 0.16346177 |
| IVNVSSM | ASSQSAWGKNTLY | 0.16296679 |
| TGIPPAPR | ASSQSAWGKNTLY | 0.22499874 |
| FDLGGGTF | ASSQSAWGKNTLY | 0.20953242 |
| FDLGGGTFDVS | ASSQSAWGKNTLY | 0.72704595 |
| NEPTAAA | ASSQSAWGKNTLY | 0.25962532 |

| | | |
|---|---|---|
| NEPTAAALA | ASSQSAWGKNTLY | 0.3417939 |
| GSIINISSI | ASSQSAWGKNTLY | 0.18571489 |
| VAPGNAGTA | ASSQSAWGKNTLY | 0.21046299 |
| TGGGASLE | ASSQSAWGKNTLY | 0.14599301 |
| VAVYNHYKRI | ASSQSAWGKNTLY | 0.19589308 |
| TDPAPATT | ASSQSAWGKNTLY | 0.162661 |
| TNRPDILDPALLRPG | ASSQSAWGKNTLY | 0.21420999 |
| LTTTTTS | ASSQSAWGKNTLY | 0.14985828 |
| PTGSGKTL | ASSQSAWGKNTLY | 0.37231332 |
| TGSGKTL | ASSQSAWGKNTLY | 0.20082982 |
| TIGHVDHGK | ASSQSAWGKNTLY | 0.27254152 |
| TIGHVDHGKTT | ASSQSAWGKNTLY | 0.17570157 |
| GHVDHGKTT | ASSQSAWGKNTLY | 0.17689288 |
| AQFEHTLL | ASSQSAWGKNTLY | 0.18976104 |
| IVNQDEL | ASSQSAWGKNTLY | 0.36018044 |
| TNGSQFF | ASSQSAWGKNTLY | 0.16034643 |
| TGKGIVE | ASSQSAWGKNTLY | 0.1440489 |
| LISNASDA | ASSQSAWGKNTLY | 0.17129296 |
| AELRTLGELTKIAKAHGAQ | ASSQSAWGKNTLY | 0.1598011 |
| APNAVAAAEPSW | ASSQSAWGKNTLY | 0.4701003 |
| APQVAGAATAVL | ASSQSAWGKNTLY | 0.3244083 |
| ARRDPAAAAAILPTDA | ASSQSAWGKNTLY | 0.25905415 |
| DAAARDVLAEAGLAEYF | ASSQSAWGKNTLY | 0.72877616 |
| DDQIVLSAKANLAKVEQ | ASSQSAWGKNTLY | 0.13606086 |
| DELLPDTGEDSDEDGH | ASSQSAWGKNTLY | 0.1898414 |
| ELPEAPANRATVPQD | ASSQSAWGKNTLY | 0.4447899 |
| FGQSKIERVTGIG | ASSQSAWGKNTLY | 0.21113032 |
| GAGGTGSTAGGAGGAGGAGGLY | ASSQSAWGKNTLY | 0.23368774 |
| GHGGTGAAGGNGGTGGAGGSGI | ASSQSAWGKNTLY | 0.31762177 |
| GNGGNGGAGGEGGPGGAGGAGA | ASSQSAWGKNTLY | 0.17101632 |
| GSTAGNGGNGGNATNGTIAGQP | ASSQSAWGKNTLY | 0.20308773 |
| GTGGDGGNGGNGGTGG | ASSQSAWGKNTLY | 0.21440476 |
| GVRRPAGIPADAPV | ASSQSAWGKNTLY | 0.23053674 |
| GYRQVTDAHLLTLARR | ASSQSAWGKNTLY | 0.17303698 |
| IAKMGQDGHDRG | ASSQSAWGKNTLY | 0.16077852 |
| KAEPSIEIQYT | ASSQSAWGKNTLY | 0.23030484 |
| KPVRIRMAADEDTVAVT | ASSQSAWGKNTLY | 0.7627574 |
| MPPRIAGMRLLVIKPEP | ASSQSAWGKNTLY | 0.23061898 |
| MKFYAVALFRSRGVPPTR | ASSQSAWGKNTLY | 0.21516041 |
| NAGGFGILGAVAHSP | ASSQSAWGKNTLY | 0.21339826 |
| NEGREITLAMAEM | ASSQSAWGKNTLY | 0.5034895 |
| NELVEWSHPLGHRWRWE | ASSQSAWGKNTLY | 0.1855486 |
| NLGAGGTGGNGGNPG | ASSQSAWGKNTLY | 0.15541267 |
| NTEWPKWETVMAADPVD | ASSQSAWGKNTLY | 0.35447174 |
| PASAAPAETQP | ASSQSAWGKNTLY | 0.15214713 |
| QQSSDGKNGVRAAEITGE | ASSQSAWGKNTLY | 0.20202725 |
| RAPDAVTTSWWKE | ASSQSAWGKNTLY | 0.19131503 |
| SGFGNVGAHTSGWFNQ | ASSQSAWGKNTLY | 0.76365083 |
| SGVAGAVGQAASVGGLKV | ASSQSAWGKNTLY | 0.28277898 |
| SNGGAGTGSAGGNGGT | ASSQSAWGKNTLY | 0.21803021 |
| TGGAGGEGGAAG | ASSQSAWGKNTLY | 0.16522102 |

| | | |
|---|---|---|
| TGGDGGAAGAGGAPGHG | ASSQSAWGKNTLY | 0.2060914 |
| TMPPLLAAIANAGRHGV | ASSQSAWGKNTLY | 0.3069866 |
| TSSTAGNSGGA | ASSQSAWGKNTLY | 0.15417917 |
| VPATVPVGILC | ASSQSAWGKNTLY | 0.32280594 |
| VQRPDSGGVVLTGRIS | ASSQSAWGKNTLY | 0.16267596 |
| WTPNPVPATKGVPAGLVQAT | ASSQSAWGKNTLY | 0.22498602 |
| YALKRALAKFSDRFDVVII | ASSQSAWGKNTLY | 0.2851378 |
| YLIGSVIV | ASSQSAWGKNTLY | 0.8064872 |
| YGQLVAVID | ASSQSAWGKNTLY | 0.20031711 |
| VTVGGMTLPA | ASSQSAWGKNTLY | 0.32459205 |
| VLALPAV | ASSQSAWGKNTLY | 0.17131084 |
| VIPPTLNYET | ASSQSAWGKNTLY | 0.21592523 |
| VGMGGMHPSSG | ASSQSAWGKNTLY | 0.54906064 |
| VGLAGGAATA | ASSQSAWGKNTLY | 0.4076117 |
| VGGAGGAGGSGP | ASSQSAWGKNTLY | 0.7812792 |
| VFAAFMV | ASSQSAWGKNTLY | 0.19795232 |
| TLYDKLR | ASSQSAWGKNTLY | 0.1526716 |
| TLAVFTVL | ASSQSAWGKNTLY | 0.18851909 |
| TGSVQPARV | ASSQSAWGKNTLY | 0.26639095 |
| TGGGAGSGGT | ASSQSAWGKNTLY | 0.40013507 |
| SHLETSL | ASSQSAWGKNTLY | 0.35266754 |
| SFTTGMT | ASSQSAWGKNTLY | 0.21931654 |
| QVEVAKRMGVRQPTVS | ASSQSAWGKNTLY | 0.19522795 |
| QGSGGPALAM | ASSQSAWGKNTLY | 0.439462 |
| MLAPSNP | ASSQSAWGKNTLY | 0.38888237 |
| LVNNAAI | ASSQSAWGKNTLY | 0.22559005 |
| LTGEINTLA | ASSQSAWGKNTLY | 0.15835805 |
| LSIFGPV | ASSQSAWGKNTLY | 0.15964165 |
| LLALLANL | ASSQSAWGKNTLY | 0.28696495 |
| LISNASDALDKLR | ASSQSAWGKNTLY | 0.30381423 |
| LINEAVP | ASSQSAWGKNTLY | 0.15793282 |
| LGYAFSGL | ASSQSAWGKNTLY | 0.25027618 |
| LDFVAVAL | ASSQSAWGKNTLY | 0.27067417 |
| LAQPGAE | ASSQSAWGKNTLY | 0.1533015 |
| ISNLGAVTGF | ASSQSAWGKNTLY | 0.21216747 |
| HQEDPGRI | ASSQSAWGKNTLY | 0.48436293 |
| GTPGEPTP | ASSQSAWGKNTLY | 0.22052512 |
| GFFNSGS | ASSQSAWGKNTLY | 0.1627969 |
| GAGGGGLTAGAN | ASSQSAWGKNTLY | 0.39089182 |
| FVDALVNM | ASSQSAWGKNTLY | 0.68649447 |
| FLTVAPEL | ASSQSAWGKNTLY | 0.40600753 |
| FETLTAL | ASSQSAWGKNTLY | 0.15189192 |
| DVINPIEL | ASSQSAWGKNTLY | 0.13709772 |
| APVSGAPV | ASSQSAWGKNTLY | 0.250215 |
| ALVGGLAW | ASSQSAWGKNTLY | 0.6832894 |
| ALAAAAATAI | ASSQSAWGKNTLY | 0.20109533 |
| AFGLGLLL | ASSQSAWGKNTLY | 0.17466336 |
| AFFAAASL | ASSQSAWGKNTLY | 0.2093176 |
| AAAALPAFSPP | ASSQSAWGKNTLY | 0.37767357 |
| VVLAPAL | ASSQSAWGKNTLY | 0.17488708 |
| TIGHVDHGKTTL | ASSQSAWGKNTLY | 0.15159751 |

| | | |
|---|---|---|
| LISNASDALDK | ASSQSAWGKNTLY | 0.23661748 |
| LISNASDAL | ASSQSAWGKNTLY | 0.30351678 |
| ISNASDALDK | ASSQSAWGKNTLY | 0.16555063 |
| NQAETLVYQTE | ASSQSAWGKNTLY | 0.18662241 |
| TGGTGGTAGAG | ASSQSAWGKNTLY | 0.3048252 |
| TGSTAGGAGGA | ASSQSAWGKNTLY | 0.21926725 |
| DPLAVPVAAGPV | ASSQSAWGKNTLY | 0.24055006 |
| GGTGGTPGNGGN | ASSQSAWGKNTLY | 0.16248451 |
| PAGLAGVDPATL | ASSQSAWGKNTLY | 0.4104363 |
| GNNLVVEAPHAGASV | ASSQSAWGKNTLY | 0.16082658 |
| KVTRIMSASTLEQPA | ASSQSAWGKNTLY | 0.1917712 |
| LGFALPIALPRRTVV | ASSQSAWGKNTLY | 0.23503606 |
| TRLGNLTARKLVVLV | ASSQSAWGKNTLY | 0.21510935 |
| WSQQRRLIYFPSAGP | ASSQSAWGKNTLY | 0.20888212 |
| GPNGNPDPMAAAVDIR | ASSQSAWGKNTLY | 0.1776643 |
| TQRCVSVADLRAAASR | ASSQSAWGKNTLY | 0.16135897 |
| QFVLDVAEHPVEVLTETV | ASSQSAWGKNTLY | 0.15238048 |
| GGAGGTGGAGGNGGTQAGDG | ASSQSAWGKNTLY | 0.378151 |
| PLLATLRAIAVDVDAKPAQV | ASSQSAWGKNTLY | 0.31112787 |
| APKFRTGGYMIDFWGVGYQVAK | ASSQSAWGKNTLY | 0.30416024 |
| GAGGAGGAGGAAGGPGGTGGTG | ASSQSAWGKNTLY | 0.24255233 |
| IGALGLLATDVAPAGSVAFGVA | ASSQSAWGKNTLY | 0.18065137 |
| AGGAGGIGGNGGAGGAGGVLMG | ASSQSAWGKNTLY | 0.23661268 |
| ADELIGLL | ASSQSAWGKNTLY | 0.15324394 |
| ADRALKA | ASSQSAWGKNTLY | 0.25084722 |
| AGAVGQVGGAGGA | ASSQSAWGKNTLY | 0.17515983 |
| AGGGGGLGGAP | ASSQSAWGKNTLY | 0.2258482 |
| AGGLLYGNGGAG | ASSQSAWGKNTLY | 0.18770178 |
| AGLESIGL | ASSQSAWGKNTLY | 0.21075915 |
| AGLKTGII | ASSQSAWGKNTLY | 0.2506186 |
| ASLAALLL | ASSQSAWGKNTLY | 0.17437811 |
| ASVINISSI | ASSQSAWGKNTLY | 0.22752133 |
| ATPPGPA | ASSQSAWGKNTLY | 0.41652775 |
| ATVASALV | ASSQSAWGKNTLY | 0.233169 |
| ATVASLIL | ASSQSAWGKNTLY | 0.16574807 |
| AVNGPVLIRH | ASSQSAWGKNTLY | 0.23467816 |
| DGAAGAVGI | ASSQSAWGKNTLY | 0.17437594 |
| DPLLHGI | ASSQSAWGKNTLY | 0.2783899 |
| ELGAALGAL | ASSQSAWGKNTLY | 0.20320353 |
| ELGRTQAV | ASSQSAWGKNTLY | 0.17909478 |
| FVAGGYLS | ASSQSAWGKNTLY | 0.2258826 |
| GASGGAGGAGGW | ASSQSAWGKNTLY | 0.15256734 |
| GGAGAGGDGGAGGI | ASSQSAWGKNTLY | 0.16774343 |
| GTGHNGSQPGMGGNG | ASSQSAWGKNTLY | 0.17770492 |
| GYAAASASAS | ASSQSAWGKNTLY | 0.24749687 |
| HQVDVAAVQQG | ASSQSAWGKNTLY | 0.23936054 |
| IGAAIAHT | ASSQSAWGKNTLY | 0.16386041 |
| IGMIEVAQL | ASSQSAWGKNTLY | 0.21700943 |
| IRLAPEHTSAMD | ASSQSAWGKNTLY | 0.19421573 |
| IYNVSVL | ASSQSAWGKNTLY | 0.1713635 |
| KQGFLGM | ASSQSAWGKNTLY | 0.19937196 |

| | | |
|---|---|---|
| LESVITQL | ASSQSAWGKNTLY | 0.18387878 |
| LGLGALVVV | ASSQSAWGKNTLY | 0.1476819 |
| LIGNGGNGGSGGI | ASSQSAWGKNTLY | 0.23614085 |
| LNAINAPTL | ASSQSAWGKNTLY | 0.1849821 |
| LQLLNTGV | ASSQSAWGKNTLY | 0.21340744 |
| LQTSIEF | ASSQSAWGKNTLY | 0.26168573 |
| MSFAVSEL | ASSQSAWGKNTLY | 0.15054902 |
| NTGALGL | ASSQSAWGKNTLY | 0.21679035 |
| PPGAGGAGGAGTT | ASSQSAWGKNTLY | 0.19818704 |
| QLVGLGI | ASSQSAWGKNTLY | 0.25889295 |
| RFLEQVHQL | ASSQSAWGKNTLY | 0.23823225 |
| RGVGYVLR | ASSQSAWGKNTLY | 0.17541993 |
| SATGVFAGVI | ASSQSAWGKNTLY | 0.3207658 |
| SLPAITLP | ASSQSAWGKNTLY | 0.23955928 |
| STALAFTV | ASSQSAWGKNTLY | 0.22752322 |
| SVLTVGLGK | ASSQSAWGKNTLY | 0.22765194 |
| TALAGAGSGI | ASSQSAWGKNTLY | 0.65033185 |
| TGLPFTGI | ASSQSAWGKNTLY | 0.2741684 |
| VAEHVTV | ASSQSAWGKNTLY | 0.16692221 |
| VDIVLNSL | ASSQSAWGKNTLY | 0.14476706 |
| VLTGGGAVL | ASSQSAWGKNTLY | 0.22644371 |
| VNVGLAYGI | ASSQSAWGKNTLY | 0.30128878 |
| VSSAAVGAV | ASSQSAWGKNTLY | 0.17985876 |
| VTSVYEAGV | ASSQSAWGKNTLY | 0.14551844 |
| WVALDRGA | ASSQSAWGKNTLY | 0.4123071 |
| YLVGSVVI | ASSQSAWGKNTLY | 0.5839072 |
| GPPGTGKTLL | ASSQSAWGKNTLY | 0.17792529 |
| GVGMTKF | ASSQSAWGKNTLY | 0.4446237 |
| IGIGGSDL | ASSQSAWGKNTLY | 0.15023983 |
| IGTLLAEL | ASSQSAWGKNTLY | 0.2588792 |
| IYTSGTT | ASSQSAWGKNTLY | 0.4437252 |
| LSFPAGV | ASSQSAWGKNTLY | 0.15397443 |
| LYTSGTT | ASSQSAWGKNTLY | 0.2725931 |
| VGAFPFGF | ASSQSAWGKNTLY | 0.19514716 |
| VGAGTVEF | ASSQSAWGKNTLY | 0.17605802 |
| VGKSTLLL | ASSQSAWGKNTLY | 0.3413632 |
| AALAHAGGGIV | ASSQSAWGKNTLY | 0.18612596 |
| AALVAVSLL | ASSQSAWGKNTLY | 0.23695688 |
| AAVSRAVAAL | ASSQSAWGKNTLY | 0.21160594 |
| AGGSSGIGGF | ASSQSAWGKNTLY | 0.16934012 |
| AGLTVANANG | ASSQSAWGKNTLY | 0.22836648 |
| AGLVGAVL | ASSQSAWGKNTLY | 0.28465256 |
| ALSQAFTV | ASSQSAWGKNTLY | 0.21326438 |
| ALTFSVSL | ASSQSAWGKNTLY | 0.29598328 |
| ALTGAALVAL | ASSQSAWGKNTLY | 0.16974473 |
| ASFSIGAGVL | ASSQSAWGKNTLY | 0.67982817 |
| DGSVYKI | ASSQSAWGKNTLY | 0.6870673 |
| EQLAEIV | ASSQSAWGKNTLY | 0.22794701 |
| FAVSELTVM | ASSQSAWGKNTLY | 0.16980386 |
| GAAPLAAGL | ASSQSAWGKNTLY | 0.5541852 |
| GADGGDGGSGGAGG | ASSQSAWGKNTLY | 0.21523853 |

| | | |
|---|---|---|
| GFGLDTGGAGG | ASSQSAWGKNTLY | 0.23086546 |
| GLASGGLL | ASSQSAWGKNTLY | 0.28599757 |
| GRIAGAAVLL | ASSQSAWGKNTLY | 0.778736 |
| GSAGSLNTSF | ASSQSAWGKNTLY | 0.8924435 |
| GTGQIGI | ASSQSAWGKNTLY | 0.8641496 |
| GTLIALIL | ASSQSAWGKNTLY | 0.85487115 |
| IAATMGFVVG | ASSQSAWGKNTLY | 0.80996466 |
| IFAQALL | ASSQSAWGKNTLY | 0.53414327 |
| IGAAVALGF | ASSQSAWGKNTLY | 0.8320836 |
| IGLTPLAL | ASSQSAWGKNTLY | 0.7717873 |
| ILGSNTSTL | ASSQSAWGKNTLY | 0.8550792 |
| IVFSLTVL | ASSQSAWGKNTLY | 0.8769752 |
| LASKATGF | ASSQSAWGKNTLY | 0.7904502 |
| LASVALIL | ASSQSAWGKNTLY | 0.89415944 |
| LFAVGLTL | ASSQSAWGKNTLY | 0.8900772 |
| LFNGGIGTYI | ASSQSAWGKNTLY | 0.65614414 |
| LTLGAGII | ASSQSAWGKNTLY | 0.8106278 |
| LVAAALGDI | ASSQSAWGKNTLY | 0.78494924 |
| NTAFVQLGI | ASSQSAWGKNTLY | 0.84847265 |
| QLFSGAEL | ASSQSAWGKNTLY | 0.37784758 |
| SAATFITL | ASSQSAWGKNTLY | 0.8842633 |
| SNYGLANLGV | ASSQSAWGKNTLY | 0.7873147 |
| SYGGAYAVM | ASSQSAWGKNTLY | 0.16665494 |
| TTGVGGLTLG | ASSQSAWGKNTLY | 0.8519714 |
| TTQTIYI | ASSQSAWGKNTLY | 0.8189585 |
| VAAGAGLGAVI | ASSQSAWGKNTLY | 0.8403454 |
| VAAVVTLL | ASSQSAWGKNTLY | 0.88192546 |
| VGFLGGLI | ASSQSAWGKNTLY | 0.4666912 |
| VGVIPILL | ASSQSAWGKNTLY | 0.7881282 |
| VIFASSLI | ASSQSAWGKNTLY | 0.87428194 |
| VKGGDGGAAAG | ASSQSAWGKNTLY | 0.8958107 |
| VTKTFTNI | ASSQSAWGKNTLY | 0.73295087 |
| VVAAAILL | ASSQSAWGKNTLY | 0.86078453 |
| YAALDVEL | ASSQSAWGKNTLY | 0.8914113 |
| LASVALLL | ASSQSAWGKNTLY | 0.8711541 |
| LDAVHRV | ASSQSAWGKNTLY | 0.88215137 |
| QGLSSSIF | ASSQSAWGKNTLY | 0.895206 |
| RGTFANIRL | ASSQSAWGKNTLY | 0.7612066 |
| RQLWWGHRI | ASSQSAWGKNTLY | 0.8887203 |
| YGPPGTGKTLL | ASSQSAWGKNTLY | 0.8945738 |
| AALVGAPANVA | ASSQSAWGKNTLY | 0.8168272 |
| AATFRRTE | ASSQSAWGKNTLY | 0.84783584 |
| AGALGGGI | ASSQSAWGKNTLY | 0.7306496 |
| AGGAGGAGGIGAG | ASSQSAWGKNTLY | 0.80428624 |
| AGSGALGL | ASSQSAWGKNTLY | 0.37752408 |
| AQIADQI | ASSQSAWGKNTLY | 0.8485081 |
| ASAVAVTYL | ASSQSAWGKNTLY | 0.88927686 |
| ASTVLGGTGG | ASSQSAWGKNTLY | 0.8080951 |
| AVGGEPAK | ASSQSAWGKNTLY | 0.7026509 |
| CEQGTFGGTC | ASSQSAWGKNTLY | 0.74492884 |
| DEGKAPAG | ASSQSAWGKNTLY | 0.8126306 |

| | | |
|---|---|---|
| DTVVFPAL | ASSQSAWGKNTLY | 0.83818805 |
| FEGITMAG | ASSQSAWGKNTLY | 0.3479152 |
| FPLVVQD | ASSQSAWGKNTLY | 0.7863572 |
| GAFTTPQ | ASSQSAWGKNTLY | 0.89035225 |
| GFGGFGTAGGVGGA | ASSQSAWGKNTLY | 0.8904122 |
| GFLAQFL | ASSQSAWGKNTLY | 0.5317884 |
| GGAGGAGGNAGVA | ASSQSAWGKNTLY | 0.88943833 |
| GGVGTAGAGGAGGAGG | ASSQSAWGKNTLY | 0.8758228 |
| GNGGGGDAGNAGSGGN | ASSQSAWGKNTLY | 0.89548075 |
| GTGGAGGAGNDAGS | ASSQSAWGKNTLY | 0.8307984 |
| GTTNTGWFNS | ASSQSAWGKNTLY | 0.7684703 |
| IFMGPLFTL | ASSQSAWGKNTLY | 0.86663616 |
| ILAHAIEAI | ASSQSAWGKNTLY | 0.7950386 |
| IRSAIREAL | ASSQSAWGKNTLY | 0.6842187 |
| ITFAGGG | ASSQSAWGKNTLY | 0.88471174 |
| ITGMSGHQEV | ASSQSAWGKNTLY | 0.83954346 |
| IVAGLACTL | ASSQSAWGKNTLY | 0.7650074 |
| IVGGGLAGA | ASSQSAWGKNTLY | 0.76865566 |
| IVTGVLNI | ASSQSAWGKNTLY | 0.8885526 |
| LAQQGCI | ASSQSAWGKNTLY | 0.6904829 |
| LCLHVEN | ASSQSAWGKNTLY | 0.7390955 |
| LDAISDGS | ASSQSAWGKNTLY | 0.6923722 |
| LGGFAAML | ASSQSAWGKNTLY | 0.84383065 |
| LGGKNPMI | ASSQSAWGKNTLY | 0.8878515 |
| LGPKGVAAIL | ASSQSAWGKNTLY | 0.8653454 |
| LMVGVSI | ASSQSAWGKNTLY | 0.7604814 |
| LVGAAFVL | ASSQSAWGKNTLY | 0.8689823 |
| LVLAVAPL | ASSQSAWGKNTLY | 0.8675965 |
| MAAIAVIDAA | ASSQSAWGKNTLY | 0.85909367 |
| NNGINAELQ | ASSQSAWGKNTLY | 0.868588 |
| RGEPLVLVHG | ASSQSAWGKNTLY | 0.7406373 |
| RGGAGGAAGLI | ASSQSAWGKNTLY | 0.88620174 |
| RSYRMHPAV | ASSQSAWGKNTLY | 0.8718424 |
| SAAHLLAGQLA | ASSQSAWGKNTLY | 0.882902 |
| SAAVGSY | ASSQSAWGKNTLY | 0.8756759 |
| SPHAVLHRI | ASSQSAWGKNTLY | 0.8563491 |
| SQIAKVL | ASSQSAWGKNTLY | 0.85063744 |
| TALGHLVP | ASSQSAWGKNTLY | 0.47411567 |
| TAPTTSGAA | ASSQSAWGKNTLY | 0.7670536 |
| TDGDDGNGGSPGT | ASSQSAWGKNTLY | 0.8512464 |
| TDGLGEN | ASSQSAWGKNTLY | 0.8412577 |
| TGGPPLPGQA | ASSQSAWGKNTLY | 0.86924946 |
| TIPAITI | ASSQSAWGKNTLY | 0.8672822 |
| TIPGFTI | ASSQSAWGKNTLY | 0.77711755 |
| TSIASGALA | ASSQSAWGKNTLY | 0.88148624 |
| TVTLPGAAI | ASSQSAWGKNTLY | 0.8754156 |
| VASGLSAALP | ASSQSAWGKNTLY | 0.597424 |
| VGAGVIGI | ASSQSAWGKNTLY | 0.8214289 |
| VGPKRFGV | ASSQSAWGKNTLY | 0.90068996 |
| VNVAFPD | ASSQSAWGKNTLY | 0.86016923 |
| VSMTNSGVS | ASSQSAWGKNTLY | 0.8223549 |

| | | |
|---|---|---|
| VTDVLGDLSV | ASSQSAWGKNTLY | 0.8214818 |
| VVVVGAGFAGL | ASSQSAWGKNTLY | 0.899053 |
| VYTVGTVV | ASSQSAWGKNTLY | 0.7695383 |
| WTTFSGP | ASSQSAWGKNTLY | 0.90006703 |
| YDELVEAF | ASSQSAWGKNTLY | 0.3322798 |
| YGTGGAGGNGG | ASSQSAWGKNTLY | 0.8091761 |
| YTNVMAV | ASSQSAWGKNTLY | 0.773316 |
| FLEQVHQL | ASSQSAWGKNTLY | 0.61976826 |
| GLSLLAVAV | ASSQSAWGKNTLY | 0.89055896 |
| LEQVHQL | ASSQSAWGKNTLY | 0.8288385 |
| LNAAGAI | ASSQSAWGKNTLY | 0.8803412 |
| PAVHASL | ASSQSAWGKNTLY | 0.8300674 |
| TELEDDA | ASSQSAWGKNTLY | 0.88571143 |
| TPPGPPN | ASSQSAWGKNTLY | 0.837257 |
| TPPPPPPP | ASSQSAWGKNTLY | 0.86210346 |
| TQSPALI | AWSLILGVHAEQF | 0.7367647 |
| EFAAAFPQ | AWSLILGVHAEQF | 0.8849695 |
| LLSGNLDVL | AWSLILGVHAEQF | 0.8891814 |
| GPANLGNYNI | AWSLILGVHAEQF | 0.87487817 |
| TLGPTLP | AWSLILGVHAEQF | 0.68167055 |
| VYAYSSCCK | AWSLILGVHAEQF | 0.79539555 |
| APAPAPAPNI | AWSLILGVHAEQF | 0.8648201 |
| PGGPVPIP | AWSLILGVHAEQF | 0.25852916 |
| FLTVALEP | AWSLILGVHAEQF | 0.27854663 |
| SQYFGANL | AWSLILGVHAEQF | 0.791189 |
| GGVKPAA | AWSLILGVHAEQF | 0.6874534 |
| GGTGGAGGAGGW | AWSLILGVHAEQF | 0.88850933 |
| GVGGVAGLGGAGG | AWSLILGVHAEQF | 0.8760223 |
| VLLVLATTI | AWSLILGVHAEQF | 0.8314739 |
| IGIFAAML | AWSLILGVHAEQF | 0.8692652 |
| AVGDLID | AWSLILGVHAEQF | 0.8483715 |
| NISPLPL | AWSLILGVHAEQF | 0.11415555 |
| VTAVFVAI | AWSLILGVHAEQF | 0.7521341 |
| SGDDNDAAL | AWSLILGVHAEQF | 0.6894565 |
| AVAALAHS | AWSLILGVHAEQF | 0.89513636 |
| GGVTVMATP | AWSLILGVHAEQF | 0.7889862 |
| AGLAANVRD | AWSLILGVHAEQF | 0.71860313 |
| TNRPDLIDPALLRPG | AWSLILGVHAEQF | 0.8346398 |
| LANLGVTAL | AWSLILGVHAEQF | 0.83947134 |
| TGERIEL | AWSLILGVHAEQF | 0.812922 |
| NGEELAA | AWSLILGVHAEQF | 0.67976534 |
| VLSLVWG | AWSLILGVHAEQF | 0.19474751 |
| IGYALWL | AWSLILGVHAEQF | 0.8856218 |
| LSLLLAPL | AWSLILGVHAEQF | 0.38922107 |
| TLDEALK | AWSLILGVHAEQF | 0.7875829 |
| TDGAGTAEGG | AWSLILGVHAEQF | 0.30420005 |
| NAPGGAGGNGG | AWSLILGVHAEQF | 0.15649337 |
| VSVFMGMGGNF | AWSLILGVHAEQF | 0.5098782 |
| LASYLAAL | AWSLILGVHAEQF | 0.88027704 |
| LVGGTPAL | AWSLILGVHAEQF | 0.79645145 |
| VRNADVL | AWSLILGVHAEQF | 0.77224576 |

| | | |
|---|---|---|
| TCSSPFLVSAM | AWSLILGVHAEQF | 0.8934495 |
| VHITDAE | AWSLILGVHAEQF | 0.8366674 |
| EYAAPSF | AWSLILGVHAEQF | 0.7871567 |
| VVDAFHGGL | AWSLILGVHAEQF | 0.794188 |
| ASVVDPQ | AWSLILGVHAEQF | 0.7356271 |
| SYNEFGGLS | AWSLILGVHAEQF | 0.78728557 |
| DPDGSVY | AWSLILGVHAEQF | 0.78317577 |
| MVSTIHATE | AWSLILGVHAEQF | 0.8982123 |
| LAAGVADPV | AWSLILGVHAEQF | 0.8264782 |
| TKEQIAA | AWSLILGVHAEQF | 0.6223465 |
| SLASPAAL | AWSLILGVHAEQF | 0.8436744 |
| IGLVFYGV | AWSLILGVHAEQF | 0.8901166 |
| FLLDASNL | AWSLILGVHAEQF | 0.904497 |
| YQVGNFP | AWSLILGVHAEQF | 0.7574804 |
| ESLLSQST | AWSLILGVHAEQF | 0.827205 |
| LFTVGFAAVG | AWSLILGVHAEQF | 0.8207841 |
| ATPPTPT | AWSLILGVHAEQF | 0.83307076 |
| TDLATLG | AWSLILGVHAEQF | 0.79218584 |
| SAGGTGR | AWSLILGVHAEQF | 0.66269183 |
| GFSNSGS | AWSLILGVHAEQF | 0.87537986 |
| GGNGGSGGTSV | AWSLILGVHAEQF | 0.90602493 |
| GTSVATGGAGN | AWSLILGVHAEQF | 0.8206681 |
| NGGNGGSGGTS | AWSLILGVHAEQF | 0.611714 |
| AVVKFPH | AWSLILGVHAEQF | 0.82690454 |
| TPPPPPP | AWSLILGVHAEQF | 0.74002147 |
| AAPAAAAAPA | AWSLILGVHAEQF | 0.8021053 |
| IALNEDL | AWSLILGVHAEQF | 0.86089754 |
| IVNVSSM | AWSLILGVHAEQF | 0.5785944 |
| TGIPPAPR | AWSLILGVHAEQF | 0.7864538 |
| FDLGGGTF | AWSLILGVHAEQF | 0.9039433 |
| FDLGGGTFDVS | AWSLILGVHAEQF | 0.8132818 |
| NEPTAAA | AWSLILGVHAEQF | 0.85741913 |
| NEPTAAALA | AWSLILGVHAEQF | 0.88442576 |
| GSIINISSI | AWSLILGVHAEQF | 0.8660829 |
| VAPGNAGTA | AWSLILGVHAEQF | 0.89523476 |
| TGGGASLE | AWSLILGVHAEQF | 0.87883604 |
| VAVYNHYKRI | AWSLILGVHAEQF | 0.83413064 |
| TDPAPATT | AWSLILGVHAEQF | 0.8285691 |
| TNRPDILDPALLRPG | AWSLILGVHAEQF | 0.81933033 |
| LTTTTTS | AWSLILGVHAEQF | 0.841696 |
| PTGSGKTL | AWSLILGVHAEQF | 0.8215536 |
| TGSGKTL | AWSLILGVHAEQF | 0.750775 |
| TIGHVDHGK | AWSLILGVHAEQF | 0.89053416 |
| TIGHVDHGKTT | AWSLILGVHAEQF | 0.8337725 |
| GHVDHGKTT | AWSLILGVHAEQF | 0.8858001 |
| AQFEHTLL | AWSLILGVHAEQF | 0.7175251 |
| IVNQDEL | AWSLILGVHAEQF | 0.73741597 |
| TNGSQFF | AWSLILGVHAEQF | 0.87724245 |
| TGKGIVE | AWSLILGVHAEQF | 0.14667328 |
| LISNASDA | AWSLILGVHAEQF | 0.35664552 |
| AELRTLGELTKIAKAHGAQ | AWSLILGVHAEQF | 0.84484756 |

| | | |
|---|---|---|
| APNAVAAAEPSW | AWSLILGVHAEQF | 0.8928593 |
| APQVAGAATAVL | AWSLILGVHAEQF | 0.16900118 |
| ARRDPAAAAAILPTDA | AWSLILGVHAEQF | 0.76328826 |
| DAAARDVLAEAGLAEYF | AWSLILGVHAEQF | 0.6058538 |
| DDQIVLSAKANLAKVEQ | AWSLILGVHAEQF | 0.86164033 |
| DELLPDTGEDSDEDGH | AWSLILGVHAEQF | 0.8439641 |
| ELPEAPANRATVPQD | AWSLILGVHAEQF | 0.5226122 |
| FGQSKIERVTGIG | AWSLILGVHAEQF | 0.879591 |
| GAGGTGSTAGGAGGAGGAGGLY | AWSLILGVHAEQF | 0.5048111 |
| GHGGTGAAGGNGGTGGAGGSGI | AWSLILGVHAEQF | 0.34553626 |
| GNGGNGGAGGEGGPGGAGGAGA | AWSLILGVHAEQF | 0.8840689 |
| GSTAGNGGNGGNATNGTIAGQP | AWSLILGVHAEQF | 0.890603 |
| GTGGDGGNGGNGGTGG | AWSLILGVHAEQF | 0.7306057 |
| GVRRPAGIPADAPV | AWSLILGVHAEQF | 0.8811908 |
| GYRQVTDAHLLTLARR | AWSLILGVHAEQF | 0.737206 |
| IAKMGQDGHDRG | AWSLILGVHAEQF | 0.9044242 |
| KAEPSIEIQYT | AWSLILGVHAEQF | 0.81101024 |
| KPVRIRMAADEDTVAVT | AWSLILGVHAEQF | 0.56379277 |
| MPPRIAGMRLLVIKPEP | AWSLILGVHAEQF | 0.88222003 |
| MKFYAVALFRSRGVPPTR | AWSLILGVHAEQF | 0.8644203 |
| NAGGFGILGAVAHSP | AWSLILGVHAEQF | 0.86036205 |
| NEGREITLAMAEM | AWSLILGVHAEQF | 0.8790878 |
| NELVEWSHPLGHRWRWE | AWSLILGVHAEQF | 0.74619484 |
| NLGAGGTGGNGGNPG | AWSLILGVHAEQF | 0.80896616 |
| NTEWPKWETVMAADPVD | AWSLILGVHAEQF | 0.88692373 |
| PASAAPAETQP | AWSLILGVHAEQF | 0.877931 |
| QQSSDGKNGVRAAEITGE | AWSLILGVHAEQF | 0.70739615 |
| RAPDAVTTSWWKE | AWSLILGVHAEQF | 0.8819603 |
| SGFGNVGAHTSGWFNQ | AWSLILGVHAEQF | 0.83252656 |
| SGVAGAVGQAASVGGLKV | AWSLILGVHAEQF | 0.773255 |
| SNGGAGTGSAGGNGGT | AWSLILGVHAEQF | 0.90238154 |
| TGGAGGEGGAAG | AWSLILGVHAEQF | 0.8117918 |
| TGGDGGAAGAGGAPGHG | AWSLILGVHAEQF | 0.875346 |
| TMPPLLAAIANAGRHGV | AWSLILGVHAEQF | 0.8868332 |
| TSSTAGNSGGA | AWSLILGVHAEQF | 0.8816985 |
| VPATVPVGILC | AWSLILGVHAEQF | 0.6471623 |
| VQRPDSGGVVLTGRIS | AWSLILGVHAEQF | 0.86991787 |
| WTPNPVPATKGVPAGLVQAT | AWSLILGVHAEQF | 0.84731185 |
| YALKRALAKFSDRFDVVII | AWSLILGVHAEQF | 0.8703886 |
| YLIGSVIV | AWSLILGVHAEQF | 0.85815215 |
| YGQLVAVID | AWSLILGVHAEQF | 0.6513269 |
| VTVGGMTLPA | AWSLILGVHAEQF | 0.86483777 |
| VLALPAV | AWSLILGVHAEQF | 0.88838553 |
| VIPPTLNYET | AWSLILGVHAEQF | 0.8505609 |
| VGMGGMHPSSG | AWSLILGVHAEQF | 0.83782065 |
| VGLAGGAATA | AWSLILGVHAEQF | 0.8992845 |
| VGGAGGAGGSGP | AWSLILGVHAEQF | 0.79013133 |
| VFAAFMV | AWSLILGVHAEQF | 0.8241393 |
| TLYDKLR | AWSLILGVHAEQF | 0.88421094 |
| TLAVFTVL | AWSLILGVHAEQF | 0.8787206 |
| TGSVQPARV | AWSLILGVHAEQF | 0.82374656 |

| | | |
|---|---|---|
| TGGGAGSGGT | AWSLILGVHAEQF | 0.8944625 |
| SHLETSL | AWSLILGVHAEQF | 0.7009914 |
| SFTTGMT | AWSLILGVHAEQF | 0.8778779 |
| QVEVAKRMGVRQPTVS | AWSLILGVHAEQF | 0.7301523 |
| QGSGGPALAM | AWSLILGVHAEQF | 0.86353385 |
| MLAPSNP | AWSLILGVHAEQF | 0.74101436 |
| LVNNAAI | AWSLILGVHAEQF | 0.8816571 |
| LTGEINTLA | AWSLILGVHAEQF | 0.87696826 |
| LSIFGPV | AWSLILGVHAEQF | 0.8805095 |
| LLALLANL | AWSLILGVHAEQF | 0.25084248 |
| LISNASDALDKLR | AWSLILGVHAEQF | 0.88148737 |
| LINEAVP | AWSLILGVHAEQF | 0.8802584 |
| LGYAFSGL | AWSLILGVHAEQF | 0.906216 |
| LDFVAVAL | AWSLILGVHAEQF | 0.82833993 |
| LAQPGAE | AWSLILGVHAEQF | 0.87512696 |
| ISNLGAVTGF | AWSLILGVHAEQF | 0.8252939 |
| HQEDPGRI | AWSLILGVHAEQF | 0.8304906 |
| GTPGEPTP | AWSLILGVHAEQF | 0.7032243 |
| GFFNSGS | AWSLILGVHAEQF | 0.81810117 |
| GAGGGGLTAGAN | AWSLILGVHAEQF | 0.7742646 |
| FVDALVNM | AWSLILGVHAEQF | 0.75182295 |
| FLTVAPEL | AWSLILGVHAEQF | 0.8863988 |
| FETLTAL | AWSLILGVHAEQF | 0.8797075 |
| DVINPIEL | AWSLILGVHAEQF | 0.8825494 |
| APVSGAPV | AWSLILGVHAEQF | 0.88396156 |
| ALVGGLAW | AWSLILGVHAEQF | 0.88388145 |
| ALAAAAATAI | AWSLILGVHAEQF | 0.88469493 |
| AFGLGLLL | AWSLILGVHAEQF | 0.8101592 |
| AFFAAASL | AWSLILGVHAEQF | 0.81456095 |
| AAAALPAFSPP | AWSLILGVHAEQF | 0.7137264 |
| VVLAPAL | AWSLILGVHAEQF | 0.78920317 |
| TIGHVDHGKTTL | AWSLILGVHAEQF | 0.80025995 |
| LISNASDALDK | AWSLILGVHAEQF | 0.8515911 |
| LISNASDAL | AWSLILGVHAEQF | 0.10567781 |
| ISNASDALDK | AWSLILGVHAEQF | 0.8228216 |
| NQAETLVYQTE | AWSLILGVHAEQF | 0.89339685 |
| TGGTGGTAGAG | AWSLILGVHAEQF | 0.7493098 |
| TGSTAGGAGGA | AWSLILGVHAEQF | 0.86749923 |
| DPLAVPVAAGPV | AWSLILGVHAEQF | 0.79225343 |
| GGTGGTPGNGGN | AWSLILGVHAEQF | 0.7729143 |
| PAGLAGVDPATL | AWSLILGVHAEQF | 0.8730843 |
| GNNLVVEAPHAGASV | AWSLILGVHAEQF | 0.72968805 |
| KVTRIMSASTLEQPA | AWSLILGVHAEQF | 0.87471294 |
| LGFALPIALPRRTVV | AWSLILGVHAEQF | 0.89887965 |
| TRLGNLTARKLVVLV | AWSLILGVHAEQF | 0.84904796 |
| WSQQRRLIYFPSAGP | AWSLILGVHAEQF | 0.8364438 |
| GPNGNPDPMAAAVDIR | AWSLILGVHAEQF | 0.8462741 |
| TQRCVSVADLRAAASR | AWSLILGVHAEQF | 0.5170294 |
| QFVLDVAEHPVEVLTETV | AWSLILGVHAEQF | 0.8464768 |
| GGAGGTGGAGGNGGTQAGDG | AWSLILGVHAEQF | 0.094339915 |
| PLLATLRAIAVDVDAKPAQV | AWSLILGVHAEQF | 0.81966734 |

| | | |
|---|---|---|
| APKFRTGGYMIDFWGVGYQVAK | AWSLILGVHAEQF | 0.7570858 |
| GAGGAGGAGGAAGGPGGTGGTG | AWSLILGVHAEQF | 0.8607639 |
| IGALGLLATDVAPAGSVAFGVA | AWSLILGVHAEQF | 0.83855104 |
| AGGAGGIGGNGGAGGAGGVLMG | AWSLILGVHAEQF | 0.8677889 |
| ADELIGLL | AWSLILGVHAEQF | 0.76035976 |
| ADRALKA | AWSLILGVHAEQF | 0.80901194 |
| AGAVGQVGGAGGA | AWSLILGVHAEQF | 0.7702316 |
| AGGGGGLGGAP | AWSLILGVHAEQF | 0.7253613 |
| AGGLLYGNGGAG | AWSLILGVHAEQF | 0.8792005 |
| AGLESIGL | AWSLILGVHAEQF | 0.84049714 |
| AGLKTGII | AWSLILGVHAEQF | 0.77146524 |
| ASLAALLL | AWSLILGVHAEQF | 0.89153755 |
| ASVINISSI | AWSLILGVHAEQF | 0.8525015 |
| ATPPGPA | AWSLILGVHAEQF | 0.49397352 |
| ATVASALV | AWSLILGVHAEQF | 0.8198498 |
| ATVASLIL | AWSLILGVHAEQF | 0.83456314 |
| AVNGPVLIRH | AWSLILGVHAEQF | 0.8336375 |
| DGAAGAVGI | AWSLILGVHAEQF | 0.10752024 |
| DPLLHGI | AWSLILGVHAEQF | 0.8610227 |
| ELGAALGAL | AWSLILGVHAEQF | 0.77541137 |
| ELGRTQAV | AWSLILGVHAEQF | 0.7188915 |
| FVAGGYLS | AWSLILGVHAEQF | 0.784696 |
| GASGGAGGAGGW | AWSLILGVHAEQF | 0.8768022 |
| GGAGAGGDGGAGGI | AWSLILGVHAEQF | 0.5642361 |
| GTGHNGSQPGMGGNG | AWSLILGVHAEQF | 0.8691747 |
| GYAAASASAS | AWSLILGVHAEQF | 0.8684033 |
| HQVDVAAVQQG | AWSLILGVHAEQF | 0.87844145 |
| IGAAIAHT | AWSLILGVHAEQF | 0.8965081 |
| IGMIEVAQL | AWSLILGVHAEQF | 0.87191653 |
| IRLAPEHTSAMD | AWSLILGVHAEQF | 0.8294903 |
| IYNVSVL | AWSLILGVHAEQF | 0.87203074 |
| KQGFLGM | AWSLILGVHAEQF | 0.83083034 |
| LESVITQL | AWSLILGVHAEQF | 0.5154043 |
| LGLGALVVV | AWSLILGVHAEQF | 0.48079577 |
| LIGNGGNGGSGGI | AWSLILGVHAEQF | 0.85624075 |
| LNAINAPTL | AWSLILGVHAEQF | 0.41486767 |
| LQLLNTGV | AWSLILGVHAEQF | 0.8537171 |
| LQTSIEF | AWSLILGVHAEQF | 0.8895968 |
| MSFAVSEL | AWSLILGVHAEQF | 0.83365035 |
| NTGALGL | AWSLILGVHAEQF | 0.64202887 |
| PPGAGGAGGAGTT | AWSLILGVHAEQF | 0.8113973 |
| QLVGLGI | AWSLILGVHAEQF | 0.7632221 |
| RFLEQVHQL | AWSLILGVHAEQF | 0.8778918 |
| RGVGYVLR | AWSLILGVHAEQF | 0.8821115 |
| SATGVFAGVI | AWSLILGVHAEQF | 0.90216136 |
| SLPAITLP | AWSLILGVHAEQF | 0.88977754 |
| STALAFTV | AWSLILGVHAEQF | 0.80333287 |
| SVLTVGLGK | AWSLILGVHAEQF | 0.86908203 |
| TALAGAGSGI | AWSLILGVHAEQF | 0.7924019 |
| TGLPFTGI | AWSLILGVHAEQF | 0.81214476 |
| VAEHVTV | AWSLILGVHAEQF | 0.83517444 |

| | | |
|---|---|---|
| VDIVLNSL | AWSLILGVHAEQF | 0.8557967 |
| VLTGGGAVL | AWSLILGVHAEQF | 0.82151806 |
| VNVGLAYGI | AWSLILGVHAEQF | 0.802306 |
| VSSAAVGAV | AWSLILGVHAEQF | 0.5673486 |
| VTSVYEAGV | AWSLILGVHAEQF | 0.8776463 |
| WVALDRGA | AWSLILGVHAEQF | 0.8478025 |
| YLVGSVVI | AWSLILGVHAEQF | 0.44231576 |
| GPPGTGKTLL | AWSLILGVHAEQF | 0.77606153 |
| GVGMTKF | AWSLILGVHAEQF | 0.9037112 |
| IGIGGSDL | AWSLILGVHAEQF | 0.8925441 |
| IGTLLAEL | AWSLILGVHAEQF | 0.74513185 |
| IYTSGTT | AWSLILGVHAEQF | 0.77185243 |
| LSFPAGV | AWSLILGVHAEQF | 0.8249854 |
| LYTSGTT | AWSLILGVHAEQF | 0.51491225 |
| VGAFPFGF | AWSLILGVHAEQF | 0.8625099 |
| VGAGTVEF | AWSLILGVHAEQF | 0.81703305 |
| VGKSTLLL | AWSLILGVHAEQF | 0.8116939 |
| AALAHAGGGIV | AWSLILGVHAEQF | 0.78353435 |
| AALVAVSLL | AWSLILGVHAEQF | 0.82757056 |
| AAVSRAVAAL | AWSLILGVHAEQF | 0.7956003 |
| AGGSSGIGGF | AWSLILGVHAEQF | 0.8672209 |
| AGLTVANANG | AWSLILGVHAEQF | 0.15641993 |
| AGLVGAVL | AWSLILGVHAEQF | 0.8720763 |
| ALSQAFTV | AWSLILGVHAEQF | 0.72033226 |
| ALTFSVSL | AWSLILGVHAEQF | 0.7291999 |
| ALTGAALVAL | AWSLILGVHAEQF | 0.54258835 |
| ASFSIGAGVL | AWSLILGVHAEQF | 0.82948166 |
| DGSVYKI | AWSLILGVHAEQF | 0.8877641 |
| EQLAEIV | AWSLILGVHAEQF | 0.8795239 |
| FAVSELTVM | AWSLILGVHAEQF | 0.6659323 |
| GAAPLAAGL | AWSLILGVHAEQF | 0.8820389 |
| GADGGDGGSGGAGG | AWSLILGVHAEQF | 0.8999468 |
| GFGLDTGGAGG | AWSLILGVHAEQF | 0.7655015 |
| GLASGGLL | AWSLILGVHAEQF | 0.48322338 |
| GRIAGAAVLL | AWSLILGVHAEQF | 0.8695291 |
| GSAGSLNTSF | AWSLILGVHAEQF | 0.8284987 |
| GTGQIGI | AWSLILGVHAEQF | 0.87003016 |
| GTLIALIL | AWSLILGVHAEQF | 0.87302315 |
| IAATMGFVVG | AWSLILGVHAEQF | 0.66758376 |
| IFAQALL | AWSLILGVHAEQF | 0.88759494 |
| IGAAVALGF | AWSLILGVHAEQF | 0.88477063 |
| IGLTPLAL | AWSLILGVHAEQF | 0.75095487 |
| ILGSNTSTL | AWSLILGVHAEQF | 0.7723732 |
| IVFSLTVL | AWSLILGVHAEQF | 0.7532116 |
| LASKATGF | AWSLILGVHAEQF | 0.49660552 |
| LASVALIL | AWSLILGVHAEQF | 0.8767117 |
| LFAVGLTL | AWSLILGVHAEQF | 0.740224 |
| LFNGGIGTYI | AWSLILGVHAEQF | 0.80411804 |
| LTLGAGII | AWSLILGVHAEQF | 0.84801036 |
| LVAAALGDI | AWSLILGVHAEQF | 0.8074334 |
| NTAFVQLGI | AWSLILGVHAEQF | 0.8136617 |

| | | |
|---|---|---|
| QLFSGAEL | AWSLILGVHAEQF | 0.8232361 |
| SAATFITL | AWSLILGVHAEQF | 0.859073 |
| SNYGLANLGV | AWSLILGVHAEQF | 0.84411603 |
| SYGGAYAVM | AWSLILGVHAEQF | 0.84129477 |
| TTGVGGLTLG | AWSLILGVHAEQF | 0.839693 |
| TTQTIYI | AWSLILGVHAEQF | 0.8897197 |
| VAAGAGLGAVI | AWSLILGVHAEQF | 0.68805337 |
| VAAVVTLL | AWSLILGVHAEQF | 0.71108866 |
| VGFLGGLI | AWSLILGVHAEQF | 0.8643106 |
| VGVIPILL | AWSLILGVHAEQF | 0.61958575 |
| VIFASSLI | AWSLILGVHAEQF | 0.8739418 |
| VKGGDGGAAAG | AWSLILGVHAEQF | 0.84100044 |
| VTKTFTNI | AWSLILGVHAEQF | 0.8496169 |
| VVAAAILL | AWSLILGVHAEQF | 0.82197857 |
| YAALDVEL | AWSLILGVHAEQF | 0.90157115 |
| LASVALLL | AWSLILGVHAEQF | 0.885875 |
| LDAVHRV | AWSLILGVHAEQF | 0.8353834 |
| QGLSSSIF | AWSLILGVHAEQF | 0.85993475 |
| RGTFANIRL | AWSLILGVHAEQF | 0.6661588 |
| RQLWWGHRI | AWSLILGVHAEQF | 0.88231504 |
| YGPPGTGKTLL | AWSLILGVHAEQF | 0.86957633 |
| AALVGAPANVA | AWSLILGVHAEQF | 0.6961841 |
| AATFRRTE | AWSLILGVHAEQF | 0.816278 |
| AGALGGGI | AWSLILGVHAEQF | 0.89221126 |
| AGGAGGAGGIGAG | AWSLILGVHAEQF | 0.88590693 |
| AGSGALGL | AWSLILGVHAEQF | 0.5386783 |
| AQIADQI | AWSLILGVHAEQF | 0.80838424 |
| ASAVAVTYL | AWSLILGVHAEQF | 0.3648081 |
| ASTVLGGTGG | AWSLILGVHAEQF | 0.7934725 |
| AVGGEPAK | AWSLILGVHAEQF | 0.6773371 |
| CEQGTFGGTC | AWSLILGVHAEQF | 0.8586699 |
| DEGKAPAG | AWSLILGVHAEQF | 0.8042954 |
| DTVVFPAL | AWSLILGVHAEQF | 0.8644028 |
| FEGITMAG | AWSLILGVHAEQF | 0.85740113 |
| FPLVVQD | AWSLILGVHAEQF | 0.8984336 |
| GAFTTPQ | AWSLILGVHAEQF | 0.7821295 |
| GFGGFGTAGGVGGA | AWSLILGVHAEQF | 0.8317261 |
| GFLAQFL | AWSLILGVHAEQF | 0.6050948 |
| GGAGGAGGNAGVA | AWSLILGVHAEQF | 0.5736704 |
| GGVGTAGAGGAGGAGG | AWSLILGVHAEQF | 0.7768106 |
| GNGGGGDAGNAGSGGN | AWSLILGVHAEQF | 0.829046 |
| GTGGAGGAGNDAGS | AWSLILGVHAEQF | 0.21957439 |
| GTTNTGWFNS | AWSLILGVHAEQF | 0.8899417 |
| IFMGPLFTL | AWSLILGVHAEQF | 0.862394 |
| ILAHAIEAI | AWSLILGVHAEQF | 0.8844702 |
| IRSAIREAL | AWSLILGVHAEQF | 0.78499573 |
| ITFAGGG | AWSLILGVHAEQF | 0.8078859 |
| ITGMSGHQEV | AWSLILGVHAEQF | 0.79276514 |
| IVAGLACTL | AWSLILGVHAEQF | 0.87737346 |
| IVGGGLAGA | AWSLILGVHAEQF | 0.7672235 |
| IVTGVLNI | AWSLILGVHAEQF | 0.6020222 |

| | | |
|---|---|---|
| LAQQGCI | AWSLILGVHAEQF | 0.7792535 |
| LCLHVEN | AWSLILGVHAEQF | 0.61637723 |
| LDAISDGS | AWSLILGVHAEQF | 0.8912985 |
| LGGFAAML | AWSLILGVHAEQF | 0.88518405 |
| LGGKNPMI | AWSLILGVHAEQF | 0.8588327 |
| LGPKGVAAIL | AWSLILGVHAEQF | 0.88433224 |
| LMVGVSI | AWSLILGVHAEQF | 0.2528297 |
| LVGAAFVL | AWSLILGVHAEQF | 0.78705394 |
| LVLAVAPL | AWSLILGVHAEQF | 0.89835775 |
| MAAIAVIDAA | AWSLILGVHAEQF | 0.8663161 |
| NNGINAELQ | AWSLILGVHAEQF | 0.81571686 |
| RGEPLVLVHG | AWSLILGVHAEQF | 0.89321107 |
| RGGAGGAAGLI | AWSLILGVHAEQF | 0.780409 |
| RSYRMHPAV | AWSLILGVHAEQF | 0.8354131 |
| SAAHLLAGQLA | AWSLILGVHAEQF | 0.847304 |
| SAAVGSY | AWSLILGVHAEQF | 0.87200856 |
| SPHAVLHRI | AWSLILGVHAEQF | 0.8605486 |
| SQIAKVL | AWSLILGVHAEQF | 0.7660796 |
| TALGHLVP | AWSLILGVHAEQF | 0.86496663 |
| TAPTTSGAA | AWSLILGVHAEQF | 0.89961815 |
| TDGDDGNGGSPGT | AWSLILGVHAEQF | 0.819427 |
| TDGLGEN | AWSLILGVHAEQF | 0.8823353 |
| TGGPPLPGQA | AWSLILGVHAEQF | 0.8655079 |
| TIPAITI | AWSLILGVHAEQF | 0.88103515 |
| TIPGFTI | AWSLILGVHAEQF | 0.75273067 |
| TSIASGALA | AWSLILGVHAEQF | 0.77803886 |
| TVTLPGAAI | AWSLILGVHAEQF | 0.82604945 |
| VASGLSAALP | AWSLILGVHAEQF | 0.693012 |
| VGAGVIGI | AWSLILGVHAEQF | 0.82983184 |
| VGPKRFGV | AWSLILGVHAEQF | 0.8024092 |
| VNVAFPD | AWSLILGVHAEQF | 0.9081012 |
| VSMTNSGVS | AWSLILGVHAEQF | 0.8670051 |
| VTDVLGDLSV | AWSLILGVHAEQF | 0.8635309 |
| VVVVGAGFAGL | AWSLILGVHAEQF | 0.75487244 |
| VYTVGTVV | AWSLILGVHAEQF | 0.85460806 |
| WTTFSGP | AWSLILGVHAEQF | 0.8562704 |
| YDELVEAF | AWSLILGVHAEQF | 0.57662225 |
| YGTGGAGGNGG | AWSLILGVHAEQF | 0.89275926 |
| YTNVMAV | AWSLILGVHAEQF | 0.7610229 |
| FLEQVHQL | AWSLILGVHAEQF | 0.772076 |
| GLSLLAVAV | AWSLILGVHAEQF | 0.8783214 |
| LEQVHQL | AWSLILGVHAEQF | 0.8978934 |
| LNAAGAI | AWSLILGVHAEQF | 0.79052794 |
| PAVHASL | AWSLILGVHAEQF | 0.80719197 |
| TELEDDA | AWSLILGVHAEQF | 0.75611746 |
| TPPGPPN | AWSLILGVHAEQF | 0.87710774 |
| TPPPPPPP | AWSLILGVHAEQF | 0.8907368 |
| TQSPALI | ATRGTGEDTQY | 0.83652633 |
| EFAAAFPQ | ATRGTGEDTQY | 0.8970226 |
| LLSGNLDVL | ATRGTGEDTQY | 0.8657544 |
| GPANLGNYNI | ATRGTGEDTQY | 0.52481437 |

| | | |
|---|---|---|
| TLGPTLP | ATRGTGEDTQY | 0.84494364 |
| VYAYSSCCK | ATRGTGEDTQY | 0.8814682 |
| APAPAPAPNI | ATRGTGEDTQY | 0.8743374 |
| PGGPVPIP | ATRGTGEDTQY | 0.90448093 |
| FLTVALEP | ATRGTGEDTQY | 0.87337196 |
| SQYFGANL | ATRGTGEDTQY | 0.7962846 |
| GGVKPAA | ATRGTGEDTQY | 0.89002025 |
| GGTGGAGGAGGW | ATRGTGEDTQY | 0.9028795 |
| GVGGVAGLGGAGG | ATRGTGEDTQY | 0.8861912 |
| VLLVLATTI | ATRGTGEDTQY | 0.7967334 |
| IGIFAAML | ATRGTGEDTQY | 0.70647144 |
| AVGDLID | ATRGTGEDTQY | 0.85449207 |
| NISPLPL | ATRGTGEDTQY | 0.8870417 |
| VTAVFVAI | ATRGTGEDTQY | 0.7254568 |
| SGDDNDAAL | ATRGTGEDTQY | 0.8289295 |
| AVAALAHS | ATRGTGEDTQY | 0.85754406 |
| GGVTVMATP | ATRGTGEDTQY | 0.6345742 |
| AGLAANVRD | ATRGTGEDTQY | 0.873658 |
| TNRPDLIDPALLRPG | ATRGTGEDTQY | 0.8885376 |
| LANLGVTAL | ATRGTGEDTQY | 0.8355782 |
| TGERIEL | ATRGTGEDTQY | 0.80077124 |
| NGEELAA | ATRGTGEDTQY | 0.883507 |
| VLSLVWG | ATRGTGEDTQY | 0.90795255 |
| IGYALWL | ATRGTGEDTQY | 0.8940675 |
| LSLLLAPL | ATRGTGEDTQY | 0.6100851 |
| TLDEALK | ATRGTGEDTQY | 0.3291981 |
| TDGAGTAEGG | ATRGTGEDTQY | 0.7608263 |
| NAPGGAGGNGG | ATRGTGEDTQY | 0.8697674 |
| VSVFMGMGGNF | ATRGTGEDTQY | 0.8258662 |
| LASYLAAL | ATRGTGEDTQY | 0.81995565 |
| LVGGTPAL | ATRGTGEDTQY | 0.85861135 |
| VRNADVL | ATRGTGEDTQY | 0.88592297 |
| TCSSPFLVSAM | ATRGTGEDTQY | 0.7774898 |
| VHITDAE | ATRGTGEDTQY | 0.635689 |
| EYAAPSF | ATRGTGEDTQY | 0.85662866 |
| VVDAFHGGL | ATRGTGEDTQY | 0.87769806 |
| ASVVDPQ | ATRGTGEDTQY | 0.7436884 |
| SYNEFGGLS | ATRGTGEDTQY | 0.8229827 |
| DPDGSVY | ATRGTGEDTQY | 0.7536271 |
| MVSTIHATE | ATRGTGEDTQY | 0.47393164 |
| LAAGVADPV | ATRGTGEDTQY | 0.8437676 |
| TKEQIAA | ATRGTGEDTQY | 0.72331643 |
| SLASPAAL | ATRGTGEDTQY | 0.8927574 |
| IGLVFYGV | ATRGTGEDTQY | 0.7380978 |
| FLLDASNL | ATRGTGEDTQY | 0.8781201 |
| YQVGNFP | ATRGTGEDTQY | 0.8587073 |
| ESLLSQST | ATRGTGEDTQY | 0.8711295 |
| LFTVGFAAVG | ATRGTGEDTQY | 0.80485666 |
| ATPPTPT | ATRGTGEDTQY | 0.75685036 |
| TDLATLG | ATRGTGEDTQY | 0.7105241 |
| SAGGTGR | ATRGTGEDTQY | 0.43215436 |

| | | |
|---|---|---|
| GFSNSGS | ATRGTGEDTQY | 0.88105303 |
| GGNGGSGGTSV | ATRGTGEDTQY | 0.16596763 |
| GTSVATGGAGN | ATRGTGEDTQY | 0.8834855 |
| NGGNGGSGGTS | ATRGTGEDTQY | 0.8834578 |
| AVVKFPH | ATRGTGEDTQY | 0.75987554 |
| TPPPPPP | ATRGTGEDTQY | 0.86105716 |
| AAPAAAAAPA | ATRGTGEDTQY | 0.7509249 |
| IALNEDL | ATRGTGEDTQY | 0.7512938 |
| IVNVSSM | ATRGTGEDTQY | 0.8822315 |
| TGIPPAPR | ATRGTGEDTQY | 0.87386864 |
| FDLGGGTF | ATRGTGEDTQY | 0.76146054 |
| FDLGGGTFDVS | ATRGTGEDTQY | 0.8542916 |
| NEPTAAA | ATRGTGEDTQY | 0.8508369 |
| NEPTAAALA | ATRGTGEDTQY | 0.78841835 |
| GSIINISSI | ATRGTGEDTQY | 0.8475505 |
| VAPGNAGTA | ATRGTGEDTQY | 0.8821479 |
| TGGGASLE | ATRGTGEDTQY | 0.86423755 |
| VAVYNHYKRI | ATRGTGEDTQY | 0.7906203 |
| TDPAPATT | ATRGTGEDTQY | 0.8714576 |
| TNRPDILDPALLRPG | ATRGTGEDTQY | 0.6736714 |
| LTTTTTS | ATRGTGEDTQY | 0.85440457 |
| PTGSGKTL | ATRGTGEDTQY | 0.8331944 |
| TGSGKTL | ATRGTGEDTQY | 0.82550806 |
| TIGHVDHGK | ATRGTGEDTQY | 0.8726633 |
| TIGHVDHGKTT | ATRGTGEDTQY | 0.7008139 |
| GHVDHGKTT | ATRGTGEDTQY | 0.7493633 |
| AQFEHTLL | ATRGTGEDTQY | 0.8182077 |
| IVNQDEL | ATRGTGEDTQY | 0.8639329 |
| TNGSQFF | ATRGTGEDTQY | 0.8798735 |
| TGKGIVE | ATRGTGEDTQY | 0.75540364 |
| LISNASDA | ATRGTGEDTQY | 0.8427074 |
| AELRTLGELTKIAKAHGAQ | ATRGTGEDTQY | 0.8085115 |
| APNAVAAAEPSW | ATRGTGEDTQY | 0.75924665 |
| APQVAGAATAVL | ATRGTGEDTQY | 0.8474269 |
| ARRDPAAAAAILPTDA | ATRGTGEDTQY | 0.823774 |
| DAAARDVLAEAGLAEYF | ATRGTGEDTQY | 0.7502667 |
| DDQIVLSAKANLAKVEQ | ATRGTGEDTQY | 0.8616327 |
| DELLPDTGEDSDEDGH | ATRGTGEDTQY | 0.7934978 |
| ELPEAPANRATVPQD | ATRGTGEDTQY | 0.86732167 |
| FGQSKIERVTGIG | ATRGTGEDTQY | 0.8999946 |
| GAGGTGSTAGGAGGAGGAGGLY | ATRGTGEDTQY | 0.7971412 |
| GHGGTGAAGGNGGTGGAGGSGI | ATRGTGEDTQY | 0.87450296 |
| GNGGNGGAGGEGGPGGAGGAGA | ATRGTGEDTQY | 0.865217 |
| GSTAGNGGNGGNATNGTIAGQP | ATRGTGEDTQY | 0.8487474 |
| GTGGDGGNGGNGGTGG | ATRGTGEDTQY | 0.83300734 |
| GVRRPAGIPADAPV | ATRGTGEDTQY | 0.7824718 |
| GYRQVTDAHLLTLARR | ATRGTGEDTQY | 0.8918906 |
| IAKMGQDGHDRG | ATRGTGEDTQY | 0.875913 |
| KAEPSIEIQYT | ATRGTGEDTQY | 0.8810836 |
| KPVRIRMAADEDTVAVT | ATRGTGEDTQY | 0.85959643 |
| MPPRIAGMRLLVIKPEP | ATRGTGEDTQY | 0.8754827 |

| | | |
|---|---|---|
| MKFYAVALFRSRGVPPTR | ATRGTGEDTQY | 0.87378466 |
| NAGGFGILGAVAHSP | ATRGTGEDTQY | 0.77670366 |
| NEGREITLAMAEM | ATRGTGEDTQY | 0.6220522 |
| NELVEWSHPLGHRWRWE | ATRGTGEDTQY | 0.81250536 |
| NLGAGGTGGNGGNPG | ATRGTGEDTQY | 0.8118149 |
| NTEWPKWETVMAADPVD | ATRGTGEDTQY | 0.8017994 |
| PASAAPAETQP | ATRGTGEDTQY | 0.39831173 |
| QQSSDGKNGVRAAEITGE | ATRGTGEDTQY | 0.88526785 |
| RAPDAVTTSWWKE | ATRGTGEDTQY | 0.9069024 |
| SGFGNVGAHTSGWFNQ | ATRGTGEDTQY | 0.7573676 |
| SGVAGAVGQAASVGGLKV | ATRGTGEDTQY | 0.8968705 |
| SNGGAGTGSAGGNGGT | ATRGTGEDTQY | 0.6558384 |
| TGGAGGEGGAAG | ATRGTGEDTQY | 0.8762716 |
| TGGDGGAAGAGGAPGHG | ATRGTGEDTQY | 0.8890708 |
| TMPPLLAAIANAGRHGV | ATRGTGEDTQY | 0.7014903 |
| TSSTAGNSGGA | ATRGTGEDTQY | 0.8749045 |
| VPATVPVGILC | ATRGTGEDTQY | 0.78792465 |
| VQRPDSGGVVLTGRIS | ATRGTGEDTQY | 0.8128662 |
| WTPNPVPATKGVPAGLVQAT | ATRGTGEDTQY | 0.89220494 |
| YALKRALAKFSDRFDVVII | ATRGTGEDTQY | 0.82802486 |
| YLIGSVIV | ATRGTGEDTQY | 0.8537394 |
| YGQLVAVID | ATRGTGEDTQY | 0.85340416 |
| VTVGGMTLPA | ATRGTGEDTQY | 0.77323854 |
| VLALPAV | ATRGTGEDTQY | 0.8557036 |
| VIPPTLNYET | ATRGTGEDTQY | 0.88880855 |
| VGMGGMHPSSG | ATRGTGEDTQY | 0.7004794 |
| VGLAGGAATA | ATRGTGEDTQY | 0.89918137 |
| VGGAGGAGGSGP | ATRGTGEDTQY | 0.77162653 |
| VFAAFMV | ATRGTGEDTQY | 0.9041449 |
| TLYDKLR | ATRGTGEDTQY | 0.866122 |
| TLAVFTVL | ATRGTGEDTQY | 0.83263135 |
| TGSVQPARV | ATRGTGEDTQY | 0.8028481 |
| TGGGAGSGGT | ATRGTGEDTQY | 0.89050066 |
| SHLETSL | ATRGTGEDTQY | 0.83065164 |
| SFTTGMT | ATRGTGEDTQY | 0.8642223 |
| QVEVAKRMGVRQPTVS | ATRGTGEDTQY | 0.89209 |
| QGSGGPALAM | ATRGTGEDTQY | 0.44931665 |
| MLAPSNP | ATRGTGEDTQY | 0.80543643 |
| LVNNAAI | ATRGTGEDTQY | 0.6628299 |
| LTGEINTLA | ATRGTGEDTQY | 0.6485486 |
| LSIFGPV | ATRGTGEDTQY | 0.5134794 |
| LLALLANL | ATRGTGEDTQY | 0.2649237 |
| LISNASDALDKLR | ATRGTGEDTQY | 0.60032094 |
| LINEAVP | ATRGTGEDTQY | 0.45779094 |
| LGYAFSGL | ATRGTGEDTQY | 0.73586965 |
| LDFVAVAL | ATRGTGEDTQY | 0.73218197 |
| LAQPGAE | ATRGTGEDTQY | 0.52341044 |
| ISNLGAVTGF | ATRGTGEDTQY | 0.77913356 |
| HQEDPGRI | ATRGTGEDTQY | 0.78115773 |
| GTPGEPTP | ATRGTGEDTQY | 0.39414057 |
| GFFNSGS | ATRGTGEDTQY | 0.57845104 |

| | | |
|---|---|---|
| GAGGGGLTAGAN | ATRGTGEDTQY | 0.44231653 |
| FVDALVNM | ATRGTGEDTQY | 0.61285186 |
| FLTVAPEL | ATRGTGEDTQY | 0.27203262 |
| FETLTAL | ATRGTGEDTQY | 0.6989971 |
| DVINPIEL | ATRGTGEDTQY | 0.42915 |
| APVSGAPV | ATRGTGEDTQY | 0.16825382 |
| ALVGGLAW | ATRGTGEDTQY | 0.6905575 |
| ALAAAAATAI | ATRGTGEDTQY | 0.60497355 |
| AFGLGLLL | ATRGTGEDTQY | 0.5385069 |
| AFFAAASL | ATRGTGEDTQY | 0.7573574 |
| AAAALPAFSPP | ATRGTGEDTQY | 0.36144245 |
| VVLAPAL | ATRGTGEDTQY | 0.37696818 |
| TIGHVDHGKTTL | ATRGTGEDTQY | 0.7118176 |
| LISNASDALDK | ATRGTGEDTQY | 0.79314554 |
| LISNASDAL | ATRGTGEDTQY | 0.3410014 |
| ISNASDALDK | ATRGTGEDTQY | 0.63358206 |
| NQAETLVYQTE | ATRGTGEDTQY | 0.794533 |
| TGGTGGTAGAG | ATRGTGEDTQY | 0.6849932 |
| TGSTAGGAGGA | ATRGTGEDTQY | 0.7608045 |
| DPLAVPVAAGPV | ATRGTGEDTQY | 0.78582054 |
| GGTGGTPGNGGN | ATRGTGEDTQY | 0.5116876 |
| PAGLAGVDPATL | ATRGTGEDTQY | 0.78093123 |
| GNNLVVEAPHAGASV | ATRGTGEDTQY | 0.77894306 |
| KVTRIMSASTLEQPA | ATRGTGEDTQY | 0.58471924 |
| LGFALPIALPRRTVV | ATRGTGEDTQY | 0.6269638 |
| TRLGNLTARKLVVLV | ATRGTGEDTQY | 0.4357322 |
| WSQQRRLIYFPSAGP | ATRGTGEDTQY | 0.5710056 |
| GPNGNPDPMAAAVDIR | ATRGTGEDTQY | 0.2510044 |
| TQRCVSVADLRAAASR | ATRGTGEDTQY | 0.6828872 |
| QFVLDVAEHPVEVLTETV | ATRGTGEDTQY | 0.7327659 |
| GGAGGTGGAGGNGGTQAGDG | ATRGTGEDTQY | 0.59967583 |
| PLLATLRAIAVDVDAKPAQV | ATRGTGEDTQY | 0.38174453 |
| APKFRTGGYMIDFWGVGYQVAK | ATRGTGEDTQY | 0.3387306 |
| GAGGAGGAGGAAGGPGGTGGTG | ATRGTGEDTQY | 0.6496698 |
| IGALGLLATDVAPAGSVAFGVA | ATRGTGEDTQY | 0.42893687 |
| AGGAGGIGGNGGAGGAGGVLMG | ATRGTGEDTQY | 0.2795461 |
| ADELIGLL | ATRGTGEDTQY | 0.5939476 |
| ADRALKA | ATRGTGEDTQY | 0.76357514 |
| AGAVGQVGGAGGA | ATRGTGEDTQY | 0.74357295 |
| AGGGGGLGGAP | ATRGTGEDTQY | 0.39864114 |
| AGGLLYGNGGAG | ATRGTGEDTQY | 0.8215765 |
| AGLESIGL | ATRGTGEDTQY | 0.70449144 |
| AGLKTGII | ATRGTGEDTQY | 0.7106255 |
| ASLAALLL | ATRGTGEDTQY | 0.56935525 |
| ASVINISSI | ATRGTGEDTQY | 0.4650343 |
| ATPPGPA | ATRGTGEDTQY | 0.7829422 |
| ATVASALV | ATRGTGEDTQY | 0.57371014 |
| ATVASLIL | ATRGTGEDTQY | 0.4798818 |
| AVNGPVLIRH | ATRGTGEDTQY | 0.7484587 |
| DGAAGAVGI | ATRGTGEDTQY | 0.52316236 |
| DPLLHGI | ATRGTGEDTQY | 0.48745966 |

| | | |
|---|---|---|
| ELGAALGAL | ATRGTGEDTQY | 0.54523635 |
| ELGRTQAV | ATRGTGEDTQY | 0.7675249 |
| FVAGGYLS | ATRGTGEDTQY | 0.42487 |
| GASGGAGGAGGW | ATRGTGEDTQY | 0.42250416 |
| GGAGAGGDGGAGGI | ATRGTGEDTQY | 0.44781435 |
| GTGHNGSQPGMGGNG | ATRGTGEDTQY | 0.6165689 |
| GYAAASASAS | ATRGTGEDTQY | 0.7915833 |
| HQVDVAAVQQG | ATRGTGEDTQY | 0.6789747 |
| IGAAIAHT | ATRGTGEDTQY | 0.51494926 |
| IGMIEVAQL | ATRGTGEDTQY | 0.6799379 |
| IRLAPEHTSAMD | ATRGTGEDTQY | 0.6845673 |
| IYNVSVL | ATRGTGEDTQY | 0.6655908 |
| KQGFLGM | ATRGTGEDTQY | 0.66842026 |
| LESVITQL | ATRGTGEDTQY | 0.50686485 |
| LGLGALVVV | ATRGTGEDTQY | 0.7741291 |
| LIGNGGNGGSGGI | ATRGTGEDTQY | 0.72109616 |
| LNAINAPTL | ATRGTGEDTQY | 0.7249056 |
| LQLLNTGV | ATRGTGEDTQY | 0.75378036 |
| LQTSIEF | ATRGTGEDTQY | 0.5948135 |
| MSFAVSEL | ATRGTGEDTQY | 0.609347 |
| NTGALGL | ATRGTGEDTQY | 0.2775699 |
| PPGAGGAGGAGTT | ATRGTGEDTQY | 0.56124437 |
| QLVGLGI | ATRGTGEDTQY | 0.62270856 |
| RFLEQVHQL | ATRGTGEDTQY | 0.6618538 |
| RGVGYVLR | ATRGTGEDTQY | 0.72979546 |
| SATGVFAGVI | ATRGTGEDTQY | 0.6470263 |
| SLPAITLP | ATRGTGEDTQY | 0.48464555 |
| STALAFTV | ATRGTGEDTQY | 0.7060493 |
| SVLTVGLGK | ATRGTGEDTQY | 0.7815844 |
| TALAGAGSGI | ATRGTGEDTQY | 0.36781436 |
| TGLPFTGI | ATRGTGEDTQY | 0.6187993 |
| VAEHVTV | ATRGTGEDTQY | 0.8433783 |
| VDIVLNSL | ATRGTGEDTQY | 0.6540804 |
| VLTGGGAVL | ATRGTGEDTQY | 0.565924 |
| VNVGLAYGI | ATRGTGEDTQY | 0.518379 |
| VSSAAVGAV | ATRGTGEDTQY | 0.7922658 |
| VTSVYEAGV | ATRGTGEDTQY | 0.43210453 |
| WVALDRGA | ATRGTGEDTQY | 0.80692947 |
| YLVGSVVI | ATRGTGEDTQY | 0.19528069 |
| GPPGTGKTLL | ATRGTGEDTQY | 0.5259219 |
| GVGMTKF | ATRGTGEDTQY | 0.43466723 |
| IGIGGSDL | ATRGTGEDTQY | 0.24052572 |
| IGTLLAEL | ATRGTGEDTQY | 0.7546878 |
| IYTSGTT | ATRGTGEDTQY | 0.5291699 |
| LSFPAGV | ATRGTGEDTQY | 0.72989076 |
| LYTSGTT | ATRGTGEDTQY | 0.592076 |
| VGAFPFGF | ATRGTGEDTQY | 0.77938783 |
| VGAGTVEF | ATRGTGEDTQY | 0.66582215 |
| VGKSTLLL | ATRGTGEDTQY | 0.62840176 |
| AALAHAGGGIV | ATRGTGEDTQY | 0.49562484 |
| AALVAVSLL | ATRGTGEDTQY | 0.7327348 |

| | | |
|---|---|---|
| AAVSRAVAAL | ATRGTGEDTQY | 0.75546724 |
| AGGSSGIGGF | ATRGTGEDTQY | 0.701386 |
| AGLTVANANG | ATRGTGEDTQY | 0.2578955 |
| AGLVGAVL | ATRGTGEDTQY | 0.59675074 |
| ALSQAFTV | ATRGTGEDTQY | 0.6430594 |
| ALTFSVSL | ATRGTGEDTQY | 0.28758478 |
| ALTGAALVAL | ATRGTGEDTQY | 0.21792397 |
| ASFSIGAGVL | ATRGTGEDTQY | 0.4158818 |
| DGSVYKI | ATRGTGEDTQY | 0.5729629 |
| EQLAEIV | ATRGTGEDTQY | 0.76203996 |
| FAVSELTVM | ATRGTGEDTQY | 0.7351959 |
| GAAPLAAGL | ATRGTGEDTQY | 0.60072815 |
| GADGGDGGSGGAGG | ATRGTGEDTQY | 0.6909083 |
| GFGLDTGGAGG | ATRGTGEDTQY | 0.6369384 |
| GLASGGLL | ATRGTGEDTQY | 0.11640029 |
| GRIAGAAVLL | ATRGTGEDTQY | 0.4993093 |
| GSAGSLNTSF | ATRGTGEDTQY | 0.36520398 |
| GTGQIGI | ATRGTGEDTQY | 0.79633224 |
| GTLIALIL | ATRGTGEDTQY | 0.52539045 |
| IAATMGFVVG | ATRGTGEDTQY | 0.5546674 |
| IFAQALL | ATRGTGEDTQY | 0.67095834 |
| IGAAVALGF | ATRGTGEDTQY | 0.6540673 |
| IGLTPLAL | ATRGTGEDTQY | 0.6122539 |
| ILGSNTSTL | ATRGTGEDTQY | 0.38583046 |
| IVFSLTVL | ATRGTGEDTQY | 0.17319623 |
| LASKATGF | ATRGTGEDTQY | 0.72117984 |
| LASVALIL | ATRGTGEDTQY | 0.34830555 |
| LFAVGLTL | ATRGTGEDTQY | 0.39094597 |
| LFNGGIGTYI | ATRGTGEDTQY | 0.28830093 |
| LTLGAGII | ATRGTGEDTQY | 0.47921813 |
| LVAAALGDI | ATRGTGEDTQY | 0.2632108 |
| NTAFVQLGI | ATRGTGEDTQY | 0.7204015 |
| QLFSGAEL | ATRGTGEDTQY | 0.52809423 |
| SAATFITL | ATRGTGEDTQY | 0.61882347 |
| SNYGLANLGV | ATRGTGEDTQY | 0.82184774 |
| SYGGAYAVM | ATRGTGEDTQY | 0.62793833 |
| TTGVGGLTLG | ATRGTGEDTQY | 0.5963803 |
| TTQTIYI | ATRGTGEDTQY | 0.46535748 |
| VAAGAGLGAVI | ATRGTGEDTQY | 0.45186117 |
| VAAVVTLL | ATRGTGEDTQY | 0.41437715 |
| VGFLGGLI | ATRGTGEDTQY | 0.41879222 |
| VGVIPILL | ATRGTGEDTQY | 0.8066462 |
| VIFASSLI | ATRGTGEDTQY | 0.59851515 |
| VKGGDGGAAAG | ATRGTGEDTQY | 0.46207517 |
| VTKTFTNI | ATRGTGEDTQY | 0.66612095 |
| VVAAAILL | ATRGTGEDTQY | 0.80381197 |
| YAALDVEL | ATRGTGEDTQY | 0.8242222 |
| LASVALLL | ATRGTGEDTQY | 0.4913451 |
| LDAVHRV | ATRGTGEDTQY | 0.46753603 |
| QGLSSSIF | ATRGTGEDTQY | 0.52750766 |
| RGTFANIRL | ATRGTGEDTQY | 0.63526976 |

| | | |
|---|---|---|
| RQLWWGHRI | ATRGTGEDTQY | 0.48334584 |
| YGPPGTGKTLL | ATRGTGEDTQY | 0.4095797 |
| AALVGAPANVA | ATRGTGEDTQY | 0.73176956 |
| AATFRRTE | ATRGTGEDTQY | 0.83026206 |
| AGALGGGI | ATRGTGEDTQY | 0.57097524 |
| AGGAGGAGGIGAG | ATRGTGEDTQY | 0.2839422 |
| AGSGALGL | ATRGTGEDTQY | 0.6388334 |
| AQIADQI | ATRGTGEDTQY | 0.3746304 |
| ASAVAVTYL | ATRGTGEDTQY | 0.5203048 |
| ASTVLGGTGG | ATRGTGEDTQY | 0.5168859 |
| AVGGEPAK | ATRGTGEDTQY | 0.35527807 |
| CEQGTFGGTC | ATRGTGEDTQY | 0.6005262 |
| DEGKAPAG | ATRGTGEDTQY | 0.7946387 |
| DTVVFPAL | ATRGTGEDTQY | 0.46439806 |
| FEGITMAG | ATRGTGEDTQY | 0.7002833 |
| FPLVVQD | ATRGTGEDTQY | 0.80074835 |
| GAFTTPQ | ATRGTGEDTQY | 0.73785114 |
| GFGGFGTAGGVGGA | ATRGTGEDTQY | 0.7931242 |
| GFLAQFL | ATRGTGEDTQY | 0.7211629 |
| GGAGGAGGNAGVA | ATRGTGEDTQY | 0.6192504 |
| GGVGTAGAGGAGGAGG | ATRGTGEDTQY | 0.5957395 |
| GNGGGGDAGNAGSGGN | ATRGTGEDTQY | 0.7822696 |
| GTGGAGGAGNDAGS | ATRGTGEDTQY | 0.6601716 |
| GTTNTGWFNS | ATRGTGEDTQY | 0.6137285 |
| IFMGPLFTL | ATRGTGEDTQY | 0.522192 |
| ILAHAIEAI | ATRGTGEDTQY | 0.7714716 |
| IRSAIREAL | ATRGTGEDTQY | 0.59104526 |
| ITFAGGG | ATRGTGEDTQY | 0.7834321 |
| ITGMSGHQEV | ATRGTGEDTQY | 0.2823227 |
| IVAGLACTL | ATRGTGEDTQY | 0.36650705 |
| IVGGGLAGA | ATRGTGEDTQY | 0.77369803 |
| IVTGVLNI | ATRGTGEDTQY | 0.13493097 |
| LAQQGCI | ATRGTGEDTQY | 0.23352344 |
| LCLHVEN | ATRGTGEDTQY | 0.61005235 |
| LDAISDGS | ATRGTGEDTQY | 0.7591243 |
| LGGFAAML | ATRGTGEDTQY | 0.151878 |
| LGGKNPMI | ATRGTGEDTQY | 0.42394 |
| LGPKGVAAIL | ATRGTGEDTQY | 0.3051815 |
| LMVGVSI | ATRGTGEDTQY | 0.6720368 |
| LVGAAFVL | ATRGTGEDTQY | 0.5761145 |
| LVLAVAPL | ATRGTGEDTQY | 0.2847383 |
| MAAIAVIDAA | ATRGTGEDTQY | 0.7237491 |
| NNGINAELQ | ATRGTGEDTQY | 0.29668692 |
| RGEPLVLVHG | ATRGTGEDTQY | 0.5528639 |
| RGGAGGAAGLI | ATRGTGEDTQY | 0.74797285 |
| RSYRMHPAV | ATRGTGEDTQY | 0.8015001 |
| SAAHLLAGQLA | ATRGTGEDTQY | 0.4491548 |
| SAAVGSY | ATRGTGEDTQY | 0.70014226 |
| SPHAVLHRI | ATRGTGEDTQY | 0.48131454 |
| SQIAKVL | ATRGTGEDTQY | 0.82766664 |
| TALGHLVP | ATRGTGEDTQY | 0.56304085 |

| | | |
|---|---|---|
| TAPTTSGAA | ATRGTGEDTQY | 0.31889778 |
| TDGDDGNGGSPGT | ATRGTGEDTQY | 0.7401975 |
| TDGLGEN | ATRGTGEDTQY | 0.76524246 |
| TGGPPLPGQA | ATRGTGEDTQY | 0.56631136 |
| TIPAITI | ATRGTGEDTQY | 0.7253757 |
| TIPGFTI | ATRGTGEDTQY | 0.5118964 |
| TSIASGALA | ATRGTGEDTQY | 0.61752975 |
| TVTLPGAAI | ATRGTGEDTQY | 0.7841244 |
| VASGLSAALP | ATRGTGEDTQY | 0.7360743 |
| VGAGVIGI | ATRGTGEDTQY | 0.42927575 |
| VGPKRFGV | ATRGTGEDTQY | 0.7641317 |
| VNVAFPD | ATRGTGEDTQY | 0.6189342 |
| VSMTNSGVS | ATRGTGEDTQY | 0.46523684 |
| VTDVLGDLSV | ATRGTGEDTQY | 0.81577206 |
| VVVVGAGFAGL | ATRGTGEDTQY | 0.5658054 |
| VYTVGTVV | ATRGTGEDTQY | 0.7504774 |
| WTTFSGP | ATRGTGEDTQY | 0.70028615 |
| YDELVEAF | ATRGTGEDTQY | 0.786042 |
| YGTGGAGGNGG | ATRGTGEDTQY | 0.34912118 |
| YTNVMAV | ATRGTGEDTQY | 0.71592796 |
| FLEQVHQL | ATRGTGEDTQY | 0.6068485 |
| GLSLLAVAV | ATRGTGEDTQY | 0.74763227 |
| LEQVHQL | ATRGTGEDTQY | 0.5915829 |
| LNAAGAI | ATRGTGEDTQY | 0.47245067 |
| PAVHASL | ATRGTGEDTQY | 0.6996542 |
| TELEDDA | ATRGTGEDTQY | 0.7963275 |
| TPPGPPN | ATRGTGEDTQY | 0.63602453 |
| TPPPPPPP | ATRGTGEDTQY | 0.6443952 |
| TQSPALI | TCSADGGGAYAEQF | 0.8145747 |
| EFAAAFPQ | TCSADGGGAYAEQF | 0.45212322 |
| LLSGNLDVL | TCSADGGGAYAEQF | 0.6260494 |
| GPANLGNYNI | TCSADGGGAYAEQF | 0.73970264 |
| TLGPTLP | TCSADGGGAYAEQF | 0.76543325 |
| VYAYSSCCK | TCSADGGGAYAEQF | 0.5976264 |
| APAPAPAPNI | TCSADGGGAYAEQF | 0.7968739 |
| PGGPVPIP | TCSADGGGAYAEQF | 0.43695176 |
| FLTVALEP | TCSADGGGAYAEQF | 0.72239166 |
| SQYFGANL | TCSADGGGAYAEQF | 0.5362388 |
| GGVKPAA | TCSADGGGAYAEQF | 0.58676434 |
| GGTGGAGGAGGW | TCSADGGGAYAEQF | 0.40957296 |
| GVGGVAGLGGAGG | TCSADGGGAYAEQF | 0.6792646 |
| VLLVLATTI | TCSADGGGAYAEQF | 0.74142385 |
| IGIFAAML | TCSADGGGAYAEQF | 0.7321782 |
| AVGDLID | TCSADGGGAYAEQF | 0.20360662 |
| NISPLPL | TCSADGGGAYAEQF | 0.74411696 |
| VTAVFVAI | TCSADGGGAYAEQF | 0.7540229 |
| SGDDNDAAL | TCSADGGGAYAEQF | 0.8261658 |
| AVAALAHS | TCSADGGGAYAEQF | 0.53793555 |
| GGVTVMATP | TCSADGGGAYAEQF | 0.72027606 |
| AGLAANVRD | TCSADGGGAYAEQF | 0.5038656 |
| TNRPDLIDPALLRPG | TCSADGGGAYAEQF | 0.5789628 |

| | | |
|---|---|---|
| LANLGVTAL | TCSADGGGAYAEQF | 0.31276086 |
| TGERIEL | TCSADGGGAYAEQF | 0.5251598 |
| NGEELAA | TCSADGGGAYAEQF | 0.5671756 |
| VLSLVWG | TCSADGGGAYAEQF | 0.49141675 |
| IGYALWL | TCSADGGGAYAEQF | 0.7401892 |
| LSLLLAPL | TCSADGGGAYAEQF | 0.743095 |
| TLDEALK | TCSADGGGAYAEQF | 0.7375697 |
| TDGAGTAEGG | TCSADGGGAYAEQF | 0.76894194 |
| NAPGGAGGNGG | TCSADGGGAYAEQF | 0.8191973 |
| VSVFMGMGGNF | TCSADGGGAYAEQF | 0.7111653 |
| LASYLAAL | TCSADGGGAYAEQF | 0.48011586 |
| LVGGTPAL | TCSADGGGAYAEQF | 0.56734836 |
| VRNADVL | TCSADGGGAYAEQF | 0.39142585 |
| TCSSPFLVSAM | TCSADGGGAYAEQF | 0.60272557 |
| VHITDAE | TCSADGGGAYAEQF | 0.49987933 |
| EYAAPSF | TCSADGGGAYAEQF | 0.6795541 |
| VVDAFHGGL | TCSADGGGAYAEQF | 0.13317476 |
| ASVVDPQ | TCSADGGGAYAEQF | 0.6553924 |
| SYNEFGGLS | TCSADGGGAYAEQF | 0.81597906 |
| DPDGSVY | TCSADGGGAYAEQF | 0.43844014 |
| MVSTIHATE | TCSADGGGAYAEQF | 0.6431974 |
| LAAGVADPV | TCSADGGGAYAEQF | 0.73647416 |
| TKEQIAA | TCSADGGGAYAEQF | 0.48257583 |
| SLASPAAL | TCSADGGGAYAEQF | 0.7042035 |
| IGLVFYGV | TCSADGGGAYAEQF | 0.4377896 |
| FLLDASNL | TCSADGGGAYAEQF | 0.69627064 |
| YQVGNFP | TCSADGGGAYAEQF | 0.8021835 |
| ESLLSQST | TCSADGGGAYAEQF | 0.65430677 |
| LFTVGFAAVG | TCSADGGGAYAEQF | 0.5075869 |
| ATPPTPT | TCSADGGGAYAEQF | 0.5633145 |
| TDLATLG | TCSADGGGAYAEQF | 0.27692232 |
| SAGGTGR | TCSADGGGAYAEQF | 0.58071697 |
| GFSNSGS | TCSADGGGAYAEQF | 0.29680905 |
| GGNGGSGGTSV | TCSADGGGAYAEQF | 0.6383196 |
| GTSVATGGAGN | TCSADGGGAYAEQF | 0.41987142 |
| NGGNGGSGGTS | TCSADGGGAYAEQF | 0.69376785 |
| AVVKFPH | TCSADGGGAYAEQF | 0.6603194 |
| TPPPPPP | TCSADGGGAYAEQF | 0.65629995 |
| AAPAAAAAPA | TCSADGGGAYAEQF | 0.3861952 |
| IALNEDL | TCSADGGGAYAEQF | 0.6080559 |
| IVNVSSM | TCSADGGGAYAEQF | 0.52732813 |
| TGIPPAPR | TCSADGGGAYAEQF | 0.31977645 |
| FDLGGGTF | TCSADGGGAYAEQF | 0.7723512 |
| FDLGGGTFDVS | TCSADGGGAYAEQF | 0.550595 |
| NEPTAAA | TCSADGGGAYAEQF | 0.6223963 |
| NEPTAAALA | TCSADGGGAYAEQF | 0.7595757 |
| GSIINISSI | TCSADGGGAYAEQF | 0.6113148 |
| VAPGNAGTA | TCSADGGGAYAEQF | 0.2891985 |
| TGGGASLE | TCSADGGGAYAEQF | 0.53699553 |
| VAVYNHYKRI | TCSADGGGAYAEQF | 0.6506249 |
| TDPAPATT | TCSADGGGAYAEQF | 0.48301455 |

| | | |
|---|---|---|
| TNRPDILDPALLRPG | TCSADGGGAYAEQF | 0.1748941 |
| LTTTTTS | TCSADGGGAYAEQF | 0.7455598 |
| PTGSGKTL | TCSADGGGAYAEQF | 0.4676476 |
| TGSGKTL | TCSADGGGAYAEQF | 0.46236825 |
| TIGHVDHGK | TCSADGGGAYAEQF | 0.58925 |
| TIGHVDHGKTT | TCSADGGGAYAEQF | 0.75413895 |
| GHVDHGKTT | TCSADGGGAYAEQF | 0.46415854 |
| AQFEHTLL | TCSADGGGAYAEQF | 0.5114963 |
| IVNQDEL | TCSADGGGAYAEQF | 0.7103509 |
| TNGSQFF | TCSADGGGAYAEQF | 0.63396233 |
| TGKGIVE | TCSADGGGAYAEQF | 0.79051775 |
| LISNASDA | TCSADGGGAYAEQF | 0.68518245 |
| AELRTLGELTKIAKAHGAQ | TCSADGGGAYAEQF | 0.56017435 |
| APNAVAAAEPSW | TCSADGGGAYAEQF | 0.70530874 |
| APQVAGAATAVL | TCSADGGGAYAEQF | 0.56007105 |
| ARRDPAAAAAILPTDA | TCSADGGGAYAEQF | 0.3430924 |
| DAAARDVLAEAGLAEYF | TCSADGGGAYAEQF | 0.24813384 |
| DDQIVLSAKANLAKVEQ | TCSADGGGAYAEQF | 0.51324487 |
| DELLPDTGEDSDEDGH | TCSADGGGAYAEQF | 0.28100222 |
| ELPEAPANRATVPQD | TCSADGGGAYAEQF | 0.669063 |
| FGQSKIERVTGIG | TCSADGGGAYAEQF | 0.7682523 |
| GAGGTGSTAGGAGGAGGAGGLY | TCSADGGGAYAEQF | 0.58096814 |
| GHGGTGAAGGNGGTGGAGGSGI | TCSADGGGAYAEQF | 0.35138148 |
| GNGGNGGAGGEGGPGGAGGAGA | TCSADGGGAYAEQF | 0.56336343 |
| GSTAGNGGNGGNATNGTIAGQP | TCSADGGGAYAEQF | 0.49625164 |
| GTGGDGGNGGNGGTGG | TCSADGGGAYAEQF | 0.7095322 |
| GVRRPAGIPADAPV | TCSADGGGAYAEQF | 0.6465008 |
| GYRQVTDAHLLTLARR | TCSADGGGAYAEQF | 0.8105284 |
| IAKMGQDGHDRG | TCSADGGGAYAEQF | 0.77772605 |
| KAEPSIEIQYT | TCSADGGGAYAEQF | 0.5262667 |
| KPVRIRMAADEDTVAVT | TCSADGGGAYAEQF | 0.7519674 |
| MPPRIAGMRLLVIKPEP | TCSADGGGAYAEQF | 0.54604423 |
| MKFYAVALFRSRGVPPTR | TCSADGGGAYAEQF | 0.65301454 |
| NAGGFGILGAVAHSP | TCSADGGGAYAEQF | 0.6129402 |
| NEGREITLAMAEM | TCSADGGGAYAEQF | 0.66773057 |
| NELVEWSHPLGHRWRWE | TCSADGGGAYAEQF | 0.49772075 |
| NLGAGGTGGNGGNPG | TCSADGGGAYAEQF | 0.5160637 |
| NTEWPKWETVMAADPVD | TCSADGGGAYAEQF | 0.27313405 |
| PASAAPAETQP | TCSADGGGAYAEQF | 0.6629887 |
| QQSSDGKNGVRAAEITGE | TCSADGGGAYAEQF | 0.3887105 |
| RAPDAVTTSWWKE | TCSADGGGAYAEQF | 0.27550524 |
| SGFGNVGAHTSGWFNQ | TCSADGGGAYAEQF | 0.5472863 |
| SGVAGAVGQAASVGGLKV | TCSADGGGAYAEQF | 0.8320936 |
| SNGGAGTGSAGGNGGT | TCSADGGGAYAEQF | 0.7739091 |
| TGGAGGEGGAAG | TCSADGGGAYAEQF | 0.47207004 |
| TGGDGGAAGAGGAPGHG | TCSADGGGAYAEQF | 0.42171937 |
| TMPPLLAAIANAGRHGV | TCSADGGGAYAEQF | 0.69337416 |
| TSSTAGNSGGA | TCSADGGGAYAEQF | 0.33870584 |
| VPATVPVGILC | TCSADGGGAYAEQF | 0.69876933 |
| VQRPDSGGVVLTGRIS | TCSADGGGAYAEQF | 0.62160957 |
| WTPNPVPATKGVPAGLVQAT | TCSADGGGAYAEQF | 0.6234628 |

| | | |
|---|---|---|
| YALKRALAKFSDRFDVVII | TCSADGGGAYAEQF | 0.42462158 |
| YLIGSVIV | TCSADGGGAYAEQF | 0.6849305 |
| YGQLVAVID | TCSADGGGAYAEQF | 0.56980824 |
| VTVGGMTLPA | TCSADGGGAYAEQF | 0.6454974 |
| VLALPAV | TCSADGGGAYAEQF | 0.14804836 |
| VIPPTLNYET | TCSADGGGAYAEQF | 0.7509904 |
| VGMGGMHPSSG | TCSADGGGAYAEQF | 0.46854052 |
| VGLAGGAATA | TCSADGGGAYAEQF | 0.43210167 |
| VGGAGGAGGSGP | TCSADGGGAYAEQF | 0.23258275 |
| VFAAFMV | TCSADGGGAYAEQF | 0.64874774 |
| TLYDKLR | TCSADGGGAYAEQF | 0.69032216 |
| TLAVFTVL | TCSADGGGAYAEQF | 0.7761502 |
| TGSVQPARV | TCSADGGGAYAEQF | 0.46183825 |
| TGGGAGSGGT | TCSADGGGAYAEQF | 0.73280513 |
| SHLETSL | TCSADGGGAYAEQF | 0.74667025 |
| SFTTGMT | TCSADGGGAYAEQF | 0.5762721 |
| QVEVAKRMGVRQPTVS | TCSADGGGAYAEQF | 0.28233725 |
| QGSGGPALAM | TCSADGGGAYAEQF | 0.6434957 |
| MLAPSNP | TCSADGGGAYAEQF | 0.60624605 |
| LVNNAAI | TCSADGGGAYAEQF | 0.75551134 |
| LTGEINTLA | TCSADGGGAYAEQF | 0.6817062 |
| LSIFGPV | TCSADGGGAYAEQF | 0.39517957 |
| LLALLANL | TCSADGGGAYAEQF | 0.67648405 |
| LISNASDALDKLR | TCSADGGGAYAEQF | 0.7611892 |
| LINEAVP | TCSADGGGAYAEQF | 0.3574598 |
| LGYAFSGL | TCSADGGGAYAEQF | 0.51607084 |
| LDFVAVAL | TCSADGGGAYAEQF | 0.566034 |
| LAQPGAE | TCSADGGGAYAEQF | 0.26924402 |
| ISNLGAVTGF | TCSADGGGAYAEQF | 0.66823983 |
| HQEDPGRI | TCSADGGGAYAEQF | 0.55273676 |
| GTPGEPTP | TCSADGGGAYAEQF | 0.40493354 |
| GFFNSGS | TCSADGGGAYAEQF | 0.6405092 |
| GAGGGGLTAGAN | TCSADGGGAYAEQF | 0.424698 |
| FVDALVNM | TCSADGGGAYAEQF | 0.4774375 |
| FLTVAPEL | TCSADGGGAYAEQF | 0.5107622 |
| FETLTAL | TCSADGGGAYAEQF | 0.62350357 |
| DVINPIEL | TCSADGGGAYAEQF | 0.7002923 |
| APVSGAPV | TCSADGGGAYAEQF | 0.5838755 |
| ALVGGLAW | TCSADGGGAYAEQF | 0.6632725 |
| ALAAAAATAI | TCSADGGGAYAEQF | 0.8101354 |
| AFGLGLLL | TCSADGGGAYAEQF | 0.29595184 |
| AFFAAASL | TCSADGGGAYAEQF | 0.46145564 |
| AAAALPAFSPP | TCSADGGGAYAEQF | 0.6695763 |
| VVLAPAL | TCSADGGGAYAEQF | 0.3750062 |
| TIGHVDHGKTTL | TCSADGGGAYAEQF | 0.73731744 |
| LISNASDALDK | TCSADGGGAYAEQF | 0.6573677 |
| LISNASDAL | TCSADGGGAYAEQF | 0.7022357 |
| ISNASDALDK | TCSADGGGAYAEQF | 0.5551195 |
| NQAETLVYQTE | TCSADGGGAYAEQF | 0.83420265 |
| TGGTGGTAGAG | TCSADGGGAYAEQF | 0.7116161 |
| TGSTAGGAGGA | TCSADGGGAYAEQF | 0.6382023 |

| | | |
|---|---|---|
| DPLAVPVAAGPV | TCSADGGGAYAEQF | 0.7277324 |
| GGTGGTPGNGGN | TCSADGGGAYAEQF | 0.44335237 |
| PAGLAGVDPATL | TCSADGGGAYAEQF | 0.73452055 |
| GNNLVVEAPHAGASV | TCSADGGGAYAEQF | 0.7114322 |
| KVTRIMSASTLEQPA | TCSADGGGAYAEQF | 0.3734669 |
| LGFALPIALPRRTVV | TCSADGGGAYAEQF | 0.62977093 |
| TRLGNLTARKLVVLV | TCSADGGGAYAEQF | 0.7743461 |
| WSQQRRLIYFPSAGP | TCSADGGGAYAEQF | 0.79664683 |
| GPNGNPDPMAAAVDIR | TCSADGGGAYAEQF | 0.31701595 |
| TQRCVSVADLRAAASR | TCSADGGGAYAEQF | 0.56477845 |
| QFVLDVAEHPVEVLTETV | TCSADGGGAYAEQF | 0.32941514 |
| GGAGGTGGAGGNGGTQAGDG | TCSADGGGAYAEQF | 0.5015372 |
| PLLATLRAIAVDVDAKPAQV | TCSADGGGAYAEQF | 0.41497996 |
| APKFRTGGYMIDFWGVGYQVAK | TCSADGGGAYAEQF | 0.69971883 |
| GAGGAGGAGGAAGGPGGTGGTG | TCSADGGGAYAEQF | 0.53681815 |
| IGALGLLATDVAPAGSVAFGVA | TCSADGGGAYAEQF | 0.7214447 |
| AGGAGGIGGNGGAGGAGGVLMG | TCSADGGGAYAEQF | 0.6703044 |
| ADELIGLL | TCSADGGGAYAEQF | 0.7815294 |
| ADRALKA | TCSADGGGAYAEQF | 0.46726978 |
| AGAVGQVGGAGGA | TCSADGGGAYAEQF | 0.58399063 |
| AGGGGGLGGAP | TCSADGGGAYAEQF | 0.28299114 |
| AGGLLYGNGGAG | TCSADGGGAYAEQF | 0.29287288 |
| AGLESIGL | TCSADGGGAYAEQF | 0.43184826 |
| AGLKTGII | TCSADGGGAYAEQF | 0.5278727 |
| ASLAALLL | TCSADGGGAYAEQF | 0.19636276 |
| ASVINISSI | TCSADGGGAYAEQF | 0.8144744 |
| ATPPGPA | TCSADGGGAYAEQF | 0.56381774 |
| ATVASALV | TCSADGGGAYAEQF | 0.72354907 |
| ATVASLIL | TCSADGGGAYAEQF | 0.46373564 |
| AVNGPVLIRH | TCSADGGGAYAEQF | 0.48038167 |
| DGAAGAVGI | TCSADGGGAYAEQF | 0.511643 |
| DPLLHGI | TCSADGGGAYAEQF | 0.707212 |
| ELGAALGAL | TCSADGGGAYAEQF | 0.47801483 |
| ELGRTQAV | TCSADGGGAYAEQF | 0.2593889 |
| FVAGGYLS | TCSADGGGAYAEQF | 0.43496117 |
| GASGGAGGAGGW | TCSADGGGAYAEQF | 0.39032227 |
| GGAGAGGDGGAGGI | TCSADGGGAYAEQF | 0.82507384 |
| GTGHNGSQPGMGGNG | TCSADGGGAYAEQF | 0.7634815 |
| GYAAASASAS | TCSADGGGAYAEQF | 0.67400014 |
| HQVDVAAVQQG | TCSADGGGAYAEQF | 0.81677914 |
| IGAAIAHT | TCSADGGGAYAEQF | 0.17242055 |
| IGMIEVAQL | TCSADGGGAYAEQF | 0.5476576 |
| IRLAPEHTSAMD | TCSADGGGAYAEQF | 0.81407964 |
| IYNVSVL | TCSADGGGAYAEQF | 0.60575545 |
| KQGFLGM | TCSADGGGAYAEQF | 0.5985856 |
| LESVITQL | TCSADGGGAYAEQF | 0.7766704 |
| LGLGALVVV | TCSADGGGAYAEQF | 0.44899553 |
| LIGNGGNGGSGGI | TCSADGGGAYAEQF | 0.48405397 |
| LNAINAPTL | TCSADGGGAYAEQF | 0.5919417 |
| LQLLNTGV | TCSADGGGAYAEQF | 0.71919817 |
| LQTSIEF | TCSADGGGAYAEQF | 0.6395669 |

| | | |
|---|---|---|
| MSFAVSEL | TCSADGGGAYAEQF | 0.36549634 |
| NTGALGL | TCSADGGGAYAEQF | 0.6425203 |
| PPGAGGAGGAGTT | TCSADGGGAYAEQF | 0.8437746 |
| QLVGLGI | TCSADGGGAYAEQF | 0.551689 |
| RFLEQVHQL | TCSADGGGAYAEQF | 0.7186413 |
| RGVGYVLR | TCSADGGGAYAEQF | 0.6309442 |
| SATGVFAGVI | TCSADGGGAYAEQF | 0.47904474 |
| SLPAITLP | TCSADGGGAYAEQF | 0.4585811 |
| STALAFTV | TCSADGGGAYAEQF | 0.4284928 |
| SVLTVGLGK | TCSADGGGAYAEQF | 0.6086663 |
| TALAGAGSGI | TCSADGGGAYAEQF | 0.40242484 |
| TGLPFTGI | TCSADGGGAYAEQF | 0.6166358 |
| VAEHVTV | TCSADGGGAYAEQF | 0.593767 |
| VDIVLNSL | TCSADGGGAYAEQF | 0.8498751 |
| VLTGGGAVL | TCSADGGGAYAEQF | 0.7029683 |
| VNVGLAYGI | TCSADGGGAYAEQF | 0.6777178 |
| VSSAAVGAV | TCSADGGGAYAEQF | 0.4521304 |
| VTSVYEAGV | TCSADGGGAYAEQF | 0.61757755 |
| WVALDRGA | TCSADGGGAYAEQF | 0.7029985 |
| YLVGSVVI | TCSADGGGAYAEQF | 0.3090687 |
| GPPGTGKTLL | TCSADGGGAYAEQF | 0.7781952 |
| GVGMTKF | TCSADGGGAYAEQF | 0.4454681 |
| IGIGGSDL | TCSADGGGAYAEQF | 0.6182595 |
| IGTLLAEL | TCSADGGGAYAEQF | 0.7101648 |
| IYTSGTT | TCSADGGGAYAEQF | 0.77930784 |
| LSFPAGV | TCSADGGGAYAEQF | 0.61517555 |
| LYTSGTT | TCSADGGGAYAEQF | 0.5824098 |
| VGAFPFGF | TCSADGGGAYAEQF | 0.44426888 |
| VGAGTVEF | TCSADGGGAYAEQF | 0.73998386 |
| VGKSTLLL | TCSADGGGAYAEQF | 0.78547275 |
| AALAHAGGGIV | TCSADGGGAYAEQF | 0.56545556 |
| AALVAVSLL | TCSADGGGAYAEQF | 0.82398397 |
| AAVSRAVAAL | TCSADGGGAYAEQF | 0.67625386 |
| AGGSSGIGGF | TCSADGGGAYAEQF | 0.34174955 |
| AGLTVANANG | TCSADGGGAYAEQF | 0.5589502 |
| AGLVGAVL | TCSADGGGAYAEQF | 0.7868416 |
| ALSQAFTV | TCSADGGGAYAEQF | 0.7765796 |
| ALTFSVSL | TCSADGGGAYAEQF | 0.7173097 |
| ALTGAALVAL | TCSADGGGAYAEQF | 0.6934707 |
| ASFSIGAGVL | TCSADGGGAYAEQF | 0.54413855 |
| DGSVYKI | TCSADGGGAYAEQF | 0.7992784 |
| EQLAEIV | TCSADGGGAYAEQF | 0.83334744 |
| FAVSELTVM | TCSADGGGAYAEQF | 0.6972929 |
| GAAPLAAGL | TCSADGGGAYAEQF | 0.4409448 |
| GADGGDGGSGGAGG | TCSADGGGAYAEQF | 0.4167815 |
| GFGLDTGGAGG | TCSADGGGAYAEQF | 0.66506463 |
| GLASGGLL | TCSADGGGAYAEQF | 0.77265656 |
| GRIAGAAVLL | TCSADGGGAYAEQF | 0.36304164 |
| GSAGSLNTSF | TCSADGGGAYAEQF | 0.6484666 |
| GTGQIGI | TCSADGGGAYAEQF | 0.64396864 |
| GTLIALIL | TCSADGGGAYAEQF | 0.2677471 |

| | | |
|---|---|---|
| IAATMGFVVG | TCSADGGGAYAEQF | 0.7130685 |
| IFAQALL | TCSADGGGAYAEQF | 0.80409026 |
| IGAAVALGF | TCSADGGGAYAEQF | 0.65179455 |
| IGLTPLAL | TCSADGGGAYAEQF | 0.368496 |
| ILGSNTSTL | TCSADGGGAYAEQF | 0.7806038 |
| IVFSLTVL | TCSADGGGAYAEQF | 0.83103263 |
| LASKATGF | TCSADGGGAYAEQF | 0.7893995 |
| LASVALIL | TCSADGGGAYAEQF | 0.2762663 |
| LFAVGLTL | TCSADGGGAYAEQF | 0.1893904 |
| LFNGGIGTYI | TCSADGGGAYAEQF | 0.56639135 |
| LTLGAGII | TCSADGGGAYAEQF | 0.6966464 |
| LVAAALGDI | TCSADGGGAYAEQF | 0.49422607 |
| NTAFVQLGI | TCSADGGGAYAEQF | 0.42071763 |
| QLFSGAEL | TCSADGGGAYAEQF | 0.695588 |
| SAATFITL | TCSADGGGAYAEQF | 0.7838385 |
| SNYGLANLGV | TCSADGGGAYAEQF | 0.4336821 |
| SYGGAYAVM | TCSADGGGAYAEQF | 0.33933097 |
| TTGVGGLTLG | TCSADGGGAYAEQF | 0.7535532 |
| TTQTIYI | TCSADGGGAYAEQF | 0.7479042 |
| VAAGAGLGAVI | TCSADGGGAYAEQF | 0.42123014 |
| VAAVVTLL | TCSADGGGAYAEQF | 0.55084985 |
| VGFLGGLI | TCSADGGGAYAEQF | 0.61431396 |
| VGVIPILL | TCSADGGGAYAEQF | 0.45053887 |
| VIFASSLI | TCSADGGGAYAEQF | 0.67584157 |
| VKGGDGGAAAG | TCSADGGGAYAEQF | 0.4095993 |
| VTKTFTNI | TCSADGGGAYAEQF | 0.7910142 |
| VVAAAILL | TCSADGGGAYAEQF | 0.4145446 |
| YAALDVEL | TCSADGGGAYAEQF | 0.72000724 |
| LASVALLL | TCSADGGGAYAEQF | 0.6426611 |
| LDAVHRV | TCSADGGGAYAEQF | 0.69688916 |
| QGLSSSIF | TCSADGGGAYAEQF | 0.58568615 |
| RGTFANIRL | TCSADGGGAYAEQF | 0.39972094 |
| RQLWWGHRI | TCSADGGGAYAEQF | 0.41553771 |
| YGPPGTGKTLL | TCSADGGGAYAEQF | 0.2296874 |
| AALVGAPANVA | TCSADGGGAYAEQF | 0.7673612 |
| AATFRRTE | TCSADGGGAYAEQF | 0.4405054 |
| AGALGGGI | TCSADGGGAYAEQF | 0.77266943 |
| AGGAGGAGGIGAG | TCSADGGGAYAEQF | 0.66805243 |
| AGSGALGL | TCSADGGGAYAEQF | 0.4789098 |
| AQIADQI | TCSADGGGAYAEQF | 0.7048047 |
| ASAVAVTYL | TCSADGGGAYAEQF | 0.32359543 |
| ASTVLGGTGG | TCSADGGGAYAEQF | 0.45758334 |
| AVGGEPAK | TCSADGGGAYAEQF | 0.62293565 |
| CEQGTFGGTC | TCSADGGGAYAEQF | 0.7219596 |
| DEGKAPAG | TCSADGGGAYAEQF | 0.4597678 |
| DTVVFPAL | TCSADGGGAYAEQF | 0.69716656 |
| FEGITMAG | TCSADGGGAYAEQF | 0.6873505 |
| FPLVVQD | TCSADGGGAYAEQF | 0.38620603 |
| GAFTTPQ | TCSADGGGAYAEQF | 0.6455456 |
| GFGGFGTAGGVGGA | TCSADGGGAYAEQF | 0.7735651 |
| GFLAQFL | TCSADGGGAYAEQF | 0.64535856 |

| | | |
|---|---|---|
| GGAGGAGGNAGVA | TCSADGGGAYAEQF | 0.39254516 |
| GGVGTAGAGGAGGAGG | TCSADGGGAYAEQF | 0.6471227 |
| GNGGGGDAGNAGSGGN | TCSADGGGAYAEQF | 0.4727444 |
| GTGGAGGAGNDAGS | TCSADGGGAYAEQF | 0.6990094 |
| GTTNTGWFNS | TCSADGGGAYAEQF | 0.6599068 |
| IFMGPLFTL | TCSADGGGAYAEQF | 0.49897736 |
| ILAHAIEAI | TCSADGGGAYAEQF | 0.70210934 |
| IRSAIREAL | TCSADGGGAYAEQF | 0.4072416 |
| ITFAGGG | TCSADGGGAYAEQF | 0.44313243 |
| ITGMSGHQEV | TCSADGGGAYAEQF | 0.5558938 |
| IVAGLACTL | TCSADGGGAYAEQF | 0.642565 |
| IVGGGLAGA | TCSADGGGAYAEQF | 0.7168799 |
| IVTGVLNI | TCSADGGGAYAEQF | 0.4167162 |
| LAQQGCI | TCSADGGGAYAEQF | 0.6195389 |
| LCLHVEN | TCSADGGGAYAEQF | 0.51219654 |
| LDAISDGS | TCSADGGGAYAEQF | 0.3882982 |
| LGGFAAML | TCSADGGGAYAEQF | 0.6372773 |
| LGGKNPMI | TCSADGGGAYAEQF | 0.5266541 |
| LGPKGVAAIL | TCSADGGGAYAEQF | 0.3118744 |
| LMVGVSI | TCSADGGGAYAEQF | 0.7094052 |
| LVGAAFVL | TCSADGGGAYAEQF | 0.4339742 |
| LVLAVAPL | TCSADGGGAYAEQF | 0.6053651 |
| MAAIAVIDAA | TCSADGGGAYAEQF | 0.72689736 |
| NNGINAELQ | TCSADGGGAYAEQF | 0.3829188 |
| RGEPLVLVHG | TCSADGGGAYAEQF | 0.712183 |
| RGGAGGAAGLI | TCSADGGGAYAEQF | 0.67215884 |
| RSYRMHPAV | TCSADGGGAYAEQF | 0.6531851 |
| SAAHLLAGQLA | TCSADGGGAYAEQF | 0.5481945 |
| SAAVGSY | TCSADGGGAYAEQF | 0.51417744 |
| SPHAVLHRI | TCSADGGGAYAEQF | 0.81556433 |
| SQIAKVL | TCSADGGGAYAEQF | 0.68366957 |
| TALGHLVP | TCSADGGGAYAEQF | 0.69931364 |
| TAPTTSGAA | TCSADGGGAYAEQF | 0.73943835 |
| TDGDDGNGGSPGT | TCSADGGGAYAEQF | 0.725875 |
| TDGLGEN | TCSADGGGAYAEQF | 0.73510486 |
| TGGPPLPGQA | TCSADGGGAYAEQF | 0.4844403 |
| TIPAITI | TCSADGGGAYAEQF | 0.28132987 |
| TIPGFTI | TCSADGGGAYAEQF | 0.55342644 |
| TSIASGALA | TCSADGGGAYAEQF | 0.5504235 |
| TVTLPGAAI | TCSADGGGAYAEQF | 0.51408696 |
| VASGLSAALP | TCSADGGGAYAEQF | 0.29149687 |
| VGAGVIGI | TCSADGGGAYAEQF | 0.77111393 |
| VGPKRFGV | TCSADGGGAYAEQF | 0.8465569 |
| VNVAFPD | TCSADGGGAYAEQF | 0.37784517 |
| VSMTNSGVS | TCSADGGGAYAEQF | 0.836205 |
| VTDVLGDLSV | TCSADGGGAYAEQF | 0.37054604 |
| VVVVGAGFAGL | TCSADGGGAYAEQF | 0.6427249 |
| VYTVGTVV | TCSADGGGAYAEQF | 0.7772908 |
| WTTFSGP | TCSADGGGAYAEQF | 0.37911445 |
| YDELVEAF | TCSADGGGAYAEQF | 0.6893997 |
| YGTGGAGGNGG | TCSADGGGAYAEQF | 0.44253486 |

| | | |
|---|---|---|
| YTNVMAV | TCSADGGGAYAEQF | 0.5114099 |
| FLEQVHQL | TCSADGGGAYAEQF | 0.78470474 |
| GLSLLAVAV | TCSADGGGAYAEQF | 0.48353872 |
| LEQVHQL | TCSADGGGAYAEQF | 0.6462924 |
| LNAAGAI | TCSADGGGAYAEQF | 0.5366104 |
| PAVHASL | TCSADGGGAYAEQF | 0.4640799 |
| TELEDDA | TCSADGGGAYAEQF | 0.69311404 |
| TPPGPPN | TCSADGGGAYAEQF | 0.7575366 |
| TPPPPPPP | TCSADGGGAYAEQF | 0.4607307 |
| TQSPALI | ASSIIWAANTGQLY | 0.8204787 |
| EFAAAFPQ | ASSIIWAANTGQLY | 0.4264477 |
| LLSGNLDVL | ASSIIWAANTGQLY | 0.82653534 |
| GPANLGNYNI | ASSIIWAANTGQLY | 0.79437876 |
| TLGPTLP | ASSIIWAANTGQLY | 0.64787716 |
| VYAYSSCCK | ASSIIWAANTGQLY | 0.52591896 |
| APAPAPAPNI | ASSIIWAANTGQLY | 0.7427093 |
| PGGPVPIP | ASSIIWAANTGQLY | 0.54200715 |
| FLTVALEP | ASSIIWAANTGQLY | 0.66138697 |
| SQYFGANL | ASSIIWAANTGQLY | 0.7672379 |
| GGVKPAA | ASSIIWAANTGQLY | 0.64361405 |
| GGTGGAGGAGGW | ASSIIWAANTGQLY | 0.79370576 |
| GVGGVAGLGGAGG | ASSIIWAANTGQLY | 0.7382301 |
| VLLVLATTI | ASSIIWAANTGQLY | 0.7180049 |
| IGIFAAML | ASSIIWAANTGQLY | 0.6304697 |
| AVGDLID | ASSIIWAANTGQLY | 0.40787175 |
| NISPLPL | ASSIIWAANTGQLY | 0.72520024 |
| VTAVFVAI | ASSIIWAANTGQLY | 0.54293036 |
| SGDDNDAAL | ASSIIWAANTGQLY | 0.7466889 |
| AVAALAHS | ASSIIWAANTGQLY | 0.7831636 |
| GGVTVMATP | ASSIIWAANTGQLY | 0.61179185 |
| AGLAANVRD | ASSIIWAANTGQLY | 0.820286 |
| TNRPDLIDPALLRPG | ASSIIWAANTGQLY | 0.8072377 |
| LANLGVTAL | ASSIIWAANTGQLY | 0.44365355 |
| TGERIEL | ASSIIWAANTGQLY | 0.6258663 |
| NGEELAA | ASSIIWAANTGQLY | 0.6062905 |
| VLSLVWG | ASSIIWAANTGQLY | 0.7137395 |
| IGYALWL | ASSIIWAANTGQLY | 0.37522247 |
| LSLLLAPL | ASSIIWAANTGQLY | 0.7765997 |
| TLDEALK | ASSIIWAANTGQLY | 0.5924323 |
| TDGAGTAEGG | ASSIIWAANTGQLY | 0.17527789 |
| NAPGGAGGNGG | ASSIIWAANTGQLY | 0.75162554 |
| VSVFMGMGGNF | ASSIIWAANTGQLY | 0.6693854 |
| LASYLAAL | ASSIIWAANTGQLY | 0.72488356 |
| LVGGTPAL | ASSIIWAANTGQLY | 0.789904 |
| VRNADVL | ASSIIWAANTGQLY | 0.5137447 |
| TCSSPFLVSAM | ASSIIWAANTGQLY | 0.5945193 |
| VHITDAE | ASSIIWAANTGQLY | 0.74961656 |
| EYAAPSF | ASSIIWAANTGQLY | 0.8167044 |
| VVDAFHGGL | ASSIIWAANTGQLY | 0.5466098 |
| ASVVDPQ | ASSIIWAANTGQLY | 0.72638583 |
| SYNEFGGLS | ASSIIWAANTGQLY | 0.8053083 |

| | | |
|---|---|---|
| DPDGSVY | ASSIIWAANTGQLY | 0.75795174 |
| MVSTIHATE | ASSIIWAANTGQLY | 0.8131216 |
| LAAGVADPV | ASSIIWAANTGQLY | 0.8337983 |
| TKEQIAA | ASSIIWAANTGQLY | 0.5603862 |
| SLASPAAL | ASSIIWAANTGQLY | 0.79934174 |
| IGLVFYGV | ASSIIWAANTGQLY | 0.81370544 |
| FLLDASNL | ASSIIWAANTGQLY | 0.70073855 |
| YQVGNFP | ASSIIWAANTGQLY | 0.68764114 |
| ESLLSQST | ASSIIWAANTGQLY | 0.6299754 |
| LFTVGFAAVG | ASSIIWAANTGQLY | 0.7132205 |
| ATPPTPT | ASSIIWAANTGQLY | 0.32169393 |
| TDLATLG | ASSIIWAANTGQLY | 0.71989673 |
| SAGGTGR | ASSIIWAANTGQLY | 0.80218655 |
| GFSNSGS | ASSIIWAANTGQLY | 0.63864964 |
| GGNGGSGGTSV | ASSIIWAANTGQLY | 0.49015564 |
| GTSVATGGAGN | ASSIIWAANTGQLY | 0.42955646 |
| NGGNGGSGGTS | ASSIIWAANTGQLY | 0.6439237 |
| AVVKFPH | ASSIIWAANTGQLY | 0.6872666 |
| TPPPPPP | ASSIIWAANTGQLY | 0.31729385 |
| AAPAAAAAPA | ASSIIWAANTGQLY | 0.6595609 |
| IALNEDL | ASSIIWAANTGQLY | 0.80368316 |
| IVNVSSM | ASSIIWAANTGQLY | 0.7935695 |
| TGIPPAPR | ASSIIWAANTGQLY | 0.3700658 |
| FDLGGGTF | ASSIIWAANTGQLY | 0.8131652 |
| FDLGGGTFDVS | ASSIIWAANTGQLY | 0.7606774 |
| NEPTAAA | ASSIIWAANTGQLY | 0.7723136 |
| NEPTAAALA | ASSIIWAANTGQLY | 0.64852405 |
| GSIINISSI | ASSIIWAANTGQLY | 0.62152314 |
| VAPGNAGTA | ASSIIWAANTGQLY | 0.76183593 |
| TGGGASLE | ASSIIWAANTGQLY | 0.6174811 |
| VAVYNHYKRI | ASSIIWAANTGQLY | 0.5994802 |
| TDPAPATT | ASSIIWAANTGQLY | 0.79772305 |
| TNRPDILDPALLRPG | ASSIIWAANTGQLY | 0.6974493 |
| LTTTTTS | ASSIIWAANTGQLY | 0.6070936 |
| PTGSGKTL | ASSIIWAANTGQLY | 0.54671746 |
| TGSGKTL | ASSIIWAANTGQLY | 0.79890776 |
| TIGHVDHGK | ASSIIWAANTGQLY | 0.46786922 |
| TIGHVDHGKTT | ASSIIWAANTGQLY | 0.6239625 |
| GHVDHGKTT | ASSIIWAANTGQLY | 0.48535573 |
| AQFEHTLL | ASSIIWAANTGQLY | 0.7029761 |
| IVNQDEL | ASSIIWAANTGQLY | 0.79775226 |
| TNGSQFF | ASSIIWAANTGQLY | 0.7352151 |
| TGKGIVE | ASSIIWAANTGQLY | 0.64792085 |
| LISNASDA | ASSIIWAANTGQLY | 0.75093764 |
| AELRTLGELTKIAKAHGAQ | ASSIIWAANTGQLY | 0.752532 |
| APNAVAAAEPSW | ASSIIWAANTGQLY | 0.7290333 |
| APQVAGAATAVL | ASSIIWAANTGQLY | 0.80953944 |
| ARRDPAAAAAILPTDA | ASSIIWAANTGQLY | 0.54585207 |
| DAAARDVLAEAGLAEYF | ASSIIWAANTGQLY | 0.7949343 |
| DDQIVLSAKANLAKVEQ | ASSIIWAANTGQLY | 0.7575417 |
| DELLPDTGEDSDEDGH | ASSIIWAANTGQLY | 0.8216232 |

| | | |
|---|---|---|
| ELPEAPANRATVPQD | ASSIIWAANTGQLY | 0.80674064 |
| FGQSKIERVTGIG | ASSIIWAANTGQLY | 0.7542232 |
| GAGGTGSTAGGAGGAGGAGGLY | ASSIIWAANTGQLY | 0.6971735 |
| GHGGTGAAGGNGGTGGAGGSGI | ASSIIWAANTGQLY | 0.29758435 |
| GNGGNGGAGGEGGPGGAGGAGA | ASSIIWAANTGQLY | 0.607105 |
| GSTAGNGGNGGNATNGTIAGQP | ASSIIWAANTGQLY | 0.7865006 |
| GTGGDGGNGGNGGTGG | ASSIIWAANTGQLY | 0.7032039 |
| GVRRPAGIPADAPV | ASSIIWAANTGQLY | 0.79771096 |
| GYRQVTDAHLLTLARR | ASSIIWAANTGQLY | 0.7795245 |
| IAKMGQDGHDRG | ASSIIWAANTGQLY | 0.5752296 |
| KAEPSIEIQYT | ASSIIWAANTGQLY | 0.82300925 |
| KPVRIRMAADEDTVAVT | ASSIIWAANTGQLY | 0.780519 |
| MPPRIAGMRLLVIKPEP | ASSIIWAANTGQLY | 0.3810957 |
| MKFYAVALFRSRGVPPTR | ASSIIWAANTGQLY | 0.6300782 |
| NAGGFGILGAVAHSP | ASSIIWAANTGQLY | 0.82277966 |
| NEGREITLAMAEM | ASSIIWAANTGQLY | 0.7118599 |
| NELVEWSHPLGHRWRWE | ASSIIWAANTGQLY | 0.66496086 |
| NLGAGGTGGNGGNPG | ASSIIWAANTGQLY | 0.67601573 |
| NTEWPKWETVMAADPVD | ASSIIWAANTGQLY | 0.817444 |
| PASAAPAETQP | ASSIIWAANTGQLY | 0.5788151 |
| QQSSDGKNGVRAAEITGE | ASSIIWAANTGQLY | 0.8203969 |
| RAPDAVTTSWWKE | ASSIIWAANTGQLY | 0.21896152 |
| SGFGNVGAHTSGWFNQ | ASSIIWAANTGQLY | 0.6145547 |
| SGVAGAVGQAASVGGLKV | ASSIIWAANTGQLY | 0.53852344 |
| SNGGAGTGSAGGNGGT | ASSIIWAANTGQLY | 0.48519123 |
| TGGAGGEGGAAG | ASSIIWAANTGQLY | 0.79995894 |
| TGGDGGAAGAGGAPGHG | ASSIIWAANTGQLY | 0.68527085 |
| TMPPLLAAIANAGRHGV | ASSIIWAANTGQLY | 0.79352325 |
| TSSTAGNSGGA | ASSIIWAANTGQLY | 0.67496645 |
| VPATVPVGILC | ASSIIWAANTGQLY | 0.8006269 |
| VQRPDSGGVVLTGRIS | ASSIIWAANTGQLY | 0.6656084 |
| WTPNPVPATKGVPAGLVQAT | ASSIIWAANTGQLY | 0.7570007 |
| YALKRALAKFSDRFDVVII | ASSIIWAANTGQLY | 0.64915574 |
| YLIGSVIV | ASSIIWAANTGQLY | 0.78790545 |
| YGQLVAVID | ASSIIWAANTGQLY | 0.83554816 |
| VTVGGMTLPA | ASSIIWAANTGQLY | 0.7555342 |
| VLALPAV | ASSIIWAANTGQLY | 0.5017482 |
| VIPPTLNYET | ASSIIWAANTGQLY | 0.6364554 |
| VGMGGMHPSSG | ASSIIWAANTGQLY | 0.7230122 |
| VGLAGGAATA | ASSIIWAANTGQLY | 0.3202337 |
| VGGAGGAGGSGP | ASSIIWAANTGQLY | 0.28428438 |
| VFAAFMV | ASSIIWAANTGQLY | 0.5873331 |
| TLYDKLR | ASSIIWAANTGQLY | 0.56649446 |
| TLAVFTVL | ASSIIWAANTGQLY | 0.8026634 |
| TGSVQPARV | ASSIIWAANTGQLY | 0.76462966 |
| TGGGAGSGGT | ASSIIWAANTGQLY | 0.6793432 |
| SHLETSL | ASSIIWAANTGQLY | 0.77598816 |
| SFTTGMT | ASSIIWAANTGQLY | 0.760013 |
| QVEVAKRMGVRQPTVS | ASSIIWAANTGQLY | 0.1545314 |
| QGSGGPALAM | ASSIIWAANTGQLY | 0.6109237 |
| MLAPSNP | ASSIIWAANTGQLY | 0.48733342 |

| | | |
|---|---|---|
| LVNNAAI | ASSIIWAANTGQLY | 0.8307055 |
| LTGEINTLA | ASSIIWAANTGQLY | 0.56723017 |
| LSIFGPV | ASSIIWAANTGQLY | 0.5286575 |
| LLALLANL | ASSIIWAANTGQLY | 0.7016103 |
| LISNASDALDKLR | ASSIIWAANTGQLY | 0.67405915 |
| LINEAVP | ASSIIWAANTGQLY | 0.6485021 |
| LGYAFSGL | ASSIIWAANTGQLY | 0.4587103 |
| LDFVAVAL | ASSIIWAANTGQLY | 0.22294647 |
| LAQPGAE | ASSIIWAANTGQLY | 0.79779744 |
| ISNLGAVTGF | ASSIIWAANTGQLY | 0.42683476 |
| HQEDPGRI | ASSIIWAANTGQLY | 0.6026486 |
| GTPGEPTP | ASSIIWAANTGQLY | 0.44228074 |
| GFFNSGS | ASSIIWAANTGQLY | 0.5709261 |
| GAGGGGLTAGAN | ASSIIWAANTGQLY | 0.38901916 |
| FVDALVNM | ASSIIWAANTGQLY | 0.81472117 |
| FLTVAPEL | ASSIIWAANTGQLY | 0.6228408 |
| FETLTAL | ASSIIWAANTGQLY | 0.6094006 |
| DVINPIEL | ASSIIWAANTGQLY | 0.81194 |
| APVSGAPV | ASSIIWAANTGQLY | 0.6753599 |
| ALVGGLAW | ASSIIWAANTGQLY | 0.62989205 |
| ALAAAAATAI | ASSIIWAANTGQLY | 0.6709188 |
| AFGLGLLL | ASSIIWAANTGQLY | 0.5563439 |
| AFFAAASL | ASSIIWAANTGQLY | 0.6274267 |
| AAAALPAFSPP | ASSIIWAANTGQLY | 0.6233218 |
| VVLAPAL | ASSIIWAANTGQLY | 0.830501 |
| TIGHVDHGKTTL | ASSIIWAANTGQLY | 0.67313445 |
| LISNASDALDK | ASSIIWAANTGQLY | 0.57279944 |
| LISNASDAL | ASSIIWAANTGQLY | 0.693562 |
| ISNASDALDK | ASSIIWAANTGQLY | 0.8133109 |
| NQAETLVYQTE | ASSIIWAANTGQLY | 0.8248061 |
| TGGTGGTAGAG | ASSIIWAANTGQLY | 0.6141958 |
| TGSTAGGAGGA | ASSIIWAANTGQLY | 0.623464 |
| DPLAVPVAAGPV | ASSIIWAANTGQLY | 0.6300114 |
| GGTGGTPGNGGN | ASSIIWAANTGQLY | 0.6652379 |
| PAGLAGVDPATL | ASSIIWAANTGQLY | 0.7038424 |
| GNNLVVEAPHAGASV | ASSIIWAANTGQLY | 0.5545047 |
| KVTRIMSASTLEQPA | ASSIIWAANTGQLY | 0.7673369 |
| LGFALPIALPRRTVV | ASSIIWAANTGQLY | 0.837306 |
| TRLGNLTARKLVVLV | ASSIIWAANTGQLY | 0.66868126 |
| WSQQRRLIYFPSAGP | ASSIIWAANTGQLY | 0.3966739 |
| GPNGNPDPMAAAVDIR | ASSIIWAANTGQLY | 0.66268593 |
| TQRCVSVADLRAAASR | ASSIIWAANTGQLY | 0.5252795 |
| QFVLDVAEHPVEVLTETV | ASSIIWAANTGQLY | 0.62920487 |
| GGAGGTGGAGGNGGTQAGDG | ASSIIWAANTGQLY | 0.754506 |
| PLLATLRAIAVDVDAKPAQV | ASSIIWAANTGQLY | 0.36095983 |
| APKFRTGGYMIDFWGVGYQVAK | ASSIIWAANTGQLY | 0.6134571 |
| GAGGAGGAGGAAGGPGGTGGTG | ASSIIWAANTGQLY | 0.82471 |
| IGALGLLATDVAPAGSVAFGVA | ASSIIWAANTGQLY | 0.59283835 |
| AGGAGGIGGNGGAGGAGGVLMG | ASSIIWAANTGQLY | 0.7074197 |
| ADELIGLL | ASSIIWAANTGQLY | 0.7909732 |
| ADRALKA | ASSIIWAANTGQLY | 0.75270677 |

| | | |
|---|---|---|
| AGAVGQVGGAGGA | ASSIIWAANTGQLY | 0.81586456 |
| AGGGGGLGGAP | ASSIIWAANTGQLY | 0.79661435 |
| AGGLLYGNGGAG | ASSIIWAANTGQLY | 0.7519787 |
| AGLESIGL | ASSIIWAANTGQLY | 0.7072059 |
| AGLKTGII | ASSIIWAANTGQLY | 0.70396245 |
| ASLAALLL | ASSIIWAANTGQLY | 0.7333727 |
| ASVINISSI | ASSIIWAANTGQLY | 0.66008306 |
| ATPPGPA | ASSIIWAANTGQLY | 0.6056945 |
| ATVASALV | ASSIIWAANTGQLY | 0.783078 |
| ATVASLIL | ASSIIWAANTGQLY | 0.67631996 |
| AVNGPVLIRH | ASSIIWAANTGQLY | 0.7894807 |
| DGAAGAVGI | ASSIIWAANTGQLY | 0.5674203 |
| DPLLHGI | ASSIIWAANTGQLY | 0.61195576 |
| ELGAALGAL | ASSIIWAANTGQLY | 0.77966 |
| ELGRTQAV | ASSIIWAANTGQLY | 0.18622091 |
| FVAGGYLS | ASSIIWAANTGQLY | 0.29397723 |
| GASGGAGGAGGW | ASSIIWAANTGQLY | 0.69281596 |
| GGAGAGGDGGAGGI | ASSIIWAANTGQLY | 0.80500495 |
| GTGHNGSQPGMGGNG | ASSIIWAANTGQLY | 0.23270307 |
| GYAAASASAS | ASSIIWAANTGQLY | 0.5769358 |
| HQVDVAAVQQG | ASSIIWAANTGQLY | 0.46898204 |
| IGAAIAHT | ASSIIWAANTGQLY | 0.78176755 |
| IGMIEVAQL | ASSIIWAANTGQLY | 0.7066932 |
| IRLAPEHTSAMD | ASSIIWAANTGQLY | 0.35614428 |
| IYNVSVL | ASSIIWAANTGQLY | 0.78473717 |
| KQGFLGM | ASSIIWAANTGQLY | 0.35554093 |
| LESVITQL | ASSIIWAANTGQLY | 0.5477131 |
| LGLGALVVV | ASSIIWAANTGQLY | 0.8007498 |
| LIGNGGNGGSGGI | ASSIIWAANTGQLY | 0.82543504 |
| LNAINAPTL | ASSIIWAANTGQLY | 0.50601965 |
| LQLLNTGV | ASSIIWAANTGQLY | 0.78906167 |
| LQTSIEF | ASSIIWAANTGQLY | 0.56819886 |
| MSFAVSEL | ASSIIWAANTGQLY | 0.82874745 |
| NTGALGL | ASSIIWAANTGQLY | 0.65353435 |
| PPGAGGAGGAGTT | ASSIIWAANTGQLY | 0.3956646 |
| QLVGLGI | ASSIIWAANTGQLY | 0.7709806 |
| RFLEQVHQL | ASSIIWAANTGQLY | 0.7620963 |
| RGVGYVLR | ASSIIWAANTGQLY | 0.76723945 |
| SATGVFAGVI | ASSIIWAANTGQLY | 0.750256 |
| SLPAITLP | ASSIIWAANTGQLY | 0.6027739 |
| STALAFTV | ASSIIWAANTGQLY | 0.6376883 |
| SVLTVGLGK | ASSIIWAANTGQLY | 0.8037321 |
| TALAGAGSGI | ASSIIWAANTGQLY | 0.8022245 |
| TGLPFTGI | ASSIIWAANTGQLY | 0.58607244 |
| VAEHVTV | ASSIIWAANTGQLY | 0.80870986 |
| VDIVLNSL | ASSIIWAANTGQLY | 0.6848129 |
| VLTGGGAVL | ASSIIWAANTGQLY | 0.64786744 |
| VNVGLAYGI | ASSIIWAANTGQLY | 0.81885207 |
| VSSAAVGAV | ASSIIWAANTGQLY | 0.75904995 |
| VTSVYEAGV | ASSIIWAANTGQLY | 0.7825006 |
| WVALDRGA | ASSIIWAANTGQLY | 0.78035104 |

| | | |
|---|---|---|
| YLVGSVVI | ASSIIWAANTGQLY | 0.8074728 |
| GPPGTGKTLL | ASSIIWAANTGQLY | 0.50317645 |
| GVGMTKF | ASSIIWAANTGQLY | 0.77681196 |
| IGIGGSDL | ASSIIWAANTGQLY | 0.67644686 |
| IGTLLAEL | ASSIIWAANTGQLY | 0.80200917 |
| IYTSGTT | ASSIIWAANTGQLY | 0.7455378 |
| LSFPAGV | ASSIIWAANTGQLY | 0.54496384 |
| LYTSGTT | ASSIIWAANTGQLY | 0.7485498 |
| VGAFPFGF | ASSIIWAANTGQLY | 0.7997278 |
| VGAGTVEF | ASSIIWAANTGQLY | 0.73533857 |
| VGKSTLLL | ASSIIWAANTGQLY | 0.7803058 |
| AALAHAGGGIV | ASSIIWAANTGQLY | 0.8247754 |
| AALVAVSLL | ASSIIWAANTGQLY | 0.63042516 |
| AAVSRAVAAL | ASSIIWAANTGQLY | 0.68491364 |
| AGGSSGIGGF | ASSIIWAANTGQLY | 0.8022306 |
| AGLTVANANG | ASSIIWAANTGQLY | 0.82284755 |
| AGLVGAVL | ASSIIWAANTGQLY | 0.6852986 |
| ALSQAFTV | ASSIIWAANTGQLY | 0.8241112 |
| ALTFSVSL | ASSIIWAANTGQLY | 0.54318935 |
| ALTGAALVAL | ASSIIWAANTGQLY | 0.7923783 |
| ASFSIGAGVL | ASSIIWAANTGQLY | 0.52746993 |
| DGSVYKI | ASSIIWAANTGQLY | 0.7416456 |
| EQLAEIV | ASSIIWAANTGQLY | 0.5508121 |
| FAVSELTVM | ASSIIWAANTGQLY | 0.7805021 |
| GAAPLAAGL | ASSIIWAANTGQLY | 0.7769114 |
| GADGGDGGSGGAGG | ASSIIWAANTGQLY | 0.79493934 |
| GFGLDTGGAGG | ASSIIWAANTGQLY | 0.3200769 |
| GLASGGLL | ASSIIWAANTGQLY | 0.79004705 |
| GRIAGAAVLL | ASSIIWAANTGQLY | 0.797729 |
| GSAGSLNTSF | ASSIIWAANTGQLY | 0.8355998 |
| GTGQIGI | ASSIIWAANTGQLY | 0.69875246 |
| GTLIALIL | ASSIIWAANTGQLY | 0.7401234 |
| IAATMGFVVG | ASSIIWAANTGQLY | 0.6905699 |
| IFAQALL | ASSIIWAANTGQLY | 0.67946434 |
| IGAAVALGF | ASSIIWAANTGQLY | 0.47843504 |
| IGLTPLAL | ASSIIWAANTGQLY | 0.6729274 |
| ILGSNTSTL | ASSIIWAANTGQLY | 0.5920893 |
| IVFSLTVL | ASSIIWAANTGQLY | 0.56775606 |
| LASKATGF | ASSIIWAANTGQLY | 0.7789853 |
| LASVALIL | ASSIIWAANTGQLY | 0.821478 |
| LFAVGLTL | ASSIIWAANTGQLY | 0.78953946 |
| LFNGGIGTYI | ASSIIWAANTGQLY | 0.77307665 |
| LTLGAGII | ASSIIWAANTGQLY | 0.80177605 |
| LVAAALGDI | ASSIIWAANTGQLY | 0.8217843 |
| NTAFVQLGI | ASSIIWAANTGQLY | 0.64050955 |
| QLFSGAEL | ASSIIWAANTGQLY | 0.6810634 |
| SAATFITL | ASSIIWAANTGQLY | 0.52921844 |
| SNYGLANLGV | ASSIIWAANTGQLY | 0.5909637 |
| SYGGAYAVM | ASSIIWAANTGQLY | 0.6383537 |
| TTGVGGLTLG | ASSIIWAANTGQLY | 0.7275034 |
| TTQTIYI | ASSIIWAANTGQLY | 0.18686022 |

| | | |
|---|---|---|
| VAAGAGLGAVI | ASSIIWAANTGQLY | 0.6652174 |
| VAAVVTLL | ASSIIWAANTGQLY | 0.8260723 |
| VGFLGGLI | ASSIIWAANTGQLY | 0.5474317 |
| VGVIPILL | ASSIIWAANTGQLY | 0.7306587 |
| VIFASSLI | ASSIIWAANTGQLY | 0.6659363 |
| VKGGDGGAAAG | ASSIIWAANTGQLY | 0.60631216 |
| VTKTFTNI | ASSIIWAANTGQLY | 0.75479805 |
| VVAAAILL | ASSIIWAANTGQLY | 0.5931948 |
| YAALDVEL | ASSIIWAANTGQLY | 0.7771609 |
| LASVALLL | ASSIIWAANTGQLY | 0.8174103 |
| LDAVHRV | ASSIIWAANTGQLY | 0.69094396 |
| QGLSSSIF | ASSIIWAANTGQLY | 0.7691785 |
| RGTFANIRL | ASSIIWAANTGQLY | 0.69634974 |
| RQLWWGHRI | ASSIIWAANTGQLY | 0.4727336 |
| YGPPGTGKTLL | ASSIIWAANTGQLY | 0.7748759 |
| AALVGAPANVA | ASSIIWAANTGQLY | 0.42449287 |
| AATFRRTE | ASSIIWAANTGQLY | 0.6550609 |
| AGALGGGI | ASSIIWAANTGQLY | 0.5595399 |
| AGGAGGAGGIGAG | ASSIIWAANTGQLY | 0.7218489 |
| AGSGALGL | ASSIIWAANTGQLY | 0.71691644 |
| AQIADQI | ASSIIWAANTGQLY | 0.7611948 |
| ASAVAVTYL | ASSIIWAANTGQLY | 0.5089412 |
| ASTVLGGTGG | ASSIIWAANTGQLY | 0.665932 |
| AVGGEPAK | ASSIIWAANTGQLY | 0.59156084 |
| CEQGTFGGTC | ASSIIWAANTGQLY | 0.49582362 |
| DEGKAPAG | ASSIIWAANTGQLY | 0.7897758 |
| DTVVFPAL | ASSIIWAANTGQLY | 0.7030422 |
| FEGITMAG | ASSIIWAANTGQLY | 0.6332434 |
| FPLVVQD | ASSIIWAANTGQLY | 0.80019116 |
| GAFTTPQ | ASSIIWAANTGQLY | 0.7623246 |
| GFGGFGTAGGVGGA | ASSIIWAANTGQLY | 0.44062668 |
| GFLAQFL | ASSIIWAANTGQLY | 0.62766814 |
| GGAGGAGGNAGVA | ASSIIWAANTGQLY | 0.6543864 |
| GGVGTAGAGGAGGAGG | ASSIIWAANTGQLY | 0.6766218 |
| GNGGGGDAGNAGSGGN | ASSIIWAANTGQLY | 0.22813763 |
| GTGGAGGAGNDAGS | ASSIIWAANTGQLY | 0.775765 |
| GTTNTGWFNS | ASSIIWAANTGQLY | 0.6582605 |
| IFMGPLFTL | ASSIIWAANTGQLY | 0.46426392 |
| ILAHAIEAI | ASSIIWAANTGQLY | 0.599085 |
| IRSAIREAL | ASSIIWAANTGQLY | 0.77245045 |
| ITFAGGG | ASSIIWAANTGQLY | 0.6089896 |
| ITGMSGHQEV | ASSIIWAANTGQLY | 0.7416966 |
| IVAGLACTL | ASSIIWAANTGQLY | 0.796288 |
| IVGGGLAGA | ASSIIWAANTGQLY | 0.81497717 |
| IVTGVLNI | ASSIIWAANTGQLY | 0.82042086 |
| LAQQGCI | ASSIIWAANTGQLY | 0.7776903 |
| LCLHVEN | ASSIIWAANTGQLY | 0.6967011 |
| LDAISDGS | ASSIIWAANTGQLY | 0.77996576 |
| LGGFAAML | ASSIIWAANTGQLY | 0.7642463 |
| LGGKNPMI | ASSIIWAANTGQLY | 0.40127838 |
| LGPKGVAAIL | ASSIIWAANTGQLY | 0.31439674 |

| | | |
|---|---|---|
| LMVGVSI | ASSIIWAANTGQLY | 0.6789708 |
| LVGAAFVL | ASSIIWAANTGQLY | 0.37652406 |
| LVLAVAPL | ASSIIWAANTGQLY | 0.7081845 |
| MAAIAVIDAA | ASSIIWAANTGQLY | 0.82773423 |
| NNGINAELQ | ASSIIWAANTGQLY | 0.607877 |
| RGEPLVLVHG | ASSIIWAANTGQLY | 0.54861265 |
| RGGAGGAAGLI | ASSIIWAANTGQLY | 0.6470613 |
| RSYRMHPAV | ASSIIWAANTGQLY | 0.6105077 |
| SAAHLLAGQLA | ASSIIWAANTGQLY | 0.79558456 |
| SAAVGSY | ASSIIWAANTGQLY | 0.76910573 |
| SPHAVLHRI | ASSIIWAANTGQLY | 0.829486 |
| SQIAKVL | ASSIIWAANTGQLY | 0.800214 |
| TALGHLVP | ASSIIWAANTGQLY | 0.62967694 |
| TAPTTSGAA | ASSIIWAANTGQLY | 0.7703604 |
| TDGDDGNGGSPGT | ASSIIWAANTGQLY | 0.6219577 |
| TDGLGEN | ASSIIWAANTGQLY | 0.6989163 |
| TGGPPLPGQA | ASSIIWAANTGQLY | 0.7290036 |
| TIPAITI | ASSIIWAANTGQLY | 0.75951636 |
| TIPGFTI | ASSIIWAANTGQLY | 0.7566384 |
| TSIASGALA | ASSIIWAANTGQLY | 0.7123395 |
| TVTLPGAAI | ASSIIWAANTGQLY | 0.41839975 |
| VASGLSAALP | ASSIIWAANTGQLY | 0.7998557 |
| VGAGVIGI | ASSIIWAANTGQLY | 0.71592915 |
| VGPKRFGV | ASSIIWAANTGQLY | 0.3403475 |
| VNVAFPD | ASSIIWAANTGQLY | 0.69963384 |
| VSMTNSGVS | ASSIIWAANTGQLY | 0.82887775 |
| VTDVLGDLSV | ASSIIWAANTGQLY | 0.8168069 |
| VVVVGAGFAGL | ASSIIWAANTGQLY | 0.5181719 |
| VYTVGTVV | ASSIIWAANTGQLY | 0.58126086 |
| WTTFSGP | ASSIIWAANTGQLY | 0.6685405 |
| YDELVEAF | ASSIIWAANTGQLY | 0.31868383 |
| YGTGGAGGNGG | ASSIIWAANTGQLY | 0.7497127 |
| YTNVMAV | ASSIIWAANTGQLY | 0.6745895 |
| FLEQVHQL | ASSIIWAANTGQLY | 0.65191394 |
| GLSLLAVAV | ASSIIWAANTGQLY | 0.5486399 |
| LEQVHQL | ASSIIWAANTGQLY | 0.68542826 |
| LNAAGAI | ASSIIWAANTGQLY | 0.6450638 |
| PAVHASL | ASSIIWAANTGQLY | 0.8082687 |
| TELEDDA | ASSIIWAANTGQLY | 0.20382476 |
| TPPGPPN | ASSIIWAANTGQLY | 0.77323794 |
| TPPPPPPP | ASSIIWAANTGQLY | 0.54524523 |
| TQSPALI | ASRRDSSYEQY | 0.5761979 |
| EFAAAFPQ | ASRRDSSYEQY | 0.4363332 |
| LLSGNLDVL | ASRRDSSYEQY | 0.73064154 |
| GPANLGNYNI | ASRRDSSYEQY | 0.7607684 |
| TLGPTLP | ASRRDSSYEQY | 0.7866251 |
| VYAYSSCCK | ASRRDSSYEQY | 0.5443552 |
| APAPAPAPNI | ASRRDSSYEQY | 0.80048907 |
| PGGPVPIP | ASRRDSSYEQY | 0.82285976 |
| FLTVALEP | ASRRDSSYEQY | 0.62157434 |
| SQYFGANL | ASRRDSSYEQY | 0.49911326 |

| | | |
|---|---|---|
| GGVKPAA | ASRRDSSYEQY | 0.76161766 |
| GGTGGAGGAGGW | ASRRDSSYEQY | 0.698945 |
| GVGGVAGLGGAGG | ASRRDSSYEQY | 0.7877487 |
| VLLVLATTI | ASRRDSSYEQY | 0.7905367 |
| IGIFAAML | ASRRDSSYEQY | 0.45459202 |
| AVGDLID | ASRRDSSYEQY | 0.7714492 |
| NISPLPL | ASRRDSSYEQY | 0.79064554 |
| VTAVFVAI | ASRRDSSYEQY | 0.54556656 |
| SGDDNDAAL | ASRRDSSYEQY | 0.6894356 |
| AVAALAHS | ASRRDSSYEQY | 0.6914376 |
| GGVTVMATP | ASRRDSSYEQY | 0.43150198 |
| AGLAANVRD | ASRRDSSYEQY | 0.802243 |
| TNRPDLIDPALLRPG | ASRRDSSYEQY | 0.54851234 |
| LANLGVTAL | ASRRDSSYEQY | 0.6193027 |
| TGERIEL | ASRRDSSYEQY | 0.70214415 |
| NGEELAA | ASRRDSSYEQY | 0.6048206 |
| VLSLVWG | ASRRDSSYEQY | 0.68339294 |
| IGYALWL | ASRRDSSYEQY | 0.6844144 |
| LSLLLAPL | ASRRDSSYEQY | 0.6959089 |
| TLDEALK | ASRRDSSYEQY | 0.7124913 |
| TDGAGTAEGG | ASRRDSSYEQY | 0.6363977 |
| NAPGGAGGNGG | ASRRDSSYEQY | 0.7090627 |
| VSVFMGMGGNF | ASRRDSSYEQY | 0.8043915 |
| LASYLAAL | ASRRDSSYEQY | 0.45925453 |
| LVGGTPAL | ASRRDSSYEQY | 0.579473 |
| VRNADVL | ASRRDSSYEQY | 0.7484019 |
| TCSSPFLVSAM | ASRRDSSYEQY | 0.4198434 |
| VHITDAE | ASRRDSSYEQY | 0.7529754 |
| EYAAPSF | ASRRDSSYEQY | 0.72883224 |
| VVDAFHGGL | ASRRDSSYEQY | 0.73090786 |
| ASVVDPQ | ASRRDSSYEQY | 0.6344078 |
| SYNEFGGLS | ASRRDSSYEQY | 0.83805287 |
| DPDGSVY | ASRRDSSYEQY | 0.7929164 |
| MVSTIHATE | ASRRDSSYEQY | 0.7104475 |
| LAAGVADPV | ASRRDSSYEQY | 0.7636105 |
| TKEQIAA | ASRRDSSYEQY | 0.6501803 |
| SLASPAAL | ASRRDSSYEQY | 0.8062759 |
| IGLVFYGV | ASRRDSSYEQY | 0.7803314 |
| FLLDASNL | ASRRDSSYEQY | 0.47217065 |
| YQVGNFP | ASRRDSSYEQY | 0.64431727 |
| ESLLSQST | ASRRDSSYEQY | 0.8095636 |
| LFTVGFAAVG | ASRRDSSYEQY | 0.8016311 |
| ATPPTPT | ASRRDSSYEQY | 0.37815252 |
| TDLATLG | ASRRDSSYEQY | 0.6270058 |
| SAGGTGR | ASRRDSSYEQY | 0.35813046 |
| GFSNSGS | ASRRDSSYEQY | 0.60315585 |
| GGNGGSGGTSV | ASRRDSSYEQY | 0.61973816 |
| GTSVATGGAGN | ASRRDSSYEQY | 0.73215103 |
| NGGNGGSGGTS | ASRRDSSYEQY | 0.6274374 |
| AVVKFPH | ASRRDSSYEQY | 0.75325626 |
| TPPPPPP | ASRRDSSYEQY | 0.7276117 |

| | | |
|---|---|---|
| AAPAAAAAPA | ASRRDSSYEQY | 0.8145958 |
| IALNEDL | ASRRDSSYEQY | 0.56522477 |
| IVNVSSM | ASRRDSSYEQY | 0.72099656 |
| TGIPPAPR | ASRRDSSYEQY | 0.51237583 |
| FDLGGGTF | ASRRDSSYEQY | 0.5431733 |
| FDLGGGTFDVS | ASRRDSSYEQY | 0.5956614 |
| NEPTAAA | ASRRDSSYEQY | 0.6719151 |
| NEPTAAALA | ASRRDSSYEQY | 0.3010285 |
| GSIINISSI | ASRRDSSYEQY | 0.7990011 |
| VAPGNAGTA | ASRRDSSYEQY | 0.7422328 |
| TGGGASLE | ASRRDSSYEQY | 0.7840818 |
| VAVYNHYKRI | ASRRDSSYEQY | 0.6733495 |
| TDPAPATT | ASRRDSSYEQY | 0.73610276 |
| TNRPDILDPALLRPG | ASRRDSSYEQY | 0.6316598 |
| LTTTTTS | ASRRDSSYEQY | 0.7549994 |
| PTGSGKTL | ASRRDSSYEQY | 0.5573398 |
| TGSGKTL | ASRRDSSYEQY | 0.41216284 |
| TIGHVDHGK | ASRRDSSYEQY | 0.62517583 |
| TIGHVDHGKTT | ASRRDSSYEQY | 0.46555227 |
| GHVDHGKTT | ASRRDSSYEQY | 0.8103821 |
| AQFEHTLL | ASRRDSSYEQY | 0.7808458 |
| IVNQDEL | ASRRDSSYEQY | 0.7263633 |
| TNGSQFF | ASRRDSSYEQY | 0.796618 |
| TGKGIVE | ASRRDSSYEQY | 0.2963777 |
| LISNASDA | ASRRDSSYEQY | 0.63963926 |
| AELRTLGELTKIAKAHGAQ | ASRRDSSYEQY | 0.8197185 |
| APNAVAAAEPSW | ASRRDSSYEQY | 0.7388031 |
| APQVAGAATAVL | ASRRDSSYEQY | 0.6546496 |
| ARRDPAAAAAILPTDA | ASRRDSSYEQY | 0.81863046 |
| DAAARDVLAEAGLAEYF | ASRRDSSYEQY | 0.62382215 |
| DDQIVLSAKANLAKVEQ | ASRRDSSYEQY | 0.67093843 |
| DELLPDTGEDSDEDGH | ASRRDSSYEQY | 0.7365643 |
| ELPEAPANRATVPQD | ASRRDSSYEQY | 0.78860396 |
| FGQSKIERVTGIG | ASRRDSSYEQY | 0.7385838 |
| GAGGTGSTAGGAGGAGGAGGLY | ASRRDSSYEQY | 0.6431214 |
| GHGGTGAAGGNGGTGGAGGSGI | ASRRDSSYEQY | 0.72950673 |
| GNGGNGGAGGEGGPGGAGGAGA | ASRRDSSYEQY | 0.8191996 |
| GSTAGNGGNGGNATNGTIAGQP | ASRRDSSYEQY | 0.6607137 |
| GTGGDGGNGGNGGTGG | ASRRDSSYEQY | 0.78026736 |
| GVRRPAGIPADAPV | ASRRDSSYEQY | 0.7539889 |
| GYRQVTDAHLLTLARR | ASRRDSSYEQY | 0.69683486 |
| IAKMGQDGHDRG | ASRRDSSYEQY | 0.53061444 |
| KAEPSIEIQYT | ASRRDSSYEQY | 0.6229655 |
| KPVRIRMAADEDTVAVT | ASRRDSSYEQY | 0.6801513 |
| MPPRIAGMRLLVIKPEP | ASRRDSSYEQY | 0.5125924 |
| MKFYAVALFRSRGVPPTR | ASRRDSSYEQY | 0.67676747 |
| NAGGFGILGAVAHSP | ASRRDSSYEQY | 0.6376327 |
| NEGREITLAMAEM | ASRRDSSYEQY | 0.8373307 |
| NELVEWSHPLGHRWRWE | ASRRDSSYEQY | 0.76123124 |
| NLGAGGTGGNGGNPG | ASRRDSSYEQY | 0.7673778 |
| NTEWPKWETVMAADPVD | ASRRDSSYEQY | 0.56157136 |

| | | |
|---|---|---|
| PASAAPAETQP | ASRRDSSYEQY | 0.7222663 |
| QQSSDGKNGVRAAEITGE | ASRRDSSYEQY | 0.7243049 |
| RAPDAVTTSWWKE | ASRRDSSYEQY | 0.45604402 |
| SGFGNVGAHTSGWFNQ | ASRRDSSYEQY | 0.81766653 |
| SGVAGAVGQAASVGGLKV | ASRRDSSYEQY | 0.6094957 |
| SNGGAGTGSAGGNGGT | ASRRDSSYEQY | 0.5848563 |
| TGGAGGEGGAAG | ASRRDSSYEQY | 0.7817888 |
| TGGDGGAAGAGGAPGHG | ASRRDSSYEQY | 0.8162076 |
| TMPPLLAAIANAGRHGV | ASRRDSSYEQY | 0.6596995 |
| TSSTAGNSGGA | ASRRDSSYEQY | 0.6400422 |
| VPATVPVGILC | ASRRDSSYEQY | 0.5801289 |
| VQRPDSGGVVLTGRIS | ASRRDSSYEQY | 0.7664498 |
| WTPNPVPATKGVPAGLVQAT | ASRRDSSYEQY | 0.80434895 |
| YALKRALAKFSDRFDVVII | ASRRDSSYEQY | 0.6755712 |
| YLIGSVIV | ASRRDSSYEQY | 0.8308124 |
| YGQLVAVID | ASRRDSSYEQY | 0.79049635 |
| VTVGGMTLPA | ASRRDSSYEQY | 0.49002188 |
| VLALPAV | ASRRDSSYEQY | 0.71861154 |
| VIPPTLNYET | ASRRDSSYEQY | 0.7873175 |
| VGMGGMHPSSG | ASRRDSSYEQY | 0.77170783 |
| VGLAGGAATA | ASRRDSSYEQY | 0.8179035 |
| VGGAGGAGGSGP | ASRRDSSYEQY | 0.8067141 |
| VFAAFMV | ASRRDSSYEQY | 0.59359825 |
| TLYDKLR | ASRRDSSYEQY | 0.79499876 |
| TLAVFTVL | ASRRDSSYEQY | 0.82081574 |
| TGSVQPARV | ASRRDSSYEQY | 0.7928337 |
| TGGGAGSGGT | ASRRDSSYEQY | 0.6302029 |
| SHLETSL | ASRRDSSYEQY | 0.5614396 |
| SFTTGMT | ASRRDSSYEQY | 0.72042435 |
| QVEVAKRMGVRQPTVS | ASRRDSSYEQY | 0.7915513 |
| QGSGGPALAM | ASRRDSSYEQY | 0.5485184 |
| MLAPSNP | ASRRDSSYEQY | 0.709143 |
| LVNNAAI | ASRRDSSYEQY | 0.7269272 |
| LTGEINTLA | ASRRDSSYEQY | 0.50519544 |
| LSIFGPV | ASRRDSSYEQY | 0.75322586 |
| LLALLANL | ASRRDSSYEQY | 0.8165708 |
| LISNASDALDKLR | ASRRDSSYEQY | 0.69164574 |
| LINEAVP | ASRRDSSYEQY | 0.5869379 |
| LGYAFSGL | ASRRDSSYEQY | 0.7994443 |
| LDFVAVAL | ASRRDSSYEQY | 0.84196454 |
| LAQPGAE | ASRRDSSYEQY | 0.81034243 |
| ISNLGAVTGF | ASRRDSSYEQY | 0.4772491 |
| HQEDPGRI | ASRRDSSYEQY | 0.24439767 |
| GTPGEPTP | ASRRDSSYEQY | 0.6141851 |
| GFFNSGS | ASRRDSSYEQY | 0.7689352 |
| GAGGGGLTAGAN | ASRRDSSYEQY | 0.6980921 |
| FVDALVNM | ASRRDSSYEQY | 0.6613445 |
| FLTVAPEL | ASRRDSSYEQY | 0.732028 |
| FETLTAL | ASRRDSSYEQY | 0.7843582 |
| DVINPIEL | ASRRDSSYEQY | 0.5991061 |
| APVSGAPV | ASRRDSSYEQY | 0.51725143 |

| | | |
|---|---|---|
| ALVGGLAW | ASRRDSSYEQY | 0.7439024 |
| ALAAAAATAI | ASRRDSSYEQY | 0.7697573 |
| AFGLGLLL | ASRRDSSYEQY | 0.5184252 |
| AFFAAASL | ASRRDSSYEQY | 0.65795565 |
| AAAALPAFSPP | ASRRDSSYEQY | 0.59005785 |
| VVLAPAL | ASRRDSSYEQY | 0.58171403 |
| TIGHVDHGKTTL | ASRRDSSYEQY | 0.6960759 |
| LISNASDALDK | ASRRDSSYEQY | 0.46132472 |
| LISNASDAL | ASRRDSSYEQY | 0.8096024 |
| ISNASDALDK | ASRRDSSYEQY | 0.5494448 |
| NQAETLVYQTE | ASRRDSSYEQY | 0.78383505 |
| TGGTGGTAGAG | ASRRDSSYEQY | 0.7783841 |
| TGSTAGGAGGA | ASRRDSSYEQY | 0.76534367 |
| DPLAVPVAAGPV | ASRRDSSYEQY | 0.6433185 |
| GGTGGTPGNGGN | ASRRDSSYEQY | 0.56086516 |
| PAGLAGVDPATL | ASRRDSSYEQY | 0.4557873 |
| GNNLVVEAPHAGASV | ASRRDSSYEQY | 0.295998 |
| KVTRIMSASTLEQPA | ASRRDSSYEQY | 0.82233584 |
| LGFALPIALPRRTVV | ASRRDSSYEQY | 0.54346275 |
| TRLGNLTARKLVVLV | ASRRDSSYEQY | 0.8217579 |
| WSQQRRLIYFPSAGP | ASRRDSSYEQY | 0.78834844 |
| GPNGNPDPMAAAVDIR | ASRRDSSYEQY | 0.6059155 |
| TQRCVSVADLRAAASR | ASRRDSSYEQY | 0.75249135 |
| QFVLDVAEHPVEVLTETV | ASRRDSSYEQY | 0.5938864 |
| GGAGGTGGAGGNGGTQAGDG | ASRRDSSYEQY | 0.5812948 |
| PLLATLRAIAVDVDAKPAQV | ASRRDSSYEQY | 0.7567869 |
| APKFRTGGYMIDFWGVGYQVAK | ASRRDSSYEQY | 0.7617959 |
| GAGGAGGAGGAAGGPGGTGGTG | ASRRDSSYEQY | 0.5191137 |
| IGALGLLATDVAPAGSVAFGVA | ASRRDSSYEQY | 0.7480703 |
| AGGAGGIGGNGGAGGAGGVLMG | ASRRDSSYEQY | 0.6980431 |
| ADELIGLL | ASRRDSSYEQY | 0.5146909 |
| ADRALKA | ASRRDSSYEQY | 0.68849 |
| AGAVGQVGGAGGA | ASRRDSSYEQY | 0.7993345 |
| AGGGGGLGGAP | ASRRDSSYEQY | 0.754465 |
| AGGLLYGNGGAG | ASRRDSSYEQY | 0.5848567 |
| AGLESIGL | ASRRDSSYEQY | 0.77066344 |
| AGLKTGII | ASRRDSSYEQY | 0.50632274 |
| ASLAALLL | ASRRDSSYEQY | 0.7170917 |
| ASVINISSI | ASRRDSSYEQY | 0.74650204 |
| ATPPGPA | ASRRDSSYEQY | 0.67330223 |
| ATVASALV | ASRRDSSYEQY | 0.76908755 |
| ATVASLIL | ASRRDSSYEQY | 0.58332896 |
| AVNGPVLIRH | ASRRDSSYEQY | 0.5379481 |
| DGAAGAVGI | ASRRDSSYEQY | 0.6773282 |
| DPLLHGI | ASRRDSSYEQY | 0.7416957 |
| ELGAALGAL | ASRRDSSYEQY | 0.7794564 |
| ELGRTQAV | ASRRDSSYEQY | 0.5401113 |
| FVAGGYLS | ASRRDSSYEQY | 0.6847778 |
| GASGGAGGAGGW | ASRRDSSYEQY | 0.63688576 |
| GGAGAGGDGGAGGI | ASRRDSSYEQY | 0.53500926 |
| GTGHNGSQPGMGGNG | ASRRDSSYEQY | 0.7225441 |

| | | |
|---|---|---|
| GYAAASASAS | ASRRDSSYEQY | 0.70501626 |
| HQVDVAAVQQG | ASRRDSSYEQY | 0.58749205 |
| IGAAIAHT | ASRRDSSYEQY | 0.793273 |
| IGMIEVAQL | ASRRDSSYEQY | 0.6521886 |
| IRLAPEHTSAMD | ASRRDSSYEQY | 0.75324416 |
| IYNVSVL | ASRRDSSYEQY | 0.81931394 |
| KQGFLGM | ASRRDSSYEQY | 0.71219075 |
| LESVITQL | ASRRDSSYEQY | 0.7744343 |
| LGLGALVVV | ASRRDSSYEQY | 0.7553165 |
| LIGNGGNGGSGGI | ASRRDSSYEQY | 0.7863765 |
| LNAINAPTL | ASRRDSSYEQY | 0.68748045 |
| LQLLNTGV | ASRRDSSYEQY | 0.5715221 |
| LQTSIEF | ASRRDSSYEQY | 0.79514694 |
| MSFAVSEL | ASRRDSSYEQY | 0.7690102 |
| NTGALGL | ASRRDSSYEQY | 0.7974303 |
| PPGAGGAGGAGTT | ASRRDSSYEQY | 0.798875 |
| QLVGLGI | ASRRDSSYEQY | 0.7694942 |
| RFLEQVHQL | ASRRDSSYEQY | 0.76898575 |
| RGVGYVLR | ASRRDSSYEQY | 0.64443123 |
| SATGVFAGVI | ASRRDSSYEQY | 0.5013146 |
| SLPAITLP | ASRRDSSYEQY | 0.67773366 |
| STALAFTV | ASRRDSSYEQY | 0.6303299 |
| SVLTVGLGK | ASRRDSSYEQY | 0.703301 |
| TALAGAGSGI | ASRRDSSYEQY | 0.37697214 |
| TGLPFTGI | ASRRDSSYEQY | 0.7933501 |
| VAEHVTV | ASRRDSSYEQY | 0.839713 |
| VDIVLNSL | ASRRDSSYEQY | 0.6082626 |
| VLTGGGAVL | ASRRDSSYEQY | 0.8168392 |
| VNVGLAYGI | ASRRDSSYEQY | 0.44956645 |
| VSSAAVGAV | ASRRDSSYEQY | 0.76276624 |
| VTSVYEAGV | ASRRDSSYEQY | 0.8038418 |
| WVALDRGA | ASRRDSSYEQY | 0.5534549 |
| YLVGSVVI | ASRRDSSYEQY | 0.7803194 |
| GPPGTGKTLL | ASRRDSSYEQY | 0.6080044 |
| GVGMTKF | ASRRDSSYEQY | 0.65239286 |
| IGIGGSDL | ASRRDSSYEQY | 0.82505965 |
| IGTLLAEL | ASRRDSSYEQY | 0.6716461 |
| IYTSGTT | ASRRDSSYEQY | 0.75689936 |
| LSFPAGV | ASRRDSSYEQY | 0.7319103 |
| LYTSGTT | ASRRDSSYEQY | 0.6357547 |
| VGAFPFGF | ASRRDSSYEQY | 0.7610454 |
| VGAGTVEF | ASRRDSSYEQY | 0.8315219 |
| VGKSTLLL | ASRRDSSYEQY | 0.54718703 |
| AALAHAGGGIV | ASRRDSSYEQY | 0.8155016 |
| AALVAVSLL | ASRRDSSYEQY | 0.68220603 |
| AAVSRAVAAL | ASRRDSSYEQY | 0.8232502 |
| AGGSSGIGGF | ASRRDSSYEQY | 0.76169 |
| AGLTVANANG | ASRRDSSYEQY | 0.6832816 |
| AGLVGAVL | ASRRDSSYEQY | 0.663678 |
| ALSQAFTV | ASRRDSSYEQY | 0.79697466 |
| ALTFSVSL | ASRRDSSYEQY | 0.6663748 |

| | | |
|---|---|---|
| ALTGAALVAL | ASRRDSSYEQY | 0.8042687 |
| ASFSIGAGVL | ASRRDSSYEQY | 0.8282738 |
| DGSVYKI | ASRRDSSYEQY | 0.33777612 |
| EQLAEIV | ASRRDSSYEQY | 0.86483777 |
| FAVSELTVM | ASRRDSSYEQY | 0.7573219 |
| GAAPLAAGL | ASRRDSSYEQY | 0.81037164 |
| GADGGDGGSGGAGG | ASRRDSSYEQY | 0.5194402 |
| GFGLDTGGAGG | ASRRDSSYEQY | 0.26047796 |
| GLASGGLL | ASRRDSSYEQY | 0.6362191 |
| GRIAGAAVLL | ASRRDSSYEQY | 0.43492556 |
| GSAGSLNTSF | ASRRDSSYEQY | 0.8381773 |
| GTGQIGI | ASRRDSSYEQY | 0.8328021 |
| GTLIALIL | ASRRDSSYEQY | 0.5224087 |
| IAATMGFVVG | ASRRDSSYEQY | 0.84491116 |
| IFAQALL | ASRRDSSYEQY | 0.8719777 |
| IGAAVALGF | ASRRDSSYEQY | 0.29569954 |
| IGLTPLAL | ASRRDSSYEQY | 0.6811501 |
| ILGSNTSTL | ASRRDSSYEQY | 0.37646455 |
| IVFSLTVL | ASRRDSSYEQY | 0.6102469 |
| LASKATGF | ASRRDSSYEQY | 0.19463447 |
| LASVALIL | ASRRDSSYEQY | 0.8131013 |
| LFAVGLTL | ASRRDSSYEQY | 0.525408 |
| LFNGGIGTYI | ASRRDSSYEQY | 0.18652898 |
| LTLGAGII | ASRRDSSYEQY | 0.7829019 |
| LVAAALGDI | ASRRDSSYEQY | 0.5834547 |
| NTAFVQLGI | ASRRDSSYEQY | 0.48911083 |
| QLFSGAEL | ASRRDSSYEQY | 0.82490253 |
| SAATFITL | ASRRDSSYEQY | 0.6655787 |
| SNYGLANLGV | ASRRDSSYEQY | 0.39952385 |
| SYGGAYAVM | ASRRDSSYEQY | 0.8256527 |
| TTGVGGLTLG | ASRRDSSYEQY | 0.8682904 |
| TTQTIYI | ASRRDSSYEQY | 0.36247087 |
| VAAGAGLGAVI | ASRRDSSYEQY | 0.74478114 |
| VAAVVTLL | ASRRDSSYEQY | 0.8798182 |
| VGFLGGLI | ASRRDSSYEQY | 0.7066676 |
| VGVIPILL | ASRRDSSYEQY | 0.8585885 |
| VIFASSLI | ASRRDSSYEQY | 0.872115 |
| VKGGDGGAAAG | ASRRDSSYEQY | 0.6994458 |
| VTKTFTNI | ASRRDSSYEQY | 0.8592027 |
| VVAAAILL | ASRRDSSYEQY | 0.8580556 |
| YAALDVEL | ASRRDSSYEQY | 0.6115184 |
| LASVALLL | ASRRDSSYEQY | 0.726491 |
| LDAVHRV | ASRRDSSYEQY | 0.3427028 |
| QGLSSSIF | ASRRDSSYEQY | 0.757836 |
| RGTFANIRL | ASRRDSSYEQY | 0.3340205 |
| RQLWWGHRI | ASRRDSSYEQY | 0.77681935 |
| YGPPGTGKTLL | ASRRDSSYEQY | 0.847556 |
| AALVGAPANVA | ASRRDSSYEQY | 0.67995685 |
| AATFRRTE | ASRRDSSYEQY | 0.38569403 |
| AGALGGGI | ASRRDSSYEQY | 0.33507875 |
| AGGAGGAGGIGAG | ASRRDSSYEQY | 0.76809716 |

| | | |
|---|---|---|
| AGSGALGL | ASRRDSSYEQY | 0.3709414 |
| AQIADQI | ASRRDSSYEQY | 0.3488709 |
| ASAVAVTYL | ASRRDSSYEQY | 0.58522105 |
| ASTVLGGTGG | ASRRDSSYEQY | 0.85380304 |
| AVGGEPAK | ASRRDSSYEQY | 0.82303387 |
| CEQGTFGGTC | ASRRDSSYEQY | 0.54355085 |
| DEGKAPAG | ASRRDSSYEQY | 0.87534404 |
| DTVVFPAL | ASRRDSSYEQY | 0.81752074 |
| FEGITMAG | ASRRDSSYEQY | 0.8581244 |
| FPLVVQD | ASRRDSSYEQY | 0.63076836 |
| GAFTTPQ | ASRRDSSYEQY | 0.33145586 |
| GFGGFGTAGGVGGA | ASRRDSSYEQY | 0.84996104 |
| GFLAQFL | ASRRDSSYEQY | 0.6817252 |
| GGAGGAGGNAGVA | ASRRDSSYEQY | 0.7999234 |
| GGVGTAGAGGAGGAGG | ASRRDSSYEQY | 0.84723794 |
| GNGGGGDAGNAGSGGN | ASRRDSSYEQY | 0.59175706 |
| GTGGAGGAGNDAGS | ASRRDSSYEQY | 0.49420267 |
| GTTNTGWFNS | ASRRDSSYEQY | 0.7276851 |
| IFMGPLFTL | ASRRDSSYEQY | 0.85212165 |
| ILAHAIEAI | ASRRDSSYEQY | 0.51987433 |
| IRSAIREAL | ASRRDSSYEQY | 0.31163603 |
| ITFAGGG | ASRRDSSYEQY | 0.6846285 |
| ITGMSGHQEV | ASRRDSSYEQY | 0.6590916 |
| IVAGLACTL | ASRRDSSYEQY | 0.86317444 |
| IVGGGLAGA | ASRRDSSYEQY | 0.73745966 |
| IVTGVLNI | ASRRDSSYEQY | 0.63527715 |
| LAQQGCI | ASRRDSSYEQY | 0.80023944 |
| LCLHVEN | ASRRDSSYEQY | 0.77946806 |
| LDAISDGS | ASRRDSSYEQY | 0.7352809 |
| LGGFAAML | ASRRDSSYEQY | 0.80559576 |
| LGGKNPMI | ASRRDSSYEQY | 0.57959825 |
| LGPKGVAAIL | ASRRDSSYEQY | 0.84020454 |
| LMVGVSI | ASRRDSSYEQY | 0.8165534 |
| LVGAAFVL | ASRRDSSYEQY | 0.7806276 |
| LVLAVAPL | ASRRDSSYEQY | 0.8543543 |
| MAAIAVIDAA | ASRRDSSYEQY | 0.68011224 |
| NNGINAELQ | ASRRDSSYEQY | 0.7692569 |
| RGEPLVLVHG | ASRRDSSYEQY | 0.34572095 |
| RGGAGGAAGLI | ASRRDSSYEQY | 0.7159921 |
| RSYRMHPAV | ASRRDSSYEQY | 0.7519449 |
| SAAHLLAGQLA | ASRRDSSYEQY | 0.7730893 |
| SAAVGSY | ASRRDSSYEQY | 0.84406817 |
| SPHAVLHRI | ASRRDSSYEQY | 0.80750036 |
| SQIAKVL | ASRRDSSYEQY | 0.5933833 |
| TALGHLVP | ASRRDSSYEQY | 0.878518 |
| TAPTTSGAA | ASRRDSSYEQY | 0.871297 |
| TDGDDGNGGSPGT | ASRRDSSYEQY | 0.37152404 |
| TDGLGEN | ASRRDSSYEQY | 0.6397511 |
| TGGPPLPGQA | ASRRDSSYEQY | 0.87078273 |
| TIPAITI | ASRRDSSYEQY | 0.7967708 |
| TIPGFTI | ASRRDSSYEQY | 0.6473875 |

| | | |
|---|---|---|
| TSIASGALA | ASRRDSSYEQY | 0.70280445 |
| TVTLPGAAI | ASRRDSSYEQY | 0.8644205 |
| VASGLSAALP | ASRRDSSYEQY | 0.75087124 |
| VGAGVIGI | ASRRDSSYEQY | 0.8830206 |
| VGPKRFGV | ASRRDSSYEQY | 0.21111995 |
| VNVAFPD | ASRRDSSYEQY | 0.6172687 |
| VSMTNSGVS | ASRRDSSYEQY | 0.5474861 |
| VTDVLGDLSV | ASRRDSSYEQY | 0.14411728 |
| VVVVGAGFAGL | ASRRDSSYEQY | 0.84252304 |
| VYTVGTVV | ASRRDSSYEQY | 0.6364484 |
| WTTFSGP | ASRRDSSYEQY | 0.81477046 |
| YDELVEAF | ASRRDSSYEQY | 0.61059284 |
| YGTGGAGGNGG | ASRRDSSYEQY | 0.84790164 |
| YTNVMAV | ASRRDSSYEQY | 0.7703313 |
| FLEQVHQL | ASRRDSSYEQY | 0.65509725 |
| GLSLLAVAV | ASRRDSSYEQY | 0.38318408 |
| LEQVHQL | ASRRDSSYEQY | 0.78391576 |
| LNAAGAI | ASRRDSSYEQY | 0.88943636 |
| PAVHASL | ASRRDSSYEQY | 0.82957774 |
| TELEDDA | ASRRDSSYEQY | 0.22852464 |
| TPPGPPN | ASRRDSSYEQY | 0.80439603 |
| TPPPPPPP | ASRRDSSYEQY | 0.7067907 |
| TQSPALI | ASSPQDNSPLY | 0.24361172 |
| EFAAAFPQ | ASSPQDNSPLY | 0.23059571 |
| LLSGNLDVL | ASSPQDNSPLY | 0.5856297 |
| GPANLGNYNI | ASSPQDNSPLY | 0.65700316 |
| TLGPTLP | ASSPQDNSPLY | 0.8081116 |
| VYAYSSCCK | ASSPQDNSPLY | 0.8177835 |
| APAPAPAPNI | ASSPQDNSPLY | 0.5732256 |
| PGGPVPIP | ASSPQDNSPLY | 0.8091027 |
| FLTVALEP | ASSPQDNSPLY | 0.7754155 |
| SQYFGANL | ASSPQDNSPLY | 0.084699094 |
| GGVKPAA | ASSPQDNSPLY | 0.5312015 |
| GGTGGAGGAGGW | ASSPQDNSPLY | 0.3107258 |
| GVGGVAGLGGAGG | ASSPQDNSPLY | 0.87203085 |
| VLLVLATTI | ASSPQDNSPLY | 0.44101906 |
| IGIFAAML | ASSPQDNSPLY | 0.60935 |
| AVGDLID | ASSPQDNSPLY | 0.6610383 |
| NISPLPL | ASSPQDNSPLY | 0.7943649 |
| VTAVFVAI | ASSPQDNSPLY | 0.738887 |
| SGDDNDAAL | ASSPQDNSPLY | 0.47100243 |
| AVAALAHS | ASSPQDNSPLY | 0.30599356 |
| GGVTVMATP | ASSPQDNSPLY | 0.78936636 |
| AGLAANVRD | ASSPQDNSPLY | 0.4683777 |
| TNRPDLIDPALLRPG | ASSPQDNSPLY | 0.46547413 |
| LANLGVTAL | ASSPQDNSPLY | 0.44943446 |
| TGERIEL | ASSPQDNSPLY | 0.8230445 |
| NGEELAA | ASSPQDNSPLY | 0.2282385 |
| VLSLVWG | ASSPQDNSPLY | 0.8318815 |
| IGYALWL | ASSPQDNSPLY | 0.47393507 |
| LSLLLAPL | ASSPQDNSPLY | 0.68438196 |

| | | |
|---|---|---|
| TLDEALK | ASSPQDNSPLY | 0.8884615 |
| TDGAGTAEGG | ASSPQDNSPLY | 0.76986456 |
| NAPGGAGGNGG | ASSPQDNSPLY | 0.68404526 |
| VSVFMGMGGNF | ASSPQDNSPLY | 0.43252647 |
| LASYLAAL | ASSPQDNSPLY | 0.3925727 |
| LVGGTPAL | ASSPQDNSPLY | 0.4289597 |
| VRNADVL | ASSPQDNSPLY | 0.45214576 |
| TCSSPFLVSAM | ASSPQDNSPLY | 0.8798489 |
| VHITDAE | ASSPQDNSPLY | 0.68403673 |
| EYAAPSF | ASSPQDNSPLY | 0.3432329 |
| VVDAFHGGL | ASSPQDNSPLY | 0.7536341 |
| ASVVDPQ | ASSPQDNSPLY | 0.8763266 |
| SYNEFGGLS | ASSPQDNSPLY | 0.88806033 |
| DPDGSVY | ASSPQDNSPLY | 0.4347713 |
| MVSTIHATE | ASSPQDNSPLY | 0.52198243 |
| LAAGVADPV | ASSPQDNSPLY | 0.69806373 |
| TKEQIAA | ASSPQDNSPLY | 0.7353123 |
| SLASPAAL | ASSPQDNSPLY | 0.43241414 |
| IGLVFYGV | ASSPQDNSPLY | 0.3604259 |
| FLLDASNL | ASSPQDNSPLY | 0.8119809 |
| YQVGNFP | ASSPQDNSPLY | 0.8873894 |
| ESLLSQST | ASSPQDNSPLY | 0.6499983 |
| LFTVGFAAVG | ASSPQDNSPLY | 0.19162296 |
| ATPPTPT | ASSPQDNSPLY | 0.68807596 |
| TDLATLG | ASSPQDNSPLY | 0.4366866 |
| SAGGTGR | ASSPQDNSPLY | 0.502639 |
| GFSNSGS | ASSPQDNSPLY | 0.62439096 |
| GGNGGSGGTSV | ASSPQDNSPLY | 0.29803807 |
| GTSVATGGAGN | ASSPQDNSPLY | 0.70234513 |
| NGGNGGSGGTS | ASSPQDNSPLY | 0.88186115 |
| AVVKFPH | ASSPQDNSPLY | 0.64654964 |
| TPPPPPP | ASSPQDNSPLY | 0.792212 |
| AAPAAAAAPA | ASSPQDNSPLY | 0.86129737 |
| IALNEDL | ASSPQDNSPLY | 0.8003615 |
| IVNVSSM | ASSPQDNSPLY | 0.8581942 |
| TGIPPAPR | ASSPQDNSPLY | 0.83069867 |
| FDLGGGTF | ASSPQDNSPLY | 0.71126163 |
| FDLGGGTFDVS | ASSPQDNSPLY | 0.73470324 |
| NEPTAAA | ASSPQDNSPLY | 0.8488294 |
| NEPTAAALA | ASSPQDNSPLY | 0.6607519 |
| GSIINISSI | ASSPQDNSPLY | 0.7227308 |
| VAPGNAGTA | ASSPQDNSPLY | 0.6994252 |
| TGGGASLE | ASSPQDNSPLY | 0.84174 |
| VAVYNHYKRI | ASSPQDNSPLY | 0.5826285 |
| TDPAPATT | ASSPQDNSPLY | 0.867666 |
| TNRPDILDPALLRPG | ASSPQDNSPLY | 0.18431175 |
| LTTTTTS | ASSPQDNSPLY | 0.325805 |
| PTGSGKTL | ASSPQDNSPLY | 0.8481975 |
| TGSGKTL | ASSPQDNSPLY | 0.11743019 |
| TIGHVDHGK | ASSPQDNSPLY | 0.22394381 |
| TIGHVDHGKTT | ASSPQDNSPLY | 0.6878247 |

| | | |
|---|---|---|
| GHVDHGKTT | ASSPQDNSPLY | 0.8608414 |
| AQFEHTLL | ASSPQDNSPLY | 0.11330231 |
| IVNQDEL | ASSPQDNSPLY | 0.5348259 |
| TNGSQFF | ASSPQDNSPLY | 0.26577228 |
| TGKGIVE | ASSPQDNSPLY | 0.7961049 |
| LISNASDA | ASSPQDNSPLY | 0.6477981 |
| AELRTLGELTKIAKAHGAQ | ASSPQDNSPLY | 0.26136252 |
| APNAVAAAEPSW | ASSPQDNSPLY | 0.7972022 |
| APQVAGAATAVL | ASSPQDNSPLY | 0.23013407 |
| ARRDPAAAAAILPTDA | ASSPQDNSPLY | 0.72815514 |
| DAAARDVLAEAGLAEYF | ASSPQDNSPLY | 0.829225 |
| DDQIVLSAKANLAKVEQ | ASSPQDNSPLY | 0.8643513 |
| DELLPDTGEDSDEDGH | ASSPQDNSPLY | 0.38510117 |
| ELPEAPANRATVPQD | ASSPQDNSPLY | 0.7918489 |
| FGQSKIERVTGIG | ASSPQDNSPLY | 0.64456713 |
| GAGGTGSTAGGAGGAGGAGGLY | ASSPQDNSPLY | 0.89759886 |
| GHGGTGAAGGNGGTGGAGGSGI | ASSPQDNSPLY | 0.60925806 |
| GNGGNGGAGGEGGPGGAGGAGA | ASSPQDNSPLY | 0.27536866 |
| GSTAGNGGNGGNATNGTIAGQP | ASSPQDNSPLY | 0.8352588 |
| GTGGDGGNGGNGGTGG | ASSPQDNSPLY | 0.81105095 |
| GVRRPAGIPADAPV | ASSPQDNSPLY | 0.6909294 |
| GYRQVTDAHLLTLARR | ASSPQDNSPLY | 0.8220961 |
| IAKMGQDGHDRG | ASSPQDNSPLY | 0.6121779 |
| KAEPSIEIQYT | ASSPQDNSPLY | 0.67053735 |
| KPVRIRMAADEDTVAVT | ASSPQDNSPLY | 0.82442296 |
| MPPRIAGMRLLVIKPEP | ASSPQDNSPLY | 0.82129264 |
| MKFYAVALFRSRGVPPTR | ASSPQDNSPLY | 0.37521297 |
| NAGGFGILGAVAHSP | ASSPQDNSPLY | 0.85060245 |
| NEGREITLAMAEM | ASSPQDNSPLY | 0.7140337 |
| NELVEWSHPLGHRWRWE | ASSPQDNSPLY | 0.42746493 |
| NLGAGGTGGNGGNPG | ASSPQDNSPLY | 0.8882896 |
| NTEWPKWETVMAADPVD | ASSPQDNSPLY | 0.59936357 |
| PASAAPAETQP | ASSPQDNSPLY | 0.8202107 |
| QQSSDGKNGVRAAEITGE | ASSPQDNSPLY | 0.8124592 |
| RAPDAVTTSWWKE | ASSPQDNSPLY | 0.86219233 |
| SGFGNVGAHTSGWFNQ | ASSPQDNSPLY | 0.3423357 |
| SGVAGAVGQAASVGGLKV | ASSPQDNSPLY | 0.78758717 |
| SNGGAGTGSAGGNGGT | ASSPQDNSPLY | 0.7391294 |
| TGGAGGEGGAAG | ASSPQDNSPLY | 0.8206397 |
| TGGDGGAAGAGGAPGHG | ASSPQDNSPLY | 0.65988934 |
| TMPPLLAAIANAGRHGV | ASSPQDNSPLY | 0.66088957 |
| TSSTAGNSGGA | ASSPQDNSPLY | 0.8101342 |
| VPATVPVGILC | ASSPQDNSPLY | 0.85454565 |
| VQRPDSGGVVLTGRIS | ASSPQDNSPLY | 0.67884576 |
| WTPNPVPATKGVPAGLVQAT | ASSPQDNSPLY | 0.8071074 |
| YALKRALAKFSDRFDVVII | ASSPQDNSPLY | 0.8716626 |
| YLIGSVIV | ASSPQDNSPLY | 0.37566096 |
| YGQLVAVID | ASSPQDNSPLY | 0.6987399 |
| VTVGGMTLPA | ASSPQDNSPLY | 0.82331043 |
| VLALPAV | ASSPQDNSPLY | 0.88124573 |
| VIPPTLNYET | ASSPQDNSPLY | 0.6577581 |

| | | |
|---|---|---|
| VGMGGMHPSSG | ASSPQDNSPLY | 0.8668517 |
| VGLAGGAATA | ASSPQDNSPLY | 0.6372231 |
| VGGAGGAGGSGP | ASSPQDNSPLY | 0.80062133 |
| VFAAFMV | ASSPQDNSPLY | 0.732395 |
| TLYDKLR | ASSPQDNSPLY | 0.6289321 |
| TLAVFTVL | ASSPQDNSPLY | 0.42649505 |
| TGSVQPARV | ASSPQDNSPLY | 0.766669 |
| TGGGAGSGGT | ASSPQDNSPLY | 0.82000107 |
| SHLETSL | ASSPQDNSPLY | 0.79964983 |
| SFTTGMT | ASSPQDNSPLY | 0.2489444 |
| QVEVAKRMGVRQPTVS | ASSPQDNSPLY | 0.76992977 |
| QGSGGPALAM | ASSPQDNSPLY | 0.80928135 |
| MLAPSNP | ASSPQDNSPLY | 0.8827462 |
| LVNNAAI | ASSPQDNSPLY | 0.6602473 |
| LTGEINTLA | ASSPQDNSPLY | 0.8549628 |
| LSIFGPV | ASSPQDNSPLY | 0.57964027 |
| LLALLANL | ASSPQDNSPLY | 0.67841244 |
| LISNASDALDKLR | ASSPQDNSPLY | 0.354773 |
| LINEAVP | ASSPQDNSPLY | 0.5546161 |
| LGYAFSGL | ASSPQDNSPLY | 0.70163035 |
| LDFVAVAL | ASSPQDNSPLY | 0.51579213 |
| LAQPGAE | ASSPQDNSPLY | 0.8350448 |
| ISNLGAVTGF | ASSPQDNSPLY | 0.8619225 |
| HQEDPGRI | ASSPQDNSPLY | 0.84834445 |
| GTPGEPTP | ASSPQDNSPLY | 0.8523333 |
| GFFNSGS | ASSPQDNSPLY | 0.87846446 |
| GAGGGGLTAGAN | ASSPQDNSPLY | 0.8012992 |
| FVDALVNM | ASSPQDNSPLY | 0.544698 |
| FLTVAPEL | ASSPQDNSPLY | 0.69158506 |
| FETLTAL | ASSPQDNSPLY | 0.26758942 |
| DVINPIEL | ASSPQDNSPLY | 0.6462831 |
| APVSGAPV | ASSPQDNSPLY | 0.55602586 |
| ALVGGLAW | ASSPQDNSPLY | 0.72277343 |
| ALAAAAATAI | ASSPQDNSPLY | 0.101887405 |
| AFGLGLLL | ASSPQDNSPLY | 0.7912607 |
| AFFAAASL | ASSPQDNSPLY | 0.8812856 |
| AAAALPAFSPP | ASSPQDNSPLY | 0.6448906 |
| VVLAPAL | ASSPQDNSPLY | 0.77924573 |
| TIGHVDHGKTTL | ASSPQDNSPLY | 0.8184997 |
| LISNASDALDK | ASSPQDNSPLY | 0.5736853 |
| LISNASDAL | ASSPQDNSPLY | 0.79919314 |
| ISNASDALDK | ASSPQDNSPLY | 0.5551868 |
| NQAETLVYQTE | ASSPQDNSPLY | 0.8268741 |
| TGGTGGTAGAG | ASSPQDNSPLY | 0.86311376 |
| TGSTAGGAGGA | ASSPQDNSPLY | 0.80989635 |
| DPLAVPVAAGPV | ASSPQDNSPLY | 0.7859812 |
| GGTGGTPGNGGN | ASSPQDNSPLY | 0.7776382 |
| PAGLAGVDPATL | ASSPQDNSPLY | 0.2771832 |
| GNNLVVEAPHAGASV | ASSPQDNSPLY | 0.8148336 |
| KVTRIMSASTLEQPA | ASSPQDNSPLY | 0.57766706 |
| LGFALPIALPRRTVV | ASSPQDNSPLY | 0.7420809 |

| | | |
|---|---|---|
| TRLGNLTARKLVVLV | ASSPQDNSPLY | 0.4674652 |
| WSQQRRLIYFPSAGP | ASSPQDNSPLY | 0.7934209 |
| GPNGNPDPMAAAVDIR | ASSPQDNSPLY | 0.748724 |
| TQRCVSVADLRAAASR | ASSPQDNSPLY | 0.7594315 |
| QFVLDVAEHPVEVLTETV | ASSPQDNSPLY | 0.47637922 |
| GGAGGTGGAGGNGGTQAGDG | ASSPQDNSPLY | 0.8031732 |
| PLLATLRAIAVDVDAKPAQV | ASSPQDNSPLY | 0.7909889 |
| APKFRTGGYMIDFWGVGYQVAK | ASSPQDNSPLY | 0.34022275 |
| GAGGAGGAGGAAGGPGGTGGTG | ASSPQDNSPLY | 0.8385299 |
| IGALGLLATDVAPAGSVAFGVA | ASSPQDNSPLY | 0.6283959 |
| AGGAGGIGGNGGAGGAGGVLMG | ASSPQDNSPLY | 0.6833291 |
| ADELIGLL | ASSPQDNSPLY | 0.86419594 |
| ADRALKA | ASSPQDNSPLY | 0.7142662 |
| AGAVGQVGGAGGA | ASSPQDNSPLY | 0.21068665 |
| AGGGGGLGGAP | ASSPQDNSPLY | 0.60379934 |
| AGGLLYGNGGAG | ASSPQDNSPLY | 0.77112913 |
| AGLESIGL | ASSPQDNSPLY | 0.6303533 |
| AGLKTGII | ASSPQDNSPLY | 0.16693947 |
| ASLAALLL | ASSPQDNSPLY | 0.8016538 |
| ASVINISSI | ASSPQDNSPLY | 0.432891 |
| ATPPGPA | ASSPQDNSPLY | 0.5938778 |
| ATVASALV | ASSPQDNSPLY | 0.5733295 |
| ATVASLIL | ASSPQDNSPLY | 0.8327403 |
| AVNGPVLIRH | ASSPQDNSPLY | 0.76203406 |
| DGAAGAVGI | ASSPQDNSPLY | 0.5105868 |
| DPLLHGI | ASSPQDNSPLY | 0.8035062 |
| ELGAALGAL | ASSPQDNSPLY | 0.7607173 |
| ELGRTQAV | ASSPQDNSPLY | 0.86460805 |
| FVAGGYLS | ASSPQDNSPLY | 0.7167525 |
| GASGGAGGAGGW | ASSPQDNSPLY | 0.69763637 |
| GGAGAGGDGGAGGI | ASSPQDNSPLY | 0.8349793 |
| GTGHNGSQPGMGGNG | ASSPQDNSPLY | 0.49810755 |
| GYAAASASAS | ASSPQDNSPLY | 0.5579339 |
| HQVDVAAVQQG | ASSPQDNSPLY | 0.38961142 |
| IGAAIAHT | ASSPQDNSPLY | 0.35730225 |
| IGMIEVAQL | ASSPQDNSPLY | 0.46583238 |
| IRLAPEHTSAMD | ASSPQDNSPLY | 0.75628734 |
| IYNVSVL | ASSPQDNSPLY | 0.8506886 |
| KQGFLGM | ASSPQDNSPLY | 0.6765419 |
| LESVITQL | ASSPQDNSPLY | 0.29228833 |
| LGLGALVVV | ASSPQDNSPLY | 0.65269464 |
| LIGNGGNGGSGGI | ASSPQDNSPLY | 0.54890597 |
| LNAINAPTL | ASSPQDNSPLY | 0.796072 |
| LQLLNTGV | ASSPQDNSPLY | 0.8170692 |
| LQTSIEF | ASSPQDNSPLY | 0.87650955 |
| MSFAVSEL | ASSPQDNSPLY | 0.83102137 |
| NTGALGL | ASSPQDNSPLY | 0.5963625 |
| PPGAGGAGGAGTT | ASSPQDNSPLY | 0.81834 |
| QLVGLGI | ASSPQDNSPLY | 0.6281228 |
| RFLEQVHQL | ASSPQDNSPLY | 0.7643899 |
| RGVGYVLR | ASSPQDNSPLY | 0.7445173 |

| | | |
|---|---|---|
| SATGVFAGVI | ASSPQDNSPLY | 0.7698195 |
| SLPAITLP | ASSPQDNSPLY | 0.6032341 |
| STALAFTV | ASSPQDNSPLY | 0.6288596 |
| SVLTVGLGK | ASSPQDNSPLY | 0.21450773 |
| TALAGAGSGI | ASSPQDNSPLY | 0.787948 |
| TGLPFTGI | ASSPQDNSPLY | 0.3540728 |
| VAEHVTV | ASSPQDNSPLY | 0.4080574 |
| VDIVLNSL | ASSPQDNSPLY | 0.65812963 |
| VLTGGGAVL | ASSPQDNSPLY | 0.8868626 |
| VNVGLAYGI | ASSPQDNSPLY | 0.8616924 |
| VSSAAVGAV | ASSPQDNSPLY | 0.64159465 |
| VTSVYEAGV | ASSPQDNSPLY | 0.44202518 |
| WVALDRGA | ASSPQDNSPLY | 0.80335784 |
| YLVGSVVI | ASSPQDNSPLY | 0.50709206 |
| GPPGTGKTLL | ASSPQDNSPLY | 0.7830837 |
| GVGMTKF | ASSPQDNSPLY | 0.71365905 |
| IGIGGSDL | ASSPQDNSPLY | 0.73136055 |
| IGTLLAEL | ASSPQDNSPLY | 0.6558142 |
| IYTSGTT | ASSPQDNSPLY | 0.8139298 |
| LSFPAGV | ASSPQDNSPLY | 0.60694206 |
| LYTSGTT | ASSPQDNSPLY | 0.72616416 |
| VGAFPFGF | ASSPQDNSPLY | 0.14509232 |
| VGAGTVEF | ASSPQDNSPLY | 0.8373116 |
| VGKSTLLL | ASSPQDNSPLY | 0.46841305 |
| AALAHAGGGIV | ASSPQDNSPLY | 0.40769958 |
| AALVAVSLL | ASSPQDNSPLY | 0.16965152 |
| AAVSRAVAAL | ASSPQDNSPLY | 0.7042706 |
| AGGSSGIGGF | ASSPQDNSPLY | 0.79345804 |
| AGLTVANANG | ASSPQDNSPLY | 0.8514732 |
| AGLVGAVL | ASSPQDNSPLY | 0.7424702 |
| ALSQAFTV | ASSPQDNSPLY | 0.83045983 |
| ALTFSVSL | ASSPQDNSPLY | 0.84174794 |
| ALTGAALVAL | ASSPQDNSPLY | 0.6285494 |
| ASFSIGAGVL | ASSPQDNSPLY | 0.20620915 |
| DGSVYKI | ASSPQDNSPLY | 0.74042904 |
| EQLAEIV | ASSPQDNSPLY | 0.7399264 |
| FAVSELTVM | ASSPQDNSPLY | 0.8509755 |
| GAAPLAAGL | ASSPQDNSPLY | 0.8167943 |
| GADGGDGGSGGAGG | ASSPQDNSPLY | 0.31704208 |
| GFGLDTGGAGG | ASSPQDNSPLY | 0.8719317 |
| GLASGGLL | ASSPQDNSPLY | 0.80467415 |
| GRIAGAAVLL | ASSPQDNSPLY | 0.37619603 |
| GSAGSLNTSF | ASSPQDNSPLY | 0.50510633 |
| GTGQIGI | ASSPQDNSPLY | 0.52756083 |
| GTLIALIL | ASSPQDNSPLY | 0.3836161 |
| IAATMGFVVG | ASSPQDNSPLY | 0.8102102 |
| IFAQALL | ASSPQDNSPLY | 0.6317381 |
| IGAAVALGF | ASSPQDNSPLY | 0.5592729 |
| IGLTPLAL | ASSPQDNSPLY | 0.68370557 |
| ILGSNTSTL | ASSPQDNSPLY | 0.49916375 |
| IVFSLTVL | ASSPQDNSPLY | 0.4998694 |

| | | |
|---|---|---|
| LASKATGF | ASSPQDNSPLY | 0.5767227 |
| LASVALIL | ASSPQDNSPLY | 0.7063948 |
| LFAVGLTL | ASSPQDNSPLY | 0.81312203 |
| LFNGGIGTYI | ASSPQDNSPLY | 0.6789017 |
| LTLGAGII | ASSPQDNSPLY | 0.75131845 |
| LVAAALGDI | ASSPQDNSPLY | 0.870037 |
| NTAFVQLGI | ASSPQDNSPLY | 0.26167864 |
| QLFSGAEL | ASSPQDNSPLY | 0.5465374 |
| SAATFITL | ASSPQDNSPLY | 0.7594774 |
| SNYGLANLGV | ASSPQDNSPLY | 0.615759 |
| SYGGAYAVM | ASSPQDNSPLY | 0.84335804 |
| TTGVGGLTLG | ASSPQDNSPLY | 0.7803272 |
| TTQTIYI | ASSPQDNSPLY | 0.7493874 |
| VAAGAGLGAVI | ASSPQDNSPLY | 0.71202874 |
| VAAVVTLL | ASSPQDNSPLY | 0.88995045 |
| VGFLGGLI | ASSPQDNSPLY | 0.7803967 |
| VGVIPILL | ASSPQDNSPLY | 0.6694127 |
| VIFASSLI | ASSPQDNSPLY | 0.82861614 |
| VKGGDGGAAAG | ASSPQDNSPLY | 0.36505038 |
| VTKTFTNI | ASSPQDNSPLY | 0.84883714 |
| VVAAAILL | ASSPQDNSPLY | 0.79946697 |
| YAALDVEL | ASSPQDNSPLY | 0.44361448 |
| LASVALLL | ASSPQDNSPLY | 0.68719673 |
| LDAVHRV | ASSPQDNSPLY | 0.8716172 |
| QGLSSSIF | ASSPQDNSPLY | 0.84574777 |
| RGTFANIRL | ASSPQDNSPLY | 0.5389477 |
| RQLWWGHRI | ASSPQDNSPLY | 0.54913044 |
| YGPPGTGKTLL | ASSPQDNSPLY | 0.46114385 |
| AALVGAPANVA | ASSPQDNSPLY | 0.55138206 |
| AATFRRTE | ASSPQDNSPLY | 0.27817035 |
| AGALGGGI | ASSPQDNSPLY | 0.79142976 |
| AGGAGGAGGIGAG | ASSPQDNSPLY | 0.63391834 |
| AGSGALGL | ASSPQDNSPLY | 0.83522666 |
| AQIADQI | ASSPQDNSPLY | 0.7515374 |
| ASAVAVTYL | ASSPQDNSPLY | 0.8524143 |
| ASTVLGGTGG | ASSPQDNSPLY | 0.6085471 |
| AVGGEPAK | ASSPQDNSPLY | 0.6504976 |
| CEQGTFGGTC | ASSPQDNSPLY | 0.22328466 |
| DEGKAPAG | ASSPQDNSPLY | 0.45294604 |
| DTVVFPAL | ASSPQDNSPLY | 0.44635695 |
| FEGITMAG | ASSPQDNSPLY | 0.6488177 |
| FPLVVQD | ASSPQDNSPLY | 0.17303318 |
| GAFTTPQ | ASSPQDNSPLY | 0.8683859 |
| GFGGFGTAGGVGGA | ASSPQDNSPLY | 0.5979966 |
| GFLAQFL | ASSPQDNSPLY | 0.7847947 |
| GGAGGAGGNAGVA | ASSPQDNSPLY | 0.6019541 |
| GGVGTAGAGGAGGAGG | ASSPQDNSPLY | 0.45955336 |
| GNGGGGDAGNAGSGGN | ASSPQDNSPLY | 0.49943435 |
| GTGGAGGAGNDAGS | ASSPQDNSPLY | 0.82394564 |
| GTTNTGWFNS | ASSPQDNSPLY | 0.41845083 |
| IFMGPLFTL | ASSPQDNSPLY | 0.16137858 |

| | | |
|---|---|---|
| ILAHAIEAI | ASSPQDNSPLY | 0.38128883 |
| IRSAIREAL | ASSPQDNSPLY | 0.39053655 |
| ITFAGGG | ASSPQDNSPLY | 0.88223064 |
| ITGMSGHQEV | ASSPQDNSPLY | 0.85614073 |
| IVAGLACTL | ASSPQDNSPLY | 0.7613458 |
| IVGGGLAGA | ASSPQDNSPLY | 0.87988883 |
| IVTGVLNI | ASSPQDNSPLY | 0.12297584 |
| LAQQGCI | ASSPQDNSPLY | 0.61975574 |
| LCLHVEN | ASSPQDNSPLY | 0.8841032 |
| LDAISDGS | ASSPQDNSPLY | 0.7842791 |
| LGGFAAML | ASSPQDNSPLY | 0.682974 |
| LGGKNPMI | ASSPQDNSPLY | 0.82625246 |
| LGPKGVAAIL | ASSPQDNSPLY | 0.5616314 |
| LMVGVSI | ASSPQDNSPLY | 0.6323362 |
| LVGAAFVL | ASSPQDNSPLY | 0.6323411 |
| LVLAVAPL | ASSPQDNSPLY | 0.8353137 |
| MAAIAVIDAA | ASSPQDNSPLY | 0.760933 |
| NNGINAELQ | ASSPQDNSPLY | 0.27349028 |
| RGEPLVLVHG | ASSPQDNSPLY | 0.82295007 |
| RGGAGGAAGLI | ASSPQDNSPLY | 0.8921933 |
| RSYRMHPAV | ASSPQDNSPLY | 0.61878276 |
| SAAHLLAGQLA | ASSPQDNSPLY | 0.74388826 |
| SAAVGSY | ASSPQDNSPLY | 0.7187855 |
| SPHAVLHRI | ASSPQDNSPLY | 0.5565141 |
| SQIAKVL | ASSPQDNSPLY | 0.5098982 |
| TALGHLVP | ASSPQDNSPLY | 0.40115994 |
| TAPTTSGAA | ASSPQDNSPLY | 0.8233861 |
| TDGDDGNGGSPGT | ASSPQDNSPLY | 0.4518709 |
| TDGLGEN | ASSPQDNSPLY | 0.70552385 |
| TGGPPLPGQA | ASSPQDNSPLY | 0.65994203 |
| TIPAITI | ASSPQDNSPLY | 0.89885163 |
| TIPGFTI | ASSPQDNSPLY | 0.78848517 |
| TSIASGALA | ASSPQDNSPLY | 0.7348728 |
| TVTLPGAAI | ASSPQDNSPLY | 0.5228011 |
| VASGLSAALP | ASSPQDNSPLY | 0.7631394 |
| VGAGVIGI | ASSPQDNSPLY | 0.8134577 |
| VGPKRFGV | ASSPQDNSPLY | 0.2395252 |
| VNVAFPD | ASSPQDNSPLY | 0.86271286 |
| VSMTNSGVS | ASSPQDNSPLY | 0.37761623 |
| VTDVLGDLSV | ASSPQDNSPLY | 0.7515038 |
| VVVVGAGFAGL | ASSPQDNSPLY | 0.74556863 |
| VYTVGTVV | ASSPQDNSPLY | 0.8518164 |
| WTTFSGP | ASSPQDNSPLY | 0.7136233 |
| YDELVEAF | ASSPQDNSPLY | 0.8188519 |
| YGTGGAGGNGG | ASSPQDNSPLY | 0.42097002 |
| YTNVMAV | ASSPQDNSPLY | 0.80209994 |
| FLEQVHQL | ASSPQDNSPLY | 0.8454751 |
| GLSLLAVAV | ASSPQDNSPLY | 0.77421945 |
| LEQVHQL | ASSPQDNSPLY | 0.8788068 |
| LNAAGAI | ASSPQDNSPLY | 0.76482356 |
| PAVHASL | ASSPQDNSPLY | 0.2071991 |

| | | |
|---|---|---|
| TELEDDA | ASSPQDNSPLY | 0.6687168 |
| TPPGPPN | ASSPQDNSPLY | 0.8639777 |
| TPPPPPPP | ASSPQDNSPLY | 0.85752106 |
| TQSPALI | ASSLSQGASSYNSPLY | 0.79428077 |
| EFAAAFPQ | ASSLSQGASSYNSPLY | 0.82391924 |
| LLSGNLDVL | ASSLSQGASSYNSPLY | 0.7306874 |
| GPANLGNYNI | ASSLSQGASSYNSPLY | 0.87506783 |
| TLGPTLP | ASSLSQGASSYNSPLY | 0.8975675 |
| VYAYSSCCK | ASSLSQGASSYNSPLY | 0.797789 |
| APAPAPAPNI | ASSLSQGASSYNSPLY | 0.4061294 |
| PGGPVPIP | ASSLSQGASSYNSPLY | 0.3665623 |
| FLTVALEP | ASSLSQGASSYNSPLY | 0.83066845 |
| SQYFGANL | ASSLSQGASSYNSPLY | 0.8386948 |
| GGVKPAA | ASSLSQGASSYNSPLY | 0.336578 |
| GGTGGAGGAGGW | ASSLSQGASSYNSPLY | 0.7084649 |
| GVGGVAGLGGAGG | ASSLSQGASSYNSPLY | 0.710654 |
| VLLVLATTI | ASSLSQGASSYNSPLY | 0.1898152 |
| IGIFAAML | ASSLSQGASSYNSPLY | 0.7998467 |
| AVGDLID | ASSLSQGASSYNSPLY | 0.873778 |
| NISPLPL | ASSLSQGASSYNSPLY | 0.77972996 |
| VTAVFVAI | ASSLSQGASSYNSPLY | 0.39761508 |
| SGDDNDAAL | ASSLSQGASSYNSPLY | 0.86532617 |
| AVAALAHS | ASSLSQGASSYNSPLY | 0.86864805 |
| GGVTVMATP | ASSLSQGASSYNSPLY | 0.8778306 |
| AGLAANVRD | ASSLSQGASSYNSPLY | 0.24191673 |
| TNRPDLIDPALLRPG | ASSLSQGASSYNSPLY | 0.13267995 |
| LANLGVTAL | ASSLSQGASSYNSPLY | 0.79049027 |
| TGERIEL | ASSLSQGASSYNSPLY | 0.8136852 |
| NGEELAA | ASSLSQGASSYNSPLY | 0.5397001 |
| VLSLVWG | ASSLSQGASSYNSPLY | 0.42388088 |
| IGYALWL | ASSLSQGASSYNSPLY | 0.7539309 |
| LSLLLAPL | ASSLSQGASSYNSPLY | 0.852445 |
| TLDEALK | ASSLSQGASSYNSPLY | 0.38292673 |
| TDGAGTAEGG | ASSLSQGASSYNSPLY | 0.25166827 |
| NAPGGAGGNGG | ASSLSQGASSYNSPLY | 0.81253654 |
| VSVFMGMGGNF | ASSLSQGASSYNSPLY | 0.81883293 |
| LASYLAAL | ASSLSQGASSYNSPLY | 0.49767607 |
| LVGGTPAL | ASSLSQGASSYNSPLY | 0.645734 |
| VRNADVL | ASSLSQGASSYNSPLY | 0.80784935 |
| TCSSPFLVSAM | ASSLSQGASSYNSPLY | 0.7916169 |
| VHITDAE | ASSLSQGASSYNSPLY | 0.76854104 |
| EYAAPSF | ASSLSQGASSYNSPLY | 0.34130234 |
| VVDAFHGGL | ASSLSQGASSYNSPLY | 0.83295333 |
| ASVVDPQ | ASSLSQGASSYNSPLY | 0.2985196 |
| SYNEFGGLS | ASSLSQGASSYNSPLY | 0.8253881 |
| DPDGSVY | ASSLSQGASSYNSPLY | 0.7772859 |
| MVSTIHATE | ASSLSQGASSYNSPLY | 0.8046194 |
| LAAGVADPV | ASSLSQGASSYNSPLY | 0.667565 |
| TKEQIAA | ASSLSQGASSYNSPLY | 0.34971356 |
| SLASPAAL | ASSLSQGASSYNSPLY | 0.39686805 |
| IGLVFYGV | ASSLSQGASSYNSPLY | 0.34979284 |

| | | |
|---|---|---|
| FLLDASNL | ASSLSQGASSYNSPLY | 0.8762033 |
| YQVGNFP | ASSLSQGASSYNSPLY | 0.79570127 |
| ESLLSQST | ASSLSQGASSYNSPLY | 0.8364557 |
| LFTVGFAAVG | ASSLSQGASSYNSPLY | 0.7041346 |
| ATPPTPT | ASSLSQGASSYNSPLY | 0.38200027 |
| TDLATLG | ASSLSQGASSYNSPLY | 0.7650633 |
| SAGGTGR | ASSLSQGASSYNSPLY | 0.25505263 |
| GFSNSGS | ASSLSQGASSYNSPLY | 0.80269045 |
| GGNGGSGGTSV | ASSLSQGASSYNSPLY | 0.74578893 |
| GTSVATGGAGN | ASSLSQGASSYNSPLY | 0.7958849 |
| NGGNGGSGGTS | ASSLSQGASSYNSPLY | 0.5427379 |
| AVVKFPH | ASSLSQGASSYNSPLY | 0.77682126 |
| TPPPPPP | ASSLSQGASSYNSPLY | 0.81451124 |
| AAPAAAAAPA | ASSLSQGASSYNSPLY | 0.4441068 |
| IALNEDL | ASSLSQGASSYNSPLY | 0.8144554 |
| IVNVSSM | ASSLSQGASSYNSPLY | 0.8356427 |
| TGIPPAPR | ASSLSQGASSYNSPLY | 0.7752131 |
| FDLGGGTF | ASSLSQGASSYNSPLY | 0.27752313 |
| FDLGGGTFDVS | ASSLSQGASSYNSPLY | 0.7554468 |
| NEPTAAA | ASSLSQGASSYNSPLY | 0.579391 |
| NEPTAAALA | ASSLSQGASSYNSPLY | 0.7782675 |
| GSIINISSI | ASSLSQGASSYNSPLY | 0.6914761 |
| VAPGNAGTA | ASSLSQGASSYNSPLY | 0.5568546 |
| TGGGASLE | ASSLSQGASSYNSPLY | 0.80844873 |
| VAVYNHYKRI | ASSLSQGASSYNSPLY | 0.29446957 |
| TDPAPATT | ASSLSQGASSYNSPLY | 0.61663073 |
| TNRPDILDPALLRPG | ASSLSQGASSYNSPLY | 0.6551106 |
| LTTTTTS | ASSLSQGASSYNSPLY | 0.74788094 |
| PTGSGKTL | ASSLSQGASSYNSPLY | 0.80037284 |
| TGSGKTL | ASSLSQGASSYNSPLY | 0.44088048 |
| TIGHVDHGK | ASSLSQGASSYNSPLY | 0.77155375 |
| TIGHVDHGKTT | ASSLSQGASSYNSPLY | 0.65837157 |
| GHVDHGKTT | ASSLSQGASSYNSPLY | 0.46193027 |
| AQFEHTLL | ASSLSQGASSYNSPLY | 0.7265184 |
| IVNQDEL | ASSLSQGASSYNSPLY | 0.5692775 |
| TNGSQFF | ASSLSQGASSYNSPLY | 0.29915673 |
| TGKGIVE | ASSLSQGASSYNSPLY | 0.8299398 |
| LISNASDA | ASSLSQGASSYNSPLY | 0.41142085 |
| AELRTLGELTKIAKAHGAQ | ASSLSQGASSYNSPLY | 0.78288317 |
| APNAVAAAEPSW | ASSLSQGASSYNSPLY | 0.8910551 |
| APQVAGAATAVL | ASSLSQGASSYNSPLY | 0.40553638 |
| ARRDPAAAAAILPTDA | ASSLSQGASSYNSPLY | 0.7947222 |
| DAAARDVLAEAGLAEYF | ASSLSQGASSYNSPLY | 0.7308545 |
| DDQIVLSAKANLAKVEQ | ASSLSQGASSYNSPLY | 0.7651986 |
| DELLPDTGEDSDEDGH | ASSLSQGASSYNSPLY | 0.64332163 |
| ELPEAPANRATVPQD | ASSLSQGASSYNSPLY | 0.48404813 |
| FGQSKIERVTGIG | ASSLSQGASSYNSPLY | 0.8623363 |
| GAGGTGSTAGGAGGAGGAGGLY | ASSLSQGASSYNSPLY | 0.7892081 |
| GHGGTGAAGGNGGTGGAGGSGI | ASSLSQGASSYNSPLY | 0.8440101 |
| GNGGNGGAGGEGGPGGAGGAGA | ASSLSQGASSYNSPLY | 0.8161979 |
| GSTAGNGGNGGNATNGTIAGQP | ASSLSQGASSYNSPLY | 0.82936996 |

| | | |
|---|---|---|
| GTGGDGGNGGNGGTGG | ASSLSQGASSYNSPLY | 0.82907534 |
| GVRRPAGIPADAPV | ASSLSQGASSYNSPLY | 0.5372902 |
| GYRQVTDAHLLTLARR | ASSLSQGASSYNSPLY | 0.23749107 |
| IAKMGQDGHDRG | ASSLSQGASSYNSPLY | 0.6388022 |
| KAEPSIEIQYT | ASSLSQGASSYNSPLY | 0.66827345 |
| KPVRIRMAADEDTVAVT | ASSLSQGASSYNSPLY | 0.42691666 |
| MPPRIAGMRLLVIKPEP | ASSLSQGASSYNSPLY | 0.23632641 |
| MKFYAVALFRSRGVPPTR | ASSLSQGASSYNSPLY | 0.84572077 |
| NAGGFGILGAVAHSP | ASSLSQGASSYNSPLY | 0.89964104 |
| NEGREITLAMAEM | ASSLSQGASSYNSPLY | 0.5023735 |
| NELVEWSHPLGHRWRWE | ASSLSQGASSYNSPLY | 0.8906563 |
| NLGAGGTGGNGGNPG | ASSLSQGASSYNSPLY | 0.3478052 |
| NTEWPKWETVMAADPVD | ASSLSQGASSYNSPLY | 0.8138325 |
| PASAAPAETQP | ASSLSQGASSYNSPLY | 0.8547622 |
| QQSSDGKNGVRAAEITGE | ASSLSQGASSYNSPLY | 0.3556185 |
| RAPDAVTTSWWKE | ASSLSQGASSYNSPLY | 0.7874696 |
| SGFGNVGAHTSGWFNQ | ASSLSQGASSYNSPLY | 0.44083396 |
| SGVAGAVGQAASVGGLKV | ASSLSQGASSYNSPLY | 0.4375077 |
| SNGGAGTGSAGGNGGT | ASSLSQGASSYNSPLY | 0.85949457 |
| TGGAGGEGGAAG | ASSLSQGASSYNSPLY | 0.6028334 |
| TGGDGGAAGAGGAPGHG | ASSLSQGASSYNSPLY | 0.83910465 |
| TMPPLLAAIANAGRHGV | ASSLSQGASSYNSPLY | 0.49665678 |
| TSSTAGNSGGA | ASSLSQGASSYNSPLY | 0.39849597 |
| VPATVPVGILC | ASSLSQGASSYNSPLY | 0.7658025 |
| VQRPDSGGVVLTGRIS | ASSLSQGASSYNSPLY | 0.833796 |
| WTPNPVPATKGVPAGLVQAT | ASSLSQGASSYNSPLY | 0.729564 |
| YALKRALAKFSDRFDVVII | ASSLSQGASSYNSPLY | 0.8922052 |
| YLIGSVIV | ASSLSQGASSYNSPLY | 0.5448667 |
| YGQLVAVID | ASSLSQGASSYNSPLY | 0.87438 |
| VTVGGMTLPA | ASSLSQGASSYNSPLY | 0.858063 |
| VLALPAV | ASSLSQGASSYNSPLY | 0.7341411 |
| VIPPTLNYET | ASSLSQGASSYNSPLY | 0.5620979 |
| VGMGGMHPSSG | ASSLSQGASSYNSPLY | 0.83202624 |
| VGLAGGAATA | ASSLSQGASSYNSPLY | 0.6069589 |
| VGGAGGAGGSGP | ASSLSQGASSYNSPLY | 0.83530915 |
| VFAAFMV | ASSLSQGASSYNSPLY | 0.85635674 |
| TLYDKLR | ASSLSQGASSYNSPLY | 0.7223785 |
| TLAVFTVL | ASSLSQGASSYNSPLY | 0.85889816 |
| TGSVQPARV | ASSLSQGASSYNSPLY | 0.8347127 |
| TGGGAGSGGT | ASSLSQGASSYNSPLY | 0.84795445 |
| SHLETSL | ASSLSQGASSYNSPLY | 0.6686505 |
| SFTTGMT | ASSLSQGASSYNSPLY | 0.51461196 |
| QVEVAKRMGVRQPTVS | ASSLSQGASSYNSPLY | 0.8128611 |
| QGSGGPALAM | ASSLSQGASSYNSPLY | 0.5763912 |
| MLAPSNP | ASSLSQGASSYNSPLY | 0.8655728 |
| LVNNAAI | ASSLSQGASSYNSPLY | 0.86872745 |
| LTGEINTLA | ASSLSQGASSYNSPLY | 0.66774106 |
| LSIFGPV | ASSLSQGASSYNSPLY | 0.82366836 |
| LLALLANL | ASSLSQGASSYNSPLY | 0.8678359 |
| LISNASDALDKLR | ASSLSQGASSYNSPLY | 0.6176315 |
| LINEAVP | ASSLSQGASSYNSPLY | 0.7950728 |

| | | |
|---|---|---|
| LGYAFSGL | ASSLSQGASSYNSPLY | 0.61310637 |
| LDFVAVAL | ASSLSQGASSYNSPLY | 0.8053689 |
| LAQPGAE | ASSLSQGASSYNSPLY | 0.40667206 |
| ISNLGAVTGF | ASSLSQGASSYNSPLY | 0.85326123 |
| HQEDPGRI | ASSLSQGASSYNSPLY | 0.72245145 |
| GTPGEPTP | ASSLSQGASSYNSPLY | 0.3394221 |
| GFFNSGS | ASSLSQGASSYNSPLY | 0.83422244 |
| GAGGGGLTAGAN | ASSLSQGASSYNSPLY | 0.71675503 |
| FVDALVNM | ASSLSQGASSYNSPLY | 0.78878087 |
| FLTVAPEL | ASSLSQGASSYNSPLY | 0.8600496 |
| FETLTAL | ASSLSQGASSYNSPLY | 0.7830018 |
| DVINPIEL | ASSLSQGASSYNSPLY | 0.5962045 |
| APVSGAPV | ASSLSQGASSYNSPLY | 0.8229707 |
| ALVGGLAW | ASSLSQGASSYNSPLY | 0.8636495 |
| ALAAAAATAI | ASSLSQGASSYNSPLY | 0.5315184 |
| AFGLGLLL | ASSLSQGASSYNSPLY | 0.8187798 |
| AFFAAASL | ASSLSQGASSYNSPLY | 0.870988 |
| AAAALPAFSPP | ASSLSQGASSYNSPLY | 0.75837064 |
| VVLAPAL | ASSLSQGASSYNSPLY | 0.88219106 |
| TIGHVDHGKTTL | ASSLSQGASSYNSPLY | 0.8876913 |
| LISNASDALDK | ASSLSQGASSYNSPLY | 0.77014005 |
| LISNASDAL | ASSLSQGASSYNSPLY | 0.8624029 |
| ISNASDALDK | ASSLSQGASSYNSPLY | 0.8606602 |
| NQAETLVYQTE | ASSLSQGASSYNSPLY | 0.7762698 |
| TGGTGGTAGAG | ASSLSQGASSYNSPLY | 0.73879653 |
| TGSTAGGAGGA | ASSLSQGASSYNSPLY | 0.714802 |
| DPLAVPVAAGPV | ASSLSQGASSYNSPLY | 0.8244076 |
| GGTGGTPGNGGN | ASSLSQGASSYNSPLY | 0.6849164 |
| PAGLAGVDPATL | ASSLSQGASSYNSPLY | 0.7795919 |
| GNNLVVEAPHAGASV | ASSLSQGASSYNSPLY | 0.87915635 |
| KVTRIMSASTLEQPA | ASSLSQGASSYNSPLY | 0.7021154 |
| LGFALPIALPRRTVV | ASSLSQGASSYNSPLY | 0.6446811 |
| TRLGNLTARKLVVLV | ASSLSQGASSYNSPLY | 0.6073034 |
| WSQQRRLIYFPSAGP | ASSLSQGASSYNSPLY | 0.8388381 |
| GPNGNPDPMAAAVDIR | ASSLSQGASSYNSPLY | 0.7160883 |
| TQRCVSVADLRAAASR | ASSLSQGASSYNSPLY | 0.5905024 |
| QFVLDVAEHPVEVLTETV | ASSLSQGASSYNSPLY | 0.777589 |
| GGAGGTGGAGGNGGTQAGDG | ASSLSQGASSYNSPLY | 0.838445 |
| PLLATLRAIAVDVDAKPAQV | ASSLSQGASSYNSPLY | 0.8534603 |
| APKFRTGGYMIDFWGVGYQVAK | ASSLSQGASSYNSPLY | 0.62741184 |
| GAGGAGGAGGAAGGPGGTGGTG | ASSLSQGASSYNSPLY | 0.87567943 |
| IGALGLLATDVAPAGSVAFGVA | ASSLSQGASSYNSPLY | 0.8488687 |
| AGGAGGIGGNGGAGGAGGVLMG | ASSLSQGASSYNSPLY | 0.8824171 |
| ADELIGLL | ASSLSQGASSYNSPLY | 0.8092096 |
| ADRALKA | ASSLSQGASSYNSPLY | 0.70313716 |
| AGAVGQVGGAGGA | ASSLSQGASSYNSPLY | 0.8437311 |
| AGGGGGLGGAP | ASSLSQGASSYNSPLY | 0.7008245 |
| AGGLLYGNGGAG | ASSLSQGASSYNSPLY | 0.7512261 |
| AGLESIGL | ASSLSQGASSYNSPLY | 0.85464704 |
| AGLKTGII | ASSLSQGASSYNSPLY | 0.6907842 |
| ASLAALLL | ASSLSQGASSYNSPLY | 0.5875624 |

| | | |
|---|---|---|
| ASVINISSI | ASSLSQGASSYNSPLY | 0.7538322 |
| ATPPGPA | ASSLSQGASSYNSPLY | 0.88471425 |
| ATVASALV | ASSLSQGASSYNSPLY | 0.64756286 |
| ATVASLIL | ASSLSQGASSYNSPLY | 0.5967648 |
| AVNGPVLIRH | ASSLSQGASSYNSPLY | 0.7839737 |
| DGAAGAVGI | ASSLSQGASSYNSPLY | 0.7668754 |
| DPLLHGI | ASSLSQGASSYNSPLY | 0.86699396 |
| ELGAALGAL | ASSLSQGASSYNSPLY | 0.8549211 |
| ELGRTQAV | ASSLSQGASSYNSPLY | 0.78219604 |
| FVAGGYLS | ASSLSQGASSYNSPLY | 0.84097064 |
| GASGGAGGAGGW | ASSLSQGASSYNSPLY | 0.8433063 |
| GGAGAGGDGGAGGI | ASSLSQGASSYNSPLY | 0.80103576 |
| GTGHNGSQPGMGGNG | ASSLSQGASSYNSPLY | 0.8724066 |
| GYAAASASAS | ASSLSQGASSYNSPLY | 0.60633886 |
| HQVDVAAVQQG | ASSLSQGASSYNSPLY | 0.8759516 |
| IGAAIAHT | ASSLSQGASSYNSPLY | 0.8290594 |
| IGMIEVAQL | ASSLSQGASSYNSPLY | 0.8705574 |
| IRLAPEHTSAMD | ASSLSQGASSYNSPLY | 0.8794167 |
| IYNVSVL | ASSLSQGASSYNSPLY | 0.8200263 |
| KQGFLGM | ASSLSQGASSYNSPLY | 0.8352975 |
| LESVITQL | ASSLSQGASSYNSPLY | 0.50835085 |
| LGLGALVVV | ASSLSQGASSYNSPLY | 0.7763871 |
| LIGNGGNGGSGGI | ASSLSQGASSYNSPLY | 0.8353076 |
| LNAINAPTL | ASSLSQGASSYNSPLY | 0.8176869 |
| LQLLNTGV | ASSLSQGASSYNSPLY | 0.86643934 |
| LQTSIEF | ASSLSQGASSYNSPLY | 0.86885786 |
| MSFAVSEL | ASSLSQGASSYNSPLY | 0.72885436 |
| NTGALGL | ASSLSQGASSYNSPLY | 0.8899946 |
| PPGAGGAGGAGTT | ASSLSQGASSYNSPLY | 0.87368107 |
| QLVGLGI | ASSLSQGASSYNSPLY | 0.48903015 |
| RFLEQVHQL | ASSLSQGASSYNSPLY | 0.756393 |
| RGVGYVLR | ASSLSQGASSYNSPLY | 0.8809002 |
| SATGVFAGVI | ASSLSQGASSYNSPLY | 0.8106567 |
| SLPAITLP | ASSLSQGASSYNSPLY | 0.7189398 |
| STALAFTV | ASSLSQGASSYNSPLY | 0.8120599 |
| SVLTVGLGK | ASSLSQGASSYNSPLY | 0.86838967 |
| TALAGAGSGI | ASSLSQGASSYNSPLY | 0.79175204 |
| TGLPFTGI | ASSLSQGASSYNSPLY | 0.884604 |
| VAEHVTV | ASSLSQGASSYNSPLY | 0.38682747 |
| VDIVLNSL | ASSLSQGASSYNSPLY | 0.82487 |
| VLTGGGAVL | ASSLSQGASSYNSPLY | 0.79610634 |
| VNVGLAYGI | ASSLSQGASSYNSPLY | 0.31899822 |
| VSSAAVGAV | ASSLSQGASSYNSPLY | 0.8180444 |
| VTSVYEAGV | ASSLSQGASSYNSPLY | 0.81384575 |
| WVALDRGA | ASSLSQGASSYNSPLY | 0.8638631 |
| YLVGSVVI | ASSLSQGASSYNSPLY | 0.77362883 |
| GPPGTGKTLL | ASSLSQGASSYNSPLY | 0.85243046 |
| GVGMTKF | ASSLSQGASSYNSPLY | 0.7586243 |
| IGIGGSDL | ASSLSQGASSYNSPLY | 0.8409182 |
| IGTLLAEL | ASSLSQGASSYNSPLY | 0.7667415 |
| IYTSGTT | ASSLSQGASSYNSPLY | 0.8008671 |

| | | |
|---|---|---|
| LSFPAGV | ASSLSQGASSYNSPLY | 0.88945806 |
| LYTSGTT | ASSLSQGASSYNSPLY | 0.8838572 |
| VGAFPFGF | ASSLSQGASSYNSPLY | 0.3733545 |
| VGAGTVEF | ASSLSQGASSYNSPLY | 0.82825106 |
| VGKSTLLL | ASSLSQGASSYNSPLY | 0.73122597 |
| AALAHAGGGIV | ASSLSQGASSYNSPLY | 0.4750753 |
| AALVAVSLL | ASSLSQGASSYNSPLY | 0.49127644 |
| AAVSRAVAAL | ASSLSQGASSYNSPLY | 0.7678311 |
| AGGSSGIGGF | ASSLSQGASSYNSPLY | 0.7539546 |
| AGLTVANANG | ASSLSQGASSYNSPLY | 0.80890894 |
| AGLVGAVL | ASSLSQGASSYNSPLY | 0.8017024 |
| ALSQAFTV | ASSLSQGASSYNSPLY | 0.6879055 |
| ALTFSVSL | ASSLSQGASSYNSPLY | 0.8412447 |
| ALTGAALVAL | ASSLSQGASSYNSPLY | 0.78186023 |
| ASFSIGAGVL | ASSLSQGASSYNSPLY | 0.12515593 |
| DGSVYKI | ASSLSQGASSYNSPLY | 0.7596334 |
| EQLAEIV | ASSLSQGASSYNSPLY | 0.65048236 |
| FAVSELTVM | ASSLSQGASSYNSPLY | 0.88414097 |
| GAAPLAAGL | ASSLSQGASSYNSPLY | 0.7139896 |
| GADGGDGGSGGAGG | ASSLSQGASSYNSPLY | 0.6462766 |
| GFGLDTGGAGG | ASSLSQGASSYNSPLY | 0.7485336 |
| GLASGGLL | ASSLSQGASSYNSPLY | 0.7970801 |
| GRIAGAAVLL | ASSLSQGASSYNSPLY | 0.76960945 |
| GSAGSLNTSF | ASSLSQGASSYNSPLY | 0.728789 |
| GTGQIGI | ASSLSQGASSYNSPLY | 0.5988939 |
| GTLIALIL | ASSLSQGASSYNSPLY | 0.8134098 |
| IAATMGFVVG | ASSLSQGASSYNSPLY | 0.7360681 |
| IFAQALL | ASSLSQGASSYNSPLY | 0.75028193 |
| IGAAVALGF | ASSLSQGASSYNSPLY | 0.72772914 |
| IGLTPLAL | ASSLSQGASSYNSPLY | 0.74405813 |
| ILGSNTSTL | ASSLSQGASSYNSPLY | 0.49048683 |
| IVFSLTVL | ASSLSQGASSYNSPLY | 0.877103 |
| LASKATGF | ASSLSQGASSYNSPLY | 0.74428654 |
| LASVALIL | ASSLSQGASSYNSPLY | 0.67639524 |
| LFAVGLTL | ASSLSQGASSYNSPLY | 0.8797395 |
| LFNGGIGTYI | ASSLSQGASSYNSPLY | 0.77275264 |
| LTLGAGII | ASSLSQGASSYNSPLY | 0.710757 |
| LVAAALGDI | ASSLSQGASSYNSPLY | 0.7307575 |
| NTAFVQLGI | ASSLSQGASSYNSPLY | 0.67161036 |
| QLFSGAEL | ASSLSQGASSYNSPLY | 0.6547833 |
| SAATFITL | ASSLSQGASSYNSPLY | 0.7030269 |
| SNYGLANLGV | ASSLSQGASSYNSPLY | 0.8740402 |
| SYGGAYAVM | ASSLSQGASSYNSPLY | 0.83942294 |
| TTGVGGLTLG | ASSLSQGASSYNSPLY | 0.6868891 |
| TTQTIYI | ASSLSQGASSYNSPLY | 0.86521614 |
| VAAGAGLGAVI | ASSLSQGASSYNSPLY | 0.87600744 |
| VAAVVTLL | ASSLSQGASSYNSPLY | 0.8837782 |
| VGFLGGLI | ASSLSQGASSYNSPLY | 0.7420335 |
| VGVIPILL | ASSLSQGASSYNSPLY | 0.81010354 |
| VIFASSLI | ASSLSQGASSYNSPLY | 0.8025774 |
| VKGGDGGAAAG | ASSLSQGASSYNSPLY | 0.69561577 |

| | | |
|---|---|---|
| VTKTFTNI | ASSLSQGASSYNSPLY | 0.80576575 |
| VVAAAILL | ASSLSQGASSYNSPLY | 0.5849986 |
| YAALDVEL | ASSLSQGASSYNSPLY | 0.871349 |
| LASVALLL | ASSLSQGASSYNSPLY | 0.89258176 |
| LDAVHRV | ASSLSQGASSYNSPLY | 0.77984285 |
| QGLSSSIF | ASSLSQGASSYNSPLY | 0.31940168 |
| RGTFANIRL | ASSLSQGASSYNSPLY | 0.7273443 |
| RQLWWGHRI | ASSLSQGASSYNSPLY | 0.6630045 |
| YGPPGTGKTLL | ASSLSQGASSYNSPLY | 0.61554337 |
| AALVGAPANVA | ASSLSQGASSYNSPLY | 0.77418256 |
| AATFRRTE | ASSLSQGASSYNSPLY | 0.49935713 |
| AGALGGGI | ASSLSQGASSYNSPLY | 0.7865244 |
| AGGAGGAGGIGAG | ASSLSQGASSYNSPLY | 0.87541145 |
| AGSGALGL | ASSLSQGASSYNSPLY | 0.7098928 |
| AQIADQI | ASSLSQGASSYNSPLY | 0.84888977 |
| ASAVAVTYL | ASSLSQGASSYNSPLY | 0.8622694 |
| ASTVLGGTGG | ASSLSQGASSYNSPLY | 0.8368641 |
| AVGGEPAK | ASSLSQGASSYNSPLY | 0.87061346 |
| CEQGTFGGTC | ASSLSQGASSYNSPLY | 0.8628905 |
| DEGKAPAG | ASSLSQGASSYNSPLY | 0.8341615 |
| DTVVFPAL | ASSLSQGASSYNSPLY | 0.8242701 |
| FEGITMAG | ASSLSQGASSYNSPLY | 0.8542407 |
| FPLVVQD | ASSLSQGASSYNSPLY | 0.7397367 |
| GAFTTPQ | ASSLSQGASSYNSPLY | 0.75859344 |
| GFGGFGTAGGVGGA | ASSLSQGASSYNSPLY | 0.76688683 |
| GFLAQFL | ASSLSQGASSYNSPLY | 0.83575845 |
| GGAGGAGGNAGVA | ASSLSQGASSYNSPLY | 0.7744663 |
| GGVGTAGAGGAGGAGG | ASSLSQGASSYNSPLY | 0.87956476 |
| GNGGGGDAGNAGSGGN | ASSLSQGASSYNSPLY | 0.40296692 |
| GTGGAGGAGNDAGS | ASSLSQGASSYNSPLY | 0.54896975 |
| GTTNTGWFNS | ASSLSQGASSYNSPLY | 0.8564745 |
| IFMGPLFTL | ASSLSQGASSYNSPLY | 0.2430093 |
| ILAHAIEAI | ASSLSQGASSYNSPLY | 0.53426635 |
| IRSAIREAL | ASSLSQGASSYNSPLY | 0.8295495 |
| ITFAGGG | ASSLSQGASSYNSPLY | 0.86650634 |
| ITGMSGHQEV | ASSLSQGASSYNSPLY | 0.30048954 |
| IVAGLACTL | ASSLSQGASSYNSPLY | 0.6870673 |
| IVGGGLAGA | ASSLSQGASSYNSPLY | 0.6366867 |
| IVTGVLNI | ASSLSQGASSYNSPLY | 0.8826921 |
| LAQQGCI | ASSLSQGASSYNSPLY | 0.73156154 |
| LCLHVEN | ASSLSQGASSYNSPLY | 0.42984644 |
| LDAISDGS | ASSLSQGASSYNSPLY | 0.8692405 |
| LGGFAAML | ASSLSQGASSYNSPLY | 0.41722035 |
| LGGKNPMI | ASSLSQGASSYNSPLY | 0.8003976 |
| LGPKGVAAIL | ASSLSQGASSYNSPLY | 0.8540468 |
| LMVGVSI | ASSLSQGASSYNSPLY | 0.8838109 |
| LVGAAFVL | ASSLSQGASSYNSPLY | 0.67433363 |
| LVLAVAPL | ASSLSQGASSYNSPLY | 0.8373941 |
| MAAIAVIDAA | ASSLSQGASSYNSPLY | 0.753628 |
| NNGINAELQ | ASSLSQGASSYNSPLY | 0.8952712 |
| RGEPLVLVHG | ASSLSQGASSYNSPLY | 0.70069546 |

| | | |
|---|---|---|
| RGGAGGAAGLI | ASSLSQGASSYNSPLY | 0.6273712 |
| RSYRMHPAV | ASSLSQGASSYNSPLY | 0.866274 |
| SAAHLLAGQLA | ASSLSQGASSYNSPLY | 0.8700429 |
| SAAVGSY | ASSLSQGASSYNSPLY | 0.83885723 |
| SPHAVLHRI | ASSLSQGASSYNSPLY | 0.8497703 |
| SQIAKVL | ASSLSQGASSYNSPLY | 0.82488453 |
| TALGHLVP | ASSLSQGASSYNSPLY | 0.7523372 |
| TAPTTSGAA | ASSLSQGASSYNSPLY | 0.83798516 |
| TDGDDGNGGSPGT | ASSLSQGASSYNSPLY | 0.8542604 |
| TDGLGEN | ASSLSQGASSYNSPLY | 0.62132037 |
| TGGPPLPGQA | ASSLSQGASSYNSPLY | 0.8805915 |
| TIPAITI | ASSLSQGASSYNSPLY | 0.75452816 |
| TIPGFTI | ASSLSQGASSYNSPLY | 0.7579937 |
| TSIASGALA | ASSLSQGASSYNSPLY | 0.8750503 |
| TVTLPGAAI | ASSLSQGASSYNSPLY | 0.8446077 |
| VASGLSAALP | ASSLSQGASSYNSPLY | 0.84802425 |
| VGAGVIGI | ASSLSQGASSYNSPLY | 0.8523202 |
| VGPKRFGV | ASSLSQGASSYNSPLY | 0.8866389 |
| VNVAFPD | ASSLSQGASSYNSPLY | 0.56899947 |
| VSMTNSGVS | ASSLSQGASSYNSPLY | 0.8543521 |
| VTDVLGDLSV | ASSLSQGASSYNSPLY | 0.79408205 |
| VVVVGAGFAGL | ASSLSQGASSYNSPLY | 0.86689395 |
| VYTVGTVV | ASSLSQGASSYNSPLY | 0.8432046 |
| WTTFSGP | ASSLSQGASSYNSPLY | 0.7645646 |
| YDELVEAF | ASSLSQGASSYNSPLY | 0.8563924 |
| YGTGGAGGNGG | ASSLSQGASSYNSPLY | 0.8693514 |
| YTNVMAV | ASSLSQGASSYNSPLY | 0.83730197 |
| FLEQVHQL | ASSLSQGASSYNSPLY | 0.85886323 |
| GLSLLAVAV | ASSLSQGASSYNSPLY | 0.8779892 |
| LEQVHQL | ASSLSQGASSYNSPLY | 0.77617025 |
| LNAAGAI | ASSLSQGASSYNSPLY | 0.7607324 |
| PAVHASL | ASSLSQGASSYNSPLY | 0.8636243 |
| TELEDDA | ASSLSQGASSYNSPLY | 0.8841187 |
| TPPGPPN | ASSLSQGASSYNSPLY | 0.81930774 |
| TPPPPPPP | ASSLSQGASSYNSPLY | 0.881742 |
| TQSPALI | ASSLSTGHQDTQY | 0.7207034 |
| EFAAAFPQ | ASSLSTGHQDTQY | 0.8179321 |
| LLSGNLDVL | ASSLSTGHQDTQY | 0.74549854 |
| GPANLGNYNI | ASSLSTGHQDTQY | 0.7472137 |
| TLGPTLP | ASSLSTGHQDTQY | 0.6365996 |
| VYAYSSCCK | ASSLSTGHQDTQY | 0.79032314 |
| APAPAPAPNI | ASSLSTGHQDTQY | 0.84032714 |
| PGGPVPIP | ASSLSTGHQDTQY | 0.8059784 |
| FLTVALEP | ASSLSTGHQDTQY | 0.47050303 |
| SQYFGANL | ASSLSTGHQDTQY | 0.78572965 |
| GGVKPAA | ASSLSTGHQDTQY | 0.8306165 |
| GGTGGAGGAGGW | ASSLSTGHQDTQY | 0.8811929 |
| GVGGVAGLGGAGG | ASSLSTGHQDTQY | 0.79759496 |
| VLLVLATTI | ASSLSTGHQDTQY | 0.8606998 |
| IGIFAAML | ASSLSTGHQDTQY | 0.69937307 |
| AVGDLID | ASSLSTGHQDTQY | 0.8136259 |

| | | |
|---|---|---|
| NISPLPL | ASSLSTGHQDTQY | 0.6185662 |
| VTAVFVAI | ASSLSTGHQDTQY | 0.7907036 |
| SGDDNDAAL | ASSLSTGHQDTQY | 0.67835855 |
| AVAALAHS | ASSLSTGHQDTQY | 0.65761495 |
| GGVTVMATP | ASSLSTGHQDTQY | 0.81831694 |
| AGLAANVRD | ASSLSTGHQDTQY | 0.8739694 |
| TNRPDLIDPALLRPG | ASSLSTGHQDTQY | 0.8377162 |
| LANLGVTAL | ASSLSTGHQDTQY | 0.8634439 |
| TGERIEL | ASSLSTGHQDTQY | 0.8703753 |
| NGEELAA | ASSLSTGHQDTQY | 0.8870264 |
| VLSLVWG | ASSLSTGHQDTQY | 0.6645466 |
| IGYALWL | ASSLSTGHQDTQY | 0.7951105 |
| LSLLLAPL | ASSLSTGHQDTQY | 0.49326825 |
| TLDEALK | ASSLSTGHQDTQY | 0.75594497 |
| TDGAGTAEGG | ASSLSTGHQDTQY | 0.7101501 |
| NAPGGAGGNGG | ASSLSTGHQDTQY | 0.76183695 |
| VSVFMGMGGNF | ASSLSTGHQDTQY | 0.19023906 |
| LASYLAAL | ASSLSTGHQDTQY | 0.79263484 |
| LVGGTPAL | ASSLSTGHQDTQY | 0.8780171 |
| VRNADVL | ASSLSTGHQDTQY | 0.7480147 |
| TCSSPFLVSAM | ASSLSTGHQDTQY | 0.84373325 |
| VHITDAE | ASSLSTGHQDTQY | 0.8234587 |
| EYAAPSF | ASSLSTGHQDTQY | 0.7356913 |
| VVDAFHGGL | ASSLSTGHQDTQY | 0.8367603 |
| ASVVDPQ | ASSLSTGHQDTQY | 0.7397293 |
| SYNEFGGLS | ASSLSTGHQDTQY | 0.86412144 |
| DPDGSVY | ASSLSTGHQDTQY | 0.8709388 |
| MVSTIHATE | ASSLSTGHQDTQY | 0.83455455 |
| LAAGVADPV | ASSLSTGHQDTQY | 0.8353377 |
| TKEQIAA | ASSLSTGHQDTQY | 0.83765054 |
| SLASPAAL | ASSLSTGHQDTQY | 0.45062497 |
| IGLVFYGV | ASSLSTGHQDTQY | 0.84374976 |
| FLLDASNL | ASSLSTGHQDTQY | 0.7381167 |
| YQVGNFP | ASSLSTGHQDTQY | 0.7686064 |
| ESLLSQST | ASSLSTGHQDTQY | 0.72357583 |
| LFTVGFAAVG | ASSLSTGHQDTQY | 0.83141255 |
| ATPPTPT | ASSLSTGHQDTQY | 0.76041687 |
| TDLATLG | ASSLSTGHQDTQY | 0.8473362 |
| SAGGTGR | ASSLSTGHQDTQY | 0.6150007 |
| GFSNSGS | ASSLSTGHQDTQY | 0.8224949 |
| GGNGGSGGTSV | ASSLSTGHQDTQY | 0.7880486 |
| GTSVATGGAGN | ASSLSTGHQDTQY | 0.58412015 |
| NGGNGGSGGTS | ASSLSTGHQDTQY | 0.8696181 |
| AVVKFPH | ASSLSTGHQDTQY | 0.83521485 |
| TPPPPPP | ASSLSTGHQDTQY | 0.71415114 |
| AAPAAAAAPA | ASSLSTGHQDTQY | 0.8608554 |
| IALNEDL | ASSLSTGHQDTQY | 0.82954955 |
| IVNVSSM | ASSLSTGHQDTQY | 0.41443932 |
| TGIPPAPR | ASSLSTGHQDTQY | 0.7400651 |
| FDLGGGTF | ASSLSTGHQDTQY | 0.80907524 |
| FDLGGGTFDVS | ASSLSTGHQDTQY | 0.73792815 |

| | | |
|---|---|---|
| NEPTAAA | ASSLSTGHQDTQY | 0.37424403 |
| NEPTAAALA | ASSLSTGHQDTQY | 0.8563403 |
| GSIINISSI | ASSLSTGHQDTQY | 0.7584926 |
| VAPGNAGTA | ASSLSTGHQDTQY | 0.59972924 |
| TGGGASLE | ASSLSTGHQDTQY | 0.73176134 |
| VAVYNHYKRI | ASSLSTGHQDTQY | 0.86069405 |
| TDPAPATT | ASSLSTGHQDTQY | 0.80064905 |
| TNRPDILDPALLRPG | ASSLSTGHQDTQY | 0.8333678 |
| LTTTTTS | ASSLSTGHQDTQY | 0.85010505 |
| PTGSGKTL | ASSLSTGHQDTQY | 0.78772235 |
| TGSGKTL | ASSLSTGHQDTQY | 0.84891784 |
| TIGHVDHGK | ASSLSTGHQDTQY | 0.8443488 |
| TIGHVDHGKTT | ASSLSTGHQDTQY | 0.8378266 |
| GHVDHGKTT | ASSLSTGHQDTQY | 0.8555676 |
| AQFEHTLL | ASSLSTGHQDTQY | 0.80686754 |
| IVNQDEL | ASSLSTGHQDTQY | 0.6898343 |
| TNGSQFF | ASSLSTGHQDTQY | 0.6630204 |
| TGKGIVE | ASSLSTGHQDTQY | 0.636906 |
| LISNASDA | ASSLSTGHQDTQY | 0.6951255 |
| AELRTLGELTKIAKAHGAQ | ASSLSTGHQDTQY | 0.8006053 |
| APNAVAAAEPSW | ASSLSTGHQDTQY | 0.8905323 |
| APQVAGAATAVL | ASSLSTGHQDTQY | 0.8200098 |
| ARRDPAAAAAILPTDA | ASSLSTGHQDTQY | 0.64158523 |
| DAAARDVLAEAGLAEYF | ASSLSTGHQDTQY | 0.7846935 |
| DDQIVLSAKANLAKVEQ | ASSLSTGHQDTQY | 0.6799406 |
| DELLPDTGEDSDEDGH | ASSLSTGHQDTQY | 0.8473039 |
| ELPEAPANRATVPQD | ASSLSTGHQDTQY | 0.8811782 |
| FGQSKIERVTGIG | ASSLSTGHQDTQY | 0.88398075 |
| GAGGTGSTAGGAGGAGGAGGLY | ASSLSTGHQDTQY | 0.82486904 |
| GHGGTGAAGGNGGTGGAGGSGI | ASSLSTGHQDTQY | 0.68202055 |
| GNGGNGGAGGEGGPGGAGGAGA | ASSLSTGHQDTQY | 0.85856175 |
| GSTAGNGGNGGNATNGTIAGQP | ASSLSTGHQDTQY | 0.7451948 |
| GTGGDGGNGGNGGTGG | ASSLSTGHQDTQY | 0.84247935 |
| GVRRPAGIPADAPV | ASSLSTGHQDTQY | 0.8323845 |
| GYRQVTDAHLLTLARR | ASSLSTGHQDTQY | 0.8331851 |
| IAKMGQDGHDRG | ASSLSTGHQDTQY | 0.8268953 |
| KAEPSIEIQYT | ASSLSTGHQDTQY | 0.8311637 |
| KPVRIRMAADEDTVAVT | ASSLSTGHQDTQY | 0.4628967 |
| MPPRIAGMRLLVIKPEP | ASSLSTGHQDTQY | 0.862586 |
| MKFYAVALFRSRGVPPTR | ASSLSTGHQDTQY | 0.5960944 |
| NAGGFGILGAVAHSP | ASSLSTGHQDTQY | 0.67664224 |
| NEGREITLAMAEM | ASSLSTGHQDTQY | 0.7312231 |
| NELVEWSHPLGHRWRWE | ASSLSTGHQDTQY | 0.87602794 |
| NLGAGGTGGNGGNPG | ASSLSTGHQDTQY | 0.87191224 |
| NTEWPKWETVMAADPVD | ASSLSTGHQDTQY | 0.6896533 |
| PASAAPAETQP | ASSLSTGHQDTQY | 0.6799805 |
| QQSSDGKNGVRAAEITGE | ASSLSTGHQDTQY | 0.8106087 |
| RAPDAVTTSWWKE | ASSLSTGHQDTQY | 0.61829215 |
| SGFGNVGAHTSGWFNQ | ASSLSTGHQDTQY | 0.8643293 |
| SGVAGAVGQAASVGGLKV | ASSLSTGHQDTQY | 0.70217454 |
| SNGGAGTGSAGGNGGT | ASSLSTGHQDTQY | 0.72286546 |

| | | |
|---|---|---|
| TGGAGGEGGAAG | ASSLSTGHQDTQY | 0.785277 |
| TGGDGGAAGAGGAPGHG | ASSLSTGHQDTQY | 0.8460737 |
| TMPPLLAAIANAGRHGV | ASSLSTGHQDTQY | 0.727842 |
| TSSTAGNSGGA | ASSLSTGHQDTQY | 0.8766979 |
| VPATVPVGILC | ASSLSTGHQDTQY | 0.34511662 |
| VQRPDSGGVVLTGRIS | ASSLSTGHQDTQY | 0.8510263 |
| WTPNPVPATKGVPAGLVQAT | ASSLSTGHQDTQY | 0.5645044 |
| YALKRALAKFSDRFDVVII | ASSLSTGHQDTQY | 0.70314384 |
| YLIGSVIV | ASSLSTGHQDTQY | 0.5065652 |
| YGQLVAVID | ASSLSTGHQDTQY | 0.8260964 |
| VTVGGMTLPA | ASSLSTGHQDTQY | 0.84064436 |
| VLALPAV | ASSLSTGHQDTQY | 0.8660458 |
| VIPPTLNYET | ASSLSTGHQDTQY | 0.80436766 |
| VGMGGMHPSSG | ASSLSTGHQDTQY | 0.85353315 |
| VGLAGGAATA | ASSLSTGHQDTQY | 0.860628 |
| VGGAGGAGGSGP | ASSLSTGHQDTQY | 0.64180994 |
| VFAAFMV | ASSLSTGHQDTQY | 0.45082068 |
| TLYDKLR | ASSLSTGHQDTQY | 0.8111541 |
| TLAVFTVL | ASSLSTGHQDTQY | 0.7924534 |
| TGSVQPARV | ASSLSTGHQDTQY | 0.86921126 |
| TGGGAGSGGT | ASSLSTGHQDTQY | 0.84279966 |
| SHLETSL | ASSLSTGHQDTQY | 0.5618174 |
| SFTTGMT | ASSLSTGHQDTQY | 0.85846436 |
| QVEVAKRMGVRQPTVS | ASSLSTGHQDTQY | 0.848048 |
| QGSGGPALAM | ASSLSTGHQDTQY | 0.6521374 |
| MLAPSNP | ASSLSTGHQDTQY | 0.76283145 |
| LVNNAAI | ASSLSTGHQDTQY | 0.7789446 |
| LTGEINTLA | ASSLSTGHQDTQY | 0.65001076 |
| LSIFGPV | ASSLSTGHQDTQY | 0.8284631 |
| LLALLANL | ASSLSTGHQDTQY | 0.72260785 |
| LISNASDALDKLR | ASSLSTGHQDTQY | 0.69499415 |
| LINEAVP | ASSLSTGHQDTQY | 0.78530216 |
| LGYAFSGL | ASSLSTGHQDTQY | 0.70838284 |
| LDFVAVAL | ASSLSTGHQDTQY | 0.70616233 |
| LAQPGAE | ASSLSTGHQDTQY | 0.70008564 |
| ISNLGAVTGF | ASSLSTGHQDTQY | 0.8128589 |
| HQEDPGRI | ASSLSTGHQDTQY | 0.8511176 |
| GTPGEPTP | ASSLSTGHQDTQY | 0.7484899 |
| GFFNSGS | ASSLSTGHQDTQY | 0.8381926 |
| GAGGGGLTAGAN | ASSLSTGHQDTQY | 0.8454312 |
| FVDALVNM | ASSLSTGHQDTQY | 0.4750965 |
| FLTVAPEL | ASSLSTGHQDTQY | 0.6684694 |
| FETLTAL | ASSLSTGHQDTQY | 0.85520256 |
| DVINPIEL | ASSLSTGHQDTQY | 0.701643 |
| APVSGAPV | ASSLSTGHQDTQY | 0.8663221 |
| ALVGGLAW | ASSLSTGHQDTQY | 0.789874 |
| ALAAAAATAI | ASSLSTGHQDTQY | 0.7825419 |
| AFGLGLLL | ASSLSTGHQDTQY | 0.79978085 |
| AFFAAASL | ASSLSTGHQDTQY | 0.8872199 |
| AAAALPAFSPP | ASSLSTGHQDTQY | 0.8425878 |
| VVLAPAL | ASSLSTGHQDTQY | 0.8170614 |

| | | |
|---|---|---|
| TIGHVDHGKTTL | ASSLSTGHQDTQY | 0.85041535 |
| LISNASDALDK | ASSLSTGHQDTQY | 0.7398572 |
| LISNASDAL | ASSLSTGHQDTQY | 0.86116666 |
| ISNASDALDK | ASSLSTGHQDTQY | 0.8629383 |
| NQAETLVYQTE | ASSLSTGHQDTQY | 0.5391217 |
| TGGTGGTAGAG | ASSLSTGHQDTQY | 0.7551404 |
| TGSTAGGAGGA | ASSLSTGHQDTQY | 0.8812673 |
| DPLAVPVAAGPV | ASSLSTGHQDTQY | 0.8746667 |
| GGTGGTPGNGGN | ASSLSTGHQDTQY | 0.7209867 |
| PAGLAGVDPATL | ASSLSTGHQDTQY | 0.7153716 |
| GNNLVVEAPHAGASV | ASSLSTGHQDTQY | 0.7296799 |
| KVTRIMSASTLEQPA | ASSLSTGHQDTQY | 0.8003994 |
| LGFALPIALPRRTVV | ASSLSTGHQDTQY | 0.6371589 |
| TRLGNLTARKLVVLV | ASSLSTGHQDTQY | 0.8368113 |
| WSQQRRLIYFPSAGP | ASSLSTGHQDTQY | 0.7109869 |
| GPNGNPDPMAAAVDIR | ASSLSTGHQDTQY | 0.8553759 |
| TQRCVSVADLRAAASR | ASSLSTGHQDTQY | 0.7867714 |
| QFVLDVAEHPVEVLTETV | ASSLSTGHQDTQY | 0.8441022 |
| GGAGGTGGAGGNGGTQAGDG | ASSLSTGHQDTQY | 0.77539444 |
| PLLATLRAIAVDVDAKPAQV | ASSLSTGHQDTQY | 0.7328627 |
| APKFRTGGYMIDFWGVGYQVAK | ASSLSTGHQDTQY | 0.6375184 |
| GAGGAGGAGGAAGGPGGTGGTG | ASSLSTGHQDTQY | 0.6911335 |
| IGALGLLATDVAPAGSVAFGVA | ASSLSTGHQDTQY | 0.7667283 |
| AGGAGGIGGNGGAGGAGGVLMG | ASSLSTGHQDTQY | 0.71090883 |
| ADELIGLL | ASSLSTGHQDTQY | 0.46285975 |
| ADRALKA | ASSLSTGHQDTQY | 0.8789227 |
| AGAVGQVGGAGGA | ASSLSTGHQDTQY | 0.79212606 |
| AGGGGGLGGAP | ASSLSTGHQDTQY | 0.81547964 |
| AGGLLYGNGGAG | ASSLSTGHQDTQY | 0.67104083 |
| AGLESIGL | ASSLSTGHQDTQY | 0.78615695 |
| AGLKTGII | ASSLSTGHQDTQY | 0.6487925 |
| ASLAALLL | ASSLSTGHQDTQY | 0.8594321 |
| ASVINISSI | ASSLSTGHQDTQY | 0.65135634 |
| ATPPGPA | ASSLSTGHQDTQY | 0.27998784 |
| ATVASALV | ASSLSTGHQDTQY | 0.764067 |
| ATVASLIL | ASSLSTGHQDTQY | 0.46710482 |
| AVNGPVLIRH | ASSLSTGHQDTQY | 0.8822228 |
| DGAAGAVGI | ASSLSTGHQDTQY | 0.8602278 |
| DPLLHGI | ASSLSTGHQDTQY | 0.812671 |
| ELGAALGAL | ASSLSTGHQDTQY | 0.89149207 |
| ELGRTQAV | ASSLSTGHQDTQY | 0.22715263 |
| FVAGGYLS | ASSLSTGHQDTQY | 0.6954166 |
| GASGGAGGAGGW | ASSLSTGHQDTQY | 0.8616271 |
| GGAGAGGDGGAGGI | ASSLSTGHQDTQY | 0.82299036 |
| GTGHNGSQPGMGGNG | ASSLSTGHQDTQY | 0.74726677 |
| GYAAASASAS | ASSLSTGHQDTQY | 0.8563976 |
| HQVDVAAVQQG | ASSLSTGHQDTQY | 0.6449878 |
| IGAAIAHT | ASSLSTGHQDTQY | 0.76336 |
| IGMIEVAQL | ASSLSTGHQDTQY | 0.80210555 |
| IRLAPEHTSAMD | ASSLSTGHQDTQY | 0.86825496 |
| IYNVSVL | ASSLSTGHQDTQY | 0.85422945 |

| | | |
|---|---|---|
| KQGFLGM | ASSLSTGHQDTQY | 0.6427222 |
| LESVITQL | ASSLSTGHQDTQY | 0.85138696 |
| LGLGALVVV | ASSLSTGHQDTQY | 0.8770859 |
| LIGNGGNGGSGGI | ASSLSTGHQDTQY | 0.7503237 |
| LNAINAPTL | ASSLSTGHQDTQY | 0.8154671 |
| LQLLNTGV | ASSLSTGHQDTQY | 0.8222827 |
| LQTSIEF | ASSLSTGHQDTQY | 0.76085925 |
| MSFAVSEL | ASSLSTGHQDTQY | 0.6314919 |
| NTGALGL | ASSLSTGHQDTQY | 0.62617475 |
| PPGAGGAGGAGTT | ASSLSTGHQDTQY | 0.8224783 |
| QLVGLGI | ASSLSTGHQDTQY | 0.5683775 |
| RFLEQVHQL | ASSLSTGHQDTQY | 0.82285047 |
| RGVGYVLR | ASSLSTGHQDTQY | 0.7551843 |
| SATGVFAGVI | ASSLSTGHQDTQY | 0.89220536 |
| SLPAITLP | ASSLSTGHQDTQY | 0.83467436 |
| STALAFTV | ASSLSTGHQDTQY | 0.8181429 |
| SVLTVGLGK | ASSLSTGHQDTQY | 0.59189034 |
| TALAGAGSGI | ASSLSTGHQDTQY | 0.8423773 |
| TGLPFTGI | ASSLSTGHQDTQY | 0.80074954 |
| VAEHVTV | ASSLSTGHQDTQY | 0.45902753 |
| VDIVLNSL | ASSLSTGHQDTQY | 0.86130095 |
| VLTGGGAVL | ASSLSTGHQDTQY | 0.55006075 |
| VNVGLAYGI | ASSLSTGHQDTQY | 0.7848625 |
| VSSAAVGAV | ASSLSTGHQDTQY | 0.84177953 |
| VTSVYEAGV | ASSLSTGHQDTQY | 0.86623144 |
| WVALDRGA | ASSLSTGHQDTQY | 0.7481846 |
| YLVGSVVI | ASSLSTGHQDTQY | 0.8019859 |
| GPPGTGKTLL | ASSLSTGHQDTQY | 0.65268624 |
| GVGMTKF | ASSLSTGHQDTQY | 0.854887 |
| IGIGGSDL | ASSLSTGHQDTQY | 0.85867506 |
| IGTLLAEL | ASSLSTGHQDTQY | 0.82961327 |
| IYTSGTT | ASSLSTGHQDTQY | 0.88865626 |
| LSFPAGV | ASSLSTGHQDTQY | 0.86759996 |
| LYTSGTT | ASSLSTGHQDTQY | 0.5850525 |
| VGAFPFGF | ASSLSTGHQDTQY | 0.8413202 |
| VGAGTVEF | ASSLSTGHQDTQY | 0.8683294 |
| VGKSTLLL | ASSLSTGHQDTQY | 0.86833835 |
| AALAHAGGGIV | ASSLSTGHQDTQY | 0.8609209 |
| AALVAVSLL | ASSLSTGHQDTQY | 0.8796552 |
| AAVSRAVAAL | ASSLSTGHQDTQY | 0.73775756 |
| AGGSSGIGGF | ASSLSTGHQDTQY | 0.8684207 |
| AGLTVANANG | ASSLSTGHQDTQY | 0.8981581 |
| AGLVGAVL | ASSLSTGHQDTQY | 0.84363 |
| ALSQAFTV | ASSLSTGHQDTQY | 0.74667907 |
| ALTFSVSL | ASSLSTGHQDTQY | 0.75284827 |
| ALTGAALVAL | ASSLSTGHQDTQY | 0.8491948 |
| ASFSIGAGVL | ASSLSTGHQDTQY | 0.8362805 |
| DGSVYKI | ASSLSTGHQDTQY | 0.7047334 |
| EQLAEIV | ASSLSTGHQDTQY | 0.76904815 |
| FAVSELTVM | ASSLSTGHQDTQY | 0.7980518 |
| GAAPLAAGL | ASSLSTGHQDTQY | 0.55352175 |

| | | |
|---|---|---|
| GADGGDGGSGGAGG | ASSLSTGHQDTQY | 0.8454083 |
| GFGLDTGGAGG | ASSLSTGHQDTQY | 0.8736898 |
| GLASGGLL | ASSLSTGHQDTQY | 0.8488235 |
| GRIAGAAVLL | ASSLSTGHQDTQY | 0.6926134 |
| GSAGSLNTSF | ASSLSTGHQDTQY | 0.8659321 |
| GTGQIGI | ASSLSTGHQDTQY | 0.86465514 |
| GTLIALIL | ASSLSTGHQDTQY | 0.8865474 |
| IAATMGFVVG | ASSLSTGHQDTQY | 0.41181803 |
| IFAQALL | ASSLSTGHQDTQY | 0.2610255 |
| IGAAVALGF | ASSLSTGHQDTQY | 0.80981296 |
| IGLTPLAL | ASSLSTGHQDTQY | 0.8275504 |
| ILGSNTSTL | ASSLSTGHQDTQY | 0.75993097 |
| IVFSLTVL | ASSLSTGHQDTQY | 0.7473104 |
| LASKATGF | ASSLSTGHQDTQY | 0.8264454 |
| LASVALIL | ASSLSTGHQDTQY | 0.84279764 |
| LFAVGLTL | ASSLSTGHQDTQY | 0.76956964 |
| LFNGGIGTYI | ASSLSTGHQDTQY | 0.6962329 |
| LTLGAGII | ASSLSTGHQDTQY | 0.8550823 |
| LVAAALGDI | ASSLSTGHQDTQY | 0.8665958 |
| NTAFVQLGI | ASSLSTGHQDTQY | 0.6638075 |
| QLFSGAEL | ASSLSTGHQDTQY | 0.7528783 |
| SAATFITL | ASSLSTGHQDTQY | 0.78530484 |
| SNYGLANLGV | ASSLSTGHQDTQY | 0.80644715 |
| SYGGAYAVM | ASSLSTGHQDTQY | 0.75698614 |
| TTGVGGLTLG | ASSLSTGHQDTQY | 0.58703625 |
| TTQTIYI | ASSLSTGHQDTQY | 0.83766526 |
| VAAGAGLGAVI | ASSLSTGHQDTQY | 0.43596384 |
| VAAVVTLL | ASSLSTGHQDTQY | 0.859332 |
| VGFLGGLI | ASSLSTGHQDTQY | 0.84096885 |
| VGVIPILL | ASSLSTGHQDTQY | 0.8450296 |
| VIFASSLI | ASSLSTGHQDTQY | 0.70400333 |
| VKGGDGGAAAG | ASSLSTGHQDTQY | 0.60047305 |
| VTKTFTNI | ASSLSTGHQDTQY | 0.6261575 |
| VVAAAILL | ASSLSTGHQDTQY | 0.6551522 |
| YAALDVEL | ASSLSTGHQDTQY | 0.8765495 |
| LASVALLL | ASSLSTGHQDTQY | 0.79081726 |
| LDAVHRV | ASSLSTGHQDTQY | 0.87092954 |
| QGLSSSIF | ASSLSTGHQDTQY | 0.77313936 |
| RGTFANIRL | ASSLSTGHQDTQY | 0.71871096 |
| RQLWWGHRI | ASSLSTGHQDTQY | 0.8501488 |
| YGPPGTGKTLL | ASSLSTGHQDTQY | 0.61368465 |
| AALVGAPANVA | ASSLSTGHQDTQY | 0.8200245 |
| AATFRRTE | ASSLSTGHQDTQY | 0.8256345 |
| AGALGGGI | ASSLSTGHQDTQY | 0.8591097 |
| AGGAGGAGGIGAG | ASSLSTGHQDTQY | 0.6556362 |
| AGSGALGL | ASSLSTGHQDTQY | 0.860781 |
| AQIADQI | ASSLSTGHQDTQY | 0.8317913 |
| ASAVAVTYL | ASSLSTGHQDTQY | 0.74058324 |
| ASTVLGGTGG | ASSLSTGHQDTQY | 0.84945226 |
| AVGGEPAK | ASSLSTGHQDTQY | 0.84750664 |
| CEQGTFGGTC | ASSLSTGHQDTQY | 0.8457205 |

| | | |
|---|---|---|
| DEGKAPAG | ASSLSTGHQDTQY | 0.61502624 |
| DTVVFPAL | ASSLSTGHQDTQY | 0.76961356 |
| FEGITMAG | ASSLSTGHQDTQY | 0.69481146 |
| FPLVVQD | ASSLSTGHQDTQY | 0.7997517 |
| GAFTTPQ | ASSLSTGHQDTQY | 0.8066182 |
| GFGGFGTAGGVGGA | ASSLSTGHQDTQY | 0.7902827 |
| GFLAQFL | ASSLSTGHQDTQY | 0.86249113 |
| GGAGGAGGNAGVA | ASSLSTGHQDTQY | 0.7281519 |
| GGVGTAGAGGAGGAGG | ASSLSTGHQDTQY | 0.7101028 |
| GNGGGGDAGNAGSGGN | ASSLSTGHQDTQY | 0.7311686 |
| GTGGAGGAGNDAGS | ASSLSTGHQDTQY | 0.78643286 |
| GTTNTGWFNS | ASSLSTGHQDTQY | 0.8338694 |
| IFMGPLFTL | ASSLSTGHQDTQY | 0.53520507 |
| ILAHAIEAI | ASSLSTGHQDTQY | 0.8135589 |
| IRSAIREAL | ASSLSTGHQDTQY | 0.79910237 |
| ITFAGGG | ASSLSTGHQDTQY | 0.60912794 |
| ITGMSGHQEV | ASSLSTGHQDTQY | 0.84427404 |
| IVAGLACTL | ASSLSTGHQDTQY | 0.8131121 |
| IVGGGLAGA | ASSLSTGHQDTQY | 0.6096133 |
| IVTGVLNI | ASSLSTGHQDTQY | 0.86068344 |
| LAQQGCI | ASSLSTGHQDTQY | 0.74391544 |
| LCLHVEN | ASSLSTGHQDTQY | 0.8410885 |
| LDAISDGS | ASSLSTGHQDTQY | 0.81722665 |
| LGGFAAML | ASSLSTGHQDTQY | 0.7223274 |
| LGGKNPMI | ASSLSTGHQDTQY | 0.8553997 |
| LGPKGVAAIL | ASSLSTGHQDTQY | 0.79559994 |
| LMVGVSI | ASSLSTGHQDTQY | 0.84248394 |
| LVGAAFVL | ASSLSTGHQDTQY | 0.7154001 |
| LVLAVAPL | ASSLSTGHQDTQY | 0.7158698 |
| MAAIAVIDAA | ASSLSTGHQDTQY | 0.8601631 |
| NNGINAELQ | ASSLSTGHQDTQY | 0.8122921 |
| RGEPLVLVHG | ASSLSTGHQDTQY | 0.86025083 |
| RGGAGGAAGLI | ASSLSTGHQDTQY | 0.86614615 |
| RSYRMHPAV | ASSLSTGHQDTQY | 0.8445933 |
| SAAHLLAGQLA | ASSLSTGHQDTQY | 0.8639332 |
| SAAVGSY | ASSLSTGHQDTQY | 0.6401465 |
| SPHAVLHRI | ASSLSTGHQDTQY | 0.58794713 |
| SQIAKVL | ASSLSTGHQDTQY | 0.7328456 |
| TALGHLVP | ASSLSTGHQDTQY | 0.7060349 |
| TAPTTSGAA | ASSLSTGHQDTQY | 0.7793848 |
| TDGDDGNGGSPGT | ASSLSTGHQDTQY | 0.59000486 |
| TDGLGEN | ASSLSTGHQDTQY | 0.84025156 |
| TGGPPLPGQA | ASSLSTGHQDTQY | 0.88898325 |
| TIPAITI | ASSLSTGHQDTQY | 0.65029806 |
| TIPGFTI | ASSLSTGHQDTQY | 0.89479125 |
| TSIASGALA | ASSLSTGHQDTQY | 0.6473854 |
| TVTLPGAAI | ASSLSTGHQDTQY | 0.8327023 |
| VASGLSAALP | ASSLSTGHQDTQY | 0.85593677 |
| VGAGVIGI | ASSLSTGHQDTQY | 0.5265815 |
| VGPKRFGV | ASSLSTGHQDTQY | 0.81959 |
| VNVAFPD | ASSLSTGHQDTQY | 0.67147374 |

| | | |
|---|---|---|
| VSMTNSGVS | ASSLSTGHQDTQY | 0.7237015 |
| VTDVLGDLSV | ASSLSTGHQDTQY | 0.86526173 |
| VVVVGAGFAGL | ASSLSTGHQDTQY | 0.70808023 |
| VYTVGTVV | ASSLSTGHQDTQY | 0.856217 |
| WTTFSGP | ASSLSTGHQDTQY | 0.8059655 |
| YDELVEAF | ASSLSTGHQDTQY | 0.65781903 |
| YGTGGAGGNGG | ASSLSTGHQDTQY | 0.8556525 |
| YTNVMAV | ASSLSTGHQDTQY | 0.87365866 |
| FLEQVHQL | ASSLSTGHQDTQY | 0.80436015 |
| GLSLLAVAV | ASSLSTGHQDTQY | 0.89634526 |
| LEQVHQL | ASSLSTGHQDTQY | 0.65476894 |
| LNAAGAI | ASSLSTGHQDTQY | 0.8668666 |
| PAVHASL | ASSLSTGHQDTQY | 0.8701552 |
| TELEDDA | ASSLSTGHQDTQY | 0.76115525 |
| TPPGPPN | ASSLSTGHQDTQY | 0.78689927 |
| TPPPPPPP | ASSLSTGHQDTQY | 0.8176067 |
| TQSPALI | AWSPALQGCGNTLY | 0.8225655 |
| EFAAAFPQ | AWSPALQGCGNTLY | 0.8649137 |
| LLSGNLDVL | AWSPALQGCGNTLY | 0.88549125 |
| GPANLGNYNI | AWSPALQGCGNTLY | 0.46595424 |
| TLGPTLP | AWSPALQGCGNTLY | 0.84299135 |
| VYAYSSCCK | AWSPALQGCGNTLY | 0.803826 |
| APAPAPAPNI | AWSPALQGCGNTLY | 0.75854266 |
| PGGPVPIP | AWSPALQGCGNTLY | 0.62418836 |
| FLTVALEP | AWSPALQGCGNTLY | 0.33196968 |
| SQYFGANL | AWSPALQGCGNTLY | 0.6532722 |
| GGVKPAA | AWSPALQGCGNTLY | 0.60421646 |
| GGTGGAGGAGGW | AWSPALQGCGNTLY | 0.82495344 |
| GVGGVAGLGGAGG | AWSPALQGCGNTLY | 0.8243764 |
| VLLVLATTI | AWSPALQGCGNTLY | 0.6455751 |
| IGIFAAML | AWSPALQGCGNTLY | 0.8360361 |
| AVGDLID | AWSPALQGCGNTLY | 0.7551842 |
| NISPLPL | AWSPALQGCGNTLY | 0.45157665 |
| VTAVFVAI | AWSPALQGCGNTLY | 0.76178014 |
| SGDDNDAAL | AWSPALQGCGNTLY | 0.5452582 |
| AVAALAHS | AWSPALQGCGNTLY | 0.75014806 |
| GGVTVMATP | AWSPALQGCGNTLY | 0.39576 |
| AGLAANVRD | AWSPALQGCGNTLY | 0.81105906 |
| TNRPDLIDPALLRPG | AWSPALQGCGNTLY | 0.69295645 |
| LANLGVTAL | AWSPALQGCGNTLY | 0.31253454 |
| TGERIEL | AWSPALQGCGNTLY | 0.8108641 |
| NGEELAA | AWSPALQGCGNTLY | 0.7150565 |
| VLSLVWG | AWSPALQGCGNTLY | 0.72601736 |
| IGYALWL | AWSPALQGCGNTLY | 0.8265474 |
| LSLLLAPL | AWSPALQGCGNTLY | 0.5128641 |
| TLDEALK | AWSPALQGCGNTLY | 0.5706637 |
| TDGAGTAEGG | AWSPALQGCGNTLY | 0.8277551 |
| NAPGGAGGNGG | AWSPALQGCGNTLY | 0.8280223 |
| VSVFMGMGGNF | AWSPALQGCGNTLY | 0.53386337 |
| LASYLAAL | AWSPALQGCGNTLY | 0.7715478 |
| LVGGTPAL | AWSPALQGCGNTLY | 0.82464314 |

| | | |
|---|---|---|
| VRNADVL | AWSPALQGCGNTLY | 0.74045026 |
| TCSSPFLVSAM | AWSPALQGCGNTLY | 0.826936 |
| VHITDAE | AWSPALQGCGNTLY | 0.8333028 |
| EYAAPSF | AWSPALQGCGNTLY | 0.5949509 |
| VVDAFHGGL | AWSPALQGCGNTLY | 0.8381307 |
| ASVVDPQ | AWSPALQGCGNTLY | 0.84851 |
| SYNEFGGLS | AWSPALQGCGNTLY | 0.68943715 |
| DPDGSVY | AWSPALQGCGNTLY | 0.7647456 |
| MVSTIHATE | AWSPALQGCGNTLY | 0.41423494 |
| LAAGVADPV | AWSPALQGCGNTLY | 0.6786548 |
| TKEQIAA | AWSPALQGCGNTLY | 0.40518582 |
| SLASPAAL | AWSPALQGCGNTLY | 0.8063878 |
| IGLVFYGV | AWSPALQGCGNTLY | 0.82902414 |
| FLLDASNL | AWSPALQGCGNTLY | 0.76777077 |
| YQVGNFP | AWSPALQGCGNTLY | 0.5754602 |
| ESLLSQST | AWSPALQGCGNTLY | 0.48909062 |
| LFTVGFAAVG | AWSPALQGCGNTLY | 0.7977699 |
| ATPPTPT | AWSPALQGCGNTLY | 0.44916168 |
| TDLATLG | AWSPALQGCGNTLY | 0.29784644 |
| SAGGTGR | AWSPALQGCGNTLY | 0.65861726 |
| GFSNSGS | AWSPALQGCGNTLY | 0.8124826 |
| GGNGGSGGTSV | AWSPALQGCGNTLY | 0.81272894 |
| GTSVATGGAGN | AWSPALQGCGNTLY | 0.6871799 |
| NGGNGGSGGTS | AWSPALQGCGNTLY | 0.8461118 |
| AVVKFPH | AWSPALQGCGNTLY | 0.817986 |
| TPPPPPP | AWSPALQGCGNTLY | 0.73818207 |
| AAPAAAAAPA | AWSPALQGCGNTLY | 0.61770886 |
| IALNEDL | AWSPALQGCGNTLY | 0.46849838 |
| IVNVSSM | AWSPALQGCGNTLY | 0.83932555 |
| TGIPPAPR | AWSPALQGCGNTLY | 0.75251764 |
| FDLGGGTF | AWSPALQGCGNTLY | 0.78459877 |
| FDLGGGTFDVS | AWSPALQGCGNTLY | 0.8223858 |
| NEPTAAA | AWSPALQGCGNTLY | 0.7430364 |
| NEPTAAALA | AWSPALQGCGNTLY | 0.6510962 |
| GSIINISSI | AWSPALQGCGNTLY | 0.7638528 |
| VAPGNAGTA | AWSPALQGCGNTLY | 0.83316296 |
| TGGGASLE | AWSPALQGCGNTLY | 0.6826889 |
| VAVYNHYKRI | AWSPALQGCGNTLY | 0.39244848 |
| TDPAPATT | AWSPALQGCGNTLY | 0.6618782 |
| TNRPDILDPALLRPG | AWSPALQGCGNTLY | 0.73782563 |
| LTTTTTS | AWSPALQGCGNTLY | 0.8334273 |
| PTGSGKTL | AWSPALQGCGNTLY | 0.73056173 |
| TGSGKTL | AWSPALQGCGNTLY | 0.63143027 |
| TIGHVDHGK | AWSPALQGCGNTLY | 0.8089392 |
| TIGHVDHGKTT | AWSPALQGCGNTLY | 0.76942694 |
| GHVDHGKTT | AWSPALQGCGNTLY | 0.7434271 |
| AQFEHTLL | AWSPALQGCGNTLY | 0.72343946 |
| IVNQDEL | AWSPALQGCGNTLY | 0.7304747 |
| TNGSQFF | AWSPALQGCGNTLY | 0.8229139 |
| TGKGIVE | AWSPALQGCGNTLY | 0.82129884 |
| LISNASDA | AWSPALQGCGNTLY | 0.7828584 |

| | | |
|---|---|---|
| AELRTLGELTKIAKAHGAQ | AWSPALQGCGNTLY | 0.8221346 |
| APNAVAAAEPSW | AWSPALQGCGNTLY | 0.6675645 |
| APQVAGAATAVL | AWSPALQGCGNTLY | 0.7524201 |
| ARRDPAAAAAILPTDA | AWSPALQGCGNTLY | 0.43209034 |
| DAAARDVLAEAGLAEYF | AWSPALQGCGNTLY | 0.44567817 |
| DDQIVLSAKANLAKVEQ | AWSPALQGCGNTLY | 0.7787144 |
| DELLPDTGEDSDEDGH | AWSPALQGCGNTLY | 0.8029907 |
| ELPEAPANRATVPQD | AWSPALQGCGNTLY | 0.8214209 |
| FGQSKIERVTGIG | AWSPALQGCGNTLY | 0.7499355 |
| GAGGTGSTAGGAGGAGGAGGLY | AWSPALQGCGNTLY | 0.7084189 |
| GHGGTGAAGGNGGTGGAGGSGI | AWSPALQGCGNTLY | 0.79795206 |
| GNGGNGGAGGEGGPGGAGGAGA | AWSPALQGCGNTLY | 0.8202569 |
| GSTAGNGGNGGNATNGTIAGQP | AWSPALQGCGNTLY | 0.5232527 |
| GTGGDGGNGGNGGTGG | AWSPALQGCGNTLY | 0.7402557 |
| GVRRPAGIPADAPV | AWSPALQGCGNTLY | 0.85493255 |
| GYRQVTDAHLLTLARR | AWSPALQGCGNTLY | 0.7997224 |
| IAKMGQDGHDRG | AWSPALQGCGNTLY | 0.7395252 |
| KAEPSIEIQYT | AWSPALQGCGNTLY | 0.7068832 |
| KPVRIRMAADEDTVAVT | AWSPALQGCGNTLY | 0.8394569 |
| MPPRIAGMRLLVIKPEP | AWSPALQGCGNTLY | 0.7124115 |
| MKFYAVALFRSRGVPPTR | AWSPALQGCGNTLY | 0.8528408 |
| NAGGFGILGAVAHSP | AWSPALQGCGNTLY | 0.36449158 |
| NEGREITLAMAEM | AWSPALQGCGNTLY | 0.64805037 |
| NELVEWSHPLGHRWRWE | AWSPALQGCGNTLY | 0.48246244 |
| NLGAGGTGGNGGNPG | AWSPALQGCGNTLY | 0.2146625 |
| NTEWPKWETVMAADPVD | AWSPALQGCGNTLY | 0.83342767 |
| PASAAPAETQP | AWSPALQGCGNTLY | 0.57376134 |
| QQSSDGKNGVRAAEITGE | AWSPALQGCGNTLY | 0.76251495 |
| RAPDAVTTSWWKE | AWSPALQGCGNTLY | 0.7166745 |
| SGFGNVGAHTSGWFNQ | AWSPALQGCGNTLY | 0.8361366 |
| SGVAGAVGQAASVGGLKV | AWSPALQGCGNTLY | 0.78903854 |
| SNGGAGTGSAGGNGGT | AWSPALQGCGNTLY | 0.6102396 |
| TGGAGGEGGAAG | AWSPALQGCGNTLY | 0.50760055 |
| TGGDGGAAGAGGAPGHG | AWSPALQGCGNTLY | 0.8058439 |
| TMPPLLAAIANAGRHGV | AWSPALQGCGNTLY | 0.82037985 |
| TSSTAGNSGGA | AWSPALQGCGNTLY | 0.7799827 |
| VPATVPVGILC | AWSPALQGCGNTLY | 0.33561498 |
| VQRPDSGGVVLTGRIS | AWSPALQGCGNTLY | 0.7997072 |
| WTPNPVPATKGVPAGLVQAT | AWSPALQGCGNTLY | 0.72629344 |
| YALKRALAKFSDRFDVVII | AWSPALQGCGNTLY | 0.4001647 |
| YLIGSVIV | AWSPALQGCGNTLY | 0.2784927 |
| YGQLVAVID | AWSPALQGCGNTLY | 0.6874636 |
| VTVGGMTLPA | AWSPALQGCGNTLY | 0.716574 |
| VLALPAV | AWSPALQGCGNTLY | 0.83111876 |
| VIPPTLNYET | AWSPALQGCGNTLY | 0.831529 |
| VGMGGMHPSSG | AWSPALQGCGNTLY | 0.7332984 |
| VGLAGGAATA | AWSPALQGCGNTLY | 0.79142714 |
| VGGAGGAGGSGP | AWSPALQGCGNTLY | 0.78535795 |
| VFAAFMV | AWSPALQGCGNTLY | 0.12182902 |
| TLYDKLR | AWSPALQGCGNTLY | 0.5476674 |
| TLAVFTVL | AWSPALQGCGNTLY | 0.48253483 |

| | | |
|---|---|---|
| TGSVQPARV | AWSPALQGCGNTLY | 0.8307402 |
| TGGGAGSGGT | AWSPALQGCGNTLY | 0.65604174 |
| SHLETSL | AWSPALQGCGNTLY | 0.7607375 |
| SFTTGMT | AWSPALQGCGNTLY | 0.7689678 |
| QVEVAKRMGVRQPTVS | AWSPALQGCGNTLY | 0.8060181 |
| QGSGGPALAM | AWSPALQGCGNTLY | 0.78143597 |
| MLAPSNP | AWSPALQGCGNTLY | 0.48897195 |
| LVNNAAI | AWSPALQGCGNTLY | 0.4478511 |
| LTGEINTLA | AWSPALQGCGNTLY | 0.79998654 |
| LSIFGPV | AWSPALQGCGNTLY | 0.64754826 |
| LLALLANL | AWSPALQGCGNTLY | 0.56329465 |
| LISNASDALDKLR | AWSPALQGCGNTLY | 0.5509985 |
| LINEAVP | AWSPALQGCGNTLY | 0.82277924 |
| LGYAFSGL | AWSPALQGCGNTLY | 0.4441729 |
| LDFVAVAL | AWSPALQGCGNTLY | 0.79224235 |
| LAQPGAE | AWSPALQGCGNTLY | 0.61482227 |
| ISNLGAVTGF | AWSPALQGCGNTLY | 0.7713532 |
| HQEDPGRI | AWSPALQGCGNTLY | 0.8487867 |
| GTPGEPTP | AWSPALQGCGNTLY | 0.7988539 |
| GFFNSGS | AWSPALQGCGNTLY | 0.7464912 |
| GAGGGGLTAGAN | AWSPALQGCGNTLY | 0.6428163 |
| FVDALVNM | AWSPALQGCGNTLY | 0.6472539 |
| FLTVAPEL | AWSPALQGCGNTLY | 0.6592117 |
| FETLTAL | AWSPALQGCGNTLY | 0.6829311 |
| DVINPIEL | AWSPALQGCGNTLY | 0.8245815 |
| APVSGAPV | AWSPALQGCGNTLY | 0.74295056 |
| ALVGGLAW | AWSPALQGCGNTLY | 0.46176198 |
| ALAAAAATAI | AWSPALQGCGNTLY | 0.7579533 |
| AFGLGLLL | AWSPALQGCGNTLY | 0.84908736 |
| AFFAAASL | AWSPALQGCGNTLY | 0.8449619 |
| AAAALPAFSPP | AWSPALQGCGNTLY | 0.6178092 |
| VVLAPAL | AWSPALQGCGNTLY | 0.524463 |
| TIGHVDHGKTTL | AWSPALQGCGNTLY | 0.5441025 |
| LISNASDALDK | AWSPALQGCGNTLY | 0.7674185 |
| LISNASDAL | AWSPALQGCGNTLY | 0.41417485 |
| ISNASDALDK | AWSPALQGCGNTLY | 0.5720294 |
| NQAETLVYQTE | AWSPALQGCGNTLY | 0.8070508 |
| TGGTGGTAGAG | AWSPALQGCGNTLY | 0.851961 |
| TGSTAGGAGGA | AWSPALQGCGNTLY | 0.6481308 |
| DPLAVPVAAGPV | AWSPALQGCGNTLY | 0.44566607 |
| GGTGGTPGNGGN | AWSPALQGCGNTLY | 0.7621017 |
| PAGLAGVDPATL | AWSPALQGCGNTLY | 0.61919147 |
| GNNLVVEAPHAGASV | AWSPALQGCGNTLY | 0.6675591 |
| KVTRIMSASTLEQPA | AWSPALQGCGNTLY | 0.526844 |
| LGFALPIALPRRTVV | AWSPALQGCGNTLY | 0.48977852 |
| TRLGNLTARKLVVLV | AWSPALQGCGNTLY | 0.76145643 |
| WSQQRRLIYFPSAGP | AWSPALQGCGNTLY | 0.84370804 |
| GPNGNPDPMAAAVDIR | AWSPALQGCGNTLY | 0.75251174 |
| TQRCVSVADLRAAASR | AWSPALQGCGNTLY | 0.77014035 |
| QFVLDVAEHPVEVLTETV | AWSPALQGCGNTLY | 0.8470442 |
| GGAGGTGGAGGNGGTQAGDG | AWSPALQGCGNTLY | 0.8173771 |

| | | |
|---|---|---|
| PLLATLRAIAVDVDAKPAQV | AWSPALQGCGNTLY | 0.8403411 |
| APKFRTGGYMIDFWGVGYQVAK | AWSPALQGCGNTLY | 0.8224654 |
| GAGGAGGAGGAAGGPGGTGGTG | AWSPALQGCGNTLY | 0.72186387 |
| IGALGLLATDVAPAGSVAFGVA | AWSPALQGCGNTLY | 0.70798516 |
| AGGAGGIGGNGGAGGAGGVLMG | AWSPALQGCGNTLY | 0.8417096 |
| ADELIGLL | AWSPALQGCGNTLY | 0.75401014 |
| ADRALKA | AWSPALQGCGNTLY | 0.7845505 |
| AGAVGQVGGAGGA | AWSPALQGCGNTLY | 0.69581836 |
| AGGGGGLGGAP | AWSPALQGCGNTLY | 0.84053564 |
| AGGLLYGNGGAG | AWSPALQGCGNTLY | 0.6678353 |
| AGLESIGL | AWSPALQGCGNTLY | 0.83903825 |
| AGLKTGII | AWSPALQGCGNTLY | 0.2211039 |
| ASLAALLL | AWSPALQGCGNTLY | 0.531412 |
| ASVINISSI | AWSPALQGCGNTLY | 0.8338127 |
| ATPPGPA | AWSPALQGCGNTLY | 0.13762508 |
| ATVASALV | AWSPALQGCGNTLY | 0.28652793 |
| ATVASLIL | AWSPALQGCGNTLY | 0.74853987 |
| AVNGPVLIRH | AWSPALQGCGNTLY | 0.82407224 |
| DGAAGAVGI | AWSPALQGCGNTLY | 0.17876536 |
| DPLLHGI | AWSPALQGCGNTLY | 0.61058694 |
| ELGAALGAL | AWSPALQGCGNTLY | 0.34323603 |
| ELGRTQAV | AWSPALQGCGNTLY | 0.73887897 |
| FVAGGYLS | AWSPALQGCGNTLY | 0.7609168 |
| GASGGAGGAGGW | AWSPALQGCGNTLY | 0.34020334 |
| GGAGAGGDGGAGGI | AWSPALQGCGNTLY | 0.80823493 |
| GTGHNGSQPGMGGNG | AWSPALQGCGNTLY | 0.4119339 |
| GYAAASASAS | AWSPALQGCGNTLY | 0.7532824 |
| HQVDVAAVQQG | AWSPALQGCGNTLY | 0.82963765 |
| IGAAIAHT | AWSPALQGCGNTLY | 0.846439 |
| IGMIEVAQL | AWSPALQGCGNTLY | 0.6055237 |
| IRLAPEHTSAMD | AWSPALQGCGNTLY | 0.7907843 |
| IYNVSVL | AWSPALQGCGNTLY | 0.68980205 |
| KQGFLGM | AWSPALQGCGNTLY | 0.85739625 |
| LESVITQL | AWSPALQGCGNTLY | 0.65717494 |
| LGLGALVVV | AWSPALQGCGNTLY | 0.3858732 |
| LIGNGGNGGSGGI | AWSPALQGCGNTLY | 0.82624745 |
| LNAINAPTL | AWSPALQGCGNTLY | 0.7800728 |
| LQLLNTGV | AWSPALQGCGNTLY | 0.70376515 |
| LQTSIEF | AWSPALQGCGNTLY | 0.82262146 |
| MSFAVSEL | AWSPALQGCGNTLY | 0.6184326 |
| NTGALGL | AWSPALQGCGNTLY | 0.7519154 |
| PPGAGGAGGAGTT | AWSPALQGCGNTLY | 0.8280996 |
| QLVGLGI | AWSPALQGCGNTLY | 0.82448184 |
| RFLEQVHQL | AWSPALQGCGNTLY | 0.61162555 |
| RGVGYVLR | AWSPALQGCGNTLY | 0.81423044 |
| SATGVFAGVI | AWSPALQGCGNTLY | 0.7758697 |
| SLPAITLP | AWSPALQGCGNTLY | 0.49249378 |
| STALAFTV | AWSPALQGCGNTLY | 0.8533428 |
| SVLTVGLGK | AWSPALQGCGNTLY | 0.51287395 |
| TALAGAGSGI | AWSPALQGCGNTLY | 0.8236089 |
| TGLPFTGI | AWSPALQGCGNTLY | 0.8017638 |

| | | |
|---|---|---|
| VAEHVTV | AWSPALQGCGNTLY | 0.81531537 |
| VDIVLNSL | AWSPALQGCGNTLY | 0.4565013 |
| VLTGGGAVL | AWSPALQGCGNTLY | 0.72201383 |
| VNVGLAYGI | AWSPALQGCGNTLY | 0.7217064 |
| VSSAAVGAV | AWSPALQGCGNTLY | 0.7955014 |
| VTSVYEAGV | AWSPALQGCGNTLY | 0.72379357 |
| WVALDRGA | AWSPALQGCGNTLY | 0.78869987 |
| YLVGSVVI | AWSPALQGCGNTLY | 0.81471246 |
| GPPGTGKTLL | AWSPALQGCGNTLY | 0.84323084 |
| GVGMTKF | AWSPALQGCGNTLY | 0.74828523 |
| IGIGGSDL | AWSPALQGCGNTLY | 0.7372617 |
| IGTLLAEL | AWSPALQGCGNTLY | 0.85034823 |
| IYTSGTT | AWSPALQGCGNTLY | 0.51859796 |
| LSFPAGV | AWSPALQGCGNTLY | 0.76288754 |
| LYTSGTT | AWSPALQGCGNTLY | 0.8193907 |
| VGAFPFGF | AWSPALQGCGNTLY | 0.8152499 |
| VGAGTVEF | AWSPALQGCGNTLY | 0.71930885 |
| VGKSTLLL | AWSPALQGCGNTLY | 0.82966363 |
| AALAHAGGGIV | AWSPALQGCGNTLY | 0.5121409 |
| AALVAVSLL | AWSPALQGCGNTLY | 0.811923 |
| AAVSRAVAAL | AWSPALQGCGNTLY | 0.7403428 |
| AGGSSGIGGF | AWSPALQGCGNTLY | 0.65264237 |
| AGLTVANANG | AWSPALQGCGNTLY | 0.6765919 |
| AGLVGAVL | AWSPALQGCGNTLY | 0.82439995 |
| ALSQAFTV | AWSPALQGCGNTLY | 0.82076335 |
| ALTFSVSL | AWSPALQGCGNTLY | 0.81623733 |
| ALTGAALVAL | AWSPALQGCGNTLY | 0.30952156 |
| ASFSIGAGVL | AWSPALQGCGNTLY | 0.81920236 |
| DGSVYKI | AWSPALQGCGNTLY | 0.81010604 |
| EQLAEIV | AWSPALQGCGNTLY | 0.85485435 |
| FAVSELTVM | AWSPALQGCGNTLY | 0.6556579 |
| GAAPLAAGL | AWSPALQGCGNTLY | 0.74697685 |
| GADGGDGGSGGAGG | AWSPALQGCGNTLY | 0.6888546 |
| GFGLDTGGAGG | AWSPALQGCGNTLY | 0.7315046 |
| GLASGGLL | AWSPALQGCGNTLY | 0.47809023 |
| GRIAGAAVLL | AWSPALQGCGNTLY | 0.5337169 |
| GSAGSLNTSF | AWSPALQGCGNTLY | 0.73972213 |
| GTGQIGI | AWSPALQGCGNTLY | 0.7029263 |
| GTLIALIL | AWSPALQGCGNTLY | 0.8342753 |
| IAATMGFVVG | AWSPALQGCGNTLY | 0.8309391 |
| IFAQALL | AWSPALQGCGNTLY | 0.8269532 |
| IGAAVALGF | AWSPALQGCGNTLY | 0.8354789 |
| IGLTPLAL | AWSPALQGCGNTLY | 0.8460788 |
| ILGSNTSTL | AWSPALQGCGNTLY | 0.77324134 |
| IVFSLTVL | AWSPALQGCGNTLY | 0.7062854 |
| LASKATGF | AWSPALQGCGNTLY | 0.6509719 |
| LASVALIL | AWSPALQGCGNTLY | 0.56191075 |
| LFAVGLTL | AWSPALQGCGNTLY | 0.7053988 |
| LFNGGIGTYI | AWSPALQGCGNTLY | 0.7119272 |
| LTLGAGII | AWSPALQGCGNTLY | 0.7780145 |
| LVAAALGDI | AWSPALQGCGNTLY | 0.13635665 |

| | | |
|---|---|---|
| NTAFVQLGI | AWSPALQGCGNTLY | 0.8044297 |
| QLFSGAEL | AWSPALQGCGNTLY | 0.84056044 |
| SAATFITL | AWSPALQGCGNTLY | 0.70563054 |
| SNYGLANLGV | AWSPALQGCGNTLY | 0.7873137 |
| SYGGAYAVM | AWSPALQGCGNTLY | 0.8169011 |
| TTGVGGLTLG | AWSPALQGCGNTLY | 0.678824 |
| TTQTIYI | AWSPALQGCGNTLY | 0.8046639 |
| VAAGAGLGAVI | AWSPALQGCGNTLY | 0.6800115 |
| VAAVVTLL | AWSPALQGCGNTLY | 0.80898225 |
| VGFLGGLI | AWSPALQGCGNTLY | 0.8369769 |
| VGVIPILL | AWSPALQGCGNTLY | 0.80314887 |
| VIFASSLI | AWSPALQGCGNTLY | 0.6087322 |
| VKGGDGGAAAG | AWSPALQGCGNTLY | 0.71919477 |
| VTKTFTNI | AWSPALQGCGNTLY | 0.30799353 |
| VVAAAILL | AWSPALQGCGNTLY | 0.6876224 |
| YAALDVEL | AWSPALQGCGNTLY | 0.60784155 |
| LASVALLL | AWSPALQGCGNTLY | 0.7851183 |
| LDAVHRV | AWSPALQGCGNTLY | 0.55490524 |
| QGLSSSIF | AWSPALQGCGNTLY | 0.8085094 |
| RGTFANIRL | AWSPALQGCGNTLY | 0.79453456 |
| RQLWWGHRI | AWSPALQGCGNTLY | 0.7717978 |
| YGPPGTGKTLL | AWSPALQGCGNTLY | 0.63796675 |
| AALVGAPANVA | AWSPALQGCGNTLY | 0.8009081 |
| AATFRRTE | AWSPALQGCGNTLY | 0.7820754 |
| AGALGGGI | AWSPALQGCGNTLY | 0.47693416 |
| AGGAGGAGGIGAG | AWSPALQGCGNTLY | 0.8104642 |
| AGSGALGL | AWSPALQGCGNTLY | 0.71958303 |
| AQIADQI | AWSPALQGCGNTLY | 0.7404975 |
| ASAVAVTYL | AWSPALQGCGNTLY | 0.82558054 |
| ASTVLGGTGG | AWSPALQGCGNTLY | 0.76728654 |
| AVGGEPAK | AWSPALQGCGNTLY | 0.34418732 |
| CEQGTFGGTC | AWSPALQGCGNTLY | 0.69165087 |
| DEGKAPAG | AWSPALQGCGNTLY | 0.7867988 |
| DTVVFPAL | AWSPALQGCGNTLY | 0.66752374 |
| FEGITMAG | AWSPALQGCGNTLY | 0.18633056 |
| FPLVVQD | AWSPALQGCGNTLY | 0.787074 |
| GAFTTPQ | AWSPALQGCGNTLY | 0.48834988 |
| GFGGFGTAGGVGGA | AWSPALQGCGNTLY | 0.63753355 |
| GFLAQFL | AWSPALQGCGNTLY | 0.7436576 |
| GGAGGAGGNAGVA | AWSPALQGCGNTLY | 0.8291895 |
| GGVGTAGAGGAGGAGG | AWSPALQGCGNTLY | 0.56652564 |
| GNGGGGDAGNAGSGGN | AWSPALQGCGNTLY | 0.5998205 |
| GTGGAGGAGNDAGS | AWSPALQGCGNTLY | 0.7862194 |
| GTTNTGWFNS | AWSPALQGCGNTLY | 0.791481 |
| IFMGPLFTL | AWSPALQGCGNTLY | 0.81219184 |
| ILAHAIEAI | AWSPALQGCGNTLY | 0.7655833 |
| IRSAIREAL | AWSPALQGCGNTLY | 0.6363784 |
| ITFAGGG | AWSPALQGCGNTLY | 0.7242231 |
| ITGMSGHQEV | AWSPALQGCGNTLY | 0.47354585 |
| IVAGLACTL | AWSPALQGCGNTLY | 0.6718533 |
| IVGGGLAGA | AWSPALQGCGNTLY | 0.5794192 |

| | | |
|---|---|---|
| IVTGVLNI | AWSPALQGCGNTLY | 0.6482165 |
| LAQQGCI | AWSPALQGCGNTLY | 0.59853095 |
| LCLHVEN | AWSPALQGCGNTLY | 0.7579119 |
| LDAISDGS | AWSPALQGCGNTLY | 0.80927694 |
| LGGFAAML | AWSPALQGCGNTLY | 0.628952 |
| LGGKNPMI | AWSPALQGCGNTLY | 0.37462026 |
| LGPKGVAAIL | AWSPALQGCGNTLY | 0.73902094 |
| LMVGVSI | AWSPALQGCGNTLY | 0.5884867 |
| LVGAAFVL | AWSPALQGCGNTLY | 0.8153229 |
| LVLAVAPL | AWSPALQGCGNTLY | 0.7692162 |
| MAAIAVIDAA | AWSPALQGCGNTLY | 0.8477094 |
| NNGINAELQ | AWSPALQGCGNTLY | 0.8358727 |
| RGEPLVLVHG | AWSPALQGCGNTLY | 0.74944896 |
| RGGAGGAAGLI | AWSPALQGCGNTLY | 0.83178806 |
| RSYRMHPAV | AWSPALQGCGNTLY | 0.6727332 |
| SAAHLLAGQLA | AWSPALQGCGNTLY | 0.7548674 |
| SAAVGSY | AWSPALQGCGNTLY | 0.7661599 |
| SPHAVLHRI | AWSPALQGCGNTLY | 0.7613785 |
| SQIAKVL | AWSPALQGCGNTLY | 0.57297707 |
| TALGHLVP | AWSPALQGCGNTLY | 0.7104139 |
| TAPTTSGAA | AWSPALQGCGNTLY | 0.38405532 |
| TDGDDGNGGSPGT | AWSPALQGCGNTLY | 0.8043829 |
| TDGLGEN | AWSPALQGCGNTLY | 0.4722017 |
| TGGPPLPGQA | AWSPALQGCGNTLY | 0.31533962 |
| TIPAITI | AWSPALQGCGNTLY | 0.7244597 |
| TIPGFTI | AWSPALQGCGNTLY | 0.8484669 |
| TSIASGALA | AWSPALQGCGNTLY | 0.82017714 |
| TVTLPGAAI | AWSPALQGCGNTLY | 0.70162594 |
| VASGLSAALP | AWSPALQGCGNTLY | 0.6415658 |
| VGAGVIGI | AWSPALQGCGNTLY | 0.8159354 |
| VGPKRFGV | AWSPALQGCGNTLY | 0.6282227 |
| VNVAFPD | AWSPALQGCGNTLY | 0.7905309 |
| VSMTNSGVS | AWSPALQGCGNTLY | 0.7666129 |
| VTDVLGDLSV | AWSPALQGCGNTLY | 0.75685906 |
| VVVVGAGFAGL | AWSPALQGCGNTLY | 0.46389058 |
| VYTVGTVV | AWSPALQGCGNTLY | 0.81476253 |
| WTTFSGP | AWSPALQGCGNTLY | 0.73289347 |
| YDELVEAF | AWSPALQGCGNTLY | 0.63752323 |
| YGTGGAGGNGG | AWSPALQGCGNTLY | 0.39758343 |
| YTNVMAV | AWSPALQGCGNTLY | 0.82598895 |
| FLEQVHQL | AWSPALQGCGNTLY | 0.6725153 |
| GLSLLAVAV | AWSPALQGCGNTLY | 0.4643476 |
| LEQVHQL | AWSPALQGCGNTLY | 0.21140133 |
| LNAAGAI | AWSPALQGCGNTLY | 0.7037064 |
| PAVHASL | AWSPALQGCGNTLY | 0.79692334 |
| TELEDDA | AWSPALQGCGNTLY | 0.8406775 |
| TPPGPPN | AWSPALQGCGNTLY | 0.74506164 |
| TPPPPPPP | AWSPALQGCGNTLY | 0.7898296 |
| TQSPALI | SSSQSEGTQDTQY | 0.823569 |
| EFAAAFPQ | SSSQSEGTQDTQY | 0.7616533 |
| LLSGNLDVL | SSSQSEGTQDTQY | 0.2854878 |

| | | |
|---|---|---|
| GPANLGNYNI | SSSQSEGTQDTQY | 0.743623 |
| TLGPTLP | SSSQSEGTQDTQY | 0.7445693 |
| VYAYSSCCK | SSSQSEGTQDTQY | 0.83080053 |
| APAPAPAPNI | SSSQSEGTQDTQY | 0.7858007 |
| PGGPVPIP | SSSQSEGTQDTQY | 0.59607244 |
| FLTVALEP | SSSQSEGTQDTQY | 0.7502574 |
| SQYFGANL | SSSQSEGTQDTQY | 0.7847956 |
| GGVKPAA | SSSQSEGTQDTQY | 0.46768904 |
| GGTGGAGGAGGW | SSSQSEGTQDTQY | 0.633921 |
| GVGGVAGLGGAGG | SSSQSEGTQDTQY | 0.57385474 |
| VLLVLATTI | SSSQSEGTQDTQY | 0.44967398 |
| IGIFAAML | SSSQSEGTQDTQY | 0.80074346 |
| AVGDLID | SSSQSEGTQDTQY | 0.762123 |
| NISPLPL | SSSQSEGTQDTQY | 0.6634309 |
| VTAVFVAI | SSSQSEGTQDTQY | 0.76008177 |
| SGDDNDAAL | SSSQSEGTQDTQY | 0.6638264 |
| AVAALAHS | SSSQSEGTQDTQY | 0.64087015 |
| GGVTVMATP | SSSQSEGTQDTQY | 0.7107599 |
| AGLAANVRD | SSSQSEGTQDTQY | 0.7854634 |
| TNRPDLIDPALLRPG | SSSQSEGTQDTQY | 0.8204224 |
| LANLGVTAL | SSSQSEGTQDTQY | 0.7413378 |
| TGERIEL | SSSQSEGTQDTQY | 0.67701715 |
| NGEELAA | SSSQSEGTQDTQY | 0.8542124 |
| VLSLVWG | SSSQSEGTQDTQY | 0.37217748 |
| IGYALWL | SSSQSEGTQDTQY | 0.60781956 |
| LSLLLAPL | SSSQSEGTQDTQY | 0.76223576 |
| TLDEALK | SSSQSEGTQDTQY | 0.69256437 |
| TDGAGTAEGG | SSSQSEGTQDTQY | 0.80945885 |
| NAPGGAGGNGG | SSSQSEGTQDTQY | 0.80320376 |
| VSVFMGMGGNF | SSSQSEGTQDTQY | 0.8003139 |
| LASYLAAL | SSSQSEGTQDTQY | 0.7357605 |
| LVGGTPAL | SSSQSEGTQDTQY | 0.84684384 |
| VRNADVL | SSSQSEGTQDTQY | 0.8148438 |
| TCSSPFLVSAM | SSSQSEGTQDTQY | 0.6840944 |
| VHITDAE | SSSQSEGTQDTQY | 0.8204198 |
| EYAAPSF | SSSQSEGTQDTQY | 0.32698807 |
| VVDAFHGGL | SSSQSEGTQDTQY | 0.78894436 |
| ASVVDPQ | SSSQSEGTQDTQY | 0.7974372 |
| SYNEFGGLS | SSSQSEGTQDTQY | 0.5849371 |
| DPDGSVY | SSSQSEGTQDTQY | 0.7563817 |
| MVSTIHATE | SSSQSEGTQDTQY | 0.8433207 |
| LAAGVADPV | SSSQSEGTQDTQY | 0.8340628 |
| TKEQIAA | SSSQSEGTQDTQY | 0.5244942 |
| SLASPAAL | SSSQSEGTQDTQY | 0.67338264 |
| IGLVFYGV | SSSQSEGTQDTQY | 0.47532284 |
| FLLDASNL | SSSQSEGTQDTQY | 0.47468546 |
| YQVGNFP | SSSQSEGTQDTQY | 0.33151695 |
| ESLLSQST | SSSQSEGTQDTQY | 0.7749485 |
| LFTVGFAAVG | SSSQSEGTQDTQY | 0.7336934 |
| ATPPTPT | SSSQSEGTQDTQY | 0.818459 |
| TDLATLG | SSSQSEGTQDTQY | 0.7778355 |

| | | |
|---|---|---|
| SAGGTGR | SSSQSEGTQDTQY | 0.8386052 |
| GFSNSGS | SSSQSEGTQDTQY | 0.6899831 |
| GGNGGSGGTSV | SSSQSEGTQDTQY | 0.77575225 |
| GTSVATGGAGN | SSSQSEGTQDTQY | 0.31467962 |
| NGGNGGSGGTS | SSSQSEGTQDTQY | 0.48160622 |
| AVVKFPH | SSSQSEGTQDTQY | 0.56844914 |
| TPPPPPP | SSSQSEGTQDTQY | 0.74465734 |
| AAPAAAAAPA | SSSQSEGTQDTQY | 0.25356302 |
| IALNEDL | SSSQSEGTQDTQY | 0.8380547 |
| IVNVSSM | SSSQSEGTQDTQY | 0.68299615 |
| TGIPPAPR | SSSQSEGTQDTQY | 0.8010427 |
| FDLGGGTF | SSSQSEGTQDTQY | 0.67481357 |
| FDLGGGTFDVS | SSSQSEGTQDTQY | 0.48401684 |
| NEPTAAA | SSSQSEGTQDTQY | 0.7013719 |
| NEPTAAALA | SSSQSEGTQDTQY | 0.81352144 |
| GSIINISSI | SSSQSEGTQDTQY | 0.6181693 |
| VAPGNAGTA | SSSQSEGTQDTQY | 0.25628233 |
| TGGGASLE | SSSQSEGTQDTQY | 0.4480307 |
| VAVYNHYKRI | SSSQSEGTQDTQY | 0.6435541 |
| TDPAPATT | SSSQSEGTQDTQY | 0.85174626 |
| TNRPDILDPALLRPG | SSSQSEGTQDTQY | 0.83703303 |
| LTTTTTS | SSSQSEGTQDTQY | 0.786841 |
| PTGSGKTL | SSSQSEGTQDTQY | 0.84625846 |
| TGSGKTL | SSSQSEGTQDTQY | 0.1484877 |
| TIGHVDHGK | SSSQSEGTQDTQY | 0.68984264 |
| TIGHVDHGKTT | SSSQSEGTQDTQY | 0.8193062 |
| GHVDHGKTT | SSSQSEGTQDTQY | 0.7345902 |
| AQFEHTLL | SSSQSEGTQDTQY | 0.74989074 |
| IVNQDEL | SSSQSEGTQDTQY | 0.8329965 |
| TNGSQFF | SSSQSEGTQDTQY | 0.68600535 |
| TGKGIVE | SSSQSEGTQDTQY | 0.66635746 |
| LISNASDA | SSSQSEGTQDTQY | 0.7580177 |
| AELRTLGELTKIAKAHGAQ | SSSQSEGTQDTQY | 0.81207985 |
| APNAVAAAEPSW | SSSQSEGTQDTQY | 0.7413177 |
| APQVAGAATAVL | SSSQSEGTQDTQY | 0.46506524 |
| ARRDPAAAAAILPTDA | SSSQSEGTQDTQY | 0.75337875 |
| DAAARDVLAEAGLAEYF | SSSQSEGTQDTQY | 0.8656697 |
| DDQIVLSAKANLAKVEQ | SSSQSEGTQDTQY | 0.74994695 |
| DELLPDTGEDSDEDGH | SSSQSEGTQDTQY | 0.80531275 |
| ELPEAPANRATVPQD | SSSQSEGTQDTQY | 0.7312722 |
| FGQSKIERVTGIG | SSSQSEGTQDTQY | 0.6985181 |
| GAGGTGSTAGGAGGAGGAGGLY | SSSQSEGTQDTQY | 0.6687205 |
| GHGGTGAAGGNGGTGGAGGSGI | SSSQSEGTQDTQY | 0.58577716 |
| GNGGNGGAGGEGGPGGAGGAGA | SSSQSEGTQDTQY | 0.8068719 |
| GSTAGNGGNGGNATNGTIAGQP | SSSQSEGTQDTQY | 0.5164668 |
| GTGGDGGNGGNGGTGG | SSSQSEGTQDTQY | 0.7248974 |
| GVRRPAGIPADAPV | SSSQSEGTQDTQY | 0.7006007 |
| GYRQVTDAHLLTLARR | SSSQSEGTQDTQY | 0.8628698 |
| IAKMGQDGHDRG | SSSQSEGTQDTQY | 0.80709875 |
| KAEPSIEIQYT | SSSQSEGTQDTQY | 0.76482254 |
| KPVRIRMAADEDTVAVT | SSSQSEGTQDTQY | 0.68929005 |

| | | |
|---|---|---|
| MPPRIAGMRLLVIKPEP | SSSQSEGTQDTQY | 0.7730409 |
| MKFYAVALFRSRGVPPTR | SSSQSEGTQDTQY | 0.81615865 |
| NAGGFGILGAVAHSP | SSSQSEGTQDTQY | 0.41917598 |
| NEGREITLAMAEM | SSSQSEGTQDTQY | 0.8286561 |
| NELVEWSHPLGHRWRWE | SSSQSEGTQDTQY | 0.5614341 |
| NLGAGGTGGNGGNPG | SSSQSEGTQDTQY | 0.7593496 |
| NTEWPKWETVMAADPVD | SSSQSEGTQDTQY | 0.6906605 |
| PASAAPAETQP | SSSQSEGTQDTQY | 0.8411431 |
| QQSSDGKNGVRAAEITGE | SSSQSEGTQDTQY | 0.7729391 |
| RAPDAVTTSWWKE | SSSQSEGTQDTQY | 0.8000216 |
| SGFGNVGAHTSGWFNQ | SSSQSEGTQDTQY | 0.6756038 |
| SGVAGAVGQAASVGGLKV | SSSQSEGTQDTQY | 0.7690036 |
| SNGGAGTGSAGGNGGT | SSSQSEGTQDTQY | 0.8426746 |
| TGGAGGEGGAAG | SSSQSEGTQDTQY | 0.67573476 |
| TGGDGGAAGAGGAPGHG | SSSQSEGTQDTQY | 0.85363704 |
| TMPPLLAAIANAGRHGV | SSSQSEGTQDTQY | 0.724509 |
| TSSTAGNSGGA | SSSQSEGTQDTQY | 0.24326764 |
| VPATVPVGILC | SSSQSEGTQDTQY | 0.66261363 |
| VQRPDSGGVVLTGRIS | SSSQSEGTQDTQY | 0.83883834 |
| WTPNPVPATKGVPAGLVQAT | SSSQSEGTQDTQY | 0.8424264 |
| YALKRALAKFSDRFDVVII | SSSQSEGTQDTQY | 0.8260945 |
| YLIGSVIV | SSSQSEGTQDTQY | 0.7841847 |
| YGQLVAVID | SSSQSEGTQDTQY | 0.7382681 |
| VTVGGMTLPA | SSSQSEGTQDTQY | 0.84474707 |
| VLALPAV | SSSQSEGTQDTQY | 0.84157413 |
| VIPPTLNYET | SSSQSEGTQDTQY | 0.75913894 |
| VGMGGMHPSSG | SSSQSEGTQDTQY | 0.46729246 |
| VGLAGGAATA | SSSQSEGTQDTQY | 0.34984857 |
| VGGAGGAGGSGP | SSSQSEGTQDTQY | 0.81892025 |
| VFAAFMV | SSSQSEGTQDTQY | 0.8328731 |
| TLYDKLR | SSSQSEGTQDTQY | 0.42956465 |
| TLAVFTVL | SSSQSEGTQDTQY | 0.768611 |
| TGSVQPARV | SSSQSEGTQDTQY | 0.78166515 |
| TGGGAGSGGT | SSSQSEGTQDTQY | 0.28162932 |
| SHLETSL | SSSQSEGTQDTQY | 0.7472662 |
| SFTTGMT | SSSQSEGTQDTQY | 0.8409326 |
| QVEVAKRMGVRQPTVS | SSSQSEGTQDTQY | 0.77791 |
| QGSGGPALAM | SSSQSEGTQDTQY | 0.46431416 |
| MLAPSNP | SSSQSEGTQDTQY | 0.83439094 |
| LVNNAAI | SSSQSEGTQDTQY | 0.85863197 |
| LTGEINTLA | SSSQSEGTQDTQY | 0.8440032 |
| LSIFGPV | SSSQSEGTQDTQY | 0.37579054 |
| LLALLANL | SSSQSEGTQDTQY | 0.1994332 |
| LISNASDALDKLR | SSSQSEGTQDTQY | 0.7818916 |
| LINEAVP | SSSQSEGTQDTQY | 0.81134796 |
| LGYAFSGL | SSSQSEGTQDTQY | 0.73042834 |
| LDFVAVAL | SSSQSEGTQDTQY | 0.61132646 |
| LAQPGAE | SSSQSEGTQDTQY | 0.7914045 |
| ISNLGAVTGF | SSSQSEGTQDTQY | 0.8365729 |
| HQEDPGRI | SSSQSEGTQDTQY | 0.5006681 |
| GTPGEPTP | SSSQSEGTQDTQY | 0.30639678 |

| | | |
|---|---|---|
| GFFNSGS | SSSQSEGTQDTQY | 0.8252227 |
| GAGGGGLTAGAN | SSSQSEGTQDTQY | 0.8156316 |
| FVDALVNM | SSSQSEGTQDTQY | 0.6160367 |
| FLTVAPEL | SSSQSEGTQDTQY | 0.74802744 |
| FETLTAL | SSSQSEGTQDTQY | 0.79031885 |
| DVINPIEL | SSSQSEGTQDTQY | 0.77233 |
| APVSGAPV | SSSQSEGTQDTQY | 0.7926233 |
| ALVGGLAW | SSSQSEGTQDTQY | 0.44656095 |
| ALAAAAATAI | SSSQSEGTQDTQY | 0.8410827 |
| AFGLGLLL | SSSQSEGTQDTQY | 0.55690545 |
| AFFAAASL | SSSQSEGTQDTQY | 0.80956066 |
| AAAALPAFSPP | SSSQSEGTQDTQY | 0.7795876 |
| VVLAPAL | SSSQSEGTQDTQY | 0.7842015 |
| TIGHVDHGKTTL | SSSQSEGTQDTQY | 0.7614875 |
| LISNASDALDK | SSSQSEGTQDTQY | 0.49239278 |
| LISNASDAL | SSSQSEGTQDTQY | 0.5793893 |
| ISNASDALDK | SSSQSEGTQDTQY | 0.48841923 |
| NQAETLVYQTE | SSSQSEGTQDTQY | 0.83545065 |
| TGGTGGTAGAG | SSSQSEGTQDTQY | 0.7461849 |
| TGSTAGGAGGA | SSSQSEGTQDTQY | 0.82342714 |
| DPLAVPVAAGPV | SSSQSEGTQDTQY | 0.78176296 |
| GGTGGTPGNGGN | SSSQSEGTQDTQY | 0.63690805 |
| PAGLAGVDPATL | SSSQSEGTQDTQY | 0.8010274 |
| GNNLVVEAPHAGASV | SSSQSEGTQDTQY | 0.4264734 |
| KVTRIMSASTLEQPA | SSSQSEGTQDTQY | 0.7465219 |
| LGFALPIALPRRTVV | SSSQSEGTQDTQY | 0.57631856 |
| TRLGNLTARKLVVLV | SSSQSEGTQDTQY | 0.7591516 |
| WSQQRRLIYFPSAGP | SSSQSEGTQDTQY | 0.6480236 |
| GPNGNPDPMAAAVDIR | SSSQSEGTQDTQY | 0.7899017 |
| TQRCVSVADLRAAASR | SSSQSEGTQDTQY | 0.7974623 |
| QFVLDVAEHPVEVLTETV | SSSQSEGTQDTQY | 0.39020944 |
| GGAGGTGGAGGNGGTQAGDG | SSSQSEGTQDTQY | 0.76897484 |
| PLLATLRAIAVDVDAKPAQV | SSSQSEGTQDTQY | 0.8367273 |
| APKFRTGGYMIDFWGVGYQVAK | SSSQSEGTQDTQY | 0.7776623 |
| GAGGAGGAGGAAGGPGGTGGTG | SSSQSEGTQDTQY | 0.5421111 |
| IGALGLLATDVAPAGSVAFGVA | SSSQSEGTQDTQY | 0.79559094 |
| AGGAGGIGGNGGAGGAGGVLMG | SSSQSEGTQDTQY | 0.64714754 |
| ADELIGLL | SSSQSEGTQDTQY | 0.80419195 |
| ADRALKA | SSSQSEGTQDTQY | 0.7807052 |
| AGAVGQVGGAGGA | SSSQSEGTQDTQY | 0.60874903 |
| AGGGGGLGGAP | SSSQSEGTQDTQY | 0.79925287 |
| AGGLLYGNGGAG | SSSQSEGTQDTQY | 0.26837966 |
| AGLESIGL | SSSQSEGTQDTQY | 0.61796266 |
| AGLKTGII | SSSQSEGTQDTQY | 0.67372555 |
| ASLAALLL | SSSQSEGTQDTQY | 0.7989489 |
| ASVINISSI | SSSQSEGTQDTQY | 0.7836891 |
| ATPPGPA | SSSQSEGTQDTQY | 0.55088115 |
| ATVASALV | SSSQSEGTQDTQY | 0.7517773 |
| ATVASLIL | SSSQSEGTQDTQY | 0.6552075 |
| AVNGPVLIRH | SSSQSEGTQDTQY | 0.5339212 |
| DGAAGAVGI | SSSQSEGTQDTQY | 0.73949873 |

| | | |
|---|---|---|
| DPLLHGI | SSSQSEGTQDTQY | 0.6103591 |
| ELGAALGAL | SSSQSEGTQDTQY | 0.29879022 |
| ELGRTQAV | SSSQSEGTQDTQY | 0.8081515 |
| FVAGGYLS | SSSQSEGTQDTQY | 0.61167014 |
| GASGGAGGAGGW | SSSQSEGTQDTQY | 0.7707952 |
| GGAGAGGDGGAGGI | SSSQSEGTQDTQY | 0.8169082 |
| GTGHNGSQPGMGGNG | SSSQSEGTQDTQY | 0.30373797 |
| GYAAASASAS | SSSQSEGTQDTQY | 0.79265845 |
| HQVDVAAVQQG | SSSQSEGTQDTQY | 0.7308368 |
| IGAAIAHT | SSSQSEGTQDTQY | 0.787431 |
| IGMIEVAQL | SSSQSEGTQDTQY | 0.76680124 |
| IRLAPEHTSAMD | SSSQSEGTQDTQY | 0.68349195 |
| IYNVSVL | SSSQSEGTQDTQY | 0.8461448 |
| KQGFLGM | SSSQSEGTQDTQY | 0.79621315 |
| LESVITQL | SSSQSEGTQDTQY | 0.6240579 |
| LGLGALVVV | SSSQSEGTQDTQY | 0.76082146 |
| LIGNGGNGGSGGI | SSSQSEGTQDTQY | 0.8215773 |
| LNAINAPTL | SSSQSEGTQDTQY | 0.8160833 |
| LQLLNTGV | SSSQSEGTQDTQY | 0.679376 |
| LQTSIEF | SSSQSEGTQDTQY | 0.31299585 |
| MSFAVSEL | SSSQSEGTQDTQY | 0.77067375 |
| NTGALGL | SSSQSEGTQDTQY | 0.7473228 |
| PPGAGGAGGAGTT | SSSQSEGTQDTQY | 0.57066315 |
| QLVGLGI | SSSQSEGTQDTQY | 0.24196592 |
| RFLEQVHQL | SSSQSEGTQDTQY | 0.811822 |
| RGVGYVLR | SSSQSEGTQDTQY | 0.8586913 |
| SATGVFAGVI | SSSQSEGTQDTQY | 0.6033548 |
| SLPAITLP | SSSQSEGTQDTQY | 0.8536658 |
| STALAFTV | SSSQSEGTQDTQY | 0.45934522 |
| SVLTVGLGK | SSSQSEGTQDTQY | 0.80887485 |
| TALAGAGSGI | SSSQSEGTQDTQY | 0.8200451 |
| TGLPFTGI | SSSQSEGTQDTQY | 0.5267772 |
| VAEHVTV | SSSQSEGTQDTQY | 0.73635787 |
| VDIVLNSL | SSSQSEGTQDTQY | 0.66223073 |
| VLTGGGAVL | SSSQSEGTQDTQY | 0.6063137 |
| VNVGLAYGI | SSSQSEGTQDTQY | 0.82661426 |
| VSSAAVGAV | SSSQSEGTQDTQY | 0.65136755 |
| VTSVYEAGV | SSSQSEGTQDTQY | 0.77232236 |
| WVALDRGA | SSSQSEGTQDTQY | 0.69625294 |
| YLVGSVVI | SSSQSEGTQDTQY | 0.6210197 |
| GPPGTGKTLL | SSSQSEGTQDTQY | 0.79483515 |
| GVGMTKF | SSSQSEGTQDTQY | 0.78167343 |
| IGIGGSDL | SSSQSEGTQDTQY | 0.69265234 |
| IGTLLAEL | SSSQSEGTQDTQY | 0.8206851 |
| IYTSGTT | SSSQSEGTQDTQY | 0.6656642 |
| LSFPAGV | SSSQSEGTQDTQY | 0.86044276 |
| LYTSGTT | SSSQSEGTQDTQY | 0.83344054 |
| VGAFPFGF | SSSQSEGTQDTQY | 0.7921018 |
| VGAGTVEF | SSSQSEGTQDTQY | 0.67890084 |
| VGKSTLLL | SSSQSEGTQDTQY | 0.8368665 |
| AALAHAGGGIV | SSSQSEGTQDTQY | 0.6898495 |

| | | |
|---|---|---|
| AALVAVSLL | SSSQSEGTQDTQY | 0.756408 |
| AAVSRAVAAL | SSSQSEGTQDTQY | 0.83132374 |
| AGGSSGIGGF | SSSQSEGTQDTQY | 0.56843543 |
| AGLTVANANG | SSSQSEGTQDTQY | 0.84204113 |
| AGLVGAVL | SSSQSEGTQDTQY | 0.75245965 |
| ALSQAFTV | SSSQSEGTQDTQY | 0.74069107 |
| ALTFSVSL | SSSQSEGTQDTQY | 0.603124 |
| ALTGAALVAL | SSSQSEGTQDTQY | 0.42559236 |
| ASFSIGAGVL | SSSQSEGTQDTQY | 0.6989922 |
| DGSVYKI | SSSQSEGTQDTQY | 0.53862685 |
| EQLAEIV | SSSQSEGTQDTQY | 0.80325115 |
| FAVSELTVM | SSSQSEGTQDTQY | 0.79857457 |
| GAAPLAAGL | SSSQSEGTQDTQY | 0.57517415 |
| GADGGDGGSGGAGG | SSSQSEGTQDTQY | 0.83157706 |
| GFGLDTGGAGG | SSSQSEGTQDTQY | 0.8314694 |
| GLASGGLL | SSSQSEGTQDTQY | 0.4222225 |
| GRIAGAAVLL | SSSQSEGTQDTQY | 0.6774358 |
| GSAGSLNTSF | SSSQSEGTQDTQY | 0.6311478 |
| GTGQIGI | SSSQSEGTQDTQY | 0.69274926 |
| GTLIALIL | SSSQSEGTQDTQY | 0.2905155 |
| IAATMGFVVG | SSSQSEGTQDTQY | 0.8054584 |
| IFAQALL | SSSQSEGTQDTQY | 0.5893944 |
| IGAAVALGF | SSSQSEGTQDTQY | 0.18262967 |
| IGLTPLAL | SSSQSEGTQDTQY | 0.7290318 |
| ILGSNTSTL | SSSQSEGTQDTQY | 0.7014352 |
| IVFSLTVL | SSSQSEGTQDTQY | 0.68150413 |
| LASKATGF | SSSQSEGTQDTQY | 0.8198709 |
| LASVALIL | SSSQSEGTQDTQY | 0.54323167 |
| LFAVGLTL | SSSQSEGTQDTQY | 0.5368094 |
| LFNGGIGTYI | SSSQSEGTQDTQY | 0.8021718 |
| LTLGAGII | SSSQSEGTQDTQY | 0.8267983 |
| LVAAALGDI | SSSQSEGTQDTQY | 0.46349645 |
| NTAFVQLGI | SSSQSEGTQDTQY | 0.769364 |
| QLFSGAEL | SSSQSEGTQDTQY | 0.8238169 |
| SAATFITL | SSSQSEGTQDTQY | 0.76969683 |
| SNYGLANLGV | SSSQSEGTQDTQY | 0.82740897 |
| SYGGAYAVM | SSSQSEGTQDTQY | 0.8377338 |
| TTGVGGLTLG | SSSQSEGTQDTQY | 0.6164936 |
| TTQTIYI | SSSQSEGTQDTQY | 0.82342386 |
| VAAGAGLGAVI | SSSQSEGTQDTQY | 0.8436233 |
| VAAVVTLL | SSSQSEGTQDTQY | 0.682808 |
| VGFLGGLI | SSSQSEGTQDTQY | 0.7206923 |
| VGVIPILL | SSSQSEGTQDTQY | 0.5180527 |
| VIFASSLI | SSSQSEGTQDTQY | 0.6787603 |
| VKGGDGGAAAG | SSSQSEGTQDTQY | 0.34375736 |
| VTKTFTNI | SSSQSEGTQDTQY | 0.7721938 |
| VVAAAILL | SSSQSEGTQDTQY | 0.8255347 |
| YAALDVEL | SSSQSEGTQDTQY | 0.67165184 |
| LASVALLL | SSSQSEGTQDTQY | 0.5134724 |
| LDAVHRV | SSSQSEGTQDTQY | 0.5036242 |
| QGLSSSIF | SSSQSEGTQDTQY | 0.7292402 |

| | | |
|---|---|---|
| RGTFANIRL | SSSQSEGTQDTQY | 0.60909545 |
| RQLWWGHRI | SSSQSEGTQDTQY | 0.38212028 |
| YGPPGTGKTLL | SSSQSEGTQDTQY | 0.6511564 |
| AALVGAPANVA | SSSQSEGTQDTQY | 0.8158506 |
| AATFRRTE | SSSQSEGTQDTQY | 0.81690836 |
| AGALGGGI | SSSQSEGTQDTQY | 0.3783183 |
| AGGAGGAGGIGAG | SSSQSEGTQDTQY | 0.8430195 |
| AGSGALGL | SSSQSEGTQDTQY | 0.79734147 |
| AQIADQI | SSSQSEGTQDTQY | 0.8387718 |
| ASAVAVTYL | SSSQSEGTQDTQY | 0.7451346 |
| ASTVLGGTGG | SSSQSEGTQDTQY | 0.57175887 |
| AVGGEPAK | SSSQSEGTQDTQY | 0.8211764 |
| CEQGTFGGTC | SSSQSEGTQDTQY | 0.64914775 |
| DEGKAPAG | SSSQSEGTQDTQY | 0.54888034 |
| DTVVFPAL | SSSQSEGTQDTQY | 0.80263174 |
| FEGITMAG | SSSQSEGTQDTQY | 0.692866 |
| FPLVVQD | SSSQSEGTQDTQY | 0.58360493 |
| GAFTTPQ | SSSQSEGTQDTQY | 0.6221069 |
| GFGGFGTAGGVGGA | SSSQSEGTQDTQY | 0.827675 |
| GFLAQFL | SSSQSEGTQDTQY | 0.49983838 |
| GGAGGAGGNAGVA | SSSQSEGTQDTQY | 0.49949914 |
| GGVGTAGAGGAGGAGG | SSSQSEGTQDTQY | 0.57131803 |
| GNGGGGDAGNAGSGGN | SSSQSEGTQDTQY | 0.7069775 |
| GTGGAGGAGNDAGS | SSSQSEGTQDTQY | 0.8174402 |
| GTTNTGWFNS | SSSQSEGTQDTQY | 0.7760441 |
| IFMGPLFTL | SSSQSEGTQDTQY | 0.64881563 |
| ILAHAIEAI | SSSQSEGTQDTQY | 0.7648536 |
| IRSAIREAL | SSSQSEGTQDTQY | 0.78404623 |
| ITFAGGG | SSSQSEGTQDTQY | 0.74584067 |
| ITGMSGHQEV | SSSQSEGTQDTQY | 0.73984253 |
| IVAGLACTL | SSSQSEGTQDTQY | 0.5946584 |
| IVGGGLAGA | SSSQSEGTQDTQY | 0.83092713 |
| IVTGVLNI | SSSQSEGTQDTQY | 0.7809087 |
| LAQQGCI | SSSQSEGTQDTQY | 0.806823 |
| LCLHVEN | SSSQSEGTQDTQY | 0.8186201 |
| LDAISDGS | SSSQSEGTQDTQY | 0.77058554 |
| LGGFAAML | SSSQSEGTQDTQY | 0.7586617 |
| LGGKNPMI | SSSQSEGTQDTQY | 0.30402392 |
| LGPKGVAAIL | SSSQSEGTQDTQY | 0.666908 |
| LMVGVSI | SSSQSEGTQDTQY | 0.7208105 |
| LVGAAFVL | SSSQSEGTQDTQY | 0.76457757 |
| LVLAVAPL | SSSQSEGTQDTQY | 0.7918558 |
| MAAIAVIDAA | SSSQSEGTQDTQY | 0.7909399 |
| NNGINAELQ | SSSQSEGTQDTQY | 0.60913634 |
| RGEPLVLVHG | SSSQSEGTQDTQY | 0.8052789 |
| RGGAGGAAGLI | SSSQSEGTQDTQY | 0.82734025 |
| RSYRMHPAV | SSSQSEGTQDTQY | 0.4247767 |
| SAAHLLAGQLA | SSSQSEGTQDTQY | 0.662659 |
| SAAVGSY | SSSQSEGTQDTQY | 0.8614455 |
| SPHAVLHRI | SSSQSEGTQDTQY | 0.76057625 |
| SQIAKVL | SSSQSEGTQDTQY | 0.64758223 |

| | | |
|---|---|---|
| TALGHLVP | SSSQSEGTQDTQY | 0.64498883 |
| TAPTTSGAA | SSSQSEGTQDTQY | 0.8481486 |
| TDGDDGNGGSPGT | SSSQSEGTQDTQY | 0.63345087 |
| TDGLGEN | SSSQSEGTQDTQY | 0.8401423 |
| TGGPPLPGQA | SSSQSEGTQDTQY | 0.23178926 |
| TIPAITI | SSSQSEGTQDTQY | 0.67821395 |
| TIPGFTI | SSSQSEGTQDTQY | 0.60138434 |
| TSIASGALA | SSSQSEGTQDTQY | 0.30299136 |
| TVTLPGAAI | SSSQSEGTQDTQY | 0.80811054 |
| VASGLSAALP | SSSQSEGTQDTQY | 0.6874189 |
| VGAGVIGI | SSSQSEGTQDTQY | 0.8029654 |
| VGPKRFGV | SSSQSEGTQDTQY | 0.72714233 |
| VNVAFPD | SSSQSEGTQDTQY | 0.8206917 |
| VSMTNSGVS | SSSQSEGTQDTQY | 0.7450095 |
| VTDVLGDLSV | SSSQSEGTQDTQY | 0.71993184 |
| VVVVGAGFAGL | SSSQSEGTQDTQY | 0.5828209 |
| VYTVGTVV | SSSQSEGTQDTQY | 0.791052 |
| WTTFSGP | SSSQSEGTQDTQY | 0.82518256 |
| YDELVEAF | SSSQSEGTQDTQY | 0.8080605 |
| YGTGGAGGNGG | SSSQSEGTQDTQY | 0.35404187 |
| YTNVMAV | SSSQSEGTQDTQY | 0.70189273 |
| FLEQVHQL | SSSQSEGTQDTQY | 0.7265836 |
| GLSLLAVAV | SSSQSEGTQDTQY | 0.30911258 |
| LEQVHQL | SSSQSEGTQDTQY | 0.27086687 |
| LNAAGAI | SSSQSEGTQDTQY | 0.6988325 |
| PAVHASL | SSSQSEGTQDTQY | 0.6198427 |
| TELEDDA | SSSQSEGTQDTQY | 0.81335753 |
| TPPGPPN | SSSQSEGTQDTQY | 0.7935066 |
| TPPPPPPP | SSSQSEGTQDTQY | 0.6682594 |
| TQSPALI | ASSLDEGGAETLY | 0.7840884 |
| EFAAAFPQ | ASSLDEGGAETLY | 0.71165705 |
| LLSGNLDVL | ASSLDEGGAETLY | 0.12671883 |
| GPANLGNYNI | ASSLDEGGAETLY | 0.67307776 |
| TLGPTLP | ASSLDEGGAETLY | 0.42177695 |
| VYAYSSCCK | ASSLDEGGAETLY | 0.8435697 |
| APAPAPAPNI | ASSLDEGGAETLY | 0.59866405 |
| PGGPVPIP | ASSLDEGGAETLY | 0.56875217 |
| FLTVALEP | ASSLDEGGAETLY | 0.69985294 |
| SQYFGANL | ASSLDEGGAETLY | 0.7445132 |
| GGVKPAA | ASSLDEGGAETLY | 0.72170347 |
| GGTGGAGGAGGW | ASSLDEGGAETLY | 0.5699526 |
| GVGGVAGLGGAGG | ASSLDEGGAETLY | 0.21228978 |
| VLLVLATTI | ASSLDEGGAETLY | 0.7977713 |
| IGIFAAML | ASSLDEGGAETLY | 0.48967284 |
| AVGDLID | ASSLDEGGAETLY | 0.5046404 |
| NISPLPL | ASSLDEGGAETLY | 0.3021925 |
| VTAVFVAI | ASSLDEGGAETLY | 0.33820948 |
| SGDDNDAAL | ASSLDEGGAETLY | 0.41089037 |
| AVAALAHS | ASSLDEGGAETLY | 0.80398893 |
| GGVTVMATP | ASSLDEGGAETLY | 0.6904577 |
| AGLAANVRD | ASSLDEGGAETLY | 0.6650578 |

| | | |
|---|---|---|
| TNRPDLIDPALLRPG | ASSLDEGGAETLY | 0.85586464 |
| LANLGVTAL | ASSLDEGGAETLY | 0.7156826 |
| TGERIEL | ASSLDEGGAETLY | 0.65944695 |
| NGEELAA | ASSLDEGGAETLY | 0.5856491 |
| VLSLVWG | ASSLDEGGAETLY | 0.5181968 |
| IGYALWL | ASSLDEGGAETLY | 0.5374565 |
| LSLLLAPL | ASSLDEGGAETLY | 0.5531336 |
| TLDEALK | ASSLDEGGAETLY | 0.8485805 |
| TDGAGTAEGG | ASSLDEGGAETLY | 0.6686175 |
| NAPGGAGGNGG | ASSLDEGGAETLY | 0.5006656 |
| VSVFMGMGGNF | ASSLDEGGAETLY | 0.753489 |
| LASYLAAL | ASSLDEGGAETLY | 0.8462602 |
| LVGGTPAL | ASSLDEGGAETLY | 0.8563829 |
| VRNADVL | ASSLDEGGAETLY | 0.5034014 |
| TCSSPFLVSAM | ASSLDEGGAETLY | 0.68091154 |
| VHITDAE | ASSLDEGGAETLY | 0.6693893 |
| EYAAPSF | ASSLDEGGAETLY | 0.7185879 |
| VVDAFHGGL | ASSLDEGGAETLY | 0.60560644 |
| ASVVDPQ | ASSLDEGGAETLY | 0.49634236 |
| SYNEFGGLS | ASSLDEGGAETLY | 0.81343234 |
| DPDGSVY | ASSLDEGGAETLY | 0.85907656 |
| MVSTIHATE | ASSLDEGGAETLY | 0.6567534 |
| LAAGVADPV | ASSLDEGGAETLY | 0.40168297 |
| TKEQIAA | ASSLDEGGAETLY | 0.6689563 |
| SLASPAAL | ASSLDEGGAETLY | 0.52601045 |
| IGLVFYGV | ASSLDEGGAETLY | 0.59373885 |
| FLLDASNL | ASSLDEGGAETLY | 0.71642673 |
| YQVGNFP | ASSLDEGGAETLY | 0.43393442 |
| ESLLSQST | ASSLDEGGAETLY | 0.7132481 |
| LFTVGFAAVG | ASSLDEGGAETLY | 0.8534943 |
| ATPPTPT | ASSLDEGGAETLY | 0.6534102 |
| TDLATLG | ASSLDEGGAETLY | 0.79445976 |
| SAGGTGR | ASSLDEGGAETLY | 0.8364363 |
| GFSNSGS | ASSLDEGGAETLY | 0.79579353 |
| GGNGGSGGTSV | ASSLDEGGAETLY | 0.8388602 |
| GTSVATGGAGN | ASSLDEGGAETLY | 0.8003348 |
| NGGNGGSGGTS | ASSLDEGGAETLY | 0.69634223 |
| AVVKFPH | ASSLDEGGAETLY | 0.74409884 |
| TPPPPPP | ASSLDEGGAETLY | 0.7879889 |
| AAPAAAAAPA | ASSLDEGGAETLY | 0.6853186 |
| IALNEDL | ASSLDEGGAETLY | 0.6837784 |
| IVNVSSM | ASSLDEGGAETLY | 0.61711514 |
| TGIPPAPR | ASSLDEGGAETLY | 0.8205403 |
| FDLGGGTF | ASSLDEGGAETLY | 0.69524765 |
| FDLGGGTFDVS | ASSLDEGGAETLY | 0.83135474 |
| NEPTAAA | ASSLDEGGAETLY | 0.40015352 |
| NEPTAAALA | ASSLDEGGAETLY | 0.45334214 |
| GSIINISSI | ASSLDEGGAETLY | 0.82062745 |
| VAPGNAGTA | ASSLDEGGAETLY | 0.1699187 |
| TGGGASLE | ASSLDEGGAETLY | 0.28460267 |
| VAVYNHYKRI | ASSLDEGGAETLY | 0.75423723 |

| | | |
|---|---|---|
| TDPAPATT | ASSLDEGGAETLY | 0.8290036 |
| TNRPDILDPALLRPG | ASSLDEGGAETLY | 0.18157189 |
| LTTTTTS | ASSLDEGGAETLY | 0.52658576 |
| PTGSGKTL | ASSLDEGGAETLY | 0.460517 |
| TGSGKTL | ASSLDEGGAETLY | 0.8057351 |
| TIGHVDHGK | ASSLDEGGAETLY | 0.7148073 |
| TIGHVDHGKTT | ASSLDEGGAETLY | 0.37340036 |
| GHVDHGKTT | ASSLDEGGAETLY | 0.8062556 |
| AQFEHTLL | ASSLDEGGAETLY | 0.33815134 |
| IVNQDEL | ASSLDEGGAETLY | 0.5816456 |
| TNGSQFF | ASSLDEGGAETLY | 0.80443585 |
| TGKGIVE | ASSLDEGGAETLY | 0.8334416 |
| LISNASDA | ASSLDEGGAETLY | 0.51134276 |
| AELRTLGELTKIAKAHGAQ | ASSLDEGGAETLY | 0.79719424 |
| APNAVAAAEPSW | ASSLDEGGAETLY | 0.64096886 |
| APQVAGAATAVL | ASSLDEGGAETLY | 0.8593801 |
| ARRDPAAAAAILPTDA | ASSLDEGGAETLY | 0.6616106 |
| DAAARDVLAEAGLAEYF | ASSLDEGGAETLY | 0.4311111 |
| DDQIVLSAKANLAKVEQ | ASSLDEGGAETLY | 0.8183447 |
| DELLPDTGEDSDEDGH | ASSLDEGGAETLY | 0.80167854 |
| ELPEAPANRATVPQD | ASSLDEGGAETLY | 0.6983192 |
| FGQSKIERVTGIG | ASSLDEGGAETLY | 0.8080923 |
| GAGGTGSTAGGAGGAGGAGGLY | ASSLDEGGAETLY | 0.6215646 |
| GHGGTGAAGGNGGTGGAGGSGI | ASSLDEGGAETLY | 0.6946152 |
| GNGGNGGAGGEGGPGGAGGAGA | ASSLDEGGAETLY | 0.8321941 |
| GSTAGNGGNGGNATNGTIAGQP | ASSLDEGGAETLY | 0.7972268 |
| GTGGDGGNGGNGGTGG | ASSLDEGGAETLY | 0.56222904 |
| GVRRPAGIPADAPV | ASSLDEGGAETLY | 0.8082324 |
| GYRQVTDAHLLTLARR | ASSLDEGGAETLY | 0.6836529 |
| IAKMGQDGHDRG | ASSLDEGGAETLY | 0.5477075 |
| KAEPSIEIQYT | ASSLDEGGAETLY | 0.8514606 |
| KPVRIRMAADEDTVAVT | ASSLDEGGAETLY | 0.72569925 |
| MPPRIAGMRLLVIKPEP | ASSLDEGGAETLY | 0.8038616 |
| MKFYAVALFRSRGVPPTR | ASSLDEGGAETLY | 0.8046442 |
| NAGGFGILGAVAHSP | ASSLDEGGAETLY | 0.8322543 |
| NEGREITLAMAEM | ASSLDEGGAETLY | 0.4751064 |
| NELVEWSHPLGHRWRWE | ASSLDEGGAETLY | 0.75154805 |
| NLGAGGTGGNGGNPG | ASSLDEGGAETLY | 0.7116852 |
| NTEWPKWETVMAADPVD | ASSLDEGGAETLY | 0.80633307 |
| PASAAPAETQP | ASSLDEGGAETLY | 0.7163342 |
| QQSSDGKNGVRAAEITGE | ASSLDEGGAETLY | 0.57088155 |
| RAPDAVTTSWWKE | ASSLDEGGAETLY | 0.7842815 |
| SGFGNVGAHTSGWFNQ | ASSLDEGGAETLY | 0.8359331 |
| SGVAGAVGQAASVGGLKV | ASSLDEGGAETLY | 0.71994233 |
| SNGGAGTGSAGGNGGT | ASSLDEGGAETLY | 0.7178646 |
| TGGAGGEGGAAG | ASSLDEGGAETLY | 0.847844 |
| TGGDGGAAGAGGAPGHG | ASSLDEGGAETLY | 0.57742745 |
| TMPPLLAAIANAGRHGV | ASSLDEGGAETLY | 0.66434157 |
| TSSTAGNSGGA | ASSLDEGGAETLY | 0.8038745 |
| VPATVPVGILC | ASSLDEGGAETLY | 0.8174063 |
| VQRPDSGGVVLTGRIS | ASSLDEGGAETLY | 0.71666 |

| | | |
|---|---|---|
| WTPNPVPATKGVPAGLVQAT | ASSLDEGGAETLY | 0.83197165 |
| YALKRALAKFSDRFDVVII | ASSLDEGGAETLY | 0.57843435 |
| YLIGSVIV | ASSLDEGGAETLY | 0.7940304 |
| YGQLVAVID | ASSLDEGGAETLY | 0.6163741 |
| VTVGGMTLPA | ASSLDEGGAETLY | 0.7650043 |
| VLALPAV | ASSLDEGGAETLY | 0.55889213 |
| VIPPTLNYET | ASSLDEGGAETLY | 0.78013647 |
| VGMGGMHPSSG | ASSLDEGGAETLY | 0.80914134 |
| VGLAGGAATA | ASSLDEGGAETLY | 0.8131075 |
| VGGAGGAGGSGP | ASSLDEGGAETLY | 0.2586622 |
| VFAAFMV | ASSLDEGGAETLY | 0.8097663 |
| TLYDKLR | ASSLDEGGAETLY | 0.811301 |
| TLAVFTVL | ASSLDEGGAETLY | 0.86248064 |
| TGSVQPARV | ASSLDEGGAETLY | 0.70074797 |
| TGGGAGSGGT | ASSLDEGGAETLY | 0.7887092 |
| SHLETSL | ASSLDEGGAETLY | 0.62874377 |
| SFTTGMT | ASSLDEGGAETLY | 0.6845644 |
| QVEVAKRMGVRQPTVS | ASSLDEGGAETLY | 0.44938186 |
| QGSGGPALAM | ASSLDEGGAETLY | 0.6418192 |
| MLAPSNP | ASSLDEGGAETLY | 0.63738984 |
| LVNNAAI | ASSLDEGGAETLY | 0.57021606 |
| LTGEINTLA | ASSLDEGGAETLY | 0.78937393 |
| LSIFGPV | ASSLDEGGAETLY | 0.80291766 |
| LLALLANL | ASSLDEGGAETLY | 0.82084405 |
| LISNASDALDKLR | ASSLDEGGAETLY | 0.8322398 |
| LINEAVP | ASSLDEGGAETLY | 0.8414788 |
| LGYAFSGL | ASSLDEGGAETLY | 0.81421995 |
| LDFVAVAL | ASSLDEGGAETLY | 0.55788434 |
| LAQPGAE | ASSLDEGGAETLY | 0.7012877 |
| ISNLGAVTGF | ASSLDEGGAETLY | 0.44753152 |
| HQEDPGRI | ASSLDEGGAETLY | 0.7086551 |
| GTPGEPTP | ASSLDEGGAETLY | 0.63203967 |
| GFFNSGS | ASSLDEGGAETLY | 0.7602123 |
| GAGGGGLTAGAN | ASSLDEGGAETLY | 0.18494391 |
| FVDALVNM | ASSLDEGGAETLY | 0.75155425 |
| FLTVAPEL | ASSLDEGGAETLY | 0.84864455 |
| FETLTAL | ASSLDEGGAETLY | 0.5660537 |
| DVINPIEL | ASSLDEGGAETLY | 0.76597905 |
| APVSGAPV | ASSLDEGGAETLY | 0.7771505 |
| ALVGGLAW | ASSLDEGGAETLY | 0.6208967 |
| ALAAAAATAI | ASSLDEGGAETLY | 0.79763603 |
| AFGLGLLL | ASSLDEGGAETLY | 0.5566418 |
| AFFAAASL | ASSLDEGGAETLY | 0.7911085 |
| AAAALPAFSPP | ASSLDEGGAETLY | 0.8477732 |
| VVLAPAL | ASSLDEGGAETLY | 0.759635 |
| TIGHVDHGKTTL | ASSLDEGGAETLY | 0.6799508 |
| LISNASDALDK | ASSLDEGGAETLY | 0.7083714 |
| LISNASDAL | ASSLDEGGAETLY | 0.40269607 |
| ISNASDALDK | ASSLDEGGAETLY | 0.72649264 |
| NQAETLVYQTE | ASSLDEGGAETLY | 0.16421013 |
| TGGTGGTAGAG | ASSLDEGGAETLY | 0.7353848 |

| | | |
|---|---|---|
| TGSTAGGAGGA | ASSLDEGGAETLY | 0.5287817 |
| DPLAVPVAAGPV | ASSLDEGGAETLY | 0.7381387 |
| GGTGGTPGNGGN | ASSLDEGGAETLY | 0.719325 |
| PAGLAGVDPATL | ASSLDEGGAETLY | 0.7579854 |
| GNNLVVEAPHAGASV | ASSLDEGGAETLY | 0.49832377 |
| KVTRIMSASTLEQPA | ASSLDEGGAETLY | 0.72693765 |
| LGFALPIALPRRTVV | ASSLDEGGAETLY | 0.6233058 |
| TRLGNLTARKLVVLV | ASSLDEGGAETLY | 0.45493495 |
| WSQQRRLIYFPSAGP | ASSLDEGGAETLY | 0.8011571 |
| GPNGNPDPMAAAVDIR | ASSLDEGGAETLY | 0.6977176 |
| TQRCVSVADLRAAASR | ASSLDEGGAETLY | 0.699374 |
| QFVLDVAEHPVEVLTETV | ASSLDEGGAETLY | 0.83039975 |
| GGAGGTGGAGGNGGTQAGDG | ASSLDEGGAETLY | 0.7382713 |
| PLLATLRAIAVDVDAKPAQV | ASSLDEGGAETLY | 0.38500425 |
| APKFRTGGYMIDFWGVGYQVAK | ASSLDEGGAETLY | 0.6841356 |
| GAGGAGGAGGAAGGPGGTGGTG | ASSLDEGGAETLY | 0.72499216 |
| IGALGLLATDVAPAGSVAFGVA | ASSLDEGGAETLY | 0.6620494 |
| AGGAGGIGGNGGAGGAGGVLMG | ASSLDEGGAETLY | 0.15247978 |
| ADELIGLL | ASSLDEGGAETLY | 0.78335106 |
| ADRALKA | ASSLDEGGAETLY | 0.54519683 |
| AGAVGQVGGAGGA | ASSLDEGGAETLY | 0.5653225 |
| AGGGGGLGGAP | ASSLDEGGAETLY | 0.6204476 |
| AGGLLYGNGGAG | ASSLDEGGAETLY | 0.8148514 |
| AGLESIGL | ASSLDEGGAETLY | 0.60585403 |
| AGLKTGII | ASSLDEGGAETLY | 0.7228664 |
| ASLAALLL | ASSLDEGGAETLY | 0.7822272 |
| ASVINISSI | ASSLDEGGAETLY | 0.75668234 |
| ATPPGPA | ASSLDEGGAETLY | 0.8364774 |
| ATVASALV | ASSLDEGGAETLY | 0.76587874 |
| ATVASLIL | ASSLDEGGAETLY | 0.71453226 |
| AVNGPVLIRH | ASSLDEGGAETLY | 0.7790489 |
| DGAAGAVGI | ASSLDEGGAETLY | 0.6852524 |
| DPLLHGI | ASSLDEGGAETLY | 0.43125904 |
| ELGAALGAL | ASSLDEGGAETLY | 0.30372703 |
| ELGRTQAV | ASSLDEGGAETLY | 0.7201644 |
| FVAGGYLS | ASSLDEGGAETLY | 0.34972405 |
| GASGGAGGAGGW | ASSLDEGGAETLY | 0.73131293 |
| GGAGAGGDGGAGGI | ASSLDEGGAETLY | 0.83249104 |
| GTGHNGSQPGMGGNG | ASSLDEGGAETLY | 0.7122438 |
| GYAAASASAS | ASSLDEGGAETLY | 0.4527877 |
| HQVDVAAVQQG | ASSLDEGGAETLY | 0.6638438 |
| IGAAIAHT | ASSLDEGGAETLY | 0.68571836 |
| IGMIEVAQL | ASSLDEGGAETLY | 0.79341906 |
| IRLAPEHTSAMD | ASSLDEGGAETLY | 0.7971407 |
| IYNVSVL | ASSLDEGGAETLY | 0.8449508 |
| KQGFLGM | ASSLDEGGAETLY | 0.80476034 |
| LESVITQL | ASSLDEGGAETLY | 0.5971603 |
| LGLGALVVV | ASSLDEGGAETLY | 0.7897649 |
| LIGNGGNGGSGGI | ASSLDEGGAETLY | 0.6894925 |
| LNAINAPTL | ASSLDEGGAETLY | 0.72551996 |
| LQLLNTGV | ASSLDEGGAETLY | 0.7169324 |

| | | |
|---|---|---|
| LQTSIEF | ASSLDEGGAETLY | 0.7506483 |
| MSFAVSEL | ASSLDEGGAETLY | 0.6585755 |
| NTGALGL | ASSLDEGGAETLY | 0.64111936 |
| PPGAGGAGGAGTT | ASSLDEGGAETLY | 0.40217793 |
| QLVGLGI | ASSLDEGGAETLY | 0.7949865 |
| RFLEQVHQL | ASSLDEGGAETLY | 0.79464084 |
| RGVGYVLR | ASSLDEGGAETLY | 0.42153707 |
| SATGVFAGVI | ASSLDEGGAETLY | 0.6275159 |
| SLPAITLP | ASSLDEGGAETLY | 0.8496735 |
| STALAFTV | ASSLDEGGAETLY | 0.8339768 |
| SVLTVGLGK | ASSLDEGGAETLY | 0.5961057 |
| TALAGAGSGI | ASSLDEGGAETLY | 0.58327293 |
| TGLPFTGI | ASSLDEGGAETLY | 0.76466566 |
| VAEHVTV | ASSLDEGGAETLY | 0.3430121 |
| VDIVLNSL | ASSLDEGGAETLY | 0.7739338 |
| VLTGGGAVL | ASSLDEGGAETLY | 0.71920216 |
| VNVGLAYGI | ASSLDEGGAETLY | 0.71402967 |
| VSSAAVGAV | ASSLDEGGAETLY | 0.6202033 |
| VTSVYEAGV | ASSLDEGGAETLY | 0.7579062 |
| WVALDRGA | ASSLDEGGAETLY | 0.6466775 |
| YLVGSVVI | ASSLDEGGAETLY | 0.78505754 |
| GPPGTGKTLL | ASSLDEGGAETLY | 0.1782267 |
| GVGMTKF | ASSLDEGGAETLY | 0.78406703 |
| IGIGGSDL | ASSLDEGGAETLY | 0.50110483 |
| IGTLLAEL | ASSLDEGGAETLY | 0.5691371 |
| IYTSGTT | ASSLDEGGAETLY | 0.32864606 |
| LSFPAGV | ASSLDEGGAETLY | 0.702695 |
| LYTSGTT | ASSLDEGGAETLY | 0.81110334 |
| VGAFPFGF | ASSLDEGGAETLY | 0.8201115 |
| VGAGTVEF | ASSLDEGGAETLY | 0.54042184 |
| VGKSTLLL | ASSLDEGGAETLY | 0.8058939 |
| AALAHAGGGIV | ASSLDEGGAETLY | 0.8417045 |
| AALVAVSLL | ASSLDEGGAETLY | 0.66456944 |
| AAVSRAVAAL | ASSLDEGGAETLY | 0.35196844 |
| AGGSSGIGGF | ASSLDEGGAETLY | 0.77298373 |
| AGLTVANANG | ASSLDEGGAETLY | 0.68919307 |
| AGLVGAVL | ASSLDEGGAETLY | 0.8069999 |
| ALSQAFTV | ASSLDEGGAETLY | 0.77842474 |
| ALTFSVSL | ASSLDEGGAETLY | 0.44698364 |
| ALTGAALVAL | ASSLDEGGAETLY | 0.8077239 |
| ASFSIGAGVL | ASSLDEGGAETLY | 0.8320029 |
| DGSVYKI | ASSLDEGGAETLY | 0.512988 |
| EQLAEIV | ASSLDEGGAETLY | 0.541029 |
| FAVSELTVM | ASSLDEGGAETLY | 0.6197591 |
| GAAPLAAGL | ASSLDEGGAETLY | 0.30251247 |
| GADGGDGGSGGAGG | ASSLDEGGAETLY | 0.7776642 |
| GFGLDTGGAGG | ASSLDEGGAETLY | 0.61997294 |
| GLASGGLL | ASSLDEGGAETLY | 0.5653479 |
| GRIAGAAVLL | ASSLDEGGAETLY | 0.73942953 |
| GSAGSLNTSF | ASSLDEGGAETLY | 0.5808564 |
| GTGQIGI | ASSLDEGGAETLY | 0.57852453 |

| | | |
|---|---|---|
| GTLIALIL | ASSLDEGGAETLY | 0.65015936 |
| IAATMGFVVG | ASSLDEGGAETLY | 0.73148906 |
| IFAQALL | ASSLDEGGAETLY | 0.77906406 |
| IGAAVALGF | ASSLDEGGAETLY | 0.72270197 |
| IGLTPLAL | ASSLDEGGAETLY | 0.7484698 |
| ILGSNTSTL | ASSLDEGGAETLY | 0.8404866 |
| IVFSLTVL | ASSLDEGGAETLY | 0.4595105 |
| LASKATGF | ASSLDEGGAETLY | 0.5334253 |
| LASVALIL | ASSLDEGGAETLY | 0.7813954 |
| LFAVGLTL | ASSLDEGGAETLY | 0.45496765 |
| LFNGGIGTYI | ASSLDEGGAETLY | 0.8044902 |
| LTLGAGII | ASSLDEGGAETLY | 0.71311647 |
| LVAAALGDI | ASSLDEGGAETLY | 0.7646534 |
| NTAFVQLGI | ASSLDEGGAETLY | 0.7206168 |
| QLFSGAEL | ASSLDEGGAETLY | 0.8546399 |
| SAATFITL | ASSLDEGGAETLY | 0.8140963 |
| SNYGLANLGV | ASSLDEGGAETLY | 0.7219446 |
| SYGGAYAVM | ASSLDEGGAETLY | 0.75118995 |
| TTGVGGLTLG | ASSLDEGGAETLY | 0.51147527 |
| TTQTIYI | ASSLDEGGAETLY | 0.8175405 |
| VAAGAGLGAVI | ASSLDEGGAETLY | 0.7900367 |
| VAAVVTLL | ASSLDEGGAETLY | 0.500875 |
| VGFLGGLI | ASSLDEGGAETLY | 0.7016267 |
| VGVIPILL | ASSLDEGGAETLY | 0.83056694 |
| VIFASSLI | ASSLDEGGAETLY | 0.83502096 |
| VKGGDGGAAAG | ASSLDEGGAETLY | 0.37384298 |
| VTKTFTNI | ASSLDEGGAETLY | 0.6472806 |
| VVAAAILL | ASSLDEGGAETLY | 0.35654753 |
| YAALDVEL | ASSLDEGGAETLY | 0.6344707 |
| LASVALLL | ASSLDEGGAETLY | 0.46813318 |
| LDAVHRV | ASSLDEGGAETLY | 0.7824638 |
| QGLSSSIF | ASSLDEGGAETLY | 0.620719 |
| RGTFANIRL | ASSLDEGGAETLY | 0.7815018 |
| RQLWWGHRI | ASSLDEGGAETLY | 0.7325027 |
| YGPPGTGKTLL | ASSLDEGGAETLY | 0.8228988 |
| AALVGAPANVA | ASSLDEGGAETLY | 0.6001515 |
| AATFRRTE | ASSLDEGGAETLY | 0.65529037 |
| AGALGGGI | ASSLDEGGAETLY | 0.3931748 |
| AGGAGGAGGIGAG | ASSLDEGGAETLY | 0.48012173 |
| AGSGALGL | ASSLDEGGAETLY | 0.57455206 |
| AQIADQI | ASSLDEGGAETLY | 0.6522583 |
| ASAVAVTYL | ASSLDEGGAETLY | 0.2585629 |
| ASTVLGGTGG | ASSLDEGGAETLY | 0.84626186 |
| AVGGEPAK | ASSLDEGGAETLY | 0.7387467 |
| CEQGTFGGTC | ASSLDEGGAETLY | 0.79527843 |
| DEGKAPAG | ASSLDEGGAETLY | 0.5823405 |
| DTVVFPAL | ASSLDEGGAETLY | 0.6049733 |
| FEGITMAG | ASSLDEGGAETLY | 0.593506 |
| FPLVVQD | ASSLDEGGAETLY | 0.7762946 |
| GAFTTPQ | ASSLDEGGAETLY | 0.56183517 |
| GFGGFGTAGGVGGA | ASSLDEGGAETLY | 0.3273351 |

| | | |
|---|---|---|
| GFLAQFL | ASSLDEGGAETLY | 0.5908465 |
| GGAGGAGGNAGVA | ASSLDEGGAETLY | 0.46246475 |
| GGVGTAGAGGAGGAGG | ASSLDEGGAETLY | 0.8431243 |
| GNGGGGDAGNAGSGGN | ASSLDEGGAETLY | 0.82236814 |
| GTGGAGGAGNDAGS | ASSLDEGGAETLY | 0.77049017 |
| GTTNTGWFNS | ASSLDEGGAETLY | 0.8405236 |
| IFMGPLFTL | ASSLDEGGAETLY | 0.26396984 |
| ILAHAIEAI | ASSLDEGGAETLY | 0.6282048 |
| IRSAIREAL | ASSLDEGGAETLY | 0.83552283 |
| ITFAGGG | ASSLDEGGAETLY | 0.77357864 |
| ITGMSGHQEV | ASSLDEGGAETLY | 0.672801 |
| IVAGLACTL | ASSLDEGGAETLY | 0.82356405 |
| IVGGGLAGA | ASSLDEGGAETLY | 0.52171963 |
| IVTGVLNI | ASSLDEGGAETLY | 0.6484833 |
| LAQQGCI | ASSLDEGGAETLY | 0.7253888 |
| LCLHVEN | ASSLDEGGAETLY | 0.79536283 |
| LDAISDGS | ASSLDEGGAETLY | 0.75275624 |
| LGGFAAML | ASSLDEGGAETLY | 0.4665822 |
| LGGKNPMI | ASSLDEGGAETLY | 0.76923263 |
| LGPKGVAAIL | ASSLDEGGAETLY | 0.860175 |
| LMVGVSI | ASSLDEGGAETLY | 0.64548635 |
| LVGAAFVL | ASSLDEGGAETLY | 0.8043163 |
| LVLAVAPL | ASSLDEGGAETLY | 0.7734794 |
| MAAIAVIDAA | ASSLDEGGAETLY | 0.7829398 |
| NNGINAELQ | ASSLDEGGAETLY | 0.56037885 |
| RGEPLVLVHG | ASSLDEGGAETLY | 0.5758356 |
| RGGAGGAAGLI | ASSLDEGGAETLY | 0.72192 |
| RSYRMHPAV | ASSLDEGGAETLY | 0.4997601 |
| SAAHLLAGQLA | ASSLDEGGAETLY | 0.71398175 |
| SAAVGSY | ASSLDEGGAETLY | 0.68330693 |
| SPHAVLHRI | ASSLDEGGAETLY | 0.8672521 |
| SQIAKVL | ASSLDEGGAETLY | 0.7751845 |
| TALGHLVP | ASSLDEGGAETLY | 0.77943575 |
| TAPTTSGAA | ASSLDEGGAETLY | 0.5185691 |
| TDGDDGNGGSPGT | ASSLDEGGAETLY | 0.7542645 |
| TDGLGEN | ASSLDEGGAETLY | 0.779119 |
| TGGPPLPGQA | ASSLDEGGAETLY | 0.45891458 |
| TIPAITI | ASSLDEGGAETLY | 0.8358189 |
| TIPGFTI | ASSLDEGGAETLY | 0.53255963 |
| TSIASGALA | ASSLDEGGAETLY | 0.7014414 |
| TVTLPGAAI | ASSLDEGGAETLY | 0.80513835 |
| VASGLSAALP | ASSLDEGGAETLY | 0.8263509 |
| VGAGVIGI | ASSLDEGGAETLY | 0.672391 |
| VGPKRFGV | ASSLDEGGAETLY | 0.6934304 |
| VNVAFPD | ASSLDEGGAETLY | 0.53516203 |
| VSMTNSGVS | ASSLDEGGAETLY | 0.8150593 |
| VTDVLGDLSV | ASSLDEGGAETLY | 0.81982243 |
| VVVVGAGFAGL | ASSLDEGGAETLY | 0.6961726 |
| VYTVGTVV | ASSLDEGGAETLY | 0.8482559 |
| WTTFSGP | ASSLDEGGAETLY | 0.7755155 |
| YDELVEAF | ASSLDEGGAETLY | 0.390858 |

| | | |
|---|---|---|
| YGTGGAGGNGG | ASSLDEGGAETLY | 0.6915545 |
| YTNVMAV | ASSLDEGGAETLY | 0.8222293 |
| FLEQVHQL | ASSLDEGGAETLY | 0.82341933 |
| GLSLLAVAV | ASSLDEGGAETLY | 0.83620435 |
| LEQVHQL | ASSLDEGGAETLY | 0.7560084 |
| LNAAGAI | ASSLDEGGAETLY | 0.64789337 |
| PAVHASL | ASSLDEGGAETLY | 0.8431852 |
| TELEDDA | ASSLDEGGAETLY | 0.8598101 |
| TPPGPPN | ASSLDEGGAETLY | 0.80423784 |
| TPPPPPPP | ASSLDEGGAETLY | 0.60919476 |
| TQSPALI | ASSQGDWGNQDTQY | 0.5936087 |
| EFAAAFPQ | ASSQGDWGNQDTQY | 0.7658283 |
| LLSGNLDVL | ASSQGDWGNQDTQY | 0.8070236 |
| GPANLGNYNI | ASSQGDWGNQDTQY | 0.5087098 |
| TLGPTLP | ASSQGDWGNQDTQY | 0.74720496 |
| VYAYSSCCK | ASSQGDWGNQDTQY | 0.74057806 |
| APAPAPAPNI | ASSQGDWGNQDTQY | 0.34179908 |
| PGGPVPIP | ASSQGDWGNQDTQY | 0.80781484 |
| FLTVALEP | ASSQGDWGNQDTQY | 0.8305805 |
| SQYFGANL | ASSQGDWGNQDTQY | 0.7418765 |
| GGVKPAA | ASSQGDWGNQDTQY | 0.6233607 |
| GGTGGAGGAGGW | ASSQGDWGNQDTQY | 0.8243336 |
| GVGGVAGLGGAGG | ASSQGDWGNQDTQY | 0.8633473 |
| VLLVLATTI | ASSQGDWGNQDTQY | 0.84557664 |
| IGIFAAML | ASSQGDWGNQDTQY | 0.34732765 |
| AVGDLID | ASSQGDWGNQDTQY | 0.2401049 |
| NISPLPL | ASSQGDWGNQDTQY | 0.6173025 |
| VTAVFVAI | ASSQGDWGNQDTQY | 0.79792625 |
| SGDDNDAAL | ASSQGDWGNQDTQY | 0.6520928 |
| AVAALAHS | ASSQGDWGNQDTQY | 0.60264087 |
| GGVTVMATP | ASSQGDWGNQDTQY | 0.7683766 |
| AGLAANVRD | ASSQGDWGNQDTQY | 0.83165383 |
| TNRPDLIDPALLRPG | ASSQGDWGNQDTQY | 0.5616368 |
| LANLGVTAL | ASSQGDWGNQDTQY | 0.44285497 |
| TGERIEL | ASSQGDWGNQDTQY | 0.79267 |
| NGEELAA | ASSQGDWGNQDTQY | 0.79721344 |
| VLSLVWG | ASSQGDWGNQDTQY | 0.55482864 |
| IGYALWL | ASSQGDWGNQDTQY | 0.6817135 |
| LSLLLAPL | ASSQGDWGNQDTQY | 0.6566637 |
| TLDEALK | ASSQGDWGNQDTQY | 0.66434395 |
| TDGAGTAEGG | ASSQGDWGNQDTQY | 0.76261914 |
| NAPGGAGGNGG | ASSQGDWGNQDTQY | 0.45639744 |
| VSVFMGMGGNF | ASSQGDWGNQDTQY | 0.832868 |
| LASYLAAL | ASSQGDWGNQDTQY | 0.48079962 |
| LVGGTPAL | ASSQGDWGNQDTQY | 0.79136837 |
| VRNADVL | ASSQGDWGNQDTQY | 0.7404288 |
| TCSSPFLVSAM | ASSQGDWGNQDTQY | 0.77224874 |
| VHITDAE | ASSQGDWGNQDTQY | 0.6303578 |
| EYAAPSF | ASSQGDWGNQDTQY | 0.59603655 |
| VVDAFHGGL | ASSQGDWGNQDTQY | 0.52363896 |
| ASVVDPQ | ASSQGDWGNQDTQY | 0.28592384 |

| | | |
|---|---|---|
| SYNEFGGLS | ASSQGDWGNQDTQY | 0.8142557 |
| DPDGSVY | ASSQGDWGNQDTQY | 0.22227252 |
| MVSTIHATE | ASSQGDWGNQDTQY | 0.8242414 |
| LAAGVADPV | ASSQGDWGNQDTQY | 0.7771991 |
| TKEQIAA | ASSQGDWGNQDTQY | 0.53923297 |
| SLASPAAL | ASSQGDWGNQDTQY | 0.7660726 |
| IGLVFYGV | ASSQGDWGNQDTQY | 0.5217521 |
| FLLDASNL | ASSQGDWGNQDTQY | 0.6027835 |
| YQVGNFP | ASSQGDWGNQDTQY | 0.79698294 |
| ESLLSQST | ASSQGDWGNQDTQY | 0.7972907 |
| LFTVGFAAVG | ASSQGDWGNQDTQY | 0.53192973 |
| ATPPTPT | ASSQGDWGNQDTQY | 0.7656133 |
| TDLATLG | ASSQGDWGNQDTQY | 0.7772918 |
| SAGGTGR | ASSQGDWGNQDTQY | 0.6070727 |
| GFSNSGS | ASSQGDWGNQDTQY | 0.7404848 |
| GGNGGSGGTSV | ASSQGDWGNQDTQY | 0.8215249 |
| GTSVATGGAGN | ASSQGDWGNQDTQY | 0.7528615 |
| NGGNGGSGGTS | ASSQGDWGNQDTQY | 0.5281912 |
| AVVKFPH | ASSQGDWGNQDTQY | 0.7904285 |
| TPPPPPP | ASSQGDWGNQDTQY | 0.5625357 |
| AAPAAAAAPA | ASSQGDWGNQDTQY | 0.775232 |
| IALNEDL | ASSQGDWGNQDTQY | 0.7010816 |
| IVNVSSM | ASSQGDWGNQDTQY | 0.64457494 |
| TGIPPAPR | ASSQGDWGNQDTQY | 0.8111246 |
| FDLGGGTF | ASSQGDWGNQDTQY | 0.49937204 |
| FDLGGGTFDVS | ASSQGDWGNQDTQY | 0.57946575 |
| NEPTAAA | ASSQGDWGNQDTQY | 0.6542374 |
| NEPTAAALA | ASSQGDWGNQDTQY | 0.72208154 |
| GSIINISSI | ASSQGDWGNQDTQY | 0.817539 |
| VAPGNAGTA | ASSQGDWGNQDTQY | 0.51154536 |
| TGGGASLE | ASSQGDWGNQDTQY | 0.7560289 |
| VAVYNHYKRI | ASSQGDWGNQDTQY | 0.6523761 |
| TDPAPATT | ASSQGDWGNQDTQY | 0.4889964 |
| TNRPDILDPALLRPG | ASSQGDWGNQDTQY | 0.72593474 |
| LTTTTTS | ASSQGDWGNQDTQY | 0.66911066 |
| PTGSGKTL | ASSQGDWGNQDTQY | 0.55818355 |
| TGSGKTL | ASSQGDWGNQDTQY | 0.7866591 |
| TIGHVDHGK | ASSQGDWGNQDTQY | 0.57855594 |
| TIGHVDHGKTT | ASSQGDWGNQDTQY | 0.76474476 |
| GHVDHGKTT | ASSQGDWGNQDTQY | 0.84953487 |
| AQFEHTLL | ASSQGDWGNQDTQY | 0.5262914 |
| IVNQDEL | ASSQGDWGNQDTQY | 0.79262286 |
| TNGSQFF | ASSQGDWGNQDTQY | 0.77451295 |
| TGKGIVE | ASSQGDWGNQDTQY | 0.75594056 |
| LISNASDA | ASSQGDWGNQDTQY | 0.6561704 |
| AELRTLGELTKIAKAHGAQ | ASSQGDWGNQDTQY | 0.5959128 |
| APNAVAAAEPSW | ASSQGDWGNQDTQY | 0.84320426 |
| APQVAGAATAVL | ASSQGDWGNQDTQY | 0.7858351 |
| ARRDPAAAAAILPTDA | ASSQGDWGNQDTQY | 0.79728734 |
| DAAARDVLAEAGLAEYF | ASSQGDWGNQDTQY | 0.797745 |
| DDQIVLSAKANLAKVEQ | ASSQGDWGNQDTQY | 0.78843856 |

| | | |
|---|---|---|
| DELLPDTGEDSDEDGH | ASSQGDWGNQDTQY | 0.80131924 |
| ELPEAPANRATVPQD | ASSQGDWGNQDTQY | 0.56710225 |
| FGQSKIERVTGIG | ASSQGDWGNQDTQY | 0.39180934 |
| GAGGTGSTAGGAGGAGGAGGLY | ASSQGDWGNQDTQY | 0.64975387 |
| GHGGTGAAGGNGGTGGAGGSGI | ASSQGDWGNQDTQY | 0.766474 |
| GNGGNGGAGGEGGPGGAGGAGA | ASSQGDWGNQDTQY | 0.6113758 |
| GSTAGNGGNGGNATNGTIAGQP | ASSQGDWGNQDTQY | 0.38665816 |
| GTGGDGGNGGNGGTGG | ASSQGDWGNQDTQY | 0.8211951 |
| GVRRPAGIPADAPV | ASSQGDWGNQDTQY | 0.8593358 |
| GYRQVTDAHLLTLARR | ASSQGDWGNQDTQY | 0.54334074 |
| IAKMGQDGHDRG | ASSQGDWGNQDTQY | 0.8565013 |
| KAEPSIEIQYT | ASSQGDWGNQDTQY | 0.47901347 |
| KPVRIRMAADEDTVAVT | ASSQGDWGNQDTQY | 0.778328 |
| MPPRIAGMRLLVIKPEP | ASSQGDWGNQDTQY | 0.8288604 |
| MKFYAVALFRSRGVPPTR | ASSQGDWGNQDTQY | 0.4893684 |
| NAGGFGILGAVAHSP | ASSQGDWGNQDTQY | 0.78582823 |
| NEGREITLAMAEM | ASSQGDWGNQDTQY | 0.58192444 |
| NELVEWSHPLGHRWRWE | ASSQGDWGNQDTQY | 0.6773093 |
| NLGAGGTGGNGGNPG | ASSQGDWGNQDTQY | 0.8327041 |
| NTEWPKWETVMAADPVD | ASSQGDWGNQDTQY | 0.60946584 |
| PASAAPAETQP | ASSQGDWGNQDTQY | 0.7681508 |
| QQSSDGKNGVRAAEITGE | ASSQGDWGNQDTQY | 0.66586256 |
| RAPDAVTTSWWKE | ASSQGDWGNQDTQY | 0.51285607 |
| SGFGNVGAHTSGWFNQ | ASSQGDWGNQDTQY | 0.74663985 |
| SGVAGAVGQAASVGGLKV | ASSQGDWGNQDTQY | 0.8214201 |
| SNGGAGTGSAGGNGGT | ASSQGDWGNQDTQY | 0.6299572 |
| TGGAGGEGGAAG | ASSQGDWGNQDTQY | 0.85507274 |
| TGGDGGAAGAGGAPGHG | ASSQGDWGNQDTQY | 0.51299703 |
| TMPPLLAAIANAGRHGV | ASSQGDWGNQDTQY | 0.85636485 |
| TSSTAGNSGGA | ASSQGDWGNQDTQY | 0.8178176 |
| VPATVPVGILC | ASSQGDWGNQDTQY | 0.7320118 |
| VQRPDSGGVVLTGRIS | ASSQGDWGNQDTQY | 0.6378541 |
| WTPNPVPATKGVPAGLVQAT | ASSQGDWGNQDTQY | 0.81237 |
| YALKRALAKFSDRFDVVII | ASSQGDWGNQDTQY | 0.62759507 |
| YLIGSVIV | ASSQGDWGNQDTQY | 0.7412166 |
| YGQLVAVID | ASSQGDWGNQDTQY | 0.8348235 |
| VTVGGMTLPA | ASSQGDWGNQDTQY | 0.122393884 |
| VLALPAV | ASSQGDWGNQDTQY | 0.40520334 |
| VIPPTLNYET | ASSQGDWGNQDTQY | 0.28392726 |
| VGMGGMHPSSG | ASSQGDWGNQDTQY | 0.3471192 |
| VGLAGGAATA | ASSQGDWGNQDTQY | 0.15104988 |
| VGGAGGAGGSGP | ASSQGDWGNQDTQY | 0.12068854 |
| VFAAFMV | ASSQGDWGNQDTQY | 0.17790055 |
| TLYDKLR | ASSQGDWGNQDTQY | 0.13921386 |
| TLAVFTVL | ASSQGDWGNQDTQY | 0.6333664 |
| TGSVQPARV | ASSQGDWGNQDTQY | 0.25526407 |
| TGGGAGSGGT | ASSQGDWGNQDTQY | 0.14625354 |
| SHLETSL | ASSQGDWGNQDTQY | 0.3016572 |
| SFTTGMT | ASSQGDWGNQDTQY | 0.40173885 |
| QVEVAKRMGVRQPTVS | ASSQGDWGNQDTQY | 0.11815621 |
| QGSGGPALAM | ASSQGDWGNQDTQY | 0.23363246 |

| | | |
|---|---|---|
| MLAPSNP | ASSQGDWGNQDTQY | 0.18206728 |
| LVNNAAI | ASSQGDWGNQDTQY | 0.16618241 |
| LTGEINTLA | ASSQGDWGNQDTQY | 0.10724606 |
| LSIFGPV | ASSQGDWGNQDTQY | 0.21552867 |
| LLALLANL | ASSQGDWGNQDTQY | 0.16841412 |
| LISNASDALDKLR | ASSQGDWGNQDTQY | 0.103214726 |
| LINEAVP | ASSQGDWGNQDTQY | 0.29943347 |
| LGYAFSGL | ASSQGDWGNQDTQY | 0.13646203 |
| LDFVAVAL | ASSQGDWGNQDTQY | 0.1601548 |
| LAQPGAE | ASSQGDWGNQDTQY | 0.33364534 |
| ISNLGAVTGF | ASSQGDWGNQDTQY | 0.21451151 |
| HQEDPGRI | ASSQGDWGNQDTQY | 0.14579009 |
| GTPGEPTP | ASSQGDWGNQDTQY | 0.26281917 |
| GFFNSGS | ASSQGDWGNQDTQY | 0.4392069 |
| GAGGGGLTAGAN | ASSQGDWGNQDTQY | 0.1357116 |
| FVDALVNM | ASSQGDWGNQDTQY | 0.20546174 |
| FLTVAPEL | ASSQGDWGNQDTQY | 0.6497536 |
| FETLTAL | ASSQGDWGNQDTQY | 0.20395514 |
| DVINPIEL | ASSQGDWGNQDTQY | 0.48731267 |
| APVSGAPV | ASSQGDWGNQDTQY | 0.4988517 |
| ALVGGLAW | ASSQGDWGNQDTQY | 0.14782672 |
| ALAAAAATAI | ASSQGDWGNQDTQY | 0.37723175 |
| AFGLGLLL | ASSQGDWGNQDTQY | 0.48614877 |
| AFFAAASL | ASSQGDWGNQDTQY | 0.19745421 |
| AAAALPAFSPP | ASSQGDWGNQDTQY | 0.18558209 |
| VVLAPAL | ASSQGDWGNQDTQY | 0.12521261 |
| TIGHVDHGKTTL | ASSQGDWGNQDTQY | 0.18371913 |
| LISNASDALDK | ASSQGDWGNQDTQY | 0.1227245 |
| LISNASDAL | ASSQGDWGNQDTQY | 0.27952185 |
| ISNASDALDK | ASSQGDWGNQDTQY | 0.40392756 |
| NQAETLVYQTE | ASSQGDWGNQDTQY | 0.19525473 |
| TGGTGGTAGAG | ASSQGDWGNQDTQY | 0.11057113 |
| TGSTAGGAGGA | ASSQGDWGNQDTQY | 0.11723406 |
| DPLAVPVAAGPV | ASSQGDWGNQDTQY | 0.2539681 |
| GGTGGTPGNGGN | ASSQGDWGNQDTQY | 0.12878627 |
| PAGLAGVDPATL | ASSQGDWGNQDTQY | 0.121636555 |
| GNNLVVEAPHAGASV | ASSQGDWGNQDTQY | 0.17234138 |
| KVTRIMSASTLEQPA | ASSQGDWGNQDTQY | 0.3828705 |
| LGFALPIALPRRTVV | ASSQGDWGNQDTQY | 0.41028482 |
| TRLGNLTARKLVVLV | ASSQGDWGNQDTQY | 0.14983417 |
| WSQQRRLIYFPSAGP | ASSQGDWGNQDTQY | 0.4606531 |
| GPNGNPDPMAAAVDIR | ASSQGDWGNQDTQY | 0.55965483 |
| TQRCVSVADLRAAASR | ASSQGDWGNQDTQY | 0.5359702 |
| QFVLDVAEHPVEVLTETV | ASSQGDWGNQDTQY | 0.2058419 |
| GGAGGTGGAGGNGGTQAGDG | ASSQGDWGNQDTQY | 0.12524705 |
| PLLATLRAIAVDVDAKPAQV | ASSQGDWGNQDTQY | 0.5565779 |
| APKFRTGGYMIDFWGVGYQVAK | ASSQGDWGNQDTQY | 0.15548714 |
| GAGGAGGAGGAAGGPGGTGGTG | ASSQGDWGNQDTQY | 0.23525538 |
| IGALGLLATDVAPAGSVAFGVA | ASSQGDWGNQDTQY | 0.3467664 |
| AGGAGGIGGNGGAGGAGGVLMG | ASSQGDWGNQDTQY | 0.20649429 |
| ADELIGLL | ASSQGDWGNQDTQY | 0.13327673 |

| | | |
|---|---|---|
| ADRALKA | ASSQGDWGNQDTQY | 0.4653873 |
| AGAVGQVGGAGGA | ASSQGDWGNQDTQY | 0.41647828 |
| AGGGGGLGGAP | ASSQGDWGNQDTQY | 0.11957662 |
| AGGLLYGNGGAG | ASSQGDWGNQDTQY | 0.12723726 |
| AGLESIGL | ASSQGDWGNQDTQY | 0.16184542 |
| AGLKTGII | ASSQGDWGNQDTQY | 0.16930428 |
| ASLAALLL | ASSQGDWGNQDTQY | 0.45412022 |
| ASVINISSI | ASSQGDWGNQDTQY | 0.224973 |
| ATPPGPA | ASSQGDWGNQDTQY | 0.22668122 |
| ATVASALV | ASSQGDWGNQDTQY | 0.32788366 |
| ATVASLIL | ASSQGDWGNQDTQY | 0.2998596 |
| AVNGPVLIRH | ASSQGDWGNQDTQY | 0.20202789 |
| DGAAGAVGI | ASSQGDWGNQDTQY | 0.3834942 |
| DPLLHGI | ASSQGDWGNQDTQY | 0.1425194 |
| ELGAALGAL | ASSQGDWGNQDTQY | 0.3810777 |
| ELGRTQAV | ASSQGDWGNQDTQY | 0.37042534 |
| FVAGGYLS | ASSQGDWGNQDTQY | 0.30287552 |
| GASGGAGGAGGW | ASSQGDWGNQDTQY | 0.43472958 |
| GGAGAGGDGGAGGI | ASSQGDWGNQDTQY | 0.1998573 |
| GTGHNGSQPGMGGNG | ASSQGDWGNQDTQY | 0.22896495 |
| GYAAASASAS | ASSQGDWGNQDTQY | 0.13756868 |
| HQVDVAAVQQG | ASSQGDWGNQDTQY | 0.17032546 |
| IGAAIAHT | ASSQGDWGNQDTQY | 0.2283215 |
| IGMIEVAQL | ASSQGDWGNQDTQY | 0.2833104 |
| IRLAPEHTSAMD | ASSQGDWGNQDTQY | 0.32913285 |
| IYNVSVL | ASSQGDWGNQDTQY | 0.3076095 |
| KQGFLGM | ASSQGDWGNQDTQY | 0.13673206 |
| LESVITQL | ASSQGDWGNQDTQY | 0.49350256 |
| LGLGALVVV | ASSQGDWGNQDTQY | 0.51533383 |
| LIGNGGNGGSGGI | ASSQGDWGNQDTQY | 0.12590203 |
| LNAINAPTL | ASSQGDWGNQDTQY | 0.18983361 |
| LQLLNTGV | ASSQGDWGNQDTQY | 0.4125879 |
| LQTSIEF | ASSQGDWGNQDTQY | 0.41170913 |
| MSFAVSEL | ASSQGDWGNQDTQY | 0.16750723 |
| NTGALGL | ASSQGDWGNQDTQY | 0.2463423 |
| PPGAGGAGGAGTT | ASSQGDWGNQDTQY | 0.69594353 |
| QLVGLGI | ASSQGDWGNQDTQY | 0.1931287 |
| RFLEQVHQL | ASSQGDWGNQDTQY | 0.50974214 |
| RGVGYVLR | ASSQGDWGNQDTQY | 0.12053439 |
| SATGVFAGVI | ASSQGDWGNQDTQY | 0.20807248 |
| SLPAITLP | ASSQGDWGNQDTQY | 0.16043621 |
| STALAFTV | ASSQGDWGNQDTQY | 0.10484919 |
| SVLTVGLGK | ASSQGDWGNQDTQY | 0.37356558 |
| TALAGAGSGI | ASSQGDWGNQDTQY | 0.15083191 |
| TGLPFTGI | ASSQGDWGNQDTQY | 0.32057315 |
| VAEHVTV | ASSQGDWGNQDTQY | 0.15091555 |
| VDIVLNSL | ASSQGDWGNQDTQY | 0.3886455 |
| VLTGGGAVL | ASSQGDWGNQDTQY | 0.24149922 |
| VNVGLAYGI | ASSQGDWGNQDTQY | 0.20446363 |
| VSSAAVGAV | ASSQGDWGNQDTQY | 0.14270571 |
| VTSVYEAGV | ASSQGDWGNQDTQY | 0.2620493 |

| | | |
|---|---|---|
| WVALDRGA | ASSQGDWGNQDTQY | 0.45355496 |
| YLVGSVVI | ASSQGDWGNQDTQY | 0.44811127 |
| GPPGTGKTLL | ASSQGDWGNQDTQY | 0.116956934 |
| GVGMTKF | ASSQGDWGNQDTQY | 0.39487505 |
| IGIGGSDL | ASSQGDWGNQDTQY | 0.21250752 |
| IGTLLAEL | ASSQGDWGNQDTQY | 0.13534407 |
| IYTSGTT | ASSQGDWGNQDTQY | 0.10946172 |
| LSFPAGV | ASSQGDWGNQDTQY | 0.14538944 |
| LYTSGTT | ASSQGDWGNQDTQY | 0.1718935 |
| VGAFPFGF | ASSQGDWGNQDTQY | 0.25079587 |
| VGAGTVEF | ASSQGDWGNQDTQY | 0.3702653 |
| VGKSTLLL | ASSQGDWGNQDTQY | 0.14321622 |
| AALAHAGGGIV | ASSQGDWGNQDTQY | 0.35063088 |
| AALVAVSLL | ASSQGDWGNQDTQY | 0.19717532 |
| AAVSRAVAAL | ASSQGDWGNQDTQY | 0.09937262 |
| AGGSSGIGGF | ASSQGDWGNQDTQY | 0.14902942 |
| AGLTVANANG | ASSQGDWGNQDTQY | 0.1154359 |
| AGLVGAVL | ASSQGDWGNQDTQY | 0.4189642 |
| ALSQAFTV | ASSQGDWGNQDTQY | 0.12851347 |
| ALTFSVSL | ASSQGDWGNQDTQY | 0.16993622 |
| ALTGAALVAL | ASSQGDWGNQDTQY | 0.1611022 |
| ASFSIGAGVL | ASSQGDWGNQDTQY | 0.57968944 |
| DGSVYKI | ASSQGDWGNQDTQY | 0.24112788 |
| EQLAEIV | ASSQGDWGNQDTQY | 0.14664388 |
| FAVSELTVM | ASSQGDWGNQDTQY | 0.120246805 |
| GAAPLAAGL | ASSQGDWGNQDTQY | 0.27088434 |
| GADGGDGGSGGAGG | ASSQGDWGNQDTQY | 0.19510326 |
| GFGLDTGGAGG | ASSQGDWGNQDTQY | 0.1574798 |
| GLASGGLL | ASSQGDWGNQDTQY | 0.15044932 |
| GRIAGAAVLL | ASSQGDWGNQDTQY | 0.5968037 |
| GSAGSLNTSF | ASSQGDWGNQDTQY | 0.11718289 |
| GTGQIGI | ASSQGDWGNQDTQY | 0.41684288 |
| GTLIALIL | ASSQGDWGNQDTQY | 0.1354265 |
| IAATMGFVVG | ASSQGDWGNQDTQY | 0.1545504 |
| IFAQALL | ASSQGDWGNQDTQY | 0.48474237 |
| IGAAVALGF | ASSQGDWGNQDTQY | 0.51358914 |
| IGLTPLAL | ASSQGDWGNQDTQY | 0.1704223 |
| ILGSNTSTL | ASSQGDWGNQDTQY | 0.1771428 |
| IVFSLTVL | ASSQGDWGNQDTQY | 0.12645198 |
| LASKATGF | ASSQGDWGNQDTQY | 0.14370155 |
| LASVALIL | ASSQGDWGNQDTQY | 0.14276525 |
| LFAVGLTL | ASSQGDWGNQDTQY | 0.4728393 |
| LFNGGIGTYI | ASSQGDWGNQDTQY | 0.2264489 |
| LTLGAGII | ASSQGDWGNQDTQY | 0.123323 |
| LVAAALGDI | ASSQGDWGNQDTQY | 0.44588163 |
| NTAFVQLGI | ASSQGDWGNQDTQY | 0.41228563 |
| QLFSGAEL | ASSQGDWGNQDTQY | 0.65253925 |
| SAATFITL | ASSQGDWGNQDTQY | 0.13100745 |
| SNYGLANLGV | ASSQGDWGNQDTQY | 0.16798373 |
| SYGGAYAVM | ASSQGDWGNQDTQY | 0.23106208 |
| TTGVGGLTLG | ASSQGDWGNQDTQY | 0.16821814 |

| | | |
|---|---|---|
| TTQTIYI | ASSQGDWGNQDTQY | 0.14585991 |
| VAAGAGLGAVI | ASSQGDWGNQDTQY | 0.14099401 |
| VAAVVTLL | ASSQGDWGNQDTQY | 0.40644336 |
| VGFLGGLI | ASSQGDWGNQDTQY | 0.78771055 |
| VGVIPILL | ASSQGDWGNQDTQY | 0.16264606 |
| VIFASSLI | ASSQGDWGNQDTQY | 0.11191418 |
| VKGGDGGAAAG | ASSQGDWGNQDTQY | 0.16642508 |
| VTKTFTNI | ASSQGDWGNQDTQY | 0.12957837 |
| VVAAAILL | ASSQGDWGNQDTQY | 0.12739582 |
| YAALDVEL | ASSQGDWGNQDTQY | 0.17408112 |
| LASVALLL | ASSQGDWGNQDTQY | 0.112968266 |
| LDAVHRV | ASSQGDWGNQDTQY | 0.15677604 |
| QGLSSSIF | ASSQGDWGNQDTQY | 0.52039653 |
| RGTFANIRL | ASSQGDWGNQDTQY | 0.189998 |
| RQLWWGHRI | ASSQGDWGNQDTQY | 0.34996927 |
| YGPPGTGKTLL | ASSQGDWGNQDTQY | 0.4494509 |
| AALVGAPANVA | ASSQGDWGNQDTQY | 0.24993715 |
| AATFRRTE | ASSQGDWGNQDTQY | 0.47951558 |
| AGALGGGI | ASSQGDWGNQDTQY | 0.6343918 |
| AGGAGGAGGIGAG | ASSQGDWGNQDTQY | 0.20687066 |
| AGSGALGL | ASSQGDWGNQDTQY | 0.20515752 |
| AQIADQI | ASSQGDWGNQDTQY | 0.45132357 |
| ASAVAVTYL | ASSQGDWGNQDTQY | 0.16412511 |
| ASTVLGGTGG | ASSQGDWGNQDTQY | 0.21253309 |
| AVGGEPAK | ASSQGDWGNQDTQY | 0.236496 |
| CEQGTFGGTC | ASSQGDWGNQDTQY | 0.29371387 |
| DEGKAPAG | ASSQGDWGNQDTQY | 0.21214372 |
| DTVVFPAL | ASSQGDWGNQDTQY | 0.55456674 |
| FEGITMAG | ASSQGDWGNQDTQY | 0.10976972 |
| FPLVVQD | ASSQGDWGNQDTQY | 0.13795336 |
| GAFTTPQ | ASSQGDWGNQDTQY | 0.5148404 |
| GFGGFGTAGGVGGA | ASSQGDWGNQDTQY | 0.10516345 |
| GFLAQFL | ASSQGDWGNQDTQY | 0.12729594 |
| GGAGGAGGNAGVA | ASSQGDWGNQDTQY | 0.23648433 |
| GGVGTAGAGGAGGAGG | ASSQGDWGNQDTQY | 0.5528453 |
| GNGGGGDAGNAGSGGN | ASSQGDWGNQDTQY | 0.10817145 |
| GTGGAGGAGNDAGS | ASSQGDWGNQDTQY | 0.19009563 |
| GTTNTGWFNS | ASSQGDWGNQDTQY | 0.12456452 |
| IFMGPLFTL | ASSQGDWGNQDTQY | 0.37098736 |
| ILAHAIEAI | ASSQGDWGNQDTQY | 0.16008988 |
| IRSAIREAL | ASSQGDWGNQDTQY | 0.113919005 |
| ITFAGGG | ASSQGDWGNQDTQY | 0.32221633 |
| ITGMSGHQEV | ASSQGDWGNQDTQY | 0.12119341 |
| IVAGLACTL | ASSQGDWGNQDTQY | 0.34056574 |
| IVGGGLAGA | ASSQGDWGNQDTQY | 0.37942535 |
| IVTGVLNI | ASSQGDWGNQDTQY | 0.75538963 |
| LAQQGCI | ASSQGDWGNQDTQY | 0.11978415 |
| LCLHVEN | ASSQGDWGNQDTQY | 0.30023855 |
| LDAISDGS | ASSQGDWGNQDTQY | 0.13484326 |
| LGGFAAML | ASSQGDWGNQDTQY | 0.75291383 |
| LGGKNPMI | ASSQGDWGNQDTQY | 0.17081001 |

| | | |
|---|---|---|
| LGPKGVAAIL | ASSQGDWGNQDTQY | 0.114683844 |
| LMVGVSI | ASSQGDWGNQDTQY | 0.31974977 |
| LVGAAFVL | ASSQGDWGNQDTQY | 0.3200158 |
| LVLAVAPL | ASSQGDWGNQDTQY | 0.20522824 |
| MAAIAVIDAA | ASSQGDWGNQDTQY | 0.5381906 |
| NNGINAELQ | ASSQGDWGNQDTQY | 0.21577466 |
| RGEPLVLVHG | ASSQGDWGNQDTQY | 0.15131363 |
| RGGAGGAAGLI | ASSQGDWGNQDTQY | 0.3233261 |
| RSYRMHPAV | ASSQGDWGNQDTQY | 0.30902204 |
| SAAHLLAGQLA | ASSQGDWGNQDTQY | 0.15721108 |
| SAAVGSY | ASSQGDWGNQDTQY | 0.39798218 |
| SPHAVLHRI | ASSQGDWGNQDTQY | 0.21626727 |
| SQIAKVL | ASSQGDWGNQDTQY | 0.1360104 |
| TALGHLVP | ASSQGDWGNQDTQY | 0.6353292 |
| TAPTTSGAA | ASSQGDWGNQDTQY | 0.22395504 |
| TDGDDGNGGSPGT | ASSQGDWGNQDTQY | 0.3136234 |
| TDGLGEN | ASSQGDWGNQDTQY | 0.25823504 |
| TGGPPLPGQA | ASSQGDWGNQDTQY | 0.77109236 |
| TIPAITI | ASSQGDWGNQDTQY | 0.147612 |
| TIPGFTI | ASSQGDWGNQDTQY | 0.30311096 |
| TSIASGALA | ASSQGDWGNQDTQY | 0.23981221 |
| TVTLPGAAI | ASSQGDWGNQDTQY | 0.4285565 |
| VASGLSAALP | ASSQGDWGNQDTQY | 0.19952023 |
| VGAGVIGI | ASSQGDWGNQDTQY | 0.350999 |
| VGPKRFGV | ASSQGDWGNQDTQY | 0.24126196 |
| VNVAFPD | ASSQGDWGNQDTQY | 0.38881564 |
| VSMTNSGVS | ASSQGDWGNQDTQY | 0.19006102 |
| VTDVLGDLSV | ASSQGDWGNQDTQY | 0.25030947 |
| VVVVGAGFAGL | ASSQGDWGNQDTQY | 0.60566545 |
| VYTVGTVV | ASSQGDWGNQDTQY | 0.13262805 |
| WTTFSGP | ASSQGDWGNQDTQY | 0.22229716 |
| YDELVEAF | ASSQGDWGNQDTQY | 0.36415827 |
| YGTGGAGGNGG | ASSQGDWGNQDTQY | 0.46664974 |
| YTNVMAV | ASSQGDWGNQDTQY | 0.2072519 |
| FLEQVHQL | ASSQGDWGNQDTQY | 0.5384852 |
| GLSLLAVAV | ASSQGDWGNQDTQY | 0.19083565 |
| LEQVHQL | ASSQGDWGNQDTQY | 0.2882623 |
| LNAAGAI | ASSQGDWGNQDTQY | 0.26749858 |
| PAVHASL | ASSQGDWGNQDTQY | 0.15856716 |
| TELEDDA | ASSQGDWGNQDTQY | 0.1316489 |
| TPPGPPN | ASSQGDWGNQDTQY | 0.19216803 |
| TPPPPPPP | ASSQGDWGNQDTQY | 0.34329587 |
| TQSPALI | ASGDGPRTNSDYT | 0.24483815 |
| EFAAAFPQ | ASGDGPRTNSDYT | 0.11604912 |
| LLSGNLDVL | ASGDGPRTNSDYT | 0.23880419 |
| GPANLGNYNI | ASGDGPRTNSDYT | 0.36285716 |
| TLGPTLP | ASGDGPRTNSDYT | 0.75571406 |
| VYAYSSCCK | ASGDGPRTNSDYT | 0.22553802 |
| APAPAPAPNI | ASGDGPRTNSDYT | 0.5120889 |
| PGGPVPIP | ASGDGPRTNSDYT | 0.21846622 |
| FLTVALEP | ASGDGPRTNSDYT | 0.2515657 |

| | | |
|---|---|---|
| SQYFGANL | ASGDGPRTNSDYT | 0.11287899 |
| GGVKPAA | ASGDGPRTNSDYT | 0.1579304 |
| GGTGGAGGAGGW | ASGDGPRTNSDYT | 0.16357426 |
| GVGGVAGLGGAGG | ASGDGPRTNSDYT | 0.13555974 |
| VLLVLATTI | ASGDGPRTNSDYT | 0.522671 |
| IGIFAAML | ASGDGPRTNSDYT | 0.5322273 |
| AVGDLID | ASGDGPRTNSDYT | 0.30211344 |
| NISPLPL | ASGDGPRTNSDYT | 0.538007 |
| VTAVFVAI | ASGDGPRTNSDYT | 0.56821704 |
| SGDDNDAAL | ASGDGPRTNSDYT | 0.35748416 |
| AVAALAHS | ASGDGPRTNSDYT | 0.1705006 |
| GGVTVMATP | ASGDGPRTNSDYT | 0.1812093 |
| AGLAANVRD | ASGDGPRTNSDYT | 0.1131664 |
| TNRPDLIDPALLRPG | ASGDGPRTNSDYT | 0.13341038 |
| LANLGVTAL | ASGDGPRTNSDYT | 0.15492892 |
| TGERIEL | ASGDGPRTNSDYT | 0.17106867 |
| NGEELAA | ASGDGPRTNSDYT | 0.10289302 |
| VLSLVWG | ASGDGPRTNSDYT | 0.46199208 |
| IGYALWL | ASGDGPRTNSDYT | 0.4588495 |
| LSLLLAPL | ASGDGPRTNSDYT | 0.23922 |
| TLDEALK | ASGDGPRTNSDYT | 0.28843334 |
| TDGAGTAEGG | ASGDGPRTNSDYT | 0.2562675 |
| NAPGGAGGNGG | ASGDGPRTNSDYT | 0.1399313 |
| VSVFMGMGGNF | ASGDGPRTNSDYT | 0.38851425 |
| LASYLAAL | ASGDGPRTNSDYT | 0.15850607 |
| LVGGTPAL | ASGDGPRTNSDYT | 0.47444624 |
| VRNADVL | ASGDGPRTNSDYT | 0.43742996 |
| TCSSPFLVSAM | ASGDGPRTNSDYT | 0.2518655 |
| VHITDAE | ASGDGPRTNSDYT | 0.20272592 |
| EYAAPSF | ASGDGPRTNSDYT | 0.29239988 |
| VVDAFHGGL | ASGDGPRTNSDYT | 0.14841576 |
| ASVVDPQ | ASGDGPRTNSDYT | 0.23258486 |
| SYNEFGGLS | ASGDGPRTNSDYT | 0.15651736 |
| DPDGSVY | ASGDGPRTNSDYT | 0.25487214 |
| MVSTIHATE | ASGDGPRTNSDYT | 0.12969331 |
| LAAGVADPV | ASGDGPRTNSDYT | 0.32797274 |
| TKEQIAA | ASGDGPRTNSDYT | 0.19820663 |
| SLASPAAL | ASGDGPRTNSDYT | 0.27959096 |
| IGLVFYGV | ASGDGPRTNSDYT | 0.13319296 |
| FLLDASNL | ASGDGPRTNSDYT | 0.2793504 |
| YQVGNFP | ASGDGPRTNSDYT | 0.32106778 |
| ESLLSQST | ASGDGPRTNSDYT | 0.124277756 |
| LFTVGFAAVG | ASGDGPRTNSDYT | 0.41172948 |
| ATPPTPT | ASGDGPRTNSDYT | 0.17862943 |
| TDLATLG | ASGDGPRTNSDYT | 0.19492304 |
| SAGGTGR | ASGDGPRTNSDYT | 0.38807952 |
| GFSNSGS | ASGDGPRTNSDYT | 0.19062215 |
| GGNGGSGGTSV | ASGDGPRTNSDYT | 0.13418058 |
| GTSVATGGAGN | ASGDGPRTNSDYT | 0.14676046 |
| NGGNGGSGGTS | ASGDGPRTNSDYT | 0.2538209 |
| AVVKFPH | ASGDGPRTNSDYT | 0.1742625 |

| | | |
|---|---|---|
| TPPPPPP | ASGDGPRTNSDYT | 0.10691209 |
| AAPAAAAAPA | ASGDGPRTNSDYT | 0.24014473 |
| IALNEDL | ASGDGPRTNSDYT | 0.13730721 |
| IVNVSSM | ASGDGPRTNSDYT | 0.14403927 |
| TGIPPAPR | ASGDGPRTNSDYT | 0.15704937 |
| FDLGGGTF | ASGDGPRTNSDYT | 0.42507312 |
| FDLGGGTFDVS | ASGDGPRTNSDYT | 0.40635514 |
| NEPTAAA | ASGDGPRTNSDYT | 0.20030567 |
| NEPTAAALA | ASGDGPRTNSDYT | 0.34959456 |
| GSIINISSI | ASGDGPRTNSDYT | 0.21429834 |
| VAPGNAGTA | ASGDGPRTNSDYT | 0.33686426 |
| TGGGASLE | ASGDGPRTNSDYT | 0.29780304 |
| VAVYNHYKRI | ASGDGPRTNSDYT | 0.21943931 |
| TDPAPATT | ASGDGPRTNSDYT | 0.29864028 |
| TNRPDILDPALLRPG | ASGDGPRTNSDYT | 0.1719763 |
| LTTTTTS | ASGDGPRTNSDYT | 0.12228343 |
| PTGSGKTL | ASGDGPRTNSDYT | 0.1482598 |
| TGSGKTL | ASGDGPRTNSDYT | 0.14284469 |
| TIGHVDHGK | ASGDGPRTNSDYT | 0.12274921 |
| TIGHVDHGKTT | ASGDGPRTNSDYT | 0.28320366 |
| GHVDHGKTT | ASGDGPRTNSDYT | 0.45621312 |
| AQFEHTLL | ASGDGPRTNSDYT | 0.257676 |
| IVNQDEL | ASGDGPRTNSDYT | 0.11333508 |
| TNGSQFF | ASGDGPRTNSDYT | 0.16533142 |
| TGKGIVE | ASGDGPRTNSDYT | 0.17780861 |
| LISNASDA | ASGDGPRTNSDYT | 0.34850624 |
| AELRTLGELTKIAKAHGAQ | ASGDGPRTNSDYT | 0.33720058 |
| APNAVAAAEPSW | ASGDGPRTNSDYT | 0.7591273 |
| APQVAGAATAVL | ASGDGPRTNSDYT | 0.3835253 |
| ARRDPAAAAAILPTDA | ASGDGPRTNSDYT | 0.15594779 |
| DAAARDVLAEAGLAEYF | ASGDGPRTNSDYT | 0.34604448 |
| DDQIVLSAKANLAKVEQ | ASGDGPRTNSDYT | 0.15868166 |
| DELLPDTGEDSDEDGH | ASGDGPRTNSDYT | 0.22687528 |
| ELPEAPANRATVPQD | ASGDGPRTNSDYT | 0.19031708 |
| FGQSKIERVTGIG | ASGDGPRTNSDYT | 0.3048414 |
| GAGGTGSTAGGAGGAGGAGGLY | ASGDGPRTNSDYT | 0.1738625 |
| GHGGTGAAGGNGGTGGAGGSGI | ASGDGPRTNSDYT | 0.18655376 |
| GNGGNGGAGGEGGPGGAGGAGA | ASGDGPRTNSDYT | 0.12604481 |
| GSTAGNGGNGGNATNGTIAGQP | ASGDGPRTNSDYT | 0.30296636 |
| GTGGDGGNGGNGGTGG | ASGDGPRTNSDYT | 0.13227586 |
| GVRRPAGIPADAPV | ASGDGPRTNSDYT | 0.13895358 |
| GYRQVTDAHLLTLARR | ASGDGPRTNSDYT | 0.17355463 |
| IAKMGQDGHDRG | ASGDGPRTNSDYT | 0.62994206 |
| KAEPSIEIQYT | ASGDGPRTNSDYT | 0.5995875 |
| KPVRIRMAADEDTVAVT | ASGDGPRTNSDYT | 0.16822273 |
| MPPRIAGMRLLVIKPEP | ASGDGPRTNSDYT | 0.15727018 |
| MKFYAVALFRSRGVPPTR | ASGDGPRTNSDYT | 0.53708947 |
| NAGGFGILGAVAHSP | ASGDGPRTNSDYT | 0.184077 |
| NEGREITLAMAEM | ASGDGPRTNSDYT | 0.28183863 |
| NELVEWSHPLGHRWRWE | ASGDGPRTNSDYT | 0.20216992 |
| NLGAGGTGGNGGNPG | ASGDGPRTNSDYT | 0.16563065 |

| | | |
|---|---|---|
| NTEWPKWETVMAADPVD | ASGDGPRTNSDYT | 0.1746129 |
| PASAAPAETQP | ASGDGPRTNSDYT | 0.3532264 |
| QQSSDGKNGVRAAEITGE | ASGDGPRTNSDYT | 0.16406867 |
| RAPDAVTTSWWKE | ASGDGPRTNSDYT | 0.3181667 |
| SGFGNVGAHTSGWFNQ | ASGDGPRTNSDYT | 0.12529095 |
| SGVAGAVGQAASVGGLKV | ASGDGPRTNSDYT | 0.32953012 |
| SNGGAGTGSAGGNGGT | ASGDGPRTNSDYT | 0.1234442 |
| TGGAGGEGGAAG | ASGDGPRTNSDYT | 0.13756737 |
| TGGDGGAAGAGGAPGHG | ASGDGPRTNSDYT | 0.109390505 |
| TMPPLLAAIANAGRHGV | ASGDGPRTNSDYT | 0.17302454 |
| TSSTAGNSGGA | ASGDGPRTNSDYT | 0.36544818 |
| VPATVPVGILC | ASGDGPRTNSDYT | 0.6181303 |
| VQRPDSGGVVLTGRIS | ASGDGPRTNSDYT | 0.22281322 |
| WTPNPVPATKGVPAGLVQAT | ASGDGPRTNSDYT | 0.25736514 |
| YALKRALAKFSDRFDVVII | ASGDGPRTNSDYT | 0.530622 |
| YLIGSVIV | ASGDGPRTNSDYT | 0.13973828 |
| YGQLVAVID | ASGDGPRTNSDYT | 0.11729615 |
| VTVGGMTLPA | ASGDGPRTNSDYT | 0.2197077 |
| VLALPAV | ASGDGPRTNSDYT | 0.22508515 |
| VIPPTLNYET | ASGDGPRTNSDYT | 0.49136412 |
| VGMGGMHPSSG | ASGDGPRTNSDYT | 0.49348927 |
| VGLAGGAATA | ASGDGPRTNSDYT | 0.108934864 |
| VGGAGGAGGSGP | ASGDGPRTNSDYT | 0.33021772 |
| VFAAFMV | ASGDGPRTNSDYT | 0.2780941 |
| TLYDKLR | ASGDGPRTNSDYT | 0.122682445 |
| TLAVFTVL | ASGDGPRTNSDYT | 0.1457186 |
| TGSVQPARV | ASGDGPRTNSDYT | 0.14197229 |
| TGGGAGSGGT | ASGDGPRTNSDYT | 0.12998757 |
| SHLETSL | ASGDGPRTNSDYT | 0.21531335 |
| SFTTGMT | ASGDGPRTNSDYT | 0.1468623 |
| QVEVAKRMGVRQPTVS | ASGDGPRTNSDYT | 0.1889839 |
| QGSGGPALAM | ASGDGPRTNSDYT | 0.15291591 |
| MLAPSNP | ASGDGPRTNSDYT | 0.14801456 |
| LVNNAAI | ASGDGPRTNSDYT | 0.17569235 |
| LTGEINTLA | ASGDGPRTNSDYT | 0.17693976 |
| LSIFGPV | ASGDGPRTNSDYT | 0.27737898 |
| LLALLANL | ASGDGPRTNSDYT | 0.4102023 |
| LISNASDALDKLR | ASGDGPRTNSDYT | 0.1484151 |
| LINEAVP | ASGDGPRTNSDYT | 0.22358747 |
| LGYAFSGL | ASGDGPRTNSDYT | 0.43693197 |
| LDFVAVAL | ASGDGPRTNSDYT | 0.12380973 |
| LAQPGAE | ASGDGPRTNSDYT | 0.13313112 |
| ISNLGAVTGF | ASGDGPRTNSDYT | 0.31504443 |
| HQEDPGRI | ASGDGPRTNSDYT | 0.15653004 |
| GTPGEPTP | ASGDGPRTNSDYT | 0.50871265 |
| GFFNSGS | ASGDGPRTNSDYT | 0.23926678 |
| GAGGGGLTAGAN | ASGDGPRTNSDYT | 0.19583173 |
| FVDALVNM | ASGDGPRTNSDYT | 0.25854304 |
| FLTVAPEL | ASGDGPRTNSDYT | 0.68476534 |
| FETLTAL | ASGDGPRTNSDYT | 0.25099722 |
| DVINPIEL | ASGDGPRTNSDYT | 0.14903185 |

| | | |
|---|---|---|
| APVSGAPV | ASGDGPRTNSDYT | 0.3053869 |
| ALVGGLAW | ASGDGPRTNSDYT | 0.13713148 |
| ALAAAAATAI | ASGDGPRTNSDYT | 0.30627912 |
| AFGLGLLL | ASGDGPRTNSDYT | 0.44500253 |
| AFFAAASL | ASGDGPRTNSDYT | 0.12718397 |
| AAAALPAFSPP | ASGDGPRTNSDYT | 0.15317553 |
| VVLAPAL | ASGDGPRTNSDYT | 0.51302695 |
| TIGHVDHGKTTL | ASGDGPRTNSDYT | 0.4157751 |
| LISNASDALDK | ASGDGPRTNSDYT | 0.14529255 |
| LISNASDAL | ASGDGPRTNSDYT | 0.13073991 |
| ISNASDALDK | ASGDGPRTNSDYT | 0.16230565 |
| NQAETLVYQTE | ASGDGPRTNSDYT | 0.20884335 |
| TGGTGGTAGAG | ASGDGPRTNSDYT | 0.123308405 |
| TGSTAGGAGGA | ASGDGPRTNSDYT | 0.30466643 |
| DPLAVPVAAGPV | ASGDGPRTNSDYT | 0.16768873 |
| GGTGGTPGNGGN | ASGDGPRTNSDYT | 0.6577041 |
| PAGLAGVDPATL | ASGDGPRTNSDYT | 0.25438124 |
| GNNLVVEAPHAGASV | ASGDGPRTNSDYT | 0.4020722 |
| KVTRIMSASTLEQPA | ASGDGPRTNSDYT | 0.1638292 |
| LGFALPIALPRRTVV | ASGDGPRTNSDYT | 0.19199565 |
| TRLGNLTARKLVVLV | ASGDGPRTNSDYT | 0.12072022 |
| WSQQRRLIYFPSAGP | ASGDGPRTNSDYT | 0.13220029 |
| GPNGNPDPMAAAVDIR | ASGDGPRTNSDYT | 0.13962093 |
| TQRCVSVADLRAAASR | ASGDGPRTNSDYT | 0.18661189 |
| QFVLDVAEHPVEVLTETV | ASGDGPRTNSDYT | 0.11872192 |
| GGAGGTGGAGGNGGTQAGDG | ASGDGPRTNSDYT | 0.487665 |
| PLLATLRAIAVDVDAKPAQV | ASGDGPRTNSDYT | 0.1565708 |
| APKFRTGGYMIDFWGVGYQVAK | ASGDGPRTNSDYT | 0.3093049 |
| GAGGAGGAGGAAGGPGGTGGTG | ASGDGPRTNSDYT | 0.15939003 |
| IGALGLLATDVAPAGSVAFGVA | ASGDGPRTNSDYT | 0.13677806 |
| AGGAGGIGGNGGAGGAGGVLMG | ASGDGPRTNSDYT | 0.15051161 |
| ADELIGLL | ASGDGPRTNSDYT | 0.5104308 |
| ADRALKA | ASGDGPRTNSDYT | 0.12043115 |
| AGAVGQVGGAGGA | ASGDGPRTNSDYT | 0.11316005 |
| AGGGGGLGGAP | ASGDGPRTNSDYT | 0.13677733 |
| AGGLLYGNGGAG | ASGDGPRTNSDYT | 0.1869813 |
| AGLESIGL | ASGDGPRTNSDYT | 0.59155846 |
| AGLKTGII | ASGDGPRTNSDYT | 0.4023946 |
| ASLAALLL | ASGDGPRTNSDYT | 0.19598672 |
| ASVINISSI | ASGDGPRTNSDYT | 0.587894 |
| ATPPGPA | ASGDGPRTNSDYT | 0.103874944 |
| ATVASALV | ASGDGPRTNSDYT | 0.15562396 |
| ATVASLIL | ASGDGPRTNSDYT | 0.6005821 |
| AVNGPVLIRH | ASGDGPRTNSDYT | 0.24752352 |
| DGAAGAVGI | ASGDGPRTNSDYT | 0.18477848 |
| DPLLHGI | ASGDGPRTNSDYT | 0.3707177 |
| ELGAALGAL | ASGDGPRTNSDYT | 0.14433533 |
| ELGRTQAV | ASGDGPRTNSDYT | 0.1847637 |
| FVAGGYLS | ASGDGPRTNSDYT | 0.21321565 |
| GASGGAGGAGGW | ASGDGPRTNSDYT | 0.26966336 |
| GGAGAGGDGGAGGI | ASGDGPRTNSDYT | 0.3733289 |

| | | |
|---|---|---|
| GTGHNGSQPGMGGNG | ASGDGPRTNSDYT | 0.11677684 |
| GYAAASASAS | ASGDGPRTNSDYT | 0.38357654 |
| HQVDVAAVQQG | ASGDGPRTNSDYT | 0.7002846 |
| IGAAIAHT | ASGDGPRTNSDYT | 0.16855401 |
| IGMIEVAQL | ASGDGPRTNSDYT | 0.41518968 |
| IRLAPEHTSAMD | ASGDGPRTNSDYT | 0.20478414 |
| IYNVSVL | ASGDGPRTNSDYT | 0.23018208 |
| KQGFLGM | ASGDGPRTNSDYT | 0.13966046 |
| LESVITQL | ASGDGPRTNSDYT | 0.11972289 |
| LGLGALVVV | ASGDGPRTNSDYT | 0.2666178 |
| LIGNGGNGGSGGI | ASGDGPRTNSDYT | 0.11912663 |
| LNAINAPTL | ASGDGPRTNSDYT | 0.27054602 |
| LQLLNTGV | ASGDGPRTNSDYT | 0.18116958 |
| LQTSIEF | ASGDGPRTNSDYT | 0.77974993 |
| MSFAVSEL | ASGDGPRTNSDYT | 0.28204465 |
| NTGALGL | ASGDGPRTNSDYT | 0.23816925 |
| PPGAGGAGGAGTT | ASGDGPRTNSDYT | 0.13959266 |
| QLVGLGI | ASGDGPRTNSDYT | 0.2090908 |
| RFLEQVHQL | ASGDGPRTNSDYT | 0.29880494 |
| RGVGYVLR | ASGDGPRTNSDYT | 0.12760767 |
| SATGVFAGVI | ASGDGPRTNSDYT | 0.44635624 |
| SLPAITLP | ASGDGPRTNSDYT | 0.13135283 |
| STALAFTV | ASGDGPRTNSDYT | 0.21535593 |
| SVLTVGLGK | ASGDGPRTNSDYT | 0.26125723 |
| TALAGAGSGI | ASGDGPRTNSDYT | 0.69360757 |
| TGLPFTGI | ASGDGPRTNSDYT | 0.17836094 |
| VAEHVTV | ASGDGPRTNSDYT | 0.31692392 |
| VDIVLNSL | ASGDGPRTNSDYT | 0.13229427 |
| VLTGGGAVL | ASGDGPRTNSDYT | 0.32059634 |
| VNVGLAYGI | ASGDGPRTNSDYT | 0.50814307 |
| VSSAAVGAV | ASGDGPRTNSDYT | 0.28994673 |
| VTSVYEAGV | ASGDGPRTNSDYT | 0.7168524 |
| WVALDRGA | ASGDGPRTNSDYT | 0.29677874 |
| YLVGSVVI | ASGDGPRTNSDYT | 0.12295619 |
| GPPGTGKTLL | ASGDGPRTNSDYT | 0.20424491 |
| GVGMTKF | ASGDGPRTNSDYT | 0.3745333 |
| IGIGGSDL | ASGDGPRTNSDYT | 0.64538467 |
| IGTLLAEL | ASGDGPRTNSDYT | 0.32315338 |
| IYTSGTT | ASGDGPRTNSDYT | 0.43039182 |
| LSFPAGV | ASGDGPRTNSDYT | 0.17918131 |
| LYTSGTT | ASGDGPRTNSDYT | 0.514258 |
| VGAFPFGF | ASGDGPRTNSDYT | 0.7758719 |
| VGAGTVEF | ASGDGPRTNSDYT | 0.29551843 |
| VGKSTLLL | ASGDGPRTNSDYT | 0.14231136 |
| AALAHAGGGIV | ASGDGPRTNSDYT | 0.15433903 |
| AALVAVSLL | ASGDGPRTNSDYT | 0.25497356 |
| AAVSRAVAAL | ASGDGPRTNSDYT | 0.3747149 |
| AGGSSGIGGF | ASGDGPRTNSDYT | 0.14512153 |
| AGLTVANANG | ASGDGPRTNSDYT | 0.15744053 |
| AGLVGAVL | ASGDGPRTNSDYT | 0.20380597 |
| ALSQAFTV | ASGDGPRTNSDYT | 0.10936932 |

| | | |
|---|---|---|
| ALTFSVSL | ASGDGPRTNSDYT | 0.36097592 |
| ALTGAALVAL | ASGDGPRTNSDYT | 0.66071945 |
| ASFSIGAGVL | ASGDGPRTNSDYT | 0.25169867 |
| DGSVYKI | ASGDGPRTNSDYT | 0.1229914 |
| EQLAEIV | ASGDGPRTNSDYT | 0.46711496 |
| FAVSELTVM | ASGDGPRTNSDYT | 0.6570462 |
| GAAPLAAGL | ASGDGPRTNSDYT | 0.451574 |
| GADGGDGGSGGAGG | ASGDGPRTNSDYT | 0.12219039 |
| GFGLDTGGAGG | ASGDGPRTNSDYT | 0.10331456 |
| GLASGGLL | ASGDGPRTNSDYT | 0.2992487 |
| GRIAGAAVLL | ASGDGPRTNSDYT | 0.350955 |
| GSAGSLNTSF | ASGDGPRTNSDYT | 0.15945928 |
| GTGQIGI | ASGDGPRTNSDYT | 0.13126096 |
| GTLIALIL | ASGDGPRTNSDYT | 0.2421391 |
| IAATMGFVVG | ASGDGPRTNSDYT | 0.35091087 |
| IFAQALL | ASGDGPRTNSDYT | 0.13122252 |
| IGAAVALGF | ASGDGPRTNSDYT | 0.13074422 |
| IGLTPLAL | ASGDGPRTNSDYT | 0.2364269 |
| ILGSNTSTL | ASGDGPRTNSDYT | 0.5120202 |
| IVFSLTVL | ASGDGPRTNSDYT | 0.11888043 |
| LASKATGF | ASGDGPRTNSDYT | 0.15636137 |
| LASVALIL | ASGDGPRTNSDYT | 0.32835984 |
| LFAVGLTL | ASGDGPRTNSDYT | 0.4196837 |
| LFNGGIGTYI | ASGDGPRTNSDYT | 0.22639361 |
| LTLGAGII | ASGDGPRTNSDYT | 0.12901844 |
| LVAAALGDI | ASGDGPRTNSDYT | 0.28151804 |
| NTAFVQLGI | ASGDGPRTNSDYT | 0.1192173 |
| QLFSGAEL | ASGDGPRTNSDYT | 0.47068474 |
| SAATFITL | ASGDGPRTNSDYT | 0.2292559 |
| SNYGLANLGV | ASGDGPRTNSDYT | 0.24228907 |
| SYGGAYAVM | ASGDGPRTNSDYT | 0.16027513 |
| TTGVGGLTLG | ASGDGPRTNSDYT | 0.1186767 |
| TTQTIYI | ASGDGPRTNSDYT | 0.13112865 |
| VAAGAGLGAVI | ASGDGPRTNSDYT | 0.13673389 |
| VAAVVTLL | ASGDGPRTNSDYT | 0.47466257 |
| VGFLGGLI | ASGDGPRTNSDYT | 0.37529546 |
| VGVIPILL | ASGDGPRTNSDYT | 0.33120024 |
| VIFASSLI | ASGDGPRTNSDYT | 0.22093055 |
| VKGGDGGAAAG | ASGDGPRTNSDYT | 0.13308084 |
| VTKTFTNI | ASGDGPRTNSDYT | 0.27878195 |
| VVAAAILL | ASGDGPRTNSDYT | 0.119032584 |
| YAALDVEL | ASGDGPRTNSDYT | 0.22324237 |
| LASVALLL | ASGDGPRTNSDYT | 0.19454679 |
| LDAVHRV | ASGDGPRTNSDYT | 0.32283637 |
| QGLSSSIF | ASGDGPRTNSDYT | 0.16289905 |
| RGTFANIRL | ASGDGPRTNSDYT | 0.31285816 |
| RQLWWGHRI | ASGDGPRTNSDYT | 0.28515887 |
| YGPPGTGKTLL | ASGDGPRTNSDYT | 0.14553632 |
| AALVGAPANVA | ASGDGPRTNSDYT | 0.226617 |
| AATFRRTE | ASGDGPRTNSDYT | 0.41046816 |
| AGALGGGI | ASGDGPRTNSDYT | 0.30650753 |

| | | |
|---|---|---|
| AGGAGGAGGIGAG | ASGDGPRTNSDYT | 0.11391402 |
| AGSGALGL | ASGDGPRTNSDYT | 0.18551956 |
| AQIADQI | ASGDGPRTNSDYT | 0.13841255 |
| ASAVAVTYL | ASGDGPRTNSDYT | 0.22090912 |
| ASTVLGGTGG | ASGDGPRTNSDYT | 0.21250826 |
| AVGGEPAK | ASGDGPRTNSDYT | 0.21154292 |
| CEQGTFGGTC | ASGDGPRTNSDYT | 0.24065593 |
| DEGKAPAG | ASGDGPRTNSDYT | 0.13291074 |
| DTVVFPAL | ASGDGPRTNSDYT | 0.1432172 |
| FEGITMAG | ASGDGPRTNSDYT | 0.16746709 |
| FPLVVQD | ASGDGPRTNSDYT | 0.28869194 |
| GAFTTPQ | ASGDGPRTNSDYT | 0.26757973 |
| GFGGFGTAGGVGGA | ASGDGPRTNSDYT | 0.12083163 |
| GFLAQFL | ASGDGPRTNSDYT | 0.26083136 |
| GGAGGAGGNAGVA | ASGDGPRTNSDYT | 0.18873245 |
| GGVGTAGAGGAGGAGG | ASGDGPRTNSDYT | 0.13520105 |
| GNGGGGDAGNAGSGGN | ASGDGPRTNSDYT | 0.21827357 |
| GTGGAGGAGNDAGS | ASGDGPRTNSDYT | 0.14585643 |
| GTTNTGWFNS | ASGDGPRTNSDYT | 0.124985695 |
| IFMGPLFTL | ASGDGPRTNSDYT | 0.31329706 |
| ILAHAIEAI | ASGDGPRTNSDYT | 0.15463562 |
| IRSAIREAL | ASGDGPRTNSDYT | 0.23545031 |
| ITFAGGG | ASGDGPRTNSDYT | 0.39241695 |
| ITGMSGHQEV | ASGDGPRTNSDYT | 0.12720884 |
| IVAGLACTL | ASGDGPRTNSDYT | 0.34687003 |
| IVGGGLAGA | ASGDGPRTNSDYT | 0.17355332 |
| IVTGVLNI | ASGDGPRTNSDYT | 0.2641253 |
| LAQQGCI | ASGDGPRTNSDYT | 0.19312103 |
| LCLHVEN | ASGDGPRTNSDYT | 0.13482195 |
| LDAISDGS | ASGDGPRTNSDYT | 0.32445872 |
| LGGFAAML | ASGDGPRTNSDYT | 0.24385598 |
| LGGKNPMI | ASGDGPRTNSDYT | 0.27610368 |
| LGPKGVAAIL | ASGDGPRTNSDYT | 0.21234532 |
| LMVGVSI | ASGDGPRTNSDYT | 0.40849298 |
| LVGAAFVL | ASGDGPRTNSDYT | 0.37395474 |
| LVLAVAPL | ASGDGPRTNSDYT | 0.12843217 |
| MAAIAVIDAA | ASGDGPRTNSDYT | 0.11513195 |
| NNGINAELQ | ASGDGPRTNSDYT | 0.18732545 |
| RGEPLVLVHG | ASGDGPRTNSDYT | 0.18293375 |
| RGGAGGAAGLI | ASGDGPRTNSDYT | 0.13875793 |
| RSYRMHPAV | ASGDGPRTNSDYT | 0.12795433 |
| SAAHLLAGQLA | ASGDGPRTNSDYT | 0.3652705 |
| SAAVGSY | ASGDGPRTNSDYT | 0.70111394 |
| SPHAVLHRI | ASGDGPRTNSDYT | 0.1686053 |
| SQIAKVL | ASGDGPRTNSDYT | 0.71990216 |
| TALGHLVP | ASGDGPRTNSDYT | 0.1169015 |
| TAPTTSGAA | ASGDGPRTNSDYT | 0.26006937 |
| TDGDDGNGGSPGT | ASGDGPRTNSDYT | 0.42034784 |
| TDGLGEN | ASGDGPRTNSDYT | 0.12712848 |
| TGGPPLPGQA | ASGDGPRTNSDYT | 0.23882294 |
| TIPAITI | ASGDGPRTNSDYT | 0.16936502 |

| | | |
|---|---|---|
| TIPGFTI | ASGDGPRTNSDYT | 0.13936663 |
| TSIASGALA | ASGDGPRTNSDYT | 0.3736713 |
| TVTLPGAAI | ASGDGPRTNSDYT | 0.19308245 |
| VASGLSAALP | ASGDGPRTNSDYT | 0.26206407 |
| VGAGVIGI | ASGDGPRTNSDYT | 0.17476931 |
| VGPKRFGV | ASGDGPRTNSDYT | 0.1378518 |
| VNVAFPD | ASGDGPRTNSDYT | 0.23687735 |
| VSMTNSGVS | ASGDGPRTNSDYT | 0.37692177 |
| VTDVLGDLSV | ASGDGPRTNSDYT | 0.21986805 |
| VVVVGAGFAGL | ASGDGPRTNSDYT | 0.7639303 |
| VYTVGTVV | ASGDGPRTNSDYT | 0.13772023 |
| WTTFSGP | ASGDGPRTNSDYT | 0.65515584 |
| YDELVEAF | ASGDGPRTNSDYT | 0.6974074 |
| YGTGGAGGNGG | ASGDGPRTNSDYT | 0.22699544 |
| YTNVMAV | ASGDGPRTNSDYT | 0.19033186 |
| FLEQVHQL | ASGDGPRTNSDYT | 0.39489526 |
| GLSLLAVAV | ASGDGPRTNSDYT | 0.24756697 |
| LEQVHQL | ASGDGPRTNSDYT | 0.31055897 |
| LNAAGAI | ASGDGPRTNSDYT | 0.37700707 |
| PAVHASL | ASGDGPRTNSDYT | 0.5694101 |
| TELEDDA | ASGDGPRTNSDYT | 0.77944696 |
| TPPGPPN | ASGDGPRTNSDYT | 0.6842866 |
| TPPPPPPP | ASGDGPRTNSDYT | 0.71168303 |
| TQSPALI | ASSLKVSAETLY | 0.6128803 |
| EFAAAFPQ | ASSLKVSAETLY | 0.4419025 |
| LLSGNLDVL | ASSLKVSAETLY | 0.7136017 |
| GPANLGNYNI | ASSLKVSAETLY | 0.5363955 |
| TLGPTLP | ASSLKVSAETLY | 0.71869814 |
| VYAYSSCCK | ASSLKVSAETLY | 0.7566643 |
| APAPAPAPNI | ASSLKVSAETLY | 0.6096664 |
| PGGPVPIP | ASSLKVSAETLY | 0.7886784 |
| FLTVALEP | ASSLKVSAETLY | 0.79308915 |
| SQYFGANL | ASSLKVSAETLY | 0.4667058 |
| GGVKPAA | ASSLKVSAETLY | 0.6239444 |
| GGTGGAGGAGGW | ASSLKVSAETLY | 0.6232785 |
| GVGGVAGLGGAGG | ASSLKVSAETLY | 0.64803326 |
| VLLVLATTI | ASSLKVSAETLY | 0.35714427 |
| IGIFAAML | ASSLKVSAETLY | 0.752787 |
| AVGDLID | ASSLKVSAETLY | 0.52748495 |
| NISPLPL | ASSLKVSAETLY | 0.2163849 |
| VTAVFVAI | ASSLKVSAETLY | 0.68970686 |
| SGDDNDAAL | ASSLKVSAETLY | 0.64926916 |
| AVAALAHS | ASSLKVSAETLY | 0.63550067 |
| GGVTVMATP | ASSLKVSAETLY | 0.75934476 |
| AGLAANVRD | ASSLKVSAETLY | 0.3613146 |
| TNRPDLIDPALLRPG | ASSLKVSAETLY | 0.61479586 |
| LANLGVTAL | ASSLKVSAETLY | 0.7543697 |
| TGERIEL | ASSLKVSAETLY | 0.7897289 |
| NGEELAA | ASSLKVSAETLY | 0.52197397 |
| VLSLVWG | ASSLKVSAETLY | 0.70233643 |
| IGYALWL | ASSLKVSAETLY | 0.7943392 |

| | | |
|---|---|---|
| LSLLLAPL | ASSLKVSAETLY | 0.73429465 |
| TLDEALK | ASSLKVSAETLY | 0.7783798 |
| TDGAGTAEGG | ASSLKVSAETLY | 0.7902182 |
| NAPGGAGGNGG | ASSLKVSAETLY | 0.6082868 |
| VSVFMGMGGNF | ASSLKVSAETLY | 0.7745179 |
| LASYLAAL | ASSLKVSAETLY | 0.7907064 |
| LVGGTPAL | ASSLKVSAETLY | 0.6809179 |
| VRNADVL | ASSLKVSAETLY | 0.69636446 |
| TCSSPFLVSAM | ASSLKVSAETLY | 0.57845426 |
| VHITDAE | ASSLKVSAETLY | 0.63833606 |
| EYAAPSF | ASSLKVSAETLY | 0.31106994 |
| VVDAFHGGL | ASSLKVSAETLY | 0.69142246 |
| ASVVDPQ | ASSLKVSAETLY | 0.7847694 |
| SYNEFGGLS | ASSLKVSAETLY | 0.6245311 |
| DPDGSVY | ASSLKVSAETLY | 0.4873303 |
| MVSTIHATE | ASSLKVSAETLY | 0.5432864 |
| LAAGVADPV | ASSLKVSAETLY | 0.60954285 |
| TKEQIAA | ASSLKVSAETLY | 0.63574564 |
| SLASPAAL | ASSLKVSAETLY | 0.35299426 |
| IGLVFYGV | ASSLKVSAETLY | 0.5400236 |
| FLLDASNL | ASSLKVSAETLY | 0.7870233 |
| YQVGNFP | ASSLKVSAETLY | 0.78420264 |
| ESLLSQST | ASSLKVSAETLY | 0.3628949 |
| LFTVGFAAVG | ASSLKVSAETLY | 0.7797775 |
| ATPPTPT | ASSLKVSAETLY | 0.72823715 |
| TDLATLG | ASSLKVSAETLY | 0.79379183 |
| SAGGTGR | ASSLKVSAETLY | 0.7041669 |
| GFSNSGS | ASSLKVSAETLY | 0.5531175 |
| GGNGGSGGTSV | ASSLKVSAETLY | 0.729692 |
| GTSVATGGAGN | ASSLKVSAETLY | 0.5985967 |
| NGGNGGSGGTS | ASSLKVSAETLY | 0.50187826 |
| AVVKFPH | ASSLKVSAETLY | 0.76889217 |
| TPPPPPP | ASSLKVSAETLY | 0.64865816 |
| AAPAAAAAPA | ASSLKVSAETLY | 0.5655414 |
| IALNEDL | ASSLKVSAETLY | 0.5491207 |
| IVNVSSM | ASSLKVSAETLY | 0.76869255 |
| TGIPPAPR | ASSLKVSAETLY | 0.48398134 |
| FDLGGGTF | ASSLKVSAETLY | 0.53929025 |
| FDLGGGTFDVS | ASSLKVSAETLY | 0.48284072 |
| NEPTAAA | ASSLKVSAETLY | 0.6765315 |
| NEPTAAALA | ASSLKVSAETLY | 0.77895427 |
| GSIINISSI | ASSLKVSAETLY | 0.73784554 |
| VAPGNAGTA | ASSLKVSAETLY | 0.5825725 |
| TGGGASLE | ASSLKVSAETLY | 0.72469974 |
| VAVYNHYKRI | ASSLKVSAETLY | 0.73014927 |
| TDPAPATT | ASSLKVSAETLY | 0.70822024 |
| TNRPDILDPALLRPG | ASSLKVSAETLY | 0.7209354 |
| LTTTTTS | ASSLKVSAETLY | 0.51913726 |
| PTGSGKTL | ASSLKVSAETLY | 0.76186 |
| TGSGKTL | ASSLKVSAETLY | 0.740092 |
| TIGHVDHGK | ASSLKVSAETLY | 0.7662678 |

| | | |
|---|---|---|
| TIGHVDHGKTT | ASSLKVSAETLY | 0.7691762 |
| GHVDHGKTT | ASSLKVSAETLY | 0.69938576 |
| AQFEHTLL | ASSLKVSAETLY | 0.6980259 |
| IVNQDEL | ASSLKVSAETLY | 0.3150354 |
| TNGSQFF | ASSLKVSAETLY | 0.6226578 |
| TGKGIVE | ASSLKVSAETLY | 0.6967313 |
| LISNASDA | ASSLKVSAETLY | 0.68422455 |
| AELRTLGELTKIAKAHGAQ | ASSLKVSAETLY | 0.73357403 |
| APNAVAAAEPSW | ASSLKVSAETLY | 0.7578446 |
| APQVAGAATAVL | ASSLKVSAETLY | 0.5949187 |
| ARRDPAAAAAILPTDA | ASSLKVSAETLY | 0.7708484 |
| DAAARDVLAEAGLAEYF | ASSLKVSAETLY | 0.7695411 |
| DDQIVLSAKANLAKVEQ | ASSLKVSAETLY | 0.41985232 |
| DELLPDTGEDSDEDGH | ASSLKVSAETLY | 0.6251526 |
| ELPEAPANRATVPQD | ASSLKVSAETLY | 0.7901816 |
| FGQSKIERVTGIG | ASSLKVSAETLY | 0.6951947 |
| GAGGTGSTAGGAGGAGGAGGLY | ASSLKVSAETLY | 0.6554088 |
| GHGGTGAAGGNGGTGGAGGSGI | ASSLKVSAETLY | 0.61128616 |
| GNGGNGGAGGEGGPGGAGGAGA | ASSLKVSAETLY | 0.79102916 |
| GSTAGNGGNGGNATNGTIAGQP | ASSLKVSAETLY | 0.5166374 |
| GTGGDGGNGGNGGTGG | ASSLKVSAETLY | 0.7965324 |
| GVRRPAGIPADAPV | ASSLKVSAETLY | 0.2644054 |
| GYRQVTDAHLLTLARR | ASSLKVSAETLY | 0.59233063 |
| IAKMGQDGHDRG | ASSLKVSAETLY | 0.5425378 |
| KAEPSIEIQYT | ASSLKVSAETLY | 0.33796278 |
| KPVRIRMAADEDTVAVT | ASSLKVSAETLY | 0.7747234 |
| MPPRIAGMRLLVIKPEP | ASSLKVSAETLY | 0.6676188 |
| MKFYAVALFRSRGVPPTR | ASSLKVSAETLY | 0.7510574 |
| NAGGFGILGAVAHSP | ASSLKVSAETLY | 0.6842153 |
| NEGREITLAMAEM | ASSLKVSAETLY | 0.77178186 |
| NELVEWSHPLGHRWRWE | ASSLKVSAETLY | 0.68543303 |
| NLGAGGTGGNGGNPG | ASSLKVSAETLY | 0.6961725 |
| NTEWPKWETVMAADPVD | ASSLKVSAETLY | 0.56180924 |
| PASAAPAETQP | ASSLKVSAETLY | 0.76487255 |
| QQSSDGKNGVRAAEITGE | ASSLKVSAETLY | 0.787318 |
| RAPDAVTTSWWKE | ASSLKVSAETLY | 0.7415011 |
| SGFGNVGAHTSGWFNQ | ASSLKVSAETLY | 0.51141053 |
| SGVAGAVGQAASVGGLKV | ASSLKVSAETLY | 0.53585714 |
| SNGGAGTGSAGGNGGT | ASSLKVSAETLY | 0.72891116 |
| TGGAGGEGGAAG | ASSLKVSAETLY | 0.31378457 |
| TGGDGGAAGAGGAPGHG | ASSLKVSAETLY | 0.24168508 |
| TMPPLLAAIANAGRHGV | ASSLKVSAETLY | 0.5575583 |
| TSSTAGNSGGA | ASSLKVSAETLY | 0.48766017 |
| VPATVPVGILC | ASSLKVSAETLY | 0.7712236 |
| VQRPDSGGVVLTGRIS | ASSLKVSAETLY | 0.76237744 |
| WTPNPVPATKGVPAGLVQAT | ASSLKVSAETLY | 0.6518824 |
| YALKRALAKFSDRFDVVII | ASSLKVSAETLY | 0.7426814 |
| YLIGSVIV | ASSLKVSAETLY | 0.71007687 |
| YGQLVAVID | ASSLKVSAETLY | 0.16822176 |
| VTVGGMTLPA | ASSLKVSAETLY | 0.63691527 |
| VLALPAV | ASSLKVSAETLY | 0.48556885 |

| | | |
|---|---|---|
| VIPPTLNYET | ASSLKVSAETLY | 0.8031948 |
| VGMGGMHPSSG | ASSLKVSAETLY | 0.5155289 |
| VGLAGGAATA | ASSLKVSAETLY | 0.5349856 |
| VGGAGGAGGSGP | ASSLKVSAETLY | 0.67580116 |
| VFAAFMV | ASSLKVSAETLY | 0.67258775 |
| TLYDKLR | ASSLKVSAETLY | 0.6537154 |
| TLAVFTVL | ASSLKVSAETLY | 0.48718634 |
| TGSVQPARV | ASSLKVSAETLY | 0.21232866 |
| TGGGAGSGGT | ASSLKVSAETLY | 0.77815497 |
| SHLETSL | ASSLKVSAETLY | 0.43121517 |
| SFTTGMT | ASSLKVSAETLY | 0.58265626 |
| QVEVAKRMGVRQPTVS | ASSLKVSAETLY | 0.30733192 |
| QGSGGPALAM | ASSLKVSAETLY | 0.23745799 |
| MLAPSNP | ASSLKVSAETLY | 0.43938014 |
| LVNNAAI | ASSLKVSAETLY | 0.7625681 |
| LTGEINTLA | ASSLKVSAETLY | 0.6264566 |
| LSIFGPV | ASSLKVSAETLY | 0.6155499 |
| LLALLANL | ASSLKVSAETLY | 0.8009583 |
| LISNASDALDKLR | ASSLKVSAETLY | 0.6582622 |
| LINEAVP | ASSLKVSAETLY | 0.60816884 |
| LGYAFSGL | ASSLKVSAETLY | 0.6077923 |
| LDFVAVAL | ASSLKVSAETLY | 0.52677697 |
| LAQPGAE | ASSLKVSAETLY | 0.5690782 |
| ISNLGAVTGF | ASSLKVSAETLY | 0.5791461 |
| HQEDPGRI | ASSLKVSAETLY | 0.8019576 |
| GTPGEPTP | ASSLKVSAETLY | 0.6105053 |
| GFFNSGS | ASSLKVSAETLY | 0.5423125 |
| GAGGGGLTAGAN | ASSLKVSAETLY | 0.6908423 |
| FVDALVNM | ASSLKVSAETLY | 0.78575146 |
| FLTVAPEL | ASSLKVSAETLY | 0.81239164 |
| FETLTAL | ASSLKVSAETLY | 0.5450865 |
| DVINPIEL | ASSLKVSAETLY | 0.69496286 |
| APVSGAPV | ASSLKVSAETLY | 0.651001 |
| ALVGGLAW | ASSLKVSAETLY | 0.65726185 |
| ALAAAAATAI | ASSLKVSAETLY | 0.60146713 |
| AFGLGLLL | ASSLKVSAETLY | 0.51080596 |
| AFFAAASL | ASSLKVSAETLY | 0.7453297 |
| AAAALPAFSPP | ASSLKVSAETLY | 0.8058896 |
| VVLAPAL | ASSLKVSAETLY | 0.6347422 |
| TIGHVDHGKTTL | ASSLKVSAETLY | 0.52540237 |
| LISNASDALDK | ASSLKVSAETLY | 0.6482082 |
| LISNASDAL | ASSLKVSAETLY | 0.53208166 |
| ISNASDALDK | ASSLKVSAETLY | 0.59705013 |
| NQAETLVYQTE | ASSLKVSAETLY | 0.7229514 |
| TGGTGGTAGAG | ASSLKVSAETLY | 0.3988551 |
| TGSTAGGAGGA | ASSLKVSAETLY | 0.63497937 |
| DPLAVPVAAGPV | ASSLKVSAETLY | 0.8069308 |
| GGTGGTPGNGGN | ASSLKVSAETLY | 0.6224448 |
| PAGLAGVDPATL | ASSLKVSAETLY | 0.7152926 |
| GNNLVVEAPHAGASV | ASSLKVSAETLY | 0.7712954 |
| KVTRIMSASTLEQPA | ASSLKVSAETLY | 0.7223573 |

| | | |
|---|---|---|
| LGFALPIALPRRTVV | ASSLKVSAETLY | 0.78016675 |
| TRLGNLTARKLVVLV | ASSLKVSAETLY | 0.7468926 |
| WSQQRRLIYFPSAGP | ASSLKVSAETLY | 0.6695974 |
| GPNGNPDPMAAAVDIR | ASSLKVSAETLY | 0.6671349 |
| TQRCVSVADLRAAASR | ASSLKVSAETLY | 0.67554945 |
| QFVLDVAEHPVEVLTETV | ASSLKVSAETLY | 0.6893387 |
| GGAGGTGGAGGNGGTQAGDG | ASSLKVSAETLY | 0.6495812 |
| PLLATLRAIAVDVDAKPAQV | ASSLKVSAETLY | 0.5786983 |
| APKFRTGGYMIDFWGVGYQVAK | ASSLKVSAETLY | 0.7800418 |
| GAGGAGGAGGAAGGPGGTGGTG | ASSLKVSAETLY | 0.65448916 |
| IGALGLLATDVAPAGSVAFGVA | ASSLKVSAETLY | 0.7487034 |
| AGGAGGIGGNGGAGGAGGVLMG | ASSLKVSAETLY | 0.44136506 |
| ADELIGLL | ASSLKVSAETLY | 0.49975246 |
| ADRALKA | ASSLKVSAETLY | 0.7470739 |
| AGAVGQVGGAGGA | ASSLKVSAETLY | 0.20386328 |
| AGGGGGLGGAP | ASSLKVSAETLY | 0.27878723 |
| AGGLLYGNGGAG | ASSLKVSAETLY | 0.68746126 |
| AGLESIGL | ASSLKVSAETLY | 0.7807588 |
| AGLKTGII | ASSLKVSAETLY | 0.22310962 |
| ASLAALLL | ASSLKVSAETLY | 0.5935492 |
| ASVINISSI | ASSLKVSAETLY | 0.44525257 |
| ATPPGPA | ASSLKVSAETLY | 0.7274768 |
| ATVASALV | ASSLKVSAETLY | 0.6880401 |
| ATVASLIL | ASSLKVSAETLY | 0.39431357 |
| AVNGPVLIRH | ASSLKVSAETLY | 0.74225867 |
| DGAAGAVGI | ASSLKVSAETLY | 0.38310805 |
| DPLLHGI | ASSLKVSAETLY | 0.37788662 |
| ELGAALGAL | ASSLKVSAETLY | 0.76747906 |
| ELGRTQAV | ASSLKVSAETLY | 0.7683294 |
| FVAGGYLS | ASSLKVSAETLY | 0.5163892 |
| GASGGAGGAGGW | ASSLKVSAETLY | 0.75961864 |
| GGAGAGGDGGAGGI | ASSLKVSAETLY | 0.56229615 |
| GTGHNGSQPGMGGNG | ASSLKVSAETLY | 0.8076165 |
| GYAAASASAS | ASSLKVSAETLY | 0.6463976 |
| HQVDVAAVQQG | ASSLKVSAETLY | 0.41212672 |
| IGAAIAHT | ASSLKVSAETLY | 0.76448023 |
| IGMIEVAQL | ASSLKVSAETLY | 0.7158766 |
| IRLAPEHTSAMD | ASSLKVSAETLY | 0.7225108 |
| IYNVSVL | ASSLKVSAETLY | 0.7213514 |
| KQGFLGM | ASSLKVSAETLY | 0.5549554 |
| LESVITQL | ASSLKVSAETLY | 0.6377964 |
| LGLGALVVV | ASSLKVSAETLY | 0.77416027 |
| LIGNGGNGGSGGI | ASSLKVSAETLY | 0.7508224 |
| LNAINAPTL | ASSLKVSAETLY | 0.49816608 |
| LQLLNTGV | ASSLKVSAETLY | 0.7684352 |
| LQTSIEF | ASSLKVSAETLY | 0.6535003 |
| MSFAVSEL | ASSLKVSAETLY | 0.60577184 |
| NTGALGL | ASSLKVSAETLY | 0.80367285 |
| PPGAGGAGGAGTT | ASSLKVSAETLY | 0.6913512 |
| QLVGLGI | ASSLKVSAETLY | 0.7467666 |
| RFLEQVHQL | ASSLKVSAETLY | 0.77029747 |

| | | |
|---|---|---|
| RGVGYVLR | ASSLKVSAETLY | 0.79683936 |
| SATGVFAGVI | ASSLKVSAETLY | 0.44700113 |
| SLPAITLP | ASSLKVSAETLY | 0.7155037 |
| STALAFTV | ASSLKVSAETLY | 0.6808224 |
| SVLTVGLGK | ASSLKVSAETLY | 0.75992596 |
| TALAGAGSGI | ASSLKVSAETLY | 0.70321715 |
| TGLPFTGI | ASSLKVSAETLY | 0.4884942 |
| VAEHVTV | ASSLKVSAETLY | 0.71799535 |
| VDIVLNSL | ASSLKVSAETLY | 0.78317153 |
| VLTGGGAVL | ASSLKVSAETLY | 0.66483694 |
| VNVGLAYGI | ASSLKVSAETLY | 0.7023305 |
| VSSAAVGAV | ASSLKVSAETLY | 0.7922356 |
| VTSVYEAGV | ASSLKVSAETLY | 0.57869256 |
| WVALDRGA | ASSLKVSAETLY | 0.67307866 |
| YLVGSVVI | ASSLKVSAETLY | 0.7714255 |
| GPPGTGKTLL | ASSLKVSAETLY | 0.77162826 |
| GVGMTKF | ASSLKVSAETLY | 0.6426187 |
| IGIGGSDL | ASSLKVSAETLY | 0.78667444 |
| IGTLLAEL | ASSLKVSAETLY | 0.5489291 |
| IYTSGTT | ASSLKVSAETLY | 0.7537977 |
| LSFPAGV | ASSLKVSAETLY | 0.55906916 |
| LYTSGTT | ASSLKVSAETLY | 0.7260754 |
| VGAFPFGF | ASSLKVSAETLY | 0.4600238 |
| VGAGTVEF | ASSLKVSAETLY | 0.7612735 |
| VGKSTLLL | ASSLKVSAETLY | 0.7563058 |
| AALAHAGGGIV | ASSLKVSAETLY | 0.75953543 |
| AALVAVSLL | ASSLKVSAETLY | 0.26743492 |
| AAVSRAVAAL | ASSLKVSAETLY | 0.7507526 |
| AGGSSGIGGF | ASSLKVSAETLY | 0.7670716 |
| AGLTVANANG | ASSLKVSAETLY | 0.8019042 |
| AGLVGAVL | ASSLKVSAETLY | 0.63389033 |
| ALSQAFTV | ASSLKVSAETLY | 0.75260913 |
| ALTFSVSL | ASSLKVSAETLY | 0.6741414 |
| ALTGAALVAL | ASSLKVSAETLY | 0.640222 |
| ASFSIGAGVL | ASSLKVSAETLY | 0.4522004 |
| DGSVYKI | ASSLKVSAETLY | 0.6303222 |
| EQLAEIV | ASSLKVSAETLY | 0.5968088 |
| FAVSELTVM | ASSLKVSAETLY | 0.52688766 |
| GAAPLAAGL | ASSLKVSAETLY | 0.73474926 |
| GADGGDGGSGGAGG | ASSLKVSAETLY | 0.76030636 |
| GFGLDTGGAGG | ASSLKVSAETLY | 0.7730687 |
| GLASGGLL | ASSLKVSAETLY | 0.7808566 |
| GRIAGAAVLL | ASSLKVSAETLY | 0.7738279 |
| GSAGSLNTSF | ASSLKVSAETLY | 0.7648532 |
| GTGQIGI | ASSLKVSAETLY | 0.5854203 |
| GTLIALIL | ASSLKVSAETLY | 0.6510037 |
| IAATMGFVVG | ASSLKVSAETLY | 0.48128048 |
| IFAQALL | ASSLKVSAETLY | 0.6419052 |
| IGAAVALGF | ASSLKVSAETLY | 0.58616245 |
| IGLTPLAL | ASSLKVSAETLY | 0.7055087 |
| ILGSNTSTL | ASSLKVSAETLY | 0.1848637 |

| | | |
|---|---|---|
| IVFSLTVL | ASSLKVSAETLY | 0.63595533 |
| LASKATGF | ASSLKVSAETLY | 0.7956917 |
| LASVALIL | ASSLKVSAETLY | 0.50543433 |
| LFAVGLTL | ASSLKVSAETLY | 0.72718036 |
| LFNGGIGTYI | ASSLKVSAETLY | 0.64417326 |
| LTLGAGII | ASSLKVSAETLY | 0.5130802 |
| LVAAALGDI | ASSLKVSAETLY | 0.7549913 |
| NTAFVQLGI | ASSLKVSAETLY | 0.5264606 |
| QLFSGAEL | ASSLKVSAETLY | 0.73461235 |
| SAATFITL | ASSLKVSAETLY | 0.8012653 |
| SNYGLANLGV | ASSLKVSAETLY | 0.7203693 |
| SYGGAYAVM | ASSLKVSAETLY | 0.68859774 |
| TTGVGGLTLG | ASSLKVSAETLY | 0.6833674 |
| TTQTIYI | ASSLKVSAETLY | 0.4211659 |
| VAAGAGLGAVI | ASSLKVSAETLY | 0.7073976 |
| VAAVVTLL | ASSLKVSAETLY | 0.1776747 |
| VGFLGGLI | ASSLKVSAETLY | 0.6670257 |
| VGVIPILL | ASSLKVSAETLY | 0.55194455 |
| VIFASSLI | ASSLKVSAETLY | 0.7239627 |
| VKGGDGGAAAG | ASSLKVSAETLY | 0.6922428 |
| VTKTFTNI | ASSLKVSAETLY | 0.7277392 |
| VVAAAILL | ASSLKVSAETLY | 0.5729907 |
| YAALDVEL | ASSLKVSAETLY | 0.59476656 |
| LASVALLL | ASSLKVSAETLY | 0.552882 |
| LDAVHRV | ASSLKVSAETLY | 0.5197449 |
| QGLSSSIF | ASSLKVSAETLY | 0.7497963 |
| RGTFANIRL | ASSLKVSAETLY | 0.66737986 |
| RQLWWGHRI | ASSLKVSAETLY | 0.60905176 |
| YGPPGTGKTLL | ASSLKVSAETLY | 0.78986675 |
| AALVGAPANVA | ASSLKVSAETLY | 0.7137473 |
| AATFRRTE | ASSLKVSAETLY | 0.4333965 |
| AGALGGGI | ASSLKVSAETLY | 0.6408448 |
| AGGAGGAGGIGAG | ASSLKVSAETLY | 0.6969231 |
| AGSGALGL | ASSLKVSAETLY | 0.6450467 |
| AQIADQI | ASSLKVSAETLY | 0.1881331 |
| ASAVAVTYL | ASSLKVSAETLY | 0.7004208 |
| ASTVLGGTGG | ASSLKVSAETLY | 0.6164518 |
| AVGGEPAK | ASSLKVSAETLY | 0.50678223 |
| CEQGTFGGTC | ASSLKVSAETLY | 0.56444126 |
| DEGKAPAG | ASSLKVSAETLY | 0.75039226 |
| DTVVFPAL | ASSLKVSAETLY | 0.36207148 |
| FEGITMAG | ASSLKVSAETLY | 0.7083435 |
| FPLVVQD | ASSLKVSAETLY | 0.7358628 |
| GAFTTPQ | ASSLKVSAETLY | 0.7831326 |
| GFGGFGTAGGVGGA | ASSLKVSAETLY | 0.78780985 |
| GFLAQFL | ASSLKVSAETLY | 0.7136861 |
| GGAGGAGGNAGVA | ASSLKVSAETLY | 0.6510601 |
| GGVGTAGAGGAGGAGG | ASSLKVSAETLY | 0.75354946 |
| GNGGGGDAGNAGSGGN | ASSLKVSAETLY | 0.7193346 |
| GTGGAGGAGNDAGS | ASSLKVSAETLY | 0.3150398 |
| GTTNTGWFNS | ASSLKVSAETLY | 0.28513175 |

| | | |
|---|---|---|
| IFMGPLFTL | ASSLKVSAETLY | 0.7152245 |
| ILAHAIEAI | ASSLKVSAETLY | 0.30183804 |
| IRSAIREAL | ASSLKVSAETLY | 0.72199786 |
| ITFAGGG | ASSLKVSAETLY | 0.78706825 |
| ITGMSGHQEV | ASSLKVSAETLY | 0.631565 |
| IVAGLACTL | ASSLKVSAETLY | 0.4923747 |
| IVGGGLAGA | ASSLKVSAETLY | 0.6295259 |
| IVTGVLNI | ASSLKVSAETLY | 0.6035996 |
| LAQQGCI | ASSLKVSAETLY | 0.7447284 |
| LCLHVEN | ASSLKVSAETLY | 0.745806 |
| LDAISDGS | ASSLKVSAETLY | 0.79139334 |
| LGGFAAML | ASSLKVSAETLY | 0.7724606 |
| LGGKNPMI | ASSLKVSAETLY | 0.6012912 |
| LGPKGVAAIL | ASSLKVSAETLY | 0.7322451 |
| LMVGVSI | ASSLKVSAETLY | 0.6183464 |
| LVGAAFVL | ASSLKVSAETLY | 0.661969 |
| LVLAVAPL | ASSLKVSAETLY | 0.66530764 |
| MAAIAVIDAA | ASSLKVSAETLY | 0.69595075 |
| NNGINAELQ | ASSLKVSAETLY | 0.618382 |
| RGEPLVLVHG | ASSLKVSAETLY | 0.6012443 |
| RGGAGGAAGLI | ASSLKVSAETLY | 0.4511181 |
| RSYRMHPAV | ASSLKVSAETLY | 0.7204812 |
| SAAHLLAGQLA | ASSLKVSAETLY | 0.7056237 |
| SAAVGSY | ASSLKVSAETLY | 0.33740157 |
| SPHAVLHRI | ASSLKVSAETLY | 0.61561495 |
| SQIAKVL | ASSLKVSAETLY | 0.7995376 |
| TALGHLVP | ASSLKVSAETLY | 0.7993715 |
| TAPTTSGAA | ASSLKVSAETLY | 0.53775376 |
| TDGDDGNGGSPGT | ASSLKVSAETLY | 0.57303417 |
| TDGLGEN | ASSLKVSAETLY | 0.6337437 |
| TGGPPLPGQA | ASSLKVSAETLY | 0.323286 |
| TIPAITI | ASSLKVSAETLY | 0.70506346 |
| TIPGFTI | ASSLKVSAETLY | 0.6476022 |
| TSIASGALA | ASSLKVSAETLY | 0.6301143 |
| TVTLPGAAI | ASSLKVSAETLY | 0.5770149 |
| VASGLSAALP | ASSLKVSAETLY | 0.6685715 |
| VGAGVIGI | ASSLKVSAETLY | 0.5950253 |
| VGPKRFGV | ASSLKVSAETLY | 0.74140036 |
| VNVAFPD | ASSLKVSAETLY | 0.21659535 |
| VSMTNSGVS | ASSLKVSAETLY | 0.728209 |
| VTDVLGDLSV | ASSLKVSAETLY | 0.5216864 |
| VVVVGAGFAGL | ASSLKVSAETLY | 0.55139196 |
| VYTVGTVV | ASSLKVSAETLY | 0.34261316 |
| WTTFSGP | ASSLKVSAETLY | 0.7002004 |
| YDELVEAF | ASSLKVSAETLY | 0.76959836 |
| YGTGGAGGNGG | ASSLKVSAETLY | 0.751208 |
| YTNVMAV | ASSLKVSAETLY | 0.38868427 |
| FLEQVHQL | ASSLKVSAETLY | 0.7648037 |
| GLSLLAVAV | ASSLKVSAETLY | 0.7940721 |
| LEQVHQL | ASSLKVSAETLY | 0.591411 |
| LNAAGAI | ASSLKVSAETLY | 0.39174777 |

| | | |
|---|---|---|
| PAVHASL | ASSLKVSAETLY | 0.74321926 |
| TELEDDA | ASSLKVSAETLY | 0.6487938 |
| TPPGPPN | ASSLKVSAETLY | 0.74446225 |
| TPPPPPPP | ASSLKVSAETLY | 0.74812835 |
| TQSPALI | ASSLGGATSAETLY | 0.42656672 |
| EFAAAFPQ | ASSLGGATSAETLY | 0.7844985 |
| LLSGNLDVL | ASSLGGATSAETLY | 0.7852277 |
| GPANLGNYNI | ASSLGGATSAETLY | 0.5183888 |
| TLGPTLP | ASSLGGATSAETLY | 0.6021293 |
| VYAYSSCCK | ASSLGGATSAETLY | 0.61319816 |
| APAPAPAPNI | ASSLGGATSAETLY | 0.28880623 |
| PGGPVPIP | ASSLGGATSAETLY | 0.7866696 |
| FLTVALEP | ASSLGGATSAETLY | 0.55263805 |
| SQYFGANL | ASSLGGATSAETLY | 0.61985135 |
| GGVKPAA | ASSLGGATSAETLY | 0.6864814 |
| GGTGGAGGAGGW | ASSLGGATSAETLY | 0.61526465 |
| GVGGVAGLGGAGG | ASSLGGATSAETLY | 0.64852476 |
| VLLVLATTI | ASSLGGATSAETLY | 0.64929265 |
| IGIFAAML | ASSLGGATSAETLY | 0.69754195 |
| AVGDLID | ASSLGGATSAETLY | 0.69543636 |
| NISPLPL | ASSLGGATSAETLY | 0.6828705 |
| VTAVFVAI | ASSLGGATSAETLY | 0.7062814 |
| SGDDNDAAL | ASSLGGATSAETLY | 0.7846274 |
| AVAALAHS | ASSLGGATSAETLY | 0.43655166 |
| GGVTVMATP | ASSLGGATSAETLY | 0.5819321 |
| AGLAANVRD | ASSLGGATSAETLY | 0.7153832 |
| TNRPDLIDPALLRPG | ASSLGGATSAETLY | 0.38590604 |
| LANLGVTAL | ASSLGGATSAETLY | 0.7496882 |
| TGERIEL | ASSLGGATSAETLY | 0.7042998 |
| NGEELAA | ASSLGGATSAETLY | 0.70299935 |
| VLSLVWG | ASSLGGATSAETLY | 0.6270579 |
| IGYALWL | ASSLGGATSAETLY | 0.8050059 |
| LSLLLAPL | ASSLGGATSAETLY | 0.76641107 |
| TLDEALK | ASSLGGATSAETLY | 0.6912196 |
| TDGAGTAEGG | ASSLGGATSAETLY | 0.68965 |
| NAPGGAGGNGG | ASSLGGATSAETLY | 0.5571576 |
| VSVFMGMGGNF | ASSLGGATSAETLY | 0.7672909 |
| LASYLAAL | ASSLGGATSAETLY | 0.73536444 |
| LVGGTPAL | ASSLGGATSAETLY | 0.46393725 |
| VRNADVL | ASSLGGATSAETLY | 0.6483578 |
| TCSSPFLVSAM | ASSLGGATSAETLY | 0.7766603 |
| VHITDAE | ASSLGGATSAETLY | 0.7694545 |
| EYAAPSF | ASSLGGATSAETLY | 0.37314484 |
| VVDAFHGGL | ASSLGGATSAETLY | 0.62960184 |
| ASVVDPQ | ASSLGGATSAETLY | 0.34688437 |
| SYNEFGGLS | ASSLGGATSAETLY | 0.5520412 |
| DPDGSVY | ASSLGGATSAETLY | 0.56200683 |
| MVSTIHATE | ASSLGGATSAETLY | 0.71992296 |
| LAAGVADPV | ASSLGGATSAETLY | 0.6423273 |
| TKEQIAA | ASSLGGATSAETLY | 0.6998651 |
| SLASPAAL | ASSLGGATSAETLY | 0.7122306 |

| | | |
|---|---|---|
| IGLVFYGV | ASSLGGATSAETLY | 0.7932153 |
| FLLDASNL | ASSLGGATSAETLY | 0.58692235 |
| YQVGNFP | ASSLGGATSAETLY | 0.69186956 |
| ESLLSQST | ASSLGGATSAETLY | 0.40894052 |
| LFTVGFAAVG | ASSLGGATSAETLY | 0.4612807 |
| ATPPTPT | ASSLGGATSAETLY | 0.55553454 |
| TDLATLG | ASSLGGATSAETLY | 0.66637313 |
| SAGGTGR | ASSLGGATSAETLY | 0.28127003 |
| GFSNSGS | ASSLGGATSAETLY | 0.7706898 |
| GGNGGSGGTSV | ASSLGGATSAETLY | 0.70865685 |
| GTSVATGGAGN | ASSLGGATSAETLY | 0.76203465 |
| NGGNGGSGGTS | ASSLGGATSAETLY | 0.6806743 |
| AVVKFPH | ASSLGGATSAETLY | 0.6272819 |
| TPPPPPP | ASSLGGATSAETLY | 0.56256884 |
| AAPAAAAAPA | ASSLGGATSAETLY | 0.73961496 |
| IALNEDL | ASSLGGATSAETLY | 0.49613273 |
| IVNVSSM | ASSLGGATSAETLY | 0.4580878 |
| TGIPPAPR | ASSLGGATSAETLY | 0.6353799 |
| FDLGGGTF | ASSLGGATSAETLY | 0.5023249 |
| FDLGGGTFDVS | ASSLGGATSAETLY | 0.780586 |
| NEPTAAA | ASSLGGATSAETLY | 0.77532303 |
| NEPTAAALA | ASSLGGATSAETLY | 0.70765674 |
| GSIINISSI | ASSLGGATSAETLY | 0.7591784 |
| VAPGNAGTA | ASSLGGATSAETLY | 0.30689985 |
| TGGGASLE | ASSLGGATSAETLY | 0.6533369 |
| VAVYNHYKRI | ASSLGGATSAETLY | 0.8041204 |
| TDPAPATT | ASSLGGATSAETLY | 0.7358057 |
| TNRPDILDPALLRPG | ASSLGGATSAETLY | 0.6590277 |
| LTTTTTS | ASSLGGATSAETLY | 0.77720594 |
| PTGSGKTL | ASSLGGATSAETLY | 0.6021749 |
| TGSGKTL | ASSLGGATSAETLY | 0.6525752 |
| TIGHVDHGK | ASSLGGATSAETLY | 0.6818242 |
| TIGHVDHGKTT | ASSLGGATSAETLY | 0.7273406 |
| GHVDHGKTT | ASSLGGATSAETLY | 0.7006899 |
| AQFEHTLL | ASSLGGATSAETLY | 0.5222332 |
| IVNQDEL | ASSLGGATSAETLY | 0.7437813 |
| TNGSQFF | ASSLGGATSAETLY | 0.8053602 |
| TGKGIVE | ASSLGGATSAETLY | 0.6300678 |
| LISNASDA | ASSLGGATSAETLY | 0.7648948 |
| AELRTLGELTKIAKAHGAQ | ASSLGGATSAETLY | 0.72670215 |
| APNAVAAAEPSW | ASSLGGATSAETLY | 0.73061454 |
| APQVAGAATAVL | ASSLGGATSAETLY | 0.53782296 |
| ARRDPAAAAAILPTDA | ASSLGGATSAETLY | 0.5328185 |
| DAAARDVLAEAGLAEYF | ASSLGGATSAETLY | 0.62108994 |
| DDQIVLSAKANLAKVEQ | ASSLGGATSAETLY | 0.51209724 |
| DELLPDTGEDSDEDGH | ASSLGGATSAETLY | 0.67903316 |
| ELPEAPANRATVPQD | ASSLGGATSAETLY | 0.6511912 |
| FGQSKIERVTGIG | ASSLGGATSAETLY | 0.8178081 |
| GAGGTGSTAGGAGGAGGAGGLY | ASSLGGATSAETLY | 0.72670865 |
| GHGGTGAAGGNGGTGGAGGSGI | ASSLGGATSAETLY | 0.71278965 |
| GNGGNGGAGGEGGPGGAGGAGA | ASSLGGATSAETLY | 0.52024454 |

| | | |
|---|---|---|
| GSTAGNGGNGGNATNGTIAGQP | ASSLGGATSAETLY | 0.6911334 |
| GTGGDGGNGGNGGTGG | ASSLGGATSAETLY | 0.70483345 |
| GVRRPAGIPADAPV | ASSLGGATSAETLY | 0.48580387 |
| GYRQVTDAHLLTLARR | ASSLGGATSAETLY | 0.7920157 |
| IAKMGQDGHDRG | ASSLGGATSAETLY | 0.5553907 |
| KAEPSIEIQYT | ASSLGGATSAETLY | 0.5837142 |
| KPVRIRMAADEDTVAVT | ASSLGGATSAETLY | 0.7454418 |
| MPPRIAGMRLLVIKPEP | ASSLGGATSAETLY | 0.77016497 |
| MKFYAVALFRSRGVPPTR | ASSLGGATSAETLY | 0.61712515 |
| NAGGFGILGAVAHSP | ASSLGGATSAETLY | 0.5882247 |
| NEGREITLAMAEM | ASSLGGATSAETLY | 0.51459086 |
| NELVEWSHPLGHRWRWE | ASSLGGATSAETLY | 0.75176895 |
| NLGAGGTGGNGGNPG | ASSLGGATSAETLY | 0.77293026 |
| NTEWPKWETVMAADPVD | ASSLGGATSAETLY | 0.65143144 |
| PASAAPAETQP | ASSLGGATSAETLY | 0.78125376 |
| QQSSDGKNGVRAAEITGE | ASSLGGATSAETLY | 0.7416724 |
| RAPDAVTTSWWKE | ASSLGGATSAETLY | 0.4331236 |
| SGFGNVGAHTSGWFNQ | ASSLGGATSAETLY | 0.6813284 |
| SGVAGAVGQAASVGGLKV | ASSLGGATSAETLY | 0.7630928 |
| SNGGAGTGSAGGNGGT | ASSLGGATSAETLY | 0.7511676 |
| TGGAGGEGGAAG | ASSLGGATSAETLY | 0.8034266 |
| TGGDGGAAGAGGAPGHG | ASSLGGATSAETLY | 0.74150145 |
| TMPPLLAAIANAGRHGV | ASSLGGATSAETLY | 0.61314607 |
| TSSTAGNSGGA | ASSLGGATSAETLY | 0.786745 |
| VPATVPVGILC | ASSLGGATSAETLY | 0.8090945 |
| VQRPDSGGVVLTGRIS | ASSLGGATSAETLY | 0.7669245 |
| WTPNPVPATKGVPAGLVQAT | ASSLGGATSAETLY | 0.59387004 |
| YALKRALAKFSDRFDVVII | ASSLGGATSAETLY | 0.51991403 |
| YLIGSVIV | ASSLGGATSAETLY | 0.7081474 |
| YGQLVAVID | ASSLGGATSAETLY | 0.7422228 |
| VTVGGMTLPA | ASSLGGATSAETLY | 0.5063824 |
| VLALPAV | ASSLGGATSAETLY | 0.6725826 |
| VIPPTLNYET | ASSLGGATSAETLY | 0.70300925 |
| VGMGGMHPSSG | ASSLGGATSAETLY | 0.43819207 |
| VGLAGGAATA | ASSLGGATSAETLY | 0.770213 |
| VGGAGGAGGSGP | ASSLGGATSAETLY | 0.7734977 |
| VFAAFMV | ASSLGGATSAETLY | 0.6575843 |
| TLYDKLR | ASSLGGATSAETLY | 0.5882782 |
| TLAVFTVL | ASSLGGATSAETLY | 0.76904345 |
| TGSVQPARV | ASSLGGATSAETLY | 0.8159479 |
| TGGGAGSGGT | ASSLGGATSAETLY | 0.7842903 |
| SHLETSL | ASSLGGATSAETLY | 0.45439702 |
| SFTTGMT | ASSLGGATSAETLY | 0.28145665 |
| QVEVAKRMGVRQPTVS | ASSLGGATSAETLY | 0.5492882 |
| QGSGGPALAM | ASSLGGATSAETLY | 0.73142654 |
| MLAPSNP | ASSLGGATSAETLY | 0.6696832 |
| LVNNAAI | ASSLGGATSAETLY | 0.6075995 |
| LTGEINTLA | ASSLGGATSAETLY | 0.69649273 |
| LSIFGPV | ASSLGGATSAETLY | 0.7774643 |
| LLALLANL | ASSLGGATSAETLY | 0.57114077 |
| LISNASDALDKLR | ASSLGGATSAETLY | 0.44495493 |

| | | |
|---|---|---|
| LINEAVP | ASSLGGATSAETLY | 0.7404502 |
| LGYAFSGL | ASSLGGATSAETLY | 0.75317585 |
| LDFVAVAL | ASSLGGATSAETLY | 0.52562904 |
| LAQPGAE | ASSLGGATSAETLY | 0.629639 |
| ISNLGAVTGF | ASSLGGATSAETLY | 0.59611183 |
| HQEDPGRI | ASSLGGATSAETLY | 0.41611424 |
| GTPGEPTP | ASSLGGATSAETLY | 0.68066853 |
| GFFNSGS | ASSLGGATSAETLY | 0.49002337 |
| GAGGGGLTAGAN | ASSLGGATSAETLY | 0.76467276 |
| FVDALVNM | ASSLGGATSAETLY | 0.50995696 |
| FLTVAPEL | ASSLGGATSAETLY | 0.7368632 |
| FETLTAL | ASSLGGATSAETLY | 0.7124695 |
| DVINPIEL | ASSLGGATSAETLY | 0.7463283 |
| APVSGAPV | ASSLGGATSAETLY | 0.6209805 |
| ALVGGLAW | ASSLGGATSAETLY | 0.58284247 |
| ALAAAAATAI | ASSLGGATSAETLY | 0.48420906 |
| AFGLGLLL | ASSLGGATSAETLY | 0.26234356 |
| AFFAAASL | ASSLGGATSAETLY | 0.7725718 |
| AAAALPAFSPP | ASSLGGATSAETLY | 0.22661798 |
| VVLAPAL | ASSLGGATSAETLY | 0.76416683 |
| TIGHVDHGKTTL | ASSLGGATSAETLY | 0.7585366 |
| LISNASDALDK | ASSLGGATSAETLY | 0.52194566 |
| LISNASDAL | ASSLGGATSAETLY | 0.6962707 |
| ISNASDALDK | ASSLGGATSAETLY | 0.4692471 |
| NQAETLVYQTE | ASSLGGATSAETLY | 0.5077514 |
| TGGTGGTAGAG | ASSLGGATSAETLY | 0.7480107 |
| TGSTAGGAGGA | ASSLGGATSAETLY | 0.7398963 |
| DPLAVPVAAGPV | ASSLGGATSAETLY | 0.5704978 |
| GGTGGTPGNGGN | ASSLGGATSAETLY | 0.6994094 |
| PAGLAGVDPATL | ASSLGGATSAETLY | 0.7164527 |
| GNNLVVEAPHAGASV | ASSLGGATSAETLY | 0.611014 |
| KVTRIMSASTLEQPA | ASSLGGATSAETLY | 0.67767906 |
| LGFALPIALPRRTVV | ASSLGGATSAETLY | 0.7767197 |
| TRLGNLTARKLVVLV | ASSLGGATSAETLY | 0.7046231 |
| WSQQRRLIYFPSAGP | ASSLGGATSAETLY | 0.53715754 |
| GPNGNPDPMAAAVDIR | ASSLGGATSAETLY | 0.75304985 |
| TQRCVSVADLRAAASR | ASSLGGATSAETLY | 0.47130078 |
| QFVLDVAEHPVEVLTETV | ASSLGGATSAETLY | 0.7078391 |
| GGAGGTGGAGGNGGTQAGDG | ASSLGGATSAETLY | 0.6521035 |
| PLLATLRAIAVDVDAKPAQV | ASSLGGATSAETLY | 0.67092645 |
| APKFRTGGYMIDFWGVGYQVAK | ASSLGGATSAETLY | 0.75341696 |
| GAGGAGGAGGAAGGPGGTGGTG | ASSLGGATSAETLY | 0.5350007 |
| IGALGLLATDVAPAGSVAFGVA | ASSLGGATSAETLY | 0.56659263 |
| AGGAGGIGGNGGAGGAGGVLMG | ASSLGGATSAETLY | 0.6156188 |
| ADELIGLL | ASSLGGATSAETLY | 0.73147434 |
| ADRALKA | ASSLGGATSAETLY | 0.76293904 |
| AGAVGQVGGAGGA | ASSLGGATSAETLY | 0.5406311 |
| AGGGGGLGGAP | ASSLGGATSAETLY | 0.7073614 |
| AGGLLYGNGGAG | ASSLGGATSAETLY | 0.6490758 |
| AGLESIGL | ASSLGGATSAETLY | 0.552662 |
| AGLKTGII | ASSLGGATSAETLY | 0.6638726 |

| | | |
|---|---|---|
| ASLAALLL | ASSLGGATSAETLY | 0.6533552 |
| ASVINISSI | ASSLGGATSAETLY | 0.56232876 |
| ATPPGPA | ASSLGGATSAETLY | 0.73560745 |
| ATVASALV | ASSLGGATSAETLY | 0.5518603 |
| ATVASLIL | ASSLGGATSAETLY | 0.7537422 |
| AVNGPVLIRH | ASSLGGATSAETLY | 0.8067041 |
| DGAAGAVGI | ASSLGGATSAETLY | 0.5943974 |
| DPLLHGI | ASSLGGATSAETLY | 0.7352915 |
| ELGAALGAL | ASSLGGATSAETLY | 0.71699184 |
| ELGRTQAV | ASSLGGATSAETLY | 0.7041153 |
| FVAGGYLS | ASSLGGATSAETLY | 0.6564021 |
| GASGGAGGAGGW | ASSLGGATSAETLY | 0.5912609 |
| GGAGAGGDGGAGGI | ASSLGGATSAETLY | 0.7821626 |
| GTGHNGSQPGMGGNG | ASSLGGATSAETLY | 0.7575187 |
| GYAAASASAS | ASSLGGATSAETLY | 0.7592673 |
| HQVDVAAVQQG | ASSLGGATSAETLY | 0.7100622 |
| IGAAIAHT | ASSLGGATSAETLY | 0.73387116 |
| IGMIEVAQL | ASSLGGATSAETLY | 0.74642146 |
| IRLAPEHTSAMD | ASSLGGATSAETLY | 0.57323503 |
| IYNVSVL | ASSLGGATSAETLY | 0.39512834 |
| KQGFLGM | ASSLGGATSAETLY | 0.63811386 |
| LESVITQL | ASSLGGATSAETLY | 0.6411241 |
| LGLGALVVV | ASSLGGATSAETLY | 0.6613411 |
| LIGNGGNGGSGGI | ASSLGGATSAETLY | 0.38216662 |
| LNAINAPTL | ASSLGGATSAETLY | 0.77121866 |
| LQLLNTGV | ASSLGGATSAETLY | 0.81297797 |
| LQTSIEF | ASSLGGATSAETLY | 0.5886364 |
| MSFAVSEL | ASSLGGATSAETLY | 0.79892516 |
| NTGALGL | ASSLGGATSAETLY | 0.45921206 |
| PPGAGGAGGAGTT | ASSLGGATSAETLY | 0.74232996 |
| QLVGLGI | ASSLGGATSAETLY | 0.77709645 |
| RFLEQVHQL | ASSLGGATSAETLY | 0.46863514 |
| RGVGYVLR | ASSLGGATSAETLY | 0.7623295 |
| SATGVFAGVI | ASSLGGATSAETLY | 0.5468564 |
| SLPAITLP | ASSLGGATSAETLY | 0.6171271 |
| STALAFTV | ASSLGGATSAETLY | 0.78327155 |
| SVLTVGLGK | ASSLGGATSAETLY | 0.6848757 |
| TALAGAGSGI | ASSLGGATSAETLY | 0.7121124 |
| TGLPFTGI | ASSLGGATSAETLY | 0.6501776 |
| VAEHVTV | ASSLGGATSAETLY | 0.52846986 |
| VDIVLNSL | ASSLGGATSAETLY | 0.69106406 |
| VLTGGGAVL | ASSLGGATSAETLY | 0.77887636 |
| VNVGLAYGI | ASSLGGATSAETLY | 0.45478916 |
| VSSAAVGAV | ASSLGGATSAETLY | 0.80709374 |
| VTSVYEAGV | ASSLGGATSAETLY | 0.6021994 |
| WVALDRGA | ASSLGGATSAETLY | 0.8094823 |
| YLVGSVVI | ASSLGGATSAETLY | 0.7375009 |
| GPPGTGKTLL | ASSLGGATSAETLY | 0.66581833 |
| GVGMTKF | ASSLGGATSAETLY | 0.6021052 |
| IGIGGSDL | ASSLGGATSAETLY | 0.7747961 |
| IGTLLAEL | ASSLGGATSAETLY | 0.6486013 |

| | | |
|---|---|---|
| IYTSGTT | ASSLGGATSAETLY | 0.7294188 |
| LSFPAGV | ASSLGGATSAETLY | 0.7872486 |
| LYTSGTT | ASSLGGATSAETLY | 0.53816366 |
| VGAFPFGF | ASSLGGATSAETLY | 0.65103096 |
| VGAGTVEF | ASSLGGATSAETLY | 0.6151712 |
| VGKSTLLL | ASSLGGATSAETLY | 0.62463236 |
| AALAHAGGGIV | ASSLGGATSAETLY | 0.5797449 |
| AALVAVSLL | ASSLGGATSAETLY | 0.46376455 |
| AAVSRAVAAL | ASSLGGATSAETLY | 0.6085134 |
| AGGSSGIGGF | ASSLGGATSAETLY | 0.53288764 |
| AGLTVANANG | ASSLGGATSAETLY | 0.6390465 |
| AGLVGAVL | ASSLGGATSAETLY | 0.6395661 |
| ALSQAFTV | ASSLGGATSAETLY | 0.58418816 |
| ALTFSVSL | ASSLGGATSAETLY | 0.65133595 |
| ALTGAALVAL | ASSLGGATSAETLY | 0.6551372 |
| ASFSIGAGVL | ASSLGGATSAETLY | 0.47104898 |
| DGSVYKI | ASSLGGATSAETLY | 0.59823513 |
| EQLAEIV | ASSLGGATSAETLY | 0.57177794 |
| FAVSELTVM | ASSLGGATSAETLY | 0.5953747 |
| GAAPLAAGL | ASSLGGATSAETLY | 0.4159165 |
| GADGGDGGSGGAGG | ASSLGGATSAETLY | 0.636855 |
| GFGLDTGGAGG | ASSLGGATSAETLY | 0.56214505 |
| GLASGGLL | ASSLGGATSAETLY | 0.32686126 |
| GRIAGAAVLL | ASSLGGATSAETLY | 0.6206118 |
| GSAGSLNTSF | ASSLGGATSAETLY | 0.6102182 |
| GTGQIGI | ASSLGGATSAETLY | 0.57893956 |
| GTLIALIL | ASSLGGATSAETLY | 0.6405593 |
| IAATMGFVVG | ASSLGGATSAETLY | 0.51908755 |
| IFAQALL | ASSLGGATSAETLY | 0.5625375 |
| IGAAVALGF | ASSLGGATSAETLY | 0.6321418 |
| IGLTPLAL | ASSLGGATSAETLY | 0.65024817 |
| ILGSNTSTL | ASSLGGATSAETLY | 0.5215198 |
| IVFSLTVL | ASSLGGATSAETLY | 0.6119344 |
| LASKATGF | ASSLGGATSAETLY | 0.6550987 |
| LASVALIL | ASSLGGATSAETLY | 0.6130375 |
| LFAVGLTL | ASSLGGATSAETLY | 0.6483423 |
| LFNGGIGTYI | ASSLGGATSAETLY | 0.6559423 |
| LTLGAGII | ASSLGGATSAETLY | 0.578028 |
| LVAAALGDI | ASSLGGATSAETLY | 0.64509434 |
| NTAFVQLGI | ASSLGGATSAETLY | 0.65583575 |
| QLFSGAEL | ASSLGGATSAETLY | 0.6027721 |
| SAATFITL | ASSLGGATSAETLY | 0.6091903 |
| SNYGLANLGV | ASSLGGATSAETLY | 0.5481971 |
| SYGGAYAVM | ASSLGGATSAETLY | 0.6097297 |
| TTGVGGLTLG | ASSLGGATSAETLY | 0.40113428 |
| TTQTIYI | ASSLGGATSAETLY | 0.6150654 |
| VAAGAGLGAVI | ASSLGGATSAETLY | 0.64924884 |
| VAAVVTLL | ASSLGGATSAETLY | 0.58244205 |
| VGFLGGLI | ASSLGGATSAETLY | 0.4937619 |
| VGVIPILL | ASSLGGATSAETLY | 0.52718854 |
| VIFASSLI | ASSLGGATSAETLY | 0.601257 |

| | | |
|---|---|---|
| VKGGDGGAAAG | ASSLGGATSAETLY | 0.5601466 |
| VTKTFTNI | ASSLGGATSAETLY | 0.40863386 |
| VVAAAILL | ASSLGGATSAETLY | 0.56968606 |
| YAALDVEL | ASSLGGATSAETLY | 0.6504215 |
| LASVALLL | ASSLGGATSAETLY | 0.64979327 |
| LDAVHRV | ASSLGGATSAETLY | 0.48014092 |
| QGLSSSIF | ASSLGGATSAETLY | 0.655495 |
| RGTFANIRL | ASSLGGATSAETLY | 0.6330438 |
| RQLWWGHRI | ASSLGGATSAETLY | 0.6579433 |
| YGPPGTGKTLL | ASSLGGATSAETLY | 0.6229491 |
| AALVGAPANVA | ASSLGGATSAETLY | 0.53955764 |
| AATFRRTE | ASSLGGATSAETLY | 0.64481646 |
| AGALGGGI | ASSLGGATSAETLY | 0.5807056 |
| AGGAGGAGGIGAG | ASSLGGATSAETLY | 0.557559 |
| AGSGALGL | ASSLGGATSAETLY | 0.64698523 |
| AQIADQI | ASSLGGATSAETLY | 0.6027931 |
| ASAVAVTYL | ASSLGGATSAETLY | 0.55823267 |
| ASTVLGGTGG | ASSLGGATSAETLY | 0.5682444 |
| AVGGEPAK | ASSLGGATSAETLY | 0.6531187 |
| CEQGTFGGTC | ASSLGGATSAETLY | 0.5222511 |
| DEGKAPAG | ASSLGGATSAETLY | 0.53681606 |
| DTVVFPAL | ASSLGGATSAETLY | 0.54894847 |
| FEGITMAG | ASSLGGATSAETLY | 0.6082229 |
| FPLVVQD | ASSLGGATSAETLY | 0.6496369 |
| GAFTTPQ | ASSLGGATSAETLY | 0.6334061 |
| GFGGFGTAGGVGGA | ASSLGGATSAETLY | 0.58552134 |
| GFLAQFL | ASSLGGATSAETLY | 0.6202059 |
| GGAGGAGGNAGVA | ASSLGGATSAETLY | 0.63463503 |
| GGVGTAGAGGAGGAGG | ASSLGGATSAETLY | 0.6226744 |
| GNGGGGDAGNAGSGGN | ASSLGGATSAETLY | 0.6419103 |
| GTGGAGGAGNDAGS | ASSLGGATSAETLY | 0.5415307 |
| GTTNTGWFNS | ASSLGGATSAETLY | 0.65106654 |
| IFMGPLFTL | ASSLGGATSAETLY | 0.6372507 |
| ILAHAIEAI | ASSLGGATSAETLY | 0.6466566 |
| IRSAIREAL | ASSLGGATSAETLY | 0.64516187 |
| ITFAGGG | ASSLGGATSAETLY | 0.62888795 |
| ITGMSGHQEV | ASSLGGATSAETLY | 0.6200315 |
| IVAGLACTL | ASSLGGATSAETLY | 0.4184466 |
| IVGGGLAGA | ASSLGGATSAETLY | 0.59251094 |
| IVTGVLNI | ASSLGGATSAETLY | 0.6176971 |
| LAQQGCI | ASSLGGATSAETLY | 0.61685556 |
| LCLHVEN | ASSLGGATSAETLY | 0.6356241 |
| LDAISDGS | ASSLGGATSAETLY | 0.6383492 |
| LGGFAAML | ASSLGGATSAETLY | 0.56324756 |
| LGGKNPMI | ASSLGGATSAETLY | 0.6532895 |
| LGPKGVAAIL | ASSLGGATSAETLY | 0.6473434 |
| LMVGVSI | ASSLGGATSAETLY | 0.45798606 |
| LVGAAFVL | ASSLGGATSAETLY | 0.58357334 |
| LVLAVAPL | ASSLGGATSAETLY | 0.65878797 |
| MAAIAVIDAA | ASSLGGATSAETLY | 0.61232275 |
| NNGINAELQ | ASSLGGATSAETLY | 0.5969477 |

| | | |
|---|---|---|
| RGEPLVLVHG | ASSLGGATSAETLY | 0.59190315 |
| RGGAGGAAGLI | ASSLGGATSAETLY | 0.6567222 |
| RSYRMHPAV | ASSLGGATSAETLY | 0.5514789 |
| SAAHLLAGQLA | ASSLGGATSAETLY | 0.6573616 |
| SAAVGSY | ASSLGGATSAETLY | 0.36401233 |
| SPHAVLHRI | ASSLGGATSAETLY | 0.5805402 |
| SQIAKVL | ASSLGGATSAETLY | 0.5704814 |
| TALGHLVP | ASSLGGATSAETLY | 0.38009265 |
| TAPTTSGAA | ASSLGGATSAETLY | 0.63580084 |
| TDGDDGNGGSPGT | ASSLGGATSAETLY | 0.590624 |
| TDGLGEN | ASSLGGATSAETLY | 0.6354973 |
| TGGPPLPGQA | ASSLGGATSAETLY | 0.6069498 |
| TIPAITI | ASSLGGATSAETLY | 0.6425193 |
| TIPGFTI | ASSLGGATSAETLY | 0.6096316 |
| TSIASGALA | ASSLGGATSAETLY | 0.62655073 |
| TVTLPGAAI | ASSLGGATSAETLY | 0.54489523 |
| VASGLSAALP | ASSLGGATSAETLY | 0.63591254 |
| VGAGVIGI | ASSLGGATSAETLY | 0.65564054 |
| VGPKRFGV | ASSLGGATSAETLY | 0.6412536 |
| VNVAFPD | ASSLGGATSAETLY | 0.47533455 |
| VSMTNSGVS | ASSLGGATSAETLY | 0.57016987 |
| VTDVLGDLSV | ASSLGGATSAETLY | 0.60248196 |
| VVVVGAGFAGL | ASSLGGATSAETLY | 0.39443752 |
| VYTVGTVV | ASSLGGATSAETLY | 0.36874688 |
| WTTFSGP | ASSLGGATSAETLY | 0.56346977 |
| YDELVEAF | ASSLGGATSAETLY | 0.5618876 |
| YGTGGAGGNGG | ASSLGGATSAETLY | 0.64612794 |
| YTNVMAV | ASSLGGATSAETLY | 0.63885045 |
| FLEQVHQL | ASSLGGATSAETLY | 0.58864814 |
| GLSLLAVAV | ASSLGGATSAETLY | 0.6394813 |
| LEQVHQL | ASSLGGATSAETLY | 0.61577517 |
| LNAAGAI | ASSLGGATSAETLY | 0.30260748 |
| PAVHASL | ASSLGGATSAETLY | 0.589526 |
| TELEDDA | ASSLGGATSAETLY | 0.48570174 |
| TPPGPPN | ASSLGGATSAETLY | 0.6578629 |
| TPPPPPPP | ASSLGGATSAETLY | 0.5330507 |
| TQSPALI | ASGDRHANSDYT | 0.55039966 |
| EFAAAFPQ | ASGDRHANSDYT | 0.61427367 |
| LLSGNLDVL | ASGDRHANSDYT | 0.6023458 |
| GPANLGNYNI | ASGDRHANSDYT | 0.60318154 |
| TLGPTLP | ASGDRHANSDYT | 0.5461329 |
| VYAYSSCCK | ASGDRHANSDYT | 0.3556397 |
| APAPAPAPNI | ASGDRHANSDYT | 0.64124876 |
| PGGPVPIP | ASGDRHANSDYT | 0.49704844 |
| FLTVALEP | ASGDRHANSDYT | 0.5344921 |
| SQYFGANL | ASGDRHANSDYT | 0.43304652 |
| GGVKPAA | ASGDRHANSDYT | 0.512532 |
| GGTGGAGGAGGW | ASGDRHANSDYT | 0.45639873 |
| GVGGVAGLGGAGG | ASGDRHANSDYT | 0.6438356 |
| VLLVLATTI | ASGDRHANSDYT | 0.5788708 |
| IGIFAAML | ASGDRHANSDYT | 0.5870007 |

| | | |
|---|---|---|
| AVGDLID | ASGDRHANSDYT | 0.6568972 |
| NISPLPL | ASGDRHANSDYT | 0.6092457 |
| VTAVFVAI | ASGDRHANSDYT | 0.58915734 |
| SGDDNDAAL | ASGDRHANSDYT | 0.57969993 |
| AVAALAHS | ASGDRHANSDYT | 0.5344258 |
| GGVTVMATP | ASGDRHANSDYT | 0.55534494 |
| AGLAANVRD | ASGDRHANSDYT | 0.56059223 |
| TNRPDLIDPALLRPG | ASGDRHANSDYT | 0.6589665 |
| LANLGVTAL | ASGDRHANSDYT | 0.5913862 |
| TGERIEL | ASGDRHANSDYT | 0.5021432 |
| NGEELAA | ASGDRHANSDYT | 0.62184715 |
| VLSLVWG | ASGDRHANSDYT | 0.65692097 |
| IGYALWL | ASGDRHANSDYT | 0.66323495 |
| LSLLLAPL | ASGDRHANSDYT | 0.54620296 |
| TLDEALK | ASGDRHANSDYT | 0.6132736 |
| TDGAGTAEGG | ASGDRHANSDYT | 0.59707385 |
| NAPGGAGGNGG | ASGDRHANSDYT | 0.59286594 |
| VSVFMGMGGNF | ASGDRHANSDYT | 0.57427794 |
| LASYLAAL | ASGDRHANSDYT | 0.50226015 |
| LVGGTPAL | ASGDRHANSDYT | 0.6425574 |
| VRNADVL | ASGDRHANSDYT | 0.6643066 |
| TCSSPFLVSAM | ASGDRHANSDYT | 0.59408116 |
| VHITDAE | ASGDRHANSDYT | 0.45503193 |
| EYAAPSF | ASGDRHANSDYT | 0.6072992 |
| VVDAFHGGL | ASGDRHANSDYT | 0.54378784 |
| ASVVDPQ | ASGDRHANSDYT | 0.57071936 |
| SYNEFGGLS | ASGDRHANSDYT | 0.60772544 |
| DPDGSVY | ASGDRHANSDYT | 0.42638275 |
| MVSTIHATE | ASGDRHANSDYT | 0.59287935 |
| LAAGVADPV | ASGDRHANSDYT | 0.65973353 |
| TKEQIAA | ASGDRHANSDYT | 0.57851636 |
| SLASPAAL | ASGDRHANSDYT | 0.6255304 |
| IGLVFYGV | ASGDRHANSDYT | 0.6453713 |
| FLLDASNL | ASGDRHANSDYT | 0.62797475 |
| YQVGNFP | ASGDRHANSDYT | 0.64774126 |
| ESLLSQST | ASGDRHANSDYT | 0.6369024 |
| LFTVGFAAVG | ASGDRHANSDYT | 0.6203397 |
| ATPPTPT | ASGDRHANSDYT | 0.6193593 |
| TDLATLG | ASGDRHANSDYT | 0.6307249 |
| SAGGTGR | ASGDRHANSDYT | 0.6153907 |
| GFSNSGS | ASGDRHANSDYT | 0.6057596 |
| GGNGGSGGTSV | ASGDRHANSDYT | 0.57186 |
| GTSVATGGAGN | ASGDRHANSDYT | 0.64456224 |
| NGGNGGSGGTS | ASGDRHANSDYT | 0.61262274 |
| AVVKFPH | ASGDRHANSDYT | 0.6504009 |
| TPPPPPP | ASGDRHANSDYT | 0.4552461 |
| AAPAAAAAPA | ASGDRHANSDYT | 0.52741873 |
| IALNEDL | ASGDRHANSDYT | 0.63840157 |
| IVNVSSM | ASGDRHANSDYT | 0.33788365 |
| TGIPPAPR | ASGDRHANSDYT | 0.38095453 |
| FDLGGGTF | ASGDRHANSDYT | 0.6160615 |

| | | |
|---|---|---|
| FDLGGGTFDVS | ASGDRHANSDYT | 0.65385675 |
| NEPTAAA | ASGDRHANSDYT | 0.32953772 |
| NEPTAAALA | ASGDRHANSDYT | 0.55504483 |
| GSIINISSI | ASGDRHANSDYT | 0.50016177 |
| VAPGNAGTA | ASGDRHANSDYT | 0.6402654 |
| TGGGASLE | ASGDRHANSDYT | 0.6034359 |
| VAVYNHYKRI | ASGDRHANSDYT | 0.45507428 |
| TDPAPATT | ASGDRHANSDYT | 0.635982 |
| TNRPDILDPALLRPG | ASGDRHANSDYT | 0.43739256 |
| LTTTTTS | ASGDRHANSDYT | 0.56327844 |
| PTGSGKTL | ASGDRHANSDYT | 0.64177114 |
| TGSGKTL | ASGDRHANSDYT | 0.6537899 |
| TIGHVDHGK | ASGDRHANSDYT | 0.5115602 |
| TIGHVDHGKTT | ASGDRHANSDYT | 0.6419752 |
| GHVDHGKTT | ASGDRHANSDYT | 0.5600103 |
| AQFEHTLL | ASGDRHANSDYT | 0.6648722 |
| IVNQDEL | ASGDRHANSDYT | 0.598557 |
| TNGSQFF | ASGDRHANSDYT | 0.4413437 |
| TGKGIVE | ASGDRHANSDYT | 0.6349567 |
| LISNASDA | ASGDRHANSDYT | 0.63394624 |
| AELRTLGELTKIAKAHGAQ | ASGDRHANSDYT | 0.6270959 |
| APNAVAAAEPSW | ASGDRHANSDYT | 0.6347495 |
| APQVAGAATAVL | ASGDRHANSDYT | 0.5747625 |
| ARRDPAAAAAILPTDA | ASGDRHANSDYT | 0.589747 |
| DAAARDVLAEAGLAEYF | ASGDRHANSDYT | 0.6470231 |
| DDQIVLSAKANLAKVEQ | ASGDRHANSDYT | 0.63791037 |
| DELLPDTGEDSDEDGH | ASGDRHANSDYT | 0.501276 |
| ELPEAPANRATVPQD | ASGDRHANSDYT | 0.6483617 |
| FGQSKIERVTGIG | ASGDRHANSDYT | 0.60451245 |
| GAGGTGSTAGGAGGAGGAGGLY | ASGDRHANSDYT | 0.5700441 |
| GHGGTGAAGGNGGTGGAGGSGI | ASGDRHANSDYT | 0.6551429 |
| GNGGNGGAGGEGGPGGAGGAGA | ASGDRHANSDYT | 0.61815464 |
| GSTAGNGGNGGNATNGTIAGQP | ASGDRHANSDYT | 0.6415506 |
| GTGGDGGNGGNGGTGG | ASGDRHANSDYT | 0.6362276 |
| GVRRPAGIPADAPV | ASGDRHANSDYT | 0.65704286 |
| GYRQVTDAHLLTLARR | ASGDRHANSDYT | 0.4880966 |
| IAKMGQDGHDRG | ASGDRHANSDYT | 0.62998235 |
| KAEPSIEIQYT | ASGDRHANSDYT | 0.60886693 |
| KPVRIRMAADEDTVAVT | ASGDRHANSDYT | 0.64565957 |
| MPPRIAGMRLLVIKPEP | ASGDRHANSDYT | 0.60939276 |
| MKFYAVALFRSRGVPPTR | ASGDRHANSDYT | 0.5520302 |
| NAGGFGILGAVAHSP | ASGDRHANSDYT | 0.6317717 |
| NEGREITLAMAEM | ASGDRHANSDYT | 0.6521987 |
| NELVEWSHPLGHRWRWE | ASGDRHANSDYT | 0.6174283 |
| NLGAGGTGGNGGNPG | ASGDRHANSDYT | 0.62710816 |
| NTEWPKWETVMAADPVD | ASGDRHANSDYT | 0.6562421 |
| PASAAPAETQP | ASGDRHANSDYT | 0.55208147 |
| QQSSDGKNGVRAAEITGE | ASGDRHANSDYT | 0.6012714 |
| RAPDAVTTSWWKE | ASGDRHANSDYT | 0.64256865 |
| SGFGNVGAHTSGWFNQ | ASGDRHANSDYT | 0.6512743 |
| SGVAGAVGQAASVGGLKV | ASGDRHANSDYT | 0.5988651 |

| | | |
|---|---|---|
| SNGGAGTGSAGGNGGT | ASGDRHANSDYT | 0.65148145 |
| TGGAGGEGGAAG | ASGDRHANSDYT | 0.5564798 |
| TGGDGGAAGAGGAPGHG | ASGDRHANSDYT | 0.6343081 |
| TMPPLLAAIANAGRHGV | ASGDRHANSDYT | 0.56017905 |
| TSSTAGNSGGA | ASGDRHANSDYT | 0.6163114 |
| VPATVPVGILC | ASGDRHANSDYT | 0.5191608 |
| VQRPDSGGVVLTGRIS | ASGDRHANSDYT | 0.6291503 |
| WTPNPVPATKGVPAGLVQAT | ASGDRHANSDYT | 0.64129084 |
| YALKRALAKFSDRFDVVII | ASGDRHANSDYT | 0.6417852 |
| YLIGSVIV | ASGDRHANSDYT | 0.36933497 |
| YGQLVAVID | ASGDRHANSDYT | 0.6332606 |
| VTVGGMTLPA | ASGDRHANSDYT | 0.6431719 |
| VLALPAV | ASGDRHANSDYT | 0.6587331 |
| VIPPTLNYET | ASGDRHANSDYT | 0.60846543 |
| VGMGGMHPSSG | ASGDRHANSDYT | 0.6377063 |
| VGLAGGAATA | ASGDRHANSDYT | 0.5889831 |
| VGGAGGAGGSGP | ASGDRHANSDYT | 0.5961007 |
| VFAAFMV | ASGDRHANSDYT | 0.5163023 |
| TLYDKLR | ASGDRHANSDYT | 0.5961196 |
| TLAVFTVL | ASGDRHANSDYT | 0.5710602 |
| TGSVQPARV | ASGDRHANSDYT | 0.5409127 |
| TGGGAGSGGT | ASGDRHANSDYT | 0.6396349 |
| SHLETSL | ASGDRHANSDYT | 0.6448518 |
| SFTTGMT | ASGDRHANSDYT | 0.64342195 |
| QVEVAKRMGVRQPTVS | ASGDRHANSDYT | 0.64931715 |
| QGSGGPALAM | ASGDRHANSDYT | 0.6552517 |
| MLAPSNP | ASGDRHANSDYT | 0.6490203 |
| LVNNAAI | ASGDRHANSDYT | 0.57259274 |
| LTGEINTLA | ASGDRHANSDYT | 0.6085399 |
| LSIFGPV | ASGDRHANSDYT | 0.4945161 |
| LLALLANL | ASGDRHANSDYT | 0.59047276 |
| LISNASDALDKLR | ASGDRHANSDYT | 0.56777406 |
| LINEAVP | ASGDRHANSDYT | 0.6222438 |
| LGYAFSGL | ASGDRHANSDYT | 0.32561025 |
| LDFVAVAL | ASGDRHANSDYT | 0.60082906 |
| LAQPGAE | ASGDRHANSDYT | 0.6582327 |
| ISNLGAVTGF | ASGDRHANSDYT | 0.56076205 |
| HQEDPGRI | ASGDRHANSDYT | 0.63340163 |
| GTPGEPTP | ASGDRHANSDYT | 0.62096643 |
| GFFNSGS | ASGDRHANSDYT | 0.5583764 |
| GAGGGGLTAGAN | ASGDRHANSDYT | 0.6408752 |
| FVDALVNM | ASGDRHANSDYT | 0.5464241 |
| FLTVAPEL | ASGDRHANSDYT | 0.63061285 |
| FETLTAL | ASGDRHANSDYT | 0.6560067 |
| DVINPIEL | ASGDRHANSDYT | 0.615998 |
| APVSGAPV | ASGDRHANSDYT | 0.61376774 |
| ALVGGLAW | ASGDRHANSDYT | 0.6093374 |
| ALAAAAATAI | ASGDRHANSDYT | 0.45365015 |
| AFGLGLLL | ASGDRHANSDYT | 0.6219909 |
| AFFAAASL | ASGDRHANSDYT | 0.46823066 |
| AAAALPAFSPP | ASGDRHANSDYT | 0.5977884 |

| | | |
|---|---|---|
| VVLAPAL | ASGDRHANSDYT | 0.5335 |
| TIGHVDHGKTTL | ASGDRHANSDYT | 0.6248361 |
| LISNASDALDK | ASGDRHANSDYT | 0.61423886 |
| LISNASDAL | ASGDRHANSDYT | 0.6344021 |
| ISNASDALDK | ASGDRHANSDYT | 0.5470716 |
| NQAETLVYQTE | ASGDRHANSDYT | 0.588781 |
| TGGTGGTAGAG | ASGDRHANSDYT | 0.55915725 |
| TGSTAGGAGGA | ASGDRHANSDYT | 0.50658935 |
| DPLAVPVAAGPV | ASGDRHANSDYT | 0.6404682 |
| GGTGGTPGNGGN | ASGDRHANSDYT | 0.60168445 |
| PAGLAGVDPATL | ASGDRHANSDYT | 0.5826179 |
| GNNLVVEAPHAGASV | ASGDRHANSDYT | 0.65055645 |
| KVTRIMSASTLEQPA | ASGDRHANSDYT | 0.6159554 |
| LGFALPIALPRRTVV | ASGDRHANSDYT | 0.44683668 |
| TRLGNLTARKLVVLV | ASGDRHANSDYT | 0.5912687 |
| WSQQRRLIYFPSAGP | ASGDRHANSDYT | 0.6121352 |
| GPNGNPDPMAAAVDIR | ASGDRHANSDYT | 0.5931065 |
| TQRCVSVADLRAAASR | ASGDRHANSDYT | 0.33438617 |
| QFVLDVAEHPVEVLTETV | ASGDRHANSDYT | 0.6325431 |
| GGAGGTGGAGGNGGTQAGDG | ASGDRHANSDYT | 0.5708006 |
| PLLATLRAIAVDVDAKPAQV | ASGDRHANSDYT | 0.5299222 |
| APKFRTGGYMIDFWGVGYQVAK | ASGDRHANSDYT | 0.5576578 |
| GAGGAGGAGGAAGGPGGTGGTG | ASGDRHANSDYT | 0.6364863 |
| IGALGLLATDVAPAGSVAFGVA | ASGDRHANSDYT | 0.5859784 |
| AGGAGGIGGNGGAGGAGGVLMG | ASGDRHANSDYT | 0.5939305 |
| ADELIGLL | ASGDRHANSDYT | 0.6339845 |
| ADRALKA | ASGDRHANSDYT | 0.6408155 |
| AGAVGQVGGAGGA | ASGDRHANSDYT | 0.65090376 |
| AGGGGGLGGAP | ASGDRHANSDYT | 0.6379241 |
| AGGLLYGNGGAG | ASGDRHANSDYT | 0.61773884 |
| AGLESIGL | ASGDRHANSDYT | 0.6438746 |
| AGLKTGII | ASGDRHANSDYT | 0.6096219 |
| ASLAALLL | ASGDRHANSDYT | 0.4191431 |
| ASVINISSI | ASGDRHANSDYT | 0.40402102 |
| ATPPGPA | ASGDRHANSDYT | 0.61102843 |
| ATVASALV | ASGDRHANSDYT | 0.42361072 |
| ATVASLIL | ASGDRHANSDYT | 0.6267568 |
| AVNGPVLIRH | ASGDRHANSDYT | 0.6555003 |
| DGAAGAVGI | ASGDRHANSDYT | 0.6067327 |
| DPLLHGI | ASGDRHANSDYT | 0.49938732 |
| ELGAALGAL | ASGDRHANSDYT | 0.59238714 |
| ELGRTQAV | ASGDRHANSDYT | 0.5790348 |
| FVAGGYLS | ASGDRHANSDYT | 0.63416684 |
| GASGGAGGAGGW | ASGDRHANSDYT | 0.6465913 |
| GGAGAGGDGGAGGI | ASGDRHANSDYT | 0.6557878 |
| GTGHNGSQPGMGGNG | ASGDRHANSDYT | 0.6426207 |
| GYAAASASAS | ASGDRHANSDYT | 0.5695057 |
| HQVDVAAVQQG | ASGDRHANSDYT | 0.6375073 |
| IGAAIAHT | ASGDRHANSDYT | 0.5670946 |
| IGMIEVAQL | ASGDRHANSDYT | 0.6110626 |
| IRLAPEHTSAMD | ASGDRHANSDYT | 0.60771215 |

| | | |
|---|---|---|
| IYNVSVL | ASGDRHANSDYT | 0.61946464 |
| KQGFLGM | ASGDRHANSDYT | 0.60176563 |
| LESVITQL | ASGDRHANSDYT | 0.59044814 |
| LGLGALVVV | ASGDRHANSDYT | 0.47485757 |
| LIGNGGNGGSGGI | ASGDRHANSDYT | 0.6411167 |
| LNAINAPTL | ASGDRHANSDYT | 0.6153229 |
| LQLLNTGV | ASGDRHANSDYT | 0.42560703 |
| LQTSIEF | ASGDRHANSDYT | 0.5833748 |
| MSFAVSEL | ASGDRHANSDYT | 0.6559544 |
| NTGALGL | ASGDRHANSDYT | 0.6582441 |
| PPGAGGAGGAGTT | ASGDRHANSDYT | 0.5432848 |
| QLVGLGI | ASGDRHANSDYT | 0.5525999 |
| RFLEQVHQL | ASGDRHANSDYT | 0.6148101 |
| RGVGYVLR | ASGDRHANSDYT | 0.46527866 |
| SATGVFAGVI | ASGDRHANSDYT | 0.6283737 |
| SLPAITLP | ASGDRHANSDYT | 0.58840036 |
| STALAFTV | ASGDRHANSDYT | 0.58498496 |
| SVLTVGLGK | ASGDRHANSDYT | 0.5753031 |
| TALAGAGSGI | ASGDRHANSDYT | 0.6157731 |
| TGLPFTGI | ASGDRHANSDYT | 0.58226836 |
| VAEHVTV | ASGDRHANSDYT | 0.64329636 |
| VDIVLNSL | ASGDRHANSDYT | 0.3408256 |
| VLTGGGAVL | ASGDRHANSDYT | 0.62092865 |
| VNVGLAYGI | ASGDRHANSDYT | 0.5458077 |
| VSSAAVGAV | ASGDRHANSDYT | 0.56803405 |
| VTSVYEAGV | ASGDRHANSDYT | 0.40862566 |
| WVALDRGA | ASGDRHANSDYT | 0.6176721 |
| YLVGSVVI | ASGDRHANSDYT | 0.6406485 |
| GPPGTGKTLL | ASGDRHANSDYT | 0.6471242 |
| GVGMTKF | ASGDRHANSDYT | 0.52658963 |
| IGIGGSDL | ASGDRHANSDYT | 0.63771766 |
| IGTLLAEL | ASGDRHANSDYT | 0.66164184 |
| IYTSGTT | ASGDRHANSDYT | 0.56851137 |
| LSFPAGV | ASGDRHANSDYT | 0.43260485 |
| LYTSGTT | ASGDRHANSDYT | 0.620869 |
| VGAFPFGF | ASGDRHANSDYT | 0.59303916 |
| VGAGTVEF | ASGDRHANSDYT | 0.63930756 |
| VGKSTLLL | ASGDRHANSDYT | 0.6443559 |
| AALAHAGGGIV | ASGDRHANSDYT | 0.49409372 |
| AALVAVSLL | ASGDRHANSDYT | 0.6459098 |
| AAVSRAVAAL | ASGDRHANSDYT | 0.64508057 |
| AGGSSGIGGF | ASGDRHANSDYT | 0.543581 |
| AGLTVANANG | ASGDRHANSDYT | 0.55670035 |
| AGLVGAVL | ASGDRHANSDYT | 0.5800729 |
| ALSQAFTV | ASGDRHANSDYT | 0.39739978 |
| ALTFSVSL | ASGDRHANSDYT | 0.6420019 |
| ALTGAALVAL | ASGDRHANSDYT | 0.5657662 |
| ASFSIGAGVL | ASGDRHANSDYT | 0.5709911 |
| DGSVYKI | ASGDRHANSDYT | 0.60596335 |
| EQLAEIV | ASGDRHANSDYT | 0.5776897 |
| FAVSELTVM | ASGDRHANSDYT | 0.5696196 |

| | | |
|---|---|---|
| GAAPLAAGL | ASGDRHANSDYT | 0.58809054 |
| GADGGDGGSGGAGG | ASGDRHANSDYT | 0.6111393 |
| GFGLDTGGAGG | ASGDRHANSDYT | 0.6253971 |
| GLASGGLL | ASGDRHANSDYT | 0.5963703 |
| GRIAGAAVLL | ASGDRHANSDYT | 0.6249757 |
| GSAGSLNTSF | ASGDRHANSDYT | 0.6510436 |
| GTGQIGI | ASGDRHANSDYT | 0.4985913 |
| GTLIALIL | ASGDRHANSDYT | 0.55412024 |
| IAATMGFVVG | ASGDRHANSDYT | 0.6282909 |
| IFAQALL | ASGDRHANSDYT | 0.47358203 |
| IGAAVALGF | ASGDRHANSDYT | 0.63145727 |
| IGLTPLAL | ASGDRHANSDYT | 0.61821204 |
| ILGSNTSTL | ASGDRHANSDYT | 0.62736 |
| IVFSLTVL | ASGDRHANSDYT | 0.61433417 |
| LASKATGF | ASGDRHANSDYT | 0.6588317 |
| LASVALIL | ASGDRHANSDYT | 0.628376 |
| LFAVGLTL | ASGDRHANSDYT | 0.60154283 |
| LFNGGIGTYI | ASGDRHANSDYT | 0.6239992 |
| LTLGAGII | ASGDRHANSDYT | 0.5600275 |
| LVAAALGDI | ASGDRHANSDYT | 0.64162326 |
| NTAFVQLGI | ASGDRHANSDYT | 0.6419816 |
| QLFSGAEL | ASGDRHANSDYT | 0.5089122 |
| SAATFITL | ASGDRHANSDYT | 0.5951577 |
| SNYGLANLGV | ASGDRHANSDYT | 0.65431476 |
| SYGGAYAVM | ASGDRHANSDYT | 0.6446133 |
| TTGVGGLTLG | ASGDRHANSDYT | 0.44446805 |
| TTQTIYI | ASGDRHANSDYT | 0.5641582 |
| VAAGAGLGAVI | ASGDRHANSDYT | 0.49483824 |
| VAAVVTLL | ASGDRHANSDYT | 0.60026395 |
| VGFLGGLI | ASGDRHANSDYT | 0.5158704 |
| VGVIPILL | ASGDRHANSDYT | 0.63167393 |
| VIFASSLI | ASGDRHANSDYT | 0.5879414 |
| VKGGDGGAAAG | ASGDRHANSDYT | 0.6333615 |
| VTKTFTNI | ASGDRHANSDYT | 0.6250666 |
| VVAAAILL | ASGDRHANSDYT | 0.64973485 |
| YAALDVEL | ASGDRHANSDYT | 0.5790917 |
| LASVALLL | ASGDRHANSDYT | 0.6027501 |
| LDAVHRV | ASGDRHANSDYT | 0.45636746 |
| QGLSSSIF | ASGDRHANSDYT | 0.52917457 |
| RGTFANIRL | ASGDRHANSDYT | 0.54782796 |
| RQLWWGHRI | ASGDRHANSDYT | 0.59320456 |
| YGPPGTGKTLL | ASGDRHANSDYT | 0.3768732 |
| AALVGAPANVA | ASGDRHANSDYT | 0.6549206 |
| AATFRRTE | ASGDRHANSDYT | 0.6003791 |
| AGALGGGI | ASGDRHANSDYT | 0.6368971 |
| AGGAGGAGGIGAG | ASGDRHANSDYT | 0.5778611 |
| AGSGALGL | ASGDRHANSDYT | 0.5754068 |
| AQIADQI | ASGDRHANSDYT | 0.5609268 |
| ASAVAVTYL | ASGDRHANSDYT | 0.6325631 |
| ASTVLGGTGG | ASGDRHANSDYT | 0.5187286 |
| AVGGEPAK | ASGDRHANSDYT | 0.43218192 |

| | | |
|---|---|---|
| CEQGTFGGTC | ASGDRHANSDYT | 0.5741686 |
| DEGKAPAG | ASGDRHANSDYT | 0.4943707 |
| DTVVFPAL | ASGDRHANSDYT | 0.6586534 |
| FEGITMAG | ASGDRHANSDYT | 0.6453296 |
| FPLVVQD | ASGDRHANSDYT | 0.61405444 |
| GAFTTPQ | ASGDRHANSDYT | 0.65500385 |
| GFGGFGTAGGVGGA | ASGDRHANSDYT | 0.36623552 |
| GFLAQFL | ASGDRHANSDYT | 0.5823358 |
| GGAGGAGGNAGVA | ASGDRHANSDYT | 0.6563794 |
| GGVGTAGAGGAGGAGG | ASGDRHANSDYT | 0.62343764 |
| GNGGGGDAGNAGSGGN | ASGDRHANSDYT | 0.6067515 |
| GTGGAGGAGNDAGS | ASGDRHANSDYT | 0.64678234 |
| GTTNTGWFNS | ASGDRHANSDYT | 0.5510527 |
| IFMGPLFTL | ASGDRHANSDYT | 0.6033035 |
| ILAHAIEAI | ASGDRHANSDYT | 0.5943481 |
| IRSAIREAL | ASGDRHANSDYT | 0.63928014 |
| ITFAGGG | ASGDRHANSDYT | 0.6244064 |
| ITGMSGHQEV | ASGDRHANSDYT | 0.5331618 |
| IVAGLACTL | ASGDRHANSDYT | 0.62928206 |
| IVGGGLAGA | ASGDRHANSDYT | 0.6611074 |
| IVTGVLNI | ASGDRHANSDYT | 0.5802218 |
| LAQQGCI | ASGDRHANSDYT | 0.6469741 |
| LCLHVEN | ASGDRHANSDYT | 0.6256297 |
| LDAISDGS | ASGDRHANSDYT | 0.6176739 |
| LGGFAAML | ASGDRHANSDYT | 0.5509934 |
| LGGKNPMI | ASGDRHANSDYT | 0.54919046 |
| LGPKGVAAIL | ASGDRHANSDYT | 0.58692706 |
| LMVGVSI | ASGDRHANSDYT | 0.523099 |
| LVGAAFVL | ASGDRHANSDYT | 0.6095751 |
| LVLAVAPL | ASGDRHANSDYT | 0.5810256 |
| MAAIAVIDAA | ASGDRHANSDYT | 0.668044 |
| NNGINAELQ | ASGDRHANSDYT | 0.632084 |
| RGEPLVLVHG | ASGDRHANSDYT | 0.622948 |
| RGGAGGAAGLI | ASGDRHANSDYT | 0.526519 |
| RSYRMHPAV | ASGDRHANSDYT | 0.6205853 |
| SAAHLLAGQLA | ASGDRHANSDYT | 0.617091 |
| SAAVGSY | ASGDRHANSDYT | 0.47465622 |
| SPHAVLHRI | ASGDRHANSDYT | 0.6553369 |
| SQIAKVL | ASGDRHANSDYT | 0.54866505 |
| TALGHLVP | ASGDRHANSDYT | 0.58514637 |
| TAPTTSGAA | ASGDRHANSDYT | 0.6370808 |
| TDGDDGNGGSPGT | ASGDRHANSDYT | 0.6467329 |
| TDGLGEN | ASGDRHANSDYT | 0.58724594 |
| TGGPPLPGQA | ASGDRHANSDYT | 0.5912318 |
| TIPAITI | ASGDRHANSDYT | 0.52502936 |
| TIPGFTI | ASGDRHANSDYT | 0.643297 |
| TSIASGALA | ASGDRHANSDYT | 0.64812356 |
| TVTLPGAAI | ASGDRHANSDYT | 0.6104993 |
| VASGLSAALP | ASGDRHANSDYT | 0.6613611 |
| VGAGVIGI | ASGDRHANSDYT | 0.6235553 |
| VGPKRFGV | ASGDRHANSDYT | 0.47020486 |

| | | |
|---|---|---|
| VNVAFPD | ASGDRHANSDYT | 0.61502004 |
| VSMTNSGVS | ASGDRHANSDYT | 0.64262784 |
| VTDVLGDLSV | ASGDRHANSDYT | 0.64624614 |
| VVVVGAGFAGL | ASGDRHANSDYT | 0.6575567 |
| VYTVGTVV | ASGDRHANSDYT | 0.6370317 |
| WTTFSGP | ASGDRHANSDYT | 0.57137823 |
| YDELVEAF | ASGDRHANSDYT | 0.6528788 |
| YGTGGAGGNGG | ASGDRHANSDYT | 0.6674385 |
| YTNVMAV | ASGDRHANSDYT | 0.64016473 |
| FLEQVHQL | ASGDRHANSDYT | 0.5780965 |
| GLSLLAVAV | ASGDRHANSDYT | 0.57779956 |
| LEQVHQL | ASGDRHANSDYT | 0.62062603 |
| LNAAGAI | ASGDRHANSDYT | 0.6301341 |
| PAVHASL | ASGDRHANSDYT | 0.52805465 |
| TELEDDA | ASGDRHANSDYT | 0.61361015 |
| TPPGPPN | ASGDRHANSDYT | 0.6258559 |
| TPPPPPPP | ASGDRHANSDYT | 0.4673953 |
| TQSPALI | ASSFLQGGAEQF | 0.6352364 |
| EFAAAFPQ | ASSFLQGGAEQF | 0.6489432 |
| LLSGNLDVL | ASSFLQGGAEQF | 0.6014919 |
| GPANLGNYNI | ASSFLQGGAEQF | 0.5446722 |
| TLGPTLP | ASSFLQGGAEQF | 0.64969707 |
| VYAYSSCCK | ASSFLQGGAEQF | 0.6645381 |
| APAPAPAPNI | ASSFLQGGAEQF | 0.6493421 |
| PGGPVPIP | ASSFLQGGAEQF | 0.44182688 |
| FLTVALEP | ASSFLQGGAEQF | 0.36859104 |
| SQYFGANL | ASSFLQGGAEQF | 0.55748326 |
| GGVKPAA | ASSFLQGGAEQF | 0.6380112 |
| GGTGGAGGAGGW | ASSFLQGGAEQF | 0.59484607 |
| GVGGVAGLGGAGG | ASSFLQGGAEQF | 0.5470048 |
| VLLVLATTI | ASSFLQGGAEQF | 0.6138249 |
| IGIFAAML | ASSFLQGGAEQF | 0.6478971 |
| AVGDLID | ASSFLQGGAEQF | 0.5527519 |
| NISPLPL | ASSFLQGGAEQF | 0.5241432 |
| VTAVFVAI | ASSFLQGGAEQF | 0.6298041 |
| SGDDNDAAL | ASSFLQGGAEQF | 0.6435697 |
| AVAALAHS | ASSFLQGGAEQF | 0.51431847 |
| GGVTVMATP | ASSFLQGGAEQF | 0.57914984 |
| AGLAANVRD | ASSFLQGGAEQF | 0.58731335 |
| TNRPDLIDPALLRPG | ASSFLQGGAEQF | 0.56067514 |
| LANLGVTAL | ASSFLQGGAEQF | 0.6134006 |
| TGERIEL | ASSFLQGGAEQF | 0.5352987 |
| NGEELAA | ASSFLQGGAEQF | 0.6469328 |
| VLSLVWG | ASSFLQGGAEQF | 0.50450337 |
| IGYALWL | ASSFLQGGAEQF | 0.63840765 |
| LSLLLAPL | ASSFLQGGAEQF | 0.623647 |
| TLDEALK | ASSFLQGGAEQF | 0.6314047 |
| TDGAGTAEGG | ASSFLQGGAEQF | 0.5792553 |
| NAPGGAGGNGG | ASSFLQGGAEQF | 0.5676234 |
| VSVFMGMGGNF | ASSFLQGGAEQF | 0.51577157 |
| LASYLAAL | ASSFLQGGAEQF | 0.42600036 |

| | | |
|---|---|---|
| LVGGTPAL | ASSFLQGGAEQF | 0.64756495 |
| VRNADVL | ASSFLQGGAEQF | 0.52774745 |
| TCSSPFLVSAM | ASSFLQGGAEQF | 0.6459248 |
| VHITDAE | ASSFLQGGAEQF | 0.62129974 |
| EYAAPSF | ASSFLQGGAEQF | 0.5442477 |
| VVDAFHGGL | ASSFLQGGAEQF | 0.6311567 |
| ASVVDPQ | ASSFLQGGAEQF | 0.49508357 |
| SYNEFGGLS | ASSFLQGGAEQF | 0.53552806 |
| DPDGSVY | ASSFLQGGAEQF | 0.63606614 |
| MVSTIHATE | ASSFLQGGAEQF | 0.63151926 |
| LAAGVADPV | ASSFLQGGAEQF | 0.5614368 |
| TKEQIAA | ASSFLQGGAEQF | 0.6305324 |
| SLASPAAL | ASSFLQGGAEQF | 0.62014973 |
| IGLVFYGV | ASSFLQGGAEQF | 0.5769433 |
| FLLDASNL | ASSFLQGGAEQF | 0.60925007 |
| YQVGNFP | ASSFLQGGAEQF | 0.64926237 |
| ESLLSQST | ASSFLQGGAEQF | 0.62157583 |
| LFTVGFAAVG | ASSFLQGGAEQF | 0.5162304 |
| ATPPTPT | ASSFLQGGAEQF | 0.62252384 |
| TDLATLG | ASSFLQGGAEQF | 0.5385604 |
| SAGGTGR | ASSFLQGGAEQF | 0.6225011 |
| GFSNSGS | ASSFLQGGAEQF | 0.6079581 |
| GGNGGSGGTSV | ASSFLQGGAEQF | 0.6046542 |
| GTSVATGGAGN | ASSFLQGGAEQF | 0.63625693 |
| NGGNGGSGGTS | ASSFLQGGAEQF | 0.50196713 |
| AVVKFPH | ASSFLQGGAEQF | 0.5443676 |
| TPPPPPP | ASSFLQGGAEQF | 0.5860561 |
| AAPAAAAAPA | ASSFLQGGAEQF | 0.61984664 |
| IALNEDL | ASSFLQGGAEQF | 0.6353336 |
| IVNVSSM | ASSFLQGGAEQF | 0.52135646 |
| TGIPPAPR | ASSFLQGGAEQF | 0.6194382 |
| FDLGGGTF | ASSFLQGGAEQF | 0.59451 |
| FDLGGGTFDVS | ASSFLQGGAEQF | 0.54808605 |
| NEPTAAA | ASSFLQGGAEQF | 0.6042415 |
| NEPTAAALA | ASSFLQGGAEQF | 0.58171296 |
| GSIINISSI | ASSFLQGGAEQF | 0.5318193 |
| VAPGNAGTA | ASSFLQGGAEQF | 0.6362961 |
| TGGGASLE | ASSFLQGGAEQF | 0.5414087 |
| VAVYNHYKRI | ASSFLQGGAEQF | 0.63375413 |
| TDPAPATT | ASSFLQGGAEQF | 0.60832936 |
| TNRPDILDPALLRPG | ASSFLQGGAEQF | 0.56827086 |
| LTTTTTS | ASSFLQGGAEQF | 0.6371654 |
| PTGSGKTL | ASSFLQGGAEQF | 0.6191658 |
| TGSGKTL | ASSFLQGGAEQF | 0.62368476 |
| TIGHVDHGK | ASSFLQGGAEQF | 0.5959823 |
| TIGHVDHGKTT | ASSFLQGGAEQF | 0.5670285 |
| GHVDHGKTT | ASSFLQGGAEQF | 0.64455897 |
| AQFEHTLL | ASSFLQGGAEQF | 0.6373943 |
| IVNQDEL | ASSFLQGGAEQF | 0.642676 |
| TNGSQFF | ASSFLQGGAEQF | 0.6327077 |
| TGKGIVE | ASSFLQGGAEQF | 0.6305072 |

| | | |
|---|---|---|
| LISNASDA | ASSFLQGGAEQF | 0.6346799 |
| AELRTLGELTKIAKAHGAQ | ASSFLQGGAEQF | 0.5656194 |
| APNAVAAAEPSW | ASSFLQGGAEQF | 0.46711802 |
| APQVAGAATAVL | ASSFLQGGAEQF | 0.58935034 |
| ARRDPAAAAAILPTDA | ASSFLQGGAEQF | 0.5922684 |
| DAAARDVLAEAGLAEYF | ASSFLQGGAEQF | 0.5764513 |
| DDQIVLSAKANLAKVEQ | ASSFLQGGAEQF | 0.45142785 |
| DELLPDTGEDSDEDGH | ASSFLQGGAEQF | 0.64534914 |
| ELPEAPANRATVPQD | ASSFLQGGAEQF | 0.66537356 |
| FGQSKIERVTGIG | ASSFLQGGAEQF | 0.5688221 |
| GAGGTGSTAGGAGGAGGAGGLY | ASSFLQGGAEQF | 0.6612644 |
| GHGGTGAAGGNGGTGGAGGSGI | ASSFLQGGAEQF | 0.46020514 |
| GNGGNGGAGGEGGPGGAGGAGA | ASSFLQGGAEQF | 0.63356304 |
| GSTAGNGGNGGNATNGTIAGQP | ASSFLQGGAEQF | 0.64657056 |
| GTGGDGGNGGNGGTGG | ASSFLQGGAEQF | 0.5310559 |
| GVRRPAGIPADAPV | ASSFLQGGAEQF | 0.63770604 |
| GYRQVTDAHLLTLARR | ASSFLQGGAEQF | 0.5507304 |
| IAKMGQDGHDRG | ASSFLQGGAEQF | 0.57601005 |
| KAEPSIEIQYT | ASSFLQGGAEQF | 0.6519411 |
| KPVRIRMAADEDTVAVT | ASSFLQGGAEQF | 0.5987925 |
| MPPRIAGMRLLVIKPEP | ASSFLQGGAEQF | 0.6334278 |
| MKFYAVALFRSRGVPPTR | ASSFLQGGAEQF | 0.58021665 |
| NAGGFGILGAVAHSP | ASSFLQGGAEQF | 0.5443029 |
| NEGREITLAMAEM | ASSFLQGGAEQF | 0.62456477 |
| NELVEWSHPLGHRWRWE | ASSFLQGGAEQF | 0.65296036 |
| NLGAGGTGGNGGNPG | ASSFLQGGAEQF | 0.55218375 |
| NTEWPKWETVMAADPVD | ASSFLQGGAEQF | 0.6661382 |
| PASAAPAETQP | ASSFLQGGAEQF | 0.56110084 |
| QQSSDGKNGVRAAEITGE | ASSFLQGGAEQF | 0.6615069 |
| RAPDAVTTSWWKE | ASSFLQGGAEQF | 0.6414367 |
| SGFGNVGAHTSGWFNQ | ASSFLQGGAEQF | 0.59992236 |
| SGVAGAVGQAASVGGLKV | ASSFLQGGAEQF | 0.58725643 |
| SNGGAGTGSAGGNGGT | ASSFLQGGAEQF | 0.64359313 |
| TGGAGGEGGAAG | ASSFLQGGAEQF | 0.5957674 |
| TGGDGGAAGAGGAPGHG | ASSFLQGGAEQF | 0.63332033 |
| TMPPLLAAIANAGRHGV | ASSFLQGGAEQF | 0.6538803 |
| TSSTAGNSGGA | ASSFLQGGAEQF | 0.39006305 |
| VPATVPVGILC | ASSFLQGGAEQF | 0.8015212 |
| VQRPDSGGVVLTGRIS | ASSFLQGGAEQF | 0.74863017 |
| WTPNPVPATKGVPAGLVQAT | ASSFLQGGAEQF | 0.58519644 |
| YALKRALAKFSDRFDVVII | ASSFLQGGAEQF | 0.6354782 |
| YLIGSVIV | ASSFLQGGAEQF | 0.2679916 |
| YGQLVAVID | ASSFLQGGAEQF | 0.5726382 |
| VTVGGMTLPA | ASSFLQGGAEQF | 0.5596572 |
| VLALPAV | ASSFLQGGAEQF | 0.8163188 |
| VIPPTLNYET | ASSFLQGGAEQF | 0.7580538 |
| VGMGGMHPSSG | ASSFLQGGAEQF | 0.603099 |
| VGLAGGAATA | ASSFLQGGAEQF | 0.74417114 |
| VGGAGGAGGSGP | ASSFLQGGAEQF | 0.553262 |
| VFAAFMV | ASSFLQGGAEQF | 0.49029702 |
| TLYDKLR | ASSFLQGGAEQF | 0.61053216 |

| | | |
|---|---|---|
| TLAVFTVL | ASSFLQGGAEQF | 0.5005059 |
| TGSVQPARV | ASSFLQGGAEQF | 0.7181056 |
| TGGGAGSGGT | ASSFLQGGAEQF | 0.36743492 |
| SHLETSL | ASSFLQGGAEQF | 0.74685764 |
| SFTTGMT | ASSFLQGGAEQF | 0.6452491 |
| QVEVAKRMGVRQPTVS | ASSFLQGGAEQF | 0.25003886 |
| QGSGGPALAM | ASSFLQGGAEQF | 0.7880189 |
| MLAPSNP | ASSFLQGGAEQF | 0.57951486 |
| LVNNAAI | ASSFLQGGAEQF | 0.67165434 |
| LTGEINTLA | ASSFLQGGAEQF | 0.7951133 |
| LSIFGPV | ASSFLQGGAEQF | 0.5976825 |
| LLALLANL | ASSFLQGGAEQF | 0.45571923 |
| LISNASDALDKLR | ASSFLQGGAEQF | 0.7781945 |
| LINEAVP | ASSFLQGGAEQF | 0.7550068 |
| LGYAFSGL | ASSFLQGGAEQF | 0.33649534 |
| LDFVAVAL | ASSFLQGGAEQF | 0.7333312 |
| LAQPGAE | ASSFLQGGAEQF | 0.7435602 |
| ISNLGAVTGF | ASSFLQGGAEQF | 0.7134762 |
| HQEDPGRI | ASSFLQGGAEQF | 0.7902327 |
| GTPGEPTP | ASSFLQGGAEQF | 0.79429686 |
| GFFNSGS | ASSFLQGGAEQF | 0.47082934 |
| GAGGGGLTAGAN | ASSFLQGGAEQF | 0.78927827 |
| FVDALVNM | ASSFLQGGAEQF | 0.7832414 |
| FLTVAPEL | ASSFLQGGAEQF | 0.5016966 |
| FETLTAL | ASSFLQGGAEQF | 0.6846653 |
| DVINPIEL | ASSFLQGGAEQF | 0.4391827 |
| APVSGAPV | ASSFLQGGAEQF | 0.6206144 |
| ALVGGLAW | ASSFLQGGAEQF | 0.49209866 |
| ALAAAAATAI | ASSFLQGGAEQF | 0.7818494 |
| AFGLGLLL | ASSFLQGGAEQF | 0.76600087 |
| AFFAAASL | ASSFLQGGAEQF | 0.74803495 |
| AAAALPAFSPP | ASSFLQGGAEQF | 0.49622628 |
| VVLAPAL | ASSFLQGGAEQF | 0.40232444 |
| TIGHVDHGKTTL | ASSFLQGGAEQF | 0.73198456 |
| LISNASDALDK | ASSFLQGGAEQF | 0.35009906 |
| LISNASDAL | ASSFLQGGAEQF | 0.3927836 |
| ISNASDALDK | ASSFLQGGAEQF | 0.59261453 |
| NQAETLVYQTE | ASSFLQGGAEQF | 0.7247424 |
| TGGTGGTAGAG | ASSFLQGGAEQF | 0.72873354 |
| TGSTAGGAGGA | ASSFLQGGAEQF | 0.59364283 |
| DPLAVPVAAGPV | ASSFLQGGAEQF | 0.81593096 |
| GGTGGTPGNGGN | ASSFLQGGAEQF | 0.7985036 |
| PAGLAGVDPATL | ASSFLQGGAEQF | 0.75690866 |
| GNNLVVEAPHAGASV | ASSFLQGGAEQF | 0.47052824 |
| KVTRIMSASTLEQPA | ASSFLQGGAEQF | 0.4229977 |
| LGFALPIALPRRTVV | ASSFLQGGAEQF | 0.8207335 |
| TRLGNLTARKLVVLV | ASSFLQGGAEQF | 0.72311056 |
| WSQQRRLIYFPSAGP | ASSFLQGGAEQF | 0.7285816 |
| GPNGNPDPMAAAVDIR | ASSFLQGGAEQF | 0.7801329 |
| TQRCVSVADLRAAASR | ASSFLQGGAEQF | 0.70787734 |
| QFVLDVAEHPVEVLTETV | ASSFLQGGAEQF | 0.5278548 |

| | | |
|---|---|---|
| GGAGGTGGAGGNGGTQAGDG | ASSFLQGGAEQF | 0.763541 |
| PLLATLRAIAVDVDAKPAQV | ASSFLQGGAEQF | 0.8115727 |
| APKFRTGGYMIDFWGVGYQVAK | ASSFLQGGAEQF | 0.5442382 |
| GAGGAGGAGGAAGGPGGTGGTG | ASSFLQGGAEQF | 0.35211885 |
| IGALGLLATDVAPAGSVAFGVA | ASSFLQGGAEQF | 0.6151298 |
| AGGAGGIGGNGGAGGAGGVLMG | ASSFLQGGAEQF | 0.6573008 |
| ADELIGLL | ASSFLQGGAEQF | 0.7678708 |
| ADRALKA | ASSFLQGGAEQF | 0.67898965 |
| AGAVGQVGGAGGA | ASSFLQGGAEQF | 0.6133046 |
| AGGGGGLGGAP | ASSFLQGGAEQF | 0.7418498 |
| AGGLLYGNGGAG | ASSFLQGGAEQF | 0.7076429 |
| AGLESIGL | ASSFLQGGAEQF | 0.71194404 |
| AGLKTGII | ASSFLQGGAEQF | 0.65860045 |
| ASLAALLL | ASSFLQGGAEQF | 0.6949406 |
| ASVINISSI | ASSFLQGGAEQF | 0.8058306 |
| ATPPGPA | ASSFLQGGAEQF | 0.78935844 |
| ATVASALV | ASSFLQGGAEQF | 0.74213505 |
| ATVASLIL | ASSFLQGGAEQF | 0.75491333 |
| AVNGPVLIRH | ASSFLQGGAEQF | 0.51824576 |
| DGAAGAVGI | ASSFLQGGAEQF | 0.70390993 |
| DPLLHGI | ASSFLQGGAEQF | 0.3482352 |
| ELGAALGAL | ASSFLQGGAEQF | 0.4016688 |
| ELGRTQAV | ASSFLQGGAEQF | 0.6878594 |
| FVAGGYLS | ASSFLQGGAEQF | 0.7654891 |
| GASGGAGGAGGW | ASSFLQGGAEQF | 0.7904364 |
| GGAGAGGDGGAGGI | ASSFLQGGAEQF | 0.5525222 |
| GTGHNGSQPGMGGNG | ASSFLQGGAEQF | 0.56659544 |
| GYAAASASAS | ASSFLQGGAEQF | 0.7223731 |
| HQVDVAAVQQG | ASSFLQGGAEQF | 0.7356726 |
| IGAAIAHT | ASSFLQGGAEQF | 0.5837605 |
| IGMIEVAQL | ASSFLQGGAEQF | 0.65122104 |
| IRLAPEHTSAMD | ASSFLQGGAEQF | 0.8208205 |
| IYNVSVL | ASSFLQGGAEQF | 0.79476947 |
| KQGFLGM | ASSFLQGGAEQF | 0.68604994 |
| LESVITQL | ASSFLQGGAEQF | 0.51498115 |
| LGLGALVVV | ASSFLQGGAEQF | 0.82034993 |
| LIGNGGNGGSGGI | ASSFLQGGAEQF | 0.7104779 |
| LNAINAPTL | ASSFLQGGAEQF | 0.8136618 |
| LQLLNTGV | ASSFLQGGAEQF | 0.25766882 |
| LQTSIEF | ASSFLQGGAEQF | 0.6068383 |
| MSFAVSEL | ASSFLQGGAEQF | 0.4293263 |
| NTGALGL | ASSFLQGGAEQF | 0.21111049 |
| PPGAGGAGGAGTT | ASSFLQGGAEQF | 0.795853 |
| QLVGLGI | ASSFLQGGAEQF | 0.38806492 |
| RFLEQVHQL | ASSFLQGGAEQF | 0.5714047 |
| RGVGYVLR | ASSFLQGGAEQF | 0.626631 |
| SATGVFAGVI | ASSFLQGGAEQF | 0.7781541 |
| SLPAITLP | ASSFLQGGAEQF | 0.79170126 |
| STALAFTV | ASSFLQGGAEQF | 0.5012246 |
| SVLTVGLGK | ASSFLQGGAEQF | 0.49170023 |
| TALAGAGSGI | ASSFLQGGAEQF | 0.72943246 |

| | | |
|---|---|---|
| TGLPFTGI | ASSFLQGGAEQF | 0.7632495 |
| VAEHVTV | ASSFLQGGAEQF | 0.7605331 |
| VDIVLNSL | ASSFLQGGAEQF | 0.23092139 |
| VLTGGGAVL | ASSFLQGGAEQF | 0.7969864 |
| VNVGLAYGI | ASSFLQGGAEQF | 0.59625196 |
| VSSAAVGAV | ASSFLQGGAEQF | 0.47156334 |
| VTSVYEAGV | ASSFLQGGAEQF | 0.34034017 |
| WVALDRGA | ASSFLQGGAEQF | 0.6086746 |
| YLVGSVVI | ASSFLQGGAEQF | 0.7094385 |
| GPPGTGKTLL | ASSFLQGGAEQF | 0.7858388 |
| GVGMTKF | ASSFLQGGAEQF | 0.79518354 |
| IGIGGSDL | ASSFLQGGAEQF | 0.6880778 |
| IGTLLAEL | ASSFLQGGAEQF | 0.70027375 |
| IYTSGTT | ASSFLQGGAEQF | 0.67547536 |
| LSFPAGV | ASSFLQGGAEQF | 0.14391221 |
| LYTSGTT | ASSFLQGGAEQF | 0.41643503 |
| VGAFPFGF | ASSFLQGGAEQF | 0.5119924 |
| VGAGTVEF | ASSFLQGGAEQF | 0.748156 |
| VGKSTLLL | ASSFLQGGAEQF | 0.52363604 |
| AALAHAGGGIV | ASSFLQGGAEQF | 0.71911955 |
| AALVAVSLL | ASSFLQGGAEQF | 0.75457907 |
| AAVSRAVAAL | ASSFLQGGAEQF | 0.7978531 |
| AGGSSGIGGF | ASSFLQGGAEQF | 0.7292886 |
| AGLTVANANG | ASSFLQGGAEQF | 0.44199833 |
| AGLVGAVL | ASSFLQGGAEQF | 0.52691793 |
| ALSQAFTV | ASSFLQGGAEQF | 0.73415446 |
| ALTFSVSL | ASSFLQGGAEQF | 0.66155326 |
| ALTGAALVAL | ASSFLQGGAEQF | 0.3359343 |
| ASFSIGAGVL | ASSFLQGGAEQF | 0.5732472 |
| DGSVYKI | ASSFLQGGAEQF | 0.81431323 |
| EQLAEIV | ASSFLQGGAEQF | 0.26185155 |
| FAVSELTVM | ASSFLQGGAEQF | 0.7796787 |
| GAAPLAAGL | ASSFLQGGAEQF | 0.56695366 |
| GADGGDGGSGGAGG | ASSFLQGGAEQF | 0.6941655 |
| GFGLDTGGAGG | ASSFLQGGAEQF | 0.8152078 |
| GLASGGLL | ASSFLQGGAEQF | 0.7843753 |
| GRIAGAAVLL | ASSFLQGGAEQF | 0.75829744 |
| GSAGSLNTSF | ASSFLQGGAEQF | 0.4133287 |
| GTGQIGI | ASSFLQGGAEQF | 0.58141613 |
| GTLIALIL | ASSFLQGGAEQF | 0.63826567 |
| IAATMGFVVG | ASSFLQGGAEQF | 0.46917367 |
| IFAQALL | ASSFLQGGAEQF | 0.7471036 |
| IGAAVALGF | ASSFLQGGAEQF | 0.75819325 |
| IGLTPLAL | ASSFLQGGAEQF | 0.40146232 |
| ILGSNTSTL | ASSFLQGGAEQF | 0.7049855 |
| IVFSLTVL | ASSFLQGGAEQF | 0.8155674 |
| LASKATGF | ASSFLQGGAEQF | 0.82847905 |
| LASVALIL | ASSFLQGGAEQF | 0.62027085 |
| LFAVGLTL | ASSFLQGGAEQF | 0.33587855 |
| LFNGGIGTYI | ASSFLQGGAEQF | 0.3432968 |
| LTLGAGII | ASSFLQGGAEQF | 0.7125647 |

| | | |
|---|---|---|
| LVAAALGDI | ASSFLQGGAEQF | 0.3194089 |
| NTAFVQLGI | ASSFLQGGAEQF | 0.44233948 |
| QLFSGAEL | ASSFLQGGAEQF | 0.7938145 |
| SAATFITL | ASSFLQGGAEQF | 0.82986605 |
| SNYGLANLGV | ASSFLQGGAEQF | 0.5843854 |
| SYGGAYAVM | ASSFLQGGAEQF | 0.33449107 |
| TTGVGGLTLG | ASSFLQGGAEQF | 0.713138 |
| TTQTIYI | ASSFLQGGAEQF | 0.4960336 |
| VAAGAGLGAVI | ASSFLQGGAEQF | 0.6701499 |
| VAAVVTLL | ASSFLQGGAEQF | 0.37776446 |
| VGFLGGLI | ASSFLQGGAEQF | 0.4586843 |
| VGVIPILL | ASSFLQGGAEQF | 0.62952614 |
| VIFASSLI | ASSFLQGGAEQF | 0.80591416 |
| VKGGDGGAAAG | ASSFLQGGAEQF | 0.7438662 |
| VTKTFTNI | ASSFLQGGAEQF | 0.7005549 |
| VVAAAILL | ASSFLQGGAEQF | 0.8261807 |
| YAALDVEL | ASSFLQGGAEQF | 0.8023205 |
| LASVALLL | ASSFLQGGAEQF | 0.82321703 |
| LDAVHRV | ASSFLQGGAEQF | 0.80125725 |
| QGLSSSIF | ASSFLQGGAEQF | 0.6395775 |
| RGTFANIRL | ASSFLQGGAEQF | 0.6274956 |
| RQLWWGHRI | ASSFLQGGAEQF | 0.82510555 |
| YGPPGTGKTLL | ASSFLQGGAEQF | 0.69960314 |
| AALVGAPANVA | ASSFLQGGAEQF | 0.77023417 |
| AATFRRTE | ASSFLQGGAEQF | 0.6967881 |
| AGALGGGI | ASSFLQGGAEQF | 0.79793173 |
| AGGAGGAGGIGAG | ASSFLQGGAEQF | 0.5509538 |
| AGSGALGL | ASSFLQGGAEQF | 0.82744193 |
| AQIADQI | ASSFLQGGAEQF | 0.1996292 |
| ASAVAVTYL | ASSFLQGGAEQF | 0.48816198 |
| ASTVLGGTGG | ASSFLQGGAEQF | 0.8191833 |
| AVGGEPAK | ASSFLQGGAEQF | 0.15855512 |
| CEQGTFGGTC | ASSFLQGGAEQF | 0.25035265 |
| DEGKAPAG | ASSFLQGGAEQF | 0.6909983 |
| DTVVFPAL | ASSFLQGGAEQF | 0.7972643 |
| FEGITMAG | ASSFLQGGAEQF | 0.20083481 |
| FPLVVQD | ASSFLQGGAEQF | 0.5314491 |
| GAFTTPQ | ASSFLQGGAEQF | 0.30691662 |
| GFGGFGTAGGVGGA | ASSFLQGGAEQF | 0.66590726 |
| GFLAQFL | ASSFLQGGAEQF | 0.52575064 |
| GGAGGAGGNAGVA | ASSFLQGGAEQF | 0.32166752 |
| GGVGTAGAGGAGGAGG | ASSFLQGGAEQF | 0.7794043 |
| GNGGGGDAGNAGSGGN | ASSFLQGGAEQF | 0.49938264 |
| GTGGAGGAGNDAGS | ASSFLQGGAEQF | 0.78690505 |
| GTTNTGWFNS | ASSFLQGGAEQF | 0.784496 |
| IFMGPLFTL | ASSFLQGGAEQF | 0.82800317 |
| ILAHAIEAI | ASSFLQGGAEQF | 0.43370664 |
| IRSAIREAL | ASSFLQGGAEQF | 0.70949346 |
| ITFAGGG | ASSFLQGGAEQF | 0.5849725 |
| ITGMSGHQEV | ASSFLQGGAEQF | 0.835373 |
| IVAGLACTL | ASSFLQGGAEQF | 0.6413811 |

| | | |
|---|---|---|
| IVGGGLAGA | ASSFLQGGAEQF | 0.35921192 |
| IVTGVLNI | ASSFLQGGAEQF | 0.7854152 |
| LAQQGCI | ASSFLQGGAEQF | 0.7992579 |
| LCLHVEN | ASSFLQGGAEQF | 0.45325536 |
| LDAISDGS | ASSFLQGGAEQF | 0.8112701 |
| LGGFAAML | ASSFLQGGAEQF | 0.6600528 |
| LGGKNPMI | ASSFLQGGAEQF | 0.726265 |
| LGPKGVAAIL | ASSFLQGGAEQF | 0.78629464 |
| LMVGVSI | ASSFLQGGAEQF | 0.75653344 |
| LVGAAFVL | ASSFLQGGAEQF | 0.583094 |
| LVLAVAPL | ASSFLQGGAEQF | 0.77225804 |
| MAAIAVIDAA | ASSFLQGGAEQF | 0.72210634 |
| NNGINAELQ | ASSFLQGGAEQF | 0.38364217 |
| RGEPLVLVHG | ASSFLQGGAEQF | 0.8261304 |
| RGGAGGAAGLI | ASSFLQGGAEQF | 0.5922924 |
| RSYRMHPAV | ASSFLQGGAEQF | 0.8010647 |
| SAAHLLAGQLA | ASSFLQGGAEQF | 0.75748193 |
| SAAVGSY | ASSFLQGGAEQF | 0.80544645 |
| SPHAVLHRI | ASSFLQGGAEQF | 0.43435538 |
| SQIAKVL | ASSFLQGGAEQF | 0.5795602 |
| TALGHLVP | ASSFLQGGAEQF | 0.62670004 |
| TAPTTSGAA | ASSFLQGGAEQF | 0.6858053 |
| TDGDDGNGGSPGT | ASSFLQGGAEQF | 0.6504414 |
| TDGLGEN | ASSFLQGGAEQF | 0.77125883 |
| TGGPPLPGQA | ASSFLQGGAEQF | 0.7898017 |
| TIPAITI | ASSFLQGGAEQF | 0.8039534 |
| TIPGFTI | ASSFLQGGAEQF | 0.6387037 |
| TSIASGALA | ASSFLQGGAEQF | 0.6227982 |
| TVTLPGAAI | ASSFLQGGAEQF | 0.8317574 |
| VASGLSAALP | ASSFLQGGAEQF | 0.52968985 |
| VGAGVIGI | ASSFLQGGAEQF | 0.7085154 |
| VGPKRFGV | ASSFLQGGAEQF | 0.7408254 |
| VNVAFPD | ASSFLQGGAEQF | 0.7565906 |
| VSMTNSGVS | ASSFLQGGAEQF | 0.6837766 |
| VTDVLGDLSV | ASSFLQGGAEQF | 0.80715317 |
| VVVVGAGFAGL | ASSFLQGGAEQF | 0.49730247 |
| VYTVGTVV | ASSFLQGGAEQF | 0.76983345 |
| WTTFSGP | ASSFLQGGAEQF | 0.69853234 |
| YDELVEAF | ASSFLQGGAEQF | 0.4849637 |
| YGTGGAGGNGG | ASSFLQGGAEQF | 0.6647291 |
| YTNVMAV | ASSFLQGGAEQF | 0.7675349 |
| FLEQVHQL | ASSFLQGGAEQF | 0.79319537 |
| GLSLLAVAV | ASSFLQGGAEQF | 0.69612986 |
| LEQVHQL | ASSFLQGGAEQF | 0.28855085 |
| LNAAGAI | ASSFLQGGAEQF | 0.78539133 |
| PAVHASL | ASSFLQGGAEQF | 0.67031425 |
| TELEDDA | ASSFLQGGAEQF | 0.834184 |
| TPPGPPN | ASSFLQGGAEQF | 0.6272828 |
| TPPPPPPP | ASSFLQGGAEQF | 0.5427538 |
| TQSPALI | GARDRDRGLDSDYT | 0.6581715 |
| EFAAAFPQ | GARDRDRGLDSDYT | 0.72656095 |

| | | |
|---|---|---|
| LLSGNLDVL | GARDRDRGLDSDYT | 0.3926804 |
| GPANLGNYNI | GARDRDRGLDSDYT | 0.5807828 |
| TLGPTLP | GARDRDRGLDSDYT | 0.72055924 |
| VYAYSSCCK | GARDRDRGLDSDYT | 0.6961503 |
| APAPAPAPNI | GARDRDRGLDSDYT | 0.81273794 |
| PGGPVPIP | GARDRDRGLDSDYT | 0.8045942 |
| FLTVALEP | GARDRDRGLDSDYT | 0.7111081 |
| SQYFGANL | GARDRDRGLDSDYT | 0.7886768 |
| GGVKPAA | GARDRDRGLDSDYT | 0.829625 |
| GGTGGAGGAGGW | GARDRDRGLDSDYT | 0.78238744 |
| GVGGVAGLGGAGG | GARDRDRGLDSDYT | 0.666316 |
| VLLVLATTI | GARDRDRGLDSDYT | 0.5248556 |
| IGIFAAML | GARDRDRGLDSDYT | 0.5202974 |
| AVGDLID | GARDRDRGLDSDYT | 0.5776996 |
| NISPLPL | GARDRDRGLDSDYT | 0.6756605 |
| VTAVFVAI | GARDRDRGLDSDYT | 0.7159107 |
| SGDDNDAAL | GARDRDRGLDSDYT | 0.16788341 |
| AVAALAHS | GARDRDRGLDSDYT | 0.79226696 |
| GGVTVMATP | GARDRDRGLDSDYT | 0.7918314 |
| AGLAANVRD | GARDRDRGLDSDYT | 0.70382524 |
| TNRPDLIDPALLRPG | GARDRDRGLDSDYT | 0.70895815 |
| LANLGVTAL | GARDRDRGLDSDYT | 0.7982094 |
| TGERIEL | GARDRDRGLDSDYT | 0.58287126 |
| NGEELAA | GARDRDRGLDSDYT | 0.73125196 |
| VLSLVWG | GARDRDRGLDSDYT | 0.6077664 |
| IGYALWL | GARDRDRGLDSDYT | 0.77723455 |
| LSLLLAPL | GARDRDRGLDSDYT | 0.7927887 |
| TLDEALK | GARDRDRGLDSDYT | 0.74559754 |
| TDGAGTAEGG | GARDRDRGLDSDYT | 0.5082343 |
| NAPGGAGGNGG | GARDRDRGLDSDYT | 0.5589402 |
| VSVFMGMGGNF | GARDRDRGLDSDYT | 0.353687 |
| LASYLAAL | GARDRDRGLDSDYT | 0.56245303 |
| LVGGTPAL | GARDRDRGLDSDYT | 0.7638382 |
| VRNADVL | GARDRDRGLDSDYT | 0.7675097 |
| TCSSPFLVSAM | GARDRDRGLDSDYT | 0.53103083 |
| VHITDAE | GARDRDRGLDSDYT | 0.74191 |
| EYAAPSF | GARDRDRGLDSDYT | 0.72672087 |
| VVDAFHGGL | GARDRDRGLDSDYT | 0.67705095 |
| ASVVDPQ | GARDRDRGLDSDYT | 0.58656204 |
| SYNEFGGLS | GARDRDRGLDSDYT | 0.80705804 |
| DPDGSVY | GARDRDRGLDSDYT | 0.7670481 |
| MVSTIHATE | GARDRDRGLDSDYT | 0.31326562 |
| LAAGVADPV | GARDRDRGLDSDYT | 0.8108369 |
| TKEQIAA | GARDRDRGLDSDYT | 0.6125583 |
| SLASPAAL | GARDRDRGLDSDYT | 0.74835527 |
| IGLVFYGV | GARDRDRGLDSDYT | 0.74265397 |
| FLLDASNL | GARDRDRGLDSDYT | 0.65112185 |
| YQVGNFP | GARDRDRGLDSDYT | 0.46406198 |
| ESLLSQST | GARDRDRGLDSDYT | 0.6103442 |
| LFTVGFAAVG | GARDRDRGLDSDYT | 0.7171734 |
| ATPPTPT | GARDRDRGLDSDYT | 0.6698874 |

| | | |
|---|---|---|
| TDLATLG | GARDRDRGLDSDYT | 0.25894493 |
| SAGGTGR | GARDRDRGLDSDYT | 0.7662525 |
| GFSNSGS | GARDRDRGLDSDYT | 0.38040745 |
| GGNGGSGGTSV | GARDRDRGLDSDYT | 0.6654383 |
| GTSVATGGAGN | GARDRDRGLDSDYT | 0.7313421 |
| NGGNGGSGGTS | GARDRDRGLDSDYT | 0.8005631 |
| AVVKFPH | GARDRDRGLDSDYT | 0.70549524 |
| TPPPPPP | GARDRDRGLDSDYT | 0.3849876 |
| AAPAAAAAPA | GARDRDRGLDSDYT | 0.784088 |
| IALNEDL | GARDRDRGLDSDYT | 0.6073298 |
| IVNVSSM | GARDRDRGLDSDYT | 0.6973474 |
| TGIPPAPR | GARDRDRGLDSDYT | 0.6862283 |
| FDLGGGTF | GARDRDRGLDSDYT | 0.52297246 |
| FDLGGGTFDVS | GARDRDRGLDSDYT | 0.59901583 |
| NEPTAAA | GARDRDRGLDSDYT | 0.37473622 |
| NEPTAAALA | GARDRDRGLDSDYT | 0.6721951 |
| GSIINISSI | GARDRDRGLDSDYT | 0.63005316 |
| VAPGNAGTA | GARDRDRGLDSDYT | 0.44459838 |
| TGGGASLE | GARDRDRGLDSDYT | 0.48026332 |
| VAVYNHYKRI | GARDRDRGLDSDYT | 0.7120212 |
| TDPAPATT | GARDRDRGLDSDYT | 0.79563016 |
| TNRPDILDPALLRPG | GARDRDRGLDSDYT | 0.52439404 |
| LTTTTTS | GARDRDRGLDSDYT | 0.30077592 |
| PTGSGKTL | GARDRDRGLDSDYT | 0.56128144 |
| TGSGKTL | GARDRDRGLDSDYT | 0.6473576 |
| TIGHVDHGK | GARDRDRGLDSDYT | 0.79312587 |
| TIGHVDHGKTT | GARDRDRGLDSDYT | 0.7088289 |
| GHVDHGKTT | GARDRDRGLDSDYT | 0.82516944 |
| AQFEHTLL | GARDRDRGLDSDYT | 0.7771776 |
| IVNQDEL | GARDRDRGLDSDYT | 0.654707 |
| TNGSQFF | GARDRDRGLDSDYT | 0.8110964 |
| TGKGIVE | GARDRDRGLDSDYT | 0.68087876 |
| LISNASDA | GARDRDRGLDSDYT | 0.7249915 |
| AELRTLGELTKIAKAHGAQ | GARDRDRGLDSDYT | 0.75146794 |
| APNAVAAAEPSW | GARDRDRGLDSDYT | 0.7642096 |
| APQVAGAATAVL | GARDRDRGLDSDYT | 0.46304047 |
| ARRDPAAAAAILPTDA | GARDRDRGLDSDYT | 0.62956864 |
| DAAARDVLAEAGLAEYF | GARDRDRGLDSDYT | 0.3438733 |
| DDQIVLSAKANLAKVEQ | GARDRDRGLDSDYT | 0.7884736 |
| DELLPDTGEDSDEDGH | GARDRDRGLDSDYT | 0.43201312 |
| ELPEAPANRATVPQD | GARDRDRGLDSDYT | 0.42709252 |
| FGQSKIERVTGIG | GARDRDRGLDSDYT | 0.647598 |
| GAGGTGSTAGGAGGAGGAGGLY | GARDRDRGLDSDYT | 0.81135833 |
| GHGGTGAAGGNGGTGGAGGSGI | GARDRDRGLDSDYT | 0.7632301 |
| GNGGNGGAGGEGGPGGAGGAGA | GARDRDRGLDSDYT | 0.6481962 |
| GSTAGNGGNGGNATNGTIAGQP | GARDRDRGLDSDYT | 0.5835006 |
| GTGGDGGNGGNGGTGG | GARDRDRGLDSDYT | 0.8010478 |
| GVRRPAGIPADAPV | GARDRDRGLDSDYT | 0.5837868 |
| GYRQVTDAHLLTLARR | GARDRDRGLDSDYT | 0.782719 |
| IAKMGQDGHDRG | GARDRDRGLDSDYT | 0.74721056 |
| KAEPSIEIQYT | GARDRDRGLDSDYT | 0.71033156 |

| | | |
|---|---|---|
| KPVRIRMAADEDTVAVT | GARDRDRGLDSDYT | 0.2745133 |
| MPPRIAGMRLLVIKPEP | GARDRDRGLDSDYT | 0.76250434 |
| MKFYAVALFRSRGVPPTR | GARDRDRGLDSDYT | 0.66657 |
| NAGGFGILGAVAHSP | GARDRDRGLDSDYT | 0.6789732 |
| NEGREITLAMAEM | GARDRDRGLDSDYT | 0.4520399 |
| NELVEWSHPLGHRWRWE | GARDRDRGLDSDYT | 0.7528907 |
| NLGAGGTGGNGGNPG | GARDRDRGLDSDYT | 0.55057347 |
| NTEWPKWETVMAADPVD | GARDRDRGLDSDYT | 0.47979093 |
| PASAAPAETQP | GARDRDRGLDSDYT | 0.20802605 |
| QQSSDGKNGVRAAEITGE | GARDRDRGLDSDYT | 0.5957457 |
| RAPDAVTTSWWKE | GARDRDRGLDSDYT | 0.7684268 |
| SGFGNVGAHTSGWFNQ | GARDRDRGLDSDYT | 0.82065606 |
| SGVAGAVGQAASVGGLKV | GARDRDRGLDSDYT | 0.7858049 |
| SNGGAGTGSAGGNGGT | GARDRDRGLDSDYT | 0.68683803 |
| TGGAGGEGGAAG | GARDRDRGLDSDYT | 0.7989959 |
| TGGDGGAAGAGGAPGHG | GARDRDRGLDSDYT | 0.6949452 |
| TMPPLLAAIANAGRHGV | GARDRDRGLDSDYT | 0.28116947 |
| TSSTAGNSGGA | GARDRDRGLDSDYT | 0.58825517 |
| VPATVPVGILC | GARDRDRGLDSDYT | 0.716593 |
| VQRPDSGGVVLTGRIS | GARDRDRGLDSDYT | 0.8010603 |
| WTPNPVPATKGVPAGLVQAT | GARDRDRGLDSDYT | 0.75937593 |
| YALKRALAKFSDRFDVVII | GARDRDRGLDSDYT | 0.5458583 |
| YLIGSVIV | GARDRDRGLDSDYT | 0.51028955 |
| YGQLVAVID | GARDRDRGLDSDYT | 0.74778926 |
| VTVGGMTLPA | GARDRDRGLDSDYT | 0.3423322 |
| VLALPAV | GARDRDRGLDSDYT | 0.55103326 |
| VIPPTLNYET | GARDRDRGLDSDYT | 0.43244043 |
| VGMGGMHPSSG | GARDRDRGLDSDYT | 0.54155767 |
| VGLAGGAATA | GARDRDRGLDSDYT | 0.6255733 |
| VGGAGGAGGSGP | GARDRDRGLDSDYT | 0.70200694 |
| VFAAFMV | GARDRDRGLDSDYT | 0.4554835 |
| TLYDKLR | GARDRDRGLDSDYT | 0.6992297 |
| TLAVFTVL | GARDRDRGLDSDYT | 0.3966462 |
| TGSVQPARV | GARDRDRGLDSDYT | 0.53061736 |
| TGGGAGSGGT | GARDRDRGLDSDYT | 0.5796772 |
| SHLETSL | GARDRDRGLDSDYT | 0.6751784 |
| SFTTGMT | GARDRDRGLDSDYT | 0.7923381 |
| QVEVAKRMGVRQPTVS | GARDRDRGLDSDYT | 0.5911206 |
| QGSGGPALAM | GARDRDRGLDSDYT | 0.61661065 |
| MLAPSNP | GARDRDRGLDSDYT | 0.8244583 |
| LVNNAAI | GARDRDRGLDSDYT | 0.2837439 |
| LTGEINTLA | GARDRDRGLDSDYT | 0.4480924 |
| LSIFGPV | GARDRDRGLDSDYT | 0.73305494 |
| LLALLANL | GARDRDRGLDSDYT | 0.72631216 |
| LISNASDALDKLR | GARDRDRGLDSDYT | 0.774928 |
| LINEAVP | GARDRDRGLDSDYT | 0.7172761 |
| LGYAFSGL | GARDRDRGLDSDYT | 0.7707158 |
| LDFVAVAL | GARDRDRGLDSDYT | 0.7175188 |
| LAQPGAE | GARDRDRGLDSDYT | 0.817841 |
| ISNLGAVTGF | GARDRDRGLDSDYT | 0.76087826 |
| HQEDPGRI | GARDRDRGLDSDYT | 0.6176557 |

| | | |
|---|---|---|
| GTPGEPTP | GARDRDRGLDSDYT | 0.80604154 |
| GFFNSGS | GARDRDRGLDSDYT | 0.27623594 |
| GAGGGGLTAGAN | GARDRDRGLDSDYT | 0.69604707 |
| FVDALVNM | GARDRDRGLDSDYT | 0.7645303 |
| FLTVAPEL | GARDRDRGLDSDYT | 0.62562823 |
| FETLTAL | GARDRDRGLDSDYT | 0.72089493 |
| DVINPIEL | GARDRDRGLDSDYT | 0.81530666 |
| APVSGAPV | GARDRDRGLDSDYT | 0.8046409 |
| ALVGGLAW | GARDRDRGLDSDYT | 0.56283164 |
| ALAAAAATAI | GARDRDRGLDSDYT | 0.64785945 |
| AFGLGLLL | GARDRDRGLDSDYT | 0.39722848 |
| AFFAAASL | GARDRDRGLDSDYT | 0.37541187 |
| AAAALPAFSPP | GARDRDRGLDSDYT | 0.26673728 |
| VVLAPAL | GARDRDRGLDSDYT | 0.7612828 |
| TIGHVDHGKTTL | GARDRDRGLDSDYT | 0.6646815 |
| LISNASDALDK | GARDRDRGLDSDYT | 0.8003756 |
| LISNASDAL | GARDRDRGLDSDYT | 0.77439713 |
| ISNASDALDK | GARDRDRGLDSDYT | 0.80436176 |
| NQAETLVYQTE | GARDRDRGLDSDYT | 0.606802 |
| TGGTGGTAGAG | GARDRDRGLDSDYT | 0.6638583 |
| TGSTAGGAGGA | GARDRDRGLDSDYT | 0.28740796 |
| DPLAVPVAAGPV | GARDRDRGLDSDYT | 0.51419604 |
| GGTGGTPGNGGN | GARDRDRGLDSDYT | 0.39980844 |
| PAGLAGVDPATL | GARDRDRGLDSDYT | 0.60216653 |
| GNNLVVEAPHAGASV | GARDRDRGLDSDYT | 0.3315556 |
| KVTRIMSASTLEQPA | GARDRDRGLDSDYT | 0.8015245 |
| LGFALPIALPRRTVV | GARDRDRGLDSDYT | 0.6623279 |
| TRLGNLTARKLVVLV | GARDRDRGLDSDYT | 0.77371055 |
| WSQQRRLIYFPSAGP | GARDRDRGLDSDYT | 0.47368163 |
| GPNGNPDPMAAAVDIR | GARDRDRGLDSDYT | 0.42226762 |
| TQRCVSVADLRAAASR | GARDRDRGLDSDYT | 0.72351074 |
| QFVLDVAEHPVEVLTETV | GARDRDRGLDSDYT | 0.73634887 |
| GGAGGTGGAGGNGGTQAGDG | GARDRDRGLDSDYT | 0.47925985 |
| PLLATLRAIAVDVDAKPAQV | GARDRDRGLDSDYT | 0.35588086 |
| APKFRTGGYMIDFWGVGYQVAK | GARDRDRGLDSDYT | 0.368873 |
| GAGGAGGAGGAAGGPGGTGGTG | GARDRDRGLDSDYT | 0.6619876 |
| IGALGLLATDVAPAGSVAFGVA | GARDRDRGLDSDYT | 0.83141 |
| AGGAGGIGGNGGAGGAGGVLMG | GARDRDRGLDSDYT | 0.8096422 |
| ADELIGLL | GARDRDRGLDSDYT | 0.7586045 |
| ADRALKA | GARDRDRGLDSDYT | 0.83427936 |
| AGAVGQVGGAGGA | GARDRDRGLDSDYT | 0.16360633 |
| AGGGGGLGGAP | GARDRDRGLDSDYT | 0.53532636 |
| AGGLLYGNGGAG | GARDRDRGLDSDYT | 0.73208225 |
| AGLESIGL | GARDRDRGLDSDYT | 0.72746515 |
| AGLKTGII | GARDRDRGLDSDYT | 0.7008205 |
| ASLAALLL | GARDRDRGLDSDYT | 0.8111175 |
| ASVINISSI | GARDRDRGLDSDYT | 0.4737577 |
| ATPPGPA | GARDRDRGLDSDYT | 0.5992301 |
| ATVASALV | GARDRDRGLDSDYT | 0.7272632 |
| ATVASLIL | GARDRDRGLDSDYT | 0.79700935 |
| AVNGPVLIRH | GARDRDRGLDSDYT | 0.7721939 |

| | | |
|---|---|---|
| DGAAGAVGI | GARDRDRGLDSDYT | 0.3753702 |
| DPLLHGI | GARDRDRGLDSDYT | 0.5935771 |
| ELGAALGAL | GARDRDRGLDSDYT | 0.8404262 |
| ELGRTQAV | GARDRDRGLDSDYT | 0.68963003 |
| FVAGGYLS | GARDRDRGLDSDYT | 0.7659414 |
| GASGGAGGAGGW | GARDRDRGLDSDYT | 0.63743776 |
| GGAGAGGDGGAGGI | GARDRDRGLDSDYT | 0.5919516 |
| GTGHNGSQPGMGGNG | GARDRDRGLDSDYT | 0.6301712 |
| GYAAASASAS | GARDRDRGLDSDYT | 0.49089092 |
| HQVDVAAVQQG | GARDRDRGLDSDYT | 0.79398257 |
| IGAAIAHT | GARDRDRGLDSDYT | 0.4985345 |
| IGMIEVAQL | GARDRDRGLDSDYT | 0.58967525 |
| IRLAPEHTSAMD | GARDRDRGLDSDYT | 0.6852713 |
| IYNVSVL | GARDRDRGLDSDYT | 0.84026194 |
| KQGFLGM | GARDRDRGLDSDYT | 0.7581714 |
| LESVITQL | GARDRDRGLDSDYT | 0.73864543 |
| LGLGALVVV | GARDRDRGLDSDYT | 0.65058845 |
| LIGNGGNGGSGGI | GARDRDRGLDSDYT | 0.70987797 |
| LNAINAPTL | GARDRDRGLDSDYT | 0.7953208 |
| LQLLNTGV | GARDRDRGLDSDYT | 0.3569223 |
| LQTSIEF | GARDRDRGLDSDYT | 0.7793241 |
| MSFAVSEL | GARDRDRGLDSDYT | 0.45525476 |
| NTGALGL | GARDRDRGLDSDYT | 0.73056424 |
| PPGAGGAGGAGTT | GARDRDRGLDSDYT | 0.48811254 |
| QLVGLGI | GARDRDRGLDSDYT | 0.8154302 |
| RFLEQVHQL | GARDRDRGLDSDYT | 0.7724421 |
| RGVGYVLR | GARDRDRGLDSDYT | 0.8002658 |
| SATGVFAGVI | GARDRDRGLDSDYT | 0.60122633 |
| SLPAITLP | GARDRDRGLDSDYT | 0.7687552 |
| STALAFTV | GARDRDRGLDSDYT | 0.82392514 |
| SVLTVGLGK | GARDRDRGLDSDYT | 0.76433337 |
| TALAGAGSGI | GARDRDRGLDSDYT | 0.8367516 |
| TGLPFTGI | GARDRDRGLDSDYT | 0.6875498 |
| VAEHVTV | GARDRDRGLDSDYT | 0.22208634 |
| VDIVLNSL | GARDRDRGLDSDYT | 0.66809285 |
| VLTGGGAVL | GARDRDRGLDSDYT | 0.78848743 |
| VNVGLAYGI | GARDRDRGLDSDYT | 0.8206463 |
| VSSAAVGAV | GARDRDRGLDSDYT | 0.6633772 |
| VTSVYEAGV | GARDRDRGLDSDYT | 0.7169664 |
| WVALDRGA | GARDRDRGLDSDYT | 0.68807614 |
| YLVGSVVI | GARDRDRGLDSDYT | 0.8203452 |
| GPPGTGKTLL | GARDRDRGLDSDYT | 0.8047645 |
| GVGMTKF | GARDRDRGLDSDYT | 0.69400525 |
| IGIGGSDL | GARDRDRGLDSDYT | 0.4024254 |
| IGTLLAEL | GARDRDRGLDSDYT | 0.2893088 |
| IYTSGTT | GARDRDRGLDSDYT | 0.7664685 |
| LSFPAGV | GARDRDRGLDSDYT | 0.815463 |
| LYTSGTT | GARDRDRGLDSDYT | 0.37344587 |
| VGAFPFGF | GARDRDRGLDSDYT | 0.7017803 |
| VGAGTVEF | GARDRDRGLDSDYT | 0.72210914 |
| VGKSTLLL | GARDRDRGLDSDYT | 0.2304604 |

| | | |
|---|---|---|
| AALAHAGGGIV | GARDRDRGLDSDYT | 0.7186563 |
| AALVAVSLL | GARDRDRGLDSDYT | 0.81201386 |
| AAVSRAVAAL | GARDRDRGLDSDYT | 0.7405918 |
| AGGSSGIGGF | GARDRDRGLDSDYT | 0.29828653 |
| AGLTVANANG | GARDRDRGLDSDYT | 0.79667854 |
| AGLVGAVL | GARDRDRGLDSDYT | 0.83316475 |
| ALSQAFTV | GARDRDRGLDSDYT | 0.8215907 |
| ALTFSVSL | GARDRDRGLDSDYT | 0.26018345 |
| ALTGAALVAL | GARDRDRGLDSDYT | 0.2007214 |
| ASFSIGAGVL | GARDRDRGLDSDYT | 0.7424475 |
| DGSVYKI | GARDRDRGLDSDYT | 0.7842922 |
| EQLAEIV | GARDRDRGLDSDYT | 0.5157163 |
| FAVSELTVM | GARDRDRGLDSDYT | 0.5128896 |
| GAAPLAAGL | GARDRDRGLDSDYT | 0.7474421 |
| GADGGDGGSGGAGG | GARDRDRGLDSDYT | 0.80705774 |
| GFGLDTGGAGG | GARDRDRGLDSDYT | 0.4043383 |
| GLASGGLL | GARDRDRGLDSDYT | 0.30759412 |
| GRIAGAAVLL | GARDRDRGLDSDYT | 0.774395 |
| GSAGSLNTSF | GARDRDRGLDSDYT | 0.79651916 |
| GTGQIGI | GARDRDRGLDSDYT | 0.5455083 |
| GTLIALIL | GARDRDRGLDSDYT | 0.7249089 |
| IAATMGFVVG | GARDRDRGLDSDYT | 0.7455856 |
| IFAQALL | GARDRDRGLDSDYT | 0.77448964 |
| IGAAVALGF | GARDRDRGLDSDYT | 0.7581721 |
| IGLTPLAL | GARDRDRGLDSDYT | 0.38074905 |
| ILGSNTSTL | GARDRDRGLDSDYT | 0.800045 |
| IVFSLTVL | GARDRDRGLDSDYT | 0.6005542 |
| LASKATGF | GARDRDRGLDSDYT | 0.7769966 |
| LASVALIL | GARDRDRGLDSDYT | 0.69770753 |
| LFAVGLTL | GARDRDRGLDSDYT | 0.77341473 |
| LFNGGIGTYI | GARDRDRGLDSDYT | 0.72016245 |
| LTLGAGII | GARDRDRGLDSDYT | 0.39320546 |
| LVAAALGDI | GARDRDRGLDSDYT | 0.4996563 |
| NTAFVQLGI | GARDRDRGLDSDYT | 0.56733155 |
| QLFSGAEL | GARDRDRGLDSDYT | 0.79527164 |
| SAATFITL | GARDRDRGLDSDYT | 0.7925433 |
| SNYGLANLGV | GARDRDRGLDSDYT | 0.8089384 |
| SYGGAYAVM | GARDRDRGLDSDYT | 0.72095346 |
| TTGVGGLTLG | GARDRDRGLDSDYT | 0.5715409 |
| TTQTIYI | GARDRDRGLDSDYT | 0.77967095 |
| VAAGAGLGAVI | GARDRDRGLDSDYT | 0.38206074 |
| VAAVVTLL | GARDRDRGLDSDYT | 0.76828027 |
| VGFLGGLI | GARDRDRGLDSDYT | 0.5489994 |
| VGVIPILL | GARDRDRGLDSDYT | 0.68861103 |
| VIFASSLI | GARDRDRGLDSDYT | 0.6311697 |
| VKGGDGGAAAG | GARDRDRGLDSDYT | 0.77589864 |
| VTKTFTNI | GARDRDRGLDSDYT | 0.76621443 |
| VVAAAILL | GARDRDRGLDSDYT | 0.30461684 |
| YAALDVEL | GARDRDRGLDSDYT | 0.64871705 |
| LASVALLL | GARDRDRGLDSDYT | 0.79635763 |
| LDAVHRV | GARDRDRGLDSDYT | 0.7209451 |

| | | |
|---|---|---|
| QGLSSSIF | GARDRDRGLDSDYT | 0.38382497 |
| RGTFANIRL | GARDRDRGLDSDYT | 0.5962453 |
| RQLWWGHRI | GARDRDRGLDSDYT | 0.6258502 |
| YGPPGTGKTLL | GARDRDRGLDSDYT | 0.78393304 |
| AALVGAPANVA | GARDRDRGLDSDYT | 0.75897473 |
| AATFRRTE | GARDRDRGLDSDYT | 0.4121494 |
| AGALGGGI | GARDRDRGLDSDYT | 0.7086309 |
| AGGAGGAGGIGAG | GARDRDRGLDSDYT | 0.23911461 |
| AGSGALGL | GARDRDRGLDSDYT | 0.6269847 |
| AQIADQI | GARDRDRGLDSDYT | 0.6934827 |
| ASAVAVTYL | GARDRDRGLDSDYT | 0.7155918 |
| ASTVLGGTGG | GARDRDRGLDSDYT | 0.71549726 |
| AVGGEPAK | GARDRDRGLDSDYT | 0.48308626 |
| CEQGTFGGTC | GARDRDRGLDSDYT | 0.72169256 |
| DEGKAPAG | GARDRDRGLDSDYT | 0.53121847 |
| DTVVFPAL | GARDRDRGLDSDYT | 0.46707278 |
| FEGITMAG | GARDRDRGLDSDYT | 0.7478751 |
| FPLVVQD | GARDRDRGLDSDYT | 0.5026199 |
| GAFTTPQ | GARDRDRGLDSDYT | 0.25742754 |
| GFGGFGTAGGVGGA | GARDRDRGLDSDYT | 0.7875761 |
| GFLAQFL | GARDRDRGLDSDYT | 0.49913985 |
| GGAGGAGGNAGVA | GARDRDRGLDSDYT | 0.54305315 |
| GGVGTAGAGGAGGAGG | GARDRDRGLDSDYT | 0.6803489 |
| GNGGGGDAGNAGSGGN | GARDRDRGLDSDYT | 0.30894122 |
| GTGGAGGAGNDAGS | GARDRDRGLDSDYT | 0.7732147 |
| GTTNTGWFNS | GARDRDRGLDSDYT | 0.6527977 |
| IFMGPLFTL | GARDRDRGLDSDYT | 0.706476 |
| ILAHAIEAI | GARDRDRGLDSDYT | 0.66220915 |
| IRSAIREAL | GARDRDRGLDSDYT | 0.49018025 |
| ITFAGGG | GARDRDRGLDSDYT | 0.80219257 |
| ITGMSGHQEV | GARDRDRGLDSDYT | 0.6586794 |
| IVAGLACTL | GARDRDRGLDSDYT | 0.42855632 |
| IVGGGLAGA | GARDRDRGLDSDYT | 0.7198597 |
| IVTGVLNI | GARDRDRGLDSDYT | 0.80415314 |
| LAQQGCI | GARDRDRGLDSDYT | 0.79502034 |
| LCLHVEN | GARDRDRGLDSDYT | 0.6565696 |
| LDAISDGS | GARDRDRGLDSDYT | 0.29824227 |
| LGGFAAML | GARDRDRGLDSDYT | 0.7057476 |
| LGGKNPMI | GARDRDRGLDSDYT | 0.7324171 |
| LGPKGVAAIL | GARDRDRGLDSDYT | 0.5610634 |
| LMVGVSI | GARDRDRGLDSDYT | 0.2462766 |
| LVGAAFVL | GARDRDRGLDSDYT | 0.6927368 |
| LVLAVAPL | GARDRDRGLDSDYT | 0.833244 |
| MAAIAVIDAA | GARDRDRGLDSDYT | 0.42503995 |
| NNGINAELQ | GARDRDRGLDSDYT | 0.83522105 |
| RGEPLVLVHG | GARDRDRGLDSDYT | 0.44130665 |
| RGGAGGAAGLI | GARDRDRGLDSDYT | 0.6460391 |
| RSYRMHPAV | GARDRDRGLDSDYT | 0.72356707 |
| SAAHLLAGQLA | GARDRDRGLDSDYT | 0.42180517 |
| SAAVGSY | GARDRDRGLDSDYT | 0.55920315 |
| SPHAVLHRI | GARDRDRGLDSDYT | 0.61155415 |

| | | |
|---|---|---|
| SQIAKVL | GARDRDRGLDSDYT | 0.47927183 |
| TALGHLVP | GARDRDRGLDSDYT | 0.78470254 |
| TAPTTSGAA | GARDRDRGLDSDYT | 0.41702348 |
| TDGDDGNGGSPGT | GARDRDRGLDSDYT | 0.66600776 |
| TDGLGEN | GARDRDRGLDSDYT | 0.5814477 |
| TGGPPLPGQA | GARDRDRGLDSDYT | 0.5505271 |
| TIPAITI | GARDRDRGLDSDYT | 0.78995955 |
| TIPGFTI | GARDRDRGLDSDYT | 0.6584934 |
| TSIASGALA | GARDRDRGLDSDYT | 0.7388047 |
| TVTLPGAAI | GARDRDRGLDSDYT | 0.7779815 |
| VASGLSAALP | GARDRDRGLDSDYT | 0.65528506 |
| VGAGVIGI | GARDRDRGLDSDYT | 0.8329052 |
| VGPKRFGV | GARDRDRGLDSDYT | 0.821158 |
| VNVAFPD | GARDRDRGLDSDYT | 0.77935183 |
| VSMTNSGVS | GARDRDRGLDSDYT | 0.6528348 |
| VTDVLGDLSV | GARDRDRGLDSDYT | 0.7967293 |
| VVVVGAGFAGL | GARDRDRGLDSDYT | 0.50748813 |
| VYTVGTVV | GARDRDRGLDSDYT | 0.6640585 |
| WTTFSGP | GARDRDRGLDSDYT | 0.79903233 |
| YDELVEAF | GARDRDRGLDSDYT | 0.60696536 |
| YGTGGAGGNGG | GARDRDRGLDSDYT | 0.72211045 |
| YTNVMAV | GARDRDRGLDSDYT | 0.6832928 |
| FLEQVHQL | GARDRDRGLDSDYT | 0.66669345 |
| GLSLLAVAV | GARDRDRGLDSDYT | 0.6160982 |
| LEQVHQL | GARDRDRGLDSDYT | 0.5389855 |
| LNAAGAI | GARDRDRGLDSDYT | 0.623311 |
| PAVHASL | GARDRDRGLDSDYT | 0.60324454 |
| TELEDDA | GARDRDRGLDSDYT | 0.69083554 |
| TPPGPPN | GARDRDRGLDSDYT | 0.71146643 |
| TPPPPPPP | GARDRDRGLDSDYT | 0.6107376 |
| TQSPALI | ASRDSNYAEQF | 0.7296141 |
| EFAAAFPQ | ASRDSNYAEQF | 0.720478 |
| LLSGNLDVL | ASRDSNYAEQF | 0.49030906 |
| GPANLGNYNI | ASRDSNYAEQF | 0.64358646 |
| TLGPTLP | ASRDSNYAEQF | 0.5850141 |
| VYAYSSCCK | ASRDSNYAEQF | 0.6570357 |
| APAPAPAPNI | ASRDSNYAEQF | 0.3774721 |
| PGGPVPIP | ASRDSNYAEQF | 0.70395386 |
| FLTVALEP | ASRDSNYAEQF | 0.56044346 |
| SQYFGANL | ASRDSNYAEQF | 0.2889704 |
| GGVKPAA | ASRDSNYAEQF | 0.69193476 |
| GGTGGAGGAGGW | ASRDSNYAEQF | 0.65839964 |
| GVGGVAGLGGAGG | ASRDSNYAEQF | 0.67891335 |
| VLLVLATTI | ASRDSNYAEQF | 0.7041131 |
| IGIFAAML | ASRDSNYAEQF | 0.5691789 |
| AVGDLID | ASRDSNYAEQF | 0.6265062 |
| NISPLPL | ASRDSNYAEQF | 0.69412756 |
| VTAVFVAI | ASRDSNYAEQF | 0.7190217 |
| SGDDNDAAL | ASRDSNYAEQF | 0.616359 |
| AVAALAHS | ASRDSNYAEQF | 0.66794324 |
| GGVTVMATP | ASRDSNYAEQF | 0.7217328 |

| | | |
|---|---|---|
| AGLAANVRD | ASRDSNYAEQF | 0.6894017 |
| TNRPDLIDPALLRPG | ASRDSNYAEQF | 0.71532595 |
| LANLGVTAL | ASRDSNYAEQF | 0.72799444 |
| TGERIEL | ASRDSNYAEQF | 0.5776833 |
| NGEELAA | ASRDSNYAEQF | 0.7173314 |
| VLSLVWG | ASRDSNYAEQF | 0.7328407 |
| IGYALWL | ASRDSNYAEQF | 0.6499586 |
| LSLLLAPL | ASRDSNYAEQF | 0.65759146 |
| TLDEALK | ASRDSNYAEQF | 0.5354246 |
| TDGAGTAEGG | ASRDSNYAEQF | 0.6638945 |
| NAPGGAGGNGG | ASRDSNYAEQF | 0.473168 |
| VSVFMGMGGNF | ASRDSNYAEQF | 0.66815275 |
| LASYLAAL | ASRDSNYAEQF | 0.72679484 |
| LVGGTPAL | ASRDSNYAEQF | 0.62687016 |
| VRNADVL | ASRDSNYAEQF | 0.5357535 |
| TCSSPFLVSAM | ASRDSNYAEQF | 0.51578265 |
| VHITDAE | ASRDSNYAEQF | 0.65875536 |
| EYAAPSF | ASRDSNYAEQF | 0.60855234 |
| VVDAFHGGL | ASRDSNYAEQF | 0.4265019 |
| ASVVDPQ | ASRDSNYAEQF | 0.63774705 |
| SYNEFGGLS | ASRDSNYAEQF | 0.71686924 |
| DPDGSVY | ASRDSNYAEQF | 0.71809727 |
| MVSTIHATE | ASRDSNYAEQF | 0.53323483 |
| LAAGVADPV | ASRDSNYAEQF | 0.7273749 |
| TKEQIAA | ASRDSNYAEQF | 0.6771436 |
| SLASPAAL | ASRDSNYAEQF | 0.7244743 |
| IGLVFYGV | ASRDSNYAEQF | 0.6335302 |
| FLLDASNL | ASRDSNYAEQF | 0.6005411 |
| YQVGNFP | ASRDSNYAEQF | 0.68830705 |
| ESLLSQST | ASRDSNYAEQF | 0.6209847 |
| LFTVGFAAVG | ASRDSNYAEQF | 0.65915453 |
| ATPPTPT | ASRDSNYAEQF | 0.7101821 |
| TDLATLG | ASRDSNYAEQF | 0.6891834 |
| SAGGTGR | ASRDSNYAEQF | 0.5973906 |
| GFSNSGS | ASRDSNYAEQF | 0.58583736 |
| GGNGGSGGTSV | ASRDSNYAEQF | 0.7244611 |
| GTSVATGGAGN | ASRDSNYAEQF | 0.54162896 |
| NGGNGGSGGTS | ASRDSNYAEQF | 0.62624437 |
| AVVKFPH | ASRDSNYAEQF | 0.6069239 |
| TPPPPPP | ASRDSNYAEQF | 0.65471613 |
| AAPAAAAAPA | ASRDSNYAEQF | 0.7149558 |
| IALNEDL | ASRDSNYAEQF | 0.70605636 |
| IVNVSSM | ASRDSNYAEQF | 0.6148957 |
| TGIPPAPR | ASRDSNYAEQF | 0.68082523 |
| FDLGGGTF | ASRDSNYAEQF | 0.7011244 |
| FDLGGGTFDVS | ASRDSNYAEQF | 0.6797035 |
| NEPTAAA | ASRDSNYAEQF | 0.7056881 |
| NEPTAAALA | ASRDSNYAEQF | 0.5920869 |
| GSIINISSI | ASRDSNYAEQF | 0.71766096 |
| VAPGNAGTA | ASRDSNYAEQF | 0.6919657 |
| TGGGASLE | ASRDSNYAEQF | 0.70897514 |

| | | |
|---|---|---|
| VAVYNHYKRI | ASRDSNYAEQF | 0.7136221 |
| TDPAPATT | ASRDSNYAEQF | 0.71021354 |
| TNRPDILDPALLRPG | ASRDSNYAEQF | 0.666436 |
| LTTTTTS | ASRDSNYAEQF | 0.4343352 |
| PTGSGKTL | ASRDSNYAEQF | 0.5095794 |
| TGSGKTL | ASRDSNYAEQF | 0.6808481 |
| TIGHVDHGK | ASRDSNYAEQF | 0.6738441 |
| TIGHVDHGKTT | ASRDSNYAEQF | 0.70514745 |
| GHVDHGKTT | ASRDSNYAEQF | 0.7070958 |
| AQFEHTLL | ASRDSNYAEQF | 0.59864515 |
| IVNQDEL | ASRDSNYAEQF | 0.72247654 |
| TNGSQFF | ASRDSNYAEQF | 0.7141296 |
| TGKGIVE | ASRDSNYAEQF | 0.44151586 |
| LISNASDA | ASRDSNYAEQF | 0.63804615 |
| AELRTLGELTKIAKAHGAQ | ASRDSNYAEQF | 0.7401958 |
| APNAVAAAEPSW | ASRDSNYAEQF | 0.65454787 |
| APQVAGAATAVL | ASRDSNYAEQF | 0.6314451 |
| ARRDPAAAAAILPTDA | ASRDSNYAEQF | 0.61860335 |
| DAAARDVLAEAGLAEYF | ASRDSNYAEQF | 0.71904415 |
| DDQIVLSAKANLAKVEQ | ASRDSNYAEQF | 0.6364134 |
| DELLPDTGEDSDEDGH | ASRDSNYAEQF | 0.7379961 |
| ELPEAPANRATVPQD | ASRDSNYAEQF | 0.37194556 |
| FGQSKIERVTGIG | ASRDSNYAEQF | 0.6135205 |
| GAGGTGSTAGGAGGAGGAGGLY | ASRDSNYAEQF | 0.59023905 |
| GHGGTGAAGGNGGTGGAGGSGI | ASRDSNYAEQF | 0.43424544 |
| GNGGNGGAGGEGGPGGAGGAGA | ASRDSNYAEQF | 0.7104518 |
| GSTAGNGGNGGNATNGTIAGQP | ASRDSNYAEQF | 0.6242649 |
| GTGGDGGNGGNGGTGG | ASRDSNYAEQF | 0.7050507 |
| GVRRPAGIPADAPV | ASRDSNYAEQF | 0.6425893 |
| GYRQVTDAHLLTLARR | ASRDSNYAEQF | 0.71992224 |
| IAKMGQDGHDRG | ASRDSNYAEQF | 0.64256287 |
| KAEPSIEIQYT | ASRDSNYAEQF | 0.6706679 |
| KPVRIRMAADEDTVAVT | ASRDSNYAEQF | 0.6329833 |
| MPPRIAGMRLLVIKPEP | ASRDSNYAEQF | 0.7058006 |
| MKFYAVALFRSRGVPPTR | ASRDSNYAEQF | 0.73029774 |
| NAGGFGILGAVAHSP | ASRDSNYAEQF | 0.7117932 |
| NEGREITLAMAEM | ASRDSNYAEQF | 0.57005095 |
| NELVEWSHPLGHRWRWE | ASRDSNYAEQF | 0.6682159 |
| NLGAGGTGGNGGNPG | ASRDSNYAEQF | 0.670338 |
| NTEWPKWETVMAADPVD | ASRDSNYAEQF | 0.43165776 |
| PASAAPAETQP | ASRDSNYAEQF | 0.38394207 |
| QQSSDGKNGVRAAEITGE | ASRDSNYAEQF | 0.65986454 |
| RAPDAVTTSWWKE | ASRDSNYAEQF | 0.59781855 |
| SGFGNVGAHTSGWFNQ | ASRDSNYAEQF | 0.7064056 |
| SGVAGAVGQAASVGGLKV | ASRDSNYAEQF | 0.69240403 |
| SNGGAGTGSAGGNGGT | ASRDSNYAEQF | 0.6333465 |
| TGGAGGEGGAAG | ASRDSNYAEQF | 0.68613946 |
| TGGDGGAAGAGGAPGHG | ASRDSNYAEQF | 0.6915801 |
| TMPPLLAAIANAGRHGV | ASRDSNYAEQF | 0.24986272 |
| TSSTAGNSGGA | ASRDSNYAEQF | 0.5522166 |
| VPATVPVGILC | ASRDSNYAEQF | 0.4984183 |

| | | |
|---|---|---|
| VQRPDSGGVVLTGRIS | ASRDSNYAEQF | 0.728504 |
| WTPNPVPATKGVPAGLVQAT | ASRDSNYAEQF | 0.6029581 |
| YALKRALAKFSDRFDVVII | ASRDSNYAEQF | 0.5518117 |
| YLIGSVIV | ASRDSNYAEQF | 0.6551484 |
| YGQLVAVID | ASRDSNYAEQF | 0.64642334 |
| VTVGGMTLPA | ASRDSNYAEQF | 0.64081335 |
| VLALPAV | ASRDSNYAEQF | 0.55866385 |
| VIPPTLNYET | ASRDSNYAEQF | 0.38794625 |
| VGMGGMHPSSG | ASRDSNYAEQF | 0.7136542 |
| VGLAGGAATA | ASRDSNYAEQF | 0.5789219 |
| VGGAGGAGGSGP | ASRDSNYAEQF | 0.63120276 |
| VFAAFMV | ASRDSNYAEQF | 0.42385653 |
| TLYDKLR | ASRDSNYAEQF | 0.64080936 |
| TLAVFTVL | ASRDSNYAEQF | 0.4965653 |
| TGSVQPARV | ASRDSNYAEQF | 0.7140219 |
| TGGGAGSGGT | ASRDSNYAEQF | 0.60169506 |
| SHLETSL | ASRDSNYAEQF | 0.6267086 |
| SFTTGMT | ASRDSNYAEQF | 0.7356789 |
| QVEVAKRMGVRQPTVS | ASRDSNYAEQF | 0.62751585 |
| QGSGGPALAM | ASRDSNYAEQF | 0.63215494 |
| MLAPSNP | ASRDSNYAEQF | 0.6760666 |
| LVNNAAI | ASRDSNYAEQF | 0.55274516 |
| LTGEINTLA | ASRDSNYAEQF | 0.6423545 |
| LSIFGPV | ASRDSNYAEQF | 0.6571589 |
| LLALLANL | ASRDSNYAEQF | 0.73406005 |
| LISNASDALDKLR | ASRDSNYAEQF | 0.6465738 |
| LINEAVP | ASRDSNYAEQF | 0.50870013 |
| LGYAFSGL | ASRDSNYAEQF | 0.67252886 |
| LDFVAVAL | ASRDSNYAEQF | 0.7289245 |
| LAQPGAE | ASRDSNYAEQF | 0.73221976 |
| ISNLGAVTGF | ASRDSNYAEQF | 0.58510077 |
| HQEDPGRI | ASRDSNYAEQF | 0.6314144 |
| GTPGEPTP | ASRDSNYAEQF | 0.60199666 |
| GFFNSGS | ASRDSNYAEQF | 0.6322334 |
| GAGGGGLTAGAN | ASRDSNYAEQF | 0.6120028 |
| FVDALVNM | ASRDSNYAEQF | 0.6117186 |
| FLTVAPEL | ASRDSNYAEQF | 0.7033536 |
| FETLTAL | ASRDSNYAEQF | 0.7240413 |
| DVINPIEL | ASRDSNYAEQF | 0.6525256 |
| APVSGAPV | ASRDSNYAEQF | 0.38750368 |
| ALVGGLAW | ASRDSNYAEQF | 0.642337 |
| ALAAAAATAI | ASRDSNYAEQF | 0.57344085 |
| AFGLGLLL | ASRDSNYAEQF | 0.60509455 |
| AFFAAASL | ASRDSNYAEQF | 0.64166766 |
| AAAALPAFSPP | ASRDSNYAEQF | 0.4339939 |
| VVLAPAL | ASRDSNYAEQF | 0.65539825 |
| TIGHVDHGKTTL | ASRDSNYAEQF | 0.73910874 |
| LISNASDALDK | ASRDSNYAEQF | 0.5971043 |
| LISNASDAL | ASRDSNYAEQF | 0.67955977 |
| ISNASDALDK | ASRDSNYAEQF | 0.7230599 |
| NQAETLVYQTE | ASRDSNYAEQF | 0.6933145 |

| | | |
|---|---|---|
| TGGTGGTAGAG | ASRDSNYAEQF | 0.71028495 |
| TGSTAGGAGGA | ASRDSNYAEQF | 0.6906072 |
| DPLAVPVAAGPV | ASRDSNYAEQF | 0.70670253 |
| GGTGGTPGNGGN | ASRDSNYAEQF | 0.6694238 |
| PAGLAGVDPATL | ASRDSNYAEQF | 0.689112 |
| GNNLVVEAPHAGASV | ASRDSNYAEQF | 0.65067524 |
| KVTRIMSASTLEQPA | ASRDSNYAEQF | 0.6421733 |
| LGFALPIALPRRTVV | ASRDSNYAEQF | 0.57611465 |
| TRLGNLTARKLVVLV | ASRDSNYAEQF | 0.71284676 |
| WSQQRRLIYFPSAGP | ASRDSNYAEQF | 0.62186706 |
| GPNGNPDPMAAAVDIR | ASRDSNYAEQF | 0.7086574 |
| TQRCVSVADLRAAASR | ASRDSNYAEQF | 0.4846504 |
| QFVLDVAEHPVEVLTETV | ASRDSNYAEQF | 0.6184229 |
| GGAGGTGGAGGNGGTQAGDG | ASRDSNYAEQF | 0.70401156 |
| PLLATLRAIAVDVDAKPAQV | ASRDSNYAEQF | 0.26905543 |
| APKFRTGGYMIDFWGVGYQVAK | ASRDSNYAEQF | 0.37504485 |
| GAGGAGGAGGAAGGPGGTGGTG | ASRDSNYAEQF | 0.6469205 |
| IGALGLLATDVAPAGSVAFGVA | ASRDSNYAEQF | 0.71666384 |
| AGGAGGIGGNGGAGGAGGVLMG | ASRDSNYAEQF | 0.2993913 |
| ADELIGLL | ASRDSNYAEQF | 0.6009611 |
| ADRALKA | ASRDSNYAEQF | 0.4549711 |
| AGAVGQVGGAGGA | ASRDSNYAEQF | 0.71811074 |
| AGGGGGLGGAP | ASRDSNYAEQF | 0.6805922 |
| AGGLLYGNGGAG | ASRDSNYAEQF | 0.42145 |
| AGLESIGL | ASRDSNYAEQF | 0.705615 |
| AGLKTGII | ASRDSNYAEQF | 0.40426093 |
| ASLAALLL | ASRDSNYAEQF | 0.62925744 |
| ASVINISSI | ASRDSNYAEQF | 0.71313286 |
| ATPPGPA | ASRDSNYAEQF | 0.7132164 |
| ATVASALV | ASRDSNYAEQF | 0.48834908 |
| ATVASLIL | ASRDSNYAEQF | 0.69252664 |
| AVNGPVLIRH | ASRDSNYAEQF | 0.59513783 |
| DGAAGAVGI | ASRDSNYAEQF | 0.7383005 |
| DPLLHGI | ASRDSNYAEQF | 0.64422333 |
| ELGAALGAL | ASRDSNYAEQF | 0.45696226 |
| ELGRTQAV | ASRDSNYAEQF | 0.71514714 |
| FVAGGYLS | ASRDSNYAEQF | 0.6859369 |
| GASGGAGGAGGW | ASRDSNYAEQF | 0.70159125 |
| GGAGAGGDGGAGGI | ASRDSNYAEQF | 0.68492794 |
| GTGHNGSQPGMGGNG | ASRDSNYAEQF | 0.62834024 |
| GYAAASASAS | ASRDSNYAEQF | 0.623469 |
| HQVDVAAVQQG | ASRDSNYAEQF | 0.7135328 |
| IGAAIAHT | ASRDSNYAEQF | 0.70899606 |
| IGMIEVAQL | ASRDSNYAEQF | 0.64415765 |
| IRLAPEHTSAMD | ASRDSNYAEQF | 0.72421825 |
| IYNVSVL | ASRDSNYAEQF | 0.65797967 |
| KQGFLGM | ASRDSNYAEQF | 0.6033262 |
| LESVITQL | ASRDSNYAEQF | 0.7312732 |
| LGLGALVVV | ASRDSNYAEQF | 0.64983463 |
| LIGNGGNGGSGGI | ASRDSNYAEQF | 0.7009685 |
| LNAINAPTL | ASRDSNYAEQF | 0.7118279 |

| | | |
|---|---|---|
| LQLLNTGV | ASRDSNYAEQF | 0.69560957 |
| LQTSIEF | ASRDSNYAEQF | 0.55362636 |
| MSFAVSEL | ASRDSNYAEQF | 0.6673112 |
| NTGALGL | ASRDSNYAEQF | 0.6504042 |
| PPGAGGAGGAGTT | ASRDSNYAEQF | 0.70489025 |
| QLVGLGI | ASRDSNYAEQF | 0.64807785 |
| RFLEQVHQL | ASRDSNYAEQF | 0.62848556 |
| RGVGYVLR | ASRDSNYAEQF | 0.70242405 |
| SATGVFAGVI | ASRDSNYAEQF | 0.7278644 |
| SLPAITLP | ASRDSNYAEQF | 0.6682497 |
| STALAFTV | ASRDSNYAEQF | 0.6894823 |
| SVLTVGLGK | ASRDSNYAEQF | 0.72093856 |
| TALAGAGSGI | ASRDSNYAEQF | 0.5962838 |
| TGLPFTGI | ASRDSNYAEQF | 0.6676389 |
| VAEHVTV | ASRDSNYAEQF | 0.7028762 |
| VDIVLNSL | ASRDSNYAEQF | 0.72236955 |
| VLTGGGAVL | ASRDSNYAEQF | 0.6477721 |
| VNVGLAYGI | ASRDSNYAEQF | 0.7245892 |
| VSSAAVGAV | ASRDSNYAEQF | 0.55761623 |
| VTSVYEAGV | ASRDSNYAEQF | 0.68769026 |
| WVALDRGA | ASRDSNYAEQF | 0.56556666 |
| YLVGSVVI | ASRDSNYAEQF | 0.66451085 |
| GPPGTGKTLL | ASRDSNYAEQF | 0.6274569 |
| GVGMTKF | ASRDSNYAEQF | 0.7065493 |
| IGIGGSDL | ASRDSNYAEQF | 0.7015257 |
| IGTLLAEL | ASRDSNYAEQF | 0.7114383 |
| IYTSGTT | ASRDSNYAEQF | 0.34771812 |
| LSFPAGV | ASRDSNYAEQF | 0.70177156 |
| LYTSGTT | ASRDSNYAEQF | 0.7079077 |
| VGAFPFGF | ASRDSNYAEQF | 0.7253817 |
| VGAGTVEF | ASRDSNYAEQF | 0.65081394 |
| VGKSTLLL | ASRDSNYAEQF | 0.6799306 |
| AALAHAGGGIV | ASRDSNYAEQF | 0.6306574 |
| AALVAVSLL | ASRDSNYAEQF | 0.650162 |
| AAVSRAVAAL | ASRDSNYAEQF | 0.5090279 |
| AGGSSGIGGF | ASRDSNYAEQF | 0.6034454 |
| AGLTVANANG | ASRDSNYAEQF | 0.57296634 |
| AGLVGAVL | ASRDSNYAEQF | 0.576613 |
| ALSQAFTV | ASRDSNYAEQF | 0.69648206 |
| ALTFSVSL | ASRDSNYAEQF | 0.7184302 |
| ALTGAALVAL | ASRDSNYAEQF | 0.7187537 |
| ASFSIGAGVL | ASRDSNYAEQF | 0.713353 |
| DGSVYKI | ASRDSNYAEQF | 0.7160361 |
| EQLAEIV | ASRDSNYAEQF | 0.7185556 |
| FAVSELTVM | ASRDSNYAEQF | 0.63946193 |
| GAAPLAAGL | ASRDSNYAEQF | 0.60038906 |
| GADGGDGGSGGAGG | ASRDSNYAEQF | 0.52092016 |
| GFGLDTGGAGG | ASRDSNYAEQF | 0.617782 |
| GLASGGLL | ASRDSNYAEQF | 0.6499268 |
| GRIAGAAVLL | ASRDSNYAEQF | 0.67181724 |
| GSAGSLNTSF | ASRDSNYAEQF | 0.27903974 |

| | | |
|---|---|---|
| GTGQIGI | ASRDSNYAEQF | 0.6340373 |
| GTLIALIL | ASRDSNYAEQF | 0.7322604 |
| IAATMGFVVG | ASRDSNYAEQF | 0.61942416 |
| IFAQALL | ASRDSNYAEQF | 0.6824758 |
| IGAAVALGF | ASRDSNYAEQF | 0.66253835 |
| IGLTPLAL | ASRDSNYAEQF | 0.6070381 |
| ILGSNTSTL | ASRDSNYAEQF | 0.6932752 |
| IVFSLTVL | ASRDSNYAEQF | 0.6071793 |
| LASKATGF | ASRDSNYAEQF | 0.6978049 |
| LASVALIL | ASRDSNYAEQF | 0.7273288 |
| LFAVGLTL | ASRDSNYAEQF | 0.6770876 |
| LFNGGIGTYI | ASRDSNYAEQF | 0.66486776 |
| LTLGAGII | ASRDSNYAEQF | 0.6510563 |
| LVAAALGDI | ASRDSNYAEQF | 0.49513155 |
| NTAFVQLGI | ASRDSNYAEQF | 0.6823535 |
| QLFSGAEL | ASRDSNYAEQF | 0.5108459 |
| SAATFITL | ASRDSNYAEQF | 0.63175094 |
| SNYGLANLGV | ASRDSNYAEQF | 0.58306944 |
| SYGGAYAVM | ASRDSNYAEQF | 0.6811615 |
| TTGVGGLTLG | ASRDSNYAEQF | 0.6786666 |
| TTQTIYI | ASRDSNYAEQF | 0.67497116 |
| VAAGAGLGAVI | ASRDSNYAEQF | 0.540051 |
| VAAVVTLL | ASRDSNYAEQF | 0.67932177 |
| VGFLGGLI | ASRDSNYAEQF | 0.6570671 |
| VGVIPILL | ASRDSNYAEQF | 0.5873879 |
| VIFASSLI | ASRDSNYAEQF | 0.70404387 |
| VKGGDGGAAAG | ASRDSNYAEQF | 0.6525238 |
| VTKTFTNI | ASRDSNYAEQF | 0.6289916 |
| VVAAAILL | ASRDSNYAEQF | 0.7198167 |
| YAALDVEL | ASRDSNYAEQF | 0.6883471 |
| LASVALLL | ASRDSNYAEQF | 0.4097039 |
| LDAVHRV | ASRDSNYAEQF | 0.6187135 |
| QGLSSSIF | ASRDSNYAEQF | 0.6476857 |
| RGTFANIRL | ASRDSNYAEQF | 0.65599763 |
| RQLWWGHRI | ASRDSNYAEQF | 0.29423708 |
| YGPPGTGKTLL | ASRDSNYAEQF | 0.70237327 |
| AALVGAPANVA | ASRDSNYAEQF | 0.6159693 |
| AATFRRTE | ASRDSNYAEQF | 0.527033 |
| AGALGGGI | ASRDSNYAEQF | 0.63644457 |
| AGGAGGAGGIGAG | ASRDSNYAEQF | 0.6958122 |
| AGSGALGL | ASRDSNYAEQF | 0.6389597 |
| AQIADQI | ASRDSNYAEQF | 0.6815325 |
| ASAVAVTYL | ASRDSNYAEQF | 0.6903566 |
| ASTVLGGTGG | ASRDSNYAEQF | 0.7204057 |
| AVGGEPAK | ASRDSNYAEQF | 0.7238248 |
| CEQGTFGGTC | ASRDSNYAEQF | 0.7166461 |
| DEGKAPAG | ASRDSNYAEQF | 0.65192664 |
| DTVVFPAL | ASRDSNYAEQF | 0.70587355 |
| FEGITMAG | ASRDSNYAEQF | 0.6264945 |
| FPLVVQD | ASRDSNYAEQF | 0.51607394 |
| GAFTTPQ | ASRDSNYAEQF | 0.46776783 |

| | | |
|---|---|---|
| GFGGFGTAGGVGGA | ASRDSNYAEQF | 0.6368833 |
| GFLAQFL | ASRDSNYAEQF | 0.4864683 |
| GGAGGAGGNAGVA | ASRDSNYAEQF | 0.6735279 |
| GGVGTAGAGGAGGAGG | ASRDSNYAEQF | 0.72509557 |
| GNGGGGDAGNAGSGGN | ASRDSNYAEQF | 0.6377327 |
| GTGGAGGAGNDAGS | ASRDSNYAEQF | 0.45915514 |
| GTTNTGWFNS | ASRDSNYAEQF | 0.62716377 |
| IFMGPLFTL | ASRDSNYAEQF | 0.58801067 |
| ILAHAIEAI | ASRDSNYAEQF | 0.6965533 |
| IRSAIREAL | ASRDSNYAEQF | 0.71460444 |
| ITFAGGG | ASRDSNYAEQF | 0.72421086 |
| ITGMSGHQEV | ASRDSNYAEQF | 0.71730953 |
| IVAGLACTL | ASRDSNYAEQF | 0.62847424 |
| IVGGGLAGA | ASRDSNYAEQF | 0.6948993 |
| IVTGVLNI | ASRDSNYAEQF | 0.6120742 |
| LAQQGCI | ASRDSNYAEQF | 0.6350119 |
| LCLHVEN | ASRDSNYAEQF | 0.67003155 |
| LDAISDGS | ASRDSNYAEQF | 0.6768402 |
| LGGFAAML | ASRDSNYAEQF | 0.6948558 |
| LGGKNPMI | ASRDSNYAEQF | 0.65589166 |
| LGPKGVAAIL | ASRDSNYAEQF | 0.44803733 |
| LMVGVSI | ASRDSNYAEQF | 0.7001356 |
| LVGAAFVL | ASRDSNYAEQF | 0.6632706 |
| LVLAVAPL | ASRDSNYAEQF | 0.44111764 |
| MAAIAVIDAA | ASRDSNYAEQF | 0.63571465 |
| NNGINAELQ | ASRDSNYAEQF | 0.7243322 |
| RGEPLVLVHG | ASRDSNYAEQF | 0.720335 |
| RGGAGGAAGLI | ASRDSNYAEQF | 0.5800078 |
| RSYRMHPAV | ASRDSNYAEQF | 0.61972034 |
| SAAHLLAGQLA | ASRDSNYAEQF | 0.6399742 |
| SAAVGSY | ASRDSNYAEQF | 0.49300057 |
| SPHAVLHRI | ASRDSNYAEQF | 0.69386435 |
| SQIAKVL | ASRDSNYAEQF | 0.6316143 |
| TALGHLVP | ASRDSNYAEQF | 0.6482554 |
| TAPTTSGAA | ASRDSNYAEQF | 0.5759668 |
| TDGDDGNGGSPGT | ASRDSNYAEQF | 0.67608905 |
| TDGLGEN | ASRDSNYAEQF | 0.6551666 |
| TGGPPLPGQA | ASRDSNYAEQF | 0.71020114 |
| TIPAITI | ASRDSNYAEQF | 0.29422784 |
| TIPGFTI | ASRDSNYAEQF | 0.7141356 |
| TSIASGALA | ASRDSNYAEQF | 0.55276537 |
| TVTLPGAAI | ASRDSNYAEQF | 0.5762544 |
| VASGLSAALP | ASRDSNYAEQF | 0.3995352 |
| VGAGVIGI | ASRDSNYAEQF | 0.64942664 |
| VGPKRFGV | ASRDSNYAEQF | 0.6950378 |
| VNVAFPD | ASRDSNYAEQF | 0.707783 |
| VSMTNSGVS | ASRDSNYAEQF | 0.6154575 |
| VTDVLGDLSV | ASRDSNYAEQF | 0.7109594 |
| VVVVGAGFAGL | ASRDSNYAEQF | 0.7162573 |
| VYTVGTVV | ASRDSNYAEQF | 0.61272514 |
| WTTFSGP | ASRDSNYAEQF | 0.47238165 |

| | | |
|---|---|---|
| YDELVEAF | ASRDSNYAEQF | 0.6971453 |
| YGTGGAGGNGG | ASRDSNYAEQF | 0.5964675 |
| YTNVMAV | ASRDSNYAEQF | 0.707912 |
| FLEQVHQL | ASRDSNYAEQF | 0.6953733 |
| GLSLLAVAV | ASRDSNYAEQF | 0.50482744 |
| LEQVHQL | ASRDSNYAEQF | 0.7077218 |
| LNAAGAI | ASRDSNYAEQF | 0.70549 |
| PAVHASL | ASRDSNYAEQF | 0.56199646 |
| TELEDDA | ASRDSNYAEQF | 0.52272606 |
| TPPGPPN | ASRDSNYAEQF | 0.611196 |
| TPPPPPPP | ASRDSNYAEQF | 0.47274834 |
| TQSPALI | ASSFWDRTSGNTLY | 0.7235424 |
| EFAAAFPQ | ASSFWDRTSGNTLY | 0.60066855 |
| LLSGNLDVL | ASSFWDRTSGNTLY | 0.65202826 |
| GPANLGNYNI | ASSFWDRTSGNTLY | 0.6584648 |
| TLGPTLP | ASSFWDRTSGNTLY | 0.67120194 |
| VYAYSSCCK | ASSFWDRTSGNTLY | 0.65685904 |
| APAPAPAPNI | ASSFWDRTSGNTLY | 0.6780213 |
| PGGPVPIP | ASSFWDRTSGNTLY | 0.7015177 |
| FLTVALEP | ASSFWDRTSGNTLY | 0.68733513 |
| SQYFGANL | ASSFWDRTSGNTLY | 0.6261976 |
| GGVKPAA | ASSFWDRTSGNTLY | 0.6296674 |
| GGTGGAGGAGGW | ASSFWDRTSGNTLY | 0.7195586 |
| GVGGVAGLGGAGG | ASSFWDRTSGNTLY | 0.6031953 |
| VLLVLATTI | ASSFWDRTSGNTLY | 0.6345253 |
| IGIFAAML | ASSFWDRTSGNTLY | 0.69113827 |
| AVGDLID | ASSFWDRTSGNTLY | 0.56654173 |
| NISPLPL | ASSFWDRTSGNTLY | 0.6999829 |
| VTAVFVAI | ASSFWDRTSGNTLY | 0.6864274 |
| SGDDNDAAL | ASSFWDRTSGNTLY | 0.67474496 |
| AVAALAHS | ASSFWDRTSGNTLY | 0.60936546 |
| GGVTVMATP | ASSFWDRTSGNTLY | 0.7308849 |
| AGLAANVRD | ASSFWDRTSGNTLY | 0.70013267 |
| TNRPDLIDPALLRPG | ASSFWDRTSGNTLY | 0.64824885 |
| LANLGVTAL | ASSFWDRTSGNTLY | 0.69753206 |
| TGERIEL | ASSFWDRTSGNTLY | 0.54209226 |
| NGEELAA | ASSFWDRTSGNTLY | 0.7154424 |
| VLSLVWG | ASSFWDRTSGNTLY | 0.69351697 |
| IGYALWL | ASSFWDRTSGNTLY | 0.5302514 |
| LSLLLAPL | ASSFWDRTSGNTLY | 0.629575 |
| TLDEALK | ASSFWDRTSGNTLY | 0.72438353 |
| TDGAGTAEGG | ASSFWDRTSGNTLY | 0.71795297 |
| NAPGGAGGNGG | ASSFWDRTSGNTLY | 0.5355271 |
| VSVFMGMGGNF | ASSFWDRTSGNTLY | 0.60796916 |
| LASYLAAL | ASSFWDRTSGNTLY | 0.47734463 |
| LVGGTPAL | ASSFWDRTSGNTLY | 0.5544763 |
| VRNADVL | ASSFWDRTSGNTLY | 0.582987 |
| TCSSPFLVSAM | ASSFWDRTSGNTLY | 0.691836 |
| VHITDAE | ASSFWDRTSGNTLY | 0.61006117 |
| EYAAPSF | ASSFWDRTSGNTLY | 0.66389716 |
| VVDAFHGGL | ASSFWDRTSGNTLY | 0.6745775 |

| | | |
|---|---|---|
| ASVVDPQ | ASSFWDRTSGNTLY | 0.7249608 |
| SYNEFGGLS | ASSFWDRTSGNTLY | 0.584898 |
| DPDGSVY | ASSFWDRTSGNTLY | 0.6842427 |
| MVSTIHATE | ASSFWDRTSGNTLY | 0.49394864 |
| LAAGVADPV | ASSFWDRTSGNTLY | 0.6361343 |
| TKEQIAA | ASSFWDRTSGNTLY | 0.5868664 |
| SLASPAAL | ASSFWDRTSGNTLY | 0.6767933 |
| IGLVFYGV | ASSFWDRTSGNTLY | 0.3904472 |
| FLLDASNL | ASSFWDRTSGNTLY | 0.73009366 |
| YQVGNFP | ASSFWDRTSGNTLY | 0.6414241 |
| ESLLSQST | ASSFWDRTSGNTLY | 0.70628047 |
| LFTVGFAAVG | ASSFWDRTSGNTLY | 0.6458315 |
| ATPPTPT | ASSFWDRTSGNTLY | 0.6477821 |
| TDLATLG | ASSFWDRTSGNTLY | 0.6394067 |
| SAGGTGR | ASSFWDRTSGNTLY | 0.68911725 |
| GFSNSGS | ASSFWDRTSGNTLY | 0.5793632 |
| GGNGGSGGTSV | ASSFWDRTSGNTLY | 0.35561007 |
| GTSVATGGAGN | ASSFWDRTSGNTLY | 0.6052306 |
| NGGNGGSGGTS | ASSFWDRTSGNTLY | 0.43118793 |
| AVVKFPH | ASSFWDRTSGNTLY | 0.72644323 |
| TPPPPPP | ASSFWDRTSGNTLY | 0.71907485 |
| AAPAAAAAPA | ASSFWDRTSGNTLY | 0.6632537 |
| IALNEDL | ASSFWDRTSGNTLY | 0.7180795 |
| IVNVSSM | ASSFWDRTSGNTLY | 0.2929057 |
| TGIPPAPR | ASSFWDRTSGNTLY | 0.6446457 |
| FDLGGGTF | ASSFWDRTSGNTLY | 0.7294994 |
| FDLGGGTFDVS | ASSFWDRTSGNTLY | 0.69256246 |
| NEPTAAA | ASSFWDRTSGNTLY | 0.6411789 |
| NEPTAAALA | ASSFWDRTSGNTLY | 0.7186913 |
| GSIINISSI | ASSFWDRTSGNTLY | 0.63671345 |
| VAPGNAGTA | ASSFWDRTSGNTLY | 0.65438706 |
| TGGGASLE | ASSFWDRTSGNTLY | 0.6900355 |
| VAVYNHYKRI | ASSFWDRTSGNTLY | 0.71836257 |
| TDPAPATT | ASSFWDRTSGNTLY | 0.67591196 |
| TNRPDILDPALLRPG | ASSFWDRTSGNTLY | 0.6036761 |
| LTTTTTS | ASSFWDRTSGNTLY | 0.6690153 |
| PTGSGKTL | ASSFWDRTSGNTLY | 0.737718 |
| TGSGKTL | ASSFWDRTSGNTLY | 0.6112977 |
| TIGHVDHGK | ASSFWDRTSGNTLY | 0.70341414 |
| TIGHVDHGKTT | ASSFWDRTSGNTLY | 0.6785179 |
| GHVDHGKTT | ASSFWDRTSGNTLY | 0.66252285 |
| AQFEHTLL | ASSFWDRTSGNTLY | 0.5659409 |
| IVNQDEL | ASSFWDRTSGNTLY | 0.57443106 |
| TNGSQFF | ASSFWDRTSGNTLY | 0.6877233 |
| TGKGIVE | ASSFWDRTSGNTLY | 0.4995417 |
| LISNASDA | ASSFWDRTSGNTLY | 0.63948596 |
| AELRTLGELTKIAKAHGAQ | ASSFWDRTSGNTLY | 0.62553227 |
| APNAVAAAEPSW | ASSFWDRTSGNTLY | 0.74404454 |
| APQVAGAATAVL | ASSFWDRTSGNTLY | 0.68630046 |
| ARRDPAAAAAILPTDA | ASSFWDRTSGNTLY | 0.7023883 |
| DAAARDVLAEAGLAEYF | ASSFWDRTSGNTLY | 0.6095545 |

| | | |
|---|---|---|
| DDQIVLSAKANLAKVEQ | ASSFWDRTSGNTLY | 0.6795467 |
| DELLPDTGEDSDEDGH | ASSFWDRTSGNTLY | 0.6883322 |
| ELPEAPANRATVPQD | ASSFWDRTSGNTLY | 0.48651674 |
| FGQSKIERVTGIG | ASSFWDRTSGNTLY | 0.7178167 |
| GAGGTGSTAGGAGGAGGAGGLY | ASSFWDRTSGNTLY | 0.5889005 |
| GHGGTGAAGGNGGTGGAGGSGI | ASSFWDRTSGNTLY | 0.63837314 |
| GNGGNGGAGGEGGPGGAGGAGA | ASSFWDRTSGNTLY | 0.7105746 |
| GSTAGNGGNGGNATNGTIAGQP | ASSFWDRTSGNTLY | 0.7100994 |
| GTGGDGGNGGNGGTGG | ASSFWDRTSGNTLY | 0.63795376 |
| GVRRPAGIPADAPV | ASSFWDRTSGNTLY | 0.663011 |
| GYRQVTDAHLLTLARR | ASSFWDRTSGNTLY | 0.6171526 |
| IAKMGQDGHDRG | ASSFWDRTSGNTLY | 0.70228606 |
| KAEPSIEIQYT | ASSFWDRTSGNTLY | 0.71054244 |
| KPVRIRMAADEDTVAVT | ASSFWDRTSGNTLY | 0.646172 |
| MPPRIAGMRLLVIKPEP | ASSFWDRTSGNTLY | 0.7313735 |
| MKFYAVALFRSRGVPPTR | ASSFWDRTSGNTLY | 0.699864 |
| NAGGFGILGAVAHSP | ASSFWDRTSGNTLY | 0.4688841 |
| NEGREITLAMAEM | ASSFWDRTSGNTLY | 0.6550598 |
| NELVEWSHPLGHRWRWE | ASSFWDRTSGNTLY | 0.71168953 |
| NLGAGGTGGNGGNPG | ASSFWDRTSGNTLY | 0.7091793 |
| NTEWPKWETVMAADPVD | ASSFWDRTSGNTLY | 0.7363882 |
| PASAAPAETQP | ASSFWDRTSGNTLY | 0.6966838 |
| QQSSDGKNGVRAAEITGE | ASSFWDRTSGNTLY | 0.61027646 |
| RAPDAVTTSWWKE | ASSFWDRTSGNTLY | 0.7219226 |
| SGFGNVGAHTSGWFNQ | ASSFWDRTSGNTLY | 0.7357609 |
| SGVAGAVGQAASVGGLKV | ASSFWDRTSGNTLY | 0.6952357 |
| SNGGAGTGSAGGNGGT | ASSFWDRTSGNTLY | 0.5917138 |
| TGGAGGEGGAAG | ASSFWDRTSGNTLY | 0.5934067 |
| TGGDGGAAGAGGAPGHG | ASSFWDRTSGNTLY | 0.6892142 |
| TMPPLLAAIANAGRHGV | ASSFWDRTSGNTLY | 0.7096802 |
| TSSTAGNSGGA | ASSFWDRTSGNTLY | 0.5214992 |
| VPATVPVGILC | ASSFWDRTSGNTLY | 0.6646442 |
| VQRPDSGGVVLTGRIS | ASSFWDRTSGNTLY | 0.67603135 |
| WTPNPVPATKGVPAGLVQAT | ASSFWDRTSGNTLY | 0.45945075 |
| YALKRALAKFSDRFDVVII | ASSFWDRTSGNTLY | 0.6793757 |
| YLIGSVIV | ASSFWDRTSGNTLY | 0.7116004 |
| YGQLVAVID | ASSFWDRTSGNTLY | 0.6744009 |
| VTVGGMTLPA | ASSFWDRTSGNTLY | 0.56365347 |
| VLALPAV | ASSFWDRTSGNTLY | 0.7146888 |
| VIPPTLNYET | ASSFWDRTSGNTLY | 0.735445 |
| VGMGGMHPSSG | ASSFWDRTSGNTLY | 0.7255559 |
| VGLAGGAATA | ASSFWDRTSGNTLY | 0.5045203 |
| VGGAGGAGGSGP | ASSFWDRTSGNTLY | 0.3057536 |
| VFAAFMV | ASSFWDRTSGNTLY | 0.6536743 |
| TLYDKLR | ASSFWDRTSGNTLY | 0.6866454 |
| TLAVFTVL | ASSFWDRTSGNTLY | 0.6763947 |
| TGSVQPARV | ASSFWDRTSGNTLY | 0.6451551 |
| TGGGAGSGGT | ASSFWDRTSGNTLY | 0.65999216 |
| SHLETSL | ASSFWDRTSGNTLY | 0.71152097 |
| SFTTGMT | ASSFWDRTSGNTLY | 0.6044129 |
| QVEVAKRMGVRQPTVS | ASSFWDRTSGNTLY | 0.53574425 |

| | | |
|---|---|---|
| QGSGGPALAM | ASSFWDRTSGNTLY | 0.6831838 |
| MLAPSNP | ASSFWDRTSGNTLY | 0.69139314 |
| LVNNAAI | ASSFWDRTSGNTLY | 0.5753665 |
| LTGEINTLA | ASSFWDRTSGNTLY | 0.62698853 |
| LSIFGPV | ASSFWDRTSGNTLY | 0.6663355 |
| LLALLANL | ASSFWDRTSGNTLY | 0.6308391 |
| LISNASDALDKLR | ASSFWDRTSGNTLY | 0.67016274 |
| LINEAVP | ASSFWDRTSGNTLY | 0.52231425 |
| LGYAFSGL | ASSFWDRTSGNTLY | 0.72549695 |
| LDFVAVAL | ASSFWDRTSGNTLY | 0.5073813 |
| LAQPGAE | ASSFWDRTSGNTLY | 0.6877663 |
| ISNLGAVTGF | ASSFWDRTSGNTLY | 0.6910973 |
| HQEDPGRI | ASSFWDRTSGNTLY | 0.6982235 |
| GTPGEPTP | ASSFWDRTSGNTLY | 0.6185061 |
| GFFNSGS | ASSFWDRTSGNTLY | 0.547258 |
| GAGGGGLTAGAN | ASSFWDRTSGNTLY | 0.5275408 |
| FVDALVNM | ASSFWDRTSGNTLY | 0.4571054 |
| FLTVAPEL | ASSFWDRTSGNTLY | 0.7182524 |
| FETLTAL | ASSFWDRTSGNTLY | 0.6367015 |
| DVINPIEL | ASSFWDRTSGNTLY | 0.722754 |
| APVSGAPV | ASSFWDRTSGNTLY | 0.6945303 |
| ALVGGLAW | ASSFWDRTSGNTLY | 0.61986077 |
| ALAAAAATAI | ASSFWDRTSGNTLY | 0.6891944 |
| AFGLGLLL | ASSFWDRTSGNTLY | 0.6178791 |
| AFFAAASL | ASSFWDRTSGNTLY | 0.6216132 |
| AAAALPAFSPP | ASSFWDRTSGNTLY | 0.6787088 |
| VVLAPAL | ASSFWDRTSGNTLY | 0.6999621 |
| TIGHVDHGKTTL | ASSFWDRTSGNTLY | 0.53144586 |
| LISNASDALDK | ASSFWDRTSGNTLY | 0.68474287 |
| LISNASDAL | ASSFWDRTSGNTLY | 0.67428267 |
| ISNASDALDK | ASSFWDRTSGNTLY | 0.52496296 |
| NQAETLVYQTE | ASSFWDRTSGNTLY | 0.675447 |
| TGGTGGTAGAG | ASSFWDRTSGNTLY | 0.7184273 |
| TGSTAGGAGGA | ASSFWDRTSGNTLY | 0.68785584 |
| DPLAVPVAAGPV | ASSFWDRTSGNTLY | 0.5254957 |
| GGTGGTPGNGGN | ASSFWDRTSGNTLY | 0.7016877 |
| PAGLAGVDPATL | ASSFWDRTSGNTLY | 0.5455137 |
| GNNLVVEAPHAGASV | ASSFWDRTSGNTLY | 0.6687776 |
| KVTRIMSASTLEQPA | ASSFWDRTSGNTLY | 0.67932504 |
| LGFALPIALPRRTVV | ASSFWDRTSGNTLY | 0.6364763 |
| TRLGNLTARKLVVLV | ASSFWDRTSGNTLY | 0.71682274 |
| WSQQRRLIYFPSAGP | ASSFWDRTSGNTLY | 0.5286033 |
| GPNGNPDPMAAAVDIR | ASSFWDRTSGNTLY | 0.5841487 |
| TQRCVSVADLRAAASR | ASSFWDRTSGNTLY | 0.6323894 |
| QFVLDVAEHPVEVLTETV | ASSFWDRTSGNTLY | 0.6960627 |
| GGAGGTGGAGGNGGTQAGDG | ASSFWDRTSGNTLY | 0.7016552 |
| PLLATLRAIAVDVDAKPAQV | ASSFWDRTSGNTLY | 0.56707615 |
| APKFRTGGYMIDFWGVGYQVAK | ASSFWDRTSGNTLY | 0.6504666 |
| GAGGAGGAGGAAGGPGGTGGTG | ASSFWDRTSGNTLY | 0.6432974 |
| IGALGLLATDVAPAGSVAFGVA | ASSFWDRTSGNTLY | 0.58166075 |
| AGGAGGIGGNGGAGGAGGVLMG | ASSFWDRTSGNTLY | 0.6683422 |

| | | |
|---|---|---|
| ADELIGLL | ASSFWDRTSGNTLY | 0.64479303 |
| ADRALKA | ASSFWDRTSGNTLY | 0.4991604 |
| AGAVGQVGGAGGA | ASSFWDRTSGNTLY | 0.7123761 |
| AGGGGGLGGAP | ASSFWDRTSGNTLY | 0.66272986 |
| AGGLLYGNGGAG | ASSFWDRTSGNTLY | 0.7116281 |
| AGLESIGL | ASSFWDRTSGNTLY | 0.71556365 |
| AGLKTGII | ASSFWDRTSGNTLY | 0.58281827 |
| ASLAALLL | ASSFWDRTSGNTLY | 0.68797624 |
| ASVINISSI | ASSFWDRTSGNTLY | 0.67594934 |
| ATPPGPA | ASSFWDRTSGNTLY | 0.6877149 |
| ATVASALV | ASSFWDRTSGNTLY | 0.67672974 |
| ATVASLIL | ASSFWDRTSGNTLY | 0.573882 |
| AVNGPVLIRH | ASSFWDRTSGNTLY | 0.71648055 |
| DGAAGAVGI | ASSFWDRTSGNTLY | 0.711958 |
| DPLLHGI | ASSFWDRTSGNTLY | 0.698155 |
| ELGAALGAL | ASSFWDRTSGNTLY | 0.70572126 |
| ELGRTQAV | ASSFWDRTSGNTLY | 0.66802996 |
| FVAGGYLS | ASSFWDRTSGNTLY | 0.6983769 |
| GASGGAGGAGGW | ASSFWDRTSGNTLY | 0.5956415 |
| GGAGAGGDGGAGGI | ASSFWDRTSGNTLY | 0.49519256 |
| GTGHNGSQPGMGGNG | ASSFWDRTSGNTLY | 0.6835941 |
| GYAAASASAS | ASSFWDRTSGNTLY | 0.63509065 |
| HQVDVAAVQQG | ASSFWDRTSGNTLY | 0.5773857 |
| IGAAIAHT | ASSFWDRTSGNTLY | 0.43546206 |
| IGMIEVAQL | ASSFWDRTSGNTLY | 0.7140665 |
| IRLAPEHTSAMD | ASSFWDRTSGNTLY | 0.73765296 |
| IYNVSVL | ASSFWDRTSGNTLY | 0.6201917 |
| KQGFLGM | ASSFWDRTSGNTLY | 0.7352873 |
| LESVITQL | ASSFWDRTSGNTLY | 0.49534208 |
| LGLGALVVV | ASSFWDRTSGNTLY | 0.7129117 |
| LIGNGGNGGSGGI | ASSFWDRTSGNTLY | 0.71901214 |
| LNAINAPTL | ASSFWDRTSGNTLY | 0.5417848 |
| LQLLNTGV | ASSFWDRTSGNTLY | 0.7179607 |
| LQTSIEF | ASSFWDRTSGNTLY | 0.57269025 |
| MSFAVSEL | ASSFWDRTSGNTLY | 0.6185295 |
| NTGALGL | ASSFWDRTSGNTLY | 0.7301353 |
| PPGAGGAGGAGTT | ASSFWDRTSGNTLY | 0.66672 |
| QLVGLGI | ASSFWDRTSGNTLY | 0.7034046 |
| RFLEQVHQL | ASSFWDRTSGNTLY | 0.68471795 |
| RGVGYVLR | ASSFWDRTSGNTLY | 0.6360424 |
| SATGVFAGVI | ASSFWDRTSGNTLY | 0.7022821 |
| SLPAITLP | ASSFWDRTSGNTLY | 0.7235935 |
| STALAFTV | ASSFWDRTSGNTLY | 0.60772717 |
| SVLTVGLGK | ASSFWDRTSGNTLY | 0.7320411 |
| TALAGAGSGI | ASSFWDRTSGNTLY | 0.59280056 |
| TGLPFTGI | ASSFWDRTSGNTLY | 0.7363646 |
| VAEHVTV | ASSFWDRTSGNTLY | 0.68939734 |
| VDIVLNSL | ASSFWDRTSGNTLY | 0.64904344 |
| VLTGGGAVL | ASSFWDRTSGNTLY | 0.6558002 |
| VNVGLAYGI | ASSFWDRTSGNTLY | 0.70018536 |
| VSSAAVGAV | ASSFWDRTSGNTLY | 0.6218376 |

| | | |
|---|---|---|
| VTSVYEAGV | ASSFWDRTSGNTLY | 0.69227785 |
| WVALDRGA | ASSFWDRTSGNTLY | 0.7288326 |
| YLVGSVVI | ASSFWDRTSGNTLY | 0.73662484 |
| GPPGTGKTLL | ASSFWDRTSGNTLY | 0.90185064 |
| GVGMTKF | ASSFWDRTSGNTLY | 0.8760059 |
| IGIGGSDL | ASSFWDRTSGNTLY | 0.8726239 |
| IGTLLAEL | ASSFWDRTSGNTLY | 0.7948095 |
| IYTSGTT | ASSFWDRTSGNTLY | 0.44714013 |
| LSFPAGV | ASSFWDRTSGNTLY | 0.82460916 |
| LYTSGTT | ASSFWDRTSGNTLY | 0.7391931 |
| VGAFPFGF | ASSFWDRTSGNTLY | 0.8692224 |
| VGAGTVEF | ASSFWDRTSGNTLY | 0.891076 |
| VGKSTLLL | ASSFWDRTSGNTLY | 0.79656225 |
| AALAHAGGGIV | ASSFWDRTSGNTLY | 0.8936619 |
| AALVAVSLL | ASSFWDRTSGNTLY | 0.90356505 |
| AAVSRAVAAL | ASSFWDRTSGNTLY | 0.52781785 |
| AGGSSGIGGF | ASSFWDRTSGNTLY | 0.8391045 |
| AGLTVANANG | ASSFWDRTSGNTLY | 0.7440753 |
| AGLVGAVL | ASSFWDRTSGNTLY | 0.83911836 |
| ALSQAFTV | ASSFWDRTSGNTLY | 0.36263546 |
| ALTFSVSL | ASSFWDRTSGNTLY | 0.89391637 |
| ALTGAALVAL | ASSFWDRTSGNTLY | 0.8018711 |
| ASFSIGAGVL | ASSFWDRTSGNTLY | 0.16326867 |
| DGSVYKI | ASSFWDRTSGNTLY | 0.8528648 |
| EQLAEIV | ASSFWDRTSGNTLY | 0.8383225 |
| FAVSELTVM | ASSFWDRTSGNTLY | 0.8302438 |
| GAAPLAAGL | ASSFWDRTSGNTLY | 0.89432824 |
| GADGGDGGSGGAGG | ASSFWDRTSGNTLY | 0.7544435 |
| GFGLDTGGAGG | ASSFWDRTSGNTLY | 0.74934447 |
| GLASGGLL | ASSFWDRTSGNTLY | 0.878837 |
| GRIAGAAVLL | ASSFWDRTSGNTLY | 0.9008881 |
| GSAGSLNTSF | ASSFWDRTSGNTLY | 0.6943661 |
| GTGQIGI | ASSFWDRTSGNTLY | 0.8709363 |
| GTLIALIL | ASSFWDRTSGNTLY | 0.9031171 |
| IAATMGFVVG | ASSFWDRTSGNTLY | 0.87054336 |
| IFAQALL | ASSFWDRTSGNTLY | 0.89986026 |
| IGAAVALGF | ASSFWDRTSGNTLY | 0.9067264 |
| IGLTPLAL | ASSFWDRTSGNTLY | 0.7671763 |
| ILGSNTSTL | ASSFWDRTSGNTLY | 0.89504397 |
| IVFSLTVL | ASSFWDRTSGNTLY | 0.89318585 |
| LASKATGF | ASSFWDRTSGNTLY | 0.8380159 |
| LASVALIL | ASSFWDRTSGNTLY | 0.85407215 |
| LFAVGLTL | ASSFWDRTSGNTLY | 0.70987445 |
| LFNGGIGTYI | ASSFWDRTSGNTLY | 0.8322227 |
| LTLGAGII | ASSFWDRTSGNTLY | 0.56017447 |
| LVAAALGDI | ASSFWDRTSGNTLY | 0.86558557 |
| NTAFVQLGI | ASSFWDRTSGNTLY | 0.8987796 |
| QLFSGAEL | ASSFWDRTSGNTLY | 0.8273338 |
| SAATFITL | ASSFWDRTSGNTLY | 0.7085289 |
| SNYGLANLGV | ASSFWDRTSGNTLY | 0.7025316 |
| SYGGAYAVM | ASSFWDRTSGNTLY | 0.855481 |

| | | |
|---|---|---|
| TTGVGGLTLG | ASSFWDRTSGNTLY | 0.8402909 |
| TTQTIYI | ASSFWDRTSGNTLY | 0.4912017 |
| VAAGAGLGAVI | ASSFWDRTSGNTLY | 0.7911037 |
| VAAVVTLL | ASSFWDRTSGNTLY | 0.89449406 |
| VGFLGGLI | ASSFWDRTSGNTLY | 0.89828134 |
| VGVIPILL | ASSFWDRTSGNTLY | 0.6326914 |
| VIFASSLI | ASSFWDRTSGNTLY | 0.8978714 |
| VKGGDGGAAAG | ASSFWDRTSGNTLY | 0.8862051 |
| VTKTFTNI | ASSFWDRTSGNTLY | 0.9035721 |
| VVAAAILL | ASSFWDRTSGNTLY | 0.86184055 |
| YAALDVEL | ASSFWDRTSGNTLY | 0.72740334 |
| LASVALLL | ASSFWDRTSGNTLY | 0.8889935 |
| LDAVHRV | ASSFWDRTSGNTLY | 0.821476 |
| QGLSSSIF | ASSFWDRTSGNTLY | 0.7782984 |
| RGTFANIRL | ASSFWDRTSGNTLY | 0.8920169 |
| RQLWWGHRI | ASSFWDRTSGNTLY | 0.83553284 |
| YGPPGTGKTLL | ASSFWDRTSGNTLY | 0.7860287 |
| AALVGAPANVA | ASSFWDRTSGNTLY | 0.8145059 |
| AATFRRTE | ASSFWDRTSGNTLY | 0.904964 |
| AGALGGGI | ASSFWDRTSGNTLY | 0.6982468 |
| AGGAGGAGGIGAG | ASSFWDRTSGNTLY | 0.7500265 |
| AGSGALGL | ASSFWDRTSGNTLY | 0.79614335 |
| AQIADQI | ASSFWDRTSGNTLY | 0.85124934 |
| ASAVAVTYL | ASSFWDRTSGNTLY | 0.89562994 |
| ASTVLGGTGG | ASSFWDRTSGNTLY | 0.8827318 |
| AVGGEPAK | ASSFWDRTSGNTLY | 0.8329754 |
| CEQGTFGGTC | ASSFWDRTSGNTLY | 0.8754213 |
| DEGKAPAG | ASSFWDRTSGNTLY | 0.8926182 |
| DTVVFPAL | ASSFWDRTSGNTLY | 0.86509806 |
| FEGITMAG | ASSFWDRTSGNTLY | 0.8884588 |
| FPLVVQD | ASSFWDRTSGNTLY | 0.7775403 |
| GAFTTPQ | ASSFWDRTSGNTLY | 0.9011169 |
| GFGGFGTAGGVGGA | ASSFWDRTSGNTLY | 0.8864083 |
| GFLAQFL | ASSFWDRTSGNTLY | 0.8980622 |
| GGAGGAGGNAGVA | ASSFWDRTSGNTLY | 0.8948478 |
| GGVGTAGAGGAGGAGG | ASSFWDRTSGNTLY | 0.8792583 |
| GNGGGGDAGNAGSGGN | ASSFWDRTSGNTLY | 0.87591183 |
| GTGGAGGAGNDAGS | ASSFWDRTSGNTLY | 0.46388513 |
| GTTNTGWFNS | ASSFWDRTSGNTLY | 0.8021908 |
| IFMGPLFTL | ASSFWDRTSGNTLY | 0.86959386 |
| ILAHAIEAI | ASSFWDRTSGNTLY | 0.87375295 |
| IRSAIREAL | ASSFWDRTSGNTLY | 0.8841816 |
| ITFAGGG | ASSFWDRTSGNTLY | 0.8895122 |
| ITGMSGHQEV | ASSFWDRTSGNTLY | 0.78331196 |
| IVAGLACTL | ASSFWDRTSGNTLY | 0.9005935 |
| IVGGGLAGA | ASSFWDRTSGNTLY | 0.89509916 |
| IVTGVLNI | ASSFWDRTSGNTLY | 0.56297123 |
| LAQQGCI | ASSFWDRTSGNTLY | 0.81473446 |
| LCLHVEN | ASSFWDRTSGNTLY | 0.905301 |
| LDAISDGS | ASSFWDRTSGNTLY | 0.8700006 |
| LGGFAAML | ASSFWDRTSGNTLY | 0.81560254 |

| | | |
|---|---|---|
| LGGKNPMI | ASSFWDRTSGNTLY | 0.83566725 |
| LGPKGVAAIL | ASSFWDRTSGNTLY | 0.8997015 |
| LMVGVSI | ASSFWDRTSGNTLY | 0.82597554 |
| LVGAAFVL | ASSFWDRTSGNTLY | 0.9049761 |
| LVLAVAPL | ASSFWDRTSGNTLY | 0.30252415 |
| MAAIAVIDAA | ASSFWDRTSGNTLY | 0.85014665 |
| NNGINAELQ | ASSFWDRTSGNTLY | 0.7829604 |
| RGEPLVLVHG | ASSFWDRTSGNTLY | 0.46312177 |
| RGGAGGAAGLI | ASSFWDRTSGNTLY | 0.89021045 |
| RSYRMHPAV | ASSFWDRTSGNTLY | 0.81828535 |
| SAAHLLAGQLA | ASSFWDRTSGNTLY | 0.8886889 |
| SAAVGSY | ASSFWDRTSGNTLY | 0.83078 |
| SPHAVLHRI | ASSFWDRTSGNTLY | 0.89284885 |
| SQIAKVL | ASSFWDRTSGNTLY | 0.8530098 |
| TALGHLVP | ASSFWDRTSGNTLY | 0.86659116 |
| TAPTTSGAA | ASSFWDRTSGNTLY | 0.7466278 |
| TDGDDGNGGSPGT | ASSFWDRTSGNTLY | 0.88548887 |
| TDGLGEN | ASSFWDRTSGNTLY | 0.9019184 |
| TGGPPLPGQA | ASSFWDRTSGNTLY | 0.89879084 |
| TIPAITI | ASSFWDRTSGNTLY | 0.54941714 |
| TIPGFTI | ASSFWDRTSGNTLY | 0.8345244 |
| TSIASGALA | ASSFWDRTSGNTLY | 0.85693836 |
| TVTLPGAAI | ASSFWDRTSGNTLY | 0.2986407 |
| VASGLSAALP | ASSFWDRTSGNTLY | 0.28239834 |
| VGAGVIGI | ASSFWDRTSGNTLY | 0.8258498 |
| VGPKRFGV | ASSFWDRTSGNTLY | 0.7811638 |
| VNVAFPD | ASSFWDRTSGNTLY | 0.8893256 |
| VSMTNSGVS | ASSFWDRTSGNTLY | 0.87785184 |
| VTDVLGDLSV | ASSFWDRTSGNTLY | 0.83562684 |
| VVVVGAGFAGL | ASSFWDRTSGNTLY | 0.88586485 |
| VYTVGTVV | ASSFWDRTSGNTLY | 0.86230636 |
| WTTFSGP | ASSFWDRTSGNTLY | 0.102629825 |
| YDELVEAF | ASSFWDRTSGNTLY | 0.8211466 |
| YGTGGAGGNGG | ASSFWDRTSGNTLY | 0.5718467 |
| YTNVMAV | ASSFWDRTSGNTLY | 0.9049963 |
| FLEQVHQL | ASSFWDRTSGNTLY | 0.7329818 |
| GLSLLAVAV | ASSFWDRTSGNTLY | 0.7467629 |
| LEQVHQL | ASSFWDRTSGNTLY | 0.85035396 |
| LNAAGAI | ASSFWDRTSGNTLY | 0.8557589 |
| PAVHASL | ASSFWDRTSGNTLY | 0.8489088 |
| TELEDDA | ASSFWDRTSGNTLY | 0.77274334 |
| TPPGPPN | ASSFWDRTSGNTLY | 0.2443916 |
| TPPPPPPP | ASSFWDRTSGNTLY | 0.8876364 |
| TQSPALI | ASSQLRGGNQAPL | 0.65567505 |
| EFAAAFPQ | ASSQLRGGNQAPL | 0.69736296 |
| LLSGNLDVL | ASSQLRGGNQAPL | 0.36949855 |
| GPANLGNYNI | ASSQLRGGNQAPL | 0.2922034 |
| TLGPTLP | ASSQLRGGNQAPL | 0.51955736 |
| VYAYSSCCK | ASSQLRGGNQAPL | 0.89849484 |
| APAPAPAPNI | ASSQLRGGNQAPL | 0.8115944 |
| PGGPVPIP | ASSQLRGGNQAPL | 0.8009294 |

| | | |
|---|---|---|
| FLTVALEP | ASSQLRGGNQAPL | 0.90055877 |
| SQYFGANL | ASSQLRGGNQAPL | 0.8543513 |
| GGVKPAA | ASSQLRGGNQAPL | 0.82576585 |
| GGTGGAGGAGGW | ASSQLRGGNQAPL | 0.7970084 |
| GVGGVAGLGGAGG | ASSQLRGGNQAPL | 0.69990623 |
| VLLVLATTI | ASSQLRGGNQAPL | 0.75113297 |
| IGIFAAML | ASSQLRGGNQAPL | 0.7587157 |
| AVGDLID | ASSQLRGGNQAPL | 0.9067601 |
| NISPLPL | ASSQLRGGNQAPL | 0.84234405 |
| VTAVFVAI | ASSQLRGGNQAPL | 0.5530284 |
| SGDDNDAAL | ASSQLRGGNQAPL | 0.87556887 |
| AVAALAHS | ASSQLRGGNQAPL | 0.9036975 |
| GGVTVMATP | ASSQLRGGNQAPL | 0.9086275 |
| AGLAANVRD | ASSQLRGGNQAPL | 0.72811496 |
| TNRPDLIDPALLRPG | ASSQLRGGNQAPL | 0.8458657 |
| LANLGVTAL | ASSQLRGGNQAPL | 0.83091927 |
| TGERIEL | ASSQLRGGNQAPL | 0.8372335 |
| NGEELAA | ASSQLRGGNQAPL | 0.8216925 |
| VLSLVWG | ASSQLRGGNQAPL | 0.61591625 |
| IGYALWL | ASSQLRGGNQAPL | 0.89760506 |
| LSLLLAPL | ASSQLRGGNQAPL | 0.9121286 |
| TLDEALK | ASSQLRGGNQAPL | 0.83483124 |
| TDGAGTAEGG | ASSQLRGGNQAPL | 0.4299044 |
| NAPGGAGGNGG | ASSQLRGGNQAPL | 0.83095884 |
| VSVFMGMGGNF | ASSQLRGGNQAPL | 0.767444 |
| LASYLAAL | ASSQLRGGNQAPL | 0.7904328 |
| LVGGTPAL | ASSQLRGGNQAPL | 0.86452734 |
| VRNADVL | ASSQLRGGNQAPL | 0.52760273 |
| TCSSPFLVSAM | ASSQLRGGNQAPL | 0.8237916 |
| VHITDAE | ASSQLRGGNQAPL | 0.9079657 |
| EYAAPSF | ASSQLRGGNQAPL | 0.81834376 |
| VVDAFHGGL | ASSQLRGGNQAPL | 0.87221605 |
| ASVVDPQ | ASSQLRGGNQAPL | 0.8958056 |
| SYNEFGGLS | ASSQLRGGNQAPL | 0.87824965 |
| DPDGSVY | ASSQLRGGNQAPL | 0.90076387 |
| MVSTIHATE | ASSQLRGGNQAPL | 0.89118123 |
| LAAGVADPV | ASSQLRGGNQAPL | 0.8492215 |
| TKEQIAA | ASSQLRGGNQAPL | 0.85818493 |
| SLASPAAL | ASSQLRGGNQAPL | 0.84473747 |
| IGLVFYGV | ASSQLRGGNQAPL | 0.84471625 |
| FLLDASNL | ASSQLRGGNQAPL | 0.84132284 |
| YQVGNFP | ASSQLRGGNQAPL | 0.8033803 |
| ESLLSQST | ASSQLRGGNQAPL | 0.89139754 |
| LFTVGFAAVG | ASSQLRGGNQAPL | 0.8633485 |
| ATPPTPT | ASSQLRGGNQAPL | 0.89989066 |
| TDLATLG | ASSQLRGGNQAPL | 0.63533914 |
| SAGGTGR | ASSQLRGGNQAPL | 0.71440625 |
| GFSNSGS | ASSQLRGGNQAPL | 0.8954286 |
| GGNGGSGGTSV | ASSQLRGGNQAPL | 0.19372828 |
| GTSVATGGAGN | ASSQLRGGNQAPL | 0.3522483 |
| NGGNGGSGGTS | ASSQLRGGNQAPL | 0.8728785 |

| | | |
|---|---|---|
| AVVKFPH | ASSQLRGGNQAPL | 0.90218997 |
| TPPPPPP | ASSQLRGGNQAPL | 0.17377846 |
| AAPAAAAAPA | ASSQLRGGNQAPL | 0.7024612 |
| IALNEDL | ASSQLRGGNQAPL | 0.6826511 |
| IVNVSSM | ASSQLRGGNQAPL | 0.8957963 |
| TGIPPAPR | ASSQLRGGNQAPL | 0.8564854 |
| FDLGGGTF | ASSQLRGGNQAPL | 0.5371175 |
| FDLGGGTFDVS | ASSQLRGGNQAPL | 0.8916208 |
| NEPTAAA | ASSQLRGGNQAPL | 0.47579885 |
| NEPTAAALA | ASSQLRGGNQAPL | 0.7108839 |
| GSIINISSI | ASSQLRGGNQAPL | 0.8886874 |
| VAPGNAGTA | ASSQLRGGNQAPL | 0.901055 |
| TGGGASLE | ASSQLRGGNQAPL | 0.61823606 |
| VAVYNHYKRI | ASSQLRGGNQAPL | 0.8922063 |
| TDPAPATT | ASSQLRGGNQAPL | 0.823965 |
| TNRPDILDPALLRPG | ASSQLRGGNQAPL | 0.90777427 |
| LTTTTTS | ASSQLRGGNQAPL | 0.8099067 |
| PTGSGKTL | ASSQLRGGNQAPL | 0.5663486 |
| TGSGKTL | ASSQLRGGNQAPL | 0.8920579 |
| TIGHVDHGK | ASSQLRGGNQAPL | 0.8808931 |
| TIGHVDHGKTT | ASSQLRGGNQAPL | 0.8660089 |
| GHVDHGKTT | ASSQLRGGNQAPL | 0.8899334 |
| AQFEHTLL | ASSQLRGGNQAPL | 0.83736145 |
| IVNQDEL | ASSQLRGGNQAPL | 0.8235527 |
| TNGSQFF | ASSQLRGGNQAPL | 0.8943552 |
| TGKGIVE | ASSQLRGGNQAPL | 0.88827884 |
| LISNASDA | ASSQLRGGNQAPL | 0.68165445 |
| AELRTLGELTKIAKAHGAQ | ASSQLRGGNQAPL | 0.8950043 |
| APNAVAAAEPSW | ASSQLRGGNQAPL | 0.8493175 |
| APQVAGAATAVL | ASSQLRGGNQAPL | 0.7235205 |
| ARRDPAAAAAILPTDA | ASSQLRGGNQAPL | 0.906576 |
| DAAARDVLAEAGLAEYF | ASSQLRGGNQAPL | 0.8754256 |
| DDQIVLSAKANLAKVEQ | ASSQLRGGNQAPL | 0.8922261 |
| DELLPDTGEDSDEDGH | ASSQLRGGNQAPL | 0.8947929 |
| ELPEAPANRATVPQD | ASSQLRGGNQAPL | 0.9050648 |
| FGQSKIERVTGIG | ASSQLRGGNQAPL | 0.5492909 |
| GAGGTGSTAGGAGGAGGAGGLY | ASSQLRGGNQAPL | 0.86199546 |
| GHGGTGAAGGNGGTGGAGGSGI | ASSQLRGGNQAPL | 0.8436732 |
| GNGGNGGAGGEGGPGGAGGAGA | ASSQLRGGNQAPL | 0.88982236 |
| GSTAGNGGNGGNATNGTIAGQP | ASSQLRGGNQAPL | 0.860152 |
| GTGGDGGNGGNGGTGG | ASSQLRGGNQAPL | 0.70980954 |
| GVRRPAGIPADAPV | ASSQLRGGNQAPL | 0.8835261 |
| GYRQVTDAHLLTLARR | ASSQLRGGNQAPL | 0.89881676 |
| IAKMGQDGHDRG | ASSQLRGGNQAPL | 0.8572835 |
| KAEPSIEIQYT | ASSQLRGGNQAPL | 0.8760773 |
| KPVRIRMAADEDTVAVT | ASSQLRGGNQAPL | 0.904572 |
| MPPRIAGMRLLVIKPEP | ASSQLRGGNQAPL | 0.76162285 |
| MKFYAVALFRSRGVPPTR | ASSQLRGGNQAPL | 0.84675854 |
| NAGGFGILGAVAHSP | ASSQLRGGNQAPL | 0.89151824 |
| NEGREITLAMAEM | ASSQLRGGNQAPL | 0.89656806 |
| NELVEWSHPLGHRWRWE | ASSQLRGGNQAPL | 0.85143405 |

| | | |
|---|---|---|
| NLGAGGTGGNGGNPG | ASSQLRGGNQAPL | 0.90310806 |
| NTEWPKWETVMAADPVD | ASSQLRGGNQAPL | 0.7479778 |
| PASAAPAETQP | ASSQLRGGNQAPL | 0.88418573 |
| QQSSDGKNGVRAAEITGE | ASSQLRGGNQAPL | 0.7593825 |
| RAPDAVTTSWWKE | ASSQLRGGNQAPL | 0.8634908 |
| SGFGNVGAHTSGWFNQ | ASSQLRGGNQAPL | 0.74225307 |
| SGVAGAVGQAASVGGLKV | ASSQLRGGNQAPL | 0.8700664 |
| SNGGAGTGSAGGNGGT | ASSQLRGGNQAPL | 0.88831264 |
| TGGAGGEGGAAG | ASSQLRGGNQAPL | 0.88145924 |
| TGGDGGAAGAGGAPGHG | ASSQLRGGNQAPL | 0.25003636 |
| TMPPLLAAIANAGRHGV | ASSQLRGGNQAPL | 0.8819903 |
| TSSTAGNSGGA | ASSQLRGGNQAPL | 0.88581854 |
| VPATVPVGILC | ASSQLRGGNQAPL | 0.910255 |
| VQRPDSGGVVLTGRIS | ASSQLRGGNQAPL | 0.8587017 |
| WTPNPVPATKGVPAGLVQAT | ASSQLRGGNQAPL | 0.88563603 |
| YALKRALAKFSDRFDVVII | ASSQLRGGNQAPL | 0.8110031 |
| YLIGSVIV | ASSQLRGGNQAPL | 0.852769 |
| YGQLVAVID | ASSQLRGGNQAPL | 0.7266775 |
| VTVGGMTLPA | ASSQLRGGNQAPL | 0.8201037 |
| VLALPAV | ASSQLRGGNQAPL | 0.8026008 |
| VIPPTLNYET | ASSQLRGGNQAPL | 0.7632927 |
| VGMGGMHPSSG | ASSQLRGGNQAPL | 0.8856554 |
| VGLAGGAATA | ASSQLRGGNQAPL | 0.8927736 |
| VGGAGGAGGSGP | ASSQLRGGNQAPL | 0.8891031 |
| VFAAFMV | ASSQLRGGNQAPL | 0.8980734 |
| TLYDKLR | ASSQLRGGNQAPL | 0.896752 |
| TLAVFTVL | ASSQLRGGNQAPL | 0.9029623 |
| TGSVQPARV | ASSQLRGGNQAPL | 0.8050902 |
| TGGGAGSGGT | ASSQLRGGNQAPL | 0.8212994 |
| SHLETSL | ASSQLRGGNQAPL | 0.63349813 |
| SFTTGMT | ASSQLRGGNQAPL | 0.8127326 |
| QVEVAKRMGVRQPTVS | ASSQLRGGNQAPL | 0.8179558 |
| QGSGGPALAM | ASSQLRGGNQAPL | 0.8604488 |
| MLAPSNP | ASSQLRGGNQAPL | 0.13020176 |
| LVNNAAI | ASSQLRGGNQAPL | 0.8542595 |
| LTGEINTLA | ASSQLRGGNQAPL | 0.90488243 |
| LSIFGPV | ASSQLRGGNQAPL | 0.7943118 |
| LLALLANL | ASSQLRGGNQAPL | 0.8786751 |
| LISNASDALDKLR | ASSQLRGGNQAPL | 0.8389089 |
| LINEAVP | ASSQLRGGNQAPL | 0.80847853 |
| LGYAFSGL | ASSQLRGGNQAPL | 0.88461804 |
| LDFVAVAL | ASSQLRGGNQAPL | 0.79154325 |
| LAQPGAE | ASSQLRGGNQAPL | 0.89344394 |
| ISNLGAVTGF | ASSQLRGGNQAPL | 0.9032913 |
| HQEDPGRI | ASSQLRGGNQAPL | 0.85979486 |
| GTPGEPTP | ASSQLRGGNQAPL | 0.8653725 |
| GFFNSGS | ASSQLRGGNQAPL | 0.86835986 |
| GAGGGGLTAGAN | ASSQLRGGNQAPL | 0.43243933 |
| FVDALVNM | ASSQLRGGNQAPL | 0.8705646 |
| FLTVAPEL | ASSQLRGGNQAPL | 0.10452569 |
| FETLTAL | ASSQLRGGNQAPL | 0.8484179 |

| | | |
|---|---|---|
| DVINPIEL | ASSQLRGGNQAPL | 0.751696 |
| APVSGAPV | ASSQLRGGNQAPL | 0.86074483 |
| ALVGGLAW | ASSQLRGGNQAPL | 0.84842813 |
| ALAAAAATAI | ASSQLRGGNQAPL | 0.8795719 |
| AFGLGLLL | ASSQLRGGNQAPL | 0.7154469 |
| AFFAAASL | ASSQLRGGNQAPL | 0.8457291 |
| AAAALPAFSPP | ASSQLRGGNQAPL | 0.7948738 |
| VVLAPAL | ASSQLRGGNQAPL | 0.6270369 |
| TIGHVDHGKTTL | ASSQLRGGNQAPL | 0.8772309 |
| LISNASDALDK | ASSQLRGGNQAPL | 0.8483635 |
| LISNASDAL | ASSQLRGGNQAPL | 0.82629335 |
| ISNASDALDK | ASSQLRGGNQAPL | 0.8967175 |
| NQAETLVYQTE | ASSQLRGGNQAPL | 0.8698477 |
| TGGTGGTAGAG | ASSQLRGGNQAPL | 0.45082343 |
| TGSTAGGAGGA | ASSQLRGGNQAPL | 0.8293304 |
| DPLAVPVAAGPV | ASSQLRGGNQAPL | 0.85367674 |
| GGTGGTPGNGGN | ASSQLRGGNQAPL | 0.8392115 |
| PAGLAGVDPATL | ASSQLRGGNQAPL | 0.13287072 |
| GNNLVVEAPHAGASV | ASSQLRGGNQAPL | 0.87345207 |
| KVTRIMSASTLEQPA | ASSQLRGGNQAPL | 0.73336273 |
| LGFALPIALPRRTVV | ASSQLRGGNQAPL | 0.7317129 |
| TRLGNLTARKLVVLV | ASSQLRGGNQAPL | 0.7887089 |
| WSQQRRLIYFPSAGP | ASSQLRGGNQAPL | 0.8885635 |
| GPNGNPDPMAAAVDIR | ASSQLRGGNQAPL | 0.8125117 |
| TQRCVSVADLRAAASR | ASSQLRGGNQAPL | 0.8596723 |
| QFVLDVAEHPVEVLTETV | ASSQLRGGNQAPL | 0.88695705 |
| GGAGGTGGAGGNGGTQAGDG | ASSQLRGGNQAPL | 0.87977916 |
| PLLATLRAIAVDVDAKPAQV | ASSQLRGGNQAPL | 0.901446 |
| APKFRTGGYMIDFWGVGYQVAK | ASSQLRGGNQAPL | 0.8756349 |
| GAGGAGGAGGAAGGPGGTGGTG | ASSQLRGGNQAPL | 0.85270715 |
| IGALGLLATDVAPAGSVAFGVA | ASSQLRGGNQAPL | 0.8791913 |
| AGGAGGIGGNGGAGGAGGVLMG | ASSQLRGGNQAPL | 0.84964144 |
| ADELIGLL | ASSQLRGGNQAPL | 0.5288905 |
| ADRALKA | ASSQLRGGNQAPL | 0.4843717 |
| AGAVGQVGGAGGA | ASSQLRGGNQAPL | 0.84284914 |
| AGGGGGLGGAP | ASSQLRGGNQAPL | 0.44195205 |
| AGGLLYGNGGAG | ASSQLRGGNQAPL | 0.85906684 |
| AGLESIGL | ASSQLRGGNQAPL | 0.906816 |
| AGLKTGII | ASSQLRGGNQAPL | 0.8577686 |
| ASLAALLL | ASSQLRGGNQAPL | 0.5735784 |
| ASVINISSI | ASSQLRGGNQAPL | 0.82581383 |
| ATPPGPA | ASSQLRGGNQAPL | 0.79236865 |
| ATVASALV | ASSQLRGGNQAPL | 0.8893702 |
| ATVASLIL | ASSQLRGGNQAPL | 0.8944972 |
| AVNGPVLIRH | ASSQLRGGNQAPL | 0.90545535 |
| DGAAGAVGI | ASSQLRGGNQAPL | 0.88719463 |
| DPLLHGI | ASSQLRGGNQAPL | 0.8085724 |
| ELGAALGAL | ASSQLRGGNQAPL | 0.8851419 |
| ELGRTQAV | ASSQLRGGNQAPL | 0.8359457 |
| FVAGGYLS | ASSQLRGGNQAPL | 0.8101498 |
| GASGGAGGAGGW | ASSQLRGGNQAPL | 0.85341763 |

| | | |
|---|---|---|
| GGAGAGGDGGAGGI | ASSQLRGGNQAPL | 0.87322193 |
| GTGHNGSQPGMGGNG | ASSQLRGGNQAPL | 0.8646928 |
| GYAAASASAS | ASSQLRGGNQAPL | 0.8092251 |
| HQVDVAAVQQG | ASSQLRGGNQAPL | 0.44361395 |
| IGAAIAHT | ASSQLRGGNQAPL | 0.89042234 |
| IGMIEVAQL | ASSQLRGGNQAPL | 0.8829211 |
| IRLAPEHTSAMD | ASSQLRGGNQAPL | 0.6788671 |
| IYNVSVL | ASSQLRGGNQAPL | 0.8134563 |
| KQGFLGM | ASSQLRGGNQAPL | 0.90576196 |
| LESVITQL | ASSQLRGGNQAPL | 0.90265006 |
| LGLGALVVV | ASSQLRGGNQAPL | 0.7832121 |
| LIGNGGNGGSGGI | ASSQLRGGNQAPL | 0.7773232 |
| LNAINAPTL | ASSQLRGGNQAPL | 0.8519473 |
| LQLLNTGV | ASSQLRGGNQAPL | 0.60863495 |
| LQTSIEF | ASSQLRGGNQAPL | 0.8861319 |
| MSFAVSEL | ASSQLRGGNQAPL | 0.84048283 |
| NTGALGL | ASSQLRGGNQAPL | 0.8342655 |
| PPGAGGAGGAGTT | ASSQLRGGNQAPL | 0.7991018 |
| QLVGLGI | ASSQLRGGNQAPL | 0.86807954 |
| RFLEQVHQL | ASSQLRGGNQAPL | 0.7996775 |
| RGVGYVLR | ASSQLRGGNQAPL | 0.897566 |
| SATGVFAGVI | ASSQLRGGNQAPL | 0.16816172 |
| SLPAITLP | ASSQLRGGNQAPL | 0.8745078 |
| STALAFTV | ASSQLRGGNQAPL | 0.74058414 |
| SVLTVGLGK | ASSQLRGGNQAPL | 0.8197466 |
| TALAGAGSGI | ASSQLRGGNQAPL | 0.5039489 |
| TGLPFTGI | ASSQLRGGNQAPL | 0.81867343 |
| VAEHVTV | ASSQLRGGNQAPL | 0.8817979 |
| VDIVLNSL | ASSQLRGGNQAPL | 0.89733183 |
| VLTGGGAVL | ASSQLRGGNQAPL | 0.7473589 |
| VNVGLAYGI | ASSQLRGGNQAPL | 0.89012027 |
| VSSAAVGAV | ASSQLRGGNQAPL | 0.90564036 |
| VTSVYEAGV | ASSQLRGGNQAPL | 0.7996428 |
| WVALDRGA | ASSQLRGGNQAPL | 0.33160844 |
| YLVGSVVI | ASSQLRGGNQAPL | 0.8820287 |
| GPPGTGKTLL | ASSQLRGGNQAPL | 0.80608237 |
| GVGMTKF | ASSQLRGGNQAPL | 0.89059675 |
| IGIGGSDL | ASSQLRGGNQAPL | 0.8875247 |
| IGTLLAEL | ASSQLRGGNQAPL | 0.65108097 |
| IYTSGTT | ASSQLRGGNQAPL | 0.8996104 |
| LSFPAGV | ASSQLRGGNQAPL | 0.8932894 |
| LYTSGTT | ASSQLRGGNQAPL | 0.77145654 |
| VGAFPFGF | ASSQLRGGNQAPL | 0.7185987 |
| VGAGTVEF | ASSQLRGGNQAPL | 0.7154776 |
| VGKSTLLL | ASSQLRGGNQAPL | 0.52060723 |
| AALAHAGGGIV | ASSQLRGGNQAPL | 0.886899 |
| AALVAVSLL | ASSQLRGGNQAPL | 0.7858529 |
| AAVSRAVAAL | ASSQLRGGNQAPL | 0.79874265 |
| AGGSSGIGGF | ASSQLRGGNQAPL | 0.8569116 |
| AGLTVANANG | ASSQLRGGNQAPL | 0.793555 |
| AGLVGAVL | ASSQLRGGNQAPL | 0.79013884 |

| | | |
|---|---|---|
| ALSQAFTV | ASSQLRGGNQAPL | 0.81140375 |
| ALTFSVSL | ASSQLRGGNQAPL | 0.8476465 |
| ALTGAALVAL | ASSQLRGGNQAPL | 0.88004243 |
| ASFSIGAGVL | ASSQLRGGNQAPL | 0.8408306 |
| DGSVYKI | ASSQLRGGNQAPL | 0.8580397 |
| EQLAEIV | ASSQLRGGNQAPL | 0.8983893 |
| FAVSELTVM | ASSQLRGGNQAPL | 0.6501161 |
| GAAPLAAGL | ASSQLRGGNQAPL | 0.76556385 |
| GADGGDGGSGGAGG | ASSQLRGGNQAPL | 0.8882688 |
| GFGLDTGGAGG | ASSQLRGGNQAPL | 0.67748266 |
| GLASGGLL | ASSQLRGGNQAPL | 0.8902551 |
| GRIAGAAVLL | ASSQLRGGNQAPL | 0.8576133 |
| GSAGSLNTSF | ASSQLRGGNQAPL | 0.8694981 |
| GTGQIGI | ASSQLRGGNQAPL | 0.8471489 |
| GTLIALIL | ASSQLRGGNQAPL | 0.9064726 |
| IAATMGFVVG | ASSQLRGGNQAPL | 0.88947415 |
| IFAQALL | ASSQLRGGNQAPL | 0.82645524 |
| IGAAVALGF | ASSQLRGGNQAPL | 0.8485697 |
| IGLTPLAL | ASSQLRGGNQAPL | 0.68759227 |
| ILGSNTSTL | ASSQLRGGNQAPL | 0.8925909 |
| IVFSLTVL | ASSQLRGGNQAPL | 0.8910723 |
| LASKATGF | ASSQLRGGNQAPL | 0.7167902 |
| LASVALIL | ASSQLRGGNQAPL | 0.82763284 |
| LFAVGLTL | ASSQLRGGNQAPL | 0.9038827 |
| LFNGGIGTYI | ASSQLRGGNQAPL | 0.8956376 |
| LTLGAGII | ASSQLRGGNQAPL | 0.57743704 |
| LVAAALGDI | ASSQLRGGNQAPL | 0.80400217 |
| NTAFVQLGI | ASSQLRGGNQAPL | 0.59357274 |
| QLFSGAEL | ASSQLRGGNQAPL | 0.8436715 |
| SAATFITL | ASSQLRGGNQAPL | 0.55758953 |
| SNYGLANLGV | ASSQLRGGNQAPL | 0.8780069 |
| SYGGAYAVM | ASSQLRGGNQAPL | 0.8011931 |
| TTGVGGLTLG | ASSQLRGGNQAPL | 0.8844416 |
| TTQTIYI | ASSQLRGGNQAPL | 0.86102605 |
| VAAGAGLGAVI | ASSQLRGGNQAPL | 0.90062404 |
| VAAVVTLL | ASSQLRGGNQAPL | 0.7990443 |
| VGFLGGLI | ASSQLRGGNQAPL | 0.8255583 |
| VGVIPILL | ASSQLRGGNQAPL | 0.5715914 |
| VIFASSLI | ASSQLRGGNQAPL | 0.6331924 |
| VKGGDGGAAAG | ASSQLRGGNQAPL | 0.78879744 |
| VTKTFTNI | ASSQLRGGNQAPL | 0.8177306 |
| VVAAAILL | ASSQLRGGNQAPL | 0.29259783 |
| YAALDVEL | ASSQLRGGNQAPL | 0.9022379 |
| LASVALLL | ASSQLRGGNQAPL | 0.868509 |
| LDAVHRV | ASSQLRGGNQAPL | 0.88798904 |
| QGLSSSIF | ASSQLRGGNQAPL | 0.8049772 |
| RGTFANIRL | ASSQLRGGNQAPL | 0.8235047 |
| RQLWWGHRI | ASSQLRGGNQAPL | 0.7928035 |
| YGPPGTGKTLL | ASSQLRGGNQAPL | 0.8841295 |
| AALVGAPANVA | ASSQLRGGNQAPL | 0.69510394 |
| AATFRRTE | ASSQLRGGNQAPL | 0.37234825 |

| | | |
|---|---|---|
| AGALGGGI | ASSQLRGGNQAPL | 0.7855197 |
| AGGAGGAGGIGAG | ASSQLRGGNQAPL | 0.55127186 |
| AGSGALGL | ASSQLRGGNQAPL | 0.90380126 |
| AQIADQI | ASSQLRGGNQAPL | 0.8963499 |
| ASAVAVTYL | ASSQLRGGNQAPL | 0.8708254 |
| ASTVLGGTGG | ASSQLRGGNQAPL | 0.9026227 |
| AVGGEPAK | ASSQLRGGNQAPL | 0.24410558 |
| CEQGTFGGTC | ASSQLRGGNQAPL | 0.8152412 |
| DEGKAPAG | ASSQLRGGNQAPL | 0.9039874 |
| DTVVFPAL | ASSQLRGGNQAPL | 0.87639546 |
| FEGITMAG | ASSQLRGGNQAPL | 0.8236613 |
| FPLVVQD | ASSQLRGGNQAPL | 0.8997792 |
| GAFTTPQ | ASSQLRGGNQAPL | 0.77670336 |
| GFGGFGTAGGVGGA | ASSQLRGGNQAPL | 0.8426155 |
| GFLAQFL | ASSQLRGGNQAPL | 0.8444572 |
| GGAGGAGGNAGVA | ASSQLRGGNQAPL | 0.8853141 |
| GGVGTAGAGGAGGAGG | ASSQLRGGNQAPL | 0.87593687 |
| GNGGGGDAGNAGSGGN | ASSQLRGGNQAPL | 0.68025434 |
| GTGGAGGAGNDAGS | ASSQLRGGNQAPL | 0.8866365 |
| GTTNTGWFNS | ASSQLRGGNQAPL | 0.9047868 |
| IFMGPLFTL | ASSQLRGGNQAPL | 0.8118379 |
| ILAHAIEAI | ASSQLRGGNQAPL | 0.89113337 |
| IRSAIREAL | ASSQLRGGNQAPL | 0.8783282 |
| ITFAGGG | ASSQLRGGNQAPL | 0.86829877 |
| ITGMSGHQEV | ASSQLRGGNQAPL | 0.7571448 |
| IVAGLACTL | ASSQLRGGNQAPL | 0.7755334 |
| IVGGGLAGA | ASSQLRGGNQAPL | 0.8485683 |
| IVTGVLNI | ASSQLRGGNQAPL | 0.74147016 |
| LAQQGCI | ASSQLRGGNQAPL | 0.84944195 |
| LCLHVEN | ASSQLRGGNQAPL | 0.81213814 |
| LDAISDGS | ASSQLRGGNQAPL | 0.9122036 |
| LGGFAAML | ASSQLRGGNQAPL | 0.8796216 |
| LGGKNPMI | ASSQLRGGNQAPL | 0.8783319 |
| LGPKGVAAIL | ASSQLRGGNQAPL | 0.7231183 |
| LMVGVSI | ASSQLRGGNQAPL | 0.87586707 |
| LVGAAFVL | ASSQLRGGNQAPL | 0.8778668 |
| LVLAVAPL | ASSQLRGGNQAPL | 0.47595847 |
| MAAIAVIDAA | ASSQLRGGNQAPL | 0.900478 |
| NNGINAELQ | ASSQLRGGNQAPL | 0.7804345 |
| RGEPLVLVHG | ASSQLRGGNQAPL | 0.8397634 |
| RGGAGGAAGLI | ASSQLRGGNQAPL | 0.89187276 |
| RSYRMHPAV | ASSQLRGGNQAPL | 0.89620847 |
| SAAHLLAGQLA | ASSQLRGGNQAPL | 0.83044636 |
| SAAVGSY | ASSQLRGGNQAPL | 0.8426559 |
| SPHAVLHRI | ASSQLRGGNQAPL | 0.7222549 |
| SQIAKVL | ASSQLRGGNQAPL | 0.8956734 |
| TALGHLVP | ASSQLRGGNQAPL | 0.8942457 |
| TAPTTSGAA | ASSQLRGGNQAPL | 0.83614814 |
| TDGDDGNGGSPGT | ASSQLRGGNQAPL | 0.905877 |
| TDGLGEN | ASSQLRGGNQAPL | 0.8859587 |
| TGGPPLPGQA | ASSQLRGGNQAPL | 0.47651035 |

| | | |
|---|---|---|
| TIPAITI | ASSQLRGGNQAPL | 0.8594947 |
| TIPGFTI | ASSQLRGGNQAPL | 0.8887782 |
| TSIASGALA | ASSQLRGGNQAPL | 0.89646447 |
| TVTLPGAAI | ASSQLRGGNQAPL | 0.90382683 |
| VASGLSAALP | ASSQLRGGNQAPL | 0.8824727 |
| VGAGVIGI | ASSQLRGGNQAPL | 0.8034527 |
| VGPKRFGV | ASSQLRGGNQAPL | 0.9042268 |
| VNVAFPD | ASSQLRGGNQAPL | 0.9126162 |
| VSMTNSGVS | ASSQLRGGNQAPL | 0.89309764 |
| VTDVLGDLSV | ASSQLRGGNQAPL | 0.83427304 |
| VVVVGAGFAGL | ASSQLRGGNQAPL | 0.83409625 |
| VYTVGTVV | ASSQLRGGNQAPL | 0.86562794 |
| WTTFSGP | ASSQLRGGNQAPL | 0.88901234 |
| YDELVEAF | ASSQLRGGNQAPL | 0.7458311 |
| YGTGGAGGNGG | ASSQLRGGNQAPL | 0.85842276 |
| YTNVMAV | ASSQLRGGNQAPL | 0.8676293 |
| FLEQVHQL | ASSQLRGGNQAPL | 0.4324373 |
| GLSLLAVAV | ASSQLRGGNQAPL | 0.8760715 |
| LEQVHQL | ASSQLRGGNQAPL | 0.8940129 |
| LNAAGAI | ASSQLRGGNQAPL | 0.86224186 |
| PAVHASL | ASSQLRGGNQAPL | 0.8150457 |
| TELEDDA | ASSQLRGGNQAPL | 0.8954713 |
| TPPGPPN | ASSQLRGGNQAPL | 0.90977836 |
| TPPPPPPP | ASSQLRGGNQAPL | 0.90467155 |
| TQSPALI | ASSMGRGRKAEQF | 0.45572528 |
| EFAAAFPQ | ASSMGRGRKAEQF | 0.32132393 |
| LLSGNLDVL | ASSMGRGRKAEQF | 0.8235423 |
| GPANLGNYNI | ASSMGRGRKAEQF | 0.8777508 |
| TLGPTLP | ASSMGRGRKAEQF | 0.82137406 |
| VYAYSSCCK | ASSMGRGRKAEQF | 0.80894166 |
| APAPAPAPNI | ASSMGRGRKAEQF | 0.8517418 |
| PGGPVPIP | ASSMGRGRKAEQF | 0.8995178 |
| FLTVALEP | ASSMGRGRKAEQF | 0.7326366 |
| SQYFGANL | ASSMGRGRKAEQF | 0.68189275 |
| GGVKPAA | ASSMGRGRKAEQF | 0.8577152 |
| GGTGGAGGAGGW | ASSMGRGRKAEQF | 0.89234626 |
| GVGGVAGLGGAGG | ASSMGRGRKAEQF | 0.75082576 |
| VLLVLATTI | ASSMGRGRKAEQF | 0.8237737 |
| IGIFAAML | ASSMGRGRKAEQF | 0.8117943 |
| AVGDLID | ASSMGRGRKAEQF | 0.8124466 |
| NISPLPL | ASSMGRGRKAEQF | 0.8660238 |
| VTAVFVAI | ASSMGRGRKAEQF | 0.74690115 |
| SGDDNDAAL | ASSMGRGRKAEQF | 0.8917948 |
| AVAALAHS | ASSMGRGRKAEQF | 0.6618657 |
| GGVTVMATP | ASSMGRGRKAEQF | 0.89278316 |
| AGLAANVRD | ASSMGRGRKAEQF | 0.8682772 |
| TNRPDLIDPALLRPG | ASSMGRGRKAEQF | 0.87913406 |
| LANLGVTAL | ASSMGRGRKAEQF | 0.8142156 |
| TGERIEL | ASSMGRGRKAEQF | 0.7698157 |
| NGEELAA | ASSMGRGRKAEQF | 0.6653955 |
| VLSLVWG | ASSMGRGRKAEQF | 0.57024133 |

| | | |
|---|---|---|
| IGYALWL | ASSMGRGRKAEQF | 0.89334214 |
| LSLLLAPL | ASSMGRGRKAEQF | 0.24211988 |
| TLDEALK | ASSMGRGRKAEQF | 0.899487 |
| TDGAGTAEGG | ASSMGRGRKAEQF | 0.8812427 |
| NAPGGAGGNGG | ASSMGRGRKAEQF | 0.7032069 |
| VSVFMGMGGNF | ASSMGRGRKAEQF | 0.88162065 |
| LASYLAAL | ASSMGRGRKAEQF | 0.72076976 |
| LVGGTPAL | ASSMGRGRKAEQF | 0.8090663 |
| VRNADVL | ASSMGRGRKAEQF | 0.88729906 |
| TCSSPFLVSAM | ASSMGRGRKAEQF | 0.8892882 |
| VHITDAE | ASSMGRGRKAEQF | 0.7306237 |
| EYAAPSF | ASSMGRGRKAEQF | 0.8829733 |
| VVDAFHGGL | ASSMGRGRKAEQF | 0.8683491 |
| ASVVDPQ | ASSMGRGRKAEQF | 0.76054454 |
| SYNEFGGLS | ASSMGRGRKAEQF | 0.8578701 |
| DPDGSVY | ASSMGRGRKAEQF | 0.894363 |
| MVSTIHATE | ASSMGRGRKAEQF | 0.87708485 |
| LAAGVADPV | ASSMGRGRKAEQF | 0.6980692 |
| TKEQIAA | ASSMGRGRKAEQF | 0.87790024 |
| SLASPAAL | ASSMGRGRKAEQF | 0.75630426 |
| IGLVFYGV | ASSMGRGRKAEQF | 0.86887515 |
| FLLDASNL | ASSMGRGRKAEQF | 0.84965384 |
| YQVGNFP | ASSMGRGRKAEQF | 0.8260428 |
| ESLLSQST | ASSMGRGRKAEQF | 0.8886551 |
| LFTVGFAAVG | ASSMGRGRKAEQF | 0.7090732 |
| ATPPTPT | ASSMGRGRKAEQF | 0.7587445 |
| TDLATLG | ASSMGRGRKAEQF | 0.84116054 |
| SAGGTGR | ASSMGRGRKAEQF | 0.855204 |
| GFSNSGS | ASSMGRGRKAEQF | 0.89142144 |
| GGNGGSGGTSV | ASSMGRGRKAEQF | 0.74757385 |
| GTSVATGGAGN | ASSMGRGRKAEQF | 0.8689445 |
| NGGNGGSGGTS | ASSMGRGRKAEQF | 0.8155341 |
| AVVKFPH | ASSMGRGRKAEQF | 0.73665303 |
| TPPPPPP | ASSMGRGRKAEQF | 0.8666757 |
| AAPAAAAAPA | ASSMGRGRKAEQF | 0.8319319 |
| IALNEDL | ASSMGRGRKAEQF | 0.7528138 |
| IVNVSSM | ASSMGRGRKAEQF | 0.88159823 |
| TGIPPAPR | ASSMGRGRKAEQF | 0.7186364 |
| FDLGGGTF | ASSMGRGRKAEQF | 0.88244087 |
| FDLGGGTFDVS | ASSMGRGRKAEQF | 0.8997514 |
| NEPTAAA | ASSMGRGRKAEQF | 0.82677877 |
| NEPTAAALA | ASSMGRGRKAEQF | 0.8910136 |
| GSIINISSI | ASSMGRGRKAEQF | 0.868694 |
| VAPGNAGTA | ASSMGRGRKAEQF | 0.8720903 |
| TGGGASLE | ASSMGRGRKAEQF | 0.8211477 |
| VAVYNHYKRI | ASSMGRGRKAEQF | 0.7422369 |
| TDPAPATT | ASSMGRGRKAEQF | 0.89607704 |
| TNRPDILDPALLRPG | ASSMGRGRKAEQF | 0.8847953 |
| LTTTTTS | ASSMGRGRKAEQF | 0.8923315 |
| PTGSGKTL | ASSMGRGRKAEQF | 0.87995076 |
| TGSGKTL | ASSMGRGRKAEQF | 0.87127227 |

| | | |
|---|---|---|
| TIGHVDHGK | ASSMGRGRKAEQF | 0.8892126 |
| TIGHVDHGKTT | ASSMGRGRKAEQF | 0.80167764 |
| GHVDHGKTT | ASSMGRGRKAEQF | 0.68186975 |
| AQFEHTLL | ASSMGRGRKAEQF | 0.8269594 |
| IVNQDEL | ASSMGRGRKAEQF | 0.8318091 |
| TNGSQFF | ASSMGRGRKAEQF | 0.78367126 |
| TGKGIVE | ASSMGRGRKAEQF | 0.5995698 |
| LISNASDA | ASSMGRGRKAEQF | 0.89361256 |
| AELRTLGELTKIAKAHGAQ | ASSMGRGRKAEQF | 0.91037023 |
| APNAVAAAEPSW | ASSMGRGRKAEQF | 0.77526677 |
| APQVAGAATAVL | ASSMGRGRKAEQF | 0.9083652 |
| ARRDPAAAAAILPTDA | ASSMGRGRKAEQF | 0.68601894 |
| DAAARDVLAEAGLAEYF | ASSMGRGRKAEQF | 0.88695955 |
| DDQIVLSAKANLAKVEQ | ASSMGRGRKAEQF | 0.8963628 |
| DELLPDTGEDSDEDGH | ASSMGRGRKAEQF | 0.7582443 |
| ELPEAPANRATVPQD | ASSMGRGRKAEQF | 0.8864917 |
| FGQSKIERVTGIG | ASSMGRGRKAEQF | 0.73784125 |
| GAGGTGSTAGGAGGAGGAGGLY | ASSMGRGRKAEQF | 0.8042544 |
| GHGGTGAAGGNGGTGGAGGSGI | ASSMGRGRKAEQF | 0.89979374 |
| GNGGNGGAGGEGGPGGAGGAGA | ASSMGRGRKAEQF | 0.8227201 |
| GSTAGNGGNGGNATNGTIAGQP | ASSMGRGRKAEQF | 0.8698099 |
| GTGGDGGNGGNGGTGG | ASSMGRGRKAEQF | 0.8393277 |
| GVRRPAGIPADAPV | ASSMGRGRKAEQF | 0.7547538 |
| GYRQVTDAHLLTLARR | ASSMGRGRKAEQF | 0.8715023 |
| IAKMGQDGHDRG | ASSMGRGRKAEQF | 0.9001126 |
| KAEPSIEIQYT | ASSMGRGRKAEQF | 0.8171859 |
| KPVRIRMAADEDTVAVT | ASSMGRGRKAEQF | 0.9117775 |
| MPPRIAGMRLLVIKPEP | ASSMGRGRKAEQF | 0.7141422 |
| MKFYAVALFRSRGVPPTR | ASSMGRGRKAEQF | 0.9034882 |
| NAGGFGILGAVAHSP | ASSMGRGRKAEQF | 0.891655 |
| NEGREITLAMAEM | ASSMGRGRKAEQF | 0.85032433 |
| NELVEWSHPLGHRWRWE | ASSMGRGRKAEQF | 0.8292185 |
| NLGAGGTGGNGGNPG | ASSMGRGRKAEQF | 0.88941276 |
| NTEWPKWETVMAADPVD | ASSMGRGRKAEQF | 0.8297732 |
| PASAAPAETQP | ASSMGRGRKAEQF | 0.88203216 |
| QQSSDGKNGVRAAEITGE | ASSMGRGRKAEQF | 0.9056561 |
| RAPDAVTTSWWKE | ASSMGRGRKAEQF | 0.22706097 |
| SGFGNVGAHTSGWFNQ | ASSMGRGRKAEQF | 0.7553572 |
| SGVAGAVGQAASVGGLKV | ASSMGRGRKAEQF | 0.63610315 |
| SNGGAGTGSAGGNGGT | ASSMGRGRKAEQF | 0.44647187 |
| TGGAGGEGGAAG | ASSMGRGRKAEQF | 0.26515132 |
| TGGDGGAAGAGGAPGHG | ASSMGRGRKAEQF | 0.17401144 |
| TMPPLLAAIANAGRHGV | ASSMGRGRKAEQF | 0.37738013 |
| TSSTAGNSGGA | ASSMGRGRKAEQF | 0.2301736 |
| VPATVPVGILC | ASSMGRGRKAEQF | 0.5141498 |
| VQRPDSGGVVLTGRIS | ASSMGRGRKAEQF | 0.51203704 |
| WTPNPVPATKGVPAGLVQAT | ASSMGRGRKAEQF | 0.3013125 |
| YALKRALAKFSDRFDVVII | ASSMGRGRKAEQF | 0.74485177 |
| YLIGSVIV | ASSMGRGRKAEQF | 0.7219262 |
| YGQLVAVID | ASSMGRGRKAEQF | 0.19464731 |
| VTVGGMTLPA | ASSMGRGRKAEQF | 0.4458879 |

| | | |
|---|---|---|
| VLALPAV | ASSMGRGRKAEQF | 0.25839812 |
| VIPPTLNYET | ASSMGRGRKAEQF | 0.36574703 |
| VGMGGMHPSSG | ASSMGRGRKAEQF | 0.16859834 |
| VGLAGGAATA | ASSMGRGRKAEQF | 0.61234534 |
| VGGAGGAGGSGP | ASSMGRGRKAEQF | 0.34576315 |
| VFAAFMV | ASSMGRGRKAEQF | 0.16198948 |
| TLYDKLR | ASSMGRGRKAEQF | 0.37305057 |
| TLAVFTVL | ASSMGRGRKAEQF | 0.32059494 |
| TGSVQPARV | ASSMGRGRKAEQF | 0.29213205 |
| TGGGAGSGGT | ASSMGRGRKAEQF | 0.7272128 |
| SHLETSL | ASSMGRGRKAEQF | 0.17256227 |
| SFTTGMT | ASSMGRGRKAEQF | 0.22700787 |
| QVEVAKRMGVRQPTVS | ASSMGRGRKAEQF | 0.7043146 |
| QGSGGPALAM | ASSMGRGRKAEQF | 0.75786465 |
| MLAPSNP | ASSMGRGRKAEQF | 0.20381202 |
| LVNNAAI | ASSMGRGRKAEQF | 0.65969676 |
| LTGEINTLA | ASSMGRGRKAEQF | 0.76800144 |
| LSIFGPV | ASSMGRGRKAEQF | 0.57101023 |
| LLALLANL | ASSMGRGRKAEQF | 0.67326856 |
| LISNASDALDKLR | ASSMGRGRKAEQF | 0.7304493 |
| LINEAVP | ASSMGRGRKAEQF | 0.22544035 |
| LGYAFSGL | ASSMGRGRKAEQF | 0.7103448 |
| LDFVAVAL | ASSMGRGRKAEQF | 0.7232859 |
| LAQPGAE | ASSMGRGRKAEQF | 0.34013686 |
| ISNLGAVTGF | ASSMGRGRKAEQF | 0.47517163 |
| HQEDPGRI | ASSMGRGRKAEQF | 0.22891468 |
| GTPGEPTP | ASSMGRGRKAEQF | 0.31716 |
| GFFNSGS | ASSMGRGRKAEQF | 0.16855954 |
| GAGGGGLTAGAN | ASSMGRGRKAEQF | 0.668482 |
| FVDALVNM | ASSMGRGRKAEQF | 0.6712681 |
| FLTVAPEL | ASSMGRGRKAEQF | 0.46371463 |
| FETLTAL | ASSMGRGRKAEQF | 0.21930511 |
| DVINPIEL | ASSMGRGRKAEQF | 0.26634085 |
| APVSGAPV | ASSMGRGRKAEQF | 0.52590454 |
| ALVGGLAW | ASSMGRGRKAEQF | 0.44086134 |
| ALAAAAATAI | ASSMGRGRKAEQF | 0.17783438 |
| AFGLGLLL | ASSMGRGRKAEQF | 0.28800365 |
| AFFAAASL | ASSMGRGRKAEQF | 0.71586037 |
| AAAALPAFSPP | ASSMGRGRKAEQF | 0.6987146 |
| VVLAPAL | ASSMGRGRKAEQF | 0.19635537 |
| TIGHVDHGKTTL | ASSMGRGRKAEQF | 0.74246436 |
| LISNASDALDK | ASSMGRGRKAEQF | 0.7384788 |
| LISNASDAL | ASSMGRGRKAEQF | 0.73817915 |
| ISNASDALDK | ASSMGRGRKAEQF | 0.37240824 |
| NQAETLVYQTE | ASSMGRGRKAEQF | 0.27272347 |
| TGGTGGTAGAG | ASSMGRGRKAEQF | 0.70189524 |
| TGSTAGGAGGA | ASSMGRGRKAEQF | 0.49901694 |
| DPLAVPVAAGPV | ASSMGRGRKAEQF | 0.21718338 |
| GGTGGTPGNGGN | ASSMGRGRKAEQF | 0.5986698 |
| PAGLAGVDPATL | ASSMGRGRKAEQF | 0.2878006 |
| GNNLVVEAPHAGASV | ASSMGRGRKAEQF | 0.31261677 |

| | | |
|---|---|---|
| KVTRIMSASTLEQPA | ASSMGRGRKAEQF | 0.63939667 |
| LGFALPIALPRRTVV | ASSMGRGRKAEQF | 0.700794 |
| TRLGNLTARKLVVLV | ASSMGRGRKAEQF | 0.25495943 |
| WSQQRRLIYFPSAGP | ASSMGRGRKAEQF | 0.23143655 |
| GPNGNPDPMAAAVDIR | ASSMGRGRKAEQF | 0.2326824 |
| TQRCVSVADLRAAASR | ASSMGRGRKAEQF | 0.597658 |
| QFVLDVAEHPVEVLTETV | ASSMGRGRKAEQF | 0.7282995 |
| GGAGGTGGAGGNGGTQAGDG | ASSMGRGRKAEQF | 0.42824587 |
| PLLATLRAIAVDVDAKPAQV | ASSMGRGRKAEQF | 0.42530006 |
| APKFRTGGYMIDFWGVGYQVAK | ASSMGRGRKAEQF | 0.7097888 |
| GAGGAGGAGGAAGGPGGTGGTG | ASSMGRGRKAEQF | 0.6267401 |
| IGALGLLATDVAPAGSVAFGVA | ASSMGRGRKAEQF | 0.4108198 |
| AGGAGGIGGNGGAGGAGGVLMG | ASSMGRGRKAEQF | 0.6487093 |
| ADELIGLL | ASSMGRGRKAEQF | 0.33871078 |
| ADRALKA | ASSMGRGRKAEQF | 0.6705958 |
| AGAVGQVGGAGGA | ASSMGRGRKAEQF | 0.5581402 |
| AGGGGGLGGAP | ASSMGRGRKAEQF | 0.6148671 |
| AGGLLYGNGGAG | ASSMGRGRKAEQF | 0.6722959 |
| AGLESIGL | ASSMGRGRKAEQF | 0.57538986 |
| AGLKTGII | ASSMGRGRKAEQF | 0.4771281 |
| ASLAALLL | ASSMGRGRKAEQF | 0.18077558 |
| ASVINISSI | ASSMGRGRKAEQF | 0.34329772 |
| ATPPGPA | ASSMGRGRKAEQF | 0.618577 |
| ATVASALV | ASSMGRGRKAEQF | 0.6236795 |
| ATVASLIL | ASSMGRGRKAEQF | 0.6375379 |
| AVNGPVLIRH | ASSMGRGRKAEQF | 0.4695507 |
| DGAAGAVGI | ASSMGRGRKAEQF | 0.27149418 |
| DPLLHGI | ASSMGRGRKAEQF | 0.69938356 |
| ELGAALGAL | ASSMGRGRKAEQF | 0.7310007 |
| ELGRTQAV | ASSMGRGRKAEQF | 0.19949028 |
| FVAGGYLS | ASSMGRGRKAEQF | 0.3127733 |
| GASGGAGGAGGW | ASSMGRGRKAEQF | 0.76343626 |
| GGAGAGGDGGAGGI | ASSMGRGRKAEQF | 0.65984875 |
| GTGHNGSQPGMGGNG | ASSMGRGRKAEQF | 0.37300384 |
| GYAAASASAS | ASSMGRGRKAEQF | 0.34878796 |
| HQVDVAAVQQG | ASSMGRGRKAEQF | 0.72295225 |
| IGAAIAHT | ASSMGRGRKAEQF | 0.27835938 |
| IGMIEVAQL | ASSMGRGRKAEQF | 0.75604177 |
| IRLAPEHTSAMD | ASSMGRGRKAEQF | 0.16906571 |
| IYNVSVL | ASSMGRGRKAEQF | 0.3420331 |
| KQGFLGM | ASSMGRGRKAEQF | 0.2638945 |
| LESVITQL | ASSMGRGRKAEQF | 0.18267982 |
| LGLGALVVV | ASSMGRGRKAEQF | 0.66470575 |
| LIGNGGNGGSGGI | ASSMGRGRKAEQF | 0.27199528 |
| LNAINAPTL | ASSMGRGRKAEQF | 0.7095605 |
| LQLLNTGV | ASSMGRGRKAEQF | 0.3719179 |
| LQTSIEF | ASSMGRGRKAEQF | 0.60934925 |
| MSFAVSEL | ASSMGRGRKAEQF | 0.63627684 |
| NTGALGL | ASSMGRGRKAEQF | 0.42286175 |
| PPGAGGAGGAGTT | ASSMGRGRKAEQF | 0.23661515 |
| QLVGLGI | ASSMGRGRKAEQF | 0.63478255 |

| | | |
|---|---|---|
| RFLEQVHQL | ASSMGRGRKAEQF | 0.6830559 |
| RGVGYVLR | ASSMGRGRKAEQF | 0.5968501 |
| SATGVFAGVI | ASSMGRGRKAEQF | 0.19515422 |
| SLPAITLP | ASSMGRGRKAEQF | 0.40919337 |
| STALAFTV | ASSMGRGRKAEQF | 0.46609265 |
| SVLTVGLGK | ASSMGRGRKAEQF | 0.16534586 |
| TALAGAGSGI | ASSMGRGRKAEQF | 0.16603422 |
| TGLPFTGI | ASSMGRGRKAEQF | 0.39388418 |
| VAEHVTV | ASSMGRGRKAEQF | 0.29350638 |
| VDIVLNSL | ASSMGRGRKAEQF | 0.737461 |
| VLTGGGAVL | ASSMGRGRKAEQF | 0.7124719 |
| VNVGLAYGI | ASSMGRGRKAEQF | 0.47184008 |
| VSSAAVGAV | ASSMGRGRKAEQF | 0.7059231 |
| VTSVYEAGV | ASSMGRGRKAEQF | 0.59777075 |
| WVALDRGA | ASSMGRGRKAEQF | 0.1601506 |
| YLVGSVVI | ASSMGRGRKAEQF | 0.30170634 |
| GPPGTGKTLL | ASSMGRGRKAEQF | 0.19559628 |
| GVGMTKF | ASSMGRGRKAEQF | 0.68600625 |
| IGIGGSDL | ASSMGRGRKAEQF | 0.20474026 |
| IGTLLAEL | ASSMGRGRKAEQF | 0.2818417 |
| IYTSGTT | ASSMGRGRKAEQF | 0.5401936 |
| LSFPAGV | ASSMGRGRKAEQF | 0.60076475 |
| LYTSGTT | ASSMGRGRKAEQF | 0.53106654 |
| VGAFPFGF | ASSMGRGRKAEQF | 0.27378535 |
| VGAGTVEF | ASSMGRGRKAEQF | 0.16276498 |
| VGKSTLLL | ASSMGRGRKAEQF | 0.6470386 |
| AALAHAGGGIV | ASSMGRGRKAEQF | 0.17285536 |
| AALVAVSLL | ASSMGRGRKAEQF | 0.22331119 |
| AAVSRAVAAL | ASSMGRGRKAEQF | 0.16902523 |
| AGGSSGIGGF | ASSMGRGRKAEQF | 0.16296136 |
| AGLTVANANG | ASSMGRGRKAEQF | 0.17788969 |
| AGLVGAVL | ASSMGRGRKAEQF | 0.62694913 |
| ALSQAFTV | ASSMGRGRKAEQF | 0.43164024 |
| ALTFSVSL | ASSMGRGRKAEQF | 0.3938411 |
| ALTGAALVAL | ASSMGRGRKAEQF | 0.7398952 |
| ASFSIGAGVL | ASSMGRGRKAEQF | 0.7017274 |
| DGSVYKI | ASSMGRGRKAEQF | 0.4060616 |
| EQLAEIV | ASSMGRGRKAEQF | 0.28951204 |
| FAVSELTVM | ASSMGRGRKAEQF | 0.22572848 |
| GAAPLAAGL | ASSMGRGRKAEQF | 0.2203139 |
| GADGGDGGSGGAGG | ASSMGRGRKAEQF | 0.21362302 |
| GFGLDTGGAGG | ASSMGRGRKAEQF | 0.7588371 |
| GLASGGLL | ASSMGRGRKAEQF | 0.3344204 |
| GRIAGAAVLL | ASSMGRGRKAEQF | 0.18507698 |
| GSAGSLNTSF | ASSMGRGRKAEQF | 0.5511731 |
| GTGQIGI | ASSMGRGRKAEQF | 0.73216575 |
| GTLIALIL | ASSMGRGRKAEQF | 0.7686786 |
| IAATMGFVVG | ASSMGRGRKAEQF | 0.25401556 |
| IFAQALL | ASSMGRGRKAEQF | 0.35846034 |
| IGAAVALGF | ASSMGRGRKAEQF | 0.35780978 |
| IGLTPLAL | ASSMGRGRKAEQF | 0.5793323 |

| | | |
|---|---|---|
| ILGSNTSTL | ASSMGRGRKAEQF | 0.34653142 |
| IVFSLTVL | ASSMGRGRKAEQF | 0.19430058 |
| LASKATGF | ASSMGRGRKAEQF | 0.67697835 |
| LASVALIL | ASSMGRGRKAEQF | 0.78905475 |
| LFAVGLTL | ASSMGRGRKAEQF | 0.2830261 |
| LFNGGIGTYI | ASSMGRGRKAEQF | 0.18455684 |
| LTLGAGII | ASSMGRGRKAEQF | 0.50966865 |
| LVAAALGDI | ASSMGRGRKAEQF | 0.32286936 |
| NTAFVQLGI | ASSMGRGRKAEQF | 0.41432044 |
| QLFSGAEL | ASSMGRGRKAEQF | 0.5854139 |
| SAATFITL | ASSMGRGRKAEQF | 0.18841872 |
| SNYGLANLGV | ASSMGRGRKAEQF | 0.26244444 |
| SYGGAYAVM | ASSMGRGRKAEQF | 0.77066886 |
| TTGVGGLTLG | ASSMGRGRKAEQF | 0.45279318 |
| TTQTIYI | ASSMGRGRKAEQF | 0.5519877 |
| VAAGAGLGAVI | ASSMGRGRKAEQF | 0.73911065 |
| VAAVVTLL | ASSMGRGRKAEQF | 0.67848736 |
| VGFLGGLI | ASSMGRGRKAEQF | 0.75785935 |
| VGVIPILL | ASSMGRGRKAEQF | 0.7213152 |
| VIFASSLI | ASSMGRGRKAEQF | 0.29005557 |
| VKGGDGGAAAG | ASSMGRGRKAEQF | 0.48924658 |
| VTKTFTNI | ASSMGRGRKAEQF | 0.46233466 |
| VVAAAILL | ASSMGRGRKAEQF | 0.5401447 |
| YAALDVEL | ASSMGRGRKAEQF | 0.4587203 |
| LASVALLL | ASSMGRGRKAEQF | 0.27151632 |
| LDAVHRV | ASSMGRGRKAEQF | 0.72694206 |
| QGLSSSIF | ASSMGRGRKAEQF | 0.5409012 |
| RGTFANIRL | ASSMGRGRKAEQF | 0.68128395 |
| RQLWWGHRI | ASSMGRGRKAEQF | 0.20675297 |
| YGPPGTGKTLL | ASSMGRGRKAEQF | 0.20745124 |
| AALVGAPANVA | ASSMGRGRKAEQF | 0.71767354 |
| AATFRRTE | ASSMGRGRKAEQF | 0.16492586 |
| AGALGGGI | ASSMGRGRKAEQF | 0.18050317 |
| AGGAGGAGGIGAG | ASSMGRGRKAEQF | 0.5638092 |
| AGSGALGL | ASSMGRGRKAEQF | 0.7523328 |
| AQIADQI | ASSMGRGRKAEQF | 0.16310224 |
| ASAVAVTYL | ASSMGRGRKAEQF | 0.2144376 |
| ASTVLGGTGG | ASSMGRGRKAEQF | 0.22291535 |
| AVGGEPAK | ASSMGRGRKAEQF | 0.63029224 |
| CEQGTFGGTC | ASSMGRGRKAEQF | 0.44797933 |
| DEGKAPAG | ASSMGRGRKAEQF | 0.18838646 |
| DTVVFPAL | ASSMGRGRKAEQF | 0.57668746 |
| FEGITMAG | ASSMGRGRKAEQF | 0.18074387 |
| FPLVVQD | ASSMGRGRKAEQF | 0.17476061 |
| GAFTTPQ | ASSMGRGRKAEQF | 0.60856485 |
| GFGGFGTAGGVGGA | ASSMGRGRKAEQF | 0.7514303 |
| GFLAQFL | ASSMGRGRKAEQF | 0.1956647 |
| GGAGGAGGNAGVA | ASSMGRGRKAEQF | 0.6997115 |
| GGVGTAGAGGAGGAGG | ASSMGRGRKAEQF | 0.3501495 |
| GNGGGGDAGNAGSGGN | ASSMGRGRKAEQF | 0.76060474 |
| GTGGAGGAGNDAGS | ASSMGRGRKAEQF | 0.29525548 |

| | | |
|---|---|---|
| GTTNTGWFNS | ASSMGRGRKAEQF | 0.18065171 |
| IFMGPLFTL | ASSMGRGRKAEQF | 0.6616627 |
| ILAHAIEAI | ASSMGRGRKAEQF | 0.5837023 |
| IRSAIREAL | ASSMGRGRKAEQF | 0.42282015 |
| ITFAGGG | ASSMGRGRKAEQF | 0.72570145 |
| ITGMSGHQEV | ASSMGRGRKAEQF | 0.3737056 |
| IVAGLACTL | ASSMGRGRKAEQF | 0.4946738 |
| IVGGGLAGA | ASSMGRGRKAEQF | 0.73032165 |
| IVTGVLNI | ASSMGRGRKAEQF | 0.66719544 |
| LAQQGCI | ASSMGRGRKAEQF | 0.20810698 |
| LCLHVEN | ASSMGRGRKAEQF | 0.5758027 |
| LDAISDGS | ASSMGRGRKAEQF | 0.541191 |
| LGGFAAML | ASSMGRGRKAEQF | 0.26819825 |
| LGGKNPMI | ASSMGRGRKAEQF | 0.74279267 |
| LGPKGVAAIL | ASSMGRGRKAEQF | 0.44749492 |
| LMVGVSI | ASSMGRGRKAEQF | 0.6960511 |
| LVGAAFVL | ASSMGRGRKAEQF | 0.53693616 |
| LVLAVAPL | ASSMGRGRKAEQF | 0.7623519 |
| MAAIAVIDAA | ASSMGRGRKAEQF | 0.1805182 |
| NNGINAELQ | ASSMGRGRKAEQF | 0.55647415 |
| RGEPLVLVHG | ASSMGRGRKAEQF | 0.51168406 |
| RGGAGGAAGLI | ASSMGRGRKAEQF | 0.6462362 |
| RSYRMHPAV | ASSMGRGRKAEQF | 0.43357366 |
| SAAHLLAGQLA | ASSMGRGRKAEQF | 0.17845352 |
| SAAVGSY | ASSMGRGRKAEQF | 0.4535842 |
| SPHAVLHRI | ASSMGRGRKAEQF | 0.7014413 |
| SQIAKVL | ASSMGRGRKAEQF | 0.42887068 |
| TALGHLVP | ASSMGRGRKAEQF | 0.53445846 |
| TAPTTSGAA | ASSMGRGRKAEQF | 0.7728815 |
| TDGDDGNGGSPGT | ASSMGRGRKAEQF | 0.24151224 |
| TDGLGEN | ASSMGRGRKAEQF | 0.40862298 |
| TGGPPLPGQA | ASSMGRGRKAEQF | 0.64921093 |
| TIPAITI | ASSMGRGRKAEQF | 0.7102145 |
| TIPGFTI | ASSMGRGRKAEQF | 0.3469954 |
| TSIASGALA | ASSMGRGRKAEQF | 0.74035597 |
| TVTLPGAAI | ASSMGRGRKAEQF | 0.25816137 |
| VASGLSAALP | ASSMGRGRKAEQF | 0.71897644 |
| VGAGVIGI | ASSMGRGRKAEQF | 0.28994596 |
| VGPKRFGV | ASSMGRGRKAEQF | 0.6318456 |
| VNVAFPD | ASSMGRGRKAEQF | 0.24740782 |
| VSMTNSGVS | ASSMGRGRKAEQF | 0.6928475 |
| VTDVLGDLSV | ASSMGRGRKAEQF | 0.7264109 |
| VVVVGAGFAGL | ASSMGRGRKAEQF | 0.71688783 |
| VYTVGTVV | ASSMGRGRKAEQF | 0.16407563 |
| WTTFSGP | ASSMGRGRKAEQF | 0.7141423 |
| YDELVEAF | ASSMGRGRKAEQF | 0.7280873 |
| YGTGGAGGNGG | ASSMGRGRKAEQF | 0.7788156 |
| YTNVMAV | ASSMGRGRKAEQF | 0.5288328 |
| FLEQVHQL | ASSMGRGRKAEQF | 0.6018729 |
| GLSLLAVAV | ASSMGRGRKAEQF | 0.39293867 |
| LEQVHQL | ASSMGRGRKAEQF | 0.42151737 |

| | | |
|---|---|---|
| LNAAGAI | ASSMGRGRKAEQF | 0.29381266 |
| PAVHASL | ASSMGRGRKAEQF | 0.44082502 |
| TELEDDA | ASSMGRGRKAEQF | 0.48806918 |
| TPPGPPN | ASSMGRGRKAEQF | 0.33828276 |
| TPPPPPPP | ASSMGRGRKAEQF | 0.6856618 |
| TQSPALI | ASSSGQTNERLF | 0.7188971 |
| EFAAAFPQ | ASSSGQTNERLF | 0.7096324 |
| LLSGNLDVL | ASSSGQTNERLF | 0.75354016 |
| GPANLGNYNI | ASSSGQTNERLF | 0.7296915 |
| TLGPTLP | ASSSGQTNERLF | 0.62534714 |
| VYAYSSCCK | ASSSGQTNERLF | 0.30504763 |
| APAPAPAPNI | ASSSGQTNERLF | 0.36428374 |
| PGGPVPIP | ASSSGQTNERLF | 0.21168831 |
| FLTVALEP | ASSSGQTNERLF | 0.27761912 |
| SQYFGANL | ASSSGQTNERLF | 0.28732407 |
| GGVKPAA | ASSSGQTNERLF | 0.49945998 |
| GGTGGAGGAGGW | ASSSGQTNERLF | 0.16203125 |
| GVGGVAGLGGAGG | ASSSGQTNERLF | 0.6439863 |
| VLLVLATTI | ASSSGQTNERLF | 0.75004 |
| IGIFAAML | ASSSGQTNERLF | 0.30682564 |
| AVGDLID | ASSSGQTNERLF | 0.56734896 |
| NISPLPL | ASSSGQTNERLF | 0.36651373 |
| VTAVFVAI | ASSSGQTNERLF | 0.3325621 |
| SGDDNDAAL | ASSSGQTNERLF | 0.7202062 |
| AVAALAHS | ASSSGQTNERLF | 0.2421172 |
| GGVTVMATP | ASSSGQTNERLF | 0.6332202 |
| AGLAANVRD | ASSSGQTNERLF | 0.73411167 |
| TNRPDLIDPALLRPG | ASSSGQTNERLF | 0.51269054 |
| LANLGVTAL | ASSSGQTNERLF | 0.5644891 |
| TGERIEL | ASSSGQTNERLF | 0.4300204 |
| NGEELAA | ASSSGQTNERLF | 0.17690131 |
| VLSLVWG | ASSSGQTNERLF | 0.6228572 |
| IGYALWL | ASSSGQTNERLF | 0.16053465 |
| LSLLLAPL | ASSSGQTNERLF | 0.5684392 |
| TLDEALK | ASSSGQTNERLF | 0.20155357 |
| TDGAGTAEGG | ASSSGQTNERLF | 0.61546516 |
| NAPGGAGGNGG | ASSSGQTNERLF | 0.5186266 |
| VSVFMGMGGNF | ASSSGQTNERLF | 0.6460525 |
| LASYLAAL | ASSSGQTNERLF | 0.26659235 |
| LVGGTPAL | ASSSGQTNERLF | 0.38004208 |
| VRNADVL | ASSSGQTNERLF | 0.281053 |
| TCSSPFLVSAM | ASSSGQTNERLF | 0.19419771 |
| VHITDAE | ASSSGQTNERLF | 0.6409745 |
| EYAAPSF | ASSSGQTNERLF | 0.34475648 |
| VVDAFHGGL | ASSSGQTNERLF | 0.4565009 |
| ASVVDPQ | ASSSGQTNERLF | 0.6927966 |
| SYNEFGGLS | ASSSGQTNERLF | 0.46687108 |
| DPDGSVY | ASSSGQTNERLF | 0.18630572 |
| MVSTIHATE | ASSSGQTNERLF | 0.6504875 |
| LAAGVADPV | ASSSGQTNERLF | 0.4730152 |
| TKEQIAA | ASSSGQTNERLF | 0.3731043 |

| | | |
|---|---|---|
| SLASPAAL | ASSSGQTNERLF | 0.16015178 |
| IGLVFYGV | ASSSGQTNERLF | 0.5016749 |
| FLLDASNL | ASSSGQTNERLF | 0.26954687 |
| YQVGNFP | ASSSGQTNERLF | 0.39951548 |
| ESLLSQST | ASSSGQTNERLF | 0.26031584 |
| LFTVGFAAVG | ASSSGQTNERLF | 0.7395679 |
| ATPPTPT | ASSSGQTNERLF | 0.18804072 |
| TDLATLG | ASSSGQTNERLF | 0.5456037 |
| SAGGTGR | ASSSGQTNERLF | 0.6654449 |
| GFSNSGS | ASSSGQTNERLF | 0.54686666 |
| GGNGGSGGTSV | ASSSGQTNERLF | 0.7649448 |
| GTSVATGGAGN | ASSSGQTNERLF | 0.5196023 |
| NGGNGGSGGTS | ASSSGQTNERLF | 0.36182785 |
| AVVKFPH | ASSSGQTNERLF | 0.41663992 |
| TPPPPPP | ASSSGQTNERLF | 0.41176915 |
| AAPAAAAAPA | ASSSGQTNERLF | 0.17272596 |
| IALNEDL | ASSSGQTNERLF | 0.17156564 |
| IVNVSSM | ASSSGQTNERLF | 0.6153761 |
| TGIPPAPR | ASSSGQTNERLF | 0.17545494 |
| FDLGGGTF | ASSSGQTNERLF | 0.46137786 |
| FDLGGGTFDVS | ASSSGQTNERLF | 0.64829886 |
| NEPTAAA | ASSSGQTNERLF | 0.4990439 |
| NEPTAAALA | ASSSGQTNERLF | 0.19444159 |
| GSIINISSI | ASSSGQTNERLF | 0.3848436 |
| VAPGNAGTA | ASSSGQTNERLF | 0.2866416 |
| TGGGASLE | ASSSGQTNERLF | 0.6947017 |
| VAVYNHYKRI | ASSSGQTNERLF | 0.54279804 |
| TDPAPATT | ASSSGQTNERLF | 0.76996547 |
| TNRPDILDPALLRPG | ASSSGQTNERLF | 0.72497714 |
| LTTTTTS | ASSSGQTNERLF | 0.43448693 |
| PTGSGKTL | ASSSGQTNERLF | 0.69483656 |
| TGSGKTL | ASSSGQTNERLF | 0.35960373 |
| TIGHVDHGK | ASSSGQTNERLF | 0.302211 |
| TIGHVDHGKTT | ASSSGQTNERLF | 0.31956625 |
| GHVDHGKTT | ASSSGQTNERLF | 0.64572346 |
| AQFEHTLL | ASSSGQTNERLF | 0.3664778 |
| IVNQDEL | ASSSGQTNERLF | 0.26593968 |
| TNGSQFF | ASSSGQTNERLF | 0.18049702 |
| TGKGIVE | ASSSGQTNERLF | 0.61471 |
| LISNASDA | ASSSGQTNERLF | 0.4483686 |
| AELRTLGELTKIAKAHGAQ | ASSSGQTNERLF | 0.19843385 |
| APNAVAAAEPSW | ASSSGQTNERLF | 0.30747595 |
| APQVAGAATAVL | ASSSGQTNERLF | 0.77378345 |
| ARRDPAAAAAILPTDA | ASSSGQTNERLF | 0.772787 |
| DAAARDVLAEAGLAEYF | ASSSGQTNERLF | 0.3171717 |
| DDQIVLSAKANLAKVEQ | ASSSGQTNERLF | 0.23590444 |
| DELLPDTGEDSDEDGH | ASSSGQTNERLF | 0.65638447 |
| ELPEAPANRATVPQD | ASSSGQTNERLF | 0.1863798 |
| FGQSKIERVTGIG | ASSSGQTNERLF | 0.4775527 |
| GAGGTGSTAGGAGGAGGAGGLY | ASSSGQTNERLF | 0.59832525 |
| GHGGTGAAGGNGGTGGAGGSGI | ASSSGQTNERLF | 0.5598299 |

| | | |
|---|---|---|
| GNGGNGGAGGEGGPGGAGGAGA | ASSSGGQTNERLF | 0.27825898 |
| GSTAGNGGNGGNATNGTIAGQP | ASSSGGQTNERLF | 0.57789636 |
| GTGGDGGNGGNGGTGG | ASSSGGQTNERLF | 0.45245737 |
| GVRRPAGIPADAPV | ASSSGGQTNERLF | 0.5915558 |
| GYRQVTDAHLLTLARR | ASSSGGQTNERLF | 0.1615148 |
| IAKMGQDGHDRG | ASSSGGQTNERLF | 0.4202604 |
| KAEPSIEIQYT | ASSSGGQTNERLF | 0.27063996 |
| KPVRIRMAADEDTVAVT | ASSSGGQTNERLF | 0.288383 |
| MPPRIAGMRLLVIKPEP | ASSSGGQTNERLF | 0.18683322 |
| MKFYAVALFRSRGVPPTR | ASSSGGQTNERLF | 0.3502869 |
| NAGGFGILGAVAHSP | ASSSGGQTNERLF | 0.72278976 |
| NEGREITLAMAEM | ASSSGGQTNERLF | 0.73726475 |
| NELVEWSHPLGHRWRWE | ASSSGGQTNERLF | 0.21140958 |
| NLGAGGTGGNGGNPG | ASSSGGQTNERLF | 0.7110225 |
| NTEWPKWETVMAADPVD | ASSSGGQTNERLF | 0.77253366 |
| PASAAPAETQP | ASSSGGQTNERLF | 0.32138467 |
| QQSSDGKNGVRAAEITGE | ASSSGGQTNERLF | 0.17395686 |
| RAPDAVTTSWWKE | ASSSGGQTNERLF | 0.5680682 |
| SGFGNVGAHTSGWFNQ | ASSSGGQTNERLF | 0.3027485 |
| SGVAGAVGQAASVGGLKV | ASSSGGQTNERLF | 0.67327493 |
| SNGGAGTGSAGGNGGT | ASSSGGQTNERLF | 0.706715 |
| TGGAGGEGGAAG | ASSSGGQTNERLF | 0.21139029 |
| TGGDGGAAGAGGAPGHG | ASSSGGQTNERLF | 0.63574517 |
| TMPPLLAAIANAGRHGV | ASSSGGQTNERLF | 0.697502 |
| TSSTAGNSGGA | ASSSGGQTNERLF | 0.27718666 |
| VPATVPVGILC | ASSSGGQTNERLF | 0.23367861 |
| VQRPDSGGVVLTGRIS | ASSSGGQTNERLF | 0.21439408 |
| WTPNPVPATKGVPAGLVQAT | ASSSGGQTNERLF | 0.16949265 |
| YALKRALAKFSDRFDVVII | ASSSGGQTNERLF | 0.5077809 |
| YLIGSVIV | ASSSGGQTNERLF | 0.33550167 |
| YGQLVAVID | ASSSGGQTNERLF | 0.26360935 |
| VTVGGMTLPA | ASSSGGQTNERLF | 0.50844777 |
| VLALPAV | ASSSGGQTNERLF | 0.22270258 |
| VIPPTLNYET | ASSSGGQTNERLF | 0.29268152 |
| VGMGGMHPSSG | ASSSGGQTNERLF | 0.27581754 |
| VGLAGGAATA | ASSSGGQTNERLF | 0.35766202 |
| VGGAGGAGGSGP | ASSSGGQTNERLF | 0.64326346 |
| VFAAFMV | ASSSGGQTNERLF | 0.4470324 |
| TLYDKLR | ASSSGGQTNERLF | 0.48023877 |
| TLAVFTVL | ASSSGGQTNERLF | 0.7558243 |
| TGSVQPARV | ASSSGGQTNERLF | 0.1977722 |
| TGGGAGSGGT | ASSSGGQTNERLF | 0.26048267 |
| SHLETSL | ASSSGGQTNERLF | 0.6820832 |
| SFTTGMT | ASSSGGQTNERLF | 0.1830855 |
| QVEVAKRMGVRQPTVS | ASSSGGQTNERLF | 0.70875406 |
| QGSGGPALAM | ASSSGGQTNERLF | 0.5486267 |
| MLAPSNP | ASSSGGQTNERLF | 0.55839276 |
| LVNNAAI | ASSSGGQTNERLF | 0.434709 |
| LTGEINTLA | ASSSGGQTNERLF | 0.76967525 |
| LSIFGPV | ASSSGGQTNERLF | 0.67292285 |
| LLALLANL | ASSSGGQTNERLF | 0.28981227 |

| | | |
|---|---|---|
| LISNASDALDKLR | ASSSGQTNERLF | 0.19759955 |
| LINEAVP | ASSSGQTNERLF | 0.21615063 |
| LGYAFSGL | ASSSGQTNERLF | 0.704066 |
| LDFVAVAL | ASSSGQTNERLF | 0.6206596 |
| LAQPGAE | ASSSGQTNERLF | 0.27543363 |
| ISNLGAVTGF | ASSSGQTNERLF | 0.5053098 |
| HQEDPGRI | ASSSGQTNERLF | 0.73481166 |
| GTPGEPTP | ASSSGQTNERLF | 0.73554134 |
| GFFNSGS | ASSSGQTNERLF | 0.17810954 |
| GAGGGGLTAGAN | ASSSGQTNERLF | 0.37246227 |
| FVDALVNM | ASSSGQTNERLF | 0.19437304 |
| FLTVAPEL | ASSSGQTNERLF | 0.43042183 |
| FETLTAL | ASSSGQTNERLF | 0.20767751 |
| DVINPIEL | ASSSGQTNERLF | 0.6876215 |
| APVSGAPV | ASSSGQTNERLF | 0.3547401 |
| ALVGGLAW | ASSSGQTNERLF | 0.7305417 |
| ALAAAAATAI | ASSSGQTNERLF | 0.46508905 |
| AFGLGLLL | ASSSGQTNERLF | 0.70498943 |
| AFFAAASL | ASSSGQTNERLF | 0.32032874 |
| AAAALPAFSPP | ASSSGQTNERLF | 0.36413747 |
| VVLAPAL | ASSSGQTNERLF | 0.19448648 |
| TIGHVDHGKTTL | ASSSGQTNERLF | 0.17815883 |
| LISNASDALDK | ASSSGQTNERLF | 0.27359626 |
| LISNASDAL | ASSSGQTNERLF | 0.29469937 |
| ISNASDALDK | ASSSGQTNERLF | 0.1639825 |
| NQAETLVYQTE | ASSSGQTNERLF | 0.7517294 |
| TGGTGGTAGAG | ASSSGQTNERLF | 0.63258755 |
| TGSTAGGAGGA | ASSSGQTNERLF | 0.65820646 |
| DPLAVPVAAGPV | ASSSGQTNERLF | 0.28219903 |
| GGTGGTPGNGGN | ASSSGQTNERLF | 0.29680896 |
| PAGLAGVDPATL | ASSSGQTNERLF | 0.27082992 |
| GNNLVVEAPHAGASV | ASSSGQTNERLF | 0.7340466 |
| KVTRIMSASTLEQPA | ASSSGQTNERLF | 0.2099882 |
| LGFALPIALPRRTVV | ASSSGQTNERLF | 0.182497 |
| TRLGNLTARKLVVLV | ASSSGQTNERLF | 0.30001366 |
| WSQQRRLIYFPSAGP | ASSSGQTNERLF | 0.19283547 |
| GPNGNPDPMAAAVDIR | ASSSGQTNERLF | 0.7412183 |
| TQRCVSVADLRAAASR | ASSSGQTNERLF | 0.7124442 |
| QFVLDVAEHPVEVLTETV | ASSSGQTNERLF | 0.6134572 |
| GGAGGTGGAGGNGGTQAGDG | ASSSGQTNERLF | 0.6845592 |
| PLLATLRAIAVDVDAKPAQV | ASSSGQTNERLF | 0.17025374 |
| APKFRTGGYMIDFWGVGYQVAK | ASSSGQTNERLF | 0.27692667 |
| GAGGAGGAGGAAGGPGGTGGTG | ASSSGQTNERLF | 0.7779057 |
| IGALGLLATDVAPAGSVAFGVA | ASSSGQTNERLF | 0.55838776 |
| AGGAGGIGGNGGAGGAGGVLMG | ASSSGQTNERLF | 0.39925998 |
| ADELIGLL | ASSSGQTNERLF | 0.729347 |
| ADRALKA | ASSSGQTNERLF | 0.33087373 |
| AGAVGQVGGAGGA | ASSSGQTNERLF | 0.38798022 |
| AGGGGGLGGAP | ASSSGQTNERLF | 0.38445833 |
| AGGLLYGNGGAG | ASSSGQTNERLF | 0.3814155 |
| AGLESIGL | ASSSGQTNERLF | 0.4813893 |

| | | |
|---|---|---|
| AGLKTGII | ASSSGQTNERLF | 0.20693538 |
| ASLAALLL | ASSSGQTNERLF | 0.54353803 |
| ASVINISSI | ASSSGQTNERLF | 0.77077556 |
| ATPPGPA | ASSSGQTNERLF | 0.5516119 |
| ATVASALV | ASSSGQTNERLF | 0.705846 |
| ATVASLIL | ASSSGQTNERLF | 0.6266662 |
| AVNGPVLIRH | ASSSGQTNERLF | 0.7034049 |
| DGAAGAVGI | ASSSGQTNERLF | 0.28445882 |
| DPLLHGI | ASSSGQTNERLF | 0.35385764 |
| ELGAALGAL | ASSSGQTNERLF | 0.399699 |
| ELGRTQAV | ASSSGQTNERLF | 0.3386706 |
| FVAGGYLS | ASSSGQTNERLF | 0.4594884 |
| GASGGAGGAGGW | ASSSGQTNERLF | 0.28618217 |
| GGAGAGGDGGAGGI | ASSSGQTNERLF | 0.7878085 |
| GTGHNGSQPGMGGNG | ASSSGQTNERLF | 0.6688644 |
| GYAAASASAS | ASSSGQTNERLF | 0.57562554 |
| HQVDVAAVQQG | ASSSGQTNERLF | 0.24452081 |
| IGAAIAHT | ASSSGQTNERLF | 0.54355025 |
| IGMIEVAQL | ASSSGQTNERLF | 0.5883826 |
| IRLAPEHTSAMD | ASSSGQTNERLF | 0.18023494 |
| IYNVSVL | ASSSGQTNERLF | 0.7518466 |
| KQGFLGM | ASSSGQTNERLF | 0.29450443 |
| LESVITQL | ASSSGQTNERLF | 0.45844948 |
| LGLGALVVV | ASSSGQTNERLF | 0.640133 |
| LIGNGGNGGSGGI | ASSSGQTNERLF | 0.7665032 |
| LNAINAPTL | ASSSGQTNERLF | 0.32353616 |
| LQLLNTGV | ASSSGQTNERLF | 0.39171895 |
| LQTSIEF | ASSSGQTNERLF | 0.22421665 |
| MSFAVSEL | ASSSGQTNERLF | 0.6968881 |
| NTGALGL | ASSSGQTNERLF | 0.7456336 |
| PPGAGGAGGAGTT | ASSSGQTNERLF | 0.2833983 |
| QLVGLGI | ASSSGQTNERLF | 0.7507515 |
| RFLEQVHQL | ASSSGQTNERLF | 0.4667235 |
| RGVGYVLR | ASSSGQTNERLF | 0.19885784 |
| SATGVFAGVI | ASSSGQTNERLF | 0.4508872 |
| SLPAITLP | ASSSGQTNERLF | 0.6497262 |
| STALAFTV | ASSSGQTNERLF | 0.74814224 |
| SVLTVGLGK | ASSSGQTNERLF | 0.70399714 |
| TALAGAGSGI | ASSSGQTNERLF | 0.59355175 |
| TGLPFTGI | ASSSGQTNERLF | 0.41296813 |
| VAEHVTV | ASSSGQTNERLF | 0.7558017 |
| VDIVLNSL | ASSSGQTNERLF | 0.77388495 |
| VLTGGGAVL | ASSSGQTNERLF | 0.7099228 |
| VNVGLAYGI | ASSSGQTNERLF | 0.48863357 |
| VSSAAVGAV | ASSSGQTNERLF | 0.32688695 |
| VTSVYEAGV | ASSSGQTNERLF | 0.41804254 |
| WVALDRGA | ASSSGQTNERLF | 0.72745574 |
| YLVGSVVI | ASSSGQTNERLF | 0.2502673 |
| GPPGTGKTLL | ASSSGQTNERLF | 0.4962061 |
| GVGMTKF | ASSSGQTNERLF | 0.61226165 |
| IGIGGSDL | ASSSGQTNERLF | 0.19045717 |

| | | |
|---|---|---|
| IGTLLAEL | ASSSGQTNERLF | 0.5860866 |
| IYTSGTT | ASSSGQTNERLF | 0.7404722 |
| LSFPAGV | ASSSGQTNERLF | 0.36019754 |
| LYTSGTT | ASSSGQTNERLF | 0.3091572 |
| VGAFPFGF | ASSSGQTNERLF | 0.74100363 |
| VGAGTVEF | ASSSGQTNERLF | 0.7890476 |
| VGKSTLLL | ASSSGQTNERLF | 0.69162756 |
| AALAHAGGGIV | ASSSGQTNERLF | 0.18646228 |
| AALVAVSLL | ASSSGQTNERLF | 0.1721939 |
| AAVSRAVAAL | ASSSGQTNERLF | 0.39629498 |
| AGGSSGIGGF | ASSSGQTNERLF | 0.6308212 |
| AGLTVANANG | ASSSGQTNERLF | 0.25170162 |
| AGLVGAVL | ASSSGQTNERLF | 0.2826524 |
| ALSQAFTV | ASSSGQTNERLF | 0.6409139 |
| ALTFSVSL | ASSSGQTNERLF | 0.71694505 |
| ALTGAALVAL | ASSSGQTNERLF | 0.23104756 |
| ASFSIGAGVL | ASSSGQTNERLF | 0.21824338 |
| DGSVYKI | ASSSGQTNERLF | 0.47084197 |
| EQLAEIV | ASSSGQTNERLF | 0.6793505 |
| FAVSELTVM | ASSSGQTNERLF | 0.2932808 |
| GAAPLAAGL | ASSSGQTNERLF | 0.5420402 |
| GADGGDGGSGGAGG | ASSSGQTNERLF | 0.2995994 |
| GFGLDTGGAGG | ASSSGQTNERLF | 0.28813168 |
| GLASGGLL | ASSSGQTNERLF | 0.5748036 |
| GRIAGAAVLL | ASSSGQTNERLF | 0.2637431 |
| GSAGSLNTSF | ASSSGQTNERLF | 0.73759365 |
| GTGQIGI | ASSSGQTNERLF | 0.25207013 |
| GTLIALIL | ASSSGQTNERLF | 0.70871276 |
| IAATMGFVVG | ASSSGQTNERLF | 0.59813786 |
| IFAQALL | ASSSGQTNERLF | 0.4395368 |
| IGAAVALGF | ASSSGQTNERLF | 0.42344132 |
| IGLTPLAL | ASSSGQTNERLF | 0.29831362 |
| ILGSNTSTL | ASSSGQTNERLF | 0.20339677 |
| IVFSLTVL | ASSSGQTNERLF | 0.17277183 |
| LASKATGF | ASSSGQTNERLF | 0.7135568 |
| LASVALIL | ASSSGQTNERLF | 0.16143778 |
| LFAVGLTL | ASSSGQTNERLF | 0.6616056 |
| LFNGGIGTYI | ASSSGQTNERLF | 0.6014686 |
| LTLGAGII | ASSSGQTNERLF | 0.22219661 |
| LVAAALGDI | ASSSGQTNERLF | 0.570081 |
| NTAFVQLGI | ASSSGQTNERLF | 0.2595122 |
| QLFSGAEL | ASSSGQTNERLF | 0.25105694 |
| SAATFITL | ASSSGQTNERLF | 0.73433614 |
| SNYGLANLGV | ASSSGQTNERLF | 0.528878 |
| SYGGAYAVM | ASSSGQTNERLF | 0.25496083 |
| TTGVGGLTLG | ASSSGQTNERLF | 0.5061188 |
| TTQTIYI | ASSSGQTNERLF | 0.56973034 |
| VAAGAGLGAVI | ASSSGQTNERLF | 0.26741943 |
| VAAVVTLL | ASSSGQTNERLF | 0.34726256 |
| VGFLGGLI | ASSSGQTNERLF | 0.7202663 |
| VGVIPILL | ASSSGQTNERLF | 0.43340698 |

| | | |
|---|---|---|
| VIFASSLI | ASSSGQTNERLF | 0.24763736 |
| VKGGDGGAAAG | ASSSGQTNERLF | 0.64271116 |
| VTKTFTNI | ASSSGQTNERLF | 0.27279738 |
| VVAAAILL | ASSSGQTNERLF | 0.6290433 |
| YAALDVEL | ASSSGQTNERLF | 0.34244224 |
| LASVALLL | ASSSGQTNERLF | 0.3864808 |
| LDAVHRV | ASSSGQTNERLF | 0.6461787 |
| QGLSSSIF | ASSSGQTNERLF | 0.21361552 |
| RGTFANIRL | ASSSGQTNERLF | 0.27209347 |
| RQLWWGHRI | ASSSGQTNERLF | 0.37737322 |
| YGPPGTGKTLL | ASSSGQTNERLF | 0.43326694 |
| AALVGAPANVA | ASSSGQTNERLF | 0.66742915 |
| AATFRRTE | ASSSGQTNERLF | 0.26092607 |
| AGALGGGI | ASSSGQTNERLF | 0.5435531 |
| AGGAGGAGGIGAG | ASSSGQTNERLF | 0.3008345 |
| AGSGALGL | ASSSGQTNERLF | 0.2319599 |
| AQIADQI | ASSSGQTNERLF | 0.357372 |
| ASAVAVTYL | ASSSGQTNERLF | 0.3211934 |
| ASTVLGGTGG | ASSSGQTNERLF | 0.31145695 |
| AVGGEPAK | ASSSGQTNERLF | 0.45483613 |
| CEQGTFGGTC | ASSSGQTNERLF | 0.21846563 |
| DEGKAPAG | ASSSGQTNERLF | 0.47640544 |
| DTVVFPAL | ASSSGQTNERLF | 0.5522 |
| FEGITMAG | ASSSGQTNERLF | 0.36070347 |
| FPLVVQD | ASSSGQTNERLF | 0.6686537 |
| GAFTTPQ | ASSSGQTNERLF | 0.65344065 |
| GFGGFGTAGGVGGA | ASSSGQTNERLF | 0.64698875 |
| GFLAQFL | ASSSGQTNERLF | 0.28482693 |
| GGAGGAGGNAGVA | ASSSGQTNERLF | 0.25959283 |
| GGVGTAGAGGAGGAGG | ASSSGQTNERLF | 0.66655934 |
| GNGGGGDAGNAGSGGN | ASSSGQTNERLF | 0.6233065 |
| GTGGAGGAGNDAGS | ASSSGQTNERLF | 0.7032956 |
| GTTNTGWFNS | ASSSGQTNERLF | 0.6447409 |
| IFMGPLFTL | ASSSGQTNERLF | 0.603587 |
| ILAHAIEAI | ASSSGQTNERLF | 0.65948766 |
| IRSAIREAL | ASSSGQTNERLF | 0.43699706 |
| ITFAGGG | ASSSGQTNERLF | 0.22057387 |
| ITGMSGHQEV | ASSSGQTNERLF | 0.29267132 |
| IVAGLACTL | ASSSGQTNERLF | 0.3338856 |
| IVGGGLAGA | ASSSGQTNERLF | 0.25513756 |
| IVTGVLNI | ASSSGQTNERLF | 0.19806434 |
| LAQQGCI | ASSSGQTNERLF | 0.7278555 |
| LCLHVEN | ASSSGQTNERLF | 0.7856978 |
| LDAISDGS | ASSSGQTNERLF | 0.25599733 |
| LGGFAAML | ASSSGQTNERLF | 0.7542106 |
| LGGKNPMI | ASSSGQTNERLF | 0.19694598 |
| LGPKGVAAIL | ASSSGQTNERLF | 0.61530364 |
| LMVGVSI | ASSSGQTNERLF | 0.73906815 |
| LVGAAFVL | ASSSGQTNERLF | 0.30166364 |
| LVLAVAPL | ASSSGQTNERLF | 0.53970575 |
| MAAIAVIDAA | ASSSGQTNERLF | 0.24529035 |

| | | |
|---|---|---|
| NNGINAELQ | ASSSGQTNERLF | 0.2648963 |
| RGEPLVLVHG | ASSSGQTNERLF | 0.7444474 |
| RGGAGGAAGLI | ASSSGQTNERLF | 0.3117771 |
| RSYRMHPAV | ASSSGQTNERLF | 0.39196277 |
| SAAHLLAGQLA | ASSSGQTNERLF | 0.31415862 |
| SAAVGSY | ASSSGQTNERLF | 0.22617233 |
| SPHAVLHRI | ASSSGQTNERLF | 0.34689254 |
| SQIAKVL | ASSSGQTNERLF | 0.70729166 |
| TALGHLVP | ASSSGQTNERLF | 0.28570437 |
| TAPTTSGAA | ASSSGQTNERLF | 0.77164125 |
| TDGDDGNGGSPGT | ASSSGQTNERLF | 0.22612764 |
| TDGLGEN | ASSSGQTNERLF | 0.7570437 |
| TGGPPLPGQA | ASSSGQTNERLF | 0.7123787 |
| TIPAITI | ASSSGQTNERLF | 0.61928654 |
| TIPGFTI | ASSSGQTNERLF | 0.30879354 |
| TSIASGALA | ASSSGQTNERLF | 0.74813485 |
| TVTLPGAAI | ASSSGQTNERLF | 0.5310413 |
| VASGLSAALP | ASSSGQTNERLF | 0.64905214 |
| VGAGVIGI | ASSSGQTNERLF | 0.66119945 |
| VGPKRFGV | ASSSGQTNERLF | 0.6429043 |
| VNVAFPD | ASSSGQTNERLF | 0.8377534 |
| VSMTNSGVS | ASSSGQTNERLF | 0.8176102 |
| VTDVLGDLSV | ASSSGQTNERLF | 0.7999654 |
| VVVVGAGFAGL | ASSSGQTNERLF | 0.7554859 |
| VYTVGTVV | ASSSGQTNERLF | 0.52090824 |
| WTTFSGP | ASSSGQTNERLF | 0.7769598 |
| YDELVEAF | ASSSGQTNERLF | 0.7167679 |
| YGTGGAGGNGG | ASSSGQTNERLF | 0.8335298 |
| YTNVMAV | ASSSGQTNERLF | 0.8261041 |
| FLEQVHQL | ASSSGQTNERLF | 0.7145351 |
| GLSLLAVAV | ASSSGQTNERLF | 0.83124655 |
| LEQVHQL | ASSSGQTNERLF | 0.83057666 |
| LNAAGAI | ASSSGQTNERLF | 0.5880945 |
| PAVHASL | ASSSGQTNERLF | 0.7886384 |
| TELEDDA | ASSSGQTNERLF | 0.7461692 |
| TPPGPPN | ASSSGQTNERLF | 0.7905052 |
| TPPPPPPP | ASSSGQTNERLF | 0.43677044 |
| TQSPALI | ASRNRGNSPLY | 0.8247667 |
| EFAAAFPQ | ASRNRGNSPLY | 0.76255107 |
| LLSGNLDVL | ASRNRGNSPLY | 0.21306479 |
| GPANLGNYNI | ASRNRGNSPLY | 0.812982 |
| TLGPTLP | ASRNRGNSPLY | 0.76604205 |
| VYAYSSCCK | ASRNRGNSPLY | 0.77833384 |
| APAPAPAPNI | ASRNRGNSPLY | 0.83184886 |
| PGGPVPIP | ASRNRGNSPLY | 0.6497868 |
| FLTVALEP | ASRNRGNSPLY | 0.66066194 |
| SQYFGANL | ASRNRGNSPLY | 0.8208908 |
| GGVKPAA | ASRNRGNSPLY | 0.83267224 |
| GGTGGAGGAGGW | ASRNRGNSPLY | 0.59988654 |
| GVGGVAGLGGAGG | ASRNRGNSPLY | 0.81484216 |
| VLLVLATTI | ASRNRGNSPLY | 0.836341 |

| | | |
|---|---|---|
| IGIFAAML | ASRNRGNSPLY | 0.80788743 |
| AVGDLID | ASRNRGNSPLY | 0.83765113 |
| NISPLPL | ASRNRGNSPLY | 0.8409961 |
| VTAVFVAI | ASRNRGNSPLY | 0.6911614 |
| SGDDNDAAL | ASRNRGNSPLY | 0.83315635 |
| AVAALAHS | ASRNRGNSPLY | 0.83831924 |
| GGVTVMATP | ASRNRGNSPLY | 0.75486463 |
| AGLAANVRD | ASRNRGNSPLY | 0.78831667 |
| TNRPDLIDPALLRPG | ASRNRGNSPLY | 0.63218254 |
| LANLGVTAL | ASRNRGNSPLY | 0.78830516 |
| TGERIEL | ASRNRGNSPLY | 0.5276136 |
| NGEELAA | ASRNRGNSPLY | 0.81457984 |
| VLSLVWG | ASRNRGNSPLY | 0.8379896 |
| IGYALWL | ASRNRGNSPLY | 0.7855371 |
| LSLLLAPL | ASRNRGNSPLY | 0.6523843 |
| TLDEALK | ASRNRGNSPLY | 0.6323459 |
| TDGAGTAEGG | ASRNRGNSPLY | 0.8060497 |
| NAPGGAGGNGG | ASRNRGNSPLY | 0.69017315 |
| VSVFMGMGGNF | ASRNRGNSPLY | 0.52404845 |
| LASYLAAL | ASRNRGNSPLY | 0.7451614 |
| LVGGTPAL | ASRNRGNSPLY | 0.8253749 |
| VRNADVL | ASRNRGNSPLY | 0.83070135 |
| TCSSPFLVSAM | ASRNRGNSPLY | 0.66018116 |
| VHITDAE | ASRNRGNSPLY | 0.84017533 |
| EYAAPSF | ASRNRGNSPLY | 0.83055645 |
| VVDAFHGGL | ASRNRGNSPLY | 0.8397363 |
| ASVVDPQ | ASRNRGNSPLY | 0.7992655 |
| SYNEFGGLS | ASRNRGNSPLY | 0.67500573 |
| DPDGSVY | ASRNRGNSPLY | 0.8333324 |
| MVSTIHATE | ASRNRGNSPLY | 0.77293766 |
| LAAGVADPV | ASRNRGNSPLY | 0.7742707 |
| TKEQIAA | ASRNRGNSPLY | 0.8259709 |
| SLASPAAL | ASRNRGNSPLY | 0.7872126 |
| IGLVFYGV | ASRNRGNSPLY | 0.71895707 |
| FLLDASNL | ASRNRGNSPLY | 0.79440796 |
| YQVGNFP | ASRNRGNSPLY | 0.8429779 |
| ESLLSQST | ASRNRGNSPLY | 0.6929396 |
| LFTVGFAAVG | ASRNRGNSPLY | 0.56874156 |
| ATPPTPT | ASRNRGNSPLY | 0.764472 |
| TDLATLG | ASRNRGNSPLY | 0.7902125 |
| SAGGTGR | ASRNRGNSPLY | 0.83287597 |
| GFSNSGS | ASRNRGNSPLY | 0.8237842 |
| GGNGGSGGTSV | ASRNRGNSPLY | 0.7685683 |
| GTSVATGGAGN | ASRNRGNSPLY | 0.81573385 |
| NGGNGGSGGTS | ASRNRGNSPLY | 0.8260846 |
| AVVKFPH | ASRNRGNSPLY | 0.8044697 |
| TPPPPPP | ASRNRGNSPLY | 0.8260902 |
| AAPAAAAAPA | ASRNRGNSPLY | 0.75356907 |
| IALNEDL | ASRNRGNSPLY | 0.839954 |
| IVNVSSM | ASRNRGNSPLY | 0.8279339 |
| TGIPPAPR | ASRNRGNSPLY | 0.8308633 |

| | | |
|---|---|---|
| FDLGGGTF | ASRNRGNSPLY | 0.83480895 |
| FDLGGGTFDVS | ASRNRGNSPLY | 0.8037131 |
| NEPTAAA | ASRNRGNSPLY | 0.8122754 |
| NEPTAAALA | ASRNRGNSPLY | 0.5271286 |
| GSIINISSI | ASRNRGNSPLY | 0.7374834 |
| VAPGNAGTA | ASRNRGNSPLY | 0.8010231 |
| TGGGASLE | ASRNRGNSPLY | 0.8170129 |
| VAVYNHYKRI | ASRNRGNSPLY | 0.8268068 |
| TDPAPATT | ASRNRGNSPLY | 0.81777394 |
| TNRPDILDPALLRPG | ASRNRGNSPLY | 0.72115785 |
| LTTTTTS | ASRNRGNSPLY | 0.83680606 |
| PTGSGKTL | ASRNRGNSPLY | 0.8353902 |
| TGSGKTL | ASRNRGNSPLY | 0.6573019 |
| TIGHVDHGK | ASRNRGNSPLY | 0.759745 |
| TIGHVDHGKTT | ASRNRGNSPLY | 0.84404767 |
| GHVDHGKTT | ASRNRGNSPLY | 0.81643915 |
| AQFEHTLL | ASRNRGNSPLY | 0.765306 |
| IVNQDEL | ASRNRGNSPLY | 0.7736118 |
| TNGSQFF | ASRNRGNSPLY | 0.8411339 |
| TGKGIVE | ASRNRGNSPLY | 0.7805301 |
| LISNASDA | ASRNRGNSPLY | 0.840249 |
| AELRTLGELTKIAKAHGAQ | ASRNRGNSPLY | 0.38904583 |
| APNAVAAAEPSW | ASRNRGNSPLY | 0.79707193 |
| APQVAGAATAVL | ASRNRGNSPLY | 0.76655734 |
| ARRDPAAAAAILPTDA | ASRNRGNSPLY | 0.29315272 |
| DAAARDVLAEAGLAEYF | ASRNRGNSPLY | 0.8281451 |
| DDQIVLSAKANLAKVEQ | ASRNRGNSPLY | 0.7192712 |
| DELLPDTGEDSDEDGH | ASRNRGNSPLY | 0.8212681 |
| ELPEAPANRATVPQD | ASRNRGNSPLY | 0.7585183 |
| FGQSKIERVTGIG | ASRNRGNSPLY | 0.8329152 |
| GAGGTGSTAGGAGGAGGAGGLY | ASRNRGNSPLY | 0.80631274 |
| GHGGTGAAGGNGGTGGAGGSGI | ASRNRGNSPLY | 0.78243417 |
| GNGGNGGAGGEGGPGGAGGAGA | ASRNRGNSPLY | 0.6868045 |
| GSTAGNGGNGGNATNGTIAGQP | ASRNRGNSPLY | 0.8174931 |
| GTGGDGGNGGNGGTGG | ASRNRGNSPLY | 0.8366631 |
| GVRRPAGIPADAPV | ASRNRGNSPLY | 0.8383024 |
| GYRQVTDAHLLTLARR | ASRNRGNSPLY | 0.39865324 |
| IAKMGQDGHDRG | ASRNRGNSPLY | 0.8128277 |
| KAEPSIEIQYT | ASRNRGNSPLY | 0.7783356 |
| KPVRIRMAADEDTVAVT | ASRNRGNSPLY | 0.29435772 |
| MPPRIAGMRLLVIKPEP | ASRNRGNSPLY | 0.3462033 |
| MKFYAVALFRSRGVPPTR | ASRNRGNSPLY | 0.78430426 |
| NAGGFGILGAVAHSP | ASRNRGNSPLY | 0.78880537 |
| NEGREITLAMAEM | ASRNRGNSPLY | 0.8237342 |
| NELVEWSHPLGHRWRWE | ASRNRGNSPLY | 0.81904596 |
| NLGAGGTGGNGGNPG | ASRNRGNSPLY | 0.7805091 |
| NTEWPKWETVMAADPVD | ASRNRGNSPLY | 0.82084763 |
| PASAAPAETQP | ASRNRGNSPLY | 0.7854743 |
| QQSSDGKNGVRAAEITGE | ASRNRGNSPLY | 0.14261174 |
| RAPDAVTTSWWKE | ASRNRGNSPLY | 0.728923 |
| SGFGNVGAHTSGWFNQ | ASRNRGNSPLY | 0.56777626 |

| | | |
|---|---|---|
| SGVAGAVGQAASVGGLKV | ASRNRGNSPLY | 0.8382657 |
| SNGGAGTGSAGGNGGT | ASRNRGNSPLY | 0.68426156 |
| TGGAGGEGGAAG | ASRNRGNSPLY | 0.7535519 |
| TGGDGGAAGAGGAPGHG | ASRNRGNSPLY | 0.794285 |
| TMPPLLAAIANAGRHGV | ASRNRGNSPLY | 0.8172135 |
| TSSTAGNSGGA | ASRNRGNSPLY | 0.796903 |
| VPATVPVGILC | ASRNRGNSPLY | 0.7488047 |
| VQRPDSGGVVLTGRIS | ASRNRGNSPLY | 0.36106864 |
| WTPNPVPATKGVPAGLVQAT | ASRNRGNSPLY | 0.8166194 |
| YALKRALAKFSDRFDVVII | ASRNRGNSPLY | 0.6555785 |
| YLIGSVIV | ASRNRGNSPLY | 0.6299777 |
| YGQLVAVID | ASRNRGNSPLY | 0.5760342 |
| VTVGGMTLPA | ASRNRGNSPLY | 0.762738 |
| VLALPAV | ASRNRGNSPLY | 0.48248637 |
| VIPPTLNYET | ASRNRGNSPLY | 0.8334204 |
| VGMGGMHPSSG | ASRNRGNSPLY | 0.73064417 |
| VGLAGGAATA | ASRNRGNSPLY | 0.7725493 |
| VGGAGGAGGSGP | ASRNRGNSPLY | 0.8411337 |
| VFAAFMV | ASRNRGNSPLY | 0.8057488 |
| TLYDKLR | ASRNRGNSPLY | 0.7844537 |
| TLAVFTVL | ASRNRGNSPLY | 0.7190417 |
| TGSVQPARV | ASRNRGNSPLY | 0.68887925 |
| TGGGAGSGGT | ASRNRGNSPLY | 0.73524976 |
| SHLETSL | ASRNRGNSPLY | 0.7265167 |
| SFTTGMT | ASRNRGNSPLY | 0.8391311 |
| QVEVAKRMGVRQPTVS | ASRNRGNSPLY | 0.80281746 |
| QGSGGPALAM | ASRNRGNSPLY | 0.617543 |
| MLAPSNP | ASRNRGNSPLY | 0.81899756 |
| LVNNAAI | ASRNRGNSPLY | 0.8402214 |
| LTGEINTLA | ASRNRGNSPLY | 0.8427865 |
| LSIFGPV | ASRNRGNSPLY | 0.70213234 |
| LLALLANL | ASRNRGNSPLY | 0.77456367 |
| LISNASDALDKLR | ASRNRGNSPLY | 0.75475 |
| LINEAVP | ASRNRGNSPLY | 0.78196335 |
| LGYAFSGL | ASRNRGNSPLY | 0.6984583 |
| LDFVAVAL | ASRNRGNSPLY | 0.66916966 |
| LAQPGAE | ASRNRGNSPLY | 0.8356578 |
| ISNLGAVTGF | ASRNRGNSPLY | 0.8450836 |
| HQEDPGRI | ASRNRGNSPLY | 0.77934164 |
| GTPGEPTP | ASRNRGNSPLY | 0.551226 |
| GFFNSGS | ASRNRGNSPLY | 0.78006524 |
| GAGGGGLTAGAN | ASRNRGNSPLY | 0.7247738 |
| FVDALVNM | ASRNRGNSPLY | 0.7504796 |
| FLTVAPEL | ASRNRGNSPLY | 0.72353506 |
| FETLTAL | ASRNRGNSPLY | 0.61676466 |
| DVINPIEL | ASRNRGNSPLY | 0.79510283 |
| APVSGAPV | ASRNRGNSPLY | 0.8419314 |
| ALVGGLAW | ASRNRGNSPLY | 0.7794853 |
| ALAAAAATAI | ASRNRGNSPLY | 0.8161241 |
| AFGLGLLL | ASRNRGNSPLY | 0.83909535 |
| AFFAAASL | ASRNRGNSPLY | 0.8278064 |

| | | |
|---|---|---|
| AAAALPAFSPP | ASRNRGNSPLY | 0.83761066 |
| VVLAPAL | ASRNRGNSPLY | 0.83287895 |
| TIGHVDHGKTTL | ASRNRGNSPLY | 0.7934512 |
| LISNASDALDK | ASRNRGNSPLY | 0.80777717 |
| LISNASDAL | ASRNRGNSPLY | 0.82853305 |
| ISNASDALDK | ASRNRGNSPLY | 0.78059953 |
| NQAETLVYQTE | ASRNRGNSPLY | 0.79637545 |
| TGGTGGTAGAG | ASRNRGNSPLY | 0.76332426 |
| TGSTAGGAGGA | ASRNRGNSPLY | 0.8266526 |
| DPLAVPVAAGPV | ASRNRGNSPLY | 0.79732794 |
| GGTGGTPGNGGN | ASRNRGNSPLY | 0.8422768 |
| PAGLAGVDPATL | ASRNRGNSPLY | 0.3082447 |
| GNNLVVEAPHAGASV | ASRNRGNSPLY | 0.65890473 |
| KVTRIMSASTLEQPA | ASRNRGNSPLY | 0.8343103 |
| LGFALPIALPRRTVV | ASRNRGNSPLY | 0.19035257 |
| TRLGNLTARKLVVLV | ASRNRGNSPLY | 0.36132756 |
| WSQQRRLIYFPSAGP | ASRNRGNSPLY | 0.81248856 |
| GPNGNPDPMAAAVDIR | ASRNRGNSPLY | 0.8349062 |
| TQRCVSVADLRAAASR | ASRNRGNSPLY | 0.19467986 |
| QFVLDVAEHPVEVLTETV | ASRNRGNSPLY | 0.6914834 |
| GGAGGTGGAGGNGGTQAGDG | ASRNRGNSPLY | 0.55261636 |
| PLLATLRAIAVDVDAKPAQV | ASRNRGNSPLY | 0.8297747 |
| APKFRTGGYMIDFWGVGYQVAK | ASRNRGNSPLY | 0.78814113 |
| GAGGAGGAGGAAGGPGGTGGTG | ASRNRGNSPLY | 0.4720051 |
| IGALGLLATDVAPAGSVAFGVA | ASRNRGNSPLY | 0.8293813 |
| AGGAGGIGGNGGAGGAGGVLMG | ASRNRGNSPLY | 0.5899302 |
| ADELIGLL | ASRNRGNSPLY | 0.77675545 |
| ADRALKA | ASRNRGNSPLY | 0.8307583 |
| AGAVGQVGGAGGA | ASRNRGNSPLY | 0.84240437 |
| AGGGGGLGGAP | ASRNRGNSPLY | 0.5706644 |
| AGGLLYGNGGAG | ASRNRGNSPLY | 0.8217355 |
| AGLESIGL | ASRNRGNSPLY | 0.76655257 |
| AGLKTGII | ASRNRGNSPLY | 0.84455085 |
| ASLAALLL | ASRNRGNSPLY | 0.7620332 |
| ASVINISSI | ASRNRGNSPLY | 0.5763977 |
| ATPPGPA | ASRNRGNSPLY | 0.8335979 |
| ATVASALV | ASRNRGNSPLY | 0.82981133 |
| ATVASLIL | ASRNRGNSPLY | 0.7672743 |
| AVNGPVLIRH | ASRNRGNSPLY | 0.83391273 |
| DGAAGAVGI | ASRNRGNSPLY | 0.78758687 |
| DPLLHGI | ASRNRGNSPLY | 0.78108966 |
| ELGAALGAL | ASRNRGNSPLY | 0.8309374 |
| ELGRTQAV | ASRNRGNSPLY | 0.82356346 |
| FVAGGYLS | ASRNRGNSPLY | 0.6838268 |
| GASGGAGGAGGW | ASRNRGNSPLY | 0.8376449 |
| GGAGAGGDGGAGGI | ASRNRGNSPLY | 0.7927251 |
| GTGHNGSQPGMGGNG | ASRNRGNSPLY | 0.65314716 |
| GYAAASASAS | ASRNRGNSPLY | 0.8429569 |
| HQVDVAAVQQG | ASRNRGNSPLY | 0.8025491 |
| IGAAIAHT | ASRNRGNSPLY | 0.8313526 |
| IGMIEVAQL | ASRNRGNSPLY | 0.82718045 |

| | | |
|---|---|---|
| IRLAPEHTSAMD | ASRNRGNSPLY | 0.8420881 |
| IYNVSVL | ASRNRGNSPLY | 0.6166721 |
| KQGFLGM | ASRNRGNSPLY | 0.80290437 |
| LESVITQL | ASRNRGNSPLY | 0.7981457 |
| LGLGALVVV | ASRNRGNSPLY | 0.8271609 |
| LIGNGGNGGSGGI | ASRNRGNSPLY | 0.78479946 |
| LNAINAPTL | ASRNRGNSPLY | 0.75457275 |
| LQLLNTGV | ASRNRGNSPLY | 0.823248 |
| LQTSIEF | ASRNRGNSPLY | 0.8387008 |
| MSFAVSEL | ASRNRGNSPLY | 0.8054441 |
| NTGALGL | ASRNRGNSPLY | 0.8099923 |
| PPGAGGAGGAGTT | ASRNRGNSPLY | 0.8412031 |
| QLVGLGI | ASRNRGNSPLY | 0.69689214 |
| RFLEQVHQL | ASRNRGNSPLY | 0.7862373 |
| RGVGYVLR | ASRNRGNSPLY | 0.8281562 |
| SATGVFAGVI | ASRNRGNSPLY | 0.8323787 |
| SLPAITLP | ASRNRGNSPLY | 0.7972547 |
| STALAFTV | ASRNRGNSPLY | 0.8382967 |
| SVLTVGLGK | ASRNRGNSPLY | 0.7263794 |
| TALAGAGSGI | ASRNRGNSPLY | 0.8132799 |
| TGLPFTGI | ASRNRGNSPLY | 0.7572906 |
| VAEHVTV | ASRNRGNSPLY | 0.7907932 |
| VDIVLNSL | ASRNRGNSPLY | 0.69842356 |
| VLTGGGAVL | ASRNRGNSPLY | 0.8091589 |
| VNVGLAYGI | ASRNRGNSPLY | 0.82976365 |
| VSSAAVGAV | ASRNRGNSPLY | 0.8210789 |
| VTSVYEAGV | ASRNRGNSPLY | 0.31154147 |
| WVALDRGA | ASRNRGNSPLY | 0.82473904 |
| YLVGSVVI | ASRNRGNSPLY | 0.82040954 |
| GPPGTGKTLL | ASRNRGNSPLY | 0.8447435 |
| GVGMTKF | ASRNRGNSPLY | 0.7943048 |
| IGIGGSDL | ASRNRGNSPLY | 0.81689984 |
| IGTLLAEL | ASRNRGNSPLY | 0.7086483 |
| IYTSGTT | ASRNRGNSPLY | 0.81331563 |
| LSFPAGV | ASRNRGNSPLY | 0.6336509 |
| LYTSGTT | ASRNRGNSPLY | 0.7151735 |
| VGAFPFGF | ASRNRGNSPLY | 0.7617252 |
| VGAGTVEF | ASRNRGNSPLY | 0.74536 |
| VGKSTLLL | ASRNRGNSPLY | 0.8293228 |
| AALAHAGGGIV | ASRNRGNSPLY | 0.83407974 |
| AALVAVSLL | ASRNRGNSPLY | 0.82917887 |
| AAVSRAVAAL | ASRNRGNSPLY | 0.8353429 |
| AGGSSGIGGF | ASRNRGNSPLY | 0.83977526 |
| AGLTVANANG | ASRNRGNSPLY | 0.83927333 |
| AGLVGAVL | ASRNRGNSPLY | 0.7732611 |
| ALSQAFTV | ASRNRGNSPLY | 0.77456766 |
| ALTFSVSL | ASRNRGNSPLY | 0.62862855 |
| ALTGAALVAL | ASRNRGNSPLY | 0.7623453 |
| ASFSIGAGVL | ASRNRGNSPLY | 0.7724132 |
| DGSVYKI | ASRNRGNSPLY | 0.80231524 |
| EQLAEIV | ASRNRGNSPLY | 0.16921286 |

| | | |
|---|---|---|
| FAVSELTVM | ASRNRGNSPLY | 0.8139419 |
| GAAPLAAGL | ASRNRGNSPLY | 0.8404561 |
| GADGGDGGSGGAGG | ASRNRGNSPLY | 0.75567895 |
| GFGLDTGGAGG | ASRNRGNSPLY | 0.81776786 |
| GLASGGLL | ASRNRGNSPLY | 0.82108617 |
| GRIAGAAVLL | ASRNRGNSPLY | 0.7726799 |
| GSAGSLNTSF | ASRNRGNSPLY | 0.8227836 |
| GTGQIGI | ASRNRGNSPLY | 0.7484027 |
| GTLIALIL | ASRNRGNSPLY | 0.82972205 |
| IAATMGFVVG | ASRNRGNSPLY | 0.84033954 |
| IFAQALL | ASRNRGNSPLY | 0.81482714 |
| IGAAVALGF | ASRNRGNSPLY | 0.7926401 |
| IGLTPLAL | ASRNRGNSPLY | 0.7988679 |
| ILGSNTSTL | ASRNRGNSPLY | 0.51727605 |
| IVFSLTVL | ASRNRGNSPLY | 0.8050487 |
| LASKATGF | ASRNRGNSPLY | 0.5768802 |
| LASVALIL | ASRNRGNSPLY | 0.80033445 |
| LFAVGLTL | ASRNRGNSPLY | 0.7203435 |
| LFNGGIGTYI | ASRNRGNSPLY | 0.80422735 |
| LTLGAGII | ASRNRGNSPLY | 0.78931344 |
| LVAAALGDI | ASRNRGNSPLY | 0.8160881 |
| NTAFVQLGI | ASRNRGNSPLY | 0.6725809 |
| QLFSGAEL | ASRNRGNSPLY | 0.8171221 |
| SAATFITL | ASRNRGNSPLY | 0.7851309 |
| SNYGLANLGV | ASRNRGNSPLY | 0.59808195 |
| SYGGAYAVM | ASRNRGNSPLY | 0.83341223 |
| TTGVGGLTLG | ASRNRGNSPLY | 0.8034386 |
| TTQTIYI | ASRNRGNSPLY | 0.792964 |
| VAAGAGLGAVI | ASRNRGNSPLY | 0.8303533 |
| VAAVVTLL | ASRNRGNSPLY | 0.80535877 |
| VGFLGGLI | ASRNRGNSPLY | 0.58058816 |
| VGVIPILL | ASRNRGNSPLY | 0.7744744 |
| VIFASSLI | ASRNRGNSPLY | 0.793988 |
| VKGGDGGAAAG | ASRNRGNSPLY | 0.7865344 |
| VTKTFTNI | ASRNRGNSPLY | 0.27502838 |
| VVAAAILL | ASRNRGNSPLY | 0.82213694 |
| YAALDVEL | ASRNRGNSPLY | 0.64530945 |
| LASVALLL | ASRNRGNSPLY | 0.73504275 |
| LDAVHRV | ASRNRGNSPLY | 0.78484195 |
| QGLSSSIF | ASRNRGNSPLY | 0.8306894 |
| RGTFANIRL | ASRNRGNSPLY | 0.7814404 |
| RQLWWGHRI | ASRNRGNSPLY | 0.75955117 |
| YGPPGTGKTLL | ASRNRGNSPLY | 0.82521737 |
| AALVGAPANVA | ASRNRGNSPLY | 0.7953948 |
| AATFRRTE | ASRNRGNSPLY | 0.83278257 |
| AGALGGGI | ASRNRGNSPLY | 0.8022918 |
| AGGAGGAGGIGAG | ASRNRGNSPLY | 0.8124063 |
| AGSGALGL | ASRNRGNSPLY | 0.8109584 |
| AQIADQI | ASRNRGNSPLY | 0.7550138 |
| ASAVAVTYL | ASRNRGNSPLY | 0.6681441 |
| ASTVLGGTGG | ASRNRGNSPLY | 0.63320404 |

| | | |
|---|---|---|
| AVGGEPAK | ASRNRGNSPLY | 0.7033185 |
| CEQGTFGGTC | ASRNRGNSPLY | 0.53736377 |
| DEGKAPAG | ASRNRGNSPLY | 0.8011428 |
| DTVVFPAL | ASRNRGNSPLY | 0.83925843 |
| FEGITMAG | ASRNRGNSPLY | 0.789449 |
| FPLVVQD | ASRNRGNSPLY | 0.5184411 |
| GAFTTPQ | ASRNRGNSPLY | 0.7718403 |
| GFGGFGTAGGVGGA | ASRNRGNSPLY | 0.75827885 |
| GFLAQFL | ASRNRGNSPLY | 0.8257804 |
| GGAGGAGGNAGVA | ASRNRGNSPLY | 0.83107185 |
| GGVGTAGAGGAGGAGG | ASRNRGNSPLY | 0.8432511 |
| GNGGGGDAGNAGSGGN | ASRNRGNSPLY | 0.8278842 |
| GTGGAGGAGNDAGS | ASRNRGNSPLY | 0.75207883 |
| GTTNTGWFNS | ASRNRGNSPLY | 0.8294462 |
| IFMGPLFTL | ASRNRGNSPLY | 0.78547734 |
| ILAHAIEAI | ASRNRGNSPLY | 0.8076388 |
| IRSAIREAL | ASRNRGNSPLY | 0.8053578 |
| ITFAGGG | ASRNRGNSPLY | 0.8193393 |
| ITGMSGHQEV | ASRNRGNSPLY | 0.76252186 |
| IVAGLACTL | ASRNRGNSPLY | 0.7694088 |
| IVGGGLAGA | ASRNRGNSPLY | 0.50408137 |
| IVTGVLNI | ASRNRGNSPLY | 0.8317495 |
| LAQQGCI | ASRNRGNSPLY | 0.6951649 |
| LCLHVEN | ASRNRGNSPLY | 0.669683 |
| LDAISDGS | ASRNRGNSPLY | 0.7575345 |
| LGGFAAML | ASRNRGNSPLY | 0.84045887 |
| LGGKNPMI | ASRNRGNSPLY | 0.8369409 |
| LGPKGVAAIL | ASRNRGNSPLY | 0.75150126 |
| LMVGVSI | ASRNRGNSPLY | 0.72969615 |
| LVGAAFVL | ASRNRGNSPLY | 0.81805897 |
| LVLAVAPL | ASRNRGNSPLY | 0.67946374 |
| MAAIAVIDAA | ASRNRGNSPLY | 0.8286936 |
| NNGINAELQ | ASRNRGNSPLY | 0.79287136 |
| RGEPLVLVHG | ASRNRGNSPLY | 0.7867447 |
| RGGAGGAAGLI | ASRNRGNSPLY | 0.7466717 |
| RSYRMHPAV | ASRNRGNSPLY | 0.822193 |
| SAAHLLAGQLA | ASRNRGNSPLY | 0.7622888 |
| SAAVGSY | ASRNRGNSPLY | 0.82923424 |
| SPHAVLHRI | ASRNRGNSPLY | 0.23593378 |
| SQIAKVL | ASRNRGNSPLY | 0.8277781 |
| TALGHLVP | ASRNRGNSPLY | 0.6774327 |
| TAPTTSGAA | ASRNRGNSPLY | 0.7509078 |
| TDGDDGNGGSPGT | ASRNRGNSPLY | 0.34482756 |
| TDGLGEN | ASRNRGNSPLY | 0.7782836 |
| TGGPPLPGQA | ASRNRGNSPLY | 0.8241289 |
| TIPAITI | ASRNRGNSPLY | 0.8366837 |
| TIPGFTI | ASRNRGNSPLY | 0.7752527 |
| TSIASGALA | ASRNRGNSPLY | 0.8273957 |
| TVTLPGAAI | ASRNRGNSPLY | 0.8365464 |
| VASGLSAALP | ASRNRGNSPLY | 0.77215075 |
| VGAGVIGI | ASRNRGNSPLY | 0.33900428 |

| | | |
|---|---|---|
| VGPKRFGV | ASRNRGNSPLY | 0.8016389 |
| VNVAFPD | ASRNRGNSPLY | 0.80011654 |
| VSMTNSGVS | ASRNRGNSPLY | 0.83518636 |
| VTDVLGDLSV | ASRNRGNSPLY | 0.80810654 |
| VVVVGAGFAGL | ASRNRGNSPLY | 0.65054137 |
| VYTVGTVV | ASRNRGNSPLY | 0.80699456 |
| WTTFSGP | ASRNRGNSPLY | 0.82242936 |
| YDELVEAF | ASRNRGNSPLY | 0.64408815 |
| YGTGGAGGNGG | ASRNRGNSPLY | 0.71075106 |
| YTNVMAV | ASRNRGNSPLY | 0.70505214 |
| FLEQVHQL | ASRNRGNSPLY | 0.5225806 |
| GLSLLAVAV | ASRNRGNSPLY | 0.80690134 |
| LEQVHQL | ASRNRGNSPLY | 0.7674966 |
| LNAAGAI | ASRNRGNSPLY | 0.7362772 |
| PAVHASL | ASRNRGNSPLY | 0.79514056 |
| TELEDDA | ASRNRGNSPLY | 0.7307198 |
| TPPGPPN | ASRNRGNSPLY | 0.71841383 |
| TPPPPPPP | ASRNRGNSPLY | 0.7495679 |
| TQSPALI | ASSPDWGTSAETLY | 0.807631 |
| EFAAAFPQ | ASSPDWGTSAETLY | 0.82933223 |
| LLSGNLDVL | ASSPDWGTSAETLY | 0.79017633 |
| GPANLGNYNI | ASSPDWGTSAETLY | 0.8013191 |
| TLGPTLP | ASSPDWGTSAETLY | 0.83675843 |
| VYAYSSCCK | ASSPDWGTSAETLY | 0.6008495 |
| APAPAPAPNI | ASSPDWGTSAETLY | 0.688196 |
| PGGPVPIP | ASSPDWGTSAETLY | 0.8256149 |
| FLTVALEP | ASSPDWGTSAETLY | 0.74322325 |
| SQYFGANL | ASSPDWGTSAETLY | 0.8288233 |
| GGVKPAA | ASSPDWGTSAETLY | 0.7963197 |
| GGTGGAGGAGGW | ASSPDWGTSAETLY | 0.8172228 |
| GVGGVAGLGGAGG | ASSPDWGTSAETLY | 0.805738 |
| VLLVLATTI | ASSPDWGTSAETLY | 0.8425326 |
| IGIFAAML | ASSPDWGTSAETLY | 0.8255595 |
| AVGDLID | ASSPDWGTSAETLY | 0.7720255 |
| NISPLPL | ASSPDWGTSAETLY | 0.82862175 |
| VTAVFVAI | ASSPDWGTSAETLY | 0.5708432 |
| SGDDNDAAL | ASSPDWGTSAETLY | 0.8306303 |
| AVAALAHS | ASSPDWGTSAETLY | 0.8255708 |
| GGVTVMATP | ASSPDWGTSAETLY | 0.7181258 |
| AGLAANVRD | ASSPDWGTSAETLY | 0.7818072 |
| TNRPDLIDPALLRPG | ASSPDWGTSAETLY | 0.841151 |
| LANLGVTAL | ASSPDWGTSAETLY | 0.8360671 |
| TGERIEL | ASSPDWGTSAETLY | 0.67691034 |
| NGEELAA | ASSPDWGTSAETLY | 0.75268763 |
| VLSLVWG | ASSPDWGTSAETLY | 0.61451674 |
| IGYALWL | ASSPDWGTSAETLY | 0.77720565 |
| LSLLLAPL | ASSPDWGTSAETLY | 0.42227453 |
| TLDEALK | ASSPDWGTSAETLY | 0.82445854 |
| TDGAGTAEGG | ASSPDWGTSAETLY | 0.74522185 |
| NAPGGAGGNGG | ASSPDWGTSAETLY | 0.829982 |
| VSVFMGMGGNF | ASSPDWGTSAETLY | 0.8094807 |

| | | |
|---|---|---|
| LASYLAAL | ASSPDWGTSAETLY | 0.83450073 |
| LVGGTPAL | ASSPDWGTSAETLY | 0.7523906 |
| VRNADVL | ASSPDWGTSAETLY | 0.7543719 |
| TCSSPFLVSAM | ASSPDWGTSAETLY | 0.50345385 |
| VHITDAE | ASSPDWGTSAETLY | 0.57758427 |
| EYAAPSF | ASSPDWGTSAETLY | 0.7243065 |
| VVDAFHGGL | ASSPDWGTSAETLY | 0.75938773 |
| ASVVDPQ | ASSPDWGTSAETLY | 0.358922 |
| SYNEFGGLS | ASSPDWGTSAETLY | 0.8416346 |
| DPDGSVY | ASSPDWGTSAETLY | 0.7818205 |
| MVSTIHATE | ASSPDWGTSAETLY | 0.8259066 |
| LAAGVADPV | ASSPDWGTSAETLY | 0.6783998 |
| TKEQIAA | ASSPDWGTSAETLY | 0.6778517 |
| SLASPAAL | ASSPDWGTSAETLY | 0.77084434 |
| IGLVFYGV | ASSPDWGTSAETLY | 0.8203526 |
| FLLDASNL | ASSPDWGTSAETLY | 0.64469993 |
| YQVGNFP | ASSPDWGTSAETLY | 0.39195356 |
| ESLLSQST | ASSPDWGTSAETLY | 0.70326054 |
| LFTVGFAAVG | ASSPDWGTSAETLY | 0.6787584 |
| ATPPTPT | ASSPDWGTSAETLY | 0.8433958 |
| TDLATLG | ASSPDWGTSAETLY | 0.83516324 |
| SAGGTGR | ASSPDWGTSAETLY | 0.80936307 |
| GFSNSGS | ASSPDWGTSAETLY | 0.84341186 |
| GGNGGSGGTSV | ASSPDWGTSAETLY | 0.19601995 |
| GTSVATGGAGN | ASSPDWGTSAETLY | 0.73929536 |
| NGGNGGSGGTS | ASSPDWGTSAETLY | 0.836106 |
| AVVKFPH | ASSPDWGTSAETLY | 0.8238952 |
| TPPPPPP | ASSPDWGTSAETLY | 0.77594376 |
| AAPAAAAAPA | ASSPDWGTSAETLY | 0.83510673 |
| IALNEDL | ASSPDWGTSAETLY | 0.6969045 |
| IVNVSSM | ASSPDWGTSAETLY | 0.77369976 |
| TGIPPAPR | ASSPDWGTSAETLY | 0.79945827 |
| FDLGGGTF | ASSPDWGTSAETLY | 0.826553 |
| FDLGGGTFDVS | ASSPDWGTSAETLY | 0.8129626 |
| NEPTAAA | ASSPDWGTSAETLY | 0.55405354 |
| NEPTAAALA | ASSPDWGTSAETLY | 0.8111255 |
| GSIINISSI | ASSPDWGTSAETLY | 0.8444941 |
| VAPGNAGTA | ASSPDWGTSAETLY | 0.7652972 |
| TGGGASLE | ASSPDWGTSAETLY | 0.8247683 |
| VAVYNHYKRI | ASSPDWGTSAETLY | 0.8080481 |
| TDPAPATT | ASSPDWGTSAETLY | 0.78745985 |
| TNRPDILDPALLRPG | ASSPDWGTSAETLY | 0.71713996 |
| LTTTTTS | ASSPDWGTSAETLY | 0.72042066 |
| PTGSGKTL | ASSPDWGTSAETLY | 0.81269914 |
| TGSGKTL | ASSPDWGTSAETLY | 0.65232736 |
| TIGHVDHGK | ASSPDWGTSAETLY | 0.7825757 |
| TIGHVDHGKTT | ASSPDWGTSAETLY | 0.7821562 |
| GHVDHGKTT | ASSPDWGTSAETLY | 0.84708214 |
| AQFEHTLL | ASSPDWGTSAETLY | 0.82194465 |
| IVNQDEL | ASSPDWGTSAETLY | 0.8234395 |
| TNGSQFF | ASSPDWGTSAETLY | 0.70265377 |

| | | |
|---|---|---|
| TGKGIVE | ASSPDWGTSAETLY | 0.8210683 |
| LISNASDA | ASSPDWGTSAETLY | 0.82244754 |
| AELRTLGELTKIAKAHGAQ | ASSPDWGTSAETLY | 0.58777833 |
| APNAVAAAEPSW | ASSPDWGTSAETLY | 0.83258194 |
| APQVAGAATAVL | ASSPDWGTSAETLY | 0.6701728 |
| ARRDPAAAAAILPTDA | ASSPDWGTSAETLY | 0.79986286 |
| DAAARDVLAEAGLAEYF | ASSPDWGTSAETLY | 0.8210854 |
| DDQIVLSAKANLAKVEQ | ASSPDWGTSAETLY | 0.83567524 |
| DELLPDTGEDSDEDGH | ASSPDWGTSAETLY | 0.78453636 |
| ELPEAPANRATVPQD | ASSPDWGTSAETLY | 0.8127765 |
| FGQSKIERVTGIG | ASSPDWGTSAETLY | 0.69401586 |
| GAGGTGSTAGGAGGAGGAGGLY | ASSPDWGTSAETLY | 0.83152723 |
| GHGGTGAAGGNGGTGGAGGSGI | ASSPDWGTSAETLY | 0.83375967 |
| GNGGNGGAGGEGGPGGAGGAGA | ASSPDWGTSAETLY | 0.8099349 |
| GSTAGNGGNGGNATNGTIAGQP | ASSPDWGTSAETLY | 0.8456157 |
| GTGGDGGNGGNGGTGG | ASSPDWGTSAETLY | 0.8091186 |
| GVRRPAGIPADAPV | ASSPDWGTSAETLY | 0.37693453 |
| GYRQVTDAHLLTLARR | ASSPDWGTSAETLY | 0.7912893 |
| IAKMGQDGHDRG | ASSPDWGTSAETLY | 0.83481836 |
| KAEPSIEIQYT | ASSPDWGTSAETLY | 0.8375342 |
| KPVRIRMAADEDTVAVT | ASSPDWGTSAETLY | 0.8324714 |
| MPPRIAGMRLLVIKPEP | ASSPDWGTSAETLY | 0.81789875 |
| MKFYAVALFRSRGVPPTR | ASSPDWGTSAETLY | 0.7743055 |
| NAGGFGILGAVAHSP | ASSPDWGTSAETLY | 0.83851904 |
| NEGREITLAMAEM | ASSPDWGTSAETLY | 0.8474953 |
| NELVEWSHPLGHRWRWE | ASSPDWGTSAETLY | 0.823476 |
| NLGAGGTGGNGGNPG | ASSPDWGTSAETLY | 0.7416289 |
| NTEWPKWETVMAADPVD | ASSPDWGTSAETLY | 0.709505 |
| PASAAPAETQP | ASSPDWGTSAETLY | 0.821443 |
| QQSSDGKNGVRAAEITGE | ASSPDWGTSAETLY | 0.83349675 |
| RAPDAVTTSWWKE | ASSPDWGTSAETLY | 0.6388993 |
| SGFGNVGAHTSGWFNQ | ASSPDWGTSAETLY | 0.7950777 |
| SGVAGAVGQAASVGGLKV | ASSPDWGTSAETLY | 0.8081834 |
| SNGGAGTGSAGGNGGT | ASSPDWGTSAETLY | 0.3866155 |
| TGGAGGEGGAAG | ASSPDWGTSAETLY | 0.8115846 |
| TGGDGGAAGAGGAPGHG | ASSPDWGTSAETLY | 0.8372065 |
| TMPPLLAAIANAGRHGV | ASSPDWGTSAETLY | 0.8203958 |
| TSSTAGNSGGA | ASSPDWGTSAETLY | 0.61847204 |
| VPATVPVGILC | ASSPDWGTSAETLY | 0.83385897 |
| VQRPDSGGVVLTGRIS | ASSPDWGTSAETLY | 0.8408021 |
| WTPNPVPATKGVPAGLVQAT | ASSPDWGTSAETLY | 0.8427861 |
| YALKRALAKFSDRFDVVII | ASSPDWGTSAETLY | 0.42291874 |
| YLIGSVIV | ASSPDWGTSAETLY | 0.3103276 |
| YGQLVAVID | ASSPDWGTSAETLY | 0.7883492 |
| VTVGGMTLPA | ASSPDWGTSAETLY | 0.8266316 |
| VLALPAV | ASSPDWGTSAETLY | 0.7542668 |
| VIPPTLNYET | ASSPDWGTSAETLY | 0.7235764 |
| VGMGGMHPSSG | ASSPDWGTSAETLY | 0.8118572 |
| VGLAGGAATA | ASSPDWGTSAETLY | 0.83364034 |
| VGGAGGAGGSGP | ASSPDWGTSAETLY | 0.65084857 |
| VFAAFMV | ASSPDWGTSAETLY | 0.6813524 |

| | | |
|---|---|---|
| TLYDKLR | ASSPDWGTSAETLY | 0.8282603 |
| TLAVFTVL | ASSPDWGTSAETLY | 0.82982767 |
| TGSVQPARV | ASSPDWGTSAETLY | 0.70286137 |
| TGGGAGSGGT | ASSPDWGTSAETLY | 0.7860932 |
| SHLETSL | ASSPDWGTSAETLY | 0.7918986 |
| SFTTGMT | ASSPDWGTSAETLY | 0.7893641 |
| QVEVAKRMGVRQPTVS | ASSPDWGTSAETLY | 0.80883193 |
| QGSGGPALAM | ASSPDWGTSAETLY | 0.6327905 |
| MLAPSNP | ASSPDWGTSAETLY | 0.8342704 |
| LVNNAAI | ASSPDWGTSAETLY | 0.66548973 |
| LTGEINTLA | ASSPDWGTSAETLY | 0.8279643 |
| LSIFGPV | ASSPDWGTSAETLY | 0.79930246 |
| LLALLANL | ASSPDWGTSAETLY | 0.81171834 |
| LISNASDALDKLR | ASSPDWGTSAETLY | 0.7627808 |
| LINEAVP | ASSPDWGTSAETLY | 0.6757063 |
| LGYAFSGL | ASSPDWGTSAETLY | 0.65113264 |
| LDFVAVAL | ASSPDWGTSAETLY | 0.66743517 |
| LAQPGAE | ASSPDWGTSAETLY | 0.8328046 |
| ISNLGAVTGF | ASSPDWGTSAETLY | 0.7510638 |
| HQEDPGRI | ASSPDWGTSAETLY | 0.83442295 |
| GTPGEPTP | ASSPDWGTSAETLY | 0.8075782 |
| GFFNSGS | ASSPDWGTSAETLY | 0.7125175 |
| GAGGGGLTAGAN | ASSPDWGTSAETLY | 0.8274721 |
| FVDALVNM | ASSPDWGTSAETLY | 0.6006273 |
| FLTVAPEL | ASSPDWGTSAETLY | 0.7877221 |
| FETLTAL | ASSPDWGTSAETLY | 0.8032031 |
| DVINPIEL | ASSPDWGTSAETLY | 0.82577336 |
| APVSGAPV | ASSPDWGTSAETLY | 0.6975989 |
| ALVGGLAW | ASSPDWGTSAETLY | 0.82549673 |
| ALAAAAATAI | ASSPDWGTSAETLY | 0.8164957 |
| AFGLGLLL | ASSPDWGTSAETLY | 0.6632011 |
| AFFAAASL | ASSPDWGTSAETLY | 0.8131182 |
| AAAALPAFSPP | ASSPDWGTSAETLY | 0.83061355 |
| VVLAPAL | ASSPDWGTSAETLY | 0.81648344 |
| TIGHVDHGKTTL | ASSPDWGTSAETLY | 0.5550459 |
| LISNASDALDK | ASSPDWGTSAETLY | 0.81152195 |
| LISNASDAL | ASSPDWGTSAETLY | 0.73737764 |
| ISNASDALDK | ASSPDWGTSAETLY | 0.811497 |
| NQAETLVYQTE | ASSPDWGTSAETLY | 0.8065516 |
| TGGTGGTAGAG | ASSPDWGTSAETLY | 0.7631766 |
| TGSTAGGAGGA | ASSPDWGTSAETLY | 0.82775253 |
| DPLAVPVAAGPV | ASSPDWGTSAETLY | 0.54494244 |
| GGTGGTPGNGGN | ASSPDWGTSAETLY | 0.7531346 |
| PAGLAGVDPATL | ASSPDWGTSAETLY | 0.7561857 |
| GNNLVVEAPHAGASV | ASSPDWGTSAETLY | 0.7874859 |
| KVTRIMSASTLEQPA | ASSPDWGTSAETLY | 0.82438713 |
| LGFALPIALPRRTVV | ASSPDWGTSAETLY | 0.6723556 |
| TRLGNLTARKLVVLV | ASSPDWGTSAETLY | 0.80402386 |
| WSQQRRLIYFPSAGP | ASSPDWGTSAETLY | 0.7649348 |
| GPNGNPDPMAAAVDIR | ASSPDWGTSAETLY | 0.6818323 |
| TQRCVSVADLRAAASR | ASSPDWGTSAETLY | 0.81351364 |

| | | |
|---|---|---|
| QFVLDVAEHPVEVLTETV | ASSPDWGTSAETLY | 0.77100354 |
| GGAGGTGGAGGNGGTQAGDG | ASSPDWGTSAETLY | 0.5566153 |
| PLLATLRAIAVDVDAKPAQV | ASSPDWGTSAETLY | 0.8231795 |
| APKFRTGGYMIDFWGVGYQVAK | ASSPDWGTSAETLY | 0.70817137 |
| GAGGAGGAGGAAGGPGGTGGTG | ASSPDWGTSAETLY | 0.8064718 |
| IGALGLLATDVAPAGSVAFGVA | ASSPDWGTSAETLY | 0.838772 |
| AGGAGGIGGNGGAGGAGGVLMG | ASSPDWGTSAETLY | 0.5830233 |
| ADELIGLL | ASSPDWGTSAETLY | 0.8291793 |
| ADRALKA | ASSPDWGTSAETLY | 0.81277543 |
| AGAVGQVGGAGGA | ASSPDWGTSAETLY | 0.80981386 |
| AGGGGGLGGAP | ASSPDWGTSAETLY | 0.7720661 |
| AGGLLYGNGGAG | ASSPDWGTSAETLY | 0.6484267 |
| AGLESIGL | ASSPDWGTSAETLY | 0.8371456 |
| AGLKTGII | ASSPDWGTSAETLY | 0.81519794 |
| ASLAALLL | ASSPDWGTSAETLY | 0.82455254 |
| ASVINISSI | ASSPDWGTSAETLY | 0.8220931 |
| ATPPGPA | ASSPDWGTSAETLY | 0.8260758 |
| ATVASALV | ASSPDWGTSAETLY | 0.8300809 |
| ATVASLIL | ASSPDWGTSAETLY | 0.73898077 |
| AVNGPVLIRH | ASSPDWGTSAETLY | 0.5154959 |
| DGAAGAVGI | ASSPDWGTSAETLY | 0.7733601 |
| DPLLHGI | ASSPDWGTSAETLY | 0.7754009 |
| ELGAALGAL | ASSPDWGTSAETLY | 0.7562483 |
| ELGRTQAV | ASSPDWGTSAETLY | 0.53445685 |
| FVAGGYLS | ASSPDWGTSAETLY | 0.8266423 |
| GASGGAGGAGGW | ASSPDWGTSAETLY | 0.84499407 |
| GGAGAGGDGGAGGI | ASSPDWGTSAETLY | 0.7153599 |
| GTGHNGSQPGMGGNG | ASSPDWGTSAETLY | 0.8454634 |
| GYAAASASAS | ASSPDWGTSAETLY | 0.60619694 |
| HQVDVAAVQQG | ASSPDWGTSAETLY | 0.82391477 |
| IGAAIAHT | ASSPDWGTSAETLY | 0.83089244 |
| IGMIEVAQL | ASSPDWGTSAETLY | 0.69595337 |
| IRLAPEHTSAMD | ASSPDWGTSAETLY | 0.8145766 |
| IYNVSVL | ASSPDWGTSAETLY | 0.69174635 |
| KQGFLGM | ASSPDWGTSAETLY | 0.7352778 |
| LESVITQL | ASSPDWGTSAETLY | 0.8324175 |
| LGLGALVVV | ASSPDWGTSAETLY | 0.69295603 |
| LIGNGGNGGSGGI | ASSPDWGTSAETLY | 0.81874865 |
| LNAINAPTL | ASSPDWGTSAETLY | 0.7772717 |
| LQLLNTGV | ASSPDWGTSAETLY | 0.7059292 |
| LQTSIEF | ASSPDWGTSAETLY | 0.81953496 |
| MSFAVSEL | ASSPDWGTSAETLY | 0.83160645 |
| NTGALGL | ASSPDWGTSAETLY | 0.79717946 |
| PPGAGGAGGAGTT | ASSPDWGTSAETLY | 0.8467545 |
| QLVGLGI | ASSPDWGTSAETLY | 0.67679274 |
| RFLEQVHQL | ASSPDWGTSAETLY | 0.8406288 |
| RGVGYVLR | ASSPDWGTSAETLY | 0.83543074 |
| SATGVFAGVI | ASSPDWGTSAETLY | 0.80391806 |
| SLPAITLP | ASSPDWGTSAETLY | 0.76876473 |
| STALAFTV | ASSPDWGTSAETLY | 0.82585114 |
| SVLTVGLGK | ASSPDWGTSAETLY | 0.7486203 |

| | | |
|---|---|---|
| TALAGAGSGI | ASSPDWGTSAETLY | 0.8165913 |
| TGLPFTGI | ASSPDWGTSAETLY | 0.8404211 |
| VAEHVTV | ASSPDWGTSAETLY | 0.69573337 |
| VDIVLNSL | ASSPDWGTSAETLY | 0.8160454 |
| VLTGGGAVL | ASSPDWGTSAETLY | 0.7481439 |
| VNVGLAYGI | ASSPDWGTSAETLY | 0.7894871 |
| VSSAAVGAV | ASSPDWGTSAETLY | 0.6832187 |
| VTSVYEAGV | ASSPDWGTSAETLY | 0.46372065 |
| WVALDRGA | ASSPDWGTSAETLY | 0.75126487 |
| YLVGSVVI | ASSPDWGTSAETLY | 0.61288494 |
| GPPGTGKTLL | ASSPDWGTSAETLY | 0.80769503 |
| GVGMTKF | ASSPDWGTSAETLY | 0.80892223 |
| IGIGGSDL | ASSPDWGTSAETLY | 0.67821765 |
| IGTLLAEL | ASSPDWGTSAETLY | 0.8231169 |
| IYTSGTT | ASSPDWGTSAETLY | 0.81116015 |
| LSFPAGV | ASSPDWGTSAETLY | 0.538743 |
| LYTSGTT | ASSPDWGTSAETLY | 0.7009394 |
| VGAFPFGF | ASSPDWGTSAETLY | 0.66736424 |
| VGAGTVEF | ASSPDWGTSAETLY | 0.7759307 |
| VGKSTLLL | ASSPDWGTSAETLY | 0.43929833 |
| AALAHAGGGIV | ASSPDWGTSAETLY | 0.8145613 |
| AALVAVSLL | ASSPDWGTSAETLY | 0.6695816 |
| AAVSRAVAAL | ASSPDWGTSAETLY | 0.28738108 |
| AGGSSGIGGF | ASSPDWGTSAETLY | 0.76562834 |
| AGLTVANANG | ASSPDWGTSAETLY | 0.74855816 |
| AGLVGAVL | ASSPDWGTSAETLY | 0.75795126 |
| ALSQAFTV | ASSPDWGTSAETLY | 0.81620294 |
| ALTFSVSL | ASSPDWGTSAETLY | 0.6298683 |
| ALTGAALVAL | ASSPDWGTSAETLY | 0.6417937 |
| ASFSIGAGVL | ASSPDWGTSAETLY | 0.77653086 |
| DGSVYKI | ASSPDWGTSAETLY | 0.8183119 |
| EQLAEIV | ASSPDWGTSAETLY | 0.63332486 |
| FAVSELTVM | ASSPDWGTSAETLY | 0.76759344 |
| GAAPLAAGL | ASSPDWGTSAETLY | 0.82620406 |
| GADGGDGGSGGAGG | ASSPDWGTSAETLY | 0.76793957 |
| GFGLDTGGAGG | ASSPDWGTSAETLY | 0.8184748 |
| GLASGGLL | ASSPDWGTSAETLY | 0.8219596 |
| GRIAGAAVLL | ASSPDWGTSAETLY | 0.6086443 |
| GSAGSLNTSF | ASSPDWGTSAETLY | 0.81990194 |
| GTGQIGI | ASSPDWGTSAETLY | 0.82145286 |
| GTLIALIL | ASSPDWGTSAETLY | 0.7279402 |
| IAATMGFVVG | ASSPDWGTSAETLY | 0.7574934 |
| IFAQALL | ASSPDWGTSAETLY | 0.64054674 |
| IGAAVALGF | ASSPDWGTSAETLY | 0.73393863 |
| IGLTPLAL | ASSPDWGTSAETLY | 0.4470747 |
| ILGSNTSTL | ASSPDWGTSAETLY | 0.7639121 |
| IVFSLTVL | ASSPDWGTSAETLY | 0.8250537 |
| LASKATGF | ASSPDWGTSAETLY | 0.7179838 |
| LASVALIL | ASSPDWGTSAETLY | 0.6211364 |
| LFAVGLTL | ASSPDWGTSAETLY | 0.552145 |
| LFNGGIGTYI | ASSPDWGTSAETLY | 0.7590038 |

| | | |
|---|---|---|
| LTLGAGII | ASSPDWGTSAETLY | 0.72524273 |
| LVAAALGDI | ASSPDWGTSAETLY | 0.48410332 |
| NTAFVQLGI | ASSPDWGTSAETLY | 0.6549969 |
| QLFSGAEL | ASSPDWGTSAETLY | 0.81294525 |
| SAATFITL | ASSPDWGTSAETLY | 0.81606525 |
| SNYGLANLGV | ASSPDWGTSAETLY | 0.5025668 |
| SYGGAYAVM | ASSPDWGTSAETLY | 0.830376 |
| TTGVGGLTLG | ASSPDWGTSAETLY | 0.799729 |
| TTQTIYI | ASSPDWGTSAETLY | 0.7790388 |
| VAAGAGLGAVI | ASSPDWGTSAETLY | 0.7559701 |
| VAAVVTLL | ASSPDWGTSAETLY | 0.65646183 |
| VGFLGGLI | ASSPDWGTSAETLY | 0.8259271 |
| VGVIPILL | ASSPDWGTSAETLY | 0.6610509 |
| VIFASSLI | ASSPDWGTSAETLY | 0.702595 |
| VKGGDGGAAAG | ASSPDWGTSAETLY | 0.8042699 |
| VTKTFTNI | ASSPDWGTSAETLY | 0.74808663 |
| VVAAAILL | ASSPDWGTSAETLY | 0.6304771 |
| YAALDVEL | ASSPDWGTSAETLY | 0.686962 |
| LASVALLL | ASSPDWGTSAETLY | 0.8215483 |
| LDAVHRV | ASSPDWGTSAETLY | 0.55391836 |
| QGLSSSIF | ASSPDWGTSAETLY | 0.6228696 |
| RGTFANIRL | ASSPDWGTSAETLY | 0.7073126 |
| RQLWWGHRI | ASSPDWGTSAETLY | 0.75326675 |
| YGPPGTGKTLL | ASSPDWGTSAETLY | 0.8174554 |
| AALVGAPANVA | ASSPDWGTSAETLY | 0.7850256 |
| AATFRRTE | ASSPDWGTSAETLY | 0.701521 |
| AGALGGGI | ASSPDWGTSAETLY | 0.79211533 |
| AGGAGGAGGIGAG | ASSPDWGTSAETLY | 0.79598826 |
| AGSGALGL | ASSPDWGTSAETLY | 0.7873124 |
| AQIADQI | ASSPDWGTSAETLY | 0.7654711 |
| ASAVAVTYL | ASSPDWGTSAETLY | 0.62031007 |
| ASTVLGGTGG | ASSPDWGTSAETLY | 0.8185869 |
| AVGGEPAK | ASSPDWGTSAETLY | 0.8074932 |
| CEQGTFGGTC | ASSPDWGTSAETLY | 0.8172789 |
| DEGKAPAG | ASSPDWGTSAETLY | 0.80466545 |
| DTVVFPAL | ASSPDWGTSAETLY | 0.764398 |
| FEGITMAG | ASSPDWGTSAETLY | 0.7756399 |
| FPLVVQD | ASSPDWGTSAETLY | 0.359357 |
| GAFTTPQ | ASSPDWGTSAETLY | 0.63329935 |
| GFGGFGTAGGVGGA | ASSPDWGTSAETLY | 0.7494397 |
| GFLAQFL | ASSPDWGTSAETLY | 0.7607494 |
| GGAGGAGGNAGVA | ASSPDWGTSAETLY | 0.7978888 |
| GGVGTAGAGGAGGAGG | ASSPDWGTSAETLY | 0.7995145 |
| GNGGGGDAGNAGSGGN | ASSPDWGTSAETLY | 0.6532528 |
| GTGGAGGAGNDAGS | ASSPDWGTSAETLY | 0.79671335 |
| GTTNTGWFNS | ASSPDWGTSAETLY | 0.8258702 |
| IFMGPLFTL | ASSPDWGTSAETLY | 0.5096601 |
| ILAHAIEAI | ASSPDWGTSAETLY | 0.6575264 |
| IRSAIREAL | ASSPDWGTSAETLY | 0.8391176 |
| ITFAGGG | ASSPDWGTSAETLY | 0.7801445 |
| ITGMSGHQEV | ASSPDWGTSAETLY | 0.71283436 |

| | | |
|---|---|---|
| IVAGLACTL | ASSPDWGTSAETLY | 0.75829935 |
| IVGGGLAGA | ASSPDWGTSAETLY | 0.8370701 |
| IVTGVLNI | ASSPDWGTSAETLY | 0.718817 |
| LAQQGCI | ASSPDWGTSAETLY | 0.8242364 |
| LCLHVEN | ASSPDWGTSAETLY | 0.32440236 |
| LDAISDGS | ASSPDWGTSAETLY | 0.742846 |
| LGGFAAML | ASSPDWGTSAETLY | 0.6275815 |
| LGGKNPMI | ASSPDWGTSAETLY | 0.43397623 |
| LGPKGVAAIL | ASSPDWGTSAETLY | 0.8055918 |
| LMVGVSI | ASSPDWGTSAETLY | 0.74914557 |
| LVGAAFVL | ASSPDWGTSAETLY | 0.80969244 |
| LVLAVAPL | ASSPDWGTSAETLY | 0.71590245 |
| MAAIAVIDAA | ASSPDWGTSAETLY | 0.8067229 |
| NNGINAELQ | ASSPDWGTSAETLY | 0.73441553 |
| RGEPLVLVHG | ASSPDWGTSAETLY | 0.7640995 |
| RGGAGGAAGLI | ASSPDWGTSAETLY | 0.71556103 |
| RSYRMHPAV | ASSPDWGTSAETLY | 0.80316067 |
| SAAHLLAGQLA | ASSPDWGTSAETLY | 0.8121465 |
| SAAVGSY | ASSPDWGTSAETLY | 0.81169665 |
| SPHAVLHRI | ASSPDWGTSAETLY | 0.48028123 |
| SQIAKVL | ASSPDWGTSAETLY | 0.7095512 |
| TALGHLVP | ASSPDWGTSAETLY | 0.7464006 |
| TAPTTSGAA | ASSPDWGTSAETLY | 0.4453649 |
| TDGDDGNGGSPGT | ASSPDWGTSAETLY | 0.3954629 |
| TDGLGEN | ASSPDWGTSAETLY | 0.64973825 |
| TGGPPLPGQA | ASSPDWGTSAETLY | 0.69300586 |
| TIPAITI | ASSPDWGTSAETLY | 0.809348 |
| TIPGFTI | ASSPDWGTSAETLY | 0.78168947 |
| TSIASGALA | ASSPDWGTSAETLY | 0.75276095 |
| TVTLPGAAI | ASSPDWGTSAETLY | 0.78604454 |
| VASGLSAALP | ASSPDWGTSAETLY | 0.7263144 |
| VGAGVIGI | ASSPDWGTSAETLY | 0.18422544 |
| VGPKRFGV | ASSPDWGTSAETLY | 0.6802957 |
| VNVAFPD | ASSPDWGTSAETLY | 0.59010303 |
| VSMTNSGVS | ASSPDWGTSAETLY | 0.8290713 |
| VTDVLGDLSV | ASSPDWGTSAETLY | 0.6294422 |
| VVVVGAGFAGL | ASSPDWGTSAETLY | 0.634422 |
| VYTVGTVV | ASSPDWGTSAETLY | 0.76374376 |
| WTTFSGP | ASSPDWGTSAETLY | 0.7623856 |
| YDELVEAF | ASSPDWGTSAETLY | 0.73818743 |
| YGTGGAGGNGG | ASSPDWGTSAETLY | 0.6316227 |
| YTNVMAV | ASSPDWGTSAETLY | 0.29770526 |
| FLEQVHQL | ASSPDWGTSAETLY | 0.8085892 |
| GLSLLAVAV | ASSPDWGTSAETLY | 0.5270758 |
| LEQVHQL | ASSPDWGTSAETLY | 0.66628397 |
| LNAAGAI | ASSPDWGTSAETLY | 0.4762191 |
| PAVHASL | ASSPDWGTSAETLY | 0.64012325 |
| TELEDDA | ASSPDWGTSAETLY | 0.4635721 |
| TPPGPPN | ASSPDWGTSAETLY | 0.8151074 |
| TPPPPPPP | ASSPDWGTSAETLY | 0.7085259 |
| TQSPALI | TCSAGQGGNTEVF | 0.6982335 |

| | | |
|---|---|---|
| EFAAAFPQ | TCSAGQGGNTEVF | 0.8346684 |
| LLSGNLDVL | TCSAGQGGNTEVF | 0.7559303 |
| GPANLGNYNI | TCSAGQGGNTEVF | 0.70043766 |
| TLGPTLP | TCSAGQGGNTEVF | 0.70719594 |
| VYAYSSCCK | TCSAGQGGNTEVF | 0.6450732 |
| APAPAPAPNI | TCSAGQGGNTEVF | 0.66141284 |
| PGGPVPIP | TCSAGQGGNTEVF | 0.7118398 |
| FLTVALEP | TCSAGQGGNTEVF | 0.8257652 |
| SQYFGANL | TCSAGQGGNTEVF | 0.770438 |
| GGVKPAA | TCSAGQGGNTEVF | 0.61532176 |
| GGTGGAGGAGGW | TCSAGQGGNTEVF | 0.7909013 |
| GVGGVAGLGGAGG | TCSAGQGGNTEVF | 0.8343433 |
| VLLVLATTI | TCSAGQGGNTEVF | 0.82320994 |
| IGIFAAML | TCSAGQGGNTEVF | 0.64485097 |
| AVGDLID | TCSAGQGGNTEVF | 0.7001457 |
| NISPLPL | TCSAGQGGNTEVF | 0.73738766 |
| VTAVFVAI | TCSAGQGGNTEVF | 0.7149199 |
| SGDDNDAAL | TCSAGQGGNTEVF | 0.7212719 |
| AVAALAHS | TCSAGQGGNTEVF | 0.6512741 |
| GGVTVMATP | TCSAGQGGNTEVF | 0.8068457 |
| AGLAANVRD | TCSAGQGGNTEVF | 0.847834 |
| TNRPDLIDPALLRPG | TCSAGQGGNTEVF | 0.7530019 |
| LANLGVTAL | TCSAGQGGNTEVF | 0.44141364 |
| TGERIEL | TCSAGQGGNTEVF | 0.7326958 |
| NGEELAA | TCSAGQGGNTEVF | 0.57426196 |
| VLSLVWG | TCSAGQGGNTEVF | 0.69392234 |
| IGYALWL | TCSAGQGGNTEVF | 0.7441458 |
| LSLLLAPL | TCSAGQGGNTEVF | 0.5295131 |
| TLDEALK | TCSAGQGGNTEVF | 0.7582378 |
| TDGAGTAEGG | TCSAGQGGNTEVF | 0.8282132 |
| NAPGGAGGNGG | TCSAGQGGNTEVF | 0.66429704 |
| VSVFMGMGGNF | TCSAGQGGNTEVF | 0.79346204 |
| LASYLAAL | TCSAGQGGNTEVF | 0.81815666 |
| LVGGTPAL | TCSAGQGGNTEVF | 0.79962504 |
| VRNADVL | TCSAGQGGNTEVF | 0.827188 |
| TCSSPFLVSAM | TCSAGQGGNTEVF | 0.79799795 |
| VHITDAE | TCSAGQGGNTEVF | 0.77111804 |
| EYAAPSF | TCSAGQGGNTEVF | 0.76536584 |
| VVDAFHGGL | TCSAGQGGNTEVF | 0.79593337 |
| ASVVDPQ | TCSAGQGGNTEVF | 0.7430889 |
| SYNEFGGLS | TCSAGQGGNTEVF | 0.73009366 |
| DPDGSVY | TCSAGQGGNTEVF | 0.6875742 |
| MVSTIHATE | TCSAGQGGNTEVF | 0.8171525 |
| LAAGVADPV | TCSAGQGGNTEVF | 0.75351286 |
| TKEQIAA | TCSAGQGGNTEVF | 0.8327349 |
| SLASPAAL | TCSAGQGGNTEVF | 0.56896764 |
| IGLVFYGV | TCSAGQGGNTEVF | 0.6661974 |
| FLLDASNL | TCSAGQGGNTEVF | 0.8111243 |
| YQVGNFP | TCSAGQGGNTEVF | 0.20225623 |
| ESLLSQST | TCSAGQGGNTEVF | 0.43143484 |
| LFTVGFAAVG | TCSAGQGGNTEVF | 0.7611619 |

| | | |
|---|---|---|
| ATPPTPT | TCSAGQGGNTEVF | 0.8147928 |
| TDLATLG | TCSAGQGGNTEVF | 0.25586447 |
| SAGGTGR | TCSAGQGGNTEVF | 0.6622873 |
| GFSNSGS | TCSAGQGGNTEVF | 0.57084465 |
| GGNGGSGGTSV | TCSAGQGGNTEVF | 0.8012871 |
| GTSVATGGAGN | TCSAGQGGNTEVF | 0.7560947 |
| NGGNGGSGGTS | TCSAGQGGNTEVF | 0.38454175 |
| AVVKFPH | TCSAGQGGNTEVF | 0.81534207 |
| TPPPPPP | TCSAGQGGNTEVF | 0.48049396 |
| AAPAAAAAPA | TCSAGQGGNTEVF | 0.68808496 |
| IALNEDL | TCSAGQGGNTEVF | 0.8032512 |
| IVNVSSM | TCSAGQGGNTEVF | 0.83130443 |
| TGIPPAPR | TCSAGQGGNTEVF | 0.62616944 |
| FDLGGGTF | TCSAGQGGNTEVF | 0.8017068 |
| FDLGGGTFDVS | TCSAGQGGNTEVF | 0.6787714 |
| NEPTAAA | TCSAGQGGNTEVF | 0.8313695 |
| NEPTAAALA | TCSAGQGGNTEVF | 0.7142531 |
| GSIINISSI | TCSAGQGGNTEVF | 0.57979035 |
| VAPGNAGTA | TCSAGQGGNTEVF | 0.81619126 |
| TGGGASLE | TCSAGQGGNTEVF | 0.80449533 |
| VAVYNHYKRI | TCSAGQGGNTEVF | 0.7723093 |
| TDPAPATT | TCSAGQGGNTEVF | 0.807981 |
| TNRPDILDPALLRPG | TCSAGQGGNTEVF | 0.729293 |
| LTTTTTS | TCSAGQGGNTEVF | 0.7483834 |
| PTGSGKTL | TCSAGQGGNTEVF | 0.8201139 |
| TGSGKTL | TCSAGQGGNTEVF | 0.7949892 |
| TIGHVDHGK | TCSAGQGGNTEVF | 0.65831196 |
| TIGHVDHGKTT | TCSAGQGGNTEVF | 0.8079389 |
| GHVDHGKTT | TCSAGQGGNTEVF | 0.7294281 |
| AQFEHTLL | TCSAGQGGNTEVF | 0.68088895 |
| IVNQDEL | TCSAGQGGNTEVF | 0.8223437 |
| TNGSQFF | TCSAGQGGNTEVF | 0.7592914 |
| TGKGIVE | TCSAGQGGNTEVF | 0.8154522 |
| LISNASDA | TCSAGQGGNTEVF | 0.811804 |
| AELRTLGELTKIAKAHGAQ | TCSAGQGGNTEVF | 0.8371803 |
| APNAVAAAEPSW | TCSAGQGGNTEVF | 0.48713997 |
| APQVAGAATAVL | TCSAGQGGNTEVF | 0.7492488 |
| ARRDPAAAAAILPTDA | TCSAGQGGNTEVF | 0.7324919 |
| DAAARDVLAEAGLAEYF | TCSAGQGGNTEVF | 0.8021022 |
| DDQIVLSAKANLAKVEQ | TCSAGQGGNTEVF | 0.7533126 |
| DELLPDTGEDSDEDGH | TCSAGQGGNTEVF | 0.5852053 |
| ELPEAPANRATVPQD | TCSAGQGGNTEVF | 0.7968776 |
| FGQSKIERVTGIG | TCSAGQGGNTEVF | 0.82752573 |
| GAGGTGSTAGGAGGAGGAGGLY | TCSAGQGGNTEVF | 0.7733141 |
| GHGGTGAAGGNGGTGGAGGSGI | TCSAGQGGNTEVF | 0.7542341 |
| GNGGNGGAGGEGGPGGAGGAGA | TCSAGQGGNTEVF | 0.83443326 |
| GSTAGNGGNGGNATNGTIAGQP | TCSAGQGGNTEVF | 0.68771696 |
| GTGGDGGNGGNGGTGG | TCSAGQGGNTEVF | 0.7338814 |
| GVRRPAGIPADAPV | TCSAGQGGNTEVF | 0.8128566 |
| GYRQVTDAHLLTLARR | TCSAGQGGNTEVF | 0.81318337 |
| IAKMGQDGHDRG | TCSAGQGGNTEVF | 0.76050746 |

| | | |
|---|---|---|
| KAEPSIEIQYT | TCSAGQGGNTEVF | 0.8082537 |
| KPVRIRMAADEDTVAVT | TCSAGQGGNTEVF | 0.5096552 |
| MPPRIAGMRLLVIKPEP | TCSAGQGGNTEVF | 0.79543555 |
| MKFYAVALFRSRGVPPTR | TCSAGQGGNTEVF | 0.6151379 |
| NAGGFGILGAVAHSP | TCSAGQGGNTEVF | 0.76749986 |
| NEGREITLAMAEM | TCSAGQGGNTEVF | 0.5846784 |
| NELVEWSHPLGHRWRWE | TCSAGQGGNTEVF | 0.7677 |
| NLGAGGTGGNGGNPG | TCSAGQGGNTEVF | 0.7985303 |
| NTEWPKWETVMAADPVD | TCSAGQGGNTEVF | 0.8046453 |
| PASAAPAETQP | TCSAGQGGNTEVF | 0.3843618 |
| QQSSDGKNGVRAAEITGE | TCSAGQGGNTEVF | 0.79569197 |
| RAPDAVTTSWWKE | TCSAGQGGNTEVF | 0.80568933 |
| SGFGNVGAHTSGWFNQ | TCSAGQGGNTEVF | 0.84683114 |
| SGVAGAVGQAASVGGLKV | TCSAGQGGNTEVF | 0.7220463 |
| SNGGAGTGSAGGNGGT | TCSAGQGGNTEVF | 0.8001493 |
| TGGAGGEGGAAG | TCSAGQGGNTEVF | 0.6787244 |
| TGGDGGAAGAGGAPGHG | TCSAGQGGNTEVF | 0.7521443 |
| TMPPLLAAIANAGRHGV | TCSAGQGGNTEVF | 0.5665758 |
| TSSTAGNSGGA | TCSAGQGGNTEVF | 0.6684192 |
| VPATVPVGILC | TCSAGQGGNTEVF | 0.6691084 |
| VQRPDSGGVVLTGRIS | TCSAGQGGNTEVF | 0.6519029 |
| WTPNPVPATKGVPAGLVQAT | TCSAGQGGNTEVF | 0.8061825 |
| YALKRALAKFSDRFDVVII | TCSAGQGGNTEVF | 0.7985974 |
| YLIGSVIV | TCSAGQGGNTEVF | 0.80983365 |
| YGQLVAVID | TCSAGQGGNTEVF | 0.8197104 |
| VTVGGMTLPA | TCSAGQGGNTEVF | 0.83348083 |
| VLALPAV | TCSAGQGGNTEVF | 0.8231595 |
| VIPPTLNYET | TCSAGQGGNTEVF | 0.6689814 |
| VGMGGMHPSSG | TCSAGQGGNTEVF | 0.6934651 |
| VGLAGGAATA | TCSAGQGGNTEVF | 0.56931233 |
| VGGAGGAGGSGP | TCSAGQGGNTEVF | 0.74114513 |
| VFAAFMV | TCSAGQGGNTEVF | 0.69982 |
| TLYDKLR | TCSAGQGGNTEVF | 0.7819663 |
| TLAVFTVL | TCSAGQGGNTEVF | 0.21442889 |
| TGSVQPARV | TCSAGQGGNTEVF | 0.7689584 |
| TGGGAGSGGT | TCSAGQGGNTEVF | 0.82783735 |
| SHLETSL | TCSAGQGGNTEVF | 0.5209711 |
| SFTTGMT | TCSAGQGGNTEVF | 0.7960599 |
| QVEVAKRMGVRQPTVS | TCSAGQGGNTEVF | 0.7719547 |
| QGSGGPALAM | TCSAGQGGNTEVF | 0.668035 |
| MLAPSNP | TCSAGQGGNTEVF | 0.8021476 |
| LVNNAAI | TCSAGQGGNTEVF | 0.68487924 |
| LTGEINTLA | TCSAGQGGNTEVF | 0.8102625 |
| LSIFGPV | TCSAGQGGNTEVF | 0.8299457 |
| LLALLANL | TCSAGQGGNTEVF | 0.7888591 |
| LISNASDALDKLR | TCSAGQGGNTEVF | 0.70556456 |
| LINEAVP | TCSAGQGGNTEVF | 0.76791346 |
| LGYAFSGL | TCSAGQGGNTEVF | 0.43080038 |
| LDFVAVAL | TCSAGQGGNTEVF | 0.7297824 |
| LAQPGAE | TCSAGQGGNTEVF | 0.46769255 |
| ISNLGAVTGF | TCSAGQGGNTEVF | 0.736564 |

| | | |
|---|---|---|
| HQEDPGRI | TCSAGQGGNTEVF | 0.6619258 |
| GTPGEPTP | TCSAGQGGNTEVF | 0.77051353 |
| GFFNSGS | TCSAGQGGNTEVF | 0.7499566 |
| GAGGGGLTAGAN | TCSAGQGGNTEVF | 0.7712035 |
| FVDALVNM | TCSAGQGGNTEVF | 0.546872 |
| FLTVAPEL | TCSAGQGGNTEVF | 0.7604188 |
| FETLTAL | TCSAGQGGNTEVF | 0.7112039 |
| DVINPIEL | TCSAGQGGNTEVF | 0.57707953 |
| APVSGAPV | TCSAGQGGNTEVF | 0.7827085 |
| ALVGGLAW | TCSAGQGGNTEVF | 0.77403724 |
| ALAAAAATAI | TCSAGQGGNTEVF | 0.70065564 |
| AFGLGLLL | TCSAGQGGNTEVF | 0.8190094 |
| AFFAAASL | TCSAGQGGNTEVF | 0.7893251 |
| AAAALPAFSPP | TCSAGQGGNTEVF | 0.4496983 |
| VVLAPAL | TCSAGQGGNTEVF | 0.6797447 |
| TIGHVDHGKTTL | TCSAGQGGNTEVF | 0.75110006 |
| LISNASDALDK | TCSAGQGGNTEVF | 0.69298667 |
| LISNASDAL | TCSAGQGGNTEVF | 0.29710427 |
| ISNASDALDK | TCSAGQGGNTEVF | 0.7856103 |
| NQAETLVYQTE | TCSAGQGGNTEVF | 0.680801 |
| TGGTGGTAGAG | TCSAGQGGNTEVF | 0.5659915 |
| TGSTAGGAGGA | TCSAGQGGNTEVF | 0.684016 |
| DPLAVPVAAGPV | TCSAGQGGNTEVF | 0.8122187 |
| GGTGGTPGNGGN | TCSAGQGGNTEVF | 0.62979865 |
| PAGLAGVDPATL | TCSAGQGGNTEVF | 0.7816708 |
| GNNLVVEAPHAGASV | TCSAGQGGNTEVF | 0.77192265 |
| KVTRIMSASTLEQPA | TCSAGQGGNTEVF | 0.78593063 |
| LGFALPIALPRRTVV | TCSAGQGGNTEVF | 0.8187466 |
| TRLGNLTARKLVVLV | TCSAGQGGNTEVF | 0.7843908 |
| WSQQRRLIYFPSAGP | TCSAGQGGNTEVF | 0.73102725 |
| GPNGNPDPMAAAVDIR | TCSAGQGGNTEVF | 0.79106146 |
| TQRCVSVADLRAAASR | TCSAGQGGNTEVF | 0.7433699 |
| QFVLDVAEHPVEVLTETV | TCSAGQGGNTEVF | 0.6582655 |
| GGAGGTGGAGGNGGTQAGDG | TCSAGQGGNTEVF | 0.487337 |
| PLLATLRAIAVDVDAKPAQV | TCSAGQGGNTEVF | 0.6795162 |
| APKFRTGGYMIDFWGVGYQVAK | TCSAGQGGNTEVF | 0.54552376 |
| GAGGAGGAGGAAGGPGGTGGTG | TCSAGQGGNTEVF | 0.75682473 |
| IGALGLLATDVAPAGSVAFGVA | TCSAGQGGNTEVF | 0.83068967 |
| AGGAGGIGGNGGAGGAGGVLMG | TCSAGQGGNTEVF | 0.691317 |
| ADELIGLL | TCSAGQGGNTEVF | 0.59321237 |
| ADRALKA | TCSAGQGGNTEVF | 0.7068096 |
| AGAVGQVGGAGGA | TCSAGQGGNTEVF | 0.59095174 |
| AGGGGGLGGAP | TCSAGQGGNTEVF | 0.80645 |
| AGGLLYGNGGAG | TCSAGQGGNTEVF | 0.7723973 |
| AGLESIGL | TCSAGQGGNTEVF | 0.84153795 |
| AGLKTGII | TCSAGQGGNTEVF | 0.7971497 |
| ASLAALLL | TCSAGQGGNTEVF | 0.68106556 |
| ASVINISSI | TCSAGQGGNTEVF | 0.8009881 |
| ATPPGPA | TCSAGQGGNTEVF | 0.72114515 |
| ATVASALV | TCSAGQGGNTEVF | 0.74431074 |
| ATVASLIL | TCSAGQGGNTEVF | 0.7516882 |

| | | |
|---|---|---|
| AVNGPVLIRH | TCSAGQGGNTEVF | 0.7697944 |
| DGAAGAVGI | TCSAGQGGNTEVF | 0.70111454 |
| DPLLHGI | TCSAGQGGNTEVF | 0.7390186 |
| ELGAALGAL | TCSAGQGGNTEVF | 0.43334338 |
| ELGRTQAV | TCSAGQGGNTEVF | 0.8101129 |
| FVAGGYLS | TCSAGQGGNTEVF | 0.6677237 |
| GASGGAGGAGGW | TCSAGQGGNTEVF | 0.5671599 |
| GGAGAGGDGGAGGI | TCSAGQGGNTEVF | 0.68304193 |
| GTGHNGSQPGMGGNG | TCSAGQGGNTEVF | 0.82419044 |
| GYAAASASAS | TCSAGQGGNTEVF | 0.82562596 |
| HQVDVAAVQQG | TCSAGQGGNTEVF | 0.651626 |
| IGAAIAHT | TCSAGQGGNTEVF | 0.6241826 |
| IGMIEVAQL | TCSAGQGGNTEVF | 0.7663173 |
| IRLAPEHTSAMD | TCSAGQGGNTEVF | 0.45277324 |
| IYNVSVL | TCSAGQGGNTEVF | 0.79809564 |
| KQGFLGM | TCSAGQGGNTEVF | 0.7174382 |
| LESVITQL | TCSAGQGGNTEVF | 0.7119427 |
| LGLGALVVV | TCSAGQGGNTEVF | 0.6806 |
| LIGNGGNGGSGGI | TCSAGQGGNTEVF | 0.767004 |
| LNAINAPTL | TCSAGQGGNTEVF | 0.6490827 |
| LQLLNTGV | TCSAGQGGNTEVF | 0.79736435 |
| LQTSIEF | TCSAGQGGNTEVF | 0.22057508 |
| MSFAVSEL | TCSAGQGGNTEVF | 0.7730167 |
| NTGALGL | TCSAGQGGNTEVF | 0.62394595 |
| PPGAGGAGGAGTT | TCSAGQGGNTEVF | 0.66093755 |
| QLVGLGI | TCSAGQGGNTEVF | 0.458767 |
| RFLEQVHQL | TCSAGQGGNTEVF | 0.7589116 |
| RGVGYVLR | TCSAGQGGNTEVF | 0.766313 |
| SATGVFAGVI | TCSAGQGGNTEVF | 0.81075275 |
| SLPAITLP | TCSAGQGGNTEVF | 0.63779086 |
| STALAFTV | TCSAGQGGNTEVF | 0.7883044 |
| SVLTVGLGK | TCSAGQGGNTEVF | 0.7967398 |
| TALAGAGSGI | TCSAGQGGNTEVF | 0.704782 |
| TGLPFTGI | TCSAGQGGNTEVF | 0.4133535 |
| VAEHVTV | TCSAGQGGNTEVF | 0.78460723 |
| VDIVLNSL | TCSAGQGGNTEVF | 0.72463745 |
| VLTGGGAVL | TCSAGQGGNTEVF | 0.80885684 |
| VNVGLAYGI | TCSAGQGGNTEVF | 0.7306429 |
| VSSAAVGAV | TCSAGQGGNTEVF | 0.56418425 |
| VTSVYEAGV | TCSAGQGGNTEVF | 0.80447364 |
| WVALDRGA | TCSAGQGGNTEVF | 0.81788826 |
| YLVGSVVI | TCSAGQGGNTEVF | 0.58555967 |
| GPPGTGKTLL | TCSAGQGGNTEVF | 0.6696898 |
| GVGMTKF | TCSAGQGGNTEVF | 0.7241901 |
| IGIGGSDL | TCSAGQGGNTEVF | 0.42209512 |
| IGTLLAEL | TCSAGQGGNTEVF | 0.78771424 |
| IYTSGTT | TCSAGQGGNTEVF | 0.6647215 |
| LSFPAGV | TCSAGQGGNTEVF | 0.6866188 |
| LYTSGTT | TCSAGQGGNTEVF | 0.7695682 |
| VGAFPFGF | TCSAGQGGNTEVF | 0.70431006 |
| VGAGTVEF | TCSAGQGGNTEVF | 0.73936194 |

| | | |
|---|---|---|
| VGKSTLLL | TCSAGQGGNTEVF | 0.74882895 |
| AALAHAGGGIV | TCSAGQGGNTEVF | 0.73580194 |
| AALVAVSLL | TCSAGQGGNTEVF | 0.77236575 |
| AAVSRAVAAL | TCSAGQGGNTEVF | 0.7227486 |
| AGGSSGIGGF | TCSAGQGGNTEVF | 0.7538439 |
| AGLTVANANG | TCSAGQGGNTEVF | 0.8269683 |
| AGLVGAVL | TCSAGQGGNTEVF | 0.54331535 |
| ALSQAFTV | TCSAGQGGNTEVF | 0.654796 |
| ALTFSVSL | TCSAGQGGNTEVF | 0.7900809 |
| ALTGAALVAL | TCSAGQGGNTEVF | 0.6096958 |
| ASFSIGAGVL | TCSAGQGGNTEVF | 0.80858934 |
| DGSVYKI | TCSAGQGGNTEVF | 0.7442138 |
| EQLAEIV | TCSAGQGGNTEVF | 0.786369 |
| FAVSELTVM | TCSAGQGGNTEVF | 0.73153746 |
| GAAPLAAGL | TCSAGQGGNTEVF | 0.82815355 |
| GADGGDGGSGGAGG | TCSAGQGGNTEVF | 0.8167292 |
| GFGLDTGGAGG | TCSAGQGGNTEVF | 0.7662117 |
| GLASGGLL | TCSAGQGGNTEVF | 0.8100174 |
| GRIAGAAVLL | TCSAGQGGNTEVF | 0.64808816 |
| GSAGSLNTSF | TCSAGQGGNTEVF | 0.79125977 |
| GTGQIGI | TCSAGQGGNTEVF | 0.7944259 |
| GTLIALIL | TCSAGQGGNTEVF | 0.5494544 |
| IAATMGFVVG | TCSAGQGGNTEVF | 0.74856305 |
| IFAQALL | TCSAGQGGNTEVF | 0.81642795 |
| IGAAVALGF | TCSAGQGGNTEVF | 0.82322085 |
| IGLTPLAL | TCSAGQGGNTEVF | 0.55576146 |
| ILGSNTSTL | TCSAGQGGNTEVF | 0.69744885 |
| IVFSLTVL | TCSAGQGGNTEVF | 0.52886665 |
| LASKATGF | TCSAGQGGNTEVF | 0.63732415 |
| LASVALIL | TCSAGQGGNTEVF | 0.61331695 |
| LFAVGLTL | TCSAGQGGNTEVF | 0.78439045 |
| LFNGGIGTYI | TCSAGQGGNTEVF | 0.6961229 |
| LTLGAGII | TCSAGQGGNTEVF | 0.8170024 |
| LVAAALGDI | TCSAGQGGNTEVF | 0.76520395 |
| NTAFVQLGI | TCSAGQGGNTEVF | 0.8202683 |
| QLFSGAEL | TCSAGQGGNTEVF | 0.6730826 |
| SAATFITL | TCSAGQGGNTEVF | 0.73444533 |
| SNYGLANLGV | TCSAGQGGNTEVF | 0.53100777 |
| SYGGAYAVM | TCSAGQGGNTEVF | 0.50128883 |
| TTGVGGLTLG | TCSAGQGGNTEVF | 0.6567059 |
| TTQTIYI | TCSAGQGGNTEVF | 0.72434664 |
| VAAGAGLGAVI | TCSAGQGGNTEVF | 0.3323613 |
| VAAVVTLL | TCSAGQGGNTEVF | 0.83148855 |
| VGFLGGLI | TCSAGQGGNTEVF | 0.7696742 |
| VGVIPILL | TCSAGQGGNTEVF | 0.8011197 |
| VIFASSLI | TCSAGQGGNTEVF | 0.695988 |
| VKGGDGGAAAG | TCSAGQGGNTEVF | 0.7300461 |
| VTKTFTNI | TCSAGQGGNTEVF | 0.6710989 |
| VVAAAILL | TCSAGQGGNTEVF | 0.79220426 |
| YAALDVEL | TCSAGQGGNTEVF | 0.56575525 |
| LASVALLL | TCSAGQGGNTEVF | 0.50754905 |

| | | |
|---|---|---|
| LDAVHRV | TCSAGQGGNTEVF | 0.659623 |
| QGLSSSIF | TCSAGQGGNTEVF | 0.53652155 |
| RGTFANIRL | TCSAGQGGNTEVF | 0.82960486 |
| RQLWWGHRI | TCSAGQGGNTEVF | 0.81066537 |
| YGPPGTGKTLL | TCSAGQGGNTEVF | 0.79102296 |
| AALVGAPANVA | TCSAGQGGNTEVF | 0.82994545 |
| AATFRRTE | TCSAGQGGNTEVF | 0.31130746 |
| AGALGGGI | TCSAGQGGNTEVF | 0.6853456 |
| AGGAGGAGGIGAG | TCSAGQGGNTEVF | 0.82638526 |
| AGSGALGL | TCSAGQGGNTEVF | 0.7722434 |
| AQIADQI | TCSAGQGGNTEVF | 0.7253349 |
| ASAVAVTYL | TCSAGQGGNTEVF | 0.8216395 |
| ASTVLGGTGG | TCSAGQGGNTEVF | 0.6449646 |
| AVGGEPAK | TCSAGQGGNTEVF | 0.6812112 |
| CEQGTFGGTC | TCSAGQGGNTEVF | 0.7656347 |
| DEGKAPAG | TCSAGQGGNTEVF | 0.79512244 |
| DTVVFPAL | TCSAGQGGNTEVF | 0.77689236 |
| FEGITMAG | TCSAGQGGNTEVF | 0.5868469 |
| FPLVVQD | TCSAGQGGNTEVF | 0.7963113 |
| GAFTTPQ | TCSAGQGGNTEVF | 0.8386252 |
| GFGGFGTAGGVGGA | TCSAGQGGNTEVF | 0.71433336 |
| GFLAQFL | TCSAGQGGNTEVF | 0.7805923 |
| GGAGGAGGNAGVA | TCSAGQGGNTEVF | 0.795414 |
| GGVGTAGAGGAGGAGG | TCSAGQGGNTEVF | 0.71142983 |
| GNGGGGDAGNAGSGGN | TCSAGQGGNTEVF | 0.63009727 |
| GTGGAGGAGNDAGS | TCSAGQGGNTEVF | 0.64449227 |
| GTTNTGWFNS | TCSAGQGGNTEVF | 0.77275 |
| IFMGPLFTL | TCSAGQGGNTEVF | 0.5745059 |
| ILAHAIEAI | TCSAGQGGNTEVF | 0.75443906 |
| IRSAIREAL | TCSAGQGGNTEVF | 0.74905014 |
| ITFAGGG | TCSAGQGGNTEVF | 0.8390425 |
| ITGMSGHQEV | TCSAGQGGNTEVF | 0.8014902 |
| IVAGLACTL | TCSAGQGGNTEVF | 0.7844695 |
| IVGGGLAGA | TCSAGQGGNTEVF | 0.5882764 |
| IVTGVLNI | TCSAGQGGNTEVF | 0.768031 |
| LAQQGCI | TCSAGQGGNTEVF | 0.7806068 |
| LCLHVEN | TCSAGQGGNTEVF | 0.47711045 |
| LDAISDGS | TCSAGQGGNTEVF | 0.82519615 |
| LGGFAAML | TCSAGQGGNTEVF | 0.665674 |
| LGGKNPMI | TCSAGQGGNTEVF | 0.7090644 |
| LGPKGVAAIL | TCSAGQGGNTEVF | 0.8025468 |
| LMVGVSI | TCSAGQGGNTEVF | 0.83832866 |
| LVGAAFVL | TCSAGQGGNTEVF | 0.7338767 |
| LVLAVAPL | TCSAGQGGNTEVF | 0.75358886 |
| MAAIAVIDAA | TCSAGQGGNTEVF | 0.63710064 |
| NNGINAELQ | TCSAGQGGNTEVF | 0.81325555 |
| RGEPLVLVHG | TCSAGQGGNTEVF | 0.8203527 |
| RGGAGGAAGLI | TCSAGQGGNTEVF | 0.724604 |
| RSYRMHPAV | TCSAGQGGNTEVF | 0.8354063 |
| SAAHLLAGQLA | TCSAGQGGNTEVF | 0.78767055 |
| SAAVGSY | TCSAGQGGNTEVF | 0.51010853 |

| | | |
|---|---|---|
| SPHAVLHRI | TCSAGQGGNTEVF | 0.76297873 |
| SQIAKVL | TCSAGQGGNTEVF | 0.7962441 |
| TALGHLVP | TCSAGQGGNTEVF | 0.80456007 |
| TAPTTSGAA | TCSAGQGGNTEVF | 0.81169957 |
| TDGDDGNGGSPGT | TCSAGQGGNTEVF | 0.7835597 |
| TDGLGEN | TCSAGQGGNTEVF | 0.6424843 |
| TGGPPLPGQA | TCSAGQGGNTEVF | 0.8251755 |
| TIPAITI | TCSAGQGGNTEVF | 0.82895494 |
| TIPGFTI | TCSAGQGGNTEVF | 0.80034125 |
| TSIASGALA | TCSAGQGGNTEVF | 0.69302046 |
| TVTLPGAAI | TCSAGQGGNTEVF | 0.6821769 |
| VASGLSAALP | TCSAGQGGNTEVF | 0.79219127 |
| VGAGVIGI | TCSAGQGGNTEVF | 0.7951818 |
| VGPKRFGV | TCSAGQGGNTEVF | 0.6302838 |
| VNVAFPD | TCSAGQGGNTEVF | 0.7602448 |
| VSMTNSGVS | TCSAGQGGNTEVF | 0.7817577 |
| VTDVLGDLSV | TCSAGQGGNTEVF | 0.50808716 |
| VVVVGAGFAGL | TCSAGQGGNTEVF | 0.7720934 |
| VYTVGTVV | TCSAGQGGNTEVF | 0.806274 |
| WTTFSGP | TCSAGQGGNTEVF | 0.78648597 |
| YDELVEAF | TCSAGQGGNTEVF | 0.6844183 |
| YGTGGAGGNGG | TCSAGQGGNTEVF | 0.8198355 |
| YTNVMAV | TCSAGQGGNTEVF | 0.8357218 |
| FLEQVHQL | TCSAGQGGNTEVF | 0.83738506 |
| GLSLLAVAV | TCSAGQGGNTEVF | 0.5517537 |
| LEQVHQL | TCSAGQGGNTEVF | 0.37035233 |
| LNAAGAI | TCSAGQGGNTEVF | 0.6716081 |
| PAVHASL | TCSAGQGGNTEVF | 0.8182146 |
| TELEDDA | TCSAGQGGNTEVF | 0.73849124 |
| TPPGPPN | TCSAGQGGNTEVF | 0.7080232 |
| TPPPPPPP | TCSAGQGGNTEVF | 0.7725757 |
| TQSPALI | ASSHRENTEVF | 0.81515396 |
| EFAAAFPQ | ASSHRENTEVF | 0.6745421 |
| LLSGNLDVL | ASSHRENTEVF | 0.5209091 |
| GPANLGNYNI | ASSHRENTEVF | 0.79172015 |
| TLGPTLP | ASSHRENTEVF | 0.8057538 |
| VYAYSSCCK | ASSHRENTEVF | 0.6448482 |
| APAPAPAPNI | ASSHRENTEVF | 0.7010486 |
| PGGPVPIP | ASSHRENTEVF | 0.7052886 |
| FLTVALEP | ASSHRENTEVF | 0.7310501 |
| SQYFGANL | ASSHRENTEVF | 0.7408984 |
| GGVKPAA | ASSHRENTEVF | 0.6075686 |
| GGTGGAGGAGGW | ASSHRENTEVF | 0.8076154 |
| GVGGVAGLGGAGG | ASSHRENTEVF | 0.65931886 |
| VLLVLATTI | ASSHRENTEVF | 0.7831585 |
| IGIFAAML | ASSHRENTEVF | 0.7714078 |
| AVGDLID | ASSHRENTEVF | 0.77865565 |
| NISPLPL | ASSHRENTEVF | 0.69285196 |
| VTAVFVAI | ASSHRENTEVF | 0.6670369 |
| SGDDNDAAL | ASSHRENTEVF | 0.50893754 |
| AVAALAHS | ASSHRENTEVF | 0.47000757 |

| | | |
|---|---|---|
| GGVTVMATP | ASSHRENTEVF | 0.8078307 |
| AGLAANVRD | ASSHRENTEVF | 0.5885305 |
| TNRPDLIDPALLRPG | ASSHRENTEVF | 0.8132932 |
| LANLGVTAL | ASSHRENTEVF | 0.801313 |
| TGERIEL | ASSHRENTEVF | 0.6586731 |
| NGEELAA | ASSHRENTEVF | 0.79505455 |
| VLSLVWG | ASSHRENTEVF | 0.5966686 |
| IGYALWL | ASSHRENTEVF | 0.7312628 |
| LSLLLAPL | ASSHRENTEVF | 0.73704153 |
| TLDEALK | ASSHRENTEVF | 0.80429715 |
| TDGAGTAEGG | ASSHRENTEVF | 0.6182146 |
| NAPGGAGGNGG | ASSHRENTEVF | 0.7965883 |
| VSVFMGMGGNF | ASSHRENTEVF | 0.7830603 |
| LASYLAAL | ASSHRENTEVF | 0.63445294 |
| LVGGTPAL | ASSHRENTEVF | 0.7866059 |
| VRNADVL | ASSHRENTEVF | 0.8106447 |
| TCSSPFLVSAM | ASSHRENTEVF | 0.7934408 |
| VHITDAE | ASSHRENTEVF | 0.6111729 |
| EYAAPSF | ASSHRENTEVF | 0.7936934 |
| VVDAFHGGL | ASSHRENTEVF | 0.6355592 |
| ASVVDPQ | ASSHRENTEVF | 0.7775678 |
| SYNEFGGLS | ASSHRENTEVF | 0.74884135 |
| DPDGSVY | ASSHRENTEVF | 0.7356385 |
| MVSTIHATE | ASSHRENTEVF | 0.8121397 |
| LAAGVADPV | ASSHRENTEVF | 0.5406541 |
| TKEQIAA | ASSHRENTEVF | 0.61835295 |
| SLASPAAL | ASSHRENTEVF | 0.7074979 |
| IGLVFYGV | ASSHRENTEVF | 0.7713327 |
| FLLDASNL | ASSHRENTEVF | 0.80412996 |
| YQVGNFP | ASSHRENTEVF | 0.61824405 |
| ESLLSQST | ASSHRENTEVF | 0.7646806 |
| LFTVGFAAVG | ASSHRENTEVF | 0.7225084 |
| ATPPTPT | ASSHRENTEVF | 0.5981999 |
| TDLATLG | ASSHRENTEVF | 0.7943052 |
| SAGGTGR | ASSHRENTEVF | 0.73507243 |
| GFSNSGS | ASSHRENTEVF | 0.64864576 |
| GGNGGSGGTSV | ASSHRENTEVF | 0.7818514 |
| GTSVATGGAGN | ASSHRENTEVF | 0.7251211 |
| NGGNGGSGGTS | ASSHRENTEVF | 0.7873621 |
| AVVKFPH | ASSHRENTEVF | 0.80541086 |
| TPPPPPP | ASSHRENTEVF | 0.61637694 |
| AAPAAAAAPA | ASSHRENTEVF | 0.7884452 |
| IALNEDL | ASSHRENTEVF | 0.75768465 |
| IVNVSSM | ASSHRENTEVF | 0.77018875 |
| TGIPPAPR | ASSHRENTEVF | 0.7432402 |
| FDLGGGTF | ASSHRENTEVF | 0.62734187 |
| FDLGGGTFDVS | ASSHRENTEVF | 0.8146824 |
| NEPTAAA | ASSHRENTEVF | 0.7998425 |
| NEPTAAALA | ASSHRENTEVF | 0.7737616 |
| GSIINISSI | ASSHRENTEVF | 0.79274654 |
| VAPGNAGTA | ASSHRENTEVF | 0.8065012 |

| | | |
|---|---|---|
| TGGGASLE | ASSHRENTEVF | 0.80938566 |
| VAVYNHYKRI | ASSHRENTEVF | 0.65289545 |
| TDPAPATT | ASSHRENTEVF | 0.52148974 |
| TNRPDILDPALLRPG | ASSHRENTEVF | 0.7247325 |
| LTTTTTS | ASSHRENTEVF | 0.7213205 |
| PTGSGKTL | ASSHRENTEVF | 0.74639606 |
| TGSGKTL | ASSHRENTEVF | 0.509168 |
| TIGHVDHGK | ASSHRENTEVF | 0.8114586 |
| TIGHVDHGKTT | ASSHRENTEVF | 0.83719856 |
| GHVDHGKTT | ASSHRENTEVF | 0.65276253 |
| AQFEHTLL | ASSHRENTEVF | 0.84173435 |
| IVNQDEL | ASSHRENTEVF | 0.61709887 |
| TNGSQFF | ASSHRENTEVF | 0.7875057 |
| TGKGIVE | ASSHRENTEVF | 0.81937796 |
| LISNASDA | ASSHRENTEVF | 0.5801318 |
| AELRTLGELTKIAKAHGAQ | ASSHRENTEVF | 0.7857456 |
| APNAVAAAEPSW | ASSHRENTEVF | 0.6174035 |
| APQVAGAATAVL | ASSHRENTEVF | 0.6783585 |
| ARRDPAAAAAILPTDA | ASSHRENTEVF | 0.818954 |
| DAAARDVLAEAGLAEYF | ASSHRENTEVF | 0.7175842 |
| DDQIVLSAKANLAKVEQ | ASSHRENTEVF | 0.77925396 |
| DELLPDTGEDSDEDGH | ASSHRENTEVF | 0.7678783 |
| ELPEAPANRATVPQD | ASSHRENTEVF | 0.65393555 |
| FGQSKIERVTGIG | ASSHRENTEVF | 0.77940166 |
| GAGGTGSTAGGAGGAGGAGGLY | ASSHRENTEVF | 0.8050702 |
| GHGGTGAAGGNGGTGGAGGSGI | ASSHRENTEVF | 0.7117068 |
| GNGGNGGAGGEGGPGGAGGAGA | ASSHRENTEVF | 0.8313731 |
| GSTAGNGGNGGNATNGTIAGQP | ASSHRENTEVF | 0.66079074 |
| GTGGDGGNGGNGGTGG | ASSHRENTEVF | 0.82589746 |
| GVRRPAGIPADAPV | ASSHRENTEVF | 0.8121315 |
| GYRQVTDAHLLTLARR | ASSHRENTEVF | 0.7367404 |
| IAKMGQDGHDRG | ASSHRENTEVF | 0.7192982 |
| KAEPSIEIQYT | ASSHRENTEVF | 0.80983216 |
| KPVRIRMAADEDTVAVT | ASSHRENTEVF | 0.7336334 |
| MPPRIAGMRLLVIKPEP | ASSHRENTEVF | 0.76439196 |
| MKFYAVALFRSRGVPPTR | ASSHRENTEVF | 0.8250201 |
| NAGGFGILGAVAHSP | ASSHRENTEVF | 0.58591837 |
| NEGREITLAMAEM | ASSHRENTEVF | 0.7769979 |
| NELVEWSHPLGHRWRWE | ASSHRENTEVF | 0.776454 |
| NLGAGGTGGNGGNPG | ASSHRENTEVF | 0.78774464 |
| NTEWPKWETVMAADPVD | ASSHRENTEVF | 0.5380987 |
| PASAAPAETQP | ASSHRENTEVF | 0.26135555 |
| QQSSDGKNGVRAAEITGE | ASSHRENTEVF | 0.74215764 |
| RAPDAVTTSWWKE | ASSHRENTEVF | 0.45584708 |
| SGFGNVGAHTSGWFNQ | ASSHRENTEVF | 0.7901723 |
| SGVAGAVGQAASVGGLKV | ASSHRENTEVF | 0.7972859 |
| SNGGAGTGSAGGNGGT | ASSHRENTEVF | 0.4906066 |
| TGGAGGEGGAAG | ASSHRENTEVF | 0.7684908 |
| TGGDGGAAGAGGAPGHG | ASSHRENTEVF | 0.78600514 |
| TMPPLLAAIANAGRHGV | ASSHRENTEVF | 0.5120408 |
| TSSTAGNSGGA | ASSHRENTEVF | 0.7280025 |

| | | |
|---|---|---|
| VPATVPVGILC | ASSHRENTEVF | 0.45334238 |
| VQRPDSGGVVLTGRIS | ASSHRENTEVF | 0.7283816 |
| WTPNPVPATKGVPAGLVQAT | ASSHRENTEVF | 0.2841674 |
| YALKRALAKFSDRFDVVII | ASSHRENTEVF | 0.75839436 |
| YLIGSVIV | ASSHRENTEVF | 0.70357186 |
| YGQLVAVID | ASSHRENTEVF | 0.23004857 |
| VTVGGMTLPA | ASSHRENTEVF | 0.7646407 |
| VLALPAV | ASSHRENTEVF | 0.56603825 |
| VIPPTLNYET | ASSHRENTEVF | 0.7166378 |
| VGMGGMHPSSG | ASSHRENTEVF | 0.8029537 |
| VGLAGGAATA | ASSHRENTEVF | 0.6340736 |
| VGGAGGAGGSGP | ASSHRENTEVF | 0.38901454 |
| VFAAFMV | ASSHRENTEVF | 0.75609684 |
| TLYDKLR | ASSHRENTEVF | 0.7989906 |
| TLAVFTVL | ASSHRENTEVF | 0.3361321 |
| TGSVQPARV | ASSHRENTEVF | 0.7780884 |
| TGGGAGSGGT | ASSHRENTEVF | 0.78923047 |
| SHLETSL | ASSHRENTEVF | 0.60532117 |
| SFTTGMT | ASSHRENTEVF | 0.812181 |
| QVEVAKRMGVRQPTVS | ASSHRENTEVF | 0.8118407 |
| QGSGGPALAM | ASSHRENTEVF | 0.67962474 |
| MLAPSNP | ASSHRENTEVF | 0.7994311 |
| LVNNAAI | ASSHRENTEVF | 0.7928975 |
| LTGEINTLA | ASSHRENTEVF | 0.63821137 |
| LSIFGPV | ASSHRENTEVF | 0.6657183 |
| LLALLANL | ASSHRENTEVF | 0.5839059 |
| LISNASDALDKLR | ASSHRENTEVF | 0.7444162 |
| LINEAVP | ASSHRENTEVF | 0.5697267 |
| LGYAFSGL | ASSHRENTEVF | 0.7335953 |
| LDFVAVAL | ASSHRENTEVF | 0.7898513 |
| LAQPGAE | ASSHRENTEVF | 0.6558088 |
| ISNLGAVTGF | ASSHRENTEVF | 0.55590963 |
| HQEDPGRI | ASSHRENTEVF | 0.3416513 |
| GTPGEPTP | ASSHRENTEVF | 0.7365625 |
| GFFNSGS | ASSHRENTEVF | 0.65046805 |
| GAGGGGLTAGAN | ASSHRENTEVF | 0.43594784 |
| FVDALVNM | ASSHRENTEVF | 0.614634 |
| FLTVAPEL | ASSHRENTEVF | 0.7227514 |
| FETLTAL | ASSHRENTEVF | 0.74660146 |
| DVINPIEL | ASSHRENTEVF | 0.43690744 |
| APVSGAPV | ASSHRENTEVF | 0.8130547 |
| ALVGGLAW | ASSHRENTEVF | 0.7760041 |
| ALAAAAATAI | ASSHRENTEVF | 0.8016659 |
| AFGLGLLL | ASSHRENTEVF | 0.70237213 |
| AFFAAASL | ASSHRENTEVF | 0.5178335 |
| AAAALPAFSPP | ASSHRENTEVF | 0.7907773 |
| VVLAPAL | ASSHRENTEVF | 0.6335809 |
| TIGHVDHGKTTL | ASSHRENTEVF | 0.6638494 |
| LISNASDALDK | ASSHRENTEVF | 0.7849927 |
| LISNASDAL | ASSHRENTEVF | 0.53114617 |
| ISNASDALDK | ASSHRENTEVF | 0.44971198 |

| | | |
|---|---|---|
| NQAETLVYQTE | ASSHRENTEVF | 0.73572975 |
| TGGTGGTAGAG | ASSHRENTEVF | 0.8012196 |
| TGSTAGGAGGA | ASSHRENTEVF | 0.5300894 |
| DPLAVPVAAGPV | ASSHRENTEVF | 0.35928535 |
| GGTGGTPGNGGN | ASSHRENTEVF | 0.69441956 |
| PAGLAGVDPATL | ASSHRENTEVF | 0.62976176 |
| GNNLVVEAPHAGASV | ASSHRENTEVF | 0.79799473 |
| KVTRIMSASTLEQPA | ASSHRENTEVF | 0.76785636 |
| LGFALPIALPRRTVV | ASSHRENTEVF | 0.7213497 |
| TRLGNLTARKLVVLV | ASSHRENTEVF | 0.795626 |
| WSQQRRLIYFPSAGP | ASSHRENTEVF | 0.7476636 |
| GPNGNPDPMAAAVDIR | ASSHRENTEVF | 0.68767846 |
| TQRCVSVADLRAAASR | ASSHRENTEVF | 0.773534 |
| QFVLDVAEHPVEVLTETV | ASSHRENTEVF | 0.55763733 |
| GGAGGTGGAGGNGGTQAGDG | ASSHRENTEVF | 0.7952502 |
| PLLATLRAIAVDVDAKPAQV | ASSHRENTEVF | 0.73589814 |
| APKFRTGGYMIDFWGVGYQVAK | ASSHRENTEVF | 0.80257195 |
| GAGGAGGAGGAAGGPGGTGGTG | ASSHRENTEVF | 0.80376583 |
| IGALGLLATDVAPAGSVAFGVA | ASSHRENTEVF | 0.595433 |
| AGGAGGIGGNGGAGGAGGVLMG | ASSHRENTEVF | 0.75767267 |
| ADELIGLL | ASSHRENTEVF | 0.3175265 |
| ADRALKA | ASSHRENTEVF | 0.66574526 |
| AGAVGQVGGAGGA | ASSHRENTEVF | 0.7702509 |
| AGGGGGLGGAP | ASSHRENTEVF | 0.7697208 |
| AGGLLYGNGGAG | ASSHRENTEVF | 0.7914806 |
| AGLESIGL | ASSHRENTEVF | 0.78462756 |
| AGLKTGII | ASSHRENTEVF | 0.6384362 |
| ASLAALLL | ASSHRENTEVF | 0.78953266 |
| ASVINISSI | ASSHRENTEVF | 0.79995227 |
| ATPPGPA | ASSHRENTEVF | 0.40132308 |
| ATVASALV | ASSHRENTEVF | 0.6857661 |
| ATVASLIL | ASSHRENTEVF | 0.8049131 |
| AVNGPVLIRH | ASSHRENTEVF | 0.78827316 |
| DGAAGAVGI | ASSHRENTEVF | 0.58756816 |
| DPLLHGI | ASSHRENTEVF | 0.77615887 |
| ELGAALGAL | ASSHRENTEVF | 0.795339 |
| ELGRTQAV | ASSHRENTEVF | 0.75455153 |
| FVAGGYLS | ASSHRENTEVF | 0.79732573 |
| GASGGAGGAGGW | ASSHRENTEVF | 0.2939128 |
| GGAGAGGDGGAGGI | ASSHRENTEVF | 0.77541804 |
| GTGHNGSQPGMGGNG | ASSHRENTEVF | 0.6357882 |
| GYAAASASAS | ASSHRENTEVF | 0.2566629 |
| HQVDVAAVQQG | ASSHRENTEVF | 0.7625106 |
| IGAAIAHT | ASSHRENTEVF | 0.71406233 |
| IGMIEVAQL | ASSHRENTEVF | 0.80532366 |
| IRLAPEHTSAMD | ASSHRENTEVF | 0.6537244 |
| IYNVSVL | ASSHRENTEVF | 0.79881275 |
| KQGFLGM | ASSHRENTEVF | 0.7289156 |
| LESVITQL | ASSHRENTEVF | 0.73527634 |
| LGLGALVVV | ASSHRENTEVF | 0.63372135 |
| LIGNGGNGGSGGI | ASSHRENTEVF | 0.6907243 |

| | | |
|---|---|---|
| LNAINAPTL | ASSHRENTEVF | 0.79797786 |
| LQLLNTGV | ASSHRENTEVF | 0.7949031 |
| LQTSIEF | ASSHRENTEVF | 0.22715849 |
| MSFAVSEL | ASSHRENTEVF | 0.76964104 |
| NTGALGL | ASSHRENTEVF | 0.55183595 |
| PPGAGGAGGAGTT | ASSHRENTEVF | 0.229277 |
| QLVGLGI | ASSHRENTEVF | 0.34598655 |
| RFLEQVHQL | ASSHRENTEVF | 0.63300717 |
| RGVGYVLR | ASSHRENTEVF | 0.6850332 |
| SATGVFAGVI | ASSHRENTEVF | 0.74957633 |
| SLPAITLP | ASSHRENTEVF | 0.74101937 |
| STALAFTV | ASSHRENTEVF | 0.5905155 |
| SVLTVGLGK | ASSHRENTEVF | 0.7690254 |
| TALAGAGSGI | ASSHRENTEVF | 0.56078744 |
| TGLPFTGI | ASSHRENTEVF | 0.1671558 |
| VAEHVTV | ASSHRENTEVF | 0.6050159 |
| VDIVLNSL | ASSHRENTEVF | 0.5432881 |
| VLTGGGAVL | ASSHRENTEVF | 0.80345625 |
| VNVGLAYGI | ASSHRENTEVF | 0.59044266 |
| VSSAAVGAV | ASSHRENTEVF | 0.619913 |
| VTSVYEAGV | ASSHRENTEVF | 0.6087042 |
| WVALDRGA | ASSHRENTEVF | 0.76094615 |
| YLVGSVVI | ASSHRENTEVF | 0.70944726 |
| GPPGTGKTLL | ASSHRENTEVF | 0.5679794 |
| GVGMTKF | ASSHRENTEVF | 0.39546406 |
| IGIGGSDL | ASSHRENTEVF | 0.6766608 |
| IGTLLAEL | ASSHRENTEVF | 0.5060598 |
| IYTSGTT | ASSHRENTEVF | 0.58144766 |
| LSFPAGV | ASSHRENTEVF | 0.6893705 |
| LYTSGTT | ASSHRENTEVF | 0.5458409 |
| VGAFPFGF | ASSHRENTEVF | 0.35798806 |
| VGAGTVEF | ASSHRENTEVF | 0.80296755 |
| VGKSTLLL | ASSHRENTEVF | 0.7285538 |
| AALAHAGGGIV | ASSHRENTEVF | 0.58320284 |
| AALVAVSLL | ASSHRENTEVF | 0.8006549 |
| AAVSRAVAAL | ASSHRENTEVF | 0.7358988 |
| AGGSSGIGGF | ASSHRENTEVF | 0.6603575 |
| AGLTVANANG | ASSHRENTEVF | 0.50213146 |
| AGLVGAVL | ASSHRENTEVF | 0.5572262 |
| ALSQAFTV | ASSHRENTEVF | 0.45005468 |
| ALTFSVSL | ASSHRENTEVF | 0.4238885 |
| ALTGAALVAL | ASSHRENTEVF | 0.80413145 |
| ASFSIGAGVL | ASSHRENTEVF | 0.79204357 |
| DGSVYKI | ASSHRENTEVF | 0.5694808 |
| EQLAEIV | ASSHRENTEVF | 0.7984079 |
| FAVSELTVM | ASSHRENTEVF | 0.79777586 |
| GAAPLAAGL | ASSHRENTEVF | 0.8022705 |
| GADGGDGGSGGAGG | ASSHRENTEVF | 0.6742143 |
| GFGLDTGGAGG | ASSHRENTEVF | 0.6693617 |
| GLASGGLL | ASSHRENTEVF | 0.7621411 |
| GRIAGAAVLL | ASSHRENTEVF | 0.63733745 |

| | | |
|---|---|---|
| GSAGSLNTSF | ASSHRENTEVF | 0.72208816 |
| GTGQIGI | ASSHRENTEVF | 0.3948355 |
| GTLIALIL | ASSHRENTEVF | 0.7934002 |
| IAATMGFVVG | ASSHRENTEVF | 0.81247735 |
| IFAQALL | ASSHRENTEVF | 0.65934825 |
| IGAAVALGF | ASSHRENTEVF | 0.31273142 |
| IGLTPLAL | ASSHRENTEVF | 0.5303843 |
| ILGSNTSTL | ASSHRENTEVF | 0.4983227 |
| IVFSLTVL | ASSHRENTEVF | 0.4768736 |
| LASKATGF | ASSHRENTEVF | 0.7129314 |
| LASVALIL | ASSHRENTEVF | 0.53017294 |
| LFAVGLTL | ASSHRENTEVF | 0.6553093 |
| LFNGGIGTYI | ASSHRENTEVF | 0.79189074 |
| LTLGAGII | ASSHRENTEVF | 0.68883705 |
| LVAAALGDI | ASSHRENTEVF | 0.7944302 |
| NTAFVQLGI | ASSHRENTEVF | 0.7962723 |
| QLFSGAEL | ASSHRENTEVF | 0.7889162 |
| SAATFITL | ASSHRENTEVF | 0.8116849 |
| SNYGLANLGV | ASSHRENTEVF | 0.79410523 |
| SYGGAYAVM | ASSHRENTEVF | 0.69108266 |
| TTGVGGLTLG | ASSHRENTEVF | 0.72741127 |
| TTQTIYI | ASSHRENTEVF | 0.7798819 |
| VAAGAGLGAVI | ASSHRENTEVF | 0.59024554 |
| VAAVVTLL | ASSHRENTEVF | 0.66457564 |
| VGFLGGLI | ASSHRENTEVF | 0.70208293 |
| VGVIPILL | ASSHRENTEVF | 0.7672199 |
| VIFASSLI | ASSHRENTEVF | 0.6633093 |
| VKGGDGGAAAG | ASSHRENTEVF | 0.8032098 |
| VTKTFTNI | ASSHRENTEVF | 0.3630398 |
| VVAAAILL | ASSHRENTEVF | 0.42566288 |
| YAALDVEL | ASSHRENTEVF | 0.80263674 |
| LASVALLL | ASSHRENTEVF | 0.177546 |
| LDAVHRV | ASSHRENTEVF | 0.50769293 |
| QGLSSSIF | ASSHRENTEVF | 0.7533448 |
| RGTFANIRL | ASSHRENTEVF | 0.78778815 |
| RQLWWGHRI | ASSHRENTEVF | 0.19155093 |
| YGPPGTGKTLL | ASSHRENTEVF | 0.48011267 |
| AALVGAPANVA | ASSHRENTEVF | 0.54461503 |
| AATFRRTE | ASSHRENTEVF | 0.7920065 |
| AGALGGGI | ASSHRENTEVF | 0.5865066 |
| AGGAGGAGGIGAG | ASSHRENTEVF | 0.24770117 |
| AGSGALGL | ASSHRENTEVF | 0.80540144 |
| AQIADQI | ASSHRENTEVF | 0.52830887 |
| ASAVAVTYL | ASSHRENTEVF | 0.70956206 |
| ASTVLGGTGG | ASSHRENTEVF | 0.80357397 |
| AVGGEPAK | ASSHRENTEVF | 0.8141545 |
| CEQGTFGGTC | ASSHRENTEVF | 0.59710157 |
| DEGKAPAG | ASSHRENTEVF | 0.7845516 |
| DTVVFPAL | ASSHRENTEVF | 0.6958645 |
| FEGITMAG | ASSHRENTEVF | 0.8148049 |
| FPLVVQD | ASSHRENTEVF | 0.5418122 |

| | | |
|---|---|---|
| GAFTTPQ | ASSHRENTEVF | 0.49726027 |
| GFGGFGTAGGVGGA | ASSHRENTEVF | 0.7974988 |
| GFLAQFL | ASSHRENTEVF | 0.804012 |
| GGAGGAGGNAGVA | ASSHRENTEVF | 0.74101603 |
| GGVGTAGAGGAGGAGG | ASSHRENTEVF | 0.7935244 |
| GNGGGGDAGNAGSGGN | ASSHRENTEVF | 0.76579744 |
| GTGGAGGAGNDAGS | ASSHRENTEVF | 0.66953456 |
| GTTNTGWFNS | ASSHRENTEVF | 0.77855635 |
| IFMGPLFTL | ASSHRENTEVF | 0.7764604 |
| ILAHAIEAI | ASSHRENTEVF | 0.44361067 |
| IRSAIREAL | ASSHRENTEVF | 0.8058286 |
| ITFAGGG | ASSHRENTEVF | 0.5820856 |
| ITGMSGHQEV | ASSHRENTEVF | 0.5659458 |
| IVAGLACTL | ASSHRENTEVF | 0.8043941 |
| IVGGGLAGA | ASSHRENTEVF | 0.7705808 |
| IVTGVLNI | ASSHRENTEVF | 0.78603494 |
| LAQQGCI | ASSHRENTEVF | 0.7391106 |
| LCLHVEN | ASSHRENTEVF | 0.8018055 |
| LDAISDGS | ASSHRENTEVF | 0.3437141 |
| LGGFAAML | ASSHRENTEVF | 0.7649901 |
| LGGKNPMI | ASSHRENTEVF | 0.7031662 |
| LGPKGVAAIL | ASSHRENTEVF | 0.79270077 |
| LMVGVSI | ASSHRENTEVF | 0.7743094 |
| LVGAAFVL | ASSHRENTEVF | 0.6742289 |
| LVLAVAPL | ASSHRENTEVF | 0.78431475 |
| MAAIAVIDAA | ASSHRENTEVF | 0.7892711 |
| NNGINAELQ | ASSHRENTEVF | 0.7473897 |
| RGEPLVLVHG | ASSHRENTEVF | 0.7478388 |
| RGGAGGAAGLI | ASSHRENTEVF | 0.8143455 |
| RSYRMHPAV | ASSHRENTEVF | 0.6253073 |
| SAAHLLAGQLA | ASSHRENTEVF | 0.6305073 |
| SAAVGSY | ASSHRENTEVF | 0.78225785 |
| SPHAVLHRI | ASSHRENTEVF | 0.8002598 |
| SQIAKVL | ASSHRENTEVF | 0.7815221 |
| TALGHLVP | ASSHRENTEVF | 0.79846686 |
| TAPTTSGAA | ASSHRENTEVF | 0.57408166 |
| TDGDDGNGGSPGT | ASSHRENTEVF | 0.74441093 |
| TDGLGEN | ASSHRENTEVF | 0.720538 |
| TGGPPLPGQA | ASSHRENTEVF | 0.6990598 |
| TIPAITI | ASSHRENTEVF | 0.48492435 |
| TIPGFTI | ASSHRENTEVF | 0.690945 |
| TSIASGALA | ASSHRENTEVF | 0.7771358 |
| TVTLPGAAI | ASSHRENTEVF | 0.74626523 |
| VASGLSAALP | ASSHRENTEVF | 0.47673607 |
| VGAGVIGI | ASSHRENTEVF | 0.72709715 |
| VGPKRFGV | ASSHRENTEVF | 0.73037624 |
| VNVAFPD | ASSHRENTEVF | 0.81734866 |
| VSMTNSGVS | ASSHRENTEVF | 0.68379676 |
| VTDVLGDLSV | ASSHRENTEVF | 0.79893684 |
| VVVVGAGFAGL | ASSHRENTEVF | 0.51405585 |
| VYTVGTVV | ASSHRENTEVF | 0.75281525 |

| | | |
|---|---|---|
| WTTFSGP | ASSHRENTEVF | 0.4502902 |
| YDELVEAF | ASSHRENTEVF | 0.6397 |
| YGTGGAGGNGG | ASSHRENTEVF | 0.63936484 |
| YTNVMAV | ASSHRENTEVF | 0.6067202 |
| FLEQVHQL | ASSHRENTEVF | 0.77312994 |
| GLSLLAVAV | ASSHRENTEVF | 0.7982718 |
| LEQVHQL | ASSHRENTEVF | 0.7203426 |
| LNAAGAI | ASSHRENTEVF | 0.78903514 |
| PAVHASL | ASSHRENTEVF | 0.80221343 |
| TELEDDA | ASSHRENTEVF | 0.8141996 |
| TPPGPPN | ASSHRENTEVF | 0.5744594 |
| TPPPPPPP | ASSHRENTEVF | 0.69953674 |
| TQSPALI | ASRGHQNTLY | 0.39850616 |
| EFAAAFPQ | ASRGHQNTLY | 0.6255263 |
| LLSGNLDVL | ASRGHQNTLY | 0.62056386 |
| GPANLGNYNI | ASRGHQNTLY | 0.645018 |
| TLGPTLP | ASRGHQNTLY | 0.18354756 |
| VYAYSSCCK | ASRGHQNTLY | 0.7650621 |
| APAPAPAPNI | ASRGHQNTLY | 0.8071956 |
| PGGPVPIP | ASRGHQNTLY | 0.54475427 |
| FLTVALEP | ASRGHQNTLY | 0.7688464 |
| SQYFGANL | ASRGHQNTLY | 0.76770097 |
| GGVKPAA | ASRGHQNTLY | 0.6839012 |
| GGTGGAGGAGGW | ASRGHQNTLY | 0.7563896 |
| GVGGVAGLGGAGG | ASRGHQNTLY | 0.63219464 |
| VLLVLATTI | ASRGHQNTLY | 0.80685544 |
| IGIFAAML | ASRGHQNTLY | 0.7872511 |
| AVGDLID | ASRGHQNTLY | 0.75670457 |
| NISPLPL | ASRGHQNTLY | 0.7052486 |
| VTAVFVAI | ASRGHQNTLY | 0.78019994 |
| SGDDNDAAL | ASRGHQNTLY | 0.30565366 |
| AVAALAHS | ASRGHQNTLY | 0.723831 |
| GGVTVMATP | ASRGHQNTLY | 0.22794369 |
| AGLAANVRD | ASRGHQNTLY | 0.745081 |
| TNRPDLIDPALLRPG | ASRGHQNTLY | 0.54173905 |
| LANLGVTAL | ASRGHQNTLY | 0.76322204 |
| TGERIEL | ASRGHQNTLY | 0.6012572 |
| NGEELAA | ASRGHQNTLY | 0.74796814 |
| VLSLVWG | ASRGHQNTLY | 0.43329608 |
| IGYALWL | ASRGHQNTLY | 0.7814254 |
| LSLLLAPL | ASRGHQNTLY | 0.7345721 |
| TLDEALK | ASRGHQNTLY | 0.2898119 |
| TDGAGTAEGG | ASRGHQNTLY | 0.7985425 |
| NAPGGAGGNGG | ASRGHQNTLY | 0.7427001 |
| VSVFMGMGGNF | ASRGHQNTLY | 0.6850755 |
| LASYLAAL | ASRGHQNTLY | 0.76988536 |
| LVGGTPAL | ASRGHQNTLY | 0.77613443 |
| VRNADVL | ASRGHQNTLY | 0.33526412 |
| TCSSPFLVSAM | ASRGHQNTLY | 0.67865586 |
| VHITDAE | ASRGHQNTLY | 0.7262269 |
| EYAAPSF | ASRGHQNTLY | 0.599208 |

| | | |
|---|---|---|
| VVDAFHGGL | ASRGHQNTLY | 0.23883785 |
| ASVVDPQ | ASRGHQNTLY | 0.77117497 |
| SYNEFGGLS | ASRGHQNTLY | 0.57608944 |
| DPDGSVY | ASRGHQNTLY | 0.54842585 |
| MVSTIHATE | ASRGHQNTLY | 0.6736988 |
| LAAGVADPV | ASRGHQNTLY | 0.80644226 |
| TKEQIAA | ASRGHQNTLY | 0.6542781 |
| SLASPAAL | ASRGHQNTLY | 0.71508336 |
| IGLVFYGV | ASRGHQNTLY | 0.7881551 |
| FLLDASNL | ASRGHQNTLY | 0.52129817 |
| YQVGNFP | ASRGHQNTLY | 0.78729594 |
| ESLLSQST | ASRGHQNTLY | 0.67024225 |
| LFTVGFAAVG | ASRGHQNTLY | 0.7244952 |
| ATPPTPT | ASRGHQNTLY | 0.7725441 |
| TDLATLG | ASRGHQNTLY | 0.6554532 |
| SAGGTGR | ASRGHQNTLY | 0.6307417 |
| GFSNSGS | ASRGHQNTLY | 0.6032821 |
| GGNGGSGGTSV | ASRGHQNTLY | 0.43631983 |
| GTSVATGGAGN | ASRGHQNTLY | 0.53661966 |
| NGGNGGSGGTS | ASRGHQNTLY | 0.7099844 |
| AVVKFPH | ASRGHQNTLY | 0.81186676 |
| TPPPPPP | ASRGHQNTLY | 0.7301406 |
| AAPAAAAAPA | ASRGHQNTLY | 0.4799931 |
| IALNEDL | ASRGHQNTLY | 0.68936545 |
| IVNVSSM | ASRGHQNTLY | 0.50918394 |
| TGIPPAPR | ASRGHQNTLY | 0.7994804 |
| FDLGGGTF | ASRGHQNTLY | 0.7906922 |
| FDLGGGTFDVS | ASRGHQNTLY | 0.8069762 |
| NEPTAAA | ASRGHQNTLY | 0.7320484 |
| NEPTAAALA | ASRGHQNTLY | 0.51473904 |
| GSIINISSI | ASRGHQNTLY | 0.8076064 |
| VAPGNAGTA | ASRGHQNTLY | 0.7103593 |
| TGGGASLE | ASRGHQNTLY | 0.7634256 |
| VAVYNHYKRI | ASRGHQNTLY | 0.76436317 |
| TDPAPATT | ASRGHQNTLY | 0.73695457 |
| TNRPDILDPALLRPG | ASRGHQNTLY | 0.63334465 |
| LTTTTTS | ASRGHQNTLY | 0.7696456 |
| PTGSGKTL | ASRGHQNTLY | 0.35247362 |
| TGSGKTL | ASRGHQNTLY | 0.76370436 |
| TIGHVDHGK | ASRGHQNTLY | 0.34677732 |
| TIGHVDHGKTT | ASRGHQNTLY | 0.6490799 |
| GHVDHGKTT | ASRGHQNTLY | 0.66387904 |
| AQFEHTLL | ASRGHQNTLY | 0.80871445 |
| IVNQDEL | ASRGHQNTLY | 0.79201066 |
| TNGSQFF | ASRGHQNTLY | 0.59161687 |
| TGKGIVE | ASRGHQNTLY | 0.53765893 |
| LISNASDA | ASRGHQNTLY | 0.7726295 |
| AELRTLGELTKIAKAHGAQ | ASRGHQNTLY | 0.5107434 |
| APNAVAAAEPSW | ASRGHQNTLY | 0.8039367 |
| APQVAGAATAVL | ASRGHQNTLY | 0.67939687 |
| ARRDPAAAAAILPTDA | ASRGHQNTLY | 0.6204785 |

| | | |
|---|---|---|
| DAAARDVLAEAGLAEYF | ASRGHQNTLY | 0.71170574 |
| DDQIVLSAKANLAKVEQ | ASRGHQNTLY | 0.78144306 |
| DELLPDTGEDSDEDGH | ASRGHQNTLY | 0.5202279 |
| ELPEAPANRATVPQD | ASRGHQNTLY | 0.7935909 |
| FGQSKIERVTGIG | ASRGHQNTLY | 0.21227415 |
| GAGGTGSTAGGAGGAGGAGGLY | ASRGHQNTLY | 0.78014797 |
| GHGGTGAAGGNGGTGGAGGSGI | ASRGHQNTLY | 0.42990625 |
| GNGGNGGAGGEGGPGGAGGAGA | ASRGHQNTLY | 0.50861436 |
| GSTAGNGGNGGNATNGTIAGQP | ASRGHQNTLY | 0.3229004 |
| GTGGDGGNGGNGGTGG | ASRGHQNTLY | 0.74992085 |
| GVRRPAGIPADAPV | ASRGHQNTLY | 0.7744045 |
| GYRQVTDAHLLTLARR | ASRGHQNTLY | 0.80175793 |
| IAKMGQDGHDRG | ASRGHQNTLY | 0.755492 |
| KAEPSIEIQYT | ASRGHQNTLY | 0.7823926 |
| KPVRIRMAADEDTVAVT | ASRGHQNTLY | 0.78813934 |
| MPPRIAGMRLLVIKPEP | ASRGHQNTLY | 0.59798527 |
| MKFYAVALFRSRGVPPTR | ASRGHQNTLY | 0.2654585 |
| NAGGFGILGAVAHSP | ASRGHQNTLY | 0.76125324 |
| NEGREITLAMAEM | ASRGHQNTLY | 0.7138335 |
| NELVEWSHPLGHRWRWE | ASRGHQNTLY | 0.80229163 |
| NLGAGGTGGNGGNPG | ASRGHQNTLY | 0.7266787 |
| NTEWPKWETVMAADPVD | ASRGHQNTLY | 0.43970668 |
| PASAAPAETQP | ASRGHQNTLY | 0.7877398 |
| QQSSDGKNGVRAAEITGE | ASRGHQNTLY | 0.7846663 |
| RAPDAVTTSWWKE | ASRGHQNTLY | 0.41524386 |
| SGFGNVGAHTSGWFNQ | ASRGHQNTLY | 0.6878271 |
| SGVAGAVGQAASVGGLKV | ASRGHQNTLY | 0.6172963 |
| SNGGAGTGSAGGNGGT | ASRGHQNTLY | 0.6016333 |
| TGGAGGEGGAAG | ASRGHQNTLY | 0.6187037 |
| TGGDGGAAGAGGAPGHG | ASRGHQNTLY | 0.6626296 |
| TMPPLLAAIANAGRHGV | ASRGHQNTLY | 0.5087307 |
| TSSTAGNSGGA | ASRGHQNTLY | 0.6825609 |
| VPATVPVGILC | ASRGHQNTLY | 0.41432256 |
| VQRPDSGGVVLTGRIS | ASRGHQNTLY | 0.51508033 |
| WTPNPVPATKGVPAGLVQAT | ASRGHQNTLY | 0.5348991 |
| YALKRALAKFSDRFDVVII | ASRGHQNTLY | 0.6819383 |
| YLIGSVIV | ASRGHQNTLY | 0.7864493 |
| YGQLVAVID | ASRGHQNTLY | 0.6586492 |
| VTVGGMTLPA | ASRGHQNTLY | 0.7767972 |
| VLALPAV | ASRGHQNTLY | 0.78942215 |
| VIPPTLNYET | ASRGHQNTLY | 0.28689152 |
| VGMGGMHPSSG | ASRGHQNTLY | 0.42594984 |
| VGLAGGAATA | ASRGHQNTLY | 0.7863016 |
| VGGAGGAGGSGP | ASRGHQNTLY | 0.6396605 |
| VFAAFMV | ASRGHQNTLY | 0.8013011 |
| TLYDKLR | ASRGHQNTLY | 0.5766773 |
| TLAVFTVL | ASRGHQNTLY | 0.68739754 |
| TGSVQPARV | ASRGHQNTLY | 0.7331814 |
| TGGGAGSGGT | ASRGHQNTLY | 0.8130851 |
| SHLETSL | ASRGHQNTLY | 0.798221 |
| SFTTGMT | ASRGHQNTLY | 0.74016786 |

| | | |
|---|---|---|
| QVEVAKRMGVRQPTVS | ASRGHQNTLY | 0.7894833 |
| QGSGGPALAM | ASRGHQNTLY | 0.41842443 |
| MLAPSNP | ASRGHQNTLY | 0.7795661 |
| LVNNAAI | ASRGHQNTLY | 0.8032895 |
| LTGEINTLA | ASRGHQNTLY | 0.4785161 |
| LSIFGPV | ASRGHQNTLY | 0.66640574 |
| LLALLANL | ASRGHQNTLY | 0.80252814 |
| LISNASDALDKLR | ASRGHQNTLY | 0.8115511 |
| LINEAVP | ASRGHQNTLY | 0.5588814 |
| LGYAFSGL | ASRGHQNTLY | 0.66984475 |
| LDFVAVAL | ASRGHQNTLY | 0.46531522 |
| LAQPGAE | ASRGHQNTLY | 0.612556 |
| ISNLGAVTGF | ASRGHQNTLY | 0.31674102 |
| HQEDPGRI | ASRGHQNTLY | 0.7802565 |
| GTPGEPTP | ASRGHQNTLY | 0.5826681 |
| GFFNSGS | ASRGHQNTLY | 0.79648614 |
| GAGGGGLTAGAN | ASRGHQNTLY | 0.7076156 |
| FVDALVNM | ASRGHQNTLY | 0.7591961 |
| FLTVAPEL | ASRGHQNTLY | 0.6483951 |
| FETLTAL | ASRGHQNTLY | 0.6082739 |
| DVINPIEL | ASRGHQNTLY | 0.60012376 |
| APVSGAPV | ASRGHQNTLY | 0.33521226 |
| ALVGGLAW | ASRGHQNTLY | 0.6083611 |
| ALAAAAATAI | ASRGHQNTLY | 0.56650126 |
| AFGLGLLL | ASRGHQNTLY | 0.25949237 |
| AFFAAASL | ASRGHQNTLY | 0.7993162 |
| AAAALPAFSPP | ASRGHQNTLY | 0.76870775 |
| VVLAPAL | ASRGHQNTLY | 0.6831279 |
| TIGHVDHGKTTL | ASRGHQNTLY | 0.46799895 |
| LISNASDALDK | ASRGHQNTLY | 0.576092 |
| LISNASDAL | ASRGHQNTLY | 0.584683 |
| ISNASDALDK | ASRGHQNTLY | 0.80075926 |
| NQAETLVYQTE | ASRGHQNTLY | 0.48706287 |
| TGGTGGTAGAG | ASRGHQNTLY | 0.2663462 |
| TGSTAGGAGGA | ASRGHQNTLY | 0.57928765 |
| DPLAVPVAAGPV | ASRGHQNTLY | 0.52119184 |
| GGTGGTPGNGGN | ASRGHQNTLY | 0.802229 |
| PAGLAGVDPATL | ASRGHQNTLY | 0.7892643 |
| GNNLVVEAPHAGASV | ASRGHQNTLY | 0.766019 |
| KVTRIMSASTLEQPA | ASRGHQNTLY | 0.81322616 |
| LGFALPIALPRRTVV | ASRGHQNTLY | 0.21267188 |
| TRLGNLTARKLVVLV | ASRGHQNTLY | 0.54322815 |
| WSQQRRLIYFPSAGP | ASRGHQNTLY | 0.788993 |
| GPNGNPDPMAAAVDIR | ASRGHQNTLY | 0.74767786 |
| TQRCVSVADLRAAASR | ASRGHQNTLY | 0.6057329 |
| QFVLDVAEHPVEVLTETV | ASRGHQNTLY | 0.80646574 |
| GGAGGTGGAGGNGGTQAGDG | ASRGHQNTLY | 0.45005542 |
| PLLATLRAIAVDVDAKPAQV | ASRGHQNTLY | 0.5865909 |
| APKFRTGGYMIDFWGVGYQVAK | ASRGHQNTLY | 0.74943554 |
| GAGGAGGAGGAAGGPGGTGGTG | ASRGHQNTLY | 0.79222107 |
| IGALGLLATDVAPAGSVAFGVA | ASRGHQNTLY | 0.76857144 |

| | | |
|---|---|---|
| AGGAGGIGGNGGAGGAGGVLMG | ASRGHQNTLY | 0.38490295 |
| ADELIGLL | ASRGHQNTLY | 0.7956736 |
| ADRALKA | ASRGHQNTLY | 0.8086325 |
| AGAVGQVGGAGGA | ASRGHQNTLY | 0.7031274 |
| AGGGGGLGGAP | ASRGHQNTLY | 0.71873605 |
| AGGLLYGNGGAG | ASRGHQNTLY | 0.76995945 |
| AGLESIGL | ASRGHQNTLY | 0.63589 |
| AGLKTGII | ASRGHQNTLY | 0.5239807 |
| ASLAALLL | ASRGHQNTLY | 0.4343741 |
| ASVINISSI | ASRGHQNTLY | 0.7772535 |
| ATPPGPA | ASRGHQNTLY | 0.43107402 |
| ATVASALV | ASRGHQNTLY | 0.7823374 |
| ATVASLIL | ASRGHQNTLY | 0.6605998 |
| AVNGPVLIRH | ASRGHQNTLY | 0.8159896 |
| DGAAGAVGI | ASRGHQNTLY | 0.7684068 |
| DPLLHGI | ASRGHQNTLY | 0.7557856 |
| ELGAALGAL | ASRGHQNTLY | 0.45884487 |
| ELGRTQAV | ASRGHQNTLY | 0.76263845 |
| FVAGGYLS | ASRGHQNTLY | 0.7566216 |
| GASGGAGGAGGW | ASRGHQNTLY | 0.3481481 |
| GGAGAGGDGGAGGI | ASRGHQNTLY | 0.7831378 |
| GTGHNGSQPGMGGNG | ASRGHQNTLY | 0.42727113 |
| GYAAASASAS | ASRGHQNTLY | 0.7235602 |
| HQVDVAAVQQG | ASRGHQNTLY | 0.7209532 |
| IGAAIAHT | ASRGHQNTLY | 0.7986776 |
| IGMIEVAQL | ASRGHQNTLY | 0.70335484 |
| IRLAPEHTSAMD | ASRGHQNTLY | 0.72649014 |
| IYNVSVL | ASRGHQNTLY | 0.47739273 |
| KQGFLGM | ASRGHQNTLY | 0.7875532 |
| LESVITQL | ASRGHQNTLY | 0.80094224 |
| LGLGALVVV | ASRGHQNTLY | 0.74920636 |
| LIGNGGNGGSGGI | ASRGHQNTLY | 0.81354547 |
| LNAINAPTL | ASRGHQNTLY | 0.7713857 |
| LQLLNTGV | ASRGHQNTLY | 0.30147192 |
| LQTSIEF | ASRGHQNTLY | 0.7778193 |
| MSFAVSEL | ASRGHQNTLY | 0.8052472 |
| NTGALGL | ASRGHQNTLY | 0.80240977 |
| PPGAGGAGGAGTT | ASRGHQNTLY | 0.74023926 |
| QLVGLGI | ASRGHQNTLY | 0.7860396 |
| RFLEQVHQL | ASRGHQNTLY | 0.6755817 |
| RGVGYVLR | ASRGHQNTLY | 0.79688907 |
| SATGVFAGVI | ASRGHQNTLY | 0.816444 |
| SLPAITLP | ASRGHQNTLY | 0.79339993 |
| STALAFTV | ASRGHQNTLY | 0.6705913 |
| SVLTVGLGK | ASRGHQNTLY | 0.58284336 |
| TALAGAGSGI | ASRGHQNTLY | 0.77364516 |
| TGLPFTGI | ASRGHQNTLY | 0.7949757 |
| VAEHVTV | ASRGHQNTLY | 0.67688227 |
| VDIVLNSL | ASRGHQNTLY | 0.6661508 |
| VLTGGGAVL | ASRGHQNTLY | 0.63789916 |
| VNVGLAYGI | ASRGHQNTLY | 0.35919476 |

| | | |
|---|---|---|
| VSSAAVGAV | ASRGHQNTLY | 0.7669101 |
| VTSVYEAGV | ASRGHQNTLY | 0.8025445 |
| WVALDRGA | ASRGHQNTLY | 0.78673416 |
| YLVGSVVI | ASRGHQNTLY | 0.5827513 |
| GPPGTGKTLL | ASRGHQNTLY | 0.79090333 |
| GVGMTKF | ASRGHQNTLY | 0.80334574 |
| IGIGGSDL | ASRGHQNTLY | 0.81357294 |
| IGTLLAEL | ASRGHQNTLY | 0.27688408 |
| IYTSGTT | ASRGHQNTLY | 0.24347723 |
| LSFPAGV | ASRGHQNTLY | 0.76378417 |
| LYTSGTT | ASRGHQNTLY | 0.76629335 |
| VGAFPFGF | ASRGHQNTLY | 0.4944812 |
| VGAGTVEF | ASRGHQNTLY | 0.6325409 |
| VGKSTLLL | ASRGHQNTLY | 0.78099793 |
| AALAHAGGGIV | ASRGHQNTLY | 0.77261746 |
| AALVAVSLL | ASRGHQNTLY | 0.5686753 |
| AAVSRAVAAL | ASRGHQNTLY | 0.3843132 |
| AGGSSGIGGF | ASRGHQNTLY | 0.7894751 |
| AGLTVANANG | ASRGHQNTLY | 0.80712706 |
| AGLVGAVL | ASRGHQNTLY | 0.5312333 |
| ALSQAFTV | ASRGHQNTLY | 0.70650977 |
| ALTFSVSL | ASRGHQNTLY | 0.6468451 |
| ALTGAALVAL | ASRGHQNTLY | 0.7462608 |
| ASFSIGAGVL | ASRGHQNTLY | 0.6931911 |
| DGSVYKI | ASRGHQNTLY | 0.32110462 |
| EQLAEIV | ASRGHQNTLY | 0.7575136 |
| FAVSELTVM | ASRGHQNTLY | 0.35475814 |
| GAAPLAAGL | ASRGHQNTLY | 0.78826416 |
| GADGGDGGSGGAGG | ASRGHQNTLY | 0.76757836 |
| GFGLDTGGAGG | ASRGHQNTLY | 0.76875836 |
| GLASGGLL | ASRGHQNTLY | 0.6282181 |
| GRIAGAAVLL | ASRGHQNTLY | 0.43585667 |
| GSAGSLNTSF | ASRGHQNTLY | 0.43903184 |
| GTGQIGI | ASRGHQNTLY | 0.49305764 |
| GTLIALIL | ASRGHQNTLY | 0.8013192 |
| IAATMGFVVG | ASRGHQNTLY | 0.35627282 |
| IFAQALL | ASRGHQNTLY | 0.80421925 |
| IGAAVALGF | ASRGHQNTLY | 0.67324996 |
| IGLTPLAL | ASRGHQNTLY | 0.5809698 |
| ILGSNTSTL | ASRGHQNTLY | 0.7777622 |
| IVFSLTVL | ASRGHQNTLY | 0.4163063 |
| LASKATGF | ASRGHQNTLY | 0.76233643 |
| LASVALIL | ASRGHQNTLY | 0.765198 |
| LFAVGLTL | ASRGHQNTLY | 0.803133 |
| LFNGGIGTYI | ASRGHQNTLY | 0.56343055 |
| LTLGAGII | ASRGHQNTLY | 0.8044819 |
| LVAAALGDI | ASRGHQNTLY | 0.7629286 |
| NTAFVQLGI | ASRGHQNTLY | 0.5398326 |
| QLFSGAEL | ASRGHQNTLY | 0.78093445 |
| SAATFITL | ASRGHQNTLY | 0.7752148 |
| SNYGLANLGV | ASRGHQNTLY | 0.7879276 |

| | | |
|---|---|---|
| SYGGAYAVM | ASRGHQNTLY | 0.45330864 |
| TTGVGGLTLG | ASRGHQNTLY | 0.6577689 |
| TTQTIYI | ASRGHQNTLY | 0.65653193 |
| VAAGAGLGAVI | ASRGHQNTLY | 0.7214303 |
| VAAVVTLL | ASRGHQNTLY | 0.73365235 |
| VGFLGGLI | ASRGHQNTLY | 0.642627 |
| VGVIPILL | ASRGHQNTLY | 0.8000729 |
| VIFASSLI | ASRGHQNTLY | 0.3969719 |
| VKGGDGGAAAG | ASRGHQNTLY | 0.6193023 |
| VTKTFTNI | ASRGHQNTLY | 0.6893753 |
| VVAAAILL | ASRGHQNTLY | 0.6575943 |
| YAALDVEL | ASRGHQNTLY | 0.7850324 |
| LASVALLL | ASRGHQNTLY | 0.4331116 |
| LDAVHRV | ASRGHQNTLY | 0.7456098 |
| QGLSSSIF | ASRGHQNTLY | 0.68346184 |
| RGTFANIRL | ASRGHQNTLY | 0.44638503 |
| RQLWWGHRI | ASRGHQNTLY | 0.7975931 |
| YGPPGTGKTLL | ASRGHQNTLY | 0.7154461 |
| AALVGAPANVA | ASRGHQNTLY | 0.57096124 |
| AATFRRTE | ASRGHQNTLY | 0.7839546 |
| AGALGGGI | ASRGHQNTLY | 0.56861407 |
| AGGAGGAGGIGAG | ASRGHQNTLY | 0.67373186 |
| AGSGALGL | ASRGHQNTLY | 0.8048427 |
| AQIADQI | ASRGHQNTLY | 0.49331433 |
| ASAVAVTYL | ASRGHQNTLY | 0.78503495 |
| ASTVLGGTGG | ASRGHQNTLY | 0.7189168 |
| AVGGEPAK | ASRGHQNTLY | 0.7766106 |
| CEQGTFGGTC | ASRGHQNTLY | 0.5518909 |
| DEGKAPAG | ASRGHQNTLY | 0.5632609 |
| DTVVFPAL | ASRGHQNTLY | 0.79995555 |
| FEGITMAG | ASRGHQNTLY | 0.6508485 |
| FPLVVQD | ASRGHQNTLY | 0.7769092 |
| GAFTTPQ | ASRGHQNTLY | 0.78071404 |
| GFGGFGTAGGVGGA | ASRGHQNTLY | 0.8007674 |
| GFLAQFL | ASRGHQNTLY | 0.804794 |
| GGAGGAGGNAGVA | ASRGHQNTLY | 0.47332835 |
| GGVGTAGAGGAGGAGG | ASRGHQNTLY | 0.55083925 |
| GNGGGGDAGNAGSGGN | ASRGHQNTLY | 0.63015234 |
| GTGGAGGAGNDAGS | ASRGHQNTLY | 0.6105089 |
| GTTNTGWFNS | ASRGHQNTLY | 0.7077087 |
| IFMGPLFTL | ASRGHQNTLY | 0.3329327 |
| ILAHAIEAI | ASRGHQNTLY | 0.77629423 |
| IRSAIREAL | ASRGHQNTLY | 0.8158723 |
| ITFAGGG | ASRGHQNTLY | 0.4409349 |
| ITGMSGHQEV | ASRGHQNTLY | 0.8166862 |
| IVAGLACTL | ASRGHQNTLY | 0.5449237 |
| IVGGGLAGA | ASRGHQNTLY | 0.69636273 |
| IVTGVLNI | ASRGHQNTLY | 0.7945894 |
| LAQQGCI | ASRGHQNTLY | 0.40699613 |
| LCLHVEN | ASRGHQNTLY | 0.6738696 |
| LDAISDGS | ASRGHQNTLY | 0.545314 |

| | | |
|---|---|---|
| LGGFAAML | ASRGHQNTLY | 0.54845387 |
| LGGKNPMI | ASRGHQNTLY | 0.802518 |
| LGPKGVAAIL | ASRGHQNTLY | 0.47211272 |
| LMVGVSI | ASRGHQNTLY | 0.7656859 |
| LVGAAFVL | ASRGHQNTLY | 0.7341909 |
| LVLAVAPL | ASRGHQNTLY | 0.48100522 |
| MAAIAVIDAA | ASRGHQNTLY | 0.78294206 |
| NNGINAELQ | ASRGHQNTLY | 0.7913615 |
| RGEPLVLVHG | ASRGHQNTLY | 0.75820756 |
| RGGAGGAAGLI | ASRGHQNTLY | 0.8158512 |
| RSYRMHPAV | ASRGHQNTLY | 0.4612492 |
| SAAHLLAGQLA | ASRGHQNTLY | 0.8001939 |
| SAAVGSY | ASRGHQNTLY | 0.81314147 |
| SPHAVLHRI | ASRGHQNTLY | 0.7160897 |
| SQIAKVL | ASRGHQNTLY | 0.68831944 |
| TALGHLVP | ASRGHQNTLY | 0.77395827 |
| TAPTTSGAA | ASRGHQNTLY | 0.7767683 |
| TDGDDGNGGSPGT | ASRGHQNTLY | 0.76555085 |
| TDGLGEN | ASRGHQNTLY | 0.8120376 |
| TGGPPLPGQA | ASRGHQNTLY | 0.5523819 |
| TIPAITI | ASRGHQNTLY | 0.7706796 |
| TIPGFTI | ASRGHQNTLY | 0.7534003 |
| TSIASGALA | ASRGHQNTLY | 0.7215508 |
| TVTLPGAAI | ASRGHQNTLY | 0.67983615 |
| VASGLSAALP | ASRGHQNTLY | 0.46804455 |
| VGAGVIGI | ASRGHQNTLY | 0.67779577 |
| VGPKRFGV | ASRGHQNTLY | 0.66690385 |
| VNVAFPD | ASRGHQNTLY | 0.7644328 |
| VSMTNSGVS | ASRGHQNTLY | 0.76003915 |
| VTDVLGDLSV | ASRGHQNTLY | 0.6448768 |
| VVVVGAGFAGL | ASRGHQNTLY | 0.76329124 |
| VYTVGTVV | ASRGHQNTLY | 0.76162434 |
| WTTFSGP | ASRGHQNTLY | 0.570645 |
| YDELVEAF | ASRGHQNTLY | 0.7172909 |
| YGTGGAGGNGG | ASRGHQNTLY | 0.69215256 |
| YTNVMAV | ASRGHQNTLY | 0.7303804 |
| FLEQVHQL | ASRGHQNTLY | 0.4698835 |
| GLSLLAVAV | ASRGHQNTLY | 0.75823057 |
| LEQVHQL | ASRGHQNTLY | 0.6755667 |
| LNAAGAI | ASRGHQNTLY | 0.28960353 |
| PAVHASL | ASRGHQNTLY | 0.7433522 |
| TELEDDA | ASRGHQNTLY | 0.6898022 |
| TPPGPPN | ASRGHQNTLY | 0.7271275 |
| TPPPPPPP | ASRGHQNTLY | 0.7671768 |
| TQSPALI | ASSPPGQGTGNTLY | 0.58187926 |
| EFAAAFPQ | ASSPPGQGTGNTLY | 0.60275865 |
| LLSGNLDVL | ASSPPGQGTGNTLY | 0.7503499 |
| GPANLGNYNI | ASSPPGQGTGNTLY | 0.76357424 |
| TLGPTLP | ASSPPGQGTGNTLY | 0.5485383 |
| VYAYSSCCK | ASSPPGQGTGNTLY | 0.74321824 |
| APAPAPAPNI | ASSPPGQGTGNTLY | 0.7589626 |

| | | |
|---|---|---|
| PGGPVPIP | ASSPPGQGTGNTLY | 0.73746604 |
| FLTVALEP | ASSPPGQGTGNTLY | 0.7714629 |
| SQYFGANL | ASSPPGQGTGNTLY | 0.7764062 |
| GGVKPAA | ASSPPGQGTGNTLY | 0.63605255 |
| GGTGGAGGAGGW | ASSPPGQGTGNTLY | 0.76944304 |
| GVGGVAGLGGAGG | ASSPPGQGTGNTLY | 0.77568936 |
| VLLVLATTI | ASSPPGQGTGNTLY | 0.6776193 |
| IGIFAAML | ASSPPGQGTGNTLY | 0.7162349 |
| AVGDLID | ASSPPGQGTGNTLY | 0.5116198 |
| NISPLPL | ASSPPGQGTGNTLY | 0.69172645 |
| VTAVFVAI | ASSPPGQGTGNTLY | 0.5306915 |
| SGDDNDAAL | ASSPPGQGTGNTLY | 0.7420429 |
| AVAALAHS | ASSPPGQGTGNTLY | 0.7681783 |
| GGVTVMATP | ASSPPGQGTGNTLY | 0.7081452 |
| AGLAANVRD | ASSPPGQGTGNTLY | 0.6188251 |
| TNRPDLIDPALLRPG | ASSPPGQGTGNTLY | 0.49687043 |
| LANLGVTAL | ASSPPGQGTGNTLY | 0.72602665 |
| TGERIEL | ASSPPGQGTGNTLY | 0.6092985 |
| NGEELAA | ASSPPGQGTGNTLY | 0.38129938 |
| VLSLVWG | ASSPPGQGTGNTLY | 0.713051 |
| IGYALWL | ASSPPGQGTGNTLY | 0.75546455 |
| LSLLLAPL | ASSPPGQGTGNTLY | 0.75626683 |
| TLDEALK | ASSPPGQGTGNTLY | 0.5732864 |
| TDGAGTAEGG | ASSPPGQGTGNTLY | 0.7782924 |
| NAPGGAGGNGG | ASSPPGQGTGNTLY | 0.7532269 |
| VSVFMGMGGNF | ASSPPGQGTGNTLY | 0.7679727 |
| LASYLAAL | ASSPPGQGTGNTLY | 0.7230004 |
| LVGGTPAL | ASSPPGQGTGNTLY | 0.59391034 |
| VRNADVL | ASSPPGQGTGNTLY | 0.7631955 |
| TCSSPFLVSAM | ASSPPGQGTGNTLY | 0.6830972 |
| VHITDAE | ASSPPGQGTGNTLY | 0.706877 |
| EYAAPSF | ASSPPGQGTGNTLY | 0.7529386 |
| VVDAFHGGL | ASSPPGQGTGNTLY | 0.7284943 |
| ASVVDPQ | ASSPPGQGTGNTLY | 0.6477742 |
| SYNEFGGLS | ASSPPGQGTGNTLY | 0.7093242 |
| DPDGSVY | ASSPPGQGTGNTLY | 0.7721131 |
| MVSTIHATE | ASSPPGQGTGNTLY | 0.6399393 |
| LAAGVADPV | ASSPPGQGTGNTLY | 0.598107 |
| TKEQIAA | ASSPPGQGTGNTLY | 0.6657436 |
| SLASPAAL | ASSPPGQGTGNTLY | 0.71630555 |
| IGLVFYGV | ASSPPGQGTGNTLY | 0.7573882 |
| FLLDASNL | ASSPPGQGTGNTLY | 0.7472316 |
| YQVGNFP | ASSPPGQGTGNTLY | 0.6606011 |
| ESLLSQST | ASSPPGQGTGNTLY | 0.7494327 |
| LFTVGFAAVG | ASSPPGQGTGNTLY | 0.7423358 |
| ATPPTPT | ASSPPGQGTGNTLY | 0.73914707 |
| TDLATLG | ASSPPGQGTGNTLY | 0.7173426 |
| SAGGTGR | ASSPPGQGTGNTLY | 0.6826123 |
| GFSNSGS | ASSPPGQGTGNTLY | 0.7705321 |
| GGNGGSGGTSV | ASSPPGQGTGNTLY | 0.7494885 |
| GTSVATGGAGN | ASSPPGQGTGNTLY | 0.7592169 |

| | | |
|---|---|---|
| NGGNGGSGGTS | ASSPPGQGTGNTLY | 0.765705 |
| AVVKFPH | ASSPPGQGTGNTLY | 0.70374036 |
| TPPPPPP | ASSPPGQGTGNTLY | 0.7345164 |
| AAPAAAAAPA | ASSPPGQGTGNTLY | 0.4765232 |
| IALNEDL | ASSPPGQGTGNTLY | 0.6469185 |
| IVNVSSM | ASSPPGQGTGNTLY | 0.71881795 |
| TGIPPAPR | ASSPPGQGTGNTLY | 0.747649 |
| FDLGGGTF | ASSPPGQGTGNTLY | 0.7572679 |
| FDLGGGTFDVS | ASSPPGQGTGNTLY | 0.74519694 |
| NEPTAAA | ASSPPGQGTGNTLY | 0.6516466 |
| NEPTAAALA | ASSPPGQGTGNTLY | 0.74390864 |
| GSIINISSI | ASSPPGQGTGNTLY | 0.7638289 |
| VAPGNAGTA | ASSPPGQGTGNTLY | 0.5833617 |
| TGGGASLE | ASSPPGQGTGNTLY | 0.71412015 |
| VAVYNHYKRI | ASSPPGQGTGNTLY | 0.78192395 |
| TDPAPATT | ASSPPGQGTGNTLY | 0.7388184 |
| TNRPDILDPALLRPG | ASSPPGQGTGNTLY | 0.6958461 |
| LTTTTTS | ASSPPGQGTGNTLY | 0.70716006 |
| PTGSGKTL | ASSPPGQGTGNTLY | 0.7746788 |
| TGSGKTL | ASSPPGQGTGNTLY | 0.71004915 |
| TIGHVDHGK | ASSPPGQGTGNTLY | 0.7713624 |
| TIGHVDHGKTT | ASSPPGQGTGNTLY | 0.39945918 |
| GHVDHGKTT | ASSPPGQGTGNTLY | 0.7162832 |
| AQFEHTLL | ASSPPGQGTGNTLY | 0.62177336 |
| IVNQDEL | ASSPPGQGTGNTLY | 0.35288036 |
| TNGSQFF | ASSPPGQGTGNTLY | 0.7671782 |
| TGKGIVE | ASSPPGQGTGNTLY | 0.6640407 |
| LISNASDA | ASSPPGQGTGNTLY | 0.75468266 |
| AELRTLGELTKIAKAHGAQ | ASSPPGQGTGNTLY | 0.71231115 |
| APNAVAAAEPSW | ASSPPGQGTGNTLY | 0.76700175 |
| APQVAGAATAVL | ASSPPGQGTGNTLY | 0.73259073 |
| ARRDPAAAAAILPTDA | ASSPPGQGTGNTLY | 0.70510626 |
| DAAARDVLAEAGLAEYF | ASSPPGQGTGNTLY | 0.59289855 |
| DDQIVLSAKANLAKVEQ | ASSPPGQGTGNTLY | 0.75518477 |
| DELLPDTGEDSDEDGH | ASSPPGQGTGNTLY | 0.75671434 |
| ELPEAPANRATVPQD | ASSPPGQGTGNTLY | 0.76280165 |
| FGQSKIERVTGIG | ASSPPGQGTGNTLY | 0.38818234 |
| GAGGTGSTAGGAGGAGGAGGLY | ASSPPGQGTGNTLY | 0.74387956 |
| GHGGTGAAGGNGGTGGAGGSGI | ASSPPGQGTGNTLY | 0.7259706 |
| GNGGNGGAGGEGGPGGAGGAGA | ASSPPGQGTGNTLY | 0.29259562 |
| GSTAGNGGNGGNATNGTIAGQP | ASSPPGQGTGNTLY | 0.31688005 |
| GTGGDGGNGGNGGTGG | ASSPPGQGTGNTLY | 0.7070233 |
| GVRRPAGIPADAPV | ASSPPGQGTGNTLY | 0.703284 |
| GYRQVTDAHLLTLARR | ASSPPGQGTGNTLY | 0.759875 |
| IAKMGQDGHDRG | ASSPPGQGTGNTLY | 0.7480942 |
| KAEPSIEIQYT | ASSPPGQGTGNTLY | 0.6856247 |
| KPVRIRMAADEDTVAVT | ASSPPGQGTGNTLY | 0.7328126 |
| MPPRIAGMRLLVIKPEP | ASSPPGQGTGNTLY | 0.70863503 |
| MKFYAVALFRSRGVPPTR | ASSPPGQGTGNTLY | 0.19758388 |
| NAGGFGILGAVAHSP | ASSPPGQGTGNTLY | 0.62385786 |
| NEGREITLAMAEM | ASSPPGQGTGNTLY | 0.56287295 |

| | | |
|---|---|---|
| NELVEWSHPLGHRWRWE | ASSPPGQGTGNTLY | 0.77433836 |
| NLGAGGTGGNGGNPG | ASSPPGQGTGNTLY | 0.7005407 |
| NTEWPKWETVMAADPVD | ASSPPGQGTGNTLY | 0.69208574 |
| PASAAPAETQP | ASSPPGQGTGNTLY | 0.70838886 |
| QQSSDGKNGVRAAEITGE | ASSPPGQGTGNTLY | 0.7466929 |
| RAPDAVTTSWWKE | ASSPPGQGTGNTLY | 0.7208706 |
| SGFGNVGAHTSGWFNQ | ASSPPGQGTGNTLY | 0.58062303 |
| SGVAGAVGQAASVGGLKV | ASSPPGQGTGNTLY | 0.36660838 |
| SNGGAGTGSAGGNGGT | ASSPPGQGTGNTLY | 0.7391969 |
| TGGAGGEGGAAG | ASSPPGQGTGNTLY | 0.5845684 |
| TGGDGGAAGAGGAPGHG | ASSPPGQGTGNTLY | 0.59981394 |
| TMPPLLAAIANAGRHGV | ASSPPGQGTGNTLY | 0.4636957 |
| TSSTAGNSGGA | ASSPPGQGTGNTLY | 0.71377873 |
| VPATVPVGILC | ASSPPGQGTGNTLY | 0.44586843 |
| VQRPDSGGVVLTGRIS | ASSPPGQGTGNTLY | 0.76612633 |
| WTPNPVPATKGVPAGLVQAT | ASSPPGQGTGNTLY | 0.6686929 |
| YALKRALAKFSDRFDVVII | ASSPPGQGTGNTLY | 0.70169836 |
| YLIGSVIV | ASSPPGQGTGNTLY | 0.7783543 |
| YGQLVAVID | ASSPPGQGTGNTLY | 0.73222697 |
| VTVGGMTLPA | ASSPPGQGTGNTLY | 0.6858319 |
| VLALPAV | ASSPPGQGTGNTLY | 0.6478987 |
| VIPPTLNYET | ASSPPGQGTGNTLY | 0.6466804 |
| VGMGGMHPSSG | ASSPPGQGTGNTLY | 0.68588877 |
| VGLAGGAATA | ASSPPGQGTGNTLY | 0.6761724 |
| VGGAGGAGGSGP | ASSPPGQGTGNTLY | 0.7724933 |
| VFAAFMV | ASSPPGQGTGNTLY | 0.74292123 |
| TLYDKLR | ASSPPGQGTGNTLY | 0.5993959 |
| TLAVFTVL | ASSPPGQGTGNTLY | 0.7623011 |
| TGSVQPARV | ASSPPGQGTGNTLY | 0.7751025 |
| TGGGAGSGGT | ASSPPGQGTGNTLY | 0.77807045 |
| SHLETSL | ASSPPGQGTGNTLY | 0.63953054 |
| SFTTGMT | ASSPPGQGTGNTLY | 0.684446 |
| QVEVAKRMGVRQPTVS | ASSPPGQGTGNTLY | 0.6755443 |
| QGSGGPALAM | ASSPPGQGTGNTLY | 0.7219371 |
| MLAPSNP | ASSPPGQGTGNTLY | 0.6038342 |
| LVNNAAI | ASSPPGQGTGNTLY | 0.61390996 |
| LTGEINTLA | ASSPPGQGTGNTLY | 0.7612401 |
| LSIFGPV | ASSPPGQGTGNTLY | 0.7807043 |
| LLALLANL | ASSPPGQGTGNTLY | 0.69916743 |
| LISNASDALDKLR | ASSPPGQGTGNTLY | 0.62129414 |
| LINEAVP | ASSPPGQGTGNTLY | 0.7024329 |
| LGYAFSGL | ASSPPGQGTGNTLY | 0.6156869 |
| LDFVAVAL | ASSPPGQGTGNTLY | 0.6632803 |
| LAQPGAE | ASSPPGQGTGNTLY | 0.6481546 |
| ISNLGAVTGF | ASSPPGQGTGNTLY | 0.56537426 |
| HQEDPGRI | ASSPPGQGTGNTLY | 0.7076868 |
| GTPGEPTP | ASSPPGQGTGNTLY | 0.7758807 |
| GFFNSGS | ASSPPGQGTGNTLY | 0.71407986 |
| GAGGGGLTAGAN | ASSPPGQGTGNTLY | 0.758151 |
| FVDALVNM | ASSPPGQGTGNTLY | 0.7747264 |
| FLTVAPEL | ASSPPGQGTGNTLY | 0.7619271 |

| | | |
|---|---|---|
| FETLTAL | ASSPPGQGTGNTLY | 0.7742063 |
| DVINPIEL | ASSPPGQGTGNTLY | 0.75498223 |
| APVSGAPV | ASSPPGQGTGNTLY | 0.70913506 |
| ALVGGLAW | ASSPPGQGTGNTLY | 0.69245934 |
| ALAAAAATAI | ASSPPGQGTGNTLY | 0.7667004 |
| AFGLGLLL | ASSPPGQGTGNTLY | 0.6976391 |
| AFFAAASL | ASSPPGQGTGNTLY | 0.70462567 |
| AAAALPAFSPP | ASSPPGQGTGNTLY | 0.68146205 |
| VVLAPAL | ASSPPGQGTGNTLY | 0.76629555 |
| TIGHVDHGKTTL | ASSPPGQGTGNTLY | 0.70555574 |
| LISNASDALDK | ASSPPGQGTGNTLY | 0.77292216 |
| LISNASDAL | ASSPPGQGTGNTLY | 0.34644747 |
| ISNASDALDK | ASSPPGQGTGNTLY | 0.5959895 |
| NQAETLVYQTE | ASSPPGQGTGNTLY | 0.7709959 |
| TGGTGGTAGAG | ASSPPGQGTGNTLY | 0.20046104 |
| TGSTAGGAGGA | ASSPPGQGTGNTLY | 0.37881917 |
| DPLAVPVAAGPV | ASSPPGQGTGNTLY | 0.723477 |
| GGTGGTPGNGGN | ASSPPGQGTGNTLY | 0.76417816 |
| PAGLAGVDPATL | ASSPPGQGTGNTLY | 0.21949421 |
| GNNLVVEAPHAGASV | ASSPPGQGTGNTLY | 0.56853354 |
| KVTRIMSASTLEQPA | ASSPPGQGTGNTLY | 0.40151465 |
| LGFALPIALPRRTVV | ASSPPGQGTGNTLY | 0.76009 |
| TRLGNLTARKLVVLV | ASSPPGQGTGNTLY | 0.7055632 |
| WSQQRRLIYFPSAGP | ASSPPGQGTGNTLY | 0.4123823 |
| GPNGNPDPMAAAVDIR | ASSPPGQGTGNTLY | 0.7605518 |
| TQRCVSVADLRAAASR | ASSPPGQGTGNTLY | 0.5312147 |
| QFVLDVAEHPVEVLTETV | ASSPPGQGTGNTLY | 0.6551124 |
| GGAGGTGGAGGNGGTQAGDG | ASSPPGQGTGNTLY | 0.76501405 |
| PLLATLRAIAVDVDAKPAQV | ASSPPGQGTGNTLY | 0.7758262 |
| APKFRTGGYMIDFWGVGYQVAK | ASSPPGQGTGNTLY | 0.56508327 |
| GAGGAGGAGGAAGGPGGTGGTG | ASSPPGQGTGNTLY | 0.7400067 |
| IGALGLLATDVAPAGSVAFGVA | ASSPPGQGTGNTLY | 0.684834 |
| AGGAGGIGGNGGAGGAGGVLMG | ASSPPGQGTGNTLY | 0.7826425 |
| ADELIGLL | ASSPPGQGTGNTLY | 0.6967144 |
| ADRALKA | ASSPPGQGTGNTLY | 0.5313732 |
| AGAVGQVGGAGGA | ASSPPGQGTGNTLY | 0.76780283 |
| AGGGGGLGGAP | ASSPPGQGTGNTLY | 0.7667775 |
| AGGLLYGNGGAG | ASSPPGQGTGNTLY | 0.693344 |
| AGLESIGL | ASSPPGQGTGNTLY | 0.7617264 |
| AGLKTGII | ASSPPGQGTGNTLY | 0.7113895 |
| ASLAALLL | ASSPPGQGTGNTLY | 0.70806193 |
| ASVINISSI | ASSPPGQGTGNTLY | 0.7629887 |
| ATPPGPA | ASSPPGQGTGNTLY | 0.754629 |
| ATVASALV | ASSPPGQGTGNTLY | 0.63741636 |
| ATVASLIL | ASSPPGQGTGNTLY | 0.7670357 |
| AVNGPVLIRH | ASSPPGQGTGNTLY | 0.7166065 |
| DGAAGAVGI | ASSPPGQGTGNTLY | 0.475997 |
| DPLLHGI | ASSPPGQGTGNTLY | 0.78174055 |
| ELGAALGAL | ASSPPGQGTGNTLY | 0.6895317 |
| ELGRTQAV | ASSPPGQGTGNTLY | 0.76032424 |
| FVAGGYLS | ASSPPGQGTGNTLY | 0.76433146 |

| | | |
|---|---|---|
| GASGGAGGAGGW | ASSPPGQGTGNTLY | 0.7676533 |
| GGAGAGGDGGAGGI | ASSPPGQGTGNTLY | 0.5542413 |
| GTGHNGSQPGMGGNG | ASSPPGQGTGNTLY | 0.7275625 |
| GYAAASASAS | ASSPPGQGTGNTLY | 0.721558 |
| HQVDVAAVQQG | ASSPPGQGTGNTLY | 0.75466657 |
| IGAAIAHT | ASSPPGQGTGNTLY | 0.7307355 |
| IGMIEVAQL | ASSPPGQGTGNTLY | 0.71876794 |
| IRLAPEHTSAMD | ASSPPGQGTGNTLY | 0.760519 |
| IYNVSVL | ASSPPGQGTGNTLY | 0.7727448 |
| KQGFLGM | ASSPPGQGTGNTLY | 0.72369915 |
| LESVITQL | ASSPPGQGTGNTLY | 0.68686813 |
| LGLGALVVV | ASSPPGQGTGNTLY | 0.7755045 |
| LIGNGGNGGSGGI | ASSPPGQGTGNTLY | 0.56786746 |
| LNAINAPTL | ASSPPGQGTGNTLY | 0.7140628 |
| LQLLNTGV | ASSPPGQGTGNTLY | 0.7586663 |
| LQTSIEF | ASSPPGQGTGNTLY | 0.7562884 |
| MSFAVSEL | ASSPPGQGTGNTLY | 0.7347196 |
| NTGALGL | ASSPPGQGTGNTLY | 0.7663591 |
| PPGAGGAGGAGTT | ASSPPGQGTGNTLY | 0.5806577 |
| QLVGLGI | ASSPPGQGTGNTLY | 0.7364452 |
| RFLEQVHQL | ASSPPGQGTGNTLY | 0.68625194 |
| RGVGYVLR | ASSPPGQGTGNTLY | 0.7153429 |
| SATGVFAGVI | ASSPPGQGTGNTLY | 0.67253345 |
| SLPAITLP | ASSPPGQGTGNTLY | 0.7402035 |
| STALAFTV | ASSPPGQGTGNTLY | 0.75635123 |
| SVLTVGLGK | ASSPPGQGTGNTLY | 0.74979925 |
| TALAGAGSGI | ASSPPGQGTGNTLY | 0.323177 |
| TGLPFTGI | ASSPPGQGTGNTLY | 0.7518377 |
| VAEHVTV | ASSPPGQGTGNTLY | 0.74453235 |
| VDIVLNSL | ASSPPGQGTGNTLY | 0.783087 |
| VLTGGGAVL | ASSPPGQGTGNTLY | 0.67589796 |
| VNVGLAYGI | ASSPPGQGTGNTLY | 0.7441132 |
| VSSAAVGAV | ASSPPGQGTGNTLY | 0.6482668 |
| VTSVYEAGV | ASSPPGQGTGNTLY | 0.732515 |
| WVALDRGA | ASSPPGQGTGNTLY | 0.5289472 |
| YLVGSVVI | ASSPPGQGTGNTLY | 0.60537463 |
| GPPGTGKTLL | ASSPPGQGTGNTLY | 0.6631547 |
| GVGMTKF | ASSPPGQGTGNTLY | 0.66496813 |
| IGIGGSDL | ASSPPGQGTGNTLY | 0.7634678 |
| IGTLLAEL | ASSPPGQGTGNTLY | 0.7605014 |
| IYTSGTT | ASSPPGQGTGNTLY | 0.75960934 |
| LSFPAGV | ASSPPGQGTGNTLY | 0.76667106 |
| LYTSGTT | ASSPPGQGTGNTLY | 0.7757307 |
| VGAFPFGF | ASSPPGQGTGNTLY | 0.7687346 |
| VGAGTVEF | ASSPPGQGTGNTLY | 0.7032345 |
| VGKSTLLL | ASSPPGQGTGNTLY | 0.70178586 |
| AALAHAGGGIV | ASSPPGQGTGNTLY | 0.6068504 |
| AALVAVSLL | ASSPPGQGTGNTLY | 0.6736194 |
| AAVSRAVAAL | ASSPPGQGTGNTLY | 0.7192027 |
| AGGSSGIGGF | ASSPPGQGTGNTLY | 0.7090596 |
| AGLTVANANG | ASSPPGQGTGNTLY | 0.20210561 |

| | | |
|---|---|---|
| AGLVGAVL | ASSPPGQGTGNTLY | 0.7374195 |
| ALSQAFTV | ASSPPGQGTGNTLY | 0.7755866 |
| ALTFSVSL | ASSPPGQGTGNTLY | 0.6814006 |
| ALTGAALVAL | ASSPPGQGTGNTLY | 0.7474339 |
| ASFSIGAGVL | ASSPPGQGTGNTLY | 0.75201666 |
| DGSVYKI | ASSPPGQGTGNTLY | 0.6954498 |
| EQLAEIV | ASSPPGQGTGNTLY | 0.7441056 |
| FAVSELTVM | ASSPPGQGTGNTLY | 0.651339 |
| GAAPLAAGL | ASSPPGQGTGNTLY | 0.7629053 |
| GADGGDGGSGGAGG | ASSPPGQGTGNTLY | 0.78021073 |
| GFGLDTGGAGG | ASSPPGQGTGNTLY | 0.744969 |
| GLASGGLL | ASSPPGQGTGNTLY | 0.63030577 |
| GRIAGAAVLL | ASSPPGQGTGNTLY | 0.7230935 |
| GSAGSLNTSF | ASSPPGQGTGNTLY | 0.45661783 |
| GTGQIGI | ASSPPGQGTGNTLY | 0.6628327 |
| GTLIALIL | ASSPPGQGTGNTLY | 0.4548782 |
| IAATMGFVVG | ASSPPGQGTGNTLY | 0.7266416 |
| IFAQALL | ASSPPGQGTGNTLY | 0.6443039 |
| IGAAVALGF | ASSPPGQGTGNTLY | 0.7352804 |
| IGLTPLAL | ASSPPGQGTGNTLY | 0.7214385 |
| ILGSNTSTL | ASSPPGQGTGNTLY | 0.71597743 |
| IVFSLTVL | ASSPPGQGTGNTLY | 0.6417796 |
| LASKATGF | ASSPPGQGTGNTLY | 0.7560127 |
| LASVALIL | ASSPPGQGTGNTLY | 0.73253417 |
| LFAVGLTL | ASSPPGQGTGNTLY | 0.55329394 |
| LFNGGIGTYI | ASSPPGQGTGNTLY | 0.7696344 |
| LTLGAGII | ASSPPGQGTGNTLY | 0.71752584 |
| LVAAALGDI | ASSPPGQGTGNTLY | 0.71787405 |
| NTAFVQLGI | ASSPPGQGTGNTLY | 0.7654057 |
| QLFSGAEL | ASSPPGQGTGNTLY | 0.7158317 |
| SAATFITL | ASSPPGQGTGNTLY | 0.5624378 |
| SNYGLANLGV | ASSPPGQGTGNTLY | 0.7025691 |
| SYGGAYAVM | ASSPPGQGTGNTLY | 0.724488 |
| TTGVGGLTLG | ASSPPGQGTGNTLY | 0.6999887 |
| TTQTIYI | ASSPPGQGTGNTLY | 0.28894627 |
| VAAGAGLGAVI | ASSPPGQGTGNTLY | 0.75109357 |
| VAAVVTLL | ASSPPGQGTGNTLY | 0.46654677 |
| VGFLGGLI | ASSPPGQGTGNTLY | 0.67307585 |
| VGVIPILL | ASSPPGQGTGNTLY | 0.72638845 |
| VIFASSLI | ASSPPGQGTGNTLY | 0.7604134 |
| VKGGDGGAAAG | ASSPPGQGTGNTLY | 0.6489729 |
| VTKTFTNI | ASSPPGQGTGNTLY | 0.7201507 |
| VVAAAILL | ASSPPGQGTGNTLY | 0.744673 |
| YAALDVEL | ASSPPGQGTGNTLY | 0.68885756 |
| LASVALLL | ASSPPGQGTGNTLY | 0.7656895 |
| LDAVHRV | ASSPPGQGTGNTLY | 0.7457631 |
| QGLSSSIF | ASSPPGQGTGNTLY | 0.6897253 |
| RGTFANIRL | ASSPPGQGTGNTLY | 0.7412118 |
| RQLWWGHRI | ASSPPGQGTGNTLY | 0.6239055 |
| YGPPGTGKTLL | ASSPPGQGTGNTLY | 0.6303005 |
| AALVGAPANVA | ASSPPGQGTGNTLY | 0.6290866 |

| | | |
|---|---|---|
| AATFRRTE | ASSPPGQGTGNTLY | 0.64562964 |
| AGALGGGI | ASSPPGQGTGNTLY | 0.49151003 |
| AGGAGGAGGIGAG | ASSPPGQGTGNTLY | 0.7363534 |
| AGSGALGL | ASSPPGQGTGNTLY | 0.7716185 |
| AQIADQI | ASSPPGQGTGNTLY | 0.72218513 |
| ASAVAVTYL | ASSPPGQGTGNTLY | 0.4116271 |
| ASTVLGGTGG | ASSPPGQGTGNTLY | 0.6946094 |
| AVGGEPAK | ASSPPGQGTGNTLY | 0.68163615 |
| CEQGTFGGTC | ASSPPGQGTGNTLY | 0.75601244 |
| DEGKAPAG | ASSPPGQGTGNTLY | 0.76386374 |
| DTVVFPAL | ASSPPGQGTGNTLY | 0.7723597 |
| FEGITMAG | ASSPPGQGTGNTLY | 0.7586901 |
| FPLVVQD | ASSPPGQGTGNTLY | 0.682155 |
| GAFTTPQ | ASSPPGQGTGNTLY | 0.75623536 |
| GFGGFGTAGGVGGA | ASSPPGQGTGNTLY | 0.6979793 |
| GFLAQFL | ASSPPGQGTGNTLY | 0.7365823 |
| GGAGGAGGNAGVA | ASSPPGQGTGNTLY | 0.72310233 |
| GGVGTAGAGGAGGAGG | ASSPPGQGTGNTLY | 0.74542683 |
| GNGGGGDAGNAGSGGN | ASSPPGQGTGNTLY | 0.6352879 |
| GTGGAGGAGNDAGS | ASSPPGQGTGNTLY | 0.6801549 |
| GTTNTGWFNS | ASSPPGQGTGNTLY | 0.46982116 |
| IFMGPLFTL | ASSPPGQGTGNTLY | 0.7445272 |
| ILAHAIEAI | ASSPPGQGTGNTLY | 0.55879617 |
| IRSAIREAL | ASSPPGQGTGNTLY | 0.46164745 |
| ITFAGGG | ASSPPGQGTGNTLY | 0.6505198 |
| ITGMSGHQEV | ASSPPGQGTGNTLY | 0.7775457 |
| IVAGLACTL | ASSPPGQGTGNTLY | 0.7623994 |
| IVGGGLAGA | ASSPPGQGTGNTLY | 0.6493362 |
| IVTGVLNI | ASSPPGQGTGNTLY | 0.66539955 |
| LAQQGCI | ASSPPGQGTGNTLY | 0.73805356 |
| LCLHVEN | ASSPPGQGTGNTLY | 0.5711452 |
| LDAISDGS | ASSPPGQGTGNTLY | 0.76119876 |
| LGGFAAML | ASSPPGQGTGNTLY | 0.7181655 |
| LGGKNPMI | ASSPPGQGTGNTLY | 0.7243106 |
| LGPKGVAAIL | ASSPPGQGTGNTLY | 0.63033795 |
| LMVGVSI | ASSPPGQGTGNTLY | 0.75339794 |
| LVGAAFVL | ASSPPGQGTGNTLY | 0.7026524 |
| LVLAVAPL | ASSPPGQGTGNTLY | 0.73747015 |
| MAAIAVIDAA | ASSPPGQGTGNTLY | 0.2393301 |
| NNGINAELQ | ASSPPGQGTGNTLY | 0.7690741 |
| RGEPLVLVHG | ASSPPGQGTGNTLY | 0.63225037 |
| RGGAGGAAGLI | ASSPPGQGTGNTLY | 0.5917592 |
| RSYRMHPAV | ASSPPGQGTGNTLY | 0.3506436 |
| SAAHLLAGQLA | ASSPPGQGTGNTLY | 0.6943291 |
| SAAVGSY | ASSPPGQGTGNTLY | 0.7565279 |
| SPHAVLHRI | ASSPPGQGTGNTLY | 0.7713864 |
| SQIAKVL | ASSPPGQGTGNTLY | 0.69889295 |
| TALGHLVP | ASSPPGQGTGNTLY | 0.75648427 |
| TAPTTSGAA | ASSPPGQGTGNTLY | 0.7633498 |
| TDGDDGNGGSPGT | ASSPPGQGTGNTLY | 0.70434725 |
| TDGLGEN | ASSPPGQGTGNTLY | 0.41627735 |

| | | |
|---|---|---|
| TGGPPLPGQA | ASSPPGQGTGNTLY | 0.73950654 |
| TIPAITI | ASSPPGQGTGNTLY | 0.7101728 |
| TIPGFTI | ASSPPGQGTGNTLY | 0.77207315 |
| TSIASGALA | ASSPPGQGTGNTLY | 0.72889626 |
| TVTLPGAAI | ASSPPGQGTGNTLY | 0.5640586 |
| VASGLSAALP | ASSPPGQGTGNTLY | 0.7487239 |
| VGAGVIGI | ASSPPGQGTGNTLY | 0.75923777 |
| VGPKRFGV | ASSPPGQGTGNTLY | 0.5305766 |
| VNVAFPD | ASSPPGQGTGNTLY | 0.614257 |
| VSMTNSGVS | ASSPPGQGTGNTLY | 0.636049 |
| VTDVLGDLSV | ASSPPGQGTGNTLY | 0.54479444 |
| VVVVGAGFAGL | ASSPPGQGTGNTLY | 0.719854 |
| VYTVGTVV | ASSPPGQGTGNTLY | 0.69794416 |
| WTTFSGP | ASSPPGQGTGNTLY | 0.64797956 |
| YDELVEAF | ASSPPGQGTGNTLY | 0.7261721 |
| YGTGGAGGNGG | ASSPPGQGTGNTLY | 0.6721612 |
| YTNVMAV | ASSPPGQGTGNTLY | 0.6296268 |
| FLEQVHQL | ASSPPGQGTGNTLY | 0.6644604 |
| GLSLLAVAV | ASSPPGQGTGNTLY | 0.7611669 |
| LEQVHQL | ASSPPGQGTGNTLY | 0.7574152 |
| LNAAGAI | ASSPPGQGTGNTLY | 0.72738916 |
| PAVHASL | ASSPPGQGTGNTLY | 0.7165189 |
| TELEDDA | ASSPPGQGTGNTLY | 0.7741254 |
| TPPGPPN | ASSPPGQGTGNTLY | 0.49954724 |
| TPPPPPPP | ASSPPGQGTGNTLY | 0.56432456 |
| TQSPALI | ASSPPTGRDTEVL | 0.75091493 |
| EFAAAFPQ | ASSPPTGRDTEVL | 0.69104564 |
| LLSGNLDVL | ASSPPTGRDTEVL | 0.7614063 |
| GPANLGNYNI | ASSPPTGRDTEVL | 0.72475094 |
| TLGPTLP | ASSPPTGRDTEVL | 0.7234067 |
| VYAYSSCCK | ASSPPTGRDTEVL | 0.69952893 |
| APAPAPAPNI | ASSPPTGRDTEVL | 0.7774708 |
| PGGPVPIP | ASSPPTGRDTEVL | 0.7601643 |
| FLTVALEP | ASSPPTGRDTEVL | 0.7275541 |
| SQYFGANL | ASSPPTGRDTEVL | 0.7673037 |
| GGVKPAA | ASSPPTGRDTEVL | 0.46750224 |
| GGTGGAGGAGGW | ASSPPTGRDTEVL | 0.7562796 |
| GVGGVAGLGGAGG | ASSPPTGRDTEVL | 0.76113874 |
| VLLVLATTI | ASSPPTGRDTEVL | 0.62493074 |
| IGIFAAML | ASSPPTGRDTEVL | 0.710562 |
| AVGDLID | ASSPPTGRDTEVL | 0.773206 |
| NISPLPL | ASSPPTGRDTEVL | 0.77343994 |
| VTAVFVAI | ASSPPTGRDTEVL | 0.6109725 |
| SGDDNDAAL | ASSPPTGRDTEVL | 0.67585516 |
| AVAALAHS | ASSPPTGRDTEVL | 0.36509436 |
| GGVTVMATP | ASSPPTGRDTEVL | 0.5711322 |
| AGLAANVRD | ASSPPTGRDTEVL | 0.38281754 |
| TNRPDLIDPALLRPG | ASSPPTGRDTEVL | 0.7414886 |
| LANLGVTAL | ASSPPTGRDTEVL | 0.67022127 |
| TGERIEL | ASSPPTGRDTEVL | 0.7549088 |
| NGEELAA | ASSPPTGRDTEVL | 0.7335695 |

| | | |
|---|---|---|
| VLSLVWG | ASSPPTGRDTEVL | 0.76937217 |
| IGYALWL | ASSPPTGRDTEVL | 0.656867 |
| LSLLLAPL | ASSPPTGRDTEVL | 0.70713174 |
| TLDEALK | ASSPPTGRDTEVL | 0.47864437 |
| TDGAGTAEGG | ASSPPTGRDTEVL | 0.51381946 |
| NAPGGAGGNGG | ASSPPTGRDTEVL | 0.6401702 |
| VSVFMGMGGNF | ASSPPTGRDTEVL | 0.7180556 |
| LASYLAAL | ASSPPTGRDTEVL | 0.28342855 |
| LVGGTPAL | ASSPPTGRDTEVL | 0.7763683 |
| VRNADVL | ASSPPTGRDTEVL | 0.7092023 |
| TCSSPFLVSAM | ASSPPTGRDTEVL | 0.74710655 |
| VHITDAE | ASSPPTGRDTEVL | 0.6099667 |
| EYAAPSF | ASSPPTGRDTEVL | 0.5754582 |
| VVDAFHGGL | ASSPPTGRDTEVL | 0.7092149 |
| ASVVDPQ | ASSPPTGRDTEVL | 0.74816006 |
| SYNEFGGLS | ASSPPTGRDTEVL | 0.6751793 |
| DPDGSVY | ASSPPTGRDTEVL | 0.44583154 |
| MVSTIHATE | ASSPPTGRDTEVL | 0.52625054 |
| LAAGVADPV | ASSPPTGRDTEVL | 0.6436551 |
| TKEQIAA | ASSPPTGRDTEVL | 0.77890205 |
| SLASPAAL | ASSPPTGRDTEVL | 0.76890194 |
| IGLVFYGV | ASSPPTGRDTEVL | 0.7400789 |
| FLLDASNL | ASSPPTGRDTEVL | 0.77547634 |
| YQVGNFP | ASSPPTGRDTEVL | 0.21730222 |
| ESLLSQST | ASSPPTGRDTEVL | 0.6464413 |
| LFTVGFAAVG | ASSPPTGRDTEVL | 0.7648026 |
| ATPPTPT | ASSPPTGRDTEVL | 0.74313927 |
| TDLATLG | ASSPPTGRDTEVL | 0.69697547 |
| SAGGTGR | ASSPPTGRDTEVL | 0.76804185 |
| GFSNSGS | ASSPPTGRDTEVL | 0.6177876 |
| GGNGGSGGTSV | ASSPPTGRDTEVL | 0.6867024 |
| GTSVATGGAGN | ASSPPTGRDTEVL | 0.74201155 |
| NGGNGGSGGTS | ASSPPTGRDTEVL | 0.7574688 |
| AVVKFPH | ASSPPTGRDTEVL | 0.7408142 |
| TPPPPPP | ASSPPTGRDTEVL | 0.54767 |
| AAPAAAAAPA | ASSPPTGRDTEVL | 0.73231435 |
| IALNEDL | ASSPPTGRDTEVL | 0.7830339 |
| IVNVSSM | ASSPPTGRDTEVL | 0.69806373 |
| TGIPPAPR | ASSPPTGRDTEVL | 0.7555225 |
| FDLGGGTF | ASSPPTGRDTEVL | 0.7164536 |
| FDLGGGTFDVS | ASSPPTGRDTEVL | 0.72086686 |
| NEPTAAA | ASSPPTGRDTEVL | 0.6480533 |
| NEPTAAALA | ASSPPTGRDTEVL | 0.64675224 |
| GSIINISSI | ASSPPTGRDTEVL | 0.7564359 |
| VAPGNAGTA | ASSPPTGRDTEVL | 0.5346451 |
| TGGGASLE | ASSPPTGRDTEVL | 0.7221573 |
| VAVYNHYKRI | ASSPPTGRDTEVL | 0.70537996 |
| TDPAPATT | ASSPPTGRDTEVL | 0.78377026 |
| TNRPDILDPALLRPG | ASSPPTGRDTEVL | 0.74528503 |
| LTTTTTS | ASSPPTGRDTEVL | 0.7430898 |
| PTGSGKTL | ASSPPTGRDTEVL | 0.6779505 |

| | | |
|---|---|---|
| TGSGKTL | ASSPPTGRDTEVL | 0.7393357 |
| TIGHVDHGK | ASSPPTGRDTEVL | 0.75069726 |
| TIGHVDHGKTT | ASSPPTGRDTEVL | 0.5198971 |
| GHVDHGKTT | ASSPPTGRDTEVL | 0.7580945 |
| AQFEHTLL | ASSPPTGRDTEVL | 0.5664828 |
| IVNQDEL | ASSPPTGRDTEVL | 0.71990466 |
| TNGSQFF | ASSPPTGRDTEVL | 0.7450592 |
| TGKGIVE | ASSPPTGRDTEVL | 0.76676774 |
| LISNASDA | ASSPPTGRDTEVL | 0.7099285 |
| AELRTLGELTKIAKAHGAQ | ASSPPTGRDTEVL | 0.73996603 |
| APNAVAAAEPSW | ASSPPTGRDTEVL | 0.659754 |
| APQVAGAATAVL | ASSPPTGRDTEVL | 0.7580024 |
| ARRDPAAAAAILPTDA | ASSPPTGRDTEVL | 0.7677928 |
| DAAARDVLAEAGLAEYF | ASSPPTGRDTEVL | 0.72570974 |
| DDQIVLSAKANLAKVEQ | ASSPPTGRDTEVL | 0.7791406 |
| DELLPDTGEDSDEDGH | ASSPPTGRDTEVL | 0.75242746 |
| ELPEAPANRATVPQD | ASSPPTGRDTEVL | 0.32338464 |
| FGQSKIERVTGIG | ASSPPTGRDTEVL | 0.7080265 |
| GAGGTGSTAGGAGGAGGAGGLY | ASSPPTGRDTEVL | 0.7646966 |
| GHGGTGAAGGNGGTGGAGGSGI | ASSPPTGRDTEVL | 0.7710004 |
| GNGGNGGAGGEGGPGGAGGAGA | ASSPPTGRDTEVL | 0.7765943 |
| GSTAGNGGNGGNATNGTIAGQP | ASSPPTGRDTEVL | 0.7455609 |
| GTGGDGGNGGNGGTGG | ASSPPTGRDTEVL | 0.6730454 |
| GVRRPAGIPADAPV | ASSPPTGRDTEVL | 0.77206695 |
| GYRQVTDAHLLTLARR | ASSPPTGRDTEVL | 0.7821702 |
| IAKMGQDGHDRG | ASSPPTGRDTEVL | 0.7392005 |
| KAEPSIEIQYT | ASSPPTGRDTEVL | 0.5679137 |
| KPVRIRMAADEDTVAVT | ASSPPTGRDTEVL | 0.54075843 |
| MPPRIAGMRLLVIKPEP | ASSPPTGRDTEVL | 0.7555702 |
| MKFYAVALFRSRGVPPTR | ASSPPTGRDTEVL | 0.7741939 |
| NAGGFGILGAVAHSP | ASSPPTGRDTEVL | 0.5685477 |
| NEGREITLAMAEM | ASSPPTGRDTEVL | 0.7124295 |
| NELVEWSHPLGHRWRWE | ASSPPTGRDTEVL | 0.7332344 |
| NLGAGGTGGNGGNPG | ASSPPTGRDTEVL | 0.3146213 |
| NTEWPKWETVMAADPVD | ASSPPTGRDTEVL | 0.73092866 |
| PASAAPAETQP | ASSPPTGRDTEVL | 0.7684946 |
| QQSSDGKNGVRAAEITGE | ASSPPTGRDTEVL | 0.7562399 |
| RAPDAVTTSWWKE | ASSPPTGRDTEVL | 0.5756933 |
| SGFGNVGAHTSGWFNQ | ASSPPTGRDTEVL | 0.76156175 |
| SGVAGAVGQAASVGGLKV | ASSPPTGRDTEVL | 0.7781551 |
| SNGGAGTGSAGGNGGT | ASSPPTGRDTEVL | 0.781718 |
| TGGAGGEGGAAG | ASSPPTGRDTEVL | 0.434909 |
| TGGDGGAAGAGGAPGHG | ASSPPTGRDTEVL | 0.2918344 |
| TMPPLLAAIANAGRHGV | ASSPPTGRDTEVL | 0.6859375 |
| TSSTAGNSGGA | ASSPPTGRDTEVL | 0.756032 |
| VPATVPVGILC | ASSPPTGRDTEVL | 0.70558274 |
| VQRPDSGGVVLTGRIS | ASSPPTGRDTEVL | 0.6785233 |
| WTPNPVPATKGVPAGLVQAT | ASSPPTGRDTEVL | 0.7533628 |
| YALKRALAKFSDRFDVVII | ASSPPTGRDTEVL | 0.76449865 |
| YLIGSVIV | ASSPPTGRDTEVL | 0.5310182 |
| YGQLVAVID | ASSPPTGRDTEVL | 0.48653638 |

| | | |
|---|---|---|
| VTVGGMTLPA | ASSPPTGRDTEVL | 0.76998186 |
| VLALPAV | ASSPPTGRDTEVL | 0.7597886 |
| VIPPTLNYET | ASSPPTGRDTEVL | 0.59970856 |
| VGMGGMHPSSG | ASSPPTGRDTEVL | 0.71328294 |
| VGLAGGAATA | ASSPPTGRDTEVL | 0.6901368 |
| VGGAGGAGGSGP | ASSPPTGRDTEVL | 0.6877004 |
| VFAAFMV | ASSPPTGRDTEVL | 0.732472 |
| TLYDKLR | ASSPPTGRDTEVL | 0.5120237 |
| TLAVFTVL | ASSPPTGRDTEVL | 0.76220095 |
| TGSVQPARV | ASSPPTGRDTEVL | 0.5670817 |
| TGGGAGSGGT | ASSPPTGRDTEVL | 0.7514893 |
| SHLETSL | ASSPPTGRDTEVL | 0.7250631 |
| SFTTGMT | ASSPPTGRDTEVL | 0.74596035 |
| QVEVAKRMGVRQPTVS | ASSPPTGRDTEVL | 0.68592703 |
| QGSGGPALAM | ASSPPTGRDTEVL | 0.5597654 |
| MLAPSNP | ASSPPTGRDTEVL | 0.6348026 |
| LVNNAAI | ASSPPTGRDTEVL | 0.52387834 |
| LTGEINTLA | ASSPPTGRDTEVL | 0.7630974 |
| LSIFGPV | ASSPPTGRDTEVL | 0.71187675 |
| LLALLANL | ASSPPTGRDTEVL | 0.7664898 |
| LISNASDALDKLR | ASSPPTGRDTEVL | 0.7450861 |
| LINEAVP | ASSPPTGRDTEVL | 0.63101757 |
| LGYAFSGL | ASSPPTGRDTEVL | 0.7536331 |
| LDFVAVAL | ASSPPTGRDTEVL | 0.58788574 |
| LAQPGAE | ASSPPTGRDTEVL | 0.70090425 |
| ISNLGAVTGF | ASSPPTGRDTEVL | 0.71783817 |
| HQEDPGRI | ASSPPTGRDTEVL | 0.76053447 |
| GTPGEPTP | ASSPPTGRDTEVL | 0.63347226 |
| GFFNSGS | ASSPPTGRDTEVL | 0.7575226 |
| GAGGGGLTAGAN | ASSPPTGRDTEVL | 0.7452961 |
| FVDALVNM | ASSPPTGRDTEVL | 0.60692877 |
| FLTVAPEL | ASSPPTGRDTEVL | 0.7495496 |
| FETLTAL | ASSPPTGRDTEVL | 0.7571221 |
| DVINPIEL | ASSPPTGRDTEVL | 0.7484312 |
| APVSGAPV | ASSPPTGRDTEVL | 0.60857826 |
| ALVGGLAW | ASSPPTGRDTEVL | 0.7408546 |
| ALAAAAATAI | ASSPPTGRDTEVL | 0.65495354 |
| AFGLGLLL | ASSPPTGRDTEVL | 0.7410579 |
| AFFAAASL | ASSPPTGRDTEVL | 0.73277277 |
| AAAALPAFSPP | ASSPPTGRDTEVL | 0.6636827 |
| VVLAPAL | ASSPPTGRDTEVL | 0.7597905 |
| TIGHVDHGKTTL | ASSPPTGRDTEVL | 0.5318692 |
| LISNASDALDK | ASSPPTGRDTEVL | 0.62870353 |
| LISNASDAL | ASSPPTGRDTEVL | 0.66938907 |
| ISNASDALDK | ASSPPTGRDTEVL | 0.71059877 |
| NQAETLVYQTE | ASSPPTGRDTEVL | 0.74936616 |
| TGGTGGTAGAG | ASSPPTGRDTEVL | 0.6103388 |
| TGSTAGGAGGA | ASSPPTGRDTEVL | 0.7234992 |
| DPLAVPVAAGPV | ASSPPTGRDTEVL | 0.7057122 |
| GGTGGTPGNGGN | ASSPPTGRDTEVL | 0.6172686 |
| PAGLAGVDPATL | ASSPPTGRDTEVL | 0.7512845 |

| | | |
|---|---|---|
| GNNLVVEAPHAGASV | ASSPPTGRDTEVL | 0.6834555 |
| KVTRIMSASTLEQPA | ASSPPTGRDTEVL | 0.47848243 |
| LGFALPIALPRRTVV | ASSPPTGRDTEVL | 0.751773 |
| TRLGNLTARKLVVLV | ASSPPTGRDTEVL | 0.68741727 |
| WSQQRRLIYFPSAGP | ASSPPTGRDTEVL | 0.7039615 |
| GPNGNPDPMAAAVDIR | ASSPPTGRDTEVL | 0.77630377 |
| TQRCVSVADLRAAASR | ASSPPTGRDTEVL | 0.42910987 |
| QFVLDVAEHPVEVLTETV | ASSPPTGRDTEVL | 0.75138474 |
| GGAGGTGGAGGNGGTQAGDG | ASSPPTGRDTEVL | 0.73901474 |
| PLLATLRAIAVDVDAKPAQV | ASSPPTGRDTEVL | 0.7297932 |
| APKFRTGGYMIDFWGVGYQVAK | ASSPPTGRDTEVL | 0.7111782 |
| GAGGAGGAGGAAGGPGGTGGTG | ASSPPTGRDTEVL | 0.6657493 |
| IGALGLLATDVAPAGSVAFGVA | ASSPPTGRDTEVL | 0.7770841 |
| AGGAGGIGGNGGAGGAGGVLMG | ASSPPTGRDTEVL | 0.7186359 |
| ADELIGLL | ASSPPTGRDTEVL | 0.7446432 |
| ADRALKA | ASSPPTGRDTEVL | 0.7550639 |
| AGAVGQVGGAGGA | ASSPPTGRDTEVL | 0.75516963 |
| AGGGGGLGGAP | ASSPPTGRDTEVL | 0.75380635 |
| AGGLLYGNGGAG | ASSPPTGRDTEVL | 0.6210135 |
| AGLESIGL | ASSPPTGRDTEVL | 0.45457262 |
| AGLKTGII | ASSPPTGRDTEVL | 0.708617 |
| ASLAALLL | ASSPPTGRDTEVL | 0.6907239 |
| ASVINISSI | ASSPPTGRDTEVL | 0.6523651 |
| ATPPGPA | ASSPPTGRDTEVL | 0.35566705 |
| ATVASALV | ASSPPTGRDTEVL | 0.7599766 |
| ATVASLIL | ASSPPTGRDTEVL | 0.7815806 |
| AVNGPVLIRH | ASSPPTGRDTEVL | 0.60085815 |
| DGAAGAVGI | ASSPPTGRDTEVL | 0.78415024 |
| DPLLHGI | ASSPPTGRDTEVL | 0.58793175 |
| ELGAALGAL | ASSPPTGRDTEVL | 0.73628336 |
| ELGRTQAV | ASSPPTGRDTEVL | 0.7637851 |
| FVAGGYLS | ASSPPTGRDTEVL | 0.54493076 |
| GASGGAGGAGGW | ASSPPTGRDTEVL | 0.7430104 |
| GGAGAGGDGGAGGI | ASSPPTGRDTEVL | 0.6828434 |
| GTGHNGSQPGMGGNG | ASSPPTGRDTEVL | 0.6709459 |
| GYAAASASAS | ASSPPTGRDTEVL | 0.7653328 |
| HQVDVAAVQQG | ASSPPTGRDTEVL | 0.65115404 |
| IGAAIAHT | ASSPPTGRDTEVL | 0.73120105 |
| IGMIEVAQL | ASSPPTGRDTEVL | 0.7190063 |
| IRLAPEHTSAMD | ASSPPTGRDTEVL | 0.6688974 |
| IYNVSVL | ASSPPTGRDTEVL | 0.74647427 |
| KQGFLGM | ASSPPTGRDTEVL | 0.7549306 |
| LESVITQL | ASSPPTGRDTEVL | 0.694654 |
| LGLGALVVV | ASSPPTGRDTEVL | 0.7807857 |
| LIGNGGNGGSGGI | ASSPPTGRDTEVL | 0.6114057 |
| LNAINAPTL | ASSPPTGRDTEVL | 0.7801541 |
| LQLLNTGV | ASSPPTGRDTEVL | 0.765342 |
| LQTSIEF | ASSPPTGRDTEVL | 0.73288214 |
| MSFAVSEL | ASSPPTGRDTEVL | 0.6896233 |
| NTGALGL | ASSPPTGRDTEVL | 0.7629899 |
| PPGAGGAGGAGTT | ASSPPTGRDTEVL | 0.68909407 |

| | | |
|---|---|---|
| QLVGLGI | ASSPPTGRDTEVL | 0.71191967 |
| RFLEQVHQL | ASSPPTGRDTEVL | 0.7753551 |
| RGVGYVLR | ASSPPTGRDTEVL | 0.66885716 |
| SATGVFAGVI | ASSPPTGRDTEVL | 0.8388318 |
| SLPAITLP | ASSPPTGRDTEVL | 0.82165945 |
| STALAFTV | ASSPPTGRDTEVL | 0.78847265 |
| SVLTVGLGK | ASSPPTGRDTEVL | 0.76664567 |
| TALAGAGSGI | ASSPPTGRDTEVL | 0.41780266 |
| TGLPFTGI | ASSPPTGRDTEVL | 0.72785985 |
| VAEHVTV | ASSPPTGRDTEVL | 0.7225257 |
| VDIVLNSL | ASSPPTGRDTEVL | 0.83081245 |
| VLTGGGAVL | ASSPPTGRDTEVL | 0.8287253 |
| VNVGLAYGI | ASSPPTGRDTEVL | 0.7275218 |
| VSSAAVGAV | ASSPPTGRDTEVL | 0.83586526 |
| VTSVYEAGV | ASSPPTGRDTEVL | 0.8128241 |
| WVALDRGA | ASSPPTGRDTEVL | 0.59072614 |
| YLVGSVVI | ASSPPTGRDTEVL | 0.78194493 |
| GPPGTGKTLL | ASSPPTGRDTEVL | 0.686609 |
| GVGMTKF | ASSPPTGRDTEVL | 0.8037606 |
| IGIGGSDL | ASSPPTGRDTEVL | 0.33345956 |
| IGTLLAEL | ASSPPTGRDTEVL | 0.83014643 |
| IYTSGTT | ASSPPTGRDTEVL | 0.7371682 |
| LSFPAGV | ASSPPTGRDTEVL | 0.2021415 |
| LYTSGTT | ASSPPTGRDTEVL | 0.8044157 |
| VGAFPFGF | ASSPPTGRDTEVL | 0.78332996 |
| VGAGTVEF | ASSPPTGRDTEVL | 0.7920445 |
| VGKSTLLL | ASSPPTGRDTEVL | 0.8329109 |
| AALAHAGGGIV | ASSPPTGRDTEVL | 0.52454066 |
| AALVAVSLL | ASSPPTGRDTEVL | 0.6228181 |
| AAVSRAVAAL | ASSPPTGRDTEVL | 0.82577693 |
| AGGSSGIGGF | ASSPPTGRDTEVL | 0.83836746 |
| AGLTVANANG | ASSPPTGRDTEVL | 0.6159022 |
| AGLVGAVL | ASSPPTGRDTEVL | 0.8189071 |
| ALSQAFTV | ASSPPTGRDTEVL | 0.8326819 |
| ALTFSVSL | ASSPPTGRDTEVL | 0.8211488 |
| ALTGAALVAL | ASSPPTGRDTEVL | 0.8334407 |
| ASFSIGAGVL | ASSPPTGRDTEVL | 0.8405484 |
| DGSVYKI | ASSPPTGRDTEVL | 0.72684133 |
| EQLAEIV | ASSPPTGRDTEVL | 0.83630705 |
| FAVSELTVM | ASSPPTGRDTEVL | 0.8365919 |
| GAAPLAAGL | ASSPPTGRDTEVL | 0.7403904 |
| GADGGDGGSGGAGG | ASSPPTGRDTEVL | 0.8001932 |
| GFGLDTGGAGG | ASSPPTGRDTEVL | 0.5730039 |
| GLASGGLL | ASSPPTGRDTEVL | 0.7706778 |
| GRIAGAAVLL | ASSPPTGRDTEVL | 0.47615743 |
| GSAGSLNTSF | ASSPPTGRDTEVL | 0.817334 |
| GTGQIGI | ASSPPTGRDTEVL | 0.8405188 |
| GTLIALIL | ASSPPTGRDTEVL | 0.79525256 |
| IAATMGFVVG | ASSPPTGRDTEVL | 0.6843275 |
| IFAQALL | ASSPPTGRDTEVL | 0.7141054 |
| IGAAVALGF | ASSPPTGRDTEVL | 0.7912803 |

| | | |
|---|---|---|
| IGLTPLAL | ASSPPTGRDTEVL | 0.76115865 |
| ILGSNTSTL | ASSPPTGRDTEVL | 0.35152584 |
| IVFSLTVL | ASSPPTGRDTEVL | 0.7682669 |
| LASKATGF | ASSPPTGRDTEVL | 0.8348646 |
| LASVALIL | ASSPPTGRDTEVL | 0.83192664 |
| LFAVGLTL | ASSPPTGRDTEVL | 0.6410674 |
| LFNGGIGTYI | ASSPPTGRDTEVL | 0.84009105 |
| LTLGAGII | ASSPPTGRDTEVL | 0.82066894 |
| LVAAALGDI | ASSPPTGRDTEVL | 0.84155726 |
| NTAFVQLGI | ASSPPTGRDTEVL | 0.7049843 |
| QLFSGAEL | ASSPPTGRDTEVL | 0.66669697 |
| SAATFITL | ASSPPTGRDTEVL | 0.826774 |
| SNYGLANLGV | ASSPPTGRDTEVL | 0.7880687 |
| SYGGAYAVM | ASSPPTGRDTEVL | 0.7667154 |
| TTGVGGLTLG | ASSPPTGRDTEVL | 0.82661605 |
| TTQTIYI | ASSPPTGRDTEVL | 0.792495 |
| VAAGAGLGAVI | ASSPPTGRDTEVL | 0.71869344 |
| VAAVVTLL | ASSPPTGRDTEVL | 0.78889024 |
| VGFLGGLI | ASSPPTGRDTEVL | 0.83944225 |
| VGVIPILL | ASSPPTGRDTEVL | 0.7011821 |
| VIFASSLI | ASSPPTGRDTEVL | 0.7521117 |
| VKGGDGGAAAG | ASSPPTGRDTEVL | 0.6585507 |
| VTKTFTNI | ASSPPTGRDTEVL | 0.7864648 |
| VVAAAILL | ASSPPTGRDTEVL | 0.83422554 |
| YAALDVEL | ASSPPTGRDTEVL | 0.8165126 |
| LASVALLL | ASSPPTGRDTEVL | 0.76457554 |
| LDAVHRV | ASSPPTGRDTEVL | 0.8211441 |
| QGLSSSIF | ASSPPTGRDTEVL | 0.82436883 |
| RGTFANIRL | ASSPPTGRDTEVL | 0.8076257 |
| RQLWWGHRI | ASSPPTGRDTEVL | 0.8079828 |
| YGPPGTGKTLL | ASSPPTGRDTEVL | 0.7586879 |
| AALVGAPANVA | ASSPPTGRDTEVL | 0.83464134 |
| AATFRRTE | ASSPPTGRDTEVL | 0.8263314 |
| AGALGGGI | ASSPPTGRDTEVL | 0.8283189 |
| AGGAGGAGGIGAG | ASSPPTGRDTEVL | 0.83198285 |
| AGSGALGL | ASSPPTGRDTEVL | 0.8138981 |
| AQIADQI | ASSPPTGRDTEVL | 0.80191207 |
| ASAVAVTYL | ASSPPTGRDTEVL | 0.5137082 |
| ASTVLGGTGG | ASSPPTGRDTEVL | 0.553675 |
| AVGGEPAK | ASSPPTGRDTEVL | 0.81648093 |
| CEQGTFGGTC | ASSPPTGRDTEVL | 0.8131926 |
| DEGKAPAG | ASSPPTGRDTEVL | 0.8282519 |
| DTVVFPAL | ASSPPTGRDTEVL | 0.7996846 |
| FEGITMAG | ASSPPTGRDTEVL | 0.7552126 |
| FPLVVQD | ASSPPTGRDTEVL | 0.8251083 |
| GAFTTPQ | ASSPPTGRDTEVL | 0.8321095 |
| GFGGFGTAGGVGGA | ASSPPTGRDTEVL | 0.597983 |
| GFLAQFL | ASSPPTGRDTEVL | 0.75846034 |
| GGAGGAGGNAGVA | ASSPPTGRDTEVL | 0.84327316 |
| GGVGTAGAGGAGGAGG | ASSPPTGRDTEVL | 0.81245226 |
| GNGGGGDAGNAGSGGN | ASSPPTGRDTEVL | 0.7716694 |

| | | |
|---|---|---|
| GTGGAGGAGNDAGS | ASSPPTGRDTEVL | 0.76519334 |
| GTTNTGWFNS | ASSPPTGRDTEVL | 0.84272647 |
| IFMGPLFTL | ASSPPTGRDTEVL | 0.7672181 |
| ILAHAIEAI | ASSPPTGRDTEVL | 0.8413569 |
| IRSAIREAL | ASSPPTGRDTEVL | 0.3910731 |
| ITFAGGG | ASSPPTGRDTEVL | 0.7637356 |
| ITGMSGHQEV | ASSPPTGRDTEVL | 0.71898705 |
| IVAGLACTL | ASSPPTGRDTEVL | 0.40905917 |
| IVGGGLAGA | ASSPPTGRDTEVL | 0.83322513 |
| IVTGVLNI | ASSPPTGRDTEVL | 0.68526584 |
| LAQQGCI | ASSPPTGRDTEVL | 0.818292 |
| LCLHVEN | ASSPPTGRDTEVL | 0.7842201 |
| LDAISDGS | ASSPPTGRDTEVL | 0.8358844 |
| LGGFAAML | ASSPPTGRDTEVL | 0.81147397 |
| LGGKNPMI | ASSPPTGRDTEVL | 0.7907002 |
| LGPKGVAAIL | ASSPPTGRDTEVL | 0.70901144 |
| LMVGVSI | ASSPPTGRDTEVL | 0.82724214 |
| LVGAAFVL | ASSPPTGRDTEVL | 0.83146363 |
| LVLAVAPL | ASSPPTGRDTEVL | 0.8312903 |
| MAAIAVIDAA | ASSPPTGRDTEVL | 0.43391076 |
| NNGINAELQ | ASSPPTGRDTEVL | 0.805902 |
| RGEPLVLVHG | ASSPPTGRDTEVL | 0.8044044 |
| RGGAGGAAGLI | ASSPPTGRDTEVL | 0.25390407 |
| RSYRMHPAV | ASSPPTGRDTEVL | 0.26672006 |
| SAAHLLAGQLA | ASSPPTGRDTEVL | 0.75679946 |
| SAAVGSY | ASSPPTGRDTEVL | 0.76147246 |
| SPHAVLHRI | ASSPPTGRDTEVL | 0.8293864 |
| SQIAKVL | ASSPPTGRDTEVL | 0.82311 |
| TALGHLVP | ASSPPTGRDTEVL | 0.7966478 |
| TAPTTSGAA | ASSPPTGRDTEVL | 0.8189738 |
| TDGDDGNGGSPGT | ASSPPTGRDTEVL | 0.81014884 |
| TDGLGEN | ASSPPTGRDTEVL | 0.14908014 |
| TGGPPLPGQA | ASSPPTGRDTEVL | 0.6797015 |
| TIPAITI | ASSPPTGRDTEVL | 0.620176 |
| TIPGFTI | ASSPPTGRDTEVL | 0.8376728 |
| TSIASGALA | ASSPPTGRDTEVL | 0.7087977 |
| TVTLPGAAI | ASSPPTGRDTEVL | 0.7212022 |
| VASGLSAALP | ASSPPTGRDTEVL | 0.80719566 |
| VGAGVIGI | ASSPPTGRDTEVL | 0.812474 |
| VGPKRFGV | ASSPPTGRDTEVL | 0.7886578 |
| VNVAFPD | ASSPPTGRDTEVL | 0.72427857 |
| VSMTNSGVS | ASSPPTGRDTEVL | 0.3318323 |
| VTDVLGDLSV | ASSPPTGRDTEVL | 0.82998544 |
| VVVVGAGFAGL | ASSPPTGRDTEVL | 0.5324808 |
| VYTVGTVV | ASSPPTGRDTEVL | 0.6402781 |
| WTTFSGP | ASSPPTGRDTEVL | 0.31571156 |
| YDELVEAF | ASSPPTGRDTEVL | 0.78691375 |
| YGTGGAGGNGG | ASSPPTGRDTEVL | 0.34499305 |
| YTNVMAV | ASSPPTGRDTEVL | 0.8292553 |
| FLEQVHQL | ASSPPTGRDTEVL | 0.7651578 |
| GLSLLAVAV | ASSPPTGRDTEVL | 0.77619934 |

| | | |
|---|---|---|
| LEQVHQL | ASSPPTGRDTEVL | 0.8431407 |
| LNAAGAI | ASSPPTGRDTEVL | 0.8102417 |
| PAVHASL | ASSPPTGRDTEVL | 0.7859877 |
| TELEDDA | ASSPPTGRDTEVL | 0.7208015 |
| TPPGPPN | ASSPPTGRDTEVL | 0.699888 |
| TPPPPPPP | ASSPPTGRDTEVL | 0.7335324 |
| TQSPALI | ASSQGTGDEQY | 0.72832227 |
| EFAAAFPQ | ASSQGTGDEQY | 0.8361687 |
| LLSGNLDVL | ASSQGTGDEQY | 0.79545355 |
| GPANLGNYNI | ASSQGTGDEQY | 0.50468457 |
| TLGPTLP | ASSQGTGDEQY | 0.7895107 |
| VYAYSSCCK | ASSQGTGDEQY | 0.84080815 |
| APAPAPAPNI | ASSQGTGDEQY | 0.8475757 |
| PGGPVPIP | ASSQGTGDEQY | 0.74177366 |
| FLTVALEP | ASSQGTGDEQY | 0.7128662 |
| SQYFGANL | ASSQGTGDEQY | 0.66721225 |
| GGVKPAA | ASSQGTGDEQY | 0.7965345 |
| GGTGGAGGAGGW | ASSQGTGDEQY | 0.67576146 |
| GVGGVAGLGGAGG | ASSQGTGDEQY | 0.6130972 |
| VLLVLATTI | ASSQGTGDEQY | 0.8303906 |
| IGIFAAML | ASSQGTGDEQY | 0.8478296 |
| AVGDLID | ASSQGTGDEQY | 0.7694856 |
| NISPLPL | ASSQGTGDEQY | 0.38277984 |
| VTAVFVAI | ASSQGTGDEQY | 0.7944599 |
| SGDDNDAAL | ASSQGTGDEQY | 0.7335608 |
| AVAALAHS | ASSQGTGDEQY | 0.7730825 |
| GGVTVMATP | ASSQGTGDEQY | 0.7725285 |
| AGLAANVRD | ASSQGTGDEQY | 0.55574447 |
| TNRPDLIDPALLRPG | ASSQGTGDEQY | 0.7950245 |
| LANLGVTAL | ASSQGTGDEQY | 0.8479348 |
| TGERIEL | ASSQGTGDEQY | 0.7892819 |
| NGEELAA | ASSQGTGDEQY | 0.81622887 |
| VLSLVWG | ASSQGTGDEQY | 0.8405881 |
| IGYALWL | ASSQGTGDEQY | 0.8291974 |
| LSLLLAPL | ASSQGTGDEQY | 0.8384757 |
| TLDEALK | ASSQGTGDEQY | 0.82639766 |
| TDGAGTAEGG | ASSQGTGDEQY | 0.78875124 |
| NAPGGAGGNGG | ASSQGTGDEQY | 0.79548335 |
| VSVFMGMGGNF | ASSQGTGDEQY | 0.8206328 |
| LASYLAAL | ASSQGTGDEQY | 0.79821026 |
| LVGGTPAL | ASSQGTGDEQY | 0.7991032 |
| VRNADVL | ASSQGTGDEQY | 0.7287462 |
| TCSSPFLVSAM | ASSQGTGDEQY | 0.83393323 |
| VHITDAE | ASSQGTGDEQY | 0.7614791 |
| EYAAPSF | ASSQGTGDEQY | 0.83820844 |
| VVDAFHGGL | ASSQGTGDEQY | 0.44496483 |
| ASVVDPQ | ASSQGTGDEQY | 0.67215884 |
| SYNEFGGLS | ASSQGTGDEQY | 0.83357906 |
| DPDGSVY | ASSQGTGDEQY | 0.16800421 |
| MVSTIHATE | ASSQGTGDEQY | 0.28068477 |
| LAAGVADPV | ASSQGTGDEQY | 0.79902 |

| | | |
|---|---|---|
| TKEQIAA | ASSQGTGDEQY | 0.84004563 |
| SLASPAAL | ASSQGTGDEQY | 0.17496338 |
| IGLVFYGV | ASSQGTGDEQY | 0.5414052 |
| FLLDASNL | ASSQGTGDEQY | 0.5856966 |
| YQVGNFP | ASSQGTGDEQY | 0.80737495 |
| ESLLSQST | ASSQGTGDEQY | 0.8003656 |
| LFTVGFAAVG | ASSQGTGDEQY | 0.42712224 |
| ATPPTPT | ASSQGTGDEQY | 0.8312801 |
| TDLATLG | ASSQGTGDEQY | 0.40962496 |
| SAGGTGR | ASSQGTGDEQY | 0.7360715 |
| GFSNSGS | ASSQGTGDEQY | 0.82940847 |
| GGNGGSGGTSV | ASSQGTGDEQY | 0.83726025 |
| GTSVATGGAGN | ASSQGTGDEQY | 0.59713763 |
| NGGNGGSGGTS | ASSQGTGDEQY | 0.82199323 |
| AVVKFPH | ASSQGTGDEQY | 0.75972664 |
| TPPPPPP | ASSQGTGDEQY | 0.8482368 |
| AAPAAAAAPA | ASSQGTGDEQY | 0.7594876 |
| IALNEDL | ASSQGTGDEQY | 0.5290035 |
| IVNVSSM | ASSQGTGDEQY | 0.83451885 |
| TGIPPAPR | ASSQGTGDEQY | 0.82572323 |
| FDLGGGTF | ASSQGTGDEQY | 0.75731343 |
| FDLGGGTFDVS | ASSQGTGDEQY | 0.82727146 |
| NEPTAAA | ASSQGTGDEQY | 0.7480053 |
| NEPTAAALA | ASSQGTGDEQY | 0.7953979 |
| GSIINISSI | ASSQGTGDEQY | 0.83392346 |
| VAPGNAGTA | ASSQGTGDEQY | 0.8282819 |
| TGGGASLE | ASSQGTGDEQY | 0.6360662 |
| VAVYNHYKRI | ASSQGTGDEQY | 0.83966225 |
| TDPAPATT | ASSQGTGDEQY | 0.80426526 |
| TNRPDILDPALLRPG | ASSQGTGDEQY | 0.62397003 |
| LTTTTTS | ASSQGTGDEQY | 0.84595287 |
| PTGSGKTL | ASSQGTGDEQY | 0.77745 |
| TGSGKTL | ASSQGTGDEQY | 0.83203286 |
| TIGHVDHGK | ASSQGTGDEQY | 0.834003 |
| TIGHVDHGKTT | ASSQGTGDEQY | 0.84034884 |
| GHVDHGKTT | ASSQGTGDEQY | 0.48009074 |
| AQFEHTLL | ASSQGTGDEQY | 0.7732837 |
| IVNQDEL | ASSQGTGDEQY | 0.7844469 |
| TNGSQFF | ASSQGTGDEQY | 0.82279056 |
| TGKGIVE | ASSQGTGDEQY | 0.78333664 |
| LISNASDA | ASSQGTGDEQY | 0.7286805 |
| AELRTLGELTKIAKAHGAQ | ASSQGTGDEQY | 0.8280205 |
| APNAVAAAEPSW | ASSQGTGDEQY | 0.8382914 |
| APQVAGAATAVL | ASSQGTGDEQY | 0.7968312 |
| ARRDPAAAAAILPTDA | ASSQGTGDEQY | 0.80127597 |
| DAAARDVLAEAGLAEYF | ASSQGTGDEQY | 0.84265864 |
| DDQIVLSAKANLAKVEQ | ASSQGTGDEQY | 0.6878325 |
| DELLPDTGEDSDEDGH | ASSQGTGDEQY | 0.7960061 |
| ELPEAPANRATVPQD | ASSQGTGDEQY | 0.8249435 |
| FGQSKIERVTGIG | ASSQGTGDEQY | 0.8277977 |
| GAGGTGSTAGGAGGAGGAGGLY | ASSQGTGDEQY | 0.7901447 |

| | | |
|---|---|---|
| GHGGTGAAGGNGGTGGAGGSGI | ASSQGTGDEQY | 0.8387575 |
| GNGGNGGAGGEGGPGGAGGAGA | ASSQGTGDEQY | 0.6757698 |
| GSTAGNGGNGGNATNGTIAGQP | ASSQGTGDEQY | 0.82916737 |
| GTGGDGGNGGNGGTGG | ASSQGTGDEQY | 0.7595486 |
| GVRRPAGIPADAPV | ASSQGTGDEQY | 0.79225826 |
| GYRQVTDAHLLTLARR | ASSQGTGDEQY | 0.7386391 |
| IAKMGQDGHDRG | ASSQGTGDEQY | 0.8244144 |
| KAEPSIEIQYT | ASSQGTGDEQY | 0.8298904 |
| KPVRIRMAADEDTVAVT | ASSQGTGDEQY | 0.82888776 |
| MPPRIAGMRLLVIKPEP | ASSQGTGDEQY | 0.24006933 |
| MKFYAVALFRSRGVPPTR | ASSQGTGDEQY | 0.8316034 |
| NAGGFGILGAVAHSP | ASSQGTGDEQY | 0.8272042 |
| NEGREITLAMAEM | ASSQGTGDEQY | 0.8447308 |
| NELVEWSHPLGHRWRWE | ASSQGTGDEQY | 0.7889999 |
| NLGAGGTGGNGGNPG | ASSQGTGDEQY | 0.7836342 |
| NTEWPKWETVMAADPVD | ASSQGTGDEQY | 0.76436156 |
| PASAAPAETQP | ASSQGTGDEQY | 0.8001785 |
| QQSSDGKNGVRAAEITGE | ASSQGTGDEQY | 0.66655153 |
| RAPDAVTTSWWKE | ASSQGTGDEQY | 0.73847425 |
| SGFGNVGAHTSGWFNQ | ASSQGTGDEQY | 0.76140904 |
| SGVAGAVGQAASVGGLKV | ASSQGTGDEQY | 0.7572311 |
| SNGGAGTGSAGGNGGT | ASSQGTGDEQY | 0.82952774 |
| TGGAGGEGGAAG | ASSQGTGDEQY | 0.83480805 |
| TGGDGGAAGAGGAPGHG | ASSQGTGDEQY | 0.8320191 |
| TMPPLLAAIANAGRHGV | ASSQGTGDEQY | 0.83009213 |
| TSSTAGNSGGA | ASSQGTGDEQY | 0.8385198 |
| VPATVPVGILC | ASSQGTGDEQY | 0.83152306 |
| VQRPDSGGVVLTGRIS | ASSQGTGDEQY | 0.78625655 |
| WTPNPVPATKGVPAGLVQAT | ASSQGTGDEQY | 0.7128252 |
| YALKRALAKFSDRFDVVII | ASSQGTGDEQY | 0.6273577 |
| YLIGSVIV | ASSQGTGDEQY | 0.7676388 |
| YGQLVAVID | ASSQGTGDEQY | 0.77899307 |
| VTVGGMTLPA | ASSQGTGDEQY | 0.8083991 |
| VLALPAV | ASSQGTGDEQY | 0.16080949 |
| VIPPTLNYET | ASSQGTGDEQY | 0.80399215 |
| VGMGGMHPSSG | ASSQGTGDEQY | 0.8381685 |
| VGLAGGAATA | ASSQGTGDEQY | 0.7743222 |
| VGGAGGAGGSGP | ASSQGTGDEQY | 0.8110227 |
| VFAAFMV | ASSQGTGDEQY | 0.80904156 |
| TLYDKLR | ASSQGTGDEQY | 0.76365817 |
| TLAVFTVL | ASSQGTGDEQY | 0.8130272 |
| TGSVQPARV | ASSQGTGDEQY | 0.7388171 |
| TGGGAGSGGT | ASSQGTGDEQY | 0.8309516 |
| SHLETSL | ASSQGTGDEQY | 0.8431872 |
| SFTTGMT | ASSQGTGDEQY | 0.81324625 |
| QVEVAKRMGVRQPTVS | ASSQGTGDEQY | 0.78090316 |
| QGSGGPALAM | ASSQGTGDEQY | 0.79704857 |
| MLAPSNP | ASSQGTGDEQY | 0.36090934 |
| LVNNAAI | ASSQGTGDEQY | 0.79543585 |
| LTGEINTLA | ASSQGTGDEQY | 0.5524608 |
| LSIFGPV | ASSQGTGDEQY | 0.79684436 |

| | | |
|---|---|---|
| LLALLANL | ASSQGTGDEQY | 0.7541477 |
| LISNASDALDKLR | ASSQGTGDEQY | 0.8160434 |
| LINEAVP | ASSQGTGDEQY | 0.79938096 |
| LGYAFSGL | ASSQGTGDEQY | 0.80827 |
| LDFVAVAL | ASSQGTGDEQY | 0.71933484 |
| LAQPGAE | ASSQGTGDEQY | 0.81587976 |
| ISNLGAVTGF | ASSQGTGDEQY | 0.8047459 |
| HQEDPGRI | ASSQGTGDEQY | 0.575057 |
| GTPGEPTP | ASSQGTGDEQY | 0.82915163 |
| GFFNSGS | ASSQGTGDEQY | 0.79033613 |
| GAGGGGLTAGAN | ASSQGTGDEQY | 0.7883091 |
| FVDALVNM | ASSQGTGDEQY | 0.83652544 |
| FLTVAPEL | ASSQGTGDEQY | 0.8041114 |
| FETLTAL | ASSQGTGDEQY | 0.33252043 |
| DVINPIEL | ASSQGTGDEQY | 0.79051965 |
| APVSGAPV | ASSQGTGDEQY | 0.7912663 |
| ALVGGLAW | ASSQGTGDEQY | 0.802866 |
| ALAAAAATAI | ASSQGTGDEQY | 0.18917088 |
| AFGLGLLL | ASSQGTGDEQY | 0.8242761 |
| AFFAAASL | ASSQGTGDEQY | 0.6212906 |
| AAAALPAFSPP | ASSQGTGDEQY | 0.7409556 |
| VVLAPAL | ASSQGTGDEQY | 0.7568792 |
| TIGHVDHGKTTL | ASSQGTGDEQY | 0.8330086 |
| LISNASDALDK | ASSQGTGDEQY | 0.77894294 |
| LISNASDAL | ASSQGTGDEQY | 0.7932664 |
| ISNASDALDK | ASSQGTGDEQY | 0.82107294 |
| NQAETLVYQTE | ASSQGTGDEQY | 0.8056519 |
| TGGTGGTAGAG | ASSQGTGDEQY | 0.83731806 |
| TGSTAGGAGGA | ASSQGTGDEQY | 0.8073239 |
| DPLAVPVAAGPV | ASSQGTGDEQY | 0.7840272 |
| GGTGGTPGNGGN | ASSQGTGDEQY | 0.78727853 |
| PAGLAGVDPATL | ASSQGTGDEQY | 0.6066179 |
| GNNLVVEAPHAGASV | ASSQGTGDEQY | 0.599016 |
| KVTRIMSASTLEQPA | ASSQGTGDEQY | 0.6160705 |
| LGFALPIALPRRTVV | ASSQGTGDEQY | 0.80231726 |
| TRLGNLTARKLVVLV | ASSQGTGDEQY | 0.4756469 |
| WSQQRRLIYFPSAGP | ASSQGTGDEQY | 0.80346066 |
| GPNGNPDPMAAAVDIR | ASSQGTGDEQY | 0.8327631 |
| TQRCVSVADLRAAASR | ASSQGTGDEQY | 0.77097166 |
| QFVLDVAEHPVEVLTETV | ASSQGTGDEQY | 0.5221799 |
| GGAGGTGGAGGNGGTQAGDG | ASSQGTGDEQY | 0.7652507 |
| PLLATLRAIAVDVDAKPAQV | ASSQGTGDEQY | 0.67123187 |
| APKFRTGGYMIDFWGVGYQVAK | ASSQGTGDEQY | 0.83045006 |
| GAGGAGGAGGAAGGPGGTGGTG | ASSQGTGDEQY | 0.8311324 |
| IGALGLLATDVAPAGSVAFGVA | ASSQGTGDEQY | 0.8461943 |
| AGGAGGIGGNGGAGGAGGVLMG | ASSQGTGDEQY | 0.832771 |
| ADELIGLL | ASSQGTGDEQY | 0.78266567 |
| ADRALKA | ASSQGTGDEQY | 0.82930887 |
| AGAVGQVGGAGGA | ASSQGTGDEQY | 0.7763641 |
| AGGGGGLGGAP | ASSQGTGDEQY | 0.7585005 |
| AGGLLYGNGGAG | ASSQGTGDEQY | 0.8036789 |

| | | |
|---|---|---|
| AGLESIGL | ASSQGTGDEQY | 0.8159983 |
| AGLKTGII | ASSQGTGDEQY | 0.77737594 |
| ASLAALLL | ASSQGTGDEQY | 0.7717819 |
| ASVINISSI | ASSQGTGDEQY | 0.42833778 |
| ATPPGPA | ASSQGTGDEQY | 0.8221252 |
| ATVASALV | ASSQGTGDEQY | 0.7839816 |
| ATVASLIL | ASSQGTGDEQY | 0.42223677 |
| AVNGPVLIRH | ASSQGTGDEQY | 0.771376 |
| DGAAGAVGI | ASSQGTGDEQY | 0.84306204 |
| DPLLHGI | ASSQGTGDEQY | 0.83187795 |
| ELGAALGAL | ASSQGTGDEQY | 0.75721157 |
| ELGRTQAV | ASSQGTGDEQY | 0.75091684 |
| FVAGGYLS | ASSQGTGDEQY | 0.79801846 |
| GASGGAGGAGGW | ASSQGTGDEQY | 0.6334282 |
| GGAGAGGDGGAGGI | ASSQGTGDEQY | 0.8193413 |
| GTGHNGSQPGMGGNG | ASSQGTGDEQY | 0.80060863 |
| GYAAASASAS | ASSQGTGDEQY | 0.8043609 |
| HQVDVAAVQQG | ASSQGTGDEQY | 0.63785434 |
| IGAAIAHT | ASSQGTGDEQY | 0.812386 |
| IGMIEVAQL | ASSQGTGDEQY | 0.7872216 |
| IRLAPEHTSAMD | ASSQGTGDEQY | 0.8159231 |
| IYNVSVL | ASSQGTGDEQY | 0.16871557 |
| KQGFLGM | ASSQGTGDEQY | 0.8203755 |
| LESVITQL | ASSQGTGDEQY | 0.72120005 |
| LGLGALVVV | ASSQGTGDEQY | 0.7093319 |
| LIGNGGNGGSGGI | ASSQGTGDEQY | 0.41866052 |
| LNAINAPTL | ASSQGTGDEQY | 0.7263948 |
| LQLLNTGV | ASSQGTGDEQY | 0.82799935 |
| LQTSIEF | ASSQGTGDEQY | 0.8321142 |
| MSFAVSEL | ASSQGTGDEQY | 0.7462437 |
| NTGALGL | ASSQGTGDEQY | 0.8235483 |
| PPGAGGAGGAGTT | ASSQGTGDEQY | 0.8400222 |
| QLVGLGI | ASSQGTGDEQY | 0.7706827 |
| RFLEQVHQL | ASSQGTGDEQY | 0.32378027 |
| RGVGYVLR | ASSQGTGDEQY | 0.81470746 |
| SATGVFAGVI | ASSQGTGDEQY | 0.75259244 |
| SLPAITLP | ASSQGTGDEQY | 0.8303542 |
| STALAFTV | ASSQGTGDEQY | 0.82729053 |
| SVLTVGLGK | ASSQGTGDEQY | 0.6630278 |
| TALAGAGSGI | ASSQGTGDEQY | 0.8106424 |
| TGLPFTGI | ASSQGTGDEQY | 0.8358059 |
| VAEHVTV | ASSQGTGDEQY | 0.7036635 |
| VDIVLNSL | ASSQGTGDEQY | 0.6090611 |
| VLTGGGAVL | ASSQGTGDEQY | 0.54963404 |
| VNVGLAYGI | ASSQGTGDEQY | 0.442253 |
| VSSAAVGAV | ASSQGTGDEQY | 0.8219488 |
| VTSVYEAGV | ASSQGTGDEQY | 0.76767445 |
| WVALDRGA | ASSQGTGDEQY | 0.68630177 |
| YLVGSVVI | ASSQGTGDEQY | 0.8107353 |
| GPPGTGKTLL | ASSQGTGDEQY | 0.7189273 |
| GVGMTKF | ASSQGTGDEQY | 0.7414237 |

| | | |
|---|---|---|
| IGIGGSDL | ASSQGTGDEQY | 0.75012934 |
| IGTLLAEL | ASSQGTGDEQY | 0.7945119 |
| IYTSGTT | ASSQGTGDEQY | 0.8211394 |
| LSFPAGV | ASSQGTGDEQY | 0.7843308 |
| LYTSGTT | ASSQGTGDEQY | 0.76600754 |
| VGAFPFGF | ASSQGTGDEQY | 0.83954 |
| VGAGTVEF | ASSQGTGDEQY | 0.5667995 |
| VGKSTLLL | ASSQGTGDEQY | 0.67312264 |
| AALAHAGGGIV | ASSQGTGDEQY | 0.822205 |
| AALVAVSLL | ASSQGTGDEQY | 0.7188169 |
| AAVSRAVAAL | ASSQGTGDEQY | 0.8326626 |
| AGGSSGIGGF | ASSQGTGDEQY | 0.8051139 |
| AGLTVANANG | ASSQGTGDEQY | 0.81800395 |
| AGLVGAVL | ASSQGTGDEQY | 0.76738185 |
| ALSQAFTV | ASSQGTGDEQY | 0.84096366 |
| ALTFSVSL | ASSQGTGDEQY | 0.83141255 |
| ALTGAALVAL | ASSQGTGDEQY | 0.76533645 |
| ASFSIGAGVL | ASSQGTGDEQY | 0.81632257 |
| DGSVYKI | ASSQGTGDEQY | 0.4152298 |
| EQLAEIV | ASSQGTGDEQY | 0.82375896 |
| FAVSELTVM | ASSQGTGDEQY | 0.8171731 |
| GAAPLAAGL | ASSQGTGDEQY | 0.7269244 |
| GADGGDGGSGGAGG | ASSQGTGDEQY | 0.7966694 |
| GFGLDTGGAGG | ASSQGTGDEQY | 0.83927643 |
| GLASGGLL | ASSQGTGDEQY | 0.8378345 |
| GRIAGAAVLL | ASSQGTGDEQY | 0.68352085 |
| GSAGSLNTSF | ASSQGTGDEQY | 0.7790873 |
| GTGQIGI | ASSQGTGDEQY | 0.4366333 |
| GTLIALIL | ASSQGTGDEQY | 0.7198861 |
| IAATMGFVVG | ASSQGTGDEQY | 0.43034458 |
| IFAQALL | ASSQGTGDEQY | 0.81921583 |
| IGAAVALGF | ASSQGTGDEQY | 0.757699 |
| IGLTPLAL | ASSQGTGDEQY | 0.82485104 |
| ILGSNTSTL | ASSQGTGDEQY | 0.8128319 |
| IVFSLTVL | ASSQGTGDEQY | 0.84127617 |
| LASKATGF | ASSQGTGDEQY | 0.7380948 |
| LASVALIL | ASSQGTGDEQY | 0.78489697 |
| LFAVGLTL | ASSQGTGDEQY | 0.60385996 |
| LFNGGIGTYI | ASSQGTGDEQY | 0.69680375 |
| LTLGAGII | ASSQGTGDEQY | 0.7430645 |
| LVAAALGDI | ASSQGTGDEQY | 0.7695838 |
| NTAFVQLGI | ASSQGTGDEQY | 0.22020024 |
| QLFSGAEL | ASSQGTGDEQY | 0.84046674 |
| SAATFITL | ASSQGTGDEQY | 0.80358964 |
| SNYGLANLGV | ASSQGTGDEQY | 0.8328705 |
| SYGGAYAVM | ASSQGTGDEQY | 0.71338004 |
| TTGVGGLTLG | ASSQGTGDEQY | 0.6911732 |
| TTQTIYI | ASSQGTGDEQY | 0.76641804 |
| VAAGAGLGAVI | ASSQGTGDEQY | 0.82224226 |
| VAAVVTLL | ASSQGTGDEQY | 0.6972711 |
| VGFLGGLI | ASSQGTGDEQY | 0.36381802 |

| | | |
|---|---|---|
| VGVIPILL | ASSQGTGDEQY | 0.5900677 |
| VIFASSLI | ASSQGTGDEQY | 0.4704721 |
| VKGGDGGAAAG | ASSQGTGDEQY | 0.8424009 |
| VTKTFTNI | ASSQGTGDEQY | 0.83887196 |
| VVAAAILL | ASSQGTGDEQY | 0.8153498 |
| YAALDVEL | ASSQGTGDEQY | 0.8363792 |
| LASVALLL | ASSQGTGDEQY | 0.18156706 |
| LDAVHRV | ASSQGTGDEQY | 0.7201158 |
| QGLSSSIF | ASSQGTGDEQY | 0.8339504 |
| RGTFANIRL | ASSQGTGDEQY | 0.82078356 |
| RQLWWGHRI | ASSQGTGDEQY | 0.7779721 |
| YGPPGTGKTLL | ASSQGTGDEQY | 0.83892083 |
| AALVGAPANVA | ASSQGTGDEQY | 0.7522266 |
| AATFRRTE | ASSQGTGDEQY | 0.7962395 |
| AGALGGGI | ASSQGTGDEQY | 0.7974604 |
| AGGAGGAGGIGAG | ASSQGTGDEQY | 0.82654226 |
| AGSGALGL | ASSQGTGDEQY | 0.8011918 |
| AQIADQI | ASSQGTGDEQY | 0.6563971 |
| ASAVAVTYL | ASSQGTGDEQY | 0.8025646 |
| ASTVLGGTGG | ASSQGTGDEQY | 0.84513986 |
| AVGGEPAK | ASSQGTGDEQY | 0.78665286 |
| CEQGTFGGTC | ASSQGTGDEQY | 0.8325041 |
| DEGKAPAG | ASSQGTGDEQY | 0.81277853 |
| DTVVFPAL | ASSQGTGDEQY | 0.8228308 |
| FEGITMAG | ASSQGTGDEQY | 0.7388347 |
| FPLVVQD | ASSQGTGDEQY | 0.72982013 |
| GAFTTPQ | ASSQGTGDEQY | 0.812922 |
| GFGGFGTAGGVGGA | ASSQGTGDEQY | 0.67651033 |
| GFLAQFL | ASSQGTGDEQY | 0.76154065 |
| GGAGGAGGNAGVA | ASSQGTGDEQY | 0.7665559 |
| GGVGTAGAGGAGGAGG | ASSQGTGDEQY | 0.851092 |
| GNGGGGDAGNAGSGGN | ASSQGTGDEQY | 0.8232254 |
| GTGGAGGAGNDAGS | ASSQGTGDEQY | 0.81749964 |
| GTTNTGWFNS | ASSQGTGDEQY | 0.7392698 |
| IFMGPLFTL | ASSQGTGDEQY | 0.81289726 |
| ILAHAIEAI | ASSQGTGDEQY | 0.8228514 |
| IRSAIREAL | ASSQGTGDEQY | 0.4530486 |
| ITFAGGG | ASSQGTGDEQY | 0.8327286 |
| ITGMSGHQEV | ASSQGTGDEQY | 0.6908339 |
| IVAGLACTL | ASSQGTGDEQY | 0.7973591 |
| IVGGGLAGA | ASSQGTGDEQY | 0.8047663 |
| IVTGVLNI | ASSQGTGDEQY | 0.83919424 |
| LAQQGCI | ASSQGTGDEQY | 0.7858725 |
| LCLHVEN | ASSQGTGDEQY | 0.8084414 |
| LDAISDGS | ASSQGTGDEQY | 0.7149837 |
| LGGFAAML | ASSQGTGDEQY | 0.8264134 |
| LGGKNPMI | ASSQGTGDEQY | 0.83800626 |
| LGPKGVAAIL | ASSQGTGDEQY | 0.7989976 |
| LMVGVSI | ASSQGTGDEQY | 0.84549606 |
| LVGAAFVL | ASSQGTGDEQY | 0.78772616 |
| LVLAVAPL | ASSQGTGDEQY | 0.3066542 |

| | | |
|---|---|---|
| MAAIAVIDAA | ASSQGTGDEQY | 0.790334 |
| NNGINAELQ | ASSQGTGDEQY | 0.83152014 |
| RGEPLVLVHG | ASSQGTGDEQY | 0.83408403 |
| RGGAGGAAGLI | ASSQGTGDEQY | 0.84329903 |
| RSYRMHPAV | ASSQGTGDEQY | 0.80149233 |
| SAAHLLAGQLA | ASSQGTGDEQY | 0.7806463 |
| SAAVGSY | ASSQGTGDEQY | 0.8397129 |
| SPHAVLHRI | ASSQGTGDEQY | 0.8462446 |
| SQIAKVL | ASSQGTGDEQY | 0.82395446 |
| TALGHLVP | ASSQGTGDEQY | 0.72760296 |
| TAPTTSGAA | ASSQGTGDEQY | 0.6077945 |
| TDGDDGNGGSPGT | ASSQGTGDEQY | 0.8189086 |
| TDGLGEN | ASSQGTGDEQY | 0.83747727 |
| TGGPPLPGQA | ASSQGTGDEQY | 0.6063604 |
| TIPAITI | ASSQGTGDEQY | 0.8103272 |
| TIPGFTI | ASSQGTGDEQY | 0.8077141 |
| TSIASGALA | ASSQGTGDEQY | 0.33840054 |
| TVTLPGAAI | ASSQGTGDEQY | 0.8145908 |
| VASGLSAALP | ASSQGTGDEQY | 0.8321913 |
| VGAGVIGI | ASSQGTGDEQY | 0.8136562 |
| VGPKRFGV | ASSQGTGDEQY | 0.71530193 |
| VNVAFPD | ASSQGTGDEQY | 0.83221793 |
| VSMTNSGVS | ASSQGTGDEQY | 0.8472706 |
| VTDVLGDLSV | ASSQGTGDEQY | 0.8435068 |
| VVVVGAGFAGL | ASSQGTGDEQY | 0.41793442 |
| VYTVGTVV | ASSQGTGDEQY | 0.27352515 |
| WTTFSGP | ASSQGTGDEQY | 0.79089457 |
| YDELVEAF | ASSQGTGDEQY | 0.8271539 |
| YGTGGAGGNGG | ASSQGTGDEQY | 0.7596217 |
| YTNVMAV | ASSQGTGDEQY | 0.7695335 |
| FLEQVHQL | ASSQGTGDEQY | 0.8133669 |
| GLSLLAVAV | ASSQGTGDEQY | 0.83797 |
| LEQVHQL | ASSQGTGDEQY | 0.6550128 |
| LNAAGAI | ASSQGTGDEQY | 0.57692397 |
| PAVHASL | ASSQGTGDEQY | 0.80417097 |
| TELEDDA | ASSQGTGDEQY | 0.82574546 |
| TPPGPPN | ASSQGTGDEQY | 0.74903804 |
| TPPPPPPP | ASSQGTGDEQY | 0.79905045 |
| TQSPALI | AWPDRVSNERLF | 0.78111625 |
| EFAAAFPQ | AWPDRVSNERLF | 0.7918805 |
| LLSGNLDVL | AWPDRVSNERLF | 0.8103099 |
| GPANLGNYNI | AWPDRVSNERLF | 0.62227905 |
| TLGPTLP | AWPDRVSNERLF | 0.8352684 |
| VYAYSSCCK | AWPDRVSNERLF | 0.6676757 |
| APAPAPAPNI | AWPDRVSNERLF | 0.8263221 |
| PGGPVPIP | AWPDRVSNERLF | 0.80960023 |
| FLTVALEP | AWPDRVSNERLF | 0.822639 |
| SQYFGANL | AWPDRVSNERLF | 0.7761451 |
| GGVKPAA | AWPDRVSNERLF | 0.7201334 |
| GGTGGAGGAGGW | AWPDRVSNERLF | 0.62048626 |
| GVGGVAGLGGAGG | AWPDRVSNERLF | 0.5234915 |

| | | |
|---|---|---|
| VLLVLATTI | AWPDRVSNERLF | 0.8324673 |
| IGIFAAML | AWPDRVSNERLF | 0.7656008 |
| AVGDLID | AWPDRVSNERLF | 0.8357193 |
| NISPLPL | AWPDRVSNERLF | 0.8253282 |
| VTAVFVAI | AWPDRVSNERLF | 0.70508313 |
| SGDDNDAAL | AWPDRVSNERLF | 0.8219848 |
| AVAALAHS | AWPDRVSNERLF | 0.658547 |
| GGVTVMATP | AWPDRVSNERLF | 0.72852206 |
| AGLAANVRD | AWPDRVSNERLF | 0.8172113 |
| TNRPDLIDPALLRPG | AWPDRVSNERLF | 0.82723653 |
| LANLGVTAL | AWPDRVSNERLF | 0.6664654 |
| TGERIEL | AWPDRVSNERLF | 0.8169954 |
| NGEELAA | AWPDRVSNERLF | 0.81330335 |
| VLSLVWG | AWPDRVSNERLF | 0.575312 |
| IGYALWL | AWPDRVSNERLF | 0.81169945 |
| LSLLLAPL | AWPDRVSNERLF | 0.8354615 |
| TLDEALK | AWPDRVSNERLF | 0.81626093 |
| TDGAGTAEGG | AWPDRVSNERLF | 0.6751576 |
| NAPGGAGGNGG | AWPDRVSNERLF | 0.8221824 |
| VSVFMGMGGNF | AWPDRVSNERLF | 0.7409682 |
| LASYLAAL | AWPDRVSNERLF | 0.81648415 |
| LVGGTPAL | AWPDRVSNERLF | 0.79685295 |
| VRNADVL | AWPDRVSNERLF | 0.7086339 |
| TCSSPFLVSAM | AWPDRVSNERLF | 0.8331032 |
| VHITDAE | AWPDRVSNERLF | 0.35960042 |
| EYAAPSF | AWPDRVSNERLF | 0.7444306 |
| VVDAFHGGL | AWPDRVSNERLF | 0.7876883 |
| ASVVDPQ | AWPDRVSNERLF | 0.8017595 |
| SYNEFGGLS | AWPDRVSNERLF | 0.8276285 |
| DPDGSVY | AWPDRVSNERLF | 0.7237668 |
| MVSTIHATE | AWPDRVSNERLF | 0.79739934 |
| LAAGVADPV | AWPDRVSNERLF | 0.77240974 |
| TKEQIAA | AWPDRVSNERLF | 0.7241762 |
| SLASPAAL | AWPDRVSNERLF | 0.8041774 |
| IGLVFYGV | AWPDRVSNERLF | 0.7498071 |
| FLLDASNL | AWPDRVSNERLF | 0.59743553 |
| YQVGNFP | AWPDRVSNERLF | 0.82291263 |
| ESLLSQST | AWPDRVSNERLF | 0.70873106 |
| LFTVGFAAVG | AWPDRVSNERLF | 0.8051113 |
| ATPPTPT | AWPDRVSNERLF | 0.836441 |
| TDLATLG | AWPDRVSNERLF | 0.6240226 |
| SAGGTGR | AWPDRVSNERLF | 0.8229176 |
| GFSNSGS | AWPDRVSNERLF | 0.81626934 |
| GGNGGSGGTSV | AWPDRVSNERLF | 0.8201629 |
| GTSVATGGAGN | AWPDRVSNERLF | 0.77097833 |
| NGGNGGSGGTS | AWPDRVSNERLF | 0.70933586 |
| AVVKFPH | AWPDRVSNERLF | 0.8375166 |
| TPPPPPP | AWPDRVSNERLF | 0.821346 |
| AAPAAAAAPA | AWPDRVSNERLF | 0.77911294 |
| IALNEDL | AWPDRVSNERLF | 0.8198112 |
| IVNVSSM | AWPDRVSNERLF | 0.81242865 |

| | | |
|---|---|---|
| TGIPPAPR | AWPDRVSNERLF | 0.82764745 |
| FDLGGGTF | AWPDRVSNERLF | 0.77304524 |
| FDLGGGTFDVS | AWPDRVSNERLF | 0.62820697 |
| NEPTAAA | AWPDRVSNERLF | 0.7927263 |
| NEPTAAALA | AWPDRVSNERLF | 0.7724056 |
| GSIINISSI | AWPDRVSNERLF | 0.6897551 |
| VAPGNAGTA | AWPDRVSNERLF | 0.3668105 |
| TGGGASLE | AWPDRVSNERLF | 0.82921684 |
| VAVYNHYKRI | AWPDRVSNERLF | 0.8487039 |
| TDPAPATT | AWPDRVSNERLF | 0.55806804 |
| TNRPDILDPALLRPG | AWPDRVSNERLF | 0.8446173 |
| LTTTTTS | AWPDRVSNERLF | 0.6399318 |
| PTGSGKTL | AWPDRVSNERLF | 0.8284323 |
| TGSGKTL | AWPDRVSNERLF | 0.8320832 |
| TIGHVDHGK | AWPDRVSNERLF | 0.6537231 |
| TIGHVDHGKTT | AWPDRVSNERLF | 0.80855656 |
| GHVDHGKTT | AWPDRVSNERLF | 0.7056421 |
| AQFEHTLL | AWPDRVSNERLF | 0.7093689 |
| IVNQDEL | AWPDRVSNERLF | 0.8358207 |
| TNGSQFF | AWPDRVSNERLF | 0.6936163 |
| TGKGIVE | AWPDRVSNERLF | 0.8095343 |
| LISNASDA | AWPDRVSNERLF | 0.79852444 |
| AELRTLGELTKIAKAHGAQ | AWPDRVSNERLF | 0.7349347 |
| APNAVAAAEPSW | AWPDRVSNERLF | 0.8214204 |
| APQVAGAATAVL | AWPDRVSNERLF | 0.8206376 |
| ARRDPAAAAAILPTDA | AWPDRVSNERLF | 0.7258397 |
| DAAARDVLAEAGLAEYF | AWPDRVSNERLF | 0.8449781 |
| DDQIVLSAKANLAKVEQ | AWPDRVSNERLF | 0.7144673 |
| DELLPDTGEDSDEDGH | AWPDRVSNERLF | 0.8416575 |
| ELPEAPANRATVPQD | AWPDRVSNERLF | 0.818377 |
| FGQSKIERVTGIG | AWPDRVSNERLF | 0.8100369 |
| GAGGTGSTAGGAGGAGGAGGLY | AWPDRVSNERLF | 0.7784895 |
| GHGGTGAAGGNGGTGGAGGSGI | AWPDRVSNERLF | 0.8329763 |
| GNGGNGGAGGEGGPGGAGGAGA | AWPDRVSNERLF | 0.7747855 |
| GSTAGNGGNGGNATNGTIAGQP | AWPDRVSNERLF | 0.80724984 |
| GTGGDGGNGGNGGTGG | AWPDRVSNERLF | 0.8395295 |
| GVRRPAGIPADAPV | AWPDRVSNERLF | 0.6199794 |
| GYRQVTDAHLLTLARR | AWPDRVSNERLF | 0.79679084 |
| IAKMGQDGHDRG | AWPDRVSNERLF | 0.7563279 |
| KAEPSIEIQYT | AWPDRVSNERLF | 0.71291286 |
| KPVRIRMAADEDTVAVT | AWPDRVSNERLF | 0.63427675 |
| MPPRIAGMRLLVIKPEP | AWPDRVSNERLF | 0.4243445 |
| MKFYAVALFRSRGVPPTR | AWPDRVSNERLF | 0.7024746 |
| NAGGFGILGAVAHSP | AWPDRVSNERLF | 0.6071595 |
| NEGREITLAMAEM | AWPDRVSNERLF | 0.7513231 |
| NELVEWSHPLGHRWRWE | AWPDRVSNERLF | 0.7659107 |
| NLGAGGTGGNGGNPG | AWPDRVSNERLF | 0.5962957 |
| NTEWPKWETVMAADPVD | AWPDRVSNERLF | 0.7960184 |
| PASAAPAETQP | AWPDRVSNERLF | 0.78145593 |
| QQSSDGKNGVRAAEITGE | AWPDRVSNERLF | 0.44738036 |
| RAPDAVTTSWWKE | AWPDRVSNERLF | 0.68907857 |

| | | |
|---|---|---|
| SGFGNVGAHTSGWFNQ | AWPDRVSNERLF | 0.66323197 |
| SGVAGAVGQAASVGGLKV | AWPDRVSNERLF | 0.6932486 |
| SNGGAGTGSAGGNGGT | AWPDRVSNERLF | 0.46918717 |
| TGGAGGEGGAAG | AWPDRVSNERLF | 0.7687839 |
| TGGDGGAAGAGGAPGHG | AWPDRVSNERLF | 0.64741784 |
| TMPPLLAAIANAGRHGV | AWPDRVSNERLF | 0.21043359 |
| TSSTAGNSGGA | AWPDRVSNERLF | 0.7223036 |
| VPATVPVGILC | AWPDRVSNERLF | 0.69876957 |
| VQRPDSGGVVLTGRIS | AWPDRVSNERLF | 0.69767797 |
| WTPNPVPATKGVPAGLVQAT | AWPDRVSNERLF | 0.78028756 |
| YALKRALAKFSDRFDVVII | AWPDRVSNERLF | 0.43773043 |
| YLIGSVIV | AWPDRVSNERLF | 0.6355555 |
| YGQLVAVID | AWPDRVSNERLF | 0.7772442 |
| VTVGGMTLPA | AWPDRVSNERLF | 0.79201657 |
| VLALPAV | AWPDRVSNERLF | 0.58290255 |
| VIPPTLNYET | AWPDRVSNERLF | 0.74643755 |
| VGMGGMHPSSG | AWPDRVSNERLF | 0.78599465 |
| VGLAGGAATA | AWPDRVSNERLF | 0.7608434 |
| VGGAGGAGGSGP | AWPDRVSNERLF | 0.7745653 |
| VFAAFMV | AWPDRVSNERLF | 0.7906992 |
| TLYDKLR | AWPDRVSNERLF | 0.62112963 |
| TLAVFTVL | AWPDRVSNERLF | 0.78319436 |
| TGSVQPARV | AWPDRVSNERLF | 0.7951791 |
| TGGGAGSGGT | AWPDRVSNERLF | 0.66981864 |
| SHLETSL | AWPDRVSNERLF | 0.7310208 |
| SFTTGMT | AWPDRVSNERLF | 0.57827914 |
| QVEVAKRMGVRQPTVS | AWPDRVSNERLF | 0.69787484 |
| QGSGGPALAM | AWPDRVSNERLF | 0.29642856 |
| MLAPSNP | AWPDRVSNERLF | 0.75277996 |
| LVNNAAI | AWPDRVSNERLF | 0.78601766 |
| LTGEINTLA | AWPDRVSNERLF | 0.6834496 |
| LSIFGPV | AWPDRVSNERLF | 0.60665286 |
| LLALLANL | AWPDRVSNERLF | 0.57039124 |
| LISNASDALDKLR | AWPDRVSNERLF | 0.7021437 |
| LINEAVP | AWPDRVSNERLF | 0.68679035 |
| LGYAFSGL | AWPDRVSNERLF | 0.30896825 |
| LDFVAVAL | AWPDRVSNERLF | 0.6727073 |
| LAQPGAE | AWPDRVSNERLF | 0.78589725 |
| ISNLGAVTGF | AWPDRVSNERLF | 0.7789139 |
| HQEDPGRI | AWPDRVSNERLF | 0.3968055 |
| GTPGEPTP | AWPDRVSNERLF | 0.789762 |
| GFFNSGS | AWPDRVSNERLF | 0.7570006 |
| GAGGGGLTAGAN | AWPDRVSNERLF | 0.7951429 |
| FVDALVNM | AWPDRVSNERLF | 0.7085856 |
| FLTVAPEL | AWPDRVSNERLF | 0.63489425 |
| FETLTAL | AWPDRVSNERLF | 0.7520945 |
| DVINPIEL | AWPDRVSNERLF | 0.69177604 |
| APVSGAPV | AWPDRVSNERLF | 0.6253128 |
| ALVGGLAW | AWPDRVSNERLF | 0.7746332 |
| ALAAAAATAI | AWPDRVSNERLF | 0.7235688 |
| AFGLGLLL | AWPDRVSNERLF | 0.6319116 |

| | | |
|---|---|---|
| AFFAAASL | AWPDRVSNERLF | 0.63486993 |
| AAAALPAFSPP | AWPDRVSNERLF | 0.7846498 |
| VVLAPAL | AWPDRVSNERLF | 0.56653744 |
| TIGHVDHGKTTL | AWPDRVSNERLF | 0.58019876 |
| LISNASDALDK | AWPDRVSNERLF | 0.5151389 |
| LISNASDAL | AWPDRVSNERLF | 0.7340586 |
| ISNASDALDK | AWPDRVSNERLF | 0.78233635 |
| NQAETLVYQTE | AWPDRVSNERLF | 0.74787056 |
| TGGTGGTAGAG | AWPDRVSNERLF | 0.6746315 |
| TGSTAGGAGGA | AWPDRVSNERLF | 0.7578168 |
| DPLAVPVAAGPV | AWPDRVSNERLF | 0.7431967 |
| GGTGGTPGNGGN | AWPDRVSNERLF | 0.7339119 |
| PAGLAGVDPATL | AWPDRVSNERLF | 0.772619 |
| GNNLVVEAPHAGASV | AWPDRVSNERLF | 0.6321284 |
| KVTRIMSASTLEQPA | AWPDRVSNERLF | 0.78240025 |
| LGFALPIALPRRTVV | AWPDRVSNERLF | 0.7488979 |
| TRLGNLTARKLVVLV | AWPDRVSNERLF | 0.7637999 |
| WSQQRRLIYFPSAGP | AWPDRVSNERLF | 0.7728797 |
| GPNGNPDPMAAAVDIR | AWPDRVSNERLF | 0.7737285 |
| TQRCVSVADLRAAASR | AWPDRVSNERLF | 0.72169673 |
| QFVLDVAEHPVEVLTETV | AWPDRVSNERLF | 0.30417475 |
| GGAGGTGGAGGNGGTQAGDG | AWPDRVSNERLF | 0.6798489 |
| PLLATLRAIAVDVDAKPAQV | AWPDRVSNERLF | 0.7574066 |
| APKFRTGGYMIDFWGVGYQVAK | AWPDRVSNERLF | 0.7447935 |
| GAGGAGGAGGAAGGPGGTGGTG | AWPDRVSNERLF | 0.7630007 |
| IGALGLLATDVAPAGSVAFGVA | AWPDRVSNERLF | 0.7363579 |
| AGGAGGIGGNGGAGGAGGVLMG | AWPDRVSNERLF | 0.6431631 |
| ADELIGLL | AWPDRVSNERLF | 0.77862704 |
| ADRALKA | AWPDRVSNERLF | 0.77544355 |
| AGAVGQVGGAGGA | AWPDRVSNERLF | 0.39203462 |
| AGGGGGLGGAP | AWPDRVSNERLF | 0.6893373 |
| AGGLLYGNGGAG | AWPDRVSNERLF | 0.79868877 |
| AGLESIGL | AWPDRVSNERLF | 0.7253628 |
| AGLKTGII | AWPDRVSNERLF | 0.69088423 |
| ASLAALLL | AWPDRVSNERLF | 0.6343857 |
| ASVINISSI | AWPDRVSNERLF | 0.78501785 |
| ATPPGPA | AWPDRVSNERLF | 0.61428124 |
| ATVASALV | AWPDRVSNERLF | 0.7995306 |
| ATVASLIL | AWPDRVSNERLF | 0.22728552 |
| AVNGPVLIRH | AWPDRVSNERLF | 0.56822157 |
| DGAAGAVGI | AWPDRVSNERLF | 0.5965369 |
| DPLLHGI | AWPDRVSNERLF | 0.3879946 |
| ELGAALGAL | AWPDRVSNERLF | 0.7837733 |
| ELGRTQAV | AWPDRVSNERLF | 0.62899953 |
| FVAGGYLS | AWPDRVSNERLF | 0.76872635 |
| GASGGAGGAGGW | AWPDRVSNERLF | 0.73343205 |
| GGAGAGGDGGAGGI | AWPDRVSNERLF | 0.7794251 |
| GTGHNGSQPGMGGNG | AWPDRVSNERLF | 0.7373677 |
| GYAAASASAS | AWPDRVSNERLF | 0.72077495 |
| HQVDVAAVQQG | AWPDRVSNERLF | 0.54296905 |
| IGAAIAHT | AWPDRVSNERLF | 0.7731004 |

| | | |
|---|---|---|
| IGMIEVAQL | AWPDRVSNERLF | 0.7883401 |
| IRLAPEHTSAMD | AWPDRVSNERLF | 0.74493873 |
| IYNVSVL | AWPDRVSNERLF | 0.530617 |
| KQGFLGM | AWPDRVSNERLF | 0.6780524 |
| LESVITQL | AWPDRVSNERLF | 0.7305481 |
| LGLGALVVV | AWPDRVSNERLF | 0.28016186 |
| LIGNGGNGGSGGI | AWPDRVSNERLF | 0.257333 |
| LNAINAPTL | AWPDRVSNERLF | 0.6866139 |
| LQLLNTGV | AWPDRVSNERLF | 0.6116693 |
| LQTSIEF | AWPDRVSNERLF | 0.78679043 |
| MSFAVSEL | AWPDRVSNERLF | 0.778543 |
| NTGALGL | AWPDRVSNERLF | 0.6971435 |
| PPGAGGAGGAGTT | AWPDRVSNERLF | 0.77692497 |
| QLVGLGI | AWPDRVSNERLF | 0.7536726 |
| RFLEQVHQL | AWPDRVSNERLF | 0.17925814 |
| RGVGYVLR | AWPDRVSNERLF | 0.6387797 |
| SATGVFAGVI | AWPDRVSNERLF | 0.4672287 |
| SLPAITLP | AWPDRVSNERLF | 0.78024656 |
| STALAFTV | AWPDRVSNERLF | 0.6219974 |
| SVLTVGLGK | AWPDRVSNERLF | 0.609692 |
| TALAGAGSGI | AWPDRVSNERLF | 0.71803594 |
| TGLPFTGI | AWPDRVSNERLF | 0.7073463 |
| VAEHVTV | AWPDRVSNERLF | 0.70461696 |
| VDIVLNSL | AWPDRVSNERLF | 0.5442622 |
| VLTGGGAVL | AWPDRVSNERLF | 0.21607277 |
| VNVGLAYGI | AWPDRVSNERLF | 0.7777525 |
| VSSAAVGAV | AWPDRVSNERLF | 0.37127563 |
| VTSVYEAGV | AWPDRVSNERLF | 0.6086278 |
| WVALDRGA | AWPDRVSNERLF | 0.27812886 |
| YLVGSVVI | AWPDRVSNERLF | 0.6396096 |
| GPPGTGKTLL | AWPDRVSNERLF | 0.38206762 |
| GVGMTKF | AWPDRVSNERLF | 0.7719101 |
| IGIGGSDL | AWPDRVSNERLF | 0.6697216 |
| IGTLLAEL | AWPDRVSNERLF | 0.67804515 |
| IYTSGTT | AWPDRVSNERLF | 0.7970464 |
| LSFPAGV | AWPDRVSNERLF | 0.70473343 |
| LYTSGTT | AWPDRVSNERLF | 0.68133473 |
| VGAFPFGF | AWPDRVSNERLF | 0.641482 |
| VGAGTVEF | AWPDRVSNERLF | 0.55399084 |
| VGKSTLLL | AWPDRVSNERLF | 0.62232226 |
| AALAHAGGGIV | AWPDRVSNERLF | 0.6276699 |
| AALVAVSLL | AWPDRVSNERLF | 0.79855233 |
| AAVSRAVAAL | AWPDRVSNERLF | 0.58887607 |
| AGGSSGIGGF | AWPDRVSNERLF | 0.46960753 |
| AGLTVANANG | AWPDRVSNERLF | 0.6631942 |
| AGLVGAVL | AWPDRVSNERLF | 0.79200304 |
| ALSQAFTV | AWPDRVSNERLF | 0.80501366 |
| ALTFSVSL | AWPDRVSNERLF | 0.6467826 |
| ALTGAALVAL | AWPDRVSNERLF | 0.69606006 |
| ASFSIGAGVL | AWPDRVSNERLF | 0.6725868 |
| DGSVYKI | AWPDRVSNERLF | 0.72394395 |

| | | |
|---|---|---|
| EQLAEIV | AWPDRVSNERLF | 0.66457367 |
| FAVSELTVM | AWPDRVSNERLF | 0.4560832 |
| GAAPLAAGL | AWPDRVSNERLF | 0.7640689 |
| GADGGDGGSGGAGG | AWPDRVSNERLF | 0.7907164 |
| GFGLDTGGAGG | AWPDRVSNERLF | 0.6690163 |
| GLASGGLL | AWPDRVSNERLF | 0.52881455 |
| GRIAGAAVLL | AWPDRVSNERLF | 0.7098516 |
| GSAGSLNTSF | AWPDRVSNERLF | 0.6050106 |
| GTGQIGI | AWPDRVSNERLF | 0.6697984 |
| GTLIALIL | AWPDRVSNERLF | 0.75161016 |
| IAATMGFVVG | AWPDRVSNERLF | 0.40603727 |
| IFAQALL | AWPDRVSNERLF | 0.6711416 |
| IGAAVALGF | AWPDRVSNERLF | 0.80655956 |
| IGLTPLAL | AWPDRVSNERLF | 0.6703519 |
| ILGSNTSTL | AWPDRVSNERLF | 0.73920286 |
| IVFSLTVL | AWPDRVSNERLF | 0.78865206 |
| LASKATGF | AWPDRVSNERLF | 0.7652496 |
| LASVALIL | AWPDRVSNERLF | 0.7896849 |
| LFAVGLTL | AWPDRVSNERLF | 0.7596202 |
| LFNGGIGTYI | AWPDRVSNERLF | 0.66900504 |
| LTLGAGII | AWPDRVSNERLF | 0.7419019 |
| LVAAALGDI | AWPDRVSNERLF | 0.72900474 |
| NTAFVQLGI | AWPDRVSNERLF | 0.73004866 |
| QLFSGAEL | AWPDRVSNERLF | 0.69041795 |
| SAATFITL | AWPDRVSNERLF | 0.5660642 |
| SNYGLANLGV | AWPDRVSNERLF | 0.781366 |
| SYGGAYAVM | AWPDRVSNERLF | 0.7245756 |
| TTGVGGLTLG | AWPDRVSNERLF | 0.7559798 |
| TTQTIYI | AWPDRVSNERLF | 0.4912167 |
| VAAGAGLGAVI | AWPDRVSNERLF | 0.5601479 |
| VAAVVTLL | AWPDRVSNERLF | 0.7728014 |
| VGFLGGLI | AWPDRVSNERLF | 0.2076717 |
| VGVIPILL | AWPDRVSNERLF | 0.25508532 |
| VIFASSLI | AWPDRVSNERLF | 0.7282816 |
| VKGGDGGAAAG | AWPDRVSNERLF | 0.79325217 |
| VTKTFTNI | AWPDRVSNERLF | 0.22507451 |
| VVAAAILL | AWPDRVSNERLF | 0.57016516 |
| YAALDVEL | AWPDRVSNERLF | 0.45800894 |
| LASVALLL | AWPDRVSNERLF | 0.75702775 |
| LDAVHRV | AWPDRVSNERLF | 0.7393367 |
| QGLSSSIF | AWPDRVSNERLF | 0.3721827 |
| RGTFANIRL | AWPDRVSNERLF | 0.7555717 |
| RQLWWGHRI | AWPDRVSNERLF | 0.4226711 |
| YGPPGTGKTLL | AWPDRVSNERLF | 0.445221 |
| AALVGAPANVA | AWPDRVSNERLF | 0.77911144 |
| AATFRRTE | AWPDRVSNERLF | 0.78114223 |
| AGALGGGI | AWPDRVSNERLF | 0.5394457 |
| AGGAGGAGGIGAG | AWPDRVSNERLF | 0.7745325 |
| AGSGALGL | AWPDRVSNERLF | 0.6081354 |
| AQIADQI | AWPDRVSNERLF | 0.8064353 |
| ASAVAVTYL | AWPDRVSNERLF | 0.67298424 |

| | | |
|---|---|---|
| ASTVLGGTGG | AWPDRVSNERLF | 0.38073277 |
| AVGGEPAK | AWPDRVSNERLF | 0.7731948 |
| CEQGTFGGTC | AWPDRVSNERLF | 0.7405559 |
| DEGKAPAG | AWPDRVSNERLF | 0.7401072 |
| DTVVFPAL | AWPDRVSNERLF | 0.76552105 |
| FEGITMAG | AWPDRVSNERLF | 0.56887645 |
| FPLVVQD | AWPDRVSNERLF | 0.6903717 |
| GAFTTPQ | AWPDRVSNERLF | 0.7811246 |
| GFGGFGTAGGVGGA | AWPDRVSNERLF | 0.7785404 |
| GFLAQFL | AWPDRVSNERLF | 0.5009043 |
| GGAGGAGGNAGVA | AWPDRVSNERLF | 0.77745795 |
| GGVGTAGAGGAGGAGG | AWPDRVSNERLF | 0.7111472 |
| GNGGGGDAGNAGSGGN | AWPDRVSNERLF | 0.6244365 |
| GTGGAGGAGNDAGS | AWPDRVSNERLF | 0.8009399 |
| GTTNTGWFNS | AWPDRVSNERLF | 0.6900313 |
| IFMGPLFTL | AWPDRVSNERLF | 0.7789071 |
| ILAHAIEAI | AWPDRVSNERLF | 0.7683973 |
| IRSAIREAL | AWPDRVSNERLF | 0.7786794 |
| ITFAGGG | AWPDRVSNERLF | 0.4867851 |
| ITGMSGHQEV | AWPDRVSNERLF | 0.75288504 |
| IVAGLACTL | AWPDRVSNERLF | 0.73996484 |
| IVGGGLAGA | AWPDRVSNERLF | 0.76173127 |
| IVTGVLNI | AWPDRVSNERLF | 0.70572096 |
| LAQQGCI | AWPDRVSNERLF | 0.53776264 |
| LCLHVEN | AWPDRVSNERLF | 0.74036306 |
| LDAISDGS | AWPDRVSNERLF | 0.79162157 |
| LGGFAAML | AWPDRVSNERLF | 0.7313897 |
| LGGKNPMI | AWPDRVSNERLF | 0.74596804 |
| LGPKGVAAIL | AWPDRVSNERLF | 0.7943706 |
| LMVGVSI | AWPDRVSNERLF | 0.58712536 |
| LVGAAFVL | AWPDRVSNERLF | 0.69149834 |
| LVLAVAPL | AWPDRVSNERLF | 0.77520716 |
| MAAIAVIDAA | AWPDRVSNERLF | 0.78285635 |
| NNGINAELQ | AWPDRVSNERLF | 0.65345657 |
| RGEPLVLVHG | AWPDRVSNERLF | 0.7900266 |
| RGGAGGAAGLI | AWPDRVSNERLF | 0.64498216 |
| RSYRMHPAV | AWPDRVSNERLF | 0.76685894 |
| SAAHLLAGQLA | AWPDRVSNERLF | 0.6686331 |
| SAAVGSY | AWPDRVSNERLF | 0.7553756 |
| SPHAVLHRI | AWPDRVSNERLF | 0.63972485 |
| SQIAKVL | AWPDRVSNERLF | 0.7787285 |
| TALGHLVP | AWPDRVSNERLF | 0.78217685 |
| TAPTTSGAA | AWPDRVSNERLF | 0.7806537 |
| TDGDDGNGGSPGT | AWPDRVSNERLF | 0.2579703 |
| TDGLGEN | AWPDRVSNERLF | 0.78126264 |
| TGGPPLPGQA | AWPDRVSNERLF | 0.7669037 |
| TIPAITI | AWPDRVSNERLF | 0.7990853 |
| TIPGFTI | AWPDRVSNERLF | 0.7335489 |
| TSIASGALA | AWPDRVSNERLF | 0.7483198 |
| TVTLPGAAI | AWPDRVSNERLF | 0.71924317 |
| VASGLSAALP | AWPDRVSNERLF | 0.6632714 |

| | | |
|---|---|---|
| VGAGVIGI | AWPDRVSNERLF | 0.5740615 |
| VGPKRFGV | AWPDRVSNERLF | 0.6945491 |
| VNVAFPD | AWPDRVSNERLF | 0.6861533 |
| VSMTNSGVS | AWPDRVSNERLF | 0.61691225 |
| VTDVLGDLSV | AWPDRVSNERLF | 0.76816374 |
| VVVVGAGFAGL | AWPDRVSNERLF | 0.7872717 |
| VYTVGTVV | AWPDRVSNERLF | 0.7893421 |
| WTTFSGP | AWPDRVSNERLF | 0.7866701 |
| YDELVEAF | AWPDRVSNERLF | 0.7896632 |
| YGTGGAGGNGG | AWPDRVSNERLF | 0.76396775 |
| YTNVMAV | AWPDRVSNERLF | 0.67965764 |
| FLEQVHQL | AWPDRVSNERLF | 0.7037777 |
| GLSLLAVAV | AWPDRVSNERLF | 0.55148447 |
| LEQVHQL | AWPDRVSNERLF | 0.6625719 |
| LNAAGAI | AWPDRVSNERLF | 0.6733494 |
| PAVHASL | AWPDRVSNERLF | 0.7412505 |
| TELEDDA | AWPDRVSNERLF | 0.19068061 |
| TPPGPPN | AWPDRVSNERLF | 0.7121544 |
| TPPPPPPP | AWPDRVSNERLF | 0.7965636 |
| TQSPALI | ASGDGSYYAEQF | 0.63887656 |
| EFAAAFPQ | ASGDGSYYAEQF | 0.74850345 |
| LLSGNLDVL | ASGDGSYYAEQF | 0.7097745 |
| GPANLGNYNI | ASGDGSYYAEQF | 0.6583048 |
| TLGPTLP | ASGDGSYYAEQF | 0.77453804 |
| VYAYSSCCK | ASGDGSYYAEQF | 0.56395626 |
| APAPAPAPNI | ASGDGSYYAEQF | 0.7529799 |
| PGGPVPIP | ASGDGSYYAEQF | 0.79596055 |
| FLTVALEP | ASGDGSYYAEQF | 0.74024534 |
| SQYFGANL | ASGDGSYYAEQF | 0.7418165 |
| GGVKPAA | ASGDGSYYAEQF | 0.7062716 |
| GGTGGAGGAGGW | ASGDGSYYAEQF | 0.42892998 |
| GVGGVAGLGGAGG | ASGDGSYYAEQF | 0.7509093 |
| VLLVLATTI | ASGDGSYYAEQF | 0.3301449 |
| IGIFAAML | ASGDGSYYAEQF | 0.6984752 |
| AVGDLID | ASGDGSYYAEQF | 0.55953443 |
| NISPLPL | ASGDGSYYAEQF | 0.74975157 |
| VTAVFVAI | ASGDGSYYAEQF | 0.74281543 |
| SGDDNDAAL | ASGDGSYYAEQF | 0.7625973 |
| AVAALAHS | ASGDGSYYAEQF | 0.57293475 |
| GGVTVMATP | ASGDGSYYAEQF | 0.7061305 |
| AGLAANVRD | ASGDGSYYAEQF | 0.6822163 |
| TNRPDLIDPALLRPG | ASGDGSYYAEQF | 0.5466666 |
| LANLGVTAL | ASGDGSYYAEQF | 0.7792135 |
| TGERIEL | ASGDGSYYAEQF | 0.6877818 |
| NGEELAA | ASGDGSYYAEQF | 0.6917289 |
| VLSLVWG | ASGDGSYYAEQF | 0.78824526 |
| IGYALWL | ASGDGSYYAEQF | 0.7497702 |
| LSLLLAPL | ASGDGSYYAEQF | 0.42694727 |
| TLDEALK | ASGDGSYYAEQF | 0.71865475 |
| TDGAGTAEGG | ASGDGSYYAEQF | 0.70846444 |
| NAPGGAGGNGG | ASGDGSYYAEQF | 0.70065176 |

| | | |
|---|---|---|
| VSVFMGMGGNF | ASGDGSYYAEQF | 0.22032058 |
| LASYLAAL | ASGDGSYYAEQF | 0.7575333 |
| LVGGTPAL | ASGDGSYYAEQF | 0.6268258 |
| VRNADVL | ASGDGSYYAEQF | 0.61640775 |
| TCSSPFLVSAM | ASGDGSYYAEQF | 0.6367413 |
| VHITDAE | ASGDGSYYAEQF | 0.7720066 |
| EYAAPSF | ASGDGSYYAEQF | 0.6894572 |
| VVDAFHGGL | ASGDGSYYAEQF | 0.7147991 |
| ASVVDPQ | ASGDGSYYAEQF | 0.7671849 |
| SYNEFGGLS | ASGDGSYYAEQF | 0.7827636 |
| DPDGSVY | ASGDGSYYAEQF | 0.7950025 |
| MVSTIHATE | ASGDGSYYAEQF | 0.75693 |
| LAAGVADPV | ASGDGSYYAEQF | 0.70539194 |
| TKEQIAA | ASGDGSYYAEQF | 0.7457655 |
| SLASPAAL | ASGDGSYYAEQF | 0.7148086 |
| IGLVFYGV | ASGDGSYYAEQF | 0.3501136 |
| FLLDASNL | ASGDGSYYAEQF | 0.33112934 |
| YQVGNFP | ASGDGSYYAEQF | 0.74664533 |
| ESLLSQST | ASGDGSYYAEQF | 0.3075535 |
| LFTVGFAAVG | ASGDGSYYAEQF | 0.72507507 |
| ATPPTPT | ASGDGSYYAEQF | 0.76273394 |
| TDLATLG | ASGDGSYYAEQF | 0.71279293 |
| SAGGTGR | ASGDGSYYAEQF | 0.50597644 |
| GFSNSGS | ASGDGSYYAEQF | 0.69636494 |
| GGNGGSGGTSV | ASGDGSYYAEQF | 0.6547015 |
| GTSVATGGAGN | ASGDGSYYAEQF | 0.77284914 |
| NGGNGGSGGTS | ASGDGSYYAEQF | 0.7790829 |
| AVVKFPH | ASGDGSYYAEQF | 0.788079 |
| TPPPPPP | ASGDGSYYAEQF | 0.7859904 |
| AAPAAAAAPA | ASGDGSYYAEQF | 0.69292915 |
| IALNEDL | ASGDGSYYAEQF | 0.7613535 |
| IVNVSSM | ASGDGSYYAEQF | 0.6367979 |
| TGIPPAPR | ASGDGSYYAEQF | 0.70618665 |
| FDLGGGTF | ASGDGSYYAEQF | 0.6982225 |
| FDLGGGTFDVS | ASGDGSYYAEQF | 0.7441702 |
| NEPTAAA | ASGDGSYYAEQF | 0.71528506 |
| NEPTAAALA | ASGDGSYYAEQF | 0.6589248 |
| GSIINISSI | ASGDGSYYAEQF | 0.37326932 |
| VAPGNAGTA | ASGDGSYYAEQF | 0.7473625 |
| TGGGASLE | ASGDGSYYAEQF | 0.75452435 |
| VAVYNHYKRI | ASGDGSYYAEQF | 0.33275706 |
| TDPAPATT | ASGDGSYYAEQF | 0.64804196 |
| TNRPDILDPALLRPG | ASGDGSYYAEQF | 0.79805005 |
| LTTTTTS | ASGDGSYYAEQF | 0.79463273 |
| PTGSGKTL | ASGDGSYYAEQF | 0.591311 |
| TGSGKTL | ASGDGSYYAEQF | 0.6238327 |
| TIGHVDHGK | ASGDGSYYAEQF | 0.71147394 |
| TIGHVDHGKTT | ASGDGSYYAEQF | 0.32667994 |
| GHVDHGKTT | ASGDGSYYAEQF | 0.7136847 |
| AQFEHTLL | ASGDGSYYAEQF | 0.7154986 |
| IVNQDEL | ASGDGSYYAEQF | 0.731986 |

| | | |
|---|---|---|
| TNGSQFF | ASGDGSYYAEQF | 0.5887644 |
| TGKGIVE | ASGDGSYYAEQF | 0.73115945 |
| LISNASDA | ASGDGSYYAEQF | 0.72240406 |
| AELRTLGELTKIAKAHGAQ | ASGDGSYYAEQF | 0.74939096 |
| APNAVAAAEPSW | ASGDGSYYAEQF | 0.20759659 |
| APQVAGAATAVL | ASGDGSYYAEQF | 0.76598763 |
| ARRDPAAAAAILPTDA | ASGDGSYYAEQF | 0.5852904 |
| DAAARDVLAEAGLAEYF | ASGDGSYYAEQF | 0.61108625 |
| DDQIVLSAKANLAKVEQ | ASGDGSYYAEQF | 0.41313097 |
| DELLPDTGEDSDEDGH | ASGDGSYYAEQF | 0.7146015 |
| ELPEAPANRATVPQD | ASGDGSYYAEQF | 0.7842947 |
| FGQSKIERVTGIG | ASGDGSYYAEQF | 0.7800556 |
| GAGGTGSTAGGAGGAGGAGGLY | ASGDGSYYAEQF | 0.5035893 |
| GHGGTGAAGGNGGTGGAGGSGI | ASGDGSYYAEQF | 0.77862024 |
| GNGGNGGAGGEGGPGGAGGAGA | ASGDGSYYAEQF | 0.8054451 |
| GSTAGNGGNGGNATNGTIAGQP | ASGDGSYYAEQF | 0.67751527 |
| GTGGDGGNGGNGGTGG | ASGDGSYYAEQF | 0.40950578 |
| GVRRPAGIPADAPV | ASGDGSYYAEQF | 0.76272833 |
| GYRQVTDAHLLTLARR | ASGDGSYYAEQF | 0.7041826 |
| IAKMGQDGHDRG | ASGDGSYYAEQF | 0.768891 |
| KAEPSIEIQYT | ASGDGSYYAEQF | 0.76394624 |
| KPVRIRMAADEDTVAVT | ASGDGSYYAEQF | 0.51955783 |
| MPPRIAGMRLLVIKPEP | ASGDGSYYAEQF | 0.765148 |
| MKFYAVALFRSRGVPPTR | ASGDGSYYAEQF | 0.78335106 |
| NAGGFGILGAVAHSP | ASGDGSYYAEQF | 0.6289711 |
| NEGREITLAMAEM | ASGDGSYYAEQF | 0.65851927 |
| NELVEWSHPLGHRWRWE | ASGDGSYYAEQF | 0.61738753 |
| NLGAGGTGGNGGNPG | ASGDGSYYAEQF | 0.34401575 |
| NTEWPKWETVMAADPVD | ASGDGSYYAEQF | 0.7826993 |
| PASAAPAETQP | ASGDGSYYAEQF | 0.6446185 |
| QQSSDGKNGVRAAEITGE | ASGDGSYYAEQF | 0.65650666 |
| RAPDAVTTSWWKE | ASGDGSYYAEQF | 0.7255868 |
| SGFGNVGAHTSGWFNQ | ASGDGSYYAEQF | 0.64209676 |
| SGVAGAVGQAASVGGLKV | ASGDGSYYAEQF | 0.63921535 |
| SNGGAGTGSAGGNGGT | ASGDGSYYAEQF | 0.69750595 |
| TGGAGGEGGAAG | ASGDGSYYAEQF | 0.7541243 |
| TGGDGGAAGAGGAPGHG | ASGDGSYYAEQF | 0.7487632 |
| TMPPLLAAIANAGRHGV | ASGDGSYYAEQF | 0.72354555 |
| TSSTAGNSGGA | ASGDGSYYAEQF | 0.6881361 |
| VPATVPVGILC | ASGDGSYYAEQF | 0.795348 |
| VQRPDSGGVVLTGRIS | ASGDGSYYAEQF | 0.5322744 |
| WTPNPVPATKGVPAGLVQAT | ASGDGSYYAEQF | 0.61768293 |
| YALKRALAKFSDRFDVVII | ASGDGSYYAEQF | 0.7502587 |
| YLIGSVIV | ASGDGSYYAEQF | 0.45936802 |
| YGQLVAVID | ASGDGSYYAEQF | 0.7603699 |
| VTVGGMTLPA | ASGDGSYYAEQF | 0.7426572 |
| VLALPAV | ASGDGSYYAEQF | 0.7428185 |
| VIPPTLNYET | ASGDGSYYAEQF | 0.710119 |
| VGMGGMHPSSG | ASGDGSYYAEQF | 0.8016148 |
| VGLAGGAATA | ASGDGSYYAEQF | 0.7786127 |
| VGGAGGAGGSGP | ASGDGSYYAEQF | 0.6834732 |

| | | |
|---|---|---|
| VFAAFMV | ASGDGSYYAEQF | 0.7534014 |
| TLYDKLR | ASGDGSYYAEQF | 0.52910644 |
| TLAVFTVL | ASGDGSYYAEQF | 0.7783644 |
| TGSVQPARV | ASGDGSYYAEQF | 0.74616534 |
| TGGGAGSGGT | ASGDGSYYAEQF | 0.56824964 |
| SHLETSL | ASGDGSYYAEQF | 0.6984377 |
| SFTTGMT | ASGDGSYYAEQF | 0.7962517 |
| QVEVAKRMGVRQPTVS | ASGDGSYYAEQF | 0.77938646 |
| QGSGGPALAM | ASGDGSYYAEQF | 0.38047397 |
| MLAPSNP | ASGDGSYYAEQF | 0.6754241 |
| LVNNAAI | ASGDGSYYAEQF | 0.34882593 |
| LTGEINTLA | ASGDGSYYAEQF | 0.65826565 |
| LSIFGPV | ASGDGSYYAEQF | 0.52419823 |
| LLALLANL | ASGDGSYYAEQF | 0.76055497 |
| LISNASDALDKLR | ASGDGSYYAEQF | 0.6683581 |
| LINEAVP | ASGDGSYYAEQF | 0.73270774 |
| LGYAFSGL | ASGDGSYYAEQF | 0.7265868 |
| LDFVAVAL | ASGDGSYYAEQF | 0.79106283 |
| LAQPGAE | ASGDGSYYAEQF | 0.61273813 |
| ISNLGAVTGF | ASGDGSYYAEQF | 0.71924675 |
| HQEDPGRI | ASGDGSYYAEQF | 0.4124079 |
| GTPGEPTP | ASGDGSYYAEQF | 0.5175203 |
| GFFNSGS | ASGDGSYYAEQF | 0.63723564 |
| GAGGGGLTAGAN | ASGDGSYYAEQF | 0.71924686 |
| FVDALVNM | ASGDGSYYAEQF | 0.27565706 |
| FLTVAPEL | ASGDGSYYAEQF | 0.78890204 |
| FETLTAL | ASGDGSYYAEQF | 0.7361103 |
| DVINPIEL | ASGDGSYYAEQF | 0.76818025 |
| APVSGAPV | ASGDGSYYAEQF | 0.68053645 |
| ALVGGLAW | ASGDGSYYAEQF | 0.69236296 |
| ALAAAAATAI | ASGDGSYYAEQF | 0.66899 |
| AFGLGLLL | ASGDGSYYAEQF | 0.76312554 |
| AFFAAASL | ASGDGSYYAEQF | 0.61494786 |
| AAAALPAFSPP | ASGDGSYYAEQF | 0.38424343 |
| VVLAPAL | ASGDGSYYAEQF | 0.6800156 |
| TIGHVDHGKTTL | ASGDGSYYAEQF | 0.54453266 |
| LISNASDALDK | ASGDGSYYAEQF | 0.79357016 |
| LISNASDAL | ASGDGSYYAEQF | 0.7851201 |
| ISNASDALDK | ASGDGSYYAEQF | 0.74541616 |
| NQAETLVYQTE | ASGDGSYYAEQF | 0.7637256 |
| TGGTGGTAGAG | ASGDGSYYAEQF | 0.27689537 |
| TGSTAGGAGGA | ASGDGSYYAEQF | 0.67961407 |
| DPLAVPVAAGPV | ASGDGSYYAEQF | 0.79364187 |
| GGTGGTPGNGGN | ASGDGSYYAEQF | 0.7586539 |
| PAGLAGVDPATL | ASGDGSYYAEQF | 0.68796456 |
| GNNLVVEAPHAGASV | ASGDGSYYAEQF | 0.7889946 |
| KVTRIMSASTLEQPA | ASGDGSYYAEQF | 0.6784862 |
| LGFALPIALPRRTVV | ASGDGSYYAEQF | 0.69569683 |
| TRLGNLTARKLVVLV | ASGDGSYYAEQF | 0.73486334 |
| WSQQRRLIYFPSAGP | ASGDGSYYAEQF | 0.74920046 |
| GPNGNPDPMAAAVDIR | ASGDGSYYAEQF | 0.7521629 |

| | | |
|---|---|---|
| TQRCVSVADLRAAASR | ASGDGSYYAEQF | 0.5940532 |
| QFVLDVAEHPVEVLTETV | ASGDGSYYAEQF | 0.7302452 |
| GGAGGTGGAGGNGGTQAGDG | ASGDGSYYAEQF | 0.8075289 |
| PLLATLRAIAVDVDAKPAQV | ASGDGSYYAEQF | 0.6991421 |
| APKFRTGGYMIDFWGVGYQVAK | ASGDGSYYAEQF | 0.7837132 |
| GAGGAGGAGGAAGGPGGTGGTG | ASGDGSYYAEQF | 0.7557765 |
| IGALGLLATDVAPAGSVAFGVA | ASGDGSYYAEQF | 0.74282444 |
| AGGAGGIGGNGGAGGAGGVLMG | ASGDGSYYAEQF | 0.582165 |
| ADELIGLL | ASGDGSYYAEQF | 0.6756214 |
| ADRALKA | ASGDGSYYAEQF | 0.71733344 |
| AGAVGQVGGAGGA | ASGDGSYYAEQF | 0.62495935 |
| AGGGGGLGGAP | ASGDGSYYAEQF | 0.67446196 |
| AGGLLYGNGGAG | ASGDGSYYAEQF | 0.633228 |
| AGLESIGL | ASGDGSYYAEQF | 0.81230044 |
| AGLKTGII | ASGDGSYYAEQF | 0.7605741 |
| ASLAALLL | ASGDGSYYAEQF | 0.7649175 |
| ASVINISSI | ASGDGSYYAEQF | 0.6148976 |
| ATPPGPA | ASGDGSYYAEQF | 0.74650645 |
| ATVASALV | ASGDGSYYAEQF | 0.75404876 |
| ATVASLIL | ASGDGSYYAEQF | 0.42671484 |
| AVNGPVLIRH | ASGDGSYYAEQF | 0.7967279 |
| DGAAGAVGI | ASGDGSYYAEQF | 0.6060699 |
| DPLLHGI | ASGDGSYYAEQF | 0.6861725 |
| ELGAALGAL | ASGDGSYYAEQF | 0.7542541 |
| ELGRTQAV | ASGDGSYYAEQF | 0.7735067 |
| FVAGGYLS | ASGDGSYYAEQF | 0.66211545 |
| GASGGAGGAGGW | ASGDGSYYAEQF | 0.69685197 |
| GGAGAGGDGGAGGI | ASGDGSYYAEQF | 0.61354303 |
| GTGHNGSQPGMGGNG | ASGDGSYYAEQF | 0.7693602 |
| GYAAASASAS | ASGDGSYYAEQF | 0.7843255 |
| HQVDVAAVQQG | ASGDGSYYAEQF | 0.71381104 |
| IGAAIAHT | ASGDGSYYAEQF | 0.7891272 |
| IGMIEVAQL | ASGDGSYYAEQF | 0.719537 |
| IRLAPEHTSAMD | ASGDGSYYAEQF | 0.41178828 |
| IYNVSVL | ASGDGSYYAEQF | 0.70248556 |
| KQGFLGM | ASGDGSYYAEQF | 0.77551836 |
| LESVITQL | ASGDGSYYAEQF | 0.7841716 |
| LGLGALVVV | ASGDGSYYAEQF | 0.80141115 |
| LIGNGGNGGSGGI | ASGDGSYYAEQF | 0.7571666 |
| LNAINAPTL | ASGDGSYYAEQF | 0.67646503 |
| LQLLNTGV | ASGDGSYYAEQF | 0.78990626 |
| LQTSIEF | ASGDGSYYAEQF | 0.8045243 |
| MSFAVSEL | ASGDGSYYAEQF | 0.7686019 |
| NTGALGL | ASGDGSYYAEQF | 0.6339253 |
| PPGAGGAGGAGTT | ASGDGSYYAEQF | 0.5656339 |
| QLVGLGI | ASGDGSYYAEQF | 0.7421576 |
| RFLEQVHQL | ASGDGSYYAEQF | 0.7859128 |
| RGVGYVLR | ASGDGSYYAEQF | 0.49410343 |
| SATGVFAGVI | ASGDGSYYAEQF | 0.73140365 |
| SLPAITLP | ASGDGSYYAEQF | 0.74706095 |
| STALAFTV | ASGDGSYYAEQF | 0.4546095 |

| | | |
|---|---|---|
| SVLTVGLGK | ASGDGSYYAEQF | 0.76565754 |
| TALAGAGSGI | ASGDGSYYAEQF | 0.78897786 |
| TGLPFTGI | ASGDGSYYAEQF | 0.6746949 |
| VAEHVTV | ASGDGSYYAEQF | 0.6439818 |
| VDIVLNSL | ASGDGSYYAEQF | 0.7901753 |
| VLTGGGAVL | ASGDGSYYAEQF | 0.8094994 |
| VNVGLAYGI | ASGDGSYYAEQF | 0.78487337 |
| VSSAAVGAV | ASGDGSYYAEQF | 0.45043612 |
| VTSVYEAGV | ASGDGSYYAEQF | 0.27613443 |
| WVALDRGA | ASGDGSYYAEQF | 0.64169043 |
| YLVGSVVI | ASGDGSYYAEQF | 0.7572874 |
| GPPGTGKTLL | ASGDGSYYAEQF | 0.6981009 |
| GVGMTKF | ASGDGSYYAEQF | 0.63904345 |
| IGIGGSDL | ASGDGSYYAEQF | 0.7504478 |
| IGTLLAEL | ASGDGSYYAEQF | 0.78143644 |
| IYTSGTT | ASGDGSYYAEQF | 0.55276173 |
| LSFPAGV | ASGDGSYYAEQF | 0.49521053 |
| LYTSGTT | ASGDGSYYAEQF | 0.7514579 |
| VGAFPFGF | ASGDGSYYAEQF | 0.7513311 |
| VGAGTVEF | ASGDGSYYAEQF | 0.6155474 |
| VGKSTLLL | ASGDGSYYAEQF | 0.706203 |
| AALAHAGGGIV | ASGDGSYYAEQF | 0.6397798 |
| AALVAVSLL | ASGDGSYYAEQF | 0.61828184 |
| AAVSRAVAAL | ASGDGSYYAEQF | 0.7420335 |
| AGGSSGIGGF | ASGDGSYYAEQF | 0.5136264 |
| AGLTVANANG | ASGDGSYYAEQF | 0.7950778 |
| AGLVGAVL | ASGDGSYYAEQF | 0.55495286 |
| ALSQAFTV | ASGDGSYYAEQF | 0.77667785 |
| ALTFSVSL | ASGDGSYYAEQF | 0.75515705 |
| ALTGAALVAL | ASGDGSYYAEQF | 0.7476344 |
| ASFSIGAGVL | ASGDGSYYAEQF | 0.674301 |
| DGSVYKI | ASGDGSYYAEQF | 0.61344117 |
| EQLAEIV | ASGDGSYYAEQF | 0.51748616 |
| FAVSELTVM | ASGDGSYYAEQF | 0.32376865 |
| GAAPLAAGL | ASGDGSYYAEQF | 0.7876395 |
| GADGGDGGSGGAGG | ASGDGSYYAEQF | 0.6534554 |
| GFGLDTGGAGG | ASGDGSYYAEQF | 0.7800879 |
| GLASGGLL | ASGDGSYYAEQF | 0.7724769 |
| GRIAGAAVLL | ASGDGSYYAEQF | 0.6072933 |
| GSAGSLNTSF | ASGDGSYYAEQF | 0.75446725 |
| GTGQIGI | ASGDGSYYAEQF | 0.57571113 |
| GTLIALIL | ASGDGSYYAEQF | 0.49611855 |
| IAATMGFVVG | ASGDGSYYAEQF | 0.77653766 |
| IFAQALL | ASGDGSYYAEQF | 0.741819 |
| IGAAVALGF | ASGDGSYYAEQF | 0.5804976 |
| IGLTPLAL | ASGDGSYYAEQF | 0.72662836 |
| ILGSNTSTL | ASGDGSYYAEQF | 0.7368807 |
| IVFSLTVL | ASGDGSYYAEQF | 0.6344347 |
| LASKATGF | ASGDGSYYAEQF | 0.6828904 |
| LASVALIL | ASGDGSYYAEQF | 0.7828604 |
| LFAVGLTL | ASGDGSYYAEQF | 0.73004884 |

| | | |
|---|---|---|
| LFNGGIGTYI | ASGDGSYYAEQF | 0.6303048 |
| LTLGAGII | ASGDGSYYAEQF | 0.7793605 |
| LVAAALGDI | ASGDGSYYAEQF | 0.5463587 |
| NTAFVQLGI | ASGDGSYYAEQF | 0.74564743 |
| QLFSGAEL | ASGDGSYYAEQF | 0.7166779 |
| SAATFITL | ASGDGSYYAEQF | 0.6949493 |
| SNYGLANLGV | ASGDGSYYAEQF | 0.77738106 |
| SYGGAYAVM | ASGDGSYYAEQF | 0.5732298 |
| TTGVGGLTLG | ASGDGSYYAEQF | 0.64040166 |
| TTQTIYI | ASGDGSYYAEQF | 0.67448807 |
| VAAGAGLGAVI | ASGDGSYYAEQF | 0.73971593 |
| VAAVVTLL | ASGDGSYYAEQF | 0.7738913 |
| VGFLGGLI | ASGDGSYYAEQF | 0.6022515 |
| VGVIPILL | ASGDGSYYAEQF | 0.7224162 |
| VIFASSLI | ASGDGSYYAEQF | 0.6710272 |
| VKGGDGGAAAG | ASGDGSYYAEQF | 0.5633023 |
| VTKTFTNI | ASGDGSYYAEQF | 0.64088166 |
| VVAAAILL | ASGDGSYYAEQF | 0.70403326 |
| YAALDVEL | ASGDGSYYAEQF | 0.6255249 |
| LASVALLL | ASGDGSYYAEQF | 0.74804205 |
| LDAVHRV | ASGDGSYYAEQF | 0.62982374 |
| QGLSSSIF | ASGDGSYYAEQF | 0.76355565 |
| RGTFANIRL | ASGDGSYYAEQF | 0.8048724 |
| RQLWWGHRI | ASGDGSYYAEQF | 0.6685266 |
| YGPPGTGKTLL | ASGDGSYYAEQF | 0.77188647 |
| AALVGAPANVA | ASGDGSYYAEQF | 0.75713736 |
| AATFRRTE | ASGDGSYYAEQF | 0.7572229 |
| AGALGGGI | ASGDGSYYAEQF | 0.70759434 |
| AGGAGGAGGIGAG | ASGDGSYYAEQF | 0.64937043 |
| AGSGALGL | ASGDGSYYAEQF | 0.78825206 |
| AQIADQI | ASGDGSYYAEQF | 0.7767483 |
| ASAVAVTYL | ASGDGSYYAEQF | 0.7732034 |
| ASTVLGGTGG | ASGDGSYYAEQF | 0.7626618 |
| AVGGEPAK | ASGDGSYYAEQF | 0.74382734 |
| CEQGTFGGTC | ASGDGSYYAEQF | 0.7600764 |
| DEGKAPAG | ASGDGSYYAEQF | 0.659358 |
| DTVVFPAL | ASGDGSYYAEQF | 0.4550569 |
| FEGITMAG | ASGDGSYYAEQF | 0.70814204 |
| FPLVVQD | ASGDGSYYAEQF | 0.6892657 |
| GAFTTPQ | ASGDGSYYAEQF | 0.664279 |
| GFGGFGTAGGVGGA | ASGDGSYYAEQF | 0.3368133 |
| GFLAQFL | ASGDGSYYAEQF | 0.78087735 |
| GGAGGAGGNAGVA | ASGDGSYYAEQF | 0.8074288 |
| GGVGTAGAGGAGGAGG | ASGDGSYYAEQF | 0.6340771 |
| GNGGGGDAGNAGSGGN | ASGDGSYYAEQF | 0.7990149 |
| GTGGAGGAGNDAGS | ASGDGSYYAEQF | 0.4638072 |
| GTTNTGWFNS | ASGDGSYYAEQF | 0.7765282 |
| IFMGPLFTL | ASGDGSYYAEQF | 0.78749347 |
| ILAHAIEAI | ASGDGSYYAEQF | 0.575395 |
| IRSAIREAL | ASGDGSYYAEQF | 0.76185876 |
| ITFAGGG | ASGDGSYYAEQF | 0.66894054 |

| | | |
|---|---|---|
| ITGMSGHQEV | ASGDGSYYAEQF | 0.69150543 |
| IVAGLACTL | ASGDGSYYAEQF | 0.7932241 |
| IVGGGLAGA | ASGDGSYYAEQF | 0.68867993 |
| IVTGVLNI | ASGDGSYYAEQF | 0.7412741 |
| LAQQGCI | ASGDGSYYAEQF | 0.6763512 |
| LCLHVEN | ASGDGSYYAEQF | 0.61187303 |
| LDAISDGS | ASGDGSYYAEQF | 0.7234484 |
| LGGFAAML | ASGDGSYYAEQF | 0.7802849 |
| LGGKNPMI | ASGDGSYYAEQF | 0.5303594 |
| LGPKGVAAIL | ASGDGSYYAEQF | 0.8029015 |
| LMVGVSI | ASGDGSYYAEQF | 0.66405565 |
| LVGAAFVL | ASGDGSYYAEQF | 0.80285054 |
| LVLAVAPL | ASGDGSYYAEQF | 0.7522919 |
| MAAIAVIDAA | ASGDGSYYAEQF | 0.722513 |
| NNGINAELQ | ASGDGSYYAEQF | 0.65439403 |
| RGEPLVLVHG | ASGDGSYYAEQF | 0.7878783 |
| RGGAGGAAGLI | ASGDGSYYAEQF | 0.67155373 |
| RSYRMHPAV | ASGDGSYYAEQF | 0.7729269 |
| SAAHLLAGQLA | ASGDGSYYAEQF | 0.78221095 |
| SAAVGSY | ASGDGSYYAEQF | 0.387385 |
| SPHAVLHRI | ASGDGSYYAEQF | 0.7741266 |
| SQIAKVL | ASGDGSYYAEQF | 0.6828589 |
| TALGHLVP | ASGDGSYYAEQF | 0.6836153 |
| TAPTTSGAA | ASGDGSYYAEQF | 0.5905577 |
| TDGDDGNGGSPGT | ASGDGSYYAEQF | 0.30875698 |
| TDGLGEN | ASGDGSYYAEQF | 0.629207 |
| TGGPPLPGQA | ASGDGSYYAEQF | 0.52010393 |
| TIPAITI | ASGDGSYYAEQF | 0.7798199 |
| TIPGFTI | ASGDGSYYAEQF | 0.7293423 |
| TSIASGALA | ASGDGSYYAEQF | 0.59263176 |
| TVTLPGAAI | ASGDGSYYAEQF | 0.7674334 |
| VASGLSAALP | ASGDGSYYAEQF | 0.77559525 |
| VGAGVIGI | ASGDGSYYAEQF | 0.29907778 |
| VGPKRFGV | ASGDGSYYAEQF | 0.5866988 |
| VNVAFPD | ASGDGSYYAEQF | 0.58561844 |
| VSMTNSGVS | ASGDGSYYAEQF | 0.644894 |
| VTDVLGDLSV | ASGDGSYYAEQF | 0.32665676 |
| VVVVGAGFAGL | ASGDGSYYAEQF | 0.70480275 |
| VYTVGTVV | ASGDGSYYAEQF | 0.46725133 |
| WTTFSGP | ASGDGSYYAEQF | 0.20771834 |
| YDELVEAF | ASGDGSYYAEQF | 0.65748864 |
| YGTGGAGGNGG | ASGDGSYYAEQF | 0.6906235 |
| YTNVMAV | ASGDGSYYAEQF | 0.5934252 |
| FLEQVHQL | ASGDGSYYAEQF | 0.7377263 |
| GLSLLAVAV | ASGDGSYYAEQF | 0.4297278 |
| LEQVHQL | ASGDGSYYAEQF | 0.4783791 |
| LNAAGAI | ASGDGSYYAEQF | 0.74117476 |
| PAVHASL | ASGDGSYYAEQF | 0.76505893 |
| TELEDDA | ASGDGSYYAEQF | 0.4080893 |
| TPPGPPN | ASGDGSYYAEQF | 0.71914995 |
| TPPPPPPP | ASGDGSYYAEQF | 0.78345597 |

| | | |
|---|---|---|
| TQSPALI | ASRRDIASGNTL | 0.6739898 |
| EFAAAFPQ | ASRRDIASGNTL | 0.7582985 |
| LLSGNLDVL | ASRRDIASGNTL | 0.77202845 |
| GPANLGNYNI | ASRRDIASGNTL | 0.59995776 |
| TLGPTLP | ASRRDIASGNTL | 0.7684387 |
| VYAYSSCCK | ASRRDIASGNTL | 0.7764362 |
| APAPAPAPNI | ASRRDIASGNTL | 0.6428872 |
| PGGPVPIP | ASRRDIASGNTL | 0.70390856 |
| FLTVALEP | ASRRDIASGNTL | 0.36019015 |
| SQYFGANL | ASRRDIASGNTL | 0.6360977 |
| GGVKPAA | ASRRDIASGNTL | 0.2807135 |
| GGTGGAGGAGGW | ASRRDIASGNTL | 0.7333856 |
| GVGGVAGLGGAGG | ASRRDIASGNTL | 0.76209 |
| VLLVLATTI | ASRRDIASGNTL | 0.64007926 |
| IGIFAAML | ASRRDIASGNTL | 0.3425095 |
| AVGDLID | ASRRDIASGNTL | 0.42196417 |
| NISPLPL | ASRRDIASGNTL | 0.66217226 |
| VTAVFVAI | ASRRDIASGNTL | 0.4216698 |
| SGDDNDAAL | ASRRDIASGNTL | 0.2806999 |
| AVAALAHS | ASRRDIASGNTL | 0.5397963 |
| GGVTVMATP | ASRRDIASGNTL | 0.7527243 |
| AGLAANVRD | ASRRDIASGNTL | 0.770774 |
| TNRPDLIDPALLRPG | ASRRDIASGNTL | 0.4168281 |
| LANLGVTAL | ASRRDIASGNTL | 0.7762272 |
| TGERIEL | ASRRDIASGNTL | 0.7482873 |
| NGEELAA | ASRRDIASGNTL | 0.7800094 |
| VLSLVWG | ASRRDIASGNTL | 0.6760305 |
| IGYALWL | ASRRDIASGNTL | 0.3706064 |
| LSLLLAPL | ASRRDIASGNTL | 0.77299076 |
| TLDEALK | ASRRDIASGNTL | 0.6552896 |
| TDGAGTAEGG | ASRRDIASGNTL | 0.69227463 |
| NAPGGAGGNGG | ASRRDIASGNTL | 0.7226677 |
| VSVFMGMGGNF | ASRRDIASGNTL | 0.55198914 |
| LASYLAAL | ASRRDIASGNTL | 0.55527174 |
| LVGGTPAL | ASRRDIASGNTL | 0.7189687 |
| VRNADVL | ASRRDIASGNTL | 0.75881296 |
| TCSSPFLVSAM | ASRRDIASGNTL | 0.47201622 |
| VHITDAE | ASRRDIASGNTL | 0.36200002 |
| EYAAPSF | ASRRDIASGNTL | 0.55886966 |
| VVDAFHGGL | ASRRDIASGNTL | 0.6812459 |
| ASVVDPQ | ASRRDIASGNTL | 0.75942373 |
| SYNEFGGLS | ASRRDIASGNTL | 0.69763345 |
| DPDGSVY | ASRRDIASGNTL | 0.6078817 |
| MVSTIHATE | ASRRDIASGNTL | 0.7308655 |
| LAAGVADPV | ASRRDIASGNTL | 0.7328062 |
| TKEQIAA | ASRRDIASGNTL | 0.7173338 |
| SLASPAAL | ASRRDIASGNTL | 0.7403827 |
| IGLVFYGV | ASRRDIASGNTL | 0.6281341 |
| FLLDASNL | ASRRDIASGNTL | 0.75023043 |
| YQVGNFP | ASRRDIASGNTL | 0.739257 |
| ESLLSQST | ASRRDIASGNTL | 0.7262254 |

| | | |
|---|---|---|
| LFTVGFAAVG | ASRRDIASGNTL | 0.7576763 |
| ATPPTPT | ASRRDIASGNTL | 0.69759583 |
| TDLATLG | ASRRDIASGNTL | 0.6957703 |
| SAGGTGR | ASRRDIASGNTL | 0.3245334 |
| GFSNSGS | ASRRDIASGNTL | 0.66728175 |
| GGNGGSGGTSV | ASRRDIASGNTL | 0.7145529 |
| GTSVATGGAGN | ASRRDIASGNTL | 0.6671086 |
| NGGNGGSGGTS | ASRRDIASGNTL | 0.75793874 |
| AVVKFPH | ASRRDIASGNTL | 0.6849059 |
| TPPPPPP | ASRRDIASGNTL | 0.52623093 |
| AAPAAAAAPA | ASRRDIASGNTL | 0.77461195 |
| IALNEDL | ASRRDIASGNTL | 0.77736545 |
| IVNVSSM | ASRRDIASGNTL | 0.4232089 |
| TGIPPAPR | ASRRDIASGNTL | 0.63204753 |
| FDLGGGTF | ASRRDIASGNTL | 0.7841127 |
| FDLGGGTFDVS | ASRRDIASGNTL | 0.7339791 |
| NEPTAAA | ASRRDIASGNTL | 0.61673343 |
| NEPTAAALA | ASRRDIASGNTL | 0.64556336 |
| GSIINISSI | ASRRDIASGNTL | 0.78919506 |
| VAPGNAGTA | ASRRDIASGNTL | 0.66284704 |
| TGGGASLE | ASRRDIASGNTL | 0.7807956 |
| VAVYNHYKRI | ASRRDIASGNTL | 0.28124145 |
| TDPAPATT | ASRRDIASGNTL | 0.5795512 |
| TNRPDILDPALLRPG | ASRRDIASGNTL | 0.45264497 |
| LTTTTTS | ASRRDIASGNTL | 0.21439028 |
| PTGSGKTL | ASRRDIASGNTL | 0.75751054 |
| TGSGKTL | ASRRDIASGNTL | 0.55166304 |
| TIGHVDHGK | ASRRDIASGNTL | 0.75052035 |
| TIGHVDHGKTT | ASRRDIASGNTL | 0.6806108 |
| GHVDHGKTT | ASRRDIASGNTL | 0.75853205 |
| AQFEHTLL | ASRRDIASGNTL | 0.73134696 |
| IVNQDEL | ASRRDIASGNTL | 0.6441237 |
| TNGSQFF | ASRRDIASGNTL | 0.39193338 |
| TGKGIVE | ASRRDIASGNTL | 0.71016425 |
| LISNASDA | ASRRDIASGNTL | 0.77406085 |
| AELRTLGELTKIAKAHGAQ | ASRRDIASGNTL | 0.71440864 |
| APNAVAAAEPSW | ASRRDIASGNTL | 0.28369427 |
| APQVAGAATAVL | ASRRDIASGNTL | 0.733241 |
| ARRDPAAAAAILPTDA | ASRRDIASGNTL | 0.67742234 |
| DAAARDVLAEAGLAEYF | ASRRDIASGNTL | 0.23245057 |
| DDQIVLSAKANLAKVEQ | ASRRDIASGNTL | 0.23856838 |
| DELLPDTGEDSDEDGH | ASRRDIASGNTL | 0.3775624 |
| ELPEAPANRATVPQD | ASRRDIASGNTL | 0.62296194 |
| FGQSKIERVTGIG | ASRRDIASGNTL | 0.75685287 |
| GAGGTGSTAGGAGGAGGAGGLY | ASRRDIASGNTL | 0.75086194 |
| GHGGTGAAGGNGGTGGAGGSGI | ASRRDIASGNTL | 0.6294186 |
| GNGGNGGAGGEGGPGGAGGAGA | ASRRDIASGNTL | 0.75225 |
| GSTAGNGGNGGNATNGTIAGQP | ASRRDIASGNTL | 0.6977152 |
| GTGGDGGNGGNGGTGG | ASRRDIASGNTL | 0.18001804 |
| GVRRPAGIPADAPV | ASRRDIASGNTL | 0.63827986 |
| GYRQVTDAHLLTLARR | ASRRDIASGNTL | 0.3171852 |

| | | |
|---|---|---|
| IAKMGQDGHDRG | ASRRDIASGNTL | 0.7697601 |
| KAEPSIEIQYT | ASRRDIASGNTL | 0.49040842 |
| KPVRIRMAADEDTVAVT | ASRRDIASGNTL | 0.59041774 |
| MPPRIAGMRLLVIKPEP | ASRRDIASGNTL | 0.7002301 |
| MKFYAVALFRSRGVPPTR | ASRRDIASGNTL | 0.7509016 |
| NAGGFGILGAVAHSP | ASRRDIASGNTL | 0.72093165 |
| NEGREITLAMAEM | ASRRDIASGNTL | 0.51209503 |
| NELVEWSHPLGHRWRWE | ASRRDIASGNTL | 0.21873507 |
| NLGAGGTGGNGGNPG | ASRRDIASGNTL | 0.7194307 |
| NTEWPKWETVMAADPVD | ASRRDIASGNTL | 0.5254213 |
| PASAAPAETQP | ASRRDIASGNTL | 0.3630244 |
| QQSSDGKNGVRAAEITGE | ASRRDIASGNTL | 0.34416246 |
| RAPDAVTTSWWKE | ASRRDIASGNTL | 0.5645829 |
| SGFGNVGAHTSGWFNQ | ASRRDIASGNTL | 0.34730554 |
| SGVAGAVGQAASVGGLKV | ASRRDIASGNTL | 0.7370042 |
| SNGGAGTGSAGGNGGT | ASRRDIASGNTL | 0.49861133 |
| TGGAGGEGGAAG | ASRRDIASGNTL | 0.62291145 |
| TGGDGGAAGAGGAPGHG | ASRRDIASGNTL | 0.77756983 |
| TMPPLLAAIANAGRHGV | ASRRDIASGNTL | 0.7539933 |
| TSSTAGNSGGA | ASRRDIASGNTL | 0.6663813 |
| VPATVPVGILC | ASRRDIASGNTL | 0.5095013 |
| VQRPDSGGVVLTGRIS | ASRRDIASGNTL | 0.44545567 |
| WTPNPVPATKGVPAGLVQAT | ASRRDIASGNTL | 0.4601025 |
| YALKRALAKFSDRFDVVII | ASRRDIASGNTL | 0.4906195 |
| YLIGSVIV | ASRRDIASGNTL | 0.7778323 |
| YGQLVAVID | ASRRDIASGNTL | 0.6383188 |
| VTVGGMTLPA | ASRRDIASGNTL | 0.43867332 |
| VLALPAV | ASRRDIASGNTL | 0.6973211 |
| VIPPTLNYET | ASRRDIASGNTL | 0.7839937 |
| VGMGGMHPSSG | ASRRDIASGNTL | 0.7905591 |
| VGLAGGAATA | ASRRDIASGNTL | 0.38826367 |
| VGGAGGAGGSGP | ASRRDIASGNTL | 0.66604173 |
| VFAAFMV | ASRRDIASGNTL | 0.6214533 |
| TLYDKLR | ASRRDIASGNTL | 0.66899765 |
| TLAVFTVL | ASRRDIASGNTL | 0.48853517 |
| TGSVQPARV | ASRRDIASGNTL | 0.4578499 |
| TGGGAGSGGT | ASRRDIASGNTL | 0.7371912 |
| SHLETSL | ASRRDIASGNTL | 0.7949246 |
| SFTTGMT | ASRRDIASGNTL | 0.6078318 |
| QVEVAKRMGVRQPTVS | ASRRDIASGNTL | 0.41609648 |
| QGSGGPALAM | ASRRDIASGNTL | 0.68461597 |
| MLAPSNP | ASRRDIASGNTL | 0.5161811 |
| LVNNAAI | ASRRDIASGNTL | 0.6227264 |
| LTGEINTLA | ASRRDIASGNTL | 0.62065995 |
| LSIFGPV | ASRRDIASGNTL | 0.34817183 |
| LLALLANL | ASRRDIASGNTL | 0.67355716 |
| LISNASDALDKLR | ASRRDIASGNTL | 0.78837204 |
| LINEAVP | ASRRDIASGNTL | 0.6892449 |
| LGYAFSGL | ASRRDIASGNTL | 0.72016895 |
| LDFVAVAL | ASRRDIASGNTL | 0.7781333 |
| LAQPGAE | ASRRDIASGNTL | 0.7185416 |

| | | |
|---|---|---|
| ISNLGAVTGF | ASRRDIASGNTL | 0.7710562 |
| HQEDPGRI | ASRRDIASGNTL | 0.7571151 |
| GTPGEPTP | ASRRDIASGNTL | 0.56886053 |
| GFFNSGS | ASRRDIASGNTL | 0.6697822 |
| GAGGGGLTAGAN | ASRRDIASGNTL | 0.7474618 |
| FVDALVNM | ASRRDIASGNTL | 0.7144549 |
| FLTVAPEL | ASRRDIASGNTL | 0.6679662 |
| FETLTAL | ASRRDIASGNTL | 0.6655112 |
| DVINPIEL | ASRRDIASGNTL | 0.7761451 |
| APVSGAPV | ASRRDIASGNTL | 0.6460972 |
| ALVGGLAW | ASRRDIASGNTL | 0.7806121 |
| ALAAAAATAI | ASRRDIASGNTL | 0.24435705 |
| AFGLGLLL | ASRRDIASGNTL | 0.40850943 |
| AFFAAASL | ASRRDIASGNTL | 0.7531238 |
| AAAALPAFSPP | ASRRDIASGNTL | 0.19412328 |
| VVLAPAL | ASRRDIASGNTL | 0.28917134 |
| TIGHVDHGKTTL | ASRRDIASGNTL | 0.6311411 |
| LISNASDALDK | ASRRDIASGNTL | 0.772704 |
| LISNASDAL | ASRRDIASGNTL | 0.18763036 |
| ISNASDALDK | ASRRDIASGNTL | 0.48518932 |
| NQAETLVYQTE | ASRRDIASGNTL | 0.25758883 |
| TGGTGGTAGAG | ASRRDIASGNTL | 0.7141838 |
| TGSTAGGAGGA | ASRRDIASGNTL | 0.69137037 |
| DPLAVPVAAGPV | ASRRDIASGNTL | 0.3229328 |
| GGTGGTPGNGGN | ASRRDIASGNTL | 0.7259941 |
| PAGLAGVDPATL | ASRRDIASGNTL | 0.4347865 |
| GNNLVVEAPHAGASV | ASRRDIASGNTL | 0.6765987 |
| KVTRIMSASTLEQPA | ASRRDIASGNTL | 0.7472925 |
| LGFALPIALPRRTVV | ASRRDIASGNTL | 0.77696085 |
| TRLGNLTARKLVVLV | ASRRDIASGNTL | 0.3764953 |
| WSQQRRLIYFPSAGP | ASRRDIASGNTL | 0.73433566 |
| GPNGNPDPMAAAVDIR | ASRRDIASGNTL | 0.6059146 |
| TQRCVSVADLRAAASR | ASRRDIASGNTL | 0.793296 |
| QFVLDVAEHPVEVLTETV | ASRRDIASGNTL | 0.607441 |
| GGAGGTGGAGGNGGTQAGDG | ASRRDIASGNTL | 0.2953396 |
| PLLATLRAIAVDVDAKPAQV | ASRRDIASGNTL | 0.7475369 |
| APKFRTGGYMIDFWGVGYQVAK | ASRRDIASGNTL | 0.73046094 |
| GAGGAGGAGGAAGGPGGTGGTG | ASRRDIASGNTL | 0.5519543 |
| IGALGLLATDVAPAGSVAFGVA | ASRRDIASGNTL | 0.75646615 |
| AGGAGGIGGNGGAGGAGGVLMG | ASRRDIASGNTL | 0.5776284 |
| ADELIGLL | ASRRDIASGNTL | 0.6940206 |
| ADRALKA | ASRRDIASGNTL | 0.7772529 |
| AGAVGQVGGAGGA | ASRRDIASGNTL | 0.74249923 |
| AGGGGGLGGAP | ASRRDIASGNTL | 0.44623893 |
| AGGLLYGNGGAG | ASRRDIASGNTL | 0.75052637 |
| AGLESIGL | ASRRDIASGNTL | 0.6652435 |
| AGLKTGII | ASRRDIASGNTL | 0.39535284 |
| ASLAALLL | ASRRDIASGNTL | 0.7890229 |
| ASVINISSI | ASRRDIASGNTL | 0.5982705 |
| ATPPGPA | ASRRDIASGNTL | 0.7643788 |
| ATVASALV | ASRRDIASGNTL | 0.7260556 |

| | | |
|---|---|---|
| ATVASLIL | ASRRDIASGNTL | 0.79050004 |
| AVNGPVLIRH | ASRRDIASGNTL | 0.33583245 |
| DGAAGAVGI | ASRRDIASGNTL | 0.708893 |
| DPLLHGI | ASRRDIASGNTL | 0.7130702 |
| ELGAALGAL | ASRRDIASGNTL | 0.7455207 |
| ELGRTQAV | ASRRDIASGNTL | 0.57807374 |
| FVAGGYLS | ASRRDIASGNTL | 0.565588 |
| GASGGAGGAGGW | ASRRDIASGNTL | 0.7238735 |
| GGAGAGGDGGAGGI | ASRRDIASGNTL | 0.7720673 |
| GTGHNGSQPGMGGNG | ASRRDIASGNTL | 0.6002935 |
| GYAAASASAS | ASRRDIASGNTL | 0.7277758 |
| HQVDVAAVQQG | ASRRDIASGNTL | 0.77685237 |
| IGAAIAHT | ASRRDIASGNTL | 0.3757605 |
| IGMIEVAQL | ASRRDIASGNTL | 0.6822763 |
| IRLAPEHTSAMD | ASRRDIASGNTL | 0.7070002 |
| IYNVSVL | ASRRDIASGNTL | 0.76692736 |
| KQGFLGM | ASRRDIASGNTL | 0.6252291 |
| LESVITQL | ASRRDIASGNTL | 0.774729 |
| LGLGALVVV | ASRRDIASGNTL | 0.5748869 |
| LIGNGGNGGSGGI | ASRRDIASGNTL | 0.74557334 |
| LNAINAPTL | ASRRDIASGNTL | 0.5858907 |
| LQLLNTGV | ASRRDIASGNTL | 0.72968274 |
| LQTSIEF | ASRRDIASGNTL | 0.3500547 |
| MSFAVSEL | ASRRDIASGNTL | 0.73689795 |
| NTGALGL | ASRRDIASGNTL | 0.75998896 |
| PPGAGGAGGAGTT | ASRRDIASGNTL | 0.75987566 |
| QLVGLGI | ASRRDIASGNTL | 0.27189487 |
| RFLEQVHQL | ASRRDIASGNTL | 0.761802 |
| RGVGYVLR | ASRRDIASGNTL | 0.7640358 |
| SATGVFAGVI | ASRRDIASGNTL | 0.7837828 |
| SLPAITLP | ASRRDIASGNTL | 0.65393937 |
| STALAFTV | ASRRDIASGNTL | 0.7374161 |
| SVLTVGLGK | ASRRDIASGNTL | 0.55355734 |
| TALAGAGSGI | ASRRDIASGNTL | 0.62175137 |
| TGLPFTGI | ASRRDIASGNTL | 0.44107544 |
| VAEHVTV | ASRRDIASGNTL | 0.47176212 |
| VDIVLNSL | ASRRDIASGNTL | 0.688167 |
| VLTGGGAVL | ASRRDIASGNTL | 0.5248426 |
| VNVGLAYGI | ASRRDIASGNTL | 0.77261895 |
| VSSAAVGAV | ASRRDIASGNTL | 0.7664448 |
| VTSVYEAGV | ASRRDIASGNTL | 0.75673306 |
| WVALDRGA | ASRRDIASGNTL | 0.77219886 |
| YLVGSVVI | ASRRDIASGNTL | 0.7794761 |
| GPPGTGKTLL | ASRRDIASGNTL | 0.74620956 |
| GVGMTKF | ASRRDIASGNTL | 0.60539573 |
| IGIGGSDL | ASRRDIASGNTL | 0.7051906 |
| IGTLLAEL | ASRRDIASGNTL | 0.3260812 |
| IYTSGTT | ASRRDIASGNTL | 0.6643153 |
| LSFPAGV | ASRRDIASGNTL | 0.56965524 |
| LYTSGTT | ASRRDIASGNTL | 0.73007786 |
| VGAFPFGF | ASRRDIASGNTL | 0.18766809 |

| | | |
|---|---|---|
| VGAGTVEF | ASRRDIASGNTL | 0.7364762 |
| VGKSTLLL | ASRRDIASGNTL | 0.7789781 |
| AALAHAGGGIV | ASRRDIASGNTL | 0.6224416 |
| AALVAVSLL | ASRRDIASGNTL | 0.7533412 |
| AAVSRAVAAL | ASRRDIASGNTL | 0.74001527 |
| AGGSSGIGGF | ASRRDIASGNTL | 0.49494576 |
| AGLTVANANG | ASRRDIASGNTL | 0.7438468 |
| AGLVGAVL | ASRRDIASGNTL | 0.48903278 |
| ALSQAFTV | ASRRDIASGNTL | 0.7384838 |
| ALTFSVSL | ASRRDIASGNTL | 0.7772336 |
| ALTGAALVAL | ASRRDIASGNTL | 0.72895384 |
| ASFSIGAGVL | ASRRDIASGNTL | 0.68597454 |
| DGSVYKI | ASRRDIASGNTL | 0.678778 |
| EQLAEIV | ASRRDIASGNTL | 0.30802464 |
| FAVSELTVM | ASRRDIASGNTL | 0.7056161 |
| GAAPLAAGL | ASRRDIASGNTL | 0.24871014 |
| GADGGDGGSGGAGG | ASRRDIASGNTL | 0.7174543 |
| GFGLDTGGAGG | ASRRDIASGNTL | 0.43417034 |
| GLASGGLL | ASRRDIASGNTL | 0.6729066 |
| GRIAGAAVLL | ASRRDIASGNTL | 0.67599535 |
| GSAGSLNTSF | ASRRDIASGNTL | 0.67486805 |
| GTGQIGI | ASRRDIASGNTL | 0.55357236 |
| GTLIALIL | ASRRDIASGNTL | 0.7361413 |
| IAATMGFVVG | ASRRDIASGNTL | 0.695452 |
| IFAQALL | ASRRDIASGNTL | 0.36188313 |
| IGAAVALGF | ASRRDIASGNTL | 0.7590172 |
| IGLTPLAL | ASRRDIASGNTL | 0.56711257 |
| ILGSNTSTL | ASRRDIASGNTL | 0.69275236 |
| IVFSLTVL | ASRRDIASGNTL | 0.7576022 |
| LASKATGF | ASRRDIASGNTL | 0.60645187 |
| LASVALIL | ASRRDIASGNTL | 0.35885808 |
| LFAVGLTL | ASRRDIASGNTL | 0.64558244 |
| LFNGGIGTYI | ASRRDIASGNTL | 0.65610194 |
| LTLGAGII | ASRRDIASGNTL | 0.61663485 |
| LVAAALGDI | ASRRDIASGNTL | 0.20769727 |
| NTAFVQLGI | ASRRDIASGNTL | 0.714707 |
| QLFSGAEL | ASRRDIASGNTL | 0.38469073 |
| SAATFITL | ASRRDIASGNTL | 0.6301743 |
| SNYGLANLGV | ASRRDIASGNTL | 0.60530806 |
| SYGGAYAVM | ASRRDIASGNTL | 0.7524638 |
| TTGVGGLTLG | ASRRDIASGNTL | 0.58728516 |
| TTQTIYI | ASRRDIASGNTL | 0.5830831 |
| VAAGAGLGAVI | ASRRDIASGNTL | 0.7342944 |
| VAAVVTLL | ASRRDIASGNTL | 0.7016635 |
| VGFLGGLI | ASRRDIASGNTL | 0.7790092 |
| VGVIPILL | ASRRDIASGNTL | 0.68065584 |
| VIFASSLI | ASRRDIASGNTL | 0.60629845 |
| VKGGDGGAAAG | ASRRDIASGNTL | 0.68632805 |
| VTKTFTNI | ASRRDIASGNTL | 0.64264333 |
| VVAAAILL | ASRRDIASGNTL | 0.40923643 |
| YAALDVEL | ASRRDIASGNTL | 0.4171371 |

| | | |
|---|---|---|
| LASVALLL | ASRRDIASGNTL | 0.6452868 |
| LDAVHRV | ASRRDIASGNTL | 0.36250198 |
| QGLSSSIF | ASRRDIASGNTL | 0.7222379 |
| RGTFANIRL | ASRRDIASGNTL | 0.7641601 |
| RQLWWGHRI | ASRRDIASGNTL | 0.6718331 |
| YGPPGTGKTLL | ASRRDIASGNTL | 0.30167583 |
| AALVGAPANVA | ASRRDIASGNTL | 0.50263137 |
| AATFRRTE | ASRRDIASGNTL | 0.5671334 |
| AGALGGGI | ASRRDIASGNTL | 0.73240006 |
| AGGAGGAGGIGAG | ASRRDIASGNTL | 0.7266623 |
| AGSGALGL | ASRRDIASGNTL | 0.7885885 |
| AQIADQI | ASRRDIASGNTL | 0.7685546 |
| ASAVAVTYL | ASRRDIASGNTL | 0.60940206 |
| ASTVLGGTGG | ASRRDIASGNTL | 0.7328806 |
| AVGGEPAK | ASRRDIASGNTL | 0.641574 |
| CEQGTFGGTC | ASRRDIASGNTL | 0.654852 |
| DEGKAPAG | ASRRDIASGNTL | 0.6077637 |
| DTVVFPAL | ASRRDIASGNTL | 0.6734393 |
| FEGITMAG | ASRRDIASGNTL | 0.6320916 |
| FPLVVQD | ASRRDIASGNTL | 0.54544675 |
| GAFTTPQ | ASRRDIASGNTL | 0.39416242 |
| GFGGFGTAGGVGGA | ASRRDIASGNTL | 0.68678594 |
| GFLAQFL | ASRRDIASGNTL | 0.47547007 |
| GGAGGAGGNAGVA | ASRRDIASGNTL | 0.3581463 |
| GGVGTAGAGGAGGAGG | ASRRDIASGNTL | 0.63956267 |
| GNGGGGDAGNAGSGGN | ASRRDIASGNTL | 0.7825178 |
| GTGGAGGAGNDAGS | ASRRDIASGNTL | 0.77792764 |
| GTTNTGWFNS | ASRRDIASGNTL | 0.5924957 |
| IFMGPLFTL | ASRRDIASGNTL | 0.49304122 |
| ILAHAIEAI | ASRRDIASGNTL | 0.7394489 |
| IRSAIREAL | ASRRDIASGNTL | 0.38247165 |
| ITFAGGG | ASRRDIASGNTL | 0.71549577 |
| ITGMSGHQEV | ASRRDIASGNTL | 0.7154487 |
| IVAGLACTL | ASRRDIASGNTL | 0.7032455 |
| IVGGGLAGA | ASRRDIASGNTL | 0.5610662 |
| IVTGVLNI | ASRRDIASGNTL | 0.6813521 |
| LAQQGCI | ASRRDIASGNTL | 0.6752847 |
| LCLHVEN | ASRRDIASGNTL | 0.7111865 |
| LDAISDGS | ASRRDIASGNTL | 0.20470124 |
| LGGFAAML | ASRRDIASGNTL | 0.7386654 |
| LGGKNPMI | ASRRDIASGNTL | 0.4677903 |
| LGPKGVAAIL | ASRRDIASGNTL | 0.4990971 |
| LMVGVSI | ASRRDIASGNTL | 0.22320832 |
| LVGAAFVL | ASRRDIASGNTL | 0.66785216 |
| LVLAVAPL | ASRRDIASGNTL | 0.7523501 |
| MAAIAVIDAA | ASRRDIASGNTL | 0.7493799 |
| NNGINAELQ | ASRRDIASGNTL | 0.60798323 |
| RGEPLVLVHG | ASRRDIASGNTL | 0.7634385 |
| RGGAGGAAGLI | ASRRDIASGNTL | 0.7705545 |
| RSYRMHPAV | ASRRDIASGNTL | 0.64411974 |
| SAAHLLAGQLA | ASRRDIASGNTL | 0.24013047 |

| | | |
|---|---|---|
| SAAVGSY | ASRRDIASGNTL | 0.7127196 |
| SPHAVLHRI | ASRRDIASGNTL | 0.69763964 |
| SQIAKVL | ASRRDIASGNTL | 0.7499414 |
| TALGHLVP | ASRRDIASGNTL | 0.7535512 |
| TAPTTSGAA | ASRRDIASGNTL | 0.3935591 |
| TDGDDGNGGSPGT | ASRRDIASGNTL | 0.7414855 |
| TDGLGEN | ASRRDIASGNTL | 0.7453228 |
| TGGPPLPGQA | ASRRDIASGNTL | 0.42401734 |
| TIPAITI | ASRRDIASGNTL | 0.5233883 |
| TIPGFTI | ASRRDIASGNTL | 0.57255304 |
| TSIASGALA | ASRRDIASGNTL | 0.30963737 |
| TVTLPGAAI | ASRRDIASGNTL | 0.71058905 |
| VASGLSAALP | ASRRDIASGNTL | 0.59609705 |
| VGAGVIGI | ASRRDIASGNTL | 0.57033604 |
| VGPKRFGV | ASRRDIASGNTL | 0.6031548 |
| VNVAFPD | ASRRDIASGNTL | 0.5008746 |
| VSMTNSGVS | ASRRDIASGNTL | 0.5678266 |
| VTDVLGDLSV | ASRRDIASGNTL | 0.55426013 |
| VVVVGAGFAGL | ASRRDIASGNTL | 0.74326265 |
| VYTVGTVV | ASRRDIASGNTL | 0.71279633 |
| WTTFSGP | ASRRDIASGNTL | 0.7036572 |
| YDELVEAF | ASRRDIASGNTL | 0.7028652 |
| YGTGGAGGNGG | ASRRDIASGNTL | 0.77762055 |
| YTNVMAV | ASRRDIASGNTL | 0.35949373 |
| FLEQVHQL | ASRRDIASGNTL | 0.5756062 |
| GLSLLAVAV | ASRRDIASGNTL | 0.7105591 |
| LEQVHQL | ASRRDIASGNTL | 0.44213855 |
| LNAAGAI | ASRRDIASGNTL | 0.7482016 |
| PAVHASL | ASRRDIASGNTL | 0.6762963 |
| TELEDDA | ASRRDIASGNTL | 0.7291605 |
| TPPGPPN | ASRRDIASGNTL | 0.67678 |
| TPPPPPPP | ASRRDIASGNTL | 0.7847794 |
| TQSPALI | ASSQEKGPNSGNTLY | 0.7237135 |
| EFAAAFPQ | ASSQEKGPNSGNTLY | 0.64327633 |
| LLSGNLDVL | ASSQEKGPNSGNTLY | 0.7079824 |
| GPANLGNYNI | ASSQEKGPNSGNTLY | 0.45952588 |
| TLGPTLP | ASSQEKGPNSGNTLY | 0.7606994 |
| VYAYSSCCK | ASSQEKGPNSGNTLY | 0.73841864 |
| APAPAPAPNI | ASSQEKGPNSGNTLY | 0.5466783 |
| PGGPVPIP | ASSQEKGPNSGNTLY | 0.6991259 |
| FLTVALEP | ASSQEKGPNSGNTLY | 0.763885 |
| SQYFGANL | ASSQEKGPNSGNTLY | 0.7667569 |
| GGVKPAA | ASSQEKGPNSGNTLY | 0.400204 |
| GGTGGAGGAGGW | ASSQEKGPNSGNTLY | 0.50535953 |
| GVGGVAGLGGAGG | ASSQEKGPNSGNTLY | 0.30191705 |
| VLLVLATTI | ASSQEKGPNSGNTLY | 0.52950907 |
| IGIFAAML | ASSQEKGPNSGNTLY | 0.2795822 |
| AVGDLID | ASSQEKGPNSGNTLY | 0.73191345 |
| NISPLPL | ASSQEKGPNSGNTLY | 0.6142789 |
| VTAVFVAI | ASSQEKGPNSGNTLY | 0.7745332 |
| SGDDNDAAL | ASSQEKGPNSGNTLY | 0.71981406 |

| | | |
|---|---|---|
| AVAALAHS | ASSQEKGPNSGNTLY | 0.76645553 |
| GGVTVMATP | ASSQEKGPNSGNTLY | 0.5692886 |
| AGLAANVRD | ASSQEKGPNSGNTLY | 0.63018596 |
| TNRPDLIDPALLRPG | ASSQEKGPNSGNTLY | 0.28678867 |
| LANLGVTAL | ASSQEKGPNSGNTLY | 0.3853035 |
| TGERIEL | ASSQEKGPNSGNTLY | 0.39443025 |
| NGEELAA | ASSQEKGPNSGNTLY | 0.6310681 |
| VLSLVWG | ASSQEKGPNSGNTLY | 0.22196501 |
| IGYALWL | ASSQEKGPNSGNTLY | 0.78598 |
| LSLLLAPL | ASSQEKGPNSGNTLY | 0.6478267 |
| TLDEALK | ASSQEKGPNSGNTLY | 0.71768475 |
| TDGAGTAEGG | ASSQEKGPNSGNTLY | 0.54119766 |
| NAPGGAGGNGG | ASSQEKGPNSGNTLY | 0.46439463 |
| VSVFMGMGGNF | ASSQEKGPNSGNTLY | 0.6072597 |
| LASYLAAL | ASSQEKGPNSGNTLY | 0.74444294 |
| LVGGTPAL | ASSQEKGPNSGNTLY | 0.40383583 |
| VRNADVL | ASSQEKGPNSGNTLY | 0.29532564 |
| TCSSPFLVSAM | ASSQEKGPNSGNTLY | 0.57612014 |
| VHITDAE | ASSQEKGPNSGNTLY | 0.4870801 |
| EYAAPSF | ASSQEKGPNSGNTLY | 0.78196275 |
| VVDAFHGGL | ASSQEKGPNSGNTLY | 0.7688117 |
| ASVVDPQ | ASSQEKGPNSGNTLY | 0.6862731 |
| SYNEFGGLS | ASSQEKGPNSGNTLY | 0.7811584 |
| DPDGSVY | ASSQEKGPNSGNTLY | 0.19617452 |
| MVSTIHATE | ASSQEKGPNSGNTLY | 0.579566 |
| LAAGVADPV | ASSQEKGPNSGNTLY | 0.78658724 |
| TKEQIAA | ASSQEKGPNSGNTLY | 0.714812 |
| SLASPAAL | ASSQEKGPNSGNTLY | 0.64331853 |
| IGLVFYGV | ASSQEKGPNSGNTLY | 0.76045144 |
| FLLDASNL | ASSQEKGPNSGNTLY | 0.5461762 |
| YQVGNFP | ASSQEKGPNSGNTLY | 0.6288097 |
| ESLLSQST | ASSQEKGPNSGNTLY | 0.6222439 |
| LFTVGFAAVG | ASSQEKGPNSGNTLY | 0.69868785 |
| ATPPTPT | ASSQEKGPNSGNTLY | 0.58597445 |
| TDLATLG | ASSQEKGPNSGNTLY | 0.33974832 |
| SAGGTGR | ASSQEKGPNSGNTLY | 0.7110599 |
| GFSNSGS | ASSQEKGPNSGNTLY | 0.78479004 |
| GGNGGSGGTSV | ASSQEKGPNSGNTLY | 0.5776094 |
| GTSVATGGAGN | ASSQEKGPNSGNTLY | 0.75147945 |
| NGGNGGSGGTS | ASSQEKGPNSGNTLY | 0.70396686 |
| AVVKFPH | ASSQEKGPNSGNTLY | 0.70905626 |
| TPPPPPP | ASSQEKGPNSGNTLY | 0.58582634 |
| AAPAAAAAPA | ASSQEKGPNSGNTLY | 0.40005386 |
| IALNEDL | ASSQEKGPNSGNTLY | 0.74240243 |
| IVNVSSM | ASSQEKGPNSGNTLY | 0.5371089 |
| TGIPPAPR | ASSQEKGPNSGNTLY | 0.613966 |
| FDLGGGTF | ASSQEKGPNSGNTLY | 0.6894136 |
| FDLGGGTFDVS | ASSQEKGPNSGNTLY | 0.79482377 |
| NEPTAAA | ASSQEKGPNSGNTLY | 0.75528 |
| NEPTAAALA | ASSQEKGPNSGNTLY | 0.6985682 |
| GSIINISSI | ASSQEKGPNSGNTLY | 0.48838216 |

| | | |
|---|---|---|
| VAPGNAGTA | ASSQEKGPNSGNTLY | 0.6799792 |
| TGGGASLE | ASSQEKGPNSGNTLY | 0.7290865 |
| VAVYNHYKRI | ASSQEKGPNSGNTLY | 0.39298865 |
| TDPAPATT | ASSQEKGPNSGNTLY | 0.77394146 |
| TNRPDILDPALLRPG | ASSQEKGPNSGNTLY | 0.4747014 |
| LTTTTTS | ASSQEKGPNSGNTLY | 0.7281623 |
| PTGSGKTL | ASSQEKGPNSGNTLY | 0.75660634 |
| TGSGKTL | ASSQEKGPNSGNTLY | 0.7827009 |
| TIGHVDHGK | ASSQEKGPNSGNTLY | 0.67327565 |
| TIGHVDHGKTT | ASSQEKGPNSGNTLY | 0.7083872 |
| GHVDHGKTT | ASSQEKGPNSGNTLY | 0.3912423 |
| AQFEHTLL | ASSQEKGPNSGNTLY | 0.7535244 |
| IVNQDEL | ASSQEKGPNSGNTLY | 0.7596224 |
| TNGSQFF | ASSQEKGPNSGNTLY | 0.7082745 |
| TGKGIVE | ASSQEKGPNSGNTLY | 0.7843746 |
| LISNASDA | ASSQEKGPNSGNTLY | 0.71078205 |
| AELRTLGELTKIAKAHGAQ | ASSQEKGPNSGNTLY | 0.2543695 |
| APNAVAAAEPSW | ASSQEKGPNSGNTLY | 0.55899006 |
| APQVAGAATAVL | ASSQEKGPNSGNTLY | 0.7493885 |
| ARRDPAAAAAILPTDA | ASSQEKGPNSGNTLY | 0.75085604 |
| DAAARDVLAEAGLAEYF | ASSQEKGPNSGNTLY | 0.7755704 |
| DDQIVLSAKANLAKVEQ | ASSQEKGPNSGNTLY | 0.7273264 |
| DELLPDTGEDSDEDGH | ASSQEKGPNSGNTLY | 0.63277006 |
| ELPEAPANRATVPQD | ASSQEKGPNSGNTLY | 0.7781409 |
| FGQSKIERVTGIG | ASSQEKGPNSGNTLY | 0.7929934 |
| GAGGTGSTAGGAGGAGGAGGLY | ASSQEKGPNSGNTLY | 0.7463968 |
| GHGGTGAAGGNGGTGGAGGSGI | ASSQEKGPNSGNTLY | 0.5847448 |
| GNGGNGGAGGEGGPGGAGGAGA | ASSQEKGPNSGNTLY | 0.51801074 |
| GSTAGNGGNGGNATNGTIAGQP | ASSQEKGPNSGNTLY | 0.7306776 |
| GTGGDGGNGGNGGTGG | ASSQEKGPNSGNTLY | 0.76696074 |
| GVRRPAGIPADAPV | ASSQEKGPNSGNTLY | 0.36048225 |
| GYRQVTDAHLLTLARR | ASSQEKGPNSGNTLY | 0.7229173 |
| IAKMGQDGHDRG | ASSQEKGPNSGNTLY | 0.6723489 |
| KAEPSIEIQYT | ASSQEKGPNSGNTLY | 0.24355522 |
| KPVRIRMAADEDTVAVT | ASSQEKGPNSGNTLY | 0.73896307 |
| MPPRIAGMRLLVIKPEP | ASSQEKGPNSGNTLY | 0.77027476 |
| MKFYAVALFRSRGVPPTR | ASSQEKGPNSGNTLY | 0.703195 |
| NAGGFGILGAVAHSP | ASSQEKGPNSGNTLY | 0.40681738 |
| NEGREITLAMAEM | ASSQEKGPNSGNTLY | 0.77503604 |
| NELVEWSHPLGHRWRWE | ASSQEKGPNSGNTLY | 0.786654 |
| NLGAGGTGGNGGNPG | ASSQEKGPNSGNTLY | 0.76940393 |
| NTEWPKWETVMAADPVD | ASSQEKGPNSGNTLY | 0.34067723 |
| PASAAPAETQP | ASSQEKGPNSGNTLY | 0.2221513 |
| QQSSDGKNGVRAAEITGE | ASSQEKGPNSGNTLY | 0.6261932 |
| RAPDAVTTSWWKE | ASSQEKGPNSGNTLY | 0.7656459 |
| SGFGNVGAHTSGWFNQ | ASSQEKGPNSGNTLY | 0.58201015 |
| SGVAGAVGQAASVGGLKV | ASSQEKGPNSGNTLY | 0.4630576 |
| SNGGAGTGSAGGNGGT | ASSQEKGPNSGNTLY | 0.66399264 |
| TGGAGGEGGAAG | ASSQEKGPNSGNTLY | 0.77204174 |
| TGGDGGAAGAGGAPGHG | ASSQEKGPNSGNTLY | 0.36656258 |
| TMPPLLAAIANAGRHGV | ASSQEKGPNSGNTLY | 0.38248324 |

| | | |
|---|---|---|
| TSSTAGNSGGA | ASSQEKGPNSGNTLY | 0.6916199 |
| VPATVPVGILC | ASSQEKGPNSGNTLY | 0.74428666 |
| VQRPDSGGVVLTGRIS | ASSQEKGPNSGNTLY | 0.49414906 |
| WTPNPVPATKGVPAGLVQAT | ASSQEKGPNSGNTLY | 0.590467 |
| YALKRALAKFSDRFDVVII | ASSQEKGPNSGNTLY | 0.5899757 |
| YLIGSVIV | ASSQEKGPNSGNTLY | 0.6061962 |
| YGQLVAVID | ASSQEKGPNSGNTLY | 0.70160604 |
| VTVGGMTLPA | ASSQEKGPNSGNTLY | 0.47071564 |
| VLALPAV | ASSQEKGPNSGNTLY | 0.76850754 |
| VIPPTLNYET | ASSQEKGPNSGNTLY | 0.4507401 |
| VGMGGMHPSSG | ASSQEKGPNSGNTLY | 0.7172277 |
| VGLAGGAATA | ASSQEKGPNSGNTLY | 0.6957512 |
| VGGAGGAGGSGP | ASSQEKGPNSGNTLY | 0.6870749 |
| VFAAFMV | ASSQEKGPNSGNTLY | 0.6278086 |
| TLYDKLR | ASSQEKGPNSGNTLY | 0.6306223 |
| TLAVFTVL | ASSQEKGPNSGNTLY | 0.41446462 |
| TGSVQPARV | ASSQEKGPNSGNTLY | 0.36488885 |
| TGGGAGSGGT | ASSQEKGPNSGNTLY | 0.7675631 |
| SHLETSL | ASSQEKGPNSGNTLY | 0.6227647 |
| SFTTGMT | ASSQEKGPNSGNTLY | 0.75981677 |
| QVEVAKRMGVRQPTVS | ASSQEKGPNSGNTLY | 0.71487975 |
| QGSGGPALAM | ASSQEKGPNSGNTLY | 0.37860096 |
| MLAPSNP | ASSQEKGPNSGNTLY | 0.7131301 |
| LVNNAAI | ASSQEKGPNSGNTLY | 0.32655 |
| LTGEINTLA | ASSQEKGPNSGNTLY | 0.6101879 |
| LSIFGPV | ASSQEKGPNSGNTLY | 0.70454746 |
| LLALLANL | ASSQEKGPNSGNTLY | 0.72642344 |
| LISNASDALDKLR | ASSQEKGPNSGNTLY | 0.5538721 |
| LINEAVP | ASSQEKGPNSGNTLY | 0.67522645 |
| LGYAFSGL | ASSQEKGPNSGNTLY | 0.7276739 |
| LDFVAVAL | ASSQEKGPNSGNTLY | 0.49407905 |
| LAQPGAE | ASSQEKGPNSGNTLY | 0.71066505 |
| ISNLGAVTGF | ASSQEKGPNSGNTLY | 0.77603066 |
| HQEDPGRI | ASSQEKGPNSGNTLY | 0.7099638 |
| GTPGEPTP | ASSQEKGPNSGNTLY | 0.38907933 |
| GFFNSGS | ASSQEKGPNSGNTLY | 0.73510283 |
| GAGGGGLTAGAN | ASSQEKGPNSGNTLY | 0.53513956 |
| FVDALVNM | ASSQEKGPNSGNTLY | 0.7329936 |
| FLTVAPEL | ASSQEKGPNSGNTLY | 0.60390294 |
| FETLTAL | ASSQEKGPNSGNTLY | 0.6932422 |
| DVINPIEL | ASSQEKGPNSGNTLY | 0.69972414 |
| APVSGAPV | ASSQEKGPNSGNTLY | 0.50140214 |
| ALVGGLAW | ASSQEKGPNSGNTLY | 0.55093366 |
| ALAAAAATAI | ASSQEKGPNSGNTLY | 0.61903155 |
| AFGLGLLL | ASSQEKGPNSGNTLY | 0.6418242 |
| AFFAAASL | ASSQEKGPNSGNTLY | 0.7460667 |
| AAAALPAFSPP | ASSQEKGPNSGNTLY | 0.4836582 |
| VVLAPAL | ASSQEKGPNSGNTLY | 0.7017154 |
| TIGHVDHGKTTL | ASSQEKGPNSGNTLY | 0.5500669 |
| LISNASDALDK | ASSQEKGPNSGNTLY | 0.50213003 |
| LISNASDAL | ASSQEKGPNSGNTLY | 0.66948926 |

| | | |
|---|---|---|
| ISNASDALDK | ASSQEKGPNSGNTLY | 0.6347321 |
| NQAETLVYQTE | ASSQEKGPNSGNTLY | 0.31433725 |
| TGGTGGTAGAG | ASSQEKGPNSGNTLY | 0.7069018 |
| TGSTAGGAGGA | ASSQEKGPNSGNTLY | 0.36953777 |
| DPLAVPVAAGPV | ASSQEKGPNSGNTLY | 0.6703889 |
| GGTGGTPGNGGN | ASSQEKGPNSGNTLY | 0.7795868 |
| PAGLAGVDPATL | ASSQEKGPNSGNTLY | 0.39861783 |
| GNNLVVEAPHAGASV | ASSQEKGPNSGNTLY | 0.7066356 |
| KVTRIMSASTLEQPA | ASSQEKGPNSGNTLY | 0.7172952 |
| LGFALPIALPRRTVV | ASSQEKGPNSGNTLY | 0.72917056 |
| TRLGNLTARKLVVLV | ASSQEKGPNSGNTLY | 0.5535903 |
| WSQQRRLIYFPSAGP | ASSQEKGPNSGNTLY | 0.5686836 |
| GPNGNPDPMAAAVDIR | ASSQEKGPNSGNTLY | 0.77394146 |
| TQRCVSVADLRAAASR | ASSQEKGPNSGNTLY | 0.7379697 |
| QFVLDVAEHPVEVLTETV | ASSQEKGPNSGNTLY | 0.7615241 |
| GGAGGTGGAGGNGGTQAGDG | ASSQEKGPNSGNTLY | 0.7424258 |
| PLLATLRAIAVDVDAKPAQV | ASSQEKGPNSGNTLY | 0.7618201 |
| APKFRTGGYMIDFWGVGYQVAK | ASSQEKGPNSGNTLY | 0.7315055 |
| GAGGAGGAGGAAGGPGGTGGTG | ASSQEKGPNSGNTLY | 0.6271759 |
| IGALGLLATDVAPAGSVAFGVA | ASSQEKGPNSGNTLY | 0.2780274 |
| AGGAGGIGGNGGAGGAGGVLMG | ASSQEKGPNSGNTLY | 0.6363376 |
| ADELIGLL | ASSQEKGPNSGNTLY | 0.6135593 |
| ADRALKA | ASSQEKGPNSGNTLY | 0.56398135 |
| AGAVGQVGGAGGA | ASSQEKGPNSGNTLY | 0.32617226 |
| AGGGGGLGGAP | ASSQEKGPNSGNTLY | 0.7775606 |
| AGGLLYGNGGAG | ASSQEKGPNSGNTLY | 0.7917543 |
| AGLESIGL | ASSQEKGPNSGNTLY | 0.49614072 |
| AGLKTGII | ASSQEKGPNSGNTLY | 0.7813207 |
| ASLAALLL | ASSQEKGPNSGNTLY | 0.41626704 |
| ASVINISSI | ASSQEKGPNSGNTLY | 0.74022764 |
| ATPPGPA | ASSQEKGPNSGNTLY | 0.7796827 |
| ATVASALV | ASSQEKGPNSGNTLY | 0.47309744 |
| ATVASLIL | ASSQEKGPNSGNTLY | 0.75054586 |
| AVNGPVLIRH | ASSQEKGPNSGNTLY | 0.5273974 |
| DGAAGAVGI | ASSQEKGPNSGNTLY | 0.5216292 |
| DPLLHGI | ASSQEKGPNSGNTLY | 0.7743211 |
| ELGAALGAL | ASSQEKGPNSGNTLY | 0.5282607 |
| ELGRTQAV | ASSQEKGPNSGNTLY | 0.692737 |
| FVAGGYLS | ASSQEKGPNSGNTLY | 0.6022093 |
| GASGGAGGAGGW | ASSQEKGPNSGNTLY | 0.41043946 |
| GGAGAGGDGGAGGI | ASSQEKGPNSGNTLY | 0.6654961 |
| GTGHNGSQPGMGGNG | ASSQEKGPNSGNTLY | 0.77298784 |
| GYAAASASAS | ASSQEKGPNSGNTLY | 0.62944984 |
| HQVDVAAVQQG | ASSQEKGPNSGNTLY | 0.79009396 |
| IGAAIAHT | ASSQEKGPNSGNTLY | 0.4883891 |
| IGMIEVAQL | ASSQEKGPNSGNTLY | 0.7921449 |
| IRLAPEHTSAMD | ASSQEKGPNSGNTLY | 0.76833284 |
| IYNVSVL | ASSQEKGPNSGNTLY | 0.7016104 |
| KQGFLGM | ASSQEKGPNSGNTLY | 0.5674775 |
| LESVITQL | ASSQEKGPNSGNTLY | 0.7749089 |
| LGLGALVVV | ASSQEKGPNSGNTLY | 0.6086987 |

| | | |
|---|---|---|
| LIGNGGNGGSGGI | ASSQEKGPNSGNTLY | 0.7399614 |
| LNAINAPTL | ASSQEKGPNSGNTLY | 0.7643875 |
| LQLLNTGV | ASSQEKGPNSGNTLY | 0.6537266 |
| LQTSIEF | ASSQEKGPNSGNTLY | 0.815969 |
| MSFAVSEL | ASSQEKGPNSGNTLY | 0.7510154 |
| NTGALGL | ASSQEKGPNSGNTLY | 0.7586014 |
| PPGAGGAGGAGTT | ASSQEKGPNSGNTLY | 0.68592817 |
| QLVGLGI | ASSQEKGPNSGNTLY | 0.57369655 |
| RFLEQVHQL | ASSQEKGPNSGNTLY | 0.72059864 |
| RGVGYVLR | ASSQEKGPNSGNTLY | 0.68104124 |
| SATGVFAGVI | ASSQEKGPNSGNTLY | 0.7451392 |
| SLPAITLP | ASSQEKGPNSGNTLY | 0.7949789 |
| STALAFTV | ASSQEKGPNSGNTLY | 0.6706069 |
| SVLTVGLGK | ASSQEKGPNSGNTLY | 0.7991475 |
| TALAGAGSGI | ASSQEKGPNSGNTLY | 0.8142462 |
| TGLPFTGI | ASSQEKGPNSGNTLY | 0.5206143 |
| VAEHVTV | ASSQEKGPNSGNTLY | 0.70141184 |
| VDIVLNSL | ASSQEKGPNSGNTLY | 0.64608085 |
| VLTGGGAVL | ASSQEKGPNSGNTLY | 0.7223249 |
| VNVGLAYGI | ASSQEKGPNSGNTLY | 0.38062853 |
| VSSAAVGAV | ASSQEKGPNSGNTLY | 0.79423296 |
| VTSVYEAGV | ASSQEKGPNSGNTLY | 0.6240716 |
| WVALDRGA | ASSQEKGPNSGNTLY | 0.18364836 |
| YLVGSVVI | ASSQEKGPNSGNTLY | 0.73366314 |
| GPPGTGKTLL | ASSQEKGPNSGNTLY | 0.70945555 |
| GVGMTKF | ASSQEKGPNSGNTLY | 0.75493336 |
| IGIGGSDL | ASSQEKGPNSGNTLY | 0.7773968 |
| IGTLLAEL | ASSQEKGPNSGNTLY | 0.4357659 |
| IYTSGTT | ASSQEKGPNSGNTLY | 0.70535827 |
| LSFPAGV | ASSQEKGPNSGNTLY | 0.7634827 |
| LYTSGTT | ASSQEKGPNSGNTLY | 0.81290543 |
| VGAFPFGF | ASSQEKGPNSGNTLY | 0.6802251 |
| VGAGTVEF | ASSQEKGPNSGNTLY | 0.7306561 |
| VGKSTLLL | ASSQEKGPNSGNTLY | 0.8181249 |
| AALAHAGGGIV | ASSQEKGPNSGNTLY | 0.77138704 |
| AALVAVSLL | ASSQEKGPNSGNTLY | 0.7995389 |
| AAVSRAVAAL | ASSQEKGPNSGNTLY | 0.822606 |
| AGGSSGIGGF | ASSQEKGPNSGNTLY | 0.62463534 |
| AGLTVANANG | ASSQEKGPNSGNTLY | 0.7984364 |
| AGLVGAVL | ASSQEKGPNSGNTLY | 0.8036729 |
| ALSQAFTV | ASSQEKGPNSGNTLY | 0.7493597 |
| ALTFSVSL | ASSQEKGPNSGNTLY | 0.71836907 |
| ALTGAALVAL | ASSQEKGPNSGNTLY | 0.62587214 |
| ASFSIGAGVL | ASSQEKGPNSGNTLY | 0.6865061 |
| DGSVYKI | ASSQEKGPNSGNTLY | 0.37750563 |
| EQLAEIV | ASSQEKGPNSGNTLY | 0.72855204 |
| FAVSELTVM | ASSQEKGPNSGNTLY | 0.8055186 |
| GAAPLAAGL | ASSQEKGPNSGNTLY | 0.67314863 |
| GADGGDGGSGGAGG | ASSQEKGPNSGNTLY | 0.5578267 |
| GFGLDTGGAGG | ASSQEKGPNSGNTLY | 0.58021694 |
| GLASGGLL | ASSQEKGPNSGNTLY | 0.6992105 |

| | | |
|---|---|---|
| GRIAGAAVLL | ASSQEKGPNSGNTLY | 0.68474907 |
| GSAGSLNTSF | ASSQEKGPNSGNTLY | 0.34733534 |
| GTGQIGI | ASSQEKGPNSGNTLY | 0.6257795 |
| GTLIALIL | ASSQEKGPNSGNTLY | 0.8084066 |
| IAATMGFVVG | ASSQEKGPNSGNTLY | 0.8180274 |
| IFAQALL | ASSQEKGPNSGNTLY | 0.44765136 |
| IGAAVALGF | ASSQEKGPNSGNTLY | 0.7962488 |
| IGLTPLAL | ASSQEKGPNSGNTLY | 0.7611741 |
| ILGSNTSTL | ASSQEKGPNSGNTLY | 0.81609225 |
| IVFSLTVL | ASSQEKGPNSGNTLY | 0.75814754 |
| LASKATGF | ASSQEKGPNSGNTLY | 0.6352575 |
| LASVALIL | ASSQEKGPNSGNTLY | 0.75287294 |
| LFAVGLTL | ASSQEKGPNSGNTLY | 0.63079095 |
| LFNGGIGTYI | ASSQEKGPNSGNTLY | 0.591795 |
| LTLGAGII | ASSQEKGPNSGNTLY | 0.79179657 |
| LVAAALGDI | ASSQEKGPNSGNTLY | 0.7559868 |
| NTAFVQLGI | ASSQEKGPNSGNTLY | 0.6371318 |
| QLFSGAEL | ASSQEKGPNSGNTLY | 0.5937597 |
| SAATFITL | ASSQEKGPNSGNTLY | 0.81777984 |
| SNYGLANLGV | ASSQEKGPNSGNTLY | 0.52477425 |
| SYGGAYAVM | ASSQEKGPNSGNTLY | 0.7072844 |
| TTGVGGLTLG | ASSQEKGPNSGNTLY | 0.5952242 |
| TTQTIYI | ASSQEKGPNSGNTLY | 0.7221937 |
| VAAGAGLGAVI | ASSQEKGPNSGNTLY | 0.8058545 |
| VAAVVTLL | ASSQEKGPNSGNTLY | 0.79113597 |
| VGFLGGLI | ASSQEKGPNSGNTLY | 0.6508049 |
| VGVIPILL | ASSQEKGPNSGNTLY | 0.7514609 |
| VIFASSLI | ASSQEKGPNSGNTLY | 0.79234064 |
| VKGGDGGAAAG | ASSQEKGPNSGNTLY | 0.7556391 |
| VTKTFTNI | ASSQEKGPNSGNTLY | 0.7757368 |
| VVAAAILL | ASSQEKGPNSGNTLY | 0.6109202 |
| YAALDVEL | ASSQEKGPNSGNTLY | 0.8097464 |
| LASVALLL | ASSQEKGPNSGNTLY | 0.773153 |
| LDAVHRV | ASSQEKGPNSGNTLY | 0.80806714 |
| QGLSSSIF | ASSQEKGPNSGNTLY | 0.7858777 |
| RGTFANIRL | ASSQEKGPNSGNTLY | 0.79257643 |
| RQLWWGHRI | ASSQEKGPNSGNTLY | 0.7521127 |
| YGPPGTGKTLL | ASSQEKGPNSGNTLY | 0.3863558 |
| AALVGAPANVA | ASSQEKGPNSGNTLY | 0.66022825 |
| AATFRRTE | ASSQEKGPNSGNTLY | 0.72626257 |
| AGALGGGI | ASSQEKGPNSGNTLY | 0.73205674 |
| AGGAGGAGGIGAG | ASSQEKGPNSGNTLY | 0.75687754 |
| AGSGALGL | ASSQEKGPNSGNTLY | 0.8054376 |
| AQIADQI | ASSQEKGPNSGNTLY | 0.64077955 |
| ASAVAVTYL | ASSQEKGPNSGNTLY | 0.80358243 |
| ASTVLGGTGG | ASSQEKGPNSGNTLY | 0.78097624 |
| AVGGEPAK | ASSQEKGPNSGNTLY | 0.48329678 |
| CEQGTFGGTC | ASSQEKGPNSGNTLY | 0.71093976 |
| DEGKAPAG | ASSQEKGPNSGNTLY | 0.8227898 |
| DTVVFPAL | ASSQEKGPNSGNTLY | 0.7007314 |
| FEGITMAG | ASSQEKGPNSGNTLY | 0.67205 |

| | | |
|---|---|---|
| FPLVVQD | ASSQEKGPNSGNTLY | 0.70140785 |
| GAFTTPQ | ASSQEKGPNSGNTLY | 0.8087904 |
| GFGGFGTAGGVGGA | ASSQEKGPNSGNTLY | 0.6213493 |
| GFLAQFL | ASSQEKGPNSGNTLY | 0.82170326 |
| GGAGGAGGNAGVA | ASSQEKGPNSGNTLY | 0.30092704 |
| GGVGTAGAGGAGGAGG | ASSQEKGPNSGNTLY | 0.7231994 |
| GNGGGGDAGNAGSGGN | ASSQEKGPNSGNTLY | 0.6981846 |
| GTGGAGGAGNDAGS | ASSQEKGPNSGNTLY | 0.49542427 |
| GTTNTGWFNS | ASSQEKGPNSGNTLY | 0.80239785 |
| IFMGPLFTL | ASSQEKGPNSGNTLY | 0.7162599 |
| ILAHAIEAI | ASSQEKGPNSGNTLY | 0.7784833 |
| IRSAIREAL | ASSQEKGPNSGNTLY | 0.6893275 |
| ITFAGGG | ASSQEKGPNSGNTLY | 0.80287755 |
| ITGMSGHQEV | ASSQEKGPNSGNTLY | 0.6909464 |
| IVAGLACTL | ASSQEKGPNSGNTLY | 0.74620324 |
| IVGGGLAGA | ASSQEKGPNSGNTLY | 0.67654693 |
| IVTGVLNI | ASSQEKGPNSGNTLY | 0.79580176 |
| LAQQGCI | ASSQEKGPNSGNTLY | 0.80626154 |
| LCLHVEN | ASSQEKGPNSGNTLY | 0.80311334 |
| LDAISDGS | ASSQEKGPNSGNTLY | 0.62672925 |
| LGGFAAML | ASSQEKGPNSGNTLY | 0.62917626 |
| LGGKNPMI | ASSQEKGPNSGNTLY | 0.7635792 |
| LGPKGVAAIL | ASSQEKGPNSGNTLY | 0.28439206 |
| LMVGVSI | ASSQEKGPNSGNTLY | 0.24927974 |
| LVGAAFVL | ASSQEKGPNSGNTLY | 0.70086396 |
| LVLAVAPL | ASSQEKGPNSGNTLY | 0.56602937 |
| MAAIAVIDAA | ASSQEKGPNSGNTLY | 0.7967127 |
| NNGINAELQ | ASSQEKGPNSGNTLY | 0.7671633 |
| RGEPLVLVHG | ASSQEKGPNSGNTLY | 0.6842571 |
| RGGAGGAAGLI | ASSQEKGPNSGNTLY | 0.7599635 |
| RSYRMHPAV | ASSQEKGPNSGNTLY | 0.7411065 |
| SAAHLLAGQLA | ASSQEKGPNSGNTLY | 0.14015673 |
| SAAVGSY | ASSQEKGPNSGNTLY | 0.6753696 |
| SPHAVLHRI | ASSQEKGPNSGNTLY | 0.5382839 |
| SQIAKVL | ASSQEKGPNSGNTLY | 0.8279824 |
| TALGHLVP | ASSQEKGPNSGNTLY | 0.6609954 |
| TAPTTSGAA | ASSQEKGPNSGNTLY | 0.56221765 |
| TDGDDGNGGSPGT | ASSQEKGPNSGNTLY | 0.7146351 |
| TDGLGEN | ASSQEKGPNSGNTLY | 0.68657637 |
| TGGPPLPGQA | ASSQEKGPNSGNTLY | 0.69296163 |
| TIPAITI | ASSQEKGPNSGNTLY | 0.6207629 |
| TIPGFTI | ASSQEKGPNSGNTLY | 0.19627519 |
| TSIASGALA | ASSQEKGPNSGNTLY | 0.8004044 |
| TVTLPGAAI | ASSQEKGPNSGNTLY | 0.4762388 |
| VASGLSAALP | ASSQEKGPNSGNTLY | 0.7070453 |
| VGAGVIGI | ASSQEKGPNSGNTLY | 0.31963834 |
| VGPKRFGV | ASSQEKGPNSGNTLY | 0.457506 |
| VNVAFPD | ASSQEKGPNSGNTLY | 0.5737121 |
| VSMTNSGVS | ASSQEKGPNSGNTLY | 0.7956484 |
| VTDVLGDLSV | ASSQEKGPNSGNTLY | 0.6266271 |
| VVVVGAGFAGL | ASSQEKGPNSGNTLY | 0.6270106 |

| | | |
|---|---|---|
| VYTVGTVV | ASSQEKGPNSGNTLY | 0.80575645 |
| WTTFSGP | ASSQEKGPNSGNTLY | 0.6974601 |
| YDELVEAF | ASSQEKGPNSGNTLY | 0.6341855 |
| YGTGGAGGNGG | ASSQEKGPNSGNTLY | 0.73563427 |
| YTNVMAV | ASSQEKGPNSGNTLY | 0.6083435 |
| FLEQVHQL | ASSQEKGPNSGNTLY | 0.7058049 |
| GLSLLAVAV | ASSQEKGPNSGNTLY | 0.7155824 |
| LEQVHQL | ASSQEKGPNSGNTLY | 0.820667 |
| LNAAGAI | ASSQEKGPNSGNTLY | 0.72250915 |
| PAVHASL | ASSQEKGPNSGNTLY | 0.58905554 |
| TELEDDA | ASSQEKGPNSGNTLY | 0.75971067 |
| TPPGPPN | ASSQEKGPNSGNTLY | 0.8161837 |
| TPPPPPPP | ASSQEKGPNSGNTLY | 0.8274614 |
| TQSPALI | ASSLESFLEQY | 0.59551597 |
| EFAAAFPQ | ASSLESFLEQY | 0.7343572 |
| LLSGNLDVL | ASSLESFLEQY | 0.703021 |
| GPANLGNYNI | ASSLESFLEQY | 0.6795587 |
| TLGPTLP | ASSLESFLEQY | 0.6800277 |
| VYAYSSCCK | ASSLESFLEQY | 0.6669829 |
| APAPAPAPNI | ASSLESFLEQY | 0.79661304 |
| PGGPVPIP | ASSLESFLEQY | 0.8337523 |
| FLTVALEP | ASSLESFLEQY | 0.7251896 |
| SQYFGANL | ASSLESFLEQY | 0.47069347 |
| GGVKPAA | ASSLESFLEQY | 0.68961096 |
| GGTGGAGGAGGW | ASSLESFLEQY | 0.5865305 |
| GVGGVAGLGGAGG | ASSLESFLEQY | 0.6515379 |
| VLLVLATTI | ASSLESFLEQY | 0.7529486 |
| IGIFAAML | ASSLESFLEQY | 0.46086392 |
| AVGDLID | ASSLESFLEQY | 0.67643297 |
| NISPLPL | ASSLESFLEQY | 0.8311026 |
| VTAVFVAI | ASSLESFLEQY | 0.6487242 |
| SGDDNDAAL | ASSLESFLEQY | 0.72310305 |
| AVAALAHS | ASSLESFLEQY | 0.7889673 |
| GGVTVMATP | ASSLESFLEQY | 0.7421521 |
| AGLAANVRD | ASSLESFLEQY | 0.80004984 |
| TNRPDLIDPALLRPG | ASSLESFLEQY | 0.77301264 |
| LANLGVTAL | ASSLESFLEQY | 0.7682791 |
| TGERIEL | ASSLESFLEQY | 0.7204309 |
| NGEELAA | ASSLESFLEQY | 0.7073103 |
| VLSLVWG | ASSLESFLEQY | 0.71108884 |
| IGYALWL | ASSLESFLEQY | 0.6852187 |
| LSLLLAPL | ASSLESFLEQY | 0.6168425 |
| TLDEALK | ASSLESFLEQY | 0.80285215 |
| TDGAGTAEGG | ASSLESFLEQY | 0.74108714 |
| NAPGGAGGNGG | ASSLESFLEQY | 0.79300135 |
| VSVFMGMGGNF | ASSLESFLEQY | 0.6087339 |
| LASYLAAL | ASSLESFLEQY | 0.6861091 |
| LVGGTPAL | ASSLESFLEQY | 0.78482074 |
| VRNADVL | ASSLESFLEQY | 0.19496995 |
| TCSSPFLVSAM | ASSLESFLEQY | 0.31722862 |
| VHITDAE | ASSLESFLEQY | 0.73574746 |

| | | |
|---|---|---|
| EYAAPSF | ASSLESFLEQY | 0.80748916 |
| VVDAFHGGL | ASSLESFLEQY | 0.197782 |
| ASVVDPQ | ASSLESFLEQY | 0.6829781 |
| SYNEFGGLS | ASSLESFLEQY | 0.5244289 |
| DPDGSVY | ASSLESFLEQY | 0.79771405 |
| MVSTIHATE | ASSLESFLEQY | 0.75033367 |
| LAAGVADPV | ASSLESFLEQY | 0.52908385 |
| TKEQIAA | ASSLESFLEQY | 0.78580827 |
| SLASPAAL | ASSLESFLEQY | 0.40074116 |
| IGLVFYGV | ASSLESFLEQY | 0.5240753 |
| FLLDASNL | ASSLESFLEQY | 0.79224515 |
| YQVGNFP | ASSLESFLEQY | 0.80474854 |
| ESLLSQST | ASSLESFLEQY | 0.58049554 |
| LFTVGFAAVG | ASSLESFLEQY | 0.79007727 |
| ATPPTPT | ASSLESFLEQY | 0.6401627 |
| TDLATLG | ASSLESFLEQY | 0.8300403 |
| SAGGTGR | ASSLESFLEQY | 0.6914587 |
| GFSNSGS | ASSLESFLEQY | 0.5209703 |
| GGNGGSGGTSV | ASSLESFLEQY | 0.7836621 |
| GTSVATGGAGN | ASSLESFLEQY | 0.7402833 |
| NGGNGGSGGTS | ASSLESFLEQY | 0.79493636 |
| AVVKFPH | ASSLESFLEQY | 0.76387274 |
| TPPPPPP | ASSLESFLEQY | 0.69534326 |
| AAPAAAAAPA | ASSLESFLEQY | 0.66583717 |
| IALNEDL | ASSLESFLEQY | 0.79809916 |
| IVNVSSM | ASSLESFLEQY | 0.7788615 |
| TGIPPAPR | ASSLESFLEQY | 0.6589551 |
| FDLGGGTF | ASSLESFLEQY | 0.79760355 |
| FDLGGGTFDVS | ASSLESFLEQY | 0.7135454 |
| NEPTAAA | ASSLESFLEQY | 0.6442789 |
| NEPTAAALA | ASSLESFLEQY | 0.8263474 |
| GSIINISSI | ASSLESFLEQY | 0.7537252 |
| VAPGNAGTA | ASSLESFLEQY | 0.77252585 |
| TGGGASLE | ASSLESFLEQY | 0.81003654 |
| VAVYNHYKRI | ASSLESFLEQY | 0.7997309 |
| TDPAPATT | ASSLESFLEQY | 0.6462074 |
| TNRPDILDPALLRPG | ASSLESFLEQY | 0.74064285 |
| LTTTTTS | ASSLESFLEQY | 0.70938593 |
| PTGSGKTL | ASSLESFLEQY | 0.7867829 |
| TGSGKTL | ASSLESFLEQY | 0.7100766 |
| TIGHVDHGK | ASSLESFLEQY | 0.5273463 |
| TIGHVDHGKTT | ASSLESFLEQY | 0.76699746 |
| GHVDHGKTT | ASSLESFLEQY | 0.80260324 |
| AQFEHTLL | ASSLESFLEQY | 0.7438012 |
| IVNQDEL | ASSLESFLEQY | 0.7420504 |
| TNGSQFF | ASSLESFLEQY | 0.8201443 |
| TGKGIVE | ASSLESFLEQY | 0.6641129 |
| LISNASDA | ASSLESFLEQY | 0.7222495 |
| AELRTLGELTKIAKAHGAQ | ASSLESFLEQY | 0.794152 |
| APNAVAAAEPSW | ASSLESFLEQY | 0.7879992 |
| APQVAGAATAVL | ASSLESFLEQY | 0.730672 |

| | | |
|---|---|---|
| ARRDPAAAAAILPTDA | ASSLESFLEQY | 0.8108952 |
| DAAARDVLAEAGLAEYF | ASSLESFLEQY | 0.616969 |
| DDQIVLSAKANLAKVEQ | ASSLESFLEQY | 0.76508343 |
| DELLPDTGEDSDEDGH | ASSLESFLEQY | 0.5506659 |
| ELPEAPANRATVPQD | ASSLESFLEQY | 0.74095666 |
| FGQSKIERVTGIG | ASSLESFLEQY | 0.6239956 |
| GAGGTGSTAGGAGGAGGAGGLY | ASSLESFLEQY | 0.75625485 |
| GHGGTGAAGGNGGTGGAGGSGI | ASSLESFLEQY | 0.7695291 |
| GNGGNGGAGGEGGPGGAGGAGA | ASSLESFLEQY | 0.7664747 |
| GSTAGNGGNGGNATNGTIAGQP | ASSLESFLEQY | 0.24239928 |
| GTGGDGGNGGNGGTGG | ASSLESFLEQY | 0.7858786 |
| GVRRPAGIPADAPV | ASSLESFLEQY | 0.79081136 |
| GYRQVTDAHLLTLARR | ASSLESFLEQY | 0.8245344 |
| IAKMGQDGHDRG | ASSLESFLEQY | 0.71862894 |
| KAEPSIEIQYT | ASSLESFLEQY | 0.78042173 |
| KPVRIRMAADEDTVAVT | ASSLESFLEQY | 0.70547104 |
| MPPRIAGMRLLVIKPEP | ASSLESFLEQY | 0.72029495 |
| MKFYAVALFRSRGVPPTR | ASSLESFLEQY | 0.5463996 |
| NAGGFGILGAVAHSP | ASSLESFLEQY | 0.67148495 |
| NEGREITLAMAEM | ASSLESFLEQY | 0.609961 |
| NELVEWSHPLGHRWRWE | ASSLESFLEQY | 0.589471 |
| NLGAGGTGGNGGNPG | ASSLESFLEQY | 0.77729493 |
| NTEWPKWETVMAADPVD | ASSLESFLEQY | 0.78288954 |
| PASAAPAETQP | ASSLESFLEQY | 0.7817025 |
| QQSSDGKNGVRAAEITGE | ASSLESFLEQY | 0.8052451 |
| RAPDAVTTSWWKE | ASSLESFLEQY | 0.7954485 |
| SGFGNVGAHTSGWFNQ | ASSLESFLEQY | 0.816398 |
| SGVAGAVGQAASVGGLKV | ASSLESFLEQY | 0.70743805 |
| SNGGAGTGSAGGNGGT | ASSLESFLEQY | 0.69093 |
| TGGAGGEGGAAG | ASSLESFLEQY | 0.5506921 |
| TGGDGGAAGAGGAPGHG | ASSLESFLEQY | 0.67207265 |
| TMPPLLAAIANAGRHGV | ASSLESFLEQY | 0.7150383 |
| TSSTAGNSGGA | ASSLESFLEQY | 0.73425937 |
| VPATVPVGILC | ASSLESFLEQY | 0.17292292 |
| VQRPDSGGVVLTGRIS | ASSLESFLEQY | 0.6701162 |
| WTPNPVPATKGVPAGLVQAT | ASSLESFLEQY | 0.8135909 |
| YALKRALAKFSDRFDVVII | ASSLESFLEQY | 0.5894698 |
| YLIGSVIV | ASSLESFLEQY | 0.7745106 |
| YGQLVAVID | ASSLESFLEQY | 0.66724205 |
| VTVGGMTLPA | ASSLESFLEQY | 0.63317615 |
| VLALPAV | ASSLESFLEQY | 0.78169024 |
| VIPPTLNYET | ASSLESFLEQY | 0.6369678 |
| VGMGGMHPSSG | ASSLESFLEQY | 0.77119076 |
| VGLAGGAATA | ASSLESFLEQY | 0.8192411 |
| VGGAGGAGGSGP | ASSLESFLEQY | 0.73018616 |
| VFAAFMV | ASSLESFLEQY | 0.70917857 |
| TLYDKLR | ASSLESFLEQY | 0.7369136 |
| TLAVFTVL | ASSLESFLEQY | 0.58443874 |
| TGSVQPARV | ASSLESFLEQY | 0.72870946 |
| TGGGAGSGGT | ASSLESFLEQY | 0.24410926 |
| SHLETSL | ASSLESFLEQY | 0.6866631 |

| | | |
|---|---|---|
| SFTTGMT | ASSLESFLEQY | 0.65720224 |
| QVEVAKRMGVRQPTVS | ASSLESFLEQY | 0.74650997 |
| QGSGGPALAM | ASSLESFLEQY | 0.7225454 |
| MLAPSNP | ASSLESFLEQY | 0.7680276 |
| LVNNAAI | ASSLESFLEQY | 0.5624744 |
| LTGEINTLA | ASSLESFLEQY | 0.6971325 |
| LSIFGPV | ASSLESFLEQY | 0.61600405 |
| LLALLANL | ASSLESFLEQY | 0.6536604 |
| LISNASDALDKLR | ASSLESFLEQY | 0.7544527 |
| LINEAVP | ASSLESFLEQY | 0.7524569 |
| LGYAFSGL | ASSLESFLEQY | 0.64008987 |
| LDFVAVAL | ASSLESFLEQY | 0.80774796 |
| LAQPGAE | ASSLESFLEQY | 0.75517255 |
| ISNLGAVTGF | ASSLESFLEQY | 0.5091085 |
| HQEDPGRI | ASSLESFLEQY | 0.6589376 |
| GTPGEPTP | ASSLESFLEQY | 0.711346 |
| GFFNSGS | ASSLESFLEQY | 0.7169374 |
| GAGGGGLTAGAN | ASSLESFLEQY | 0.18449943 |
| FVDALVNM | ASSLESFLEQY | 0.75310856 |
| FLTVAPEL | ASSLESFLEQY | 0.64574593 |
| FETLTAL | ASSLESFLEQY | 0.54141474 |
| DVINPIEL | ASSLESFLEQY | 0.70231783 |
| APVSGAPV | ASSLESFLEQY | 0.7762517 |
| ALVGGLAW | ASSLESFLEQY | 0.6083743 |
| ALAAAAATAI | ASSLESFLEQY | 0.7883171 |
| AFGLGLLL | ASSLESFLEQY | 0.7636892 |
| AFFAAASL | ASSLESFLEQY | 0.79824543 |
| AAAALPAFSPP | ASSLESFLEQY | 0.81111664 |
| VVLAPAL | ASSLESFLEQY | 0.8020991 |
| TIGHVDHGKTTL | ASSLESFLEQY | 0.735945 |
| LISNASDALDK | ASSLESFLEQY | 0.78265965 |
| LISNASDAL | ASSLESFLEQY | 0.74559706 |
| ISNASDALDK | ASSLESFLEQY | 0.4107899 |
| NQAETLVYQTE | ASSLESFLEQY | 0.40603358 |
| TGGTGGTAGAG | ASSLESFLEQY | 0.710227 |
| TGSTAGGAGGA | ASSLESFLEQY | 0.36217 |
| DPLAVPVAAGPV | ASSLESFLEQY | 0.75227857 |
| GGTGGTPGNGGN | ASSLESFLEQY | 0.82405615 |
| PAGLAGVDPATL | ASSLESFLEQY | 0.7231556 |
| GNNLVVEAPHAGASV | ASSLESFLEQY | 0.56039083 |
| KVTRIMSASTLEQPA | ASSLESFLEQY | 0.6910461 |
| LGFALPIALPRRTVV | ASSLESFLEQY | 0.6844319 |
| TRLGNLTARKLVVLV | ASSLESFLEQY | 0.77112854 |
| WSQQRRLIYFPSAGP | ASSLESFLEQY | 0.8017862 |
| GPNGNPDPMAAAVDIR | ASSLESFLEQY | 0.8079879 |
| TQRCVSVADLRAAASR | ASSLESFLEQY | 0.8036558 |
| QFVLDVAEHPVEVLTETV | ASSLESFLEQY | 0.6603135 |
| GGAGGTGGAGGNGGTQAGDG | ASSLESFLEQY | 0.76072943 |
| PLLATLRAIAVDVDAKPAQV | ASSLESFLEQY | 0.6751604 |
| APKFRTGGYMIDFWGVGYQVAK | ASSLESFLEQY | 0.6458621 |
| GAGGAGGAGGAAGGPGGTGGTG | ASSLESFLEQY | 0.7229957 |

| | | |
|---|---|---|
| IGALGLLATDVAPAGSVAFGVA | ASSLESFLEQY | 0.7308246 |
| AGGAGGIGGNGGAGGAGGVLMG | ASSLESFLEQY | 0.75444824 |
| ADELIGLL | ASSLESFLEQY | 0.6775194 |
| ADRALKA | ASSLESFLEQY | 0.5208692 |
| AGAVGQVGGAGGA | ASSLESFLEQY | 0.7775913 |
| AGGGGGLGGAP | ASSLESFLEQY | 0.738634 |
| AGGLLYGNGGAG | ASSLESFLEQY | 0.41492417 |
| AGLESIGL | ASSLESFLEQY | 0.6656058 |
| AGLKTGII | ASSLESFLEQY | 0.8247545 |
| ASLAALLL | ASSLESFLEQY | 0.8177248 |
| ASVINISSI | ASSLESFLEQY | 0.5961113 |
| ATPPGPA | ASSLESFLEQY | 0.70451576 |
| ATVASALV | ASSLESFLEQY | 0.6810815 |
| ATVASLIL | ASSLESFLEQY | 0.4375436 |
| AVNGPVLIRH | ASSLESFLEQY | 0.77592003 |
| DGAAGAVGI | ASSLESFLEQY | 0.6865108 |
| DPLLHGI | ASSLESFLEQY | 0.6901624 |
| ELGAALGAL | ASSLESFLEQY | 0.670014 |
| ELGRTQAV | ASSLESFLEQY | 0.71144706 |
| FVAGGYLS | ASSLESFLEQY | 0.6812602 |
| GASGGAGGAGGW | ASSLESFLEQY | 0.7989842 |
| GGAGAGGDGGAGGI | ASSLESFLEQY | 0.18687394 |
| GTGHNGSQPGMGGNG | ASSLESFLEQY | 0.791221 |
| GYAAASASAS | ASSLESFLEQY | 0.5639739 |
| HQVDVAAVQQG | ASSLESFLEQY | 0.6901978 |
| IGAAIAHT | ASSLESFLEQY | 0.45905387 |
| IGMIEVAQL | ASSLESFLEQY | 0.7034441 |
| IRLAPEHTSAMD | ASSLESFLEQY | 0.77640367 |
| IYNVSVL | ASSLESFLEQY | 0.79061306 |
| KQGFLGM | ASSLESFLEQY | 0.49270684 |
| LESVITQL | ASSLESFLEQY | 0.7796652 |
| LGLGALVVV | ASSLESFLEQY | 0.81670725 |
| LIGNGGNGGSGGI | ASSLESFLEQY | 0.6151696 |
| LNAINAPTL | ASSLESFLEQY | 0.5392014 |
| LQLLNTGV | ASSLESFLEQY | 0.78983265 |
| LQTSIEF | ASSLESFLEQY | 0.62682056 |
| MSFAVSEL | ASSLESFLEQY | 0.7863523 |
| NTGALGL | ASSLESFLEQY | 0.7698358 |
| PPGAGGAGGAGTT | ASSLESFLEQY | 0.5077684 |
| QLVGLGI | ASSLESFLEQY | 0.8181782 |
| RFLEQVHQL | ASSLESFLEQY | 0.79310185 |
| RGVGYVLR | ASSLESFLEQY | 0.60712796 |
| SATGVFAGVI | ASSLESFLEQY | 0.62923056 |
| SLPAITLP | ASSLESFLEQY | 0.67079014 |
| STALAFTV | ASSLESFLEQY | 0.30396056 |
| SVLTVGLGK | ASSLESFLEQY | 0.8058876 |
| TALAGAGSGI | ASSLESFLEQY | 0.5897424 |
| TGLPFTGI | ASSLESFLEQY | 0.72462505 |
| VAEHVTV | ASSLESFLEQY | 0.7361198 |
| VDIVLNSL | ASSLESFLEQY | 0.75037897 |
| VLTGGGAVL | ASSLESFLEQY | 0.7372641 |

| | | |
|---|---|---|
| VNVGLAYGI | ASSLESFLEQY | 0.7413314 |
| VSSAAVGAV | ASSLESFLEQY | 0.77707875 |
| VTSVYEAGV | ASSLESFLEQY | 0.7375348 |
| WVALDRGA | ASSLESFLEQY | 0.70051616 |
| YLVGSVVI | ASSLESFLEQY | 0.7183158 |
| GPPGTGKTLL | ASSLESFLEQY | 0.8013455 |
| GVGMTKF | ASSLESFLEQY | 0.61445516 |
| IGIGGSDL | ASSLESFLEQY | 0.6774631 |
| IGTLLAEL | ASSLESFLEQY | 0.77416545 |
| IYTSGTT | ASSLESFLEQY | 0.5291583 |
| LSFPAGV | ASSLESFLEQY | 0.7626348 |
| LYTSGTT | ASSLESFLEQY | 0.73157024 |
| VGAFPFGF | ASSLESFLEQY | 0.7339124 |
| VGAGTVEF | ASSLESFLEQY | 0.70147544 |
| VGKSTLLL | ASSLESFLEQY | 0.8119425 |
| AALAHAGGGIV | ASSLESFLEQY | 0.78642964 |
| AALVAVSLL | ASSLESFLEQY | 0.7239355 |
| AAVSRAVAAL | ASSLESFLEQY | 0.71358955 |
| AGGSSGIGGF | ASSLESFLEQY | 0.6592915 |
| AGLTVANANG | ASSLESFLEQY | 0.78329545 |
| AGLVGAVL | ASSLESFLEQY | 0.7841176 |
| ALSQAFTV | ASSLESFLEQY | 0.5448836 |
| ALTFSVSL | ASSLESFLEQY | 0.6752535 |
| ALTGAALVAL | ASSLESFLEQY | 0.8124137 |
| ASFSIGAGVL | ASSLESFLEQY | 0.7842701 |
| DGSVYKI | ASSLESFLEQY | 0.47382393 |
| EQLAEIV | ASSLESFLEQY | 0.6569719 |
| FAVSELTVM | ASSLESFLEQY | 0.42699158 |
| GAAPLAAGL | ASSLESFLEQY | 0.7186235 |
| GADGGDGGSGGAGG | ASSLESFLEQY | 0.6081799 |
| GFGLDTGGAGG | ASSLESFLEQY | 0.7450329 |
| GLASGGLL | ASSLESFLEQY | 0.64812905 |
| GRIAGAAVLL | ASSLESFLEQY | 0.7369775 |
| GSAGSLNTSF | ASSLESFLEQY | 0.75075155 |
| GTGQIGI | ASSLESFLEQY | 0.8245185 |
| GTLIALIL | ASSLESFLEQY | 0.65004057 |
| IAATMGFVVG | ASSLESFLEQY | 0.7422535 |
| IFAQALL | ASSLESFLEQY | 0.5172862 |
| IGAAVALGF | ASSLESFLEQY | 0.5905364 |
| IGLTPLAL | ASSLESFLEQY | 0.6440764 |
| ILGSNTSTL | ASSLESFLEQY | 0.7530913 |
| IVFSLTVL | ASSLESFLEQY | 0.28014007 |
| LASKATGF | ASSLESFLEQY | 0.8120529 |
| LASVALIL | ASSLESFLEQY | 0.7164407 |
| LFAVGLTL | ASSLESFLEQY | 0.79422563 |
| LFNGGIGTYI | ASSLESFLEQY | 0.7182709 |
| LTLGAGII | ASSLESFLEQY | 0.693856 |
| LVAAALGDI | ASSLESFLEQY | 0.69448984 |
| NTAFVQLGI | ASSLESFLEQY | 0.77847433 |
| QLFSGAEL | ASSLESFLEQY | 0.6354096 |
| SAATFITL | ASSLESFLEQY | 0.42899096 |

| | | |
|---|---|---|
| SNYGLANLGV | ASSLESFLEQY | 0.6788945 |
| SYGGAYAVM | ASSLESFLEQY | 0.48306865 |
| TTGVGGLTLG | ASSLESFLEQY | 0.809456 |
| TTQTIYI | ASSLESFLEQY | 0.79248583 |
| VAAGAGLGAVI | ASSLESFLEQY | 0.7411498 |
| VAAVVTLL | ASSLESFLEQY | 0.80178976 |
| VGFLGGLI | ASSLESFLEQY | 0.279333 |
| VGVIPILL | ASSLESFLEQY | 0.7192364 |
| VIFASSLI | ASSLESFLEQY | 0.82423496 |
| VKGGDGGAAAG | ASSLESFLEQY | 0.7631777 |
| VTKTFTNI | ASSLESFLEQY | 0.67314583 |
| VVAAAILL | ASSLESFLEQY | 0.80684525 |
| YAALDVEL | ASSLESFLEQY | 0.6617286 |
| LASVALLL | ASSLESFLEQY | 0.726729 |
| LDAVHRV | ASSLESFLEQY | 0.76208365 |
| QGLSSSIF | ASSLESFLEQY | 0.7790781 |
| RGTFANIRL | ASSLESFLEQY | 0.730313 |
| RQLWWGHRI | ASSLESFLEQY | 0.6207613 |
| YGPPGTGKTLL | ASSLESFLEQY | 0.77607584 |
| AALVGAPANVA | ASSLESFLEQY | 0.8173574 |
| AATFRRTE | ASSLESFLEQY | 0.66450393 |
| AGALGGGI | ASSLESFLEQY | 0.78665984 |
| AGGAGGAGGIGAG | ASSLESFLEQY | 0.7531829 |
| AGSGALGL | ASSLESFLEQY | 0.7255039 |
| AQIADQI | ASSLESFLEQY | 0.5940924 |
| ASAVAVTYL | ASSLESFLEQY | 0.61816853 |
| ASTVLGGTGG | ASSLESFLEQY | 0.70219636 |
| AVGGEPAK | ASSLESFLEQY | 0.5365503 |
| CEQGTFGGTC | ASSLESFLEQY | 0.72103095 |
| DEGKAPAG | ASSLESFLEQY | 0.7009553 |
| DTVVFPAL | ASSLESFLEQY | 0.8412391 |
| FEGITMAG | ASSLESFLEQY | 0.7524673 |
| FPLVVQD | ASSLESFLEQY | 0.7803303 |
| GAFTTPQ | ASSLESFLEQY | 0.6506269 |
| GFGGFGTAGGVGGA | ASSLESFLEQY | 0.7493067 |
| GFLAQFL | ASSLESFLEQY | 0.7474226 |
| GGAGGAGGNAGVA | ASSLESFLEQY | 0.53421336 |
| GGVGTAGAGGAGGAGG | ASSLESFLEQY | 0.8048998 |
| GNGGGGDAGNAGSGGN | ASSLESFLEQY | 0.63324577 |
| GTGGAGGAGNDAGS | ASSLESFLEQY | 0.6619363 |
| GTTNTGWFNS | ASSLESFLEQY | 0.80153275 |
| IFMGPLFTL | ASSLESFLEQY | 0.79077965 |
| ILAHAIEAI | ASSLESFLEQY | 0.6726087 |
| IRSAIREAL | ASSLESFLEQY | 0.6371342 |
| ITFAGGG | ASSLESFLEQY | 0.6701342 |
| ITGMSGHQEV | ASSLESFLEQY | 0.7898525 |
| IVAGLACTL | ASSLESFLEQY | 0.79946065 |
| IVGGGLAGA | ASSLESFLEQY | 0.68094444 |
| IVTGVLNI | ASSLESFLEQY | 0.81578827 |
| LAQQGCI | ASSLESFLEQY | 0.7957441 |
| LCLHVEN | ASSLESFLEQY | 0.54156464 |

| | | |
|---|---|---|
| LDAISDGS | ASSLESFLEQY | 0.73317087 |
| LGGFAAML | ASSLESFLEQY | 0.7843781 |
| LGGKNPMI | ASSLESFLEQY | 0.7774977 |
| LGPKGVAAIL | ASSLESFLEQY | 0.8357097 |
| LMVGVSI | ASSLESFLEQY | 0.74164504 |
| LVGAAFVL | ASSLESFLEQY | 0.62986386 |
| LVLAVAPL | ASSLESFLEQY | 0.81016076 |
| MAAIAVIDAA | ASSLESFLEQY | 0.8352115 |
| NNGINAELQ | ASSLESFLEQY | 0.79256445 |
| RGEPLVLVHG | ASSLESFLEQY | 0.6817254 |
| RGGAGGAAGLI | ASSLESFLEQY | 0.7113938 |
| RSYRMHPAV | ASSLESFLEQY | 0.7427732 |
| SAAHLLAGQLA | ASSLESFLEQY | 0.7896173 |
| SAAVGSY | ASSLESFLEQY | 0.61356056 |
| SPHAVLHRI | ASSLESFLEQY | 0.70624053 |
| SQIAKVL | ASSLESFLEQY | 0.74518096 |
| TALGHLVP | ASSLESFLEQY | 0.49099493 |
| TAPTTSGAA | ASSLESFLEQY | 0.75263256 |
| TDGDDGNGGSPGT | ASSLESFLEQY | 0.7838576 |
| TDGLGEN | ASSLESFLEQY | 0.73002094 |
| TGGPPLPGQA | ASSLESFLEQY | 0.66813946 |
| TIPAITI | ASSLESFLEQY | 0.78503895 |
| TIPGFTI | ASSLESFLEQY | 0.83399665 |
| TSIASGALA | ASSLESFLEQY | 0.8143929 |
| TVTLPGAAI | ASSLESFLEQY | 0.5991659 |
| VASGLSAALP | ASSLESFLEQY | 0.31605276 |
| VGAGVIGI | ASSLESFLEQY | 0.6058847 |
| VGPKRFGV | ASSLESFLEQY | 0.75403345 |
| VNVAFPD | ASSLESFLEQY | 0.7498591 |
| VSMTNSGVS | ASSLESFLEQY | 0.7203026 |
| VTDVLGDLSV | ASSLESFLEQY | 0.70723987 |
| VVVVGAGFAGL | ASSLESFLEQY | 0.802403 |
| VYTVGTVV | ASSLESFLEQY | 0.6637038 |
| WTTFSGP | ASSLESFLEQY | 0.6536474 |
| YDELVEAF | ASSLESFLEQY | 0.71373725 |
| YGTGGAGGNGG | ASSLESFLEQY | 0.78952634 |
| YTNVMAV | ASSLESFLEQY | 0.57627153 |
| FLEQVHQL | ASSLESFLEQY | 0.65714544 |
| GLSLLAVAV | ASSLESFLEQY | 0.618102 |
| LEQVHQL | ASSLESFLEQY | 0.55333763 |
| LNAAGAI | ASSLESFLEQY | 0.72957706 |
| PAVHASL | ASSLESFLEQY | 0.6279017 |
| TELEDDA | ASSLESFLEQY | 0.7905365 |
| TPPGPPN | ASSLESFLEQY | 0.5781115 |
| TPPPPPPP | ASSLESFLEQY | 0.7738699 |
| TQSPALI | ASGVQNSDYT | 0.7420285 |
| EFAAAFPQ | ASGVQNSDYT | 0.7769993 |
| LLSGNLDVL | ASGVQNSDYT | 0.64703816 |
| GPANLGNYNI | ASGVQNSDYT | 0.60658973 |
| TLGPTLP | ASGVQNSDYT | 0.58736914 |
| VYAYSSCCK | ASGVQNSDYT | 0.42837524 |

| | | |
|---|---|---|
| APAPAPAPNI | ASGVQNSDYT | 0.7821916 |
| PGGPVPIP | ASGVQNSDYT | 0.52243155 |
| FLTVALEP | ASGVQNSDYT | 0.7956407 |
| SQYFGANL | ASGVQNSDYT | 0.78842777 |
| GGVKPAA | ASGVQNSDYT | 0.6203208 |
| GGTGGAGGAGGW | ASGVQNSDYT | 0.75857574 |
| GVGGVAGLGGAGG | ASGVQNSDYT | 0.64628756 |
| VLLVLATTI | ASGVQNSDYT | 0.60107076 |
| IGIFAAML | ASGVQNSDYT | 0.76775014 |
| AVGDLID | ASGVQNSDYT | 0.78230804 |
| NISPLPL | ASGVQNSDYT | 0.58874613 |
| VTAVFVAI | ASGVQNSDYT | 0.7554269 |
| SGDDNDAAL | ASGVQNSDYT | 0.7292502 |
| AVAALAHS | ASGVQNSDYT | 0.6632799 |
| GGVTVMATP | ASGVQNSDYT | 0.748303 |
| AGLAANVRD | ASGVQNSDYT | 0.7986193 |
| TNRPDLIDPALLRPG | ASGVQNSDYT | 0.76380366 |
| LANLGVTAL | ASGVQNSDYT | 0.62286484 |
| TGERIEL | ASGVQNSDYT | 0.7635429 |
| NGEELAA | ASGVQNSDYT | 0.5482353 |
| VLSLVWG | ASGVQNSDYT | 0.74471885 |
| IGYALWL | ASGVQNSDYT | 0.71948904 |
| LSLLLAPL | ASGVQNSDYT | 0.714874 |
| TLDEALK | ASGVQNSDYT | 0.7732271 |
| TDGAGTAEGG | ASGVQNSDYT | 0.6073691 |
| NAPGGAGGNGG | ASGVQNSDYT | 0.6031864 |
| VSVFMGMGGNF | ASGVQNSDYT | 0.66705436 |
| LASYLAAL | ASGVQNSDYT | 0.77275926 |
| LVGGTPAL | ASGVQNSDYT | 0.78982294 |
| VRNADVL | ASGVQNSDYT | 0.60079664 |
| TCSSPFLVSAM | ASGVQNSDYT | 0.73955834 |
| VHITDAE | ASGVQNSDYT | 0.6957203 |
| EYAAPSF | ASGVQNSDYT | 0.63999975 |
| VVDAFHGGL | ASGVQNSDYT | 0.75986683 |
| ASVVDPQ | ASGVQNSDYT | 0.7076503 |
| SYNEFGGLS | ASGVQNSDYT | 0.6202463 |
| DPDGSVY | ASGVQNSDYT | 0.7619022 |
| MVSTIHATE | ASGVQNSDYT | 0.723402 |
| LAAGVADPV | ASGVQNSDYT | 0.7916878 |
| TKEQIAA | ASGVQNSDYT | 0.7802386 |
| SLASPAAL | ASGVQNSDYT | 0.66399276 |
| IGLVFYGV | ASGVQNSDYT | 0.772595 |
| FLLDASNL | ASGVQNSDYT | 0.7540895 |
| YQVGNFP | ASGVQNSDYT | 0.7258087 |
| ESLLSQST | ASGVQNSDYT | 0.75446844 |
| LFTVGFAAVG | ASGVQNSDYT | 0.6464299 |
| ATPPTPT | ASGVQNSDYT | 0.80868304 |
| TDLATLG | ASGVQNSDYT | 0.78960836 |
| SAGGTGR | ASGVQNSDYT | 0.78065324 |
| GFSNSGS | ASGVQNSDYT | 0.7489647 |
| GGNGGSGGTSV | ASGVQNSDYT | 0.737272 |

| | | |
|---|---|---|
| GTSVATGGAGN | ASGVQNSDYT | 0.7746797 |
| NGGNGGSGGTS | ASGVQNSDYT | 0.62202644 |
| AVVKFPH | ASGVQNSDYT | 0.5315945 |
| TPPPPPP | ASGVQNSDYT | 0.73690003 |
| AAPAAAAAPA | ASGVQNSDYT | 0.6511346 |
| IALNEDL | ASGVQNSDYT | 0.70047694 |
| IVNVSSM | ASGVQNSDYT | 0.5496643 |
| TGIPPAPR | ASGVQNSDYT | 0.79462415 |
| FDLGGGTF | ASGVQNSDYT | 0.83694965 |
| FDLGGGTFDVS | ASGVQNSDYT | 0.7078413 |
| NEPTAAA | ASGVQNSDYT | 0.8185105 |
| NEPTAAALA | ASGVQNSDYT | 0.546989 |
| GSIINISSI | ASGVQNSDYT | 0.7872968 |
| VAPGNAGTA | ASGVQNSDYT | 0.80368274 |
| TGGGASLE | ASGVQNSDYT | 0.57578695 |
| VAVYNHYKRI | ASGVQNSDYT | 0.80557007 |
| TDPAPATT | ASGVQNSDYT | 0.63868564 |
| TNRPDILDPALLRPG | ASGVQNSDYT | 0.717003 |
| LTTTTTS | ASGVQNSDYT | 0.80519444 |
| PTGSGKTL | ASGVQNSDYT | 0.75475484 |
| TGSGKTL | ASGVQNSDYT | 0.7460095 |
| TIGHVDHGK | ASGVQNSDYT | 0.766057 |
| TIGHVDHGKTT | ASGVQNSDYT | 0.7009786 |
| GHVDHGKTT | ASGVQNSDYT | 0.7531731 |
| AQFEHTLL | ASGVQNSDYT | 0.8095088 |
| IVNQDEL | ASGVQNSDYT | 0.60794693 |
| TNGSQFF | ASGVQNSDYT | 0.83181727 |
| TGKGIVE | ASGVQNSDYT | 0.60746956 |
| LISNASDA | ASGVQNSDYT | 0.8262589 |
| AELRTLGELTKIAKAHGAQ | ASGVQNSDYT | 0.7586907 |
| APNAVAAAEPSW | ASGVQNSDYT | 0.7077376 |
| APQVAGAATAVL | ASGVQNSDYT | 0.6898319 |
| ARRDPAAAAAILPTDA | ASGVQNSDYT | 0.7942321 |
| DAAARDVLAEAGLAEYF | ASGVQNSDYT | 0.707295 |
| DDQIVLSAKANLAKVEQ | ASGVQNSDYT | 0.7527988 |
| DELLPDTGEDSDEDGH | ASGVQNSDYT | 0.82484716 |
| ELPEAPANRATVPQD | ASGVQNSDYT | 0.5778049 |
| FGQSKIERVTGIG | ASGVQNSDYT | 0.84257436 |
| GAGGTGSTAGGAGGAGGAGGLY | ASGVQNSDYT | 0.78627497 |
| GHGGTGAAGGNGGTGGAGGSGI | ASGVQNSDYT | 0.82022345 |
| GNGGNGGAGGEGGPGGAGGAGA | ASGVQNSDYT | 0.69422346 |
| GSTAGNGGNGGNATNGTIAGQP | ASGVQNSDYT | 0.39497405 |
| GTGGDGGNGGNGGTGG | ASGVQNSDYT | 0.78575087 |
| GVRRPAGIPADAPV | ASGVQNSDYT | 0.61283565 |
| GYRQVTDAHLLTLARR | ASGVQNSDYT | 0.8142461 |
| IAKMGQDGHDRG | ASGVQNSDYT | 0.8243818 |
| KAEPSIEIQYT | ASGVQNSDYT | 0.7155291 |
| KPVRIRMAADEDTVAVT | ASGVQNSDYT | 0.83054245 |
| MPPRIAGMRLLVIKPEP | ASGVQNSDYT | 0.8387522 |
| MKFYAVALFRSRGVPPTR | ASGVQNSDYT | 0.4963253 |
| NAGGFGILGAVAHSP | ASGVQNSDYT | 0.7467351 |

| | | |
|---|---|---|
| NEGREITLAMAEM | ASGVQNSDYT | 0.7213758 |
| NELVEWSHPLGHRWRWE | ASGVQNSDYT | 0.76191163 |
| NLGAGGTGGNGGNPG | ASGVQNSDYT | 0.42411596 |
| NTEWPKWETVMAADPVD | ASGVQNSDYT | 0.82171106 |
| PASAAPAETQP | ASGVQNSDYT | 0.71708566 |
| QQSSDGKNGVRAAEITGE | ASGVQNSDYT | 0.18303868 |
| RAPDAVTTSWWKE | ASGVQNSDYT | 0.79896426 |
| SGFGNVGAHTSGWFNQ | ASGVQNSDYT | 0.72678757 |
| SGVAGAVGQAASVGGLKV | ASGVQNSDYT | 0.69746596 |
| SNGGAGTGSAGGNGGT | ASGVQNSDYT | 0.83032805 |
| TGGAGGEGGAAG | ASGVQNSDYT | 0.538523 |
| TGGDGGAAGAGGAPGHG | ASGVQNSDYT | 0.6054005 |
| TMPPLLAAIANAGRHGV | ASGVQNSDYT | 0.80890936 |
| TSSTAGNSGGA | ASGVQNSDYT | 0.8394207 |
| VPATVPVGILC | ASGVQNSDYT | 0.50356346 |
| VQRPDSGGVVLTGRIS | ASGVQNSDYT | 0.79406357 |
| WTPNPVPATKGVPAGLVQAT | ASGVQNSDYT | 0.85004103 |
| YALKRALAKFSDRFDVVII | ASGVQNSDYT | 0.7986134 |
| YLIGSVIV | ASGVQNSDYT | 0.84411323 |
| YGQLVAVID | ASGVQNSDYT | 0.848518 |
| VTVGGMTLPA | ASGVQNSDYT | 0.72264266 |
| VLALPAV | ASGVQNSDYT | 0.8315822 |
| VIPPTLNYET | ASGVQNSDYT | 0.8411974 |
| VGMGGMHPSSG | ASGVQNSDYT | 0.7319851 |
| VGLAGGAATA | ASGVQNSDYT | 0.7843657 |
| VGGAGGAGGSGP | ASGVQNSDYT | 0.6201234 |
| VFAAFMV | ASGVQNSDYT | 0.77639353 |
| TLYDKLR | ASGVQNSDYT | 0.3197313 |
| TLAVFTVL | ASGVQNSDYT | 0.7933382 |
| TGSVQPARV | ASGVQNSDYT | 0.8409851 |
| TGGGAGSGGT | ASGVQNSDYT | 0.7657331 |
| SHLETSL | ASGVQNSDYT | 0.5607939 |
| SFTTGMT | ASGVQNSDYT | 0.5986891 |
| QVEVAKRMGVRQPTVS | ASGVQNSDYT | 0.79438066 |
| QGSGGPALAM | ASGVQNSDYT | 0.65806794 |
| MLAPSNP | ASGVQNSDYT | 0.4334339 |
| LVNNAAI | ASGVQNSDYT | 0.6873348 |
| LTGEINTLA | ASGVQNSDYT | 0.8390155 |
| LSIFGPV | ASGVQNSDYT | 0.8418454 |
| LLALLANL | ASGVQNSDYT | 0.60653985 |
| LISNASDALDKLR | ASGVQNSDYT | 0.8450731 |
| LINEAVP | ASGVQNSDYT | 0.81824476 |
| LGYAFSGL | ASGVQNSDYT | 0.85052884 |
| LDFVAVAL | ASGVQNSDYT | 0.7980087 |
| LAQPGAE | ASGVQNSDYT | 0.6043712 |
| ISNLGAVTGF | ASGVQNSDYT | 0.832284 |
| HQEDPGRI | ASGVQNSDYT | 0.7478272 |
| GTPGEPTP | ASGVQNSDYT | 0.7602611 |
| GFFNSGS | ASGVQNSDYT | 0.83611035 |
| GAGGGGLTAGAN | ASGVQNSDYT | 0.7330116 |
| FVDALVNM | ASGVQNSDYT | 0.6532062 |

| | | |
|---|---|---|
| FLTVAPEL | ASGVQNSDYT | 0.758204 |
| FETLTAL | ASGVQNSDYT | 0.8470767 |
| DVINPIEL | ASGVQNSDYT | 0.6536908 |
| APVSGAPV | ASGVQNSDYT | 0.56443906 |
| ALVGGLAW | ASGVQNSDYT | 0.71387607 |
| ALAAAAATAI | ASGVQNSDYT | 0.78228176 |
| AFGLGLLL | ASGVQNSDYT | 0.8380157 |
| AFFAAASL | ASGVQNSDYT | 0.81040156 |
| AAAALPAFSPP | ASGVQNSDYT | 0.7595706 |
| VVLAPAL | ASGVQNSDYT | 0.8083844 |
| TIGHVDHGKTTL | ASGVQNSDYT | 0.82967013 |
| LISNASDALDK | ASGVQNSDYT | 0.8032396 |
| LISNASDAL | ASGVQNSDYT | 0.81212026 |
| ISNASDALDK | ASGVQNSDYT | 0.6822958 |
| NQAETLVYQTE | ASGVQNSDYT | 0.8423869 |
| TGGTGGTAGAG | ASGVQNSDYT | 0.83618486 |
| TGSTAGGAGGA | ASGVQNSDYT | 0.83253944 |
| DPLAVPVAAGPV | ASGVQNSDYT | 0.83380556 |
| GGTGGTPGNGGN | ASGVQNSDYT | 0.7971889 |
| PAGLAGVDPATL | ASGVQNSDYT | 0.80957043 |
| GNNLVVEAPHAGASV | ASGVQNSDYT | 0.51966536 |
| KVTRIMSASTLEQPA | ASGVQNSDYT | 0.7341619 |
| LGFALPIALPRRTVV | ASGVQNSDYT | 0.7756034 |
| TRLGNLTARKLVVLV | ASGVQNSDYT | 0.7960058 |
| WSQQRRLIYFPSAGP | ASGVQNSDYT | 0.8082148 |
| GPNGNPDPMAAAVDIR | ASGVQNSDYT | 0.81471914 |
| TQRCVSVADLRAAASR | ASGVQNSDYT | 0.7110461 |
| QFVLDVAEHPVEVLTETV | ASGVQNSDYT | 0.8353329 |
| GGAGGTGGAGGNGGTQAGDG | ASGVQNSDYT | 0.8383195 |
| PLLATLRAIAVDVDAKPAQV | ASGVQNSDYT | 0.5582522 |
| APKFRTGGYMIDFWGVGYQVAK | ASGVQNSDYT | 0.7633191 |
| GAGGAGGAGGAAGGPGGTGGTG | ASGVQNSDYT | 0.84535766 |
| IGALGLLATDVAPAGSVAFGVA | ASGVQNSDYT | 0.80265975 |
| AGGAGGIGGNGGAGGAGGVLMG | ASGVQNSDYT | 0.75899357 |
| ADELIGLL | ASGVQNSDYT | 0.7793857 |
| ADRALKA | ASGVQNSDYT | 0.84392977 |
| AGAVGQVGGAGGA | ASGVQNSDYT | 0.7312838 |
| AGGGGGLGGAP | ASGVQNSDYT | 0.8462084 |
| AGGLLYGNGGAG | ASGVQNSDYT | 0.3361164 |
| AGLESIGL | ASGVQNSDYT | 0.7603141 |
| AGLKTGII | ASGVQNSDYT | 0.7434442 |
| ASLAALLL | ASGVQNSDYT | 0.23257375 |
| ASVINISSI | ASGVQNSDYT | 0.83065236 |
| ATPPGPA | ASGVQNSDYT | 0.71131444 |
| ATVASALV | ASGVQNSDYT | 0.8261085 |
| ATVASLIL | ASGVQNSDYT | 0.750306 |
| AVNGPVLIRH | ASGVQNSDYT | 0.8323092 |
| DGAAGAVGI | ASGVQNSDYT | 0.79945046 |
| DPLLHGI | ASGVQNSDYT | 0.7878752 |
| ELGAALGAL | ASGVQNSDYT | 0.6680815 |
| ELGRTQAV | ASGVQNSDYT | 0.82000387 |

| | | |
|---|---|---|
| FVAGGYLS | ASGVQNSDYT | 0.83367103 |
| GASGGAGGAGGW | ASGVQNSDYT | 0.8379979 |
| GGAGAGGDGGAGGI | ASGVQNSDYT | 0.34908152 |
| GTGHNGSQPGMGGNG | ASGVQNSDYT | 0.7615922 |
| GYAAASASAS | ASGVQNSDYT | 0.7667633 |
| HQVDVAAVQQG | ASGVQNSDYT | 0.2878159 |
| IGAAIAHT | ASGVQNSDYT | 0.2528402 |
| IGMIEVAQL | ASGVQNSDYT | 0.694986 |
| IRLAPEHTSAMD | ASGVQNSDYT | 0.7386166 |
| IYNVSVL | ASGVQNSDYT | 0.82083344 |
| KQGFLGM | ASGVQNSDYT | 0.8269876 |
| LESVITQL | ASGVQNSDYT | 0.75004005 |
| LGLGALVVV | ASGVQNSDYT | 0.8245386 |
| LIGNGGNGGSGGI | ASGVQNSDYT | 0.78471416 |
| LNAINAPTL | ASGVQNSDYT | 0.12033504 |
| LQLLNTGV | ASGVQNSDYT | 0.7536887 |
| LQTSIEF | ASGVQNSDYT | 0.49777567 |
| MSFAVSEL | ASGVQNSDYT | 0.84289813 |
| NTGALGL | ASGVQNSDYT | 0.63957703 |
| PPGAGGAGGAGTT | ASGVQNSDYT | 0.71857613 |
| QLVGLGI | ASGVQNSDYT | 0.78258455 |
| RFLEQVHQL | ASGVQNSDYT | 0.79106355 |
| RGVGYVLR | ASGVQNSDYT | 0.7887345 |
| SATGVFAGVI | ASGVQNSDYT | 0.65320945 |
| SLPAITLP | ASGVQNSDYT | 0.21496652 |
| STALAFTV | ASGVQNSDYT | 0.82896125 |
| SVLTVGLGK | ASGVQNSDYT | 0.59002995 |
| TALAGAGSGI | ASGVQNSDYT | 0.5261916 |
| TGLPFTGI | ASGVQNSDYT | 0.5711947 |
| VAEHVTV | ASGVQNSDYT | 0.5279974 |
| VDIVLNSL | ASGVQNSDYT | 0.43051338 |
| VLTGGGAVL | ASGVQNSDYT | 0.8380054 |
| VNVGLAYGI | ASGVQNSDYT | 0.7234293 |
| VSSAAVGAV | ASGVQNSDYT | 0.7479763 |
| VTSVYEAGV | ASGVQNSDYT | 0.8434125 |
| WVALDRGA | ASGVQNSDYT | 0.78295434 |
| YLVGSVVI | ASGVQNSDYT | 0.767112 |
| GPPGTGKTLL | ASGVQNSDYT | 0.65432227 |
| GVGMTKF | ASGVQNSDYT | 0.6459591 |
| IGIGGSDL | ASGVQNSDYT | 0.67308295 |
| IGTLLAEL | ASGVQNSDYT | 0.6350056 |
| IYTSGTT | ASGVQNSDYT | 0.84627503 |
| LSFPAGV | ASGVQNSDYT | 0.7771106 |
| LYTSGTT | ASGVQNSDYT | 0.5767412 |
| VGAFPFGF | ASGVQNSDYT | 0.8271897 |
| VGAGTVEF | ASGVQNSDYT | 0.8455806 |
| VGKSTLLL | ASGVQNSDYT | 0.85133624 |
| AALAHAGGGIV | ASGVQNSDYT | 0.6444266 |
| AALVAVSLL | ASGVQNSDYT | 0.7745508 |
| AAVSRAVAAL | ASGVQNSDYT | 0.7711055 |
| AGGSSGIGGF | ASGVQNSDYT | 0.7676165 |

| | | |
|---|---|---|
| AGLTVANANG | ASGVQNSDYT | 0.6890644 |
| AGLVGAVL | ASGVQNSDYT | 0.5415486 |
| ALSQAFTV | ASGVQNSDYT | 0.8375709 |
| ALTFSVSL | ASGVQNSDYT | 0.8547296 |
| ALTGAALVAL | ASGVQNSDYT | 0.7761174 |
| ASFSIGAGVL | ASGVQNSDYT | 0.61507964 |
| DGSVYKI | ASGVQNSDYT | 0.7614219 |
| EQLAEIV | ASGVQNSDYT | 0.6369804 |
| FAVSELTVM | ASGVQNSDYT | 0.70501435 |
| GAAPLAAGL | ASGVQNSDYT | 0.779449 |
| GADGGDGGSGGAGG | ASGVQNSDYT | 0.49807736 |
| GFGLDTGGAGG | ASGVQNSDYT | 0.757563 |
| GLASGGLL | ASGVQNSDYT | 0.8543015 |
| GRIAGAAVLL | ASGVQNSDYT | 0.7446302 |
| GSAGSLNTSF | ASGVQNSDYT | 0.8180009 |
| GTGQIGI | ASGVQNSDYT | 0.8369498 |
| GTLIALIL | ASGVQNSDYT | 0.8151878 |
| IAATMGFVVG | ASGVQNSDYT | 0.8439807 |
| IFAQALL | ASGVQNSDYT | 0.8182547 |
| IGAAVALGF | ASGVQNSDYT | 0.78304124 |
| IGLTPLAL | ASGVQNSDYT | 0.7928771 |
| ILGSNTSTL | ASGVQNSDYT | 0.80064857 |
| IVFSLTVL | ASGVQNSDYT | 0.76439583 |
| LASKATGF | ASGVQNSDYT | 0.79200137 |
| LASVALIL | ASGVQNSDYT | 0.7698114 |
| LFAVGLTL | ASGVQNSDYT | 0.8202139 |
| LFNGGIGTYI | ASGVQNSDYT | 0.79802406 |
| LTLGAGII | ASGVQNSDYT | 0.84060687 |
| LVAAALGDI | ASGVQNSDYT | 0.30284745 |
| NTAFVQLGI | ASGVQNSDYT | 0.54759216 |
| QLFSGAEL | ASGVQNSDYT | 0.83233297 |
| SAATFITL | ASGVQNSDYT | 0.19519496 |
| SNYGLANLGV | ASGVQNSDYT | 0.3237778 |
| SYGGAYAVM | ASGVQNSDYT | 0.7957325 |
| TTGVGGLTLG | ASGVQNSDYT | 0.8442011 |
| TTQTIYI | ASGVQNSDYT | 0.16650316 |
| VAAGAGLGAVI | ASGVQNSDYT | 0.6225083 |
| VAAVVTLL | ASGVQNSDYT | 0.5321873 |
| VGFLGGLI | ASGVQNSDYT | 0.8258065 |
| VGVIPILL | ASGVQNSDYT | 0.76109385 |
| VIFASSLI | ASGVQNSDYT | 0.39399934 |
| VKGGDGGAAAG | ASGVQNSDYT | 0.8249974 |
| VTKTFTNI | ASGVQNSDYT | 0.61672056 |
| VVAAAILL | ASGVQNSDYT | 0.70414746 |
| YAALDVEL | ASGVQNSDYT | 0.83436954 |
| LASVALLL | ASGVQNSDYT | 0.841135 |
| LDAVHRV | ASGVQNSDYT | 0.5931772 |
| QGLSSSIF | ASGVQNSDYT | 0.8197148 |
| RGTFANIRL | ASGVQNSDYT | 0.72929144 |
| RQLWWGHRI | ASGVQNSDYT | 0.8551596 |
| YGPPGTGKTLL | ASGVQNSDYT | 0.7324562 |

| | | |
|---|---|---|
| AALVGAPANVA | ASGVQNSDYT | 0.4026938 |
| AATFRRTE | ASGVQNSDYT | 0.8279253 |
| AGALGGGI | ASGVQNSDYT | 0.821771 |
| AGGAGGAGGIGAG | ASGVQNSDYT | 0.7733026 |
| AGSGALGL | ASGVQNSDYT | 0.81751794 |
| AQIADQI | ASGVQNSDYT | 0.7559694 |
| ASAVAVTYL | ASGVQNSDYT | 0.7461829 |
| ASTVLGGTGG | ASGVQNSDYT | 0.8260257 |
| AVGGEPAK | ASGVQNSDYT | 0.8136766 |
| CEQGTFGGTC | ASGVQNSDYT | 0.57182384 |
| DEGKAPAG | ASGVQNSDYT | 0.837855 |
| DTVVFPAL | ASGVQNSDYT | 0.784448 |
| FEGITMAG | ASGVQNSDYT | 0.6593278 |
| FPLVVQD | ASGVQNSDYT | 0.8538427 |
| GAFTTPQ | ASGVQNSDYT | 0.7859463 |
| GFGGFGTAGGVGGA | ASGVQNSDYT | 0.8304417 |
| GFLAQFL | ASGVQNSDYT | 0.8218167 |
| GGAGGAGGNAGVA | ASGVQNSDYT | 0.8388163 |
| GGVGTAGAGGAGGAGG | ASGVQNSDYT | 0.44108868 |
| GNGGGGDAGNAGSGGN | ASGVQNSDYT | 0.794043 |
| GTGGAGGAGNDAGS | ASGVQNSDYT | 0.7957095 |
| GTTNTGWFNS | ASGVQNSDYT | 0.8218359 |
| IFMGPLFTL | ASGVQNSDYT | 0.74365973 |
| ILAHAIEAI | ASGVQNSDYT | 0.6818269 |
| IRSAIREAL | ASGVQNSDYT | 0.82288194 |
| ITFAGGG | ASGVQNSDYT | 0.84434354 |
| ITGMSGHQEV | ASGVQNSDYT | 0.7998532 |
| IVAGLACTL | ASGVQNSDYT | 0.79103875 |
| IVGGGLAGA | ASGVQNSDYT | 0.85068214 |
| IVTGVLNI | ASGVQNSDYT | 0.5744566 |
| LAQQGCI | ASGVQNSDYT | 0.7785132 |
| LCLHVEN | ASGVQNSDYT | 0.83765864 |
| LDAISDGS | ASGVQNSDYT | 0.83297205 |
| LGGFAAML | ASGVQNSDYT | 0.7822267 |
| LGGKNPMI | ASGVQNSDYT | 0.839341 |
| LGPKGVAAIL | ASGVQNSDYT | 0.70460737 |
| LMVGVSI | ASGVQNSDYT | 0.810287 |
| LVGAAFVL | ASGVQNSDYT | 0.69962287 |
| LVLAVAPL | ASGVQNSDYT | 0.790113 |
| MAAIAVIDAA | ASGVQNSDYT | 0.59157634 |
| NNGINAELQ | ASGVQNSDYT | 0.7999897 |
| RGEPLVLVHG | ASGVQNSDYT | 0.82991844 |
| RGGAGGAAGLI | ASGVQNSDYT | 0.81159556 |
| RSYRMHPAV | ASGVQNSDYT | 0.33369806 |
| SAAHLLAGQLA | ASGVQNSDYT | 0.81355494 |
| SAAVGSY | ASGVQNSDYT | 0.82134295 |
| SPHAVLHRI | ASGVQNSDYT | 0.8515503 |
| SQIAKVL | ASGVQNSDYT | 0.78440166 |
| TALGHLVP | ASGVQNSDYT | 0.8300928 |
| TAPTTSGAA | ASGVQNSDYT | 0.73173016 |
| TDGDDGNGGSPGT | ASGVQNSDYT | 0.7960876 |

| | | |
|---|---|---|
| TDGLGEN | ASGVQNSDYT | 0.61561584 |
| TGGPPLPGQA | ASGVQNSDYT | 0.7031528 |
| TIPAITI | ASGVQNSDYT | 0.734292 |
| TIPGFTI | ASGVQNSDYT | 0.66103387 |
| TSIASGALA | ASGVQNSDYT | 0.8284652 |
| TVTLPGAAI | ASGVQNSDYT | 0.82849526 |
| VASGLSAALP | ASGVQNSDYT | 0.8287202 |
| VGAGVIGI | ASGVQNSDYT | 0.83869135 |
| VGPKRFGV | ASGVQNSDYT | 0.8421271 |
| VNVAFPD | ASGVQNSDYT | 0.8341258 |
| VSMTNSGVS | ASGVQNSDYT | 0.71476483 |
| VTDVLGDLSV | ASGVQNSDYT | 0.76229745 |
| VVVVGAGFAGL | ASGVQNSDYT | 0.53722525 |
| VYTVGTVV | ASGVQNSDYT | 0.7349084 |
| WTTFSGP | ASGVQNSDYT | 0.71456337 |
| YDELVEAF | ASGVQNSDYT | 0.7950272 |
| YGTGGAGGNGG | ASGVQNSDYT | 0.15017734 |
| YTNVMAV | ASGVQNSDYT | 0.7766124 |
| FLEQVHQL | ASGVQNSDYT | 0.8453558 |
| GLSLLAVAV | ASGVQNSDYT | 0.7229908 |
| LEQVHQL | ASGVQNSDYT | 0.82253134 |
| LNAAGAI | ASGVQNSDYT | 0.8033198 |
| PAVHASL | ASGVQNSDYT | 0.7173549 |
| TELEDDA | ASGVQNSDYT | 0.8215979 |
| TPPGPPN | ASGVQNSDYT | 0.6636597 |
| TPPPPPPP | ASGVQNSDYT | 0.8323863 |
| TQSPALI | ASSIAGRVFNQDTQY | 0.85009253 |
| EFAAAFPQ | ASSIAGRVFNQDTQY | 0.80328536 |
| LLSGNLDVL | ASSIAGRVFNQDTQY | 0.75971997 |
| GPANLGNYNI | ASSIAGRVFNQDTQY | 0.7941619 |
| TLGPTLP | ASSIAGRVFNQDTQY | 0.37547863 |
| VYAYSSCCK | ASSIAGRVFNQDTQY | 0.7678236 |
| APAPAPAPNI | ASSIAGRVFNQDTQY | 0.16198023 |
| PGGPVPIP | ASSIAGRVFNQDTQY | 0.7786869 |
| FLTVALEP | ASSIAGRVFNQDTQY | 0.6385858 |
| SQYFGANL | ASSIAGRVFNQDTQY | 0.79361475 |
| GGVKPAA | ASSIAGRVFNQDTQY | 0.78086257 |
| GGTGGAGGAGGW | ASSIAGRVFNQDTQY | 0.8222768 |
| GVGGVAGLGGAGG | ASSIAGRVFNQDTQY | 0.68249667 |
| VLLVLATTI | ASSIAGRVFNQDTQY | 0.77529454 |
| IGIFAAML | ASSIAGRVFNQDTQY | 0.7594288 |
| AVGDLID | ASSIAGRVFNQDTQY | 0.47136414 |
| NISPLPL | ASSIAGRVFNQDTQY | 0.82349634 |
| VTAVFVAI | ASSIAGRVFNQDTQY | 0.77885735 |
| SGDDNDAAL | ASSIAGRVFNQDTQY | 0.7545028 |
| AVAALAHS | ASSIAGRVFNQDTQY | 0.8348352 |
| GGVTVMATP | ASSIAGRVFNQDTQY | 0.7837236 |
| AGLAANVRD | ASSIAGRVFNQDTQY | 0.49321163 |
| TNRPDLIDPALLRPG | ASSIAGRVFNQDTQY | 0.7531364 |
| LANLGVTAL | ASSIAGRVFNQDTQY | 0.7818742 |
| TGERIEL | ASSIAGRVFNQDTQY | 0.7461338 |

| | | |
|---|---|---|
| NGEELAA | ASSIAGRVFNQDTQY | 0.1528169 |
| VLSLVWG | ASSIAGRVFNQDTQY | 0.8171408 |
| IGYALWL | ASSIAGRVFNQDTQY | 0.6582719 |
| LSLLLAPL | ASSIAGRVFNQDTQY | 0.6842158 |
| TLDEALK | ASSIAGRVFNQDTQY | 0.72878814 |
| TDGAGTAEGG | ASSIAGRVFNQDTQY | 0.8319204 |
| NAPGGAGGNGG | ASSIAGRVFNQDTQY | 0.67460907 |
| VSVFMGMGGNF | ASSIAGRVFNQDTQY | 0.7404543 |
| LASYLAAL | ASSIAGRVFNQDTQY | 0.81382865 |
| LVGGTPAL | ASSIAGRVFNQDTQY | 0.79540133 |
| VRNADVL | ASSIAGRVFNQDTQY | 0.8399848 |
| TCSSPFLVSAM | ASSIAGRVFNQDTQY | 0.8013138 |
| VHITDAE | ASSIAGRVFNQDTQY | 0.800495 |
| EYAAPSF | ASSIAGRVFNQDTQY | 0.8159453 |
| VVDAFHGGL | ASSIAGRVFNQDTQY | 0.78544307 |
| ASVVDPQ | ASSIAGRVFNQDTQY | 0.48744148 |
| SYNEFGGLS | ASSIAGRVFNQDTQY | 0.5258481 |
| DPDGSVY | ASSIAGRVFNQDTQY | 0.773724 |
| MVSTIHATE | ASSIAGRVFNQDTQY | 0.4701658 |
| LAAGVADPV | ASSIAGRVFNQDTQY | 0.8009486 |
| TKEQIAA | ASSIAGRVFNQDTQY | 0.8418291 |
| SLASPAAL | ASSIAGRVFNQDTQY | 0.7737538 |
| IGLVFYGV | ASSIAGRVFNQDTQY | 0.5049592 |
| FLLDASNL | ASSIAGRVFNQDTQY | 0.7450454 |
| YQVGNFP | ASSIAGRVFNQDTQY | 0.6745355 |
| ESLLSQST | ASSIAGRVFNQDTQY | 0.82123876 |
| LFTVGFAAVG | ASSIAGRVFNQDTQY | 0.8205072 |
| ATPPTPT | ASSIAGRVFNQDTQY | 0.8406522 |
| TDLATLG | ASSIAGRVFNQDTQY | 0.82837856 |
| SAGGTGR | ASSIAGRVFNQDTQY | 0.72318053 |
| GFSNSGS | ASSIAGRVFNQDTQY | 0.83113533 |
| GGNGGSGGTSV | ASSIAGRVFNQDTQY | 0.74305046 |
| GTSVATGGAGN | ASSIAGRVFNQDTQY | 0.78813845 |
| NGGNGGSGGTS | ASSIAGRVFNQDTQY | 0.78845215 |
| AVVKFPH | ASSIAGRVFNQDTQY | 0.8064499 |
| TPPPPPP | ASSIAGRVFNQDTQY | 0.708933 |
| AAPAAAAAPA | ASSIAGRVFNQDTQY | 0.7460731 |
| IALNEDL | ASSIAGRVFNQDTQY | 0.58293295 |
| IVNVSSM | ASSIAGRVFNQDTQY | 0.81808853 |
| TGIPPAPR | ASSIAGRVFNQDTQY | 0.7294909 |
| FDLGGGTF | ASSIAGRVFNQDTQY | 0.5386264 |
| FDLGGGTFDVS | ASSIAGRVFNQDTQY | 0.7386962 |
| NEPTAAA | ASSIAGRVFNQDTQY | 0.8448099 |
| NEPTAAALA | ASSIAGRVFNQDTQY | 0.8480607 |
| GSIINISSI | ASSIAGRVFNQDTQY | 0.68039334 |
| VAPGNAGTA | ASSIAGRVFNQDTQY | 0.6602755 |
| TGGGASLE | ASSIAGRVFNQDTQY | 0.79166794 |
| VAVYNHYKRI | ASSIAGRVFNQDTQY | 0.5900921 |
| TDPAPATT | ASSIAGRVFNQDTQY | 0.8184568 |
| TNRPDILDPALLRPG | ASSIAGRVFNQDTQY | 0.7652935 |
| LTTTTTS | ASSIAGRVFNQDTQY | 0.75861764 |

| | | |
|---|---|---|
| PTGSGKTL | ASSIAGRVFNQDTQY | 0.7044443 |
| TGSGKTL | ASSIAGRVFNQDTQY | 0.80598223 |
| TIGHVDHGK | ASSIAGRVFNQDTQY | 0.74528337 |
| TIGHVDHGKTT | ASSIAGRVFNQDTQY | 0.8247422 |
| GHVDHGKTT | ASSIAGRVFNQDTQY | 0.25549543 |
| AQFEHTLL | ASSIAGRVFNQDTQY | 0.81203926 |
| IVNQDEL | ASSIAGRVFNQDTQY | 0.6586793 |
| TNGSQFF | ASSIAGRVFNQDTQY | 0.6989448 |
| TGKGIVE | ASSIAGRVFNQDTQY | 0.2977316 |
| LISNASDA | ASSIAGRVFNQDTQY | 0.80267495 |
| AELRTLGELTKIAKAHGAQ | ASSIAGRVFNQDTQY | 0.8247905 |
| APNAVAAAEPSW | ASSIAGRVFNQDTQY | 0.83925205 |
| APQVAGAATAVL | ASSIAGRVFNQDTQY | 0.7121099 |
| ARRDPAAAAAILPTDA | ASSIAGRVFNQDTQY | 0.81433463 |
| DAAARDVLAEAGLAEYF | ASSIAGRVFNQDTQY | 0.8484362 |
| DDQIVLSAKANLAKVEQ | ASSIAGRVFNQDTQY | 0.73966527 |
| DELLPDTGEDSDEDGH | ASSIAGRVFNQDTQY | 0.29055148 |
| ELPEAPANRATVPQD | ASSIAGRVFNQDTQY | 0.7968995 |
| FGQSKIERVTGIG | ASSIAGRVFNQDTQY | 0.7584375 |
| GAGGTGSTAGGAGGAGGAGGLY | ASSIAGRVFNQDTQY | 0.829592 |
| GHGGTGAAGGNGGTGGAGGSGI | ASSIAGRVFNQDTQY | 0.82128537 |
| GNGGNGGAGGEGGPGGAGGAGA | ASSIAGRVFNQDTQY | 0.56074905 |
| GSTAGNGGNGGNATNGTIAGQP | ASSIAGRVFNQDTQY | 0.81920516 |
| GTGGDGGNGGNGGTGG | ASSIAGRVFNQDTQY | 0.82430166 |
| GVRRPAGIPADAPV | ASSIAGRVFNQDTQY | 0.63995856 |
| GYRQVTDAHLLTLARR | ASSIAGRVFNQDTQY | 0.7359799 |
| IAKMGQDGHDRG | ASSIAGRVFNQDTQY | 0.73100007 |
| KAEPSIEIQYT | ASSIAGRVFNQDTQY | 0.33450818 |
| KPVRIRMAADEDTVAVT | ASSIAGRVFNQDTQY | 0.798612 |
| MPPRIAGMRLLVIKPEP | ASSIAGRVFNQDTQY | 0.72472763 |
| MKFYAVALFRSRGVPPTR | ASSIAGRVFNQDTQY | 0.66877794 |
| NAGGFGILGAVAHSP | ASSIAGRVFNQDTQY | 0.77660024 |
| NEGREITLAMAEM | ASSIAGRVFNQDTQY | 0.6244253 |
| NELVEWSHPLGHRWRWE | ASSIAGRVFNQDTQY | 0.665759 |
| NLGAGGTGGNGGNPG | ASSIAGRVFNQDTQY | 0.6929298 |
| NTEWPKWETVMAADPVD | ASSIAGRVFNQDTQY | 0.78617823 |
| PASAAPAETQP | ASSIAGRVFNQDTQY | 0.81558514 |
| QQSSDGKNGVRAAEITGE | ASSIAGRVFNQDTQY | 0.7642231 |
| RAPDAVTTSWWKE | ASSIAGRVFNQDTQY | 0.8116248 |
| SGFGNVGAHTSGWFNQ | ASSIAGRVFNQDTQY | 0.84183806 |
| SGVAGAVGQAASVGGLKV | ASSIAGRVFNQDTQY | 0.49235117 |
| SNGGAGTGSAGGNGGT | ASSIAGRVFNQDTQY | 0.6276529 |
| TGGAGGEGGAAG | ASSIAGRVFNQDTQY | 0.8188832 |
| TGGDGGAAGAGGAPGHG | ASSIAGRVFNQDTQY | 0.64261305 |
| TMPPLLAAIANAGRHGV | ASSIAGRVFNQDTQY | 0.83169794 |
| TSSTAGNSGGA | ASSIAGRVFNQDTQY | 0.7892192 |
| VPATVPVGILC | ASSIAGRVFNQDTQY | 0.81035423 |
| VQRPDSGGVVLTGRIS | ASSIAGRVFNQDTQY | 0.77935743 |
| WTPNPVPATKGVPAGLVQAT | ASSIAGRVFNQDTQY | 0.84902185 |
| YALKRALAKFSDRFDVVII | ASSIAGRVFNQDTQY | 0.81846344 |
| YLIGSVIV | ASSIAGRVFNQDTQY | 0.778992 |

| | | |
|---|---|---|
| YGQLVAVID | ASSIAGRVFNQDTQY | 0.7858468 |
| VTVGGMTLPA | ASSIAGRVFNQDTQY | 0.6111028 |
| VLALPAV | ASSIAGRVFNQDTQY | 0.8224598 |
| VIPPTLNYET | ASSIAGRVFNQDTQY | 0.8297374 |
| VGMGGMHPSSG | ASSIAGRVFNQDTQY | 0.6331037 |
| VGLAGGAATA | ASSIAGRVFNQDTQY | 0.7499532 |
| VGGAGGAGGSGP | ASSIAGRVFNQDTQY | 0.8508754 |
| VFAAFMV | ASSIAGRVFNQDTQY | 0.8352649 |
| TLYDKLR | ASSIAGRVFNQDTQY | 0.5351427 |
| TLAVFTVL | ASSIAGRVFNQDTQY | 0.71722263 |
| TGSVQPARV | ASSIAGRVFNQDTQY | 0.5919111 |
| TGGGAGSGGT | ASSIAGRVFNQDTQY | 0.75916845 |
| SHLETSL | ASSIAGRVFNQDTQY | 0.43550476 |
| SFTTGMT | ASSIAGRVFNQDTQY | 0.81489694 |
| QVEVAKRMGVRQPTVS | ASSIAGRVFNQDTQY | 0.7477515 |
| QGSGGPALAM | ASSIAGRVFNQDTQY | 0.821725 |
| MLAPSNP | ASSIAGRVFNQDTQY | 0.8007262 |
| LVNNAAI | ASSIAGRVFNQDTQY | 0.84329677 |
| LTGEINTLA | ASSIAGRVFNQDTQY | 0.7204683 |
| LSIFGPV | ASSIAGRVFNQDTQY | 0.71699387 |
| LLALLANL | ASSIAGRVFNQDTQY | 0.4520051 |
| LISNASDALDKLR | ASSIAGRVFNQDTQY | 0.5859201 |
| LINEAVP | ASSIAGRVFNQDTQY | 0.6784235 |
| LGYAFSGL | ASSIAGRVFNQDTQY | 0.7090845 |
| LDFVAVAL | ASSIAGRVFNQDTQY | 0.30080467 |
| LAQPGAE | ASSIAGRVFNQDTQY | 0.84661436 |
| ISNLGAVTGF | ASSIAGRVFNQDTQY | 0.77312267 |
| HQEDPGRI | ASSIAGRVFNQDTQY | 0.82801473 |
| GTPGEPTP | ASSIAGRVFNQDTQY | 0.68517244 |
| GFFNSGS | ASSIAGRVFNQDTQY | 0.5877484 |
| GAGGGGLTAGAN | ASSIAGRVFNQDTQY | 0.70910895 |
| FVDALVNM | ASSIAGRVFNQDTQY | 0.81891465 |
| FLTVAPEL | ASSIAGRVFNQDTQY | 0.5935954 |
| FETLTAL | ASSIAGRVFNQDTQY | 0.49653798 |
| DVINPIEL | ASSIAGRVFNQDTQY | 0.7340312 |
| APVSGAPV | ASSIAGRVFNQDTQY | 0.7030854 |
| ALVGGLAW | ASSIAGRVFNQDTQY | 0.8509946 |
| ALAAAAATAI | ASSIAGRVFNQDTQY | 0.8363111 |
| AFGLGLLL | ASSIAGRVFNQDTQY | 0.78884757 |
| AFFAAASL | ASSIAGRVFNQDTQY | 0.8487138 |
| AAAALPAFSPP | ASSIAGRVFNQDTQY | 0.19929126 |
| VVLAPAL | ASSIAGRVFNQDTQY | 0.70054007 |
| TIGHVDHGKTTL | ASSIAGRVFNQDTQY | 0.85024333 |
| LISNASDALDK | ASSIAGRVFNQDTQY | 0.8080251 |
| LISNASDAL | ASSIAGRVFNQDTQY | 0.7726373 |
| ISNASDALDK | ASSIAGRVFNQDTQY | 0.8393316 |
| NQAETLVYQTE | ASSIAGRVFNQDTQY | 0.65714574 |
| TGGTGGTAGAG | ASSIAGRVFNQDTQY | 0.7545384 |
| TGSTAGGAGGA | ASSIAGRVFNQDTQY | 0.7657475 |
| DPLAVPVAAGPV | ASSIAGRVFNQDTQY | 0.8208461 |
| GGTGGTPGNGGN | ASSIAGRVFNQDTQY | 0.79287845 |

| | | |
|---|---|---|
| PAGLAGVDPATL | ASSIAGRVFNQDTQY | 0.5206175 |
| GNNLVVEAPHAGASV | ASSIAGRVFNQDTQY | 0.8208344 |
| KVTRIMSASTLEQPA | ASSIAGRVFNQDTQY | 0.8522791 |
| LGFALPIALPRRTVV | ASSIAGRVFNQDTQY | 0.7298771 |
| TRLGNLTARKLVVLV | ASSIAGRVFNQDTQY | 0.82475656 |
| WSQQRRLIYFPSAGP | ASSIAGRVFNQDTQY | 0.8037303 |
| GPNGNPDPMAAAVDIR | ASSIAGRVFNQDTQY | 0.78791845 |
| TQRCVSVADLRAAASR | ASSIAGRVFNQDTQY | 0.6938406 |
| QFVLDVAEHPVEVLTETV | ASSIAGRVFNQDTQY | 0.6471157 |
| GGAGGTGGAGGNGGTQAGDG | ASSIAGRVFNQDTQY | 0.78607976 |
| PLLATLRAIAVDVDAKPAQV | ASSIAGRVFNQDTQY | 0.62664866 |
| APKFRTGGYMIDFWGVGYQVAK | ASSIAGRVFNQDTQY | 0.7939652 |
| GAGGAGGAGGAAGGPGGTGGTG | ASSIAGRVFNQDTQY | 0.7641846 |
| IGALGLLATDVAPAGSVAFGVA | ASSIAGRVFNQDTQY | 0.8605623 |
| AGGAGGIGGNGGAGGAGGVLMG | ASSIAGRVFNQDTQY | 0.8198043 |
| ADELIGLL | ASSIAGRVFNQDTQY | 0.8082979 |
| ADRALKA | ASSIAGRVFNQDTQY | 0.6121734 |
| AGAVGQVGGAGGA | ASSIAGRVFNQDTQY | 0.8091787 |
| AGGGGGLGGAP | ASSIAGRVFNQDTQY | 0.799896 |
| AGGLLYGNGGAG | ASSIAGRVFNQDTQY | 0.5876002 |
| AGLESIGL | ASSIAGRVFNQDTQY | 0.84557104 |
| AGLKTGII | ASSIAGRVFNQDTQY | 0.6101279 |
| ASLAALLL | ASSIAGRVFNQDTQY | 0.7772901 |
| ASVINISSI | ASSIAGRVFNQDTQY | 0.8242352 |
| ATPPGPA | ASSIAGRVFNQDTQY | 0.8283961 |
| ATVASALV | ASSIAGRVFNQDTQY | 0.76669145 |
| ATVASLIL | ASSIAGRVFNQDTQY | 0.80286896 |
| AVNGPVLIRH | ASSIAGRVFNQDTQY | 0.5929158 |
| DGAAGAVGI | ASSIAGRVFNQDTQY | 0.8304813 |
| DPLLHGI | ASSIAGRVFNQDTQY | 0.8356165 |
| ELGAALGAL | ASSIAGRVFNQDTQY | 0.74604404 |
| ELGRTQAV | ASSIAGRVFNQDTQY | 0.8455167 |
| FVAGGYLS | ASSIAGRVFNQDTQY | 0.7901157 |
| GASGGAGGAGGW | ASSIAGRVFNQDTQY | 0.35974044 |
| GGAGAGGDGGAGGI | ASSIAGRVFNQDTQY | 0.7831018 |
| GTGHNGSQPGMGGNG | ASSIAGRVFNQDTQY | 0.83438796 |
| GYAAASASAS | ASSIAGRVFNQDTQY | 0.83699256 |
| HQVDVAAVQQG | ASSIAGRVFNQDTQY | 0.8384117 |
| IGAAIAHT | ASSIAGRVFNQDTQY | 0.77989364 |
| IGMIEVAQL | ASSIAGRVFNQDTQY | 0.7365215 |
| IRLAPEHTSAMD | ASSIAGRVFNQDTQY | 0.8447641 |
| IYNVSVL | ASSIAGRVFNQDTQY | 0.86424065 |
| KQGFLGM | ASSIAGRVFNQDTQY | 0.827999 |
| LESVITQL | ASSIAGRVFNQDTQY | 0.72475964 |
| LGLGALVVV | ASSIAGRVFNQDTQY | 0.7119469 |
| LIGNGGNGGSGGI | ASSIAGRVFNQDTQY | 0.8157822 |
| LNAINAPTL | ASSIAGRVFNQDTQY | 0.82506776 |
| LQLLNTGV | ASSIAGRVFNQDTQY | 0.6267831 |
| LQTSIEF | ASSIAGRVFNQDTQY | 0.7892319 |
| MSFAVSEL | ASSIAGRVFNQDTQY | 0.8040421 |
| NTGALGL | ASSIAGRVFNQDTQY | 0.36202157 |

| | | |
|---|---|---|
| PPGAGGAGGAGTT | ASSIAGRVFNQDTQY | 0.8084747 |
| QLVGLGI | ASSIAGRVFNQDTQY | 0.83536035 |
| RFLEQVHQL | ASSIAGRVFNQDTQY | 0.8035413 |
| RGVGYVLR | ASSIAGRVFNQDTQY | 0.6214477 |
| SATGVFAGVI | ASSIAGRVFNQDTQY | 0.83907634 |
| SLPAITLP | ASSIAGRVFNQDTQY | 0.8502725 |
| STALAFTV | ASSIAGRVFNQDTQY | 0.84537756 |
| SVLTVGLGK | ASSIAGRVFNQDTQY | 0.3543741 |
| TALAGAGSGI | ASSIAGRVFNQDTQY | 0.28217795 |
| TGLPFTGI | ASSIAGRVFNQDTQY | 0.7527784 |
| VAEHVTV | ASSIAGRVFNQDTQY | 0.8147855 |
| VDIVLNSL | ASSIAGRVFNQDTQY | 0.68995106 |
| VLTGGGAVL | ASSIAGRVFNQDTQY | 0.6216141 |
| VNVGLAYGI | ASSIAGRVFNQDTQY | 0.7707233 |
| VSSAAVGAV | ASSIAGRVFNQDTQY | 0.8378931 |
| VTSVYEAGV | ASSIAGRVFNQDTQY | 0.55899084 |
| WVALDRGA | ASSIAGRVFNQDTQY | 0.62215775 |
| YLVGSVVI | ASSIAGRVFNQDTQY | 0.82014567 |
| GPPGTGKTLL | ASSIAGRVFNQDTQY | 0.83876956 |
| GVGMTKF | ASSIAGRVFNQDTQY | 0.6170062 |
| IGIGGSDL | ASSIAGRVFNQDTQY | 0.7356945 |
| IGTLLAEL | ASSIAGRVFNQDTQY | 0.77655065 |
| IYTSGTT | ASSIAGRVFNQDTQY | 0.7632685 |
| LSFPAGV | ASSIAGRVFNQDTQY | 0.78523713 |
| LYTSGTT | ASSIAGRVFNQDTQY | 0.53779626 |
| VGAFPFGF | ASSIAGRVFNQDTQY | 0.8279668 |
| VGAGTVEF | ASSIAGRVFNQDTQY | 0.5883975 |
| VGKSTLLL | ASSIAGRVFNQDTQY | 0.83550644 |
| AALAHAGGGIV | ASSIAGRVFNQDTQY | 0.7803861 |
| AALVAVSLL | ASSIAGRVFNQDTQY | 0.801608 |
| AAVSRAVAAL | ASSIAGRVFNQDTQY | 0.7469145 |
| AGGSSGIGGF | ASSIAGRVFNQDTQY | 0.6776989 |
| AGLTVANANG | ASSIAGRVFNQDTQY | 0.5913577 |
| AGLVGAVL | ASSIAGRVFNQDTQY | 0.53326404 |
| ALSQAFTV | ASSIAGRVFNQDTQY | 0.8298802 |
| ALTFSVSL | ASSIAGRVFNQDTQY | 0.33268094 |
| ALTGAALVAL | ASSIAGRVFNQDTQY | 0.8238201 |
| ASFSIGAGVL | ASSIAGRVFNQDTQY | 0.80734515 |
| DGSVYKI | ASSIAGRVFNQDTQY | 0.6233762 |
| EQLAEIV | ASSIAGRVFNQDTQY | 0.81965 |
| FAVSELTVM | ASSIAGRVFNQDTQY | 0.49973026 |
| GAAPLAAGL | ASSIAGRVFNQDTQY | 0.75924027 |
| GADGGDGGSGGAGG | ASSIAGRVFNQDTQY | 0.8090788 |
| GFGLDTGGAGG | ASSIAGRVFNQDTQY | 0.820763 |
| GLASGGLL | ASSIAGRVFNQDTQY | 0.69552284 |
| GRIAGAAVLL | ASSIAGRVFNQDTQY | 0.82548213 |
| GSAGSLNTSF | ASSIAGRVFNQDTQY | 0.8114923 |
| GTGQIGI | ASSIAGRVFNQDTQY | 0.7101751 |
| GTLIALIL | ASSIAGRVFNQDTQY | 0.7967624 |
| IAATMGFVVG | ASSIAGRVFNQDTQY | 0.8287593 |
| IFAQALL | ASSIAGRVFNQDTQY | 0.81609964 |

| | | |
|---|---|---|
| IGAAVALGF | ASSIAGRVFNQDTQY | 0.5868698 |
| IGLTPLAL | ASSIAGRVFNQDTQY | 0.80251837 |
| ILGSNTSTL | ASSIAGRVFNQDTQY | 0.6850867 |
| IVFSLTVL | ASSIAGRVFNQDTQY | 0.801561 |
| LASKATGF | ASSIAGRVFNQDTQY | 0.7839179 |
| LASVALIL | ASSIAGRVFNQDTQY | 0.76346374 |
| LFAVGLTL | ASSIAGRVFNQDTQY | 0.8242104 |
| LFNGGIGTYI | ASSIAGRVFNQDTQY | 0.53126264 |
| LTLGAGII | ASSIAGRVFNQDTQY | 0.6993406 |
| LVAAALGDI | ASSIAGRVFNQDTQY | 0.744933 |
| NTAFVQLGI | ASSIAGRVFNQDTQY | 0.7620983 |
| QLFSGAEL | ASSIAGRVFNQDTQY | 0.8207326 |
| SAATFITL | ASSIAGRVFNQDTQY | 0.62964153 |
| SNYGLANLGV | ASSIAGRVFNQDTQY | 0.79662114 |
| SYGGAYAVM | ASSIAGRVFNQDTQY | 0.74594986 |
| TTGVGGLTLG | ASSIAGRVFNQDTQY | 0.6223681 |
| TTQTIYI | ASSIAGRVFNQDTQY | 0.80024874 |
| VAAGAGLGAVI | ASSIAGRVFNQDTQY | 0.7347243 |
| VAAVVTLL | ASSIAGRVFNQDTQY | 0.6000311 |
| VGFLGGLI | ASSIAGRVFNQDTQY | 0.8203262 |
| VGVIPILL | ASSIAGRVFNQDTQY | 0.6222853 |
| VIFASSLI | ASSIAGRVFNQDTQY | 0.7939108 |
| VKGGDGGAAAG | ASSIAGRVFNQDTQY | 0.83134085 |
| VTKTFTNI | ASSIAGRVFNQDTQY | 0.61534715 |
| VVAAAILL | ASSIAGRVFNQDTQY | 0.83383226 |
| YAALDVEL | ASSIAGRVFNQDTQY | 0.793838 |
| LASVALLL | ASSIAGRVFNQDTQY | 0.7868081 |
| LDAVHRV | ASSIAGRVFNQDTQY | 0.7279764 |
| QGLSSSIF | ASSIAGRVFNQDTQY | 0.7029179 |
| RGTFANIRL | ASSIAGRVFNQDTQY | 0.8368981 |
| RQLWWGHRI | ASSIAGRVFNQDTQY | 0.8182287 |
| YGPPGTGKTLL | ASSIAGRVFNQDTQY | 0.8244301 |
| AALVGAPANVA | ASSIAGRVFNQDTQY | 0.8059982 |
| AATFRRTE | ASSIAGRVFNQDTQY | 0.8206471 |
| AGALGGGI | ASSIAGRVFNQDTQY | 0.82983947 |
| AGGAGGAGGIGAG | ASSIAGRVFNQDTQY | 0.6894686 |
| AGSGALGL | ASSIAGRVFNQDTQY | 0.4543532 |
| AQIADQI | ASSIAGRVFNQDTQY | 0.72381645 |
| ASAVAVTYL | ASSIAGRVFNQDTQY | 0.7116789 |
| ASTVLGGTGG | ASSIAGRVFNQDTQY | 0.7222085 |
| AVGGEPAK | ASSIAGRVFNQDTQY | 0.4409724 |
| CEQGTFGGTC | ASSIAGRVFNQDTQY | 0.82822967 |
| DEGKAPAG | ASSIAGRVFNQDTQY | 0.8552772 |
| DTVVFPAL | ASSIAGRVFNQDTQY | 0.6081334 |
| FEGITMAG | ASSIAGRVFNQDTQY | 0.8484029 |
| FPLVVQD | ASSIAGRVFNQDTQY | 0.57596415 |
| GAFTTPQ | ASSIAGRVFNQDTQY | 0.81811666 |
| GFGGFGTAGGVGGA | ASSIAGRVFNQDTQY | 0.83755106 |
| GFLAQFL | ASSIAGRVFNQDTQY | 0.5769753 |
| GGAGGAGGNAGVA | ASSIAGRVFNQDTQY | 0.8061311 |
| GGVGTAGAGGAGGAGG | ASSIAGRVFNQDTQY | 0.67755944 |

| | | |
|---|---|---|
| GNGGGGDAGNAGSGGN | ASSIAGRVFNQDTQY | 0.7048038 |
| GTGGAGGAGNDAGS | ASSIAGRVFNQDTQY | 0.83573735 |
| GTTNTGWFNS | ASSIAGRVFNQDTQY | 0.68121123 |
| IFMGPLFTL | ASSIAGRVFNQDTQY | 0.8114325 |
| ILAHAIEAI | ASSIAGRVFNQDTQY | 0.70606077 |
| IRSAIREAL | ASSIAGRVFNQDTQY | 0.6479887 |
| ITFAGGG | ASSIAGRVFNQDTQY | 0.7977698 |
| ITGMSGHQEV | ASSIAGRVFNQDTQY | 0.8323909 |
| IVAGLACTL | ASSIAGRVFNQDTQY | 0.74095386 |
| IVGGGLAGA | ASSIAGRVFNQDTQY | 0.8630591 |
| IVTGVLNI | ASSIAGRVFNQDTQY | 0.6921337 |
| LAQQGCI | ASSIAGRVFNQDTQY | 0.84364927 |
| LCLHVEN | ASSIAGRVFNQDTQY | 0.8362354 |
| LDAISDGS | ASSIAGRVFNQDTQY | 0.7823186 |
| LGGFAAML | ASSIAGRVFNQDTQY | 0.7542701 |
| LGGKNPMI | ASSIAGRVFNQDTQY | 0.8290497 |
| LGPKGVAAIL | ASSIAGRVFNQDTQY | 0.7474745 |
| LMVGVSI | ASSIAGRVFNQDTQY | 0.8115612 |
| LVGAAFVL | ASSIAGRVFNQDTQY | 0.8393705 |
| LVLAVAPL | ASSIAGRVFNQDTQY | 0.84055686 |
| MAAIAVIDAA | ASSIAGRVFNQDTQY | 0.8753358 |
| NNGINAELQ | ASSIAGRVFNQDTQY | 0.860493 |
| RGEPLVLVHG | ASSIAGRVFNQDTQY | 0.8599358 |
| RGGAGGAAGLI | ASSIAGRVFNQDTQY | 0.82153785 |
| RSYRMHPAV | ASSIAGRVFNQDTQY | 0.6049151 |
| SAAHLLAGQLA | ASSIAGRVFNQDTQY | 0.8555087 |
| SAAVGSY | ASSIAGRVFNQDTQY | 0.81048703 |
| SPHAVLHRI | ASSIAGRVFNQDTQY | 0.8560494 |
| SQIAKVL | ASSIAGRVFNQDTQY | 0.8724084 |
| TALGHLVP | ASSIAGRVFNQDTQY | 0.82248086 |
| TAPTTSGAA | ASSIAGRVFNQDTQY | 0.87473446 |
| TDGDDGNGGSPGT | ASSIAGRVFNQDTQY | 0.86886734 |
| TDGLGEN | ASSIAGRVFNQDTQY | 0.7528577 |
| TGGPPLPGQA | ASSIAGRVFNQDTQY | 0.8540839 |
| TIPAITI | ASSIAGRVFNQDTQY | 0.81046414 |
| TIPGFTI | ASSIAGRVFNQDTQY | 0.8583092 |
| TSIASGALA | ASSIAGRVFNQDTQY | 0.743276 |
| TVTLPGAAI | ASSIAGRVFNQDTQY | 0.870265 |
| VASGLSAALP | ASSIAGRVFNQDTQY | 0.8264314 |
| VGAGVIGI | ASSIAGRVFNQDTQY | 0.4701699 |
| VGPKRFGV | ASSIAGRVFNQDTQY | 0.8649038 |
| VNVAFPD | ASSIAGRVFNQDTQY | 0.854998 |
| VSMTNSGVS | ASSIAGRVFNQDTQY | 0.8547795 |
| VTDVLGDLSV | ASSIAGRVFNQDTQY | 0.8699775 |
| VVVVGAGFAGL | ASSIAGRVFNQDTQY | 0.7630775 |
| VYTVGTVV | ASSIAGRVFNQDTQY | 0.820464 |
| WTTFSGP | ASSIAGRVFNQDTQY | 0.8660809 |
| YDELVEAF | ASSIAGRVFNQDTQY | 0.87351215 |
| YGTGGAGGNGG | ASSIAGRVFNQDTQY | 0.80096895 |
| YTNVMAV | ASSIAGRVFNQDTQY | 0.85665035 |
| FLEQVHQL | ASSIAGRVFNQDTQY | 0.8724643 |

| | | |
|---|---|---|
| GLSLLAVAV | ASSIAGRVFNQDTQY | 0.8567006 |
| LEQVHQL | ASSIAGRVFNQDTQY | 0.8754759 |
| LNAAGAI | ASSIAGRVFNQDTQY | 0.8768639 |
| PAVHASL | ASSIAGRVFNQDTQY | 0.7674128 |
| TELEDDA | ASSIAGRVFNQDTQY | 0.87349385 |
| TPPGPPN | ASSIAGRVFNQDTQY | 0.87676066 |
| TPPPPPPP | ASSIAGRVFNQDTQY | 0.8427315 |
| TQSPALI | ASSGDRNNERLF | 0.8572131 |
| EFAAAFPQ | ASSGDRNNERLF | 0.7834679 |
| LLSGNLDVL | ASSGDRNNERLF | 0.8286672 |
| GPANLGNYNI | ASSGDRNNERLF | 0.5601933 |
| TLGPTLP | ASSGDRNNERLF | 0.8566995 |
| VYAYSSCCK | ASSGDRNNERLF | 0.8750975 |
| APAPAPAPNI | ASSGDRNNERLF | 0.8440866 |
| PGGPVPIP | ASSGDRNNERLF | 0.8131675 |
| FLTVALEP | ASSGDRNNERLF | 0.79964656 |
| SQYFGANL | ASSGDRNNERLF | 0.8528642 |
| GGVKPAA | ASSGDRNNERLF | 0.8306276 |
| GGTGGAGGAGGW | ASSGDRNNERLF | 0.70950764 |
| GVGGVAGLGGAGG | ASSGDRNNERLF | 0.82210875 |
| VLLVLATTI | ASSGDRNNERLF | 0.86865187 |
| IGIFAAML | ASSGDRNNERLF | 0.8696338 |
| AVGDLID | ASSGDRNNERLF | 0.7784437 |
| NISPLPL | ASSGDRNNERLF | 0.87608045 |
| VTAVFVAI | ASSGDRNNERLF | 0.86495715 |
| SGDDNDAAL | ASSGDRNNERLF | 0.86836684 |
| AVAALAHS | ASSGDRNNERLF | 0.8583809 |
| GGVTVMATP | ASSGDRNNERLF | 0.8258969 |
| AGLAANVRD | ASSGDRNNERLF | 0.8680786 |
| TNRPDLIDPALLRPG | ASSGDRNNERLF | 0.82632273 |
| LANLGVTAL | ASSGDRNNERLF | 0.83206576 |
| TGERIEL | ASSGDRNNERLF | 0.86506665 |
| NGEELAA | ASSGDRNNERLF | 0.8439332 |
| VLSLVWG | ASSGDRNNERLF | 0.8221839 |
| IGYALWL | ASSGDRNNERLF | 0.8176155 |
| LSLLLAPL | ASSGDRNNERLF | 0.87617624 |
| TLDEALK | ASSGDRNNERLF | 0.7952148 |
| TDGAGTAEGG | ASSGDRNNERLF | 0.72584105 |
| NAPGGAGGNGG | ASSGDRNNERLF | 0.8144714 |
| VSVFMGMGGNF | ASSGDRNNERLF | 0.8492409 |
| LASYLAAL | ASSGDRNNERLF | 0.8738581 |
| LVGGTPAL | ASSGDRNNERLF | 0.869812 |
| VRNADVL | ASSGDRNNERLF | 0.82739764 |
| TCSSPFLVSAM | ASSGDRNNERLF | 0.8635793 |
| VHITDAE | ASSGDRNNERLF | 0.8672492 |
| EYAAPSF | ASSGDRNNERLF | 0.8555236 |
| VVDAFHGGL | ASSGDRNNERLF | 0.8572792 |
| ASVVDPQ | ASSGDRNNERLF | 0.81935835 |
| SYNEFGGLS | ASSGDRNNERLF | 0.87620515 |
| DPDGSVY | ASSGDRNNERLF | 0.8680276 |
| MVSTIHATE | ASSGDRNNERLF | 0.872825 |

| | | |
|---|---|---|
| LAAGVADPV | ASSGDRNNERLF | 0.8745605 |
| TKEQIAA | ASSGDRNNERLF | 0.85172737 |
| SLASPAAL | ASSGDRNNERLF | 0.85202163 |
| IGLVFYGV | ASSGDRNNERLF | 0.5993984 |
| FLLDASNL | ASSGDRNNERLF | 0.70612955 |
| YQVGNFP | ASSGDRNNERLF | 0.8563826 |
| ESLLSQST | ASSGDRNNERLF | 0.8596041 |
| LFTVGFAAVG | ASSGDRNNERLF | 0.8661839 |
| ATPPTPT | ASSGDRNNERLF | 0.86987114 |
| TDLATLG | ASSGDRNNERLF | 0.81723493 |
| SAGGTGR | ASSGDRNNERLF | 0.86860955 |
| GFSNSGS | ASSGDRNNERLF | 0.87191564 |
| GGNGGSGGTSV | ASSGDRNNERLF | 0.746057 |
| GTSVATGGAGN | ASSGDRNNERLF | 0.83585465 |
| NGGNGGSGGTS | ASSGDRNNERLF | 0.8799982 |
| AVVKFPH | ASSGDRNNERLF | 0.8543064 |
| TPPPPPP | ASSGDRNNERLF | 0.8387961 |
| AAPAAAAAPA | ASSGDRNNERLF | 0.8421802 |
| IALNEDL | ASSGDRNNERLF | 0.87345254 |
| IVNVSSM | ASSGDRNNERLF | 0.83225214 |
| TGIPPAPR | ASSGDRNNERLF | 0.87935495 |
| FDLGGGTF | ASSGDRNNERLF | 0.5371814 |
| FDLGGGTFDVS | ASSGDRNNERLF | 0.8543911 |
| NEPTAAA | ASSGDRNNERLF | 0.8130769 |
| NEPTAAALA | ASSGDRNNERLF | 0.6613067 |
| GSIINISSI | ASSGDRNNERLF | 0.8695054 |
| VAPGNAGTA | ASSGDRNNERLF | 0.8487756 |
| TGGGASLE | ASSGDRNNERLF | 0.8624453 |
| VAVYNHYKRI | ASSGDRNNERLF | 0.8490102 |
| TDPAPATT | ASSGDRNNERLF | 0.8759981 |
| TNRPDILDPALLRPG | ASSGDRNNERLF | 0.85181296 |
| LTTTTTS | ASSGDRNNERLF | 0.8322959 |
| PTGSGKTL | ASSGDRNNERLF | 0.82304287 |
| TGSGKTL | ASSGDRNNERLF | 0.8664251 |
| TIGHVDHGK | ASSGDRNNERLF | 0.87242275 |
| TIGHVDHGKTT | ASSGDRNNERLF | 0.8750116 |
| GHVDHGKTT | ASSGDRNNERLF | 0.6941532 |
| AQFEHTLL | ASSGDRNNERLF | 0.8505511 |
| IVNQDEL | ASSGDRNNERLF | 0.8505026 |
| TNGSQFF | ASSGDRNNERLF | 0.4143866 |
| TGKGIVE | ASSGDRNNERLF | 0.50786746 |
| LISNASDA | ASSGDRNNERLF | 0.8318223 |
| AELRTLGELTKIAKAHGAQ | ASSGDRNNERLF | 0.82821155 |
| APNAVAAAEPSW | ASSGDRNNERLF | 0.87046874 |
| APQVAGAATAVL | ASSGDRNNERLF | 0.8666713 |
| ARRDPAAAAAILPTDA | ASSGDRNNERLF | 0.8422203 |
| DAAARDVLAEAGLAEYF | ASSGDRNNERLF | 0.85900474 |
| DDQIVLSAKANLAKVEQ | ASSGDRNNERLF | 0.8537475 |
| DELLPDTGEDSDEDGH | ASSGDRNNERLF | 0.13955131 |
| ELPEAPANRATVPQD | ASSGDRNNERLF | 0.8161681 |
| FGQSKIERVTGIG | ASSGDRNNERLF | 0.75571454 |

| | | |
|---|---|---|
| GAGGTGSTAGGAGGAGGAGGLY | ASSGDRNNERLF | 0.87627167 |
| GHGGTGAAGGNGGTGGAGGSGI | ASSGDRNNERLF | 0.82831824 |
| GNGGNGGAGGEGGPGGAGGAGA | ASSGDRNNERLF | 0.8233727 |
| GSTAGNGGNGGNATNGTIAGQP | ASSGDRNNERLF | 0.8523395 |
| GTGGDGGNGGNGGTGG | ASSGDRNNERLF | 0.8514334 |
| GVRRPAGIPADAPV | ASSGDRNNERLF | 0.83383316 |
| GYRQVTDAHLLTLARR | ASSGDRNNERLF | 0.8173584 |
| IAKMGQDGHDRG | ASSGDRNNERLF | 0.31488135 |
| KAEPSIEIQYT | ASSGDRNNERLF | 0.8666929 |
| KPVRIRMAADEDTVAVT | ASSGDRNNERLF | 0.75471103 |
| MPPRIAGMRLLVIKPEP | ASSGDRNNERLF | 0.8097221 |
| MKFYAVALFRSRGVPPTR | ASSGDRNNERLF | 0.73913515 |
| NAGGFGILGAVAHSP | ASSGDRNNERLF | 0.7882639 |
| NEGREITLAMAEM | ASSGDRNNERLF | 0.7419246 |
| NELVEWSHPLGHRWRWE | ASSGDRNNERLF | 0.8747988 |
| NLGAGGTGGNGGNPG | ASSGDRNNERLF | 0.81544673 |
| NTEWPKWETVMAADPVD | ASSGDRNNERLF | 0.84110016 |
| PASAAPAETQP | ASSGDRNNERLF | 0.877277 |
| QQSSDGKNGVRAAEITGE | ASSGDRNNERLF | 0.84291464 |
| RAPDAVTTSWWKE | ASSGDRNNERLF | 0.8369238 |
| SGFGNVGAHTSGWFNQ | ASSGDRNNERLF | 0.8386117 |
| SGVAGAVGQAASVGGLKV | ASSGDRNNERLF | 0.8209679 |
| SNGGAGTGSAGGNGGT | ASSGDRNNERLF | 0.8259171 |
| TGGAGGEGGAAG | ASSGDRNNERLF | 0.834116 |
| TGGDGGAAGAGGAPGHG | ASSGDRNNERLF | 0.8773522 |
| TMPPLLAAIANAGRHGV | ASSGDRNNERLF | 0.8563098 |
| TSSTAGNSGGA | ASSGDRNNERLF | 0.81239384 |
| VPATVPVGILC | ASSGDRNNERLF | 0.86809725 |
| VQRPDSGGVVLTGRIS | ASSGDRNNERLF | 0.8764129 |
| WTPNPVPATKGVPAGLVQAT | ASSGDRNNERLF | 0.87651813 |
| YALKRALAKFSDRFDVVII | ASSGDRNNERLF | 0.7971297 |
| YLIGSVIV | ASSGDRNNERLF | 0.8403293 |
| YGQLVAVID | ASSGDRNNERLF | 0.84193933 |
| VTVGGMTLPA | ASSGDRNNERLF | 0.8425929 |
| VLALPAV | ASSGDRNNERLF | 0.8451095 |
| VIPPTLNYET | ASSGDRNNERLF | 0.8091516 |
| VGMGGMHPSSG | ASSGDRNNERLF | 0.87126535 |
| VGLAGGAATA | ASSGDRNNERLF | 0.8760521 |
| VGGAGGAGGSGP | ASSGDRNNERLF | 0.8448862 |
| VFAAFMV | ASSGDRNNERLF | 0.7585858 |
| TLYDKLR | ASSGDRNNERLF | 0.8432487 |
| TLAVFTVL | ASSGDRNNERLF | 0.8120569 |
| TGSVQPARV | ASSGDRNNERLF | 0.8279753 |
| TGGGAGSGGT | ASSGDRNNERLF | 0.82480955 |
| SHLETSL | ASSGDRNNERLF | 0.78670275 |
| SFTTGMT | ASSGDRNNERLF | 0.8508341 |
| QVEVAKRMGVRQPTVS | ASSGDRNNERLF | 0.8756657 |
| QGSGGPALAM | ASSGDRNNERLF | 0.81964 |
| MLAPSNP | ASSGDRNNERLF | 0.86225694 |
| LVNNAAI | ASSGDRNNERLF | 0.87676215 |
| LTGEINTLA | ASSGDRNNERLF | 0.86746144 |

| | | |
|---|---|---|
| LSIFGPV | ASSGDRNNERLF | 0.87456477 |
| LLALLANL | ASSGDRNNERLF | 0.86906517 |
| LISNASDALDKLR | ASSGDRNNERLF | 0.85918856 |
| LINEAVP | ASSGDRNNERLF | 0.85637015 |
| LGYAFSGL | ASSGDRNNERLF | 0.85640574 |
| LDFVAVAL | ASSGDRNNERLF | 0.8483499 |
| LAQPGAE | ASSGDRNNERLF | 0.84817535 |
| ISNLGAVTGF | ASSGDRNNERLF | 0.83369356 |
| HQEDPGRI | ASSGDRNNERLF | 0.87208796 |
| GTPGEPTP | ASSGDRNNERLF | 0.8551119 |
| GFFNSGS | ASSGDRNNERLF | 0.8693398 |
| GAGGGGLTAGAN | ASSGDRNNERLF | 0.7443727 |
| FVDALVNM | ASSGDRNNERLF | 0.80206966 |
| FLTVAPEL | ASSGDRNNERLF | 0.8699224 |
| FETLTAL | ASSGDRNNERLF | 0.15917099 |
| DVINPIEL | ASSGDRNNERLF | 0.60336614 |
| APVSGAPV | ASSGDRNNERLF | 0.8629528 |
| ALVGGLAW | ASSGDRNNERLF | 0.8689248 |
| ALAAAAATAI | ASSGDRNNERLF | 0.20203528 |
| AFGLGLLL | ASSGDRNNERLF | 0.81285137 |
| AFFAAASL | ASSGDRNNERLF | 0.74403304 |
| AAAALPAFSPP | ASSGDRNNERLF | 0.87111104 |
| VVLAPAL | ASSGDRNNERLF | 0.85451657 |
| TIGHVDHGKTTL | ASSGDRNNERLF | 0.5217683 |
| LISNASDALDK | ASSGDRNNERLF | 0.871376 |
| LISNASDAL | ASSGDRNNERLF | 0.7644607 |
| ISNASDALDK | ASSGDRNNERLF | 0.7750815 |
| NQAETLVYQTE | ASSGDRNNERLF | 0.8745997 |
| TGGTGGTAGAG | ASSGDRNNERLF | 0.8747233 |
| TGSTAGGAGGA | ASSGDRNNERLF | 0.8099539 |
| DPLAVPVAAGPV | ASSGDRNNERLF | 0.8670738 |
| GGTGGTPGNGGN | ASSGDRNNERLF | 0.8305631 |
| PAGLAGVDPATL | ASSGDRNNERLF | 0.8763532 |
| GNNLVVEAPHAGASV | ASSGDRNNERLF | 0.82122314 |
| KVTRIMSASTLEQPA | ASSGDRNNERLF | 0.798497 |
| LGFALPIALPRRTVV | ASSGDRNNERLF | 0.8727323 |
| TRLGNLTARKLVVLV | ASSGDRNNERLF | 0.8595903 |
| WSQQRRLIYFPSAGP | ASSGDRNNERLF | 0.85304636 |
| GPNGNPDPMAAAVDIR | ASSGDRNNERLF | 0.8691825 |
| TQRCVSVADLRAAASR | ASSGDRNNERLF | 0.8393632 |
| QFVLDVAEHPVEVLTETV | ASSGDRNNERLF | 0.84192616 |
| GGAGGTGGAGGNGGTQAGDG | ASSGDRNNERLF | 0.8730215 |
| PLLATLRAIAVDVDAKPAQV | ASSGDRNNERLF | 0.8721951 |
| APKFRTGGYMIDFWGVGYQVAK | ASSGDRNNERLF | 0.74420804 |
| GAGGAGGAGGAAGGPGGTGGTG | ASSGDRNNERLF | 0.87539166 |
| IGALGLLATDVAPAGSVAFGVA | ASSGDRNNERLF | 0.84769547 |
| AGGAGGIGGNGGAGGAGGVLMG | ASSGDRNNERLF | 0.81071687 |
| ADELIGLL | ASSGDRNNERLF | 0.87724066 |
| ADRALKA | ASSGDRNNERLF | 0.85070014 |
| AGAVGQVGGAGGA | ASSGDRNNERLF | 0.8685732 |
| AGGGGGLGGAP | ASSGDRNNERLF | 0.86942905 |

| | | |
|---|---|---|
| AGGLLYGNGGAG | ASSGDRNNERLF | 0.8564894 |
| AGLESIGL | ASSGDRNNERLF | 0.5945879 |
| AGLKTGII | ASSGDRNNERLF | 0.8474557 |
| ASLAALLL | ASSGDRNNERLF | 0.8346709 |
| ASVINISSI | ASSGDRNNERLF | 0.87147474 |
| ATPPGPA | ASSGDRNNERLF | 0.8534105 |
| ATVASALV | ASSGDRNNERLF | 0.7656574 |
| ATVASLIL | ASSGDRNNERLF | 0.86065614 |
| AVNGPVLIRH | ASSGDRNNERLF | 0.87654847 |
| DGAAGAVGI | ASSGDRNNERLF | 0.863664 |
| DPLLHGI | ASSGDRNNERLF | 0.8599661 |
| ELGAALGAL | ASSGDRNNERLF | 0.8763676 |
| ELGRTQAV | ASSGDRNNERLF | 0.8223876 |
| FVAGGYLS | ASSGDRNNERLF | 0.85349905 |
| GASGGAGGAGGW | ASSGDRNNERLF | 0.8734941 |
| GGAGAGGDGGAGGI | ASSGDRNNERLF | 0.87057483 |
| GTGHNGSQPGMGGNG | ASSGDRNNERLF | 0.85785306 |
| GYAAASASAS | ASSGDRNNERLF | 0.87237006 |
| HQVDVAAVQQG | ASSGDRNNERLF | 0.7390858 |
| IGAAIAHT | ASSGDRNNERLF | 0.862606 |
| IGMIEVAQL | ASSGDRNNERLF | 0.7905051 |
| IRLAPEHTSAMD | ASSGDRNNERLF | 0.85249734 |
| IYNVSVL | ASSGDRNNERLF | 0.7854609 |
| KQGFLGM | ASSGDRNNERLF | 0.86375785 |
| LESVITQL | ASSGDRNNERLF | 0.86820143 |
| LGLGALVVV | ASSGDRNNERLF | 0.86778116 |
| LIGNGGNGGSGGI | ASSGDRNNERLF | 0.6112578 |
| LNAINAPTL | ASSGDRNNERLF | 0.86844134 |
| LQLLNTGV | ASSGDRNNERLF | 0.86422896 |
| LQTSIEF | ASSGDRNNERLF | 0.8792353 |
| MSFAVSEL | ASSGDRNNERLF | 0.841628 |
| NTGALGL | ASSGDRNNERLF | 0.86393905 |
| PPGAGGAGGAGTT | ASSGDRNNERLF | 0.81821436 |
| QLVGLGI | ASSGDRNNERLF | 0.8546207 |
| RFLEQVHQL | ASSGDRNNERLF | 0.78032 |
| RGVGYVLR | ASSGDRNNERLF | 0.7887298 |
| SATGVFAGVI | ASSGDRNNERLF | 0.81736827 |
| SLPAITLP | ASSGDRNNERLF | 0.8165035 |
| STALAFTV | ASSGDRNNERLF | 0.8682146 |
| SVLTVGLGK | ASSGDRNNERLF | 0.86895716 |
| TALAGAGSGI | ASSGDRNNERLF | 0.8677737 |
| TGLPFTGI | ASSGDRNNERLF | 0.87434345 |
| VAEHVTV | ASSGDRNNERLF | 0.8707942 |
| VDIVLNSL | ASSGDRNNERLF | 0.8748326 |
| VLTGGGAVL | ASSGDRNNERLF | 0.82968664 |
| VNVGLAYGI | ASSGDRNNERLF | 0.8074769 |
| VSSAAVGAV | ASSGDRNNERLF | 0.81221074 |
| VTSVYEAGV | ASSGDRNNERLF | 0.8496057 |
| WVALDRGA | ASSGDRNNERLF | 0.836288 |
| YLVGSVVI | ASSGDRNNERLF | 0.8624202 |
| GPPGTGKTLL | ASSGDRNNERLF | 0.14017504 |

| | | |
|---|---|---|
| GVGMTKF | ASSGDRNNERLF | 0.8455706 |
| IGIGGSDL | ASSGDRNNERLF | 0.87763876 |
| IGTLLAEL | ASSGDRNNERLF | 0.7754754 |
| IYTSGTT | ASSGDRNNERLF | 0.85507095 |
| LSFPAGV | ASSGDRNNERLF | 0.85102046 |
| LYTSGTT | ASSGDRNNERLF | 0.83822286 |
| VGAFPFGF | ASSGDRNNERLF | 0.86786467 |
| VGAGTVEF | ASSGDRNNERLF | 0.8288876 |
| VGKSTLLL | ASSGDRNNERLF | 0.868279 |
| AALAHAGGGIV | ASSGDRNNERLF | 0.87466717 |
| AALVAVSLL | ASSGDRNNERLF | 0.85785127 |
| AAVSRAVAAL | ASSGDRNNERLF | 0.8407949 |
| AGGSSGIGGF | ASSGDRNNERLF | 0.8529084 |
| AGLTVANANG | ASSGDRNNERLF | 0.5027177 |
| AGLVGAVL | ASSGDRNNERLF | 0.85023856 |
| ALSQAFTV | ASSGDRNNERLF | 0.2790594 |
| ALTFSVSL | ASSGDRNNERLF | 0.8396936 |
| ALTGAALVAL | ASSGDRNNERLF | 0.8068136 |
| ASFSIGAGVL | ASSGDRNNERLF | 0.8593796 |
| DGSVYKI | ASSGDRNNERLF | 0.8531351 |
| EQLAEIV | ASSGDRNNERLF | 0.86646974 |
| FAVSELTVM | ASSGDRNNERLF | 0.77959657 |
| GAAPLAAGL | ASSGDRNNERLF | 0.8509022 |
| GADGGDGGSGGAGG | ASSGDRNNERLF | 0.83268476 |
| GFGLDTGGAGG | ASSGDRNNERLF | 0.7781863 |
| GLASGGLL | ASSGDRNNERLF | 0.86943 |
| GRIAGAAVLL | ASSGDRNNERLF | 0.8629706 |
| GSAGSLNTSF | ASSGDRNNERLF | 0.825528 |
| GTGQIGI | ASSGDRNNERLF | 0.8698073 |
| GTLIALIL | ASSGDRNNERLF | 0.85935694 |
| IAATMGFVVG | ASSGDRNNERLF | 0.6833257 |
| IFAQALL | ASSGDRNNERLF | 0.82249665 |
| IGAAVALGF | ASSGDRNNERLF | 0.8534663 |
| IGLTPLAL | ASSGDRNNERLF | 0.8180505 |
| ILGSNTSTL | ASSGDRNNERLF | 0.2742696 |
| IVFSLTVL | ASSGDRNNERLF | 0.85730505 |
| LASKATGF | ASSGDRNNERLF | 0.81025994 |
| LASVALIL | ASSGDRNNERLF | 0.7814035 |
| LFAVGLTL | ASSGDRNNERLF | 0.84513414 |
| LFNGGIGTYI | ASSGDRNNERLF | 0.8701965 |
| LTLGAGII | ASSGDRNNERLF | 0.7188295 |
| LVAAALGDI | ASSGDRNNERLF | 0.81201476 |
| NTAFVQLGI | ASSGDRNNERLF | 0.85990787 |
| QLFSGAEL | ASSGDRNNERLF | 0.8670417 |
| SAATFITL | ASSGDRNNERLF | 0.8716902 |
| SNYGLANLGV | ASSGDRNNERLF | 0.86008644 |
| SYGGAYAVM | ASSGDRNNERLF | 0.84957826 |
| TTGVGGLTLG | ASSGDRNNERLF | 0.8583919 |
| TTQTIYI | ASSGDRNNERLF | 0.8556093 |
| VAAGAGLGAVI | ASSGDRNNERLF | 0.7955755 |
| VAAVVTLL | ASSGDRNNERLF | 0.7199703 |

| | | |
|---|---|---|
| VGFLGGLI | ASSGDRNNERLF | 0.841635 |
| VGVIPILL | ASSGDRNNERLF | 0.7539378 |
| VIFASSLI | ASSGDRNNERLF | 0.84988827 |
| VKGGDGGAAAG | ASSGDRNNERLF | 0.8776917 |
| VTKTFTNI | ASSGDRNNERLF | 0.8285353 |
| VVAAAILL | ASSGDRNNERLF | 0.7158721 |
| YAALDVEL | ASSGDRNNERLF | 0.85214615 |
| LASVALLL | ASSGDRNNERLF | 0.7707056 |
| LDAVHRV | ASSGDRNNERLF | 0.85973334 |
| QGLSSSIF | ASSGDRNNERLF | 0.86212754 |
| RGTFANIRL | ASSGDRNNERLF | 0.8772917 |
| RQLWWGHRI | ASSGDRNNERLF | 0.87063265 |
| YGPPGTGKTLL | ASSGDRNNERLF | 0.83162296 |
| AALVGAPANVA | ASSGDRNNERLF | 0.8702832 |
| AATFRRTE | ASSGDRNNERLF | 0.8245878 |
| AGALGGGI | ASSGDRNNERLF | 0.8483387 |
| AGGAGGAGGIGAG | ASSGDRNNERLF | 0.8528416 |
| AGSGALGL | ASSGDRNNERLF | 0.86062264 |
| AQIADQI | ASSGDRNNERLF | 0.847371 |
| ASAVAVTYL | ASSGDRNNERLF | 0.8407636 |
| ASTVLGGTGG | ASSGDRNNERLF | 0.654551 |
| AVGGEPAK | ASSGDRNNERLF | 0.8723766 |
| CEQGTFGGTC | ASSGDRNNERLF | 0.7816141 |
| DEGKAPAG | ASSGDRNNERLF | 0.7646891 |
| DTVVFPAL | ASSGDRNNERLF | 0.81713384 |
| FEGITMAG | ASSGDRNNERLF | 0.876227 |
| FPLVVQD | ASSGDRNNERLF | 0.87264925 |
| GAFTTPQ | ASSGDRNNERLF | 0.8167403 |
| GFGGFGTAGGVGGA | ASSGDRNNERLF | 0.827953 |
| GFLAQFL | ASSGDRNNERLF | 0.8550985 |
| GGAGGAGGNAGVA | ASSGDRNNERLF | 0.75336695 |
| GGVGTAGAGGAGGAGG | ASSGDRNNERLF | 0.8687118 |
| GNGGGGDAGNAGSGGN | ASSGDRNNERLF | 0.8468194 |
| GTGGAGGAGNDAGS | ASSGDRNNERLF | 0.84194756 |
| GTTNTGWFNS | ASSGDRNNERLF | 0.8317712 |
| IFMGPLFTL | ASSGDRNNERLF | 0.8628442 |
| ILAHAIEAI | ASSGDRNNERLF | 0.8295469 |
| IRSAIREAL | ASSGDRNNERLF | 0.86624694 |
| ITFAGGG | ASSGDRNNERLF | 0.2929964 |
| ITGMSGHQEV | ASSGDRNNERLF | 0.87050605 |
| IVAGLACTL | ASSGDRNNERLF | 0.8067386 |
| IVGGGLAGA | ASSGDRNNERLF | 0.8207102 |
| IVTGVLNI | ASSGDRNNERLF | 0.7195003 |
| LAQQGCI | ASSGDRNNERLF | 0.8611269 |
| LCLHVEN | ASSGDRNNERLF | 0.86320543 |
| LDAISDGS | ASSGDRNNERLF | 0.8724926 |
| LGGFAAML | ASSGDRNNERLF | 0.7935123 |
| LGGKNPMI | ASSGDRNNERLF | 0.86913437 |
| LGPKGVAAIL | ASSGDRNNERLF | 0.86718905 |
| LMVGVSI | ASSGDRNNERLF | 0.8376256 |
| LVGAAFVL | ASSGDRNNERLF | 0.4779393 |

| | | |
|---|---|---|
| LVLAVAPL | ASSGDRNNERLF | 0.8626571 |
| MAAIAVIDAA | ASSGDRNNERLF | 0.8444927 |
| NNGINAELQ | ASSGDRNNERLF | 0.87020457 |
| RGEPLVLVHG | ASSGDRNNERLF | 0.84388095 |
| RGGAGGAAGLI | ASSGDRNNERLF | 0.7849981 |
| RSYRMHPAV | ASSGDRNNERLF | 0.86600786 |
| SAAHLLAGQLA | ASSGDRNNERLF | 0.86732835 |
| SAAVGSY | ASSGDRNNERLF | 0.7874774 |
| SPHAVLHRI | ASSGDRNNERLF | 0.83809036 |
| SQIAKVL | ASSGDRNNERLF | 0.8483639 |
| TALGHLVP | ASSGDRNNERLF | 0.72486323 |
| TAPTTSGAA | ASSGDRNNERLF | 0.86829937 |
| TDGDDGNGGSPGT | ASSGDRNNERLF | 0.8307527 |
| TDGLGEN | ASSGDRNNERLF | 0.827 |
| TGGPPLPGQA | ASSGDRNNERLF | 0.85961235 |
| TIPAITI | ASSGDRNNERLF | 0.8448302 |
| TIPGFTI | ASSGDRNNERLF | 0.8346416 |
| TSIASGALA | ASSGDRNNERLF | 0.8429041 |
| TVTLPGAAI | ASSGDRNNERLF | 0.8493706 |
| VASGLSAALP | ASSGDRNNERLF | 0.8660364 |
| VGAGVIGI | ASSGDRNNERLF | 0.8458625 |
| VGPKRFGV | ASSGDRNNERLF | 0.8541744 |
| VNVAFPD | ASSGDRNNERLF | 0.87596506 |
| VSMTNSGVS | ASSGDRNNERLF | 0.73729944 |
| VTDVLGDLSV | ASSGDRNNERLF | 0.8187688 |
| VVVVGAGFAGL | ASSGDRNNERLF | 0.8633158 |
| VYTVGTVV | ASSGDRNNERLF | 0.79048145 |
| WTTFSGP | ASSGDRNNERLF | 0.8683785 |
| YDELVEAF | ASSGDRNNERLF | 0.853033 |
| YGTGGAGGNGG | ASSGDRNNERLF | 0.86398005 |
| YTNVMAV | ASSGDRNNERLF | 0.8397225 |
| FLEQVHQL | ASSGDRNNERLF | 0.87723094 |
| GLSLLAVAV | ASSGDRNNERLF | 0.86954576 |
| LEQVHQL | ASSGDRNNERLF | 0.8628055 |
| LNAAGAI | ASSGDRNNERLF | 0.8661497 |
| PAVHASL | ASSGDRNNERLF | 0.8061712 |
| TELEDDA | ASSGDRNNERLF | 0.8701713 |
| TPPGPPN | ASSGDRNNERLF | 0.8686462 |
| TPPPPPPP | ASSGDRNNERLF | 0.76832116 |
| TQSPALI | ASSDGTDNERLF | 0.84338635 |
| EFAAAFPQ | ASSDGTDNERLF | 0.87612784 |
| LLSGNLDVL | ASSDGTDNERLF | 0.87431633 |
| GPANLGNYNI | ASSDGTDNERLF | 0.702976 |
| TLGPTLP | ASSDGTDNERLF | 0.83690125 |
| VYAYSSCCK | ASSDGTDNERLF | 0.69950926 |
| APAPAPAPNI | ASSDGTDNERLF | 0.8290739 |
| PGGPVPIP | ASSDGTDNERLF | 0.75458395 |
| FLTVALEP | ASSDGTDNERLF | 0.8587025 |
| SQYFGANL | ASSDGTDNERLF | 0.83202034 |
| GGVKPAA | ASSDGTDNERLF | 0.85427 |
| GGTGGAGGAGGW | ASSDGTDNERLF | 0.85586286 |

| | | |
|---|---|---|
| GVGGVAGLGGAGG | ASSDGTDNERLF | 0.87519574 |
| VLLVLATTI | ASSDGTDNERLF | 0.83572423 |
| IGIFAAML | ASSDGTDNERLF | 0.8463278 |
| AVGDLID | ASSDGTDNERLF | 0.7406944 |
| NISPLPL | ASSDGTDNERLF | 0.40035576 |
| VTAVFVAI | ASSDGTDNERLF | 0.83819234 |
| SGDDNDAAL | ASSDGTDNERLF | 0.8379718 |
| AVAALAHS | ASSDGTDNERLF | 0.3520692 |
| GGVTVMATP | ASSDGTDNERLF | 0.8751419 |
| AGLAANVRD | ASSDGTDNERLF | 0.8550431 |
| TNRPDLIDPALLRPG | ASSDGTDNERLF | 0.8659977 |
| LANLGVTAL | ASSDGTDNERLF | 0.83546436 |
| TGERIEL | ASSDGTDNERLF | 0.8542697 |
| NGEELAA | ASSDGTDNERLF | 0.8310331 |
| VLSLVWG | ASSDGTDNERLF | 0.8642592 |
| IGYALWL | ASSDGTDNERLF | 0.787035 |
| LSLLLAPL | ASSDGTDNERLF | 0.74956167 |
| TLDEALK | ASSDGTDNERLF | 0.84840333 |
| TDGAGTAEGG | ASSDGTDNERLF | 0.8006706 |
| NAPGGAGGNGG | ASSDGTDNERLF | 0.8712052 |
| VSVFMGMGGNF | ASSDGTDNERLF | 0.87332255 |
| LASYLAAL | ASSDGTDNERLF | 0.860628 |
| LVGGTPAL | ASSDGTDNERLF | 0.8748674 |
| VRNADVL | ASSDGTDNERLF | 0.28644016 |
| TCSSPFLVSAM | ASSDGTDNERLF | 0.83861405 |
| VHITDAE | ASSDGTDNERLF | 0.874808 |
| EYAAPSF | ASSDGTDNERLF | 0.86268365 |
| VVDAFHGGL | ASSDGTDNERLF | 0.8420678 |
| ASVVDPQ | ASSDGTDNERLF | 0.875669 |
| SYNEFGGLS | ASSDGTDNERLF | 0.82146436 |
| DPDGSVY | ASSDGTDNERLF | 0.8088124 |
| MVSTIHATE | ASSDGTDNERLF | 0.85663927 |
| LAAGVADPV | ASSDGTDNERLF | 0.8686764 |
| TKEQIAA | ASSDGTDNERLF | 0.8632206 |
| SLASPAAL | ASSDGTDNERLF | 0.79920757 |
| IGLVFYGV | ASSDGTDNERLF | 0.86162674 |
| FLLDASNL | ASSDGTDNERLF | 0.8810205 |
| YQVGNFP | ASSDGTDNERLF | 0.83889997 |
| ESLLSQST | ASSDGTDNERLF | 0.8644545 |
| LFTVGFAAVG | ASSDGTDNERLF | 0.8616189 |
| ATPPTPT | ASSDGTDNERLF | 0.79704726 |
| TDLATLG | ASSDGTDNERLF | 0.81596863 |
| SAGGTGR | ASSDGTDNERLF | 0.82237554 |
| GFSNSGS | ASSDGTDNERLF | 0.8507297 |
| GGNGGSGGTSV | ASSDGTDNERLF | 0.76874876 |
| GTSVATGGAGN | ASSDGTDNERLF | 0.85607874 |
| NGGNGGSGGTS | ASSDGTDNERLF | 0.84533787 |
| AVVKFPH | ASSDGTDNERLF | 0.8796327 |
| TPPPPPP | ASSDGTDNERLF | 0.8595038 |
| AAPAAAAAPA | ASSDGTDNERLF | 0.8667094 |
| IALNEDL | ASSDGTDNERLF | 0.804925 |

| | | |
|---|---|---|
| IVNVSSM | ASSDGTDNERLF | 0.86518264 |
| TGIPPAPR | ASSDGTDNERLF | 0.863222 |
| FDLGGGTF | ASSDGTDNERLF | 0.68551326 |
| FDLGGGTFDVS | ASSDGTDNERLF | 0.87417614 |
| NEPTAAA | ASSDGTDNERLF | 0.81458974 |
| NEPTAAALA | ASSDGTDNERLF | 0.8384793 |
| GSIINISSI | ASSDGTDNERLF | 0.8665559 |
| VAPGNAGTA | ASSDGTDNERLF | 0.87094593 |
| TGGGASLE | ASSDGTDNERLF | 0.85123295 |
| VAVYNHYKRI | ASSDGTDNERLF | 0.84733695 |
| TDPAPATT | ASSDGTDNERLF | 0.82466567 |
| TNRPDILDPALLRPG | ASSDGTDNERLF | 0.8657401 |
| LTTTTTS | ASSDGTDNERLF | 0.87244296 |
| PTGSGKTL | ASSDGTDNERLF | 0.8155299 |
| TGSGKTL | ASSDGTDNERLF | 0.8741714 |
| TIGHVDHGK | ASSDGTDNERLF | 0.8664116 |
| TIGHVDHGKTT | ASSDGTDNERLF | 0.61170447 |
| GHVDHGKTT | ASSDGTDNERLF | 0.82970965 |
| AQFEHTLL | ASSDGTDNERLF | 0.87307304 |
| IVNQDEL | ASSDGTDNERLF | 0.86976415 |
| TNGSQFF | ASSDGTDNERLF | 0.8741601 |
| TGKGIVE | ASSDGTDNERLF | 0.86051977 |
| LISNASDA | ASSDGTDNERLF | 0.80202675 |
| AELRTLGELTKIAKAHGAQ | ASSDGTDNERLF | 0.87562066 |
| APNAVAAAEPSW | ASSDGTDNERLF | 0.8792196 |
| APQVAGAATAVL | ASSDGTDNERLF | 0.8684461 |
| ARRDPAAAAAILPTDA | ASSDGTDNERLF | 0.8200281 |
| DAAARDVLAEAGLAEYF | ASSDGTDNERLF | 0.7988355 |
| DDQIVLSAKANLAKVEQ | ASSDGTDNERLF | 0.8621734 |
| DELLPDTGEDSDEDGH | ASSDGTDNERLF | 0.86948097 |
| ELPEAPANRATVPQD | ASSDGTDNERLF | 0.81752104 |
| FGQSKIERVTGIG | ASSDGTDNERLF | 0.8524319 |
| GAGGTGSTAGGAGGAGGAGGLY | ASSDGTDNERLF | 0.8560654 |
| GHGGTGAAGGNGGTGGAGGSGI | ASSDGTDNERLF | 0.635653 |
| GNGGNGGAGGEGGPGGAGGAGA | ASSDGTDNERLF | 0.85771286 |
| GSTAGNGGNGGNATNGTIAGQP | ASSDGTDNERLF | 0.8717949 |
| GTGGDGGNGGNGGTGG | ASSDGTDNERLF | 0.8637859 |
| GVRRPAGIPADAPV | ASSDGTDNERLF | 0.8164667 |
| GYRQVTDAHLLTLARR | ASSDGTDNERLF | 0.87299764 |
| IAKMGQDGHDRG | ASSDGTDNERLF | 0.8775838 |
| KAEPSIEIQYT | ASSDGTDNERLF | 0.8772558 |
| KPVRIRMAADEDTVAVT | ASSDGTDNERLF | 0.7319808 |
| MPPRIAGMRLLVIKPEP | ASSDGTDNERLF | 0.63033944 |
| MKFYAVALFRSRGVPPTR | ASSDGTDNERLF | 0.8361786 |
| NAGGFGILGAVAHSP | ASSDGTDNERLF | 0.8594466 |
| NEGREITLAMAEM | ASSDGTDNERLF | 0.84543765 |
| NELVEWSHPLGHRWRWE | ASSDGTDNERLF | 0.8320956 |
| NLGAGGTGGNGGNPG | ASSDGTDNERLF | 0.8555473 |
| NTEWPKWETVMAADPVD | ASSDGTDNERLF | 0.87331784 |
| PASAAPAETQP | ASSDGTDNERLF | 0.80824417 |
| QQSSDGKNGVRAAEITGE | ASSDGTDNERLF | 0.7387874 |

| | | |
|---|---|---|
| RAPDAVTTSWWKE | ASSDGTDNERLF | 0.86874664 |
| SGFGNVGAHTSGWFNQ | ASSDGTDNERLF | 0.8707808 |
| SGVAGAVGQAASVGGLKV | ASSDGTDNERLF | 0.81530285 |
| SNGGAGTGSAGGNGGT | ASSDGTDNERLF | 0.83989894 |
| TGGAGGEGGAAG | ASSDGTDNERLF | 0.83650523 |
| TGGDGGAAGAGGAPGHG | ASSDGTDNERLF | 0.82042843 |
| TMPPLLAAIANAGRHGV | ASSDGTDNERLF | 0.8547729 |
| TSSTAGNSGGA | ASSDGTDNERLF | 0.7785721 |
| VPATVPVGILC | ASSDGTDNERLF | 0.8759439 |
| VQRPDSGGVVLTGRIS | ASSDGTDNERLF | 0.80688643 |
| WTPNPVPATKGVPAGLVQAT | ASSDGTDNERLF | 0.864269 |
| YALKRALAKFSDRFDVVII | ASSDGTDNERLF | 0.86216646 |
| YLIGSVIV | ASSDGTDNERLF | 0.8613987 |
| YGQLVAVID | ASSDGTDNERLF | 0.839169 |
| VTVGGMTLPA | ASSDGTDNERLF | 0.8095604 |
| VLALPAV | ASSDGTDNERLF | 0.7639327 |
| VIPPTLNYET | ASSDGTDNERLF | 0.7375946 |
| VGMGGMHPSSG | ASSDGTDNERLF | 0.87066567 |
| VGLAGGAATA | ASSDGTDNERLF | 0.59003377 |
| VGGAGGAGGSGP | ASSDGTDNERLF | 0.8743236 |
| VFAAFMV | ASSDGTDNERLF | 0.8643559 |
| TLYDKLR | ASSDGTDNERLF | 0.815714 |
| TLAVFTVL | ASSDGTDNERLF | 0.86932236 |
| TGSVQPARV | ASSDGTDNERLF | 0.7233299 |
| TGGGAGSGGT | ASSDGTDNERLF | 0.8155688 |
| SHLETSL | ASSDGTDNERLF | 0.8362733 |
| SFTTGMT | ASSDGTDNERLF | 0.8704645 |
| QVEVAKRMGVRQPTVS | ASSDGTDNERLF | 0.81249046 |
| QGSGGPALAM | ASSDGTDNERLF | 0.86790574 |
| MLAPSNP | ASSDGTDNERLF | 0.85477436 |
| LVNNAAI | ASSDGTDNERLF | 0.78964436 |
| LTGEINTLA | ASSDGTDNERLF | 0.8629862 |
| LSIFGPV | ASSDGTDNERLF | 0.8727249 |
| LLALLANL | ASSDGTDNERLF | 0.8667127 |
| LISNASDALDKLR | ASSDGTDNERLF | 0.78540295 |
| LINEAVP | ASSDGTDNERLF | 0.8647063 |
| LGYAFSGL | ASSDGTDNERLF | 0.7934762 |
| LDFVAVAL | ASSDGTDNERLF | 0.861322 |
| LAQPGAE | ASSDGTDNERLF | 0.85873806 |
| ISNLGAVTGF | ASSDGTDNERLF | 0.8299129 |
| HQEDPGRI | ASSDGTDNERLF | 0.86760753 |
| GTPGEPTP | ASSDGTDNERLF | 0.7476882 |
| GFFNSGS | ASSDGTDNERLF | 0.8078426 |
| GAGGGGLTAGAN | ASSDGTDNERLF | 0.8325541 |
| FVDALVNM | ASSDGTDNERLF | 0.85815597 |
| FLTVAPEL | ASSDGTDNERLF | 0.8696526 |
| FETLTAL | ASSDGTDNERLF | 0.79380465 |
| DVINPIEL | ASSDGTDNERLF | 0.844432 |
| APVSGAPV | ASSDGTDNERLF | 0.8297952 |
| ALVGGLAW | ASSDGTDNERLF | 0.7533728 |
| ALAAAAATAI | ASSDGTDNERLF | 0.8665952 |

| | | |
|---|---|---|
| AFGLGLLL | ASSDGTDNERLF | 0.8591851 |
| AFFAAASL | ASSDGTDNERLF | 0.78071135 |
| AAAALPAFSPP | ASSDGTDNERLF | 0.8657322 |
| VVLAPAL | ASSDGTDNERLF | 0.8140101 |
| TIGHVDHGKTTL | ASSDGTDNERLF | 0.86640465 |
| LISNASDALDK | ASSDGTDNERLF | 0.8776599 |
| LISNASDAL | ASSDGTDNERLF | 0.7919032 |
| ISNASDALDK | ASSDGTDNERLF | 0.8692313 |
| NQAETLVYQTE | ASSDGTDNERLF | 0.85878694 |
| TGGTGGTAGAG | ASSDGTDNERLF | 0.8602622 |
| TGSTAGGAGGA | ASSDGTDNERLF | 0.84494203 |
| DPLAVPVAAGPV | ASSDGTDNERLF | 0.80705863 |
| GGTGGTPGNGGN | ASSDGTDNERLF | 0.8761819 |
| PAGLAGVDPATL | ASSDGTDNERLF | 0.86896515 |
| GNNLVVEAPHAGASV | ASSDGTDNERLF | 0.8573149 |
| KVTRIMSASTLEQPA | ASSDGTDNERLF | 0.8622238 |
| LGFALPIALPRRTVV | ASSDGTDNERLF | 0.8686041 |
| TRLGNLTARKLVVLV | ASSDGTDNERLF | 0.8701486 |
| WSQQRRLIYFPSAGP | ASSDGTDNERLF | 0.8146362 |
| GPNGNPDPMAAAVDIR | ASSDGTDNERLF | 0.71944106 |
| TQRCVSVADLRAAASR | ASSDGTDNERLF | 0.8451805 |
| QFVLDVAEHPVEVLTETV | ASSDGTDNERLF | 0.8301912 |
| GGAGGTGGAGGNGGTQAGDG | ASSDGTDNERLF | 0.8528581 |
| PLLATLRAIAVDVDAKPAQV | ASSDGTDNERLF | 0.6898402 |
| APKFRTGGYMIDFWGVGYQVAK | ASSDGTDNERLF | 0.86998034 |
| GAGGAGGAGGAAGGPGGTGGTG | ASSDGTDNERLF | 0.87954783 |
| IGALGLLATDVAPAGSVAFGVA | ASSDGTDNERLF | 0.82358277 |
| AGGAGGIGGNGGAGGAGGVLMG | ASSDGTDNERLF | 0.877609 |
| ADELIGLL | ASSDGTDNERLF | 0.79898787 |
| ADRALKA | ASSDGTDNERLF | 0.8658846 |
| AGAVGQVGGAGGA | ASSDGTDNERLF | 0.8715576 |
| AGGGGGLGGAP | ASSDGTDNERLF | 0.8030895 |
| AGGLLYGNGGAG | ASSDGTDNERLF | 0.8628473 |
| AGLESIGL | ASSDGTDNERLF | 0.7868223 |
| AGLKTGII | ASSDGTDNERLF | 0.83830124 |
| ASLAALLL | ASSDGTDNERLF | 0.8756569 |
| ASVINISSI | ASSDGTDNERLF | 0.8281605 |
| ATPPGPA | ASSDGTDNERLF | 0.8577993 |
| ATVASALV | ASSDGTDNERLF | 0.8552722 |
| ATVASLIL | ASSDGTDNERLF | 0.8201617 |
| AVNGPVLIRH | ASSDGTDNERLF | 0.86548513 |
| DGAAGAVGI | ASSDGTDNERLF | 0.87265635 |
| DPLLHGI | ASSDGTDNERLF | 0.81736124 |
| ELGAALGAL | ASSDGTDNERLF | 0.8788059 |
| ELGRTQAV | ASSDGTDNERLF | 0.8136406 |
| FVAGGYLS | ASSDGTDNERLF | 0.87913865 |
| GASGGAGGAGGW | ASSDGTDNERLF | 0.87195545 |
| GGAGAGGDGGAGGI | ASSDGTDNERLF | 0.8495947 |
| GTGHNGSQPGMGGNG | ASSDGTDNERLF | 0.8407912 |
| GYAAASASAS | ASSDGTDNERLF | 0.8676688 |
| HQVDVAAVQQG | ASSDGTDNERLF | 0.83463174 |

| | | |
|---|---|---|
| IGAAIAHT | ASSDGTDNERLF | 0.86128545 |
| IGMIEVAQL | ASSDGTDNERLF | 0.87791383 |
| IRLAPEHTSAMD | ASSDGTDNERLF | 0.3713132 |
| IYNVSVL | ASSDGTDNERLF | 0.67618847 |
| KQGFLGM | ASSDGTDNERLF | 0.6287751 |
| LESVITQL | ASSDGTDNERLF | 0.5254499 |
| LGLGALVVV | ASSDGTDNERLF | 0.5066341 |
| LIGNGGNGGSGGI | ASSDGTDNERLF | 0.36523926 |
| LNAINAPTL | ASSDGTDNERLF | 0.47235712 |
| LQLLNTGV | ASSDGTDNERLF | 0.46986812 |
| LQTSIEF | ASSDGTDNERLF | 0.69772416 |
| MSFAVSEL | ASSDGTDNERLF | 0.6133477 |
| NTGALGL | ASSDGTDNERLF | 0.459003 |
| PPGAGGAGGAGTT | ASSDGTDNERLF | 0.6626855 |
| QLVGLGI | ASSDGTDNERLF | 0.5983694 |
| RFLEQVHQL | ASSDGTDNERLF | 0.3864332 |
| RGVGYVLR | ASSDGTDNERLF | 0.5252274 |
| SATGVFAGVI | ASSDGTDNERLF | 0.42063463 |
| SLPAITLP | ASSDGTDNERLF | 0.5325568 |
| STALAFTV | ASSDGTDNERLF | 0.30994207 |
| SVLTVGLGK | ASSDGTDNERLF | 0.61623305 |
| TALAGAGSGI | ASSDGTDNERLF | 0.5346334 |
| TGLPFTGI | ASSDGTDNERLF | 0.3071304 |
| VAEHVTV | ASSDGTDNERLF | 0.61454827 |
| VDIVLNSL | ASSDGTDNERLF | 0.54294235 |
| VLTGGGAVL | ASSDGTDNERLF | 0.5737696 |
| VNVGLAYGI | ASSDGTDNERLF | 0.66631985 |
| VSSAAVGAV | ASSDGTDNERLF | 0.47486782 |
| VTSVYEAGV | ASSDGTDNERLF | 0.4637763 |
| WVALDRGA | ASSDGTDNERLF | 0.6471213 |
| YLVGSVVI | ASSDGTDNERLF | 0.65552014 |
| GPPGTGKTLL | ASSDGTDNERLF | 0.4246248 |
| GVGMTKF | ASSDGTDNERLF | 0.6205611 |
| IGIGGSDL | ASSDGTDNERLF | 0.6522398 |
| IGTLLAEL | ASSDGTDNERLF | 0.59341085 |
| IYTSGTT | ASSDGTDNERLF | 0.6486478 |
| LSFPAGV | ASSDGTDNERLF | 0.6611378 |
| LYTSGTT | ASSDGTDNERLF | 0.44977546 |
| VGAFPFGF | ASSDGTDNERLF | 0.6642276 |
| VGAGTVEF | ASSDGTDNERLF | 0.6753725 |
| VGKSTLLL | ASSDGTDNERLF | 0.4805386 |
| AALAHAGGGIV | ASSDGTDNERLF | 0.52638054 |
| AALVAVSLL | ASSDGTDNERLF | 0.35767394 |
| AAVSRAVAAL | ASSDGTDNERLF | 0.53692824 |
| AGGSSGIGGF | ASSDGTDNERLF | 0.46753293 |
| AGLTVANANG | ASSDGTDNERLF | 0.65304375 |
| AGLVGAVL | ASSDGTDNERLF | 0.6713443 |
| ALSQAFTV | ASSDGTDNERLF | 0.56431186 |
| ALTFSVSL | ASSDGTDNERLF | 0.3791294 |
| ALTGAALVAL | ASSDGTDNERLF | 0.35580125 |
| ASFSIGAGVL | ASSDGTDNERLF | 0.56976557 |

| | | |
|---|---|---|
| DGSVYKI | ASSDGTDNERLF | 0.36872852 |
| EQLAEIV | ASSDGTDNERLF | 0.35547027 |
| FAVSELTVM | ASSDGTDNERLF | 0.4828729 |
| GAAPLAAGL | ASSDGTDNERLF | 0.6569363 |
| GADGGDGGSGGAGG | ASSDGTDNERLF | 0.64643335 |
| GFGLDTGGAGG | ASSDGTDNERLF | 0.52386 |
| GLASGGLL | ASSDGTDNERLF | 0.6724057 |
| GRIAGAAVLL | ASSDGTDNERLF | 0.6758829 |
| GSAGSLNTSF | ASSDGTDNERLF | 0.6693611 |
| GTGQIGI | ASSDGTDNERLF | 0.48482674 |
| GTLIALIL | ASSDGTDNERLF | 0.38485909 |
| IAATMGFVVG | ASSDGTDNERLF | 0.6930561 |
| IFAQALL | ASSDGTDNERLF | 0.58496106 |
| IGAAVALGF | ASSDGTDNERLF | 0.64093864 |
| IGLTPLAL | ASSDGTDNERLF | 0.6308292 |
| ILGSNTSTL | ASSDGTDNERLF | 0.5957608 |
| IVFSLTVL | ASSDGTDNERLF | 0.46042418 |
| LASKATGF | ASSDGTDNERLF | 0.64473456 |
| LASVALIL | ASSDGTDNERLF | 0.6468885 |
| LFAVGLTL | ASSDGTDNERLF | 0.44653884 |
| LFNGGIGTYI | ASSDGTDNERLF | 0.3961453 |
| LTLGAGII | ASSDGTDNERLF | 0.5721985 |
| LVAAALGDI | ASSDGTDNERLF | 0.5886966 |
| NTAFVQLGI | ASSDGTDNERLF | 0.65083766 |
| QLFSGAEL | ASSDGTDNERLF | 0.57018787 |
| SAATFITL | ASSDGTDNERLF | 0.46550354 |
| SNYGLANLGV | ASSDGTDNERLF | 0.63695836 |
| SYGGAYAVM | ASSDGTDNERLF | 0.591451 |
| TTGVGGLTLG | ASSDGTDNERLF | 0.5884268 |
| TTQTIYI | ASSDGTDNERLF | 0.5442531 |
| VAAGAGLGAVI | ASSDGTDNERLF | 0.52930206 |
| VAAVVTLL | ASSDGTDNERLF | 0.63362026 |
| VGFLGGLI | ASSDGTDNERLF | 0.62947667 |
| VGVIPILL | ASSDGTDNERLF | 0.55647147 |
| VIFASSLI | ASSDGTDNERLF | 0.63892937 |
| VKGGDGGAAAG | ASSDGTDNERLF | 0.58977795 |
| VTKTFTNI | ASSDGTDNERLF | 0.52933884 |
| VVAAAILL | ASSDGTDNERLF | 0.37657076 |
| YAALDVEL | ASSDGTDNERLF | 0.50194764 |
| LASVALLL | ASSDGTDNERLF | 0.59468997 |
| LDAVHRV | ASSDGTDNERLF | 0.6121037 |
| QGLSSSIF | ASSDGTDNERLF | 0.65906686 |
| RGTFANIRL | ASSDGTDNERLF | 0.5292598 |
| RQLWWGHRI | ASSDGTDNERLF | 0.492857 |
| YGPPGTGKTLL | ASSDGTDNERLF | 0.63086426 |
| AALVGAPANVA | ASSDGTDNERLF | 0.6533637 |
| AATFRRTE | ASSDGTDNERLF | 0.3839953 |
| AGALGGGI | ASSDGTDNERLF | 0.5150934 |
| AGGAGGAGGIGAG | ASSDGTDNERLF | 0.7000275 |
| AGSGALGL | ASSDGTDNERLF | 0.6549875 |
| AQIADQI | ASSDGTDNERLF | 0.52942574 |

| | | |
|---|---|---|
| ASAVAVTYL | ASSDGTDNERLF | 0.4619316 |
| ASTVLGGTGG | ASSDGTDNERLF | 0.70548636 |
| AVGGEPAK | ASSDGTDNERLF | 0.57599163 |
| CEQGTFGGTC | ASSDGTDNERLF | 0.68344754 |
| DEGKAPAG | ASSDGTDNERLF | 0.3211347 |
| DTVVFPAL | ASSDGTDNERLF | 0.45064917 |
| FEGITMAG | ASSDGTDNERLF | 0.41374615 |
| FPLVVQD | ASSDGTDNERLF | 0.28818306 |
| GAFTTPQ | ASSDGTDNERLF | 0.6443298 |
| GFGGFGTAGGVGGA | ASSDGTDNERLF | 0.42742413 |
| GFLAQFL | ASSDGTDNERLF | 0.60453117 |
| GGAGGAGGNAGVA | ASSDGTDNERLF | 0.525017 |
| GGVGTAGAGGAGGAGG | ASSDGTDNERLF | 0.65826786 |
| GNGGGGDAGNAGSGGN | ASSDGTDNERLF | 0.5969093 |
| GTGGAGGAGNDAGS | ASSDGTDNERLF | 0.54101753 |
| GTTNTGWFNS | ASSDGTDNERLF | 0.41497445 |
| IFMGPLFTL | ASSDGTDNERLF | 0.60817397 |
| ILAHAIEAI | ASSDGTDNERLF | 0.65494573 |
| IRSAIREAL | ASSDGTDNERLF | 0.5751432 |
| ITFAGGG | ASSDGTDNERLF | 0.35330507 |
| ITGMSGHQEV | ASSDGTDNERLF | 0.6396828 |
| IVAGLACTL | ASSDGTDNERLF | 0.5194874 |
| IVGGGLAGA | ASSDGTDNERLF | 0.36241305 |
| IVTGVLNI | ASSDGTDNERLF | 0.3248932 |
| LAQQGCI | ASSDGTDNERLF | 0.5129483 |
| LCLHVEN | ASSDGTDNERLF | 0.54168373 |
| LDAISDGS | ASSDGTDNERLF | 0.63865644 |
| LGGFAAML | ASSDGTDNERLF | 0.6511908 |
| LGGKNPMI | ASSDGTDNERLF | 0.5333752 |
| LGPKGVAAIL | ASSDGTDNERLF | 0.6172034 |
| LMVGVSI | ASSDGTDNERLF | 0.6078205 |
| LVGAAFVL | ASSDGTDNERLF | 0.25952068 |
| LVLAVAPL | ASSDGTDNERLF | 0.41077626 |
| MAAIAVIDAA | ASSDGTDNERLF | 0.3730766 |
| NNGINAELQ | ASSDGTDNERLF | 0.65085053 |
| RGEPLVLVHG | ASSDGTDNERLF | 0.46694672 |
| RGGAGGAAGLI | ASSDGTDNERLF | 0.5299915 |
| RSYRMHPAV | ASSDGTDNERLF | 0.5830095 |
| SAAHLLAGQLA | ASSDGTDNERLF | 0.666785 |
| SAAVGSY | ASSDGTDNERLF | 0.56492424 |
| SPHAVLHRI | ASSDGTDNERLF | 0.4039744 |
| SQIAKVL | ASSDGTDNERLF | 0.37876657 |
| TALGHLVP | ASSDGTDNERLF | 0.60808396 |
| TAPTTSGAA | ASSDGTDNERLF | 0.50961995 |
| TDGDDGNGGSPGT | ASSDGTDNERLF | 0.44613367 |
| TDGLGEN | ASSDGTDNERLF | 0.3846374 |
| TGGPPLPGQA | ASSDGTDNERLF | 0.62762576 |
| TIPAITI | ASSDGTDNERLF | 0.31195217 |
| TIPGFTI | ASSDGTDNERLF | 0.6039034 |
| TSIASGALA | ASSDGTDNERLF | 0.39771828 |
| TVTLPGAAI | ASSDGTDNERLF | 0.56621796 |

| | | |
|---|---|---|
| VASGLSAALP | ASSDGTDNERLF | 0.6946837 |
| VGAGVIGI | ASSDGTDNERLF | 0.66919315 |
| VGPKRFGV | ASSDGTDNERLF | 0.5370686 |
| VNVAFPD | ASSDGTDNERLF | 0.48256204 |
| VSMTNSGVS | ASSDGTDNERLF | 0.43437865 |
| VTDVLGDLSV | ASSDGTDNERLF | 0.5057212 |
| VVVVGAGFAGL | ASSDGTDNERLF | 0.46890855 |
| VYTVGTVV | ASSDGTDNERLF | 0.6648226 |
| WTTFSGP | ASSDGTDNERLF | 0.5017174 |
| YDELVEAF | ASSDGTDNERLF | 0.3671412 |
| YGTGGAGGNGG | ASSDGTDNERLF | 0.4899877 |
| YTNVMAV | ASSDGTDNERLF | 0.68404907 |
| FLEQVHQL | ASSDGTDNERLF | 0.6944971 |
| GLSLLAVAV | ASSDGTDNERLF | 0.46826637 |
| LEQVHQL | ASSDGTDNERLF | 0.39122665 |
| LNAAGAI | ASSDGTDNERLF | 0.4133555 |
| PAVHASL | ASSDGTDNERLF | 0.57893705 |
| TELEDDA | ASSDGTDNERLF | 0.3770124 |
| TPPGPPN | ASSDGTDNERLF | 0.43274227 |
| TPPPPPPP | ASSDGTDNERLF | 0.60308295 |
| TQSPALI | GARRTGGSERLF | 0.7090559 |
| EFAAAFPQ | GARRTGGSERLF | 0.5288879 |
| LLSGNLDVL | GARRTGGSERLF | 0.30497813 |
| GPANLGNYNI | GARRTGGSERLF | 0.5759593 |
| TLGPTLP | GARRTGGSERLF | 0.431275 |
| VYAYSSCCK | GARRTGGSERLF | 0.51614404 |
| APAPAPAPNI | GARRTGGSERLF | 0.46334594 |
| PGGPVPIP | GARRTGGSERLF | 0.34409702 |
| FLTVALEP | GARRTGGSERLF | 0.57772785 |
| SQYFGANL | GARRTGGSERLF | 0.6858139 |
| GGVKPAA | GARRTGGSERLF | 0.6190239 |
| GGTGGAGGAGGW | GARRTGGSERLF | 0.5391108 |
| GVGGVAGLGGAGG | GARRTGGSERLF | 0.6909741 |
| VLLVLATTI | GARRTGGSERLF | 0.64967775 |
| IGIFAAML | GARRTGGSERLF | 0.68294954 |
| AVGDLID | GARRTGGSERLF | 0.6824022 |
| NISPLPL | GARRTGGSERLF | 0.52248716 |
| VTAVFVAI | GARRTGGSERLF | 0.5177926 |
| SGDDNDAAL | GARRTGGSERLF | 0.68462706 |
| AVAALAHS | GARRTGGSERLF | 0.5447078 |
| GGVTVMATP | GARRTGGSERLF | 0.603896 |
| AGLAANVRD | GARRTGGSERLF | 0.43409508 |
| TNRPDLIDPALLRPG | GARRTGGSERLF | 0.66355234 |
| LANLGVTAL | GARRTGGSERLF | 0.44299224 |
| TGERIEL | GARRTGGSERLF | 0.69918895 |
| NGEELAA | GARRTGGSERLF | 0.29968762 |
| VLSLVWG | GARRTGGSERLF | 0.45383358 |
| IGYALWL | GARRTGGSERLF | 0.67386043 |
| LSLLLAPL | GARRTGGSERLF | 0.2664892 |
| TLDEALK | GARRTGGSERLF | 0.3087344 |
| TDGAGTAEGG | GARRTGGSERLF | 0.5220246 |

| | | |
|---|---|---|
| NAPGGAGGNGG | GARRTGGSERLF | 0.66261244 |
| VSVFMGMGGNF | GARRTGGSERLF | 0.2722073 |
| LASYLAAL | GARRTGGSERLF | 0.52160895 |
| LVGGTPAL | GARRTGGSERLF | 0.333644 |
| VRNADVL | GARRTGGSERLF | 0.53378457 |
| TCSSPFLVSAM | GARRTGGSERLF | 0.5579275 |
| VHITDAE | GARRTGGSERLF | 0.36076778 |
| EYAAPSF | GARRTGGSERLF | 0.6287899 |
| VVDAFHGGL | GARRTGGSERLF | 0.32064354 |
| ASVVDPQ | GARRTGGSERLF | 0.60763717 |
| SYNEFGGLS | GARRTGGSERLF | 0.62927216 |
| DPDGSVY | GARRTGGSERLF | 0.70092857 |
| MVSTIHATE | GARRTGGSERLF | 0.38498217 |
| LAAGVADPV | GARRTGGSERLF | 0.59863394 |
| TKEQIAA | GARRTGGSERLF | 0.44428682 |
| SLASPAAL | GARRTGGSERLF | 0.70893586 |
| IGLVFYGV | GARRTGGSERLF | 0.525811 |
| FLLDASNL | GARRTGGSERLF | 0.33728045 |
| YQVGNFP | GARRTGGSERLF | 0.6655 |
| ESLLSQST | GARRTGGSERLF | 0.5774467 |
| LFTVGFAAVG | GARRTGGSERLF | 0.47657508 |
| ATPPTPT | GARRTGGSERLF | 0.6830431 |
| TDLATLG | GARRTGGSERLF | 0.4228359 |
| SAGGTGR | GARRTGGSERLF | 0.5775757 |
| GFSNSGS | GARRTGGSERLF | 0.66308594 |
| GGNGGSGGTSV | GARRTGGSERLF | 0.6495856 |
| GTSVATGGAGN | GARRTGGSERLF | 0.45077276 |
| NGGNGGSGGTS | GARRTGGSERLF | 0.63036394 |
| AVVKFPH | GARRTGGSERLF | 0.5999807 |
| TPPPPPP | GARRTGGSERLF | 0.3966239 |
| AAPAAAAAPA | GARRTGGSERLF | 0.68755287 |
| IALNEDL | GARRTGGSERLF | 0.47997642 |
| IVNVSSM | GARRTGGSERLF | 0.64554363 |
| TGIPPAPR | GARRTGGSERLF | 0.6127149 |
| FDLGGGTF | GARRTGGSERLF | 0.69877875 |
| FDLGGGTFDVS | GARRTGGSERLF | 0.45809588 |
| NEPTAAA | GARRTGGSERLF | 0.5616918 |
| NEPTAAALA | GARRTGGSERLF | 0.5759208 |
| GSIINISSI | GARRTGGSERLF | 0.61893 |
| VAPGNAGTA | GARRTGGSERLF | 0.4662608 |
| TGGGASLE | GARRTGGSERLF | 0.610669 |
| VAVYNHYKRI | GARRTGGSERLF | 0.61841667 |
| TDPAPATT | GARRTGGSERLF | 0.6696414 |
| TNRPDILDPALLRPG | GARRTGGSERLF | 0.5220885 |
| LTTTTTS | GARRTGGSERLF | 0.544324 |
| PTGSGKTL | GARRTGGSERLF | 0.69652075 |
| TGSGKTL | GARRTGGSERLF | 0.43420696 |
| TIGHVDHGK | GARRTGGSERLF | 0.5532601 |
| TIGHVDHGKTT | GARRTGGSERLF | 0.6164171 |
| GHVDHGKTT | GARRTGGSERLF | 0.6510608 |
| AQFEHTLL | GARRTGGSERLF | 0.49751142 |

| | | |
|---|---|---|
| IVNQDEL | GARRTGGSERLF | 0.6647489 |
| TNGSQFF | GARRTGGSERLF | 0.4901814 |
| TGKGIVE | GARRTGGSERLF | 0.6446799 |
| LISNASDA | GARRTGGSERLF | 0.5416832 |
| AELRTLGELTKIAKAHGAQ | GARRTGGSERLF | 0.48499516 |
| APNAVAAAEPSW | GARRTGGSERLF | 0.5094929 |
| APQVAGAATAVL | GARRTGGSERLF | 0.6213483 |
| ARRDPAAAAAILPTDA | GARRTGGSERLF | 0.6670297 |
| DAAARDVLAEAGLAEYF | GARRTGGSERLF | 0.64401186 |
| DDQIVLSAKANLAKVEQ | GARRTGGSERLF | 0.3191282 |
| DELLPDTGEDSDEDGH | GARRTGGSERLF | 0.6550905 |
| ELPEAPANRATVPQD | GARRTGGSERLF | 0.6308988 |
| FGQSKIERVTGIG | GARRTGGSERLF | 0.7061508 |
| GAGGTGSTAGGAGGAGGAGGLY | GARRTGGSERLF | 0.5093056 |
| GHGGTGAAGGNGGTGGAGGSGI | GARRTGGSERLF | 0.54228413 |
| GNGGNGGAGGEGGPGGAGGAGA | GARRTGGSERLF | 0.54806757 |
| GSTAGNGGNGGNATNGTIAGQP | GARRTGGSERLF | 0.53895974 |
| GTGGDGGNGGNGGTGG | GARRTGGSERLF | 0.35381958 |
| GVRRPAGIPADAPV | GARRTGGSERLF | 0.39617053 |
| GYRQVTDAHLLTLARR | GARRTGGSERLF | 0.5703447 |
| IAKMGQDGHDRG | GARRTGGSERLF | 0.52227056 |
| KAEPSIEIQYT | GARRTGGSERLF | 0.6861012 |
| KPVRIRMAADEDTVAVT | GARRTGGSERLF | 0.67056966 |
| MPPRIAGMRLLVIKPEP | GARRTGGSERLF | 0.65324974 |
| MKFYAVALFRSRGVPPTR | GARRTGGSERLF | 0.67031777 |
| NAGGFGILGAVAHSP | GARRTGGSERLF | 0.6795756 |
| NEGREITLAMAEM | GARRTGGSERLF | 0.616604 |
| NELVEWSHPLGHRWRWE | GARRTGGSERLF | 0.5059862 |
| NLGAGGTGGNGGNPG | GARRTGGSERLF | 0.5610356 |
| NTEWPKWETVMAADPVD | GARRTGGSERLF | 0.38121474 |
| PASAAPAETQP | GARRTGGSERLF | 0.5135956 |
| QQSSDGKNGVRAAEITGE | GARRTGGSERLF | 0.52008057 |
| RAPDAVTTSWWKE | GARRTGGSERLF | 0.5877906 |
| SGFGNVGAHTSGWFNQ | GARRTGGSERLF | 0.26771522 |
| SGVAGAVGQAASVGGLKV | GARRTGGSERLF | 0.6669798 |
| SNGGAGTGSAGGNGGT | GARRTGGSERLF | 0.67825407 |
| TGGAGGEGGAAG | GARRTGGSERLF | 0.60676265 |
| TGGDGGAAGAGGAPGHG | GARRTGGSERLF | 0.6208786 |
| TMPPLLAAIANAGRHGV | GARRTGGSERLF | 0.64799744 |
| TSSTAGNSGGA | GARRTGGSERLF | 0.43838033 |
| VPATVPVGILC | GARRTGGSERLF | 0.6109638 |
| VQRPDSGGVVLTGRIS | GARRTGGSERLF | 0.44508523 |
| WTPNPVPATKGVPAGLVQAT | GARRTGGSERLF | 0.63143855 |
| YALKRALAKFSDRFDVVII | GARRTGGSERLF | 0.6619513 |
| YLIGSVIV | GARRTGGSERLF | 0.6183524 |
| YGQLVAVID | GARRTGGSERLF | 0.49217376 |
| VTVGGMTLPA | GARRTGGSERLF | 0.5103845 |
| VLALPAV | GARRTGGSERLF | 0.39998525 |
| VIPPTLNYET | GARRTGGSERLF | 0.542171 |
| VGMGGMHPSSG | GARRTGGSERLF | 0.52645105 |
| VGLAGGAATA | GARRTGGSERLF | 0.58923614 |

| | | |
|---|---|---|
| VGGAGGAGGSGP | GARRTGGSERLF | 0.4472549 |
| VFAAFMV | GARRTGGSERLF | 0.62563246 |
| TLYDKLR | GARRTGGSERLF | 0.61282694 |
| TLAVFTVL | GARRTGGSERLF | 0.54676485 |
| TGSVQPARV | GARRTGGSERLF | 0.4385941 |
| TGGGAGSGGT | GARRTGGSERLF | 0.662536 |
| SHLETSL | GARRTGGSERLF | 0.65738034 |
| SFTTGMT | GARRTGGSERLF | 0.38324708 |
| QVEVAKRMGVRQPTVS | GARRTGGSERLF | 0.6283747 |
| QGSGGPALAM | GARRTGGSERLF | 0.463246 |
| MLAPSNP | GARRTGGSERLF | 0.5490492 |
| LVNNAAI | GARRTGGSERLF | 0.663452 |
| LTGEINTLA | GARRTGGSERLF | 0.5701418 |
| LSIFGPV | GARRTGGSERLF | 0.31998476 |
| LLALLANL | GARRTGGSERLF | 0.5725262 |
| LISNASDALDKLR | GARRTGGSERLF | 0.5826003 |
| LINEAVP | GARRTGGSERLF | 0.5726029 |
| LGYAFSGL | GARRTGGSERLF | 0.28346744 |
| LDFVAVAL | GARRTGGSERLF | 0.6229776 |
| LAQPGAE | GARRTGGSERLF | 0.39026183 |
| ISNLGAVTGF | GARRTGGSERLF | 0.46591228 |
| HQEDPGRI | GARRTGGSERLF | 0.51545876 |
| GTPGEPTP | GARRTGGSERLF | 0.6697918 |
| GFFNSGS | GARRTGGSERLF | 0.60273504 |
| GAGGGGLTAGAN | GARRTGGSERLF | 0.4806779 |
| FVDALVNM | GARRTGGSERLF | 0.6264392 |
| FLTVAPEL | GARRTGGSERLF | 0.5705331 |
| FETLTAL | GARRTGGSERLF | 0.67109 |
| DVINPIEL | GARRTGGSERLF | 0.61465925 |
| APVSGAPV | GARRTGGSERLF | 0.47845024 |
| ALVGGLAW | GARRTGGSERLF | 0.5327223 |
| ALAAAAATAI | GARRTGGSERLF | 0.39263192 |
| AFGLGLLL | GARRTGGSERLF | 0.5152375 |
| AFFAAASL | GARRTGGSERLF | 0.45500207 |
| AAAALPAFSPP | GARRTGGSERLF | 0.39934808 |
| VVLAPAL | GARRTGGSERLF | 0.46392465 |
| TIGHVDHGKTTL | GARRTGGSERLF | 0.6055001 |
| LISNASDALDK | GARRTGGSERLF | 0.6281557 |
| LISNASDAL | GARRTGGSERLF | 0.51731586 |
| ISNASDALDK | GARRTGGSERLF | 0.33242992 |
| NQAETLVYQTE | GARRTGGSERLF | 0.45750695 |
| TGGTGGTAGAG | GARRTGGSERLF | 0.5265076 |
| TGSTAGGAGGA | GARRTGGSERLF | 0.6588553 |
| DPLAVPVAAGPV | GARRTGGSERLF | 0.63732266 |
| GGTGGTPGNGGN | GARRTGGSERLF | 0.70823646 |
| PAGLAGVDPATL | GARRTGGSERLF | 0.6609015 |
| GNNLVVEAPHAGASV | GARRTGGSERLF | 0.5356029 |
| KVTRIMSASTLEQPA | GARRTGGSERLF | 0.6573383 |
| LGFALPIALPRRTVV | GARRTGGSERLF | 0.4563419 |
| TRLGNLTARKLVVLV | GARRTGGSERLF | 0.5221613 |
| WSQQRRLIYFPSAGP | GARRTGGSERLF | 0.5225976 |

| | | |
|---|---|---|
| GPNGNPDPMAAAVDIR | GARRTGGSERLF | 0.5671327 |
| TQRCVSVADLRAAASR | GARRTGGSERLF | 0.50076956 |
| QFVLDVAEHPVEVLTETV | GARRTGGSERLF | 0.46910536 |
| GGAGGTGGAGGNGGTQAGDG | GARRTGGSERLF | 0.31230277 |
| PLLATLRAIAVDVDAKPAQV | GARRTGGSERLF | 0.65032625 |
| APKFRTGGYMIDFWGVGYQVAK | GARRTGGSERLF | 0.42006186 |
| GAGGAGGAGGAAGGPGGTGGTG | GARRTGGSERLF | 0.36190397 |
| IGALGLLATDVAPAGSVAFGVA | GARRTGGSERLF | 0.46558833 |
| AGGAGGIGGNGGAGGAGGVLMG | GARRTGGSERLF | 0.6896709 |
| ADELIGLL | GARRTGGSERLF | 0.66300166 |
| ADRALKA | GARRTGGSERLF | 0.51082563 |
| AGAVGQVGGAGGA | GARRTGGSERLF | 0.44087225 |
| AGGGGGLGGAP | GARRTGGSERLF | 0.6652819 |
| AGGLLYGNGGAG | GARRTGGSERLF | 0.48573413 |
| AGLESIGL | GARRTGGSERLF | 0.5862611 |
| AGLKTGII | GARRTGGSERLF | 0.59979546 |
| ASLAALLL | GARRTGGSERLF | 0.6065247 |
| ASVINISSI | GARRTGGSERLF | 0.36839375 |
| ATPPGPA | GARRTGGSERLF | 0.62238395 |
| ATVASALV | GARRTGGSERLF | 0.56866115 |
| ATVASLIL | GARRTGGSERLF | 0.5411803 |
| AVNGPVLIRH | GARRTGGSERLF | 0.28771806 |
| DGAAGAVGI | GARRTGGSERLF | 0.6163566 |
| DPLLHGI | GARRTGGSERLF | 0.45269868 |
| ELGAALGAL | GARRTGGSERLF | 0.3895886 |
| ELGRTQAV | GARRTGGSERLF | 0.29500055 |
| FVAGGYLS | GARRTGGSERLF | 0.5063026 |
| GASGGAGGAGGW | GARRTGGSERLF | 0.6521487 |
| GGAGAGGDGGAGGI | GARRTGGSERLF | 0.6716738 |
| GTGHNGSQPGMGGNG | GARRTGGSERLF | 0.60474443 |
| GYAAASASAS | GARRTGGSERLF | 0.647973 |
| HQVDVAAVQQG | GARRTGGSERLF | 0.64876235 |
| IGAAIAHT | GARRTGGSERLF | 0.5154481 |
| IGMIEVAQL | GARRTGGSERLF | 0.33413255 |
| IRLAPEHTSAMD | GARRTGGSERLF | 0.55427974 |
| IYNVSVL | GARRTGGSERLF | 0.63606584 |
| KQGFLGM | GARRTGGSERLF | 0.63024837 |
| LESVITQL | GARRTGGSERLF | 0.6518696 |
| LGLGALVVV | GARRTGGSERLF | 0.42671552 |
| LIGNGGNGGSGGI | GARRTGGSERLF | 0.5589421 |
| LNAINAPTL | GARRTGGSERLF | 0.583264 |
| LQLLNTGV | GARRTGGSERLF | 0.36103526 |
| LQTSIEF | GARRTGGSERLF | 0.37773842 |
| MSFAVSEL | GARRTGGSERLF | 0.4341156 |
| NTGALGL | GARRTGGSERLF | 0.40897352 |
| PPGAGGAGGAGTT | GARRTGGSERLF | 0.59240925 |
| QLVGLGI | GARRTGGSERLF | 0.54301775 |
| RFLEQVHQL | GARRTGGSERLF | 0.5457008 |
| RGVGYVLR | GARRTGGSERLF | 0.54583544 |
| SATGVFAGVI | GARRTGGSERLF | 0.4687285 |
| SLPAITLP | GARRTGGSERLF | 0.49049518 |

| | | |
|---|---|---|
| STALAFTV | GARRTGGSERLF | 0.57387805 |
| SVLTVGLGK | GARRTGGSERLF | 0.6153308 |
| TALAGAGSGI | GARRTGGSERLF | 0.6481325 |
| TGLPFTGI | GARRTGGSERLF | 0.5607813 |
| VAEHVTV | GARRTGGSERLF | 0.48441714 |
| VDIVLNSL | GARRTGGSERLF | 0.68921614 |
| VLTGGGAVL | GARRTGGSERLF | 0.4082337 |
| VNVGLAYGI | GARRTGGSERLF | 0.45457354 |
| VSSAAVGAV | GARRTGGSERLF | 0.6358548 |
| VTSVYEAGV | GARRTGGSERLF | 0.53966236 |
| WVALDRGA | GARRTGGSERLF | 0.657022 |
| YLVGSVVI | GARRTGGSERLF | 0.62090635 |
| GPPGTGKTLL | GARRTGGSERLF | 0.6267884 |
| GVGMTKF | GARRTGGSERLF | 0.6101377 |
| IGIGGSDL | GARRTGGSERLF | 0.69432104 |
| IGTLLAEL | GARRTGGSERLF | 0.63207215 |
| IYTSGTT | GARRTGGSERLF | 0.47342223 |
| LSFPAGV | GARRTGGSERLF | 0.6889795 |
| LYTSGTT | GARRTGGSERLF | 0.34573576 |
| VGAFPFGF | GARRTGGSERLF | 0.6514025 |
| VGAGTVEF | GARRTGGSERLF | 0.6079561 |
| VGKSTLLL | GARRTGGSERLF | 0.46109164 |
| AALAHAGGGIV | GARRTGGSERLF | 0.57771206 |
| AALVAVSLL | GARRTGGSERLF | 0.6692127 |
| AAVSRAVAAL | GARRTGGSERLF | 0.67950743 |
| AGGSSGIGGF | GARRTGGSERLF | 0.5091497 |
| AGLTVANANG | GARRTGGSERLF | 0.511263 |
| AGLVGAVL | GARRTGGSERLF | 0.41375005 |
| ALSQAFTV | GARRTGGSERLF | 0.3501016 |
| ALTFSVSL | GARRTGGSERLF | 0.338139 |
| ALTGAALVAL | GARRTGGSERLF | 0.6242976 |
| ASFSIGAGVL | GARRTGGSERLF | 0.5208626 |
| DGSVYKI | GARRTGGSERLF | 0.6823951 |
| EQLAEIV | GARRTGGSERLF | 0.6216351 |
| FAVSELTVM | GARRTGGSERLF | 0.6600957 |
| GAAPLAAGL | GARRTGGSERLF | 0.4537052 |
| GADGGDGGSGGAGG | GARRTGGSERLF | 0.5756308 |
| GFGLDTGGAGG | GARRTGGSERLF | 0.3174407 |
| GLASGGLL | GARRTGGSERLF | 0.48994723 |
| GRIAGAAVLL | GARRTGGSERLF | 0.38801336 |
| GSAGSLNTSF | GARRTGGSERLF | 0.57624066 |
| GTGQIGI | GARRTGGSERLF | 0.32480675 |
| GTLIALIL | GARRTGGSERLF | 0.6814108 |
| IAATMGFVVG | GARRTGGSERLF | 0.4885765 |
| IFAQALL | GARRTGGSERLF | 0.6333668 |
| IGAAVALGF | GARRTGGSERLF | 0.44552186 |
| IGLTPLAL | GARRTGGSERLF | 0.3897618 |
| ILGSNTSTL | GARRTGGSERLF | 0.5334931 |
| IVFSLTVL | GARRTGGSERLF | 0.64241856 |
| LASKATGF | GARRTGGSERLF | 0.45670438 |
| LASVALIL | GARRTGGSERLF | 0.28691122 |

| | | |
|---|---|---|
| LFAVGLTL | GARRTGGSERLF | 0.37617952 |
| LFNGGIGTYI | GARRTGGSERLF | 0.3568506 |
| LTLGAGII | GARRTGGSERLF | 0.6966269 |
| LVAAALGDI | GARRTGGSERLF | 0.66639423 |
| NTAFVQLGI | GARRTGGSERLF | 0.58888686 |
| QLFSGAEL | GARRTGGSERLF | 0.69899106 |
| SAATFITL | GARRTGGSERLF | 0.27385882 |
| SNYGLANLGV | GARRTGGSERLF | 0.4884066 |
| SYGGAYAVM | GARRTGGSERLF | 0.6698648 |
| TTGVGGLTLG | GARRTGGSERLF | 0.6240107 |
| TTQTIYI | GARRTGGSERLF | 0.55911857 |
| VAAGAGLGAVI | GARRTGGSERLF | 0.65353173 |
| VAAVVTLL | GARRTGGSERLF | 0.4934523 |
| VGFLGGLI | GARRTGGSERLF | 0.5549938 |
| VGVIPILL | GARRTGGSERLF | 0.59875256 |
| VIFASSLI | GARRTGGSERLF | 0.5810908 |
| VKGGDGGAAAG | GARRTGGSERLF | 0.5148788 |
| VTKTFTNI | GARRTGGSERLF | 0.35422194 |
| VVAAAILL | GARRTGGSERLF | 0.52946866 |
| YAALDVEL | GARRTGGSERLF | 0.7089282 |
| LASVALLL | GARRTGGSERLF | 0.56257296 |
| LDAVHRV | GARRTGGSERLF | 0.6000405 |
| QGLSSSIF | GARRTGGSERLF | 0.55594534 |
| RGTFANIRL | GARRTGGSERLF | 0.62097776 |
| RQLWWGHRI | GARRTGGSERLF | 0.4912961 |
| YGPPGTGKTLL | GARRTGGSERLF | 0.42108163 |
| AALVGAPANVA | GARRTGGSERLF | 0.66234696 |
| AATFRRTE | GARRTGGSERLF | 0.42081457 |
| AGALGGGI | GARRTGGSERLF | 0.49265626 |
| AGGAGGAGGIGAG | GARRTGGSERLF | 0.51659596 |
| AGSGALGL | GARRTGGSERLF | 0.71015835 |
| AQIADQI | GARRTGGSERLF | 0.60513985 |
| ASAVAVTYL | GARRTGGSERLF | 0.6177074 |
| ASTVLGGTGG | GARRTGGSERLF | 0.54792285 |
| AVGGEPAK | GARRTGGSERLF | 0.5491712 |
| CEQGTFGGTC | GARRTGGSERLF | 0.6445744 |
| DEGKAPAG | GARRTGGSERLF | 0.32929647 |
| DTVVFPAL | GARRTGGSERLF | 0.65349746 |
| FEGITMAG | GARRTGGSERLF | 0.39106697 |
| FPLVVQD | GARRTGGSERLF | 0.561684 |
| GAFTTPQ | GARRTGGSERLF | 0.5855151 |
| GFGGFGTAGGVGGA | GARRTGGSERLF | 0.6998757 |
| GFLAQFL | GARRTGGSERLF | 0.55777365 |
| GGAGGAGGNAGVA | GARRTGGSERLF | 0.6700053 |
| GGVGTAGAGGAGGAGG | GARRTGGSERLF | 0.50352526 |
| GNGGGDAGNAGSGGN | GARRTGGSERLF | 0.6193664 |
| GTGGAGGAGNDAGS | GARRTGGSERLF | 0.68451935 |
| GTTNTGWFNS | GARRTGGSERLF | 0.63919824 |
| IFMGPLFTL | GARRTGGSERLF | 0.70864654 |
| ILAHAIEAI | GARRTGGSERLF | 0.5213134 |
| IRSAIREAL | GARRTGGSERLF | 0.31120998 |

| | | |
|---|---|---|
| ITFAGGG | GARRTGGSERLF | 0.5093634 |
| ITGMSGHQEV | GARRTGGSERLF | 0.64967537 |
| IVAGLACTL | GARRTGGSERLF | 0.68682516 |
| IVGGGLAGA | GARRTGGSERLF | 0.6516864 |
| IVTGVLNI | GARRTGGSERLF | 0.61839086 |
| LAQQGCI | GARRTGGSERLF | 0.5073179 |
| LCLHVEN | GARRTGGSERLF | 0.6756685 |
| LDAISDGS | GARRTGGSERLF | 0.692476 |
| LGGFAAML | GARRTGGSERLF | 0.59277594 |
| LGGKNPMI | GARRTGGSERLF | 0.38481364 |
| LGPKGVAAIL | GARRTGGSERLF | 0.3670378 |
| LMVGVSI | GARRTGGSERLF | 0.6348958 |
| LVGAAFVL | GARRTGGSERLF | 0.66368544 |
| LVLAVAPL | GARRTGGSERLF | 0.33901563 |
| MAAIAVIDAA | GARRTGGSERLF | 0.5986765 |
| NNGINAELQ | GARRTGGSERLF | 0.61502033 |
| RGEPLVLVHG | GARRTGGSERLF | 0.30715168 |
| RGGAGGAAGLI | GARRTGGSERLF | 0.58033895 |
| RSYRMHPAV | GARRTGGSERLF | 0.6761488 |
| SAAHLLAGQLA | GARRTGGSERLF | 0.55537546 |
| SAAVGSY | GARRTGGSERLF | 0.377243 |
| SPHAVLHRI | GARRTGGSERLF | 0.67311484 |
| SQIAKVL | GARRTGGSERLF | 0.68873477 |
| TALGHLVP | GARRTGGSERLF | 0.68394256 |
| TAPTTSGAA | GARRTGGSERLF | 0.33761272 |
| TDGDDGNGGSPGT | GARRTGGSERLF | 0.27826288 |
| TDGLGEN | GARRTGGSERLF | 0.6240689 |
| TGGPPLPGQA | GARRTGGSERLF | 0.6780484 |
| TIPAITI | GARRTGGSERLF | 0.52227664 |
| TIPGFTI | GARRTGGSERLF | 0.46458066 |
| TSIASGALA | GARRTGGSERLF | 0.60319597 |
| TVTLPGAAI | GARRTGGSERLF | 0.6578975 |
| VASGLSAALP | GARRTGGSERLF | 0.3942184 |
| VGAGVIGI | GARRTGGSERLF | 0.36839265 |
| VGPKRFGV | GARRTGGSERLF | 0.59997165 |
| VNVAFPD | GARRTGGSERLF | 0.6270874 |
| VSMTNSGVS | GARRTGGSERLF | 0.44510773 |
| VTDVLGDLSV | GARRTGGSERLF | 0.5607299 |
| VVVVGAGFAGL | GARRTGGSERLF | 0.6464324 |
| VYTVGTVV | GARRTGGSERLF | 0.5818169 |
| WTTFSGP | GARRTGGSERLF | 0.60471505 |
| YDELVEAF | GARRTGGSERLF | 0.37606716 |
| YGTGGAGGNGG | GARRTGGSERLF | 0.6716664 |
| YTNVMAV | GARRTGGSERLF | 0.3840136 |
| FLEQVHQL | GARRTGGSERLF | 0.6103634 |
| GLSLLAVAV | GARRTGGSERLF | 0.5932081 |
| LEQVHQL | GARRTGGSERLF | 0.57907456 |
| LNAAGAI | GARRTGGSERLF | 0.55545056 |
| PAVHASL | GARRTGGSERLF | 0.38395 |
| TELEDDA | GARRTGGSERLF | 0.42739975 |
| TPPGPPN | GARRTGGSERLF | 0.42691684 |

| | | |
|---|---|---|
| TPPPPPPP | GARRTGGSERLF | 0.67283636 |
| TQSPALI | ASSLDIQDTQY | 0.6202998 |
| EFAAAFPQ | ASSLDIQDTQY | 0.6551049 |
| LLSGNLDVL | ASSLDIQDTQY | 0.5670155 |
| GPANLGNYNI | ASSLDIQDTQY | 0.48219508 |
| TLGPTLP | ASSLDIQDTQY | 0.6085266 |
| VYAYSSCCK | ASSLDIQDTQY | 0.39506912 |
| APAPAPAPNI | ASSLDIQDTQY | 0.6291913 |
| PGGPVPIP | ASSLDIQDTQY | 0.5027085 |
| FLTVALEP | ASSLDIQDTQY | 0.60048926 |
| SQYFGANL | ASSLDIQDTQY | 0.43043602 |
| GGVKPAA | ASSLDIQDTQY | 0.5687382 |
| GGTGGAGGAGGW | ASSLDIQDTQY | 0.61043644 |
| GVGGVAGLGGAGG | ASSLDIQDTQY | 0.35348606 |
| VLLVLATTI | ASSLDIQDTQY | 0.55602944 |
| IGIFAAML | ASSLDIQDTQY | 0.6627271 |
| AVGDLID | ASSLDIQDTQY | 0.5482617 |
| NISPLPL | ASSLDIQDTQY | 0.37698203 |
| VTAVFVAI | ASSLDIQDTQY | 0.58799493 |
| SGDDNDAAL | ASSLDIQDTQY | 0.4797448 |
| AVAALAHS | ASSLDIQDTQY | 0.6381083 |
| GGVTVMATP | ASSLDIQDTQY | 0.5907607 |
| AGLAANVRD | ASSLDIQDTQY | 0.41031462 |
| TNRPDLIDPALLRPG | ASSLDIQDTQY | 0.59636796 |
| LANLGVTAL | ASSLDIQDTQY | 0.32769912 |
| TGERIEL | ASSLDIQDTQY | 0.4794313 |
| NGEELAA | ASSLDIQDTQY | 0.49200696 |
| VLSLVWG | ASSLDIQDTQY | 0.6255183 |
| IGYALWL | ASSLDIQDTQY | 0.62339824 |
| LSLLLAPL | ASSLDIQDTQY | 0.40619844 |
| TLDEALK | ASSLDIQDTQY | 0.5688213 |
| TDGAGTAEGG | ASSLDIQDTQY | 0.522714 |
| NAPGGAGGNGG | ASSLDIQDTQY | 0.42749226 |
| VSVFMGMGGNF | ASSLDIQDTQY | 0.49344343 |
| LASYLAAL | ASSLDIQDTQY | 0.43266356 |
| LVGGTPAL | ASSLDIQDTQY | 0.3356349 |
| VRNADVL | ASSLDIQDTQY | 0.5884922 |
| TCSSPFLVSAM | ASSLDIQDTQY | 0.43985164 |
| VHITDAE | ASSLDIQDTQY | 0.53802454 |
| EYAAPSF | ASSLDIQDTQY | 0.6289017 |
| VVDAFHGGL | ASSLDIQDTQY | 0.3448453 |
| ASVVDPQ | ASSLDIQDTQY | 0.5903683 |
| SYNEFGGLS | ASSLDIQDTQY | 0.5538816 |
| DPDGSVY | ASSLDIQDTQY | 0.6180519 |
| MVSTIHATE | ASSLDIQDTQY | 0.5941487 |
| LAAGVADPV | ASSLDIQDTQY | 0.4528796 |
| TKEQIAA | ASSLDIQDTQY | 0.66900027 |
| SLASPAAL | ASSLDIQDTQY | 0.61847043 |
| IGLVFYGV | ASSLDIQDTQY | 0.56986576 |
| FLLDASNL | ASSLDIQDTQY | 0.59372604 |
| YQVGNFP | ASSLDIQDTQY | 0.6724962 |

| | | |
|---|---|---|
| ESLLSQST | ASSLDIQDTQY | 0.649305 |
| LFTVGFAAVG | ASSLDIQDTQY | 0.51476 |
| ATPPTPT | ASSLDIQDTQY | 0.3094617 |
| TDLATLG | ASSLDIQDTQY | 0.5906458 |
| SAGGTGR | ASSLDIQDTQY | 0.60757124 |
| GFSNSGS | ASSLDIQDTQY | 0.41126502 |
| GGNGGSGGTSV | ASSLDIQDTQY | 0.31442708 |
| GTSVATGGAGN | ASSLDIQDTQY | 0.65906596 |
| NGGNGGSGGTS | ASSLDIQDTQY | 0.7026427 |
| AVVKFPH | ASSLDIQDTQY | 0.50135696 |
| TPPPPPP | ASSLDIQDTQY | 0.70393026 |
| AAPAAAAAPA | ASSLDIQDTQY | 0.33839378 |
| IALNEDL | ASSLDIQDTQY | 0.6402403 |
| IVNVSSM | ASSLDIQDTQY | 0.67202175 |
| TGIPPAPR | ASSLDIQDTQY | 0.39981997 |
| FDLGGGTF | ASSLDIQDTQY | 0.60626054 |
| FDLGGGTFDVS | ASSLDIQDTQY | 0.458655 |
| NEPTAAA | ASSLDIQDTQY | 0.465537 |
| NEPTAAALA | ASSLDIQDTQY | 0.6674794 |
| GSIINISSI | ASSLDIQDTQY | 0.517354 |
| VAPGNAGTA | ASSLDIQDTQY | 0.57965267 |
| TGGGASLE | ASSLDIQDTQY | 0.5358424 |
| VAVYNHYKRI | ASSLDIQDTQY | 0.44451156 |
| TDPAPATT | ASSLDIQDTQY | 0.57654977 |
| TNRPDILDPALLRPG | ASSLDIQDTQY | 0.61865 |
| LTTTTTS | ASSLDIQDTQY | 0.56872165 |
| PTGSGKTL | ASSLDIQDTQY | 0.6823585 |
| TGSGKTL | ASSLDIQDTQY | 0.42797622 |
| TIGHVDHGK | ASSLDIQDTQY | 0.7016485 |
| TIGHVDHGKTT | ASSLDIQDTQY | 0.68673754 |
| GHVDHGKTT | ASSLDIQDTQY | 0.6347938 |
| AQFEHTLL | ASSLDIQDTQY | 0.45688972 |
| IVNQDEL | ASSLDIQDTQY | 0.669904 |
| TNGSQFF | ASSLDIQDTQY | 0.47992435 |
| TGKGIVE | ASSLDIQDTQY | 0.5682393 |
| LISNASDA | ASSLDIQDTQY | 0.6355976 |
| AELRTLGELTKIAKAHGAQ | ASSLDIQDTQY | 0.5543048 |
| APNAVAAAEPSW | ASSLDIQDTQY | 0.88454527 |
| APQVAGAATAVL | ASSLDIQDTQY | 0.8462039 |
| ARRDPAAAAAILPTDA | ASSLDIQDTQY | 0.80388725 |
| DAAARDVLAEAGLAEYF | ASSLDIQDTQY | 0.63468754 |
| DDQIVLSAKANLAKVEQ | ASSLDIQDTQY | 0.32370847 |
| DELLPDTGEDSDEDGH | ASSLDIQDTQY | 0.79013014 |
| ELPEAPANRATVPQD | ASSLDIQDTQY | 0.54033196 |
| FGQSKIERVTGIG | ASSLDIQDTQY | 0.86849517 |
| GAGGTGSTAGGAGGAGGAGGLY | ASSLDIQDTQY | 0.8580922 |
| GHGGTGAAGGNGGTGGAGGSGI | ASSLDIQDTQY | 0.6357399 |
| GNGGNGGAGGEGGPGGAGGAGA | ASSLDIQDTQY | 0.8716167 |
| GSTAGNGGNGGNATNGTIAGQP | ASSLDIQDTQY | 0.87335503 |
| GTGGDGGNGGNGGTGG | ASSLDIQDTQY | 0.3970798 |
| GVRRPAGIPADAPV | ASSLDIQDTQY | 0.7858433 |

| | | |
|---|---|---|
| GYRQVTDAHLLTLARR | ASSLDIQDTQY | 0.7941972 |
| IAKMGQDGHDRG | ASSLDIQDTQY | 0.74999386 |
| KAEPSIEIQYT | ASSLDIQDTQY | 0.5074812 |
| KPVRIRMAADEDTVAVT | ASSLDIQDTQY | 0.8472934 |
| MPPRIAGMRLLVIKPEP | ASSLDIQDTQY | 0.7902205 |
| MKFYAVALFRSRGVPPTR | ASSLDIQDTQY | 0.15117455 |
| NAGGFGILGAVAHSP | ASSLDIQDTQY | 0.8076838 |
| NEGREITLAMAEM | ASSLDIQDTQY | 0.7771293 |
| NELVEWSHPLGHRWRWE | ASSLDIQDTQY | 0.6991837 |
| NLGAGGTGGNGGNPG | ASSLDIQDTQY | 0.87427473 |
| NTEWPKWETVMAADPVD | ASSLDIQDTQY | 0.69307065 |
| PASAAPAETQP | ASSLDIQDTQY | 0.61003196 |
| QQSSDGKNGVRAAEITGE | ASSLDIQDTQY | 0.86353445 |
| RAPDAVTTSWWKE | ASSLDIQDTQY | 0.87343985 |
| SGFGNVGAHTSGWFNQ | ASSLDIQDTQY | 0.39961368 |
| SGVAGAVGQAASVGGLKV | ASSLDIQDTQY | 0.85475445 |
| SNGGAGTGSAGGNGGT | ASSLDIQDTQY | 0.87886834 |
| TGGAGGEGGAAG | ASSLDIQDTQY | 0.7789217 |
| TGGDGGAAGAGGAPGHG | ASSLDIQDTQY | 0.88101876 |
| TMPPLLAAIANAGRHGV | ASSLDIQDTQY | 0.8836979 |
| TSSTAGNSGGA | ASSLDIQDTQY | 0.6907656 |
| VPATVPVGILC | ASSLDIQDTQY | 0.8729373 |
| VQRPDSGGVVLTGRIS | ASSLDIQDTQY | 0.8831214 |
| WTPNPVPATKGVPAGLVQAT | ASSLDIQDTQY | 0.6916066 |
| YALKRALAKFSDRFDVVII | ASSLDIQDTQY | 0.7919791 |
| YLIGSVIV | ASSLDIQDTQY | 0.5617837 |
| YGQLVAVID | ASSLDIQDTQY | 0.7880391 |
| VTVGGMTLPA | ASSLDIQDTQY | 0.3228625 |
| VLALPAV | ASSLDIQDTQY | 0.85629797 |
| VIPPTLNYET | ASSLDIQDTQY | 0.8696371 |
| VGMGGMHPSSG | ASSLDIQDTQY | 0.75969315 |
| VGLAGGAATA | ASSLDIQDTQY | 0.6073748 |
| VGGAGGAGGSGP | ASSLDIQDTQY | 0.6113657 |
| VFAAFMV | ASSLDIQDTQY | 0.828454 |
| TLYDKLR | ASSLDIQDTQY | 0.6432335 |
| TLAVFTVL | ASSLDIQDTQY | 0.37370694 |
| TGSVQPARV | ASSLDIQDTQY | 0.7926078 |
| TGGGAGSGGT | ASSLDIQDTQY | 0.8535423 |
| SHLETSL | ASSLDIQDTQY | 0.85651505 |
| SFTTGMT | ASSLDIQDTQY | 0.46230674 |
| QVEVAKRMGVRQPTVS | ASSLDIQDTQY | 0.88654923 |
| QGSGGPALAM | ASSLDIQDTQY | 0.876109 |
| MLAPSNP | ASSLDIQDTQY | 0.881482 |
| LVNNAAI | ASSLDIQDTQY | 0.8089437 |
| LTGEINTLA | ASSLDIQDTQY | 0.6324861 |
| LSIFGPV | ASSLDIQDTQY | 0.87897855 |
| LLALLANL | ASSLDIQDTQY | 0.74962395 |
| LISNASDALDKLR | ASSLDIQDTQY | 0.71115875 |
| LINEAVP | ASSLDIQDTQY | 0.8653167 |
| LGYAFSGL | ASSLDIQDTQY | 0.68527615 |
| LDFVAVAL | ASSLDIQDTQY | 0.6894097 |

| | | |
|---|---|---|
| LAQPGAE | ASSLDIQDTQY | 0.8174925 |
| ISNLGAVTGF | ASSLDIQDTQY | 0.8830857 |
| HQEDPGRI | ASSLDIQDTQY | 0.6313667 |
| GTPGEPTP | ASSLDIQDTQY | 0.41785556 |
| GFFNSGS | ASSLDIQDTQY | 0.6708251 |
| GAGGGGLTAGAN | ASSLDIQDTQY | 0.7818786 |
| FVDALVNM | ASSLDIQDTQY | 0.87279904 |
| FLTVAPEL | ASSLDIQDTQY | 0.8209861 |
| FETLTAL | ASSLDIQDTQY | 0.8131921 |
| DVINPIEL | ASSLDIQDTQY | 0.8532603 |
| APVSGAPV | ASSLDIQDTQY | 0.8413082 |
| ALVGGLAW | ASSLDIQDTQY | 0.80556077 |
| ALAAAAATAI | ASSLDIQDTQY | 0.8757149 |
| AFGLGLLL | ASSLDIQDTQY | 0.7385512 |
| AFFAAASL | ASSLDIQDTQY | 0.88316405 |
| AAAALPAFSPP | ASSLDIQDTQY | 0.85725653 |
| VVLAPAL | ASSLDIQDTQY | 0.8681803 |
| TIGHVDHGKTTL | ASSLDIQDTQY | 0.88095033 |
| LISNASDALDK | ASSLDIQDTQY | 0.8249445 |
| LISNASDAL | ASSLDIQDTQY | 0.8255764 |
| ISNASDALDK | ASSLDIQDTQY | 0.2896142 |
| NQAETLVYQTE | ASSLDIQDTQY | 0.75985587 |
| TGGTGGTAGAG | ASSLDIQDTQY | 0.8463266 |
| TGSTAGGAGGA | ASSLDIQDTQY | 0.85417354 |
| DPLAVPVAAGPV | ASSLDIQDTQY | 0.8649167 |
| GGTGGTPGNGGN | ASSLDIQDTQY | 0.84624237 |
| PAGLAGVDPATL | ASSLDIQDTQY | 0.6408758 |
| GNNLVVEAPHAGASV | ASSLDIQDTQY | 0.87818074 |
| KVTRIMSASTLEQPA | ASSLDIQDTQY | 0.87652206 |
| LGFALPIALPRRTVV | ASSLDIQDTQY | 0.46383882 |
| TRLGNLTARKLVVLV | ASSLDIQDTQY | 0.7633908 |
| WSQQRRLIYFPSAGP | ASSLDIQDTQY | 0.8897394 |
| GPNGNPDPMAAAVDIR | ASSLDIQDTQY | 0.85889345 |
| TQRCVSVADLRAAASR | ASSLDIQDTQY | 0.7349514 |
| QFVLDVAEHPVEVLTETV | ASSLDIQDTQY | 0.6959038 |
| GGAGGTGGAGGNGGTQAGDG | ASSLDIQDTQY | 0.88338864 |
| PLLATLRAIAVDVDAKPAQV | ASSLDIQDTQY | 0.76683384 |
| APKFRTGGYMIDFWGVGYQVAK | ASSLDIQDTQY | 0.8833271 |
| GAGGAGGAGGAAGGPGGTGGTG | ASSLDIQDTQY | 0.20516571 |
| IGALGLLATDVAPAGSVAFGVA | ASSLDIQDTQY | 0.7579385 |
| AGGAGGIGGNGGAGGAGGVLMG | ASSLDIQDTQY | 0.630567 |
| ADELIGLL | ASSLDIQDTQY | 0.21171129 |
| ADRALKA | ASSLDIQDTQY | 0.84334123 |
| AGAVGQVGGAGGA | ASSLDIQDTQY | 0.6299044 |
| AGGGGGLGGAP | ASSLDIQDTQY | 0.8526951 |
| AGGLLYGNGGAG | ASSLDIQDTQY | 0.81196344 |
| AGLESIGL | ASSLDIQDTQY | 0.8679448 |
| AGLKTGII | ASSLDIQDTQY | 0.85278726 |
| ASLAALLL | ASSLDIQDTQY | 0.79266864 |
| ASVINISSI | ASSLDIQDTQY | 0.5555508 |
| ATPPGPA | ASSLDIQDTQY | 0.8172228 |

| | | |
|---|---|---|
| ATVASALV | ASSLDIQDTQY | 0.8782194 |
| ATVASLIL | ASSLDIQDTQY | 0.8706837 |
| AVNGPVLIRH | ASSLDIQDTQY | 0.2781775 |
| DGAAGAVGI | ASSLDIQDTQY | 0.84331405 |
| DPLLHGI | ASSLDIQDTQY | 0.72438383 |
| ELGAALGAL | ASSLDIQDTQY | 0.2669832 |
| ELGRTQAV | ASSLDIQDTQY | 0.23030725 |
| FVAGGYLS | ASSLDIQDTQY | 0.77280927 |
| GASGGAGGAGGW | ASSLDIQDTQY | 0.8021841 |
| GGAGAGGDGGAGGI | ASSLDIQDTQY | 0.8649138 |
| GTGHNGSQPGMGGNG | ASSLDIQDTQY | 0.8707012 |
| GYAAASASAS | ASSLDIQDTQY | 0.74736404 |
| HQVDVAAVQQG | ASSLDIQDTQY | 0.8653754 |
| IGAAIAHT | ASSLDIQDTQY | 0.7816435 |
| IGMIEVAQL | ASSLDIQDTQY | 0.105090246 |
| IRLAPEHTSAMD | ASSLDIQDTQY | 0.76459205 |
| IYNVSVL | ASSLDIQDTQY | 0.41137502 |
| KQGFLGM | ASSLDIQDTQY | 0.8822653 |
| LESVITQL | ASSLDIQDTQY | 0.6259756 |
| LGLGALVVV | ASSLDIQDTQY | 0.74808156 |
| LIGNGGNGGSGGI | ASSLDIQDTQY | 0.79300153 |
| LNAINAPTL | ASSLDIQDTQY | 0.8528017 |
| LQLLNTGV | ASSLDIQDTQY | 0.8120191 |
| LQTSIEF | ASSLDIQDTQY | 0.71686053 |
| MSFAVSEL | ASSLDIQDTQY | 0.20395246 |
| NTGALGL | ASSLDIQDTQY | 0.8263359 |
| PPGAGGAGGAGTT | ASSLDIQDTQY | 0.6839292 |
| QLVGLGI | ASSLDIQDTQY | 0.43860334 |
| RFLEQVHQL | ASSLDIQDTQY | 0.4918192 |
| RGVGYVLR | ASSLDIQDTQY | 0.4703773 |
| SATGVFAGVI | ASSLDIQDTQY | 0.38759634 |
| SLPAITLP | ASSLDIQDTQY | 0.87379754 |
| STALAFTV | ASSLDIQDTQY | 0.7944329 |
| SVLTVGLGK | ASSLDIQDTQY | 0.76832277 |
| TALAGAGSGI | ASSLDIQDTQY | 0.8853556 |
| TGLPFTGI | ASSLDIQDTQY | 0.83848643 |
| VAEHVTV | ASSLDIQDTQY | 0.79575825 |
| VDIVLNSL | ASSLDIQDTQY | 0.5690637 |
| VLTGGGAVL | ASSLDIQDTQY | 0.60981095 |
| VNVGLAYGI | ASSLDIQDTQY | 0.54785216 |
| VSSAAVGAV | ASSLDIQDTQY | 0.51371723 |
| VTSVYEAGV | ASSLDIQDTQY | 0.88553214 |
| WVALDRGA | ASSLDIQDTQY | 0.77015775 |
| YLVGSVVI | ASSLDIQDTQY | 0.5298598 |
| GPPGTGKTLL | ASSLDIQDTQY | 0.8494157 |
| GVGMTKF | ASSLDIQDTQY | 0.88496596 |
| IGIGGSDL | ASSLDIQDTQY | 0.8904568 |
| IGTLLAEL | ASSLDIQDTQY | 0.59619534 |
| IYTSGTT | ASSLDIQDTQY | 0.7704849 |
| LSFPAGV | ASSLDIQDTQY | 0.7355983 |
| LYTSGTT | ASSLDIQDTQY | 0.8061521 |

| | | |
|---|---|---|
| VGAFPFGF | ASSLDIQDTQY | 0.75615036 |
| VGAGTVEF | ASSLDIQDTQY | 0.42068285 |
| VGKSTLLL | ASSLDIQDTQY | 0.87785834 |
| AALAHAGGGIV | ASSLDIQDTQY | 0.8897041 |
| AALVAVSLL | ASSLDIQDTQY | 0.6961527 |
| AAVSRAVAAL | ASSLDIQDTQY | 0.6952481 |
| AGGSSGIGGF | ASSLDIQDTQY | 0.78323233 |
| AGLTVANANG | ASSLDIQDTQY | 0.6839364 |
| AGLVGAVL | ASSLDIQDTQY | 0.71747935 |
| ALSQAFTV | ASSLDIQDTQY | 0.75519437 |
| ALTFSVSL | ASSLDIQDTQY | 0.47276592 |
| ALTGAALVAL | ASSLDIQDTQY | 0.75234723 |
| ASFSIGAGVL | ASSLDIQDTQY | 0.88509953 |
| DGSVYKI | ASSLDIQDTQY | 0.7797326 |
| EQLAEIV | ASSLDIQDTQY | 0.853639 |
| FAVSELTVM | ASSLDIQDTQY | 0.8801716 |
| GAAPLAAGL | ASSLDIQDTQY | 0.86183715 |
| GADGGDGGSGGAGG | ASSLDIQDTQY | 0.8831024 |
| GFGLDTGGAGG | ASSLDIQDTQY | 0.88000894 |
| GLASGGLL | ASSLDIQDTQY | 0.74439526 |
| GRIAGAAVLL | ASSLDIQDTQY | 0.8220161 |
| GSAGSLNTSF | ASSLDIQDTQY | 0.8605539 |
| GTGQIGI | ASSLDIQDTQY | 0.7852782 |
| GTLIALIL | ASSLDIQDTQY | 0.80021155 |
| IAATMGFVVG | ASSLDIQDTQY | 0.77088547 |
| IFAQALL | ASSLDIQDTQY | 0.86466944 |
| IGAAVALGF | ASSLDIQDTQY | 0.8235387 |
| IGLTPLAL | ASSLDIQDTQY | 0.8786073 |
| ILGSNTSTL | ASSLDIQDTQY | 0.30542946 |
| IVFSLTVL | ASSLDIQDTQY | 0.43682045 |
| LASKATGF | ASSLDIQDTQY | 0.87802035 |
| LASVALIL | ASSLDIQDTQY | 0.14966331 |
| LFAVGLTL | ASSLDIQDTQY | 0.24799988 |
| LFNGGIGTYI | ASSLDIQDTQY | 0.8428581 |
| LTLGAGII | ASSLDIQDTQY | 0.87962306 |
| LVAAALGDI | ASSLDIQDTQY | 0.16889696 |
| NTAFVQLGI | ASSLDIQDTQY | 0.6176261 |
| QLFSGAEL | ASSLDIQDTQY | 0.46378458 |
| SAATFITL | ASSLDIQDTQY | 0.87074447 |
| SNYGLANLGV | ASSLDIQDTQY | 0.7751275 |
| SYGGAYAVM | ASSLDIQDTQY | 0.35210246 |
| TTGVGGLTLG | ASSLDIQDTQY | 0.85874104 |
| TTQTIYI | ASSLDIQDTQY | 0.59756887 |
| VAAGAGLGAVI | ASSLDIQDTQY | 0.7437322 |
| VAAVVTLL | ASSLDIQDTQY | 0.87038994 |
| VGFLGGLI | ASSLDIQDTQY | 0.88744736 |
| VGVIPILL | ASSLDIQDTQY | 0.5430052 |
| VIFASSLI | ASSLDIQDTQY | 0.86149514 |
| VKGGDGGAAAG | ASSLDIQDTQY | 0.7964069 |
| VTKTFTNI | ASSLDIQDTQY | 0.89066255 |
| VVAAAILL | ASSLDIQDTQY | 0.67653334 |

| | | |
|---|---|---|
| YAALDVEL | ASSLDIQDTQY | 0.41672623 |
| LASVALLL | ASSLDIQDTQY | 0.8681042 |
| LDAVHRV | ASSLDIQDTQY | 0.86828446 |
| QGLSSSIF | ASSLDIQDTQY | 0.7013182 |
| RGTFANIRL | ASSLDIQDTQY | 0.87686384 |
| RQLWWGHRI | ASSLDIQDTQY | 0.7861004 |
| YGPPGTGKTLL | ASSLDIQDTQY | 0.7865497 |
| AALVGAPANVA | ASSLDIQDTQY | 0.87303746 |
| AATFRRTE | ASSLDIQDTQY | 0.8749357 |
| AGALGGGI | ASSLDIQDTQY | 0.42544326 |
| AGGAGGAGGIGAG | ASSLDIQDTQY | 0.86077666 |
| AGSGALGL | ASSLDIQDTQY | 0.78177273 |
| AQIADQI | ASSLDIQDTQY | 0.59855103 |
| ASAVAVTYL | ASSLDIQDTQY | 0.8841336 |
| ASTVLGGTGG | ASSLDIQDTQY | 0.83332664 |
| AVGGEPAK | ASSLDIQDTQY | 0.8678051 |
| CEQGTFGGTC | ASSLDIQDTQY | 0.8089379 |
| DEGKAPAG | ASSLDIQDTQY | 0.8822246 |
| DTVVFPAL | ASSLDIQDTQY | 0.40281564 |
| FEGITMAG | ASSLDIQDTQY | 0.84935856 |
| FPLVVQD | ASSLDIQDTQY | 0.8084521 |
| GAFTTPQ | ASSLDIQDTQY | 0.8749026 |
| GFGGFGTAGGVGGA | ASSLDIQDTQY | 0.82980967 |
| GFLAQFL | ASSLDIQDTQY | 0.7349154 |
| GGAGGAGGNAGVA | ASSLDIQDTQY | 0.833501 |
| GGVGTAGAGGAGGAGG | ASSLDIQDTQY | 0.87561774 |
| GNGGGGDAGNAGSGGN | ASSLDIQDTQY | 0.83643484 |
| GTGGAGGAGNDAGS | ASSLDIQDTQY | 0.8513112 |
| GTTNTGWFNS | ASSLDIQDTQY | 0.886665 |
| IFMGPLFTL | ASSLDIQDTQY | 0.5354242 |
| ILAHAIEAI | ASSLDIQDTQY | 0.74751717 |
| IRSAIREAL | ASSLDIQDTQY | 0.8652098 |
| ITFAGGG | ASSLDIQDTQY | 0.8758384 |
| ITGMSGHQEV | ASSLDIQDTQY | 0.8012508 |
| IVAGLACTL | ASSLDIQDTQY | 0.88649726 |
| IVGGGLAGA | ASSLDIQDTQY | 0.7256668 |
| IVTGVLNI | ASSLDIQDTQY | 0.85281986 |
| LAQQGCI | ASSLDIQDTQY | 0.7690996 |
| LCLHVEN | ASSLDIQDTQY | 0.8220201 |
| LDAISDGS | ASSLDIQDTQY | 0.5455367 |
| LGGFAAML | ASSLDIQDTQY | 0.84700674 |
| LGGKNPMI | ASSLDIQDTQY | 0.8743356 |
| LGPKGVAAIL | ASSLDIQDTQY | 0.86243594 |
| LMVGVSI | ASSLDIQDTQY | 0.21853687 |
| LVGAAFVL | ASSLDIQDTQY | 0.8616794 |
| LVLAVAPL | ASSLDIQDTQY | 0.8597625 |
| MAAIAVIDAA | ASSLDIQDTQY | 0.8917316 |
| NNGINAELQ | ASSLDIQDTQY | 0.8037359 |
| RGEPLVLVHG | ASSLDIQDTQY | 0.84131557 |
| RGGAGGAAGLI | ASSLDIQDTQY | 0.6779215 |
| RSYRMHPAV | ASSLDIQDTQY | 0.7909887 |

| | | |
|---|---|---|
| SAAHLLAGQLA | ASSLDIQDTQY | 0.603068 |
| SAAVGSY | ASSLDIQDTQY | 0.71472174 |
| SPHAVLHRI | ASSLDIQDTQY | 0.7808517 |
| SQIAKVL | ASSLDIQDTQY | 0.6800375 |
| TALGHLVP | ASSLDIQDTQY | 0.8668523 |
| TAPTTSGAA | ASSLDIQDTQY | 0.8810817 |
| TDGDDGNGGSPGT | ASSLDIQDTQY | 0.8647225 |
| TDGLGEN | ASSLDIQDTQY | 0.88123786 |
| TGGPPLPGQA | ASSLDIQDTQY | 0.88483924 |
| TIPAITI | ASSLDIQDTQY | 0.8734087 |
| TIPGFTI | ASSLDIQDTQY | 0.62465173 |
| TSIASGALA | ASSLDIQDTQY | 0.79797673 |
| TVTLPGAAI | ASSLDIQDTQY | 0.48753774 |
| VASGLSAALP | ASSLDIQDTQY | 0.7836824 |
| VGAGVIGI | ASSLDIQDTQY | 0.6862598 |
| VGPKRFGV | ASSLDIQDTQY | 0.82814014 |
| VNVAFPD | ASSLDIQDTQY | 0.1318955 |
| VSMTNSGVS | ASSLDIQDTQY | 0.8536236 |
| VTDVLGDLSV | ASSLDIQDTQY | 0.88571304 |
| VVVVGAGFAGL | ASSLDIQDTQY | 0.76263237 |
| VYTVGTVV | ASSLDIQDTQY | 0.823333 |
| WTTFSGP | ASSLDIQDTQY | 0.8565952 |
| YDELVEAF | ASSLDIQDTQY | 0.7760171 |
| YGTGGAGGNGG | ASSLDIQDTQY | 0.86579716 |
| YTNVMAV | ASSLDIQDTQY | 0.60894793 |
| FLEQVHQL | ASSLDIQDTQY | 0.8566859 |
| GLSLLAVAV | ASSLDIQDTQY | 0.8779589 |
| LEQVHQL | ASSLDIQDTQY | 0.8212389 |
| LNAAGAI | ASSLDIQDTQY | 0.829253 |
| PAVHASL | ASSLDIQDTQY | 0.76427263 |
| TELEDDA | ASSLDIQDTQY | 0.33547157 |
| TPPGPPN | ASSLDIQDTQY | 0.8437258 |
| TPPPPPPP | ASSLDIQDTQY | 0.14086299 |
| TQSPALI | ASSLGTGNSGNTLY | 0.83036697 |
| EFAAAFPQ | ASSLGTGNSGNTLY | 0.46887666 |
| LLSGNLDVL | ASSLGTGNSGNTLY | 0.79970723 |
| GPANLGNYNI | ASSLGTGNSGNTLY | 0.78583705 |
| TLGPTLP | ASSLGTGNSGNTLY | 0.86333174 |
| VYAYSSCCK | ASSLGTGNSGNTLY | 0.6319842 |
| APAPAPAPNI | ASSLGTGNSGNTLY | 0.8439132 |
| PGGPVPIP | ASSLGTGNSGNTLY | 0.7542672 |
| FLTVALEP | ASSLGTGNSGNTLY | 0.39101756 |
| SQYFGANL | ASSLGTGNSGNTLY | 0.87746346 |
| GGVKPAA | ASSLGTGNSGNTLY | 0.7638458 |
| GGTGGAGGAGGW | ASSLGTGNSGNTLY | 0.8231932 |
| GVGGVAGLGGAGG | ASSLGTGNSGNTLY | 0.8516217 |
| VLLVLATTI | ASSLGTGNSGNTLY | 0.7858728 |
| IGIFAAML | ASSLGTGNSGNTLY | 0.50453675 |
| AVGDLID | ASSLGTGNSGNTLY | 0.8016269 |
| NISPLPL | ASSLGTGNSGNTLY | 0.8019713 |
| VTAVFVAI | ASSLGTGNSGNTLY | 0.67701954 |

| | | |
|---|---|---|
| SGDDNDAAL | ASSLGTGNSGNTLY | 0.14613894 |
| AVAALAHS | ASSLGTGNSGNTLY | 0.85920626 |
| GGVTVMATP | ASSLGTGNSGNTLY | 0.58762383 |
| AGLAANVRD | ASSLGTGNSGNTLY | 0.75964594 |
| TNRPDLIDPALLRPG | ASSLGTGNSGNTLY | 0.7484538 |
| LANLGVTAL | ASSLGTGNSGNTLY | 0.871385 |
| TGERIEL | ASSLGTGNSGNTLY | 0.7477466 |
| NGEELAA | ASSLGTGNSGNTLY | 0.55508804 |
| VLSLVWG | ASSLGTGNSGNTLY | 0.87269914 |
| IGYALWL | ASSLGTGNSGNTLY | 0.8141725 |
| LSLLLAPL | ASSLGTGNSGNTLY | 0.8708934 |
| TLDEALK | ASSLGTGNSGNTLY | 0.7886116 |
| TDGAGTAEGG | ASSLGTGNSGNTLY | 0.7961554 |
| NAPGGAGGNGG | ASSLGTGNSGNTLY | 0.823175 |
| VSVFMGMGGNF | ASSLGTGNSGNTLY | 0.8188917 |
| LASYLAAL | ASSLGTGNSGNTLY | 0.48224363 |
| LVGGTPAL | ASSLGTGNSGNTLY | 0.39189136 |
| VRNADVL | ASSLGTGNSGNTLY | 0.83358973 |
| TCSSPFLVSAM | ASSLGTGNSGNTLY | 0.34179467 |
| VHITDAE | ASSLGTGNSGNTLY | 0.77822506 |
| EYAAPSF | ASSLGTGNSGNTLY | 0.88252336 |
| VVDAFHGGL | ASSLGTGNSGNTLY | 0.8402769 |
| ASVVDPQ | ASSLGTGNSGNTLY | 0.42817992 |
| SYNEFGGLS | ASSLGTGNSGNTLY | 0.76683664 |
| DPDGSVY | ASSLGTGNSGNTLY | 0.75098044 |
| MVSTIHATE | ASSLGTGNSGNTLY | 0.8617562 |
| LAAGVADPV | ASSLGTGNSGNTLY | 0.8561032 |
| TKEQIAA | ASSLGTGNSGNTLY | 0.88391894 |
| SLASPAAL | ASSLGTGNSGNTLY | 0.86554486 |
| IGLVFYGV | ASSLGTGNSGNTLY | 0.6703734 |
| FLLDASNL | ASSLGTGNSGNTLY | 0.8707482 |
| YQVGNFP | ASSLGTGNSGNTLY | 0.76419646 |
| ESLLSQST | ASSLGTGNSGNTLY | 0.8419451 |
| LFTVGFAAVG | ASSLGTGNSGNTLY | 0.8044607 |
| ATPPTPT | ASSLGTGNSGNTLY | 0.8577653 |
| TDLATLG | ASSLGTGNSGNTLY | 0.748492 |
| SAGGTGR | ASSLGTGNSGNTLY | 0.72962207 |
| GFSNSGS | ASSLGTGNSGNTLY | 0.47577733 |
| GGNGGSGGTSV | ASSLGTGNSGNTLY | 0.8673215 |
| GTSVATGGAGN | ASSLGTGNSGNTLY | 0.80926895 |
| NGGNGGSGGTS | ASSLGTGNSGNTLY | 0.60354155 |
| AVVKFPH | ASSLGTGNSGNTLY | 0.68528885 |
| TPPPPPP | ASSLGTGNSGNTLY | 0.8873245 |
| AAPAAAAAPA | ASSLGTGNSGNTLY | 0.88250095 |
| IALNEDL | ASSLGTGNSGNTLY | 0.69645923 |
| IVNVSSM | ASSLGTGNSGNTLY | 0.57838726 |
| TGIPPAPR | ASSLGTGNSGNTLY | 0.8638314 |
| FDLGGGTF | ASSLGTGNSGNTLY | 0.43449286 |
| FDLGGGTFDVS | ASSLGTGNSGNTLY | 0.85336304 |
| NEPTAAA | ASSLGTGNSGNTLY | 0.82538223 |
| NEPTAAALA | ASSLGTGNSGNTLY | 0.81463146 |

| | | |
|---|---|---|
| GSIINISSI | ASSLGTGNSGNTLY | 0.67841434 |
| VAPGNAGTA | ASSLGTGNSGNTLY | 0.85606647 |
| TGGGASLE | ASSLGTGNSGNTLY | 0.7450143 |
| VAVYNHYKRI | ASSLGTGNSGNTLY | 0.86691093 |
| TDPAPATT | ASSLGTGNSGNTLY | 0.2199702 |
| TNRPDILDPALLRPG | ASSLGTGNSGNTLY | 0.852937 |
| LTTTTTS | ASSLGTGNSGNTLY | 0.61551 |
| PTGSGKTL | ASSLGTGNSGNTLY | 0.6820812 |
| TGSGKTL | ASSLGTGNSGNTLY | 0.340483 |
| TIGHVDHGK | ASSLGTGNSGNTLY | 0.81171644 |
| TIGHVDHGKTT | ASSLGTGNSGNTLY | 0.87327963 |
| GHVDHGKTT | ASSLGTGNSGNTLY | 0.88138103 |
| AQFEHTLL | ASSLGTGNSGNTLY | 0.75462085 |
| IVNQDEL | ASSLGTGNSGNTLY | 0.8727297 |
| TNGSQFF | ASSLGTGNSGNTLY | 0.88669133 |
| TGKGIVE | ASSLGTGNSGNTLY | 0.77220666 |
| LISNASDA | ASSLGTGNSGNTLY | 0.23870374 |
| AELRTLGELTKIAKAHGAQ | ASSLGTGNSGNTLY | 0.80772436 |
| APNAVAAAEPSW | ASSLGTGNSGNTLY | 0.84231365 |
| APQVAGAATAVL | ASSLGTGNSGNTLY | 0.87908876 |
| ARRDPAAAAAILPTDA | ASSLGTGNSGNTLY | 0.8592742 |
| DAAARDVLAEAGLAEYF | ASSLGTGNSGNTLY | 0.56614304 |
| DDQIVLSAKANLAKVEQ | ASSLGTGNSGNTLY | 0.832977 |
| DELLPDTGEDSDEDGH | ASSLGTGNSGNTLY | 0.8623864 |
| ELPEAPANRATVPQD | ASSLGTGNSGNTLY | 0.60596496 |
| FGQSKIERVTGIG | ASSLGTGNSGNTLY | 0.73167896 |
| GAGGTGSTAGGAGGAGGAGGLY | ASSLGTGNSGNTLY | 0.7013719 |
| GHGGTGAAGGNGGTGGAGGSGI | ASSLGTGNSGNTLY | 0.43721157 |
| GNGGNGGAGGEGGPGGAGGAGA | ASSLGTGNSGNTLY | 0.825297 |
| GSTAGNGGNGGNATNGTIAGQP | ASSLGTGNSGNTLY | 0.7461313 |
| GTGGDGGNGGNGGTGG | ASSLGTGNSGNTLY | 0.61434174 |
| GVRRPAGIPADAPV | ASSLGTGNSGNTLY | 0.7958127 |
| GYRQVTDAHLLTLARR | ASSLGTGNSGNTLY | 0.5304795 |
| IAKMGQDGHDRG | ASSLGTGNSGNTLY | 0.6092246 |
| KAEPSIEIQYT | ASSLGTGNSGNTLY | 0.648595 |
| KPVRIRMAADEDTVAVT | ASSLGTGNSGNTLY | 0.81071055 |
| MPPRIAGMRLLVIKPEP | ASSLGTGNSGNTLY | 0.8690984 |
| MKFYAVALFRSRGVPPTR | ASSLGTGNSGNTLY | 0.8399174 |
| NAGGFGILGAVAHSP | ASSLGTGNSGNTLY | 0.85462505 |
| NEGREITLAMAEM | ASSLGTGNSGNTLY | 0.8806199 |
| NELVEWSHPLGHRWRWE | ASSLGTGNSGNTLY | 0.39766037 |
| NLGAGGTGGNGGNPG | ASSLGTGNSGNTLY | 0.600134 |
| NTEWPKWETVMAADPVD | ASSLGTGNSGNTLY | 0.86294085 |
| PASAAPAETQP | ASSLGTGNSGNTLY | 0.5458824 |
| QQSSDGKNGVRAAEITGE | ASSLGTGNSGNTLY | 0.872686 |
| RAPDAVTTSWWKE | ASSLGTGNSGNTLY | 0.79335046 |
| SGFGNVGAHTSGWFNQ | ASSLGTGNSGNTLY | 0.84734017 |
| SGVAGAVGQAASVGGLKV | ASSLGTGNSGNTLY | 0.84802866 |
| SNGGAGTGSAGGNGGT | ASSLGTGNSGNTLY | 0.89023864 |
| TGGAGGEGGAAG | ASSLGTGNSGNTLY | 0.85243684 |
| TGGDGGAAGAGGAPGHG | ASSLGTGNSGNTLY | 0.790216 |

| | | |
|---|---|---|
| TMPPLLAAIANAGRHGV | ASSLGTGNSGNTLY | 0.8637068 |
| TSSTAGNSGGA | ASSLGTGNSGNTLY | 0.5510963 |
| VPATVPVGILC | ASSLGTGNSGNTLY | 0.87429506 |
| VQRPDSGGVVLTGRIS | ASSLGTGNSGNTLY | 0.85812634 |
| WTPNPVPATKGVPAGLVQAT | ASSLGTGNSGNTLY | 0.63282096 |
| YALKRALAKFSDRFDVVII | ASSLGTGNSGNTLY | 0.79132795 |
| YLIGSVIV | ASSLGTGNSGNTLY | 0.885558 |
| YGQLVAVID | ASSLGTGNSGNTLY | 0.88021946 |
| VTVGGMTLPA | ASSLGTGNSGNTLY | 0.33923018 |
| VLALPAV | ASSLGTGNSGNTLY | 0.7217141 |
| VIPPTLNYET | ASSLGTGNSGNTLY | 0.43415493 |
| VGMGGMHPSSG | ASSLGTGNSGNTLY | 0.76691 |
| VGLAGGAATA | ASSLGTGNSGNTLY | 0.3574106 |
| VGGAGGAGGSGP | ASSLGTGNSGNTLY | 0.86139745 |
| VFAAFMV | ASSLGTGNSGNTLY | 0.7053611 |
| TLYDKLR | ASSLGTGNSGNTLY | 0.8630837 |
| TLAVFTVL | ASSLGTGNSGNTLY | 0.81396246 |
| TGSVQPARV | ASSLGTGNSGNTLY | 0.8621954 |
| TGGGAGSGGT | ASSLGTGNSGNTLY | 0.69924426 |
| SHLETSL | ASSLGTGNSGNTLY | 0.6703615 |
| SFTTGMT | ASSLGTGNSGNTLY | 0.36114806 |
| QVEVAKRMGVRQPTVS | ASSLGTGNSGNTLY | 0.41178316 |
| QGSGGPALAM | ASSLGTGNSGNTLY | 0.7293336 |
| MLAPSNP | ASSLGTGNSGNTLY | 0.73261285 |
| LVNNAAI | ASSLGTGNSGNTLY | 0.3515092 |
| LTGEINTLA | ASSLGTGNSGNTLY | 0.884405 |
| LSIFGPV | ASSLGTGNSGNTLY | 0.8061007 |
| LLALLANL | ASSLGTGNSGNTLY | 0.83667725 |
| LISNASDALDKLR | ASSLGTGNSGNTLY | 0.6192362 |
| LINEAVP | ASSLGTGNSGNTLY | 0.71527386 |
| LGYAFSGL | ASSLGTGNSGNTLY | 0.67186046 |
| LDFVAVAL | ASSLGTGNSGNTLY | 0.85414857 |
| LAQPGAE | ASSLGTGNSGNTLY | 0.5827897 |
| ISNLGAVTGF | ASSLGTGNSGNTLY | 0.43179083 |
| HQEDPGRI | ASSLGTGNSGNTLY | 0.7589711 |
| GTPGEPTP | ASSLGTGNSGNTLY | 0.7789593 |
| GFFNSGS | ASSLGTGNSGNTLY | 0.88566196 |
| GAGGGGLTAGAN | ASSLGTGNSGNTLY | 0.86855745 |
| FVDALVNM | ASSLGTGNSGNTLY | 0.835738 |
| FLTVAPEL | ASSLGTGNSGNTLY | 0.8829763 |
| FETLTAL | ASSLGTGNSGNTLY | 0.19046496 |
| DVINPIEL | ASSLGTGNSGNTLY | 0.70022416 |
| APVSGAPV | ASSLGTGNSGNTLY | 0.87999356 |
| ALVGGLAW | ASSLGTGNSGNTLY | 0.8401568 |
| ALAAAAATAI | ASSLGTGNSGNTLY | 0.7481595 |
| AFGLGLLL | ASSLGTGNSGNTLY | 0.8765594 |
| AFFAAASL | ASSLGTGNSGNTLY | 0.55620337 |
| AAAALPAFSPP | ASSLGTGNSGNTLY | 0.6527286 |
| VVLAPAL | ASSLGTGNSGNTLY | 0.7880499 |
| TIGHVDHGKTTL | ASSLGTGNSGNTLY | 0.8432127 |
| LISNASDALDK | ASSLGTGNSGNTLY | 0.8418981 |

| | | |
|---|---|---|
| LISNASDAL | ASSLGTGNSGNTLY | 0.48699296 |
| ISNASDALDK | ASSLGTGNSGNTLY | 0.81838673 |
| NQAETLVYQTE | ASSLGTGNSGNTLY | 0.89223576 |
| TGGTGGTAGAG | ASSLGTGNSGNTLY | 0.72987735 |
| TGSTAGGAGGA | ASSLGTGNSGNTLY | 0.8781117 |
| DPLAVPVAAGPV | ASSLGTGNSGNTLY | 0.82346535 |
| GGTGGTPGNGGN | ASSLGTGNSGNTLY | 0.83361685 |
| PAGLAGVDPATL | ASSLGTGNSGNTLY | 0.63638365 |
| GNNLVVEAPHAGASV | ASSLGTGNSGNTLY | 0.690835 |
| KVTRIMSASTLEQPA | ASSLGTGNSGNTLY | 0.83944684 |
| LGFALPIALPRRTVV | ASSLGTGNSGNTLY | 0.6303056 |
| TRLGNLTARKLVVLV | ASSLGTGNSGNTLY | 0.79576814 |
| WSQQRRLIYFPSAGP | ASSLGTGNSGNTLY | 0.7558392 |
| GPNGNPDPMAAAVDIR | ASSLGTGNSGNTLY | 0.89160264 |
| TQRCVSVADLRAAASR | ASSLGTGNSGNTLY | 0.8493387 |
| QFVLDVAEHPVEVLTETV | ASSLGTGNSGNTLY | 0.8479289 |
| GGAGGTGGAGGNGGTQAGDG | ASSLGTGNSGNTLY | 0.57161236 |
| PLLATLRAIAVDVDAKPAQV | ASSLGTGNSGNTLY | 0.8490079 |
| APKFRTGGYMIDFWGVGYQVAK | ASSLGTGNSGNTLY | 0.8564255 |
| GAGGAGGAGGAAGGPGGTGGTG | ASSLGTGNSGNTLY | 0.5340771 |
| IGALGLLATDVAPAGSVAFGVA | ASSLGTGNSGNTLY | 0.8812758 |
| AGGAGGIGGNGGAGGAGGVLMG | ASSLGTGNSGNTLY | 0.57013804 |
| ADELIGLL | ASSLGTGNSGNTLY | 0.8252585 |
| ADRALKA | ASSLGTGNSGNTLY | 0.8471639 |
| AGAVGQVGGAGGA | ASSLGTGNSGNTLY | 0.8758912 |
| AGGGGGLGGAP | ASSLGTGNSGNTLY | 0.7321124 |
| AGGLLYGNGGAG | ASSLGTGNSGNTLY | 0.8117645 |
| AGLESIGL | ASSLGTGNSGNTLY | 0.53971666 |
| AGLKTGII | ASSLGTGNSGNTLY | 0.8733435 |
| ASLAALLL | ASSLGTGNSGNTLY | 0.86973804 |
| ASVINISSI | ASSLGTGNSGNTLY | 0.8289648 |
| ATPPGPA | ASSLGTGNSGNTLY | 0.8900403 |
| ATVASALV | ASSLGTGNSGNTLY | 0.85876715 |
| ATVASLIL | ASSLGTGNSGNTLY | 0.30313516 |
| AVNGPVLIRH | ASSLGTGNSGNTLY | 0.7554896 |
| DGAAGAVGI | ASSLGTGNSGNTLY | 0.87797046 |
| DPLLHGI | ASSLGTGNSGNTLY | 0.88376606 |
| ELGAALGAL | ASSLGTGNSGNTLY | 0.8698487 |
| ELGRTQAV | ASSLGTGNSGNTLY | 0.855729 |
| FVAGGYLS | ASSLGTGNSGNTLY | 0.76721305 |
| GASGGAGGAGGW | ASSLGTGNSGNTLY | 0.8832451 |
| GGAGAGGDGGAGGI | ASSLGTGNSGNTLY | 0.8930541 |
| GTGHNGSQPGMGGNG | ASSLGTGNSGNTLY | 0.85560566 |
| GYAAASASAS | ASSLGTGNSGNTLY | 0.7521125 |
| HQVDVAAVQQG | ASSLGTGNSGNTLY | 0.650401 |
| IGAAIAHT | ASSLGTGNSGNTLY | 0.8259771 |
| IGMIEVAQL | ASSLGTGNSGNTLY | 0.8710078 |
| IRLAPEHTSAMD | ASSLGTGNSGNTLY | 0.5242988 |
| IYNVSVL | ASSLGTGNSGNTLY | 0.8060384 |
| KQGFLGM | ASSLGTGNSGNTLY | 0.81095165 |
| LESVITQL | ASSLGTGNSGNTLY | 0.30625165 |

| | | |
|---|---|---|
| LGLGALVVV | ASSLGTGNSGNTLY | 0.86273706 |
| LIGNGGNGGSGGI | ASSLGTGNSGNTLY | 0.8860056 |
| LNAINAPTL | ASSLGTGNSGNTLY | 0.81338024 |
| LQLLNTGV | ASSLGTGNSGNTLY | 0.6918386 |
| LQTSIEF | ASSLGTGNSGNTLY | 0.8773633 |
| MSFAVSEL | ASSLGTGNSGNTLY | 0.88993233 |
| NTGALGL | ASSLGTGNSGNTLY | 0.8814584 |
| PPGAGGAGGAGTT | ASSLGTGNSGNTLY | 0.2867351 |
| QLVGLGI | ASSLGTGNSGNTLY | 0.23998556 |
| RFLEQVHQL | ASSLGTGNSGNTLY | 0.7393554 |
| RGVGYVLR | ASSLGTGNSGNTLY | 0.86507076 |
| SATGVFAGVI | ASSLGTGNSGNTLY | 0.6567837 |
| SLPAITLP | ASSLGTGNSGNTLY | 0.58637404 |
| STALAFTV | ASSLGTGNSGNTLY | 0.85759234 |
| SVLTVGLGK | ASSLGTGNSGNTLY | 0.87203276 |
| TALAGAGSGI | ASSLGTGNSGNTLY | 0.47151464 |
| TGLPFTGI | ASSLGTGNSGNTLY | 0.48392332 |
| VAEHVTV | ASSLGTGNSGNTLY | 0.8692994 |
| VDIVLNSL | ASSLGTGNSGNTLY | 0.868488 |
| VLTGGGAVL | ASSLGTGNSGNTLY | 0.6437259 |
| VNVGLAYGI | ASSLGTGNSGNTLY | 0.7633359 |
| VSSAAVGAV | ASSLGTGNSGNTLY | 0.733248 |
| VTSVYEAGV | ASSLGTGNSGNTLY | 0.8103937 |
| WVALDRGA | ASSLGTGNSGNTLY | 0.8470299 |
| YLVGSVVI | ASSLGTGNSGNTLY | 0.5255647 |
| GPPGTGKTLL | ASSLGTGNSGNTLY | 0.8829111 |
| GVGMTKF | ASSLGTGNSGNTLY | 0.5523045 |
| IGIGGSDL | ASSLGTGNSGNTLY | 0.87689406 |
| IGTLLAEL | ASSLGTGNSGNTLY | 0.81615674 |
| IYTSGTT | ASSLGTGNSGNTLY | 0.8223157 |
| LSFPAGV | ASSLGTGNSGNTLY | 0.7248808 |
| LYTSGTT | ASSLGTGNSGNTLY | 0.69814813 |
| VGAFPFGF | ASSLGTGNSGNTLY | 0.5155909 |
| VGAGTVEF | ASSLGTGNSGNTLY | 0.4932875 |
| VGKSTLLL | ASSLGTGNSGNTLY | 0.87911105 |
| AALAHAGGGIV | ASSLGTGNSGNTLY | 0.3658955 |
| AALVAVSLL | ASSLGTGNSGNTLY | 0.8780634 |
| AAVSRAVAAL | ASSLGTGNSGNTLY | 0.77739096 |
| AGGSSGIGGF | ASSLGTGNSGNTLY | 0.548627 |
| AGLTVANANG | ASSLGTGNSGNTLY | 0.8640172 |
| AGLVGAVL | ASSLGTGNSGNTLY | 0.4453879 |
| ALSQAFTV | ASSLGTGNSGNTLY | 0.72406816 |
| ALTFSVSL | ASSLGTGNSGNTLY | 0.85644865 |
| ALTGAALVAL | ASSLGTGNSGNTLY | 0.854764 |
| ASFSIGAGVL | ASSLGTGNSGNTLY | 0.64183265 |
| DGSVYKI | ASSLGTGNSGNTLY | 0.8480121 |
| EQLAEIV | ASSLGTGNSGNTLY | 0.84800625 |
| FAVSELTVM | ASSLGTGNSGNTLY | 0.69347084 |
| GAAPLAAGL | ASSLGTGNSGNTLY | 0.7761357 |
| GADGGDGGSGGAGG | ASSLGTGNSGNTLY | 0.8705326 |
| GFGLDTGGAGG | ASSLGTGNSGNTLY | 0.8089712 |

| | | |
|---|---|---|
| GLASGGLL | ASSLGTGNSGNTLY | 0.5494189 |
| GRIAGAAVLL | ASSLGTGNSGNTLY | 0.83663625 |
| GSAGSLNTSF | ASSLGTGNSGNTLY | 0.6848843 |
| GTGQIGI | ASSLGTGNSGNTLY | 0.85072327 |
| GTLIALIL | ASSLGTGNSGNTLY | 0.81184995 |
| IAATMGFVVG | ASSLGTGNSGNTLY | 0.6923995 |
| IFAQALL | ASSLGTGNSGNTLY | 0.8523598 |
| IGAAVALGF | ASSLGTGNSGNTLY | 0.63560784 |
| IGLTPLAL | ASSLGTGNSGNTLY | 0.7011794 |
| ILGSNTSTL | ASSLGTGNSGNTLY | 0.68490034 |
| IVFSLTVL | ASSLGTGNSGNTLY | 0.72707736 |
| LASKATGF | ASSLGTGNSGNTLY | 0.8589653 |
| LASVALIL | ASSLGTGNSGNTLY | 0.54819363 |
| LFAVGLTL | ASSLGTGNSGNTLY | 0.8388931 |
| LFNGGIGTYI | ASSLGTGNSGNTLY | 0.6340407 |
| LTLGAGII | ASSLGTGNSGNTLY | 0.4335898 |
| LVAAALGDI | ASSLGTGNSGNTLY | 0.7939838 |
| NTAFVQLGI | ASSLGTGNSGNTLY | 0.7800009 |
| QLFSGAEL | ASSLGTGNSGNTLY | 0.62314844 |
| SAATFITL | ASSLGTGNSGNTLY | 0.8562089 |
| SNYGLANLGV | ASSLGTGNSGNTLY | 0.4449782 |
| SYGGAYAVM | ASSLGTGNSGNTLY | 0.7917447 |
| TTGVGGLTLG | ASSLGTGNSGNTLY | 0.8879067 |
| TTQTIYI | ASSLGTGNSGNTLY | 0.70465446 |
| VAAGAGLGAVI | ASSLGTGNSGNTLY | 0.87354875 |
| VAAVVTLL | ASSLGTGNSGNTLY | 0.8495135 |
| VGFLGGLI | ASSLGTGNSGNTLY | 0.8594919 |
| VGVIPILL | ASSLGTGNSGNTLY | 0.684657 |
| VIFASSLI | ASSLGTGNSGNTLY | 0.6867889 |
| VKGGDGGAAAG | ASSLGTGNSGNTLY | 0.8832598 |
| VTKTFTNI | ASSLGTGNSGNTLY | 0.8398342 |
| VVAAAILL | ASSLGTGNSGNTLY | 0.86497986 |
| YAALDVEL | ASSLGTGNSGNTLY | 0.86149144 |
| LASVALLL | ASSLGTGNSGNTLY | 0.871402 |
| LDAVHRV | ASSLGTGNSGNTLY | 0.8660629 |
| QGLSSSIF | ASSLGTGNSGNTLY | 0.7354069 |
| RGTFANIRL | ASSLGTGNSGNTLY | 0.37151438 |
| RQLWWGHRI | ASSLGTGNSGNTLY | 0.66461134 |
| YGPPGTGKTLL | ASSLGTGNSGNTLY | 0.7419536 |
| AALVGAPANVA | ASSLGTGNSGNTLY | 0.73536235 |
| AATFRRTE | ASSLGTGNSGNTLY | 0.4172386 |
| AGALGGGI | ASSLGTGNSGNTLY | 0.86840045 |
| AGGAGGAGGIGAG | ASSLGTGNSGNTLY | 0.89153 |
| AGSGALGL | ASSLGTGNSGNTLY | 0.6441701 |
| AQIADQI | ASSLGTGNSGNTLY | 0.89053786 |
| ASAVAVTYL | ASSLGTGNSGNTLY | 0.51592743 |
| ASTVLGGTGG | ASSLGTGNSGNTLY | 0.8242184 |
| AVGGEPAK | ASSLGTGNSGNTLY | 0.86863947 |
| CEQGTFGGTC | ASSLGTGNSGNTLY | 0.6244614 |
| DEGKAPAG | ASSLGTGNSGNTLY | 0.8213613 |
| DTVVFPAL | ASSLGTGNSGNTLY | 0.7183313 |

| | | |
|---|---|---|
| FEGITMAG | ASSLGTGNSGNTLY | 0.7242575 |
| FPLVVQD | ASSLGTGNSGNTLY | 0.8754264 |
| GAFTTPQ | ASSLGTGNSGNTLY | 0.59669507 |
| GFGGFGTAGGVGGA | ASSLGTGNSGNTLY | 0.8344367 |
| GFLAQFL | ASSLGTGNSGNTLY | 0.66865605 |
| GGAGGAGGNAGVA | ASSLGTGNSGNTLY | 0.51268935 |
| GGVGTAGAGGAGGAGG | ASSLGTGNSGNTLY | 0.81988394 |
| GNGGGGDAGNAGSGGN | ASSLGTGNSGNTLY | 0.8755609 |
| GTGGAGGAGNDAGS | ASSLGTGNSGNTLY | 0.80753857 |
| GTTNTGWFNS | ASSLGTGNSGNTLY | 0.892231 |
| IFMGPLFTL | ASSLGTGNSGNTLY | 0.63817596 |
| ILAHAIEAI | ASSLGTGNSGNTLY | 0.8910654 |
| IRSAIREAL | ASSLGTGNSGNTLY | 0.88316613 |
| ITFAGGG | ASSLGTGNSGNTLY | 0.8291848 |
| ITGMSGHQEV | ASSLGTGNSGNTLY | 0.7358193 |
| IVAGLACTL | ASSLGTGNSGNTLY | 0.8688227 |
| IVGGGLAGA | ASSLGTGNSGNTLY | 0.64305526 |
| IVTGVLNI | ASSLGTGNSGNTLY | 0.86504304 |
| LAQQGCI | ASSLGTGNSGNTLY | 0.8829304 |
| LCLHVEN | ASSLGTGNSGNTLY | 0.36314496 |
| LDAISDGS | ASSLGTGNSGNTLY | 0.8154459 |
| LGGFAAML | ASSLGTGNSGNTLY | 0.747316 |
| LGGKNPMI | ASSLGTGNSGNTLY | 0.7487135 |
| LGPKGVAAIL | ASSLGTGNSGNTLY | 0.55041856 |
| LMVGVSI | ASSLGTGNSGNTLY | 0.32060015 |
| LVGAAFVL | ASSLGTGNSGNTLY | 0.66502213 |
| LVLAVAPL | ASSLGTGNSGNTLY | 0.4780991 |
| MAAIAVIDAA | ASSLGTGNSGNTLY | 0.8047602 |
| NNGINAELQ | ASSLGTGNSGNTLY | 0.79687774 |
| RGEPLVLVHG | ASSLGTGNSGNTLY | 0.5170084 |
| RGGAGGAAGLI | ASSLGTGNSGNTLY | 0.79021955 |
| RSYRMHPAV | ASSLGTGNSGNTLY | 0.81103027 |
| SAAHLLAGQLA | ASSLGTGNSGNTLY | 0.30436426 |
| SAAVGSY | ASSLGTGNSGNTLY | 0.61928785 |
| SPHAVLHRI | ASSLGTGNSGNTLY | 0.4906783 |
| SQIAKVL | ASSLGTGNSGNTLY | 0.5917732 |
| TALGHLVP | ASSLGTGNSGNTLY | 0.18052736 |
| TAPTTSGAA | ASSLGTGNSGNTLY | 0.7887999 |
| TDGDDGNGGSPGT | ASSLGTGNSGNTLY | 0.55954826 |
| TDGLGEN | ASSLGTGNSGNTLY | 0.15858328 |
| TGGPPLPGQA | ASSLGTGNSGNTLY | 0.705516 |
| TIPAITI | ASSLGTGNSGNTLY | 0.594746 |
| TIPGFTI | ASSLGTGNSGNTLY | 0.58399343 |
| TSIASGALA | ASSLGTGNSGNTLY | 0.79751384 |
| TVTLPGAAI | ASSLGTGNSGNTLY | 0.50454235 |
| VASGLSAALP | ASSLGTGNSGNTLY | 0.47444844 |
| VGAGVIGI | ASSLGTGNSGNTLY | 0.76969624 |
| VGPKRFGV | ASSLGTGNSGNTLY | 0.8181046 |
| VNVAFPD | ASSLGTGNSGNTLY | 0.39685732 |
| VSMTNSGVS | ASSLGTGNSGNTLY | 0.73877305 |
| VTDVLGDLSV | ASSLGTGNSGNTLY | 0.833827 |

| | | |
|---|---|---|
| VVVVGAGFAGL | ASSLGTGNSGNTLY | 0.68689215 |
| VYTVGTVV | ASSLGTGNSGNTLY | 0.82535285 |
| WTTFSGP | ASSLGTGNSGNTLY | 0.8363054 |
| YDELVEAF | ASSLGTGNSGNTLY | 0.61747056 |
| YGTGGAGGNGG | ASSLGTGNSGNTLY | 0.8090757 |
| YTNVMAV | ASSLGTGNSGNTLY | 0.81725144 |
| FLEQVHQL | ASSLGTGNSGNTLY | 0.5997292 |
| GLSLLAVAV | ASSLGTGNSGNTLY | 0.6576817 |
| LEQVHQL | ASSLGTGNSGNTLY | 0.40394408 |
| LNAAGAI | ASSLGTGNSGNTLY | 0.6468731 |
| PAVHASL | ASSLGTGNSGNTLY | 0.36149085 |
| TELEDDA | ASSLGTGNSGNTLY | 0.7294253 |
| TPPGPPN | ASSLGTGNSGNTLY | 0.81616074 |
| TPPPPPPP | ASSLGTGNSGNTLY | 0.6370375 |
| TQSPALI | ASSGQGAQDTQY | 0.3476599 |
| EFAAAFPQ | ASSGQGAQDTQY | 0.3230805 |
| LLSGNLDVL | ASSGQGAQDTQY | 0.7088531 |
| GPANLGNYNI | ASSGQGAQDTQY | 0.52994597 |
| TLGPTLP | ASSGQGAQDTQY | 0.3598409 |
| VYAYSSCCK | ASSGQGAQDTQY | 0.45318556 |
| APAPAPAPNI | ASSGQGAQDTQY | 0.79863644 |
| PGGPVPIP | ASSGQGAQDTQY | 0.8036339 |
| FLTVALEP | ASSGQGAQDTQY | 0.4192486 |
| SQYFGANL | ASSGQGAQDTQY | 0.83526146 |
| GGVKPAA | ASSGQGAQDTQY | 0.8032078 |
| GGTGGAGGAGGW | ASSGQGAQDTQY | 0.83920205 |
| GVGGVAGLGGAGG | ASSGQGAQDTQY | 0.7012458 |
| VLLVLATTI | ASSGQGAQDTQY | 0.3538515 |
| IGIFAAML | ASSGQGAQDTQY | 0.8196997 |
| AVGDLID | ASSGQGAQDTQY | 0.6147932 |
| NISPLPL | ASSGQGAQDTQY | 0.63974965 |
| VTAVFVAI | ASSGQGAQDTQY | 0.7581013 |
| SGDDNDAAL | ASSGQGAQDTQY | 0.63436025 |
| AVAALAHS | ASSGQGAQDTQY | 0.51908934 |
| GGVTVMATP | ASSGQGAQDTQY | 0.71544147 |
| AGLAANVRD | ASSGQGAQDTQY | 0.8230201 |
| TNRPDLIDPALLRPG | ASSGQGAQDTQY | 0.45729414 |
| LANLGVTAL | ASSGQGAQDTQY | 0.35927105 |
| TGERIEL | ASSGQGAQDTQY | 0.6591177 |
| NGEELAA | ASSGQGAQDTQY | 0.6768968 |
| VLSLVWG | ASSGQGAQDTQY | 0.8128616 |
| IGYALWL | ASSGQGAQDTQY | 0.73745894 |
| LSLLLAPL | ASSGQGAQDTQY | 0.6007465 |
| TLDEALK | ASSGQGAQDTQY | 0.76295376 |
| TDGAGTAEGG | ASSGQGAQDTQY | 0.7686945 |
| NAPGGAGGNGG | ASSGQGAQDTQY | 0.703747 |
| VSVFMGMGGNF | ASSGQGAQDTQY | 0.76038945 |
| LASYLAAL | ASSGQGAQDTQY | 0.50634027 |
| LVGGTPAL | ASSGQGAQDTQY | 0.81443524 |
| VRNADVL | ASSGQGAQDTQY | 0.7829725 |
| TCSSPFLVSAM | ASSGQGAQDTQY | 0.8063283 |

| | | |
|---|---|---|
| VHITDAE | ASSGQGAQDTQY | 0.82082355 |
| EYAAPSF | ASSGQGAQDTQY | 0.7146453 |
| VVDAFHGGL | ASSGQGAQDTQY | 0.7358724 |
| ASVVDPQ | ASSGQGAQDTQY | 0.31444845 |
| SYNEFGGLS | ASSGQGAQDTQY | 0.64875484 |
| DPDGSVY | ASSGQGAQDTQY | 0.69865656 |
| MVSTIHATE | ASSGQGAQDTQY | 0.74516237 |
| LAAGVADPV | ASSGQGAQDTQY | 0.7803136 |
| TKEQIAA | ASSGQGAQDTQY | 0.80258036 |
| SLASPAAL | ASSGQGAQDTQY | 0.53420025 |
| IGLVFYGV | ASSGQGAQDTQY | 0.8194165 |
| FLLDASNL | ASSGQGAQDTQY | 0.82007414 |
| YQVGNFP | ASSGQGAQDTQY | 0.3704782 |
| ESLLSQST | ASSGQGAQDTQY | 0.5666532 |
| LFTVGFAAVG | ASSGQGAQDTQY | 0.8295045 |
| ATPPTPT | ASSGQGAQDTQY | 0.7790005 |
| TDLATLG | ASSGQGAQDTQY | 0.5748674 |
| SAGGTGR | ASSGQGAQDTQY | 0.63491327 |
| GFSNSGS | ASSGQGAQDTQY | 0.84007657 |
| GGNGGSGGTSV | ASSGQGAQDTQY | 0.69172406 |
| GTSVATGGAGN | ASSGQGAQDTQY | 0.8279388 |
| NGGNGGSGGTS | ASSGQGAQDTQY | 0.24187702 |
| AVVKFPH | ASSGQGAQDTQY | 0.66861844 |
| TPPPPPP | ASSGQGAQDTQY | 0.5587107 |
| AAPAAAAAPA | ASSGQGAQDTQY | 0.1821325 |
| IALNEDL | ASSGQGAQDTQY | 0.7955456 |
| IVNVSSM | ASSGQGAQDTQY | 0.58556736 |
| TGIPPAPR | ASSGQGAQDTQY | 0.77269745 |
| FDLGGGTF | ASSGQGAQDTQY | 0.58766377 |
| FDLGGGTFDVS | ASSGQGAQDTQY | 0.7999506 |
| NEPTAAA | ASSGQGAQDTQY | 0.71616054 |
| NEPTAAALA | ASSGQGAQDTQY | 0.58740664 |
| GSIINISSI | ASSGQGAQDTQY | 0.40435487 |
| VAPGNAGTA | ASSGQGAQDTQY | 0.7619419 |
| TGGGASLE | ASSGQGAQDTQY | 0.8205085 |
| VAVYNHYKRI | ASSGQGAQDTQY | 0.8014351 |
| TDPAPATT | ASSGQGAQDTQY | 0.27732667 |
| TNRPDILDPALLRPG | ASSGQGAQDTQY | 0.7095914 |
| LTTTTTS | ASSGQGAQDTQY | 0.6808456 |
| PTGSGKTL | ASSGQGAQDTQY | 0.21431199 |
| TGSGKTL | ASSGQGAQDTQY | 0.20876014 |
| TIGHVDHGK | ASSGQGAQDTQY | 0.63231707 |
| TIGHVDHGKTT | ASSGQGAQDTQY | 0.5785443 |
| GHVDHGKTT | ASSGQGAQDTQY | 0.7773895 |
| AQFEHTLL | ASSGQGAQDTQY | 0.76748633 |
| IVNQDEL | ASSGQGAQDTQY | 0.57638067 |
| TNGSQFF | ASSGQGAQDTQY | 0.7692142 |
| TGKGIVE | ASSGQGAQDTQY | 0.7098293 |
| LISNASDA | ASSGQGAQDTQY | 0.12177592 |
| AELRTLGELTKIAKAHGAQ | ASSGQGAQDTQY | 0.5104056 |
| APNAVAAAEPSW | ASSGQGAQDTQY | 0.3122392 |

| | | |
|---|---|---|
| APQVAGAATAVL | ASSGQGAQDTQY | 0.8366461 |
| ARRDPAAAAAILPTDA | ASSGQGAQDTQY | 0.37325206 |
| DAAARDVLAEAGLAEYF | ASSGQGAQDTQY | 0.5544622 |
| DDQIVLSAKANLAKVEQ | ASSGQGAQDTQY | 0.6409718 |
| DELLPDTGEDSDEDGH | ASSGQGAQDTQY | 0.7833676 |
| ELPEAPANRATVPQD | ASSGQGAQDTQY | 0.695027 |
| FGQSKIERVTGIG | ASSGQGAQDTQY | 0.4697336 |
| GAGGTGSTAGGAGGAGGAGGLY | ASSGQGAQDTQY | 0.20926869 |
| GHGGTGAAGGNGGTGGAGGSGI | ASSGQGAQDTQY | 0.75245667 |
| GNGGNGGAGGEGGPGGAGGAGA | ASSGQGAQDTQY | 0.44340974 |
| GSTAGNGGNGGNATNGTIAGQP | ASSGQGAQDTQY | 0.4373566 |
| GTGGDGGNGGNGGTGG | ASSGQGAQDTQY | 0.3320297 |
| GVRRPAGIPADAPV | ASSGQGAQDTQY | 0.24716058 |
| GYRQVTDAHLLTLARR | ASSGQGAQDTQY | 0.3541282 |
| IAKMGQDGHDRG | ASSGQGAQDTQY | 0.80942863 |
| KAEPSIEIQYT | ASSGQGAQDTQY | 0.4995581 |
| KPVRIRMAADEDTVAVT | ASSGQGAQDTQY | 0.6192504 |
| MPPRIAGMRLLVIKPEP | ASSGQGAQDTQY | 0.84347 |
| MKFYAVALFRSRGVPPTR | ASSGQGAQDTQY | 0.75052667 |
| NAGGFGILGAVAHSP | ASSGQGAQDTQY | 0.60383296 |
| NEGREITLAMAEM | ASSGQGAQDTQY | 0.46135932 |
| NELVEWSHPLGHRWRWE | ASSGQGAQDTQY | 0.36912388 |
| NLGAGGTGGNGGNPG | ASSGQGAQDTQY | 0.45692843 |
| NTEWPKWETVMAADPVD | ASSGQGAQDTQY | 0.49968284 |
| PASAAPAETQP | ASSGQGAQDTQY | 0.8220147 |
| QQSSDGKNGVRAAEITGE | ASSGQGAQDTQY | 0.6892471 |
| RAPDAVTTSWWKE | ASSGQGAQDTQY | 0.38194495 |
| SGFGNVGAHTSGWFNQ | ASSGQGAQDTQY | 0.7807455 |
| SGVAGAVGQAASVGGLKV | ASSGQGAQDTQY | 0.83148015 |
| SNGGAGTGSAGGNGGT | ASSGQGAQDTQY | 0.8425437 |
| TGGAGGEGGAAG | ASSGQGAQDTQY | 0.37537044 |
| TGGDGGAAGAGGAPGHG | ASSGQGAQDTQY | 0.64905226 |
| TMPPLLAAIANAGRHGV | ASSGQGAQDTQY | 0.6637089 |
| TSSTAGNSGGA | ASSGQGAQDTQY | 0.6641082 |
| VPATVPVGILC | ASSGQGAQDTQY | 0.47655153 |
| VQRPDSGGVVLTGRIS | ASSGQGAQDTQY | 0.41775882 |
| WTPNPVPATKGVPAGLVQAT | ASSGQGAQDTQY | 0.7959995 |
| YALKRALAKFSDRFDVVII | ASSGQGAQDTQY | 0.85726625 |
| YLIGSVIV | ASSGQGAQDTQY | 0.62693584 |
| YGQLVAVID | ASSGQGAQDTQY | 0.27450597 |
| VTVGGMTLPA | ASSGQGAQDTQY | 0.61404073 |
| VLALPAV | ASSGQGAQDTQY | 0.51294297 |
| VIPPTLNYET | ASSGQGAQDTQY | 0.4691201 |
| VGMGGMHPSSG | ASSGQGAQDTQY | 0.6468444 |
| VGLAGGAATA | ASSGQGAQDTQY | 0.3147465 |
| VGGAGGAGGSGP | ASSGQGAQDTQY | 0.64559877 |
| VFAAFMV | ASSGQGAQDTQY | 0.8415375 |
| TLYDKLR | ASSGQGAQDTQY | 0.659191 |
| TLAVFTVL | ASSGQGAQDTQY | 0.7666863 |
| TGSVQPARV | ASSGQGAQDTQY | 0.8146305 |
| TGGGAGSGGT | ASSGQGAQDTQY | 0.7639702 |

| | | |
|---|---|---|
| SHLETSL | ASSGQGAQDTQY | 0.819717 |
| SFTTGMT | ASSGQGAQDTQY | 0.8129895 |
| QVEVAKRMGVRQPTVS | ASSGQGAQDTQY | 0.6000825 |
| QGSGGPALAM | ASSGQGAQDTQY | 0.7017029 |
| MLAPSNP | ASSGQGAQDTQY | 0.794707 |
| LVNNAAI | ASSGQGAQDTQY | 0.6258868 |
| LTGEINTLA | ASSGQGAQDTQY | 0.65162814 |
| LSIFGPV | ASSGQGAQDTQY | 0.61553717 |
| LLALLANL | ASSGQGAQDTQY | 0.79644006 |
| LISNASDALDKLR | ASSGQGAQDTQY | 0.6287643 |
| LINEAVP | ASSGQGAQDTQY | 0.8405236 |
| LGYAFSGL | ASSGQGAQDTQY | 0.2534229 |
| LDFVAVAL | ASSGQGAQDTQY | 0.39119473 |
| LAQPGAE | ASSGQGAQDTQY | 0.8121835 |
| ISNLGAVTGF | ASSGQGAQDTQY | 0.15064324 |
| HQEDPGRI | ASSGQGAQDTQY | 0.25238872 |
| GTPGEPTP | ASSGQGAQDTQY | 0.6930252 |
| GFFNSGS | ASSGQGAQDTQY | 0.81952786 |
| GAGGGGLTAGAN | ASSGQGAQDTQY | 0.15467566 |
| FVDALVNM | ASSGQGAQDTQY | 0.5612663 |
| FLTVAPEL | ASSGQGAQDTQY | 0.32342482 |
| FETLTAL | ASSGQGAQDTQY | 0.79995024 |
| DVINPIEL | ASSGQGAQDTQY | 0.65037155 |
| APVSGAPV | ASSGQGAQDTQY | 0.28294516 |
| ALVGGLAW | ASSGQGAQDTQY | 0.7945791 |
| ALAAAAATAI | ASSGQGAQDTQY | 0.2760893 |
| AFGLGLLL | ASSGQGAQDTQY | 0.46910754 |
| AFFAAASL | ASSGQGAQDTQY | 0.79686534 |
| AAAALPAFSPP | ASSGQGAQDTQY | 0.83844125 |
| VVLAPAL | ASSGQGAQDTQY | 0.32461175 |
| TIGHVDHGKTTL | ASSGQGAQDTQY | 0.7739388 |
| LISNASDALDK | ASSGQGAQDTQY | 0.65301967 |
| LISNASDAL | ASSGQGAQDTQY | 0.8500101 |
| ISNASDALDK | ASSGQGAQDTQY | 0.58731115 |
| NQAETLVYQTE | ASSGQGAQDTQY | 0.31525016 |
| TGGTGGTAGAG | ASSGQGAQDTQY | 0.8098334 |
| TGSTAGGAGGA | ASSGQGAQDTQY | 0.7966408 |
| DPLAVPVAAGPV | ASSGQGAQDTQY | 0.7049096 |
| GGTGGTPGNGGN | ASSGQGAQDTQY | 0.8058226 |
| PAGLAGVDPATL | ASSGQGAQDTQY | 0.6436733 |
| GNNLVVEAPHAGASV | ASSGQGAQDTQY | 0.61810005 |
| KVTRIMSASTLEQPA | ASSGQGAQDTQY | 0.7879175 |
| LGFALPIALPRRTVV | ASSGQGAQDTQY | 0.7516454 |
| TRLGNLTARKLVVLV | ASSGQGAQDTQY | 0.4696312 |
| WSQQRRLIYFPSAGP | ASSGQGAQDTQY | 0.82298756 |
| GPNGNPDPMAAAVDIR | ASSGQGAQDTQY | 0.6880983 |
| TQRCVSVADLRAAASR | ASSGQGAQDTQY | 0.41765696 |
| QFVLDVAEHPVEVLTETV | ASSGQGAQDTQY | 0.84830296 |
| GGAGGTGGAGGNGGTQAGDG | ASSGQGAQDTQY | 0.7163627 |
| PLLATLRAIAVDVDAKPAQV | ASSGQGAQDTQY | 0.7814008 |
| APKFRTGGYMIDFWGVGYQVAK | ASSGQGAQDTQY | 0.78375757 |

| | | |
|---|---|---|
| GAGGAGGAGGAAGGPGGTGGTG | ASSGQGAQDTQY | 0.85009253 |
| IGALGLLATDVAPAGSVAFGVA | ASSGQGAQDTQY | 0.38884324 |
| AGGAGGIGGNGGAGGAGGVLMG | ASSGQGAQDTQY | 0.77234244 |
| ADELIGLL | ASSGQGAQDTQY | 0.70357823 |
| ADRALKA | ASSGQGAQDTQY | 0.7933606 |
| AGAVGQVGGAGGA | ASSGQGAQDTQY | 0.71234775 |
| AGGGGGLGGAP | ASSGQGAQDTQY | 0.6115037 |
| AGGLLYGNGGAG | ASSGQGAQDTQY | 0.7680323 |
| AGLESIGL | ASSGQGAQDTQY | 0.8177146 |
| AGLKTGII | ASSGQGAQDTQY | 0.6543225 |
| ASLAALLL | ASSGQGAQDTQY | 0.7328055 |
| ASVINISSI | ASSGQGAQDTQY | 0.8309458 |
| ATPPGPA | ASSGQGAQDTQY | 0.40653592 |
| ATVASALV | ASSGQGAQDTQY | 0.6693815 |
| ATVASLIL | ASSGQGAQDTQY | 0.78998697 |
| AVNGPVLIRH | ASSGQGAQDTQY | 0.813081 |
| DGAAGAVGI | ASSGQGAQDTQY | 0.67174673 |
| DPLLHGI | ASSGQGAQDTQY | 0.8050176 |
| ELGAALGAL | ASSGQGAQDTQY | 0.5853307 |
| ELGRTQAV | ASSGQGAQDTQY | 0.7368146 |
| FVAGGYLS | ASSGQGAQDTQY | 0.6447277 |
| GASGGAGGAGGW | ASSGQGAQDTQY | 0.66450834 |
| GGAGAGGDGGAGGI | ASSGQGAQDTQY | 0.42365962 |
| GTGHNGSQPGMGGNG | ASSGQGAQDTQY | 0.7006765 |
| GYAAASASAS | ASSGQGAQDTQY | 0.78793144 |
| HQVDVAAVQQG | ASSGQGAQDTQY | 0.7480863 |
| IGAAIAHT | ASSGQGAQDTQY | 0.2159126 |
| IGMIEVAQL | ASSGQGAQDTQY | 0.75033605 |
| IRLAPEHTSAMD | ASSGQGAQDTQY | 0.75394726 |
| IYNVSVL | ASSGQGAQDTQY | 0.85430276 |
| KQGFLGM | ASSGQGAQDTQY | 0.65423965 |
| LESVITQL | ASSGQGAQDTQY | 0.8095201 |
| LGLGALVVV | ASSGQGAQDTQY | 0.52068233 |
| LIGNGGNGGSGGI | ASSGQGAQDTQY | 0.6991239 |
| LNAINAPTL | ASSGQGAQDTQY | 0.34107396 |
| LQLLNTGV | ASSGQGAQDTQY | 0.5924393 |
| LQTSIEF | ASSGQGAQDTQY | 0.59455585 |
| MSFAVSEL | ASSGQGAQDTQY | 0.46059805 |
| NTGALGL | ASSGQGAQDTQY | 0.80864 |
| PPGAGGAGGAGTT | ASSGQGAQDTQY | 0.8028867 |
| QLVGLGI | ASSGQGAQDTQY | 0.8124359 |
| RFLEQVHQL | ASSGQGAQDTQY | 0.816965 |
| RGVGYVLR | ASSGQGAQDTQY | 0.835287 |
| SATGVFAGVI | ASSGQGAQDTQY | 0.81779087 |
| SLPAITLP | ASSGQGAQDTQY | 0.5676861 |
| STALAFTV | ASSGQGAQDTQY | 0.6385661 |
| SVLTVGLGK | ASSGQGAQDTQY | 0.3126017 |
| TALAGAGSGI | ASSGQGAQDTQY | 0.62131894 |
| TGLPFTGI | ASSGQGAQDTQY | 0.6038625 |
| VAEHVTV | ASSGQGAQDTQY | 0.68910825 |
| VDIVLNSL | ASSGQGAQDTQY | 0.14608368 |

| | | |
|---|---|---|
| VLTGGGAVL | ASSGQGAQDTQY | 0.76076007 |
| VNVGLAYGI | ASSGQGAQDTQY | 0.82403874 |
| VSSAAVGAV | ASSGQGAQDTQY | 0.58043844 |
| VTSVYEAGV | ASSGQGAQDTQY | 0.7570054 |
| WVALDRGA | ASSGQGAQDTQY | 0.78263843 |
| YLVGSVVI | ASSGQGAQDTQY | 0.55575454 |
| GPPGTGKTLL | ASSGQGAQDTQY | 0.7793639 |
| GVGMTKF | ASSGQGAQDTQY | 0.55217683 |
| IGIGGSDL | ASSGQGAQDTQY | 0.79626 |
| IGTLLAEL | ASSGQGAQDTQY | 0.832902 |
| IYTSGTT | ASSGQGAQDTQY | 0.730178 |
| LSFPAGV | ASSGQGAQDTQY | 0.71843386 |
| LYTSGTT | ASSGQGAQDTQY | 0.72297573 |
| VGAFPFGF | ASSGQGAQDTQY | 0.33278507 |
| VGAGTVEF | ASSGQGAQDTQY | 0.72883415 |
| VGKSTLLL | ASSGQGAQDTQY | 0.14566946 |
| AALAHAGGGIV | ASSGQGAQDTQY | 0.71373594 |
| AALVAVSLL | ASSGQGAQDTQY | 0.4651361 |
| AAVSRAVAAL | ASSGQGAQDTQY | 0.71831006 |
| AGGSSGIGGF | ASSGQGAQDTQY | 0.6831708 |
| AGLTVANANG | ASSGQGAQDTQY | 0.7236648 |
| AGLVGAVL | ASSGQGAQDTQY | 0.4360927 |
| ALSQAFTV | ASSGQGAQDTQY | 0.7459189 |
| ALTFSVSL | ASSGQGAQDTQY | 0.68203485 |
| ALTGAALVAL | ASSGQGAQDTQY | 0.37927112 |
| ASFSIGAGVL | ASSGQGAQDTQY | 0.8053535 |
| DGSVYKI | ASSGQGAQDTQY | 0.6474147 |
| EQLAEIV | ASSGQGAQDTQY | 0.63755786 |
| FAVSELTVM | ASSGQGAQDTQY | 0.8107235 |
| GAAPLAAGL | ASSGQGAQDTQY | 0.6959276 |
| GADGGDGGSGGAGG | ASSGQGAQDTQY | 0.2904854 |
| GFGLDTGGAGG | ASSGQGAQDTQY | 0.6386632 |
| GLASGGLL | ASSGQGAQDTQY | 0.7062495 |
| GRIAGAAVLL | ASSGQGAQDTQY | 0.6099057 |
| GSAGSLNTSF | ASSGQGAQDTQY | 0.16517185 |
| GTGQIGI | ASSGQGAQDTQY | 0.72718257 |
| GTLIALIL | ASSGQGAQDTQY | 0.42819732 |
| IAATMGFVVG | ASSGQGAQDTQY | 0.46990228 |
| IFAQALL | ASSGQGAQDTQY | 0.57155806 |
| IGAAVALGF | ASSGQGAQDTQY | 0.79834783 |
| IGLTPLAL | ASSGQGAQDTQY | 0.63853544 |
| ILGSNTSTL | ASSGQGAQDTQY | 0.68220526 |
| IVFSLTVL | ASSGQGAQDTQY | 0.76741254 |
| LASKATGF | ASSGQGAQDTQY | 0.73879135 |
| LASVALIL | ASSGQGAQDTQY | 0.8193829 |
| LFAVGLTL | ASSGQGAQDTQY | 0.7324907 |
| LFNGGIGTYI | ASSGQGAQDTQY | 0.7069634 |
| LTLGAGII | ASSGQGAQDTQY | 0.73703706 |
| LVAAALGDI | ASSGQGAQDTQY | 0.57997614 |
| NTAFVQLGI | ASSGQGAQDTQY | 0.3857587 |
| QLFSGAEL | ASSGQGAQDTQY | 0.37778157 |

| | | |
|---|---|---|
| SAATFITL | ASSGQGAQDTQY | 0.47762477 |
| SNYGLANLGV | ASSGQGAQDTQY | 0.2864073 |
| SYGGAYAVM | ASSGQGAQDTQY | 0.73517287 |
| TTGVGGLTLG | ASSGQGAQDTQY | 0.8374909 |
| TTQTIYI | ASSGQGAQDTQY | 0.66113544 |
| VAAGAGLGAVI | ASSGQGAQDTQY | 0.3174048 |
| VAAVVTLL | ASSGQGAQDTQY | 0.61099565 |
| VGFLGGLI | ASSGQGAQDTQY | 0.5311499 |
| VGVIPILL | ASSGQGAQDTQY | 0.7606628 |
| VIFASSLI | ASSGQGAQDTQY | 0.79890096 |
| VKGGDGGAAAG | ASSGQGAQDTQY | 0.84363425 |
| VTKTFTNI | ASSGQGAQDTQY | 0.78457993 |
| VVAAAILL | ASSGQGAQDTQY | 0.5637786 |
| YAALDVEL | ASSGQGAQDTQY | 0.7746682 |
| LASVALLL | ASSGQGAQDTQY | 0.6437011 |
| LDAVHRV | ASSGQGAQDTQY | 0.742265 |
| QGLSSSIF | ASSGQGAQDTQY | 0.67026895 |
| RGTFANIRL | ASSGQGAQDTQY | 0.733484 |
| RQLWWGHRI | ASSGQGAQDTQY | 0.5806409 |
| YGPPGTGKTLL | ASSGQGAQDTQY | 0.50194156 |
| AALVGAPANVA | ASSGQGAQDTQY | 0.29051602 |
| AATFRRTE | ASSGQGAQDTQY | 0.788661 |
| AGALGGGI | ASSGQGAQDTQY | 0.59656703 |
| AGGAGGAGGIGAG | ASSGQGAQDTQY | 0.44032317 |
| AGSGALGL | ASSGQGAQDTQY | 0.57938206 |
| AQIADQI | ASSGQGAQDTQY | 0.8349422 |
| ASAVAVTYL | ASSGQGAQDTQY | 0.8337586 |
| ASTVLGGTGG | ASSGQGAQDTQY | 0.5882897 |
| AVGGEPAK | ASSGQGAQDTQY | 0.509851 |
| CEQGTFGGTC | ASSGQGAQDTQY | 0.77393514 |
| DEGKAPAG | ASSGQGAQDTQY | 0.44270653 |
| DTVVFPAL | ASSGQGAQDTQY | 0.784179 |
| FEGITMAG | ASSGQGAQDTQY | 0.6913644 |
| FPLVVQD | ASSGQGAQDTQY | 0.5911765 |
| GAFTTPQ | ASSGQGAQDTQY | 0.59547836 |
| GFGGFGTAGGVGGA | ASSGQGAQDTQY | 0.7549627 |
| GFLAQFL | ASSGQGAQDTQY | 0.5839248 |
| GGAGGAGGNAGVA | ASSGQGAQDTQY | 0.796817 |
| GGVGTAGAGGAGGAGG | ASSGQGAQDTQY | 0.15806733 |
| GNGGGGDAGNAGSGGN | ASSGQGAQDTQY | 0.7823659 |
| GTGGAGGAGNDAGS | ASSGQGAQDTQY | 0.43613377 |
| GTTNTGWFNS | ASSGQGAQDTQY | 0.43380773 |
| IFMGPLFTL | ASSGQGAQDTQY | 0.213638 |
| ILAHAIEAI | ASSGQGAQDTQY | 0.66750824 |
| IRSAIREAL | ASSGQGAQDTQY | 0.7849329 |
| ITFAGGG | ASSGQGAQDTQY | 0.8179256 |
| ITGMSGHQEV | ASSGQGAQDTQY | 0.599662 |
| IVAGLACTL | ASSGQGAQDTQY | 0.7695928 |
| IVGGGLAGA | ASSGQGAQDTQY | 0.82075155 |
| IVTGVLNI | ASSGQGAQDTQY | 0.55411386 |
| LAQQGCI | ASSGQGAQDTQY | 0.23474002 |

| | | |
|---|---|---|
| LCLHVEN | ASSGQGAQDTQY | 0.73731303 |
| LDAISDGS | ASSGQGAQDTQY | 0.71242857 |
| LGGFAAML | ASSGQGAQDTQY | 0.80955267 |
| LGGKNPMI | ASSGQGAQDTQY | 0.77805257 |
| LGPKGVAAIL | ASSGQGAQDTQY | 0.3018373 |
| LMVGVSI | ASSGQGAQDTQY | 0.8193182 |
| LVGAAFVL | ASSGQGAQDTQY | 0.7876227 |
| LVLAVAPL | ASSGQGAQDTQY | 0.39534307 |
| MAAIAVIDAA | ASSGQGAQDTQY | 0.49347523 |
| NNGINAELQ | ASSGQGAQDTQY | 0.4976259 |
| RGEPLVLVHG | ASSGQGAQDTQY | 0.28604752 |
| RGGAGGAAGLI | ASSGQGAQDTQY | 0.77130973 |
| RSYRMHPAV | ASSGQGAQDTQY | 0.58299565 |
| SAAHLLAGQLA | ASSGQGAQDTQY | 0.59521735 |
| SAAVGSY | ASSGQGAQDTQY | 0.65937006 |
| SPHAVLHRI | ASSGQGAQDTQY | 0.53250545 |
| SQIAKVL | ASSGQGAQDTQY | 0.5249302 |
| TALGHLVP | ASSGQGAQDTQY | 0.59440935 |
| TAPTTSGAA | ASSGQGAQDTQY | 0.7044333 |
| TDGDDGNGGSPGT | ASSGQGAQDTQY | 0.78066695 |
| TDGLGEN | ASSGQGAQDTQY | 0.65368825 |
| TGGPPLPGQA | ASSGQGAQDTQY | 0.7473501 |
| TIPAITI | ASSGQGAQDTQY | 0.8171364 |
| TIPGFTI | ASSGQGAQDTQY | 0.36633694 |
| TSIASGALA | ASSGQGAQDTQY | 0.48228937 |
| TVTLPGAAI | ASSGQGAQDTQY | 0.76943743 |
| VASGLSAALP | ASSGQGAQDTQY | 0.55231094 |
| VGAGVIGI | ASSGQGAQDTQY | 0.8047727 |
| VGPKRFGV | ASSGQGAQDTQY | 0.7168302 |
| VNVAFPD | ASSGQGAQDTQY | 0.7245658 |
| VSMTNSGVS | ASSGQGAQDTQY | 0.70456725 |
| VTDVLGDLSV | ASSGQGAQDTQY | 0.83702433 |
| VVVVGAGFAGL | ASSGQGAQDTQY | 0.77547127 |
| VYTVGTVV | ASSGQGAQDTQY | 0.67494327 |
| WTTFSGP | ASSGQGAQDTQY | 0.785159 |
| YDELVEAF | ASSGQGAQDTQY | 0.40542722 |
| YGTGGAGGNGG | ASSGQGAQDTQY | 0.77546775 |
| YTNVMAV | ASSGQGAQDTQY | 0.78626907 |
| FLEQVHQL | ASSGQGAQDTQY | 0.41309267 |
| GLSLLAVAV | ASSGQGAQDTQY | 0.6293516 |
| LEQVHQL | ASSGQGAQDTQY | 0.8229559 |
| LNAAGAI | ASSGQGAQDTQY | 0.8040118 |
| PAVHASL | ASSGQGAQDTQY | 0.40198588 |
| TELEDDA | ASSGQGAQDTQY | 0.49782997 |
| TPPGPPN | ASSGQGAQDTQY | 0.35644782 |
| TPPPPPPP | ASSGQGAQDTQY | 0.57462883 |
| TQSPALI | ASSLEGTGWSDYT | 0.30059174 |
| EFAAAFPQ | ASSLEGTGWSDYT | 0.75845265 |
| LLSGNLDVL | ASSLEGTGWSDYT | 0.54817325 |
| GPANLGNYNI | ASSLEGTGWSDYT | 0.8128418 |
| TLGPTLP | ASSLEGTGWSDYT | 0.7268865 |

| | | |
|---|---|---|
| VYAYSSCCK | ASSLEGTGWSDYT | 0.8104576 |
| APAPAPAPNI | ASSLEGTGWSDYT | 0.54072696 |
| PGGPVPIP | ASSLEGTGWSDYT | 0.5666119 |
| FLTVALEP | ASSLEGTGWSDYT | 0.29437846 |
| SQYFGANL | ASSLEGTGWSDYT | 0.37778258 |
| GGVKPAA | ASSLEGTGWSDYT | 0.46293694 |
| GGTGGAGGAGGW | ASSLEGTGWSDYT | 0.62912405 |
| GVGGVAGLGGAGG | ASSLEGTGWSDYT | 0.2136853 |
| VLLVLATTI | ASSLEGTGWSDYT | 0.829903 |
| IGIFAAML | ASSLEGTGWSDYT | 0.6730113 |
| AVGDLID | ASSLEGTGWSDYT | 0.76534796 |
| NISPLPL | ASSLEGTGWSDYT | 0.5727223 |
| VTAVFVAI | ASSLEGTGWSDYT | 0.45071143 |
| SGDDNDAAL | ASSLEGTGWSDYT | 0.5644258 |
| AVAALAHS | ASSLEGTGWSDYT | 0.7877155 |
| GGVTVMATP | ASSLEGTGWSDYT | 0.31349623 |
| AGLAANVRD | ASSLEGTGWSDYT | 0.23506773 |
| TNRPDLIDPALLRPG | ASSLEGTGWSDYT | 0.47451144 |
| LANLGVTAL | ASSLEGTGWSDYT | 0.37747157 |
| TGERIEL | ASSLEGTGWSDYT | 0.8486506 |
| NGEELAA | ASSLEGTGWSDYT | 0.8084999 |
| VLSLVWG | ASSLEGTGWSDYT | 0.7218019 |
| IGYALWL | ASSLEGTGWSDYT | 0.8425664 |
| LSLLLAPL | ASSLEGTGWSDYT | 0.17700851 |
| TLDEALK | ASSLEGTGWSDYT | 0.5861623 |
| TDGAGTAEGG | ASSLEGTGWSDYT | 0.83539927 |
| NAPGGAGGNGG | ASSLEGTGWSDYT | 0.7539923 |
| VSVFMGMGGNF | ASSLEGTGWSDYT | 0.59921944 |
| LASYLAAL | ASSLEGTGWSDYT | 0.7958353 |
| LVGGTPAL | ASSLEGTGWSDYT | 0.49852812 |
| VRNADVL | ASSLEGTGWSDYT | 0.5989455 |
| TCSSPFLVSAM | ASSLEGTGWSDYT | 0.65917575 |
| VHITDAE | ASSLEGTGWSDYT | 0.7591451 |
| EYAAPSF | ASSLEGTGWSDYT | 0.72055626 |
| VVDAFHGGL | ASSLEGTGWSDYT | 0.29315948 |
| ASVVDPQ | ASSLEGTGWSDYT | 0.7813333 |
| SYNEFGGLS | ASSLEGTGWSDYT | 0.84945303 |
| DPDGSVY | ASSLEGTGWSDYT | 0.5913389 |
| MVSTIHATE | ASSLEGTGWSDYT | 0.7592483 |
| LAAGVADPV | ASSLEGTGWSDYT | 0.73593867 |
| TKEQIAA | ASSLEGTGWSDYT | 0.783535 |
| SLASPAAL | ASSLEGTGWSDYT | 0.46069044 |
| IGLVFYGV | ASSLEGTGWSDYT | 0.40505558 |
| FLLDASNL | ASSLEGTGWSDYT | 0.7446872 |
| YQVGNFP | ASSLEGTGWSDYT | 0.39721075 |
| ESLLSQST | ASSLEGTGWSDYT | 0.68874186 |
| LFTVGFAAVG | ASSLEGTGWSDYT | 0.6511407 |
| ATPPTPT | ASSLEGTGWSDYT | 0.8582138 |
| TDLATLG | ASSLEGTGWSDYT | 0.75303745 |
| SAGGTGR | ASSLEGTGWSDYT | 0.7332263 |
| GFSNSGS | ASSLEGTGWSDYT | 0.4670263 |

| | | |
|---|---|---|
| GGNGGSGGTSV | ASSLEGTGWSDYT | 0.7285938 |
| GTSVATGGAGN | ASSLEGTGWSDYT | 0.77155924 |
| NGGNGGSGGTS | ASSLEGTGWSDYT | 0.32086286 |
| AVVKFPH | ASSLEGTGWSDYT | 0.8155334 |
| TPPPPPP | ASSLEGTGWSDYT | 0.4353215 |
| AAPAAAAAPA | ASSLEGTGWSDYT | 0.69803685 |
| IALNEDL | ASSLEGTGWSDYT | 0.7583827 |
| IVNVSSM | ASSLEGTGWSDYT | 0.82626337 |
| TGIPPAPR | ASSLEGTGWSDYT | 0.6213134 |
| FDLGGGTF | ASSLEGTGWSDYT | 0.727275 |
| FDLGGGTFDVS | ASSLEGTGWSDYT | 0.3855401 |
| NEPTAAA | ASSLEGTGWSDYT | 0.7940906 |
| NEPTAAALA | ASSLEGTGWSDYT | 0.8087089 |
| GSIINISSI | ASSLEGTGWSDYT | 0.7495732 |
| VAPGNAGTA | ASSLEGTGWSDYT | 0.8442174 |
| TGGGASLE | ASSLEGTGWSDYT | 0.7709129 |
| VAVYNHYKRI | ASSLEGTGWSDYT | 0.25482455 |
| TDPAPATT | ASSLEGTGWSDYT | 0.64141154 |
| TNRPDILDPALLRPG | ASSLEGTGWSDYT | 0.814802 |
| LTTTTTS | ASSLEGTGWSDYT | 0.8211735 |
| PTGSGKTL | ASSLEGTGWSDYT | 0.8095942 |
| TGSGKTL | ASSLEGTGWSDYT | 0.7738483 |
| TIGHVDHGK | ASSLEGTGWSDYT | 0.6219045 |
| TIGHVDHGKTT | ASSLEGTGWSDYT | 0.82774305 |
| GHVDHGKTT | ASSLEGTGWSDYT | 0.86190975 |
| AQFEHTLL | ASSLEGTGWSDYT | 0.79358786 |
| IVNQDEL | ASSLEGTGWSDYT | 0.5115855 |
| TNGSQFF | ASSLEGTGWSDYT | 0.48849398 |
| TGKGIVE | ASSLEGTGWSDYT | 0.7561031 |
| LISNASDA | ASSLEGTGWSDYT | 0.7961693 |
| AELRTLGELTKIAKAHGAQ | ASSLEGTGWSDYT | 0.4046855 |
| APNAVAAAEPSW | ASSLEGTGWSDYT | 0.688079 |
| APQVAGAATAVL | ASSLEGTGWSDYT | 0.7148497 |
| ARRDPAAAAAILPTDA | ASSLEGTGWSDYT | 0.24045824 |
| DAAARDVLAEAGLAEYF | ASSLEGTGWSDYT | 0.7683241 |
| DDQIVLSAKANLAKVEQ | ASSLEGTGWSDYT | 0.81871545 |
| DELLPDTGEDSDEDGH | ASSLEGTGWSDYT | 0.74165994 |
| ELPEAPANRATVPQD | ASSLEGTGWSDYT | 0.44901416 |
| FGQSKIERVTGIG | ASSLEGTGWSDYT | 0.8168086 |
| GAGGTGSTAGGAGGAGGAGGLY | ASSLEGTGWSDYT | 0.8404794 |
| GHGGTGAAGGNGGTGGAGGSGI | ASSLEGTGWSDYT | 0.84254473 |
| GNGGNGGAGGEGGPGGAGGAGA | ASSLEGTGWSDYT | 0.27602035 |
| GSTAGNGGNGGNATNGTIAGQP | ASSLEGTGWSDYT | 0.17646787 |
| GTGGDGGNGGNGGTGG | ASSLEGTGWSDYT | 0.6332481 |
| GVRRPAGIPADAPV | ASSLEGTGWSDYT | 0.8051081 |
| GYRQVTDAHLLTLARR | ASSLEGTGWSDYT | 0.6247646 |
| IAKMGQDGHDRG | ASSLEGTGWSDYT | 0.55959535 |
| KAEPSIEIQYT | ASSLEGTGWSDYT | 0.7407617 |
| KPVRIRMAADEDTVAVT | ASSLEGTGWSDYT | 0.8029636 |
| MPPRIAGMRLLVIKPEP | ASSLEGTGWSDYT | 0.39665496 |
| MKFYAVALFRSRGVPPTR | ASSLEGTGWSDYT | 0.3600033 |

| | | |
|---|---|---|
| NAGGFGILGAVAHSP | ASSLEGTGWSDYT | 0.79080164 |
| NEGREITLAMAEM | ASSLEGTGWSDYT | 0.800043 |
| NELVEWSHPLGHRWRWE | ASSLEGTGWSDYT | 0.40259206 |
| NLGAGGTGGNGGNPG | ASSLEGTGWSDYT | 0.599052 |
| NTEWPKWETVMAADPVD | ASSLEGTGWSDYT | 0.6908921 |
| PASAAPAETQP | ASSLEGTGWSDYT | 0.7120116 |
| QQSSDGKNGVRAAEITGE | ASSLEGTGWSDYT | 0.7372569 |
| RAPDAVTTSWWKE | ASSLEGTGWSDYT | 0.3759211 |
| SGFGNVGAHTSGWFNQ | ASSLEGTGWSDYT | 0.78578156 |
| SGVAGAVGQAASVGGLKV | ASSLEGTGWSDYT | 0.32208696 |
| SNGGAGTGSAGGNGGT | ASSLEGTGWSDYT | 0.7779345 |
| TGGAGGEGGAAG | ASSLEGTGWSDYT | 0.6985464 |
| TGGDGGAAGAGGAPGHG | ASSLEGTGWSDYT | 0.7385989 |
| TMPPLLAAIANAGRHGV | ASSLEGTGWSDYT | 0.5744461 |
| TSSTAGNSGGA | ASSLEGTGWSDYT | 0.44966245 |
| VPATVPVGILC | ASSLEGTGWSDYT | 0.39986184 |
| VQRPDSGGVVLTGRIS | ASSLEGTGWSDYT | 0.35920772 |
| WTPNPVPATKGVPAGLVQAT | ASSLEGTGWSDYT | 0.806538 |
| YALKRALAKFSDRFDVVII | ASSLEGTGWSDYT | 0.17303656 |
| YLIGSVIV | ASSLEGTGWSDYT | 0.79591894 |
| YGQLVAVID | ASSLEGTGWSDYT | 0.70789796 |
| VTVGGMTLPA | ASSLEGTGWSDYT | 0.38089374 |
| VLALPAV | ASSLEGTGWSDYT | 0.7335719 |
| VIPPTLNYET | ASSLEGTGWSDYT | 0.36939067 |
| VGMGGMHPSSG | ASSLEGTGWSDYT | 0.65783966 |
| VGLAGGAATA | ASSLEGTGWSDYT | 0.78021383 |
| VGGAGGAGGSGP | ASSLEGTGWSDYT | 0.78791535 |
| VFAAFMV | ASSLEGTGWSDYT | 0.45184284 |
| TLYDKLR | ASSLEGTGWSDYT | 0.766289 |
| TLAVFTVL | ASSLEGTGWSDYT | 0.7552638 |
| TGSVQPARV | ASSLEGTGWSDYT | 0.45865393 |
| TGGGAGSGGT | ASSLEGTGWSDYT | 0.75387096 |
| SHLETSL | ASSLEGTGWSDYT | 0.78569555 |
| SFTTGMT | ASSLEGTGWSDYT | 0.74331516 |
| QVEVAKRMGVRQPTVS | ASSLEGTGWSDYT | 0.2689393 |
| QGSGGPALAM | ASSLEGTGWSDYT | 0.75246537 |
| MLAPSNP | ASSLEGTGWSDYT | 0.48915076 |
| LVNNAAI | ASSLEGTGWSDYT | 0.7313633 |
| LTGEINTLA | ASSLEGTGWSDYT | 0.61491 |
| LSIFGPV | ASSLEGTGWSDYT | 0.58661085 |
| LLALLANL | ASSLEGTGWSDYT | 0.7825395 |
| LISNASDALDKLR | ASSLEGTGWSDYT | 0.34674755 |
| LINEAVP | ASSLEGTGWSDYT | 0.5525948 |
| LGYAFSGL | ASSLEGTGWSDYT | 0.61720574 |
| LDFVAVAL | ASSLEGTGWSDYT | 0.67550683 |
| LAQPGAE | ASSLEGTGWSDYT | 0.7769519 |
| ISNLGAVTGF | ASSLEGTGWSDYT | 0.42707255 |
| HQEDPGRI | ASSLEGTGWSDYT | 0.7226918 |
| GTPGEPTP | ASSLEGTGWSDYT | 0.5990397 |
| GFFNSGS | ASSLEGTGWSDYT | 0.4221266 |
| GAGGGGLTAGAN | ASSLEGTGWSDYT | 0.74139917 |

| | | |
|---|---|---|
| FVDALVNM | ASSLEGTGWSDYT | 0.59017414 |
| FLTVAPEL | ASSLEGTGWSDYT | 0.3924811 |
| FETLTAL | ASSLEGTGWSDYT | 0.7591126 |
| DVINPIEL | ASSLEGTGWSDYT | 0.4569791 |
| APVSGAPV | ASSLEGTGWSDYT | 0.7565056 |
| ALVGGLAW | ASSLEGTGWSDYT | 0.8438964 |
| ALAAAAATAI | ASSLEGTGWSDYT | 0.40330255 |
| AFGLGLLL | ASSLEGTGWSDYT | 0.76761675 |
| AFFAAASL | ASSLEGTGWSDYT | 0.7257026 |
| AAAALPAFSPP | ASSLEGTGWSDYT | 0.722389 |
| VVLAPAL | ASSLEGTGWSDYT | 0.618801 |
| TIGHVDHGKTTL | ASSLEGTGWSDYT | 0.4347608 |
| LISNASDALDK | ASSLEGTGWSDYT | 0.8142891 |
| LISNASDAL | ASSLEGTGWSDYT | 0.7608743 |
| ISNASDALDK | ASSLEGTGWSDYT | 0.7787436 |
| NQAETLVYQTE | ASSLEGTGWSDYT | 0.7711771 |
| TGGTGGTAGAG | ASSLEGTGWSDYT | 0.79949445 |
| TGSTAGGAGGA | ASSLEGTGWSDYT | 0.79390085 |
| DPLAVPVAAGPV | ASSLEGTGWSDYT | 0.5168081 |
| GGTGGTPGNGGN | ASSLEGTGWSDYT | 0.3117513 |
| PAGLAGVDPATL | ASSLEGTGWSDYT | 0.5583153 |
| GNNLVVEAPHAGASV | ASSLEGTGWSDYT | 0.6509249 |
| KVTRIMSASTLEQPA | ASSLEGTGWSDYT | 0.5314958 |
| LGFALPIALPRRTVV | ASSLEGTGWSDYT | 0.29650703 |
| TRLGNLTARKLVVLV | ASSLEGTGWSDYT | 0.79222107 |
| WSQQRRLIYFPSAGP | ASSLEGTGWSDYT | 0.8482529 |
| GPNGNPDPMAAAVDIR | ASSLEGTGWSDYT | 0.5445943 |
| TQRCVSVADLRAAASR | ASSLEGTGWSDYT | 0.8544177 |
| QFVLDVAEHPVEVLTETV | ASSLEGTGWSDYT | 0.40150708 |
| GGAGGTGGAGGNGGTQAGDG | ASSLEGTGWSDYT | 0.7761972 |
| PLLATLRAIAVDVDAKPAQV | ASSLEGTGWSDYT | 0.8020791 |
| APKFRTGGYMIDFWGVGYQVAK | ASSLEGTGWSDYT | 0.4492966 |
| GAGGAGGAGGAAGGPGGTGGTG | ASSLEGTGWSDYT | 0.74784803 |
| IGALGLLATDVAPAGSVAFGVA | ASSLEGTGWSDYT | 0.40739787 |
| AGGAGGIGGNGGAGGAGGVLMG | ASSLEGTGWSDYT | 0.46341786 |
| ADELIGLL | ASSLEGTGWSDYT | 0.8006254 |
| ADRALKA | ASSLEGTGWSDYT | 0.6199797 |
| AGAVGQVGGAGGA | ASSLEGTGWSDYT | 0.733701 |
| AGGGGGLGGAP | ASSLEGTGWSDYT | 0.6293353 |
| AGGLLYGNGGAG | ASSLEGTGWSDYT | 0.46350724 |
| AGLESIGL | ASSLEGTGWSDYT | 0.6818913 |
| AGLKTGII | ASSLEGTGWSDYT | 0.7861115 |
| ASLAALLL | ASSLEGTGWSDYT | 0.68164164 |
| ASVINISSI | ASSLEGTGWSDYT | 0.8608248 |
| ATPPGPA | ASSLEGTGWSDYT | 0.43177187 |
| ATVASALV | ASSLEGTGWSDYT | 0.8313762 |
| ATVASLIL | ASSLEGTGWSDYT | 0.82684326 |
| AVNGPVLIRH | ASSLEGTGWSDYT | 0.70305824 |
| DGAAGAVGI | ASSLEGTGWSDYT | 0.54907125 |
| DPLLHGI | ASSLEGTGWSDYT | 0.7981931 |
| ELGAALGAL | ASSLEGTGWSDYT | 0.60615385 |

| | | |
|---|---|---|
| ELGRTQAV | ASSLEGTGWSDYT | 0.76905984 |
| FVAGGYLS | ASSLEGTGWSDYT | 0.8305484 |
| GASGGAGGAGGW | ASSLEGTGWSDYT | 0.45173883 |
| GGAGAGGDGGAGGI | ASSLEGTGWSDYT | 0.6531024 |
| GTGHNGSQPGMGGNG | ASSLEGTGWSDYT | 0.59607345 |
| GYAAASASAS | ASSLEGTGWSDYT | 0.55627966 |
| HQVDVAAVQQG | ASSLEGTGWSDYT | 0.50291765 |
| IGAAIAHT | ASSLEGTGWSDYT | 0.40813452 |
| IGMIEVAQL | ASSLEGTGWSDYT | 0.49480647 |
| IRLAPEHTSAMD | ASSLEGTGWSDYT | 0.5088246 |
| IYNVSVL | ASSLEGTGWSDYT | 0.63719547 |
| KQGFLGM | ASSLEGTGWSDYT | 0.6092879 |
| LESVITQL | ASSLEGTGWSDYT | 0.48194912 |
| LGLGALVVV | ASSLEGTGWSDYT | 0.6533333 |
| LIGNGGNGGSGGI | ASSLEGTGWSDYT | 0.65901357 |
| LNAINAPTL | ASSLEGTGWSDYT | 0.38494846 |
| LQLLNTGV | ASSLEGTGWSDYT | 0.5351762 |
| LQTSIEF | ASSLEGTGWSDYT | 0.44555598 |
| MSFAVSEL | ASSLEGTGWSDYT | 0.5309787 |
| NTGALGL | ASSLEGTGWSDYT | 0.32494122 |
| PPGAGGAGGAGTT | ASSLEGTGWSDYT | 0.62420106 |
| QLVGLGI | ASSLEGTGWSDYT | 0.45051944 |
| RFLEQVHQL | ASSLEGTGWSDYT | 0.3002863 |
| RGVGYVLR | ASSLEGTGWSDYT | 0.5520895 |
| SATGVFAGVI | ASSLEGTGWSDYT | 0.56576496 |
| SLPAITLP | ASSLEGTGWSDYT | 0.59638983 |
| STALAFTV | ASSLEGTGWSDYT | 0.638504 |
| SVLTVGLGK | ASSLEGTGWSDYT | 0.4313375 |
| TALAGAGSGI | ASSLEGTGWSDYT | 0.5036822 |
| TGLPFTGI | ASSLEGTGWSDYT | 0.6217251 |
| VAEHVTV | ASSLEGTGWSDYT | 0.65417284 |
| VDIVLNSL | ASSLEGTGWSDYT | 0.47720158 |
| VLTGGGAVL | ASSLEGTGWSDYT | 0.5916853 |
| VNVGLAYGI | ASSLEGTGWSDYT | 0.65063906 |
| VSSAAVGAV | ASSLEGTGWSDYT | 0.6011191 |
| VTSVYEAGV | ASSLEGTGWSDYT | 0.6284125 |
| WVALDRGA | ASSLEGTGWSDYT | 0.65037954 |
| YLVGSVVI | ASSLEGTGWSDYT | 0.42099607 |
| GPPGTGKTLL | ASSLEGTGWSDYT | 0.64375967 |
| GVGMTKF | ASSLEGTGWSDYT | 0.6541769 |
| IGIGGSDL | ASSLEGTGWSDYT | 0.5356925 |
| IGTLLAEL | ASSLEGTGWSDYT | 0.5482531 |
| IYTSGTT | ASSLEGTGWSDYT | 0.42624715 |
| LSFPAGV | ASSLEGTGWSDYT | 0.53404427 |
| LYTSGTT | ASSLEGTGWSDYT | 0.37192276 |
| VGAFPFGF | ASSLEGTGWSDYT | 0.6072394 |
| VGAGTVEF | ASSLEGTGWSDYT | 0.6489864 |
| VGKSTLLL | ASSLEGTGWSDYT | 0.54339415 |
| AALAHAGGGIV | ASSLEGTGWSDYT | 0.40966257 |
| AALVAVSLL | ASSLEGTGWSDYT | 0.3994707 |
| AAVSRAVAAL | ASSLEGTGWSDYT | 0.55782485 |

| | | |
|---|---|---|
| AGGSSGIGGF | ASSLEGTGWSDYT | 0.473299 |
| AGLTVANANG | ASSLEGTGWSDYT | 0.34396783 |
| AGLVGAVL | ASSLEGTGWSDYT | 0.49237025 |
| ALSQAFTV | ASSLEGTGWSDYT | 0.6534644 |
| ALTFSVSL | ASSLEGTGWSDYT | 0.651332 |
| ALTGAALVAL | ASSLEGTGWSDYT | 0.41457316 |
| ASFSIGAGVL | ASSLEGTGWSDYT | 0.6629859 |
| DGSVYKI | ASSLEGTGWSDYT | 0.6078398 |
| EQLAEIV | ASSLEGTGWSDYT | 0.629773 |
| FAVSELTVM | ASSLEGTGWSDYT | 0.5285628 |
| GAAPLAAGL | ASSLEGTGWSDYT | 0.45944092 |
| GADGGDGGSGGAGG | ASSLEGTGWSDYT | 0.63980216 |
| GFGLDTGGAGG | ASSLEGTGWSDYT | 0.54150295 |
| GLASGGLL | ASSLEGTGWSDYT | 0.46645063 |
| GRIAGAAVLL | ASSLEGTGWSDYT | 0.6211107 |
| GSAGSLNTSF | ASSLEGTGWSDYT | 0.59859216 |
| GTGQIGI | ASSLEGTGWSDYT | 0.5011696 |
| GTLIALIL | ASSLEGTGWSDYT | 0.5336703 |
| IAATMGFVVG | ASSLEGTGWSDYT | 0.6208043 |
| IFAQALL | ASSLEGTGWSDYT | 0.43481055 |
| IGAAVALGF | ASSLEGTGWSDYT | 0.49432015 |
| IGLTPLAL | ASSLEGTGWSDYT | 0.4740162 |
| ILGSNTSTL | ASSLEGTGWSDYT | 0.581033 |
| IVFSLTVL | ASSLEGTGWSDYT | 0.649982 |
| LASKATGF | ASSLEGTGWSDYT | 0.5928583 |
| LASVALIL | ASSLEGTGWSDYT | 0.47250304 |
| LFAVGLTL | ASSLEGTGWSDYT | 0.60847104 |
| LFNGGIGTYI | ASSLEGTGWSDYT | 0.6023047 |
| LTLGAGII | ASSLEGTGWSDYT | 0.57200575 |
| LVAAALGDI | ASSLEGTGWSDYT | 0.625666 |
| NTAFVQLGI | ASSLEGTGWSDYT | 0.5228471 |
| QLFSGAEL | ASSLEGTGWSDYT | 0.6133697 |
| SAATFITL | ASSLEGTGWSDYT | 0.59598637 |
| SNYGLANLGV | ASSLEGTGWSDYT | 0.61193615 |
| SYGGAYAVM | ASSLEGTGWSDYT | 0.6246542 |
| TTGVGGLTLG | ASSLEGTGWSDYT | 0.6179886 |
| TTQTIYI | ASSLEGTGWSDYT | 0.5374152 |
| VAAGAGLGAVI | ASSLEGTGWSDYT | 0.36224633 |
| VAAVVTLL | ASSLEGTGWSDYT | 0.44184074 |
| VGFLGGLI | ASSLEGTGWSDYT | 0.594285 |
| VGVIPILL | ASSLEGTGWSDYT | 0.587544 |
| VIFASSLI | ASSLEGTGWSDYT | 0.6305758 |
| VKGGDGGAAAG | ASSLEGTGWSDYT | 0.6091589 |
| VTKTFTNI | ASSLEGTGWSDYT | 0.48524228 |
| VVAAAILL | ASSLEGTGWSDYT | 0.6516384 |
| YAALDVEL | ASSLEGTGWSDYT | 0.641627 |
| LASVALLL | ASSLEGTGWSDYT | 0.3715071 |
| LDAVHRV | ASSLEGTGWSDYT | 0.5182995 |
| QGLSSSIF | ASSLEGTGWSDYT | 0.6721194 |
| RGTFANIRL | ASSLEGTGWSDYT | 0.5888099 |
| RQLWWGHRI | ASSLEGTGWSDYT | 0.5221192 |

| | | |
|---|---|---|
| YGPPGTGKTLL | ASSLEGTGWSDYT | 0.4877078 |
| AALVGAPANVA | ASSLEGTGWSDYT | 0.66448426 |
| AATFRRTE | ASSLEGTGWSDYT | 0.46737576 |
| AGALGGGI | ASSLEGTGWSDYT | 0.6546352 |
| AGGAGGAGGIGAG | ASSLEGTGWSDYT | 0.32486147 |
| AGSGALGL | ASSLEGTGWSDYT | 0.4709442 |
| AQIADQI | ASSLEGTGWSDYT | 0.45386392 |
| ASAVAVTYL | ASSLEGTGWSDYT | 0.3623864 |
| ASTVLGGTGG | ASSLEGTGWSDYT | 0.63324535 |
| AVGGEPAK | ASSLEGTGWSDYT | 0.48949713 |
| CEQGTFGGTC | ASSLEGTGWSDYT | 0.6186557 |
| DEGKAPAG | ASSLEGTGWSDYT | 0.5192474 |
| DTVVFPAL | ASSLEGTGWSDYT | 0.6380877 |
| FEGITMAG | ASSLEGTGWSDYT | 0.55892694 |
| FPLVVQD | ASSLEGTGWSDYT | 0.5769378 |
| GAFTTPQ | ASSLEGTGWSDYT | 0.46832845 |
| GFGGFGTAGGVGGA | ASSLEGTGWSDYT | 0.6240897 |
| GFLAQFL | ASSLEGTGWSDYT | 0.6561147 |
| GGAGGAGGNAGVA | ASSLEGTGWSDYT | 0.5700268 |
| GGVGTAGAGGAGGAGG | ASSLEGTGWSDYT | 0.41517046 |
| GNGGGGDAGNAGSGGN | ASSLEGTGWSDYT | 0.5101853 |
| GTGGAGGAGNDAGS | ASSLEGTGWSDYT | 0.5917209 |
| GTTNTGWFNS | ASSLEGTGWSDYT | 0.33961377 |
| IFMGPLFTL | ASSLEGTGWSDYT | 0.3318827 |
| ILAHAIEAI | ASSLEGTGWSDYT | 0.53968173 |
| IRSAIREAL | ASSLEGTGWSDYT | 0.52410066 |
| ITFAGGG | ASSLEGTGWSDYT | 0.6420196 |
| ITGMSGHQEV | ASSLEGTGWSDYT | 0.6276303 |
| IVAGLACTL | ASSLEGTGWSDYT | 0.52639925 |
| IVGGGLAGA | ASSLEGTGWSDYT | 0.6050174 |
| IVTGVLNI | ASSLEGTGWSDYT | 0.602603 |
| LAQQGCI | ASSLEGTGWSDYT | 0.28749156 |
| LCLHVEN | ASSLEGTGWSDYT | 0.46244726 |
| LDAISDGS | ASSLEGTGWSDYT | 0.38829997 |
| LGGFAAML | ASSLEGTGWSDYT | 0.64861715 |
| LGGKNPMI | ASSLEGTGWSDYT | 0.47552833 |
| LGPKGVAAIL | ASSLEGTGWSDYT | 0.4707583 |
| LMVGVSI | ASSLEGTGWSDYT | 0.57001543 |
| LVGAAFVL | ASSLEGTGWSDYT | 0.59737563 |
| LVLAVAPL | ASSLEGTGWSDYT | 0.55539715 |
| MAAIAVIDAA | ASSLEGTGWSDYT | 0.4411241 |
| NNGINAELQ | ASSLEGTGWSDYT | 0.3259561 |
| RGEPLVLVHG | ASSLEGTGWSDYT | 0.63420177 |
| RGGAGGAAGLI | ASSLEGTGWSDYT | 0.42970526 |
| RSYRMHPAV | ASSLEGTGWSDYT | 0.49002966 |
| SAAHLLAGQLA | ASSLEGTGWSDYT | 0.33482242 |
| SAAVGSY | ASSLEGTGWSDYT | 0.53814626 |
| SPHAVLHRI | ASSLEGTGWSDYT | 0.39734325 |
| SQIAKVL | ASSLEGTGWSDYT | 0.6109907 |
| TALGHLVP | ASSLEGTGWSDYT | 0.5015514 |
| TAPTTSGAA | ASSLEGTGWSDYT | 0.5429155 |

| | | |
|---|---|---|
| TDGDDGNGGSPGT | ASSLEGTGWSDYT | 0.66846037 |
| TDGLGEN | ASSLEGTGWSDYT | 0.5996317 |
| TGGPPLPGQA | ASSLEGTGWSDYT | 0.52569205 |
| TIPAITI | ASSLEGTGWSDYT | 0.5339271 |
| TIPGFTI | ASSLEGTGWSDYT | 0.42215288 |
| TSIASGALA | ASSLEGTGWSDYT | 0.50219584 |
| TVTLPGAAI | ASSLEGTGWSDYT | 0.52121913 |
| VASGLSAALP | ASSLEGTGWSDYT | 0.6642084 |
| VGAGVIGI | ASSLEGTGWSDYT | 0.5002252 |
| VGPKRFGV | ASSLEGTGWSDYT | 0.39920917 |
| VNVAFPD | ASSLEGTGWSDYT | 0.5377234 |
| VSMTNSGVS | ASSLEGTGWSDYT | 0.6569722 |
| VTDVLGDLSV | ASSLEGTGWSDYT | 0.6627649 |
| VVVVGAGFAGL | ASSLEGTGWSDYT | 0.4544356 |
| VYTVGTVV | ASSLEGTGWSDYT | 0.5057063 |
| WTTFSGP | ASSLEGTGWSDYT | 0.49070752 |
| YDELVEAF | ASSLEGTGWSDYT | 0.57026124 |
| YGTGGAGGNGG | ASSLEGTGWSDYT | 0.45267892 |
| YTNVMAV | ASSLEGTGWSDYT | 0.47449228 |
| FLEQVHQL | ASSLEGTGWSDYT | 0.5888006 |
| GLSLLAVAV | ASSLEGTGWSDYT | 0.6723175 |
| LEQVHQL | ASSLEGTGWSDYT | 0.51390606 |
| LNAAGAI | ASSLEGTGWSDYT | 0.34613946 |
| PAVHASL | ASSLEGTGWSDYT | 0.55721134 |
| TELEDDA | ASSLEGTGWSDYT | 0.4526662 |
| TPPGPPN | ASSLEGTGWSDYT | 0.5025244 |
| TPPPPPPP | ASSLEGTGWSDYT | 0.5688988 |
| TQSPALI | ASSLVGVQDTQY | 0.36749646 |
| EFAAAFPQ | ASSLVGVQDTQY | 0.5623194 |
| LLSGNLDVL | ASSLVGVQDTQY | 0.67080903 |
| GPANLGNYNI | ASSLVGVQDTQY | 0.5311382 |
| TLGPTLP | ASSLVGVQDTQY | 0.5781157 |
| VYAYSSCCK | ASSLVGVQDTQY | 0.64780647 |
| APAPAPAPNI | ASSLVGVQDTQY | 0.6172483 |
| PGGPVPIP | ASSLVGVQDTQY | 0.6487944 |
| FLTVALEP | ASSLVGVQDTQY | 0.6226445 |
| SQYFGANL | ASSLVGVQDTQY | 0.55933565 |
| GGVKPAA | ASSLVGVQDTQY | 0.54349667 |
| GGTGGAGGAGGW | ASSLVGVQDTQY | 0.63187194 |
| GVGGVAGLGGAGG | ASSLVGVQDTQY | 0.56956786 |
| VLLVLATTI | ASSLVGVQDTQY | 0.541843 |
| IGIFAAML | ASSLVGVQDTQY | 0.41835985 |
| AVGDLID | ASSLVGVQDTQY | 0.6496243 |
| NISPLPL | ASSLVGVQDTQY | 0.5203037 |
| VTAVFVAI | ASSLVGVQDTQY | 0.6404371 |
| SGDDNDAAL | ASSLVGVQDTQY | 0.3861699 |
| AVAALAHS | ASSLVGVQDTQY | 0.47766647 |
| GGVTVMATP | ASSLVGVQDTQY | 0.6338245 |
| AGLAANVRD | ASSLVGVQDTQY | 0.2987797 |
| TNRPDLIDPALLRPG | ASSLVGVQDTQY | 0.3254209 |
| LANLGVTAL | ASSLVGVQDTQY | 0.51458657 |

| | | |
|---|---|---|
| TGERIEL | ASSLVGVQDTQY | 0.6456638 |
| NGEELAA | ASSLVGVQDTQY | 0.2987767 |
| VLSLVWG | ASSLVGVQDTQY | 0.48203874 |
| IGYALWL | ASSLVGVQDTQY | 0.38070115 |
| LSLLLAPL | ASSLVGVQDTQY | 0.6100643 |
| TLDEALK | ASSLVGVQDTQY | 0.6042011 |
| TDGAGTAEGG | ASSLVGVQDTQY | 0.36430496 |
| NAPGGAGGNGG | ASSLVGVQDTQY | 0.6192869 |
| VSVFMGMGGNF | ASSLVGVQDTQY | 0.34273657 |
| LASYLAAL | ASSLVGVQDTQY | 0.5026385 |
| LVGGTPAL | ASSLVGVQDTQY | 0.61533844 |
| VRNADVL | ASSLVGVQDTQY | 0.6520629 |
| TCSSPFLVSAM | ASSLVGVQDTQY | 0.39278293 |
| VHITDAE | ASSLVGVQDTQY | 0.6133984 |
| EYAAPSF | ASSLVGVQDTQY | 0.479732 |
| VVDAFHGGL | ASSLVGVQDTQY | 0.66843617 |
| ASVVDPQ | ASSLVGVQDTQY | 0.5295784 |
| SYNEFGGLS | ASSLVGVQDTQY | 0.37735984 |
| DPDGSVY | ASSLVGVQDTQY | 0.6386346 |
| MVSTIHATE | ASSLVGVQDTQY | 0.6000126 |
| LAAGVADPV | ASSLVGVQDTQY | 0.59297156 |
| TKEQIAA | ASSLVGVQDTQY | 0.61750543 |
| SLASPAAL | ASSLVGVQDTQY | 0.49060768 |
| IGLVFYGV | ASSLVGVQDTQY | 0.5458171 |
| FLLDASNL | ASSLVGVQDTQY | 0.6437998 |
| YQVGNFP | ASSLVGVQDTQY | 0.625813 |
| ESLLSQST | ASSLVGVQDTQY | 0.5166073 |
| LFTVGFAAVG | ASSLVGVQDTQY | 0.6334493 |
| ATPPTPT | ASSLVGVQDTQY | 0.58208853 |
| TDLATLG | ASSLVGVQDTQY | 0.47081146 |
| SAGGTGR | ASSLVGVQDTQY | 0.66034836 |
| GFSNSGS | ASSLVGVQDTQY | 0.5134888 |
| GGNGGSGGTSV | ASSLVGVQDTQY | 0.62209636 |
| GTSVATGGAGN | ASSLVGVQDTQY | 0.6230503 |
| NGGNGGSGGTS | ASSLVGVQDTQY | 0.65609246 |
| AVVKFPH | ASSLVGVQDTQY | 0.42396823 |
| TPPPPPP | ASSLVGVQDTQY | 0.5725836 |
| AAPAAAAAPA | ASSLVGVQDTQY | 0.5619844 |
| IALNEDL | ASSLVGVQDTQY | 0.6263165 |
| IVNVSSM | ASSLVGVQDTQY | 0.51821136 |
| TGIPPAPR | ASSLVGVQDTQY | 0.5311001 |
| FDLGGGTF | ASSLVGVQDTQY | 0.5910064 |
| FDLGGGTFDVS | ASSLVGVQDTQY | 0.6532329 |
| NEPTAAA | ASSLVGVQDTQY | 0.5261388 |
| NEPTAAALA | ASSLVGVQDTQY | 0.6013115 |
| GSIINISSI | ASSLVGVQDTQY | 0.66208816 |
| VAPGNAGTA | ASSLVGVQDTQY | 0.46562356 |
| TGGGASLE | ASSLVGVQDTQY | 0.5607567 |
| VAVYNHYKRI | ASSLVGVQDTQY | 0.5966166 |
| TDPAPATT | ASSLVGVQDTQY | 0.6500375 |
| TNRPDILDPALLRPG | ASSLVGVQDTQY | 0.52383727 |

| | | |
|---|---|---|
| LTTTTTS | ASSLVGVQDTQY | 0.64254594 |
| PTGSGKTL | ASSLVGVQDTQY | 0.4517029 |
| TGSGKTL | ASSLVGVQDTQY | 0.61804557 |
| TIGHVDHGK | ASSLVGVQDTQY | 0.48231938 |
| TIGHVDHGKTT | ASSLVGVQDTQY | 0.5837433 |
| GHVDHGKTT | ASSLVGVQDTQY | 0.5369185 |
| AQFEHTLL | ASSLVGVQDTQY | 0.61542374 |
| IVNQDEL | ASSLVGVQDTQY | 0.62885994 |
| TNGSQFF | ASSLVGVQDTQY | 0.6367663 |
| TGKGIVE | ASSLVGVQDTQY | 0.31575692 |
| LISNASDA | ASSLVGVQDTQY | 0.63892573 |
| AELRTLGELTKIAKAHGAQ | ASSLVGVQDTQY | 0.635909 |
| APNAVAAAEPSW | ASSLVGVQDTQY | 0.6694064 |
| APQVAGAATAVL | ASSLVGVQDTQY | 0.527225 |
| ARRDPAAAAAILPTDA | ASSLVGVQDTQY | 0.59912604 |
| DAAARDVLAEAGLAEYF | ASSLVGVQDTQY | 0.5396139 |
| DDQIVLSAKANLAKVEQ | ASSLVGVQDTQY | 0.53359795 |
| DELLPDTGEDSDEDGH | ASSLVGVQDTQY | 0.4165576 |
| ELPEAPANRATVPQD | ASSLVGVQDTQY | 0.5337544 |
| FGQSKIERVTGIG | ASSLVGVQDTQY | 0.50769866 |
| GAGGTGSTAGGAGGAGGAGGLY | ASSLVGVQDTQY | 0.49600142 |
| GHGGTGAAGGNGGTGGAGGSGI | ASSLVGVQDTQY | 0.6374403 |
| GNGGNGGAGGEGGPGGAGGAGA | ASSLVGVQDTQY | 0.6387405 |
| GSTAGNGGNGGNATNGTIAGQP | ASSLVGVQDTQY | 0.6472392 |
| GTGGDGGNGGNGGTGG | ASSLVGVQDTQY | 0.6455765 |
| GVRRPAGIPADAPV | ASSLVGVQDTQY | 0.6522639 |
| GYRQVTDAHLLTLARR | ASSLVGVQDTQY | 0.61852396 |
| IAKMGQDGHDRG | ASSLVGVQDTQY | 0.53315616 |
| KAEPSIEIQYT | ASSLVGVQDTQY | 0.50944084 |
| KPVRIRMAADEDTVAVT | ASSLVGVQDTQY | 0.4212028 |
| MPPRIAGMRLLVIKPEP | ASSLVGVQDTQY | 0.5138274 |
| MKFYAVALFRSRGVPPTR | ASSLVGVQDTQY | 0.5481698 |
| NAGGFGILGAVAHSP | ASSLVGVQDTQY | 0.5928596 |
| NEGREITLAMAEM | ASSLVGVQDTQY | 0.30615047 |
| NELVEWSHPLGHRWRWE | ASSLVGVQDTQY | 0.5772158 |
| NLGAGGTGGNGGNPG | ASSLVGVQDTQY | 0.6651214 |
| NTEWPKWETVMAADPVD | ASSLVGVQDTQY | 0.5117875 |
| PASAAPAETQP | ASSLVGVQDTQY | 0.5997466 |
| QQSSDGKNGVRAAEITGE | ASSLVGVQDTQY | 0.60309005 |
| RAPDAVTTSWWKE | ASSLVGVQDTQY | 0.48212057 |
| SGFGNVGAHTSGWFNQ | ASSLVGVQDTQY | 0.62504226 |
| SGVAGAVGQAASVGGLKV | ASSLVGVQDTQY | 0.46110013 |
| SNGGAGTGSAGGNGGT | ASSLVGVQDTQY | 0.5960578 |
| TGGAGGEGGAAG | ASSLVGVQDTQY | 0.6572354 |
| TGGDGGAAGAGGAPGHG | ASSLVGVQDTQY | 0.5745288 |
| TMPPLLAAIANAGRHGV | ASSLVGVQDTQY | 0.5763273 |
| TSSTAGNSGGA | ASSLVGVQDTQY | 0.5211617 |
| VPATVPVGILC | ASSLVGVQDTQY | 0.40357277 |
| VQRPDSGGVVLTGRIS | ASSLVGVQDTQY | 0.6005887 |
| WTPNPVPATKGVPAGLVQAT | ASSLVGVQDTQY | 0.38162798 |
| YALKRALAKFSDRFDVVII | ASSLVGVQDTQY | 0.54954076 |

| | | |
|---|---|---|
| YLIGSVIV | ASSLVGVQDTQY | 0.4448728 |
| YGQLVAVID | ASSLVGVQDTQY | 0.6018463 |
| VTVGGMTLPA | ASSLVGVQDTQY | 0.60246193 |
| VLALPAV | ASSLVGVQDTQY | 0.55448365 |
| VIPPTLNYET | ASSLVGVQDTQY | 0.43393248 |
| VGMGGMHPSSG | ASSLVGVQDTQY | 0.54478616 |
| VGLAGGAATA | ASSLVGVQDTQY | 0.492836 |
| VGGAGGAGGSGP | ASSLVGVQDTQY | 0.4516366 |
| VFAAFMV | ASSLVGVQDTQY | 0.61511475 |
| TLYDKLR | ASSLVGVQDTQY | 0.50220823 |
| TLAVFTVL | ASSLVGVQDTQY | 0.5503322 |
| TGSVQPARV | ASSLVGVQDTQY | 0.6521604 |
| TGGGAGSGGT | ASSLVGVQDTQY | 0.59533834 |
| SHLETSL | ASSLVGVQDTQY | 0.3655803 |
| SFTTGMT | ASSLVGVQDTQY | 0.5500804 |
| QVEVAKRMGVRQPTVS | ASSLVGVQDTQY | 0.5473187 |
| QGSGGPALAM | ASSLVGVQDTQY | 0.5410472 |
| MLAPSNP | ASSLVGVQDTQY | 0.29695535 |
| LVNNAAI | ASSLVGVQDTQY | 0.60006756 |
| LTGEINTLA | ASSLVGVQDTQY | 0.46832576 |
| LSIFGPV | ASSLVGVQDTQY | 0.43591386 |
| LLALLANL | ASSLVGVQDTQY | 0.5110718 |
| LISNASDALDKLR | ASSLVGVQDTQY | 0.63535285 |
| LINEAVP | ASSLVGVQDTQY | 0.528366 |
| LGYAFSGL | ASSLVGVQDTQY | 0.5920946 |
| LDFVAVAL | ASSLVGVQDTQY | 0.6148416 |
| LAQPGAE | ASSLVGVQDTQY | 0.6240312 |
| ISNLGAVTGF | ASSLVGVQDTQY | 0.6547401 |
| HQEDPGRI | ASSLVGVQDTQY | 0.6425698 |
| GTPGEPTP | ASSLVGVQDTQY | 0.54105586 |
| GFFNSGS | ASSLVGVQDTQY | 0.5997866 |
| GAGGGGLTAGAN | ASSLVGVQDTQY | 0.48901758 |
| FVDALVNM | ASSLVGVQDTQY | 0.36954916 |
| FLTVAPEL | ASSLVGVQDTQY | 0.3416808 |
| FETLTAL | ASSLVGVQDTQY | 0.518311 |
| DVINPIEL | ASSLVGVQDTQY | 0.3480718 |
| APVSGAPV | ASSLVGVQDTQY | 0.58477604 |
| ALVGGLAW | ASSLVGVQDTQY | 0.6281303 |
| ALAAAAATAI | ASSLVGVQDTQY | 0.5146495 |
| AFGLGLLL | ASSLVGVQDTQY | 0.3948591 |
| AFFAAASL | ASSLVGVQDTQY | 0.49022657 |
| AAAALPAFSPP | ASSLVGVQDTQY | 0.4777978 |
| VVLAPAL | ASSLVGVQDTQY | 0.6258655 |
| TIGHVDHGKTTL | ASSLVGVQDTQY | 0.6091728 |
| LISNASDALDK | ASSLVGVQDTQY | 0.6688994 |
| LISNASDAL | ASSLVGVQDTQY | 0.65042686 |
| ISNASDALDK | ASSLVGVQDTQY | 0.53459156 |
| NQAETLVYQTE | ASSLVGVQDTQY | 0.6164784 |
| TGGTGGTAGAG | ASSLVGVQDTQY | 0.4888128 |
| TGSTAGGAGGA | ASSLVGVQDTQY | 0.5367352 |
| DPLAVPVAAGPV | ASSLVGVQDTQY | 0.51649266 |

| | | |
|---|---|---|
| GGTGGTPGNGGN | ASSLVGVQDTQY | 0.56161296 |
| PAGLAGVDPATL | ASSLVGVQDTQY | 0.58229214 |
| GNNLVVEAPHAGASV | ASSLVGVQDTQY | 0.48224133 |
| KVTRIMSASTLEQPA | ASSLVGVQDTQY | 0.36807853 |
| LGFALPIALPRRTVV | ASSLVGVQDTQY | 0.5807303 |
| TRLGNLTARKLVVLV | ASSLVGVQDTQY | 0.5185982 |
| WSQQRRLIYFPSAGP | ASSLVGVQDTQY | 0.3560844 |
| GPNGNPDPMAAAVDIR | ASSLVGVQDTQY | 0.47754228 |
| TQRCVSVADLRAAASR | ASSLVGVQDTQY | 0.66392344 |
| QFVLDVAEHPVEVLTETV | ASSLVGVQDTQY | 0.655627 |
| GGAGGTGGAGGNGGTQAGDG | ASSLVGVQDTQY | 0.4896929 |
| PLLATLRAIAVDVDAKPAQV | ASSLVGVQDTQY | 0.49091348 |
| APKFRTGGYMIDFWGVGYQVAK | ASSLVGVQDTQY | 0.58964056 |
| GAGGAGGAGGAAGGPGGTGGTG | ASSLVGVQDTQY | 0.38619047 |
| IGALGLLATDVAPAGSVAFGVA | ASSLVGVQDTQY | 0.57353175 |
| AGGAGGIGGNGGAGGAGGVLMG | ASSLVGVQDTQY | 0.57954895 |
| ADELIGLL | ASSLVGVQDTQY | 0.5787853 |
| ADRALKA | ASSLVGVQDTQY | 0.46132916 |
| AGAVGQVGGAGGA | ASSLVGVQDTQY | 0.5830834 |
| AGGGGGLGGAP | ASSLVGVQDTQY | 0.56306463 |
| AGGLLYGNGGAG | ASSLVGVQDTQY | 0.58594847 |
| AGLESIGL | ASSLVGVQDTQY | 0.3043532 |
| AGLKTGII | ASSLVGVQDTQY | 0.6147182 |
| ASLAALLL | ASSLVGVQDTQY | 0.46131477 |
| ASVINISSI | ASSLVGVQDTQY | 0.45774642 |
| ATPPGPA | ASSLVGVQDTQY | 0.33234608 |
| ATVASALV | ASSLVGVQDTQY | 0.52048874 |
| ATVASLIL | ASSLVGVQDTQY | 0.6173452 |
| AVNGPVLIRH | ASSLVGVQDTQY | 0.63470894 |
| DGAAGAVGI | ASSLVGVQDTQY | 0.42381364 |
| DPLLHGI | ASSLVGVQDTQY | 0.63630867 |
| ELGAALGAL | ASSLVGVQDTQY | 0.64245087 |
| ELGRTQAV | ASSLVGVQDTQY | 0.5249272 |
| FVAGGYLS | ASSLVGVQDTQY | 0.38173783 |
| GASGGAGGAGGW | ASSLVGVQDTQY | 0.6255908 |
| GGAGAGGDGGAGGI | ASSLVGVQDTQY | 0.5186724 |
| GTGHNGSQPGMGGNG | ASSLVGVQDTQY | 0.63547385 |
| GYAAASASAS | ASSLVGVQDTQY | 0.6106962 |
| HQVDVAAVQQG | ASSLVGVQDTQY | 0.41200322 |
| IGAAIAHT | ASSLVGVQDTQY | 0.63517755 |
| IGMIEVAQL | ASSLVGVQDTQY | 0.61521494 |
| IRLAPEHTSAMD | ASSLVGVQDTQY | 0.41778067 |
| IYNVSVL | ASSLVGVQDTQY | 0.4014312 |
| KQGFLGM | ASSLVGVQDTQY | 0.47707844 |
| LESVITQL | ASSLVGVQDTQY | 0.3506208 |
| LGLGALVVV | ASSLVGVQDTQY | 0.6250495 |
| LIGNGGNGGSGGI | ASSLVGVQDTQY | 0.5116704 |
| LNAINAPTL | ASSLVGVQDTQY | 0.5328667 |
| LQLLNTGV | ASSLVGVQDTQY | 0.57060313 |
| LQTSIEF | ASSLVGVQDTQY | 0.5351307 |
| MSFAVSEL | ASSLVGVQDTQY | 0.53038156 |

| | | |
|---|---|---|
| NTGALGL | ASSLVGVQDTQY | 0.5651023 |
| PPGAGGAGGAGTT | ASSLVGVQDTQY | 0.6074246 |
| QLVGLGI | ASSLVGVQDTQY | 0.6015047 |
| RFLEQVHQL | ASSLVGVQDTQY | 0.5390039 |
| RGVGYVLR | ASSLVGVQDTQY | 0.533432 |
| SATGVFAGVI | ASSLVGVQDTQY | 0.65599954 |
| SLPAITLP | ASSLVGVQDTQY | 0.4692046 |
| STALAFTV | ASSLVGVQDTQY | 0.49008662 |
| SVLTVGLGK | ASSLVGVQDTQY | 0.61252767 |
| TALAGAGSGI | ASSLVGVQDTQY | 0.41440895 |
| TGLPFTGI | ASSLVGVQDTQY | 0.6151243 |
| VAEHVTV | ASSLVGVQDTQY | 0.5954694 |
| VDIVLNSL | ASSLVGVQDTQY | 0.6005515 |
| VLTGGGAVL | ASSLVGVQDTQY | 0.50802463 |
| VNVGLAYGI | ASSLVGVQDTQY | 0.6641114 |
| VSSAAVGAV | ASSLVGVQDTQY | 0.6333382 |
| VTSVYEAGV | ASSLVGVQDTQY | 0.49938366 |
| WVALDRGA | ASSLVGVQDTQY | 0.6348493 |
| YLVGSVVI | ASSLVGVQDTQY | 0.4382442 |
| GPPGTGKTLL | ASSLVGVQDTQY | 0.6349204 |
| GVGMTKF | ASSLVGVQDTQY | 0.6022901 |
| IGIGGSDL | ASSLVGVQDTQY | 0.44503683 |
| IGTLLAEL | ASSLVGVQDTQY | 0.55159664 |
| IYTSGTT | ASSLVGVQDTQY | 0.6345976 |
| LSFPAGV | ASSLVGVQDTQY | 0.6338956 |
| LYTSGTT | ASSLVGVQDTQY | 0.3674022 |
| VGAFPFGF | ASSLVGVQDTQY | 0.5089799 |
| VGAGTVEF | ASSLVGVQDTQY | 0.37800595 |
| VGKSTLLL | ASSLVGVQDTQY | 0.46182823 |
| AALAHAGGGIV | ASSLVGVQDTQY | 0.42570952 |
| AALVAVSLL | ASSLVGVQDTQY | 0.60814863 |
| AAVSRAVAAL | ASSLVGVQDTQY | 0.501264 |
| AGGSSGIGGF | ASSLVGVQDTQY | 0.61782014 |
| AGLTVANANG | ASSLVGVQDTQY | 0.58663404 |
| AGLVGAVL | ASSLVGVQDTQY | 0.65459365 |
| ALSQAFTV | ASSLVGVQDTQY | 0.45642728 |
| ALTFSVSL | ASSLVGVQDTQY | 0.5927852 |
| ALTGAALVAL | ASSLVGVQDTQY | 0.3612522 |
| ASFSIGAGVL | ASSLVGVQDTQY | 0.49277434 |
| DGSVYKI | ASSLVGVQDTQY | 0.41965365 |
| EQLAEIV | ASSLVGVQDTQY | 0.58548766 |
| FAVSELTVM | ASSLVGVQDTQY | 0.3426824 |
| GAAPLAAGL | ASSLVGVQDTQY | 0.6591939 |
| GADGGDGGSGGAGG | ASSLVGVQDTQY | 0.5579762 |
| GFGLDTGGAGG | ASSLVGVQDTQY | 0.6253349 |
| GLASGGLL | ASSLVGVQDTQY | 0.4927074 |
| GRIAGAAVLL | ASSLVGVQDTQY | 0.49242797 |
| GSAGSLNTSF | ASSLVGVQDTQY | 0.5497498 |
| GTGQIGI | ASSLVGVQDTQY | 0.62231725 |
| GTLIALIL | ASSLVGVQDTQY | 0.45622867 |
| IAATMGFVVG | ASSLVGVQDTQY | 0.3291354 |

| | | |
|---|---|---|
| IFAQALL | ASSLVGVQDTQY | 0.43487915 |
| IGAAVALGF | ASSLVGVQDTQY | 0.36929345 |
| IGLTPLAL | ASSLVGVQDTQY | 0.6490028 |
| ILGSNTSTL | ASSLVGVQDTQY | 0.63526124 |
| IVFSLTVL | ASSLVGVQDTQY | 0.5818818 |
| LASKATGF | ASSLVGVQDTQY | 0.63201505 |
| LASVALIL | ASSLVGVQDTQY | 0.314555 |
| LFAVGLTL | ASSLVGVQDTQY | 0.5185721 |
| LFNGGIGTYI | ASSLVGVQDTQY | 0.6635813 |
| LTLGAGII | ASSLVGVQDTQY | 0.5984497 |
| LVAAALGDI | ASSLVGVQDTQY | 0.5386164 |
| NTAFVQLGI | ASSLVGVQDTQY | 0.6477215 |
| QLFSGAEL | ASSLVGVQDTQY | 0.51769 |
| SAATFITL | ASSLVGVQDTQY | 0.5467187 |
| SNYGLANLGV | ASSLVGVQDTQY | 0.6063179 |
| SYGGAYAVM | ASSLVGVQDTQY | 0.58367246 |
| TTGVGGLTLG | ASSLVGVQDTQY | 0.5423304 |
| TTQTIYI | ASSLVGVQDTQY | 0.41322693 |
| VAAGAGLGAVI | ASSLVGVQDTQY | 0.5693739 |
| VAAVVTLL | ASSLVGVQDTQY | 0.6716641 |
| VGFLGGLI | ASSLVGVQDTQY | 0.5407703 |
| VGVIPILL | ASSLVGVQDTQY | 0.63027483 |
| VIFASSLI | ASSLVGVQDTQY | 0.60346884 |
| VKGGDGGAAAG | ASSLVGVQDTQY | 0.6002986 |
| VTKTFTNI | ASSLVGVQDTQY | 0.4588129 |
| VVAAAILL | ASSLVGVQDTQY | 0.49285382 |
| YAALDVEL | ASSLVGVQDTQY | 0.5435494 |
| LASVALLL | ASSLVGVQDTQY | 0.4154132 |
| LDAVHRV | ASSLVGVQDTQY | 0.51499295 |
| QGLSSSIF | ASSLVGVQDTQY | 0.49027607 |
| RGTFANIRL | ASSLVGVQDTQY | 0.6774202 |
| RQLWWGHRI | ASSLVGVQDTQY | 0.6065193 |
| YGPPGTGKTLL | ASSLVGVQDTQY | 0.61661005 |
| AALVGAPANVA | ASSLVGVQDTQY | 0.51851505 |
| AATFRRTE | ASSLVGVQDTQY | 0.54105884 |
| AGALGGGI | ASSLVGVQDTQY | 0.60923845 |
| AGGAGGAGGIGAG | ASSLVGVQDTQY | 0.38883483 |
| AGSGALGL | ASSLVGVQDTQY | 0.64650893 |
| AQIADQI | ASSLVGVQDTQY | 0.47225556 |
| ASAVAVTYL | ASSLVGVQDTQY | 0.54118454 |
| ASTVLGGTGG | ASSLVGVQDTQY | 0.62458926 |
| AVGGEPAK | ASSLVGVQDTQY | 0.6523169 |
| CEQGTFGGTC | ASSLVGVQDTQY | 0.49975386 |
| DEGKAPAG | ASSLVGVQDTQY | 0.5085828 |
| DTVVFPAL | ASSLVGVQDTQY | 0.5243058 |
| FEGITMAG | ASSLVGVQDTQY | 0.6147464 |
| FPLVVQD | ASSLVGVQDTQY | 0.64680314 |
| GAFTTPQ | ASSLVGVQDTQY | 0.56980306 |
| GFGGFGTAGGVGGA | ASSLVGVQDTQY | 0.6650666 |
| GFLAQFL | ASSLVGVQDTQY | 0.59396446 |
| GGAGGAGGNAGVA | ASSLVGVQDTQY | 0.3884353 |

| | | |
|---|---|---|
| GGVGTAGAGGAGGAGG | ASSLVGVQDTQY | 0.5313649 |
| GNGGGGDAGNAGSGGN | ASSLVGVQDTQY | 0.6235964 |
| GTGGAGGAGNDAGS | ASSLVGVQDTQY | 0.6422075 |
| GTTNTGWFNS | ASSLVGVQDTQY | 0.66004425 |
| IFMGPLFTL | ASSLVGVQDTQY | 0.6083745 |
| ILAHAIEAI | ASSLVGVQDTQY | 0.48768836 |
| IRSAIREAL | ASSLVGVQDTQY | 0.6476188 |
| ITFAGGG | ASSLVGVQDTQY | 0.66501343 |
| ITGMSGHQEV | ASSLVGVQDTQY | 0.62106127 |
| IVAGLACTL | ASSLVGVQDTQY | 0.4656367 |
| IVGGGLAGA | ASSLVGVQDTQY | 0.46640202 |
| IVTGVLNI | ASSLVGVQDTQY | 0.57019794 |
| LAQQGCI | ASSLVGVQDTQY | 0.64621323 |
| LCLHVEN | ASSLVGVQDTQY | 0.40104792 |
| LDAISDGS | ASSLVGVQDTQY | 0.58662176 |
| LGGFAAML | ASSLVGVQDTQY | 0.5997206 |
| LGGKNPMI | ASSLVGVQDTQY | 0.36967576 |
| LGPKGVAAIL | ASSLVGVQDTQY | 0.54338455 |
| LMVGVSI | ASSLVGVQDTQY | 0.6477596 |
| LVGAAFVL | ASSLVGVQDTQY | 0.5389102 |
| LVLAVAPL | ASSLVGVQDTQY | 0.45452997 |
| MAAIAVIDAA | ASSLVGVQDTQY | 0.6429141 |
| NNGINAELQ | ASSLVGVQDTQY | 0.6749046 |
| RGEPLVLVHG | ASSLVGVQDTQY | 0.64640546 |
| RGGAGGAAGLI | ASSLVGVQDTQY | 0.39226827 |
| RSYRMHPAV | ASSLVGVQDTQY | 0.30754977 |
| SAAHLLAGQLA | ASSLVGVQDTQY | 0.5312021 |
| SAAVGSY | ASSLVGVQDTQY | 0.62995523 |
| SPHAVLHRI | ASSLVGVQDTQY | 0.5714069 |
| SQIAKVL | ASSLVGVQDTQY | 0.5361851 |
| TALGHLVP | ASSLVGVQDTQY | 0.5983613 |
| TAPTTSGAA | ASSLVGVQDTQY | 0.6419313 |
| TDGDDGNGGSPGT | ASSLVGVQDTQY | 0.46284136 |
| TDGLGEN | ASSLVGVQDTQY | 0.43237975 |
| TGGPPLPGQA | ASSLVGVQDTQY | 0.5918835 |
| TIPAITI | ASSLVGVQDTQY | 0.60691965 |
| TIPGFTI | ASSLVGVQDTQY | 0.4469279 |
| TSIASGALA | ASSLVGVQDTQY | 0.5468755 |
| TVTLPGAAI | ASSLVGVQDTQY | 0.5429157 |
| VASGLSAALP | ASSLVGVQDTQY | 0.48869085 |
| VGAGVIGI | ASSLVGVQDTQY | 0.59473705 |
| VGPKRFGV | ASSLVGVQDTQY | 0.4332563 |
| VNVAFPD | ASSLVGVQDTQY | 0.6473133 |
| VSMTNSGVS | ASSLVGVQDTQY | 0.40393007 |
| VTDVLGDLSV | ASSLVGVQDTQY | 0.584909 |
| VVVVGAGFAGL | ASSLVGVQDTQY | 0.60414064 |
| VYTVGTVV | ASSLVGVQDTQY | 0.5743879 |
| WTTFSGP | ASSLVGVQDTQY | 0.5187661 |
| YDELVEAF | ASSLVGVQDTQY | 0.46273524 |
| YGTGGAGGNGG | ASSLVGVQDTQY | 0.40664047 |
| YTNVMAV | ASSLVGVQDTQY | 0.34211507 |

| | | |
|---|---|---|
| FLEQVHQL | ASSLVGVQDTQY | 0.6446674 |
| GLSLLAVAV | ASSLVGVQDTQY | 0.51919025 |
| LEQVHQL | ASSLVGVQDTQY | 0.6380698 |
| LNAAGAI | ASSLVGVQDTQY | 0.6137601 |
| PAVHASL | ASSLVGVQDTQY | 0.47432348 |
| TELEDDA | ASSLVGVQDTQY | 0.58636934 |
| TPPGPPN | ASSLVGVQDTQY | 0.5271869 |
| TPPPPPPP | ASSLVGVQDTQY | 0.44993114 |
| TQSPALI | ASSSGTGEDTQY | 0.5897716 |
| EFAAAFPQ | ASSSGTGEDTQY | 0.61215276 |
| LLSGNLDVL | ASSSGTGEDTQY | 0.4267053 |
| GPANLGNYNI | ASSSGTGEDTQY | 0.56746805 |
| TLGPTLP | ASSSGTGEDTQY | 0.5802061 |
| VYAYSSCCK | ASSSGTGEDTQY | 0.42285293 |
| APAPAPAPNI | ASSSGTGEDTQY | 0.5483458 |
| PGGPVPIP | ASSSGTGEDTQY | 0.6404039 |
| FLTVALEP | ASSSGTGEDTQY | 0.5543213 |
| SQYFGANL | ASSSGTGEDTQY | 0.42079154 |
| GGVKPAA | ASSSGTGEDTQY | 0.6204057 |
| GGTGGAGGAGGW | ASSSGTGEDTQY | 0.46670443 |
| GVGGVAGLGGAGG | ASSSGTGEDTQY | 0.6007517 |
| VLLVLATTI | ASSSGTGEDTQY | 0.56686485 |
| IGIFAAML | ASSSGTGEDTQY | 0.5179794 |
| AVGDLID | ASSSGTGEDTQY | 0.6338843 |
| NISPLPL | ASSSGTGEDTQY | 0.39660248 |
| VTAVFVAI | ASSSGTGEDTQY | 0.44465113 |
| SGDDNDAAL | ASSSGTGEDTQY | 0.49432054 |
| AVAALAHS | ASSSGTGEDTQY | 0.602921 |
| GGVTVMATP | ASSSGTGEDTQY | 0.63447374 |
| AGLAANVRD | ASSSGTGEDTQY | 0.4682529 |
| TNRPDLIDPALLRPG | ASSSGTGEDTQY | 0.5496184 |
| LANLGVTAL | ASSSGTGEDTQY | 0.50790757 |
| TGERIEL | ASSSGTGEDTQY | 0.4562129 |
| NGEELAA | ASSSGTGEDTQY | 0.5186689 |
| VLSLVWG | ASSSGTGEDTQY | 0.4829145 |
| IGYALWL | ASSSGTGEDTQY | 0.42309022 |
| LSLLLAPL | ASSSGTGEDTQY | 0.5872299 |
| TLDEALK | ASSSGTGEDTQY | 0.53761494 |
| TDGAGTAEGG | ASSSGTGEDTQY | 0.59606576 |
| NAPGGAGGNGG | ASSSGTGEDTQY | 0.6455309 |
| VSVFMGMGGNF | ASSSGTGEDTQY | 0.46172017 |
| LASYLAAL | ASSSGTGEDTQY | 0.5695102 |
| LVGGTPAL | ASSSGTGEDTQY | 0.60676485 |
| VRNADVL | ASSSGTGEDTQY | 0.6069534 |
| TCSSPFLVSAM | ASSSGTGEDTQY | 0.57145214 |
| VHITDAE | ASSSGTGEDTQY | 0.42521888 |
| EYAAPSF | ASSSGTGEDTQY | 0.64904165 |
| VVDAFHGGL | ASSSGTGEDTQY | 0.63358504 |
| ASVVDPQ | ASSSGTGEDTQY | 0.6286307 |
| SYNEFGGLS | ASSSGTGEDTQY | 0.6335484 |
| DPDGSVY | ASSSGTGEDTQY | 0.5986601 |

| | | |
|---|---|---|
| MVSTIHATE | ASSSGTGEDTQY | 0.6185562 |
| LAAGVADPV | ASSSGTGEDTQY | 0.5138716 |
| TKEQIAA | ASSSGTGEDTQY | 0.374971 |
| SLASPAAL | ASSSGTGEDTQY | 0.58334917 |
| IGLVFYGV | ASSSGTGEDTQY | 0.549279 |
| FLLDASNL | ASSSGTGEDTQY | 0.45289826 |
| YQVGNFP | ASSSGTGEDTQY | 0.37359798 |
| ESLLSQST | ASSSGTGEDTQY | 0.6466614 |
| LFTVGFAAVG | ASSSGTGEDTQY | 0.6755053 |
| ATPPTPT | ASSSGTGEDTQY | 0.50294536 |
| TDLATLG | ASSSGTGEDTQY | 0.6623699 |
| SAGGTGR | ASSSGTGEDTQY | 0.37816817 |
| GFSNSGS | ASSSGTGEDTQY | 0.6327758 |
| GGNGGSGGTSV | ASSSGTGEDTQY | 0.64228845 |
| GTSVATGGAGN | ASSSGTGEDTQY | 0.4476174 |
| NGGNGGSGGTS | ASSSGTGEDTQY | 0.64430934 |
| AVVKFPH | ASSSGTGEDTQY | 0.412431 |
| TPPPPPP | ASSSGTGEDTQY | 0.4986028 |
| AAPAAAAAPA | ASSSGTGEDTQY | 0.6532627 |
| IALNEDL | ASSSGTGEDTQY | 0.5620562 |
| IVNVSSM | ASSSGTGEDTQY | 0.58051175 |
| TGIPPAPR | ASSSGTGEDTQY | 0.581609 |
| FDLGGGTF | ASSSGTGEDTQY | 0.5105023 |
| FDLGGGTFDVS | ASSSGTGEDTQY | 0.57069844 |
| NEPTAAA | ASSSGTGEDTQY | 0.6436407 |
| NEPTAAALA | ASSSGTGEDTQY | 0.44109863 |
| GSIINISSI | ASSSGTGEDTQY | 0.6572852 |
| VAPGNAGTA | ASSSGTGEDTQY | 0.42989472 |
| TGGGASLE | ASSSGTGEDTQY | 0.66301066 |
| VAVYNHYKRI | ASSSGTGEDTQY | 0.6203203 |
| TDPAPATT | ASSSGTGEDTQY | 0.5852399 |
| TNRPDILDPALLRPG | ASSSGTGEDTQY | 0.49929163 |
| LTTTTTS | ASSSGTGEDTQY | 0.6429152 |
| PTGSGKTL | ASSSGTGEDTQY | 0.5268374 |
| TGSGKTL | ASSSGTGEDTQY | 0.6076492 |
| TIGHVDHGK | ASSSGTGEDTQY | 0.640061 |
| TIGHVDHGKTT | ASSSGTGEDTQY | 0.22214894 |
| GHVDHGKTT | ASSSGTGEDTQY | 0.746494 |
| AQFEHTLL | ASSSGTGEDTQY | 0.62065136 |
| IVNQDEL | ASSSGTGEDTQY | 0.46239445 |
| TNGSQFF | ASSSGTGEDTQY | 0.28215766 |
| TGKGIVE | ASSSGTGEDTQY | 0.19894303 |
| LISNASDA | ASSSGTGEDTQY | 0.34275597 |
| AELRTLGELTKIAKAHGAQ | ASSSGTGEDTQY | 0.25535214 |
| APNAVAAAEPSW | ASSSGTGEDTQY | 0.79013586 |
| APQVAGAATAVL | ASSSGTGEDTQY | 0.59594405 |
| ARRDPAAAAAILPTDA | ASSSGTGEDTQY | 0.2964475 |
| DAAARDVLAEAGLAEYF | ASSSGTGEDTQY | 0.71070004 |
| DDQIVLSAKANLAKVEQ | ASSSGTGEDTQY | 0.6752658 |
| DELLPDTGEDSDEDGH | ASSSGTGEDTQY | 0.2159288 |
| ELPEAPANRATVPQD | ASSSGTGEDTQY | 0.432789 |

| | | |
|---|---|---|
| FGQSKIERVTGIG | ASSSGTGEDTQY | 0.24400094 |
| GAGGTGSTAGGAGGAGGAGGLY | ASSSGTGEDTQY | 0.3600844 |
| GHGGTGAAGGNGGTGGAGGSGI | ASSSGTGEDTQY | 0.17778094 |
| GNGGNGGAGGEGGPGGAGGAGA | ASSSGTGEDTQY | 0.5043495 |
| GSTAGNGGNGGNATNGTIAGQP | ASSSGTGEDTQY | 0.3732484 |
| GTGGDGGNGGNGGTGG | ASSSGTGEDTQY | 0.1525708 |
| GVRRPAGIPADAPV | ASSSGTGEDTQY | 0.5696541 |
| GYRQVTDAHLLTLARR | ASSSGTGEDTQY | 0.34163845 |
| IAKMGQDGHDRG | ASSSGTGEDTQY | 0.38728917 |
| KAEPSIEIQYT | ASSSGTGEDTQY | 0.6492176 |
| KPVRIRMAADEDTVAVT | ASSSGTGEDTQY | 0.34686732 |
| MPPRIAGMRLLVIKPEP | ASSSGTGEDTQY | 0.2695007 |
| MKFYAVALFRSRGVPPTR | ASSSGTGEDTQY | 0.6112425 |
| NAGGFGILGAVAHSP | ASSSGTGEDTQY | 0.66035986 |
| NEGREITLAMAEM | ASSSGTGEDTQY | 0.2368499 |
| NELVEWSHPLGHRWRWE | ASSSGTGEDTQY | 0.46050483 |
| NLGAGGTGGNGGNPG | ASSSGTGEDTQY | 0.75400853 |
| NTEWPKWETVMAADPVD | ASSSGTGEDTQY | 0.4569255 |
| PASAAPAETQP | ASSSGTGEDTQY | 0.6427562 |
| QQSSDGKNGVRAAEITGE | ASSSGTGEDTQY | 0.6719806 |
| RAPDAVTTSWWKE | ASSSGTGEDTQY | 0.3210585 |
| SGFGNVGAHTSGWFNQ | ASSSGTGEDTQY | 0.67643553 |
| SGVAGAVGQAASVGGLKV | ASSSGTGEDTQY | 0.7915641 |
| SNGGAGTGSAGGNGGT | ASSSGTGEDTQY | 0.33697602 |
| TGGAGGEGGAAG | ASSSGTGEDTQY | 0.3996607 |
| TGGDGGAAGAGGAPGHG | ASSSGTGEDTQY | 0.23074912 |
| TMPPLLAAIANAGRHGV | ASSSGTGEDTQY | 0.44237733 |
| TSSTAGNSGGA | ASSSGTGEDTQY | 0.19814584 |
| VPATVPVGILC | ASSSGTGEDTQY | 0.68235755 |
| VQRPDSGGVVLTGRIS | ASSSGTGEDTQY | 0.6291554 |
| WTPNPVPATKGVPAGLVQAT | ASSSGTGEDTQY | 0.44768566 |
| YALKRALAKFSDRFDVVII | ASSSGTGEDTQY | 0.20735952 |
| YLIGSVIV | ASSSGTGEDTQY | 0.16816357 |
| YGQLVAVID | ASSSGTGEDTQY | 0.5571272 |
| VTVGGMTLPA | ASSSGTGEDTQY | 0.22206762 |
| VLALPAV | ASSSGTGEDTQY | 0.1982951 |
| VIPPTLNYET | ASSSGTGEDTQY | 0.3779261 |
| VGMGGMHPSSG | ASSSGTGEDTQY | 0.6291468 |
| VGLAGGAATA | ASSSGTGEDTQY | 0.6420058 |
| VGGAGGAGGSGP | ASSSGTGEDTQY | 0.34380525 |
| VFAAFMV | ASSSGTGEDTQY | 0.7556151 |
| TLYDKLR | ASSSGTGEDTQY | 0.7365922 |
| TLAVFTVL | ASSSGTGEDTQY | 0.58150136 |
| TGSVQPARV | ASSSGTGEDTQY | 0.37969995 |
| TGGGAGSGGT | ASSSGTGEDTQY | 0.23347269 |
| SHLETSL | ASSSGTGEDTQY | 0.80235463 |
| SFTTGMT | ASSSGTGEDTQY | 0.44522578 |
| QVEVAKRMGVRQPTVS | ASSSGTGEDTQY | 0.53567564 |
| QGSGGPALAM | ASSSGTGEDTQY | 0.6313236 |
| MLAPSNP | ASSSGTGEDTQY | 0.46005517 |
| LVNNAAI | ASSSGTGEDTQY | 0.295124 |

| | | |
|---|---|---|
| LTGEINTLA | ASSSGTGEDTQY | 0.6484097 |
| LSIFGPV | ASSSGTGEDTQY | 0.62396955 |
| LLALLANL | ASSSGTGEDTQY | 0.261459 |
| LISNASDALDKLR | ASSSGTGEDTQY | 0.21175921 |
| LINEAVP | ASSSGTGEDTQY | 0.44790196 |
| LGYAFSGL | ASSSGTGEDTQY | 0.50052834 |
| LDFVAVAL | ASSSGTGEDTQY | 0.6565603 |
| LAQPGAE | ASSSGTGEDTQY | 0.47989488 |
| ISNLGAVTGF | ASSSGTGEDTQY | 0.3663234 |
| HQEDPGRI | ASSSGTGEDTQY | 0.6147921 |
| GTPGEPTP | ASSSGTGEDTQY | 0.51346785 |
| GFFNSGS | ASSSGTGEDTQY | 0.49205726 |
| GAGGGGLTAGAN | ASSSGTGEDTQY | 0.5326413 |
| FVDALVNM | ASSSGTGEDTQY | 0.4044975 |
| FLTVAPEL | ASSSGTGEDTQY | 0.59568053 |
| FETLTAL | ASSSGTGEDTQY | 0.6256826 |
| DVINPIEL | ASSSGTGEDTQY | 0.46774614 |
| APVSGAPV | ASSSGTGEDTQY | 0.6308419 |
| ALVGGLAW | ASSSGTGEDTQY | 0.44364145 |
| ALAAAAATAI | ASSSGTGEDTQY | 0.42681494 |
| AFGLGLLL | ASSSGTGEDTQY | 0.19163519 |
| AFFAAASL | ASSSGTGEDTQY | 0.43827495 |
| AAAALPAFSPP | ASSSGTGEDTQY | 0.43663526 |
| VVLAPAL | ASSSGTGEDTQY | 0.6122006 |
| TIGHVDHGKTTL | ASSSGTGEDTQY | 0.66532934 |
| LISNASDALDK | ASSSGTGEDTQY | 0.47881216 |
| LISNASDAL | ASSSGTGEDTQY | 0.32549927 |
| ISNASDALDK | ASSSGTGEDTQY | 0.60393476 |
| NQAETLVYQTE | ASSSGTGEDTQY | 0.71912587 |
| TGGTGGTAGAG | ASSSGTGEDTQY | 0.22014868 |
| TGSTAGGAGGA | ASSSGTGEDTQY | 0.45079255 |
| DPLAVPVAAGPV | ASSSGTGEDTQY | 0.7897167 |
| GGTGGTPGNGGN | ASSSGTGEDTQY | 0.6493842 |
| PAGLAGVDPATL | ASSSGTGEDTQY | 0.37461537 |
| GNNLVVEAPHAGASV | ASSSGTGEDTQY | 0.37742025 |
| KVTRIMSASTLEQPA | ASSSGTGEDTQY | 0.80760884 |
| LGFALPIALPRRTVV | ASSSGTGEDTQY | 0.39743513 |
| TRLGNLTARKLVVLV | ASSSGTGEDTQY | 0.7724679 |
| WSQQRRLIYFPSAGP | ASSSGTGEDTQY | 0.16537492 |
| GPNGNPDPMAAAVDIR | ASSSGTGEDTQY | 0.31332025 |
| TQRCVSVADLRAAASR | ASSSGTGEDTQY | 0.24869862 |
| QFVLDVAEHPVEVLTETV | ASSSGTGEDTQY | 0.15534873 |
| GGAGGTGGAGGNGGTQAGDG | ASSSGTGEDTQY | 0.69575983 |
| PLLATLRAIAVDVDAKPAQV | ASSSGTGEDTQY | 0.29731688 |
| APKFRTGGYMIDFWGVGYQVAK | ASSSGTGEDTQY | 0.6179347 |
| GAGGAGGAGGAAGGPGGTGGTG | ASSSGTGEDTQY | 0.3557706 |
| IGALGLLATDVAPAGSVAFGVA | ASSSGTGEDTQY | 0.664566 |
| AGGAGGIGGNGGAGGAGGVLMG | ASSSGTGEDTQY | 0.5392596 |
| ADELIGLL | ASSSGTGEDTQY | 0.40736306 |
| ADRALKA | ASSSGTGEDTQY | 0.24590157 |
| AGAVGQVGGAGGA | ASSSGTGEDTQY | 0.55409265 |

| | | |
|---|---|---|
| AGGGGGLGGAP | ASSSGTGEDTQY | 0.67149067 |
| AGGLLYGNGGAG | ASSSGTGEDTQY | 0.5234988 |
| AGLESIGL | ASSSGTGEDTQY | 0.18088564 |
| AGLKTGII | ASSSGTGEDTQY | 0.6001095 |
| ASLAALLL | ASSSGTGEDTQY | 0.3768764 |
| ASVINISSI | ASSSGTGEDTQY | 0.2066072 |
| ATPPGPA | ASSSGTGEDTQY | 0.17198817 |
| ATVASALV | ASSSGTGEDTQY | 0.33163816 |
| ATVASLIL | ASSSGTGEDTQY | 0.5076951 |
| AVNGPVLIRH | ASSSGTGEDTQY | 0.67090064 |
| DGAAGAVGI | ASSSGTGEDTQY | 0.68623114 |
| DPLLHGI | ASSSGTGEDTQY | 0.31877056 |
| ELGAALGAL | ASSSGTGEDTQY | 0.65338075 |
| ELGRTQAV | ASSSGTGEDTQY | 0.5528264 |
| FVAGGYLS | ASSSGTGEDTQY | 0.12812811 |
| GASGGAGGAGGW | ASSSGTGEDTQY | 0.32092473 |
| GGAGAGGDGGAGGI | ASSSGTGEDTQY | 0.20868349 |
| GTGHNGSQPGMGGNG | ASSSGTGEDTQY | 0.6261189 |
| GYAAASASAS | ASSSGTGEDTQY | 0.30816406 |
| HQVDVAAVQQG | ASSSGTGEDTQY | 0.444001 |
| IGAAIAHT | ASSSGTGEDTQY | 0.41301653 |
| IGMIEVAQL | ASSSGTGEDTQY | 0.76513773 |
| IRLAPEHTSAMD | ASSSGTGEDTQY | 0.5192951 |
| IYNVSVL | ASSSGTGEDTQY | 0.22463132 |
| KQGFLGM | ASSSGTGEDTQY | 0.19825548 |
| LESVITQL | ASSSGTGEDTQY | 0.50780606 |
| LGLGALVVV | ASSSGTGEDTQY | 0.32590088 |
| LIGNGGNGGSGGI | ASSSGTGEDTQY | 0.25472382 |
| LNAINAPTL | ASSSGTGEDTQY | 0.23262683 |
| LQLLNTGV | ASSSGTGEDTQY | 0.7417189 |
| LQTSIEF | ASSSGTGEDTQY | 0.18784164 |
| MSFAVSEL | ASSSGTGEDTQY | 0.555399 |
| NTGALGL | ASSSGTGEDTQY | 0.24651954 |
| PPGAGGAGGAGTT | ASSSGTGEDTQY | 0.44921082 |
| QLVGLGI | ASSSGTGEDTQY | 0.755081 |
| RFLEQVHQL | ASSSGTGEDTQY | 0.7572366 |
| RGVGYVLR | ASSSGTGEDTQY | 0.39112473 |
| SATGVFAGVI | ASSSGTGEDTQY | 0.2992581 |
| SLPAITLP | ASSSGTGEDTQY | 0.24522254 |
| STALAFTV | ASSSGTGEDTQY | 0.28406113 |
| SVLTVGLGK | ASSSGTGEDTQY | 0.31280607 |
| TALAGAGSGI | ASSSGTGEDTQY | 0.71928304 |
| TGLPFTGI | ASSSGTGEDTQY | 0.38092673 |
| VAEHVTV | ASSSGTGEDTQY | 0.22281699 |
| VDIVLNSL | ASSSGTGEDTQY | 0.52498186 |
| VLTGGGAVL | ASSSGTGEDTQY | 0.75298226 |
| VNVGLAYGI | ASSSGTGEDTQY | 0.7957537 |
| VSSAAVGAV | ASSSGTGEDTQY | 0.301656 |
| VTSVYEAGV | ASSSGTGEDTQY | 0.302548 |
| WVALDRGA | ASSSGTGEDTQY | 0.3397086 |
| YLVGSVVI | ASSSGTGEDTQY | 0.5057739 |

| | | |
|---|---|---|
| GPPGTGKTLL | ASSSGTGEDTQY | 0.2622151 |
| GVGMTKF | ASSSGTGEDTQY | 0.26701403 |
| IGIGGSDL | ASSSGTGEDTQY | 0.55921876 |
| IGTLLAEL | ASSSGTGEDTQY | 0.8328036 |
| IYTSGTT | ASSSGTGEDTQY | 0.39943498 |
| LSFPAGV | ASSSGTGEDTQY | 0.17941365 |
| LYTSGTT | ASSSGTGEDTQY | 0.4026076 |
| VGAFPFGF | ASSSGTGEDTQY | 0.23078232 |
| VGAGTVEF | ASSSGTGEDTQY | 0.2742707 |
| VGKSTLLL | ASSSGTGEDTQY | 0.35539338 |
| AALAHAGGGIV | ASSSGTGEDTQY | 0.19766797 |
| AALVAVSLL | ASSSGTGEDTQY | 0.41719773 |
| AAVSRAVAAL | ASSSGTGEDTQY | 0.7257132 |
| AGGSSGIGGF | ASSSGTGEDTQY | 0.38652897 |
| AGLTVANANG | ASSSGTGEDTQY | 0.4975052 |
| AGLVGAVL | ASSSGTGEDTQY | 0.7697481 |
| ALSQAFTV | ASSSGTGEDTQY | 0.6015277 |
| ALTFSVSL | ASSSGTGEDTQY | 0.75124407 |
| ALTGAALVAL | ASSSGTGEDTQY | 0.7480601 |
| ASFSIGAGVL | ASSSGTGEDTQY | 0.3825525 |
| DGSVYKI | ASSSGTGEDTQY | 0.36883533 |
| EQLAEIV | ASSSGTGEDTQY | 0.7934477 |
| FAVSELTVM | ASSSGTGEDTQY | 0.37760085 |
| GAAPLAAGL | ASSSGTGEDTQY | 0.4827517 |
| GADGGDGGSGGAGG | ASSSGTGEDTQY | 0.369146 |
| GFGLDTGGAGG | ASSSGTGEDTQY | 0.71552825 |
| GLASGGLL | ASSSGTGEDTQY | 0.38565797 |
| GRIAGAAVLL | ASSSGTGEDTQY | 0.7766759 |
| GSAGSLNTSF | ASSSGTGEDTQY | 0.16956381 |
| GTGQIGI | ASSSGTGEDTQY | 0.25003242 |
| GTLIALIL | ASSSGTGEDTQY | 0.7406606 |
| IAATMGFVVG | ASSSGTGEDTQY | 0.13802204 |
| IFAQALL | ASSSGTGEDTQY | 0.17629443 |
| IGAAVALGF | ASSSGTGEDTQY | 0.40767103 |
| IGLTPLAL | ASSSGTGEDTQY | 0.7181912 |
| ILGSNTSTL | ASSSGTGEDTQY | 0.14428796 |
| IVFSLTVL | ASSSGTGEDTQY | 0.32269728 |
| LASKATGF | ASSSGTGEDTQY | 0.18371548 |
| LASVALIL | ASSSGTGEDTQY | 0.5410631 |
| LFAVGLTL | ASSSGTGEDTQY | 0.41458204 |
| LFNGGIGTYI | ASSSGTGEDTQY | 0.18904264 |
| LTLGAGII | ASSSGTGEDTQY | 0.5883324 |
| LVAAALGDI | ASSSGTGEDTQY | 0.18496937 |
| NTAFVQLGI | ASSSGTGEDTQY | 0.5534277 |
| QLFSGAEL | ASSSGTGEDTQY | 0.6312574 |
| SAATFITL | ASSSGTGEDTQY | 0.8153517 |
| SNYGLANLGV | ASSSGTGEDTQY | 0.2285823 |
| SYGGAYAVM | ASSSGTGEDTQY | 0.5293853 |
| TTGVGGLTLG | ASSSGTGEDTQY | 0.3285002 |
| TTQTIYI | ASSSGTGEDTQY | 0.8300448 |
| VAAGAGLGAVI | ASSSGTGEDTQY | 0.3234678 |

| | | |
|---|---|---|
| VAAVVTLL | ASSSGTGEDTQY | 0.18205152 |
| VGFLGGLI | ASSSGTGEDTQY | 0.65598804 |
| VGVIPILL | ASSSGTGEDTQY | 0.52753705 |
| VIFASSLI | ASSSGTGEDTQY | 0.36205155 |
| VKGGDGGAAAG | ASSSGTGEDTQY | 0.7440295 |
| VTKTFTNI | ASSSGTGEDTQY | 0.3379576 |
| VVAAAILL | ASSSGTGEDTQY | 0.37983003 |
| YAALDVEL | ASSSGTGEDTQY | 0.70675814 |
| LASVALLL | ASSSGTGEDTQY | 0.60415316 |
| LDAVHRV | ASSSGTGEDTQY | 0.31010115 |
| QGLSSSIF | ASSSGTGEDTQY | 0.6520668 |
| RGTFANIRL | ASSSGTGEDTQY | 0.54286736 |
| RQLWWGHRI | ASSSGTGEDTQY | 0.25076798 |
| YGPPGTGKTLL | ASSSGTGEDTQY | 0.7978256 |
| AALVGAPANVA | ASSSGTGEDTQY | 0.31082714 |
| AATFRRTE | ASSSGTGEDTQY | 0.6641915 |
| AGALGGGI | ASSSGTGEDTQY | 0.5187111 |
| AGGAGGAGGIGAG | ASSSGTGEDTQY | 0.8097442 |
| AGSGALGL | ASSSGTGEDTQY | 0.25885594 |
| AQIADQI | ASSSGTGEDTQY | 0.60983384 |
| ASAVAVTYL | ASSSGTGEDTQY | 0.52669823 |
| ASTVLGGTGG | ASSSGTGEDTQY | 0.5329035 |
| AVGGEPAK | ASSSGTGEDTQY | 0.36438307 |
| CEQGTFGGTC | ASSSGTGEDTQY | 0.5379587 |
| DEGKAPAG | ASSSGTGEDTQY | 0.5206147 |
| DTVVFPAL | ASSSGTGEDTQY | 0.72747463 |
| FEGITMAG | ASSSGTGEDTQY | 0.39426422 |
| FPLVVQD | ASSSGTGEDTQY | 0.42599326 |
| GAFTTPQ | ASSSGTGEDTQY | 0.7992189 |
| GFGGFGTAGGVGGA | ASSSGTGEDTQY | 0.2438735 |
| GFLAQFL | ASSSGTGEDTQY | 0.4644464 |
| GGAGGAGGNAGVA | ASSSGTGEDTQY | 0.59267175 |
| GGVGTAGAGGAGGAGG | ASSSGTGEDTQY | 0.67488414 |
| GNGGGGDAGNAGSGGN | ASSSGTGEDTQY | 0.35507828 |
| GTGGAGGAGNDAGS | ASSSGTGEDTQY | 0.70501167 |
| GTTNTGWFNS | ASSSGTGEDTQY | 0.4192598 |
| IFMGPLFTL | ASSSGTGEDTQY | 0.6375332 |
| ILAHAIEAI | ASSSGTGEDTQY | 0.5446985 |
| IRSAIREAL | ASSSGTGEDTQY | 0.39073178 |
| ITFAGGG | ASSSGTGEDTQY | 0.29768974 |
| ITGMSGHQEV | ASSSGTGEDTQY | 0.5545865 |
| IVAGLACTL | ASSSGTGEDTQY | 0.70629406 |
| IVGGGLAGA | ASSSGTGEDTQY | 0.63976204 |
| IVTGVLNI | ASSSGTGEDTQY | 0.17570479 |
| LAQQGCI | ASSSGTGEDTQY | 0.6244724 |
| LCLHVEN | ASSSGTGEDTQY | 0.62452734 |
| LDAISDGS | ASSSGTGEDTQY | 0.82981336 |
| LGGFAAML | ASSSGTGEDTQY | 0.33476573 |
| LGGKNPMI | ASSSGTGEDTQY | 0.5846945 |
| LGPKGVAAIL | ASSSGTGEDTQY | 0.40760517 |
| LMVGVSI | ASSSGTGEDTQY | 0.41856313 |

| | | |
|---|---|---|
| LVGAAFVL | ASSSGTGEDTQY | 0.19447534 |
| LVLAVAPL | ASSSGTGEDTQY | 0.30129087 |
| MAAIAVIDAA | ASSSGTGEDTQY | 0.46500933 |
| NNGINAELQ | ASSSGTGEDTQY | 0.33113855 |
| RGEPLVLVHG | ASSSGTGEDTQY | 0.77798146 |
| RGGAGGAAGLI | ASSSGTGEDTQY | 0.69216776 |
| RSYRMHPAV | ASSSGTGEDTQY | 0.65853715 |
| SAAHLLAGQLA | ASSSGTGEDTQY | 0.75360316 |
| SAAVGSY | ASSSGTGEDTQY | 0.7802923 |
| SPHAVLHRI | ASSSGTGEDTQY | 0.53086436 |
| SQIAKVL | ASSSGTGEDTQY | 0.29598504 |
| TALGHLVP | ASSSGTGEDTQY | 0.45513332 |
| TAPTTSGAA | ASSSGTGEDTQY | 0.22225222 |
| TDGDDGNGGSPGT | ASSSGTGEDTQY | 0.32192776 |
| TDGLGEN | ASSSGTGEDTQY | 0.32481104 |
| TGGPPLPGQA | ASSSGTGEDTQY | 0.45045564 |
| TIPAITI | ASSSGTGEDTQY | 0.14068994 |
| TIPGFTI | ASSSGTGEDTQY | 0.7550679 |
| TSIASGALA | ASSSGTGEDTQY | 0.7416419 |
| TVTLPGAAI | ASSSGTGEDTQY | 0.4697088 |
| VASGLSAALP | ASSSGTGEDTQY | 0.5985082 |
| VGAGVIGI | ASSSGTGEDTQY | 0.6183165 |
| VGPKRFGV | ASSSGTGEDTQY | 0.3274059 |
| VNVAFPD | ASSSGTGEDTQY | 0.64688104 |
| VSMTNSGVS | ASSSGTGEDTQY | 0.3055398 |
| VTDVLGDLSV | ASSSGTGEDTQY | 0.6253704 |
| VVVVGAGFAGL | ASSSGTGEDTQY | 0.7039418 |
| VYTVGTVV | ASSSGTGEDTQY | 0.5295929 |
| WTTFSGP | ASSSGTGEDTQY | 0.34205234 |
| YDELVEAF | ASSSGTGEDTQY | 0.42886126 |
| YGTGGAGGNGG | ASSSGTGEDTQY | 0.21251778 |
| YTNVMAV | ASSSGTGEDTQY | 0.37378007 |
| FLEQVHQL | ASSSGTGEDTQY | 0.2606122 |
| GLSLLAVAV | ASSSGTGEDTQY | 0.53518623 |
| LEQVHQL | ASSSGTGEDTQY | 0.216661 |
| LNAAGAI | ASSSGTGEDTQY | 0.5650079 |
| PAVHASL | ASSSGTGEDTQY | 0.5449363 |
| TELEDDA | ASSSGTGEDTQY | 0.44663936 |
| TPPGPPN | ASSSGTGEDTQY | 0.22911075 |
| TPPPPPPP | ASSSGTGEDTQY | 0.6282363 |
| TQSPALI | ASSPGGPQDTQY | 0.5809829 |
| EFAAAFPQ | ASSPGGPQDTQY | 0.20676923 |
| LLSGNLDVL | ASSPGGPQDTQY | 0.5847865 |
| GPANLGNYNI | ASSPGGPQDTQY | 0.32687756 |
| TLGPTLP | ASSPGGPQDTQY | 0.49473622 |
| VYAYSSCCK | ASSPGGPQDTQY | 0.65834904 |
| APAPAPAPNI | ASSPGGPQDTQY | 0.4404832 |
| PGGPVPIP | ASSPGGPQDTQY | 0.1829301 |
| FLTVALEP | ASSPGGPQDTQY | 0.4499488 |
| SQYFGANL | ASSPGGPQDTQY | 0.4807307 |
| GGVKPAA | ASSPGGPQDTQY | 0.2938535 |

| | | |
|---|---|---|
| GGTGGAGGAGGW | ASSPGGPQDTQY | 0.14439467 |
| GVGGVAGLGGAGG | ASSPGGPQDTQY | 0.58128977 |
| VLLVLATTI | ASSPGGPQDTQY | 0.25144595 |
| IGIFAAML | ASSPGGPQDTQY | 0.31928897 |
| AVGDLID | ASSPGGPQDTQY | 0.41722926 |
| NISPLPL | ASSPGGPQDTQY | 0.68264157 |
| VTAVFVAI | ASSPGGPQDTQY | 0.44086242 |
| SGDDNDAAL | ASSPGGPQDTQY | 0.3515306 |
| AVAALAHS | ASSPGGPQDTQY | 0.5999317 |
| GGVTVMATP | ASSPGGPQDTQY | 0.45920712 |
| AGLAANVRD | ASSPGGPQDTQY | 0.726681 |
| TNRPDLIDPALLRPG | ASSPGGPQDTQY | 0.48941314 |
| LANLGVTAL | ASSPGGPQDTQY | 0.31731123 |
| TGERIEL | ASSPGGPQDTQY | 0.56962997 |
| NGEELAA | ASSPGGPQDTQY | 0.33074906 |
| VLSLVWG | ASSPGGPQDTQY | 0.27500978 |
| IGYALWL | ASSPGGPQDTQY | 0.24482076 |
| LSLLLAPL | ASSPGGPQDTQY | 0.21233723 |
| TLDEALK | ASSPGGPQDTQY | 0.22389026 |
| TDGAGTAEGG | ASSPGGPQDTQY | 0.48627836 |
| NAPGGAGGNGG | ASSPGGPQDTQY | 0.61088026 |
| VSVFMGMGGNF | ASSPGGPQDTQY | 0.4175756 |
| LASYLAAL | ASSPGGPQDTQY | 0.19457285 |
| LVGGTPAL | ASSPGGPQDTQY | 0.3629852 |
| VRNADVL | ASSPGGPQDTQY | 0.37250936 |
| TCSSPFLVSAM | ASSPGGPQDTQY | 0.6656518 |
| VHITDAE | ASSPGGPQDTQY | 0.45690548 |
| EYAAPSF | ASSPGGPQDTQY | 0.8263552 |
| VVDAFHGGL | ASSPGGPQDTQY | 0.7007791 |
| ASVVDPQ | ASSPGGPQDTQY | 0.4022077 |
| SYNEFGGLS | ASSPGGPQDTQY | 0.64553076 |
| DPDGSVY | ASSPGGPQDTQY | 0.31031892 |
| MVSTIHATE | ASSPGGPQDTQY | 0.43967447 |
| LAAGVADPV | ASSPGGPQDTQY | 0.43657723 |
| TKEQIAA | ASSPGGPQDTQY | 0.55427945 |
| SLASPAAL | ASSPGGPQDTQY | 0.30302262 |
| IGLVFYGV | ASSPGGPQDTQY | 0.34154627 |
| FLLDASNL | ASSPGGPQDTQY | 0.17497009 |
| YQVGNFP | ASSPGGPQDTQY | 0.60473937 |
| ESLLSQST | ASSPGGPQDTQY | 0.21703702 |
| LFTVGFAAVG | ASSPGGPQDTQY | 0.20932779 |
| ATPPTPT | ASSPGGPQDTQY | 0.32272625 |
| TDLATLG | ASSPGGPQDTQY | 0.800776 |
| SAGGTGR | ASSPGGPQDTQY | 0.7648711 |
| GFSNSGS | ASSPGGPQDTQY | 0.31461036 |
| GGNGGSGGTSV | ASSPGGPQDTQY | 0.24528378 |
| GTSVATGGAGN | ASSPGGPQDTQY | 0.76495564 |
| NGGNGGSGGTS | ASSPGGPQDTQY | 0.26501286 |
| AVVKFPH | ASSPGGPQDTQY | 0.48883796 |
| TPPPPPP | ASSPGGPQDTQY | 0.53937805 |
| AAPAAAAAPA | ASSPGGPQDTQY | 0.49832082 |

| | | |
|---|---|---|
| IALNEDL | ASSPGGPQDTQY | 0.25829613 |
| IVNVSSM | ASSPGGPQDTQY | 0.6523653 |
| TGIPPAPR | ASSPGGPQDTQY | 0.40639508 |
| FDLGGGTF | ASSPGGPQDTQY | 0.40573874 |
| FDLGGGTFDVS | ASSPGGPQDTQY | 0.20837146 |
| NEPTAAA | ASSPGGPQDTQY | 0.6558678 |
| NEPTAAALA | ASSPGGPQDTQY | 0.2793575 |
| GSIINISSI | ASSPGGPQDTQY | 0.23739348 |
| VAPGNAGTA | ASSPGGPQDTQY | 0.15868169 |
| TGGGASLE | ASSPGGPQDTQY | 0.36101282 |
| VAVYNHYKRI | ASSPGGPQDTQY | 0.6827061 |
| TDPAPATT | ASSPGGPQDTQY | 0.7850901 |
| TNRPDILDPALLRPG | ASSPGGPQDTQY | 0.47375962 |
| LTTTTTS | ASSPGGPQDTQY | 0.6164584 |
| PTGSGKTL | ASSPGGPQDTQY | 0.69671357 |
| TGSGKTL | ASSPGGPQDTQY | 0.39940876 |
| TIGHVDHGK | ASSPGGPQDTQY | 0.18491384 |
| TIGHVDHGKTT | ASSPGGPQDTQY | 0.4778421 |
| GHVDHGKTT | ASSPGGPQDTQY | 0.49577123 |
| AQFEHTLL | ASSPGGPQDTQY | 0.71378565 |
| IVNQDEL | ASSPGGPQDTQY | 0.7009846 |
| TNGSQFF | ASSPGGPQDTQY | 0.20903139 |
| TGKGIVE | ASSPGGPQDTQY | 0.48277122 |
| LISNASDA | ASSPGGPQDTQY | 0.52334297 |
| AELRTLGELTKIAKAHGAQ | ASSPGGPQDTQY | 0.20650306 |
| APNAVAAAEPSW | ASSPGGPQDTQY | 0.31073135 |
| APQVAGAATAVL | ASSPGGPQDTQY | 0.27760893 |
| ARRDPAAAAAILPTDA | ASSPGGPQDTQY | 0.22858605 |
| DAAARDVLAEAGLAEYF | ASSPGGPQDTQY | 0.4835526 |
| DDQIVLSAKANLAKVEQ | ASSPGGPQDTQY | 0.41943514 |
| DELLPDTGEDSDEDGH | ASSPGGPQDTQY | 0.36618835 |
| ELPEAPANRATVPQD | ASSPGGPQDTQY | 0.3608325 |
| FGQSKIERVTGIG | ASSPGGPQDTQY | 0.30441976 |
| GAGGTGSTAGGAGGAGGAGGLY | ASSPGGPQDTQY | 0.305582 |
| GHGGTGAAGGNGGTGGAGGSGI | ASSPGGPQDTQY | 0.38418418 |
| GNGGNGGAGGEGGPGGAGGAGA | ASSPGGPQDTQY | 0.50282145 |
| GSTAGNGGNGGNATNGTIAGQP | ASSPGGPQDTQY | 0.68481445 |
| GTGGDGGNGGNGGTGG | ASSPGGPQDTQY | 0.4875397 |
| GVRRPAGIPADAPV | ASSPGGPQDTQY | 0.43330628 |
| GYRQVTDAHLLTLARR | ASSPGGPQDTQY | 0.7736089 |
| IAKMGQDGHDRG | ASSPGGPQDTQY | 0.2090007 |
| KAEPSIEIQYT | ASSPGGPQDTQY | 0.27326354 |
| KPVRIRMAADEDTVAVT | ASSPGGPQDTQY | 0.5816338 |
| MPPRIAGMRLLVIKPEP | ASSPGGPQDTQY | 0.34474677 |
| MKFYAVALFRSRGVPPTR | ASSPGGPQDTQY | 0.6309255 |
| NAGGFGILGAVAHSP | ASSPGGPQDTQY | 0.58960766 |
| NEGREITLAMAEM | ASSPGGPQDTQY | 0.5267402 |
| NELVEWSHPLGHRWRWE | ASSPGGPQDTQY | 0.48375985 |
| NLGAGGTGGNGGNPG | ASSPGGPQDTQY | 0.8176819 |
| NTEWPKWETVMAADPVD | ASSPGGPQDTQY | 0.59233063 |
| PASAAPAETQP | ASSPGGPQDTQY | 0.31764847 |

| | | |
|---|---|---|
| QQSSDGKNGVRAAEITGE | ASSPGGPQDTQY | 0.73877287 |
| RAPDAVTTSWWKE | ASSPGGPQDTQY | 0.21811284 |
| SGFGNVGAHTSGWFNQ | ASSPGGPQDTQY | 0.6078288 |
| SGVAGAVGQAASVGGLKV | ASSPGGPQDTQY | 0.5884172 |
| SNGGAGTGSAGGNGGT | ASSPGGPQDTQY | 0.2601787 |
| TGGAGGEGGAAG | ASSPGGPQDTQY | 0.38763964 |
| TGGDGGAAGAGGAPGHG | ASSPGGPQDTQY | 0.6898399 |
| TMPPLLAAIANAGRHGV | ASSPGGPQDTQY | 0.71360564 |
| TSSTAGNSGGA | ASSPGGPQDTQY | 0.21108142 |
| VPATVPVGILC | ASSPGGPQDTQY | 0.32672626 |
| VQRPDSGGVVLTGRIS | ASSPGGPQDTQY | 0.24024945 |
| WTPNPVPATKGVPAGLVQAT | ASSPGGPQDTQY | 0.2521785 |
| YALKRALAKFSDRFDVVII | ASSPGGPQDTQY | 0.21365663 |
| YLIGSVIV | ASSPGGPQDTQY | 0.5798904 |
| YGQLVAVID | ASSPGGPQDTQY | 0.3674559 |
| VTVGGMTLPA | ASSPGGPQDTQY | 0.78426504 |
| VLALPAV | ASSPGGPQDTQY | 0.48081672 |
| VIPPTLNYET | ASSPGGPQDTQY | 0.69481206 |
| VGMGGMHPSSG | ASSPGGPQDTQY | 0.2929014 |
| VGLAGGAATA | ASSPGGPQDTQY | 0.44906622 |
| VGGAGGAGGSGP | ASSPGGPQDTQY | 0.17758118 |
| VFAAFMV | ASSPGGPQDTQY | 0.23371188 |
| TLYDKLR | ASSPGGPQDTQY | 0.23206924 |
| TLAVFTVL | ASSPGGPQDTQY | 0.45743233 |
| TGSVQPARV | ASSPGGPQDTQY | 0.16867092 |
| TGGGAGSGGT | ASSPGGPQDTQY | 0.75569063 |
| SHLETSL | ASSPGGPQDTQY | 0.34349376 |
| SFTTGMT | ASSPGGPQDTQY | 0.5503036 |
| QVEVAKRMGVRQPTVS | ASSPGGPQDTQY | 0.29849324 |
| QGSGGPALAM | ASSPGGPQDTQY | 0.22792527 |
| MLAPSNP | ASSPGGPQDTQY | 0.35659826 |
| LVNNAAI | ASSPGGPQDTQY | 0.6552502 |
| LTGEINTLA | ASSPGGPQDTQY | 0.28126127 |
| LSIFGPV | ASSPGGPQDTQY | 0.16202073 |
| LLALLANL | ASSPGGPQDTQY | 0.26919484 |
| LISNASDALDKLR | ASSPGGPQDTQY | 0.24850999 |
| LINEAVP | ASSPGGPQDTQY | 0.79739046 |
| LGYAFSGL | ASSPGGPQDTQY | 0.6900338 |
| LDFVAVAL | ASSPGGPQDTQY | 0.4613755 |
| LAQPGAE | ASSPGGPQDTQY | 0.7824549 |
| ISNLGAVTGF | ASSPGGPQDTQY | 0.14233743 |
| HQEDPGRI | ASSPGGPQDTQY | 0.36800334 |
| GTPGEPTP | ASSPGGPQDTQY | 0.7718787 |
| GFFNSGS | ASSPGGPQDTQY | 0.4163788 |
| GAGGGGLTAGAN | ASSPGGPQDTQY | 0.40195227 |
| FVDALVNM | ASSPGGPQDTQY | 0.6557564 |
| FLTVAPEL | ASSPGGPQDTQY | 0.3182053 |
| FETLTAL | ASSPGGPQDTQY | 0.29530188 |
| DVINPIEL | ASSPGGPQDTQY | 0.48664021 |
| APVSGAPV | ASSPGGPQDTQY | 0.5590236 |
| ALVGGLAW | ASSPGGPQDTQY | 0.52070713 |

| | | |
|---|---|---|
| ALAAAAATAI | ASSPGGPQDTQY | 0.19967358 |
| AFGLGLLL | ASSPGGPQDTQY | 0.46664357 |
| AFFAAASL | ASSPGGPQDTQY | 0.83796734 |
| AAAALPAFSPP | ASSPGGPQDTQY | 0.40105382 |
| VVLAPAL | ASSPGGPQDTQY | 0.5491129 |
| TIGHVDHGKTTL | ASSPGGPQDTQY | 0.44190687 |
| LISNASDALDK | ASSPGGPQDTQY | 0.30428806 |
| LISNASDAL | ASSPGGPQDTQY | 0.28169006 |
| ISNASDALDK | ASSPGGPQDTQY | 0.2570108 |
| NQAETLVYQTE | ASSPGGPQDTQY | 0.6268074 |
| TGGTGGTAGAG | ASSPGGPQDTQY | 0.23729968 |
| TGSTAGGAGGA | ASSPGGPQDTQY | 0.3879192 |
| DPLAVPVAAGPV | ASSPGGPQDTQY | 0.41037452 |
| GGTGGTPGNGGN | ASSPGGPQDTQY | 0.8355979 |
| PAGLAGVDPATL | ASSPGGPQDTQY | 0.5998837 |
| GNNLVVEAPHAGASV | ASSPGGPQDTQY | 0.5523983 |
| KVTRIMSASTLEQPA | ASSPGGPQDTQY | 0.32039958 |
| LGFALPIALPRRTVV | ASSPGGPQDTQY | 0.40969783 |
| TRLGNLTARKLVVLV | ASSPGGPQDTQY | 0.66146016 |
| WSQQRRLIYFPSAGP | ASSPGGPQDTQY | 0.18359257 |
| GPNGNPDPMAAAVDIR | ASSPGGPQDTQY | 0.72657174 |
| TQRCVSVADLRAAASR | ASSPGGPQDTQY | 0.24254915 |
| QFVLDVAEHPVEVLTETV | ASSPGGPQDTQY | 0.5024234 |
| GGAGGTGGAGGNGGTQAGDG | ASSPGGPQDTQY | 0.50417304 |
| PLLATLRAIAVDVDAKPAQV | ASSPGGPQDTQY | 0.8074114 |
| APKFRTGGYMIDFWGVGYQVAK | ASSPGGPQDTQY | 0.4235233 |
| GAGGAGGAGGAAGGPGGTGGTG | ASSPGGPQDTQY | 0.5670532 |
| IGALGLLATDVAPAGSVAFGVA | ASSPGGPQDTQY | 0.29096824 |
| AGGAGGIGGNGGAGGAGGVLMG | ASSPGGPQDTQY | 0.5917375 |
| ADELIGLL | ASSPGGPQDTQY | 0.76064956 |
| ADRALKA | ASSPGGPQDTQY | 0.4985669 |
| AGAVGQVGGAGGA | ASSPGGPQDTQY | 0.8247444 |
| AGGGGGLGGAP | ASSPGGPQDTQY | 0.45351726 |
| AGGLLYGNGGAG | ASSPGGPQDTQY | 0.19009122 |
| AGLESIGL | ASSPGGPQDTQY | 0.35367906 |
| AGLKTGII | ASSPGGPQDTQY | 0.6927006 |
| ASLAALLL | ASSPGGPQDTQY | 0.7860616 |
| ASVINISSI | ASSPGGPQDTQY | 0.5243861 |
| ATPPGPA | ASSPGGPQDTQY | 0.60510004 |
| ATVASALV | ASSPGGPQDTQY | 0.35952812 |
| ATVASLIL | ASSPGGPQDTQY | 0.7312757 |
| AVNGPVLIRH | ASSPGGPQDTQY | 0.8234978 |
| DGAAGAVGI | ASSPGGPQDTQY | 0.5549752 |
| DPLLHGI | ASSPGGPQDTQY | 0.24305739 |
| ELGAALGAL | ASSPGGPQDTQY | 0.24803369 |
| ELGRTQAV | ASSPGGPQDTQY | 0.56642044 |
| FVAGGYLS | ASSPGGPQDTQY | 0.7490425 |
| GASGGAGGAGGW | ASSPGGPQDTQY | 0.20782885 |
| GGAGAGGDGGAGGI | ASSPGGPQDTQY | 0.44058776 |
| GTGHNGSQPGMGGNG | ASSPGGPQDTQY | 0.5372833 |
| GYAAASASAS | ASSPGGPQDTQY | 0.17712879 |

| | | |
|---|---|---|
| HQVDVAAVQQG | ASSPGGPQDTQY | 0.54315734 |
| IGAAIAHT | ASSPGGPQDTQY | 0.7937385 |
| IGMIEVAQL | ASSPGGPQDTQY | 0.44845748 |
| IRLAPEHTSAMD | ASSPGGPQDTQY | 0.21224785 |
| IYNVSVL | ASSPGGPQDTQY | 0.75444734 |
| KQGFLGM | ASSPGGPQDTQY | 0.8214764 |
| LESVITQL | ASSPGGPQDTQY | 0.7244147 |
| LGLGALVVV | ASSPGGPQDTQY | 0.19293526 |
| LIGNGGNGGSGGI | ASSPGGPQDTQY | 0.14458573 |
| LNAINAPTL | ASSPGGPQDTQY | 0.5412442 |
| LQLLNTGV | ASSPGGPQDTQY | 0.72210544 |
| LQTSIEF | ASSPGGPQDTQY | 0.38949543 |
| MSFAVSEL | ASSPGGPQDTQY | 0.2668463 |
| NTGALGL | ASSPGGPQDTQY | 0.54166716 |
| PPGAGGAGGAGTT | ASSPGGPQDTQY | 0.6922145 |
| QLVGLGI | ASSPGGPQDTQY | 0.23583116 |
| RFLEQVHQL | ASSPGGPQDTQY | 0.19085078 |
| RGVGYVLR | ASSPGGPQDTQY | 0.6356355 |
| SATGVFAGVI | ASSPGGPQDTQY | 0.62224865 |
| SLPAITLP | ASSPGGPQDTQY | 0.24571943 |
| STALAFTV | ASSPGGPQDTQY | 0.35824674 |
| SVLTVGLGK | ASSPGGPQDTQY | 0.5704845 |
| TALAGAGSGI | ASSPGGPQDTQY | 0.46272185 |
| TGLPFTGI | ASSPGGPQDTQY | 0.534523 |
| VAEHVTV | ASSPGGPQDTQY | 0.20997512 |
| VDIVLNSL | ASSPGGPQDTQY | 0.72543484 |
| VLTGGGAVL | ASSPGGPQDTQY | 0.20374463 |
| VNVGLAYGI | ASSPGGPQDTQY | 0.6272794 |
| VSSAAVGAV | ASSPGGPQDTQY | 0.48560786 |
| VTSVYEAGV | ASSPGGPQDTQY | 0.4977569 |
| WVALDRGA | ASSPGGPQDTQY | 0.41822764 |
| YLVGSVVI | ASSPGGPQDTQY | 0.2069213 |
| GPPGTGKTLL | ASSPGGPQDTQY | 0.24957219 |
| GVGMTKF | ASSPGGPQDTQY | 0.22859606 |
| IGIGGSDL | ASSPGGPQDTQY | 0.7241777 |
| IGTLLAEL | ASSPGGPQDTQY | 0.6356795 |
| IYTSGTT | ASSPGGPQDTQY | 0.61053586 |
| LSFPAGV | ASSPGGPQDTQY | 0.46071482 |
| LYTSGTT | ASSPGGPQDTQY | 0.26433372 |
| VGAFPFGF | ASSPGGPQDTQY | 0.53824496 |
| VGAGTVEF | ASSPGGPQDTQY | 0.22649133 |
| VGKSTLLL | ASSPGGPQDTQY | 0.45426178 |
| AALAHAGGGIV | ASSPGGPQDTQY | 0.3768634 |
| AALVAVSLL | ASSPGGPQDTQY | 0.54586947 |
| AAVSRAVAAL | ASSPGGPQDTQY | 0.2757682 |
| AGGSSGIGGF | ASSPGGPQDTQY | 0.4683646 |
| AGLTVANANG | ASSPGGPQDTQY | 0.566216 |
| AGLVGAVL | ASSPGGPQDTQY | 0.25414008 |
| ALSQAFTV | ASSPGGPQDTQY | 0.43278676 |
| ALTFSVSL | ASSPGGPQDTQY | 0.72899425 |
| ALTGAALVAL | ASSPGGPQDTQY | 0.4881232 |

| | | |
|---|---|---|
| ASFSIGAGVL | ASSPGGPQDTQY | 0.19942696 |
| DGSVYKI | ASSPGGPQDTQY | 0.5364752 |
| EQLAEIV | ASSPGGPQDTQY | 0.29185385 |
| FAVSELTVM | ASSPGGPQDTQY | 0.55726117 |
| GAAPLAAGL | ASSPGGPQDTQY | 0.49874997 |
| GADGGDGGSGGAGG | ASSPGGPQDTQY | 0.35195267 |
| GFGLDTGGAGG | ASSPGGPQDTQY | 0.5297204 |
| GLASGGLL | ASSPGGPQDTQY | 0.23258905 |
| GRIAGAAVLL | ASSPGGPQDTQY | 0.32004857 |
| GSAGSLNTSF | ASSPGGPQDTQY | 0.32799786 |
| GTGQIGI | ASSPGGPQDTQY | 0.5365006 |
| GTLIALIL | ASSPGGPQDTQY | 0.58057 |
| IAATMGFVVG | ASSPGGPQDTQY | 0.2483747 |
| IFAQALL | ASSPGGPQDTQY | 0.4866042 |
| IGAAVALGF | ASSPGGPQDTQY | 0.29914087 |
| IGLTPLAL | ASSPGGPQDTQY | 0.22909167 |
| ILGSNTSTL | ASSPGGPQDTQY | 0.38702276 |
| IVFSLTVL | ASSPGGPQDTQY | 0.31979704 |
| LASKATGF | ASSPGGPQDTQY | 0.19924967 |
| LASVALIL | ASSPGGPQDTQY | 0.57002974 |
| LFAVGLTL | ASSPGGPQDTQY | 0.28536484 |
| LFNGGIGTYI | ASSPGGPQDTQY | 0.4018712 |
| LTLGAGII | ASSPGGPQDTQY | 0.7294238 |
| LVAAALGDI | ASSPGGPQDTQY | 0.20356578 |
| NTAFVQLGI | ASSPGGPQDTQY | 0.5227839 |
| QLFSGAEL | ASSPGGPQDTQY | 0.43820184 |
| SAATFITL | ASSPGGPQDTQY | 0.4802887 |
| SNYGLANLGV | ASSPGGPQDTQY | 0.46296468 |
| SYGGAYAVM | ASSPGGPQDTQY | 0.25646353 |
| TTGVGGLTLG | ASSPGGPQDTQY | 0.73486173 |
| TTQTIYI | ASSPGGPQDTQY | 0.5148955 |
| VAAGAGLGAVI | ASSPGGPQDTQY | 0.60009074 |
| VAAVVTLL | ASSPGGPQDTQY | 0.548376 |
| VGFLGGLI | ASSPGGPQDTQY | 0.7304479 |
| VGVIPILL | ASSPGGPQDTQY | 0.61405146 |
| VIFASSLI | ASSPGGPQDTQY | 0.28648326 |
| VKGGDGGAAAG | ASSPGGPQDTQY | 0.17821746 |
| VTKTFTNI | ASSPGGPQDTQY | 0.45414132 |
| VVAAAILL | ASSPGGPQDTQY | 0.46835244 |
| YAALDVEL | ASSPGGPQDTQY | 0.24801129 |
| LASVALLL | ASSPGGPQDTQY | 0.18512039 |
| LDAVHRV | ASSPGGPQDTQY | 0.72656333 |
| QGLSSSIF | ASSPGGPQDTQY | 0.81535435 |
| RGTFANIRL | ASSPGGPQDTQY | 0.34899682 |
| RQLWWGHRI | ASSPGGPQDTQY | 0.81916475 |
| YGPPGTGKTLL | ASSPGGPQDTQY | 0.19838317 |
| AALVGAPANVA | ASSPGGPQDTQY | 0.5400582 |
| AATFRRTE | ASSPGGPQDTQY | 0.7433211 |
| AGALGGGI | ASSPGGPQDTQY | 0.24585268 |
| AGGAGGAGGIGAG | ASSPGGPQDTQY | 0.5069135 |
| AGSGALGL | ASSPGGPQDTQY | 0.28742018 |

| | | |
|---|---|---|
| AQIADQI | ASSPGGPQDTQY | 0.28386322 |
| ASAVAVTYL | ASSPGGPQDTQY | 0.70874596 |
| ASTVLGGTGG | ASSPGGPQDTQY | 0.3611064 |
| AVGGEPAK | ASSPGGPQDTQY | 0.5314685 |
| CEQGTFGGTC | ASSPGGPQDTQY | 0.35846904 |
| DEGKAPAG | ASSPGGPQDTQY | 0.25782782 |
| DTVVFPAL | ASSPGGPQDTQY | 0.5163791 |
| FEGITMAG | ASSPGGPQDTQY | 0.60742885 |
| FPLVVQD | ASSPGGPQDTQY | 0.45568678 |
| GAFTTPQ | ASSPGGPQDTQY | 0.8032361 |
| GFGGFGTAGGVGGA | ASSPGGPQDTQY | 0.29307485 |
| GFLAQFL | ASSPGGPQDTQY | 0.82562184 |
| GGAGGAGGNAGVA | ASSPGGPQDTQY | 0.80704474 |
| GGVGTAGAGGAGGAGG | ASSPGGPQDTQY | 0.5811747 |
| GNGGGGDAGNAGSGGN | ASSPGGPQDTQY | 0.3164034 |
| GTGGAGGAGNDAGS | ASSPGGPQDTQY | 0.7557209 |
| GTTNTGWFNS | ASSPGGPQDTQY | 0.3325321 |
| IFMGPLFTL | ASSPGGPQDTQY | 0.4744898 |
| ILAHAIEAI | ASSPGGPQDTQY | 0.61265135 |
| IRSAIREAL | ASSPGGPQDTQY | 0.69792235 |
| ITFAGGG | ASSPGGPQDTQY | 0.78281105 |
| ITGMSGHQEV | ASSPGGPQDTQY | 0.75237286 |
| IVAGLACTL | ASSPGGPQDTQY | 0.75963193 |
| IVGGGLAGA | ASSPGGPQDTQY | 0.7018862 |
| IVTGVLNI | ASSPGGPQDTQY | 0.52577436 |
| LAQQGCI | ASSPGGPQDTQY | 0.75860757 |
| LCLHVEN | ASSPGGPQDTQY | 0.6464803 |
| LDAISDGS | ASSPGGPQDTQY | 0.7812655 |
| LGGFAAML | ASSPGGPQDTQY | 0.7792407 |
| LGGKNPMI | ASSPGGPQDTQY | 0.7174638 |
| LGPKGVAAIL | ASSPGGPQDTQY | 0.7974401 |
| LMVGVSI | ASSPGGPQDTQY | 0.79944587 |
| LVGAAFVL | ASSPGGPQDTQY | 0.5414209 |
| LVLAVAPL | ASSPGGPQDTQY | 0.7145002 |
| MAAIAVIDAA | ASSPGGPQDTQY | 0.7179866 |
| NNGINAELQ | ASSPGGPQDTQY | 0.7397515 |
| RGEPLVLVHG | ASSPGGPQDTQY | 0.5341216 |
| RGGAGGAAGLI | ASSPGGPQDTQY | 0.7843496 |
| RSYRMHPAV | ASSPGGPQDTQY | 0.70190936 |
| SAAHLLAGQLA | ASSPGGPQDTQY | 0.24792635 |
| SAAVGSY | ASSPGGPQDTQY | 0.7582247 |
| SPHAVLHRI | ASSPGGPQDTQY | 0.74362355 |
| SQIAKVL | ASSPGGPQDTQY | 0.6979854 |
| TALGHLVP | ASSPGGPQDTQY | 0.7824116 |
| TAPTTSGAA | ASSPGGPQDTQY | 0.4676083 |
| TDGDDGNGGSPGT | ASSPGGPQDTQY | 0.6756813 |
| TDGLGEN | ASSPGGPQDTQY | 0.7734159 |
| TGGPPLPGQA | ASSPGGPQDTQY | 0.7937262 |
| TIPAITI | ASSPGGPQDTQY | 0.6226613 |
| TIPGFTI | ASSPGGPQDTQY | 0.753111 |
| TSIASGALA | ASSPGGPQDTQY | 0.7956989 |

| | | |
|---|---|---|
| TVTLPGAAI | ASSPGGPQDTQY | 0.76753044 |
| VASGLSAALP | ASSPGGPQDTQY | 0.7926425 |
| VGAGVIGI | ASSPGGPQDTQY | 0.8002539 |
| VGPKRFGV | ASSPGGPQDTQY | 0.72782004 |
| VNVAFPD | ASSPGGPQDTQY | 0.7818452 |
| VSMTNSGVS | ASSPGGPQDTQY | 0.80117095 |
| VTDVLGDLSV | ASSPGGPQDTQY | 0.71362954 |
| VVVVGAGFAGL | ASSPGGPQDTQY | 0.75511533 |
| VYTVGTVV | ASSPGGPQDTQY | 0.70792365 |
| WTTFSGP | ASSPGGPQDTQY | 0.7464924 |
| YDELVEAF | ASSPGGPQDTQY | 0.36891267 |
| YGTGGAGGNGG | ASSPGGPQDTQY | 0.7535595 |
| YTNVMAV | ASSPGGPQDTQY | 0.78395116 |
| FLEQVHQL | ASSPGGPQDTQY | 0.7015197 |
| GLSLLAVAV | ASSPGGPQDTQY | 0.63336957 |
| LEQVHQL | ASSPGGPQDTQY | 0.6361828 |
| LNAAGAI | ASSPGGPQDTQY | 0.69220686 |
| PAVHASL | ASSPGGPQDTQY | 0.7339193 |
| TELEDDA | ASSPGGPQDTQY | 0.40779614 |
| TPPGPPN | ASSPGGPQDTQY | 0.68735933 |
| TPPPPPPP | ASSPGGPQDTQY | 0.7875047 |
| TQSPALI | ASRDNYAEQF | 0.78925085 |
| EFAAAFPQ | ASRDNYAEQF | 0.47889674 |
| LLSGNLDVL | ASRDNYAEQF | 0.78875494 |
| GPANLGNYNI | ASRDNYAEQF | 0.7927624 |
| TLGPTLP | ASRDNYAEQF | 0.795256 |
| VYAYSSCCK | ASRDNYAEQF | 0.7499394 |
| APAPAPAPNI | ASRDNYAEQF | 0.6478764 |
| PGGPVPIP | ASRDNYAEQF | 0.7866856 |
| FLTVALEP | ASRDNYAEQF | 0.7035295 |
| SQYFGANL | ASRDNYAEQF | 0.657063 |
| GGVKPAA | ASRDNYAEQF | 0.7927524 |
| GGTGGAGGAGGW | ASRDNYAEQF | 0.71561956 |
| GVGGVAGLGGAGG | ASRDNYAEQF | 0.67949665 |
| VLLVLATTI | ASRDNYAEQF | 0.70264363 |
| IGIFAAML | ASRDNYAEQF | 0.7908125 |
| AVGDLID | ASRDNYAEQF | 0.6141746 |
| NISPLPL | ASRDNYAEQF | 0.6287241 |
| VTAVFVAI | ASRDNYAEQF | 0.6297865 |
| SGDDNDAAL | ASRDNYAEQF | 0.7469586 |
| AVAALAHS | ASRDNYAEQF | 0.78331196 |
| GGVTVMATP | ASRDNYAEQF | 0.768788 |
| AGLAANVRD | ASRDNYAEQF | 0.71330756 |
| TNRPDLIDPALLRPG | ASRDNYAEQF | 0.7684543 |
| LANLGVTAL | ASRDNYAEQF | 0.76682585 |
| TGERIEL | ASRDNYAEQF | 0.7610267 |
| NGEELAA | ASRDNYAEQF | 0.7883433 |
| VLSLVWG | ASRDNYAEQF | 0.64696145 |
| IGYALWL | ASRDNYAEQF | 0.7892785 |
| LSLLLAPL | ASRDNYAEQF | 0.77387947 |
| TLDEALK | ASRDNYAEQF | 0.7917654 |

| | | |
|---|---|---|
| TDGAGTAEGG | ASRDNYAEQF | 0.78387964 |
| NAPGGAGGNGG | ASRDNYAEQF | 0.7474077 |
| VSVFMGMGGNF | ASRDNYAEQF | 0.75565815 |
| LASYLAAL | ASRDNYAEQF | 0.38258436 |
| LVGGTPAL | ASRDNYAEQF | 0.7458615 |
| VRNADVL | ASRDNYAEQF | 0.77027094 |
| TCSSPFLVSAM | ASRDNYAEQF | 0.74497867 |
| VHITDAE | ASRDNYAEQF | 0.77809834 |
| EYAAPSF | ASRDNYAEQF | 0.78506184 |
| VVDAFHGGL | ASRDNYAEQF | 0.6962203 |
| ASVVDPQ | ASRDNYAEQF | 0.7979488 |
| SYNEFGGLS | ASRDNYAEQF | 0.7777834 |
| DPDGSVY | ASRDNYAEQF | 0.51706535 |
| MVSTIHATE | ASRDNYAEQF | 0.7159834 |
| LAAGVADPV | ASRDNYAEQF | 0.79129094 |
| TKEQIAA | ASRDNYAEQF | 0.7775177 |
| SLASPAAL | ASRDNYAEQF | 0.74164236 |
| IGLVFYGV | ASRDNYAEQF | 0.74002236 |
| FLLDASNL | ASRDNYAEQF | 0.7999418 |
| YQVGNFP | ASRDNYAEQF | 0.67178446 |
| ESLLSQST | ASRDNYAEQF | 0.79936814 |
| LFTVGFAAVG | ASRDNYAEQF | 0.30093914 |
| ATPPTPT | ASRDNYAEQF | 0.7139063 |
| TDLATLG | ASRDNYAEQF | 0.6197886 |
| SAGGTGR | ASRDNYAEQF | 0.46156734 |
| GFSNSGS | ASRDNYAEQF | 0.78401256 |
| GGNGGSGGTSV | ASRDNYAEQF | 0.7409128 |
| GTSVATGGAGN | ASRDNYAEQF | 0.78529185 |
| NGGNGGSGGTS | ASRDNYAEQF | 0.75511324 |
| AVVKFPH | ASRDNYAEQF | 0.7786776 |
| TPPPPPP | ASRDNYAEQF | 0.7406687 |
| AAPAAAAAPA | ASRDNYAEQF | 0.7746104 |
| IALNEDL | ASRDNYAEQF | 0.6101535 |
| IVNVSSM | ASRDNYAEQF | 0.78351367 |
| TGIPPAPR | ASRDNYAEQF | 0.7999863 |
| FDLGGGTF | ASRDNYAEQF | 0.77229345 |
| FDLGGGTFDVS | ASRDNYAEQF | 0.6268845 |
| NEPTAAA | ASRDNYAEQF | 0.65856266 |
| NEPTAAALA | ASRDNYAEQF | 0.74898803 |
| GSIINISSI | ASRDNYAEQF | 0.32389504 |
| VAPGNAGTA | ASRDNYAEQF | 0.32822138 |
| TGGGASLE | ASRDNYAEQF | 0.648569 |
| VAVYNHYKRI | ASRDNYAEQF | 0.6272695 |
| TDPAPATT | ASRDNYAEQF | 0.7904413 |
| TNRPDILDPALLRPG | ASRDNYAEQF | 0.7756177 |
| LTTTTTS | ASRDNYAEQF | 0.727644 |
| PTGSGKTL | ASRDNYAEQF | 0.7818963 |
| TGSGKTL | ASRDNYAEQF | 0.75808954 |
| TIGHVDHGK | ASRDNYAEQF | 0.19127482 |
| TIGHVDHGKTT | ASRDNYAEQF | 0.7268548 |
| GHVDHGKTT | ASRDNYAEQF | 0.5869053 |

| | | |
|---|---|---|
| AQFEHTLL | ASRDNYAEQF | 0.8026449 |
| IVNQDEL | ASRDNYAEQF | 0.65361094 |
| TNGSQFF | ASRDNYAEQF | 0.65393454 |
| TGKGIVE | ASRDNYAEQF | 0.73699284 |
| LISNASDA | ASRDNYAEQF | 0.7452066 |
| AELRTLGELTKIAKAHGAQ | ASRDNYAEQF | 0.74516284 |
| APNAVAAAEPSW | ASRDNYAEQF | 0.6366902 |
| APQVAGAATAVL | ASRDNYAEQF | 0.24908288 |
| ARRDPAAAAAILPTDA | ASRDNYAEQF | 0.78369683 |
| DAAARDVLAEAGLAEYF | ASRDNYAEQF | 0.513957 |
| DDQIVLSAKANLAKVEQ | ASRDNYAEQF | 0.6383926 |
| DELLPDTGEDSDEDGH | ASRDNYAEQF | 0.452562 |
| ELPEAPANRATVPQD | ASRDNYAEQF | 0.42186618 |
| FGQSKIERVTGIG | ASRDNYAEQF | 0.5940684 |
| GAGGTGSTAGGAGGAGGAGGLY | ASRDNYAEQF | 0.79384243 |
| GHGGTGAAGGNGGTGGAGGSGI | ASRDNYAEQF | 0.6890065 |
| GNGGNGGAGGEGGPGGAGGAGA | ASRDNYAEQF | 0.69379616 |
| GSTAGNGGNGGNATNGTIAGQP | ASRDNYAEQF | 0.79166305 |
| GTGGDGGNGGNGGTGG | ASRDNYAEQF | 0.7553282 |
| GVRRPAGIPADAPV | ASRDNYAEQF | 0.70486856 |
| GYRQVTDAHLLTLARR | ASRDNYAEQF | 0.69007605 |
| IAKMGQDGHDRG | ASRDNYAEQF | 0.66039723 |
| KAEPSIEIQYT | ASRDNYAEQF | 0.6494841 |
| KPVRIRMAADEDTVAVT | ASRDNYAEQF | 0.6594547 |
| MPPRIAGMRLLVIKPEP | ASRDNYAEQF | 0.7999873 |
| MKFYAVALFRSRGVPPTR | ASRDNYAEQF | 0.71737015 |
| NAGGFGILGAVAHSP | ASRDNYAEQF | 0.6067332 |
| NEGREITLAMAEM | ASRDNYAEQF | 0.75009143 |
| NELVEWSHPLGHRWRWE | ASRDNYAEQF | 0.79971504 |
| NLGAGGTGGNGGNPG | ASRDNYAEQF | 0.8046894 |
| NTEWPKWETVMAADPVD | ASRDNYAEQF | 0.66422355 |
| PASAAPAETQP | ASRDNYAEQF | 0.75359595 |
| QQSSDGKNGVRAAEITGE | ASRDNYAEQF | 0.72502 |
| RAPDAVTTSWWKE | ASRDNYAEQF | 0.73900187 |
| SGFGNVGAHTSGWFNQ | ASRDNYAEQF | 0.72335124 |
| SGVAGAVGQAASVGGLKV | ASRDNYAEQF | 0.5385792 |
| SNGGAGTGSAGGNGGT | ASRDNYAEQF | 0.77704936 |
| TGGAGGEGGAAG | ASRDNYAEQF | 0.81087375 |
| TGGDGGAAGAGGAPGHG | ASRDNYAEQF | 0.7077217 |
| TMPPLLAAIANAGRHGV | ASRDNYAEQF | 0.64076567 |
| TSSTAGNSGGA | ASRDNYAEQF | 0.7320757 |
| VPATVPVGILC | ASRDNYAEQF | 0.64747727 |
| VQRPDSGGVVLTGRIS | ASRDNYAEQF | 0.68846667 |
| WTPNPVPATKGVPAGLVQAT | ASRDNYAEQF | 0.763556 |
| YALKRALAKFSDRFDVVII | ASRDNYAEQF | 0.5175584 |
| YLIGSVIV | ASRDNYAEQF | 0.6958001 |
| YGQLVAVID | ASRDNYAEQF | 0.8014024 |
| VTVGGMTLPA | ASRDNYAEQF | 0.70730543 |
| VLALPAV | ASRDNYAEQF | 0.74391353 |
| VIPPTLNYET | ASRDNYAEQF | 0.7793815 |
| VGMGGMHPSSG | ASRDNYAEQF | 0.7612668 |

| | | |
|---|---|---|
| VGLAGGAATA | ASRDNYAEQF | 0.79184127 |
| VGGAGGAGGSGP | ASRDNYAEQF | 0.78979945 |
| VFAAFMV | ASRDNYAEQF | 0.7479714 |
| TLYDKLR | ASRDNYAEQF | 0.7505472 |
| TLAVFTVL | ASRDNYAEQF | 0.73152626 |
| TGSVQPARV | ASRDNYAEQF | 0.75582975 |
| TGGGAGSGGT | ASRDNYAEQF | 0.7297434 |
| SHLETSL | ASRDNYAEQF | 0.70134866 |
| SFTTGMT | ASRDNYAEQF | 0.79036605 |
| QVEVAKRMGVRQPTVS | ASRDNYAEQF | 0.7500963 |
| QGSGGPALAM | ASRDNYAEQF | 0.790153 |
| MLAPSNP | ASRDNYAEQF | 0.6782144 |
| LVNNAAI | ASRDNYAEQF | 0.6263999 |
| LTGEINTLA | ASRDNYAEQF | 0.7744137 |
| LSIFGPV | ASRDNYAEQF | 0.2554032 |
| LLALLANL | ASRDNYAEQF | 0.3604877 |
| LISNASDALDKLR | ASRDNYAEQF | 0.7517686 |
| LINEAVP | ASRDNYAEQF | 0.7980939 |
| LGYAFSGL | ASRDNYAEQF | 0.24023032 |
| LDFVAVAL | ASRDNYAEQF | 0.64861023 |
| LAQPGAE | ASRDNYAEQF | 0.6084806 |
| ISNLGAVTGF | ASRDNYAEQF | 0.76523185 |
| HQEDPGRI | ASRDNYAEQF | 0.7526535 |
| GTPGEPTP | ASRDNYAEQF | 0.5110632 |
| GFFNSGS | ASRDNYAEQF | 0.76987743 |
| GAGGGGLTAGAN | ASRDNYAEQF | 0.5446518 |
| FVDALVNM | ASRDNYAEQF | 0.4282236 |
| FLTVAPEL | ASRDNYAEQF | 0.7834912 |
| FETLTAL | ASRDNYAEQF | 0.7926543 |
| DVINPIEL | ASRDNYAEQF | 0.6371228 |
| APVSGAPV | ASRDNYAEQF | 0.78259444 |
| ALVGGLAW | ASRDNYAEQF | 0.6955938 |
| ALAAAAATAI | ASRDNYAEQF | 0.8048627 |
| AFGLGLLL | ASRDNYAEQF | 0.72475183 |
| AFFAAASL | ASRDNYAEQF | 0.51603913 |
| AAAALPAFSPP | ASRDNYAEQF | 0.7718679 |
| VVLAPAL | ASRDNYAEQF | 0.75730634 |
| TIGHVDHGKTTL | ASRDNYAEQF | 0.76860344 |
| LISNASDALDK | ASRDNYAEQF | 0.7828119 |
| LISNASDAL | ASRDNYAEQF | 0.6566236 |
| ISNASDALDK | ASRDNYAEQF | 0.7017716 |
| NQAETLVYQTE | ASRDNYAEQF | 0.7892797 |
| TGGTGGTAGAG | ASRDNYAEQF | 0.78850484 |
| TGSTAGGAGGA | ASRDNYAEQF | 0.5569239 |
| DPLAVPVAAGPV | ASRDNYAEQF | 0.7790322 |
| GGTGGTPGNGGN | ASRDNYAEQF | 0.7318525 |
| PAGLAGVDPATL | ASRDNYAEQF | 0.70717776 |
| GNNLVVEAPHAGASV | ASRDNYAEQF | 0.7963731 |
| KVTRIMSASTLEQPA | ASRDNYAEQF | 0.7569845 |
| LGFALPIALPRRTVV | ASRDNYAEQF | 0.7823131 |
| TRLGNLTARKLVVLV | ASRDNYAEQF | 0.7806102 |

| | | |
|---|---|---|
| WSQQRRLIYFPSAGP | ASRDNYAEQF | 0.8031306 |
| GPNGNPDPMAAAVDIR | ASRDNYAEQF | 0.5751872 |
| TQRCVSVADLRAAASR | ASRDNYAEQF | 0.781083 |
| QFVLDVAEHPVEVLTETV | ASRDNYAEQF | 0.75219953 |
| GGAGGTGGAGGNGGTQAGDG | ASRDNYAEQF | 0.7790072 |
| PLLATLRAIAVDVDAKPAQV | ASRDNYAEQF | 0.7601254 |
| APKFRTGGYMIDFWGVGYQVAK | ASRDNYAEQF | 0.55313134 |
| GAGGAGGAGGAAGGPGGTGGTG | ASRDNYAEQF | 0.7668295 |
| IGALGLLATDVAPAGSVAFGVA | ASRDNYAEQF | 0.7916701 |
| AGGAGGIGGNGGAGGAGGVLMG | ASRDNYAEQF | 0.7537092 |
| ADELIGLL | ASRDNYAEQF | 0.77323616 |
| ADRALKA | ASRDNYAEQF | 0.79385257 |
| AGAVGQVGGAGGA | ASRDNYAEQF | 0.70590377 |
| AGGGGGLGGAP | ASRDNYAEQF | 0.73505497 |
| AGGLLYGNGGAG | ASRDNYAEQF | 0.7816921 |
| AGLESIGL | ASRDNYAEQF | 0.7911389 |
| AGLKTGII | ASRDNYAEQF | 0.7320372 |
| ASLAALLL | ASRDNYAEQF | 0.79335976 |
| ASVINISSI | ASRDNYAEQF | 0.6546179 |
| ATPPGPA | ASRDNYAEQF | 0.7734878 |
| ATVASALV | ASRDNYAEQF | 0.66987544 |
| ATVASLIL | ASRDNYAEQF | 0.7760479 |
| AVNGPVLIRH | ASRDNYAEQF | 0.58232987 |
| DGAAGAVGI | ASRDNYAEQF | 0.776358 |
| DPLLHGI | ASRDNYAEQF | 0.7815186 |
| ELGAALGAL | ASRDNYAEQF | 0.783701 |
| ELGRTQAV | ASRDNYAEQF | 0.39076826 |
| FVAGGYLS | ASRDNYAEQF | 0.7740245 |
| GASGGAGGAGGW | ASRDNYAEQF | 0.7845994 |
| GGAGAGGDGGAGGI | ASRDNYAEQF | 0.80651253 |
| GTGHNGSQPGMGGNG | ASRDNYAEQF | 0.7534046 |
| GYAAASASAS | ASRDNYAEQF | 0.7744747 |
| HQVDVAAVQQG | ASRDNYAEQF | 0.7307943 |
| IGAAIAHT | ASRDNYAEQF | 0.7190585 |
| IGMIEVAQL | ASRDNYAEQF | 0.6464635 |
| IRLAPEHTSAMD | ASRDNYAEQF | 0.7265799 |
| IYNVSVL | ASRDNYAEQF | 0.70446634 |
| KQGFLGM | ASRDNYAEQF | 0.64052 |
| LESVITQL | ASRDNYAEQF | 0.7868087 |
| LGLGALVVV | ASRDNYAEQF | 0.7918905 |
| LIGNGGNGGSGGI | ASRDNYAEQF | 0.7826344 |
| LNAINAPTL | ASRDNYAEQF | 0.7872362 |
| LQLLNTGV | ASRDNYAEQF | 0.78893113 |
| LQTSIEF | ASRDNYAEQF | 0.7884623 |
| MSFAVSEL | ASRDNYAEQF | 0.688171 |
| NTGALGL | ASRDNYAEQF | 0.75491655 |
| PPGAGGAGGAGTT | ASRDNYAEQF | 0.6201712 |
| QLVGLGI | ASRDNYAEQF | 0.7107419 |
| RFLEQVHQL | ASRDNYAEQF | 0.6919104 |
| RGVGYVLR | ASRDNYAEQF | 0.7549926 |
| SATGVFAGVI | ASRDNYAEQF | 0.2074006 |

| | | |
|---|---|---|
| SLPAITLP | ASRDNYAEQF | 0.7483332 |
| STALAFTV | ASRDNYAEQF | 0.7985947 |
| SVLTVGLGK | ASRDNYAEQF | 0.6490277 |
| TALAGAGSGI | ASRDNYAEQF | 0.781821 |
| TGLPFTGI | ASRDNYAEQF | 0.7283462 |
| VAEHVTV | ASRDNYAEQF | 0.68415713 |
| VDIVLNSL | ASRDNYAEQF | 0.7803807 |
| VLTGGGAVL | ASRDNYAEQF | 0.6544808 |
| VNVGLAYGI | ASRDNYAEQF | 0.7757826 |
| VSSAAVGAV | ASRDNYAEQF | 0.7973331 |
| VTSVYEAGV | ASRDNYAEQF | 0.75021327 |
| WVALDRGA | ASRDNYAEQF | 0.76808476 |
| YLVGSVVI | ASRDNYAEQF | 0.7491104 |
| GPPGTGKTLL | ASRDNYAEQF | 0.5610879 |
| GVGMTKF | ASRDNYAEQF | 0.7693917 |
| IGIGGSDL | ASRDNYAEQF | 0.23717134 |
| IGTLLAEL | ASRDNYAEQF | 0.72687626 |
| IYTSGTT | ASRDNYAEQF | 0.6355106 |
| LSFPAGV | ASRDNYAEQF | 0.75582343 |
| LYTSGTT | ASRDNYAEQF | 0.7399608 |
| VGAFPFGF | ASRDNYAEQF | 0.7719939 |
| VGAGTVEF | ASRDNYAEQF | 0.6784347 |
| VGKSTLLL | ASRDNYAEQF | 0.6943091 |
| AALAHAGGGIV | ASRDNYAEQF | 0.6512208 |
| AALVAVSLL | ASRDNYAEQF | 0.5729786 |
| AAVSRAVAAL | ASRDNYAEQF | 0.77882975 |
| AGGSSGIGGF | ASRDNYAEQF | 0.73671246 |
| AGLTVANANG | ASRDNYAEQF | 0.697261 |
| AGLVGAVL | ASRDNYAEQF | 0.7867805 |
| ALSQAFTV | ASRDNYAEQF | 0.749166 |
| ALTFSVSL | ASRDNYAEQF | 0.5600654 |
| ALTGAALVAL | ASRDNYAEQF | 0.71825945 |
| ASFSIGAGVL | ASRDNYAEQF | 0.74940604 |
| DGSVYKI | ASRDNYAEQF | 0.729432 |
| EQLAEIV | ASRDNYAEQF | 0.21485932 |
| FAVSELTVM | ASRDNYAEQF | 0.75344646 |
| GAAPLAAGL | ASRDNYAEQF | 0.7147909 |
| GADGGDGGSGGAGG | ASRDNYAEQF | 0.6084132 |
| GFGLDTGGAGG | ASRDNYAEQF | 0.66856277 |
| GLASGGLL | ASRDNYAEQF | 0.7698667 |
| GRIAGAAVLL | ASRDNYAEQF | 0.63412845 |
| GSAGSLNTSF | ASRDNYAEQF | 0.75329554 |
| GTGQIGI | ASRDNYAEQF | 0.77853 |
| GTLIALIL | ASRDNYAEQF | 0.79076505 |
| IAATMGFVVG | ASRDNYAEQF | 0.7987827 |
| IFAQALL | ASRDNYAEQF | 0.7616199 |
| IGAAVALGF | ASRDNYAEQF | 0.7212624 |
| IGLTPLAL | ASRDNYAEQF | 0.78786576 |
| ILGSNTSTL | ASRDNYAEQF | 0.77359945 |
| IVFSLTVL | ASRDNYAEQF | 0.3950565 |
| LASKATGF | ASRDNYAEQF | 0.35467458 |

| | | |
|---|---|---|
| LASVALIL | ASRDNYAEQF | 0.7660384 |
| LFAVGLTL | ASRDNYAEQF | 0.4352999 |
| LFNGGIGTYI | ASRDNYAEQF | 0.76096755 |
| LTLGAGII | ASRDNYAEQF | 0.7977048 |
| LVAAALGDI | ASRDNYAEQF | 0.7378641 |
| NTAFVQLGI | ASRDNYAEQF | 0.63508046 |
| QLFSGAEL | ASRDNYAEQF | 0.71263754 |
| SAATFITL | ASRDNYAEQF | 0.73270833 |
| SNYGLANLGV | ASRDNYAEQF | 0.7837826 |
| SYGGAYAVM | ASRDNYAEQF | 0.78431773 |
| TTGVGGLTLG | ASRDNYAEQF | 0.801086 |
| TTQTIYI | ASRDNYAEQF | 0.7789097 |
| VAAGAGLGAVI | ASRDNYAEQF | 0.6952764 |
| VAAVVTLL | ASRDNYAEQF | 0.7675879 |
| VGFLGGLI | ASRDNYAEQF | 0.7091763 |
| VGVIPILL | ASRDNYAEQF | 0.7493155 |
| VIFASSLI | ASRDNYAEQF | 0.7521076 |
| VKGGDGGAAAG | ASRDNYAEQF | 0.7499066 |
| VTKTFTNI | ASRDNYAEQF | 0.7356428 |
| VVAAAILL | ASRDNYAEQF | 0.7263311 |
| YAALDVEL | ASRDNYAEQF | 0.57048666 |
| LASVALLL | ASRDNYAEQF | 0.7776019 |
| LDAVHRV | ASRDNYAEQF | 0.75582767 |
| QGLSSSIF | ASRDNYAEQF | 0.4847116 |
| RGTFANIRL | ASRDNYAEQF | 0.7259756 |
| RQLWWGHRI | ASRDNYAEQF | 0.7978827 |
| YGPPGTGKTLL | ASRDNYAEQF | 0.7956625 |
| AALVGAPANVA | ASRDNYAEQF | 0.6368644 |
| AATFRRTE | ASRDNYAEQF | 0.6751571 |
| AGALGGGI | ASRDNYAEQF | 0.7424414 |
| AGGAGGAGGIGAG | ASRDNYAEQF | 0.43645123 |
| AGSGALGL | ASRDNYAEQF | 0.76686597 |
| AQIADQI | ASRDNYAEQF | 0.7186129 |
| ASAVAVTYL | ASRDNYAEQF | 0.7025197 |
| ASTVLGGTGG | ASRDNYAEQF | 0.70173776 |
| AVGGEPAK | ASRDNYAEQF | 0.7303339 |
| CEQGTFGGTC | ASRDNYAEQF | 0.67091334 |
| DEGKAPAG | ASRDNYAEQF | 0.7872347 |
| DTVVFPAL | ASRDNYAEQF | 0.24268733 |
| FEGITMAG | ASRDNYAEQF | 0.755497 |
| FPLVVQD | ASRDNYAEQF | 0.6611252 |
| GAFTTPQ | ASRDNYAEQF | 0.70177853 |
| GFGGFGTAGGVGGA | ASRDNYAEQF | 0.53113866 |
| GFLAQFL | ASRDNYAEQF | 0.75566816 |
| GGAGGAGGNAGVA | ASRDNYAEQF | 0.78483325 |
| GGVGTAGAGGAGGAGG | ASRDNYAEQF | 0.77798134 |
| GNGGGGDAGNAGSGGN | ASRDNYAEQF | 0.571314 |
| GTGGAGGAGNDAGS | ASRDNYAEQF | 0.78218186 |
| GTTNTGWFNS | ASRDNYAEQF | 0.80447686 |
| IFMGPLFTL | ASRDNYAEQF | 0.7053188 |
| ILAHAIEAI | ASRDNYAEQF | 0.5131454 |

| | | |
|---|---|---|
| IRSAIREAL | ASRDNYAEQF | 0.77149075 |
| ITFAGGG | ASRDNYAEQF | 0.7484148 |
| ITGMSGHQEV | ASRDNYAEQF | 0.7693832 |
| IVAGLACTL | ASRDNYAEQF | 0.786513 |
| IVGGGLAGA | ASRDNYAEQF | 0.5890873 |
| IVTGVLNI | ASRDNYAEQF | 0.7937574 |
| LAQQGCI | ASRDNYAEQF | 0.7801988 |
| LCLHVEN | ASRDNYAEQF | 0.6769077 |
| LDAISDGS | ASRDNYAEQF | 0.7188406 |
| LGGFAAML | ASRDNYAEQF | 0.70102227 |
| LGGKNPMI | ASRDNYAEQF | 0.41328275 |
| LGPKGVAAIL | ASRDNYAEQF | 0.790604 |
| LMVGVSI | ASRDNYAEQF | 0.65936685 |
| LVGAAFVL | ASRDNYAEQF | 0.7000618 |
| LVLAVAPL | ASRDNYAEQF | 0.74141884 |
| MAAIAVIDAA | ASRDNYAEQF | 0.7020923 |
| NNGINAELQ | ASRDNYAEQF | 0.7153592 |
| RGEPLVLVHG | ASRDNYAEQF | 0.70533067 |
| RGGAGGAAGLI | ASRDNYAEQF | 0.7492425 |
| RSYRMHPAV | ASRDNYAEQF | 0.7400628 |
| SAAHLLAGQLA | ASRDNYAEQF | 0.75439024 |
| SAAVGSY | ASRDNYAEQF | 0.7639576 |
| SPHAVLHRI | ASRDNYAEQF | 0.79178846 |
| SQIAKVL | ASRDNYAEQF | 0.54469186 |
| TALGHLVP | ASRDNYAEQF | 0.6757132 |
| TAPTTSGAA | ASRDNYAEQF | 0.75780684 |
| TDGDDGNGGSPGT | ASRDNYAEQF | 0.48551118 |
| TDGLGEN | ASRDNYAEQF | 0.7722735 |
| TGGPPLPGQA | ASRDNYAEQF | 0.74828064 |
| TIPAITI | ASRDNYAEQF | 0.75672114 |
| TIPGFTI | ASRDNYAEQF | 0.7495275 |
| TSIASGALA | ASRDNYAEQF | 0.8006558 |
| TVTLPGAAI | ASRDNYAEQF | 0.77940464 |
| VASGLSAALP | ASRDNYAEQF | 0.7527614 |
| VGAGVIGI | ASRDNYAEQF | 0.7675872 |
| VGPKRFGV | ASRDNYAEQF | 0.69019485 |
| VNVAFPD | ASRDNYAEQF | 0.7858338 |
| VSMTNSGVS | ASRDNYAEQF | 0.7811532 |
| VTDVLGDLSV | ASRDNYAEQF | 0.5924498 |
| VVVVGAGFAGL | ASRDNYAEQF | 0.70532787 |
| VYTVGTVV | ASRDNYAEQF | 0.79702556 |
| WTTFSGP | ASRDNYAEQF | 0.775715 |
| YDELVEAF | ASRDNYAEQF | 0.41804278 |
| YGTGGAGGNGG | ASRDNYAEQF | 0.70606494 |
| YTNVMAV | ASRDNYAEQF | 0.47908077 |
| FLEQVHQL | ASRDNYAEQF | 0.7006811 |
| GLSLLAVAV | ASRDNYAEQF | 0.67868656 |
| LEQVHQL | ASRDNYAEQF | 0.77032125 |
| LNAAGAI | ASRDNYAEQF | 0.73096794 |
| PAVHASL | ASRDNYAEQF | 0.78255755 |
| TELEDDA | ASRDNYAEQF | 0.7508017 |

| | | |
|---|---|---|
| TPPGPPN | ASRDNYAEQF | 0.79287946 |
| TPPPPPPP | ASRDNYAEQF | 0.70311564 |
| TQSPALI | ASSLGLGGSAETLY | 0.7464808 |
| EFAAAFPQ | ASSLGLGGSAETLY | 0.527626 |
| LLSGNLDVL | ASSLGLGGSAETLY | 0.49234045 |
| GPANLGNYNI | ASSLGLGGSAETLY | 0.6905634 |
| TLGPTLP | ASSLGLGGSAETLY | 0.7227307 |
| VYAYSSCCK | ASSLGLGGSAETLY | 0.31370205 |
| APAPAPAPNI | ASSLGLGGSAETLY | 0.78497684 |
| PGGPVPIP | ASSLGLGGSAETLY | 0.7602463 |
| FLTVALEP | ASSLGLGGSAETLY | 0.77890587 |
| SQYFGANL | ASSLGLGGSAETLY | 0.73169947 |
| GGVKPAA | ASSLGLGGSAETLY | 0.75068164 |
| GGTGGAGGAGGW | ASSLGLGGSAETLY | 0.6747221 |
| GVGGVAGLGGAGG | ASSLGLGGSAETLY | 0.771772 |
| VLLVLATTI | ASSLGLGGSAETLY | 0.64495677 |
| IGIFAAML | ASSLGLGGSAETLY | 0.5641422 |
| AVGDLID | ASSLGLGGSAETLY | 0.73987937 |
| NISPLPL | ASSLGLGGSAETLY | 0.62791973 |
| VTAVFVAI | ASSLGLGGSAETLY | 0.793869 |
| SGDDNDAAL | ASSLGLGGSAETLY | 0.78279376 |
| AVAALAHS | ASSLGLGGSAETLY | 0.7512571 |
| GGVTVMATP | ASSLGLGGSAETLY | 0.7859012 |
| AGLAANVRD | ASSLGLGGSAETLY | 0.3773094 |
| TNRPDLIDPALLRPG | ASSLGLGGSAETLY | 0.7293186 |
| LANLGVTAL | ASSLGLGGSAETLY | 0.8004459 |
| TGERIEL | ASSLGLGGSAETLY | 0.76243305 |
| NGEELAA | ASSLGLGGSAETLY | 0.7416769 |
| VLSLVWG | ASSLGLGGSAETLY | 0.79463387 |
| IGYALWL | ASSLGLGGSAETLY | 0.68458253 |
| LSLLLAPL | ASSLGLGGSAETLY | 0.74541473 |
| TLDEALK | ASSLGLGGSAETLY | 0.73679906 |
| TDGAGTAEGG | ASSLGLGGSAETLY | 0.7801547 |
| NAPGGAGGNGG | ASSLGLGGSAETLY | 0.77343 |
| VSVFMGMGGNF | ASSLGLGGSAETLY | 0.71543586 |
| LASYLAAL | ASSLGLGGSAETLY | 0.77226543 |
| LVGGTPAL | ASSLGLGGSAETLY | 0.7985942 |
| VRNADVL | ASSLGLGGSAETLY | 0.7001096 |
| TCSSPFLVSAM | ASSLGLGGSAETLY | 0.7891383 |
| VHITDAE | ASSLGLGGSAETLY | 0.7683817 |
| EYAAPSF | ASSLGLGGSAETLY | 0.77472603 |
| VVDAFHGGL | ASSLGLGGSAETLY | 0.67392933 |
| ASVVDPQ | ASSLGLGGSAETLY | 0.6918359 |
| SYNEFGGLS | ASSLGLGGSAETLY | 0.7104616 |
| DPDGSVY | ASSLGLGGSAETLY | 0.6592887 |
| MVSTIHATE | ASSLGLGGSAETLY | 0.7540026 |
| LAAGVADPV | ASSLGLGGSAETLY | 0.7013967 |
| TKEQIAA | ASSLGLGGSAETLY | 0.80592775 |
| SLASPAAL | ASSLGLGGSAETLY | 0.7776377 |
| IGLVFYGV | ASSLGLGGSAETLY | 0.7524276 |
| FLLDASNL | ASSLGLGGSAETLY | 0.62477595 |

| | | |
|---|---|---|
| YQVGNFP | ASSLGLGGSAETLY | 0.759948 |
| ESLLSQST | ASSLGLGGSAETLY | 0.7612114 |
| LFTVGFAAVG | ASSLGLGGSAETLY | 0.58006835 |
| ATPPTPT | ASSLGLGGSAETLY | 0.79842484 |
| TDLATLG | ASSLGLGGSAETLY | 0.69065595 |
| SAGGTGR | ASSLGLGGSAETLY | 0.66099215 |
| GFSNSGS | ASSLGLGGSAETLY | 0.77621806 |
| GGNGGSGGTSV | ASSLGLGGSAETLY | 0.80284536 |
| GTSVATGGAGN | ASSLGLGGSAETLY | 0.72355056 |
| NGGNGGSGGTS | ASSLGLGGSAETLY | 0.6971879 |
| AVVKFPH | ASSLGLGGSAETLY | 0.63139534 |
| TPPPPPP | ASSLGLGGSAETLY | 0.78596634 |
| AAPAAAAAPA | ASSLGLGGSAETLY | 0.7831284 |
| IALNEDL | ASSLGLGGSAETLY | 0.77007496 |
| IVNVSSM | ASSLGLGGSAETLY | 0.79715526 |
| TGIPPAPR | ASSLGLGGSAETLY | 0.7709334 |
| FDLGGGTF | ASSLGLGGSAETLY | 0.56348205 |
| FDLGGGTFDVS | ASSLGLGGSAETLY | 0.7483164 |
| NEPTAAA | ASSLGLGGSAETLY | 0.77843916 |
| NEPTAAALA | ASSLGLGGSAETLY | 0.775823 |
| GSIINISSI | ASSLGLGGSAETLY | 0.8052919 |
| VAPGNAGTA | ASSLGLGGSAETLY | 0.77993715 |
| TGGGASLE | ASSLGLGGSAETLY | 0.682572 |
| VAVYNHYKRI | ASSLGLGGSAETLY | 0.7906604 |
| TDPAPATT | ASSLGLGGSAETLY | 0.80341315 |
| TNRPDILDPALLRPG | ASSLGLGGSAETLY | 0.78225005 |
| LTTTTTS | ASSLGLGGSAETLY | 0.72406757 |
| PTGSGKTL | ASSLGLGGSAETLY | 0.6846514 |
| TGSGKTL | ASSLGLGGSAETLY | 0.755795 |
| TIGHVDHGK | ASSLGLGGSAETLY | 0.7706492 |
| TIGHVDHGKTT | ASSLGLGGSAETLY | 0.67855173 |
| GHVDHGKTT | ASSLGLGGSAETLY | 0.75699997 |
| AQFEHTLL | ASSLGLGGSAETLY | 0.7625215 |
| IVNQDEL | ASSLGLGGSAETLY | 0.6293375 |
| TNGSQFF | ASSLGLGGSAETLY | 0.7674873 |
| TGKGIVE | ASSLGLGGSAETLY | 0.7885033 |
| LISNASDA | ASSLGLGGSAETLY | 0.72394156 |
| AELRTLGELTKIAKAHGAQ | ASSLGLGGSAETLY | 0.7110338 |
| APNAVAAAEPSW | ASSLGLGGSAETLY | 0.7886243 |
| APQVAGAATAVL | ASSLGLGGSAETLY | 0.80483437 |
| ARRDPAAAAAILPTDA | ASSLGLGGSAETLY | 0.79086673 |
| DAAARDVLAEAGLAEYF | ASSLGLGGSAETLY | 0.56267357 |
| DDQIVLSAKANLAKVEQ | ASSLGLGGSAETLY | 0.4143848 |
| DELLPDTGEDSDEDGH | ASSLGLGGSAETLY | 0.6499511 |
| ELPEAPANRATVPQD | ASSLGLGGSAETLY | 0.7866174 |
| FGQSKIERVTGIG | ASSLGLGGSAETLY | 0.71143645 |
| GAGGTGSTAGGAGGAGGAGGLY | ASSLGLGGSAETLY | 0.678968 |
| GHGGTGAAGGNGGTGGAGGSGI | ASSLGLGGSAETLY | 0.757781 |
| GNGGNGGAGGEGGPGGAGGAGA | ASSLGLGGSAETLY | 0.7887962 |
| GSTAGNGGNGGNATNGTIAGQP | ASSLGLGGSAETLY | 0.7106173 |
| GTGGDGGNGGNGGTGG | ASSLGLGGSAETLY | 0.53398377 |

| | | |
|---|---|---|
| GVRRPAGIPADAPV | ASSLGLGGSAETLY | 0.7732071 |
| GYRQVTDAHLLTLARR | ASSLGLGGSAETLY | 0.7770897 |
| IAKMGQDGHDRG | ASSLGLGGSAETLY | 0.65576005 |
| KAEPSIEIQYT | ASSLGLGGSAETLY | 0.70510745 |
| KPVRIRMAADEDTVAVT | ASSLGLGGSAETLY | 0.65821487 |
| MPPRIAGMRLLVIKPEP | ASSLGLGGSAETLY | 0.5280462 |
| MKFYAVALFRSRGVPPTR | ASSLGLGGSAETLY | 0.7487122 |
| NAGGFGILGAVAHSP | ASSLGLGGSAETLY | 0.62417185 |
| NEGREITLAMAEM | ASSLGLGGSAETLY | 0.7817137 |
| NELVEWSHPLGHRWRWE | ASSLGLGGSAETLY | 0.63722086 |
| NLGAGGTGGNGGNPG | ASSLGLGGSAETLY | 0.78628385 |
| NTEWPKWETVMAADPVD | ASSLGLGGSAETLY | 0.7644756 |
| PASAAPAETQP | ASSLGLGGSAETLY | 0.7775 |
| QQSSDGKNGVRAAEITGE | ASSLGLGGSAETLY | 0.7245079 |
| RAPDAVTTSWWKE | ASSLGLGGSAETLY | 0.71210253 |
| SGFGNVGAHTSGWFNQ | ASSLGLGGSAETLY | 0.61484474 |
| SGVAGAVGQAASVGGLKV | ASSLGLGGSAETLY | 0.3842374 |
| SNGGAGTGSAGGNGGT | ASSLGLGGSAETLY | 0.79373544 |
| TGGAGGEGGAAG | ASSLGLGGSAETLY | 0.37694973 |
| TGGDGGAAGAGGAPGHG | ASSLGLGGSAETLY | 0.78941005 |
| TMPPLLAAIANAGRHGV | ASSLGLGGSAETLY | 0.7796545 |
| TSSTAGNSGGA | ASSLGLGGSAETLY | 0.62144166 |
| VPATVPVGILC | ASSLGLGGSAETLY | 0.7698519 |
| VQRPDSGGVVLTGRIS | ASSLGLGGSAETLY | 0.5954949 |
| WTPNPVPATKGVPAGLVQAT | ASSLGLGGSAETLY | 0.6282643 |
| YALKRALAKFSDRFDVVII | ASSLGLGGSAETLY | 0.79051447 |
| YLIGSVIV | ASSLGLGGSAETLY | 0.77099717 |
| YGQLVAVID | ASSLGLGGSAETLY | 0.6929872 |
| VTVGGMTLPA | ASSLGLGGSAETLY | 0.7669325 |
| VLALPAV | ASSLGLGGSAETLY | 0.7486613 |
| VIPPTLNYET | ASSLGLGGSAETLY | 0.72275805 |
| VGMGGMHPSSG | ASSLGLGGSAETLY | 0.73685485 |
| VGLAGGAATA | ASSLGLGGSAETLY | 0.7829801 |
| VGGAGGAGGSGP | ASSLGLGGSAETLY | 0.7661791 |
| VFAAFMV | ASSLGLGGSAETLY | 0.68754685 |
| TLYDKLR | ASSLGLGGSAETLY | 0.7741362 |
| TLAVFTVL | ASSLGLGGSAETLY | 0.63090825 |
| TGSVQPARV | ASSLGLGGSAETLY | 0.7566712 |
| TGGGAGSGGT | ASSLGLGGSAETLY | 0.73234653 |
| SHLETSL | ASSLGLGGSAETLY | 0.7622744 |
| SFTTGMT | ASSLGLGGSAETLY | 0.76729274 |
| QVEVAKRMGVRQPTVS | ASSLGLGGSAETLY | 0.6817073 |
| QGSGGPALAM | ASSLGLGGSAETLY | 0.6529824 |
| MLAPSNP | ASSLGLGGSAETLY | 0.74427825 |
| LVNNAAI | ASSLGLGGSAETLY | 0.7565955 |
| LTGEINTLA | ASSLGLGGSAETLY | 0.7789588 |
| LSIFGPV | ASSLGLGGSAETLY | 0.6559901 |
| LLALLANL | ASSLGLGGSAETLY | 0.7574365 |
| LISNASDALDKLR | ASSLGLGGSAETLY | 0.73146784 |
| LINEAVP | ASSLGLGGSAETLY | 0.68687344 |
| LGYAFSGL | ASSLGLGGSAETLY | 0.74421185 |

| | | |
|---|---|---|
| LDFVAVAL | ASSLGLGGSAETLY | 0.7503326 |
| LAQPGAE | ASSLGLGGSAETLY | 0.70270777 |
| ISNLGAVTGF | ASSLGLGGSAETLY | 0.7768017 |
| HQEDPGRI | ASSLGLGGSAETLY | 0.62640655 |
| GTPGEPTP | ASSLGLGGSAETLY | 0.77157193 |
| GFFNSGS | ASSLGLGGSAETLY | 0.7884825 |
| GAGGGGLTAGAN | ASSLGLGGSAETLY | 0.7343947 |
| FVDALVNM | ASSLGLGGSAETLY | 0.78002447 |
| FLTVAPEL | ASSLGLGGSAETLY | 0.7486756 |
| FETLTAL | ASSLGLGGSAETLY | 0.7673554 |
| DVINPIEL | ASSLGLGGSAETLY | 0.7235162 |
| APVSGAPV | ASSLGLGGSAETLY | 0.71635276 |
| ALVGGLAW | ASSLGLGGSAETLY | 0.78413063 |
| ALAAAAATAI | ASSLGLGGSAETLY | 0.77652323 |
| AFGLGLLL | ASSLGLGGSAETLY | 0.788455 |
| AFFAAASL | ASSLGLGGSAETLY | 0.7575338 |
| AAAALPAFSPP | ASSLGLGGSAETLY | 0.7882676 |
| VVLAPAL | ASSLGLGGSAETLY | 0.7697307 |
| TIGHVDHGKTTL | ASSLGLGGSAETLY | 0.68713886 |
| LISNASDALDK | ASSLGLGGSAETLY | 0.54957706 |
| LISNASDAL | ASSLGLGGSAETLY | 0.7158848 |
| ISNASDALDK | ASSLGLGGSAETLY | 0.7020741 |
| NQAETLVYQTE | ASSLGLGGSAETLY | 0.7443658 |
| TGGTGGTAGAG | ASSLGLGGSAETLY | 0.46639693 |
| TGSTAGGAGGA | ASSLGLGGSAETLY | 0.790228 |
| DPLAVPVAAGPV | ASSLGLGGSAETLY | 0.80395156 |
| GGTGGTPGNGGN | ASSLGLGGSAETLY | 0.69748473 |
| PAGLAGVDPATL | ASSLGLGGSAETLY | 0.799085 |
| GNNLVVEAPHAGASV | ASSLGLGGSAETLY | 0.5960597 |
| KVTRIMSASTLEQPA | ASSLGLGGSAETLY | 0.77635086 |
| LGFALPIALPRRTVV | ASSLGLGGSAETLY | 0.79494536 |
| TRLGNLTARKLVVLV | ASSLGLGGSAETLY | 0.6378578 |
| WSQQRRLIYFPSAGP | ASSLGLGGSAETLY | 0.78060246 |
| GPNGNPDPMAAAVDIR | ASSLGLGGSAETLY | 0.72945595 |
| TQRCVSVADLRAAASR | ASSLGLGGSAETLY | 0.73401964 |
| QFVLDVAEHPVEVLTETV | ASSLGLGGSAETLY | 0.7961049 |
| GGAGGTGGAGGNGGTQAGDG | ASSLGLGGSAETLY | 0.7411052 |
| PLLATLRAIAVDVDAKPAQV | ASSLGLGGSAETLY | 0.7764015 |
| APKFRTGGYMIDFWGVGYQVAK | ASSLGLGGSAETLY | 0.7216027 |
| GAGGAGGAGGAAGGPGGTGGTG | ASSLGLGGSAETLY | 0.6543425 |
| IGALGLLATDVAPAGSVAFGVA | ASSLGLGGSAETLY | 0.7572602 |
| AGGAGGIGGNGGAGGAGGVLMG | ASSLGLGGSAETLY | 0.7975181 |
| ADELIGLL | ASSLGLGGSAETLY | 0.65043795 |
| ADRALKA | ASSLGLGGSAETLY | 0.8019022 |
| AGAVGQVGGAGGA | ASSLGLGGSAETLY | 0.71119326 |
| AGGGGGLGGAP | ASSLGLGGSAETLY | 0.8022716 |
| AGGLLYGNGGAG | ASSLGLGGSAETLY | 0.77335835 |
| AGLESIGL | ASSLGLGGSAETLY | 0.7208667 |
| AGLKTGII | ASSLGLGGSAETLY | 0.7266516 |
| ASLAALLL | ASSLGLGGSAETLY | 0.79148436 |
| ASVINISSI | ASSLGLGGSAETLY | 0.7261269 |

| | | |
|---|---|---|
| ATPPGPA | ASSLGLGGSAETLY | 0.78689444 |
| ATVASALV | ASSLGLGGSAETLY | 0.79672945 |
| ATVASLIL | ASSLGLGGSAETLY | 0.4644631 |
| AVNGPVLIRH | ASSLGLGGSAETLY | 0.74352574 |
| DGAAGAVGI | ASSLGLGGSAETLY | 0.7113197 |
| DPLLHGI | ASSLGLGGSAETLY | 0.71503663 |
| ELGAALGAL | ASSLGLGGSAETLY | 0.5963394 |
| ELGRTQAV | ASSLGLGGSAETLY | 0.45156163 |
| FVAGGYLS | ASSLGLGGSAETLY | 0.6834627 |
| GASGGAGGAGGW | ASSLGLGGSAETLY | 0.5319258 |
| GGAGAGGDGGAGGI | ASSLGLGGSAETLY | 0.744557 |
| GTGHNGSQPGMGGNG | ASSLGLGGSAETLY | 0.73437977 |
| GYAAASASAS | ASSLGLGGSAETLY | 0.5793151 |
| HQVDVAAVQQG | ASSLGLGGSAETLY | 0.73918223 |
| IGAAIAHT | ASSLGLGGSAETLY | 0.7470064 |
| IGMIEVAQL | ASSLGLGGSAETLY | 0.406812 |
| IRLAPEHTSAMD | ASSLGLGGSAETLY | 0.6723275 |
| IYNVSVL | ASSLGLGGSAETLY | 0.6298487 |
| KQGFLGM | ASSLGLGGSAETLY | 0.63859326 |
| LESVITQL | ASSLGLGGSAETLY | 0.37556708 |
| LGLGALVVV | ASSLGLGGSAETLY | 0.7287134 |
| LIGNGGNGGSGGI | ASSLGLGGSAETLY | 0.65549845 |
| LNAINAPTL | ASSLGLGGSAETLY | 0.29783237 |
| LQLLNTGV | ASSLGLGGSAETLY | 0.71312046 |
| LQTSIEF | ASSLGLGGSAETLY | 0.6404378 |
| MSFAVSEL | ASSLGLGGSAETLY | 0.5865207 |
| NTGALGL | ASSLGLGGSAETLY | 0.73782516 |
| PPGAGGAGGAGTT | ASSLGLGGSAETLY | 0.66357124 |
| QLVGLGI | ASSLGLGGSAETLY | 0.56790674 |
| RFLEQVHQL | ASSLGLGGSAETLY | 0.7285018 |
| RGVGYVLR | ASSLGLGGSAETLY | 0.7428314 |
| SATGVFAGVI | ASSLGLGGSAETLY | 0.48097125 |
| SLPAITLP | ASSLGLGGSAETLY | 0.7130901 |
| STALAFTV | ASSLGLGGSAETLY | 0.74901676 |
| SVLTVGLGK | ASSLGLGGSAETLY | 0.68434584 |
| TALAGAGSGI | ASSLGLGGSAETLY | 0.74804175 |
| TGLPFTGI | ASSLGLGGSAETLY | 0.75073385 |
| VAEHVTV | ASSLGLGGSAETLY | 0.6603126 |
| VDIVLNSL | ASSLGLGGSAETLY | 0.7433976 |
| VLTGGGAVL | ASSLGLGGSAETLY | 0.7528386 |
| VNVGLAYGI | ASSLGLGGSAETLY | 0.60825086 |
| VSSAAVGAV | ASSLGLGGSAETLY | 0.68567634 |
| VTSVYEAGV | ASSLGLGGSAETLY | 0.501678 |
| WVALDRGA | ASSLGLGGSAETLY | 0.7016145 |
| YLVGSVVI | ASSLGLGGSAETLY | 0.4139846 |
| GPPGTGKTLL | ASSLGLGGSAETLY | 0.7208239 |
| GVGMTKF | ASSLGLGGSAETLY | 0.73968613 |
| IGIGGSDL | ASSLGLGGSAETLY | 0.66689456 |
| IGTLLAEL | ASSLGLGGSAETLY | 0.5128886 |
| IYTSGTT | ASSLGLGGSAETLY | 0.49664715 |
| LSFPAGV | ASSLGLGGSAETLY | 0.7106655 |

| | | |
|---|---|---|
| LYTSGTT | ASSLGLGGSAETLY | 0.53502095 |
| VGAFPFGF | ASSLGLGGSAETLY | 0.47932547 |
| VGAGTVEF | ASSLGLGGSAETLY | 0.57651556 |
| VGKSTLLL | ASSLGLGGSAETLY | 0.7331464 |
| AALAHAGGGIV | ASSLGLGGSAETLY | 0.72908974 |
| AALVAVSLL | ASSLGLGGSAETLY | 0.5569678 |
| AAVSRAVAAL | ASSLGLGGSAETLY | 0.7506836 |
| AGGSSGIGGF | ASSLGLGGSAETLY | 0.7419051 |
| AGLTVANANG | ASSLGLGGSAETLY | 0.74967754 |
| AGLVGAVL | ASSLGLGGSAETLY | 0.6885112 |
| ALSQAFTV | ASSLGLGGSAETLY | 0.45670524 |
| ALTFSVSL | ASSLGLGGSAETLY | 0.74546266 |
| ALTGAALVAL | ASSLGLGGSAETLY | 0.664369 |
| ASFSIGAGVL | ASSLGLGGSAETLY | 0.7075079 |
| DGSVYKI | ASSLGLGGSAETLY | 0.7336525 |
| EQLAEIV | ASSLGLGGSAETLY | 0.6665914 |
| FAVSELTVM | ASSLGLGGSAETLY | 0.5797472 |
| GAAPLAAGL | ASSLGLGGSAETLY | 0.70509696 |
| GADGGDGGSGGAGG | ASSLGLGGSAETLY | 0.7456949 |
| GFGLDTGGAGG | ASSLGLGGSAETLY | 0.5874522 |
| GLASGGLL | ASSLGLGGSAETLY | 0.4362965 |
| GRIAGAAVLL | ASSLGLGGSAETLY | 0.6667616 |
| GSAGSLNTSF | ASSLGLGGSAETLY | 0.6717376 |
| GTGQIGI | ASSLGLGGSAETLY | 0.7402855 |
| GTLIALIL | ASSLGLGGSAETLY | 0.70319426 |
| IAATMGFVVG | ASSLGLGGSAETLY | 0.65014297 |
| IFAQALL | ASSLGLGGSAETLY | 0.7243985 |
| IGAAVALGF | ASSLGLGGSAETLY | 0.71545434 |
| IGLTPLAL | ASSLGLGGSAETLY | 0.6958647 |
| ILGSNTSTL | ASSLGLGGSAETLY | 0.73903584 |
| IVFSLTVL | ASSLGLGGSAETLY | 0.6292257 |
| LASKATGF | ASSLGLGGSAETLY | 0.7441342 |
| LASVALIL | ASSLGLGGSAETLY | 0.7305256 |
| LFAVGLTL | ASSLGLGGSAETLY | 0.7355056 |
| LFNGGIGTYI | ASSLGLGGSAETLY | 0.746449 |
| LTLGAGII | ASSLGLGGSAETLY | 0.6771165 |
| LVAAALGDI | ASSLGLGGSAETLY | 0.7086291 |
| NTAFVQLGI | ASSLGLGGSAETLY | 0.4575769 |
| QLFSGAEL | ASSLGLGGSAETLY | 0.67140365 |
| SAATFITL | ASSLGLGGSAETLY | 0.7209182 |
| SNYGLANLGV | ASSLGLGGSAETLY | 0.7186271 |
| SYGGAYAVM | ASSLGLGGSAETLY | 0.74306846 |
| TTGVGGLTLG | ASSLGLGGSAETLY | 0.7334256 |
| TTQTIYI | ASSLGLGGSAETLY | 0.6221193 |
| VAAGAGLGAVI | ASSLGLGGSAETLY | 0.75119674 |
| VAAVVTLL | ASSLGLGGSAETLY | 0.7450304 |
| VGFLGGLI | ASSLGLGGSAETLY | 0.4902686 |
| VGVIPILL | ASSLGLGGSAETLY | 0.652469 |
| VIFASSLI | ASSLGLGGSAETLY | 0.7493136 |
| VKGGDGGAAAG | ASSLGLGGSAETLY | 0.73448575 |
| VTKTFTNI | ASSLGLGGSAETLY | 0.64623547 |

| | | |
|---|---|---|
| VVAAAILL | ASSLGLGGSAETLY | 0.69094205 |
| YAALDVEL | ASSLGLGGSAETLY | 0.7509067 |
| LASVALLL | ASSLGLGGSAETLY | 0.7139813 |
| LDAVHRV | ASSLGLGGSAETLY | 0.7487682 |
| QGLSSSIF | ASSLGLGGSAETLY | 0.36006302 |
| RGTFANIRL | ASSLGLGGSAETLY | 0.6655917 |
| RQLWWGHRI | ASSLGLGGSAETLY | 0.641742 |
| YGPPGTGKTLL | ASSLGLGGSAETLY | 0.278321 |
| AALVGAPANVA | ASSLGLGGSAETLY | 0.73698604 |
| AATFRRTE | ASSLGLGGSAETLY | 0.6539161 |
| AGALGGGI | ASSLGLGGSAETLY | 0.73046714 |
| AGGAGGAGGIGAG | ASSLGLGGSAETLY | 0.6639051 |
| AGSGALGL | ASSLGLGGSAETLY | 0.7360276 |
| AQIADQI | ASSLGLGGSAETLY | 0.7191444 |
| ASAVAVTYL | ASSLGLGGSAETLY | 0.6543488 |
| ASTVLGGTGG | ASSLGLGGSAETLY | 0.4631215 |
| AVGGEPAK | ASSLGLGGSAETLY | 0.7233149 |
| CEQGTFGGTC | ASSLGLGGSAETLY | 0.754007 |
| DEGKAPAG | ASSLGLGGSAETLY | 0.738206 |
| DTVVFPAL | ASSLGLGGSAETLY | 0.3517881 |
| FEGITMAG | ASSLGLGGSAETLY | 0.7239784 |
| FPLVVQD | ASSLGLGGSAETLY | 0.6788324 |
| GAFTTPQ | ASSLGLGGSAETLY | 0.34871647 |
| GFGGFGTAGGVGGA | ASSLGLGGSAETLY | 0.3933001 |
| GFLAQFL | ASSLGLGGSAETLY | 0.667392 |
| GGAGGAGGNAGVA | ASSLGLGGSAETLY | 0.68473816 |
| GGVGTAGAGGAGGAGG | ASSLGLGGSAETLY | 0.7289083 |
| GNGGGGDAGNAGSGGN | ASSLGLGGSAETLY | 0.7279326 |
| GTGGAGGAGNDAGS | ASSLGLGGSAETLY | 0.6353767 |
| GTTNTGWFNS | ASSLGLGGSAETLY | 0.73119664 |
| IFMGPLFTL | ASSLGLGGSAETLY | 0.67996067 |
| ILAHAIEAI | ASSLGLGGSAETLY | 0.22198151 |
| IRSAIREAL | ASSLGLGGSAETLY | 0.60286045 |
| ITFAGGG | ASSLGLGGSAETLY | 0.440868 |
| ITGMSGHQEV | ASSLGLGGSAETLY | 0.7496227 |
| IVAGLACTL | ASSLGLGGSAETLY | 0.5397016 |
| IVGGGLAGA | ASSLGLGGSAETLY | 0.6560383 |
| IVTGVLNI | ASSLGLGGSAETLY | 0.66297865 |
| LAQQGCI | ASSLGLGGSAETLY | 0.72830594 |
| LCLHVEN | ASSLGLGGSAETLY | 0.70039356 |
| LDAISDGS | ASSLGLGGSAETLY | 0.5689031 |
| LGGFAAML | ASSLGLGGSAETLY | 0.36599982 |
| LGGKNPMI | ASSLGLGGSAETLY | 0.715639 |
| LGPKGVAAIL | ASSLGLGGSAETLY | 0.58920956 |
| LMVGVSI | ASSLGLGGSAETLY | 0.509402 |
| LVGAAFVL | ASSLGLGGSAETLY | 0.55986154 |
| LVLAVAPL | ASSLGLGGSAETLY | 0.73387927 |
| MAAIAVIDAA | ASSLGLGGSAETLY | 0.43504548 |
| NNGINAELQ | ASSLGLGGSAETLY | 0.74354255 |
| RGEPLVLVHG | ASSLGLGGSAETLY | 0.6096766 |
| RGGAGGAAGLI | ASSLGLGGSAETLY | 0.67111546 |

| | | |
|---|---|---|
| RSYRMHPAV | ASSLGLGGSAETLY | 0.752038 |
| SAAHLLAGQLA | ASSLGLGGSAETLY | 0.7180486 |
| SAAVGSY | ASSLGLGGSAETLY | 0.67361194 |
| SPHAVLHRI | ASSLGLGGSAETLY | 0.5097178 |
| SQIAKVL | ASSLGLGGSAETLY | 0.5168566 |
| TALGHLVP | ASSLGLGGSAETLY | 0.5456208 |
| TAPTTSGAA | ASSLGLGGSAETLY | 0.58208704 |
| TDGDDGNGGSPGT | ASSLGLGGSAETLY | 0.7522541 |
| TDGLGEN | ASSLGLGGSAETLY | 0.6979239 |
| TGGPPLPGQA | ASSLGLGGSAETLY | 0.47019464 |
| TIPAITI | ASSLGLGGSAETLY | 0.71812963 |
| TIPGFTI | ASSLGLGGSAETLY | 0.75110495 |
| TSIASGALA | ASSLGLGGSAETLY | 0.75548697 |
| TVTLPGAAI | ASSLGLGGSAETLY | 0.5031308 |
| VASGLSAALP | ASSLGLGGSAETLY | 0.6456196 |
| VGAGVIGI | ASSLGLGGSAETLY | 0.68718255 |
| VGPKRFGV | ASSLGLGGSAETLY | 0.6894915 |
| VNVAFPD | ASSLGLGGSAETLY | 0.5471989 |
| VSMTNSGVS | ASSLGLGGSAETLY | 0.47625482 |
| VTDVLGDLSV | ASSLGLGGSAETLY | 0.73426163 |
| VVVVGAGFAGL | ASSLGLGGSAETLY | 0.756548 |
| VYTVGTVV | ASSLGLGGSAETLY | 0.64775646 |
| WTTFSGP | ASSLGLGGSAETLY | 0.46602774 |
| YDELVEAF | ASSLGLGGSAETLY | 0.6681286 |
| YGTGGAGGNGG | ASSLGLGGSAETLY | 0.5754891 |
| YTNVMAV | ASSLGLGGSAETLY | 0.5889051 |
| FLEQVHQL | ASSLGLGGSAETLY | 0.6682426 |
| GLSLLAVAV | ASSLGLGGSAETLY | 0.4431974 |
| LEQVHQL | ASSLGLGGSAETLY | 0.6803603 |
| LNAAGAI | ASSLGLGGSAETLY | 0.7505163 |
| PAVHASL | ASSLGLGGSAETLY | 0.6856639 |
| TELEDDA | ASSLGLGGSAETLY | 0.7247248 |
| TPPGPPN | ASSLGLGGSAETLY | 0.74574274 |
| TPPPPPPP | ASSLGLGGSAETLY | 0.72835195 |
| TQSPALI | ASRQGRVTGQLY | 0.74637884 |
| EFAAAFPQ | ASRQGRVTGQLY | 0.745069 |
| LLSGNLDVL | ASRQGRVTGQLY | 0.66711694 |
| GPANLGNYNI | ASRQGRVTGQLY | 0.70960855 |
| TLGPTLP | ASRQGRVTGQLY | 0.73982215 |
| VYAYSSCCK | ASRQGRVTGQLY | 0.65556836 |
| APAPAPAPNI | ASRQGRVTGQLY | 0.68413615 |
| PGGPVPIP | ASRQGRVTGQLY | 0.6793754 |
| FLTVALEP | ASRQGRVTGQLY | 0.7369254 |
| SQYFGANL | ASRQGRVTGQLY | 0.66955805 |
| GGVKPAA | ASRQGRVTGQLY | 0.7514849 |
| GGTGGAGGAGGW | ASRQGRVTGQLY | 0.32375452 |
| GVGGVAGLGGAGG | ASRQGRVTGQLY | 0.47705773 |
| VLLVLATTI | ASRQGRVTGQLY | 0.7433176 |
| IGIFAAML | ASRQGRVTGQLY | 0.257403 |
| AVGDLID | ASRQGRVTGQLY | 0.35252944 |
| NISPLPL | ASRQGRVTGQLY | 0.69888735 |

| | | |
|---|---|---|
| VTAVFVAI | ASRQGRVTGQLY | 0.74408495 |
| SGDDNDAAL | ASRQGRVTGQLY | 0.24710776 |
| AVAALAHS | ASRQGRVTGQLY | 0.5937586 |
| GGVTVMATP | ASRQGRVTGQLY | 0.43636245 |
| AGLAANVRD | ASRQGRVTGQLY | 0.7315725 |
| TNRPDLIDPALLRPG | ASRQGRVTGQLY | 0.661532 |
| LANLGVTAL | ASRQGRVTGQLY | 0.38106245 |
| TGERIEL | ASRQGRVTGQLY | 0.7265522 |
| NGEELAA | ASRQGRVTGQLY | 0.51153785 |
| VLSLVWG | ASRQGRVTGQLY | 0.69932175 |
| IGYALWL | ASRQGRVTGQLY | 0.73900217 |
| LSLLLAPL | ASRQGRVTGQLY | 0.75252336 |
| TLDEALK | ASRQGRVTGQLY | 0.48723865 |
| TDGAGTAEGG | ASRQGRVTGQLY | 0.726213 |
| NAPGGAGGNGG | ASRQGRVTGQLY | 0.6690115 |
| VSVFMGMGGNF | ASRQGRVTGQLY | 0.75637436 |
| LASYLAAL | ASRQGRVTGQLY | 0.6397418 |
| LVGGTPAL | ASRQGRVTGQLY | 0.41306168 |
| VRNADVL | ASRQGRVTGQLY | 0.73604417 |
| TCSSPFLVSAM | ASRQGRVTGQLY | 0.73061156 |
| VHITDAE | ASRQGRVTGQLY | 0.6651712 |
| EYAAPSF | ASRQGRVTGQLY | 0.7436038 |
| VVDAFHGGL | ASRQGRVTGQLY | 0.65859985 |
| ASVVDPQ | ASRQGRVTGQLY | 0.66557556 |
| SYNEFGGLS | ASRQGRVTGQLY | 0.734635 |
| DPDGSVY | ASRQGRVTGQLY | 0.7358484 |
| MVSTIHATE | ASRQGRVTGQLY | 0.5101682 |
| LAAGVADPV | ASRQGRVTGQLY | 0.7455149 |
| TKEQIAA | ASRQGRVTGQLY | 0.6853646 |
| SLASPAAL | ASRQGRVTGQLY | 0.5154138 |
| IGLVFYGV | ASRQGRVTGQLY | 0.75381136 |
| FLLDASNL | ASRQGRVTGQLY | 0.6669495 |
| YQVGNFP | ASRQGRVTGQLY | 0.73462176 |
| ESLLSQST | ASRQGRVTGQLY | 0.7305467 |
| LFTVGFAAVG | ASRQGRVTGQLY | 0.75094926 |
| ATPPTPT | ASRQGRVTGQLY | 0.45547175 |
| TDLATLG | ASRQGRVTGQLY | 0.71553934 |
| SAGGTGR | ASRQGRVTGQLY | 0.7002772 |
| GFSNSGS | ASRQGRVTGQLY | 0.7339543 |
| GGNGGSGGTSV | ASRQGRVTGQLY | 0.68942446 |
| GTSVATGGAGN | ASRQGRVTGQLY | 0.6875142 |
| NGGNGGSGGTS | ASRQGRVTGQLY | 0.7250274 |
| AVVKFPH | ASRQGRVTGQLY | 0.74401486 |
| TPPPPPP | ASRQGRVTGQLY | 0.6809806 |
| AAPAAAAAPA | ASRQGRVTGQLY | 0.732571 |
| IALNEDL | ASRQGRVTGQLY | 0.7502818 |
| IVNVSSM | ASRQGRVTGQLY | 0.5202565 |
| TGIPPAPR | ASRQGRVTGQLY | 0.66880035 |
| FDLGGGTF | ASRQGRVTGQLY | 0.73093987 |
| FDLGGGTFDVS | ASRQGRVTGQLY | 0.7518369 |
| NEPTAAA | ASRQGRVTGQLY | 0.6764082 |

| | | |
|---|---|---|
| NEPTAAALA | ASRQGRVTGQLY | 0.7445512 |
| GSIINISSI | ASRQGRVTGQLY | 0.65556484 |
| VAPGNAGTA | ASRQGRVTGQLY | 0.721544 |
| TGGGASLE | ASRQGRVTGQLY | 0.67504525 |
| VAVYNHYKRI | ASRQGRVTGQLY | 0.69339037 |
| TDPAPATT | ASRQGRVTGQLY | 0.5490196 |
| TNRPDILDPALLRPG | ASRQGRVTGQLY | 0.70760834 |
| LTTTTTS | ASRQGRVTGQLY | 0.7361854 |
| PTGSGKTL | ASRQGRVTGQLY | 0.7236174 |
| TGSGKTL | ASRQGRVTGQLY | 0.41459745 |
| TIGHVDHGK | ASRQGRVTGQLY | 0.7155646 |
| TIGHVDHGKTT | ASRQGRVTGQLY | 0.72136927 |
| GHVDHGKTT | ASRQGRVTGQLY | 0.75665855 |
| AQFEHTLL | ASRQGRVTGQLY | 0.6926104 |
| IVNQDEL | ASRQGRVTGQLY | 0.73952734 |
| TNGSQFF | ASRQGRVTGQLY | 0.6198027 |
| TGKGIVE | ASRQGRVTGQLY | 0.68142486 |
| LISNASDA | ASRQGRVTGQLY | 0.49359035 |
| AELRTLGELTKIAKAHGAQ | ASRQGRVTGQLY | 0.63793933 |
| APNAVAAAEPSW | ASRQGRVTGQLY | 0.68082786 |
| APQVAGAATAVL | ASRQGRVTGQLY | 0.59596896 |
| ARRDPAAAAAILPTDA | ASRQGRVTGQLY | 0.7403223 |
| DAAARDVLAEAGLAEYF | ASRQGRVTGQLY | 0.74820966 |
| DDQIVLSAKANLAKVEQ | ASRQGRVTGQLY | 0.73793495 |
| DELLPDTGEDSDEDGH | ASRQGRVTGQLY | 0.74249226 |
| ELPEAPANRATVPQD | ASRQGRVTGQLY | 0.7512283 |
| FGQSKIERVTGIG | ASRQGRVTGQLY | 0.7377364 |
| GAGGTGSTAGGAGGAGGAGGLY | ASRQGRVTGQLY | 0.6107068 |
| GHGGTGAAGGNGGTGGAGGSGI | ASRQGRVTGQLY | 0.6858448 |
| GNGGNGGAGGEGGPGGAGGAGA | ASRQGRVTGQLY | 0.4343334 |
| GSTAGNGGNGGNATNGTIAGQP | ASRQGRVTGQLY | 0.65436697 |
| GTGGDGGNGGNGGTGG | ASRQGRVTGQLY | 0.6304101 |
| GVRRPAGIPADAPV | ASRQGRVTGQLY | 0.69134015 |
| GYRQVTDAHLLTLARR | ASRQGRVTGQLY | 0.24161327 |
| IAKMGQDGHDRG | ASRQGRVTGQLY | 0.72874594 |
| KAEPSIEIQYT | ASRQGRVTGQLY | 0.7528061 |
| KPVRIRMAADEDTVAVT | ASRQGRVTGQLY | 0.6514324 |
| MPPRIAGMRLLVIKPEP | ASRQGRVTGQLY | 0.7206092 |
| MKFYAVALFRSRGVPPTR | ASRQGRVTGQLY | 0.7313006 |
| NAGGFGILGAVAHSP | ASRQGRVTGQLY | 0.6481381 |
| NEGREITLAMAEM | ASRQGRVTGQLY | 0.7259208 |
| NELVEWSHPLGHRWRWE | ASRQGRVTGQLY | 0.6190796 |
| NLGAGGTGGNGGNPG | ASRQGRVTGQLY | 0.7355388 |
| NTEWPKWETVMAADPVD | ASRQGRVTGQLY | 0.74965453 |
| PASAAPAETQP | ASRQGRVTGQLY | 0.72166073 |
| QQSSDGKNGVRAAEITGE | ASRQGRVTGQLY | 0.7240491 |
| RAPDAVTTSWWKE | ASRQGRVTGQLY | 0.71062076 |
| SGFGNVGAHTSGWFNQ | ASRQGRVTGQLY | 0.43757132 |
| SGVAGAVGQAASVGGLKV | ASRQGRVTGQLY | 0.73069865 |
| SNGGAGTGSAGGNGGT | ASRQGRVTGQLY | 0.6400666 |
| TGGAGGEGGAAG | ASRQGRVTGQLY | 0.6996645 |

| | | |
|---|---|---|
| TGGDGGAAGAGGAPGHG | ASRQGRVTGQLY | 0.54500455 |
| TMPPLLAAIANAGRHGV | ASRQGRVTGQLY | 0.6995019 |
| TSSTAGNSGGA | ASRQGRVTGQLY | 0.6874792 |
| VPATVPVGILC | ASRQGRVTGQLY | 0.71128345 |
| VQRPDSGGVVLTGRIS | ASRQGRVTGQLY | 0.5633812 |
| WTPNPVPATKGVPAGLVQAT | ASRQGRVTGQLY | 0.72817004 |
| YALKRALAKFSDRFDVVII | ASRQGRVTGQLY | 0.7132802 |
| YLIGSVIV | ASRQGRVTGQLY | 0.48490605 |
| YGQLVAVID | ASRQGRVTGQLY | 0.7430768 |
| VTVGGMTLPA | ASRQGRVTGQLY | 0.6575406 |
| VLALPAV | ASRQGRVTGQLY | 0.6923164 |
| VIPPTLNYET | ASRQGRVTGQLY | 0.7408767 |
| VGMGGMHPSSG | ASRQGRVTGQLY | 0.6847223 |
| VGLAGGAATA | ASRQGRVTGQLY | 0.45375043 |
| VGGAGGAGGSGP | ASRQGRVTGQLY | 0.66997665 |
| VFAAFMV | ASRQGRVTGQLY | 0.6982219 |
| TLYDKLR | ASRQGRVTGQLY | 0.64550406 |
| TLAVFTVL | ASRQGRVTGQLY | 0.3174494 |
| TGSVQPARV | ASRQGRVTGQLY | 0.72094905 |
| TGGGAGSGGT | ASRQGRVTGQLY | 0.510232 |
| SHLETSL | ASRQGRVTGQLY | 0.6398438 |
| SFTTGMT | ASRQGRVTGQLY | 0.63175416 |
| QVEVAKRMGVRQPTVS | ASRQGRVTGQLY | 0.73389125 |
| QGSGGPALAM | ASRQGRVTGQLY | 0.70308137 |
| MLAPSNP | ASRQGRVTGQLY | 0.6287452 |
| LVNNAAI | ASRQGRVTGQLY | 0.7341304 |
| LTGEINTLA | ASRQGRVTGQLY | 0.69829047 |
| LSIFGPV | ASRQGRVTGQLY | 0.744269 |
| LLALLANL | ASRQGRVTGQLY | 0.6883694 |
| LISNASDALDKLR | ASRQGRVTGQLY | 0.70057887 |
| LINEAVP | ASRQGRVTGQLY | 0.7224963 |
| LGYAFSGL | ASRQGRVTGQLY | 0.6172042 |
| LDFVAVAL | ASRQGRVTGQLY | 0.5933808 |
| LAQPGAE | ASRQGRVTGQLY | 0.5133326 |
| ISNLGAVTGF | ASRQGRVTGQLY | 0.6150718 |
| HQEDPGRI | ASRQGRVTGQLY | 0.5071547 |
| GTPGEPTP | ASRQGRVTGQLY | 0.7027532 |
| GFFNSGS | ASRQGRVTGQLY | 0.7491468 |
| GAGGGGLTAGAN | ASRQGRVTGQLY | 0.6639304 |
| FVDALVNM | ASRQGRVTGQLY | 0.4391464 |
| FLTVAPEL | ASRQGRVTGQLY | 0.65523887 |
| FETLTAL | ASRQGRVTGQLY | 0.6382855 |
| DVINPIEL | ASRQGRVTGQLY | 0.7298397 |
| APVSGAPV | ASRQGRVTGQLY | 0.7292494 |
| ALVGGLAW | ASRQGRVTGQLY | 0.7549406 |
| ALAAAAATAI | ASRQGRVTGQLY | 0.7316507 |
| AFGLGLLL | ASRQGRVTGQLY | 0.6265403 |
| AFFAAASL | ASRQGRVTGQLY | 0.73099256 |
| AAAALPAFSPP | ASRQGRVTGQLY | 0.6652156 |
| VVLAPAL | ASRQGRVTGQLY | 0.7280148 |
| TIGHVDHGKTTL | ASRQGRVTGQLY | 0.7069037 |

| | | |
|---|---|---|
| LISNASDALDK | ASRQGRVTGQLY | 0.7180998 |
| LISNASDAL | ASRQGRVTGQLY | 0.6557356 |
| ISNASDALDK | ASRQGRVTGQLY | 0.6398717 |
| NQAETLVYQTE | ASRQGRVTGQLY | 0.42698014 |
| TGGTGGTAGAG | ASRQGRVTGQLY | 0.7345956 |
| TGSTAGGAGGA | ASRQGRVTGQLY | 0.6148196 |
| DPLAVPVAAGPV | ASRQGRVTGQLY | 0.57969105 |
| GGTGGTPGNGGN | ASRQGRVTGQLY | 0.6532337 |
| PAGLAGVDPATL | ASRQGRVTGQLY | 0.7524991 |
| GNNLVVEAPHAGASV | ASRQGRVTGQLY | 0.7436186 |
| KVTRIMSASTLEQPA | ASRQGRVTGQLY | 0.63785446 |
| LGFALPIALPRRTVV | ASRQGRVTGQLY | 0.5957098 |
| TRLGNLTARKLVVLV | ASRQGRVTGQLY | 0.73190236 |
| WSQQRRLIYFPSAGP | ASRQGRVTGQLY | 0.53385717 |
| GPNGNPDPMAAAVDIR | ASRQGRVTGQLY | 0.72580534 |
| TQRCVSVADLRAAASR | ASRQGRVTGQLY | 0.69333243 |
| QFVLDVAEHPVEVLTETV | ASRQGRVTGQLY | 0.6885434 |
| GGAGGTGGAGGNGGTQAGDG | ASRQGRVTGQLY | 0.66267574 |
| PLLATLRAIAVDVDAKPAQV | ASRQGRVTGQLY | 0.72791076 |
| APKFRTGGYMIDFWGVGYQVAK | ASRQGRVTGQLY | 0.645247 |
| GAGGAGGAGGAAGGPGGTGGTG | ASRQGRVTGQLY | 0.72012913 |
| IGALGLLATDVAPAGSVAFGVA | ASRQGRVTGQLY | 0.2931349 |
| AGGAGGIGGNGGAGGAGGVLMG | ASRQGRVTGQLY | 0.73609734 |
| ADELIGLL | ASRQGRVTGQLY | 0.5589379 |
| ADRALKA | ASRQGRVTGQLY | 0.5578183 |
| AGAVGQVGGAGGA | ASRQGRVTGQLY | 0.2954627 |
| AGGGGGLGGAP | ASRQGRVTGQLY | 0.69332546 |
| AGGLLYGNGGAG | ASRQGRVTGQLY | 0.7308471 |
| AGLESIGL | ASRQGRVTGQLY | 0.74378896 |
| AGLKTGII | ASRQGRVTGQLY | 0.70176744 |
| ASLAALLL | ASRQGRVTGQLY | 0.73823875 |
| ASVINISSI | ASRQGRVTGQLY | 0.7469933 |
| ATPPGPA | ASRQGRVTGQLY | 0.66324604 |
| ATVASALV | ASRQGRVTGQLY | 0.3198328 |
| ATVASLIL | ASRQGRVTGQLY | 0.7036444 |
| AVNGPVLIRH | ASRQGRVTGQLY | 0.7183461 |
| DGAAGAVGI | ASRQGRVTGQLY | 0.744406 |
| DPLLHGI | ASRQGRVTGQLY | 0.72389376 |
| ELGAALGAL | ASRQGRVTGQLY | 0.48521543 |
| ELGRTQAV | ASRQGRVTGQLY | 0.74684614 |
| FVAGGYLS | ASRQGRVTGQLY | 0.7233149 |
| GASGGAGGAGGW | ASRQGRVTGQLY | 0.5172893 |
| GGAGAGGDGGAGGI | ASRQGRVTGQLY | 0.6334043 |
| GTGHNGSQPGMGGNG | ASRQGRVTGQLY | 0.58557695 |
| GYAAASASAS | ASRQGRVTGQLY | 0.4385755 |
| HQVDVAAVQQG | ASRQGRVTGQLY | 0.7232624 |
| IGAAIAHT | ASRQGRVTGQLY | 0.65989035 |
| IGMIEVAQL | ASRQGRVTGQLY | 0.61453176 |
| IRLAPEHTSAMD | ASRQGRVTGQLY | 0.6769809 |
| IYNVSVL | ASRQGRVTGQLY | 0.6142107 |
| KQGFLGM | ASRQGRVTGQLY | 0.57477295 |

| | | |
|---|---|---|
| LESVITQL | ASRQGRVTGQLY | 0.62180835 |
| LGLGALVVV | ASRQGRVTGQLY | 0.7138964 |
| LIGNGGNGGSGGI | ASRQGRVTGQLY | 0.7331268 |
| LNAINAPTL | ASRQGRVTGQLY | 0.69211406 |
| LQLLNTGV | ASRQGRVTGQLY | 0.71984655 |
| LQTSIEF | ASRQGRVTGQLY | 0.74710727 |
| MSFAVSEL | ASRQGRVTGQLY | 0.45653093 |
| NTGALGL | ASRQGRVTGQLY | 0.58286357 |
| PPGAGGAGGAGTT | ASRQGRVTGQLY | 0.71460223 |
| QLVGLGI | ASRQGRVTGQLY | 0.63034236 |
| RFLEQVHQL | ASRQGRVTGQLY | 0.73935235 |
| RGVGYVLR | ASRQGRVTGQLY | 0.6988248 |
| SATGVFAGVI | ASRQGRVTGQLY | 0.70862347 |
| SLPAITLP | ASRQGRVTGQLY | 0.7150485 |
| STALAFTV | ASRQGRVTGQLY | 0.7545773 |
| SVLTVGLGK | ASRQGRVTGQLY | 0.7188757 |
| TALAGAGSGI | ASRQGRVTGQLY | 0.6867564 |
| TGLPFTGI | ASRQGRVTGQLY | 0.74695873 |
| VAEHVTV | ASRQGRVTGQLY | 0.49336886 |
| VDIVLNSL | ASRQGRVTGQLY | 0.74160266 |
| VLTGGGAVL | ASRQGRVTGQLY | 0.72834 |
| VNVGLAYGI | ASRQGRVTGQLY | 0.5546615 |
| VSSAAVGAV | ASRQGRVTGQLY | 0.6701261 |
| VTSVYEAGV | ASRQGRVTGQLY | 0.7510483 |
| WVALDRGA | ASRQGRVTGQLY | 0.7430644 |
| YLVGSVVI | ASRQGRVTGQLY | 0.5729215 |
| GPPGTGKTLL | ASRQGRVTGQLY | 0.59436774 |
| GVGMTKF | ASRQGRVTGQLY | 0.52144605 |
| IGIGGSDL | ASRQGRVTGQLY | 0.6308273 |
| IGTLLAEL | ASRQGRVTGQLY | 0.36783043 |
| IYTSGTT | ASRQGRVTGQLY | 0.7266752 |
| LSFPAGV | ASRQGRVTGQLY | 0.62798023 |
| LYTSGTT | ASRQGRVTGQLY | 0.74105525 |
| VGAFPFGF | ASRQGRVTGQLY | 0.7011214 |
| VGAGTVEF | ASRQGRVTGQLY | 0.7418299 |
| VGKSTLLL | ASRQGRVTGQLY | 0.62692267 |
| AALAHAGGGIV | ASRQGRVTGQLY | 0.6135117 |
| AALVAVSLL | ASRQGRVTGQLY | 0.3643335 |
| AAVSRAVAAL | ASRQGRVTGQLY | 0.57894015 |
| AGGSSGIGGF | ASRQGRVTGQLY | 0.5883908 |
| AGLTVANANG | ASRQGRVTGQLY | 0.65867776 |
| AGLVGAVL | ASRQGRVTGQLY | 0.33179867 |
| ALSQAFTV | ASRQGRVTGQLY | 0.7455886 |
| ALTFSVSL | ASRQGRVTGQLY | 0.66360784 |
| ALTGAALVAL | ASRQGRVTGQLY | 0.7211179 |
| ASFSIGAGVL | ASRQGRVTGQLY | 0.60115826 |
| DGSVYKI | ASRQGRVTGQLY | 0.6016077 |
| EQLAEIV | ASRQGRVTGQLY | 0.59961677 |
| FAVSELTVM | ASRQGRVTGQLY | 0.72785497 |
| GAAPLAAGL | ASRQGRVTGQLY | 0.5071863 |
| GADGGDGGSGGAGG | ASRQGRVTGQLY | 0.3678229 |

| | | |
|---|---|---|
| GFGLDTGGAGG | ASRQGRVTGQLY | 0.5592046 |
| GLASGGLL | ASRQGRVTGQLY | 0.63844883 |
| GRIAGAAVLL | ASRQGRVTGQLY | 0.75198424 |
| GSAGSLNTSF | ASRQGRVTGQLY | 0.7416064 |
| GTGQIGI | ASRQGRVTGQLY | 0.70474386 |
| GTLIALIL | ASRQGRVTGQLY | 0.7518755 |
| IAATMGFVVG | ASRQGRVTGQLY | 0.26270998 |
| IFAQALL | ASRQGRVTGQLY | 0.6255708 |
| IGAAVALGF | ASRQGRVTGQLY | 0.74987537 |
| IGLTPLAL | ASRQGRVTGQLY | 0.7168505 |
| ILGSNTSTL | ASRQGRVTGQLY | 0.6617332 |
| IVFSLTVL | ASRQGRVTGQLY | 0.7427212 |
| LASKATGF | ASRQGRVTGQLY | 0.55923307 |
| LASVALIL | ASRQGRVTGQLY | 0.63037187 |
| LFAVGLTL | ASRQGRVTGQLY | 0.6684344 |
| LFNGGIGTYI | ASRQGRVTGQLY | 0.73349965 |
| LTLGAGII | ASRQGRVTGQLY | 0.71022904 |
| LVAAALGDI | ASRQGRVTGQLY | 0.4378701 |
| NTAFVQLGI | ASRQGRVTGQLY | 0.7270118 |
| QLFSGAEL | ASRQGRVTGQLY | 0.7555114 |
| SAATFITL | ASRQGRVTGQLY | 0.6249464 |
| SNYGLANLGV | ASRQGRVTGQLY | 0.7375653 |
| SYGGAYAVM | ASRQGRVTGQLY | 0.7110288 |
| TTGVGGLTLG | ASRQGRVTGQLY | 0.7001149 |
| TTQTIYI | ASRQGRVTGQLY | 0.5875163 |
| VAAGAGLGAVI | ASRQGRVTGQLY | 0.55683756 |
| VAAVVTLL | ASRQGRVTGQLY | 0.73115516 |
| VGFLGGLI | ASRQGRVTGQLY | 0.5416491 |
| VGVIPILL | ASRQGRVTGQLY | 0.68332064 |
| VIFASSLI | ASRQGRVTGQLY | 0.6640694 |
| VKGGDGGAAAG | ASRQGRVTGQLY | 0.7570207 |
| VTKTFTNI | ASRQGRVTGQLY | 0.72616804 |
| VVAAAILL | ASRQGRVTGQLY | 0.70615876 |
| YAALDVEL | ASRQGRVTGQLY | 0.57157826 |
| LASVALLL | ASRQGRVTGQLY | 0.7181162 |
| LDAVHRV | ASRQGRVTGQLY | 0.73085296 |
| QGLSSSIF | ASRQGRVTGQLY | 0.4534989 |
| RGTFANIRL | ASRQGRVTGQLY | 0.74457884 |
| RQLWWGHRI | ASRQGRVTGQLY | 0.5106602 |
| YGPPGTGKTLL | ASRQGRVTGQLY | 0.6953375 |
| AALVGAPANVA | ASRQGRVTGQLY | 0.7102386 |
| AATFRRTE | ASRQGRVTGQLY | 0.7465079 |
| AGALGGGI | ASRQGRVTGQLY | 0.6818147 |
| AGGAGGAGGIGAG | ASRQGRVTGQLY | 0.7299792 |
| AGSGALGL | ASRQGRVTGQLY | 0.49825484 |
| AQIADQI | ASRQGRVTGQLY | 0.73264325 |
| ASAVAVTYL | ASRQGRVTGQLY | 0.7375026 |
| ASTVLGGTGG | ASRQGRVTGQLY | 0.73126936 |
| AVGGEPAK | ASRQGRVTGQLY | 0.7554251 |
| CEQGTFGGTC | ASRQGRVTGQLY | 0.7247335 |
| DEGKAPAG | ASRQGRVTGQLY | 0.31836382 |

| | | |
|---|---|---|
| DTVVFPAL | ASRQGRVTGQLY | 0.6584873 |
| FEGITMAG | ASRQGRVTGQLY | 0.74484074 |
| FPLVVQD | ASRQGRVTGQLY | 0.74737275 |
| GAFTTPQ | ASRQGRVTGQLY | 0.74640834 |
| GFGGFGTAGGVGGA | ASRQGRVTGQLY | 0.74201894 |
| GFLAQFL | ASRQGRVTGQLY | 0.6854105 |
| GGAGGAGGNAGVA | ASRQGRVTGQLY | 0.74822044 |
| GGVGTAGAGGAGGAGG | ASRQGRVTGQLY | 0.75819844 |
| GNGGGGDAGNAGSGGN | ASRQGRVTGQLY | 0.7283697 |
| GTGGAGGAGNDAGS | ASRQGRVTGQLY | 0.5516403 |
| GTTNTGWFNS | ASRQGRVTGQLY | 0.5196291 |
| IFMGPLFTL | ASRQGRVTGQLY | 0.73084426 |
| ILAHAIEAI | ASRQGRVTGQLY | 0.73982215 |
| IRSAIREAL | ASRQGRVTGQLY | 0.49653268 |
| ITFAGGG | ASRQGRVTGQLY | 0.6823462 |
| ITGMSGHQEV | ASRQGRVTGQLY | 0.6972834 |
| IVAGLACTL | ASRQGRVTGQLY | 0.3286763 |
| IVGGGLAGA | ASRQGRVTGQLY | 0.71465576 |
| IVTGVLNI | ASRQGRVTGQLY | 0.75073034 |
| LAQQGCI | ASRQGRVTGQLY | 0.7179676 |
| LCLHVEN | ASRQGRVTGQLY | 0.5619667 |
| LDAISDGS | ASRQGRVTGQLY | 0.74579287 |
| LGGFAAML | ASRQGRVTGQLY | 0.7511337 |
| LGGKNPMI | ASRQGRVTGQLY | 0.75123715 |
| LGPKGVAAIL | ASRQGRVTGQLY | 0.36837628 |
| LMVGVSI | ASRQGRVTGQLY | 0.28395632 |
| LVGAAFVL | ASRQGRVTGQLY | 0.70195246 |
| LVLAVAPL | ASRQGRVTGQLY | 0.7357809 |
| MAAIAVIDAA | ASRQGRVTGQLY | 0.6320868 |
| NNGINAELQ | ASRQGRVTGQLY | 0.54223615 |
| RGEPLVLVHG | ASRQGRVTGQLY | 0.71752703 |
| RGGAGGAAGLI | ASRQGRVTGQLY | 0.73972744 |
| RSYRMHPAV | ASRQGRVTGQLY | 0.5113946 |
| SAAHLLAGQLA | ASRQGRVTGQLY | 0.41202962 |
| SAAVGSY | ASRQGRVTGQLY | 0.74314 |
| SPHAVLHRI | ASRQGRVTGQLY | 0.7317344 |
| SQIAKVL | ASRQGRVTGQLY | 0.5880288 |
| TALGHLVP | ASRQGRVTGQLY | 0.6541215 |
| TAPTTSGAA | ASRQGRVTGQLY | 0.7047291 |
| TDGDDGNGGSPGT | ASRQGRVTGQLY | 0.71784186 |
| TDGLGEN | ASRQGRVTGQLY | 0.71575546 |
| TGGPPLPGQA | ASRQGRVTGQLY | 0.46505767 |
| TIPAITI | ASRQGRVTGQLY | 0.73746705 |
| TIPGFTI | ASRQGRVTGQLY | 0.48344976 |
| TSIASGALA | ASRQGRVTGQLY | 0.7377229 |
| TVTLPGAAI | ASRQGRVTGQLY | 0.7178967 |
| VASGLSAALP | ASRQGRVTGQLY | 0.72374475 |
| VGAGVIGI | ASRQGRVTGQLY | 0.6519728 |
| VGPKRFGV | ASRQGRVTGQLY | 0.5371747 |
| VNVAFPD | ASRQGRVTGQLY | 0.52206314 |
| VSMTNSGVS | ASRQGRVTGQLY | 0.5019238 |

| | | |
|---|---|---|
| VTDVLGDLSV | ASRQGRVTGQLY | 0.7508842 |
| VVVVGAGFAGL | ASRQGRVTGQLY | 0.7094977 |
| VYTVGTVV | ASRQGRVTGQLY | 0.742015 |
| WTTFSGP | ASRQGRVTGQLY | 0.7029631 |
| YDELVEAF | ASRQGRVTGQLY | 0.4850209 |
| YGTGGAGGNGG | ASRQGRVTGQLY | 0.72297865 |
| YTNVMAV | ASRQGRVTGQLY | 0.4156316 |
| FLEQVHQL | ASRQGRVTGQLY | 0.7162992 |
| GLSLLAVAV | ASRQGRVTGQLY | 0.6986263 |
| LEQVHQL | ASRQGRVTGQLY | 0.72225034 |
| LNAAGAI | ASRQGRVTGQLY | 0.59358 |
| PAVHASL | ASRQGRVTGQLY | 0.7250519 |
| TELEDDA | ASRQGRVTGQLY | 0.72530913 |
| TPPGPPN | ASRQGRVTGQLY | 0.5755079 |
| TPPPPPPP | ASRQGRVTGQLY | 0.7229752 |
| TQSPALI | ASGSQGEVERLF | 0.73185885 |
| EFAAAFPQ | ASGSQGEVERLF | 0.72389305 |
| LLSGNLDVL | ASGSQGEVERLF | 0.41078436 |
| GPANLGNYNI | ASGSQGEVERLF | 0.7074859 |
| TLGPTLP | ASGSQGEVERLF | 0.610925 |
| VYAYSSCCK | ASGSQGEVERLF | 0.7125199 |
| APAPAPAPNI | ASGSQGEVERLF | 0.6741471 |
| PGGPVPIP | ASGSQGEVERLF | 0.62658817 |
| FLTVALEP | ASGSQGEVERLF | 0.72742844 |
| SQYFGANL | ASGSQGEVERLF | 0.40501818 |
| GGVKPAA | ASGSQGEVERLF | 0.6481788 |
| GGTGGAGGAGGW | ASGSQGEVERLF | 0.64686406 |
| GVGGVAGLGGAGG | ASGSQGEVERLF | 0.6797656 |
| VLLVLATTI | ASGSQGEVERLF | 0.72466147 |
| IGIFAAML | ASGSQGEVERLF | 0.5324352 |
| AVGDLID | ASGSQGEVERLF | 0.708187 |
| NISPLPL | ASGSQGEVERLF | 0.63948935 |
| VTAVFVAI | ASGSQGEVERLF | 0.5274464 |
| SGDDNDAAL | ASGSQGEVERLF | 0.7078613 |
| AVAALAHS | ASGSQGEVERLF | 0.63915145 |
| GGVTVMATP | ASGSQGEVERLF | 0.48408973 |
| AGLAANVRD | ASGSQGEVERLF | 0.73096645 |
| TNRPDLIDPALLRPG | ASGSQGEVERLF | 0.54302275 |
| LANLGVTAL | ASGSQGEVERLF | 0.7068109 |
| TGERIEL | ASGSQGEVERLF | 0.75659394 |
| NGEELAA | ASGSQGEVERLF | 0.57159185 |
| VLSLVWG | ASGSQGEVERLF | 0.730932 |
| IGYALWL | ASGSQGEVERLF | 0.706788 |
| LSLLLAPL | ASGSQGEVERLF | 0.72555363 |
| TLDEALK | ASGSQGEVERLF | 0.65598285 |
| TDGAGTAEGG | ASGSQGEVERLF | 0.5826442 |
| NAPGGAGGNGG | ASGSQGEVERLF | 0.74434006 |
| VSVFMGMGGNF | ASGSQGEVERLF | 0.71004623 |
| LASYLAAL | ASGSQGEVERLF | 0.7436917 |
| LVGGTPAL | ASGSQGEVERLF | 0.7334215 |
| VRNADVL | ASGSQGEVERLF | 0.74457705 |

| | | |
|---|---|---|
| TCSSPFLVSAM | ASGSQGEVERLF | 0.7326518 |
| VHITDAE | ASGSQGEVERLF | 0.6136558 |
| EYAAPSF | ASGSQGEVERLF | 0.3850518 |
| VVDAFHGGL | ASGSQGEVERLF | 0.6316418 |
| ASVVDPQ | ASGSQGEVERLF | 0.6653813 |
| SYNEFGGLS | ASGSQGEVERLF | 0.6010009 |
| DPDGSVY | ASGSQGEVERLF | 0.37408656 |
| MVSTIHATE | ASGSQGEVERLF | 0.7360811 |
| LAAGVADPV | ASGSQGEVERLF | 0.7572691 |
| TKEQIAA | ASGSQGEVERLF | 0.5926204 |
| SLASPAAL | ASGSQGEVERLF | 0.7549628 |
| IGLVFYGV | ASGSQGEVERLF | 0.49490947 |
| FLLDASNL | ASGSQGEVERLF | 0.7219422 |
| YQVGNFP | ASGSQGEVERLF | 0.7417842 |
| ESLLSQST | ASGSQGEVERLF | 0.5059657 |
| LFTVGFAAVG | ASGSQGEVERLF | 0.71913576 |
| ATPPTPT | ASGSQGEVERLF | 0.5765879 |
| TDLATLG | ASGSQGEVERLF | 0.5861343 |
| SAGGTGR | ASGSQGEVERLF | 0.74598515 |
| GFSNSGS | ASGSQGEVERLF | 0.620515 |
| GGNGGSGGTSV | ASGSQGEVERLF | 0.72925186 |
| GTSVATGGAGN | ASGSQGEVERLF | 0.5989906 |
| NGGNGGSGGTS | ASGSQGEVERLF | 0.50553215 |
| AVVKFPH | ASGSQGEVERLF | 0.7036686 |
| TPPPPPP | ASGSQGEVERLF | 0.74215996 |
| AAPAAAAAPA | ASGSQGEVERLF | 0.7110742 |
| IALNEDL | ASGSQGEVERLF | 0.75682026 |
| IVNVSSM | ASGSQGEVERLF | 0.59902555 |
| TGIPPAPR | ASGSQGEVERLF | 0.7520956 |
| FDLGGGTF | ASGSQGEVERLF | 0.7460401 |
| FDLGGGTFDVS | ASGSQGEVERLF | 0.69840205 |
| NEPTAAA | ASGSQGEVERLF | 0.6231879 |
| NEPTAAALA | ASGSQGEVERLF | 0.7344258 |
| GSIINISSI | ASGSQGEVERLF | 0.60584486 |
| VAPGNAGTA | ASGSQGEVERLF | 0.7426118 |
| TGGGASLE | ASGSQGEVERLF | 0.74855685 |
| VAVYNHYKRI | ASGSQGEVERLF | 0.4784582 |
| TDPAPATT | ASGSQGEVERLF | 0.8276891 |
| TNRPDILDPALLRPG | ASGSQGEVERLF | 0.7555999 |
| LTTTTTS | ASGSQGEVERLF | 0.76426136 |
| PTGSGKTL | ASGSQGEVERLF | 0.5416372 |
| TGSGKTL | ASGSQGEVERLF | 0.28436267 |
| TIGHVDHGK | ASGSQGEVERLF | 0.6854235 |
| TIGHVDHGKTT | ASGSQGEVERLF | 0.4339621 |
| GHVDHGKTT | ASGSQGEVERLF | 0.81608725 |
| AQFEHTLL | ASGSQGEVERLF | 0.79146326 |
| IVNQDEL | ASGSQGEVERLF | 0.57712615 |
| TNGSQFF | ASGSQGEVERLF | 0.8064805 |
| TGKGIVE | ASGSQGEVERLF | 0.8185544 |
| LISNASDA | ASGSQGEVERLF | 0.36953205 |
| AELRTLGELTKIAKAHGAQ | ASGSQGEVERLF | 0.7119074 |

| | | |
|---|---|---|
| APNAVAAAEPSW | ASGSQGEVERLF | 0.53377855 |
| APQVAGAATAVL | ASGSQGEVERLF | 0.7089766 |
| ARRDPAAAAAILPTDA | ASGSQGEVERLF | 0.21635358 |
| DAAARDVLAEAGLAEYF | ASGSQGEVERLF | 0.80302876 |
| DDQIVLSAKANLAKVEQ | ASGSQGEVERLF | 0.69167054 |
| DELLPDTGEDSDEDGH | ASGSQGEVERLF | 0.17882434 |
| ELPEAPANRATVPQD | ASGSQGEVERLF | 0.68243366 |
| FGQSKIERVTGIG | ASGSQGEVERLF | 0.6610831 |
| GAGGTGSTAGGAGGAGGAGGLY | ASGSQGEVERLF | 0.67062956 |
| GHGGTGAAGGNGGTGGAGGSGI | ASGSQGEVERLF | 0.81285155 |
| GNGGNGGAGGEGGPGGAGGAGA | ASGSQGEVERLF | 0.641189 |
| GSTAGNGGNGGNATNGTIAGQP | ASGSQGEVERLF | 0.43046314 |
| GTGGDGGNGGNGGTGG | ASGSQGEVERLF | 0.79783547 |
| GVRRPAGIPADAPV | ASGSQGEVERLF | 0.8160795 |
| GYRQVTDAHLLTLARR | ASGSQGEVERLF | 0.37154567 |
| IAKMGQDGHDRG | ASGSQGEVERLF | 0.78597057 |
| KAEPSIEIQYT | ASGSQGEVERLF | 0.82797015 |
| KPVRIRMAADEDTVAVT | ASGSQGEVERLF | 0.71664333 |
| MPPRIAGMRLLVIKPEP | ASGSQGEVERLF | 0.8184306 |
| MKFYAVALFRSRGVPPTR | ASGSQGEVERLF | 0.8246609 |
| NAGGFGILGAVAHSP | ASGSQGEVERLF | 0.57705605 |
| NEGREITLAMAEM | ASGSQGEVERLF | 0.8152774 |
| NELVEWSHPLGHRWRWE | ASGSQGEVERLF | 0.8094367 |
| NLGAGGTGGNGGNPG | ASGSQGEVERLF | 0.6661284 |
| NTEWPKWETVMAADPVD | ASGSQGEVERLF | 0.706782 |
| PASAAPAETQP | ASGSQGEVERLF | 0.48247814 |
| QQSSDGKNGVRAAEITGE | ASGSQGEVERLF | 0.72131777 |
| RAPDAVTTSWWKE | ASGSQGEVERLF | 0.2701213 |
| SGFGNVGAHTSGWFNQ | ASGSQGEVERLF | 0.78713024 |
| SGVAGAVGQAASVGGLKV | ASGSQGEVERLF | 0.8184921 |
| SNGGAGTGSAGGNGGT | ASGSQGEVERLF | 0.70986927 |
| TGGAGGEGGAAG | ASGSQGEVERLF | 0.4334197 |
| TGGDGGAAGAGGAPGHG | ASGSQGEVERLF | 0.50114614 |
| TMPPLLAAIANAGRHGV | ASGSQGEVERLF | 0.7512465 |
| TSSTAGNSGGA | ASGSQGEVERLF | 0.6460372 |
| VPATVPVGILC | ASGSQGEVERLF | 0.3039032 |
| VQRPDSGGVVLTGRIS | ASGSQGEVERLF | 0.7008312 |
| WTPNPVPATKGVPAGLVQAT | ASGSQGEVERLF | 0.80758446 |
| YALKRALAKFSDRFDVVII | ASGSQGEVERLF | 0.8115059 |
| YLIGSVIV | ASGSQGEVERLF | 0.3886871 |
| YGQLVAVID | ASGSQGEVERLF | 0.82424843 |
| VTVGGMTLPA | ASGSQGEVERLF | 0.8158217 |
| VLALPAV | ASGSQGEVERLF | 0.8216678 |
| VIPPTLNYET | ASGSQGEVERLF | 0.7314801 |
| VGMGGMHPSSG | ASGSQGEVERLF | 0.5133112 |
| VGLAGGAATA | ASGSQGEVERLF | 0.8209117 |
| VGGAGGAGGSGP | ASGSQGEVERLF | 0.6674446 |
| VFAAFMV | ASGSQGEVERLF | 0.6137302 |
| TLYDKLR | ASGSQGEVERLF | 0.80344343 |
| TLAVFTVL | ASGSQGEVERLF | 0.576718 |
| TGSVQPARV | ASGSQGEVERLF | 0.5471699 |

| | | |
|---|---|---|
| TGGGAGSGGT | ASGSQGEVERLF | 0.7690608 |
| SHLETSL | ASGSQGEVERLF | 0.8184054 |
| SFTTGMT | ASGSQGEVERLF | 0.53070056 |
| QVEVAKRMGVRQPTVS | ASGSQGEVERLF | 0.48090702 |
| QGSGGPALAM | ASGSQGEVERLF | 0.6196005 |
| MLAPSNP | ASGSQGEVERLF | 0.7204866 |
| LVNNAAI | ASGSQGEVERLF | 0.81740415 |
| LTGEINTLA | ASGSQGEVERLF | 0.7700176 |
| LSIFGPV | ASGSQGEVERLF | 0.73416173 |
| LLALLANL | ASGSQGEVERLF | 0.7917329 |
| LISNASDALDKLR | ASGSQGEVERLF | 0.80051935 |
| LINEAVP | ASGSQGEVERLF | 0.73501503 |
| LGYAFSGL | ASGSQGEVERLF | 0.7977767 |
| LDFVAVAL | ASGSQGEVERLF | 0.59833086 |
| LAQPGAE | ASGSQGEVERLF | 0.814661 |
| ISNLGAVTGF | ASGSQGEVERLF | 0.7944268 |
| HQEDPGRI | ASGSQGEVERLF | 0.8134531 |
| GTPGEPTP | ASGSQGEVERLF | 0.8167409 |
| GFFNSGS | ASGSQGEVERLF | 0.7788588 |
| GAGGGGLTAGAN | ASGSQGEVERLF | 0.7671783 |
| FVDALVNM | ASGSQGEVERLF | 0.3018248 |
| FLTVAPEL | ASGSQGEVERLF | 0.62681687 |
| FETLTAL | ASGSQGEVERLF | 0.7719567 |
| DVINPIEL | ASGSQGEVERLF | 0.78204197 |
| APVSGAPV | ASGSQGEVERLF | 0.8010533 |
| ALVGGLAW | ASGSQGEVERLF | 0.796311 |
| ALAAAAATAI | ASGSQGEVERLF | 0.5620507 |
| AFGLGLLL | ASGSQGEVERLF | 0.818746 |
| AFFAAASL | ASGSQGEVERLF | 0.818383 |
| AAAALPAFSPP | ASGSQGEVERLF | 0.3937987 |
| VVLAPAL | ASGSQGEVERLF | 0.627828 |
| TIGHVDHGKTTL | ASGSQGEVERLF | 0.8302115 |
| LISNASDALDK | ASGSQGEVERLF | 0.79844075 |
| LISNASDAL | ASGSQGEVERLF | 0.66533667 |
| ISNASDALDK | ASGSQGEVERLF | 0.6925235 |
| NQAETLVYQTE | ASGSQGEVERLF | 0.82249284 |
| TGGTGGTAGAG | ASGSQGEVERLF | 0.6732209 |
| TGSTAGGAGGA | ASGSQGEVERLF | 0.82258177 |
| DPLAVPVAAGPV | ASGSQGEVERLF | 0.2381101 |
| GGTGGTPGNGGN | ASGSQGEVERLF | 0.7089443 |
| PAGLAGVDPATL | ASGSQGEVERLF | 0.56526804 |
| GNNLVVEAPHAGASV | ASGSQGEVERLF | 0.2172154 |
| KVTRIMSASTLEQPA | ASGSQGEVERLF | 0.7819302 |
| LGFALPIALPRRTVV | ASGSQGEVERLF | 0.57964206 |
| TRLGNLTARKLVVLV | ASGSQGEVERLF | 0.8062608 |
| WSQQRRLIYFPSAGP | ASGSQGEVERLF | 0.68658924 |
| GPNGNPDPMAAAVDIR | ASGSQGEVERLF | 0.79959965 |
| TQRCVSVADLRAAASR | ASGSQGEVERLF | 0.7838393 |
| QFVLDVAEHPVEVLTETV | ASGSQGEVERLF | 0.730284 |
| GGAGGTGGAGGNGGTQAGDG | ASGSQGEVERLF | 0.5372363 |
| PLLATLRAIAVDVDAKPAQV | ASGSQGEVERLF | 0.76276666 |

| | | |
|---|---|---|
| APKFRTGGYMIDFWGVGYQVAK | ASGSQGEVERLF | 0.81367934 |
| GAGGAGGAGGAAGGPGGTGGTG | ASGSQGEVERLF | 0.8010472 |
| IGALGLLATDVAPAGSVAFGVA | ASGSQGEVERLF | 0.26686853 |
| AGGAGGIGGNGGAGGAGGVLMG | ASGSQGEVERLF | 0.7204907 |
| ADELIGLL | ASGSQGEVERLF | 0.6911288 |
| ADRALKA | ASGSQGEVERLF | 0.26616105 |
| AGAVGQVGGAGGA | ASGSQGEVERLF | 0.20518841 |
| AGGGGGLGGAP | ASGSQGEVERLF | 0.551527 |
| AGGLLYGNGGAG | ASGSQGEVERLF | 0.7004858 |
| AGLESIGL | ASGSQGEVERLF | 0.8037309 |
| AGLKTGII | ASGSQGEVERLF | 0.7997743 |
| ASLAALLL | ASGSQGEVERLF | 0.68703014 |
| ASVINISSI | ASGSQGEVERLF | 0.8037375 |
| ATPPGPA | ASGSQGEVERLF | 0.68204665 |
| ATVASALV | ASGSQGEVERLF | 0.15136886 |
| ATVASLIL | ASGSQGEVERLF | 0.66055906 |
| AVNGPVLIRH | ASGSQGEVERLF | 0.38485208 |
| DGAAGAVGI | ASGSQGEVERLF | 0.8234489 |
| DPLLHGI | ASGSQGEVERLF | 0.43777025 |
| ELGAALGAL | ASGSQGEVERLF | 0.5823574 |
| ELGRTQAV | ASGSQGEVERLF | 0.7401986 |
| FVAGGYLS | ASGSQGEVERLF | 0.7886586 |
| GASGGAGGAGGW | ASGSQGEVERLF | 0.75751394 |
| GGAGAGGDGGAGGI | ASGSQGEVERLF | 0.59405965 |
| GTGHNGSQPGMGGNG | ASGSQGEVERLF | 0.19799888 |
| GYAAASASAS | ASGSQGEVERLF | 0.7843889 |
| HQVDVAAVQQG | ASGSQGEVERLF | 0.56315863 |
| IGAAIAHT | ASGSQGEVERLF | 0.44587705 |
| IGMIEVAQL | ASGSQGEVERLF | 0.287891 |
| IRLAPEHTSAMD | ASGSQGEVERLF | 0.4337837 |
| IYNVSVL | ASGSQGEVERLF | 0.31627545 |
| KQGFLGM | ASGSQGEVERLF | 0.80768627 |
| LESVITQL | ASGSQGEVERLF | 0.7315305 |
| LGLGALVVV | ASGSQGEVERLF | 0.657864 |
| LIGNGGNGGSGGI | ASGSQGEVERLF | 0.8234341 |
| LNAINAPTL | ASGSQGEVERLF | 0.7877102 |
| LQLLNTGV | ASGSQGEVERLF | 0.7163998 |
| LQTSIEF | ASGSQGEVERLF | 0.5246068 |
| MSFAVSEL | ASGSQGEVERLF | 0.51031303 |
| NTGALGL | ASGSQGEVERLF | 0.46339473 |
| PPGAGGAGGAGTT | ASGSQGEVERLF | 0.4805847 |
| QLVGLGI | ASGSQGEVERLF | 0.8210475 |
| RFLEQVHQL | ASGSQGEVERLF | 0.70869875 |
| RGVGYVLR | ASGSQGEVERLF | 0.3976035 |
| SATGVFAGVI | ASGSQGEVERLF | 0.79631793 |
| SLPAITLP | ASGSQGEVERLF | 0.82623625 |
| STALAFTV | ASGSQGEVERLF | 0.83268136 |
| SVLTVGLGK | ASGSQGEVERLF | 0.50857955 |
| TALAGAGSGI | ASGSQGEVERLF | 0.70316327 |
| TGLPFTGI | ASGSQGEVERLF | 0.66942763 |
| VAEHVTV | ASGSQGEVERLF | 0.7277192 |

| | | |
|---|---|---|
| VDIVLNSL | ASGSQGEVERLF | 0.614022 |
| VLTGGGAVL | ASGSQGEVERLF | 0.3517826 |
| VNVGLAYGI | ASGSQGEVERLF | 0.8154006 |
| VSSAAVGAV | ASGSQGEVERLF | 0.83664626 |
| VTSVYEAGV | ASGSQGEVERLF | 0.6618479 |
| WVALDRGA | ASGSQGEVERLF | 0.3204506 |
| YLVGSVVI | ASGSQGEVERLF | 0.69975984 |
| GPPGTGKTLL | ASGSQGEVERLF | 0.60550374 |
| GVGMTKF | ASGSQGEVERLF | 0.6307616 |
| IGIGGSDL | ASGSQGEVERLF | 0.734646 |
| IGTLLAEL | ASGSQGEVERLF | 0.39641052 |
| IYTSGTT | ASGSQGEVERLF | 0.69015825 |
| LSFPAGV | ASGSQGEVERLF | 0.8316945 |
| LYTSGTT | ASGSQGEVERLF | 0.72148263 |
| VGAFPFGF | ASGSQGEVERLF | 0.78848934 |
| VGAGTVEF | ASGSQGEVERLF | 0.81722915 |
| VGKSTLLL | ASGSQGEVERLF | 0.79976404 |
| AALAHAGGGIV | ASGSQGEVERLF | 0.8192142 |
| AALVAVSLL | ASGSQGEVERLF | 0.81995106 |
| AAVSRAVAAL | ASGSQGEVERLF | 0.67876476 |
| AGGSSGIGGF | ASGSQGEVERLF | 0.7582889 |
| AGLTVANANG | ASGSQGEVERLF | 0.79568577 |
| AGLVGAVL | ASGSQGEVERLF | 0.7027842 |
| ALSQAFTV | ASGSQGEVERLF | 0.73681325 |
| ALTFSVSL | ASGSQGEVERLF | 0.6485806 |
| ALTGAALVAL | ASGSQGEVERLF | 0.81015813 |
| ASFSIGAGVL | ASGSQGEVERLF | 0.7582512 |
| DGSVYKI | ASGSQGEVERLF | 0.826169 |
| EQLAEIV | ASGSQGEVERLF | 0.31166205 |
| FAVSELTVM | ASGSQGEVERLF | 0.41287738 |
| GAAPLAAGL | ASGSQGEVERLF | 0.81874883 |
| GADGGDGGSGGAGG | ASGSQGEVERLF | 0.19406326 |
| GFGLDTGGAGG | ASGSQGEVERLF | 0.25719655 |
| GLASGGLL | ASGSQGEVERLF | 0.76946753 |
| GRIAGAAVLL | ASGSQGEVERLF | 0.8269184 |
| GSAGSLNTSF | ASGSQGEVERLF | 0.20305072 |
| GTGQIGI | ASGSQGEVERLF | 0.46384954 |
| GTLIALIL | ASGSQGEVERLF | 0.48855165 |
| IAATMGFVVG | ASGSQGEVERLF | 0.812 |
| IFAQALL | ASGSQGEVERLF | 0.70431983 |
| IGAAVALGF | ASGSQGEVERLF | 0.29611826 |
| IGLTPLAL | ASGSQGEVERLF | 0.80544996 |
| ILGSNTSTL | ASGSQGEVERLF | 0.32069656 |
| IVFSLTVL | ASGSQGEVERLF | 0.666172 |
| LASKATGF | ASGSQGEVERLF | 0.7941567 |
| LASVALIL | ASGSQGEVERLF | 0.8293622 |
| LFAVGLTL | ASGSQGEVERLF | 0.39826846 |
| LFNGGIGTYI | ASGSQGEVERLF | 0.80595934 |
| LTLGAGII | ASGSQGEVERLF | 0.71462095 |
| LVAAALGDI | ASGSQGEVERLF | 0.83402354 |
| NTAFVQLGI | ASGSQGEVERLF | 0.58197093 |

| | | |
|---|---|---|
| QLFSGAEL | ASGSQGEVERLF | 0.32495117 |
| SAATFITL | ASGSQGEVERLF | 0.8104405 |
| SNYGLANLGV | ASGSQGEVERLF | 0.81084275 |
| SYGGAYAVM | ASGSQGEVERLF | 0.70853853 |
| TTGVGGLTLG | ASGSQGEVERLF | 0.8168055 |
| TTQTIYI | ASGSQGEVERLF | 0.72688806 |
| VAAGAGLGAVI | ASGSQGEVERLF | 0.7342255 |
| VAAVVTLL | ASGSQGEVERLF | 0.8089417 |
| VGFLGGLI | ASGSQGEVERLF | 0.7975908 |
| VGVIPILL | ASGSQGEVERLF | 0.37824994 |
| VIFASSLI | ASGSQGEVERLF | 0.80030525 |
| VKGGDGGAAAG | ASGSQGEVERLF | 0.70952165 |
| VTKTFTNI | ASGSQGEVERLF | 0.56898254 |
| VVAAAILL | ASGSQGEVERLF | 0.8253946 |
| YAALDVEL | ASGSQGEVERLF | 0.775824 |
| LASVALLL | ASGSQGEVERLF | 0.8109154 |
| LDAVHRV | ASGSQGEVERLF | 0.77796984 |
| QGLSSSIF | ASGSQGEVERLF | 0.83730686 |
| RGTFANIRL | ASGSQGEVERLF | 0.28652525 |
| RQLWWGHRI | ASGSQGEVERLF | 0.732461 |
| YGPPGTGKTLL | ASGSQGEVERLF | 0.7292801 |
| AALVGAPANVA | ASGSQGEVERLF | 0.8124303 |
| AATFRRTE | ASGSQGEVERLF | 0.7080716 |
| AGALGGGI | ASGSQGEVERLF | 0.4803874 |
| AGGAGGAGGIGAG | ASGSQGEVERLF | 0.78639084 |
| AGSGALGL | ASGSQGEVERLF | 0.8166188 |
| AQIADQI | ASGSQGEVERLF | 0.7476266 |
| ASAVAVTYL | ASGSQGEVERLF | 0.783329 |
| ASTVLGGTGG | ASGSQGEVERLF | 0.8289252 |
| AVGGEPAK | ASGSQGEVERLF | 0.5099741 |
| CEQGTFGGTC | ASGSQGEVERLF | 0.6922467 |
| DEGKAPAG | ASGSQGEVERLF | 0.8052492 |
| DTVVFPAL | ASGSQGEVERLF | 0.81370986 |
| FEGITMAG | ASGSQGEVERLF | 0.7394032 |
| FPLVVQD | ASGSQGEVERLF | 0.8195298 |
| GAFTTPQ | ASGSQGEVERLF | 0.53947973 |
| GFGGFGTAGGVGGA | ASGSQGEVERLF | 0.788625 |
| GFLAQFL | ASGSQGEVERLF | 0.6741015 |
| GGAGGAGGNAGVA | ASGSQGEVERLF | 0.76006544 |
| GGVGTAGAGGAGGAGG | ASGSQGEVERLF | 0.4461809 |
| GNGGGGDAGNAGSGGN | ASGSQGEVERLF | 0.77562916 |
| GTGGAGGAGNDAGS | ASGSQGEVERLF | 0.8112794 |
| GTTNTGWFNS | ASGSQGEVERLF | 0.7952552 |
| IFMGPLFTL | ASGSQGEVERLF | 0.21552727 |
| ILAHAIEAI | ASGSQGEVERLF | 0.7927654 |
| IRSAIREAL | ASGSQGEVERLF | 0.8008106 |
| ITFAGGG | ASGSQGEVERLF | 0.8344382 |
| ITGMSGHQEV | ASGSQGEVERLF | 0.732141 |
| IVAGLACTL | ASGSQGEVERLF | 0.79319286 |
| IVGGGLAGA | ASGSQGEVERLF | 0.53871953 |
| IVTGVLNI | ASGSQGEVERLF | 0.7489056 |

| | | |
|---|---|---|
| LAQQGCI | ASGSQGEVERLF | 0.5510802 |
| LCLHVEN | ASGSQGEVERLF | 0.5826056 |
| LDAISDGS | ASGSQGEVERLF | 0.6840944 |
| LGGFAAML | ASGSQGEVERLF | 0.5850961 |
| LGGKNPMI | ASGSQGEVERLF | 0.80609065 |
| LGPKGVAAIL | ASGSQGEVERLF | 0.80809915 |
| LMVGVSI | ASGSQGEVERLF | 0.8088167 |
| LVGAAFVL | ASGSQGEVERLF | 0.8226248 |
| LVLAVAPL | ASGSQGEVERLF | 0.82783955 |
| MAAIAVIDAA | ASGSQGEVERLF | 0.8180351 |
| NNGINAELQ | ASGSQGEVERLF | 0.58193994 |
| RGEPLVLVHG | ASGSQGEVERLF | 0.69390666 |
| RGGAGGAAGLI | ASGSQGEVERLF | 0.35587296 |
| RSYRMHPAV | ASGSQGEVERLF | 0.67075145 |
| SAAHLLAGQLA | ASGSQGEVERLF | 0.6346291 |
| SAAVGSY | ASGSQGEVERLF | 0.75197196 |
| SPHAVLHRI | ASGSQGEVERLF | 0.17798607 |
| SQIAKVL | ASGSQGEVERLF | 0.79539937 |
| TALGHLVP | ASGSQGEVERLF | 0.82189244 |
| TAPTTSGAA | ASGSQGEVERLF | 0.63265157 |
| TDGDDGNGGSPGT | ASGSQGEVERLF | 0.79189605 |
| TDGLGEN | ASGSQGEVERLF | 0.78275335 |
| TGGPPLPGQA | ASGSQGEVERLF | 0.6581249 |
| TIPAITI | ASGSQGEVERLF | 0.8077151 |
| TIPGFTI | ASGSQGEVERLF | 0.60050434 |
| TSIASGALA | ASGSQGEVERLF | 0.8116819 |
| TVTLPGAAI | ASGSQGEVERLF | 0.8218128 |
| VASGLSAALP | ASGSQGEVERLF | 0.7680762 |
| VGAGVIGI | ASGSQGEVERLF | 0.7611162 |
| VGPKRFGV | ASGSQGEVERLF | 0.7465238 |
| VNVAFPD | ASGSQGEVERLF | 0.25561094 |
| VSMTNSGVS | ASGSQGEVERLF | 0.77094686 |
| VTDVLGDLSV | ASGSQGEVERLF | 0.21550626 |
| VVVVGAGFAGL | ASGSQGEVERLF | 0.76307344 |
| VYTVGTVV | ASGSQGEVERLF | 0.48610926 |
| WTTFSGP | ASGSQGEVERLF | 0.7625388 |
| YDELVEAF | ASGSQGEVERLF | 0.6851727 |
| YGTGGAGGNGG | ASGSQGEVERLF | 0.7911093 |
| YTNVMAV | ASGSQGEVERLF | 0.51979774 |
| FLEQVHQL | ASGSQGEVERLF | 0.76374054 |
| GLSLLAVAV | ASGSQGEVERLF | 0.6904174 |
| LEQVHQL | ASGSQGEVERLF | 0.3512469 |
| LNAAGAI | ASGSQGEVERLF | 0.7993272 |
| PAVHASL | ASGSQGEVERLF | 0.70489997 |
| TELEDDA | ASGSQGEVERLF | 0.745296 |
| TPPGPPN | ASGSQGEVERLF | 0.8101994 |
| TPPPPPPP | ASGSQGEVERLF | 0.7526313 |
| TQSPALI | ASSPGGTLY | 0.26983997 |
| EFAAAFPQ | ASSPGGTLY | 0.746316 |
| LLSGNLDVL | ASSPGGTLY | 0.7416607 |
| GPANLGNYNI | ASSPGGTLY | 0.6249292 |

| | | |
|---|---|---|
| TLGPTLP | ASSPGGTLY | 0.18098807 |
| VYAYSSCCK | ASSPGGTLY | 0.7964276 |
| APAPAPAPNI | ASSPGGTLY | 0.5720672 |
| PGGPVPIP | ASSPGGTLY | 0.62164366 |
| FLTVALEP | ASSPGGTLY | 0.5991746 |
| SQYFGANL | ASSPGGTLY | 0.8159977 |
| GGVKPAA | ASSPGGTLY | 0.7682928 |
| GGTGGAGGAGGW | ASSPGGTLY | 0.6619898 |
| GVGGVAGLGGAGG | ASSPGGTLY | 0.7998529 |
| VLLVLATTI | ASSPGGTLY | 0.71714365 |
| IGIFAAML | ASSPGGTLY | 0.8167342 |
| AVGDLID | ASSPGGTLY | 0.74559116 |
| NISPLPL | ASSPGGTLY | 0.70959723 |
| VTAVFVAI | ASSPGGTLY | 0.7523507 |
| SGDDNDAAL | ASSPGGTLY | 0.6875684 |
| AVAALAHS | ASSPGGTLY | 0.33699128 |
| GGVTVMATP | ASSPGGTLY | 0.3396997 |
| AGLAANVRD | ASSPGGTLY | 0.731848 |
| TNRPDLIDPALLRPG | ASSPGGTLY | 0.33379975 |
| LANLGVTAL | ASSPGGTLY | 0.71962345 |
| TGERIEL | ASSPGGTLY | 0.82089 |
| NGEELAA | ASSPGGTLY | 0.7742056 |
| VLSLVWG | ASSPGGTLY | 0.36938238 |
| IGYALWL | ASSPGGTLY | 0.6783644 |
| LSLLLAPL | ASSPGGTLY | 0.5632822 |
| TLDEALK | ASSPGGTLY | 0.80379397 |
| TDGAGTAEGG | ASSPGGTLY | 0.7667327 |
| NAPGGAGGNGG | ASSPGGTLY | 0.8294921 |
| VSVFMGMGGNF | ASSPGGTLY | 0.8046829 |
| LASYLAAL | ASSPGGTLY | 0.6045855 |
| LVGGTPAL | ASSPGGTLY | 0.8110254 |
| VRNADVL | ASSPGGTLY | 0.69384813 |
| TCSSPFLVSAM | ASSPGGTLY | 0.7062293 |
| VHITDAE | ASSPGGTLY | 0.7267357 |
| EYAAPSF | ASSPGGTLY | 0.7887586 |
| VVDAFHGGL | ASSPGGTLY | 0.7070061 |
| ASVVDPQ | ASSPGGTLY | 0.6609429 |
| SYNEFGGLS | ASSPGGTLY | 0.32576966 |
| DPDGSVY | ASSPGGTLY | 0.7907487 |
| MVSTIHATE | ASSPGGTLY | 0.7363272 |
| LAAGVADPV | ASSPGGTLY | 0.42024672 |
| TKEQIAA | ASSPGGTLY | 0.61457366 |
| SLASPAAL | ASSPGGTLY | 0.82856226 |
| IGLVFYGV | ASSPGGTLY | 0.8278682 |
| FLLDASNL | ASSPGGTLY | 0.6186031 |
| YQVGNFP | ASSPGGTLY | 0.49568513 |
| ESLLSQST | ASSPGGTLY | 0.7960697 |
| LFTVGFAAVG | ASSPGGTLY | 0.40100205 |
| ATPPTPT | ASSPGGTLY | 0.78247654 |
| TDLATLG | ASSPGGTLY | 0.7465024 |
| SAGGTGR | ASSPGGTLY | 0.74225134 |

| | | |
|---|---|---|
| GFSNSGS | ASSPGGTLY | 0.6044734 |
| GGNGGSGGTSV | ASSPGGTLY | 0.7823479 |
| GTSVATGGAGN | ASSPGGTLY | 0.6478162 |
| NGGNGGSGGTS | ASSPGGTLY | 0.8051712 |
| AVVKFPH | ASSPGGTLY | 0.19244461 |
| TPPPPPP | ASSPGGTLY | 0.7684821 |
| AAPAAAAAPA | ASSPGGTLY | 0.48672214 |
| IALNEDL | ASSPGGTLY | 0.64119005 |
| IVNVSSM | ASSPGGTLY | 0.26755533 |
| TGIPPAPR | ASSPGGTLY | 0.70210385 |
| FDLGGGTF | ASSPGGTLY | 0.8030769 |
| FDLGGGTFDVS | ASSPGGTLY | 0.8187837 |
| NEPTAAA | ASSPGGTLY | 0.6129794 |
| NEPTAAALA | ASSPGGTLY | 0.79756427 |
| GSIINISSI | ASSPGGTLY | 0.8304505 |
| VAPGNAGTA | ASSPGGTLY | 0.661608 |
| TGGGASLE | ASSPGGTLY | 0.22533152 |
| VAVYNHYKRI | ASSPGGTLY | 0.77557665 |
| TDPAPATT | ASSPGGTLY | 0.77187264 |
| TNRPDILDPALLRPG | ASSPGGTLY | 0.81911355 |
| LTTTTTS | ASSPGGTLY | 0.7864472 |
| PTGSGKTL | ASSPGGTLY | 0.43827963 |
| TGSGKTL | ASSPGGTLY | 0.8056878 |
| TIGHVDHGK | ASSPGGTLY | 0.81198996 |
| TIGHVDHGKTT | ASSPGGTLY | 0.51547813 |
| GHVDHGKTT | ASSPGGTLY | 0.49050754 |
| AQFEHTLL | ASSPGGTLY | 0.5080812 |
| IVNQDEL | ASSPGGTLY | 0.261186 |
| TNGSQFF | ASSPGGTLY | 0.7492378 |
| TGKGIVE | ASSPGGTLY | 0.66029036 |
| LISNASDA | ASSPGGTLY | 0.5524487 |
| AELRTLGELTKIAKAHGAQ | ASSPGGTLY | 0.7488446 |
| APNAVAAAEPSW | ASSPGGTLY | 0.4839552 |
| APQVAGAATAVL | ASSPGGTLY | 0.6027685 |
| ARRDPAAAAAILPTDA | ASSPGGTLY | 0.5491172 |
| DAAARDVLAEAGLAEYF | ASSPGGTLY | 0.6311555 |
| DDQIVLSAKANLAKVEQ | ASSPGGTLY | 0.79567456 |
| DELLPDTGEDSDEDGH | ASSPGGTLY | 0.7452477 |
| ELPEAPANRATVPQD | ASSPGGTLY | 0.78534144 |
| FGQSKIERVTGIG | ASSPGGTLY | 0.8211431 |
| GAGGTGSTAGGAGGAGGAGGLY | ASSPGGTLY | 0.3479388 |
| GHGGTGAAGGNGGTGGAGGSGI | ASSPGGTLY | 0.50947756 |
| GNGGNGGAGGEGGPGGAGGAGA | ASSPGGTLY | 0.8093494 |
| GSTAGNGGNGGNATNGTIAGQP | ASSPGGTLY | 0.5046103 |
| GTGGDGGNGGNGGTGG | ASSPGGTLY | 0.8146157 |
| GVRRPAGIPADAPV | ASSPGGTLY | 0.68311846 |
| GYRQVTDAHLLTLARR | ASSPGGTLY | 0.77668333 |
| IAKMGQDGHDRG | ASSPGGTLY | 0.7739152 |
| KAEPSIEIQYT | ASSPGGTLY | 0.8305516 |
| KPVRIRMAADEDTVAVT | ASSPGGTLY | 0.80369216 |
| MPPRIAGMRLLVIKPEP | ASSPGGTLY | 0.6726204 |

| | | |
|---|---|---|
| MKFYAVALFRSRGVPPTR | ASSPGGTLY | 0.76414865 |
| NAGGFGILGAVAHSP | ASSPGGTLY | 0.43895757 |
| NEGREITLAMAEM | ASSPGGTLY | 0.80129075 |
| NELVEWSHPLGHRWRWE | ASSPGGTLY | 0.8049528 |
| NLGAGGTGGNGGNPG | ASSPGGTLY | 0.52166605 |
| NTEWPKWETVMAADPVD | ASSPGGTLY | 0.73822296 |
| PASAAPAETQP | ASSPGGTLY | 0.82279414 |
| QQSSDGKNGVRAAEITGE | ASSPGGTLY | 0.8122375 |
| RAPDAVTTSWWKE | ASSPGGTLY | 0.36151254 |
| SGFGNVGAHTSGWFNQ | ASSPGGTLY | 0.6657001 |
| SGVAGAVGQAASVGGLKV | ASSPGGTLY | 0.4222405 |
| SNGGAGTGSAGGNGGT | ASSPGGTLY | 0.6774141 |
| TGGAGGEGGAAG | ASSPGGTLY | 0.3450075 |
| TGGDGGAAGAGGAPGHG | ASSPGGTLY | 0.8017706 |
| TMPPLLAAIANAGRHGV | ASSPGGTLY | 0.64470917 |
| TSSTAGNSGGA | ASSPGGTLY | 0.8191446 |
| VPATVPVGILC | ASSPGGTLY | 0.7428471 |
| VQRPDSGGVVLTGRIS | ASSPGGTLY | 0.8084593 |
| WTPNPVPATKGVPAGLVQAT | ASSPGGTLY | 0.6286912 |
| YALKRALAKFSDRFDVVII | ASSPGGTLY | 0.54882276 |
| YLIGSVIV | ASSPGGTLY | 0.36698884 |
| YGQLVAVID | ASSPGGTLY | 0.336101 |
| VTVGGMTLPA | ASSPGGTLY | 0.57460093 |
| VLALPAV | ASSPGGTLY | 0.6476328 |
| VIPPTLNYET | ASSPGGTLY | 0.24371223 |
| VGMGGMHPSSG | ASSPGGTLY | 0.8290956 |
| VGLAGGAATA | ASSPGGTLY | 0.7583934 |
| VGGAGGAGGSGP | ASSPGGTLY | 0.7884654 |
| VFAAFMV | ASSPGGTLY | 0.54247034 |
| TLYDKLR | ASSPGGTLY | 0.5306822 |
| TLAVFTVL | ASSPGGTLY | 0.56562376 |
| TGSVQPARV | ASSPGGTLY | 0.80712134 |
| TGGGAGSGGT | ASSPGGTLY | 0.41917187 |
| SHLETSL | ASSPGGTLY | 0.26072913 |
| SFTTGMT | ASSPGGTLY | 0.59157765 |
| QVEVAKRMGVRQPTVS | ASSPGGTLY | 0.3787944 |
| QGSGGPALAM | ASSPGGTLY | 0.8311449 |
| MLAPSNP | ASSPGGTLY | 0.81577086 |
| LVNNAAI | ASSPGGTLY | 0.7816083 |
| LTGEINTLA | ASSPGGTLY | 0.8267973 |
| LSIFGPV | ASSPGGTLY | 0.20065725 |
| LLALLANL | ASSPGGTLY | 0.603128 |
| LISNASDALDKLR | ASSPGGTLY | 0.8260453 |
| LINEAVP | ASSPGGTLY | 0.7785328 |
| LGYAFSGL | ASSPGGTLY | 0.69456077 |
| LDFVAVAL | ASSPGGTLY | 0.80846536 |
| LAQPGAE | ASSPGGTLY | 0.49978983 |
| ISNLGAVTGF | ASSPGGTLY | 0.6233541 |
| HQEDPGRI | ASSPGGTLY | 0.69726425 |
| GTPGEPTP | ASSPGGTLY | 0.76087093 |
| GFFNSGS | ASSPGGTLY | 0.71991205 |

| | | |
|---|---|---|
| GAGGGGLTAGAN | ASSPGGTLY | 0.40392432 |
| FVDALVNM | ASSPGGTLY | 0.78951585 |
| FLTVAPEL | ASSPGGTLY | 0.831032 |
| FETLTAL | ASSPGGTLY | 0.70054567 |
| DVINPIEL | ASSPGGTLY | 0.81496227 |
| APVSGAPV | ASSPGGTLY | 0.7764402 |
| ALVGGLAW | ASSPGGTLY | 0.80162334 |
| ALAAAAATAI | ASSPGGTLY | 0.56847566 |
| AFGLGLLL | ASSPGGTLY | 0.56628746 |
| AFFAAASL | ASSPGGTLY | 0.75369674 |
| AAAALPAFSPP | ASSPGGTLY | 0.51133776 |
| VVLAPAL | ASSPGGTLY | 0.7416344 |
| TIGHVDHGKTTL | ASSPGGTLY | 0.66375625 |
| LISNASDALDK | ASSPGGTLY | 0.8372423 |
| LISNASDAL | ASSPGGTLY | 0.7992531 |
| ISNASDALDK | ASSPGGTLY | 0.77847505 |
| NQAETLVYQTE | ASSPGGTLY | 0.44935226 |
| TGGTGGTAGAG | ASSPGGTLY | 0.7658959 |
| TGSTAGGAGGA | ASSPGGTLY | 0.7920547 |
| DPLAVPVAAGPV | ASSPGGTLY | 0.30924332 |
| GGTGGTPGNGGN | ASSPGGTLY | 0.8212223 |
| PAGLAGVDPATL | ASSPGGTLY | 0.51942897 |
| GNNLVVEAPHAGASV | ASSPGGTLY | 0.75888574 |
| KVTRIMSASTLEQPA | ASSPGGTLY | 0.7989081 |
| LGFALPIALPRRTVV | ASSPGGTLY | 0.8265846 |
| TRLGNLTARKLVVLV | ASSPGGTLY | 0.6391355 |
| WSQQRRLIYFPSAGP | ASSPGGTLY | 0.74501884 |
| GPNGNPDPMAAAVDIR | ASSPGGTLY | 0.45114237 |
| TQRCVSVADLRAAASR | ASSPGGTLY | 0.81573564 |
| QFVLDVAEHPVEVLTETV | ASSPGGTLY | 0.81611085 |
| GGAGGTGGAGGNGGTQAGDG | ASSPGGTLY | 0.78148043 |
| PLLATLRAIAVDVDAKPAQV | ASSPGGTLY | 0.8322808 |
| APKFRTGGYMIDFWGVGYQVAK | ASSPGGTLY | 0.78667325 |
| GAGGAGGAGGAAGGPGGTGGTG | ASSPGGTLY | 0.31005728 |
| IGALGLLATDVAPAGSVAFGVA | ASSPGGTLY | 0.75330174 |
| AGGAGGIGGNGGAGGAGGVLMG | ASSPGGTLY | 0.8110435 |
| ADELIGLL | ASSPGGTLY | 0.8205878 |
| ADRALKA | ASSPGGTLY | 0.77588457 |
| AGAVGQVGGAGGA | ASSPGGTLY | 0.75176585 |
| AGGGGGLGGAP | ASSPGGTLY | 0.68492794 |
| AGGLLYGNGGAG | ASSPGGTLY | 0.8252156 |
| AGLESIGL | ASSPGGTLY | 0.8365519 |
| AGLKTGII | ASSPGGTLY | 0.8075217 |
| ASLAALLL | ASSPGGTLY | 0.71422064 |
| ASVINISSI | ASSPGGTLY | 0.65262246 |
| ATPPGPA | ASSPGGTLY | 0.77041775 |
| ATVASALV | ASSPGGTLY | 0.80956054 |
| ATVASLIL | ASSPGGTLY | 0.51774055 |
| AVNGPVLIRH | ASSPGGTLY | 0.74715143 |
| DGAAGAVGI | ASSPGGTLY | 0.759876 |
| DPLLHGI | ASSPGGTLY | 0.29187012 |

| | | |
|---|---|---|
| ELGAALGAL | ASSPGGTLY | 0.7725854 |
| ELGRTQAV | ASSPGGTLY | 0.82035553 |
| FVAGGYLS | ASSPGGTLY | 0.74490976 |
| GASGGAGGAGGW | ASSPGGTLY | 0.60201323 |
| GGAGAGGDGGAGGI | ASSPGGTLY | 0.8189881 |
| GTGHNGSQPGMGGNG | ASSPGGTLY | 0.83304524 |
| GYAAASASAS | ASSPGGTLY | 0.82496977 |
| HQVDVAAVQQG | ASSPGGTLY | 0.27399403 |
| IGAAIAHT | ASSPGGTLY | 0.21685144 |
| IGMIEVAQL | ASSPGGTLY | 0.66637605 |
| IRLAPEHTSAMD | ASSPGGTLY | 0.8131336 |
| IYNVSVL | ASSPGGTLY | 0.5998772 |
| KQGFLGM | ASSPGGTLY | 0.61253285 |
| LESVITQL | ASSPGGTLY | 0.78264654 |
| LGLGALVVV | ASSPGGTLY | 0.81396294 |
| LIGNGGNGGSGGI | ASSPGGTLY | 0.46496552 |
| LNAINAPTL | ASSPGGTLY | 0.43956164 |
| LQLLNTGV | ASSPGGTLY | 0.7697262 |
| LQTSIEF | ASSPGGTLY | 0.817162 |
| MSFAVSEL | ASSPGGTLY | 0.5663498 |
| NTGALGL | ASSPGGTLY | 0.71294844 |
| PPGAGGAGGAGTT | ASSPGGTLY | 0.6807234 |
| QLVGLGI | ASSPGGTLY | 0.7889617 |
| RFLEQVHQL | ASSPGGTLY | 0.7626853 |
| RGVGYVLR | ASSPGGTLY | 0.48465455 |
| SATGVFAGVI | ASSPGGTLY | 0.8170368 |
| SLPAITLP | ASSPGGTLY | 0.44538924 |
| STALAFTV | ASSPGGTLY | 0.80813307 |
| SVLTVGLGK | ASSPGGTLY | 0.7615278 |
| TALAGAGSGI | ASSPGGTLY | 0.76319927 |
| TGLPFTGI | ASSPGGTLY | 0.6417378 |
| VAEHVTV | ASSPGGTLY | 0.6212162 |
| VDIVLNSL | ASSPGGTLY | 0.38163203 |
| VLTGGGAVL | ASSPGGTLY | 0.38161725 |
| VNVGLAYGI | ASSPGGTLY | 0.8089324 |
| VSSAAVGAV | ASSPGGTLY | 0.33891928 |
| VTSVYEAGV | ASSPGGTLY | 0.8083041 |
| WVALDRGA | ASSPGGTLY | 0.74578136 |
| YLVGSVVI | ASSPGGTLY | 0.4600815 |
| GPPGTGKTLL | ASSPGGTLY | 0.79831517 |
| GVGMTKF | ASSPGGTLY | 0.43684024 |
| IGIGGSDL | ASSPGGTLY | 0.65170074 |
| IGTLLAEL | ASSPGGTLY | 0.8003501 |
| IYTSGTT | ASSPGGTLY | 0.7969197 |
| LSFPAGV | ASSPGGTLY | 0.50054175 |
| LYTSGTT | ASSPGGTLY | 0.7937893 |
| VGAFPFGF | ASSPGGTLY | 0.7846984 |
| VGAGTVEF | ASSPGGTLY | 0.57710534 |
| VGKSTLLL | ASSPGGTLY | 0.7183515 |
| AALAHAGGGIV | ASSPGGTLY | 0.8112112 |
| AALVAVSLL | ASSPGGTLY | 0.7500334 |

| | | |
|---|---|---|
| AAVSRAVAAL | ASSPGGTLY | 0.4601642 |
| AGGSSGIGGF | ASSPGGTLY | 0.7793615 |
| AGLTVANANG | ASSPGGTLY | 0.6321626 |
| AGLVGAVL | ASSPGGTLY | 0.78350663 |
| ALSQAFTV | ASSPGGTLY | 0.72041166 |
| ALTFSVSL | ASSPGGTLY | 0.65964067 |
| ALTGAALVAL | ASSPGGTLY | 0.80877244 |
| ASFSIGAGVL | ASSPGGTLY | 0.45427778 |
| DGSVYKI | ASSPGGTLY | 0.5997212 |
| EQLAEIV | ASSPGGTLY | 0.64454466 |
| FAVSELTVM | ASSPGGTLY | 0.63640356 |
| GAAPLAAGL | ASSPGGTLY | 0.8013585 |
| GADGGDGGSGGAGG | ASSPGGTLY | 0.46346146 |
| GFGLDTGGAGG | ASSPGGTLY | 0.781083 |
| GLASGGLL | ASSPGGTLY | 0.6571115 |
| GRIAGAAVLL | ASSPGGTLY | 0.43016195 |
| GSAGSLNTSF | ASSPGGTLY | 0.75398153 |
| GTGQIGI | ASSPGGTLY | 0.6481278 |
| GTLIALIL | ASSPGGTLY | 0.5251649 |
| IAATMGFVVG | ASSPGGTLY | 0.7854277 |
| IFAQALL | ASSPGGTLY | 0.45433414 |
| IGAAVALGF | ASSPGGTLY | 0.7526867 |
| IGLTPLAL | ASSPGGTLY | 0.82492316 |
| ILGSNTSTL | ASSPGGTLY | 0.59362257 |
| IVFSLTVL | ASSPGGTLY | 0.8038848 |
| LASKATGF | ASSPGGTLY | 0.79265916 |
| LASVALIL | ASSPGGTLY | 0.7829609 |
| LFAVGLTL | ASSPGGTLY | 0.57443774 |
| LFNGGIGTYI | ASSPGGTLY | 0.54747415 |
| LTLGAGII | ASSPGGTLY | 0.8171246 |
| LVAAALGDI | ASSPGGTLY | 0.7832153 |
| NTAFVQLGI | ASSPGGTLY | 0.7890377 |
| QLFSGAEL | ASSPGGTLY | 0.80688155 |
| SAATFITL | ASSPGGTLY | 0.79683197 |
| SNYGLANLGV | ASSPGGTLY | 0.8102874 |
| SYGGAYAVM | ASSPGGTLY | 0.66671896 |
| TTGVGGLTLG | ASSPGGTLY | 0.44391045 |
| TTQTIYI | ASSPGGTLY | 0.5919894 |
| VAAGAGLGAVI | ASSPGGTLY | 0.63513136 |
| VAAVVTLL | ASSPGGTLY | 0.58072656 |
| VGFLGGLI | ASSPGGTLY | 0.36252013 |
| VGVIPILL | ASSPGGTLY | 0.80964094 |
| VIFASSLI | ASSPGGTLY | 0.8348992 |
| VKGGDGGAAAG | ASSPGGTLY | 0.48135197 |
| VTKTFTNI | ASSPGGTLY | 0.8348312 |
| VVAAAILL | ASSPGGTLY | 0.40076914 |
| YAALDVEL | ASSPGGTLY | 0.78183526 |
| LASVALLL | ASSPGGTLY | 0.8104634 |
| LDAVHRV | ASSPGGTLY | 0.5121755 |
| QGLSSSIF | ASSPGGTLY | 0.7643317 |
| RGTFANIRL | ASSPGGTLY | 0.49514544 |

| | | |
|---|---|---|
| RQLWWGHRI | ASSPGGTLY | 0.5575501 |
| YGPPGTGKTLL | ASSPGGTLY | 0.80870354 |
| AALVGAPANVA | ASSPGGTLY | 0.5555934 |
| AATFRRTE | ASSPGGTLY | 0.76914036 |
| AGALGGGI | ASSPGGTLY | 0.6877357 |
| AGGAGGAGGIGAG | ASSPGGTLY | 0.48265618 |
| AGSGALGL | ASSPGGTLY | 0.7505052 |
| AQIADQI | ASSPGGTLY | 0.80783105 |
| ASAVAVTYL | ASSPGGTLY | 0.6858909 |
| ASTVLGGTGG | ASSPGGTLY | 0.8363246 |
| AVGGEPAK | ASSPGGTLY | 0.4287096 |
| CEQGTFGGTC | ASSPGGTLY | 0.8291811 |
| DEGKAPAG | ASSPGGTLY | 0.8207966 |
| DTVVFPAL | ASSPGGTLY | 0.7637952 |
| FEGITMAG | ASSPGGTLY | 0.6858032 |
| FPLVVQD | ASSPGGTLY | 0.81286955 |
| GAFTTPQ | ASSPGGTLY | 0.71802545 |
| GFGGFGTAGGVGGA | ASSPGGTLY | 0.7804119 |
| GFLAQFL | ASSPGGTLY | 0.82091534 |
| GGAGGAGGNAGVA | ASSPGGTLY | 0.49231973 |
| GGVGTAGAGGAGGAGG | ASSPGGTLY | 0.8372679 |
| GNGGGGDAGNAGSGGN | ASSPGGTLY | 0.7221534 |
| GTGGAGGAGNDAGS | ASSPGGTLY | 0.815537 |
| GTTNTGWFNS | ASSPGGTLY | 0.6245144 |
| IFMGPLFTL | ASSPGGTLY | 0.41012952 |
| ILAHAIEAI | ASSPGGTLY | 0.7305164 |
| IRSAIREAL | ASSPGGTLY | 0.55032384 |
| ITFAGGG | ASSPGGTLY | 0.8227761 |
| ITGMSGHQEV | ASSPGGTLY | 0.8255333 |
| IVAGLACTL | ASSPGGTLY | 0.6252971 |
| IVGGGLAGA | ASSPGGTLY | 0.8106393 |
| IVTGVLNI | ASSPGGTLY | 0.85262144 |
| LAQQGCI | ASSPGGTLY | 0.42402315 |
| LCLHVEN | ASSPGGTLY | 0.6780515 |
| LDAISDGS | ASSPGGTLY | 0.6547525 |
| LGGFAAML | ASSPGGTLY | 0.7085136 |
| LGGKNPMI | ASSPGGTLY | 0.28729585 |
| LGPKGVAAIL | ASSPGGTLY | 0.78910697 |
| LMVGVSI | ASSPGGTLY | 0.53549063 |
| LVGAAFVL | ASSPGGTLY | 0.17584972 |
| LVLAVAPL | ASSPGGTLY | 0.7855191 |
| MAAIAVIDAA | ASSPGGTLY | 0.609027 |
| NNGINAELQ | ASSPGGTLY | 0.64785624 |
| RGEPLVLVHG | ASSPGGTLY | 0.8041347 |
| RGGAGGAAGLI | ASSPGGTLY | 0.6646298 |
| RSYRMHPAV | ASSPGGTLY | 0.54594004 |
| SAAHLLAGQLA | ASSPGGTLY | 0.7453313 |
| SAAVGSY | ASSPGGTLY | 0.82919395 |
| SPHAVLHRI | ASSPGGTLY | 0.50027853 |
| SQIAKVL | ASSPGGTLY | 0.6997047 |
| TALGHLVP | ASSPGGTLY | 0.8616308 |

| | | |
|---|---|---|
| TAPTTSGAA | ASSPGGTLY | 0.7182081 |
| TDGDDGNGGSPGT | ASSPGGTLY | 0.8562933 |
| TDGLGEN | ASSPGGTLY | 0.86367667 |
| TGGPPLPGQA | ASSPGGTLY | 0.65941733 |
| TIPAITI | ASSPGGTLY | 0.8146914 |
| TIPGFTI | ASSPGGTLY | 0.8340962 |
| TSIASGALA | ASSPGGTLY | 0.70131034 |
| TVTLPGAAI | ASSPGGTLY | 0.71963894 |
| VASGLSAALP | ASSPGGTLY | 0.47182143 |
| VGAGVIGI | ASSPGGTLY | 0.7156384 |
| VGPKRFGV | ASSPGGTLY | 0.42151615 |
| VNVAFPD | ASSPGGTLY | 0.7028773 |
| VSMTNSGVS | ASSPGGTLY | 0.8356483 |
| VTDVLGDLSV | ASSPGGTLY | 0.674401 |
| VVVVGAGFAGL | ASSPGGTLY | 0.3597022 |
| VYTVGTVV | ASSPGGTLY | 0.39357334 |
| WTTFSGP | ASSPGGTLY | 0.7852675 |
| YDELVEAF | ASSPGGTLY | 0.5226637 |
| YGTGGAGGNGG | ASSPGGTLY | 0.42177606 |
| YTNVMAV | ASSPGGTLY | 0.6089617 |
| FLEQVHQL | ASSPGGTLY | 0.83371234 |
| GLSLLAVAV | ASSPGGTLY | 0.839502 |
| LEQVHQL | ASSPGGTLY | 0.5780139 |
| LNAAGAI | ASSPGGTLY | 0.8605227 |
| PAVHASL | ASSPGGTLY | 0.7947716 |
| TELEDDA | ASSPGGTLY | 0.8581267 |
| TPPGPPN | ASSPGGTLY | 0.7340803 |
| TPPPPPPP | ASSPGGTLY | 0.5045809 |
| TQSPALI | ASGDKGLQDTQY | 0.8303068 |
| EFAAAFPQ | ASGDKGLQDTQY | 0.6011356 |
| LLSGNLDVL | ASGDKGLQDTQY | 0.7888304 |
| GPANLGNYNI | ASGDKGLQDTQY | 0.8369597 |
| TLGPTLP | ASGDKGLQDTQY | 0.64972454 |
| VYAYSSCCK | ASGDKGLQDTQY | 0.56237686 |
| APAPAPAPNI | ASGDKGLQDTQY | 0.65881574 |
| PGGPVPIP | ASGDKGLQDTQY | 0.8518547 |
| FLTVALEP | ASGDKGLQDTQY | 0.46684664 |
| SQYFGANL | ASGDKGLQDTQY | 0.4553256 |
| GGVKPAA | ASGDKGLQDTQY | 0.6463772 |
| GGTGGAGGAGGW | ASGDKGLQDTQY | 0.6632526 |
| GVGGVAGLGGAGG | ASGDKGLQDTQY | 0.8459228 |
| VLLVLATTI | ASGDKGLQDTQY | 0.7936461 |
| IGIFAAML | ASGDKGLQDTQY | 0.6753708 |
| AVGDLID | ASGDKGLQDTQY | 0.7616389 |
| NISPLPL | ASGDKGLQDTQY | 0.8120067 |
| VTAVFVAI | ASGDKGLQDTQY | 0.74753106 |
| SGDDNDAAL | ASGDKGLQDTQY | 0.77752686 |
| AVAALAHS | ASGDKGLQDTQY | 0.5727172 |
| GGVTVMATP | ASGDKGLQDTQY | 0.8442919 |
| AGLAANVRD | ASGDKGLQDTQY | 0.8291817 |
| TNRPDLIDPALLRPG | ASGDKGLQDTQY | 0.83132756 |

| | | |
|---|---|---|
| LANLGVTAL | ASGDKGLQDTQY | 0.8395169 |
| TGERIEL | ASGDKGLQDTQY | 0.75253093 |
| NGEELAA | ASGDKGLQDTQY | 0.7470256 |
| VLSLVWG | ASGDKGLQDTQY | 0.46452105 |
| IGYALWL | ASGDKGLQDTQY | 0.62170196 |
| LSLLLAPL | ASGDKGLQDTQY | 0.67165524 |
| TLDEALK | ASGDKGLQDTQY | 0.70817745 |
| TDGAGTAEGG | ASGDKGLQDTQY | 0.78732216 |
| NAPGGAGGNGG | ASGDKGLQDTQY | 0.83104396 |
| VSVFMGMGGNF | ASGDKGLQDTQY | 0.56257224 |
| LASYLAAL | ASGDKGLQDTQY | 0.843477 |
| LVGGTPAL | ASGDKGLQDTQY | 0.83662164 |
| VRNADVL | ASGDKGLQDTQY | 0.45621482 |
| TCSSPFLVSAM | ASGDKGLQDTQY | 0.66984177 |
| VHITDAE | ASGDKGLQDTQY | 0.87136924 |
| EYAAPSF | ASGDKGLQDTQY | 0.77992374 |
| VVDAFHGGL | ASGDKGLQDTQY | 0.6333836 |
| ASVVDPQ | ASGDKGLQDTQY | 0.7457228 |
| SYNEFGGLS | ASGDKGLQDTQY | 0.8583522 |
| DPDGSVY | ASGDKGLQDTQY | 0.7021139 |
| MVSTIHATE | ASGDKGLQDTQY | 0.8478764 |
| LAAGVADPV | ASGDKGLQDTQY | 0.33084413 |
| TKEQIAA | ASGDKGLQDTQY | 0.7106712 |
| SLASPAAL | ASGDKGLQDTQY | 0.66628754 |
| IGLVFYGV | ASGDKGLQDTQY | 0.21199113 |
| FLLDASNL | ASGDKGLQDTQY | 0.82126904 |
| YQVGNFP | ASGDKGLQDTQY | 0.7025138 |
| ESLLSQST | ASGDKGLQDTQY | 0.81773674 |
| LFTVGFAAVG | ASGDKGLQDTQY | 0.6202645 |
| ATPPTPT | ASGDKGLQDTQY | 0.8458396 |
| TDLATLG | ASGDKGLQDTQY | 0.7467975 |
| SAGGTGR | ASGDKGLQDTQY | 0.7327231 |
| GFSNSGS | ASGDKGLQDTQY | 0.6489488 |
| GGNGGSGGTSV | ASGDKGLQDTQY | 0.7866606 |
| GTSVATGGAGN | ASGDKGLQDTQY | 0.8451287 |
| NGGNGGSGGTS | ASGDKGLQDTQY | 0.8606545 |
| AVVKFPH | ASGDKGLQDTQY | 0.3640154 |
| TPPPPPP | ASGDKGLQDTQY | 0.74939 |
| AAPAAAAAPA | ASGDKGLQDTQY | 0.7136941 |
| IALNEDL | ASGDKGLQDTQY | 0.35067266 |
| IVNVSSM | ASGDKGLQDTQY | 0.28565896 |
| TGIPPAPR | ASGDKGLQDTQY | 0.5788392 |
| FDLGGGTF | ASGDKGLQDTQY | 0.6675439 |
| FDLGGGTFDVS | ASGDKGLQDTQY | 0.7736878 |
| NEPTAAA | ASGDKGLQDTQY | 0.79546535 |
| NEPTAAALA | ASGDKGLQDTQY | 0.64106005 |
| GSIINISSI | ASGDKGLQDTQY | 0.76490545 |
| VAPGNAGTA | ASGDKGLQDTQY | 0.73666847 |
| TGGGASLE | ASGDKGLQDTQY | 0.088249564 |
| VAVYNHYKRI | ASGDKGLQDTQY | 0.6339536 |
| TDPAPATT | ASGDKGLQDTQY | 0.40023717 |

| | | |
|---|---|---|
| TNRPDILDPALLRPG | ASGDKGLQDTQY | 0.8647812 |
| LTTTTTS | ASGDKGLQDTQY | 0.47337693 |
| PTGSGKTL | ASGDKGLQDTQY | 0.64178145 |
| TGSGKTL | ASGDKGLQDTQY | 0.69886565 |
| TIGHVDHGK | ASGDKGLQDTQY | 0.73640954 |
| TIGHVDHGKTT | ASGDKGLQDTQY | 0.70065266 |
| GHVDHGKTT | ASGDKGLQDTQY | 0.5609057 |
| AQFEHTLL | ASGDKGLQDTQY | 0.29108804 |
| IVNQDEL | ASGDKGLQDTQY | 0.80879605 |
| TNGSQFF | ASGDKGLQDTQY | 0.6436459 |
| TGKGIVE | ASGDKGLQDTQY | 0.5545956 |
| LISNASDA | ASGDKGLQDTQY | 0.59168327 |
| AELRTLGELTKIAKAHGAQ | ASGDKGLQDTQY | 0.5759398 |
| APNAVAAAEPSW | ASGDKGLQDTQY | 0.4077788 |
| APQVAGAATAVL | ASGDKGLQDTQY | 0.8474258 |
| ARRDPAAAAAILPTDA | ASGDKGLQDTQY | 0.4999956 |
| DAAARDVLAEAGLAEYF | ASGDKGLQDTQY | 0.62654865 |
| DDQIVLSAKANLAKVEQ | ASGDKGLQDTQY | 0.84879565 |
| DELLPDTGEDSDEDGH | ASGDKGLQDTQY | 0.71911323 |
| ELPEAPANRATVPQD | ASGDKGLQDTQY | 0.6866921 |
| FGQSKIERVTGIG | ASGDKGLQDTQY | 0.6406847 |
| GAGGTGSTAGGAGGAGGAGGLY | ASGDKGLQDTQY | 0.51714385 |
| GHGGTGAAGGNGGTGGAGGSGI | ASGDKGLQDTQY | 0.53320295 |
| GNGGNGGAGGEGGPGGAGGAGA | ASGDKGLQDTQY | 0.5592659 |
| GSTAGNGGNGGNATNGTIAGQP | ASGDKGLQDTQY | 0.8566171 |
| GTGGDGGNGGNGGTGG | ASGDKGLQDTQY | 0.7716043 |
| GVRRPAGIPADAPV | ASGDKGLQDTQY | 0.51775134 |
| GYRQVTDAHLLTLARR | ASGDKGLQDTQY | 0.8469142 |
| IAKMGQDGHDRG | ASGDKGLQDTQY | 0.84163976 |
| KAEPSIEIQYT | ASGDKGLQDTQY | 0.86305064 |
| KPVRIRMAADEDTVAVT | ASGDKGLQDTQY | 0.48867756 |
| MPPRIAGMRLLVIKPEP | ASGDKGLQDTQY | 0.69643664 |
| MKFYAVALFRSRGVPPTR | ASGDKGLQDTQY | 0.72327507 |
| NAGGFGILGAVAHSP | ASGDKGLQDTQY | 0.6791832 |
| NEGREITLAMAEM | ASGDKGLQDTQY | 0.61004686 |
| NELVEWSHPLGHRWRWE | ASGDKGLQDTQY | 0.54511714 |
| NLGAGGTGGNGGNPG | ASGDKGLQDTQY | 0.8282337 |
| NTEWPKWETVMAADPVD | ASGDKGLQDTQY | 0.88010335 |
| PASAAPAETQP | ASGDKGLQDTQY | 0.7141514 |
| QQSSDGKNGVRAAEITGE | ASGDKGLQDTQY | 0.3250563 |
| RAPDAVTTSWWKE | ASGDKGLQDTQY | 0.6449128 |
| SGFGNVGAHTSGWFNQ | ASGDKGLQDTQY | 0.4919061 |
| SGVAGAVGQAASVGGLKV | ASGDKGLQDTQY | 0.53292036 |
| SNGGAGTGSAGGNGGT | ASGDKGLQDTQY | 0.62051743 |
| TGGAGGEGGAAG | ASGDKGLQDTQY | 0.36466873 |
| TGGDGGAAGAGGAPGHG | ASGDKGLQDTQY | 0.67966664 |
| TMPPLLAAIANAGRHGV | ASGDKGLQDTQY | 0.858282 |
| TSSTAGNSGGA | ASGDKGLQDTQY | 0.62075436 |
| VPATVPVGILC | ASGDKGLQDTQY | 0.77735186 |
| VQRPDSGGVVLTGRIS | ASGDKGLQDTQY | 0.8178321 |
| WTPNPVPATKGVPAGLVQAT | ASGDKGLQDTQY | 0.7502476 |

| | | |
|---|---|---|
| YALKRALAKFSDRFDVVII | ASGDKGLQDTQY | 0.83547986 |
| YLIGSVIV | ASGDKGLQDTQY | 0.8134905 |
| YGQLVAVID | ASGDKGLQDTQY | 0.8021542 |
| VTVGGMTLPA | ASGDKGLQDTQY | 0.7425252 |
| VLALPAV | ASGDKGLQDTQY | 0.7517481 |
| VIPPTLNYET | ASGDKGLQDTQY | 0.6836314 |
| VGMGGMHPSSG | ASGDKGLQDTQY | 0.7236633 |
| VGLAGGAATA | ASGDKGLQDTQY | 0.7198769 |
| VGGAGGAGGSGP | ASGDKGLQDTQY | 0.76381326 |
| VFAAFMV | ASGDKGLQDTQY | 0.6886801 |
| TLYDKLR | ASGDKGLQDTQY | 0.85373163 |
| TLAVFTVL | ASGDKGLQDTQY | 0.2762965 |
| TGSVQPARV | ASGDKGLQDTQY | 0.5210989 |
| TGGGAGSGGT | ASGDKGLQDTQY | 0.8142126 |
| SHLETSL | ASGDKGLQDTQY | 0.1978215 |
| SFTTGMT | ASGDKGLQDTQY | 0.37746072 |
| QVEVAKRMGVRQPTVS | ASGDKGLQDTQY | 0.72856 |
| QGSGGPALAM | ASGDKGLQDTQY | 0.8282044 |
| MLAPSNP | ASGDKGLQDTQY | 0.18537605 |
| LVNNAAI | ASGDKGLQDTQY | 0.68294775 |
| LTGEINTLA | ASGDKGLQDTQY | 0.40496042 |
| LSIFGPV | ASGDKGLQDTQY | 0.86515546 |
| LLALLANL | ASGDKGLQDTQY | 0.7065579 |
| LISNASDALDKLR | ASGDKGLQDTQY | 0.36554727 |
| LINEAVP | ASGDKGLQDTQY | 0.8304863 |
| LGYAFSGL | ASGDKGLQDTQY | 0.5066251 |
| LDFVAVAL | ASGDKGLQDTQY | 0.70649445 |
| LAQPGAE | ASGDKGLQDTQY | 0.8399755 |
| ISNLGAVTGF | ASGDKGLQDTQY | 0.86564934 |
| HQEDPGRI | ASGDKGLQDTQY | 0.42220157 |
| GTPGEPTP | ASGDKGLQDTQY | 0.8035084 |
| GFFNSGS | ASGDKGLQDTQY | 0.61798525 |
| GAGGGGLTAGAN | ASGDKGLQDTQY | 0.87202907 |
| FVDALVNM | ASGDKGLQDTQY | 0.6431827 |
| FLTVAPEL | ASGDKGLQDTQY | 0.37086326 |
| FETLTAL | ASGDKGLQDTQY | 0.80319804 |
| DVINPIEL | ASGDKGLQDTQY | 0.82246923 |
| APVSGAPV | ASGDKGLQDTQY | 0.75817716 |
| ALVGGLAW | ASGDKGLQDTQY | 0.7845546 |
| ALAAAAATAI | ASGDKGLQDTQY | 0.74799323 |
| AFGLGLLL | ASGDKGLQDTQY | 0.60418934 |
| AFFAAASL | ASGDKGLQDTQY | 0.8223529 |
| AAAALPAFSPP | ASGDKGLQDTQY | 0.77352315 |
| VVLAPAL | ASGDKGLQDTQY | 0.56725603 |
| TIGHVDHGKTTL | ASGDKGLQDTQY | 0.8601903 |
| LISNASDALDK | ASGDKGLQDTQY | 0.749475 |
| LISNASDAL | ASGDKGLQDTQY | 0.57683635 |
| ISNASDALDK | ASGDKGLQDTQY | 0.86078507 |
| NQAETLVYQTE | ASGDKGLQDTQY | 0.75251323 |
| TGGTGGTAGAG | ASGDKGLQDTQY | 0.7851393 |
| TGSTAGGAGGA | ASGDKGLQDTQY | 0.788958 |

| | | |
|---|---|---|
| DPLAVPVAAGPV | ASGDKGLQDTQY | 0.8701081 |
| GGTGGTPGNGGN | ASGDKGLQDTQY | 0.47064433 |
| PAGLAGVDPATL | ASGDKGLQDTQY | 0.7789808 |
| GNNLVVEAPHAGASV | ASGDKGLQDTQY | 0.7203306 |
| KVTRIMSASTLEQPA | ASGDKGLQDTQY | 0.82441163 |
| LGFALPIALPRRTVV | ASGDKGLQDTQY | 0.7500818 |
| TRLGNLTARKLVVLV | ASGDKGLQDTQY | 0.6541396 |
| WSQQRRLIYFPSAGP | ASGDKGLQDTQY | 0.79030037 |
| GPNGNPDPMAAAVDIR | ASGDKGLQDTQY | 0.8345969 |
| TQRCVSVADLRAAASR | ASGDKGLQDTQY | 0.74022317 |
| QFVLDVAEHPVEVLTETV | ASGDKGLQDTQY | 0.7676702 |
| GGAGGTGGAGGNGGTQAGDG | ASGDKGLQDTQY | 0.85920274 |
| PLLATLRAIAVDVDAKPAQV | ASGDKGLQDTQY | 0.50679725 |
| APKFRTGGYMIDFWGVGYQVAK | ASGDKGLQDTQY | 0.7459732 |
| GAGGAGGAGGAAGGPGGTGGTG | ASGDKGLQDTQY | 0.8207023 |
| IGALGLLATDVAPAGSVAFGVA | ASGDKGLQDTQY | 0.8408014 |
| AGGAGGIGGNGGAGGAGGVLMG | ASGDKGLQDTQY | 0.72731614 |
| ADELIGLL | ASGDKGLQDTQY | 0.8335128 |
| ADRALKA | ASGDKGLQDTQY | 0.65898836 |
| AGAVGQVGGAGGA | ASGDKGLQDTQY | 0.75071347 |
| AGGGGGLGGAP | ASGDKGLQDTQY | 0.60262096 |
| AGGLLYGNGGAG | ASGDKGLQDTQY | 0.70779055 |
| AGLESIGL | ASGDKGLQDTQY | 0.4610061 |
| AGLKTGII | ASGDKGLQDTQY | 0.72963494 |
| ASLAALLL | ASGDKGLQDTQY | 0.77220964 |
| ASVINISSI | ASGDKGLQDTQY | 0.77397513 |
| ATPPGPA | ASGDKGLQDTQY | 0.33260557 |
| ATVASALV | ASGDKGLQDTQY | 0.7689626 |
| ATVASLIL | ASGDKGLQDTQY | 0.80175215 |
| AVNGPVLIRH | ASGDKGLQDTQY | 0.87238544 |
| DGAAGAVGI | ASGDKGLQDTQY | 0.71728814 |
| DPLLHGI | ASGDKGLQDTQY | 0.828865 |
| ELGAALGAL | ASGDKGLQDTQY | 0.6528723 |
| ELGRTQAV | ASGDKGLQDTQY | 0.7555016 |
| FVAGGYLS | ASGDKGLQDTQY | 0.38445434 |
| GASGGAGGAGGW | ASGDKGLQDTQY | 0.5616074 |
| GGAGAGGDGGAGGI | ASGDKGLQDTQY | 0.64543784 |
| GTGHNGSQPGMGGNG | ASGDKGLQDTQY | 0.5238944 |
| GYAAASASAS | ASGDKGLQDTQY | 0.79667574 |
| HQVDVAAVQQG | ASGDKGLQDTQY | 0.81745756 |
| IGAAIAHT | ASGDKGLQDTQY | 0.82215196 |
| IGMIEVAQL | ASGDKGLQDTQY | 0.8046076 |
| IRLAPEHTSAMD | ASGDKGLQDTQY | 0.81973183 |
| IYNVSVL | ASGDKGLQDTQY | 0.8524217 |
| KQGFLGM | ASGDKGLQDTQY | 0.61728376 |
| LESVITQL | ASGDKGLQDTQY | 0.6788781 |
| LGLGALVVV | ASGDKGLQDTQY | 0.38592887 |
| LIGNGGNGGSGGI | ASGDKGLQDTQY | 0.60469073 |
| LNAINAPTL | ASGDKGLQDTQY | 0.6076213 |
| LQLLNTGV | ASGDKGLQDTQY | 0.7126773 |
| LQTSIEF | ASGDKGLQDTQY | 0.16444856 |

| | | |
|---|---|---|
| MSFAVSEL | ASGDKGLQDTQY | 0.6984192 |
| NTGALGL | ASGDKGLQDTQY | 0.85536265 |
| PPGAGGAGGAGTT | ASGDKGLQDTQY | 0.60154784 |
| QLVGLGI | ASGDKGLQDTQY | 0.7846641 |
| RFLEQVHQL | ASGDKGLQDTQY | 0.76674366 |
| RGVGYVLR | ASGDKGLQDTQY | 0.58572274 |
| SATGVFAGVI | ASGDKGLQDTQY | 0.76181245 |
| SLPAITLP | ASGDKGLQDTQY | 0.5917777 |
| STALAFTV | ASGDKGLQDTQY | 0.8252286 |
| SVLTVGLGK | ASGDKGLQDTQY | 0.84593093 |
| TALAGAGSGI | ASGDKGLQDTQY | 0.7422383 |
| TGLPFTGI | ASGDKGLQDTQY | 0.66626084 |
| VAEHVTV | ASGDKGLQDTQY | 0.78058267 |
| VDIVLNSL | ASGDKGLQDTQY | 0.49990556 |
| VLTGGGAVL | ASGDKGLQDTQY | 0.705026 |
| VNVGLAYGI | ASGDKGLQDTQY | 0.24802957 |
| VSSAAVGAV | ASGDKGLQDTQY | 0.6819259 |
| VTSVYEAGV | ASGDKGLQDTQY | 0.52643824 |
| WVALDRGA | ASGDKGLQDTQY | 0.7599267 |
| YLVGSVVI | ASGDKGLQDTQY | 0.73099583 |
| GPPGTGKTLL | ASGDKGLQDTQY | 0.7799447 |
| GVGMTKF | ASGDKGLQDTQY | 0.49408418 |
| IGIGGSDL | ASGDKGLQDTQY | 0.7521597 |
| IGTLLAEL | ASGDKGLQDTQY | 0.6915929 |
| IYTSGTT | ASGDKGLQDTQY | 0.42121536 |
| LSFPAGV | ASGDKGLQDTQY | 0.8302671 |
| LYTSGTT | ASGDKGLQDTQY | 0.71531504 |
| VGAFPFGF | ASGDKGLQDTQY | 0.6783646 |
| VGAGTVEF | ASGDKGLQDTQY | 0.8233379 |
| VGKSTLLL | ASGDKGLQDTQY | 0.75742114 |
| AALAHAGGGIV | ASGDKGLQDTQY | 0.5427935 |
| AALVAVSLL | ASGDKGLQDTQY | 0.56962645 |
| AAVSRAVAAL | ASGDKGLQDTQY | 0.70472515 |
| AGGSSGIGGF | ASGDKGLQDTQY | 0.64147025 |
| AGLTVANANG | ASGDKGLQDTQY | 0.2078145 |
| AGLVGAVL | ASGDKGLQDTQY | 0.8146623 |
| ALSQAFTV | ASGDKGLQDTQY | 0.585363 |
| ALTFSVSL | ASGDKGLQDTQY | 0.579441 |
| ALTGAALVAL | ASGDKGLQDTQY | 0.6761813 |
| ASFSIGAGVL | ASGDKGLQDTQY | 0.8163546 |
| DGSVYKI | ASGDKGLQDTQY | 0.7317476 |
| EQLAEIV | ASGDKGLQDTQY | 0.7050737 |
| FAVSELTVM | ASGDKGLQDTQY | 0.7747122 |
| GAAPLAAGL | ASGDKGLQDTQY | 0.7887989 |
| GADGGDGGSGGAGG | ASGDKGLQDTQY | 0.83880603 |
| GFGLDTGGAGG | ASGDKGLQDTQY | 0.78969204 |
| GLASGGLL | ASGDKGLQDTQY | 0.75347304 |
| GRIAGAAVLL | ASGDKGLQDTQY | 0.8296405 |
| GSAGSLNTSF | ASGDKGLQDTQY | 0.729316 |
| GTGQIGI | ASGDKGLQDTQY | 0.4889186 |
| GTLIALIL | ASGDKGLQDTQY | 0.44837865 |

| | | |
|---|---|---|
| IAATMGFVVG | ASGDKGLQDTQY | 0.621798 |
| IFAQALL | ASGDKGLQDTQY | 0.4594501 |
| IGAAVALGF | ASGDKGLQDTQY | 0.7701594 |
| IGLTPLAL | ASGDKGLQDTQY | 0.8778931 |
| ILGSNTSTL | ASGDKGLQDTQY | 0.73804677 |
| IVFSLTVL | ASGDKGLQDTQY | 0.37521666 |
| LASKATGF | ASGDKGLQDTQY | 0.62035346 |
| LASVALIL | ASGDKGLQDTQY | 0.6668595 |
| LFAVGLTL | ASGDKGLQDTQY | 0.7818912 |
| LFNGGIGTYI | ASGDKGLQDTQY | 0.8257296 |
| LTLGAGII | ASGDKGLQDTQY | 0.8563361 |
| LVAAALGDI | ASGDKGLQDTQY | 0.81757927 |
| NTAFVQLGI | ASGDKGLQDTQY | 0.61941624 |
| QLFSGAEL | ASGDKGLQDTQY | 0.80993605 |
| SAATFITL | ASGDKGLQDTQY | 0.655169 |
| SNYGLANLGV | ASGDKGLQDTQY | 0.76554304 |
| SYGGAYAVM | ASGDKGLQDTQY | 0.7466136 |
| TTGVGGLTLG | ASGDKGLQDTQY | 0.79785424 |
| TTQTIYI | ASGDKGLQDTQY | 0.6427667 |
| VAAGAGLGAVI | ASGDKGLQDTQY | 0.7079716 |
| VAAVVTLL | ASGDKGLQDTQY | 0.46348608 |
| VGFLGGLI | ASGDKGLQDTQY | 0.82287 |
| VGVIPILL | ASGDKGLQDTQY | 0.5380722 |
| VIFASSLI | ASGDKGLQDTQY | 0.5090717 |
| VKGGDGGAAAG | ASGDKGLQDTQY | 0.69636226 |
| VTKTFTNI | ASGDKGLQDTQY | 0.8419206 |
| VVAAAILL | ASGDKGLQDTQY | 0.8400407 |
| YAALDVEL | ASGDKGLQDTQY | 0.5849575 |
| LASVALLL | ASGDKGLQDTQY | 0.5797069 |
| LDAVHRV | ASGDKGLQDTQY | 0.75023234 |
| QGLSSSIF | ASGDKGLQDTQY | 0.60659844 |
| RGTFANIRL | ASGDKGLQDTQY | 0.81413496 |
| RQLWWGHRI | ASGDKGLQDTQY | 0.6883918 |
| YGPPGTGKTLL | ASGDKGLQDTQY | 0.68041563 |
| AALVGAPANVA | ASGDKGLQDTQY | 0.6712836 |
| AATFRRTE | ASGDKGLQDTQY | 0.772408 |
| AGALGGGI | ASGDKGLQDTQY | 0.6837635 |
| AGGAGGAGGIGAG | ASGDKGLQDTQY | 0.8217117 |
| AGSGALGL | ASGDKGLQDTQY | 0.2256985 |
| AQIADQI | ASGDKGLQDTQY | 0.830176 |
| ASAVAVTYL | ASGDKGLQDTQY | 0.48701337 |
| ASTVLGGTGG | ASGDKGLQDTQY | 0.57359695 |
| AVGGEPAK | ASGDKGLQDTQY | 0.2653778 |
| CEQGTFGGTC | ASGDKGLQDTQY | 0.7737551 |
| DEGKAPAG | ASGDKGLQDTQY | 0.7681508 |
| DTVVFPAL | ASGDKGLQDTQY | 0.82499367 |
| FEGITMAG | ASGDKGLQDTQY | 0.6957477 |
| FPLVVQD | ASGDKGLQDTQY | 0.79593 |
| GAFTTPQ | ASGDKGLQDTQY | 0.82573265 |
| GFGGFGTAGGVGGA | ASGDKGLQDTQY | 0.59916705 |
| GFLAQFL | ASGDKGLQDTQY | 0.29637495 |

| | | |
|---|---|---|
| GGAGGAGGNAGVA | ASGDKGLQDTQY | 0.75653696 |
| GGVGTAGAGGAGGAGG | ASGDKGLQDTQY | 0.75056136 |
| GNGGGGDAGNAGSGGN | ASGDKGLQDTQY | 0.84607065 |
| GTGGAGGAGNDAGS | ASGDKGLQDTQY | 0.7961556 |
| GTTNTGWFNS | ASGDKGLQDTQY | 0.3887331 |
| IFMGPLFTL | ASGDKGLQDTQY | 0.82327855 |
| ILAHAIEAI | ASGDKGLQDTQY | 0.7906492 |
| IRSAIREAL | ASGDKGLQDTQY | 0.44683662 |
| ITFAGGG | ASGDKGLQDTQY | 0.6452028 |
| ITGMSGHQEV | ASGDKGLQDTQY | 0.69451165 |
| IVAGLACTL | ASGDKGLQDTQY | 0.38157603 |
| IVGGGLAGA | ASGDKGLQDTQY | 0.7979469 |
| IVTGVLNI | ASGDKGLQDTQY | 0.5801606 |
| LAQQGCI | ASGDKGLQDTQY | 0.65253866 |
| LCLHVEN | ASGDKGLQDTQY | 0.6937811 |
| LDAISDGS | ASGDKGLQDTQY | 0.5901756 |
| LGGFAAML | ASGDKGLQDTQY | 0.68035793 |
| LGGKNPMI | ASGDKGLQDTQY | 0.65532243 |
| LGPKGVAAIL | ASGDKGLQDTQY | 0.7848077 |
| LMVGVSI | ASGDKGLQDTQY | 0.7736335 |
| LVGAAFVL | ASGDKGLQDTQY | 0.6819169 |
| LVLAVAPL | ASGDKGLQDTQY | 0.8082447 |
| MAAIAVIDAA | ASGDKGLQDTQY | 0.8183727 |
| NNGINAELQ | ASGDKGLQDTQY | 0.43522617 |
| RGEPLVLVHG | ASGDKGLQDTQY | 0.6151234 |
| RGGAGGAAGLI | ASGDKGLQDTQY | 0.788831 |
| RSYRMHPAV | ASGDKGLQDTQY | 0.56351984 |
| SAAHLLAGQLA | ASGDKGLQDTQY | 0.81183535 |
| SAAVGSY | ASGDKGLQDTQY | 0.75720936 |
| SPHAVLHRI | ASGDKGLQDTQY | 0.748998 |
| SQIAKVL | ASGDKGLQDTQY | 0.69084376 |
| TALGHLVP | ASGDKGLQDTQY | 0.87134176 |
| TAPTTSGAA | ASGDKGLQDTQY | 0.77753174 |
| TDGDDGNGGSPGT | ASGDKGLQDTQY | 0.7386523 |
| TDGLGEN | ASGDKGLQDTQY | 0.79816484 |
| TGGPPLPGQA | ASGDKGLQDTQY | 0.5833869 |
| TIPAITI | ASGDKGLQDTQY | 0.8053612 |
| TIPGFTI | ASGDKGLQDTQY | 0.8183218 |
| TSIASGALA | ASGDKGLQDTQY | 0.4802835 |
| TVTLPGAAI | ASGDKGLQDTQY | 0.6087934 |
| VASGLSAALP | ASGDKGLQDTQY | 0.8480465 |
| VGAGVIGI | ASGDKGLQDTQY | 0.8373821 |
| VGPKRFGV | ASGDKGLQDTQY | 0.53129035 |
| VNVAFPD | ASGDKGLQDTQY | 0.6212837 |
| VSMTNSGVS | ASGDKGLQDTQY | 0.5788733 |
| VTDVLGDLSV | ASGDKGLQDTQY | 0.67817026 |
| VVVVGAGFAGL | ASGDKGLQDTQY | 0.46963322 |
| VYTVGTVV | ASGDKGLQDTQY | 0.75685704 |
| WTTFSGP | ASGDKGLQDTQY | 0.6289543 |
| YDELVEAF | ASGDKGLQDTQY | 0.81677604 |
| YGTGGAGGNGG | ASGDKGLQDTQY | 0.7674589 |

| | | |
|---|---|---|
| YTNVMAV | ASGDKGLQDTQY | 0.83851457 |
| FLEQVHQL | ASGDKGLQDTQY | 0.5857323 |
| GLSLLAVAV | ASGDKGLQDTQY | 0.69619024 |
| LEQVHQL | ASGDKGLQDTQY | 0.3865414 |
| LNAAGAI | ASGDKGLQDTQY | 0.38581026 |
| PAVHASL | ASGDKGLQDTQY | 0.56014264 |
| TELEDDA | ASGDKGLQDTQY | 0.68235326 |
| TPPGPPN | ASGDKGLQDTQY | 0.29388946 |
| TPPPPPPP | ASGDKGLQDTQY | 0.86264026 |
| TQSPALI | ASSDMTAANSDYT | 0.66266495 |
| EFAAAFPQ | ASSDMTAANSDYT | 0.81606686 |
| LLSGNLDVL | ASSDMTAANSDYT | 0.6903029 |
| GPANLGNYNI | ASSDMTAANSDYT | 0.4845261 |
| TLGPTLP | ASSDMTAANSDYT | 0.6118067 |
| VYAYSSCCK | ASSDMTAANSDYT | 0.78469914 |
| APAPAPAPNI | ASSDMTAANSDYT | 0.4014666 |
| PGGPVPIP | ASSDMTAANSDYT | 0.37813646 |
| FLTVALEP | ASSDMTAANSDYT | 0.6486782 |
| SQYFGANL | ASSDMTAANSDYT | 0.6788575 |
| GGVKPAA | ASSDMTAANSDYT | 0.85606545 |
| GGTGGAGGAGGW | ASSDMTAANSDYT | 0.8129735 |
| GVGGVAGLGGAGG | ASSDMTAANSDYT | 0.69561327 |
| VLLVLATTI | ASSDMTAANSDYT | 0.86914957 |
| IGIFAAML | ASSDMTAANSDYT | 0.18456747 |
| AVGDLID | ASSDMTAANSDYT | 0.7094969 |
| NISPLPL | ASSDMTAANSDYT | 0.8643942 |
| VTAVFVAI | ASSDMTAANSDYT | 0.7495044 |
| SGDDNDAAL | ASSDMTAANSDYT | 0.67133796 |
| AVAALAHS | ASSDMTAANSDYT | 0.82884926 |
| GGVTVMATP | ASSDMTAANSDYT | 0.58528703 |
| AGLAANVRD | ASSDMTAANSDYT | 0.6649569 |
| TNRPDLIDPALLRPG | ASSDMTAANSDYT | 0.68218845 |
| LANLGVTAL | ASSDMTAANSDYT | 0.82117563 |
| TGERIEL | ASSDMTAANSDYT | 0.7773349 |
| NGEELAA | ASSDMTAANSDYT | 0.408315 |
| VLSLVWG | ASSDMTAANSDYT | 0.80774796 |
| IGYALWL | ASSDMTAANSDYT | 0.85893 |
| LSLLLAPL | ASSDMTAANSDYT | 0.59768665 |
| TLDEALK | ASSDMTAANSDYT | 0.82136667 |
| TDGAGTAEGG | ASSDMTAANSDYT | 0.7558465 |
| NAPGGAGGNGG | ASSDMTAANSDYT | 0.5856142 |
| VSVFMGMGGNF | ASSDMTAANSDYT | 0.55364245 |
| LASYLAAL | ASSDMTAANSDYT | 0.51137435 |
| LVGGTPAL | ASSDMTAANSDYT | 0.75600946 |
| VRNADVL | ASSDMTAANSDYT | 0.46074212 |
| TCSSPFLVSAM | ASSDMTAANSDYT | 0.7340946 |
| VHITDAE | ASSDMTAANSDYT | 0.72606486 |
| EYAAPSF | ASSDMTAANSDYT | 0.8710438 |
| VVDAFHGGL | ASSDMTAANSDYT | 0.76697916 |
| ASVVDPQ | ASSDMTAANSDYT | 0.7803786 |
| SYNEFGGLS | ASSDMTAANSDYT | 0.5589032 |

| | | |
|---|---|---|
| DPDGSVY | ASSDMTAANSDYT | 0.7607355 |
| MVSTIHATE | ASSDMTAANSDYT | 0.74769807 |
| LAAGVADPV | ASSDMTAANSDYT | 0.49266204 |
| TKEQIAA | ASSDMTAANSDYT | 0.8253747 |
| SLASPAAL | ASSDMTAANSDYT | 0.5287786 |
| IGLVFYGV | ASSDMTAANSDYT | 0.7405597 |
| FLLDASNL | ASSDMTAANSDYT | 0.8171822 |
| YQVGNFP | ASSDMTAANSDYT | 0.8263233 |
| ESLLSQST | ASSDMTAANSDYT | 0.69198775 |
| LFTVGFAAVG | ASSDMTAANSDYT | 0.7806914 |
| ATPPTPT | ASSDMTAANSDYT | 0.5521772 |
| TDLATLG | ASSDMTAANSDYT | 0.8053436 |
| SAGGTGR | ASSDMTAANSDYT | 0.8253366 |
| GFSNSGS | ASSDMTAANSDYT | 0.72789794 |
| GGNGGSGGTSV | ASSDMTAANSDYT | 0.8708546 |
| GTSVATGGAGN | ASSDMTAANSDYT | 0.8333863 |
| NGGNGGSGGTS | ASSDMTAANSDYT | 0.39486548 |
| AVVKFPH | ASSDMTAANSDYT | 0.7129723 |
| TPPPPPP | ASSDMTAANSDYT | 0.8430142 |
| AAPAAAAAPA | ASSDMTAANSDYT | 0.809539 |
| IALNEDL | ASSDMTAANSDYT | 0.81211954 |
| IVNVSSM | ASSDMTAANSDYT | 0.7828489 |
| TGIPPAPR | ASSDMTAANSDYT | 0.6202738 |
| FDLGGGTF | ASSDMTAANSDYT | 0.8343786 |
| FDLGGGTFDVS | ASSDMTAANSDYT | 0.89206433 |
| NEPTAAA | ASSDMTAANSDYT | 0.82368696 |
| NEPTAAALA | ASSDMTAANSDYT | 0.58576006 |
| GSIINISSI | ASSDMTAANSDYT | 0.6146653 |
| VAPGNAGTA | ASSDMTAANSDYT | 0.77811444 |
| TGGGASLE | ASSDMTAANSDYT | 0.8152512 |
| VAVYNHYKRI | ASSDMTAANSDYT | 0.49901927 |
| TDPAPATT | ASSDMTAANSDYT | 0.67766786 |
| TNRPDILDPALLRPG | ASSDMTAANSDYT | 0.7103354 |
| LTTTTTS | ASSDMTAANSDYT | 0.28757817 |
| PTGSGKTL | ASSDMTAANSDYT | 0.805832 |
| TGSGKTL | ASSDMTAANSDYT | 0.8450032 |
| TIGHVDHGK | ASSDMTAANSDYT | 0.79388237 |
| TIGHVDHGKTT | ASSDMTAANSDYT | 0.43260098 |
| GHVDHGKTT | ASSDMTAANSDYT | 0.81193626 |
| AQFEHTLL | ASSDMTAANSDYT | 0.8398758 |
| IVNQDEL | ASSDMTAANSDYT | 0.851591 |
| TNGSQFF | ASSDMTAANSDYT | 0.41511032 |
| TGKGIVE | ASSDMTAANSDYT | 0.19498552 |
| LISNASDA | ASSDMTAANSDYT | 0.7407454 |
| AELRTLGELTKIAKAHGAQ | ASSDMTAANSDYT | 0.8121769 |
| APNAVAAAEPSW | ASSDMTAANSDYT | 0.64297175 |
| APQVAGAATAVL | ASSDMTAANSDYT | 0.5052793 |
| ARRDPAAAAAILPTDA | ASSDMTAANSDYT | 0.6889015 |
| DAAARDVLAEAGLAEYF | ASSDMTAANSDYT | 0.8227047 |
| DDQIVLSAKANLAKVEQ | ASSDMTAANSDYT | 0.51558876 |
| DELLPDTGEDSDEDGH | ASSDMTAANSDYT | 0.4823258 |

| | | |
|---|---|---|
| ELPEAPANRATVPQD | ASSDMTAANSDYT | 0.8087518 |
| FGQSKIERVTGIG | ASSDMTAANSDYT | 0.8322338 |
| GAGGTGSTAGGAGGAGGAGGLY | ASSDMTAANSDYT | 0.50191915 |
| GHGGTGAAGGNGGTGGAGGSGI | ASSDMTAANSDYT | 0.56700075 |
| GNGGNGGAGGEGGPGGAGGAGA | ASSDMTAANSDYT | 0.7647032 |
| GSTAGNGGNGGNATNGTIAGQP | ASSDMTAANSDYT | 0.77399015 |
| GTGGDGGNGGNGGTGG | ASSDMTAANSDYT | 0.7185036 |
| GVRRPAGIPADAPV | ASSDMTAANSDYT | 0.4380806 |
| GYRQVTDAHLLTLARR | ASSDMTAANSDYT | 0.82667416 |
| IAKMGQDGHDRG | ASSDMTAANSDYT | 0.4309118 |
| KAEPSIEIQYT | ASSDMTAANSDYT | 0.80692977 |
| KPVRIRMAADEDTVAVT | ASSDMTAANSDYT | 0.72741926 |
| MPPRIAGMRLLVIKPEP | ASSDMTAANSDYT | 0.7819613 |
| MKFYAVALFRSRGVPPTR | ASSDMTAANSDYT | 0.60344076 |
| NAGGFGILGAVAHSP | ASSDMTAANSDYT | 0.46451858 |
| NEGREITLAMAEM | ASSDMTAANSDYT | 0.5010629 |
| NELVEWSHPLGHRWRWE | ASSDMTAANSDYT | 0.4016939 |
| NLGAGGTGGNGGNPG | ASSDMTAANSDYT | 0.82234037 |
| NTEWPKWETVMAADPVD | ASSDMTAANSDYT | 0.43778652 |
| PASAAPAETQP | ASSDMTAANSDYT | 0.833246 |
| QQSSDGKNGVRAAEITGE | ASSDMTAANSDYT | 0.7428818 |
| RAPDAVTTSWWKE | ASSDMTAANSDYT | 0.60263944 |
| SGFGNVGAHTSGWFNQ | ASSDMTAANSDYT | 0.7815588 |
| SGVAGAVGQAASVGGLKV | ASSDMTAANSDYT | 0.37684092 |
| SNGGAGTGSAGGNGGT | ASSDMTAANSDYT | 0.74671406 |
| TGGAGGEGGAAG | ASSDMTAANSDYT | 0.7159903 |
| TGGDGGAAGAGGAPGHG | ASSDMTAANSDYT | 0.82711494 |
| TMPPLLAAIANAGRHGV | ASSDMTAANSDYT | 0.6028538 |
| TSSTAGNSGGA | ASSDMTAANSDYT | 0.79734635 |
| VPATVPVGILC | ASSDMTAANSDYT | 0.7469875 |
| VQRPDSGGVVLTGRIS | ASSDMTAANSDYT | 0.54488856 |
| WTPNPVPATKGVPAGLVQAT | ASSDMTAANSDYT | 0.79839504 |
| YALKRALAKFSDRFDVVII | ASSDMTAANSDYT | 0.81378907 |
| YLIGSVIV | ASSDMTAANSDYT | 0.8052865 |
| YGQLVAVID | ASSDMTAANSDYT | 0.31063566 |
| VTVGGMTLPA | ASSDMTAANSDYT | 0.7645457 |
| VLALPAV | ASSDMTAANSDYT | 0.58031344 |
| VIPPTLNYET | ASSDMTAANSDYT | 0.7324374 |
| VGMGGMHPSSG | ASSDMTAANSDYT | 0.75761724 |
| VGLAGGAATA | ASSDMTAANSDYT | 0.6926886 |
| VGGAGGAGGSGP | ASSDMTAANSDYT | 0.80232394 |
| VFAAFMV | ASSDMTAANSDYT | 0.5320856 |
| TLYDKLR | ASSDMTAANSDYT | 0.58323383 |
| TLAVFTVL | ASSDMTAANSDYT | 0.70204186 |
| TGSVQPARV | ASSDMTAANSDYT | 0.7634323 |
| TGGGAGSGGT | ASSDMTAANSDYT | 0.7976358 |
| SHLETSL | ASSDMTAANSDYT | 0.5122986 |
| SFTTGMT | ASSDMTAANSDYT | 0.72587454 |
| QVEVAKRMGVRQPTVS | ASSDMTAANSDYT | 0.67507434 |
| QGSGGPALAM | ASSDMTAANSDYT | 0.5428275 |
| MLAPSNP | ASSDMTAANSDYT | 0.773309 |

| | | |
|---|---|---|
| LVNNAAI | ASSDMTAANSDYT | 0.69493663 |
| LTGEINTLA | ASSDMTAANSDYT | 0.3681548 |
| LSIFGPV | ASSDMTAANSDYT | 0.8164884 |
| LLALLANL | ASSDMTAANSDYT | 0.51353556 |
| LISNASDALDKLR | ASSDMTAANSDYT | 0.776162 |
| LINEAVP | ASSDMTAANSDYT | 0.86013335 |
| LGYAFSGL | ASSDMTAANSDYT | 0.4016818 |
| LDFVAVAL | ASSDMTAANSDYT | 0.7998991 |
| LAQPGAE | ASSDMTAANSDYT | 0.7430454 |
| ISNLGAVTGF | ASSDMTAANSDYT | 0.7044444 |
| HQEDPGRI | ASSDMTAANSDYT | 0.6926278 |
| GTPGEPTP | ASSDMTAANSDYT | 0.5673562 |
| GFFNSGS | ASSDMTAANSDYT | 0.835698 |
| GAGGGGLTAGAN | ASSDMTAANSDYT | 0.8028727 |
| FVDALVNM | ASSDMTAANSDYT | 0.8086164 |
| FLTVAPEL | ASSDMTAANSDYT | 0.8110271 |
| FETLTAL | ASSDMTAANSDYT | 0.81621146 |
| DVINPIEL | ASSDMTAANSDYT | 0.8284503 |
| APVSGAPV | ASSDMTAANSDYT | 0.5843079 |
| ALVGGLAW | ASSDMTAANSDYT | 0.36623335 |
| ALAAAAATAI | ASSDMTAANSDYT | 0.678154 |
| AFGLGLLL | ASSDMTAANSDYT | 0.6301796 |
| AFFAAASL | ASSDMTAANSDYT | 0.6368916 |
| AAAALPAFSPP | ASSDMTAANSDYT | 0.4352759 |
| VVLAPAL | ASSDMTAANSDYT | 0.821221 |
| TIGHVDHGKTTL | ASSDMTAANSDYT | 0.87724257 |
| LISNASDALDK | ASSDMTAANSDYT | 0.60627294 |
| LISNASDAL | ASSDMTAANSDYT | 0.8617728 |
| ISNASDALDK | ASSDMTAANSDYT | 0.45893574 |
| NQAETLVYQTE | ASSDMTAANSDYT | 0.79522717 |
| TGGTGGTAGAG | ASSDMTAANSDYT | 0.8376177 |
| TGSTAGGAGGA | ASSDMTAANSDYT | 0.5197601 |
| DPLAVPVAAGPV | ASSDMTAANSDYT | 0.802477 |
| GGTGGTPGNGGN | ASSDMTAANSDYT | 0.5122099 |
| PAGLAGVDPATL | ASSDMTAANSDYT | 0.5341219 |
| GNNLVVEAPHAGASV | ASSDMTAANSDYT | 0.8307539 |
| KVTRIMSASTLEQPA | ASSDMTAANSDYT | 0.67878973 |
| LGFALPIALPRRTVV | ASSDMTAANSDYT | 0.8053513 |
| TRLGNLTARKLVVLV | ASSDMTAANSDYT | 0.6842284 |
| WSQQRRLIYFPSAGP | ASSDMTAANSDYT | 0.5731158 |
| GPNGNPDPMAAAVDIR | ASSDMTAANSDYT | 0.80303514 |
| TQRCVSVADLRAAASR | ASSDMTAANSDYT | 0.83973 |
| QFVLDVAEHPVEVLTETV | ASSDMTAANSDYT | 0.7070228 |
| GGAGGTGGAGGNGGTQAGDG | ASSDMTAANSDYT | 0.8881918 |
| PLLATLRAIAVDVDAKPAQV | ASSDMTAANSDYT | 0.5960671 |
| APKFRTGGYMIDFWGVGYQVAK | ASSDMTAANSDYT | 0.858378 |
| GAGGAGGAGGAAGGPGGTGGTG | ASSDMTAANSDYT | 0.8532264 |
| IGALGLLATDVAPAGSVAFGVA | ASSDMTAANSDYT | 0.68543684 |
| AGGAGGIGGNGGAGGAGGVLMG | ASSDMTAANSDYT | 0.6920345 |
| ADELIGLL | ASSDMTAANSDYT | 0.78930557 |
| ADRALKA | ASSDMTAANSDYT | 0.6782533 |

| | | |
|---|---|---|
| AGAVGQVGGAGGA | ASSDMTAANSDYT | 0.77321315 |
| AGGGGGLGGAP | ASSDMTAANSDYT | 0.85477316 |
| AGGLLYGNGGAG | ASSDMTAANSDYT | 0.43498054 |
| AGLESIGL | ASSDMTAANSDYT | 0.77656865 |
| AGLKTGII | ASSDMTAANSDYT | 0.69600797 |
| ASLAALLL | ASSDMTAANSDYT | 0.71899277 |
| ASVINISSI | ASSDMTAANSDYT | 0.5337938 |
| ATPPGPA | ASSDMTAANSDYT | 0.43457127 |
| ATVASALV | ASSDMTAANSDYT | 0.6439465 |
| ATVASLIL | ASSDMTAANSDYT | 0.4759147 |
| AVNGPVLIRH | ASSDMTAANSDYT | 0.7540325 |
| DGAAGAVGI | ASSDMTAANSDYT | 0.75852853 |
| DPLLHGI | ASSDMTAANSDYT | 0.52853274 |
| ELGAALGAL | ASSDMTAANSDYT | 0.7318421 |
| ELGRTQAV | ASSDMTAANSDYT | 0.75237733 |
| FVAGGYLS | ASSDMTAANSDYT | 0.3792593 |
| GASGGAGGAGGW | ASSDMTAANSDYT | 0.62144935 |
| GGAGAGGDGGAGGI | ASSDMTAANSDYT | 0.6758616 |
| GTGHNGSQPGMGGNG | ASSDMTAANSDYT | 0.63178235 |
| GYAAASASAS | ASSDMTAANSDYT | 0.35877597 |
| HQVDVAAVQQG | ASSDMTAANSDYT | 0.72824645 |
| IGAAIAHT | ASSDMTAANSDYT | 0.51052797 |
| IGMIEVAQL | ASSDMTAANSDYT | 0.2692707 |
| IRLAPEHTSAMD | ASSDMTAANSDYT | 0.7037197 |
| IYNVSVL | ASSDMTAANSDYT | 0.5652412 |
| KQGFLGM | ASSDMTAANSDYT | 0.55832815 |
| LESVITQL | ASSDMTAANSDYT | 0.7519995 |
| LGLGALVVV | ASSDMTAANSDYT | 0.60456294 |
| LIGNGGNGGSGGI | ASSDMTAANSDYT | 0.49758613 |
| LNAINAPTL | ASSDMTAANSDYT | 0.69741267 |
| LQLLNTGV | ASSDMTAANSDYT | 0.76522034 |
| LQTSIEF | ASSDMTAANSDYT | 0.43575466 |
| MSFAVSEL | ASSDMTAANSDYT | 0.66944367 |
| NTGALGL | ASSDMTAANSDYT | 0.78429455 |
| PPGAGGAGGAGTT | ASSDMTAANSDYT | 0.6760507 |
| QLVGLGI | ASSDMTAANSDYT | 0.7995548 |
| RFLEQVHQL | ASSDMTAANSDYT | 0.8058714 |
| RGVGYVLR | ASSDMTAANSDYT | 0.5874881 |
| SATGVFAGVI | ASSDMTAANSDYT | 0.7663405 |
| SLPAITLP | ASSDMTAANSDYT | 0.7560232 |
| STALAFTV | ASSDMTAANSDYT | 0.6374834 |
| SVLTVGLGK | ASSDMTAANSDYT | 0.6379291 |
| TALAGAGSGI | ASSDMTAANSDYT | 0.42544118 |
| TGLPFTGI | ASSDMTAANSDYT | 0.6443714 |
| VAEHVTV | ASSDMTAANSDYT | 0.44492993 |
| VDIVLNSL | ASSDMTAANSDYT | 0.6706403 |
| VLTGGGAVL | ASSDMTAANSDYT | 0.78910804 |
| VNVGLAYGI | ASSDMTAANSDYT | 0.5966162 |
| VSSAAVGAV | ASSDMTAANSDYT | 0.3966785 |
| VTSVYEAGV | ASSDMTAANSDYT | 0.41600168 |
| WVALDRGA | ASSDMTAANSDYT | 0.703344 |

| | | |
|---|---|---|
| YLVGSVVI | ASSDMTAANSDYT | 0.4641394 |
| GPPGTGKTLL | ASSDMTAANSDYT | 0.4691018 |
| GVGMTKF | ASSDMTAANSDYT | 0.5903467 |
| IGIGGSDL | ASSDMTAANSDYT | 0.7492446 |
| IGTLLAEL | ASSDMTAANSDYT | 0.7555487 |
| IYTSGTT | ASSDMTAANSDYT | 0.49102575 |
| LSFPAGV | ASSDMTAANSDYT | 0.7747124 |
| LYTSGTT | ASSDMTAANSDYT | 0.7262173 |
| VGAFPFGF | ASSDMTAANSDYT | 0.7966752 |
| VGAGTVEF | ASSDMTAANSDYT | 0.6858763 |
| VGKSTLLL | ASSDMTAANSDYT | 0.4392487 |
| AALAHAGGGIV | ASSDMTAANSDYT | 0.73529696 |
| AALVAVSLL | ASSDMTAANSDYT | 0.54847085 |
| AAVSRAVAAL | ASSDMTAANSDYT | 0.7411481 |
| AGGSSGIGGF | ASSDMTAANSDYT | 0.7450646 |
| AGLTVANANG | ASSDMTAANSDYT | 0.5912091 |
| AGLVGAVL | ASSDMTAANSDYT | 0.5019673 |
| ALSQAFTV | ASSDMTAANSDYT | 0.61125934 |
| ALTFSVSL | ASSDMTAANSDYT | 0.8038665 |
| ALTGAALVAL | ASSDMTAANSDYT | 0.4390011 |
| ASFSIGAGVL | ASSDMTAANSDYT | 0.48311815 |
| DGSVYKI | ASSDMTAANSDYT | 0.6233541 |
| EQLAEIV | ASSDMTAANSDYT | 0.62782764 |
| FAVSELTVM | ASSDMTAANSDYT | 0.7715951 |
| GAAPLAAGL | ASSDMTAANSDYT | 0.737216 |
| GADGGDGGSGGAGG | ASSDMTAANSDYT | 0.64385957 |
| GFGLDTGGAGG | ASSDMTAANSDYT | 0.7053221 |
| GLASGGLL | ASSDMTAANSDYT | 0.75901985 |
| GRIAGAAVLL | ASSDMTAANSDYT | 0.6776429 |
| GSAGSLNTSF | ASSDMTAANSDYT | 0.69432235 |
| GTGQIGI | ASSDMTAANSDYT | 0.50011086 |
| GTLIALIL | ASSDMTAANSDYT | 0.79869384 |
| IAATMGFVVG | ASSDMTAANSDYT | 0.74588174 |
| IFAQALL | ASSDMTAANSDYT | 0.7597579 |
| IGAAVALGF | ASSDMTAANSDYT | 0.7839294 |
| IGLTPLAL | ASSDMTAANSDYT | 0.6958388 |
| ILGSNTSTL | ASSDMTAANSDYT | 0.705793 |
| IVFSLTVL | ASSDMTAANSDYT | 0.43312016 |
| LASKATGF | ASSDMTAANSDYT | 0.63032854 |
| LASVALIL | ASSDMTAANSDYT | 0.62024426 |
| LFAVGLTL | ASSDMTAANSDYT | 0.6912371 |
| LFNGGIGTYI | ASSDMTAANSDYT | 0.73284197 |
| LTLGAGII | ASSDMTAANSDYT | 0.74887663 |
| LVAAALGDI | ASSDMTAANSDYT | 0.5374392 |
| NTAFVQLGI | ASSDMTAANSDYT | 0.7782624 |
| QLFSGAEL | ASSDMTAANSDYT | 0.77934337 |
| SAATFITL | ASSDMTAANSDYT | 0.42469406 |
| SNYGLANLGV | ASSDMTAANSDYT | 0.61590254 |
| SYGGAYAVM | ASSDMTAANSDYT | 0.7884181 |
| TTGVGGLTLG | ASSDMTAANSDYT | 0.6856072 |
| TTQTIYI | ASSDMTAANSDYT | 0.58291346 |

| | | |
|---|---|---|
| VAAGAGLGAVI | ASSDMTAANSDYT | 0.6637716 |
| VAAVVTLL | ASSDMTAANSDYT | 0.7715911 |
| VGFLGGLI | ASSDMTAANSDYT | 0.697193 |
| VGVIPILL | ASSDMTAANSDYT | 0.78580964 |
| VIFASSLI | ASSDMTAANSDYT | 0.3205051 |
| VKGGDGGAAAG | ASSDMTAANSDYT | 0.6683241 |
| VTKTFTNI | ASSDMTAANSDYT | 0.61702466 |
| VVAAAILL | ASSDMTAANSDYT | 0.29234618 |
| YAALDVEL | ASSDMTAANSDYT | 0.75298685 |
| LASVALLL | ASSDMTAANSDYT | 0.5842416 |
| LDAVHRV | ASSDMTAANSDYT | 0.7394137 |
| QGLSSSIF | ASSDMTAANSDYT | 0.5962888 |
| RGTFANIRL | ASSDMTAANSDYT | 0.7687861 |
| RQLWWGHRI | ASSDMTAANSDYT | 0.6780096 |
| YGPPGTGKTLL | ASSDMTAANSDYT | 0.65441525 |
| AALVGAPANVA | ASSDMTAANSDYT | 0.5864355 |
| AATFRRTE | ASSDMTAANSDYT | 0.6974022 |
| AGALGGGI | ASSDMTAANSDYT | 0.7858561 |
| AGGAGGAGGIGAG | ASSDMTAANSDYT | 0.8015428 |
| AGSGALGL | ASSDMTAANSDYT | 0.34335807 |
| AQIADQI | ASSDMTAANSDYT | 0.71868134 |
| ASAVAVTYL | ASSDMTAANSDYT | 0.6352452 |
| ASTVLGGTGG | ASSDMTAANSDYT | 0.45573294 |
| AVGGEPAK | ASSDMTAANSDYT | 0.3504824 |
| CEQGTFGGTC | ASSDMTAANSDYT | 0.5639512 |
| DEGKAPAG | ASSDMTAANSDYT | 0.59829307 |
| DTVVFPAL | ASSDMTAANSDYT | 0.6811414 |
| FEGITMAG | ASSDMTAANSDYT | 0.72365654 |
| FPLVVQD | ASSDMTAANSDYT | 0.56527185 |
| GAFTTPQ | ASSDMTAANSDYT | 0.7265727 |
| GFGGFGTAGGVGGA | ASSDMTAANSDYT | 0.62649184 |
| GFLAQFL | ASSDMTAANSDYT | 0.15449008 |
| GGAGGAGGNAGVA | ASSDMTAANSDYT | 0.5872976 |
| GGVGTAGAGGAGGAGG | ASSDMTAANSDYT | 0.3619501 |
| GNGGGGDAGNAGSGGN | ASSDMTAANSDYT | 0.7994281 |
| GTGGAGGAGNDAGS | ASSDMTAANSDYT | 0.49563208 |
| GTTNTGWFNS | ASSDMTAANSDYT | 0.5621948 |
| IFMGPLFTL | ASSDMTAANSDYT | 0.6213778 |
| ILAHAIEAI | ASSDMTAANSDYT | 0.66681427 |
| IRSAIREAL | ASSDMTAANSDYT | 0.6334388 |
| ITFAGGG | ASSDMTAANSDYT | 0.5498209 |
| ITGMSGHQEV | ASSDMTAANSDYT | 0.3929466 |
| IVAGLACTL | ASSDMTAANSDYT | 0.72058636 |
| IVGGGLAGA | ASSDMTAANSDYT | 0.65806127 |
| IVTGVLNI | ASSDMTAANSDYT | 0.49287164 |
| LAQQGCI | ASSDMTAANSDYT | 0.50239795 |
| LCLHVEN | ASSDMTAANSDYT | 0.6928028 |
| LDAISDGS | ASSDMTAANSDYT | 0.35804477 |
| LGGFAAML | ASSDMTAANSDYT | 0.78012806 |
| LGGKNPMI | ASSDMTAANSDYT | 0.5047023 |
| LGPKGVAAIL | ASSDMTAANSDYT | 0.55635834 |

| | | |
|---|---|---|
| LMVGVSI | ASSDMTAANSDYT | 0.7843885 |
| LVGAAFVL | ASSDMTAANSDYT | 0.62770176 |
| LVLAVAPL | ASSDMTAANSDYT | 0.60426676 |
| MAAIAVIDAA | ASSDMTAANSDYT | 0.5553307 |
| NNGINAELQ | ASSDMTAANSDYT | 0.48098397 |
| RGEPLVLVHG | ASSDMTAANSDYT | 0.49831733 |
| RGGAGGAAGLI | ASSDMTAANSDYT | 0.50956887 |
| RSYRMHPAV | ASSDMTAANSDYT | 0.786354 |
| SAAHLLAGQLA | ASSDMTAANSDYT | 0.7120347 |
| SAAVGSY | ASSDMTAANSDYT | 0.46500134 |
| SPHAVLHRI | ASSDMTAANSDYT | 0.7559977 |
| SQIAKVL | ASSDMTAANSDYT | 0.78632915 |
| TALGHLVP | ASSDMTAANSDYT | 0.80423373 |
| TAPTTSGAA | ASSDMTAANSDYT | 0.48780864 |
| TDGDDGNGGSPGT | ASSDMTAANSDYT | 0.62118423 |
| TDGLGEN | ASSDMTAANSDYT | 0.6354646 |
| TGGPPLPGQA | ASSDMTAANSDYT | 0.59694755 |
| TIPAITI | ASSDMTAANSDYT | 0.59297466 |
| TIPGFTI | ASSDMTAANSDYT | 0.48964083 |
| TSIASGALA | ASSDMTAANSDYT | 0.7838201 |
| TVTLPGAAI | ASSDMTAANSDYT | 0.80856377 |
| VASGLSAALP | ASSDMTAANSDYT | 0.6453375 |
| VGAGVIGI | ASSDMTAANSDYT | 0.3766939 |
| VGPKRFGV | ASSDMTAANSDYT | 0.5878707 |
| VNVAFPD | ASSDMTAANSDYT | 0.45398325 |
| VSMTNSGVS | ASSDMTAANSDYT | 0.49392575 |
| VTDVLGDLSV | ASSDMTAANSDYT | 0.5982299 |
| VVVVGAGFAGL | ASSDMTAANSDYT | 0.35199714 |
| VYTVGTVV | ASSDMTAANSDYT | 0.6383121 |
| WTTFSGP | ASSDMTAANSDYT | 0.7900199 |
| YDELVEAF | ASSDMTAANSDYT | 0.53477377 |
| YGTGGAGGNGG | ASSDMTAANSDYT | 0.70802045 |
| YTNVMAV | ASSDMTAANSDYT | 0.7743598 |
| FLEQVHQL | ASSDMTAANSDYT | 0.7248142 |
| GLSLLAVAV | ASSDMTAANSDYT | 0.7728122 |
| LEQVHQL | ASSDMTAANSDYT | 0.7382541 |
| LNAAGAI | ASSDMTAANSDYT | 0.69893396 |
| PAVHASL | ASSDMTAANSDYT | 0.6801184 |
| TELEDDA | ASSDMTAANSDYT | 0.7291622 |
| TPPGPPN | ASSDMTAANSDYT | 0.5800374 |
| TPPPPPPP | ASSDMTAANSDYT | 0.64509875 |
| TQSPALI | ASGDALGGSEQY | 0.64745736 |
| EFAAAFPQ | ASGDALGGSEQY | 0.69242936 |
| LLSGNLDVL | ASGDALGGSEQY | 0.65501463 |
| GPANLGNYNI | ASGDALGGSEQY | 0.7898896 |
| TLGPTLP | ASGDALGGSEQY | 0.32338512 |
| VYAYSSCCK | ASGDALGGSEQY | 0.46496743 |
| APAPAPAPNI | ASGDALGGSEQY | 0.7656374 |
| PGGPVPIP | ASGDALGGSEQY | 0.24623819 |
| FLTVALEP | ASGDALGGSEQY | 0.43186966 |
| SQYFGANL | ASGDALGGSEQY | 0.68443125 |

| | | |
|---|---|---|
| GGVKPAA | ASGDALGGSEQY | 0.76803243 |
| GGTGGAGGAGGW | ASGDALGGSEQY | 0.26025862 |
| GVGGVAGLGGAGG | ASGDALGGSEQY | 0.57792586 |
| VLLVLATTI | ASGDALGGSEQY | 0.40538824 |
| IGIFAAML | ASGDALGGSEQY | 0.7839508 |
| AVGDLID | ASGDALGGSEQY | 0.58765864 |
| NISPLPL | ASGDALGGSEQY | 0.42547303 |
| VTAVFVAI | ASGDALGGSEQY | 0.7498107 |
| SGDDNDAAL | ASGDALGGSEQY | 0.4679702 |
| AVAALAHS | ASGDALGGSEQY | 0.70175713 |
| GGVTVMATP | ASGDALGGSEQY | 0.7623763 |
| AGLAANVRD | ASGDALGGSEQY | 0.79383063 |
| TNRPDLIDPALLRPG | ASGDALGGSEQY | 0.4566172 |
| LANLGVTAL | ASGDALGGSEQY | 0.7211704 |
| TGERIEL | ASGDALGGSEQY | 0.5776936 |
| NGEELAA | ASGDALGGSEQY | 0.8237191 |
| VLSLVWG | ASGDALGGSEQY | 0.56709015 |
| IGYALWL | ASGDALGGSEQY | 0.42967132 |
| LSLLLAPL | ASGDALGGSEQY | 0.7491579 |
| TLDEALK | ASGDALGGSEQY | 0.7522755 |
| TDGAGTAEGG | ASGDALGGSEQY | 0.63994104 |
| NAPGGAGGNGG | ASGDALGGSEQY | 0.7231059 |
| VSVFMGMGGNF | ASGDALGGSEQY | 0.6682772 |
| LASYLAAL | ASGDALGGSEQY | 0.5889896 |
| LVGGTPAL | ASGDALGGSEQY | 0.7303222 |
| VRNADVL | ASGDALGGSEQY | 0.71409816 |
| TCSSPFLVSAM | ASGDALGGSEQY | 0.5291549 |
| VHITDAE | ASGDALGGSEQY | 0.78201306 |
| EYAAPSF | ASGDALGGSEQY | 0.63720804 |
| VVDAFHGGL | ASGDALGGSEQY | 0.4794955 |
| ASVVDPQ | ASGDALGGSEQY | 0.7999439 |
| SYNEFGGLS | ASGDALGGSEQY | 0.66781276 |
| DPDGSVY | ASGDALGGSEQY | 0.74208325 |
| MVSTIHATE | ASGDALGGSEQY | 0.73684007 |
| LAAGVADPV | ASGDALGGSEQY | 0.7824705 |
| TKEQIAA | ASGDALGGSEQY | 0.5442395 |
| SLASPAAL | ASGDALGGSEQY | 0.7076044 |
| IGLVFYGV | ASGDALGGSEQY | 0.65071523 |
| FLLDASNL | ASGDALGGSEQY | 0.7559611 |
| YQVGNFP | ASGDALGGSEQY | 0.66535836 |
| ESLLSQST | ASGDALGGSEQY | 0.667721 |
| LFTVGFAAVG | ASGDALGGSEQY | 0.7434664 |
| ATPPTPT | ASGDALGGSEQY | 0.778304 |
| TDLATLG | ASGDALGGSEQY | 0.6968611 |
| SAGGTGR | ASGDALGGSEQY | 0.6612773 |
| GFSNSGS | ASGDALGGSEQY | 0.7846035 |
| GGNGGSGGTSV | ASGDALGGSEQY | 0.50240463 |
| GTSVATGGAGN | ASGDALGGSEQY | 0.6453582 |
| NGGNGGSGGTS | ASGDALGGSEQY | 0.7640521 |
| AVVKFPH | ASGDALGGSEQY | 0.7696773 |
| TPPPPPP | ASGDALGGSEQY | 0.667161 |

| | | |
|---|---|---|
| AAPAAAAAPA | ASGDALGGSEQY | 0.782773 |
| IALNEDL | ASGDALGGSEQY | 0.60622454 |
| IVNVSSM | ASGDALGGSEQY | 0.6874465 |
| TGIPPAPR | ASGDALGGSEQY | 0.5550332 |
| FDLGGGTF | ASGDALGGSEQY | 0.61368537 |
| FDLGGGTFDVS | ASGDALGGSEQY | 0.4384897 |
| NEPTAAA | ASGDALGGSEQY | 0.610883 |
| NEPTAAALA | ASGDALGGSEQY | 0.7416099 |
| GSIINISSI | ASGDALGGSEQY | 0.67895234 |
| VAPGNAGTA | ASGDALGGSEQY | 0.37661698 |
| TGGGASLE | ASGDALGGSEQY | 0.70946383 |
| VAVYNHYKRI | ASGDALGGSEQY | 0.7158637 |
| TDPAPATT | ASGDALGGSEQY | 0.7932856 |
| TNRPDILDPALLRPG | ASGDALGGSEQY | 0.6605725 |
| LTTTTTS | ASGDALGGSEQY | 0.7499875 |
| PTGSGKTL | ASGDALGGSEQY | 0.5901524 |
| TGSGKTL | ASGDALGGSEQY | 0.70800966 |
| TIGHVDHGK | ASGDALGGSEQY | 0.37758458 |
| TIGHVDHGKTT | ASGDALGGSEQY | 0.5365814 |
| GHVDHGKTT | ASGDALGGSEQY | 0.60996336 |
| AQFEHTLL | ASGDALGGSEQY | 0.47090182 |
| IVNQDEL | ASGDALGGSEQY | 0.7050375 |
| TNGSQFF | ASGDALGGSEQY | 0.756507 |
| TGKGIVE | ASGDALGGSEQY | 0.73072696 |
| LISNASDA | ASGDALGGSEQY | 0.7666174 |
| AELRTLGELTKIAKAHGAQ | ASGDALGGSEQY | 0.77545357 |
| APNAVAAAEPSW | ASGDALGGSEQY | 0.78601146 |
| APQVAGAATAVL | ASGDALGGSEQY | 0.5215138 |
| ARRDPAAAAAILPTDA | ASGDALGGSEQY | 0.648295 |
| DAAARDVLAEAGLAEYF | ASGDALGGSEQY | 0.36269578 |
| DDQIVLSAKANLAKVEQ | ASGDALGGSEQY | 0.5840355 |
| DELLPDTGEDSDEDGH | ASGDALGGSEQY | 0.54914284 |
| ELPEAPANRATVPQD | ASGDALGGSEQY | 0.6591426 |
| FGQSKIERVTGIG | ASGDALGGSEQY | 0.21263957 |
| GAGGTGSTAGGAGGAGGAGGLY | ASGDALGGSEQY | 0.6454663 |
| GHGGTGAAGGNGGTGGAGGSGI | ASGDALGGSEQY | 0.78935784 |
| GNGGNGGAGGEGGPGGAGGAGA | ASGDALGGSEQY | 0.62033767 |
| GSTAGNGGNGGNATNGTIAGQP | ASGDALGGSEQY | 0.70806813 |
| GTGGDGGNGGNGGTGG | ASGDALGGSEQY | 0.71812063 |
| GVRRPAGIPADAPV | ASGDALGGSEQY | 0.56041163 |
| GYRQVTDAHLLTLARR | ASGDALGGSEQY | 0.717667 |
| IAKMGQDGHDRG | ASGDALGGSEQY | 0.5521805 |
| KAEPSIEIQYT | ASGDALGGSEQY | 0.7676166 |
| KPVRIRMAADEDTVAVT | ASGDALGGSEQY | 0.77591705 |
| MPPRIAGMRLLVIKPEP | ASGDALGGSEQY | 0.6916185 |
| MKFYAVALFRSRGVPPTR | ASGDALGGSEQY | 0.6046986 |
| NAGGFGILGAVAHSP | ASGDALGGSEQY | 0.7122371 |
| NEGREITLAMAEM | ASGDALGGSEQY | 0.5083847 |
| NELVEWSHPLGHRWRWE | ASGDALGGSEQY | 0.64689386 |
| NLGAGGTGGNGGNPG | ASGDALGGSEQY | 0.52404684 |
| NTEWPKWETVMAADPVD | ASGDALGGSEQY | 0.6352521 |

| | | |
|---|---|---|
| PASAAPAETQP | ASGDALGGSEQY | 0.5227962 |
| QQSSDGKNGVRAAEITGE | ASGDALGGSEQY | 0.6862159 |
| RAPDAVTTSWWKE | ASGDALGGSEQY | 0.64206797 |
| SGFGNVGAHTSGWFNQ | ASGDALGGSEQY | 0.721432 |
| SGVAGAVGQAASVGGLKV | ASGDALGGSEQY | 0.43808675 |
| SNGGAGTGSAGGNGGT | ASGDALGGSEQY | 0.7581041 |
| TGGAGGEGGAAG | ASGDALGGSEQY | 0.7083078 |
| TGGDGGAAGAGGAPGHG | ASGDALGGSEQY | 0.40926883 |
| TMPPLLAAIANAGRHGV | ASGDALGGSEQY | 0.76697755 |
| TSSTAGNSGGA | ASGDALGGSEQY | 0.6736361 |
| VPATVPVGILC | ASGDALGGSEQY | 0.5778897 |
| VQRPDSGGVVLTGRIS | ASGDALGGSEQY | 0.75808156 |
| WTPNPVPATKGVPAGLVQAT | ASGDALGGSEQY | 0.641891 |
| YALKRALAKFSDRFDVVII | ASGDALGGSEQY | 0.52577555 |
| YLIGSVIV | ASGDALGGSEQY | 0.5355009 |
| YGQLVAVID | ASGDALGGSEQY | 0.66192937 |
| VTVGGMTLPA | ASGDALGGSEQY | 0.5959387 |
| VLALPAV | ASGDALGGSEQY | 0.33574915 |
| VIPPTLNYET | ASGDALGGSEQY | 0.7356745 |
| VGMGGMHPSSG | ASGDALGGSEQY | 0.4846198 |
| VGLAGGAATA | ASGDALGGSEQY | 0.5141425 |
| VGGAGGAGGSGP | ASGDALGGSEQY | 0.57330245 |
| VFAAFMV | ASGDALGGSEQY | 0.74908423 |
| TLYDKLR | ASGDALGGSEQY | 0.68587554 |
| TLAVFTVL | ASGDALGGSEQY | 0.61103505 |
| TGSVQPARV | ASGDALGGSEQY | 0.6784393 |
| TGGGAGSGGT | ASGDALGGSEQY | 0.6772414 |
| SHLETSL | ASGDALGGSEQY | 0.7658783 |
| SFTTGMT | ASGDALGGSEQY | 0.6836984 |
| QVEVAKRMGVRQPTVS | ASGDALGGSEQY | 0.70079076 |
| QGSGGPALAM | ASGDALGGSEQY | 0.7156305 |
| MLAPSNP | ASGDALGGSEQY | 0.66695654 |
| LVNNAAI | ASGDALGGSEQY | 0.5609733 |
| LTGEINTLA | ASGDALGGSEQY | 0.5392801 |
| LSIFGPV | ASGDALGGSEQY | 0.5969522 |
| LLALLANL | ASGDALGGSEQY | 0.52690136 |
| LISNASDALDKLR | ASGDALGGSEQY | 0.668977 |
| LINEAVP | ASGDALGGSEQY | 0.80433124 |
| LGYAFSGL | ASGDALGGSEQY | 0.6873433 |
| LDFVAVAL | ASGDALGGSEQY | 0.4043041 |
| LAQPGAE | ASGDALGGSEQY | 0.584695 |
| ISNLGAVTGF | ASGDALGGSEQY | 0.6458895 |
| HQEDPGRI | ASGDALGGSEQY | 0.7388681 |
| GTPGEPTP | ASGDALGGSEQY | 0.7587775 |
| GFFNSGS | ASGDALGGSEQY | 0.7878514 |
| GAGGGGLTAGAN | ASGDALGGSEQY | 0.73520416 |
| FVDALVNM | ASGDALGGSEQY | 0.518938 |
| FLTVAPEL | ASGDALGGSEQY | 0.73842704 |
| FETLTAL | ASGDALGGSEQY | 0.6121429 |
| DVINPIEL | ASGDALGGSEQY | 0.6894221 |
| APVSGAPV | ASGDALGGSEQY | 0.6909763 |

| | | |
|---|---|---|
| ALVGGLAW | ASGDALGGSEQY | 0.70277506 |
| ALAAAAATAI | ASGDALGGSEQY | 0.59786487 |
| AFGLGLLL | ASGDALGGSEQY | 0.6338403 |
| AFFAAASL | ASGDALGGSEQY | 0.43517047 |
| AAAALPAFSPP | ASGDALGGSEQY | 0.73358625 |
| VVLAPAL | ASGDALGGSEQY | 0.5287434 |
| TIGHVDHGKTTL | ASGDALGGSEQY | 0.5579792 |
| LISNASDALDK | ASGDALGGSEQY | 0.5746297 |
| LISNASDAL | ASGDALGGSEQY | 0.78123707 |
| ISNASDALDK | ASGDALGGSEQY | 0.784892 |
| NQAETLVYQTE | ASGDALGGSEQY | 0.5387323 |
| TGGTGGTAGAG | ASGDALGGSEQY | 0.52218586 |
| TGSTAGGAGGA | ASGDALGGSEQY | 0.68007994 |
| DPLAVPVAAGPV | ASGDALGGSEQY | 0.50879085 |
| GGTGGTPGNGGN | ASGDALGGSEQY | 0.7589762 |
| PAGLAGVDPATL | ASGDALGGSEQY | 0.6115008 |
| GNNLVVEAPHAGASV | ASGDALGGSEQY | 0.63235897 |
| KVTRIMSASTLEQPA | ASGDALGGSEQY | 0.5926284 |
| LGFALPIALPRRTVV | ASGDALGGSEQY | 0.7330532 |
| TRLGNLTARKLVVLV | ASGDALGGSEQY | 0.5790205 |
| WSQQRRLIYFPSAGP | ASGDALGGSEQY | 0.75044227 |
| GPNGNPDPMAAAVDIR | ASGDALGGSEQY | 0.31297857 |
| TQRCVSVADLRAAASR | ASGDALGGSEQY | 0.7658264 |
| QFVLDVAEHPVEVLTETV | ASGDALGGSEQY | 0.4242704 |
| GGAGGTGGAGGNGGTQAGDG | ASGDALGGSEQY | 0.5532776 |
| PLLATLRAIAVDVDAKPAQV | ASGDALGGSEQY | 0.32809067 |
| APKFRTGGYMIDFWGVGYQVAK | ASGDALGGSEQY | 0.6766411 |
| GAGGAGGAGGAAGGPGGTGGTG | ASGDALGGSEQY | 0.7363277 |
| IGALGLLATDVAPAGSVAFGVA | ASGDALGGSEQY | 0.77653694 |
| AGGAGGIGGNGGAGGAGGVLMG | ASGDALGGSEQY | 0.6988454 |
| ADELIGLL | ASGDALGGSEQY | 0.709491 |
| ADRALKA | ASGDALGGSEQY | 0.7881626 |
| AGAVGQVGGAGGA | ASGDALGGSEQY | 0.5559067 |
| AGGGGGLGGAP | ASGDALGGSEQY | 0.36410862 |
| AGGLLYGNGGAG | ASGDALGGSEQY | 0.671456 |
| AGLESIGL | ASGDALGGSEQY | 0.62224025 |
| AGLKTGII | ASGDALGGSEQY | 0.7707678 |
| ASLAALLL | ASGDALGGSEQY | 0.7218914 |
| ASVINISSI | ASGDALGGSEQY | 0.35348865 |
| ATPPGPA | ASGDALGGSEQY | 0.72759765 |
| ATVASALV | ASGDALGGSEQY | 0.73082846 |
| ATVASLIL | ASGDALGGSEQY | 0.42030483 |
| AVNGPVLIRH | ASGDALGGSEQY | 0.5619381 |
| DGAAGAVGI | ASGDALGGSEQY | 0.5986786 |
| DPLLHGI | ASGDALGGSEQY | 0.44446713 |
| ELGAALGAL | ASGDALGGSEQY | 0.68600595 |
| ELGRTQAV | ASGDALGGSEQY | 0.5463971 |
| FVAGGYLS | ASGDALGGSEQY | 0.57568824 |
| GASGGAGGAGGW | ASGDALGGSEQY | 0.63918704 |
| GGAGAGGDGGAGGI | ASGDALGGSEQY | 0.5177543 |
| GTGHNGSQPGMGGNG | ASGDALGGSEQY | 0.5855552 |

| | | |
|---|---|---|
| GYAAASASAS | ASGDALGGSEQY | 0.57260734 |
| HQVDVAAVQQG | ASGDALGGSEQY | 0.7372132 |
| IGAAIAHT | ASGDALGGSEQY | 0.74893516 |
| IGMIEVAQL | ASGDALGGSEQY | 0.6166156 |
| IRLAPEHTSAMD | ASGDALGGSEQY | 0.70113945 |
| IYNVSVL | ASGDALGGSEQY | 0.76431334 |
| KQGFLGM | ASGDALGGSEQY | 0.41332507 |
| LESVITQL | ASGDALGGSEQY | 0.5206584 |
| LGLGALVVV | ASGDALGGSEQY | 0.7281039 |
| LIGNGGNGGSGGI | ASGDALGGSEQY | 0.6008711 |
| LNAINAPTL | ASGDALGGSEQY | 0.7730458 |
| LQLLNTGV | ASGDALGGSEQY | 0.6596739 |
| LQTSIEF | ASGDALGGSEQY | 0.6809398 |
| MSFAVSEL | ASGDALGGSEQY | 0.64851147 |
| NTGALGL | ASGDALGGSEQY | 0.8015287 |
| PPGAGGAGGAGTT | ASGDALGGSEQY | 0.7130771 |
| QLVGLGI | ASGDALGGSEQY | 0.646467 |
| RFLEQVHQL | ASGDALGGSEQY | 0.7236401 |
| RGVGYVLR | ASGDALGGSEQY | 0.4923213 |
| SATGVFAGVI | ASGDALGGSEQY | 0.7458744 |
| SLPAITLP | ASGDALGGSEQY | 0.7443976 |
| STALAFTV | ASGDALGGSEQY | 0.43707982 |
| SVLTVGLGK | ASGDALGGSEQY | 0.5985385 |
| TALAGAGSGI | ASGDALGGSEQY | 0.803121 |
| TGLPFTGI | ASGDALGGSEQY | 0.76915014 |
| VAEHVTV | ASGDALGGSEQY | 0.6054218 |
| VDIVLNSL | ASGDALGGSEQY | 0.54060626 |
| VLTGGGAVL | ASGDALGGSEQY | 0.5500045 |
| VNVGLAYGI | ASGDALGGSEQY | 0.64205897 |
| VSSAAVGAV | ASGDALGGSEQY | 0.4268881 |
| VTSVYEAGV | ASGDALGGSEQY | 0.71915674 |
| WVALDRGA | ASGDALGGSEQY | 0.564281 |
| YLVGSVVI | ASGDALGGSEQY | 0.7318858 |
| GPPGTGKTLL | ASGDALGGSEQY | 0.6710958 |
| GVGMTKF | ASGDALGGSEQY | 0.75953114 |
| IGIGGSDL | ASGDALGGSEQY | 0.5784209 |
| IGTLLAEL | ASGDALGGSEQY | 0.5837494 |
| IYTSGTT | ASGDALGGSEQY | 0.3894866 |
| LSFPAGV | ASGDALGGSEQY | 0.5340216 |
| LYTSGTT | ASGDALGGSEQY | 0.59092206 |
| VGAFPFGF | ASGDALGGSEQY | 0.5986293 |
| VGAGTVEF | ASGDALGGSEQY | 0.40513432 |
| VGKSTLLL | ASGDALGGSEQY | 0.7951287 |
| AALAHAGGGIV | ASGDALGGSEQY | 0.58776987 |
| AALVAVSLL | ASGDALGGSEQY | 0.72672266 |
| AAVSRAVAAL | ASGDALGGSEQY | 0.5715124 |
| AGGSSGIGGF | ASGDALGGSEQY | 0.49815664 |
| AGLTVANANG | ASGDALGGSEQY | 0.5343424 |
| AGLVGAVL | ASGDALGGSEQY | 0.7269601 |
| ALSQAFTV | ASGDALGGSEQY | 0.4552583 |
| ALTFSVSL | ASGDALGGSEQY | 0.35989723 |

| | | |
|---|---|---|
| ALTGAALVAL | ASGDALGGSEQY | 0.5762337 |
| ASFSIGAGVL | ASGDALGGSEQY | 0.64455986 |
| DGSVYKI | ASGDALGGSEQY | 0.7921488 |
| EQLAEIV | ASGDALGGSEQY | 0.77058804 |
| FAVSELTVM | ASGDALGGSEQY | 0.6652008 |
| GAAPLAAGL | ASGDALGGSEQY | 0.8014916 |
| GADGGDGGSGGAGG | ASGDALGGSEQY | 0.23926546 |
| GFGLDTGGAGG | ASGDALGGSEQY | 0.59802914 |
| GLASGGLL | ASGDALGGSEQY | 0.7744924 |
| GRIAGAAVLL | ASGDALGGSEQY | 0.71181184 |
| GSAGSLNTSF | ASGDALGGSEQY | 0.6027777 |
| GTGQIGI | ASGDALGGSEQY | 0.7666379 |
| GTLIALIL | ASGDALGGSEQY | 0.45653418 |
| IAATMGFVVG | ASGDALGGSEQY | 0.59815794 |
| IFAQALL | ASGDALGGSEQY | 0.6392262 |
| IGAAVALGF | ASGDALGGSEQY | 0.76521075 |
| IGLTPLAL | ASGDALGGSEQY | 0.7395836 |
| ILGSNTSTL | ASGDALGGSEQY | 0.41422778 |
| IVFSLTVL | ASGDALGGSEQY | 0.7342167 |
| LASKATGF | ASGDALGGSEQY | 0.80093586 |
| LASVALIL | ASGDALGGSEQY | 0.50335675 |
| LFAVGLTL | ASGDALGGSEQY | 0.76625144 |
| LFNGGIGTYI | ASGDALGGSEQY | 0.6556337 |
| LTLGAGII | ASGDALGGSEQY | 0.6197753 |
| LVAAALGDI | ASGDALGGSEQY | 0.48789763 |
| NTAFVQLGI | ASGDALGGSEQY | 0.43398803 |
| QLFSGAEL | ASGDALGGSEQY | 0.7605385 |
| SAATFITL | ASGDALGGSEQY | 0.50183976 |
| SNYGLANLGV | ASGDALGGSEQY | 0.6779343 |
| SYGGAYAVM | ASGDALGGSEQY | 0.6501678 |
| TTGVGGLTLG | ASGDALGGSEQY | 0.8118949 |
| TTQTIYI | ASGDALGGSEQY | 0.70629805 |
| VAAGAGLGAVI | ASGDALGGSEQY | 0.720924 |
| VAAVVTLL | ASGDALGGSEQY | 0.487733 |
| VGFLGGLI | ASGDALGGSEQY | 0.7229861 |
| VGVIPILL | ASGDALGGSEQY | 0.7059841 |
| VIFASSLI | ASGDALGGSEQY | 0.48220903 |
| VKGGDGGAAAG | ASGDALGGSEQY | 0.7710138 |
| VTKTFTNI | ASGDALGGSEQY | 0.45191985 |
| VVAAAILL | ASGDALGGSEQY | 0.65500414 |
| YAALDVEL | ASGDALGGSEQY | 0.7399481 |
| LASVALLL | ASGDALGGSEQY | 0.75245833 |
| LDAVHRV | ASGDALGGSEQY | 0.65059936 |
| QGLSSSIF | ASGDALGGSEQY | 0.7580414 |
| RGTFANIRL | ASGDALGGSEQY | 0.47400382 |
| RQLWWGHRI | ASGDALGGSEQY | 0.75630176 |
| YGPPGTGKTLL | ASGDALGGSEQY | 0.75049376 |
| AALVGAPANVA | ASGDALGGSEQY | 0.68437743 |
| AATFRRTE | ASGDALGGSEQY | 0.80497116 |
| AGALGGGI | ASGDALGGSEQY | 0.7558561 |
| AGGAGGAGGIGAG | ASGDALGGSEQY | 0.398091 |

| | | |
|---|---|---|
| AGSGALGL | ASGDALGGSEQY | 0.6820971 |
| AQIADQI | ASGDALGGSEQY | 0.7657853 |
| ASAVAVTYL | ASGDALGGSEQY | 0.77982914 |
| ASTVLGGTGG | ASGDALGGSEQY | 0.76386505 |
| AVGGEPAK | ASGDALGGSEQY | 0.7313793 |
| CEQGTFGGTC | ASGDALGGSEQY | 0.59332687 |
| DEGKAPAG | ASGDALGGSEQY | 0.7886099 |
| DTVVFPAL | ASGDALGGSEQY | 0.83171624 |
| FEGITMAG | ASGDALGGSEQY | 0.71280515 |
| FPLVVQD | ASGDALGGSEQY | 0.5438321 |
| GAFTTPQ | ASGDALGGSEQY | 0.5644275 |
| GFGGFGTAGGVGGA | ASGDALGGSEQY | 0.7310864 |
| GFLAQFL | ASGDALGGSEQY | 0.74883425 |
| GGAGGAGGNAGVA | ASGDALGGSEQY | 0.47940782 |
| GGVGTAGAGGAGGAGG | ASGDALGGSEQY | 0.63972914 |
| GNGGGGDAGNAGSGGN | ASGDALGGSEQY | 0.6599261 |
| GTGGAGGAGNDAGS | ASGDALGGSEQY | 0.2998283 |
| GTTNTGWFNS | ASGDALGGSEQY | 0.74114823 |
| IFMGPLFTL | ASGDALGGSEQY | 0.7719766 |
| ILAHAIEAI | ASGDALGGSEQY | 0.73818505 |
| IRSAIREAL | ASGDALGGSEQY | 0.46375352 |
| ITFAGGG | ASGDALGGSEQY | 0.7645062 |
| ITGMSGHQEV | ASGDALGGSEQY | 0.7598173 |
| IVAGLACTL | ASGDALGGSEQY | 0.8026855 |
| IVGGGLAGA | ASGDALGGSEQY | 0.38790303 |
| IVTGVLNI | ASGDALGGSEQY | 0.2505282 |
| LAQQGCI | ASGDALGGSEQY | 0.69225264 |
| LCLHVEN | ASGDALGGSEQY | 0.7057644 |
| LDAISDGS | ASGDALGGSEQY | 0.5570492 |
| LGGFAAML | ASGDALGGSEQY | 0.46082526 |
| LGGKNPMI | ASGDALGGSEQY | 0.6762259 |
| LGPKGVAAIL | ASGDALGGSEQY | 0.7605762 |
| LMVGVSI | ASGDALGGSEQY | 0.4753418 |
| LVGAAFVL | ASGDALGGSEQY | 0.4908442 |
| LVLAVAPL | ASGDALGGSEQY | 0.74083894 |
| MAAIAVIDAA | ASGDALGGSEQY | 0.7602141 |
| NNGINAELQ | ASGDALGGSEQY | 0.46466652 |
| RGEPLVLVHG | ASGDALGGSEQY | 0.5626804 |
| RGGAGGAAGLI | ASGDALGGSEQY | 0.6886733 |
| RSYRMHPAV | ASGDALGGSEQY | 0.70715904 |
| SAAHLLAGQLA | ASGDALGGSEQY | 0.6648787 |
| SAAVGSY | ASGDALGGSEQY | 0.43098542 |
| SPHAVLHRI | ASGDALGGSEQY | 0.7346799 |
| SQIAKVL | ASGDALGGSEQY | 0.42957026 |
| TALGHLVP | ASGDALGGSEQY | 0.76785815 |
| TAPTTSGAA | ASGDALGGSEQY | 0.6266222 |
| TDGDDGNGGSPGT | ASGDALGGSEQY | 0.7099918 |
| TDGLGEN | ASGDALGGSEQY | 0.57016563 |
| TGGPPLPGQA | ASGDALGGSEQY | 0.43870682 |
| TIPAITI | ASGDALGGSEQY | 0.48238564 |
| TIPGFTI | ASGDALGGSEQY | 0.46197867 |

| | | |
|---|---|---|
| TSIASGALA | ASGDALGGSEQY | 0.76395696 |
| TVTLPGAAI | ASGDALGGSEQY | 0.68252516 |
| VASGLSAALP | ASGDALGGSEQY | 0.7364987 |
| VGAGVIGI | ASGDALGGSEQY | 0.6654525 |
| VGPKRFGV | ASGDALGGSEQY | 0.52076316 |
| VNVAFPD | ASGDALGGSEQY | 0.74041635 |
| VSMTNSGVS | ASGDALGGSEQY | 0.38193637 |
| VTDVLGDLSV | ASGDALGGSEQY | 0.7261951 |
| VVVVGAGFAGL | ASGDALGGSEQY | 0.6670135 |
| VYTVGTVV | ASGDALGGSEQY | 0.7394524 |
| WTTFSGP | ASGDALGGSEQY | 0.5347543 |
| YDELVEAF | ASGDALGGSEQY | 0.74939847 |
| YGTGGAGGNGG | ASGDALGGSEQY | 0.7100868 |
| YTNVMAV | ASGDALGGSEQY | 0.53778154 |
| FLEQVHQL | ASGDALGGSEQY | 0.72402334 |
| GLSLLAVAV | ASGDALGGSEQY | 0.7424793 |
| LEQVHQL | ASGDALGGSEQY | 0.7427802 |
| LNAAGAI | ASGDALGGSEQY | 0.37275025 |
| PAVHASL | ASGDALGGSEQY | 0.66600245 |
| TELEDDA | ASGDALGGSEQY | 0.5056443 |
| TPPGPPN | ASGDALGGSEQY | 0.67654014 |
| TPPPPPPP | ASGDALGGSEQY | 0.6805785 |
| TQSPALI | TCSADGGDQDTQY | 0.58604056 |
| EFAAAFPQ | TCSADGGDQDTQY | 0.7098255 |
| LLSGNLDVL | TCSADGGDQDTQY | 0.5444516 |
| GPANLGNYNI | TCSADGGDQDTQY | 0.5309812 |
| TLGPTLP | TCSADGGDQDTQY | 0.59317243 |
| VYAYSSCCK | TCSADGGDQDTQY | 0.67568386 |
| APAPAPAPNI | TCSADGGDQDTQY | 0.7230925 |
| PGGPVPIP | TCSADGGDQDTQY | 0.41485798 |
| FLTVALEP | TCSADGGDQDTQY | 0.6883458 |
| SQYFGANL | TCSADGGDQDTQY | 0.61257565 |
| GGVKPAA | TCSADGGDQDTQY | 0.45058736 |
| GGTGGAGGAGGW | TCSADGGDQDTQY | 0.7063155 |
| GVGGVAGLGGAGG | TCSADGGDQDTQY | 0.6501996 |
| VLLVLATTI | TCSADGGDQDTQY | 0.40298176 |
| IGIFAAML | TCSADGGDQDTQY | 0.74653256 |
| AVGDLID | TCSADGGDQDTQY | 0.5020807 |
| NISPLPL | TCSADGGDQDTQY | 0.70900947 |
| VTAVFVAI | TCSADGGDQDTQY | 0.80229545 |
| SGDDNDAAL | TCSADGGDQDTQY | 0.39221257 |
| AVAALAHS | TCSADGGDQDTQY | 0.7641241 |
| GGVTVMATP | TCSADGGDQDTQY | 0.6434363 |
| AGLAANVRD | TCSADGGDQDTQY | 0.6545073 |
| TNRPDLIDPALLRPG | TCSADGGDQDTQY | 0.6079791 |
| LANLGVTAL | TCSADGGDQDTQY | 0.5742521 |
| TGERIEL | TCSADGGDQDTQY | 0.77186006 |
| NGEELAA | TCSADGGDQDTQY | 0.7119223 |
| VLSLVWG | TCSADGGDQDTQY | 0.7209809 |
| IGYALWL | TCSADGGDQDTQY | 0.70757145 |
| LSLLLAPL | TCSADGGDQDTQY | 0.7281367 |

| | | |
|---|---|---|
| TLDEALK | TCSADGGDQDTQY | 0.754259 |
| TDGAGTAEGG | TCSADGGDQDTQY | 0.48940223 |
| NAPGGAGGNGG | TCSADGGDQDTQY | 0.34388685 |
| VSVFMGMGGNF | TCSADGGDQDTQY | 0.5995367 |
| LASYLAAL | TCSADGGDQDTQY | 0.57990986 |
| LVGGTPAL | TCSADGGDQDTQY | 0.579943 |
| VRNADVL | TCSADGGDQDTQY | 0.4641031 |
| TCSSPFLVSAM | TCSADGGDQDTQY | 0.7314296 |
| VHITDAE | TCSADGGDQDTQY | 0.8097422 |
| EYAAPSF | TCSADGGDQDTQY | 0.5345516 |
| VVDAFHGGL | TCSADGGDQDTQY | 0.8100219 |
| ASVVDPQ | TCSADGGDQDTQY | 0.4499734 |
| SYNEFGGLS | TCSADGGDQDTQY | 0.7207677 |
| DPDGSVY | TCSADGGDQDTQY | 0.7691811 |
| MVSTIHATE | TCSADGGDQDTQY | 0.42662823 |
| LAAGVADPV | TCSADGGDQDTQY | 0.69927096 |
| TKEQIAA | TCSADGGDQDTQY | 0.5885704 |
| SLASPAAL | TCSADGGDQDTQY | 0.5659214 |
| IGLVFYGV | TCSADGGDQDTQY | 0.7546685 |
| FLLDASNL | TCSADGGDQDTQY | 0.5596971 |
| YQVGNFP | TCSADGGDQDTQY | 0.73656285 |
| ESLLSQST | TCSADGGDQDTQY | 0.5840687 |
| LFTVGFAAVG | TCSADGGDQDTQY | 0.5057527 |
| ATPPTPT | TCSADGGDQDTQY | 0.720382 |
| TDLATLG | TCSADGGDQDTQY | 0.74795467 |
| SAGGTGR | TCSADGGDQDTQY | 0.69380677 |
| GFSNSGS | TCSADGGDQDTQY | 0.82836956 |
| GGNGGSGGTSV | TCSADGGDQDTQY | 0.47881255 |
| GTSVATGGAGN | TCSADGGDQDTQY | 0.7905579 |
| NGGNGGSGGTS | TCSADGGDQDTQY | 0.7621575 |
| AVVKFPH | TCSADGGDQDTQY | 0.6393626 |
| TPPPPPP | TCSADGGDQDTQY | 0.6214424 |
| AAPAAAAAPA | TCSADGGDQDTQY | 0.72036767 |
| IALNEDL | TCSADGGDQDTQY | 0.5875129 |
| IVNVSSM | TCSADGGDQDTQY | 0.7018595 |
| TGIPPAPR | TCSADGGDQDTQY | 0.80025476 |
| FDLGGGTF | TCSADGGDQDTQY | 0.6176085 |
| FDLGGGTFDVS | TCSADGGDQDTQY | 0.84553176 |
| NEPTAAA | TCSADGGDQDTQY | 0.77624047 |
| NEPTAAALA | TCSADGGDQDTQY | 0.77514803 |
| GSIINISSI | TCSADGGDQDTQY | 0.598418 |
| VAPGNAGTA | TCSADGGDQDTQY | 0.42902064 |
| TGGGASLE | TCSADGGDQDTQY | 0.73143965 |
| VAVYNHYKRI | TCSADGGDQDTQY | 0.5128827 |
| TDPAPATT | TCSADGGDQDTQY | 0.7659322 |
| TNRPDILDPALLRPG | TCSADGGDQDTQY | 0.8111557 |
| LTTTTTS | TCSADGGDQDTQY | 0.6386641 |
| PTGSGKTL | TCSADGGDQDTQY | 0.8459426 |
| TGSGKTL | TCSADGGDQDTQY | 0.84288806 |
| TIGHVDHGK | TCSADGGDQDTQY | 0.4647954 |
| TIGHVDHGKTT | TCSADGGDQDTQY | 0.6977703 |

| | | |
|---|---|---|
| GHVDHGKTT | TCSADGGDQDTQY | 0.6417714 |
| AQFEHTLL | TCSADGGDQDTQY | 0.7330904 |
| IVNQDEL | TCSADGGDQDTQY | 0.34809119 |
| TNGSQFF | TCSADGGDQDTQY | 0.8106601 |
| TGKGIVE | TCSADGGDQDTQY | 0.6178637 |
| LISNASDA | TCSADGGDQDTQY | 0.17282541 |
| AELRTLGELTKIAKAHGAQ | TCSADGGDQDTQY | 0.7822962 |
| APNAVAAAEPSW | TCSADGGDQDTQY | 0.7197605 |
| APQVAGAATAVL | TCSADGGDQDTQY | 0.6933007 |
| ARRDPAAAAAILPTDA | TCSADGGDQDTQY | 0.8302429 |
| DAAARDVLAEAGLAEYF | TCSADGGDQDTQY | 0.558373 |
| DDQIVLSAKANLAKVEQ | TCSADGGDQDTQY | 0.5896648 |
| DELLPDTGEDSDEDGH | TCSADGGDQDTQY | 0.80388147 |
| ELPEAPANRATVPQD | TCSADGGDQDTQY | 0.8377466 |
| FGQSKIERVTGIG | TCSADGGDQDTQY | 0.52540135 |
| GAGGTGSTAGGAGGAGGAGGLY | TCSADGGDQDTQY | 0.7680622 |
| GHGGTGAAGGNGGTGGAGGSGI | TCSADGGDQDTQY | 0.8331945 |
| GNGGNGGAGGEGGPGGAGGAGA | TCSADGGDQDTQY | 0.77748597 |
| GSTAGNGGNGGNATNGTIAGQP | TCSADGGDQDTQY | 0.8268892 |
| GTGGDGGNGGNGGTGG | TCSADGGDQDTQY | 0.8401195 |
| GVRRPAGIPADAPV | TCSADGGDQDTQY | 0.6596335 |
| GYRQVTDAHLLTLARR | TCSADGGDQDTQY | 0.8277999 |
| IAKMGQDGHDRG | TCSADGGDQDTQY | 0.8503672 |
| KAEPSIEIQYT | TCSADGGDQDTQY | 0.6952275 |
| KPVRIRMAADEDTVAVT | TCSADGGDQDTQY | 0.7626896 |
| MPPRIAGMRLLVIKPEP | TCSADGGDQDTQY | 0.58149177 |
| MKFYAVALFRSRGVPPTR | TCSADGGDQDTQY | 0.70836294 |
| NAGGFGILGAVAHSP | TCSADGGDQDTQY | 0.3190654 |
| NEGREITLAMAEM | TCSADGGDQDTQY | 0.7669778 |
| NELVEWSHPLGHRWRWE | TCSADGGDQDTQY | 0.83473337 |
| NLGAGGTGGNGGNPG | TCSADGGDQDTQY | 0.70487154 |
| NTEWPKWETVMAADPVD | TCSADGGDQDTQY | 0.5383623 |
| PASAAPAETQP | TCSADGGDQDTQY | 0.47689927 |
| QQSSDGKNGVRAAEITGE | TCSADGGDQDTQY | 0.75984156 |
| RAPDAVTTSWWKE | TCSADGGDQDTQY | 0.6724696 |
| SGFGNVGAHTSGWFNQ | TCSADGGDQDTQY | 0.33936232 |
| SGVAGAVGQAASVGGLKV | TCSADGGDQDTQY | 0.7444477 |
| SNGGAGTGSAGGNGGT | TCSADGGDQDTQY | 0.8310043 |
| TGGAGGEGGAAG | TCSADGGDQDTQY | 0.8258735 |
| TGGDGGAAGAGGAPGHG | TCSADGGDQDTQY | 0.44475168 |
| TMPPLLAAIANAGRHGV | TCSADGGDQDTQY | 0.8522241 |
| TSSTAGNSGGA | TCSADGGDQDTQY | 0.7755814 |
| VPATVPVGILC | TCSADGGDQDTQY | 0.84380007 |
| VQRPDSGGVVLTGRIS | TCSADGGDQDTQY | 0.73208994 |
| WTPNPVPATKGVPAGLVQAT | TCSADGGDQDTQY | 0.63308144 |
| YALKRALAKFSDRFDVVII | TCSADGGDQDTQY | 0.81673235 |
| YLIGSVIV | TCSADGGDQDTQY | 0.6936364 |
| YGQLVAVID | TCSADGGDQDTQY | 0.6250048 |
| VTVGGMTLPA | TCSADGGDQDTQY | 0.8331363 |
| VLALPAV | TCSADGGDQDTQY | 0.73825574 |
| VIPPTLNYET | TCSADGGDQDTQY | 0.6188072 |

| | | |
|---|---|---|
| VGMGGMHPSSG | TCSADGGDQDTQY | 0.62882364 |
| VGLAGGAATA | TCSADGGDQDTQY | 0.8307437 |
| VGGAGGAGGSGP | TCSADGGDQDTQY | 0.5690823 |
| VFAAFMV | TCSADGGDQDTQY | 0.5577424 |
| TLYDKLR | TCSADGGDQDTQY | 0.5901967 |
| TLAVFTVL | TCSADGGDQDTQY | 0.72472 |
| TGSVQPARV | TCSADGGDQDTQY | 0.8334829 |
| TGGGAGSGGT | TCSADGGDQDTQY | 0.78751796 |
| SHLETSL | TCSADGGDQDTQY | 0.6912212 |
| SFTTGMT | TCSADGGDQDTQY | 0.78028804 |
| QVEVAKRMGVRQPTVS | TCSADGGDQDTQY | 0.79265773 |
| QGSGGPALAM | TCSADGGDQDTQY | 0.77273756 |
| MLAPSNP | TCSADGGDQDTQY | 0.77015346 |
| LVNNAAI | TCSADGGDQDTQY | 0.6255162 |
| LTGEINTLA | TCSADGGDQDTQY | 0.82976 |
| LSIFGPV | TCSADGGDQDTQY | 0.8087023 |
| LLALLANL | TCSADGGDQDTQY | 0.81962407 |
| LISNASDALDKLR | TCSADGGDQDTQY | 0.8179483 |
| LINEAVP | TCSADGGDQDTQY | 0.79545075 |
| LGYAFSGL | TCSADGGDQDTQY | 0.76594466 |
| LDFVAVAL | TCSADGGDQDTQY | 0.30481222 |
| LAQPGAE | TCSADGGDQDTQY | 0.6896888 |
| ISNLGAVTGF | TCSADGGDQDTQY | 0.75323206 |
| HQEDPGRI | TCSADGGDQDTQY | 0.7760234 |
| GTPGEPTP | TCSADGGDQDTQY | 0.7918481 |
| GFFNSGS | TCSADGGDQDTQY | 0.8122252 |
| GAGGGGLTAGAN | TCSADGGDQDTQY | 0.6343068 |
| FVDALVNM | TCSADGGDQDTQY | 0.80344415 |
| FLTVAPEL | TCSADGGDQDTQY | 0.82838 |
| FETLTAL | TCSADGGDQDTQY | 0.4251746 |
| DVINPIEL | TCSADGGDQDTQY | 0.72615653 |
| APVSGAPV | TCSADGGDQDTQY | 0.86201763 |
| ALVGGLAW | TCSADGGDQDTQY | 0.76317275 |
| ALAAAAATAI | TCSADGGDQDTQY | 0.7115897 |
| AFGLGLLL | TCSADGGDQDTQY | 0.6938169 |
| AFFAAASL | TCSADGGDQDTQY | 0.8316965 |
| AAAALPAFSPP | TCSADGGDQDTQY | 0.6278478 |
| VVLAPAL | TCSADGGDQDTQY | 0.8448411 |
| TIGHVDHGKTTL | TCSADGGDQDTQY | 0.26088375 |
| LISNASDALDK | TCSADGGDQDTQY | 0.6707687 |
| LISNASDAL | TCSADGGDQDTQY | 0.5809075 |
| ISNASDALDK | TCSADGGDQDTQY | 0.32936776 |
| NQAETLVYQTE | TCSADGGDQDTQY | 0.8191123 |
| TGGTGGTAGAG | TCSADGGDQDTQY | 0.71997416 |
| TGSTAGGAGGA | TCSADGGDQDTQY | 0.8179157 |
| DPLAVPVAAGPV | TCSADGGDQDTQY | 0.7281661 |
| GGTGGTPGNGGN | TCSADGGDQDTQY | 0.8303939 |
| PAGLAGVDPATL | TCSADGGDQDTQY | 0.7692883 |
| GNNLVVEAPHAGASV | TCSADGGDQDTQY | 0.7641704 |
| KVTRIMSASTLEQPA | TCSADGGDQDTQY | 0.6324706 |
| LGFALPIALPRRTVV | TCSADGGDQDTQY | 0.80978584 |

| | | |
|---|---|---|
| TRLGNLTARKLVVLV | TCSADGGDQDTQY | 0.82726276 |
| WSQQRRLIYFPSAGP | TCSADGGDQDTQY | 0.80921507 |
| GPNGNPDPMAAAVDIR | TCSADGGDQDTQY | 0.44252545 |
| TQRCVSVADLRAAASR | TCSADGGDQDTQY | 0.6852433 |
| QFVLDVAEHPVEVLTETV | TCSADGGDQDTQY | 0.72633946 |
| GGAGGTGGAGGNGGTQAGDG | TCSADGGDQDTQY | 0.3042418 |
| PLLATLRAIAVDVDAKPAQV | TCSADGGDQDTQY | 0.24107416 |
| APKFRTGGYMIDFWGVGYQVAK | TCSADGGDQDTQY | 0.70626837 |
| GAGGAGGAGGAAGGPGGTGGTG | TCSADGGDQDTQY | 0.6758889 |
| IGALGLLATDVAPAGSVAFGVA | TCSADGGDQDTQY | 0.8242737 |
| AGGAGGIGGNGGAGGAGGVLMG | TCSADGGDQDTQY | 0.8115764 |
| ADELIGLL | TCSADGGDQDTQY | 0.71332145 |
| ADRALKA | TCSADGGDQDTQY | 0.8082285 |
| AGAVGQVGGAGGA | TCSADGGDQDTQY | 0.7726946 |
| AGGGGGLGGAP | TCSADGGDQDTQY | 0.1304222 |
| AGGLLYGNGGAG | TCSADGGDQDTQY | 0.68714476 |
| AGLESIGL | TCSADGGDQDTQY | 0.47927842 |
| AGLKTGII | TCSADGGDQDTQY | 0.8423135 |
| ASLAALLL | TCSADGGDQDTQY | 0.64401585 |
| ASVINISSI | TCSADGGDQDTQY | 0.630811 |
| ATPPGPA | TCSADGGDQDTQY | 0.74524033 |
| ATVASALV | TCSADGGDQDTQY | 0.7365515 |
| ATVASLIL | TCSADGGDQDTQY | 0.7401345 |
| AVNGPVLIRH | TCSADGGDQDTQY | 0.5434716 |
| DGAAGAVGI | TCSADGGDQDTQY | 0.21343625 |
| DPLLHGI | TCSADGGDQDTQY | 0.81437266 |
| ELGAALGAL | TCSADGGDQDTQY | 0.4885499 |
| ELGRTQAV | TCSADGGDQDTQY | 0.5954627 |
| FVAGGYLS | TCSADGGDQDTQY | 0.3282118 |
| GASGGAGGAGGW | TCSADGGDQDTQY | 0.23756485 |
| GGAGAGGDGGAGGI | TCSADGGDQDTQY | 0.42484465 |
| GTGHNGSQPGMGGNG | TCSADGGDQDTQY | 0.8274765 |
| GYAAASASAS | TCSADGGDQDTQY | 0.71156573 |
| HQVDVAAVQQG | TCSADGGDQDTQY | 0.69987977 |
| IGAAIAHT | TCSADGGDQDTQY | 0.85471284 |
| IGMIEVAQL | TCSADGGDQDTQY | 0.7103369 |
| IRLAPEHTSAMD | TCSADGGDQDTQY | 0.7265446 |
| IYNVSVL | TCSADGGDQDTQY | 0.6288985 |
| KQGFLGM | TCSADGGDQDTQY | 0.5802165 |
| LESVITQL | TCSADGGDQDTQY | 0.6075895 |
| LGLGALVVV | TCSADGGDQDTQY | 0.6115135 |
| LIGNGGNGGSGGI | TCSADGGDQDTQY | 0.8537477 |
| LNAINAPTL | TCSADGGDQDTQY | 0.69099593 |
| LQLLNTGV | TCSADGGDQDTQY | 0.5397879 |
| LQTSIEF | TCSADGGDQDTQY | 0.76324296 |
| MSFAVSEL | TCSADGGDQDTQY | 0.844385 |
| NTGALGL | TCSADGGDQDTQY | 0.86319125 |
| PPGAGGAGGAGTT | TCSADGGDQDTQY | 0.6053368 |
| QLVGLGI | TCSADGGDQDTQY | 0.7019578 |
| RFLEQVHQL | TCSADGGDQDTQY | 0.71535045 |
| RGVGYVLR | TCSADGGDQDTQY | 0.73213625 |

| | | |
|---|---|---|
| SATGVFAGVI | TCSADGGDQDTQY | 0.6656872 |
| SLPAITLP | TCSADGGDQDTQY | 0.5169982 |
| STALAFTV | TCSADGGDQDTQY | 0.8108196 |
| SVLTVGLGK | TCSADGGDQDTQY | 0.8412779 |
| TALAGAGSGI | TCSADGGDQDTQY | 0.7230978 |
| TGLPFTGI | TCSADGGDQDTQY | 0.40603107 |
| VAEHVTV | TCSADGGDQDTQY | 0.7088156 |
| VDIVLNSL | TCSADGGDQDTQY | 0.5737834 |
| VLTGGGAVL | TCSADGGDQDTQY | 0.6432792 |
| VNVGLAYGI | TCSADGGDQDTQY | 0.75182533 |
| VSSAAVGAV | TCSADGGDQDTQY | 0.44619402 |
| VTSVYEAGV | TCSADGGDQDTQY | 0.7226161 |
| WVALDRGA | TCSADGGDQDTQY | 0.8606728 |
| YLVGSVVI | TCSADGGDQDTQY | 0.6570581 |
| GPPGTGKTLL | TCSADGGDQDTQY | 0.8086101 |
| GVGMTKF | TCSADGGDQDTQY | 0.83222413 |
| IGIGGSDL | TCSADGGDQDTQY | 0.803241 |
| IGTLLAEL | TCSADGGDQDTQY | 0.8422563 |
| IYTSGTT | TCSADGGDQDTQY | 0.7777381 |
| LSFPAGV | TCSADGGDQDTQY | 0.7681074 |
| LYTSGTT | TCSADGGDQDTQY | 0.74100035 |
| VGAFPFGF | TCSADGGDQDTQY | 0.7749982 |
| VGAGTVEF | TCSADGGDQDTQY | 0.73774934 |
| VGKSTLLL | TCSADGGDQDTQY | 0.73214704 |
| AALAHAGGGIV | TCSADGGDQDTQY | 0.6653339 |
| AALVAVSLL | TCSADGGDQDTQY | 0.82080746 |
| AAVSRAVAAL | TCSADGGDQDTQY | 0.7223612 |
| AGGSSGIGGF | TCSADGGDQDTQY | 0.8118756 |
| AGLTVANANG | TCSADGGDQDTQY | 0.41698653 |
| AGLVGAVL | TCSADGGDQDTQY | 0.5340098 |
| ALSQAFTV | TCSADGGDQDTQY | 0.82372713 |
| ALTFSVSL | TCSADGGDQDTQY | 0.16853872 |
| ALTGAALVAL | TCSADGGDQDTQY | 0.24832946 |
| ASFSIGAGVL | TCSADGGDQDTQY | 0.7566401 |
| DGSVYKI | TCSADGGDQDTQY | 0.84286183 |
| EQLAEIV | TCSADGGDQDTQY | 0.18619059 |
| FAVSELTVM | TCSADGGDQDTQY | 0.5808445 |
| GAAPLAAGL | TCSADGGDQDTQY | 0.4727583 |
| GADGGDGGSGGAGG | TCSADGGDQDTQY | 0.81637216 |
| GFGLDTGGAGG | TCSADGGDQDTQY | 0.7451581 |
| GLASGGLL | TCSADGGDQDTQY | 0.41192734 |
| GRIAGAAVLL | TCSADGGDQDTQY | 0.8110031 |
| GSAGSLNTSF | TCSADGGDQDTQY | 0.40628344 |
| GTGQIGI | TCSADGGDQDTQY | 0.4789383 |
| GTLIALIL | TCSADGGDQDTQY | 0.82570887 |
| IAATMGFVVG | TCSADGGDQDTQY | 0.8234899 |
| IFAQALL | TCSADGGDQDTQY | 0.5372525 |
| IGAAVALGF | TCSADGGDQDTQY | 0.811901 |
| IGLTPLAL | TCSADGGDQDTQY | 0.6356565 |
| ILGSNTSTL | TCSADGGDQDTQY | 0.86606115 |
| IVFSLTVL | TCSADGGDQDTQY | 0.67661005 |

| | | |
|---|---|---|
| LASKATGF | TCSADGGDQDTQY | 0.41483638 |
| LASVALIL | TCSADGGDQDTQY | 0.8188325 |
| LFAVGLTL | TCSADGGDQDTQY | 0.8170852 |
| LFNGGIGTYI | TCSADGGDQDTQY | 0.770713 |
| LTLGAGII | TCSADGGDQDTQY | 0.7923512 |
| LVAAALGDI | TCSADGGDQDTQY | 0.64841616 |
| NTAFVQLGI | TCSADGGDQDTQY | 0.69634795 |
| QLFSGAEL | TCSADGGDQDTQY | 0.83460236 |
| SAATFITL | TCSADGGDQDTQY | 0.8150712 |
| SNYGLANLGV | TCSADGGDQDTQY | 0.59033644 |
| SYGGAYAVM | TCSADGGDQDTQY | 0.83278143 |
| TTGVGGLTLG | TCSADGGDQDTQY | 0.7472787 |
| TTQTIYI | TCSADGGDQDTQY | 0.6319684 |
| VAAGAGLGAVI | TCSADGGDQDTQY | 0.85344315 |
| VAAVVTLL | TCSADGGDQDTQY | 0.74735975 |
| VGFLGGLI | TCSADGGDQDTQY | 0.81443614 |
| VGVIPILL | TCSADGGDQDTQY | 0.8035497 |
| VIFASSLI | TCSADGGDQDTQY | 0.7752893 |
| VKGGDGGAAAG | TCSADGGDQDTQY | 0.50467145 |
| VTKTFTNI | TCSADGGDQDTQY | 0.8023445 |
| VVAAAILL | TCSADGGDQDTQY | 0.745734 |
| YAALDVEL | TCSADGGDQDTQY | 0.8066417 |
| LASVALLL | TCSADGGDQDTQY | 0.74661565 |
| LDAVHRV | TCSADGGDQDTQY | 0.5872604 |
| QGLSSSIF | TCSADGGDQDTQY | 0.7969254 |
| RGTFANIRL | TCSADGGDQDTQY | 0.84653604 |
| RQLWWGHRI | TCSADGGDQDTQY | 0.7562624 |
| YGPPGTGKTLL | TCSADGGDQDTQY | 0.7421998 |
| AALVGAPANVA | TCSADGGDQDTQY | 0.8502779 |
| AATFRRTE | TCSADGGDQDTQY | 0.61054206 |
| AGALGGGI | TCSADGGDQDTQY | 0.7432213 |
| AGGAGGAGGIGAG | TCSADGGDQDTQY | 0.8171093 |
| AGSGALGL | TCSADGGDQDTQY | 0.8253551 |
| AQIADQI | TCSADGGDQDTQY | 0.7235135 |
| ASAVAVTYL | TCSADGGDQDTQY | 0.83095133 |
| ASTVLGGTGG | TCSADGGDQDTQY | 0.64136183 |
| AVGGEPAK | TCSADGGDQDTQY | 0.81087714 |
| CEQGTFGGTC | TCSADGGDQDTQY | 0.64672184 |
| DEGKAPAG | TCSADGGDQDTQY | 0.7846759 |
| DTVVFPAL | TCSADGGDQDTQY | 0.61508626 |
| FEGITMAG | TCSADGGDQDTQY | 0.8030561 |
| FPLVVQD | TCSADGGDQDTQY | 0.8205527 |
| GAFTTPQ | TCSADGGDQDTQY | 0.8250449 |
| GFGGFGTAGGVGGA | TCSADGGDQDTQY | 0.2604535 |
| GFLAQFL | TCSADGGDQDTQY | 0.8168187 |
| GGAGGAGGNAGVA | TCSADGGDQDTQY | 0.81380713 |
| GGVGTAGAGGAGGAGG | TCSADGGDQDTQY | 0.8583566 |
| GNGGGGDAGNAGSGGN | TCSADGGDQDTQY | 0.7164531 |
| GTGGAGGAGNDAGS | TCSADGGDQDTQY | 0.8067918 |
| GTTNTGWFNS | TCSADGGDQDTQY | 0.68432486 |
| IFMGPLFTL | TCSADGGDQDTQY | 0.71002614 |

| | | |
|---|---|---|
| ILAHAIEAI | TCSADGGDQDTQY | 0.5122541 |
| IRSAIREAL | TCSADGGDQDTQY | 0.68126464 |
| ITFAGGG | TCSADGGDQDTQY | 0.6639948 |
| ITGMSGHQEV | TCSADGGDQDTQY | 0.59797573 |
| IVAGLACTL | TCSADGGDQDTQY | 0.8034292 |
| IVGGGLAGA | TCSADGGDQDTQY | 0.8154187 |
| IVTGVLNI | TCSADGGDQDTQY | 0.8387531 |
| LAQQGCI | TCSADGGDQDTQY | 0.825698 |
| LCLHVEN | TCSADGGDQDTQY | 0.844141 |
| LDAISDGS | TCSADGGDQDTQY | 0.8185038 |
| LGGFAAML | TCSADGGDQDTQY | 0.61839247 |
| LGGKNPMI | TCSADGGDQDTQY | 0.7207063 |
| LGPKGVAAIL | TCSADGGDQDTQY | 0.48970324 |
| LMVGVSI | TCSADGGDQDTQY | 0.7105272 |
| LVGAAFVL | TCSADGGDQDTQY | 0.660362 |
| LVLAVAPL | TCSADGGDQDTQY | 0.77251804 |
| MAAIAVIDAA | TCSADGGDQDTQY | 0.15281424 |
| NNGINAELQ | TCSADGGDQDTQY | 0.72631323 |
| RGEPLVLVHG | TCSADGGDQDTQY | 0.8545077 |
| RGGAGGAAGLI | TCSADGGDQDTQY | 0.6646923 |
| RSYRMHPAV | TCSADGGDQDTQY | 0.78261197 |
| SAAHLLAGQLA | TCSADGGDQDTQY | 0.78028727 |
| SAAVGSY | TCSADGGDQDTQY | 0.6659182 |
| SPHAVLHRI | TCSADGGDQDTQY | 0.8046802 |
| SQIAKVL | TCSADGGDQDTQY | 0.6049229 |
| TALGHLVP | TCSADGGDQDTQY | 0.80251217 |
| TAPTTSGAA | TCSADGGDQDTQY | 0.8478002 |
| TDGDDGNGGSPGT | TCSADGGDQDTQY | 0.7679634 |
| TDGLGEN | TCSADGGDQDTQY | 0.7273503 |
| TGGPPLPGQA | TCSADGGDQDTQY | 0.7487558 |
| TIPAITI | TCSADGGDQDTQY | 0.42486593 |
| TIPGFTI | TCSADGGDQDTQY | 0.7389715 |
| TSIASGALA | TCSADGGDQDTQY | 0.13787565 |
| TVTLPGAAI | TCSADGGDQDTQY | 0.73585165 |
| VASGLSAALP | TCSADGGDQDTQY | 0.5509666 |
| VGAGVIGI | TCSADGGDQDTQY | 0.7692587 |
| VGPKRFGV | TCSADGGDQDTQY | 0.7686345 |
| VNVAFPD | TCSADGGDQDTQY | 0.77934813 |
| VSMTNSGVS | TCSADGGDQDTQY | 0.5723474 |
| VTDVLGDLSV | TCSADGGDQDTQY | 0.72257495 |
| VVVVGAGFAGL | TCSADGGDQDTQY | 0.656346 |
| VYTVGTVV | TCSADGGDQDTQY | 0.49593675 |
| WTTFSGP | TCSADGGDQDTQY | 0.8313868 |
| YDELVEAF | TCSADGGDQDTQY | 0.7219538 |
| YGTGGAGGNGG | TCSADGGDQDTQY | 0.730546 |
| YTNVMAV | TCSADGGDQDTQY | 0.83543754 |
| FLEQVHQL | TCSADGGDQDTQY | 0.77784014 |
| GLSLLAVAV | TCSADGGDQDTQY | 0.42278087 |
| LEQVHQL | TCSADGGDQDTQY | 0.71012586 |
| LNAAGAI | TCSADGGDQDTQY | 0.74385923 |
| PAVHASL | TCSADGGDQDTQY | 0.65506023 |

| | | |
|---|---|---|
| TELEDDA | TCSADGGDQDTQY | 0.16090599 |
| TPPGPPN | TCSADGGDQDTQY | 0.8113242 |
| TPPPPPPP | TCSADGGDQDTQY | 0.64327383 |
| TQSPALI | ASSLGDRGLEQY | 0.5912655 |
| EFAAAFPQ | ASSLGDRGLEQY | 0.6792431 |
| LLSGNLDVL | ASSLGDRGLEQY | 0.8176297 |
| GPANLGNYNI | ASSLGDRGLEQY | 0.67358875 |
| TLGPTLP | ASSLGDRGLEQY | 0.7378887 |
| VYAYSSCCK | ASSLGDRGLEQY | 0.8017427 |
| APAPAPAPNI | ASSLGDRGLEQY | 0.8077291 |
| PGGPVPIP | ASSLGDRGLEQY | 0.8464136 |
| FLTVALEP | ASSLGDRGLEQY | 0.7881643 |
| SQYFGANL | ASSLGDRGLEQY | 0.7047472 |
| GGVKPAA | ASSLGDRGLEQY | 0.8079016 |
| GGTGGAGGAGGW | ASSLGDRGLEQY | 0.73146313 |
| GVGGVAGLGGAGG | ASSLGDRGLEQY | 0.41456422 |
| VLLVLATTI | ASSLGDRGLEQY | 0.3442589 |
| IGIFAAML | ASSLGDRGLEQY | 0.739771 |
| AVGDLID | ASSLGDRGLEQY | 0.36585206 |
| NISPLPL | ASSLGDRGLEQY | 0.75071764 |
| VTAVFVAI | ASSLGDRGLEQY | 0.83719444 |
| SGDDNDAAL | ASSLGDRGLEQY | 0.7458876 |
| AVAALAHS | ASSLGDRGLEQY | 0.4550988 |
| GGVTVMATP | ASSLGDRGLEQY | 0.6914474 |
| AGLAANVRD | ASSLGDRGLEQY | 0.68172824 |
| TNRPDLIDPALLRPG | ASSLGDRGLEQY | 0.8046188 |
| LANLGVTAL | ASSLGDRGLEQY | 0.80510664 |
| TGERIEL | ASSLGDRGLEQY | 0.8315588 |
| NGEELAA | ASSLGDRGLEQY | 0.8120805 |
| VLSLVWG | ASSLGDRGLEQY | 0.6666982 |
| IGYALWL | ASSLGDRGLEQY | 0.8112489 |
| LSLLLAPL | ASSLGDRGLEQY | 0.6977303 |
| TLDEALK | ASSLGDRGLEQY | 0.756523 |
| TDGAGTAEGG | ASSLGDRGLEQY | 0.7580483 |
| NAPGGAGGNGG | ASSLGDRGLEQY | 0.7833983 |
| VSVFMGMGGNF | ASSLGDRGLEQY | 0.7111912 |
| LASYLAAL | ASSLGDRGLEQY | 0.7029397 |
| LVGGTPAL | ASSLGDRGLEQY | 0.38884705 |
| VRNADVL | ASSLGDRGLEQY | 0.7720934 |
| TCSSPFLVSAM | ASSLGDRGLEQY | 0.81758195 |
| VHITDAE | ASSLGDRGLEQY | 0.3683497 |
| EYAAPSF | ASSLGDRGLEQY | 0.6906456 |
| VVDAFHGGL | ASSLGDRGLEQY | 0.85693926 |
| ASVVDPQ | ASSLGDRGLEQY | 0.8400545 |
| SYNEFGGLS | ASSLGDRGLEQY | 0.59724474 |
| DPDGSVY | ASSLGDRGLEQY | 0.56880975 |
| MVSTIHATE | ASSLGDRGLEQY | 0.7399563 |
| LAAGVADPV | ASSLGDRGLEQY | 0.38079393 |
| TKEQIAA | ASSLGDRGLEQY | 0.78692526 |
| SLASPAAL | ASSLGDRGLEQY | 0.7329096 |
| IGLVFYGV | ASSLGDRGLEQY | 0.7553358 |

| | | |
|---|---|---|
| FLLDASNL | ASSLGDRGLEQY | 0.6579423 |
| YQVGNFP | ASSLGDRGLEQY | 0.76303315 |
| ESLLSQST | ASSLGDRGLEQY | 0.7035556 |
| LFTVGFAAVG | ASSLGDRGLEQY | 0.79764724 |
| ATPPTPT | ASSLGDRGLEQY | 0.18175203 |
| TDLATLG | ASSLGDRGLEQY | 0.821158 |
| SAGGTGR | ASSLGDRGLEQY | 0.5852008 |
| GFSNSGS | ASSLGDRGLEQY | 0.5895488 |
| GGNGGSGGTSV | ASSLGDRGLEQY | 0.34377766 |
| GTSVATGGAGN | ASSLGDRGLEQY | 0.7541012 |
| NGGNGGSGGTS | ASSLGDRGLEQY | 0.82686174 |
| AVVKFPH | ASSLGDRGLEQY | 0.82752854 |
| TPPPPPP | ASSLGDRGLEQY | 0.59311366 |
| AAPAAAAAPA | ASSLGDRGLEQY | 0.81693375 |
| IALNEDL | ASSLGDRGLEQY | 0.8524873 |
| IVNVSSM | ASSLGDRGLEQY | 0.70010406 |
| TGIPPAPR | ASSLGDRGLEQY | 0.3683382 |
| FDLGGGTF | ASSLGDRGLEQY | 0.7902219 |
| FDLGGGTFDVS | ASSLGDRGLEQY | 0.6944689 |
| NEPTAAA | ASSLGDRGLEQY | 0.8155212 |
| NEPTAAALA | ASSLGDRGLEQY | 0.8189043 |
| GSIINISSI | ASSLGDRGLEQY | 0.4890083 |
| VAPGNAGTA | ASSLGDRGLEQY | 0.81032443 |
| TGGGASLE | ASSLGDRGLEQY | 0.8350351 |
| VAVYNHYKRI | ASSLGDRGLEQY | 0.58997786 |
| TDPAPATT | ASSLGDRGLEQY | 0.6142159 |
| TNRPDILDPALLRPG | ASSLGDRGLEQY | 0.68719816 |
| LTTTTTS | ASSLGDRGLEQY | 0.40651298 |
| PTGSGKTL | ASSLGDRGLEQY | 0.8170495 |
| TGSGKTL | ASSLGDRGLEQY | 0.6681596 |
| TIGHVDHGK | ASSLGDRGLEQY | 0.6264059 |
| TIGHVDHGKTT | ASSLGDRGLEQY | 0.7586892 |
| GHVDHGKTT | ASSLGDRGLEQY | 0.64980066 |
| AQFEHTLL | ASSLGDRGLEQY | 0.6623757 |
| IVNQDEL | ASSLGDRGLEQY | 0.7066436 |
| TNGSQFF | ASSLGDRGLEQY | 0.77801454 |
| TGKGIVE | ASSLGDRGLEQY | 0.782574 |
| LISNASDA | ASSLGDRGLEQY | 0.7559956 |
| AELRTLGELTKIAKAHGAQ | ASSLGDRGLEQY | 0.7811914 |
| APNAVAAAEPSW | ASSLGDRGLEQY | 0.8464196 |
| APQVAGAATAVL | ASSLGDRGLEQY | 0.5094928 |
| ARRDPAAAAAILPTDA | ASSLGDRGLEQY | 0.6258596 |
| DAAARDVLAEAGLAEYF | ASSLGDRGLEQY | 0.7960585 |
| DDQIVLSAKANLAKVEQ | ASSLGDRGLEQY | 0.44914034 |
| DELLPDTGEDSDEDGH | ASSLGDRGLEQY | 0.81063414 |
| ELPEAPANRATVPQD | ASSLGDRGLEQY | 0.76831293 |
| FGQSKIERVTGIG | ASSLGDRGLEQY | 0.7868554 |
| GAGGTGSTAGGAGGAGGAGGLY | ASSLGDRGLEQY | 0.6849135 |
| GHGGTGAAGGNGGTGGAGGSGI | ASSLGDRGLEQY | 0.86031103 |
| GNGGNGGAGGEGGPGGAGGAGA | ASSLGDRGLEQY | 0.8168198 |
| GSTAGNGGNGGNATNGTIAGQP | ASSLGDRGLEQY | 0.7592714 |

| | | |
|---|---|---|
| GTGGDGGNGGNGGTGG | ASSLGDRGLEQY | 0.7787988 |
| GVRRPAGIPADAPV | ASSLGDRGLEQY | 0.5587947 |
| GYRQVTDAHLLTLARR | ASSLGDRGLEQY | 0.82187235 |
| IAKMGQDGHDRG | ASSLGDRGLEQY | 0.79874796 |
| KAEPSIEIQYT | ASSLGDRGLEQY | 0.52714205 |
| KPVRIRMAADEDTVAVT | ASSLGDRGLEQY | 0.7033789 |
| MPPRIAGMRLLVIKPEP | ASSLGDRGLEQY | 0.8363768 |
| MKFYAVALFRSRGVPPTR | ASSLGDRGLEQY | 0.84011805 |
| NAGGFGILGAVAHSP | ASSLGDRGLEQY | 0.35595852 |
| NEGREITLAMAEM | ASSLGDRGLEQY | 0.69423723 |
| NELVEWSHPLGHRWRWE | ASSLGDRGLEQY | 0.390085 |
| NLGAGGTGGNGGNPG | ASSLGDRGLEQY | 0.60853326 |
| NTEWPKWETVMAADPVD | ASSLGDRGLEQY | 0.5385277 |
| PASAAPAETQP | ASSLGDRGLEQY | 0.79171383 |
| QQSSDGKNGVRAAEITGE | ASSLGDRGLEQY | 0.6877732 |
| RAPDAVTTSWWKE | ASSLGDRGLEQY | 0.75737107 |
| SGFGNVGAHTSGWFNQ | ASSLGDRGLEQY | 0.7475016 |
| SGVAGAVGQAASVGGLKV | ASSLGDRGLEQY | 0.8356578 |
| SNGGAGTGSAGGNGGT | ASSLGDRGLEQY | 0.6108302 |
| TGGAGGEGGAAG | ASSLGDRGLEQY | 0.71500313 |
| TGGDGGAAGAGGAPGHG | ASSLGDRGLEQY | 0.48063114 |
| TMPPLLAAIANAGRHGV | ASSLGDRGLEQY | 0.44419804 |
| TSSTAGNSGGA | ASSLGDRGLEQY | 0.5878571 |
| VPATVPVGILC | ASSLGDRGLEQY | 0.7023071 |
| VQRPDSGGVVLTGRIS | ASSLGDRGLEQY | 0.2802388 |
| WTPNPVPATKGVPAGLVQAT | ASSLGDRGLEQY | 0.84698105 |
| YALKRALAKFSDRFDVVII | ASSLGDRGLEQY | 0.758623 |
| YLIGSVIV | ASSLGDRGLEQY | 0.8096349 |
| YGQLVAVID | ASSLGDRGLEQY | 0.6702637 |
| VTVGGMTLPA | ASSLGDRGLEQY | 0.6444039 |
| VLALPAV | ASSLGDRGLEQY | 0.6243907 |
| VIPPTLNYET | ASSLGDRGLEQY | 0.7821841 |
| VGMGGMHPSSG | ASSLGDRGLEQY | 0.60453093 |
| VGLAGGAATA | ASSLGDRGLEQY | 0.37773126 |
| VGGAGGAGGSGP | ASSLGDRGLEQY | 0.63677436 |
| VFAAFMV | ASSLGDRGLEQY | 0.50355875 |
| TLYDKLR | ASSLGDRGLEQY | 0.84675074 |
| TLAVFTVL | ASSLGDRGLEQY | 0.8226133 |
| TGSVQPARV | ASSLGDRGLEQY | 0.7663049 |
| TGGGAGSGGT | ASSLGDRGLEQY | 0.8314048 |
| SHLETSL | ASSLGDRGLEQY | 0.24556975 |
| SFTTGMT | ASSLGDRGLEQY | 0.6810318 |
| QVEVAKRMGVRQPTVS | ASSLGDRGLEQY | 0.849475 |
| QGSGGPALAM | ASSLGDRGLEQY | 0.7817215 |
| MLAPSNP | ASSLGDRGLEQY | 0.73376644 |
| LVNNAAI | ASSLGDRGLEQY | 0.83739364 |
| LTGEINTLA | ASSLGDRGLEQY | 0.629197 |
| LSIFGPV | ASSLGDRGLEQY | 0.6579925 |
| LLALLANL | ASSLGDRGLEQY | 0.75021917 |
| LISNASDALDKLR | ASSLGDRGLEQY | 0.81346536 |
| LINEAVP | ASSLGDRGLEQY | 0.78630817 |

| | | |
|---|---|---|
| LGYAFSGL | ASSLGDRGLEQY | 0.5177773 |
| LDFVAVAL | ASSLGDRGLEQY | 0.7960764 |
| LAQPGAE | ASSLGDRGLEQY | 0.85786724 |
| ISNLGAVTGF | ASSLGDRGLEQY | 0.6753356 |
| HQEDPGRI | ASSLGDRGLEQY | 0.82954395 |
| GTPGEPTP | ASSLGDRGLEQY | 0.78823507 |
| GFFNSGS | ASSLGDRGLEQY | 0.78564215 |
| GAGGGGLTAGAN | ASSLGDRGLEQY | 0.58754516 |
| FVDALVNM | ASSLGDRGLEQY | 0.6430354 |
| FLTVAPEL | ASSLGDRGLEQY | 0.72757006 |
| FETLTAL | ASSLGDRGLEQY | 0.56176805 |
| DVINPIEL | ASSLGDRGLEQY | 0.73307514 |
| APVSGAPV | ASSLGDRGLEQY | 0.7221638 |
| ALVGGLAW | ASSLGDRGLEQY | 0.86723214 |
| ALAAAAATAI | ASSLGDRGLEQY | 0.7921529 |
| AFGLGLLL | ASSLGDRGLEQY | 0.8071288 |
| AFFAAASL | ASSLGDRGLEQY | 0.54307896 |
| AAAALPAFSPP | ASSLGDRGLEQY | 0.7697097 |
| VVLAPAL | ASSLGDRGLEQY | 0.7863574 |
| TIGHVDHGKTTL | ASSLGDRGLEQY | 0.46764666 |
| LISNASDALDK | ASSLGDRGLEQY | 0.8471361 |
| LISNASDAL | ASSLGDRGLEQY | 0.60319996 |
| ISNASDALDK | ASSLGDRGLEQY | 0.70987314 |
| NQAETLVYQTE | ASSLGDRGLEQY | 0.8156572 |
| TGGTGGTAGAG | ASSLGDRGLEQY | 0.80394566 |
| TGSTAGGAGGA | ASSLGDRGLEQY | 0.7090233 |
| DPLAVPVAAGPV | ASSLGDRGLEQY | 0.7238641 |
| GGTGGTPGNGGN | ASSLGDRGLEQY | 0.592927 |
| PAGLAGVDPATL | ASSLGDRGLEQY | 0.81865084 |
| GNNLVVEAPHAGASV | ASSLGDRGLEQY | 0.8301858 |
| KVTRIMSASTLEQPA | ASSLGDRGLEQY | 0.6887958 |
| LGFALPIALPRRTVV | ASSLGDRGLEQY | 0.8431712 |
| TRLGNLTARKLVVLV | ASSLGDRGLEQY | 0.8011961 |
| WSQQRRLIYFPSAGP | ASSLGDRGLEQY | 0.4091821 |
| GPNGNPDPMAAAVDIR | ASSLGDRGLEQY | 0.741492 |
| TQRCVSVADLRAAASR | ASSLGDRGLEQY | 0.837759 |
| QFVLDVAEHPVEVLTETV | ASSLGDRGLEQY | 0.8290273 |
| GGAGGTGGAGGNGGTQAGDG | ASSLGDRGLEQY | 0.8432915 |
| PLLATLRAIAVDVDAKPAQV | ASSLGDRGLEQY | 0.7841306 |
| APKFRTGGYMIDFWGVGYQVAK | ASSLGDRGLEQY | 0.6680578 |
| GAGGAGGAGGAAGGPGGTGGTG | ASSLGDRGLEQY | 0.8394538 |
| IGALGLLATDVAPAGSVAFGVA | ASSLGDRGLEQY | 0.8586674 |
| AGGAGGIGGNGGAGGAGGVLMG | ASSLGDRGLEQY | 0.82240087 |
| ADELIGLL | ASSLGDRGLEQY | 0.6582753 |
| ADRALKA | ASSLGDRGLEQY | 0.61438996 |
| AGAVGQVGGAGGA | ASSLGDRGLEQY | 0.774147 |
| AGGGGGLGGAP | ASSLGDRGLEQY | 0.8264093 |
| AGGLLYGNGGAG | ASSLGDRGLEQY | 0.5469794 |
| AGLESIGL | ASSLGDRGLEQY | 0.7712542 |
| AGLKTGII | ASSLGDRGLEQY | 0.76715946 |
| ASLAALLL | ASSLGDRGLEQY | 0.41459286 |

| | | |
|---|---|---|
| ASVINISSI | ASSLGDRGLEQY | 0.8210582 |
| ATPPGPA | ASSLGDRGLEQY | 0.84233445 |
| ATVASALV | ASSLGDRGLEQY | 0.7583958 |
| ATVASLIL | ASSLGDRGLEQY | 0.6120214 |
| AVNGPVLIRH | ASSLGDRGLEQY | 0.8328621 |
| DGAAGAVGI | ASSLGDRGLEQY | 0.86593366 |
| DPLLHGI | ASSLGDRGLEQY | 0.8408871 |
| ELGAALGAL | ASSLGDRGLEQY | 0.40362397 |
| ELGRTQAV | ASSLGDRGLEQY | 0.24102487 |
| FVAGGYLS | ASSLGDRGLEQY | 0.6727649 |
| GASGGAGGAGGW | ASSLGDRGLEQY | 0.78062 |
| GGAGAGGDGGAGGI | ASSLGDRGLEQY | 0.6967655 |
| GTGHNGSQPGMGGNG | ASSLGDRGLEQY | 0.6382964 |
| GYAAASASAS | ASSLGDRGLEQY | 0.788586 |
| HQVDVAAVQQG | ASSLGDRGLEQY | 0.8312812 |
| IGAAIAHT | ASSLGDRGLEQY | 0.5981622 |
| IGMIEVAQL | ASSLGDRGLEQY | 0.45662904 |
| IRLAPEHTSAMD | ASSLGDRGLEQY | 0.8129691 |
| IYNVSVL | ASSLGDRGLEQY | 0.806273 |
| KQGFLGM | ASSLGDRGLEQY | 0.60275257 |
| LESVITQL | ASSLGDRGLEQY | 0.69055593 |
| LGLGALVVV | ASSLGDRGLEQY | 0.70062435 |
| LIGNGGNGGSGGI | ASSLGDRGLEQY | 0.6119753 |
| LNAINAPTL | ASSLGDRGLEQY | 0.77456665 |
| LQLLNTGV | ASSLGDRGLEQY | 0.4718274 |
| LQTSIEF | ASSLGDRGLEQY | 0.84672344 |
| MSFAVSEL | ASSLGDRGLEQY | 0.49963504 |
| NTGALGL | ASSLGDRGLEQY | 0.8133723 |
| PPGAGGAGGAGTT | ASSLGDRGLEQY | 0.7654828 |
| QLVGLGI | ASSLGDRGLEQY | 0.7965065 |
| RFLEQVHQL | ASSLGDRGLEQY | 0.6836317 |
| RGVGYVLR | ASSLGDRGLEQY | 0.6204688 |
| SATGVFAGVI | ASSLGDRGLEQY | 0.5544112 |
| SLPAITLP | ASSLGDRGLEQY | 0.3540392 |
| STALAFTV | ASSLGDRGLEQY | 0.8265914 |
| SVLTVGLGK | ASSLGDRGLEQY | 0.30787414 |
| TALAGAGSGI | ASSLGDRGLEQY | 0.8351517 |
| TGLPFTGI | ASSLGDRGLEQY | 0.7888735 |
| VAEHVTV | ASSLGDRGLEQY | 0.6283871 |
| VDIVLNSL | ASSLGDRGLEQY | 0.78496945 |
| VLTGGGAVL | ASSLGDRGLEQY | 0.5615597 |
| VNVGLAYGI | ASSLGDRGLEQY | 0.63579965 |
| VSSAAVGAV | ASSLGDRGLEQY | 0.8147956 |
| VTSVYEAGV | ASSLGDRGLEQY | 0.8090819 |
| WVALDRGA | ASSLGDRGLEQY | 0.5954617 |
| YLVGSVVI | ASSLGDRGLEQY | 0.79437804 |
| GPPGTGKTLL | ASSLGDRGLEQY | 0.7798193 |
| GVGMTKF | ASSLGDRGLEQY | 0.5986746 |
| IGIGGSDL | ASSLGDRGLEQY | 0.758569 |
| IGTLLAEL | ASSLGDRGLEQY | 0.8207076 |
| IYTSGTT | ASSLGDRGLEQY | 0.7879275 |

| | | |
|---|---|---|
| LSFPAGV | ASSLGDRGLEQY | 0.51487255 |
| LYTSGTT | ASSLGDRGLEQY | 0.81042403 |
| VGAFPFGF | ASSLGDRGLEQY | 0.59745693 |
| VGAGTVEF | ASSLGDRGLEQY | 0.7820055 |
| VGKSTLLL | ASSLGDRGLEQY | 0.7722198 |
| AALAHAGGGIV | ASSLGDRGLEQY | 0.7008904 |
| AALVAVSLL | ASSLGDRGLEQY | 0.8317478 |
| AAVSRAVAAL | ASSLGDRGLEQY | 0.6195103 |
| AGGSSGIGGF | ASSLGDRGLEQY | 0.61773705 |
| AGLTVANANG | ASSLGDRGLEQY | 0.716593 |
| AGLVGAVL | ASSLGDRGLEQY | 0.7604487 |
| ALSQAFTV | ASSLGDRGLEQY | 0.822066 |
| ALTFSVSL | ASSLGDRGLEQY | 0.542481 |
| ALTGAALVAL | ASSLGDRGLEQY | 0.759432 |
| ASFSIGAGVL | ASSLGDRGLEQY | 0.67433107 |
| DGSVYKI | ASSLGDRGLEQY | 0.510976 |
| EQLAEIV | ASSLGDRGLEQY | 0.7443894 |
| FAVSELTVM | ASSLGDRGLEQY | 0.7071574 |
| GAAPLAAGL | ASSLGDRGLEQY | 0.59526795 |
| GADGGDGGSGGAGG | ASSLGDRGLEQY | 0.80437297 |
| GFGLDTGGAGG | ASSLGDRGLEQY | 0.6423025 |
| GLASGGLL | ASSLGDRGLEQY | 0.78767854 |
| GRIAGAAVLL | ASSLGDRGLEQY | 0.85927093 |
| GSAGSLNTSF | ASSLGDRGLEQY | 0.5705423 |
| GTGQIGI | ASSLGDRGLEQY | 0.8064398 |
| GTLIALIL | ASSLGDRGLEQY | 0.79701567 |
| IAATMGFVVG | ASSLGDRGLEQY | 0.7754431 |
| IFAQALL | ASSLGDRGLEQY | 0.7056309 |
| IGAAVALGF | ASSLGDRGLEQY | 0.62571305 |
| IGLTPLAL | ASSLGDRGLEQY | 0.8503735 |
| ILGSNTSTL | ASSLGDRGLEQY | 0.81170535 |
| IVFSLTVL | ASSLGDRGLEQY | 0.81769264 |
| LASKATGF | ASSLGDRGLEQY | 0.81236124 |
| LASVALIL | ASSLGDRGLEQY | 0.79076344 |
| LFAVGLTL | ASSLGDRGLEQY | 0.8126863 |
| LFNGGIGTYI | ASSLGDRGLEQY | 0.6387019 |
| LTLGAGII | ASSLGDRGLEQY | 0.4634863 |
| LVAAALGDI | ASSLGDRGLEQY | 0.6879029 |
| NTAFVQLGI | ASSLGDRGLEQY | 0.7627194 |
| QLFSGAEL | ASSLGDRGLEQY | 0.6722969 |
| SAATFITL | ASSLGDRGLEQY | 0.3653685 |
| SNYGLANLGV | ASSLGDRGLEQY | 0.8307146 |
| SYGGAYAVM | ASSLGDRGLEQY | 0.86351883 |
| TTGVGGLTLG | ASSLGDRGLEQY | 0.5981083 |
| TTQTIYI | ASSLGDRGLEQY | 0.85613906 |
| VAAGAGLGAVI | ASSLGDRGLEQY | 0.48873332 |
| VAAVVTLL | ASSLGDRGLEQY | 0.7997085 |
| VGFLGGLI | ASSLGDRGLEQY | 0.8372031 |
| VGVIPILL | ASSLGDRGLEQY | 0.5380266 |
| VIFASSLI | ASSLGDRGLEQY | 0.7979607 |
| VKGGDGGAAAG | ASSLGDRGLEQY | 0.5888097 |

| | | |
|---|---|---|
| VTKTFTNI | ASSLGDRGLEQY | 0.7073823 |
| VVAAAILL | ASSLGDRGLEQY | 0.8397485 |
| YAALDVEL | ASSLGDRGLEQY | 0.69771767 |
| LASVALLL | ASSLGDRGLEQY | 0.79285765 |
| LDAVHRV | ASSLGDRGLEQY | 0.7018316 |
| QGLSSSIF | ASSLGDRGLEQY | 0.5664557 |
| RGTFANIRL | ASSLGDRGLEQY | 0.78941655 |
| RQLWWGHRI | ASSLGDRGLEQY | 0.8293301 |
| YGPPGTGKTLL | ASSLGDRGLEQY | 0.6332256 |
| AALVGAPANVA | ASSLGDRGLEQY | 0.8526102 |
| AATFRRTE | ASSLGDRGLEQY | 0.61774075 |
| AGALGGGI | ASSLGDRGLEQY | 0.8638162 |
| AGGAGGAGGIGAG | ASSLGDRGLEQY | 0.82167876 |
| AGSGALGL | ASSLGDRGLEQY | 0.7425654 |
| AQIADQI | ASSLGDRGLEQY | 0.69930327 |
| ASAVAVTYL | ASSLGDRGLEQY | 0.8196343 |
| ASTVLGGTGG | ASSLGDRGLEQY | 0.66448927 |
| AVGGEPAK | ASSLGDRGLEQY | 0.7724941 |
| CEQGTFGGTC | ASSLGDRGLEQY | 0.83715856 |
| DEGKAPAG | ASSLGDRGLEQY | 0.3234965 |
| DTVVFPAL | ASSLGDRGLEQY | 0.86756784 |
| FEGITMAG | ASSLGDRGLEQY | 0.80750924 |
| FPLVVQD | ASSLGDRGLEQY | 0.84066844 |
| GAFTTPQ | ASSLGDRGLEQY | 0.58983755 |
| GFGGFGTAGGVGGA | ASSLGDRGLEQY | 0.34149566 |
| GFLAQFL | ASSLGDRGLEQY | 0.7004986 |
| GGAGGAGGNAGVA | ASSLGDRGLEQY | 0.5244437 |
| GGVGTAGAGGAGGAGG | ASSLGDRGLEQY | 0.837337 |
| GNGGGGDAGNAGSGGN | ASSLGDRGLEQY | 0.83913517 |
| GTGGAGGAGNDAGS | ASSLGDRGLEQY | 0.5894968 |
| GTTNTGWFNS | ASSLGDRGLEQY | 0.83216286 |
| IFMGPLFTL | ASSLGDRGLEQY | 0.8564131 |
| ILAHAIEAI | ASSLGDRGLEQY | 0.34384245 |
| IRSAIREAL | ASSLGDRGLEQY | 0.730543 |
| ITFAGGG | ASSLGDRGLEQY | 0.7308168 |
| ITGMSGHQEV | ASSLGDRGLEQY | 0.6511502 |
| IVAGLACTL | ASSLGDRGLEQY | 0.25448862 |
| IVGGGLAGA | ASSLGDRGLEQY | 0.80969745 |
| IVTGVLNI | ASSLGDRGLEQY | 0.64091736 |
| LAQQGCI | ASSLGDRGLEQY | 0.16396484 |
| LCLHVEN | ASSLGDRGLEQY | 0.8059717 |
| LDAISDGS | ASSLGDRGLEQY | 0.5846066 |
| LGGFAAML | ASSLGDRGLEQY | 0.5826413 |
| LGGKNPMI | ASSLGDRGLEQY | 0.83631086 |
| LGPKGVAAIL | ASSLGDRGLEQY | 0.6694313 |
| LMVGVSI | ASSLGDRGLEQY | 0.62580365 |
| LVGAAFVL | ASSLGDRGLEQY | 0.82058567 |
| LVLAVAPL | ASSLGDRGLEQY | 0.8532882 |
| MAAIAVIDAA | ASSLGDRGLEQY | 0.49558598 |
| NNGINAELQ | ASSLGDRGLEQY | 0.7522085 |
| RGEPLVLVHG | ASSLGDRGLEQY | 0.8829973 |

| | | |
|---|---|---|
| RGGAGGAAGLI | ASSLGDRGLEQY | 0.7625209 |
| RSYRMHPAV | ASSLGDRGLEQY | 0.8803562 |
| SAAHLLAGQLA | ASSLGDRGLEQY | 0.885954 |
| SAAVGSY | ASSLGDRGLEQY | 0.67701876 |
| SPHAVLHRI | ASSLGDRGLEQY | 0.85421216 |
| SQIAKVL | ASSLGDRGLEQY | 0.8563562 |
| TALGHLVP | ASSLGDRGLEQY | 0.6628186 |
| TAPTTSGAA | ASSLGDRGLEQY | 0.73772085 |
| TDGDDGNGGSPGT | ASSLGDRGLEQY | 0.40816188 |
| TDGLGEN | ASSLGDRGLEQY | 0.7486777 |
| TGGPPLPGQA | ASSLGDRGLEQY | 0.39652264 |
| TIPAITI | ASSLGDRGLEQY | 0.7880572 |
| TIPGFTI | ASSLGDRGLEQY | 0.8689498 |
| TSIASGALA | ASSLGDRGLEQY | 0.7341453 |
| TVTLPGAAI | ASSLGDRGLEQY | 0.36248565 |
| VASGLSAALP | ASSLGDRGLEQY | 0.40058476 |
| VGAGVIGI | ASSLGDRGLEQY | 0.8221371 |
| VGPKRFGV | ASSLGDRGLEQY | 0.3838628 |
| VNVAFPD | ASSLGDRGLEQY | 0.4020875 |
| VSMTNSGVS | ASSLGDRGLEQY | 0.68782985 |
| VTDVLGDLSV | ASSLGDRGLEQY | 0.85762346 |
| VVVVGAGFAGL | ASSLGDRGLEQY | 0.8534747 |
| VYTVGTVV | ASSLGDRGLEQY | 0.56595993 |
| WTTFSGP | ASSLGDRGLEQY | 0.8729934 |
| YDELVEAF | ASSLGDRGLEQY | 0.8250519 |
| YGTGGAGGNGG | ASSLGDRGLEQY | 0.87764996 |
| YTNVMAV | ASSLGDRGLEQY | 0.7591238 |
| FLEQVHQL | ASSLGDRGLEQY | 0.37626135 |
| GLSLLAVAV | ASSLGDRGLEQY | 0.83823055 |
| LEQVHQL | ASSLGDRGLEQY | 0.6708742 |
| LNAAGAI | ASSLGDRGLEQY | 0.827493 |
| PAVHASL | ASSLGDRGLEQY | 0.8515866 |
| TELEDDA | ASSLGDRGLEQY | 0.7567279 |
| TPPGPPN | ASSLGDRGLEQY | 0.5482451 |
| TPPPPPPP | ASSLGDRGLEQY | 0.7276688 |
| TQSPALI | ASSPTVANSDYT | 0.880965 |
| EFAAAFPQ | ASSPTVANSDYT | 0.49973062 |
| LLSGNLDVL | ASSPTVANSDYT | 0.39572465 |
| GPANLGNYNI | ASSPTVANSDYT | 0.6500393 |
| TLGPTLP | ASSPTVANSDYT | 0.6905308 |
| VYAYSSCCK | ASSPTVANSDYT | 0.86942637 |
| APAPAPAPNI | ASSPTVANSDYT | 0.76370186 |
| PGGPVPIP | ASSPTVANSDYT | 0.7153832 |
| FLTVALEP | ASSPTVANSDYT | 0.8067272 |
| SQYFGANL | ASSPTVANSDYT | 0.8235466 |
| GGVKPAA | ASSPTVANSDYT | 0.7663044 |
| GGTGGAGGAGGW | ASSPTVANSDYT | 0.8262439 |
| GVGGVAGLGGAGG | ASSPTVANSDYT | 0.59559184 |
| VLLVLATTI | ASSPTVANSDYT | 0.87428844 |
| IGIFAAML | ASSPTVANSDYT | 0.8545977 |
| AVGDLID | ASSPTVANSDYT | 0.8245692 |

| | | |
|---|---|---|
| NISPLPL | ASSPTVANSDYT | 0.863075 |
| VTAVFVAI | ASSPTVANSDYT | 0.7765996 |
| SGDDNDAAL | ASSPTVANSDYT | 0.7744271 |
| AVAALAHS | ASSPTVANSDYT | 0.43796694 |
| GGVTVMATP | ASSPTVANSDYT | 0.7121446 |
| AGLAANVRD | ASSPTVANSDYT | 0.7578307 |
| TNRPDLIDPALLRPG | ASSPTVANSDYT | 0.7888737 |
| LANLGVTAL | ASSPTVANSDYT | 0.8336178 |
| TGERIEL | ASSPTVANSDYT | 0.84177583 |
| NGEELAA | ASSPTVANSDYT | 0.61114705 |
| VLSLVWG | ASSPTVANSDYT | 0.876546 |
| IGYALWL | ASSPTVANSDYT | 0.86265844 |
| LSLLLAPL | ASSPTVANSDYT | 0.47906455 |
| TLDEALK | ASSPTVANSDYT | 0.72009426 |
| TDGAGTAEGG | ASSPTVANSDYT | 0.8586092 |
| NAPGGAGGNGG | ASSPTVANSDYT | 0.8124477 |
| VSVFMGMGGNF | ASSPTVANSDYT | 0.6926607 |
| LASYLAAL | ASSPTVANSDYT | 0.7480427 |
| LVGGTPAL | ASSPTVANSDYT | 0.85836077 |
| VRNADVL | ASSPTVANSDYT | 0.7339777 |
| TCSSPFLVSAM | ASSPTVANSDYT | 0.87693745 |
| VHITDAE | ASSPTVANSDYT | 0.23503569 |
| EYAAPSF | ASSPTVANSDYT | 0.67300206 |
| VVDAFHGGL | ASSPTVANSDYT | 0.7037506 |
| ASVVDPQ | ASSPTVANSDYT | 0.15243284 |
| SYNEFGGLS | ASSPTVANSDYT | 0.853782 |
| DPDGSVY | ASSPTVANSDYT | 0.6017817 |
| MVSTIHATE | ASSPTVANSDYT | 0.79696274 |
| LAAGVADPV | ASSPTVANSDYT | 0.66096836 |
| TKEQIAA | ASSPTVANSDYT | 0.86076564 |
| SLASPAAL | ASSPTVANSDYT | 0.8060545 |
| IGLVFYGV | ASSPTVANSDYT | 0.64283377 |
| FLLDASNL | ASSPTVANSDYT | 0.4436292 |
| YQVGNFP | ASSPTVANSDYT | 0.80626136 |
| ESLLSQST | ASSPTVANSDYT | 0.88291556 |
| LFTVGFAAVG | ASSPTVANSDYT | 0.8801813 |
| ATPPTPT | ASSPTVANSDYT | 0.31867656 |
| TDLATLG | ASSPTVANSDYT | 0.80735195 |
| SAGGTGR | ASSPTVANSDYT | 0.7272779 |
| GFSNSGS | ASSPTVANSDYT | 0.51102453 |
| GGNGGSGGTSV | ASSPTVANSDYT | 0.21980709 |
| GTSVATGGAGN | ASSPTVANSDYT | 0.6904369 |
| NGGNGGSGGTS | ASSPTVANSDYT | 0.73105663 |
| AVVKFPH | ASSPTVANSDYT | 0.8037529 |
| TPPPPPP | ASSPTVANSDYT | 0.8592154 |
| AAPAAAAAPA | ASSPTVANSDYT | 0.60863435 |
| IALNEDL | ASSPTVANSDYT | 0.832134 |
| IVNVSSM | ASSPTVANSDYT | 0.75972915 |
| TGIPPAPR | ASSPTVANSDYT | 0.081542164 |
| FDLGGGTF | ASSPTVANSDYT | 0.634045 |
| FDLGGGTFDVS | ASSPTVANSDYT | 0.3184591 |

| | | |
|---|---|---|
| NEPTAAA | ASSPTVANSDYT | 0.8664892 |
| NEPTAAALA | ASSPTVANSDYT | 0.48157343 |
| GSIINISSI | ASSPTVANSDYT | 0.68934745 |
| VAPGNAGTA | ASSPTVANSDYT | 0.7111163 |
| TGGGASLE | ASSPTVANSDYT | 0.7950306 |
| VAVYNHYKRI | ASSPTVANSDYT | 0.7559054 |
| TDPAPATT | ASSPTVANSDYT | 0.6071496 |
| TNRPDILDPALLRPG | ASSPTVANSDYT | 0.48726428 |
| LTTTTTS | ASSPTVANSDYT | 0.82898587 |
| PTGSGKTL | ASSPTVANSDYT | 0.74661356 |
| TGSGKTL | ASSPTVANSDYT | 0.5192029 |
| TIGHVDHGK | ASSPTVANSDYT | 0.56966245 |
| TIGHVDHGKTT | ASSPTVANSDYT | 0.864123 |
| GHVDHGKTT | ASSPTVANSDYT | 0.2846074 |
| AQFEHTLL | ASSPTVANSDYT | 0.8690696 |
| IVNQDEL | ASSPTVANSDYT | 0.5496109 |
| TNGSQFF | ASSPTVANSDYT | 0.66738796 |
| TGKGIVE | ASSPTVANSDYT | 0.8792909 |
| LISNASDA | ASSPTVANSDYT | 0.75972503 |
| AELRTLGELTKIAKAHGAQ | ASSPTVANSDYT | 0.745739 |
| APNAVAAAEPSW | ASSPTVANSDYT | 0.56191736 |
| APQVAGAATAVL | ASSPTVANSDYT | 0.4838993 |
| ARRDPAAAAAILPTDA | ASSPTVANSDYT | 0.5873548 |
| DAAARDVLAEAGLAEYF | ASSPTVANSDYT | 0.56683683 |
| DDQIVLSAKANLAKVEQ | ASSPTVANSDYT | 0.8753266 |
| DELLPDTGEDSDEDGH | ASSPTVANSDYT | 0.7376462 |
| ELPEAPANRATVPQD | ASSPTVANSDYT | 0.44187933 |
| FGQSKIERVTGIG | ASSPTVANSDYT | 0.82783806 |
| GAGGTGSTAGGAGGAGGAGGLY | ASSPTVANSDYT | 0.87224317 |
| GHGGTGAAGGNGGTGGAGGSGI | ASSPTVANSDYT | 0.8800355 |
| GNGGNGGAGGEGGPGGAGGAGA | ASSPTVANSDYT | 0.42864186 |
| GSTAGNGGNGGNATNGTIAGQP | ASSPTVANSDYT | 0.61249816 |
| GTGGDGGNGGNGGTGG | ASSPTVANSDYT | 0.7396146 |
| GVRRPAGIPADAPV | ASSPTVANSDYT | 0.7084162 |
| GYRQVTDAHLLTLARR | ASSPTVANSDYT | 0.5178162 |
| IAKMGQDGHDRG | ASSPTVANSDYT | 0.4921122 |
| KAEPSIEIQYT | ASSPTVANSDYT | 0.8627137 |
| KPVRIRMAADEDTVAVT | ASSPTVANSDYT | 0.87421894 |
| MPPRIAGMRLLVIKPEP | ASSPTVANSDYT | 0.70720506 |
| MKFYAVALFRSRGVPPTR | ASSPTVANSDYT | 0.3036911 |
| NAGGFGILGAVAHSP | ASSPTVANSDYT | 0.6726601 |
| NEGREITLAMAEM | ASSPTVANSDYT | 0.53962713 |
| NELVEWSHPLGHRWRWE | ASSPTVANSDYT | 0.55494046 |
| NLGAGGTGGNGGNPG | ASSPTVANSDYT | 0.7322004 |
| NTEWPKWETVMAADPVD | ASSPTVANSDYT | 0.32213002 |
| PASAAPAETQP | ASSPTVANSDYT | 0.70978165 |
| QQSSDGKNGVRAAEITGE | ASSPTVANSDYT | 0.89013505 |
| RAPDAVTTSWWKE | ASSPTVANSDYT | 0.6579709 |
| SGFGNVGAHTSGWFNQ | ASSPTVANSDYT | 0.8358729 |
| SGVAGAVGQAASVGGLKV | ASSPTVANSDYT | 0.86610043 |
| SNGGAGTGSAGGNGGT | ASSPTVANSDYT | 0.8102284 |

| | | |
|---|---|---|
| TGGAGGEGGAAG | ASSPTVANSDYT | 0.8602327 |
| TGGDGGAAGAGGAPGHG | ASSPTVANSDYT | 0.82738876 |
| TMPPLLAAIANAGRHGV | ASSPTVANSDYT | 0.7346678 |
| TSSTAGNSGGA | ASSPTVANSDYT | 0.7618973 |
| VPATVPVGILC | ASSPTVANSDYT | 0.8343841 |
| VQRPDSGGVVLTGRIS | ASSPTVANSDYT | 0.64921486 |
| WTPNPVPATKGVPAGLVQAT | ASSPTVANSDYT | 0.77408487 |
| YALKRALAKFSDRFDVVII | ASSPTVANSDYT | 0.7619593 |
| YLIGSVIV | ASSPTVANSDYT | 0.8082526 |
| YGQLVAVID | ASSPTVANSDYT | 0.75958574 |
| VTVGGMTLPA | ASSPTVANSDYT | 0.8632003 |
| VLALPAV | ASSPTVANSDYT | 0.19768418 |
| VIPPTLNYET | ASSPTVANSDYT | 0.4838795 |
| VGMGGMHPSSG | ASSPTVANSDYT | 0.85453033 |
| VGLAGGAATA | ASSPTVANSDYT | 0.17196739 |
| VGGAGGAGGSGP | ASSPTVANSDYT | 0.257048 |
| VFAAFMV | ASSPTVANSDYT | 0.7934511 |
| TLYDKLR | ASSPTVANSDYT | 0.86613595 |
| TLAVFTVL | ASSPTVANSDYT | 0.23536423 |
| TGSVQPARV | ASSPTVANSDYT | 0.71352196 |
| TGGGAGSGGT | ASSPTVANSDYT | 0.3399083 |
| SHLETSL | ASSPTVANSDYT | 0.8548769 |
| SFTTGMT | ASSPTVANSDYT | 0.6970335 |
| QVEVAKRMGVRQPTVS | ASSPTVANSDYT | 0.32979807 |
| QGSGGPALAM | ASSPTVANSDYT | 0.8230539 |
| MLAPSNP | ASSPTVANSDYT | 0.53495127 |
| LVNNAAI | ASSPTVANSDYT | 0.7885647 |
| LTGEINTLA | ASSPTVANSDYT | 0.8619945 |
| LSIFGPV | ASSPTVANSDYT | 0.8716156 |
| LLALLANL | ASSPTVANSDYT | 0.418135 |
| LISNASDALDKLR | ASSPTVANSDYT | 0.8252145 |
| LINEAVP | ASSPTVANSDYT | 0.62901103 |
| LGYAFSGL | ASSPTVANSDYT | 0.89402974 |
| LDFVAVAL | ASSPTVANSDYT | 0.6564725 |
| LAQPGAE | ASSPTVANSDYT | 0.29089022 |
| ISNLGAVTGF | ASSPTVANSDYT | 0.84341234 |
| HQEDPGRI | ASSPTVANSDYT | 0.8296143 |
| GTPGEPTP | ASSPTVANSDYT | 0.7166225 |
| GFFNSGS | ASSPTVANSDYT | 0.82045174 |
| GAGGGGLTAGAN | ASSPTVANSDYT | 0.7322587 |
| FVDALVNM | ASSPTVANSDYT | 0.63706225 |
| FLTVAPEL | ASSPTVANSDYT | 0.8158404 |
| FETLTAL | ASSPTVANSDYT | 0.8258316 |
| DVINPIEL | ASSPTVANSDYT | 0.60226023 |
| APVSGAPV | ASSPTVANSDYT | 0.8767015 |
| ALVGGLAW | ASSPTVANSDYT | 0.75152063 |
| ALAAAAATAI | ASSPTVANSDYT | 0.45450038 |
| AFGLGLLL | ASSPTVANSDYT | 0.8876349 |
| AFFAAASL | ASSPTVANSDYT | 0.74360245 |
| AAAALPAFSPP | ASSPTVANSDYT | 0.8384449 |
| VVLAPAL | ASSPTVANSDYT | 0.82207376 |

| | | |
|---|---|---|
| TIGHVDHGKTTL | ASSPTVANSDYT | 0.85048795 |
| LISNASDALDK | ASSPTVANSDYT | 0.5935291 |
| LISNASDAL | ASSPTVANSDYT | 0.7843882 |
| ISNASDALDK | ASSPTVANSDYT | 0.77060735 |
| NQAETLVYQTE | ASSPTVANSDYT | 0.83386266 |
| TGGTGGTAGAG | ASSPTVANSDYT | 0.73412216 |
| TGSTAGGAGGA | ASSPTVANSDYT | 0.8086042 |
| DPLAVPVAAGPV | ASSPTVANSDYT | 0.81990933 |
| GGTGGTPGNGGN | ASSPTVANSDYT | 0.87791544 |
| PAGLAGVDPATL | ASSPTVANSDYT | 0.76358634 |
| GNNLVVEAPHAGASV | ASSPTVANSDYT | 0.8067315 |
| KVTRIMSASTLEQPA | ASSPTVANSDYT | 0.87728417 |
| LGFALPIALPRRTVV | ASSPTVANSDYT | 0.42612568 |
| TRLGNLTARKLVVLV | ASSPTVANSDYT | 0.77000237 |
| WSQQRRLIYFPSAGP | ASSPTVANSDYT | 0.86687434 |
| GPNGNPDPMAAAVDIR | ASSPTVANSDYT | 0.87699854 |
| TQRCVSVADLRAAASR | ASSPTVANSDYT | 0.71418756 |
| QFVLDVAEHPVEVLTETV | ASSPTVANSDYT | 0.8686838 |
| GGAGGTGGAGGNGGTQAGDG | ASSPTVANSDYT | 0.7385526 |
| PLLATLRAIAVDVDAKPAQV | ASSPTVANSDYT | 0.80256784 |
| APKFRTGGYMIDFWGVGYQVAK | ASSPTVANSDYT | 0.7245971 |
| GAGGAGGAGGAAGGPGGTGGTG | ASSPTVANSDYT | 0.6417276 |
| IGALGLLATDVAPAGSVAFGVA | ASSPTVANSDYT | 0.5455141 |
| AGGAGGIGGNGGAGGAGGVLMG | ASSPTVANSDYT | 0.7467231 |
| ADELIGLL | ASSPTVANSDYT | 0.83901155 |
| ADRALKA | ASSPTVANSDYT | 0.7749529 |
| AGAVGQVGGAGGA | ASSPTVANSDYT | 0.32761943 |
| AGGGGGLGGAP | ASSPTVANSDYT | 0.80297315 |
| AGGLLYGNGGAG | ASSPTVANSDYT | 0.7968612 |
| AGLESIGL | ASSPTVANSDYT | 0.8787023 |
| AGLKTGII | ASSPTVANSDYT | 0.76603246 |
| ASLAALLL | ASSPTVANSDYT | 0.8615136 |
| ASVINISSI | ASSPTVANSDYT | 0.7239076 |
| ATPPGPA | ASSPTVANSDYT | 0.76445985 |
| ATVASALV | ASSPTVANSDYT | 0.36406475 |
| ATVASLIL | ASSPTVANSDYT | 0.6401588 |
| AVNGPVLIRH | ASSPTVANSDYT | 0.7375628 |
| DGAAGAVGI | ASSPTVANSDYT | 0.5276584 |
| DPLLHGI | ASSPTVANSDYT | 0.8207456 |
| ELGAALGAL | ASSPTVANSDYT | 0.8664993 |
| ELGRTQAV | ASSPTVANSDYT | 0.8596453 |
| FVAGGYLS | ASSPTVANSDYT | 0.86089325 |
| GASGGAGGAGGW | ASSPTVANSDYT | 0.8724947 |
| GGAGAGGDGGAGGI | ASSPTVANSDYT | 0.8442838 |
| GTGHNGSQPGMGGNG | ASSPTVANSDYT | 0.63885605 |
| GYAAASASAS | ASSPTVANSDYT | 0.6815265 |
| HQVDVAAVQQG | ASSPTVANSDYT | 0.30794942 |
| IGAAIAHT | ASSPTVANSDYT | 0.6154157 |
| IGMIEVAQL | ASSPTVANSDYT | 0.6543802 |
| IRLAPEHTSAMD | ASSPTVANSDYT | 0.74915314 |
| IYNVSVL | ASSPTVANSDYT | 0.1315414 |

| | | |
|---|---|---|
| KQGFLGM | ASSPTVANSDYT | 0.7644745 |
| LESVITQL | ASSPTVANSDYT | 0.8779426 |
| LGLGALVVV | ASSPTVANSDYT | 0.7404006 |
| LIGNGGNGGSGGI | ASSPTVANSDYT | 0.80974984 |
| LNAINAPTL | ASSPTVANSDYT | 0.8214432 |
| LQLLNTGV | ASSPTVANSDYT | 0.6319702 |
| LQTSIEF | ASSPTVANSDYT | 0.83141184 |
| MSFAVSEL | ASSPTVANSDYT | 0.6486299 |
| NTGALGL | ASSPTVANSDYT | 0.85627556 |
| PPGAGGAGGAGTT | ASSPTVANSDYT | 0.87301314 |
| QLVGLGI | ASSPTVANSDYT | 0.80901587 |
| RFLEQVHQL | ASSPTVANSDYT | 0.78132355 |
| RGVGYVLR | ASSPTVANSDYT | 0.8074912 |
| SATGVFAGVI | ASSPTVANSDYT | 0.57234526 |
| SLPAITLP | ASSPTVANSDYT | 0.79477817 |
| STALAFTV | ASSPTVANSDYT | 0.6580594 |
| SVLTVGLGK | ASSPTVANSDYT | 0.7493753 |
| TALAGAGSGI | ASSPTVANSDYT | 0.5388093 |
| TGLPFTGI | ASSPTVANSDYT | 0.79116285 |
| VAEHVTV | ASSPTVANSDYT | 0.7509285 |
| VDIVLNSL | ASSPTVANSDYT | 0.8401196 |
| VLTGGGAVL | ASSPTVANSDYT | 0.5033195 |
| VNVGLAYGI | ASSPTVANSDYT | 0.80369616 |
| VSSAAVGAV | ASSPTVANSDYT | 0.77524114 |
| VTSVYEAGV | ASSPTVANSDYT | 0.41460872 |
| WVALDRGA | ASSPTVANSDYT | 0.85718876 |
| YLVGSVVI | ASSPTVANSDYT | 0.76382446 |
| GPPGTGKTLL | ASSPTVANSDYT | 0.7358979 |
| GVGMTKF | ASSPTVANSDYT | 0.8562752 |
| IGIGGSDL | ASSPTVANSDYT | 0.7500135 |
| IGTLLAEL | ASSPTVANSDYT | 0.5208348 |
| IYTSGTT | ASSPTVANSDYT | 0.63692814 |
| LSFPAGV | ASSPTVANSDYT | 0.7644735 |
| LYTSGTT | ASSPTVANSDYT | 0.69839066 |
| VGAFPFGF | ASSPTVANSDYT | 0.21939033 |
| VGAGTVEF | ASSPTVANSDYT | 0.8217868 |
| VGKSTLLL | ASSPTVANSDYT | 0.4583597 |
| AALAHAGGGIV | ASSPTVANSDYT | 0.65550905 |
| AALVAVSLL | ASSPTVANSDYT | 0.64155823 |
| AAVSRAVAAL | ASSPTVANSDYT | 0.84701204 |
| AGGSSGIGGF | ASSPTVANSDYT | 0.7821171 |
| AGLTVANANG | ASSPTVANSDYT | 0.62242246 |
| AGLVGAVL | ASSPTVANSDYT | 0.8252293 |
| ALSQAFTV | ASSPTVANSDYT | 0.81171787 |
| ALTFSVSL | ASSPTVANSDYT | 0.84118515 |
| ALTGAALVAL | ASSPTVANSDYT | 0.76750934 |
| ASFSIGAGVL | ASSPTVANSDYT | 0.79295206 |
| DGSVYKI | ASSPTVANSDYT | 0.8094808 |
| EQLAEIV | ASSPTVANSDYT | 0.6663249 |
| FAVSELTVM | ASSPTVANSDYT | 0.49038297 |
| GAAPLAAGL | ASSPTVANSDYT | 0.5464729 |

| | | |
|---|---|---|
| GADGGDGGSGGAGG | ASSPTVANSDYT | 0.6285803 |
| GFGLDTGGAGG | ASSPTVANSDYT | 0.41955864 |
| GLASGGLL | ASSPTVANSDYT | 0.7755206 |
| GRIAGAAVLL | ASSPTVANSDYT | 0.8778311 |
| GSAGSLNTSF | ASSPTVANSDYT | 0.78780437 |
| GTGQIGI | ASSPTVANSDYT | 0.30807102 |
| GTLIALIL | ASSPTVANSDYT | 0.69776785 |
| IAATMGFVVG | ASSPTVANSDYT | 0.7483567 |
| IFAQALL | ASSPTVANSDYT | 0.8313358 |
| IGAAVALGF | ASSPTVANSDYT | 0.8500005 |
| IGLTPLAL | ASSPTVANSDYT | 0.85417444 |
| ILGSNTSTL | ASSPTVANSDYT | 0.8313967 |
| IVFSLTVL | ASSPTVANSDYT | 0.62689847 |
| LASKATGF | ASSPTVANSDYT | 0.84126747 |
| LASVALIL | ASSPTVANSDYT | 0.6531297 |
| LFAVGLTL | ASSPTVANSDYT | 0.8184714 |
| LFNGGIGTYI | ASSPTVANSDYT | 0.76984775 |
| LTLGAGII | ASSPTVANSDYT | 0.82489294 |
| LVAAALGDI | ASSPTVANSDYT | 0.66560006 |
| NTAFVQLGI | ASSPTVANSDYT | 0.66382515 |
| QLFSGAEL | ASSPTVANSDYT | 0.37709174 |
| SAATFITL | ASSPTVANSDYT | 0.83456886 |
| SNYGLANLGV | ASSPTVANSDYT | 0.6635363 |
| SYGGAYAVM | ASSPTVANSDYT | 0.5140047 |
| TTGVGGLTLG | ASSPTVANSDYT | 0.65565103 |
| TTQTIYI | ASSPTVANSDYT | 0.87315774 |
| VAAGAGLGAVI | ASSPTVANSDYT | 0.87935185 |
| VAAVVTLL | ASSPTVANSDYT | 0.64799136 |
| VGFLGGLI | ASSPTVANSDYT | 0.5915649 |
| VGVIPILL | ASSPTVANSDYT | 0.80837584 |
| VIFASSLI | ASSPTVANSDYT | 0.6006446 |
| VKGGDGGAAAG | ASSPTVANSDYT | 0.8157823 |
| VTKTFTNI | ASSPTVANSDYT | 0.73904836 |
| VVAAAILL | ASSPTVANSDYT | 0.73475504 |
| YAALDVEL | ASSPTVANSDYT | 0.6467068 |
| LASVALLL | ASSPTVANSDYT | 0.8265406 |
| LDAVHRV | ASSPTVANSDYT | 0.66884196 |
| QGLSSSIF | ASSPTVANSDYT | 0.81083834 |
| RGTFANIRL | ASSPTVANSDYT | 0.49601102 |
| RQLWWGHRI | ASSPTVANSDYT | 0.8482981 |
| YGPPGTGKTLL | ASSPTVANSDYT | 0.46935147 |
| AALVGAPANVA | ASSPTVANSDYT | 0.55706567 |
| AATFRRTE | ASSPTVANSDYT | 0.23717296 |
| AGALGGGI | ASSPTVANSDYT | 0.7412816 |
| AGGAGGAGGIGAG | ASSPTVANSDYT | 0.82521886 |
| AGSGALGL | ASSPTVANSDYT | 0.8712931 |
| AQIADQI | ASSPTVANSDYT | 0.7616783 |
| ASAVAVTYL | ASSPTVANSDYT | 0.8005184 |
| ASTVLGGTGG | ASSPTVANSDYT | 0.8693616 |
| AVGGEPAK | ASSPTVANSDYT | 0.65173686 |
| CEQGTFGGTC | ASSPTVANSDYT | 0.2741887 |

| | | |
|---|---|---|
| DEGKAPAG | ASSPTVANSDYT | 0.79709315 |
| DTVVFPAL | ASSPTVANSDYT | 0.7341638 |
| FEGITMAG | ASSPTVANSDYT | 0.86697584 |
| FPLVVQD | ASSPTVANSDYT | 0.8331655 |
| GAFTTPQ | ASSPTVANSDYT | 0.3450307 |
| GFGGFGTAGGVGGA | ASSPTVANSDYT | 0.84980315 |
| GFLAQFL | ASSPTVANSDYT | 0.82652545 |
| GGAGGAGGNAGVA | ASSPTVANSDYT | 0.4638427 |
| GGVGTAGAGGAGGAGG | ASSPTVANSDYT | 0.654824 |
| GNGGGGDAGNAGSGGN | ASSPTVANSDYT | 0.5758221 |
| GTGGAGGAGNDAGS | ASSPTVANSDYT | 0.48769265 |
| GTTNTGWFNS | ASSPTVANSDYT | 0.82891273 |
| IFMGPLFTL | ASSPTVANSDYT | 0.6465753 |
| ILAHAIEAI | ASSPTVANSDYT | 0.69347024 |
| IRSAIREAL | ASSPTVANSDYT | 0.68108535 |
| ITFAGGG | ASSPTVANSDYT | 0.6006665 |
| ITGMSGHQEV | ASSPTVANSDYT | 0.6690881 |
| IVAGLACTL | ASSPTVANSDYT | 0.69076633 |
| IVGGGLAGA | ASSPTVANSDYT | 0.7996483 |
| IVTGVLNI | ASSPTVANSDYT | 0.83935773 |
| LAQQGCI | ASSPTVANSDYT | 0.69262916 |
| LCLHVEN | ASSPTVANSDYT | 0.8085755 |
| LDAISDGS | ASSPTVANSDYT | 0.871529 |
| LGGFAAML | ASSPTVANSDYT | 0.45952582 |
| LGGKNPMI | ASSPTVANSDYT | 0.5512698 |
| LGPKGVAAIL | ASSPTVANSDYT | 0.81520253 |
| LMVGVSI | ASSPTVANSDYT | 0.6603142 |
| LVGAAFVL | ASSPTVANSDYT | 0.851137 |
| LVLAVAPL | ASSPTVANSDYT | 0.77505493 |
| MAAIAVIDAA | ASSPTVANSDYT | 0.78571796 |
| NNGINAELQ | ASSPTVANSDYT | 0.74681777 |
| RGEPLVLVHG | ASSPTVANSDYT | 0.8886156 |
| RGGAGGAAGLI | ASSPTVANSDYT | 0.8084307 |
| RSYRMHPAV | ASSPTVANSDYT | 0.6837088 |
| SAAHLLAGQLA | ASSPTVANSDYT | 0.79732805 |
| SAAVGSY | ASSPTVANSDYT | 0.42594367 |
| SPHAVLHRI | ASSPTVANSDYT | 0.8344213 |
| SQIAKVL | ASSPTVANSDYT | 0.8264817 |
| TALGHLVP | ASSPTVANSDYT | 0.52815384 |
| TAPTTSGAA | ASSPTVANSDYT | 0.64227736 |
| TDGDDGNGGSPGT | ASSPTVANSDYT | 0.89306617 |
| TDGLGEN | ASSPTVANSDYT | 0.84323925 |
| TGGPPLPGQA | ASSPTVANSDYT | 0.60395193 |
| TIPAITI | ASSPTVANSDYT | 0.5583262 |
| TIPGFTI | ASSPTVANSDYT | 0.62096983 |
| TSIASGALA | ASSPTVANSDYT | 0.6987605 |
| TVTLPGAAI | ASSPTVANSDYT | 0.2878149 |
| VASGLSAALP | ASSPTVANSDYT | 0.8230351 |
| VGAGVIGI | ASSPTVANSDYT | 0.6805143 |
| VGPKRFGV | ASSPTVANSDYT | 0.8235719 |
| VNVAFPD | ASSPTVANSDYT | 0.7966876 |

| | | |
|---|---|---|
| VSMTNSGVS | ASSPTVANSDYT | 0.86983204 |
| VTDVLGDLSV | ASSPTVANSDYT | 0.6395886 |
| VVVVGAGFAGL | ASSPTVANSDYT | 0.6635027 |
| VYTVGTVV | ASSPTVANSDYT | 0.27344507 |
| WTTFSGP | ASSPTVANSDYT | 0.54650486 |
| YDELVEAF | ASSPTVANSDYT | 0.61672705 |
| YGTGGAGGNGG | ASSPTVANSDYT | 0.73484945 |
| YTNVMAV | ASSPTVANSDYT | 0.43051398 |
| FLEQVHQL | ASSPTVANSDYT | 0.86876595 |
| GLSLLAVAV | ASSPTVANSDYT | 0.64315104 |
| LEQVHQL | ASSPTVANSDYT | 0.83601177 |
| LNAAGAI | ASSPTVANSDYT | 0.7190965 |
| PAVHASL | ASSPTVANSDYT | 0.5074533 |
| TELEDDA | ASSPTVANSDYT | 0.6215178 |
| TPPGPPN | ASSPTVANSDYT | 0.83123803 |
| TPPPPPPP | ASSPTVANSDYT | 0.44885302 |
| TQSPALI | ASSSTGGNTGQLY | 0.30429333 |
| EFAAAFPQ | ASSSTGGNTGQLY | 0.57433563 |
| LLSGNLDVL | ASSSTGGNTGQLY | 0.776285 |
| GPANLGNYNI | ASSSTGGNTGQLY | 0.87982666 |
| TLGPTLP | ASSSTGGNTGQLY | 0.8609638 |
| VYAYSSCCK | ASSSTGGNTGQLY | 0.747711 |
| APAPAPAPNI | ASSSTGGNTGQLY | 0.88221204 |
| PGGPVPIP | ASSSTGGNTGQLY | 0.13867496 |
| FLTVALEP | ASSSTGGNTGQLY | 0.68401134 |
| SQYFGANL | ASSSTGGNTGQLY | 0.8768733 |
| GGVKPAA | ASSSTGGNTGQLY | 0.81586474 |
| GGTGGAGGAGGW | ASSSTGGNTGQLY | 0.718734 |
| GVGGVAGLGGAGG | ASSSTGGNTGQLY | 0.8559931 |
| VLLVLATTI | ASSSTGGNTGQLY | 0.56529814 |
| IGIFAAML | ASSSTGGNTGQLY | 0.6927006 |
| AVGDLID | ASSSTGGNTGQLY | 0.7258331 |
| NISPLPL | ASSSTGGNTGQLY | 0.8405681 |
| VTAVFVAI | ASSSTGGNTGQLY | 0.8543749 |
| SGDDNDAAL | ASSSTGGNTGQLY | 0.30810702 |
| AVAALAHS | ASSSTGGNTGQLY | 0.8385712 |
| GGVTVMATP | ASSSTGGNTGQLY | 0.89046943 |
| AGLAANVRD | ASSSTGGNTGQLY | 0.60278606 |
| TNRPDLIDPALLRPG | ASSSTGGNTGQLY | 0.8591175 |
| LANLGVTAL | ASSSTGGNTGQLY | 0.7700324 |
| TGERIEL | ASSSTGGNTGQLY | 0.73720676 |
| NGEELAA | ASSSTGGNTGQLY | 0.55532694 |
| VLSLVWG | ASSSTGGNTGQLY | 0.4802624 |
| IGYALWL | ASSSTGGNTGQLY | 0.8013859 |
| LSLLLAPL | ASSSTGGNTGQLY | 0.52381 |
| TLDEALK | ASSSTGGNTGQLY | 0.75524235 |
| TDGAGTAEGG | ASSSTGGNTGQLY | 0.74953985 |
| NAPGGAGGNGG | ASSSTGGNTGQLY | 0.8997803 |
| VSVFMGMGGNF | ASSSTGGNTGQLY | 0.8070735 |
| LASYLAAL | ASSSTGGNTGQLY | 0.81097996 |
| LVGGTPAL | ASSSTGGNTGQLY | 0.57314533 |

| | | |
|---|---|---|
| VRNADVL | ASSSTGGNTGQLY | 0.8155134 |
| TCSSPFLVSAM | ASSSTGGNTGQLY | 0.8156673 |
| VHITDAE | ASSSTGGNTGQLY | 0.4364729 |
| EYAAPSF | ASSSTGGNTGQLY | 0.8672595 |
| VVDAFHGGL | ASSSTGGNTGQLY | 0.48851633 |
| ASVVDPQ | ASSSTGGNTGQLY | 0.78686935 |
| SYNEFGGLS | ASSSTGGNTGQLY | 0.7600329 |
| DPDGSVY | ASSSTGGNTGQLY | 0.82296884 |
| MVSTIHATE | ASSSTGGNTGQLY | 0.7441882 |
| LAAGVADPV | ASSSTGGNTGQLY | 0.83632374 |
| TKEQIAA | ASSSTGGNTGQLY | 0.49802464 |
| SLASPAAL | ASSSTGGNTGQLY | 0.8367302 |
| IGLVFYGV | ASSSTGGNTGQLY | 0.85147977 |
| FLLDASNL | ASSSTGGNTGQLY | 0.78466713 |
| YQVGNFP | ASSSTGGNTGQLY | 0.8748566 |
| ESLLSQST | ASSSTGGNTGQLY | 0.82638013 |
| LFTVGFAAVG | ASSSTGGNTGQLY | 0.24491514 |
| ATPPTPT | ASSSTGGNTGQLY | 0.6988108 |
| TDLATLG | ASSSTGGNTGQLY | 0.86127377 |
| SAGGTGR | ASSSTGGNTGQLY | 0.87021446 |
| GFSNSGS | ASSSTGGNTGQLY | 0.84627867 |
| GGNGGSGGTSV | ASSSTGGNTGQLY | 0.84108746 |
| GTSVATGGAGN | ASSSTGGNTGQLY | 0.70491934 |
| NGGNGGSGGTS | ASSSTGGNTGQLY | 0.87277246 |
| AVVKFPH | ASSSTGGNTGQLY | 0.9107211 |
| TPPPPPP | ASSSTGGNTGQLY | 0.8403201 |
| AAPAAAAAPA | ASSSTGGNTGQLY | 0.5639661 |
| IALNEDL | ASSSTGGNTGQLY | 0.52970994 |
| IVNVSSM | ASSSTGGNTGQLY | 0.8336064 |
| TGIPPAPR | ASSSTGGNTGQLY | 0.83818793 |
| FDLGGGTF | ASSSTGGNTGQLY | 0.44572705 |
| FDLGGGTFDVS | ASSSTGGNTGQLY | 0.6997305 |
| NEPTAAA | ASSSTGGNTGQLY | 0.75020593 |
| NEPTAAALA | ASSSTGGNTGQLY | 0.20311403 |
| GSIINISSI | ASSSTGGNTGQLY | 0.8301443 |
| VAPGNAGTA | ASSSTGGNTGQLY | 0.8786144 |
| TGGGASLE | ASSSTGGNTGQLY | 0.8076122 |
| VAVYNHYKRI | ASSSTGGNTGQLY | 0.42051265 |
| TDPAPATT | ASSSTGGNTGQLY | 0.8646661 |
| TNRPDILDPALLRPG | ASSSTGGNTGQLY | 0.878603 |
| LTTTTTS | ASSSTGGNTGQLY | 0.8745103 |
| PTGSGKTL | ASSSTGGNTGQLY | 0.35075772 |
| TGSGKTL | ASSSTGGNTGQLY | 0.14941284 |
| TIGHVDHGK | ASSSTGGNTGQLY | 0.79798746 |
| TIGHVDHGKTT | ASSSTGGNTGQLY | 0.8105314 |
| GHVDHGKTT | ASSSTGGNTGQLY | 0.6365263 |
| AQFEHTLL | ASSSTGGNTGQLY | 0.46948832 |
| IVNQDEL | ASSSTGGNTGQLY | 0.7630177 |
| TNGSQFF | ASSSTGGNTGQLY | 0.85531706 |
| TGKGIVE | ASSSTGGNTGQLY | 0.4041432 |
| LISNASDA | ASSSTGGNTGQLY | 0.46819878 |

| | | |
|---|---|---|
| AELRTLGELTKIAKAHGAQ | ASSSTGGNTGQLY | 0.82386863 |
| APNAVAAAEPSW | ASSSTGGNTGQLY | 0.85303766 |
| APQVAGAATAVL | ASSSTGGNTGQLY | 0.5079508 |
| ARRDPAAAAAILPTDA | ASSSTGGNTGQLY | 0.6304602 |
| DAAARDVLAEAGLAEYF | ASSSTGGNTGQLY | 0.8113984 |
| DDQIVLSAKANLAKVEQ | ASSSTGGNTGQLY | 0.79960334 |
| DELLPDTGEDSDEDGH | ASSSTGGNTGQLY | 0.7875931 |
| ELPEAPANRATVPQD | ASSSTGGNTGQLY | 0.3698334 |
| FGQSKIERVTGIG | ASSSTGGNTGQLY | 0.84246325 |
| GAGGTGSTAGGAGGAGGAGGLY | ASSSTGGNTGQLY | 0.43983468 |
| GHGGTGAAGGNGGTGGAGGSGI | ASSSTGGNTGQLY | 0.8641644 |
| GNGGNGGAGGEGGPGGAGGAGA | ASSSTGGNTGQLY | 0.7679329 |
| GSTAGNGGNGGNATNGTIAGQP | ASSSTGGNTGQLY | 0.8145169 |
| GTGGDGGNGGNGGTGG | ASSSTGGNTGQLY | 0.6814699 |
| GVRRPAGIPADAPV | ASSSTGGNTGQLY | 0.39912605 |
| GYRQVTDAHLLTLARR | ASSSTGGNTGQLY | 0.48016468 |
| IAKMGQDGHDRG | ASSSTGGNTGQLY | 0.5576667 |
| KAEPSIEIQYT | ASSSTGGNTGQLY | 0.87011266 |
| KPVRIRMAADEDTVAVT | ASSSTGGNTGQLY | 0.7962874 |
| MPPRIAGMRLLVIKPEP | ASSSTGGNTGQLY | 0.84169734 |
| MKFYAVALFRSRGVPPTR | ASSSTGGNTGQLY | 0.7755065 |
| NAGGFGILGAVAHSP | ASSSTGGNTGQLY | 0.44571006 |
| NEGREITLAMAEM | ASSSTGGNTGQLY | 0.83108807 |
| NELVEWSHPLGHRWRWE | ASSSTGGNTGQLY | 0.36317244 |
| NLGAGGTGGNGGNPG | ASSSTGGNTGQLY | 0.7754693 |
| NTEWPKWETVMAADPVD | ASSSTGGNTGQLY | 0.73643243 |
| PASAAPAETQP | ASSSTGGNTGQLY | 0.81870776 |
| QQSSDGKNGVRAAEITGE | ASSSTGGNTGQLY | 0.6485354 |
| RAPDAVTTSWWKE | ASSSTGGNTGQLY | 0.82722116 |
| SGFGNVGAHTSGWFNQ | ASSSTGGNTGQLY | 0.80816174 |
| SGVAGAVGQAASVGGLKV | ASSSTGGNTGQLY | 0.526444 |
| SNGGAGTGSAGGNGGT | ASSSTGGNTGQLY | 0.7960835 |
| TGGAGGEGGAAG | ASSSTGGNTGQLY | 0.83123696 |
| TGGDGGAAGAGGAPGHG | ASSSTGGNTGQLY | 0.82649374 |
| TMPPLLAAIANAGRHGV | ASSSTGGNTGQLY | 0.3236406 |
| TSSTAGNSGGA | ASSSTGGNTGQLY | 0.799626 |
| VPATVPVGILC | ASSSTGGNTGQLY | 0.5197538 |
| VQRPDSGGVVLTGRIS | ASSSTGGNTGQLY | 0.78188777 |
| WTPNPVPATKGVPAGLVQAT | ASSSTGGNTGQLY | 0.7474451 |
| YALKRALAKFSDRFDVVII | ASSSTGGNTGQLY | 0.684989 |
| YLIGSVIV | ASSSTGGNTGQLY | 0.842174 |
| YGQLVAVID | ASSSTGGNTGQLY | 0.41761518 |
| VTVGGMTLPA | ASSSTGGNTGQLY | 0.67051935 |
| VLALPAV | ASSSTGGNTGQLY | 0.6764276 |
| VIPPTLNYET | ASSSTGGNTGQLY | 0.7887244 |
| VGMGGMHPSSG | ASSSTGGNTGQLY | 0.8265984 |
| VGLAGGAATA | ASSSTGGNTGQLY | 0.42488867 |
| VGGAGGAGGSGP | ASSSTGGNTGQLY | 0.7890197 |
| VFAAFMV | ASSSTGGNTGQLY | 0.69995517 |
| TLYDKLR | ASSSTGGNTGQLY | 0.47755632 |
| TLAVFTVL | ASSSTGGNTGQLY | 0.7369587 |

| | | |
|---|---|---|
| TGSVQPARV | ASSSTGGNTGQLY | 0.66023964 |
| TGGGAGSGGT | ASSSTGGNTGQLY | 0.38876992 |
| SHLETSL | ASSSTGGNTGQLY | 0.8402609 |
| SFTTGMT | ASSSTGGNTGQLY | 0.58393204 |
| QVEVAKRMGVRQPTVS | ASSSTGGNTGQLY | 0.8072703 |
| QGSGGPALAM | ASSSTGGNTGQLY | 0.8781159 |
| MLAPSNP | ASSSTGGNTGQLY | 0.4885239 |
| LVNNAAI | ASSSTGGNTGQLY | 0.8600125 |
| LTGEINTLA | ASSSTGGNTGQLY | 0.74414325 |
| LSIFGPV | ASSSTGGNTGQLY | 0.7711898 |
| LLALLANL | ASSSTGGNTGQLY | 0.72470677 |
| LISNASDALDKLR | ASSSTGGNTGQLY | 0.56631446 |
| LINEAVP | ASSSTGGNTGQLY | 0.8621058 |
| LGYAFSGL | ASSSTGGNTGQLY | 0.837456 |
| LDFVAVAL | ASSSTGGNTGQLY | 0.83011436 |
| LAQPGAE | ASSSTGGNTGQLY | 0.80958116 |
| ISNLGAVTGF | ASSSTGGNTGQLY | 0.8168612 |
| HQEDPGRI | ASSSTGGNTGQLY | 0.84875035 |
| GTPGEPTP | ASSSTGGNTGQLY | 0.5918495 |
| GFFNSGS | ASSSTGGNTGQLY | 0.32669878 |
| GAGGGGLTAGAN | ASSSTGGNTGQLY | 0.680849 |
| FVDALVNM | ASSSTGGNTGQLY | 0.7084917 |
| FLTVAPEL | ASSSTGGNTGQLY | 0.54116017 |
| FETLTAL | ASSSTGGNTGQLY | 0.3583606 |
| DVINPIEL | ASSSTGGNTGQLY | 0.8178878 |
| APVSGAPV | ASSSTGGNTGQLY | 0.89435756 |
| ALVGGLAW | ASSSTGGNTGQLY | 0.6897311 |
| ALAAAAATAI | ASSSTGGNTGQLY | 0.8835533 |
| AFGLGLLL | ASSSTGGNTGQLY | 0.3760383 |
| AFFAAASL | ASSSTGGNTGQLY | 0.84133077 |
| AAAALPAFSPP | ASSSTGGNTGQLY | 0.85001326 |
| VVLAPAL | ASSSTGGNTGQLY | 0.4877541 |
| TIGHVDHGKTTL | ASSSTGGNTGQLY | 0.76157284 |
| LISNASDALDK | ASSSTGGNTGQLY | 0.6825658 |
| LISNASDAL | ASSSTGGNTGQLY | 0.59089553 |
| ISNASDALDK | ASSSTGGNTGQLY | 0.862206 |
| NQAETLVYQTE | ASSSTGGNTGQLY | 0.6757897 |
| TGGTGGTAGAG | ASSSTGGNTGQLY | 0.82055557 |
| TGSTAGGAGGA | ASSSTGGNTGQLY | 0.5812753 |
| DPLAVPVAAGPV | ASSSTGGNTGQLY | 0.5182973 |
| GGTGGTPGNGGN | ASSSTGGNTGQLY | 0.788334 |
| PAGLAGVDPATL | ASSSTGGNTGQLY | 0.83815897 |
| GNNLVVEAPHAGASV | ASSSTGGNTGQLY | 0.75105643 |
| KVTRIMSASTLEQPA | ASSSTGGNTGQLY | 0.90818346 |
| LGFALPIALPRRTVV | ASSSTGGNTGQLY | 0.651736 |
| TRLGNLTARKLVVLV | ASSSTGGNTGQLY | 0.88194525 |
| WSQQRRLIYFPSAGP | ASSSTGGNTGQLY | 0.86525667 |
| GPNGNPDPMAAAVDIR | ASSSTGGNTGQLY | 0.750401 |
| TQRCVSVADLRAAASR | ASSSTGGNTGQLY | 0.66275215 |
| QFVLDVAEHPVEVLTETV | ASSSTGGNTGQLY | 0.8398912 |
| GGAGGTGGAGGNGGTQAGDG | ASSSTGGNTGQLY | 0.59766066 |

| | | |
|---|---|---|
| PLLATLRAIAVDVDAKPAQV | ASSSTGGNTGQLY | 0.8441534 |
| APKFRTGGYMIDFWGVGYQVAK | ASSSTGGNTGQLY | 0.868678 |
| GAGGAGGAGGAAGGPGGTGGTG | ASSSTGGNTGQLY | 0.4949066 |
| IGALGLLATDVAPAGSVAFGVA | ASSSTGGNTGQLY | 0.7592992 |
| AGGAGGIGGNGGAGGAGGVLMG | ASSSTGGNTGQLY | 0.7323191 |
| ADELIGLL | ASSSTGGNTGQLY | 0.693131 |
| ADRALKA | ASSSTGGNTGQLY | 0.65109795 |
| AGAVGQVGGAGGA | ASSSTGGNTGQLY | 0.44889045 |
| AGGGGGLGGAP | ASSSTGGNTGQLY | 0.6105548 |
| AGGLLYGNGGAG | ASSSTGGNTGQLY | 0.6207609 |
| AGLESIGL | ASSSTGGNTGQLY | 0.75625527 |
| AGLKTGII | ASSSTGGNTGQLY | 0.7387465 |
| ASLAALLL | ASSSTGGNTGQLY | 0.6250202 |
| ASVINISSI | ASSSTGGNTGQLY | 0.74220634 |
| ATPPGPA | ASSSTGGNTGQLY | 0.74908125 |
| ATVASALV | ASSSTGGNTGQLY | 0.4436683 |
| ATVASLIL | ASSSTGGNTGQLY | 0.6896416 |
| AVNGPVLIRH | ASSSTGGNTGQLY | 0.54889786 |
| DGAAGAVGI | ASSSTGGNTGQLY | 0.6729914 |
| DPLLHGI | ASSSTGGNTGQLY | 0.33027512 |
| ELGAALGAL | ASSSTGGNTGQLY | 0.73904026 |
| ELGRTQAV | ASSSTGGNTGQLY | 0.63108766 |
| FVAGGYLS | ASSSTGGNTGQLY | 0.26889867 |
| GASGGAGGAGGW | ASSSTGGNTGQLY | 0.7161608 |
| GGAGAGGDGGAGGI | ASSSTGGNTGQLY | 0.68418676 |
| GTGHNGSQPGMGGNG | ASSSTGGNTGQLY | 0.6938261 |
| GYAAASASAS | ASSSTGGNTGQLY | 0.74624157 |
| HQVDVAAVQQG | ASSSTGGNTGQLY | 0.6579088 |
| IGAAIAHT | ASSSTGGNTGQLY | 0.5771099 |
| IGMIEVAQL | ASSSTGGNTGQLY | 0.73603046 |
| IRLAPEHTSAMD | ASSSTGGNTGQLY | 0.752154 |
| IYNVSVL | ASSSTGGNTGQLY | 0.5467166 |
| KQGFLGM | ASSSTGGNTGQLY | 0.72582996 |
| LESVITQL | ASSSTGGNTGQLY | 0.7551702 |
| LGLGALVVV | ASSSTGGNTGQLY | 0.7185874 |
| LIGNGGNGGSGGI | ASSSTGGNTGQLY | 0.7528764 |
| LNAINAPTL | ASSSTGGNTGQLY | 0.75901973 |
| LQLLNTGV | ASSSTGGNTGQLY | 0.5984173 |
| LQTSIEF | ASSSTGGNTGQLY | 0.7515065 |
| MSFAVSEL | ASSSTGGNTGQLY | 0.74669427 |
| NTGALGL | ASSSTGGNTGQLY | 0.6549917 |
| PPGAGGAGGAGTT | ASSSTGGNTGQLY | 0.6889572 |
| QLVGLGI | ASSSTGGNTGQLY | 0.43921387 |
| RFLEQVHQL | ASSSTGGNTGQLY | 0.6749535 |
| RGVGYVLR | ASSSTGGNTGQLY | 0.57321703 |
| SATGVFAGVI | ASSSTGGNTGQLY | 0.7348444 |
| SLPAITLP | ASSSTGGNTGQLY | 0.75801885 |
| STALAFTV | ASSSTGGNTGQLY | 0.694123 |
| SVLTVGLGK | ASSSTGGNTGQLY | 0.5420069 |
| TALAGAGSGI | ASSSTGGNTGQLY | 0.5505142 |
| TGLPFTGI | ASSSTGGNTGQLY | 0.73122936 |

| | | |
|---|---|---|
| VAEHVTV | ASSSTGGNTGQLY | 0.51838243 |
| VDIVLNSL | ASSSTGGNTGQLY | 0.37829316 |
| VLTGGGAVL | ASSSTGGNTGQLY | 0.68267953 |
| VNVGLAYGI | ASSSTGGNTGQLY | 0.7462426 |
| VSSAAVGAV | ASSSTGGNTGQLY | 0.7517026 |
| VTSVYEAGV | ASSSTGGNTGQLY | 0.59055185 |
| WVALDRGA | ASSSTGGNTGQLY | 0.7568837 |
| YLVGSVVI | ASSSTGGNTGQLY | 0.74522674 |
| GPPGTGKTLL | ASSSTGGNTGQLY | 0.7549972 |
| GVGMTKF | ASSSTGGNTGQLY | 0.6844161 |
| IGIGGSDL | ASSSTGGNTGQLY | 0.53807735 |
| IGTLLAEL | ASSSTGGNTGQLY | 0.75299454 |
| IYTSGTT | ASSSTGGNTGQLY | 0.7005547 |
| LSFPAGV | ASSSTGGNTGQLY | 0.69798017 |
| LYTSGTT | ASSSTGGNTGQLY | 0.7382146 |
| VGAFPFGF | ASSSTGGNTGQLY | 0.6949142 |
| VGAGTVEF | ASSSTGGNTGQLY | 0.635746 |
| VGKSTLLL | ASSSTGGNTGQLY | 0.71414614 |
| AALAHAGGGIV | ASSSTGGNTGQLY | 0.75467265 |
| AALVAVSLL | ASSSTGGNTGQLY | 0.61257285 |
| AAVSRAVAAL | ASSSTGGNTGQLY | 0.5652833 |
| AGGSSGIGGF | ASSSTGGNTGQLY | 0.6850718 |
| AGLTVANANG | ASSSTGGNTGQLY | 0.7085482 |
| AGLVGAVL | ASSSTGGNTGQLY | 0.75336635 |
| ALSQAFTV | ASSSTGGNTGQLY | 0.7254945 |
| ALTFSVSL | ASSSTGGNTGQLY | 0.6783534 |
| ALTGAALVAL | ASSSTGGNTGQLY | 0.72308934 |
| ASFSIGAGVL | ASSSTGGNTGQLY | 0.7387105 |
| DGSVYKI | ASSSTGGNTGQLY | 0.71524763 |
| EQLAEIV | ASSSTGGNTGQLY | 0.7458658 |
| FAVSELTVM | ASSSTGGNTGQLY | 0.6701264 |
| GAAPLAAGL | ASSSTGGNTGQLY | 0.74984646 |
| GADGGDGGSGGAGG | ASSSTGGNTGQLY | 0.74278986 |
| GFGLDTGGAGG | ASSSTGGNTGQLY | 0.73462397 |
| GLASGGLL | ASSSTGGNTGQLY | 0.75146425 |
| GRIAGAAVLL | ASSSTGGNTGQLY | 0.7333098 |
| GSAGSLNTSF | ASSSTGGNTGQLY | 0.69132984 |
| GTGQIGI | ASSSTGGNTGQLY | 0.46258056 |
| GTLIALIL | ASSSTGGNTGQLY | 0.6406102 |
| IAATMGFVVG | ASSSTGGNTGQLY | 0.71194553 |
| IFAQALL | ASSSTGGNTGQLY | 0.7288104 |
| IGAAVALGF | ASSSTGGNTGQLY | 0.7419815 |
| IGLTPLAL | ASSSTGGNTGQLY | 0.71071327 |
| ILGSNTSTL | ASSSTGGNTGQLY | 0.6410259 |
| IVFSLTVL | ASSSTGGNTGQLY | 0.7548928 |
| LASKATGF | ASSSTGGNTGQLY | 0.7476206 |
| LASVALIL | ASSSTGGNTGQLY | 0.48649502 |
| LFAVGLTL | ASSSTGGNTGQLY | 0.68780184 |
| LFNGGIGTYI | ASSSTGGNTGQLY | 0.7667382 |
| LTLGAGII | ASSSTGGNTGQLY | 0.7230463 |
| LVAAALGDI | ASSSTGGNTGQLY | 0.6523918 |

| | | |
|---|---|---|
| NTAFVQLGI | ASSSTGGNTGQLY | 0.6113901 |
| QLFSGAEL | ASSSTGGNTGQLY | 0.76165795 |
| SAATFITL | ASSSTGGNTGQLY | 0.6580037 |
| SNYGLANLGV | ASSSTGGNTGQLY | 0.75618243 |
| SYGGAYAVM | ASSSTGGNTGQLY | 0.34596613 |
| TTGVGGLTLG | ASSSTGGNTGQLY | 0.62554747 |
| TTQTIYI | ASSSTGGNTGQLY | 0.6204126 |
| VAAGAGLGAVI | ASSSTGGNTGQLY | 0.29259026 |
| VAAVVTLL | ASSSTGGNTGQLY | 0.7458559 |
| VGFLGGLI | ASSSTGGNTGQLY | 0.49035606 |
| VGVIPILL | ASSSTGGNTGQLY | 0.7225559 |
| VIFASSLI | ASSSTGGNTGQLY | 0.64056206 |
| VKGGDGGAAAG | ASSSTGGNTGQLY | 0.7529857 |
| VTKTFTNI | ASSSTGGNTGQLY | 0.7101883 |
| VVAAAILL | ASSSTGGNTGQLY | 0.70054543 |
| YAALDVEL | ASSSTGGNTGQLY | 0.5690651 |
| LASVALLL | ASSSTGGNTGQLY | 0.73043966 |
| LDAVHRV | ASSSTGGNTGQLY | 0.754217 |
| QGLSSSIF | ASSSTGGNTGQLY | 0.7413993 |
| RGTFANIRL | ASSSTGGNTGQLY | 0.36996725 |
| RQLWWGHRI | ASSSTGGNTGQLY | 0.7251756 |
| YGPPGTGKTLL | ASSSTGGNTGQLY | 0.6911532 |
| AALVGAPANVA | ASSSTGGNTGQLY | 0.4646652 |
| AATFRRTE | ASSSTGGNTGQLY | 0.29530227 |
| AGALGGGI | ASSSTGGNTGQLY | 0.6901518 |
| AGGAGGAGGIGAG | ASSSTGGNTGQLY | 0.70581627 |
| AGSGALGL | ASSSTGGNTGQLY | 0.7337693 |
| AQIADQI | ASSSTGGNTGQLY | 0.7382836 |
| ASAVAVTYL | ASSSTGGNTGQLY | 0.6692078 |
| ASTVLGGTGG | ASSSTGGNTGQLY | 0.73522556 |
| AVGGEPAK | ASSSTGGNTGQLY | 0.71808755 |
| CEQGTFGGTC | ASSSTGGNTGQLY | 0.21254532 |
| DEGKAPAG | ASSSTGGNTGQLY | 0.6256043 |
| DTVVFPAL | ASSSTGGNTGQLY | 0.43099567 |
| FEGITMAG | ASSSTGGNTGQLY | 0.7542788 |
| FPLVVQD | ASSSTGGNTGQLY | 0.62805474 |
| GAFTTPQ | ASSSTGGNTGQLY | 0.6470682 |
| GFGGFGTAGGVGGA | ASSSTGGNTGQLY | 0.70555747 |
| GFLAQFL | ASSSTGGNTGQLY | 0.7350121 |
| GGAGGAGGNAGVA | ASSSTGGNTGQLY | 0.7091725 |
| GGVGTAGAGGAGGAGG | ASSSTGGNTGQLY | 0.6321558 |
| GNGGGGDAGNAGSGGN | ASSSTGGNTGQLY | 0.44282293 |
| GTGGAGGAGNDAGS | ASSSTGGNTGQLY | 0.7328831 |
| GTTNTGWFNS | ASSSTGGNTGQLY | 0.6380743 |
| IFMGPLFTL | ASSSTGGNTGQLY | 0.54948276 |
| ILAHAIEAI | ASSSTGGNTGQLY | 0.45236316 |
| IRSAIREAL | ASSSTGGNTGQLY | 0.72188234 |
| ITFAGGG | ASSSTGGNTGQLY | 0.38015956 |
| ITGMSGHQEV | ASSSTGGNTGQLY | 0.73812467 |
| IVAGLACTL | ASSSTGGNTGQLY | 0.5172885 |
| IVGGGLAGA | ASSSTGGNTGQLY | 0.6897339 |

| | | |
|---|---|---|
| IVTGVLNI | ASSSTGGNTGQLY | 0.76338875 |
| LAQQGCI | ASSSTGGNTGQLY | 0.7300443 |
| LCLHVEN | ASSSTGGNTGQLY | 0.6943382 |
| LDAISDGS | ASSSTGGNTGQLY | 0.6064211 |
| LGGFAAML | ASSSTGGNTGQLY | 0.5529533 |
| LGGKNPMI | ASSSTGGNTGQLY | 0.62083673 |
| LGPKGVAAIL | ASSSTGGNTGQLY | 0.6341953 |
| LMVGVSI | ASSSTGGNTGQLY | 0.7572526 |
| LVGAAFVL | ASSSTGGNTGQLY | 0.6749888 |
| LVLAVAPL | ASSSTGGNTGQLY | 0.41577303 |
| MAAIAVIDAA | ASSSTGGNTGQLY | 0.7166381 |
| NNGINAELQ | ASSSTGGNTGQLY | 0.76013947 |
| RGEPLVLVHG | ASSSTGGNTGQLY | 0.7627585 |
| RGGAGGAAGLI | ASSSTGGNTGQLY | 0.59811485 |
| RSYRMHPAV | ASSSTGGNTGQLY | 0.64277714 |
| SAAHLLAGQLA | ASSSTGGNTGQLY | 0.57453156 |
| SAAVGSY | ASSSTGGNTGQLY | 0.6982194 |
| SPHAVLHRI | ASSSTGGNTGQLY | 0.5627482 |
| SQIAKVL | ASSSTGGNTGQLY | 0.5908158 |
| TALGHLVP | ASSSTGGNTGQLY | 0.73968726 |
| TAPTTSGAA | ASSSTGGNTGQLY | 0.76749754 |
| TDGDDGNGGSPGT | ASSSTGGNTGQLY | 0.67845464 |
| TDGLGEN | ASSSTGGNTGQLY | 0.31944513 |
| TGGPPLPGQA | ASSSTGGNTGQLY | 0.69669783 |
| TIPAITI | ASSSTGGNTGQLY | 0.6380307 |
| TIPGFTI | ASSSTGGNTGQLY | 0.65501773 |
| TSIASGALA | ASSSTGGNTGQLY | 0.6128332 |
| TVTLPGAAI | ASSSTGGNTGQLY | 0.44813138 |
| VASGLSAALP | ASSSTGGNTGQLY | 0.69777554 |
| VGAGVIGI | ASSSTGGNTGQLY | 0.7638789 |
| VGPKRFGV | ASSSTGGNTGQLY | 0.6812241 |
| VNVAFPD | ASSSTGGNTGQLY | 0.64323 |
| VSMTNSGVS | ASSSTGGNTGQLY | 0.7567705 |
| VTDVLGDLSV | ASSSTGGNTGQLY | 0.73737234 |
| VVVVGAGFAGL | ASSSTGGNTGQLY | 0.7539542 |
| VYTVGTVV | ASSSTGGNTGQLY | 0.7531382 |
| WTTFSGP | ASSSTGGNTGQLY | 0.691992 |
| YDELVEAF | ASSSTGGNTGQLY | 0.7021341 |
| YGTGGAGGNGG | ASSSTGGNTGQLY | 0.7470443 |
| YTNVMAV | ASSSTGGNTGQLY | 0.688259 |
| FLEQVHQL | ASSSTGGNTGQLY | 0.7068744 |
| GLSLLAVAV | ASSSTGGNTGQLY | 0.61381984 |
| LEQVHQL | ASSSTGGNTGQLY | 0.7441596 |
| LNAAGAI | ASSSTGGNTGQLY | 0.68269527 |
| PAVHASL | ASSSTGGNTGQLY | 0.76029426 |
| TELEDDA | ASSSTGGNTGQLY | 0.29471973 |
| TPPGPPN | ASSSTGGNTGQLY | 0.55809474 |
| TPPPPPPP | ASSSTGGNTGQLY | 0.75296867 |
| TQSPALI | AWSLPGQRNQAPL | 0.23989674 |
| EFAAAFPQ | AWSLPGQRNQAPL | 0.2958735 |
| LLSGNLDVL | AWSLPGQRNQAPL | 0.66255903 |

| | | |
|---|---|---|
| GPANLGNYNI | AWSLPGQRNQAPL | 0.7545717 |
| TLGPTLP | AWSLPGQRNQAPL | 0.26779833 |
| VYAYSSCCK | AWSLPGQRNQAPL | 0.58913016 |
| APAPAPAPNI | AWSLPGQRNQAPL | 0.39866114 |
| PGGPVPIP | AWSLPGQRNQAPL | 0.75342697 |
| FLTVALEP | AWSLPGQRNQAPL | 0.7038374 |
| SQYFGANL | AWSLPGQRNQAPL | 0.47225207 |
| GGVKPAA | AWSLPGQRNQAPL | 0.74335 |
| GGTGGAGGAGGW | AWSLPGQRNQAPL | 0.37253273 |
| GVGGVAGLGGAGG | AWSLPGQRNQAPL | 0.72705734 |
| VLLVLATTI | AWSLPGQRNQAPL | 0.74298173 |
| IGIFAAML | AWSLPGQRNQAPL | 0.7634438 |
| AVGDLID | AWSLPGQRNQAPL | 0.50828993 |
| NISPLPL | AWSLPGQRNQAPL | 0.7343607 |
| VTAVFVAI | AWSLPGQRNQAPL | 0.66462934 |
| SGDDNDAAL | AWSLPGQRNQAPL | 0.7681546 |
| AVAALAHS | AWSLPGQRNQAPL | 0.6563967 |
| GGVTVMATP | AWSLPGQRNQAPL | 0.4396534 |
| AGLAANVRD | AWSLPGQRNQAPL | 0.7497635 |
| TNRPDLIDPALLRPG | AWSLPGQRNQAPL | 0.7238729 |
| LANLGVTAL | AWSLPGQRNQAPL | 0.657264 |
| TGERIEL | AWSLPGQRNQAPL | 0.748374 |
| NGEELAA | AWSLPGQRNQAPL | 0.70097435 |
| VLSLVWG | AWSLPGQRNQAPL | 0.69103587 |
| IGYALWL | AWSLPGQRNQAPL | 0.74588084 |
| LSLLLAPL | AWSLPGQRNQAPL | 0.7443955 |
| TLDEALK | AWSLPGQRNQAPL | 0.6328816 |
| TDGAGTAEGG | AWSLPGQRNQAPL | 0.74820685 |
| NAPGGAGGNGG | AWSLPGQRNQAPL | 0.7089071 |
| VSVFMGMGGNF | AWSLPGQRNQAPL | 0.47727907 |
| LASYLAAL | AWSLPGQRNQAPL | 0.7651701 |
| LVGGTPAL | AWSLPGQRNQAPL | 0.678465 |
| VRNADVL | AWSLPGQRNQAPL | 0.74316096 |
| TCSSPFLVSAM | AWSLPGQRNQAPL | 0.7379638 |
| VHITDAE | AWSLPGQRNQAPL | 0.7599957 |
| EYAAPSF | AWSLPGQRNQAPL | 0.58942115 |
| VVDAFHGGL | AWSLPGQRNQAPL | 0.6948463 |
| ASVVDPQ | AWSLPGQRNQAPL | 0.7065067 |
| SYNEFGGLS | AWSLPGQRNQAPL | 0.7420682 |
| DPDGSVY | AWSLPGQRNQAPL | 0.67062986 |
| MVSTIHATE | AWSLPGQRNQAPL | 0.69879794 |
| LAAGVADPV | AWSLPGQRNQAPL | 0.7271848 |
| TKEQIAA | AWSLPGQRNQAPL | 0.7557609 |
| SLASPAAL | AWSLPGQRNQAPL | 0.6815895 |
| IGLVFYGV | AWSLPGQRNQAPL | 0.7298389 |
| FLLDASNL | AWSLPGQRNQAPL | 0.7616457 |
| YQVGNFP | AWSLPGQRNQAPL | 0.4945336 |
| ESLLSQST | AWSLPGQRNQAPL | 0.6954281 |
| LFTVGFAAVG | AWSLPGQRNQAPL | 0.72804147 |
| ATPPTPT | AWSLPGQRNQAPL | 0.7490288 |
| TDLATLG | AWSLPGQRNQAPL | 0.6899071 |

| | | |
|---|---|---|
| SAGGTGR | AWSLPGQRNQAPL | 0.75740194 |
| GFSNSGS | AWSLPGQRNQAPL | 0.6611177 |
| GGNGGSGGTSV | AWSLPGQRNQAPL | 0.73191893 |
| GTSVATGGAGN | AWSLPGQRNQAPL | 0.6735121 |
| NGGNGGSGGTS | AWSLPGQRNQAPL | 0.66078043 |
| AVVKFPH | AWSLPGQRNQAPL | 0.6350026 |
| TPPPPPP | AWSLPGQRNQAPL | 0.70691633 |
| AAPAAAAAPA | AWSLPGQRNQAPL | 0.7436574 |
| IALNEDL | AWSLPGQRNQAPL | 0.7284379 |
| IVNVSSM | AWSLPGQRNQAPL | 0.27555397 |
| TGIPPAPR | AWSLPGQRNQAPL | 0.7411769 |
| FDLGGGTF | AWSLPGQRNQAPL | 0.7324297 |
| FDLGGGTFDVS | AWSLPGQRNQAPL | 0.76462716 |
| NEPTAAA | AWSLPGQRNQAPL | 0.6963424 |
| NEPTAAALA | AWSLPGQRNQAPL | 0.7048203 |
| GSIINISSI | AWSLPGQRNQAPL | 0.60842013 |
| VAPGNAGTA | AWSLPGQRNQAPL | 0.7057835 |
| TGGGASLE | AWSLPGQRNQAPL | 0.5444422 |
| VAVYNHYKRI | AWSLPGQRNQAPL | 0.5162035 |
| TDPAPATT | AWSLPGQRNQAPL | 0.68577147 |
| TNRPDILDPALLRPG | AWSLPGQRNQAPL | 0.64272153 |
| LTTTTTS | AWSLPGQRNQAPL | 0.7498205 |
| PTGSGKTL | AWSLPGQRNQAPL | 0.751935 |
| TGSGKTL | AWSLPGQRNQAPL | 0.7443034 |
| TIGHVDHGK | AWSLPGQRNQAPL | 0.7528205 |
| TIGHVDHGKTT | AWSLPGQRNQAPL | 0.75553846 |
| GHVDHGKTT | AWSLPGQRNQAPL | 0.7495436 |
| AQFEHTLL | AWSLPGQRNQAPL | 0.6756093 |
| IVNQDEL | AWSLPGQRNQAPL | 0.6707541 |
| TNGSQFF | AWSLPGQRNQAPL | 0.4896769 |
| TGKGIVE | AWSLPGQRNQAPL | 0.6653017 |
| LISNASDA | AWSLPGQRNQAPL | 0.6830375 |
| AELRTLGELTKIAKAHGAQ | AWSLPGQRNQAPL | 0.7171453 |
| APNAVAAAEPSW | AWSLPGQRNQAPL | 0.23557465 |
| APQVAGAATAVL | AWSLPGQRNQAPL | 0.73086166 |
| ARRDPAAAAAILPTDA | AWSLPGQRNQAPL | 0.7587485 |
| DAAARDVLAEAGLAEYF | AWSLPGQRNQAPL | 0.7049554 |
| DDQIVLSAKANLAKVEQ | AWSLPGQRNQAPL | 0.73015016 |
| DELLPDTGEDSDEDGH | AWSLPGQRNQAPL | 0.7280028 |
| ELPEAPANRATVPQD | AWSLPGQRNQAPL | 0.6621902 |
| FGQSKIERVTGIG | AWSLPGQRNQAPL | 0.7327762 |
| GAGGTGSTAGGAGGAGGAGGLY | AWSLPGQRNQAPL | 0.6533024 |
| GHGGTGAAGGNGGTGGAGGSGI | AWSLPGQRNQAPL | 0.738935 |
| GNGGNGGAGGEGGPGGAGGAGA | AWSLPGQRNQAPL | 0.75766325 |
| GSTAGNGGNGGNATNGTIAGQP | AWSLPGQRNQAPL | 0.7153662 |
| GTGGDGGNGGNGGTGG | AWSLPGQRNQAPL | 0.69376767 |
| GVRRPAGIPADAPV | AWSLPGQRNQAPL | 0.66375566 |
| GYRQVTDAHLLTLARR | AWSLPGQRNQAPL | 0.45966637 |
| IAKMGQDGHDRG | AWSLPGQRNQAPL | 0.70988154 |
| KAEPSIEIQYT | AWSLPGQRNQAPL | 0.5658705 |
| KPVRIRMAADEDTVAVT | AWSLPGQRNQAPL | 0.70478714 |

| | | |
|---|---|---|
| MPPRIAGMRLLVIKPEP | AWSLPGQRNQAPL | 0.6149676 |
| MKFYAVALFRSRGVPPTR | AWSLPGQRNQAPL | 0.71747446 |
| NAGGFGILGAVAHSP | AWSLPGQRNQAPL | 0.7081076 |
| NEGREITLAMAEM | AWSLPGQRNQAPL | 0.70800143 |
| NELVEWSHPLGHRWRWE | AWSLPGQRNQAPL | 0.5417 |
| NLGAGGTGGNGGNPG | AWSLPGQRNQAPL | 0.7352339 |
| NTEWPKWETVMAADPVD | AWSLPGQRNQAPL | 0.7052685 |
| PASAAPAETQP | AWSLPGQRNQAPL | 0.5074115 |
| QQSSDGKNGVRAAEITGE | AWSLPGQRNQAPL | 0.7397512 |
| RAPDAVTTSWWKE | AWSLPGQRNQAPL | 0.62900674 |
| SGFGNVGAHTSGWFNQ | AWSLPGQRNQAPL | 0.710063 |
| SGVAGAVGQAASVGGLKV | AWSLPGQRNQAPL | 0.7477342 |
| SNGGAGTGSAGGNGGT | AWSLPGQRNQAPL | 0.70635605 |
| TGGAGGEGGAAG | AWSLPGQRNQAPL | 0.35078415 |
| TGGDGGAAGAGGAPGHG | AWSLPGQRNQAPL | 0.68030566 |
| TMPPLLAAIANAGRHGV | AWSLPGQRNQAPL | 0.690165 |
| TSSTAGNSGGA | AWSLPGQRNQAPL | 0.7013777 |
| VPATVPVGILC | AWSLPGQRNQAPL | 0.25635204 |
| VQRPDSGGVVLTGRIS | AWSLPGQRNQAPL | 0.7374852 |
| WTPNPVPATKGVPAGLVQAT | AWSLPGQRNQAPL | 0.4646404 |
| YALKRALAKFSDRFDVVII | AWSLPGQRNQAPL | 0.627187 |
| YLIGSVIV | AWSLPGQRNQAPL | 0.67660046 |
| YGQLVAVID | AWSLPGQRNQAPL | 0.74748254 |
| VTVGGMTLPA | AWSLPGQRNQAPL | 0.73052454 |
| VLALPAV | AWSLPGQRNQAPL | 0.6281292 |
| VIPPTLNYET | AWSLPGQRNQAPL | 0.7335197 |
| VGMGGMHPSSG | AWSLPGQRNQAPL | 0.70453465 |
| VGLAGGAATA | AWSLPGQRNQAPL | 0.7512709 |
| VGGAGGAGGSGP | AWSLPGQRNQAPL | 0.72313374 |
| VFAAFMV | AWSLPGQRNQAPL | 0.7000254 |
| TLYDKLR | AWSLPGQRNQAPL | 0.7190047 |
| TLAVFTVL | AWSLPGQRNQAPL | 0.6253363 |
| TGSVQPARV | AWSLPGQRNQAPL | 0.55142903 |
| TGGGAGSGGT | AWSLPGQRNQAPL | 0.48788804 |
| SHLETSL | AWSLPGQRNQAPL | 0.58371925 |
| SFTTGMT | AWSLPGQRNQAPL | 0.42882153 |
| QVEVAKRMGVRQPTVS | AWSLPGQRNQAPL | 0.7126244 |
| QGSGGPALAM | AWSLPGQRNQAPL | 0.75260293 |
| MLAPSNP | AWSLPGQRNQAPL | 0.7161726 |
| LVNNAAI | AWSLPGQRNQAPL | 0.3656656 |
| LTGEINTLA | AWSLPGQRNQAPL | 0.650958 |
| LSIFGPV | AWSLPGQRNQAPL | 0.68189144 |
| LLALLANL | AWSLPGQRNQAPL | 0.73662317 |
| LISNASDALDKLR | AWSLPGQRNQAPL | 0.7538471 |
| LINEAVP | AWSLPGQRNQAPL | 0.7666441 |
| LGYAFSGL | AWSLPGQRNQAPL | 0.7494445 |
| LDFVAVAL | AWSLPGQRNQAPL | 0.65777445 |
| LAQPGAE | AWSLPGQRNQAPL | 0.7450077 |
| ISNLGAVTGF | AWSLPGQRNQAPL | 0.6676471 |
| HQEDPGRI | AWSLPGQRNQAPL | 0.63867104 |
| GTPGEPTP | AWSLPGQRNQAPL | 0.6772125 |

| | | |
|---|---|---|
| GFFNSGS | AWSLPGQRNQAPL | 0.724751 |
| GAGGGGLTAGAN | AWSLPGQRNQAPL | 0.6801808 |
| FVDALVNM | AWSLPGQRNQAPL | 0.6268889 |
| FLTVAPEL | AWSLPGQRNQAPL | 0.40677536 |
| FETLTAL | AWSLPGQRNQAPL | 0.73348176 |
| DVINPIEL | AWSLPGQRNQAPL | 0.5154687 |
| APVSGAPV | AWSLPGQRNQAPL | 0.45408517 |
| ALVGGLAW | AWSLPGQRNQAPL | 0.6512108 |
| ALAAAAATAI | AWSLPGQRNQAPL | 0.7599043 |
| AFGLGLLL | AWSLPGQRNQAPL | 0.7565411 |
| AFFAAASL | AWSLPGQRNQAPL | 0.65979826 |
| AAAALPAFSPP | AWSLPGQRNQAPL | 0.6503991 |
| VVLAPAL | AWSLPGQRNQAPL | 0.73715365 |
| TIGHVDHGKTTL | AWSLPGQRNQAPL | 0.5932714 |
| LISNASDALDK | AWSLPGQRNQAPL | 0.73809505 |
| LISNASDAL | AWSLPGQRNQAPL | 0.71378946 |
| ISNASDALDK | AWSLPGQRNQAPL | 0.71576494 |
| NQAETLVYQTE | AWSLPGQRNQAPL | 0.5186522 |
| TGGTGGTAGAG | AWSLPGQRNQAPL | 0.7365872 |
| TGSTAGGAGGA | AWSLPGQRNQAPL | 0.68534434 |
| DPLAVPVAAGPV | AWSLPGQRNQAPL | 0.73165214 |
| GGTGGTPGNGGN | AWSLPGQRNQAPL | 0.3579113 |
| PAGLAGVDPATL | AWSLPGQRNQAPL | 0.7522183 |
| GNNLVVEAPHAGASV | AWSLPGQRNQAPL | 0.5973544 |
| KVTRIMSASTLEQPA | AWSLPGQRNQAPL | 0.60967505 |
| LGFALPIALPRRTVV | AWSLPGQRNQAPL | 0.28951368 |
| TRLGNLTARKLVVLV | AWSLPGQRNQAPL | 0.65436786 |
| WSQQRRLIYFPSAGP | AWSLPGQRNQAPL | 0.7312659 |
| GPNGNPDPMAAAVDIR | AWSLPGQRNQAPL | 0.7551874 |
| TQRCVSVADLRAAASR | AWSLPGQRNQAPL | 0.6646448 |
| QFVLDVAEHPVEVLTETV | AWSLPGQRNQAPL | 0.741477 |
| GGAGGTGGAGGNGGTQAGDG | AWSLPGQRNQAPL | 0.75113106 |
| PLLATLRAIAVDVDAKPAQV | AWSLPGQRNQAPL | 0.66772306 |
| APKFRTGGYMIDFWGVGYQVAK | AWSLPGQRNQAPL | 0.33245742 |
| GAGGAGGAGGAAGGPGGTGGTG | AWSLPGQRNQAPL | 0.7099174 |
| IGALGLLATDVAPAGSVAFGVA | AWSLPGQRNQAPL | 0.6899078 |
| AGGAGGIGGNGGAGGAGGVLMG | AWSLPGQRNQAPL | 0.75611514 |
| ADELIGLL | AWSLPGQRNQAPL | 0.73342896 |
| ADRALKA | AWSLPGQRNQAPL | 0.52840126 |
| AGAVGQVGGAGGA | AWSLPGQRNQAPL | 0.70459485 |
| AGGGGGLGGAP | AWSLPGQRNQAPL | 0.7127899 |
| AGGLLYGNGGAG | AWSLPGQRNQAPL | 0.5576376 |
| AGLESIGL | AWSLPGQRNQAPL | 0.513895 |
| AGLKTGII | AWSLPGQRNQAPL | 0.545588 |
| ASLAALLL | AWSLPGQRNQAPL | 0.4836946 |
| ASVINISSI | AWSLPGQRNQAPL | 0.7196376 |
| ATPPGPA | AWSLPGQRNQAPL | 0.6826894 |
| ATVASALV | AWSLPGQRNQAPL | 0.66766655 |
| ATVASLIL | AWSLPGQRNQAPL | 0.6942053 |
| AVNGPVLIRH | AWSLPGQRNQAPL | 0.62531865 |
| DGAAGAVGI | AWSLPGQRNQAPL | 0.6205829 |

| | | |
|---|---|---|
| DPLLHGI | AWSLPGQRNQAPL | 0.67487514 |
| ELGAALGAL | AWSLPGQRNQAPL | 0.7265673 |
| ELGRTQAV | AWSLPGQRNQAPL | 0.7454354 |
| FVAGGYLS | AWSLPGQRNQAPL | 0.69069624 |
| GASGGAGGAGGW | AWSLPGQRNQAPL | 0.63763165 |
| GGAGAGGDGGAGGI | AWSLPGQRNQAPL | 0.75586665 |
| GTGHNGSQPGMGGNG | AWSLPGQRNQAPL | 0.5283402 |
| GYAAASASAS | AWSLPGQRNQAPL | 0.6218884 |
| HQVDVAAVQQG | AWSLPGQRNQAPL | 0.7427001 |
| IGAAIAHT | AWSLPGQRNQAPL | 0.64004076 |
| IGMIEVAQL | AWSLPGQRNQAPL | 0.7488129 |
| IRLAPEHTSAMD | AWSLPGQRNQAPL | 0.7153454 |
| IYNVSVL | AWSLPGQRNQAPL | 0.7319356 |
| KQGFLGM | AWSLPGQRNQAPL | 0.70118314 |
| LESVITQL | AWSLPGQRNQAPL | 0.76078045 |
| LGLGALVVV | AWSLPGQRNQAPL | 0.7346953 |
| LIGNGGNGGSGGI | AWSLPGQRNQAPL | 0.6390902 |
| LNAINAPTL | AWSLPGQRNQAPL | 0.74827087 |
| LQLLNTGV | AWSLPGQRNQAPL | 0.44359437 |
| LQTSIEF | AWSLPGQRNQAPL | 0.7454954 |
| MSFAVSEL | AWSLPGQRNQAPL | 0.72764933 |
| NTGALGL | AWSLPGQRNQAPL | 0.617905 |
| PPGAGGAGGAGTT | AWSLPGQRNQAPL | 0.69464517 |
| QLVGLGI | AWSLPGQRNQAPL | 0.75553894 |
| RFLEQVHQL | AWSLPGQRNQAPL | 0.74965787 |
| RGVGYVLR | AWSLPGQRNQAPL | 0.58590454 |
| SATGVFAGVI | AWSLPGQRNQAPL | 0.6447474 |
| SLPAITLP | AWSLPGQRNQAPL | 0.5119732 |
| STALAFTV | AWSLPGQRNQAPL | 0.6118913 |
| SVLTVGLGK | AWSLPGQRNQAPL | 0.35160545 |
| TALAGAGSGI | AWSLPGQRNQAPL | 0.73574287 |
| TGLPFTGI | AWSLPGQRNQAPL | 0.6219425 |
| VAEHVTV | AWSLPGQRNQAPL | 0.7464913 |
| VDIVLNSL | AWSLPGQRNQAPL | 0.7257862 |
| VLTGGGAVL | AWSLPGQRNQAPL | 0.75456285 |
| VNVGLAYGI | AWSLPGQRNQAPL | 0.620425 |
| VSSAAVGAV | AWSLPGQRNQAPL | 0.66756785 |
| VTSVYEAGV | AWSLPGQRNQAPL | 0.37627894 |
| WVALDRGA | AWSLPGQRNQAPL | 0.6382611 |
| YLVGSVVI | AWSLPGQRNQAPL | 0.54686797 |
| GPPGTGKTLL | AWSLPGQRNQAPL | 0.69456285 |
| GVGMTKF | AWSLPGQRNQAPL | 0.47546363 |
| IGIGGSDL | AWSLPGQRNQAPL | 0.7646628 |
| IGTLLAEL | AWSLPGQRNQAPL | 0.6623318 |
| IYTSGTT | AWSLPGQRNQAPL | 0.7399417 |
| LSFPAGV | AWSLPGQRNQAPL | 0.5551235 |
| LYTSGTT | AWSLPGQRNQAPL | 0.55072016 |
| VGAFPFGF | AWSLPGQRNQAPL | 0.67872703 |
| VGAGTVEF | AWSLPGQRNQAPL | 0.730453 |
| VGKSTLLL | AWSLPGQRNQAPL | 0.5828414 |
| AALAHAGGGIV | AWSLPGQRNQAPL | 0.25365904 |

| | | |
|---|---|---|
| AALVAVSLL | AWSLPGQRNQAPL | 0.56844854 |
| AAVSRAVAAL | AWSLPGQRNQAPL | 0.43445468 |
| AGGSSGIGGF | AWSLPGQRNQAPL | 0.76019806 |
| AGLTVANANG | AWSLPGQRNQAPL | 0.7534777 |
| AGLVGAVL | AWSLPGQRNQAPL | 0.713286 |
| ALSQAFTV | AWSLPGQRNQAPL | 0.76106536 |
| ALTFSVSL | AWSLPGQRNQAPL | 0.23396063 |
| ALTGAALVAL | AWSLPGQRNQAPL | 0.6333546 |
| ASFSIGAGVL | AWSLPGQRNQAPL | 0.7560332 |
| DGSVYKI | AWSLPGQRNQAPL | 0.7351935 |
| EQLAEIV | AWSLPGQRNQAPL | 0.6578268 |
| FAVSELTVM | AWSLPGQRNQAPL | 0.74968207 |
| GAAPLAAGL | AWSLPGQRNQAPL | 0.6063341 |
| GADGGDGGSGGAGG | AWSLPGQRNQAPL | 0.6942495 |
| GFGLDTGGAGG | AWSLPGQRNQAPL | 0.71644354 |
| GLASGGLL | AWSLPGQRNQAPL | 0.72540736 |
| GRIAGAAVLL | AWSLPGQRNQAPL | 0.703328 |
| GSAGSLNTSF | AWSLPGQRNQAPL | 0.44481605 |
| GTGQIGI | AWSLPGQRNQAPL | 0.6789006 |
| GTLIALIL | AWSLPGQRNQAPL | 0.7654098 |
| IAATMGFVVG | AWSLPGQRNQAPL | 0.66686547 |
| IFAQALL | AWSLPGQRNQAPL | 0.74082184 |
| IGAAVALGF | AWSLPGQRNQAPL | 0.7076912 |
| IGLTPLAL | AWSLPGQRNQAPL | 0.7088846 |
| ILGSNTSTL | AWSLPGQRNQAPL | 0.61206365 |
| IVFSLTVL | AWSLPGQRNQAPL | 0.6025799 |
| LASKATGF | AWSLPGQRNQAPL | 0.72932255 |
| LASVALIL | AWSLPGQRNQAPL | 0.58905804 |
| LFAVGLTL | AWSLPGQRNQAPL | 0.6388123 |
| LFNGGIGTYI | AWSLPGQRNQAPL | 0.6751081 |
| LTLGAGII | AWSLPGQRNQAPL | 0.76954556 |
| LVAAALGDI | AWSLPGQRNQAPL | 0.7263743 |
| NTAFVQLGI | AWSLPGQRNQAPL | 0.7402822 |
| QLFSGAEL | AWSLPGQRNQAPL | 0.6326156 |
| SAATFITL | AWSLPGQRNQAPL | 0.7066955 |
| SNYGLANLGV | AWSLPGQRNQAPL | 0.74081445 |
| SYGGAYAVM | AWSLPGQRNQAPL | 0.4120344 |
| TTGVGGLTLG | AWSLPGQRNQAPL | 0.751637 |
| TTQTIYI | AWSLPGQRNQAPL | 0.5595243 |
| VAAGAGLGAVI | AWSLPGQRNQAPL | 0.722479 |
| VAAVVTLL | AWSLPGQRNQAPL | 0.72885776 |
| VGFLGGLI | AWSLPGQRNQAPL | 0.7553569 |
| VGVIPILL | AWSLPGQRNQAPL | 0.6914368 |
| VIFASSLI | AWSLPGQRNQAPL | 0.72638506 |
| VKGGDGGAAAG | AWSLPGQRNQAPL | 0.5998074 |
| VTKTFTNI | AWSLPGQRNQAPL | 0.74443376 |
| VVAAAILL | AWSLPGQRNQAPL | 0.752796 |
| YAALDVEL | AWSLPGQRNQAPL | 0.7272771 |
| LASVALLL | AWSLPGQRNQAPL | 0.7669301 |
| LDAVHRV | AWSLPGQRNQAPL | 0.744912 |
| QGLSSSIF | AWSLPGQRNQAPL | 0.32085383 |

| | | |
|---|---|---|
| RGTFANIRL | AWSLPGQRNQAPL | 0.6630637 |
| RQLWWGHRI | AWSLPGQRNQAPL | 0.7538821 |
| YGPPGTGKTLL | AWSLPGQRNQAPL | 0.7569183 |
| AALVGAPANVA | AWSLPGQRNQAPL | 0.74537617 |
| AATFRRTE | AWSLPGQRNQAPL | 0.72031933 |
| AGALGGGI | AWSLPGQRNQAPL | 0.67241395 |
| AGGAGGAGGIGAG | AWSLPGQRNQAPL | 0.7589861 |
| AGSGALGL | AWSLPGQRNQAPL | 0.7699287 |
| AQIADQI | AWSLPGQRNQAPL | 0.7306601 |
| ASAVAVTYL | AWSLPGQRNQAPL | 0.62424594 |
| ASTVLGGTGG | AWSLPGQRNQAPL | 0.57366824 |
| AVGGEPAK | AWSLPGQRNQAPL | 0.7268935 |
| CEQGTFGGTC | AWSLPGQRNQAPL | 0.75348055 |
| DEGKAPAG | AWSLPGQRNQAPL | 0.4164734 |
| DTVVFPAL | AWSLPGQRNQAPL | 0.7158329 |
| FEGITMAG | AWSLPGQRNQAPL | 0.7211391 |
| FPLVVQD | AWSLPGQRNQAPL | 0.28711522 |
| GAFTTPQ | AWSLPGQRNQAPL | 0.7120483 |
| GFGGFGTAGGVGGA | AWSLPGQRNQAPL | 0.7518948 |
| GFLAQFL | AWSLPGQRNQAPL | 0.7087612 |
| GGAGGAGGNAGVA | AWSLPGQRNQAPL | 0.54015577 |
| GGVGTAGAGGAGGAGG | AWSLPGQRNQAPL | 0.75190896 |
| GNGGGGDAGNAGSGGN | AWSLPGQRNQAPL | 0.75607574 |
| GTGGAGGAGNDAGS | AWSLPGQRNQAPL | 0.76096964 |
| GTTNTGWFNS | AWSLPGQRNQAPL | 0.34126604 |
| IFMGPLFTL | AWSLPGQRNQAPL | 0.26249844 |
| ILAHAIEAI | AWSLPGQRNQAPL | 0.7116575 |
| IRSAIREAL | AWSLPGQRNQAPL | 0.7393322 |
| ITFAGGG | AWSLPGQRNQAPL | 0.67091656 |
| ITGMSGHQEV | AWSLPGQRNQAPL | 0.6302042 |
| IVAGLACTL | AWSLPGQRNQAPL | 0.71083057 |
| IVGGGLAGA | AWSLPGQRNQAPL | 0.7537497 |
| IVTGVLNI | AWSLPGQRNQAPL | 0.451455 |
| LAQQGCI | AWSLPGQRNQAPL | 0.5410912 |
| LCLHVEN | AWSLPGQRNQAPL | 0.7121371 |
| LDAISDGS | AWSLPGQRNQAPL | 0.74037796 |
| LGGFAAML | AWSLPGQRNQAPL | 0.5999061 |
| LGGKNPMI | AWSLPGQRNQAPL | 0.68877894 |
| LGPKGVAAIL | AWSLPGQRNQAPL | 0.7155781 |
| LMVGVSI | AWSLPGQRNQAPL | 0.70739114 |
| LVGAAFVL | AWSLPGQRNQAPL | 0.72884303 |
| LVLAVAPL | AWSLPGQRNQAPL | 0.55599207 |
| MAAIAVIDAA | AWSLPGQRNQAPL | 0.75162345 |
| NNGINAELQ | AWSLPGQRNQAPL | 0.47286725 |
| RGEPLVLVHG | AWSLPGQRNQAPL | 0.7360089 |
| RGGAGGAAGLI | AWSLPGQRNQAPL | 0.70902705 |
| RSYRMHPAV | AWSLPGQRNQAPL | 0.7093089 |
| SAAHLLAGQLA | AWSLPGQRNQAPL | 0.6703953 |
| SAAVGSY | AWSLPGQRNQAPL | 0.54432404 |
| SPHAVLHRI | AWSLPGQRNQAPL | 0.571723 |
| SQIAKVL | AWSLPGQRNQAPL | 0.5378854 |

| | | |
|---|---|---|
| TALGHLVP | AWSLPGQRNQAPL | 0.74947625 |
| TAPTTSGAA | AWSLPGQRNQAPL | 0.7267631 |
| TDGDDGNGGSPGT | AWSLPGQRNQAPL | 0.7476214 |
| TDGLGEN | AWSLPGQRNQAPL | 0.71228445 |
| TGGPPLPGQA | AWSLPGQRNQAPL | 0.58795 |
| TIPAITI | AWSLPGQRNQAPL | 0.7314973 |
| TIPGFTI | AWSLPGQRNQAPL | 0.58715844 |
| TSIASGALA | AWSLPGQRNQAPL | 0.66391855 |
| TVTLPGAAI | AWSLPGQRNQAPL | 0.66647893 |
| VASGLSAALP | AWSLPGQRNQAPL | 0.7328296 |
| VGAGVIGI | AWSLPGQRNQAPL | 0.52268505 |
| VGPKRFGV | AWSLPGQRNQAPL | 0.7202621 |
| VNVAFPD | AWSLPGQRNQAPL | 0.7141764 |
| VSMTNSGVS | AWSLPGQRNQAPL | 0.51710856 |
| VTDVLGDLSV | AWSLPGQRNQAPL | 0.68320847 |
| VVVVGAGFAGL | AWSLPGQRNQAPL | 0.7480713 |
| VYTVGTVV | AWSLPGQRNQAPL | 0.69386005 |
| WTTFSGP | AWSLPGQRNQAPL | 0.44923666 |
| YDELVEAF | AWSLPGQRNQAPL | 0.7172258 |
| YGTGGAGGNGG | AWSLPGQRNQAPL | 0.6372764 |
| YTNVMAV | AWSLPGQRNQAPL | 0.72781134 |
| FLEQVHQL | AWSLPGQRNQAPL | 0.7081297 |
| GLSLLAVAV | AWSLPGQRNQAPL | 0.6302574 |
| LEQVHQL | AWSLPGQRNQAPL | 0.73547894 |
| LNAAGAI | AWSLPGQRNQAPL | 0.50749654 |
| PAVHASL | AWSLPGQRNQAPL | 0.64750695 |
| TELEDDA | AWSLPGQRNQAPL | 0.67034334 |
| TPPGPPN | AWSLPGQRNQAPL | 0.69517374 |
| TPPPPPPP | AWSLPGQRNQAPL | 0.7397758 |
| TQSPALI | ASTGDKNTEVF | 0.5721349 |
| EFAAAFPQ | ASTGDKNTEVF | 0.701157 |
| LLSGNLDVL | ASTGDKNTEVF | 0.65999967 |
| GPANLGNYNI | ASTGDKNTEVF | 0.5726541 |
| TLGPTLP | ASTGDKNTEVF | 0.6777231 |
| VYAYSSCCK | ASTGDKNTEVF | 0.6410856 |
| APAPAPAPNI | ASTGDKNTEVF | 0.45113826 |
| PGGPVPIP | ASTGDKNTEVF | 0.7281649 |
| FLTVALEP | ASTGDKNTEVF | 0.56674814 |
| SQYFGANL | ASTGDKNTEVF | 0.70668757 |
| GGVKPAA | ASTGDKNTEVF | 0.74984217 |
| GGTGGAGGAGGW | ASTGDKNTEVF | 0.5405431 |
| GVGGVAGLGGAGG | ASTGDKNTEVF | 0.72764605 |
| VLLVLATTI | ASTGDKNTEVF | 0.72231495 |
| IGIFAAML | ASTGDKNTEVF | 0.72319317 |
| AVGDLID | ASTGDKNTEVF | 0.69833827 |
| NISPLPL | ASTGDKNTEVF | 0.5744381 |
| VTAVFVAI | ASTGDKNTEVF | 0.75732076 |
| SGDDNDAAL | ASTGDKNTEVF | 0.73585224 |
| AVAALAHS | ASTGDKNTEVF | 0.72933364 |
| GGVTVMATP | ASTGDKNTEVF | 0.73908484 |
| AGLAANVRD | ASTGDKNTEVF | 0.7308443 |

| | | |
|---|---|---|
| TNRPDLIDPALLRPG | ASTGDKNTEVF | 0.74869764 |
| LANLGVTAL | ASTGDKNTEVF | 0.6643414 |
| TGERIEL | ASTGDKNTEVF | 0.43108132 |
| NGEELAA | ASTGDKNTEVF | 0.68534684 |
| VLSLVWG | ASTGDKNTEVF | 0.6708244 |
| IGYALWL | ASTGDKNTEVF | 0.5366516 |
| LSLLLAPL | ASTGDKNTEVF | 0.41797 |
| TLDEALK | ASTGDKNTEVF | 0.74500453 |
| TDGAGTAEGG | ASTGDKNTEVF | 0.7666116 |
| NAPGGAGGNGG | ASTGDKNTEVF | 0.6419204 |
| VSVFMGMGGNF | ASTGDKNTEVF | 0.76744723 |
| LASYLAAL | ASTGDKNTEVF | 0.49616104 |
| LVGGTPAL | ASTGDKNTEVF | 0.74477935 |
| VRNADVL | ASTGDKNTEVF | 0.74457216 |
| TCSSPFLVSAM | ASTGDKNTEVF | 0.5980887 |
| VHITDAE | ASTGDKNTEVF | 0.73948616 |
| EYAAPSF | ASTGDKNTEVF | 0.5069896 |
| VVDAFHGGL | ASTGDKNTEVF | 0.6358118 |
| ASVVDPQ | ASTGDKNTEVF | 0.7468444 |
| SYNEFGGLS | ASTGDKNTEVF | 0.6104497 |
| DPDGSVY | ASTGDKNTEVF | 0.7082269 |
| MVSTIHATE | ASTGDKNTEVF | 0.67562664 |
| LAAGVADPV | ASTGDKNTEVF | 0.6157006 |
| TKEQIAA | ASTGDKNTEVF | 0.7212045 |
| SLASPAAL | ASTGDKNTEVF | 0.7431015 |
| IGLVFYGV | ASTGDKNTEVF | 0.6904502 |
| FLLDASNL | ASTGDKNTEVF | 0.7678688 |
| YQVGNFP | ASTGDKNTEVF | 0.5304339 |
| ESLLSQST | ASTGDKNTEVF | 0.75742555 |
| LFTVGFAAVG | ASTGDKNTEVF | 0.7534783 |
| ATPPTPT | ASTGDKNTEVF | 0.7235797 |
| TDLATLG | ASTGDKNTEVF | 0.66846114 |
| SAGGTGR | ASTGDKNTEVF | 0.74382925 |
| GFSNSGS | ASTGDKNTEVF | 0.62182474 |
| GGNGGSGGTSV | ASTGDKNTEVF | 0.728145 |
| GTSVATGGAGN | ASTGDKNTEVF | 0.7566526 |
| NGGNGGSGGTS | ASTGDKNTEVF | 0.33427116 |
| AVVKFPH | ASTGDKNTEVF | 0.8783728 |
| TPPPPPP | ASTGDKNTEVF | 0.8258511 |
| AAPAAAAAPA | ASTGDKNTEVF | 0.84551203 |
| IALNEDL | ASTGDKNTEVF | 0.61294544 |
| IVNVSSM | ASTGDKNTEVF | 0.37099567 |
| TGIPPAPR | ASTGDKNTEVF | 0.73236716 |
| FDLGGGTF | ASTGDKNTEVF | 0.52782184 |
| FDLGGGTFDVS | ASTGDKNTEVF | 0.8578771 |
| NEPTAAA | ASTGDKNTEVF | 0.85053176 |
| NEPTAAALA | ASTGDKNTEVF | 0.6142317 |
| GSIINISSI | ASTGDKNTEVF | 0.84899366 |
| VAPGNAGTA | ASTGDKNTEVF | 0.86946064 |
| TGGGASLE | ASTGDKNTEVF | 0.36352646 |
| VAVYNHYKRI | ASTGDKNTEVF | 0.75200987 |

| | | |
|---|---|---|
| TDPAPATT | ASTGDKNTEVF | 0.7770545 |
| TNRPDILDPALLRPG | ASTGDKNTEVF | 0.6594492 |
| LTTTTTS | ASTGDKNTEVF | 0.2704649 |
| PTGSGKTL | ASTGDKNTEVF | 0.82978463 |
| TGSGKTL | ASTGDKNTEVF | 0.6970842 |
| TIGHVDHGK | ASTGDKNTEVF | 0.18582991 |
| TIGHVDHGKTT | ASTGDKNTEVF | 0.8196769 |
| GHVDHGKTT | ASTGDKNTEVF | 0.6448616 |
| AQFEHTLL | ASTGDKNTEVF | 0.6091419 |
| IVNQDEL | ASTGDKNTEVF | 0.854633 |
| TNGSQFF | ASTGDKNTEVF | 0.73044 |
| TGKGIVE | ASTGDKNTEVF | 0.6655061 |
| LISNASDA | ASTGDKNTEVF | 0.83871424 |
| AELRTLGELTKIAKAHGAQ | ASTGDKNTEVF | 0.8667798 |
| APNAVAAAEPSW | ASTGDKNTEVF | 0.5208637 |
| APQVAGAATAVL | ASTGDKNTEVF | 0.7796341 |
| ARRDPAAAAAILPTDA | ASTGDKNTEVF | 0.88673276 |
| DAAARDVLAEAGLAEYF | ASTGDKNTEVF | 0.7816508 |
| DDQIVLSAKANLAKVEQ | ASTGDKNTEVF | 0.8869273 |
| DELLPDTGEDSDEDGH | ASTGDKNTEVF | 0.8913227 |
| ELPEAPANRATVPQD | ASTGDKNTEVF | 0.70908505 |
| FGQSKIERVTGIG | ASTGDKNTEVF | 0.8661556 |
| GAGGTGSTAGGAGGAGGAGGLY | ASTGDKNTEVF | 0.8710926 |
| GHGGTGAAGGNGGTGGAGGSGI | ASTGDKNTEVF | 0.68421006 |
| GNGGNGGAGGEGGPGGAGGAGA | ASTGDKNTEVF | 0.7615974 |
| GSTAGNGGNGGNATNGTIAGQP | ASTGDKNTEVF | 0.4218924 |
| GTGGDGGNGGNGGTGG | ASTGDKNTEVF | 0.7793019 |
| GVRRPAGIPADAPV | ASTGDKNTEVF | 0.4427651 |
| GYRQVTDAHLLTLARR | ASTGDKNTEVF | 0.8147857 |
| IAKMGQDGHDRG | ASTGDKNTEVF | 0.87882185 |
| KAEPSIEIQYT | ASTGDKNTEVF | 0.7561573 |
| KPVRIRMAADEDTVAVT | ASTGDKNTEVF | 0.40727308 |
| MPPRIAGMRLLVIKPEP | ASTGDKNTEVF | 0.4630193 |
| MKFYAVALFRSRGVPPTR | ASTGDKNTEVF | 0.8358942 |
| NAGGFGILGAVAHSP | ASTGDKNTEVF | 0.41800737 |
| NEGREITLAMAEM | ASTGDKNTEVF | 0.46290338 |
| NELVEWSHPLGHRWRWE | ASTGDKNTEVF | 0.7056378 |
| NLGAGGTGGNGGNPG | ASTGDKNTEVF | 0.86664283 |
| NTEWPKWETVMAADPVD | ASTGDKNTEVF | 0.86186063 |
| PASAAPAETQP | ASTGDKNTEVF | 0.58137226 |
| QQSSDGKNGVRAAEITGE | ASTGDKNTEVF | 0.8820838 |
| RAPDAVTTSWWKE | ASTGDKNTEVF | 0.8432372 |
| SGFGNVGAHTSGWFNQ | ASTGDKNTEVF | 0.8873503 |
| SGVAGAVGQAASVGGLKV | ASTGDKNTEVF | 0.7925824 |
| SNGGAGTGSAGGNGGT | ASTGDKNTEVF | 0.38842815 |
| TGGAGGEGGAAG | ASTGDKNTEVF | 0.85776925 |
| TGGDGGAAGAGGAPGHG | ASTGDKNTEVF | 0.7058762 |
| TMPPLLAAIANAGRHGV | ASTGDKNTEVF | 0.8354749 |
| TSSTAGNSGGA | ASTGDKNTEVF | 0.86371636 |
| VPATVPVGILC | ASTGDKNTEVF | 0.78569525 |
| VQRPDSGGVVLTGRIS | ASTGDKNTEVF | 0.5873829 |

| | | |
|---|---|---|
| WTPNPVPATKGVPAGLVQAT | ASTGDKNTEVF | 0.7617127 |
| YALKRALAKFSDRFDVVII | ASTGDKNTEVF | 0.8869306 |
| YLIGSVIV | ASTGDKNTEVF | 0.54128057 |
| YGQLVAVID | ASTGDKNTEVF | 0.42517203 |
| VTVGGMTLPA | ASTGDKNTEVF | 0.7063041 |
| VLALPAV | ASTGDKNTEVF | 0.73476994 |
| VIPPTLNYET | ASTGDKNTEVF | 0.8779096 |
| VGMGGMHPSSG | ASTGDKNTEVF | 0.7889739 |
| VGLAGGAATA | ASTGDKNTEVF | 0.74803823 |
| VGGAGGAGGSGP | ASTGDKNTEVF | 0.82769454 |
| VFAAFMV | ASTGDKNTEVF | 0.8398737 |
| TLYDKLR | ASTGDKNTEVF | 0.78218126 |
| TLAVFTVL | ASTGDKNTEVF | 0.8326365 |
| TGSVQPARV | ASTGDKNTEVF | 0.6317247 |
| TGGGAGSGGT | ASTGDKNTEVF | 0.8796125 |
| SHLETSL | ASTGDKNTEVF | 0.8647743 |
| SFTTGMT | ASTGDKNTEVF | 0.84305274 |
| QVEVAKRMGVRQPTVS | ASTGDKNTEVF | 0.87446296 |
| QGSGGPALAM | ASTGDKNTEVF | 0.8036637 |
| MLAPSNP | ASTGDKNTEVF | 0.8035365 |
| LVNNAAI | ASTGDKNTEVF | 0.42859524 |
| LTGEINTLA | ASTGDKNTEVF | 0.7558868 |
| LSIFGPV | ASTGDKNTEVF | 0.7849562 |
| LLALLANL | ASTGDKNTEVF | 0.81626344 |
| LISNASDALDKLR | ASTGDKNTEVF | 0.84866154 |
| LINEAVP | ASTGDKNTEVF | 0.85617757 |
| LGYAFSGL | ASTGDKNTEVF | 0.6526842 |
| LDFVAVAL | ASTGDKNTEVF | 0.88244045 |
| LAQPGAE | ASTGDKNTEVF | 0.8716582 |
| ISNLGAVTGF | ASTGDKNTEVF | 0.49637777 |
| HQEDPGRI | ASTGDKNTEVF | 0.7199724 |
| GTPGEPTP | ASTGDKNTEVF | 0.8725487 |
| GFFNSGS | ASTGDKNTEVF | 0.82585466 |
| GAGGGGLTAGAN | ASTGDKNTEVF | 0.719102 |
| FVDALVNM | ASTGDKNTEVF | 0.7593026 |
| FLTVAPEL | ASTGDKNTEVF | 0.8723954 |
| FETLTAL | ASTGDKNTEVF | 0.76633227 |
| DVINPIEL | ASTGDKNTEVF | 0.88337564 |
| APVSGAPV | ASTGDKNTEVF | 0.24251404 |
| ALVGGLAW | ASTGDKNTEVF | 0.7030342 |
| ALAAAAATAI | ASTGDKNTEVF | 0.7166537 |
| AFGLGLLL | ASTGDKNTEVF | 0.14868096 |
| AFFAAASL | ASTGDKNTEVF | 0.86193216 |
| AAAALPAFSPP | ASTGDKNTEVF | 0.64310086 |
| VVLAPAL | ASTGDKNTEVF | 0.81575793 |
| TIGHVDHGKTTL | ASTGDKNTEVF | 0.7204395 |
| LISNASDALDK | ASTGDKNTEVF | 0.867508 |
| LISNASDAL | ASTGDKNTEVF | 0.8249427 |
| ISNASDALDK | ASTGDKNTEVF | 0.67014724 |
| NQAETLVYQTE | ASTGDKNTEVF | 0.46932638 |
| TGGTGGTAGAG | ASTGDKNTEVF | 0.81904507 |

| | | |
|---|---|---|
| TGSTAGGAGGA | ASTGDKNTEVF | 0.8889462 |
| DPLAVPVAAGPV | ASTGDKNTEVF | 0.8859477 |
| GGTGGTPGNGGN | ASTGDKNTEVF | 0.33424735 |
| PAGLAGVDPATL | ASTGDKNTEVF | 0.82330066 |
| GNNLVVEAPHAGASV | ASTGDKNTEVF | 0.7255198 |
| KVTRIMSASTLEQPA | ASTGDKNTEVF | 0.51825094 |
| LGFALPIALPRRTVV | ASTGDKNTEVF | 0.2329651 |
| TRLGNLTARKLVVLV | ASTGDKNTEVF | 0.72724587 |
| WSQQRRLIYFPSAGP | ASTGDKNTEVF | 0.7661132 |
| GPNGNPDPMAAAVDIR | ASTGDKNTEVF | 0.8246526 |
| TQRCVSVADLRAAASR | ASTGDKNTEVF | 0.8705562 |
| QFVLDVAEHPVEVLTETV | ASTGDKNTEVF | 0.65267164 |
| GGAGGTGGAGGNGGTQAGDG | ASTGDKNTEVF | 0.85149515 |
| PLLATLRAIAVDVDAKPAQV | ASTGDKNTEVF | 0.7713155 |
| APKFRTGGYMIDFWGVGYQVAK | ASTGDKNTEVF | 0.08113016 |
| GAGGAGGAGGAAGGPGGTGGTG | ASTGDKNTEVF | 0.67499495 |
| IGALGLLATDVAPAGSVAFGVA | ASTGDKNTEVF | 0.3459813 |
| AGGAGGIGGNGGAGGAGGVLMG | ASTGDKNTEVF | 0.875966 |
| ADELIGLL | ASTGDKNTEVF | 0.4751364 |
| ADRALKA | ASTGDKNTEVF | 0.7168 |
| AGAVGQVGGAGGA | ASTGDKNTEVF | 0.7427799 |
| AGGGGGLGGAP | ASTGDKNTEVF | 0.82044506 |
| AGGLLYGNGGAG | ASTGDKNTEVF | 0.7875881 |
| AGLESIGL | ASTGDKNTEVF | 0.6215181 |
| AGLKTGII | ASTGDKNTEVF | 0.5228269 |
| ASLAALLL | ASTGDKNTEVF | 0.8361689 |
| ASVINISSI | ASTGDKNTEVF | 0.7789711 |
| ATPPGPA | ASTGDKNTEVF | 0.52960706 |
| ATVASALV | ASTGDKNTEVF | 0.6022488 |
| ATVASLIL | ASTGDKNTEVF | 0.8738319 |
| AVNGPVLIRH | ASTGDKNTEVF | 0.31142026 |
| DGAAGAVGI | ASTGDKNTEVF | 0.8784952 |
| DPLLHGI | ASTGDKNTEVF | 0.6258517 |
| ELGAALGAL | ASTGDKNTEVF | 0.69795924 |
| ELGRTQAV | ASTGDKNTEVF | 0.8890163 |
| FVAGGYLS | ASTGDKNTEVF | 0.7900638 |
| GASGGAGGAGGW | ASTGDKNTEVF | 0.7723194 |
| GGAGAGGDGGAGGI | ASTGDKNTEVF | 0.5665175 |
| GTGHNGSQPGMGGNG | ASTGDKNTEVF | 0.50596595 |
| GYAAASASAS | ASTGDKNTEVF | 0.60069895 |
| HQVDVAAVQQG | ASTGDKNTEVF | 0.59764016 |
| IGAAIAHT | ASTGDKNTEVF | 0.8854909 |
| IGMIEVAQL | ASTGDKNTEVF | 0.750705 |
| IRLAPEHTSAMD | ASTGDKNTEVF | 0.44501764 |
| IYNVSVL | ASTGDKNTEVF | 0.8366114 |
| KQGFLGM | ASTGDKNTEVF | 0.88376117 |
| LESVITQL | ASTGDKNTEVF | 0.8910071 |
| LGLGALVVV | ASTGDKNTEVF | 0.4519804 |
| LIGNGGNGGSGGI | ASTGDKNTEVF | 0.66005075 |
| LNAINAPTL | ASTGDKNTEVF | 0.75451005 |
| LQLLNTGV | ASTGDKNTEVF | 0.73234165 |

| | | |
|---|---|---|
| LQTSIEF | ASTGDKNTEVF | 0.5695812 |
| MSFAVSEL | ASTGDKNTEVF | 0.5005548 |
| NTGALGL | ASTGDKNTEVF | 0.8717631 |
| PPGAGGAGGAGTT | ASTGDKNTEVF | 0.8853185 |
| QLVGLGI | ASTGDKNTEVF | 0.72609514 |
| RFLEQVHQL | ASTGDKNTEVF | 0.3446117 |
| RGVGYVLR | ASTGDKNTEVF | 0.7095316 |
| SATGVFAGVI | ASTGDKNTEVF | 0.5982672 |
| SLPAITLP | ASTGDKNTEVF | 0.5949311 |
| STALAFTV | ASTGDKNTEVF | 0.7640393 |
| SVLTVGLGK | ASTGDKNTEVF | 0.34908378 |
| TALAGAGSGI | ASTGDKNTEVF | 0.7442564 |
| TGLPFTGI | ASTGDKNTEVF | 0.8943783 |
| VAEHVTV | ASTGDKNTEVF | 0.70899105 |
| VDIVLNSL | ASTGDKNTEVF | 0.8469299 |
| VLTGGGAVL | ASTGDKNTEVF | 0.87904155 |
| VNVGLAYGI | ASTGDKNTEVF | 0.8291229 |
| VSSAAVGAV | ASTGDKNTEVF | 0.87169117 |
| VTSVYEAGV | ASTGDKNTEVF | 0.8457247 |
| WVALDRGA | ASTGDKNTEVF | 0.7444085 |
| YLVGSVVI | ASTGDKNTEVF | 0.79007196 |
| GPPGTGKTLL | ASTGDKNTEVF | 0.8553762 |
| GVGMTKF | ASTGDKNTEVF | 0.68670344 |
| IGIGGSDL | ASTGDKNTEVF | 0.7931413 |
| IGTLLAEL | ASTGDKNTEVF | 0.7735149 |
| IYTSGTT | ASTGDKNTEVF | 0.8238841 |
| LSFPAGV | ASTGDKNTEVF | 0.7798759 |
| LYTSGTT | ASTGDKNTEVF | 0.8755157 |
| VGAFPFGF | ASTGDKNTEVF | 0.20875564 |
| VGAGTVEF | ASTGDKNTEVF | 0.5254785 |
| VGKSTLLL | ASTGDKNTEVF | 0.8674869 |
| AALAHAGGGIV | ASTGDKNTEVF | 0.17416309 |
| AALVAVSLL | ASTGDKNTEVF | 0.2742768 |
| AAVSRAVAAL | ASTGDKNTEVF | 0.8111208 |
| AGGSSGIGGF | ASTGDKNTEVF | 0.87634987 |
| AGLTVANANG | ASTGDKNTEVF | 0.22058384 |
| AGLVGAVL | ASTGDKNTEVF | 0.74245834 |
| ALSQAFTV | ASTGDKNTEVF | 0.39320162 |
| ALTFSVSL | ASTGDKNTEVF | 0.86588186 |
| ALTGAALVAL | ASTGDKNTEVF | 0.7182652 |
| ASFSIGAGVL | ASTGDKNTEVF | 0.364589 |
| DGSVYKI | ASTGDKNTEVF | 0.83919394 |
| EQLAEIV | ASTGDKNTEVF | 0.5884399 |
| FAVSELTVM | ASTGDKNTEVF | 0.81056285 |
| GAAPLAAGL | ASTGDKNTEVF | 0.87099385 |
| GADGGDGGSGGAGG | ASTGDKNTEVF | 0.88226867 |
| GFGLDTGGAGG | ASTGDKNTEVF | 0.44226867 |
| GLASGGLL | ASTGDKNTEVF | 0.84582335 |
| GRIAGAAVLL | ASTGDKNTEVF | 0.6784296 |
| GSAGSLNTSF | ASTGDKNTEVF | 0.9006808 |
| GTGQIGI | ASTGDKNTEVF | 0.68394613 |

| | | |
|---|---|---|
| GTLIALIL | ASTGDKNTEVF | 0.32101747 |
| IAATMGFVVG | ASTGDKNTEVF | 0.85706997 |
| IFAQALL | ASTGDKNTEVF | 0.84172094 |
| IGAAVALGF | ASTGDKNTEVF | 0.7248155 |
| IGLTPLAL | ASTGDKNTEVF | 0.84019494 |
| ILGSNTSTL | ASTGDKNTEVF | 0.75110334 |
| IVFSLTVL | ASTGDKNTEVF | 0.6841368 |
| LASKATGF | ASTGDKNTEVF | 0.838218 |
| LASVALIL | ASTGDKNTEVF | 0.8375372 |
| LFAVGLTL | ASTGDKNTEVF | 0.63956684 |
| LFNGGIGTYI | ASTGDKNTEVF | 0.8811333 |
| LTLGAGII | ASTGDKNTEVF | 0.772409 |
| LVAAALGDI | ASTGDKNTEVF | 0.46948507 |
| NTAFVQLGI | ASTGDKNTEVF | 0.8938486 |
| QLFSGAEL | ASTGDKNTEVF | 0.7739643 |
| SAATFITL | ASTGDKNTEVF | 0.85589045 |
| SNYGLANLGV | ASTGDKNTEVF | 0.82588243 |
| SYGGAYAVM | ASTGDKNTEVF | 0.86133933 |
| TTGVGGLTLG | ASTGDKNTEVF | 0.6147891 |
| TTQTIYI | ASTGDKNTEVF | 0.8046137 |
| VAAGAGLGAVI | ASTGDKNTEVF | 0.79443073 |
| VAAVVTLL | ASTGDKNTEVF | 0.8476759 |
| VGFLGGLI | ASTGDKNTEVF | 0.7712387 |
| VGVIPILL | ASTGDKNTEVF | 0.8300674 |
| VIFASSLI | ASTGDKNTEVF | 0.837398 |
| VKGGDGGAAAG | ASTGDKNTEVF | 0.88410807 |
| VTKTFTNI | ASTGDKNTEVF | 0.77036977 |
| VVAAAILL | ASTGDKNTEVF | 0.8202394 |
| YAALDVEL | ASTGDKNTEVF | 0.8854421 |
| LASVALLL | ASTGDKNTEVF | 0.44517028 |
| LDAVHRV | ASTGDKNTEVF | 0.7824171 |
| QGLSSSIF | ASTGDKNTEVF | 0.87289524 |
| RGTFANIRL | ASTGDKNTEVF | 0.8837923 |
| RQLWWGHRI | ASTGDKNTEVF | 0.72872317 |
| YGPPGTGKTLL | ASTGDKNTEVF | 0.8754426 |
| AALVGAPANVA | ASTGDKNTEVF | 0.76587826 |
| AATFRRTE | ASTGDKNTEVF | 0.82235646 |
| AGALGGGI | ASTGDKNTEVF | 0.75766516 |
| AGGAGGAGGIGAG | ASTGDKNTEVF | 0.70251155 |
| AGSGALGL | ASTGDKNTEVF | 0.59270394 |
| AQIADQI | ASTGDKNTEVF | 0.77922714 |
| ASAVAVTYL | ASTGDKNTEVF | 0.8577068 |
| ASTVLGGTGG | ASTGDKNTEVF | 0.8054904 |
| AVGGEPAK | ASTGDKNTEVF | 0.34602636 |
| CEQGTFGGTC | ASTGDKNTEVF | 0.8287989 |
| DEGKAPAG | ASTGDKNTEVF | 0.81879437 |
| DTVVFPAL | ASTGDKNTEVF | 0.88812643 |
| FEGITMAG | ASTGDKNTEVF | 0.7864306 |
| FPLVVQD | ASTGDKNTEVF | 0.86685246 |
| GAFTTPQ | ASTGDKNTEVF | 0.7457231 |
| GFGGFGTAGGVGGA | ASTGDKNTEVF | 0.7835575 |

| | | |
|---|---|---|
| GFLAQFL | ASTGDKNTEVF | 0.4084013 |
| GGAGGAGGNAGVA | ASTGDKNTEVF | 0.66823953 |
| GGVGTAGAGGAGGAGG | ASTGDKNTEVF | 0.7729272 |
| GNGGGGDAGNAGSGGN | ASTGDKNTEVF | 0.5571449 |
| GTGGAGGAGNDAGS | ASTGDKNTEVF | 0.8462038 |
| GTTNTGWFNS | ASTGDKNTEVF | 0.87392473 |
| IFMGPLFTL | ASTGDKNTEVF | 0.8687605 |
| ILAHAIEAI | ASTGDKNTEVF | 0.87414604 |
| IRSAIREAL | ASTGDKNTEVF | 0.8820452 |
| ITFAGGG | ASTGDKNTEVF | 0.8590261 |
| ITGMSGHQEV | ASTGDKNTEVF | 0.6572624 |
| IVAGLACTL | ASTGDKNTEVF | 0.7433742 |
| IVGGGLAGA | ASTGDKNTEVF | 0.31877607 |
| IVTGVLNI | ASTGDKNTEVF | 0.66833806 |
| LAQQGCI | ASTGDKNTEVF | 0.68531233 |
| LCLHVEN | ASTGDKNTEVF | 0.7834833 |
| LDAISDGS | ASTGDKNTEVF | 0.12896378 |
| LGGFAAML | ASTGDKNTEVF | 0.7963373 |
| LGGKNPMI | ASTGDKNTEVF | 0.88758343 |
| LGPKGVAAIL | ASTGDKNTEVF | 0.7550861 |
| LMVGVSI | ASTGDKNTEVF | 0.825969 |
| LVGAAFVL | ASTGDKNTEVF | 0.8439458 |
| LVLAVAPL | ASTGDKNTEVF | 0.6622959 |
| MAAIAVIDAA | ASTGDKNTEVF | 0.84954655 |
| NNGINAELQ | ASTGDKNTEVF | 0.6945305 |
| RGEPLVLVHG | ASTGDKNTEVF | 0.866137 |
| RGGAGGAAGLI | ASTGDKNTEVF | 0.87781096 |
| RSYRMHPAV | ASTGDKNTEVF | 0.825042 |
| SAAHLLAGQLA | ASTGDKNTEVF | 0.79938895 |
| SAAVGSY | ASTGDKNTEVF | 0.8176423 |
| SPHAVLHRI | ASTGDKNTEVF | 0.60971665 |
| SQIAKVL | ASTGDKNTEVF | 0.8112037 |
| TALGHLVP | ASTGDKNTEVF | 0.7068647 |
| TAPTTSGAA | ASTGDKNTEVF | 0.77878916 |
| TDGDDGNGGSPGT | ASTGDKNTEVF | 0.5638984 |
| TDGLGEN | ASTGDKNTEVF | 0.80561674 |
| TGGPPLPGQA | ASTGDKNTEVF | 0.7691293 |
| TIPAITI | ASTGDKNTEVF | 0.84742206 |
| TIPGFTI | ASTGDKNTEVF | 0.5451888 |
| TSIASGALA | ASTGDKNTEVF | 0.8271056 |
| TVTLPGAAI | ASTGDKNTEVF | 0.8035864 |
| VASGLSAALP | ASTGDKNTEVF | 0.4348095 |
| VGAGVIGI | ASTGDKNTEVF | 0.86504406 |
| VGPKRFGV | ASTGDKNTEVF | 0.7727324 |
| VNVAFPD | ASTGDKNTEVF | 0.7682234 |
| VSMTNSGVS | ASTGDKNTEVF | 0.86663276 |
| VTDVLGDLSV | ASTGDKNTEVF | 0.77208525 |
| VVVVGAGFAGL | ASTGDKNTEVF | 0.55650914 |
| VYTVGTVV | ASTGDKNTEVF | 0.69531083 |
| WTTFSGP | ASTGDKNTEVF | 0.78966177 |
| YDELVEAF | ASTGDKNTEVF | 0.7217386 |

| | | |
|---|---|---|
| YGTGGAGGNGG | ASTGDKNTEVF | 0.22956048 |
| YTNVMAV | ASTGDKNTEVF | 0.8342553 |
| FLEQVHQL | ASTGDKNTEVF | 0.47136545 |
| GLSLLAVAV | ASTGDKNTEVF | 0.69070363 |
| LEQVHQL | ASTGDKNTEVF | 0.64938164 |
| LNAAGAI | ASTGDKNTEVF | 0.85907423 |
| PAVHASL | ASTGDKNTEVF | 0.8136169 |
| TELEDDA | ASTGDKNTEVF | 0.64648426 |
| TPPGPPN | ASTGDKNTEVF | 0.8395685 |
| TPPPPPPP | ASTGDKNTEVF | 0.8250842 |
| TQSPALI | ASSPPGGDYEQY | 0.8577653 |
| EFAAAFPQ | ASSPPGGDYEQY | 0.7778202 |
| LLSGNLDVL | ASSPPGGDYEQY | 0.80895513 |
| GPANLGNYNI | ASSPPGGDYEQY | 0.8196632 |
| TLGPTLP | ASSPPGGDYEQY | 0.7279252 |
| VYAYSSCCK | ASSPPGGDYEQY | 0.53962636 |
| APAPAPAPNI | ASSPPGGDYEQY | 0.5710417 |
| PGGPVPIP | ASSPPGGDYEQY | 0.6860473 |
| FLTVALEP | ASSPPGGDYEQY | 0.45404667 |
| SQYFGANL | ASSPPGGDYEQY | 0.79564965 |
| GGVKPAA | ASSPPGGDYEQY | 0.8851645 |
| GGTGGAGGAGGW | ASSPPGGDYEQY | 0.7999175 |
| GVGGVAGLGGAGG | ASSPPGGDYEQY | 0.32322228 |
| VLLVLATTI | ASSPPGGDYEQY | 0.72895247 |
| IGIFAAML | ASSPPGGDYEQY | 0.77768975 |
| AVGDLID | ASSPPGGDYEQY | 0.84910893 |
| NISPLPL | ASSPPGGDYEQY | 0.8657973 |
| VTAVFVAI | ASSPPGGDYEQY | 0.87223864 |
| SGDDNDAAL | ASSPPGGDYEQY | 0.84381723 |
| AVAALAHS | ASSPPGGDYEQY | 0.64981604 |
| GGVTVMATP | ASSPPGGDYEQY | 0.85315824 |
| AGLAANVRD | ASSPPGGDYEQY | 0.68813086 |
| TNRPDLIDPALLRPG | ASSPPGGDYEQY | 0.83550763 |
| LANLGVTAL | ASSPPGGDYEQY | 0.7928989 |
| TGERIEL | ASSPPGGDYEQY | 0.83903515 |
| NGEELAA | ASSPPGGDYEQY | 0.6921742 |
| VLSLVWG | ASSPPGGDYEQY | 0.67838687 |
| IGYALWL | ASSPPGGDYEQY | 0.40143424 |
| LSLLLAPL | ASSPPGGDYEQY | 0.84994555 |
| TLDEALK | ASSPPGGDYEQY | 0.70119166 |
| TDGAGTAEGG | ASSPPGGDYEQY | 0.5903082 |
| NAPGGAGGNGG | ASSPPGGDYEQY | 0.68916976 |
| VSVFMGMGGNF | ASSPPGGDYEQY | 0.88171303 |
| LASYLAAL | ASSPPGGDYEQY | 0.8885337 |
| LVGGTPAL | ASSPPGGDYEQY | 0.6750075 |
| VRNADVL | ASSPPGGDYEQY | 0.6278546 |
| TCSSPFLVSAM | ASSPPGGDYEQY | 0.83188224 |
| VHITDAE | ASSPPGGDYEQY | 0.60590947 |
| EYAAPSF | ASSPPGGDYEQY | 0.8255631 |
| VVDAFHGGL | ASSPPGGDYEQY | 0.7714462 |
| ASVVDPQ | ASSPPGGDYEQY | 0.76163125 |

| | | |
|---|---|---|
| SYNEFGGLS | ASSPPGGDYEQY | 0.6706406 |
| DPDGSVY | ASSPPGGDYEQY | 0.8441682 |
| MVSTIHATE | ASSPPGGDYEQY | 0.7006127 |
| LAAGVADPV | ASSPPGGDYEQY | 0.8292811 |
| TKEQIAA | ASSPPGGDYEQY | 0.48995692 |
| SLASPAAL | ASSPPGGDYEQY | 0.8560938 |
| IGLVFYGV | ASSPPGGDYEQY | 0.5152389 |
| FLLDASNL | ASSPPGGDYEQY | 0.57726187 |
| YQVGNFP | ASSPPGGDYEQY | 0.2518304 |
| ESLLSQST | ASSPPGGDYEQY | 0.76349914 |
| LFTVGFAAVG | ASSPPGGDYEQY | 0.85085416 |
| ATPPTPT | ASSPPGGDYEQY | 0.8809153 |
| TDLATLG | ASSPPGGDYEQY | 0.78921884 |
| SAGGTGR | ASSPPGGDYEQY | 0.8283138 |
| GFSNSGS | ASSPPGGDYEQY | 0.8840877 |
| GGNGGSGGTSV | ASSPPGGDYEQY | 0.68864816 |
| GTSVATGGAGN | ASSPPGGDYEQY | 0.27540174 |
| NGGNGGSGGTS | ASSPPGGDYEQY | 0.8120237 |
| AVVKFPH | ASSPPGGDYEQY | 0.7550887 |
| TPPPPPP | ASSPPGGDYEQY | 0.87581277 |
| AAPAAAAAPA | ASSPPGGDYEQY | 0.8463191 |
| IALNEDL | ASSPPGGDYEQY | 0.37606686 |
| IVNVSSM | ASSPPGGDYEQY | 0.86208344 |
| TGIPPAPR | ASSPPGGDYEQY | 0.8453901 |
| FDLGGGTF | ASSPPGGDYEQY | 0.52551746 |
| FDLGGGTFDVS | ASSPPGGDYEQY | 0.66418624 |
| NEPTAAA | ASSPPGGDYEQY | 0.5906562 |
| NEPTAAALA | ASSPPGGDYEQY | 0.5274759 |
| GSIINISSI | ASSPPGGDYEQY | 0.84090924 |
| VAPGNAGTA | ASSPPGGDYEQY | 0.69164324 |
| TGGGASLE | ASSPPGGDYEQY | 0.72671294 |
| VAVYNHYKRI | ASSPPGGDYEQY | 0.70893383 |
| TDPAPATT | ASSPPGGDYEQY | 0.6348904 |
| TNRPDILDPALLRPG | ASSPPGGDYEQY | 0.6860994 |
| LTTTTTS | ASSPPGGDYEQY | 0.7198266 |
| PTGSGKTL | ASSPPGGDYEQY | 0.8230227 |
| TGSGKTL | ASSPPGGDYEQY | 0.8576926 |
| TIGHVDHGK | ASSPPGGDYEQY | 0.7134663 |
| TIGHVDHGKTT | ASSPPGGDYEQY | 0.8259469 |
| GHVDHGKTT | ASSPPGGDYEQY | 0.8809217 |
| AQFEHTLL | ASSPPGGDYEQY | 0.5161438 |
| IVNQDEL | ASSPPGGDYEQY | 0.5774812 |
| TNGSQFF | ASSPPGGDYEQY | 0.83351076 |
| TGKGIVE | ASSPPGGDYEQY | 0.6881119 |
| LISNASDA | ASSPPGGDYEQY | 0.866323 |
| AELRTLGELTKIAKAHGAQ | ASSPPGGDYEQY | 0.7909167 |
| APNAVAAAEPSW | ASSPPGGDYEQY | 0.8148171 |
| APQVAGAATAVL | ASSPPGGDYEQY | 0.7926053 |
| ARRDPAAAAAILPTDA | ASSPPGGDYEQY | 0.893962 |
| DAAARDVLAEAGLAEYF | ASSPPGGDYEQY | 0.82339764 |
| DDQIVLSAKANLAKVEQ | ASSPPGGDYEQY | 0.7005491 |

| | | |
|---|---|---|
| DELLPDTGEDSDEDGH | ASSPPGGDYEQY | 0.82630044 |
| ELPEAPANRATVPQD | ASSPPGGDYEQY | 0.42374748 |
| FGQSKIERVTGIG | ASSPPGGDYEQY | 0.8530122 |
| GAGGTGSTAGGAGGAGGAGGLY | ASSPPGGDYEQY | 0.84199125 |
| GHGGTGAAGGNGGTGGAGGSGI | ASSPPGGDYEQY | 0.5629476 |
| GNGGNGGAGGEGGPGGAGGAGA | ASSPPGGDYEQY | 0.6879053 |
| GSTAGNGGNGGNATNGTIAGQP | ASSPPGGDYEQY | 0.8963608 |
| GTGGDGGNGGNGGTGG | ASSPPGGDYEQY | 0.85598135 |
| GVRRPAGIPADAPV | ASSPPGGDYEQY | 0.6283251 |
| GYRQVTDAHLLTLARR | ASSPPGGDYEQY | 0.5846036 |
| IAKMGQDGHDRG | ASSPPGGDYEQY | 0.6295736 |
| KAEPSIEIQYT | ASSPPGGDYEQY | 0.717183 |
| KPVRIRMAADEDTVAVT | ASSPPGGDYEQY | 0.27725986 |
| MPPRIAGMRLLVIKPEP | ASSPPGGDYEQY | 0.8440659 |
| MKFYAVALFRSRGVPPTR | ASSPPGGDYEQY | 0.70463586 |
| NAGGFGILGAVAHSP | ASSPPGGDYEQY | 0.8379444 |
| NEGREITLAMAEM | ASSPPGGDYEQY | 0.8127163 |
| NELVEWSHPLGHRWRWE | ASSPPGGDYEQY | 0.875166 |
| NLGAGGTGGNGGNPG | ASSPPGGDYEQY | 0.66706765 |
| NTEWPKWETVMAADPVD | ASSPPGGDYEQY | 0.67863595 |
| PASAAPAETQP | ASSPPGGDYEQY | 0.27731124 |
| QQSSDGKNGVRAAEITGE | ASSPPGGDYEQY | 0.5945156 |
| RAPDAVTTSWWKE | ASSPPGGDYEQY | 0.65872955 |
| SGFGNVGAHTSGWFNQ | ASSPPGGDYEQY | 0.76170635 |
| SGVAGAVGQAASVGGLKV | ASSPPGGDYEQY | 0.48803234 |
| SNGGAGTGSAGGNGGT | ASSPPGGDYEQY | 0.879174 |
| TGGAGGEGGAAG | ASSPPGGDYEQY | 0.6881268 |
| TGGDGGAAGAGGAPGHG | ASSPPGGDYEQY | 0.8432384 |
| TMPPLLAAIANAGRHGV | ASSPPGGDYEQY | 0.74401027 |
| TSSTAGNSGGA | ASSPPGGDYEQY | 0.55971277 |
| VPATVPVGILC | ASSPPGGDYEQY | 0.6396111 |
| VQRPDSGGVVLTGRIS | ASSPPGGDYEQY | 0.8458106 |
| WTPNPVPATKGVPAGLVQAT | ASSPPGGDYEQY | 0.44469854 |
| YALKRALAKFSDRFDVVII | ASSPPGGDYEQY | 0.3298394 |
| YLIGSVIV | ASSPPGGDYEQY | 0.63135564 |
| YGQLVAVID | ASSPPGGDYEQY | 0.79781175 |
| VTVGGMTLPA | ASSPPGGDYEQY | 0.8881287 |
| VLALPAV | ASSPPGGDYEQY | 0.8736699 |
| VIPPTLNYET | ASSPPGGDYEQY | 0.77901304 |
| VGMGGMHPSSG | ASSPPGGDYEQY | 0.88958573 |
| VGLAGGAATA | ASSPPGGDYEQY | 0.13777842 |
| VGGAGGAGGSGP | ASSPPGGDYEQY | 0.7219962 |
| VFAAFMV | ASSPPGGDYEQY | 0.88403225 |
| TLYDKLR | ASSPPGGDYEQY | 0.83373296 |
| TLAVFTVL | ASSPPGGDYEQY | 0.7434666 |
| TGSVQPARV | ASSPPGGDYEQY | 0.86924 |
| TGGGAGSGGT | ASSPPGGDYEQY | 0.5805912 |
| SHLETSL | ASSPPGGDYEQY | 0.7128233 |
| SFTTGMT | ASSPPGGDYEQY | 0.74979126 |
| QVEVAKRMGVRQPTVS | ASSPPGGDYEQY | 0.8530413 |
| QGSGGPALAM | ASSPPGGDYEQY | 0.86789465 |

| | | |
|---|---|---|
| MLAPSNP | ASSPPGGDYEQY | 0.31981307 |
| LVNNAAI | ASSPPGGDYEQY | 0.84802663 |
| LTGEINTLA | ASSPPGGDYEQY | 0.89732945 |
| LSIFGPV | ASSPPGGDYEQY | 0.64470875 |
| LLALLANL | ASSPPGGDYEQY | 0.8740549 |
| LISNASDALDKLR | ASSPPGGDYEQY | 0.7957171 |
| LINEAVP | ASSPPGGDYEQY | 0.7716156 |
| LGYAFSGL | ASSPPGGDYEQY | 0.5856873 |
| LDFVAVAL | ASSPPGGDYEQY | 0.5055821 |
| LAQPGAE | ASSPPGGDYEQY | 0.8243414 |
| ISNLGAVTGF | ASSPPGGDYEQY | 0.59188473 |
| HQEDPGRI | ASSPPGGDYEQY | 0.7718233 |
| GTPGEPTP | ASSPPGGDYEQY | 0.7712712 |
| GFFNSGS | ASSPPGGDYEQY | 0.90406317 |
| GAGGGGLTAGAN | ASSPPGGDYEQY | 0.8245161 |
| FVDALVNM | ASSPPGGDYEQY | 0.8320378 |
| FLTVAPEL | ASSPPGGDYEQY | 0.59118783 |
| FETLTAL | ASSPPGGDYEQY | 0.8306638 |
| DVINPIEL | ASSPPGGDYEQY | 0.83712316 |
| APVSGAPV | ASSPPGGDYEQY | 0.46338302 |
| ALVGGLAW | ASSPPGGDYEQY | 0.87934893 |
| ALAAAAATAI | ASSPPGGDYEQY | 0.531148 |
| AFGLGLLL | ASSPPGGDYEQY | 0.80706537 |
| AFFAAASL | ASSPPGGDYEQY | 0.78238106 |
| AAAALPAFSPP | ASSPPGGDYEQY | 0.84315026 |
| VVLAPAL | ASSPPGGDYEQY | 0.76685613 |
| TIGHVDHGKTTL | ASSPPGGDYEQY | 0.85121447 |
| LISNASDALDK | ASSPPGGDYEQY | 0.5230495 |
| LISNASDAL | ASSPPGGDYEQY | 0.8544674 |
| ISNASDALDK | ASSPPGGDYEQY | 0.8629595 |
| NQAETLVYQTE | ASSPPGGDYEQY | 0.80999994 |
| TGGTGGTAGAG | ASSPPGGDYEQY | 0.8852302 |
| TGSTAGGAGGA | ASSPPGGDYEQY | 0.8403651 |
| DPLAVPVAAGPV | ASSPPGGDYEQY | 0.23032889 |
| GGTGGTPGNGGN | ASSPPGGDYEQY | 0.72576284 |
| PAGLAGVDPATL | ASSPPGGDYEQY | 0.8708619 |
| GNNLVVEAPHAGASV | ASSPPGGDYEQY | 0.88142717 |
| KVTRIMSASTLEQPA | ASSPPGGDYEQY | 0.86376506 |
| LGFALPIALPRRTVV | ASSPPGGDYEQY | 0.8521932 |
| TRLGNLTARKLVVLV | ASSPPGGDYEQY | 0.7345854 |
| WSQQRRLIYFPSAGP | ASSPPGGDYEQY | 0.88362694 |
| GPNGNPDPMAAAVDIR | ASSPPGGDYEQY | 0.91138923 |
| TQRCVSVADLRAAASR | ASSPPGGDYEQY | 0.8549236 |
| QFVLDVAEHPVEVLTETV | ASSPPGGDYEQY | 0.6113728 |
| GGAGGTGGAGGNGGTQAGDG | ASSPPGGDYEQY | 0.5750376 |
| PLLATLRAIAVDVDAKPAQV | ASSPPGGDYEQY | 0.8456878 |
| APKFRTGGYMIDFWGVGYQVAK | ASSPPGGDYEQY | 0.84781706 |
| GAGGAGGAGGAAGGPGGTGGTG | ASSPPGGDYEQY | 0.4644087 |
| IGALGLLATDVAPAGSVAFGVA | ASSPPGGDYEQY | 0.7498043 |
| AGGAGGIGGNGGAGGAGGVLMG | ASSPPGGDYEQY | 0.78110296 |
| ADELIGLL | ASSPPGGDYEQY | 0.20601292 |

| | | |
|---|---|---|
| ADRALKA | ASSPPGGDYEQY | 0.8468648 |
| AGAVGQVGGAGGA | ASSPPGGDYEQY | 0.8832262 |
| AGGGGGLGGAP | ASSPPGGDYEQY | 0.81984156 |
| AGGLLYGNGGAG | ASSPPGGDYEQY | 0.50014055 |
| AGLESIGL | ASSPPGGDYEQY | 0.8761501 |
| AGLKTGII | ASSPPGGDYEQY | 0.8882896 |
| ASLAALLL | ASSPPGGDYEQY | 0.88390243 |
| ASVINISSI | ASSPPGGDYEQY | 0.36578208 |
| ATPPGPA | ASSPPGGDYEQY | 0.15755874 |
| ATVASALV | ASSPPGGDYEQY | 0.8132844 |
| ATVASLIL | ASSPPGGDYEQY | 0.8303844 |
| AVNGPVLIRH | ASSPPGGDYEQY | 0.66964644 |
| DGAAGAVGI | ASSPPGGDYEQY | 0.5075694 |
| DPLLHGI | ASSPPGGDYEQY | 0.80144566 |
| ELGAALGAL | ASSPPGGDYEQY | 0.8646176 |
| ELGRTQAV | ASSPPGGDYEQY | 0.41604337 |
| FVAGGYLS | ASSPPGGDYEQY | 0.4813191 |
| GASGGAGGAGGW | ASSPPGGDYEQY | 0.84541845 |
| GGAGAGGDGGAGGI | ASSPPGGDYEQY | 0.8624345 |
| GTGHNGSQPGMGGNG | ASSPPGGDYEQY | 0.55957484 |
| GYAAASASAS | ASSPPGGDYEQY | 0.65899 |
| HQVDVAAVQQG | ASSPPGGDYEQY | 0.8209506 |
| IGAAIAHT | ASSPPGGDYEQY | 0.8220617 |
| IGMIEVAQL | ASSPPGGDYEQY | 0.8126059 |
| IRLAPEHTSAMD | ASSPPGGDYEQY | 0.3972019 |
| IYNVSVL | ASSPPGGDYEQY | 0.8560204 |
| KQGFLGM | ASSPPGGDYEQY | 0.4862711 |
| LESVITQL | ASSPPGGDYEQY | 0.87015694 |
| LGLGALVVV | ASSPPGGDYEQY | 0.7952823 |
| LIGNGGNGGSGGI | ASSPPGGDYEQY | 0.82822883 |
| LNAINAPTL | ASSPPGGDYEQY | 0.7144983 |
| LQLLNTGV | ASSPPGGDYEQY | 0.48291618 |
| LQTSIEF | ASSPPGGDYEQY | 0.49062413 |
| MSFAVSEL | ASSPPGGDYEQY | 0.58493966 |
| NTGALGL | ASSPPGGDYEQY | 0.8788762 |
| PPGAGGAGGAGTT | ASSPPGGDYEQY | 0.82083845 |
| QLVGLGI | ASSPPGGDYEQY | 0.8558209 |
| RFLEQVHQL | ASSPPGGDYEQY | 0.7913914 |
| RGVGYVLR | ASSPPGGDYEQY | 0.46340364 |
| SATGVFAGVI | ASSPPGGDYEQY | 0.84144187 |
| SLPAITLP | ASSPPGGDYEQY | 0.39930293 |
| STALAFTV | ASSPPGGDYEQY | 0.7980957 |
| SVLTVGLGK | ASSPPGGDYEQY | 0.7717731 |
| TALAGAGSGI | ASSPPGGDYEQY | 0.8306601 |
| TGLPFTGI | ASSPPGGDYEQY | 0.67454845 |
| VAEHVTV | ASSPPGGDYEQY | 0.84090436 |
| VDIVLNSL | ASSPPGGDYEQY | 0.8312218 |
| VLTGGGAVL | ASSPPGGDYEQY | 0.60113335 |
| VNVGLAYGI | ASSPPGGDYEQY | 0.8087136 |
| VSSAAVGAV | ASSPPGGDYEQY | 0.84901804 |
| VTSVYEAGV | ASSPPGGDYEQY | 0.8348824 |

| | | |
|---|---|---|
| WVALDRGA | ASSPPGGDYEQY | 0.34998024 |
| YLVGSVVI | ASSPPGGDYEQY | 0.8222111 |
| GPPGTGKTLL | ASSPPGGDYEQY | 0.5494065 |
| GVGMTKF | ASSPPGGDYEQY | 0.80456585 |
| IGIGGSDL | ASSPPGGDYEQY | 0.76797986 |
| IGTLLAEL | ASSPPGGDYEQY | 0.7035834 |
| IYTSGTT | ASSPPGGDYEQY | 0.8561238 |
| LSFPAGV | ASSPPGGDYEQY | 0.42965832 |
| LYTSGTT | ASSPPGGDYEQY | 0.7134824 |
| VGAFPFGF | ASSPPGGDYEQY | 0.7031033 |
| VGAGTVEF | ASSPPGGDYEQY | 0.7983422 |
| VGKSTLLL | ASSPPGGDYEQY | 0.8429389 |
| AALAHAGGGIV | ASSPPGGDYEQY | 0.44813806 |
| AALVAVSLL | ASSPPGGDYEQY | 0.81654024 |
| AAVSRAVAAL | ASSPPGGDYEQY | 0.7190024 |
| AGGSSGIGGF | ASSPPGGDYEQY | 0.50272834 |
| AGLTVANANG | ASSPPGGDYEQY | 0.7606857 |
| AGLVGAVL | ASSPPGGDYEQY | 0.71531093 |
| ALSQAFTV | ASSPPGGDYEQY | 0.44923827 |
| ALTFSVSL | ASSPPGGDYEQY | 0.8540603 |
| ALTGAALVAL | ASSPPGGDYEQY | 0.5904983 |
| ASFSIGAGVL | ASSPPGGDYEQY | 0.82172006 |
| DGSVYKI | ASSPPGGDYEQY | 0.88721865 |
| EQLAEIV | ASSPPGGDYEQY | 0.5282588 |
| FAVSELTVM | ASSPPGGDYEQY | 0.8669529 |
| GAAPLAAGL | ASSPPGGDYEQY | 0.77937067 |
| GADGGDGGSGGAGG | ASSPPGGDYEQY | 0.7990699 |
| GFGLDTGGAGG | ASSPPGGDYEQY | 0.7511269 |
| GLASGGLL | ASSPPGGDYEQY | 0.6030135 |
| GRIAGAAVLL | ASSPPGGDYEQY | 0.8706714 |
| GSAGSLNTSF | ASSPPGGDYEQY | 0.8509498 |
| GTGQIGI | ASSPPGGDYEQY | 0.8513138 |
| GTLIALIL | ASSPPGGDYEQY | 0.8312526 |
| IAATMGFVVG | ASSPPGGDYEQY | 0.84018433 |
| IFAQALL | ASSPPGGDYEQY | 0.8600343 |
| IGAAVALGF | ASSPPGGDYEQY | 0.659106 |
| IGLTPLAL | ASSPPGGDYEQY | 0.34933653 |
| ILGSNTSTL | ASSPPGGDYEQY | 0.6990053 |
| IVFSLTVL | ASSPPGGDYEQY | 0.7436465 |
| LASKATGF | ASSPPGGDYEQY | 0.587795 |
| LASVALIL | ASSPPGGDYEQY | 0.3871331 |
| LFAVGLTL | ASSPPGGDYEQY | 0.8382044 |
| LFNGGIGTYI | ASSPPGGDYEQY | 0.8974366 |
| LTLGAGII | ASSPPGGDYEQY | 0.7250606 |
| LVAAALGDI | ASSPPGGDYEQY | 0.892218 |
| NTAFVQLGI | ASSPPGGDYEQY | 0.4213068 |
| QLFSGAEL | ASSPPGGDYEQY | 0.85142505 |
| SAATFITL | ASSPPGGDYEQY | 0.8658035 |
| SNYGLANLGV | ASSPPGGDYEQY | 0.5342185 |
| SYGGAYAVM | ASSPPGGDYEQY | 0.7913232 |
| TTGVGGLTLG | ASSPPGGDYEQY | 0.7325362 |

| | | |
|---|---|---|
| TTQTIYI | ASSPPGGDYEQY | 0.6065711 |
| VAAGAGLGAVI | ASSPPGGDYEQY | 0.8740735 |
| VAAVVTLL | ASSPPGGDYEQY | 0.705163 |
| VGFLGGLI | ASSPPGGDYEQY | 0.83826697 |
| VGVIPILL | ASSPPGGDYEQY | 0.60917294 |
| VIFASSLI | ASSPPGGDYEQY | 0.5205461 |
| VKGGDGGAAAG | ASSPPGGDYEQY | 0.80263805 |
| VTKTFTNI | ASSPPGGDYEQY | 0.8536936 |
| VVAAAILL | ASSPPGGDYEQY | 0.7849342 |
| YAALDVEL | ASSPPGGDYEQY | 0.9096515 |
| LASVALLL | ASSPPGGDYEQY | 0.67484295 |
| LDAVHRV | ASSPPGGDYEQY | 0.8911471 |
| QGLSSSIF | ASSPPGGDYEQY | 0.87721133 |
| RGTFANIRL | ASSPPGGDYEQY | 0.77783084 |
| RQLWWGHRI | ASSPPGGDYEQY | 0.69037986 |
| YGPPGTGKTLL | ASSPPGGDYEQY | 0.8560642 |
| AALVGAPANVA | ASSPPGGDYEQY | 0.6200411 |
| AATFRRTE | ASSPPGGDYEQY | 0.85261285 |
| AGALGGGI | ASSPPGGDYEQY | 0.87894845 |
| AGGAGGAGGIGAG | ASSPPGGDYEQY | 0.5678531 |
| AGSGALGL | ASSPPGGDYEQY | 0.81327176 |
| AQIADQI | ASSPPGGDYEQY | 0.7165831 |
| ASAVAVTYL | ASSPPGGDYEQY | 0.77095175 |
| ASTVLGGTGG | ASSPPGGDYEQY | 0.61121106 |
| AVGGEPAK | ASSPPGGDYEQY | 0.4038705 |
| CEQGTFGGTC | ASSPPGGDYEQY | 0.7253407 |
| DEGKAPAG | ASSPPGGDYEQY | 0.53915083 |
| DTVVFPAL | ASSPPGGDYEQY | 0.73032314 |
| FEGITMAG | ASSPPGGDYEQY | 0.7799701 |
| FPLVVQD | ASSPPGGDYEQY | 0.6011078 |
| GAFTTPQ | ASSPPGGDYEQY | 0.80242693 |
| GFGGFGTAGGVGGA | ASSPPGGDYEQY | 0.821602 |
| GFLAQFL | ASSPPGGDYEQY | 0.3911028 |
| GGAGGAGGNAGVA | ASSPPGGDYEQY | 0.6460263 |
| GGVGTAGAGGAGGAGG | ASSPPGGDYEQY | 0.6007814 |
| GNGGGGDAGNAGSGGN | ASSPPGGDYEQY | 0.67688346 |
| GTGGAGGAGNDAGS | ASSPPGGDYEQY | 0.28492957 |
| GTTNTGWFNS | ASSPPGGDYEQY | 0.7770059 |
| IFMGPLFTL | ASSPPGGDYEQY | 0.5336354 |
| ILAHAIEAI | ASSPPGGDYEQY | 0.15376396 |
| IRSAIREAL | ASSPPGGDYEQY | 0.7365519 |
| ITFAGGG | ASSPPGGDYEQY | 0.62866175 |
| ITGMSGHQEV | ASSPPGGDYEQY | 0.6432875 |
| IVAGLACTL | ASSPPGGDYEQY | 0.77358305 |
| IVGGGLAGA | ASSPPGGDYEQY | 0.4574586 |
| IVTGVLNI | ASSPPGGDYEQY | 0.61517644 |
| LAQQGCI | ASSPPGGDYEQY | 0.7462071 |
| LCLHVEN | ASSPPGGDYEQY | 0.8102033 |
| LDAISDGS | ASSPPGGDYEQY | 0.541827 |
| LGGFAAML | ASSPPGGDYEQY | 0.7030624 |
| LGGKNPMI | ASSPPGGDYEQY | 0.8367624 |

| | | |
|---|---|---|
| LGPKGVAAIL | ASSPPGGDYEQY | 0.7511536 |
| LMVGVSI | ASSPPGGDYEQY | 0.81882125 |
| LVGAAFVL | ASSPPGGDYEQY | 0.8312316 |
| LVLAVAPL | ASSPPGGDYEQY | 0.6320901 |
| MAAIAVIDAA | ASSPPGGDYEQY | 0.78195727 |
| NNGINAELQ | ASSPPGGDYEQY | 0.83011234 |
| RGEPLVLVHG | ASSPPGGDYEQY | 0.67164946 |
| RGGAGGAAGLI | ASSPPGGDYEQY | 0.7171086 |
| RSYRMHPAV | ASSPPGGDYEQY | 0.57690704 |
| SAAHLLAGQLA | ASSPPGGDYEQY | 0.6718005 |
| SAAVGSY | ASSPPGGDYEQY | 0.27630368 |
| SPHAVLHRI | ASSPPGGDYEQY | 0.6850712 |
| SQIAKVL | ASSPPGGDYEQY | 0.8037878 |
| TALGHLVP | ASSPPGGDYEQY | 0.6373439 |
| TAPTTSGAA | ASSPPGGDYEQY | 0.4356792 |
| TDGDDGNGGSPGT | ASSPPGGDYEQY | 0.48262346 |
| TDGLGEN | ASSPPGGDYEQY | 0.68656576 |
| TGGPPLPGQA | ASSPPGGDYEQY | 0.6289126 |
| TIPAITI | ASSPPGGDYEQY | 0.32313472 |
| TIPGFTI | ASSPPGGDYEQY | 0.5525625 |
| TSIASGALA | ASSPPGGDYEQY | 0.8102511 |
| TVTLPGAAI | ASSPPGGDYEQY | 0.8205241 |
| VASGLSAALP | ASSPPGGDYEQY | 0.43948615 |
| VGAGVIGI | ASSPPGGDYEQY | 0.7929311 |
| VGPKRFGV | ASSPPGGDYEQY | 0.7550002 |
| VNVAFPD | ASSPPGGDYEQY | 0.8500452 |
| VSMTNSGVS | ASSPPGGDYEQY | 0.7507219 |
| VTDVLGDLSV | ASSPPGGDYEQY | 0.5767612 |
| VVVVGAGFAGL | ASSPPGGDYEQY | 0.7581325 |
| VYTVGTVV | ASSPPGGDYEQY | 0.6013781 |
| WTTFSGP | ASSPPGGDYEQY | 0.6026591 |
| YDELVEAF | ASSPPGGDYEQY | 0.8003784 |
| YGTGGAGGNGG | ASSPPGGDYEQY | 0.691196 |
| YTNVMAV | ASSPPGGDYEQY | 0.5658898 |
| FLEQVHQL | ASSPPGGDYEQY | 0.5953079 |
| GLSLLAVAV | ASSPPGGDYEQY | 0.81914544 |
| LEQVHQL | ASSPPGGDYEQY | 0.46164912 |
| LNAAGAI | ASSPPGGDYEQY | 0.5006293 |
| PAVHASL | ASSPPGGDYEQY | 0.5627206 |
| TELEDDA | ASSPPGGDYEQY | 0.6991942 |
| TPPGPPN | ASSPPGGDYEQY | 0.7978883 |
| TPPPPPPP | ASSPPGGDYEQY | 0.7704307 |
| TQSPALI | ASSLDGTNERLF | 0.63979185 |
| EFAAAFPQ | ASSLDGTNERLF | 0.7436501 |
| LLSGNLDVL | ASSLDGTNERLF | 0.7851334 |
| GPANLGNYNI | ASSLDGTNERLF | 0.73333234 |
| TLGPTLP | ASSLDGTNERLF | 0.76854146 |
| VYAYSSCCK | ASSLDGTNERLF | 0.51876175 |
| APAPAPAPNI | ASSLDGTNERLF | 0.81308794 |
| PGGPVPIP | ASSLDGTNERLF | 0.7842331 |
| FLTVALEP | ASSLDGTNERLF | 0.78913975 |

| | | |
|---|---|---|
| SQYFGANL | ASSLDGTNERLF | 0.7919908 |
| GGVKPAA | ASSLDGTNERLF | 0.7573297 |
| GGTGGAGGAGGW | ASSLDGTNERLF | 0.74245083 |
| GVGGVAGLGGAGG | ASSLDGTNERLF | 0.3834932 |
| VLLVLATTI | ASSLDGTNERLF | 0.63444364 |
| IGIFAAML | ASSLDGTNERLF | 0.6839926 |
| AVGDLID | ASSLDGTNERLF | 0.6907606 |
| NISPLPL | ASSLDGTNERLF | 0.7386825 |
| VTAVFVAI | ASSLDGTNERLF | 0.8057866 |
| SGDDNDAAL | ASSLDGTNERLF | 0.55810446 |
| AVAALAHS | ASSLDGTNERLF | 0.81785774 |
| GGVTVMATP | ASSLDGTNERLF | 0.79862785 |
| AGLAANVRD | ASSLDGTNERLF | 0.40253526 |
| TNRPDLIDPALLRPG | ASSLDGTNERLF | 0.63833404 |
| LANLGVTAL | ASSLDGTNERLF | 0.8246778 |
| TGERIEL | ASSLDGTNERLF | 0.70648277 |
| NGEELAA | ASSLDGTNERLF | 0.6334001 |
| VLSLVWG | ASSLDGTNERLF | 0.7122957 |
| IGYALWL | ASSLDGTNERLF | 0.80805767 |
| LSLLLAPL | ASSLDGTNERLF | 0.5687876 |
| TLDEALK | ASSLDGTNERLF | 0.8292917 |
| TDGAGTAEGG | ASSLDGTNERLF | 0.25848734 |
| NAPGGAGGNGG | ASSLDGTNERLF | 0.66996 |
| VSVFMGMGGNF | ASSLDGTNERLF | 0.67556185 |
| LASYLAAL | ASSLDGTNERLF | 0.2769071 |
| LVGGTPAL | ASSLDGTNERLF | 0.7864973 |
| VRNADVL | ASSLDGTNERLF | 0.71521115 |
| TCSSPFLVSAM | ASSLDGTNERLF | 0.78922105 |
| VHITDAE | ASSLDGTNERLF | 0.6600009 |
| EYAAPSF | ASSLDGTNERLF | 0.80099094 |
| VVDAFHGGL | ASSLDGTNERLF | 0.69586754 |
| ASVVDPQ | ASSLDGTNERLF | 0.7329178 |
| SYNEFGGLS | ASSLDGTNERLF | 0.5899336 |
| DPDGSVY | ASSLDGTNERLF | 0.77720565 |
| MVSTIHATE | ASSLDGTNERLF | 0.8160914 |
| LAAGVADPV | ASSLDGTNERLF | 0.8105074 |
| TKEQIAA | ASSLDGTNERLF | 0.45619783 |
| SLASPAAL | ASSLDGTNERLF | 0.6086036 |
| IGLVFYGV | ASSLDGTNERLF | 0.721812 |
| FLLDASNL | ASSLDGTNERLF | 0.27867606 |
| YQVGNFP | ASSLDGTNERLF | 0.21097942 |
| ESLLSQST | ASSLDGTNERLF | 0.54899 |
| LFTVGFAAVG | ASSLDGTNERLF | 0.53437537 |
| ATPPTPT | ASSLDGTNERLF | 0.76838756 |
| TDLATLG | ASSLDGTNERLF | 0.769068 |
| SAGGTGR | ASSLDGTNERLF | 0.63497776 |
| GFSNSGS | ASSLDGTNERLF | 0.76362646 |
| GGNGGSGGTSV | ASSLDGTNERLF | 0.75269544 |
| GTSVATGGAGN | ASSLDGTNERLF | 0.112608604 |
| NGGNGGSGGTS | ASSLDGTNERLF | 0.65532184 |
| AVVKFPH | ASSLDGTNERLF | 0.3998763 |

| | | |
|---|---|---|
| TPPPPPP | ASSLDGTNERLF | 0.8329656 |
| AAPAAAAAPA | ASSLDGTNERLF | 0.5328468 |
| IALNEDL | ASSLDGTNERLF | 0.52481115 |
| IVNVSSM | ASSLDGTNERLF | 0.6830272 |
| TGIPPAPR | ASSLDGTNERLF | 0.66079664 |
| FDLGGGTF | ASSLDGTNERLF | 0.678839 |
| FDLGGGTFDVS | ASSLDGTNERLF | 0.55100405 |
| NEPTAAA | ASSLDGTNERLF | 0.18746927 |
| NEPTAAALA | ASSLDGTNERLF | 0.80199945 |
| GSIINISSI | ASSLDGTNERLF | 0.4059471 |
| VAPGNAGTA | ASSLDGTNERLF | 0.57244694 |
| TGGGASLE | ASSLDGTNERLF | 0.35040307 |
| VAVYNHYKRI | ASSLDGTNERLF | 0.24132273 |
| TDPAPATT | ASSLDGTNERLF | 0.44267058 |
| TNRPDILDPALLRPG | ASSLDGTNERLF | 0.8123076 |
| LTTTTTS | ASSLDGTNERLF | 0.55524784 |
| PTGSGKTL | ASSLDGTNERLF | 0.60460716 |
| TGSGKTL | ASSLDGTNERLF | 0.8096069 |
| TIGHVDHGK | ASSLDGTNERLF | 0.66790396 |
| TIGHVDHGKTT | ASSLDGTNERLF | 0.6051881 |
| GHVDHGKTT | ASSLDGTNERLF | 0.65559286 |
| AQFEHTLL | ASSLDGTNERLF | 0.4650284 |
| IVNQDEL | ASSLDGTNERLF | 0.593897 |
| TNGSQFF | ASSLDGTNERLF | 0.6110985 |
| TGKGIVE | ASSLDGTNERLF | 0.8282622 |
| LISNASDA | ASSLDGTNERLF | 0.67800164 |
| AELRTLGELTKIAKAHGAQ | ASSLDGTNERLF | 0.5163867 |
| APNAVAAAEPSW | ASSLDGTNERLF | 0.78400004 |
| APQVAGAATAVL | ASSLDGTNERLF | 0.8182681 |
| ARRDPAAAAAILPTDA | ASSLDGTNERLF | 0.84815586 |
| DAAARDVLAEAGLAEYF | ASSLDGTNERLF | 0.5194154 |
| DDQIVLSAKANLAKVEQ | ASSLDGTNERLF | 0.7420086 |
| DELLPDTGEDSDEDGH | ASSLDGTNERLF | 0.709951 |
| ELPEAPANRATVPQD | ASSLDGTNERLF | 0.6624045 |
| FGQSKIERVTGIG | ASSLDGTNERLF | 0.6692387 |
| GAGGTGSTAGGAGGAGGAGGLY | ASSLDGTNERLF | 0.5292276 |
| GHGGTGAAGGNGGTGGAGGSGI | ASSLDGTNERLF | 0.79370284 |
| GNGGNGGAGGEGGPGGAGGAGA | ASSLDGTNERLF | 0.8462281 |
| GSTAGNGGNGGNATNGTIAGQP | ASSLDGTNERLF | 0.687983 |
| GTGGDGGNGGNGGTGG | ASSLDGTNERLF | 0.457191 |
| GVRRPAGIPADAPV | ASSLDGTNERLF | 0.630149 |
| GYRQVTDAHLLTLARR | ASSLDGTNERLF | 0.49364677 |
| IAKMGQDGHDRG | ASSLDGTNERLF | 0.570104 |
| KAEPSIEIQYT | ASSLDGTNERLF | 0.72723037 |
| KPVRIRMAADEDTVAVT | ASSLDGTNERLF | 0.3385458 |
| MPPRIAGMRLLVIKPEP | ASSLDGTNERLF | 0.6500411 |
| MKFYAVALFRSRGVPPTR | ASSLDGTNERLF | 0.8473719 |
| NAGGFGILGAVAHSP | ASSLDGTNERLF | 0.58653724 |
| NEGREITLAMAEM | ASSLDGTNERLF | 0.72509944 |
| NELVEWSHPLGHRWRWE | ASSLDGTNERLF | 0.781841 |
| NLGAGGTGGNGGNPG | ASSLDGTNERLF | 0.7168918 |

| | | |
|---|---|---|
| NTEWPKWETVMAADPVD | ASSLDGTNERLF | 0.8057058 |
| PASAAPAETQP | ASSLDGTNERLF | 0.7626195 |
| QQSSDGKNGVRAAEITGE | ASSLDGTNERLF | 0.7327434 |
| RAPDAVTTSWWKE | ASSLDGTNERLF | 0.71516407 |
| SGFGNVGAHTSGWFNQ | ASSLDGTNERLF | 0.6595199 |
| SGVAGAVGQAASVGGLKV | ASSLDGTNERLF | 0.6829309 |
| SNGGAGTGSAGGNGGT | ASSLDGTNERLF | 0.6672038 |
| TGGAGGEGGAAG | ASSLDGTNERLF | 0.6295831 |
| TGGDGGAAGAGGAPGHG | ASSLDGTNERLF | 0.7752004 |
| TMPPLLAAIANAGRHGV | ASSLDGTNERLF | 0.7079601 |
| TSSTAGNSGGA | ASSLDGTNERLF | 0.78855395 |
| VPATVPVGILC | ASSLDGTNERLF | 0.35665917 |
| VQRPDSGGVVLTGRIS | ASSLDGTNERLF | 0.5156187 |
| WTPNPVPATKGVPAGLVQAT | ASSLDGTNERLF | 0.76863694 |
| YALKRALAKFSDRFDVVII | ASSLDGTNERLF | 0.19599073 |
| YLIGSVIV | ASSLDGTNERLF | 0.267569 |
| YGQLVAVID | ASSLDGTNERLF | 0.72153556 |
| VTVGGMTLPA | ASSLDGTNERLF | 0.8223283 |
| VLALPAV | ASSLDGTNERLF | 0.18896948 |
| VIPPTLNYET | ASSLDGTNERLF | 0.5915725 |
| VGMGGMHPSSG | ASSLDGTNERLF | 0.3902411 |
| VGLAGGAATA | ASSLDGTNERLF | 0.81684685 |
| VGGAGGAGGSGP | ASSLDGTNERLF | 0.70509404 |
| VFAAFMV | ASSLDGTNERLF | 0.28886583 |
| TLYDKLR | ASSLDGTNERLF | 0.7771417 |
| TLAVFTVL | ASSLDGTNERLF | 0.3621356 |
| TGSVQPARV | ASSLDGTNERLF | 0.3788733 |
| TGGGAGSGGT | ASSLDGTNERLF | 0.7957934 |
| SHLETSL | ASSLDGTNERLF | 0.8020443 |
| SFTTGMT | ASSLDGTNERLF | 0.44953996 |
| QVEVAKRMGVRQPTVS | ASSLDGTNERLF | 0.77177835 |
| QGSGGPALAM | ASSLDGTNERLF | 0.5905691 |
| MLAPSNP | ASSLDGTNERLF | 0.85261464 |
| LVNNAAI | ASSLDGTNERLF | 0.6485918 |
| LTGEINTLA | ASSLDGTNERLF | 0.36484578 |
| LSIFGPV | ASSLDGTNERLF | 0.7795268 |
| LLALLANL | ASSLDGTNERLF | 0.73695827 |
| LISNASDALDKLR | ASSLDGTNERLF | 0.7749557 |
| LINEAVP | ASSLDGTNERLF | 0.73819506 |
| LGYAFSGL | ASSLDGTNERLF | 0.6465539 |
| LDFVAVAL | ASSLDGTNERLF | 0.6032712 |
| LAQPGAE | ASSLDGTNERLF | 0.78151727 |
| ISNLGAVTGF | ASSLDGTNERLF | 0.7634059 |
| HQEDPGRI | ASSLDGTNERLF | 0.5085401 |
| GTPGEPTP | ASSLDGTNERLF | 0.80405676 |
| GFFNSGS | ASSLDGTNERLF | 0.70326805 |
| GAGGGGLTAGAN | ASSLDGTNERLF | 0.64667845 |
| FVDALVNM | ASSLDGTNERLF | 0.8379867 |
| FLTVAPEL | ASSLDGTNERLF | 0.7249439 |
| FETLTAL | ASSLDGTNERLF | 0.7709797 |
| DVINPIEL | ASSLDGTNERLF | 0.79535526 |

| | | |
|---|---|---|
| APVSGAPV | ASSLDGTNERLF | 0.81426376 |
| ALVGGLAW | ASSLDGTNERLF | 0.40966922 |
| ALAAAAATAI | ASSLDGTNERLF | 0.78798914 |
| AFGLGLLL | ASSLDGTNERLF | 0.71754754 |
| AFFAAASL | ASSLDGTNERLF | 0.78756094 |
| AAAALPAFSPP | ASSLDGTNERLF | 0.72788715 |
| VVLAPAL | ASSLDGTNERLF | 0.53085166 |
| TIGHVDHGKTTL | ASSLDGTNERLF | 0.7505467 |
| LISNASDALDK | ASSLDGTNERLF | 0.81082845 |
| LISNASDAL | ASSLDGTNERLF | 0.720706 |
| ISNASDALDK | ASSLDGTNERLF | 0.74833727 |
| NQAETLVYQTE | ASSLDGTNERLF | 0.8214454 |
| TGGTGGTAGAG | ASSLDGTNERLF | 0.5581583 |
| TGSTAGGAGGA | ASSLDGTNERLF | 0.7101854 |
| DPLAVPVAAGPV | ASSLDGTNERLF | 0.80564165 |
| GGTGGTPGNGGN | ASSLDGTNERLF | 0.79925156 |
| PAGLAGVDPATL | ASSLDGTNERLF | 0.6801027 |
| GNNLVVEAPHAGASV | ASSLDGTNERLF | 0.8113054 |
| KVTRIMSASTLEQPA | ASSLDGTNERLF | 0.5251075 |
| LGFALPIALPRRTVV | ASSLDGTNERLF | 0.75402683 |
| TRLGNLTARKLVVLV | ASSLDGTNERLF | 0.54939437 |
| WSQQRRLIYFPSAGP | ASSLDGTNERLF | 0.727895 |
| GPNGNPDPMAAAVDIR | ASSLDGTNERLF | 0.422194 |
| TQRCVSVADLRAAASR | ASSLDGTNERLF | 0.748062 |
| QFVLDVAEHPVEVLTETV | ASSLDGTNERLF | 0.7676555 |
| GGAGGTGGAGGNGGTQAGDG | ASSLDGTNERLF | 0.7681104 |
| PLLATLRAIAVDVDAKPAQV | ASSLDGTNERLF | 0.24598116 |
| APKFRTGGYMIDFWGVGYQVAK | ASSLDGTNERLF | 0.7402878 |
| GAGGAGGAGGAAGGPGGTGGTG | ASSLDGTNERLF | 0.7841325 |
| IGALGLLATDVAPAGSVAFGVA | ASSLDGTNERLF | 0.8325341 |
| AGGAGGIGGNGGAGGAGGVLMG | ASSLDGTNERLF | 0.6992709 |
| ADELIGLL | ASSLDGTNERLF | 0.8006515 |
| ADRALKA | ASSLDGTNERLF | 0.6617584 |
| AGAVGQVGGAGGA | ASSLDGTNERLF | 0.68837017 |
| AGGGGGLGGAP | ASSLDGTNERLF | 0.47712028 |
| AGGLLYGNGGAG | ASSLDGTNERLF | 0.6883919 |
| AGLESIGL | ASSLDGTNERLF | 0.5767484 |
| AGLKTGII | ASSLDGTNERLF | 0.48060387 |
| ASLAALLL | ASSLDGTNERLF | 0.76372385 |
| ASVINISSI | ASSLDGTNERLF | 0.796955 |
| ATPPGPA | ASSLDGTNERLF | 0.7987054 |
| ATVASALV | ASSLDGTNERLF | 0.80262995 |
| ATVASLIL | ASSLDGTNERLF | 0.79460394 |
| AVNGPVLIRH | ASSLDGTNERLF | 0.8022604 |
| DGAAGAVGI | ASSLDGTNERLF | 0.61286646 |
| DPLLHGI | ASSLDGTNERLF | 0.63284206 |
| ELGAALGAL | ASSLDGTNERLF | 0.4511341 |
| ELGRTQAV | ASSLDGTNERLF | 0.6223385 |
| FVAGGYLS | ASSLDGTNERLF | 0.6081491 |
| GASGGAGGAGGW | ASSLDGTNERLF | 0.7104381 |
| GGAGAGGDGGAGGI | ASSLDGTNERLF | 0.14626686 |

| | | |
|---|---|---|
| GTGHNGSQPGMGGNG | ASSLDGTNERLF | 0.6314299 |
| GYAAASASAS | ASSLDGTNERLF | 0.81760764 |
| HQVDVAAVQQG | ASSLDGTNERLF | 0.6033623 |
| IGAAIAHT | ASSLDGTNERLF | 0.7764938 |
| IGMIEVAQL | ASSLDGTNERLF | 0.65905213 |
| IRLAPEHTSAMD | ASSLDGTNERLF | 0.5648603 |
| IYNVSVL | ASSLDGTNERLF | 0.7686671 |
| KQGFLGM | ASSLDGTNERLF | 0.54149604 |
| LESVITQL | ASSLDGTNERLF | 0.79101115 |
| LGLGALVVV | ASSLDGTNERLF | 0.8301519 |
| LIGNGGNGGSGGI | ASSLDGTNERLF | 0.72230196 |
| LNAINAPTL | ASSLDGTNERLF | 0.6962782 |
| LQLLNTGV | ASSLDGTNERLF | 0.7502252 |
| LQTSIEF | ASSLDGTNERLF | 0.36357808 |
| MSFAVSEL | ASSLDGTNERLF | 0.71325517 |
| NTGALGL | ASSLDGTNERLF | 0.12956466 |
| PPGAGGAGGAGTT | ASSLDGTNERLF | 0.6519343 |
| QLVGLGI | ASSLDGTNERLF | 0.5213379 |
| RFLEQVHQL | ASSLDGTNERLF | 0.7252457 |
| RGVGYVLR | ASSLDGTNERLF | 0.71926945 |
| SATGVFAGVI | ASSLDGTNERLF | 0.7736975 |
| SLPAITLP | ASSLDGTNERLF | 0.536236 |
| STALAFTV | ASSLDGTNERLF | 0.62535965 |
| SVLTVGLGK | ASSLDGTNERLF | 0.59796166 |
| TALAGAGSGI | ASSLDGTNERLF | 0.51319945 |
| TGLPFTGI | ASSLDGTNERLF | 0.7863815 |
| VAEHVTV | ASSLDGTNERLF | 0.7170067 |
| VDIVLNSL | ASSLDGTNERLF | 0.5894193 |
| VLTGGGAVL | ASSLDGTNERLF | 0.80926055 |
| VNVGLAYGI | ASSLDGTNERLF | 0.72878945 |
| VSSAAVGAV | ASSLDGTNERLF | 0.3875947 |
| VTSVYEAGV | ASSLDGTNERLF | 0.6281563 |
| WVALDRGA | ASSLDGTNERLF | 0.701132 |
| YLVGSVVI | ASSLDGTNERLF | 0.6622575 |
| GPPGTGKTLL | ASSLDGTNERLF | 0.14626646 |
| GVGMTKF | ASSLDGTNERLF | 0.74331975 |
| IGIGGSDL | ASSLDGTNERLF | 0.6588781 |
| IGTLLAEL | ASSLDGTNERLF | 0.48439893 |
| IYTSGTT | ASSLDGTNERLF | 0.61593235 |
| LSFPAGV | ASSLDGTNERLF | 0.7707169 |
| LYTSGTT | ASSLDGTNERLF | 0.52816784 |
| VGAFPFGF | ASSLDGTNERLF | 0.74727994 |
| VGAGTVEF | ASSLDGTNERLF | 0.75127006 |
| VGKSTLLL | ASSLDGTNERLF | 0.7992478 |
| AALAHAGGGIV | ASSLDGTNERLF | 0.8186519 |
| AALVAVSLL | ASSLDGTNERLF | 0.7630526 |
| AAVSRAVAAL | ASSLDGTNERLF | 0.73195785 |
| AGGSSGIGGF | ASSLDGTNERLF | 0.78492004 |
| AGLTVANANG | ASSLDGTNERLF | 0.7639701 |
| AGLVGAVL | ASSLDGTNERLF | 0.27576244 |
| ALSQAFTV | ASSLDGTNERLF | 0.26031682 |

| | | |
|---|---|---|
| ALTFSVSL | ASSLDGTNERLF | 0.70179486 |
| ALTGAALVAL | ASSLDGTNERLF | 0.31880304 |
| ASFSIGAGVL | ASSLDGTNERLF | 0.74150753 |
| DGSVYKI | ASSLDGTNERLF | 0.83348846 |
| EQLAEIV | ASSLDGTNERLF | 0.71618164 |
| FAVSELTVM | ASSLDGTNERLF | 0.47266552 |
| GAAPLAAGL | ASSLDGTNERLF | 0.6297675 |
| GADGGDGGSGGAGG | ASSLDGTNERLF | 0.4995133 |
| GFGLDTGGAGG | ASSLDGTNERLF | 0.7550515 |
| GLASGGLL | ASSLDGTNERLF | 0.79773074 |
| GRIAGAAVLL | ASSLDGTNERLF | 0.8078128 |
| GSAGSLNTSF | ASSLDGTNERLF | 0.7925609 |
| GTGQIGI | ASSLDGTNERLF | 0.62409616 |
| GTLIALIL | ASSLDGTNERLF | 0.75246143 |
| IAATMGFVVG | ASSLDGTNERLF | 0.6085696 |
| IFAQALL | ASSLDGTNERLF | 0.700575 |
| IGAAVALGF | ASSLDGTNERLF | 0.69255894 |
| IGLTPLAL | ASSLDGTNERLF | 0.74264103 |
| ILGSNTSTL | ASSLDGTNERLF | 0.6758107 |
| IVFSLTVL | ASSLDGTNERLF | 0.64814544 |
| LASKATGF | ASSLDGTNERLF | 0.43010798 |
| LASVALIL | ASSLDGTNERLF | 0.7590955 |
| LFAVGLTL | ASSLDGTNERLF | 0.7287353 |
| LFNGGIGTYI | ASSLDGTNERLF | 0.3630132 |
| LTLGAGII | ASSLDGTNERLF | 0.61297333 |
| LVAAALGDI | ASSLDGTNERLF | 0.82502484 |
| NTAFVQLGI | ASSLDGTNERLF | 0.8235947 |
| QLFSGAEL | ASSLDGTNERLF | 0.55851513 |
| SAATFITL | ASSLDGTNERLF | 0.5912936 |
| SNYGLANLGV | ASSLDGTNERLF | 0.6467153 |
| SYGGAYAVM | ASSLDGTNERLF | 0.406003 |
| TTGVGGLTLG | ASSLDGTNERLF | 0.7505262 |
| TTQTIYI | ASSLDGTNERLF | 0.62372756 |
| VAAGAGLGAVI | ASSLDGTNERLF | 0.6305564 |
| VAAVVTLL | ASSLDGTNERLF | 0.65135986 |
| VGFLGGLI | ASSLDGTNERLF | 0.7056064 |
| VGVIPILL | ASSLDGTNERLF | 0.6395434 |
| VIFASSLI | ASSLDGTNERLF | 0.796677 |
| VKGGDGGAAAG | ASSLDGTNERLF | 0.16533704 |
| VTKTFTNI | ASSLDGTNERLF | 0.76529163 |
| VVAAAILL | ASSLDGTNERLF | 0.4991416 |
| YAALDVEL | ASSLDGTNERLF | 0.5822916 |
| LASVALLL | ASSLDGTNERLF | 0.3153888 |
| LDAVHRV | ASSLDGTNERLF | 0.74764717 |
| QGLSSSIF | ASSLDGTNERLF | 0.7947297 |
| RGTFANIRL | ASSLDGTNERLF | 0.78765845 |
| RQLWWGHRI | ASSLDGTNERLF | 0.4392101 |
| YGPPGTGKTLL | ASSLDGTNERLF | 0.76509094 |
| AALVGAPANVA | ASSLDGTNERLF | 0.84031105 |
| AATFRRTE | ASSLDGTNERLF | 0.56264603 |
| AGALGGGI | ASSLDGTNERLF | 0.29858983 |

| | | |
|---|---|---|
| AGGAGGAGGIGAG | ASSLDGTNERLF | 0.7537436 |
| AGSGALGL | ASSLDGTNERLF | 0.6820419 |
| AQIADQI | ASSLDGTNERLF | 0.7675508 |
| ASAVAVTYL | ASSLDGTNERLF | 0.8192376 |
| ASTVLGGTGG | ASSLDGTNERLF | 0.3723633 |
| AVGGEPAK | ASSLDGTNERLF | 0.8154713 |
| CEQGTFGGTC | ASSLDGTNERLF | 0.77517104 |
| DEGKAPAG | ASSLDGTNERLF | 0.50036967 |
| DTVVFPAL | ASSLDGTNERLF | 0.65401226 |
| FEGITMAG | ASSLDGTNERLF | 0.68727297 |
| FPLVVQD | ASSLDGTNERLF | 0.28704762 |
| GAFTTPQ | ASSLDGTNERLF | 0.80619866 |
| GFGGFGTAGGVGGA | ASSLDGTNERLF | 0.5353166 |
| GFLAQFL | ASSLDGTNERLF | 0.65366185 |
| GGAGGAGGNAGVA | ASSLDGTNERLF | 0.699113 |
| GGVGTAGAGGAGGAGG | ASSLDGTNERLF | 0.6522372 |
| GNGGGGDAGNAGSGGN | ASSLDGTNERLF | 0.65983826 |
| GTGGAGGAGNDAGS | ASSLDGTNERLF | 0.65918094 |
| GTTNTGWFNS | ASSLDGTNERLF | 0.7423789 |
| IFMGPLFTL | ASSLDGTNERLF | 0.7116881 |
| ILAHAIEAI | ASSLDGTNERLF | 0.68681765 |
| IRSAIREAL | ASSLDGTNERLF | 0.74759763 |
| ITFAGGG | ASSLDGTNERLF | 0.7967086 |
| ITGMSGHQEV | ASSLDGTNERLF | 0.49001166 |
| IVAGLACTL | ASSLDGTNERLF | 0.59816486 |
| IVGGGLAGA | ASSLDGTNERLF | 0.7670121 |
| IVTGVLNI | ASSLDGTNERLF | 0.41173494 |
| LAQQGCI | ASSLDGTNERLF | 0.77341765 |
| LCLHVEN | ASSLDGTNERLF | 0.73350453 |
| LDAISDGS | ASSLDGTNERLF | 0.71464443 |
| LGGFAAML | ASSLDGTNERLF | 0.66142696 |
| LGGKNPMI | ASSLDGTNERLF | 0.83698773 |
| LGPKGVAAIL | ASSLDGTNERLF | 0.7694352 |
| LMVGVSI | ASSLDGTNERLF | 0.72482455 |
| LVGAAFVL | ASSLDGTNERLF | 0.70139706 |
| LVLAVAPL | ASSLDGTNERLF | 0.64853376 |
| MAAIAVIDAA | ASSLDGTNERLF | 0.77806634 |
| NNGINAELQ | ASSLDGTNERLF | 0.77990127 |
| RGEPLVLVHG | ASSLDGTNERLF | 0.427319 |
| RGGAGGAAGLI | ASSLDGTNERLF | 0.6130893 |
| RSYRMHPAV | ASSLDGTNERLF | 0.8242327 |
| SAAHLLAGQLA | ASSLDGTNERLF | 0.7933874 |
| SAAVGSY | ASSLDGTNERLF | 0.37876156 |
| SPHAVLHRI | ASSLDGTNERLF | 0.6064307 |
| SQIAKVL | ASSLDGTNERLF | 0.41119727 |
| TALGHLVP | ASSLDGTNERLF | 0.69426227 |
| TAPTTSGAA | ASSLDGTNERLF | 0.56956017 |
| TDGDDGNGGSPGT | ASSLDGTNERLF | 0.7307101 |
| TDGLGEN | ASSLDGTNERLF | 0.62831455 |
| TGGPPLPGQA | ASSLDGTNERLF | 0.7488521 |
| TIPAITI | ASSLDGTNERLF | 0.72223115 |

| | | |
|---|---|---|
| TIPGFTI | ASSLDGTNERLF | 0.8155983 |
| TSIASGALA | ASSLDGTNERLF | 0.6109632 |
| TVTLPGAAI | ASSLDGTNERLF | 0.68463624 |
| VASGLSAALP | ASSLDGTNERLF | 0.40436286 |
| VGAGVIGI | ASSLDGTNERLF | 0.46715635 |
| VGPKRFGV | ASSLDGTNERLF | 0.5569157 |
| VNVAFPD | ASSLDGTNERLF | 0.66942096 |
| VSMTNSGVS | ASSLDGTNERLF | 0.2372038 |
| VTDVLGDLSV | ASSLDGTNERLF | 0.8190159 |
| VVVVGAGFAGL | ASSLDGTNERLF | 0.6918222 |
| VYTVGTVV | ASSLDGTNERLF | 0.78951955 |
| WTTFSGP | ASSLDGTNERLF | 0.6570072 |
| YDELVEAF | ASSLDGTNERLF | 0.6352912 |
| YGTGGAGGNGG | ASSLDGTNERLF | 0.58706933 |
| YTNVMAV | ASSLDGTNERLF | 0.77364874 |
| FLEQVHQL | ASSLDGTNERLF | 0.47040325 |
| GLSLLAVAV | ASSLDGTNERLF | 0.3862701 |
| LEQVHQL | ASSLDGTNERLF | 0.71632487 |
| LNAAGAI | ASSLDGTNERLF | 0.45375246 |
| PAVHASL | ASSLDGTNERLF | 0.82407653 |
| TELEDDA | ASSLDGTNERLF | 0.7940071 |
| TPPGPPN | ASSLDGTNERLF | 0.69328827 |
| TPPPPPPP | ASSLDGTNERLF | 0.8091631 |
| TQSPALI | AWKTGTGGGEQY | 0.23109992 |
| EFAAAFPQ | AWKTGTGGGEQY | 0.6333955 |
| LLSGNLDVL | AWKTGTGGGEQY | 0.8352728 |
| GPANLGNYNI | AWKTGTGGGEQY | 0.7476448 |
| TLGPTLP | AWKTGTGGGEQY | 0.6606791 |
| VYAYSSCCK | AWKTGTGGGEQY | 0.80026734 |
| APAPAPAPNI | AWKTGTGGGEQY | 0.603437 |
| PGGPVPIP | AWKTGTGGGEQY | 0.68876565 |
| FLTVALEP | AWKTGTGGGEQY | 0.7050537 |
| SQYFGANL | AWKTGTGGGEQY | 0.76693904 |
| GGVKPAA | AWKTGTGGGEQY | 0.76631206 |
| GGTGGAGGAGGW | AWKTGTGGGEQY | 0.4931 |
| GVGGVAGLGGAGG | AWKTGTGGGEQY | 0.77742445 |
| VLLVLATTI | AWKTGTGGGEQY | 0.83273965 |
| IGIFAAML | AWKTGTGGGEQY | 0.6067366 |
| AVGDLID | AWKTGTGGGEQY | 0.77640885 |
| NISPLPL | AWKTGTGGGEQY | 0.74056077 |
| VTAVFVAI | AWKTGTGGGEQY | 0.74539334 |
| SGDDNDAAL | AWKTGTGGGEQY | 0.53591746 |
| AVAALAHS | AWKTGTGGGEQY | 0.53140235 |
| GGVTVMATP | AWKTGTGGGEQY | 0.6639199 |
| AGLAANVRD | AWKTGTGGGEQY | 0.45991322 |
| TNRPDLIDPALLRPG | AWKTGTGGGEQY | 0.73004127 |
| LANLGVTAL | AWKTGTGGGEQY | 0.66484296 |
| TGERIEL | AWKTGTGGGEQY | 0.8566464 |
| NGEELAA | AWKTGTGGGEQY | 0.7610191 |
| VLSLVWG | AWKTGTGGGEQY | 0.73665273 |
| IGYALWL | AWKTGTGGGEQY | 0.52751684 |

| | | |
|---|---|---|
| LSLLLAPL | AWKTGTGGGEQY | 0.7346188 |
| TLDEALK | AWKTGTGGGEQY | 0.70281845 |
| TDGAGTAEGG | AWKTGTGGGEQY | 0.44180986 |
| NAPGGAGGNGG | AWKTGTGGGEQY | 0.8193291 |
| VSVFMGMGGNF | AWKTGTGGGEQY | 0.5331297 |
| LASYLAAL | AWKTGTGGGEQY | 0.65336144 |
| LVGGTPAL | AWKTGTGGGEQY | 0.7976939 |
| VRNADVL | AWKTGTGGGEQY | 0.7928344 |
| TCSSPFLVSAM | AWKTGTGGGEQY | 0.66070104 |
| VHITDAE | AWKTGTGGGEQY | 0.6664585 |
| EYAAPSF | AWKTGTGGGEQY | 0.5445102 |
| VVDAFHGGL | AWKTGTGGGEQY | 0.7860599 |
| ASVVDPQ | AWKTGTGGGEQY | 0.786618 |
| SYNEFGGLS | AWKTGTGGGEQY | 0.70703447 |
| DPDGSVY | AWKTGTGGGEQY | 0.80565006 |
| MVSTIHATE | AWKTGTGGGEQY | 0.78899455 |
| LAAGVADPV | AWKTGTGGGEQY | 0.4699264 |
| TKEQIAA | AWKTGTGGGEQY | 0.69507617 |
| SLASPAAL | AWKTGTGGGEQY | 0.7913678 |
| IGLVFYGV | AWKTGTGGGEQY | 0.7753638 |
| FLLDASNL | AWKTGTGGGEQY | 0.83140767 |
| YQVGNFP | AWKTGTGGGEQY | 0.7565641 |
| ESLLSQST | AWKTGTGGGEQY | 0.5994048 |
| LFTVGFAAVG | AWKTGTGGGEQY | 0.8101975 |
| ATPPTPT | AWKTGTGGGEQY | 0.86704916 |
| TDLATLG | AWKTGTGGGEQY | 0.7874809 |
| SAGGTGR | AWKTGTGGGEQY | 0.6445949 |
| GFSNSGS | AWKTGTGGGEQY | 0.6409336 |
| GGNGGSGGTSV | AWKTGTGGGEQY | 0.7563184 |
| GTSVATGGAGN | AWKTGTGGGEQY | 0.7622201 |
| NGGNGGSGGTS | AWKTGTGGGEQY | 0.55762386 |
| AVVKFPH | AWKTGTGGGEQY | 0.6921326 |
| TPPPPPP | AWKTGTGGGEQY | 0.72478724 |
| AAPAAAAAPA | AWKTGTGGGEQY | 0.43079713 |
| IALNEDL | AWKTGTGGGEQY | 0.7803737 |
| IVNVSSM | AWKTGTGGGEQY | 0.7952746 |
| TGIPPAPR | AWKTGTGGGEQY | 0.7123419 |
| FDLGGGTF | AWKTGTGGGEQY | 0.5578316 |
| FDLGGGTFDVS | AWKTGTGGGEQY | 0.79201955 |
| NEPTAAA | AWKTGTGGGEQY | 0.83096313 |
| NEPTAAALA | AWKTGTGGGEQY | 0.81500274 |
| GSIINISSI | AWKTGTGGGEQY | 0.42047775 |
| VAPGNAGTA | AWKTGTGGGEQY | 0.23623115 |
| TGGGASLE | AWKTGTGGGEQY | 0.6028912 |
| VAVYNHYKRI | AWKTGTGGGEQY | 0.7717772 |
| TDPAPATT | AWKTGTGGGEQY | 0.6865027 |
| TNRPDILDPALLRPG | AWKTGTGGGEQY | 0.58705664 |
| LTTTTTS | AWKTGTGGGEQY | 0.6822723 |
| PTGSGKTL | AWKTGTGGGEQY | 0.7956457 |
| TGSGKTL | AWKTGTGGGEQY | 0.5914375 |
| TIGHVDHGK | AWKTGTGGGEQY | 0.49647683 |

| | | |
|---|---|---|
| TIGHVDHGKTT | AWKTGTGGGEQY | 0.7744938 |
| GHVDHGKTT | AWKTGTGGGEQY | 0.78871787 |
| AQFEHTLL | AWKTGTGGGEQY | 0.5065681 |
| IVNQDEL | AWKTGTGGGEQY | 0.6197612 |
| TNGSQFF | AWKTGTGGGEQY | 0.6570815 |
| TGKGIVE | AWKTGTGGGEQY | 0.5147219 |
| LISNASDA | AWKTGTGGGEQY | 0.6922091 |
| AELRTLGELTKIAKAHGAQ | AWKTGTGGGEQY | 0.47279394 |
| APNAVAAAEPSW | AWKTGTGGGEQY | 0.7966634 |
| APQVAGAATAVL | AWKTGTGGGEQY | 0.42924017 |
| ARRDPAAAAAILPTDA | AWKTGTGGGEQY | 0.7885734 |
| DAAARDVLAEAGLAEYF | AWKTGTGGGEQY | 0.7205149 |
| DDQIVLSAKANLAKVEQ | AWKTGTGGGEQY | 0.75283754 |
| DELLPDTGEDSDEDGH | AWKTGTGGGEQY | 0.6107459 |
| ELPEAPANRATVPQD | AWKTGTGGGEQY | 0.5488678 |
| FGQSKIERVTGIG | AWKTGTGGGEQY | 0.44983706 |
| GAGGTGSTAGGAGGAGGAGGLY | AWKTGTGGGEQY | 0.29909965 |
| GHGGTGAAGGNGGTGGAGGSGI | AWKTGTGGGEQY | 0.79843163 |
| GNGGNGGAGGEGGPGGAGGAGA | AWKTGTGGGEQY | 0.22691551 |
| GSTAGNGGNGGNATNGTIAGQP | AWKTGTGGGEQY | 0.77590275 |
| GTGGDGGNGGNGGTGG | AWKTGTGGGEQY | 0.76059294 |
| GVRRPAGIPADAPV | AWKTGTGGGEQY | 0.5471319 |
| GYRQVTDAHLLTLARR | AWKTGTGGGEQY | 0.74105906 |
| IAKMGQDGHDRG | AWKTGTGGGEQY | 0.45231587 |
| KAEPSIEIQYT | AWKTGTGGGEQY | 0.5889862 |
| KPVRIRMAADEDTVAVT | AWKTGTGGGEQY | 0.7852346 |
| MPPRIAGMRLLVIKPEP | AWKTGTGGGEQY | 0.7847693 |
| MKFYAVALFRSRGVPPTR | AWKTGTGGGEQY | 0.5646776 |
| NAGGFGILGAVAHSP | AWKTGTGGGEQY | 0.74175715 |
| NEGREITLAMAEM | AWKTGTGGGEQY | 0.72637296 |
| NELVEWSHPLGHRWRWE | AWKTGTGGGEQY | 0.62848806 |
| NLGAGGTGGNGGNPG | AWKTGTGGGEQY | 0.7271971 |
| NTEWPKWETVMAADPVD | AWKTGTGGGEQY | 0.79005414 |
| PASAAPAETQP | AWKTGTGGGEQY | 0.7645078 |
| QQSSDGKNGVRAAEITGE | AWKTGTGGGEQY | 0.46277732 |
| RAPDAVTTSWWKE | AWKTGTGGGEQY | 0.7703853 |
| SGFGNVGAHTSGWFNQ | AWKTGTGGGEQY | 0.49140006 |
| SGVAGAVGQAASVGGLKV | AWKTGTGGGEQY | 0.71575505 |
| SNGGAGTGSAGGNGGT | AWKTGTGGGEQY | 0.6772064 |
| TGGAGGEGGAAG | AWKTGTGGGEQY | 0.71226215 |
| TGGDGGAAGAGGAPGHG | AWKTGTGGGEQY | 0.7707025 |
| TMPPLLAAIANAGRHGV | AWKTGTGGGEQY | 0.63202244 |
| TSSTAGNSGGA | AWKTGTGGGEQY | 0.56040615 |
| VPATVPVGILC | AWKTGTGGGEQY | 0.6594795 |
| VQRPDSGGVVLTGRIS | AWKTGTGGGEQY | 0.7260754 |
| WTPNPVPATKGVPAGLVQAT | AWKTGTGGGEQY | 0.7751959 |
| YALKRALAKFSDRFDVVII | AWKTGTGGGEQY | 0.510518 |
| YLIGSVIV | AWKTGTGGGEQY | 0.71662563 |
| YGQLVAVID | AWKTGTGGGEQY | 0.67476237 |
| VTVGGMTLPA | AWKTGTGGGEQY | 0.5463319 |
| VLALPAV | AWKTGTGGGEQY | 0.70088017 |

| | | |
|---|---|---|
| VIPPTLNYET | AWKTGTGGGEQY | 0.69126976 |
| VGMGGMHPSSG | AWKTGTGGGEQY | 0.53117394 |
| VGLAGGAATA | AWKTGTGGGEQY | 0.7674469 |
| VGGAGGAGGSGP | AWKTGTGGGEQY | 0.587795 |
| VFAAFMV | AWKTGTGGGEQY | 0.79095364 |
| TLYDKLR | AWKTGTGGGEQY | 0.84275407 |
| TLAVFTVL | AWKTGTGGGEQY | 0.57458675 |
| TGSVQPARV | AWKTGTGGGEQY | 0.7816639 |
| TGGGAGSGGT | AWKTGTGGGEQY | 0.709014 |
| SHLETSL | AWKTGTGGGEQY | 0.6941479 |
| SFTTGMT | AWKTGTGGGEQY | 0.67371875 |
| QVEVAKRMGVRQPTVS | AWKTGTGGGEQY | 0.56258523 |
| QGSGGPALAM | AWKTGTGGGEQY | 0.78756845 |
| MLAPSNP | AWKTGTGGGEQY | 0.779825 |
| LVNNAAI | AWKTGTGGGEQY | 0.7933237 |
| LTGEINTLA | AWKTGTGGGEQY | 0.7446661 |
| LSIFGPV | AWKTGTGGGEQY | 0.76402456 |
| LLALLANL | AWKTGTGGGEQY | 0.7774044 |
| LISNASDALDKLR | AWKTGTGGGEQY | 0.5253306 |
| LINEAVP | AWKTGTGGGEQY | 0.3577653 |
| LGYAFSGL | AWKTGTGGGEQY | 0.6478483 |
| LDFVAVAL | AWKTGTGGGEQY | 0.5814286 |
| LAQPGAE | AWKTGTGGGEQY | 0.64225316 |
| ISNLGAVTGF | AWKTGTGGGEQY | 0.39464185 |
| HQEDPGRI | AWKTGTGGGEQY | 0.7919421 |
| GTPGEPTP | AWKTGTGGGEQY | 0.8418368 |
| GFFNSGS | AWKTGTGGGEQY | 0.6086576 |
| GAGGGGLTAGAN | AWKTGTGGGEQY | 0.8202157 |
| FVDALVNM | AWKTGTGGGEQY | 0.43017465 |
| FLTVAPEL | AWKTGTGGGEQY | 0.7861438 |
| FETLTAL | AWKTGTGGGEQY | 0.81602657 |
| DVINPIEL | AWKTGTGGGEQY | 0.48428863 |
| APVSGAPV | AWKTGTGGGEQY | 0.7762365 |
| ALVGGLAW | AWKTGTGGGEQY | 0.5192385 |
| ALAAAAATAI | AWKTGTGGGEQY | 0.6256762 |
| AFGLGLLL | AWKTGTGGGEQY | 0.80755544 |
| AFFAAASL | AWKTGTGGGEQY | 0.6891653 |
| AAAALPAFSPP | AWKTGTGGGEQY | 0.76040995 |
| VVLAPAL | AWKTGTGGGEQY | 0.67738026 |
| TIGHVDHGKTTL | AWKTGTGGGEQY | 0.5560158 |
| LISNASDALDK | AWKTGTGGGEQY | 0.7177626 |
| LISNASDAL | AWKTGTGGGEQY | 0.8098691 |
| ISNASDALDK | AWKTGTGGGEQY | 0.55350465 |
| NQAETLVYQTE | AWKTGTGGGEQY | 0.86308837 |
| TGGTGGTAGAG | AWKTGTGGGEQY | 0.57121885 |
| TGSTAGGAGGA | AWKTGTGGGEQY | 0.8267416 |
| DPLAVPVAAGPV | AWKTGTGGGEQY | 0.7621256 |
| GGTGGTPGNGGN | AWKTGTGGGEQY | 0.6616086 |
| PAGLAGVDPATL | AWKTGTGGGEQY | 0.65762836 |
| GNNLVVEAPHAGASV | AWKTGTGGGEQY | 0.78211105 |
| KVTRIMSASTLEQPA | AWKTGTGGGEQY | 0.6695157 |

| | | |
|---|---|---|
| LGFALPIALPRRTVV | AWKTGTGGGEQY | 0.77104056 |
| TRLGNLTARKLVVLV | AWKTGTGGGEQY | 0.81305015 |
| WSQQRRLIYFPSAGP | AWKTGTGGGEQY | 0.6693574 |
| GPNGNPDPMAAAVDIR | AWKTGTGGGEQY | 0.8540664 |
| TQRCVSVADLRAAASR | AWKTGTGGGEQY | 0.8158737 |
| QFVLDVAEHPVEVLTETV | AWKTGTGGGEQY | 0.80492604 |
| GGAGGTGGAGGNGGTQAGDG | AWKTGTGGGEQY | 0.6733233 |
| PLLATLRAIAVDVDAKPAQV | AWKTGTGGGEQY | 0.3183232 |
| APKFRTGGYMIDFWGVGYQVAK | AWKTGTGGGEQY | 0.7679109 |
| GAGGAGGAGGAAGGPGGTGGTG | AWKTGTGGGEQY | 0.6287682 |
| IGALGLLATDVAPAGSVAFGVA | AWKTGTGGGEQY | 0.833308 |
| AGGAGGIGGNGGAGGAGGVLMG | AWKTGTGGGEQY | 0.8309992 |
| ADELIGLL | AWKTGTGGGEQY | 0.67779106 |
| ADRALKA | AWKTGTGGGEQY | 0.8497833 |
| AGAVGQVGGAGGA | AWKTGTGGGEQY | 0.84660655 |
| AGGGGGLGGAP | AWKTGTGGGEQY | 0.4852811 |
| AGGLLYGNGGAG | AWKTGTGGGEQY | 0.78475934 |
| AGLESIGL | AWKTGTGGGEQY | 0.6561282 |
| AGLKTGII | AWKTGTGGGEQY | 0.7975832 |
| ASLAALLL | AWKTGTGGGEQY | 0.40459532 |
| ASVINISSI | AWKTGTGGGEQY | 0.838698 |
| ATPPGPA | AWKTGTGGGEQY | 0.7585577 |
| ATVASALV | AWKTGTGGGEQY | 0.1582668 |
| ATVASLIL | AWKTGTGGGEQY | 0.80126685 |
| AVNGPVLIRH | AWKTGTGGGEQY | 0.77174073 |
| DGAAGAVGI | AWKTGTGGGEQY | 0.76997375 |
| DPLLHGI | AWKTGTGGGEQY | 0.84941983 |
| ELGAALGAL | AWKTGTGGGEQY | 0.4495428 |
| ELGRTQAV | AWKTGTGGGEQY | 0.6475359 |
| FVAGGYLS | AWKTGTGGGEQY | 0.83941823 |
| GASGGAGGAGGW | AWKTGTGGGEQY | 0.8467458 |
| GGAGAGGDGGAGGI | AWKTGTGGGEQY | 0.54938626 |
| GTGHNGSQPGMGGNG | AWKTGTGGGEQY | 0.83177745 |
| GYAAASASAS | AWKTGTGGGEQY | 0.8481747 |
| HQVDVAAVQQG | AWKTGTGGGEQY | 0.8087084 |
| IGAAIAHT | AWKTGTGGGEQY | 0.8447269 |
| IGMIEVAQL | AWKTGTGGGEQY | 0.8474293 |
| IRLAPEHTSAMD | AWKTGTGGGEQY | 0.6950789 |
| IYNVSVL | AWKTGTGGGEQY | 0.85008943 |
| KQGFLGM | AWKTGTGGGEQY | 0.8568449 |
| LESVITQL | AWKTGTGGGEQY | 0.70991004 |
| LGLGALVVV | AWKTGTGGGEQY | 0.804077 |
| LIGNGGNGGSGGI | AWKTGTGGGEQY | 0.6185244 |
| LNAINAPTL | AWKTGTGGGEQY | 0.78357863 |
| LQLLNTGV | AWKTGTGGGEQY | 0.2865529 |
| LQTSIEF | AWKTGTGGGEQY | 0.8339186 |
| MSFAVSEL | AWKTGTGGGEQY | 0.84598297 |
| NTGALGL | AWKTGTGGGEQY | 0.79677737 |
| PPGAGGAGGAGTT | AWKTGTGGGEQY | 0.63201296 |
| QLVGLGI | AWKTGTGGGEQY | 0.5933492 |
| RFLEQVHQL | AWKTGTGGGEQY | 0.8139746 |

| | | |
|---|---|---|
| RGVGYVLR | AWKTGTGGGEQY | 0.72502136 |
| SATGVFAGVI | AWKTGTGGGEQY | 0.40270278 |
| SLPAITLP | AWKTGTGGGEQY | 0.7293087 |
| STALAFTV | AWKTGTGGGEQY | 0.8392807 |
| SVLTVGLGK | AWKTGTGGGEQY | 0.84201014 |
| TALAGAGSGI | AWKTGTGGGEQY | 0.5131552 |
| TGLPFTGI | AWKTGTGGGEQY | 0.85683084 |
| VAEHVTV | AWKTGTGGGEQY | 0.8427787 |
| VDIVLNSL | AWKTGTGGGEQY | 0.85608107 |
| VLTGGGAVL | AWKTGTGGGEQY | 0.7902013 |
| VNVGLAYGI | AWKTGTGGGEQY | 0.6897307 |
| VSSAAVGAV | AWKTGTGGGEQY | 0.84864324 |
| VTSVYEAGV | AWKTGTGGGEQY | 0.7744236 |
| WVALDRGA | AWKTGTGGGEQY | 0.69774306 |
| YLVGSVVI | AWKTGTGGGEQY | 0.84177184 |
| GPPGTGKTLL | AWKTGTGGGEQY | 0.7725599 |
| GVGMTKF | AWKTGTGGGEQY | 0.7092191 |
| IGIGGSDL | AWKTGTGGGEQY | 0.8000729 |
| IGTLLAEL | AWKTGTGGGEQY | 0.85227096 |
| IYTSGTT | AWKTGTGGGEQY | 0.6611795 |
| LSFPAGV | AWKTGTGGGEQY | 0.4658571 |
| LYTSGTT | AWKTGTGGGEQY | 0.732808 |
| VGAFPFGF | AWKTGTGGGEQY | 0.7818477 |
| VGAGTVEF | AWKTGTGGGEQY | 0.8456013 |
| VGKSTLLL | AWKTGTGGGEQY | 0.81636167 |
| AALAHAGGGIV | AWKTGTGGGEQY | 0.7779853 |
| AALVAVSLL | AWKTGTGGGEQY | 0.825856 |
| AAVSRAVAAL | AWKTGTGGGEQY | 0.82699305 |
| AGGSSGIGGF | AWKTGTGGGEQY | 0.81143636 |
| AGLTVANANG | AWKTGTGGGEQY | 0.78680634 |
| AGLVGAVL | AWKTGTGGGEQY | 0.7692345 |
| ALSQAFTV | AWKTGTGGGEQY | 0.8519461 |
| ALTFSVSL | AWKTGTGGGEQY | 0.8390634 |
| ALTGAALVAL | AWKTGTGGGEQY | 0.8401209 |
| ASFSIGAGVL | AWKTGTGGGEQY | 0.8399216 |
| DGSVYKI | AWKTGTGGGEQY | 0.7972729 |
| EQLAEIV | AWKTGTGGGEQY | 0.8198425 |
| FAVSELTVM | AWKTGTGGGEQY | 0.3234684 |
| GAAPLAAGL | AWKTGTGGGEQY | 0.65904534 |
| GADGGDGGSGGAGG | AWKTGTGGGEQY | 0.8107712 |
| GFGLDTGGAGG | AWKTGTGGGEQY | 0.8353565 |
| GLASGGLL | AWKTGTGGGEQY | 0.8353113 |
| GRIAGAAVLL | AWKTGTGGGEQY | 0.8164226 |
| GSAGSLNTSF | AWKTGTGGGEQY | 0.7117187 |
| GTGQIGI | AWKTGTGGGEQY | 0.82355595 |
| GTLIALIL | AWKTGTGGGEQY | 0.84795463 |
| IAATMGFVVG | AWKTGTGGGEQY | 0.49632174 |
| IFAQALL | AWKTGTGGGEQY | 0.71321315 |
| IGAAVALGF | AWKTGTGGGEQY | 0.8601385 |
| IGLTPLAL | AWKTGTGGGEQY | 0.83469033 |
| ILGSNTSTL | AWKTGTGGGEQY | 0.7553913 |

| | | |
|---|---|---|
| IVFSLTVL | AWKTGTGGGEQY | 0.74967766 |
| LASKATGF | AWKTGTGGGEQY | 0.8527731 |
| LASVALIL | AWKTGTGGGEQY | 0.75283825 |
| LFAVGLTL | AWKTGTGGGEQY | 0.8555847 |
| LFNGGIGTYI | AWKTGTGGGEQY | 0.2393817 |
| LTLGAGII | AWKTGTGGGEQY | 0.7634686 |
| LVAAALGDI | AWKTGTGGGEQY | 0.6829125 |
| NTAFVQLGI | AWKTGTGGGEQY | 0.258958 |
| QLFSGAEL | AWKTGTGGGEQY | 0.8444162 |
| SAATFITL | AWKTGTGGGEQY | 0.73646873 |
| SNYGLANLGV | AWKTGTGGGEQY | 0.8414871 |
| SYGGAYAVM | AWKTGTGGGEQY | 0.80496645 |
| TTGVGGLTLG | AWKTGTGGGEQY | 0.8412405 |
| TTQTIYI | AWKTGTGGGEQY | 0.831354 |
| VAAGAGLGAVI | AWKTGTGGGEQY | 0.7896088 |
| VAAVVTLL | AWKTGTGGGEQY | 0.6247157 |
| VGFLGGLI | AWKTGTGGGEQY | 0.83347315 |
| VGVIPILL | AWKTGTGGGEQY | 0.83831716 |
| VIFASSLI | AWKTGTGGGEQY | 0.84379494 |
| VKGGDGGAAAG | AWKTGTGGGEQY | 0.36604744 |
| VTKTFTNI | AWKTGTGGGEQY | 0.8140337 |
| VVAAAILL | AWKTGTGGGEQY | 0.7893722 |
| YAALDVEL | AWKTGTGGGEQY | 0.24062783 |
| LASVALLL | AWKTGTGGGEQY | 0.2652383 |
| LDAVHRV | AWKTGTGGGEQY | 0.7628667 |
| QGLSSSIF | AWKTGTGGGEQY | 0.76702344 |
| RGTFANIRL | AWKTGTGGGEQY | 0.84702367 |
| RQLWWGHRI | AWKTGTGGGEQY | 0.84076667 |
| YGPPGTGKTLL | AWKTGTGGGEQY | 0.7924329 |
| AALVGAPANVA | AWKTGTGGGEQY | 0.83906025 |
| AATFRRTE | AWKTGTGGGEQY | 0.79900044 |
| AGALGGGI | AWKTGTGGGEQY | 0.13488302 |
| AGGAGGAGGIGAG | AWKTGTGGGEQY | 0.7070148 |
| AGSGALGL | AWKTGTGGGEQY | 0.5394126 |
| AQIADQI | AWKTGTGGGEQY | 0.84945714 |
| ASAVAVTYL | AWKTGTGGGEQY | 0.5817015 |
| ASTVLGGTGG | AWKTGTGGGEQY | 0.71439046 |
| AVGGEPAK | AWKTGTGGGEQY | 0.81418043 |
| CEQGTFGGTC | AWKTGTGGGEQY | 0.8285953 |
| DEGKAPAG | AWKTGTGGGEQY | 0.81400406 |
| DTVVFPAL | AWKTGTGGGEQY | 0.6595273 |
| FEGITMAG | AWKTGTGGGEQY | 0.18362467 |
| FPLVVQD | AWKTGTGGGEQY | 0.83051383 |
| GAFTTPQ | AWKTGTGGGEQY | 0.39092007 |
| GFGGFGTAGGVGGA | AWKTGTGGGEQY | 0.5549418 |
| GFLAQFL | AWKTGTGGGEQY | 0.32748014 |
| GGAGGAGGNAGVA | AWKTGTGGGEQY | 0.23486954 |
| GGVGTAGAGGAGGAGG | AWKTGTGGGEQY | 0.38357705 |
| GNGGGGDAGNAGSGGN | AWKTGTGGGEQY | 0.8410147 |
| GTGGAGGAGNDAGS | AWKTGTGGGEQY | 0.78648233 |
| GTTNTGWFNS | AWKTGTGGGEQY | 0.7715411 |

| | | |
|---|---|---|
| IFMGPLFTL | AWKTGTGGGEQY | 0.85727656 |
| ILAHAIEAI | AWKTGTGGGEQY | 0.8212975 |
| IRSAIREAL | AWKTGTGGGEQY | 0.796419 |
| ITFAGGG | AWKTGTGGGEQY | 0.6699546 |
| ITGMSGHQEV | AWKTGTGGGEQY | 0.63671243 |
| IVAGLACTL | AWKTGTGGGEQY | 0.6919825 |
| IVGGGLAGA | AWKTGTGGGEQY | 0.67116976 |
| IVTGVLNI | AWKTGTGGGEQY | 0.85604584 |
| LAQQGCI | AWKTGTGGGEQY | 0.79427934 |
| LCLHVEN | AWKTGTGGGEQY | 0.5016314 |
| LDAISDGS | AWKTGTGGGEQY | 0.8325106 |
| LGGFAAML | AWKTGTGGGEQY | 0.85604084 |
| LGGKNPMI | AWKTGTGGGEQY | 0.8596157 |
| LGPKGVAAIL | AWKTGTGGGEQY | 0.6610542 |
| LMVGVSI | AWKTGTGGGEQY | 0.754045 |
| LVGAAFVL | AWKTGTGGGEQY | 0.7526441 |
| LVLAVAPL | AWKTGTGGGEQY | 0.8131733 |
| MAAIAVIDAA | AWKTGTGGGEQY | 0.6461488 |
| NNGINAELQ | AWKTGTGGGEQY | 0.5347348 |
| RGEPLVLVHG | AWKTGTGGGEQY | 0.84909433 |
| RGGAGGAAGLI | AWKTGTGGGEQY | 0.8593627 |
| RSYRMHPAV | AWKTGTGGGEQY | 0.73591506 |
| SAAHLLAGQLA | AWKTGTGGGEQY | 0.5587845 |
| SAAVGSY | AWKTGTGGGEQY | 0.78685695 |
| SPHAVLHRI | AWKTGTGGGEQY | 0.70068234 |
| SQIAKVL | AWKTGTGGGEQY | 0.7696619 |
| TALGHLVP | AWKTGTGGGEQY | 0.7801361 |
| TAPTTSGAA | AWKTGTGGGEQY | 0.4751482 |
| TDGDDGNGGSPGT | AWKTGTGGGEQY | 0.7851483 |
| TDGLGEN | AWKTGTGGGEQY | 0.8582871 |
| TGGPPLPGQA | AWKTGTGGGEQY | 0.775403 |
| TIPAITI | AWKTGTGGGEQY | 0.83499026 |
| TIPGFTI | AWKTGTGGGEQY | 0.8558767 |
| TSIASGALA | AWKTGTGGGEQY | 0.8464989 |
| TVTLPGAAI | AWKTGTGGGEQY | 0.8553187 |
| VASGLSAALP | AWKTGTGGGEQY | 0.8387109 |
| VGAGVIGI | AWKTGTGGGEQY | 0.7348081 |
| VGPKRFGV | AWKTGTGGGEQY | 0.81082106 |
| VNVAFPD | AWKTGTGGGEQY | 0.8277868 |
| VSMTNSGVS | AWKTGTGGGEQY | 0.7984561 |
| VTDVLGDLSV | AWKTGTGGGEQY | 0.7942701 |
| VVVVGAGFAGL | AWKTGTGGGEQY | 0.77663517 |
| VYTVGTVV | AWKTGTGGGEQY | 0.84597814 |
| WTTFSGP | AWKTGTGGGEQY | 0.782792 |
| YDELVEAF | AWKTGTGGGEQY | 0.8498173 |
| YGTGGAGGNGG | AWKTGTGGGEQY | 0.32532454 |
| YTNVMAV | AWKTGTGGGEQY | 0.603296 |
| FLEQVHQL | AWKTGTGGGEQY | 0.85238904 |
| GLSLLAVAV | AWKTGTGGGEQY | 0.15020734 |
| LEQVHQL | AWKTGTGGGEQY | 0.2643803 |
| LNAAGAI | AWKTGTGGGEQY | 0.82648075 |

| | | |
|---|---|---|
| PAVHASL | AWKTGTGGGEQY | 0.8518882 |
| TELEDDA | AWKTGTGGGEQY | 0.1627685 |
| TPPGPPN | AWKTGTGGGEQY | 0.5750125 |
| TPPPPPPP | AWKTGTGGGEQY | 0.47918022 |
| TQSPALI | ASSDAFSNERLF | 0.8370391 |
| EFAAAFPQ | ASSDAFSNERLF | 0.7846225 |
| LLSGNLDVL | ASSDAFSNERLF | 0.2476397 |
| GPANLGNYNI | ASSDAFSNERLF | 0.84063303 |
| TLGPTLP | ASSDAFSNERLF | 0.5115603 |
| VYAYSSCCK | ASSDAFSNERLF | 0.59641707 |
| APAPAPAPNI | ASSDAFSNERLF | 0.8408454 |
| PGGPVPIP | ASSDAFSNERLF | 0.8538978 |
| FLTVALEP | ASSDAFSNERLF | 0.44016895 |
| SQYFGANL | ASSDAFSNERLF | 0.84284735 |
| GGVKPAA | ASSDAFSNERLF | 0.7814182 |
| GGTGGAGGAGGW | ASSDAFSNERLF | 0.8594202 |
| GVGGVAGLGGAGG | ASSDAFSNERLF | 0.7176833 |
| VLLVLATTI | ASSDAFSNERLF | 0.35831666 |
| IGIFAAML | ASSDAFSNERLF | 0.8484665 |
| AVGDLID | ASSDAFSNERLF | 0.8403832 |
| NISPLPL | ASSDAFSNERLF | 0.7397232 |
| VTAVFVAI | ASSDAFSNERLF | 0.84558666 |
| SGDDNDAAL | ASSDAFSNERLF | 0.7577783 |
| AVAALAHS | ASSDAFSNERLF | 0.8047637 |
| GGVTVMATP | ASSDAFSNERLF | 0.8508838 |
| AGLAANVRD | ASSDAFSNERLF | 0.8344015 |
| TNRPDLIDPALLRPG | ASSDAFSNERLF | 0.5012304 |
| LANLGVTAL | ASSDAFSNERLF | 0.8447998 |
| TGERIEL | ASSDAFSNERLF | 0.8018131 |
| NGEELAA | ASSDAFSNERLF | 0.6404539 |
| VLSLVWG | ASSDAFSNERLF | 0.8563929 |
| IGYALWL | ASSDAFSNERLF | 0.77691793 |
| LSLLLAPL | ASSDAFSNERLF | 0.8481333 |
| TLDEALK | ASSDAFSNERLF | 0.83958685 |
| TDGAGTAEGG | ASSDAFSNERLF | 0.8459376 |
| NAPGGAGGNGG | ASSDAFSNERLF | 0.31405288 |
| VSVFMGMGGNF | ASSDAFSNERLF | 0.7694664 |
| LASYLAAL | ASSDAFSNERLF | 0.78369 |
| LVGGTPAL | ASSDAFSNERLF | 0.8351599 |
| VRNADVL | ASSDAFSNERLF | 0.77074116 |
| TCSSPFLVSAM | ASSDAFSNERLF | 0.6327677 |
| VHITDAE | ASSDAFSNERLF | 0.82911706 |
| EYAAPSF | ASSDAFSNERLF | 0.85322136 |
| VVDAFHGGL | ASSDAFSNERLF | 0.78145397 |
| ASVVDPQ | ASSDAFSNERLF | 0.7677966 |
| SYNEFGGLS | ASSDAFSNERLF | 0.85940826 |
| DPDGSVY | ASSDAFSNERLF | 0.6513859 |
| MVSTIHATE | ASSDAFSNERLF | 0.7804046 |
| LAAGVADPV | ASSDAFSNERLF | 0.8382541 |
| TKEQIAA | ASSDAFSNERLF | 0.8415868 |
| SLASPAAL | ASSDAFSNERLF | 0.7925746 |

| | | |
|---|---|---|
| IGLVFYGV | ASSDAFSNERLF | 0.8510033 |
| FLLDASNL | ASSDAFSNERLF | 0.61481774 |
| YQVGNFP | ASSDAFSNERLF | 0.8425659 |
| ESLLSQST | ASSDAFSNERLF | 0.7797044 |
| LFTVGFAAVG | ASSDAFSNERLF | 0.82703996 |
| ATPPTPT | ASSDAFSNERLF | 0.63968 |
| TDLATLG | ASSDAFSNERLF | 0.83657074 |
| SAGGTGR | ASSDAFSNERLF | 0.8480275 |
| GFSNSGS | ASSDAFSNERLF | 0.84271526 |
| GGNGGSGGTSV | ASSDAFSNERLF | 0.23192774 |
| GTSVATGGAGN | ASSDAFSNERLF | 0.8424876 |
| NGGNGGSGGTS | ASSDAFSNERLF | 0.84155315 |
| AVVKFPH | ASSDAFSNERLF | 0.86215854 |
| TPPPPPP | ASSDAFSNERLF | 0.7854037 |
| AAPAAAAAPA | ASSDAFSNERLF | 0.82711554 |
| IALNEDL | ASSDAFSNERLF | 0.7535727 |
| IVNVSSM | ASSDAFSNERLF | 0.7973276 |
| TGIPPAPR | ASSDAFSNERLF | 0.6404235 |
| FDLGGGTF | ASSDAFSNERLF | 0.74804527 |
| FDLGGGTFDVS | ASSDAFSNERLF | 0.73255754 |
| NEPTAAA | ASSDAFSNERLF | 0.74542016 |
| NEPTAAALA | ASSDAFSNERLF | 0.84471726 |
| GSIINISSI | ASSDAFSNERLF | 0.8417347 |
| VAPGNAGTA | ASSDAFSNERLF | 0.84221023 |
| TGGGASLE | ASSDAFSNERLF | 0.85101444 |
| VAVYNHYKRI | ASSDAFSNERLF | 0.8543116 |
| TDPAPATT | ASSDAFSNERLF | 0.839527 |
| TNRPDILDPALLRPG | ASSDAFSNERLF | 0.732576 |
| LTTTTTS | ASSDAFSNERLF | 0.7617513 |
| PTGSGKTL | ASSDAFSNERLF | 0.6131166 |
| TGSGKTL | ASSDAFSNERLF | 0.757033 |
| TIGHVDHGK | ASSDAFSNERLF | 0.75343907 |
| TIGHVDHGKTT | ASSDAFSNERLF | 0.8177678 |
| GHVDHGKTT | ASSDAFSNERLF | 0.14240041 |
| AQFEHTLL | ASSDAFSNERLF | 0.82216215 |
| IVNQDEL | ASSDAFSNERLF | 0.8566566 |
| TNGSQFF | ASSDAFSNERLF | 0.74093515 |
| TGKGIVE | ASSDAFSNERLF | 0.82640326 |
| LISNASDA | ASSDAFSNERLF | 0.81480813 |
| AELRTLGELTKIAKAHGAQ | ASSDAFSNERLF | 0.76455927 |
| APNAVAAAEPSW | ASSDAFSNERLF | 0.83832467 |
| APQVAGAATAVL | ASSDAFSNERLF | 0.713485 |
| ARRDPAAAAAILPTDA | ASSDAFSNERLF | 0.84209734 |
| DAAARDVLAEAGLAEYF | ASSDAFSNERLF | 0.85780907 |
| DDQIVLSAKANLAKVEQ | ASSDAFSNERLF | 0.8275312 |
| DELLPDTGEDSDEDGH | ASSDAFSNERLF | 0.75257826 |
| ELPEAPANRATVPQD | ASSDAFSNERLF | 0.778365 |
| FGQSKIERVTGIG | ASSDAFSNERLF | 0.30013028 |
| GAGGTGSTAGGAGGAGGAGGLY | ASSDAFSNERLF | 0.7814179 |
| GHGGTGAAGGNGGTGGAGGSGI | ASSDAFSNERLF | 0.13465813 |
| GNGGNGGAGGEGGPGGAGGAGA | ASSDAFSNERLF | 0.82155395 |

| | | |
|---|---|---|
| GSTAGNGGNGGNATNGTIAGQP | ASSDAFSNERLF | 0.5532832 |
| GTGGDGGNGGNGGTGG | ASSDAFSNERLF | 0.8026657 |
| GVRRPAGIPADAPV | ASSDAFSNERLF | 0.80686843 |
| GYRQVTDAHLLTLARR | ASSDAFSNERLF | 0.82664806 |
| IAKMGQDGHDRG | ASSDAFSNERLF | 0.65985847 |
| KAEPSIEIQYT | ASSDAFSNERLF | 0.82018197 |
| KPVRIRMAADEDTVAVT | ASSDAFSNERLF | 0.75616497 |
| MPPRIAGMRLLVIKPEP | ASSDAFSNERLF | 0.5141645 |
| MKFYAVALFRSRGVPPTR | ASSDAFSNERLF | 0.8434517 |
| NAGGFGILGAVAHSP | ASSDAFSNERLF | 0.7800369 |
| NEGREITLAMAEM | ASSDAFSNERLF | 0.8075272 |
| NELVEWSHPLGHRWRWE | ASSDAFSNERLF | 0.846638 |
| NLGAGGTGGNGGNPG | ASSDAFSNERLF | 0.8062111 |
| NTEWPKWETVMAADPVD | ASSDAFSNERLF | 0.39712924 |
| PASAAPAETQP | ASSDAFSNERLF | 0.7997305 |
| QQSSDGKNGVRAAEITGE | ASSDAFSNERLF | 0.8057649 |
| RAPDAVTTSWWKE | ASSDAFSNERLF | 0.6917127 |
| SGFGNVGAHTSGWFNQ | ASSDAFSNERLF | 0.14367062 |
| SGVAGAVGQAASVGGLKV | ASSDAFSNERLF | 0.82927024 |
| SNGGAGTGSAGGNGGT | ASSDAFSNERLF | 0.6376107 |
| TGGAGGEGGAAG | ASSDAFSNERLF | 0.7283727 |
| TGGDGGAAGAGGAPGHG | ASSDAFSNERLF | 0.75307906 |
| TMPPLLAAIANAGRHGV | ASSDAFSNERLF | 0.84649837 |
| TSSTAGNSGGA | ASSDAFSNERLF | 0.57542944 |
| VPATVPVGILC | ASSDAFSNERLF | 0.77207196 |
| VQRPDSGGVVLTGRIS | ASSDAFSNERLF | 0.83710337 |
| WTPNPVPATKGVPAGLVQAT | ASSDAFSNERLF | 0.79753494 |
| YALKRALAKFSDRFDVVII | ASSDAFSNERLF | 0.85094166 |
| YLIGSVIV | ASSDAFSNERLF | 0.79461217 |
| YGQLVAVID | ASSDAFSNERLF | 0.7765174 |
| VTVGGMTLPA | ASSDAFSNERLF | 0.8230373 |
| VLALPAV | ASSDAFSNERLF | 0.7189865 |
| VIPPTLNYET | ASSDAFSNERLF | 0.5355623 |
| VGMGGMHPSSG | ASSDAFSNERLF | 0.4580884 |
| VGLAGGAATA | ASSDAFSNERLF | 0.80276597 |
| VGGAGGAGGSGP | ASSDAFSNERLF | 0.38425815 |
| VFAAFMV | ASSDAFSNERLF | 0.7918398 |
| TLYDKLR | ASSDAFSNERLF | 0.85015136 |
| TLAVFTVL | ASSDAFSNERLF | 0.77749777 |
| TGSVQPARV | ASSDAFSNERLF | 0.46372107 |
| TGGGAGSGGT | ASSDAFSNERLF | 0.78004694 |
| SHLETSL | ASSDAFSNERLF | 0.5930042 |
| SFTTGMT | ASSDAFSNERLF | 0.8433537 |
| QVEVAKRMGVRQPTVS | ASSDAFSNERLF | 0.81854105 |
| QGSGGPALAM | ASSDAFSNERLF | 0.8547004 |
| MLAPSNP | ASSDAFSNERLF | 0.8410263 |
| LVNNAAI | ASSDAFSNERLF | 0.7657435 |
| LTGEINTLA | ASSDAFSNERLF | 0.84441483 |
| LSIFGPV | ASSDAFSNERLF | 0.7892531 |
| LLALLANL | ASSDAFSNERLF | 0.7888051 |
| LISNASDALDKLR | ASSDAFSNERLF | 0.79788953 |

| | | |
|---|---|---|
| LINEAVP | ASSDAFSNERLF | 0.83747095 |
| LGYAFSGL | ASSDAFSNERLF | 0.6872418 |
| LDFVAVAL | ASSDAFSNERLF | 0.74946845 |
| LAQPGAE | ASSDAFSNERLF | 0.4678303 |
| ISNLGAVTGF | ASSDAFSNERLF | 0.8393942 |
| HQEDPGRI | ASSDAFSNERLF | 0.785174 |
| GTPGEPTP | ASSDAFSNERLF | 0.54511315 |
| GFFNSGS | ASSDAFSNERLF | 0.73860586 |
| GAGGGGLTAGAN | ASSDAFSNERLF | 0.85355365 |
| FVDALVNM | ASSDAFSNERLF | 0.8509414 |
| FLTVAPEL | ASSDAFSNERLF | 0.74222404 |
| FETLTAL | ASSDAFSNERLF | 0.69549227 |
| DVINPIEL | ASSDAFSNERLF | 0.82829744 |
| APVSGAPV | ASSDAFSNERLF | 0.41759545 |
| ALVGGLAW | ASSDAFSNERLF | 0.82948315 |
| ALAAAAATAI | ASSDAFSNERLF | 0.81327826 |
| AFGLGLLL | ASSDAFSNERLF | 0.80911946 |
| AFFAAASL | ASSDAFSNERLF | 0.6636034 |
| AAAALPAFSPP | ASSDAFSNERLF | 0.83435774 |
| VVLAPAL | ASSDAFSNERLF | 0.78911185 |
| TIGHVDHGKTTL | ASSDAFSNERLF | 0.82740724 |
| LISNASDALDK | ASSDAFSNERLF | 0.17508665 |
| LISNASDAL | ASSDAFSNERLF | 0.7702452 |
| ISNASDALDK | ASSDAFSNERLF | 0.6581515 |
| NQAETLVYQTE | ASSDAFSNERLF | 0.67030716 |
| TGGTGGTAGAG | ASSDAFSNERLF | 0.29922318 |
| TGSTAGGAGGA | ASSDAFSNERLF | 0.7708584 |
| DPLAVPVAAGPV | ASSDAFSNERLF | 0.847167 |
| GGTGGTPGNGGN | ASSDAFSNERLF | 0.85425067 |
| PAGLAGVDPATL | ASSDAFSNERLF | 0.720761 |
| GNNLVVEAPHAGASV | ASSDAFSNERLF | 0.8435254 |
| KVTRIMSASTLEQPA | ASSDAFSNERLF | 0.8569567 |
| LGFALPIALPRRTVV | ASSDAFSNERLF | 0.7790618 |
| TRLGNLTARKLVVLV | ASSDAFSNERLF | 0.24005362 |
| WSQQRRLIYFPSAGP | ASSDAFSNERLF | 0.8215872 |
| GPNGNPDPMAAAVDIR | ASSDAFSNERLF | 0.7968246 |
| TQRCVSVADLRAAASR | ASSDAFSNERLF | 0.84506243 |
| QFVLDVAEHPVEVLTETV | ASSDAFSNERLF | 0.83818734 |
| GGAGGTGGAGGNGGTQAGDG | ASSDAFSNERLF | 0.61557925 |
| PLLATLRAIAVDVDAKPAQV | ASSDAFSNERLF | 0.8246044 |
| APKFRTGGYMIDFWGVGYQVAK | ASSDAFSNERLF | 0.8470877 |
| GAGGAGGAGGAAGGPGGTGGTG | ASSDAFSNERLF | 0.6517171 |
| IGALGLLATDVAPAGSVAFGVA | ASSDAFSNERLF | 0.66131264 |
| AGGAGGIGGNGGAGGAGGVLMG | ASSDAFSNERLF | 0.626335 |
| ADELIGLL | ASSDAFSNERLF | 0.34122142 |
| ADRALKA | ASSDAFSNERLF | 0.8158112 |
| AGAVGQVGGAGGA | ASSDAFSNERLF | 0.77056265 |
| AGGGGGLGGAP | ASSDAFSNERLF | 0.68990374 |
| AGGLLYGNGGAG | ASSDAFSNERLF | 0.82346797 |
| AGLESIGL | ASSDAFSNERLF | 0.6685593 |
| AGLKTGII | ASSDAFSNERLF | 0.6828295 |

| | | |
|---|---|---|
| ASLAALLL | ASSDAFSNERLF | 0.74179447 |
| ASVINISSI | ASSDAFSNERLF | 0.7633481 |
| ATPPGPA | ASSDAFSNERLF | 0.84120595 |
| ATVASALV | ASSDAFSNERLF | 0.79066616 |
| ATVASLIL | ASSDAFSNERLF | 0.8202933 |
| AVNGPVLIRH | ASSDAFSNERLF | 0.8559345 |
| DGAAGAVGI | ASSDAFSNERLF | 0.4726624 |
| DPLLHGI | ASSDAFSNERLF | 0.6267511 |
| ELGAALGAL | ASSDAFSNERLF | 0.8346894 |
| ELGRTQAV | ASSDAFSNERLF | 0.6506996 |
| FVAGGYLS | ASSDAFSNERLF | 0.84391904 |
| GASGGAGGAGGW | ASSDAFSNERLF | 0.80132437 |
| GGAGAGGDGGAGGI | ASSDAFSNERLF | 0.83011854 |
| GTGHNGSQPGMGGNG | ASSDAFSNERLF | 0.79295 |
| GYAAASASAS | ASSDAFSNERLF | 0.8607924 |
| HQVDVAAVQQG | ASSDAFSNERLF | 0.84415424 |
| IGAAIAHT | ASSDAFSNERLF | 0.7627052 |
| IGMIEVAQL | ASSDAFSNERLF | 0.8249147 |
| IRLAPEHTSAMD | ASSDAFSNERLF | 0.4273584 |
| IYNVSVL | ASSDAFSNERLF | 0.84429216 |
| KQGFLGM | ASSDAFSNERLF | 0.8381541 |
| LESVITQL | ASSDAFSNERLF | 0.6793345 |
| LGLGALVVV | ASSDAFSNERLF | 0.8043725 |
| LIGNGGNGGSGGI | ASSDAFSNERLF | 0.8511459 |
| LNAINAPTL | ASSDAFSNERLF | 0.85552955 |
| LQLLNTGV | ASSDAFSNERLF | 0.40486622 |
| LQTSIEF | ASSDAFSNERLF | 0.7775011 |
| MSFAVSEL | ASSDAFSNERLF | 0.3508531 |
| NTGALGL | ASSDAFSNERLF | 0.72378814 |
| PPGAGGAGGAGTT | ASSDAFSNERLF | 0.4067157 |
| QLVGLGI | ASSDAFSNERLF | 0.8376368 |
| RFLEQVHQL | ASSDAFSNERLF | 0.74872965 |
| RGVGYVLR | ASSDAFSNERLF | 0.83640134 |
| SATGVFAGVI | ASSDAFSNERLF | 0.7958221 |
| SLPAITLP | ASSDAFSNERLF | 0.8499488 |
| STALAFTV | ASSDAFSNERLF | 0.73407364 |
| SVLTVGLGK | ASSDAFSNERLF | 0.7501031 |
| TALAGAGSGI | ASSDAFSNERLF | 0.43365952 |
| TGLPFTGI | ASSDAFSNERLF | 0.3959474 |
| VAEHVTV | ASSDAFSNERLF | 0.6844286 |
| VDIVLNSL | ASSDAFSNERLF | 0.7615928 |
| VLTGGGAVL | ASSDAFSNERLF | 0.21330963 |
| VNVGLAYGI | ASSDAFSNERLF | 0.8558573 |
| VSSAAVGAV | ASSDAFSNERLF | 0.81972504 |
| VTSVYEAGV | ASSDAFSNERLF | 0.8394565 |
| WVALDRGA | ASSDAFSNERLF | 0.6575708 |
| YLVGSVVI | ASSDAFSNERLF | 0.65941465 |
| GPPGTGKTLL | ASSDAFSNERLF | 0.759724 |
| GVGMTKF | ASSDAFSNERLF | 0.82346046 |
| IGIGGSDL | ASSDAFSNERLF | 0.5428533 |
| IGTLLAEL | ASSDAFSNERLF | 0.48014572 |

| | | |
|---|---|---|
| IYTSGTT | ASSDAFSNERLF | 0.6947401 |
| LSFPAGV | ASSDAFSNERLF | 0.58716625 |
| LYTSGTT | ASSDAFSNERLF | 0.85655355 |
| VGAFPFGF | ASSDAFSNERLF | 0.8506011 |
| VGAGTVEF | ASSDAFSNERLF | 0.8339474 |
| VGKSTLLL | ASSDAFSNERLF | 0.855687 |
| AALAHAGGGIV | ASSDAFSNERLF | 0.2090932 |
| AALVAVSLL | ASSDAFSNERLF | 0.69046175 |
| AAVSRAVAAL | ASSDAFSNERLF | 0.8528736 |
| AGGSSGIGGF | ASSDAFSNERLF | 0.8225061 |
| AGLTVANANG | ASSDAFSNERLF | 0.7674984 |
| AGLVGAVL | ASSDAFSNERLF | 0.85083646 |
| ALSQAFTV | ASSDAFSNERLF | 0.7020537 |
| ALTFSVSL | ASSDAFSNERLF | 0.66012466 |
| ALTGAALVAL | ASSDAFSNERLF | 0.80010957 |
| ASFSIGAGVL | ASSDAFSNERLF | 0.8330635 |
| DGSVYKI | ASSDAFSNERLF | 0.82339007 |
| EQLAEIV | ASSDAFSNERLF | 0.5190941 |
| FAVSELTVM | ASSDAFSNERLF | 0.8133185 |
| GAAPLAAGL | ASSDAFSNERLF | 0.86160374 |
| GADGGDGGSGGAGG | ASSDAFSNERLF | 0.77906764 |
| GFGLDTGGAGG | ASSDAFSNERLF | 0.8425691 |
| GLASGGLL | ASSDAFSNERLF | 0.8279094 |
| GRIAGAAVLL | ASSDAFSNERLF | 0.81843626 |
| GSAGSLNTSF | ASSDAFSNERLF | 0.7231674 |
| GTGQIGI | ASSDAFSNERLF | 0.74263096 |
| GTLIALIL | ASSDAFSNERLF | 0.82126737 |
| IAATMGFVVG | ASSDAFSNERLF | 0.5831972 |
| IFAQALL | ASSDAFSNERLF | 0.7774607 |
| IGAAVALGF | ASSDAFSNERLF | 0.74842954 |
| IGLTPLAL | ASSDAFSNERLF | 0.8624048 |
| ILGSNTSTL | ASSDAFSNERLF | 0.836581 |
| IVFSLTVL | ASSDAFSNERLF | 0.83636624 |
| LASKATGF | ASSDAFSNERLF | 0.691862 |
| LASVALIL | ASSDAFSNERLF | 0.8226835 |
| LFAVGLTL | ASSDAFSNERLF | 0.83527637 |
| LFNGGIGTYI | ASSDAFSNERLF | 0.4871644 |
| LTLGAGII | ASSDAFSNERLF | 0.8496623 |
| LVAAALGDI | ASSDAFSNERLF | 0.65307736 |
| NTAFVQLGI | ASSDAFSNERLF | 0.81079185 |
| QLFSGAEL | ASSDAFSNERLF | 0.8323949 |
| SAATFITL | ASSDAFSNERLF | 0.849136 |
| SNYGLANLGV | ASSDAFSNERLF | 0.7623072 |
| SYGGAYAVM | ASSDAFSNERLF | 0.80807674 |
| TTGVGGLTLG | ASSDAFSNERLF | 0.67979246 |
| TTQTIYI | ASSDAFSNERLF | 0.84970164 |
| VAAGAGLGAVI | ASSDAFSNERLF | 0.85154176 |
| VAAVVTLL | ASSDAFSNERLF | 0.8080921 |
| VGFLGGLI | ASSDAFSNERLF | 0.8567616 |
| VGVIPILL | ASSDAFSNERLF | 0.7958659 |
| VIFASSLI | ASSDAFSNERLF | 0.32767284 |

| | | |
|---|---|---|
| VKGGDGGAAAG | ASSDAFSNERLF | 0.76591367 |
| VTKTFTNI | ASSDAFSNERLF | 0.8325895 |
| VVAAAILL | ASSDAFSNERLF | 0.8525838 |
| YAALDVEL | ASSDAFSNERLF | 0.84534365 |
| LASVALLL | ASSDAFSNERLF | 0.8052447 |
| LDAVHRV | ASSDAFSNERLF | 0.78617716 |
| QGLSSSIF | ASSDAFSNERLF | 0.8564731 |
| RGTFANIRL | ASSDAFSNERLF | 0.8616172 |
| RQLWWGHRI | ASSDAFSNERLF | 0.8402935 |
| YGPPGTGKTLL | ASSDAFSNERLF | 0.73220026 |
| AALVGAPANVA | ASSDAFSNERLF | 0.65606093 |
| AATFRRTE | ASSDAFSNERLF | 0.8256209 |
| AGALGGGI | ASSDAFSNERLF | 0.84811074 |
| AGGAGGAGGIGAG | ASSDAFSNERLF | 0.60734296 |
| AGSGALGL | ASSDAFSNERLF | 0.7961592 |
| AQIADQI | ASSDAFSNERLF | 0.8152587 |
| ASAVAVTYL | ASSDAFSNERLF | 0.3704679 |
| ASTVLGGTGG | ASSDAFSNERLF | 0.83702016 |
| AVGGEPAK | ASSDAFSNERLF | 0.8527091 |
| CEQGTFGGTC | ASSDAFSNERLF | 0.8127823 |
| DEGKAPAG | ASSDAFSNERLF | 0.6474403 |
| DTVVFPAL | ASSDAFSNERLF | 0.848087 |
| FEGITMAG | ASSDAFSNERLF | 0.86159873 |
| FPLVVQD | ASSDAFSNERLF | 0.8558084 |
| GAFTTPQ | ASSDAFSNERLF | 0.37395 |
| GFGGFGTAGGVGGA | ASSDAFSNERLF | 0.27352563 |
| GFLAQFL | ASSDAFSNERLF | 0.6970358 |
| GGAGGAGGNAGVA | ASSDAFSNERLF | 0.837651 |
| GGVGTAGAGGAGGAGG | ASSDAFSNERLF | 0.7470008 |
| GNGGGGDAGNAGSGGN | ASSDAFSNERLF | 0.70355797 |
| GTGGAGGAGNDAGS | ASSDAFSNERLF | 0.82668155 |
| GTTNTGWFNS | ASSDAFSNERLF | 0.85190314 |
| IFMGPLFTL | ASSDAFSNERLF | 0.6161293 |
| ILAHAIEAI | ASSDAFSNERLF | 0.43431717 |
| IRSAIREAL | ASSDAFSNERLF | 0.837172 |
| ITFAGGG | ASSDAFSNERLF | 0.8428266 |
| ITGMSGHQEV | ASSDAFSNERLF | 0.7295134 |
| IVAGLACTL | ASSDAFSNERLF | 0.8017999 |
| IVGGGLAGA | ASSDAFSNERLF | 0.71221435 |
| IVTGVLNI | ASSDAFSNERLF | 0.7015457 |
| LAQQGCI | ASSDAFSNERLF | 0.82753474 |
| LCLHVEN | ASSDAFSNERLF | 0.58330935 |
| LDAISDGS | ASSDAFSNERLF | 0.8545011 |
| LGGFAAML | ASSDAFSNERLF | 0.63228583 |
| LGGKNPMI | ASSDAFSNERLF | 0.83489805 |
| LGPKGVAAIL | ASSDAFSNERLF | 0.817072 |
| LMVGVSI | ASSDAFSNERLF | 0.82395464 |
| LVGAAFVL | ASSDAFSNERLF | 0.73501813 |
| LVLAVAPL | ASSDAFSNERLF | 0.676922 |
| MAAIAVIDAA | ASSDAFSNERLF | 0.5082263 |
| NNGINAELQ | ASSDAFSNERLF | 0.50966084 |

| | | |
|---|---|---|
| RGEPLVLVHG | ASSDAFSNERLF | 0.84371346 |
| RGGAGGAAGLI | ASSDAFSNERLF | 0.22118141 |
| RSYRMHPAV | ASSDAFSNERLF | 0.8514836 |
| SAAHLLAGQLA | ASSDAFSNERLF | 0.83227146 |
| SAAVGSY | ASSDAFSNERLF | 0.649506 |
| SPHAVLHRI | ASSDAFSNERLF | 0.8362582 |
| SQIAKVL | ASSDAFSNERLF | 0.52427626 |
| TALGHLVP | ASSDAFSNERLF | 0.7600913 |
| TAPTTSGAA | ASSDAFSNERLF | 0.84111845 |
| TDGDDGNGGSPGT | ASSDAFSNERLF | 0.832187 |
| TDGLGEN | ASSDAFSNERLF | 0.63402545 |
| TGGPPLPGQA | ASSDAFSNERLF | 0.82493865 |
| TIPAITI | ASSDAFSNERLF | 0.83430815 |
| TIPGFTI | ASSDAFSNERLF | 0.6447351 |
| TSIASGALA | ASSDAFSNERLF | 0.79467046 |
| TVTLPGAAI | ASSDAFSNERLF | 0.8490224 |
| VASGLSAALP | ASSDAFSNERLF | 0.82083523 |
| VGAGVIGI | ASSDAFSNERLF | 0.55832523 |
| VGPKRFGV | ASSDAFSNERLF | 0.83505 |
| VNVAFPD | ASSDAFSNERLF | 0.6856367 |
| VSMTNSGVS | ASSDAFSNERLF | 0.83380806 |
| VTDVLGDLSV | ASSDAFSNERLF | 0.80748296 |
| VVVVGAGFAGL | ASSDAFSNERLF | 0.75377476 |
| VYTVGTVV | ASSDAFSNERLF | 0.8356092 |
| WTTFSGP | ASSDAFSNERLF | 0.5014239 |
| YDELVEAF | ASSDAFSNERLF | 0.76030624 |
| YGTGGAGGNGG | ASSDAFSNERLF | 0.7731416 |
| YTNVMAV | ASSDAFSNERLF | 0.7832906 |
| FLEQVHQL | ASSDAFSNERLF | 0.84395635 |
| GLSLLAVAV | ASSDAFSNERLF | 0.68589747 |
| LEQVHQL | ASSDAFSNERLF | 0.8264603 |
| LNAAGAI | ASSDAFSNERLF | 0.70925796 |
| PAVHASL | ASSDAFSNERLF | 0.5323386 |
| TELEDDA | ASSDAFSNERLF | 0.82139707 |
| TPPGPPN | ASSDAFSNERLF | 0.72813666 |
| TPPPPPPP | ASSDAFSNERLF | 0.6299288 |
| TQSPALI | ASSLRVNTEVF | 0.8342869 |
| EFAAAFPQ | ASSLRVNTEVF | 0.6107048 |
| LLSGNLDVL | ASSLRVNTEVF | 0.7951244 |
| GPANLGNYNI | ASSLRVNTEVF | 0.8569617 |
| TLGPTLP | ASSLRVNTEVF | 0.57436645 |
| VYAYSSCCK | ASSLRVNTEVF | 0.83209634 |
| APAPAPAPNI | ASSLRVNTEVF | 0.8299438 |
| PGGPVPIP | ASSLRVNTEVF | 0.82275176 |
| FLTVALEP | ASSLRVNTEVF | 0.74117863 |
| SQYFGANL | ASSLRVNTEVF | 0.6426954 |
| GGVKPAA | ASSLRVNTEVF | 0.8442657 |
| GGTGGAGGAGGW | ASSLRVNTEVF | 0.8228971 |
| GVGGVAGLGGAGG | ASSLRVNTEVF | 0.8328676 |
| VLLVLATTI | ASSLRVNTEVF | 0.8335589 |
| IGIFAAML | ASSLRVNTEVF | 0.83732396 |

| | | |
|---|---|---|
| AVGDLID | ASSLRVNTEVF | 0.84214216 |
| NISPLPL | ASSLRVNTEVF | 0.7524299 |
| VTAVFVAI | ASSLRVNTEVF | 0.45185828 |
| SGDDNDAAL | ASSLRVNTEVF | 0.7827135 |
| AVAALAHS | ASSLRVNTEVF | 0.7669672 |
| GGVTVMATP | ASSLRVNTEVF | 0.69205403 |
| AGLAANVRD | ASSLRVNTEVF | 0.351913 |
| TNRPDLIDPALLRPG | ASSLRVNTEVF | 0.8464217 |
| LANLGVTAL | ASSLRVNTEVF | 0.86076653 |
| TGERIEL | ASSLRVNTEVF | 0.6645746 |
| NGEELAA | ASSLRVNTEVF | 0.85924906 |
| VLSLVWG | ASSLRVNTEVF | 0.52783984 |
| IGYALWL | ASSLRVNTEVF | 0.8137255 |
| LSLLLAPL | ASSLRVNTEVF | 0.849522 |
| TLDEALK | ASSLRVNTEVF | 0.6698768 |
| TDGAGTAEGG | ASSLRVNTEVF | 0.82070124 |
| NAPGGAGGNGG | ASSLRVNTEVF | 0.5903758 |
| VSVFMGMGGNF | ASSLRVNTEVF | 0.6675577 |
| LASYLAAL | ASSLRVNTEVF | 0.85111606 |
| LVGGTPAL | ASSLRVNTEVF | 0.68085027 |
| VRNADVL | ASSLRVNTEVF | 0.81317735 |
| TCSSPFLVSAM | ASSLRVNTEVF | 0.74916303 |
| VHITDAE | ASSLRVNTEVF | 0.6231915 |
| EYAAPSF | ASSLRVNTEVF | 0.8158486 |
| VVDAFHGGL | ASSLRVNTEVF | 0.8477481 |
| ASVVDPQ | ASSLRVNTEVF | 0.77518314 |
| SYNEFGGLS | ASSLRVNTEVF | 0.86108804 |
| DPDGSVY | ASSLRVNTEVF | 0.69142234 |
| MVSTIHATE | ASSLRVNTEVF | 0.85853374 |
| LAAGVADPV | ASSLRVNTEVF | 0.85215706 |
| TKEQIAA | ASSLRVNTEVF | 0.82632697 |
| SLASPAAL | ASSLRVNTEVF | 0.7658595 |
| IGLVFYGV | ASSLRVNTEVF | 0.84556466 |
| FLLDASNL | ASSLRVNTEVF | 0.7332113 |
| YQVGNFP | ASSLRVNTEVF | 0.78314173 |
| ESLLSQST | ASSLRVNTEVF | 0.84961337 |
| LFTVGFAAVG | ASSLRVNTEVF | 0.3235371 |
| ATPPTPT | ASSLRVNTEVF | 0.7637123 |
| TDLATLG | ASSLRVNTEVF | 0.6094384 |
| SAGGTGR | ASSLRVNTEVF | 0.61588305 |
| GFSNSGS | ASSLRVNTEVF | 0.45876914 |
| GGNGGSGGTSV | ASSLRVNTEVF | 0.28430787 |
| GTSVATGGAGN | ASSLRVNTEVF | 0.5512933 |
| NGGNGGSGGTS | ASSLRVNTEVF | 0.38281578 |
| AVVKFPH | ASSLRVNTEVF | 0.77783215 |
| TPPPPPP | ASSLRVNTEVF | 0.713292 |
| AAPAAAAAPA | ASSLRVNTEVF | 0.44961697 |
| IALNEDL | ASSLRVNTEVF | 0.7537756 |
| IVNVSSM | ASSLRVNTEVF | 0.73255134 |
| TGIPPAPR | ASSLRVNTEVF | 0.3354436 |
| FDLGGGTF | ASSLRVNTEVF | 0.53592145 |

| | | |
|---|---|---|
| FDLGGGTFDVS | ASSLRVNTEVF | 0.56540835 |
| NEPTAAA | ASSLRVNTEVF | 0.5519273 |
| NEPTAAALA | ASSLRVNTEVF | 0.27428767 |
| GSIINISSI | ASSLRVNTEVF | 0.64882374 |
| VAPGNAGTA | ASSLRVNTEVF | 0.42238808 |
| TGGGASLE | ASSLRVNTEVF | 0.19344036 |
| VAVYNHYKRI | ASSLRVNTEVF | 0.6768607 |
| TDPAPATT | ASSLRVNTEVF | 0.57370114 |
| TNRPDILDPALLRPG | ASSLRVNTEVF | 0.45016426 |
| LTTTTTS | ASSLRVNTEVF | 0.7213534 |
| PTGSGKTL | ASSLRVNTEVF | 0.45796752 |
| TGSGKTL | ASSLRVNTEVF | 0.39757138 |
| TIGHVDHGK | ASSLRVNTEVF | 0.7235361 |
| TIGHVDHGKTT | ASSLRVNTEVF | 0.7591232 |
| GHVDHGKTT | ASSLRVNTEVF | 0.30989516 |
| AQFEHTLL | ASSLRVNTEVF | 0.6371535 |
| IVNQDEL | ASSLRVNTEVF | 0.76267946 |
| TNGSQFF | ASSLRVNTEVF | 0.600979 |
| TGKGIVE | ASSLRVNTEVF | 0.7553704 |
| LISNASDA | ASSLRVNTEVF | 0.75943905 |
| AELRTLGELTKIAKAHGAQ | ASSLRVNTEVF | 0.5344859 |
| APNAVAAAEPSW | ASSLRVNTEVF | 0.7688835 |
| APQVAGAATAVL | ASSLRVNTEVF | 0.7902894 |
| ARRDPAAAAAILPTDA | ASSLRVNTEVF | 0.55997914 |
| DAAARDVLAEAGLAEYF | ASSLRVNTEVF | 0.6333916 |
| DDQIVLSAKANLAKVEQ | ASSLRVNTEVF | 0.32559475 |
| DELLPDTGEDSDEDGH | ASSLRVNTEVF | 0.6436203 |
| ELPEAPANRATVPQD | ASSLRVNTEVF | 0.26221985 |
| FGQSKIERVTGIG | ASSLRVNTEVF | 0.69717526 |
| GAGGTGSTAGGAGGAGGAGGLY | ASSLRVNTEVF | 0.74061817 |
| GHGGTGAAGGNGGTGGAGGSGI | ASSLRVNTEVF | 0.5884279 |
| GNGGNGGAGGEGGPGGAGGAGA | ASSLRVNTEVF | 0.3006158 |
| GSTAGNGGNGGNATNGTIAGQP | ASSLRVNTEVF | 0.33205456 |
| GTGGDGGNGGNGGTGG | ASSLRVNTEVF | 0.6682888 |
| GVRRPAGIPADAPV | ASSLRVNTEVF | 0.2998195 |
| GYRQVTDAHLLTLARR | ASSLRVNTEVF | 0.2683878 |
| IAKMGQDGHDRG | ASSLRVNTEVF | 0.5750969 |
| KAEPSIEIQYT | ASSLRVNTEVF | 0.73739064 |
| KPVRIRMAADEDTVAVT | ASSLRVNTEVF | 0.7408432 |
| MPPRIAGMRLLVIKPEP | ASSLRVNTEVF | 0.37275162 |
| MKFYAVALFRSRGVPPTR | ASSLRVNTEVF | 0.7917925 |
| NAGGFGILGAVAHSP | ASSLRVNTEVF | 0.72413087 |
| NEGREITLAMAEM | ASSLRVNTEVF | 0.7603153 |
| NELVEWSHPLGHRWRWE | ASSLRVNTEVF | 0.55423397 |
| NLGAGGTGGNGGNPG | ASSLRVNTEVF | 0.3200601 |
| NTEWPKWETVMAADPVD | ASSLRVNTEVF | 0.7806847 |
| PASAAPAETQP | ASSLRVNTEVF | 0.58362275 |
| QQSSDGKNGVRAAEITGE | ASSLRVNTEVF | 0.6511445 |
| RAPDAVTTSWWKE | ASSLRVNTEVF | 0.73428774 |
| SGFGNVGAHTSGWFNQ | ASSLRVNTEVF | 0.48855942 |
| SGVAGAVGQAASVGGLKV | ASSLRVNTEVF | 0.41666073 |

| | | |
|---|---|---|
| SNGGAGTGSAGGNGGT | ASSLRVNTEVF | 0.65292263 |
| TGGAGGEGGAAG | ASSLRVNTEVF | 0.74163973 |
| TGGDGGAAGAGGAPGHG | ASSLRVNTEVF | 0.37137336 |
| TMPPLLAAIANAGRHGV | ASSLRVNTEVF | 0.32500446 |
| TSSTAGNSGGA | ASSLRVNTEVF | 0.4403728 |
| VPATVPVGILC | ASSLRVNTEVF | 0.5659869 |
| VQRPDSGGVVLTGRIS | ASSLRVNTEVF | 0.76830983 |
| WTPNPVPATKGVPAGLVQAT | ASSLRVNTEVF | 0.64623415 |
| YALKRALAKFSDRFDVVII | ASSLRVNTEVF | 0.50802773 |
| YLIGSVIV | ASSLRVNTEVF | 0.7138186 |
| YGQLVAVID | ASSLRVNTEVF | 0.64178973 |
| VTVGGMTLPA | ASSLRVNTEVF | 0.6246355 |
| VLALPAV | ASSLRVNTEVF | 0.53187746 |
| VIPPTLNYET | ASSLRVNTEVF | 0.49440452 |
| VGMGGMHPSSG | ASSLRVNTEVF | 0.73795176 |
| VGLAGGAATA | ASSLRVNTEVF | 0.68784815 |
| VGGAGGAGGSGP | ASSLRVNTEVF | 0.6627914 |
| VFAAFMV | ASSLRVNTEVF | 0.73702335 |
| TLYDKLR | ASSLRVNTEVF | 0.60887384 |
| TLAVFTVL | ASSLRVNTEVF | 0.6283794 |
| TGSVQPARV | ASSLRVNTEVF | 0.29507488 |
| TGGGAGSGGT | ASSLRVNTEVF | 0.5641273 |
| SHLETSL | ASSLRVNTEVF | 0.6098954 |
| SFTTGMT | ASSLRVNTEVF | 0.6183564 |
| QVEVAKRMGVRQPTVS | ASSLRVNTEVF | 0.73441255 |
| QGSGGPALAM | ASSLRVNTEVF | 0.6597475 |
| MLAPSNP | ASSLRVNTEVF | 0.49652386 |
| LVNNAAI | ASSLRVNTEVF | 0.6649611 |
| LTGEINTLA | ASSLRVNTEVF | 0.7705182 |
| LSIFGPV | ASSLRVNTEVF | 0.3383511 |
| LLALLANL | ASSLRVNTEVF | 0.5719122 |
| LISNASDALDKLR | ASSLRVNTEVF | 0.80084103 |
| LINEAVP | ASSLRVNTEVF | 0.713405 |
| LGYAFSGL | ASSLRVNTEVF | 0.566102 |
| LDFVAVAL | ASSLRVNTEVF | 0.5200428 |
| LAQPGAE | ASSLRVNTEVF | 0.7883415 |
| ISNLGAVTGF | ASSLRVNTEVF | 0.5203707 |
| HQEDPGRI | ASSLRVNTEVF | 0.78192145 |
| GTPGEPTP | ASSLRVNTEVF | 0.2305052 |
| GFFNSGS | ASSLRVNTEVF | 0.5102343 |
| GAGGGGLTAGAN | ASSLRVNTEVF | 0.3941341 |
| FVDALVNM | ASSLRVNTEVF | 0.19450112 |
| FLTVAPEL | ASSLRVNTEVF | 0.74534726 |
| FETLTAL | ASSLRVNTEVF | 0.48245347 |
| DVINPIEL | ASSLRVNTEVF | 0.69717574 |
| APVSGAPV | ASSLRVNTEVF | 0.57853365 |
| ALVGGLAW | ASSLRVNTEVF | 0.7476269 |
| ALAAAAATAI | ASSLRVNTEVF | 0.71128654 |
| AFGLGLLL | ASSLRVNTEVF | 0.5529481 |
| AFFAAASL | ASSLRVNTEVF | 0.4099448 |
| AAAALPAFSPP | ASSLRVNTEVF | 0.6448501 |

| | | |
|---|---|---|
| VVLAPAL | ASSLRVNTEVF | 0.7258517 |
| TIGHVDHGKTTL | ASSLRVNTEVF | 0.7150413 |
| LISNASDALDK | ASSLRVNTEVF | 0.23519278 |
| LISNASDAL | ASSLRVNTEVF | 0.6497779 |
| ISNASDALDK | ASSLRVNTEVF | 0.56313807 |
| NQAETLVYQTE | ASSLRVNTEVF | 0.29845276 |
| TGGTGGTAGAG | ASSLRVNTEVF | 0.21509297 |
| TGSTAGGAGGA | ASSLRVNTEVF | 0.39331847 |
| DPLAVPVAAGPV | ASSLRVNTEVF | 0.5927995 |
| GGTGGTPGNGGN | ASSLRVNTEVF | 0.7225625 |
| PAGLAGVDPATL | ASSLRVNTEVF | 0.7483699 |
| GNNLVVEAPHAGASV | ASSLRVNTEVF | 0.49279022 |
| KVTRIMSASTLEQPA | ASSLRVNTEVF | 0.7268301 |
| LGFALPIALPRRTVV | ASSLRVNTEVF | 0.5612709 |
| TRLGNLTARKLVVLV | ASSLRVNTEVF | 0.15825266 |
| WSQQRRLIYFPSAGP | ASSLRVNTEVF | 0.5125648 |
| GPNGNPDPMAAAVDIR | ASSLRVNTEVF | 0.33438796 |
| TQRCVSVADLRAAASR | ASSLRVNTEVF | 0.74354935 |
| QFVLDVAEHPVEVLTETV | ASSLRVNTEVF | 0.38853213 |
| GGAGGTGGAGGNGGTQAGDG | ASSLRVNTEVF | 0.5668051 |
| PLLATLRAIAVDVDAKPAQV | ASSLRVNTEVF | 0.6063411 |
| APKFRTGGYMIDFWGVGYQVAK | ASSLRVNTEVF | 0.7277721 |
| GAGGAGGAGGAAGGPGGTGGTG | ASSLRVNTEVF | 0.6649499 |
| IGALGLLATDVAPAGSVAFGVA | ASSLRVNTEVF | 0.38665944 |
| AGGAGGIGGNGGAGGAGGVLMG | ASSLRVNTEVF | 0.25894123 |
| ADELIGLL | ASSLRVNTEVF | 0.6784884 |
| ADRALKA | ASSLRVNTEVF | 0.5747222 |
| AGAVGQVGGAGGA | ASSLRVNTEVF | 0.33112562 |
| AGGGGGLGGAP | ASSLRVNTEVF | 0.37613806 |
| AGGLLYGNGGAG | ASSLRVNTEVF | 0.7601471 |
| AGLESIGL | ASSLRVNTEVF | 0.2597757 |
| AGLKTGII | ASSLRVNTEVF | 0.70559675 |
| ASLAALLL | ASSLRVNTEVF | 0.44349995 |
| ASVINISSI | ASSLRVNTEVF | 0.55746144 |
| ATPPGPA | ASSLRVNTEVF | 0.7888263 |
| ATVASALV | ASSLRVNTEVF | 0.7131791 |
| ATVASLIL | ASSLRVNTEVF | 0.6272578 |
| AVNGPVLIRH | ASSLRVNTEVF | 0.41497564 |
| DGAAGAVGI | ASSLRVNTEVF | 0.4220479 |
| DPLLHGI | ASSLRVNTEVF | 0.37269327 |
| ELGAALGAL | ASSLRVNTEVF | 0.4005127 |
| ELGRTQAV | ASSLRVNTEVF | 0.76354736 |
| FVAGGYLS | ASSLRVNTEVF | 0.567296 |
| GASGGAGGAGGW | ASSLRVNTEVF | 0.3794045 |
| GGAGAGGDGGAGGI | ASSLRVNTEVF | 0.67378473 |
| GTGHNGSQPGMGGNG | ASSLRVNTEVF | 0.7763926 |
| GYAAASASAS | ASSLRVNTEVF | 0.7951604 |
| HQVDVAAVQQG | ASSLRVNTEVF | 0.39683282 |
| IGAAIAHT | ASSLRVNTEVF | 0.44698972 |
| IGMIEVAQL | ASSLRVNTEVF | 0.50326306 |
| IRLAPEHTSAMD | ASSLRVNTEVF | 0.5883808 |

| | | |
|---|---|---|
| IYNVSVL | ASSLRVNTEVF | 0.34295207 |
| KQGFLGM | ASSLRVNTEVF | 0.35376838 |
| LESVITQL | ASSLRVNTEVF | 0.7115056 |
| LGLGALVVV | ASSLRVNTEVF | 0.8014329 |
| LIGNGGNGGSGGI | ASSLRVNTEVF | 0.4922828 |
| LNAINAPTL | ASSLRVNTEVF | 0.31028432 |
| LQLLNTGV | ASSLRVNTEVF | 0.5611399 |
| LQTSIEF | ASSLRVNTEVF | 0.3958009 |
| MSFAVSEL | ASSLRVNTEVF | 0.45194072 |
| NTGALGL | ASSLRVNTEVF | 0.548084 |
| PPGAGGAGGAGTT | ASSLRVNTEVF | 0.2859861 |
| QLVGLGI | ASSLRVNTEVF | 0.6167371 |
| RFLEQVHQL | ASSLRVNTEVF | 0.7908489 |
| RGVGYVLR | ASSLRVNTEVF | 0.6535204 |
| SATGVFAGVI | ASSLRVNTEVF | 0.70352054 |
| SLPAITLP | ASSLRVNTEVF | 0.7797581 |
| STALAFTV | ASSLRVNTEVF | 0.68352056 |
| SVLTVGLGK | ASSLRVNTEVF | 0.76765406 |
| TALAGAGSGI | ASSLRVNTEVF | 0.746261 |
| TGLPFTGI | ASSLRVNTEVF | 0.56157804 |
| VAEHVTV | ASSLRVNTEVF | 0.5921556 |
| VDIVLNSL | ASSLRVNTEVF | 0.7730667 |
| VLTGGGAVL | ASSLRVNTEVF | 0.62502366 |
| VNVGLAYGI | ASSLRVNTEVF | 0.6063467 |
| VSSAAVGAV | ASSLRVNTEVF | 0.64465386 |
| VTSVYEAGV | ASSLRVNTEVF | 0.7539119 |
| WVALDRGA | ASSLRVNTEVF | 0.50393164 |
| YLVGSVVI | ASSLRVNTEVF | 0.77358997 |
| GPPGTGKTLL | ASSLRVNTEVF | 0.22160053 |
| GVGMTKF | ASSLRVNTEVF | 0.31398612 |
| IGIGGSDL | ASSLRVNTEVF | 0.74398065 |
| IGTLLAEL | ASSLRVNTEVF | 0.17531621 |
| IYTSGTT | ASSLRVNTEVF | 0.24035054 |
| LSFPAGV | ASSLRVNTEVF | 0.62997186 |
| LYTSGTT | ASSLRVNTEVF | 0.7597065 |
| VGAFPFGF | ASSLRVNTEVF | 0.18113528 |
| VGAGTVEF | ASSLRVNTEVF | 0.472382 |
| VGKSTLLL | ASSLRVNTEVF | 0.2704119 |
| AALAHAGGGIV | ASSLRVNTEVF | 0.6731111 |
| AALVAVSLL | ASSLRVNTEVF | 0.6080148 |
| AAVSRAVAAL | ASSLRVNTEVF | 0.2758018 |
| AGGSSGIGGF | ASSLRVNTEVF | 0.667925 |
| AGLTVANANG | ASSLRVNTEVF | 0.39432815 |
| AGLVGAVL | ASSLRVNTEVF | 0.6679081 |
| ALSQAFTV | ASSLRVNTEVF | 0.7485291 |
| ALTFSVSL | ASSLRVNTEVF | 0.7895347 |
| ALTGAALVAL | ASSLRVNTEVF | 0.3346763 |
| ASFSIGAGVL | ASSLRVNTEVF | 0.7061657 |
| DGSVYKI | ASSLRVNTEVF | 0.45013487 |
| EQLAEIV | ASSLRVNTEVF | 0.81057703 |
| FAVSELTVM | ASSLRVNTEVF | 0.57293177 |

| | | |
|---|---|---|
| GAAPLAAGL | ASSLRVNTEVF | 0.26536083 |
| GADGGDGGSGGAGG | ASSLRVNTEVF | 0.7293296 |
| GFGLDTGGAGG | ASSLRVNTEVF | 0.70748633 |
| GLASGGLL | ASSLRVNTEVF | 0.48638928 |
| GRIAGAAVLL | ASSLRVNTEVF | 0.7396916 |
| GSAGSLNTSF | ASSLRVNTEVF | 0.46898422 |
| GTGQIGI | ASSLRVNTEVF | 0.5866101 |
| GTLIALIL | ASSLRVNTEVF | 0.7795696 |
| IAATMGFVVG | ASSLRVNTEVF | 0.67876875 |
| IFAQALL | ASSLRVNTEVF | 0.39194542 |
| IGAAVALGF | ASSLRVNTEVF | 0.7411355 |
| IGLTPLAL | ASSLRVNTEVF | 0.55793893 |
| ILGSNTSTL | ASSLRVNTEVF | 0.35605252 |
| IVFSLTVL | ASSLRVNTEVF | 0.78982806 |
| LASKATGF | ASSLRVNTEVF | 0.48864052 |
| LASVALIL | ASSLRVNTEVF | 0.73425686 |
| LFAVGLTL | ASSLRVNTEVF | 0.66127396 |
| LFNGGIGTYI | ASSLRVNTEVF | 0.7695929 |
| LTLGAGII | ASSLRVNTEVF | 0.37142247 |
| LVAAALGDI | ASSLRVNTEVF | 0.61567116 |
| NTAFVQLGI | ASSLRVNTEVF | 0.63865554 |
| QLFSGAEL | ASSLRVNTEVF | 0.70430934 |
| SAATFITL | ASSLRVNTEVF | 0.51655555 |
| SNYGLANLGV | ASSLRVNTEVF | 0.67223084 |
| SYGGAYAVM | ASSLRVNTEVF | 0.7133585 |
| TTGVGGLTLG | ASSLRVNTEVF | 0.7682762 |
| TTQTIYI | ASSLRVNTEVF | 0.5175508 |
| VAAGAGLGAVI | ASSLRVNTEVF | 0.5272064 |
| VAAVVTLL | ASSLRVNTEVF | 0.7883531 |
| VGFLGGLI | ASSLRVNTEVF | 0.36156964 |
| VGVIPILL | ASSLRVNTEVF | 0.58766115 |
| VIFASSLI | ASSLRVNTEVF | 0.7019723 |
| VKGGDGGAAAG | ASSLRVNTEVF | 0.73974633 |
| VTKTFTNI | ASSLRVNTEVF | 0.5294566 |
| VVAAAILL | ASSLRVNTEVF | 0.752561 |
| YAALDVEL | ASSLRVNTEVF | 0.51364326 |
| LASVALLL | ASSLRVNTEVF | 0.7466328 |
| LDAVHRV | ASSLRVNTEVF | 0.6080309 |
| QGLSSSIF | ASSLRVNTEVF | 0.63314116 |
| RGTFANIRL | ASSLRVNTEVF | 0.36188388 |
| RQLWWGHRI | ASSLRVNTEVF | 0.71217024 |
| YGPPGTGKTLL | ASSLRVNTEVF | 0.72957593 |
| AALVGAPANVA | ASSLRVNTEVF | 0.73043245 |
| AATFRRTE | ASSLRVNTEVF | 0.2583947 |
| AGALGGGI | ASSLRVNTEVF | 0.72183335 |
| AGGAGGAGGIGAG | ASSLRVNTEVF | 0.74561405 |
| AGSGALGL | ASSLRVNTEVF | 0.80374736 |
| AQIADQI | ASSLRVNTEVF | 0.5491091 |
| ASAVAVTYL | ASSLRVNTEVF | 0.69672704 |
| ASTVLGGTGG | ASSLRVNTEVF | 0.52991295 |
| AVGGEPAK | ASSLRVNTEVF | 0.55557764 |

| | | |
|---|---|---|
| CEQGTFGGTC | ASSLRVNTEVF | 0.27940696 |
| DEGKAPAG | ASSLRVNTEVF | 0.4340306 |
| DTVVFPAL | ASSLRVNTEVF | 0.6399662 |
| FEGITMAG | ASSLRVNTEVF | 0.47731617 |
| FPLVVQD | ASSLRVNTEVF | 0.75073594 |
| GAFTTPQ | ASSLRVNTEVF | 0.7230375 |
| GFGGFGTAGGVGGA | ASSLRVNTEVF | 0.7441727 |
| GFLAQFL | ASSLRVNTEVF | 0.7592546 |
| GGAGGAGGNAGVA | ASSLRVNTEVF | 0.790709 |
| GGVGTAGAGGAGGAGG | ASSLRVNTEVF | 0.67037046 |
| GNGGGGDAGNAGSGGN | ASSLRVNTEVF | 0.46819645 |
| GTGGAGGAGNDAGS | ASSLRVNTEVF | 0.6295985 |
| GTTNTGWFNS | ASSLRVNTEVF | 0.26292655 |
| IFMGPLFTL | ASSLRVNTEVF | 0.5580633 |
| ILAHAIEAI | ASSLRVNTEVF | 0.4405442 |
| IRSAIREAL | ASSLRVNTEVF | 0.68675995 |
| ITFAGGG | ASSLRVNTEVF | 0.17733264 |
| ITGMSGHQEV | ASSLRVNTEVF | 0.70475155 |
| IVAGLACTL | ASSLRVNTEVF | 0.77705663 |
| IVGGGLAGA | ASSLRVNTEVF | 0.5406458 |
| IVTGVLNI | ASSLRVNTEVF | 0.69776046 |
| LAQQGCI | ASSLRVNTEVF | 0.76444924 |
| LCLHVEN | ASSLRVNTEVF | 0.39922303 |
| LDAISDGS | ASSLRVNTEVF | 0.69800115 |
| LGGFAAML | ASSLRVNTEVF | 0.43476808 |
| LGGKNPMI | ASSLRVNTEVF | 0.7143383 |
| LGPKGVAAIL | ASSLRVNTEVF | 0.73086226 |
| LMVGVSI | ASSLRVNTEVF | 0.7027509 |
| LVGAAFVL | ASSLRVNTEVF | 0.42644325 |
| LVLAVAPL | ASSLRVNTEVF | 0.6204996 |
| MAAIAVIDAA | ASSLRVNTEVF | 0.3434758 |
| NNGINAELQ | ASSLRVNTEVF | 0.4772174 |
| RGEPLVLVHG | ASSLRVNTEVF | 0.5096311 |
| RGGAGGAAGLI | ASSLRVNTEVF | 0.67495644 |
| RSYRMHPAV | ASSLRVNTEVF | 0.39267302 |
| SAAHLLAGQLA | ASSLRVNTEVF | 0.67071784 |
| SAAVGSY | ASSLRVNTEVF | 0.6066855 |
| SPHAVLHRI | ASSLRVNTEVF | 0.6268621 |
| SQIAKVL | ASSLRVNTEVF | 0.460213 |
| TALGHLVP | ASSLRVNTEVF | 0.6837389 |
| TAPTTSGAA | ASSLRVNTEVF | 0.625191 |
| TDGDDGNGGSPGT | ASSLRVNTEVF | 0.29097667 |
| TDGLGEN | ASSLRVNTEVF | 0.75533617 |
| TGGPPLPGQA | ASSLRVNTEVF | 0.54055905 |
| TIPAITI | ASSLRVNTEVF | 0.6370574 |
| TIPGFTI | ASSLRVNTEVF | 0.7519716 |
| TSIASGALA | ASSLRVNTEVF | 0.5837244 |
| TVTLPGAAI | ASSLRVNTEVF | 0.3696358 |
| VASGLSAALP | ASSLRVNTEVF | 0.51790065 |
| VGAGVIGI | ASSLRVNTEVF | 0.6218834 |
| VGPKRFGV | ASSLRVNTEVF | 0.54683816 |

| | | |
|---|---|---|
| VNVAFPD | ASSLRVNTEVF | 0.21902344 |
| VSMTNSGVS | ASSLRVNTEVF | 0.7207216 |
| VTDVLGDLSV | ASSLRVNTEVF | 0.34940723 |
| VVVVGAGFAGL | ASSLRVNTEVF | 0.55556315 |
| VYTVGTVV | ASSLRVNTEVF | 0.4963625 |
| WTTFSGP | ASSLRVNTEVF | 0.75234437 |
| YDELVEAF | ASSLRVNTEVF | 0.5658881 |
| YGTGGAGGNGG | ASSLRVNTEVF | 0.49852931 |
| YTNVMAV | ASSLRVNTEVF | 0.6906379 |
| FLEQVHQL | ASSLRVNTEVF | 0.63988733 |
| GLSLLAVAV | ASSLRVNTEVF | 0.7537551 |
| LEQVHQL | ASSLRVNTEVF | 0.6053263 |
| LNAAGAI | ASSLRVNTEVF | 0.5511965 |
| PAVHASL | ASSLRVNTEVF | 0.69768393 |
| TELEDDA | ASSLRVNTEVF | 0.4302248 |
| TPPGPPN | ASSLRVNTEVF | 0.3823176 |
| TPPPPPPP | ASSLRVNTEVF | 0.33969462 |
| TQSPALI | ASSPILGSAETLY | 0.5539704 |
| EFAAAFPQ | ASSPILGSAETLY | 0.31287587 |
| LLSGNLDVL | ASSPILGSAETLY | 0.6897611 |
| GPANLGNYNI | ASSPILGSAETLY | 0.738106 |
| TLGPTLP | ASSPILGSAETLY | 0.5649767 |
| VYAYSSCCK | ASSPILGSAETLY | 0.28532004 |
| APAPAPAPNI | ASSPILGSAETLY | 0.46661416 |
| PGGPVPIP | ASSPILGSAETLY | 0.6232754 |
| FLTVALEP | ASSPILGSAETLY | 0.7221881 |
| SQYFGANL | ASSPILGSAETLY | 0.64424884 |
| GGVKPAA | ASSPILGSAETLY | 0.7950864 |
| GGTGGAGGAGGW | ASSPILGSAETLY | 0.7448407 |
| GVGGVAGLGGAGG | ASSPILGSAETLY | 0.48153168 |
| VLLVLATTI | ASSPILGSAETLY | 0.7124015 |
| IGIFAAML | ASSPILGSAETLY | 0.5399292 |
| AVGDLID | ASSPILGSAETLY | 0.6717216 |
| NISPLPL | ASSPILGSAETLY | 0.61269873 |
| VTAVFVAI | ASSPILGSAETLY | 0.646494 |
| SGDDNDAAL | ASSPILGSAETLY | 0.45040423 |
| AVAALAHS | ASSPILGSAETLY | 0.5312039 |
| GGVTVMATP | ASSPILGSAETLY | 0.3164478 |
| AGLAANVRD | ASSPILGSAETLY | 0.66092765 |
| TNRPDLIDPALLRPG | ASSPILGSAETLY | 0.37466106 |
| LANLGVTAL | ASSPILGSAETLY | 0.29039577 |
| TGERIEL | ASSPILGSAETLY | 0.5559157 |
| NGEELAA | ASSPILGSAETLY | 0.7889626 |
| VLSLVWG | ASSPILGSAETLY | 0.7688784 |
| IGYALWL | ASSPILGSAETLY | 0.51457846 |
| LSLLLAPL | ASSPILGSAETLY | 0.40433246 |
| TLDEALK | ASSPILGSAETLY | 0.7275828 |
| TDGAGTAEGG | ASSPILGSAETLY | 0.3577731 |
| NAPGGAGGNGG | ASSPILGSAETLY | 0.6371275 |
| VSVFMGMGGNF | ASSPILGSAETLY | 0.65252596 |
| LASYLAAL | ASSPILGSAETLY | 0.6202827 |

| | | |
|---|---|---|
| LVGGTPAL | ASSPILGSAETLY | 0.4294123 |
| VRNADVL | ASSPILGSAETLY | 0.6726198 |
| TCSSPFLVSAM | ASSPILGSAETLY | 0.562346 |
| VHITDAE | ASSPILGSAETLY | 0.5848979 |
| EYAAPSF | ASSPILGSAETLY | 0.2431674 |
| VVDAFHGGL | ASSPILGSAETLY | 0.7079245 |
| ASVVDPQ | ASSPILGSAETLY | 0.37905997 |
| SYNEFGGLS | ASSPILGSAETLY | 0.36549926 |
| DPDGSVY | ASSPILGSAETLY | 0.20969693 |
| MVSTIHATE | ASSPILGSAETLY | 0.55994844 |
| LAAGVADPV | ASSPILGSAETLY | 0.7164536 |
| TKEQIAA | ASSPILGSAETLY | 0.7495719 |
| SLASPAAL | ASSPILGSAETLY | 0.5945193 |
| IGLVFYGV | ASSPILGSAETLY | 0.7006277 |
| FLLDASNL | ASSPILGSAETLY | 0.7663365 |
| YQVGNFP | ASSPILGSAETLY | 0.55972314 |
| ESLLSQST | ASSPILGSAETLY | 0.24663287 |
| LFTVGFAAVG | ASSPILGSAETLY | 0.65368533 |
| ATPPTPT | ASSPILGSAETLY | 0.65988183 |
| TDLATLG | ASSPILGSAETLY | 0.7470796 |
| SAGGTGR | ASSPILGSAETLY | 0.71843827 |
| GFSNSGS | ASSPILGSAETLY | 0.3552919 |
| GGNGGSGGTSV | ASSPILGSAETLY | 0.6307408 |
| GTSVATGGAGN | ASSPILGSAETLY | 0.7511885 |
| NGGNGGSGGTS | ASSPILGSAETLY | 0.31742436 |
| AVVKFPH | ASSPILGSAETLY | 0.4498396 |
| TPPPPPP | ASSPILGSAETLY | 0.4599855 |
| AAPAAAAAPA | ASSPILGSAETLY | 0.41944617 |
| IALNEDL | ASSPILGSAETLY | 0.63738847 |
| IVNVSSM | ASSPILGSAETLY | 0.5064857 |
| TGIPPAPR | ASSPILGSAETLY | 0.4606378 |
| FDLGGGTF | ASSPILGSAETLY | 0.5021859 |
| FDLGGGTFDVS | ASSPILGSAETLY | 0.3981085 |
| NEPTAAA | ASSPILGSAETLY | 0.44143796 |
| NEPTAAALA | ASSPILGSAETLY | 0.5002431 |
| GSIINISSI | ASSPILGSAETLY | 0.70920545 |
| VAPGNAGTA | ASSPILGSAETLY | 0.690794 |
| TGGGASLE | ASSPILGSAETLY | 0.56195396 |
| VAVYNHYKRI | ASSPILGSAETLY | 0.67057073 |
| TDPAPATT | ASSPILGSAETLY | 0.78415763 |
| TNRPDILDPALLRPG | ASSPILGSAETLY | 0.29960278 |
| LTTTTTS | ASSPILGSAETLY | 0.39827335 |
| PTGSGKTL | ASSPILGSAETLY | 0.6346364 |
| TGSGKTL | ASSPILGSAETLY | 0.37292427 |
| TIGHVDHGK | ASSPILGSAETLY | 0.7391659 |
| TIGHVDHGKTT | ASSPILGSAETLY | 0.5963038 |
| GHVDHGKTT | ASSPILGSAETLY | 0.6752497 |
| AQFEHTLL | ASSPILGSAETLY | 0.60051835 |
| IVNQDEL | ASSPILGSAETLY | 0.80291307 |
| TNGSQFF | ASSPILGSAETLY | 0.70683193 |
| TGKGIVE | ASSPILGSAETLY | 0.5311022 |

| | | |
|---|---|---|
| LISNASDA | ASSPILGSAETLY | 0.71577895 |
| AELRTLGELTKIAKAHGAQ | ASSPILGSAETLY | 0.31506446 |
| APNAVAAAEPSW | ASSPILGSAETLY | 0.7192246 |
| APQVAGAATAVL | ASSPILGSAETLY | 0.7106903 |
| ARRDPAAAAAILPTDA | ASSPILGSAETLY | 0.41798925 |
| DAAARDVLAEAGLAEYF | ASSPILGSAETLY | 0.59366775 |
| DDQIVLSAKANLAKVEQ | ASSPILGSAETLY | 0.7641425 |
| DELLPDTGEDSDEDGH | ASSPILGSAETLY | 0.7742132 |
| ELPEAPANRATVPQD | ASSPILGSAETLY | 0.34100643 |
| FGQSKIERVTGIG | ASSPILGSAETLY | 0.46359652 |
| GAGGTGSTAGGAGGAGGAGGLY | ASSPILGSAETLY | 0.32078093 |
| GHGGTGAAGGNGGTGGAGGSGI | ASSPILGSAETLY | 0.4003136 |
| GNGGNGGAGGEGGPGGAGGAGA | ASSPILGSAETLY | 0.27503043 |
| GSTAGNGGNGGNATNGTIAGQP | ASSPILGSAETLY | 0.71256864 |
| GTGGDGGNGGNGGTGG | ASSPILGSAETLY | 0.57759666 |
| GVRRPAGIPADAPV | ASSPILGSAETLY | 0.7499378 |
| GYRQVTDAHLLTLARR | ASSPILGSAETLY | 0.6635034 |
| IAKMGQDGHDRG | ASSPILGSAETLY | 0.7407694 |
| KAEPSIEIQYT | ASSPILGSAETLY | 0.49271864 |
| KPVRIRMAADEDTVAVT | ASSPILGSAETLY | 0.5468508 |
| MPPRIAGMRLLVIKPEP | ASSPILGSAETLY | 0.26342303 |
| MKFYAVALFRSRGVPPTR | ASSPILGSAETLY | 0.34475425 |
| NAGGFGILGAVAHSP | ASSPILGSAETLY | 0.35246283 |
| NEGREITLAMAEM | ASSPILGSAETLY | 0.47455055 |
| NELVEWSHPLGHRWRWE | ASSPILGSAETLY | 0.2678368 |
| NLGAGGTGGNGGNPG | ASSPILGSAETLY | 0.77898574 |
| NTEWPKWETVMAADPVD | ASSPILGSAETLY | 0.5394999 |
| PASAAPAETQP | ASSPILGSAETLY | 0.67050767 |
| QQSSDGKNGVRAAEITGE | ASSPILGSAETLY | 0.54725975 |
| RAPDAVTTSWWKE | ASSPILGSAETLY | 0.33933032 |
| SGFGNVGAHTSGWFNQ | ASSPILGSAETLY | 0.42542043 |
| SGVAGAVGQAASVGGLKV | ASSPILGSAETLY | 0.7258698 |
| SNGGAGTGSAGGNGGT | ASSPILGSAETLY | 0.35662472 |
| TGGAGGEGGAAG | ASSPILGSAETLY | 0.2834856 |
| TGGDGGAAGAGGAPGHG | ASSPILGSAETLY | 0.3700034 |
| TMPPLLAAIANAGRHGV | ASSPILGSAETLY | 0.58854043 |
| TSSTAGNSGGA | ASSPILGSAETLY | 0.7915283 |
| VPATVPVGILC | ASSPILGSAETLY | 0.7521888 |
| VQRPDSGGVVLTGRIS | ASSPILGSAETLY | 0.64176905 |
| WTPNPVPATKGVPAGLVQAT | ASSPILGSAETLY | 0.78106254 |
| YALKRALAKFSDRFDVVII | ASSPILGSAETLY | 0.18687722 |
| YLIGSVIV | ASSPILGSAETLY | 0.5767392 |
| YGQLVAVID | ASSPILGSAETLY | 0.785102 |
| VTVGGMTLPA | ASSPILGSAETLY | 0.6581923 |
| VLALPAV | ASSPILGSAETLY | 0.61321175 |
| VIPPTLNYET | ASSPILGSAETLY | 0.7652619 |
| VGMGGMHPSSG | ASSPILGSAETLY | 0.41873324 |
| VGLAGGAATA | ASSPILGSAETLY | 0.5737681 |
| VGGAGGAGGSGP | ASSPILGSAETLY | 0.5726639 |
| VFAAFMV | ASSPILGSAETLY | 0.71620256 |
| TLYDKLR | ASSPILGSAETLY | 0.6814627 |

| | | |
|---|---|---|
| TLAVFTVL | ASSPILGSAETLY | 0.29715908 |
| TGSVQPARV | ASSPILGSAETLY | 0.6710774 |
| TGGGAGSGGT | ASSPILGSAETLY | 0.8025274 |
| SHLETSL | ASSPILGSAETLY | 0.5125769 |
| SFTTGMT | ASSPILGSAETLY | 0.75009465 |
| QVEVAKRMGVRQPTVS | ASSPILGSAETLY | 0.64193887 |
| QGSGGPALAM | ASSPILGSAETLY | 0.566809 |
| MLAPSNP | ASSPILGSAETLY | 0.49101686 |
| LVNNAAI | ASSPILGSAETLY | 0.36137465 |
| LTGEINTLA | ASSPILGSAETLY | 0.67684805 |
| LSIFGPV | ASSPILGSAETLY | 0.43585765 |
| LLALLANL | ASSPILGSAETLY | 0.57425034 |
| LISNASDALDKLR | ASSPILGSAETLY | 0.60549927 |
| LINEAVP | ASSPILGSAETLY | 0.8101021 |
| LGYAFSGL | ASSPILGSAETLY | 0.69932777 |
| LDFVAVAL | ASSPILGSAETLY | 0.6902106 |
| LAQPGAE | ASSPILGSAETLY | 0.37668198 |
| ISNLGAVTGF | ASSPILGSAETLY | 0.6374272 |
| HQEDPGRI | ASSPILGSAETLY | 0.6973589 |
| GTPGEPTP | ASSPILGSAETLY | 0.35613626 |
| GFFNSGS | ASSPILGSAETLY | 0.7627028 |
| GAGGGGLTAGAN | ASSPILGSAETLY | 0.3940084 |
| FVDALVNM | ASSPILGSAETLY | 0.67293227 |
| FLTVAPEL | ASSPILGSAETLY | 0.66958594 |
| FETLTAL | ASSPILGSAETLY | 0.7718246 |
| DVINPIEL | ASSPILGSAETLY | 0.613335 |
| APVSGAPV | ASSPILGSAETLY | 0.6694179 |
| ALVGGLAW | ASSPILGSAETLY | 0.33815405 |
| ALAAAAATAI | ASSPILGSAETLY | 0.7109524 |
| AFGLGLLL | ASSPILGSAETLY | 0.7644607 |
| AFFAAASL | ASSPILGSAETLY | 0.6715013 |
| AAAALPAFSPP | ASSPILGSAETLY | 0.8003093 |
| VVLAPAL | ASSPILGSAETLY | 0.6476646 |
| TIGHVDHGKTTL | ASSPILGSAETLY | 0.24893194 |
| LISNASDALDK | ASSPILGSAETLY | 0.5215502 |
| LISNASDAL | ASSPILGSAETLY | 0.74924356 |
| ISNASDALDK | ASSPILGSAETLY | 0.7609049 |
| NQAETLVYQTE | ASSPILGSAETLY | 0.7251178 |
| TGGTGGTAGAG | ASSPILGSAETLY | 0.6351826 |
| TGSTAGGAGGA | ASSPILGSAETLY | 0.5919306 |
| DPLAVPVAAGPV | ASSPILGSAETLY | 0.7733871 |
| GGTGGTPGNGGN | ASSPILGSAETLY | 0.8070665 |
| PAGLAGVDPATL | ASSPILGSAETLY | 0.7117328 |
| GNNLVVEAPHAGASV | ASSPILGSAETLY | 0.4123922 |
| KVTRIMSASTLEQPA | ASSPILGSAETLY | 0.3746235 |
| LGFALPIALPRRTVV | ASSPILGSAETLY | 0.7086779 |
| TRLGNLTARKLVVLV | ASSPILGSAETLY | 0.74030524 |
| WSQQRRLIYFPSAGP | ASSPILGSAETLY | 0.32443228 |
| GPNGNPDPMAAAVDIR | ASSPILGSAETLY | 0.637885 |
| TQRCVSVADLRAAASR | ASSPILGSAETLY | 0.5780507 |
| QFVLDVAEHPVEVLTETV | ASSPILGSAETLY | 0.2266422 |

| | | |
|---|---|---|
| GGAGGTGGAGGNGGTQAGDG | ASSPILGSAETLY | 0.73249304 |
| PLLATLRAIAVDVDAKPAQV | ASSPILGSAETLY | 0.787924 |
| APKFRTGGYMIDFWGVGYQVAK | ASSPILGSAETLY | 0.63942593 |
| GAGGAGGAGGAAGGPGGTGGTG | ASSPILGSAETLY | 0.30583063 |
| IGALGLLATDVAPAGSVAFGVA | ASSPILGSAETLY | 0.7745522 |
| AGGAGGIGGNGGAGGAGGVLMG | ASSPILGSAETLY | 0.8055494 |
| ADELIGLL | ASSPILGSAETLY | 0.7622744 |
| ADRALKA | ASSPILGSAETLY | 0.28441077 |
| AGAVGQVGGAGGA | ASSPILGSAETLY | 0.18493327 |
| AGGGGGLGGAP | ASSPILGSAETLY | 0.59241134 |
| AGGLLYGNGGAG | ASSPILGSAETLY | 0.74443626 |
| AGLESIGL | ASSPILGSAETLY | 0.46469706 |
| AGLKTGII | ASSPILGSAETLY | 0.36254838 |
| ASLAALLL | ASSPILGSAETLY | 0.6426581 |
| ASVINISSI | ASSPILGSAETLY | 0.74488354 |
| ATPPGPA | ASSPILGSAETLY | 0.36655456 |
| ATVASALV | ASSPILGSAETLY | 0.2909013 |
| ATVASLIL | ASSPILGSAETLY | 0.6365409 |
| AVNGPVLIRH | ASSPILGSAETLY | 0.7207329 |
| DGAAGAVGI | ASSPILGSAETLY | 0.4131173 |
| DPLLHGI | ASSPILGSAETLY | 0.48853573 |
| ELGAALGAL | ASSPILGSAETLY | 0.6472622 |
| ELGRTQAV | ASSPILGSAETLY | 0.666525 |
| FVAGGYLS | ASSPILGSAETLY | 0.6411951 |
| GASGGAGGAGGW | ASSPILGSAETLY | 0.31301978 |
| GGAGAGGDGGAGGI | ASSPILGSAETLY | 0.77000374 |
| GTGHNGSQPGMGGNG | ASSPILGSAETLY | 0.36520767 |
| GYAAASASAS | ASSPILGSAETLY | 0.6827367 |
| HQVDVAAVQQG | ASSPILGSAETLY | 0.6063603 |
| IGAAIAHT | ASSPILGSAETLY | 0.68913174 |
| IGMIEVAQL | ASSPILGSAETLY | 0.5666355 |
| IRLAPEHTSAMD | ASSPILGSAETLY | 0.46596724 |
| IYNVSVL | ASSPILGSAETLY | 0.34277102 |
| KQGFLGM | ASSPILGSAETLY | 0.27961326 |
| LESVITQL | ASSPILGSAETLY | 0.7446087 |
| LGLGALVVV | ASSPILGSAETLY | 0.7193917 |
| LIGNGGNGGSGGI | ASSPILGSAETLY | 0.730053 |
| LNAINAPTL | ASSPILGSAETLY | 0.6676992 |
| LQLLNTGV | ASSPILGSAETLY | 0.40858945 |
| LQTSIEF | ASSPILGSAETLY | 0.6674559 |
| MSFAVSEL | ASSPILGSAETLY | 0.2742146 |
| NTGALGL | ASSPILGSAETLY | 0.63627404 |
| PPGAGGAGGAGTT | ASSPILGSAETLY | 0.591457 |
| QLVGLGI | ASSPILGSAETLY | 0.66020906 |
| RFLEQVHQL | ASSPILGSAETLY | 0.56917644 |
| RGVGYVLR | ASSPILGSAETLY | 0.6292286 |
| SATGVFAGVI | ASSPILGSAETLY | 0.7058171 |
| SLPAITLP | ASSPILGSAETLY | 0.37623447 |
| STALAFTV | ASSPILGSAETLY | 0.6543337 |
| SVLTVGLGK | ASSPILGSAETLY | 0.7734935 |
| TALAGAGSGI | ASSPILGSAETLY | 0.6580315 |

| | | |
|---|---|---|
| TGLPFTGI | ASSPILGSAETLY | 0.3118491 |
| VAEHVTV | ASSPILGSAETLY | 0.69551814 |
| VDIVLNSL | ASSPILGSAETLY | 0.4709912 |
| VLTGGGAVL | ASSPILGSAETLY | 0.68067163 |
| VNVGLAYGI | ASSPILGSAETLY | 0.62669134 |
| VSSAAVGAV | ASSPILGSAETLY | 0.5264475 |
| VTSVYEAGV | ASSPILGSAETLY | 0.69483775 |
| WVALDRGA | ASSPILGSAETLY | 0.3112964 |
| YLVGSVVI | ASSPILGSAETLY | 0.47935003 |
| GPPGTGKTLL | ASSPILGSAETLY | 0.5641624 |
| GVGMTKF | ASSPILGSAETLY | 0.6316329 |
| IGIGGSDL | ASSPILGSAETLY | 0.73365426 |
| IGTLLAEL | ASSPILGSAETLY | 0.37822917 |
| IYTSGTT | ASSPILGSAETLY | 0.67002195 |
| LSFPAGV | ASSPILGSAETLY | 0.5008423 |
| LYTSGTT | ASSPILGSAETLY | 0.45268282 |
| VGAFPFGF | ASSPILGSAETLY | 0.57219034 |
| VGAGTVEF | ASSPILGSAETLY | 0.4555252 |
| VGKSTLLL | ASSPILGSAETLY | 0.29583335 |
| AALAHAGGGIV | ASSPILGSAETLY | 0.7028265 |
| AALVAVSLL | ASSPILGSAETLY | 0.3721978 |
| AAVSRAVAAL | ASSPILGSAETLY | 0.6208195 |
| AGGSSGIGGF | ASSPILGSAETLY | 0.78463995 |
| AGLTVANANG | ASSPILGSAETLY | 0.26892126 |
| AGLVGAVL | ASSPILGSAETLY | 0.67854714 |
| ALSQAFTV | ASSPILGSAETLY | 0.6687777 |
| ALTFSVSL | ASSPILGSAETLY | 0.6660583 |
| ALTGAALVAL | ASSPILGSAETLY | 0.45867282 |
| ASFSIGAGVL | ASSPILGSAETLY | 0.42206022 |
| DGSVYKI | ASSPILGSAETLY | 0.7707535 |
| EQLAEIV | ASSPILGSAETLY | 0.7134284 |
| FAVSELTVM | ASSPILGSAETLY | 0.64107823 |
| GAAPLAAGL | ASSPILGSAETLY | 0.6903068 |
| GADGGDGGSGGAGG | ASSPILGSAETLY | 0.7546499 |
| GFGLDTGGAGG | ASSPILGSAETLY | 0.7242351 |
| GLASGGLL | ASSPILGSAETLY | 0.49377364 |
| GRIAGAAVLL | ASSPILGSAETLY | 0.25253248 |
| GSAGSLNTSF | ASSPILGSAETLY | 0.5660838 |
| GTGQIGI | ASSPILGSAETLY | 0.59414655 |
| GTLIALIL | ASSPILGSAETLY | 0.44505808 |
| IAATMGFVVG | ASSPILGSAETLY | 0.2602052 |
| IFAQALL | ASSPILGSAETLY | 0.7679541 |
| IGAAVALGF | ASSPILGSAETLY | 0.8110333 |
| IGLTPLAL | ASSPILGSAETLY | 0.42876446 |
| ILGSNTSTL | ASSPILGSAETLY | 0.80207264 |
| IVFSLTVL | ASSPILGSAETLY | 0.29197764 |
| LASKATGF | ASSPILGSAETLY | 0.69240105 |
| LASVALIL | ASSPILGSAETLY | 0.76840657 |
| LFAVGLTL | ASSPILGSAETLY | 0.35148484 |
| LFNGGIGTYI | ASSPILGSAETLY | 0.6832069 |
| LTLGAGII | ASSPILGSAETLY | 0.525065 |

| | | |
|---|---|---|
| LVAAALGDI | ASSPILGSAETLY | 0.41585338 |
| NTAFVQLGI | ASSPILGSAETLY | 0.77075195 |
| QLFSGAEL | ASSPILGSAETLY | 0.41515014 |
| SAATFITL | ASSPILGSAETLY | 0.6681305 |
| SNYGLANLGV | ASSPILGSAETLY | 0.45088208 |
| SYGGAYAVM | ASSPILGSAETLY | 0.35462472 |
| TTGVGGLTLG | ASSPILGSAETLY | 0.6717702 |
| TTQTIYI | ASSPILGSAETLY | 0.7195394 |
| VAAGAGLGAVI | ASSPILGSAETLY | 0.521838 |
| VAAVVTLL | ASSPILGSAETLY | 0.8019203 |
| VGFLGGLI | ASSPILGSAETLY | 0.4931058 |
| VGVIPILL | ASSPILGSAETLY | 0.8106209 |
| VIFASSLI | ASSPILGSAETLY | 0.7805652 |
| VKGGDGGAAAG | ASSPILGSAETLY | 0.6136643 |
| VTKTFTNI | ASSPILGSAETLY | 0.49921006 |
| VVAAAILL | ASSPILGSAETLY | 0.7684312 |
| YAALDVEL | ASSPILGSAETLY | 0.50431806 |
| LASVALLL | ASSPILGSAETLY | 0.5537023 |
| LDAVHRV | ASSPILGSAETLY | 0.74328196 |
| QGLSSSIF | ASSPILGSAETLY | 0.63482726 |
| RGTFANIRL | ASSPILGSAETLY | 0.7059772 |
| RQLWWGHRI | ASSPILGSAETLY | 0.6738098 |
| YGPPGTGKTLL | ASSPILGSAETLY | 0.68959355 |
| AALVGAPANVA | ASSPILGSAETLY | 0.64892507 |
| AATFRRTE | ASSPILGSAETLY | 0.4967448 |
| AGALGGGI | ASSPILGSAETLY | 0.6682441 |
| AGGAGGAGGIGAG | ASSPILGSAETLY | 0.6215397 |
| AGSGALGL | ASSPILGSAETLY | 0.6834203 |
| AQIADQI | ASSPILGSAETLY | 0.69123733 |
| ASAVAVTYL | ASSPILGSAETLY | 0.637574 |
| ASTVLGGTGG | ASSPILGSAETLY | 0.7076786 |
| AVGGEPAK | ASSPILGSAETLY | 0.7091111 |
| CEQGTFGGTC | ASSPILGSAETLY | 0.537007 |
| DEGKAPAG | ASSPILGSAETLY | 0.66103065 |
| DTVVFPAL | ASSPILGSAETLY | 0.6140089 |
| FEGITMAG | ASSPILGSAETLY | 0.6707933 |
| FPLVVQD | ASSPILGSAETLY | 0.38124016 |
| GAFTTPQ | ASSPILGSAETLY | 0.7012385 |
| GFGGFGTAGGVGGA | ASSPILGSAETLY | 0.6253675 |
| GFLAQFL | ASSPILGSAETLY | 0.27738285 |
| GGAGGAGGNAGVA | ASSPILGSAETLY | 0.6715196 |
| GGVGTAGAGGAGGAGG | ASSPILGSAETLY | 0.67129445 |
| GNGGGGDAGNAGSGGN | ASSPILGSAETLY | 0.66994053 |
| GTGGAGGAGNDAGS | ASSPILGSAETLY | 0.69455725 |
| GTTNTGWFNS | ASSPILGSAETLY | 0.38913822 |
| IFMGPLFTL | ASSPILGSAETLY | 0.62005055 |
| ILAHAIEAI | ASSPILGSAETLY | 0.6895286 |
| IRSAIREAL | ASSPILGSAETLY | 0.7054192 |
| ITFAGGG | ASSPILGSAETLY | 0.613373 |
| ITGMSGHQEV | ASSPILGSAETLY | 0.6788266 |
| IVAGLACTL | ASSPILGSAETLY | 0.7100121 |

| | | |
|---|---|---|
| IVGGGLAGA | ASSPILGSAETLY | 0.68265164 |
| IVTGVLNI | ASSPILGSAETLY | 0.70294833 |
| LAQQGCI | ASSPILGSAETLY | 0.7101958 |
| LCLHVEN | ASSPILGSAETLY | 0.6006446 |
| LDAISDGS | ASSPILGSAETLY | 0.70103925 |
| LGGFAAML | ASSPILGSAETLY | 0.70347303 |
| LGGKNPMI | ASSPILGSAETLY | 0.66196 |
| LGPKGVAAIL | ASSPILGSAETLY | 0.67966354 |
| LMVGVSI | ASSPILGSAETLY | 0.5996734 |
| LVGAAFVL | ASSPILGSAETLY | 0.6441163 |
| LVLAVAPL | ASSPILGSAETLY | 0.38294327 |
| MAAIAVIDAA | ASSPILGSAETLY | 0.67646897 |
| NNGINAELQ | ASSPILGSAETLY | 0.6995135 |
| RGEPLVLVHG | ASSPILGSAETLY | 0.6499239 |
| RGGAGGAAGLI | ASSPILGSAETLY | 0.57405704 |
| RSYRMHPAV | ASSPILGSAETLY | 0.59159875 |
| SAAHLLAGQLA | ASSPILGSAETLY | 0.64686894 |
| SAAVGSY | ASSPILGSAETLY | 0.66039145 |
| SPHAVLHRI | ASSPILGSAETLY | 0.42464587 |
| SQIAKVL | ASSPILGSAETLY | 0.6176172 |
| TALGHLVP | ASSPILGSAETLY | 0.70319617 |
| TAPTTSGAA | ASSPILGSAETLY | 0.7092006 |
| TDGDDGNGGSPGT | ASSPILGSAETLY | 0.5093039 |
| TDGLGEN | ASSPILGSAETLY | 0.7043339 |
| TGGPPLPGQA | ASSPILGSAETLY | 0.6921073 |
| TIPAITI | ASSPILGSAETLY | 0.70704234 |
| TIPGFTI | ASSPILGSAETLY | 0.6728322 |
| TSIASGALA | ASSPILGSAETLY | 0.60823417 |
| TVTLPGAAI | ASSPILGSAETLY | 0.69507223 |
| VASGLSAALP | ASSPILGSAETLY | 0.6383826 |
| VGAGVIGI | ASSPILGSAETLY | 0.5694079 |
| VGPKRFGV | ASSPILGSAETLY | 0.7019861 |
| VNVAFPD | ASSPILGSAETLY | 0.6683247 |
| VSMTNSGVS | ASSPILGSAETLY | 0.6154866 |
| VTDVLGDLSV | ASSPILGSAETLY | 0.6243466 |
| VVVVGAGFAGL | ASSPILGSAETLY | 0.7014871 |
| VYTVGTVV | ASSPILGSAETLY | 0.5829742 |
| WTTFSGP | ASSPILGSAETLY | 0.5561754 |
| YDELVEAF | ASSPILGSAETLY | 0.61461496 |
| YGTGGAGGNGG | ASSPILGSAETLY | 0.67139363 |
| YTNVMAV | ASSPILGSAETLY | 0.700715 |
| FLEQVHQL | ASSPILGSAETLY | 0.6944072 |
| GLSLLAVAV | ASSPILGSAETLY | 0.6066689 |
| LEQVHQL | ASSPILGSAETLY | 0.68869436 |
| LNAAGAI | ASSPILGSAETLY | 0.6951611 |
| PAVHASL | ASSPILGSAETLY | 0.6802064 |
| TELEDDA | ASSPILGSAETLY | 0.6854826 |
| TPPGPPN | ASSPILGSAETLY | 0.6193732 |
| TPPPPPPP | ASSPILGSAETLY | 0.69930303 |
| TQSPALI | ASSDGGGSYNSPLY | 0.69334584 |
| EFAAAFPQ | ASSDGGGSYNSPLY | 0.7035525 |

| | | |
|---|---|---|
| LLSGNLDVL | ASSDGGGSYNSPLY | 0.6963873 |
| GPANLGNYNI | ASSDGGGSYNSPLY | 0.6731175 |
| TLGPTLP | ASSDGGGSYNSPLY | 0.6753199 |
| VYAYSSCCK | ASSDGGGSYNSPLY | 0.40867612 |
| APAPAPAPNI | ASSDGGGSYNSPLY | 0.6219268 |
| PGGPVPIP | ASSDGGGSYNSPLY | 0.6750368 |
| FLTVALEP | ASSDGGGSYNSPLY | 0.67152464 |
| SQYFGANL | ASSDGGGSYNSPLY | 0.68737054 |
| GGVKPAA | ASSDGGGSYNSPLY | 0.7006577 |
| GGTGGAGGAGGW | ASSDGGGSYNSPLY | 0.61951387 |
| GVGGVAGLGGAGG | ASSDGGGSYNSPLY | 0.70239437 |
| VLLVLATTI | ASSDGGGSYNSPLY | 0.70037735 |
| IGIFAAML | ASSDGGGSYNSPLY | 0.52662575 |
| AVGDLID | ASSDGGGSYNSPLY | 0.6444447 |
| NISPLPL | ASSDGGGSYNSPLY | 0.71345603 |
| VTAVFVAI | ASSDGGGSYNSPLY | 0.672136 |
| SGDDNDAAL | ASSDGGGSYNSPLY | 0.65189326 |
| AVAALAHS | ASSDGGGSYNSPLY | 0.66677165 |
| GGVTVMATP | ASSDGGGSYNSPLY | 0.7127133 |
| AGLAANVRD | ASSDGGGSYNSPLY | 0.64084864 |
| TNRPDLIDPALLRPG | ASSDGGGSYNSPLY | 0.7127551 |
| LANLGVTAL | ASSDGGGSYNSPLY | 0.35893738 |
| TGERIEL | ASSDGGGSYNSPLY | 0.66151327 |
| NGEELAA | ASSDGGGSYNSPLY | 0.6185601 |
| VLSLVWG | ASSDGGGSYNSPLY | 0.4385484 |
| IGYALWL | ASSDGGGSYNSPLY | 0.7013008 |
| LSLLLAPL | ASSDGGGSYNSPLY | 0.6662463 |
| TLDEALK | ASSDGGGSYNSPLY | 0.7009218 |
| TDGAGTAEGG | ASSDGGGSYNSPLY | 0.6607537 |
| NAPGGAGGNGG | ASSDGGGSYNSPLY | 0.70316875 |
| VSVFMGMGGNF | ASSDGGGSYNSPLY | 0.66172564 |
| LASYLAAL | ASSDGGGSYNSPLY | 0.6755794 |
| LVGGTPAL | ASSDGGGSYNSPLY | 0.561159 |
| VRNADVL | ASSDGGGSYNSPLY | 0.69906014 |
| TCSSPFLVSAM | ASSDGGGSYNSPLY | 0.70308316 |
| VHITDAE | ASSDGGGSYNSPLY | 0.69624174 |
| EYAAPSF | ASSDGGGSYNSPLY | 0.55229694 |
| VVDAFHGGL | ASSDGGGSYNSPLY | 0.6344286 |
| ASVVDPQ | ASSDGGGSYNSPLY | 0.68629235 |
| SYNEFGGLS | ASSDGGGSYNSPLY | 0.30096358 |
| DPDGSVY | ASSDGGGSYNSPLY | 0.37519592 |
| MVSTIHATE | ASSDGGGSYNSPLY | 0.63575006 |
| LAAGVADPV | ASSDGGGSYNSPLY | 0.5841662 |
| TKEQIAA | ASSDGGGSYNSPLY | 0.702747 |
| SLASPAAL | ASSDGGGSYNSPLY | 0.6897181 |
| IGLVFYGV | ASSDGGGSYNSPLY | 0.6640294 |
| FLLDASNL | ASSDGGGSYNSPLY | 0.68340254 |
| YQVGNFP | ASSDGGGSYNSPLY | 0.6751267 |
| ESLLSQST | ASSDGGGSYNSPLY | 0.25805363 |
| LFTVGFAAVG | ASSDGGGSYNSPLY | 0.6406911 |
| ATPPTPT | ASSDGGGSYNSPLY | 0.5563494 |

| | | |
|---|---|---|
| TDLATLG | ASSDGGGSYNSPLY | 0.71212244 |
| SAGGTGR | ASSDGGGSYNSPLY | 0.60369354 |
| GFSNSGS | ASSDGGGSYNSPLY | 0.59427416 |
| GGNGGSGGTSV | ASSDGGGSYNSPLY | 0.668362 |
| GTSVATGGAGN | ASSDGGGSYNSPLY | 0.6731279 |
| NGGNGGSGGTS | ASSDGGGSYNSPLY | 0.67219687 |
| AVVKFPH | ASSDGGGSYNSPLY | 0.57670236 |
| TPPPPPP | ASSDGGGSYNSPLY | 0.27668744 |
| AAPAAAAAPA | ASSDGGGSYNSPLY | 0.7011824 |
| IALNEDL | ASSDGGGSYNSPLY | 0.42773473 |
| IVNVSSM | ASSDGGGSYNSPLY | 0.6137977 |
| TGIPPAPR | ASSDGGGSYNSPLY | 0.35830337 |
| FDLGGGTF | ASSDGGGSYNSPLY | 0.29319456 |
| FDLGGGTFDVS | ASSDGGGSYNSPLY | 0.52656096 |
| NEPTAAA | ASSDGGGSYNSPLY | 0.7012573 |
| NEPTAAALA | ASSDGGGSYNSPLY | 0.6314919 |
| GSIINISSI | ASSDGGGSYNSPLY | 0.6284934 |
| VAPGNAGTA | ASSDGGGSYNSPLY | 0.7064622 |
| TGGGASLE | ASSDGGGSYNSPLY | 0.6784704 |
| VAVYNHYKRI | ASSDGGGSYNSPLY | 0.6360528 |
| TDPAPATT | ASSDGGGSYNSPLY | 0.65491533 |
| TNRPDILDPALLRPG | ASSDGGGSYNSPLY | 0.6017567 |
| LTTTTTS | ASSDGGGSYNSPLY | 0.60280514 |
| PTGSGKTL | ASSDGGGSYNSPLY | 0.6345767 |
| TGSGKTL | ASSDGGGSYNSPLY | 0.70937705 |
| TIGHVDHGK | ASSDGGGSYNSPLY | 0.6649172 |
| TIGHVDHGKTT | ASSDGGGSYNSPLY | 0.5308573 |
| GHVDHGKTT | ASSDGGGSYNSPLY | 0.6848016 |
| AQFEHTLL | ASSDGGGSYNSPLY | 0.7114155 |
| IVNQDEL | ASSDGGGSYNSPLY | 0.7099457 |
| TNGSQFF | ASSDGGGSYNSPLY | 0.58781624 |
| TGKGIVE | ASSDGGGSYNSPLY | 0.6743766 |
| LISNASDA | ASSDGGGSYNSPLY | 0.6584621 |
| AELRTLGELTKIAKAHGAQ | ASSDGGGSYNSPLY | 0.6601164 |
| APNAVAAAEPSW | ASSDGGGSYNSPLY | 0.6276834 |
| APQVAGAATAVL | ASSDGGGSYNSPLY | 0.55422556 |
| ARRDPAAAAAILPTDA | ASSDGGGSYNSPLY | 0.6919061 |
| DAAARDVLAEAGLAEYF | ASSDGGGSYNSPLY | 0.71676135 |
| DDQIVLSAKANLAKVEQ | ASSDGGGSYNSPLY | 0.64551365 |
| DELLPDTGEDSDEDGH | ASSDGGGSYNSPLY | 0.46657974 |
| ELPEAPANRATVPQD | ASSDGGGSYNSPLY | 0.6597841 |
| FGQSKIERVTGIG | ASSDGGGSYNSPLY | 0.60418034 |
| GAGGTGSTAGGAGGAGGAGGLY | ASSDGGGSYNSPLY | 0.63643 |
| GHGGTGAAGGNGGTGGAGGSGI | ASSDGGGSYNSPLY | 0.6656954 |
| GNGGNGGAGGEGGPGGAGGAGA | ASSDGGGSYNSPLY | 0.53443366 |
| GSTAGNGGNGGNATNGTIAGQP | ASSDGGGSYNSPLY | 0.6578983 |
| GTGGDGGNGGNGGTGG | ASSDGGGSYNSPLY | 0.7131424 |
| GVRRPAGIPADAPV | ASSDGGGSYNSPLY | 0.65485424 |
| GYRQVTDAHLLTLARR | ASSDGGGSYNSPLY | 0.678843 |
| IAKMGQDGHDRG | ASSDGGGSYNSPLY | 0.70345193 |
| KAEPSIEIQYT | ASSDGGGSYNSPLY | 0.68340605 |

| | | |
|---|---|---|
| KPVRIRMAADEDTVAVT | ASSDGGGSYNSPLY | 0.7044668 |
| MPPRIAGMRLLVIKPEP | ASSDGGGSYNSPLY | 0.69431895 |
| MKFYAVALFRSRGVPPTR | ASSDGGGSYNSPLY | 0.67792165 |
| NAGGFGILGAVAHSP | ASSDGGGSYNSPLY | 0.67382944 |
| NEGREITLAMAEM | ASSDGGGSYNSPLY | 0.6705226 |
| NELVEWSHPLGHRWRWE | ASSDGGGSYNSPLY | 0.6734701 |
| NLGAGGTGGNGGNPG | ASSDGGGSYNSPLY | 0.6550264 |
| NTEWPKWETVMAADPVD | ASSDGGGSYNSPLY | 0.63077754 |
| PASAAPAETQP | ASSDGGGSYNSPLY | 0.70704114 |
| QQSSDGKNGVRAAEITGE | ASSDGGGSYNSPLY | 0.6664934 |
| RAPDAVTTSWWKE | ASSDGGGSYNSPLY | 0.7043855 |
| SGFGNVGAHTSGWFNQ | ASSDGGGSYNSPLY | 0.58953357 |
| SGVAGAVGQAASVGGLKV | ASSDGGGSYNSPLY | 0.5942459 |
| SNGGAGTGSAGGNGGT | ASSDGGGSYNSPLY | 0.69658667 |
| TGGAGGEGGAAG | ASSDGGGSYNSPLY | 0.2706573 |
| TGGDGGAAGAGGAPGHG | ASSDGGGSYNSPLY | 0.3841892 |
| TMPPLLAAIANAGRHGV | ASSDGGGSYNSPLY | 0.66446924 |
| TSSTAGNSGGA | ASSDGGGSYNSPLY | 0.7053535 |
| VPATVPVGILC | ASSDGGGSYNSPLY | 0.27039102 |
| VQRPDSGGVVLTGRIS | ASSDGGGSYNSPLY | 0.608622 |
| WTPNPVPATKGVPAGLVQAT | ASSDGGGSYNSPLY | 0.53474116 |
| YALKRALAKFSDRFDVVII | ASSDGGGSYNSPLY | 0.6978141 |
| YLIGSVIV | ASSDGGGSYNSPLY | 0.6808857 |
| YGQLVAVID | ASSDGGGSYNSPLY | 0.4994552 |
| VTVGGMTLPA | ASSDGGGSYNSPLY | 0.6971746 |
| VLALPAV | ASSDGGGSYNSPLY | 0.4716162 |
| VIPPTLNYET | ASSDGGGSYNSPLY | 0.47782865 |
| VGMGGMHPSSG | ASSDGGGSYNSPLY | 0.696025 |
| VGLAGGAATA | ASSDGGGSYNSPLY | 0.7030566 |
| VGGAGGAGGSGP | ASSDGGGSYNSPLY | 0.5519702 |
| VFAAFMV | ASSDGGGSYNSPLY | 0.69817555 |
| TLYDKLR | ASSDGGGSYNSPLY | 0.6280141 |
| TLAVFTVL | ASSDGGGSYNSPLY | 0.711058 |
| TGSVQPARV | ASSDGGGSYNSPLY | 0.65184045 |
| TGGGAGSGGT | ASSDGGGSYNSPLY | 0.4892461 |
| SHLETSL | ASSDGGGSYNSPLY | 0.6969876 |
| SFTTGMT | ASSDGGGSYNSPLY | 0.6826143 |
| QVEVAKRMGVRQPTVS | ASSDGGGSYNSPLY | 0.67624784 |
| QGSGGPALAM | ASSDGGGSYNSPLY | 0.69505537 |
| MLAPSNP | ASSDGGGSYNSPLY | 0.6202027 |
| LVNNAAI | ASSDGGGSYNSPLY | 0.6580499 |
| LTGEINTLA | ASSDGGGSYNSPLY | 0.70716643 |
| LSIFGPV | ASSDGGGSYNSPLY | 0.6917498 |
| LLALLANL | ASSDGGGSYNSPLY | 0.6064024 |
| LISNASDALDKLR | ASSDGGGSYNSPLY | 0.6981407 |
| LINEAVP | ASSDGGGSYNSPLY | 0.6627163 |
| LGYAFSGL | ASSDGGGSYNSPLY | 0.64141417 |
| LDFVAVAL | ASSDGGGSYNSPLY | 0.7102933 |
| LAQPGAE | ASSDGGGSYNSPLY | 0.6676839 |
| ISNLGAVTGF | ASSDGGGSYNSPLY | 0.6957048 |
| HQEDPGRI | ASSDGGGSYNSPLY | 0.70582926 |

| | | |
|---|---|---|
| GTPGEPTP | ASSDGGGSYNSPLY | 0.6980515 |
| GFFNSGS | ASSDGGGSYNSPLY | 0.5245878 |
| GAGGGGLTAGAN | ASSDGGGSYNSPLY | 0.6728735 |
| FVDALVNM | ASSDGGGSYNSPLY | 0.66810274 |
| FLTVAPEL | ASSDGGGSYNSPLY | 0.69555885 |
| FETLTAL | ASSDGGGSYNSPLY | 0.66035175 |
| DVINPIEL | ASSDGGGSYNSPLY | 0.4499849 |
| APVSGAPV | ASSDGGGSYNSPLY | 0.6874699 |
| ALVGGLAW | ASSDGGGSYNSPLY | 0.7043537 |
| ALAAAAATAI | ASSDGGGSYNSPLY | 0.6637484 |
| AFGLGLLL | ASSDGGGSYNSPLY | 0.67600703 |
| AFFAAASL | ASSDGGGSYNSPLY | 0.70881104 |
| AAAALPAFSPP | ASSDGGGSYNSPLY | 0.6299083 |
| VVLAPAL | ASSDGGGSYNSPLY | 0.66620785 |
| TIGHVDHGKTTL | ASSDGGGSYNSPLY | 0.69629014 |
| LISNASDALDK | ASSDGGGSYNSPLY | 0.6954881 |
| LISNASDAL | ASSDGGGSYNSPLY | 0.6684264 |
| ISNASDALDK | ASSDGGGSYNSPLY | 0.7017237 |
| NQAETLVYQTE | ASSDGGGSYNSPLY | 0.59754443 |
| TGGTGGTAGAG | ASSDGGGSYNSPLY | 0.68863666 |
| TGSTAGGAGGA | ASSDGGGSYNSPLY | 0.5515455 |
| DPLAVPVAAGPV | ASSDGGGSYNSPLY | 0.68925285 |
| GGTGGTPGNGGN | ASSDGGGSYNSPLY | 0.61539197 |
| PAGLAGVDPATL | ASSDGGGSYNSPLY | 0.6958742 |
| GNNLVVEAPHAGASV | ASSDGGGSYNSPLY | 0.68951356 |
| KVTRIMSASTLEQPA | ASSDGGGSYNSPLY | 0.6982614 |
| LGFALPIALPRRTVV | ASSDGGGSYNSPLY | 0.35486525 |
| TRLGNLTARKLVVLV | ASSDGGGSYNSPLY | 0.6971104 |
| WSQQRRLIYFPSAGP | ASSDGGGSYNSPLY | 0.7039101 |
| GPNGNPDPMAAAVDIR | ASSDGGGSYNSPLY | 0.71544415 |
| TQRCVSVADLRAAASR | ASSDGGGSYNSPLY | 0.670079 |
| QFVLDVAEHPVEVLTETV | ASSDGGGSYNSPLY | 0.6949525 |
| GGAGGTGGAGGNGGTQAGDG | ASSDGGGSYNSPLY | 0.64289176 |
| PLLATLRAIAVDVDAKPAQV | ASSDGGGSYNSPLY | 0.65310323 |
| APKFRTGGYMIDFWGVGYQVAK | ASSDGGGSYNSPLY | 0.59645593 |
| GAGGAGGAGGAAGGPGGTGGTG | ASSDGGGSYNSPLY | 0.6381852 |
| IGALGLLATDVAPAGSVAFGVA | ASSDGGGSYNSPLY | 0.622704 |
| AGGAGGIGGNGGAGGAGGVLMG | ASSDGGGSYNSPLY | 0.61790884 |
| ADELIGLL | ASSDGGGSYNSPLY | 0.6982167 |
| ADRALKA | ASSDGGGSYNSPLY | 0.69265825 |
| AGAVGQVGGAGGA | ASSDGGGSYNSPLY | 0.6986948 |
| AGGGGGLGGAP | ASSDGGGSYNSPLY | 0.7060045 |
| AGGLLYGNGGAG | ASSDGGGSYNSPLY | 0.6967043 |
| AGLESIGL | ASSDGGGSYNSPLY | 0.7038593 |
| AGLKTGII | ASSDGGGSYNSPLY | 0.6446824 |
| ASLAALLL | ASSDGGGSYNSPLY | 0.65073395 |
| ASVINISSI | ASSDGGGSYNSPLY | 0.5768976 |
| ATPPGPA | ASSDGGGSYNSPLY | 0.6499078 |
| ATVASALV | ASSDGGGSYNSPLY | 0.6463003 |
| ATVASLIL | ASSDGGGSYNSPLY | 0.6820554 |
| AVNGPVLIRH | ASSDGGGSYNSPLY | 0.26486588 |

| | | |
|---|---|---|
| DGAAGAVGI | ASSDGGGSYNSPLY | 0.6597944 |
| DPLLHGI | ASSDGGGSYNSPLY | 0.7073885 |
| ELGAALGAL | ASSDGGGSYNSPLY | 0.59645844 |
| ELGRTQAV | ASSDGGGSYNSPLY | 0.6968454 |
| FVAGGYLS | ASSDGGGSYNSPLY | 0.65508723 |
| GASGGAGGAGGW | ASSDGGGSYNSPLY | 0.6232024 |
| GGAGAGGDGGAGGI | ASSDGGGSYNSPLY | 0.69803977 |
| GTGHNGSQPGMGGNG | ASSDGGGSYNSPLY | 0.616136 |
| GYAAASASAS | ASSDGGGSYNSPLY | 0.69416976 |
| HQVDVAAVQQG | ASSDGGGSYNSPLY | 0.7098778 |
| IGAAIAHT | ASSDGGGSYNSPLY | 0.6811792 |
| IGMIEVAQL | ASSDGGGSYNSPLY | 0.6651529 |
| IRLAPEHTSAMD | ASSDGGGSYNSPLY | 0.6816768 |
| IYNVSVL | ASSDGGGSYNSPLY | 0.4840574 |
| KQGFLGM | ASSDGGGSYNSPLY | 0.6717436 |
| LESVITQL | ASSDGGGSYNSPLY | 0.25724447 |
| LGLGALVVV | ASSDGGGSYNSPLY | 0.66263247 |
| LIGNGGNGGSGGI | ASSDGGGSYNSPLY | 0.5775311 |
| LNAINAPTL | ASSDGGGSYNSPLY | 0.6801803 |
| LQLLNTGV | ASSDGGGSYNSPLY | 0.669392 |
| LQTSIEF | ASSDGGGSYNSPLY | 0.6825855 |
| MSFAVSEL | ASSDGGGSYNSPLY | 0.6158283 |
| NTGALGL | ASSDGGGSYNSPLY | 0.6534933 |
| PPGAGGAGGAGTT | ASSDGGGSYNSPLY | 0.6135858 |
| QLVGLGI | ASSDGGGSYNSPLY | 0.56408906 |
| RFLEQVHQL | ASSDGGGSYNSPLY | 0.6839615 |
| RGVGYVLR | ASSDGGGSYNSPLY | 0.64807373 |
| SATGVFAGVI | ASSDGGGSYNSPLY | 0.63436115 |
| SLPAITLP | ASSDGGGSYNSPLY | 0.7043406 |
| STALAFTV | ASSDGGGSYNSPLY | 0.6816833 |
| SVLTVGLGK | ASSDGGGSYNSPLY | 0.4097071 |
| TALAGAGSGI | ASSDGGGSYNSPLY | 0.64947146 |
| TGLPFTGI | ASSDGGGSYNSPLY | 0.6689228 |
| VAEHVTV | ASSDGGGSYNSPLY | 0.66080046 |
| VDIVLNSL | ASSDGGGSYNSPLY | 0.25875357 |
| VLTGGGAVL | ASSDGGGSYNSPLY | 0.6738118 |
| VNVGLAYGI | ASSDGGGSYNSPLY | 0.64098614 |
| VSSAAVGAV | ASSDGGGSYNSPLY | 0.5886581 |
| VTSVYEAGV | ASSDGGGSYNSPLY | 0.6298317 |
| WVALDRGA | ASSDGGGSYNSPLY | 0.69526327 |
| YLVGSVVI | ASSDGGGSYNSPLY | 0.40243107 |
| GPPGTGKTLL | ASSDGGGSYNSPLY | 0.68855774 |
| GVGMTKF | ASSDGGGSYNSPLY | 0.6858555 |
| IGIGGSDL | ASSDGGGSYNSPLY | 0.6975734 |
| IGTLLAEL | ASSDGGGSYNSPLY | 0.70970315 |
| IYTSGTT | ASSDGGGSYNSPLY | 0.6848304 |
| LSFPAGV | ASSDGGGSYNSPLY | 0.65853107 |
| LYTSGTT | ASSDGGGSYNSPLY | 0.6960767 |
| VGAFPFGF | ASSDGGGSYNSPLY | 0.6686106 |
| VGAGTVEF | ASSDGGGSYNSPLY | 0.46071142 |
| VGKSTLLL | ASSDGGGSYNSPLY | 0.45446467 |

| | | |
|---|---|---|
| AALAHAGGGIV | ASSDGGGSYNSPLY | 0.6677959 |
| AALVAVSLL | ASSDGGGSYNSPLY | 0.41294312 |
| AAVSRAVAAL | ASSDGGGSYNSPLY | 0.6838933 |
| AGGSSGIGGF | ASSDGGGSYNSPLY | 0.71007633 |
| AGLTVANANG | ASSDGGGSYNSPLY | 0.6541682 |
| AGLVGAVL | ASSDGGGSYNSPLY | 0.5299427 |
| ALSQAFTV | ASSDGGGSYNSPLY | 0.64335424 |
| ALTFSVSL | ASSDGGGSYNSPLY | 0.64064807 |
| ALTGAALVAL | ASSDGGGSYNSPLY | 0.6927711 |
| ASFSIGAGVL | ASSDGGGSYNSPLY | 0.69399846 |
| DGSVYKI | ASSDGGGSYNSPLY | 0.7102585 |
| EQLAEIV | ASSDGGGSYNSPLY | 0.7031896 |
| FAVSELTVM | ASSDGGGSYNSPLY | 0.65206695 |
| GAAPLAAGL | ASSDGGGSYNSPLY | 0.685904 |
| GADGGDGGSGGAGG | ASSDGGGSYNSPLY | 0.65153277 |
| GFGLDTGGAGG | ASSDGGGSYNSPLY | 0.6313101 |
| GLASGGLL | ASSDGGGSYNSPLY | 0.6633886 |
| GRIAGAAVLL | ASSDGGGSYNSPLY | 0.6626965 |
| GSAGSLNTSF | ASSDGGGSYNSPLY | 0.66509974 |
| GTGQIGI | ASSDGGGSYNSPLY | 0.64833933 |
| GTLIALIL | ASSDGGGSYNSPLY | 0.48780784 |
| IAATMGFVVG | ASSDGGGSYNSPLY | 0.6919333 |
| IFAQALL | ASSDGGGSYNSPLY | 0.67657197 |
| IGAAVALGF | ASSDGGGSYNSPLY | 0.47794378 |
| IGLTPLAL | ASSDGGGSYNSPLY | 0.6512896 |
| ILGSNTSTL | ASSDGGGSYNSPLY | 0.7097628 |
| IVFSLTVL | ASSDGGGSYNSPLY | 0.7065438 |
| LASKATGF | ASSDGGGSYNSPLY | 0.61350185 |
| LASVALIL | ASSDGGGSYNSPLY | 0.62185645 |
| LFAVGLTL | ASSDGGGSYNSPLY | 0.6605069 |
| LFNGGIGTYI | ASSDGGGSYNSPLY | 0.45097905 |
| LTLGAGII | ASSDGGGSYNSPLY | 0.68560183 |
| LVAAALGDI | ASSDGGGSYNSPLY | 0.6593597 |
| NTAFVQLGI | ASSDGGGSYNSPLY | 0.6483091 |
| QLFSGAEL | ASSDGGGSYNSPLY | 0.6371317 |
| SAATFITL | ASSDGGGSYNSPLY | 0.66948915 |
| SNYGLANLGV | ASSDGGGSYNSPLY | 0.629977 |
| SYGGAYAVM | ASSDGGGSYNSPLY | 0.69408303 |
| TTGVGGLTLG | ASSDGGGSYNSPLY | 0.26816043 |
| TTQTIYI | ASSDGGGSYNSPLY | 0.6773881 |
| VAAGAGLGAVI | ASSDGGGSYNSPLY | 0.5943836 |
| VAAVVTLL | ASSDGGGSYNSPLY | 0.5979553 |
| VGFLGGLI | ASSDGGGSYNSPLY | 0.38496137 |
| VGVIPILL | ASSDGGGSYNSPLY | 0.6648398 |
| VIFASSLI | ASSDGGGSYNSPLY | 0.69317377 |
| VKGGDGGAAAG | ASSDGGGSYNSPLY | 0.6933984 |
| VTKTFTNI | ASSDGGGSYNSPLY | 0.56070495 |
| VVAAAILL | ASSDGGGSYNSPLY | 0.69673765 |
| YAALDVEL | ASSDGGGSYNSPLY | 0.7003541 |
| LASVALLL | ASSDGGGSYNSPLY | 0.63100445 |
| LDAVHRV | ASSDGGGSYNSPLY | 0.39318013 |

| | | |
|---|---|---|
| QGLSSSIF | ASSDGGGSYNSPLY | 0.69339347 |
| RGTFANIRL | ASSDGGGSYNSPLY | 0.63979733 |
| RQLWWGHRI | ASSDGGGSYNSPLY | 0.69337 |
| YGPPGTGKTLL | ASSDGGGSYNSPLY | 0.68532896 |
| AALVGAPANVA | ASSDGGGSYNSPLY | 0.58025706 |
| AATFRRTE | ASSDGGGSYNSPLY | 0.7092954 |
| AGALGGGI | ASSDGGGSYNSPLY | 0.6995206 |
| AGGAGGAGGIGAG | ASSDGGGSYNSPLY | 0.60481036 |
| AGSGALGL | ASSDGGGSYNSPLY | 0.5944141 |
| AQIADQI | ASSDGGGSYNSPLY | 0.61366576 |
| ASAVAVTYL | ASSDGGGSYNSPLY | 0.32965913 |
| ASTVLGGTGG | ASSDGGGSYNSPLY | 0.6954648 |
| AVGGEPAK | ASSDGGGSYNSPLY | 0.61856234 |
| CEQGTFGGTC | ASSDGGGSYNSPLY | 0.65162003 |
| DEGKAPAG | ASSDGGGSYNSPLY | 0.6824975 |
| DTVVFPAL | ASSDGGGSYNSPLY | 0.6539568 |
| FEGITMAG | ASSDGGGSYNSPLY | 0.65541196 |
| FPLVVQD | ASSDGGGSYNSPLY | 0.66281813 |
| GAFTTPQ | ASSDGGGSYNSPLY | 0.6763898 |
| GFGGFGTAGGVGGA | ASSDGGGSYNSPLY | 0.6694279 |
| GFLAQFL | ASSDGGGSYNSPLY | 0.6711116 |
| GGAGGAGGNAGVA | ASSDGGGSYNSPLY | 0.67618346 |
| GGVGTAGAGGAGGAGG | ASSDGGGSYNSPLY | 0.7066103 |
| GNGGGGDAGNAGSGGN | ASSDGGGSYNSPLY | 0.5792866 |
| GTGGAGGAGNDAGS | ASSDGGGSYNSPLY | 0.6302924 |
| GTTNTGWFNS | ASSDGGGSYNSPLY | 0.68384826 |
| IFMGPLFTL | ASSDGGGSYNSPLY | 0.55840546 |
| ILAHAIEAI | ASSDGGGSYNSPLY | 0.6873835 |
| IRSAIREAL | ASSDGGGSYNSPLY | 0.6725186 |
| ITFAGGG | ASSDGGGSYNSPLY | 0.6813599 |
| ITGMSGHQEV | ASSDGGGSYNSPLY | 0.66621244 |
| IVAGLACTL | ASSDGGGSYNSPLY | 0.7081591 |
| IVGGGLAGA | ASSDGGGSYNSPLY | 0.7006871 |
| IVTGVLNI | ASSDGGGSYNSPLY | 0.6726109 |
| LAQQGCI | ASSDGGGSYNSPLY | 0.66129756 |
| LCLHVEN | ASSDGGGSYNSPLY | 0.58750457 |
| LDAISDGS | ASSDGGGSYNSPLY | 0.69694245 |
| LGGFAAML | ASSDGGGSYNSPLY | 0.6944863 |
| LGGKNPMI | ASSDGGGSYNSPLY | 0.5895802 |
| LGPKGVAAIL | ASSDGGGSYNSPLY | 0.658937 |
| LMVGVSI | ASSDGGGSYNSPLY | 0.7057265 |
| LVGAAFVL | ASSDGGGSYNSPLY | 0.6996405 |
| LVLAVAPL | ASSDGGGSYNSPLY | 0.44797087 |
| MAAIAVIDAA | ASSDGGGSYNSPLY | 0.6479852 |
| NNGINAELQ | ASSDGGGSYNSPLY | 0.40503493 |
| RGEPLVLVHG | ASSDGGGSYNSPLY | 0.6343664 |
| RGGAGGAAGLI | ASSDGGGSYNSPLY | 0.57863575 |
| RSYRMHPAV | ASSDGGGSYNSPLY | 0.67987585 |
| SAAHLLAGQLA | ASSDGGGSYNSPLY | 0.64329344 |
| SAAVGSY | ASSDGGGSYNSPLY | 0.69333375 |
| SPHAVLHRI | ASSDGGGSYNSPLY | 0.67853856 |

| | | |
|---|---|---|
| SQIAKVL | ASSDGGGSYNSPLY | 0.71032774 |
| TALGHLVP | ASSDGGGSYNSPLY | 0.63025534 |
| TAPTTSGAA | ASSDGGGSYNSPLY | 0.66905344 |
| TDGDDGNGGSPGT | ASSDGGGSYNSPLY | 0.4595502 |
| TDGLGEN | ASSDGGGSYNSPLY | 0.42355707 |
| TGGPPLPGQA | ASSDGGGSYNSPLY | 0.5900061 |
| TIPAITI | ASSDGGGSYNSPLY | 0.67092246 |
| TIPGFTI | ASSDGGGSYNSPLY | 0.30386147 |
| TSIASGALA | ASSDGGGSYNSPLY | 0.70865965 |
| TVTLPGAAI | ASSDGGGSYNSPLY | 0.6767334 |
| VASGLSAALP | ASSDGGGSYNSPLY | 0.69960153 |
| VGAGVIGI | ASSDGGGSYNSPLY | 0.6173396 |
| VGPKRFGV | ASSDGGGSYNSPLY | 0.6477771 |
| VNVAFPD | ASSDGGGSYNSPLY | 0.6325586 |
| VSMTNSGVS | ASSDGGGSYNSPLY | 0.69384664 |
| VTDVLGDLSV | ASSDGGGSYNSPLY | 0.59381294 |
| VVVVGAGFAGL | ASSDGGGSYNSPLY | 0.4406343 |
| VYTVGTVV | ASSDGGGSYNSPLY | 0.6375851 |
| WTTFSGP | ASSDGGGSYNSPLY | 0.51898754 |
| YDELVEAF | ASSDGGGSYNSPLY | 0.7022919 |
| YGTGGAGGNGG | ASSDGGGSYNSPLY | 0.69898254 |
| YTNVMAV | ASSDGGGSYNSPLY | 0.6838916 |
| FLEQVHQL | ASSDGGGSYNSPLY | 0.7049596 |
| GLSLLAVAV | ASSDGGGSYNSPLY | 0.30710515 |
| LEQVHQL | ASSDGGGSYNSPLY | 0.6241397 |
| LNAAGAI | ASSDGGGSYNSPLY | 0.7129501 |
| PAVHASL | ASSDGGGSYNSPLY | 0.6789308 |
| TELEDDA | ASSDGGGSYNSPLY | 0.65572596 |
| TPPGPPN | ASSDGGGSYNSPLY | 0.70507777 |
| TPPPPPPP | ASSDGGGSYNSPLY | 0.63617134 |
| TQSPALI | ASSLGNSYNSPLY | 0.6692138 |
| EFAAAFPQ | ASSLGNSYNSPLY | 0.6660279 |
| LLSGNLDVL | ASSLGNSYNSPLY | 0.6894574 |
| GPANLGNYNI | ASSLGNSYNSPLY | 0.6407437 |
| TLGPTLP | ASSLGNSYNSPLY | 0.5916425 |
| VYAYSSCCK | ASSLGNSYNSPLY | 0.6943291 |
| APAPAPAPNI | ASSLGNSYNSPLY | 0.7102294 |
| PGGPVPIP | ASSLGNSYNSPLY | 0.6526998 |
| FLTVALEP | ASSLGNSYNSPLY | 0.6925633 |
| SQYFGANL | ASSLGNSYNSPLY | 0.68892527 |
| GGVKPAA | ASSLGNSYNSPLY | 0.6906028 |
| GGTGGAGGAGGW | ASSLGNSYNSPLY | 0.6250756 |
| GVGGVAGLGGAGG | ASSLGNSYNSPLY | 0.6229453 |
| VLLVLATTI | ASSLGNSYNSPLY | 0.6595652 |
| IGIFAAML | ASSLGNSYNSPLY | 0.58318233 |
| AVGDLID | ASSLGNSYNSPLY | 0.67271584 |
| NISPLPL | ASSLGNSYNSPLY | 0.6534262 |
| VTAVFVAI | ASSLGNSYNSPLY | 0.71677256 |
| SGDDNDAAL | ASSLGNSYNSPLY | 0.6923678 |
| AVAALAHS | ASSLGNSYNSPLY | 0.68101484 |
| GGVTVMATP | ASSLGNSYNSPLY | 0.6015668 |

| | | |
|---|---|---|
| AGLAANVRD | ASSLGNSYNSPLY | 0.6795249 |
| TNRPDLIDPALLRPG | ASSLGNSYNSPLY | 0.6854902 |
| LANLGVTAL | ASSLGNSYNSPLY | 0.48944643 |
| TGERIEL | ASSLGNSYNSPLY | 0.7062775 |
| NGEELAA | ASSLGNSYNSPLY | 0.62428725 |
| VLSLVWG | ASSLGNSYNSPLY | 0.6408664 |
| IGYALWL | ASSLGNSYNSPLY | 0.704244 |
| LSLLLAPL | ASSLGNSYNSPLY | 0.70882004 |
| TLDEALK | ASSLGNSYNSPLY | 0.6421741 |
| TDGAGTAEGG | ASSLGNSYNSPLY | 0.6367171 |
| NAPGGAGGNGG | ASSLGNSYNSPLY | 0.6253507 |
| VSVFMGMGGNF | ASSLGNSYNSPLY | 0.69595814 |
| LASYLAAL | ASSLGNSYNSPLY | 0.7025448 |
| LVGGTPAL | ASSLGNSYNSPLY | 0.632084 |
| VRNADVL | ASSLGNSYNSPLY | 0.70644796 |
| TCSSPFLVSAM | ASSLGNSYNSPLY | 0.69015765 |
| VHITDAE | ASSLGNSYNSPLY | 0.50230074 |
| EYAAPSF | ASSLGNSYNSPLY | 0.661779 |
| VVDAFHGGL | ASSLGNSYNSPLY | 0.6949153 |
| ASVVDPQ | ASSLGNSYNSPLY | 0.68806803 |
| SYNEFGGLS | ASSLGNSYNSPLY | 0.71243817 |
| DPDGSVY | ASSLGNSYNSPLY | 0.6696323 |
| MVSTIHATE | ASSLGNSYNSPLY | 0.6274513 |
| LAAGVADPV | ASSLGNSYNSPLY | 0.7074799 |
| TKEQIAA | ASSLGNSYNSPLY | 0.71255314 |
| SLASPAAL | ASSLGNSYNSPLY | 0.69852483 |
| IGLVFYGV | ASSLGNSYNSPLY | 0.6438595 |
| FLLDASNL | ASSLGNSYNSPLY | 0.63618827 |
| YQVGNFP | ASSLGNSYNSPLY | 0.6866553 |
| ESLLSQST | ASSLGNSYNSPLY | 0.69417876 |
| LFTVGFAAVG | ASSLGNSYNSPLY | 0.5991752 |
| ATPPTPT | ASSLGNSYNSPLY | 0.6736884 |
| TDLATLG | ASSLGNSYNSPLY | 0.68227994 |
| SAGGTGR | ASSLGNSYNSPLY | 0.5348033 |
| GFSNSGS | ASSLGNSYNSPLY | 0.67585045 |
| GGNGGSGGTSV | ASSLGNSYNSPLY | 0.69454604 |
| GTSVATGGAGN | ASSLGNSYNSPLY | 0.6761581 |
| NGGNGGSGGTS | ASSLGNSYNSPLY | 0.6514833 |
| AVVKFPH | ASSLGNSYNSPLY | 0.69815445 |
| TPPPPPP | ASSLGNSYNSPLY | 0.71494806 |
| AAPAAAAAPA | ASSLGNSYNSPLY | 0.70926654 |
| IALNEDL | ASSLGNSYNSPLY | 0.512395 |
| IVNVSSM | ASSLGNSYNSPLY | 0.36199254 |
| TGIPPAPR | ASSLGNSYNSPLY | 0.60384715 |
| FDLGGGTF | ASSLGNSYNSPLY | 0.6951952 |
| FDLGGGTFDVS | ASSLGNSYNSPLY | 0.66161466 |
| NEPTAAA | ASSLGNSYNSPLY | 0.64611965 |
| NEPTAAALA | ASSLGNSYNSPLY | 0.67272943 |
| GSIINISSI | ASSLGNSYNSPLY | 0.70549595 |
| VAPGNAGTA | ASSLGNSYNSPLY | 0.63650656 |
| TGGGASLE | ASSLGNSYNSPLY | 0.4807953 |

| | | |
|---|---|---|
| VAVYNHYKRI | ASSLGNSYNSPLY | 0.67572105 |
| TDPAPATT | ASSLGNSYNSPLY | 0.6978256 |
| TNRPDILDPALLRPG | ASSLGNSYNSPLY | 0.61784786 |
| LTTTTTS | ASSLGNSYNSPLY | 0.645067 |
| PTGSGKTL | ASSLGNSYNSPLY | 0.59927 |
| TGSGKTL | ASSLGNSYNSPLY | 0.4676688 |
| TIGHVDHGK | ASSLGNSYNSPLY | 0.6738434 |
| TIGHVDHGKTT | ASSLGNSYNSPLY | 0.60757226 |
| GHVDHGKTT | ASSLGNSYNSPLY | 0.7027932 |
| AQFEHTLL | ASSLGNSYNSPLY | 0.601624 |
| IVNQDEL | ASSLGNSYNSPLY | 0.68081856 |
| TNGSQFF | ASSLGNSYNSPLY | 0.6789621 |
| TGKGIVE | ASSLGNSYNSPLY | 0.687534 |
| LISNASDA | ASSLGNSYNSPLY | 0.64261854 |
| AELRTLGELTKIAKAHGAQ | ASSLGNSYNSPLY | 0.62945557 |
| APNAVAAAEPSW | ASSLGNSYNSPLY | 0.57440376 |
| APQVAGAATAVL | ASSLGNSYNSPLY | 0.47105786 |
| ARRDPAAAAAILPTDA | ASSLGNSYNSPLY | 0.6949241 |
| DAAARDVLAEAGLAEYF | ASSLGNSYNSPLY | 0.27687156 |
| DDQIVLSAKANLAKVEQ | ASSLGNSYNSPLY | 0.7033045 |
| DELLPDTGEDSDEDGH | ASSLGNSYNSPLY | 0.69991076 |
| ELPEAPANRATVPQD | ASSLGNSYNSPLY | 0.5968343 |
| FGQSKIERVTGIG | ASSLGNSYNSPLY | 0.6838411 |
| GAGGTGSTAGGAGGAGGAGGLY | ASSLGNSYNSPLY | 0.60813063 |
| GHGGTGAAGGNGGTGGAGGSGI | ASSLGNSYNSPLY | 0.5982137 |
| GNGGNGGAGGEGGPGGAGGAGA | ASSLGNSYNSPLY | 0.6911886 |
| GSTAGNGGNGGNATNGTIAGQP | ASSLGNSYNSPLY | 0.69630754 |
| GTGGDGGNGGNGGTGG | ASSLGNSYNSPLY | 0.60807765 |
| GVRRPAGIPADAPV | ASSLGNSYNSPLY | 0.6810174 |
| GYRQVTDAHLLTLARR | ASSLGNSYNSPLY | 0.67411375 |
| IAKMGQDGHDRG | ASSLGNSYNSPLY | 0.63031113 |
| KAEPSIEIQYT | ASSLGNSYNSPLY | 0.6868527 |
| KPVRIRMAADEDTVAVT | ASSLGNSYNSPLY | 0.7071582 |
| MPPRIAGMRLLVIKPEP | ASSLGNSYNSPLY | 0.686164 |
| MKFYAVALFRSRGVPPTR | ASSLGNSYNSPLY | 0.6232667 |
| NAGGFGILGAVAHSP | ASSLGNSYNSPLY | 0.6892446 |
| NEGREITLAMAEM | ASSLGNSYNSPLY | 0.5810019 |
| NELVEWSHPLGHRWRWE | ASSLGNSYNSPLY | 0.6825756 |
| NLGAGGTGGNGGNPG | ASSLGNSYNSPLY | 0.6565057 |
| NTEWPKWETVMAADPVD | ASSLGNSYNSPLY | 0.67415494 |
| PASAAPAETQP | ASSLGNSYNSPLY | 0.6923133 |
| QQSSDGKNGVRAAEITGE | ASSLGNSYNSPLY | 0.50872666 |
| RAPDAVTTSWWKE | ASSLGNSYNSPLY | 0.5976585 |
| SGFGNVGAHTSGWFNQ | ASSLGNSYNSPLY | 0.649327 |
| SGVAGAVGQAASVGGLKV | ASSLGNSYNSPLY | 0.68527204 |
| SNGGAGTGSAGGNGGT | ASSLGNSYNSPLY | 0.6997973 |
| TGGAGGEGGAAG | ASSLGNSYNSPLY | 0.6216831 |
| TGGDGGAAGAGGAPGHG | ASSLGNSYNSPLY | 0.67155266 |
| TMPPLLAAIANAGRHGV | ASSLGNSYNSPLY | 0.660457 |
| TSSTAGNSGGA | ASSLGNSYNSPLY | 0.62676585 |
| VPATVPVGILC | ASSLGNSYNSPLY | 0.6847261 |

| | | |
|---|---|---|
| VQRPDSGGVVLTGRIS | ASSLGNSYNSPLY | 0.65538013 |
| WTPNPVPATKGVPAGLVQAT | ASSLGNSYNSPLY | 0.5805958 |
| YALKRALAKFSDRFDVVII | ASSLGNSYNSPLY | 0.68851143 |
| YLIGSVIV | ASSLGNSYNSPLY | 0.64113605 |
| YGQLVAVID | ASSLGNSYNSPLY | 0.6841665 |
| VTVGGMTLPA | ASSLGNSYNSPLY | 0.70055455 |
| VLALPAV | ASSLGNSYNSPLY | 0.6409731 |
| VIPPTLNYET | ASSLGNSYNSPLY | 0.6834487 |
| VGMGGMHPSSG | ASSLGNSYNSPLY | 0.6788081 |
| VGLAGGAATA | ASSLGNSYNSPLY | 0.67461354 |
| VGGAGGAGGSGP | ASSLGNSYNSPLY | 0.6704431 |
| VFAAFMV | ASSLGNSYNSPLY | 0.6387635 |
| TLYDKLR | ASSLGNSYNSPLY | 0.70604634 |
| TLAVFTVL | ASSLGNSYNSPLY | 0.6954789 |
| TGSVQPARV | ASSLGNSYNSPLY | 0.6970639 |
| TGGGAGSGGT | ASSLGNSYNSPLY | 0.68604755 |
| SHLETSL | ASSLGNSYNSPLY | 0.6891942 |
| SFTTGMT | ASSLGNSYNSPLY | 0.69221854 |
| QVEVAKRMGVRQPTVS | ASSLGNSYNSPLY | 0.62723756 |
| QGSGGPALAM | ASSLGNSYNSPLY | 0.51959294 |
| MLAPSNP | ASSLGNSYNSPLY | 0.664008 |
| LVNNAAI | ASSLGNSYNSPLY | 0.672897 |
| LTGEINTLA | ASSLGNSYNSPLY | 0.6437011 |
| LSIFGPV | ASSLGNSYNSPLY | 0.45757684 |
| LLALLANL | ASSLGNSYNSPLY | 0.70723724 |
| LISNASDALDKLR | ASSLGNSYNSPLY | 0.71353775 |
| LINEAVP | ASSLGNSYNSPLY | 0.5964276 |
| LGYAFSGL | ASSLGNSYNSPLY | 0.7066937 |
| LDFVAVAL | ASSLGNSYNSPLY | 0.54971784 |
| LAQPGAE | ASSLGNSYNSPLY | 0.6952181 |
| ISNLGAVTGF | ASSLGNSYNSPLY | 0.70838046 |
| HQEDPGRI | ASSLGNSYNSPLY | 0.58884025 |
| GTPGEPTP | ASSLGNSYNSPLY | 0.7004615 |
| GFFNSGS | ASSLGNSYNSPLY | 0.62317985 |
| GAGGGGLTAGAN | ASSLGNSYNSPLY | 0.6462597 |
| FVDALVNM | ASSLGNSYNSPLY | 0.7048038 |
| FLTVAPEL | ASSLGNSYNSPLY | 0.66214323 |
| FETLTAL | ASSLGNSYNSPLY | 0.686125 |
| DVINPIEL | ASSLGNSYNSPLY | 0.6758066 |
| APVSGAPV | ASSLGNSYNSPLY | 0.6110648 |
| ALVGGLAW | ASSLGNSYNSPLY | 0.6828873 |
| ALAAAAATAI | ASSLGNSYNSPLY | 0.7051796 |
| AFGLGLLL | ASSLGNSYNSPLY | 0.60840213 |
| AFFAAASL | ASSLGNSYNSPLY | 0.71152794 |
| AAAALPAFSPP | ASSLGNSYNSPLY | 0.6086627 |
| VVLAPAL | ASSLGNSYNSPLY | 0.71426713 |
| TIGHVDHGKTTL | ASSLGNSYNSPLY | 0.6892488 |
| LISNASDALDK | ASSLGNSYNSPLY | 0.66089547 |
| LISNASDAL | ASSLGNSYNSPLY | 0.63880944 |
| ISNASDALDK | ASSLGNSYNSPLY | 0.7036816 |
| NQAETLVYQTE | ASSLGNSYNSPLY | 0.663733 |

| | | |
|---|---|---|
| TGGTGGTAGAG | ASSLGNSYNSPLY | 0.6830791 |
| TGSTAGGAGGA | ASSLGNSYNSPLY | 0.70928913 |
| DPLAVPVAAGPV | ASSLGNSYNSPLY | 0.59060526 |
| GGTGGTPGNGGN | ASSLGNSYNSPLY | 0.7096755 |
| PAGLAGVDPATL | ASSLGNSYNSPLY | 0.65686846 |
| GNNLVVEAPHAGASV | ASSLGNSYNSPLY | 0.638193 |
| KVTRIMSASTLEQPA | ASSLGNSYNSPLY | 0.591874 |
| LGFALPIALPRRTVV | ASSLGNSYNSPLY | 0.43382618 |
| TRLGNLTARKLVVLV | ASSLGNSYNSPLY | 0.6462624 |
| WSQQRRLIYFPSAGP | ASSLGNSYNSPLY | 0.5591599 |
| GPNGNPDPMAAAVDIR | ASSLGNSYNSPLY | 0.6905328 |
| TQRCVSVADLRAAASR | ASSLGNSYNSPLY | 0.67978406 |
| QFVLDVAEHPVEVLTETV | ASSLGNSYNSPLY | 0.5685068 |
| GGAGGTGGAGGNGGTQAGDG | ASSLGNSYNSPLY | 0.71816516 |
| PLLATLRAIAVDVDAKPAQV | ASSLGNSYNSPLY | 0.69982713 |
| APKFRTGGYMIDFWGVGYQVAK | ASSLGNSYNSPLY | 0.495484 |
| GAGGAGGAGGAAGGPGGTGGTG | ASSLGNSYNSPLY | 0.6265995 |
| IGALGLLATDVAPAGSVAFGVA | ASSLGNSYNSPLY | 0.6345802 |
| AGGAGGIGGNGGAGGAGGVLMG | ASSLGNSYNSPLY | 0.64191 |
| ADELIGLL | ASSLGNSYNSPLY | 0.44706684 |
| ADRALKA | ASSLGNSYNSPLY | 0.69364566 |
| AGAVGQVGGAGGA | ASSLGNSYNSPLY | 0.5837062 |
| AGGGGGLGGAP | ASSLGNSYNSPLY | 0.2827962 |
| AGGLLYGNGGAG | ASSLGNSYNSPLY | 0.6613338 |
| AGLESIGL | ASSLGNSYNSPLY | 0.6661295 |
| AGLKTGII | ASSLGNSYNSPLY | 0.63681406 |
| ASLAALLL | ASSLGNSYNSPLY | 0.7022823 |
| ASVINISSI | ASSLGNSYNSPLY | 0.42158005 |
| ATPPGPA | ASSLGNSYNSPLY | 0.5589887 |
| ATVASALV | ASSLGNSYNSPLY | 0.7028314 |
| ATVASLIL | ASSLGNSYNSPLY | 0.7087672 |
| AVNGPVLIRH | ASSLGNSYNSPLY | 0.51200944 |
| DGAAGAVGI | ASSLGNSYNSPLY | 0.67579603 |
| DPLLHGI | ASSLGNSYNSPLY | 0.6953206 |
| ELGAALGAL | ASSLGNSYNSPLY | 0.6758378 |
| ELGRTQAV | ASSLGNSYNSPLY | 0.69647276 |
| FVAGGYLS | ASSLGNSYNSPLY | 0.70534337 |
| GASGGAGGAGGW | ASSLGNSYNSPLY | 0.53170305 |
| GGAGAGGDGGAGGI | ASSLGNSYNSPLY | 0.7072786 |
| GTGHNGSQPGMGGNG | ASSLGNSYNSPLY | 0.71639097 |
| GYAAASASAS | ASSLGNSYNSPLY | 0.61375237 |
| HQVDVAAVQQG | ASSLGNSYNSPLY | 0.6743827 |
| IGAAIAHT | ASSLGNSYNSPLY | 0.5347545 |
| IGMIEVAQL | ASSLGNSYNSPLY | 0.6069702 |
| IRLAPEHTSAMD | ASSLGNSYNSPLY | 0.33264866 |
| IYNVSVL | ASSLGNSYNSPLY | 0.67958045 |
| KQGFLGM | ASSLGNSYNSPLY | 0.69589216 |
| LESVITQL | ASSLGNSYNSPLY | 0.6318608 |
| LGLGALVVV | ASSLGNSYNSPLY | 0.58098626 |
| LIGNGGNGGSGGI | ASSLGNSYNSPLY | 0.5256808 |
| LNAINAPTL | ASSLGNSYNSPLY | 0.5803261 |

| | | |
|---|---|---|
| LQLLNTGV | ASSLGNSYNSPLY | 0.6088446 |
| LQTSIEF | ASSLGNSYNSPLY | 0.3308836 |
| MSFAVSEL | ASSLGNSYNSPLY | 0.6330192 |
| NTGALGL | ASSLGNSYNSPLY | 0.7078171 |
| PPGAGGAGGAGTT | ASSLGNSYNSPLY | 0.69507813 |
| QLVGLGI | ASSLGNSYNSPLY | 0.4506594 |
| RFLEQVHQL | ASSLGNSYNSPLY | 0.70978874 |
| RGVGYVLR | ASSLGNSYNSPLY | 0.6841817 |
| SATGVFAGVI | ASSLGNSYNSPLY | 0.69096756 |
| SLPAITLP | ASSLGNSYNSPLY | 0.59930366 |
| STALAFTV | ASSLGNSYNSPLY | 0.58556753 |
| SVLTVGLGK | ASSLGNSYNSPLY | 0.6938889 |
| TALAGAGSGI | ASSLGNSYNSPLY | 0.6275681 |
| TGLPFTGI | ASSLGNSYNSPLY | 0.5084732 |
| VAEHVTV | ASSLGNSYNSPLY | 0.7051671 |
| VDIVLNSL | ASSLGNSYNSPLY | 0.6190025 |
| VLTGGGAVL | ASSLGNSYNSPLY | 0.6079993 |
| VNVGLAYGI | ASSLGNSYNSPLY | 0.5940244 |
| VSSAAVGAV | ASSLGNSYNSPLY | 0.68108696 |
| VTSVYEAGV | ASSLGNSYNSPLY | 0.57212925 |
| WVALDRGA | ASSLGNSYNSPLY | 0.4916018 |
| YLVGSVVI | ASSLGNSYNSPLY | 0.46584165 |
| GPPGTGKTLL | ASSLGNSYNSPLY | 0.6553192 |
| GVGMTKF | ASSLGNSYNSPLY | 0.7042702 |
| IGIGGSDL | ASSLGNSYNSPLY | 0.6543753 |
| IGTLLAEL | ASSLGNSYNSPLY | 0.5542694 |
| IYTSGTT | ASSLGNSYNSPLY | 0.68530643 |
| LSFPAGV | ASSLGNSYNSPLY | 0.6540757 |
| LYTSGTT | ASSLGNSYNSPLY | 0.6580041 |
| VGAFPFGF | ASSLGNSYNSPLY | 0.66182 |
| VGAGTVEF | ASSLGNSYNSPLY | 0.6149999 |
| VGKSTLLL | ASSLGNSYNSPLY | 0.6830771 |
| AALAHAGGGIV | ASSLGNSYNSPLY | 0.67428434 |
| AALVAVSLL | ASSLGNSYNSPLY | 0.683968 |
| AAVSRAVAAL | ASSLGNSYNSPLY | 0.69331884 |
| AGGSSGIGGF | ASSLGNSYNSPLY | 0.63987666 |
| AGLTVANANG | ASSLGNSYNSPLY | 0.6369463 |
| AGLVGAVL | ASSLGNSYNSPLY | 0.32950377 |
| ALSQAFTV | ASSLGNSYNSPLY | 0.59899336 |
| ALTFSVSL | ASSLGNSYNSPLY | 0.68100905 |
| ALTGAALVAL | ASSLGNSYNSPLY | 0.6695215 |
| ASFSIGAGVL | ASSLGNSYNSPLY | 0.69466305 |
| DGSVYKI | ASSLGNSYNSPLY | 0.6452657 |
| EQLAEIV | ASSLGNSYNSPLY | 0.5795331 |
| FAVSELTVM | ASSLGNSYNSPLY | 0.67880666 |
| GAAPLAAGL | ASSLGNSYNSPLY | 0.69889605 |
| GADGGDGGSGGAGG | ASSLGNSYNSPLY | 0.47416162 |
| GFGLDTGGAGG | ASSLGNSYNSPLY | 0.62962306 |
| GLASGGLL | ASSLGNSYNSPLY | 0.72275305 |
| GRIAGAAVLL | ASSLGNSYNSPLY | 0.6567914 |
| GSAGSLNTSF | ASSLGNSYNSPLY | 0.63476163 |

| | | |
|---|---|---|
| GTGQIGI | ASSLGNSYNSPLY | 0.5755949 |
| GTLIALIL | ASSLGNSYNSPLY | 0.71465653 |
| IAATMGFVVG | ASSLGNSYNSPLY | 0.5430646 |
| IFAQALL | ASSLGNSYNSPLY | 0.71558464 |
| IGAAVALGF | ASSLGNSYNSPLY | 0.29989517 |
| IGLTPLAL | ASSLGNSYNSPLY | 0.51418227 |
| ILGSNTSTL | ASSLGNSYNSPLY | 0.45061314 |
| IVFSLTVL | ASSLGNSYNSPLY | 0.38364905 |
| LASKATGF | ASSLGNSYNSPLY | 0.70038027 |
| LASVALIL | ASSLGNSYNSPLY | 0.5644232 |
| LFAVGLTL | ASSLGNSYNSPLY | 0.6924023 |
| LFNGGIGTYI | ASSLGNSYNSPLY | 0.67669296 |
| LTLGAGII | ASSLGNSYNSPLY | 0.70478153 |
| LVAAALGDI | ASSLGNSYNSPLY | 0.6594462 |
| NTAFVQLGI | ASSLGNSYNSPLY | 0.65621364 |
| QLFSGAEL | ASSLGNSYNSPLY | 0.5293218 |
| SAATFITL | ASSLGNSYNSPLY | 0.69091296 |
| SNYGLANLGV | ASSLGNSYNSPLY | 0.6966568 |
| SYGGAYAVM | ASSLGNSYNSPLY | 0.62383103 |
| TTGVGGLTLG | ASSLGNSYNSPLY | 0.47685456 |
| TTQTIYI | ASSLGNSYNSPLY | 0.600908 |
| VAAGAGLGAVI | ASSLGNSYNSPLY | 0.66584027 |
| VAAVVTLL | ASSLGNSYNSPLY | 0.315064 |
| VGFLGGLI | ASSLGNSYNSPLY | 0.31463024 |
| VGVIPILL | ASSLGNSYNSPLY | 0.5760496 |
| VIFASSLI | ASSLGNSYNSPLY | 0.5738035 |
| VKGGDGGAAAG | ASSLGNSYNSPLY | 0.70717144 |
| VTKTFTNI | ASSLGNSYNSPLY | 0.70018697 |
| VVAAAILL | ASSLGNSYNSPLY | 0.64184964 |
| YAALDVEL | ASSLGNSYNSPLY | 0.6761811 |
| LASVALLL | ASSLGNSYNSPLY | 0.6689545 |
| LDAVHRV | ASSLGNSYNSPLY | 0.2573496 |
| QGLSSSIF | ASSLGNSYNSPLY | 0.56738114 |
| RGTFANIRL | ASSLGNSYNSPLY | 0.5258366 |
| RQLWWGHRI | ASSLGNSYNSPLY | 0.7090583 |
| YGPPGTGKTLL | ASSLGNSYNSPLY | 0.615257 |
| AALVGAPANVA | ASSLGNSYNSPLY | 0.59582967 |
| AATFRRTE | ASSLGNSYNSPLY | 0.6556094 |
| AGALGGGI | ASSLGNSYNSPLY | 0.65491503 |
| AGGAGGAGGIGAG | ASSLGNSYNSPLY | 0.6458592 |
| AGSGALGL | ASSLGNSYNSPLY | 0.5057286 |
| AQIADQI | ASSLGNSYNSPLY | 0.27277982 |
| ASAVAVTYL | ASSLGNSYNSPLY | 0.6994889 |
| ASTVLGGTGG | ASSLGNSYNSPLY | 0.388893 |
| AVGGEPAK | ASSLGNSYNSPLY | 0.55429006 |
| CEQGTFGGTC | ASSLGNSYNSPLY | 0.32146633 |
| DEGKAPAG | ASSLGNSYNSPLY | 0.5530784 |
| DTVVFPAL | ASSLGNSYNSPLY | 0.41675538 |
| FEGITMAG | ASSLGNSYNSPLY | 0.68465495 |
| FPLVVQD | ASSLGNSYNSPLY | 0.63856304 |
| GAFTTPQ | ASSLGNSYNSPLY | 0.6350324 |

| | | |
|---|---|---|
| GFGGFGTAGGVGGA | ASSLGNSYNSPLY | 0.71758306 |
| GFLAQFL | ASSLGNSYNSPLY | 0.6560147 |
| GGAGGAGGNAGVA | ASSLGNSYNSPLY | 0.6292099 |
| GGVGTAGAGGAGGAGG | ASSLGNSYNSPLY | 0.5795391 |
| GNGGGGDAGNAGSGGN | ASSLGNSYNSPLY | 0.5764687 |
| GTGGAGGAGNDAGS | ASSLGNSYNSPLY | 0.5312861 |
| GTTNTGWFNS | ASSLGNSYNSPLY | 0.5566009 |
| IFMGPLFTL | ASSLGNSYNSPLY | 0.7174569 |
| ILAHAIEAI | ASSLGNSYNSPLY | 0.54098594 |
| IRSAIREAL | ASSLGNSYNSPLY | 0.5004237 |
| ITFAGGG | ASSLGNSYNSPLY | 0.54228944 |
| ITGMSGHQEV | ASSLGNSYNSPLY | 0.7141605 |
| IVAGLACTL | ASSLGNSYNSPLY | 0.71479166 |
| IVGGGLAGA | ASSLGNSYNSPLY | 0.59151345 |
| IVTGVLNI | ASSLGNSYNSPLY | 0.54033834 |
| LAQQGCI | ASSLGNSYNSPLY | 0.57538605 |
| LCLHVEN | ASSLGNSYNSPLY | 0.66326916 |
| LDAISDGS | ASSLGNSYNSPLY | 0.5334821 |
| LGGFAAML | ASSLGNSYNSPLY | 0.4631502 |
| LGGKNPMI | ASSLGNSYNSPLY | 0.6630211 |
| LGPKGVAAIL | ASSLGNSYNSPLY | 0.71664345 |
| LMVGVSI | ASSLGNSYNSPLY | 0.5739212 |
| LVGAAFVL | ASSLGNSYNSPLY | 0.53641015 |
| LVLAVAPL | ASSLGNSYNSPLY | 0.6489627 |
| MAAIAVIDAA | ASSLGNSYNSPLY | 0.5495223 |
| NNGINAELQ | ASSLGNSYNSPLY | 0.621897 |
| RGEPLVLVHG | ASSLGNSYNSPLY | 0.6214872 |
| RGGAGGAAGLI | ASSLGNSYNSPLY | 0.50096166 |
| RSYRMHPAV | ASSLGNSYNSPLY | 0.6381852 |
| SAAHLLAGQLA | ASSLGNSYNSPLY | 0.71466315 |
| SAAVGSY | ASSLGNSYNSPLY | 0.626073 |
| SPHAVLHRI | ASSLGNSYNSPLY | 0.6694075 |
| SQIAKVL | ASSLGNSYNSPLY | 0.70887077 |
| TALGHLVP | ASSLGNSYNSPLY | 0.6902908 |
| TAPTTSGAA | ASSLGNSYNSPLY | 0.7087265 |
| TDGDDGNGGSPGT | ASSLGNSYNSPLY | 0.6904831 |
| TDGLGEN | ASSLGNSYNSPLY | 0.63747126 |
| TGGPPLPGQA | ASSLGNSYNSPLY | 0.6249856 |
| TIPAITI | ASSLGNSYNSPLY | 0.68108976 |
| TIPGFTI | ASSLGNSYNSPLY | 0.66665995 |
| TSIASGALA | ASSLGNSYNSPLY | 0.63176334 |
| TVTLPGAAI | ASSLGNSYNSPLY | 0.5323254 |
| VASGLSAALP | ASSLGNSYNSPLY | 0.7072189 |
| VGAGVIGI | ASSLGNSYNSPLY | 0.5939585 |
| VGPKRFGV | ASSLGNSYNSPLY | 0.6893117 |
| VNVAFPD | ASSLGNSYNSPLY | 0.45396274 |
| VSMTNSGVS | ASSLGNSYNSPLY | 0.50572 |
| VTDVLGDLSV | ASSLGNSYNSPLY | 0.6935755 |
| VVVVGAGFAGL | ASSLGNSYNSPLY | 0.25988176 |
| VYTVGTVV | ASSLGNSYNSPLY | 0.29915404 |
| WTTFSGP | ASSLGNSYNSPLY | 0.6251592 |

| | | |
|---|---|---|
| YDELVEAF | ASSLGNSYNSPLY | 0.70041263 |
| YGTGGAGGNGG | ASSLGNSYNSPLY | 0.26753807 |
| YTNVMAV | ASSLGNSYNSPLY | 0.53016293 |
| FLEQVHQL | ASSLGNSYNSPLY | 0.44545934 |
| GLSLLAVAV | ASSLGNSYNSPLY | 0.6402052 |
| LEQVHQL | ASSLGNSYNSPLY | 0.6776239 |
| LNAAGAI | ASSLGNSYNSPLY | 0.43286717 |
| PAVHASL | ASSLGNSYNSPLY | 0.6826915 |
| TELEDDA | ASSLGNSYNSPLY | 0.4910801 |
| TPPGPPN | ASSLGNSYNSPLY | 0.46927845 |
| TPPPPPPP | ASSLGNSYNSPLY | 0.69406366 |
| TQSPALI | ASSRLGVNYAEQF | 0.70207775 |
| EFAAAFPQ | ASSRLGVNYAEQF | 0.5655612 |
| LLSGNLDVL | ASSRLGVNYAEQF | 0.6893443 |
| GPANLGNYNI | ASSRLGVNYAEQF | 0.5621126 |
| TLGPTLP | ASSRLGVNYAEQF | 0.717162 |
| VYAYSSCCK | ASSRLGVNYAEQF | 0.613561 |
| APAPAPAPNI | ASSRLGVNYAEQF | 0.4231851 |
| PGGPVPIP | ASSRLGVNYAEQF | 0.6906574 |
| FLTVALEP | ASSRLGVNYAEQF | 0.6635419 |
| SQYFGANL | ASSRLGVNYAEQF | 0.62529075 |
| GGVKPAA | ASSRLGVNYAEQF | 0.6904713 |
| GGTGGAGGAGGW | ASSRLGVNYAEQF | 0.46535343 |
| GVGGVAGLGGAGG | ASSRLGVNYAEQF | 0.6522577 |
| VLLVLATTI | ASSRLGVNYAEQF | 0.7083021 |
| IGIFAAML | ASSRLGVNYAEQF | 0.7029232 |
| AVGDLID | ASSRLGVNYAEQF | 0.48174205 |
| NISPLPL | ASSRLGVNYAEQF | 0.6744148 |
| VTAVFVAI | ASSRLGVNYAEQF | 0.6466477 |
| SGDDNDAAL | ASSRLGVNYAEQF | 0.59398437 |
| AVAALAHS | ASSRLGVNYAEQF | 0.7063233 |
| GGVTVMATP | ASSRLGVNYAEQF | 0.59826607 |
| AGLAANVRD | ASSRLGVNYAEQF | 0.6923946 |
| TNRPDLIDPALLRPG | ASSRLGVNYAEQF | 0.6793806 |
| LANLGVTAL | ASSRLGVNYAEQF | 0.69855636 |
| TGERIEL | ASSRLGVNYAEQF | 0.5006248 |
| NGEELAA | ASSRLGVNYAEQF | 0.6445556 |
| VLSLVWG | ASSRLGVNYAEQF | 0.6486322 |
| IGYALWL | ASSRLGVNYAEQF | 0.69698346 |
| LSLLLAPL | ASSRLGVNYAEQF | 0.6150465 |
| TLDEALK | ASSRLGVNYAEQF | 0.47826177 |
| TDGAGTAEGG | ASSRLGVNYAEQF | 0.66937447 |
| NAPGGAGGNGG | ASSRLGVNYAEQF | 0.70493954 |
| VSVFMGMGGNF | ASSRLGVNYAEQF | 0.6467889 |
| LASYLAAL | ASSRLGVNYAEQF | 0.6540766 |
| LVGGTPAL | ASSRLGVNYAEQF | 0.71233207 |
| VRNADVL | ASSRLGVNYAEQF | 0.5635341 |
| TCSSPFLVSAM | ASSRLGVNYAEQF | 0.6379675 |
| VHITDAE | ASSRLGVNYAEQF | 0.67200416 |
| EYAAPSF | ASSRLGVNYAEQF | 0.69790536 |
| VVDAFHGGL | ASSRLGVNYAEQF | 0.6081248 |

| | | |
|---|---|---|
| ASVVDPQ | ASSRLGVNYAEQF | 0.7040353 |
| SYNEFGGLS | ASSRLGVNYAEQF | 0.5525895 |
| DPDGSVY | ASSRLGVNYAEQF | 0.69964004 |
| MVSTIHATE | ASSRLGVNYAEQF | 0.59603345 |
| LAAGVADPV | ASSRLGVNYAEQF | 0.6920512 |
| TKEQIAA | ASSRLGVNYAEQF | 0.5692479 |
| SLASPAAL | ASSRLGVNYAEQF | 0.7060177 |
| IGLVFYGV | ASSRLGVNYAEQF | 0.696069 |
| FLLDASNL | ASSRLGVNYAEQF | 0.71038043 |
| YQVGNFP | ASSRLGVNYAEQF | 0.3005246 |
| ESLLSQST | ASSRLGVNYAEQF | 0.7040377 |
| LFTVGFAAVG | ASSRLGVNYAEQF | 0.7045538 |
| ATPPTPT | ASSRLGVNYAEQF | 0.7193912 |
| TDLATLG | ASSRLGVNYAEQF | 0.60367393 |
| SAGGTGR | ASSRLGVNYAEQF | 0.65832305 |
| GFSNSGS | ASSRLGVNYAEQF | 0.6167712 |
| GGNGGSGGTSV | ASSRLGVNYAEQF | 0.586923 |
| GTSVATGGAGN | ASSRLGVNYAEQF | 0.55627286 |
| NGGNGGSGGTS | ASSRLGVNYAEQF | 0.5892049 |
| AVVKFPH | ASSRLGVNYAEQF | 0.63186073 |
| TPPPPPP | ASSRLGVNYAEQF | 0.60376245 |
| AAPAAAAAPA | ASSRLGVNYAEQF | 0.6917188 |
| IALNEDL | ASSRLGVNYAEQF | 0.7024753 |
| IVNVSSM | ASSRLGVNYAEQF | 0.701141 |
| TGIPPAPR | ASSRLGVNYAEQF | 0.7023845 |
| FDLGGGTF | ASSRLGVNYAEQF | 0.7047907 |
| FDLGGGTFDVS | ASSRLGVNYAEQF | 0.67644656 |
| NEPTAAA | ASSRLGVNYAEQF | 0.57169724 |
| NEPTAAALA | ASSRLGVNYAEQF | 0.6574706 |
| GSIINISSI | ASSRLGVNYAEQF | 0.55429494 |
| VAPGNAGTA | ASSRLGVNYAEQF | 0.6489477 |
| TGGGASLE | ASSRLGVNYAEQF | 0.60342205 |
| VAVYNHYKRI | ASSRLGVNYAEQF | 0.68144256 |
| TDPAPATT | ASSRLGVNYAEQF | 0.26032147 |
| TNRPDILDPALLRPG | ASSRLGVNYAEQF | 0.665661 |
| LTTTTTS | ASSRLGVNYAEQF | 0.71517956 |
| PTGSGKTL | ASSRLGVNYAEQF | 0.5297608 |
| TGSGKTL | ASSRLGVNYAEQF | 0.6639778 |
| TIGHVDHGK | ASSRLGVNYAEQF | 0.66265434 |
| TIGHVDHGKTT | ASSRLGVNYAEQF | 0.58775806 |
| GHVDHGKTT | ASSRLGVNYAEQF | 0.686587 |
| AQFEHTLL | ASSRLGVNYAEQF | 0.52412677 |
| IVNQDEL | ASSRLGVNYAEQF | 0.6683178 |
| TNGSQFF | ASSRLGVNYAEQF | 0.7072698 |
| TGKGIVE | ASSRLGVNYAEQF | 0.67204434 |
| LISNASDA | ASSRLGVNYAEQF | 0.60175884 |
| AELRTLGELTKIAKAHGAQ | ASSRLGVNYAEQF | 0.6112381 |
| APNAVAAAEPSW | ASSRLGVNYAEQF | 0.43572432 |
| APQVAGAATAVL | ASSRLGVNYAEQF | 0.6676269 |
| ARRDPAAAAAILPTDA | ASSRLGVNYAEQF | 0.29989862 |
| DAAARDVLAEAGLAEYF | ASSRLGVNYAEQF | 0.647992 |

| | | |
|---|---|---|
| DDQIVLSAKANLAKVEQ | ASSRLGVNYAEQF | 0.5031531 |
| DELLPDTGEDSDEDGH | ASSRLGVNYAEQF | 0.67934805 |
| ELPEAPANRATVPQD | ASSRLGVNYAEQF | 0.67454547 |
| FGQSKIERVTGIG | ASSRLGVNYAEQF | 0.6566669 |
| GAGGTGSTAGGAGGAGGAGGLY | ASSRLGVNYAEQF | 0.5818573 |
| GHGGTGAAGGNGGTGGAGGSGI | ASSRLGVNYAEQF | 0.6394433 |
| GNGGNGGAGGEGGPGGAGGAGA | ASSRLGVNYAEQF | 0.5824095 |
| GSTAGNGGNGGNATNGTIAGQP | ASSRLGVNYAEQF | 0.48720992 |
| GTGGDGGNGGNGGTGG | ASSRLGVNYAEQF | 0.69130766 |
| GVRRPAGIPADAPV | ASSRLGVNYAEQF | 0.5609609 |
| GYRQVTDAHLLTLARR | ASSRLGVNYAEQF | 0.6305166 |
| IAKMGQDGHDRG | ASSRLGVNYAEQF | 0.70875704 |
| KAEPSIEIQYT | ASSRLGVNYAEQF | 0.6607299 |
| KPVRIRMAADEDTVAVT | ASSRLGVNYAEQF | 0.4256915 |
| MPPRIAGMRLLVIKPEP | ASSRLGVNYAEQF | 0.6356621 |
| MKFYAVALFRSRGVPPTR | ASSRLGVNYAEQF | 0.6497346 |
| NAGGFGILGAVAHSP | ASSRLGVNYAEQF | 0.6209502 |
| NEGREITLAMAEM | ASSRLGVNYAEQF | 0.26660004 |
| NELVEWSHPLGHRWRWE | ASSRLGVNYAEQF | 0.6714374 |
| NLGAGGTGGNGGNPG | ASSRLGVNYAEQF | 0.5950943 |
| NTEWPKWETVMAADPVD | ASSRLGVNYAEQF | 0.5819437 |
| PASAAPAETQP | ASSRLGVNYAEQF | 0.59685063 |
| QQSSDGKNGVRAAEITGE | ASSRLGVNYAEQF | 0.6960056 |
| RAPDAVTTSWWKE | ASSRLGVNYAEQF | 0.5072408 |
| SGFGNVGAHTSGWFNQ | ASSRLGVNYAEQF | 0.61455464 |
| SGVAGAVGQAASVGGLKV | ASSRLGVNYAEQF | 0.6879158 |
| SNGGAGTGSAGGNGGT | ASSRLGVNYAEQF | 0.69531435 |
| TGGAGGEGGAAG | ASSRLGVNYAEQF | 0.71306205 |
| TGGDGGAAGAGGAPGHG | ASSRLGVNYAEQF | 0.65714455 |
| TMPPLLAAIANAGRHGV | ASSRLGVNYAEQF | 0.5416532 |
| TSSTAGNSGGA | ASSRLGVNYAEQF | 0.67081916 |
| VPATVPVGILC | ASSRLGVNYAEQF | 0.5967561 |
| VQRPDSGGVVLTGRIS | ASSRLGVNYAEQF | 0.43650633 |
| WTPNPVPATKGVPAGLVQAT | ASSRLGVNYAEQF | 0.34595463 |
| YALKRALAKFSDRFDVVII | ASSRLGVNYAEQF | 0.656574 |
| YLIGSVIV | ASSRLGVNYAEQF | 0.3787345 |
| YGQLVAVID | ASSRLGVNYAEQF | 0.64842033 |
| VTVGGMTLPA | ASSRLGVNYAEQF | 0.6766078 |
| VLALPAV | ASSRLGVNYAEQF | 0.57721806 |
| VIPPTLNYET | ASSRLGVNYAEQF | 0.45794368 |
| VGMGGMHPSSG | ASSRLGVNYAEQF | 0.60652125 |
| VGLAGGAATA | ASSRLGVNYAEQF | 0.65583044 |
| VGGAGGAGGSGP | ASSRLGVNYAEQF | 0.6937107 |
| VFAAFMV | ASSRLGVNYAEQF | 0.6510178 |
| TLYDKLR | ASSRLGVNYAEQF | 0.7184051 |
| TLAVFTVL | ASSRLGVNYAEQF | 0.7076119 |
| TGSVQPARV | ASSRLGVNYAEQF | 0.6250456 |
| TGGGAGSGGT | ASSRLGVNYAEQF | 0.6837465 |
| SHLETSL | ASSRLGVNYAEQF | 0.61986744 |
| SFTTGMT | ASSRLGVNYAEQF | 0.6315548 |
| QVEVAKRMGVRQPTVS | ASSRLGVNYAEQF | 0.6347727 |

| | | |
|---|---|---|
| QGSGGPALAM | ASSRLGVNYAEQF | 0.6069258 |
| MLAPSNP | ASSRLGVNYAEQF | 0.59434783 |
| LVNNAAI | ASSRLGVNYAEQF | 0.6177877 |
| LTGEINTLA | ASSRLGVNYAEQF | 0.39933145 |
| LSIFGPV | ASSRLGVNYAEQF | 0.6356143 |
| LLALLANL | ASSRLGVNYAEQF | 0.65100384 |
| LISNASDALDKLR | ASSRLGVNYAEQF | 0.35432196 |
| LINEAVP | ASSRLGVNYAEQF | 0.6064484 |
| LGYAFSGL | ASSRLGVNYAEQF | 0.71628416 |
| LDFVAVAL | ASSRLGVNYAEQF | 0.6979535 |
| LAQPGAE | ASSRLGVNYAEQF | 0.5815984 |
| ISNLGAVTGF | ASSRLGVNYAEQF | 0.52236974 |
| HQEDPGRI | ASSRLGVNYAEQF | 0.65098214 |
| GTPGEPTP | ASSRLGVNYAEQF | 0.3589541 |
| GFFNSGS | ASSRLGVNYAEQF | 0.6383005 |
| GAGGGGLTAGAN | ASSRLGVNYAEQF | 0.66042864 |
| FVDALVNM | ASSRLGVNYAEQF | 0.66737735 |
| FLTVAPEL | ASSRLGVNYAEQF | 0.50511163 |
| FETLTAL | ASSRLGVNYAEQF | 0.66105694 |
| DVINPIEL | ASSRLGVNYAEQF | 0.62419283 |
| APVSGAPV | ASSRLGVNYAEQF | 0.6366014 |
| ALVGGLAW | ASSRLGVNYAEQF | 0.26679674 |
| ALAAAAATAI | ASSRLGVNYAEQF | 0.66916394 |
| AFGLGLLL | ASSRLGVNYAEQF | 0.58114564 |
| AFFAAASL | ASSRLGVNYAEQF | 0.5242518 |
| AAAALPAFSPP | ASSRLGVNYAEQF | 0.3607032 |
| VVLAPAL | ASSRLGVNYAEQF | 0.66291076 |
| TIGHVDHGKTTL | ASSRLGVNYAEQF | 0.69175994 |
| LISNASDALDK | ASSRLGVNYAEQF | 0.6904218 |
| LISNASDAL | ASSRLGVNYAEQF | 0.49924302 |
| ISNASDALDK | ASSRLGVNYAEQF | 0.7050507 |
| NQAETLVYQTE | ASSRLGVNYAEQF | 0.7043802 |
| TGGTGGTAGAG | ASSRLGVNYAEQF | 0.63791656 |
| TGSTAGGAGGA | ASSRLGVNYAEQF | 0.4194039 |
| DPLAVPVAAGPV | ASSRLGVNYAEQF | 0.679014 |
| GGTGGTPGNGGN | ASSRLGVNYAEQF | 0.62835014 |
| PAGLAGVDPATL | ASSRLGVNYAEQF | 0.6836514 |
| GNNLVVEAPHAGASV | ASSRLGVNYAEQF | 0.66186035 |
| KVTRIMSASTLEQPA | ASSRLGVNYAEQF | 0.5635217 |
| LGFALPIALPRRTVV | ASSRLGVNYAEQF | 0.67370313 |
| TRLGNLTARKLVVLV | ASSRLGVNYAEQF | 0.7013031 |
| WSQQRRLIYFPSAGP | ASSRLGVNYAEQF | 0.5545014 |
| GPNGNPDPMAAAVDIR | ASSRLGVNYAEQF | 0.55874205 |
| TQRCVSVADLRAAASR | ASSRLGVNYAEQF | 0.5978462 |
| QFVLDVAEHPVEVLTETV | ASSRLGVNYAEQF | 0.3577519 |
| GGAGGTGGAGGNGGTQAGDG | ASSRLGVNYAEQF | 0.6904119 |
| PLLATLRAIAVDVDAKPAQV | ASSRLGVNYAEQF | 0.61279 |
| APKFRTGGYMIDFWGVGYQVAK | ASSRLGVNYAEQF | 0.5708371 |
| GAGGAGGAGGAAGGPGGTGGTG | ASSRLGVNYAEQF | 0.67773616 |
| IGALGLLATDVAPAGSVAFGVA | ASSRLGVNYAEQF | 0.568144 |
| AGGAGGIGGNGGAGGAGGVLMG | ASSRLGVNYAEQF | 0.60333043 |

| | | |
|---|---|---|
| ADELIGLL | ASSRLGVNYAEQF | 0.6240626 |
| ADRALKA | ASSRLGVNYAEQF | 0.6705669 |
| AGAVGQVGGAGGA | ASSRLGVNYAEQF | 0.6658494 |
| AGGGGGLGGAP | ASSRLGVNYAEQF | 0.6622127 |
| AGGLLYGNGGAG | ASSRLGVNYAEQF | 0.62903905 |
| AGLESIGL | ASSRLGVNYAEQF | 0.71484673 |
| AGLKTGII | ASSRLGVNYAEQF | 0.47259277 |
| ASLAALLL | ASSRLGVNYAEQF | 0.61499727 |
| ASVINISSI | ASSRLGVNYAEQF | 0.66110355 |
| ATPPGPA | ASSRLGVNYAEQF | 0.41360635 |
| ATVASALV | ASSRLGVNYAEQF | 0.682897 |
| ATVASLIL | ASSRLGVNYAEQF | 0.66528803 |
| AVNGPVLIRH | ASSRLGVNYAEQF | 0.6785526 |
| DGAAGAVGI | ASSRLGVNYAEQF | 0.63017696 |
| DPLLHGI | ASSRLGVNYAEQF | 0.7166134 |
| ELGAALGAL | ASSRLGVNYAEQF | 0.6978964 |
| ELGRTQAV | ASSRLGVNYAEQF | 0.6440753 |
| FVAGGYLS | ASSRLGVNYAEQF | 0.6847043 |
| GASGGAGGAGGW | ASSRLGVNYAEQF | 0.45929545 |
| GGAGAGGDGGAGGI | ASSRLGVNYAEQF | 0.7018541 |
| GTGHNGSQPGMGGNG | ASSRLGVNYAEQF | 0.6611481 |
| GYAAASASAS | ASSRLGVNYAEQF | 0.56812596 |
| HQVDVAAVQQG | ASSRLGVNYAEQF | 0.65462565 |
| IGAAIAHT | ASSRLGVNYAEQF | 0.70044374 |
| IGMIEVAQL | ASSRLGVNYAEQF | 0.70279515 |
| IRLAPEHTSAMD | ASSRLGVNYAEQF | 0.36931837 |
| IYNVSVL | ASSRLGVNYAEQF | 0.6432796 |
| KQGFLGM | ASSRLGVNYAEQF | 0.33365798 |
| LESVITQL | ASSRLGVNYAEQF | 0.4702691 |
| LGLGALVVV | ASSRLGVNYAEQF | 0.523244 |
| LIGNGGNGGSGGI | ASSRLGVNYAEQF | 0.67495775 |
| LNAINAPTL | ASSRLGVNYAEQF | 0.62661576 |
| LQLLNTGV | ASSRLGVNYAEQF | 0.6686722 |
| LQTSIEF | ASSRLGVNYAEQF | 0.6548595 |
| MSFAVSEL | ASSRLGVNYAEQF | 0.7077199 |
| NTGALGL | ASSRLGVNYAEQF | 0.55329585 |
| PPGAGGAGGAGTT | ASSRLGVNYAEQF | 0.6655755 |
| QLVGLGI | ASSRLGVNYAEQF | 0.38898382 |
| RFLEQVHQL | ASSRLGVNYAEQF | 0.41830528 |
| RGVGYVLR | ASSRLGVNYAEQF | 0.5303607 |
| SATGVFAGVI | ASSRLGVNYAEQF | 0.64127785 |
| SLPAITLP | ASSRLGVNYAEQF | 0.30207202 |
| STALAFTV | ASSRLGVNYAEQF | 0.7016591 |
| SVLTVGLGK | ASSRLGVNYAEQF | 0.6627242 |
| TALAGAGSGI | ASSRLGVNYAEQF | 0.6854317 |
| TGLPFTGI | ASSRLGVNYAEQF | 0.59067047 |
| VAEHVTV | ASSRLGVNYAEQF | 0.6285337 |
| VDIVLNSL | ASSRLGVNYAEQF | 0.5978066 |
| VLTGGGAVL | ASSRLGVNYAEQF | 0.6807113 |
| VNVGLAYGI | ASSRLGVNYAEQF | 0.61685175 |
| VSSAAVGAV | ASSRLGVNYAEQF | 0.39839563 |

| | | |
|---|---|---|
| VTSVYEAGV | ASSRLGVNYAEQF | 0.5659528 |
| WVALDRGA | ASSRLGVNYAEQF | 0.55759436 |
| YLVGSVVI | ASSRLGVNYAEQF | 0.7008244 |
| GPPGTGKTLL | ASSRLGVNYAEQF | 0.69692075 |
| GVGMTKF | ASSRLGVNYAEQF | 0.67974627 |
| IGIGGSDL | ASSRLGVNYAEQF | 0.68415236 |
| IGTLLAEL | ASSRLGVNYAEQF | 0.30844587 |
| IYTSGTT | ASSRLGVNYAEQF | 0.59909517 |
| LSFPAGV | ASSRLGVNYAEQF | 0.7144674 |
| LYTSGTT | ASSRLGVNYAEQF | 0.66009223 |
| VGAFPFGF | ASSRLGVNYAEQF | 0.6438009 |
| VGAGTVEF | ASSRLGVNYAEQF | 0.7054454 |
| VGKSTLLL | ASSRLGVNYAEQF | 0.6026199 |
| AALAHAGGGIV | ASSRLGVNYAEQF | 0.6171179 |
| AALVAVSLL | ASSRLGVNYAEQF | 0.6577265 |
| AAVSRAVAAL | ASSRLGVNYAEQF | 0.66990554 |
| AGGSSGIGGF | ASSRLGVNYAEQF | 0.61754423 |
| AGLTVANANG | ASSRLGVNYAEQF | 0.55963314 |
| AGLVGAVL | ASSRLGVNYAEQF | 0.6562884 |
| ALSQAFTV | ASSRLGVNYAEQF | 0.72192645 |
| ALTFSVSL | ASSRLGVNYAEQF | 0.63823915 |
| ALTGAALVAL | ASSRLGVNYAEQF | 0.69774574 |
| ASFSIGAGVL | ASSRLGVNYAEQF | 0.6734947 |
| DGSVYKI | ASSRLGVNYAEQF | 0.67289174 |
| EQLAEIV | ASSRLGVNYAEQF | 0.57244986 |
| FAVSELTVM | ASSRLGVNYAEQF | 0.6178159 |
| GAAPLAAGL | ASSRLGVNYAEQF | 0.6487462 |
| GADGGDGGSGGAGG | ASSRLGVNYAEQF | 0.5794531 |
| GFGLDTGGAGG | ASSRLGVNYAEQF | 0.62350714 |
| GLASGGLL | ASSRLGVNYAEQF | 0.61673427 |
| GRIAGAAVLL | ASSRLGVNYAEQF | 0.7226926 |
| GSAGSLNTSF | ASSRLGVNYAEQF | 0.6792593 |
| GTGQIGI | ASSRLGVNYAEQF | 0.6839479 |
| GTLIALIL | ASSRLGVNYAEQF | 0.5789385 |
| IAATMGFVVG | ASSRLGVNYAEQF | 0.643983 |
| IFAQALL | ASSRLGVNYAEQF | 0.6798512 |
| IGAAVALGF | ASSRLGVNYAEQF | 0.46873063 |
| IGLTPLAL | ASSRLGVNYAEQF | 0.7095628 |
| ILGSNTSTL | ASSRLGVNYAEQF | 0.5807198 |
| IVFSLTVL | ASSRLGVNYAEQF | 0.57133514 |
| LASKATGF | ASSRLGVNYAEQF | 0.6921538 |
| LASVALIL | ASSRLGVNYAEQF | 0.71256363 |
| LFAVGLTL | ASSRLGVNYAEQF | 0.6118826 |
| LFNGGIGTYI | ASSRLGVNYAEQF | 0.61828554 |
| LTLGAGII | ASSRLGVNYAEQF | 0.6120347 |
| LVAAALGDI | ASSRLGVNYAEQF | 0.67644083 |
| NTAFVQLGI | ASSRLGVNYAEQF | 0.7012181 |
| QLFSGAEL | ASSRLGVNYAEQF | 0.61927027 |
| SAATFITL | ASSRLGVNYAEQF | 0.7138776 |
| SNYGLANLGV | ASSRLGVNYAEQF | 0.6238875 |
| SYGGAYAVM | ASSRLGVNYAEQF | 0.4071525 |

| | | |
|---|---|---|
| TTGVGGLTLG | ASSRLGVNYAEQF | 0.6311229 |
| TTQTIYI | ASSRLGVNYAEQF | 0.6977578 |
| VAAGAGLGAVI | ASSRLGVNYAEQF | 0.6957721 |
| VAAVVTLL | ASSRLGVNYAEQF | 0.7115943 |
| VGFLGGLI | ASSRLGVNYAEQF | 0.6506656 |
| VGVIPILL | ASSRLGVNYAEQF | 0.60802484 |
| VIFASSLI | ASSRLGVNYAEQF | 0.7096575 |
| VKGGDGGAAAG | ASSRLGVNYAEQF | 0.7049544 |
| VTKTFTNI | ASSRLGVNYAEQF | 0.6838608 |
| VVAAAILL | ASSRLGVNYAEQF | 0.5397804 |
| YAALDVEL | ASSRLGVNYAEQF | 0.45255283 |
| LASVALLL | ASSRLGVNYAEQF | 0.6575528 |
| LDAVHRV | ASSRLGVNYAEQF | 0.69649947 |
| QGLSSSIF | ASSRLGVNYAEQF | 0.46352795 |
| RGTFANIRL | ASSRLGVNYAEQF | 0.67652303 |
| RQLWWGHRI | ASSRLGVNYAEQF | 0.66376257 |
| YGPPGTGKTLL | ASSRLGVNYAEQF | 0.46716952 |
| AALVGAPANVA | ASSRLGVNYAEQF | 0.67824835 |
| AATFRRTE | ASSRLGVNYAEQF | 0.70776117 |
| AGALGGGI | ASSRLGVNYAEQF | 0.6219615 |
| AGGAGGAGGIGAG | ASSRLGVNYAEQF | 0.6015865 |
| AGSGALGL | ASSRLGVNYAEQF | 0.7021972 |
| AQIADQI | ASSRLGVNYAEQF | 0.7213441 |
| ASAVAVTYL | ASSRLGVNYAEQF | 0.7051314 |
| ASTVLGGTGG | ASSRLGVNYAEQF | 0.43654782 |
| AVGGEPAK | ASSRLGVNYAEQF | 0.34340432 |
| CEQGTFGGTC | ASSRLGVNYAEQF | 0.58118796 |
| DEGKAPAG | ASSRLGVNYAEQF | 0.68348527 |
| DTVVFPAL | ASSRLGVNYAEQF | 0.61789155 |
| FEGITMAG | ASSRLGVNYAEQF | 0.55352336 |
| FPLVVQD | ASSRLGVNYAEQF | 0.6801665 |
| GAFTTPQ | ASSRLGVNYAEQF | 0.70469546 |
| GFGGFGTAGGVGGA | ASSRLGVNYAEQF | 0.55678123 |
| GFLAQFL | ASSRLGVNYAEQF | 0.39698496 |
| GGAGGAGGNAGVA | ASSRLGVNYAEQF | 0.6820748 |
| GGVGTAGAGGAGGAGG | ASSRLGVNYAEQF | 0.6724705 |
| GNGGGGDAGNAGSGGN | ASSRLGVNYAEQF | 0.6086864 |
| GTGGAGGAGNDAGS | ASSRLGVNYAEQF | 0.64464784 |
| GTTNTGWFNS | ASSRLGVNYAEQF | 0.57709205 |
| IFMGPLFTL | ASSRLGVNYAEQF | 0.49964863 |
| ILAHAIEAI | ASSRLGVNYAEQF | 0.6714324 |
| IRSAIREAL | ASSRLGVNYAEQF | 0.49736995 |
| ITFAGGG | ASSRLGVNYAEQF | 0.7152684 |
| ITGMSGHQEV | ASSRLGVNYAEQF | 0.56085086 |
| IVAGLACTL | ASSRLGVNYAEQF | 0.6689154 |
| IVGGGLAGA | ASSRLGVNYAEQF | 0.6834037 |
| IVTGVLNI | ASSRLGVNYAEQF | 0.672328 |
| LAQQGCI | ASSRLGVNYAEQF | 0.62597984 |
| LCLHVEN | ASSRLGVNYAEQF | 0.6135847 |
| LDAISDGS | ASSRLGVNYAEQF | 0.5374933 |
| LGGFAAML | ASSRLGVNYAEQF | 0.32792833 |

| | | |
|---|---|---|
| LGGKNPMI | ASSRLGVNYAEQF | 0.70742834 |
| LGPKGVAAIL | ASSRLGVNYAEQF | 0.4906605 |
| LMVGVSI | ASSRLGVNYAEQF | 0.70454884 |
| LVGAAFVL | ASSRLGVNYAEQF | 0.6904255 |
| LVLAVAPL | ASSRLGVNYAEQF | 0.6011505 |
| MAAIAVIDAA | ASSRLGVNYAEQF | 0.6689274 |
| NNGINAELQ | ASSRLGVNYAEQF | 0.5051532 |
| RGEPLVLVHG | ASSRLGVNYAEQF | 0.54676133 |
| RGGAGGAAGLI | ASSRLGVNYAEQF | 0.66638696 |
| RSYRMHPAV | ASSRLGVNYAEQF | 0.6778683 |
| SAAHLLAGQLA | ASSRLGVNYAEQF | 0.56915563 |
| SAAVGSY | ASSRLGVNYAEQF | 0.6527187 |
| SPHAVLHRI | ASSRLGVNYAEQF | 0.672626 |
| SQIAKVL | ASSRLGVNYAEQF | 0.54449606 |
| TALGHLVP | ASSRLGVNYAEQF | 0.6372752 |
| TAPTTSGAA | ASSRLGVNYAEQF | 0.7013393 |
| TDGDDGNGGSPGT | ASSRLGVNYAEQF | 0.64477605 |
| TDGLGEN | ASSRLGVNYAEQF | 0.6161576 |
| TGGPPLPGQA | ASSRLGVNYAEQF | 0.6886195 |
| TIPAITI | ASSRLGVNYAEQF | 0.5748002 |
| TIPGFTI | ASSRLGVNYAEQF | 0.6800562 |
| TSIASGALA | ASSRLGVNYAEQF | 0.6599289 |
| TVTLPGAAI | ASSRLGVNYAEQF | 0.6096362 |
| VASGLSAALP | ASSRLGVNYAEQF | 0.6847606 |
| VGAGVIGI | ASSRLGVNYAEQF | 0.4304892 |
| VGPKRFGV | ASSRLGVNYAEQF | 0.5480235 |
| VNVAFPD | ASSRLGVNYAEQF | 0.59985876 |
| VSMTNSGVS | ASSRLGVNYAEQF | 0.66777366 |
| VTDVLGDLSV | ASSRLGVNYAEQF | 0.69790465 |
| VVVVGAGFAGL | ASSRLGVNYAEQF | 0.5938761 |
| VYTVGTVV | ASSRLGVNYAEQF | 0.6508523 |
| WTTFSGP | ASSRLGVNYAEQF | 0.617805 |
| YDELVEAF | ASSRLGVNYAEQF | 0.532272 |
| YGTGGAGGNGG | ASSRLGVNYAEQF | 0.5896507 |
| YTNVMAV | ASSRLGVNYAEQF | 0.59275436 |
| FLEQVHQL | ASSRLGVNYAEQF | 0.53094965 |
| GLSLLAVAV | ASSRLGVNYAEQF | 0.6793067 |
| LEQVHQL | ASSRLGVNYAEQF | 0.5368045 |
| LNAAGAI | ASSRLGVNYAEQF | 0.6444496 |
| PAVHASL | ASSRLGVNYAEQF | 0.7186644 |
| TELEDDA | ASSRLGVNYAEQF | 0.58735776 |
| TPPGPPN | ASSRLGVNYAEQF | 0.66587627 |
| TPPPPPPP | ASSRLGVNYAEQF | 0.66992706 |
| TQSPALI | ASSEGQGEVF | 0.6857376 |
| EFAAAFPQ | ASSEGQGEVF | 0.6376497 |
| LLSGNLDVL | ASSEGQGEVF | 0.6312957 |
| GPANLGNYNI | ASSEGQGEVF | 0.7096098 |
| TLGPTLP | ASSEGQGEVF | 0.69644964 |
| VYAYSSCCK | ASSEGQGEVF | 0.6658382 |
| APAPAPAPNI | ASSEGQGEVF | 0.6837494 |
| PGGPVPIP | ASSEGQGEVF | 0.6873821 |

| | | |
|---|---|---|
| FLTVALEP | ASSEGQGEVF | 0.6846014 |
| SQYFGANL | ASSEGQGEVF | 0.61238337 |
| GGVKPAA | ASSEGQGEVF | 0.40711156 |
| GGTGGAGGAGGW | ASSEGQGEVF | 0.6262263 |
| GVGGVAGLGGAGG | ASSEGQGEVF | 0.6624324 |
| VLLVLATTI | ASSEGQGEVF | 0.61283845 |
| IGIFAAML | ASSEGQGEVF | 0.36477664 |
| AVGDLID | ASSEGQGEVF | 0.71118724 |
| NISPLPL | ASSEGQGEVF | 0.71940446 |
| VTAVFVAI | ASSEGQGEVF | 0.5305886 |
| SGDDNDAAL | ASSEGQGEVF | 0.7147505 |
| AVAALAHS | ASSEGQGEVF | 0.47197405 |
| GGVTVMATP | ASSEGQGEVF | 0.6819967 |
| AGLAANVRD | ASSEGQGEVF | 0.70955825 |
| TNRPDLIDPALLRPG | ASSEGQGEVF | 0.54334706 |
| LANLGVTAL | ASSEGQGEVF | 0.6924235 |
| TGERIEL | ASSEGQGEVF | 0.630931 |
| NGEELAA | ASSEGQGEVF | 0.6484787 |
| VLSLVWG | ASSEGQGEVF | 0.7076849 |
| IGYALWL | ASSEGQGEVF | 0.6056007 |
| LSLLLAPL | ASSEGQGEVF | 0.68019354 |
| TLDEALK | ASSEGQGEVF | 0.6095371 |
| TDGAGTAEGG | ASSEGQGEVF | 0.54261076 |
| NAPGGAGGNGG | ASSEGQGEVF | 0.66581935 |
| VSVFMGMGGNF | ASSEGQGEVF | 0.69584596 |
| LASYLAAL | ASSEGQGEVF | 0.47224745 |
| LVGGTPAL | ASSEGQGEVF | 0.70001984 |
| VRNADVL | ASSEGQGEVF | 0.57745427 |
| TCSSPFLVSAM | ASSEGQGEVF | 0.72306085 |
| VHITDAE | ASSEGQGEVF | 0.67735344 |
| EYAAPSF | ASSEGQGEVF | 0.6597532 |
| VVDAFHGGL | ASSEGQGEVF | 0.5884689 |
| ASVVDPQ | ASSEGQGEVF | 0.70613885 |
| SYNEFGGLS | ASSEGQGEVF | 0.61950564 |
| DPDGSVY | ASSEGQGEVF | 0.68015176 |
| MVSTIHATE | ASSEGQGEVF | 0.70135295 |
| LAAGVADPV | ASSEGQGEVF | 0.55094725 |
| TKEQIAA | ASSEGQGEVF | 0.69323146 |
| SLASPAAL | ASSEGQGEVF | 0.6621446 |
| IGLVFYGV | ASSEGQGEVF | 0.6316346 |
| FLLDASNL | ASSEGQGEVF | 0.605071 |
| YQVGNFP | ASSEGQGEVF | 0.41006953 |
| ESLLSQST | ASSEGQGEVF | 0.64437765 |
| LFTVGFAAVG | ASSEGQGEVF | 0.56481004 |
| ATPPTPT | ASSEGQGEVF | 0.67714536 |
| TDLATLG | ASSEGQGEVF | 0.6650468 |
| SAGGTGR | ASSEGQGEVF | 0.5833116 |
| GFSNSGS | ASSEGQGEVF | 0.6936064 |
| GGNGGSGGTSV | ASSEGQGEVF | 0.6704791 |
| GTSVATGGAGN | ASSEGQGEVF | 0.45288062 |
| NGGNGGSGGTS | ASSEGQGEVF | 0.6423877 |

| | | |
|---|---|---|
| AVVKFPH | ASSEGQGEVF | 0.61879236 |
| TPPPPPP | ASSEGQGEVF | 0.63335943 |
| AAPAAAAAPA | ASSEGQGEVF | 0.4070554 |
| IALNEDL | ASSEGQGEVF | 0.68012 |
| IVNVSSM | ASSEGQGEVF | 0.6154692 |
| TGIPPAPR | ASSEGQGEVF | 0.30457067 |
| FDLGGGTF | ASSEGQGEVF | 0.6413862 |
| FDLGGGTFDVS | ASSEGQGEVF | 0.65759164 |
| NEPTAAA | ASSEGQGEVF | 0.61517847 |
| NEPTAAALA | ASSEGQGEVF | 0.687607 |
| GSIINISSI | ASSEGQGEVF | 0.54129577 |
| VAPGNAGTA | ASSEGQGEVF | 0.5644762 |
| TGGGASLE | ASSEGQGEVF | 0.6899882 |
| VAVYNHYKRI | ASSEGQGEVF | 0.69137 |
| TDPAPATT | ASSEGQGEVF | 0.50777066 |
| TNRPDILDPALLRPG | ASSEGQGEVF | 0.67480093 |
| LTTTTTS | ASSEGQGEVF | 0.6751837 |
| PTGSGKTL | ASSEGQGEVF | 0.6487366 |
| TGSGKTL | ASSEGQGEVF | 0.68543935 |
| TIGHVDHGK | ASSEGQGEVF | 0.6885513 |
| TIGHVDHGKTT | ASSEGQGEVF | 0.4726866 |
| GHVDHGKTT | ASSEGQGEVF | 0.6931547 |
| AQFEHTLL | ASSEGQGEVF | 0.6937845 |
| IVNQDEL | ASSEGQGEVF | 0.6034 |
| TNGSQFF | ASSEGQGEVF | 0.6660421 |
| TGKGIVE | ASSEGQGEVF | 0.5342975 |
| LISNASDA | ASSEGQGEVF | 0.587894 |
| AELRTLGELTKIAKAHGAQ | ASSEGQGEVF | 0.3823107 |
| APNAVAAAEPSW | ASSEGQGEVF | 0.6793697 |
| APQVAGAATAVL | ASSEGQGEVF | 0.68528605 |
| ARRDPAAAAAILPTDA | ASSEGQGEVF | 0.64532554 |
| DAAARDVLAEAGLAEYF | ASSEGQGEVF | 0.58315474 |
| DDQIVLSAKANLAKVEQ | ASSEGQGEVF | 0.5669917 |
| DELLPDTGEDSDEDGH | ASSEGQGEVF | 0.627716 |
| ELPEAPANRATVPQD | ASSEGQGEVF | 0.60058427 |
| FGQSKIERVTGIG | ASSEGQGEVF | 0.43470258 |
| GAGGTGSTAGGAGGAGGAGGLY | ASSEGQGEVF | 0.56787235 |
| GHGGTGAAGGNGGTGGAGGSGI | ASSEGQGEVF | 0.68375033 |
| GNGGNGGAGGEGGPGGAGGAGA | ASSEGQGEVF | 0.67959344 |
| GSTAGNGGNGGNATNGTIAGQP | ASSEGQGEVF | 0.41729963 |
| GTGGDGGNGGNGGTGG | ASSEGQGEVF | 0.69580007 |
| GVRRPAGIPADAPV | ASSEGQGEVF | 0.6807674 |
| GYRQVTDAHLLTLARR | ASSEGQGEVF | 0.68020487 |
| IAKMGQDGHDRG | ASSEGQGEVF | 0.6263241 |
| KAEPSIEIQYT | ASSEGQGEVF | 0.5637862 |
| KPVRIRMAADEDTVAVT | ASSEGQGEVF | 0.68805444 |
| MPPRIAGMRLLVIKPEP | ASSEGQGEVF | 0.64468455 |
| MKFYAVALFRSRGVPPTR | ASSEGQGEVF | 0.5629292 |
| NAGGFGILGAVAHSP | ASSEGQGEVF | 0.6839106 |
| NEGREITLAMAEM | ASSEGQGEVF | 0.64147055 |
| NELVEWSHPLGHRWRWE | ASSEGQGEVF | 0.62525547 |

| | | |
|---|---|---|
| NLGAGGTGGNGGNPG | ASSEGQGEVF | 0.6235112 |
| NTEWPKWETVMAADPVD | ASSEGQGEVF | 0.67920256 |
| PASAAPAETQP | ASSEGQGEVF | 0.5970028 |
| QQSSDGKNGVRAAEITGE | ASSEGQGEVF | 0.4349935 |
| RAPDAVTTSWWKE | ASSEGQGEVF | 0.5588797 |
| SGFGNVGAHTSGWFNQ | ASSEGQGEVF | 0.65383446 |
| SGVAGAVGQAASVGGLKV | ASSEGQGEVF | 0.68955827 |
| SNGGAGTGSAGGNGGT | ASSEGQGEVF | 0.65206826 |
| TGGAGGEGGAAG | ASSEGQGEVF | 0.59525216 |
| TGGDGGAAGAGGAPGHG | ASSEGQGEVF | 0.67664003 |
| TMPPLLAAIANAGRHGV | ASSEGQGEVF | 0.6528126 |
| TSSTAGNSGGA | ASSEGQGEVF | 0.6505312 |
| VPATVPVGILC | ASSEGQGEVF | 0.6080011 |
| VQRPDSGGVVLTGRIS | ASSEGQGEVF | 0.6284579 |
| WTPNPVPATKGVPAGLVQAT | ASSEGQGEVF | 0.6849241 |
| YALKRALAKFSDRFDVVII | ASSEGQGEVF | 0.67565495 |
| YLIGSVIV | ASSEGQGEVF | 0.67977107 |
| YGQLVAVID | ASSEGQGEVF | 0.68338716 |
| VTVGGMTLPA | ASSEGQGEVF | 0.6350344 |
| VLALPAV | ASSEGQGEVF | 0.65591997 |
| VIPPTLNYET | ASSEGQGEVF | 0.33580935 |
| VGMGGMHPSSG | ASSEGQGEVF | 0.47390306 |
| VGLAGGAATA | ASSEGQGEVF | 0.6624856 |
| VGGAGGAGGSGP | ASSEGQGEVF | 0.67397034 |
| VFAAFMV | ASSEGQGEVF | 0.6793906 |
| TLYDKLR | ASSEGQGEVF | 0.6524974 |
| TLAVFTVL | ASSEGQGEVF | 0.57603455 |
| TGSVQPARV | ASSEGQGEVF | 0.6472054 |
| TGGGAGSGGT | ASSEGQGEVF | 0.6861265 |
| SHLETSL | ASSEGQGEVF | 0.47513792 |
| SFTTGMT | ASSEGQGEVF | 0.5840664 |
| QVEVAKRMGVRQPTVS | ASSEGQGEVF | 0.697193 |
| QGSGGPALAM | ASSEGQGEVF | 0.66745377 |
| MLAPSNP | ASSEGQGEVF | 0.6474627 |
| LVNNAAI | ASSEGQGEVF | 0.5515766 |
| LTGEINTLA | ASSEGQGEVF | 0.69075686 |
| LSIFGPV | ASSEGQGEVF | 0.61187756 |
| LLALLANL | ASSEGQGEVF | 0.6916933 |
| LISNASDALDKLR | ASSEGQGEVF | 0.3164469 |
| LINEAVP | ASSEGQGEVF | 0.5901139 |
| LGYAFSGL | ASSEGQGEVF | 0.47028938 |
| LDFVAVAL | ASSEGQGEVF | 0.35710302 |
| LAQPGAE | ASSEGQGEVF | 0.68297166 |
| ISNLGAVTGF | ASSEGQGEVF | 0.5589586 |
| HQEDPGRI | ASSEGQGEVF | 0.6603217 |
| GTPGEPTP | ASSEGQGEVF | 0.6662441 |
| GFFNSGS | ASSEGQGEVF | 0.68500376 |
| GAGGGGLTAGAN | ASSEGQGEVF | 0.6690749 |
| FVDALVNM | ASSEGQGEVF | 0.5919147 |
| FLTVAPEL | ASSEGQGEVF | 0.47407344 |
| FETLTAL | ASSEGQGEVF | 0.6789011 |

| | | |
|---|---|---|
| DVINPIEL | ASSEGQGEVF | 0.66946846 |
| APVSGAPV | ASSEGQGEVF | 0.66361165 |
| ALVGGLAW | ASSEGQGEVF | 0.43641204 |
| ALAAAAATAI | ASSEGQGEVF | 0.64904606 |
| AFGLGLLL | ASSEGQGEVF | 0.65082246 |
| AFFAAASL | ASSEGQGEVF | 0.3824186 |
| AAAALPAFSPP | ASSEGQGEVF | 0.35486037 |
| VVLAPAL | ASSEGQGEVF | 0.62615365 |
| TIGHVDHGKTTL | ASSEGQGEVF | 0.601301 |
| LISNASDALDK | ASSEGQGEVF | 0.6916484 |
| LISNASDAL | ASSEGQGEVF | 0.68919444 |
| ISNASDALDK | ASSEGQGEVF | 0.6541393 |
| NQAETLVYQTE | ASSEGQGEVF | 0.66597354 |
| TGGTGGTAGAG | ASSEGQGEVF | 0.6430646 |
| TGSTAGGAGGA | ASSEGQGEVF | 0.28055087 |
| DPLAVPVAAGPV | ASSEGQGEVF | 0.5496882 |
| GGTGGTPGNGGN | ASSEGQGEVF | 0.50926256 |
| PAGLAGVDPATL | ASSEGQGEVF | 0.6898583 |
| GNNLVVEAPHAGASV | ASSEGQGEVF | 0.49308866 |
| KVTRIMSASTLEQPA | ASSEGQGEVF | 0.616645 |
| LGFALPIALPRRTVV | ASSEGQGEVF | 0.66703975 |
| TRLGNLTARKLVVLV | ASSEGQGEVF | 0.6700095 |
| WSQQRRLIYFPSAGP | ASSEGQGEVF | 0.65595365 |
| GPNGNPDPMAAAVDIR | ASSEGQGEVF | 0.54921144 |
| TQRCVSVADLRAAASR | ASSEGQGEVF | 0.3167506 |
| QFVLDVAEHPVEVLTETV | ASSEGQGEVF | 0.6820172 |
| GGAGGTGGAGGNGGTQAGDG | ASSEGQGEVF | 0.52021 |
| PLLATLRAIAVDVDAKPAQV | ASSEGQGEVF | 0.49120387 |
| APKFRTGGYMIDFWGVGYQVAK | ASSEGQGEVF | 0.3954149 |
| GAGGAGGAGGAAGGPGGTGGTG | ASSEGQGEVF | 0.669542 |
| IGALGLLATDVAPAGSVAFGVA | ASSEGQGEVF | 0.42433718 |
| AGGAGGIGGNGGAGGAGGVLMG | ASSEGQGEVF | 0.679235 |
| ADELIGLL | ASSEGQGEVF | 0.648777 |
| ADRALKA | ASSEGQGEVF | 0.6499999 |
| AGAVGQVGGAGGA | ASSEGQGEVF | 0.6976304 |
| AGGGGGLGGAP | ASSEGQGEVF | 0.66198313 |
| AGGLLYGNGGAG | ASSEGQGEVF | 0.65529805 |
| AGLESIGL | ASSEGQGEVF | 0.5736539 |
| AGLKTGII | ASSEGQGEVF | 0.5496822 |
| ASLAALLL | ASSEGQGEVF | 0.5558256 |
| ASVINISSI | ASSEGQGEVF | 0.57360566 |
| ATPPGPA | ASSEGQGEVF | 0.6949515 |
| ATVASALV | ASSEGQGEVF | 0.5878553 |
| ATVASLIL | ASSEGQGEVF | 0.49214065 |
| AVNGPVLIRH | ASSEGQGEVF | 0.60363716 |
| DGAAGAVGI | ASSEGQGEVF | 0.6950307 |
| DPLLHGI | ASSEGQGEVF | 0.6966671 |
| ELGAALGAL | ASSEGQGEVF | 0.52584183 |
| ELGRTQAV | ASSEGQGEVF | 0.5169065 |
| FVAGGYLS | ASSEGQGEVF | 0.56211144 |
| GASGGAGGAGGW | ASSEGQGEVF | 0.6639023 |

| | | |
|---|---|---|
| GGAGAGGDGGAGGI | ASSEGQGEVF | 0.47984046 |
| GTGHNGSQPGMGGNG | ASSEGQGEVF | 0.47418466 |
| GYAAASASAS | ASSEGQGEVF | 0.67512214 |
| HQVDVAAVQQG | ASSEGQGEVF | 0.69185704 |
| IGAAIAHT | ASSEGQGEVF | 0.5699284 |
| IGMIEVAQL | ASSEGQGEVF | 0.5449176 |
| IRLAPEHTSAMD | ASSEGQGEVF | 0.65958196 |
| IYNVSVL | ASSEGQGEVF | 0.60648143 |
| KQGFLGM | ASSEGQGEVF | 0.6396379 |
| LESVITQL | ASSEGQGEVF | 0.5883166 |
| LGLGALVVV | ASSEGQGEVF | 0.50964266 |
| LIGNGGNGGSGGI | ASSEGQGEVF | 0.64247775 |
| LNAINAPTL | ASSEGQGEVF | 0.6954224 |
| LQLLNTGV | ASSEGQGEVF | 0.640846 |
| LQTSIEF | ASSEGQGEVF | 0.67175007 |
| MSFAVSEL | ASSEGQGEVF | 0.69541717 |
| NTGALGL | ASSEGQGEVF | 0.6832571 |
| PPGAGGAGGAGTT | ASSEGQGEVF | 0.69276893 |
| QLVGLGI | ASSEGQGEVF | 0.6838076 |
| RFLEQVHQL | ASSEGQGEVF | 0.5775924 |
| RGVGYVLR | ASSEGQGEVF | 0.64642346 |
| SATGVFAGVI | ASSEGQGEVF | 0.68457603 |
| SLPAITLP | ASSEGQGEVF | 0.6692673 |
| STALAFTV | ASSEGQGEVF | 0.64563346 |
| SVLTVGLGK | ASSEGQGEVF | 0.58256525 |
| TALAGAGSGI | ASSEGQGEVF | 0.6911194 |
| TGLPFTGI | ASSEGQGEVF | 0.6136068 |
| VAEHVTV | ASSEGQGEVF | 0.6842377 |
| VDIVLNSL | ASSEGQGEVF | 0.42699778 |
| VLTGGGAVL | ASSEGQGEVF | 0.5286033 |
| VNVGLAYGI | ASSEGQGEVF | 0.6871677 |
| VSSAAVGAV | ASSEGQGEVF | 0.27916202 |
| VTSVYEAGV | ASSEGQGEVF | 0.3321652 |
| WVALDRGA | ASSEGQGEVF | 0.6567759 |
| YLVGSVVI | ASSEGQGEVF | 0.6872968 |
| GPPGTGKTLL | ASSEGQGEVF | 0.29569784 |
| GVGMTKF | ASSEGQGEVF | 0.5311362 |
| IGIGGSDL | ASSEGQGEVF | 0.4795778 |
| IGTLLAEL | ASSEGQGEVF | 0.64670134 |
| IYTSGTT | ASSEGQGEVF | 0.6546315 |
| LSFPAGV | ASSEGQGEVF | 0.38455516 |
| LYTSGTT | ASSEGQGEVF | 0.6713135 |
| VGAFPFGF | ASSEGQGEVF | 0.53713965 |
| VGAGTVEF | ASSEGQGEVF | 0.62329435 |
| VGKSTLLL | ASSEGQGEVF | 0.6824831 |
| AALAHAGGGIV | ASSEGQGEVF | 0.6920166 |
| AALVAVSLL | ASSEGQGEVF | 0.4873982 |
| AAVSRAVAAL | ASSEGQGEVF | 0.6836221 |
| AGGSSGIGGF | ASSEGQGEVF | 0.6032565 |
| AGLTVANANG | ASSEGQGEVF | 0.69561434 |
| AGLVGAVL | ASSEGQGEVF | 0.6284518 |

| | | |
|---|---|---|
| ALSQAFTV | ASSEGQGEVF | 0.44744492 |
| ALTFSVSL | ASSEGQGEVF | 0.6829325 |
| ALTGAALVAL | ASSEGQGEVF | 0.66410285 |
| ASFSIGAGVL | ASSEGQGEVF | 0.5524633 |
| DGSVYKI | ASSEGQGEVF | 0.68661267 |
| EQLAEIV | ASSEGQGEVF | 0.5314637 |
| FAVSELTVM | ASSEGQGEVF | 0.66653556 |
| GAAPLAAGL | ASSEGQGEVF | 0.69363916 |
| GADGGDGGSGGAGG | ASSEGQGEVF | 0.6831318 |
| GFGLDTGGAGG | ASSEGQGEVF | 0.5095664 |
| GLASGGLL | ASSEGQGEVF | 0.67105985 |
| GRIAGAAVLL | ASSEGQGEVF | 0.6436749 |
| GSAGSLNTSF | ASSEGQGEVF | 0.55530053 |
| GTGQIGI | ASSEGQGEVF | 0.6876445 |
| GTLIALIL | ASSEGQGEVF | 0.612769 |
| IAATMGFVVG | ASSEGQGEVF | 0.6867195 |
| IFAQALL | ASSEGQGEVF | 0.66812515 |
| IGAAVALGF | ASSEGQGEVF | 0.6691366 |
| IGLTPLAL | ASSEGQGEVF | 0.4202154 |
| ILGSNTSTL | ASSEGQGEVF | 0.6128353 |
| IVFSLTVL | ASSEGQGEVF | 0.6420914 |
| LASKATGF | ASSEGQGEVF | 0.6826973 |
| LASVALIL | ASSEGQGEVF | 0.6268138 |
| LFAVGLTL | ASSEGQGEVF | 0.6301699 |
| LFNGGIGTYI | ASSEGQGEVF | 0.66348267 |
| LTLGAGII | ASSEGQGEVF | 0.69302315 |
| LVAAALGDI | ASSEGQGEVF | 0.63249403 |
| NTAFVQLGI | ASSEGQGEVF | 0.59611356 |
| QLFSGAEL | ASSEGQGEVF | 0.6954714 |
| SAATFITL | ASSEGQGEVF | 0.5698247 |
| SNYGLANLGV | ASSEGQGEVF | 0.63436764 |
| SYGGAYAVM | ASSEGQGEVF | 0.6657288 |
| TTGVGGLTLG | ASSEGQGEVF | 0.6772589 |
| TTQTIYI | ASSEGQGEVF | 0.6191442 |
| VAAGAGLGAVI | ASSEGQGEVF | 0.68401766 |
| VAAVVTLL | ASSEGQGEVF | 0.4771117 |
| VGFLGGLI | ASSEGQGEVF | 0.68153465 |
| VGVIPILL | ASSEGQGEVF | 0.60627997 |
| VIFASSLI | ASSEGQGEVF | 0.67507 |
| VKGGDGGAAAG | ASSEGQGEVF | 0.5859833 |
| VTKTFTNI | ASSEGQGEVF | 0.68228084 |
| VVAAAILL | ASSEGQGEVF | 0.68763715 |
| YAALDVEL | ASSEGQGEVF | 0.68842596 |
| LASVALLL | ASSEGQGEVF | 0.33716077 |
| LDAVHRV | ASSEGQGEVF | 0.68947816 |
| QGLSSSIF | ASSEGQGEVF | 0.67960346 |
| RGTFANIRL | ASSEGQGEVF | 0.6982256 |
| RQLWWGHRI | ASSEGQGEVF | 0.6236244 |
| YGPPGTGKTLL | ASSEGQGEVF | 0.65177494 |
| AALVGAPANVA | ASSEGQGEVF | 0.599171 |
| AATFRRTE | ASSEGQGEVF | 0.6100826 |

| | | |
|---|---|---|
| AGALGGGI | ASSEGQGEVF | 0.5699265 |
| AGGAGGAGGIGAG | ASSEGQGEVF | 0.5702367 |
| AGSGALGL | ASSEGQGEVF | 0.64206403 |
| AQIADQI | ASSEGQGEVF | 0.6236179 |
| ASAVAVTYL | ASSEGQGEVF | 0.68542933 |
| ASTVLGGTGG | ASSEGQGEVF | 0.68159825 |
| AVGGEPAK | ASSEGQGEVF | 0.6842023 |
| CEQGTFGGTC | ASSEGQGEVF | 0.693456 |
| DEGKAPAG | ASSEGQGEVF | 0.6939825 |
| DTVVFPAL | ASSEGQGEVF | 0.6761455 |
| FEGITMAG | ASSEGQGEVF | 0.58810025 |
| FPLVVQD | ASSEGQGEVF | 0.6223009 |
| GAFTTPQ | ASSEGQGEVF | 0.5735948 |
| GFGGFGTAGGVGGA | ASSEGQGEVF | 0.64608896 |
| GFLAQFL | ASSEGQGEVF | 0.6159766 |
| GGAGGAGGNAGVA | ASSEGQGEVF | 0.6797013 |
| GGVGTAGAGGAGGAGG | ASSEGQGEVF | 0.28911787 |
| GNGGGGDAGNAGSGGN | ASSEGQGEVF | 0.6713065 |
| GTGGAGGAGNDAGS | ASSEGQGEVF | 0.6952517 |
| GTTNTGWFNS | ASSEGQGEVF | 0.5724522 |
| IFMGPLFTL | ASSEGQGEVF | 0.6596319 |
| ILAHAIEAI | ASSEGQGEVF | 0.6714675 |
| IRSAIREAL | ASSEGQGEVF | 0.62012196 |
| ITFAGGG | ASSEGQGEVF | 0.68526745 |
| ITGMSGHQEV | ASSEGQGEVF | 0.57652014 |
| IVAGLACTL | ASSEGQGEVF | 0.66587037 |
| IVGGGLAGA | ASSEGQGEVF | 0.68919724 |
| IVTGVLNI | ASSEGQGEVF | 0.6720464 |
| LAQQGCI | ASSEGQGEVF | 0.54915977 |
| LCLHVEN | ASSEGQGEVF | 0.59488463 |
| LDAISDGS | ASSEGQGEVF | 0.40388405 |
| LGGFAAML | ASSEGQGEVF | 0.6360673 |
| LGGKNPMI | ASSEGQGEVF | 0.5182446 |
| LGPKGVAAIL | ASSEGQGEVF | 0.6646665 |
| LMVGVSI | ASSEGQGEVF | 0.5220021 |
| LVGAAFVL | ASSEGQGEVF | 0.66264737 |
| LVLAVAPL | ASSEGQGEVF | 0.6652148 |
| MAAIAVIDAA | ASSEGQGEVF | 0.6543193 |
| NNGINAELQ | ASSEGQGEVF | 0.58501244 |
| RGEPLVLVHG | ASSEGQGEVF | 0.66557336 |
| RGGAGGAAGLI | ASSEGQGEVF | 0.6011528 |
| RSYRMHPAV | ASSEGQGEVF | 0.48082063 |
| SAAHLLAGQLA | ASSEGQGEVF | 0.6811063 |
| SAAVGSY | ASSEGQGEVF | 0.61608106 |
| SPHAVLHRI | ASSEGQGEVF | 0.6435032 |
| SQIAKVL | ASSEGQGEVF | 0.68361104 |
| TALGHLVP | ASSEGQGEVF | 0.64129025 |
| TAPTTSGAA | ASSEGQGEVF | 0.42401737 |
| TDGDDGNGGSPGT | ASSEGQGEVF | 0.6448712 |
| TDGLGEN | ASSEGQGEVF | 0.66300577 |
| TGGPPLPGQA | ASSEGQGEVF | 0.6317773 |

| | | |
|---|---|---|
| TIPAITI | ASSEGQGEVF | 0.3167276 |
| TIPGFTI | ASSEGQGEVF | 0.65037346 |
| TSIASGALA | ASSEGQGEVF | 0.56032777 |
| TVTLPGAAI | ASSEGQGEVF | 0.59692824 |
| VASGLSAALP | ASSEGQGEVF | 0.59068114 |
| VGAGVIGI | ASSEGQGEVF | 0.68593884 |
| VGPKRFGV | ASSEGQGEVF | 0.60136616 |
| VNVAFPD | ASSEGQGEVF | 0.5665859 |
| VSMTNSGVS | ASSEGQGEVF | 0.67591625 |
| VTDVLGDLSV | ASSEGQGEVF | 0.6687995 |
| VVVVGAGFAGL | ASSEGQGEVF | 0.68920696 |
| VYTVGTVV | ASSEGQGEVF | 0.6206086 |
| WTTFSGP | ASSEGQGEVF | 0.5699322 |
| YDELVEAF | ASSEGQGEVF | 0.6125396 |
| YGTGGAGGNGG | ASSEGQGEVF | 0.57005465 |
| YTNVMAV | ASSEGQGEVF | 0.5079651 |
| FLEQVHQL | ASSEGQGEVF | 0.44028345 |
| GLSLLAVAV | ASSEGQGEVF | 0.6626778 |
| LEQVHQL | ASSEGQGEVF | 0.39091542 |
| LNAAGAI | ASSEGQGEVF | 0.64618945 |
| PAVHASL | ASSEGQGEVF | 0.6820165 |
| TELEDDA | ASSEGQGEVF | 0.5394603 |
| TPPGPPN | ASSEGQGEVF | 0.43954408 |
| TPPPPPPP | ASSEGQGEVF | 0.6361357 |
| TQSPALI | TCSADLGSQNTLY | 0.61448175 |
| EFAAAFPQ | TCSADLGSQNTLY | 0.68178284 |
| LLSGNLDVL | TCSADLGSQNTLY | 0.6472377 |
| GPANLGNYNI | TCSADLGSQNTLY | 0.6933044 |
| TLGPTLP | TCSADLGSQNTLY | 0.67966974 |
| VYAYSSCCK | TCSADLGSQNTLY | 0.62546027 |
| APAPAPAPNI | TCSADLGSQNTLY | 0.6786212 |
| PGGPVPIP | TCSADLGSQNTLY | 0.63571227 |
| FLTVALEP | TCSADLGSQNTLY | 0.65452385 |
| SQYFGANL | TCSADLGSQNTLY | 0.63108456 |
| GGVKPAA | TCSADLGSQNTLY | 0.6327312 |
| GGTGGAGGAGGW | TCSADLGSQNTLY | 0.5734321 |
| GVGGVAGLGGAGG | TCSADLGSQNTLY | 0.5809926 |
| VLLVLATTI | TCSADLGSQNTLY | 0.4371019 |
| IGIFAAML | TCSADLGSQNTLY | 0.6696679 |
| AVGDLID | TCSADLGSQNTLY | 0.6401983 |
| NISPLPL | TCSADLGSQNTLY | 0.48998976 |
| VTAVFVAI | TCSADLGSQNTLY | 0.5802475 |
| SGDDNDAAL | TCSADLGSQNTLY | 0.6955368 |
| AVAALAHS | TCSADLGSQNTLY | 0.6896608 |
| GGVTVMATP | TCSADLGSQNTLY | 0.61451125 |
| AGLAANVRD | TCSADLGSQNTLY | 0.5763681 |
| TNRPDLIDPALLRPG | TCSADLGSQNTLY | 0.67286766 |
| LANLGVTAL | TCSADLGSQNTLY | 0.3666998 |
| TGERIEL | TCSADLGSQNTLY | 0.6541568 |
| NGEELAA | TCSADLGSQNTLY | 0.6695014 |
| VLSLVWG | TCSADLGSQNTLY | 0.6512401 |

| | | |
|---|---|---|
| IGYALWL | TCSADLGSQNTLY | 0.4936108 |
| LSLLLAPL | TCSADLGSQNTLY | 0.66914475 |
| TLDEALK | TCSADLGSQNTLY | 0.61225677 |
| TDGAGTAEGG | TCSADLGSQNTLY | 0.6476444 |
| NAPGGAGGNGG | TCSADLGSQNTLY | 0.33237898 |
| VSVFMGMGGNF | TCSADLGSQNTLY | 0.62997925 |
| LASYLAAL | TCSADLGSQNTLY | 0.59192646 |
| LVGGTPAL | TCSADLGSQNTLY | 0.5223638 |
| VRNADVL | TCSADLGSQNTLY | 0.3548519 |
| TCSSPFLVSAM | TCSADLGSQNTLY | 0.6456361 |
| VHITDAE | TCSADLGSQNTLY | 0.68506193 |
| EYAAPSF | TCSADLGSQNTLY | 0.68863094 |
| VVDAFHGGL | TCSADLGSQNTLY | 0.5842422 |
| ASVVDPQ | TCSADLGSQNTLY | 0.6857069 |
| SYNEFGGLS | TCSADLGSQNTLY | 0.6909771 |
| DPDGSVY | TCSADLGSQNTLY | 0.6542134 |
| MVSTIHATE | TCSADLGSQNTLY | 0.3659374 |
| LAAGVADPV | TCSADLGSQNTLY | 0.66905195 |
| TKEQIAA | TCSADLGSQNTLY | 0.6379223 |
| SLASPAAL | TCSADLGSQNTLY | 0.68278587 |
| IGLVFYGV | TCSADLGSQNTLY | 0.6408117 |
| FLLDASNL | TCSADLGSQNTLY | 0.58291864 |
| YQVGNFP | TCSADLGSQNTLY | 0.63873994 |
| ESLLSQST | TCSADLGSQNTLY | 0.6901877 |
| LFTVGFAAVG | TCSADLGSQNTLY | 0.5802855 |
| ATPPTPT | TCSADLGSQNTLY | 0.542019 |
| TDLATLG | TCSADLGSQNTLY | 0.5581769 |
| SAGGTGR | TCSADLGSQNTLY | 0.44639736 |
| GFSNSGS | TCSADLGSQNTLY | 0.66079056 |
| GGNGGSGGTSV | TCSADLGSQNTLY | 0.63923955 |
| GTSVATGGAGN | TCSADLGSQNTLY | 0.57526606 |
| NGGNGGSGGTS | TCSADLGSQNTLY | 0.6681796 |
| AVVKFPH | TCSADLGSQNTLY | 0.56904 |
| TPPPPPP | TCSADLGSQNTLY | 0.6104888 |
| AAPAAAAAPA | TCSADLGSQNTLY | 0.6290406 |
| IALNEDL | TCSADLGSQNTLY | 0.62099886 |
| IVNVSSM | TCSADLGSQNTLY | 0.67500365 |
| TGIPPAPR | TCSADLGSQNTLY | 0.6302333 |
| FDLGGGTF | TCSADLGSQNTLY | 0.6439798 |
| FDLGGGTFDVS | TCSADLGSQNTLY | 0.69684434 |
| NEPTAAA | TCSADLGSQNTLY | 0.49565572 |
| NEPTAAALA | TCSADLGSQNTLY | 0.5463915 |
| GSIINISSI | TCSADLGSQNTLY | 0.66231203 |
| VAPGNAGTA | TCSADLGSQNTLY | 0.5301657 |
| TGGGASLE | TCSADLGSQNTLY | 0.6749506 |
| VAVYNHYKRI | TCSADLGSQNTLY | 0.6498015 |
| TDPAPATT | TCSADLGSQNTLY | 0.67950964 |
| TNRPDILDPALLRPG | TCSADLGSQNTLY | 0.64070743 |
| LTTTTTS | TCSADLGSQNTLY | 0.6965462 |
| PTGSGKTL | TCSADLGSQNTLY | 0.6871631 |
| TGSGKTL | TCSADLGSQNTLY | 0.64297163 |

| | | |
|---|---|---|
| TIGHVDHGK | TCSADLGSQNTLY | 0.68574613 |
| TIGHVDHGKTT | TCSADLGSQNTLY | 0.4485308 |
| GHVDHGKTT | TCSADLGSQNTLY | 0.6837086 |
| AQFEHTLL | TCSADLGSQNTLY | 0.66863084 |
| IVNQDEL | TCSADLGSQNTLY | 0.5926773 |
| TNGSQFF | TCSADLGSQNTLY | 0.66857475 |
| TGKGIVE | TCSADLGSQNTLY | 0.6868528 |
| LISNASDA | TCSADLGSQNTLY | 0.68896055 |
| AELRTLGELTKIAKAHGAQ | TCSADLGSQNTLY | 0.39328942 |
| APNAVAAAEPSW | TCSADLGSQNTLY | 0.6337841 |
| APQVAGAATAVL | TCSADLGSQNTLY | 0.3597569 |
| ARRDPAAAAAILPTDA | TCSADLGSQNTLY | 0.45396322 |
| DAAARDVLAEAGLAEYF | TCSADLGSQNTLY | 0.44736803 |
| DDQIVLSAKANLAKVEQ | TCSADLGSQNTLY | 0.6750209 |
| DELLPDTGEDSDEDGH | TCSADLGSQNTLY | 0.63942474 |
| ELPEAPANRATVPQD | TCSADLGSQNTLY | 0.6629224 |
| FGQSKIERVTGIG | TCSADLGSQNTLY | 0.6614883 |
| GAGGTGSTAGGAGGAGGAGGLY | TCSADLGSQNTLY | 0.6910907 |
| GHGGTGAAGGNGGTGGAGGSGI | TCSADLGSQNTLY | 0.6004317 |
| GNGGNGGAGGEGGPGGAGGAGA | TCSADLGSQNTLY | 0.6416149 |
| GSTAGNGGNGGNATNGTIAGQP | TCSADLGSQNTLY | 0.3935602 |
| GTGGDGGNGGNGGTGG | TCSADLGSQNTLY | 0.38694683 |
| GVRRPAGIPADAPV | TCSADLGSQNTLY | 0.5338527 |
| GYRQVTDAHLLTLARR | TCSADLGSQNTLY | 0.63242984 |
| IAKMGQDGHDRG | TCSADLGSQNTLY | 0.36773437 |
| KAEPSIEIQYT | TCSADLGSQNTLY | 0.68124473 |
| KPVRIRMAADEDTVAVT | TCSADLGSQNTLY | 0.666732 |
| MPPRIAGMRLLVIKPEP | TCSADLGSQNTLY | 0.67786926 |
| MKFYAVALFRSRGVPPTR | TCSADLGSQNTLY | 0.53398335 |
| NAGGFGILGAVAHSP | TCSADLGSQNTLY | 0.6021461 |
| NEGREITLAMAEM | TCSADLGSQNTLY | 0.60460925 |
| NELVEWSHPLGHRWRWE | TCSADLGSQNTLY | 0.6727325 |
| NLGAGGTGGNGGNPG | TCSADLGSQNTLY | 0.48300844 |
| NTEWPKWETVMAADPVD | TCSADLGSQNTLY | 0.45261765 |
| PASAAPAETQP | TCSADLGSQNTLY | 0.5419316 |
| QQSSDGKNGVRAAEITGE | TCSADLGSQNTLY | 0.59632134 |
| RAPDAVTTSWWKE | TCSADLGSQNTLY | 0.69076407 |
| SGFGNVGAHTSGWFNQ | TCSADLGSQNTLY | 0.6886403 |
| SGVAGAVGQAASVGGLKV | TCSADLGSQNTLY | 0.6786679 |
| SNGGAGTGSAGGNGGT | TCSADLGSQNTLY | 0.6873324 |
| TGGAGGEGGAAG | TCSADLGSQNTLY | 0.32491094 |
| TGGDGGAAGAGGAPGHG | TCSADLGSQNTLY | 0.5322368 |
| TMPPLLAAIANAGRHGV | TCSADLGSQNTLY | 0.6911864 |
| TSSTAGNSGGA | TCSADLGSQNTLY | 0.66350347 |
| VPATVPVGILC | TCSADLGSQNTLY | 0.6506628 |
| VQRPDSGGVVLTGRIS | TCSADLGSQNTLY | 0.6930612 |
| WTPNPVPATKGVPAGLVQAT | TCSADLGSQNTLY | 0.58936125 |
| YALKRALAKFSDRFDVVII | TCSADLGSQNTLY | 0.6190274 |
| YLIGSVIV | TCSADLGSQNTLY | 0.6489894 |
| YGQLVAVID | TCSADLGSQNTLY | 0.66076297 |
| VTVGGMTLPA | TCSADLGSQNTLY | 0.6499183 |

| | | |
|---|---|---|
| VLALPAV | TCSADLGSQNTLY | 0.49030834 |
| VIPPTLNYET | TCSADLGSQNTLY | 0.6539256 |
| VGMGGMHPSSG | TCSADLGSQNTLY | 0.6999724 |
| VGLAGGAATA | TCSADLGSQNTLY | 0.6451471 |
| VGGAGGAGGSGP | TCSADLGSQNTLY | 0.68451214 |
| VFAAFMV | TCSADLGSQNTLY | 0.6674999 |
| TLYDKLR | TCSADLGSQNTLY | 0.65575075 |
| TLAVFTVL | TCSADLGSQNTLY | 0.60388047 |
| TGSVQPARV | TCSADLGSQNTLY | 0.62976587 |
| TGGGAGSGGT | TCSADLGSQNTLY | 0.66186315 |
| SHLETSL | TCSADLGSQNTLY | 0.58729774 |
| SFTTGMT | TCSADLGSQNTLY | 0.6216975 |
| QVEVAKRMGVRQPTVS | TCSADLGSQNTLY | 0.61058563 |
| QGSGGPALAM | TCSADLGSQNTLY | 0.6986968 |
| MLAPSNP | TCSADLGSQNTLY | 0.67062503 |
| LVNNAAI | TCSADLGSQNTLY | 0.6747049 |
| LTGEINTLA | TCSADLGSQNTLY | 0.57363856 |
| LSIFGPV | TCSADLGSQNTLY | 0.6521595 |
| LLALLANL | TCSADLGSQNTLY | 0.68288624 |
| LISNASDALDKLR | TCSADLGSQNTLY | 0.48604095 |
| LINEAVP | TCSADLGSQNTLY | 0.691857 |
| LGYAFSGL | TCSADLGSQNTLY | 0.6183795 |
| LDFVAVAL | TCSADLGSQNTLY | 0.6084468 |
| LAQPGAE | TCSADLGSQNTLY | 0.6636427 |
| ISNLGAVTGF | TCSADLGSQNTLY | 0.68567526 |
| HQEDPGRI | TCSADLGSQNTLY | 0.61093616 |
| GTPGEPTP | TCSADLGSQNTLY | 0.64131796 |
| GFFNSGS | TCSADLGSQNTLY | 0.5891971 |
| GAGGGGLTAGAN | TCSADLGSQNTLY | 0.6771356 |
| FVDALVNM | TCSADLGSQNTLY | 0.6898458 |
| FLTVAPEL | TCSADLGSQNTLY | 0.6480781 |
| FETLTAL | TCSADLGSQNTLY | 0.695345 |
| DVINPIEL | TCSADLGSQNTLY | 0.6051811 |
| APVSGAPV | TCSADLGSQNTLY | 0.38752648 |
| ALVGGLAW | TCSADLGSQNTLY | 0.6209103 |
| ALAAAAATAI | TCSADLGSQNTLY | 0.6796715 |
| AFGLGLLL | TCSADLGSQNTLY | 0.6873494 |
| AFFAAASL | TCSADLGSQNTLY | 0.68555295 |
| AAAALPAFSPP | TCSADLGSQNTLY | 0.61648893 |
| VVLAPAL | TCSADLGSQNTLY | 0.620589 |
| TIGHVDHGKTTL | TCSADLGSQNTLY | 0.6946629 |
| LISNASDALDK | TCSADLGSQNTLY | 0.6887299 |
| LISNASDAL | TCSADLGSQNTLY | 0.67905986 |
| ISNASDALDK | TCSADLGSQNTLY | 0.5849245 |
| NQAETLVYQTE | TCSADLGSQNTLY | 0.49437958 |
| TGGTGGTAGAG | TCSADLGSQNTLY | 0.65858305 |
| TGSTAGGAGGA | TCSADLGSQNTLY | 0.6818545 |
| DPLAVPVAAGPV | TCSADLGSQNTLY | 0.5152745 |
| GGTGGTPGNGGN | TCSADLGSQNTLY | 0.6695121 |
| PAGLAGVDPATL | TCSADLGSQNTLY | 0.6694201 |
| GNNLVVEAPHAGASV | TCSADLGSQNTLY | 0.4466461 |

| | | |
|---|---|---|
| KVTRIMSASTLEQPA | TCSADLGSQNTLY | 0.6488819 |
| LGFALPIALPRRTVV | TCSADLGSQNTLY | 0.6922182 |
| TRLGNLTARKLVVLV | TCSADLGSQNTLY | 0.637151 |
| WSQQRRLIYFPSAGP | TCSADLGSQNTLY | 0.5826676 |
| GPNGNPDPMAAAVDIR | TCSADLGSQNTLY | 0.6888218 |
| TQRCVSVADLRAAASR | TCSADLGSQNTLY | 0.69996524 |
| QFVLDVAEHPVEVLTETV | TCSADLGSQNTLY | 0.69196165 |
| GGAGGTGGAGGNGGTQAGDG | TCSADLGSQNTLY | 0.40893015 |
| PLLATLRAIAVDVDAKPAQV | TCSADLGSQNTLY | 0.37486422 |
| APKFRTGGYMIDFWGVGYQVAK | TCSADLGSQNTLY | 0.5820539 |
| GAGGAGGAGGAAGGPGGTGGTG | TCSADLGSQNTLY | 0.6776798 |
| IGALGLLATDVAPAGSVAFGVA | TCSADLGSQNTLY | 0.6192931 |
| AGGAGGIGGNGGAGGAGGVLMG | TCSADLGSQNTLY | 0.5500003 |
| ADELIGLL | TCSADLGSQNTLY | 0.6736859 |
| ADRALKA | TCSADLGSQNTLY | 0.68987954 |
| AGAVGQVGGAGGA | TCSADLGSQNTLY | 0.540202 |
| AGGGGGLGGAP | TCSADLGSQNTLY | 0.40186653 |
| AGGLLYGNGGAG | TCSADLGSQNTLY | 0.67844814 |
| AGLESIGL | TCSADLGSQNTLY | 0.6737099 |
| AGLKTGII | TCSADLGSQNTLY | 0.6133865 |
| ASLAALLL | TCSADLGSQNTLY | 0.6535425 |
| ASVINISSI | TCSADLGSQNTLY | 0.55895776 |
| ATPPGPA | TCSADLGSQNTLY | 0.63810575 |
| ATVASALV | TCSADLGSQNTLY | 0.6787008 |
| ATVASLIL | TCSADLGSQNTLY | 0.5563351 |
| AVNGPVLIRH | TCSADLGSQNTLY | 0.6921173 |
| DGAAGAVGI | TCSADLGSQNTLY | 0.59486014 |
| DPLLHGI | TCSADLGSQNTLY | 0.6602675 |
| ELGAALGAL | TCSADLGSQNTLY | 0.6701168 |
| ELGRTQAV | TCSADLGSQNTLY | 0.6649898 |
| FVAGGYLS | TCSADLGSQNTLY | 0.63660824 |
| GASGGAGGAGGW | TCSADLGSQNTLY | 0.62549496 |
| GGAGAGGDGGAGGI | TCSADLGSQNTLY | 0.48896402 |
| GTGHNGSQPGMGGNG | TCSADLGSQNTLY | 0.4611533 |
| GYAAASASAS | TCSADLGSQNTLY | 0.6868243 |
| HQVDVAAVQQG | TCSADLGSQNTLY | 0.6245053 |
| IGAAIAHT | TCSADLGSQNTLY | 0.6908054 |
| IGMIEVAQL | TCSADLGSQNTLY | 0.6811629 |
| IRLAPEHTSAMD | TCSADLGSQNTLY | 0.56724226 |
| IYNVSVL | TCSADLGSQNTLY | 0.6711744 |
| KQGFLGM | TCSADLGSQNTLY | 0.52094185 |
| LESVITQL | TCSADLGSQNTLY | 0.577505 |
| LGLGALVVV | TCSADLGSQNTLY | 0.6621183 |
| LIGNGGNGGSGGI | TCSADLGSQNTLY | 0.6674198 |
| LNAINAPTL | TCSADLGSQNTLY | 0.58526 |
| LQLLNTGV | TCSADLGSQNTLY | 0.6590692 |
| LQTSIEF | TCSADLGSQNTLY | 0.67605627 |
| MSFAVSEL | TCSADLGSQNTLY | 0.5441892 |
| NTGALGL | TCSADLGSQNTLY | 0.6205304 |
| PPGAGGAGGAGTT | TCSADLGSQNTLY | 0.6886404 |
| QLVGLGI | TCSADLGSQNTLY | 0.6416651 |

| | | |
|---|---|---|
| RFLEQVHQL | TCSADLGSQNTLY | 0.5496931 |
| RGVGYVLR | TCSADLGSQNTLY | 0.67745245 |
| SATGVFAGVI | TCSADLGSQNTLY | 0.5669572 |
| SLPAITLP | TCSADLGSQNTLY | 0.6816759 |
| STALAFTV | TCSADLGSQNTLY | 0.64431727 |
| SVLTVGLGK | TCSADLGSQNTLY | 0.6061463 |
| TALAGAGSGI | TCSADLGSQNTLY | 0.67167 |
| TGLPFTGI | TCSADLGSQNTLY | 0.3883602 |
| VAEHVTV | TCSADLGSQNTLY | 0.57087135 |
| VDIVLNSL | TCSADLGSQNTLY | 0.6016334 |
| VLTGGGAVL | TCSADLGSQNTLY | 0.65598464 |
| VNVGLAYGI | TCSADLGSQNTLY | 0.6855639 |
| VSSAAVGAV | TCSADLGSQNTLY | 0.6012034 |
| VTSVYEAGV | TCSADLGSQNTLY | 0.66942245 |
| WVALDRGA | TCSADLGSQNTLY | 0.60946155 |
| YLVGSVVI | TCSADLGSQNTLY | 0.544822 |
| GPPGTGKTLL | TCSADLGSQNTLY | 0.62403774 |
| GVGMTKF | TCSADLGSQNTLY | 0.59287626 |
| IGIGGSDL | TCSADLGSQNTLY | 0.52909803 |
| IGTLLAEL | TCSADLGSQNTLY | 0.67107326 |
| IYTSGTT | TCSADLGSQNTLY | 0.5300478 |
| LSFPAGV | TCSADLGSQNTLY | 0.6177873 |
| LYTSGTT | TCSADLGSQNTLY | 0.6950697 |
| VGAFPFGF | TCSADLGSQNTLY | 0.5428666 |
| VGAGTVEF | TCSADLGSQNTLY | 0.66623193 |
| VGKSTLLL | TCSADLGSQNTLY | 0.6717031 |
| AALAHAGGGIV | TCSADLGSQNTLY | 0.6712483 |
| AALVAVSLL | TCSADLGSQNTLY | 0.63566506 |
| AAVSRAVAAL | TCSADLGSQNTLY | 0.6069115 |
| AGGSSGIGGF | TCSADLGSQNTLY | 0.69039315 |
| AGLTVANANG | TCSADLGSQNTLY | 0.68185127 |
| AGLVGAVL | TCSADLGSQNTLY | 0.60640717 |
| ALSQAFTV | TCSADLGSQNTLY | 0.6679473 |
| ALTFSVSL | TCSADLGSQNTLY | 0.6819755 |
| ALTGAALVAL | TCSADLGSQNTLY | 0.6764832 |
| ASFSIGAGVL | TCSADLGSQNTLY | 0.64022744 |
| DGSVYKI | TCSADLGSQNTLY | 0.43461466 |
| EQLAEIV | TCSADLGSQNTLY | 0.62448674 |
| FAVSELTVM | TCSADLGSQNTLY | 0.6635777 |
| GAAPLAAGL | TCSADLGSQNTLY | 0.6360828 |
| GADGGDGGSGGAGG | TCSADLGSQNTLY | 0.40341166 |
| GFGLDTGGAGG | TCSADLGSQNTLY | 0.68861663 |
| GLASGGLL | TCSADLGSQNTLY | 0.6985539 |
| GRIAGAAVLL | TCSADLGSQNTLY | 0.56932956 |
| GSAGSLNTSF | TCSADLGSQNTLY | 0.69625396 |
| GTGQIGI | TCSADLGSQNTLY | 0.4794316 |
| GTLIALIL | TCSADLGSQNTLY | 0.6661353 |
| IAATMGFVVG | TCSADLGSQNTLY | 0.68486905 |
| IFAQALL | TCSADLGSQNTLY | 0.6006365 |
| IGAAVALGF | TCSADLGSQNTLY | 0.6546839 |
| IGLTPLAL | TCSADLGSQNTLY | 0.5748292 |

| | | |
|---|---|---|
| ILGSNTSTL | TCSADLGSQNTLY | 0.6067705 |
| IVFSLTVL | TCSADLGSQNTLY | 0.69098675 |
| LASKATGF | TCSADLGSQNTLY | 0.5851613 |
| LASVALIL | TCSADLGSQNTLY | 0.6461047 |
| LFAVGLTL | TCSADLGSQNTLY | 0.5931122 |
| LFNGGIGTYI | TCSADLGSQNTLY | 0.52870107 |
| LTLGAGII | TCSADLGSQNTLY | 0.6471773 |
| LVAAALGDI | TCSADLGSQNTLY | 0.6722945 |
| NTAFVQLGI | TCSADLGSQNTLY | 0.60993725 |
| QLFSGAEL | TCSADLGSQNTLY | 0.69001234 |
| SAATFITL | TCSADLGSQNTLY | 0.5938756 |
| SNYGLANLGV | TCSADLGSQNTLY | 0.6972388 |
| SYGGAYAVM | TCSADLGSQNTLY | 0.6869296 |
| TTGVGGLTLG | TCSADLGSQNTLY | 0.6671195 |
| TTQTIYI | TCSADLGSQNTLY | 0.60205317 |
| VAAGAGLGAVI | TCSADLGSQNTLY | 0.69209224 |
| VAAVVTLL | TCSADLGSQNTLY | 0.59942496 |
| VGFLGGLI | TCSADLGSQNTLY | 0.62688744 |
| VGVIPILL | TCSADLGSQNTLY | 0.691293 |
| VIFASSLI | TCSADLGSQNTLY | 0.24801055 |
| VKGGDGGAAAG | TCSADLGSQNTLY | 0.7343794 |
| VTKTFTNI | TCSADLGSQNTLY | 0.65536296 |
| VVAAAILL | TCSADLGSQNTLY | 0.7146019 |
| YAALDVEL | TCSADLGSQNTLY | 0.42789912 |
| LASVALLL | TCSADLGSQNTLY | 0.26625323 |
| LDAVHRV | TCSADLGSQNTLY | 0.51206064 |
| QGLSSSIF | TCSADLGSQNTLY | 0.39378798 |
| RGTFANIRL | TCSADLGSQNTLY | 0.7425216 |
| RQLWWGHRI | TCSADLGSQNTLY | 0.6450851 |
| YGPPGTGKTLL | TCSADLGSQNTLY | 0.3879966 |
| AALVGAPANVA | TCSADLGSQNTLY | 0.72129977 |
| AATFRRTE | TCSADLGSQNTLY | 0.7283448 |
| AGALGGGI | TCSADLGSQNTLY | 0.28139836 |
| AGGAGGAGGIGAG | TCSADLGSQNTLY | 0.57198817 |
| AGSGALGL | TCSADLGSQNTLY | 0.5437534 |
| AQIADQI | TCSADLGSQNTLY | 0.5111629 |
| ASAVAVTYL | TCSADLGSQNTLY | 0.19741076 |
| ASTVLGGTGG | TCSADLGSQNTLY | 0.61964035 |
| AVGGEPAK | TCSADLGSQNTLY | 0.4892174 |
| CEQGTFGGTC | TCSADLGSQNTLY | 0.19419895 |
| DEGKAPAG | TCSADLGSQNTLY | 0.6804936 |
| DTVVFPAL | TCSADLGSQNTLY | 0.40956292 |
| FEGITMAG | TCSADLGSQNTLY | 0.46083814 |
| FPLVVQD | TCSADLGSQNTLY | 0.7237096 |
| GAFTTPQ | TCSADLGSQNTLY | 0.55453104 |
| GFGGFGTAGGVGGA | TCSADLGSQNTLY | 0.4361707 |
| GFLAQFL | TCSADLGSQNTLY | 0.68057394 |
| GGAGGAGGNAGVA | TCSADLGSQNTLY | 0.73763883 |
| GGVGTAGAGGAGGAGG | TCSADLGSQNTLY | 0.36866173 |
| GNGGGGDAGNAGSGGN | TCSADLGSQNTLY | 0.5973326 |
| GTGGAGGAGNDAGS | TCSADLGSQNTLY | 0.78444815 |

| | | |
|---|---|---|
| GTTNTGWFNS | TCSADLGSQNTLY | 0.6066421 |
| IFMGPLFTL | TCSADLGSQNTLY | 0.7785708 |
| ILAHAIEAI | TCSADLGSQNTLY | 0.78219044 |
| IRSAIREAL | TCSADLGSQNTLY | 0.383107 |
| ITFAGGG | TCSADLGSQNTLY | 0.7178681 |
| ITGMSGHQEV | TCSADLGSQNTLY | 0.7442572 |
| IVAGLACTL | TCSADLGSQNTLY | 0.54452467 |
| IVGGGLAGA | TCSADLGSQNTLY | 0.5938197 |
| IVTGVLNI | TCSADLGSQNTLY | 0.2685401 |
| LAQQGCI | TCSADLGSQNTLY | 0.5778731 |
| LCLHVEN | TCSADLGSQNTLY | 0.31936556 |
| LDAISDGS | TCSADLGSQNTLY | 0.67176914 |
| LGGFAAML | TCSADLGSQNTLY | 0.75145245 |
| LGGKNPMI | TCSADLGSQNTLY | 0.59280235 |
| LGPKGVAAIL | TCSADLGSQNTLY | 0.25982863 |
| LMVGVSI | TCSADLGSQNTLY | 0.27928907 |
| LVGAAFVL | TCSADLGSQNTLY | 0.64592147 |
| LVLAVAPL | TCSADLGSQNTLY | 0.28812024 |
| MAAIAVIDAA | TCSADLGSQNTLY | 0.33769888 |
| NNGINAELQ | TCSADLGSQNTLY | 0.3425383 |
| RGEPLVLVHG | TCSADLGSQNTLY | 0.7516198 |
| RGGAGGAAGLI | TCSADLGSQNTLY | 0.76244956 |
| RSYRMHPAV | TCSADLGSQNTLY | 0.41928244 |
| SAAHLLAGQLA | TCSADLGSQNTLY | 0.7488327 |
| SAAVGSY | TCSADLGSQNTLY | 0.72579336 |
| SPHAVLHRI | TCSADLGSQNTLY | 0.7603593 |
| SQIAKVL | TCSADLGSQNTLY | 0.6439526 |
| TALGHLVP | TCSADLGSQNTLY | 0.25481272 |
| TAPTTSGAA | TCSADLGSQNTLY | 0.7439462 |
| TDGDDGNGGSPGT | TCSADLGSQNTLY | 0.5444269 |
| TDGLGEN | TCSADLGSQNTLY | 0.6853548 |
| TGGPPLPGQA | TCSADLGSQNTLY | 0.72955275 |
| TIPAITI | TCSADLGSQNTLY | 0.59843445 |
| TIPGFTI | TCSADLGSQNTLY | 0.3852811 |
| TSIASGALA | TCSADLGSQNTLY | 0.69207597 |
| TVTLPGAAI | TCSADLGSQNTLY | 0.73796284 |
| VASGLSAALP | TCSADLGSQNTLY | 0.34764183 |
| VGAGVIGI | TCSADLGSQNTLY | 0.25081617 |
| VGPKRFGV | TCSADLGSQNTLY | 0.6048336 |
| VNVAFPD | TCSADLGSQNTLY | 0.6016497 |
| VSMTNSGVS | TCSADLGSQNTLY | 0.7466792 |
| VTDVLGDLSV | TCSADLGSQNTLY | 0.6429013 |
| VVVVGAGFAGL | TCSADLGSQNTLY | 0.5851134 |
| VYTVGTVV | TCSADLGSQNTLY | 0.70324934 |
| WTTFSGP | TCSADLGSQNTLY | 0.7139354 |
| YDELVEAF | TCSADLGSQNTLY | 0.6138065 |
| YGTGGAGGNGG | TCSADLGSQNTLY | 0.61858255 |
| YTNVMAV | TCSADLGSQNTLY | 0.46468028 |
| FLEQVHQL | TCSADLGSQNTLY | 0.73143035 |
| GLSLLAVAV | TCSADLGSQNTLY | 0.7365093 |
| LEQVHQL | TCSADLGSQNTLY | 0.67868674 |

| | | |
|---|---|---|
| LNAAGAI | TCSADLGSQNTLY | 0.7355163 |
| PAVHASL | TCSADLGSQNTLY | 0.5152695 |
| TELEDDA | TCSADLGSQNTLY | 0.6466294 |
| TPPGPPN | TCSADLGSQNTLY | 0.35787022 |
| TPPPPPPP | TCSADLGSQNTLY | 0.5271989 |
| TQSPALI | ASSQDGTGGSERLF | 0.52742463 |
| EFAAAFPQ | ASSQDGTGGSERLF | 0.6690425 |
| LLSGNLDVL | ASSQDGTGGSERLF | 0.68124056 |
| GPANLGNYNI | ASSQDGTGGSERLF | 0.6680896 |
| TLGPTLP | ASSQDGTGGSERLF | 0.44113258 |
| VYAYSSCCK | ASSQDGTGGSERLF | 0.7357368 |
| APAPAPAPNI | ASSQDGTGGSERLF | 0.7664882 |
| PGGPVPIP | ASSQDGTGGSERLF | 0.33998564 |
| FLTVALEP | ASSQDGTGGSERLF | 0.48065287 |
| SQYFGANL | ASSQDGTGGSERLF | 0.7363584 |
| GGVKPAA | ASSQDGTGGSERLF | 0.71756035 |
| GGTGGAGGAGGW | ASSQDGTGGSERLF | 0.5254097 |
| GVGGVAGLGGAGG | ASSQDGTGGSERLF | 0.63493216 |
| VLLVLATTI | ASSQDGTGGSERLF | 0.7693694 |
| IGIFAAML | ASSQDGTGGSERLF | 0.6214273 |
| AVGDLID | ASSQDGTGGSERLF | 0.7671486 |
| NISPLPL | ASSQDGTGGSERLF | 0.2575194 |
| VTAVFVAI | ASSQDGTGGSERLF | 0.56469405 |
| SGDDNDAAL | ASSQDGTGGSERLF | 0.5019111 |
| AVAALAHS | ASSQDGTGGSERLF | 0.16866241 |
| GGVTVMATP | ASSQDGTGGSERLF | 0.7454136 |
| AGLAANVRD | ASSQDGTGGSERLF | 0.4275903 |
| TNRPDLIDPALLRPG | ASSQDGTGGSERLF | 0.7119517 |
| LANLGVTAL | ASSQDGTGGSERLF | 0.44092792 |
| TGERIEL | ASSQDGTGGSERLF | 0.72808796 |
| NGEELAA | ASSQDGTGGSERLF | 0.6626745 |
| VLSLVWG | ASSQDGTGGSERLF | 0.5245418 |
| IGYALWL | ASSQDGTGGSERLF | 0.3701194 |
| LSLLLAPL | ASSQDGTGGSERLF | 0.68265533 |
| TLDEALK | ASSQDGTGGSERLF | 0.73386705 |
| TDGAGTAEGG | ASSQDGTGGSERLF | 0.75576746 |
| NAPGGAGGNGG | ASSQDGTGGSERLF | 0.25154617 |
| VSVFMGMGGNF | ASSQDGTGGSERLF | 0.71710306 |
| LASYLAAL | ASSQDGTGGSERLF | 0.5963632 |
| LVGGTPAL | ASSQDGTGGSERLF | 0.2831458 |
| VRNADVL | ASSQDGTGGSERLF | 0.21681716 |
| TCSSPFLVSAM | ASSQDGTGGSERLF | 0.49797273 |
| VHITDAE | ASSQDGTGGSERLF | 0.52790546 |
| EYAAPSF | ASSQDGTGGSERLF | 0.6720326 |
| VVDAFHGGL | ASSQDGTGGSERLF | 0.7315179 |
| ASVVDPQ | ASSQDGTGGSERLF | 0.42605096 |
| SYNEFGGLS | ASSQDGTGGSERLF | 0.70323944 |
| DPDGSVY | ASSQDGTGGSERLF | 0.5626992 |
| MVSTIHATE | ASSQDGTGGSERLF | 0.1499103 |
| LAAGVADPV | ASSQDGTGGSERLF | 0.48564354 |
| TKEQIAA | ASSQDGTGGSERLF | 0.26488978 |

| | | |
|---|---|---|
| SLASPAAL | ASSQDGTGGSERLF | 0.7314236 |
| IGLVFYGV | ASSQDGTGGSERLF | 0.27307975 |
| FLLDASNL | ASSQDGTGGSERLF | 0.5338055 |
| YQVGNFP | ASSQDGTGGSERLF | 0.543396 |
| ESLLSQST | ASSQDGTGGSERLF | 0.7258946 |
| LFTVGFAAVG | ASSQDGTGGSERLF | 0.67501915 |
| ATPPTPT | ASSQDGTGGSERLF | 0.39970922 |
| TDLATLG | ASSQDGTGGSERLF | 0.29631928 |
| SAGGTGR | ASSQDGTGGSERLF | 0.70699173 |
| GFSNSGS | ASSQDGTGGSERLF | 0.5546195 |
| GGNGGSGGTSV | ASSQDGTGGSERLF | 0.38930845 |
| GTSVATGGAGN | ASSQDGTGGSERLF | 0.4454109 |
| NGGNGGSGGTS | ASSQDGTGGSERLF | 0.76031965 |
| AVVKFPH | ASSQDGTGGSERLF | 0.24289882 |
| TPPPPPP | ASSQDGTGGSERLF | 0.75403696 |
| AAPAAAAAPA | ASSQDGTGGSERLF | 0.37689948 |
| IALNEDL | ASSQDGTGGSERLF | 0.5160456 |
| IVNVSSM | ASSQDGTGGSERLF | 0.77022535 |
| TGIPPAPR | ASSQDGTGGSERLF | 0.7114322 |
| FDLGGGTF | ASSQDGTGGSERLF | 0.59288645 |
| FDLGGGTFDVS | ASSQDGTGGSERLF | 0.49437863 |
| NEPTAAA | ASSQDGTGGSERLF | 0.3228856 |
| NEPTAAALA | ASSQDGTGGSERLF | 0.42982262 |
| GSIINISSI | ASSQDGTGGSERLF | 0.40570247 |
| VAPGNAGTA | ASSQDGTGGSERLF | 0.76620203 |
| TGGGASLE | ASSQDGTGGSERLF | 0.5997275 |
| VAVYNHYKRI | ASSQDGTGGSERLF | 0.28938696 |
| TDPAPATT | ASSQDGTGGSERLF | 0.70191497 |
| TNRPDILDPALLRPG | ASSQDGTGGSERLF | 0.75949335 |
| LTTTTTS | ASSQDGTGGSERLF | 0.7577925 |
| PTGSGKTL | ASSQDGTGGSERLF | 0.32938582 |
| TGSGKTL | ASSQDGTGGSERLF | 0.53828514 |
| TIGHVDHGK | ASSQDGTGGSERLF | 0.6021548 |
| TIGHVDHGKTT | ASSQDGTGGSERLF | 0.5419068 |
| GHVDHGKTT | ASSQDGTGGSERLF | 0.3993828 |
| AQFEHTLL | ASSQDGTGGSERLF | 0.37467822 |
| IVNQDEL | ASSQDGTGGSERLF | 0.7342794 |
| TNGSQFF | ASSQDGTGGSERLF | 0.7921902 |
| TGKGIVE | ASSQDGTGGSERLF | 0.50014013 |
| LISNASDA | ASSQDGTGGSERLF | 0.22003095 |
| AELRTLGELTKIAKAHGAQ | ASSQDGTGGSERLF | 0.5094545 |
| APNAVAAAEPSW | ASSQDGTGGSERLF | 0.35164815 |
| APQVAGAATAVL | ASSQDGTGGSERLF | 0.383803 |
| ARRDPAAAAAILPTDA | ASSQDGTGGSERLF | 0.47759646 |
| DAAARDVLAEAGLAEYF | ASSQDGTGGSERLF | 0.24732 |
| DDQIVLSAKANLAKVEQ | ASSQDGTGGSERLF | 0.5147614 |
| DELLPDTGEDSDEDGH | ASSQDGTGGSERLF | 0.7757498 |
| ELPEAPANRATVPQD | ASSQDGTGGSERLF | 0.58148134 |
| FGQSKIERVTGIG | ASSQDGTGGSERLF | 0.6972012 |
| GAGGTGSTAGGAGGAGGAGGLY | ASSQDGTGGSERLF | 0.76146805 |
| GHGGTGAAGGNGGTGGAGGSGI | ASSQDGTGGSERLF | 0.6791818 |

| | | |
|---|---|---|
| GNGGNGGAGGEGGPGGAGGAGA | ASSQDGTGGSERLF | 0.7537979 |
| GSTAGNGGNGGNATNGTIAGQP | ASSQDGTGGSERLF | 0.7376934 |
| GTGGDGGNGGNGGTGG | ASSQDGTGGSERLF | 0.57952726 |
| GVRRPAGIPADAPV | ASSQDGTGGSERLF | 0.58073485 |
| GYRQVTDAHLLTLARR | ASSQDGTGGSERLF | 0.7293457 |
| IAKMGQDGHDRG | ASSQDGTGGSERLF | 0.5147573 |
| KAEPSIEIQYT | ASSQDGTGGSERLF | 0.6208594 |
| KPVRIRMAADEDTVAVT | ASSQDGTGGSERLF | 0.62485915 |
| MPPRIAGMRLLVIKPEP | ASSQDGTGGSERLF | 0.69804007 |
| MKFYAVALFRSRGVPPTR | ASSQDGTGGSERLF | 0.6286714 |
| NAGGFGILGAVAHSP | ASSQDGTGGSERLF | 0.7659276 |
| NEGREITLAMAEM | ASSQDGTGGSERLF | 0.19255728 |
| NELVEWSHPLGHRWRWE | ASSQDGTGGSERLF | 0.3930142 |
| NLGAGGTGGNGGNPG | ASSQDGTGGSERLF | 0.75257385 |
| NTEWPKWETVMAADPVD | ASSQDGTGGSERLF | 0.17242026 |
| PASAAPAETQP | ASSQDGTGGSERLF | 0.30335414 |
| QQSSDGKNGVRAAEITGE | ASSQDGTGGSERLF | 0.6022127 |
| RAPDAVTTSWWKE | ASSQDGTGGSERLF | 0.74859303 |
| SGFGNVGAHTSGWFNQ | ASSQDGTGGSERLF | 0.18094137 |
| SGVAGAVGQAASVGGLKV | ASSQDGTGGSERLF | 0.5694054 |
| SNGGAGTGSAGGNGGT | ASSQDGTGGSERLF | 0.26365873 |
| TGGAGGEGGAAG | ASSQDGTGGSERLF | 0.73781514 |
| TGGDGGAAGAGGAPGHG | ASSQDGTGGSERLF | 0.49381453 |
| TMPPLLAAIANAGRHGV | ASSQDGTGGSERLF | 0.22919448 |
| TSSTAGNSGGA | ASSQDGTGGSERLF | 0.6978716 |
| VPATVPVGILC | ASSQDGTGGSERLF | 0.35020643 |
| VQRPDSGGVVLTGRIS | ASSQDGTGGSERLF | 0.69928944 |
| WTPNPVPATKGVPAGLVQAT | ASSQDGTGGSERLF | 0.7322717 |
| YALKRALAKFSDRFDVVII | ASSQDGTGGSERLF | 0.7814563 |
| YLIGSVIV | ASSQDGTGGSERLF | 0.2536773 |
| YGQLVAVID | ASSQDGTGGSERLF | 0.65999943 |
| VTVGGMTLPA | ASSQDGTGGSERLF | 0.46355677 |
| VLALPAV | ASSQDGTGGSERLF | 0.785362 |
| VIPPTLNYET | ASSQDGTGGSERLF | 0.5063201 |
| VGMGGMHPSSG | ASSQDGTGGSERLF | 0.23906517 |
| VGLAGGAATA | ASSQDGTGGSERLF | 0.7091752 |
| VGGAGGAGGSGP | ASSQDGTGGSERLF | 0.67957985 |
| VFAAFMV | ASSQDGTGGSERLF | 0.49954748 |
| TLYDKLR | ASSQDGTGGSERLF | 0.72342646 |
| TLAVFTVL | ASSQDGTGGSERLF | 0.55459034 |
| TGSVQPARV | ASSQDGTGGSERLF | 0.49637857 |
| TGGGAGSGGT | ASSQDGTGGSERLF | 0.7251354 |
| SHLETSL | ASSQDGTGGSERLF | 0.6105261 |
| SFTTGMT | ASSQDGTGGSERLF | 0.49145284 |
| QVEVAKRMGVRQPTVS | ASSQDGTGGSERLF | 0.72561115 |
| QGSGGPALAM | ASSQDGTGGSERLF | 0.6113936 |
| MLAPSNP | ASSQDGTGGSERLF | 0.32748982 |
| LVNNAAI | ASSQDGTGGSERLF | 0.781564 |
| LTGEINTLA | ASSQDGTGGSERLF | 0.55519027 |
| LSIFGPV | ASSQDGTGGSERLF | 0.72144544 |
| LLALLANL | ASSQDGTGGSERLF | 0.6656585 |

| | | |
|---|---|---|
| LISNASDALDKLR | ASSQDGTGGSERLF | 0.7731494 |
| LINEAVP | ASSQDGTGGSERLF | 0.4277543 |
| LGYAFSGL | ASSQDGTGGSERLF | 0.5755507 |
| LDFVAVAL | ASSQDGTGGSERLF | 0.6660075 |
| LAQPGAE | ASSQDGTGGSERLF | 0.7081286 |
| ISNLGAVTGF | ASSQDGTGGSERLF | 0.45415387 |
| HQEDPGRI | ASSQDGTGGSERLF | 0.70089066 |
| GTPGEPTP | ASSQDGTGGSERLF | 0.67594707 |
| GFFNSGS | ASSQDGTGGSERLF | 0.7624196 |
| GAGGGGLTAGAN | ASSQDGTGGSERLF | 0.5306862 |
| FVDALVNM | ASSQDGTGGSERLF | 0.57853556 |
| FLTVAPEL | ASSQDGTGGSERLF | 0.77744305 |
| FETLTAL | ASSQDGTGGSERLF | 0.2982546 |
| DVINPIEL | ASSQDGTGGSERLF | 0.62862146 |
| APVSGAPV | ASSQDGTGGSERLF | 0.72109294 |
| ALVGGLAW | ASSQDGTGGSERLF | 0.73337317 |
| ALAAAAATAI | ASSQDGTGGSERLF | 0.5662222 |
| AFGLGLLL | ASSQDGTGGSERLF | 0.73773134 |
| AFFAAASL | ASSQDGTGGSERLF | 0.5087182 |
| AAAALPAFSPP | ASSQDGTGGSERLF | 0.67962974 |
| VVLAPAL | ASSQDGTGGSERLF | 0.5349712 |
| TIGHVDHGKTTL | ASSQDGTGGSERLF | 0.5635333 |
| LISNASDALDK | ASSQDGTGGSERLF | 0.42961705 |
| LISNASDAL | ASSQDGTGGSERLF | 0.6117964 |
| ISNASDALDK | ASSQDGTGGSERLF | 0.72354764 |
| NQAETLVYQTE | ASSQDGTGGSERLF | 0.6695927 |
| TGGTGGTAGAG | ASSQDGTGGSERLF | 0.29062632 |
| TGSTAGGAGGA | ASSQDGTGGSERLF | 0.66805816 |
| DPLAVPVAAGPV | ASSQDGTGGSERLF | 0.72227246 |
| GGTGGTPGNGGN | ASSQDGTGGSERLF | 0.78530073 |
| PAGLAGVDPATL | ASSQDGTGGSERLF | 0.61026555 |
| GNNLVVEAPHAGASV | ASSQDGTGGSERLF | 0.7483083 |
| KVTRIMSASTLEQPA | ASSQDGTGGSERLF | 0.5926359 |
| LGFALPIALPRRTVV | ASSQDGTGGSERLF | 0.6287631 |
| TRLGNLTARKLVVLV | ASSQDGTGGSERLF | 0.3058331 |
| WSQQRRLIYFPSAGP | ASSQDGTGGSERLF | 0.40156937 |
| GPNGNPDPMAAAVDIR | ASSQDGTGGSERLF | 0.5863353 |
| TQRCVSVADLRAAASR | ASSQDGTGGSERLF | 0.4095834 |
| QFVLDVAEHPVEVLTETV | ASSQDGTGGSERLF | 0.7476087 |
| GGAGGTGGAGGNGGTQAGDG | ASSQDGTGGSERLF | 0.73916924 |
| PLLATLRAIAVDVDAKPAQV | ASSQDGTGGSERLF | 0.7075393 |
| APKFRTGGYMIDFWGVGYQVAK | ASSQDGTGGSERLF | 0.76205516 |
| GAGGAGGAGGAAGGPGGTGGTG | ASSQDGTGGSERLF | 0.75528634 |
| IGALGLLATDVAPAGSVAFGVA | ASSQDGTGGSERLF | 0.7146907 |
| AGGAGGIGGNGGAGGAGGVLMG | ASSQDGTGGSERLF | 0.4927385 |
| ADELIGLL | ASSQDGTGGSERLF | 0.544718 |
| ADRALKA | ASSQDGTGGSERLF | 0.24060068 |
| AGAVGQVGGAGGA | ASSQDGTGGSERLF | 0.40848923 |
| AGGGGGLGGAP | ASSQDGTGGSERLF | 0.4792976 |
| AGGLLYGNGGAG | ASSQDGTGGSERLF | 0.5823324 |
| AGLESIGL | ASSQDGTGGSERLF | 0.16633098 |

| | | |
|---|---|---|
| AGLKTGII | ASSQDGTGGSERLF | 0.69446355 |
| ASLAALLL | ASSQDGTGGSERLF | 0.76763135 |
| ASVINISSI | ASSQDGTGGSERLF | 0.5816082 |
| ATPPGPA | ASSQDGTGGSERLF | 0.71902 |
| ATVASALV | ASSQDGTGGSERLF | 0.68815947 |
| ATVASLIL | ASSQDGTGGSERLF | 0.45911855 |
| AVNGPVLIRH | ASSQDGTGGSERLF | 0.7276 |
| DGAAGAVGI | ASSQDGTGGSERLF | 0.48662573 |
| DPLLHGI | ASSQDGTGGSERLF | 0.7614925 |
| ELGAALGAL | ASSQDGTGGSERLF | 0.7661934 |
| ELGRTQAV | ASSQDGTGGSERLF | 0.6650144 |
| FVAGGYLS | ASSQDGTGGSERLF | 0.48648283 |
| GASGGAGGAGGW | ASSQDGTGGSERLF | 0.6893655 |
| GGAGAGGDGGAGGI | ASSQDGTGGSERLF | 0.43443212 |
| GTGHNGSQPGMGGNG | ASSQDGTGGSERLF | 0.5336212 |
| GYAAASASAS | ASSQDGTGGSERLF | 0.48545626 |
| HQVDVAAVQQG | ASSQDGTGGSERLF | 0.6464269 |
| IGAAIAHT | ASSQDGTGGSERLF | 0.34107572 |
| IGMIEVAQL | ASSQDGTGGSERLF | 0.68157923 |
| IRLAPEHTSAMD | ASSQDGTGGSERLF | 0.59299344 |
| IYNVSVL | ASSQDGTGGSERLF | 0.65445626 |
| KQGFLGM | ASSQDGTGGSERLF | 0.37929994 |
| LESVITQL | ASSQDGTGGSERLF | 0.7084134 |
| LGLGALVVV | ASSQDGTGGSERLF | 0.71312964 |
| LIGNGGNGGSGGI | ASSQDGTGGSERLF | 0.29809633 |
| LNAINAPTL | ASSQDGTGGSERLF | 0.7108874 |
| LQLLNTGV | ASSQDGTGGSERLF | 0.49590617 |
| LQTSIEF | ASSQDGTGGSERLF | 0.54392326 |
| MSFAVSEL | ASSQDGTGGSERLF | 0.7352853 |
| NTGALGL | ASSQDGTGGSERLF | 0.51502466 |
| PPGAGGAGGAGTT | ASSQDGTGGSERLF | 0.32660118 |
| QLVGLGI | ASSQDGTGGSERLF | 0.49940658 |
| RFLEQVHQL | ASSQDGTGGSERLF | 0.6326003 |
| RGVGYVLR | ASSQDGTGGSERLF | 0.54891855 |
| SATGVFAGVI | ASSQDGTGGSERLF | 0.18859039 |
| SLPAITLP | ASSQDGTGGSERLF | 0.60358036 |
| STALAFTV | ASSQDGTGGSERLF | 0.33216345 |
| SVLTVGLGK | ASSQDGTGGSERLF | 0.40165764 |
| TALAGAGSGI | ASSQDGTGGSERLF | 0.43900162 |
| TGLPFTGI | ASSQDGTGGSERLF | 0.74134946 |
| VAEHVTV | ASSQDGTGGSERLF | 0.6892944 |
| VDIVLNSL | ASSQDGTGGSERLF | 0.5405607 |
| VLTGGGAVL | ASSQDGTGGSERLF | 0.6465811 |
| VNVGLAYGI | ASSQDGTGGSERLF | 0.5874436 |
| VSSAAVGAV | ASSQDGTGGSERLF | 0.7544136 |
| VTSVYEAGV | ASSQDGTGGSERLF | 0.64853704 |
| WVALDRGA | ASSQDGTGGSERLF | 0.6208829 |
| YLVGSVVI | ASSQDGTGGSERLF | 0.6914359 |
| GPPGTGKTLL | ASSQDGTGGSERLF | 0.54249 |
| GVGMTKF | ASSQDGTGGSERLF | 0.39724207 |
| IGIGGSDL | ASSQDGTGGSERLF | 0.4862502 |

| | | |
|---|---|---|
| IGTLLAEL | ASSQDGTGGSERLF | 0.38263136 |
| IYTSGTT | ASSQDGTGGSERLF | 0.35537288 |
| LSFPAGV | ASSQDGTGGSERLF | 0.6580665 |
| LYTSGTT | ASSQDGTGGSERLF | 0.746516 |
| VGAFPFGF | ASSQDGTGGSERLF | 0.5336 |
| VGAGTVEF | ASSQDGTGGSERLF | 0.22436748 |
| VGKSTLLL | ASSQDGTGGSERLF | 0.47410935 |
| AALAHAGGGIV | ASSQDGTGGSERLF | 0.49969074 |
| AALVAVSLL | ASSQDGTGGSERLF | 0.7243309 |
| AAVSRAVAAL | ASSQDGTGGSERLF | 0.7166395 |
| AGGSSGIGGF | ASSQDGTGGSERLF | 0.77005273 |
| AGLTVANANG | ASSQDGTGGSERLF | 0.72713125 |
| AGLVGAVL | ASSQDGTGGSERLF | 0.47995722 |
| ALSQAFTV | ASSQDGTGGSERLF | 0.7166718 |
| ALTFSVSL | ASSQDGTGGSERLF | 0.5136653 |
| ALTGAALVAL | ASSQDGTGGSERLF | 0.6491411 |
| ASFSIGAGVL | ASSQDGTGGSERLF | 0.595055 |
| DGSVYKI | ASSQDGTGGSERLF | 0.6147287 |
| EQLAEIV | ASSQDGTGGSERLF | 0.3925169 |
| FAVSELTVM | ASSQDGTGGSERLF | 0.51443267 |
| GAAPLAAGL | ASSQDGTGGSERLF | 0.2889047 |
| GADGGDGGSGGAGG | ASSQDGTGGSERLF | 0.65802264 |
| GFGLDTGGAGG | ASSQDGTGGSERLF | 0.39607373 |
| GLASGGLL | ASSQDGTGGSERLF | 0.46660626 |
| GRIAGAAVLL | ASSQDGTGGSERLF | 0.556211 |
| GSAGSLNTSF | ASSQDGTGGSERLF | 0.74851525 |
| GTGQIGI | ASSQDGTGGSERLF | 0.784673 |
| GTLIALIL | ASSQDGTGGSERLF | 0.4901302 |
| IAATMGFVVG | ASSQDGTGGSERLF | 0.4264341 |
| IFAQALL | ASSQDGTGGSERLF | 0.7177735 |
| IGAAVALGF | ASSQDGTGGSERLF | 0.4780475 |
| IGLTPLAL | ASSQDGTGGSERLF | 0.6659287 |
| ILGSNTSTL | ASSQDGTGGSERLF | 0.617999 |
| IVFSLTVL | ASSQDGTGGSERLF | 0.48995215 |
| LASKATGF | ASSQDGTGGSERLF | 0.47802693 |
| LASVALIL | ASSQDGTGGSERLF | 0.7127849 |
| LFAVGLTL | ASSQDGTGGSERLF | 0.47068268 |
| LFNGGIGTYI | ASSQDGTGGSERLF | 0.65735924 |
| LTLGAGII | ASSQDGTGGSERLF | 0.25812235 |
| LVAAALGDI | ASSQDGTGGSERLF | 0.64709413 |
| NTAFVQLGI | ASSQDGTGGSERLF | 0.34244013 |
| QLFSGAEL | ASSQDGTGGSERLF | 0.39656174 |
| SAATFITL | ASSQDGTGGSERLF | 0.18981478 |
| SNYGLANLGV | ASSQDGTGGSERLF | 0.5581355 |
| SYGGAYAVM | ASSQDGTGGSERLF | 0.6682346 |
| TTGVGGLTLG | ASSQDGTGGSERLF | 0.77122813 |
| TTQTIYI | ASSQDGTGGSERLF | 0.6644166 |
| VAAGAGLGAVI | ASSQDGTGGSERLF | 0.6548393 |
| VAAVVTLL | ASSQDGTGGSERLF | 0.7508029 |
| VGFLGGLI | ASSQDGTGGSERLF | 0.47474787 |
| VGVIPILL | ASSQDGTGGSERLF | 0.23438577 |

| | | |
|---|---|---|
| VIFASSLI | ASSQDGTGGSERLF | 0.6107561 |
| VKGGDGGAAAG | ASSQDGTGGSERLF | 0.5430165 |
| VTKTFTNI | ASSQDGTGGSERLF | 0.7470286 |
| VVAAAILL | ASSQDGTGGSERLF | 0.6582539 |
| YAALDVEL | ASSQDGTGGSERLF | 0.25644958 |
| LASVALLL | ASSQDGTGGSERLF | 0.6669246 |
| LDAVHRV | ASSQDGTGGSERLF | 0.7110127 |
| QGLSSSIF | ASSQDGTGGSERLF | 0.2987556 |
| RGTFANIRL | ASSQDGTGGSERLF | 0.4083991 |
| RQLWWGHRI | ASSQDGTGGSERLF | 0.37877095 |
| YGPPGTGKTLL | ASSQDGTGGSERLF | 0.31628913 |
| AALVGAPANVA | ASSQDGTGGSERLF | 0.5430144 |
| AATFRRTE | ASSQDGTGGSERLF | 0.48817283 |
| AGALGGGI | ASSQDGTGGSERLF | 0.56098557 |
| AGGAGGAGGIGAG | ASSQDGTGGSERLF | 0.4818052 |
| AGSGALGL | ASSQDGTGGSERLF | 0.40873188 |
| AQIADQI | ASSQDGTGGSERLF | 0.5390332 |
| ASAVAVTYL | ASSQDGTGGSERLF | 0.5626387 |
| ASTVLGGTGG | ASSQDGTGGSERLF | 0.62124634 |
| AVGGEPAK | ASSQDGTGGSERLF | 0.7231524 |
| CEQGTFGGTC | ASSQDGTGGSERLF | 0.4113112 |
| DEGKAPAG | ASSQDGTGGSERLF | 0.64793235 |
| DTVVFPAL | ASSQDGTGGSERLF | 0.7706078 |
| FEGITMAG | ASSQDGTGGSERLF | 0.31761444 |
| FPLVVQD | ASSQDGTGGSERLF | 0.38954547 |
| GAFTTPQ | ASSQDGTGGSERLF | 0.6622175 |
| GFGGFGTAGGVGGA | ASSQDGTGGSERLF | 0.5484781 |
| GFLAQFL | ASSQDGTGGSERLF | 0.7463334 |
| GGAGGAGGNAGVA | ASSQDGTGGSERLF | 0.6254233 |
| GGVGTAGAGGAGGAGG | ASSQDGTGGSERLF | 0.62880147 |
| GNGGGGDAGNAGSGGN | ASSQDGTGGSERLF | 0.6184056 |
| GTGGAGGAGNDAGS | ASSQDGTGGSERLF | 0.77145725 |
| GTTNTGWFNS | ASSQDGTGGSERLF | 0.6713253 |
| IFMGPLFTL | ASSQDGTGGSERLF | 0.45891386 |
| ILAHAIEAI | ASSQDGTGGSERLF | 0.63905287 |
| IRSAIREAL | ASSQDGTGGSERLF | 0.2883572 |
| ITFAGGG | ASSQDGTGGSERLF | 0.69472814 |
| ITGMSGHQEV | ASSQDGTGGSERLF | 0.72838354 |
| IVAGLACTL | ASSQDGTGGSERLF | 0.36984587 |
| IVGGGLAGA | ASSQDGTGGSERLF | 0.5015193 |
| IVTGVLNI | ASSQDGTGGSERLF | 0.7786982 |
| LAQQGCI | ASSQDGTGGSERLF | 0.74544036 |
| LCLHVEN | ASSQDGTGGSERLF | 0.46009007 |
| LDAISDGS | ASSQDGTGGSERLF | 0.40453136 |
| LGGFAAML | ASSQDGTGGSERLF | 0.4608072 |
| LGGKNPMI | ASSQDGTGGSERLF | 0.54383373 |
| LGPKGVAAIL | ASSQDGTGGSERLF | 0.22675265 |
| LMVGVSI | ASSQDGTGGSERLF | 0.7116108 |
| LVGAAFVL | ASSQDGTGGSERLF | 0.51119757 |
| LVLAVAPL | ASSQDGTGGSERLF | 0.7512759 |
| MAAIAVIDAA | ASSQDGTGGSERLF | 0.6654564 |

| | | |
|---|---|---|
| NNGINAELQ | ASSQDGTGGSERLF | 0.7622051 |
| RGEPLVLVHG | ASSQDGTGGSERLF | 0.46868 |
| RGGAGGAAGLI | ASSQDGTGGSERLF | 0.5736954 |
| RSYRMHPAV | ASSQDGTGGSERLF | 0.23808384 |
| SAAHLLAGQLA | ASSQDGTGGSERLF | 0.5380906 |
| SAAVGSY | ASSQDGTGGSERLF | 0.34749353 |
| SPHAVLHRI | ASSQDGTGGSERLF | 0.6055651 |
| SQIAKVL | ASSQDGTGGSERLF | 0.30867332 |
| TALGHLVP | ASSQDGTGGSERLF | 0.7632761 |
| TAPTTSGAA | ASSQDGTGGSERLF | 0.48924524 |
| TDGDDGNGGSPGT | ASSQDGTGGSERLF | 0.720778 |
| TDGLGEN | ASSQDGTGGSERLF | 0.45949137 |
| TGGPPLPGQA | ASSQDGTGGSERLF | 0.30135855 |
| TIPAITI | ASSQDGTGGSERLF | 0.46785504 |
| TIPGFTI | ASSQDGTGGSERLF | 0.74311125 |
| TSIASGALA | ASSQDGTGGSERLF | 0.2526496 |
| TVTLPGAAI | ASSQDGTGGSERLF | 0.22070725 |
| VASGLSAALP | ASSQDGTGGSERLF | 0.40455735 |
| VGAGVIGI | ASSQDGTGGSERLF | 0.5910549 |
| VGPKRFGV | ASSQDGTGGSERLF | 0.7619952 |
| VNVAFPD | ASSQDGTGGSERLF | 0.7545736 |
| VSMTNSGVS | ASSQDGTGGSERLF | 0.6052641 |
| VTDVLGDLSV | ASSQDGTGGSERLF | 0.7788728 |
| VVVVGAGFAGL | ASSQDGTGGSERLF | 0.16629687 |
| VYTVGTVV | ASSQDGTGGSERLF | 0.46778896 |
| WTTFSGP | ASSQDGTGGSERLF | 0.78151256 |
| YDELVEAF | ASSQDGTGGSERLF | 0.65495205 |
| YGTGGAGGNGG | ASSQDGTGGSERLF | 0.57142437 |
| YTNVMAV | ASSQDGTGGSERLF | 0.73515296 |
| FLEQVHQL | ASSQDGTGGSERLF | 0.41767633 |
| GLSLLAVAV | ASSQDGTGGSERLF | 0.5659732 |
| LEQVHQL | ASSQDGTGGSERLF | 0.62007 |
| LNAAGAI | ASSQDGTGGSERLF | 0.7005613 |
| PAVHASL | ASSQDGTGGSERLF | 0.65953314 |
| TELEDDA | ASSQDGTGGSERLF | 0.21856269 |
| TPPGPPN | ASSQDGTGGSERLF | 0.72255313 |
| TPPPPPPP | ASSQDGTGGSERLF | 0.779446 |
| TQSPALI | SSSPGTDNNQAPL | 0.48923126 |
| EFAAAFPQ | SSSPGTDNNQAPL | 0.70948064 |
| LLSGNLDVL | SSSPGTDNNQAPL | 0.5697746 |
| GPANLGNYNI | SSSPGTDNNQAPL | 0.57071686 |
| TLGPTLP | SSSPGTDNNQAPL | 0.4049316 |
| VYAYSSCCK | SSSPGTDNNQAPL | 0.2855528 |
| APAPAPAPNI | SSSPGTDNNQAPL | 0.69839144 |
| PGGPVPIP | SSSPGTDNNQAPL | 0.36483845 |
| FLTVALEP | SSSPGTDNNQAPL | 0.6500256 |
| SQYFGANL | SSSPGTDNNQAPL | 0.5715716 |
| GGVKPAA | SSSPGTDNNQAPL | 0.7948873 |
| GGTGGAGGAGGW | SSSPGTDNNQAPL | 0.692973 |
| GVGGVAGLGGAGG | SSSPGTDNNQAPL | 0.6095303 |
| VLLVLATTI | SSSPGTDNNQAPL | 0.44577974 |

| | | |
|---|---|---|
| IGIFAAML | SSSPGTDNNQAPL | 0.61614275 |
| AVGDLID | SSSPGTDNNQAPL | 0.69636774 |
| NISPLPL | SSSPGTDNNQAPL | 0.28110301 |
| VTAVFVAI | SSSPGTDNNQAPL | 0.75577176 |
| SGDDNDAAL | SSSPGTDNNQAPL | 0.34826076 |
| AVAALAHS | SSSPGTDNNQAPL | 0.6211928 |
| GGVTVMATP | SSSPGTDNNQAPL | 0.68501407 |
| AGLAANVRD | SSSPGTDNNQAPL | 0.7537899 |
| TNRPDLIDPALLRPG | SSSPGTDNNQAPL | 0.60264194 |
| LANLGVTAL | SSSPGTDNNQAPL | 0.73824775 |
| TGERIEL | SSSPGTDNNQAPL | 0.398397 |
| NGEELAA | SSSPGTDNNQAPL | 0.7198976 |
| VLSLVWG | SSSPGTDNNQAPL | 0.7552966 |
| IGYALWL | SSSPGTDNNQAPL | 0.635488 |
| LSLLLAPL | SSSPGTDNNQAPL | 0.7743875 |
| TLDEALK | SSSPGTDNNQAPL | 0.6015551 |
| TDGAGTAEGG | SSSPGTDNNQAPL | 0.22233304 |
| NAPGGAGGNGG | SSSPGTDNNQAPL | 0.57878816 |
| VSVFMGMGGNF | SSSPGTDNNQAPL | 0.7047609 |
| LASYLAAL | SSSPGTDNNQAPL | 0.7628046 |
| LVGGTPAL | SSSPGTDNNQAPL | 0.7074052 |
| VRNADVL | SSSPGTDNNQAPL | 0.61055005 |
| TCSSPFLVSAM | SSSPGTDNNQAPL | 0.5690945 |
| VHITDAE | SSSPGTDNNQAPL | 0.7743207 |
| EYAAPSF | SSSPGTDNNQAPL | 0.81031466 |
| VVDAFHGGL | SSSPGTDNNQAPL | 0.69112676 |
| ASVVDPQ | SSSPGTDNNQAPL | 0.45632935 |
| SYNEFGGLS | SSSPGTDNNQAPL | 0.4551906 |
| DPDGSVY | SSSPGTDNNQAPL | 0.70139575 |
| MVSTIHATE | SSSPGTDNNQAPL | 0.68242925 |
| LAAGVADPV | SSSPGTDNNQAPL | 0.35853186 |
| TKEQIAA | SSSPGTDNNQAPL | 0.55375713 |
| SLASPAAL | SSSPGTDNNQAPL | 0.64009154 |
| IGLVFYGV | SSSPGTDNNQAPL | 0.198953 |
| FLLDASNL | SSSPGTDNNQAPL | 0.66869706 |
| YQVGNFP | SSSPGTDNNQAPL | 0.74943954 |
| ESLLSQST | SSSPGTDNNQAPL | 0.6655921 |
| LFTVGFAAVG | SSSPGTDNNQAPL | 0.29460406 |
| ATPPTPT | SSSPGTDNNQAPL | 0.75669557 |
| TDLATLG | SSSPGTDNNQAPL | 0.76930904 |
| SAGGTGR | SSSPGTDNNQAPL | 0.7726925 |
| GFSNSGS | SSSPGTDNNQAPL | 0.28207466 |
| GGNGGSGGTSV | SSSPGTDNNQAPL | 0.17672937 |
| GTSVATGGAGN | SSSPGTDNNQAPL | 0.6699604 |
| NGGNGGSGGTS | SSSPGTDNNQAPL | 0.70327294 |
| AVVKFPH | SSSPGTDNNQAPL | 0.4746074 |
| TPPPPPP | SSSPGTDNNQAPL | 0.32546902 |
| AAPAAAAAPA | SSSPGTDNNQAPL | 0.5728885 |
| IALNEDL | SSSPGTDNNQAPL | 0.7414612 |
| IVNVSSM | SSSPGTDNNQAPL | 0.28652188 |
| TGIPPAPR | SSSPGTDNNQAPL | 0.31417215 |

| | | |
|---|---|---|
| FDLGGGTF | SSSPGTDNNQAPL | 0.6536177 |
| FDLGGGTFDVS | SSSPGTDNNQAPL | 0.75340116 |
| NEPTAAA | SSSPGTDNNQAPL | 0.34288394 |
| NEPTAAALA | SSSPGTDNNQAPL | 0.47093266 |
| GSIINISSI | SSSPGTDNNQAPL | 0.7041824 |
| VAPGNAGTA | SSSPGTDNNQAPL | 0.73469746 |
| TGGGASLE | SSSPGTDNNQAPL | 0.63749456 |
| VAVYNHYKRI | SSSPGTDNNQAPL | 0.2981066 |
| TDPAPATT | SSSPGTDNNQAPL | 0.6671639 |
| TNRPDILDPALLRPG | SSSPGTDNNQAPL | 0.29686856 |
| LTTTTTS | SSSPGTDNNQAPL | 0.72767776 |
| PTGSGKTL | SSSPGTDNNQAPL | 0.5710912 |
| TGSGKTL | SSSPGTDNNQAPL | 0.6417521 |
| TIGHVDHGK | SSSPGTDNNQAPL | 0.52997947 |
| TIGHVDHGKTT | SSSPGTDNNQAPL | 0.31513894 |
| GHVDHGKTT | SSSPGTDNNQAPL | 0.31104603 |
| AQFEHTLL | SSSPGTDNNQAPL | 0.4741232 |
| IVNQDEL | SSSPGTDNNQAPL | 0.7364348 |
| TNGSQFF | SSSPGTDNNQAPL | 0.6603686 |
| TGKGIVE | SSSPGTDNNQAPL | 0.6801928 |
| LISNASDA | SSSPGTDNNQAPL | 0.6558585 |
| AELRTLGELTKIAKAHGAQ | SSSPGTDNNQAPL | 0.35515237 |
| APNAVAAAEPSW | SSSPGTDNNQAPL | 0.66770124 |
| APQVAGAATAVL | SSSPGTDNNQAPL | 0.28032148 |
| ARRDPAAAAAILPTDA | SSSPGTDNNQAPL | 0.69773495 |
| DAAARDVLAEAGLAEYF | SSSPGTDNNQAPL | 0.5522875 |
| DDQIVLSAKANLAKVEQ | SSSPGTDNNQAPL | 0.6940179 |
| DELLPDTGEDSDEDGH | SSSPGTDNNQAPL | 0.49039868 |
| ELPEAPANRATVPQD | SSSPGTDNNQAPL | 0.7058866 |
| FGQSKIERVTGIG | SSSPGTDNNQAPL | 0.69287777 |
| GAGGTGSTAGGAGGAGGAGGLY | SSSPGTDNNQAPL | 0.42248666 |
| GHGGTGAAGGNGGTGGAGGSGI | SSSPGTDNNQAPL | 0.6281473 |
| GNGGNGGAGGEGGPGGAGGAGA | SSSPGTDNNQAPL | 0.7420732 |
| GSTAGNGGNGGNATNGTIAGQP | SSSPGTDNNQAPL | 0.6969606 |
| GTGGDGGNGGNGGTGG | SSSPGTDNNQAPL | 0.22591987 |
| GVRRPAGIPADAPV | SSSPGTDNNQAPL | 0.56619984 |
| GYRQVTDAHLLTLARR | SSSPGTDNNQAPL | 0.41338065 |
| IAKMGQDGHDRG | SSSPGTDNNQAPL | 0.64008725 |
| KAEPSIEIQYT | SSSPGTDNNQAPL | 0.5758999 |
| KPVRIRMAADEDTVAVT | SSSPGTDNNQAPL | 0.61499727 |
| MPPRIAGMRLLVIKPEP | SSSPGTDNNQAPL | 0.6334753 |
| MKFYAVALFRSRGVPPTR | SSSPGTDNNQAPL | 0.2533925 |
| NAGGFGILGAVAHSP | SSSPGTDNNQAPL | 0.48903814 |
| NEGREITLAMAEM | SSSPGTDNNQAPL | 0.5176121 |
| NELVEWSHPLGHRWRWE | SSSPGTDNNQAPL | 0.67568964 |
| NLGAGGTGGNGGNPG | SSSPGTDNNQAPL | 0.6647509 |
| NTEWPKWETVMAADPVD | SSSPGTDNNQAPL | 0.3301236 |
| PASAAPAETQP | SSSPGTDNNQAPL | 0.6571048 |
| QQSSDGKNGVRAAEITGE | SSSPGTDNNQAPL | 0.5579126 |
| RAPDAVTTSWWKE | SSSPGTDNNQAPL | 0.37146503 |
| SGFGNVGAHTSGWFNQ | SSSPGTDNNQAPL | 0.57486194 |

| | | |
|---|---|---|
| SGVAGAVGQAASVGGLKV | SSSPGTDNNQAPL | 0.4788111 |
| SNGGAGTGSAGGNGGT | SSSPGTDNNQAPL | 0.25303885 |
| TGGAGGEGGAAG | SSSPGTDNNQAPL | 0.70804524 |
| TGGDGGAAGAGGAPGHG | SSSPGTDNNQAPL | 0.416129 |
| TMPPLLAAIANAGRHGV | SSSPGTDNNQAPL | 0.59830356 |
| TSSTAGNSGGA | SSSPGTDNNQAPL | 0.7478571 |
| VPATVPVGILC | SSSPGTDNNQAPL | 0.23638114 |
| VQRPDSGGVVLTGRIS | SSSPGTDNNQAPL | 0.7135593 |
| WTPNPVPATKGVPAGLVQAT | SSSPGTDNNQAPL | 0.4749392 |
| YALKRALAKFSDRFDVVII | SSSPGTDNNQAPL | 0.5816126 |
| YLIGSVIV | SSSPGTDNNQAPL | 0.58647025 |
| YGQLVAVID | SSSPGTDNNQAPL | 0.36840856 |
| VTVGGMTLPA | SSSPGTDNNQAPL | 0.6604775 |
| VLALPAV | SSSPGTDNNQAPL | 0.6754944 |
| VIPPTLNYET | SSSPGTDNNQAPL | 0.693142 |
| VGMGGMHPSSG | SSSPGTDNNQAPL | 0.5577354 |
| VGLAGGAATA | SSSPGTDNNQAPL | 0.7273511 |
| VGGAGGAGGSGP | SSSPGTDNNQAPL | 0.7352602 |
| VFAAFMV | SSSPGTDNNQAPL | 0.42041007 |
| TLYDKLR | SSSPGTDNNQAPL | 0.23385352 |
| TLAVFTVL | SSSPGTDNNQAPL | 0.5984971 |
| TGSVQPARV | SSSPGTDNNQAPL | 0.60762346 |
| TGGGAGSGGT | SSSPGTDNNQAPL | 0.36533937 |
| SHLETSL | SSSPGTDNNQAPL | 0.30709207 |
| SFTTGMT | SSSPGTDNNQAPL | 0.73261786 |
| QVEVAKRMGVRQPTVS | SSSPGTDNNQAPL | 0.77749807 |
| QGSGGPALAM | SSSPGTDNNQAPL | 0.48173553 |
| MLAPSNP | SSSPGTDNNQAPL | 0.77854633 |
| LVNNAAI | SSSPGTDNNQAPL | 0.28686193 |
| LTGEINTLA | SSSPGTDNNQAPL | 0.68984675 |
| LSIFGPV | SSSPGTDNNQAPL | 0.7699611 |
| LLALLANL | SSSPGTDNNQAPL | 0.3399046 |
| LISNASDALDKLR | SSSPGTDNNQAPL | 0.6721374 |
| LINEAVP | SSSPGTDNNQAPL | 0.50006574 |
| LGYAFSGL | SSSPGTDNNQAPL | 0.30829954 |
| LDFVAVAL | SSSPGTDNNQAPL | 0.7397802 |
| LAQPGAE | SSSPGTDNNQAPL | 0.54346454 |
| ISNLGAVTGF | SSSPGTDNNQAPL | 0.68801105 |
| HQEDPGRI | SSSPGTDNNQAPL | 0.47749978 |
| GTPGEPTP | SSSPGTDNNQAPL | 0.36369747 |
| GFFNSGS | SSSPGTDNNQAPL | 0.642207 |
| GAGGGGLTAGAN | SSSPGTDNNQAPL | 0.71871763 |
| FVDALVNM | SSSPGTDNNQAPL | 0.6645256 |
| FLTVAPEL | SSSPGTDNNQAPL | 0.8091542 |
| FETLTAL | SSSPGTDNNQAPL | 0.4142791 |
| DVINPIEL | SSSPGTDNNQAPL | 0.7594532 |
| APVSGAPV | SSSPGTDNNQAPL | 0.7675413 |
| ALVGGLAW | SSSPGTDNNQAPL | 0.6222907 |
| ALAAAAATAI | SSSPGTDNNQAPL | 0.57364774 |
| AFGLGLLL | SSSPGTDNNQAPL | 0.7446714 |
| AFFAAASL | SSSPGTDNNQAPL | 0.57365036 |

| | | |
|---|---|---|
| AAAALPAFSPP | SSSPGTDNNQAPL | 0.6531968 |
| VVLAPAL | SSSPGTDNNQAPL | 0.73649704 |
| TIGHVDHGKTTL | SSSPGTDNNQAPL | 0.30214748 |
| LISNASDALDK | SSSPGTDNNQAPL | 0.7550785 |
| LISNASDAL | SSSPGTDNNQAPL | 0.47464246 |
| ISNASDALDK | SSSPGTDNNQAPL | 0.5884818 |
| NQAETLVYQTE | SSSPGTDNNQAPL | 0.38205034 |
| TGGTGGTAGAG | SSSPGTDNNQAPL | 0.27426374 |
| TGSTAGGAGGA | SSSPGTDNNQAPL | 0.4452357 |
| DPLAVPVAAGPV | SSSPGTDNNQAPL | 0.32238698 |
| GGTGGTPGNGGN | SSSPGTDNNQAPL | 0.6583302 |
| PAGLAGVDPATL | SSSPGTDNNQAPL | 0.5924996 |
| GNNLVVEAPHAGASV | SSSPGTDNNQAPL | 0.35682118 |
| KVTRIMSASTLEQPA | SSSPGTDNNQAPL | 0.70681095 |
| LGFALPIALPRRTVV | SSSPGTDNNQAPL | 0.7574499 |
| TRLGNLTARKLVVLV | SSSPGTDNNQAPL | 0.24962439 |
| WSQQRRLIYFPSAGP | SSSPGTDNNQAPL | 0.42590165 |
| GPNGNPDPMAAAVDIR | SSSPGTDNNQAPL | 0.4286148 |
| TQRCVSVADLRAAASR | SSSPGTDNNQAPL | 0.45409673 |
| QFVLDVAEHPVEVLTETV | SSSPGTDNNQAPL | 0.17620137 |
| GGAGGTGGAGGNGGTQAGDG | SSSPGTDNNQAPL | 0.6420095 |
| PLLATLRAIAVDVDAKPAQV | SSSPGTDNNQAPL | 0.28309643 |
| APKFRTGGYMIDFWGVGYQVAK | SSSPGTDNNQAPL | 0.12541081 |
| GAGGAGGAGGAAGGPGGTGGTG | SSSPGTDNNQAPL | 0.47462732 |
| IGALGLLATDVAPAGSVAFGVA | SSSPGTDNNQAPL | 0.4752895 |
| AGGAGGIGGNGGAGGAGGVLMG | SSSPGTDNNQAPL | 0.44022688 |
| ADELIGLL | SSSPGTDNNQAPL | 0.679881 |
| ADRALKA | SSSPGTDNNQAPL | 0.21874318 |
| AGAVGQVGGAGGA | SSSPGTDNNQAPL | 0.30931228 |
| AGGGGGLGGAP | SSSPGTDNNQAPL | 0.614988 |
| AGGLLYGNGGAG | SSSPGTDNNQAPL | 0.70813036 |
| AGLESIGL | SSSPGTDNNQAPL | 0.30262658 |
| AGLKTGII | SSSPGTDNNQAPL | 0.53652155 |
| ASLAALLL | SSSPGTDNNQAPL | 0.7357303 |
| ASVINISSI | SSSPGTDNNQAPL | 0.5265737 |
| ATPPGPA | SSSPGTDNNQAPL | 0.66822445 |
| ATVASALV | SSSPGTDNNQAPL | 0.68876624 |
| ATVASLIL | SSSPGTDNNQAPL | 0.31524187 |
| AVNGPVLIRH | SSSPGTDNNQAPL | 0.7025484 |
| DGAAGAVGI | SSSPGTDNNQAPL | 0.7502588 |
| DPLLHGI | SSSPGTDNNQAPL | 0.4361216 |
| ELGAALGAL | SSSPGTDNNQAPL | 0.5018923 |
| ELGRTQAV | SSSPGTDNNQAPL | 0.31071696 |
| FVAGGYLS | SSSPGTDNNQAPL | 0.3957753 |
| GASGGAGGAGGW | SSSPGTDNNQAPL | 0.18480055 |
| GGAGAGGDGGAGGI | SSSPGTDNNQAPL | 0.5773494 |
| GTGHNGSQPGMGGNG | SSSPGTDNNQAPL | 0.6348513 |
| GYAAASASAS | SSSPGTDNNQAPL | 0.43305758 |
| HQVDVAAVQQG | SSSPGTDNNQAPL | 0.23262766 |
| IGAAIAHT | SSSPGTDNNQAPL | 0.23879255 |
| IGMIEVAQL | SSSPGTDNNQAPL | 0.40276247 |

| | | |
|---|---|---|
| IRLAPEHTSAMD | SSSPGTDNNQAPL | 0.36108232 |
| IYNVSVL | SSSPGTDNNQAPL | 0.1904497 |
| KQGFLGM | SSSPGTDNNQAPL | 0.39296675 |
| LESVITQL | SSSPGTDNNQAPL | 0.6787141 |
| LGLGALVVV | SSSPGTDNNQAPL | 0.69619393 |
| LIGNGGNGGSGGI | SSSPGTDNNQAPL | 0.20099445 |
| LNAINAPTL | SSSPGTDNNQAPL | 0.77479446 |
| LQLLNTGV | SSSPGTDNNQAPL | 0.5886284 |
| LQTSIEF | SSSPGTDNNQAPL | 0.7137681 |
| MSFAVSEL | SSSPGTDNNQAPL | 0.45944434 |
| NTGALGL | SSSPGTDNNQAPL | 0.29947364 |
| PPGAGGAGGAGTT | SSSPGTDNNQAPL | 0.73252153 |
| QLVGLGI | SSSPGTDNNQAPL | 0.39764562 |
| RFLEQVHQL | SSSPGTDNNQAPL | 0.3017543 |
| RGVGYVLR | SSSPGTDNNQAPL | 0.6775903 |
| SATGVFAGVI | SSSPGTDNNQAPL | 0.43320838 |
| SLPAITLP | SSSPGTDNNQAPL | 0.30350667 |
| STALAFTV | SSSPGTDNNQAPL | 0.48101145 |
| SVLTVGLGK | SSSPGTDNNQAPL | 0.5564165 |
| TALAGAGSGI | SSSPGTDNNQAPL | 0.24317737 |
| TGLPFTGI | SSSPGTDNNQAPL | 0.33146504 |
| VAEHVTV | SSSPGTDNNQAPL | 0.27332774 |
| VDIVLNSL | SSSPGTDNNQAPL | 0.42969358 |
| VLTGGGAVL | SSSPGTDNNQAPL | 0.70713246 |
| VNVGLAYGI | SSSPGTDNNQAPL | 0.56331056 |
| VSSAAVGAV | SSSPGTDNNQAPL | 0.31746852 |
| VTSVYEAGV | SSSPGTDNNQAPL | 0.61896265 |
| WVALDRGA | SSSPGTDNNQAPL | 0.62440836 |
| YLVGSVVI | SSSPGTDNNQAPL | 0.5188513 |
| GPPGTGKTLL | SSSPGTDNNQAPL | 0.515972 |
| GVGMTKF | SSSPGTDNNQAPL | 0.3611335 |
| IGIGGSDL | SSSPGTDNNQAPL | 0.5949489 |
| IGTLLAEL | SSSPGTDNNQAPL | 0.64211744 |
| IYTSGTT | SSSPGTDNNQAPL | 0.6140575 |
| LSFPAGV | SSSPGTDNNQAPL | 0.6094092 |
| LYTSGTT | SSSPGTDNNQAPL | 0.5064279 |
| VGAFPFGF | SSSPGTDNNQAPL | 0.46530855 |
| VGAGTVEF | SSSPGTDNNQAPL | 0.16978827 |
| VGKSTLLL | SSSPGTDNNQAPL | 0.36441427 |
| AALAHAGGGIV | SSSPGTDNNQAPL | 0.47688866 |
| AALVAVSLL | SSSPGTDNNQAPL | 0.5177063 |
| AAVSRAVAAL | SSSPGTDNNQAPL | 0.5886698 |
| AGGSSGIGGF | SSSPGTDNNQAPL | 0.5931138 |
| AGLTVANANG | SSSPGTDNNQAPL | 0.3132545 |
| AGLVGAVL | SSSPGTDNNQAPL | 0.53914225 |
| ALSQAFTV | SSSPGTDNNQAPL | 0.69094396 |
| ALTFSVSL | SSSPGTDNNQAPL | 0.24764171 |
| ALTGAALVAL | SSSPGTDNNQAPL | 0.40956423 |
| ASFSIGAGVL | SSSPGTDNNQAPL | 0.79924226 |
| DGSVYKI | SSSPGTDNNQAPL | 0.593644 |
| EQLAEIV | SSSPGTDNNQAPL | 0.37337428 |

| | | |
|---|---|---|
| FAVSELTVM | SSSPGTDNNQAPL | 0.40223742 |
| GAAPLAAGL | SSSPGTDNNQAPL | 0.7568177 |
| GADGGDGGSGGAGG | SSSPGTDNNQAPL | 0.3442487 |
| GFGLDTGGAGG | SSSPGTDNNQAPL | 0.72968817 |
| GLASGGLL | SSSPGTDNNQAPL | 0.1642719 |
| GRIAGAAVLL | SSSPGTDNNQAPL | 0.37460417 |
| GSAGSLNTSF | SSSPGTDNNQAPL | 0.29310083 |
| GTGQIGI | SSSPGTDNNQAPL | 0.16837232 |
| GTLIALIL | SSSPGTDNNQAPL | 0.65888894 |
| IAATMGFVVG | SSSPGTDNNQAPL | 0.40221277 |
| IFAQALL | SSSPGTDNNQAPL | 0.6395031 |
| IGAAVALGF | SSSPGTDNNQAPL | 0.4651466 |
| IGLTPLAL | SSSPGTDNNQAPL | 0.6864264 |
| ILGSNTSTL | SSSPGTDNNQAPL | 0.58051085 |
| IVFSLTVL | SSSPGTDNNQAPL | 0.47073466 |
| LASKATGF | SSSPGTDNNQAPL | 0.3432449 |
| LASVALIL | SSSPGTDNNQAPL | 0.60299003 |
| LFAVGLTL | SSSPGTDNNQAPL | 0.59425443 |
| LFNGGIGTYI | SSSPGTDNNQAPL | 0.55969274 |
| LTLGAGII | SSSPGTDNNQAPL | 0.23433049 |
| LVAAALGDI | SSSPGTDNNQAPL | 0.35428166 |
| NTAFVQLGI | SSSPGTDNNQAPL | 0.49254823 |
| QLFSGAEL | SSSPGTDNNQAPL | 0.18701924 |
| SAATFITL | SSSPGTDNNQAPL | 0.14280538 |
| SNYGLANLGV | SSSPGTDNNQAPL | 0.29134786 |
| SYGGAYAVM | SSSPGTDNNQAPL | 0.35278016 |
| TTGVGGLTLG | SSSPGTDNNQAPL | 0.6757846 |
| TTQTIYI | SSSPGTDNNQAPL | 0.6466122 |
| VAAGAGLGAVI | SSSPGTDNNQAPL | 0.44160172 |
| VAAVVTLL | SSSPGTDNNQAPL | 0.5457859 |
| VGFLGGLI | SSSPGTDNNQAPL | 0.46614677 |
| VGVIPILL | SSSPGTDNNQAPL | 0.11146722 |
| VIFASSLI | SSSPGTDNNQAPL | 0.3899343 |
| VKGGDGGAAAG | SSSPGTDNNQAPL | 0.26131207 |
| VTKTFTNI | SSSPGTDNNQAPL | 0.69763494 |
| VVAAAILL | SSSPGTDNNQAPL | 0.30105036 |
| YAALDVEL | SSSPGTDNNQAPL | 0.30675817 |
| LASVALLL | SSSPGTDNNQAPL | 0.490327 |
| LDAVHRV | SSSPGTDNNQAPL | 0.63416386 |
| QGLSSSIF | SSSPGTDNNQAPL | 0.52358305 |
| RGTFANIRL | SSSPGTDNNQAPL | 0.25337118 |
| RQLWWGHRI | SSSPGTDNNQAPL | 0.12948121 |
| YGPPGTGKTLL | SSSPGTDNNQAPL | 0.6027597 |
| AALVGAPANVA | SSSPGTDNNQAPL | 0.28185427 |
| AATFRRTE | SSSPGTDNNQAPL | 0.32044053 |
| AGALGGGI | SSSPGTDNNQAPL | 0.1620235 |
| AGGAGGAGGIGAG | SSSPGTDNNQAPL | 0.15099065 |
| AGSGALGL | SSSPGTDNNQAPL | 0.26618007 |
| AQIADQI | SSSPGTDNNQAPL | 0.5907758 |
| ASAVAVTYL | SSSPGTDNNQAPL | 0.39236075 |
| ASTVLGGTGG | SSSPGTDNNQAPL | 0.38139153 |

| | | |
|---|---|---|
| AVGGEPAK | SSSPGTDNNQAPL | 0.7816795 |
| CEQGTFGGTC | SSSPGTDNNQAPL | 0.6166785 |
| DEGKAPAG | SSSPGTDNNQAPL | 0.42869538 |
| DTVVFPAL | SSSPGTDNNQAPL | 0.3773633 |
| FEGITMAG | SSSPGTDNNQAPL | 0.30857286 |
| FPLVVQD | SSSPGTDNNQAPL | 0.30393454 |
| GAFTTPQ | SSSPGTDNNQAPL | 0.35160992 |
| GFGGFGTAGGVGGA | SSSPGTDNNQAPL | 0.7443527 |
| GFLAQFL | SSSPGTDNNQAPL | 0.39392352 |
| GGAGGAGGNAGVA | SSSPGTDNNQAPL | 0.3150105 |
| GGVGTAGAGGAGGAGG | SSSPGTDNNQAPL | 0.55458814 |
| GNGGGGDAGNAGSGGN | SSSPGTDNNQAPL | 0.7320802 |
| GTGGAGGAGNDAGS | SSSPGTDNNQAPL | 0.7870554 |
| GTTNTGWFNS | SSSPGTDNNQAPL | 0.26006386 |
| IFMGPLFTL | SSSPGTDNNQAPL | 0.39753452 |
| ILAHAIEAI | SSSPGTDNNQAPL | 0.4026386 |
| IRSAIREAL | SSSPGTDNNQAPL | 0.4714173 |
| ITFAGGG | SSSPGTDNNQAPL | 0.30719328 |
| ITGMSGHQEV | SSSPGTDNNQAPL | 0.32967898 |
| IVAGLACTL | SSSPGTDNNQAPL | 0.5372571 |
| IVGGGLAGA | SSSPGTDNNQAPL | 0.79384553 |
| IVTGVLNI | SSSPGTDNNQAPL | 0.4261195 |
| LAQQGCI | SSSPGTDNNQAPL | 0.29796872 |
| LCLHVEN | SSSPGTDNNQAPL | 0.41117382 |
| LDAISDGS | SSSPGTDNNQAPL | 0.2708578 |
| LGGFAAML | SSSPGTDNNQAPL | 0.32612574 |
| LGGKNPMI | SSSPGTDNNQAPL | 0.45172587 |
| LGPKGVAAIL | SSSPGTDNNQAPL | 0.24187975 |
| LMVGVSI | SSSPGTDNNQAPL | 0.48148912 |
| LVGAAFVL | SSSPGTDNNQAPL | 0.7801043 |
| LVLAVAPL | SSSPGTDNNQAPL | 0.47476256 |
| MAAIAVIDAA | SSSPGTDNNQAPL | 0.66773105 |
| NNGINAELQ | SSSPGTDNNQAPL | 0.7309388 |
| RGEPLVLVHG | SSSPGTDNNQAPL | 0.6175543 |
| RGGAGGAAGLI | SSSPGTDNNQAPL | 0.75184697 |
| RSYRMHPAV | SSSPGTDNNQAPL | 0.63029265 |
| SAAHLLAGQLA | SSSPGTDNNQAPL | 0.5124494 |
| SAAVGSY | SSSPGTDNNQAPL | 0.43448073 |
| SPHAVLHRI | SSSPGTDNNQAPL | 0.572651 |
| SQIAKVL | SSSPGTDNNQAPL | 0.45922577 |
| TALGHLVP | SSSPGTDNNQAPL | 0.4488912 |
| TAPTTSGAA | SSSPGTDNNQAPL | 0.40126956 |
| TDGDDGNGGSPGT | SSSPGTDNNQAPL | 0.7146395 |
| TDGLGEN | SSSPGTDNNQAPL | 0.43139222 |
| TGGPPLPGQA | SSSPGTDNNQAPL | 0.71280694 |
| TIPAITI | SSSPGTDNNQAPL | 0.23268555 |
| TIPGFTI | SSSPGTDNNQAPL | 0.29502535 |
| TSIASGALA | SSSPGTDNNQAPL | 0.6880463 |
| TVTLPGAAI | SSSPGTDNNQAPL | 0.12210183 |
| VASGLSAALP | SSSPGTDNNQAPL | 0.16273567 |
| VGAGVIGI | SSSPGTDNNQAPL | 0.39263862 |

| | | |
|---|---|---|
| VGPKRFGV | SSSPGTDNNQAPL | 0.72452044 |
| VNVAFPD | SSSPGTDNNQAPL | 0.12597856 |
| VSMTNSGVS | SSSPGTDNNQAPL | 0.32128912 |
| VTDVLGDLSV | SSSPGTDNNQAPL | 0.2368169 |
| VVVVGAGFAGL | SSSPGTDNNQAPL | 0.6155218 |
| VYTVGTVV | SSSPGTDNNQAPL | 0.5075671 |
| WTTFSGP | SSSPGTDNNQAPL | 0.2410905 |
| YDELVEAF | SSSPGTDNNQAPL | 0.67240477 |
| YGTGGAGGNGG | SSSPGTDNNQAPL | 0.21158357 |
| YTNVMAV | SSSPGTDNNQAPL | 0.24894556 |
| FLEQVHQL | SSSPGTDNNQAPL | 0.63753563 |
| GLSLLAVAV | SSSPGTDNNQAPL | 0.72126913 |
| LEQVHQL | SSSPGTDNNQAPL | 0.27506056 |
| LNAAGAI | SSSPGTDNNQAPL | 0.6214209 |
| PAVHASL | SSSPGTDNNQAPL | 0.2780305 |
| TELEDDA | SSSPGTDNNQAPL | 0.7845972 |
| TPPGPPN | SSSPGTDNNQAPL | 0.3890695 |
| TPPPPPPP | SSSPGTDNNQAPL | 0.23077706 |
| TQSPALI | ASSLADSNNQAPL | 0.6866225 |
| EFAAAFPQ | ASSLADSNNQAPL | 0.65704674 |
| LLSGNLDVL | ASSLADSNNQAPL | 0.4790695 |
| GPANLGNYNI | ASSLADSNNQAPL | 0.64684224 |
| TLGPTLP | ASSLADSNNQAPL | 0.37827122 |
| VYAYSSCCK | ASSLADSNNQAPL | 0.4696269 |
| APAPAPAPNI | ASSLADSNNQAPL | 0.7326299 |
| PGGPVPIP | ASSLADSNNQAPL | 0.5978148 |
| FLTVALEP | ASSLADSNNQAPL | 0.3360839 |
| SQYFGANL | ASSLADSNNQAPL | 0.61618 |
| GGVKPAA | ASSLADSNNQAPL | 0.4668566 |
| GGTGGAGGAGGW | ASSLADSNNQAPL | 0.31815764 |
| GVGGVAGLGGAGG | ASSLADSNNQAPL | 0.76251 |
| VLLVLATTI | ASSLADSNNQAPL | 0.45438376 |
| IGIFAAML | ASSLADSNNQAPL | 0.6316585 |
| AVGDLID | ASSLADSNNQAPL | 0.637974 |
| NISPLPL | ASSLADSNNQAPL | 0.7353791 |
| VTAVFVAI | ASSLADSNNQAPL | 0.28481683 |
| SGDDNDAAL | ASSLADSNNQAPL | 0.4449301 |
| AVAALAHS | ASSLADSNNQAPL | 0.4421271 |
| GGVTVMATP | ASSLADSNNQAPL | 0.63572085 |
| AGLAANVRD | ASSLADSNNQAPL | 0.4042696 |
| TNRPDLIDPALLRPG | ASSLADSNNQAPL | 0.28829312 |
| LANLGVTAL | ASSLADSNNQAPL | 0.5659503 |
| TGERIEL | ASSLADSNNQAPL | 0.7083353 |
| NGEELAA | ASSLADSNNQAPL | 0.40180153 |
| VLSLVWG | ASSLADSNNQAPL | 0.4811896 |
| IGYALWL | ASSLADSNNQAPL | 0.78451836 |
| LSLLLAPL | ASSLADSNNQAPL | 0.3052607 |
| TLDEALK | ASSLADSNNQAPL | 0.46238327 |
| TDGAGTAEGG | ASSLADSNNQAPL | 0.5782535 |
| NAPGGAGGNGG | ASSLADSNNQAPL | 0.6358418 |
| VSVFMGMGGNF | ASSLADSNNQAPL | 0.4358883 |

| | | |
|---|---|---|
| LASYLAAL | ASSLADSNNQAPL | 0.6562016 |
| LVGGTPAL | ASSLADSNNQAPL | 0.32600957 |
| VRNADVL | ASSLADSNNQAPL | 0.6205358 |
| TCSSPFLVSAM | ASSLADSNNQAPL | 0.23693493 |
| VHITDAE | ASSLADSNNQAPL | 0.55045104 |
| EYAAPSF | ASSLADSNNQAPL | 0.27550822 |
| VVDAFHGGL | ASSLADSNNQAPL | 0.61888194 |
| ASVVDPQ | ASSLADSNNQAPL | 0.6478699 |
| SYNEFGGLS | ASSLADSNNQAPL | 0.6753465 |
| DPDGSVY | ASSLADSNNQAPL | 0.15084034 |
| MVSTIHATE | ASSLADSNNQAPL | 0.67406356 |
| LAAGVADPV | ASSLADSNNQAPL | 0.6660501 |
| TKEQIAA | ASSLADSNNQAPL | 0.80302805 |
| SLASPAAL | ASSLADSNNQAPL | 0.41405287 |
| IGLVFYGV | ASSLADSNNQAPL | 0.6224511 |
| FLLDASNL | ASSLADSNNQAPL | 0.33789086 |
| YQVGNFP | ASSLADSNNQAPL | 0.4516077 |
| ESLLSQST | ASSLADSNNQAPL | 0.21927656 |
| LFTVGFAAVG | ASSLADSNNQAPL | 0.3144643 |
| ATPPTPT | ASSLADSNNQAPL | 0.37273896 |
| TDLATLG | ASSLADSNNQAPL | 0.31918058 |
| SAGGTGR | ASSLADSNNQAPL | 0.698863 |
| GFSNSGS | ASSLADSNNQAPL | 0.60846037 |
| GGNGGSGGTSV | ASSLADSNNQAPL | 0.70040894 |
| GTSVATGGAGN | ASSLADSNNQAPL | 0.7048776 |
| NGGNGGSGGTS | ASSLADSNNQAPL | 0.72582996 |
| AVVKFPH | ASSLADSNNQAPL | 0.61218035 |
| TPPPPPP | ASSLADSNNQAPL | 0.3662645 |
| AAPAAAAAPA | ASSLADSNNQAPL | 0.41535515 |
| IALNEDL | ASSLADSNNQAPL | 0.25482765 |
| IVNVSSM | ASSLADSNNQAPL | 0.43515193 |
| TGIPPAPR | ASSLADSNNQAPL | 0.39091235 |
| FDLGGGTF | ASSLADSNNQAPL | 0.56036216 |
| FDLGGGTFDVS | ASSLADSNNQAPL | 0.12053143 |
| NEPTAAA | ASSLADSNNQAPL | 0.55948687 |
| NEPTAAALA | ASSLADSNNQAPL | 0.75474405 |
| GSIINISSI | ASSLADSNNQAPL | 0.35219753 |
| VAPGNAGTA | ASSLADSNNQAPL | 0.61164623 |
| TGGGASLE | ASSLADSNNQAPL | 0.6028264 |
| VAVYNHYKRI | ASSLADSNNQAPL | 0.28974265 |
| TDPAPATT | ASSLADSNNQAPL | 0.6126586 |
| TNRPDILDPALLRPG | ASSLADSNNQAPL | 0.31503138 |
| LTTTTTS | ASSLADSNNQAPL | 0.6035964 |
| PTGSGKTL | ASSLADSNNQAPL | 0.7724811 |
| TGSGKTL | ASSLADSNNQAPL | 0.56019604 |
| TIGHVDHGK | ASSLADSNNQAPL | 0.3343392 |
| TIGHVDHGKTT | ASSLADSNNQAPL | 0.4844814 |
| GHVDHGKTT | ASSLADSNNQAPL | 0.2620393 |
| AQFEHTLL | ASSLADSNNQAPL | 0.3825934 |
| IVNQDEL | ASSLADSNNQAPL | 0.117775075 |
| TNGSQFF | ASSLADSNNQAPL | 0.5266207 |

| | | |
|---|---|---|
| TGKGIVE | ASSLADSNNQAPL | 0.30815944 |
| LISNASDA | ASSLADSNNQAPL | 0.5331103 |
| AELRTLGELTKIAKAHGAQ | ASSLADSNNQAPL | 0.48552886 |
| APNAVAAAEPSW | ASSLADSNNQAPL | 0.47593492 |
| APQVAGAATAVL | ASSLADSNNQAPL | 0.29118723 |
| ARRDPAAAAAILPTDA | ASSLADSNNQAPL | 0.47761697 |
| DAAARDVLAEAGLAEYF | ASSLADSNNQAPL | 0.30405068 |
| DDQIVLSAKANLAKVEQ | ASSLADSNNQAPL | 0.2639186 |
| DELLPDTGEDSDEDGH | ASSLADSNNQAPL | 0.60471326 |
| ELPEAPANRATVPQD | ASSLADSNNQAPL | 0.39029488 |
| FGQSKIERVTGIG | ASSLADSNNQAPL | 0.49850133 |
| GAGGTGSTAGGAGGAGGAGGLY | ASSLADSNNQAPL | 0.69860226 |
| GHGGTGAAGGNGGTGGAGGSGI | ASSLADSNNQAPL | 0.54092336 |
| GNGGNGGAGGEGGPGGAGGAGA | ASSLADSNNQAPL | 0.26815695 |
| GSTAGNGGNGGNATNGTIAGQP | ASSLADSNNQAPL | 0.44488618 |
| GTGGDGGNGGNGGTGG | ASSLADSNNQAPL | 0.4425171 |
| GVRRPAGIPADAPV | ASSLADSNNQAPL | 0.38879412 |
| GYRQVTDAHLLTLARR | ASSLADSNNQAPL | 0.119661465 |
| IAKMGQDGHDRG | ASSLADSNNQAPL | 0.5934864 |
| KAEPSIEIQYT | ASSLADSNNQAPL | 0.32485965 |
| KPVRIRMAADEDTVAVT | ASSLADSNNQAPL | 0.34086454 |
| MPPRIAGMRLLVIKPEP | ASSLADSNNQAPL | 0.4048152 |
| MKFYAVALFRSRGVPPTR | ASSLADSNNQAPL | 0.7095956 |
| NAGGFGILGAVAHSP | ASSLADSNNQAPL | 0.22336964 |
| NEGREITLAMAEM | ASSLADSNNQAPL | 0.4787858 |
| NELVEWSHPLGHRWRWE | ASSLADSNNQAPL | 0.6026574 |
| NLGAGGTGGNGGNPG | ASSLADSNNQAPL | 0.52667683 |
| NTEWPKWETVMAADPVD | ASSLADSNNQAPL | 0.7248532 |
| PASAAPAETQP | ASSLADSNNQAPL | 0.53360665 |
| QQSSDGKNGVRAAEITGE | ASSLADSNNQAPL | 0.3656331 |
| RAPDAVTTSWWKE | ASSLADSNNQAPL | 0.60289735 |
| SGFGNVGAHTSGWFNQ | ASSLADSNNQAPL | 0.42552748 |
| SGVAGAVGQAASVGGLKV | ASSLADSNNQAPL | 0.19511588 |
| SNGGAGTGSAGGNGGT | ASSLADSNNQAPL | 0.171824 |
| TGGAGGEGGAAG | ASSLADSNNQAPL | 0.4984056 |
| TGGDGGAAGAGGAPGHG | ASSLADSNNQAPL | 0.17700334 |
| TMPPLLAAIANAGRHGV | ASSLADSNNQAPL | 0.5452107 |
| TSSTAGNSGGA | ASSLADSNNQAPL | 0.68141806 |
| VPATVPVGILC | ASSLADSNNQAPL | 0.4382415 |
| VQRPDSGGVVLTGRIS | ASSLADSNNQAPL | 0.23614085 |
| WTPNPVPATKGVPAGLVQAT | ASSLADSNNQAPL | 0.3433344 |
| YALKRALAKFSDRFDVVII | ASSLADSNNQAPL | 0.42397943 |
| YLIGSVIV | ASSLADSNNQAPL | 0.61894506 |
| YGQLVAVID | ASSLADSNNQAPL | 0.48012114 |
| VTVGGMTLPA | ASSLADSNNQAPL | 0.76039606 |
| VLALPAV | ASSLADSNNQAPL | 0.6655657 |
| VIPPTLNYET | ASSLADSNNQAPL | 0.38268858 |
| VGMGGMHPSSG | ASSLADSNNQAPL | 0.6308572 |
| VGLAGGAATA | ASSLADSNNQAPL | 0.44710985 |
| VGGAGGAGGSGP | ASSLADSNNQAPL | 0.38391566 |
| VFAAFMV | ASSLADSNNQAPL | 0.4451252 |

| | | |
|---|---|---|
| TLYDKLR | ASSLADSNNQAPL | 0.46455818 |
| TLAVFTVL | ASSLADSNNQAPL | 0.38424382 |
| TGSVQPARV | ASSLADSNNQAPL | 0.3906481 |
| TGGGAGSGGT | ASSLADSNNQAPL | 0.25090414 |
| SHLETSL | ASSLADSNNQAPL | 0.47933525 |
| SFTTGMT | ASSLADSNNQAPL | 0.58890045 |
| QVEVAKRMGVRQPTVS | ASSLADSNNQAPL | 0.19873673 |
| QGSGGPALAM | ASSLADSNNQAPL | 0.39925045 |
| MLAPSNP | ASSLADSNNQAPL | 0.78464 |
| LVNNAAI | ASSLADSNNQAPL | 0.7198566 |
| LTGEINTLA | ASSLADSNNQAPL | 0.310234 |
| LSIFGPV | ASSLADSNNQAPL | 0.3483883 |
| LLALLANL | ASSLADSNNQAPL | 0.5905845 |
| LISNASDALDKLR | ASSLADSNNQAPL | 0.20201729 |
| LINEAVP | ASSLADSNNQAPL | 0.5133983 |
| LGYAFSGL | ASSLADSNNQAPL | 0.53590643 |
| LDFVAVAL | ASSLADSNNQAPL | 0.49889314 |
| LAQPGAE | ASSLADSNNQAPL | 0.32316625 |
| ISNLGAVTGF | ASSLADSNNQAPL | 0.52464795 |
| HQEDPGRI | ASSLADSNNQAPL | 0.4107344 |
| GTPGEPTP | ASSLADSNNQAPL | 0.5164612 |
| GFFNSGS | ASSLADSNNQAPL | 0.1387229 |
| GAGGGGLTAGAN | ASSLADSNNQAPL | 0.60128796 |
| FVDALVNM | ASSLADSNNQAPL | 0.24582687 |
| FLTVAPEL | ASSLADSNNQAPL | 0.33097163 |
| FETLTAL | ASSLADSNNQAPL | 0.16521497 |
| DVINPIEL | ASSLADSNNQAPL | 0.43194956 |
| APVSGAPV | ASSLADSNNQAPL | 0.64691156 |
| ALVGGLAW | ASSLADSNNQAPL | 0.6949615 |
| ALAAAAATAI | ASSLADSNNQAPL | 0.23433702 |
| AFGLGLLL | ASSLADSNNQAPL | 0.6491915 |
| AFFAAASL | ASSLADSNNQAPL | 0.67106754 |
| AAAALPAFSPP | ASSLADSNNQAPL | 0.3727193 |
| VVLAPAL | ASSLADSNNQAPL | 0.21900061 |
| TIGHVDHGKTTL | ASSLADSNNQAPL | 0.6002235 |
| LISNASDALDK | ASSLADSNNQAPL | 0.39857638 |
| LISNASDAL | ASSLADSNNQAPL | 0.67317605 |
| ISNASDALDK | ASSLADSNNQAPL | 0.5963295 |
| NQAETLVYQTE | ASSLADSNNQAPL | 0.22916415 |
| TGGTGGTAGAG | ASSLADSNNQAPL | 0.6815868 |
| TGSTAGGAGGA | ASSLADSNNQAPL | 0.7276455 |
| DPLAVPVAAGPV | ASSLADSNNQAPL | 0.26573214 |
| GGTGGTPGNGGN | ASSLADSNNQAPL | 0.3048562 |
| PAGLAGVDPATL | ASSLADSNNQAPL | 0.36542276 |
| GNNLVVEAPHAGASV | ASSLADSNNQAPL | 0.14992619 |
| KVTRIMSASTLEQPA | ASSLADSNNQAPL | 0.56347996 |
| LGFALPIALPRRTVV | ASSLADSNNQAPL | 0.353906 |
| TRLGNLTARKLVVLV | ASSLADSNNQAPL | 0.37727752 |
| WSQQRRLIYFPSAGP | ASSLADSNNQAPL | 0.52138436 |
| GPNGNPDPMAAAVDIR | ASSLADSNNQAPL | 0.39677435 |
| TQRCVSVADLRAAASR | ASSLADSNNQAPL | 0.40518048 |

| | | |
|---|---|---|
| QFVLDVAEHPVEVLTETV | ASSLADSNNQAPL | 0.4262804 |
| GGAGGTGGAGGNGGTQAGDG | ASSLADSNNQAPL | 0.54576427 |
| PLLATLRAIAVDVDAKPAQV | ASSLADSNNQAPL | 0.5391184 |
| APKFRTGGYMIDFWGVGYQVAK | ASSLADSNNQAPL | 0.5237931 |
| GAGGAGGAGGAAGGPGGTGGTG | ASSLADSNNQAPL | 0.5702014 |
| IGALGLLATDVAPAGSVAFGVA | ASSLADSNNQAPL | 0.7600112 |
| AGGAGGIGGNGGAGGAGGVLMG | ASSLADSNNQAPL | 0.26502466 |
| ADELIGLL | ASSLADSNNQAPL | 0.3348234 |
| ADRALKA | ASSLADSNNQAPL | 0.6145405 |
| AGAVGQVGGAGGA | ASSLADSNNQAPL | 0.2568523 |
| AGGGGGLGGAP | ASSLADSNNQAPL | 0.6371038 |
| AGGLLYGNGGAG | ASSLADSNNQAPL | 0.48701686 |
| AGLESIGL | ASSLADSNNQAPL | 0.59177345 |
| AGLKTGII | ASSLADSNNQAPL | 0.44929373 |
| ASLAALLL | ASSLADSNNQAPL | 0.78163624 |
| ASVINISSI | ASSLADSNNQAPL | 0.67738235 |
| ATPPGPA | ASSLADSNNQAPL | 0.45516598 |
| ATVASALV | ASSLADSNNQAPL | 0.44393295 |
| ATVASLIL | ASSLADSNNQAPL | 0.27966458 |
| AVNGPVLIRH | ASSLADSNNQAPL | 0.5991808 |
| DGAAGAVGI | ASSLADSNNQAPL | 0.58520955 |
| DPLLHGI | ASSLADSNNQAPL | 0.2706942 |
| ELGAALGAL | ASSLADSNNQAPL | 0.46059984 |
| ELGRTQAV | ASSLADSNNQAPL | 0.63568664 |
| FVAGGYLS | ASSLADSNNQAPL | 0.7116165 |
| GASGGAGGAGGW | ASSLADSNNQAPL | 0.20383283 |
| GGAGAGGDGGAGGI | ASSLADSNNQAPL | 0.3958168 |
| GTGHNGSQPGMGGNG | ASSLADSNNQAPL | 0.18717402 |
| GYAAASASAS | ASSLADSNNQAPL | 0.31141618 |
| HQVDVAAVQQG | ASSLADSNNQAPL | 0.2939263 |
| IGAAIAHT | ASSLADSNNQAPL | 0.56992596 |
| IGMIEVAQL | ASSLADSNNQAPL | 0.37320855 |
| IRLAPEHTSAMD | ASSLADSNNQAPL | 0.6473444 |
| IYNVSVL | ASSLADSNNQAPL | 0.5448122 |
| KQGFLGM | ASSLADSNNQAPL | 0.7330352 |
| LESVITQL | ASSLADSNNQAPL | 0.29806858 |
| LGLGALVVV | ASSLADSNNQAPL | 0.4065948 |
| LIGNGGNGGSGGI | ASSLADSNNQAPL | 0.20272484 |
| LNAINAPTL | ASSLADSNNQAPL | 0.22421227 |
| LQLLNTGV | ASSLADSNNQAPL | 0.26222882 |
| LQTSIEF | ASSLADSNNQAPL | 0.45124933 |
| MSFAVSEL | ASSLADSNNQAPL | 0.16059864 |
| NTGALGL | ASSLADSNNQAPL | 0.7646092 |
| PPGAGGAGGAGTT | ASSLADSNNQAPL | 0.49800858 |
| QLVGLGI | ASSLADSNNQAPL | 0.61412454 |
| RFLEQVHQL | ASSLADSNNQAPL | 0.39414135 |
| RGVGYVLR | ASSLADSNNQAPL | 0.33055353 |
| SATGVFAGVI | ASSLADSNNQAPL | 0.3620007 |
| SLPAITLP | ASSLADSNNQAPL | 0.61855376 |
| STALAFTV | ASSLADSNNQAPL | 0.27880764 |
| SVLTVGLGK | ASSLADSNNQAPL | 0.22281808 |

| | | |
|---|---|---|
| TALAGAGSGI | ASSLADSNNQAPL | 0.34744766 |
| TGLPFTGI | ASSLADSNNQAPL | 0.27801043 |
| VAEHVTV | ASSLADSNNQAPL | 0.7505538 |
| VDIVLNSL | ASSLADSNNQAPL | 0.67496 |
| VLTGGGAVL | ASSLADSNNQAPL | 0.6028237 |
| VNVGLAYGI | ASSLADSNNQAPL | 0.73596084 |
| VSSAAVGAV | ASSLADSNNQAPL | 0.14707339 |
| VTSVYEAGV | ASSLADSNNQAPL | 0.40247238 |
| WVALDRGA | ASSLADSNNQAPL | 0.77764887 |
| YLVGSVVI | ASSLADSNNQAPL | 0.5122991 |
| GPPGTGKTLL | ASSLADSNNQAPL | 0.4254027 |
| GVGMTKF | ASSLADSNNQAPL | 0.6950743 |
| IGIGGSDL | ASSLADSNNQAPL | 0.32786623 |
| IGTLLAEL | ASSLADSNNQAPL | 0.40418172 |
| IYTSGTT | ASSLADSNNQAPL | 0.46368045 |
| LSFPAGV | ASSLADSNNQAPL | 0.56940615 |
| LYTSGTT | ASSLADSNNQAPL | 0.38746092 |
| VGAFPFGF | ASSLADSNNQAPL | 0.2742332 |
| VGAGTVEF | ASSLADSNNQAPL | 0.6011784 |
| VGKSTLLL | ASSLADSNNQAPL | 0.7963174 |
| AALAHAGGGIV | ASSLADSNNQAPL | 0.45163783 |
| AALVAVSLL | ASSLADSNNQAPL | 0.699525 |
| AAVSRAVAAL | ASSLADSNNQAPL | 0.56663907 |
| AGGSSGIGGF | ASSLADSNNQAPL | 0.64088815 |
| AGLTVANANG | ASSLADSNNQAPL | 0.3067895 |
| AGLVGAVL | ASSLADSNNQAPL | 0.27477986 |
| ALSQAFTV | ASSLADSNNQAPL | 0.4523335 |
| ALTFSVSL | ASSLADSNNQAPL | 0.25026682 |
| ALTGAALVAL | ASSLADSNNQAPL | 0.44775194 |
| ASFSIGAGVL | ASSLADSNNQAPL | 0.4908905 |
| DGSVYKI | ASSLADSNNQAPL | 0.81119776 |
| EQLAEIV | ASSLADSNNQAPL | 0.5859977 |
| FAVSELTVM | ASSLADSNNQAPL | 0.5697912 |
| GAAPLAAGL | ASSLADSNNQAPL | 0.25649452 |
| GADGGDGGSGGAGG | ASSLADSNNQAPL | 0.41128048 |
| GFGLDTGGAGG | ASSLADSNNQAPL | 0.58741474 |
| GLASGGLL | ASSLADSNNQAPL | 0.29262495 |
| GRIAGAAVLL | ASSLADSNNQAPL | 0.6954453 |
| GSAGSLNTSF | ASSLADSNNQAPL | 0.3106752 |
| GTGQIGI | ASSLADSNNQAPL | 0.49393904 |
| GTLIALIL | ASSLADSNNQAPL | 0.6441977 |
| IAATMGFVVG | ASSLADSNNQAPL | 0.7181324 |
| IFAQALL | ASSLADSNNQAPL | 0.38950095 |
| IGAAVALGF | ASSLADSNNQAPL | 0.42568815 |
| IGLTPLAL | ASSLADSNNQAPL | 0.30359697 |
| ILGSNTSTL | ASSLADSNNQAPL | 0.6317395 |
| IVFSLTVL | ASSLADSNNQAPL | 0.7410751 |
| LASKATGF | ASSLADSNNQAPL | 0.40434495 |
| LASVALIL | ASSLADSNNQAPL | 0.7526777 |
| LFAVGLTL | ASSLADSNNQAPL | 0.589201 |
| LFNGGIGTYI | ASSLADSNNQAPL | 0.23101145 |

| | | |
|---|---|---|
| LTLGAGII | ASSLADSNNQAPL | 0.42345428 |
| LVAAALGDI | ASSLADSNNQAPL | 0.6887102 |
| NTAFVQLGI | ASSLADSNNQAPL | 0.6707634 |
| QLFSGAEL | ASSLADSNNQAPL | 0.7210369 |
| SAATFITL | ASSLADSNNQAPL | 0.4094876 |
| SNYGLANLGV | ASSLADSNNQAPL | 0.34609464 |
| SYGGAYAVM | ASSLADSNNQAPL | 0.7433957 |
| TTGVGGLTLG | ASSLADSNNQAPL | 0.76390886 |
| TTQTIYI | ASSLADSNNQAPL | 0.6778815 |
| VAAGAGLGAVI | ASSLADSNNQAPL | 0.3487951 |
| VAAVVTLL | ASSLADSNNQAPL | 0.3512128 |
| VGFLGGLI | ASSLADSNNQAPL | 0.51837504 |
| VGVIPILL | ASSLADSNNQAPL | 0.6975752 |
| VIFASSLI | ASSLADSNNQAPL | 0.26319945 |
| VKGGDGGAAAG | ASSLADSNNQAPL | 0.52304965 |
| VTKTFTNI | ASSLADSNNQAPL | 0.51552665 |
| VVAAAILL | ASSLADSNNQAPL | 0.19804278 |
| YAALDVEL | ASSLADSNNQAPL | 0.5365373 |
| LASVALLL | ASSLADSNNQAPL | 0.7188007 |
| LDAVHRV | ASSLADSNNQAPL | 0.47178233 |
| QGLSSSIF | ASSLADSNNQAPL | 0.3232025 |
| RGTFANIRL | ASSLADSNNQAPL | 0.67886275 |
| RQLWWGHRI | ASSLADSNNQAPL | 0.80022156 |
| YGPPGTGKTLL | ASSLADSNNQAPL | 0.7224058 |
| AALVGAPANVA | ASSLADSNNQAPL | 0.2485016 |
| AATFRRTE | ASSLADSNNQAPL | 0.1486827 |
| AGALGGGI | ASSLADSNNQAPL | 0.29271686 |
| AGGAGGAGGIGAG | ASSLADSNNQAPL | 0.6742604 |
| AGSGALGL | ASSLADSNNQAPL | 0.41769087 |
| AQIADQI | ASSLADSNNQAPL | 0.34513694 |
| ASAVAVTYL | ASSLADSNNQAPL | 0.5308197 |
| ASTVLGGTGG | ASSLADSNNQAPL | 0.7370252 |
| AVGGEPAK | ASSLADSNNQAPL | 0.30780858 |
| CEQGTFGGTC | ASSLADSNNQAPL | 0.24061832 |
| DEGKAPAG | ASSLADSNNQAPL | 0.5906111 |
| DTVVFPAL | ASSLADSNNQAPL | 0.6604039 |
| FEGITMAG | ASSLADSNNQAPL | 0.28658855 |
| FPLVVQD | ASSLADSNNQAPL | 0.39139786 |
| GAFTTPQ | ASSLADSNNQAPL | 0.32423118 |
| GFGGFGTAGGVGGA | ASSLADSNNQAPL | 0.31406423 |
| GFLAQFL | ASSLADSNNQAPL | 0.56833065 |
| GGAGGAGGNAGVA | ASSLADSNNQAPL | 0.2712884 |
| GGVGTAGAGGAGGAGG | ASSLADSNNQAPL | 0.7288092 |
| GNGGGGDAGNAGSGGN | ASSLADSNNQAPL | 0.2701156 |
| GTGGAGGAGNDAGS | ASSLADSNNQAPL | 0.5042176 |
| GTTNTGWFNS | ASSLADSNNQAPL | 0.51127243 |
| IFMGPLFTL | ASSLADSNNQAPL | 0.578941 |
| ILAHAIEAI | ASSLADSNNQAPL | 0.36174214 |
| IRSAIREAL | ASSLADSNNQAPL | 0.3331372 |
| ITFAGGG | ASSLADSNNQAPL | 0.29231375 |
| ITGMSGHQEV | ASSLADSNNQAPL | 0.15736978 |

| | | |
|---|---|---|
| IVAGLACTL | ASSLADSNNQAPL | 0.6410911 |
| IVGGGLAGA | ASSLADSNNQAPL | 0.13879524 |
| IVTGVLNI | ASSLADSNNQAPL | 0.7101857 |
| LAQQGCI | ASSLADSNNQAPL | 0.59017473 |
| LCLHVEN | ASSLADSNNQAPL | 0.30231276 |
| LDAISDGS | ASSLADSNNQAPL | 0.513951 |
| LGGFAAML | ASSLADSNNQAPL | 0.24907577 |
| LGGKNPMI | ASSLADSNNQAPL | 0.2696925 |
| LGPKGVAAIL | ASSLADSNNQAPL | 0.561537 |
| LMVGVSI | ASSLADSNNQAPL | 0.63832015 |
| LVGAAFVL | ASSLADSNNQAPL | 0.30827802 |
| LVLAVAPL | ASSLADSNNQAPL | 0.48607934 |
| MAAIAVIDAA | ASSLADSNNQAPL | 0.5449271 |
| NNGINAELQ | ASSLADSNNQAPL | 0.3080015 |
| RGEPLVLVHG | ASSLADSNNQAPL | 0.508439 |
| RGGAGGAAGLI | ASSLADSNNQAPL | 0.7223309 |
| RSYRMHPAV | ASSLADSNNQAPL | 0.5045696 |
| SAAHLLAGQLA | ASSLADSNNQAPL | 0.30011755 |
| SAAVGSY | ASSLADSNNQAPL | 0.60588646 |
| SPHAVLHRI | ASSLADSNNQAPL | 0.28076583 |
| SQIAKVL | ASSLADSNNQAPL | 0.598557 |
| TALGHLVP | ASSLADSNNQAPL | 0.45951954 |
| TAPTTSGAA | ASSLADSNNQAPL | 0.39936113 |
| TDGDDGNGGSPGT | ASSLADSNNQAPL | 0.65029466 |
| TDGLGEN | ASSLADSNNQAPL | 0.30524826 |
| TGGPPLPGQA | ASSLADSNNQAPL | 0.2853835 |
| TIPAITI | ASSLADSNNQAPL | 0.3763568 |
| TIPGFTI | ASSLADSNNQAPL | 0.49684387 |
| TSIASGALA | ASSLADSNNQAPL | 0.6675081 |
| TVTLPGAAI | ASSLADSNNQAPL | 0.29075986 |
| VASGLSAALP | ASSLADSNNQAPL | 0.4881906 |
| VGAGVIGI | ASSLADSNNQAPL | 0.38272938 |
| VGPKRFGV | ASSLADSNNQAPL | 0.30308127 |
| VNVAFPD | ASSLADSNNQAPL | 0.48353365 |
| VSMTNSGVS | ASSLADSNNQAPL | 0.361726 |
| VTDVLGDLSV | ASSLADSNNQAPL | 0.2704363 |
| VVVVGAGFAGL | ASSLADSNNQAPL | 0.6023169 |
| VYTVGTVV | ASSLADSNNQAPL | 0.36083338 |
| WTTFSGP | ASSLADSNNQAPL | 0.5090605 |
| YDELVEAF | ASSLADSNNQAPL | 0.741089 |
| YGTGGAGGNGG | ASSLADSNNQAPL | 0.28952834 |
| YTNVMAV | ASSLADSNNQAPL | 0.5150869 |
| FLEQVHQL | ASSLADSNNQAPL | 0.5913493 |
| GLSLLAVAV | ASSLADSNNQAPL | 0.533584 |
| LEQVHQL | ASSLADSNNQAPL | 0.44360578 |
| LNAAGAI | ASSLADSNNQAPL | 0.36423063 |
| PAVHASL | ASSLADSNNQAPL | 0.7673237 |
| TELEDDA | ASSLADSNNQAPL | 0.6317625 |
| TPPGPPN | ASSLADSNNQAPL | 0.6040675 |
| TPPPPPPP | ASSLADSNNQAPL | 0.63229525 |
| TQSPALI | ASSDDTISNERLF | 0.6222627 |

| | | |
|---|---|---|
| EFAAAFPQ | ASSDDTISNERLF | 0.6559602 |
| LLSGNLDVL | ASSDDTISNERLF | 0.33042616 |
| GPANLGNYNI | ASSDDTISNERLF | 0.21162915 |
| TLGPTLP | ASSDDTISNERLF | 0.41731983 |
| VYAYSSCCK | ASSDDTISNERLF | 0.4752246 |
| APAPAPAPNI | ASSDDTISNERLF | 0.39130998 |
| PGGPVPIP | ASSDDTISNERLF | 0.2378305 |
| FLTVALEP | ASSDDTISNERLF | 0.7223778 |
| SQYFGANL | ASSDDTISNERLF | 0.80446184 |
| GGVKPAA | ASSDDTISNERLF | 0.29828495 |
| GGTGGAGGAGGW | ASSDDTISNERLF | 0.76044273 |
| GVGGVAGLGGAGG | ASSDDTISNERLF | 0.2526179 |
| VLLVLATTI | ASSDDTISNERLF | 0.5878065 |
| IGIFAAML | ASSDDTISNERLF | 0.71869063 |
| AVGDLID | ASSDDTISNERLF | 0.25507462 |
| NISPLPL | ASSDDTISNERLF | 0.58355176 |
| VTAVFVAI | ASSDDTISNERLF | 0.35102385 |
| SGDDNDAAL | ASSDDTISNERLF | 0.39238 |
| AVAALAHS | ASSDDTISNERLF | 0.7221576 |
| GGVTVMATP | ASSDDTISNERLF | 0.39685088 |
| AGLAANVRD | ASSDDTISNERLF | 0.5321613 |
| TNRPDLIDPALLRPG | ASSDDTISNERLF | 0.4513181 |
| LANLGVTAL | ASSDDTISNERLF | 0.32535207 |
| TGERIEL | ASSDDTISNERLF | 0.56204015 |
| NGEELAA | ASSDDTISNERLF | 0.6707951 |
| VLSLVWG | ASSDDTISNERLF | 0.28739038 |
| IGYALWL | ASSDDTISNERLF | 0.7442604 |
| LSLLLAPL | ASSDDTISNERLF | 0.2809249 |
| TLDEALK | ASSDDTISNERLF | 0.80971265 |
| TDGAGTAEGG | ASSDDTISNERLF | 0.7189456 |
| NAPGGAGGNGG | ASSDDTISNERLF | 0.5304568 |
| VSVFMGMGGNF | ASSDDTISNERLF | 0.37519312 |
| LASYLAAL | ASSDDTISNERLF | 0.7151035 |
| LVGGTPAL | ASSDDTISNERLF | 0.40777817 |
| VRNADVL | ASSDDTISNERLF | 0.49490294 |
| TCSSPFLVSAM | ASSDDTISNERLF | 0.67173874 |
| VHITDAE | ASSDDTISNERLF | 0.61811733 |
| EYAAPSF | ASSDDTISNERLF | 0.8639909 |
| VVDAFHGGL | ASSDDTISNERLF | 0.74753344 |
| ASVVDPQ | ASSDDTISNERLF | 0.7734077 |
| SYNEFGGLS | ASSDDTISNERLF | 0.6382208 |
| DPDGSVY | ASSDDTISNERLF | 0.4449044 |
| MVSTIHATE | ASSDDTISNERLF | 0.7461628 |
| LAAGVADPV | ASSDDTISNERLF | 0.55564976 |
| TKEQIAA | ASSDDTISNERLF | 0.80821395 |
| SLASPAAL | ASSDDTISNERLF | 0.8241997 |
| IGLVFYGV | ASSDDTISNERLF | 0.6218793 |
| FLLDASNL | ASSDDTISNERLF | 0.8520701 |
| YQVGNFP | ASSDDTISNERLF | 0.8544879 |
| ESLLSQST | ASSDDTISNERLF | 0.4775173 |
| LFTVGFAAVG | ASSDDTISNERLF | 0.680348 |

| | | |
|---|---|---|
| ATPPTPT | ASSDDTISNERLF | 0.7146979 |
| TDLATLG | ASSDDTISNERLF | 0.7224498 |
| SAGGTGR | ASSDDTISNERLF | 0.3885107 |
| GFSNSGS | ASSDDTISNERLF | 0.823659 |
| GGNGGSGGTSV | ASSDDTISNERLF | 0.61837506 |
| GTSVATGGAGN | ASSDDTISNERLF | 0.18027453 |
| NGGNGGSGGTS | ASSDDTISNERLF | 0.76677585 |
| AVVKFPH | ASSDDTISNERLF | 0.7287139 |
| TPPPPPP | ASSDDTISNERLF | 0.68325675 |
| AAPAAAAAPA | ASSDDTISNERLF | 0.8295493 |
| IALNEDL | ASSDDTISNERLF | 0.56589323 |
| IVNVSSM | ASSDDTISNERLF | 0.5680293 |
| TGIPPAPR | ASSDDTISNERLF | 0.81202126 |
| FDLGGGTF | ASSDDTISNERLF | 0.8435142 |
| FDLGGGTFDVS | ASSDDTISNERLF | 0.48348224 |
| NEPTAAA | ASSDDTISNERLF | 0.77083784 |
| NEPTAAALA | ASSDDTISNERLF | 0.8488798 |
| GSIINISSI | ASSDDTISNERLF | 0.7812178 |
| VAPGNAGTA | ASSDDTISNERLF | 0.8433942 |
| TGGGASLE | ASSDDTISNERLF | 0.8576662 |
| VAVYNHYKRI | ASSDDTISNERLF | 0.65837556 |
| TDPAPATT | ASSDDTISNERLF | 0.83942664 |
| TNRPDILDPALLRPG | ASSDDTISNERLF | 0.8558701 |
| LTTTTTS | ASSDDTISNERLF | 0.7213671 |
| PTGSGKTL | ASSDDTISNERLF | 0.7465527 |
| TGSGKTL | ASSDDTISNERLF | 0.5948667 |
| TIGHVDHGK | ASSDDTISNERLF | 0.71575683 |
| TIGHVDHGKTT | ASSDDTISNERLF | 0.35782647 |
| GHVDHGKTT | ASSDDTISNERLF | 0.7682924 |
| AQFEHTLL | ASSDDTISNERLF | 0.8468899 |
| IVNQDEL | ASSDDTISNERLF | 0.68655026 |
| TNGSQFF | ASSDDTISNERLF | 0.542748 |
| TGKGIVE | ASSDDTISNERLF | 0.5067454 |
| LISNASDA | ASSDDTISNERLF | 0.7835025 |
| AELRTLGELTKIAKAHGAQ | ASSDDTISNERLF | 0.6331457 |
| APNAVAAAEPSW | ASSDDTISNERLF | 0.39485273 |
| APQVAGAATAVL | ASSDDTISNERLF | 0.71804816 |
| ARRDPAAAAAILPTDA | ASSDDTISNERLF | 0.845442 |
| DAAARDVLAEAGLAEYF | ASSDDTISNERLF | 0.84283906 |
| DDQIVLSAKANLAKVEQ | ASSDDTISNERLF | 0.40320772 |
| DELLPDTGEDSDEDGH | ASSDDTISNERLF | 0.8623897 |
| ELPEAPANRATVPQD | ASSDDTISNERLF | 0.815291 |
| FGQSKIERVTGIG | ASSDDTISNERLF | 0.8507078 |
| GAGGTGSTAGGAGGAGGAGGLY | ASSDDTISNERLF | 0.7634238 |
| GHGGTGAAGGNGGTGGAGGSGI | ASSDDTISNERLF | 0.6037001 |
| GNGGNGGAGGEGGPGGAGGAGA | ASSDDTISNERLF | 0.83811116 |
| GSTAGNGGNGGNATNGTIAGQP | ASSDDTISNERLF | 0.6514298 |
| GTGGDGGNGGNGGTGG | ASSDDTISNERLF | 0.60138136 |
| GVRRPAGIPADAPV | ASSDDTISNERLF | 0.83816206 |
| GYRQVTDAHLLTLARR | ASSDDTISNERLF | 0.6925211 |
| IAKMGQDGHDRG | ASSDDTISNERLF | 0.608404 |

| | | |
|---|---|---|
| KAEPSIEIQYT | ASSDDTISNERLF | 0.6184936 |
| KPVRIRMAADEDTVAVT | ASSDDTISNERLF | 0.8572222 |
| MPPRIAGMRLLVIKPEP | ASSDDTISNERLF | 0.52263224 |
| MKFYAVALFRSRGVPPTR | ASSDDTISNERLF | 0.5601842 |
| NAGGFGILGAVAHSP | ASSDDTISNERLF | 0.58352524 |
| NEGREITLAMAEM | ASSDDTISNERLF | 0.7285223 |
| NELVEWSHPLGHRWRWE | ASSDDTISNERLF | 0.83596104 |
| NLGAGGTGGNGGNPG | ASSDDTISNERLF | 0.8020063 |
| NTEWPKWETVMAADPVD | ASSDDTISNERLF | 0.67048013 |
| PASAAPAETQP | ASSDDTISNERLF | 0.77043676 |
| QQSSDGKNGVRAAEITGE | ASSDDTISNERLF | 0.81168044 |
| RAPDAVTTSWWKE | ASSDDTISNERLF | 0.7738011 |
| SGFGNVGAHTSGWFNQ | ASSDDTISNERLF | 0.8039998 |
| SGVAGAVGQAASVGGLKV | ASSDDTISNERLF | 0.5968945 |
| SNGGAGTGSAGGNGGT | ASSDDTISNERLF | 0.8542188 |
| TGGAGGEGGAAG | ASSDDTISNERLF | 0.8108969 |
| TGGDGGAAGAGGAPGHG | ASSDDTISNERLF | 0.83104986 |
| TMPPLLAAIANAGRHGV | ASSDDTISNERLF | 0.8402634 |
| TSSTAGNSGGA | ASSDDTISNERLF | 0.7929786 |
| VPATVPVGILC | ASSDDTISNERLF | 0.7777835 |
| VQRPDSGGVVLTGRIS | ASSDDTISNERLF | 0.29508892 |
| WTPNPVPATKGVPAGLVQAT | ASSDDTISNERLF | 0.73914415 |
| YALKRALAKFSDRFDVVII | ASSDDTISNERLF | 0.7516996 |
| YLIGSVIV | ASSDDTISNERLF | 0.7691461 |
| YGQLVAVID | ASSDDTISNERLF | 0.8042281 |
| VTVGGMTLPA | ASSDDTISNERLF | 0.8109715 |
| VLALPAV | ASSDDTISNERLF | 0.64352006 |
| VIPPTLNYET | ASSDDTISNERLF | 0.8414383 |
| VGMGGMHPSSG | ASSDDTISNERLF | 0.84743726 |
| VGLAGGAATA | ASSDDTISNERLF | 0.44866765 |
| VGGAGGAGGSGP | ASSDDTISNERLF | 0.6847505 |
| VFAAFMV | ASSDDTISNERLF | 0.8829045 |
| TLYDKLR | ASSDDTISNERLF | 0.752558 |
| TLAVFTVL | ASSDDTISNERLF | 0.68626344 |
| TGSVQPARV | ASSDDTISNERLF | 0.67709315 |
| TGGGAGSGGT | ASSDDTISNERLF | 0.86386377 |
| SHLETSL | ASSDDTISNERLF | 0.6676636 |
| SFTTGMT | ASSDDTISNERLF | 0.8611702 |
| QVEVAKRMGVRQPTVS | ASSDDTISNERLF | 0.22062625 |
| QGSGGPALAM | ASSDDTISNERLF | 0.69736123 |
| MLAPSNP | ASSDDTISNERLF | 0.632843 |
| LVNNAAI | ASSDDTISNERLF | 0.29081824 |
| LTGEINTLA | ASSDDTISNERLF | 0.82491577 |
| LSIFGPV | ASSDDTISNERLF | 0.7212581 |
| LLALLANL | ASSDDTISNERLF | 0.8138961 |
| LISNASDALDKLR | ASSDDTISNERLF | 0.7606871 |
| LINEAVP | ASSDDTISNERLF | 0.8405164 |
| LGYAFSGL | ASSDDTISNERLF | 0.7318153 |
| LDFVAVAL | ASSDDTISNERLF | 0.7683101 |
| LAQPGAE | ASSDDTISNERLF | 0.6330791 |
| ISNLGAVTGF | ASSDDTISNERLF | 0.8083678 |

| | | |
|---|---|---|
| HQEDPGRI | ASSDDTISNERLF | 0.85385513 |
| GTPGEPTP | ASSDDTISNERLF | 0.8331 |
| GFFNSGS | ASSDDTISNERLF | 0.3734275 |
| GAGGGGLTAGAN | ASSDDTISNERLF | 0.6983942 |
| FVDALVNM | ASSDDTISNERLF | 0.7410397 |
| FLTVAPEL | ASSDDTISNERLF | 0.34879258 |
| FETLTAL | ASSDDTISNERLF | 0.2650052 |
| DVINPIEL | ASSDDTISNERLF | 0.70807177 |
| APVSGAPV | ASSDDTISNERLF | 0.67682785 |
| ALVGGLAW | ASSDDTISNERLF | 0.82898784 |
| ALAAAAATAI | ASSDDTISNERLF | 0.8111454 |
| AFGLGLLL | ASSDDTISNERLF | 0.7014124 |
| AFFAAASL | ASSDDTISNERLF | 0.8091303 |
| AAAALPAFSPP | ASSDDTISNERLF | 0.71655166 |
| VVLAPAL | ASSDDTISNERLF | 0.112408064 |
| TIGHVDHGKTTL | ASSDDTISNERLF | 0.7251389 |
| LISNASDALDK | ASSDDTISNERLF | 0.50050855 |
| LISNASDAL | ASSDDTISNERLF | 0.86593235 |
| ISNASDALDK | ASSDDTISNERLF | 0.62443405 |
| NQAETLVYQTE | ASSDDTISNERLF | 0.6262779 |
| TGGTGGTAGAG | ASSDDTISNERLF | 0.72999376 |
| TGSTAGGAGGA | ASSDDTISNERLF | 0.75137913 |
| DPLAVPVAAGPV | ASSDDTISNERLF | 0.7288011 |
| GGTGGTPGNGGN | ASSDDTISNERLF | 0.5827625 |
| PAGLAGVDPATL | ASSDDTISNERLF | 0.20270745 |
| GNNLVVEAPHAGASV | ASSDDTISNERLF | 0.8300449 |
| KVTRIMSASTLEQPA | ASSDDTISNERLF | 0.5407554 |
| LGFALPIALPRRTVV | ASSDDTISNERLF | 0.522219 |
| TRLGNLTARKLVVLV | ASSDDTISNERLF | 0.3594548 |
| WSQQRRLIYFPSAGP | ASSDDTISNERLF | 0.33842042 |
| GPNGNPDPMAAAVDIR | ASSDDTISNERLF | 0.42422244 |
| TQRCVSVADLRAAASR | ASSDDTISNERLF | 0.8270946 |
| QFVLDVAEHPVEVLTETV | ASSDDTISNERLF | 0.67351615 |
| GGAGGTGGAGGNGGTQAGDG | ASSDDTISNERLF | 0.68500865 |
| PLLATLRAIAVDVDAKPAQV | ASSDDTISNERLF | 0.87742233 |
| APKFRTGGYMIDFWGVGYQVAK | ASSDDTISNERLF | 0.7182934 |
| GAGGAGGAGGAAGGPGGTGGTG | ASSDDTISNERLF | 0.6921547 |
| IGALGLLATDVAPAGSVAFGVA | ASSDDTISNERLF | 0.6281982 |
| AGGAGGIGGNGGAGGAGGVLMG | ASSDDTISNERLF | 0.585346 |
| ADELIGLL | ASSDDTISNERLF | 0.5683205 |
| ADRALKA | ASSDDTISNERLF | 0.58287203 |
| AGAVGQVGGAGGA | ASSDDTISNERLF | 0.8693701 |
| AGGGGGLGGAP | ASSDDTISNERLF | 0.70201135 |
| AGGLLYGNGGAG | ASSDDTISNERLF | 0.5845213 |
| AGLESIGL | ASSDDTISNERLF | 0.7887076 |
| AGLKTGII | ASSDDTISNERLF | 0.863112 |
| ASLAALLL | ASSDDTISNERLF | 0.8774915 |
| ASVINISSI | ASSDDTISNERLF | 0.58276206 |
| ATPPGPA | ASSDDTISNERLF | 0.7193396 |
| ATVASALV | ASSDDTISNERLF | 0.7010163 |
| ATVASLIL | ASSDDTISNERLF | 0.71104276 |

| | | |
|---|---|---|
| AVNGPVLIRH | ASSDDTISNERLF | 0.66814476 |
| DGAAGAVGI | ASSDDTISNERLF | 0.52022 |
| DPLLHGI | ASSDDTISNERLF | 0.8341466 |
| ELGAALGAL | ASSDDTISNERLF | 0.882999 |
| ELGRTQAV | ASSDDTISNERLF | 0.71420896 |
| FVAGGYLS | ASSDDTISNERLF | 0.52502644 |
| GASGGAGGAGGW | ASSDDTISNERLF | 0.6910783 |
| GGAGAGGDGGAGGI | ASSDDTISNERLF | 0.5477377 |
| GTGHNGSQPGMGGNG | ASSDDTISNERLF | 0.62659836 |
| GYAAASASAS | ASSDDTISNERLF | 0.74821997 |
| HQVDVAAVQQG | ASSDDTISNERLF | 0.43669128 |
| IGAAIAHT | ASSDDTISNERLF | 0.72173893 |
| IGMIEVAQL | ASSDDTISNERLF | 0.87056947 |
| IRLAPEHTSAMD | ASSDDTISNERLF | 0.65261555 |
| IYNVSVL | ASSDDTISNERLF | 0.7919724 |
| KQGFLGM | ASSDDTISNERLF | 0.84520805 |
| LESVITQL | ASSDDTISNERLF | 0.8035412 |
| LGLGALVVV | ASSDDTISNERLF | 0.8466738 |
| LIGNGGNGGSGGI | ASSDDTISNERLF | 0.82189155 |
| LNAINAPTL | ASSDDTISNERLF | 0.7644192 |
| LQLLNTGV | ASSDDTISNERLF | 0.7696018 |
| LQTSIEF | ASSDDTISNERLF | 0.8016805 |
| MSFAVSEL | ASSDDTISNERLF | 0.7146225 |
| NTGALGL | ASSDDTISNERLF | 0.70710766 |
| PPGAGGAGGAGTT | ASSDDTISNERLF | 0.6424087 |
| QLVGLGI | ASSDDTISNERLF | 0.8379773 |
| RFLEQVHQL | ASSDDTISNERLF | 0.74681383 |
| RGVGYVLR | ASSDDTISNERLF | 0.85056823 |
| SATGVFAGVI | ASSDDTISNERLF | 0.43000296 |
| SLPAITLP | ASSDDTISNERLF | 0.5089281 |
| STALAFTV | ASSDDTISNERLF | 0.82897377 |
| SVLTVGLGK | ASSDDTISNERLF | 0.152563 |
| TALAGAGSGI | ASSDDTISNERLF | 0.2755085 |
| TGLPFTGI | ASSDDTISNERLF | 0.76197034 |
| VAEHVTV | ASSDDTISNERLF | 0.8502878 |
| VDIVLNSL | ASSDDTISNERLF | 0.18589038 |
| VLTGGGAVL | ASSDDTISNERLF | 0.5762672 |
| VNVGLAYGI | ASSDDTISNERLF | 0.4451697 |
| VSSAAVGAV | ASSDDTISNERLF | 0.84238195 |
| VTSVYEAGV | ASSDDTISNERLF | 0.7365681 |
| WVALDRGA | ASSDDTISNERLF | 0.41547012 |
| YLVGSVVI | ASSDDTISNERLF | 0.81444424 |
| GPPGTGKTLL | ASSDDTISNERLF | 0.42274076 |
| GVGMTKF | ASSDDTISNERLF | 0.63285327 |
| IGIGGSDL | ASSDDTISNERLF | 0.82582855 |
| IGTLLAEL | ASSDDTISNERLF | 0.8520919 |
| IYTSGTT | ASSDDTISNERLF | 0.58772165 |
| LSFPAGV | ASSDDTISNERLF | 0.80996895 |
| LYTSGTT | ASSDDTISNERLF | 0.66358364 |
| VGAFPFGF | ASSDDTISNERLF | 0.8810193 |
| VGAGTVEF | ASSDDTISNERLF | 0.6813891 |

| | | |
|---|---|---|
| VGKSTLLL | ASSDDTISNERLF | 0.46467015 |
| AALAHAGGGIV | ASSDDTISNERLF | 0.8268282 |
| AALVAVSLL | ASSDDTISNERLF | 0.80771273 |
| AAVSRAVAAL | ASSDDTISNERLF | 0.73945 |
| AGGSSGIGGF | ASSDDTISNERLF | 0.82529044 |
| AGLTVANANG | ASSDDTISNERLF | 0.7014783 |
| AGLVGAVL | ASSDDTISNERLF | 0.69715655 |
| ALSQAFTV | ASSDDTISNERLF | 0.8475497 |
| ALTFSVSL | ASSDDTISNERLF | 0.8162356 |
| ALTGAALVAL | ASSDDTISNERLF | 0.56047356 |
| ASFSIGAGVL | ASSDDTISNERLF | 0.8306564 |
| DGSVYKI | ASSDDTISNERLF | 0.7014364 |
| EQLAEIV | ASSDDTISNERLF | 0.62550634 |
| FAVSELTVM | ASSDDTISNERLF | 0.8698776 |
| GAAPLAAGL | ASSDDTISNERLF | 0.7638057 |
| GADGGDGGSGGAGG | ASSDDTISNERLF | 0.8167614 |
| GFGLDTGGAGG | ASSDDTISNERLF | 0.8212904 |
| GLASGGLL | ASSDDTISNERLF | 0.8466727 |
| GRIAGAAVLL | ASSDDTISNERLF | 0.5024711 |
| GSAGSLNTSF | ASSDDTISNERLF | 0.78951883 |
| GTGQIGI | ASSDDTISNERLF | 0.7258817 |
| GTLIALIL | ASSDDTISNERLF | 0.8285052 |
| IAATMGFVVG | ASSDDTISNERLF | 0.74741757 |
| IFAQALL | ASSDDTISNERLF | 0.59171957 |
| IGAAVALGF | ASSDDTISNERLF | 0.80580413 |
| IGLTPLAL | ASSDDTISNERLF | 0.8520082 |
| ILGSNTSTL | ASSDDTISNERLF | 0.7662614 |
| IVFSLTVL | ASSDDTISNERLF | 0.7803726 |
| LASKATGF | ASSDDTISNERLF | 0.86100495 |
| LASVALIL | ASSDDTISNERLF | 0.6190551 |
| LFAVGLTL | ASSDDTISNERLF | 0.7015297 |
| LFNGGIGTYI | ASSDDTISNERLF | 0.82078743 |
| LTLGAGII | ASSDDTISNERLF | 0.84148455 |
| LVAAALGDI | ASSDDTISNERLF | 0.73707724 |
| NTAFVQLGI | ASSDDTISNERLF | 0.859566 |
| QLFSGAEL | ASSDDTISNERLF | 0.6376601 |
| SAATFITL | ASSDDTISNERLF | 0.81320506 |
| SNYGLANLGV | ASSDDTISNERLF | 0.64764273 |
| SYGGAYAVM | ASSDDTISNERLF | 0.77800953 |
| TTGVGGLTLG | ASSDDTISNERLF | 0.55458456 |
| TTQTIYI | ASSDDTISNERLF | 0.795024 |
| VAAGAGLGAVI | ASSDDTISNERLF | 0.8232676 |
| VAAVVTLL | ASSDDTISNERLF | 0.8282384 |
| VGFLGGLI | ASSDDTISNERLF | 0.26436383 |
| VGVIPILL | ASSDDTISNERLF | 0.82121253 |
| VIFASSLI | ASSDDTISNERLF | 0.8311778 |
| VKGGDGGAAAG | ASSDDTISNERLF | 0.8825038 |
| VTKTFTNI | ASSDDTISNERLF | 0.72274613 |
| VVAAAILL | ASSDDTISNERLF | 0.8050612 |
| YAALDVEL | ASSDDTISNERLF | 0.6504992 |
| LASVALLL | ASSDDTISNERLF | 0.7226153 |

| | | |
|---|---|---|
| LDAVHRV | ASSDDTISNERLF | 0.50582993 |
| QGLSSSIF | ASSDDTISNERLF | 0.6643325 |
| RGTFANIRL | ASSDDTISNERLF | 0.6637998 |
| RQLWWGHRI | ASSDDTISNERLF | 0.5762901 |
| YGPPGTGKTLL | ASSDDTISNERLF | 0.8015169 |
| AALVGAPANVA | ASSDDTISNERLF | 0.8257654 |
| AATFRRTE | ASSDDTISNERLF | 0.83578724 |
| AGALGGGI | ASSDDTISNERLF | 0.8444884 |
| AGGAGGAGGIGAG | ASSDDTISNERLF | 0.86170435 |
| AGSGALGL | ASSDDTISNERLF | 0.83806837 |
| AQIADQI | ASSDDTISNERLF | 0.6001288 |
| ASAVAVTYL | ASSDDTISNERLF | 0.7538522 |
| ASTVLGGTGG | ASSDDTISNERLF | 0.4960869 |
| AVGGEPAK | ASSDDTISNERLF | 0.73371 |
| CEQGTFGGTC | ASSDDTISNERLF | 0.65145564 |
| DEGKAPAG | ASSDDTISNERLF | 0.78223777 |
| DTVVFPAL | ASSDDTISNERLF | 0.15023586 |
| FEGITMAG | ASSDDTISNERLF | 0.76957506 |
| FPLVVQD | ASSDDTISNERLF | 0.86769956 |
| GAFTTPQ | ASSDDTISNERLF | 0.5565294 |
| GFGGFGTAGGVGGA | ASSDDTISNERLF | 0.7843448 |
| GFLAQFL | ASSDDTISNERLF | 0.7866192 |
| GGAGGAGGNAGVA | ASSDDTISNERLF | 0.630628 |
| GGVGTAGAGGAGGAGG | ASSDDTISNERLF | 0.8058306 |
| GNGGGDAGNAGSGGN | ASSDDTISNERLF | 0.5785976 |
| GTGGAGGAGNDAGS | ASSDDTISNERLF | 0.796859 |
| GTTNTGWFNS | ASSDDTISNERLF | 0.8630811 |
| IFMGPLFTL | ASSDDTISNERLF | 0.77038217 |
| ILAHAIEAI | ASSDDTISNERLF | 0.7371692 |
| IRSAIREAL | ASSDDTISNERLF | 0.7321849 |
| ITFAGGG | ASSDDTISNERLF | 0.4321046 |
| ITGMSGHQEV | ASSDDTISNERLF | 0.7644629 |
| IVAGLACTL | ASSDDTISNERLF | 0.14438747 |
| IVGGGLAGA | ASSDDTISNERLF | 0.7288702 |
| IVTGVLNI | ASSDDTISNERLF | 0.5812951 |
| LAQQGCI | ASSDDTISNERLF | 0.7584087 |
| LCLHVEN | ASSDDTISNERLF | 0.7425018 |
| LDAISDGS | ASSDDTISNERLF | 0.78385866 |
| LGGFAAML | ASSDDTISNERLF | 0.54555 |
| LGGKNPMI | ASSDDTISNERLF | 0.73474574 |
| LGPKGVAAIL | ASSDDTISNERLF | 0.64681053 |
| LMVGVSI | ASSDDTISNERLF | 0.4722841 |
| LVGAAFVL | ASSDDTISNERLF | 0.821635 |
| LVLAVAPL | ASSDDTISNERLF | 0.7344094 |
| MAAIAVIDAA | ASSDDTISNERLF | 0.7287455 |
| NNGINAELQ | ASSDDTISNERLF | 0.85066074 |
| RGEPLVLVHG | ASSDDTISNERLF | 0.7558813 |
| RGGAGGAAGLI | ASSDDTISNERLF | 0.44652176 |
| RSYRMHPAV | ASSDDTISNERLF | 0.6865882 |
| SAAHLLAGQLA | ASSDDTISNERLF | 0.7242462 |
| SAAVGSY | ASSDDTISNERLF | 0.6703099 |

| | | |
|---|---|---|
| SPHAVLHRI | ASSDDTISNERLF | 0.14618549 |
| SQIAKVL | ASSDDTISNERLF | 0.82463723 |
| TALGHLVP | ASSDDTISNERLF | 0.627414 |
| TAPTTSGAA | ASSDDTISNERLF | 0.5694249 |
| TDGDDGNGGSPGT | ASSDDTISNERLF | 0.64894474 |
| TDGLGEN | ASSDDTISNERLF | 0.81721467 |
| TGGPPLPGQA | ASSDDTISNERLF | 0.5944109 |
| TIPAITI | ASSDDTISNERLF | 0.72617203 |
| TIPGFTI | ASSDDTISNERLF | 0.8026487 |
| TSIASGALA | ASSDDTISNERLF | 0.7914766 |
| TVTLPGAAI | ASSDDTISNERLF | 0.8515278 |
| VASGLSAALP | ASSDDTISNERLF | 0.79317385 |
| VGAGVIGI | ASSDDTISNERLF | 0.74898994 |
| VGPKRFGV | ASSDDTISNERLF | 0.8096791 |
| VNVAFPD | ASSDDTISNERLF | 0.75455993 |
| VSMTNSGVS | ASSDDTISNERLF | 0.45109183 |
| VTDVLGDLSV | ASSDDTISNERLF | 0.352882 |
| VVVVGAGFAGL | ASSDDTISNERLF | 0.70417154 |
| VYTVGTVV | ASSDDTISNERLF | 0.3458278 |
| WTTFSGP | ASSDDTISNERLF | 0.74967533 |
| YDELVEAF | ASSDDTISNERLF | 0.8593514 |
| YGTGGAGGNGG | ASSDDTISNERLF | 0.76623845 |
| YTNVMAV | ASSDDTISNERLF | 0.5223288 |
| FLEQVHQL | ASSDDTISNERLF | 0.68548626 |
| GLSLLAVAV | ASSDDTISNERLF | 0.7250935 |
| LEQVHQL | ASSDDTISNERLF | 0.8041706 |
| LNAAGAI | ASSDDTISNERLF | 0.84036934 |
| PAVHASL | ASSDDTISNERLF | 0.87068224 |
| TELEDDA | ASSDDTISNERLF | 0.8291229 |
| TPPGPPN | ASSDDTISNERLF | 0.61566246 |
| TPPPPPPP | ASSDDTISNERLF | 0.8059257 |
| TQSPALI | ASSPGLGGNYAEQF | 0.71420497 |
| EFAAAFPQ | ASSPGLGGNYAEQF | 0.75691056 |
| LLSGNLDVL | ASSPGLGGNYAEQF | 0.7499479 |
| GPANLGNYNI | ASSPGLGGNYAEQF | 0.7837543 |
| TLGPTLP | ASSPGLGGNYAEQF | 0.7135199 |
| VYAYSSCCK | ASSPGLGGNYAEQF | 0.66879594 |
| APAPAPAPNI | ASSPGLGGNYAEQF | 0.4195935 |
| PGGPVPIP | ASSPGLGGNYAEQF | 0.80558777 |
| FLTVALEP | ASSPGLGGNYAEQF | 0.79071724 |
| SQYFGANL | ASSPGLGGNYAEQF | 0.4609971 |
| GGVKPAA | ASSPGLGGNYAEQF | 0.6593903 |
| GGTGGAGGAGGW | ASSPGLGGNYAEQF | 0.86788106 |
| GVGGVAGLGGAGG | ASSPGLGGNYAEQF | 0.852566 |
| VLLVLATTI | ASSPGLGGNYAEQF | 0.5920143 |
| IGIFAAML | ASSPGLGGNYAEQF | 0.5844395 |
| AVGDLID | ASSPGLGGNYAEQF | 0.76424026 |
| NISPLPL | ASSPGLGGNYAEQF | 0.35179734 |
| VTAVFVAI | ASSPGLGGNYAEQF | 0.80312926 |
| SGDDNDAAL | ASSPGLGGNYAEQF | 0.7133662 |
| AVAALAHS | ASSPGLGGNYAEQF | 0.7145444 |

| | | |
|---|---|---|
| GGVTVMATP | ASSPGLGGNYAEQF | 0.6430329 |
| AGLAANVRD | ASSPGLGGNYAEQF | 0.7568254 |
| TNRPDLIDPALLRPG | ASSPGLGGNYAEQF | 0.6558889 |
| LANLGVTAL | ASSPGLGGNYAEQF | 0.81857866 |
| TGERIEL | ASSPGLGGNYAEQF | 0.18102427 |
| NGEELAA | ASSPGLGGNYAEQF | 0.81230193 |
| VLSLVWG | ASSPGLGGNYAEQF | 0.55337155 |
| IGYALWL | ASSPGLGGNYAEQF | 0.65946364 |
| LSLLLAPL | ASSPGLGGNYAEQF | 0.3512669 |
| TLDEALK | ASSPGLGGNYAEQF | 0.7651725 |
| TDGAGTAEGG | ASSPGLGGNYAEQF | 0.8170017 |
| NAPGGAGGNGG | ASSPGLGGNYAEQF | 0.82704926 |
| VSVFMGMGGNF | ASSPGLGGNYAEQF | 0.5957682 |
| LASYLAAL | ASSPGLGGNYAEQF | 0.82752156 |
| LVGGTPAL | ASSPGLGGNYAEQF | 0.8554579 |
| VRNADVL | ASSPGLGGNYAEQF | 0.67789114 |
| TCSSPFLVSAM | ASSPGLGGNYAEQF | 0.38897195 |
| VHITDAE | ASSPGLGGNYAEQF | 0.7902271 |
| EYAAPSF | ASSPGLGGNYAEQF | 0.71098256 |
| VVDAFHGGL | ASSPGLGGNYAEQF | 0.83009475 |
| ASVVDPQ | ASSPGLGGNYAEQF | 0.7722374 |
| SYNEFGGLS | ASSPGLGGNYAEQF | 0.48864537 |
| DPDGSVY | ASSPGLGGNYAEQF | 0.83142096 |
| MVSTIHATE | ASSPGLGGNYAEQF | 0.8442015 |
| LAAGVADPV | ASSPGLGGNYAEQF | 0.5517952 |
| TKEQIAA | ASSPGLGGNYAEQF | 0.6526972 |
| SLASPAAL | ASSPGLGGNYAEQF | 0.7044823 |
| IGLVFYGV | ASSPGLGGNYAEQF | 0.35324195 |
| FLLDASNL | ASSPGLGGNYAEQF | 0.80291116 |
| YQVGNFP | ASSPGLGGNYAEQF | 0.6366782 |
| ESLLSQST | ASSPGLGGNYAEQF | 0.59587646 |
| LFTVGFAAVG | ASSPGLGGNYAEQF | 0.7518174 |
| ATPPTPT | ASSPGLGGNYAEQF | 0.61134994 |
| TDLATLG | ASSPGLGGNYAEQF | 0.6281333 |
| SAGGTGR | ASSPGLGGNYAEQF | 0.6743942 |
| GFSNSGS | ASSPGLGGNYAEQF | 0.7558218 |
| GGNGGSGGTSV | ASSPGLGGNYAEQF | 0.7845799 |
| GTSVATGGAGN | ASSPGLGGNYAEQF | 0.75233907 |
| NGGNGGSGGTS | ASSPGLGGNYAEQF | 0.7754992 |
| AVVKFPH | ASSPGLGGNYAEQF | 0.85519624 |
| TPPPPPP | ASSPGLGGNYAEQF | 0.4433857 |
| AAPAAAAPA | ASSPGLGGNYAEQF | 0.58668995 |
| IALNEDL | ASSPGLGGNYAEQF | 0.7930534 |
| IVNVSSM | ASSPGLGGNYAEQF | 0.48138496 |
| TGIPPAPR | ASSPGLGGNYAEQF | 0.81899214 |
| FDLGGGTF | ASSPGLGGNYAEQF | 0.73300093 |
| FDLGGGTFDVS | ASSPGLGGNYAEQF | 0.7887759 |
| NEPTAAA | ASSPGLGGNYAEQF | 0.7640584 |
| NEPTAAALA | ASSPGLGGNYAEQF | 0.8780633 |
| GSIINISSI | ASSPGLGGNYAEQF | 0.8197127 |
| VAPGNAGTA | ASSPGLGGNYAEQF | 0.7668734 |

| | | |
|---|---|---|
| TGGGASLE | ASSPGLGGNYAEQF | 0.76909214 |
| VAVYNHYKRI | ASSPGLGGNYAEQF | 0.60349536 |
| TDPAPATT | ASSPGLGGNYAEQF | 0.83526456 |
| TNRPDILDPALLRPG | ASSPGLGGNYAEQF | 0.79841876 |
| LTTTTTS | ASSPGLGGNYAEQF | 0.50659394 |
| PTGSGKTL | ASSPGLGGNYAEQF | 0.7077285 |
| TGSGKTL | ASSPGLGGNYAEQF | 0.85069895 |
| TIGHVDHGK | ASSPGLGGNYAEQF | 0.843861 |
| TIGHVDHGKTT | ASSPGLGGNYAEQF | 0.3790283 |
| GHVDHGKTT | ASSPGLGGNYAEQF | 0.6828977 |
| AQFEHTLL | ASSPGLGGNYAEQF | 0.39394063 |
| IVNQDEL | ASSPGLGGNYAEQF | 0.6830024 |
| TNGSQFF | ASSPGLGGNYAEQF | 0.52097285 |
| TGKGIVE | ASSPGLGGNYAEQF | 0.7953483 |
| LISNASDA | ASSPGLGGNYAEQF | 0.65587187 |
| AELRTLGELTKIAKAHGAQ | ASSPGLGGNYAEQF | 0.78834444 |
| APNAVAAAEPSW | ASSPGLGGNYAEQF | 0.75477076 |
| APQVAGAATAVL | ASSPGLGGNYAEQF | 0.8492912 |
| ARRDPAAAAAILPTDA | ASSPGLGGNYAEQF | 0.61740136 |
| DAAARDVLAEAGLAEYF | ASSPGLGGNYAEQF | 0.693683 |
| DDQIVLSAKANLAKVEQ | ASSPGLGGNYAEQF | 0.3977402 |
| DELLPDTGEDSDEDGH | ASSPGLGGNYAEQF | 0.4901514 |
| ELPEAPANRATVPQD | ASSPGLGGNYAEQF | 0.6274224 |
| FGQSKIERVTGIG | ASSPGLGGNYAEQF | 0.6720418 |
| GAGGTGSTAGGAGGAGGAGGLY | ASSPGLGGNYAEQF | 0.2934328 |
| GHGGTGAAGGNGGTGGAGGSGI | ASSPGLGGNYAEQF | 0.8703857 |
| GNGGNGGAGGEGGPGGAGGAGA | ASSPGLGGNYAEQF | 0.7341546 |
| GSTAGNGGNGGNATNGTIAGQP | ASSPGLGGNYAEQF | 0.81828165 |
| GTGGDGGNGGNGGTGG | ASSPGLGGNYAEQF | 0.64499027 |
| GVRRPAGIPADAPV | ASSPGLGGNYAEQF | 0.6875977 |
| GYRQVTDAHLLTLARR | ASSPGLGGNYAEQF | 0.6129363 |
| IAKMGQDGHDRG | ASSPGLGGNYAEQF | 0.78860927 |
| KAEPSIEIQYT | ASSPGLGGNYAEQF | 0.6015798 |
| KPVRIRMAADEDTVAVT | ASSPGLGGNYAEQF | 0.37333554 |
| MPPRIAGMRLLVIKPEP | ASSPGLGGNYAEQF | 0.65853494 |
| MKFYAVALFRSRGVPPTR | ASSPGLGGNYAEQF | 0.58064073 |
| NAGGFGILGAVAHSP | ASSPGLGGNYAEQF | 0.86429083 |
| NEGREITLAMAEM | ASSPGLGGNYAEQF | 0.8343587 |
| NELVEWSHPLGHRWRWE | ASSPGLGGNYAEQF | 0.7685243 |
| NLGAGGTGGNGGNPG | ASSPGLGGNYAEQF | 0.8604604 |
| NTEWPKWETVMAADPVD | ASSPGLGGNYAEQF | 0.26406816 |
| PASAAPAETQP | ASSPGLGGNYAEQF | 0.68632656 |
| QQSSDGKNGVRAAEITGE | ASSPGLGGNYAEQF | 0.8638775 |
| RAPDAVTTSWWKE | ASSPGLGGNYAEQF | 0.7897395 |
| SGFGNVGAHTSGWFNQ | ASSPGLGGNYAEQF | 0.7063326 |
| SGVAGAVGQAASVGGLKV | ASSPGLGGNYAEQF | 0.83515155 |
| SNGGAGTGSAGGNGGT | ASSPGLGGNYAEQF | 0.5615592 |
| TGGAGGEGGAAG | ASSPGLGGNYAEQF | 0.66221446 |
| TGGDGGAAGAGGAPGHG | ASSPGLGGNYAEQF | 0.71725476 |
| TMPPLLAAIANAGRHGV | ASSPGLGGNYAEQF | 0.81554484 |
| TSSTAGNSGGA | ASSPGLGGNYAEQF | 0.7834951 |

| | | |
|---|---|---|
| VPATVPVGILC | ASSPGLGGNYAEQF | 0.532726 |
| VQRPDSGGVVLTGRIS | ASSPGLGGNYAEQF | 0.79062366 |
| WTPNPVPATKGVPAGLVQAT | ASSPGLGGNYAEQF | 0.88129437 |
| YALKRALAKFSDRFDVVII | ASSPGLGGNYAEQF | 0.6557273 |
| YLIGSVIV | ASSPGLGGNYAEQF | 0.8173426 |
| YGQLVAVID | ASSPGLGGNYAEQF | 0.7850403 |
| VTVGGMTLPA | ASSPGLGGNYAEQF | 0.75386316 |
| VLALPAV | ASSPGLGGNYAEQF | 0.57223845 |
| VIPPTLNYET | ASSPGLGGNYAEQF | 0.5953731 |
| VGMGGMHPSSG | ASSPGLGGNYAEQF | 0.7386912 |
| VGLAGGAATA | ASSPGLGGNYAEQF | 0.56134725 |
| VGGAGGAGGSGP | ASSPGLGGNYAEQF | 0.7281797 |
| VFAAFMV | ASSPGLGGNYAEQF | 0.7244644 |
| TLYDKLR | ASSPGLGGNYAEQF | 0.8906156 |
| TLAVFTVL | ASSPGLGGNYAEQF | 0.7875358 |
| TGSVQPARV | ASSPGLGGNYAEQF | 0.79039997 |
| TGGGAGSGGT | ASSPGLGGNYAEQF | 0.52691376 |
| SHLETSL | ASSPGLGGNYAEQF | 0.7798846 |
| SFTTGMT | ASSPGLGGNYAEQF | 0.78314406 |
| QVEVAKRMGVRQPTVS | ASSPGLGGNYAEQF | 0.4782399 |
| QGSGGPALAM | ASSPGLGGNYAEQF | 0.85898167 |
| MLAPSNP | ASSPGLGGNYAEQF | 0.5807278 |
| LVNNAAI | ASSPGLGGNYAEQF | 0.7207867 |
| LTGEINTLA | ASSPGLGGNYAEQF | 0.82768047 |
| LSIFGPV | ASSPGLGGNYAEQF | 0.84449023 |
| LLALLANL | ASSPGLGGNYAEQF | 0.70134395 |
| LISNASDALDKLR | ASSPGLGGNYAEQF | 0.716319 |
| LINEAVP | ASSPGLGGNYAEQF | 0.5522888 |
| LGYAFSGL | ASSPGLGGNYAEQF | 0.8327571 |
| LDFVAVAL | ASSPGLGGNYAEQF | 0.83356565 |
| LAQPGAE | ASSPGLGGNYAEQF | 0.70291525 |
| ISNLGAVTGF | ASSPGLGGNYAEQF | 0.8726233 |
| HQEDPGRI | ASSPGLGGNYAEQF | 0.8166464 |
| GTPGEPTP | ASSPGLGGNYAEQF | 0.4290126 |
| GFFNSGS | ASSPGLGGNYAEQF | 0.72569394 |
| GAGGGGLTAGAN | ASSPGLGGNYAEQF | 0.8364812 |
| FVDALVNM | ASSPGLGGNYAEQF | 0.84005916 |
| FLTVAPEL | ASSPGLGGNYAEQF | 0.85389876 |
| FETLTAL | ASSPGLGGNYAEQF | 0.79427415 |
| DVINPIEL | ASSPGLGGNYAEQF | 0.6451031 |
| APVSGAPV | ASSPGLGGNYAEQF | 0.86036503 |
| ALVGGLAW | ASSPGLGGNYAEQF | 0.88671935 |
| ALAAAAATAI | ASSPGLGGNYAEQF | 0.82153535 |
| AFGLGLLL | ASSPGLGGNYAEQF | 0.6605717 |
| AFFAAASL | ASSPGLGGNYAEQF | 0.65033484 |
| AAAALPAFSPP | ASSPGLGGNYAEQF | 0.76980674 |
| VVLAPAL | ASSPGLGGNYAEQF | 0.8164519 |
| TIGHVDHGKTTL | ASSPGLGGNYAEQF | 0.5206795 |
| LISNASDALDK | ASSPGLGGNYAEQF | 0.7515566 |
| LISNASDAL | ASSPGLGGNYAEQF | 0.7641847 |
| ISNASDALDK | ASSPGLGGNYAEQF | 0.39578348 |

| | | |
|---|---|---|
| NQAETLVYQTE | ASSPGLGGNYAEQF | 0.8365144 |
| TGGTGGTAGAG | ASSPGLGGNYAEQF | 0.8542204 |
| TGSTAGGAGGA | ASSPGLGGNYAEQF | 0.7604728 |
| DPLAVPVAAGPV | ASSPGLGGNYAEQF | 0.63371366 |
| GGTGGTPGNGGN | ASSPGLGGNYAEQF | 0.84396756 |
| PAGLAGVDPATL | ASSPGLGGNYAEQF | 0.88345706 |
| GNNLVVEAPHAGASV | ASSPGLGGNYAEQF | 0.86105096 |
| KVTRIMSASTLEQPA | ASSPGLGGNYAEQF | 0.37660524 |
| LGFALPIALPRRTVV | ASSPGLGGNYAEQF | 0.26032975 |
| TRLGNLTARKLVVLV | ASSPGLGGNYAEQF | 0.62498605 |
| WSQQRRLIYFPSAGP | ASSPGLGGNYAEQF | 0.80442333 |
| GPNGNPDPMAAAVDIR | ASSPGLGGNYAEQF | 0.67144454 |
| TQRCVSVADLRAAASR | ASSPGLGGNYAEQF | 0.59808356 |
| QFVLDVAEHPVEVLTETV | ASSPGLGGNYAEQF | 0.77172315 |
| GGAGGTGGAGGNGGTQAGDG | ASSPGLGGNYAEQF | 0.8471239 |
| PLLATLRAIAVDVDAKPAQV | ASSPGLGGNYAEQF | 0.60566247 |
| APKFRTGGYMIDFWGVGYQVAK | ASSPGLGGNYAEQF | 0.47412297 |
| GAGGAGGAGGAAGGPGGTGGTG | ASSPGLGGNYAEQF | 0.8146087 |
| IGALGLLATDVAPAGSVAFGVA | ASSPGLGGNYAEQF | 0.8091724 |
| AGGAGGIGGNGGAGGAGGVLMG | ASSPGLGGNYAEQF | 0.56820256 |
| ADELIGLL | ASSPGLGGNYAEQF | 0.6740021 |
| ADRALKA | ASSPGLGGNYAEQF | 0.66913927 |
| AGAVGQVGGAGGA | ASSPGLGGNYAEQF | 0.67030156 |
| AGGGGGLGGAP | ASSPGLGGNYAEQF | 0.7638229 |
| AGGLLYGNGGAG | ASSPGLGGNYAEQF | 0.50252193 |
| AGLESIGL | ASSPGLGGNYAEQF | 0.85428166 |
| AGLKTGII | ASSPGLGGNYAEQF | 0.54247224 |
| ASLAALLL | ASSPGLGGNYAEQF | 0.81755286 |
| ASVINISSI | ASSPGLGGNYAEQF | 0.7669056 |
| ATPPGPA | ASSPGLGGNYAEQF | 0.79221815 |
| ATVASALV | ASSPGLGGNYAEQF | 0.66636837 |
| ATVASLIL | ASSPGLGGNYAEQF | 0.62567097 |
| AVNGPVLIRH | ASSPGLGGNYAEQF | 0.5569358 |
| DGAAGAVGI | ASSPGLGGNYAEQF | 0.3279665 |
| DPLLHGI | ASSPGLGGNYAEQF | 0.83833873 |
| ELGAALGAL | ASSPGLGGNYAEQF | 0.27217987 |
| ELGRTQAV | ASSPGLGGNYAEQF | 0.8395071 |
| FVAGGYLS | ASSPGLGGNYAEQF | 0.79225 |
| GASGGAGGAGGW | ASSPGLGGNYAEQF | 0.5925361 |
| GGAGAGGDGGAGGI | ASSPGLGGNYAEQF | 0.7900395 |
| GTGHNGSQPGMGGNG | ASSPGLGGNYAEQF | 0.49836946 |
| GYAAASASAS | ASSPGLGGNYAEQF | 0.6533967 |
| HQVDVAAVQQG | ASSPGLGGNYAEQF | 0.7986525 |
| IGAAIAHT | ASSPGLGGNYAEQF | 0.80992115 |
| IGMIEVAQL | ASSPGLGGNYAEQF | 0.5918937 |
| IRLAPEHTSAMD | ASSPGLGGNYAEQF | 0.791497 |
| IYNVSVL | ASSPGLGGNYAEQF | 0.78489226 |
| KQGFLGM | ASSPGLGGNYAEQF | 0.6528107 |
| LESVITQL | ASSPGLGGNYAEQF | 0.75970817 |
| LGLGALVVV | ASSPGLGGNYAEQF | 0.83987135 |
| LIGNGGNGGSGGI | ASSPGLGGNYAEQF | 0.7755173 |

| | | |
|---|---|---|
| LNAINAPTL | ASSPGLGGNYAEQF | 0.57426 |
| LQLLNTGV | ASSPGLGGNYAEQF | 0.8041472 |
| LQTSIEF | ASSPGLGGNYAEQF | 0.59215355 |
| MSFAVSEL | ASSPGLGGNYAEQF | 0.77473027 |
| NTGALGL | ASSPGLGGNYAEQF | 0.7508699 |
| PPGAGGAGGAGTT | ASSPGLGGNYAEQF | 0.68177265 |
| QLVGLGI | ASSPGLGGNYAEQF | 0.82250565 |
| RFLEQVHQL | ASSPGLGGNYAEQF | 0.5980808 |
| RGVGYVLR | ASSPGLGGNYAEQF | 0.58765304 |
| SATGVFAGVI | ASSPGLGGNYAEQF | 0.6620425 |
| SLPAITLP | ASSPGLGGNYAEQF | 0.7420169 |
| STALAFTV | ASSPGLGGNYAEQF | 0.82848626 |
| SVLTVGLGK | ASSPGLGGNYAEQF | 0.540661 |
| TALAGAGSGI | ASSPGLGGNYAEQF | 0.75881857 |
| TGLPFTGI | ASSPGLGGNYAEQF | 0.66723967 |
| VAEHVTV | ASSPGLGGNYAEQF | 0.5231397 |
| VDIVLNSL | ASSPGLGGNYAEQF | 0.76005054 |
| VLTGGGAVL | ASSPGLGGNYAEQF | 0.72340167 |
| VNVGLAYGI | ASSPGLGGNYAEQF | 0.57416135 |
| VSSAAVGAV | ASSPGLGGNYAEQF | 0.79825723 |
| VTSVYEAGV | ASSPGLGGNYAEQF | 0.58163863 |
| WVALDRGA | ASSPGLGGNYAEQF | 0.79275525 |
| YLVGSVVI | ASSPGLGGNYAEQF | 0.8760127 |
| GPPGTGKTLL | ASSPGLGGNYAEQF | 0.6142733 |
| GVGMTKF | ASSPGLGGNYAEQF | 0.8138653 |
| IGIGGSDL | ASSPGLGGNYAEQF | 0.78554314 |
| IGTLLAEL | ASSPGLGGNYAEQF | 0.78460264 |
| IYTSGTT | ASSPGLGGNYAEQF | 0.6795626 |
| LSFPAGV | ASSPGLGGNYAEQF | 0.6546628 |
| LYTSGTT | ASSPGLGGNYAEQF | 0.8610512 |
| VGAFPFGF | ASSPGLGGNYAEQF | 0.80962634 |
| VGAGTVEF | ASSPGLGGNYAEQF | 0.8248168 |
| VGKSTLLL | ASSPGLGGNYAEQF | 0.8178384 |
| AALAHAGGGIV | ASSPGLGGNYAEQF | 0.80818784 |
| AALVAVSLL | ASSPGLGGNYAEQF | 0.8106145 |
| AAVSRAVAAL | ASSPGLGGNYAEQF | 0.59917676 |
| AGGSSGIGGF | ASSPGLGGNYAEQF | 0.39472017 |
| AGLTVANANG | ASSPGLGGNYAEQF | 0.67459565 |
| AGLVGAVL | ASSPGLGGNYAEQF | 0.76598084 |
| ALSQAFTV | ASSPGLGGNYAEQF | 0.70277923 |
| ALTFSVSL | ASSPGLGGNYAEQF | 0.41551566 |
| ALTGAALVAL | ASSPGLGGNYAEQF | 0.83888227 |
| ASFSIGAGVL | ASSPGLGGNYAEQF | 0.88153756 |
| DGSVYKI | ASSPGLGGNYAEQF | 0.57567847 |
| EQLAEIV | ASSPGLGGNYAEQF | 0.8807603 |
| FAVSELTVM | ASSPGLGGNYAEQF | 0.51122147 |
| GAAPLAAGL | ASSPGLGGNYAEQF | 0.79504347 |
| GADGGDGGSGGAGG | ASSPGLGGNYAEQF | 0.8485141 |
| GFGLDTGGAGG | ASSPGLGGNYAEQF | 0.5115846 |
| GLASGGLL | ASSPGLGGNYAEQF | 0.81514573 |
| GRIAGAAVLL | ASSPGLGGNYAEQF | 0.6618163 |

| | | |
|---|---|---|
| GSAGSLNTSF | ASSPGLGGNYAEQF | 0.7060832 |
| GTGQIGI | ASSPGLGGNYAEQF | 0.8438699 |
| GTLIALIL | ASSPGLGGNYAEQF | 0.6479856 |
| IAATMGFVVG | ASSPGLGGNYAEQF | 0.7936004 |
| IFAQALL | ASSPGLGGNYAEQF | 0.6799414 |
| IGAAVALGF | ASSPGLGGNYAEQF | 0.5623174 |
| IGLTPLAL | ASSPGLGGNYAEQF | 0.7798373 |
| ILGSNTSTL | ASSPGLGGNYAEQF | 0.83860767 |
| IVFSLTVL | ASSPGLGGNYAEQF | 0.6682648 |
| LASKATGF | ASSPGLGGNYAEQF | 0.8832283 |
| LASVALIL | ASSPGLGGNYAEQF | 0.59320474 |
| LFAVGLTL | ASSPGLGGNYAEQF | 0.88084143 |
| LFNGGIGTYI | ASSPGLGGNYAEQF | 0.83055264 |
| LTLGAGII | ASSPGLGGNYAEQF | 0.7281989 |
| LVAAALGDI | ASSPGLGGNYAEQF | 0.6845708 |
| NTAFVQLGI | ASSPGLGGNYAEQF | 0.82826984 |
| QLFSGAEL | ASSPGLGGNYAEQF | 0.6586006 |
| SAATFITL | ASSPGLGGNYAEQF | 0.79588544 |
| SNYGLANLGV | ASSPGLGGNYAEQF | 0.86134416 |
| SYGGAYAVM | ASSPGLGGNYAEQF | 0.29077494 |
| TTGVGGLTLG | ASSPGLGGNYAEQF | 0.8020216 |
| TTQTIYI | ASSPGLGGNYAEQF | 0.7273849 |
| VAAGAGLGAVI | ASSPGLGGNYAEQF | 0.5272925 |
| VAAVVTLL | ASSPGLGGNYAEQF | 0.45758954 |
| VGFLGGLI | ASSPGLGGNYAEQF | 0.25750297 |
| VGVIPILL | ASSPGLGGNYAEQF | 0.44079325 |
| VIFASSLI | ASSPGLGGNYAEQF | 0.39070362 |
| VKGGDGGAAAG | ASSPGLGGNYAEQF | 0.7986321 |
| VTKTFTNI | ASSPGLGGNYAEQF | 0.7125779 |
| VVAAAILL | ASSPGLGGNYAEQF | 0.40160242 |
| YAALDVEL | ASSPGLGGNYAEQF | 0.7900946 |
| LASVALLL | ASSPGLGGNYAEQF | 0.66007686 |
| LDAVHRV | ASSPGLGGNYAEQF | 0.27941826 |
| QGLSSSIF | ASSPGLGGNYAEQF | 0.54198265 |
| RGTFANIRL | ASSPGLGGNYAEQF | 0.33639422 |
| RQLWWGHRI | ASSPGLGGNYAEQF | 0.46032557 |
| YGPPGTGKTLL | ASSPGLGGNYAEQF | 0.24311267 |
| AALVGAPANVA | ASSPGLGGNYAEQF | 0.6393224 |
| AATFRRTE | ASSPGLGGNYAEQF | 0.55916476 |
| AGALGGGI | ASSPGLGGNYAEQF | 0.18069662 |
| AGGAGGAGGIGAG | ASSPGLGGNYAEQF | 0.6836092 |
| AGSGALGL | ASSPGLGGNYAEQF | 0.47216803 |
| AQIADQI | ASSPGLGGNYAEQF | 0.4450469 |
| ASAVAVTYL | ASSPGLGGNYAEQF | 0.74972343 |
| ASTVLGGTGG | ASSPGLGGNYAEQF | 0.44384336 |
| AVGGEPAK | ASSPGLGGNYAEQF | 0.34437928 |
| CEQGTFGGTC | ASSPGLGGNYAEQF | 0.78191185 |
| DEGKAPAG | ASSPGLGGNYAEQF | 0.78911465 |
| DTVVFPAL | ASSPGLGGNYAEQF | 0.285971 |
| FEGITMAG | ASSPGLGGNYAEQF | 0.6048136 |
| FPLVVQD | ASSPGLGGNYAEQF | 0.7840353 |

| | | |
|---|---|---|
| GAFTTPQ | ASSPGLGGNYAEQF | 0.67633253 |
| GFGGFGTAGGVGGA | ASSPGLGGNYAEQF | 0.7168602 |
| GFLAQFL | ASSPGLGGNYAEQF | 0.74501145 |
| GGAGGAGGNAGVA | ASSPGLGGNYAEQF | 0.47991055 |
| GGVGTAGAGGAGGAGG | ASSPGLGGNYAEQF | 0.77922785 |
| GNGGGGDAGNAGSGGN | ASSPGLGGNYAEQF | 0.816975 |
| GTGGAGGAGNDAGS | ASSPGLGGNYAEQF | 0.47028118 |
| GTTNTGWFNS | ASSPGLGGNYAEQF | 0.56690645 |
| IFMGPLFTL | ASSPGLGGNYAEQF | 0.30567107 |
| ILAHAIEAI | ASSPGLGGNYAEQF | 0.60465 |
| IRSAIREAL | ASSPGLGGNYAEQF | 0.3191264 |
| ITFAGGG | ASSPGLGGNYAEQF | 0.7379725 |
| ITGMSGHQEV | ASSPGLGGNYAEQF | 0.7544719 |
| IVAGLACTL | ASSPGLGGNYAEQF | 0.57818824 |
| IVGGGLAGA | ASSPGLGGNYAEQF | 0.31263006 |
| IVTGVLNI | ASSPGLGGNYAEQF | 0.32908314 |
| LAQQGCI | ASSPGLGGNYAEQF | 0.66200423 |
| LCLHVEN | ASSPGLGGNYAEQF | 0.2976991 |
| LDAISDGS | ASSPGLGGNYAEQF | 0.2533219 |
| LGGFAAML | ASSPGLGGNYAEQF | 0.5610683 |
| LGGKNPMI | ASSPGLGGNYAEQF | 0.7559718 |
| LGPKGVAAIL | ASSPGLGGNYAEQF | 0.7381097 |
| LMVGVSI | ASSPGLGGNYAEQF | 0.3718295 |
| LVGAAFVL | ASSPGLGGNYAEQF | 0.8071977 |
| LVLAVAPL | ASSPGLGGNYAEQF | 0.7763492 |
| MAAIAVIDAA | ASSPGLGGNYAEQF | 0.7248056 |
| NNGINAELQ | ASSPGLGGNYAEQF | 0.4851517 |
| RGEPLVLVHG | ASSPGLGGNYAEQF | 0.30018172 |
| RGGAGGAAGLI | ASSPGLGGNYAEQF | 0.8062723 |
| RSYRMHPAV | ASSPGLGGNYAEQF | 0.6781457 |
| SAAHLLAGQLA | ASSPGLGGNYAEQF | 0.6245483 |
| SAAVGSY | ASSPGLGGNYAEQF | 0.7298931 |
| SPHAVLHRI | ASSPGLGGNYAEQF | 0.60066444 |
| SQIAKVL | ASSPGLGGNYAEQF | 0.42241055 |
| TALGHLVP | ASSPGLGGNYAEQF | 0.6918148 |
| TAPTTSGAA | ASSPGLGGNYAEQF | 0.7347847 |
| TDGDDGNGGSPGT | ASSPGLGGNYAEQF | 0.42535743 |
| TDGLGEN | ASSPGLGGNYAEQF | 0.29570127 |
| TGGPPLPGQA | ASSPGLGGNYAEQF | 0.44243908 |
| TIPAITI | ASSPGLGGNYAEQF | 0.5832739 |
| TIPGFTI | ASSPGLGGNYAEQF | 0.7630367 |
| TSIASGALA | ASSPGLGGNYAEQF | 0.58505976 |
| TVTLPGAAI | ASSPGLGGNYAEQF | 0.5609181 |
| VASGLSAALP | ASSPGLGGNYAEQF | 0.71435994 |
| VGAGVIGI | ASSPGLGGNYAEQF | 0.60465884 |
| VGPKRFGV | ASSPGLGGNYAEQF | 0.5622278 |
| VNVAFPD | ASSPGLGGNYAEQF | 0.7172123 |
| VSMTNSGVS | ASSPGLGGNYAEQF | 0.5166078 |
| VTDVLGDLSV | ASSPGLGGNYAEQF | 0.7172549 |
| VVVVGAGFAGL | ASSPGLGGNYAEQF | 0.6896295 |
| VYTVGTVV | ASSPGLGGNYAEQF | 0.57537746 |

| | | |
|---|---|---|
| WTTFSGP | ASSPGLGGNYAEQF | 0.73042023 |
| YDELVEAF | ASSPGLGGNYAEQF | 0.6757431 |
| YGTGGAGGNGG | ASSPGLGGNYAEQF | 0.5883998 |
| YTNVMAV | ASSPGLGGNYAEQF | 0.26041365 |
| FLEQVHQL | ASSPGLGGNYAEQF | 0.5601727 |
| GLSLLAVAV | ASSPGLGGNYAEQF | 0.7107208 |
| LEQVHQL | ASSPGLGGNYAEQF | 0.68805516 |
| LNAAGAI | ASSPGLGGNYAEQF | 0.7581179 |
| PAVHASL | ASSPGLGGNYAEQF | 0.5525685 |
| TELEDDA | ASSPGLGGNYAEQF | 0.49016386 |
| TPPGPPN | ASSPGLGGNYAEQF | 0.7551469 |
| TPPPPPPP | ASSPGLGGNYAEQF | 0.7740403 |
| TQSPALI | ASSNGTDSGNTLY | 0.28090438 |
| EFAAAFPQ | ASSNGTDSGNTLY | 0.5895852 |
| LLSGNLDVL | ASSNGTDSGNTLY | 0.8150487 |
| GPANLGNYNI | ASSNGTDSGNTLY | 0.74184567 |
| TLGPTLP | ASSNGTDSGNTLY | 0.59083474 |
| VYAYSSCCK | ASSNGTDSGNTLY | 0.3994275 |
| APAPAPAPNI | ASSNGTDSGNTLY | 0.81249225 |
| PGGPVPIP | ASSNGTDSGNTLY | 0.50527716 |
| FLTVALEP | ASSNGTDSGNTLY | 0.82106036 |
| SQYFGANL | ASSNGTDSGNTLY | 0.18421467 |
| GGVKPAA | ASSNGTDSGNTLY | 0.38529497 |
| GGTGGAGGAGGW | ASSNGTDSGNTLY | 0.38952234 |
| GVGGVAGLGGAGG | ASSNGTDSGNTLY | 0.1831966 |
| VLLVLATTI | ASSNGTDSGNTLY | 0.7576555 |
| IGIFAAML | ASSNGTDSGNTLY | 0.35363334 |
| AVGDLID | ASSNGTDSGNTLY | 0.6629538 |
| NISPLPL | ASSNGTDSGNTLY | 0.61569726 |
| VTAVFVAI | ASSNGTDSGNTLY | 0.7411645 |
| SGDDNDAAL | ASSNGTDSGNTLY | 0.6435317 |
| AVAALAHS | ASSNGTDSGNTLY | 0.5113398 |
| GGVTVMATP | ASSNGTDSGNTLY | 0.32556826 |
| AGLAANVRD | ASSNGTDSGNTLY | 0.7249469 |
| TNRPDLIDPALLRPG | ASSNGTDSGNTLY | 0.78617036 |
| LANLGVTAL | ASSNGTDSGNTLY | 0.6287688 |
| TGERIEL | ASSNGTDSGNTLY | 0.2381494 |
| NGEELAA | ASSNGTDSGNTLY | 0.71503675 |
| VLSLVWG | ASSNGTDSGNTLY | 0.5590169 |
| IGYALWL | ASSNGTDSGNTLY | 0.23705891 |
| LSLLLAPL | ASSNGTDSGNTLY | 0.2059899 |
| TLDEALK | ASSNGTDSGNTLY | 0.4813714 |
| TDGAGTAEGG | ASSNGTDSGNTLY | 0.6123969 |
| NAPGGAGGNGG | ASSNGTDSGNTLY | 0.7708894 |
| VSVFMGMGGNF | ASSNGTDSGNTLY | 0.7875141 |
| LASYLAAL | ASSNGTDSGNTLY | 0.49256307 |
| LVGGTPAL | ASSNGTDSGNTLY | 0.7604809 |
| VRNADVL | ASSNGTDSGNTLY | 0.7101541 |
| TCSSPFLVSAM | ASSNGTDSGNTLY | 0.14503649 |
| VHITDAE | ASSNGTDSGNTLY | 0.43167165 |
| EYAAPSF | ASSNGTDSGNTLY | 0.27974132 |

| | | |
|---|---|---|
| VVDAFHGGL | ASSNGTDSGNTLY | 0.6837635 |
| ASVVDPQ | ASSNGTDSGNTLY | 0.41675457 |
| SYNEFGGLS | ASSNGTDSGNTLY | 0.5674697 |
| DPDGSVY | ASSNGTDSGNTLY | 0.5829884 |
| MVSTIHATE | ASSNGTDSGNTLY | 0.7692647 |
| LAAGVADPV | ASSNGTDSGNTLY | 0.6062341 |
| TKEQIAA | ASSNGTDSGNTLY | 0.36033863 |
| SLASPAAL | ASSNGTDSGNTLY | 0.2494755 |
| IGLVFYGV | ASSNGTDSGNTLY | 0.7108004 |
| FLLDASNL | ASSNGTDSGNTLY | 0.43434814 |
| YQVGNFP | ASSNGTDSGNTLY | 0.31996274 |
| ESLLSQST | ASSNGTDSGNTLY | 0.2776661 |
| LFTVGFAAVG | ASSNGTDSGNTLY | 0.76629627 |
| ATPPTPT | ASSNGTDSGNTLY | 0.21441756 |
| TDLATLG | ASSNGTDSGNTLY | 0.68687284 |
| SAGGTGR | ASSNGTDSGNTLY | 0.4769242 |
| GFSNSGS | ASSNGTDSGNTLY | 0.63131076 |
| GGNGGSGGTSV | ASSNGTDSGNTLY | 0.8229253 |
| GTSVATGGAGN | ASSNGTDSGNTLY | 0.7567059 |
| NGGNGGSGGTS | ASSNGTDSGNTLY | 0.59612393 |
| AVVKFPH | ASSNGTDSGNTLY | 0.40505627 |
| TPPPPPP | ASSNGTDSGNTLY | 0.33972758 |
| AAPAAAAAPA | ASSNGTDSGNTLY | 0.39123505 |
| IALNEDL | ASSNGTDSGNTLY | 0.40401405 |
| IVNVSSM | ASSNGTDSGNTLY | 0.7649361 |
| TGIPPAPR | ASSNGTDSGNTLY | 0.46543476 |
| FDLGGGTF | ASSNGTDSGNTLY | 0.25516897 |
| FDLGGGTFDVS | ASSNGTDSGNTLY | 0.47549617 |
| NEPTAAA | ASSNGTDSGNTLY | 0.78982073 |
| NEPTAAALA | ASSNGTDSGNTLY | 0.8214632 |
| GSIINISSI | ASSNGTDSGNTLY | 0.5186933 |
| VAPGNAGTA | ASSNGTDSGNTLY | 0.38687384 |
| TGGGASLE | ASSNGTDSGNTLY | 0.36962688 |
| VAVYNHYKRI | ASSNGTDSGNTLY | 0.6410782 |
| TDPAPATT | ASSNGTDSGNTLY | 0.33575112 |
| TNRPDILDPALLRPG | ASSNGTDSGNTLY | 0.28740454 |
| LTTTTTS | ASSNGTDSGNTLY | 0.68095386 |
| PTGSGKTL | ASSNGTDSGNTLY | 0.8184619 |
| TGSGKTL | ASSNGTDSGNTLY | 0.5223794 |
| TIGHVDHGK | ASSNGTDSGNTLY | 0.21487692 |
| TIGHVDHGKTT | ASSNGTDSGNTLY | 0.58961093 |
| GHVDHGKTT | ASSNGTDSGNTLY | 0.38127163 |
| AQFEHTLL | ASSNGTDSGNTLY | 0.4288891 |
| IVNQDEL | ASSNGTDSGNTLY | 0.48836043 |
| TNGSQFF | ASSNGTDSGNTLY | 0.25549805 |
| TGKGIVE | ASSNGTDSGNTLY | 0.5262127 |
| LISNASDA | ASSNGTDSGNTLY | 0.81616414 |
| AELRTLGELTKIAKAHGAQ | ASSNGTDSGNTLY | 0.6607472 |
| APNAVAAAEPSW | ASSNGTDSGNTLY | 0.65028095 |
| APQVAGAATAVL | ASSNGTDSGNTLY | 0.8030368 |
| ARRDPAAAAAILPTDA | ASSNGTDSGNTLY | 0.72249055 |

| | | |
|---|---|---|
| DAAARDVLAEAGLAEYF | ASSNGTDSGNTLY | 0.7875844 |
| DDQIVLSAKANLAKVEQ | ASSNGTDSGNTLY | 0.7910558 |
| DELLPDTGEDSDEDGH | ASSNGTDSGNTLY | 0.4881944 |
| ELPEAPANRATVPQD | ASSNGTDSGNTLY | 0.6190921 |
| FGQSKIERVTGIG | ASSNGTDSGNTLY | 0.8088758 |
| GAGGTGSTAGGAGGAGGAGGLY | ASSNGTDSGNTLY | 0.602641 |
| GHGGTGAAGGNGGTGGAGGSGI | ASSNGTDSGNTLY | 0.652097 |
| GNGGNGGAGGEGGPGGAGGAGA | ASSNGTDSGNTLY | 0.44058973 |
| GSTAGNGGNGGNATNGTIAGQP | ASSNGTDSGNTLY | 0.77216065 |
| GTGGDGGNGGNGGTGG | ASSNGTDSGNTLY | 0.52923936 |
| GVRRPAGIPADAPV | ASSNGTDSGNTLY | 0.78645223 |
| GYRQVTDAHLLTLARR | ASSNGTDSGNTLY | 0.2085898 |
| IAKMGQDGHDRG | ASSNGTDSGNTLY | 0.2963639 |
| KAEPSIEIQYT | ASSNGTDSGNTLY | 0.7759605 |
| KPVRIRMAADEDTVAVT | ASSNGTDSGNTLY | 0.1623197 |
| MPPRIAGMRLLVIKPEP | ASSNGTDSGNTLY | 0.19404408 |
| MKFYAVALFRSRGVPPTR | ASSNGTDSGNTLY | 0.5831648 |
| NAGGFGILGAVAHSP | ASSNGTDSGNTLY | 0.7892102 |
| NEGREITLAMAEM | ASSNGTDSGNTLY | 0.1821992 |
| NELVEWSHPLGHRWRWE | ASSNGTDSGNTLY | 0.33903298 |
| NLGAGGTGGNGGNPG | ASSNGTDSGNTLY | 0.25825343 |
| NTEWPKWETVMAADPVD | ASSNGTDSGNTLY | 0.5676494 |
| PASAAPAETQP | ASSNGTDSGNTLY | 0.5830841 |
| QQSSDGKNGVRAAEITGE | ASSNGTDSGNTLY | 0.22867848 |
| RAPDAVTTSWWKE | ASSNGTDSGNTLY | 0.6124181 |
| SGFGNVGAHTSGWFNQ | ASSNGTDSGNTLY | 0.22424017 |
| SGVAGAVGQAASVGGLKV | ASSNGTDSGNTLY | 0.6309353 |
| SNGGAGTGSAGGNGGT | ASSNGTDSGNTLY | 0.73524714 |
| TGGAGGEGGAAG | ASSNGTDSGNTLY | 0.81412977 |
| TGGDGGAAGAGGAPGHG | ASSNGTDSGNTLY | 0.37893993 |
| TMPPLLAAIANAGRHGV | ASSNGTDSGNTLY | 0.694641 |
| TSSTAGNSGGA | ASSNGTDSGNTLY | 0.41297215 |
| VPATVPVGILC | ASSNGTDSGNTLY | 0.83540297 |
| VQRPDSGGVVLTGRIS | ASSNGTDSGNTLY | 0.53659433 |
| WTPNPVPATKGVPAGLVQAT | ASSNGTDSGNTLY | 0.22977012 |
| YALKRALAKFSDRFDVVII | ASSNGTDSGNTLY | 0.7280615 |
| YLIGSVIV | ASSNGTDSGNTLY | 0.6384767 |
| YGQLVAVID | ASSNGTDSGNTLY | 0.5185315 |
| VTVGGMTLPA | ASSNGTDSGNTLY | 0.77985835 |
| VLALPAV | ASSNGTDSGNTLY | 0.36893383 |
| VIPPTLNYET | ASSNGTDSGNTLY | 0.52352697 |
| VGMGGMHPSSG | ASSNGTDSGNTLY | 0.75911593 |
| VGLAGGAATA | ASSNGTDSGNTLY | 0.7394113 |
| VGGAGGAGGSGP | ASSNGTDSGNTLY | 0.33913493 |
| VFAAFMV | ASSNGTDSGNTLY | 0.72865677 |
| TLYDKLR | ASSNGTDSGNTLY | 0.631414 |
| TLAVFTVL | ASSNGTDSGNTLY | 0.34875864 |
| TGSVQPARV | ASSNGTDSGNTLY | 0.8200749 |
| TGGGAGSGGT | ASSNGTDSGNTLY | 0.43621486 |
| SHLETSL | ASSNGTDSGNTLY | 0.756721 |
| SFTTGMT | ASSNGTDSGNTLY | 0.6637437 |

| | | |
|---|---|---|
| QVEVAKRMGVRQPTVS | ASSNGTDSGNTLY | 0.80554104 |
| QGSGGPALAM | ASSNGTDSGNTLY | 0.357667 |
| MLAPSNP | ASSNGTDSGNTLY | 0.68948144 |
| LVNNAAI | ASSNGTDSGNTLY | 0.6846875 |
| LTGEINTLA | ASSNGTDSGNTLY | 0.63279563 |
| LSIFGPV | ASSNGTDSGNTLY | 0.45999867 |
| LLALLANL | ASSNGTDSGNTLY | 0.69824696 |
| LISNASDALDKLR | ASSNGTDSGNTLY | 0.6471368 |
| LINEAVP | ASSNGTDSGNTLY | 0.78717744 |
| LGYAFSGL | ASSNGTDSGNTLY | 0.5529791 |
| LDFVAVAL | ASSNGTDSGNTLY | 0.6514022 |
| LAQPGAE | ASSNGTDSGNTLY | 0.80649203 |
| ISNLGAVTGF | ASSNGTDSGNTLY | 0.34470946 |
| HQEDPGRI | ASSNGTDSGNTLY | 0.5607479 |
| GTPGEPTP | ASSNGTDSGNTLY | 0.6924448 |
| GFFNSGS | ASSNGTDSGNTLY | 0.77090967 |
| GAGGGGLTAGAN | ASSNGTDSGNTLY | 0.42573196 |
| FVDALVNM | ASSNGTDSGNTLY | 0.7994647 |
| FLTVAPEL | ASSNGTDSGNTLY | 0.58163786 |
| FETLTAL | ASSNGTDSGNTLY | 0.76963097 |
| DVINPIEL | ASSNGTDSGNTLY | 0.70352924 |
| APVSGAPV | ASSNGTDSGNTLY | 0.54638135 |
| ALVGGLAW | ASSNGTDSGNTLY | 0.4555462 |
| ALAAAAATAI | ASSNGTDSGNTLY | 0.7809753 |
| AFGLGLLL | ASSNGTDSGNTLY | 0.77664596 |
| AFFAAASL | ASSNGTDSGNTLY | 0.75142723 |
| AAAALPAFSPP | ASSNGTDSGNTLY | 0.19388878 |
| VVLAPAL | ASSNGTDSGNTLY | 0.72423035 |
| TIGHVDHGKTTL | ASSNGTDSGNTLY | 0.729921 |
| LISNASDALDK | ASSNGTDSGNTLY | 0.822301 |
| LISNASDAL | ASSNGTDSGNTLY | 0.5861746 |
| ISNASDALDK | ASSNGTDSGNTLY | 0.6156792 |
| NQAETLVYQTE | ASSNGTDSGNTLY | 0.59516156 |
| TGGTGGTAGAG | ASSNGTDSGNTLY | 0.47074723 |
| TGSTAGGAGGA | ASSNGTDSGNTLY | 0.29823515 |
| DPLAVPVAAGPV | ASSNGTDSGNTLY | 0.5501412 |
| GGTGGTPGNGGN | ASSNGTDSGNTLY | 0.62454224 |
| PAGLAGVDPATL | ASSNGTDSGNTLY | 0.44262993 |
| GNNLVVEAPHAGASV | ASSNGTDSGNTLY | 0.8018944 |
| KVTRIMSASTLEQPA | ASSNGTDSGNTLY | 0.79388547 |
| LGFALPIALPRRTVV | ASSNGTDSGNTLY | 0.7864026 |
| TRLGNLTARKLVVLV | ASSNGTDSGNTLY | 0.79143405 |
| WSQQRRLIYFPSAGP | ASSNGTDSGNTLY | 0.8103456 |
| GPNGNPDPMAAAVDIR | ASSNGTDSGNTLY | 0.6105909 |
| TQRCVSVADLRAAASR | ASSNGTDSGNTLY | 0.52217346 |
| QFVLDVAEHPVEVLTETV | ASSNGTDSGNTLY | 0.60924625 |
| GGAGGTGGAGGNGGTQAGDG | ASSNGTDSGNTLY | 0.2773808 |
| PLLATLRAIAVDVDAKPAQV | ASSNGTDSGNTLY | 0.48085135 |
| APKFRTGGYMIDFWGVGYQVAK | ASSNGTDSGNTLY | 0.50429875 |
| GAGGAGGAGGAAGGPGGTGGTG | ASSNGTDSGNTLY | 0.6144899 |
| IGALGLLATDVAPAGSVAFGVA | ASSNGTDSGNTLY | 0.15430671 |

| | | |
|---|---|---|
| AGGAGGIGGNGGAGGAGGVLMG | ASSNGTDSGNTLY | 0.7696367 |
| ADELIGLL | ASSNGTDSGNTLY | 0.7902827 |
| ADRALKA | ASSNGTDSGNTLY | 0.6830977 |
| AGAVGQVGGAGGA | ASSNGTDSGNTLY | 0.6635099 |
| AGGGGGLGGAP | ASSNGTDSGNTLY | 0.74028623 |
| AGGLLYGNGGAG | ASSNGTDSGNTLY | 0.46853992 |
| AGLESIGL | ASSNGTDSGNTLY | 0.74234176 |
| AGLKTGII | ASSNGTDSGNTLY | 0.41031277 |
| ASLAALLL | ASSNGTDSGNTLY | 0.66967607 |
| ASVINISSI | ASSNGTDSGNTLY | 0.7891402 |
| ATPPGPA | ASSNGTDSGNTLY | 0.67548645 |
| ATVASALV | ASSNGTDSGNTLY | 0.6342163 |
| ATVASLIL | ASSNGTDSGNTLY | 0.4599208 |
| AVNGPVLIRH | ASSNGTDSGNTLY | 0.2711011 |
| DGAAGAVGI | ASSNGTDSGNTLY | 0.6547148 |
| DPLLHGI | ASSNGTDSGNTLY | 0.47718397 |
| ELGAALGAL | ASSNGTDSGNTLY | 0.65510046 |
| ELGRTQAV | ASSNGTDSGNTLY | 0.3669659 |
| FVAGGYLS | ASSNGTDSGNTLY | 0.69804406 |
| GASGGAGGAGGW | ASSNGTDSGNTLY | 0.6685984 |
| GGAGAGGDGGAGGI | ASSNGTDSGNTLY | 0.6394962 |
| GTGHNGSQPGMGGNG | ASSNGTDSGNTLY | 0.39981222 |
| GYAAASASAS | ASSNGTDSGNTLY | 0.6943838 |
| HQVDVAAVQQG | ASSNGTDSGNTLY | 0.66473085 |
| IGAAIAHT | ASSNGTDSGNTLY | 0.27106145 |
| IGMIEVAQL | ASSNGTDSGNTLY | 0.7622192 |
| IRLAPEHTSAMD | ASSNGTDSGNTLY | 0.44019353 |
| IYNVSVL | ASSNGTDSGNTLY | 0.64109015 |
| KQGFLGM | ASSNGTDSGNTLY | 0.76752526 |
| LESVITQL | ASSNGTDSGNTLY | 0.60003483 |
| LGLGALVVV | ASSNGTDSGNTLY | 0.2205202 |
| LIGNGGNGGSGGI | ASSNGTDSGNTLY | 0.607559 |
| LNAINAPTL | ASSNGTDSGNTLY | 0.6557226 |
| LQLLNTGV | ASSNGTDSGNTLY | 0.5894794 |
| LQTSIEF | ASSNGTDSGNTLY | 0.17642257 |
| MSFAVSEL | ASSNGTDSGNTLY | 0.7202174 |
| NTGALGL | ASSNGTDSGNTLY | 0.34754527 |
| PPGAGGAGGAGTT | ASSNGTDSGNTLY | 0.44939232 |
| QLVGLGI | ASSNGTDSGNTLY | 0.4821047 |
| RFLEQVHQL | ASSNGTDSGNTLY | 0.75579816 |
| RGVGYVLR | ASSNGTDSGNTLY | 0.70961475 |
| SATGVFAGVI | ASSNGTDSGNTLY | 0.37501222 |
| SLPAITLP | ASSNGTDSGNTLY | 0.7521136 |
| STALAFTV | ASSNGTDSGNTLY | 0.7035996 |
| SVLTVGLGK | ASSNGTDSGNTLY | 0.78685844 |
| TALAGAGSGI | ASSNGTDSGNTLY | 0.6063503 |
| TGLPFTGI | ASSNGTDSGNTLY | 0.5465057 |
| VAEHVTV | ASSNGTDSGNTLY | 0.56240374 |
| VDIVLNSL | ASSNGTDSGNTLY | 0.40145427 |
| VLTGGGAVL | ASSNGTDSGNTLY | 0.36409867 |
| VNVGLAYGI | ASSNGTDSGNTLY | 0.26616687 |

| | | |
|---|---|---|
| VSSAAVGAV | ASSNGTDSGNTLY | 0.47695303 |
| VTSVYEAGV | ASSNGTDSGNTLY | 0.2937131 |
| WVALDRGA | ASSNGTDSGNTLY | 0.67055744 |
| YLVGSVVI | ASSNGTDSGNTLY | 0.68018603 |
| GPPGTGKTLL | ASSNGTDSGNTLY | 0.5899089 |
| GVGMTKF | ASSNGTDSGNTLY | 0.27017394 |
| IGIGGSDL | ASSNGTDSGNTLY | 0.5216256 |
| IGTLLAEL | ASSNGTDSGNTLY | 0.51627135 |
| IYTSGTT | ASSNGTDSGNTLY | 0.73825717 |
| LSFPAGV | ASSNGTDSGNTLY | 0.7051171 |
| LYTSGTT | ASSNGTDSGNTLY | 0.81965566 |
| VGAFPFGF | ASSNGTDSGNTLY | 0.77863586 |
| VGAGTVEF | ASSNGTDSGNTLY | 0.5112135 |
| VGKSTLLL | ASSNGTDSGNTLY | 0.7359053 |
| AALAHAGGGIV | ASSNGTDSGNTLY | 0.41640133 |
| AALVAVSLL | ASSNGTDSGNTLY | 0.605997 |
| AAVSRAVAAL | ASSNGTDSGNTLY | 0.58473575 |
| AGGSSGIGGF | ASSNGTDSGNTLY | 0.64045143 |
| AGLTVANANG | ASSNGTDSGNTLY | 0.49879938 |
| AGLVGAVL | ASSNGTDSGNTLY | 0.5135938 |
| ALSQAFTV | ASSNGTDSGNTLY | 0.22950476 |
| ALTFSVSL | ASSNGTDSGNTLY | 0.71360236 |
| ALTGAALVAL | ASSNGTDSGNTLY | 0.3095676 |
| ASFSIGAGVL | ASSNGTDSGNTLY | 0.24036291 |
| DGSVYKI | ASSNGTDSGNTLY | 0.44634587 |
| EQLAEIV | ASSNGTDSGNTLY | 0.8248639 |
| FAVSELTVM | ASSNGTDSGNTLY | 0.7988958 |
| GAAPLAAGL | ASSNGTDSGNTLY | 0.5088893 |
| GADGGDGGSGGAGG | ASSNGTDSGNTLY | 0.42958397 |
| GFGLDTGGAGG | ASSNGTDSGNTLY | 0.7834342 |
| GLASGGLL | ASSNGTDSGNTLY | 0.35231155 |
| GRIAGAAVLL | ASSNGTDSGNTLY | 0.5792509 |
| GSAGSLNTSF | ASSNGTDSGNTLY | 0.6787464 |
| GTGQIGI | ASSNGTDSGNTLY | 0.64097524 |
| GTLIALIL | ASSNGTDSGNTLY | 0.30274242 |
| IAATMGFVVG | ASSNGTDSGNTLY | 0.7099042 |
| IFAQALL | ASSNGTDSGNTLY | 0.5798662 |
| IGAAVALGF | ASSNGTDSGNTLY | 0.52935004 |
| IGLTPLAL | ASSNGTDSGNTLY | 0.26703608 |
| ILGSNTSTL | ASSNGTDSGNTLY | 0.7403561 |
| IVFSLTVL | ASSNGTDSGNTLY | 0.43663722 |
| LASKATGF | ASSNGTDSGNTLY | 0.34118187 |
| LASVALIL | ASSNGTDSGNTLY | 0.20994301 |
| LFAVGLTL | ASSNGTDSGNTLY | 0.4565999 |
| LFNGGIGTYI | ASSNGTDSGNTLY | 0.7747289 |
| LTLGAGII | ASSNGTDSGNTLY | 0.7972465 |
| LVAAALGDI | ASSNGTDSGNTLY | 0.56910086 |
| NTAFVQLGI | ASSNGTDSGNTLY | 0.71559894 |
| QLFSGAEL | ASSNGTDSGNTLY | 0.7859627 |
| SAATFITL | ASSNGTDSGNTLY | 0.6039548 |
| SNYGLANLGV | ASSNGTDSGNTLY | 0.25536263 |

| | | |
|---|---|---|
| SYGGAYAVM | ASSNGTDSGNTLY | 0.57205987 |
| TTGVGGLTLG | ASSNGTDSGNTLY | 0.6820091 |
| TTQTIYI | ASSNGTDSGNTLY | 0.731051 |
| VAAGAGLGAVI | ASSNGTDSGNTLY | 0.7893913 |
| VAAVVTLL | ASSNGTDSGNTLY | 0.3267684 |
| VGFLGGLI | ASSNGTDSGNTLY | 0.5678072 |
| VGVIPILL | ASSNGTDSGNTLY | 0.7082529 |
| VIFASSLI | ASSNGTDSGNTLY | 0.31129083 |
| VKGGDGGAAAG | ASSNGTDSGNTLY | 0.4042415 |
| VTKTFTNI | ASSNGTDSGNTLY | 0.3417784 |
| VVAAAILL | ASSNGTDSGNTLY | 0.33102697 |
| YAALDVEL | ASSNGTDSGNTLY | 0.7014216 |
| LASVALLL | ASSNGTDSGNTLY | 0.5810162 |
| LDAVHRV | ASSNGTDSGNTLY | 0.4104811 |
| QGLSSSIF | ASSNGTDSGNTLY | 0.46798044 |
| RGTFANIRL | ASSNGTDSGNTLY | 0.38498116 |
| RQLWWGHRI | ASSNGTDSGNTLY | 0.36865795 |
| YGPPGTGKTLL | ASSNGTDSGNTLY | 0.46308595 |
| AALVGAPANVA | ASSNGTDSGNTLY | 0.61981696 |
| AATFRRTE | ASSNGTDSGNTLY | 0.7377523 |
| AGALGGGI | ASSNGTDSGNTLY | 0.5278233 |
| AGGAGGAGGIGAG | ASSNGTDSGNTLY | 0.44990146 |
| AGSGALGL | ASSNGTDSGNTLY | 0.81413805 |
| AQIADQI | ASSNGTDSGNTLY | 0.27089113 |
| ASAVAVTYL | ASSNGTDSGNTLY | 0.3571869 |
| ASTVLGGTGG | ASSNGTDSGNTLY | 0.6342169 |
| AVGGEPAK | ASSNGTDSGNTLY | 0.43065196 |
| CEQGTFGGTC | ASSNGTDSGNTLY | 0.7156242 |
| DEGKAPAG | ASSNGTDSGNTLY | 0.68818307 |
| DTVVFPAL | ASSNGTDSGNTLY | 0.6836221 |
| FEGITMAG | ASSNGTDSGNTLY | 0.71977293 |
| FPLVVQD | ASSNGTDSGNTLY | 0.8233915 |
| GAFTTPQ | ASSNGTDSGNTLY | 0.6893773 |
| GFGGFGTAGGVGGA | ASSNGTDSGNTLY | 0.39421365 |
| GFLAQFL | ASSNGTDSGNTLY | 0.78830147 |
| GGAGGAGGNAGVA | ASSNGTDSGNTLY | 0.2743422 |
| GGVGTAGAGGAGGAGG | ASSNGTDSGNTLY | 0.7225392 |
| GNGGGGDAGNAGSGGN | ASSNGTDSGNTLY | 0.6806779 |
| GTGGAGGAGNDAGS | ASSNGTDSGNTLY | 0.4135864 |
| GTTNTGWFNS | ASSNGTDSGNTLY | 0.54465485 |
| IFMGPLFTL | ASSNGTDSGNTLY | 0.7818247 |
| ILAHAIEAI | ASSNGTDSGNTLY | 0.7729847 |
| IRSAIREAL | ASSNGTDSGNTLY | 0.309301 |
| ITFAGGG | ASSNGTDSGNTLY | 0.45705786 |
| ITGMSGHQEV | ASSNGTDSGNTLY | 0.33856022 |
| IVAGLACTL | ASSNGTDSGNTLY | 0.41733113 |
| IVGGGLAGA | ASSNGTDSGNTLY | 0.25842112 |
| IVTGVLNI | ASSNGTDSGNTLY | 0.6926768 |
| LAQQGCI | ASSNGTDSGNTLY | 0.5626699 |
| LCLHVEN | ASSNGTDSGNTLY | 0.7847797 |
| LDAISDGS | ASSNGTDSGNTLY | 0.68535954 |

| | | |
|---|---|---|
| LGGFAAML | ASSNGTDSGNTLY | 0.776718 |
| LGGKNPMI | ASSNGTDSGNTLY | 0.47775888 |
| LGPKGVAAIL | ASSNGTDSGNTLY | 0.5911282 |
| LMVGVSI | ASSNGTDSGNTLY | 0.22634089 |
| LVGAAFVL | ASSNGTDSGNTLY | 0.4101597 |
| LVLAVAPL | ASSNGTDSGNTLY | 0.36429715 |
| MAAIAVIDAA | ASSNGTDSGNTLY | 0.51116157 |
| NNGINAELQ | ASSNGTDSGNTLY | 0.31744704 |
| RGEPLVLVHG | ASSNGTDSGNTLY | 0.770754 |
| RGGAGGAAGLI | ASSNGTDSGNTLY | 0.49074292 |
| RSYRMHPAV | ASSNGTDSGNTLY | 0.7148959 |
| SAAHLLAGQLA | ASSNGTDSGNTLY | 0.4465225 |
| SAAVGSY | ASSNGTDSGNTLY | 0.37245125 |
| SPHAVLHRI | ASSNGTDSGNTLY | 0.50718343 |
| SQIAKVL | ASSNGTDSGNTLY | 0.7443744 |
| TALGHLVP | ASSNGTDSGNTLY | 0.36591333 |
| TAPTTSGAA | ASSNGTDSGNTLY | 0.18007594 |
| TDGDDGNGGSPGT | ASSNGTDSGNTLY | 0.35755685 |
| TDGLGEN | ASSNGTDSGNTLY | 0.29459813 |
| TGGPPLPGQA | ASSNGTDSGNTLY | 0.81257594 |
| TIPAITI | ASSNGTDSGNTLY | 0.7772081 |
| TIPGFTI | ASSNGTDSGNTLY | 0.6251632 |
| TSIASGALA | ASSNGTDSGNTLY | 0.7966667 |
| TVTLPGAAI | ASSNGTDSGNTLY | 0.17021553 |
| VASGLSAALP | ASSNGTDSGNTLY | 0.5038767 |
| VGAGVIGI | ASSNGTDSGNTLY | 0.7776705 |
| VGPKRFGV | ASSNGTDSGNTLY | 0.6970632 |
| VNVAFPD | ASSNGTDSGNTLY | 0.6108277 |
| VSMTNSGVS | ASSNGTDSGNTLY | 0.7634927 |
| VTDVLGDLSV | ASSNGTDSGNTLY | 0.44974405 |
| VVVVGAGFAGL | ASSNGTDSGNTLY | 0.57528764 |
| VYTVGTVV | ASSNGTDSGNTLY | 0.5879595 |
| WTTFSGP | ASSNGTDSGNTLY | 0.6801491 |
| YDELVEAF | ASSNGTDSGNTLY | 0.6386697 |
| YGTGGAGGNGG | ASSNGTDSGNTLY | 0.2946418 |
| YTNVMAV | ASSNGTDSGNTLY | 0.5392394 |
| FLEQVHQL | ASSNGTDSGNTLY | 0.8373759 |
| GLSLLAVAV | ASSNGTDSGNTLY | 0.5636301 |
| LEQVHQL | ASSNGTDSGNTLY | 0.7293209 |
| LNAAGAI | ASSNGTDSGNTLY | 0.59999174 |
| PAVHASL | ASSNGTDSGNTLY | 0.6918588 |
| TELEDDA | ASSNGTDSGNTLY | 0.42718476 |
| TPPGPPN | ASSNGTDSGNTLY | 0.42311704 |
| TPPPPPPP | ASSNGTDSGNTLY | 0.7025416 |
| TQSPALI | AWSWTGAETLY | 0.40536466 |
| EFAAAFPQ | AWSWTGAETLY | 0.49501163 |
| LLSGNLDVL | AWSWTGAETLY | 0.40292257 |
| GPANLGNYNI | AWSWTGAETLY | 0.8363496 |
| TLGPTLP | AWSWTGAETLY | 0.71410996 |
| VYAYSSCCK | AWSWTGAETLY | 0.6767137 |
| APAPAPAPNI | AWSWTGAETLY | 0.38038647 |

| | | |
|---|---|---|
| PGGPVPIP | AWSWTGAETLY | 0.592904 |
| FLTVALEP | AWSWTGAETLY | 0.7529872 |
| SQYFGANL | AWSWTGAETLY | 0.24369463 |
| GGVKPAA | AWSWTGAETLY | 0.79339147 |
| GGTGGAGGAGGW | AWSWTGAETLY | 0.35467678 |
| GVGGVAGLGGAGG | AWSWTGAETLY | 0.64412326 |
| VLLVLATTI | AWSWTGAETLY | 0.5545045 |
| IGIFAAML | AWSWTGAETLY | 0.80664897 |
| AVGDLID | AWSWTGAETLY | 0.548382 |
| NISPLPL | AWSWTGAETLY | 0.64355874 |
| VTAVFVAI | AWSWTGAETLY | 0.36626494 |
| SGDDNDAAL | AWSWTGAETLY | 0.6632904 |
| AVAALAHS | AWSWTGAETLY | 0.7958087 |
| GGVTVMATP | AWSWTGAETLY | 0.659652 |
| AGLAANVRD | AWSWTGAETLY | 0.82432157 |
| TNRPDLIDPALLRPG | AWSWTGAETLY | 0.47292042 |
| LANLGVTAL | AWSWTGAETLY | 0.22364616 |
| TGERIEL | AWSWTGAETLY | 0.5054041 |
| NGEELAA | AWSWTGAETLY | 0.7672982 |
| VLSLVWG | AWSWTGAETLY | 0.80913436 |
| IGYALWL | AWSWTGAETLY | 0.70250344 |
| LSLLLAPL | AWSWTGAETLY | 0.74985886 |
| TLDEALK | AWSWTGAETLY | 0.5901612 |
| TDGAGTAEGG | AWSWTGAETLY | 0.78248525 |
| NAPGGAGGNGG | AWSWTGAETLY | 0.8158555 |
| VSVFMGMGGNF | AWSWTGAETLY | 0.6373311 |
| LASYLAAL | AWSWTGAETLY | 0.3324579 |
| LVGGTPAL | AWSWTGAETLY | 0.3063628 |
| VRNADVL | AWSWTGAETLY | 0.68937266 |
| TCSSPFLVSAM | AWSWTGAETLY | 0.78012896 |
| VHITDAE | AWSWTGAETLY | 0.26206258 |
| EYAAPSF | AWSWTGAETLY | 0.64796793 |
| VVDAFHGGL | AWSWTGAETLY | 0.65123224 |
| ASVVDPQ | AWSWTGAETLY | 0.23073041 |
| SYNEFGGLS | AWSWTGAETLY | 0.70567167 |
| DPDGSVY | AWSWTGAETLY | 0.81299114 |
| MVSTIHATE | AWSWTGAETLY | 0.514942 |
| LAAGVADPV | AWSWTGAETLY | 0.28761137 |
| TKEQIAA | AWSWTGAETLY | 0.804242 |
| SLASPAAL | AWSWTGAETLY | 0.83154976 |
| IGLVFYGV | AWSWTGAETLY | 0.75253725 |
| FLLDASNL | AWSWTGAETLY | 0.21965988 |
| YQVGNFP | AWSWTGAETLY | 0.16546153 |
| ESLLSQST | AWSWTGAETLY | 0.68201816 |
| LFTVGFAAVG | AWSWTGAETLY | 0.7784159 |
| ATPPTPT | AWSWTGAETLY | 0.4393863 |
| TDLATLG | AWSWTGAETLY | 0.35294926 |
| SAGGTGR | AWSWTGAETLY | 0.6940384 |
| GFSNSGS | AWSWTGAETLY | 0.7427356 |
| GGNGGSGGTSV | AWSWTGAETLY | 0.31593573 |
| GTSVATGGAGN | AWSWTGAETLY | 0.26509687 |

| | | |
|---|---|---|
| NGGNGGSGGTS | AWSWTGAETLY | 0.72088087 |
| AVVKFPH | AWSWTGAETLY | 0.7009984 |
| TPPPPPP | AWSWTGAETLY | 0.41230384 |
| AAPAAAAAPA | AWSWTGAETLY | 0.5499822 |
| IALNEDL | AWSWTGAETLY | 0.70957184 |
| IVNVSSM | AWSWTGAETLY | 0.6299948 |
| TGIPPAPR | AWSWTGAETLY | 0.703857 |
| FDLGGGTF | AWSWTGAETLY | 0.2829873 |
| FDLGGGTFDVS | AWSWTGAETLY | 0.80767655 |
| NEPTAAA | AWSWTGAETLY | 0.27736038 |
| NEPTAAALA | AWSWTGAETLY | 0.7615683 |
| GSIINISSI | AWSWTGAETLY | 0.670287 |
| VAPGNAGTA | AWSWTGAETLY | 0.6501386 |
| TGGGASLE | AWSWTGAETLY | 0.58960396 |
| VAVYNHYKRI | AWSWTGAETLY | 0.3200917 |
| TDPAPATT | AWSWTGAETLY | 0.33707744 |
| TNRPDILDPALLRPG | AWSWTGAETLY | 0.27114305 |
| LTTTTTS | AWSWTGAETLY | 0.7964537 |
| PTGSGKTL | AWSWTGAETLY | 0.7422217 |
| TGSGKTL | AWSWTGAETLY | 0.7503107 |
| TIGHVDHGK | AWSWTGAETLY | 0.66242105 |
| TIGHVDHGKTT | AWSWTGAETLY | 0.39544398 |
| GHVDHGKTT | AWSWTGAETLY | 0.66246855 |
| AQFEHTLL | AWSWTGAETLY | 0.296058 |
| IVNQDEL | AWSWTGAETLY | 0.5354522 |
| TNGSQFF | AWSWTGAETLY | 0.5953034 |
| TGKGIVE | AWSWTGAETLY | 0.56362367 |
| LISNASDA | AWSWTGAETLY | 0.43324143 |
| AELRTLGELTKIAKAHGAQ | AWSWTGAETLY | 0.58958423 |
| APNAVAAAEPSW | AWSWTGAETLY | 0.6837022 |
| APQVAGAATAVL | AWSWTGAETLY | 0.36806667 |
| ARRDPAAAAAILPTDA | AWSWTGAETLY | 0.48741224 |
| DAAARDVLAEAGLAEYF | AWSWTGAETLY | 0.78742653 |
| DDQIVLSAKANLAKVEQ | AWSWTGAETLY | 0.52381563 |
| DELLPDTGEDSDEDGH | AWSWTGAETLY | 0.33616498 |
| ELPEAPANRATVPQD | AWSWTGAETLY | 0.72016436 |
| FGQSKIERVTGIG | AWSWTGAETLY | 0.40914553 |
| GAGGTGSTAGGAGGAGGAGGLY | AWSWTGAETLY | 0.70465136 |
| GHGGTGAAGGNGGTGGAGGSGI | AWSWTGAETLY | 0.63454187 |
| GNGGNGGAGGEGGPGGAGGAGA | AWSWTGAETLY | 0.4144534 |
| GSTAGNGGNGGNATNGTIAGQP | AWSWTGAETLY | 0.7230032 |
| GTGGDGGNGGNGGTGG | AWSWTGAETLY | 0.26536074 |
| GVRRPAGIPADAPV | AWSWTGAETLY | 0.52991617 |
| GYRQVTDAHLLTLARR | AWSWTGAETLY | 0.52522165 |
| IAKMGQDGHDRG | AWSWTGAETLY | 0.62800515 |
| KAEPSIEIQYT | AWSWTGAETLY | 0.6802808 |
| KPVRIRMAADEDTVAVT | AWSWTGAETLY | 0.36649764 |
| MPPRIAGMRLLVIKPEP | AWSWTGAETLY | 0.65563893 |
| MKFYAVALFRSRGVPPTR | AWSWTGAETLY | 0.5015547 |
| NAGGFGILGAVAHSP | AWSWTGAETLY | 0.37013257 |
| NEGREITLAMAEM | AWSWTGAETLY | 0.39967778 |

| | | |
|---|---|---|
| NELVEWSHPLGHRWRWE | AWSWTGAETLY | 0.45276448 |
| NLGAGGTGGNGGNPG | AWSWTGAETLY | 0.28440034 |
| NTEWPKWETVMAADPVD | AWSWTGAETLY | 0.69428986 |
| PASAAPAETQP | AWSWTGAETLY | 0.3984539 |
| QQSSDGKNGVRAAEITGE | AWSWTGAETLY | 0.6628232 |
| RAPDAVTTSWWKE | AWSWTGAETLY | 0.78778356 |
| SGFGNVGAHTSGWFNQ | AWSWTGAETLY | 0.33046985 |
| SGVAGAVGQAASVGGLKV | AWSWTGAETLY | 0.6886726 |
| SNGGAGTGSAGGNGGT | AWSWTGAETLY | 0.6231785 |
| TGGAGGEGGAAG | AWSWTGAETLY | 0.72298455 |
| TGGDGGAAGAGGAPGHG | AWSWTGAETLY | 0.5452473 |
| TMPPLLAAIANAGRHGV | AWSWTGAETLY | 0.43698388 |
| TSSTAGNSGGA | AWSWTGAETLY | 0.7822156 |
| VPATVPVGILC | AWSWTGAETLY | 0.7025057 |
| VQRPDSGGVVLTGRIS | AWSWTGAETLY | 0.61535525 |
| WTPNPVPATKGVPAGLVQAT | AWSWTGAETLY | 0.6812358 |
| YALKRALAKFSDRFDVVII | AWSWTGAETLY | 0.768605 |
| YLIGSVIV | AWSWTGAETLY | 0.69311786 |
| YGQLVAVID | AWSWTGAETLY | 0.4894159 |
| VTVGGMTLPA | AWSWTGAETLY | 0.22191657 |
| VLALPAV | AWSWTGAETLY | 0.55936193 |
| VIPPTLNYET | AWSWTGAETLY | 0.56770414 |
| VGMGGMHPSSG | AWSWTGAETLY | 0.3448951 |
| VGLAGGAATA | AWSWTGAETLY | 0.2032991 |
| VGGAGGAGGSGP | AWSWTGAETLY | 0.742321 |
| VFAAFMV | AWSWTGAETLY | 0.8308498 |
| TLYDKLR | AWSWTGAETLY | 0.39832833 |
| TLAVFTVL | AWSWTGAETLY | 0.8202214 |
| TGSVQPARV | AWSWTGAETLY | 0.25053364 |
| TGGGAGSGGT | AWSWTGAETLY | 0.7376505 |
| SHLETSL | AWSWTGAETLY | 0.75506234 |
| SFTTGMT | AWSWTGAETLY | 0.35373986 |
| QVEVAKRMGVRQPTVS | AWSWTGAETLY | 0.616982 |
| QGSGGPALAM | AWSWTGAETLY | 0.57731205 |
| MLAPSNP | AWSWTGAETLY | 0.4636891 |
| LVNNAAI | AWSWTGAETLY | 0.78901005 |
| LTGEINTLA | AWSWTGAETLY | 0.42770547 |
| LSIFGPV | AWSWTGAETLY | 0.64121634 |
| LLALLANL | AWSWTGAETLY | 0.40419745 |
| LISNASDALDKLR | AWSWTGAETLY | 0.38121605 |
| LINEAVP | AWSWTGAETLY | 0.6288263 |
| LGYAFSGL | AWSWTGAETLY | 0.673324 |
| LDFVAVAL | AWSWTGAETLY | 0.5130653 |
| LAQPGAE | AWSWTGAETLY | 0.7918829 |
| ISNLGAVTGF | AWSWTGAETLY | 0.5275167 |
| HQEDPGRI | AWSWTGAETLY | 0.825117 |
| GTPGEPTP | AWSWTGAETLY | 0.8040471 |
| GFFNSGS | AWSWTGAETLY | 0.6641098 |
| GAGGGGLTAGAN | AWSWTGAETLY | 0.44117048 |
| FVDALVNM | AWSWTGAETLY | 0.7963234 |
| FLTVAPEL | AWSWTGAETLY | 0.43563104 |

| | | |
|---|---|---|
| FETLTAL | AWSWTGAETLY | 0.71727735 |
| DVINPIEL | AWSWTGAETLY | 0.71748984 |
| APVSGAPV | AWSWTGAETLY | 0.48484206 |
| ALVGGLAW | AWSWTGAETLY | 0.697034 |
| ALAAAAATAI | AWSWTGAETLY | 0.6300505 |
| AFGLGLLL | AWSWTGAETLY | 0.62032527 |
| AFFAAASL | AWSWTGAETLY | 0.5291331 |
| AAAALPAFSPP | AWSWTGAETLY | 0.42279202 |
| VVLAPAL | AWSWTGAETLY | 0.5987255 |
| TIGHVDHGKTTL | AWSWTGAETLY | 0.47058827 |
| LISNASDALDK | AWSWTGAETLY | 0.67840433 |
| LISNASDAL | AWSWTGAETLY | 0.67631495 |
| ISNASDALDK | AWSWTGAETLY | 0.5041626 |
| NQAETLVYQTE | AWSWTGAETLY | 0.69232005 |
| TGGTGGTAGAG | AWSWTGAETLY | 0.6520194 |
| TGSTAGGAGGA | AWSWTGAETLY | 0.42464447 |
| DPLAVPVAAGPV | AWSWTGAETLY | 0.59157085 |
| GGTGGTPGNGGN | AWSWTGAETLY | 0.6120904 |
| PAGLAGVDPATL | AWSWTGAETLY | 0.60333896 |
| GNNLVVEAPHAGASV | AWSWTGAETLY | 0.4862863 |
| KVTRIMSASTLEQPA | AWSWTGAETLY | 0.6606601 |
| LGFALPIALPRRTVV | AWSWTGAETLY | 0.5121349 |
| TRLGNLTARKLVVLV | AWSWTGAETLY | 0.3293084 |
| WSQQRRLIYFPSAGP | AWSWTGAETLY | 0.6529269 |
| GPNGNPDPMAAAVDIR | AWSWTGAETLY | 0.5880953 |
| TQRCVSVADLRAAASR | AWSWTGAETLY | 0.5497315 |
| QFVLDVAEHPVEVLTETV | AWSWTGAETLY | 0.6736047 |
| GGAGGTGGAGGNGGTQAGDG | AWSWTGAETLY | 0.5679313 |
| PLLATLRAIAVDVDAKPAQV | AWSWTGAETLY | 0.4955693 |
| APKFRTGGYMIDFWGVGYQVAK | AWSWTGAETLY | 0.67675483 |
| GAGGAGGAGGAAGGPGGTGGTG | AWSWTGAETLY | 0.6838856 |
| IGALGLLATDVAPAGSVAFGVA | AWSWTGAETLY | 0.4301716 |
| AGGAGGIGGNGGAGGAGGVLMG | AWSWTGAETLY | 0.62611973 |
| ADELIGLL | AWSWTGAETLY | 0.6837541 |
| ADRALKA | AWSWTGAETLY | 0.6054859 |
| AGAVGQVGGAGGA | AWSWTGAETLY | 0.6916714 |
| AGGGGGLGGAP | AWSWTGAETLY | 0.6950954 |
| AGGLLYGNGGAG | AWSWTGAETLY | 0.5900919 |
| AGLESIGL | AWSWTGAETLY | 0.68853354 |
| AGLKTGII | AWSWTGAETLY | 0.69899786 |
| ASLAALLL | AWSWTGAETLY | 0.5830802 |
| ASVINISSI | AWSWTGAETLY | 0.6282137 |
| ATPPGPA | AWSWTGAETLY | 0.4244878 |
| ATVASALV | AWSWTGAETLY | 0.63036615 |
| ATVASLIL | AWSWTGAETLY | 0.40455776 |
| AVNGPVLIRH | AWSWTGAETLY | 0.6453831 |
| DGAAGAVGI | AWSWTGAETLY | 0.6875431 |
| DPLLHGI | AWSWTGAETLY | 0.57754004 |
| ELGAALGAL | AWSWTGAETLY | 0.4686614 |
| ELGRTQAV | AWSWTGAETLY | 0.4754792 |
| FVAGGYLS | AWSWTGAETLY | 0.63299453 |

| | | |
|---|---|---|
| GASGGAGGAGGW | AWSWTGAETLY | 0.4637444 |
| GGAGAGGDGGAGGI | AWSWTGAETLY | 0.40290162 |
| GTGHNGSQPGMGGNG | AWSWTGAETLY | 0.6091789 |
| GYAAASASAS | AWSWTGAETLY | 0.6688836 |
| HQVDVAAVQQG | AWSWTGAETLY | 0.66470283 |
| IGAAIAHT | AWSWTGAETLY | 0.4816338 |
| IGMIEVAQL | AWSWTGAETLY | 0.6945267 |
| IRLAPEHTSAMD | AWSWTGAETLY | 0.6705706 |
| IYNVSVL | AWSWTGAETLY | 0.6876074 |
| KQGFLGM | AWSWTGAETLY | 0.590999 |
| LESVITQL | AWSWTGAETLY | 0.46531758 |
| LGLGALVVV | AWSWTGAETLY | 0.6735043 |
| LIGNGGNGGSGGI | AWSWTGAETLY | 0.57287484 |
| LNAINAPTL | AWSWTGAETLY | 0.655079 |
| LQLLNTGV | AWSWTGAETLY | 0.6819221 |
| LQTSIEF | AWSWTGAETLY | 0.5612793 |
| MSFAVSEL | AWSWTGAETLY | 0.51708364 |
| NTGALGL | AWSWTGAETLY | 0.59017915 |
| PPGAGGAGGAGTT | AWSWTGAETLY | 0.6813767 |
| QLVGLGI | AWSWTGAETLY | 0.5290493 |
| RFLEQVHQL | AWSWTGAETLY | 0.4319735 |
| RGVGYVLR | AWSWTGAETLY | 0.5787655 |
| SATGVFAGVI | AWSWTGAETLY | 0.60227454 |
| SLPAITLP | AWSWTGAETLY | 0.68465793 |
| STALAFTV | AWSWTGAETLY | 0.648042 |
| SVLTVGLGK | AWSWTGAETLY | 0.6130597 |
| TALAGAGSGI | AWSWTGAETLY | 0.65681744 |
| TGLPFTGI | AWSWTGAETLY | 0.6472414 |
| VAEHVTV | AWSWTGAETLY | 0.63636214 |
| VDIVLNSL | AWSWTGAETLY | 0.6439622 |
| VLTGGGAVL | AWSWTGAETLY | 0.54834044 |
| VNVGLAYGI | AWSWTGAETLY | 0.68072325 |
| VSSAAVGAV | AWSWTGAETLY | 0.66122055 |
| VTSVYEAGV | AWSWTGAETLY | 0.6639383 |
| WVALDRGA | AWSWTGAETLY | 0.69031227 |
| YLVGSVVI | AWSWTGAETLY | 0.63460106 |
| GPPGTGKTLL | AWSWTGAETLY | 0.6372194 |
| GVGMTKF | AWSWTGAETLY | 0.40364727 |
| IGIGGSDL | AWSWTGAETLY | 0.56409264 |
| IGTLLAEL | AWSWTGAETLY | 0.64560586 |
| IYTSGTT | AWSWTGAETLY | 0.63859093 |
| LSFPAGV | AWSWTGAETLY | 0.67421454 |
| LYTSGTT | AWSWTGAETLY | 0.65160054 |
| VGAFPFGF | AWSWTGAETLY | 0.5472081 |
| VGAGTVEF | AWSWTGAETLY | 0.68266857 |
| VGKSTLLL | AWSWTGAETLY | 0.68718064 |
| AALAHAGGGIV | AWSWTGAETLY | 0.42805755 |
| AALVAVSLL | AWSWTGAETLY | 0.579072 |
| AAVSRAVAAL | AWSWTGAETLY | 0.7044511 |
| AGGSSGIGGF | AWSWTGAETLY | 0.63746625 |
| AGLTVANANG | AWSWTGAETLY | 0.5878389 |

| | | |
|---|---|---|
| AGLVGAVL | AWSWTGAETLY | 0.5683097 |
| ALSQAFTV | AWSWTGAETLY | 0.6900573 |
| ALTFSVSL | AWSWTGAETLY | 0.6263644 |
| ALTGAALVAL | AWSWTGAETLY | 0.7031554 |
| ASFSIGAGVL | AWSWTGAETLY | 0.3572949 |
| DGSVYKI | AWSWTGAETLY | 0.5446203 |
| EQLAEIV | AWSWTGAETLY | 0.52521 |
| FAVSELTVM | AWSWTGAETLY | 0.316807 |
| GAAPLAAGL | AWSWTGAETLY | 0.67453945 |
| GADGGDGGSGGAGG | AWSWTGAETLY | 0.5175634 |
| GFGLDTGGAGG | AWSWTGAETLY | 0.657902 |
| GLASGGLL | AWSWTGAETLY | 0.6153307 |
| GRIAGAAVLL | AWSWTGAETLY | 0.68219197 |
| GSAGSLNTSF | AWSWTGAETLY | 0.64966524 |
| GTGQIGI | AWSWTGAETLY | 0.6130689 |
| GTLIALIL | AWSWTGAETLY | 0.5286299 |
| IAATMGFVVG | AWSWTGAETLY | 0.6513557 |
| IFAQALL | AWSWTGAETLY | 0.6924809 |
| IGAAVALGF | AWSWTGAETLY | 0.67302096 |
| IGLTPLAL | AWSWTGAETLY | 0.39507934 |
| ILGSNTSTL | AWSWTGAETLY | 0.65106493 |
| IVFSLTVL | AWSWTGAETLY | 0.59232146 |
| LASKATGF | AWSWTGAETLY | 0.37530664 |
| LASVALIL | AWSWTGAETLY | 0.37401465 |
| LFAVGLTL | AWSWTGAETLY | 0.5481297 |
| LFNGGIGTYI | AWSWTGAETLY | 0.56981 |
| LTLGAGII | AWSWTGAETLY | 0.66978705 |
| LVAAALGDI | AWSWTGAETLY | 0.67584765 |
| NTAFVQLGI | AWSWTGAETLY | 0.5593989 |
| QLFSGAEL | AWSWTGAETLY | 0.6680298 |
| SAATFITL | AWSWTGAETLY | 0.6215259 |
| SNYGLANLGV | AWSWTGAETLY | 0.2604559 |
| SYGGAYAVM | AWSWTGAETLY | 0.5617446 |
| TTGVGGLTLG | AWSWTGAETLY | 0.44586414 |
| TTQTIYI | AWSWTGAETLY | 0.6999706 |
| VAAGAGLGAVI | AWSWTGAETLY | 0.527288 |
| VAAVVTLL | AWSWTGAETLY | 0.58420604 |
| VGFLGGLI | AWSWTGAETLY | 0.5966593 |
| VGVIPILL | AWSWTGAETLY | 0.6313518 |
| VIFASSLI | AWSWTGAETLY | 0.61920154 |
| VKGGDGGAAAG | AWSWTGAETLY | 0.49016416 |
| VTKTFTNI | AWSWTGAETLY | 0.37180898 |
| VVAAAILL | AWSWTGAETLY | 0.6578256 |
| YAALDVEL | AWSWTGAETLY | 0.56447566 |
| LASVALLL | AWSWTGAETLY | 0.4634173 |
| LDAVHRV | AWSWTGAETLY | 0.4851684 |
| QGLSSSIF | AWSWTGAETLY | 0.6533158 |
| RGTFANIRL | AWSWTGAETLY | 0.38286266 |
| RQLWWGHRI | AWSWTGAETLY | 0.6777544 |
| YGPPGTGKTLL | AWSWTGAETLY | 0.5519162 |
| AALVGAPANVA | AWSWTGAETLY | 0.605633 |

| | | |
|---|---|---|
| AATFRRTE | AWSWTGAETLY | 0.70425105 |
| AGALGGGI | AWSWTGAETLY | 0.6297113 |
| AGGAGGAGGIGAG | AWSWTGAETLY | 0.5991647 |
| AGSGALGL | AWSWTGAETLY | 0.5099691 |
| AQIADQI | AWSWTGAETLY | 0.48993725 |
| ASAVAVTYL | AWSWTGAETLY | 0.48580337 |
| ASTVLGGTGG | AWSWTGAETLY | 0.51015323 |
| AVGGEPAK | AWSWTGAETLY | 0.6996778 |
| CEQGTFGGTC | AWSWTGAETLY | 0.61002314 |
| DEGKAPAG | AWSWTGAETLY | 0.46809855 |
| DTVVFPAL | AWSWTGAETLY | 0.6362887 |
| FEGITMAG | AWSWTGAETLY | 0.7009022 |
| FPLVVQD | AWSWTGAETLY | 0.70246804 |
| GAFTTPQ | AWSWTGAETLY | 0.5315138 |
| GFGGFGTAGGVGGA | AWSWTGAETLY | 0.58954024 |
| GFLAQFL | AWSWTGAETLY | 0.5515182 |
| GGAGGAGGNAGVA | AWSWTGAETLY | 0.6132714 |
| GGVGTAGAGGAGGAGG | AWSWTGAETLY | 0.52318275 |
| GNGGGGDAGNAGSGGN | AWSWTGAETLY | 0.4526557 |
| GTGGAGGAGNDAGS | AWSWTGAETLY | 0.6612312 |
| GTTNTGWFNS | AWSWTGAETLY | 0.70214546 |
| IFMGPLFTL | AWSWTGAETLY | 0.58748436 |
| ILAHAIEAI | AWSWTGAETLY | 0.47188985 |
| IRSAIREAL | AWSWTGAETLY | 0.6043287 |
| ITFAGGG | AWSWTGAETLY | 0.48100343 |
| ITGMSGHQEV | AWSWTGAETLY | 0.52059513 |
| IVAGLACTL | AWSWTGAETLY | 0.53448933 |
| IVGGGLAGA | AWSWTGAETLY | 0.39667836 |
| IVTGVLNI | AWSWTGAETLY | 0.60469466 |
| LAQQGCI | AWSWTGAETLY | 0.69757587 |
| LCLHVEN | AWSWTGAETLY | 0.6025879 |
| LDAISDGS | AWSWTGAETLY | 0.64031506 |
| LGGFAAML | AWSWTGAETLY | 0.6899537 |
| LGGKNPMI | AWSWTGAETLY | 0.6576301 |
| LGPKGVAAIL | AWSWTGAETLY | 0.678707 |
| LMVGVSI | AWSWTGAETLY | 0.6761279 |
| LVGAAFVL | AWSWTGAETLY | 0.63631916 |
| LVLAVAPL | AWSWTGAETLY | 0.6344891 |
| MAAIAVIDAA | AWSWTGAETLY | 0.6770797 |
| NNGINAELQ | AWSWTGAETLY | 0.6116027 |
| RGEPLVLVHG | AWSWTGAETLY | 0.6032653 |
| RGGAGGAAGLI | AWSWTGAETLY | 0.58425796 |
| RSYRMHPAV | AWSWTGAETLY | 0.6755033 |
| SAAHLLAGQLA | AWSWTGAETLY | 0.5630621 |
| SAAVGSY | AWSWTGAETLY | 0.6821051 |
| SPHAVLHRI | AWSWTGAETLY | 0.34431127 |
| SQIAKVL | AWSWTGAETLY | 0.4248408 |
| TALGHLVP | AWSWTGAETLY | 0.67271376 |
| TAPTTSGAA | AWSWTGAETLY | 0.28606611 |
| TDGDDGNGGSPGT | AWSWTGAETLY | 0.3712954 |
| TDGLGEN | AWSWTGAETLY | 0.60353106 |

| | | |
|---|---|---|
| TGGPPLPGQA | AWSWTGAETLY | 0.6828756 |
| TIPAITI | AWSWTGAETLY | 0.290231 |
| TIPGFTI | AWSWTGAETLY | 0.5141243 |
| TSIASGALA | AWSWTGAETLY | 0.39772147 |
| TVTLPGAAI | AWSWTGAETLY | 0.67390215 |
| VASGLSAALP | AWSWTGAETLY | 0.5992254 |
| VGAGVIGI | AWSWTGAETLY | 0.39634684 |
| VGPKRFGV | AWSWTGAETLY | 0.6621568 |
| VNVAFPD | AWSWTGAETLY | 0.47412413 |
| VSMTNSGVS | AWSWTGAETLY | 0.62338734 |
| VTDVLGDLSV | AWSWTGAETLY | 0.6797831 |
| VVVVGAGFAGL | AWSWTGAETLY | 0.6925454 |
| VYTVGTVV | AWSWTGAETLY | 0.4687233 |
| WTTFSGP | AWSWTGAETLY | 0.6516699 |
| YDELVEAF | AWSWTGAETLY | 0.55905974 |
| YGTGGAGGNGG | AWSWTGAETLY | 0.70835537 |
| YTNVMAV | AWSWTGAETLY | 0.57688636 |
| FLEQVHQL | AWSWTGAETLY | 0.40055606 |
| GLSLLAVAV | AWSWTGAETLY | 0.67086613 |
| LEQVHQL | AWSWTGAETLY | 0.65140235 |
| LNAAGAI | AWSWTGAETLY | 0.573822 |
| PAVHASL | AWSWTGAETLY | 0.66083986 |
| TELEDDA | AWSWTGAETLY | 0.54511297 |
| TPPGPPN | AWSWTGAETLY | 0.58975154 |
| TPPPPPPP | AWSWTGAETLY | 0.680762 |
| TQSPALI | ASSSDWGSQNTLY | 0.6761992 |
| EFAAAFPQ | ASSSDWGSQNTLY | 0.4559423 |
| LLSGNLDVL | ASSSDWGSQNTLY | 0.6622729 |
| GPANLGNYNI | ASSSDWGSQNTLY | 0.5931348 |
| TLGPTLP | ASSSDWGSQNTLY | 0.512312 |
| VYAYSSCCK | ASSSDWGSQNTLY | 0.6962787 |
| APAPAPAPNI | ASSSDWGSQNTLY | 0.55155414 |
| PGGPVPIP | ASSSDWGSQNTLY | 0.6666324 |
| FLTVALEP | ASSSDWGSQNTLY | 0.6517005 |
| SQYFGANL | ASSSDWGSQNTLY | 0.6809372 |
| GGVKPAA | ASSSDWGSQNTLY | 0.4654438 |
| GGTGGAGGAGGW | ASSSDWGSQNTLY | 0.6287691 |
| GVGGVAGLGGAGG | ASSSDWGSQNTLY | 0.61556536 |
| VLLVLATTI | ASSSDWGSQNTLY | 0.6809274 |
| IGIFAAML | ASSSDWGSQNTLY | 0.6096524 |
| AVGDLID | ASSSDWGSQNTLY | 0.63334924 |
| NISPLPL | ASSSDWGSQNTLY | 0.6674665 |
| VTAVFVAI | ASSSDWGSQNTLY | 0.69142705 |
| SGDDNDAAL | ASSSDWGSQNTLY | 0.6433338 |
| AVAALAHS | ASSSDWGSQNTLY | 0.64501226 |
| GGVTVMATP | ASSSDWGSQNTLY | 0.6871809 |
| AGLAANVRD | ASSSDWGSQNTLY | 0.46656695 |
| TNRPDLIDPALLRPG | ASSSDWGSQNTLY | 0.5884524 |
| LANLGVTAL | ASSSDWGSQNTLY | 0.6735072 |
| TGERIEL | ASSSDWGSQNTLY | 0.6885725 |
| NGEELAA | ASSSDWGSQNTLY | 0.5943706 |

| | | |
|---|---|---|
| VLSLVWG | ASSSDWGSQNTLY | 0.6923791 |
| IGYALWL | ASSSDWGSQNTLY | 0.56000316 |
| LSLLLAPL | ASSSDWGSQNTLY | 0.67180574 |
| TLDEALK | ASSSDWGSQNTLY | 0.5994217 |
| TDGAGTAEGG | ASSSDWGSQNTLY | 0.6306117 |
| NAPGGAGGNGG | ASSSDWGSQNTLY | 0.47688884 |
| VSVFMGMGGNF | ASSSDWGSQNTLY | 0.67795837 |
| LASYLAAL | ASSSDWGSQNTLY | 0.67403895 |
| LVGGTPAL | ASSSDWGSQNTLY | 0.6735958 |
| VRNADVL | ASSSDWGSQNTLY | 0.37977248 |
| TCSSPFLVSAM | ASSSDWGSQNTLY | 0.6491991 |
| VHITDAE | ASSSDWGSQNTLY | 0.67588574 |
| EYAAPSF | ASSSDWGSQNTLY | 0.69500643 |
| VVDAFHGGL | ASSSDWGSQNTLY | 0.5966736 |
| ASVVDPQ | ASSSDWGSQNTLY | 0.6398135 |
| SYNEFGGLS | ASSSDWGSQNTLY | 0.5848457 |
| DPDGSVY | ASSSDWGSQNTLY | 0.5837705 |
| MVSTIHATE | ASSSDWGSQNTLY | 0.45645443 |
| LAAGVADPV | ASSSDWGSQNTLY | 0.55938965 |
| TKEQIAA | ASSSDWGSQNTLY | 0.60164654 |
| SLASPAAL | ASSSDWGSQNTLY | 0.5148194 |
| IGLVFYGV | ASSSDWGSQNTLY | 0.6675411 |
| FLLDASNL | ASSSDWGSQNTLY | 0.6908508 |
| YQVGNFP | ASSSDWGSQNTLY | 0.6804501 |
| ESLLSQST | ASSSDWGSQNTLY | 0.6846652 |
| LFTVGFAAVG | ASSSDWGSQNTLY | 0.6930277 |
| ATPPTPT | ASSSDWGSQNTLY | 0.663894 |
| TDLATLG | ASSSDWGSQNTLY | 0.5258972 |
| SAGGTGR | ASSSDWGSQNTLY | 0.6276842 |
| GFSNSGS | ASSSDWGSQNTLY | 0.4017855 |
| GGNGGSGGTSV | ASSSDWGSQNTLY | 0.59212965 |
| GTSVATGGAGN | ASSSDWGSQNTLY | 0.5313034 |
| NGGNGGSGGTS | ASSSDWGSQNTLY | 0.6299969 |
| AVVKFPH | ASSSDWGSQNTLY | 0.27409944 |
| TPPPPPP | ASSSDWGSQNTLY | 0.63176227 |
| AAPAAAAAPA | ASSSDWGSQNTLY | 0.7017058 |
| IALNEDL | ASSSDWGSQNTLY | 0.57938725 |
| IVNVSSM | ASSSDWGSQNTLY | 0.6429453 |
| TGIPPAPR | ASSSDWGSQNTLY | 0.6587546 |
| FDLGGGTF | ASSSDWGSQNTLY | 0.5555343 |
| FDLGGGTFDVS | ASSSDWGSQNTLY | 0.65473723 |
| NEPTAAA | ASSSDWGSQNTLY | 0.5096313 |
| NEPTAAALA | ASSSDWGSQNTLY | 0.66417897 |
| GSIINISSI | ASSSDWGSQNTLY | 0.688242 |
| VAPGNAGTA | ASSSDWGSQNTLY | 0.6544384 |
| TGGGASLE | ASSSDWGSQNTLY | 0.5979204 |
| VAVYNHYKRI | ASSSDWGSQNTLY | 0.6044974 |
| TDPAPATT | ASSSDWGSQNTLY | 0.4263184 |
| TNRPDILDPALLRPG | ASSSDWGSQNTLY | 0.64468604 |
| LTTTTTS | ASSSDWGSQNTLY | 0.5235349 |
| PTGSGKTL | ASSSDWGSQNTLY | 0.60930353 |

| | | |
|---|---|---|
| TGSGKTL | ASSSDWGSQNTLY | 0.48789376 |
| TIGHVDHGK | ASSSDWGSQNTLY | 0.64394164 |
| TIGHVDHGKTT | ASSSDWGSQNTLY | 0.6308523 |
| GHVDHGKTT | ASSSDWGSQNTLY | 0.64982545 |
| AQFEHTLL | ASSSDWGSQNTLY | 0.54534435 |
| IVNQDEL | ASSSDWGSQNTLY | 0.66106564 |
| TNGSQFF | ASSSDWGSQNTLY | 0.6605637 |
| TGKGIVE | ASSSDWGSQNTLY | 0.43337086 |
| LISNASDA | ASSSDWGSQNTLY | 0.6805286 |
| AELRTLGELTKIAKAHGAQ | ASSSDWGSQNTLY | 0.5863372 |
| APNAVAAAEPSW | ASSSDWGSQNTLY | 0.5906236 |
| APQVAGAATAVL | ASSSDWGSQNTLY | 0.67953074 |
| ARRDPAAAAAILPTDA | ASSSDWGSQNTLY | 0.60301733 |
| DAAARDVLAEAGLAEYF | ASSSDWGSQNTLY | 0.44055817 |
| DDQIVLSAKANLAKVEQ | ASSSDWGSQNTLY | 0.5809962 |
| DELLPDTGEDSDEDGH | ASSSDWGSQNTLY | 0.61937463 |
| ELPEAPANRATVPQD | ASSSDWGSQNTLY | 0.57488745 |
| FGQSKIERVTGIG | ASSSDWGSQNTLY | 0.32081392 |
| GAGGTGSTAGGAGGAGGAGGLY | ASSSDWGSQNTLY | 0.67991644 |
| GHGGTGAAGGNGGTGGAGGSGI | ASSSDWGSQNTLY | 0.51509815 |
| GNGGNGGAGGEGGPGGAGGAGA | ASSSDWGSQNTLY | 0.5697188 |
| GSTAGNGGNGGNATNGTIAGQP | ASSSDWGSQNTLY | 0.5583431 |
| GTGGDGGNGGNGGTGG | ASSSDWGSQNTLY | 0.66712785 |
| GVRRPAGIPADAPV | ASSSDWGSQNTLY | 0.59403473 |
| GYRQVTDAHLLTLARR | ASSSDWGSQNTLY | 0.49259803 |
| IAKMGQDGHDRG | ASSSDWGSQNTLY | 0.6428593 |
| KAEPSIEIQYT | ASSSDWGSQNTLY | 0.64907426 |
| KPVRIRMAADEDTVAVT | ASSSDWGSQNTLY | 0.679433 |
| MPPRIAGMRLLVIKPEP | ASSSDWGSQNTLY | 0.6113648 |
| MKFYAVALFRSRGVPPTR | ASSSDWGSQNTLY | 0.61635596 |
| NAGGFGILGAVAHSP | ASSSDWGSQNTLY | 0.649741 |
| NEGREITLAMAEM | ASSSDWGSQNTLY | 0.5589238 |
| NELVEWSHPLGHRWRWE | ASSSDWGSQNTLY | 0.5546761 |
| NLGAGGTGGNGGNPG | ASSSDWGSQNTLY | 0.47917235 |
| NTEWPKWETVMAADPVD | ASSSDWGSQNTLY | 0.59761226 |
| PASAAPAETQP | ASSSDWGSQNTLY | 0.4982983 |
| QQSSDGKNGVRAAEITGE | ASSSDWGSQNTLY | 0.62526345 |
| RAPDAVTTSWWKE | ASSSDWGSQNTLY | 0.68458354 |
| SGFGNVGAHTSGWFNQ | ASSSDWGSQNTLY | 0.5917332 |
| SGVAGAVGQAASVGGLKV | ASSSDWGSQNTLY | 0.43868375 |
| SNGGAGTGSAGGNGGT | ASSSDWGSQNTLY | 0.5847223 |
| TGGAGGEGGAAG | ASSSDWGSQNTLY | 0.5905738 |
| TGGDGGAAGAGGAPGHG | ASSSDWGSQNTLY | 0.65768194 |
| TMPPLLAAIANAGRHGV | ASSSDWGSQNTLY | 0.6730703 |
| TSSTAGNSGGA | ASSSDWGSQNTLY | 0.7011899 |
| VPATVPVGILC | ASSSDWGSQNTLY | 0.6822585 |
| VQRPDSGGVVLTGRIS | ASSSDWGSQNTLY | 0.54817647 |
| WTPNPVPATKGVPAGLVQAT | ASSSDWGSQNTLY | 0.65484595 |
| YALKRALAKFSDRFDVVII | ASSSDWGSQNTLY | 0.56847465 |
| YLIGSVIV | ASSSDWGSQNTLY | 0.6593696 |
| YGQLVAVID | ASSSDWGSQNTLY | 0.6421773 |

| | | |
|---|---|---|
| VTVGGMTLPA | ASSSDWGSQNTLY | 0.6446477 |
| VLALPAV | ASSSDWGSQNTLY | 0.55972356 |
| VIPPTLNYET | ASSSDWGSQNTLY | 0.61800194 |
| VGMGGMHPSSG | ASSSDWGSQNTLY | 0.4124666 |
| VGLAGGAATA | ASSSDWGSQNTLY | 0.66346776 |
| VGGAGGAGGSGP | ASSSDWGSQNTLY | 0.56390154 |
| VFAAFMV | ASSSDWGSQNTLY | 0.4427418 |
| TLYDKLR | ASSSDWGSQNTLY | 0.59970456 |
| TLAVFTVL | ASSSDWGSQNTLY | 0.69602 |
| TGSVQPARV | ASSSDWGSQNTLY | 0.6837549 |
| TGGGAGSGGT | ASSSDWGSQNTLY | 0.5479635 |
| SHLETSL | ASSSDWGSQNTLY | 0.50246364 |
| SFTTGMT | ASSSDWGSQNTLY | 0.6441997 |
| QVEVAKRMGVRQPTVS | ASSSDWGSQNTLY | 0.44008136 |
| QGSGGPALAM | ASSSDWGSQNTLY | 0.6402004 |
| MLAPSNP | ASSSDWGSQNTLY | 0.6060092 |
| LVNNAAI | ASSSDWGSQNTLY | 0.6326826 |
| LTGEINTLA | ASSSDWGSQNTLY | 0.5027014 |
| LSIFGPV | ASSSDWGSQNTLY | 0.65461326 |
| LLALLANL | ASSSDWGSQNTLY | 0.60442317 |
| LISNASDALDKLR | ASSSDWGSQNTLY | 0.62433857 |
| LINEAVP | ASSSDWGSQNTLY | 0.30468106 |
| LGYAFSGL | ASSSDWGSQNTLY | 0.6717245 |
| LDFVAVAL | ASSSDWGSQNTLY | 0.50886744 |
| LAQPGAE | ASSSDWGSQNTLY | 0.46868318 |
| ISNLGAVTGF | ASSSDWGSQNTLY | 0.33696213 |
| HQEDPGRI | ASSSDWGSQNTLY | 0.63493884 |
| GTPGEPTP | ASSSDWGSQNTLY | 0.6680957 |
| GFFNSGS | ASSSDWGSQNTLY | 0.676525 |
| GAGGGGLTAGAN | ASSSDWGSQNTLY | 0.62432235 |
| FVDALVNM | ASSSDWGSQNTLY | 0.67939794 |
| FLTVAPEL | ASSSDWGSQNTLY | 0.69113445 |
| FETLTAL | ASSSDWGSQNTLY | 0.5938052 |
| DVINPIEL | ASSSDWGSQNTLY | 0.38815472 |
| APVSGAPV | ASSSDWGSQNTLY | 0.606334 |
| ALVGGLAW | ASSSDWGSQNTLY | 0.64284873 |
| ALAAAAATAI | ASSSDWGSQNTLY | 0.6819804 |
| AFGLGLLL | ASSSDWGSQNTLY | 0.6564499 |
| AFFAAASL | ASSSDWGSQNTLY | 0.44871342 |
| AAAALPAFSPP | ASSSDWGSQNTLY | 0.64818394 |
| VVLAPAL | ASSSDWGSQNTLY | 0.6787387 |
| TIGHVDHGKTTL | ASSSDWGSQNTLY | 0.4536192 |
| LISNASDALDK | ASSSDWGSQNTLY | 0.5499219 |
| LISNASDAL | ASSSDWGSQNTLY | 0.56733054 |
| ISNASDALDK | ASSSDWGSQNTLY | 0.42590287 |
| NQAETLVYQTE | ASSSDWGSQNTLY | 0.66042465 |
| TGGTGGTAGAG | ASSSDWGSQNTLY | 0.575931 |
| TGSTAGGAGGA | ASSSDWGSQNTLY | 0.51183057 |
| DPLAVPVAAGPV | ASSSDWGSQNTLY | 0.614519 |
| GGTGGTPGNGGN | ASSSDWGSQNTLY | 0.5051842 |
| PAGLAGVDPATL | ASSSDWGSQNTLY | 0.50495315 |

| | | |
|---|---|---|
| GNNLVVEAPHAGASV | ASSSDWGSQNTLY | 0.5487109 |
| KVTRIMSASTLEQPA | ASSSDWGSQNTLY | 0.66315174 |
| LGFALPIALPRRTVV | ASSSDWGSQNTLY | 0.6547586 |
| TRLGNLTARKLVVLV | ASSSDWGSQNTLY | 0.6257896 |
| WSQQRRLIYFPSAGP | ASSSDWGSQNTLY | 0.61070013 |
| GPNGNPDPMAAAVDIR | ASSSDWGSQNTLY | 0.6974584 |
| TQRCVSVADLRAAASR | ASSSDWGSQNTLY | 0.41272354 |
| QFVLDVAEHPVEVLTETV | ASSSDWGSQNTLY | 0.5457251 |
| GGAGGTGGAGGNGGTQAGDG | ASSSDWGSQNTLY | 0.6312932 |
| PLLATLRAIAVDVDAKPAQV | ASSSDWGSQNTLY | 0.55080116 |
| APKFRTGGYMIDFWGVGYQVAK | ASSSDWGSQNTLY | 0.6748346 |
| GAGGAGGAGGAAGGPGGTGGTG | ASSSDWGSQNTLY | 0.6391629 |
| IGALGLLATDVAPAGSVAFGVA | ASSSDWGSQNTLY | 0.631691 |
| AGGAGGIGGNGGAGGAGGVLMG | ASSSDWGSQNTLY | 0.61930835 |
| ADELIGLL | ASSSDWGSQNTLY | 0.70275414 |
| ADRALKA | ASSSDWGSQNTLY | 0.6545435 |
| AGAVGQVGGAGGA | ASSSDWGSQNTLY | 0.6105252 |
| AGGGGGLGGAP | ASSSDWGSQNTLY | 0.66428006 |
| AGGLLYGNGGAG | ASSSDWGSQNTLY | 0.46310788 |
| AGLESIGL | ASSSDWGSQNTLY | 0.6860331 |
| AGLKTGII | ASSSDWGSQNTLY | 0.6496046 |
| ASLAALLL | ASSSDWGSQNTLY | 0.47701544 |
| ASVINISSI | ASSSDWGSQNTLY | 0.5985649 |
| ATPPGPA | ASSSDWGSQNTLY | 0.6957902 |
| ATVASALV | ASSSDWGSQNTLY | 0.6801529 |
| ATVASLIL | ASSSDWGSQNTLY | 0.52772486 |
| AVNGPVLIRH | ASSSDWGSQNTLY | 0.5591631 |
| DGAAGAVGI | ASSSDWGSQNTLY | 0.45344976 |
| DPLLHGI | ASSSDWGSQNTLY | 0.5441613 |
| ELGAALGAL | ASSSDWGSQNTLY | 0.43714526 |
| ELGRTQAV | ASSSDWGSQNTLY | 0.6500785 |
| FVAGGYLS | ASSSDWGSQNTLY | 0.57376707 |
| GASGGAGGAGGW | ASSSDWGSQNTLY | 0.6596497 |
| GGAGAGGDGGAGGI | ASSSDWGSQNTLY | 0.6235343 |
| GTGHNGSQPGMGGNG | ASSSDWGSQNTLY | 0.67741776 |
| GYAAASASAS | ASSSDWGSQNTLY | 0.5399573 |
| HQVDVAAVQQG | ASSSDWGSQNTLY | 0.5979728 |
| IGAAIAHT | ASSSDWGSQNTLY | 0.37837428 |
| IGMIEVAQL | ASSSDWGSQNTLY | 0.5595762 |
| IRLAPEHTSAMD | ASSSDWGSQNTLY | 0.5268061 |
| IYNVSVL | ASSSDWGSQNTLY | 0.5807574 |
| KQGFLGM | ASSSDWGSQNTLY | 0.3565783 |
| LESVITQL | ASSSDWGSQNTLY | 0.6941482 |
| LGLGALVVV | ASSSDWGSQNTLY | 0.58844644 |
| LIGNGGNGGSGGI | ASSSDWGSQNTLY | 0.64027786 |
| LNAINAPTL | ASSSDWGSQNTLY | 0.566322 |
| LQLLNTGV | ASSSDWGSQNTLY | 0.50741285 |
| LQTSIEF | ASSSDWGSQNTLY | 0.50351334 |
| MSFAVSEL | ASSSDWGSQNTLY | 0.65329117 |
| NTGALGL | ASSSDWGSQNTLY | 0.50550485 |
| PPGAGGAGGAGTT | ASSSDWGSQNTLY | 0.42605987 |

| | | |
|---|---|---|
| QLVGLGI | ASSSDWGSQNTLY | 0.49885577 |
| RFLEQVHQL | ASSSDWGSQNTLY | 0.61726284 |
| RGVGYVLR | ASSSDWGSQNTLY | 0.69272006 |
| SATGVFAGVI | ASSSDWGSQNTLY | 0.6873102 |
| SLPAITLP | ASSSDWGSQNTLY | 0.6313701 |
| STALAFTV | ASSSDWGSQNTLY | 0.68573207 |
| SVLTVGLGK | ASSSDWGSQNTLY | 0.30150047 |
| TALAGAGSGI | ASSSDWGSQNTLY | 0.58193374 |
| TGLPFTGI | ASSSDWGSQNTLY | 0.68581426 |
| VAEHVTV | ASSSDWGSQNTLY | 0.6472225 |
| VDIVLNSL | ASSSDWGSQNTLY | 0.60562617 |
| VLTGGGAVL | ASSSDWGSQNTLY | 0.6790182 |
| VNVGLAYGI | ASSSDWGSQNTLY | 0.5121226 |
| VSSAAVGAV | ASSSDWGSQNTLY | 0.57029647 |
| VTSVYEAGV | ASSSDWGSQNTLY | 0.59067416 |
| WVALDRGA | ASSSDWGSQNTLY | 0.6791113 |
| YLVGSVVI | ASSSDWGSQNTLY | 0.63900936 |
| GPPGTGKTLL | ASSSDWGSQNTLY | 0.45103037 |
| GVGMTKF | ASSSDWGSQNTLY | 0.6313972 |
| IGIGGSDL | ASSSDWGSQNTLY | 0.71094 |
| IGTLLAEL | ASSSDWGSQNTLY | 0.55693066 |
| IYTSGTT | ASSSDWGSQNTLY | 0.6828674 |
| LSFPAGV | ASSSDWGSQNTLY | 0.6019067 |
| LYTSGTT | ASSSDWGSQNTLY | 0.64247346 |
| VGAFPFGF | ASSSDWGSQNTLY | 0.51861596 |
| VGAGTVEF | ASSSDWGSQNTLY | 0.4986384 |
| VGKSTLLL | ASSSDWGSQNTLY | 0.67306197 |
| AALAHAGGGIV | ASSSDWGSQNTLY | 0.5124645 |
| AALVAVSLL | ASSSDWGSQNTLY | 0.6001313 |
| AAVSRAVAAL | ASSSDWGSQNTLY | 0.5902525 |
| AGGSSGIGGF | ASSSDWGSQNTLY | 0.70785636 |
| AGLTVANANG | ASSSDWGSQNTLY | 0.65417016 |
| AGLVGAVL | ASSSDWGSQNTLY | 0.6199019 |
| ALSQAFTV | ASSSDWGSQNTLY | 0.5072465 |
| ALTFSVSL | ASSSDWGSQNTLY | 0.6486257 |
| ALTGAALVAL | ASSSDWGSQNTLY | 0.6541448 |
| ASFSIGAGVL | ASSSDWGSQNTLY | 0.45583165 |
| DGSVYKI | ASSSDWGSQNTLY | 0.68834794 |
| EQLAEIV | ASSSDWGSQNTLY | 0.4670063 |
| FAVSELTVM | ASSSDWGSQNTLY | 0.617306 |
| GAAPLAAGL | ASSSDWGSQNTLY | 0.64965665 |
| GADGGDGGSGGAGG | ASSSDWGSQNTLY | 0.69210625 |
| GFGLDTGGAGG | ASSSDWGSQNTLY | 0.6013806 |
| GLASGGLL | ASSSDWGSQNTLY | 0.66396815 |
| GRIAGAAVLL | ASSSDWGSQNTLY | 0.48561788 |
| GSAGSLNTSF | ASSSDWGSQNTLY | 0.65254265 |
| GTGQIGI | ASSSDWGSQNTLY | 0.67365175 |
| GTLIALIL | ASSSDWGSQNTLY | 0.6256894 |
| IAATMGFVVG | ASSSDWGSQNTLY | 0.70017356 |
| IFAQALL | ASSSDWGSQNTLY | 0.6509379 |
| IGAAVALGF | ASSSDWGSQNTLY | 0.37818798 |

| | | |
|---|---|---|
| IGLTPLAL | ASSSDWGSQNTLY | 0.60797703 |
| ILGSNTSTL | ASSSDWGSQNTLY | 0.6788729 |
| IVFSLTVL | ASSSDWGSQNTLY | 0.6824083 |
| LASKATGF | ASSSDWGSQNTLY | 0.68838966 |
| LASVALIL | ASSSDWGSQNTLY | 0.6567778 |
| LFAVGLTL | ASSSDWGSQNTLY | 0.60997045 |
| LFNGGIGTYI | ASSSDWGSQNTLY | 0.69417423 |
| LTLGAGII | ASSSDWGSQNTLY | 0.70792353 |
| LVAAALGDI | ASSSDWGSQNTLY | 0.61833024 |
| NTAFVQLGI | ASSSDWGSQNTLY | 0.49167204 |
| QLFSGAEL | ASSSDWGSQNTLY | 0.47273633 |
| SAATFITL | ASSSDWGSQNTLY | 0.6663654 |
| SNYGLANLGV | ASSSDWGSQNTLY | 0.66470397 |
| SYGGAYAVM | ASSSDWGSQNTLY | 0.43093094 |
| TTGVGGLTLG | ASSSDWGSQNTLY | 0.6148112 |
| TTQTIYI | ASSSDWGSQNTLY | 0.62531954 |
| VAAGAGLGAVI | ASSSDWGSQNTLY | 0.3784151 |
| VAAVVTLL | ASSSDWGSQNTLY | 0.68476987 |
| VGFLGGLI | ASSSDWGSQNTLY | 0.6955997 |
| VGVIPILL | ASSSDWGSQNTLY | 0.6411736 |
| VIFASSLI | ASSSDWGSQNTLY | 0.4854666 |
| VKGGDGGAAAG | ASSSDWGSQNTLY | 0.68892497 |
| VTKTFTNI | ASSSDWGSQNTLY | 0.697019 |
| VVAAAILL | ASSSDWGSQNTLY | 0.69389296 |
| YAALDVEL | ASSSDWGSQNTLY | 0.3921777 |
| LASVALLL | ASSSDWGSQNTLY | 0.30674186 |
| LDAVHRV | ASSSDWGSQNTLY | 0.6213496 |
| QGLSSSIF | ASSSDWGSQNTLY | 0.6622754 |
| RGTFANIRL | ASSSDWGSQNTLY | 0.55651176 |
| RQLWWGHRI | ASSSDWGSQNTLY | 0.48666903 |
| YGPPGTGKTLL | ASSSDWGSQNTLY | 0.6327683 |
| AALVGAPANVA | ASSSDWGSQNTLY | 0.6828266 |
| AATFRRTE | ASSSDWGSQNTLY | 0.48318118 |
| AGALGGGI | ASSSDWGSQNTLY | 0.4057566 |
| AGGAGGAGGIGAG | ASSSDWGSQNTLY | 0.67969835 |
| AGSGALGL | ASSSDWGSQNTLY | 0.65402246 |
| AQIADQI | ASSSDWGSQNTLY | 0.5401839 |
| ASAVAVTYL | ASSSDWGSQNTLY | 0.5629735 |
| ASTVLGGTGG | ASSSDWGSQNTLY | 0.64127874 |
| AVGGEPAK | ASSSDWGSQNTLY | 0.64163923 |
| CEQGTFGGTC | ASSSDWGSQNTLY | 0.64001906 |
| DEGKAPAG | ASSSDWGSQNTLY | 0.41874295 |
| DTVVFPAL | ASSSDWGSQNTLY | 0.6977209 |
| FEGITMAG | ASSSDWGSQNTLY | 0.450844 |
| FPLVVQD | ASSSDWGSQNTLY | 0.67142045 |
| GAFTTPQ | ASSSDWGSQNTLY | 0.6358161 |
| GFGGFGTAGGVGGA | ASSSDWGSQNTLY | 0.6597306 |
| GFLAQFL | ASSSDWGSQNTLY | 0.5735065 |
| GGAGGAGGNAGVA | ASSSDWGSQNTLY | 0.4964473 |
| GGVGTAGAGGAGGAGG | ASSSDWGSQNTLY | 0.45173144 |
| GNGGGGDAGNAGSGGN | ASSSDWGSQNTLY | 0.4251668 |

| | | |
|---|---|---|
| GTGGAGGAGNDAGS | ASSSDWGSQNTLY | 0.6930945 |
| GTTNTGWFNS | ASSSDWGSQNTLY | 0.63880086 |
| IFMGPLFTL | ASSSDWGSQNTLY | 0.67666936 |
| ILAHAIEAI | ASSSDWGSQNTLY | 0.62752485 |
| IRSAIREAL | ASSSDWGSQNTLY | 0.53269374 |
| ITFAGGG | ASSSDWGSQNTLY | 0.65173984 |
| ITGMSGHQEV | ASSSDWGSQNTLY | 0.38064867 |
| IVAGLACTL | ASSSDWGSQNTLY | 0.63945067 |
| IVGGGLAGA | ASSSDWGSQNTLY | 0.6075619 |
| IVTGVLNI | ASSSDWGSQNTLY | 0.6419264 |
| LAQQGCI | ASSSDWGSQNTLY | 0.545642 |
| LCLHVEN | ASSSDWGSQNTLY | 0.6433823 |
| LDAISDGS | ASSSDWGSQNTLY | 0.66170835 |
| LGGFAAML | ASSSDWGSQNTLY | 0.505097 |
| LGGKNPMI | ASSSDWGSQNTLY | 0.64017373 |
| LGPKGVAAIL | ASSSDWGSQNTLY | 0.6873238 |
| LMVGVSI | ASSSDWGSQNTLY | 0.6394632 |
| LVGAAFVL | ASSSDWGSQNTLY | 0.4397698 |
| LVLAVAPL | ASSSDWGSQNTLY | 0.6449461 |
| MAAIAVIDAA | ASSSDWGSQNTLY | 0.54744375 |
| NNGINAELQ | ASSSDWGSQNTLY | 0.64527214 |
| RGEPLVLVHG | ASSSDWGSQNTLY | 0.6314702 |
| RGGAGGAAGLI | ASSSDWGSQNTLY | 0.53369224 |
| RSYRMHPAV | ASSSDWGSQNTLY | 0.66094565 |
| SAAHLLAGQLA | ASSSDWGSQNTLY | 0.45027804 |
| SAAVGSY | ASSSDWGSQNTLY | 0.52792525 |
| SPHAVLHRI | ASSSDWGSQNTLY | 0.5750562 |
| SQIAKVL | ASSSDWGSQNTLY | 0.62115127 |
| TALGHLVP | ASSSDWGSQNTLY | 0.6466072 |
| TAPTTSGAA | ASSSDWGSQNTLY | 0.48538786 |
| TDGDDGNGGSPGT | ASSSDWGSQNTLY | 0.6354097 |
| TDGLGEN | ASSSDWGSQNTLY | 0.5807712 |
| TGGPPLPGQA | ASSSDWGSQNTLY | 0.49890578 |
| TIPAITI | ASSSDWGSQNTLY | 0.6091621 |
| TIPGFTI | ASSSDWGSQNTLY | 0.58164704 |
| TSIASGALA | ASSSDWGSQNTLY | 0.40900162 |
| TVTLPGAAI | ASSSDWGSQNTLY | 0.6809732 |
| VASGLSAALP | ASSSDWGSQNTLY | 0.478345 |
| VGAGVIGI | ASSSDWGSQNTLY | 0.6369397 |
| VGPKRFGV | ASSSDWGSQNTLY | 0.7044858 |
| VNVAFPD | ASSSDWGSQNTLY | 0.39986813 |
| VSMTNSGVS | ASSSDWGSQNTLY | 0.65680504 |
| VTDVLGDLSV | ASSSDWGSQNTLY | 0.63231486 |
| VVVVGAGFAGL | ASSSDWGSQNTLY | 0.65003455 |
| VYTVGTVV | ASSSDWGSQNTLY | 0.57735753 |
| WTTFSGP | ASSSDWGSQNTLY | 0.559077 |
| YDELVEAF | ASSSDWGSQNTLY | 0.68654317 |
| YGTGGAGGNGG | ASSSDWGSQNTLY | 0.64352137 |
| YTNVMAV | ASSSDWGSQNTLY | 0.59065247 |
| FLEQVHQL | ASSSDWGSQNTLY | 0.6669965 |
| GLSLLAVAV | ASSSDWGSQNTLY | 0.6769941 |

| | | |
|---|---|---|
| LEQVHQL | ASSSDWGSQNTLY | 0.6667177 |
| LNAAGAI | ASSSDWGSQNTLY | 0.5351073 |
| PAVHASL | ASSSDWGSQNTLY | 0.36386105 |
| TELEDDA | ASSSDWGSQNTLY | 0.58422387 |
| TPPGPPN | ASSSDWGSQNTLY | 0.6055246 |
| TPPPPPPP | ASSSDWGSQNTLY | 0.53710854 |
| TQSPALI | ASSSVGYEQY | 0.37361678 |
| EFAAAFPQ | ASSSVGYEQY | 0.680025 |
| LLSGNLDVL | ASSSVGYEQY | 0.7068299 |
| GPANLGNYNI | ASSSVGYEQY | 0.4810372 |
| TLGPTLP | ASSSVGYEQY | 0.7003727 |
| VYAYSSCCK | ASSSVGYEQY | 0.41844958 |
| APAPAPAPNI | ASSSVGYEQY | 0.6472293 |
| PGGPVPIP | ASSSVGYEQY | 0.68549967 |
| FLTVALEP | ASSSVGYEQY | 0.42989343 |
| SQYFGANL | ASSSVGYEQY | 0.64807457 |
| GGVKPAA | ASSSVGYEQY | 0.58850074 |
| GGTGGAGGAGGW | ASSSVGYEQY | 0.5152442 |
| GVGGVAGLGGAGG | ASSSVGYEQY | 0.6874547 |
| VLLVLATTI | ASSSVGYEQY | 0.50279987 |
| IGIFAAML | ASSSVGYEQY | 0.67330515 |
| AVGDLID | ASSSVGYEQY | 0.53443015 |
| NISPLPL | ASSSVGYEQY | 0.46951434 |
| VTAVFVAI | ASSSVGYEQY | 0.65562284 |
| SGDDNDAAL | ASSSVGYEQY | 0.6707338 |
| AVAALAHS | ASSSVGYEQY | 0.60470426 |
| GGVTVMATP | ASSSVGYEQY | 0.7050079 |
| AGLAANVRD | ASSSVGYEQY | 0.56442285 |
| TNRPDLIDPALLRPG | ASSSVGYEQY | 0.70889527 |
| LANLGVTAL | ASSSVGYEQY | 0.6665826 |
| TGERIEL | ASSSVGYEQY | 0.6104736 |
| NGEELAA | ASSSVGYEQY | 0.57744354 |
| VLSLVWG | ASSSVGYEQY | 0.6742843 |
| IGYALWL | ASSSVGYEQY | 0.52511 |
| LSLLLAPL | ASSSVGYEQY | 0.6588143 |
| TLDEALK | ASSSVGYEQY | 0.69239616 |
| TDGAGTAEGG | ASSSVGYEQY | 0.518586 |
| NAPGGAGGNGG | ASSSVGYEQY | 0.85062444 |
| VSVFMGMGGNF | ASSSVGYEQY | 0.7693759 |
| LASYLAAL | ASSSVGYEQY | 0.65277576 |
| LVGGTPAL | ASSSVGYEQY | 0.5992511 |
| VRNADVL | ASSSVGYEQY | 0.29447508 |
| TCSSPFLVSAM | ASSSVGYEQY | 0.685373 |
| VHITDAE | ASSSVGYEQY | 0.50481844 |
| EYAAPSF | ASSSVGYEQY | 0.76428974 |
| VVDAFHGGL | ASSSVGYEQY | 0.7499014 |
| ASVVDPQ | ASSSVGYEQY | 0.5655609 |
| SYNEFGGLS | ASSSVGYEQY | 0.85065126 |
| DPDGSVY | ASSSVGYEQY | 0.80367815 |
| MVSTIHATE | ASSSVGYEQY | 0.3538661 |
| LAAGVADPV | ASSSVGYEQY | 0.6786963 |

| | | |
|---|---|---|
| TKEQIAA | ASSSVGYEQY | 0.6346214 |
| SLASPAAL | ASSSVGYEQY | 0.64548385 |
| IGLVFYGV | ASSSVGYEQY | 0.27649105 |
| FLLDASNL | ASSSVGYEQY | 0.80301464 |
| YQVGNFP | ASSSVGYEQY | 0.60646725 |
| ESLLSQST | ASSSVGYEQY | 0.13552861 |
| LFTVGFAAVG | ASSSVGYEQY | 0.70250285 |
| ATPPTPT | ASSSVGYEQY | 0.71875024 |
| TDLATLG | ASSSVGYEQY | 0.63579535 |
| SAGGTGR | ASSSVGYEQY | 0.8462242 |
| GFSNSGS | ASSSVGYEQY | 0.23412567 |
| GGNGGSGGTSV | ASSSVGYEQY | 0.49571955 |
| GTSVATGGAGN | ASSSVGYEQY | 0.8327998 |
| NGGNGGSGGTS | ASSSVGYEQY | 0.8504989 |
| AVVKFPH | ASSSVGYEQY | 0.37304264 |
| TPPPPPP | ASSSVGYEQY | 0.8081169 |
| AAPAAAAAPA | ASSSVGYEQY | 0.79677975 |
| IALNEDL | ASSSVGYEQY | 0.7342752 |
| IVNVSSM | ASSSVGYEQY | 0.82398176 |
| TGIPPAPR | ASSSVGYEQY | 0.83901 |
| FDLGGGTF | ASSSVGYEQY | 0.43907815 |
| FDLGGGTFDVS | ASSSVGYEQY | 0.8437511 |
| NEPTAAA | ASSSVGYEQY | 0.848228 |
| NEPTAAALA | ASSSVGYEQY | 0.57893384 |
| GSIINISSI | ASSSVGYEQY | 0.76055604 |
| VAPGNAGTA | ASSSVGYEQY | 0.480444 |
| TGGGASLE | ASSSVGYEQY | 0.51401556 |
| VAVYNHYKRI | ASSSVGYEQY | 0.23235679 |
| TDPAPATT | ASSSVGYEQY | 0.81377125 |
| TNRPDILDPALLRPG | ASSSVGYEQY | 0.80451465 |
| LTTTTTS | ASSSVGYEQY | 0.69123334 |
| PTGSGKTL | ASSSVGYEQY | 0.55751455 |
| TGSGKTL | ASSSVGYEQY | 0.47862065 |
| TIGHVDHGK | ASSSVGYEQY | 0.6806638 |
| TIGHVDHGKTT | ASSSVGYEQY | 0.645803 |
| GHVDHGKTT | ASSSVGYEQY | 0.23576462 |
| AQFEHTLL | ASSSVGYEQY | 0.59646684 |
| IVNQDEL | ASSSVGYEQY | 0.81815994 |
| TNGSQFF | ASSSVGYEQY | 0.7945106 |
| TGKGIVE | ASSSVGYEQY | 0.27124146 |
| LISNASDA | ASSSVGYEQY | 0.86016774 |
| AELRTLGELTKIAKAHGAQ | ASSSVGYEQY | 0.8090378 |
| APNAVAAAEPSW | ASSSVGYEQY | 0.82276213 |
| APQVAGAATAVL | ASSSVGYEQY | 0.6263666 |
| ARRDPAAAAAILPTDA | ASSSVGYEQY | 0.57727224 |
| DAAARDVLAEAGLAEYF | ASSSVGYEQY | 0.82631445 |
| DDQIVLSAKANLAKVEQ | ASSSVGYEQY | 0.7230861 |
| DELLPDTGEDSDEDGH | ASSSVGYEQY | 0.38434193 |
| ELPEAPANRATVPQD | ASSSVGYEQY | 0.814541 |
| FGQSKIERVTGIG | ASSSVGYEQY | 0.6800475 |
| GAGGTGSTAGGAGGAGGAGGLY | ASSSVGYEQY | 0.6607959 |

| | | |
|---|---|---|
| GHGGTGAAGGNGGTGGAGGSGI | ASSSVGYEQY | 0.6365177 |
| GNGGNGGAGGEGGPGGAGGAGA | ASSSVGYEQY | 0.78801244 |
| GSTAGNGGNGGNATNGTIAGQP | ASSSVGYEQY | 0.54005694 |
| GTGGDGGNGGNGGTGG | ASSSVGYEQY | 0.42025614 |
| GVRRPAGIPADAPV | ASSSVGYEQY | 0.40242743 |
| GYRQVTDAHLLTLARR | ASSSVGYEQY | 0.74425185 |
| IAKMGQDGHDRG | ASSSVGYEQY | 0.8414241 |
| KAEPSIEIQYT | ASSSVGYEQY | 0.6759149 |
| KPVRIRMAADEDTVAVT | ASSSVGYEQY | 0.5975438 |
| MPPRIAGMRLLVIKPEP | ASSSVGYEQY | 0.8032918 |
| MKFYAVALFRSRGVPPTR | ASSSVGYEQY | 0.69852984 |
| NAGGFGILGAVAHSP | ASSSVGYEQY | 0.71216756 |
| NEGREITLAMAEM | ASSSVGYEQY | 0.70735514 |
| NELVEWSHPLGHRWRWE | ASSSVGYEQY | 0.6646161 |
| NLGAGGTGGNGGNPG | ASSSVGYEQY | 0.810161 |
| NTEWPKWETVMAADPVD | ASSSVGYEQY | 0.7765213 |
| PASAAPAETQP | ASSSVGYEQY | 0.80433 |
| QQSSDGKNGVRAAEITGE | ASSSVGYEQY | 0.81473196 |
| RAPDAVTTSWWKE | ASSSVGYEQY | 0.722232 |
| SGFGNVGAHTSGWFNQ | ASSSVGYEQY | 0.67466295 |
| SGVAGAVGQAASVGGLKV | ASSSVGYEQY | 0.19770962 |
| SNGGAGTGSAGGNGGT | ASSSVGYEQY | 0.51556754 |
| TGGAGGEGGAAG | ASSSVGYEQY | 0.77516395 |
| TGGDGGAAGAGGAPGHG | ASSSVGYEQY | 0.789133 |
| TMPPLLAAIANAGRHGV | ASSSVGYEQY | 0.78643215 |
| TSSTAGNSGGA | ASSSVGYEQY | 0.7020071 |
| VPATVPVGILC | ASSSVGYEQY | 0.5355544 |
| VQRPDSGGVVLTGRIS | ASSSVGYEQY | 0.7434393 |
| WTPNPVPATKGVPAGLVQAT | ASSSVGYEQY | 0.8107704 |
| YALKRALAKFSDRFDVVII | ASSSVGYEQY | 0.3922024 |
| YLIGSVIV | ASSSVGYEQY | 0.6305208 |
| YGQLVAVID | ASSSVGYEQY | 0.86705875 |
| VTVGGMTLPA | ASSSVGYEQY | 0.776935 |
| VLALPAV | ASSSVGYEQY | 0.6969701 |
| VIPPTLNYET | ASSSVGYEQY | 0.4694845 |
| VGMGGMHPSSG | ASSSVGYEQY | 0.83883417 |
| VGLAGGAATA | ASSSVGYEQY | 0.56083655 |
| VGGAGGAGGSGP | ASSSVGYEQY | 0.84490967 |
| VFAAFMV | ASSSVGYEQY | 0.16506088 |
| TLYDKLR | ASSSVGYEQY | 0.49774322 |
| TLAVFTVL | ASSSVGYEQY | 0.3744113 |
| TGSVQPARV | ASSSVGYEQY | 0.28462556 |
| TGGGAGSGGT | ASSSVGYEQY | 0.8046384 |
| SHLETSL | ASSSVGYEQY | 0.49754062 |
| SFTTGMT | ASSSVGYEQY | 0.75849324 |
| QVEVAKRMGVRQPTVS | ASSSVGYEQY | 0.74857783 |
| QGSGGPALAM | ASSSVGYEQY | 0.82191205 |
| MLAPSNP | ASSSVGYEQY | 0.7922338 |
| LVNNAAI | ASSSVGYEQY | 0.6478542 |
| LTGEINTLA | ASSSVGYEQY | 0.43084258 |
| LSIFGPV | ASSSVGYEQY | 0.7979461 |

| | | |
|---|---|---|
| LLALLANL | ASSSVGYEQY | 0.78920704 |
| LISNASDALDKLR | ASSSVGYEQY | 0.7278125 |
| LINEAVP | ASSSVGYEQY | 0.33180857 |
| LGYAFSGL | ASSSVGYEQY | 0.7032689 |
| LDFVAVAL | ASSSVGYEQY | 0.7227258 |
| LAQPGAE | ASSSVGYEQY | 0.21295312 |
| ISNLGAVTGF | ASSSVGYEQY | 0.19193271 |
| HQEDPGRI | ASSSVGYEQY | 0.71457195 |
| GTPGEPTP | ASSSVGYEQY | 0.5845822 |
| GFFNSGS | ASSSVGYEQY | 0.8448446 |
| GAGGGGLTAGAN | ASSSVGYEQY | 0.8377222 |
| FVDALVNM | ASSSVGYEQY | 0.71658635 |
| FLTVAPEL | ASSSVGYEQY | 0.77568996 |
| FETLTAL | ASSSVGYEQY | 0.74043345 |
| DVINPIEL | ASSSVGYEQY | 0.11725081 |
| APVSGAPV | ASSSVGYEQY | 0.5381003 |
| ALVGGLAW | ASSSVGYEQY | 0.41378945 |
| ALAAAAATAI | ASSSVGYEQY | 0.8416084 |
| AFGLGLLL | ASSSVGYEQY | 0.49445212 |
| AFFAAASL | ASSSVGYEQY | 0.6226635 |
| AAAALPAFSPP | ASSSVGYEQY | 0.75849605 |
| VVLAPAL | ASSSVGYEQY | 0.763453 |
| TIGHVDHGKTTL | ASSSVGYEQY | 0.70566833 |
| LISNASDALDK | ASSSVGYEQY | 0.45212966 |
| LISNASDAL | ASSSVGYEQY | 0.13945526 |
| ISNASDALDK | ASSSVGYEQY | 0.80285054 |
| NQAETLVYQTE | ASSSVGYEQY | 0.28530517 |
| TGGTGGTAGAG | ASSSVGYEQY | 0.4152041 |
| TGSTAGGAGGA | ASSSVGYEQY | 0.21941143 |
| DPLAVPVAAGPV | ASSSVGYEQY | 0.17973864 |
| GGTGGTPGNGGN | ASSSVGYEQY | 0.30712265 |
| PAGLAGVDPATL | ASSSVGYEQY | 0.79740185 |
| GNNLVVEAPHAGASV | ASSSVGYEQY | 0.7192981 |
| KVTRIMSASTLEQPA | ASSSVGYEQY | 0.7030814 |
| LGFALPIALPRRTVV | ASSSVGYEQY | 0.8604 |
| TRLGNLTARKLVVLV | ASSSVGYEQY | 0.7617227 |
| WSQQRRLIYFPSAGP | ASSSVGYEQY | 0.7247008 |
| GPNGNPDPMAAAVDIR | ASSSVGYEQY | 0.5113608 |
| TQRCVSVADLRAAASR | ASSSVGYEQY | 0.51052624 |
| QFVLDVAEHPVEVLTETV | ASSSVGYEQY | 0.48042953 |
| GGAGGTGGAGGNGGTQAGDG | ASSSVGYEQY | 0.44877276 |
| PLLATLRAIAVDVDAKPAQV | ASSSVGYEQY | 0.8573512 |
| APKFRTGGYMIDFWGVGYQVAK | ASSSVGYEQY | 0.52706885 |
| GAGGAGGAGGAAGGPGGTGGTG | ASSSVGYEQY | 0.39132813 |
| IGALGLLATDVAPAGSVAFGVA | ASSSVGYEQY | 0.62434113 |
| AGGAGGIGGNGGAGGAGGVLMG | ASSSVGYEQY | 0.85080206 |
| ADELIGLL | ASSSVGYEQY | 0.8617773 |
| ADRALKA | ASSSVGYEQY | 0.5022363 |
| AGAVGQVGGAGGA | ASSSVGYEQY | 0.46693385 |
| AGGGGGLGGAP | ASSSVGYEQY | 0.53810775 |
| AGGLLYGNGGAG | ASSSVGYEQY | 0.7785108 |

| | | |
|---|---|---|
| AGLESIGL | ASSSVGYEQY | 0.48047936 |
| AGLKTGII | ASSSVGYEQY | 0.38829598 |
| ASLAALLL | ASSSVGYEQY | 0.77701116 |
| ASVINISSI | ASSSVGYEQY | 0.85011184 |
| ATPPGPA | ASSSVGYEQY | 0.5382807 |
| ATVASALV | ASSSVGYEQY | 0.44100165 |
| ATVASLIL | ASSSVGYEQY | 0.73891884 |
| AVNGPVLIRH | ASSSVGYEQY | 0.5862926 |
| DGAAGAVGI | ASSSVGYEQY | 0.6981622 |
| DPLLHGI | ASSSVGYEQY | 0.6472199 |
| ELGAALGAL | ASSSVGYEQY | 0.4202171 |
| ELGRTQAV | ASSSVGYEQY | 0.60611945 |
| FVAGGYLS | ASSSVGYEQY | 0.8565755 |
| GASGGAGGAGGW | ASSSVGYEQY | 0.67470753 |
| GGAGAGGDGGAGGI | ASSSVGYEQY | 0.7658794 |
| GTGHNGSQPGMGGNG | ASSSVGYEQY | 0.85254216 |
| GYAAASASAS | ASSSVGYEQY | 0.8181449 |
| HQVDVAAVQQG | ASSSVGYEQY | 0.8576875 |
| IGAAIAHT | ASSSVGYEQY | 0.81894934 |
| IGMIEVAQL | ASSSVGYEQY | 0.5598124 |
| IRLAPEHTSAMD | ASSSVGYEQY | 0.6319476 |
| IYNVSVL | ASSSVGYEQY | 0.7838413 |
| KQGFLGM | ASSSVGYEQY | 0.77400255 |
| LESVITQL | ASSSVGYEQY | 0.6935104 |
| LGLGALVVV | ASSSVGYEQY | 0.50850487 |
| LIGNGGNGGSGGI | ASSSVGYEQY | 0.83911383 |
| LNAINAPTL | ASSSVGYEQY | 0.612841 |
| LQLLNTGV | ASSSVGYEQY | 0.8128115 |
| LQTSIEF | ASSSVGYEQY | 0.37675637 |
| MSFAVSEL | ASSSVGYEQY | 0.40838888 |
| NTGALGL | ASSSVGYEQY | 0.8355051 |
| PPGAGGAGGAGTT | ASSSVGYEQY | 0.124168865 |
| QLVGLGI | ASSSVGYEQY | 0.17940608 |
| RFLEQVHQL | ASSSVGYEQY | 0.6980696 |
| RGVGYVLR | ASSSVGYEQY | 0.84243655 |
| SATGVFAGVI | ASSSVGYEQY | 0.1397817 |
| SLPAITLP | ASSSVGYEQY | 0.44909734 |
| STALAFTV | ASSSVGYEQY | 0.37030846 |
| SVLTVGLGK | ASSSVGYEQY | 0.7588029 |
| TALAGAGSGI | ASSSVGYEQY | 0.70422035 |
| TGLPFTGI | ASSSVGYEQY | 0.31226218 |
| VAEHVTV | ASSSVGYEQY | 0.79396653 |
| VDIVLNSL | ASSSVGYEQY | 0.39216316 |
| VLTGGGAVL | ASSSVGYEQY | 0.42451215 |
| VNVGLAYGI | ASSSVGYEQY | 0.8120903 |
| VSSAAVGAV | ASSSVGYEQY | 0.84604007 |
| VTSVYEAGV | ASSSVGYEQY | 0.44788983 |
| WVALDRGA | ASSSVGYEQY | 0.8201521 |
| YLVGSVVI | ASSSVGYEQY | 0.5539044 |
| GPPGTGKTLL | ASSSVGYEQY | 0.86169046 |
| GVGMTKF | ASSSVGYEQY | 0.62255496 |

| | | |
|---|---|---|
| IGIGGSDL | ASSSVGYEQY | 0.33399138 |
| IGTLLAEL | ASSSVGYEQY | 0.8135058 |
| IYTSGTT | ASSSVGYEQY | 0.78772116 |
| LSFPAGV | ASSSVGYEQY | 0.54635125 |
| LYTSGTT | ASSSVGYEQY | 0.82916427 |
| VGAFPFGF | ASSSVGYEQY | 0.4973597 |
| VGAGTVEF | ASSSVGYEQY | 0.74962926 |
| VGKSTLLL | ASSSVGYEQY | 0.8481188 |
| AALAHAGGGIV | ASSSVGYEQY | 0.83548963 |
| AALVAVSLL | ASSSVGYEQY | 0.48204398 |
| AAVSRAVAAL | ASSSVGYEQY | 0.76118606 |
| AGGSSGIGGF | ASSSVGYEQY | 0.717841 |
| AGLTVANANG | ASSSVGYEQY | 0.50582606 |
| AGLVGAVL | ASSSVGYEQY | 0.84756017 |
| ALSQAFTV | ASSSVGYEQY | 0.6778642 |
| ALTFSVSL | ASSSVGYEQY | 0.8158115 |
| ALTGAALVAL | ASSSVGYEQY | 0.7186012 |
| ASFSIGAGVL | ASSSVGYEQY | 0.79497397 |
| DGSVYKI | ASSSVGYEQY | 0.35323033 |
| EQLAEIV | ASSSVGYEQY | 0.70509684 |
| FAVSELTVM | ASSSVGYEQY | 0.71589386 |
| GAAPLAAGL | ASSSVGYEQY | 0.8170334 |
| GADGGDGGSGGAGG | ASSSVGYEQY | 0.71845925 |
| GFGLDTGGAGG | ASSSVGYEQY | 0.35360438 |
| GLASGGLL | ASSSVGYEQY | 0.71972024 |
| GRIAGAAVLL | ASSSVGYEQY | 0.83925885 |
| GSAGSLNTSF | ASSSVGYEQY | 0.66952884 |
| GTGQIGI | ASSSVGYEQY | 0.64890903 |
| GTLIALIL | ASSSVGYEQY | 0.86350846 |
| IAATMGFVVG | ASSSVGYEQY | 0.50172085 |
| IFAQALL | ASSSVGYEQY | 0.66729057 |
| IGAAVALGF | ASSSVGYEQY | 0.7602922 |
| IGLTPLAL | ASSSVGYEQY | 0.8011526 |
| ILGSNTSTL | ASSSVGYEQY | 0.63185227 |
| IVFSLTVL | ASSSVGYEQY | 0.8314079 |
| LASKATGF | ASSSVGYEQY | 0.41831917 |
| LASVALIL | ASSSVGYEQY | 0.81539345 |
| LFAVGLTL | ASSSVGYEQY | 0.57895875 |
| LFNGGIGTYI | ASSSVGYEQY | 0.7867684 |
| LTLGAGII | ASSSVGYEQY | 0.62841856 |
| LVAAALGDI | ASSSVGYEQY | 0.8229557 |
| NTAFVQLGI | ASSSVGYEQY | 0.83771324 |
| QLFSGAEL | ASSSVGYEQY | 0.8344153 |
| SAATFITL | ASSSVGYEQY | 0.17113616 |
| SNYGLANLGV | ASSSVGYEQY | 0.843671 |
| SYGGAYAVM | ASSSVGYEQY | 0.804847 |
| TTGVGGLTLG | ASSSVGYEQY | 0.8715036 |
| TTQTIYI | ASSSVGYEQY | 0.6113577 |
| VAAGAGLGAVI | ASSSVGYEQY | 0.7327956 |
| VAAVVTLL | ASSSVGYEQY | 0.6272184 |
| VGFLGGLI | ASSSVGYEQY | 0.6017856 |

| | | |
|---|---|---|
| VGVIPILL | ASSSVGYEQY | 0.51596284 |
| VIFASSLI | ASSSVGYEQY | 0.53854454 |
| VKGGDGGAAAG | ASSSVGYEQY | 0.72808003 |
| VTKTFTNI | ASSSVGYEQY | 0.6384187 |
| VVAAAILL | ASSSVGYEQY | 0.81478643 |
| YAALDVEL | ASSSVGYEQY | 0.81743276 |
| LASVALLL | ASSSVGYEQY | 0.8248588 |
| LDAVHRV | ASSSVGYEQY | 0.8499479 |
| QGLSSSIF | ASSSVGYEQY | 0.8466623 |
| RGTFANIRL | ASSSVGYEQY | 0.8020864 |
| RQLWWGHRI | ASSSVGYEQY | 0.5093086 |
| YGPPGTGKTLL | ASSSVGYEQY | 0.66131586 |
| AALVGAPANVA | ASSSVGYEQY | 0.5307562 |
| AATFRRTE | ASSSVGYEQY | 0.6691613 |
| AGALGGGI | ASSSVGYEQY | 0.59827805 |
| AGGAGGAGGIGAG | ASSSVGYEQY | 0.79899824 |
| AGSGALGL | ASSSVGYEQY | 0.12671553 |
| AQIADQI | ASSSVGYEQY | 0.7743001 |
| ASAVAVTYL | ASSSVGYEQY | 0.85552084 |
| ASTVLGGTGG | ASSSVGYEQY | 0.5736369 |
| AVGGEPAK | ASSSVGYEQY | 0.7117894 |
| CEQGTFGGTC | ASSSVGYEQY | 0.7477553 |
| DEGKAPAG | ASSSVGYEQY | 0.6144625 |
| DTVVFPAL | ASSSVGYEQY | 0.82725054 |
| FEGITMAG | ASSSVGYEQY | 0.46352673 |
| FPLVVQD | ASSSVGYEQY | 0.76515365 |
| GAFTTPQ | ASSSVGYEQY | 0.84062326 |
| GFGGFGTAGGVGGA | ASSSVGYEQY | 0.73528516 |
| GFLAQFL | ASSSVGYEQY | 0.59648544 |
| GGAGGAGGNAGVA | ASSSVGYEQY | 0.59373534 |
| GGVGTAGAGGAGGAGG | ASSSVGYEQY | 0.29910433 |
| GNGGGGDAGNAGSGGN | ASSSVGYEQY | 0.719679 |
| GTGGAGGAGNDAGS | ASSSVGYEQY | 0.12417486 |
| GTTNTGWFNS | ASSSVGYEQY | 0.75131154 |
| IFMGPLFTL | ASSSVGYEQY | 0.43065286 |
| ILAHAIEAI | ASSSVGYEQY | 0.76830536 |
| IRSAIREAL | ASSSVGYEQY | 0.7710577 |
| ITFAGGG | ASSSVGYEQY | 0.7322214 |
| ITGMSGHQEV | ASSSVGYEQY | 0.5574689 |
| IVAGLACTL | ASSSVGYEQY | 0.75228333 |
| IVGGGLAGA | ASSSVGYEQY | 0.4730476 |
| IVTGVLNI | ASSSVGYEQY | 0.36744934 |
| LAQQGCI | ASSSVGYEQY | 0.8211321 |
| LCLHVEN | ASSSVGYEQY | 0.57416123 |
| LDAISDGS | ASSSVGYEQY | 0.71169627 |
| LGGFAAML | ASSSVGYEQY | 0.80419743 |
| LGGKNPMI | ASSSVGYEQY | 0.70269156 |
| LGPKGVAAIL | ASSSVGYEQY | 0.3417761 |
| LMVGVSI | ASSSVGYEQY | 0.7452328 |
| LVGAAFVL | ASSSVGYEQY | 0.7314439 |
| LVLAVAPL | ASSSVGYEQY | 0.6405044 |

| | | |
|---|---|---|
| MAAIAVIDAA | ASSSVGYEQY | 0.1175738 |
| NNGINAELQ | ASSSVGYEQY | 0.7419946 |
| RGEPLVLVHG | ASSSVGYEQY | 0.52076375 |
| RGGAGGAAGLI | ASSSVGYEQY | 0.59484255 |
| RSYRMHPAV | ASSSVGYEQY | 0.582121 |
| SAAHLLAGQLA | ASSSVGYEQY | 0.83645487 |
| SAAVGSY | ASSSVGYEQY | 0.29988205 |
| SPHAVLHRI | ASSSVGYEQY | 0.61929345 |
| SQIAKVL | ASSSVGYEQY | 0.8086808 |
| TALGHLVP | ASSSVGYEQY | 0.7964065 |
| TAPTTSGAA | ASSSVGYEQY | 0.84558165 |
| TDGDDGNGGSPGT | ASSSVGYEQY | 0.67034733 |
| TDGLGEN | ASSSVGYEQY | 0.47937843 |
| TGGPPLPGQA | ASSSVGYEQY | 0.64307934 |
| TIPAITI | ASSSVGYEQY | 0.5808485 |
| TIPGFTI | ASSSVGYEQY | 0.2904529 |
| TSIASGALA | ASSSVGYEQY | 0.21767057 |
| TVTLPGAAI | ASSSVGYEQY | 0.76573616 |
| VASGLSAALP | ASSSVGYEQY | 0.2199895 |
| VGAGVIGI | ASSSVGYEQY | 0.70998585 |
| VGPKRFGV | ASSSVGYEQY | 0.810024 |
| VNVAFPD | ASSSVGYEQY | 0.57795185 |
| VSMTNSGVS | ASSSVGYEQY | 0.34900725 |
| VTDVLGDLSV | ASSSVGYEQY | 0.64046866 |
| VVVVGAGFAGL | ASSSVGYEQY | 0.7057986 |
| VYTVGTVV | ASSSVGYEQY | 0.82069135 |
| WTTFSGP | ASSSVGYEQY | 0.71638036 |
| YDELVEAF | ASSSVGYEQY | 0.84472686 |
| YGTGGAGGNGG | ASSSVGYEQY | 0.8168089 |
| YTNVMAV | ASSSVGYEQY | 0.6493559 |
| FLEQVHQL | ASSSVGYEQY | 0.81395566 |
| GLSLLAVAV | ASSSVGYEQY | 0.63430923 |
| LEQVHQL | ASSSVGYEQY | 0.68060225 |
| LNAAGAI | ASSSVGYEQY | 0.62667763 |
| PAVHASL | ASSSVGYEQY | 0.7097249 |
| TELEDDA | ASSSVGYEQY | 0.5683837 |
| TPPGPPN | ASSSVGYEQY | 0.531369 |
| TPPPPPPP | ASSSVGYEQY | 0.3198684 |
| TQSPALI | ASSLAQNSGNTLY | 0.77062595 |
| EFAAAFPQ | ASSLAQNSGNTLY | 0.7552217 |
| LLSGNLDVL | ASSLAQNSGNTLY | 0.30437356 |
| GPANLGNYNI | ASSLAQNSGNTLY | 0.55916685 |
| TLGPTLP | ASSLAQNSGNTLY | 0.8619115 |
| VYAYSSCCK | ASSLAQNSGNTLY | 0.8423221 |
| APAPAPAPNI | ASSLAQNSGNTLY | 0.59504557 |
| PGGPVPIP | ASSLAQNSGNTLY | 0.51405865 |
| FLTVALEP | ASSLAQNSGNTLY | 0.7727673 |
| SQYFGANL | ASSLAQNSGNTLY | 0.22773 |
| GGVKPAA | ASSLAQNSGNTLY | 0.72543883 |
| GGTGGAGGAGGW | ASSLAQNSGNTLY | 0.78269005 |
| GVGGVAGLGGAGG | ASSLAQNSGNTLY | 0.76081383 |

| | | |
|---|---|---|
| VLLVLATTI | ASSLAQNSGNTLY | 0.42634094 |
| IGIFAAML | ASSLAQNSGNTLY | 0.7463943 |
| AVGDLID | ASSLAQNSGNTLY | 0.6371944 |
| NISPLPL | ASSLAQNSGNTLY | 0.7388218 |
| VTAVFVAI | ASSLAQNSGNTLY | 0.14747843 |
| SGDDNDAAL | ASSLAQNSGNTLY | 0.73747396 |
| AVAALAHS | ASSLAQNSGNTLY | 0.5751575 |
| GGVTVMATP | ASSLAQNSGNTLY | 0.43464094 |
| AGLAANVRD | ASSLAQNSGNTLY | 0.27603808 |
| TNRPDLIDPALLRPG | ASSLAQNSGNTLY | 0.6823979 |
| LANLGVTAL | ASSLAQNSGNTLY | 0.8293518 |
| TGERIEL | ASSLAQNSGNTLY | 0.8342907 |
| NGEELAA | ASSLAQNSGNTLY | 0.45012218 |
| VLSLVWG | ASSLAQNSGNTLY | 0.8328454 |
| IGYALWL | ASSLAQNSGNTLY | 0.85582286 |
| LSLLLAPL | ASSLAQNSGNTLY | 0.7100247 |
| TLDEALK | ASSLAQNSGNTLY | 0.28694144 |
| TDGAGTAEGG | ASSLAQNSGNTLY | 0.7582619 |
| NAPGGAGGNGG | ASSLAQNSGNTLY | 0.6618588 |
| VSVFMGMGGNF | ASSLAQNSGNTLY | 0.8210573 |
| LASYLAAL | ASSLAQNSGNTLY | 0.7604315 |
| LVGGTPAL | ASSLAQNSGNTLY | 0.5419642 |
| VRNADVL | ASSLAQNSGNTLY | 0.7173478 |
| TCSSPFLVSAM | ASSLAQNSGNTLY | 0.82994425 |
| VHITDAE | ASSLAQNSGNTLY | 0.5493303 |
| EYAAPSF | ASSLAQNSGNTLY | 0.4920966 |
| VVDAFHGGL | ASSLAQNSGNTLY | 0.4917927 |
| ASVVDPQ | ASSLAQNSGNTLY | 0.24049369 |
| SYNEFGGLS | ASSLAQNSGNTLY | 0.76858157 |
| DPDGSVY | ASSLAQNSGNTLY | 0.6639662 |
| MVSTIHATE | ASSLAQNSGNTLY | 0.48632365 |
| LAAGVADPV | ASSLAQNSGNTLY | 0.77900904 |
| TKEQIAA | ASSLAQNSGNTLY | 0.46876308 |
| SLASPAAL | ASSLAQNSGNTLY | 0.5607288 |
| IGLVFYGV | ASSLAQNSGNTLY | 0.5968045 |
| FLLDASNL | ASSLAQNSGNTLY | 0.7051647 |
| YQVGNFP | ASSLAQNSGNTLY | 0.7832098 |
| ESLLSQST | ASSLAQNSGNTLY | 0.73517513 |
| LFTVGFAAVG | ASSLAQNSGNTLY | 0.6784768 |
| ATPPTPT | ASSLAQNSGNTLY | 0.8602756 |
| TDLATLG | ASSLAQNSGNTLY | 0.42288935 |
| SAGGTGR | ASSLAQNSGNTLY | 0.51272035 |
| GFSNSGS | ASSLAQNSGNTLY | 0.78865266 |
| GGNGGSGGTSV | ASSLAQNSGNTLY | 0.3258354 |
| GTSVATGGAGN | ASSLAQNSGNTLY | 0.8042122 |
| NGGNGGSGGTS | ASSLAQNSGNTLY | 0.7378528 |
| AVVKFPH | ASSLAQNSGNTLY | 0.80085933 |
| TPPPPPP | ASSLAQNSGNTLY | 0.64123 |
| AAPAAAAAPA | ASSLAQNSGNTLY | 0.86516476 |
| IALNEDL | ASSLAQNSGNTLY | 0.8343255 |
| IVNVSSM | ASSLAQNSGNTLY | 0.6690117 |

| | | |
|---|---|---|
| TGIPPAPR | ASSLAQNSGNTLY | 0.75969183 |
| FDLGGGTF | ASSLAQNSGNTLY | 0.3863683 |
| FDLGGGTFDVS | ASSLAQNSGNTLY | 0.8182459 |
| NEPTAAA | ASSLAQNSGNTLY | 0.7605296 |
| NEPTAAALA | ASSLAQNSGNTLY | 0.55830795 |
| GSIINISSI | ASSLAQNSGNTLY | 0.7522911 |
| VAPGNAGTA | ASSLAQNSGNTLY | 0.8202312 |
| TGGGASLE | ASSLAQNSGNTLY | 0.8386457 |
| VAVYNHYKRI | ASSLAQNSGNTLY | 0.22633232 |
| TDPAPATT | ASSLAQNSGNTLY | 0.7071382 |
| TNRPDILDPALLRPG | ASSLAQNSGNTLY | 0.21337365 |
| LTTTTTS | ASSLAQNSGNTLY | 0.3739787 |
| PTGSGKTL | ASSLAQNSGNTLY | 0.40241838 |
| TGSGKTL | ASSLAQNSGNTLY | 0.7850082 |
| TIGHVDHGK | ASSLAQNSGNTLY | 0.65616274 |
| TIGHVDHGKTT | ASSLAQNSGNTLY | 0.77001715 |
| GHVDHGKTT | ASSLAQNSGNTLY | 0.74246556 |
| AQFEHTLL | ASSLAQNSGNTLY | 0.83553374 |
| IVNQDEL | ASSLAQNSGNTLY | 0.52665454 |
| TNGSQFF | ASSLAQNSGNTLY | 0.6529919 |
| TGKGIVE | ASSLAQNSGNTLY | 0.29902798 |
| LISNASDA | ASSLAQNSGNTLY | 0.24783042 |
| AELRTLGELTKIAKAHGAQ | ASSLAQNSGNTLY | 0.4880531 |
| APNAVAAAEPSW | ASSLAQNSGNTLY | 0.6654979 |
| APQVAGAATAVL | ASSLAQNSGNTLY | 0.19214779 |
| ARRDPAAAAAILPTDA | ASSLAQNSGNTLY | 0.82892114 |
| DAAARDVLAEAGLAEYF | ASSLAQNSGNTLY | 0.782544 |
| DDQIVLSAKANLAKVEQ | ASSLAQNSGNTLY | 0.791875 |
| DELLPDTGEDSDEDGH | ASSLAQNSGNTLY | 0.47012782 |
| ELPEAPANRATVPQD | ASSLAQNSGNTLY | 0.6288123 |
| FGQSKIERVTGIG | ASSLAQNSGNTLY | 0.5851745 |
| GAGGTGSTAGGAGGAGGAGGLY | ASSLAQNSGNTLY | 0.7975747 |
| GHGGTGAAGGNGGTGGAGGSGI | ASSLAQNSGNTLY | 0.5056749 |
| GNGGNGGAGGEGGPGGAGGAGA | ASSLAQNSGNTLY | 0.31902635 |
| GSTAGNGGNGGNATNGTIAGQP | ASSLAQNSGNTLY | 0.5072749 |
| GTGGDGGNGGNGGTGG | ASSLAQNSGNTLY | 0.55035293 |
| GVRRPAGIPADAPV | ASSLAQNSGNTLY | 0.84068507 |
| GYRQVTDAHLLTLARR | ASSLAQNSGNTLY | 0.817392 |
| IAKMGQDGHDRG | ASSLAQNSGNTLY | 0.80271757 |
| KAEPSIEIQYT | ASSLAQNSGNTLY | 0.82120705 |
| KPVRIRMAADEDTVAVT | ASSLAQNSGNTLY | 0.21883856 |
| MPPRIAGMRLLVIKPEP | ASSLAQNSGNTLY | 0.5142838 |
| MKFYAVALFRSRGVPPTR | ASSLAQNSGNTLY | 0.84551156 |
| NAGGFGILGAVAHSP | ASSLAQNSGNTLY | 0.7365117 |
| NEGREITLAMAEM | ASSLAQNSGNTLY | 0.6934977 |
| NELVEWSHPLGHRWRWE | ASSLAQNSGNTLY | 0.8459869 |
| NLGAGGTGGNGGNPG | ASSLAQNSGNTLY | 0.5489033 |
| NTEWPKWETVMAADPVD | ASSLAQNSGNTLY | 0.613251 |
| PASAAPAETQP | ASSLAQNSGNTLY | 0.71881294 |
| QQSSDGKNGVRAAEITGE | ASSLAQNSGNTLY | 0.7227999 |
| RAPDAVTTSWWKE | ASSLAQNSGNTLY | 0.7033607 |

| | | |
|---|---|---|
| SGFGNVGAHTSGWFNQ | ASSLAQNSGNTLY | 0.45390254 |
| SGVAGAVGQAASVGGLKV | ASSLAQNSGNTLY | 0.7148591 |
| SNGGAGTGSAGGNGGT | ASSLAQNSGNTLY | 0.87024105 |
| TGGAGGEGGAAG | ASSLAQNSGNTLY | 0.71508336 |
| TGGDGGAAGAGGAPGHG | ASSLAQNSGNTLY | 0.8374418 |
| TMPPLLAAIANAGRHGV | ASSLAQNSGNTLY | 0.76422364 |
| TSSTAGNSGGA | ASSLAQNSGNTLY | 0.74676585 |
| VPATVPVGILC | ASSLAQNSGNTLY | 0.57615066 |
| VQRPDSGGVVLTGRIS | ASSLAQNSGNTLY | 0.6831644 |
| WTPNPVPATKGVPAGLVQAT | ASSLAQNSGNTLY | 0.73381424 |
| YALKRALAKFSDRFDVVII | ASSLAQNSGNTLY | 0.6273792 |
| YLIGSVIV | ASSLAQNSGNTLY | 0.63783026 |
| YGQLVAVID | ASSLAQNSGNTLY | 0.61418456 |
| VTVGGMTLPA | ASSLAQNSGNTLY | 0.87085867 |
| VLALPAV | ASSLAQNSGNTLY | 0.76345074 |
| VIPPTLNYET | ASSLAQNSGNTLY | 0.79971766 |
| VGMGGMHPSSG | ASSLAQNSGNTLY | 0.5334166 |
| VGLAGGAATA | ASSLAQNSGNTLY | 0.6936216 |
| VGGAGGAGGSGP | ASSLAQNSGNTLY | 0.80309224 |
| VFAAFMV | ASSLAQNSGNTLY | 0.3794563 |
| TLYDKLR | ASSLAQNSGNTLY | 0.8478205 |
| TLAVFTVL | ASSLAQNSGNTLY | 0.5967001 |
| TGSVQPARV | ASSLAQNSGNTLY | 0.5700392 |
| TGGGAGSGGT | ASSLAQNSGNTLY | 0.754668 |
| SHLETSL | ASSLAQNSGNTLY | 0.8301312 |
| SFTTGMT | ASSLAQNSGNTLY | 0.6140135 |
| QVEVAKRMGVRQPTVS | ASSLAQNSGNTLY | 0.6919285 |
| QGSGGPALAM | ASSLAQNSGNTLY | 0.5584574 |
| MLAPSNP | ASSLAQNSGNTLY | 0.7925433 |
| LVNNAAI | ASSLAQNSGNTLY | 0.84444267 |
| LTGEINTLA | ASSLAQNSGNTLY | 0.5586281 |
| LSIFGPV | ASSLAQNSGNTLY | 0.85458493 |
| LLALLANL | ASSLAQNSGNTLY | 0.69570076 |
| LISNASDALDKLR | ASSLAQNSGNTLY | 0.30138996 |
| LINEAVP | ASSLAQNSGNTLY | 0.6469606 |
| LGYAFSGL | ASSLAQNSGNTLY | 0.8222681 |
| LDFVAVAL | ASSLAQNSGNTLY | 0.83717275 |
| LAQPGAE | ASSLAQNSGNTLY | 0.8437773 |
| ISNLGAVTGF | ASSLAQNSGNTLY | 0.7065301 |
| HQEDPGRI | ASSLAQNSGNTLY | 0.5851151 |
| GTPGEPTP | ASSLAQNSGNTLY | 0.85081244 |
| GFFNSGS | ASSLAQNSGNTLY | 0.8441937 |
| GAGGGGLTAGAN | ASSLAQNSGNTLY | 0.81203926 |
| FVDALVNM | ASSLAQNSGNTLY | 0.53871363 |
| FLTVAPEL | ASSLAQNSGNTLY | 0.40448445 |
| FETLTAL | ASSLAQNSGNTLY | 0.66250837 |
| DVINPIEL | ASSLAQNSGNTLY | 0.8342129 |
| APVSGAPV | ASSLAQNSGNTLY | 0.41604182 |
| ALVGGLAW | ASSLAQNSGNTLY | 0.76460254 |
| ALAAAAATAI | ASSLAQNSGNTLY | 0.7744818 |
| AFGLGLLL | ASSLAQNSGNTLY | 0.33135197 |

| | | |
|---|---|---|
| AFFAAASL | ASSLAQNSGNTLY | 0.7608217 |
| AAAALPAFSPP | ASSLAQNSGNTLY | 0.84835 |
| VVLAPAL | ASSLAQNSGNTLY | 0.63463414 |
| TIGHVDHGKTTL | ASSLAQNSGNTLY | 0.6256467 |
| LISNASDALDK | ASSLAQNSGNTLY | 0.8335042 |
| LISNASDAL | ASSLAQNSGNTLY | 0.87238884 |
| ISNASDALDK | ASSLAQNSGNTLY | 0.8442615 |
| NQAETLVYQTE | ASSLAQNSGNTLY | 0.2908461 |
| TGGTGGTAGAG | ASSLAQNSGNTLY | 0.22435571 |
| TGSTAGGAGGA | ASSLAQNSGNTLY | 0.5399587 |
| DPLAVPVAAGPV | ASSLAQNSGNTLY | 0.78720796 |
| GGTGGTPGNGGN | ASSLAQNSGNTLY | 0.5712918 |
| PAGLAGVDPATL | ASSLAQNSGNTLY | 0.5173637 |
| GNNLVVEAPHAGASV | ASSLAQNSGNTLY | 0.8128672 |
| KVTRIMSASTLEQPA | ASSLAQNSGNTLY | 0.8365128 |
| LGFALPIALPRRTVV | ASSLAQNSGNTLY | 0.44587862 |
| TRLGNLTARKLVVLV | ASSLAQNSGNTLY | 0.32679296 |
| WSQQRRLIYFPSAGP | ASSLAQNSGNTLY | 0.7937238 |
| GPNGNPDPMAAAVDIR | ASSLAQNSGNTLY | 0.78759015 |
| TQRCVSVADLRAAASR | ASSLAQNSGNTLY | 0.6086024 |
| QFVLDVAEHPVEVLTETV | ASSLAQNSGNTLY | 0.7405156 |
| GGAGGTGGAGGNGGTQAGDG | ASSLAQNSGNTLY | 0.46951556 |
| PLLATLRAIAVDVDAKPAQV | ASSLAQNSGNTLY | 0.3323188 |
| APKFRTGGYMIDFWGVGYQVAK | ASSLAQNSGNTLY | 0.8071713 |
| GAGGAGGAGGAAGGPGGTGGTG | ASSLAQNSGNTLY | 0.4661061 |
| IGALGLLATDVAPAGSVAFGVA | ASSLAQNSGNTLY | 0.8550724 |
| AGGAGGIGGNGGAGGAGGVLMG | ASSLAQNSGNTLY | 0.5565444 |
| ADELIGLL | ASSLAQNSGNTLY | 0.77819204 |
| ADRALKA | ASSLAQNSGNTLY | 0.77784026 |
| AGAVGQVGGAGGA | ASSLAQNSGNTLY | 0.73622006 |
| AGGGGGLGGAP | ASSLAQNSGNTLY | 0.67744064 |
| AGGLLYGNGGAG | ASSLAQNSGNTLY | 0.5950111 |
| AGLESIGL | ASSLAQNSGNTLY | 0.44780397 |
| AGLKTGII | ASSLAQNSGNTLY | 0.23181956 |
| ASLAALLL | ASSLAQNSGNTLY | 0.83010775 |
| ASVINISSI | ASSLAQNSGNTLY | 0.14566031 |
| ATPPGPA | ASSLAQNSGNTLY | 0.8394202 |
| ATVASALV | ASSLAQNSGNTLY | 0.78344643 |
| ATVASLIL | ASSLAQNSGNTLY | 0.53276 |
| AVNGPVLIRH | ASSLAQNSGNTLY | 0.7535014 |
| DGAAGAVGI | ASSLAQNSGNTLY | 0.578176 |
| DPLLHGI | ASSLAQNSGNTLY | 0.43752086 |
| ELGAALGAL | ASSLAQNSGNTLY | 0.7944381 |
| ELGRTQAV | ASSLAQNSGNTLY | 0.7813701 |
| FVAGGYLS | ASSLAQNSGNTLY | 0.5449711 |
| GASGGAGGAGGW | ASSLAQNSGNTLY | 0.73152804 |
| GGAGAGGDGGAGGI | ASSLAQNSGNTLY | 0.77382547 |
| GTGHNGSQPGMGGNG | ASSLAQNSGNTLY | 0.50239414 |
| GYAAASASAS | ASSLAQNSGNTLY | 0.6094083 |
| HQVDVAAVQQG | ASSLAQNSGNTLY | 0.8287902 |
| IGAAIAHT | ASSLAQNSGNTLY | 0.66347516 |

| | | |
|---|---|---|
| IGMIEVAQL | ASSLAQNSGNTLY | 0.5608455 |
| IRLAPEHTSAMD | ASSLAQNSGNTLY | 0.79138064 |
| IYNVSVL | ASSLAQNSGNTLY | 0.48847702 |
| KQGFLGM | ASSLAQNSGNTLY | 0.82117057 |
| LESVITQL | ASSLAQNSGNTLY | 0.6984106 |
| LGLGALVVV | ASSLAQNSGNTLY | 0.59038615 |
| LIGNGGNGGSGGI | ASSLAQNSGNTLY | 0.7949915 |
| LNAINAPTL | ASSLAQNSGNTLY | 0.35061574 |
| LQLLNTGV | ASSLAQNSGNTLY | 0.4762047 |
| LQTSIEF | ASSLAQNSGNTLY | 0.6127765 |
| MSFAVSEL | ASSLAQNSGNTLY | 0.71330535 |
| NTGALGL | ASSLAQNSGNTLY | 0.8276532 |
| PPGAGGAGGAGTT | ASSLAQNSGNTLY | 0.55721855 |
| QLVGLGI | ASSLAQNSGNTLY | 0.7582557 |
| RFLEQVHQL | ASSLAQNSGNTLY | 0.54872316 |
| RGVGYVLR | ASSLAQNSGNTLY | 0.43404216 |
| SATGVFAGVI | ASSLAQNSGNTLY | 0.5930644 |
| SLPAITLP | ASSLAQNSGNTLY | 0.5597892 |
| STALAFTV | ASSLAQNSGNTLY | 0.5484619 |
| SVLTVGLGK | ASSLAQNSGNTLY | 0.7793773 |
| TALAGAGSGI | ASSLAQNSGNTLY | 0.515805 |
| TGLPFTGI | ASSLAQNSGNTLY | 0.6314724 |
| VAEHVTV | ASSLAQNSGNTLY | 0.8458369 |
| VDIVLNSL | ASSLAQNSGNTLY | 0.594406 |
| VLTGGGAVL | ASSLAQNSGNTLY | 0.7761271 |
| VNVGLAYGI | ASSLAQNSGNTLY | 0.8021209 |
| VSSAAVGAV | ASSLAQNSGNTLY | 0.78895116 |
| VTSVYEAGV | ASSLAQNSGNTLY | 0.619547 |
| WVALDRGA | ASSLAQNSGNTLY | 0.6351752 |
| YLVGSVVI | ASSLAQNSGNTLY | 0.8540768 |
| GPPGTGKTLL | ASSLAQNSGNTLY | 0.8095379 |
| GVGMTKF | ASSLAQNSGNTLY | 0.7220303 |
| IGIGGSDL | ASSLAQNSGNTLY | 0.7968961 |
| IGTLLAEL | ASSLAQNSGNTLY | 0.801741 |
| IYTSGTT | ASSLAQNSGNTLY | 0.81205916 |
| LSFPAGV | ASSLAQNSGNTLY | 0.7041569 |
| LYTSGTT | ASSLAQNSGNTLY | 0.3441536 |
| VGAFPFGF | ASSLAQNSGNTLY | 0.57819086 |
| VGAGTVEF | ASSLAQNSGNTLY | 0.7853062 |
| VGKSTLLL | ASSLAQNSGNTLY | 0.60351616 |
| AALAHAGGGIV | ASSLAQNSGNTLY | 0.30257666 |
| AALVAVSLL | ASSLAQNSGNTLY | 0.83353627 |
| AAVSRAVAAL | ASSLAQNSGNTLY | 0.8697578 |
| AGGSSGIGGF | ASSLAQNSGNTLY | 0.46791065 |
| AGLTVANANG | ASSLAQNSGNTLY | 0.86148083 |
| AGLVGAVL | ASSLAQNSGNTLY | 0.3762269 |
| ALSQAFTV | ASSLAQNSGNTLY | 0.7613892 |
| ALTFSVSL | ASSLAQNSGNTLY | 0.82455236 |
| ALTGAALVAL | ASSLAQNSGNTLY | 0.568511 |
| ASFSIGAGVL | ASSLAQNSGNTLY | 0.7329324 |
| DGSVYKI | ASSLAQNSGNTLY | 0.56547093 |

| | | |
|---|---|---|
| EQLAEIV | ASSLAQNSGNTLY | 0.6850935 |
| FAVSELTVM | ASSLAQNSGNTLY | 0.8437421 |
| GAAPLAAGL | ASSLAQNSGNTLY | 0.55043286 |
| GADGGDGGSGGAGG | ASSLAQNSGNTLY | 0.6803749 |
| GFGLDTGGAGG | ASSLAQNSGNTLY | 0.57639563 |
| GLASGGLL | ASSLAQNSGNTLY | 0.4611417 |
| GRIAGAAVLL | ASSLAQNSGNTLY | 0.69931245 |
| GSAGSLNTSF | ASSLAQNSGNTLY | 0.8112752 |
| GTGQIGI | ASSLAQNSGNTLY | 0.4990821 |
| GTLIALIL | ASSLAQNSGNTLY | 0.83861476 |
| IAATMGFVVG | ASSLAQNSGNTLY | 0.56754476 |
| IFAQALL | ASSLAQNSGNTLY | 0.8685544 |
| IGAAVALGF | ASSLAQNSGNTLY | 0.8319572 |
| IGLTPLAL | ASSLAQNSGNTLY | 0.7851223 |
| ILGSNTSTL | ASSLAQNSGNTLY | 0.5803661 |
| IVFSLTVL | ASSLAQNSGNTLY | 0.8457215 |
| LASKATGF | ASSLAQNSGNTLY | 0.57470953 |
| LASVALIL | ASSLAQNSGNTLY | 0.7425648 |
| LFAVGLTL | ASSLAQNSGNTLY | 0.8364687 |
| LFNGGIGTYI | ASSLAQNSGNTLY | 0.39864853 |
| LTLGAGII | ASSLAQNSGNTLY | 0.74806404 |
| LVAAALGDI | ASSLAQNSGNTLY | 0.69421375 |
| NTAFVQLGI | ASSLAQNSGNTLY | 0.6608616 |
| QLFSGAEL | ASSLAQNSGNTLY | 0.55947447 |
| SAATFITL | ASSLAQNSGNTLY | 0.34403756 |
| SNYGLANLGV | ASSLAQNSGNTLY | 0.5558378 |
| SYGGAYAVM | ASSLAQNSGNTLY | 0.5565402 |
| TTGVGGLTLG | ASSLAQNSGNTLY | 0.74025995 |
| TTQTIYI | ASSLAQNSGNTLY | 0.66444767 |
| VAAGAGLGAVI | ASSLAQNSGNTLY | 0.50011015 |
| VAAVVTLL | ASSLAQNSGNTLY | 0.7399659 |
| VGFLGGLI | ASSLAQNSGNTLY | 0.74270004 |
| VGVIPILL | ASSLAQNSGNTLY | 0.3595479 |
| VIFASSLI | ASSLAQNSGNTLY | 0.59289205 |
| VKGGDGGAAAG | ASSLAQNSGNTLY | 0.52991813 |
| VTKTFTNI | ASSLAQNSGNTLY | 0.6062565 |
| VVAAAILL | ASSLAQNSGNTLY | 0.26630962 |
| YAALDVEL | ASSLAQNSGNTLY | 0.69537055 |
| LASVALLL | ASSLAQNSGNTLY | 0.501823 |
| LDAVHRV | ASSLAQNSGNTLY | 0.23233779 |
| QGLSSSIF | ASSLAQNSGNTLY | 0.66123366 |
| RGTFANIRL | ASSLAQNSGNTLY | 0.5538246 |
| RQLWWGHRI | ASSLAQNSGNTLY | 0.63593066 |
| YGPPGTGKTLL | ASSLAQNSGNTLY | 0.73643154 |
| AALVGAPANVA | ASSLAQNSGNTLY | 0.5264649 |
| AATFRRTE | ASSLAQNSGNTLY | 0.4620502 |
| AGALGGGI | ASSLAQNSGNTLY | 0.72190255 |
| AGGAGGAGGIGAG | ASSLAQNSGNTLY | 0.73287284 |
| AGSGALGL | ASSLAQNSGNTLY | 0.4320418 |
| AQIADQI | ASSLAQNSGNTLY | 0.68440515 |
| ASAVAVTYL | ASSLAQNSGNTLY | 0.7417462 |

| | | |
|---|---|---|
| ASTVLGGTGG | ASSLAQNSGNTLY | 0.67159116 |
| AVGGEPAK | ASSLAQNSGNTLY | 0.73334765 |
| CEQGTFGGTC | ASSLAQNSGNTLY | 0.7398639 |
| DEGKAPAG | ASSLAQNSGNTLY | 0.41710567 |
| DTVVFPAL | ASSLAQNSGNTLY | 0.73281264 |
| FEGITMAG | ASSLAQNSGNTLY | 0.74800384 |
| FPLVVQD | ASSLAQNSGNTLY | 0.51291955 |
| GAFTTPQ | ASSLAQNSGNTLY | 0.66759384 |
| GFGGFGTAGGVGGA | ASSLAQNSGNTLY | 0.3835209 |
| GFLAQFL | ASSLAQNSGNTLY | 0.5746515 |
| GGAGGAGGNAGVA | ASSLAQNSGNTLY | 0.33780476 |
| GGVGTAGAGGAGGAGG | ASSLAQNSGNTLY | 0.7122065 |
| GNGGGGDAGNAGSGGN | ASSLAQNSGNTLY | 0.72657335 |
| GTGGAGGAGNDAGS | ASSLAQNSGNTLY | 0.6682659 |
| GTTNTGWFNS | ASSLAQNSGNTLY | 0.42313945 |
| IFMGPLFTL | ASSLAQNSGNTLY | 0.3881238 |
| ILAHAIEAI | ASSLAQNSGNTLY | 0.6704793 |
| IRSAIREAL | ASSLAQNSGNTLY | 0.4395995 |
| ITFAGGG | ASSLAQNSGNTLY | 0.28643584 |
| ITGMSGHQEV | ASSLAQNSGNTLY | 0.6286496 |
| IVAGLACTL | ASSLAQNSGNTLY | 0.7421484 |
| IVGGGLAGA | ASSLAQNSGNTLY | 0.7371077 |
| IVTGVLNI | ASSLAQNSGNTLY | 0.5055484 |
| LAQQGCI | ASSLAQNSGNTLY | 0.75063676 |
| LCLHVEN | ASSLAQNSGNTLY | 0.7062706 |
| LDAISDGS | ASSLAQNSGNTLY | 0.71917665 |
| LGGFAAML | ASSLAQNSGNTLY | 0.57822376 |
| LGGKNPMI | ASSLAQNSGNTLY | 0.43941808 |
| LGPKGVAAIL | ASSLAQNSGNTLY | 0.7421801 |
| LMVGVSI | ASSLAQNSGNTLY | 0.6475266 |
| LVGAAFVL | ASSLAQNSGNTLY | 0.53591084 |
| LVLAVAPL | ASSLAQNSGNTLY | 0.72619504 |
| MAAIAVIDAA | ASSLAQNSGNTLY | 0.6676354 |
| NNGINAELQ | ASSLAQNSGNTLY | 0.51514953 |
| RGEPLVLVHG | ASSLAQNSGNTLY | 0.662421 |
| RGGAGGAAGLI | ASSLAQNSGNTLY | 0.70304716 |
| RSYRMHPAV | ASSLAQNSGNTLY | 0.5035879 |
| SAAHLLAGQLA | ASSLAQNSGNTLY | 0.4318242 |
| SAAVGSY | ASSLAQNSGNTLY | 0.53549993 |
| SPHAVLHRI | ASSLAQNSGNTLY | 0.66066086 |
| SQIAKVL | ASSLAQNSGNTLY | 0.7374246 |
| TALGHLVP | ASSLAQNSGNTLY | 0.6447822 |
| TAPTTSGAA | ASSLAQNSGNTLY | 0.4828372 |
| TDGDDGNGGSPGT | ASSLAQNSGNTLY | 0.70789945 |
| TDGLGEN | ASSLAQNSGNTLY | 0.6838752 |
| TGGPPLPGQA | ASSLAQNSGNTLY | 0.65420675 |
| TIPAITI | ASSLAQNSGNTLY | 0.688871 |
| TIPGFTI | ASSLAQNSGNTLY | 0.62360656 |
| TSIASGALA | ASSLAQNSGNTLY | 0.6896486 |
| TVTLPGAAI | ASSLAQNSGNTLY | 0.725484 |
| VASGLSAALP | ASSLAQNSGNTLY | 0.6838141 |

| | | |
|---|---|---|
| VGAGVIGI | ASSLAQNSGNTLY | 0.7145668 |
| VGPKRFGV | ASSLAQNSGNTLY | 0.6021488 |
| VNVAFPD | ASSLAQNSGNTLY | 0.6013811 |
| VSMTNSGVS | ASSLAQNSGNTLY | 0.36187664 |
| VTDVLGDLSV | ASSLAQNSGNTLY | 0.4757827 |
| VVVVGAGFAGL | ASSLAQNSGNTLY | 0.6763551 |
| VYTVGTVV | ASSLAQNSGNTLY | 0.6877228 |
| WTTFSGP | ASSLAQNSGNTLY | 0.72131443 |
| YDELVEAF | ASSLAQNSGNTLY | 0.65800685 |
| YGTGGAGGNGG | ASSLAQNSGNTLY | 0.4579016 |
| YTNVMAV | ASSLAQNSGNTLY | 0.7225575 |
| FLEQVHQL | ASSLAQNSGNTLY | 0.7326325 |
| GLSLLAVAV | ASSLAQNSGNTLY | 0.42317843 |
| LEQVHQL | ASSLAQNSGNTLY | 0.60913175 |
| LNAAGAI | ASSLAQNSGNTLY | 0.7607254 |
| PAVHASL | ASSLAQNSGNTLY | 0.6967826 |
| TELEDDA | ASSLAQNSGNTLY | 0.575816 |
| TPPGPPN | ASSLAQNSGNTLY | 0.5612266 |
| TPPPPPPP | ASSLAQNSGNTLY | 0.74967706 |
| TQSPALI | ASSQTTNSDYT | 0.49096468 |
| EFAAAFPQ | ASSQTTNSDYT | 0.74301624 |
| LLSGNLDVL | ASSQTTNSDYT | 0.31936324 |
| GPANLGNYNI | ASSQTTNSDYT | 0.5171446 |
| TLGPTLP | ASSQTTNSDYT | 0.43931156 |
| VYAYSSCCK | ASSQTTNSDYT | 0.26524115 |
| APAPAPAPNI | ASSQTTNSDYT | 0.73003745 |
| PGGPVPIP | ASSQTTNSDYT | 0.46787512 |
| FLTVALEP | ASSQTTNSDYT | 0.71951103 |
| SQYFGANL | ASSQTTNSDYT | 0.5869674 |
| GGVKPAA | ASSQTTNSDYT | 0.72728634 |
| GGTGGAGGAGGW | ASSQTTNSDYT | 0.69242513 |
| GVGGVAGLGGAGG | ASSQTTNSDYT | 0.59375846 |
| VLLVLATTI | ASSQTTNSDYT | 0.41178498 |
| IGIFAAML | ASSQTTNSDYT | 0.7091851 |
| AVGDLID | ASSQTTNSDYT | 0.71886873 |
| NISPLPL | ASSQTTNSDYT | 0.67324317 |
| VTAVFVAI | ASSQTTNSDYT | 0.321568 |
| SGDDNDAAL | ASSQTTNSDYT | 0.62977254 |
| AVAALAHS | ASSQTTNSDYT | 0.6540503 |
| GGVTVMATP | ASSQTTNSDYT | 0.3403928 |
| AGLAANVRD | ASSQTTNSDYT | 0.26371577 |
| TNRPDLIDPALLRPG | ASSQTTNSDYT | 0.5601343 |
| LANLGVTAL | ASSQTTNSDYT | 0.64420223 |
| TGERIEL | ASSQTTNSDYT | 0.7307774 |
| NGEELAA | ASSQTTNSDYT | 0.7312269 |
| VLSLVWG | ASSQTTNSDYT | 0.6084325 |
| IGYALWL | ASSQTTNSDYT | 0.68521935 |
| LSLLLAPL | ASSQTTNSDYT | 0.6647772 |
| TLDEALK | ASSQTTNSDYT | 0.20986685 |
| TDGAGTAEGG | ASSQTTNSDYT | 0.4687208 |
| NAPGGAGGNGG | ASSQTTNSDYT | 0.38185078 |

| | | |
|---|---|---|
| VSVFMGMGGNF | ASSQTTNSDYT | 0.7285768 |
| LASYLAAL | ASSQTTNSDYT | 0.5154139 |
| LVGGTPAL | ASSQTTNSDYT | 0.56306076 |
| VRNADVL | ASSQTTNSDYT | 0.6695336 |
| TCSSPFLVSAM | ASSQTTNSDYT | 0.70536005 |
| VHITDAE | ASSQTTNSDYT | 0.66284627 |
| EYAAPSF | ASSQTTNSDYT | 0.41177332 |
| VVDAFHGGL | ASSQTTNSDYT | 0.34163207 |
| ASVVDPQ | ASSQTTNSDYT | 0.7169777 |
| SYNEFGGLS | ASSQTTNSDYT | 0.44210452 |
| DPDGSVY | ASSQTTNSDYT | 0.44871816 |
| MVSTIHATE | ASSQTTNSDYT | 0.31255957 |
| LAAGVADPV | ASSQTTNSDYT | 0.6833768 |
| TKEQIAA | ASSQTTNSDYT | 0.3344465 |
| SLASPAAL | ASSQTTNSDYT | 0.7088732 |
| IGLVFYGV | ASSQTTNSDYT | 0.5375299 |
| FLLDASNL | ASSQTTNSDYT | 0.63965136 |
| YQVGNFP | ASSQTTNSDYT | 0.75830024 |
| ESLLSQST | ASSQTTNSDYT | 0.7119498 |
| LFTVGFAAVG | ASSQTTNSDYT | 0.6368692 |
| ATPPTPT | ASSQTTNSDYT | 0.5086498 |
| TDLATLG | ASSQTTNSDYT | 0.42584825 |
| SAGGTGR | ASSQTTNSDYT | 0.51199186 |
| GFSNSGS | ASSQTTNSDYT | 0.53243184 |
| GGNGGSGGTSV | ASSQTTNSDYT | 0.75194204 |
| GTSVATGGAGN | ASSQTTNSDYT | 0.60728055 |
| NGGNGGSGGTS | ASSQTTNSDYT | 0.3811176 |
| AVVKFPH | ASSQTTNSDYT | 0.65739083 |
| TPPPPPP | ASSQTTNSDYT | 0.75024295 |
| AAPAAAAAPA | ASSQTTNSDYT | 0.75766194 |
| IALNEDL | ASSQTTNSDYT | 0.43854123 |
| IVNVSSM | ASSQTTNSDYT | 0.6007104 |
| TGIPPAPR | ASSQTTNSDYT | 0.5492147 |
| FDLGGGTF | ASSQTTNSDYT | 0.6693573 |
| FDLGGGTFDVS | ASSQTTNSDYT | 0.4042563 |
| NEPTAAA | ASSQTTNSDYT | 0.47434396 |
| NEPTAAALA | ASSQTTNSDYT | 0.67810464 |
| GSIINISSI | ASSQTTNSDYT | 0.764207 |
| VAPGNAGTA | ASSQTTNSDYT | 0.5379666 |
| TGGGASLE | ASSQTTNSDYT | 0.3699497 |
| VAVYNHYKRI | ASSQTTNSDYT | 0.6338187 |
| TDPAPATT | ASSQTTNSDYT | 0.45747578 |
| TNRPDILDPALLRPG | ASSQTTNSDYT | 0.5589242 |
| LTTTTTS | ASSQTTNSDYT | 0.49890944 |
| PTGSGKTL | ASSQTTNSDYT | 0.39283296 |
| TGSGKTL | ASSQTTNSDYT | 0.62763023 |
| TIGHVDHGK | ASSQTTNSDYT | 0.7586416 |
| TIGHVDHGKTT | ASSQTTNSDYT | 0.66382885 |
| GHVDHGKTT | ASSQTTNSDYT | 0.68945336 |
| AQFEHTLL | ASSQTTNSDYT | 0.7502707 |
| IVNQDEL | ASSQTTNSDYT | 0.71986645 |

| | | |
|---|---|---|
| TNGSQFF | ASSQTTNSDYT | 0.7480327 |
| TGKGIVE | ASSQTTNSDYT | 0.7228248 |
| LISNASDA | ASSQTTNSDYT | 0.6022618 |
| AELRTLGELTKIAKAHGAQ | ASSQTTNSDYT | 0.5488256 |
| APNAVAAAEPSW | ASSQTTNSDYT | 0.739643 |
| APQVAGAATAVL | ASSQTTNSDYT | 0.6315254 |
| ARRDPAAAAAILPTDA | ASSQTTNSDYT | 0.6553222 |
| DAAARDVLAEAGLAEYF | ASSQTTNSDYT | 0.4887805 |
| DDQIVLSAKANLAKVEQ | ASSQTTNSDYT | 0.7420653 |
| DELLPDTGEDSDEDGH | ASSQTTNSDYT | 0.5373285 |
| ELPEAPANRATVPQD | ASSQTTNSDYT | 0.73745036 |
| FGQSKIERVTGIG | ASSQTTNSDYT | 0.27750838 |
| GAGGTGSTAGGAGGAGGAGGLY | ASSQTTNSDYT | 0.44815004 |
| GHGGTGAAGGNGGTGGAGGSGI | ASSQTTNSDYT | 0.739028 |
| GNGGNGGAGGEGGPGGAGGAGA | ASSQTTNSDYT | 0.21589588 |
| GSTAGNGGNGGNATNGTIAGQP | ASSQTTNSDYT | 0.26171106 |
| GTGGDGGNGGNGGTGG | ASSQTTNSDYT | 0.56953907 |
| GVRRPAGIPADAPV | ASSQTTNSDYT | 0.73896825 |
| GYRQVTDAHLLTLARR | ASSQTTNSDYT | 0.2262594 |
| IAKMGQDGHDRG | ASSQTTNSDYT | 0.4446006 |
| KAEPSIEIQYT | ASSQTTNSDYT | 0.3382471 |
| KPVRIRMAADEDTVAVT | ASSQTTNSDYT | 0.7045727 |
| MPPRIAGMRLLVIKPEP | ASSQTTNSDYT | 0.6397414 |
| MKFYAVALFRSRGVPPTR | ASSQTTNSDYT | 0.3113669 |
| NAGGFGILGAVAHSP | ASSQTTNSDYT | 0.7105604 |
| NEGREITLAMAEM | ASSQTTNSDYT | 0.36341566 |
| NELVEWSHPLGHRWRWE | ASSQTTNSDYT | 0.64978015 |
| NLGAGGTGGNGGNPG | ASSQTTNSDYT | 0.71061516 |
| NTEWPKWETVMAADPVD | ASSQTTNSDYT | 0.7526565 |
| PASAAPAETQP | ASSQTTNSDYT | 0.36691722 |
| QQSSDGKNGVRAAEITGE | ASSQTTNSDYT | 0.6782412 |
| RAPDAVTTSWWKE | ASSQTTNSDYT | 0.5172489 |
| SGFGNVGAHTSGWFNQ | ASSQTTNSDYT | 0.7645744 |
| SGVAGAVGQAASVGGLKV | ASSQTTNSDYT | 0.61691046 |
| SNGGAGTGSAGGNGGT | ASSQTTNSDYT | 0.3456083 |
| TGGAGGEGGAAG | ASSQTTNSDYT | 0.7269319 |
| TGGDGGAAGAGGAPGHG | ASSQTTNSDYT | 0.70663404 |
| TMPPLLAAIANAGRHGV | ASSQTTNSDYT | 0.5515386 |
| TSSTAGNSGGA | ASSQTTNSDYT | 0.72608083 |
| VPATVPVGILC | ASSQTTNSDYT | 0.568655 |
| VQRPDSGGVVLTGRIS | ASSQTTNSDYT | 0.6387692 |
| WTPNPVPATKGVPAGLVQAT | ASSQTTNSDYT | 0.7389812 |
| YALKRALAKFSDRFDVVII | ASSQTTNSDYT | 0.71462107 |
| YLIGSVIV | ASSQTTNSDYT | 0.55467606 |
| YGQLVAVID | ASSQTTNSDYT | 0.7008021 |
| VTVGGMTLPA | ASSQTTNSDYT | 0.68431294 |
| VLALPAV | ASSQTTNSDYT | 0.4557037 |
| VIPPTLNYET | ASSQTTNSDYT | 0.75802684 |
| VGMGGMHPSSG | ASSQTTNSDYT | 0.529621 |
| VGLAGGAATA | ASSQTTNSDYT | 0.7192246 |
| VGGAGGAGGSGP | ASSQTTNSDYT | 0.7043552 |

| | | |
|---|---|---|
| VFAAFMV | ASSQTTNSDYT | 0.75058866 |
| TLYDKLR | ASSQTTNSDYT | 0.4117216 |
| TLAVFTVL | ASSQTTNSDYT | 0.6577859 |
| TGSVQPARV | ASSQTTNSDYT | 0.6675303 |
| TGGGAGSGGT | ASSQTTNSDYT | 0.7067673 |
| SHLETSL | ASSQTTNSDYT | 0.61532503 |
| SFTTGMT | ASSQTTNSDYT | 0.6664635 |
| QVEVAKRMGVRQPTVS | ASSQTTNSDYT | 0.6830697 |
| QGSGGPALAM | ASSQTTNSDYT | 0.7454963 |
| MLAPSNP | ASSQTTNSDYT | 0.5629894 |
| LVNNAAI | ASSQTTNSDYT | 0.66687465 |
| LTGEINTLA | ASSQTTNSDYT | 0.75827456 |
| LSIFGPV | ASSQTTNSDYT | 0.41940242 |
| LLALLANL | ASSQTTNSDYT | 0.668326 |
| LISNASDALDKLR | ASSQTTNSDYT | 0.6635175 |
| LINEAVP | ASSQTTNSDYT | 0.73416615 |
| LGYAFSGL | ASSQTTNSDYT | 0.611185 |
| LDFVAVAL | ASSQTTNSDYT | 0.7274282 |
| LAQPGAE | ASSQTTNSDYT | 0.4266659 |
| ISNLGAVTGF | ASSQTTNSDYT | 0.7060014 |
| HQEDPGRI | ASSQTTNSDYT | 0.59763527 |
| GTPGEPTP | ASSQTTNSDYT | 0.6738365 |
| GFFNSGS | ASSQTTNSDYT | 0.5564214 |
| GAGGGGLTAGAN | ASSQTTNSDYT | 0.7238736 |
| FVDALVNM | ASSQTTNSDYT | 0.7273721 |
| FLTVAPEL | ASSQTTNSDYT | 0.72381085 |
| FETLTAL | ASSQTTNSDYT | 0.2547683 |
| DVINPIEL | ASSQTTNSDYT | 0.7183792 |
| APVSGAPV | ASSQTTNSDYT | 0.7175862 |
| ALVGGLAW | ASSQTTNSDYT | 0.76246 |
| ALAAAAATAI | ASSQTTNSDYT | 0.53520286 |
| AFGLGLLL | ASSQTTNSDYT | 0.6885234 |
| AFFAAASL | ASSQTTNSDYT | 0.52326846 |
| AAAALPAFSPP | ASSQTTNSDYT | 0.628554 |
| VVLAPAL | ASSQTTNSDYT | 0.43347675 |
| TIGHVDHGKTTL | ASSQTTNSDYT | 0.52525145 |
| LISNASDALDK | ASSQTTNSDYT | 0.5979787 |
| LISNASDAL | ASSQTTNSDYT | 0.57984596 |
| ISNASDALDK | ASSQTTNSDYT | 0.7372815 |
| NQAETLVYQTE | ASSQTTNSDYT | 0.73545754 |
| TGGTGGTAGAG | ASSQTTNSDYT | 0.7258729 |
| TGSTAGGAGGA | ASSQTTNSDYT | 0.7397398 |
| DPLAVPVAAGPV | ASSQTTNSDYT | 0.75099075 |
| GGTGGTPGNGGN | ASSQTTNSDYT | 0.6967143 |
| PAGLAGVDPATL | ASSQTTNSDYT | 0.62546 |
| GNNLVVEAPHAGASV | ASSQTTNSDYT | 0.5865127 |
| KVTRIMSASTLEQPA | ASSQTTNSDYT | 0.43971097 |
| LGFALPIALPRRTVV | ASSQTTNSDYT | 0.5332379 |
| TRLGNLTARKLVVLV | ASSQTTNSDYT | 0.63115764 |
| WSQQRRLIYFPSAGP | ASSQTTNSDYT | 0.6817777 |
| GPNGNPDPMAAAVDIR | ASSQTTNSDYT | 0.21213162 |

| | | |
|---|---|---|
| TQRCVSVADLRAAASR | ASSQTTNSDYT | 0.69996697 |
| QFVLDVAEHPVEVLTETV | ASSQTTNSDYT | 0.7562736 |
| GGAGGTGGAGGNGGTQAGDG | ASSQTTNSDYT | 0.65167606 |
| PLLATLRAIAVDVDAKPAQV | ASSQTTNSDYT | 0.69452477 |
| APKFRTGGYMIDFWGVGYQVAK | ASSQTTNSDYT | 0.7181095 |
| GAGGAGGAGGAAGGPGGTGGTG | ASSQTTNSDYT | 0.5272262 |
| IGALGLLATDVAPAGSVAFGVA | ASSQTTNSDYT | 0.7075459 |
| AGGAGGIGGNGGAGGAGGVLMG | ASSQTTNSDYT | 0.5086456 |
| ADELIGLL | ASSQTTNSDYT | 0.7117399 |
| ADRALKA | ASSQTTNSDYT | 0.75034076 |
| AGAVGQVGGAGGA | ASSQTTNSDYT | 0.6852948 |
| AGGGGGLGGAP | ASSQTTNSDYT | 0.59626985 |
| AGGLLYGNGGAG | ASSQTTNSDYT | 0.6160158 |
| AGLESIGL | ASSQTTNSDYT | 0.348917 |
| AGLKTGII | ASSQTTNSDYT | 0.64197063 |
| ASLAALLL | ASSQTTNSDYT | 0.468797 |
| ASVINISSI | ASSQTTNSDYT | 0.65320444 |
| ATPPGPA | ASSQTTNSDYT | 0.43043077 |
| ATVASALV | ASSQTTNSDYT | 0.69736135 |
| ATVASLIL | ASSQTTNSDYT | 0.68644 |
| AVNGPVLIRH | ASSQTTNSDYT | 0.60092044 |
| DGAAGAVGI | ASSQTTNSDYT | 0.514385 |
| DPLLHGI | ASSQTTNSDYT | 0.6697048 |
| ELGAALGAL | ASSQTTNSDYT | 0.6038172 |
| ELGRTQAV | ASSQTTNSDYT | 0.4060092 |
| FVAGGYLS | ASSQTTNSDYT | 0.7248498 |
| GASGGAGGAGGW | ASSQTTNSDYT | 0.50376916 |
| GGAGAGGDGGAGGI | ASSQTTNSDYT | 0.66979086 |
| GTGHNGSQPGMGGNG | ASSQTTNSDYT | 0.73910904 |
| GYAAASASAS | ASSQTTNSDYT | 0.6250292 |
| HQVDVAAVQQG | ASSQTTNSDYT | 0.30371317 |
| IGAAIAHT | ASSQTTNSDYT | 0.6464845 |
| IGMIEVAQL | ASSQTTNSDYT | 0.61880773 |
| IRLAPEHTSAMD | ASSQTTNSDYT | 0.6237222 |
| IYNVSVL | ASSQTTNSDYT | 0.23571274 |
| KQGFLGM | ASSQTTNSDYT | 0.6752076 |
| LESVITQL | ASSQTTNSDYT | 0.44610676 |
| LGLGALVVV | ASSQTTNSDYT | 0.5330541 |
| LIGNGGNGGSGGI | ASSQTTNSDYT | 0.6042516 |
| LNAINAPTL | ASSQTTNSDYT | 0.74242944 |
| LQLLNTGV | ASSQTTNSDYT | 0.6281364 |
| LQTSIEF | ASSQTTNSDYT | 0.60518587 |
| MSFAVSEL | ASSQTTNSDYT | 0.69924134 |
| NTGALGL | ASSQTTNSDYT | 0.6123539 |
| PPGAGGAGGAGTT | ASSQTTNSDYT | 0.74881446 |
| QLVGLGI | ASSQTTNSDYT | 0.67938435 |
| RFLEQVHQL | ASSQTTNSDYT | 0.5389733 |
| RGVGYVLR | ASSQTTNSDYT | 0.66140795 |
| SATGVFAGVI | ASSQTTNSDYT | 0.45343214 |
| SLPAITLP | ASSQTTNSDYT | 0.3861379 |
| STALAFTV | ASSQTTNSDYT | 0.3083126 |

| | | |
|---|---|---|
| SVLTVGLGK | ASSQTTNSDYT | 0.5369998 |
| TALAGAGSGI | ASSQTTNSDYT | 0.30974105 |
| TGLPFTGI | ASSQTTNSDYT | 0.67474604 |
| VAEHVTV | ASSQTTNSDYT | 0.7041874 |
| VDIVLNSL | ASSQTTNSDYT | 0.5760447 |
| VLTGGGAVL | ASSQTTNSDYT | 0.31494093 |
| VNVGLAYGI | ASSQTTNSDYT | 0.520584 |
| VSSAAVGAV | ASSQTTNSDYT | 0.55639625 |
| VTSVYEAGV | ASSQTTNSDYT | 0.72278273 |
| WVALDRGA | ASSQTTNSDYT | 0.69575655 |
| YLVGSVVI | ASSQTTNSDYT | 0.7557826 |
| GPPGTGKTLL | ASSQTTNSDYT | 0.73383415 |
| GVGMTKF | ASSQTTNSDYT | 0.6345526 |
| IGIGGSDL | ASSQTTNSDYT | 0.7197212 |
| IGTLLAEL | ASSQTTNSDYT | 0.5622704 |
| IYTSGTT | ASSQTTNSDYT | 0.66983366 |
| LSFPAGV | ASSQTTNSDYT | 0.593753 |
| LYTSGTT | ASSQTTNSDYT | 0.6422982 |
| VGAFPFGF | ASSQTTNSDYT | 0.53239566 |
| VGAGTVEF | ASSQTTNSDYT | 0.53560644 |
| VGKSTLLL | ASSQTTNSDYT | 0.3207444 |
| AALAHAGGGIV | ASSQTTNSDYT | 0.5919099 |
| AALVAVSLL | ASSQTTNSDYT | 0.4300418 |
| AAVSRAVAAL | ASSQTTNSDYT | 0.29897785 |
| AGGSSGIGGF | ASSQTTNSDYT | 0.5192278 |
| AGLTVANANG | ASSQTTNSDYT | 0.75329924 |
| AGLVGAVL | ASSQTTNSDYT | 0.73919094 |
| ALSQAFTV | ASSQTTNSDYT | 0.5635268 |
| ALTFSVSL | ASSQTTNSDYT | 0.53159654 |
| ALTGAALVAL | ASSQTTNSDYT | 0.7024883 |
| ASFSIGAGVL | ASSQTTNSDYT | 0.44570732 |
| DGSVYKI | ASSQTTNSDYT | 0.6641737 |
| EQLAEIV | ASSQTTNSDYT | 0.6647369 |
| FAVSELTVM | ASSQTTNSDYT | 0.6931902 |
| GAAPLAAGL | ASSQTTNSDYT | 0.45722312 |
| GADGGDGGSGGAGG | ASSQTTNSDYT | 0.6966363 |
| GFGLDTGGAGG | ASSQTTNSDYT | 0.63610697 |
| GLASGGLL | ASSQTTNSDYT | 0.6179186 |
| GRIAGAAVLL | ASSQTTNSDYT | 0.32960695 |
| GSAGSLNTSF | ASSQTTNSDYT | 0.7097033 |
| GTGQIGI | ASSQTTNSDYT | 0.491481 |
| GTLIALIL | ASSQTTNSDYT | 0.40620428 |
| IAATMGFVVG | ASSQTTNSDYT | 0.25915092 |
| IFAQALL | ASSQTTNSDYT | 0.6280972 |
| IGAAVALGF | ASSQTTNSDYT | 0.7204627 |
| IGLTPLAL | ASSQTTNSDYT | 0.7401604 |
| ILGSNTSTL | ASSQTTNSDYT | 0.4964401 |
| IVFSLTVL | ASSQTTNSDYT | 0.73329926 |
| LASKATGF | ASSQTTNSDYT | 0.72183836 |
| LASVALIL | ASSQTTNSDYT | 0.6139095 |
| LFAVGLTL | ASSQTTNSDYT | 0.2924921 |

| | | |
|---|---|---|
| LFNGGIGTYI | ASSQTTNSDYT | 0.6808287 |
| LTLGAGII | ASSQTTNSDYT | 0.5952749 |
| LVAAALGDI | ASSQTTNSDYT | 0.7440504 |
| NTAFVQLGI | ASSQTTNSDYT | 0.6920319 |
| QLFSGAEL | ASSQTTNSDYT | 0.4418303 |
| SAATFITL | ASSQTTNSDYT | 0.68407273 |
| SNYGLANLGV | ASSQTTNSDYT | 0.6939267 |
| SYGGAYAVM | ASSQTTNSDYT | 0.3822634 |
| TTGVGGLTLG | ASSQTTNSDYT | 0.40150976 |
| TTQTIYI | ASSQTTNSDYT | 0.4953478 |
| VAAGAGLGAVI | ASSQTTNSDYT | 0.42211646 |
| VAAVVTLL | ASSQTTNSDYT | 0.6220323 |
| VGFLGGLI | ASSQTTNSDYT | 0.60512197 |
| VGVIPILL | ASSQTTNSDYT | 0.54634887 |
| VIFASSLI | ASSQTTNSDYT | 0.66751885 |
| VKGGDGGAAAG | ASSQTTNSDYT | 0.49876785 |
| VTKTFTNI | ASSQTTNSDYT | 0.5563312 |
| VVAAAILL | ASSQTTNSDYT | 0.61991596 |
| YAALDVEL | ASSQTTNSDYT | 0.6878029 |
| LASVALLL | ASSQTTNSDYT | 0.70203674 |
| LDAVHRV | ASSQTTNSDYT | 0.6289401 |
| QGLSSSIF | ASSQTTNSDYT | 0.6377939 |
| RGTFANIRL | ASSQTTNSDYT | 0.75749665 |
| RQLWWGHRI | ASSQTTNSDYT | 0.41168207 |
| YGPPGTGKTLL | ASSQTTNSDYT | 0.48262364 |
| AALVGAPANVA | ASSQTTNSDYT | 0.690604 |
| AATFRRTE | ASSQTTNSDYT | 0.52020544 |
| AGALGGGI | ASSQTTNSDYT | 0.71749157 |
| AGGAGGAGGIGAG | ASSQTTNSDYT | 0.66660154 |
| AGSGALGL | ASSQTTNSDYT | 0.69974554 |
| AQIADQI | ASSQTTNSDYT | 0.5590009 |
| ASAVAVTYL | ASSQTTNSDYT | 0.75633353 |
| ASTVLGGTGG | ASSQTTNSDYT | 0.7231488 |
| AVGGEPAK | ASSQTTNSDYT | 0.57508683 |
| CEQGTFGGTC | ASSQTTNSDYT | 0.7238963 |
| DEGKAPAG | ASSQTTNSDYT | 0.35585487 |
| DTVVFPAL | ASSQTTNSDYT | 0.72023404 |
| FEGITMAG | ASSQTTNSDYT | 0.6859695 |
| FPLVVQD | ASSQTTNSDYT | 0.53250897 |
| GAFTTPQ | ASSQTTNSDYT | 0.63847387 |
| GFGGFGTAGGVGGA | ASSQTTNSDYT | 0.72859484 |
| GFLAQFL | ASSQTTNSDYT | 0.74284387 |
| GGAGGAGGNAGVA | ASSQTTNSDYT | 0.34819466 |
| GGVGTAGAGGAGGAGG | ASSQTTNSDYT | 0.5703969 |
| GNGGGGDAGNAGSGGN | ASSQTTNSDYT | 0.32566506 |
| GTGGAGGAGNDAGS | ASSQTTNSDYT | 0.37251213 |
| GTTNTGWFNS | ASSQTTNSDYT | 0.33465594 |
| IFMGPLFTL | ASSQTTNSDYT | 0.6812151 |
| ILAHAIEAI | ASSQTTNSDYT | 0.5577149 |
| IRSAIREAL | ASSQTTNSDYT | 0.72869444 |
| ITFAGGG | ASSQTTNSDYT | 0.69297147 |

| | | |
|---|---|---|
| ITGMSGHQEV | ASSQTTNSDYT | 0.74578106 |
| IVAGLACTL | ASSQTTNSDYT | 0.44484544 |
| IVGGGLAGA | ASSQTTNSDYT | 0.6236276 |
| IVTGVLNI | ASSQTTNSDYT | 0.28593257 |
| LAQQGCI | ASSQTTNSDYT | 0.56962067 |
| LCLHVEN | ASSQTTNSDYT | 0.39633498 |
| LDAISDGS | ASSQTTNSDYT | 0.64346254 |
| LGGFAAML | ASSQTTNSDYT | 0.33081716 |
| LGGKNPMI | ASSQTTNSDYT | 0.75460076 |
| LGPKGVAAIL | ASSQTTNSDYT | 0.61152804 |
| LMVGVSI | ASSQTTNSDYT | 0.7219677 |
| LVGAAFVL | ASSQTTNSDYT | 0.40059805 |
| LVLAVAPL | ASSQTTNSDYT | 0.48502547 |
| MAAIAVIDAA | ASSQTTNSDYT | 0.6269878 |
| NNGINAELQ | ASSQTTNSDYT | 0.70234644 |
| RGEPLVLVHG | ASSQTTNSDYT | 0.48845577 |
| RGGAGGAAGLI | ASSQTTNSDYT | 0.26887918 |
| RSYRMHPAV | ASSQTTNSDYT | 0.41332793 |
| SAAHLLAGQLA | ASSQTTNSDYT | 0.41992384 |
| SAAVGSY | ASSQTTNSDYT | 0.74659896 |
| SPHAVLHRI | ASSQTTNSDYT | 0.7395363 |
| SQIAKVL | ASSQTTNSDYT | 0.67986774 |
| TALGHLVP | ASSQTTNSDYT | 0.7307856 |
| TAPTTSGAA | ASSQTTNSDYT | 0.22337915 |
| TDGDDGNGGSPGT | ASSQTTNSDYT | 0.48455036 |
| TDGLGEN | ASSQTTNSDYT | 0.75448793 |
| TGGPPLPGQA | ASSQTTNSDYT | 0.69379365 |
| TIPAITI | ASSQTTNSDYT | 0.6119095 |
| TIPGFTI | ASSQTTNSDYT | 0.7312833 |
| TSIASGALA | ASSQTTNSDYT | 0.5418494 |
| TVTLPGAAI | ASSQTTNSDYT | 0.6180616 |
| VASGLSAALP | ASSQTTNSDYT | 0.679111 |
| VGAGVIGI | ASSQTTNSDYT | 0.65026057 |
| VGPKRFGV | ASSQTTNSDYT | 0.6388048 |
| VNVAFPD | ASSQTTNSDYT | 0.33973503 |
| VSMTNSGVS | ASSQTTNSDYT | 0.66488314 |
| VTDVLGDLSV | ASSQTTNSDYT | 0.76305556 |
| VVVVGAGFAGL | ASSQTTNSDYT | 0.6506007 |
| VYTVGTVV | ASSQTTNSDYT | 0.7227943 |
| WTTFSGP | ASSQTTNSDYT | 0.6504911 |
| YDELVEAF | ASSQTTNSDYT | 0.69249344 |
| YGTGGAGGNGG | ASSQTTNSDYT | 0.5225794 |
| YTNVMAV | ASSQTTNSDYT | 0.4569984 |
| FLEQVHQL | ASSQTTNSDYT | 0.6417916 |
| GLSLLAVAV | ASSQTTNSDYT | 0.42740607 |
| LEQVHQL | ASSQTTNSDYT | 0.5980249 |
| LNAAGAI | ASSQTTNSDYT | 0.59004825 |
| PAVHASL | ASSQTTNSDYT | 0.76730824 |
| TELEDDA | ASSQTTNSDYT | 0.7084744 |
| TPPGPPN | ASSQTTNSDYT | 0.6875733 |
| TPPPPPPP | ASSQTTNSDYT | 0.5458057 |

| | | |
|---|---|---|
| TQSPALI | ASSERRGQANTEVF | 0.6036584 |
| EFAAAFPQ | ASSERRGQANTEVF | 0.7098955 |
| LLSGNLDVL | ASSERRGQANTEVF | 0.36460856 |
| GPANLGNYNI | ASSERRGQANTEVF | 0.7337941 |
| TLGPTLP | ASSERRGQANTEVF | 0.47486633 |
| VYAYSSCCK | ASSERRGQANTEVF | 0.6399703 |
| APAPAPAPNI | ASSERRGQANTEVF | 0.68253744 |
| PGGPVPIP | ASSERRGQANTEVF | 0.74535406 |
| FLTVALEP | ASSERRGQANTEVF | 0.6051794 |
| SQYFGANL | ASSERRGQANTEVF | 0.6560893 |
| GGVKPAA | ASSERRGQANTEVF | 0.5675148 |
| GGTGGAGGAGGW | ASSERRGQANTEVF | 0.67886865 |
| GVGGVAGLGGAGG | ASSERRGQANTEVF | 0.74816513 |
| VLLVLATTI | ASSERRGQANTEVF | 0.6626853 |
| IGIFAAML | ASSERRGQANTEVF | 0.7556447 |
| AVGDLID | ASSERRGQANTEVF | 0.6915144 |
| NISPLPL | ASSERRGQANTEVF | 0.2874165 |
| VTAVFVAI | ASSERRGQANTEVF | 0.5783434 |
| SGDDNDAAL | ASSERRGQANTEVF | 0.7286553 |
| AVAALAHS | ASSERRGQANTEVF | 0.74483776 |
| GGVTVMATP | ASSERRGQANTEVF | 0.7319925 |
| AGLAANVRD | ASSERRGQANTEVF | 0.6736114 |
| TNRPDLIDPALLRPG | ASSERRGQANTEVF | 0.5637139 |
| LANLGVTAL | ASSERRGQANTEVF | 0.74968684 |
| TGERIEL | ASSERRGQANTEVF | 0.75869095 |
| NGEELAA | ASSERRGQANTEVF | 0.7037257 |
| VLSLVWG | ASSERRGQANTEVF | 0.41325226 |
| IGYALWL | ASSERRGQANTEVF | 0.38224617 |
| LSLLLAPL | ASSERRGQANTEVF | 0.6847985 |
| TLDEALK | ASSERRGQANTEVF | 0.7452202 |
| TDGAGTAEGG | ASSERRGQANTEVF | 0.36650696 |
| NAPGGAGGNGG | ASSERRGQANTEVF | 0.6704899 |
| VSVFMGMGGNF | ASSERRGQANTEVF | 0.6886771 |
| LASYLAAL | ASSERRGQANTEVF | 0.28932637 |
| LVGGTPAL | ASSERRGQANTEVF | 0.59875643 |
| VRNADVL | ASSERRGQANTEVF | 0.74453926 |
| TCSSPFLVSAM | ASSERRGQANTEVF | 0.664374 |
| VHITDAE | ASSERRGQANTEVF | 0.429281 |
| EYAAPSF | ASSERRGQANTEVF | 0.73735595 |
| VVDAFHGGL | ASSERRGQANTEVF | 0.76212704 |
| ASVVDPQ | ASSERRGQANTEVF | 0.7499862 |
| SYNEFGGLS | ASSERRGQANTEVF | 0.3021027 |
| DPDGSVY | ASSERRGQANTEVF | 0.23784442 |
| MVSTIHATE | ASSERRGQANTEVF | 0.6447093 |
| LAAGVADPV | ASSERRGQANTEVF | 0.7040651 |
| TKEQIAA | ASSERRGQANTEVF | 0.5800158 |
| SLASPAAL | ASSERRGQANTEVF | 0.52069795 |
| IGLVFYGV | ASSERRGQANTEVF | 0.7050695 |
| FLLDASNL | ASSERRGQANTEVF | 0.74277794 |
| YQVGNFP | ASSERRGQANTEVF | 0.4019509 |
| ESLLSQST | ASSERRGQANTEVF | 0.330225 |

| | | |
|---|---|---|
| LFTVGFAAVG | ASSERRGQANTEVF | 0.71923447 |
| ATPPTPT | ASSERRGQANTEVF | 0.7192389 |
| TDLATLG | ASSERRGQANTEVF | 0.48552316 |
| SAGGTGR | ASSERRGQANTEVF | 0.6437865 |
| GFSNSGS | ASSERRGQANTEVF | 0.67085254 |
| GGNGGSGGTSV | ASSERRGQANTEVF | 0.625342 |
| GTSVATGGAGN | ASSERRGQANTEVF | 0.68124044 |
| NGGNGGSGGTS | ASSERRGQANTEVF | 0.3859294 |
| AVVKFPH | ASSERRGQANTEVF | 0.7410045 |
| TPPPPPP | ASSERRGQANTEVF | 0.39433777 |
| AAPAAAAAPA | ASSERRGQANTEVF | 0.6782963 |
| IALNEDL | ASSERRGQANTEVF | 0.6818536 |
| IVNVSSM | ASSERRGQANTEVF | 0.638718 |
| TGIPPAPR | ASSERRGQANTEVF | 0.5773221 |
| FDLGGGTF | ASSERRGQANTEVF | 0.44878274 |
| FDLGGGTFDVS | ASSERRGQANTEVF | 0.44447613 |
| NEPTAAA | ASSERRGQANTEVF | 0.3110084 |
| NEPTAAALA | ASSERRGQANTEVF | 0.7411304 |
| GSIINISSI | ASSERRGQANTEVF | 0.6989025 |
| VAPGNAGTA | ASSERRGQANTEVF | 0.72517526 |
| TGGGASLE | ASSERRGQANTEVF | 0.6991413 |
| VAVYNHYKRI | ASSERRGQANTEVF | 0.49376357 |
| TDPAPATT | ASSERRGQANTEVF | 0.6742251 |
| TNRPDILDPALLRPG | ASSERRGQANTEVF | 0.46303165 |
| LTTTTTS | ASSERRGQANTEVF | 0.5102099 |
| PTGSGKTL | ASSERRGQANTEVF | 0.57745814 |
| TGSGKTL | ASSERRGQANTEVF | 0.708341 |
| TIGHVDHGK | ASSERRGQANTEVF | 0.41711292 |
| TIGHVDHGKTT | ASSERRGQANTEVF | 0.6781425 |
| GHVDHGKTT | ASSERRGQANTEVF | 0.680243 |
| AQFEHTLL | ASSERRGQANTEVF | 0.42953473 |
| IVNQDEL | ASSERRGQANTEVF | 0.6566898 |
| TNGSQFF | ASSERRGQANTEVF | 0.732036 |
| TGKGIVE | ASSERRGQANTEVF | 0.64479136 |
| LISNASDA | ASSERRGQANTEVF | 0.4189471 |
| AELRTLGELTKIAKAHGAQ | ASSERRGQANTEVF | 0.6499768 |
| APNAVAAAEPSW | ASSERRGQANTEVF | 0.5282783 |
| APQVAGAATAVL | ASSERRGQANTEVF | 0.70374936 |
| ARRDPAAAAAILPTDA | ASSERRGQANTEVF | 0.638492 |
| DAAARDVLAEAGLAEYF | ASSERRGQANTEVF | 0.53173286 |
| DDQIVLSAKANLAKVEQ | ASSERRGQANTEVF | 0.71041685 |
| DELLPDTGEDSDEDGH | ASSERRGQANTEVF | 0.3356525 |
| ELPEAPANRATVPQD | ASSERRGQANTEVF | 0.4880218 |
| FGQSKIERVTGIG | ASSERRGQANTEVF | 0.527465 |
| GAGGTGSTAGGAGGAGGAGGLY | ASSERRGQANTEVF | 0.6615087 |
| GHGGTGAAGGNGGTGGAGGSGI | ASSERRGQANTEVF | 0.7143415 |
| GNGGNGGAGGEGGPGGAGGAGA | ASSERRGQANTEVF | 0.48079982 |
| GSTAGNGGNGGNATNGTIAGQP | ASSERRGQANTEVF | 0.6346611 |
| GTGGDGGNGGNGGTGG | ASSERRGQANTEVF | 0.57527477 |
| GVRRPAGIPADAPV | ASSERRGQANTEVF | 0.50272566 |
| GYRQVTDAHLLTLARR | ASSERRGQANTEVF | 0.6294968 |

| | | |
|---|---|---|
| IAKMGQDGHDRG | ASSERRGQANTEVF | 0.51546586 |
| KAEPSIEIQYT | ASSERRGQANTEVF | 0.34417364 |
| KPVRIRMAADEDTVAVT | ASSERRGQANTEVF | 0.67357373 |
| MPPRIAGMRLLVIKPEP | ASSERRGQANTEVF | 0.54401696 |
| MKFYAVALFRSRGVPPTR | ASSERRGQANTEVF | 0.56226766 |
| NAGGFGILGAVAHSP | ASSERRGQANTEVF | 0.74040097 |
| NEGREITLAMAEM | ASSERRGQANTEVF | 0.41152334 |
| NELVEWSHPLGHRWRWE | ASSERRGQANTEVF | 0.66030306 |
| NLGAGGTGGNGGNPG | ASSERRGQANTEVF | 0.6733376 |
| NTEWPKWETVMAADPVD | ASSERRGQANTEVF | 0.6899151 |
| PASAAPAETQP | ASSERRGQANTEVF | 0.6504392 |
| QQSSDGKNGVRAAEITGE | ASSERRGQANTEVF | 0.4689591 |
| RAPDAVTTSWWKE | ASSERRGQANTEVF | 0.74379516 |
| SGFGNVGAHTSGWFNQ | ASSERRGQANTEVF | 0.6988324 |
| SGVAGAVGQAASVGGLKV | ASSERRGQANTEVF | 0.7189481 |
| SNGGAGTGSAGGNGGT | ASSERRGQANTEVF | 0.7144221 |
| TGGAGGEGGAAG | ASSERRGQANTEVF | 0.70872104 |
| TGGDGGAAGAGGAPGHG | ASSERRGQANTEVF | 0.7285021 |
| TMPPLLAAIANAGRHGV | ASSERRGQANTEVF | 0.57052815 |
| TSSTAGNSGGA | ASSERRGQANTEVF | 0.31875023 |
| VPATVPVGILC | ASSERRGQANTEVF | 0.6384631 |
| VQRPDSGGVVLTGRIS | ASSERRGQANTEVF | 0.60614634 |
| WTPNPVPATKGVPAGLVQAT | ASSERRGQANTEVF | 0.47588152 |
| YALKRALAKFSDRFDVVII | ASSERRGQANTEVF | 0.29060787 |
| YLIGSVIV | ASSERRGQANTEVF | 0.7403448 |
| YGQLVAVID | ASSERRGQANTEVF | 0.763703 |
| VTVGGMTLPA | ASSERRGQANTEVF | 0.43153003 |
| VLALPAV | ASSERRGQANTEVF | 0.7580166 |
| VIPPTLNYET | ASSERRGQANTEVF | 0.36650348 |
| VGMGGMHPSSG | ASSERRGQANTEVF | 0.68529963 |
| VGLAGGAATA | ASSERRGQANTEVF | 0.7410432 |
| VGGAGGAGGSGP | ASSERRGQANTEVF | 0.41906118 |
| VFAAFMV | ASSERRGQANTEVF | 0.7061286 |
| TLYDKLR | ASSERRGQANTEVF | 0.48685098 |
| TLAVFTVL | ASSERRGQANTEVF | 0.5359421 |
| TGSVQPARV | ASSERRGQANTEVF | 0.74024683 |
| TGGGAGSGGT | ASSERRGQANTEVF | 0.54648364 |
| SHLETSL | ASSERRGQANTEVF | 0.65702593 |
| SFTTGMT | ASSERRGQANTEVF | 0.60611296 |
| QVEVAKRMGVRQPTVS | ASSERRGQANTEVF | 0.5142696 |
| QGSGGPALAM | ASSERRGQANTEVF | 0.661864 |
| MLAPSNP | ASSERRGQANTEVF | 0.7111432 |
| LVNNAAI | ASSERRGQANTEVF | 0.4678486 |
| LTGEINTLA | ASSERRGQANTEVF | 0.75049376 |
| LSIFGPV | ASSERRGQANTEVF | 0.4040947 |
| LLALLANL | ASSERRGQANTEVF | 0.75647104 |
| LISNASDALDKLR | ASSERRGQANTEVF | 0.7274682 |
| LINEAVP | ASSERRGQANTEVF | 0.6923878 |
| LGYAFSGL | ASSERRGQANTEVF | 0.5473558 |
| LDFVAVAL | ASSERRGQANTEVF | 0.74105126 |
| LAQPGAE | ASSERRGQANTEVF | 0.5700933 |

| | | |
|---|---|---|
| ISNLGAVTGF | ASSERRGQANTEVF | 0.67506254 |
| HQEDPGRI | ASSERRGQANTEVF | 0.72191316 |
| GTPGEPTP | ASSERRGQANTEVF | 0.36252198 |
| GFFNSGS | ASSERRGQANTEVF | 0.7263625 |
| GAGGGGLTAGAN | ASSERRGQANTEVF | 0.6164789 |
| FVDALVNM | ASSERRGQANTEVF | 0.64473224 |
| FLTVAPEL | ASSERRGQANTEVF | 0.50446767 |
| FETLTAL | ASSERRGQANTEVF | 0.3820093 |
| DVINPIEL | ASSERRGQANTEVF | 0.5211861 |
| APVSGAPV | ASSERRGQANTEVF | 0.46858215 |
| ALVGGLAW | ASSERRGQANTEVF | 0.6938649 |
| ALAAAAATAI | ASSERRGQANTEVF | 0.685801 |
| AFGLGLLL | ASSERRGQANTEVF | 0.49380094 |
| AFFAAASL | ASSERRGQANTEVF | 0.67929786 |
| AAAALPAFSPP | ASSERRGQANTEVF | 0.646133 |
| VVLAPAL | ASSERRGQANTEVF | 0.3357579 |
| TIGHVDHGKTTL | ASSERRGQANTEVF | 0.55294526 |
| LISNASDALDK | ASSERRGQANTEVF | 0.40863374 |
| LISNASDAL | ASSERRGQANTEVF | 0.55101323 |
| ISNASDALDK | ASSERRGQANTEVF | 0.2507143 |
| NQAETLVYQTE | ASSERRGQANTEVF | 0.66122514 |
| TGGTGGTAGAG | ASSERRGQANTEVF | 0.43192232 |
| TGSTAGGAGGA | ASSERRGQANTEVF | 0.23966402 |
| DPLAVPVAAGPV | ASSERRGQANTEVF | 0.6197325 |
| GGTGGTPGNGGN | ASSERRGQANTEVF | 0.48208606 |
| PAGLAGVDPATL | ASSERRGQANTEVF | 0.49157685 |
| GNNLVVEAPHAGASV | ASSERRGQANTEVF | 0.6610961 |
| KVTRIMSASTLEQPA | ASSERRGQANTEVF | 0.5244962 |
| LGFALPIALPRRTVV | ASSERRGQANTEVF | 0.42173174 |
| TRLGNLTARKLVVLV | ASSERRGQANTEVF | 0.63209736 |
| WSQQRRLIYFPSAGP | ASSERRGQANTEVF | 0.6977827 |
| GPNGNPDPMAAAVDIR | ASSERRGQANTEVF | 0.4342043 |
| TQRCVSVADLRAAASR | ASSERRGQANTEVF | 0.5556903 |
| QFVLDVAEHPVEVLTETV | ASSERRGQANTEVF | 0.74283266 |
| GGAGGTGGAGGNGGTQAGDG | ASSERRGQANTEVF | 0.59887266 |
| PLLATLRAIAVDVDAKPAQV | ASSERRGQANTEVF | 0.7080122 |
| APKFRTGGYMIDFWGVGYQVAK | ASSERRGQANTEVF | 0.7290742 |
| GAGGAGGAGGAAGGPGGTGGTG | ASSERRGQANTEVF | 0.4715097 |
| IGALGLLATDVAPAGSVAFGVA | ASSERRGQANTEVF | 0.70350325 |
| AGGAGGIGGNGGAGGAGGVLMG | ASSERRGQANTEVF | 0.69239306 |
| ADELIGLL | ASSERRGQANTEVF | 0.5780378 |
| ADRALKA | ASSERRGQANTEVF | 0.5751496 |
| AGAVGQVGGAGGA | ASSERRGQANTEVF | 0.37823254 |
| AGGGGGLGGAP | ASSERRGQANTEVF | 0.5399293 |
| AGGLLYGNGGAG | ASSERRGQANTEVF | 0.43491828 |
| AGLESIGL | ASSERRGQANTEVF | 0.6118213 |
| AGLKTGII | ASSERRGQANTEVF | 0.7391778 |
| ASLAALLL | ASSERRGQANTEVF | 0.5427422 |
| ASVINISSI | ASSERRGQANTEVF | 0.32054004 |
| ATPPGPA | ASSERRGQANTEVF | 0.31888145 |
| ATVASALV | ASSERRGQANTEVF | 0.670649 |

| | | |
|---|---|---|
| ATVASLIL | ASSERRGQANTEVF | 0.36883205 |
| AVNGPVLIRH | ASSERRGQANTEVF | 0.38898262 |
| DGAAGAVGI | ASSERRGQANTEVF | 0.4551482 |
| DPLLHGI | ASSERRGQANTEVF | 0.7039386 |
| ELGAALGAL | ASSERRGQANTEVF | 0.71242106 |
| ELGRTQAV | ASSERRGQANTEVF | 0.48099887 |
| FVAGGYLS | ASSERRGQANTEVF | 0.7181325 |
| GASGGAGGAGGW | ASSERRGQANTEVF | 0.6416866 |
| GGAGAGGDGGAGGI | ASSERRGQANTEVF | 0.7054005 |
| GTGHNGSQPGMGGNG | ASSERRGQANTEVF | 0.59984344 |
| GYAAASASAS | ASSERRGQANTEVF | 0.3968017 |
| HQVDVAAVQQG | ASSERRGQANTEVF | 0.67816645 |
| IGAAIAHT | ASSERRGQANTEVF | 0.53071165 |
| IGMIEVAQL | ASSERRGQANTEVF | 0.6543475 |
| IRLAPEHTSAMD | ASSERRGQANTEVF | 0.66389287 |
| IYNVSVL | ASSERRGQANTEVF | 0.57778805 |
| KQGFLGM | ASSERRGQANTEVF | 0.4491952 |
| LESVITQL | ASSERRGQANTEVF | 0.5623543 |
| LGLGALVVV | ASSERRGQANTEVF | 0.72404927 |
| LIGNGGNGGSGGI | ASSERRGQANTEVF | 0.39370543 |
| LNAINAPTL | ASSERRGQANTEVF | 0.42231798 |
| LQLLNTGV | ASSERRGQANTEVF | 0.55102646 |
| LQTSIEF | ASSERRGQANTEVF | 0.58720577 |
| MSFAVSEL | ASSERRGQANTEVF | 0.7071742 |
| NTGALGL | ASSERRGQANTEVF | 0.6434018 |
| PPGAGGAGGAGTT | ASSERRGQANTEVF | 0.53237456 |
| QLVGLGI | ASSERRGQANTEVF | 0.60841966 |
| RFLEQVHQL | ASSERRGQANTEVF | 0.69632876 |
| RGVGYVLR | ASSERRGQANTEVF | 0.6302637 |
| SATGVFAGVI | ASSERRGQANTEVF | 0.58478326 |
| SLPAITLP | ASSERRGQANTEVF | 0.467148 |
| STALAFTV | ASSERRGQANTEVF | 0.7028481 |
| SVLTVGLGK | ASSERRGQANTEVF | 0.68960404 |
| TALAGAGSGI | ASSERRGQANTEVF | 0.6328588 |
| TGLPFTGI | ASSERRGQANTEVF | 0.687736 |
| VAEHVTV | ASSERRGQANTEVF | 0.63302255 |
| VDIVLNSL | ASSERRGQANTEVF | 0.621299 |
| VLTGGGAVL | ASSERRGQANTEVF | 0.35921055 |
| VNVGLAYGI | ASSERRGQANTEVF | 0.46414548 |
| VSSAAVGAV | ASSERRGQANTEVF | 0.52061385 |
| VTSVYEAGV | ASSERRGQANTEVF | 0.61314374 |
| WVALDRGA | ASSERRGQANTEVF | 0.6651569 |
| YLVGSVVI | ASSERRGQANTEVF | 0.6384883 |
| GPPGTGKTLL | ASSERRGQANTEVF | 0.49060917 |
| GVGMTKF | ASSERRGQANTEVF | 0.71200156 |
| IGIGGSDL | ASSERRGQANTEVF | 0.7116987 |
| IGTLLAEL | ASSERRGQANTEVF | 0.38570696 |
| IYTSGTT | ASSERRGQANTEVF | 0.54404724 |
| LSFPAGV | ASSERRGQANTEVF | 0.7294123 |
| LYTSGTT | ASSERRGQANTEVF | 0.61790663 |
| VGAFPFGF | ASSERRGQANTEVF | 0.54437095 |

| | | |
|---|---|---|
| VGAGTVEF | ASSERRGQANTEVF | 0.5854675 |
| VGKSTLLL | ASSERRGQANTEVF | 0.7152059 |
| AALAHAGGGIV | ASSERRGQANTEVF | 0.5870913 |
| AALVAVSLL | ASSERRGQANTEVF | 0.73472625 |
| AAVSRAVAAL | ASSERRGQANTEVF | 0.2710321 |
| AGGSSGIGGF | ASSERRGQANTEVF | 0.53209054 |
| AGLTVANANG | ASSERRGQANTEVF | 0.50573766 |
| AGLVGAVL | ASSERRGQANTEVF | 0.25431243 |
| ALSQAFTV | ASSERRGQANTEVF | 0.7054621 |
| ALTFSVSL | ASSERRGQANTEVF | 0.47378117 |
| ALTGAALVAL | ASSERRGQANTEVF | 0.63216466 |
| ASFSIGAGVL | ASSERRGQANTEVF | 0.47954395 |
| DGSVYKI | ASSERRGQANTEVF | 0.70514727 |
| EQLAEIV | ASSERRGQANTEVF | 0.5819429 |
| FAVSELTVM | ASSERRGQANTEVF | 0.52695215 |
| GAAPLAAGL | ASSERRGQANTEVF | 0.47207433 |
| GADGGDGGSGGAGG | ASSERRGQANTEVF | 0.6590067 |
| GFGLDTGGAGG | ASSERRGQANTEVF | 0.72292566 |
| GLASGGLL | ASSERRGQANTEVF | 0.71766895 |
| GRIAGAAVLL | ASSERRGQANTEVF | 0.31514287 |
| GSAGSLNTSF | ASSERRGQANTEVF | 0.6479574 |
| GTGQIGI | ASSERRGQANTEVF | 0.5706805 |
| GTLIALIL | ASSERRGQANTEVF | 0.39937764 |
| IAATMGFVVG | ASSERRGQANTEVF | 0.3231529 |
| IFAQALL | ASSERRGQANTEVF | 0.52463204 |
| IGAAVALGF | ASSERRGQANTEVF | 0.57748294 |
| IGLTPLAL | ASSERRGQANTEVF | 0.6434729 |
| ILGSNTSTL | ASSERRGQANTEVF | 0.6601825 |
| IVFSLTVL | ASSERRGQANTEVF | 0.48441085 |
| LASKATGF | ASSERRGQANTEVF | 0.6463412 |
| LASVALIL | ASSERRGQANTEVF | 0.62653214 |
| LFAVGLTL | ASSERRGQANTEVF | 0.17331137 |
| LFNGGIGTYI | ASSERRGQANTEVF | 0.48345757 |
| LTLGAGII | ASSERRGQANTEVF | 0.33107287 |
| LVAAALGDI | ASSERRGQANTEVF | 0.70477545 |
| NTAFVQLGI | ASSERRGQANTEVF | 0.3966721 |
| QLFSGAEL | ASSERRGQANTEVF | 0.48618352 |
| SAATFITL | ASSERRGQANTEVF | 0.5794945 |
| SNYGLANLGV | ASSERRGQANTEVF | 0.61229694 |
| SYGGAYAVM | ASSERRGQANTEVF | 0.59596103 |
| TTGVGGLTLG | ASSERRGQANTEVF | 0.4368133 |
| TTQTIYI | ASSERRGQANTEVF | 0.3796806 |
| VAAGAGLGAVI | ASSERRGQANTEVF | 0.6715437 |
| VAAVVTLL | ASSERRGQANTEVF | 0.5386664 |
| VGFLGGLI | ASSERRGQANTEVF | 0.42339563 |
| VGVIPILL | ASSERRGQANTEVF | 0.44107965 |
| VIFASSLI | ASSERRGQANTEVF | 0.67067623 |
| VKGGDGGAAAG | ASSERRGQANTEVF | 0.3152315 |
| VTKTFTNI | ASSERRGQANTEVF | 0.6701178 |
| VVAAAILL | ASSERRGQANTEVF | 0.38077128 |
| YAALDVEL | ASSERRGQANTEVF | 0.5230137 |

| | | |
|---|---|---|
| LASVALLL | ASSERRGQANTEVF | 0.73412085 |
| LDAVHRV | ASSERRGQANTEVF | 0.6050786 |
| QGLSSSIF | ASSERRGQANTEVF | 0.55678695 |
| RGTFANIRL | ASSERRGQANTEVF | 0.5423369 |
| RQLWWGHRI | ASSERRGQANTEVF | 0.4181824 |
| YGPPGTGKTLL | ASSERRGQANTEVF | 0.495269 |
| AALVGAPANVA | ASSERRGQANTEVF | 0.4899476 |
| AATFRRTE | ASSERRGQANTEVF | 0.69685525 |
| AGALGGGI | ASSERRGQANTEVF | 0.59439594 |
| AGGAGGAGGIGAG | ASSERRGQANTEVF | 0.38078216 |
| AGSGALGL | ASSERRGQANTEVF | 0.6753025 |
| AQIADQI | ASSERRGQANTEVF | 0.7174549 |
| ASAVAVTYL | ASSERRGQANTEVF | 0.74006325 |
| ASTVLGGTGG | ASSERRGQANTEVF | 0.4136438 |
| AVGGEPAK | ASSERRGQANTEVF | 0.4897103 |
| CEQGTFGGTC | ASSERRGQANTEVF | 0.53294253 |
| DEGKAPAG | ASSERRGQANTEVF | 0.52488744 |
| DTVVFPAL | ASSERRGQANTEVF | 0.43623295 |
| FEGITMAG | ASSERRGQANTEVF | 0.4543949 |
| FPLVVQD | ASSERRGQANTEVF | 0.6905967 |
| GAFTTPQ | ASSERRGQANTEVF | 0.75415075 |
| GFGGFGTAGGVGGA | ASSERRGQANTEVF | 0.6122407 |
| GFLAQFL | ASSERRGQANTEVF | 0.2577836 |
| GGAGGAGGNAGVA | ASSERRGQANTEVF | 0.5412464 |
| GGVGTAGAGGAGGAGG | ASSERRGQANTEVF | 0.3976335 |
| GNGGGGDAGNAGSGGN | ASSERRGQANTEVF | 0.44401485 |
| GTGGAGGAGNDAGS | ASSERRGQANTEVF | 0.47493532 |
| GTTNTGWFNS | ASSERRGQANTEVF | 0.30203962 |
| IFMGPLFTL | ASSERRGQANTEVF | 0.5677721 |
| ILAHAIEAI | ASSERRGQANTEVF | 0.75153637 |
| IRSAIREAL | ASSERRGQANTEVF | 0.48837924 |
| ITFAGGG | ASSERRGQANTEVF | 0.6060676 |
| ITGMSGHQEV | ASSERRGQANTEVF | 0.7140846 |
| IVAGLACTL | ASSERRGQANTEVF | 0.6441184 |
| IVGGGLAGA | ASSERRGQANTEVF | 0.69915766 |
| IVTGVLNI | ASSERRGQANTEVF | 0.659935 |
| LAQQGCI | ASSERRGQANTEVF | 0.61323464 |
| LCLHVEN | ASSERRGQANTEVF | 0.589314 |
| LDAISDGS | ASSERRGQANTEVF | 0.6906492 |
| LGGFAAML | ASSERRGQANTEVF | 0.54274195 |
| LGGKNPMI | ASSERRGQANTEVF | 0.6060594 |
| LGPKGVAAIL | ASSERRGQANTEVF | 0.5631019 |
| LMVGVSI | ASSERRGQANTEVF | 0.66646713 |
| LVGAAFVL | ASSERRGQANTEVF | 0.58279413 |
| LVLAVAPL | ASSERRGQANTEVF | 0.7170517 |
| MAAIAVIDAA | ASSERRGQANTEVF | 0.27797756 |
| NNGINAELQ | ASSERRGQANTEVF | 0.43813342 |
| RGEPLVLVHG | ASSERRGQANTEVF | 0.6929763 |
| RGGAGGAAGLI | ASSERRGQANTEVF | 0.2394583 |
| RSYRMHPAV | ASSERRGQANTEVF | 0.34489036 |
| SAAHLLAGQLA | ASSERRGQANTEVF | 0.57282174 |

| | | |
|---|---|---|
| SAAVGSY | ASSERRGQANTEVF | 0.7312745 |
| SPHAVLHRI | ASSERRGQANTEVF | 0.23706612 |
| SQIAKVL | ASSERRGQANTEVF | 0.5203419 |
| TALGHLVP | ASSERRGQANTEVF | 0.349796 |
| TAPTTSGAA | ASSERRGQANTEVF | 0.71099097 |
| TDGDDGNGGSPGT | ASSERRGQANTEVF | 0.54213893 |
| TDGLGEN | ASSERRGQANTEVF | 0.36174738 |
| TGGPPLPGQA | ASSERRGQANTEVF | 0.6706933 |
| TIPAITI | ASSERRGQANTEVF | 0.30223516 |
| TIPGFTI | ASSERRGQANTEVF | 0.6522373 |
| TSIASGALA | ASSERRGQANTEVF | 0.693353 |
| TVTLPGAAI | ASSERRGQANTEVF | 0.7242607 |
| VASGLSAALP | ASSERRGQANTEVF | 0.36826095 |
| VGAGVIGI | ASSERRGQANTEVF | 0.639082 |
| VGPKRFGV | ASSERRGQANTEVF | 0.463674 |
| VNVAFPD | ASSERRGQANTEVF | 0.753025 |
| VSMTNSGVS | ASSERRGQANTEVF | 0.5411432 |
| VTDVLGDLSV | ASSERRGQANTEVF | 0.34126458 |
| VVVVGAGFAGL | ASSERRGQANTEVF | 0.66369295 |
| VYTVGTVV | ASSERRGQANTEVF | 0.64918375 |
| WTTFSGP | ASSERRGQANTEVF | 0.5396286 |
| YDELVEAF | ASSERRGQANTEVF | 0.6402476 |
| YGTGGAGGNGG | ASSERRGQANTEVF | 0.57459706 |
| YTNVMAV | ASSERRGQANTEVF | 0.5098763 |
| FLEQVHQL | ASSERRGQANTEVF | 0.65615594 |
| GLSLLAVAV | ASSERRGQANTEVF | 0.6162234 |
| LEQVHQL | ASSERRGQANTEVF | 0.5285248 |
| LNAAGAI | ASSERRGQANTEVF | 0.7306833 |
| PAVHASL | ASSERRGQANTEVF | 0.60963184 |
| TELEDDA | ASSERRGQANTEVF | 0.41441205 |
| TPPGPPN | ASSERRGQANTEVF | 0.7543893 |
| TPPPPPPP | ASSERRGQANTEVF | 0.51088583 |
| TQSPALI | AARGGEDTQY | 0.6846768 |
| EFAAAFPQ | AARGGEDTQY | 0.68407863 |
| LLSGNLDVL | AARGGEDTQY | 0.7375163 |
| GPANLGNYNI | AARGGEDTQY | 0.44749862 |
| TLGPTLP | AARGGEDTQY | 0.60242254 |
| VYAYSSCCK | AARGGEDTQY | 0.61164415 |
| APAPAPAPNI | AARGGEDTQY | 0.6362928 |
| PGGPVPIP | AARGGEDTQY | 0.4917742 |
| FLTVALEP | AARGGEDTQY | 0.5906049 |
| SQYFGANL | AARGGEDTQY | 0.669694 |
| GGVKPAA | AARGGEDTQY | 0.73111206 |
| GGTGGAGGAGGW | AARGGEDTQY | 0.5893317 |
| GVGGVAGLGGAGG | AARGGEDTQY | 0.5537988 |
| VLLVLATTI | AARGGEDTQY | 0.72997177 |
| IGIFAAML | AARGGEDTQY | 0.4138958 |
| AVGDLID | AARGGEDTQY | 0.63055944 |
| NISPLPL | AARGGEDTQY | 0.6853128 |
| VTAVFVAI | AARGGEDTQY | 0.6989423 |
| SGDDNDAAL | AARGGEDTQY | 0.53166336 |

| | | |
|---|---|---|
| AVAALAHS | AARGGEDTQY | 0.6937441 |
| GGVTVMATP | AARGGEDTQY | 0.5347858 |
| AGLAANVRD | AARGGEDTQY | 0.6203331 |
| TNRPDLIDPALLRPG | AARGGEDTQY | 0.5029078 |
| LANLGVTAL | AARGGEDTQY | 0.4666265 |
| TGERIEL | AARGGEDTQY | 0.43579453 |
| NGEELAA | AARGGEDTQY | 0.57467234 |
| VLSLVWG | AARGGEDTQY | 0.67191964 |
| IGYALWL | AARGGEDTQY | 0.6204754 |
| LSLLLAPL | AARGGEDTQY | 0.30896282 |
| TLDEALK | AARGGEDTQY | 0.6615858 |
| TDGAGTAEGG | AARGGEDTQY | 0.64773124 |
| NAPGGAGGNGG | AARGGEDTQY | 0.7296297 |
| VSVFMGMGGNF | AARGGEDTQY | 0.5830446 |
| LASYLAAL | AARGGEDTQY | 0.6846777 |
| LVGGTPAL | AARGGEDTQY | 0.51065624 |
| VRNADVL | AARGGEDTQY | 0.6285894 |
| TCSSPFLVSAM | AARGGEDTQY | 0.34059912 |
| VHITDAE | AARGGEDTQY | 0.4322245 |
| EYAAPSF | AARGGEDTQY | 0.5194472 |
| VVDAFHGGL | AARGGEDTQY | 0.42291516 |
| ASVVDPQ | AARGGEDTQY | 0.67769104 |
| SYNEFGGLS | AARGGEDTQY | 0.67771053 |
| DPDGSVY | AARGGEDTQY | 0.70000315 |
| MVSTIHATE | AARGGEDTQY | 0.7011563 |
| LAAGVADPV | AARGGEDTQY | 0.7162216 |
| TKEQIAA | AARGGEDTQY | 0.67237115 |
| SLASPAAL | AARGGEDTQY | 0.52747625 |
| IGLVFYGV | AARGGEDTQY | 0.5274244 |
| FLLDASNL | AARGGEDTQY | 0.3274472 |
| YQVGNFP | AARGGEDTQY | 0.47886848 |
| ESLLSQST | AARGGEDTQY | 0.5373174 |
| LFTVGFAAVG | AARGGEDTQY | 0.58459014 |
| ATPPTPT | AARGGEDTQY | 0.23530312 |
| TDLATLG | AARGGEDTQY | 0.6034631 |
| SAGGTGR | AARGGEDTQY | 0.7159118 |
| GFSNSGS | AARGGEDTQY | 0.5452597 |
| GGNGGSGGTSV | AARGGEDTQY | 0.6689612 |
| GTSVATGGAGN | AARGGEDTQY | 0.6501982 |
| NGGNGGSGGTS | AARGGEDTQY | 0.4714977 |
| AVVKFPH | AARGGEDTQY | 0.6604566 |
| TPPPPPP | AARGGEDTQY | 0.5210261 |
| AAPAAAAAPA | AARGGEDTQY | 0.6972028 |
| IALNEDL | AARGGEDTQY | 0.737569 |
| IVNVSSM | AARGGEDTQY | 0.6354491 |
| TGIPPAPR | AARGGEDTQY | 0.49936998 |
| FDLGGGTF | AARGGEDTQY | 0.62983894 |
| FDLGGGTFDVS | AARGGEDTQY | 0.41551137 |
| NEPTAAA | AARGGEDTQY | 0.53064245 |
| NEPTAAALA | AARGGEDTQY | 0.5546795 |
| GSIINISSI | AARGGEDTQY | 0.57383 |

| | | |
|---|---|---|
| VAPGNAGTA | AARGGEDTQY | 0.4903775 |
| TGGGASLE | AARGGEDTQY | 0.62961304 |
| VAVYNHYKRI | AARGGEDTQY | 0.6269191 |
| TDPAPATT | AARGGEDTQY | 0.62676793 |
| TNRPDILDPALLRPG | AARGGEDTQY | 0.36997098 |
| LTTTTTS | AARGGEDTQY | 0.6553281 |
| PTGSGKTL | AARGGEDTQY | 0.6099045 |
| TGSGKTL | AARGGEDTQY | 0.37470746 |
| TIGHVDHGK | AARGGEDTQY | 0.6479354 |
| TIGHVDHGKTT | AARGGEDTQY | 0.5611064 |
| GHVDHGKTT | AARGGEDTQY | 0.57177466 |
| AQFEHTLL | AARGGEDTQY | 0.7102226 |
| IVNQDEL | AARGGEDTQY | 0.56897384 |
| TNGSQFF | AARGGEDTQY | 0.34591544 |
| TGKGIVE | AARGGEDTQY | 0.46775585 |
| LISNASDA | AARGGEDTQY | 0.60726774 |
| AELRTLGELTKIAKAHGAQ | AARGGEDTQY | 0.57018733 |
| APNAVAAAEPSW | AARGGEDTQY | 0.25954983 |
| APQVAGAATAVL | AARGGEDTQY | 0.6708785 |
| ARRDPAAAAAILPTDA | AARGGEDTQY | 0.41791287 |
| DAAARDVLAEAGLAEYF | AARGGEDTQY | 0.45138717 |
| DDQIVLSAKANLAKVEQ | AARGGEDTQY | 0.5068064 |
| DELLPDTGEDSDEDGH | AARGGEDTQY | 0.6726767 |
| ELPEAPANRATVPQD | AARGGEDTQY | 0.6442236 |
| FGQSKIERVTGIG | AARGGEDTQY | 0.5546046 |
| GAGGTGSTAGGAGGAGGAGGLY | AARGGEDTQY | 0.59159225 |
| GHGGTGAAGGNGGTGGAGGSGI | AARGGEDTQY | 0.6337118 |
| GNGGNGGAGGEGGPGGAGGAGA | AARGGEDTQY | 0.69883454 |
| GSTAGNGGNGGNATNGTIAGQP | AARGGEDTQY | 0.63368475 |
| GTGGDGGNGGNGGTGG | AARGGEDTQY | 0.5984517 |
| GVRRPAGIPADAPV | AARGGEDTQY | 0.63259387 |
| GYRQVTDAHLLTLARR | AARGGEDTQY | 0.5259974 |
| IAKMGQDGHDRG | AARGGEDTQY | 0.46119544 |
| KAEPSIEIQYT | AARGGEDTQY | 0.44504142 |
| KPVRIRMAADEDTVAVT | AARGGEDTQY | 0.36325642 |
| MPPRIAGMRLLVIKPEP | AARGGEDTQY | 0.43629107 |
| MKFYAVALFRSRGVPPTR | AARGGEDTQY | 0.6434418 |
| NAGGFGILGAVAHSP | AARGGEDTQY | 0.7018405 |
| NEGREITLAMAEM | AARGGEDTQY | 0.54810405 |
| NELVEWSHPLGHRWRWE | AARGGEDTQY | 0.33456242 |
| NLGAGGTGGNGGNPG | AARGGEDTQY | 0.5143456 |
| NTEWPKWETVMAADPVD | AARGGEDTQY | 0.5035147 |
| PASAAPAETQP | AARGGEDTQY | 0.6626056 |
| QQSSDGKNGVRAAEITGE | AARGGEDTQY | 0.69786674 |
| RAPDAVTTSWWKE | AARGGEDTQY | 0.7187714 |
| SGFGNVGAHTSGWFNQ | AARGGEDTQY | 0.68817914 |
| SGVAGAVGQAASVGGLKV | AARGGEDTQY | 0.53427225 |
| SNGGAGTGSAGGNGGT | AARGGEDTQY | 0.67462695 |
| TGGAGGEGGAAG | AARGGEDTQY | 0.49926433 |
| TGGDGGAAGAGGAPGHG | AARGGEDTQY | 0.592448 |
| TMPPLLAAIANAGRHGV | AARGGEDTQY | 0.6029228 |

| | | |
|---|---|---|
| TSSTAGNSGGA | AARGGEDTQY | 0.6385443 |
| VPATVPVGILC | AARGGEDTQY | 0.48708028 |
| VQRPDSGGVVLTGRIS | AARGGEDTQY | 0.5344513 |
| WTPNPVPATKGVPAGLVQAT | AARGGEDTQY | 0.33265403 |
| YALKRALAKFSDRFDVVII | AARGGEDTQY | 0.64106786 |
| YLIGSVIV | AARGGEDTQY | 0.3352874 |
| YGQLVAVID | AARGGEDTQY | 0.4080848 |
| VTVGGMTLPA | AARGGEDTQY | 0.5340217 |
| VLALPAV | AARGGEDTQY | 0.73012537 |
| VIPPTLNYET | AARGGEDTQY | 0.7346484 |
| VGMGGMHPSSG | AARGGEDTQY | 0.482776 |
| VGLAGGAATA | AARGGEDTQY | 0.49002674 |
| VGGAGGAGGSGP | AARGGEDTQY | 0.61854553 |
| VFAAFMV | AARGGEDTQY | 0.47315466 |
| TLYDKLR | AARGGEDTQY | 0.65054166 |
| TLAVFTVL | AARGGEDTQY | 0.56267244 |
| TGSVQPARV | AARGGEDTQY | 0.5739594 |
| TGGGAGSGGT | AARGGEDTQY | 0.47955817 |
| SHLETSL | AARGGEDTQY | 0.66614354 |
| SFTTGMT | AARGGEDTQY | 0.5348136 |
| QVEVAKRMGVRQPTVS | AARGGEDTQY | 0.6069706 |
| QGSGGPALAM | AARGGEDTQY | 0.29891026 |
| MLAPSNP | AARGGEDTQY | 0.70917726 |
| LVNNAAI | AARGGEDTQY | 0.40037403 |
| LTGEINTLA | AARGGEDTQY | 0.465736 |
| LSIFGPV | AARGGEDTQY | 0.261508 |
| LLALLANL | AARGGEDTQY | 0.5656844 |
| LISNASDALDKLR | AARGGEDTQY | 0.6115916 |
| LINEAVP | AARGGEDTQY | 0.7172817 |
| LGYAFSGL | AARGGEDTQY | 0.6047349 |
| LDFVAVAL | AARGGEDTQY | 0.5886969 |
| LAQPGAE | AARGGEDTQY | 0.67888176 |
| ISNLGAVTGF | AARGGEDTQY | 0.49431908 |
| HQEDPGRI | AARGGEDTQY | 0.3259499 |
| GTPGEPTP | AARGGEDTQY | 0.57635653 |
| GFFNSGS | AARGGEDTQY | 0.5338431 |
| GAGGGGLTAGAN | AARGGEDTQY | 0.71820545 |
| FVDALVNM | AARGGEDTQY | 0.63803315 |
| FLTVAPEL | AARGGEDTQY | 0.31033444 |
| FETLTAL | AARGGEDTQY | 0.61498183 |
| DVINPIEL | AARGGEDTQY | 0.65094036 |
| APVSGAPV | AARGGEDTQY | 0.34916216 |
| ALVGGLAW | AARGGEDTQY | 0.47331378 |
| ALAAAAATAI | AARGGEDTQY | 0.49670714 |
| AFGLGLLL | AARGGEDTQY | 0.36212167 |
| AFFAAASL | AARGGEDTQY | 0.64505583 |
| AAAALPAFSPP | AARGGEDTQY | 0.4999687 |
| VVLAPAL | AARGGEDTQY | 0.5476539 |
| TIGHVDHGKTTL | AARGGEDTQY | 0.5422818 |
| LISNASDALDK | AARGGEDTQY | 0.4983111 |
| LISNASDAL | AARGGEDTQY | 0.5159466 |

| | | |
|---|---|---|
| ISNASDALDK | AARGGEDTQY | 0.54732907 |
| NQAETLVYQTE | AARGGEDTQY | 0.6098304 |
| TGGTGGTAGAG | AARGGEDTQY | 0.67444676 |
| TGSTAGGAGGA | AARGGEDTQY | 0.5399789 |
| DPLAVPVAAGPV | AARGGEDTQY | 0.56168956 |
| GGTGGTPGNGGN | AARGGEDTQY | 0.71738666 |
| PAGLAGVDPATL | AARGGEDTQY | 0.36949292 |
| GNNLVVEAPHAGASV | AARGGEDTQY | 0.48832226 |
| KVTRIMSASTLEQPA | AARGGEDTQY | 0.65422755 |
| LGFALPIALPRRTVV | AARGGEDTQY | 0.52601445 |
| TRLGNLTARKLVVLV | AARGGEDTQY | 0.6949403 |
| WSQQRRLIYFPSAGP | AARGGEDTQY | 0.6493719 |
| GPNGNPDPMAAAVDIR | AARGGEDTQY | 0.62208724 |
| TQRCVSVADLRAAASR | AARGGEDTQY | 0.5469359 |
| QFVLDVAEHPVEVLTETV | AARGGEDTQY | 0.7370188 |
| GGAGGTGGAGGNGGTQAGDG | AARGGEDTQY | 0.6245979 |
| PLLATLRAIAVDVDAKPAQV | AARGGEDTQY | 0.53333807 |
| APKFRTGGYMIDFWGVGYQVAK | AARGGEDTQY | 0.6286237 |
| GAGGAGGAGGAAGGPGGTGGTG | AARGGEDTQY | 0.43502676 |
| IGALGLLATDVAPAGSVAFGVA | AARGGEDTQY | 0.6416999 |
| AGGAGGIGGNGGAGGAGGVLMG | AARGGEDTQY | 0.6511068 |
| ADELIGLL | AARGGEDTQY | 0.42234403 |
| ADRALKA | AARGGEDTQY | 0.51913965 |
| AGAVGQVGGAGGA | AARGGEDTQY | 0.7378652 |
| AGGGGGLGGAP | AARGGEDTQY | 0.6780271 |
| AGGLLYGNGGAG | AARGGEDTQY | 0.48581704 |
| AGLESIGL | AARGGEDTQY | 0.43254936 |
| AGLKTGII | AARGGEDTQY | 0.5107602 |
| ASLAALLL | AARGGEDTQY | 0.49871147 |
| ASVINISSI | AARGGEDTQY | 0.39120537 |
| ATPPGPA | AARGGEDTQY | 0.6493392 |
| ATVASALV | AARGGEDTQY | 0.53699756 |
| ATVASLIL | AARGGEDTQY | 0.6818151 |
| AVNGPVLIRH | AARGGEDTQY | 0.6404496 |
| DGAAGAVGI | AARGGEDTQY | 0.7155345 |
| DPLLHGI | AARGGEDTQY | 0.49406064 |
| ELGAALGAL | AARGGEDTQY | 0.57814217 |
| ELGRTQAV | AARGGEDTQY | 0.32273877 |
| FVAGGYLS | AARGGEDTQY | 0.5180011 |
| GASGGAGGAGGW | AARGGEDTQY | 0.4376536 |
| GGAGAGGDGGAGGI | AARGGEDTQY | 0.5634717 |
| GTGHNGSQPGMGGNG | AARGGEDTQY | 0.35262728 |
| GYAAASASAS | AARGGEDTQY | 0.755524 |
| HQVDVAAVQQG | AARGGEDTQY | 0.465648 |
| IGAAIAHT | AARGGEDTQY | 0.67461765 |
| IGMIEVAQL | AARGGEDTQY | 0.4956177 |
| IRLAPEHTSAMD | AARGGEDTQY | 0.39215064 |
| IYNVSVL | AARGGEDTQY | 0.53087157 |
| KQGFLGM | AARGGEDTQY | 0.66298175 |
| LESVITQL | AARGGEDTQY | 0.3527819 |
| LGLGALVVV | AARGGEDTQY | 0.25682396 |

| | | |
|---|---|---|
| LIGNGGNGGSGGI | AARGGEDTQY | 0.45840603 |
| LNAINAPTL | AARGGEDTQY | 0.35728028 |
| LQLLNTGV | AARGGEDTQY | 0.7400701 |
| LQTSIEF | AARGGEDTQY | 0.71537006 |
| MSFAVSEL | AARGGEDTQY | 0.5638852 |
| NTGALGL | AARGGEDTQY | 0.7362232 |
| PPGAGGAGGAGTT | AARGGEDTQY | 0.21332756 |
| QLVGLGI | AARGGEDTQY | 0.5024216 |
| RFLEQVHQL | AARGGEDTQY | 0.7181862 |
| RGVGYVLR | AARGGEDTQY | 0.6413651 |
| SATGVFAGVI | AARGGEDTQY | 0.57270765 |
| SLPAITLP | AARGGEDTQY | 0.68625057 |
| STALAFTV | AARGGEDTQY | 0.4881969 |
| SVLTVGLGK | AARGGEDTQY | 0.5827951 |
| TALAGAGSGI | AARGGEDTQY | 0.5514307 |
| TGLPFTGI | AARGGEDTQY | 0.680767 |
| VAEHVTV | AARGGEDTQY | 0.6448248 |
| VDIVLNSL | AARGGEDTQY | 0.34491733 |
| VLTGGGAVL | AARGGEDTQY | 0.6496255 |
| VNVGLAYGI | AARGGEDTQY | 0.74505 |
| VSSAAVGAV | AARGGEDTQY | 0.46622127 |
| VTSVYEAGV | AARGGEDTQY | 0.61899984 |
| WVALDRGA | AARGGEDTQY | 0.5737824 |
| YLVGSVVI | AARGGEDTQY | 0.46380562 |
| GPPGTGKTLL | AARGGEDTQY | 0.45695794 |
| GVGMTKF | AARGGEDTQY | 0.36383197 |
| IGIGGSDL | AARGGEDTQY | 0.658505 |
| IGTLLAEL | AARGGEDTQY | 0.3618939 |
| IYTSGTT | AARGGEDTQY | 0.54638404 |
| LSFPAGV | AARGGEDTQY | 0.5645431 |
| LYTSGTT | AARGGEDTQY | 0.77266675 |
| VGAFPFGF | AARGGEDTQY | 0.6564411 |
| VGAGTVEF | AARGGEDTQY | 0.63770807 |
| VGKSTLLL | AARGGEDTQY | 0.44395953 |
| AALAHAGGGIV | AARGGEDTQY | 0.6120087 |
| AALVAVSLL | AARGGEDTQY | 0.64842176 |
| AAVSRAVAAL | AARGGEDTQY | 0.3526253 |
| AGGSSGIGGF | AARGGEDTQY | 0.7212607 |
| AGLTVANANG | AARGGEDTQY | 0.4081605 |
| AGLVGAVL | AARGGEDTQY | 0.6109538 |
| ALSQAFTV | AARGGEDTQY | 0.6415936 |
| ALTFSVSL | AARGGEDTQY | 0.6869599 |
| ALTGAALVAL | AARGGEDTQY | 0.5866178 |
| ASFSIGAGVL | AARGGEDTQY | 0.6649116 |
| DGSVYKI | AARGGEDTQY | 0.43483466 |
| EQLAEIV | AARGGEDTQY | 0.6640231 |
| FAVSELTVM | AARGGEDTQY | 0.7048216 |
| GAAPLAAGL | AARGGEDTQY | 0.5767182 |
| GADGGDGGSGGAGG | AARGGEDTQY | 0.73038 |
| GFGLDTGGAGG | AARGGEDTQY | 0.62742233 |
| GLASGGLL | AARGGEDTQY | 0.36263657 |

| | | |
|---|---|---|
| GRIAGAAVLL | AARGGEDTQY | 0.5947022 |
| GSAGSLNTSF | AARGGEDTQY | 0.7222401 |
| GTGQIGI | AARGGEDTQY | 0.7185545 |
| GTLIALIL | AARGGEDTQY | 0.64794344 |
| IAATMGFVVG | AARGGEDTQY | 0.6211703 |
| IFAQALL | AARGGEDTQY | 0.50403553 |
| IGAAVALGF | AARGGEDTQY | 0.725018 |
| IGLTPLAL | AARGGEDTQY | 0.78500843 |
| ILGSNTSTL | AARGGEDTQY | 0.6298093 |
| IVFSLTVL | AARGGEDTQY | 0.42074233 |
| LASKATGF | AARGGEDTQY | 0.44616455 |
| LASVALIL | AARGGEDTQY | 0.6816995 |
| LFAVGLTL | AARGGEDTQY | 0.7020673 |
| LFNGGIGTYI | AARGGEDTQY | 0.3722003 |
| LTLGAGII | AARGGEDTQY | 0.5704738 |
| LVAAALGDI | AARGGEDTQY | 0.62675375 |
| NTAFVQLGI | AARGGEDTQY | 0.27910882 |
| QLFSGAEL | AARGGEDTQY | 0.6235894 |
| SAATFITL | AARGGEDTQY | 0.7019234 |
| SNYGLANLGV | AARGGEDTQY | 0.64751697 |
| SYGGAYAVM | AARGGEDTQY | 0.34900674 |
| TTGVGGLTLG | AARGGEDTQY | 0.71760035 |
| TTQTIYI | AARGGEDTQY | 0.7202237 |
| VAAGAGLGAVI | AARGGEDTQY | 0.73799276 |
| VAAVVTLL | AARGGEDTQY | 0.34396 |
| VGFLGGLI | AARGGEDTQY | 0.23451044 |
| VGVIPILL | AARGGEDTQY | 0.60791636 |
| VIFASSLI | AARGGEDTQY | 0.65705293 |
| VKGGDGGAAAG | AARGGEDTQY | 0.5420903 |
| VTKTFTNI | AARGGEDTQY | 0.41356325 |
| VVAAAILL | AARGGEDTQY | 0.52220833 |
| YAALDVEL | AARGGEDTQY | 0.7105062 |
| LASVALLL | AARGGEDTQY | 0.38765988 |
| LDAVHRV | AARGGEDTQY | 0.40206462 |
| QGLSSSIF | AARGGEDTQY | 0.6176448 |
| RGTFANIRL | AARGGEDTQY | 0.67667496 |
| RQLWWGHRI | AARGGEDTQY | 0.40772858 |
| YGPPGTGKTLL | AARGGEDTQY | 0.48874223 |
| AALVGAPANVA | AARGGEDTQY | 0.6639427 |
| AATFRRTE | AARGGEDTQY | 0.6749972 |
| AGALGGGI | AARGGEDTQY | 0.60037404 |
| AGGAGGAGGIGAG | AARGGEDTQY | 0.38758433 |
| AGSGALGL | AARGGEDTQY | 0.68264043 |
| AQIADQI | AARGGEDTQY | 0.35873926 |
| ASAVAVTYL | AARGGEDTQY | 0.6656833 |
| ASTVLGGTGG | AARGGEDTQY | 0.5608494 |
| AVGGEPAK | AARGGEDTQY | 0.6246867 |
| CEQGTFGGTC | AARGGEDTQY | 0.53791344 |
| DEGKAPAG | AARGGEDTQY | 0.33348498 |
| DTVVFPAL | AARGGEDTQY | 0.42682797 |
| FEGITMAG | AARGGEDTQY | 0.39907235 |

| | | |
|---|---|---|
| FPLVVQD | AARGGEDTQY | 0.69439936 |
| GAFTTPQ | AARGGEDTQY | 0.6284874 |
| GFGGFGTAGGVGGA | AARGGEDTQY | 0.6399505 |
| GFLAQFL | AARGGEDTQY | 0.6327064 |
| GGAGGAGGNAGVA | AARGGEDTQY | 0.45598552 |
| GGVGTAGAGGAGGAGG | AARGGEDTQY | 0.6541431 |
| GNGGGGDAGNAGSGGN | AARGGEDTQY | 0.36490127 |
| GTGGAGGAGNDAGS | AARGGEDTQY | 0.6391936 |
| GTTNTGWFNS | AARGGEDTQY | 0.4890788 |
| IFMGPLFTL | AARGGEDTQY | 0.6158418 |
| ILAHAIEAI | AARGGEDTQY | 0.46549895 |
| IRSAIREAL | AARGGEDTQY | 0.6397917 |
| ITFAGGG | AARGGEDTQY | 0.6422263 |
| ITGMSGHQEV | AARGGEDTQY | 0.4245844 |
| IVAGLACTL | AARGGEDTQY | 0.6290829 |
| IVGGGLAGA | AARGGEDTQY | 0.6942142 |
| IVTGVLNI | AARGGEDTQY | 0.65851384 |
| LAQQGCI | AARGGEDTQY | 0.30866894 |
| LCLHVEN | AARGGEDTQY | 0.6500681 |
| LDAISDGS | AARGGEDTQY | 0.420191 |
| LGGFAAML | AARGGEDTQY | 0.6227131 |
| LGGKNPMI | AARGGEDTQY | 0.6107986 |
| LGPKGVAAIL | AARGGEDTQY | 0.5517827 |
| LMVGVSI | AARGGEDTQY | 0.6406837 |
| LVGAAFVL | AARGGEDTQY | 0.3898075 |
| LVLAVAPL | AARGGEDTQY | 0.48000437 |
| MAAIAVIDAA | AARGGEDTQY | 0.49893188 |
| NNGINAELQ | AARGGEDTQY | 0.6467837 |
| RGEPLVLVHG | AARGGEDTQY | 0.6208731 |
| RGGAGGAAGLI | AARGGEDTQY | 0.37428585 |
| RSYRMHPAV | AARGGEDTQY | 0.5990168 |
| SAAHLLAGQLA | AARGGEDTQY | 0.58492714 |
| SAAVGSY | AARGGEDTQY | 0.45196116 |
| SPHAVLHRI | AARGGEDTQY | 0.5693884 |
| SQIAKVL | AARGGEDTQY | 0.525488 |
| TALGHLVP | AARGGEDTQY | 0.30764368 |
| TAPTTSGAA | AARGGEDTQY | 0.6659068 |
| TDGDDGNGGSPGT | AARGGEDTQY | 0.47106168 |
| TDGLGEN | AARGGEDTQY | 0.6472166 |
| TGGPPLPGQA | AARGGEDTQY | 0.67145973 |
| TIPAITI | AARGGEDTQY | 0.31055084 |
| TIPGFTI | AARGGEDTQY | 0.66434145 |
| TSIASGALA | AARGGEDTQY | 0.55658454 |
| TVTLPGAAI | AARGGEDTQY | 0.5351321 |
| VASGLSAALP | AARGGEDTQY | 0.57907856 |
| VGAGVIGI | AARGGEDTQY | 0.47078487 |
| VGPKRFGV | AARGGEDTQY | 0.6940088 |
| VNVAFPD | AARGGEDTQY | 0.6722224 |
| VSMTNSGVS | AARGGEDTQY | 0.59610397 |
| VTDVLGDLSV | AARGGEDTQY | 0.63574016 |
| VVVVGAGFAGL | AARGGEDTQY | 0.64943445 |

| | | |
|---|---|---|
| VYTVGTVV | AARGGEDTQY | 0.6762372 |
| WTTFSGP | AARGGEDTQY | 0.44078657 |
| YDELVEAF | AARGGEDTQY | 0.3229149 |
| YGTGGAGGNGG | AARGGEDTQY | 0.59157467 |
| YTNVMAV | AARGGEDTQY | 0.56143844 |
| FLEQVHQL | AARGGEDTQY | 0.45854768 |
| GLSLLAVAV | AARGGEDTQY | 0.37737226 |
| LEQVHQL | AARGGEDTQY | 0.66065186 |
| LNAAGAI | AARGGEDTQY | 0.7627865 |
| PAVHASL | AARGGEDTQY | 0.5085104 |
| TELEDDA | AARGGEDTQY | 0.7597371 |
| TPPGPPN | AARGGEDTQY | 0.37446058 |
| TPPPPPPP | AARGGEDTQY | 0.6997237 |
| TQSPALI | ASSLGLGRYAEQF | 0.70250726 |
| EFAAAFPQ | ASSLGLGRYAEQF | 0.4123326 |
| LLSGNLDVL | ASSLGLGRYAEQF | 0.6013702 |
| GPANLGNYNI | ASSLGLGRYAEQF | 0.4521386 |
| TLGPTLP | ASSLGLGRYAEQF | 0.45008332 |
| VYAYSSCCK | ASSLGLGRYAEQF | 0.6708724 |
| APAPAPAPNI | ASSLGLGRYAEQF | 0.5189596 |
| PGGPVPIP | ASSLGLGRYAEQF | 0.6457991 |
| FLTVALEP | ASSLGLGRYAEQF | 0.5057374 |
| SQYFGANL | ASSLGLGRYAEQF | 0.4728452 |
| GGVKPAA | ASSLGLGRYAEQF | 0.65041006 |
| GGTGGAGGAGGW | ASSLGLGRYAEQF | 0.64923847 |
| GVGGVAGLGGAGG | ASSLGLGRYAEQF | 0.5829969 |
| VLLVLATTI | ASSLGLGRYAEQF | 0.77837515 |
| IGIFAAML | ASSLGLGRYAEQF | 0.45187068 |
| AVGDLID | ASSLGLGRYAEQF | 0.70774364 |
| NISPLPL | ASSLGLGRYAEQF | 0.70658195 |
| VTAVFVAI | ASSLGLGRYAEQF | 0.5863718 |
| SGDDNDAAL | ASSLGLGRYAEQF | 0.5418694 |
| AVAALAHS | ASSLGLGRYAEQF | 0.67984146 |
| GGVTVMATP | ASSLGLGRYAEQF | 0.51616836 |
| AGLAANVRD | ASSLGLGRYAEQF | 0.5820938 |
| TNRPDLIDPALLRPG | ASSLGLGRYAEQF | 0.7027166 |
| LANLGVTAL | ASSLGLGRYAEQF | 0.6851386 |
| TGERIEL | ASSLGLGRYAEQF | 0.7755357 |
| NGEELAA | ASSLGLGRYAEQF | 0.7548777 |
| VLSLVWG | ASSLGLGRYAEQF | 0.7122233 |
| IGYALWL | ASSLGLGRYAEQF | 0.66537404 |
| LSLLLAPL | ASSLGLGRYAEQF | 0.3811376 |
| TLDEALK | ASSLGLGRYAEQF | 0.7043118 |
| TDGAGTAEGG | ASSLGLGRYAEQF | 0.62874854 |
| NAPGGAGGNGG | ASSLGLGRYAEQF | 0.751001 |
| VSVFMGMGGNF | ASSLGLGRYAEQF | 0.76277596 |
| LASYLAAL | ASSLGLGRYAEQF | 0.6756008 |
| LVGGTPAL | ASSLGLGRYAEQF | 0.77380335 |
| VRNADVL | ASSLGLGRYAEQF | 0.7434931 |
| TCSSPFLVSAM | ASSLGLGRYAEQF | 0.61612463 |
| VHITDAE | ASSLGLGRYAEQF | 0.72449416 |

| | | |
|---|---|---|
| EYAAPSF | ASSLGLGRYAEQF | 0.6695049 |
| VVDAFHGGL | ASSLGLGRYAEQF | 0.7354392 |
| ASVVDPQ | ASSLGLGRYAEQF | 0.6114458 |
| SYNEFGGLS | ASSLGLGRYAEQF | 0.75410426 |
| DPDGSVY | ASSLGLGRYAEQF | 0.6998551 |
| MVSTIHATE | ASSLGLGRYAEQF | 0.2729627 |
| LAAGVADPV | ASSLGLGRYAEQF | 0.7464493 |
| TKEQIAA | ASSLGLGRYAEQF | 0.7321653 |
| SLASPAAL | ASSLGLGRYAEQF | 0.7166971 |
| IGLVFYGV | ASSLGLGRYAEQF | 0.76798964 |
| FLLDASNL | ASSLGLGRYAEQF | 0.62556446 |
| YQVGNFP | ASSLGLGRYAEQF | 0.6516794 |
| ESLLSQST | ASSLGLGRYAEQF | 0.76332504 |
| LFTVGFAAVG | ASSLGLGRYAEQF | 0.77358484 |
| ATPPTPT | ASSLGLGRYAEQF | 0.58781874 |
| TDLATLG | ASSLGLGRYAEQF | 0.75337625 |
| SAGGTGR | ASSLGLGRYAEQF | 0.76648164 |
| GFSNSGS | ASSLGLGRYAEQF | 0.75376254 |
| GGNGGSGGTSV | ASSLGLGRYAEQF | 0.7524116 |
| GTSVATGGAGN | ASSLGLGRYAEQF | 0.76715964 |
| NGGNGGSGGTS | ASSLGLGRYAEQF | 0.6924064 |
| AVVKFPH | ASSLGLGRYAEQF | 0.7715528 |
| TPPPPPP | ASSLGLGRYAEQF | 0.7759186 |
| AAPAAAAAPA | ASSLGLGRYAEQF | 0.7057677 |
| IALNEDL | ASSLGLGRYAEQF | 0.72701013 |
| IVNVSSM | ASSLGLGRYAEQF | 0.68077993 |
| TGIPPAPR | ASSLGLGRYAEQF | 0.6998969 |
| FDLGGGTF | ASSLGLGRYAEQF | 0.35078952 |
| FDLGGGTFDVS | ASSLGLGRYAEQF | 0.7531427 |
| NEPTAAA | ASSLGLGRYAEQF | 0.7588371 |
| NEPTAAALA | ASSLGLGRYAEQF | 0.71595013 |
| GSIINISSI | ASSLGLGRYAEQF | 0.67199904 |
| VAPGNAGTA | ASSLGLGRYAEQF | 0.6502886 |
| TGGGASLE | ASSLGLGRYAEQF | 0.74124634 |
| VAVYNHYKRI | ASSLGLGRYAEQF | 0.68492645 |
| TDPAPATT | ASSLGLGRYAEQF | 0.35702378 |
| TNRPDILDPALLRPG | ASSLGLGRYAEQF | 0.7281998 |
| LTTTTTS | ASSLGLGRYAEQF | 0.7624548 |
| PTGSGKTL | ASSLGLGRYAEQF | 0.75918365 |
| TGSGKTL | ASSLGLGRYAEQF | 0.52613896 |
| TIGHVDHGK | ASSLGLGRYAEQF | 0.7778208 |
| TIGHVDHGKTT | ASSLGLGRYAEQF | 0.7541114 |
| GHVDHGKTT | ASSLGLGRYAEQF | 0.7676844 |
| AQFEHTLL | ASSLGLGRYAEQF | 0.6772909 |
| IVNQDEL | ASSLGLGRYAEQF | 0.69696045 |
| TNGSQFF | ASSLGLGRYAEQF | 0.7612616 |
| TGKGIVE | ASSLGLGRYAEQF | 0.71786034 |
| LISNASDA | ASSLGLGRYAEQF | 0.55024946 |
| AELRTLGELTKIAKAHGAQ | ASSLGLGRYAEQF | 0.7583932 |
| APNAVAAAEPSW | ASSLGLGRYAEQF | 0.71711004 |
| APQVAGAATAVL | ASSLGLGRYAEQF | 0.68356395 |

| | | |
|---|---|---|
| ARRDPAAAAAILPTDA | ASSLGLGRYAEQF | 0.6879297 |
| DAAARDVLAEAGLAEYF | ASSLGLGRYAEQF | 0.77492726 |
| DDQIVLSAKANLAKVEQ | ASSLGLGRYAEQF | 0.6401396 |
| DELLPDTGEDSDEDGH | ASSLGLGRYAEQF | 0.63321424 |
| ELPEAPANRATVPQD | ASSLGLGRYAEQF | 0.5048541 |
| FGQSKIERVTGIG | ASSLGLGRYAEQF | 0.72950816 |
| GAGGTGSTAGGAGGAGGAGGLY | ASSLGLGRYAEQF | 0.7695815 |
| GHGGTGAAGGNGGTGGAGGSGI | ASSLGLGRYAEQF | 0.7315922 |
| GNGGNGGAGGEGGPGGAGGAGA | ASSLGLGRYAEQF | 0.71117747 |
| GSTAGNGGNGGNATNGTIAGQP | ASSLGLGRYAEQF | 0.7517408 |
| GTGGDGGNGGNGGTGG | ASSLGLGRYAEQF | 0.71705174 |
| GVRRPAGIPADAPV | ASSLGLGRYAEQF | 0.7435214 |
| GYRQVTDAHLLTLARR | ASSLGLGRYAEQF | 0.7655316 |
| IAKMGQDGHDRG | ASSLGLGRYAEQF | 0.69014406 |
| KAEPSIEIQYT | ASSLGLGRYAEQF | 0.7711345 |
| KPVRIRMAADEDTVAVT | ASSLGLGRYAEQF | 0.7398393 |
| MPPRIAGMRLLVIKPEP | ASSLGLGRYAEQF | 0.7465936 |
| MKFYAVALFRSRGVPPTR | ASSLGLGRYAEQF | 0.7613538 |
| NAGGFGILGAVAHSP | ASSLGLGRYAEQF | 0.7401986 |
| NEGREITLAMAEM | ASSLGLGRYAEQF | 0.73563695 |
| NELVEWSHPLGHRWRWE | ASSLGLGRYAEQF | 0.42829323 |
| NLGAGGTGGNGGNPG | ASSLGLGRYAEQF | 0.65427864 |
| NTEWPKWETVMAADPVD | ASSLGLGRYAEQF | 0.7580197 |
| PASAAPAETQP | ASSLGLGRYAEQF | 0.74738264 |
| QQSSDGKNGVRAAEITGE | ASSLGLGRYAEQF | 0.75976425 |
| RAPDAVTTSWWKE | ASSLGLGRYAEQF | 0.6980602 |
| SGFGNVGAHTSGWFNQ | ASSLGLGRYAEQF | 0.7164302 |
| SGVAGAVGQAASVGGLKV | ASSLGLGRYAEQF | 0.7523844 |
| SNGGAGTGSAGGNGGT | ASSLGLGRYAEQF | 0.7654804 |
| TGGAGGEGGAAG | ASSLGLGRYAEQF | 0.510102 |
| TGGDGGAAGAGGAPGHG | ASSLGLGRYAEQF | 0.7367099 |
| TMPPLLAAIANAGRHGV | ASSLGLGRYAEQF | 0.78232217 |
| TSSTAGNSGGA | ASSLGLGRYAEQF | 0.73788095 |
| VPATVPVGILC | ASSLGLGRYAEQF | 0.7164866 |
| VQRPDSGGVVLTGRIS | ASSLGLGRYAEQF | 0.65591896 |
| WTPNPVPATKGVPAGLVQAT | ASSLGLGRYAEQF | 0.7714315 |
| YALKRALAKFSDRFDVVII | ASSLGLGRYAEQF | 0.60448194 |
| YLIGSVIV | ASSLGLGRYAEQF | 0.78179973 |
| YGQLVAVID | ASSLGLGRYAEQF | 0.29061437 |
| VTVGGMTLPA | ASSLGLGRYAEQF | 0.6483177 |
| VLALPAV | ASSLGLGRYAEQF | 0.56868607 |
| VIPPTLNYET | ASSLGLGRYAEQF | 0.58292407 |
| VGMGGMHPSSG | ASSLGLGRYAEQF | 0.76429677 |
| VGLAGGAATA | ASSLGLGRYAEQF | 0.65552855 |
| VGGAGGAGGSGP | ASSLGLGRYAEQF | 0.7511153 |
| VFAAFMV | ASSLGLGRYAEQF | 0.7512269 |
| TLYDKLR | ASSLGLGRYAEQF | 0.76597303 |
| TLAVFTVL | ASSLGLGRYAEQF | 0.7440174 |
| TGSVQPARV | ASSLGLGRYAEQF | 0.7428125 |
| TGGGAGSGGT | ASSLGLGRYAEQF | 0.61685205 |
| SHLETSL | ASSLGLGRYAEQF | 0.76252073 |

| | | |
|---|---|---|
| SFTTGMT | ASSLGLGRYAEQF | 0.7642956 |
| QVEVAKRMGVRQPTVS | ASSLGLGRYAEQF | 0.73184246 |
| QGSGGPALAM | ASSLGLGRYAEQF | 0.5331894 |
| MLAPSNP | ASSLGLGRYAEQF | 0.7133648 |
| LVNNAAI | ASSLGLGRYAEQF | 0.7397765 |
| LTGEINTLA | ASSLGLGRYAEQF | 0.38125777 |
| LSIFGPV | ASSLGLGRYAEQF | 0.29801524 |
| LLALLANL | ASSLGLGRYAEQF | 0.7217642 |
| LISNASDALDKLR | ASSLGLGRYAEQF | 0.7054627 |
| LINEAVP | ASSLGLGRYAEQF | 0.7743521 |
| LGYAFSGL | ASSLGLGRYAEQF | 0.7639065 |
| LDFVAVAL | ASSLGLGRYAEQF | 0.73298836 |
| LAQPGAE | ASSLGLGRYAEQF | 0.7637988 |
| ISNLGAVTGF | ASSLGLGRYAEQF | 0.7336913 |
| HQEDPGRI | ASSLGLGRYAEQF | 0.23752561 |
| GTPGEPTP | ASSLGLGRYAEQF | 0.65536934 |
| GFFNSGS | ASSLGLGRYAEQF | 0.59708333 |
| GAGGGGLTAGAN | ASSLGLGRYAEQF | 0.75546604 |
| FVDALVNM | ASSLGLGRYAEQF | 0.73144853 |
| FLTVAPEL | ASSLGLGRYAEQF | 0.69010735 |
| FETLTAL | ASSLGLGRYAEQF | 0.742116 |
| DVINPIEL | ASSLGLGRYAEQF | 0.7361584 |
| APVSGAPV | ASSLGLGRYAEQF | 0.71782243 |
| ALVGGLAW | ASSLGLGRYAEQF | 0.61484283 |
| ALAAAAATAI | ASSLGLGRYAEQF | 0.2972691 |
| AFGLGLLL | ASSLGLGRYAEQF | 0.7530513 |
| AFFAAASL | ASSLGLGRYAEQF | 0.4331101 |
| AAAALPAFSPP | ASSLGLGRYAEQF | 0.66169095 |
| VVLAPAL | ASSLGLGRYAEQF | 0.3679936 |
| TIGHVDHGKTTL | ASSLGLGRYAEQF | 0.657048 |
| LISNASDALDK | ASSLGLGRYAEQF | 0.3986608 |
| LISNASDAL | ASSLGLGRYAEQF | 0.74565196 |
| ISNASDALDK | ASSLGLGRYAEQF | 0.71660334 |
| NQAETLVYQTE | ASSLGLGRYAEQF | 0.71895117 |
| TGGTGGTAGAG | ASSLGLGRYAEQF | 0.7798463 |
| TGSTAGGAGGA | ASSLGLGRYAEQF | 0.7224966 |
| DPLAVPVAAGPV | ASSLGLGRYAEQF | 0.7181932 |
| GGTGGTPGNGGN | ASSLGLGRYAEQF | 0.66570437 |
| PAGLAGVDPATL | ASSLGLGRYAEQF | 0.68837523 |
| GNNLVVEAPHAGASV | ASSLGLGRYAEQF | 0.6887409 |
| KVTRIMSASTLEQPA | ASSLGLGRYAEQF | 0.6419852 |
| LGFALPIALPRRTVV | ASSLGLGRYAEQF | 0.7702789 |
| TRLGNLTARKLVVLV | ASSLGLGRYAEQF | 0.6837375 |
| WSQQRRLIYFPSAGP | ASSLGLGRYAEQF | 0.5756823 |
| GPNGNPDPMAAAVDIR | ASSLGLGRYAEQF | 0.65657836 |
| TQRCVSVADLRAAASR | ASSLGLGRYAEQF | 0.7722101 |
| QFVLDVAEHPVEVLTETV | ASSLGLGRYAEQF | 0.78172064 |
| GGAGGTGGAGGNGGTQAGDG | ASSLGLGRYAEQF | 0.7270088 |
| PLLATLRAIAVDVDAKPAQV | ASSLGLGRYAEQF | 0.59588605 |
| APKFRTGGYMIDFWGVGYQVAK | ASSLGLGRYAEQF | 0.67140484 |
| GAGGAGGAGGAAGGPGGTGGTG | ASSLGLGRYAEQF | 0.7350323 |

| | | |
|---|---|---|
| IGALGLLATDVAPAGSVAFGVA | ASSLGLGRYAEQF | 0.6892141 |
| AGGAGGIGGNGGAGGAGGVLMG | ASSLGLGRYAEQF | 0.51051104 |
| ADELIGLL | ASSLGLGRYAEQF | 0.76488185 |
| ADRALKA | ASSLGLGRYAEQF | 0.77254665 |
| AGAVGQVGGAGGA | ASSLGLGRYAEQF | 0.7143028 |
| AGGGGGLGGAP | ASSLGLGRYAEQF | 0.5784867 |
| AGGLLYGNGGAG | ASSLGLGRYAEQF | 0.7376326 |
| AGLESIGL | ASSLGLGRYAEQF | 0.66570187 |
| AGLKTGII | ASSLGLGRYAEQF | 0.72599494 |
| ASLAALLL | ASSLGLGRYAEQF | 0.7172291 |
| ASVINISSI | ASSLGLGRYAEQF | 0.55740863 |
| ATPPGPA | ASSLGLGRYAEQF | 0.7191752 |
| ATVASALV | ASSLGLGRYAEQF | 0.779673 |
| ATVASLIL | ASSLGLGRYAEQF | 0.7057172 |
| AVNGPVLIRH | ASSLGLGRYAEQF | 0.7492492 |
| DGAAGAVGI | ASSLGLGRYAEQF | 0.77623165 |
| DPLLHGI | ASSLGLGRYAEQF | 0.763741 |
| ELGAALGAL | ASSLGLGRYAEQF | 0.7746646 |
| ELGRTQAV | ASSLGLGRYAEQF | 0.75960296 |
| FVAGGYLS | ASSLGLGRYAEQF | 0.675434 |
| GASGGAGGAGGW | ASSLGLGRYAEQF | 0.7192226 |
| GGAGAGGDGGAGGI | ASSLGLGRYAEQF | 0.75968647 |
| GTGHNGSQPGMGGNG | ASSLGLGRYAEQF | 0.7432533 |
| GYAAASASAS | ASSLGLGRYAEQF | 0.72896445 |
| HQVDVAAVQQG | ASSLGLGRYAEQF | 0.6002517 |
| IGAAIAHT | ASSLGLGRYAEQF | 0.77189493 |
| IGMIEVAQL | ASSLGLGRYAEQF | 0.7298132 |
| IRLAPEHTSAMD | ASSLGLGRYAEQF | 0.7657398 |
| IYNVSVL | ASSLGLGRYAEQF | 0.54009354 |
| KQGFLGM | ASSLGLGRYAEQF | 0.60493326 |
| LESVITQL | ASSLGLGRYAEQF | 0.76606846 |
| LGLGALVVV | ASSLGLGRYAEQF | 0.22353476 |
| LIGNGGNGGSGGI | ASSLGLGRYAEQF | 0.2688566 |
| LNAINAPTL | ASSLGLGRYAEQF | 0.74361277 |
| LQLLNTGV | ASSLGLGRYAEQF | 0.7724102 |
| LQTSIEF | ASSLGLGRYAEQF | 0.26783922 |
| MSFAVSEL | ASSLGLGRYAEQF | 0.59887075 |
| NTGALGL | ASSLGLGRYAEQF | 0.53330094 |
| PPGAGGAGGAGTT | ASSLGLGRYAEQF | 0.7033868 |
| QLVGLGI | ASSLGLGRYAEQF | 0.73014355 |
| RFLEQVHQL | ASSLGLGRYAEQF | 0.5484326 |
| RGVGYVLR | ASSLGLGRYAEQF | 0.75041103 |
| SATGVFAGVI | ASSLGLGRYAEQF | 0.49535915 |
| SLPAITLP | ASSLGLGRYAEQF | 0.678874 |
| STALAFTV | ASSLGLGRYAEQF | 0.76347697 |
| SVLTVGLGK | ASSLGLGRYAEQF | 0.7717147 |
| TALAGAGSGI | ASSLGLGRYAEQF | 0.7062447 |
| TGLPFTGI | ASSLGLGRYAEQF | 0.76158965 |
| VAEHVTV | ASSLGLGRYAEQF | 0.6870973 |
| VDIVLNSL | ASSLGLGRYAEQF | 0.7812731 |
| VLTGGGAVL | ASSLGLGRYAEQF | 0.6935712 |

| | | |
|---|---|---|
| VNVGLAYGI | ASSLGLGRYAEQF | 0.486395 |
| VSSAAVGAV | ASSLGLGRYAEQF | 0.76480806 |
| VTSVYEAGV | ASSLGLGRYAEQF | 0.7607374 |
| WVALDRGA | ASSLGLGRYAEQF | 0.731746 |
| YLVGSVVI | ASSLGLGRYAEQF | 0.76361704 |
| GPPGTGKTLL | ASSLGLGRYAEQF | 0.65420514 |
| GVGMTKF | ASSLGLGRYAEQF | 0.7323247 |
| IGIGGSDL | ASSLGLGRYAEQF | 0.7713741 |
| IGTLLAEL | ASSLGLGRYAEQF | 0.77282697 |
| IYTSGTT | ASSLGLGRYAEQF | 0.49953976 |
| LSFPAGV | ASSLGLGRYAEQF | 0.76557446 |
| LYTSGTT | ASSLGLGRYAEQF | 0.7223698 |
| VGAFPFGF | ASSLGLGRYAEQF | 0.65694904 |
| VGAGTVEF | ASSLGLGRYAEQF | 0.7767999 |
| VGKSTLLL | ASSLGLGRYAEQF | 0.71912575 |
| AALAHAGGGIV | ASSLGLGRYAEQF | 0.7670651 |
| AALVAVSLL | ASSLGLGRYAEQF | 0.7592949 |
| AAVSRAVAAL | ASSLGLGRYAEQF | 0.7608035 |
| AGGSSGIGGF | ASSLGLGRYAEQF | 0.5379334 |
| AGLTVANANG | ASSLGLGRYAEQF | 0.7506532 |
| AGLVGAVL | ASSLGLGRYAEQF | 0.7349757 |
| ALSQAFTV | ASSLGLGRYAEQF | 0.7535201 |
| ALTFSVSL | ASSLGLGRYAEQF | 0.73260117 |
| ALTGAALVAL | ASSLGLGRYAEQF | 0.6761622 |
| ASFSIGAGVL | ASSLGLGRYAEQF | 0.7476498 |
| DGSVYKI | ASSLGLGRYAEQF | 0.77359384 |
| EQLAEIV | ASSLGLGRYAEQF | 0.74234664 |
| FAVSELTVM | ASSLGLGRYAEQF | 0.7376249 |
| GAAPLAAGL | ASSLGLGRYAEQF | 0.77843475 |
| GADGGDGGSGGAGG | ASSLGLGRYAEQF | 0.70072305 |
| GFGLDTGGAGG | ASSLGLGRYAEQF | 0.6998254 |
| GLASGGLL | ASSLGLGRYAEQF | 0.7609638 |
| GRIAGAAVLL | ASSLGLGRYAEQF | 0.7605691 |
| GSAGSLNTSF | ASSLGLGRYAEQF | 0.69606924 |
| GTGQIGI | ASSLGLGRYAEQF | 0.7802229 |
| GTLIALIL | ASSLGLGRYAEQF | 0.6804007 |
| IAATMGFVVG | ASSLGLGRYAEQF | 0.7647573 |
| IFAQALL | ASSLGLGRYAEQF | 0.7198009 |
| IGAAVALGF | ASSLGLGRYAEQF | 0.7338108 |
| IGLTPLAL | ASSLGLGRYAEQF | 0.6828345 |
| ILGSNTSTL | ASSLGLGRYAEQF | 0.7677518 |
| IVFSLTVL | ASSLGLGRYAEQF | 0.7693555 |
| LASKATGF | ASSLGLGRYAEQF | 0.7669774 |
| LASVALIL | ASSLGLGRYAEQF | 0.25928968 |
| LFAVGLTL | ASSLGLGRYAEQF | 0.769469 |
| LFNGGIGTYI | ASSLGLGRYAEQF | 0.7608826 |
| LTLGAGII | ASSLGLGRYAEQF | 0.7789196 |
| LVAAALGDI | ASSLGLGRYAEQF | 0.71559894 |
| NTAFVQLGI | ASSLGLGRYAEQF | 0.7410346 |
| QLFSGAEL | ASSLGLGRYAEQF | 0.722591 |
| SAATFITL | ASSLGLGRYAEQF | 0.69221044 |

| | | |
|---|---|---|
| SNYGLANLGV | ASSLGLGRYAEQF | 0.6168864 |
| SYGGAYAVM | ASSLGLGRYAEQF | 0.72789407 |
| TTGVGGLTLG | ASSLGLGRYAEQF | 0.7076826 |
| TTQTIYI | ASSLGLGRYAEQF | 0.69879574 |
| VAAGAGLGAVI | ASSLGLGRYAEQF | 0.7615576 |
| VAAVVTLL | ASSLGLGRYAEQF | 0.76963437 |
| VGFLGGLI | ASSLGLGRYAEQF | 0.7664805 |
| VGVIPILL | ASSLGLGRYAEQF | 0.76753926 |
| VIFASSLI | ASSLGLGRYAEQF | 0.77613044 |
| VKGGDGGAAAG | ASSLGLGRYAEQF | 0.7545211 |
| VTKTFTNI | ASSLGLGRYAEQF | 0.6895039 |
| VVAAAILL | ASSLGLGRYAEQF | 0.71044564 |
| YAALDVEL | ASSLGLGRYAEQF | 0.66304636 |
| LASVALLL | ASSLGLGRYAEQF | 0.7233416 |
| LDAVHRV | ASSLGLGRYAEQF | 0.67260194 |
| QGLSSSIF | ASSLGLGRYAEQF | 0.7480104 |
| RGTFANIRL | ASSLGLGRYAEQF | 0.213363 |
| RQLWWGHRI | ASSLGLGRYAEQF | 0.7395622 |
| YGPPGTGKTLL | ASSLGLGRYAEQF | 0.77359766 |
| AALVGAPANVA | ASSLGLGRYAEQF | 0.67985857 |
| AATFRRTE | ASSLGLGRYAEQF | 0.74195397 |
| AGALGGGI | ASSLGLGRYAEQF | 0.74497104 |
| AGGAGGAGGIGAG | ASSLGLGRYAEQF | 0.6873108 |
| AGSGALGL | ASSLGLGRYAEQF | 0.75691986 |
| AQIADQI | ASSLGLGRYAEQF | 0.6133681 |
| ASAVAVTYL | ASSLGLGRYAEQF | 0.7350613 |
| ASTVLGGTGG | ASSLGLGRYAEQF | 0.77622527 |
| AVGGEPAK | ASSLGLGRYAEQF | 0.74512225 |
| CEQGTFGGTC | ASSLGLGRYAEQF | 0.72530353 |
| DEGKAPAG | ASSLGLGRYAEQF | 0.68439496 |
| DTVVFPAL | ASSLGLGRYAEQF | 0.38204092 |
| FEGITMAG | ASSLGLGRYAEQF | 0.73826635 |
| FPLVVQD | ASSLGLGRYAEQF | 0.5671493 |
| GAFTTPQ | ASSLGLGRYAEQF | 0.7327153 |
| GFGGFGTAGGVGGA | ASSLGLGRYAEQF | 0.61781716 |
| GFLAQFL | ASSLGLGRYAEQF | 0.74943876 |
| GGAGGAGGNAGVA | ASSLGLGRYAEQF | 0.7355579 |
| GGVGTAGAGGAGGAGG | ASSLGLGRYAEQF | 0.7512651 |
| GNGGGGDAGNAGSGGN | ASSLGLGRYAEQF | 0.65370035 |
| GTGGAGGAGNDAGS | ASSLGLGRYAEQF | 0.70740294 |
| GTTNTGWFNS | ASSLGLGRYAEQF | 0.68291086 |
| IFMGPLFTL | ASSLGLGRYAEQF | 0.57845926 |
| ILAHAIEAI | ASSLGLGRYAEQF | 0.77409756 |
| IRSAIREAL | ASSLGLGRYAEQF | 0.70843995 |
| ITFAGGG | ASSLGLGRYAEQF | 0.7267163 |
| ITGMSGHQEV | ASSLGLGRYAEQF | 0.76454735 |
| IVAGLACTL | ASSLGLGRYAEQF | 0.7390388 |
| IVGGGLAGA | ASSLGLGRYAEQF | 0.4787041 |
| IVTGVLNI | ASSLGLGRYAEQF | 0.7367116 |
| LAQQGCI | ASSLGLGRYAEQF | 0.7360393 |
| LCLHVEN | ASSLGLGRYAEQF | 0.7075634 |

| | | |
|---|---|---|
| LDAISDGS | ASSLGLGRYAEQF | 0.29165763 |
| LGGFAAML | ASSLGLGRYAEQF | 0.7648141 |
| LGGKNPMI | ASSLGLGRYAEQF | 0.6497548 |
| LGPKGVAAIL | ASSLGLGRYAEQF | 0.67348784 |
| LMVGVSI | ASSLGLGRYAEQF | 0.71344006 |
| LVGAAFVL | ASSLGLGRYAEQF | 0.7621942 |
| LVLAVAPL | ASSLGLGRYAEQF | 0.7065122 |
| MAAIAVIDAA | ASSLGLGRYAEQF | 0.68694407 |
| NNGINAELQ | ASSLGLGRYAEQF | 0.7696388 |
| RGEPLVLVHG | ASSLGLGRYAEQF | 0.75418794 |
| RGGAGGAAGLI | ASSLGLGRYAEQF | 0.7698858 |
| RSYRMHPAV | ASSLGLGRYAEQF | 0.74998367 |
| SAAHLLAGQLA | ASSLGLGRYAEQF | 0.659323 |
| SAAVGSY | ASSLGLGRYAEQF | 0.7187347 |
| SPHAVLHRI | ASSLGLGRYAEQF | 0.6743304 |
| SQIAKVL | ASSLGLGRYAEQF | 0.4318124 |
| TALGHLVP | ASSLGLGRYAEQF | 0.40290964 |
| TAPTTSGAA | ASSLGLGRYAEQF | 0.73053706 |
| TDGDDGNGGSPGT | ASSLGLGRYAEQF | 0.3490462 |
| TDGLGEN | ASSLGLGRYAEQF | 0.7280259 |
| TGGPPLPGQA | ASSLGLGRYAEQF | 0.7558441 |
| TIPAITI | ASSLGLGRYAEQF | 0.7423421 |
| TIPGFTI | ASSLGLGRYAEQF | 0.5554304 |
| TSIASGALA | ASSLGLGRYAEQF | 0.72922313 |
| TVTLPGAAI | ASSLGLGRYAEQF | 0.6930975 |
| VASGLSAALP | ASSLGLGRYAEQF | 0.7626001 |
| VGAGVIGI | ASSLGLGRYAEQF | 0.74891806 |
| VGPKRFGV | ASSLGLGRYAEQF | 0.77793384 |
| VNVAFPD | ASSLGLGRYAEQF | 0.76916987 |
| VSMTNSGVS | ASSLGLGRYAEQF | 0.6988149 |
| VTDVLGDLSV | ASSLGLGRYAEQF | 0.7674391 |
| VVVVGAGFAGL | ASSLGLGRYAEQF | 0.68727887 |
| VYTVGTVV | ASSLGLGRYAEQF | 0.7293955 |
| WTTFSGP | ASSLGLGRYAEQF | 0.71610373 |
| YDELVEAF | ASSLGLGRYAEQF | 0.74780047 |
| YGTGGAGGNGG | ASSLGLGRYAEQF | 0.7192781 |
| YTNVMAV | ASSLGLGRYAEQF | 0.6873739 |
| FLEQVHQL | ASSLGLGRYAEQF | 0.43227386 |
| GLSLLAVAV | ASSLGLGRYAEQF | 0.75509465 |
| LEQVHQL | ASSLGLGRYAEQF | 0.69266236 |
| LNAAGAI | ASSLGLGRYAEQF | 0.35494897 |
| PAVHASL | ASSLGLGRYAEQF | 0.66492033 |
| TELEDDA | ASSLGLGRYAEQF | 0.78023326 |
| TPPGPPN | ASSLGLGRYAEQF | 0.76718307 |
| TPPPPPPP | ASSLGLGRYAEQF | 0.6493945 |
| TQSPALI | ASFSPTYQDTQY | 0.62324274 |
| EFAAAFPQ | ASFSPTYQDTQY | 0.74120975 |
| LLSGNLDVL | ASFSPTYQDTQY | 0.45890158 |
| GPANLGNYNI | ASFSPTYQDTQY | 0.7493673 |
| TLGPTLP | ASFSPTYQDTQY | 0.7303411 |
| VYAYSSCCK | ASFSPTYQDTQY | 0.7373253 |

| | | |
|---|---|---|
| APAPAPAPNI | ASFSPTYQDTQY | 0.5896034 |
| PGGPVPIP | ASFSPTYQDTQY | 0.742852 |
| FLTVALEP | ASFSPTYQDTQY | 0.71812546 |
| SQYFGANL | ASFSPTYQDTQY | 0.7494976 |
| GGVKPAA | ASFSPTYQDTQY | 0.3232887 |
| GGTGGAGGAGGW | ASFSPTYQDTQY | 0.7472007 |
| GVGGVAGLGGAGG | ASFSPTYQDTQY | 0.68805826 |
| VLLVLATTI | ASFSPTYQDTQY | 0.6949339 |
| IGIFAAML | ASFSPTYQDTQY | 0.55272716 |
| AVGDLID | ASFSPTYQDTQY | 0.70828795 |
| NISPLPL | ASFSPTYQDTQY | 0.76278055 |
| VTAVFVAI | ASFSPTYQDTQY | 0.7684684 |
| SGDDNDAAL | ASFSPTYQDTQY | 0.6250335 |
| AVAALAHS | ASFSPTYQDTQY | 0.76004386 |
| GGVTVMATP | ASFSPTYQDTQY | 0.7744503 |
| AGLAANVRD | ASFSPTYQDTQY | 0.707078 |
| TNRPDLIDPALLRPG | ASFSPTYQDTQY | 0.50517946 |
| LANLGVTAL | ASFSPTYQDTQY | 0.73000777 |
| TGERIEL | ASFSPTYQDTQY | 0.72045934 |
| NGEELAA | ASFSPTYQDTQY | 0.7513398 |
| VLSLVWG | ASFSPTYQDTQY | 0.75295645 |
| IGYALWL | ASFSPTYQDTQY | 0.65303206 |
| LSLLLAPL | ASFSPTYQDTQY | 0.7453699 |
| TLDEALK | ASFSPTYQDTQY | 0.7685119 |
| TDGAGTAEGG | ASFSPTYQDTQY | 0.6793821 |
| NAPGGAGGNGG | ASFSPTYQDTQY | 0.6792253 |
| VSVFMGMGGNF | ASFSPTYQDTQY | 0.59756356 |
| LASYLAAL | ASFSPTYQDTQY | 0.5436423 |
| LVGGTPAL | ASFSPTYQDTQY | 0.7529187 |
| VRNADVL | ASFSPTYQDTQY | 0.6990981 |
| TCSSPFLVSAM | ASFSPTYQDTQY | 0.6323402 |
| VHITDAE | ASFSPTYQDTQY | 0.7375808 |
| EYAAPSF | ASFSPTYQDTQY | 0.65915656 |
| VVDAFHGGL | ASFSPTYQDTQY | 0.6630113 |
| ASVVDPQ | ASFSPTYQDTQY | 0.7039169 |
| SYNEFGGLS | ASFSPTYQDTQY | 0.73265415 |
| DPDGSVY | ASFSPTYQDTQY | 0.75650907 |
| MVSTIHATE | ASFSPTYQDTQY | 0.71493936 |
| LAAGVADPV | ASFSPTYQDTQY | 0.7172105 |
| TKEQIAA | ASFSPTYQDTQY | 0.77646625 |
| SLASPAAL | ASFSPTYQDTQY | 0.46449804 |
| IGLVFYGV | ASFSPTYQDTQY | 0.6115495 |
| FLLDASNL | ASFSPTYQDTQY | 0.7554008 |
| YQVGNFP | ASFSPTYQDTQY | 0.4800075 |
| ESLLSQST | ASFSPTYQDTQY | 0.7658837 |
| LFTVGFAAVG | ASFSPTYQDTQY | 0.73365754 |
| ATPPTPT | ASFSPTYQDTQY | 0.75451237 |
| TDLATLG | ASFSPTYQDTQY | 0.73386884 |
| SAGGTGR | ASFSPTYQDTQY | 0.7795792 |
| GFSNSGS | ASFSPTYQDTQY | 0.7611328 |
| GGNGGSGGTSV | ASFSPTYQDTQY | 0.69706345 |

| | | |
|---|---|---|
| GTSVATGGAGN | ASFSPTYQDTQY | 0.7603854 |
| NGGNGGSGGTS | ASFSPTYQDTQY | 0.5380864 |
| AVVKFPH | ASFSPTYQDTQY | 0.75482666 |
| TPPPPPP | ASFSPTYQDTQY | 0.7475096 |
| AAPAAAAAPA | ASFSPTYQDTQY | 0.630975 |
| IALNEDL | ASFSPTYQDTQY | 0.7353473 |
| IVNVSSM | ASFSPTYQDTQY | 0.7729864 |
| TGIPPAPR | ASFSPTYQDTQY | 0.7758335 |
| FDLGGGTF | ASFSPTYQDTQY | 0.34603542 |
| FDLGGGTFDVS | ASFSPTYQDTQY | 0.7189139 |
| NEPTAAA | ASFSPTYQDTQY | 0.4010767 |
| NEPTAAALA | ASFSPTYQDTQY | 0.6760432 |
| GSIINISSI | ASFSPTYQDTQY | 0.54992914 |
| VAPGNAGTA | ASFSPTYQDTQY | 0.761315 |
| TGGGASLE | ASFSPTYQDTQY | 0.70609623 |
| VAVYNHYKRI | ASFSPTYQDTQY | 0.7443216 |
| TDPAPATT | ASFSPTYQDTQY | 0.73417866 |
| TNRPDILDPALLRPG | ASFSPTYQDTQY | 0.7730504 |
| LTTTTTS | ASFSPTYQDTQY | 0.69959366 |
| PTGSGKTL | ASFSPTYQDTQY | 0.7213294 |
| TGSGKTL | ASFSPTYQDTQY | 0.545374 |
| TIGHVDHGK | ASFSPTYQDTQY | 0.38184035 |
| TIGHVDHGKTT | ASFSPTYQDTQY | 0.70414925 |
| GHVDHGKTT | ASFSPTYQDTQY | 0.6987012 |
| AQFEHTLL | ASFSPTYQDTQY | 0.27118245 |
| IVNQDEL | ASFSPTYQDTQY | 0.7706767 |
| TNGSQFF | ASFSPTYQDTQY | 0.73827326 |
| TGKGIVE | ASFSPTYQDTQY | 0.75681865 |
| LISNASDA | ASFSPTYQDTQY | 0.6666323 |
| AELRTLGELTKIAKAHGAQ | ASFSPTYQDTQY | 0.71989906 |
| APNAVAAAEPSW | ASFSPTYQDTQY | 0.6982801 |
| APQVAGAATAVL | ASFSPTYQDTQY | 0.7577295 |
| ARRDPAAAAAILPTDA | ASFSPTYQDTQY | 0.7267519 |
| DAAARDVLAEAGLAEYF | ASFSPTYQDTQY | 0.44005516 |
| DDQIVLSAKANLAKVEQ | ASFSPTYQDTQY | 0.6601851 |
| DELLPDTGEDSDEDGH | ASFSPTYQDTQY | 0.61946934 |
| ELPEAPANRATVPQD | ASFSPTYQDTQY | 0.7772802 |
| FGQSKIERVTGIG | ASFSPTYQDTQY | 0.76708436 |
| GAGGTGSTAGGAGGAGGAGGLY | ASFSPTYQDTQY | 0.7501172 |
| GHGGTGAAGGNGGTGGAGGSGI | ASFSPTYQDTQY | 0.7737891 |
| GNGGNGGAGGEGGPGGAGGAGA | ASFSPTYQDTQY | 0.31573272 |
| GSTAGNGGNGGNATNGTIAGQP | ASFSPTYQDTQY | 0.67496985 |
| GTGGDGGNGGNGGTGG | ASFSPTYQDTQY | 0.76805747 |
| GVRRPAGIPADAPV | ASFSPTYQDTQY | 0.7421587 |
| GYRQVTDAHLLTLARR | ASFSPTYQDTQY | 0.71932566 |
| IAKMGQDGHDRG | ASFSPTYQDTQY | 0.7725049 |
| KAEPSIEIQYT | ASFSPTYQDTQY | 0.6863879 |
| KPVRIRMAADEDTVAVT | ASFSPTYQDTQY | 0.70164764 |
| MPPRIAGMRLLVIKPEP | ASFSPTYQDTQY | 0.7360501 |
| MKFYAVALFRSRGVPPTR | ASFSPTYQDTQY | 0.75176287 |
| NAGGFGILGAVAHSP | ASFSPTYQDTQY | 0.75698495 |

| | | |
|---|---|---|
| NEGREITLAMAEM | ASFSPTYQDTQY | 0.70358 |
| NELVEWSHPLGHRWRWE | ASFSPTYQDTQY | 0.7259207 |
| NLGAGGTGGNGGNPG | ASFSPTYQDTQY | 0.7845082 |
| NTEWPKWETVMAADPVD | ASFSPTYQDTQY | 0.7255938 |
| PASAAPAETQP | ASFSPTYQDTQY | 0.7623529 |
| QQSSDGKNGVRAAEITGE | ASFSPTYQDTQY | 0.7419743 |
| RAPDAVTTSWWKE | ASFSPTYQDTQY | 0.7294084 |
| SGFGNVGAHTSGWFNQ | ASFSPTYQDTQY | 0.67595136 |
| SGVAGAVGQAASVGGLKV | ASFSPTYQDTQY | 0.6994419 |
| SNGGAGTGSAGGNGGT | ASFSPTYQDTQY | 0.6945528 |
| TGGAGGEGGAAG | ASFSPTYQDTQY | 0.6647866 |
| TGGDGGAAGAGGAPGHG | ASFSPTYQDTQY | 0.708341 |
| TMPPLLAAIANAGRHGV | ASFSPTYQDTQY | 0.6706535 |
| TSSTAGNSGGA | ASFSPTYQDTQY | 0.786496 |
| VPATVPVGILC | ASFSPTYQDTQY | 0.75419694 |
| VQRPDSGGVVLTGRIS | ASFSPTYQDTQY | 0.7487931 |
| WTPNPVPATKGVPAGLVQAT | ASFSPTYQDTQY | 0.6908052 |
| YALKRALAKFSDRFDVVII | ASFSPTYQDTQY | 0.7374455 |
| YLIGSVIV | ASFSPTYQDTQY | 0.7497748 |
| YGQLVAVID | ASFSPTYQDTQY | 0.46358347 |
| VTVGGMTLPA | ASFSPTYQDTQY | 0.77160406 |
| VLALPAV | ASFSPTYQDTQY | 0.6481442 |
| VIPPTLNYET | ASFSPTYQDTQY | 0.6988188 |
| VGMGGMHPSSG | ASFSPTYQDTQY | 0.72335386 |
| VGLAGGAATA | ASFSPTYQDTQY | 0.7669126 |
| VGGAGGAGGSGP | ASFSPTYQDTQY | 0.70929444 |
| VFAAFMV | ASFSPTYQDTQY | 0.6545465 |
| TLYDKLR | ASFSPTYQDTQY | 0.67657596 |
| TLAVFTVL | ASFSPTYQDTQY | 0.7649177 |
| TGSVQPARV | ASFSPTYQDTQY | 0.773529 |
| TGGGAGSGGT | ASFSPTYQDTQY | 0.7318783 |
| SHLETSL | ASFSPTYQDTQY | 0.77986 |
| SFTTGMT | ASFSPTYQDTQY | 0.6830901 |
| QVEVAKRMGVRQPTVS | ASFSPTYQDTQY | 0.448691 |
| QGSGGPALAM | ASFSPTYQDTQY | 0.72523683 |
| MLAPSNP | ASFSPTYQDTQY | 0.7641624 |
| LVNNAAI | ASFSPTYQDTQY | 0.77076054 |
| LTGEINTLA | ASFSPTYQDTQY | 0.7737199 |
| LSIFGPV | ASFSPTYQDTQY | 0.76514375 |
| LLALLANL | ASFSPTYQDTQY | 0.7223739 |
| LISNASDALDKLR | ASFSPTYQDTQY | 0.7712728 |
| LINEAVP | ASFSPTYQDTQY | 0.77362764 |
| LGYAFSGL | ASFSPTYQDTQY | 0.75014985 |
| LDFVAVAL | ASFSPTYQDTQY | 0.6705742 |
| LAQPGAE | ASFSPTYQDTQY | 0.5319427 |
| ISNLGAVTGF | ASFSPTYQDTQY | 0.7383175 |
| HQEDPGRI | ASFSPTYQDTQY | 0.76974905 |
| GTPGEPTP | ASFSPTYQDTQY | 0.5597952 |
| GFFNSGS | ASFSPTYQDTQY | 0.7385706 |
| GAGGGGLTAGAN | ASFSPTYQDTQY | 0.74110246 |
| FVDALVNM | ASFSPTYQDTQY | 0.5003569 |

| | | |
|---|---|---|
| FLTVAPEL | ASFSPTYQDTQY | 0.7569107 |
| FETLTAL | ASFSPTYQDTQY | 0.7754954 |
| DVINPIEL | ASFSPTYQDTQY | 0.7061402 |
| APVSGAPV | ASFSPTYQDTQY | 0.6902778 |
| ALVGGLAW | ASFSPTYQDTQY | 0.77165836 |
| ALAAAAATAI | ASFSPTYQDTQY | 0.7855038 |
| AFGLGLLL | ASFSPTYQDTQY | 0.7640406 |
| AFFAAASL | ASFSPTYQDTQY | 0.45594215 |
| AAAALPAFSPP | ASFSPTYQDTQY | 0.3298935 |
| VVLAPAL | ASFSPTYQDTQY | 0.6878817 |
| TIGHVDHGKTTL | ASFSPTYQDTQY | 0.75220025 |
| LISNASDALDK | ASFSPTYQDTQY | 0.6867654 |
| LISNASDAL | ASFSPTYQDTQY | 0.67862785 |
| ISNASDALDK | ASFSPTYQDTQY | 0.7549454 |
| NQAETLVYQTE | ASFSPTYQDTQY | 0.7663431 |
| TGGTGGTAGAG | ASFSPTYQDTQY | 0.6846061 |
| TGSTAGGAGGA | ASFSPTYQDTQY | 0.52535367 |
| DPLAVPVAAGPV | ASFSPTYQDTQY | 0.7562474 |
| GGTGGTPGNGGN | ASFSPTYQDTQY | 0.75632936 |
| PAGLAGVDPATL | ASFSPTYQDTQY | 0.66616917 |
| GNNLVVEAPHAGASV | ASFSPTYQDTQY | 0.73809266 |
| KVTRIMSASTLEQPA | ASFSPTYQDTQY | 0.6648186 |
| LGFALPIALPRRTVV | ASFSPTYQDTQY | 0.59728265 |
| TRLGNLTARKLVVLV | ASFSPTYQDTQY | 0.7453399 |
| WSQQRRLIYFPSAGP | ASFSPTYQDTQY | 0.65447557 |
| GPNGNPDPMAAAVDIR | ASFSPTYQDTQY | 0.7782092 |
| TQRCVSVADLRAAASR | ASFSPTYQDTQY | 0.66637635 |
| QFVLDVAEHPVEVLTETV | ASFSPTYQDTQY | 0.76890385 |
| GGAGGTGGAGGNGGTQAGDG | ASFSPTYQDTQY | 0.7464912 |
| PLLATLRAIAVDVDAKPAQV | ASFSPTYQDTQY | 0.75387806 |
| APKFRTGGYMIDFWGVGYQVAK | ASFSPTYQDTQY | 0.7187345 |
| GAGGAGGAGGAAGGPGGTGGTG | ASFSPTYQDTQY | 0.6660348 |
| IGALGLLATDVAPAGSVAFGVA | ASFSPTYQDTQY | 0.6527127 |
| AGGAGGIGGNGGAGGAGGVLMG | ASFSPTYQDTQY | 0.4578169 |
| ADELIGLL | ASFSPTYQDTQY | 0.7667318 |
| ADRALKA | ASFSPTYQDTQY | 0.6666244 |
| AGAVGQVGGAGGA | ASFSPTYQDTQY | 0.77331674 |
| AGGGGGLGGAP | ASFSPTYQDTQY | 0.75827855 |
| AGGLLYGNGGAG | ASFSPTYQDTQY | 0.6599613 |
| AGLESIGL | ASFSPTYQDTQY | 0.75558877 |
| AGLKTGII | ASFSPTYQDTQY | 0.61509264 |
| ASLAALLL | ASFSPTYQDTQY | 0.5757339 |
| ASVINISSI | ASFSPTYQDTQY | 0.755893 |
| ATPPGPA | ASFSPTYQDTQY | 0.7447099 |
| ATVASALV | ASFSPTYQDTQY | 0.66838133 |
| ATVASLIL | ASFSPTYQDTQY | 0.7387727 |
| AVNGPVLIRH | ASFSPTYQDTQY | 0.7482799 |
| DGAAGAVGI | ASFSPTYQDTQY | 0.6718278 |
| DPLLHGI | ASFSPTYQDTQY | 0.71473885 |
| ELGAALGAL | ASFSPTYQDTQY | 0.77191114 |
| ELGRTQAV | ASFSPTYQDTQY | 0.727684 |

| | | |
|---|---|---|
| FVAGGYLS | ASFSPTYQDTQY | 0.7072358 |
| GASGGAGGAGGW | ASFSPTYQDTQY | 0.7484004 |
| GGAGAGGDGGAGGI | ASFSPTYQDTQY | 0.65654576 |
| GTGHNGSQPGMGGNG | ASFSPTYQDTQY | 0.753198 |
| GYAAASASAS | ASFSPTYQDTQY | 0.74165523 |
| HQVDVAAVQQG | ASFSPTYQDTQY | 0.6969684 |
| IGAAIAHT | ASFSPTYQDTQY | 0.75864327 |
| IGMIEVAQL | ASFSPTYQDTQY | 0.49663186 |
| IRLAPEHTSAMD | ASFSPTYQDTQY | 0.7026235 |
| IYNVSVL | ASFSPTYQDTQY | 0.7037138 |
| KQGFLGM | ASFSPTYQDTQY | 0.7436794 |
| LESVITQL | ASFSPTYQDTQY | 0.756338 |
| LGLGALVVV | ASFSPTYQDTQY | 0.6205454 |
| LIGNGGNGGSGGI | ASFSPTYQDTQY | 0.73753834 |
| LNAINAPTL | ASFSPTYQDTQY | 0.65930545 |
| LQLLNTGV | ASFSPTYQDTQY | 0.5804278 |
| LQTSIEF | ASFSPTYQDTQY | 0.69545376 |
| MSFAVSEL | ASFSPTYQDTQY | 0.68904775 |
| NTGALGL | ASFSPTYQDTQY | 0.6512333 |
| PPGAGGAGGAGTT | ASFSPTYQDTQY | 0.7469928 |
| QLVGLGI | ASFSPTYQDTQY | 0.62661433 |
| RFLEQVHQL | ASFSPTYQDTQY | 0.74425507 |
| RGVGYVLR | ASFSPTYQDTQY | 0.7421118 |
| SATGVFAGVI | ASFSPTYQDTQY | 0.7336813 |
| SLPAITLP | ASFSPTYQDTQY | 0.7653425 |
| STALAFTV | ASFSPTYQDTQY | 0.75021154 |
| SVLTVGLGK | ASFSPTYQDTQY | 0.7621949 |
| TALAGAGSGI | ASFSPTYQDTQY | 0.71246576 |
| TGLPFTGI | ASFSPTYQDTQY | 0.70475245 |
| VAEHVTV | ASFSPTYQDTQY | 0.7724291 |
| VDIVLNSL | ASFSPTYQDTQY | 0.7540038 |
| VLTGGGAVL | ASFSPTYQDTQY | 0.71244377 |
| VNVGLAYGI | ASFSPTYQDTQY | 0.7508652 |
| VSSAAVGAV | ASFSPTYQDTQY | 0.7508037 |
| VTSVYEAGV | ASFSPTYQDTQY | 0.7548634 |
| WVALDRGA | ASFSPTYQDTQY | 0.70581126 |
| YLVGSVVI | ASFSPTYQDTQY | 0.52134556 |
| GPPGTGKTLL | ASFSPTYQDTQY | 0.71540344 |
| GVGMTKF | ASFSPTYQDTQY | 0.72284967 |
| IGIGGSDL | ASFSPTYQDTQY | 0.65367925 |
| IGTLLAEL | ASFSPTYQDTQY | 0.3477677 |
| IYTSGTT | ASFSPTYQDTQY | 0.7656889 |
| LSFPAGV | ASFSPTYQDTQY | 0.7837591 |
| LYTSGTT | ASFSPTYQDTQY | 0.6110256 |
| VGAFPFGF | ASFSPTYQDTQY | 0.7792221 |
| VGAGTVEF | ASFSPTYQDTQY | 0.5515411 |
| VGKSTLLL | ASFSPTYQDTQY | 0.7516371 |
| AALAHAGGGIV | ASFSPTYQDTQY | 0.76667714 |
| AALVAVSLL | ASFSPTYQDTQY | 0.62991047 |
| AAVSRAVAAL | ASFSPTYQDTQY | 0.72940755 |
| AGGSSGIGGF | ASFSPTYQDTQY | 0.6834984 |

| | | |
|---|---|---|
| AGLTVANANG | ASFSPTYQDTQY | 0.7230376 |
| AGLVGAVL | ASFSPTYQDTQY | 0.77258646 |
| ALSQAFTV | ASFSPTYQDTQY | 0.69613004 |
| ALTFSVSL | ASFSPTYQDTQY | 0.7209085 |
| ALTGAALVAL | ASFSPTYQDTQY | 0.70936537 |
| ASFSIGAGVL | ASFSPTYQDTQY | 0.6475855 |
| DGSVYKI | ASFSPTYQDTQY | 0.74826336 |
| EQLAEIV | ASFSPTYQDTQY | 0.75698376 |
| FAVSELTVM | ASFSPTYQDTQY | 0.5963036 |
| GAAPLAAGL | ASFSPTYQDTQY | 0.76841104 |
| GADGGDGGSGGAGG | ASFSPTYQDTQY | 0.67508656 |
| GFGLDTGGAGG | ASFSPTYQDTQY | 0.7836101 |
| GLASGGLL | ASFSPTYQDTQY | 0.7674295 |
| GRIAGAAVLL | ASFSPTYQDTQY | 0.74321246 |
| GSAGSLNTSF | ASFSPTYQDTQY | 0.69885194 |
| GTGQIGI | ASFSPTYQDTQY | 0.7701128 |
| GTLIALIL | ASFSPTYQDTQY | 0.71836895 |
| IAATMGFVVG | ASFSPTYQDTQY | 0.75407135 |
| IFAQALL | ASFSPTYQDTQY | 0.7647165 |
| IGAAVALGF | ASFSPTYQDTQY | 0.4694466 |
| IGLTPLAL | ASFSPTYQDTQY | 0.77132976 |
| ILGSNTSTL | ASFSPTYQDTQY | 0.72312105 |
| IVFSLTVL | ASFSPTYQDTQY | 0.6439316 |
| LASKATGF | ASFSPTYQDTQY | 0.5173997 |
| LASVALIL | ASFSPTYQDTQY | 0.34831566 |
| LFAVGLTL | ASFSPTYQDTQY | 0.60123175 |
| LFNGGIGTYI | ASFSPTYQDTQY | 0.48045784 |
| LTLGAGII | ASFSPTYQDTQY | 0.71198785 |
| LVAAALGDI | ASFSPTYQDTQY | 0.74637437 |
| NTAFVQLGI | ASFSPTYQDTQY | 0.53510064 |
| QLFSGAEL | ASFSPTYQDTQY | 0.75236106 |
| SAATFITL | ASFSPTYQDTQY | 0.7107505 |
| SNYGLANLGV | ASFSPTYQDTQY | 0.41673583 |
| SYGGAYAVM | ASFSPTYQDTQY | 0.6641121 |
| TTGVGGLTLG | ASFSPTYQDTQY | 0.502609 |
| TTQTIYI | ASFSPTYQDTQY | 0.63219094 |
| VAAGAGLGAVI | ASFSPTYQDTQY | 0.348407 |
| VAAVVTLL | ASFSPTYQDTQY | 0.7152854 |
| VGFLGGLI | ASFSPTYQDTQY | 0.59578645 |
| VGVIPILL | ASFSPTYQDTQY | 0.24477378 |
| VIFASSLI | ASFSPTYQDTQY | 0.70500064 |
| VKGGDGGAAAG | ASFSPTYQDTQY | 0.66250575 |
| VTKTFTNI | ASFSPTYQDTQY | 0.5801296 |
| VVAAAILL | ASFSPTYQDTQY | 0.7536319 |
| YAALDVEL | ASFSPTYQDTQY | 0.5327544 |
| LASVALLL | ASFSPTYQDTQY | 0.543534 |
| LDAVHRV | ASFSPTYQDTQY | 0.7357605 |
| QGLSSSIF | ASFSPTYQDTQY | 0.76212597 |
| RGTFANIRL | ASFSPTYQDTQY | 0.425278 |
| RQLWWGHRI | ASFSPTYQDTQY | 0.6940893 |
| YGPPGTGKTLL | ASFSPTYQDTQY | 0.7523899 |

| | | |
|---|---|---|
| AALVGAPANVA | ASFSPTYQDTQY | 0.7001934 |
| AATFRRTE | ASFSPTYQDTQY | 0.75857735 |
| AGALGGGI | ASFSPTYQDTQY | 0.7674986 |
| AGGAGGAGGIGAG | ASFSPTYQDTQY | 0.54420364 |
| AGSGALGL | ASFSPTYQDTQY | 0.7538278 |
| AQIADQI | ASFSPTYQDTQY | 0.7703148 |
| ASAVAVTYL | ASFSPTYQDTQY | 0.60236216 |
| ASTVLGGTGG | ASFSPTYQDTQY | 0.6577027 |
| AVGGEPAK | ASFSPTYQDTQY | 0.49058226 |
| CEQGTFGGTC | ASFSPTYQDTQY | 0.64719707 |
| DEGKAPAG | ASFSPTYQDTQY | 0.29184064 |
| DTVVFPAL | ASFSPTYQDTQY | 0.71836126 |
| FEGITMAG | ASFSPTYQDTQY | 0.74811596 |
| FPLVVQD | ASFSPTYQDTQY | 0.64899284 |
| GAFTTPQ | ASFSPTYQDTQY | 0.51285464 |
| GFGGFGTAGGVGGA | ASFSPTYQDTQY | 0.40510356 |
| GFLAQFL | ASFSPTYQDTQY | 0.70194525 |
| GGAGGAGGNAGVA | ASFSPTYQDTQY | 0.4511099 |
| GGVGTAGAGGAGGAGG | ASFSPTYQDTQY | 0.36973202 |
| GNGGGGDAGNAGSGGN | ASFSPTYQDTQY | 0.62786806 |
| GTGGAGGAGNDAGS | ASFSPTYQDTQY | 0.71950996 |
| GTTNTGWFNS | ASFSPTYQDTQY | 0.73919183 |
| IFMGPLFTL | ASFSPTYQDTQY | 0.40685874 |
| ILAHAIEAI | ASFSPTYQDTQY | 0.771338 |
| IRSAIREAL | ASFSPTYQDTQY | 0.7071552 |
| ITFAGGG | ASFSPTYQDTQY | 0.7528485 |
| ITGMSGHQEV | ASFSPTYQDTQY | 0.70677304 |
| IVAGLACTL | ASFSPTYQDTQY | 0.46587062 |
| IVGGGLAGA | ASFSPTYQDTQY | 0.74190956 |
| IVTGVLNI | ASFSPTYQDTQY | 0.6296232 |
| LAQQGCI | ASFSPTYQDTQY | 0.58578086 |
| LCLHVEN | ASFSPTYQDTQY | 0.7366163 |
| LDAISDGS | ASFSPTYQDTQY | 0.6462116 |
| LGGFAAML | ASFSPTYQDTQY | 0.54024017 |
| LGGKNPMI | ASFSPTYQDTQY | 0.6014449 |
| LGPKGVAAIL | ASFSPTYQDTQY | 0.7607644 |
| LMVGVSI | ASFSPTYQDTQY | 0.5138351 |
| LVGAAFVL | ASFSPTYQDTQY | 0.38329965 |
| LVLAVAPL | ASFSPTYQDTQY | 0.5447437 |
| MAAIAVIDAA | ASFSPTYQDTQY | 0.69795716 |
| NNGINAELQ | ASFSPTYQDTQY | 0.7604562 |
| RGEPLVLVHG | ASFSPTYQDTQY | 0.7233597 |
| RGGAGGAAGLI | ASFSPTYQDTQY | 0.58633643 |
| RSYRMHPAV | ASFSPTYQDTQY | 0.7279928 |
| SAAHLLAGQLA | ASFSPTYQDTQY | 0.6860746 |
| SAAVGSY | ASFSPTYQDTQY | 0.6836946 |
| SPHAVLHRI | ASFSPTYQDTQY | 0.7087242 |
| SQIAKVL | ASFSPTYQDTQY | 0.570781 |
| TALGHLVP | ASFSPTYQDTQY | 0.7577226 |
| TAPTTSGAA | ASFSPTYQDTQY | 0.7121866 |
| TDGDDGNGGSPGT | ASFSPTYQDTQY | 0.6998096 |

| | | |
|---|---|---|
| TDGLGEN | ASFSPTYQDTQY | 0.7534788 |
| TGGPPLPGQA | ASFSPTYQDTQY | 0.68222797 |
| TIPAITI | ASFSPTYQDTQY | 0.69210017 |
| TIPGFTI | ASFSPTYQDTQY | 0.32030183 |
| TSIASGALA | ASFSPTYQDTQY | 0.6320317 |
| TVTLPGAAI | ASFSPTYQDTQY | 0.69081867 |
| VASGLSAALP | ASFSPTYQDTQY | 0.70919466 |
| VGAGVIGI | ASFSPTYQDTQY | 0.72491455 |
| VGPKRFGV | ASFSPTYQDTQY | 0.6914451 |
| VNVAFPD | ASFSPTYQDTQY | 0.58495206 |
| VSMTNSGVS | ASFSPTYQDTQY | 0.7462491 |
| VTDVLGDLSV | ASFSPTYQDTQY | 0.7526485 |
| VVVVGAGFAGL | ASFSPTYQDTQY | 0.43690038 |
| VYTVGTVV | ASFSPTYQDTQY | 0.65359604 |
| WTTFSGP | ASFSPTYQDTQY | 0.77612793 |
| YDELVEAF | ASFSPTYQDTQY | 0.6899586 |
| YGTGGAGGNGG | ASFSPTYQDTQY | 0.6304058 |
| YTNVMAV | ASFSPTYQDTQY | 0.5480338 |
| FLEQVHQL | ASFSPTYQDTQY | 0.73656034 |
| GLSLLAVAV | ASFSPTYQDTQY | 0.5240513 |
| LEQVHQL | ASFSPTYQDTQY | 0.7715497 |
| LNAAGAI | ASFSPTYQDTQY | 0.23524944 |
| PAVHASL | ASFSPTYQDTQY | 0.6148428 |
| TELEDDA | ASFSPTYQDTQY | 0.53263366 |
| TPPGPPN | ASFSPTYQDTQY | 0.27457148 |
| TPPPPPPP | ASFSPTYQDTQY | 0.755281 |
| TQSPALI | GARDKNTGQLY | 0.5400483 |
| EFAAAFPQ | GARDKNTGQLY | 0.70637435 |
| LLSGNLDVL | GARDKNTGQLY | 0.6795584 |
| GPANLGNYNI | GARDKNTGQLY | 0.7488467 |
| TLGPTLP | GARDKNTGQLY | 0.6930816 |
| VYAYSSCCK | GARDKNTGQLY | 0.59821236 |
| APAPAPAPNI | GARDKNTGQLY | 0.4439813 |
| PGGPVPIP | GARDKNTGQLY | 0.72713685 |
| FLTVALEP | GARDKNTGQLY | 0.75819623 |
| SQYFGANL | GARDKNTGQLY | 0.717199 |
| GGVKPAA | GARDKNTGQLY | 0.30081806 |
| GGTGGAGGAGGW | GARDKNTGQLY | 0.6728925 |
| GVGGVAGLGGAGG | GARDKNTGQLY | 0.66886884 |
| VLLVLATTI | GARDKNTGQLY | 0.37017062 |
| IGIFAAML | GARDKNTGQLY | 0.25548214 |
| AVGDLID | GARDKNTGQLY | 0.61109614 |
| NISPLPL | GARDKNTGQLY | 0.6630454 |
| VTAVFVAI | GARDKNTGQLY | 0.7450731 |
| SGDDNDAAL | GARDKNTGQLY | 0.7462878 |
| AVAALAHS | GARDKNTGQLY | 0.6203289 |
| GGVTVMATP | GARDKNTGQLY | 0.7421088 |
| AGLAANVRD | GARDKNTGQLY | 0.62314475 |
| TNRPDLIDPALLRPG | GARDKNTGQLY | 0.18878332 |
| LANLGVTAL | GARDKNTGQLY | 0.57045317 |
| TGERIEL | GARDKNTGQLY | 0.38891447 |

| | | |
|---|---|---|
| NGEELAA | GARDKNTGQLY | 0.73785615 |
| VLSLVWG | GARDKNTGQLY | 0.61175686 |
| IGYALWL | GARDKNTGQLY | 0.6237665 |
| LSLLLAPL | GARDKNTGQLY | 0.65888715 |
| TLDEALK | GARDKNTGQLY | 0.6841291 |
| TDGAGTAEGG | GARDKNTGQLY | 0.670625 |
| NAPGGAGGNGG | GARDKNTGQLY | 0.49646485 |
| VSVFMGMGGNF | GARDKNTGQLY | 0.29986286 |
| LASYLAAL | GARDKNTGQLY | 0.71577936 |
| LVGGTPAL | GARDKNTGQLY | 0.47636256 |
| VRNADVL | GARDKNTGQLY | 0.48644707 |
| TCSSPFLVSAM | GARDKNTGQLY | 0.29809567 |
| VHITDAE | GARDKNTGQLY | 0.74608797 |
| EYAAPSF | GARDKNTGQLY | 0.29705685 |
| VVDAFHGGL | GARDKNTGQLY | 0.7368412 |
| ASVVDPQ | GARDKNTGQLY | 0.6170542 |
| SYNEFGGLS | GARDKNTGQLY | 0.66871554 |
| DPDGSVY | GARDKNTGQLY | 0.77879155 |
| MVSTIHATE | GARDKNTGQLY | 0.65417695 |
| LAAGVADPV | GARDKNTGQLY | 0.6240488 |
| TKEQIAA | GARDKNTGQLY | 0.4693298 |
| SLASPAAL | GARDKNTGQLY | 0.48920685 |
| IGLVFYGV | GARDKNTGQLY | 0.5112577 |
| FLLDASNL | GARDKNTGQLY | 0.5169396 |
| YQVGNFP | GARDKNTGQLY | 0.76497716 |
| ESLLSQST | GARDKNTGQLY | 0.591107 |
| LFTVGFAAVG | GARDKNTGQLY | 0.4014181 |
| ATPPTPT | GARDKNTGQLY | 0.6535663 |
| TDLATLG | GARDKNTGQLY | 0.7650482 |
| SAGGTGR | GARDKNTGQLY | 0.7817788 |
| GFSNSGS | GARDKNTGQLY | 0.5119577 |
| GGNGGSGGTSV | GARDKNTGQLY | 0.5562479 |
| GTSVATGGAGN | GARDKNTGQLY | 0.60084355 |
| NGGNGGSGGTS | GARDKNTGQLY | 0.669041 |
| AVVKFPH | GARDKNTGQLY | 0.5421325 |
| TPPPPPP | GARDKNTGQLY | 0.41264412 |
| AAPAAAAAPA | GARDKNTGQLY | 0.7370135 |
| IALNEDL | GARDKNTGQLY | 0.73637605 |
| IVNVSSM | GARDKNTGQLY | 0.627928 |
| TGIPPAPR | GARDKNTGQLY | 0.3400842 |
| FDLGGGTF | GARDKNTGQLY | 0.6562902 |
| FDLGGGTFDVS | GARDKNTGQLY | 0.483922 |
| NEPTAAA | GARDKNTGQLY | 0.57832897 |
| NEPTAAALA | GARDKNTGQLY | 0.5857574 |
| GSIINISSI | GARDKNTGQLY | 0.45375407 |
| VAPGNAGTA | GARDKNTGQLY | 0.6781461 |
| TGGGASLE | GARDKNTGQLY | 0.7789175 |
| VAVYNHYKRI | GARDKNTGQLY | 0.55518746 |
| TDPAPATT | GARDKNTGQLY | 0.715194 |
| TNRPDILDPALLRPG | GARDKNTGQLY | 0.7693728 |
| LTTTTTS | GARDKNTGQLY | 0.728261 |

| | | |
|---|---|---|
| PTGSGKTL | GARDKNTGQLY | 0.76369023 |
| TGSGKTL | GARDKNTGQLY | 0.709522 |
| TIGHVDHGK | GARDKNTGQLY | 0.6173575 |
| TIGHVDHGKTT | GARDKNTGQLY | 0.6809425 |
| GHVDHGKTT | GARDKNTGQLY | 0.75198984 |
| AQFEHTLL | GARDKNTGQLY | 0.6278616 |
| IVNQDEL | GARDKNTGQLY | 0.65125716 |
| TNGSQFF | GARDKNTGQLY | 0.5661131 |
| TGKGIVE | GARDKNTGQLY | 0.7561469 |
| LISNASDA | GARDKNTGQLY | 0.70307493 |
| AELRTLGELTKIAKAHGAQ | GARDKNTGQLY | 0.7291863 |
| APNAVAAAEPSW | GARDKNTGQLY | 0.31624407 |
| APQVAGAATAVL | GARDKNTGQLY | 0.37297124 |
| ARRDPAAAAAILPTDA | GARDKNTGQLY | 0.74480975 |
| DAAARDVLAEAGLAEYF | GARDKNTGQLY | 0.20180929 |
| DDQIVLSAKANLAKVEQ | GARDKNTGQLY | 0.2537985 |
| DELLPDTGEDSDEDGH | GARDKNTGQLY | 0.6978699 |
| ELPEAPANRATVPQD | GARDKNTGQLY | 0.7581805 |
| FGQSKIERVTGIG | GARDKNTGQLY | 0.22770213 |
| GAGGTGSTAGGAGGAGGAGGLY | GARDKNTGQLY | 0.51370627 |
| GHGGTGAAGGNGGTGGAGGSGI | GARDKNTGQLY | 0.37413603 |
| GNGGNGGAGGEGGPGGAGGAGA | GARDKNTGQLY | 0.68398917 |
| GSTAGNGGNGGNATNGTIAGQP | GARDKNTGQLY | 0.6409919 |
| GTGGDGGNGGNGGTGG | GARDKNTGQLY | 0.34726393 |
| GVRRPAGIPADAPV | GARDKNTGQLY | 0.7205523 |
| GYRQVTDAHLLTLARR | GARDKNTGQLY | 0.33895212 |
| IAKMGQDGHDRG | GARDKNTGQLY | 0.63463557 |
| KAEPSIEIQYT | GARDKNTGQLY | 0.74973667 |
| KPVRIRMAADEDTVAVT | GARDKNTGQLY | 0.74930835 |
| MPPRIAGMRLLVIKPEP | GARDKNTGQLY | 0.5793563 |
| MKFYAVALFRSRGVPPTR | GARDKNTGQLY | 0.7272303 |
| NAGGFGILGAVAHSP | GARDKNTGQLY | 0.60043275 |
| NEGREITLAMAEM | GARDKNTGQLY | 0.78911006 |
| NELVEWSHPLGHRWRWE | GARDKNTGQLY | 0.60505056 |
| NLGAGGTGGNGGNPG | GARDKNTGQLY | 0.40507448 |
| NTEWPKWETVMAADPVD | GARDKNTGQLY | 0.748192 |
| PASAAPAETQP | GARDKNTGQLY | 0.72631323 |
| QQSSDGKNGVRAAEITGE | GARDKNTGQLY | 0.6317767 |
| RAPDAVTTSWWKE | GARDKNTGQLY | 0.725348 |
| SGFGNVGAHTSGWFNQ | GARDKNTGQLY | 0.5627088 |
| SGVAGAVGQAASVGGLKV | GARDKNTGQLY | 0.6422304 |
| SNGGAGTGSAGGNGGT | GARDKNTGQLY | 0.74814165 |
| TGGAGGEGGAAG | GARDKNTGQLY | 0.74385476 |
| TGGDGGAAGAGGAPGHG | GARDKNTGQLY | 0.41801894 |
| TMPPLLAAIANAGRHGV | GARDKNTGQLY | 0.740164 |
| TSSTAGNSGGA | GARDKNTGQLY | 0.60709393 |
| VPATVPVGILC | GARDKNTGQLY | 0.5131612 |
| VQRPDSGGVVLTGRIS | GARDKNTGQLY | 0.7804572 |
| WTPNPVPATKGVPAGLVQAT | GARDKNTGQLY | 0.572697 |
| YALKRALAKFSDRFDVVII | GARDKNTGQLY | 0.742306 |
| YLIGSVIV | GARDKNTGQLY | 0.7240902 |

| | | |
|---|---|---|
| YGQLVAVID | GARDKNTGQLY | 0.71561706 |
| VTVGGMTLPA | GARDKNTGQLY | 0.4360453 |
| VLALPAV | GARDKNTGQLY | 0.66942316 |
| VIPPTLNYET | GARDKNTGQLY | 0.70379126 |
| VGMGGMHPSSG | GARDKNTGQLY | 0.7192714 |
| VGLAGGAATA | GARDKNTGQLY | 0.6227952 |
| VGGAGGAGGSGP | GARDKNTGQLY | 0.6349951 |
| VFAAFMV | GARDKNTGQLY | 0.7090244 |
| TLYDKLR | GARDKNTGQLY | 0.7718837 |
| TLAVFTVL | GARDKNTGQLY | 0.6893407 |
| TGSVQPARV | GARDKNTGQLY | 0.6910843 |
| TGGGAGSGGT | GARDKNTGQLY | 0.7685715 |
| SHLETSL | GARDKNTGQLY | 0.5249777 |
| SFTTGMT | GARDKNTGQLY | 0.61205804 |
| QVEVAKRMGVRQPTVS | GARDKNTGQLY | 0.74466383 |
| QGSGGPALAM | GARDKNTGQLY | 0.75540364 |
| MLAPSNP | GARDKNTGQLY | 0.63346773 |
| LVNNAAI | GARDKNTGQLY | 0.75871396 |
| LTGEINTLA | GARDKNTGQLY | 0.6096756 |
| LSIFGPV | GARDKNTGQLY | 0.71243334 |
| LLALLANL | GARDKNTGQLY | 0.6594357 |
| LISNASDALDKLR | GARDKNTGQLY | 0.661605 |
| LINEAVP | GARDKNTGQLY | 0.5279585 |
| LGYAFSGL | GARDKNTGQLY | 0.7050409 |
| LDFVAVAL | GARDKNTGQLY | 0.7491008 |
| LAQPGAE | GARDKNTGQLY | 0.7135935 |
| ISNLGAVTGF | GARDKNTGQLY | 0.25463426 |
| HQEDPGRI | GARDKNTGQLY | 0.7380943 |
| GTPGEPTP | GARDKNTGQLY | 0.6700476 |
| GFFNSGS | GARDKNTGQLY | 0.77377176 |
| GAGGGGLTAGAN | GARDKNTGQLY | 0.651304 |
| FVDALVNM | GARDKNTGQLY | 0.7136409 |
| FLTVAPEL | GARDKNTGQLY | 0.5696956 |
| FETLTAL | GARDKNTGQLY | 0.60438883 |
| DVINPIEL | GARDKNTGQLY | 0.40089694 |
| APVSGAPV | GARDKNTGQLY | 0.6178768 |
| ALVGGLAW | GARDKNTGQLY | 0.66557324 |
| ALAAAAATAI | GARDKNTGQLY | 0.57044506 |
| AFGLGLLL | GARDKNTGQLY | 0.72454286 |
| AFFAAASL | GARDKNTGQLY | 0.75846326 |
| AAAALPAFSPP | GARDKNTGQLY | 0.75420105 |
| VVLAPAL | GARDKNTGQLY | 0.7610838 |
| TIGHVDHGKTTL | GARDKNTGQLY | 0.77277374 |
| LISNASDALDK | GARDKNTGQLY | 0.7236334 |
| LISNASDAL | GARDKNTGQLY | 0.48918813 |
| ISNASDALDK | GARDKNTGQLY | 0.647099 |
| NQAETLVYQTE | GARDKNTGQLY | 0.41644803 |
| TGGTGGTAGAG | GARDKNTGQLY | 0.6693373 |
| TGSTAGGAGGA | GARDKNTGQLY | 0.52847624 |
| DPLAVPVAAGPV | GARDKNTGQLY | 0.7133848 |
| GGTGGTPGNGGN | GARDKNTGQLY | 0.20333537 |

| | | |
|---|---|---|
| PAGLAGVDPATL | GARDKNTGQLY | 0.68891084 |
| GNNLVVEAPHAGASV | GARDKNTGQLY | 0.77360165 |
| KVTRIMSASTLEQPA | GARDKNTGQLY | 0.4739778 |
| LGFALPIALPRRTVV | GARDKNTGQLY | 0.710764 |
| TRLGNLTARKLVVLV | GARDKNTGQLY | 0.74028337 |
| WSQQRRLIYFPSAGP | GARDKNTGQLY | 0.58629566 |
| GPNGNPDPMAAAVDIR | GARDKNTGQLY | 0.738794 |
| TQRCVSVADLRAAASR | GARDKNTGQLY | 0.5266857 |
| QFVLDVAEHPVEVLTETV | GARDKNTGQLY | 0.72428656 |
| GGAGGTGGAGGNGGTQAGDG | GARDKNTGQLY | 0.76729786 |
| PLLATLRAIAVDVDAKPAQV | GARDKNTGQLY | 0.7127065 |
| APKFRTGGYMIDFWGVGYQVAK | GARDKNTGQLY | 0.66221964 |
| GAGGAGGAGGAAGGPGGTGGTG | GARDKNTGQLY | 0.6439043 |
| IGALGLLATDVAPAGSVAFGVA | GARDKNTGQLY | 0.32767892 |
| AGGAGGIGGNGGAGGAGGVLMG | GARDKNTGQLY | 0.6756865 |
| ADELIGLL | GARDKNTGQLY | 0.41462427 |
| ADRALKA | GARDKNTGQLY | 0.6762072 |
| AGAVGQVGGAGGA | GARDKNTGQLY | 0.4245715 |
| AGGGGGLGGAP | GARDKNTGQLY | 0.7042645 |
| AGGLLYGNGGAG | GARDKNTGQLY | 0.6610483 |
| AGLESIGL | GARDKNTGQLY | 0.7115373 |
| AGLKTGII | GARDKNTGQLY | 0.42071798 |
| ASLAALLL | GARDKNTGQLY | 0.67181945 |
| ASVINISSI | GARDKNTGQLY | 0.633427 |
| ATPPGPA | GARDKNTGQLY | 0.4023011 |
| ATVASALV | GARDKNTGQLY | 0.7594117 |
| ATVASLIL | GARDKNTGQLY | 0.63839805 |
| AVNGPVLIRH | GARDKNTGQLY | 0.6655676 |
| DGAAGAVGI | GARDKNTGQLY | 0.73062515 |
| DPLLHGI | GARDKNTGQLY | 0.6629964 |
| ELGAALGAL | GARDKNTGQLY | 0.29063132 |
| ELGRTQAV | GARDKNTGQLY | 0.66046 |
| FVAGGYLS | GARDKNTGQLY | 0.6809988 |
| GASGGAGGAGGW | GARDKNTGQLY | 0.5713986 |
| GGAGAGGDGGAGGI | GARDKNTGQLY | 0.24463253 |
| GTGHNGSQPGMGGNG | GARDKNTGQLY | 0.7204989 |
| GYAAASASAS | GARDKNTGQLY | 0.49762446 |
| HQVDVAAVQQG | GARDKNTGQLY | 0.58044505 |
| IGAAIAHT | GARDKNTGQLY | 0.62580824 |
| IGMIEVAQL | GARDKNTGQLY | 0.7483438 |
| IRLAPEHTSAMD | GARDKNTGQLY | 0.5940496 |
| IYNVSVL | GARDKNTGQLY | 0.53977823 |
| KQGFLGM | GARDKNTGQLY | 0.7273241 |
| LESVITQL | GARDKNTGQLY | 0.6709278 |
| LGLGALVVV | GARDKNTGQLY | 0.7593948 |
| LIGNGGNGGSGGI | GARDKNTGQLY | 0.64815855 |
| LNAINAPTL | GARDKNTGQLY | 0.64038146 |
| LQLLNTGV | GARDKNTGQLY | 0.6664145 |
| LQTSIEF | GARDKNTGQLY | 0.57919776 |
| MSFAVSEL | GARDKNTGQLY | 0.4507993 |
| NTGALGL | GARDKNTGQLY | 0.34978864 |

| | | |
|---|---|---|
| PPGAGGAGGAGTT | GARDKNTGQLY | 0.57830644 |
| QLVGLGI | GARDKNTGQLY | 0.31150988 |
| RFLEQVHQL | GARDKNTGQLY | 0.67291176 |
| RGVGYVLR | GARDKNTGQLY | 0.73718274 |
| SATGVFAGVI | GARDKNTGQLY | 0.6903968 |
| SLPAITLP | GARDKNTGQLY | 0.4131162 |
| STALAFTV | GARDKNTGQLY | 0.66383487 |
| SVLTVGLGK | GARDKNTGQLY | 0.56739885 |
| TALAGAGSGI | GARDKNTGQLY | 0.7348662 |
| TGLPFTGI | GARDKNTGQLY | 0.6898181 |
| VAEHVTV | GARDKNTGQLY | 0.7521198 |
| VDIVLNSL | GARDKNTGQLY | 0.74424255 |
| VLTGGGAVL | GARDKNTGQLY | 0.54295695 |
| VNVGLAYGI | GARDKNTGQLY | 0.74269116 |
| VSSAAVGAV | GARDKNTGQLY | 0.5897869 |
| VTSVYEAGV | GARDKNTGQLY | 0.717226 |
| WVALDRGA | GARDKNTGQLY | 0.6690725 |
| YLVGSVVI | GARDKNTGQLY | 0.67533463 |
| GPPGTGKTLL | GARDKNTGQLY | 0.5826341 |
| GVGMTKF | GARDKNTGQLY | 0.5938376 |
| IGIGGSDL | GARDKNTGQLY | 0.2825814 |
| IGTLLAEL | GARDKNTGQLY | 0.639736 |
| IYTSGTT | GARDKNTGQLY | 0.42577672 |
| LSFPAGV | GARDKNTGQLY | 0.36042044 |
| LYTSGTT | GARDKNTGQLY | 0.4891519 |
| VGAFPFGF | GARDKNTGQLY | 0.7800635 |
| VGAGTVEF | GARDKNTGQLY | 0.7670835 |
| VGKSTLLL | GARDKNTGQLY | 0.5436441 |
| AALAHAGGGIV | GARDKNTGQLY | 0.46046156 |
| AALVAVSLL | GARDKNTGQLY | 0.71350783 |
| AAVSRAVAAL | GARDKNTGQLY | 0.3653577 |
| AGGSSGIGGF | GARDKNTGQLY | 0.6979659 |
| AGLTVANANG | GARDKNTGQLY | 0.658671 |
| AGLVGAVL | GARDKNTGQLY | 0.6589577 |
| ALSQAFTV | GARDKNTGQLY | 0.51795137 |
| ALTFSVSL | GARDKNTGQLY | 0.7177689 |
| ALTGAALVAL | GARDKNTGQLY | 0.5351034 |
| ASFSIGAGVL | GARDKNTGQLY | 0.66659844 |
| DGSVYKI | GARDKNTGQLY | 0.23642458 |
| EQLAEIV | GARDKNTGQLY | 0.74386805 |
| FAVSELTVM | GARDKNTGQLY | 0.5267651 |
| GAAPLAAGL | GARDKNTGQLY | 0.5171352 |
| GADGGDGGSGGAGG | GARDKNTGQLY | 0.31325704 |
| GFGLDTGGAGG | GARDKNTGQLY | 0.62467897 |
| GLASGGLL | GARDKNTGQLY | 0.7389957 |
| GRIAGAAVLL | GARDKNTGQLY | 0.75502217 |
| GSAGSLNTSF | GARDKNTGQLY | 0.53926885 |
| GTGQIGI | GARDKNTGQLY | 0.7275976 |
| GTLIALIL | GARDKNTGQLY | 0.7612283 |
| IAATMGFVVG | GARDKNTGQLY | 0.65248466 |
| IFAQALL | GARDKNTGQLY | 0.32996207 |

| | | |
|---|---|---|
| IGAAVALGF | GARDKNTGQLY | 0.6553267 |
| IGLTPLAL | GARDKNTGQLY | 0.6087589 |
| ILGSNTSTL | GARDKNTGQLY | 0.7434527 |
| IVFSLTVL | GARDKNTGQLY | 0.65965265 |
| LASKATGF | GARDKNTGQLY | 0.46517706 |
| LASVALIL | GARDKNTGQLY | 0.67615455 |
| LFAVGLTL | GARDKNTGQLY | 0.7459171 |
| LFNGGIGTYI | GARDKNTGQLY | 0.46033886 |
| LTLGAGII | GARDKNTGQLY | 0.57855874 |
| LVAAALGDI | GARDKNTGQLY | 0.47724557 |
| NTAFVQLGI | GARDKNTGQLY | 0.4177155 |
| QLFSGAEL | GARDKNTGQLY | 0.680183 |
| SAATFITL | GARDKNTGQLY | 0.6144902 |
| SNYGLANLGV | GARDKNTGQLY | 0.57728595 |
| SYGGAYAVM | GARDKNTGQLY | 0.6071536 |
| TTGVGGLTLG | GARDKNTGQLY | 0.5735426 |
| TTQTIYI | GARDKNTGQLY | 0.513042 |
| VAAGAGLGAVI | GARDKNTGQLY | 0.6055931 |
| VAAVVTLL | GARDKNTGQLY | 0.69051516 |
| VGFLGGLI | GARDKNTGQLY | 0.7306124 |
| VGVIPILL | GARDKNTGQLY | 0.67617995 |
| VIFASSLI | GARDKNTGQLY | 0.63791835 |
| VKGGDGGAAAG | GARDKNTGQLY | 0.7778216 |
| VTKTFTNI | GARDKNTGQLY | 0.3712047 |
| VVAAAILL | GARDKNTGQLY | 0.47195306 |
| YAALDVEL | GARDKNTGQLY | 0.7161093 |
| LASVALLL | GARDKNTGQLY | 0.446779 |
| LDAVHRV | GARDKNTGQLY | 0.7442235 |
| QGLSSSIF | GARDKNTGQLY | 0.6564654 |
| RGTFANIRL | GARDKNTGQLY | 0.7104651 |
| RQLWWGHRI | GARDKNTGQLY | 0.6713037 |
| YGPPGTGKTLL | GARDKNTGQLY | 0.7803786 |
| AALVGAPANVA | GARDKNTGQLY | 0.7385132 |
| AATFRRTE | GARDKNTGQLY | 0.61369467 |
| AGALGGGI | GARDKNTGQLY | 0.7506702 |
| AGGAGGAGGIGAG | GARDKNTGQLY | 0.36769328 |
| AGSGALGL | GARDKNTGQLY | 0.7357688 |
| AQIADQI | GARDKNTGQLY | 0.70022494 |
| ASAVAVTYL | GARDKNTGQLY | 0.48776257 |
| ASTVLGGTGG | GARDKNTGQLY | 0.6529366 |
| AVGGEPAK | GARDKNTGQLY | 0.777038 |
| CEQGTFGGTC | GARDKNTGQLY | 0.7572334 |
| DEGKAPAG | GARDKNTGQLY | 0.32045668 |
| DTVVFPAL | GARDKNTGQLY | 0.58439666 |
| FEGITMAG | GARDKNTGQLY | 0.41801625 |
| FPLVVQD | GARDKNTGQLY | 0.5846633 |
| GAFTTPQ | GARDKNTGQLY | 0.35431093 |
| GFGGFGTAGGVGGA | GARDKNTGQLY | 0.7362536 |
| GFLAQFL | GARDKNTGQLY | 0.6169014 |
| GGAGGAGGNAGVA | GARDKNTGQLY | 0.6749886 |
| GGVGTAGAGGAGGAGG | GARDKNTGQLY | 0.67185795 |

| | | |
|---|---|---|
| GNGGGGDAGNAGSGGN | GARDKNTGQLY | 0.7608763 |
| GTGGAGGAGNDAGS | GARDKNTGQLY | 0.58657455 |
| GTTNTGWFNS | GARDKNTGQLY | 0.58395094 |
| IFMGPLFTL | GARDKNTGQLY | 0.3157444 |
| ILAHAIEAI | GARDKNTGQLY | 0.32605252 |
| IRSAIREAL | GARDKNTGQLY | 0.57171565 |
| ITFAGGG | GARDKNTGQLY | 0.6121681 |
| ITGMSGHQEV | GARDKNTGQLY | 0.2570551 |
| IVAGLACTL | GARDKNTGQLY | 0.75792944 |
| IVGGGLAGA | GARDKNTGQLY | 0.6545768 |
| IVTGVLNI | GARDKNTGQLY | 0.70976734 |
| LAQQGCI | GARDKNTGQLY | 0.52508754 |
| LCLHVEN | GARDKNTGQLY | 0.5691967 |
| LDAISDGS | GARDKNTGQLY | 0.52114767 |
| LGGFAAML | GARDKNTGQLY | 0.7411213 |
| LGGKNPMI | GARDKNTGQLY | 0.5470074 |
| LGPKGVAAIL | GARDKNTGQLY | 0.27523604 |
| LMVGVSI | GARDKNTGQLY | 0.5256735 |
| LVGAAFVL | GARDKNTGQLY | 0.48559606 |
| LVLAVAPL | GARDKNTGQLY | 0.7724451 |
| MAAIAVIDAA | GARDKNTGQLY | 0.7476512 |
| NNGINAELQ | GARDKNTGQLY | 0.68394923 |
| RGEPLVLVHG | GARDKNTGQLY | 0.74652064 |
| RGGAGGAAGLI | GARDKNTGQLY | 0.22794095 |
| RSYRMHPAV | GARDKNTGQLY | 0.5885756 |
| SAAHLLAGQLA | GARDKNTGQLY | 0.7568867 |
| SAAVGSY | GARDKNTGQLY | 0.66999036 |
| SPHAVLHRI | GARDKNTGQLY | 0.6523589 |
| SQIAKVL | GARDKNTGQLY | 0.7602361 |
| TALGHLVP | GARDKNTGQLY | 0.4999047 |
| TAPTTSGAA | GARDKNTGQLY | 0.56869096 |
| TDGDDGNGGSPGT | GARDKNTGQLY | 0.67944217 |
| TDGLGEN | GARDKNTGQLY | 0.7217921 |
| TGGPPLPGQA | GARDKNTGQLY | 0.7270545 |
| TIPAITI | GARDKNTGQLY | 0.40239292 |
| TIPGFTI | GARDKNTGQLY | 0.68805563 |
| TSIASGALA | GARDKNTGQLY | 0.79004174 |
| TVTLPGAAI | GARDKNTGQLY | 0.6301104 |
| VASGLSAALP | GARDKNTGQLY | 0.71834135 |
| VGAGVIGI | GARDKNTGQLY | 0.6709392 |
| VGPKRFGV | GARDKNTGQLY | 0.49349433 |
| VNVAFPD | GARDKNTGQLY | 0.52058643 |
| VSMTNSGVS | GARDKNTGQLY | 0.51660496 |
| VTDVLGDLSV | GARDKNTGQLY | 0.6704152 |
| VVVVGAGFAGL | GARDKNTGQLY | 0.5146264 |
| VYTVGTVV | GARDKNTGQLY | 0.68024874 |
| WTTFSGP | GARDKNTGQLY | 0.6231267 |
| YDELVEAF | GARDKNTGQLY | 0.7905977 |
| YGTGGAGGNGG | GARDKNTGQLY | 0.7043036 |
| YTNVMAV | GARDKNTGQLY | 0.7021736 |
| FLEQVHQL | GARDKNTGQLY | 0.50674725 |

| | | |
|---|---|---|
| GLSLLAVAV | GARDKNTGQLY | 0.70402986 |
| LEQVHQL | GARDKNTGQLY | 0.72241426 |
| LNAAGAI | GARDKNTGQLY | 0.30566296 |
| PAVHASL | GARDKNTGQLY | 0.7683563 |
| TELEDDA | GARDKNTGQLY | 0.49689513 |
| TPPGPPN | GARDKNTGQLY | 0.67697823 |
| TPPPPPPP | GARDKNTGQLY | 0.69682014 |
| TQSPALI | ASGDLGETLY | 0.70866245 |
| EFAAAFPQ | ASGDLGETLY | 0.67973816 |
| LLSGNLDVL | ASGDLGETLY | 0.65499103 |
| GPANLGNYNI | ASGDLGETLY | 0.45585245 |
| TLGPTLP | ASGDLGETLY | 0.73407185 |
| VYAYSSCCK | ASGDLGETLY | 0.75130093 |
| APAPAPAPNI | ASGDLGETLY | 0.6148467 |
| PGGPVPIP | ASGDLGETLY | 0.7627152 |
| FLTVALEP | ASGDLGETLY | 0.66378474 |
| SQYFGANL | ASGDLGETLY | 0.31774327 |
| GGVKPAA | ASGDLGETLY | 0.63755286 |
| GGTGGAGGAGGW | ASGDLGETLY | 0.758539 |
| GVGGVAGLGGAGG | ASGDLGETLY | 0.7661027 |
| VLLVLATTI | ASGDLGETLY | 0.7524757 |
| IGIFAAML | ASGDLGETLY | 0.73833436 |
| AVGDLID | ASGDLGETLY | 0.62608874 |
| NISPLPL | ASGDLGETLY | 0.7626489 |
| VTAVFVAI | ASGDLGETLY | 0.78883815 |
| SGDDNDAAL | ASGDLGETLY | 0.69678843 |
| AVAALAHS | ASGDLGETLY | 0.52834207 |
| GGVTVMATP | ASGDLGETLY | 0.45213056 |
| AGLAANVRD | ASGDLGETLY | 0.72959566 |
| TNRPDLIDPALLRPG | ASGDLGETLY | 0.7497634 |
| LANLGVTAL | ASGDLGETLY | 0.45143062 |
| TGERIEL | ASGDLGETLY | 0.70506644 |
| NGEELAA | ASGDLGETLY | 0.6964162 |
| VLSLVWG | ASGDLGETLY | 0.32895008 |
| IGYALWL | ASGDLGETLY | 0.72145927 |
| LSLLLAPL | ASGDLGETLY | 0.77223295 |
| TLDEALK | ASGDLGETLY | 0.6798687 |
| TDGAGTAEGG | ASGDLGETLY | 0.52472997 |
| NAPGGAGGNGG | ASGDLGETLY | 0.77390385 |
| VSVFMGMGGNF | ASGDLGETLY | 0.7839445 |
| LASYLAAL | ASGDLGETLY | 0.765298 |
| LVGGTPAL | ASGDLGETLY | 0.3361726 |
| VRNADVL | ASGDLGETLY | 0.2361279 |
| TCSSPFLVSAM | ASGDLGETLY | 0.651098 |
| VHITDAE | ASGDLGETLY | 0.6954345 |
| EYAAPSF | ASGDLGETLY | 0.56874645 |
| VVDAFHGGL | ASGDLGETLY | 0.50331247 |
| ASVVDPQ | ASGDLGETLY | 0.7364044 |
| SYNEFGGLS | ASGDLGETLY | 0.7486242 |
| DPDGSVY | ASGDLGETLY | 0.48411575 |
| MVSTIHATE | ASGDLGETLY | 0.35573694 |

| | | |
|---|---|---|
| LAAGVADPV | ASGDLGETLY | 0.7539333 |
| TKEQIAA | ASGDLGETLY | 0.72721136 |
| SLASPAAL | ASGDLGETLY | 0.5169208 |
| IGLVFYGV | ASGDLGETLY | 0.61925983 |
| FLLDASNL | ASGDLGETLY | 0.65487486 |
| YQVGNFP | ASGDLGETLY | 0.62899864 |
| ESLLSQST | ASGDLGETLY | 0.69870293 |
| LFTVGFAAVG | ASGDLGETLY | 0.46466345 |
| ATPPTPT | ASGDLGETLY | 0.7584215 |
| TDLATLG | ASGDLGETLY | 0.4854023 |
| SAGGTGR | ASGDLGETLY | 0.744303 |
| GFSNSGS | ASGDLGETLY | 0.67689395 |
| GGNGGSGGTSV | ASGDLGETLY | 0.6997961 |
| GTSVATGGAGN | ASGDLGETLY | 0.66673195 |
| NGGNGGSGGTS | ASGDLGETLY | 0.50232184 |
| AVVKFPH | ASGDLGETLY | 0.46220422 |
| TPPPPPP | ASGDLGETLY | 0.3741114 |
| AAPAAAAAPA | ASGDLGETLY | 0.7615365 |
| IALNEDL | ASGDLGETLY | 0.6580621 |
| IVNVSSM | ASGDLGETLY | 0.73524636 |
| TGIPPAPR | ASGDLGETLY | 0.7146901 |
| FDLGGGTF | ASGDLGETLY | 0.4708812 |
| FDLGGGTFDVS | ASGDLGETLY | 0.71574104 |
| NEPTAAA | ASGDLGETLY | 0.38330302 |
| NEPTAAALA | ASGDLGETLY | 0.5801171 |
| GSIINISSI | ASGDLGETLY | 0.6815909 |
| VAPGNAGTA | ASGDLGETLY | 0.7135522 |
| TGGGASLE | ASGDLGETLY | 0.5710442 |
| VAVYNHYKRI | ASGDLGETLY | 0.7093453 |
| TDPAPATT | ASGDLGETLY | 0.7199054 |
| TNRPDILDPALLRPG | ASGDLGETLY | 0.5444556 |
| LTTTTTS | ASGDLGETLY | 0.65795857 |
| PTGSGKTL | ASGDLGETLY | 0.75146437 |
| TGSGKTL | ASGDLGETLY | 0.6878538 |
| TIGHVDHGK | ASGDLGETLY | 0.4634323 |
| TIGHVDHGKTT | ASGDLGETLY | 0.72799194 |
| GHVDHGKTT | ASGDLGETLY | 0.49482706 |
| AQFEHTLL | ASGDLGETLY | 0.69957185 |
| IVNQDEL | ASGDLGETLY | 0.6377678 |
| TNGSQFF | ASGDLGETLY | 0.6135353 |
| TGKGIVE | ASGDLGETLY | 0.7452433 |
| LISNASDA | ASGDLGETLY | 0.3798491 |
| AELRTLGELTKIAKAHGAQ | ASGDLGETLY | 0.601773 |
| APNAVAAAEPSW | ASGDLGETLY | 0.54426634 |
| APQVAGAATAVL | ASGDLGETLY | 0.6621547 |
| ARRDPAAAAAILPTDA | ASGDLGETLY | 0.71637046 |
| DAAARDVLAEAGLAEYF | ASGDLGETLY | 0.43284285 |
| DDQIVLSAKANLAKVEQ | ASGDLGETLY | 0.6900574 |
| DELLPDTGEDSDEDGH | ASGDLGETLY | 0.55791694 |
| ELPEAPANRATVPQD | ASGDLGETLY | 0.418979 |
| FGQSKIERVTGIG | ASGDLGETLY | 0.6589918 |

| | | |
|---|---|---|
| GAGGTGSTAGGAGGAGGAGGLY | ASGDLGETLY | 0.6242649 |
| GHGGTGAAGGNGGTGGAGGSGI | ASGDLGETLY | 0.42729035 |
| GNGGNGGAGGEGGPGGAGGAGA | ASGDLGETLY | 0.68913364 |
| GSTAGNGGNGGNATNGTIAGQP | ASGDLGETLY | 0.54563403 |
| GTGGDGGNGGNGGTGG | ASGDLGETLY | 0.6985348 |
| GVRRPAGIPADAPV | ASGDLGETLY | 0.78336024 |
| GYRQVTDAHLLTLARR | ASGDLGETLY | 0.438955 |
| IAKMGQDGHDRG | ASGDLGETLY | 0.7471864 |
| KAEPSIEIQYT | ASGDLGETLY | 0.689278 |
| KPVRIRMAADEDTVAVT | ASGDLGETLY | 0.7091563 |
| MPPRIAGMRLLVIKPEP | ASGDLGETLY | 0.6254218 |
| MKFYAVALFRSRGVPPTR | ASGDLGETLY | 0.5572188 |
| NAGGFGILGAVAHSP | ASGDLGETLY | 0.7739194 |
| NEGREITLAMAEM | ASGDLGETLY | 0.7197157 |
| NELVEWSHPLGHRWRWE | ASGDLGETLY | 0.71248615 |
| NLGAGGTGGNGGNPG | ASGDLGETLY | 0.69064724 |
| NTEWPKWETVMAADPVD | ASGDLGETLY | 0.7143173 |
| PASAAPAETQP | ASGDLGETLY | 0.7369861 |
| QQSSDGKNGVRAAEITGE | ASGDLGETLY | 0.51220155 |
| RAPDAVTTSWWKE | ASGDLGETLY | 0.3430396 |
| SGFGNVGAHTSGWFNQ | ASGDLGETLY | 0.5957264 |
| SGVAGAVGQAASVGGLKV | ASGDLGETLY | 0.6069815 |
| SNGGAGTGSAGGNGGT | ASGDLGETLY | 0.53723 |
| TGGAGGEGGAAG | ASGDLGETLY | 0.31797695 |
| TGGDGGAAGAGGAPGHG | ASGDLGETLY | 0.7374687 |
| TMPPLLAAIANAGRHGV | ASGDLGETLY | 0.7877259 |
| TSSTAGNSGGA | ASGDLGETLY | 0.5846604 |
| VPATVPVGILC | ASGDLGETLY | 0.77697474 |
| VQRPDSGGVVLTGRIS | ASGDLGETLY | 0.44045654 |
| WTPNPVPATKGVPAGLVQAT | ASGDLGETLY | 0.7299919 |
| YALKRALAKFSDRFDVVII | ASGDLGETLY | 0.75927305 |
| YLIGSVIV | ASGDLGETLY | 0.4579764 |
| YGQLVAVID | ASGDLGETLY | 0.67902005 |
| VTVGGMTLPA | ASGDLGETLY | 0.5927266 |
| VLALPAV | ASGDLGETLY | 0.59182197 |
| VIPPTLNYET | ASGDLGETLY | 0.7631241 |
| VGMGGMHPSSG | ASGDLGETLY | 0.5637598 |
| VGLAGGAATA | ASGDLGETLY | 0.6984674 |
| VGGAGGAGGSGP | ASGDLGETLY | 0.58115566 |
| VFAAFMV | ASGDLGETLY | 0.4301184 |
| TLYDKLR | ASGDLGETLY | 0.6894308 |
| TLAVFTVL | ASGDLGETLY | 0.70944166 |
| TGSVQPARV | ASGDLGETLY | 0.62355113 |
| TGGGAGSGGT | ASGDLGETLY | 0.78621453 |
| SHLETSL | ASGDLGETLY | 0.5118648 |
| SFTTGMT | ASGDLGETLY | 0.7821041 |
| QVEVAKRMGVRQPTVS | ASGDLGETLY | 0.7591524 |
| QGSGGPALAM | ASGDLGETLY | 0.67323554 |
| MLAPSNP | ASGDLGETLY | 0.56646246 |
| LVNNAAI | ASGDLGETLY | 0.75686383 |
| LTGEINTLA | ASGDLGETLY | 0.5796149 |

| | | |
|---|---|---|
| LSIFGPV | ASGDLGETLY | 0.7307745 |
| LLALLANL | ASGDLGETLY | 0.75097036 |
| LISNASDALDKLR | ASGDLGETLY | 0.6400516 |
| LINEAVP | ASGDLGETLY | 0.7397902 |
| LGYAFSGL | ASGDLGETLY | 0.71196055 |
| LDFVAVAL | ASGDLGETLY | 0.6887183 |
| LAQPGAE | ASGDLGETLY | 0.6585094 |
| ISNLGAVTGF | ASGDLGETLY | 0.3670755 |
| HQEDPGRI | ASGDLGETLY | 0.7070134 |
| GTPGEPTP | ASGDLGETLY | 0.61013234 |
| GFFNSGS | ASGDLGETLY | 0.7260765 |
| GAGGGGLTAGAN | ASGDLGETLY | 0.7181191 |
| FVDALVNM | ASGDLGETLY | 0.66130054 |
| FLTVAPEL | ASGDLGETLY | 0.74152106 |
| FETLTAL | ASGDLGETLY | 0.7408868 |
| DVINPIEL | ASGDLGETLY | 0.57983696 |
| APVSGAPV | ASGDLGETLY | 0.6834927 |
| ALVGGLAW | ASGDLGETLY | 0.66151273 |
| ALAAAAATAI | ASGDLGETLY | 0.67326784 |
| AFGLGLLL | ASGDLGETLY | 0.4574669 |
| AFFAAASL | ASGDLGETLY | 0.73035735 |
| AAAALPAFSPP | ASGDLGETLY | 0.66716766 |
| VVLAPAL | ASGDLGETLY | 0.25284395 |
| TIGHVDHGKTTL | ASGDLGETLY | 0.69258714 |
| LISNASDALDK | ASGDLGETLY | 0.6887026 |
| LISNASDAL | ASGDLGETLY | 0.68600893 |
| ISNASDALDK | ASGDLGETLY | 0.73561513 |
| NQAETLVYQTE | ASGDLGETLY | 0.32893223 |
| TGGTGGTAGAG | ASGDLGETLY | 0.61790603 |
| TGSTAGGAGGA | ASGDLGETLY | 0.7318449 |
| DPLAVPVAAGPV | ASGDLGETLY | 0.74333954 |
| GGTGGTPGNGGN | ASGDLGETLY | 0.5265497 |
| PAGLAGVDPATL | ASGDLGETLY | 0.7191849 |
| GNNLVVEAPHAGASV | ASGDLGETLY | 0.74146354 |
| KVTRIMSASTLEQPA | ASGDLGETLY | 0.7163062 |
| LGFALPIALPRRTVV | ASGDLGETLY | 0.7346047 |
| TRLGNLTARKLVVLV | ASGDLGETLY | 0.7388393 |
| WSQQRRLIYFPSAGP | ASGDLGETLY | 0.6253953 |
| GPNGNPDPMAAAVDIR | ASGDLGETLY | 0.7378092 |
| TQRCVSVADLRAAASR | ASGDLGETLY | 0.7429926 |
| QFVLDVAEHPVEVLTETV | ASGDLGETLY | 0.66437304 |
| GGAGGTGGAGGNGGTQAGDG | ASGDLGETLY | 0.70777565 |
| PLLATLRAIAVDVDAKPAQV | ASGDLGETLY | 0.63145816 |
| APKFRTGGYMIDFWGVGYQVAK | ASGDLGETLY | 0.6551591 |
| GAGGAGGAGGAAGGPGGTGGTG | ASGDLGETLY | 0.32419246 |
| IGALGLLATDVAPAGSVAFGVA | ASGDLGETLY | 0.7202225 |
| AGGAGGIGGNGGAGGAGGVLMG | ASGDLGETLY | 0.73277664 |
| ADELIGLL | ASGDLGETLY | 0.69115937 |
| ADRALKA | ASGDLGETLY | 0.6095008 |
| AGAVGQVGGAGGA | ASGDLGETLY | 0.6359512 |
| AGGGGGLGGAP | ASGDLGETLY | 0.6525841 |

| | | |
|---|---|---|
| AGGLLYGNGGAG | ASGDLGETLY | 0.68489563 |
| AGLESIGL | ASGDLGETLY | 0.34242436 |
| AGLKTGII | ASGDLGETLY | 0.59603035 |
| ASLAALLL | ASGDLGETLY | 0.73857355 |
| ASVINISSI | ASGDLGETLY | 0.7402196 |
| ATPPGPA | ASGDLGETLY | 0.4495228 |
| ATVASALV | ASGDLGETLY | 0.73837405 |
| ATVASLIL | ASGDLGETLY | 0.72812045 |
| AVNGPVLIRH | ASGDLGETLY | 0.7402736 |
| DGAAGAVGI | ASGDLGETLY | 0.69824964 |
| DPLLHGI | ASGDLGETLY | 0.6332922 |
| ELGAALGAL | ASGDLGETLY | 0.73029137 |
| ELGRTQAV | ASGDLGETLY | 0.68863714 |
| FVAGGYLS | ASGDLGETLY | 0.54296815 |
| GASGGAGGAGGW | ASGDLGETLY | 0.72859883 |
| GGAGAGGDGGAGGI | ASGDLGETLY | 0.66427076 |
| GTGHNGSQPGMGGNG | ASGDLGETLY | 0.6669018 |
| GYAAASASAS | ASGDLGETLY | 0.6752449 |
| HQVDVAAVQQG | ASGDLGETLY | 0.7232413 |
| IGAAIAHT | ASGDLGETLY | 0.621925 |
| IGMIEVAQL | ASGDLGETLY | 0.5016956 |
| IRLAPEHTSAMD | ASGDLGETLY | 0.61878073 |
| IYNVSVL | ASGDLGETLY | 0.7136408 |
| KQGFLGM | ASGDLGETLY | 0.73897517 |
| LESVITQL | ASGDLGETLY | 0.7061972 |
| LGLGALVVV | ASGDLGETLY | 0.6520301 |
| LIGNGGNGGSGGI | ASGDLGETLY | 0.7273085 |
| LNAINAPTL | ASGDLGETLY | 0.7181973 |
| LQLLNTGV | ASGDLGETLY | 0.6962764 |
| LQTSIEF | ASGDLGETLY | 0.7214062 |
| MSFAVSEL | ASGDLGETLY | 0.6596264 |
| NTGALGL | ASGDLGETLY | 0.72609985 |
| PPGAGGAGGAGTT | ASGDLGETLY | 0.7170205 |
| QLVGLGI | ASGDLGETLY | 0.7295166 |
| RFLEQVHQL | ASGDLGETLY | 0.7292974 |
| RGVGYVLR | ASGDLGETLY | 0.70690405 |
| SATGVFAGVI | ASGDLGETLY | 0.6939791 |
| SLPAITLP | ASGDLGETLY | 0.33311868 |
| STALAFTV | ASGDLGETLY | 0.6412809 |
| SVLTVGLGK | ASGDLGETLY | 0.70764893 |
| TALAGAGSGI | ASGDLGETLY | 0.71550584 |
| TGLPFTGI | ASGDLGETLY | 0.7151201 |
| VAEHVTV | ASGDLGETLY | 0.713075 |
| VDIVLNSL | ASGDLGETLY | 0.6424439 |
| VLTGGGAVL | ASGDLGETLY | 0.7258451 |
| VNVGLAYGI | ASGDLGETLY | 0.733994 |
| VSSAAVGAV | ASGDLGETLY | 0.51319396 |
| VTSVYEAGV | ASGDLGETLY | 0.6775379 |
| WVALDRGA | ASGDLGETLY | 0.7437946 |
| YLVGSVVI | ASGDLGETLY | 0.71615577 |
| GPPGTGKTLL | ASGDLGETLY | 0.69621146 |

| | | |
|---|---|---|
| GVGMTKF | ASGDLGETLY | 0.6118803 |
| IGIGGSDL | ASGDLGETLY | 0.7440545 |
| IGTLLAEL | ASGDLGETLY | 0.6451895 |
| IYTSGTT | ASGDLGETLY | 0.74290013 |
| LSFPAGV | ASGDLGETLY | 0.28609103 |
| LYTSGTT | ASGDLGETLY | 0.6136261 |
| VGAFPFGF | ASGDLGETLY | 0.5059331 |
| VGAGTVEF | ASGDLGETLY | 0.4040823 |
| VGKSTLLL | ASGDLGETLY | 0.73341215 |
| AALAHAGGGIV | ASGDLGETLY | 0.6598551 |
| AALVAVSLL | ASGDLGETLY | 0.73331964 |
| AAVSRAVAAL | ASGDLGETLY | 0.7048011 |
| AGGSSGIGGF | ASGDLGETLY | 0.72999 |
| AGLTVANANG | ASGDLGETLY | 0.7086489 |
| AGLVGAVL | ASGDLGETLY | 0.70033437 |
| ALSQAFTV | ASGDLGETLY | 0.54091454 |
| ALTFSVSL | ASGDLGETLY | 0.7317326 |
| ALTGAALVAL | ASGDLGETLY | 0.72813284 |
| ASFSIGAGVL | ASGDLGETLY | 0.7018101 |
| DGSVYKI | ASGDLGETLY | 0.4817647 |
| EQLAEIV | ASGDLGETLY | 0.6504411 |
| FAVSELTVM | ASGDLGETLY | 0.714832 |
| GAAPLAAGL | ASGDLGETLY | 0.27841294 |
| GADGGDGGSGGAGG | ASGDLGETLY | 0.28261912 |
| GFGLDTGGAGG | ASGDLGETLY | 0.6338196 |
| GLASGGLL | ASGDLGETLY | 0.58092886 |
| GRIAGAAVLL | ASGDLGETLY | 0.7385131 |
| GSAGSLNTSF | ASGDLGETLY | 0.7340226 |
| GTGQIGI | ASGDLGETLY | 0.6956208 |
| GTLIALIL | ASGDLGETLY | 0.7291089 |
| IAATMGFVVG | ASGDLGETLY | 0.71232533 |
| IFAQALL | ASGDLGETLY | 0.2330204 |
| IGAAVALGF | ASGDLGETLY | 0.6854886 |
| IGLTPLAL | ASGDLGETLY | 0.5996994 |
| ILGSNTSTL | ASGDLGETLY | 0.7398329 |
| IVFSLTVL | ASGDLGETLY | 0.6091114 |
| LASKATGF | ASGDLGETLY | 0.66187227 |
| LASVALIL | ASGDLGETLY | 0.7063817 |
| LFAVGLTL | ASGDLGETLY | 0.7145112 |
| LFNGGIGTYI | ASGDLGETLY | 0.7053087 |
| LTLGAGII | ASGDLGETLY | 0.56117857 |
| LVAAALGDI | ASGDLGETLY | 0.24377343 |
| NTAFVQLGI | ASGDLGETLY | 0.733574 |
| QLFSGAEL | ASGDLGETLY | 0.4464919 |
| SAATFITL | ASGDLGETLY | 0.5674463 |
| SNYGLANLGV | ASGDLGETLY | 0.3296983 |
| SYGGAYAVM | ASGDLGETLY | 0.26515204 |
| TTGVGGLTLG | ASGDLGETLY | 0.46107337 |
| TTQTIYI | ASGDLGETLY | 0.7281028 |
| VAAGAGLGAVI | ASGDLGETLY | 0.6835904 |
| VAAVVTLL | ASGDLGETLY | 0.68320465 |

| | | |
|---|---|---|
| VGFLGGLI | ASGDLGETLY | 0.7419764 |
| VGVIPILL | ASGDLGETLY | 0.71671236 |
| VIFASSLI | ASGDLGETLY | 0.67814195 |
| VKGGDGGAAAG | ASGDLGETLY | 0.6171528 |
| VTKTFTNI | ASGDLGETLY | 0.58764005 |
| VVAAAILL | ASGDLGETLY | 0.57290596 |
| YAALDVEL | ASGDLGETLY | 0.5328752 |
| LASVALLL | ASGDLGETLY | 0.74229383 |
| LDAVHRV | ASGDLGETLY | 0.6158894 |
| QGLSSSIF | ASGDLGETLY | 0.55399716 |
| RGTFANIRL | ASGDLGETLY | 0.6660321 |
| RQLWWGHRI | ASGDLGETLY | 0.7407304 |
| YGPPGTGKTLL | ASGDLGETLY | 0.7458401 |
| AALVGAPANVA | ASGDLGETLY | 0.6528989 |
| AATFRRTE | ASGDLGETLY | 0.68771887 |
| AGALGGGI | ASGDLGETLY | 0.6660812 |
| AGGAGGAGGIGAG | ASGDLGETLY | 0.7088268 |
| AGSGALGL | ASGDLGETLY | 0.6495465 |
| AQIADQI | ASGDLGETLY | 0.42593196 |
| ASAVAVTYL | ASGDLGETLY | 0.71869385 |
| ASTVLGGTGG | ASGDLGETLY | 0.7479717 |
| AVGGEPAK | ASGDLGETLY | 0.6357392 |
| CEQGTFGGTC | ASGDLGETLY | 0.63921833 |
| DEGKAPAG | ASGDLGETLY | 0.702857 |
| DTVVFPAL | ASGDLGETLY | 0.6458299 |
| FEGITMAG | ASGDLGETLY | 0.6845348 |
| FPLVVQD | ASGDLGETLY | 0.7194252 |
| GAFTTPQ | ASGDLGETLY | 0.51308274 |
| GFGGFGTAGGVGGA | ASGDLGETLY | 0.6328263 |
| GFLAQFL | ASGDLGETLY | 0.7449477 |
| GGAGGAGGNAGVA | ASGDLGETLY | 0.6937684 |
| GGVGTAGAGGAGGAGG | ASGDLGETLY | 0.70810986 |
| GNGGGGDAGNAGSGGN | ASGDLGETLY | 0.7349553 |
| GTGGAGGAGNDAGS | ASGDLGETLY | 0.7221365 |
| GTTNTGWFNS | ASGDLGETLY | 0.740689 |
| IFMGPLFTL | ASGDLGETLY | 0.7332455 |
| ILAHAIEAI | ASGDLGETLY | 0.6174659 |
| IRSAIREAL | ASGDLGETLY | 0.6860057 |
| ITFAGGG | ASGDLGETLY | 0.6997992 |
| ITGMSGHQEV | ASGDLGETLY | 0.71421623 |
| IVAGLACTL | ASGDLGETLY | 0.6950525 |
| IVGGGLAGA | ASGDLGETLY | 0.616947 |
| IVTGVLNI | ASGDLGETLY | 0.7403642 |
| LAQQGCI | ASGDLGETLY | 0.7029058 |
| LCLHVEN | ASGDLGETLY | 0.73308015 |
| LDAISDGS | ASGDLGETLY | 0.54319227 |
| LGGFAAML | ASGDLGETLY | 0.5212042 |
| LGGKNPMI | ASGDLGETLY | 0.73220176 |
| LGPKGVAAIL | ASGDLGETLY | 0.24475606 |
| LMVGVSI | ASGDLGETLY | 0.2813643 |
| LVGAAFVL | ASGDLGETLY | 0.69006985 |

| | | |
|---|---|---|
| LVLAVAPL | ASGDLGETLY | 0.7394614 |
| MAAIAVIDAA | ASGDLGETLY | 0.24525458 |
| NNGINAELQ | ASGDLGETLY | 0.61134547 |
| RGEPLVLVHG | ASGDLGETLY | 0.5281508 |
| RGGAGGAAGLI | ASGDLGETLY | 0.70825213 |
| RSYRMHPAV | ASGDLGETLY | 0.70936227 |
| SAAHLLAGQLA | ASGDLGETLY | 0.46648788 |
| SAAVGSY | ASGDLGETLY | 0.72653836 |
| SPHAVLHRI | ASGDLGETLY | 0.52629024 |
| SQIAKVL | ASGDLGETLY | 0.41556895 |
| TALGHLVP | ASGDLGETLY | 0.7306912 |
| TAPTTSGAA | ASGDLGETLY | 0.73792434 |
| TDGDDGNGGSPGT | ASGDLGETLY | 0.56847787 |
| TDGLGEN | ASGDLGETLY | 0.7340069 |
| TGGPPLPGQA | ASGDLGETLY | 0.6615261 |
| TIPAITI | ASGDLGETLY | 0.7454473 |
| TIPGFTI | ASGDLGETLY | 0.68284214 |
| TSIASGALA | ASGDLGETLY | 0.4232619 |
| TVTLPGAAI | ASGDLGETLY | 0.7295986 |
| VASGLSAALP | ASGDLGETLY | 0.7109164 |
| VGAGVIGI | ASGDLGETLY | 0.6867061 |
| VGPKRFGV | ASGDLGETLY | 0.7312269 |
| VNVAFPD | ASGDLGETLY | 0.5694635 |
| VSMTNSGVS | ASGDLGETLY | 0.6966092 |
| VTDVLGDLSV | ASGDLGETLY | 0.739975 |
| VVVVGAGFAGL | ASGDLGETLY | 0.72587276 |
| VYTVGTVV | ASGDLGETLY | 0.48372912 |
| WTTFSGP | ASGDLGETLY | 0.71664166 |
| YDELVEAF | ASGDLGETLY | 0.705552 |
| YGTGGAGGNGG | ASGDLGETLY | 0.66225713 |
| YTNVMAV | ASGDLGETLY | 0.74173164 |
| FLEQVHQL | ASGDLGETLY | 0.70563644 |
| GLSLLAVAV | ASGDLGETLY | 0.72935915 |
| LEQVHQL | ASGDLGETLY | 0.7201624 |
| LNAAGAI | ASGDLGETLY | 0.738912 |
| PAVHASL | ASGDLGETLY | 0.4205616 |
| TELEDDA | ASGDLGETLY | 0.7203002 |
| TPPGPPN | ASGDLGETLY | 0.70523775 |
| TPPPPPPP | ASGDLGETLY | 0.7299919 |
| TQSPALI | ASSNRDWGYEQY | 0.69942194 |
| EFAAAFPQ | ASSNRDWGYEQY | 0.4533496 |
| LLSGNLDVL | ASSNRDWGYEQY | 0.70032346 |
| GPANLGNYNI | ASSNRDWGYEQY | 0.73821795 |
| TLGPTLP | ASSNRDWGYEQY | 0.6597568 |
| VYAYSSCCK | ASSNRDWGYEQY | 0.70759284 |
| APAPAPAPNI | ASSNRDWGYEQY | 0.743611 |
| PGGPVPIP | ASSNRDWGYEQY | 0.61514294 |
| FLTVALEP | ASSNRDWGYEQY | 0.6920912 |
| SQYFGANL | ASSNRDWGYEQY | 0.71897614 |
| GGVKPAA | ASSNRDWGYEQY | 0.7274592 |
| GGTGGAGGAGGW | ASSNRDWGYEQY | 0.667007 |

| | | |
|---|---|---|
| GVGGVAGLGGAGG | ASSNRDWGYEQY | 0.7368429 |
| VLLVLATTI | ASSNRDWGYEQY | 0.60318494 |
| IGIFAAML | ASSNRDWGYEQY | 0.72794604 |
| AVGDLID | ASSNRDWGYEQY | 0.6295657 |
| NISPLPL | ASSNRDWGYEQY | 0.72759336 |
| VTAVFVAI | ASSNRDWGYEQY | 0.62427104 |
| SGDDNDAAL | ASSNRDWGYEQY | 0.73153555 |
| AVAALAHS | ASSNRDWGYEQY | 0.73374975 |
| GGVTVMATP | ASSNRDWGYEQY | 0.73196936 |
| AGLAANVRD | ASSNRDWGYEQY | 0.29827124 |
| TNRPDLIDPALLRPG | ASSNRDWGYEQY | 0.7338052 |
| LANLGVTAL | ASSNRDWGYEQY | 0.7360693 |
| TGERIEL | ASSNRDWGYEQY | 0.74768674 |
| NGEELAA | ASSNRDWGYEQY | 0.6897456 |
| VLSLVWG | ASSNRDWGYEQY | 0.7216446 |
| IGYALWL | ASSNRDWGYEQY | 0.69397295 |
| LSLLLAPL | ASSNRDWGYEQY | 0.6724513 |
| TLDEALK | ASSNRDWGYEQY | 0.6310426 |
| TDGAGTAEGG | ASSNRDWGYEQY | 0.6849309 |
| NAPGGAGGNGG | ASSNRDWGYEQY | 0.6877104 |
| VSVFMGMGGNF | ASSNRDWGYEQY | 0.66388863 |
| LASYLAAL | ASSNRDWGYEQY | 0.73341537 |
| LVGGTPAL | ASSNRDWGYEQY | 0.72953165 |
| VRNADVL | ASSNRDWGYEQY | 0.7341261 |
| TCSSPFLVSAM | ASSNRDWGYEQY | 0.7404717 |
| VHITDAE | ASSNRDWGYEQY | 0.7356501 |
| EYAAPSF | ASSNRDWGYEQY | 0.72824514 |
| VVDAFHGGL | ASSNRDWGYEQY | 0.6336356 |
| ASVVDPQ | ASSNRDWGYEQY | 0.6956581 |
| SYNEFGGLS | ASSNRDWGYEQY | 0.6358055 |
| DPDGSVY | ASSNRDWGYEQY | 0.6687801 |
| MVSTIHATE | ASSNRDWGYEQY | 0.65642214 |
| LAAGVADPV | ASSNRDWGYEQY | 0.7181531 |
| TKEQIAA | ASSNRDWGYEQY | 0.23723109 |
| SLASPAAL | ASSNRDWGYEQY | 0.7076067 |
| IGLVFYGV | ASSNRDWGYEQY | 0.74075675 |
| FLLDASNL | ASSNRDWGYEQY | 0.63739675 |
| YQVGNFP | ASSNRDWGYEQY | 0.7235954 |
| ESLLSQST | ASSNRDWGYEQY | 0.6868818 |
| LFTVGFAAVG | ASSNRDWGYEQY | 0.62630844 |
| ATPPTPT | ASSNRDWGYEQY | 0.73498404 |
| TDLATLG | ASSNRDWGYEQY | 0.56038916 |
| SAGGTGR | ASSNRDWGYEQY | 0.71944726 |
| GFSNSGS | ASSNRDWGYEQY | 0.74202126 |
| GGNGGSGGTSV | ASSNRDWGYEQY | 0.70089674 |
| GTSVATGGAGN | ASSNRDWGYEQY | 0.69273293 |
| NGGNGGSGGTS | ASSNRDWGYEQY | 0.67120016 |
| AVVKFPH | ASSNRDWGYEQY | 0.38851747 |
| TPPPPPP | ASSNRDWGYEQY | 0.70442843 |
| AAPAAAAPA | ASSNRDWGYEQY | 0.23191251 |
| IALNEDL | ASSNRDWGYEQY | 0.7049743 |

| | | |
|---|---|---|
| IVNVSSM | ASSNRDWGYEQY | 0.5185702 |
| TGIPPAPR | ASSNRDWGYEQY | 0.7210884 |
| FDLGGGTF | ASSNRDWGYEQY | 0.7109916 |
| FDLGGGTFDVS | ASSNRDWGYEQY | 0.7090368 |
| NEPTAAA | ASSNRDWGYEQY | 0.656021 |
| NEPTAAALA | ASSNRDWGYEQY | 0.6923292 |
| GSIINISSI | ASSNRDWGYEQY | 0.6134956 |
| VAPGNAGTA | ASSNRDWGYEQY | 0.48503876 |
| TGGGASLE | ASSNRDWGYEQY | 0.7270117 |
| VAVYNHYKRI | ASSNRDWGYEQY | 0.6053371 |
| TDPAPATT | ASSNRDWGYEQY | 0.66121274 |
| TNRPDILDPALLRPG | ASSNRDWGYEQY | 0.7374625 |
| LTTTTTS | ASSNRDWGYEQY | 0.70428693 |
| PTGSGKTL | ASSNRDWGYEQY | 0.46773624 |
| TGSGKTL | ASSNRDWGYEQY | 0.7023033 |
| TIGHVDHGK | ASSNRDWGYEQY | 0.7013919 |
| TIGHVDHGKTT | ASSNRDWGYEQY | 0.67800134 |
| GHVDHGKTT | ASSNRDWGYEQY | 0.23316039 |
| AQFEHTLL | ASSNRDWGYEQY | 0.69777083 |
| IVNQDEL | ASSNRDWGYEQY | 0.65988344 |
| TNGSQFF | ASSNRDWGYEQY | 0.63809717 |
| TGKGIVE | ASSNRDWGYEQY | 0.6127063 |
| LISNASDA | ASSNRDWGYEQY | 0.73363507 |
| AELRTLGELTKIAKAHGAQ | ASSNRDWGYEQY | 0.37102705 |
| APNAVAAAEPSW | ASSNRDWGYEQY | 0.7153752 |
| APQVAGAATAVL | ASSNRDWGYEQY | 0.72539353 |
| ARRDPAAAAAILPTDA | ASSNRDWGYEQY | 0.731035 |
| DAAARDVLAEAGLAEYF | ASSNRDWGYEQY | 0.74223495 |
| DDQIVLSAKANLAKVEQ | ASSNRDWGYEQY | 0.7014232 |
| DELLPDTGEDSDEDGH | ASSNRDWGYEQY | 0.63826656 |
| ELPEAPANRATVPQD | ASSNRDWGYEQY | 0.71529055 |
| FGQSKIERVTGIG | ASSNRDWGYEQY | 0.7180321 |
| GAGGTGSTAGGAGGAGGAGGLY | ASSNRDWGYEQY | 0.35938022 |
| GHGGTGAAGGNGGTGGAGGSGI | ASSNRDWGYEQY | 0.35557285 |
| GNGGNGGAGGEGGPGGAGGAGA | ASSNRDWGYEQY | 0.7194055 |
| GSTAGNGGNGGNATNGTIAGQP | ASSNRDWGYEQY | 0.29843506 |
| GTGGDGGNGGNGGTGG | ASSNRDWGYEQY | 0.71399754 |
| GVRRPAGIPADAPV | ASSNRDWGYEQY | 0.73645234 |
| GYRQVTDAHLLTLARR | ASSNRDWGYEQY | 0.69130445 |
| IAKMGQDGHDRG | ASSNRDWGYEQY | 0.5128263 |
| KAEPSIEIQYT | ASSNRDWGYEQY | 0.66937906 |
| KPVRIRMAADEDTVAVT | ASSNRDWGYEQY | 0.6906204 |
| MPPRIAGMRLLVIKPEP | ASSNRDWGYEQY | 0.7313342 |
| MKFYAVALFRSRGVPPTR | ASSNRDWGYEQY | 0.71304905 |
| NAGGFGILGAVAHSP | ASSNRDWGYEQY | 0.74367607 |
| NEGREITLAMAEM | ASSNRDWGYEQY | 0.73769104 |
| NELVEWSHPLGHRWRWE | ASSNRDWGYEQY | 0.6820443 |
| NLGAGGTGGNGGNPG | ASSNRDWGYEQY | 0.72623557 |
| NTEWPKWETVMAADPVD | ASSNRDWGYEQY | 0.66392326 |
| PASAAPAETQP | ASSNRDWGYEQY | 0.68815655 |
| QQSSDGKNGVRAAEITGE | ASSNRDWGYEQY | 0.6532569 |

| | | |
|---|---|---|
| RAPDAVTTSWWKE | ASSNRDWGYEQY | 0.68040144 |
| SGFGNVGAHTSGWFNQ | ASSNRDWGYEQY | 0.658097 |
| SGVAGAVGQAASVGGLKV | ASSNRDWGYEQY | 0.59776604 |
| SNGGAGTGSAGGNGGT | ASSNRDWGYEQY | 0.47355452 |
| TGGAGGEGGAAG | ASSNRDWGYEQY | 0.7086601 |
| TGGDGGAAGAGGAPGHG | ASSNRDWGYEQY | 0.7117723 |
| TMPPLLAAIANAGRHGV | ASSNRDWGYEQY | 0.4265799 |
| TSSTAGNSGGA | ASSNRDWGYEQY | 0.6380516 |
| VPATVPVGILC | ASSNRDWGYEQY | 0.7451809 |
| VQRPDSGGVVLTGRIS | ASSNRDWGYEQY | 0.74268585 |
| WTPNPVPATKGVPAGLVQAT | ASSNRDWGYEQY | 0.65838563 |
| YALKRALAKFSDRFDVVII | ASSNRDWGYEQY | 0.5932229 |
| YLIGSVIV | ASSNRDWGYEQY | 0.7099626 |
| YGQLVAVID | ASSNRDWGYEQY | 0.36991742 |
| VTVGGMTLPA | ASSNRDWGYEQY | 0.6985835 |
| VLALPAV | ASSNRDWGYEQY | 0.7083664 |
| VIPPTLNYET | ASSNRDWGYEQY | 0.68983614 |
| VGMGGMHPSSG | ASSNRDWGYEQY | 0.5735334 |
| VGLAGGAATA | ASSNRDWGYEQY | 0.69891703 |
| VGGAGGAGGSGP | ASSNRDWGYEQY | 0.6648709 |
| VFAAFMV | ASSNRDWGYEQY | 0.7234355 |
| TLYDKLR | ASSNRDWGYEQY | 0.25513858 |
| TLAVFTVL | ASSNRDWGYEQY | 0.7043669 |
| TGSVQPARV | ASSNRDWGYEQY | 0.6445278 |
| TGGGAGSGGT | ASSNRDWGYEQY | 0.63067573 |
| SHLETSL | ASSNRDWGYEQY | 0.36151966 |
| SFTTGMT | ASSNRDWGYEQY | 0.7002022 |
| QVEVAKRMGVRQPTVS | ASSNRDWGYEQY | 0.7363714 |
| QGSGGPALAM | ASSNRDWGYEQY | 0.73262525 |
| MLAPSNP | ASSNRDWGYEQY | 0.49879557 |
| LVNNAAI | ASSNRDWGYEQY | 0.7323001 |
| LTGEINTLA | ASSNRDWGYEQY | 0.7428715 |
| LSIFGPV | ASSNRDWGYEQY | 0.67082477 |
| LLALLANL | ASSNRDWGYEQY | 0.33724266 |
| LISNASDALDKLR | ASSNRDWGYEQY | 0.7290386 |
| LINEAVP | ASSNRDWGYEQY | 0.69179237 |
| LGYAFSGL | ASSNRDWGYEQY | 0.72257733 |
| LDFVAVAL | ASSNRDWGYEQY | 0.72619176 |
| LAQPGAE | ASSNRDWGYEQY | 0.6056267 |
| ISNLGAVTGF | ASSNRDWGYEQY | 0.7386354 |
| HQEDPGRI | ASSNRDWGYEQY | 0.73444515 |
| GTPGEPTP | ASSNRDWGYEQY | 0.60940266 |
| GFFNSGS | ASSNRDWGYEQY | 0.64875734 |
| GAGGGGLTAGAN | ASSNRDWGYEQY | 0.6102207 |
| FVDALVNM | ASSNRDWGYEQY | 0.33470052 |
| FLTVAPEL | ASSNRDWGYEQY | 0.71985066 |
| FETLTAL | ASSNRDWGYEQY | 0.6588634 |
| DVINPIEL | ASSNRDWGYEQY | 0.64556694 |
| APVSGAPV | ASSNRDWGYEQY | 0.7077486 |
| ALVGGLAW | ASSNRDWGYEQY | 0.5738149 |
| ALAAAAATAI | ASSNRDWGYEQY | 0.64308614 |

| | | |
|---|---|---|
| AFGLGLLL | ASSNRDWGYEQY | 0.65361947 |
| AFFAAASL | ASSNRDWGYEQY | 0.7108582 |
| AAAALPAFSPP | ASSNRDWGYEQY | 0.7107018 |
| VVLAPAL | ASSNRDWGYEQY | 0.7138126 |
| TIGHVDHGKTTL | ASSNRDWGYEQY | 0.71235615 |
| LISNASDALDK | ASSNRDWGYEQY | 0.74058294 |
| LISNASDAL | ASSNRDWGYEQY | 0.51076436 |
| ISNASDALDK | ASSNRDWGYEQY | 0.5979668 |
| NQAETLVYQTE | ASSNRDWGYEQY | 0.7210529 |
| TGGTGGTAGAG | ASSNRDWGYEQY | 0.4583031 |
| TGSTAGGAGGA | ASSNRDWGYEQY | 0.728811 |
| DPLAVPVAAGPV | ASSNRDWGYEQY | 0.7079338 |
| GGTGGTPGNGGN | ASSNRDWGYEQY | 0.7170335 |
| PAGLAGVDPATL | ASSNRDWGYEQY | 0.71250975 |
| GNNLVVEAPHAGASV | ASSNRDWGYEQY | 0.7421224 |
| KVTRIMSASTLEQPA | ASSNRDWGYEQY | 0.73597026 |
| LGFALPIALPRRTVV | ASSNRDWGYEQY | 0.68192387 |
| TRLGNLTARKLVVLV | ASSNRDWGYEQY | 0.71086895 |
| WSQQRRLIYFPSAGP | ASSNRDWGYEQY | 0.5669304 |
| GPNGNPDPMAAAVDIR | ASSNRDWGYEQY | 0.7290906 |
| TQRCVSVADLRAAASR | ASSNRDWGYEQY | 0.7245307 |
| QFVLDVAEHPVEVLTETV | ASSNRDWGYEQY | 0.619654 |
| GGAGGTGGAGGNGGTQAGDG | ASSNRDWGYEQY | 0.6991956 |
| PLLATLRAIAVDVDAKPAQV | ASSNRDWGYEQY | 0.7326119 |
| APKFRTGGYMIDFWGVGYQVAK | ASSNRDWGYEQY | 0.7336538 |
| GAGGAGGAGGAAGGPGGTGGTG | ASSNRDWGYEQY | 0.3273225 |
| IGALGLLATDVAPAGSVAFGVA | ASSNRDWGYEQY | 0.6905538 |
| AGGAGGIGGNGGAGGAGGVLMG | ASSNRDWGYEQY | 0.35704577 |
| ADELIGLL | ASSNRDWGYEQY | 0.59836197 |
| ADRALKA | ASSNRDWGYEQY | 0.5352497 |
| AGAVGQVGGAGGA | ASSNRDWGYEQY | 0.7222632 |
| AGGGGGLGGAP | ASSNRDWGYEQY | 0.6872622 |
| AGGLLYGNGGAG | ASSNRDWGYEQY | 0.72368056 |
| AGLESIGL | ASSNRDWGYEQY | 0.71414614 |
| AGLKTGII | ASSNRDWGYEQY | 0.74151695 |
| ASLAALLL | ASSNRDWGYEQY | 0.6366265 |
| ASVINISSI | ASSNRDWGYEQY | 0.6907513 |
| ATPPGPA | ASSNRDWGYEQY | 0.40791684 |
| ATVASALV | ASSNRDWGYEQY | 0.3895218 |
| ATVASLIL | ASSNRDWGYEQY | 0.55637884 |
| AVNGPVLIRH | ASSNRDWGYEQY | 0.66188544 |
| DGAAGAVGI | ASSNRDWGYEQY | 0.30392766 |
| DPLLHGI | ASSNRDWGYEQY | 0.7374772 |
| ELGAALGAL | ASSNRDWGYEQY | 0.7208733 |
| ELGRTQAV | ASSNRDWGYEQY | 0.7327057 |
| FVAGGYLS | ASSNRDWGYEQY | 0.65084624 |
| GASGGAGGAGGW | ASSNRDWGYEQY | 0.6614773 |
| GGAGAGGDGGAGGI | ASSNRDWGYEQY | 0.6535765 |
| GTGHNGSQPGMGGNG | ASSNRDWGYEQY | 0.7309108 |
| GYAAASASAS | ASSNRDWGYEQY | 0.599463 |
| HQVDVAAVQQG | ASSNRDWGYEQY | 0.4970912 |

| | | |
|---|---|---|
| IGAAIAHT | ASSNRDWGYEQY | 0.691975 |
| IGMIEVAQL | ASSNRDWGYEQY | 0.64108247 |
| IRLAPEHTSAMD | ASSNRDWGYEQY | 0.7359659 |
| IYNVSVL | ASSNRDWGYEQY | 0.7314653 |
| KQGFLGM | ASSNRDWGYEQY | 0.7232374 |
| LESVITQL | ASSNRDWGYEQY | 0.73192394 |
| LGLGALVVV | ASSNRDWGYEQY | 0.3033939 |
| LIGNGGNGGSGGI | ASSNRDWGYEQY | 0.6726968 |
| LNAINAPTL | ASSNRDWGYEQY | 0.744776 |
| LQLLNTGV | ASSNRDWGYEQY | 0.71750635 |
| LQTSIEF | ASSNRDWGYEQY | 0.693195 |
| MSFAVSEL | ASSNRDWGYEQY | 0.7392825 |
| NTGALGL | ASSNRDWGYEQY | 0.660166 |
| PPGAGGAGGAGTT | ASSNRDWGYEQY | 0.68400705 |
| QLVGLGI | ASSNRDWGYEQY | 0.7064939 |
| RFLEQVHQL | ASSNRDWGYEQY | 0.69510055 |
| RGVGYVLR | ASSNRDWGYEQY | 0.69282806 |
| SATGVFAGVI | ASSNRDWGYEQY | 0.568779 |
| SLPAITLP | ASSNRDWGYEQY | 0.71466 |
| STALAFTV | ASSNRDWGYEQY | 0.7443025 |
| SVLTVGLGK | ASSNRDWGYEQY | 0.7027494 |
| TALAGAGSGI | ASSNRDWGYEQY | 0.73570466 |
| TGLPFTGI | ASSNRDWGYEQY | 0.72393835 |
| VAEHVTV | ASSNRDWGYEQY | 0.73304594 |
| VDIVLNSL | ASSNRDWGYEQY | 0.64959663 |
| VLTGGGAVL | ASSNRDWGYEQY | 0.66077924 |
| VNVGLAYGI | ASSNRDWGYEQY | 0.69183886 |
| VSSAAVGAV | ASSNRDWGYEQY | 0.64996785 |
| VTSVYEAGV | ASSNRDWGYEQY | 0.69246733 |
| WVALDRGA | ASSNRDWGYEQY | 0.6475439 |
| YLVGSVVI | ASSNRDWGYEQY | 0.7477008 |
| GPPGTGKTLL | ASSNRDWGYEQY | 0.72163224 |
| GVGMTKF | ASSNRDWGYEQY | 0.720361 |
| IGIGGSDL | ASSNRDWGYEQY | 0.62692726 |
| IGTLLAEL | ASSNRDWGYEQY | 0.681315 |
| IYTSGTT | ASSNRDWGYEQY | 0.7202374 |
| LSFPAGV | ASSNRDWGYEQY | 0.46868634 |
| LYTSGTT | ASSNRDWGYEQY | 0.7409431 |
| VGAFPFGF | ASSNRDWGYEQY | 0.6512487 |
| VGAGTVEF | ASSNRDWGYEQY | 0.64290965 |
| VGKSTLLL | ASSNRDWGYEQY | 0.73060447 |
| AALAHAGGGIV | ASSNRDWGYEQY | 0.7409199 |
| AALVAVSLL | ASSNRDWGYEQY | 0.63339514 |
| AAVSRAVAAL | ASSNRDWGYEQY | 0.6686636 |
| AGGSSGIGGF | ASSNRDWGYEQY | 0.6238671 |
| AGLTVANANG | ASSNRDWGYEQY | 0.7301751 |
| AGLVGAVL | ASSNRDWGYEQY | 0.73965067 |
| ALSQAFTV | ASSNRDWGYEQY | 0.6903149 |
| ALTFSVSL | ASSNRDWGYEQY | 0.7409521 |
| ALTGAALVAL | ASSNRDWGYEQY | 0.7071153 |
| ASFSIGAGVL | ASSNRDWGYEQY | 0.44606853 |

| | | |
|---|---|---|
| DGSVYKI | ASSNRDWGYEQY | 0.69113296 |
| EQLAEIV | ASSNRDWGYEQY | 0.72911716 |
| FAVSELTVM | ASSNRDWGYEQY | 0.7319015 |
| GAAPLAAGL | ASSNRDWGYEQY | 0.7450358 |
| GADGGDGGSGGAGG | ASSNRDWGYEQY | 0.7043979 |
| GFGLDTGGAGG | ASSNRDWGYEQY | 0.6605063 |
| GLASGGLL | ASSNRDWGYEQY | 0.7405634 |
| GRIAGAAVLL | ASSNRDWGYEQY | 0.7455995 |
| GSAGSLNTSF | ASSNRDWGYEQY | 0.7362587 |
| GTGQIGI | ASSNRDWGYEQY | 0.6696825 |
| GTLIALIL | ASSNRDWGYEQY | 0.6063254 |
| IAATMGFVVG | ASSNRDWGYEQY | 0.68133104 |
| IFAQALL | ASSNRDWGYEQY | 0.7271584 |
| IGAAVALGF | ASSNRDWGYEQY | 0.55014807 |
| IGLTPLAL | ASSNRDWGYEQY | 0.70017725 |
| ILGSNTSTL | ASSNRDWGYEQY | 0.72293055 |
| IVFSLTVL | ASSNRDWGYEQY | 0.52854764 |
| LASKATGF | ASSNRDWGYEQY | 0.723473 |
| LASVALIL | ASSNRDWGYEQY | 0.73367906 |
| LFAVGLTL | ASSNRDWGYEQY | 0.6371424 |
| LFNGGIGTYI | ASSNRDWGYEQY | 0.6830869 |
| LTLGAGII | ASSNRDWGYEQY | 0.7358966 |
| LVAAALGDI | ASSNRDWGYEQY | 0.74736345 |
| NTAFVQLGI | ASSNRDWGYEQY | 0.7369262 |
| QLFSGAEL | ASSNRDWGYEQY | 0.43217847 |
| SAATFITL | ASSNRDWGYEQY | 0.36423644 |
| SNYGLANLGV | ASSNRDWGYEQY | 0.5892434 |
| SYGGAYAVM | ASSNRDWGYEQY | 0.72857094 |
| TTGVGGLTLG | ASSNRDWGYEQY | 0.64029425 |
| TTQTIYI | ASSNRDWGYEQY | 0.6450295 |
| VAAGAGLGAVI | ASSNRDWGYEQY | 0.71782184 |
| VAAVVTLL | ASSNRDWGYEQY | 0.7384213 |
| VGFLGGLI | ASSNRDWGYEQY | 0.6038829 |
| VGVIPILL | ASSNRDWGYEQY | 0.44610012 |
| VIFASSLI | ASSNRDWGYEQY | 0.7202373 |
| VKGGDGGAAAG | ASSNRDWGYEQY | 0.7311475 |
| VTKTFTNI | ASSNRDWGYEQY | 0.63694656 |
| VVAAAILL | ASSNRDWGYEQY | 0.69630337 |
| YAALDVEL | ASSNRDWGYEQY | 0.61485374 |
| LASVALLL | ASSNRDWGYEQY | 0.360966 |
| LDAVHRV | ASSNRDWGYEQY | 0.71509445 |
| QGLSSSIF | ASSNRDWGYEQY | 0.6208484 |
| RGTFANIRL | ASSNRDWGYEQY | 0.7383894 |
| RQLWWGHRI | ASSNRDWGYEQY | 0.64067054 |
| YGPPGTGKTLL | ASSNRDWGYEQY | 0.71859485 |
| AALVGAPANVA | ASSNRDWGYEQY | 0.7119912 |
| AATFRRTE | ASSNRDWGYEQY | 0.705999 |
| AGALGGGI | ASSNRDWGYEQY | 0.6853559 |
| AGGAGGAGGIGAG | ASSNRDWGYEQY | 0.6654403 |
| AGSGALGL | ASSNRDWGYEQY | 0.5511176 |
| AQIADQI | ASSNRDWGYEQY | 0.40069816 |

| | | |
|---|---|---|
| ASAVAVTYL | ASSNRDWGYEQY | 0.729814 |
| ASTVLGGTGG | ASSNRDWGYEQY | 0.25463036 |
| AVGGEPAK | ASSNRDWGYEQY | 0.731198 |
| CEQGTFGGTC | ASSNRDWGYEQY | 0.7293029 |
| DEGKAPAG | ASSNRDWGYEQY | 0.59186125 |
| DTVVFPAL | ASSNRDWGYEQY | 0.7043287 |
| FEGITMAG | ASSNRDWGYEQY | 0.605687 |
| FPLVVQD | ASSNRDWGYEQY | 0.5459614 |
| GAFTTPQ | ASSNRDWGYEQY | 0.7330849 |
| GFGGFGTAGGVGGA | ASSNRDWGYEQY | 0.7235178 |
| GFLAQFL | ASSNRDWGYEQY | 0.6509367 |
| GGAGGAGGNAGVA | ASSNRDWGYEQY | 0.70239496 |
| GGVGTAGAGGAGGAGG | ASSNRDWGYEQY | 0.7126179 |
| GNGGGGDAGNAGSGGN | ASSNRDWGYEQY | 0.659228 |
| GTGGAGGAGNDAGS | ASSNRDWGYEQY | 0.6395476 |
| GTTNTGWFNS | ASSNRDWGYEQY | 0.73787284 |
| IFMGPLFTL | ASSNRDWGYEQY | 0.6776284 |
| ILAHAIEAI | ASSNRDWGYEQY | 0.65207213 |
| IRSAIREAL | ASSNRDWGYEQY | 0.72643656 |
| ITFAGGG | ASSNRDWGYEQY | 0.5730291 |
| ITGMSGHQEV | ASSNRDWGYEQY | 0.72344846 |
| IVAGLACTL | ASSNRDWGYEQY | 0.65687096 |
| IVGGGLAGA | ASSNRDWGYEQY | 0.6892127 |
| IVTGVLNI | ASSNRDWGYEQY | 0.7203658 |
| LAQQGCI | ASSNRDWGYEQY | 0.61710036 |
| LCLHVEN | ASSNRDWGYEQY | 0.6150106 |
| LDAISDGS | ASSNRDWGYEQY | 0.65681875 |
| LGGFAAML | ASSNRDWGYEQY | 0.7141532 |
| LGGKNPMI | ASSNRDWGYEQY | 0.7362126 |
| LGPKGVAAIL | ASSNRDWGYEQY | 0.64290154 |
| LMVGVSI | ASSNRDWGYEQY | 0.71248686 |
| LVGAAFVL | ASSNRDWGYEQY | 0.6401977 |
| LVLAVAPL | ASSNRDWGYEQY | 0.60631406 |
| MAAIAVIDAA | ASSNRDWGYEQY | 0.6521423 |
| NNGINAELQ | ASSNRDWGYEQY | 0.68270445 |
| RGEPLVLVHG | ASSNRDWGYEQY | 0.6631634 |
| RGGAGGAAGLI | ASSNRDWGYEQY | 0.7082881 |
| RSYRMHPAV | ASSNRDWGYEQY | 0.5515869 |
| SAAHLLAGQLA | ASSNRDWGYEQY | 0.7024665 |
| SAAVGSY | ASSNRDWGYEQY | 0.73428416 |
| SPHAVLHRI | ASSNRDWGYEQY | 0.6740276 |
| SQIAKVL | ASSNRDWGYEQY | 0.71630144 |
| TALGHLVP | ASSNRDWGYEQY | 0.72108203 |
| TAPTTSGAA | ASSNRDWGYEQY | 0.70929706 |
| TDGDDGNGGSPGT | ASSNRDWGYEQY | 0.6653071 |
| TDGLGEN | ASSNRDWGYEQY | 0.6581453 |
| TGGPPLPGQA | ASSNRDWGYEQY | 0.74017483 |
| TIPAITI | ASSNRDWGYEQY | 0.73129267 |
| TIPGFTI | ASSNRDWGYEQY | 0.73109496 |
| TSIASGALA | ASSNRDWGYEQY | 0.71374536 |
| TVTLPGAAI | ASSNRDWGYEQY | 0.72939813 |

| | | |
|---|---|---|
| VASGLSAALP | ASSNRDWGYEQY | 0.7278782 |
| VGAGVIGI | ASSNRDWGYEQY | 0.68305326 |
| VGPKRFGV | ASSNRDWGYEQY | 0.47178113 |
| VNVAFPD | ASSNRDWGYEQY | 0.66604614 |
| VSMTNSGVS | ASSNRDWGYEQY | 0.6930933 |
| VTDVLGDLSV | ASSNRDWGYEQY | 0.67983663 |
| VVVVGAGFAGL | ASSNRDWGYEQY | 0.40241733 |
| VYTVGTVV | ASSNRDWGYEQY | 0.7394855 |
| WTTFSGP | ASSNRDWGYEQY | 0.74713445 |
| YDELVEAF | ASSNRDWGYEQY | 0.6045617 |
| YGTGGAGGNGG | ASSNRDWGYEQY | 0.7416451 |
| YTNVMAV | ASSNRDWGYEQY | 0.5260556 |
| FLEQVHQL | ASSNRDWGYEQY | 0.7259682 |
| GLSLLAVAV | ASSNRDWGYEQY | 0.7394069 |
| LEQVHQL | ASSNRDWGYEQY | 0.63362205 |
| LNAAGAI | ASSNRDWGYEQY | 0.7291384 |
| PAVHASL | ASSNRDWGYEQY | 0.61508036 |
| TELEDDA | ASSNRDWGYEQY | 0.68019617 |
| TPPGPPN | ASSNRDWGYEQY | 0.73791987 |
| TPPPPPPP | ASSNRDWGYEQY | 0.6739011 |
| TQSPALI | ASSSTGGEQY | 0.6865933 |
| EFAAAFPQ | ASSSTGGEQY | 0.6895875 |
| LLSGNLDVL | ASSSTGGEQY | 0.56772727 |
| GPANLGNYNI | ASSSTGGEQY | 0.6795938 |
| TLGPTLP | ASSSTGGEQY | 0.73414713 |
| VYAYSSCCK | ASSSTGGEQY | 0.5909094 |
| APAPAPAPNI | ASSSTGGEQY | 0.7449868 |
| PGGPVPIP | ASSSTGGEQY | 0.64109576 |
| FLTVALEP | ASSSTGGEQY | 0.74712384 |
| SQYFGANL | ASSSTGGEQY | 0.7300881 |
| GGVKPAA | ASSSTGGEQY | 0.7109598 |
| GGTGGAGGAGGW | ASSSTGGEQY | 0.63365406 |
| GVGGVAGLGGAGG | ASSSTGGEQY | 0.73999834 |
| VLLVLATTI | ASSSTGGEQY | 0.69846743 |
| IGIFAAML | ASSSTGGEQY | 0.7184931 |
| AVGDLID | ASSSTGGEQY | 0.7353225 |
| NISPLPL | ASSSTGGEQY | 0.1336371 |
| VTAVFVAI | ASSSTGGEQY | 0.15895757 |
| SGDDNDAAL | ASSSTGGEQY | 0.14325449 |
| AVAALAHS | ASSSTGGEQY | 0.1852391 |
| GGVTVMATP | ASSSTGGEQY | 0.13955137 |
| AGLAANVRD | ASSSTGGEQY | 0.13465852 |
| TNRPDLIDPALLRPG | ASSSTGGEQY | 0.16286182 |
| LANLGVTAL | ASSSTGGEQY | 0.1308339 |
| TGERIEL | ASSSTGGEQY | 0.19090134 |
| NGEELAA | ASSSTGGEQY | 0.16685441 |
| VLSLVWG | ASSSTGGEQY | 0.14069197 |
| IGYALWL | ASSSTGGEQY | 0.18356459 |
| LSLLLAPL | ASSSTGGEQY | 0.17026897 |
| TLDEALK | ASSSTGGEQY | 0.13273306 |
| TDGAGTAEGG | ASSSTGGEQY | 0.15556912 |

| | | |
|---|---|---|
| NAPGGAGGNGG | ASSSTGGEQY | 0.15304872 |
| VSVFMGMGGNF | ASSSTGGEQY | 0.14495194 |
| LASYLAAL | ASSSTGGEQY | 0.12398642 |
| LVGGTPAL | ASSSTGGEQY | 0.15610407 |
| VRNADVL | ASSSTGGEQY | 0.13842052 |
| TCSSPFLVSAM | ASSSTGGEQY | 0.12518324 |
| VHITDAE | ASSSTGGEQY | 0.16477938 |
| EYAAPSF | ASSSTGGEQY | 0.1359813 |
| VVDAFHGGL | ASSSTGGEQY | 0.13794473 |
| ASVVDPQ | ASSSTGGEQY | 0.17195138 |
| SYNEFGGLS | ASSSTGGEQY | 0.13202742 |
| DPDGSVY | ASSSTGGEQY | 0.1359356 |
| MVSTIHATE | ASSSTGGEQY | 0.15712751 |
| LAAGVADPV | ASSSTGGEQY | 0.18890053 |
| TKEQIAA | ASSSTGGEQY | 0.1361901 |
| SLASPAAL | ASSSTGGEQY | 0.14212954 |
| IGLVFYGV | ASSSTGGEQY | 0.23536994 |
| FLLDASNL | ASSSTGGEQY | 0.15166147 |
| YQVGNFP | ASSSTGGEQY | 0.18785723 |
| ESLLSQST | ASSSTGGEQY | 0.20700519 |
| LFTVGFAAVG | ASSSTGGEQY | 0.1418374 |
| ATPPTPT | ASSSTGGEQY | 0.17000924 |
| TDLATLG | ASSSTGGEQY | 0.1662486 |
| SAGGTGR | ASSSTGGEQY | 0.1549285 |
| GFSNSGS | ASSSTGGEQY | 0.15000719 |
| GGNGGSGGTSV | ASSSTGGEQY | 0.13987492 |
| GTSVATGGAGN | ASSSTGGEQY | 0.14266707 |
| NGGNGGSGGTS | ASSSTGGEQY | 0.12861957 |
| AVVKFPH | ASSSTGGEQY | 0.1376007 |
| TPPPPPP | ASSSTGGEQY | 0.17553446 |
| AAPAAAAAPA | ASSSTGGEQY | 0.14484702 |
| IALNEDL | ASSSTGGEQY | 0.12694596 |
| IVNVSSM | ASSSTGGEQY | 0.13180333 |
| TGIPPAPR | ASSSTGGEQY | 0.12948637 |
| FDLGGGTF | ASSSTGGEQY | 0.13600354 |
| FDLGGGTFDVS | ASSSTGGEQY | 0.1368285 |
| NEPTAAA | ASSSTGGEQY | 0.12657116 |
| NEPTAAALA | ASSSTGGEQY | 0.18810378 |
| GSIINISSI | ASSSTGGEQY | 0.19798991 |
| VAPGNAGTA | ASSSTGGEQY | 0.13259633 |
| TGGGASLE | ASSSTGGEQY | 0.14858432 |
| VAVYNHYKRI | ASSSTGGEQY | 0.21901704 |
| TDPAPATT | ASSSTGGEQY | 0.17283885 |
| TNRPDILDPALLRPG | ASSSTGGEQY | 0.16701819 |
| LTTTTTS | ASSSTGGEQY | 0.13079952 |
| PTGSGKTL | ASSSTGGEQY | 0.23909687 |
| TGSGKTL | ASSSTGGEQY | 0.13800907 |
| TIGHVDHGK | ASSSTGGEQY | 0.16524489 |
| TIGHVDHGKTT | ASSSTGGEQY | 0.1698879 |
| GHVDHGKTT | ASSSTGGEQY | 0.14035209 |
| AQFEHTLL | ASSSTGGEQY | 0.13110222 |

| | | |
|---|---|---|
| IVNQDEL | ASSSTGGEQY | 0.18657097 |
| TNGSQFF | ASSSTGGEQY | 0.17581148 |
| TGKGIVE | ASSSTGGEQY | 0.1280006 |
| LISNASDA | ASSSTGGEQY | 0.12798378 |
| AELRTLGELTKIAKAHGAQ | ASSSTGGEQY | 0.15448542 |
| APNAVAAAEPSW | ASSSTGGEQY | 0.15000464 |
| APQVAGAATAVL | ASSSTGGEQY | 0.19434404 |
| ARRDPAAAAAILPTDA | ASSSTGGEQY | 0.16616443 |
| DAAARDVLAEAGLAEYF | ASSSTGGEQY | 0.13759391 |
| DDQIVLSAKANLAKVEQ | ASSSTGGEQY | 0.18028298 |
| DELLPDTGEDSDEDGH | ASSSTGGEQY | 0.15940285 |
| ELPEAPANRATVPQD | ASSSTGGEQY | 0.14720698 |
| FGQSKIERVTGIG | ASSSTGGEQY | 0.15181468 |
| GAGGTGSTAGGAGGAGGAGGLY | ASSSTGGEQY | 0.12630884 |
| GHGGTGAAGGNGGTGGAGGSGI | ASSSTGGEQY | 0.16811283 |
| GNGGNGGAGGEGGPGGAGGAGA | ASSSTGGEQY | 0.17664589 |
| GSTAGNGGNGGNATNGTIAGQP | ASSSTGGEQY | 0.17173375 |
| GTGGDGGNGGNGGTGG | ASSSTGGEQY | 0.17282337 |
| GVRRPAGIPADAPV | ASSSTGGEQY | 0.14478417 |
| GYRQVTDAHLLTLARR | ASSSTGGEQY | 0.16990995 |
| IAKMGQDGHDRG | ASSSTGGEQY | 0.13329075 |
| KAEPSIEIQYT | ASSSTGGEQY | 0.15268113 |
| KPVRIRMAADEDTVAVT | ASSSTGGEQY | 0.14354576 |
| MPPRIAGMRLLVIKPEP | ASSSTGGEQY | 0.15091215 |
| MKFYAVALFRSRGVPPTR | ASSSTGGEQY | 0.15598026 |
| NAGGFGILGAVAHSP | ASSSTGGEQY | 0.19489904 |
| NEGREITLAMAEM | ASSSTGGEQY | 0.13619389 |
| NELVEWSHPLGHRWRWE | ASSSTGGEQY | 0.18490744 |
| NLGAGGTGGNGGNPG | ASSSTGGEQY | 0.20546928 |
| NTEWPKWETVMAADPVD | ASSSTGGEQY | 0.12995453 |
| PASAAPAETQP | ASSSTGGEQY | 0.14013766 |
| QQSSDGKNGVRAAEITGE | ASSSTGGEQY | 0.15172583 |
| RAPDAVTTSWWKE | ASSSTGGEQY | 0.1854701 |
| SGFGNVGAHTSGWFNQ | ASSSTGGEQY | 0.15881693 |
| SGVAGAVGQAASVGGLKV | ASSSTGGEQY | 0.16655546 |
| SNGGAGTGSAGGNGGT | ASSSTGGEQY | 0.20770109 |
| TGGAGGEGGAAG | ASSSTGGEQY | 0.14485958 |
| TGGDGGAAGAGGAPGHG | ASSSTGGEQY | 0.2105308 |
| TMPPLLAAIANAGRHGV | ASSSTGGEQY | 0.1286928 |
| TSSTAGNSGGA | ASSSTGGEQY | 0.17876002 |
| VPATVPVGILC | ASSSTGGEQY | 0.1408529 |
| VQRPDSGGVVLTGRIS | ASSSTGGEQY | 0.121774524 |
| WTPNPVPATKGVPAGLVQAT | ASSSTGGEQY | 0.17084445 |
| YALKRALAKFSDRFDVVII | ASSSTGGEQY | 0.15889381 |
| YLIGSVIV | ASSSTGGEQY | 0.17701757 |
| YGQLVAVID | ASSSTGGEQY | 0.15141568 |
| VTVGGMTLPA | ASSSTGGEQY | 0.15458737 |
| VLALPAV | ASSSTGGEQY | 0.13579871 |
| VIPPTLNYET | ASSSTGGEQY | 0.14198717 |
| VGMGGMHPSSG | ASSSTGGEQY | 0.13504398 |
| VGLAGGAATA | ASSSTGGEQY | 0.16674976 |

| | | |
|---|---|---|
| VGGAGGAGGSGP | ASSSTGGEQY | 0.17808221 |
| VFAAFMV | ASSSTGGEQY | 0.19188665 |
| TLYDKLR | ASSSTGGEQY | 0.13001797 |
| TLAVFTVL | ASSSTGGEQY | 0.14805655 |
| TGSVQPARV | ASSSTGGEQY | 0.15003166 |
| TGGGAGSGGT | ASSSTGGEQY | 0.12622438 |
| SHLETSL | ASSSTGGEQY | 0.1289591 |
| SFTTGMT | ASSSTGGEQY | 0.12606633 |
| QVEVAKRMGVRQPTVS | ASSSTGGEQY | 0.12572245 |
| QGSGGPALAM | ASSSTGGEQY | 0.16068573 |
| MLAPSNP | ASSSTGGEQY | 0.1708128 |
| LVNNAAI | ASSSTGGEQY | 0.14211665 |
| LTGEINTLA | ASSSTGGEQY | 0.16107287 |
| LSIFGPV | ASSSTGGEQY | 0.14263996 |
| LLALLANL | ASSSTGGEQY | 0.12022493 |
| LISNASDALDKLR | ASSSTGGEQY | 0.14020155 |
| LINEAVP | ASSSTGGEQY | 0.1342876 |
| LGYAFSGL | ASSSTGGEQY | 0.21195515 |
| LDFVAVAL | ASSSTGGEQY | 0.12748608 |
| LAQPGAE | ASSSTGGEQY | 0.13779078 |
| ISNLGAVTGF | ASSSTGGEQY | 0.14795609 |
| HQEDPGRI | ASSSTGGEQY | 0.1616908 |
| GTPGEPTP | ASSSTGGEQY | 0.14771275 |
| GFFNSGS | ASSSTGGEQY | 0.13355023 |
| GAGGGGLTAGAN | ASSSTGGEQY | 0.12635855 |
| FVDALVNM | ASSSTGGEQY | 0.1712156 |
| FLTVAPEL | ASSSTGGEQY | 0.13663325 |
| FETLTAL | ASSSTGGEQY | 0.14138183 |
| DVINPIEL | ASSSTGGEQY | 0.13371868 |
| APVSGAPV | ASSSTGGEQY | 0.14176694 |
| ALVGGLAW | ASSSTGGEQY | 0.13260156 |
| ALAAAAATAI | ASSSTGGEQY | 0.1983485 |
| AFGLGLLL | ASSSTGGEQY | 0.1341411 |
| AFFAAASL | ASSSTGGEQY | 0.14102425 |
| AAAALPAFSPP | ASSSTGGEQY | 0.16286865 |
| VVLAPAL | ASSSTGGEQY | 0.17347981 |
| TIGHVDHGKTTL | ASSSTGGEQY | 0.13867438 |
| LISNASDALDK | ASSSTGGEQY | 0.14485332 |
| LISNASDAL | ASSSTGGEQY | 0.13259463 |
| ISNASDALDK | ASSSTGGEQY | 0.1353097 |
| NQAETLVYQTE | ASSSTGGEQY | 0.13884 |
| TGGTGGTAGAG | ASSSTGGEQY | 0.18399753 |
| TGSTAGGAGGA | ASSSTGGEQY | 0.1586789 |
| DPLAVPVAAGPV | ASSSTGGEQY | 0.13354544 |
| GGTGGTPGNGGN | ASSSTGGEQY | 0.1673422 |
| PAGLAGVDPATL | ASSSTGGEQY | 0.1905026 |
| GNNLVVEAPHAGASV | ASSSTGGEQY | 0.18014826 |
| KVTRIMSASTLEQPA | ASSSTGGEQY | 0.13650735 |
| LGFALPIALPRRTVV | ASSSTGGEQY | 0.15759225 |
| TRLGNLTARKLVVLV | ASSSTGGEQY | 0.16224289 |
| WSQQRRLIYFPSAGP | ASSSTGGEQY | 0.13546489 |

| | | |
|---|---|---|
| GPNGNPDPMAAAVDIR | ASSSTGGEQY | 0.14430459 |
| TQRCVSVADLRAAASR | ASSSTGGEQY | 0.1365918 |
| QFVLDVAEHPVEVLTETV | ASSSTGGEQY | 0.17531101 |
| GGAGGTGGAGGNGGTQAGDG | ASSSTGGEQY | 0.5305527 |
| PLLATLRAIAVDVDAKPAQV | ASSSTGGEQY | 0.13841526 |
| APKFRTGGYMIDFWGVGYQVAK | ASSSTGGEQY | 0.12974077 |
| GAGGAGGAGGAAGGPGGTGGTG | ASSSTGGEQY | 0.14833017 |
| IGALGLLATDVAPAGSVAFGVA | ASSSTGGEQY | 0.12831846 |
| AGGAGGIGGNGGAGGAGGVLMG | ASSSTGGEQY | 0.13433748 |
| ADELIGLL | ASSSTGGEQY | 0.14777437 |
| ADRALKA | ASSSTGGEQY | 0.12585156 |
| AGAVGQVGGAGGA | ASSSTGGEQY | 0.14384502 |
| AGGGGGLGGAP | ASSSTGGEQY | 0.1711875 |
| AGGLLYGNGGAG | ASSSTGGEQY | 0.14366366 |
| AGLESIGL | ASSSTGGEQY | 0.17164396 |
| AGLKTGII | ASSSTGGEQY | 0.17623897 |
| ASLAALLL | ASSSTGGEQY | 0.147992 |
| ASVINISSI | ASSSTGGEQY | 0.17874111 |
| ATPPGPA | ASSSTGGEQY | 0.22220966 |
| ATVASALV | ASSSTGGEQY | 0.14325862 |
| ATVASLIL | ASSSTGGEQY | 0.14338331 |
| AVNGPVLIRH | ASSSTGGEQY | 0.15732016 |
| DGAAGAVGI | ASSSTGGEQY | 0.14810713 |
| DPLLHGI | ASSSTGGEQY | 0.1675081 |
| ELGAALGAL | ASSSTGGEQY | 0.14488763 |
| ELGRTQAV | ASSSTGGEQY | 0.17529482 |
| FVAGGYLS | ASSSTGGEQY | 0.14496177 |
| GASGGAGGAGGW | ASSSTGGEQY | 0.21229666 |
| GGAGAGGDGGAGGI | ASSSTGGEQY | 0.12859784 |
| GTGHNGSQPGMGGNG | ASSSTGGEQY | 0.13482223 |
| GYAAASASAS | ASSSTGGEQY | 0.1561203 |
| HQVDVAAVQQG | ASSSTGGEQY | 0.12591799 |
| IGAAIAHT | ASSSTGGEQY | 0.13773961 |
| IGMIEVAQL | ASSSTGGEQY | 0.16004588 |
| IRLAPEHTSAMD | ASSSTGGEQY | 0.15331174 |
| IYNVSVL | ASSSTGGEQY | 0.12147236 |
| KQGFLGM | ASSSTGGEQY | 0.14396513 |
| LESVITQL | ASSSTGGEQY | 0.13310738 |
| LGLGALVVV | ASSSTGGEQY | 0.15719964 |
| LIGNGGNGGSGGI | ASSSTGGEQY | 0.14735274 |
| LNAINAPTL | ASSSTGGEQY | 0.12904409 |
| LQLLNTGV | ASSSTGGEQY | 0.16265334 |
| LQTSIEF | ASSSTGGEQY | 0.12906474 |
| MSFAVSEL | ASSSTGGEQY | 0.1334364 |
| NTGALGL | ASSSTGGEQY | 0.15837893 |
| PPGAGGAGGAGTT | ASSSTGGEQY | 0.26612306 |
| QLVGLGI | ASSSTGGEQY | 0.13742793 |
| RFLEQVHQL | ASSSTGGEQY | 0.1618282 |
| RGVGYVLR | ASSSTGGEQY | 0.13359188 |
| SATGVFAGVI | ASSSTGGEQY | 0.20317505 |
| SLPAITLP | ASSSTGGEQY | 0.15032193 |

| | | |
|---|---|---|
| STALAFTV | ASSSTGGEQY | 0.13269676 |
| SVLTVGLGK | ASSSTGGEQY | 0.16187541 |
| TALAGAGSGI | ASSSTGGEQY | 0.15276368 |
| TGLPFTGI | ASSSTGGEQY | 0.17003183 |
| VAEHVTV | ASSSTGGEQY | 0.18847273 |
| VDIVLNSL | ASSSTGGEQY | 0.1410298 |
| VLTGGGAVL | ASSSTGGEQY | 0.14145017 |
| VNVGLAYGI | ASSSTGGEQY | 0.16037364 |
| VSSAAVGAV | ASSSTGGEQY | 0.14657147 |
| VTSVYEAGV | ASSSTGGEQY | 0.1299768 |
| WVALDRGA | ASSSTGGEQY | 0.16901751 |
| YLVGSVVI | ASSSTGGEQY | 0.14900136 |
| GPPGTGKTLL | ASSSTGGEQY | 0.15486255 |
| GVGMTKF | ASSSTGGEQY | 0.16533071 |
| IGIGGSDL | ASSSTGGEQY | 0.1539749 |
| IGTLLAEL | ASSSTGGEQY | 0.16168422 |
| IYTSGTT | ASSSTGGEQY | 0.16663858 |
| LSFPAGV | ASSSTGGEQY | 0.3969438 |
| LYTSGTT | ASSSTGGEQY | 0.13624729 |
| VGAFPFGF | ASSSTGGEQY | 0.15327732 |
| VGAGTVEF | ASSSTGGEQY | 0.15862122 |
| VGKSTLLL | ASSSTGGEQY | 0.16949226 |
| AALAHAGGGIV | ASSSTGGEQY | 0.1359217 |
| AALVAVSLL | ASSSTGGEQY | 0.14695773 |
| AAVSRAVAAL | ASSSTGGEQY | 0.16391711 |
| AGGSSGIGGF | ASSSTGGEQY | 0.1960496 |
| AGLTVANANG | ASSSTGGEQY | 0.14449936 |
| AGLVGAVL | ASSSTGGEQY | 0.14584036 |
| ALSQAFTV | ASSSTGGEQY | 0.20688513 |
| ALTFSVSL | ASSSTGGEQY | 0.14832419 |
| ALTGAALVAL | ASSSTGGEQY | 0.15030023 |
| ASFSIGAGVL | ASSSTGGEQY | 0.17920925 |
| DGSVYKI | ASSSTGGEQY | 0.1805455 |
| EQLAEIV | ASSSTGGEQY | 0.15284081 |
| FAVSELTVM | ASSSTGGEQY | 0.17812768 |
| GAAPLAAGL | ASSSTGGEQY | 0.13102229 |
| GADGGDGGSGGAGG | ASSSTGGEQY | 0.15650858 |
| GFGLDTGGAGG | ASSSTGGEQY | 0.12793277 |
| GLASGGLL | ASSSTGGEQY | 0.14803834 |
| GRIAGAAVLL | ASSSTGGEQY | 0.12423848 |
| GSAGSLNTSF | ASSSTGGEQY | 0.15184133 |
| GTGQIGI | ASSSTGGEQY | 0.15830222 |
| GTLIALIL | ASSSTGGEQY | 0.1627189 |
| IAATMGFVVG | ASSSTGGEQY | 0.12959377 |
| IFAQALL | ASSSTGGEQY | 0.14435935 |
| IGAAVALGF | ASSSTGGEQY | 0.17872143 |
| IGLTPLAL | ASSSTGGEQY | 0.3890803 |
| ILGSNTSTL | ASSSTGGEQY | 0.14477092 |
| IVFSLTVL | ASSSTGGEQY | 0.20448041 |
| LASKATGF | ASSSTGGEQY | 0.14522788 |
| LASVALIL | ASSSTGGEQY | 0.14124106 |

| | | |
|---|---|---|
| LFAVGLTL | ASSSTGGEQY | 0.1269191 |
| LFNGGIGTYI | ASSSTGGEQY | 0.15869482 |
| LTLGAGII | ASSSTGGEQY | 0.13601959 |
| LVAAALGDI | ASSSTGGEQY | 0.13016151 |
| NTAFVQLGI | ASSSTGGEQY | 0.17415643 |
| QLFSGAEL | ASSSTGGEQY | 0.17658462 |
| SAATFITL | ASSSTGGEQY | 0.16173437 |
| SNYGLANLGV | ASSSTGGEQY | 0.20476699 |
| SYGGAYAVM | ASSSTGGEQY | 0.16132174 |
| TTGVGGLTLG | ASSSTGGEQY | 0.1753956 |
| TTQTIYI | ASSSTGGEQY | 0.13257097 |
| VAAGAGLGAVI | ASSSTGGEQY | 0.15654929 |
| VAAVVTLL | ASSSTGGEQY | 0.12751935 |
| VGFLGGLI | ASSSTGGEQY | 0.14073445 |
| VGVIPILL | ASSSTGGEQY | 0.13192165 |
| VIFASSLI | ASSSTGGEQY | 0.15189016 |
| VKGGDGGAAAG | ASSSTGGEQY | 0.12178782 |
| VTKTFTNI | ASSSTGGEQY | 0.18245284 |
| VVAAAILL | ASSSTGGEQY | 0.18054166 |
| YAALDVEL | ASSSTGGEQY | 0.13321118 |
| LASVALLL | ASSSTGGEQY | 0.17692357 |
| LDAVHRV | ASSSTGGEQY | 0.1428538 |
| QGLSSSIF | ASSSTGGEQY | 0.12790193 |
| RGTFANIRL | ASSSTGGEQY | 0.18197924 |
| RQLWWGHRI | ASSSTGGEQY | 0.13470782 |
| YGPPGTGKTLL | ASSSTGGEQY | 0.17707463 |
| AALVGAPANVA | ASSSTGGEQY | 0.17372417 |
| AATFRRTE | ASSSTGGEQY | 0.16563654 |
| AGALGGGI | ASSSTGGEQY | 0.13846578 |
| AGGAGGAGGIGAG | ASSSTGGEQY | 0.17021033 |
| AGSGALGL | ASSSTGGEQY | 0.13620141 |
| AQIADQI | ASSSTGGEQY | 0.14543669 |
| ASAVAVTYL | ASSSTGGEQY | 0.12963389 |
| ASTVLGGTGG | ASSSTGGEQY | 0.14961974 |
| AVGGEPAK | ASSSTGGEQY | 0.1342463 |
| CEQGTFGGTC | ASSSTGGEQY | 0.1890432 |
| DEGKAPAG | ASSSTGGEQY | 0.14359407 |
| DTVVFPAL | ASSSTGGEQY | 0.15754 |
| FEGITMAG | ASSSTGGEQY | 0.14079763 |
| FPLVVQD | ASSSTGGEQY | 0.14956056 |
| GAFTTPQ | ASSSTGGEQY | 0.14717306 |
| GFGGFGTAGGVGGA | ASSSTGGEQY | 0.13138245 |
| GFLAQFL | ASSSTGGEQY | 0.15348369 |
| GGAGGAGGNAGVA | ASSSTGGEQY | 0.15030392 |
| GGVGTAGAGGAGGAGG | ASSSTGGEQY | 0.12660606 |
| GNGGGGDAGNAGSGGN | ASSSTGGEQY | 0.17505881 |
| GTGGAGGAGNDAGS | ASSSTGGEQY | 0.15066896 |
| GTTNTGWFNS | ASSSTGGEQY | 0.13006106 |
| IFMGPLFTL | ASSSTGGEQY | 0.13558315 |
| ILAHAIEAI | ASSSTGGEQY | 0.17621213 |
| IRSAIREAL | ASSSTGGEQY | 0.14816141 |

| | | |
|---|---|---|
| ITFAGGG | ASSSTGGEQY | 0.12256133 |
| ITGMSGHQEV | ASSSTGGEQY | 0.13495423 |
| IVAGLACTL | ASSSTGGEQY | 0.15595858 |
| IVGGGLAGA | ASSSTGGEQY | 0.12570575 |
| IVTGVLNI | ASSSTGGEQY | 0.13197112 |
| LAQQGCI | ASSSTGGEQY | 0.19155002 |
| LCLHVEN | ASSSTGGEQY | 0.14913486 |
| LDAISDGS | ASSSTGGEQY | 0.15257438 |
| LGGFAAML | ASSSTGGEQY | 0.16216096 |
| LGGKNPMI | ASSSTGGEQY | 0.17676559 |
| LGPKGVAAIL | ASSSTGGEQY | 0.18395732 |
| LMVGVSI | ASSSTGGEQY | 0.1438306 |
| LVGAAFVL | ASSSTGGEQY | 0.1461458 |
| LVLAVAPL | ASSSTGGEQY | 0.1555368 |
| MAAIAVIDAA | ASSSTGGEQY | 0.16367438 |
| NNGINAELQ | ASSSTGGEQY | 0.13693897 |
| RGEPLVLVHG | ASSSTGGEQY | 0.1312737 |
| RGGAGGAAGLI | ASSSTGGEQY | 0.13862747 |
| RSYRMHPAV | ASSSTGGEQY | 0.14061943 |
| SAAHLLAGQLA | ASSSTGGEQY | 0.16803867 |
| SAAVGSY | ASSSTGGEQY | 0.208313 |
| SPHAVLHRI | ASSSTGGEQY | 0.13581996 |
| SQIAKVL | ASSSTGGEQY | 0.1333801 |
| TALGHLVP | ASSSTGGEQY | 0.14697762 |
| TAPTTSGAA | ASSSTGGEQY | 0.1385162 |
| TDGDDGNGGSPGT | ASSSTGGEQY | 0.1658879 |
| TDGLGEN | ASSSTGGEQY | 0.14258735 |
| TGGPPLPGQA | ASSSTGGEQY | 0.43930858 |
| TIPAITI | ASSSTGGEQY | 0.16603027 |
| TIPGFTI | ASSSTGGEQY | 0.1335372 |
| TSIASGALA | ASSSTGGEQY | 0.17208135 |
| TVTLPGAAI | ASSSTGGEQY | 0.13690695 |
| VASGLSAALP | ASSSTGGEQY | 0.14420715 |
| VGAGVIGI | ASSSTGGEQY | 0.14680946 |
| VGPKRFGV | ASSSTGGEQY | 0.1316821 |
| VNVAFPD | ASSSTGGEQY | 0.13941552 |
| VSMTNSGVS | ASSSTGGEQY | 0.15440588 |
| VTDVLGDLSV | ASSSTGGEQY | 0.13270733 |
| VVVVGAGFAGL | ASSSTGGEQY | 0.18098462 |
| VYTVGTVV | ASSSTGGEQY | 0.13173409 |
| WTTFSGP | ASSSTGGEQY | 0.13818751 |
| YDELVEAF | ASSSTGGEQY | 0.13872668 |
| YGTGGAGGNGG | ASSSTGGEQY | 0.17708166 |
| YTNVMAV | ASSSTGGEQY | 0.23872347 |
| FLEQVHQL | ASSSTGGEQY | 0.13718347 |
| GLSLLAVAV | ASSSTGGEQY | 0.13356729 |
| LEQVHQL | ASSSTGGEQY | 0.20985174 |
| LNAAGAI | ASSSTGGEQY | 0.13223867 |
| PAVHASL | ASSSTGGEQY | 0.1654915 |
| TELEDDA | ASSSTGGEQY | 0.13171923 |
| TPPGPPN | ASSSTGGEQY | 0.12892398 |

| | | |
|---|---|---|
| TPPPPPPP | ASSSTGGEQY | 0.14984444 |
| TQSPALI | ASSIDRADTQY | 0.1573268 |
| EFAAAFPQ | ASSIDRADTQY | 0.12781294 |
| LLSGNLDVL | ASSIDRADTQY | 0.17551842 |
| GPANLGNYNI | ASSIDRADTQY | 0.12126559 |
| TLGPTLP | ASSIDRADTQY | 0.13988477 |
| VYAYSSCCK | ASSIDRADTQY | 0.13285789 |
| APAPAPAPNI | ASSIDRADTQY | 0.13474636 |
| PGGPVPIP | ASSIDRADTQY | 0.12683627 |
| FLTVALEP | ASSIDRADTQY | 0.15260886 |
| SQYFGANL | ASSIDRADTQY | 0.15865418 |
| GGVKPAA | ASSIDRADTQY | 0.1905925 |
| GGTGGAGGAGGW | ASSIDRADTQY | 0.13933542 |
| GVGGVAGLGGAGG | ASSIDRADTQY | 0.15051784 |
| VLLVLATTI | ASSIDRADTQY | 0.17163682 |
| IGIFAAML | ASSIDRADTQY | 0.1367828 |
| AVGDLID | ASSIDRADTQY | 0.12949555 |
| NISPLPL | ASSIDRADTQY | 0.15167962 |
| VTAVFVAI | ASSIDRADTQY | 0.13104284 |
| SGDDNDAAL | ASSIDRADTQY | 0.14287442 |
| AVAALAHS | ASSIDRADTQY | 0.18087712 |
| GGVTVMATP | ASSIDRADTQY | 0.124659345 |
| AGLAANVRD | ASSIDRADTQY | 0.1921814 |
| TNRPDLIDPALLRPG | ASSIDRADTQY | 0.16015969 |
| LANLGVTAL | ASSIDRADTQY | 0.13168624 |
| TGERIEL | ASSIDRADTQY | 0.13795453 |
| NGEELAA | ASSIDRADTQY | 0.1350507 |
| VLSLVWG | ASSIDRADTQY | 0.12605116 |
| IGYALWL | ASSIDRADTQY | 0.16591568 |
| LSLLLAPL | ASSIDRADTQY | 0.1345102 |
| TLDEALK | ASSIDRADTQY | 0.14892796 |
| TDGAGTAEGG | ASSIDRADTQY | 0.14058416 |
| NAPGGAGGNGG | ASSIDRADTQY | 0.14454347 |
| VSVFMGMGGNF | ASSIDRADTQY | 0.1484775 |
| LASYLAAL | ASSIDRADTQY | 0.14471434 |
| LVGGTPAL | ASSIDRADTQY | 0.14000805 |
| VRNADVL | ASSIDRADTQY | 0.15811187 |
| TCSSPFLVSAM | ASSIDRADTQY | 0.12901913 |
| VHITDAE | ASSIDRADTQY | 0.15549201 |
| EYAAPSF | ASSIDRADTQY | 0.19303863 |
| VVDAFHGGL | ASSIDRADTQY | 0.12639862 |
| ASVVDPQ | ASSIDRADTQY | 0.13456097 |
| SYNEFGGLS | ASSIDRADTQY | 0.1511965 |
| DPDGSVY | ASSIDRADTQY | 0.13305858 |
| MVSTIHATE | ASSIDRADTQY | 0.16720186 |
| LAAGVADPV | ASSIDRADTQY | 0.1480484 |
| TKEQIAA | ASSIDRADTQY | 0.153082 |
| SLASPAAL | ASSIDRADTQY | 0.16227351 |
| IGLVFYGV | ASSIDRADTQY | 0.1640139 |
| FLLDASNL | ASSIDRADTQY | 0.16361837 |
| YQVGNFP | ASSIDRADTQY | 0.14425933 |

| | | |
|---|---|---|
| ESLLSQST | ASSIDRADTQY | 0.14698538 |
| LFTVGFAAVG | ASSIDRADTQY | 0.13585488 |
| ATPPTPT | ASSIDRADTQY | 0.149275 |
| TDLATLG | ASSIDRADTQY | 0.21527942 |
| SAGGTGR | ASSIDRADTQY | 0.12692909 |
| GFSNSGS | ASSIDRADTQY | 0.14271161 |
| GGNGGSGGTSV | ASSIDRADTQY | 0.18279195 |
| GTSVATGGAGN | ASSIDRADTQY | 0.17141622 |
| NGGNGGSGGTS | ASSIDRADTQY | 0.13914233 |
| AVVKFPH | ASSIDRADTQY | 0.13464406 |
| TPPPPPP | ASSIDRADTQY | 0.14727412 |
| AAPAAAAAPA | ASSIDRADTQY | 0.15722655 |
| IALNEDL | ASSIDRADTQY | 0.13567047 |
| IVNVSSM | ASSIDRADTQY | 0.16442734 |
| TGIPPAPR | ASSIDRADTQY | 0.16293328 |
| FDLGGGTF | ASSIDRADTQY | 0.26307902 |
| FDLGGGTFDVS | ASSIDRADTQY | 0.1652237 |
| NEPTAAA | ASSIDRADTQY | 0.18850017 |
| NEPTAAALA | ASSIDRADTQY | 0.15861939 |
| GSIINISSI | ASSIDRADTQY | 0.14474785 |
| VAPGNAGTA | ASSIDRADTQY | 0.13473329 |
| TGGGASLE | ASSIDRADTQY | 0.13419119 |
| VAVYNHYKRI | ASSIDRADTQY | 0.13739592 |
| TDPAPATT | ASSIDRADTQY | 0.13930553 |
| TNRPDILDPALLRPG | ASSIDRADTQY | 0.12394997 |
| LTTTTTS | ASSIDRADTQY | 0.1523846 |
| PTGSGKTL | ASSIDRADTQY | 0.14336433 |
| TGSGKTL | ASSIDRADTQY | 0.1764999 |
| TIGHVDHGK | ASSIDRADTQY | 0.14998038 |
| TIGHVDHGKTT | ASSIDRADTQY | 0.14115055 |
| GHVDHGKTT | ASSIDRADTQY | 0.12998167 |
| AQFEHTLL | ASSIDRADTQY | 0.21299139 |
| IVNQDEL | ASSIDRADTQY | 0.12671405 |
| TNGSQFF | ASSIDRADTQY | 0.12672716 |
| TGKGIVE | ASSIDRADTQY | 0.15040675 |
| LISNASDA | ASSIDRADTQY | 0.13283984 |
| AELRTLGELTKIAKAHGAQ | ASSIDRADTQY | 0.16111667 |
| APNAVAAAEPSW | ASSIDRADTQY | 0.17933103 |
| APQVAGAATAVL | ASSIDRADTQY | 0.1476293 |
| ARRDPAAAAAILPTDA | ASSIDRADTQY | 0.22920005 |
| DAAARDVLAEAGLAEYF | ASSIDRADTQY | 0.12509765 |
| DDQIVLSAKANLAKVEQ | ASSIDRADTQY | 0.14262769 |
| DELLPDTGEDSDEDGH | ASSIDRADTQY | 0.22248508 |
| ELPEAPANRATVPQD | ASSIDRADTQY | 0.14424428 |
| FGQSKIERVTGIG | ASSIDRADTQY | 0.16714758 |
| GAGGTGSTAGGAGGAGGAGGLY | ASSIDRADTQY | 0.16227652 |
| GHGGTGAAGGNGGTGGAGGSGI | ASSIDRADTQY | 0.13604666 |
| GNGGNGGAGGEGGPGGAGGAGA | ASSIDRADTQY | 0.15181993 |
| GSTAGNGGNGGNATNGTIAGQP | ASSIDRADTQY | 0.143088 |
| GTGGDGGNGGNGGTGG | ASSIDRADTQY | 0.16237855 |
| GVRRPAGIPADAPV | ASSIDRADTQY | 0.1417469 |

| | | |
|---|---|---|
| GYRQVTDAHLLTLARR | ASSIDRADTQY | 0.12879695 |
| IAKMGQDGHDRG | ASSIDRADTQY | 0.2035296 |
| KAEPSIEIQYT | ASSIDRADTQY | 0.18469326 |
| KPVRIRMAADEDTVAVT | ASSIDRADTQY | 0.14839903 |
| MPPRIAGMRLLVIKPEP | ASSIDRADTQY | 0.15214969 |
| MKFYAVALFRSRGVPPTR | ASSIDRADTQY | 0.14782128 |
| NAGGFGILGAVAHSP | ASSIDRADTQY | 0.15958677 |
| NEGREITLAMAEM | ASSIDRADTQY | 0.14031126 |
| NELVEWSHPLGHRWRWE | ASSIDRADTQY | 0.12788275 |
| NLGAGGTGGNGGNPG | ASSIDRADTQY | 0.15305467 |
| NTEWPKWETVMAADPVD | ASSIDRADTQY | 0.12797675 |
| PASAAPAETQP | ASSIDRADTQY | 0.1891432 |
| QQSSDGKNGVRAAEITGE | ASSIDRADTQY | 0.14327557 |
| RAPDAVTTSWWKE | ASSIDRADTQY | 0.20569426 |
| SGFGNVGAHTSGWFNQ | ASSIDRADTQY | 0.16265826 |
| SGVAGAVGQAASVGGLKV | ASSIDRADTQY | 0.14075406 |
| SNGGAGTGSAGGNGGT | ASSIDRADTQY | 0.12956293 |
| TGGAGGEGGAAG | ASSIDRADTQY | 0.14362033 |
| TGGDGGAAGAGGAPGHG | ASSIDRADTQY | 0.14108731 |
| TMPPLLAAIANAGRHGV | ASSIDRADTQY | 0.13214046 |
| TSSTAGNSGGA | ASSIDRADTQY | 0.18623856 |
| VPATVPVGILC | ASSIDRADTQY | 0.13463846 |
| VQRPDSGGVVLTGRIS | ASSIDRADTQY | 0.13221626 |
| WTPNPVPATKGVPAGLVQAT | ASSIDRADTQY | 0.15882255 |
| YALKRALAKFSDRFDVVII | ASSIDRADTQY | 0.36683777 |
| YLIGSVIV | ASSIDRADTQY | 0.14437822 |
| YGQLVAVID | ASSIDRADTQY | 0.15687697 |
| VTVGGMTLPA | ASSIDRADTQY | 0.13151434 |
| VLALPAV | ASSIDRADTQY | 0.16549416 |
| VIPPTLNYET | ASSIDRADTQY | 0.20051928 |
| VGMGGMHPSSG | ASSIDRADTQY | 0.14216913 |
| VGLAGGAATA | ASSIDRADTQY | 0.26991832 |
| VGGAGGAGGSGP | ASSIDRADTQY | 0.14192481 |
| VFAAFMV | ASSIDRADTQY | 0.13591199 |
| TLYDKLR | ASSIDRADTQY | 0.16608375 |
| TLAVFTVL | ASSIDRADTQY | 0.15228571 |
| TGSVQPARV | ASSIDRADTQY | 0.16108465 |
| TGGGAGSGGT | ASSIDRADTQY | 0.1612454 |
| SHLETSL | ASSIDRADTQY | 0.15142335 |
| SFTTGMT | ASSIDRADTQY | 0.13503519 |
| QVEVAKRMGVRQPTVS | ASSIDRADTQY | 0.1841532 |
| QGSGGPALAM | ASSIDRADTQY | 0.20450713 |
| MLAPSNP | ASSIDRADTQY | 0.1667263 |
| LVNNAAI | ASSIDRADTQY | 0.14132053 |
| LTGEINTLA | ASSIDRADTQY | 0.14184667 |
| LSIFGPV | ASSIDRADTQY | 0.16455236 |
| LLALLANL | ASSIDRADTQY | 0.13926585 |
| LISNASDALDKLR | ASSIDRADTQY | 0.15260966 |
| LINEAVP | ASSIDRADTQY | 0.14526972 |
| LGYAFSGL | ASSIDRADTQY | 0.16200384 |
| LDFVAVAL | ASSIDRADTQY | 0.13453494 |

| | | |
|---|---|---|
| LAQPGAE | ASSIDRADTQY | 0.14593445 |
| ISNLGAVTGF | ASSIDRADTQY | 0.16009854 |
| HQEDPGRI | ASSIDRADTQY | 0.16054204 |
| GTPGEPTP | ASSIDRADTQY | 0.13585895 |
| GFFNSGS | ASSIDRADTQY | 0.1826261 |
| GAGGGGLTAGAN | ASSIDRADTQY | 0.17384917 |
| FVDALVNM | ASSIDRADTQY | 0.19193994 |
| FLTVAPEL | ASSIDRADTQY | 0.13086899 |
| FETLTAL | ASSIDRADTQY | 0.124199875 |
| DVINPIEL | ASSIDRADTQY | 0.15590693 |
| APVSGAPV | ASSIDRADTQY | 0.18627824 |
| ALVGGLAW | ASSIDRADTQY | 0.14286174 |
| ALAAAAATAI | ASSIDRADTQY | 0.1325493 |
| AFGLGLLL | ASSIDRADTQY | 0.13626435 |
| AFFAAASL | ASSIDRADTQY | 0.1641754 |
| AAAALPAFSPP | ASSIDRADTQY | 0.15133663 |
| VVLAPAL | ASSIDRADTQY | 0.1291556 |
| TIGHVDHGKTTL | ASSIDRADTQY | 0.14518692 |
| LISNASDALDK | ASSIDRADTQY | 0.2230593 |
| LISNASDAL | ASSIDRADTQY | 0.12979347 |
| ISNASDALDK | ASSIDRADTQY | 0.13594335 |
| NQAETLVYQTE | ASSIDRADTQY | 0.17317754 |
| TGGTGGTAGAG | ASSIDRADTQY | 0.14000438 |
| TGSTAGGAGGA | ASSIDRADTQY | 0.13882044 |
| DPLAVPVAAGPV | ASSIDRADTQY | 0.1349979 |
| GGTGGTPGNGGN | ASSIDRADTQY | 0.1392795 |
| PAGLAGVDPATL | ASSIDRADTQY | 0.13152288 |
| GNNLVVEAPHAGASV | ASSIDRADTQY | 0.16485186 |
| KVTRIMSASTLEQPA | ASSIDRADTQY | 0.14851776 |
| LGFALPIALPRRTVV | ASSIDRADTQY | 0.1593208 |
| TRLGNLTARKLVVLV | ASSIDRADTQY | 0.15121523 |
| WSQQRRLIYFPSAGP | ASSIDRADTQY | 0.13367057 |
| GPNGNPDPMAAAVDIR | ASSIDRADTQY | 0.13301624 |
| TQRCVSVADLRAAASR | ASSIDRADTQY | 0.12812828 |
| QFVLDVAEHPVEVLTETV | ASSIDRADTQY | 0.16719565 |
| GGAGGTGGAGGNGGTQAGDG | ASSIDRADTQY | 0.13378797 |
| PLLATLRAIAVDVDAKPAQV | ASSIDRADTQY | 0.16105445 |
| APKFRTGGYMIDFWGVGYQVAK | ASSIDRADTQY | 0.16039068 |
| GAGGAGGAGGAAGGPGGTGGTG | ASSIDRADTQY | 0.13333794 |
| IGALGLLATDVAPAGSVAFGVA | ASSIDRADTQY | 0.14574517 |
| AGGAGGIGGNGGAGGAGGVLMG | ASSIDRADTQY | 0.12470055 |
| ADELIGLL | ASSIDRADTQY | 0.1574484 |
| ADRALKA | ASSIDRADTQY | 0.14336598 |
| AGAVGQVGGAGGA | ASSIDRADTQY | 0.16313398 |
| AGGGGGLGGAP | ASSIDRADTQY | 0.15396644 |
| AGGLLYGNGGAG | ASSIDRADTQY | 0.17646818 |
| AGLESIGL | ASSIDRADTQY | 0.16587326 |
| AGLKTGII | ASSIDRADTQY | 0.15693167 |
| ASLAALLL | ASSIDRADTQY | 0.16791792 |
| ASVINISSI | ASSIDRADTQY | 0.16839142 |
| ATPPGPA | ASSIDRADTQY | 0.17151305 |

| | | |
|---|---|---|
| ATVASALV | ASSIDRADTQY | 0.12993513 |
| ATVASLIL | ASSIDRADTQY | 0.14709263 |
| AVNGPVLIRH | ASSIDRADTQY | 0.13188055 |
| DGAAGAVGI | ASSIDRADTQY | 0.14737247 |
| DPLLHGI | ASSIDRADTQY | 0.13578284 |
| ELGAALGAL | ASSIDRADTQY | 0.16830829 |
| ELGRTQAV | ASSIDRADTQY | 0.1488162 |
| FVAGGYLS | ASSIDRADTQY | 0.13004078 |
| GASGGAGGAGGW | ASSIDRADTQY | 0.13341402 |
| GGAGAGGDGGAGGI | ASSIDRADTQY | 0.14598624 |
| GTGHNGSQPGMGGNG | ASSIDRADTQY | 0.16840625 |
| GYAAASASAS | ASSIDRADTQY | 0.1527671 |
| HQVDVAAVQQG | ASSIDRADTQY | 0.12984855 |
| IGAAIAHT | ASSIDRADTQY | 0.16886158 |
| IGMIEVAQL | ASSIDRADTQY | 0.1501075 |
| IRLAPEHTSAMD | ASSIDRADTQY | 0.14078881 |
| IYNVSVL | ASSIDRADTQY | 0.16789731 |
| KQGFLGM | ASSIDRADTQY | 0.14667247 |
| LESVITQL | ASSIDRADTQY | 0.13926758 |
| LGLGALVVV | ASSIDRADTQY | 0.16167079 |
| LIGNGGNGGSGGI | ASSIDRADTQY | 0.13331018 |
| LNAINAPTL | ASSIDRADTQY | 0.17859225 |
| LQLLNTGV | ASSIDRADTQY | 0.12968457 |
| LQTSIEF | ASSIDRADTQY | 0.13627201 |
| MSFAVSEL | ASSIDRADTQY | 0.15808332 |
| NTGALGL | ASSIDRADTQY | 0.1300396 |
| PPGAGGAGGAGTT | ASSIDRADTQY | 0.14565098 |
| QLVGLGI | ASSIDRADTQY | 0.14806701 |
| RFLEQVHQL | ASSIDRADTQY | 0.13835129 |
| RGVGYVLR | ASSIDRADTQY | 0.14492027 |
| SATGVFAGVI | ASSIDRADTQY | 0.17542076 |
| SLPAITLP | ASSIDRADTQY | 0.16256839 |
| STALAFTV | ASSIDRADTQY | 0.13221288 |
| SVLTVGLGK | ASSIDRADTQY | 0.18280786 |
| TALAGAGSGI | ASSIDRADTQY | 0.1772602 |
| TGLPFTGI | ASSIDRADTQY | 0.13114735 |
| VAEHVTV | ASSIDRADTQY | 0.12460191 |
| VDIVLNSL | ASSIDRADTQY | 0.13991965 |
| VLTGGGAVL | ASSIDRADTQY | 0.1312949 |
| VNVGLAYGI | ASSIDRADTQY | 0.14287606 |
| VSSAAVGAV | ASSIDRADTQY | 0.13279253 |
| VTSVYEAGV | ASSIDRADTQY | 0.16603766 |
| WVALDRGA | ASSIDRADTQY | 0.18964776 |
| YLVGSVVI | ASSIDRADTQY | 0.14276308 |
| GPPGTGKTLL | ASSIDRADTQY | 0.17465962 |
| GVGMTKF | ASSIDRADTQY | 0.13127533 |
| IGIGGSDL | ASSIDRADTQY | 0.1698663 |
| IGTLLAEL | ASSIDRADTQY | 0.18553911 |
| IYTSGTT | ASSIDRADTQY | 0.12907007 |
| LSFPAGV | ASSIDRADTQY | 0.1567303 |
| LYTSGTT | ASSIDRADTQY | 0.1440294 |

| | | |
|---|---|---|
| VGAFPFGF | ASSIDRADTQY | 0.13528241 |
| VGAGTVEF | ASSIDRADTQY | 0.16828384 |
| VGKSTLLL | ASSIDRADTQY | 0.15297349 |
| AALAHAGGGIV | ASSIDRADTQY | 0.16263193 |
| AALVAVSLL | ASSIDRADTQY | 0.14264335 |
| AAVSRAVAAL | ASSIDRADTQY | 0.12994748 |
| AGGSSGIGGF | ASSIDRADTQY | 0.14767568 |
| AGLTVANANG | ASSIDRADTQY | 0.17237265 |
| AGLVGAVL | ASSIDRADTQY | 0.14777318 |
| ALSQAFTV | ASSIDRADTQY | 0.19943242 |
| ALTFSVSL | ASSIDRADTQY | 0.13834251 |
| ALTGAALVAL | ASSIDRADTQY | 0.16410175 |
| ASFSIGAGVL | ASSIDRADTQY | 0.33298644 |
| DGSVYKI | ASSIDRADTQY | 0.14305656 |
| EQLAEIV | ASSIDRADTQY | 0.13889442 |
| FAVSELTVM | ASSIDRADTQY | 0.19905095 |
| GAAPLAAGL | ASSIDRADTQY | 0.16874519 |
| GADGGDGGSGGAGG | ASSIDRADTQY | 0.16067383 |
| GFGLDTGGAGG | ASSIDRADTQY | 0.18136522 |
| GLASGGLL | ASSIDRADTQY | 0.22682384 |
| GRIAGAAVLL | ASSIDRADTQY | 0.7752436 |
| GSAGSLNTSF | ASSIDRADTQY | 0.5725252 |
| GTGQIGI | ASSIDRADTQY | 0.57013214 |
| GTLIALIL | ASSIDRADTQY | 0.41120958 |
| IAATMGFVVG | ASSIDRADTQY | 0.18724878 |
| IFAQALL | ASSIDRADTQY | 0.43383032 |
| IGAAVALGF | ASSIDRADTQY | 0.3118605 |
| IGLTPLAL | ASSIDRADTQY | 0.7055385 |
| ILGSNTSTL | ASSIDRADTQY | 0.59897757 |
| IVFSLTVL | ASSIDRADTQY | 0.4429748 |
| LASKATGF | ASSIDRADTQY | 0.76667184 |
| LASVALIL | ASSIDRADTQY | 0.73948014 |
| LFAVGLTL | ASSIDRADTQY | 0.16746126 |
| LFNGGIGTYI | ASSIDRADTQY | 0.40218067 |
| LTLGAGII | ASSIDRADTQY | 0.391185 |
| LVAAALGDI | ASSIDRADTQY | 0.44189453 |
| NTAFVQLGI | ASSIDRADTQY | 0.1206576 |
| QLFSGAEL | ASSIDRADTQY | 0.64181584 |
| SAATFITL | ASSIDRADTQY | 0.37334198 |
| SNYGLANLGV | ASSIDRADTQY | 0.090063564 |
| SYGGAYAVM | ASSIDRADTQY | 0.47111237 |
| TTGVGGLTLG | ASSIDRADTQY | 0.45753944 |
| TTQTIYI | ASSIDRADTQY | 0.3812378 |
| VAAGAGLGAVI | ASSIDRADTQY | 0.7367399 |
| VAAVVTLL | ASSIDRADTQY | 0.18041894 |
| VGFLGGLI | ASSIDRADTQY | 0.33012152 |
| VGVIPILL | ASSIDRADTQY | 0.66702545 |
| VIFASSLI | ASSIDRADTQY | 0.7930479 |
| VKGGDGGAAAG | ASSIDRADTQY | 0.28704193 |
| VTKTFTNI | ASSIDRADTQY | 0.58758384 |
| VVAAAILL | ASSIDRADTQY | 0.8168771 |

| | | |
|---|---|---|
| YAALDVEL | ASSIDRADTQY | 0.61876 |
| LASVALLL | ASSIDRADTQY | 0.74559224 |
| LDAVHRV | ASSIDRADTQY | 0.78228253 |
| QGLSSSIF | ASSIDRADTQY | 0.2588905 |
| RGTFANIRL | ASSIDRADTQY | 0.723397 |
| RQLWWGHRI | ASSIDRADTQY | 0.7896342 |
| YGPPGTGKTLL | ASSIDRADTQY | 0.43821037 |
| AALVGAPANVA | ASSIDRADTQY | 0.5603455 |
| AATFRRTE | ASSIDRADTQY | 0.28109163 |
| AGALGGGI | ASSIDRADTQY | 0.3572977 |
| AGGAGGAGGIGAG | ASSIDRADTQY | 0.14305694 |
| AGSGALGL | ASSIDRADTQY | 0.6331031 |
| AQIADQI | ASSIDRADTQY | 0.69863033 |
| ASAVAVTYL | ASSIDRADTQY | 0.48852813 |
| ASTVLGGTGG | ASSIDRADTQY | 0.19948173 |
| AVGGEPAK | ASSIDRADTQY | 0.22873662 |
| CEQGTFGGTC | ASSIDRADTQY | 0.45833176 |
| DEGKAPAG | ASSIDRADTQY | 0.42305946 |
| DTVVFPAL | ASSIDRADTQY | 0.1572415 |
| FEGITMAG | ASSIDRADTQY | 0.2847041 |
| FPLVVQD | ASSIDRADTQY | 0.74551684 |
| GAFTTPQ | ASSIDRADTQY | 0.7622893 |
| GFGGFGTAGGVGGA | ASSIDRADTQY | 0.17991196 |
| GFLAQFL | ASSIDRADTQY | 0.75432485 |
| GGAGGAGGNAGVA | ASSIDRADTQY | 0.71073097 |
| GGVGTAGAGGAGGAGG | ASSIDRADTQY | 0.8028243 |
| GNGGGGDAGNAGSGGN | ASSIDRADTQY | 0.5206691 |
| GTGGAGGAGNDAGS | ASSIDRADTQY | 0.23606999 |
| GTTNTGWFNS | ASSIDRADTQY | 0.74969393 |
| IFMGPLFTL | ASSIDRADTQY | 0.5191286 |
| ILAHAIEAI | ASSIDRADTQY | 0.3577898 |
| IRSAIREAL | ASSIDRADTQY | 0.6634557 |
| ITFAGGG | ASSIDRADTQY | 0.4634894 |
| ITGMSGHQEV | ASSIDRADTQY | 0.37200537 |
| IVAGLACTL | ASSIDRADTQY | 0.5777644 |
| IVGGGLAGA | ASSIDRADTQY | 0.66322464 |
| IVTGVLNI | ASSIDRADTQY | 0.22972657 |
| LAQQGCI | ASSIDRADTQY | 0.27302182 |
| LCLHVEN | ASSIDRADTQY | 0.2527921 |
| LDAISDGS | ASSIDRADTQY | 0.5955925 |
| LGGFAAML | ASSIDRADTQY | 0.755245 |
| LGGKNPMI | ASSIDRADTQY | 0.4910378 |
| LGPKGVAAIL | ASSIDRADTQY | 0.41654688 |
| LMVGVSI | ASSIDRADTQY | 0.70579875 |
| LVGAAFVL | ASSIDRADTQY | 0.68396366 |
| LVLAVAPL | ASSIDRADTQY | 0.52532345 |
| MAAIAVIDAA | ASSIDRADTQY | 0.59092164 |
| NNGINAELQ | ASSIDRADTQY | 0.38260758 |
| RGEPLVLVHG | ASSIDRADTQY | 0.64819336 |
| RGGAGGAAGLI | ASSIDRADTQY | 0.6945715 |
| RSYRMHPAV | ASSIDRADTQY | 0.686407 |

| | | |
|---|---|---|
| SAAHLLAGQLA | ASSIDRADTQY | 0.6696274 |
| SAAVGSY | ASSIDRADTQY | 0.610384 |
| SPHAVLHRI | ASSIDRADTQY | 0.51877356 |
| SQIAKVL | ASSIDRADTQY | 0.16507393 |
| TALGHLVP | ASSIDRADTQY | 0.42689097 |
| TAPTTSGAA | ASSIDRADTQY | 0.53304815 |
| TDGDDGNGGSPGT | ASSIDRADTQY | 0.5732343 |
| TDGLGEN | ASSIDRADTQY | 0.54235804 |
| TGGPPLPGQA | ASSIDRADTQY | 0.55883193 |
| TIPAITI | ASSIDRADTQY | 0.3153716 |
| TIPGFTI | ASSIDRADTQY | 0.6432911 |
| TSIASGALA | ASSIDRADTQY | 0.73562306 |
| TVTLPGAAI | ASSIDRADTQY | 0.20609526 |
| VASGLSAALP | ASSIDRADTQY | 0.43674123 |
| VGAGVIGI | ASSIDRADTQY | 0.76975065 |
| VGPKRFGV | ASSIDRADTQY | 0.6673757 |
| VNVAFPD | ASSIDRADTQY | 0.5068804 |
| VSMTNSGVS | ASSIDRADTQY | 0.47405186 |
| VTDVLGDLSV | ASSIDRADTQY | 0.7869394 |
| VVVVGAGFAGL | ASSIDRADTQY | 0.32071856 |
| VYTVGTVV | ASSIDRADTQY | 0.7779063 |
| WTTFSGP | ASSIDRADTQY | 0.11994484 |
| YDELVEAF | ASSIDRADTQY | 0.42088985 |
| YGTGGAGGNGG | ASSIDRADTQY | 0.29762965 |
| YTNVMAV | ASSIDRADTQY | 0.12379577 |
| FLEQVHQL | ASSIDRADTQY | 0.7519155 |
| GLSLLAVAV | ASSIDRADTQY | 0.29746366 |
| LEQVHQL | ASSIDRADTQY | 0.71160525 |
| LNAAGAI | ASSIDRADTQY | 0.44109792 |
| PAVHASL | ASSIDRADTQY | 0.68983173 |
| TELEDDA | ASSIDRADTQY | 0.61005855 |
| TPPGPPN | ASSIDRADTQY | 0.4261656 |
| TPPPPPPP | ASSIDRADTQY | 0.24175784 |
| TQSPALI | ASSPGTGEREVF | 0.740307 |
| EFAAAFPQ | ASSPGTGEREVF | 0.6389072 |
| LLSGNLDVL | ASSPGTGEREVF | 0.5878437 |
| GPANLGNYNI | ASSPGTGEREVF | 0.21430278 |
| TLGPTLP | ASSPGTGEREVF | 0.39983144 |
| VYAYSSCCK | ASSPGTGEREVF | 0.5632946 |
| APAPAPAPNI | ASSPGTGEREVF | 0.13215594 |
| PGGPVPIP | ASSPGTGEREVF | 0.10366149 |
| FLTVALEP | ASSPGTGEREVF | 0.3278368 |
| SQYFGANL | ASSPGTGEREVF | 0.35163522 |
| GGVKPAA | ASSPGTGEREVF | 0.7365067 |
| GGTGGAGGAGGW | ASSPGTGEREVF | 0.71828926 |
| GVGGVAGLGGAGG | ASSPGTGEREVF | 0.481413 |
| VLLVLATTI | ASSPGTGEREVF | 0.70499057 |
| IGIFAAML | ASSPGTGEREVF | 0.6216266 |
| AVGDLID | ASSPGTGEREVF | 0.0775265 |
| NISPLPL | ASSPGTGEREVF | 0.39704707 |
| VTAVFVAI | ASSPGTGEREVF | 0.17124204 |

| | | |
|---|---|---|
| SGDDNDAAL | ASSPGTGEREVF | 0.7329523 |
| AVAALAHS | ASSPGTGEREVF | 0.22118759 |
| GGVTVMATP | ASSPGTGEREVF | 0.36915505 |
| AGLAANVRD | ASSPGTGEREVF | 0.5497258 |
| TNRPDLIDPALLRPG | ASSPGTGEREVF | 0.6589998 |
| LANLGVTAL | ASSPGTGEREVF | 0.6089107 |
| TGERIEL | ASSPGTGEREVF | 0.2448508 |
| NGEELAA | ASSPGTGEREVF | 0.09905821 |
| VLSLVWG | ASSPGTGEREVF | 0.7052683 |
| IGYALWL | ASSPGTGEREVF | 0.21331121 |
| LSLLLAPL | ASSPGTGEREVF | 0.26497895 |
| TLDEALK | ASSPGTGEREVF | 0.13873924 |
| TDGAGTAEGG | ASSPGTGEREVF | 0.2109608 |
| NAPGGAGGNGG | ASSPGTGEREVF | 0.2244936 |
| VSVFMGMGGNF | ASSPGTGEREVF | 0.6978427 |
| LASYLAAL | ASSPGTGEREVF | 0.40793008 |
| LVGGTPAL | ASSPGTGEREVF | 0.45537505 |
| VRNADVL | ASSPGTGEREVF | 0.75958776 |
| TCSSPFLVSAM | ASSPGTGEREVF | 0.68686473 |
| VHITDAE | ASSPGTGEREVF | 0.45581436 |
| EYAAPSF | ASSPGTGEREVF | 0.3663336 |
| VVDAFHGGL | ASSPGTGEREVF | 0.24981923 |
| ASVVDPQ | ASSPGTGEREVF | 0.312302 |
| SYNEFGGLS | ASSPGTGEREVF | 0.30360654 |
| DPDGSVY | ASSPGTGEREVF | 0.7712375 |
| MVSTIHATE | ASSPGTGEREVF | 0.45011848 |
| LAAGVADPV | ASSPGTGEREVF | 0.20766649 |
| TKEQIAA | ASSPGTGEREVF | 0.67227983 |
| SLASPAAL | ASSPGTGEREVF | 0.78100806 |
| IGLVFYGV | ASSPGTGEREVF | 0.82174397 |
| FLLDASNL | ASSPGTGEREVF | 0.2057731 |
| YQVGNFP | ASSPGTGEREVF | 0.40405348 |
| ESLLSQST | ASSPGTGEREVF | 0.4214525 |
| LFTVGFAAVG | ASSPGTGEREVF | 0.56078434 |
| ATPPTPT | ASSPGTGEREVF | 0.32891503 |
| TDLATLG | ASSPGTGEREVF | 0.26038554 |
| SAGGTGR | ASSPGTGEREVF | 0.63688046 |
| GFSNSGS | ASSPGTGEREVF | 0.8360938 |
| GGNGGSGGTSV | ASSPGTGEREVF | 0.36179078 |
| GTSVATGGAGN | ASSPGTGEREVF | 0.22461705 |
| NGGNGGSGGTS | ASSPGTGEREVF | 0.52400506 |
| AVVKFPH | ASSPGTGEREVF | 0.30143476 |
| TPPPPPP | ASSPGTGEREVF | 0.3868202 |
| AAPAAAAAPA | ASSPGTGEREVF | 0.5560794 |
| IALNEDL | ASSPGTGEREVF | 0.1729877 |
| IVNVSSM | ASSPGTGEREVF | 0.39548534 |
| TGIPPAPR | ASSPGTGEREVF | 0.8267677 |
| FDLGGGTF | ASSPGTGEREVF | 0.5397697 |
| FDLGGGTFDVS | ASSPGTGEREVF | 0.6331612 |
| NEPTAAA | ASSPGTGEREVF | 0.73766124 |
| NEPTAAALA | ASSPGTGEREVF | 0.6539532 |

| | | |
|---|---|---|
| GSIINISSI | ASSPGTGEREVF | 0.8042408 |
| VAPGNAGTA | ASSPGTGEREVF | 0.722893 |
| TGGGASLE | ASSPGTGEREVF | 0.4131263 |
| VAVYNHYKRI | ASSPGTGEREVF | 0.4927729 |
| TDPAPATT | ASSPGTGEREVF | 0.556383 |
| TNRPDILDPALLRPG | ASSPGTGEREVF | 0.5733094 |
| LTTTTTS | ASSPGTGEREVF | 0.5427169 |
| PTGSGKTL | ASSPGTGEREVF | 0.43400824 |
| TGSGKTL | ASSPGTGEREVF | 0.748443 |
| TIGHVDHGK | ASSPGTGEREVF | 0.6401844 |
| TIGHVDHGKTT | ASSPGTGEREVF | 0.7367064 |
| GHVDHGKTT | ASSPGTGEREVF | 0.19979183 |
| AQFEHTLL | ASSPGTGEREVF | 0.28912404 |
| IVNQDEL | ASSPGTGEREVF | 0.7415297 |
| TNGSQFF | ASSPGTGEREVF | 0.10167934 |
| TGKGIVE | ASSPGTGEREVF | 0.14189745 |
| LISNASDA | ASSPGTGEREVF | 0.49492276 |
| AELRTLGELTKIAKAHGAQ | ASSPGTGEREVF | 0.7967962 |
| APNAVAAAEPSW | ASSPGTGEREVF | 0.098954454 |
| APQVAGAATAVL | ASSPGTGEREVF | 0.29786414 |
| ARRDPAAAAAILPTDA | ASSPGTGEREVF | 0.20831676 |
| DAAARDVLAEAGLAEYF | ASSPGTGEREVF | 0.6353282 |
| DDQIVLSAKANLAKVEQ | ASSPGTGEREVF | 0.52693176 |
| DELLPDTGEDSDEDGH | ASSPGTGEREVF | 0.17971216 |
| ELPEAPANRATVPQD | ASSPGTGEREVF | 0.6761604 |
| FGQSKIERVTGIG | ASSPGTGEREVF | 0.17250597 |
| GAGGTGSTAGGAGGAGGAGGLY | ASSPGTGEREVF | 0.1940502 |
| GHGGTGAAGGNGGTGGAGGSGI | ASSPGTGEREVF | 0.704896 |
| GNGGNGGAGGEGGPGGAGGAGA | ASSPGTGEREVF | 0.7705606 |
| GSTAGNGGNGGNATNGTIAGQP | ASSPGTGEREVF | 0.21730505 |
| GTGGDGGNGGNGGTGG | ASSPGTGEREVF | 0.7477927 |
| GVRRPAGIPADAPV | ASSPGTGEREVF | 0.3144448 |
| GYRQVTDAHLLTLARR | ASSPGTGEREVF | 0.8313004 |
| IAKMGQDGHDRG | ASSPGTGEREVF | 0.43977803 |
| KAEPSIEIQYT | ASSPGTGEREVF | 0.18447311 |
| KPVRIRMAADEDTVAVT | ASSPGTGEREVF | 0.6628615 |
| MPPRIAGMRLLVIKPEP | ASSPGTGEREVF | 0.6598675 |
| MKFYAVALFRSRGVPPTR | ASSPGTGEREVF | 0.47390115 |
| NAGGFGILGAVAHSP | ASSPGTGEREVF | 0.6824904 |
| NEGREITLAMAEM | ASSPGTGEREVF | 0.43494982 |
| NELVEWSHPLGHRWRWE | ASSPGTGEREVF | 0.44726983 |
| NLGAGGTGGNGGNPG | ASSPGTGEREVF | 0.7412331 |
| NTEWPKWETVMAADPVD | ASSPGTGEREVF | 0.63881505 |
| PASAAPAETQP | ASSPGTGEREVF | 0.2634974 |
| QQSSDGKNGVRAAEITGE | ASSPGTGEREVF | 0.60288763 |
| RAPDAVTTSWWKE | ASSPGTGEREVF | 0.5566571 |
| SGFGNVGAHTSGWFNQ | ASSPGTGEREVF | 0.28389913 |
| SGVAGAVGQAASVGGLKV | ASSPGTGEREVF | 0.81443566 |
| SNGGAGTGSAGGNGGT | ASSPGTGEREVF | 0.48258728 |
| TGGAGGEGGAAG | ASSPGTGEREVF | 0.72471607 |
| TGGDGGAAGAGGAPGHG | ASSPGTGEREVF | 0.6562768 |

| | | |
|---|---|---|
| TMPPLLAAIANAGRHGV | ASSPGTGEREVF | 0.8074465 |
| TSSTAGNSGGA | ASSPGTGEREVF | 0.22956586 |
| VPATVPVGILC | ASSPGTGEREVF | 0.6185719 |
| VQRPDSGGVVLTGRIS | ASSPGTGEREVF | 0.55848575 |
| WTPNPVPATKGVPAGLVQAT | ASSPGTGEREVF | 0.67249155 |
| YALKRALAKFSDRFDVVII | ASSPGTGEREVF | 0.5131774 |
| YLIGSVIV | ASSPGTGEREVF | 0.2429888 |
| YGQLVAVID | ASSPGTGEREVF | 0.52673393 |
| VTVGGMTLPA | ASSPGTGEREVF | 0.7715254 |
| VLALPAV | ASSPGTGEREVF | 0.40744087 |
| VIPPTLNYET | ASSPGTGEREVF | 0.51802385 |
| VGMGGMHPSSG | ASSPGTGEREVF | 0.81546676 |
| VGLAGGAATA | ASSPGTGEREVF | 0.24116382 |
| VGGAGGAGGSGP | ASSPGTGEREVF | 0.5081658 |
| VFAAFMV | ASSPGTGEREVF | 0.6825939 |
| TLYDKLR | ASSPGTGEREVF | 0.683486 |
| TLAVFTVL | ASSPGTGEREVF | 0.46731275 |
| TGSVQPARV | ASSPGTGEREVF | 0.6940433 |
| TGGGAGSGGT | ASSPGTGEREVF | 0.31129608 |
| SHLETSL | ASSPGTGEREVF | 0.66143215 |
| SFTTGMT | ASSPGTGEREVF | 0.3296956 |
| QVEVAKRMGVRQPTVS | ASSPGTGEREVF | 0.65475667 |
| QGSGGPALAM | ASSPGTGEREVF | 0.24681447 |
| MLAPSNP | ASSPGTGEREVF | 0.663165 |
| LVNNAAI | ASSPGTGEREVF | 0.7210859 |
| LTGEINTLA | ASSPGTGEREVF | 0.708887 |
| LSIFGPV | ASSPGTGEREVF | 0.119895004 |
| LLALLANL | ASSPGTGEREVF | 0.710312 |
| LISNASDALDKLR | ASSPGTGEREVF | 0.7254733 |
| LINEAVP | ASSPGTGEREVF | 0.83167744 |
| LGYAFSGL | ASSPGTGEREVF | 0.5769636 |
| LDFVAVAL | ASSPGTGEREVF | 0.66379094 |
| LAQPGAE | ASSPGTGEREVF | 0.43143678 |
| ISNLGAVTGF | ASSPGTGEREVF | 0.48709106 |
| HQEDPGRI | ASSPGTGEREVF | 0.25089353 |
| GTPGEPTP | ASSPGTGEREVF | 0.39466995 |
| GFFNSGS | ASSPGTGEREVF | 0.4873951 |
| GAGGGGLTAGAN | ASSPGTGEREVF | 0.29982406 |
| FVDALVNM | ASSPGTGEREVF | 0.78080153 |
| FLTVAPEL | ASSPGTGEREVF | 0.7156352 |
| FETLTAL | ASSPGTGEREVF | 0.7305026 |
| DVINPIEL | ASSPGTGEREVF | 0.77232283 |
| APVSGAPV | ASSPGTGEREVF | 0.77103984 |
| ALVGGLAW | ASSPGTGEREVF | 0.64904875 |
| ALAAAAATAI | ASSPGTGEREVF | 0.45429322 |
| AFGLGLLL | ASSPGTGEREVF | 0.5016128 |
| AFFAAASL | ASSPGTGEREVF | 0.23988356 |
| AAAALPAFSPP | ASSPGTGEREVF | 0.38342226 |
| VVLAPAL | ASSPGTGEREVF | 0.4262246 |
| TIGHVDHGKTTL | ASSPGTGEREVF | 0.6138607 |
| LISNASDALDK | ASSPGTGEREVF | 0.092603065 |

| | | |
|---|---|---|
| LISNASDAL | ASSPGTGEREVF | 0.61743796 |
| ISNASDALDK | ASSPGTGEREVF | 0.7702756 |
| NQAETLVYQTE | ASSPGTGEREVF | 0.39785424 |
| TGGTGGTAGAG | ASSPGTGEREVF | 0.6863295 |
| TGSTAGGAGGA | ASSPGTGEREVF | 0.6004398 |
| DPLAVPVAAGPV | ASSPGTGEREVF | 0.31172216 |
| GGTGGTPGNGGN | ASSPGTGEREVF | 0.7424522 |
| PAGLAGVDPATL | ASSPGTGEREVF | 0.2825413 |
| GNNLVVEAPHAGASV | ASSPGTGEREVF | 0.701967 |
| KVTRIMSASTLEQPA | ASSPGTGEREVF | 0.815079 |
| LGFALPIALPRRTVV | ASSPGTGEREVF | 0.51186156 |
| TRLGNLTARKLVVLV | ASSPGTGEREVF | 0.38743132 |
| WSQQRRLIYFPSAGP | ASSPGTGEREVF | 0.54093665 |
| GPNGNPDPMAAAVDIR | ASSPGTGEREVF | 0.210355 |
| TQRCVSVADLRAAASR | ASSPGTGEREVF | 0.42720726 |
| QFVLDVAEHPVEVLTETV | ASSPGTGEREVF | 0.10089226 |
| GGAGGTGGAGGNGGTQAGDG | ASSPGTGEREVF | 0.57748324 |
| PLLATLRAIAVDVDAKPAQV | ASSPGTGEREVF | 0.24586998 |
| APKFRTGGYMIDFWGVGYQVAK | ASSPGTGEREVF | 0.62357616 |
| GAGGAGGAGGAAGGPGGTGGTG | ASSPGTGEREVF | 0.5553132 |
| IGALGLLATDVAPAGSVAFGVA | ASSPGTGEREVF | 0.56047845 |
| AGGAGGIGGNGGAGGAGGVLMG | ASSPGTGEREVF | 0.30283937 |
| ADELIGLL | ASSPGTGEREVF | 0.46476048 |
| ADRALKA | ASSPGTGEREVF | 0.35351017 |
| AGAVGQVGGAGGA | ASSPGTGEREVF | 0.21512465 |
| AGGGGGLGGAP | ASSPGTGEREVF | 0.6810403 |
| AGGLLYGNGGAG | ASSPGTGEREVF | 0.39061213 |
| AGLESIGL | ASSPGTGEREVF | 0.50413096 |
| AGLKTGII | ASSPGTGEREVF | 0.7260503 |
| ASLAALLL | ASSPGTGEREVF | 0.56392795 |
| ASVINISSI | ASSPGTGEREVF | 0.21599916 |
| ATPPGPA | ASSPGTGEREVF | 0.5477929 |
| ATVASALV | ASSPGTGEREVF | 0.50903016 |
| ATVASLIL | ASSPGTGEREVF | 0.4882571 |
| AVNGPVLIRH | ASSPGTGEREVF | 0.08826735 |
| DGAAGAVGI | ASSPGTGEREVF | 0.50411403 |
| DPLLHGI | ASSPGTGEREVF | 0.286591 |
| ELGAALGAL | ASSPGTGEREVF | 0.3320873 |
| ELGRTQAV | ASSPGTGEREVF | 0.3504212 |
| FVAGGYLS | ASSPGTGEREVF | 0.73164946 |
| GASGGAGGAGGW | ASSPGTGEREVF | 0.43442518 |
| GGAGAGGDGGAGGI | ASSPGTGEREVF | 0.56871223 |
| GTGHNGSQPGMGGNG | ASSPGTGEREVF | 0.64198387 |
| GYAAASASAS | ASSPGTGEREVF | 0.6522915 |
| HQVDVAAVQQG | ASSPGTGEREVF | 0.8073117 |
| IGAAIAHT | ASSPGTGEREVF | 0.59295976 |
| IGMIEVAQL | ASSPGTGEREVF | 0.43016338 |
| IRLAPEHTSAMD | ASSPGTGEREVF | 0.5441079 |
| IYNVSVL | ASSPGTGEREVF | 0.548223 |
| KQGFLGM | ASSPGTGEREVF | 0.13304538 |
| LESVITQL | ASSPGTGEREVF | 0.13937198 |

| | | |
|---|---|---|
| LGLGALVVV | ASSPGTGEREVF | 0.5793333 |
| LIGNGGNGGSGGI | ASSPGTGEREVF | 0.14657177 |
| LNAINAPTL | ASSPGTGEREVF | 0.64947057 |
| LQLLNTGV | ASSPGTGEREVF | 0.7319256 |
| LQTSIEF | ASSPGTGEREVF | 0.54044765 |
| MSFAVSEL | ASSPGTGEREVF | 0.18203764 |
| NTGALGL | ASSPGTGEREVF | 0.3395512 |
| PPGAGGAGGAGTT | ASSPGTGEREVF | 0.38803744 |
| QLVGLGI | ASSPGTGEREVF | 0.71544814 |
| RFLEQVHQL | ASSPGTGEREVF | 0.6109715 |
| RGVGYVLR | ASSPGTGEREVF | 0.7736403 |
| SATGVFAGVI | ASSPGTGEREVF | 0.74071395 |
| SLPAITLP | ASSPGTGEREVF | 0.42868543 |
| STALAFTV | ASSPGTGEREVF | 0.70387876 |
| SVLTVGLGK | ASSPGTGEREVF | 0.39189005 |
| TALAGAGSGI | ASSPGTGEREVF | 0.49557585 |
| TGLPFTGI | ASSPGTGEREVF | 0.3704186 |
| VAEHVTV | ASSPGTGEREVF | 0.59832764 |
| VDIVLNSL | ASSPGTGEREVF | 0.34084246 |
| VLTGGGAVL | ASSPGTGEREVF | 0.31569543 |
| VNVGLAYGI | ASSPGTGEREVF | 0.24181387 |
| VSSAAVGAV | ASSPGTGEREVF | 0.55908495 |
| VTSVYEAGV | ASSPGTGEREVF | 0.6184194 |
| WVALDRGA | ASSPGTGEREVF | 0.18050846 |
| YLVGSVVI | ASSPGTGEREVF | 0.4374752 |
| GPPGTGKTLL | ASSPGTGEREVF | 0.8135952 |
| GVGMTKF | ASSPGTGEREVF | 0.81029975 |
| IGIGGSDL | ASSPGTGEREVF | 0.29413846 |
| IGTLLAEL | ASSPGTGEREVF | 0.32994983 |
| IYTSGTT | ASSPGTGEREVF | 0.64796287 |
| LSFPAGV | ASSPGTGEREVF | 0.19451094 |
| LYTSGTT | ASSPGTGEREVF | 0.55860627 |
| VGAFPFGF | ASSPGTGEREVF | 0.629283 |
| VGAGTVEF | ASSPGTGEREVF | 0.5095632 |
| VGKSTLLL | ASSPGTGEREVF | 0.30291954 |
| AALAHAGGGIV | ASSPGTGEREVF | 0.5434002 |
| AALVAVSLL | ASSPGTGEREVF | 0.44769627 |
| AAVSRAVAAL | ASSPGTGEREVF | 0.61111575 |
| AGGSSGIGGF | ASSPGTGEREVF | 0.10159178 |
| AGLTVANANG | ASSPGTGEREVF | 0.54088247 |
| AGLVGAVL | ASSPGTGEREVF | 0.3166847 |
| ALSQAFTV | ASSPGTGEREVF | 0.3572478 |
| ALTFSVSL | ASSPGTGEREVF | 0.14800303 |
| ALTGAALVAL | ASSPGTGEREVF | 0.4310702 |
| ASFSIGAGVL | ASSPGTGEREVF | 0.7359632 |
| DGSVYKI | ASSPGTGEREVF | 0.7663399 |
| EQLAEIV | ASSPGTGEREVF | 0.1951686 |
| FAVSELTVM | ASSPGTGEREVF | 0.66716623 |
| GAAPLAAGL | ASSPGTGEREVF | 0.8388957 |
| GADGGDGGSGGAGG | ASSPGTGEREVF | 0.45918733 |
| GFGLDTGGAGG | ASSPGTGEREVF | 0.1641297 |

| | | |
|---|---|---|
| GLASGGLL | ASSPGTGEREVF | 0.6386292 |
| GRIAGAAVLL | ASSPGTGEREVF | 0.53364825 |
| GSAGSLNTSF | ASSPGTGEREVF | 0.6957901 |
| GTGQIGI | ASSPGTGEREVF | 0.7480936 |
| GTLIALIL | ASSPGTGEREVF | 0.17608702 |
| IAATMGFVVG | ASSPGTGEREVF | 0.7111058 |
| IFAQALL | ASSPGTGEREVF | 0.7270489 |
| IGAAVALGF | ASSPGTGEREVF | 0.27077436 |
| IGLTPLAL | ASSPGTGEREVF | 0.3057774 |
| ILGSNTSTL | ASSPGTGEREVF | 0.28596905 |
| IVFSLTVL | ASSPGTGEREVF | 0.13192709 |
| LASKATGF | ASSPGTGEREVF | 0.5898471 |
| LASVALIL | ASSPGTGEREVF | 0.3567824 |
| LFAVGLTL | ASSPGTGEREVF | 0.4216503 |
| LFNGGIGTYI | ASSPGTGEREVF | 0.49331123 |
| LTLGAGII | ASSPGTGEREVF | 0.33474 |
| LVAAALGDI | ASSPGTGEREVF | 0.4106452 |
| NTAFVQLGI | ASSPGTGEREVF | 0.42568305 |
| QLFSGAEL | ASSPGTGEREVF | 0.53879094 |
| SAATFITL | ASSPGTGEREVF | 0.59528273 |
| SNYGLANLGV | ASSPGTGEREVF | 0.509593 |
| SYGGAYAVM | ASSPGTGEREVF | 0.5691161 |
| TTGVGGLTLG | ASSPGTGEREVF | 0.792877 |
| TTQTIYI | ASSPGTGEREVF | 0.22314173 |
| VAAGAGLGAVI | ASSPGTGEREVF | 0.32170773 |
| VAAVVTLL | ASSPGTGEREVF | 0.6665573 |
| VGFLGGLI | ASSPGTGEREVF | 0.22600475 |
| VGVIPILL | ASSPGTGEREVF | 0.6930454 |
| VIFASSLI | ASSPGTGEREVF | 0.5724801 |
| VKGGDGGAAAG | ASSPGTGEREVF | 0.64567494 |
| VTKTFTNI | ASSPGTGEREVF | 0.60956764 |
| VVAAAILL | ASSPGTGEREVF | 0.7903433 |
| YAALDVEL | ASSPGTGEREVF | 0.7012781 |
| LASVALLL | ASSPGTGEREVF | 0.35400692 |
| LDAVHRV | ASSPGTGEREVF | 0.6113919 |
| QGLSSSIF | ASSPGTGEREVF | 0.26438773 |
| RGTFANIRL | ASSPGTGEREVF | 0.63959605 |
| RQLWWGHRI | ASSPGTGEREVF | 0.72318596 |
| YGPPGTGKTLL | ASSPGTGEREVF | 0.27474296 |
| AALVGAPANVA | ASSPGTGEREVF | 0.46925417 |
| AATFRRTE | ASSPGTGEREVF | 0.75304383 |
| AGALGGGI | ASSPGTGEREVF | 0.7132008 |
| AGGAGGAGGIGAG | ASSPGTGEREVF | 0.16137393 |
| AGSGALGL | ASSPGTGEREVF | 0.32358357 |
| AQIADQI | ASSPGTGEREVF | 0.18586732 |
| ASAVAVTYL | ASSPGTGEREVF | 0.42176333 |
| ASTVLGGTGG | ASSPGTGEREVF | 0.2388762 |
| AVGGEPAK | ASSPGTGEREVF | 0.66651845 |
| CEQGTFGGTC | ASSPGTGEREVF | 0.47914547 |
| DEGKAPAG | ASSPGTGEREVF | 0.7273741 |
| DTVVFPAL | ASSPGTGEREVF | 0.6383337 |

| | | |
|---|---|---|
| FEGITMAG | ASSPGTGEREVF | 0.7928896 |
| FPLVVQD | ASSPGTGEREVF | 0.35911888 |
| GAFTTPQ | ASSPGTGEREVF | 0.4503765 |
| GFGGFGTAGGVGGA | ASSPGTGEREVF | 0.18149911 |
| GFLAQFL | ASSPGTGEREVF | 0.22627671 |
| GGAGGAGGNAGVA | ASSPGTGEREVF | 0.2553163 |
| GGVGTAGAGGAGGAGG | ASSPGTGEREVF | 0.4999672 |
| GNGGGGDAGNAGSGGN | ASSPGTGEREVF | 0.13029309 |
| GTGGAGGAGNDAGS | ASSPGTGEREVF | 0.77055156 |
| GTTNTGWFNS | ASSPGTGEREVF | 0.619708 |
| IFMGPLFTL | ASSPGTGEREVF | 0.7138561 |
| ILAHAIEAI | ASSPGTGEREVF | 0.3833405 |
| IRSAIREAL | ASSPGTGEREVF | 0.3032624 |
| ITFAGGG | ASSPGTGEREVF | 0.4295693 |
| ITGMSGHQEV | ASSPGTGEREVF | 0.7337638 |
| IVAGLACTL | ASSPGTGEREVF | 0.21293338 |
| IVGGGLAGA | ASSPGTGEREVF | 0.17463371 |
| IVTGVLNI | ASSPGTGEREVF | 0.38624492 |
| LAQQGCI | ASSPGTGEREVF | 0.23925763 |
| LCLHVEN | ASSPGTGEREVF | 0.77670366 |
| LDAISDGS | ASSPGTGEREVF | 0.7195811 |
| LGGFAAML | ASSPGTGEREVF | 0.5945287 |
| LGGKNPMI | ASSPGTGEREVF | 0.76040614 |
| LGPKGVAAIL | ASSPGTGEREVF | 0.115715474 |
| LMVGVSI | ASSPGTGEREVF | 0.36533722 |
| LVGAAFVL | ASSPGTGEREVF | 0.817639 |
| LVLAVAPL | ASSPGTGEREVF | 0.5689906 |
| MAAIAVIDAA | ASSPGTGEREVF | 0.5228677 |
| NNGINAELQ | ASSPGTGEREVF | 0.7874614 |
| RGEPLVLVHG | ASSPGTGEREVF | 0.36505777 |
| RGGAGGAAGLI | ASSPGTGEREVF | 0.50429726 |
| RSYRMHPAV | ASSPGTGEREVF | 0.50490236 |
| SAAHLLAGQLA | ASSPGTGEREVF | 0.47667244 |
| SAAVGSY | ASSPGTGEREVF | 0.5631497 |
| SPHAVLHRI | ASSPGTGEREVF | 0.20719722 |
| SQIAKVL | ASSPGTGEREVF | 0.64526737 |
| TALGHLVP | ASSPGTGEREVF | 0.8079511 |
| TAPTTSGAA | ASSPGTGEREVF | 0.50644314 |
| TDGDDGNGGSPGT | ASSPGTGEREVF | 0.73364186 |
| TDGLGEN | ASSPGTGEREVF | 0.6440108 |
| TGGPPLPGQA | ASSPGTGEREVF | 0.7009608 |
| TIPAITI | ASSPGTGEREVF | 0.33908975 |
| TIPGFTI | ASSPGTGEREVF | 0.2893815 |
| TSIASGALA | ASSPGTGEREVF | 0.4885792 |
| TVTLPGAAI | ASSPGTGEREVF | 0.30654472 |
| VASGLSAALP | ASSPGTGEREVF | 0.57827747 |
| VGAGVIGI | ASSPGTGEREVF | 0.4196048 |
| VGPKRFGV | ASSPGTGEREVF | 0.849313 |
| VNVAFPD | ASSPGTGEREVF | 0.6882733 |
| VSMTNSGVS | ASSPGTGEREVF | 0.6151786 |
| VTDVLGDLSV | ASSPGTGEREVF | 0.26557449 |

| | | |
|---|---|---|
| VVVVGAGFAGL | ASSPGTGEREVF | 0.4552804 |
| VYTVGTVV | ASSPGTGEREVF | 0.6225822 |
| WTTFSGP | ASSPGTGEREVF | 0.23572963 |
| YDELVEAF | ASSPGTGEREVF | 0.78388035 |
| YGTGGAGGNGG | ASSPGTGEREVF | 0.35302097 |
| YTNVMAV | ASSPGTGEREVF | 0.4447097 |
| FLEQVHQL | ASSPGTGEREVF | 0.6873354 |
| GLSLLAVAV | ASSPGTGEREVF | 0.76029503 |
| LEQVHQL | ASSPGTGEREVF | 0.45518428 |
| LNAAGAI | ASSPGTGEREVF | 0.518134 |
| PAVHASL | ASSPGTGEREVF | 0.26320815 |
| TELEDDA | ASSPGTGEREVF | 0.72027254 |
| TPPGPPN | ASSPGTGEREVF | 0.78655994 |
| TPPPPPPP | ASSPGTGEREVF | 0.5950071 |
| TQSPALI | ASSQDWGGQDTQY | 0.77060276 |
| EFAAAFPQ | ASSQDWGGQDTQY | 0.59431076 |
| LLSGNLDVL | ASSQDWGGQDTQY | 0.21217467 |
| GPANLGNYNI | ASSQDWGGQDTQY | 0.46536142 |
| TLGPTLP | ASSQDWGGQDTQY | 0.673065 |
| VYAYSSCCK | ASSQDWGGQDTQY | 0.7284211 |
| APAPAPAPNI | ASSQDWGGQDTQY | 0.7989464 |
| PGGPVPIP | ASSQDWGGQDTQY | 0.5559303 |
| FLTVALEP | ASSQDWGGQDTQY | 0.3671565 |
| SQYFGANL | ASSQDWGGQDTQY | 0.77733576 |
| GGVKPAA | ASSQDWGGQDTQY | 0.85305536 |
| GGTGGAGGAGGW | ASSQDWGGQDTQY | 0.7512375 |
| GVGGVAGLGGAGG | ASSQDWGGQDTQY | 0.41716915 |
| VLLVLATTI | ASSQDWGGQDTQY | 0.3705715 |
| IGIFAAML | ASSQDWGGQDTQY | 0.49626958 |
| AVGDLID | ASSQDWGGQDTQY | 0.73103696 |
| NISPLPL | ASSQDWGGQDTQY | 0.31896624 |
| VTAVFVAI | ASSQDWGGQDTQY | 0.6251364 |
| SGDDNDAAL | ASSQDWGGQDTQY | 0.6455779 |
| AVAALAHS | ASSQDWGGQDTQY | 0.17285343 |
| GGVTVMATP | ASSQDWGGQDTQY | 0.6348669 |
| AGLAANVRD | ASSQDWGGQDTQY | 0.7388649 |
| TNRPDLIDPALLRPG | ASSQDWGGQDTQY | 0.4186927 |
| LANLGVTAL | ASSQDWGGQDTQY | 0.32866812 |
| TGERIEL | ASSQDWGGQDTQY | 0.7522578 |
| NGEELAA | ASSQDWGGQDTQY | 0.8251964 |
| VLSLVWG | ASSQDWGGQDTQY | 0.74189734 |
| IGYALWL | ASSQDWGGQDTQY | 0.20477328 |
| LSLLLAPL | ASSQDWGGQDTQY | 0.12666237 |
| TLDEALK | ASSQDWGGQDTQY | 0.30333385 |
| TDGAGTAEGG | ASSQDWGGQDTQY | 0.7433989 |
| NAPGGAGGNGG | ASSQDWGGQDTQY | 0.40426806 |
| VSVFMGMGGNF | ASSQDWGGQDTQY | 0.30943358 |
| LASYLAAL | ASSQDWGGQDTQY | 0.6217267 |
| LVGGTPAL | ASSQDWGGQDTQY | 0.7608906 |
| VRNADVL | ASSQDWGGQDTQY | 0.23505628 |
| TCSSPFLVSAM | ASSQDWGGQDTQY | 0.21303399 |

| | | |
|---|---|---|
| VHITDAE | ASSQDWGGQDTQY | 0.62929356 |
| EYAAPSF | ASSQDWGGQDTQY | 0.75950265 |
| VVDAFHGGL | ASSQDWGGQDTQY | 0.21874058 |
| ASVVDPQ | ASSQDWGGQDTQY | 0.438501 |
| SYNEFGGLS | ASSQDWGGQDTQY | 0.38820273 |
| DPDGSVY | ASSQDWGGQDTQY | 0.381925 |
| MVSTIHATE | ASSQDWGGQDTQY | 0.6053003 |
| LAAGVADPV | ASSQDWGGQDTQY | 0.28868613 |
| TKEQIAA | ASSQDWGGQDTQY | 0.72649527 |
| SLASPAAL | ASSQDWGGQDTQY | 0.26815033 |
| IGLVFYGV | ASSQDWGGQDTQY | 0.6842009 |
| FLLDASNL | ASSQDWGGQDTQY | 0.49829713 |
| YQVGNFP | ASSQDWGGQDTQY | 0.44982937 |
| ESLLSQST | ASSQDWGGQDTQY | 0.5056343 |
| LFTVGFAAVG | ASSQDWGGQDTQY | 0.3964614 |
| ATPPTPT | ASSQDWGGQDTQY | 0.23614007 |
| TDLATLG | ASSQDWGGQDTQY | 0.1393681 |
| SAGGTGR | ASSQDWGGQDTQY | 0.71457046 |
| GFSNSGS | ASSQDWGGQDTQY | 0.1608535 |
| GGNGGSGGTSV | ASSQDWGGQDTQY | 0.6765735 |
| GTSVATGGAGN | ASSQDWGGQDTQY | 0.71064943 |
| NGGNGGSGGTS | ASSQDWGGQDTQY | 0.2257502 |
| AVVKFPH | ASSQDWGGQDTQY | 0.5592958 |
| TPPPPPP | ASSQDWGGQDTQY | 0.23382281 |
| AAPAAAAAPA | ASSQDWGGQDTQY | 0.25622505 |
| IALNEDL | ASSQDWGGQDTQY | 0.73364174 |
| IVNVSSM | ASSQDWGGQDTQY | 0.62326145 |
| TGIPPAPR | ASSQDWGGQDTQY | 0.34147382 |
| FDLGGGTF | ASSQDWGGQDTQY | 0.6221812 |
| FDLGGGTFDVS | ASSQDWGGQDTQY | 0.55577123 |
| NEPTAAA | ASSQDWGGQDTQY | 0.31330237 |
| NEPTAAALA | ASSQDWGGQDTQY | 0.44014123 |
| GSIINISSI | ASSQDWGGQDTQY | 0.75002503 |
| VAPGNAGTA | ASSQDWGGQDTQY | 0.5789685 |
| TGGGASLE | ASSQDWGGQDTQY | 0.2529929 |
| VAVYNHYKRI | ASSQDWGGQDTQY | 0.6720945 |
| TDPAPATT | ASSQDWGGQDTQY | 0.27231705 |
| TNRPDILDPALLRPG | ASSQDWGGQDTQY | 0.6011162 |
| LTTTTTS | ASSQDWGGQDTQY | 0.4198982 |
| PTGSGKTL | ASSQDWGGQDTQY | 0.4830191 |
| TGSGKTL | ASSQDWGGQDTQY | 0.6488607 |
| TIGHVDHGK | ASSQDWGGQDTQY | 0.2528801 |
| TIGHVDHGKTT | ASSQDWGGQDTQY | 0.2669594 |
| GHVDHGKTT | ASSQDWGGQDTQY | 0.4311129 |
| AQFEHTLL | ASSQDWGGQDTQY | 0.5340657 |
| IVNQDEL | ASSQDWGGQDTQY | 0.6983479 |
| TNGSQFF | ASSQDWGGQDTQY | 0.30095148 |
| TGKGIVE | ASSQDWGGQDTQY | 0.54720306 |
| LISNASDA | ASSQDWGGQDTQY | 0.40494573 |
| AELRTLGELTKIAKAHGAQ | ASSQDWGGQDTQY | 0.30829814 |
| APNAVAAAEPSW | ASSQDWGGQDTQY | 0.4990796 |

| | | |
|---|---|---|
| APQVAGAATAVL | ASSQDWGGQDTQY | 0.37284017 |
| ARRDPAAAAAILPTDA | ASSQDWGGQDTQY | 0.2892325 |
| DAAARDVLAEAGLAEYF | ASSQDWGGQDTQY | 0.5398385 |
| DDQIVLSAKANLAKVEQ | ASSQDWGGQDTQY | 0.2733398 |
| DELLPDTGEDSDEDGH | ASSQDWGGQDTQY | 0.56806105 |
| ELPEAPANRATVPQD | ASSQDWGGQDTQY | 0.73179144 |
| FGQSKIERVTGIG | ASSQDWGGQDTQY | 0.30232787 |
| GAGGTGSTAGGAGGAGGAGGLY | ASSQDWGGQDTQY | 0.64516485 |
| GHGGTGAAGGNGGTGGAGGSGI | ASSQDWGGQDTQY | 0.6298735 |
| GNGGNGGAGGEGGPGGAGGAGA | ASSQDWGGQDTQY | 0.54460865 |
| GSTAGNGGNGGNATNGTIAGQP | ASSQDWGGQDTQY | 0.5060209 |
| GTGGDGGNGGNGGTGG | ASSQDWGGQDTQY | 0.35544223 |
| GVRRPAGIPADAPV | ASSQDWGGQDTQY | 0.7208841 |
| GYRQVTDAHLLTLARR | ASSQDWGGQDTQY | 0.69234836 |
| IAKMGQDGHDRG | ASSQDWGGQDTQY | 0.6865504 |
| KAEPSIEIQYT | ASSQDWGGQDTQY | 0.54056364 |
| KPVRIRMAADEDTVAVT | ASSQDWGGQDTQY | 0.6915833 |
| MPPRIAGMRLLVIKPEP | ASSQDWGGQDTQY | 0.68726075 |
| MKFYAVALFRSRGVPPTR | ASSQDWGGQDTQY | 0.4113133 |
| NAGGFGILGAVAHSP | ASSQDWGGQDTQY | 0.18348151 |
| NEGREITLAMAEM | ASSQDWGGQDTQY | 0.42353028 |
| NELVEWSHPLGHRWRWE | ASSQDWGGQDTQY | 0.45771334 |
| NLGAGGTGGNGGNPG | ASSQDWGGQDTQY | 0.34622043 |
| NTEWPKWETVMAADPVD | ASSQDWGGQDTQY | 0.20269367 |
| PASAAPAETQP | ASSQDWGGQDTQY | 0.75364494 |
| QQSSDGKNGVRAAEITGE | ASSQDWGGQDTQY | 0.8357738 |
| RAPDAVTTSWWKE | ASSQDWGGQDTQY | 0.3709401 |
| SGFGNVGAHTSGWFNQ | ASSQDWGGQDTQY | 0.79960585 |
| SGVAGAVGQAASVGGLKV | ASSQDWGGQDTQY | 0.20547351 |
| SNGGAGTGSAGGNGGT | ASSQDWGGQDTQY | 0.7058534 |
| TGGAGGEGGAAG | ASSQDWGGQDTQY | 0.7820513 |
| TGGDGGAAGAGGAPGHG | ASSQDWGGQDTQY | 0.36140788 |
| TMPPLLAAIANAGRHGV | ASSQDWGGQDTQY | 0.63195765 |
| TSSTAGNSGGA | ASSQDWGGQDTQY | 0.3544415 |
| VPATVPVGILC | ASSQDWGGQDTQY | 0.39123392 |
| VQRPDSGGVVLTGRIS | ASSQDWGGQDTQY | 0.7427794 |
| WTPNPVPATKGVPAGLVQAT | ASSQDWGGQDTQY | 0.46119028 |
| YALKRALAKFSDRFDVVII | ASSQDWGGQDTQY | 0.4753549 |
| YLIGSVIV | ASSQDWGGQDTQY | 0.40808964 |
| YGQLVAVID | ASSQDWGGQDTQY | 0.31513512 |
| VTVGGMTLPA | ASSQDWGGQDTQY | 0.439243 |
| VLALPAV | ASSQDWGGQDTQY | 0.7334025 |
| VIPPTLNYET | ASSQDWGGQDTQY | 0.34679654 |
| VGMGGMHPSSG | ASSQDWGGQDTQY | 0.8425807 |
| VGLAGGAATA | ASSQDWGGQDTQY | 0.3129943 |
| VGGAGGAGGSGP | ASSQDWGGQDTQY | 0.8012784 |
| VFAAFMV | ASSQDWGGQDTQY | 0.75782746 |
| TLYDKLR | ASSQDWGGQDTQY | 0.58603525 |
| TLAVFTVL | ASSQDWGGQDTQY | 0.4032387 |
| TGSVQPARV | ASSQDWGGQDTQY | 0.783792 |
| TGGGAGSGGT | ASSQDWGGQDTQY | 0.4963025 |

| | | |
|---|---|---|
| SHLETSL | ASSQDWGGQDTQY | 0.6156505 |
| SFTTGMT | ASSQDWGGQDTQY | 0.71028656 |
| QVEVAKRMGVRQPTVS | ASSQDWGGQDTQY | 0.61927867 |
| QGSGGPALAM | ASSQDWGGQDTQY | 0.76157874 |
| MLAPSNP | ASSQDWGGQDTQY | 0.7396487 |
| LVNNAAI | ASSQDWGGQDTQY | 0.7254869 |
| LTGEINTLA | ASSQDWGGQDTQY | 0.6350918 |
| LSIFGPV | ASSQDWGGQDTQY | 0.4109891 |
| LLALLANL | ASSQDWGGQDTQY | 0.7259633 |
| LISNASDALDKLR | ASSQDWGGQDTQY | 0.5806375 |
| LINEAVP | ASSQDWGGQDTQY | 0.75654304 |
| LGYAFSGL | ASSQDWGGQDTQY | 0.7504654 |
| LDFVAVAL | ASSQDWGGQDTQY | 0.649287 |
| LAQPGAE | ASSQDWGGQDTQY | 0.7620234 |
| ISNLGAVTGF | ASSQDWGGQDTQY | 0.7617831 |
| HQEDPGRI | ASSQDWGGQDTQY | 0.46532542 |
| GTPGEPTP | ASSQDWGGQDTQY | 0.7041565 |
| GFFNSGS | ASSQDWGGQDTQY | 0.6772253 |
| GAGGGGLTAGAN | ASSQDWGGQDTQY | 0.6916281 |
| FVDALVNM | ASSQDWGGQDTQY | 0.45027223 |
| FLTVAPEL | ASSQDWGGQDTQY | 0.7496084 |
| FETLTAL | ASSQDWGGQDTQY | 0.71061075 |
| DVINPIEL | ASSQDWGGQDTQY | 0.25765455 |
| APVSGAPV | ASSQDWGGQDTQY | 0.7199474 |
| ALVGGLAW | ASSQDWGGQDTQY | 0.7118849 |
| ALAAAAATAI | ASSQDWGGQDTQY | 0.68265283 |
| AFGLGLLL | ASSQDWGGQDTQY | 0.7585355 |
| AFFAAASL | ASSQDWGGQDTQY | 0.6932436 |
| AAAALPAFSPP | ASSQDWGGQDTQY | 0.6478 |
| VVLAPAL | ASSQDWGGQDTQY | 0.75312626 |
| TIGHVDHGKTTL | ASSQDWGGQDTQY | 0.7625457 |
| LISNASDALDK | ASSQDWGGQDTQY | 0.5652486 |
| LISNASDAL | ASSQDWGGQDTQY | 0.7414838 |
| ISNASDALDK | ASSQDWGGQDTQY | 0.7631755 |
| NQAETLVYQTE | ASSQDWGGQDTQY | 0.73923373 |
| TGGTGGTAGAG | ASSQDWGGQDTQY | 0.7637594 |
| TGSTAGGAGGA | ASSQDWGGQDTQY | 0.76567066 |
| DPLAVPVAAGPV | ASSQDWGGQDTQY | 0.65542865 |
| GGTGGTPGNGGN | ASSQDWGGQDTQY | 0.7577987 |
| PAGLAGVDPATL | ASSQDWGGQDTQY | 0.76466924 |
| GNNLVVEAPHAGASV | ASSQDWGGQDTQY | 0.6787207 |
| KVTRIMSASTLEQPA | ASSQDWGGQDTQY | 0.72883445 |
| LGFALPIALPRRTVV | ASSQDWGGQDTQY | 0.6460484 |
| TRLGNLTARKLVVLV | ASSQDWGGQDTQY | 0.7204932 |
| WSQQRRLIYFPSAGP | ASSQDWGGQDTQY | 0.39103353 |
| GPNGNPDPMAAAVDIR | ASSQDWGGQDTQY | 0.7458411 |
| TQRCVSVADLRAAASR | ASSQDWGGQDTQY | 0.7594023 |
| QFVLDVAEHPVEVLTETV | ASSQDWGGQDTQY | 0.71241724 |
| GGAGGTGGAGGNGGTQAGDG | ASSQDWGGQDTQY | 0.6263409 |
| PLLATLRAIAVDVDAKPAQV | ASSQDWGGQDTQY | 0.6255722 |
| APKFRTGGYMIDFWGVGYQVAK | ASSQDWGGQDTQY | 0.7197801 |

| | | |
|---|---|---|
| GAGGAGGAGGAAGGPGGTGGTG | ASSQDWGGQDTQY | 0.690313 |
| IGALGLLATDVAPAGSVAFGVA | ASSQDWGGQDTQY | 0.43966353 |
| AGGAGGIGGNGGAGGAGGVLMG | ASSQDWGGQDTQY | 0.67244655 |
| ADELIGLL | ASSQDWGGQDTQY | 0.75895333 |
| ADRALKA | ASSQDWGGQDTQY | 0.7596746 |
| AGAVGQVGGAGGA | ASSQDWGGQDTQY | 0.46774447 |
| AGGGGGLGGAP | ASSQDWGGQDTQY | 0.76084876 |
| AGGLLYGNGGAG | ASSQDWGGQDTQY | 0.75543135 |
| AGLESIGL | ASSQDWGGQDTQY | 0.76293486 |
| AGLKTGII | ASSQDWGGQDTQY | 0.7319933 |
| ASLAALLL | ASSQDWGGQDTQY | 0.61478364 |
| ASVINISSI | ASSQDWGGQDTQY | 0.76038706 |
| ATPPGPA | ASSQDWGGQDTQY | 0.7133159 |
| ATVASALV | ASSQDWGGQDTQY | 0.68431973 |
| ATVASLIL | ASSQDWGGQDTQY | 0.75561905 |
| AVNGPVLIRH | ASSQDWGGQDTQY | 0.7183094 |
| DGAAGAVGI | ASSQDWGGQDTQY | 0.6590616 |
| DPLLHGI | ASSQDWGGQDTQY | 0.7197365 |
| ELGAALGAL | ASSQDWGGQDTQY | 0.7612201 |
| ELGRTQAV | ASSQDWGGQDTQY | 0.62239647 |
| FVAGGYLS | ASSQDWGGQDTQY | 0.52393496 |
| GASGGAGGAGGW | ASSQDWGGQDTQY | 0.6561955 |
| GGAGAGGDGGAGGI | ASSQDWGGQDTQY | 0.7309189 |
| GTGHNGSQPGMGGNG | ASSQDWGGQDTQY | 0.75940216 |
| GYAAASASAS | ASSQDWGGQDTQY | 0.74109244 |
| HQVDVAAVQQG | ASSQDWGGQDTQY | 0.7140844 |
| IGAAIAHT | ASSQDWGGQDTQY | 0.7497715 |
| IGMIEVAQL | ASSQDWGGQDTQY | 0.7501738 |
| IRLAPEHTSAMD | ASSQDWGGQDTQY | 0.7202616 |
| IYNVSVL | ASSQDWGGQDTQY | 0.7567579 |
| KQGFLGM | ASSQDWGGQDTQY | 0.68343824 |
| LESVITQL | ASSQDWGGQDTQY | 0.7597406 |
| LGLGALVVV | ASSQDWGGQDTQY | 0.7474925 |
| LIGNGGNGGSGGI | ASSQDWGGQDTQY | 0.7576641 |
| LNAINAPTL | ASSQDWGGQDTQY | 0.7589098 |
| LQLLNTGV | ASSQDWGGQDTQY | 0.7399839 |
| LQTSIEF | ASSQDWGGQDTQY | 0.7348237 |
| MSFAVSEL | ASSQDWGGQDTQY | 0.36332884 |
| NTGALGL | ASSQDWGGQDTQY | 0.70491594 |
| PPGAGGAGGAGTT | ASSQDWGGQDTQY | 0.7433736 |
| QLVGLGI | ASSQDWGGQDTQY | 0.740687 |
| RFLEQVHQL | ASSQDWGGQDTQY | 0.75297165 |
| RGVGYVLR | ASSQDWGGQDTQY | 0.7532168 |
| SATGVFAGVI | ASSQDWGGQDTQY | 0.6699902 |
| SLPAITLP | ASSQDWGGQDTQY | 0.76327974 |
| STALAFTV | ASSQDWGGQDTQY | 0.7587693 |
| SVLTVGLGK | ASSQDWGGQDTQY | 0.49341446 |
| TALAGAGSGI | ASSQDWGGQDTQY | 0.68471843 |
| TGLPFTGI | ASSQDWGGQDTQY | 0.7634543 |
| VAEHVTV | ASSQDWGGQDTQY | 0.74725616 |
| VDIVLNSL | ASSQDWGGQDTQY | 0.71435976 |

| | | |
|---|---|---|
| VLTGGGAVL | ASSQDWGGQDTQY | 0.7023748 |
| VNVGLAYGI | ASSQDWGGQDTQY | 0.7657112 |
| VSSAAVGAV | ASSQDWGGQDTQY | 0.7008426 |
| VTSVYEAGV | ASSQDWGGQDTQY | 0.765158 |
| WVALDRGA | ASSQDWGGQDTQY | 0.28797746 |
| YLVGSVVI | ASSQDWGGQDTQY | 0.7027118 |
| GPPGTGKTLL | ASSQDWGGQDTQY | 0.63688016 |
| GVGMTKF | ASSQDWGGQDTQY | 0.34908974 |
| IGIGGSDL | ASSQDWGGQDTQY | 0.7551737 |
| IGTLLAEL | ASSQDWGGQDTQY | 0.68804675 |
| IYTSGTT | ASSQDWGGQDTQY | 0.75380516 |
| LSFPAGV | ASSQDWGGQDTQY | 0.7322247 |
| LYTSGTT | ASSQDWGGQDTQY | 0.7540724 |
| VGAFPFGF | ASSQDWGGQDTQY | 0.7380533 |
| VGAGTVEF | ASSQDWGGQDTQY | 0.7273599 |
| VGKSTLLL | ASSQDWGGQDTQY | 0.59077317 |
| AALAHAGGGIV | ASSQDWGGQDTQY | 0.7503338 |
| AALVAVSLL | ASSQDWGGQDTQY | 0.7633922 |
| AAVSRAVAAL | ASSQDWGGQDTQY | 0.75300455 |
| AGGSSGIGGF | ASSQDWGGQDTQY | 0.45506454 |
| AGLTVANANG | ASSQDWGGQDTQY | 0.72275573 |
| AGLVGAVL | ASSQDWGGQDTQY | 0.7124684 |
| ALSQAFTV | ASSQDWGGQDTQY | 0.49164385 |
| ALTFSVSL | ASSQDWGGQDTQY | 0.35948232 |
| ALTGAALVAL | ASSQDWGGQDTQY | 0.69264877 |
| ASFSIGAGVL | ASSQDWGGQDTQY | 0.6983856 |
| DGSVYKI | ASSQDWGGQDTQY | 0.7577723 |
| EQLAEIV | ASSQDWGGQDTQY | 0.7558278 |
| FAVSELTVM | ASSQDWGGQDTQY | 0.7107893 |
| GAAPLAAGL | ASSQDWGGQDTQY | 0.7557467 |
| GADGGDGGSGGAGG | ASSQDWGGQDTQY | 0.7285638 |
| GFGLDTGGAGG | ASSQDWGGQDTQY | 0.23218116 |
| GLASGGLL | ASSQDWGGQDTQY | 0.7156311 |
| GRIAGAAVLL | ASSQDWGGQDTQY | 0.49694663 |
| GSAGSLNTSF | ASSQDWGGQDTQY | 0.7631706 |
| GTGQIGI | ASSQDWGGQDTQY | 0.59866565 |
| GTLIALIL | ASSQDWGGQDTQY | 0.6739578 |
| IAATMGFVVG | ASSQDWGGQDTQY | 0.7268513 |
| IFAQALL | ASSQDWGGQDTQY | 0.7404039 |
| IGAAVALGF | ASSQDWGGQDTQY | 0.7405334 |
| IGLTPLAL | ASSQDWGGQDTQY | 0.673339 |
| ILGSNTSTL | ASSQDWGGQDTQY | 0.3559983 |
| IVFSLTVL | ASSQDWGGQDTQY | 0.75164604 |
| LASKATGF | ASSQDWGGQDTQY | 0.6790961 |
| LASVALIL | ASSQDWGGQDTQY | 0.5320044 |
| LFAVGLTL | ASSQDWGGQDTQY | 0.4421418 |
| LFNGGIGTYI | ASSQDWGGQDTQY | 0.74505615 |
| LTLGAGII | ASSQDWGGQDTQY | 0.48748457 |
| LVAAALGDI | ASSQDWGGQDTQY | 0.76125365 |
| NTAFVQLGI | ASSQDWGGQDTQY | 0.7111857 |
| QLFSGAEL | ASSQDWGGQDTQY | 0.70755434 |

| | | |
|---|---|---|
| SAATFITL | ASSQDWGGQDTQY | 0.76269835 |
| SNYGLANLGV | ASSQDWGGQDTQY | 0.74002886 |
| SYGGAYAVM | ASSQDWGGQDTQY | 0.71221066 |
| TTGVGGLTLG | ASSQDWGGQDTQY | 0.63343734 |
| TTQTIYI | ASSQDWGGQDTQY | 0.5675532 |
| VAAGAGLGAVI | ASSQDWGGQDTQY | 0.625921 |
| VAAVVTLL | ASSQDWGGQDTQY | 0.6133451 |
| VGFLGGLI | ASSQDWGGQDTQY | 0.7652184 |
| VGVIPILL | ASSQDWGGQDTQY | 0.6948479 |
| VIFASSLI | ASSQDWGGQDTQY | 0.5367111 |
| VKGGDGGAAAG | ASSQDWGGQDTQY | 0.7164277 |
| VTKTFTNI | ASSQDWGGQDTQY | 0.7641377 |
| VVAAAILL | ASSQDWGGQDTQY | 0.7673514 |
| YAALDVEL | ASSQDWGGQDTQY | 0.61228967 |
| LASVALLL | ASSQDWGGQDTQY | 0.7201841 |
| LDAVHRV | ASSQDWGGQDTQY | 0.6902215 |
| QGLSSSIF | ASSQDWGGQDTQY | 0.72578025 |
| RGTFANIRL | ASSQDWGGQDTQY | 0.7015526 |
| RQLWWGHRI | ASSQDWGGQDTQY | 0.4784134 |
| YGPPGTGKTLL | ASSQDWGGQDTQY | 0.75673217 |
| AALVGAPANVA | ASSQDWGGQDTQY | 0.7691453 |
| AATFRRTE | ASSQDWGGQDTQY | 0.6552881 |
| AGALGGGI | ASSQDWGGQDTQY | 0.60581684 |
| AGGAGGAGGIGAG | ASSQDWGGQDTQY | 0.7217709 |
| AGSGALGL | ASSQDWGGQDTQY | 0.6630287 |
| AQIADQI | ASSQDWGGQDTQY | 0.6881938 |
| ASAVAVTYL | ASSQDWGGQDTQY | 0.7380085 |
| ASTVLGGTGG | ASSQDWGGQDTQY | 0.48211378 |
| AVGGEPAK | ASSQDWGGQDTQY | 0.7114917 |
| CEQGTFGGTC | ASSQDWGGQDTQY | 0.7672983 |
| DEGKAPAG | ASSQDWGGQDTQY | 0.71095014 |
| DTVVFPAL | ASSQDWGGQDTQY | 0.7402859 |
| FEGITMAG | ASSQDWGGQDTQY | 0.7599528 |
| FPLVVQD | ASSQDWGGQDTQY | 0.74859977 |
| GAFTTPQ | ASSQDWGGQDTQY | 0.76239663 |
| GFGGFGTAGGVGGA | ASSQDWGGQDTQY | 0.7581478 |
| GFLAQFL | ASSQDWGGQDTQY | 0.6965796 |
| GGAGGAGGNAGVA | ASSQDWGGQDTQY | 0.7381637 |
| GGVGTAGAGGAGGAGG | ASSQDWGGQDTQY | 0.74499655 |
| GNGGGGDAGNAGSGGN | ASSQDWGGQDTQY | 0.73004794 |
| GTGGAGGAGNDAGS | ASSQDWGGQDTQY | 0.7220293 |
| GTTNTGWFNS | ASSQDWGGQDTQY | 0.6926431 |
| IFMGPLFTL | ASSQDWGGQDTQY | 0.757719 |
| ILAHAIEAI | ASSQDWGGQDTQY | 0.74421334 |
| IRSAIREAL | ASSQDWGGQDTQY | 0.75930583 |
| ITFAGGG | ASSQDWGGQDTQY | 0.4857711 |
| ITGMSGHQEV | ASSQDWGGQDTQY | 0.5585214 |
| IVAGLACTL | ASSQDWGGQDTQY | 0.75523615 |
| IVGGGLAGA | ASSQDWGGQDTQY | 0.26210967 |
| IVTGVLNI | ASSQDWGGQDTQY | 0.34655455 |
| LAQQGCI | ASSQDWGGQDTQY | 0.73955464 |

| | | |
|---|---|---|
| LCLHVEN | ASSQDWGGQDTQY | 0.7628566 |
| LDAISDGS | ASSQDWGGQDTQY | 0.28216332 |
| LGGFAAML | ASSQDWGGQDTQY | 0.61606646 |
| LGGKNPMI | ASSQDWGGQDTQY | 0.58679795 |
| LGPKGVAAIL | ASSQDWGGQDTQY | 0.7516725 |
| LMVGVSI | ASSQDWGGQDTQY | 0.7228706 |
| LVGAAFVL | ASSQDWGGQDTQY | 0.39455712 |
| LVLAVAPL | ASSQDWGGQDTQY | 0.74876416 |
| MAAIAVIDAA | ASSQDWGGQDTQY | 0.5387724 |
| NNGINAELQ | ASSQDWGGQDTQY | 0.7072375 |
| RGEPLVLVHG | ASSQDWGGQDTQY | 0.756642 |
| RGGAGGAAGLI | ASSQDWGGQDTQY | 0.7641029 |
| RSYRMHPAV | ASSQDWGGQDTQY | 0.5727844 |
| SAAHLLAGQLA | ASSQDWGGQDTQY | 0.7569892 |
| SAAVGSY | ASSQDWGGQDTQY | 0.70996475 |
| SPHAVLHRI | ASSQDWGGQDTQY | 0.76905954 |
| SQIAKVL | ASSQDWGGQDTQY | 0.69125634 |
| TALGHLVP | ASSQDWGGQDTQY | 0.5025844 |
| TAPTTSGAA | ASSQDWGGQDTQY | 0.7537399 |
| TDGDDGNGGSPGT | ASSQDWGGQDTQY | 0.7420287 |
| TDGLGEN | ASSQDWGGQDTQY | 0.70081127 |
| TGGPPLPGQA | ASSQDWGGQDTQY | 0.7552804 |
| TIPAITI | ASSQDWGGQDTQY | 0.6694883 |
| TIPGFTI | ASSQDWGGQDTQY | 0.7117356 |
| TSIASGALA | ASSQDWGGQDTQY | 0.7607869 |
| TVTLPGAAI | ASSQDWGGQDTQY | 0.7533325 |
| VASGLSAALP | ASSQDWGGQDTQY | 0.5066776 |
| VGAGVIGI | ASSQDWGGQDTQY | 0.7512919 |
| VGPKRFGV | ASSQDWGGQDTQY | 0.7201097 |
| VNVAFPD | ASSQDWGGQDTQY | 0.658525 |
| VSMTNSGVS | ASSQDWGGQDTQY | 0.7645942 |
| VTDVLGDLSV | ASSQDWGGQDTQY | 0.7370236 |
| VVVVGAGFAGL | ASSQDWGGQDTQY | 0.756829 |
| VYTVGTVV | ASSQDWGGQDTQY | 0.7466835 |
| WTTFSGP | ASSQDWGGQDTQY | 0.7638046 |
| YDELVEAF | ASSQDWGGQDTQY | 0.42710167 |
| YGTGGAGGNGG | ASSQDWGGQDTQY | 0.7456597 |
| YTNVMAV | ASSQDWGGQDTQY | 0.7251126 |
| FLEQVHQL | ASSQDWGGQDTQY | 0.75734085 |
| GLSLLAVAV | ASSQDWGGQDTQY | 0.73005164 |
| LEQVHQL | ASSQDWGGQDTQY | 0.7197056 |
| LNAAGAI | ASSQDWGGQDTQY | 0.74085176 |
| PAVHASL | ASSQDWGGQDTQY | 0.7633921 |
| TELEDDA | ASSQDWGGQDTQY | 0.72352886 |
| TPPGPPN | ASSQDWGGQDTQY | 0.7486775 |
| TPPPPPPP | ASSQDWGGQDTQY | 0.76443815 |
| TQSPALI | ASSRLASSYEQY | 0.63794595 |
| EFAAAFPQ | ASSRLASSYEQY | 0.70710397 |
| LLSGNLDVL | ASSRLASSYEQY | 0.752687 |
| GPANLGNYNI | ASSRLASSYEQY | 0.76197463 |
| TLGPTLP | ASSRLASSYEQY | 0.7060118 |

| | | |
|---|---|---|
| VYAYSSCCK | ASSRLASSYEQY | 0.7624817 |
| APAPAPAPNI | ASSRLASSYEQY | 0.64902747 |
| PGGPVPIP | ASSRLASSYEQY | 0.74885106 |
| FLTVALEP | ASSRLASSYEQY | 0.7043337 |
| SQYFGANL | ASSRLASSYEQY | 0.7478492 |
| GGVKPAA | ASSRLASSYEQY | 0.617923 |
| GGTGGAGGAGGW | ASSRLASSYEQY | 0.74759024 |
| GVGGVAGLGGAGG | ASSRLASSYEQY | 0.75830936 |
| VLLVLATTI | ASSRLASSYEQY | 0.755165 |
| IGIFAAML | ASSRLASSYEQY | 0.3560642 |
| AVGDLID | ASSRLASSYEQY | 0.7535253 |
| NISPLPL | ASSRLASSYEQY | 0.7572336 |
| VTAVFVAI | ASSRLASSYEQY | 0.7686113 |
| SGDDNDAAL | ASSRLASSYEQY | 0.7365242 |
| AVAALAHS | ASSRLASSYEQY | 0.74406075 |
| GGVTVMATP | ASSRLASSYEQY | 0.69672465 |
| AGLAANVRD | ASSRLASSYEQY | 0.71511084 |
| TNRPDLIDPALLRPG | ASSRLASSYEQY | 0.64838946 |
| LANLGVTAL | ASSRLASSYEQY | 0.7171849 |
| TGERIEL | ASSRLASSYEQY | 0.71755475 |
| NGEELAA | ASSRLASSYEQY | 0.66135097 |
| VLSLVWG | ASSRLASSYEQY | 0.7574981 |
| IGYALWL | ASSRLASSYEQY | 0.761775 |
| LSLLLAPL | ASSRLASSYEQY | 0.75743353 |
| TLDEALK | ASSRLASSYEQY | 0.7618736 |
| TDGAGTAEGG | ASSRLASSYEQY | 0.7621956 |
| NAPGGAGGNGG | ASSRLASSYEQY | 0.75769866 |
| VSVFMGMGGNF | ASSRLASSYEQY | 0.66561973 |
| LASYLAAL | ASSRLASSYEQY | 0.7238233 |
| LVGGTPAL | ASSRLASSYEQY | 0.5580252 |
| VRNADVL | ASSRLASSYEQY | 0.7069538 |
| TCSSPFLVSAM | ASSRLASSYEQY | 0.68389165 |
| VHITDAE | ASSRLASSYEQY | 0.7421669 |
| EYAAPSF | ASSRLASSYEQY | 0.232186 |
| VVDAFHGGL | ASSRLASSYEQY | 0.7411646 |
| ASVVDPQ | ASSRLASSYEQY | 0.76536906 |
| SYNEFGGLS | ASSRLASSYEQY | 0.6919342 |
| DPDGSVY | ASSRLASSYEQY | 0.74928147 |
| MVSTIHATE | ASSRLASSYEQY | 0.7449126 |
| LAAGVADPV | ASSRLASSYEQY | 0.68596065 |
| TKEQIAA | ASSRLASSYEQY | 0.7566925 |
| SLASPAAL | ASSRLASSYEQY | 0.6463239 |
| IGLVFYGV | ASSRLASSYEQY | 0.75309885 |
| FLLDASNL | ASSRLASSYEQY | 0.7656679 |
| YQVGNFP | ASSRLASSYEQY | 0.73362553 |
| ESLLSQST | ASSRLASSYEQY | 0.7407798 |
| LFTVGFAAVG | ASSRLASSYEQY | 0.7291429 |
| ATPPTPT | ASSRLASSYEQY | 0.39896253 |
| TDLATLG | ASSRLASSYEQY | 0.7437961 |
| SAGGTGR | ASSRLASSYEQY | 0.5780461 |
| GFSNSGS | ASSRLASSYEQY | 0.7307738 |

| | | |
|---|---|---|
| GGNGGSGGTSV | ASSRLASSYEQY | 0.5831678 |
| GTSVATGGAGN | ASSRLASSYEQY | 0.7378328 |
| NGGNGGSGGTS | ASSRLASSYEQY | 0.7235726 |
| AVVKFPH | ASSRLASSYEQY | 0.7474238 |
| TPPPPPP | ASSRLASSYEQY | 0.6359141 |
| AAPAAAAAPA | ASSRLASSYEQY | 0.7306858 |
| IALNEDL | ASSRLASSYEQY | 0.69492555 |
| IVNVSSM | ASSRLASSYEQY | 0.5263195 |
| TGIPPAPR | ASSRLASSYEQY | 0.7538732 |
| FDLGGGTF | ASSRLASSYEQY | 0.70864534 |
| FDLGGGTFDVS | ASSRLASSYEQY | 0.7215563 |
| NEPTAAA | ASSRLASSYEQY | 0.7598298 |
| NEPTAAALA | ASSRLASSYEQY | 0.7258525 |
| GSIINISSI | ASSRLASSYEQY | 0.4484778 |
| VAPGNAGTA | ASSRLASSYEQY | 0.7259475 |
| TGGGASLE | ASSRLASSYEQY | 0.7245363 |
| VAVYNHYKRI | ASSRLASSYEQY | 0.6968181 |
| TDPAPATT | ASSRLASSYEQY | 0.32218575 |
| TNRPDILDPALLRPG | ASSRLASSYEQY | 0.73515975 |
| LTTTTTS | ASSRLASSYEQY | 0.6559026 |
| PTGSGKTL | ASSRLASSYEQY | 0.66269624 |
| TGSGKTL | ASSRLASSYEQY | 0.6482968 |
| TIGHVDHGK | ASSRLASSYEQY | 0.75569403 |
| TIGHVDHGKTT | ASSRLASSYEQY | 0.73329306 |
| GHVDHGKTT | ASSRLASSYEQY | 0.7296984 |
| AQFEHTLL | ASSRLASSYEQY | 0.7536004 |
| IVNQDEL | ASSRLASSYEQY | 0.744527 |
| TNGSQFF | ASSRLASSYEQY | 0.76391095 |
| TGKGIVE | ASSRLASSYEQY | 0.73419535 |
| LISNASDA | ASSRLASSYEQY | 0.72759706 |
| AELRTLGELTKIAKAHGAQ | ASSRLASSYEQY | 0.74131846 |
| APNAVAAAEPSW | ASSRLASSYEQY | 0.7432299 |
| APQVAGAATAVL | ASSRLASSYEQY | 0.49602515 |
| ARRDPAAAAAILPTDA | ASSRLASSYEQY | 0.47624174 |
| DAAARDVLAEAGLAEYF | ASSRLASSYEQY | 0.7365051 |
| DDQIVLSAKANLAKVEQ | ASSRLASSYEQY | 0.41752714 |
| DELLPDTGEDSDEDGH | ASSRLASSYEQY | 0.7311474 |
| ELPEAPANRATVPQD | ASSRLASSYEQY | 0.76309955 |
| FGQSKIERVTGIG | ASSRLASSYEQY | 0.7313231 |
| GAGGTGSTAGGAGGAGGAGGLY | ASSRLASSYEQY | 0.5735189 |
| GHGGTGAAGGNGGTGGAGGSGI | ASSRLASSYEQY | 0.7064031 |
| GNGGNGGAGGEGGPGGAGGAGA | ASSRLASSYEQY | 0.69152737 |
| GSTAGNGGNGGNATNGTIAGQP | ASSRLASSYEQY | 0.7566619 |
| GTGGDGGNGGNGGTGG | ASSRLASSYEQY | 0.751728 |
| GVRRPAGIPADAPV | ASSRLASSYEQY | 0.7651805 |
| GYRQVTDAHLLTLARR | ASSRLASSYEQY | 0.75606406 |
| IAKMGQDGHDRG | ASSRLASSYEQY | 0.68010867 |
| KAEPSIEIQYT | ASSRLASSYEQY | 0.7522305 |
| KPVRIRMAADEDTVAVT | ASSRLASSYEQY | 0.70663077 |
| MPPRIAGMRLLVIKPEP | ASSRLASSYEQY | 0.74088067 |
| MKFYAVALFRSRGVPPTR | ASSRLASSYEQY | 0.71992385 |

| | | |
|---|---|---|
| NAGGFGILGAVAHSP | ASSRLASSYEQY | 0.73594 |
| NEGREITLAMAEM | ASSRLASSYEQY | 0.7205348 |
| NELVEWSHPLGHRWRWE | ASSRLASSYEQY | 0.6855929 |
| NLGAGGTGGNGGNPG | ASSRLASSYEQY | 0.44721866 |
| NTEWPKWETVMAADPVD | ASSRLASSYEQY | 0.7448479 |
| PASAAPAETQP | ASSRLASSYEQY | 0.7322796 |
| QQSSDGKNGVRAAEITGE | ASSRLASSYEQY | 0.5936584 |
| RAPDAVTTSWWKE | ASSRLASSYEQY | 0.6858207 |
| SGFGNVGAHTSGWFNQ | ASSRLASSYEQY | 0.76430005 |
| SGVAGAVGQAASVGGLKV | ASSRLASSYEQY | 0.76529515 |
| SNGGAGTGSAGGNGGT | ASSRLASSYEQY | 0.6766101 |
| TGGAGGEGGAAG | ASSRLASSYEQY | 0.62284184 |
| TGGDGGAAGAGGAPGHG | ASSRLASSYEQY | 0.7476233 |
| TMPPLLAAIANAGRHGV | ASSRLASSYEQY | 0.4262874 |
| TSSTAGNSGGA | ASSRLASSYEQY | 0.7367105 |
| VPATVPVGILC | ASSRLASSYEQY | 0.73001987 |
| VQRPDSGGVVLTGRIS | ASSRLASSYEQY | 0.7184504 |
| WTPNPVPATKGVPAGLVQAT | ASSRLASSYEQY | 0.6536796 |
| YALKRALAKFSDRFDVVII | ASSRLASSYEQY | 0.73950076 |
| YLIGSVIV | ASSRLASSYEQY | 0.6841042 |
| YGQLVAVID | ASSRLASSYEQY | 0.7554687 |
| VTVGGMTLPA | ASSRLASSYEQY | 0.34500873 |
| VLALPAV | ASSRLASSYEQY | 0.732769 |
| VIPPTLNYET | ASSRLASSYEQY | 0.6367036 |
| VGMGGMHPSSG | ASSRLASSYEQY | 0.70793056 |
| VGLAGGAATA | ASSRLASSYEQY | 0.40591693 |
| VGGAGGAGGSGP | ASSRLASSYEQY | 0.72748125 |
| VFAAFMV | ASSRLASSYEQY | 0.7591506 |
| TLYDKLR | ASSRLASSYEQY | 0.75859517 |
| TLAVFTVL | ASSRLASSYEQY | 0.6550827 |
| TGSVQPARV | ASSRLASSYEQY | 0.75420165 |
| TGGGAGSGGT | ASSRLASSYEQY | 0.76636904 |
| SHLETSL | ASSRLASSYEQY | 0.715292 |
| SFTTGMT | ASSRLASSYEQY | 0.32929367 |
| QVEVAKRMGVRQPTVS | ASSRLASSYEQY | 0.74456155 |
| QGSGGPALAM | ASSRLASSYEQY | 0.7251526 |
| MLAPSNP | ASSRLASSYEQY | 0.7539461 |
| LVNNAAI | ASSRLASSYEQY | 0.7521603 |
| LTGEINTLA | ASSRLASSYEQY | 0.5895016 |
| LSIFGPV | ASSRLASSYEQY | 0.75371885 |
| LLALLANL | ASSRLASSYEQY | 0.7545125 |
| LISNASDALDKLR | ASSRLASSYEQY | 0.64052397 |
| LINEAVP | ASSRLASSYEQY | 0.6774535 |
| LGYAFSGL | ASSRLASSYEQY | 0.65877604 |
| LDFVAVAL | ASSRLASSYEQY | 0.5616789 |
| LAQPGAE | ASSRLASSYEQY | 0.742986 |
| ISNLGAVTGF | ASSRLASSYEQY | 0.6886909 |
| HQEDPGRI | ASSRLASSYEQY | 0.68524575 |
| GTPGEPTP | ASSRLASSYEQY | 0.71523833 |
| GFFNSGS | ASSRLASSYEQY | 0.64866704 |
| GAGGGGLTAGAN | ASSRLASSYEQY | 0.6585311 |

| | | |
|---|---|---|
| FVDALVNM | ASSRLASSYEQY | 0.6754434 |
| FLTVAPEL | ASSRLASSYEQY | 0.7207469 |
| FETLTAL | ASSRLASSYEQY | 0.7458241 |
| DVINPIEL | ASSRLASSYEQY | 0.7228439 |
| APVSGAPV | ASSRLASSYEQY | 0.74712497 |
| ALVGGLAW | ASSRLASSYEQY | 0.7642107 |
| ALAAAAATAI | ASSRLASSYEQY | 0.52615666 |
| AFGLGLLL | ASSRLASSYEQY | 0.623764 |
| AFFAAASL | ASSRLASSYEQY | 0.7470906 |
| AAAALPAFSPP | ASSRLASSYEQY | 0.55240273 |
| VVLAPAL | ASSRLASSYEQY | 0.75342 |
| TIGHVDHGKTTL | ASSRLASSYEQY | 0.72342384 |
| LISNASDALDK | ASSRLASSYEQY | 0.7445477 |
| LISNASDAL | ASSRLASSYEQY | 0.74347603 |
| ISNASDALDK | ASSRLASSYEQY | 0.765453 |
| NQAETLVYQTE | ASSRLASSYEQY | 0.7545945 |
| TGGTGGTAGAG | ASSRLASSYEQY | 0.71205914 |
| TGSTAGGAGGA | ASSRLASSYEQY | 0.7567469 |
| DPLAVPVAAGPV | ASSRLASSYEQY | 0.5936542 |
| GGTGGTPGNGGN | ASSRLASSYEQY | 0.75528145 |
| PAGLAGVDPATL | ASSRLASSYEQY | 0.75271714 |
| GNNLVVEAPHAGASV | ASSRLASSYEQY | 0.6301174 |
| KVTRIMSASTLEQPA | ASSRLASSYEQY | 0.7158972 |
| LGFALPIALPRRTVV | ASSRLASSYEQY | 0.76453674 |
| TRLGNLTARKLVVLV | ASSRLASSYEQY | 0.7574404 |
| WSQQRRLIYFPSAGP | ASSRLASSYEQY | 0.42907798 |
| GPNGNPDPMAAAVDIR | ASSRLASSYEQY | 0.67872643 |
| TQRCVSVADLRAAASR | ASSRLASSYEQY | 0.53819394 |
| QFVLDVAEHPVEVLTETV | ASSRLASSYEQY | 0.72089124 |
| GGAGGTGGAGGNGGTQAGDG | ASSRLASSYEQY | 0.48317188 |
| PLLATLRAIAVDVDAKPAQV | ASSRLASSYEQY | 0.7533134 |
| APKFRTGGYMIDFWGVGYQVAK | ASSRLASSYEQY | 0.7066203 |
| GAGGAGGAGGAAGGPGGTGGTG | ASSRLASSYEQY | 0.7531915 |
| IGALGLLATDVAPAGSVAFGVA | ASSRLASSYEQY | 0.7301327 |
| AGGAGGIGGNGGAGGAGGVLMG | ASSRLASSYEQY | 0.75976795 |
| ADELIGLL | ASSRLASSYEQY | 0.6880279 |
| ADRALKA | ASSRLASSYEQY | 0.6955027 |
| AGAVGQVGGAGGA | ASSRLASSYEQY | 0.42240453 |
| AGGGGGLGGAP | ASSRLASSYEQY | 0.62992287 |
| AGGLLYGNGGAG | ASSRLASSYEQY | 0.67562664 |
| AGLESIGL | ASSRLASSYEQY | 0.699352 |
| AGLKTGII | ASSRLASSYEQY | 0.52324605 |
| ASLAALLL | ASSRLASSYEQY | 0.76106846 |
| ASVINISSI | ASSRLASSYEQY | 0.7368723 |
| ATPPGPA | ASSRLASSYEQY | 0.75071245 |
| ATVASALV | ASSRLASSYEQY | 0.6504222 |
| ATVASLIL | ASSRLASSYEQY | 0.7134455 |
| AVNGPVLIRH | ASSRLASSYEQY | 0.65649045 |
| DGAAGAVGI | ASSRLASSYEQY | 0.7532109 |
| DPLLHGI | ASSRLASSYEQY | 0.5568628 |
| ELGAALGAL | ASSRLASSYEQY | 0.4200945 |

| | | |
|---|---|---|
| ELGRTQAV | ASSRLASSYEQY | 0.7144474 |
| FVAGGYLS | ASSRLASSYEQY | 0.632876 |
| GASGGAGGAGGW | ASSRLASSYEQY | 0.7621181 |
| GGAGAGGDGGAGGI | ASSRLASSYEQY | 0.7586255 |
| GTGHNGSQPGMGGNG | ASSRLASSYEQY | 0.7400366 |
| GYAAASASAS | ASSRLASSYEQY | 0.75890493 |
| HQVDVAAVQQG | ASSRLASSYEQY | 0.28682104 |
| IGAAIAHT | ASSRLASSYEQY | 0.68951905 |
| IGMIEVAQL | ASSRLASSYEQY | 0.7642858 |
| IRLAPEHTSAMD | ASSRLASSYEQY | 0.7473812 |
| IYNVSVL | ASSRLASSYEQY | 0.7188675 |
| KQGFLGM | ASSRLASSYEQY | 0.7627033 |
| LESVITQL | ASSRLASSYEQY | 0.63192415 |
| LGLGALVVV | ASSRLASSYEQY | 0.69991636 |
| LIGNGGNGGSGGI | ASSRLASSYEQY | 0.729317 |
| LNAINAPTL | ASSRLASSYEQY | 0.7444335 |
| LQLLNTGV | ASSRLASSYEQY | 0.75140995 |
| LQTSIEF | ASSRLASSYEQY | 0.5673634 |
| MSFAVSEL | ASSRLASSYEQY | 0.7415152 |
| NTGALGL | ASSRLASSYEQY | 0.76736313 |
| PPGAGGAGGAGTT | ASSRLASSYEQY | 0.7085911 |
| QLVGLGI | ASSRLASSYEQY | 0.7590124 |
| RFLEQVHQL | ASSRLASSYEQY | 0.7452333 |
| RGVGYVLR | ASSRLASSYEQY | 0.7482725 |
| SATGVFAGVI | ASSRLASSYEQY | 0.6420835 |
| SLPAITLP | ASSRLASSYEQY | 0.65724826 |
| STALAFTV | ASSRLASSYEQY | 0.73082274 |
| SVLTVGLGK | ASSRLASSYEQY | 0.6698033 |
| TALAGAGSGI | ASSRLASSYEQY | 0.7288616 |
| TGLPFTGI | ASSRLASSYEQY | 0.6652992 |
| VAEHVTV | ASSRLASSYEQY | 0.76966643 |
| VDIVLNSL | ASSRLASSYEQY | 0.75075006 |
| VLTGGGAVL | ASSRLASSYEQY | 0.74498534 |
| VNVGLAYGI | ASSRLASSYEQY | 0.605489 |
| VSSAAVGAV | ASSRLASSYEQY | 0.73962265 |
| VTSVYEAGV | ASSRLASSYEQY | 0.7473914 |
| WVALDRGA | ASSRLASSYEQY | 0.44306388 |
| YLVGSVVI | ASSRLASSYEQY | 0.763237 |
| GPPGTGKTLL | ASSRLASSYEQY | 0.6517368 |
| GVGMTKF | ASSRLASSYEQY | 0.71413344 |
| IGIGGSDL | ASSRLASSYEQY | 0.75310075 |
| IGTLLAEL | ASSRLASSYEQY | 0.761176 |
| IYTSGTT | ASSRLASSYEQY | 0.7010577 |
| LSFPAGV | ASSRLASSYEQY | 0.728668 |
| LYTSGTT | ASSRLASSYEQY | 0.590024 |
| VGAFPFGF | ASSRLASSYEQY | 0.75801444 |
| VGAGTVEF | ASSRLASSYEQY | 0.7593386 |
| VGKSTLLL | ASSRLASSYEQY | 0.7442585 |
| AALAHAGGGIV | ASSRLASSYEQY | 0.7649964 |
| AALVAVSLL | ASSRLASSYEQY | 0.744205 |
| AAVSRAVAAL | ASSRLASSYEQY | 0.4381488 |

| | | |
|---|---|---|
| AGGSSGIGGF | ASSRLASSYEQY | 0.70961195 |
| AGLTVANANG | ASSRLASSYEQY | 0.7549303 |
| AGLVGAVL | ASSRLASSYEQY | 0.7589979 |
| ALSQAFTV | ASSRLASSYEQY | 0.7637154 |
| ALTFSVSL | ASSRLASSYEQY | 0.7501175 |
| ALTGAALVAL | ASSRLASSYEQY | 0.7023873 |
| ASFSIGAGVL | ASSRLASSYEQY | 0.7629371 |
| DGSVYKI | ASSRLASSYEQY | 0.76863635 |
| EQLAEIV | ASSRLASSYEQY | 0.75561845 |
| FAVSELTVM | ASSRLASSYEQY | 0.7112115 |
| GAAPLAAGL | ASSRLASSYEQY | 0.6860787 |
| GADGGDGGSGGAGG | ASSRLASSYEQY | 0.73640954 |
| GFGLDTGGAGG | ASSRLASSYEQY | 0.7499893 |
| GLASGGLL | ASSRLASSYEQY | 0.6246164 |
| GRIAGAAVLL | ASSRLASSYEQY | 0.7395876 |
| GSAGSLNTSF | ASSRLASSYEQY | 0.74360204 |
| GTGQIGI | ASSRLASSYEQY | 0.47748023 |
| GTLIALIL | ASSRLASSYEQY | 0.7463635 |
| IAATMGFVVG | ASSRLASSYEQY | 0.7601454 |
| IFAQALL | ASSRLASSYEQY | 0.7144977 |
| IGAAVALGF | ASSRLASSYEQY | 0.6871464 |
| IGLTPLAL | ASSRLASSYEQY | 0.76092386 |
| ILGSNTSTL | ASSRLASSYEQY | 0.768171 |
| IVFSLTVL | ASSRLASSYEQY | 0.76153183 |
| LASKATGF | ASSRLASSYEQY | 0.43161047 |
| LASVALIL | ASSRLASSYEQY | 0.35927552 |
| LFAVGLTL | ASSRLASSYEQY | 0.68870455 |
| LFNGGIGTYI | ASSRLASSYEQY | 0.7535455 |
| LTLGAGII | ASSRLASSYEQY | 0.68547404 |
| LVAAALGDI | ASSRLASSYEQY | 0.6602926 |
| NTAFVQLGI | ASSRLASSYEQY | 0.74558944 |
| QLFSGAEL | ASSRLASSYEQY | 0.76056427 |
| SAATFITL | ASSRLASSYEQY | 0.63056195 |
| SNYGLANLGV | ASSRLASSYEQY | 0.55152315 |
| SYGGAYAVM | ASSRLASSYEQY | 0.75396633 |
| TTGVGGLTLG | ASSRLASSYEQY | 0.753989 |
| TTQTIYI | ASSRLASSYEQY | 0.638201 |
| VAAGAGLGAVI | ASSRLASSYEQY | 0.70813686 |
| VAAVVTLL | ASSRLASSYEQY | 0.6942208 |
| VGFLGGLI | ASSRLASSYEQY | 0.7212173 |
| VGVIPILL | ASSRLASSYEQY | 0.74018186 |
| VIFASSLI | ASSRLASSYEQY | 0.5865543 |
| VKGGDGGAAAG | ASSRLASSYEQY | 0.7571324 |
| VTKTFTNI | ASSRLASSYEQY | 0.59690684 |
| VVAAAILL | ASSRLASSYEQY | 0.75626564 |
| YAALDVEL | ASSRLASSYEQY | 0.74064064 |
| LASVALLL | ASSRLASSYEQY | 0.7329105 |
| LDAVHRV | ASSRLASSYEQY | 0.709237 |
| QGLSSSIF | ASSRLASSYEQY | 0.6951561 |
| RGTFANIRL | ASSRLASSYEQY | 0.49337798 |
| RQLWWGHRI | ASSRLASSYEQY | 0.55871814 |

| | | |
|---|---|---|
| YGPPGTGKTLL | ASSRLASSYEQY | 0.761963 |
| AALVGAPANVA | ASSRLASSYEQY | 0.7315224 |
| AATFRRTE | ASSRLASSYEQY | 0.75828004 |
| AGALGGGI | ASSRLASSYEQY | 0.7481231 |
| AGGAGGAGGIGAG | ASSRLASSYEQY | 0.5713991 |
| AGSGALGL | ASSRLASSYEQY | 0.74805486 |
| AQIADQI | ASSRLASSYEQY | 0.5777041 |
| ASAVAVTYL | ASSRLASSYEQY | 0.6637888 |
| ASTVLGGTGG | ASSRLASSYEQY | 0.7524247 |
| AVGGEPAK | ASSRLASSYEQY | 0.747126 |
| CEQGTFGGTC | ASSRLASSYEQY | 0.6563759 |
| DEGKAPAG | ASSRLASSYEQY | 0.74443185 |
| DTVVFPAL | ASSRLASSYEQY | 0.73983824 |
| FEGITMAG | ASSRLASSYEQY | 0.6747122 |
| FPLVVQD | ASSRLASSYEQY | 0.7014914 |
| GAFTTPQ | ASSRLASSYEQY | 0.7608949 |
| GFGGFGTAGGVGGA | ASSRLASSYEQY | 0.73427916 |
| GFLAQFL | ASSRLASSYEQY | 0.6019686 |
| GGAGGAGGNAGVA | ASSRLASSYEQY | 0.75290143 |
| GGVGTAGAGGAGGAGG | ASSRLASSYEQY | 0.6481509 |
| GNGGGGDAGNAGSGGN | ASSRLASSYEQY | 0.74154544 |
| GTGGAGGAGNDAGS | ASSRLASSYEQY | 0.708114 |
| GTTNTGWFNS | ASSRLASSYEQY | 0.7160104 |
| IFMGPLFTL | ASSRLASSYEQY | 0.7509974 |
| ILAHAIEAI | ASSRLASSYEQY | 0.6504681 |
| IRSAIREAL | ASSRLASSYEQY | 0.6755791 |
| ITFAGGG | ASSRLASSYEQY | 0.6965092 |
| ITGMSGHQEV | ASSRLASSYEQY | 0.7216978 |
| IVAGLACTL | ASSRLASSYEQY | 0.75557745 |
| IVGGGLAGA | ASSRLASSYEQY | 0.61788714 |
| IVTGVLNI | ASSRLASSYEQY | 0.7380127 |
| LAQQGCI | ASSRLASSYEQY | 0.6851286 |
| LCLHVEN | ASSRLASSYEQY | 0.617581 |
| LDAISDGS | ASSRLASSYEQY | 0.7181381 |
| LGGFAAML | ASSRLASSYEQY | 0.72406006 |
| LGGKNPMI | ASSRLASSYEQY | 0.6643913 |
| LGPKGVAAIL | ASSRLASSYEQY | 0.74315965 |
| LMVGVSI | ASSRLASSYEQY | 0.60161924 |
| LVGAAFVL | ASSRLASSYEQY | 0.7432243 |
| LVLAVAPL | ASSRLASSYEQY | 0.764037 |
| MAAIAVIDAA | ASSRLASSYEQY | 0.6996356 |
| NNGINAELQ | ASSRLASSYEQY | 0.7544875 |
| RGEPLVLVHG | ASSRLASSYEQY | 0.73863053 |
| RGGAGGAAGLI | ASSRLASSYEQY | 0.7480458 |
| RSYRMHPAV | ASSRLASSYEQY | 0.68396246 |
| SAAHLLAGQLA | ASSRLASSYEQY | 0.6466186 |
| SAAVGSY | ASSRLASSYEQY | 0.7563361 |
| SPHAVLHRI | ASSRLASSYEQY | 0.7539495 |
| SQIAKVL | ASSRLASSYEQY | 0.7569995 |
| TALGHLVP | ASSRLASSYEQY | 0.74573946 |
| TAPTTSGAA | ASSRLASSYEQY | 0.7566897 |

| | | |
|---|---|---|
| TDGDDGNGGSPGT | ASSRLASSYEQY | 0.7524483 |
| TDGLGEN | ASSRLASSYEQY | 0.70122355 |
| TGGPPLPGQA | ASSRLASSYEQY | 0.5079349 |
| TIPAITI | ASSRLASSYEQY | 0.69750094 |
| TIPGFTI | ASSRLASSYEQY | 0.7032182 |
| TSIASGALA | ASSRLASSYEQY | 0.69583964 |
| TVTLPGAAI | ASSRLASSYEQY | 0.49763542 |
| VASGLSAALP | ASSRLASSYEQY | 0.7587656 |
| VGAGVIGI | ASSRLASSYEQY | 0.767612 |
| VGPKRFGV | ASSRLASSYEQY | 0.68181443 |
| VNVAFPD | ASSRLASSYEQY | 0.7647387 |
| VSMTNSGVS | ASSRLASSYEQY | 0.5691266 |
| VTDVLGDLSV | ASSRLASSYEQY | 0.7515898 |
| VVVVGAGFAGL | ASSRLASSYEQY | 0.7626641 |
| VYTVGTVV | ASSRLASSYEQY | 0.6326417 |
| WTTFSGP | ASSRLASSYEQY | 0.7499246 |
| YDELVEAF | ASSRLASSYEQY | 0.638749 |
| YGTGGAGGNGG | ASSRLASSYEQY | 0.68703383 |
| YTNVMAV | ASSRLASSYEQY | 0.76244736 |
| FLEQVHQL | ASSRLASSYEQY | 0.690171 |
| GLSLLAVAV | ASSRLASSYEQY | 0.7422943 |
| LEQVHQL | ASSRLASSYEQY | 0.6887179 |
| LNAAGAI | ASSRLASSYEQY | 0.5649703 |
| PAVHASL | ASSRLASSYEQY | 0.72578096 |
| TELEDDA | ASSRLASSYEQY | 0.76143324 |
| TPPGPPN | ASSRLASSYEQY | 0.7091678 |
| TPPPPPPP | ASSRLASSYEQY | 0.7683744 |
| TQSPALI | ASSQERDRGDEQY | 0.6582289 |
| EFAAAFPQ | ASSQERDRGDEQY | 0.7636336 |
| LLSGNLDVL | ASSQERDRGDEQY | 0.75660574 |
| GPANLGNYNI | ASSQERDRGDEQY | 0.72696006 |
| TLGPTLP | ASSQERDRGDEQY | 0.7051257 |
| VYAYSSCCK | ASSQERDRGDEQY | 0.7613513 |
| APAPAPAPNI | ASSQERDRGDEQY | 0.70177877 |
| PGGPVPIP | ASSQERDRGDEQY | 0.7554186 |
| FLTVALEP | ASSQERDRGDEQY | 0.7626379 |
| SQYFGANL | ASSQERDRGDEQY | 0.7317037 |
| GGVKPAA | ASSQERDRGDEQY | 0.87906957 |
| GGTGGAGGAGGW | ASSQERDRGDEQY | 0.8267845 |
| GVGGVAGLGGAGG | ASSQERDRGDEQY | 0.8270067 |
| VLLVLATTI | ASSQERDRGDEQY | 0.78895015 |
| IGIFAAML | ASSQERDRGDEQY | 0.43415883 |
| AVGDLID | ASSQERDRGDEQY | 0.8031827 |
| NISPLPL | ASSQERDRGDEQY | 0.7389236 |
| VTAVFVAI | ASSQERDRGDEQY | 0.79026276 |
| SGDDNDAAL | ASSQERDRGDEQY | 0.8548347 |
| AVAALAHS | ASSQERDRGDEQY | 0.7858634 |
| GGVTVMATP | ASSQERDRGDEQY | 0.87533605 |
| AGLAANVRD | ASSQERDRGDEQY | 0.8796262 |
| TNRPDLIDPALLRPG | ASSQERDRGDEQY | 0.6102509 |
| LANLGVTAL | ASSQERDRGDEQY | 0.78106594 |

| | | |
|---|---|---|
| TGERIEL | ASSQERDRGDEQY | 0.7433766 |
| NGEELAA | ASSQERDRGDEQY | 0.8247783 |
| VLSLVWG | ASSQERDRGDEQY | 0.37809536 |
| IGYALWL | ASSQERDRGDEQY | 0.87502086 |
| LSLLLAPL | ASSQERDRGDEQY | 0.70121646 |
| TLDEALK | ASSQERDRGDEQY | 0.17974041 |
| TDGAGTAEGG | ASSQERDRGDEQY | 0.8130213 |
| NAPGGAGGNGG | ASSQERDRGDEQY | 0.80943775 |
| VSVFMGMGGNF | ASSQERDRGDEQY | 0.8069701 |
| LASYLAAL | ASSQERDRGDEQY | 0.864632 |
| LVGGTPAL | ASSQERDRGDEQY | 0.50212353 |
| VRNADVL | ASSQERDRGDEQY | 0.7084068 |
| TCSSPFLVSAM | ASSQERDRGDEQY | 0.8596362 |
| VHITDAE | ASSQERDRGDEQY | 0.8875334 |
| EYAAPSF | ASSQERDRGDEQY | 0.63470036 |
| VVDAFHGGL | ASSQERDRGDEQY | 0.83993244 |
| ASVVDPQ | ASSQERDRGDEQY | 0.8832145 |
| SYNEFGGLS | ASSQERDRGDEQY | 0.84710336 |
| DPDGSVY | ASSQERDRGDEQY | 0.8649983 |
| MVSTIHATE | ASSQERDRGDEQY | 0.88084805 |
| LAAGVADPV | ASSQERDRGDEQY | 0.6712052 |
| TKEQIAA | ASSQERDRGDEQY | 0.8800044 |
| SLASPAAL | ASSQERDRGDEQY | 0.8611498 |
| IGLVFYGV | ASSQERDRGDEQY | 0.80907536 |
| FLLDASNL | ASSQERDRGDEQY | 0.83379865 |
| YQVGNFP | ASSQERDRGDEQY | 0.7126487 |
| ESLLSQST | ASSQERDRGDEQY | 0.74653214 |
| LFTVGFAAVG | ASSQERDRGDEQY | 0.36951792 |
| ATPPTPT | ASSQERDRGDEQY | 0.84345365 |
| TDLATLG | ASSQERDRGDEQY | 0.87281036 |
| SAGGTGR | ASSQERDRGDEQY | 0.7907456 |
| GFSNSGS | ASSQERDRGDEQY | 0.6873909 |
| GGNGGSGGTSV | ASSQERDRGDEQY | 0.7022537 |
| GTSVATGGAGN | ASSQERDRGDEQY | 0.7848464 |
| NGGNGGSGGTS | ASSQERDRGDEQY | 0.79062617 |
| AVVKFPH | ASSQERDRGDEQY | 0.3899718 |
| TPPPPPP | ASSQERDRGDEQY | 0.6797426 |
| AAPAAAAAPA | ASSQERDRGDEQY | 0.87728137 |
| IALNEDL | ASSQERDRGDEQY | 0.8838868 |
| IVNVSSM | ASSQERDRGDEQY | 0.49380368 |
| TGIPPAPR | ASSQERDRGDEQY | 0.87917626 |
| FDLGGGTF | ASSQERDRGDEQY | 0.8417942 |
| FDLGGGTFDVS | ASSQERDRGDEQY | 0.8677743 |
| NEPTAAA | ASSQERDRGDEQY | 0.81649965 |
| NEPTAAALA | ASSQERDRGDEQY | 0.7158256 |
| GSIINISSI | ASSQERDRGDEQY | 0.85811335 |
| VAPGNAGTA | ASSQERDRGDEQY | 0.76826274 |
| TGGGASLE | ASSQERDRGDEQY | 0.64743745 |
| VAVYNHYKRI | ASSQERDRGDEQY | 0.8712175 |
| TDPAPATT | ASSQERDRGDEQY | 0.7895762 |
| TNRPDILDPALLRPG | ASSQERDRGDEQY | 0.7458653 |

| | | |
|---|---|---|
| LTTTTTS | ASSQERDRGDEQY | 0.67643106 |
| PTGSGKTL | ASSQERDRGDEQY | 0.878774 |
| TGSGKTL | ASSQERDRGDEQY | 0.6200272 |
| TIGHVDHGK | ASSQERDRGDEQY | 0.6970051 |
| TIGHVDHGKTT | ASSQERDRGDEQY | 0.6748387 |
| GHVDHGKTT | ASSQERDRGDEQY | 0.8198343 |
| AQFEHTLL | ASSQERDRGDEQY | 0.8807665 |
| IVNQDEL | ASSQERDRGDEQY | 0.8467207 |
| TNGSQFF | ASSQERDRGDEQY | 0.7434262 |
| TGKGIVE | ASSQERDRGDEQY | 0.8454921 |
| LISNASDA | ASSQERDRGDEQY | 0.85717165 |
| AELRTLGELTKIAKAHGAQ | ASSQERDRGDEQY | 0.84296846 |
| APNAVAAAEPSW | ASSQERDRGDEQY | 0.8452227 |
| APQVAGAATAVL | ASSQERDRGDEQY | 0.7335305 |
| ARRDPAAAAAILPTDA | ASSQERDRGDEQY | 0.8808615 |
| DAAARDVLAEAGLAEYF | ASSQERDRGDEQY | 0.8624998 |
| DDQIVLSAKANLAKVEQ | ASSQERDRGDEQY | 0.87644506 |
| DELLPDTGEDSDEDGH | ASSQERDRGDEQY | 0.8614483 |
| ELPEAPANRATVPQD | ASSQERDRGDEQY | 0.832403 |
| FGQSKIERVTGIG | ASSQERDRGDEQY | 0.8346294 |
| GAGGTGSTAGGAGGAGGAGGLY | ASSQERDRGDEQY | 0.3861915 |
| GHGGTGAAGGNGGTGGAGGSGI | ASSQERDRGDEQY | 0.7109433 |
| GNGGNGGAGGEGGPGGAGGAGA | ASSQERDRGDEQY | 0.8198397 |
| GSTAGNGGNGGNATNGTIAGQP | ASSQERDRGDEQY | 0.8255793 |
| GTGGDGGNGGNGGTGG | ASSQERDRGDEQY | 0.8407701 |
| GVRRPAGIPADAPV | ASSQERDRGDEQY | 0.85952497 |
| GYRQVTDAHLLTLARR | ASSQERDRGDEQY | 0.7391975 |
| IAKMGQDGHDRG | ASSQERDRGDEQY | 0.8570967 |
| KAEPSIEIQYT | ASSQERDRGDEQY | 0.87040627 |
| KPVRIRMAADEDTVAVT | ASSQERDRGDEQY | 0.6059427 |
| MPPRIAGMRLLVIKPEP | ASSQERDRGDEQY | 0.78005135 |
| MKFYAVALFRSRGVPPTR | ASSQERDRGDEQY | 0.88936406 |
| NAGGFGILGAVAHSP | ASSQERDRGDEQY | 0.80401534 |
| NEGREITLAMAEM | ASSQERDRGDEQY | 0.79304147 |
| NELVEWSHPLGHRWRWE | ASSQERDRGDEQY | 0.7638465 |
| NLGAGGTGGNGGNPG | ASSQERDRGDEQY | 0.86421335 |
| NTEWPKWETVMAADPVD | ASSQERDRGDEQY | 0.709191 |
| PASAAPAETQP | ASSQERDRGDEQY | 0.87900704 |
| QQSSDGKNGVRAAEITGE | ASSQERDRGDEQY | 0.27069902 |
| RAPDAVTTSWWKE | ASSQERDRGDEQY | 0.78881156 |
| SGFGNVGAHTSGWFNQ | ASSQERDRGDEQY | 0.68847287 |
| SGVAGAVGQAASVGGLKV | ASSQERDRGDEQY | 0.5098599 |
| SNGGAGTGSAGGNGGT | ASSQERDRGDEQY | 0.8752129 |
| TGGAGGEGGAAG | ASSQERDRGDEQY | 0.76370955 |
| TGGDGGAAGAGGAPGHG | ASSQERDRGDEQY | 0.85514975 |
| TMPPLLAAIANAGRHGV | ASSQERDRGDEQY | 0.7817599 |
| TSSTAGNSGGA | ASSQERDRGDEQY | 0.8670049 |
| VPATVPVGILC | ASSQERDRGDEQY | 0.81478405 |
| VQRPDSGGVVLTGRIS | ASSQERDRGDEQY | 0.81293255 |
| WTPNPVPATKGVPAGLVQAT | ASSQERDRGDEQY | 0.70865536 |
| YALKRALAKFSDRFDVVII | ASSQERDRGDEQY | 0.8661839 |

| | | |
|---|---|---|
| YLIGSVIV | ASSQERDRGDEQY | 0.86248565 |
| YGQLVAVID | ASSQERDRGDEQY | 0.85803795 |
| VTVGGMTLPA | ASSQERDRGDEQY | 0.5187218 |
| VLALPAV | ASSQERDRGDEQY | 0.7086372 |
| VIPPTLNYET | ASSQERDRGDEQY | 0.8500222 |
| VGMGGMHPSSG | ASSQERDRGDEQY | 0.30129763 |
| VGLAGGAATA | ASSQERDRGDEQY | 0.26196548 |
| VGGAGGAGGSGP | ASSQERDRGDEQY | 0.736243 |
| VFAAFMV | ASSQERDRGDEQY | 0.68135387 |
| TLYDKLR | ASSQERDRGDEQY | 0.87910193 |
| TLAVFTVL | ASSQERDRGDEQY | 0.863281 |
| TGSVQPARV | ASSQERDRGDEQY | 0.8172529 |
| TGGGAGSGGT | ASSQERDRGDEQY | 0.8573674 |
| SHLETSL | ASSQERDRGDEQY | 0.80888534 |
| SFTTGMT | ASSQERDRGDEQY | 0.1156092 |
| QVEVAKRMGVRQPTVS | ASSQERDRGDEQY | 0.7806701 |
| QGSGGPALAM | ASSQERDRGDEQY | 0.6413023 |
| MLAPSNP | ASSQERDRGDEQY | 0.88525355 |
| LVNNAAI | ASSQERDRGDEQY | 0.6997973 |
| LTGEINTLA | ASSQERDRGDEQY | 0.6746287 |
| LSIFGPV | ASSQERDRGDEQY | 0.8313779 |
| LLALLANL | ASSQERDRGDEQY | 0.78219974 |
| LISNASDALDKLR | ASSQERDRGDEQY | 0.79437435 |
| LINEAVP | ASSQERDRGDEQY | 0.6772585 |
| LGYAFSGL | ASSQERDRGDEQY | 0.1583512 |
| LDFVAVAL | ASSQERDRGDEQY | 0.8685337 |
| LAQPGAE | ASSQERDRGDEQY | 0.45105597 |
| ISNLGAVTGF | ASSQERDRGDEQY | 0.72035444 |
| HQEDPGRI | ASSQERDRGDEQY | 0.31388605 |
| GTPGEPTP | ASSQERDRGDEQY | 0.22500138 |
| GFFNSGS | ASSQERDRGDEQY | 0.5131775 |
| GAGGGGLTAGAN | ASSQERDRGDEQY | 0.8640858 |
| FVDALVNM | ASSQERDRGDEQY | 0.7569549 |
| FLTVAPEL | ASSQERDRGDEQY | 0.7427691 |
| FETLTAL | ASSQERDRGDEQY | 0.8833232 |
| DVINPIEL | ASSQERDRGDEQY | 0.79137987 |
| APVSGAPV | ASSQERDRGDEQY | 0.7782384 |
| ALVGGLAW | ASSQERDRGDEQY | 0.75227594 |
| ALAAAAATAI | ASSQERDRGDEQY | 0.73677695 |
| AFGLGLLL | ASSQERDRGDEQY | 0.7162848 |
| AFFAAASL | ASSQERDRGDEQY | 0.7130496 |
| AAAALPAFSPP | ASSQERDRGDEQY | 0.88568556 |
| VVLAPAL | ASSQERDRGDEQY | 0.78149545 |
| TIGHVDHGKTTL | ASSQERDRGDEQY | 0.59318745 |
| LISNASDALDK | ASSQERDRGDEQY | 0.8239663 |
| LISNASDAL | ASSQERDRGDEQY | 0.88765395 |
| ISNASDALDK | ASSQERDRGDEQY | 0.8934307 |
| NQAETLVYQTE | ASSQERDRGDEQY | 0.69348174 |
| TGGTGGTAGAG | ASSQERDRGDEQY | 0.7541246 |
| TGSTAGGAGGA | ASSQERDRGDEQY | 0.7731935 |
| DPLAVPVAAGPV | ASSQERDRGDEQY | 0.8193939 |

| | | |
|---|---|---|
| GGTGGTPGNGGN | ASSQERDRGDEQY | 0.721649 |
| PAGLAGVDPATL | ASSQERDRGDEQY | 0.63403594 |
| GNNLVVEAPHAGASV | ASSQERDRGDEQY | 0.8691129 |
| KVTRIMSASTLEQPA | ASSQERDRGDEQY | 0.8914328 |
| LGFALPIALPRRTVV | ASSQERDRGDEQY | 0.78912866 |
| TRLGNLTARKLVVLV | ASSQERDRGDEQY | 0.5815824 |
| WSQQRRLIYFPSAGP | ASSQERDRGDEQY | 0.8184545 |
| GPNGNPDPMAAAVDIR | ASSQERDRGDEQY | 0.68875873 |
| TQRCVSVADLRAAASR | ASSQERDRGDEQY | 0.798483 |
| QFVLDVAEHPVEVLTETV | ASSQERDRGDEQY | 0.8300903 |
| GGAGGTGGAGGNGGTQAGDG | ASSQERDRGDEQY | 0.5823925 |
| PLLATLRAIAVDVDAKPAQV | ASSQERDRGDEQY | 0.76658195 |
| APKFRTGGYMIDFWGVGYQVAK | ASSQERDRGDEQY | 0.8925605 |
| GAGGAGGAGGAAGGPGGTGGTG | ASSQERDRGDEQY | 0.7619863 |
| IGALGLLATDVAPAGSVAFGVA | ASSQERDRGDEQY | 0.8203783 |
| AGGAGGIGGNGGAGGAGGVLMG | ASSQERDRGDEQY | 0.87390697 |
| ADELIGLL | ASSQERDRGDEQY | 0.84989536 |
| ADRALKA | ASSQERDRGDEQY | 0.8856473 |
| AGAVGQVGGAGGA | ASSQERDRGDEQY | 0.8444486 |
| AGGGGGLGGAP | ASSQERDRGDEQY | 0.79686415 |
| AGGLLYGNGGAG | ASSQERDRGDEQY | 0.8060558 |
| AGLESIGL | ASSQERDRGDEQY | 0.82575333 |
| AGLKTGII | ASSQERDRGDEQY | 0.8388506 |
| ASLAALLL | ASSQERDRGDEQY | 0.8092503 |
| ASVINISSI | ASSQERDRGDEQY | 0.7138639 |
| ATPPGPA | ASSQERDRGDEQY | 0.88362813 |
| ATVASALV | ASSQERDRGDEQY | 0.8229289 |
| ATVASLIL | ASSQERDRGDEQY | 0.86494124 |
| AVNGPVLIRH | ASSQERDRGDEQY | 0.62946045 |
| DGAAGAVGI | ASSQERDRGDEQY | 0.6910494 |
| DPLLHGI | ASSQERDRGDEQY | 0.865247 |
| ELGAALGAL | ASSQERDRGDEQY | 0.16310187 |
| ELGRTQAV | ASSQERDRGDEQY | 0.30044216 |
| FVAGGYLS | ASSQERDRGDEQY | 0.8282322 |
| GASGGAGGAGGW | ASSQERDRGDEQY | 0.8751198 |
| GGAGAGGDGGAGGI | ASSQERDRGDEQY | 0.17532077 |
| GTGHNGSQPGMGGNG | ASSQERDRGDEQY | 0.7028427 |
| GYAAASASAS | ASSQERDRGDEQY | 0.5847908 |
| HQVDVAAVQQG | ASSQERDRGDEQY | 0.8447131 |
| IGAAIAHT | ASSQERDRGDEQY | 0.82671463 |
| IGMIEVAQL | ASSQERDRGDEQY | 0.55528927 |
| IRLAPEHTSAMD | ASSQERDRGDEQY | 0.8656558 |
| IYNVSVL | ASSQERDRGDEQY | 0.52815783 |
| KQGFLGM | ASSQERDRGDEQY | 0.60368055 |
| LESVITQL | ASSQERDRGDEQY | 0.85983026 |
| LGLGALVVV | ASSQERDRGDEQY | 0.8701911 |
| LIGNGGNGGSGGI | ASSQERDRGDEQY | 0.6535895 |
| LNAINAPTL | ASSQERDRGDEQY | 0.8682896 |
| LQLLNTGV | ASSQERDRGDEQY | 0.75077903 |
| LQTSIEF | ASSQERDRGDEQY | 0.8924457 |
| MSFAVSEL | ASSQERDRGDEQY | 0.7927204 |

| | | |
|---|---|---|
| NTGALGL | ASSQERDRGDEQY | 0.54335034 |
| PPGAGGAGGAGTT | ASSQERDRGDEQY | 0.8637323 |
| QLVGLGI | ASSQERDRGDEQY | 0.8407424 |
| RFLEQVHQL | ASSQERDRGDEQY | 0.8428678 |
| RGVGYVLR | ASSQERDRGDEQY | 0.857828 |
| SATGVFAGVI | ASSQERDRGDEQY | 0.75863475 |
| SLPAITLP | ASSQERDRGDEQY | 0.81584466 |
| STALAFTV | ASSQERDRGDEQY | 0.8808723 |
| SVLTVGLGK | ASSQERDRGDEQY | 0.8589734 |
| TALAGAGSGI | ASSQERDRGDEQY | 0.6634213 |
| TGLPFTGI | ASSQERDRGDEQY | 0.85153544 |
| VAEHVTV | ASSQERDRGDEQY | 0.80386275 |
| VDIVLNSL | ASSQERDRGDEQY | 0.7230152 |
| VLTGGGAVL | ASSQERDRGDEQY | 0.88555634 |
| VNVGLAYGI | ASSQERDRGDEQY | 0.8297925 |
| VSSAAVGAV | ASSQERDRGDEQY | 0.8690927 |
| VTSVYEAGV | ASSQERDRGDEQY | 0.8720716 |
| WVALDRGA | ASSQERDRGDEQY | 0.88003504 |
| YLVGSVVI | ASSQERDRGDEQY | 0.5643989 |
| GPPGTGKTLL | ASSQERDRGDEQY | 0.8146308 |
| GVGMTKF | ASSQERDRGDEQY | 0.81058216 |
| IGIGGSDL | ASSQERDRGDEQY | 0.8613571 |
| IGTLLAEL | ASSQERDRGDEQY | 0.7883625 |
| IYTSGTT | ASSQERDRGDEQY | 0.4900248 |
| LSFPAGV | ASSQERDRGDEQY | 0.8439271 |
| LYTSGTT | ASSQERDRGDEQY | 0.8769652 |
| VGAFPFGF | ASSQERDRGDEQY | 0.8173605 |
| VGAGTVEF | ASSQERDRGDEQY | 0.81051975 |
| VGKSTLLL | ASSQERDRGDEQY | 0.8947369 |
| AALAHAGGGIV | ASSQERDRGDEQY | 0.73802865 |
| AALVAVSLL | ASSQERDRGDEQY | 0.80400336 |
| AAVSRAVAAL | ASSQERDRGDEQY | 0.8677654 |
| AGGSSGIGGF | ASSQERDRGDEQY | 0.8573246 |
| AGLTVANANG | ASSQERDRGDEQY | 0.8180226 |
| AGLVGAVL | ASSQERDRGDEQY | 0.87507 |
| ALSQAFTV | ASSQERDRGDEQY | 0.63871974 |
| ALTFSVSL | ASSQERDRGDEQY | 0.8514576 |
| ALTGAALVAL | ASSQERDRGDEQY | 0.6395674 |
| ASFSIGAGVL | ASSQERDRGDEQY | 0.84137833 |
| DGSVYKI | ASSQERDRGDEQY | 0.66563594 |
| EQLAEIV | ASSQERDRGDEQY | 0.8441404 |
| FAVSELTVM | ASSQERDRGDEQY | 0.8622569 |
| GAAPLAAGL | ASSQERDRGDEQY | 0.8545358 |
| GADGGDGGSGGAGG | ASSQERDRGDEQY | 0.22663145 |
| GFGLDTGGAGG | ASSQERDRGDEQY | 0.8763267 |
| GLASGGLL | ASSQERDRGDEQY | 0.8586152 |
| GRIAGAAVLL | ASSQERDRGDEQY | 0.8961551 |
| GSAGSLNTSF | ASSQERDRGDEQY | 0.8078435 |
| GTGQIGI | ASSQERDRGDEQY | 0.8519311 |
| GTLIALIL | ASSQERDRGDEQY | 0.77138114 |
| IAATMGFVVG | ASSQERDRGDEQY | 0.8033704 |

| | | |
|---|---|---|
| IFAQALL | ASSQERDRGDEQY | 0.67645514 |
| IGAAVALGF | ASSQERDRGDEQY | 0.74496406 |
| IGLTPLAL | ASSQERDRGDEQY | 0.7708007 |
| ILGSNTSTL | ASSQERDRGDEQY | 0.7496302 |
| IVFSLTVL | ASSQERDRGDEQY | 0.845914 |
| LASKATGF | ASSQERDRGDEQY | 0.852396 |
| LASVALIL | ASSQERDRGDEQY | 0.8631875 |
| LFAVGLTL | ASSQERDRGDEQY | 0.8825611 |
| LFNGGIGTYI | ASSQERDRGDEQY | 0.8822088 |
| LTLGAGII | ASSQERDRGDEQY | 0.8775852 |
| LVAAALGDI | ASSQERDRGDEQY | 0.74236834 |
| NTAFVQLGI | ASSQERDRGDEQY | 0.7773473 |
| QLFSGAEL | ASSQERDRGDEQY | 0.683045 |
| SAATFITL | ASSQERDRGDEQY | 0.79335284 |
| SNYGLANLGV | ASSQERDRGDEQY | 0.7765302 |
| SYGGAYAVM | ASSQERDRGDEQY | 0.85376537 |
| TTGVGGLTLG | ASSQERDRGDEQY | 0.11925427 |
| TTQTIYI | ASSQERDRGDEQY | 0.790973 |
| VAAGAGLGAVI | ASSQERDRGDEQY | 0.8805176 |
| VAAVVTLL | ASSQERDRGDEQY | 0.5946305 |
| VGFLGGLI | ASSQERDRGDEQY | 0.8514209 |
| VGVIPILL | ASSQERDRGDEQY | 0.7835738 |
| VIFASSLI | ASSQERDRGDEQY | 0.72438866 |
| VKGGDGGAAAG | ASSQERDRGDEQY | 0.8682139 |
| VTKTFTNI | ASSQERDRGDEQY | 0.7086739 |
| VVAAAILL | ASSQERDRGDEQY | 0.8525607 |
| YAALDVEL | ASSQERDRGDEQY | 0.87920964 |
| LASVALLL | ASSQERDRGDEQY | 0.83039427 |
| LDAVHRV | ASSQERDRGDEQY | 0.79308 |
| QGLSSSIF | ASSQERDRGDEQY | 0.7980811 |
| RGTFANIRL | ASSQERDRGDEQY | 0.50176066 |
| RQLWWGHRI | ASSQERDRGDEQY | 0.81353426 |
| YGPPGTGKTLL | ASSQERDRGDEQY | 0.13300095 |
| AALVGAPANVA | ASSQERDRGDEQY | 0.81256366 |
| AATFRRTE | ASSQERDRGDEQY | 0.70321196 |
| AGALGGGI | ASSQERDRGDEQY | 0.84146595 |
| AGGAGGAGGIGAG | ASSQERDRGDEQY | 0.816997 |
| AGSGALGL | ASSQERDRGDEQY | 0.8355472 |
| AQIADQI | ASSQERDRGDEQY | 0.6869112 |
| ASAVAVTYL | ASSQERDRGDEQY | 0.7797862 |
| ASTVLGGTGG | ASSQERDRGDEQY | 0.6348198 |
| AVGGEPAK | ASSQERDRGDEQY | 0.618616 |
| CEQGTFGGTC | ASSQERDRGDEQY | 0.8304281 |
| DEGKAPAG | ASSQERDRGDEQY | 0.81911105 |
| DTVVFPAL | ASSQERDRGDEQY | 0.75355995 |
| FEGITMAG | ASSQERDRGDEQY | 0.8747908 |
| FPLVVQD | ASSQERDRGDEQY | 0.82958275 |
| GAFTTPQ | ASSQERDRGDEQY | 0.54365623 |
| GFGGFGTAGGVGGA | ASSQERDRGDEQY | 0.7799746 |
| GFLAQFL | ASSQERDRGDEQY | 0.8196453 |
| GGAGGAGGNAGVA | ASSQERDRGDEQY | 0.80478036 |

| | | |
|---|---|---|
| GGVGTAGAGGAGGAGG | ASSQERDRGDEQY | 0.115161866 |
| GNGGGGDAGNAGSGGN | ASSQERDRGDEQY | 0.8235666 |
| GTGGAGGAGNDAGS | ASSQERDRGDEQY | 0.719056 |
| GTTNTGWFNS | ASSQERDRGDEQY | 0.6548878 |
| IFMGPLFTL | ASSQERDRGDEQY | 0.73655385 |
| ILAHAIEAI | ASSQERDRGDEQY | 0.8685256 |
| IRSAIREAL | ASSQERDRGDEQY | 0.59350544 |
| ITFAGGG | ASSQERDRGDEQY | 0.8196319 |
| ITGMSGHQEV | ASSQERDRGDEQY | 0.8552081 |
| IVAGLACTL | ASSQERDRGDEQY | 0.8653512 |
| IVGGGLAGA | ASSQERDRGDEQY | 0.8822621 |
| IVTGVLNI | ASSQERDRGDEQY | 0.8454207 |
| LAQQGCI | ASSQERDRGDEQY | 0.7815088 |
| LCLHVEN | ASSQERDRGDEQY | 0.84019756 |
| LDAISDGS | ASSQERDRGDEQY | 0.8028722 |
| LGGFAAML | ASSQERDRGDEQY | 0.44265044 |
| LGGKNPMI | ASSQERDRGDEQY | 0.37372845 |
| LGPKGVAAIL | ASSQERDRGDEQY | 0.81420135 |
| LMVGVSI | ASSQERDRGDEQY | 0.35082027 |
| LVGAAFVL | ASSQERDRGDEQY | 0.83474994 |
| LVLAVAPL | ASSQERDRGDEQY | 0.882262 |
| MAAIAVIDAA | ASSQERDRGDEQY | 0.7790552 |
| NNGINAELQ | ASSQERDRGDEQY | 0.581789 |
| RGEPLVLVHG | ASSQERDRGDEQY | 0.76938605 |
| RGGAGGAAGLI | ASSQERDRGDEQY | 0.80819684 |
| RSYRMHPAV | ASSQERDRGDEQY | 0.8609434 |
| SAAHLLAGQLA | ASSQERDRGDEQY | 0.85231334 |
| SAAVGSY | ASSQERDRGDEQY | 0.8741005 |
| SPHAVLHRI | ASSQERDRGDEQY | 0.8667096 |
| SQIAKVL | ASSQERDRGDEQY | 0.7499763 |
| TALGHLVP | ASSQERDRGDEQY | 0.860076 |
| TAPTTSGAA | ASSQERDRGDEQY | 0.786624 |
| TDGDDGNGGSPGT | ASSQERDRGDEQY | 0.73111475 |
| TDGLGEN | ASSQERDRGDEQY | 0.7997737 |
| TGGPPLPGQA | ASSQERDRGDEQY | 0.81124854 |
| TIPAITI | ASSQERDRGDEQY | 0.79465365 |
| TIPGFTI | ASSQERDRGDEQY | 0.7374773 |
| TSIASGALA | ASSQERDRGDEQY | 0.5554261 |
| TVTLPGAAI | ASSQERDRGDEQY | 0.83495885 |
| VASGLSAALP | ASSQERDRGDEQY | 0.8016869 |
| VGAGVIGI | ASSQERDRGDEQY | 0.5063995 |
| VGPKRFGV | ASSQERDRGDEQY | 0.7299158 |
| VNVAFPD | ASSQERDRGDEQY | 0.8940247 |
| VSMTNSGVS | ASSQERDRGDEQY | 0.8856627 |
| VTDVLGDLSV | ASSQERDRGDEQY | 0.7216592 |
| VVVVGAGFAGL | ASSQERDRGDEQY | 0.7350846 |
| VYTVGTVV | ASSQERDRGDEQY | 0.7899624 |
| WTTFSGP | ASSQERDRGDEQY | 0.4644261 |
| YDELVEAF | ASSQERDRGDEQY | 0.85164654 |
| YGTGGAGGNGG | ASSQERDRGDEQY | 0.80632186 |
| YTNVMAV | ASSQERDRGDEQY | 0.7874409 |

| | | |
|---|---|---|
| FLEQVHQL | ASSQERDRGDEQY | 0.7032512 |
| GLSLLAVAV | ASSQERDRGDEQY | 0.806491 |
| LEQVHQL | ASSQERDRGDEQY | 0.712489 |
| LNAAGAI | ASSQERDRGDEQY | 0.8660308 |
| PAVHASL | ASSQERDRGDEQY | 0.17402333 |
| TELEDDA | ASSQERDRGDEQY | 0.82423043 |
| TPPGPPN | ASSQERDRGDEQY | 0.6807022 |
| TPPPPPPP | ASSQERDRGDEQY | 0.7456877 |
| TQSPALI | ASSFRDWGVNRY | 0.46053112 |
| EFAAAFPQ | ASSFRDWGVNRY | 0.79124826 |
| LLSGNLDVL | ASSFRDWGVNRY | 0.8604586 |
| GPANLGNYNI | ASSFRDWGVNRY | 0.87161183 |
| TLGPTLP | ASSFRDWGVNRY | 0.5691523 |
| VYAYSSCCK | ASSFRDWGVNRY | 0.8532175 |
| APAPAPAPNI | ASSFRDWGVNRY | 0.88841844 |
| PGGPVPIP | ASSFRDWGVNRY | 0.7544212 |
| FLTVALEP | ASSFRDWGVNRY | 0.42104578 |
| SQYFGANL | ASSFRDWGVNRY | 0.85377896 |
| GGVKPAA | ASSFRDWGVNRY | 0.7039019 |
| GGTGGAGGAGGW | ASSFRDWGVNRY | 0.854228 |
| GVGGVAGLGGAGG | ASSFRDWGVNRY | 0.82645243 |
| VLLVLATTI | ASSFRDWGVNRY | 0.65690964 |
| IGIFAAML | ASSFRDWGVNRY | 0.87357736 |
| AVGDLID | ASSFRDWGVNRY | 0.87343407 |
| NISPLPL | ASSFRDWGVNRY | 0.71100676 |
| VTAVFVAI | ASSFRDWGVNRY | 0.6877197 |
| SGDDNDAAL | ASSFRDWGVNRY | 0.6947155 |
| AVAALAHS | ASSFRDWGVNRY | 0.3119406 |
| GGVTVMATP | ASSFRDWGVNRY | 0.8552836 |
| AGLAANVRD | ASSFRDWGVNRY | 0.6895295 |
| TNRPDLIDPALLRPG | ASSFRDWGVNRY | 0.7404696 |
| LANLGVTAL | ASSFRDWGVNRY | 0.84280527 |
| TGERIEL | ASSFRDWGVNRY | 0.7163379 |
| NGEELAA | ASSFRDWGVNRY | 0.78603005 |
| VLSLVWG | ASSFRDWGVNRY | 0.784243 |
| IGYALWL | ASSFRDWGVNRY | 0.77636176 |
| LSLLLAPL | ASSFRDWGVNRY | 0.82654077 |
| TLDEALK | ASSFRDWGVNRY | 0.7942443 |
| TDGAGTAEGG | ASSFRDWGVNRY | 0.7866491 |
| NAPGGAGGNGG | ASSFRDWGVNRY | 0.88324714 |
| VSVFMGMGGNF | ASSFRDWGVNRY | 0.597626 |
| LASYLAAL | ASSFRDWGVNRY | 0.7152975 |
| LVGGTPAL | ASSFRDWGVNRY | 0.8490823 |
| VRNADVL | ASSFRDWGVNRY | 0.5634552 |
| TCSSPFLVSAM | ASSFRDWGVNRY | 0.85006464 |
| VHITDAE | ASSFRDWGVNRY | 0.80182683 |
| EYAAPSF | ASSFRDWGVNRY | 0.85016024 |
| VVDAFHGGL | ASSFRDWGVNRY | 0.759781 |
| ASVVDPQ | ASSFRDWGVNRY | 0.884871 |
| SYNEFGGLS | ASSFRDWGVNRY | 0.87664986 |
| DPDGSVY | ASSFRDWGVNRY | 0.7998885 |

| | | |
|---|---|---|
| MVSTIHATE | ASSFRDWGVNRY | 0.772982 |
| LAAGVADPV | ASSFRDWGVNRY | 0.6548045 |
| TKEQIAA | ASSFRDWGVNRY | 0.8542329 |
| SLASPAAL | ASSFRDWGVNRY | 0.8608328 |
| IGLVFYGV | ASSFRDWGVNRY | 0.6780482 |
| FLLDASNL | ASSFRDWGVNRY | 0.8190341 |
| YQVGNFP | ASSFRDWGVNRY | 0.87638366 |
| ESLLSQST | ASSFRDWGVNRY | 0.8706962 |
| LFTVGFAAVG | ASSFRDWGVNRY | 0.42880157 |
| ATPPTPT | ASSFRDWGVNRY | 0.7835071 |
| TDLATLG | ASSFRDWGVNRY | 0.42433307 |
| SAGGTGR | ASSFRDWGVNRY | 0.72263217 |
| GFSNSGS | ASSFRDWGVNRY | 0.6113388 |
| GGNGGSGGTSV | ASSFRDWGVNRY | 0.8441577 |
| GTSVATGGAGN | ASSFRDWGVNRY | 0.77024204 |
| NGGNGGSGGTS | ASSFRDWGVNRY | 0.83185506 |
| AVVKFPH | ASSFRDWGVNRY | 0.8463028 |
| TPPPPPP | ASSFRDWGVNRY | 0.88938427 |
| AAPAAAAAPA | ASSFRDWGVNRY | 0.7617219 |
| IALNEDL | ASSFRDWGVNRY | 0.80498934 |
| IVNVSSM | ASSFRDWGVNRY | 0.5386487 |
| TGIPPAPR | ASSFRDWGVNRY | 0.36894807 |
| FDLGGGTF | ASSFRDWGVNRY | 0.6766173 |
| FDLGGGTFDVS | ASSFRDWGVNRY | 0.77646196 |
| NEPTAAA | ASSFRDWGVNRY | 0.24121277 |
| NEPTAAALA | ASSFRDWGVNRY | 0.87949574 |
| GSIINISSI | ASSFRDWGVNRY | 0.8430208 |
| VAPGNAGTA | ASSFRDWGVNRY | 0.8698669 |
| TGGGASLE | ASSFRDWGVNRY | 0.7455779 |
| VAVYNHYKRI | ASSFRDWGVNRY | 0.7597396 |
| TDPAPATT | ASSFRDWGVNRY | 0.7668369 |
| TNRPDILDPALLRPG | ASSFRDWGVNRY | 0.8540453 |
| LTTTTTS | ASSFRDWGVNRY | 0.72167164 |
| PTGSGKTL | ASSFRDWGVNRY | 0.49055105 |
| TGSGKTL | ASSFRDWGVNRY | 0.7132119 |
| TIGHVDHGK | ASSFRDWGVNRY | 0.6298961 |
| TIGHVDHGKTT | ASSFRDWGVNRY | 0.8775928 |
| GHVDHGKTT | ASSFRDWGVNRY | 0.8655406 |
| AQFEHTLL | ASSFRDWGVNRY | 0.8593204 |
| IVNQDEL | ASSFRDWGVNRY | 0.87342685 |
| TNGSQFF | ASSFRDWGVNRY | 0.32171118 |
| TGKGIVE | ASSFRDWGVNRY | 0.73988265 |
| LISNASDA | ASSFRDWGVNRY | 0.89113706 |
| AELRTLGELTKIAKAHGAQ | ASSFRDWGVNRY | 0.84822243 |
| APNAVAAAEPSW | ASSFRDWGVNRY | 0.78987813 |
| APQVAGAATAVL | ASSFRDWGVNRY | 0.8835213 |
| ARRDPAAAAAILPTDA | ASSFRDWGVNRY | 0.709146 |
| DAAARDVLAEAGLAEYF | ASSFRDWGVNRY | 0.80160296 |
| DDQIVLSAKANLAKVEQ | ASSFRDWGVNRY | 0.81892544 |
| DELLPDTGEDSDEDGH | ASSFRDWGVNRY | 0.8439494 |
| ELPEAPANRATVPQD | ASSFRDWGVNRY | 0.80950034 |

| | | |
|---|---|---|
| FGQSKIERVTGIG | ASSFRDWGVNRY | 0.66295755 |
| GAGGTGSTAGGAGGAGGAGGLY | ASSFRDWGVNRY | 0.8461746 |
| GHGGTGAAGGNGGTGGAGGSGI | ASSFRDWGVNRY | 0.8850197 |
| GNGGNGGAGGEGGPGGAGGAGA | ASSFRDWGVNRY | 0.7981534 |
| GSTAGNGGNGGNATNGTIAGQP | ASSFRDWGVNRY | 0.8678033 |
| GTGGDGGNGGNGGTGG | ASSFRDWGVNRY | 0.8425032 |
| GVRRPAGIPADAPV | ASSFRDWGVNRY | 0.8278614 |
| GYRQVTDAHLLTLARR | ASSFRDWGVNRY | 0.74325204 |
| IAKMGQDGHDRG | ASSFRDWGVNRY | 0.7705679 |
| KAEPSIEIQYT | ASSFRDWGVNRY | 0.77711797 |
| KPVRIRMAADEDTVAVT | ASSFRDWGVNRY | 0.69509673 |
| MPPRIAGMRLLVIKPEP | ASSFRDWGVNRY | 0.79962456 |
| MKFYAVALFRSRGVPPTR | ASSFRDWGVNRY | 0.8028308 |
| NAGGFGILGAVAHSP | ASSFRDWGVNRY | 0.89886904 |
| NEGREITLAMAEM | ASSFRDWGVNRY | 0.84801495 |
| NELVEWSHPLGHRWRWE | ASSFRDWGVNRY | 0.8486231 |
| NLGAGGTGGNGGNPG | ASSFRDWGVNRY | 0.71343184 |
| NTEWPKWETVMAADPVD | ASSFRDWGVNRY | 0.82420087 |
| PASAAPAETQP | ASSFRDWGVNRY | 0.84226644 |
| QQSSDGKNGVRAAEITGE | ASSFRDWGVNRY | 0.5978545 |
| RAPDAVTTSWWKE | ASSFRDWGVNRY | 0.8829974 |
| SGFGNVGAHTSGWFNQ | ASSFRDWGVNRY | 0.7279267 |
| SGVAGAVGQAASVGGLKV | ASSFRDWGVNRY | 0.7243231 |
| SNGGAGTGSAGGNGGT | ASSFRDWGVNRY | 0.8643768 |
| TGGAGGEGGAAG | ASSFRDWGVNRY | 0.86753464 |
| TGGDGGAAGAGGAPGHG | ASSFRDWGVNRY | 0.789235 |
| TMPPLLAAIANAGRHGV | ASSFRDWGVNRY | 0.74741346 |
| TSSTAGNSGGA | ASSFRDWGVNRY | 0.7415822 |
| VPATVPVGILC | ASSFRDWGVNRY | 0.8734845 |
| VQRPDSGGVVLTGRIS | ASSFRDWGVNRY | 0.8850313 |
| WTPNPVPATKGVPAGLVQAT | ASSFRDWGVNRY | 0.7458925 |
| YALKRALAKFSDRFDVVII | ASSFRDWGVNRY | 0.8819945 |
| YLIGSVIV | ASSFRDWGVNRY | 0.83026904 |
| YGQLVAVID | ASSFRDWGVNRY | 0.53053397 |
| VTVGGMTLPA | ASSFRDWGVNRY | 0.82282615 |
| VLALPAV | ASSFRDWGVNRY | 0.8532983 |
| VIPPTLNYET | ASSFRDWGVNRY | 0.849688 |
| VGMGGMHPSSG | ASSFRDWGVNRY | 0.87572384 |
| VGLAGGAATA | ASSFRDWGVNRY | 0.7955137 |
| VGGAGGAGGSGP | ASSFRDWGVNRY | 0.7426721 |
| VFAAFMV | ASSFRDWGVNRY | 0.88713336 |
| TLYDKLR | ASSFRDWGVNRY | 0.8860061 |
| TLAVFTVL | ASSFRDWGVNRY | 0.87215614 |
| TGSVQPARV | ASSFRDWGVNRY | 0.7861841 |
| TGGGAGSGGT | ASSFRDWGVNRY | 0.767308 |
| SHLETSL | ASSFRDWGVNRY | 0.81943953 |
| SFTTGMT | ASSFRDWGVNRY | 0.86623085 |
| QVEVAKRMGVRQPTVS | ASSFRDWGVNRY | 0.6722727 |
| QGSGGPALAM | ASSFRDWGVNRY | 0.8237239 |
| MLAPSNP | ASSFRDWGVNRY | 0.8390598 |
| LVNNAAI | ASSFRDWGVNRY | 0.52572995 |

| | | |
|---|---|---|
| LTGEINTLA | ASSFRDWGVNRY | 0.8223932 |
| LSIFGPV | ASSFRDWGVNRY | 0.864339 |
| LLALLANL | ASSFRDWGVNRY | 0.7958004 |
| LISNASDALDKLR | ASSFRDWGVNRY | 0.76265574 |
| LINEAVP | ASSFRDWGVNRY | 0.8738352 |
| LGYAFSGL | ASSFRDWGVNRY | 0.90011585 |
| LDFVAVAL | ASSFRDWGVNRY | 0.8842813 |
| LAQPGAE | ASSFRDWGVNRY | 0.50477856 |
| ISNLGAVTGF | ASSFRDWGVNRY | 0.3727029 |
| HQEDPGRI | ASSFRDWGVNRY | 0.6769279 |
| GTPGEPTP | ASSFRDWGVNRY | 0.8397732 |
| GFFNSGS | ASSFRDWGVNRY | 0.7709962 |
| GAGGGGLTAGAN | ASSFRDWGVNRY | 0.76891214 |
| FVDALVNM | ASSFRDWGVNRY | 0.8091552 |
| FLTVAPEL | ASSFRDWGVNRY | 0.8825401 |
| FETLTAL | ASSFRDWGVNRY | 0.727082 |
| DVINPIEL | ASSFRDWGVNRY | 0.5780929 |
| APVSGAPV | ASSFRDWGVNRY | 0.7983979 |
| ALVGGLAW | ASSFRDWGVNRY | 0.86891913 |
| ALAAAAATAI | ASSFRDWGVNRY | 0.6911787 |
| AFGLGLLL | ASSFRDWGVNRY | 0.8107024 |
| AFFAAASL | ASSFRDWGVNRY | 0.6907903 |
| AAAALPAFSPP | ASSFRDWGVNRY | 0.65217185 |
| VVLAPAL | ASSFRDWGVNRY | 0.81977075 |
| TIGHVDHGKTTL | ASSFRDWGVNRY | 0.73982656 |
| LISNASDALDK | ASSFRDWGVNRY | 0.86722827 |
| LISNASDAL | ASSFRDWGVNRY | 0.73687947 |
| ISNASDALDK | ASSFRDWGVNRY | 0.8608259 |
| NQAETLVYQTE | ASSFRDWGVNRY | 0.830351 |
| TGGTGGTAGAG | ASSFRDWGVNRY | 0.85189164 |
| TGSTAGGAGGA | ASSFRDWGVNRY | 0.78126365 |
| DPLAVPVAAGPV | ASSFRDWGVNRY | 0.75261676 |
| GGTGGTPGNGGN | ASSFRDWGVNRY | 0.6583473 |
| PAGLAGVDPATL | ASSFRDWGVNRY | 0.41609848 |
| GNNLVVEAPHAGASV | ASSFRDWGVNRY | 0.8588569 |
| KVTRIMSASTLEQPA | ASSFRDWGVNRY | 0.2416683 |
| LGFALPIALPRRTVV | ASSFRDWGVNRY | 0.87732756 |
| TRLGNLTARKLVVLV | ASSFRDWGVNRY | 0.8679839 |
| WSQQRRLIYFPSAGP | ASSFRDWGVNRY | 0.68901527 |
| GPNGNPDPMAAAVDIR | ASSFRDWGVNRY | 0.8504889 |
| TQRCVSVADLRAAASR | ASSFRDWGVNRY | 0.6788248 |
| QFVLDVAEHPVEVLTETV | ASSFRDWGVNRY | 0.67425275 |
| GGAGGTGGAGGNGGTQAGDG | ASSFRDWGVNRY | 0.8545921 |
| PLLATLRAIAVDVDAKPAQV | ASSFRDWGVNRY | 0.8550234 |
| APKFRTGGYMIDFWGVGYQVAK | ASSFRDWGVNRY | 0.7054686 |
| GAGGAGGAGGAAGGPGGTGGTG | ASSFRDWGVNRY | 0.8377272 |
| IGALGLLATDVAPAGSVAFGVA | ASSFRDWGVNRY | 0.8370337 |
| AGGAGGIGGNGGAGGAGGVLMG | ASSFRDWGVNRY | 0.7208872 |
| ADELIGLL | ASSFRDWGVNRY | 0.7988358 |
| ADRALKA | ASSFRDWGVNRY | 0.8698444 |
| AGAVGQVGGAGGA | ASSFRDWGVNRY | 0.8236494 |

| | | |
|---|---|---|
| AGGGGGLGGAP | ASSFRDWGVNRY | 0.75940716 |
| AGGLLYGNGGAG | ASSFRDWGVNRY | 0.84409 |
| AGLESIGL | ASSFRDWGVNRY | 0.685741 |
| AGLKTGII | ASSFRDWGVNRY | 0.8534217 |
| ASLAALLL | ASSFRDWGVNRY | 0.81070966 |
| ASVINISSI | ASSFRDWGVNRY | 0.7980422 |
| ATPPGPA | ASSFRDWGVNRY | 0.8540946 |
| ATVASALV | ASSFRDWGVNRY | 0.5443738 |
| ATVASLIL | ASSFRDWGVNRY | 0.7003137 |
| AVNGPVLIRH | ASSFRDWGVNRY | 0.775256 |
| DGAAGAVGI | ASSFRDWGVNRY | 0.8409684 |
| DPLLHGI | ASSFRDWGVNRY | 0.87288254 |
| ELGAALGAL | ASSFRDWGVNRY | 0.71516603 |
| ELGRTQAV | ASSFRDWGVNRY | 0.8413793 |
| FVAGGYLS | ASSFRDWGVNRY | 0.7705518 |
| GASGGAGGAGGW | ASSFRDWGVNRY | 0.70953745 |
| GGAGAGGDGGAGGI | ASSFRDWGVNRY | 0.8300051 |
| GTGHNGSQPGMGGNG | ASSFRDWGVNRY | 0.75883293 |
| GYAAASASAS | ASSFRDWGVNRY | 0.7454742 |
| HQVDVAAVQQG | ASSFRDWGVNRY | 0.8371849 |
| IGAAIAHT | ASSFRDWGVNRY | 0.7326013 |
| IGMIEVAQL | ASSFRDWGVNRY | 0.8380376 |
| IRLAPEHTSAMD | ASSFRDWGVNRY | 0.86815953 |
| IYNVSVL | ASSFRDWGVNRY | 0.7671677 |
| KQGFLGM | ASSFRDWGVNRY | 0.8601657 |
| LESVITQL | ASSFRDWGVNRY | 0.84771276 |
| LGLGALVVV | ASSFRDWGVNRY | 0.82268673 |
| LIGNGGNGGSGGI | ASSFRDWGVNRY | 0.79165447 |
| LNAINAPTL | ASSFRDWGVNRY | 0.73780084 |
| LQLLNTGV | ASSFRDWGVNRY | 0.8837576 |
| LQTSIEF | ASSFRDWGVNRY | 0.8647001 |
| MSFAVSEL | ASSFRDWGVNRY | 0.8441746 |
| NTGALGL | ASSFRDWGVNRY | 0.8349719 |
| PPGAGGAGGAGTT | ASSFRDWGVNRY | 0.8294718 |
| QLVGLGI | ASSFRDWGVNRY | 0.8630972 |
| RFLEQVHQL | ASSFRDWGVNRY | 0.7647451 |
| RGVGYVLR | ASSFRDWGVNRY | 0.5779211 |
| SATGVFAGVI | ASSFRDWGVNRY | 0.7732033 |
| SLPAITLP | ASSFRDWGVNRY | 0.79618996 |
| STALAFTV | ASSFRDWGVNRY | 0.77914196 |
| SVLTVGLGK | ASSFRDWGVNRY | 0.5090016 |
| TALAGAGSGI | ASSFRDWGVNRY | 0.8760286 |
| TGLPFTGI | ASSFRDWGVNRY | 0.89968336 |
| VAEHVTV | ASSFRDWGVNRY | 0.7152882 |
| VDIVLNSL | ASSFRDWGVNRY | 0.8934958 |
| VLTGGGAVL | ASSFRDWGVNRY | 0.5962563 |
| VNVGLAYGI | ASSFRDWGVNRY | 0.8490361 |
| VSSAAVGAV | ASSFRDWGVNRY | 0.87251693 |
| VTSVYEAGV | ASSFRDWGVNRY | 0.6992489 |
| WVALDRGA | ASSFRDWGVNRY | 0.84992886 |
| YLVGSVVI | ASSFRDWGVNRY | 0.7203259 |

| | | |
|---|---|---|
| GPPGTGKTLL | ASSFRDWGVNRY | 0.79054695 |
| GVGMTKF | ASSFRDWGVNRY | 0.8755467 |
| IGIGGSDL | ASSFRDWGVNRY | 0.76710385 |
| IGTLLAEL | ASSFRDWGVNRY | 0.7939849 |
| IYTSGTT | ASSFRDWGVNRY | 0.8127824 |
| LSFPAGV | ASSFRDWGVNRY | 0.7246555 |
| LYTSGTT | ASSFRDWGVNRY | 0.82963943 |
| VGAFPFGF | ASSFRDWGVNRY | 0.8693007 |
| VGAGTVEF | ASSFRDWGVNRY | 0.70286936 |
| VGKSTLLL | ASSFRDWGVNRY | 0.88419807 |
| AALAHAGGGIV | ASSFRDWGVNRY | 0.69326794 |
| AALVAVSLL | ASSFRDWGVNRY | 0.89291584 |
| AAVSRAVAAL | ASSFRDWGVNRY | 0.8568804 |
| AGGSSGIGGF | ASSFRDWGVNRY | 0.82890177 |
| AGLTVANANG | ASSFRDWGVNRY | 0.76745063 |
| AGLVGAVL | ASSFRDWGVNRY | 0.88073105 |
| ALSQAFTV | ASSFRDWGVNRY | 0.7951687 |
| ALTFSVSL | ASSFRDWGVNRY | 0.8234987 |
| ALTGAALVAL | ASSFRDWGVNRY | 0.8851039 |
| ASFSIGAGVL | ASSFRDWGVNRY | 0.45560756 |
| DGSVYKI | ASSFRDWGVNRY | 0.6373959 |
| EQLAEIV | ASSFRDWGVNRY | 0.5973495 |
| FAVSELTVM | ASSFRDWGVNRY | 0.60116315 |
| GAAPLAAGL | ASSFRDWGVNRY | 0.51879245 |
| GADGGDGGSGGAGG | ASSFRDWGVNRY | 0.4559146 |
| GFGLDTGGAGG | ASSFRDWGVNRY | 0.5632825 |
| GLASGGLL | ASSFRDWGVNRY | 0.50871164 |
| GRIAGAAVLL | ASSFRDWGVNRY | 0.6322621 |
| GSAGSLNTSF | ASSFRDWGVNRY | 0.61855733 |
| GTGQIGI | ASSFRDWGVNRY | 0.50688195 |
| GTLIALIL | ASSFRDWGVNRY | 0.6270863 |
| IAATMGFVVG | ASSFRDWGVNRY | 0.63621044 |
| IFAQALL | ASSFRDWGVNRY | 0.38926822 |
| IGAAVALGF | ASSFRDWGVNRY | 0.5582115 |
| IGLTPLAL | ASSFRDWGVNRY | 0.5037343 |
| ILGSNTSTL | ASSFRDWGVNRY | 0.5297464 |
| IVFSLTVL | ASSFRDWGVNRY | 0.33563817 |
| LASKATGF | ASSFRDWGVNRY | 0.61373055 |
| LASVALIL | ASSFRDWGVNRY | 0.4961992 |
| LFAVGLTL | ASSFRDWGVNRY | 0.32476267 |
| LFNGGIGTYI | ASSFRDWGVNRY | 0.5780159 |
| LTLGAGII | ASSFRDWGVNRY | 0.5359296 |
| LVAAALGDI | ASSFRDWGVNRY | 0.57928616 |
| NTAFVQLGI | ASSFRDWGVNRY | 0.6238979 |
| QLFSGAEL | ASSFRDWGVNRY | 0.5432781 |
| SAATFITL | ASSFRDWGVNRY | 0.54627717 |
| SNYGLANLGV | ASSFRDWGVNRY | 0.59154844 |
| SYGGAYAVM | ASSFRDWGVNRY | 0.63835734 |
| TTGVGGLTLG | ASSFRDWGVNRY | 0.52766323 |
| TTQTIYI | ASSFRDWGVNRY | 0.5665934 |
| VAAGAGLGAVI | ASSFRDWGVNRY | 0.649892 |

| | | |
|---|---|---|
| VAAVVTLL | ASSFRDWGVNRY | 0.57181865 |
| VGFLGGLI | ASSFRDWGVNRY | 0.64844936 |
| VGVIPILL | ASSFRDWGVNRY | 0.65433216 |
| VIFASSLI | ASSFRDWGVNRY | 0.4993544 |
| VKGGDGGAAAG | ASSFRDWGVNRY | 0.6297201 |
| VTKTFTNI | ASSFRDWGVNRY | 0.642606 |
| VVAAAILL | ASSFRDWGVNRY | 0.53853244 |
| YAALDVEL | ASSFRDWGVNRY | 0.56150186 |
| LASVALLL | ASSFRDWGVNRY | 0.46092412 |
| LDAVHRV | ASSFRDWGVNRY | 0.5604876 |
| QGLSSSIF | ASSFRDWGVNRY | 0.45046648 |
| RGTFANIRL | ASSFRDWGVNRY | 0.56953126 |
| RQLWWGHRI | ASSFRDWGVNRY | 0.6450072 |
| YGPPGTGKTLL | ASSFRDWGVNRY | 0.53971404 |
| AALVGAPANVA | ASSFRDWGVNRY | 0.41330865 |
| AATFRRTE | ASSFRDWGVNRY | 0.42345548 |
| AGALGGGI | ASSFRDWGVNRY | 0.5762061 |
| AGGAGGAGGIGAG | ASSFRDWGVNRY | 0.48195985 |
| AGSGALGL | ASSFRDWGVNRY | 0.3963111 |
| AQIADQI | ASSFRDWGVNRY | 0.4927097 |
| ASAVAVTYL | ASSFRDWGVNRY | 0.63234866 |
| ASTVLGGTGG | ASSFRDWGVNRY | 0.6379768 |
| AVGGEPAK | ASSFRDWGVNRY | 0.44644636 |
| CEQGTFGGTC | ASSFRDWGVNRY | 0.6457769 |
| DEGKAPAG | ASSFRDWGVNRY | 0.60637546 |
| DTVVFPAL | ASSFRDWGVNRY | 0.64949316 |
| FEGITMAG | ASSFRDWGVNRY | 0.5869906 |
| FPLVVQD | ASSFRDWGVNRY | 0.47740462 |
| GAFTTPQ | ASSFRDWGVNRY | 0.6137884 |
| GFGGFGTAGGVGGA | ASSFRDWGVNRY | 0.5255801 |
| GFLAQFL | ASSFRDWGVNRY | 0.57155997 |
| GGAGGAGGNAGVA | ASSFRDWGVNRY | 0.61518204 |
| GGVGTAGAGGAGGAGG | ASSFRDWGVNRY | 0.6050662 |
| GNGGGGDAGNAGSGGN | ASSFRDWGVNRY | 0.47959888 |
| GTGGAGGAGNDAGS | ASSFRDWGVNRY | 0.52940583 |
| GTTNTGWFNS | ASSFRDWGVNRY | 0.6378335 |
| IFMGPLFTL | ASSFRDWGVNRY | 0.44301283 |
| ILAHAIEAI | ASSFRDWGVNRY | 0.51221997 |
| IRSAIREAL | ASSFRDWGVNRY | 0.5334519 |
| ITFAGGG | ASSFRDWGVNRY | 0.5722825 |
| ITGMSGHQEV | ASSFRDWGVNRY | 0.6408421 |
| IVAGLACTL | ASSFRDWGVNRY | 0.6074326 |
| IVGGGLAGA | ASSFRDWGVNRY | 0.53015965 |
| IVTGVLNI | ASSFRDWGVNRY | 0.6079187 |
| LAQQGCI | ASSFRDWGVNRY | 0.62065554 |
| LCLHVEN | ASSFRDWGVNRY | 0.57676685 |
| LDAISDGS | ASSFRDWGVNRY | 0.5850564 |
| LGGFAAML | ASSFRDWGVNRY | 0.48874578 |
| LGGKNPMI | ASSFRDWGVNRY | 0.63193136 |
| LGPKGVAAIL | ASSFRDWGVNRY | 0.6115285 |
| LMVGVSI | ASSFRDWGVNRY | 0.63495517 |

| | | |
|---|---|---|
| LVGAAFVL | ASSFRDWGVNRY | 0.63702905 |
| LVLAVAPL | ASSFRDWGVNRY | 0.6207154 |
| MAAIAVIDAA | ASSFRDWGVNRY | 0.5788316 |
| NNGINAELQ | ASSFRDWGVNRY | 0.4101186 |
| RGEPLVLVHG | ASSFRDWGVNRY | 0.47187415 |
| RGGAGGAAGLI | ASSFRDWGVNRY | 0.5589632 |
| RSYRMHPAV | ASSFRDWGVNRY | 0.57782376 |
| SAAHLLAGQLA | ASSFRDWGVNRY | 0.6079135 |
| SAAVGSY | ASSFRDWGVNRY | 0.64081895 |
| SPHAVLHRI | ASSFRDWGVNRY | 0.46946716 |
| SQIAKVL | ASSFRDWGVNRY | 0.6323298 |
| TALGHLVP | ASSFRDWGVNRY | 0.6393716 |
| TAPTTSGAA | ASSFRDWGVNRY | 0.4023467 |
| TDGDDGNGGSPGT | ASSFRDWGVNRY | 0.52847034 |
| TDGLGEN | ASSFRDWGVNRY | 0.645904 |
| TGGPPLPGQA | ASSFRDWGVNRY | 0.5842079 |
| TIPAITI | ASSFRDWGVNRY | 0.5370995 |
| TIPGFTI | ASSFRDWGVNRY | 0.56821096 |
| TSIASGALA | ASSFRDWGVNRY | 0.64394647 |
| TVTLPGAAI | ASSFRDWGVNRY | 0.5497284 |
| VASGLSAALP | ASSFRDWGVNRY | 0.64284986 |
| VGAGVIGI | ASSFRDWGVNRY | 0.35365745 |
| VGPKRFGV | ASSFRDWGVNRY | 0.55294424 |
| VNVAFPD | ASSFRDWGVNRY | 0.53230524 |
| VSMTNSGVS | ASSFRDWGVNRY | 0.34837043 |
| VTDVLGDLSV | ASSFRDWGVNRY | 0.62399817 |
| VVVVGAGFAGL | ASSFRDWGVNRY | 0.5418565 |
| VYTVGTVV | ASSFRDWGVNRY | 0.613948 |
| WTTFSGP | ASSFRDWGVNRY | 0.54562634 |
| YDELVEAF | ASSFRDWGVNRY | 0.6308212 |
| YGTGGAGGNGG | ASSFRDWGVNRY | 0.5626933 |
| YTNVMAV | ASSFRDWGVNRY | 0.5549031 |
| FLEQVHQL | ASSFRDWGVNRY | 0.48220307 |
| GLSLLAVAV | ASSFRDWGVNRY | 0.6180079 |
| LEQVHQL | ASSFRDWGVNRY | 0.6322472 |
| LNAAGAI | ASSFRDWGVNRY | 0.6450907 |
| PAVHASL | ASSFRDWGVNRY | 0.4407832 |
| TELEDDA | ASSFRDWGVNRY | 0.56780654 |
| TPPGPPN | ASSFRDWGVNRY | 0.588612 |
| TPPPPPPP | ASSFRDWGVNRY | 0.4524069 |
| TQSPALI | ASSSRGLERLF | 0.3727944 |
| EFAAAFPQ | ASSSRGLERLF | 0.55995107 |
| LLSGNLDVL | ASSSRGLERLF | 0.52136195 |
| GPANLGNYNI | ASSSRGLERLF | 0.6075742 |
| TLGPTLP | ASSSRGLERLF | 0.605956 |
| VYAYSSCCK | ASSSRGLERLF | 0.5145885 |
| APAPAPAPNI | ASSSRGLERLF | 0.5998417 |
| PGGPVPIP | ASSSRGLERLF | 0.58956 |
| FLTVALEP | ASSSRGLERLF | 0.29815507 |
| SQYFGANL | ASSSRGLERLF | 0.51930106 |
| GGVKPAA | ASSSRGLERLF | 0.39796472 |

| | | |
|---|---|---|
| GGTGGAGGAGGW | ASSSRGLERLF | 0.6516173 |
| GVGGVAGLGGAGG | ASSSRGLERLF | 0.45030746 |
| VLLVLATTI | ASSSRGLERLF | 0.4609406 |
| IGIFAAML | ASSSRGLERLF | 0.55500525 |
| AVGDLID | ASSSRGLERLF | 0.5748229 |
| NISPLPL | ASSSRGLERLF | 0.56935674 |
| VTAVFVAI | ASSSRGLERLF | 0.4576909 |
| SGDDNDAAL | ASSSRGLERLF | 0.4172385 |
| AVAALAHS | ASSSRGLERLF | 0.6263842 |
| GGVTVMATP | ASSSRGLERLF | 0.53431964 |
| AGLAANVRD | ASSSRGLERLF | 0.5318981 |
| TNRPDLIDPALLRPG | ASSSRGLERLF | 0.40717214 |
| LANLGVTAL | ASSSRGLERLF | 0.5756521 |
| TGERIEL | ASSSRGLERLF | 0.45514852 |
| NGEELAA | ASSSRGLERLF | 0.63973767 |
| VLSLVWG | ASSSRGLERLF | 0.48930824 |
| IGYALWL | ASSSRGLERLF | 0.5279833 |
| LSLLLAPL | ASSSRGLERLF | 0.64389485 |
| TLDEALK | ASSSRGLERLF | 0.5766893 |
| TDGAGTAEGG | ASSSRGLERLF | 0.52388895 |
| NAPGGAGGNGG | ASSSRGLERLF | 0.56153655 |
| VSVFMGMGGNF | ASSSRGLERLF | 0.42685708 |
| LASYLAAL | ASSSRGLERLF | 0.5303305 |
| LVGGTPAL | ASSSRGLERLF | 0.55582505 |
| VRNADVL | ASSSRGLERLF | 0.6440629 |
| TCSSPFLVSAM | ASSSRGLERLF | 0.5785069 |
| VHITDAE | ASSSRGLERLF | 0.44011417 |
| EYAAPSF | ASSSRGLERLF | 0.63486975 |
| VVDAFHGGL | ASSSRGLERLF | 0.64466983 |
| ASVVDPQ | ASSSRGLERLF | 0.65189713 |
| SYNEFGGLS | ASSSRGLERLF | 0.41612422 |
| DPDGSVY | ASSSRGLERLF | 0.5834535 |
| MVSTIHATE | ASSSRGLERLF | 0.55809146 |
| LAAGVADPV | ASSSRGLERLF | 0.5281669 |
| TKEQIAA | ASSSRGLERLF | 0.53533393 |
| SLASPAAL | ASSSRGLERLF | 0.5049522 |
| IGLVFYGV | ASSSRGLERLF | 0.62149465 |
| FLLDASNL | ASSSRGLERLF | 0.65932614 |
| YQVGNFP | ASSSRGLERLF | 0.5385362 |
| ESLLSQST | ASSSRGLERLF | 0.40109387 |
| LFTVGFAAVG | ASSSRGLERLF | 0.5348371 |
| ATPPTPT | ASSSRGLERLF | 0.46820134 |
| TDLATLG | ASSSRGLERLF | 0.4819045 |
| SAGGTGR | ASSSRGLERLF | 0.5838921 |
| GFSNSGS | ASSSRGLERLF | 0.3755984 |
| GGNGGSGGTSV | ASSSRGLERLF | 0.5689508 |
| GTSVATGGAGN | ASSSRGLERLF | 0.6575658 |
| NGGNGGSGGTS | ASSSRGLERLF | 0.52363443 |
| AVVKFPH | ASSSRGLERLF | 0.60719573 |
| TPPPPPP | ASSSRGLERLF | 0.62806743 |
| AAPAAAAPA | ASSSRGLERLF | 0.58869016 |

| | | |
|---|---|---|
| IALNEDL | ASSSRGLERLF | 0.63086724 |
| IVNVSSM | ASSSRGLERLF | 0.6148574 |
| TGIPPAPR | ASSSRGLERLF | 0.5815302 |
| FDLGGGTF | ASSSRGLERLF | 0.57709897 |
| FDLGGGTFDVS | ASSSRGLERLF | 0.6144192 |
| NEPTAAA | ASSSRGLERLF | 0.54262495 |
| NEPTAAALA | ASSSRGLERLF | 0.5505837 |
| GSIINISSI | ASSSRGLERLF | 0.53071654 |
| VAPGNAGTA | ASSSRGLERLF | 0.618864 |
| TGGGASLE | ASSSRGLERLF | 0.5617322 |
| VAVYNHYKRI | ASSSRGLERLF | 0.6380986 |
| TDPAPATT | ASSSRGLERLF | 0.38383153 |
| TNRPDILDPALLRPG | ASSSRGLERLF | 0.50146323 |
| LTTTTTS | ASSSRGLERLF | 0.6233247 |
| PTGSGKTL | ASSSRGLERLF | 0.3276046 |
| TGSGKTL | ASSSRGLERLF | 0.40229064 |
| TIGHVDHGK | ASSSRGLERLF | 0.5721984 |
| TIGHVDHGKTT | ASSSRGLERLF | 0.6371292 |
| GHVDHGKTT | ASSSRGLERLF | 0.34205475 |
| AQFEHTLL | ASSSRGLERLF | 0.5272598 |
| IVNQDEL | ASSSRGLERLF | 0.43397227 |
| TNGSQFF | ASSSRGLERLF | 0.6490445 |
| TGKGIVE | ASSSRGLERLF | 0.5925426 |
| LISNASDA | ASSSRGLERLF | 0.3783296 |
| AELRTLGELTKIAKAHGAQ | ASSSRGLERLF | 0.62137806 |
| APNAVAAAEPSW | ASSSRGLERLF | 0.37888235 |
| APQVAGAATAVL | ASSSRGLERLF | 0.5597275 |
| ARRDPAAAAAILPTDA | ASSSRGLERLF | 0.62328845 |
| DAAARDVLAEAGLAEYF | ASSSRGLERLF | 0.6457782 |
| DDQIVLSAKANLAKVEQ | ASSSRGLERLF | 0.40986532 |
| DELLPDTGEDSDEDGH | ASSSRGLERLF | 0.6231166 |
| ELPEAPANRATVPQD | ASSSRGLERLF | 0.50912106 |
| FGQSKIERVTGIG | ASSSRGLERLF | 0.6610687 |
| GAGGTGSTAGGAGGAGGAGGLY | ASSSRGLERLF | 0.54733324 |
| GHGGTGAAGGNGGTGGAGGSGI | ASSSRGLERLF | 0.40963873 |
| GNGGNGGAGGEGGPGGAGGAGA | ASSSRGLERLF | 0.63158435 |
| GSTAGNGGNGGNATNGTIAGQP | ASSSRGLERLF | 0.604962 |
| GTGGDGGNGGNGGTGG | ASSSRGLERLF | 0.6048248 |
| GVRRPAGIPADAPV | ASSSRGLERLF | 0.60218906 |
| GYRQVTDAHLLTLARR | ASSSRGLERLF | 0.5728729 |
| IAKMGQDGHDRG | ASSSRGLERLF | 0.53019446 |
| KAEPSIEIQYT | ASSSRGLERLF | 0.6169592 |
| KPVRIRMAADEDTVAVT | ASSSRGLERLF | 0.58632916 |
| MPPRIAGMRLLVIKPEP | ASSSRGLERLF | 0.55430454 |
| MKFYAVALFRSRGVPPTR | ASSSRGLERLF | 0.6390443 |
| NAGGFGILGAVAHSP | ASSSRGLERLF | 0.57554466 |
| NEGREITLAMAEM | ASSSRGLERLF | 0.49702436 |
| NELVEWSHPLGHRWRWE | ASSSRGLERLF | 0.6499232 |
| NLGAGGTGGNGGNPG | ASSSRGLERLF | 0.5725826 |
| NTEWPKWETVMAADPVD | ASSSRGLERLF | 0.6107262 |
| PASAAPAETQP | ASSSRGLERLF | 0.62739694 |

| | | |
|---|---|---|
| QQSSDGKNGVRAAEITGE | ASSSRGLERLF | 0.6495411 |
| RAPDAVTTSWWKE | ASSSRGLERLF | 0.44791132 |
| SGFGNVGAHTSGWFNQ | ASSSRGLERLF | 0.5815788 |
| SGVAGAVGQAASVGGLKV | ASSSRGLERLF | 0.57089376 |
| SNGGAGTGSAGGNGGT | ASSSRGLERLF | 0.6271243 |
| TGGAGGEGGAAG | ASSSRGLERLF | 0.58455056 |
| TGGDGGAAGAGGAPGHG | ASSSRGLERLF | 0.5755327 |
| TMPPLLAAIANAGRHGV | ASSSRGLERLF | 0.6093101 |
| TSSTAGNSGGA | ASSSRGLERLF | 0.64451605 |
| VPATVPVGILC | ASSSRGLERLF | 0.5585739 |
| VQRPDSGGVVLTGRIS | ASSSRGLERLF | 0.57820475 |
| WTPNPVPATKGVPAGLVQAT | ASSSRGLERLF | 0.6436953 |
| YALKRALAKFSDRFDVVII | ASSSRGLERLF | 0.49068618 |
| YLIGSVIV | ASSSRGLERLF | 0.5820807 |
| YGQLVAVID | ASSSRGLERLF | 0.62714624 |
| VTVGGMTLPA | ASSSRGLERLF | 0.63040805 |
| VLALPAV | ASSSRGLERLF | 0.56793416 |
| VIPPTLNYET | ASSSRGLERLF | 0.61755216 |
| VGMGGMHPSSG | ASSSRGLERLF | 0.45808125 |
| VGLAGGAATA | ASSSRGLERLF | 0.59252876 |
| VGGAGGAGGSGP | ASSSRGLERLF | 0.5107345 |
| VFAAFMV | ASSSRGLERLF | 0.5705125 |
| TLYDKLR | ASSSRGLERLF | 0.51301384 |
| TLAVFTVL | ASSSRGLERLF | 0.5830965 |
| TGSVQPARV | ASSSRGLERLF | 0.61371565 |
| TGGGAGSGGT | ASSSRGLERLF | 0.6047027 |
| SHLETSL | ASSSRGLERLF | 0.3671524 |
| SFTTGMT | ASSSRGLERLF | 0.5922084 |
| QVEVAKRMGVRQPTVS | ASSSRGLERLF | 0.614386 |
| QGSGGPALAM | ASSSRGLERLF | 0.6485695 |
| MLAPSNP | ASSSRGLERLF | 0.56389594 |
| LVNNAAI | ASSSRGLERLF | 0.62383175 |
| LTGEINTLA | ASSSRGLERLF | 0.5364389 |
| LSIFGPV | ASSSRGLERLF | 0.5751168 |
| LLALLANL | ASSSRGLERLF | 0.417233 |
| LISNASDALDKLR | ASSSRGLERLF | 0.5528352 |
| LINEAVP | ASSSRGLERLF | 0.49761343 |
| LGYAFSGL | ASSSRGLERLF | 0.45874912 |
| LDFVAVAL | ASSSRGLERLF | 0.6147942 |
| LAQPGAE | ASSSRGLERLF | 0.6192483 |
| ISNLGAVTGF | ASSSRGLERLF | 0.63620454 |
| HQEDPGRI | ASSSRGLERLF | 0.6377114 |
| GTPGEPTP | ASSSRGLERLF | 0.6344782 |
| GFFNSGS | ASSSRGLERLF | 0.63944954 |
| GAGGGGLTAGAN | ASSSRGLERLF | 0.55036974 |
| FVDALVNM | ASSSRGLERLF | 0.52828205 |
| FLTVAPEL | ASSSRGLERLF | 0.4139042 |
| FETLTAL | ASSSRGLERLF | 0.5156413 |
| DVINPIEL | ASSSRGLERLF | 0.5541272 |
| APVSGAPV | ASSSRGLERLF | 0.57685506 |
| ALVGGLAW | ASSSRGLERLF | 0.3363157 |

| | | |
|---|---|---|
| ALAAAAATAI | ASSSRGLERLF | 0.5502238 |
| AFGLGLLL | ASSSRGLERLF | 0.64715415 |
| AFFAAASL | ASSSRGLERLF | 0.55821115 |
| AAAALPAFSPP | ASSSRGLERLF | 0.61755025 |
| VVLAPAL | ASSSRGLERLF | 0.5841853 |
| TIGHVDHGKTTL | ASSSRGLERLF | 0.49334013 |
| LISNASDALDK | ASSSRGLERLF | 0.61771226 |
| LISNASDAL | ASSSRGLERLF | 0.5104267 |
| ISNASDALDK | ASSSRGLERLF | 0.6255652 |
| NQAETLVYQTE | ASSSRGLERLF | 0.6497841 |
| TGGTGGTAGAG | ASSSRGLERLF | 0.59612644 |
| TGSTAGGAGGA | ASSSRGLERLF | 0.5384542 |
| DPLAVPVAAGPV | ASSSRGLERLF | 0.596668 |
| GGTGGTPGNGGN | ASSSRGLERLF | 0.4287346 |
| PAGLAGVDPATL | ASSSRGLERLF | 0.5607978 |
| GNNLVVEAPHAGASV | ASSSRGLERLF | 0.46585616 |
| KVTRIMSASTLEQPA | ASSSRGLERLF | 0.5499047 |
| LGFALPIALPRRTVV | ASSSRGLERLF | 0.46733117 |
| TRLGNLTARKLVVLV | ASSSRGLERLF | 0.59704703 |
| WSQQRRLIYFPSAGP | ASSSRGLERLF | 0.5883132 |
| GPNGNPDPMAAAVDIR | ASSSRGLERLF | 0.5998086 |
| TQRCVSVADLRAAASR | ASSSRGLERLF | 0.45812 |
| QFVLDVAEHPVEVLTETV | ASSSRGLERLF | 0.58892184 |
| GGAGGTGGAGGNGGTQAGDG | ASSSRGLERLF | 0.55207485 |
| PLLATLRAIAVDVDAKPAQV | ASSSRGLERLF | 0.49379158 |
| APKFRTGGYMIDFWGVGYQVAK | ASSSRGLERLF | 0.6129619 |
| GAGGAGGAGGAAGGPGGTGGTG | ASSSRGLERLF | 0.569952 |
| IGALGLLATDVAPAGSVAFGVA | ASSSRGLERLF | 0.5335258 |
| AGGAGGIGGNGGAGGAGGVLMG | ASSSRGLERLF | 0.63327837 |
| ADELIGLL | ASSSRGLERLF | 0.6060591 |
| ADRALKA | ASSSRGLERLF | 0.3987615 |
| AGAVGQVGGAGGA | ASSSRGLERLF | 0.5426581 |
| AGGGGGLGGAP | ASSSRGLERLF | 0.5666448 |
| AGGLLYGNGGAG | ASSSRGLERLF | 0.5513603 |
| AGLESIGL | ASSSRGLERLF | 0.35025293 |
| AGLKTGII | ASSSRGLERLF | 0.5979266 |
| ASLAALLL | ASSSRGLERLF | 0.49628633 |
| ASVINISSI | ASSSRGLERLF | 0.44212693 |
| ATPPGPA | ASSSRGLERLF | 0.51929593 |
| ATVASALV | ASSSRGLERLF | 0.62447906 |
| ATVASLIL | ASSSRGLERLF | 0.5912562 |
| AVNGPVLIRH | ASSSRGLERLF | 0.627532 |
| DGAAGAVGI | ASSSRGLERLF | 0.5856157 |
| DPLLHGI | ASSSRGLERLF | 0.6167271 |
| ELGAALGAL | ASSSRGLERLF | 0.6326875 |
| ELGRTQAV | ASSSRGLERLF | 0.61393154 |
| FVAGGYLS | ASSSRGLERLF | 0.58429223 |
| GASGGAGGAGGW | ASSSRGLERLF | 0.6024061 |
| GGAGAGGDGGAGGI | ASSSRGLERLF | 0.57574546 |
| GTGHNGSQPGMGGNG | ASSSRGLERLF | 0.41660115 |
| GYAAASASAS | ASSSRGLERLF | 0.38813135 |

| | | |
|---|---|---|
| HQVDVAAVQQG | ASSSRGLERLF | 0.5478132 |
| IGAAIAHT | ASSSRGLERLF | 0.4186125 |
| IGMIEVAQL | ASSSRGLERLF | 0.59552187 |
| IRLAPEHTSAMD | ASSSRGLERLF | 0.65145147 |
| IYNVSVL | ASSSRGLERLF | 0.56515527 |
| KQGFLGM | ASSSRGLERLF | 0.4146548 |
| LESVITQL | ASSSRGLERLF | 0.51953304 |
| LGLGALVVV | ASSSRGLERLF | 0.47677058 |
| LIGNGGNGGSGGI | ASSSRGLERLF | 0.6159532 |
| LNAINAPTL | ASSSRGLERLF | 0.63426536 |
| LQLLNTGV | ASSSRGLERLF | 0.64463747 |
| LQTSIEF | ASSSRGLERLF | 0.6151859 |
| MSFAVSEL | ASSSRGLERLF | 0.5409267 |
| NTGALGL | ASSSRGLERLF | 0.60210496 |
| PPGAGGAGGAGTT | ASSSRGLERLF | 0.4994866 |
| QLVGLGI | ASSSRGLERLF | 0.5724839 |
| RFLEQVHQL | ASSSRGLERLF | 0.55592084 |
| RGVGYVLR | ASSSRGLERLF | 0.59294915 |
| SATGVFAGVI | ASSSRGLERLF | 0.5573339 |
| SLPAITLP | ASSSRGLERLF | 0.5380469 |
| STALAFTV | ASSSRGLERLF | 0.4145966 |
| SVLTVGLGK | ASSSRGLERLF | 0.6158469 |
| TALAGAGSGI | ASSSRGLERLF | 0.5480122 |
| TGLPFTGI | ASSSRGLERLF | 0.43022323 |
| VAEHVTV | ASSSRGLERLF | 0.5259461 |
| VDIVLNSL | ASSSRGLERLF | 0.6442487 |
| VLTGGGAVL | ASSSRGLERLF | 0.647541 |
| VNVGLAYGI | ASSSRGLERLF | 0.49617812 |
| VSSAAVGAV | ASSSRGLERLF | 0.5284858 |
| VTSVYEAGV | ASSSRGLERLF | 0.5734309 |
| WVALDRGA | ASSSRGLERLF | 0.44090793 |
| YLVGSVVI | ASSSRGLERLF | 0.6005428 |
| GPPGTGKTLL | ASSSRGLERLF | 0.54617846 |
| GVGMTKF | ASSSRGLERLF | 0.5612432 |
| IGIGGSDL | ASSSRGLERLF | 0.5132204 |
| IGTLLAEL | ASSSRGLERLF | 0.6090641 |
| IYTSGTT | ASSSRGLERLF | 0.5610071 |
| LSFPAGV | ASSSRGLERLF | 0.6224878 |
| LYTSGTT | ASSSRGLERLF | 0.3711512 |
| VGAFPFGF | ASSSRGLERLF | 0.6270444 |
| VGAGTVEF | ASSSRGLERLF | 0.453379 |
| VGKSTLLL | ASSSRGLERLF | 0.46060848 |
| AALAHAGGGIV | ASSSRGLERLF | 0.36718386 |
| AALVAVSLL | ASSSRGLERLF | 0.56069136 |
| AAVSRAVAAL | ASSSRGLERLF | 0.6193286 |
| AGGSSGIGGF | ASSSRGLERLF | 0.6333164 |
| AGLTVANANG | ASSSRGLERLF | 0.51793015 |
| AGLVGAVL | ASSSRGLERLF | 0.6038086 |
| ALSQAFTV | ASSSRGLERLF | 0.6357189 |
| ALTFSVSL | ASSSRGLERLF | 0.50972784 |
| ALTGAALVAL | ASSSRGLERLF | 0.38451836 |

| | | |
|---|---|---|
| ASFSIGAGVL | ASSSRGLERLF | 0.6215004 |
| DGSVYKI | ASSSRGLERLF | 0.5356482 |
| EQLAEIV | ASSSRGLERLF | 0.6429063 |
| FAVSELTVM | ASSSRGLERLF | 0.61637866 |
| GAAPLAAGL | ASSSRGLERLF | 0.3865486 |
| GADGGDGGSGGAGG | ASSSRGLERLF | 0.644071 |
| GFGLDTGGAGG | ASSSRGLERLF | 0.60290813 |
| GLASGGLL | ASSSRGLERLF | 0.44078606 |
| GRIAGAAVLL | ASSSRGLERLF | 0.4668953 |
| GSAGSLNTSF | ASSSRGLERLF | 0.509512 |
| GTGQIGI | ASSSRGLERLF | 0.43700862 |
| GTLIALIL | ASSSRGLERLF | 0.61643845 |
| IAATMGFVVG | ASSSRGLERLF | 0.504053 |
| IFAQALL | ASSSRGLERLF | 0.59056693 |
| IGAAVALGF | ASSSRGLERLF | 0.545118 |
| IGLTPLAL | ASSSRGLERLF | 0.5728285 |
| ILGSNTSTL | ASSSRGLERLF | 0.5540158 |
| IVFSLTVL | ASSSRGLERLF | 0.5824558 |
| LASKATGF | ASSSRGLERLF | 0.618857 |
| LASVALIL | ASSSRGLERLF | 0.60163796 |
| LFAVGLTL | ASSSRGLERLF | 0.55549276 |
| LFNGGIGTYI | ASSSRGLERLF | 0.58960485 |
| LTLGAGII | ASSSRGLERLF | 0.63634485 |
| LVAAALGDI | ASSSRGLERLF | 0.51435673 |
| NTAFVQLGI | ASSSRGLERLF | 0.5015016 |
| QLFSGAEL | ASSSRGLERLF | 0.61708426 |
| SAATFITL | ASSSRGLERLF | 0.48393992 |
| SNYGLANLGV | ASSSRGLERLF | 0.62089163 |
| SYGGAYAVM | ASSSRGLERLF | 0.5922269 |
| TTGVGGLTLG | ASSSRGLERLF | 0.57815784 |
| TTQTIYI | ASSSRGLERLF | 0.56411487 |
| VAAGAGLGAVI | ASSSRGLERLF | 0.64478886 |
| VAAVVTLL | ASSSRGLERLF | 0.6188746 |
| VGFLGGLI | ASSSRGLERLF | 0.54524684 |
| VGVIPILL | ASSSRGLERLF | 0.60576504 |
| VIFASSLI | ASSSRGLERLF | 0.45774388 |
| VKGGDGGAAAG | ASSSRGLERLF | 0.6123478 |
| VTKTFTNI | ASSSRGLERLF | 0.6231301 |
| VVAAAILL | ASSSRGLERLF | 0.45002168 |
| YAALDVEL | ASSSRGLERLF | 0.5296075 |
| LASVALLL | ASSSRGLERLF | 0.6455797 |
| LDAVHRV | ASSSRGLERLF | 0.6159369 |
| QGLSSSIF | ASSSRGLERLF | 0.45608145 |
| RGTFANIRL | ASSSRGLERLF | 0.48911858 |
| RQLWWGHRI | ASSSRGLERLF | 0.45527977 |
| YGPPGTGKTLL | ASSSRGLERLF | 0.49380794 |
| AALVGAPANVA | ASSSRGLERLF | 0.43196824 |
| AATFRRTE | ASSSRGLERLF | 0.59299445 |
| AGALGGGI | ASSSRGLERLF | 0.52938026 |
| AGGAGGAGGIGAG | ASSSRGLERLF | 0.61675215 |
| AGSGALGL | ASSSRGLERLF | 0.5910745 |

| | | |
|---|---|---|
| AQIADQI | ASSSRGLERLF | 0.6402191 |
| ASAVAVTYL | ASSSRGLERLF | 0.496042 |
| ASTVLGGTGG | ASSSRGLERLF | 0.5728325 |
| AVGGEPAK | ASSSRGLERLF | 0.39667997 |
| CEQGTFGGTC | ASSSRGLERLF | 0.58571625 |
| DEGKAPAG | ASSSRGLERLF | 0.51181614 |
| DTVVFPAL | ASSSRGLERLF | 0.59304136 |
| FEGITMAG | ASSSRGLERLF | 0.4324443 |
| FPLVVQD | ASSSRGLERLF | 0.6449138 |
| GAFTTPQ | ASSSRGLERLF | 0.55458915 |
| GFGGFGTAGGVGGA | ASSSRGLERLF | 0.6163968 |
| GFLAQFL | ASSSRGLERLF | 0.54091483 |
| GGAGGAGGNAGVA | ASSSRGLERLF | 0.5011572 |
| GGVGTAGAGGAGGAGG | ASSSRGLERLF | 0.53327626 |
| GNGGGGDAGNAGSGGN | ASSSRGLERLF | 0.61696434 |
| GTGGAGGAGNDAGS | ASSSRGLERLF | 0.416987 |
| GTTNTGWFNS | ASSSRGLERLF | 0.36039 |
| IFMGPLFTL | ASSSRGLERLF | 0.5472709 |
| ILAHAIEAI | ASSSRGLERLF | 0.40104714 |
| IRSAIREAL | ASSSRGLERLF | 0.6437617 |
| ITFAGGG | ASSSRGLERLF | 0.6295287 |
| ITGMSGHQEV | ASSSRGLERLF | 0.5739511 |
| IVAGLACTL | ASSSRGLERLF | 0.64198273 |
| IVGGGLAGA | ASSSRGLERLF | 0.32653534 |
| IVTGVLNI | ASSSRGLERLF | 0.5175026 |
| LAQQGCI | ASSSRGLERLF | 0.6436801 |
| LCLHVEN | ASSSRGLERLF | 0.5876503 |
| LDAISDGS | ASSSRGLERLF | 0.557189 |
| LGGFAAML | ASSSRGLERLF | 0.6329061 |
| LGGKNPMI | ASSSRGLERLF | 0.534796 |
| LGPKGVAAIL | ASSSRGLERLF | 0.5636867 |
| LMVGVSI | ASSSRGLERLF | 0.6054883 |
| LVGAAFVL | ASSSRGLERLF | 0.6050974 |
| LVLAVAPL | ASSSRGLERLF | 0.61686593 |
| MAAIAVIDAA | ASSSRGLERLF | 0.4167652 |
| NNGINAELQ | ASSSRGLERLF | 0.6207302 |
| RGEPLVLVHG | ASSSRGLERLF | 0.6465811 |
| RGGAGGAAGLI | ASSSRGLERLF | 0.5382515 |
| RSYRMHPAV | ASSSRGLERLF | 0.6234201 |
| SAAHLLAGQLA | ASSSRGLERLF | 0.60712427 |
| SAAVGSY | ASSSRGLERLF | 0.5901147 |
| SPHAVLHRI | ASSSRGLERLF | 0.4847424 |
| SQIAKVL | ASSSRGLERLF | 0.4477715 |
| TALGHLVP | ASSSRGLERLF | 0.59359187 |
| TAPTTSGAA | ASSSRGLERLF | 0.4132061 |
| TDGDDGNGGSPGT | ASSSRGLERLF | 0.5849981 |
| TDGLGEN | ASSSRGLERLF | 0.536048 |
| TGGPPLPGQA | ASSSRGLERLF | 0.6618096 |
| TIPAITI | ASSSRGLERLF | 0.600739 |
| TIPGFTI | ASSSRGLERLF | 0.613307 |
| TSIASGALA | ASSSRGLERLF | 0.47304118 |

| | | |
|---|---|---|
| TVTLPGAAI | ASSSRGLERLF | 0.5913914 |
| VASGLSAALP | ASSSRGLERLF | 0.5893008 |
| VGAGVIGI | ASSSRGLERLF | 0.41421872 |
| VGPKRFGV | ASSSRGLERLF | 0.6346421 |
| VNVAFPD | ASSSRGLERLF | 0.48681542 |
| VSMTNSGVS | ASSSRGLERLF | 0.5628582 |
| VTDVLGDLSV | ASSSRGLERLF | 0.6157355 |
| VVVVGAGFAGL | ASSSRGLERLF | 0.63238823 |
| VYTVGTVV | ASSSRGLERLF | 0.53712744 |
| WTTFSGP | ASSSRGLERLF | 0.58351886 |
| YDELVEAF | ASSSRGLERLF | 0.49567267 |
| YGTGGAGGNGG | ASSSRGLERLF | 0.61388886 |
| YTNVMAV | ASSSRGLERLF | 0.6274532 |
| FLEQVHQL | ASSSRGLERLF | 0.5738892 |
| GLSLLAVAV | ASSSRGLERLF | 0.64772856 |
| LEQVHQL | ASSSRGLERLF | 0.6449236 |
| LNAAGAI | ASSSRGLERLF | 0.412965 |
| PAVHASL | ASSSRGLERLF | 0.57361555 |
| TELEDDA | ASSSRGLERLF | 0.62873423 |
| TPPGPPN | ASSSRGLERLF | 0.6290544 |
| TPPPPPPP | ASSSRGLERLF | 0.64995044 |
| TQSPALI | ASSSPRLGGRDTQY | 0.60770273 |
| EFAAAFPQ | ASSSPRLGGRDTQY | 0.48732117 |
| LLSGNLDVL | ASSSPRLGGRDTQY | 0.6360014 |
| GPANLGNYNI | ASSSPRLGGRDTQY | 0.66489285 |
| TLGPTLP | ASSSPRLGGRDTQY | 0.623902 |
| VYAYSSCCK | ASSSPRLGGRDTQY | 0.49594492 |
| APAPAPAPNI | ASSSPRLGGRDTQY | 0.52961236 |
| PGGPVPIP | ASSSPRLGGRDTQY | 0.61276704 |
| FLTVALEP | ASSSPRLGGRDTQY | 0.6243644 |
| SQYFGANL | ASSSPRLGGRDTQY | 0.46780363 |
| GGVKPAA | ASSSPRLGGRDTQY | 0.57151854 |
| GGTGGAGGAGGW | ASSSPRLGGRDTQY | 0.5920022 |
| GVGGVAGLGGAGG | ASSSPRLGGRDTQY | 0.39200014 |
| VLLVLATTI | ASSSPRLGGRDTQY | 0.599805 |
| IGIFAAML | ASSSPRLGGRDTQY | 0.62243724 |
| AVGDLID | ASSSPRLGGRDTQY | 0.59115386 |
| NISPLPL | ASSSPRLGGRDTQY | 0.492559 |
| VTAVFVAI | ASSSPRLGGRDTQY | 0.6293782 |
| SGDDNDAAL | ASSSPRLGGRDTQY | 0.6420363 |
| AVAALAHS | ASSSPRLGGRDTQY | 0.6477827 |
| GGVTVMATP | ASSSPRLGGRDTQY | 0.43157113 |
| AGLAANVRD | ASSSPRLGGRDTQY | 0.32393914 |
| TNRPDLIDPALLRPG | ASSSPRLGGRDTQY | 0.5507079 |
| LANLGVTAL | ASSSPRLGGRDTQY | 0.6118684 |
| TGERIEL | ASSSPRLGGRDTQY | 0.58882844 |
| NGEELAA | ASSSPRLGGRDTQY | 0.53981894 |
| VLSLVWG | ASSSPRLGGRDTQY | 0.58775306 |
| IGYALWL | ASSSPRLGGRDTQY | 0.62524223 |
| LSLLLAPL | ASSSPRLGGRDTQY | 0.48679146 |
| TLDEALK | ASSSPRLGGRDTQY | 0.45023355 |

| | | |
|---|---|---|
| TDGAGTAEGG | ASSSPRLGGRDTQY | 0.6191093 |
| NAPGGAGGNGG | ASSSPRLGGRDTQY | 0.63193613 |
| VSVFMGMGGNF | ASSSPRLGGRDTQY | 0.45757776 |
| LASYLAAL | ASSSPRLGGRDTQY | 0.50670004 |
| LVGGTPAL | ASSSPRLGGRDTQY | 0.5723472 |
| VRNADVL | ASSSPRLGGRDTQY | 0.58555925 |
| TCSSPFLVSAM | ASSSPRLGGRDTQY | 0.5723962 |
| VHITDAE | ASSSPRLGGRDTQY | 0.4485259 |
| EYAAPSF | ASSSPRLGGRDTQY | 0.62047344 |
| VVDAFHGGL | ASSSPRLGGRDTQY | 0.40615705 |
| ASVVDPQ | ASSSPRLGGRDTQY | 0.6004604 |
| SYNEFGGLS | ASSSPRLGGRDTQY | 0.5751047 |
| DPDGSVY | ASSSPRLGGRDTQY | 0.59611785 |
| MVSTIHATE | ASSSPRLGGRDTQY | 0.52494204 |
| LAAGVADPV | ASSSPRLGGRDTQY | 0.47725078 |
| TKEQIAA | ASSSPRLGGRDTQY | 0.4231931 |
| SLASPAAL | ASSSPRLGGRDTQY | 0.38291654 |
| IGLVFYGV | ASSSPRLGGRDTQY | 0.6214025 |
| FLLDASNL | ASSSPRLGGRDTQY | 0.5812069 |
| YQVGNFP | ASSSPRLGGRDTQY | 0.6146025 |
| ESLLSQST | ASSSPRLGGRDTQY | 0.6146576 |
| LFTVGFAAVG | ASSSPRLGGRDTQY | 0.47699124 |
| ATPPTPT | ASSSPRLGGRDTQY | 0.60155773 |
| TDLATLG | ASSSPRLGGRDTQY | 0.5196175 |
| SAGGTGR | ASSSPRLGGRDTQY | 0.54295623 |
| GFSNSGS | ASSSPRLGGRDTQY | 0.5893626 |
| GGNGGSGGTSV | ASSSPRLGGRDTQY | 0.6169096 |
| GTSVATGGAGN | ASSSPRLGGRDTQY | 0.46700454 |
| NGGNGGSGGTS | ASSSPRLGGRDTQY | 0.6086079 |
| AVVKFPH | ASSSPRLGGRDTQY | 0.59328604 |
| TPPPPPP | ASSSPRLGGRDTQY | 0.5070105 |
| AAPAAAAAPA | ASSSPRLGGRDTQY | 0.5942143 |
| IALNEDL | ASSSPRLGGRDTQY | 0.62550503 |
| IVNVSSM | ASSSPRLGGRDTQY | 0.60126776 |
| TGIPPAPR | ASSSPRLGGRDTQY | 0.41421646 |
| FDLGGGTF | ASSSPRLGGRDTQY | 0.6133752 |
| FDLGGGTFDVS | ASSSPRLGGRDTQY | 0.4347454 |
| NEPTAAA | ASSSPRLGGRDTQY | 0.5816176 |
| NEPTAAALA | ASSSPRLGGRDTQY | 0.5575299 |
| GSIINISSI | ASSSPRLGGRDTQY | 0.56437635 |
| VAPGNAGTA | ASSSPRLGGRDTQY | 0.6361496 |
| TGGGASLE | ASSSPRLGGRDTQY | 0.52316743 |
| VAVYNHYKRI | ASSSPRLGGRDTQY | 0.46639282 |
| TDPAPATT | ASSSPRLGGRDTQY | 0.5233013 |
| TNRPDILDPALLRPG | ASSSPRLGGRDTQY | 0.60347646 |
| LTTTTTS | ASSSPRLGGRDTQY | 0.6204602 |
| PTGSGKTL | ASSSPRLGGRDTQY | 0.45120648 |
| TGSGKTL | ASSSPRLGGRDTQY | 0.58174956 |
| TIGHVDHGK | ASSSPRLGGRDTQY | 0.55717295 |
| TIGHVDHGKTT | ASSSPRLGGRDTQY | 0.4971465 |
| GHVDHGKTT | ASSSPRLGGRDTQY | 0.5829364 |

| | | |
|---|---|---|
| AQFEHTLL | ASSSPRLGGRDTQY | 0.56965065 |
| IVNQDEL | ASSSPRLGGRDTQY | 0.4529147 |
| TNGSQFF | ASSSPRLGGRDTQY | 0.60550284 |
| TGKGIVE | ASSSPRLGGRDTQY | 0.577011 |
| LISNASDA | ASSSPRLGGRDTQY | 0.61597127 |
| AELRTLGELTKIAKAHGAQ | ASSSPRLGGRDTQY | 0.62745714 |
| APNAVAAAEPSW | ASSSPRLGGRDTQY | 0.44979018 |
| APQVAGAATAVL | ASSSPRLGGRDTQY | 0.60136026 |
| ARRDPAAAAAILPTDA | ASSSPRLGGRDTQY | 0.5902642 |
| DAAARDVLAEAGLAEYF | ASSSPRLGGRDTQY | 0.57693493 |
| DDQIVLSAKANLAKVEQ | ASSSPRLGGRDTQY | 0.5849906 |
| DELLPDTGEDSDEDGH | ASSSPRLGGRDTQY | 0.4637168 |
| ELPEAPANRATVPQD | ASSSPRLGGRDTQY | 0.62992203 |
| FGQSKIERVTGIG | ASSSPRLGGRDTQY | 0.63183373 |
| GAGGTGSTAGGAGGAGGAGGLY | ASSSPRLGGRDTQY | 0.61187375 |
| GHGGTGAAGGNGGTGGAGGSGI | ASSSPRLGGRDTQY | 0.61212885 |
| GNGGNGGAGGEGGPGGAGGAGA | ASSSPRLGGRDTQY | 0.60034525 |
| GSTAGNGGNGGNATNGTIAGQP | ASSSPRLGGRDTQY | 0.62202567 |
| GTGGDGGNGGNGGTGG | ASSSPRLGGRDTQY | 0.4955353 |
| GVRRPAGIPADAPV | ASSSPRLGGRDTQY | 0.40139735 |
| GYRQVTDAHLLTLARR | ASSSPRLGGRDTQY | 0.58602357 |
| IAKMGQDGHDRG | ASSSPRLGGRDTQY | 0.54427224 |
| KAEPSIEIQYT | ASSSPRLGGRDTQY | 0.5095893 |
| KPVRIRMAADEDTVAVT | ASSSPRLGGRDTQY | 0.42901787 |
| MPPRIAGMRLLVIKPEP | ASSSPRLGGRDTQY | 0.62317777 |
| MKFYAVALFRSRGVPPTR | ASSSPRLGGRDTQY | 0.65541065 |
| NAGGFGILGAVAHSP | ASSSPRLGGRDTQY | 0.5620463 |
| NEGREITLAMAEM | ASSSPRLGGRDTQY | 0.65249467 |
| NELVEWSHPLGHRWRWE | ASSSPRLGGRDTQY | 0.42023635 |
| NLGAGGTGGNGGNPG | ASSSPRLGGRDTQY | 0.6297902 |
| NTEWPKWETVMAADPVD | ASSSPRLGGRDTQY | 0.63298595 |
| PASAAPAETQP | ASSSPRLGGRDTQY | 0.46122724 |
| QQSSDGKNGVRAAEITGE | ASSSPRLGGRDTQY | 0.62230676 |
| RAPDAVTTSWWKE | ASSSPRLGGRDTQY | 0.45211336 |
| SGFGNVGAHTSGWFNQ | ASSSPRLGGRDTQY | 0.5037227 |
| SGVAGAVGQAASVGGLKV | ASSSPRLGGRDTQY | 0.62813 |
| SNGGAGTGSAGGNGGT | ASSSPRLGGRDTQY | 0.58157825 |
| TGGAGGEGGAAG | ASSSPRLGGRDTQY | 0.6044159 |
| TGGDGGAAGAGGAPGHG | ASSSPRLGGRDTQY | 0.58532363 |
| TMPPLLAAIANAGRHGV | ASSSPRLGGRDTQY | 0.50885606 |
| TSSTAGNSGGA | ASSSPRLGGRDTQY | 0.5824315 |
| VPATVPVGILC | ASSSPRLGGRDTQY | 0.6282593 |
| VQRPDSGGVVLTGRIS | ASSSPRLGGRDTQY | 0.53101915 |
| WTPNPVPATKGVPAGLVQAT | ASSSPRLGGRDTQY | 0.6646068 |
| YALKRALAKFSDRFDVVII | ASSSPRLGGRDTQY | 0.45882192 |
| YLIGSVIV | ASSSPRLGGRDTQY | 0.63891804 |
| YGQLVAVID | ASSSPRLGGRDTQY | 0.6122967 |
| VTVGGMTLPA | ASSSPRLGGRDTQY | 0.5495627 |
| VLALPAV | ASSSPRLGGRDTQY | 0.5342859 |
| VIPPTLNYET | ASSSPRLGGRDTQY | 0.6273499 |
| VGMGGMHPSSG | ASSSPRLGGRDTQY | 0.5525207 |

| | | |
|---|---|---|
| VGLAGGAATA | ASSSPRLGGRDTQY | 0.58838445 |
| VGGAGGAGGSGP | ASSSPRLGGRDTQY | 0.6471367 |
| VFAAFMV | ASSSPRLGGRDTQY | 0.5437869 |
| TLYDKLR | ASSSPRLGGRDTQY | 0.8578776 |
| TLAVFTVL | ASSSPRLGGRDTQY | 0.8043792 |
| TGSVQPARV | ASSSPRLGGRDTQY | 0.85972834 |
| TGGGAGSGGT | ASSSPRLGGRDTQY | 0.6971024 |
| SHLETSL | ASSSPRLGGRDTQY | 0.4084097 |
| SFTTGMT | ASSSPRLGGRDTQY | 0.80118537 |
| QVEVAKRMGVRQPTVS | ASSSPRLGGRDTQY | 0.6301712 |
| QGSGGPALAM | ASSSPRLGGRDTQY | 0.86551154 |
| MLAPSNP | ASSSPRLGGRDTQY | 0.8619176 |
| LVNNAAI | ASSSPRLGGRDTQY | 0.7025704 |
| LTGEINTLA | ASSSPRLGGRDTQY | 0.8491657 |
| LSIFGPV | ASSSPRLGGRDTQY | 0.8711435 |
| LLALLANL | ASSSPRLGGRDTQY | 0.50827545 |
| LISNASDALDKLR | ASSSPRLGGRDTQY | 0.76149416 |
| LINEAVP | ASSSPRLGGRDTQY | 0.7702975 |
| LGYAFSGL | ASSSPRLGGRDTQY | 0.7734226 |
| LDFVAVAL | ASSSPRLGGRDTQY | 0.3477 |
| LAQPGAE | ASSSPRLGGRDTQY | 0.84079874 |
| ISNLGAVTGF | ASSSPRLGGRDTQY | 0.68767 |
| HQEDPGRI | ASSSPRLGGRDTQY | 0.25504088 |
| GTPGEPTP | ASSSPRLGGRDTQY | 0.83510107 |
| GFFNSGS | ASSSPRLGGRDTQY | 0.6903232 |
| GAGGGGLTAGAN | ASSSPRLGGRDTQY | 0.67714155 |
| FVDALVNM | ASSSPRLGGRDTQY | 0.84345734 |
| FLTVAPEL | ASSSPRLGGRDTQY | 0.68144566 |
| FETLTAL | ASSSPRLGGRDTQY | 0.62106025 |
| DVINPIEL | ASSSPRLGGRDTQY | 0.81585383 |
| APVSGAPV | ASSSPRLGGRDTQY | 0.8604505 |
| ALVGGLAW | ASSSPRLGGRDTQY | 0.53215647 |
| ALAAAAATAI | ASSSPRLGGRDTQY | 0.79028857 |
| AFGLGLLL | ASSSPRLGGRDTQY | 0.88752866 |
| AFFAAASL | ASSSPRLGGRDTQY | 0.79291415 |
| AAAALPAFSPP | ASSSPRLGGRDTQY | 0.8855686 |
| VVLAPAL | ASSSPRLGGRDTQY | 0.89179325 |
| TIGHVDHGKTTL | ASSSPRLGGRDTQY | 0.76525986 |
| LISNASDALDK | ASSSPRLGGRDTQY | 0.8589519 |
| LISNASDAL | ASSSPRLGGRDTQY | 0.8668798 |
| ISNASDALDK | ASSSPRLGGRDTQY | 0.7471051 |
| NQAETLVYQTE | ASSSPRLGGRDTQY | 0.79439443 |
| TGGTGGTAGAG | ASSSPRLGGRDTQY | 0.5675626 |
| TGSTAGGAGGA | ASSSPRLGGRDTQY | 0.8074881 |
| DPLAVPVAAGPV | ASSSPRLGGRDTQY | 0.4439739 |
| GGTGGTPGNGGN | ASSSPRLGGRDTQY | 0.79275775 |
| PAGLAGVDPATL | ASSSPRLGGRDTQY | 0.8718204 |
| GNNLVVEAPHAGASV | ASSSPRLGGRDTQY | 0.76092315 |
| KVTRIMSASTLEQPA | ASSSPRLGGRDTQY | 0.4697659 |
| LGFALPIALPRRTVV | ASSSPRLGGRDTQY | 0.5923993 |
| TRLGNLTARKLVVLV | ASSSPRLGGRDTQY | 0.8176593 |

| | | |
|---|---|---|
| WSQQRRLIYFPSAGP | ASSSPRLGGRDTQY | 0.5258358 |
| GPNGNPDPMAAAVDIR | ASSSPRLGGRDTQY | 0.5245502 |
| TQRCVSVADLRAAASR | ASSSPRLGGRDTQY | 0.6964731 |
| QFVLDVAEHPVEVLTETV | ASSSPRLGGRDTQY | 0.86314964 |
| GGAGGTGGAGGNGGTQAGDG | ASSSPRLGGRDTQY | 0.86935675 |
| PLLATLRAIAVDVDAKPAQV | ASSSPRLGGRDTQY | 0.6557235 |
| APKFRTGGYMIDFWGVGYQVAK | ASSSPRLGGRDTQY | 0.8707982 |
| GAGGAGGAGGAAGGPGGTGGTG | ASSSPRLGGRDTQY | 0.8508333 |
| IGALGLLATDVAPAGSVAFGVA | ASSSPRLGGRDTQY | 0.88668805 |
| AGGAGGIGGNGGAGGAGGVLMG | ASSSPRLGGRDTQY | 0.81805956 |
| ADELIGLL | ASSSPRLGGRDTQY | 0.5265562 |
| ADRALKA | ASSSPRLGGRDTQY | 0.8627069 |
| AGAVGQVGGAGGA | ASSSPRLGGRDTQY | 0.71924543 |
| AGGGGGLGGAP | ASSSPRLGGRDTQY | 0.82097924 |
| AGGLLYGNGGAG | ASSSPRLGGRDTQY | 0.86430526 |
| AGLESIGL | ASSSPRLGGRDTQY | 0.7258188 |
| AGLKTGII | ASSSPRLGGRDTQY | 0.6030124 |
| ASLAALLL | ASSSPRLGGRDTQY | 0.7726778 |
| ASVINISSI | ASSSPRLGGRDTQY | 0.8864298 |
| ATPPGPA | ASSSPRLGGRDTQY | 0.59781504 |
| ATVASALV | ASSSPRLGGRDTQY | 0.45844573 |
| ATVASLIL | ASSSPRLGGRDTQY | 0.74889493 |
| AVNGPVLIRH | ASSSPRLGGRDTQY | 0.76500607 |
| DGAAGAVGI | ASSSPRLGGRDTQY | 0.8672771 |
| DPLLHGI | ASSSPRLGGRDTQY | 0.8404192 |
| ELGAALGAL | ASSSPRLGGRDTQY | 0.75531995 |
| ELGRTQAV | ASSSPRLGGRDTQY | 0.8237559 |
| FVAGGYLS | ASSSPRLGGRDTQY | 0.8455484 |
| GASGGAGGAGGW | ASSSPRLGGRDTQY | 0.8157948 |
| GGAGAGGDGGAGGI | ASSSPRLGGRDTQY | 0.85041785 |
| GTGHNGSQPGMGGNG | ASSSPRLGGRDTQY | 0.6516564 |
| GYAAASASAS | ASSSPRLGGRDTQY | 0.87516606 |
| HQVDVAAVQQG | ASSSPRLGGRDTQY | 0.86386514 |
| IGAAIAHT | ASSSPRLGGRDTQY | 0.858272 |
| IGMIEVAQL | ASSSPRLGGRDTQY | 0.87803155 |
| IRLAPEHTSAMD | ASSSPRLGGRDTQY | 0.7734916 |
| IYNVSVL | ASSSPRLGGRDTQY | 0.82926315 |
| KQGFLGM | ASSSPRLGGRDTQY | 0.5531198 |
| LESVITQL | ASSSPRLGGRDTQY | 0.7821354 |
| LGLGALVVV | ASSSPRLGGRDTQY | 0.79320514 |
| LIGNGGNGGSGGI | ASSSPRLGGRDTQY | 0.7917329 |
| LNAINAPTL | ASSSPRLGGRDTQY | 0.8488013 |
| LQLLNTGV | ASSSPRLGGRDTQY | 0.8555651 |
| LQTSIEF | ASSSPRLGGRDTQY | 0.6957352 |
| MSFAVSEL | ASSSPRLGGRDTQY | 0.88699126 |
| NTGALGL | ASSSPRLGGRDTQY | 0.8741335 |
| PPGAGGAGGAGTT | ASSSPRLGGRDTQY | 0.6001744 |
| QLVGLGI | ASSSPRLGGRDTQY | 0.7675847 |
| RFLEQVHQL | ASSSPRLGGRDTQY | 0.8731489 |
| RGVGYVLR | ASSSPRLGGRDTQY | 0.8276514 |
| SATGVFAGVI | ASSSPRLGGRDTQY | 0.7565206 |

| | | |
|---|---|---|
| SLPAITLP | ASSSPRLGGRDTQY | 0.8214567 |
| STALAFTV | ASSSPRLGGRDTQY | 0.87635773 |
| SVLTVGLGK | ASSSPRLGGRDTQY | 0.78244877 |
| TALAGAGSGI | ASSSPRLGGRDTQY | 0.88459635 |
| TGLPFTGI | ASSSPRLGGRDTQY | 0.41142046 |
| VAEHVTV | ASSSPRLGGRDTQY | 0.80489653 |
| VDIVLNSL | ASSSPRLGGRDTQY | 0.75888574 |
| VLTGGGAVL | ASSSPRLGGRDTQY | 0.18297033 |
| VNVGLAYGI | ASSSPRLGGRDTQY | 0.8529036 |
| VSSAAVGAV | ASSSPRLGGRDTQY | 0.7714815 |
| VTSVYEAGV | ASSSPRLGGRDTQY | 0.85373276 |
| WVALDRGA | ASSSPRLGGRDTQY | 0.75990534 |
| YLVGSVVI | ASSSPRLGGRDTQY | 0.86097693 |
| GPPGTGKTLL | ASSSPRLGGRDTQY | 0.7996342 |
| GVGMTKF | ASSSPRLGGRDTQY | 0.814334 |
| IGIGGSDL | ASSSPRLGGRDTQY | 0.6680165 |
| IGTLLAEL | ASSSPRLGGRDTQY | 0.82747555 |
| IYTSGTT | ASSSPRLGGRDTQY | 0.8830341 |
| LSFPAGV | ASSSPRLGGRDTQY | 0.88511395 |
| LYTSGTT | ASSSPRLGGRDTQY | 0.3694229 |
| VGAFPFGF | ASSSPRLGGRDTQY | 0.82718265 |
| VGAGTVEF | ASSSPRLGGRDTQY | 0.76917326 |
| VGKSTLLL | ASSSPRLGGRDTQY | 0.3500253 |
| AALAHAGGGIV | ASSSPRLGGRDTQY | 0.3779335 |
| AALVAVSLL | ASSSPRLGGRDTQY | 0.65020853 |
| AAVSRAVAAL | ASSSPRLGGRDTQY | 0.7062209 |
| AGGSSGIGGF | ASSSPRLGGRDTQY | 0.8267793 |
| AGLTVANANG | ASSSPRLGGRDTQY | 0.8368499 |
| AGLVGAVL | ASSSPRLGGRDTQY | 0.7105419 |
| ALSQAFTV | ASSSPRLGGRDTQY | 0.83158016 |
| ALTFSVSL | ASSSPRLGGRDTQY | 0.79393804 |
| ALTGAALVAL | ASSSPRLGGRDTQY | 0.08429589 |
| ASFSIGAGVL | ASSSPRLGGRDTQY | 0.7501712 |
| DGSVYKI | ASSSPRLGGRDTQY | 0.5193386 |
| EQLAEIV | ASSSPRLGGRDTQY | 0.88773024 |
| FAVSELTVM | ASSSPRLGGRDTQY | 0.6348794 |
| GAAPLAAGL | ASSSPRLGGRDTQY | 0.73164326 |
| GADGGDGGSGGAGG | ASSSPRLGGRDTQY | 0.77136075 |
| GFGLDTGGAGG | ASSSPRLGGRDTQY | 0.8125056 |
| GLASGGLL | ASSSPRLGGRDTQY | 0.7949195 |
| GRIAGAAVLL | ASSSPRLGGRDTQY | 0.63988674 |
| GSAGSLNTSF | ASSSPRLGGRDTQY | 0.42305347 |
| GTGQIGI | ASSSPRLGGRDTQY | 0.84037995 |
| GTLIALIL | ASSSPRLGGRDTQY | 0.72742 |
| IAATMGFVVG | ASSSPRLGGRDTQY | 0.5852995 |
| IFAQALL | ASSSPRLGGRDTQY | 0.6980232 |
| IGAAVALGF | ASSSPRLGGRDTQY | 0.8172605 |
| IGLTPLAL | ASSSPRLGGRDTQY | 0.43941522 |
| ILGSNTSTL | ASSSPRLGGRDTQY | 0.8775225 |
| IVFSLTVL | ASSSPRLGGRDTQY | 0.6382545 |
| LASKATGF | ASSSPRLGGRDTQY | 0.71875805 |

| | | |
|---|---|---|
| LASVALIL | ASSSPRLGGRDTQY | 0.87661207 |
| LFAVGLTL | ASSSPRLGGRDTQY | 0.79481065 |
| LFNGGIGTYI | ASSSPRLGGRDTQY | 0.7527074 |
| LTLGAGII | ASSSPRLGGRDTQY | 0.70647657 |
| LVAAALGDI | ASSSPRLGGRDTQY | 0.63674533 |
| NTAFVQLGI | ASSSPRLGGRDTQY | 0.6538543 |
| QLFSGAEL | ASSSPRLGGRDTQY | 0.6382058 |
| SAATFITL | ASSSPRLGGRDTQY | 0.8779001 |
| SNYGLANLGV | ASSSPRLGGRDTQY | 0.83283716 |
| SYGGAYAVM | ASSSPRLGGRDTQY | 0.596838 |
| TTGVGGLTLG | ASSSPRLGGRDTQY | 0.86773777 |
| TTQTIYI | ASSSPRLGGRDTQY | 0.88180274 |
| VAAGAGLGAVI | ASSSPRLGGRDTQY | 0.8828144 |
| VAAVVTLL | ASSSPRLGGRDTQY | 0.6284356 |
| VGFLGGLI | ASSSPRLGGRDTQY | 0.81367314 |
| VGVIPILL | ASSSPRLGGRDTQY | 0.80567956 |
| VIFASSLI | ASSSPRLGGRDTQY | 0.75067186 |
| VKGGDGGAAAG | ASSSPRLGGRDTQY | 0.67999995 |
| VTKTFTNI | ASSSPRLGGRDTQY | 0.587157 |
| VVAAAILL | ASSSPRLGGRDTQY | 0.87265754 |
| YAALDVEL | ASSSPRLGGRDTQY | 0.901242 |
| LASVALLL | ASSSPRLGGRDTQY | 0.785781 |
| LDAVHRV | ASSSPRLGGRDTQY | 0.4131685 |
| QGLSSSIF | ASSSPRLGGRDTQY | 0.75225914 |
| RGTFANIRL | ASSSPRLGGRDTQY | 0.60566235 |
| RQLWWGHRI | ASSSPRLGGRDTQY | 0.647717 |
| YGPPGTGKTLL | ASSSPRLGGRDTQY | 0.7324047 |
| AALVGAPANVA | ASSSPRLGGRDTQY | 0.44977486 |
| AATFRRTE | ASSSPRLGGRDTQY | 0.7552431 |
| AGALGGGI | ASSSPRLGGRDTQY | 0.8840204 |
| AGGAGGAGGIGAG | ASSSPRLGGRDTQY | 0.7535942 |
| AGSGALGL | ASSSPRLGGRDTQY | 0.8414345 |
| AQIADQI | ASSSPRLGGRDTQY | 0.8661952 |
| ASAVAVTYL | ASSSPRLGGRDTQY | 0.8249335 |
| ASTVLGGTGG | ASSSPRLGGRDTQY | 0.86503935 |
| AVGGEPAK | ASSSPRLGGRDTQY | 0.8628703 |
| CEQGTFGGTC | ASSSPRLGGRDTQY | 0.82465714 |
| DEGKAPAG | ASSSPRLGGRDTQY | 0.79974097 |
| DTVVFPAL | ASSSPRLGGRDTQY | 0.8400899 |
| FEGITMAG | ASSSPRLGGRDTQY | 0.7555283 |
| FPLVVQD | ASSSPRLGGRDTQY | 0.8041216 |
| GAFTTPQ | ASSSPRLGGRDTQY | 0.8035644 |
| GFGGFGTAGGVGGA | ASSSPRLGGRDTQY | 0.84287584 |
| GFLAQFL | ASSSPRLGGRDTQY | 0.7833309 |
| GGAGGAGGNAGVA | ASSSPRLGGRDTQY | 0.88825333 |
| GGVGTAGAGGAGGAGG | ASSSPRLGGRDTQY | 0.23884402 |
| GNGGGGDAGNAGSGGN | ASSSPRLGGRDTQY | 0.5534948 |
| GTGGAGGAGNDAGS | ASSSPRLGGRDTQY | 0.8575899 |
| GTTNTGWFNS | ASSSPRLGGRDTQY | 0.18699673 |
| IFMGPLFTL | ASSSPRLGGRDTQY | 0.51942205 |
| ILAHAIEAI | ASSSPRLGGRDTQY | 0.81346697 |

| | | |
|---|---|---|
| IRSAIREAL | ASSSPRLGGRDTQY | 0.8707666 |
| ITFAGGG | ASSSPRLGGRDTQY | 0.16110064 |
| ITGMSGHQEV | ASSSPRLGGRDTQY | 0.7363515 |
| IVAGLACTL | ASSSPRLGGRDTQY | 0.44630533 |
| IVGGGLAGA | ASSSPRLGGRDTQY | 0.86623555 |
| IVTGVLNI | ASSSPRLGGRDTQY | 0.7409414 |
| LAQQGCI | ASSSPRLGGRDTQY | 0.41443148 |
| LCLHVEN | ASSSPRLGGRDTQY | 0.85133994 |
| LDAISDGS | ASSSPRLGGRDTQY | 0.60241723 |
| LGGFAAML | ASSSPRLGGRDTQY | 0.77699333 |
| LGGKNPMI | ASSSPRLGGRDTQY | 0.8666842 |
| LGPKGVAAIL | ASSSPRLGGRDTQY | 0.8867476 |
| LMVGVSI | ASSSPRLGGRDTQY | 0.5636492 |
| LVGAAFVL | ASSSPRLGGRDTQY | 0.8275353 |
| LVLAVAPL | ASSSPRLGGRDTQY | 0.724208 |
| MAAIAVIDAA | ASSSPRLGGRDTQY | 0.897634 |
| NNGINAELQ | ASSSPRLGGRDTQY | 0.7245718 |
| RGEPLVLVHG | ASSSPRLGGRDTQY | 0.42509788 |
| RGGAGGAAGLI | ASSSPRLGGRDTQY | 0.8454113 |
| RSYRMHPAV | ASSSPRLGGRDTQY | 0.84284914 |
| SAAHLLAGQLA | ASSSPRLGGRDTQY | 0.7832009 |
| SAAVGSY | ASSSPRLGGRDTQY | 0.8483755 |
| SPHAVLHRI | ASSSPRLGGRDTQY | 0.78002137 |
| SQIAKVL | ASSSPRLGGRDTQY | 0.7221501 |
| TALGHLVP | ASSSPRLGGRDTQY | 0.84714615 |
| TAPTTSGAA | ASSSPRLGGRDTQY | 0.83807236 |
| TDGDDGNGGSPGT | ASSSPRLGGRDTQY | 0.62840605 |
| TDGLGEN | ASSSPRLGGRDTQY | 0.87767386 |
| TGGPPLPGQA | ASSSPRLGGRDTQY | 0.792232 |
| TIPAITI | ASSSPRLGGRDTQY | 0.6685653 |
| TIPGFTI | ASSSPRLGGRDTQY | 0.8835584 |
| TSIASGALA | ASSSPRLGGRDTQY | 0.79651934 |
| TVTLPGAAI | ASSSPRLGGRDTQY | 0.8504608 |
| VASGLSAALP | ASSSPRLGGRDTQY | 0.84654593 |
| VGAGVIGI | ASSSPRLGGRDTQY | 0.88977545 |
| VGPKRFGV | ASSSPRLGGRDTQY | 0.58468664 |
| VNVAFPD | ASSSPRLGGRDTQY | 0.85255206 |
| VSMTNSGVS | ASSSPRLGGRDTQY | 0.8189285 |
| VTDVLGDLSV | ASSSPRLGGRDTQY | 0.85439605 |
| VVVVGAGFAGL | ASSSPRLGGRDTQY | 0.8063642 |
| VYTVGTVV | ASSSPRLGGRDTQY | 0.798007 |
| WTTFSGP | ASSSPRLGGRDTQY | 0.85632074 |
| YDELVEAF | ASSSPRLGGRDTQY | 0.87280744 |
| YGTGGAGGNGG | ASSSPRLGGRDTQY | 0.8094883 |
| YTNVMAV | ASSSPRLGGRDTQY | 0.8323915 |
| FLEQVHQL | ASSSPRLGGRDTQY | 0.8798152 |
| GLSLLAVAV | ASSSPRLGGRDTQY | 0.6087804 |
| LEQVHQL | ASSSPRLGGRDTQY | 0.80154794 |
| LNAAGAI | ASSSPRLGGRDTQY | 0.861293 |
| PAVHASL | ASSSPRLGGRDTQY | 0.87492216 |
| TELEDDA | ASSSPRLGGRDTQY | 0.7949419 |

| | | |
|---|---|---|
| TPPGPPN | ASSSPRLGGRDTQY | 0.8823389 |
| TPPPPPPP | ASSSPRLGGRDTQY | 0.7251078 |
| TQSPALI | ASFLQGPGNTLY | 0.83019286 |
| EFAAAFPQ | ASFLQGPGNTLY | 0.71758604 |
| LLSGNLDVL | ASFLQGPGNTLY | 0.76583433 |
| GPANLGNYNI | ASFLQGPGNTLY | 0.54448867 |
| TLGPTLP | ASFLQGPGNTLY | 0.8166468 |
| VYAYSSCCK | ASFLQGPGNTLY | 0.8387625 |
| APAPAPAPNI | ASFLQGPGNTLY | 0.82019186 |
| PGGPVPIP | ASFLQGPGNTLY | 0.44101822 |
| FLTVALEP | ASFLQGPGNTLY | 0.8126794 |
| SQYFGANL | ASFLQGPGNTLY | 0.84908545 |
| GGVKPAA | ASFLQGPGNTLY | 0.8891455 |
| GGTGGAGGAGGW | ASFLQGPGNTLY | 0.7911876 |
| GVGGVAGLGGAGG | ASFLQGPGNTLY | 0.87153363 |
| VLLVLATTI | ASFLQGPGNTLY | 0.7579862 |
| IGIFAAML | ASFLQGPGNTLY | 0.8103546 |
| AVGDLID | ASFLQGPGNTLY | 0.54153097 |
| NISPLPL | ASFLQGPGNTLY | 0.7358848 |
| VTAVFVAI | ASFLQGPGNTLY | 0.74634933 |
| SGDDNDAAL | ASFLQGPGNTLY | 0.6239042 |
| AVAALAHS | ASFLQGPGNTLY | 0.8442274 |
| GGVTVMATP | ASFLQGPGNTLY | 0.86823255 |
| AGLAANVRD | ASFLQGPGNTLY | 0.8410156 |
| TNRPDLIDPALLRPG | ASFLQGPGNTLY | 0.8620792 |
| LANLGVTAL | ASFLQGPGNTLY | 0.8680823 |
| TGERIEL | ASFLQGPGNTLY | 0.875609 |
| NGEELAA | ASFLQGPGNTLY | 0.7156782 |
| VLSLVWG | ASFLQGPGNTLY | 0.78639543 |
| IGYALWL | ASFLQGPGNTLY | 0.4396615 |
| LSLLLAPL | ASFLQGPGNTLY | 0.7079822 |
| TLDEALK | ASFLQGPGNTLY | 0.70443106 |
| TDGAGTAEGG | ASFLQGPGNTLY | 0.78155226 |
| NAPGGAGGNGG | ASFLQGPGNTLY | 0.11116508 |
| VSVFMGMGGNF | ASFLQGPGNTLY | 0.8016019 |
| LASYLAAL | ASFLQGPGNTLY | 0.87782276 |
| LVGGTPAL | ASFLQGPGNTLY | 0.7346285 |
| VRNADVL | ASFLQGPGNTLY | 0.8488073 |
| TCSSPFLVSAM | ASFLQGPGNTLY | 0.820269 |
| VHITDAE | ASFLQGPGNTLY | 0.6856735 |
| EYAAPSF | ASFLQGPGNTLY | 0.83634543 |
| VVDAFHGGL | ASFLQGPGNTLY | 0.6994789 |
| ASVVDPQ | ASFLQGPGNTLY | 0.867283 |
| SYNEFGGLS | ASFLQGPGNTLY | 0.88095224 |
| DPDGSVY | ASFLQGPGNTLY | 0.8215444 |
| MVSTIHATE | ASFLQGPGNTLY | 0.7898444 |
| LAAGVADPV | ASFLQGPGNTLY | 0.84031 |
| TKEQIAA | ASFLQGPGNTLY | 0.60685194 |
| SLASPAAL | ASFLQGPGNTLY | 0.799851 |
| IGLVFYGV | ASFLQGPGNTLY | 0.49752274 |
| FLLDASNL | ASFLQGPGNTLY | 0.7809771 |

| | | |
|---|---|---|
| YQVGNFP | ASFLQGPGNTLY | 0.65662265 |
| ESLLSQST | ASFLQGPGNTLY | 0.82421136 |
| LFTVGFAAVG | ASFLQGPGNTLY | 0.779071 |
| ATPPTPT | ASFLQGPGNTLY | 0.8390783 |
| TDLATLG | ASFLQGPGNTLY | 0.6976873 |
| SAGGTGR | ASFLQGPGNTLY | 0.8166364 |
| GFSNSGS | ASFLQGPGNTLY | 0.79801404 |
| GGNGGSGGTSV | ASFLQGPGNTLY | 0.4801335 |
| GTSVATGGAGN | ASFLQGPGNTLY | 0.86624193 |
| NGGNGGSGGTS | ASFLQGPGNTLY | 0.7875949 |
| AVVKFPH | ASFLQGPGNTLY | 0.73052 |
| TPPPPPP | ASFLQGPGNTLY | 0.86217093 |
| AAPAAAAAPA | ASFLQGPGNTLY | 0.77292216 |
| IALNEDL | ASFLQGPGNTLY | 0.57089126 |
| IVNVSSM | ASFLQGPGNTLY | 0.6967191 |
| TGIPPAPR | ASFLQGPGNTLY | 0.80597776 |
| FDLGGGTF | ASFLQGPGNTLY | 0.7495884 |
| FDLGGGTFDVS | ASFLQGPGNTLY | 0.19610097 |
| NEPTAAA | ASFLQGPGNTLY | 0.84191644 |
| NEPTAAALA | ASFLQGPGNTLY | 0.66485614 |
| GSIINISSI | ASFLQGPGNTLY | 0.66597074 |
| VAPGNAGTA | ASFLQGPGNTLY | 0.72683007 |
| TGGGASLE | ASFLQGPGNTLY | 0.8496257 |
| VAVYNHYKRI | ASFLQGPGNTLY | 0.7636035 |
| TDPAPATT | ASFLQGPGNTLY | 0.7617545 |
| TNRPDILDPALLRPG | ASFLQGPGNTLY | 0.83730626 |
| LTTTTTS | ASFLQGPGNTLY | 0.81067246 |
| PTGSGKTL | ASFLQGPGNTLY | 0.8696795 |
| TGSGKTL | ASFLQGPGNTLY | 0.82211614 |
| TIGHVDHGK | ASFLQGPGNTLY | 0.8097756 |
| TIGHVDHGKTT | ASFLQGPGNTLY | 0.8604449 |
| GHVDHGKTT | ASFLQGPGNTLY | 0.76989555 |
| AQFEHTLL | ASFLQGPGNTLY | 0.66755664 |
| IVNQDEL | ASFLQGPGNTLY | 0.6375504 |
| TNGSQFF | ASFLQGPGNTLY | 0.75808513 |
| TGKGIVE | ASFLQGPGNTLY | 0.65071964 |
| LISNASDA | ASFLQGPGNTLY | 0.826513 |
| AELRTLGELTKIAKAHGAQ | ASFLQGPGNTLY | 0.8913945 |
| APNAVAAAEPSW | ASFLQGPGNTLY | 0.7899555 |
| APQVAGAATAVL | ASFLQGPGNTLY | 0.49185723 |
| ARRDPAAAAAILPTDA | ASFLQGPGNTLY | 0.7379353 |
| DAAARDVLAEAGLAEYF | ASFLQGPGNTLY | 0.7341868 |
| DDQIVLSAKANLAKVEQ | ASFLQGPGNTLY | 0.84258336 |
| DELLPDTGEDSDEDGH | ASFLQGPGNTLY | 0.85375375 |
| ELPEAPANRATVPQD | ASFLQGPGNTLY | 0.88671863 |
| FGQSKIERVTGIG | ASFLQGPGNTLY | 0.8497094 |
| GAGGTGSTAGGAGGAGGAGGLY | ASFLQGPGNTLY | 0.7127218 |
| GHGGTGAAGGNGGTGGAGGSGI | ASFLQGPGNTLY | 0.85330236 |
| GNGGNGGAGGEGGPGGAGGAGA | ASFLQGPGNTLY | 0.74305916 |
| GSTAGNGGNGGNATNGTIAGQP | ASFLQGPGNTLY | 0.8355677 |
| GTGGDGGNGGNGGTGG | ASFLQGPGNTLY | 0.8198245 |

| | | |
|---|---|---|
| GVRRPAGIPADAPV | ASFLQGPGNTLY | 0.81800085 |
| GYRQVTDAHLLTLARR | ASFLQGPGNTLY | 0.73342437 |
| IAKMGQDGHDRG | ASFLQGPGNTLY | 0.79092854 |
| KAEPSIEIQYT | ASFLQGPGNTLY | 0.53876567 |
| KPVRIRMAADEDTVAVT | ASFLQGPGNTLY | 0.8565704 |
| MPPRIAGMRLLVIKPEP | ASFLQGPGNTLY | 0.61563826 |
| MKFYAVALFRSRGVPPTR | ASFLQGPGNTLY | 0.58699423 |
| NAGGFGILGAVAHSP | ASFLQGPGNTLY | 0.7544405 |
| NEGREITLAMAEM | ASFLQGPGNTLY | 0.8793681 |
| NELVEWSHPLGHRWRWE | ASFLQGPGNTLY | 0.87761295 |
| NLGAGGTGGNGGNPG | ASFLQGPGNTLY | 0.6959102 |
| NTEWPKWETVMAADPVD | ASFLQGPGNTLY | 0.6848894 |
| PASAAPAETQP | ASFLQGPGNTLY | 0.8219762 |
| QQSSDGKNGVRAAEITGE | ASFLQGPGNTLY | 0.6773336 |
| RAPDAVTTSWWKE | ASFLQGPGNTLY | 0.8578328 |
| SGFGNVGAHTSGWFNQ | ASFLQGPGNTLY | 0.74035394 |
| SGVAGAVGQAASVGGLKV | ASFLQGPGNTLY | 0.72888833 |
| SNGGAGTGSAGGNGGT | ASFLQGPGNTLY | 0.7633927 |
| TGGAGGEGGAAG | ASFLQGPGNTLY | 0.8282504 |
| TGGDGGAAGAGGAPGHG | ASFLQGPGNTLY | 0.72771966 |
| TMPPLLAAIANAGRHGV | ASFLQGPGNTLY | 0.8553691 |
| TSSTAGNSGGA | ASFLQGPGNTLY | 0.29322368 |
| VPATVPVGILC | ASFLQGPGNTLY | 0.8514048 |
| VQRPDSGGVVLTGRIS | ASFLQGPGNTLY | 0.6010835 |
| WTPNPVPATKGVPAGLVQAT | ASFLQGPGNTLY | 0.6426224 |
| YALKRALAKFSDRFDVVII | ASFLQGPGNTLY | 0.28800994 |
| YLIGSVIV | ASFLQGPGNTLY | 0.82650805 |
| YGQLVAVID | ASFLQGPGNTLY | 0.833495 |
| VTVGGMTLPA | ASFLQGPGNTLY | 0.8628652 |
| VLALPAV | ASFLQGPGNTLY | 0.80171293 |
| VIPPTLNYET | ASFLQGPGNTLY | 0.84166706 |
| VGMGGMHPSSG | ASFLQGPGNTLY | 0.86142445 |
| VGLAGGAATA | ASFLQGPGNTLY | 0.6889611 |
| VGGAGGAGGSGP | ASFLQGPGNTLY | 0.33332494 |
| VFAAFMV | ASFLQGPGNTLY | 0.7942474 |
| TLYDKLR | ASFLQGPGNTLY | 0.7925847 |
| TLAVFTVL | ASFLQGPGNTLY | 0.8625395 |
| TGSVQPARV | ASFLQGPGNTLY | 0.85603243 |
| TGGGAGSGGT | ASFLQGPGNTLY | 0.49172634 |
| SHLETSL | ASFLQGPGNTLY | 0.86304605 |
| SFTTGMT | ASFLQGPGNTLY | 0.83472717 |
| QVEVAKRMGVRQPTVS | ASFLQGPGNTLY | 0.53034043 |
| QGSGGPALAM | ASFLQGPGNTLY | 0.7541956 |
| MLAPSNP | ASFLQGPGNTLY | 0.7583237 |
| LVNNAAI | ASFLQGPGNTLY | 0.5609515 |
| LTGEINTLA | ASFLQGPGNTLY | 0.8241589 |
| LSIFGPV | ASFLQGPGNTLY | 0.6930574 |
| LLALLANL | ASFLQGPGNTLY | 0.7142066 |
| LISNASDALDKLR | ASFLQGPGNTLY | 0.77023375 |
| LINEAVP | ASFLQGPGNTLY | 0.65602076 |
| LGYAFSGL | ASFLQGPGNTLY | 0.69741344 |

| | | |
|---|---|---|
| LDFVAVAL | ASFLQGPGNTLY | 0.70868087 |
| LAQPGAE | ASFLQGPGNTLY | 0.83664334 |
| ISNLGAVTGF | ASFLQGPGNTLY | 0.838143 |
| HQEDPGRI | ASFLQGPGNTLY | 0.7545421 |
| GTPGEPTP | ASFLQGPGNTLY | 0.843235 |
| GFFNSGS | ASFLQGPGNTLY | 0.8669379 |
| GAGGGGLTAGAN | ASFLQGPGNTLY | 0.39214247 |
| FVDALVNM | ASFLQGPGNTLY | 0.6566988 |
| FLTVAPEL | ASFLQGPGNTLY | 0.82125854 |
| FETLTAL | ASFLQGPGNTLY | 0.69445914 |
| DVINPIEL | ASFLQGPGNTLY | 0.86807865 |
| APVSGAPV | ASFLQGPGNTLY | 0.80253434 |
| ALVGGLAW | ASFLQGPGNTLY | 0.80820936 |
| ALAAAAATAI | ASFLQGPGNTLY | 0.79205036 |
| AFGLGLLL | ASFLQGPGNTLY | 0.887227 |
| AFFAAASL | ASFLQGPGNTLY | 0.815588 |
| AAAALPAFSPP | ASFLQGPGNTLY | 0.79630977 |
| VVLAPAL | ASFLQGPGNTLY | 0.830801 |
| TIGHVDHGKTTL | ASFLQGPGNTLY | 0.65156126 |
| LISNASDALDK | ASFLQGPGNTLY | 0.8525399 |
| LISNASDAL | ASFLQGPGNTLY | 0.86431974 |
| ISNASDALDK | ASFLQGPGNTLY | 0.57307804 |
| NQAETLVYQTE | ASFLQGPGNTLY | 0.72817063 |
| TGGTGGTAGAG | ASFLQGPGNTLY | 0.887766 |
| TGSTAGGAGGA | ASFLQGPGNTLY | 0.86362076 |
| DPLAVPVAAGPV | ASFLQGPGNTLY | 0.67576104 |
| GGTGGTPGNGGN | ASFLQGPGNTLY | 0.6766709 |
| PAGLAGVDPATL | ASFLQGPGNTLY | 0.64365476 |
| GNNLVVEAPHAGASV | ASFLQGPGNTLY | 0.7609209 |
| KVTRIMSASTLEQPA | ASFLQGPGNTLY | 0.45823652 |
| LGFALPIALPRRTVV | ASFLQGPGNTLY | 0.8243751 |
| TRLGNLTARKLVVLV | ASFLQGPGNTLY | 0.76163244 |
| WSQQRRLIYFPSAGP | ASFLQGPGNTLY | 0.86435217 |
| GPNGNPDPMAAAVDIR | ASFLQGPGNTLY | 0.8182621 |
| TQRCVSVADLRAAASR | ASFLQGPGNTLY | 0.86838806 |
| QFVLDVAEHPVEVLTETV | ASFLQGPGNTLY | 0.74589324 |
| GGAGGTGGAGGNGGTQAGDG | ASFLQGPGNTLY | 0.7522229 |
| PLLATLRAIAVDVDAKPAQV | ASFLQGPGNTLY | 0.42536542 |
| APKFRTGGYMIDFWGVGYQVAK | ASFLQGPGNTLY | 0.5063814 |
| GAGGAGGAGGAAGGPGGTGGTG | ASFLQGPGNTLY | 0.6805531 |
| IGALGLLATDVAPAGSVAFGVA | ASFLQGPGNTLY | 0.72726536 |
| AGGAGGIGGNGGAGGAGGVLMG | ASFLQGPGNTLY | 0.30496806 |
| ADELIGLL | ASFLQGPGNTLY | 0.87782025 |
| ADRALKA | ASFLQGPGNTLY | 0.72791946 |
| AGAVGQVGGAGGA | ASFLQGPGNTLY | 0.8451587 |
| AGGGGGLGGAP | ASFLQGPGNTLY | 0.7151854 |
| AGGLLYGNGGAG | ASFLQGPGNTLY | 0.5988394 |
| AGLESIGL | ASFLQGPGNTLY | 0.69326454 |
| AGLKTGII | ASFLQGPGNTLY | 0.85026264 |
| ASLAALLL | ASFLQGPGNTLY | 0.57740057 |
| ASVINISSI | ASFLQGPGNTLY | 0.38329753 |

| | | |
|---|---|---|
| ATPPGPA | ASFLQGPGNTLY | 0.7176326 |
| ATVASALV | ASFLQGPGNTLY | 0.75093406 |
| ATVASLIL | ASFLQGPGNTLY | 0.8835928 |
| AVNGPVLIRH | ASFLQGPGNTLY | 0.86507285 |
| DGAAGAVGI | ASFLQGPGNTLY | 0.79115546 |
| DPLLHGI | ASFLQGPGNTLY | 0.8948076 |
| ELGAALGAL | ASFLQGPGNTLY | 0.15476014 |
| ELGRTQAV | ASFLQGPGNTLY | 0.725423 |
| FVAGGYLS | ASFLQGPGNTLY | 0.88481 |
| GASGGAGGAGGW | ASFLQGPGNTLY | 0.82231253 |
| GGAGAGGDGGAGGI | ASFLQGPGNTLY | 0.7812036 |
| GTGHNGSQPGMGGNG | ASFLQGPGNTLY | 0.86196566 |
| GYAAASASAS | ASFLQGPGNTLY | 0.6651639 |
| HQVDVAAVQQG | ASFLQGPGNTLY | 0.77221584 |
| IGAAIAHT | ASFLQGPGNTLY | 0.74749464 |
| IGMIEVAQL | ASFLQGPGNTLY | 0.86200106 |
| IRLAPEHTSAMD | ASFLQGPGNTLY | 0.83900905 |
| IYNVSVL | ASFLQGPGNTLY | 0.50319767 |
| KQGFLGM | ASFLQGPGNTLY | 0.86008334 |
| LESVITQL | ASFLQGPGNTLY | 0.88760376 |
| LGLGALVVV | ASFLQGPGNTLY | 0.70039 |
| LIGNGGNGGSGGI | ASFLQGPGNTLY | 0.8610972 |
| LNAINAPTL | ASFLQGPGNTLY | 0.80056226 |
| LQLLNTGV | ASFLQGPGNTLY | 0.75255156 |
| LQTSIEF | ASFLQGPGNTLY | 0.674527 |
| MSFAVSEL | ASFLQGPGNTLY | 0.5218191 |
| NTGALGL | ASFLQGPGNTLY | 0.8309952 |
| PPGAGGAGGAGTT | ASFLQGPGNTLY | 0.58903307 |
| QLVGLGI | ASFLQGPGNTLY | 0.82312655 |
| RFLEQVHQL | ASFLQGPGNTLY | 0.7849394 |
| RGVGYVLR | ASFLQGPGNTLY | 0.8987683 |
| SATGVFAGVI | ASFLQGPGNTLY | 0.83869386 |
| SLPAITLP | ASFLQGPGNTLY | 0.80513346 |
| STALAFTV | ASFLQGPGNTLY | 0.6525192 |
| SVLTVGLGK | ASFLQGPGNTLY | 0.82945997 |
| TALAGAGSGI | ASFLQGPGNTLY | 0.80940044 |
| TGLPFTGI | ASFLQGPGNTLY | 0.523039 |
| VAEHVTV | ASFLQGPGNTLY | 0.87246627 |
| VDIVLNSL | ASFLQGPGNTLY | 0.5512327 |
| VLTGGGAVL | ASFLQGPGNTLY | 0.77784276 |
| VNVGLAYGI | ASFLQGPGNTLY | 0.8390807 |
| VSSAAVGAV | ASFLQGPGNTLY | 0.8670732 |
| VTSVYEAGV | ASFLQGPGNTLY | 0.78892857 |
| WVALDRGA | ASFLQGPGNTLY | 0.8396088 |
| YLVGSVVI | ASFLQGPGNTLY | 0.6208087 |
| GPPGTGKTLL | ASFLQGPGNTLY | 0.8478611 |
| GVGMTKF | ASFLQGPGNTLY | 0.8620046 |
| IGIGGSDL | ASFLQGPGNTLY | 0.80373013 |
| IGTLLAEL | ASFLQGPGNTLY | 0.89223576 |
| IYTSGTT | ASFLQGPGNTLY | 0.8382919 |
| LSFPAGV | ASFLQGPGNTLY | 0.37644204 |

| | | |
|---|---|---|
| LYTSGTT | ASFLQGPGNTLY | 0.7983794 |
| VGAFPFGF | ASFLQGPGNTLY | 0.86931884 |
| VGAGTVEF | ASFLQGPGNTLY | 0.86155057 |
| VGKSTLLL | ASFLQGPGNTLY | 0.8679315 |
| AALAHAGGGIV | ASFLQGPGNTLY | 0.85970306 |
| AALVAVSLL | ASFLQGPGNTLY | 0.7266011 |
| AAVSRAVAAL | ASFLQGPGNTLY | 0.87447554 |
| AGGSSGIGGF | ASFLQGPGNTLY | 0.9082716 |
| AGLTVANANG | ASFLQGPGNTLY | 0.8444843 |
| AGLVGAVL | ASFLQGPGNTLY | 0.64035267 |
| ALSQAFTV | ASFLQGPGNTLY | 0.64541256 |
| ALTFSVSL | ASFLQGPGNTLY | 0.84499466 |
| ALTGAALVAL | ASFLQGPGNTLY | 0.82988644 |
| ASFSIGAGVL | ASFLQGPGNTLY | 0.63215566 |
| DGSVYKI | ASFLQGPGNTLY | 0.7645619 |
| EQLAEIV | ASFLQGPGNTLY | 0.7975496 |
| FAVSELTVM | ASFLQGPGNTLY | 0.339513 |
| GAAPLAAGL | ASFLQGPGNTLY | 0.8458725 |
| GADGGDGGSGGAGG | ASFLQGPGNTLY | 0.8729868 |
| GFGLDTGGAGG | ASFLQGPGNTLY | 0.8473372 |
| GLASGGLL | ASFLQGPGNTLY | 0.55226123 |
| GRIAGAAVLL | ASFLQGPGNTLY | 0.8660495 |
| GSAGSLNTSF | ASFLQGPGNTLY | 0.8765349 |
| GTGQIGI | ASFLQGPGNTLY | 0.88599193 |
| GTLIALIL | ASFLQGPGNTLY | 0.40973207 |
| IAATMGFVVG | ASFLQGPGNTLY | 0.19328143 |
| IFAQALL | ASFLQGPGNTLY | 0.80914235 |
| IGAAVALGF | ASFLQGPGNTLY | 0.8366157 |
| IGLTPLAL | ASFLQGPGNTLY | 0.711783 |
| ILGSNTSTL | ASFLQGPGNTLY | 0.58801126 |
| IVFSLTVL | ASFLQGPGNTLY | 0.77118134 |
| LASKATGF | ASFLQGPGNTLY | 0.8616455 |
| LASVALIL | ASFLQGPGNTLY | 0.6300732 |
| LFAVGLTL | ASFLQGPGNTLY | 0.54517007 |
| LFNGGIGTYI | ASFLQGPGNTLY | 0.85360086 |
| LTLGAGII | ASFLQGPGNTLY | 0.87209755 |
| LVAAALGDI | ASFLQGPGNTLY | 0.60500526 |
| NTAFVQLGI | ASFLQGPGNTLY | 0.71288955 |
| QLFSGAEL | ASFLQGPGNTLY | 0.8160249 |
| SAATFITL | ASFLQGPGNTLY | 0.80393195 |
| SNYGLANLGV | ASFLQGPGNTLY | 0.7848757 |
| SYGGAYAVM | ASFLQGPGNTLY | 0.49875522 |
| TTGVGGLTLG | ASFLQGPGNTLY | 0.84211326 |
| TTQTIYI | ASFLQGPGNTLY | 0.55059266 |
| VAAGAGLGAVI | ASFLQGPGNTLY | 0.8636643 |
| VAAVVTLL | ASFLQGPGNTLY | 0.79288816 |
| VGFLGGLI | ASFLQGPGNTLY | 0.8370414 |
| VGVIPILL | ASFLQGPGNTLY | 0.7542184 |
| VIFASSLI | ASFLQGPGNTLY | 0.5534029 |
| VKGGDGGAAAG | ASFLQGPGNTLY | 0.6041861 |
| VTKTFTNI | ASFLQGPGNTLY | 0.5612561 |

| | | |
|---|---|---|
| VVAAAILL | ASFLQGPGNTLY | 0.8646138 |
| YAALDVEL | ASFLQGPGNTLY | 0.7563137 |
| LASVALLL | ASFLQGPGNTLY | 0.8520062 |
| LDAVHRV | ASFLQGPGNTLY | 0.80763704 |
| QGLSSSIF | ASFLQGPGNTLY | 0.625841 |
| RGTFANIRL | ASFLQGPGNTLY | 0.83703285 |
| RQLWWGHRI | ASFLQGPGNTLY | 0.3981048 |
| YGPPGTGKTLL | ASFLQGPGNTLY | 0.8460601 |
| AALVGAPANVA | ASFLQGPGNTLY | 0.7823808 |
| AATFRRTE | ASFLQGPGNTLY | 0.8457783 |
| AGALGGGI | ASFLQGPGNTLY | 0.7542691 |
| AGGAGGAGGIGAG | ASFLQGPGNTLY | 0.85308397 |
| AGSGALGL | ASFLQGPGNTLY | 0.82606876 |
| AQIADQI | ASFLQGPGNTLY | 0.7420316 |
| ASAVAVTYL | ASFLQGPGNTLY | 0.84598875 |
| ASTVLGGTGG | ASFLQGPGNTLY | 0.85559785 |
| AVGGEPAK | ASFLQGPGNTLY | 0.8551987 |
| CEQGTFGGTC | ASFLQGPGNTLY | 0.5174324 |
| DEGKAPAG | ASFLQGPGNTLY | 0.78851616 |
| DTVVFPAL | ASFLQGPGNTLY | 0.6749368 |
| FEGITMAG | ASFLQGPGNTLY | 0.80749285 |
| FPLVVQD | ASFLQGPGNTLY | 0.7960421 |
| GAFTTPQ | ASFLQGPGNTLY | 0.79125875 |
| GFGGFGTAGGVGGA | ASFLQGPGNTLY | 0.820152 |
| GFLAQFL | ASFLQGPGNTLY | 0.5743785 |
| GGAGGAGGNAGVA | ASFLQGPGNTLY | 0.68011594 |
| GGVGTAGAGGAGGAGG | ASFLQGPGNTLY | 0.76069903 |
| GNGGGGDAGNAGSGGN | ASFLQGPGNTLY | 0.81450367 |
| GTGGAGGAGNDAGS | ASFLQGPGNTLY | 0.8288653 |
| GTTNTGWFNS | ASFLQGPGNTLY | 0.5708426 |
| IFMGPLFTL | ASFLQGPGNTLY | 0.8062377 |
| ILAHAIEAI | ASFLQGPGNTLY | 0.7703465 |
| IRSAIREAL | ASFLQGPGNTLY | 0.65164804 |
| ITFAGGG | ASFLQGPGNTLY | 0.8334145 |
| ITGMSGHQEV | ASFLQGPGNTLY | 0.7469895 |
| IVAGLACTL | ASFLQGPGNTLY | 0.40379822 |
| IVGGGLAGA | ASFLQGPGNTLY | 0.8580189 |
| IVTGVLNI | ASFLQGPGNTLY | 0.59131277 |
| LAQQGCI | ASFLQGPGNTLY | 0.8166516 |
| LCLHVEN | ASFLQGPGNTLY | 0.824344 |
| LDAISDGS | ASFLQGPGNTLY | 0.5482346 |
| LGGFAAML | ASFLQGPGNTLY | 0.8563052 |
| LGGKNPMI | ASFLQGPGNTLY | 0.7499075 |
| LGPKGVAAIL | ASFLQGPGNTLY | 0.8021906 |
| LMVGVSI | ASFLQGPGNTLY | 0.74229413 |
| LVGAAFVL | ASFLQGPGNTLY | 0.7318094 |
| LVLAVAPL | ASFLQGPGNTLY | 0.8590133 |
| MAAIAVIDAA | ASFLQGPGNTLY | 0.82939273 |
| NNGINAELQ | ASFLQGPGNTLY | 0.8579042 |
| RGEPLVLVHG | ASFLQGPGNTLY | 0.8207507 |
| RGGAGGAAGLI | ASFLQGPGNTLY | 0.8614551 |

| | | |
|---|---|---|
| RSYRMHPAV | ASFLQGPGNTLY | 0.8557379 |
| SAAHLLAGQLA | ASFLQGPGNTLY | 0.64504325 |
| SAAVGSY | ASFLQGPGNTLY | 0.38948327 |
| SPHAVLHRI | ASFLQGPGNTLY | 0.7506075 |
| SQIAKVL | ASFLQGPGNTLY | 0.7246168 |
| TALGHLVP | ASFLQGPGNTLY | 0.6918479 |
| TAPTTSGAA | ASFLQGPGNTLY | 0.42143276 |
| TDGDDGNGGSPGT | ASFLQGPGNTLY | 0.85074174 |
| TDGLGEN | ASFLQGPGNTLY | 0.89344835 |
| TGGPPLPGQA | ASFLQGPGNTLY | 0.62241304 |
| TIPAITI | ASFLQGPGNTLY | 0.88607585 |
| TIPGFTI | ASFLQGPGNTLY | 0.56787115 |
| TSIASGALA | ASFLQGPGNTLY | 0.84103966 |
| TVTLPGAAI | ASFLQGPGNTLY | 0.87243056 |
| VASGLSAALP | ASFLQGPGNTLY | 0.4851941 |
| VGAGVIGI | ASFLQGPGNTLY | 0.8244842 |
| VGPKRFGV | ASFLQGPGNTLY | 0.7692245 |
| VNVAFPD | ASFLQGPGNTLY | 0.6444292 |
| VSMTNSGVS | ASFLQGPGNTLY | 0.8669578 |
| VTDVLGDLSV | ASFLQGPGNTLY | 0.7313948 |
| VVVVGAGFAGL | ASFLQGPGNTLY | 0.85575444 |
| VYTVGTVV | ASFLQGPGNTLY | 0.70119154 |
| WTTFSGP | ASFLQGPGNTLY | 0.63335943 |
| YDELVEAF | ASFLQGPGNTLY | 0.8411902 |
| YGTGGAGGNGG | ASFLQGPGNTLY | 0.8665573 |
| YTNVMAV | ASFLQGPGNTLY | 0.7908282 |
| FLEQVHQL | ASFLQGPGNTLY | 0.9061832 |
| GLSLLAVAV | ASFLQGPGNTLY | 0.716563 |
| LEQVHQL | ASFLQGPGNTLY | 0.88839585 |
| LNAAGAI | ASFLQGPGNTLY | 0.87282395 |
| PAVHASL | ASFLQGPGNTLY | 0.7731552 |
| TELEDDA | ASFLQGPGNTLY | 0.77676356 |
| TPPGPPN | ASFLQGPGNTLY | 0.84847945 |
| TPPPPPPP | ASFLQGPGNTLY | 0.7832605 |
| TQSPALI | ASSLNTGQLY | 0.85013354 |
| EFAAAFPQ | ASSLNTGQLY | 0.88282305 |
| LLSGNLDVL | ASSLNTGQLY | 0.61227584 |
| GPANLGNYNI | ASSLNTGQLY | 0.84958476 |
| TLGPTLP | ASSLNTGQLY | 0.739426 |
| VYAYSSCCK | ASSLNTGQLY | 0.7757417 |
| APAPAPAPNI | ASSLNTGQLY | 0.63505566 |
| PGGPVPIP | ASSLNTGQLY | 0.4351966 |
| FLTVALEP | ASSLNTGQLY | 0.7463682 |
| SQYFGANL | ASSLNTGQLY | 0.5441164 |
| GGVKPAA | ASSLNTGQLY | 0.7947197 |
| GGTGGAGGAGGW | ASSLNTGQLY | 0.80790764 |
| GVGGVAGLGGAGG | ASSLNTGQLY | 0.6521565 |
| VLLVLATTI | ASSLNTGQLY | 0.85184604 |
| IGIFAAML | ASSLNTGQLY | 0.8516544 |
| AVGDLID | ASSLNTGQLY | 0.4582125 |
| NISPLPL | ASSLNTGQLY | 0.66391706 |

| | | |
|---|---|---|
| VTAVFVAI | ASSLNTGQLY | 0.68125916 |
| SGDDNDAAL | ASSLNTGQLY | 0.71269107 |
| AVAALAHS | ASSLNTGQLY | 0.3027513 |
| GGVTVMATP | ASSLNTGQLY | 0.82294893 |
| AGLAANVRD | ASSLNTGQLY | 0.6022064 |
| TNRPDLIDPALLRPG | ASSLNTGQLY | 0.16314104 |
| LANLGVTAL | ASSLNTGQLY | 0.7664094 |
| TGERIEL | ASSLNTGQLY | 0.72316515 |
| NGEELAA | ASSLNTGQLY | 0.6709745 |
| VLSLVWG | ASSLNTGQLY | 0.83683395 |
| IGYALWL | ASSLNTGQLY | 0.43932617 |
| LSLLLAPL | ASSLNTGQLY | 0.57778764 |
| TLDEALK | ASSLNTGQLY | 0.7995882 |
| TDGAGTAEGG | ASSLNTGQLY | 0.8446545 |
| NAPGGAGGNGG | ASSLNTGQLY | 0.49969766 |
| VSVFMGMGGNF | ASSLNTGQLY | 0.770295 |
| LASYLAAL | ASSLNTGQLY | 0.84469175 |
| LVGGTPAL | ASSLNTGQLY | 0.78933835 |
| VRNADVL | ASSLNTGQLY | 0.82934093 |
| TCSSPFLVSAM | ASSLNTGQLY | 0.84818757 |
| VHITDAE | ASSLNTGQLY | 0.65763736 |
| EYAAPSF | ASSLNTGQLY | 0.8307475 |
| VVDAFHGGL | ASSLNTGQLY | 0.84766936 |
| ASVVDPQ | ASSLNTGQLY | 0.68264884 |
| SYNEFGGLS | ASSLNTGQLY | 0.7638631 |
| DPDGSVY | ASSLNTGQLY | 0.6123814 |
| MVSTIHATE | ASSLNTGQLY | 0.72290397 |
| LAAGVADPV | ASSLNTGQLY | 0.27959806 |
| TKEQIAA | ASSLNTGQLY | 0.7718142 |
| SLASPAAL | ASSLNTGQLY | 0.8408613 |
| IGLVFYGV | ASSLNTGQLY | 0.7045009 |
| FLLDASNL | ASSLNTGQLY | 0.5283085 |
| YQVGNFP | ASSLNTGQLY | 0.52047837 |
| ESLLSQST | ASSLNTGQLY | 0.74840105 |
| LFTVGFAAVG | ASSLNTGQLY | 0.6553173 |
| ATPPTPT | ASSLNTGQLY | 0.3453128 |
| TDLATLG | ASSLNTGQLY | 0.6972965 |
| SAGGTGR | ASSLNTGQLY | 0.8421978 |
| GFSNSGS | ASSLNTGQLY | 0.8315736 |
| GGNGGSGGTSV | ASSLNTGQLY | 0.41644034 |
| GTSVATGGAGN | ASSLNTGQLY | 0.8586042 |
| NGGNGGSGGTS | ASSLNTGQLY | 0.8051717 |
| AVVKFPH | ASSLNTGQLY | 0.8396913 |
| TPPPPPP | ASSLNTGQLY | 0.74731666 |
| AAPAAAAAPA | ASSLNTGQLY | 0.5924424 |
| IALNEDL | ASSLNTGQLY | 0.83927333 |
| IVNVSSM | ASSLNTGQLY | 0.6857206 |
| TGIPPAPR | ASSLNTGQLY | 0.59690833 |
| FDLGGGTF | ASSLNTGQLY | 0.8381176 |
| FDLGGGTFDVS | ASSLNTGQLY | 0.71982634 |
| NEPTAAA | ASSLNTGQLY | 0.5947057 |

| | | |
|---|---|---|
| NEPTAAALA | ASSLNTGQLY | 0.63197714 |
| GSIINISSI | ASSLNTGQLY | 0.83945274 |
| VAPGNAGTA | ASSLNTGQLY | 0.5444449 |
| TGGGASLE | ASSLNTGQLY | 0.5490391 |
| VAVYNHYKRI | ASSLNTGQLY | 0.5891255 |
| TDPAPATT | ASSLNTGQLY | 0.7436218 |
| TNRPDILDPALLRPG | ASSLNTGQLY | 0.83642447 |
| LTTTTTS | ASSLNTGQLY | 0.783527 |
| PTGSGKTL | ASSLNTGQLY | 0.6361488 |
| TGSGKTL | ASSLNTGQLY | 0.7750886 |
| TIGHVDHGK | ASSLNTGQLY | 0.78654027 |
| TIGHVDHGKTT | ASSLNTGQLY | 0.7787698 |
| GHVDHGKTT | ASSLNTGQLY | 0.80878127 |
| AQFEHTLL | ASSLNTGQLY | 0.5951145 |
| IVNQDEL | ASSLNTGQLY | 0.8312179 |
| TNGSQFF | ASSLNTGQLY | 0.8094927 |
| TGKGIVE | ASSLNTGQLY | 0.8186101 |
| LISNASDA | ASSLNTGQLY | 0.8219929 |
| AELRTLGELTKIAKAHGAQ | ASSLNTGQLY | 0.76312745 |
| APNAVAAAEPSW | ASSLNTGQLY | 0.775032 |
| APQVAGAATAVL | ASSLNTGQLY | 0.3165447 |
| ARRDPAAAAAILPTDA | ASSLNTGQLY | 0.7051275 |
| DAAARDVLAEAGLAEYF | ASSLNTGQLY | 0.7619641 |
| DDQIVLSAKANLAKVEQ | ASSLNTGQLY | 0.76154363 |
| DELLPDTGEDSDEDGH | ASSLNTGQLY | 0.80293024 |
| ELPEAPANRATVPQD | ASSLNTGQLY | 0.79155713 |
| FGQSKIERVTGIG | ASSLNTGQLY | 0.63505286 |
| GAGGTGSTAGGAGGAGGAGGLY | ASSLNTGQLY | 0.8371063 |
| GHGGTGAAGGNGGTGGAGGSGI | ASSLNTGQLY | 0.83332795 |
| GNGGNGGAGGEGGPGGAGGAGA | ASSLNTGQLY | 0.43899432 |
| GSTAGNGGNGGNATNGTIAGQP | ASSLNTGQLY | 0.7101235 |
| GTGGDGGNGGNGGTGG | ASSLNTGQLY | 0.8699862 |
| GVRRPAGIPADAPV | ASSLNTGQLY | 0.77207184 |
| GYRQVTDAHLLTLARR | ASSLNTGQLY | 0.71514916 |
| IAKMGQDGHDRG | ASSLNTGQLY | 0.6904011 |
| KAEPSIEIQYT | ASSLNTGQLY | 0.85608494 |
| KPVRIRMAADEDTVAVT | ASSLNTGQLY | 0.62373686 |
| MPPRIAGMRLLVIKPEP | ASSLNTGQLY | 0.856194 |
| MKFYAVALFRSRGVPPTR | ASSLNTGQLY | 0.22597541 |
| NAGGFGILGAVAHSP | ASSLNTGQLY | 0.6420127 |
| NEGREITLAMAEM | ASSLNTGQLY | 0.616561 |
| NELVEWSHPLGHRWRWE | ASSLNTGQLY | 0.31446537 |
| NLGAGGTGGNGGNPG | ASSLNTGQLY | 0.82874477 |
| NTEWPKWETVMAADPVD | ASSLNTGQLY | 0.69847697 |
| PASAAPAETQP | ASSLNTGQLY | 0.8170526 |
| QQSSDGKNGVRAAEITGE | ASSLNTGQLY | 0.7427691 |
| RAPDAVTTSWWKE | ASSLNTGQLY | 0.83397925 |
| SGFGNVGAHTSGWFNQ | ASSLNTGQLY | 0.75303745 |
| SGVAGAVGQAASVGGLKV | ASSLNTGQLY | 0.76625943 |
| SNGGAGTGSAGGNGGT | ASSLNTGQLY | 0.6085973 |
| TGGAGGEGGAAG | ASSLNTGQLY | 0.8189668 |

| | | |
|---|---|---|
| TGGDGGAAGAGGAPGHG | ASSLNTGQLY | 0.8458607 |
| TMPPLLAAIANAGRHGV | ASSLNTGQLY | 0.78776157 |
| TSSTAGNSGGA | ASSLNTGQLY | 0.4174722 |
| VPATVPVGILC | ASSLNTGQLY | 0.669139 |
| VQRPDSGGVVLTGRIS | ASSLNTGQLY | 0.7419634 |
| WTPNPVPATKGVPAGLVQAT | ASSLNTGQLY | 0.313703 |
| YALKRALAKFSDRFDVVII | ASSLNTGQLY | 0.2519648 |
| YLIGSVIV | ASSLNTGQLY | 0.66145366 |
| YGQLVAVID | ASSLNTGQLY | 0.65513164 |
| VTVGGMTLPA | ASSLNTGQLY | 0.8336493 |
| VLALPAV | ASSLNTGQLY | 0.80560046 |
| VIPPTLNYET | ASSLNTGQLY | 0.7151646 |
| VGMGGMHPSSG | ASSLNTGQLY | 0.81524163 |
| VGLAGGAATA | ASSLNTGQLY | 0.7624328 |
| VGGAGGAGGSGP | ASSLNTGQLY | 0.1111521 |
| VFAAFMV | ASSLNTGQLY | 0.7005857 |
| TLYDKLR | ASSLNTGQLY | 0.46763846 |
| TLAVFTVL | ASSLNTGQLY | 0.8533052 |
| TGSVQPARV | ASSLNTGQLY | 0.6189606 |
| TGGGAGSGGT | ASSLNTGQLY | 0.6245446 |
| SHLETSL | ASSLNTGQLY | 0.74675715 |
| SFTTGMT | ASSLNTGQLY | 0.7489184 |
| QVEVAKRMGVRQPTVS | ASSLNTGQLY | 0.74446976 |
| QGSGGPALAM | ASSLNTGQLY | 0.5002551 |
| MLAPSNP | ASSLNTGQLY | 0.18220538 |
| LVNNAAI | ASSLNTGQLY | 0.82874084 |
| LTGEINTLA | ASSLNTGQLY | 0.4242561 |
| LSIFGPV | ASSLNTGQLY | 0.5582274 |
| LLALLANL | ASSLNTGQLY | 0.33693504 |
| LISNASDALDKLR | ASSLNTGQLY | 0.36444348 |
| LINEAVP | ASSLNTGQLY | 0.39041677 |
| LGYAFSGL | ASSLNTGQLY | 0.8153545 |
| LDFVAVAL | ASSLNTGQLY | 0.6937303 |
| LAQPGAE | ASSLNTGQLY | 0.7000665 |
| ISNLGAVTGF | ASSLNTGQLY | 0.86310416 |
| HQEDPGRI | ASSLNTGQLY | 0.72245646 |
| GTPGEPTP | ASSLNTGQLY | 0.7117935 |
| GFFNSGS | ASSLNTGQLY | 0.6049878 |
| GAGGGGLTAGAN | ASSLNTGQLY | 0.5877172 |
| FVDALVNM | ASSLNTGQLY | 0.5960713 |
| FLTVAPEL | ASSLNTGQLY | 0.5966387 |
| FETLTAL | ASSLNTGQLY | 0.86514795 |
| DVINPIEL | ASSLNTGQLY | 0.6763182 |
| APVSGAPV | ASSLNTGQLY | 0.5293926 |
| ALVGGLAW | ASSLNTGQLY | 0.73953074 |
| ALAAAAATAI | ASSLNTGQLY | 0.8616662 |
| AFGLGLLL | ASSLNTGQLY | 0.87208545 |
| AFFAAASL | ASSLNTGQLY | 0.5949791 |
| AAAALPAFSPP | ASSLNTGQLY | 0.7189375 |
| VVLAPAL | ASSLNTGQLY | 0.7147106 |
| TIGHVDHGKTTL | ASSLNTGQLY | 0.7241931 |

| | | |
|---|---|---|
| LISNASDALDK | ASSLNTGQLY | 0.6444834 |
| LISNASDAL | ASSLNTGQLY | 0.5074738 |
| ISNASDALDK | ASSLNTGQLY | 0.80608946 |
| NQAETLVYQTE | ASSLNTGQLY | 0.8720849 |
| TGGTGGTAGAG | ASSLNTGQLY | 0.7166031 |
| TGSTAGGAGGA | ASSLNTGQLY | 0.42986682 |
| DPLAVPVAAGPV | ASSLNTGQLY | 0.71255994 |
| GGTGGTPGNGGN | ASSLNTGQLY | 0.55264413 |
| PAGLAGVDPATL | ASSLNTGQLY | 0.64886576 |
| GNNLVVEAPHAGASV | ASSLNTGQLY | 0.7434597 |
| KVTRIMSASTLEQPA | ASSLNTGQLY | 0.43164963 |
| LGFALPIALPRRTVV | ASSLNTGQLY | 0.72651696 |
| TRLGNLTARKLVVLV | ASSLNTGQLY | 0.87058485 |
| WSQQRRLIYFPSAGP | ASSLNTGQLY | 0.6624204 |
| GPNGNPDPMAAAVDIR | ASSLNTGQLY | 0.7982321 |
| TQRCVSVADLRAAASR | ASSLNTGQLY | 0.84325457 |
| QFVLDVAEHPVEVLTETV | ASSLNTGQLY | 0.81288135 |
| GGAGGTGGAGGNGGTQAGDG | ASSLNTGQLY | 0.84922683 |
| PLLATLRAIAVDVDAKPAQV | ASSLNTGQLY | 0.80443984 |
| APKFRTGGYMIDFWGVGYQVAK | ASSLNTGQLY | 0.75531834 |
| GAGGAGGAGGAAGGPGGTGGTG | ASSLNTGQLY | 0.75696886 |
| IGALGLLATDVAPAGSVAFGVA | ASSLNTGQLY | 0.7931998 |
| AGGAGGIGGNGGAGGAGGVLMG | ASSLNTGQLY | 0.74888337 |
| ADELIGLL | ASSLNTGQLY | 0.72394294 |
| ADRALKA | ASSLNTGQLY | 0.64494973 |
| AGAVGQVGGAGGA | ASSLNTGQLY | 0.8326042 |
| AGGGGGLGGAP | ASSLNTGQLY | 0.7365366 |
| AGGLLYGNGGAG | ASSLNTGQLY | 0.8411093 |
| AGLESIGL | ASSLNTGQLY | 0.4223692 |
| AGLKTGII | ASSLNTGQLY | 0.52007186 |
| ASLAALLL | ASSLNTGQLY | 0.8239745 |
| ASVINISSI | ASSLNTGQLY | 0.15695068 |
| ATPPGPA | ASSLNTGQLY | 0.2511123 |
| ATVASALV | ASSLNTGQLY | 0.7416645 |
| ATVASLIL | ASSLNTGQLY | 0.8537009 |
| AVNGPVLIRH | ASSLNTGQLY | 0.16063899 |
| DGAAGAVGI | ASSLNTGQLY | 0.5781102 |
| DPLLHGI | ASSLNTGQLY | 0.4574713 |
| ELGAALGAL | ASSLNTGQLY | 0.78913265 |
| ELGRTQAV | ASSLNTGQLY | 0.7380689 |
| FVAGGYLS | ASSLNTGQLY | 0.41000912 |
| GASGGAGGAGGW | ASSLNTGQLY | 0.8067666 |
| GGAGAGGDGGAGGI | ASSLNTGQLY | 0.3868625 |
| GTGHNGSQPGMGGNG | ASSLNTGQLY | 0.56154966 |
| GYAAASASAS | ASSLNTGQLY | 0.8283274 |
| HQVDVAAVQQG | ASSLNTGQLY | 0.8473022 |
| IGAAIAHT | ASSLNTGQLY | 0.5723878 |
| IGMIEVAQL | ASSLNTGQLY | 0.81425 |
| IRLAPEHTSAMD | ASSLNTGQLY | 0.612826 |
| IYNVSVL | ASSLNTGQLY | 0.8699403 |
| KQGFLGM | ASSLNTGQLY | 0.70062983 |

| | | |
|---|---|---|
| LESVITQL | ASSLNTGQLY | 0.40759447 |
| LGLGALVVV | ASSLNTGQLY | 0.8235408 |
| LIGNGGNGGSGGI | ASSLNTGQLY | 0.81310785 |
| LNAINAPTL | ASSLNTGQLY | 0.7705193 |
| LQLLNTGV | ASSLNTGQLY | 0.8091562 |
| LQTSIEF | ASSLNTGQLY | 0.6176629 |
| MSFAVSEL | ASSLNTGQLY | 0.7018906 |
| NTGALGL | ASSLNTGQLY | 0.83625597 |
| PPGAGGAGGAGTT | ASSLNTGQLY | 0.8214525 |
| QLVGLGI | ASSLNTGQLY | 0.5612726 |
| RFLEQVHQL | ASSLNTGQLY | 0.8296703 |
| RGVGYVLR | ASSLNTGQLY | 0.72038436 |
| SATGVFAGVI | ASSLNTGQLY | 0.64663243 |
| SLPAITLP | ASSLNTGQLY | 0.8635049 |
| STALAFTV | ASSLNTGQLY | 0.7210847 |
| SVLTVGLGK | ASSLNTGQLY | 0.8294215 |
| TALAGAGSGI | ASSLNTGQLY | 0.8190892 |
| TGLPFTGI | ASSLNTGQLY | 0.83514655 |
| VAEHVTV | ASSLNTGQLY | 0.52024865 |
| VDIVLNSL | ASSLNTGQLY | 0.79435647 |
| VLTGGGAVL | ASSLNTGQLY | 0.7462298 |
| VNVGLAYGI | ASSLNTGQLY | 0.8062124 |
| VSSAAVGAV | ASSLNTGQLY | 0.7116247 |
| VTSVYEAGV | ASSLNTGQLY | 0.5204841 |
| WVALDRGA | ASSLNTGQLY | 0.8050628 |
| YLVGSVVI | ASSLNTGQLY | 0.8551606 |
| GPPGTGKTLL | ASSLNTGQLY | 0.7517042 |
| GVGMTKF | ASSLNTGQLY | 0.772176 |
| IGIGGSDL | ASSLNTGQLY | 0.85986125 |
| IGTLLAEL | ASSLNTGQLY | 0.60025644 |
| IYTSGTT | ASSLNTGQLY | 0.7214205 |
| LSFPAGV | ASSLNTGQLY | 0.8180547 |
| LYTSGTT | ASSLNTGQLY | 0.8410314 |
| VGAFPFGF | ASSLNTGQLY | 0.70213664 |
| VGAGTVEF | ASSLNTGQLY | 0.8492886 |
| VGKSTLLL | ASSLNTGQLY | 0.62992096 |
| AALAHAGGGIV | ASSLNTGQLY | 0.8042328 |
| AALVAVSLL | ASSLNTGQLY | 0.6202494 |
| AAVSRAVAAL | ASSLNTGQLY | 0.78487885 |
| AGGSSGIGGF | ASSLNTGQLY | 0.57769895 |
| AGLTVANANG | ASSLNTGQLY | 0.7986908 |
| AGLVGAVL | ASSLNTGQLY | 0.82942855 |
| ALSQAFTV | ASSLNTGQLY | 0.82495785 |
| ALTFSVSL | ASSLNTGQLY | 0.244542 |
| ALTGAALVAL | ASSLNTGQLY | 0.824873 |
| ASFSIGAGVL | ASSLNTGQLY | 0.80867434 |
| DGSVYKI | ASSLNTGQLY | 0.87634736 |
| EQLAEIV | ASSLNTGQLY | 0.7291165 |
| FAVSELTVM | ASSLNTGQLY | 0.8139055 |
| GAAPLAAGL | ASSLNTGQLY | 0.6714402 |
| GADGGDGGSGGAGG | ASSLNTGQLY | 0.69832265 |

| | | |
|---|---|---|
| GFGLDTGGAGG | ASSLNTGQLY | 0.49179548 |
| GLASGGLL | ASSLNTGQLY | 0.66731614 |
| GRIAGAAVLL | ASSLNTGQLY | 0.6637944 |
| GSAGSLNTSF | ASSLNTGQLY | 0.5859988 |
| GTGQIGI | ASSLNTGQLY | 0.8139338 |
| GTLIALIL | ASSLNTGQLY | 0.82459223 |
| IAATMGFVVG | ASSLNTGQLY | 0.8415221 |
| IFAQALL | ASSLNTGQLY | 0.838933 |
| IGAAVALGF | ASSLNTGQLY | 0.85879207 |
| IGLTPLAL | ASSLNTGQLY | 0.8093166 |
| ILGSNTSTL | ASSLNTGQLY | 0.6414247 |
| IVFSLTVL | ASSLNTGQLY | 0.72962916 |
| LASKATGF | ASSLNTGQLY | 0.47148058 |
| LASVALIL | ASSLNTGQLY | 0.71543485 |
| LFAVGLTL | ASSLNTGQLY | 0.6590175 |
| LFNGGIGTYI | ASSLNTGQLY | 0.78609836 |
| LTLGAGII | ASSLNTGQLY | 0.13554999 |
| LVAAALGDI | ASSLNTGQLY | 0.755581 |
| NTAFVQLGI | ASSLNTGQLY | 0.8657018 |
| QLFSGAEL | ASSLNTGQLY | 0.58710265 |
| SAATFITL | ASSLNTGQLY | 0.7906871 |
| SNYGLANLGV | ASSLNTGQLY | 0.7899564 |
| SYGGAYAVM | ASSLNTGQLY | 0.6143792 |
| TTGVGGLTLG | ASSLNTGQLY | 0.8170333 |
| TTQTIYI | ASSLNTGQLY | 0.5728817 |
| VAAGAGLGAVI | ASSLNTGQLY | 0.7992135 |
| VAAVVTLL | ASSLNTGQLY | 0.8615277 |
| VGFLGGLI | ASSLNTGQLY | 0.76929235 |
| VGVIPILL | ASSLNTGQLY | 0.74378246 |
| VIFASSLI | ASSLNTGQLY | 0.744926 |
| VKGGDGGAAAG | ASSLNTGQLY | 0.43307889 |
| VTKTFTNI | ASSLNTGQLY | 0.75828004 |
| VVAAAILL | ASSLNTGQLY | 0.16051382 |
| YAALDVEL | ASSLNTGQLY | 0.7300042 |
| LASVALLL | ASSLNTGQLY | 0.59527004 |
| LDAVHRV | ASSLNTGQLY | 0.7693327 |
| QGLSSSIF | ASSLNTGQLY | 0.7620522 |
| RGTFANIRL | ASSLNTGQLY | 0.7879238 |
| RQLWWGHRI | ASSLNTGQLY | 0.5714455 |
| YGPPGTGKTLL | ASSLNTGQLY | 0.7104042 |
| AALVGAPANVA | ASSLNTGQLY | 0.63233405 |
| AATFRRTE | ASSLNTGQLY | 0.49572104 |
| AGALGGGI | ASSLNTGQLY | 0.81608784 |
| AGGAGGAGGIGAG | ASSLNTGQLY | 0.69874454 |
| AGSGALGL | ASSLNTGQLY | 0.73470855 |
| AQIADQI | ASSLNTGQLY | 0.84274316 |
| ASAVAVTYL | ASSLNTGQLY | 0.7761359 |
| ASTVLGGTGG | ASSLNTGQLY | 0.42621672 |
| AVGGEPAK | ASSLNTGQLY | 0.6988908 |
| CEQGTFGGTC | ASSLNTGQLY | 0.73795384 |
| DEGKAPAG | ASSLNTGQLY | 0.6966541 |

| | | |
|---|---|---|
| DTVVFPAL | ASSLNTGQLY | 0.14472547 |
| FEGITMAG | ASSLNTGQLY | 0.8043883 |
| FPLVVQD | ASSLNTGQLY | 0.65150774 |
| GAFTTPQ | ASSLNTGQLY | 0.5714807 |
| GFGGFGTAGGVGGA | ASSLNTGQLY | 0.6511126 |
| GFLAQFL | ASSLNTGQLY | 0.81969595 |
| GGAGGAGGNAGVA | ASSLNTGQLY | 0.5594284 |
| GGVGTAGAGGAGGAGG | ASSLNTGQLY | 0.7197906 |
| GNGGGGDAGNAGSGGN | ASSLNTGQLY | 0.8124417 |
| GTGGAGGAGNDAGS | ASSLNTGQLY | 0.81620735 |
| GTTNTGWFNS | ASSLNTGQLY | 0.85210174 |
| IFMGPLFTL | ASSLNTGQLY | 0.7914982 |
| ILAHAIEAI | ASSLNTGQLY | 0.731998 |
| IRSAIREAL | ASSLNTGQLY | 0.81358176 |
| ITFAGGG | ASSLNTGQLY | 0.7409066 |
| ITGMSGHQEV | ASSLNTGQLY | 0.3984937 |
| IVAGLACTL | ASSLNTGQLY | 0.2986187 |
| IVGGGLAGA | ASSLNTGQLY | 0.7409452 |
| IVTGVLNI | ASSLNTGQLY | 0.35373086 |
| LAQQGCI | ASSLNTGQLY | 0.76691824 |
| LCLHVEN | ASSLNTGQLY | 0.8284104 |
| LDAISDGS | ASSLNTGQLY | 0.70722234 |
| LGGFAAML | ASSLNTGQLY | 0.51121056 |
| LGGKNPMI | ASSLNTGQLY | 0.68346477 |
| LGPKGVAAIL | ASSLNTGQLY | 0.6936193 |
| LMVGVSI | ASSLNTGQLY | 0.8170097 |
| LVGAAFVL | ASSLNTGQLY | 0.8089777 |
| LVLAVAPL | ASSLNTGQLY | 0.85707337 |
| MAAIAVIDAA | ASSLNTGQLY | 0.82665634 |
| NNGINAELQ | ASSLNTGQLY | 0.65585536 |
| RGEPLVLVHG | ASSLNTGQLY | 0.8178917 |
| RGGAGGAAGLI | ASSLNTGQLY | 0.69687766 |
| RSYRMHPAV | ASSLNTGQLY | 0.7503066 |
| SAAHLLAGQLA | ASSLNTGQLY | 0.74842 |
| SAAVGSY | ASSLNTGQLY | 0.7681617 |
| SPHAVLHRI | ASSLNTGQLY | 0.66030455 |
| SQIAKVL | ASSLNTGQLY | 0.68794465 |
| TALGHLVP | ASSLNTGQLY | 0.38659674 |
| TAPTTSGAA | ASSLNTGQLY | 0.78530234 |
| TDGDDGNGGSPGT | ASSLNTGQLY | 0.79649353 |
| TDGLGEN | ASSLNTGQLY | 0.36484963 |
| TGGPPLPGQA | ASSLNTGQLY | 0.6889789 |
| TIPAITI | ASSLNTGQLY | 0.8589357 |
| TIPGFTI | ASSLNTGQLY | 0.85319835 |
| TSIASGALA | ASSLNTGQLY | 0.5746125 |
| TVTLPGAAI | ASSLNTGQLY | 0.5917317 |
| VASGLSAALP | ASSLNTGQLY | 0.7615528 |
| VGAGVIGI | ASSLNTGQLY | 0.35212836 |
| VGPKRFGV | ASSLNTGQLY | 0.77214336 |
| VNVAFPD | ASSLNTGQLY | 0.72135663 |
| VSMTNSGVS | ASSLNTGQLY | 0.7354628 |

| | | |
|---|---|---|
| VTDVLGDLSV | ASSLNTGQLY | 0.6460234 |
| VVVVGAGFAGL | ASSLNTGQLY | 0.74985373 |
| VYTVGTVV | ASSLNTGQLY | 0.6725428 |
| WTTFSGP | ASSLNTGQLY | 0.78784436 |
| YDELVEAF | ASSLNTGQLY | 0.16282608 |
| YGTGGAGGNGG | ASSLNTGQLY | 0.7996062 |
| YTNVMAV | ASSLNTGQLY | 0.5788515 |
| FLEQVHQL | ASSLNTGQLY | 0.6141271 |
| GLSLLAVAV | ASSLNTGQLY | 0.3037094 |
| LEQVHQL | ASSLNTGQLY | 0.77012223 |
| LNAAGAI | ASSLNTGQLY | 0.81151927 |
| PAVHASL | ASSLNTGQLY | 0.8261845 |
| TELEDDA | ASSLNTGQLY | 0.57922447 |
| TPPGPPN | ASSLNTGQLY | 0.82213736 |
| TPPPPPPP | ASSLNTGQLY | 0.8472824 |
| TQSPALI | ASSLSGDWGYEQY | 0.7029586 |
| EFAAAFPQ | ASSLSGDWGYEQY | 0.39055356 |
| LLSGNLDVL | ASSLSGDWGYEQY | 0.7849891 |
| GPANLGNYNI | ASSLSGDWGYEQY | 0.71723616 |
| TLGPTLP | ASSLSGDWGYEQY | 0.81233513 |
| VYAYSSCCK | ASSLSGDWGYEQY | 0.8042021 |
| APAPAPAPNI | ASSLSGDWGYEQY | 0.48388147 |
| PGGPVPIP | ASSLSGDWGYEQY | 0.82222456 |
| FLTVALEP | ASSLSGDWGYEQY | 0.82806027 |
| SQYFGANL | ASSLSGDWGYEQY | 0.5654159 |
| GGVKPAA | ASSLSGDWGYEQY | 0.6771436 |
| GGTGGAGGAGGW | ASSLSGDWGYEQY | 0.69633675 |
| GVGGVAGLGGAGG | ASSLSGDWGYEQY | 0.2888591 |
| VLLVLATTI | ASSLSGDWGYEQY | 0.820122 |
| IGIFAAML | ASSLSGDWGYEQY | 0.65596235 |
| AVGDLID | ASSLSGDWGYEQY | 0.60667735 |
| NISPLPL | ASSLSGDWGYEQY | 0.75829124 |
| VTAVFVAI | ASSLSGDWGYEQY | 0.6176722 |
| SGDDNDAAL | ASSLSGDWGYEQY | 0.63232374 |
| AVAALAHS | ASSLSGDWGYEQY | 0.6979366 |
| GGVTVMATP | ASSLSGDWGYEQY | 0.7445061 |
| AGLAANVRD | ASSLSGDWGYEQY | 0.7763723 |
| TNRPDLIDPALLRPG | ASSLSGDWGYEQY | 0.72853565 |
| LANLGVTAL | ASSLSGDWGYEQY | 0.77049065 |
| TGERIEL | ASSLSGDWGYEQY | 0.85631794 |
| NGEELAA | ASSLSGDWGYEQY | 0.43388718 |
| VLSLVWG | ASSLSGDWGYEQY | 0.6126149 |
| IGYALWL | ASSLSGDWGYEQY | 0.78431857 |
| LSLLLAPL | ASSLSGDWGYEQY | 0.4439504 |
| TLDEALK | ASSLSGDWGYEQY | 0.8090142 |
| TDGAGTAEGG | ASSLSGDWGYEQY | 0.7597767 |
| NAPGGAGGNGG | ASSLSGDWGYEQY | 0.7937007 |
| VSVFMGMGGNF | ASSLSGDWGYEQY | 0.7126786 |
| LASYLAAL | ASSLSGDWGYEQY | 0.8723911 |
| LVGGTPAL | ASSLSGDWGYEQY | 0.8236835 |
| VRNADVL | ASSLSGDWGYEQY | 0.747687 |

| | | |
|---|---|---|
| TCSSPFLVSAM | ASSLSGDWGYEQY | 0.78504556 |
| VHITDAE | ASSLSGDWGYEQY | 0.58845925 |
| EYAAPSF | ASSLSGDWGYEQY | 0.8295092 |
| VVDAFHGGL | ASSLSGDWGYEQY | 0.8033924 |
| ASVVDPQ | ASSLSGDWGYEQY | 0.53021544 |
| SYNEFGGLS | ASSLSGDWGYEQY | 0.70882463 |
| DPDGSVY | ASSLSGDWGYEQY | 0.84768957 |
| MVSTIHATE | ASSLSGDWGYEQY | 0.8422266 |
| LAAGVADPV | ASSLSGDWGYEQY | 0.3659095 |
| TKEQIAA | ASSLSGDWGYEQY | 0.6783032 |
| SLASPAAL | ASSLSGDWGYEQY | 0.3998853 |
| IGLVFYGV | ASSLSGDWGYEQY | 0.65698916 |
| FLLDASNL | ASSLSGDWGYEQY | 0.5480124 |
| YQVGNFP | ASSLSGDWGYEQY | 0.8041221 |
| ESLLSQST | ASSLSGDWGYEQY | 0.68124676 |
| LFTVGFAAVG | ASSLSGDWGYEQY | 0.79361385 |
| ATPPTPT | ASSLSGDWGYEQY | 0.76248896 |
| TDLATLG | ASSLSGDWGYEQY | 0.8449433 |
| SAGGTGR | ASSLSGDWGYEQY | 0.6168553 |
| GFSNSGS | ASSLSGDWGYEQY | 0.7138197 |
| GGNGGSGGTSV | ASSLSGDWGYEQY | 0.40467927 |
| GTSVATGGAGN | ASSLSGDWGYEQY | 0.44016415 |
| NGGNGGSGGTS | ASSLSGDWGYEQY | 0.57365453 |
| AVVKFPH | ASSLSGDWGYEQY | 0.68521714 |
| TPPPPPP | ASSLSGDWGYEQY | 0.23999684 |
| AAPAAAAAPA | ASSLSGDWGYEQY | 0.85509264 |
| IALNEDL | ASSLSGDWGYEQY | 0.753845 |
| IVNVSSM | ASSLSGDWGYEQY | 0.8121743 |
| TGIPPAPR | ASSLSGDWGYEQY | 0.65186596 |
| FDLGGGTF | ASSLSGDWGYEQY | 0.64355123 |
| FDLGGGTFDVS | ASSLSGDWGYEQY | 0.6249858 |
| NEPTAAA | ASSLSGDWGYEQY | 0.7912196 |
| NEPTAAALA | ASSLSGDWGYEQY | 0.58596694 |
| GSIINISSI | ASSLSGDWGYEQY | 0.35936427 |
| VAPGNAGTA | ASSLSGDWGYEQY | 0.6233471 |
| TGGGASLE | ASSLSGDWGYEQY | 0.50528383 |
| VAVYNHYKRI | ASSLSGDWGYEQY | 0.8637953 |
| TDPAPATT | ASSLSGDWGYEQY | 0.8313348 |
| TNRPDILDPALLRPG | ASSLSGDWGYEQY | 0.76617914 |
| LTTTTTS | ASSLSGDWGYEQY | 0.8501325 |
| PTGSGKTL | ASSLSGDWGYEQY | 0.24342139 |
| TGSGKTL | ASSLSGDWGYEQY | 0.6621141 |
| TIGHVDHGK | ASSLSGDWGYEQY | 0.8576654 |
| TIGHVDHGKTT | ASSLSGDWGYEQY | 0.79154986 |
| GHVDHGKTT | ASSLSGDWGYEQY | 0.72580945 |
| AQFEHTLL | ASSLSGDWGYEQY | 0.8416234 |
| IVNQDEL | ASSLSGDWGYEQY | 0.5974546 |
| TNGSQFF | ASSLSGDWGYEQY | 0.6968881 |
| TGKGIVE | ASSLSGDWGYEQY | 0.7299311 |
| LISNASDA | ASSLSGDWGYEQY | 0.81058264 |
| AELRTLGELTKIAKAHGAQ | ASSLSGDWGYEQY | 0.74016595 |

| | | |
|---|---|---|
| APNAVAAAEPSW | ASSLSGDWGYEQY | 0.5326811 |
| APQVAGAATAVL | ASSLSGDWGYEQY | 0.7927441 |
| ARRDPAAAAAILPTDA | ASSLSGDWGYEQY | 0.8727471 |
| DAAARDVLAEAGLAEYF | ASSLSGDWGYEQY | 0.6644428 |
| DDQIVLSAKANLAKVEQ | ASSLSGDWGYEQY | 0.82791543 |
| DELLPDTGEDSDEDGH | ASSLSGDWGYEQY | 0.7965387 |
| ELPEAPANRATVPQD | ASSLSGDWGYEQY | 0.74956703 |
| FGQSKIERVTGIG | ASSLSGDWGYEQY | 0.59052247 |
| GAGGTGSTAGGAGGAGGAGGLY | ASSLSGDWGYEQY | 0.6079086 |
| GHGGTGAAGGNGGTGGAGGSGI | ASSLSGDWGYEQY | 0.72594607 |
| GNGGNGGAGGEGGPGGAGGAGA | ASSLSGDWGYEQY | 0.5515567 |
| GSTAGNGGNGGNATNGTIAGQP | ASSLSGDWGYEQY | 0.725316 |
| GTGGDGGNGGNGGTGG | ASSLSGDWGYEQY | 0.7115046 |
| GVRRPAGIPADAPV | ASSLSGDWGYEQY | 0.88224393 |
| GYRQVTDAHLLTLARR | ASSLSGDWGYEQY | 0.8117701 |
| IAKMGQDGHDRG | ASSLSGDWGYEQY | 0.79977065 |
| KAEPSIEIQYT | ASSLSGDWGYEQY | 0.53075224 |
| KPVRIRMAADEDTVAVT | ASSLSGDWGYEQY | 0.7618496 |
| MPPRIAGMRLLVIKPEP | ASSLSGDWGYEQY | 0.7814753 |
| MKFYAVALFRSRGVPPTR | ASSLSGDWGYEQY | 0.44619548 |
| NAGGFGILGAVAHSP | ASSLSGDWGYEQY | 0.85928035 |
| NEGREITLAMAEM | ASSLSGDWGYEQY | 0.58668935 |
| NELVEWSHPLGHRWRWE | ASSLSGDWGYEQY | 0.6862019 |
| NLGAGGTGGNGGNPG | ASSLSGDWGYEQY | 0.8104426 |
| NTEWPKWETVMAADPVD | ASSLSGDWGYEQY | 0.8423119 |
| PASAAPAETQP | ASSLSGDWGYEQY | 0.71552306 |
| QQSSDGKNGVRAAEITGE | ASSLSGDWGYEQY | 0.7302975 |
| RAPDAVTTSWWKE | ASSLSGDWGYEQY | 0.55541015 |
| SGFGNVGAHTSGWFNQ | ASSLSGDWGYEQY | 0.82268083 |
| SGVAGAVGQAASVGGLKV | ASSLSGDWGYEQY | 0.83719397 |
| SNGGAGTGSAGGNGGT | ASSLSGDWGYEQY | 0.68423027 |
| TGGAGGEGGAAG | ASSLSGDWGYEQY | 0.86367136 |
| TGGDGGAAGAGGAPGHG | ASSLSGDWGYEQY | 0.75728965 |
| TMPPLLAAIANAGRHGV | ASSLSGDWGYEQY | 0.421069 |
| TSSTAGNSGGA | ASSLSGDWGYEQY | 0.7415702 |
| VPATVPVGILC | ASSLSGDWGYEQY | 0.8327562 |
| VQRPDSGGVVLTGRIS | ASSLSGDWGYEQY | 0.8359983 |
| WTPNPVPATKGVPAGLVQAT | ASSLSGDWGYEQY | 0.84827715 |
| YALKRALAKFSDRFDVVII | ASSLSGDWGYEQY | 0.7853943 |
| YLIGSVIV | ASSLSGDWGYEQY | 0.659313 |
| YGQLVAVID | ASSLSGDWGYEQY | 0.848504 |
| VTVGGMTLPA | ASSLSGDWGYEQY | 0.87142146 |
| VLALPAV | ASSLSGDWGYEQY | 0.8214811 |
| VIPPTLNYET | ASSLSGDWGYEQY | 0.6098095 |
| VGMGGMHPSSG | ASSLSGDWGYEQY | 0.59479964 |
| VGLAGGAATA | ASSLSGDWGYEQY | 0.7833036 |
| VGGAGGAGGSGP | ASSLSGDWGYEQY | 0.823172 |
| VFAAFMV | ASSLSGDWGYEQY | 0.5326279 |
| TLYDKLR | ASSLSGDWGYEQY | 0.7732898 |
| TLAVFTVL | ASSLSGDWGYEQY | 0.7823335 |
| TGSVQPARV | ASSLSGDWGYEQY | 0.41710156 |

| | | |
|---|---|---|
| TGGGAGSGGT | ASSLSGDWGYEQY | 0.79597926 |
| SHLETSL | ASSLSGDWGYEQY | 0.8525294 |
| SFTTGMT | ASSLSGDWGYEQY | 0.7481719 |
| QVEVAKRMGVRQPTVS | ASSLSGDWGYEQY | 0.6010592 |
| QGSGGPALAM | ASSLSGDWGYEQY | 0.84773064 |
| MLAPSNP | ASSLSGDWGYEQY | 0.87851846 |
| LVNNAAI | ASSLSGDWGYEQY | 0.8505881 |
| LTGEINTLA | ASSLSGDWGYEQY | 0.38450468 |
| LSIFGPV | ASSLSGDWGYEQY | 0.24147964 |
| LLALLANL | ASSLSGDWGYEQY | 0.6455903 |
| LISNASDALDKLR | ASSLSGDWGYEQY | 0.8027102 |
| LINEAVP | ASSLSGDWGYEQY | 0.68854725 |
| LGYAFSGL | ASSLSGDWGYEQY | 0.60890025 |
| LDFVAVAL | ASSLSGDWGYEQY | 0.7594992 |
| LAQPGAE | ASSLSGDWGYEQY | 0.83988506 |
| ISNLGAVTGF | ASSLSGDWGYEQY | 0.5982405 |
| HQEDPGRI | ASSLSGDWGYEQY | 0.4575918 |
| GTPGEPTP | ASSLSGDWGYEQY | 0.79467106 |
| GFFNSGS | ASSLSGDWGYEQY | 0.80656683 |
| GAGGGGLTAGAN | ASSLSGDWGYEQY | 0.5416985 |
| FVDALVNM | ASSLSGDWGYEQY | 0.68695414 |
| FLTVAPEL | ASSLSGDWGYEQY | 0.67452985 |
| FETLTAL | ASSLSGDWGYEQY | 0.6199827 |
| DVINPIEL | ASSLSGDWGYEQY | 0.7620015 |
| APVSGAPV | ASSLSGDWGYEQY | 0.48864207 |
| ALVGGLAW | ASSLSGDWGYEQY | 0.84319055 |
| ALAAAAATAI | ASSLSGDWGYEQY | 0.539227 |
| AFGLGLLL | ASSLSGDWGYEQY | 0.81294453 |
| AFFAAASL | ASSLSGDWGYEQY | 0.76181245 |
| AAAALPAFSPP | ASSLSGDWGYEQY | 0.78659236 |
| VVLAPAL | ASSLSGDWGYEQY | 0.683064 |
| TIGHVDHGKTTL | ASSLSGDWGYEQY | 0.62693375 |
| LISNASDALDK | ASSLSGDWGYEQY | 0.5580642 |
| LISNASDAL | ASSLSGDWGYEQY | 0.27507597 |
| ISNASDALDK | ASSLSGDWGYEQY | 0.8358307 |
| NQAETLVYQTE | ASSLSGDWGYEQY | 0.2595403 |
| TGGTGGTAGAG | ASSLSGDWGYEQY | 0.83254415 |
| TGSTAGGAGGA | ASSLSGDWGYEQY | 0.8026254 |
| DPLAVPVAAGPV | ASSLSGDWGYEQY | 0.590018 |
| GGTGGTPGNGGN | ASSLSGDWGYEQY | 0.7825089 |
| PAGLAGVDPATL | ASSLSGDWGYEQY | 0.49518517 |
| GNNLVVEAPHAGASV | ASSLSGDWGYEQY | 0.6358321 |
| KVTRIMSASTLEQPA | ASSLSGDWGYEQY | 0.7906158 |
| LGFALPIALPRRTVV | ASSLSGDWGYEQY | 0.7991706 |
| TRLGNLTARKLVVLV | ASSLSGDWGYEQY | 0.60531926 |
| WSQQRRLIYFPSAGP | ASSLSGDWGYEQY | 0.7924312 |
| GPNGNPDPMAAAVDIR | ASSLSGDWGYEQY | 0.7835999 |
| TQRCVSVADLRAAASR | ASSLSGDWGYEQY | 0.63377583 |
| QFVLDVAEHPVEVLTETV | ASSLSGDWGYEQY | 0.7549268 |
| GGAGGTGGAGGNGGTQAGDG | ASSLSGDWGYEQY | 0.82642126 |
| PLLATLRAIAVDVDAKPAQV | ASSLSGDWGYEQY | 0.7857497 |

| | | |
|---|---|---|
| APKFRTGGYMIDFWGVGYQVAK | ASSLSGDWGYEQY | 0.5560912 |
| GAGGAGGAGGAAGGPGGTGGTG | ASSLSGDWGYEQY | 0.8150681 |
| IGALGLLATDVAPAGSVAFGVA | ASSLSGDWGYEQY | 0.5892444 |
| AGGAGGIGGNGGAGGAGGVLMG | ASSLSGDWGYEQY | 0.7780944 |
| ADELIGLL | ASSLSGDWGYEQY | 0.7574133 |
| ADRALKA | ASSLSGDWGYEQY | 0.70258623 |
| AGAVGQVGGAGGA | ASSLSGDWGYEQY | 0.82937646 |
| AGGGGGLGGAP | ASSLSGDWGYEQY | 0.50103444 |
| AGGLLYGNGGAG | ASSLSGDWGYEQY | 0.595124 |
| AGLESIGL | ASSLSGDWGYEQY | 0.6975907 |
| AGLKTGII | ASSLSGDWGYEQY | 0.7527087 |
| ASLAALLL | ASSLSGDWGYEQY | 0.82251763 |
| ASVINISSI | ASSLSGDWGYEQY | 0.553527 |
| ATPPGPA | ASSLSGDWGYEQY | 0.751559 |
| ATVASALV | ASSLSGDWGYEQY | 0.6948187 |
| ATVASLIL | ASSLSGDWGYEQY | 0.553696 |
| AVNGPVLIRH | ASSLSGDWGYEQY | 0.72793424 |
| DGAAGAVGI | ASSLSGDWGYEQY | 0.68905216 |
| DPLLHGI | ASSLSGDWGYEQY | 0.5401195 |
| ELGAALGAL | ASSLSGDWGYEQY | 0.8046696 |
| ELGRTQAV | ASSLSGDWGYEQY | 0.6035855 |
| FVAGGYLS | ASSLSGDWGYEQY | 0.78510636 |
| GASGGAGGAGGW | ASSLSGDWGYEQY | 0.86443937 |
| GGAGAGGDGGAGGI | ASSLSGDWGYEQY | 0.59243214 |
| GTGHNGSQPGMGGNG | ASSLSGDWGYEQY | 0.80838484 |
| GYAAASASAS | ASSLSGDWGYEQY | 0.7843926 |
| HQVDVAAVQQG | ASSLSGDWGYEQY | 0.7810533 |
| IGAAIAHT | ASSLSGDWGYEQY | 0.6932111 |
| IGMIEVAQL | ASSLSGDWGYEQY | 0.6478436 |
| IRLAPEHTSAMD | ASSLSGDWGYEQY | 0.8464973 |
| IYNVSVL | ASSLSGDWGYEQY | 0.8129735 |
| KQGFLGM | ASSLSGDWGYEQY | 0.82611173 |
| LESVITQL | ASSLSGDWGYEQY | 0.8094344 |
| LGLGALVVV | ASSLSGDWGYEQY | 0.8017689 |
| LIGNGGNGGSGGI | ASSLSGDWGYEQY | 0.8119017 |
| LNAINAPTL | ASSLSGDWGYEQY | 0.61575377 |
| LQLLNTGV | ASSLSGDWGYEQY | 0.42087355 |
| LQTSIEF | ASSLSGDWGYEQY | 0.68227255 |
| MSFAVSEL | ASSLSGDWGYEQY | 0.753187 |
| NTGALGL | ASSLSGDWGYEQY | 0.6885709 |
| PPGAGGAGGAGTT | ASSLSGDWGYEQY | 0.36318627 |
| QLVGLGI | ASSLSGDWGYEQY | 0.8308532 |
| RFLEQVHQL | ASSLSGDWGYEQY | 0.87273335 |
| RGVGYVLR | ASSLSGDWGYEQY | 0.5707623 |
| SATGVFAGVI | ASSLSGDWGYEQY | 0.8676031 |
| SLPAITLP | ASSLSGDWGYEQY | 0.47632658 |
| STALAFTV | ASSLSGDWGYEQY | 0.80185366 |
| SVLTVGLGK | ASSLSGDWGYEQY | 0.8392757 |
| TALAGAGSGI | ASSLSGDWGYEQY | 0.48850256 |
| TGLPFTGI | ASSLSGDWGYEQY | 0.80257607 |
| VAEHVTV | ASSLSGDWGYEQY | 0.6523624 |

| | | |
|---|---|---|
| VDIVLNSL | ASSLSGDWGYEQY | 0.69600713 |
| VLTGGGAVL | ASSLSGDWGYEQY | 0.8463377 |
| VNVGLAYGI | ASSLSGDWGYEQY | 0.67241883 |
| VSSAAVGAV | ASSLSGDWGYEQY | 0.8022553 |
| VTSVYEAGV | ASSLSGDWGYEQY | 0.65853196 |
| WVALDRGA | ASSLSGDWGYEQY | 0.5695201 |
| YLVGSVVI | ASSLSGDWGYEQY | 0.7779196 |
| GPPGTGKTLL | ASSLSGDWGYEQY | 0.8450822 |
| GVGMTKF | ASSLSGDWGYEQY | 0.60322964 |
| IGIGGSDL | ASSLSGDWGYEQY | 0.86802673 |
| IGTLLAEL | ASSLSGDWGYEQY | 0.62290746 |
| IYTSGTT | ASSLSGDWGYEQY | 0.86416495 |
| LSFPAGV | ASSLSGDWGYEQY | 0.8303639 |
| LYTSGTT | ASSLSGDWGYEQY | 0.73088133 |
| VGAFPFGF | ASSLSGDWGYEQY | 0.6824734 |
| VGAGTVEF | ASSLSGDWGYEQY | 0.8295064 |
| VGKSTLLL | ASSLSGDWGYEQY | 0.66167027 |
| AALAHAGGGIV | ASSLSGDWGYEQY | 0.79835033 |
| AALVAVSLL | ASSLSGDWGYEQY | 0.83669394 |
| AAVSRAVAAL | ASSLSGDWGYEQY | 0.62284833 |
| AGGSSGIGGF | ASSLSGDWGYEQY | 0.8406695 |
| AGLTVANANG | ASSLSGDWGYEQY | 0.7674212 |
| AGLVGAVL | ASSLSGDWGYEQY | 0.7107719 |
| ALSQAFTV | ASSLSGDWGYEQY | 0.5949856 |
| ALTFSVSL | ASSLSGDWGYEQY | 0.3097178 |
| ALTGAALVAL | ASSLSGDWGYEQY | 0.71177316 |
| ASFSIGAGVL | ASSLSGDWGYEQY | 0.503939 |
| DGSVYKI | ASSLSGDWGYEQY | 0.70773256 |
| EQLAEIV | ASSLSGDWGYEQY | 0.7880383 |
| FAVSELTVM | ASSLSGDWGYEQY | 0.55929875 |
| GAAPLAAGL | ASSLSGDWGYEQY | 0.83071876 |
| GADGGDGGSGGAGG | ASSLSGDWGYEQY | 0.8344022 |
| GFGLDTGGAGG | ASSLSGDWGYEQY | 0.38185817 |
| GLASGGLL | ASSLSGDWGYEQY | 0.69773936 |
| GRIAGAAVLL | ASSLSGDWGYEQY | 0.6463863 |
| GSAGSLNTSF | ASSLSGDWGYEQY | 0.69304824 |
| GTGQIGI | ASSLSGDWGYEQY | 0.33295345 |
| GTLIALIL | ASSLSGDWGYEQY | 0.78841025 |
| IAATMGFVVG | ASSLSGDWGYEQY | 0.5790453 |
| IFAQALL | ASSLSGDWGYEQY | 0.100349635 |
| IGAAVALGF | ASSLSGDWGYEQY | 0.76225746 |
| IGLTPLAL | ASSLSGDWGYEQY | 0.6631303 |
| ILGSNTSTL | ASSLSGDWGYEQY | 0.63897777 |
| IVFSLTVL | ASSLSGDWGYEQY | 0.80710924 |
| LASKATGF | ASSLSGDWGYEQY | 0.19743708 |
| LASVALIL | ASSLSGDWGYEQY | 0.5807788 |
| LFAVGLTL | ASSLSGDWGYEQY | 0.7961225 |
| LFNGGIGTYI | ASSLSGDWGYEQY | 0.83852756 |
| LTLGAGII | ASSLSGDWGYEQY | 0.41023955 |
| LVAAALGDI | ASSLSGDWGYEQY | 0.7523298 |
| NTAFVQLGI | ASSLSGDWGYEQY | 0.8319478 |

| | | |
|---|---|---|
| QLFSGAEL | ASSLSGDWGYEQY | 0.7911917 |
| SAATFITL | ASSLSGDWGYEQY | 0.8209229 |
| SNYGLANLGV | ASSLSGDWGYEQY | 0.8396073 |
| SYGGAYAVM | ASSLSGDWGYEQY | 0.51909363 |
| TTGVGGLTLG | ASSLSGDWGYEQY | 0.8061235 |
| TTQTIYI | ASSLSGDWGYEQY | 0.85746074 |
| VAAGAGLGAVI | ASSLSGDWGYEQY | 0.6497668 |
| VAAVVTLL | ASSLSGDWGYEQY | 0.70655113 |
| VGFLGGLI | ASSLSGDWGYEQY | 0.60642207 |
| VGVIPILL | ASSLSGDWGYEQY | 0.6400292 |
| VIFASSLI | ASSLSGDWGYEQY | 0.1673327 |
| VKGGDGGAAAG | ASSLSGDWGYEQY | 0.7300838 |
| VTKTFTNI | ASSLSGDWGYEQY | 0.81609124 |
| VVAAAILL | ASSLSGDWGYEQY | 0.62296367 |
| YAALDVEL | ASSLSGDWGYEQY | 0.5039805 |
| LASVALLL | ASSLSGDWGYEQY | 0.5028187 |
| LDAVHRV | ASSLSGDWGYEQY | 0.58652437 |
| QGLSSSIF | ASSLSGDWGYEQY | 0.60873866 |
| RGTFANIRL | ASSLSGDWGYEQY | 0.18135561 |
| RQLWWGHRI | ASSLSGDWGYEQY | 0.5424407 |
| YGPPGTGKTLL | ASSLSGDWGYEQY | 0.818522 |
| AALVGAPANVA | ASSLSGDWGYEQY | 0.83205223 |
| AATFRRTE | ASSLSGDWGYEQY | 0.21695495 |
| AGALGGGI | ASSLSGDWGYEQY | 0.8192365 |
| AGGAGGAGGIGAG | ASSLSGDWGYEQY | 0.7471862 |
| AGSGALGL | ASSLSGDWGYEQY | 0.85276073 |
| AQIADQI | ASSLSGDWGYEQY | 0.73388946 |
| ASAVAVTYL | ASSLSGDWGYEQY | 0.5917054 |
| ASTVLGGTGG | ASSLSGDWGYEQY | 0.7455223 |
| AVGGEPAK | ASSLSGDWGYEQY | 0.6404778 |
| CEQGTFGGTC | ASSLSGDWGYEQY | 0.3485636 |
| DEGKAPAG | ASSLSGDWGYEQY | 0.8158039 |
| DTVVFPAL | ASSLSGDWGYEQY | 0.60924554 |
| FEGITMAG | ASSLSGDWGYEQY | 0.5382388 |
| FPLVVQD | ASSLSGDWGYEQY | 0.56971467 |
| GAFTTPQ | ASSLSGDWGYEQY | 0.8419055 |
| GFGGFGTAGGVGGA | ASSLSGDWGYEQY | 0.46208844 |
| GFLAQFL | ASSLSGDWGYEQY | 0.41278696 |
| GGAGGAGGNAGVA | ASSLSGDWGYEQY | 0.34817064 |
| GGVGTAGAGGAGGAGG | ASSLSGDWGYEQY | 0.71469724 |
| GNGGGGDAGNAGSGGN | ASSLSGDWGYEQY | 0.82035947 |
| GTGGAGGAGNDAGS | ASSLSGDWGYEQY | 0.7845975 |
| GTTNTGWFNS | ASSLSGDWGYEQY | 0.5671485 |
| IFMGPLFTL | ASSLSGDWGYEQY | 0.7761737 |
| ILAHAIEAI | ASSLSGDWGYEQY | 0.7416737 |
| IRSAIREAL | ASSLSGDWGYEQY | 0.7281623 |
| ITFAGGG | ASSLSGDWGYEQY | 0.84058106 |
| ITGMSGHQEV | ASSLSGDWGYEQY | 0.5269301 |
| IVAGLACTL | ASSLSGDWGYEQY | 0.8332219 |
| IVGGGLAGA | ASSLSGDWGYEQY | 0.7679309 |
| IVTGVLNI | ASSLSGDWGYEQY | 0.7716012 |

| | | |
|---|---|---|
| LAQQGCI | ASSLSGDWGYEQY | 0.8040598 |
| LCLHVEN | ASSLSGDWGYEQY | 0.71816266 |
| LDAISDGS | ASSLSGDWGYEQY | 0.7212715 |
| LGGFAAML | ASSLSGDWGYEQY | 0.21960413 |
| LGGKNPMI | ASSLSGDWGYEQY | 0.63098454 |
| LGPKGVAAIL | ASSLSGDWGYEQY | 0.76437056 |
| LMVGVSI | ASSLSGDWGYEQY | 0.72938037 |
| LVGAAFVL | ASSLSGDWGYEQY | 0.7637416 |
| LVLAVAPL | ASSLSGDWGYEQY | 0.7435945 |
| MAAIAVIDAA | ASSLSGDWGYEQY | 0.5477705 |
| NNGINAELQ | ASSLSGDWGYEQY | 0.82445335 |
| RGEPLVLVHG | ASSLSGDWGYEQY | 0.79193044 |
| RGGAGGAAGLI | ASSLSGDWGYEQY | 0.33826458 |
| RSYRMHPAV | ASSLSGDWGYEQY | 0.69709396 |
| SAAHLLAGQLA | ASSLSGDWGYEQY | 0.8605301 |
| SAAVGSY | ASSLSGDWGYEQY | 0.70929575 |
| SPHAVLHRI | ASSLSGDWGYEQY | 0.6477283 |
| SQIAKVL | ASSLSGDWGYEQY | 0.5586083 |
| TALGHLVP | ASSLSGDWGYEQY | 0.8112935 |
| TAPTTSGAA | ASSLSGDWGYEQY | 0.3929018 |
| TDGDDGNGGSPGT | ASSLSGDWGYEQY | 0.8550503 |
| TDGLGEN | ASSLSGDWGYEQY | 0.1348709 |
| TGGPPLPGQA | ASSLSGDWGYEQY | 0.61354125 |
| TIPAITI | ASSLSGDWGYEQY | 0.5495328 |
| TIPGFTI | ASSLSGDWGYEQY | 0.3039134 |
| TSIASGALA | ASSLSGDWGYEQY | 0.81919694 |
| TVTLPGAAI | ASSLSGDWGYEQY | 0.5916921 |
| VASGLSAALP | ASSLSGDWGYEQY | 0.7966113 |
| VGAGVIGI | ASSLSGDWGYEQY | 0.7212036 |
| VGPKRFGV | ASSLSGDWGYEQY | 0.820906 |
| VNVAFPD | ASSLSGDWGYEQY | 0.7183653 |
| VSMTNSGVS | ASSLSGDWGYEQY | 0.75897515 |
| VTDVLGDLSV | ASSLSGDWGYEQY | 0.47270513 |
| VVVVGAGFAGL | ASSLSGDWGYEQY | 0.79629093 |
| VYTVGTVV | ASSLSGDWGYEQY | 0.8298639 |
| WTTFSGP | ASSLSGDWGYEQY | 0.77227175 |
| YDELVEAF | ASSLSGDWGYEQY | 0.32594225 |
| YGTGGAGGNGG | ASSLSGDWGYEQY | 0.5788985 |
| YTNVMAV | ASSLSGDWGYEQY | 0.7283753 |
| FLEQVHQL | ASSLSGDWGYEQY | 0.18739548 |
| GLSLLAVAV | ASSLSGDWGYEQY | 0.13300636 |
| LEQVHQL | ASSLSGDWGYEQY | 0.5145385 |
| LNAAGAI | ASSLSGDWGYEQY | 0.44553694 |
| PAVHASL | ASSLSGDWGYEQY | 0.8152126 |
| TELEDDA | ASSLSGDWGYEQY | 0.8140518 |
| TPPGPPN | ASSLSGDWGYEQY | 0.6587936 |
| TPPPPPPP | ASSLSGDWGYEQY | 0.7934122 |
| TQSPALI | AWSLAWGENTLY | 0.72151744 |
| EFAAAFPQ | AWSLAWGENTLY | 0.08799012 |
| LLSGNLDVL | AWSLAWGENTLY | 0.58337295 |
| GPANLGNYNI | AWSLAWGENTLY | 0.37830955 |

| | | |
|---|---|---|
| TLGPTLP | AWSLAWGENTLY | 0.8150741 |
| VYAYSSCCK | AWSLAWGENTLY | 0.65058106 |
| APAPAPAPNI | AWSLAWGENTLY | 0.54708123 |
| PGGPVPIP | AWSLAWGENTLY | 0.6855386 |
| FLTVALEP | AWSLAWGENTLY | 0.6653317 |
| SQYFGANL | AWSLAWGENTLY | 0.67151636 |
| GGVKPAA | AWSLAWGENTLY | 0.47852257 |
| GGTGGAGGAGGW | AWSLAWGENTLY | 0.11524359 |
| GVGGVAGLGGAGG | AWSLAWGENTLY | 0.8067222 |
| VLLVLATTI | AWSLAWGENTLY | 0.26261362 |
| IGIFAAML | AWSLAWGENTLY | 0.51270586 |
| AVGDLID | AWSLAWGENTLY | 0.18611312 |
| NISPLPL | AWSLAWGENTLY | 0.10623979 |
| VTAVFVAI | AWSLAWGENTLY | 0.31726813 |
| SGDDNDAAL | AWSLAWGENTLY | 0.8078866 |
| AVAALAHS | AWSLAWGENTLY | 0.57546484 |
| GGVTVMATP | AWSLAWGENTLY | 0.61667216 |
| AGLAANVRD | AWSLAWGENTLY | 0.83816546 |
| TNRPDLIDPALLRPG | AWSLAWGENTLY | 0.6427629 |
| LANLGVTAL | AWSLAWGENTLY | 0.6179526 |
| TGERIEL | AWSLAWGENTLY | 0.558411 |
| NGEELAA | AWSLAWGENTLY | 0.51650786 |
| VLSLVWG | AWSLAWGENTLY | 0.5493505 |
| IGYALWL | AWSLAWGENTLY | 0.49107543 |
| LSLLLAPL | AWSLAWGENTLY | 0.8566841 |
| TLDEALK | AWSLAWGENTLY | 0.5577799 |
| TDGAGTAEGG | AWSLAWGENTLY | 0.42980734 |
| NAPGGAGGNGG | AWSLAWGENTLY | 0.6562804 |
| VSVFMGMGGNF | AWSLAWGENTLY | 0.81948376 |
| LASYLAAL | AWSLAWGENTLY | 0.8700434 |
| LVGGTPAL | AWSLAWGENTLY | 0.59869564 |
| VRNADVL | AWSLAWGENTLY | 0.6474121 |
| TCSSPFLVSAM | AWSLAWGENTLY | 0.65831053 |
| VHITDAE | AWSLAWGENTLY | 0.67977977 |
| EYAAPSF | AWSLAWGENTLY | 0.6695032 |
| VVDAFHGGL | AWSLAWGENTLY | 0.35303473 |
| ASVVDPQ | AWSLAWGENTLY | 0.80557966 |
| SYNEFGGLS | AWSLAWGENTLY | 0.8128009 |
| DPDGSVY | AWSLAWGENTLY | 0.6430189 |
| MVSTIHATE | AWSLAWGENTLY | 0.55475193 |
| LAAGVADPV | AWSLAWGENTLY | 0.6603035 |
| TKEQIAA | AWSLAWGENTLY | 0.5259199 |
| SLASPAAL | AWSLAWGENTLY | 0.6141439 |
| IGLVFYGV | AWSLAWGENTLY | 0.73140466 |
| FLLDASNL | AWSLAWGENTLY | 0.38047895 |
| YQVGNFP | AWSLAWGENTLY | 0.59678113 |
| ESLLSQST | AWSLAWGENTLY | 0.86920846 |
| LFTVGFAAVG | AWSLAWGENTLY | 0.6136356 |
| ATPPTPT | AWSLAWGENTLY | 0.7186161 |
| TDLATLG | AWSLAWGENTLY | 0.82177186 |
| SAGGTGR | AWSLAWGENTLY | 0.75815016 |

| | | |
|---|---|---|
| GFSNSGS | AWSLAWGENTLY | 0.8405474 |
| GGNGGSGGTSV | AWSLAWGENTLY | 0.7660538 |
| GTSVATGGAGN | AWSLAWGENTLY | 0.67086685 |
| NGGNGGSGGTS | AWSLAWGENTLY | 0.69564235 |
| AVVKFPH | AWSLAWGENTLY | 0.6501866 |
| TPPPPPP | AWSLAWGENTLY | 0.6844172 |
| AAPAAAAAPA | AWSLAWGENTLY | 0.6677158 |
| IALNEDL | AWSLAWGENTLY | 0.50713146 |
| IVNVSSM | AWSLAWGENTLY | 0.8259387 |
| TGIPPAPR | AWSLAWGENTLY | 0.74605846 |
| FDLGGGTF | AWSLAWGENTLY | 0.7509173 |
| FDLGGGTFDVS | AWSLAWGENTLY | 0.38937923 |
| NEPTAAA | AWSLAWGENTLY | 0.39101446 |
| NEPTAAALA | AWSLAWGENTLY | 0.7750201 |
| GSIINISSI | AWSLAWGENTLY | 0.12667689 |
| VAPGNAGTA | AWSLAWGENTLY | 0.14387509 |
| TGGGASLE | AWSLAWGENTLY | 0.751075 |
| VAVYNHYKRI | AWSLAWGENTLY | 0.83224607 |
| TDPAPATT | AWSLAWGENTLY | 0.13118398 |
| TNRPDILDPALLRPG | AWSLAWGENTLY | 0.48595792 |
| LTTTTTS | AWSLAWGENTLY | 0.32672188 |
| PTGSGKTL | AWSLAWGENTLY | 0.7192725 |
| TGSGKTL | AWSLAWGENTLY | 0.7061473 |
| TIGHVDHGK | AWSLAWGENTLY | 0.26945984 |
| TIGHVDHGKTT | AWSLAWGENTLY | 0.7780045 |
| GHVDHGKTT | AWSLAWGENTLY | 0.33008176 |
| AQFEHTLL | AWSLAWGENTLY | 0.20614982 |
| IVNQDEL | AWSLAWGENTLY | 0.8240718 |
| TNGSQFF | AWSLAWGENTLY | 0.7932931 |
| TGKGIVE | AWSLAWGENTLY | 0.5661119 |
| LISNASDA | AWSLAWGENTLY | 0.79243827 |
| AELRTLGELTKIAKAHGAQ | AWSLAWGENTLY | 0.5283208 |
| APNAVAAAEPSW | AWSLAWGENTLY | 0.8732196 |
| APQVAGAATAVL | AWSLAWGENTLY | 0.6361908 |
| ARRDPAAAAAILPTDA | AWSLAWGENTLY | 0.33576646 |
| DAAARDVLAEAGLAEYF | AWSLAWGENTLY | 0.8086082 |
| DDQIVLSAKANLAKVEQ | AWSLAWGENTLY | 0.734143 |
| DELLPDTGEDSDEDGH | AWSLAWGENTLY | 0.75230134 |
| ELPEAPANRATVPQD | AWSLAWGENTLY | 0.7597394 |
| FGQSKIERVTGIG | AWSLAWGENTLY | 0.49692914 |
| GAGGTGSTAGGAGGAGGAGGLY | AWSLAWGENTLY | 0.6490715 |
| GHGGTGAAGGNGGTGGAGGSGI | AWSLAWGENTLY | 0.8170781 |
| GNGGNGGAGGEGGPGGAGGAGA | AWSLAWGENTLY | 0.8281899 |
| GSTAGNGGNGGNATNGTIAGQP | AWSLAWGENTLY | 0.34303257 |
| GTGGDGGNGGNGGTGG | AWSLAWGENTLY | 0.815567 |
| GVRRPAGIPADAPV | AWSLAWGENTLY | 0.66265833 |
| GYRQVTDAHLLTLARR | AWSLAWGENTLY | 0.6162608 |
| IAKMGQDGHDRG | AWSLAWGENTLY | 0.86799836 |
| KAEPSIEIQYT | AWSLAWGENTLY | 0.6766926 |
| KPVRIRMAADEDTVAVT | AWSLAWGENTLY | 0.7819761 |
| MPPRIAGMRLLVIKPEP | AWSLAWGENTLY | 0.8027675 |

| | | |
|---|---|---|
| MKFYAVALFRSRGVPPTR | AWSLAWGENTLY | 0.76661706 |
| NAGGFGILGAVAHSP | AWSLAWGENTLY | 0.3610373 |
| NEGREITLAMAEM | AWSLAWGENTLY | 0.7789867 |
| NELVEWSHPLGHRWRWE | AWSLAWGENTLY | 0.7406398 |
| NLGAGGTGGNGGNPG | AWSLAWGENTLY | 0.7942947 |
| NTEWPKWETVMAADPVD | AWSLAWGENTLY | 0.6881527 |
| PASAAPAETQP | AWSLAWGENTLY | 0.3492428 |
| QQSSDGKNGVRAAEITGE | AWSLAWGENTLY | 0.73597705 |
| RAPDAVTTSWWKE | AWSLAWGENTLY | 0.8257904 |
| SGFGNVGAHTSGWFNQ | AWSLAWGENTLY | 0.7436756 |
| SGVAGAVGQAASVGGLKV | AWSLAWGENTLY | 0.75987554 |
| SNGGAGTGSAGGNGGT | AWSLAWGENTLY | 0.84951746 |
| TGGAGGEGGAAG | AWSLAWGENTLY | 0.61539876 |
| TGGDGGAAGAGGAPGHG | AWSLAWGENTLY | 0.6277935 |
| TMPPLLAAIANAGRHGV | AWSLAWGENTLY | 0.80679727 |
| TSSTAGNSGGA | AWSLAWGENTLY | 0.81853986 |
| VPATVPVGILC | AWSLAWGENTLY | 0.6493281 |
| VQRPDSGGVVLTGRIS | AWSLAWGENTLY | 0.8540215 |
| WTPNPVPATKGVPAGLVQAT | AWSLAWGENTLY | 0.49921536 |
| YALKRALAKFSDRFDVVII | AWSLAWGENTLY | 0.79230106 |
| YLIGSVIV | AWSLAWGENTLY | 0.48929116 |
| YGQLVAVID | AWSLAWGENTLY | 0.7624103 |
| VTVGGMTLPA | AWSLAWGENTLY | 0.39491776 |
| VLALPAV | AWSLAWGENTLY | 0.77612734 |
| VIPPTLNYET | AWSLAWGENTLY | 0.8016707 |
| VGMGGMHPSSG | AWSLAWGENTLY | 0.7919004 |
| VGLAGGAATA | AWSLAWGENTLY | 0.14261402 |
| VGGAGGAGGSGP | AWSLAWGENTLY | 0.7944931 |
| VFAAFMV | AWSLAWGENTLY | 0.77118784 |
| TLYDKLR | AWSLAWGENTLY | 0.85186374 |
| TLAVFTVL | AWSLAWGENTLY | 0.6976098 |
| TGSVQPARV | AWSLAWGENTLY | 0.7767848 |
| TGGGAGSGGT | AWSLAWGENTLY | 0.67759216 |
| SHLETSL | AWSLAWGENTLY | 0.6220887 |
| SFTTGMT | AWSLAWGENTLY | 0.4683789 |
| QVEVAKRMGVRQPTVS | AWSLAWGENTLY | 0.66174495 |
| QGSGGPALAM | AWSLAWGENTLY | 0.6037662 |
| MLAPSNP | AWSLAWGENTLY | 0.5473833 |
| LVNNAAI | AWSLAWGENTLY | 0.7746395 |
| LTGEINTLA | AWSLAWGENTLY | 0.8348901 |
| LSIFGPV | AWSLAWGENTLY | 0.7991359 |
| LLALLANL | AWSLAWGENTLY | 0.8206894 |
| LISNASDALDKLR | AWSLAWGENTLY | 0.795172 |
| LINEAVP | AWSLAWGENTLY | 0.81011194 |
| LGYAFSGL | AWSLAWGENTLY | 0.55860424 |
| LDFVAVAL | AWSLAWGENTLY | 0.6539719 |
| LAQPGAE | AWSLAWGENTLY | 0.5109223 |
| ISNLGAVTGF | AWSLAWGENTLY | 0.6211387 |
| HQEDPGRI | AWSLAWGENTLY | 0.5542844 |
| GTPGEPTP | AWSLAWGENTLY | 0.7373738 |
| GFFNSGS | AWSLAWGENTLY | 0.10006284 |

| | | |
|---|---|---|
| GAGGGGLTAGAN | AWSLAWGENTLY | 0.643988 |
| FVDALVNM | AWSLAWGENTLY | 0.84649044 |
| FLTVAPEL | AWSLAWGENTLY | 0.5118803 |
| FETLTAL | AWSLAWGENTLY | 0.7775178 |
| DVINPIEL | AWSLAWGENTLY | 0.6109315 |
| APVSGAPV | AWSLAWGENTLY | 0.55920196 |
| ALVGGLAW | AWSLAWGENTLY | 0.7894448 |
| ALAAAAATAI | AWSLAWGENTLY | 0.42324582 |
| AFGLGLLL | AWSLAWGENTLY | 0.7567019 |
| AFFAAASL | AWSLAWGENTLY | 0.8536538 |
| AAAALPAFSPP | AWSLAWGENTLY | 0.7445167 |
| VVLAPAL | AWSLAWGENTLY | 0.7304417 |
| TIGHVDHGKTTL | AWSLAWGENTLY | 0.66485596 |
| LISNASDALDK | AWSLAWGENTLY | 0.29488558 |
| LISNASDAL | AWSLAWGENTLY | 0.7488529 |
| ISNASDALDK | AWSLAWGENTLY | 0.086560614 |
| NQAETLVYQTE | AWSLAWGENTLY | 0.68012786 |
| TGGTGGTAGAG | AWSLAWGENTLY | 0.4292397 |
| TGSTAGGAGGA | AWSLAWGENTLY | 0.7407751 |
| DPLAVPVAAGPV | AWSLAWGENTLY | 0.690695 |
| GGTGGTPGNGGN | AWSLAWGENTLY | 0.797377 |
| PAGLAGVDPATL | AWSLAWGENTLY | 0.50229734 |
| GNNLVVEAPHAGASV | AWSLAWGENTLY | 0.529868 |
| KVTRIMSASTLEQPA | AWSLAWGENTLY | 0.4134566 |
| LGFALPIALPRRTVV | AWSLAWGENTLY | 0.3945974 |
| TRLGNLTARKLVVLV | AWSLAWGENTLY | 0.8090197 |
| WSQQRRLIYFPSAGP | AWSLAWGENTLY | 0.6744597 |
| GPNGNPDPMAAAVDIR | AWSLAWGENTLY | 0.5575936 |
| TQRCVSVADLRAAASR | AWSLAWGENTLY | 0.81762624 |
| QFVLDVAEHPVEVLTETV | AWSLAWGENTLY | 0.73341644 |
| GGAGGTGGAGGNGGTQAGDG | AWSLAWGENTLY | 0.35927984 |
| PLLATLRAIAVDVDAKPAQV | AWSLAWGENTLY | 0.68963546 |
| APKFRTGGYMIDFWGVGYQVAK | AWSLAWGENTLY | 0.6941447 |
| GAGGAGGAGGAAGGPGGTGGTG | AWSLAWGENTLY | 0.6720227 |
| IGALGLLATDVAPAGSVAFGVA | AWSLAWGENTLY | 0.096307576 |
| AGGAGGIGGNGGAGGAGGVLMG | AWSLAWGENTLY | 0.7299088 |
| ADELIGLL | AWSLAWGENTLY | 0.60943156 |
| ADRALKA | AWSLAWGENTLY | 0.5114359 |
| AGAVGQVGGAGGA | AWSLAWGENTLY | 0.6302388 |
| AGGGGGLGGAP | AWSLAWGENTLY | 0.80200744 |
| AGGLLYGNGGAG | AWSLAWGENTLY | 0.2843763 |
| AGLESIGL | AWSLAWGENTLY | 0.6237484 |
| AGLKTGII | AWSLAWGENTLY | 0.8149987 |
| ASLAALLL | AWSLAWGENTLY | 0.7837944 |
| ASVINISSI | AWSLAWGENTLY | 0.8379509 |
| ATPPGPA | AWSLAWGENTLY | 0.74022853 |
| ATVASALV | AWSLAWGENTLY | 0.6686709 |
| ATVASLIL | AWSLAWGENTLY | 0.77185714 |
| AVNGPVLIRH | AWSLAWGENTLY | 0.77336085 |
| DGAAGAVGI | AWSLAWGENTLY | 0.16602726 |
| DPLLHGI | AWSLAWGENTLY | 0.1534155 |

| | | |
|---|---|---|
| ELGAALGAL | AWSLAWGENTLY | 0.74988633 |
| ELGRTQAV | AWSLAWGENTLY | 0.1550332 |
| FVAGGYLS | AWSLAWGENTLY | 0.728131 |
| GASGGAGGAGGW | AWSLAWGENTLY | 0.8223932 |
| GGAGAGGDGGAGGI | AWSLAWGENTLY | 0.70450175 |
| GTGHNGSQPGMGGNG | AWSLAWGENTLY | 0.41939694 |
| GYAAASASAS | AWSLAWGENTLY | 0.6746792 |
| HQVDVAAVQQG | AWSLAWGENTLY | 0.5612445 |
| IGAAIAHT | AWSLAWGENTLY | 0.7860613 |
| IGMIEVAQL | AWSLAWGENTLY | 0.7665975 |
| IRLAPEHTSAMD | AWSLAWGENTLY | 0.81238896 |
| IYNVSVL | AWSLAWGENTLY | 0.81832623 |
| KQGFLGM | AWSLAWGENTLY | 0.5971634 |
| LESVITQL | AWSLAWGENTLY | 0.8003799 |
| LGLGALVVV | AWSLAWGENTLY | 0.5665301 |
| LIGNGGNGGSGGI | AWSLAWGENTLY | 0.63331574 |
| LNAINAPTL | AWSLAWGENTLY | 0.6739881 |
| LQLLNTGV | AWSLAWGENTLY | 0.7074752 |
| LQTSIEF | AWSLAWGENTLY | 0.68972766 |
| MSFAVSEL | AWSLAWGENTLY | 0.6392975 |
| NTGALGL | AWSLAWGENTLY | 0.3929759 |
| PPGAGGAGGAGTT | AWSLAWGENTLY | 0.72989964 |
| QLVGLGI | AWSLAWGENTLY | 0.6755314 |
| RFLEQVHQL | AWSLAWGENTLY | 0.19369026 |
| RGVGYVLR | AWSLAWGENTLY | 0.48281085 |
| SATGVFAGVI | AWSLAWGENTLY | 0.8663317 |
| SLPAITLP | AWSLAWGENTLY | 0.83508843 |
| STALAFTV | AWSLAWGENTLY | 0.46585587 |
| SVLTVGLGK | AWSLAWGENTLY | 0.49301997 |
| TALAGAGSGI | AWSLAWGENTLY | 0.6688664 |
| TGLPFTGI | AWSLAWGENTLY | 0.2120195 |
| VAEHVTV | AWSLAWGENTLY | 0.74824786 |
| VDIVLNSL | AWSLAWGENTLY | 0.6501559 |
| VLTGGGAVL | AWSLAWGENTLY | 0.65176487 |
| VNVGLAYGI | AWSLAWGENTLY | 0.5354477 |
| VSSAAVGAV | AWSLAWGENTLY | 0.72118616 |
| VTSVYEAGV | AWSLAWGENTLY | 0.5485423 |
| WVALDRGA | AWSLAWGENTLY | 0.7876637 |
| YLVGSVVI | AWSLAWGENTLY | 0.1193354 |
| GPPGTGKTLL | AWSLAWGENTLY | 0.7648919 |
| GVGMTKF | AWSLAWGENTLY | 0.48761362 |
| IGIGGSDL | AWSLAWGENTLY | 0.5613084 |
| IGTLLAEL | AWSLAWGENTLY | 0.31781483 |
| IYTSGTT | AWSLAWGENTLY | 0.7335862 |
| LSFPAGV | AWSLAWGENTLY | 0.82814837 |
| LYTSGTT | AWSLAWGENTLY | 0.8111286 |
| VGAFPFGF | AWSLAWGENTLY | 0.28965104 |
| VGAGTVEF | AWSLAWGENTLY | 0.81443477 |
| VGKSTLLL | AWSLAWGENTLY | 0.8615448 |
| AALAHAGGGIV | AWSLAWGENTLY | 0.5912529 |
| AALVAVSLL | AWSLAWGENTLY | 0.28372335 |

| | | |
|---|---|---|
| AAVSRAVAAL | AWSLAWGENTLY | 0.7398396 |
| AGGSSGIGGF | AWSLAWGENTLY | 0.6630527 |
| AGLTVANANG | AWSLAWGENTLY | 0.74923503 |
| AGLVGAVL | AWSLAWGENTLY | 0.79609 |
| ALSQAFTV | AWSLAWGENTLY | 0.441452 |
| ALTFSVSL | AWSLAWGENTLY | 0.7513548 |
| ALTGAALVAL | AWSLAWGENTLY | 0.79155904 |
| ASFSIGAGVL | AWSLAWGENTLY | 0.5509026 |
| DGSVYKI | AWSLAWGENTLY | 0.66887856 |
| EQLAEIV | AWSLAWGENTLY | 0.5458905 |
| FAVSELTVM | AWSLAWGENTLY | 0.20253864 |
| GAAPLAAGL | AWSLAWGENTLY | 0.77957046 |
| GADGGDGGSGGAGG | AWSLAWGENTLY | 0.55911523 |
| GFGLDTGGAGG | AWSLAWGENTLY | 0.5566921 |
| GLASGGLL | AWSLAWGENTLY | 0.69485587 |
| GRIAGAAVLL | AWSLAWGENTLY | 0.5715527 |
| GSAGSLNTSF | AWSLAWGENTLY | 0.56990904 |
| GTGQIGI | AWSLAWGENTLY | 0.614213 |
| GTLIALIL | AWSLAWGENTLY | 0.74171245 |
| IAATMGFVVG | AWSLAWGENTLY | 0.7518903 |
| IFAQALL | AWSLAWGENTLY | 0.7206121 |
| IGAAVALGF | AWSLAWGENTLY | 0.68820894 |
| IGLTPLAL | AWSLAWGENTLY | 0.83465886 |
| ILGSNTSTL | AWSLAWGENTLY | 0.33135995 |
| IVFSLTVL | AWSLAWGENTLY | 0.4525066 |
| LASKATGF | AWSLAWGENTLY | 0.7832515 |
| LASVALIL | AWSLAWGENTLY | 0.255121 |
| LFAVGLTL | AWSLAWGENTLY | 0.7860061 |
| LFNGGIGTYI | AWSLAWGENTLY | 0.70176554 |
| LTLGAGII | AWSLAWGENTLY | 0.73844826 |
| LVAAALGDI | AWSLAWGENTLY | 0.66462165 |
| NTAFVQLGI | AWSLAWGENTLY | 0.8649622 |
| QLFSGAEL | AWSLAWGENTLY | 0.8083085 |
| SAATFITL | AWSLAWGENTLY | 0.65808177 |
| SNYGLANLGV | AWSLAWGENTLY | 0.682398 |
| SYGGAYAVM | AWSLAWGENTLY | 0.48149532 |
| TTGVGGLTLG | AWSLAWGENTLY | 0.8093899 |
| TTQTIYI | AWSLAWGENTLY | 0.7693955 |
| VAAGAGLGAVI | AWSLAWGENTLY | 0.4574175 |
| VAAVVTLL | AWSLAWGENTLY | 0.65527284 |
| VGFLGGLI | AWSLAWGENTLY | 0.850953 |
| VGVIPILL | AWSLAWGENTLY | 0.83307874 |
| VIFASSLI | AWSLAWGENTLY | 0.19279057 |
| VKGGDGGAAAG | AWSLAWGENTLY | 0.6409417 |
| VTKTFTNI | AWSLAWGENTLY | 0.23618338 |
| VVAAAILL | AWSLAWGENTLY | 0.66317916 |
| YAALDVEL | AWSLAWGENTLY | 0.4703137 |
| LASVALLL | AWSLAWGENTLY | 0.77192694 |
| LDAVHRV | AWSLAWGENTLY | 0.6306323 |
| QGLSSSIF | AWSLAWGENTLY | 0.6863357 |
| RGTFANIRL | AWSLAWGENTLY | 0.7293062 |

| | | |
|---|---|---|
| RQLWWGHRI | AWSLAWGENTLY | 0.8483365 |
| YGPPGTGKTLL | AWSLAWGENTLY | 0.6133748 |
| AALVGAPANVA | AWSLAWGENTLY | 0.638803 |
| AATFRRTE | AWSLAWGENTLY | 0.30300492 |
| AGALGGGI | AWSLAWGENTLY | 0.2546264 |
| AGGAGGAGGIGAG | AWSLAWGENTLY | 0.5837786 |
| AGSGALGL | AWSLAWGENTLY | 0.5840092 |
| AQIADQI | AWSLAWGENTLY | 0.15120766 |
| ASAVAVTYL | AWSLAWGENTLY | 0.8187195 |
| ASTVLGGTGG | AWSLAWGENTLY | 0.7273408 |
| AVGGEPAK | AWSLAWGENTLY | 0.80341625 |
| CEQGTFGGTC | AWSLAWGENTLY | 0.5622966 |
| DEGKAPAG | AWSLAWGENTLY | 0.7003556 |
| DTVVFPAL | AWSLAWGENTLY | 0.5593183 |
| FEGITMAG | AWSLAWGENTLY | 0.7949867 |
| FPLVVQD | AWSLAWGENTLY | 0.5702133 |
| GAFTTPQ | AWSLAWGENTLY | 0.34148133 |
| GFGGFGTAGGVGGA | AWSLAWGENTLY | 0.7250746 |
| GFLAQFL | AWSLAWGENTLY | 0.50043344 |
| GGAGGAGGNAGVA | AWSLAWGENTLY | 0.8195897 |
| GGVGTAGAGGAGGAGG | AWSLAWGENTLY | 0.7993703 |
| GNGGGGDAGNAGSGGN | AWSLAWGENTLY | 0.73457015 |
| GTGGAGGAGNDAGS | AWSLAWGENTLY | 0.78810483 |
| GTTNTGWFNS | AWSLAWGENTLY | 0.20196648 |
| IFMGPLFTL | AWSLAWGENTLY | 0.58439946 |
| ILAHAIEAI | AWSLAWGENTLY | 0.85057604 |
| IRSAIREAL | AWSLAWGENTLY | 0.74820346 |
| ITFAGGG | AWSLAWGENTLY | 0.6557393 |
| ITGMSGHQEV | AWSLAWGENTLY | 0.83474064 |
| IVAGLACTL | AWSLAWGENTLY | 0.5352646 |
| IVGGGLAGA | AWSLAWGENTLY | 0.661672 |
| IVTGVLNI | AWSLAWGENTLY | 0.7308397 |
| LAQQGCI | AWSLAWGENTLY | 0.7648958 |
| LCLHVEN | AWSLAWGENTLY | 0.7438869 |
| LDAISDGS | AWSLAWGENTLY | 0.59780395 |
| LGGFAAML | AWSLAWGENTLY | 0.73948854 |
| LGGKNPMI | AWSLAWGENTLY | 0.86645454 |
| LGPKGVAAIL | AWSLAWGENTLY | 0.67400473 |
| LMVGVSI | AWSLAWGENTLY | 0.81040406 |
| LVGAAFVL | AWSLAWGENTLY | 0.75303555 |
| LVLAVAPL | AWSLAWGENTLY | 0.70498675 |
| MAAIAVIDAA | AWSLAWGENTLY | 0.52103394 |
| NNGINAELQ | AWSLAWGENTLY | 0.60342443 |
| RGEPLVLVHG | AWSLAWGENTLY | 0.52415544 |
| RGGAGGAAGLI | AWSLAWGENTLY | 0.46763015 |
| RSYRMHPAV | AWSLAWGENTLY | 0.70604366 |
| SAAHLLAGQLA | AWSLAWGENTLY | 0.6073243 |
| SAAVGSY | AWSLAWGENTLY | 0.87861884 |
| SPHAVLHRI | AWSLAWGENTLY | 0.7510823 |
| SQIAKVL | AWSLAWGENTLY | 0.7540816 |
| TALGHLVP | AWSLAWGENTLY | 0.4799827 |

| | | |
|---|---|---|
| TAPTTSGAA | AWSLAWGENTLY | 0.7429005 |
| TDGDDGNGGSPGT | AWSLAWGENTLY | 0.71756655 |
| TDGLGEN | AWSLAWGENTLY | 0.42963725 |
| TGGPPLPGQA | AWSLAWGENTLY | 0.84285164 |
| TIPAITI | AWSLAWGENTLY | 0.49016175 |
| TIPGFTI | AWSLAWGENTLY | 0.56050146 |
| TSIASGALA | AWSLAWGENTLY | 0.79645646 |
| TVTLPGAAI | AWSLAWGENTLY | 0.764186 |
| VASGLSAALP | AWSLAWGENTLY | 0.6098958 |
| VGAGVIGI | AWSLAWGENTLY | 0.49128535 |
| VGPKRFGV | AWSLAWGENTLY | 0.49893314 |
| VNVAFPD | AWSLAWGENTLY | 0.79617 |
| VSMTNSGVS | AWSLAWGENTLY | 0.8214073 |
| VTDVLGDLSV | AWSLAWGENTLY | 0.6205605 |
| VVVVGAGFAGL | AWSLAWGENTLY | 0.8211086 |
| VYTVGTVV | AWSLAWGENTLY | 0.70970213 |
| WTTFSGP | AWSLAWGENTLY | 0.4013489 |
| YDELVEAF | AWSLAWGENTLY | 0.70185214 |
| YGTGGAGGNGG | AWSLAWGENTLY | 0.8251412 |
| YTNVMAV | AWSLAWGENTLY | 0.80198896 |
| FLEQVHQL | AWSLAWGENTLY | 0.86294043 |
| GLSLLAVAV | AWSLAWGENTLY | 0.78649795 |
| LEQVHQL | AWSLAWGENTLY | 0.6200158 |
| LNAAGAI | AWSLAWGENTLY | 0.82287526 |
| PAVHASL | AWSLAWGENTLY | 0.87093663 |
| TELEDDA | AWSLAWGENTLY | 0.7985817 |
| TPPGPPN | AWSLAWGENTLY | 0.6390344 |
| TPPPPPPP | AWSLAWGENTLY | 0.5261684 |
| TQSPALI | ASSLRDRGHERLF | 0.7303668 |
| EFAAAFPQ | ASSLRDRGHERLF | 0.80257416 |
| LLSGNLDVL | ASSLRDRGHERLF | 0.47284085 |
| GPANLGNYNI | ASSLRDRGHERLF | 0.69318664 |
| TLGPTLP | ASSLRDRGHERLF | 0.73425794 |
| VYAYSSCCK | ASSLRDRGHERLF | 0.40817314 |
| APAPAPAPNI | ASSLRDRGHERLF | 0.76365113 |
| PGGPVPIP | ASSLRDRGHERLF | 0.834983 |
| FLTVALEP | ASSLRDRGHERLF | 0.63982123 |
| SQYFGANL | ASSLRDRGHERLF | 0.5821877 |
| GGVKPAA | ASSLRDRGHERLF | 0.81984913 |
| GGTGGAGGAGGW | ASSLRDRGHERLF | 0.86906374 |
| GVGGVAGLGGAGG | ASSLRDRGHERLF | 0.81282455 |
| VLLVLATTI | ASSLRDRGHERLF | 0.30982563 |
| IGIFAAML | ASSLRDRGHERLF | 0.18335171 |
| AVGDLID | ASSLRDRGHERLF | 0.5320192 |
| NISPLPL | ASSLRDRGHERLF | 0.71289235 |
| VTAVFVAI | ASSLRDRGHERLF | 0.60611284 |
| SGDDNDAAL | ASSLRDRGHERLF | 0.5174791 |
| AVAALAHS | ASSLRDRGHERLF | 0.7433297 |
| GGVTVMATP | ASSLRDRGHERLF | 0.80973303 |
| AGLAANVRD | ASSLRDRGHERLF | 0.585666 |
| TNRPDLIDPALLRPG | ASSLRDRGHERLF | 0.40627533 |

| | | |
|---|---|---|
| LANLGVTAL | ASSLRDRGHERLF | 0.7798871 |
| TGERIEL | ASSLRDRGHERLF | 0.7979154 |
| NGEELAA | ASSLRDRGHERLF | 0.54128295 |
| VLSLVWG | ASSLRDRGHERLF | 0.689424 |
| IGYALWL | ASSLRDRGHERLF | 0.4951092 |
| LSLLLAPL | ASSLRDRGHERLF | 0.19444785 |
| TLDEALK | ASSLRDRGHERLF | 0.70976394 |
| TDGAGTAEGG | ASSLRDRGHERLF | 0.45290846 |
| NAPGGAGGNGG | ASSLRDRGHERLF | 0.83577085 |
| VSVFMGMGGNF | ASSLRDRGHERLF | 0.47228953 |
| LASYLAAL | ASSLRDRGHERLF | 0.81711084 |
| LVGGTPAL | ASSLRDRGHERLF | 0.75626576 |
| VRNADVL | ASSLRDRGHERLF | 0.773916 |
| TCSSPFLVSAM | ASSLRDRGHERLF | 0.6446636 |
| VHITDAE | ASSLRDRGHERLF | 0.55978715 |
| EYAAPSF | ASSLRDRGHERLF | 0.4507723 |
| VVDAFHGGL | ASSLRDRGHERLF | 0.18065959 |
| ASVVDPQ | ASSLRDRGHERLF | 0.83099127 |
| SYNEFGGLS | ASSLRDRGHERLF | 0.11041321 |
| DPDGSVY | ASSLRDRGHERLF | 0.81577456 |
| MVSTIHATE | ASSLRDRGHERLF | 0.79216886 |
| LAAGVADPV | ASSLRDRGHERLF | 0.525072 |
| TKEQIAA | ASSLRDRGHERLF | 0.76562774 |
| SLASPAAL | ASSLRDRGHERLF | 0.39645413 |
| IGLVFYGV | ASSLRDRGHERLF | 0.35441762 |
| FLLDASNL | ASSLRDRGHERLF | 0.7972489 |
| YQVGNFP | ASSLRDRGHERLF | 0.7808336 |
| ESLLSQST | ASSLRDRGHERLF | 0.5817497 |
| LFTVGFAAVG | ASSLRDRGHERLF | 0.71428883 |
| ATPPTPT | ASSLRDRGHERLF | 0.74244237 |
| TDLATLG | ASSLRDRGHERLF | 0.6350461 |
| SAGGTGR | ASSLRDRGHERLF | 0.62967587 |
| GFSNSGS | ASSLRDRGHERLF | 0.81355214 |
| GGNGGSGGTSV | ASSLRDRGHERLF | 0.7270524 |
| GTSVATGGAGN | ASSLRDRGHERLF | 0.57556725 |
| NGGNGGSGGTS | ASSLRDRGHERLF | 0.77380455 |
| AVVKFPH | ASSLRDRGHERLF | 0.42966878 |
| TPPPPPP | ASSLRDRGHERLF | 0.7422596 |
| AAPAAAAAPA | ASSLRDRGHERLF | 0.689548 |
| IALNEDL | ASSLRDRGHERLF | 0.6993859 |
| IVNVSSM | ASSLRDRGHERLF | 0.7801199 |
| TGIPPAPR | ASSLRDRGHERLF | 0.5511141 |
| FDLGGGTF | ASSLRDRGHERLF | 0.5738612 |
| FDLGGGTFDVS | ASSLRDRGHERLF | 0.5411557 |
| NEPTAAA | ASSLRDRGHERLF | 0.7161211 |
| NEPTAAALA | ASSLRDRGHERLF | 0.8023498 |
| GSIINISSI | ASSLRDRGHERLF | 0.46589303 |
| VAPGNAGTA | ASSLRDRGHERLF | 0.7190778 |
| TGGGASLE | ASSLRDRGHERLF | 0.5812006 |
| VAVYNHYKRI | ASSLRDRGHERLF | 0.44773802 |
| TDPAPATT | ASSLRDRGHERLF | 0.6442749 |

| | | |
|---|---|---|
| TNRPDILDPALLRPG | ASSLRDRGHERLF | 0.6767528 |
| LTTTTTS | ASSLRDRGHERLF | 0.599551 |
| PTGSGKTL | ASSLRDRGHERLF | 0.749519 |
| TGSGKTL | ASSLRDRGHERLF | 0.4674527 |
| TIGHVDHGK | ASSLRDRGHERLF | 0.756623 |
| TIGHVDHGKTT | ASSLRDRGHERLF | 0.86198694 |
| GHVDHGKTT | ASSLRDRGHERLF | 0.7045193 |
| AQFEHTLL | ASSLRDRGHERLF | 0.8080437 |
| IVNQDEL | ASSLRDRGHERLF | 0.7374899 |
| TNGSQFF | ASSLRDRGHERLF | 0.77001846 |
| TGKGIVE | ASSLRDRGHERLF | 0.6329559 |
| LISNASDA | ASSLRDRGHERLF | 0.56545424 |
| AELRTLGELTKIAKAHGAQ | ASSLRDRGHERLF | 0.83823955 |
| APNAVAAAEPSW | ASSLRDRGHERLF | 0.7926872 |
| APQVAGAATAVL | ASSLRDRGHERLF | 0.8144094 |
| ARRDPAAAAAILPTDA | ASSLRDRGHERLF | 0.73073363 |
| DAAARDVLAEAGLAEYF | ASSLRDRGHERLF | 0.7665223 |
| DDQIVLSAKANLAKVEQ | ASSLRDRGHERLF | 0.78973675 |
| DELLPDTGEDSDEDGH | ASSLRDRGHERLF | 0.53138554 |
| ELPEAPANRATVPQD | ASSLRDRGHERLF | 0.27916873 |
| FGQSKIERVTGIG | ASSLRDRGHERLF | 0.5992096 |
| GAGGTGSTAGGAGGAGGAGGLY | ASSLRDRGHERLF | 0.514547 |
| GHGGTGAAGGNGGTGGAGGSGI | ASSLRDRGHERLF | 0.6203407 |
| GNGGNGGAGGEGGPGGAGGAGA | ASSLRDRGHERLF | 0.28276974 |
| GSTAGNGGNGGNATNGTIAGQP | ASSLRDRGHERLF | 0.81592405 |
| GTGGDGGNGGNGGTGG | ASSLRDRGHERLF | 0.8736478 |
| GVRRPAGIPADAPV | ASSLRDRGHERLF | 0.56487185 |
| GYRQVTDAHLLTLARR | ASSLRDRGHERLF | 0.82383406 |
| IAKMGQDGHDRG | ASSLRDRGHERLF | 0.36643568 |
| KAEPSIEIQYT | ASSLRDRGHERLF | 0.78430355 |
| KPVRIRMAADEDTVAVT | ASSLRDRGHERLF | 0.830765 |
| MPPRIAGMRLLVIKPEP | ASSLRDRGHERLF | 0.45313984 |
| MKFYAVALFRSRGVPPTR | ASSLRDRGHERLF | 0.7725264 |
| NAGGFGILGAVAHSP | ASSLRDRGHERLF | 0.561852 |
| NEGREITLAMAEM | ASSLRDRGHERLF | 0.6949046 |
| NELVEWSHPLGHRWRWE | ASSLRDRGHERLF | 0.8432224 |
| NLGAGGTGGNGGNPG | ASSLRDRGHERLF | 0.6018503 |
| NTEWPKWETVMAADPVD | ASSLRDRGHERLF | 0.72269714 |
| PASAAPAETQP | ASSLRDRGHERLF | 0.6526717 |
| QQSSDGKNGVRAAEITGE | ASSLRDRGHERLF | 0.4546209 |
| RAPDAVTTSWWKE | ASSLRDRGHERLF | 0.7473399 |
| SGFGNVGAHTSGWFNQ | ASSLRDRGHERLF | 0.8284681 |
| SGVAGAVGQAASVGGLKV | ASSLRDRGHERLF | 0.36930507 |
| SNGGAGTGSAGGNGGT | ASSLRDRGHERLF | 0.8603363 |
| TGGAGGEGGAAG | ASSLRDRGHERLF | 0.5554869 |
| TGGDGGAAGAGGAPGHG | ASSLRDRGHERLF | 0.8728845 |
| TMPPLLAAIANAGRHGV | ASSLRDRGHERLF | 0.78299296 |
| TSSTAGNSGGA | ASSLRDRGHERLF | 0.68445706 |
| VPATVPVGILC | ASSLRDRGHERLF | 0.6267264 |
| VQRPDSGGVVLTGRIS | ASSLRDRGHERLF | 0.8236568 |
| WTPNPVPATKGVPAGLVQAT | ASSLRDRGHERLF | 0.6687082 |

| | | |
|---|---|---|
| YALKRALAKFSDRFDVVII | ASSLRDRGHERLF | 0.8160485 |
| YLIGSVIV | ASSLRDRGHERLF | 0.82004666 |
| YGQLVAVID | ASSLRDRGHERLF | 0.7951488 |
| VTVGGMTLPA | ASSLRDRGHERLF | 0.8931643 |
| VLALPAV | ASSLRDRGHERLF | 0.8687126 |
| VIPPTLNYET | ASSLRDRGHERLF | 0.85878134 |
| VGMGGMHPSSG | ASSLRDRGHERLF | 0.8155225 |
| VGLAGGAATA | ASSLRDRGHERLF | 0.4433403 |
| VGGAGGAGGSGP | ASSLRDRGHERLF | 0.84945536 |
| VFAAFMV | ASSLRDRGHERLF | 0.7443508 |
| TLYDKLR | ASSLRDRGHERLF | 0.87815917 |
| TLAVFTVL | ASSLRDRGHERLF | 0.89015603 |
| TGSVQPARV | ASSLRDRGHERLF | 0.78834116 |
| TGGGAGSGGT | ASSLRDRGHERLF | 0.900564 |
| SHLETSL | ASSLRDRGHERLF | 0.88868093 |
| SFTTGMT | ASSLRDRGHERLF | 0.6621266 |
| QVEVAKRMGVRQPTVS | ASSLRDRGHERLF | 0.84576195 |
| QGSGGPALAM | ASSLRDRGHERLF | 0.8034336 |
| MLAPSNP | ASSLRDRGHERLF | 0.8438547 |
| LVNNAAI | ASSLRDRGHERLF | 0.59159076 |
| LTGEINTLA | ASSLRDRGHERLF | 0.88583726 |
| LSIFGPV | ASSLRDRGHERLF | 0.81136703 |
| LLALLANL | ASSLRDRGHERLF | 0.17662889 |
| LISNASDALDKLR | ASSLRDRGHERLF | 0.8749381 |
| LINEAVP | ASSLRDRGHERLF | 0.8596738 |
| LGYAFSGL | ASSLRDRGHERLF | 0.8354405 |
| LDFVAVAL | ASSLRDRGHERLF | 0.8936665 |
| LAQPGAE | ASSLRDRGHERLF | 0.43315327 |
| ISNLGAVTGF | ASSLRDRGHERLF | 0.7771791 |
| HQEDPGRI | ASSLRDRGHERLF | 0.8899036 |
| GTPGEPTP | ASSLRDRGHERLF | 0.898973 |
| GFFNSGS | ASSLRDRGHERLF | 0.7284259 |
| GAGGGGLTAGAN | ASSLRDRGHERLF | 0.8764958 |
| FVDALVNM | ASSLRDRGHERLF | 0.8954471 |
| FLTVAPEL | ASSLRDRGHERLF | 0.87245667 |
| FETLTAL | ASSLRDRGHERLF | 0.894475 |
| DVINPIEL | ASSLRDRGHERLF | 0.89850533 |
| APVSGAPV | ASSLRDRGHERLF | 0.7810874 |
| ALVGGLAW | ASSLRDRGHERLF | 0.89640343 |
| ALAAAAATAI | ASSLRDRGHERLF | 0.9044371 |
| AFGLGLLL | ASSLRDRGHERLF | 0.7906918 |
| AFFAAASL | ASSLRDRGHERLF | 0.869006 |
| AAAALPAFSPP | ASSLRDRGHERLF | 0.77857876 |
| VVLAPAL | ASSLRDRGHERLF | 0.83891386 |
| TIGHVDHGKTTL | ASSLRDRGHERLF | 0.28033003 |
| LISNASDALDK | ASSLRDRGHERLF | 0.8770608 |
| LISNASDAL | ASSLRDRGHERLF | 0.8936218 |
| ISNASDALDK | ASSLRDRGHERLF | 0.84821904 |
| NQAETLVYQTE | ASSLRDRGHERLF | 0.74452823 |
| TGGTGGTAGAG | ASSLRDRGHERLF | 0.7500274 |
| TGSTAGGAGGA | ASSLRDRGHERLF | 0.8128773 |

| | | |
|---|---|---|
| DPLAVPVAAGPV | ASSLRDRGHERLF | 0.8311037 |
| GGTGGTPGNGGN | ASSLRDRGHERLF | 0.3320167 |
| PAGLAGVDPATL | ASSLRDRGHERLF | 0.7938745 |
| GNNLVVEAPHAGASV | ASSLRDRGHERLF | 0.8934114 |
| KVTRIMSASTLEQPA | ASSLRDRGHERLF | 0.8893937 |
| LGFALPIALPRRTVV | ASSLRDRGHERLF | 0.6048249 |
| TRLGNLTARKLVVLV | ASSLRDRGHERLF | 0.90137947 |
| WSQQRRLIYFPSAGP | ASSLRDRGHERLF | 0.8806871 |
| GPNGNPDPMAAAVDIR | ASSLRDRGHERLF | 0.8941067 |
| TQRCVSVADLRAAASR | ASSLRDRGHERLF | 0.79553586 |
| QFVLDVAEHPVEVLTETV | ASSLRDRGHERLF | 0.7783002 |
| GGAGGTGGAGGNGGTQAGDG | ASSLRDRGHERLF | 0.8880874 |
| PLLATLRAIAVDVDAKPAQV | ASSLRDRGHERLF | 0.8378307 |
| APKFRTGGYMIDFWGVGYQVAK | ASSLRDRGHERLF | 0.773746 |
| GAGGAGGAGGAAGGPGGTGGTG | ASSLRDRGHERLF | 0.8934959 |
| IGALGLLATDVAPAGSVAFGVA | ASSLRDRGHERLF | 0.8333266 |
| AGGAGGIGGNGGAGGAGGVLMG | ASSLRDRGHERLF | 0.79864305 |
| ADELIGLL | ASSLRDRGHERLF | 0.8078753 |
| ADRALKA | ASSLRDRGHERLF | 0.8905409 |
| AGAVGQVGGAGGA | ASSLRDRGHERLF | 0.755628 |
| AGGGGGLGGAP | ASSLRDRGHERLF | 0.58548987 |
| AGGLLYGNGGAG | ASSLRDRGHERLF | 0.6682354 |
| AGLESIGL | ASSLRDRGHERLF | 0.8453684 |
| AGLKTGII | ASSLRDRGHERLF | 0.896168 |
| ASLAALLL | ASSLRDRGHERLF | 0.8621664 |
| ASVINISSI | ASSLRDRGHERLF | 0.7845968 |
| ATPPGPA | ASSLRDRGHERLF | 0.881456 |
| ATVASALV | ASSLRDRGHERLF | 0.85639554 |
| ATVASLIL | ASSLRDRGHERLF | 0.8511246 |
| AVNGPVLIRH | ASSLRDRGHERLF | 0.88448906 |
| DGAAGAVGI | ASSLRDRGHERLF | 0.7955986 |
| DPLLHGI | ASSLRDRGHERLF | 0.8880809 |
| ELGAALGAL | ASSLRDRGHERLF | 0.88090485 |
| ELGRTQAV | ASSLRDRGHERLF | 0.8871747 |
| FVAGGYLS | ASSLRDRGHERLF | 0.89165056 |
| GASGGAGGAGGW | ASSLRDRGHERLF | 0.8446061 |
| GGAGAGGDGGAGGI | ASSLRDRGHERLF | 0.8447435 |
| GTGHNGSQPGMGGNG | ASSLRDRGHERLF | 0.4596387 |
| GYAAASASAS | ASSLRDRGHERLF | 0.789942 |
| HQVDVAAVQQG | ASSLRDRGHERLF | 0.8833782 |
| IGAAIAHT | ASSLRDRGHERLF | 0.8713696 |
| IGMIEVAQL | ASSLRDRGHERLF | 0.89167947 |
| IRLAPEHTSAMD | ASSLRDRGHERLF | 0.8675897 |
| IYNVSVL | ASSLRDRGHERLF | 0.7884609 |
| KQGFLGM | ASSLRDRGHERLF | 0.89013815 |
| LESVITQL | ASSLRDRGHERLF | 0.89295876 |
| LGLGALVVV | ASSLRDRGHERLF | 0.58286136 |
| LIGNGGNGGSGGI | ASSLRDRGHERLF | 0.8217809 |
| LNAINAPTL | ASSLRDRGHERLF | 0.9038174 |
| LQLLNTGV | ASSLRDRGHERLF | 0.86561346 |
| LQTSIEF | ASSLRDRGHERLF | 0.84552354 |

| | | |
|---|---|---|
| MSFAVSEL | ASSLRDRGHERLF | 0.8023069 |
| NTGALGL | ASSLRDRGHERLF | 0.8983 |
| PPGAGGAGGAGTT | ASSLRDRGHERLF | 0.7995579 |
| QLVGLGI | ASSLRDRGHERLF | 0.9041061 |
| RFLEQVHQL | ASSLRDRGHERLF | 0.26210877 |
| RGVGYVLR | ASSLRDRGHERLF | 0.7574256 |
| SATGVFAGVI | ASSLRDRGHERLF | 0.5642358 |
| SLPAITLP | ASSLRDRGHERLF | 0.41771722 |
| STALAFTV | ASSLRDRGHERLF | 0.89340025 |
| SVLTVGLGK | ASSLRDRGHERLF | 0.80161756 |
| TALAGAGSGI | ASSLRDRGHERLF | 0.88720423 |
| TGLPFTGI | ASSLRDRGHERLF | 0.87035286 |
| VAEHVTV | ASSLRDRGHERLF | 0.8952483 |
| VDIVLNSL | ASSLRDRGHERLF | 0.8650017 |
| VLTGGGAVL | ASSLRDRGHERLF | 0.8529191 |
| VNVGLAYGI | ASSLRDRGHERLF | 0.6976299 |
| VSSAAVGAV | ASSLRDRGHERLF | 0.89057213 |
| VTSVYEAGV | ASSLRDRGHERLF | 0.8970555 |
| WVALDRGA | ASSLRDRGHERLF | 0.86853004 |
| YLVGSVVI | ASSLRDRGHERLF | 0.65196973 |
| GPPGTGKTLL | ASSLRDRGHERLF | 0.84298176 |
| GVGMTKF | ASSLRDRGHERLF | 0.8678522 |
| IGIGGSDL | ASSLRDRGHERLF | 0.23614323 |
| IGTLLAEL | ASSLRDRGHERLF | 0.32821566 |
| IYTSGTT | ASSLRDRGHERLF | 0.81246734 |
| LSFPAGV | ASSLRDRGHERLF | 0.77488565 |
| LYTSGTT | ASSLRDRGHERLF | 0.8967817 |
| VGAFPFGF | ASSLRDRGHERLF | 0.8898314 |
| VGAGTVEF | ASSLRDRGHERLF | 0.8495853 |
| VGKSTLLL | ASSLRDRGHERLF | 0.88099563 |
| AALAHAGGGIV | ASSLRDRGHERLF | 0.8667418 |
| AALVAVSLL | ASSLRDRGHERLF | 0.10326121 |
| AAVSRAVAAL | ASSLRDRGHERLF | 0.7522782 |
| AGGSSGIGGF | ASSLRDRGHERLF | 0.7122493 |
| AGLTVANANG | ASSLRDRGHERLF | 0.89643824 |
| AGLVGAVL | ASSLRDRGHERLF | 0.78027904 |
| ALSQAFTV | ASSLRDRGHERLF | 0.8018894 |
| ALTFSVSL | ASSLRDRGHERLF | 0.8614924 |
| ALTGAALVAL | ASSLRDRGHERLF | 0.86255056 |
| ASFSIGAGVL | ASSLRDRGHERLF | 0.8544185 |
| DGSVYKI | ASSLRDRGHERLF | 0.6877434 |
| EQLAEIV | ASSLRDRGHERLF | 0.21574342 |
| FAVSELTVM | ASSLRDRGHERLF | 0.8911365 |
| GAAPLAAGL | ASSLRDRGHERLF | 0.4110782 |
| GADGGDGGSGGAGG | ASSLRDRGHERLF | 0.75922817 |
| GFGLDTGGAGG | ASSLRDRGHERLF | 0.37727708 |
| GLASGGLL | ASSLRDRGHERLF | 0.7959677 |
| GRIAGAAVLL | ASSLRDRGHERLF | 0.4567786 |
| GSAGSLNTSF | ASSLRDRGHERLF | 0.88821065 |
| GTGQIGI | ASSLRDRGHERLF | 0.83823836 |
| GTLIALIL | ASSLRDRGHERLF | 0.8355325 |

| | | |
|---|---|---|
| IAATMGFVVG | ASSLRDRGHERLF | 0.90249157 |
| IFAQALL | ASSLRDRGHERLF | 0.8526077 |
| IGAAVALGF | ASSLRDRGHERLF | 0.8427228 |
| IGLTPLAL | ASSLRDRGHERLF | 0.7928121 |
| ILGSNTSTL | ASSLRDRGHERLF | 0.7466012 |
| IVFSLTVL | ASSLRDRGHERLF | 0.7408571 |
| LASKATGF | ASSLRDRGHERLF | 0.75955296 |
| LASVALIL | ASSLRDRGHERLF | 0.90200317 |
| LFAVGLTL | ASSLRDRGHERLF | 0.7869796 |
| LFNGGIGTYI | ASSLRDRGHERLF | 0.5665685 |
| LTLGAGII | ASSLRDRGHERLF | 0.7586249 |
| LVAAALGDI | ASSLRDRGHERLF | 0.90104353 |
| NTAFVQLGI | ASSLRDRGHERLF | 0.90358686 |
| QLFSGAEL | ASSLRDRGHERLF | 0.8117144 |
| SAATFITL | ASSLRDRGHERLF | 0.7165996 |
| SNYGLANLGV | ASSLRDRGHERLF | 0.8159542 |
| SYGGAYAVM | ASSLRDRGHERLF | 0.8600477 |
| TTGVGGLTLG | ASSLRDRGHERLF | 0.78101355 |
| TTQTIYI | ASSLRDRGHERLF | 0.5481452 |
| VAAGAGLGAVI | ASSLRDRGHERLF | 0.8808795 |
| VAAVVTLL | ASSLRDRGHERLF | 0.90045524 |
| VGFLGGLI | ASSLRDRGHERLF | 0.7741616 |
| VGVIPILL | ASSLRDRGHERLF | 0.69943964 |
| VIFASSLI | ASSLRDRGHERLF | 0.852177 |
| VKGGDGGAAAG | ASSLRDRGHERLF | 0.7882252 |
| VTKTFTNI | ASSLRDRGHERLF | 0.8251047 |
| VVAAAILL | ASSLRDRGHERLF | 0.8466753 |
| YAALDVEL | ASSLRDRGHERLF | 0.6022688 |
| LASVALLL | ASSLRDRGHERLF | 0.84138167 |
| LDAVHRV | ASSLRDRGHERLF | 0.9027072 |
| QGLSSSIF | ASSLRDRGHERLF | 0.8290632 |
| RGTFANIRL | ASSLRDRGHERLF | 0.8746729 |
| RQLWWGHRI | ASSLRDRGHERLF | 0.89880013 |
| YGPPGTGKTLL | ASSLRDRGHERLF | 0.88682175 |
| AALVGAPANVA | ASSLRDRGHERLF | 0.898669 |
| AATFRRTE | ASSLRDRGHERLF | 0.884286 |
| AGALGGGI | ASSLRDRGHERLF | 0.8441868 |
| AGGAGGAGGIGAG | ASSLRDRGHERLF | 0.8450895 |
| AGSGALGL | ASSLRDRGHERLF | 0.8787192 |
| AQIADQI | ASSLRDRGHERLF | 0.8659232 |
| ASAVAVTYL | ASSLRDRGHERLF | 0.8513984 |
| ASTVLGGTGG | ASSLRDRGHERLF | 0.7985693 |
| AVGGEPAK | ASSLRDRGHERLF | 0.89366436 |
| CEQGTFGGTC | ASSLRDRGHERLF | 0.8059917 |
| DEGKAPAG | ASSLRDRGHERLF | 0.8850365 |
| DTVVFPAL | ASSLRDRGHERLF | 0.5339782 |
| FEGITMAG | ASSLRDRGHERLF | 0.70201296 |
| FPLVVQD | ASSLRDRGHERLF | 0.88734406 |
| GAFTTPQ | ASSLRDRGHERLF | 0.11748372 |
| GFGGFGTAGGVGGA | ASSLRDRGHERLF | 0.26249388 |
| GFLAQFL | ASSLRDRGHERLF | 0.8635832 |

| | | |
|---|---|---|
| GGAGGAGGNAGVA | ASSLRDRGHERLF | 0.89637375 |
| GGVGTAGAGGAGGAGG | ASSLRDRGHERLF | 0.13511156 |
| GNGGGGDAGNAGSGGN | ASSLRDRGHERLF | 0.7330852 |
| GTGGAGGAGNDAGS | ASSLRDRGHERLF | 0.6739591 |
| GTTNTGWFNS | ASSLRDRGHERLF | 0.8583039 |
| IFMGPLFTL | ASSLRDRGHERLF | 0.86708117 |
| ILAHAIEAI | ASSLRDRGHERLF | 0.38613594 |
| IRSAIREAL | ASSLRDRGHERLF | 0.88374615 |
| ITFAGGG | ASSLRDRGHERLF | 0.57707477 |
| ITGMSGHQEV | ASSLRDRGHERLF | 0.58412695 |
| IVAGLACTL | ASSLRDRGHERLF | 0.8950963 |
| IVGGGLAGA | ASSLRDRGHERLF | 0.8945285 |
| IVTGVLNI | ASSLRDRGHERLF | 0.74978405 |
| LAQQGCI | ASSLRDRGHERLF | 0.89288795 |
| LCLHVEN | ASSLRDRGHERLF | 0.7776963 |
| LDAISDGS | ASSLRDRGHERLF | 0.90503883 |
| LGGFAAML | ASSLRDRGHERLF | 0.82908404 |
| LGGKNPMI | ASSLRDRGHERLF | 0.46769875 |
| LGPKGVAAIL | ASSLRDRGHERLF | 0.8866272 |
| LMVGVSI | ASSLRDRGHERLF | 0.8785259 |
| LVGAAFVL | ASSLRDRGHERLF | 0.8600256 |
| LVLAVAPL | ASSLRDRGHERLF | 0.8824941 |
| MAAIAVIDAA | ASSLRDRGHERLF | 0.63756937 |
| NNGINAELQ | ASSLRDRGHERLF | 0.84487355 |
| RGEPLVLVHG | ASSLRDRGHERLF | 0.8963387 |
| RGGAGGAAGLI | ASSLRDRGHERLF | 0.89402044 |
| RSYRMHPAV | ASSLRDRGHERLF | 0.5754616 |
| SAAHLLAGQLA | ASSLRDRGHERLF | 0.89646757 |
| SAAVGSY | ASSLRDRGHERLF | 0.8526751 |
| SPHAVLHRI | ASSLRDRGHERLF | 0.79968053 |
| SQIAKVL | ASSLRDRGHERLF | 0.90003824 |
| TALGHLVP | ASSLRDRGHERLF | 0.82964087 |
| TAPTTSGAA | ASSLRDRGHERLF | 0.89107776 |
| TDGDDGNGGSPGT | ASSLRDRGHERLF | 0.8858805 |
| TDGLGEN | ASSLRDRGHERLF | 0.8824831 |
| TGGPPLPGQA | ASSLRDRGHERLF | 0.48908654 |
| TIPAITI | ASSLRDRGHERLF | 0.8694944 |
| TIPGFTI | ASSLRDRGHERLF | 0.85661185 |
| TSIASGALA | ASSLRDRGHERLF | 0.8878791 |
| TVTLPGAAI | ASSLRDRGHERLF | 0.83988607 |
| VASGLSAALP | ASSLRDRGHERLF | 0.7360431 |
| VGAGVIGI | ASSLRDRGHERLF | 0.87263525 |
| VGPKRFGV | ASSLRDRGHERLF | 0.89778554 |
| VNVAFPD | ASSLRDRGHERLF | 0.85178715 |
| VSMTNSGVS | ASSLRDRGHERLF | 0.8716388 |
| VTDVLGDLSV | ASSLRDRGHERLF | 0.9016752 |
| VVVVGAGFAGL | ASSLRDRGHERLF | 0.8146082 |
| VYTVGTVV | ASSLRDRGHERLF | 0.8479947 |
| WTTFSGP | ASSLRDRGHERLF | 0.88183177 |
| YDELVEAF | ASSLRDRGHERLF | 0.89473283 |
| YGTGGAGGNGG | ASSLRDRGHERLF | 0.79975057 |

| | | |
|---|---|---|
| YTNVMAV | ASSLRDRGHERLF | 0.896533 |
| FLEQVHQL | ASSLRDRGHERLF | 0.7439184 |
| GLSLLAVAV | ASSLRDRGHERLF | 0.8877398 |
| LEQVHQL | ASSLRDRGHERLF | 0.81085104 |
| LNAAGAI | ASSLRDRGHERLF | 0.88269365 |
| PAVHASL | ASSLRDRGHERLF | 0.77965844 |
| TELEDDA | ASSLRDRGHERLF | 0.89159644 |
| TPPGPPN | ASSLRDRGHERLF | 0.89038986 |
| TPPPPPPP | ASSLRDRGHERLF | 0.8942284 |
| TQSPALI | ASSQDREDTQD | 0.2373515 |
| EFAAAFPQ | ASSQDREDTQD | 0.8884219 |
| LLSGNLDVL | ASSQDREDTQD | 0.8824551 |
| GPANLGNYNI | ASSQDREDTQD | 0.9034907 |
| TLGPTLP | ASSQDREDTQD | 0.8292968 |
| VYAYSSCCK | ASSQDREDTQD | 0.8740439 |
| APAPAPAPNI | ASSQDREDTQD | 0.8371041 |
| PGGPVPIP | ASSQDREDTQD | 0.81301284 |
| FLTVALEP | ASSQDREDTQD | 0.7389313 |
| SQYFGANL | ASSQDREDTQD | 0.8417742 |
| GGVKPAA | ASSQDREDTQD | 0.8311835 |
| GGTGGAGGAGGW | ASSQDREDTQD | 0.80017483 |
| GVGGVAGLGGAGG | ASSQDREDTQD | 0.8868716 |
| VLLVLATTI | ASSQDREDTQD | 0.89798784 |
| IGIFAAML | ASSQDREDTQD | 0.89339775 |
| AVGDLID | ASSQDREDTQD | 0.8928437 |
| NISPLPL | ASSQDREDTQD | 0.89836544 |
| VTAVFVAI | ASSQDREDTQD | 0.8852987 |
| SGDDNDAAL | ASSQDREDTQD | 0.8189141 |
| AVAALAHS | ASSQDREDTQD | 0.83890617 |
| GGVTVMATP | ASSQDREDTQD | 0.7373973 |
| AGLAANVRD | ASSQDREDTQD | 0.83744985 |
| TNRPDLIDPALLRPG | ASSQDREDTQD | 0.79894805 |
| LANLGVTAL | ASSQDREDTQD | 0.87397754 |
| TGERIEL | ASSQDREDTQD | 0.11641634 |
| NGEELAA | ASSQDREDTQD | 0.8591255 |
| VLSLVWG | ASSQDREDTQD | 0.90201956 |
| IGYALWL | ASSQDREDTQD | 0.7884344 |
| LSLLLAPL | ASSQDREDTQD | 0.86994106 |
| TLDEALK | ASSQDREDTQD | 0.8733353 |
| TDGAGTAEGG | ASSQDREDTQD | 0.8069717 |
| NAPGGAGGNGG | ASSQDREDTQD | 0.88643104 |
| VSVFMGMGGNF | ASSQDREDTQD | 0.75904 |
| LASYLAAL | ASSQDREDTQD | 0.8798975 |
| LVGGTPAL | ASSQDREDTQD | 0.903037 |
| VRNADVL | ASSQDREDTQD | 0.8685485 |
| TCSSPFLVSAM | ASSQDREDTQD | 0.8580253 |
| VHITDAE | ASSQDREDTQD | 0.84826666 |
| EYAAPSF | ASSQDREDTQD | 0.43323758 |
| VVDAFHGGL | ASSQDREDTQD | 0.8737926 |
| ASVVDPQ | ASSQDREDTQD | 0.40426987 |
| SYNEFGGLS | ASSQDREDTQD | 0.8526716 |

| | | |
|---|---|---|
| DPDGSVY | ASSQDREDTQD | 0.6777922 |
| MVSTIHATE | ASSQDREDTQD | 0.87867355 |
| LAAGVADPV | ASSQDREDTQD | 0.8637366 |
| TKEQIAA | ASSQDREDTQD | 0.87814647 |
| SLASPAAL | ASSQDREDTQD | 0.7802831 |
| IGLVFYGV | ASSQDREDTQD | 0.8490997 |
| FLLDASNL | ASSQDREDTQD | 0.7671233 |
| YQVGNFP | ASSQDREDTQD | 0.650504 |
| ESLLSQST | ASSQDREDTQD | 0.8925545 |
| LFTVGFAAVG | ASSQDREDTQD | 0.7948353 |
| ATPPTPT | ASSQDREDTQD | 0.8292682 |
| TDLATLG | ASSQDREDTQD | 0.8950865 |
| SAGGTGR | ASSQDREDTQD | 0.8771644 |
| GFSNSGS | ASSQDREDTQD | 0.50273514 |
| GGNGGSGGTSV | ASSQDREDTQD | 0.8398541 |
| GTSVATGGAGN | ASSQDREDTQD | 0.86167383 |
| NGGNGGSGGTS | ASSQDREDTQD | 0.84407413 |
| AVVKFPH | ASSQDREDTQD | 0.14398593 |
| TPPPPPP | ASSQDREDTQD | 0.86697495 |
| AAPAAAAAPA | ASSQDREDTQD | 0.80236804 |
| IALNEDL | ASSQDREDTQD | 0.7799662 |
| IVNVSSM | ASSQDREDTQD | 0.79890096 |
| TGIPPAPR | ASSQDREDTQD | 0.89035827 |
| FDLGGGTF | ASSQDREDTQD | 0.7427113 |
| FDLGGGTFDVS | ASSQDREDTQD | 0.83184195 |
| NEPTAAA | ASSQDREDTQD | 0.8913263 |
| NEPTAAALA | ASSQDREDTQD | 0.88729775 |
| GSIINISSI | ASSQDREDTQD | 0.899355 |
| VAPGNAGTA | ASSQDREDTQD | 0.8629867 |
| TGGGASLE | ASSQDREDTQD | 0.7677216 |
| VAVYNHYKRI | ASSQDREDTQD | 0.8717208 |
| TDPAPATT | ASSQDREDTQD | 0.8255455 |
| TNRPDILDPALLRPG | ASSQDREDTQD | 0.5921458 |
| LTTTTTS | ASSQDREDTQD | 0.41478217 |
| PTGSGKTL | ASSQDREDTQD | 0.86887306 |
| TGSGKTL | ASSQDREDTQD | 0.44260603 |
| TIGHVDHGK | ASSQDREDTQD | 0.86714685 |
| TIGHVDHGKTT | ASSQDREDTQD | 0.8921781 |
| GHVDHGKTT | ASSQDREDTQD | 0.8029691 |
| AQFEHTLL | ASSQDREDTQD | 0.6456499 |
| IVNQDEL | ASSQDREDTQD | 0.832078 |
| TNGSQFF | ASSQDREDTQD | 0.78772837 |
| TGKGIVE | ASSQDREDTQD | 0.8902835 |
| LISNASDA | ASSQDREDTQD | 0.8502538 |
| AELRTLGELTKIAKAHGAQ | ASSQDREDTQD | 0.89939487 |
| APNAVAAAEPSW | ASSQDREDTQD | 0.8905548 |
| APQVAGAATAVL | ASSQDREDTQD | 0.82963383 |
| ARRDPAAAAAILPTDA | ASSQDREDTQD | 0.88876325 |
| DAAARDVLAEAGLAEYF | ASSQDREDTQD | 0.8050917 |
| DDQIVLSAKANLAKVEQ | ASSQDREDTQD | 0.86197895 |
| DELLPDTGEDSDEDGH | ASSQDREDTQD | 0.8582826 |

| | | |
|---|---|---|
| ELPEAPANRATVPQD | ASSQDREDTQD | 0.8579105 |
| FGQSKIERVTGIG | ASSQDREDTQD | 0.8262179 |
| GAGGTGSTAGGAGGAGGAGGLY | ASSQDREDTQD | 0.84882534 |
| GHGGTGAAGGNGGTGGAGGSGI | ASSQDREDTQD | 0.46923035 |
| GNGGNGGAGGEGGPGGAGGAGA | ASSQDREDTQD | 0.8736157 |
| GSTAGNGGNGGNATNGTIAGQP | ASSQDREDTQD | 0.8809625 |
| GTGGDGGNGGNGGTGG | ASSQDREDTQD | 0.42763495 |
| GVRRPAGIPADAPV | ASSQDREDTQD | 0.8294264 |
| GYRQVTDAHLLTLARR | ASSQDREDTQD | 0.901638 |
| IAKMGQDGHDRG | ASSQDREDTQD | 0.8935343 |
| KAEPSIEIQYT | ASSQDREDTQD | 0.7940558 |
| KPVRIRMAADEDTVAVT | ASSQDREDTQD | 0.75943726 |
| MPPRIAGMRLLVIKPEP | ASSQDREDTQD | 0.8583572 |
| MKFYAVALFRSRGVPPTR | ASSQDREDTQD | 0.55100155 |
| NAGGFGILGAVAHSP | ASSQDREDTQD | 0.855884 |
| NEGREITLAMAEM | ASSQDREDTQD | 0.8512888 |
| NELVEWSHPLGHRWRWE | ASSQDREDTQD | 0.8406595 |
| NLGAGGTGGNGGNPG | ASSQDREDTQD | 0.766281 |
| NTEWPKWETVMAADPVD | ASSQDREDTQD | 0.8723843 |
| PASAAPAETQP | ASSQDREDTQD | 0.82150847 |
| QQSSDGKNGVRAAEITGE | ASSQDREDTQD | 0.87061495 |
| RAPDAVTTSWWKE | ASSQDREDTQD | 0.14393008 |
| SGFGNVGAHTSGWFNQ | ASSQDREDTQD | 0.8640251 |
| SGVAGAVGQAASVGGLKV | ASSQDREDTQD | 0.7901093 |
| SNGGAGTGSAGGNGGT | ASSQDREDTQD | 0.6666905 |
| TGGAGGEGGAAG | ASSQDREDTQD | 0.37869883 |
| TGGDGGAAGAGGAPGHG | ASSQDREDTQD | 0.8529429 |
| TMPPLLAAIANAGRHGV | ASSQDREDTQD | 0.887759 |
| TSSTAGNSGGA | ASSQDREDTQD | 0.89191216 |
| VPATVPVGILC | ASSQDREDTQD | 0.7456921 |
| VQRPDSGGVVLTGRIS | ASSQDREDTQD | 0.89349365 |
| WTPNPVPATKGVPAGLVQAT | ASSQDREDTQD | 0.9016803 |
| YALKRALAKFSDRFDVVII | ASSQDREDTQD | 0.8385138 |
| YLIGSVIV | ASSQDREDTQD | 0.39771998 |
| YGQLVAVID | ASSQDREDTQD | 0.8712615 |
| VTVGGMTLPA | ASSQDREDTQD | 0.8373209 |
| VLALPAV | ASSQDREDTQD | 0.88202286 |
| VIPPTLNYET | ASSQDREDTQD | 0.87892735 |
| VGMGGMHPSSG | ASSQDREDTQD | 0.7387699 |
| VGLAGGAATA | ASSQDREDTQD | 0.88246727 |
| VGGAGGAGGSGP | ASSQDREDTQD | 0.8894043 |
| VFAAFMV | ASSQDREDTQD | 0.7715296 |
| TLYDKLR | ASSQDREDTQD | 0.8040497 |
| TLAVFTVL | ASSQDREDTQD | 0.74622655 |
| TGSVQPARV | ASSQDREDTQD | 0.46501157 |
| TGGGAGSGGT | ASSQDREDTQD | 0.8852922 |
| SHLETSL | ASSQDREDTQD | 0.81584966 |
| SFTTGMT | ASSQDREDTQD | 0.8018312 |
| QVEVAKRMGVRQPTVS | ASSQDREDTQD | 0.8668546 |
| QGSGGPALAM | ASSQDREDTQD | 0.79865545 |
| MLAPSNP | ASSQDREDTQD | 0.81454587 |

| | | |
|---|---|---|
| LVNNAAI | ASSQDREDTQD | 0.82835203 |
| LTGEINTLA | ASSQDREDTQD | 0.85691094 |
| LSIFGPV | ASSQDREDTQD | 0.874298 |
| LLALLANL | ASSQDREDTQD | 0.8299546 |
| LISNASDALDKLR | ASSQDREDTQD | 0.85186523 |
| LINEAVP | ASSQDREDTQD | 0.9013218 |
| LGYAFSGL | ASSQDREDTQD | 0.6118952 |
| LDFVAVAL | ASSQDREDTQD | 0.7918269 |
| LAQPGAE | ASSQDREDTQD | 0.86718017 |
| ISNLGAVTGF | ASSQDREDTQD | 0.5855293 |
| HQEDPGRI | ASSQDREDTQD | 0.88058805 |
| GTPGEPTP | ASSQDREDTQD | 0.86608917 |
| GFFNSGS | ASSQDREDTQD | 0.87839586 |
| GAGGGGLTAGAN | ASSQDREDTQD | 0.83598614 |
| FVDALVNM | ASSQDREDTQD | 0.90463585 |
| FLTVAPEL | ASSQDREDTQD | 0.8890137 |
| FETLTAL | ASSQDREDTQD | 0.82427716 |
| DVINPIEL | ASSQDREDTQD | 0.8904834 |
| APVSGAPV | ASSQDREDTQD | 0.5991049 |
| ALVGGLAW | ASSQDREDTQD | 0.8932527 |
| ALAAAAATAI | ASSQDREDTQD | 0.877398 |
| AFGLGLLL | ASSQDREDTQD | 0.75590414 |
| AFFAAASL | ASSQDREDTQD | 0.84581614 |
| AAAALPAFSPP | ASSQDREDTQD | 0.8972019 |
| VVLAPAL | ASSQDREDTQD | 0.8970108 |
| TIGHVDHGKTTL | ASSQDREDTQD | 0.43887085 |
| LISNASDALDK | ASSQDREDTQD | 0.8353247 |
| LISNASDAL | ASSQDREDTQD | 0.37322718 |
| ISNASDALDK | ASSQDREDTQD | 0.6730919 |
| NQAETLVYQTE | ASSQDREDTQD | 0.6513144 |
| TGGTGGTAGAG | ASSQDREDTQD | 0.8816628 |
| TGSTAGGAGGA | ASSQDREDTQD | 0.83786666 |
| DPLAVPVAAGPV | ASSQDREDTQD | 0.8733578 |
| GGTGGTPGNGGN | ASSQDREDTQD | 0.8682699 |
| PAGLAGVDPATL | ASSQDREDTQD | 0.89889795 |
| GNNLVVEAPHAGASV | ASSQDREDTQD | 0.7932716 |
| KVTRIMSASTLEQPA | ASSQDREDTQD | 0.8505828 |
| LGFALPIALPRRTVV | ASSQDREDTQD | 0.5069663 |
| TRLGNLTARKLVVLV | ASSQDREDTQD | 0.6243439 |
| WSQQRRLIYFPSAGP | ASSQDREDTQD | 0.7533206 |
| GPNGNPDPMAAAVDIR | ASSQDREDTQD | 0.82922447 |
| TQRCVSVADLRAAASR | ASSQDREDTQD | 0.24814355 |
| QFVLDVAEHPVEVLTETV | ASSQDREDTQD | 0.8929062 |
| GGAGGTGGAGGNGGTQAGDG | ASSQDREDTQD | 0.87127846 |
| PLLATLRAIAVDVDAKPAQV | ASSQDREDTQD | 0.8879131 |
| APKFRTGGYMIDFWGVGYQVAK | ASSQDREDTQD | 0.8133112 |
| GAGGAGGAGGAAGGPGGTGGTG | ASSQDREDTQD | 0.83680534 |
| IGALGLLATDVAPAGSVAFGVA | ASSQDREDTQD | 0.8030788 |
| AGGAGGIGGNGGAGGAGGVLMG | ASSQDREDTQD | 0.8827884 |
| ADELIGLL | ASSQDREDTQD | 0.7594539 |
| ADRALKA | ASSQDREDTQD | 0.5222025 |

| | | |
|---|---|---|
| AGAVGQVGGAGGA | ASSQDREDTQD | 0.81390166 |
| AGGGGGLGGAP | ASSQDREDTQD | 0.6992721 |
| AGGLLYGNGGAG | ASSQDREDTQD | 0.8981933 |
| AGLESIGL | ASSQDREDTQD | 0.8929436 |
| AGLKTGII | ASSQDREDTQD | 0.8774716 |
| ASLAALLL | ASSQDREDTQD | 0.89096576 |
| ASVINISSI | ASSQDREDTQD | 0.21567762 |
| ATPPGPA | ASSQDREDTQD | 0.8068314 |
| ATVASALV | ASSQDREDTQD | 0.89923334 |
| ATVASLIL | ASSQDREDTQD | 0.8723784 |
| AVNGPVLIRH | ASSQDREDTQD | 0.85234094 |
| DGAAGAVGI | ASSQDREDTQD | 0.898349 |
| DPLLHGI | ASSQDREDTQD | 0.8134537 |
| ELGAALGAL | ASSQDREDTQD | 0.84564996 |
| ELGRTQAV | ASSQDREDTQD | 0.8523981 |
| FVAGGYLS | ASSQDREDTQD | 0.88329214 |
| GASGGAGGAGGW | ASSQDREDTQD | 0.8673342 |
| GGAGAGGDGGAGGI | ASSQDREDTQD | 0.76845574 |
| GTGHNGSQPGMGGNG | ASSQDREDTQD | 0.8676236 |
| GYAAASASAS | ASSQDREDTQD | 0.90709126 |
| HQVDVAAVQQG | ASSQDREDTQD | 0.84897953 |
| IGAAIAHT | ASSQDREDTQD | 0.8929517 |
| IGMIEVAQL | ASSQDREDTQD | 0.88275033 |
| IRLAPEHTSAMD | ASSQDREDTQD | 0.8666544 |
| IYNVSVL | ASSQDREDTQD | 0.78245044 |
| KQGFLGM | ASSQDREDTQD | 0.83662796 |
| LESVITQL | ASSQDREDTQD | 0.8513333 |
| LGLGALVVV | ASSQDREDTQD | 0.7646822 |
| LIGNGGNGGSGGI | ASSQDREDTQD | 0.8411609 |
| LNAINAPTL | ASSQDREDTQD | 0.7862847 |
| LQLLNTGV | ASSQDREDTQD | 0.9071816 |
| LQTSIEF | ASSQDREDTQD | 0.88503945 |
| MSFAVSEL | ASSQDREDTQD | 0.879648 |
| NTGALGL | ASSQDREDTQD | 0.7783625 |
| PPGAGGAGGAGTT | ASSQDREDTQD | 0.86550343 |
| QLVGLGI | ASSQDREDTQD | 0.87655574 |
| RFLEQVHQL | ASSQDREDTQD | 0.5025208 |
| RGVGYVLR | ASSQDREDTQD | 0.89853936 |
| SATGVFAGVI | ASSQDREDTQD | 0.8042281 |
| SLPAITLP | ASSQDREDTQD | 0.7860234 |
| STALAFTV | ASSQDREDTQD | 0.8754275 |
| SVLTVGLGK | ASSQDREDTQD | 0.89428914 |
| TALAGAGSGI | ASSQDREDTQD | 0.82063717 |
| TGLPFTGI | ASSQDREDTQD | 0.8396424 |
| VAEHVTV | ASSQDREDTQD | 0.7814053 |
| VDIVLNSL | ASSQDREDTQD | 0.8835263 |
| VLTGGGAVL | ASSQDREDTQD | 0.8960084 |
| VNVGLAYGI | ASSQDREDTQD | 0.85500294 |
| VSSAAVGAV | ASSQDREDTQD | 0.9008206 |
| VTSVYEAGV | ASSQDREDTQD | 0.80731225 |
| WVALDRGA | ASSQDREDTQD | 0.46397895 |

| | | |
|---|---|---|
| YLVGSVVI | ASSQDREDTQD | 0.8506964 |
| GPPGTGKTLL | ASSQDREDTQD | 0.89331305 |
| GVGMTKF | ASSQDREDTQD | 0.89227575 |
| IGIGGSDL | ASSQDREDTQD | 0.9023899 |
| IGTLLAEL | ASSQDREDTQD | 0.88466346 |
| IYTSGTT | ASSQDREDTQD | 0.8186883 |
| LSFPAGV | ASSQDREDTQD | 0.8973061 |
| LYTSGTT | ASSQDREDTQD | 0.8981484 |
| VGAFPFGF | ASSQDREDTQD | 0.8898854 |
| VGAGTVEF | ASSQDREDTQD | 0.78556836 |
| VGKSTLLL | ASSQDREDTQD | 0.59936035 |
| AALAHAGGGIV | ASSQDREDTQD | 0.8595489 |
| AALVAVSLL | ASSQDREDTQD | 0.88682765 |
| AAVSRAVAAL | ASSQDREDTQD | 0.693478 |
| AGGSSGIGGF | ASSQDREDTQD | 0.87201875 |
| AGLTVANANG | ASSQDREDTQD | 0.8660515 |
| AGLVGAVL | ASSQDREDTQD | 0.62809265 |
| ALSQAFTV | ASSQDREDTQD | 0.87157536 |
| ALTFSVSL | ASSQDREDTQD | 0.8996653 |
| ALTGAALVAL | ASSQDREDTQD | 0.8348111 |
| ASFSIGAGVL | ASSQDREDTQD | 0.7945117 |
| DGSVYKI | ASSQDREDTQD | 0.8954308 |
| EQLAEIV | ASSQDREDTQD | 0.90691257 |
| FAVSELTVM | ASSQDREDTQD | 0.89689434 |
| GAAPLAAGL | ASSQDREDTQD | 0.6003828 |
| GADGGDGGSGGAGG | ASSQDREDTQD | 0.28457692 |
| GFGLDTGGAGG | ASSQDREDTQD | 0.81810987 |
| GLASGGLL | ASSQDREDTQD | 0.87768614 |
| GRIAGAAVLL | ASSQDREDTQD | 0.82025 |
| GSAGSLNTSF | ASSQDREDTQD | 0.77736527 |
| GTGQIGI | ASSQDREDTQD | 0.8780651 |
| GTLIALIL | ASSQDREDTQD | 0.89292604 |
| IAATMGFVVG | ASSQDREDTQD | 0.7962549 |
| IFAQALL | ASSQDREDTQD | 0.48719054 |
| IGAAVALGF | ASSQDREDTQD | 0.88175213 |
| IGLTPLAL | ASSQDREDTQD | 0.8820318 |
| ILGSNTSTL | ASSQDREDTQD | 0.8146528 |
| IVFSLTVL | ASSQDREDTQD | 0.85134447 |
| LASKATGF | ASSQDREDTQD | 0.81368834 |
| LASVALIL | ASSQDREDTQD | 0.53310055 |
| LFAVGLTL | ASSQDREDTQD | 0.8713967 |
| LFNGGIGTYI | ASSQDREDTQD | 0.67765266 |
| LTLGAGII | ASSQDREDTQD | 0.9004822 |
| LVAAALGDI | ASSQDREDTQD | 0.73191285 |
| NTAFVQLGI | ASSQDREDTQD | 0.88188493 |
| QLFSGAEL | ASSQDREDTQD | 0.88111997 |
| SAATFITL | ASSQDREDTQD | 0.87916285 |
| SNYGLANLGV | ASSQDREDTQD | 0.8353431 |
| SYGGAYAVM | ASSQDREDTQD | 0.8164837 |
| TTGVGGLTLG | ASSQDREDTQD | 0.6575378 |
| TTQTIYI | ASSQDREDTQD | 0.45163003 |

| | | |
|---|---|---|
| VAAGAGLGAVI | ASSQDREDTQD | 0.8969407 |
| VAAVVTLL | ASSQDREDTQD | 0.7475963 |
| VGFLGGLI | ASSQDREDTQD | 0.89763206 |
| VGVIPILL | ASSQDREDTQD | 0.8893055 |
| VIFASSLI | ASSQDREDTQD | 0.7980537 |
| VKGGDGGAAAG | ASSQDREDTQD | 0.87441754 |
| VTKTFTNI | ASSQDREDTQD | 0.6797972 |
| VVAAAILL | ASSQDREDTQD | 0.76751304 |
| YAALDVEL | ASSQDREDTQD | 0.8840196 |
| LASVALLL | ASSQDREDTQD | 0.87743485 |
| LDAVHRV | ASSQDREDTQD | 0.8005146 |
| QGLSSSIF | ASSQDREDTQD | 0.8610297 |
| RGTFANIRL | ASSQDREDTQD | 0.86954635 |
| RQLWWGHRI | ASSQDREDTQD | 0.77090806 |
| YGPPGTGKTLL | ASSQDREDTQD | 0.85072887 |
| AALVGAPANVA | ASSQDREDTQD | 0.8925561 |
| AATFRRTE | ASSQDREDTQD | 0.8663533 |
| AGALGGGI | ASSQDREDTQD | 0.78979826 |
| AGGAGGAGGIGAG | ASSQDREDTQD | 0.8852412 |
| AGSGALGL | ASSQDREDTQD | 0.7883724 |
| AQIADQI | ASSQDREDTQD | 0.87687266 |
| ASAVAVTYL | ASSQDREDTQD | 0.86077094 |
| ASTVLGGTGG | ASSQDREDTQD | 0.82232356 |
| AVGGEPAK | ASSQDREDTQD | 0.88721895 |
| CEQGTFGGTC | ASSQDREDTQD | 0.5416518 |
| DEGKAPAG | ASSQDREDTQD | 0.781556 |
| DTVVFPAL | ASSQDREDTQD | 0.8362602 |
| FEGITMAG | ASSQDREDTQD | 0.86598635 |
| FPLVVQD | ASSQDREDTQD | 0.88851297 |
| GAFTTPQ | ASSQDREDTQD | 0.77074033 |
| GFGGFGTAGGVGGA | ASSQDREDTQD | 0.8587214 |
| GFLAQFL | ASSQDREDTQD | 0.8321457 |
| GGAGGAGGNAGVA | ASSQDREDTQD | 0.77279294 |
| GGVGTAGAGGAGGAGG | ASSQDREDTQD | 0.804044 |
| GNGGGGDAGNAGSGGN | ASSQDREDTQD | 0.8169061 |
| GTGGAGGAGNDAGS | ASSQDREDTQD | 0.74705917 |
| GTTNTGWFNS | ASSQDREDTQD | 0.88490295 |
| IFMGPLFTL | ASSQDREDTQD | 0.69807655 |
| ILAHAIEAI | ASSQDREDTQD | 0.867203 |
| IRSAIREAL | ASSQDREDTQD | 0.89561814 |
| ITFAGGG | ASSQDREDTQD | 0.8127856 |
| ITGMSGHQEV | ASSQDREDTQD | 0.8758484 |
| IVAGLACTL | ASSQDREDTQD | 0.8710274 |
| IVGGGLAGA | ASSQDREDTQD | 0.8875602 |
| IVTGVLNI | ASSQDREDTQD | 0.8389132 |
| LAQQGCI | ASSQDREDTQD | 0.799811 |
| LCLHVEN | ASSQDREDTQD | 0.89185923 |
| LDAISDGS | ASSQDREDTQD | 0.8874364 |
| LGGFAAML | ASSQDREDTQD | 0.8809657 |
| LGGKNPMI | ASSQDREDTQD | 0.87877846 |
| LGPKGVAAIL | ASSQDREDTQD | 0.8801287 |

| | | |
|---|---|---|
| LMVGVSI | ASSQDREDTQD | 0.8850528 |
| LVGAAFVL | ASSQDREDTQD | 0.8206483 |
| LVLAVAPL | ASSQDREDTQD | 0.60131526 |
| MAAIAVIDAA | ASSQDREDTQD | 0.83376896 |
| NNGINAELQ | ASSQDREDTQD | 0.8477692 |
| RGEPLVLVHG | ASSQDREDTQD | 0.80324906 |
| RGGAGGAAGLI | ASSQDREDTQD | 0.32955337 |
| RSYRMHPAV | ASSQDREDTQD | 0.8942704 |
| SAAHLLAGQLA | ASSQDREDTQD | 0.90610516 |
| SAAVGSY | ASSQDREDTQD | 0.7971591 |
| SPHAVLHRI | ASSQDREDTQD | 0.9018259 |
| SQIAKVL | ASSQDREDTQD | 0.58341897 |
| TALGHLVP | ASSQDREDTQD | 0.88605684 |
| TAPTTSGAA | ASSQDREDTQD | 0.89546275 |
| TDGDDGNGGSPGT | ASSQDREDTQD | 0.7746775 |
| TDGLGEN | ASSQDREDTQD | 0.87789226 |
| TGGPPLPGQA | ASSQDREDTQD | 0.7852129 |
| TIPAITI | ASSQDREDTQD | 0.820482 |
| TIPGFTI | ASSQDREDTQD | 0.8991041 |
| TSIASGALA | ASSQDREDTQD | 0.82775843 |
| TVTLPGAAI | ASSQDREDTQD | 0.8799312 |
| VASGLSAALP | ASSQDREDTQD | 0.8308122 |
| VGAGVIGI | ASSQDREDTQD | 0.72599614 |
| VGPKRFGV | ASSQDREDTQD | 0.87583435 |
| VNVAFPD | ASSQDREDTQD | 0.8952754 |
| VSMTNSGVS | ASSQDREDTQD | 0.75708175 |
| VTDVLGDLSV | ASSQDREDTQD | 0.8973596 |
| VVVVGAGFAGL | ASSQDREDTQD | 0.8223765 |
| VYTVGTVV | ASSQDREDTQD | 0.904477 |
| WTTFSGP | ASSQDREDTQD | 0.887795 |
| YDELVEAF | ASSQDREDTQD | 0.86128765 |
| YGTGGAGGNGG | ASSQDREDTQD | 0.811792 |
| YTNVMAV | ASSQDREDTQD | 0.89627075 |
| FLEQVHQL | ASSQDREDTQD | 0.83908135 |
| GLSLLAVAV | ASSQDREDTQD | 0.88605917 |
| LEQVHQL | ASSQDREDTQD | 0.89640874 |
| LNAAGAI | ASSQDREDTQD | 0.69994533 |
| PAVHASL | ASSQDREDTQD | 0.8469579 |
| TELEDDA | ASSQDREDTQD | 0.8228489 |
| TPPGPPN | ASSQDREDTQD | 0.80306375 |
| TPPPPPPP | ASSQDREDTQD | 0.77217007 |
| TQSPALI | ASSSDSTEVF | 0.39832282 |
| EFAAAFPQ | ASSSDSTEVF | 0.775036 |
| LLSGNLDVL | ASSSDSTEVF | 0.713518 |
| GPANLGNYNI | ASSSDSTEVF | 0.7998475 |
| TLGPTLP | ASSSDSTEVF | 0.8154059 |
| VYAYSSCCK | ASSSDSTEVF | 0.7119461 |
| APAPAPAPNI | ASSSDSTEVF | 0.8525838 |
| PGGPVPIP | ASSSDSTEVF | 0.8510591 |
| FLTVALEP | ASSSDSTEVF | 0.56854355 |
| SQYFGANL | ASSSDSTEVF | 0.7860323 |

| | | |
|---|---|---|
| GGVKPAA | ASSSDSTEVF | 0.66338015 |
| GGTGGAGGAGGW | ASSSDSTEVF | 0.7788234 |
| GVGGVAGLGGAGG | ASSSDSTEVF | 0.27372968 |
| VLLVLATTI | ASSSDSTEVF | 0.84167445 |
| IGIFAAML | ASSSDSTEVF | 0.7095758 |
| AVGDLID | ASSSDSTEVF | 0.14463806 |
| NISPLPL | ASSSDSTEVF | 0.80084836 |
| VTAVFVAI | ASSSDSTEVF | 0.7717562 |
| SGDDNDAAL | ASSSDSTEVF | 0.79876757 |
| AVAALAHS | ASSSDSTEVF | 0.84199435 |
| GGVTVMATP | ASSSDSTEVF | 0.3010801 |
| AGLAANVRD | ASSSDSTEVF | 0.6920794 |
| TNRPDLIDPALLRPG | ASSSDSTEVF | 0.8313217 |
| LANLGVTAL | ASSSDSTEVF | 0.85222554 |
| TGERIEL | ASSSDSTEVF | 0.6493503 |
| NGEELAA | ASSSDSTEVF | 0.81692815 |
| VLSLVWG | ASSSDSTEVF | 0.85138226 |
| IGYALWL | ASSSDSTEVF | 0.820763 |
| LSLLLAPL | ASSSDSTEVF | 0.8369461 |
| TLDEALK | ASSSDSTEVF | 0.84914076 |
| TDGAGTAEGG | ASSSDSTEVF | 0.6295904 |
| NAPGGAGGNGG | ASSSDSTEVF | 0.8448335 |
| VSVFMGMGGNF | ASSSDSTEVF | 0.8519379 |
| LASYLAAL | ASSSDSTEVF | 0.7117375 |
| LVGGTPAL | ASSSDSTEVF | 0.81083906 |
| VRNADVL | ASSSDSTEVF | 0.61152947 |
| TCSSPFLVSAM | ASSSDSTEVF | 0.7253988 |
| VHITDAE | ASSSDSTEVF | 0.28838372 |
| EYAAPSF | ASSSDSTEVF | 0.81543374 |
| VVDAFHGGL | ASSSDSTEVF | 0.8405869 |
| ASVVDPQ | ASSSDSTEVF | 0.78287685 |
| SYNEFGGLS | ASSSDSTEVF | 0.58554304 |
| DPDGSVY | ASSSDSTEVF | 0.6519958 |
| MVSTIHATE | ASSSDSTEVF | 0.72149235 |
| LAAGVADPV | ASSSDSTEVF | 0.77743894 |
| TKEQIAA | ASSSDSTEVF | 0.2609551 |
| SLASPAAL | ASSSDSTEVF | 0.6824008 |
| IGLVFYGV | ASSSDSTEVF | 0.8491057 |
| FLLDASNL | ASSSDSTEVF | 0.8549955 |
| YQVGNFP | ASSSDSTEVF | 0.47052687 |
| ESLLSQST | ASSSDSTEVF | 0.84830195 |
| LFTVGFAAVG | ASSSDSTEVF | 0.8128356 |
| ATPPTPT | ASSSDSTEVF | 0.8554826 |
| TDLATLG | ASSSDSTEVF | 0.7662662 |
| SAGGTGR | ASSSDSTEVF | 0.67241013 |
| GFSNSGS | ASSSDSTEVF | 0.82207656 |
| GGNGGSGGTSV | ASSSDSTEVF | 0.7676978 |
| GTSVATGGAGN | ASSSDSTEVF | 0.5481367 |
| NGGNGGSGGTS | ASSSDSTEVF | 0.84093463 |
| AVVKFPH | ASSSDSTEVF | 0.7978205 |
| TPPPPPP | ASSSDSTEVF | 0.7060876 |

| | | |
|---|---|---|
| AAPAAAAAPA | ASSSDSTEVF | 0.7228683 |
| IALNEDL | ASSSDSTEVF | 0.8315277 |
| IVNVSSM | ASSSDSTEVF | 0.62538797 |
| TGIPPAPR | ASSSDSTEVF | 0.51214355 |
| FDLGGGTF | ASSSDSTEVF | 0.662174 |
| FDLGGGTFDVS | ASSSDSTEVF | 0.8062954 |
| NEPTAAA | ASSSDSTEVF | 0.8455516 |
| NEPTAAALA | ASSSDSTEVF | 0.8159045 |
| GSIINISSI | ASSSDSTEVF | 0.6518609 |
| VAPGNAGTA | ASSSDSTEVF | 0.83300567 |
| TGGGASLE | ASSSDSTEVF | 0.817775 |
| VAVYNHYKRI | ASSSDSTEVF | 0.78944004 |
| TDPAPATT | ASSSDSTEVF | 0.80899394 |
| TNRPDILDPALLRPG | ASSSDSTEVF | 0.714621 |
| LTTTTTS | ASSSDSTEVF | 0.83253276 |
| PTGSGKTL | ASSSDSTEVF | 0.8268221 |
| TGSGKTL | ASSSDSTEVF | 0.83374375 |
| TIGHVDHGK | ASSSDSTEVF | 0.82218736 |
| TIGHVDHGKTT | ASSSDSTEVF | 0.7940353 |
| GHVDHGKTT | ASSSDSTEVF | 0.78647184 |
| AQFEHTLL | ASSSDSTEVF | 0.3377589 |
| IVNQDEL | ASSSDSTEVF | 0.7075192 |
| TNGSQFF | ASSSDSTEVF | 0.8024243 |
| TGKGIVE | ASSSDSTEVF | 0.81206954 |
| LISNASDA | ASSSDSTEVF | 0.82068026 |
| AELRTLGELTKIAKAHGAQ | ASSSDSTEVF | 0.81460595 |
| APNAVAAAEPSW | ASSSDSTEVF | 0.6768968 |
| APQVAGAATAVL | ASSSDSTEVF | 0.819128 |
| ARRDPAAAAAILPTDA | ASSSDSTEVF | 0.83928907 |
| DAAARDVLAEAGLAEYF | ASSSDSTEVF | 0.47542572 |
| DDQIVLSAKANLAKVEQ | ASSSDSTEVF | 0.74615294 |
| DELLPDTGEDSDEDGH | ASSSDSTEVF | 0.8603545 |
| ELPEAPANRATVPQD | ASSSDSTEVF | 0.7995424 |
| FGQSKIERVTGIG | ASSSDSTEVF | 0.77440774 |
| GAGGTGSTAGGAGGAGGAGGLY | ASSSDSTEVF | 0.74171865 |
| GHGGTGAAGGNGGTGGAGGSGI | ASSSDSTEVF | 0.8516764 |
| GNGGNGGAGGEGGPGGAGGAGA | ASSSDSTEVF | 0.6893672 |
| GSTAGNGGNGGNATNGTIAGQP | ASSSDSTEVF | 0.85610384 |
| GTGGDGGNGGNGGTGG | ASSSDSTEVF | 0.23164783 |
| GVRRPAGIPADAPV | ASSSDSTEVF | 0.74629176 |
| GYRQVTDAHLLTLARR | ASSSDSTEVF | 0.6417414 |
| IAKMGQDGHDRG | ASSSDSTEVF | 0.423535 |
| KAEPSIEIQYT | ASSSDSTEVF | 0.84853923 |
| KPVRIRMAADEDTVAVT | ASSSDSTEVF | 0.7641239 |
| MPPRIAGMRLLVIKPEP | ASSSDSTEVF | 0.8395479 |
| MKFYAVALFRSRGVPPTR | ASSSDSTEVF | 0.7848333 |
| NAGGFGILGAVAHSP | ASSSDSTEVF | 0.8411697 |
| NEGREITLAMAEM | ASSSDSTEVF | 0.79418266 |
| NELVEWSHPLGHRWRWE | ASSSDSTEVF | 0.7948518 |
| NLGAGGTGGNGGNPG | ASSSDSTEVF | 0.57205343 |
| NTEWPKWETVMAADPVD | ASSSDSTEVF | 0.84510696 |

| | | |
|---|---|---|
| PASAAPAETQP | ASSSDSTEVF | 0.83003813 |
| QQSSDGKNGVRAAEITGE | ASSSDSTEVF | 0.8231363 |
| RAPDAVTTSWWKE | ASSSDSTEVF | 0.5151871 |
| SGFGNVGAHTSGWFNQ | ASSSDSTEVF | 0.7139965 |
| SGVAGAVGQAASVGGLKV | ASSSDSTEVF | 0.82303 |
| SNGGAGTGSAGGNGGT | ASSSDSTEVF | 0.19242887 |
| TGGAGGEGGAAG | ASSSDSTEVF | 0.20089231 |
| TGGDGGAAGAGGAPGHG | ASSSDSTEVF | 0.74639183 |
| TMPPLLAAIANAGRHGV | ASSSDSTEVF | 0.57848096 |
| TSSTAGNSGGA | ASSSDSTEVF | 0.8487145 |
| VPATVPVGILC | ASSSDSTEVF | 0.8321985 |
| VQRPDSGGVVLTGRIS | ASSSDSTEVF | 0.7953528 |
| WTPNPVPATKGVPAGLVQAT | ASSSDSTEVF | 0.8223413 |
| YALKRALAKFSDRFDVVII | ASSSDSTEVF | 0.8109434 |
| YLIGSVIV | ASSSDSTEVF | 0.12996717 |
| YGQLVAVID | ASSSDSTEVF | 0.67083216 |
| VTVGGMTLPA | ASSSDSTEVF | 0.61455166 |
| VLALPAV | ASSSDSTEVF | 0.84534067 |
| VIPPTLNYET | ASSSDSTEVF | 0.65664303 |
| VGMGGMHPSSG | ASSSDSTEVF | 0.65836084 |
| VGLAGGAATA | ASSSDSTEVF | 0.803308 |
| VGGAGGAGGSGP | ASSSDSTEVF | 0.7944639 |
| VFAAFMV | ASSSDSTEVF | 0.78751516 |
| TLYDKLR | ASSSDSTEVF | 0.5385866 |
| TLAVFTVL | ASSSDSTEVF | 0.16042838 |
| TGSVQPARV | ASSSDSTEVF | 0.84359765 |
| TGGGAGSGGT | ASSSDSTEVF | 0.2939212 |
| SHLETSL | ASSSDSTEVF | 0.68745357 |
| SFTTGMT | ASSSDSTEVF | 0.21660748 |
| QVEVAKRMGVRQPTVS | ASSSDSTEVF | 0.14844577 |
| QGSGGPALAM | ASSSDSTEVF | 0.4357224 |
| MLAPSNP | ASSSDSTEVF | 0.83143294 |
| LVNNAAI | ASSSDSTEVF | 0.75394356 |
| LTGEINTLA | ASSSDSTEVF | 0.738224 |
| LSIFGPV | ASSSDSTEVF | 0.8536832 |
| LLALLANL | ASSSDSTEVF | 0.80016017 |
| LISNASDALDKLR | ASSSDSTEVF | 0.75248075 |
| LINEAVP | ASSSDSTEVF | 0.7491171 |
| LGYAFSGL | ASSSDSTEVF | 0.63207495 |
| LDFVAVAL | ASSSDSTEVF | 0.6567896 |
| LAQPGAE | ASSSDSTEVF | 0.70234424 |
| ISNLGAVTGF | ASSSDSTEVF | 0.856727 |
| HQEDPGRI | ASSSDSTEVF | 0.73425764 |
| GTPGEPTP | ASSSDSTEVF | 0.48530024 |
| GFFNSGS | ASSSDSTEVF | 0.7918496 |
| GAGGGGLTAGAN | ASSSDSTEVF | 0.8523768 |
| FVDALVNM | ASSSDSTEVF | 0.8612114 |
| FLTVAPEL | ASSSDSTEVF | 0.6872393 |
| FETLTAL | ASSSDSTEVF | 0.757182 |
| DVINPIEL | ASSSDSTEVF | 0.7509747 |
| APVSGAPV | ASSSDSTEVF | 0.79616135 |

| | | |
|---|---|---|
| ALVGGLAW | ASSSDSTEVF | 0.6847005 |
| ALAAAAATAI | ASSSDSTEVF | 0.5011449 |
| AFGLGLLL | ASSSDSTEVF | 0.8204551 |
| AFFAAASL | ASSSDSTEVF | 0.85565656 |
| AAAALPAFSPP | ASSSDSTEVF | 0.68048346 |
| VVLAPAL | ASSSDSTEVF | 0.4474138 |
| TIGHVDHGKTTL | ASSSDSTEVF | 0.79042196 |
| LISNASDALDK | ASSSDSTEVF | 0.6876298 |
| LISNASDAL | ASSSDSTEVF | 0.7600827 |
| ISNASDALDK | ASSSDSTEVF | 0.8044973 |
| NQAETLVYQTE | ASSSDSTEVF | 0.45935374 |
| TGGTGGTAGAG | ASSSDSTEVF | 0.7453014 |
| TGSTAGGAGGA | ASSSDSTEVF | 0.8619861 |
| DPLAVPVAAGPV | ASSSDSTEVF | 0.77549917 |
| GGTGGTPGNGGN | ASSSDSTEVF | 0.8187448 |
| PAGLAGVDPATL | ASSSDSTEVF | 0.84843826 |
| GNNLVVEAPHAGASV | ASSSDSTEVF | 0.82878315 |
| KVTRIMSASTLEQPA | ASSSDSTEVF | 0.8537071 |
| LGFALPIALPRRTVV | ASSSDSTEVF | 0.81156546 |
| TRLGNLTARKLVVLV | ASSSDSTEVF | 0.7356335 |
| WSQQRRLIYFPSAGP | ASSSDSTEVF | 0.7844211 |
| GPNGNPDPMAAAVDIR | ASSSDSTEVF | 0.78428626 |
| TQRCVSVADLRAAASR | ASSSDSTEVF | 0.80931723 |
| QFVLDVAEHPVEVLTETV | ASSSDSTEVF | 0.7876295 |
| GGAGGTGGAGGNGGTQAGDG | ASSSDSTEVF | 0.6781807 |
| PLLATLRAIAVDVDAKPAQV | ASSSDSTEVF | 0.8528887 |
| APKFRTGGYMIDFWGVGYQVAK | ASSSDSTEVF | 0.76425874 |
| GAGGAGGAGGAAGGPGGTGGTG | ASSSDSTEVF | 0.8318727 |
| IGALGLLATDVAPAGSVAFGVA | ASSSDSTEVF | 0.5773833 |
| AGGAGGIGGNGGAGGAGGVLMG | ASSSDSTEVF | 0.62976134 |
| ADELIGLL | ASSSDSTEVF | 0.8401657 |
| ADRALKA | ASSSDSTEVF | 0.14938396 |
| AGAVGQVGGAGGA | ASSSDSTEVF | 0.21518128 |
| AGGGGGLGGAP | ASSSDSTEVF | 0.79004943 |
| AGGLLYGNGGAG | ASSSDSTEVF | 0.84844387 |
| AGLESIGL | ASSSDSTEVF | 0.15158455 |
| AGLKTGII | ASSSDSTEVF | 0.65654504 |
| ASLAALLL | ASSSDSTEVF | 0.4771727 |
| ASVINISSI | ASSSDSTEVF | 0.8043614 |
| ATPPGPA | ASSSDSTEVF | 0.80954623 |
| ATVASALV | ASSSDSTEVF | 0.3671273 |
| ATVASLIL | ASSSDSTEVF | 0.835946 |
| AVNGPVLIRH | ASSSDSTEVF | 0.3879395 |
| DGAAGAVGI | ASSSDSTEVF | 0.28388947 |
| DPLLHGI | ASSSDSTEVF | 0.8329887 |
| ELGAALGAL | ASSSDSTEVF | 0.8365788 |
| ELGRTQAV | ASSSDSTEVF | 0.51027644 |
| FVAGGYLS | ASSSDSTEVF | 0.840821 |
| GASGGAGGAGGW | ASSSDSTEVF | 0.66213626 |
| GGAGAGGDGGAGGI | ASSSDSTEVF | 0.8620224 |
| GTGHNGSQPGMGGNG | ASSSDSTEVF | 0.7851765 |

| | | |
|---|---|---|
| GYAAASASAS | ASSSDSTEVF | 0.39754438 |
| HQVDVAAVQQG | ASSSDSTEVF | 0.84129727 |
| IGAAIAHT | ASSSDSTEVF | 0.8162477 |
| IGMIEVAQL | ASSSDSTEVF | 0.79586613 |
| IRLAPEHTSAMD | ASSSDSTEVF | 0.82731575 |
| IYNVSVL | ASSSDSTEVF | 0.6096921 |
| KQGFLGM | ASSSDSTEVF | 0.78372806 |
| LESVITQL | ASSSDSTEVF | 0.85380465 |
| LGLGALVVV | ASSSDSTEVF | 0.8250919 |
| LIGNGGNGGSGGI | ASSSDSTEVF | 0.59137833 |
| LNAINAPTL | ASSSDSTEVF | 0.83364344 |
| LQLLNTGV | ASSSDSTEVF | 0.8030667 |
| LQTSIEF | ASSSDSTEVF | 0.7002096 |
| MSFAVSEL | ASSSDSTEVF | 0.85734165 |
| NTGALGL | ASSSDSTEVF | 0.73349965 |
| PPGAGGAGGAGTT | ASSSDSTEVF | 0.83687735 |
| QLVGLGI | ASSSDSTEVF | 0.84671646 |
| RFLEQVHQL | ASSSDSTEVF | 0.82621145 |
| RGVGYVLR | ASSSDSTEVF | 0.45633778 |
| SATGVFAGVI | ASSSDSTEVF | 0.814425 |
| SLPAITLP | ASSSDSTEVF | 0.80167127 |
| STALAFTV | ASSSDSTEVF | 0.82127297 |
| SVLTVGLGK | ASSSDSTEVF | 0.7792336 |
| TALAGAGSGI | ASSSDSTEVF | 0.48354137 |
| TGLPFTGI | ASSSDSTEVF | 0.8107888 |
| VAEHVTV | ASSSDSTEVF | 0.8502103 |
| VDIVLNSL | ASSSDSTEVF | 0.739743 |
| VLTGGGAVL | ASSSDSTEVF | 0.7627762 |
| VNVGLAYGI | ASSSDSTEVF | 0.85817885 |
| VSSAAVGAV | ASSSDSTEVF | 0.7060214 |
| VTSVYEAGV | ASSSDSTEVF | 0.7877412 |
| WVALDRGA | ASSSDSTEVF | 0.83453935 |
| YLVGSVVI | ASSSDSTEVF | 0.8293981 |
| GPPGTGKTLL | ASSSDSTEVF | 0.7554977 |
| GVGMTKF | ASSSDSTEVF | 0.8447789 |
| IGIGGSDL | ASSSDSTEVF | 0.59887135 |
| IGTLLAEL | ASSSDSTEVF | 0.83556485 |
| IYTSGTT | ASSSDSTEVF | 0.66847897 |
| LSFPAGV | ASSSDSTEVF | 0.8246231 |
| LYTSGTT | ASSSDSTEVF | 0.70508313 |
| VGAFPFGF | ASSSDSTEVF | 0.83205163 |
| VGAGTVEF | ASSSDSTEVF | 0.83730745 |
| VGKSTLLL | ASSSDSTEVF | 0.83825654 |
| AALAHAGGGIV | ASSSDSTEVF | 0.2011562 |
| AALVAVSLL | ASSSDSTEVF | 0.83840084 |
| AAVSRAVAAL | ASSSDSTEVF | 0.8413017 |
| AGGSSGIGGF | ASSSDSTEVF | 0.8621127 |
| AGLTVANANG | ASSSDSTEVF | 0.7870467 |
| AGLVGAVL | ASSSDSTEVF | 0.8296027 |
| ALSQAFTV | ASSSDSTEVF | 0.77882487 |
| ALTFSVSL | ASSSDSTEVF | 0.7538275 |

| | | |
|---|---|---|
| ALTGAALVAL | ASSSDSTEVF | 0.6363845 |
| ASFSIGAGVL | ASSSDSTEVF | 0.7662848 |
| DGSVYKI | ASSSDSTEVF | 0.7272483 |
| EQLAEIV | ASSSDSTEVF | 0.7288412 |
| FAVSELTVM | ASSSDSTEVF | 0.83744156 |
| GAAPLAAGL | ASSSDSTEVF | 0.83137876 |
| GADGGDGGSGGAGG | ASSSDSTEVF | 0.8433724 |
| GFGLDTGGAGG | ASSSDSTEVF | 0.8499657 |
| GLASGGLL | ASSSDSTEVF | 0.84316236 |
| GRIAGAAVLL | ASSSDSTEVF | 0.83435893 |
| GSAGSLNTSF | ASSSDSTEVF | 0.7546582 |
| GTGQIGI | ASSSDSTEVF | 0.7579727 |
| GTLIALIL | ASSSDSTEVF | 0.64717925 |
| IAATMGFVVG | ASSSDSTEVF | 0.7000412 |
| IFAQALL | ASSSDSTEVF | 0.7508863 |
| IGAAVALGF | ASSSDSTEVF | 0.8134532 |
| IGLTPLAL | ASSSDSTEVF | 0.13355865 |
| ILGSNTSTL | ASSSDSTEVF | 0.76351786 |
| IVFSLTVL | ASSSDSTEVF | 0.8538273 |
| LASKATGF | ASSSDSTEVF | 0.6954917 |
| LASVALIL | ASSSDSTEVF | 0.8300539 |
| LFAVGLTL | ASSSDSTEVF | 0.7564477 |
| LFNGGIGTYI | ASSSDSTEVF | 0.69224584 |
| LTLGAGII | ASSSDSTEVF | 0.8414315 |
| LVAAALGDI | ASSSDSTEVF | 0.6452532 |
| NTAFVQLGI | ASSSDSTEVF | 0.8327873 |
| QLFSGAEL | ASSSDSTEVF | 0.8572427 |
| SAATFITL | ASSSDSTEVF | 0.81302047 |
| SNYGLANLGV | ASSSDSTEVF | 0.75310695 |
| SYGGAYAVM | ASSSDSTEVF | 0.7889294 |
| TTGVGGLTLG | ASSSDSTEVF | 0.39394253 |
| TTQTIYI | ASSSDSTEVF | 0.7791244 |
| VAAGAGLGAVI | ASSSDSTEVF | 0.12203887 |
| VAAVVTLL | ASSSDSTEVF | 0.7942138 |
| VGFLGGLI | ASSSDSTEVF | 0.5653183 |
| VGVIPILL | ASSSDSTEVF | 0.82515585 |
| VIFASSLI | ASSSDSTEVF | 0.80808353 |
| VKGGDGGAAAG | ASSSDSTEVF | 0.8005921 |
| VTKTFTNI | ASSSDSTEVF | 0.700027 |
| VVAAAILL | ASSSDSTEVF | 0.74996245 |
| YAALDVEL | ASSSDSTEVF | 0.66649723 |
| LASVALLL | ASSSDSTEVF | 0.57096624 |
| LDAVHRV | ASSSDSTEVF | 0.8323636 |
| QGLSSSIF | ASSSDSTEVF | 0.72929597 |
| RGTFANIRL | ASSSDSTEVF | 0.68906546 |
| RQLWWGHRI | ASSSDSTEVF | 0.84980553 |
| YGPPGTGKTLL | ASSSDSTEVF | 0.8137939 |
| AALVGAPANVA | ASSSDSTEVF | 0.3814333 |
| AATFRRTE | ASSSDSTEVF | 0.78205645 |
| AGALGGGI | ASSSDSTEVF | 0.7985156 |
| AGGAGGAGGIGAG | ASSSDSTEVF | 0.79408324 |

| | | |
|---|---|---|
| AGSGALGL | ASSSDSTEVF | 0.1270316 |
| AQIADQI | ASSSDSTEVF | 0.78413075 |
| ASAVAVTYL | ASSSDSTEVF | 0.7013321 |
| ASTVLGGTGG | ASSSDSTEVF | 0.64626396 |
| AVGGEPAK | ASSSDSTEVF | 0.7013987 |
| CEQGTFGGTC | ASSSDSTEVF | 0.8422064 |
| DEGKAPAG | ASSSDSTEVF | 0.373445 |
| DTVVFPAL | ASSSDSTEVF | 0.826578 |
| FEGITMAG | ASSSDSTEVF | 0.82452285 |
| FPLVVQD | ASSSDSTEVF | 0.8328155 |
| GAFTTPQ | ASSSDSTEVF | 0.8556895 |
| GFGGFGTAGGVGGA | ASSSDSTEVF | 0.81865287 |
| GFLAQFL | ASSSDSTEVF | 0.7352178 |
| GGAGGAGGNAGVA | ASSSDSTEVF | 0.8180113 |
| GGVGTAGAGGAGGAGG | ASSSDSTEVF | 0.7521243 |
| GNGGGGDAGNAGSGGN | ASSSDSTEVF | 0.29237553 |
| GTGGAGGAGNDAGS | ASSSDSTEVF | 0.28326374 |
| GTTNTGWFNS | ASSSDSTEVF | 0.80722344 |
| IFMGPLFTL | ASSSDSTEVF | 0.23465036 |
| ILAHAIEAI | ASSSDSTEVF | 0.8231522 |
| IRSAIREAL | ASSSDSTEVF | 0.84236413 |
| ITFAGGG | ASSSDSTEVF | 0.75670755 |
| ITGMSGHQEV | ASSSDSTEVF | 0.45107943 |
| IVAGLACTL | ASSSDSTEVF | 0.7494159 |
| IVGGGLAGA | ASSSDSTEVF | 0.6879742 |
| IVTGVLNI | ASSSDSTEVF | 0.83793443 |
| LAQQGCI | ASSSDSTEVF | 0.82916003 |
| LCLHVEN | ASSSDSTEVF | 0.8485287 |
| LDAISDGS | ASSSDSTEVF | 0.8451761 |
| LGGFAAML | ASSSDSTEVF | 0.780586 |
| LGGKNPMI | ASSSDSTEVF | 0.82855415 |
| LGPKGVAAIL | ASSSDSTEVF | 0.7233939 |
| LMVGVSI | ASSSDSTEVF | 0.7473577 |
| LVGAAFVL | ASSSDSTEVF | 0.7567852 |
| LVLAVAPL | ASSSDSTEVF | 0.7895827 |
| MAAIAVIDAA | ASSSDSTEVF | 0.73510814 |
| NNGINAELQ | ASSSDSTEVF | 0.7098243 |
| RGEPLVLVHG | ASSSDSTEVF | 0.40123346 |
| RGGAGGAAGLI | ASSSDSTEVF | 0.8020689 |
| RSYRMHPAV | ASSSDSTEVF | 0.8156144 |
| SAAHLLAGQLA | ASSSDSTEVF | 0.31461793 |
| SAAVGSY | ASSSDSTEVF | 0.7159574 |
| SPHAVLHRI | ASSSDSTEVF | 0.8584994 |
| SQIAKVL | ASSSDSTEVF | 0.8531792 |
| TALGHLVP | ASSSDSTEVF | 0.71454966 |
| TAPTTSGAA | ASSSDSTEVF | 0.68107593 |
| TDGDDGNGGSPGT | ASSSDSTEVF | 0.76970017 |
| TDGLGEN | ASSSDSTEVF | 0.3884077 |
| TGGPPLPGQA | ASSSDSTEVF | 0.8001892 |
| TIPAITI | ASSSDSTEVF | 0.78737104 |
| TIPGFTI | ASSSDSTEVF | 0.76663506 |

| | | |
|---|---|---|
| TSIASGALA | ASSSDSTEVF | 0.66711533 |
| TVTLPGAAI | ASSSDSTEVF | 0.80203295 |
| VASGLSAALP | ASSSDSTEVF | 0.7455747 |
| VGAGVIGI | ASSSDSTEVF | 0.80968016 |
| VGPKRFGV | ASSSDSTEVF | 0.14628153 |
| VNVAFPD | ASSSDSTEVF | 0.7999126 |
| VSMTNSGVS | ASSSDSTEVF | 0.65951174 |
| VTDVLGDLSV | ASSSDSTEVF | 0.6022279 |
| VVVVGAGFAGL | ASSSDSTEVF | 0.31172585 |
| VYTVGTVV | ASSSDSTEVF | 0.76176924 |
| WTTFSGP | ASSSDSTEVF | 0.84645224 |
| YDELVEAF | ASSSDSTEVF | 0.8430823 |
| YGTGGAGGNGG | ASSSDSTEVF | 0.4863968 |
| YTNVMAV | ASSSDSTEVF | 0.8395719 |
| FLEQVHQL | ASSSDSTEVF | 0.8562858 |
| GLSLLAVAV | ASSSDSTEVF | 0.71695304 |
| LEQVHQL | ASSSDSTEVF | 0.3082439 |
| LNAAGAI | ASSSDSTEVF | 0.8327668 |
| PAVHASL | ASSSDSTEVF | 0.73633546 |
| TELEDDA | ASSSDSTEVF | 0.81591886 |
| TPPGPPN | ASSSDSTEVF | 0.8310617 |
| TPPPPPPP | ASSSDSTEVF | 0.6026646 |
| TQSPALI | ASSLRDGGQLY | 0.84626323 |
| EFAAAFPQ | ASSLRDGGQLY | 0.84342796 |
| LLSGNLDVL | ASSLRDGGQLY | 0.6647674 |
| GPANLGNYNI | ASSLRDGGQLY | 0.60581607 |
| TLGPTLP | ASSLRDGGQLY | 0.6027248 |
| VYAYSSCCK | ASSLRDGGQLY | 0.32874435 |
| APAPAPAPNI | ASSLRDGGQLY | 0.82085645 |
| PGGPVPIP | ASSLRDGGQLY | 0.71531504 |
| FLTVALEP | ASSLRDGGQLY | 0.7570005 |
| SQYFGANL | ASSLRDGGQLY | 0.81413305 |
| GGVKPAA | ASSLRDGGQLY | 0.7303553 |
| GGTGGAGGAGGW | ASSLRDGGQLY | 0.74121416 |
| GVGGVAGLGGAGG | ASSLRDGGQLY | 0.78023994 |
| VLLVLATTI | ASSLRDGGQLY | 0.8055229 |
| IGIFAAML | ASSLRDGGQLY | 0.80565226 |
| AVGDLID | ASSLRDGGQLY | 0.78662986 |
| NISPLPL | ASSLRDGGQLY | 0.78297555 |
| VTAVFVAI | ASSLRDGGQLY | 0.8528049 |
| SGDDNDAAL | ASSLRDGGQLY | 0.60399806 |
| AVAALAHS | ASSLRDGGQLY | 0.6370292 |
| GGVTVMATP | ASSLRDGGQLY | 0.8207911 |
| AGLAANVRD | ASSLRDGGQLY | 0.5291518 |
| TNRPDLIDPALLRPG | ASSLRDGGQLY | 0.8243511 |
| LANLGVTAL | ASSLRDGGQLY | 0.8070715 |
| TGERIEL | ASSLRDGGQLY | 0.8149272 |
| NGEELAA | ASSLRDGGQLY | 0.758827 |
| VLSLVWG | ASSLRDGGQLY | 0.85858893 |
| IGYALWL | ASSLRDGGQLY | 0.84768075 |
| LSLLLAPL | ASSLRDGGQLY | 0.7663256 |

| | | |
|---|---|---|
| TLDEALK | ASSLRDGGQLY | 0.795249 |
| TDGAGTAEGG | ASSLRDGGQLY | 0.5330478 |
| NAPGGAGGNGG | ASSLRDGGQLY | 0.8383494 |
| VSVFMGMGGNF | ASSLRDGGQLY | 0.83359843 |
| LASYLAAL | ASSLRDGGQLY | 0.64941 |
| LVGGTPAL | ASSLRDGGQLY | 0.7829883 |
| VRNADVL | ASSLRDGGQLY | 0.84650195 |
| TCSSPFLVSAM | ASSLRDGGQLY | 0.8483765 |
| VHITDAE | ASSLRDGGQLY | 0.31552216 |
| EYAAPSF | ASSLRDGGQLY | 0.76828885 |
| VVDAFHGGL | ASSLRDGGQLY | 0.2969128 |
| ASVVDPQ | ASSLRDGGQLY | 0.6685915 |
| SYNEFGGLS | ASSLRDGGQLY | 0.50169694 |
| DPDGSVY | ASSLRDGGQLY | 0.8194294 |
| MVSTIHATE | ASSLRDGGQLY | 0.764481 |
| LAAGVADPV | ASSLRDGGQLY | 0.81439537 |
| TKEQIAA | ASSLRDGGQLY | 0.82333905 |
| SLASPAAL | ASSLRDGGQLY | 0.8585124 |
| IGLVFYGV | ASSLRDGGQLY | 0.7072109 |
| FLLDASNL | ASSLRDGGQLY | 0.7931629 |
| YQVGNFP | ASSLRDGGQLY | 0.42350364 |
| ESLLSQST | ASSLRDGGQLY | 0.33607846 |
| LFTVGFAAVG | ASSLRDGGQLY | 0.59620154 |
| ATPPTPT | ASSLRDGGQLY | 0.77947485 |
| TDLATLG | ASSLRDGGQLY | 0.18565571 |
| SAGGTGR | ASSLRDGGQLY | 0.85006523 |
| GFSNSGS | ASSLRDGGQLY | 0.8184056 |
| GGNGGSGGTSV | ASSLRDGGQLY | 0.8461324 |
| GTSVATGGAGN | ASSLRDGGQLY | 0.68207234 |
| NGGNGGSGGTS | ASSLRDGGQLY | 0.715654 |
| AVVKFPH | ASSLRDGGQLY | 0.744719 |
| TPPPPPP | ASSLRDGGQLY | 0.837726 |
| AAPAAAAAPA | ASSLRDGGQLY | 0.63819706 |
| IALNEDL | ASSLRDGGQLY | 0.40360034 |
| IVNVSSM | ASSLRDGGQLY | 0.7022649 |
| TGIPPAPR | ASSLRDGGQLY | 0.47436756 |
| FDLGGGTF | ASSLRDGGQLY | 0.8475187 |
| FDLGGGTFDVS | ASSLRDGGQLY | 0.84289026 |
| NEPTAAA | ASSLRDGGQLY | 0.82618004 |
| NEPTAAALA | ASSLRDGGQLY | 0.8391925 |
| GSIINISSI | ASSLRDGGQLY | 0.20440304 |
| VAPGNAGTA | ASSLRDGGQLY | 0.68666506 |
| TGGGASLE | ASSLRDGGQLY | 0.85917133 |
| VAVYNHYKRI | ASSLRDGGQLY | 0.81209785 |
| TDPAPATT | ASSLRDGGQLY | 0.7792288 |
| TNRPDILDPALLRPG | ASSLRDGGQLY | 0.8509784 |
| LTTTTTS | ASSLRDGGQLY | 0.75552666 |
| PTGSGKTL | ASSLRDGGQLY | 0.80075896 |
| TGSGKTL | ASSLRDGGQLY | 0.7997104 |
| TIGHVDHGK | ASSLRDGGQLY | 0.813059 |
| TIGHVDHGKTT | ASSLRDGGQLY | 0.8009044 |

| | | |
|---|---|---|
| GHVDHGKTT | ASSLRDGGQLY | 0.5964976 |
| AQFEHTLL | ASSLRDGGQLY | 0.8197943 |
| IVNQDEL | ASSLRDGGQLY | 0.858197 |
| TNGSQFF | ASSLRDGGQLY | 0.80049455 |
| TGKGIVE | ASSLRDGGQLY | 0.8388486 |
| LISNASDA | ASSLRDGGQLY | 0.82863283 |
| AELRTLGELTKIAKAHGAQ | ASSLRDGGQLY | 0.8382821 |
| APNAVAAAEPSW | ASSLRDGGQLY | 0.7271491 |
| APQVAGAATAVL | ASSLRDGGQLY | 0.7389761 |
| ARRDPAAAAAILPTDA | ASSLRDGGQLY | 0.7517179 |
| DAAARDVLAEAGLAEYF | ASSLRDGGQLY | 0.6294755 |
| DDQIVLSAKANLAKVEQ | ASSLRDGGQLY | 0.78370655 |
| DELLPDTGEDSDEDGH | ASSLRDGGQLY | 0.7055937 |
| ELPEAPANRATVPQD | ASSLRDGGQLY | 0.86511827 |
| FGQSKIERVTGIG | ASSLRDGGQLY | 0.8281971 |
| GAGGTGSTAGGAGGAGGAGGLY | ASSLRDGGQLY | 0.816833 |
| GHGGTGAAGGNGGTGGAGGSGI | ASSLRDGGQLY | 0.7002363 |
| GNGGNGGAGGEGGPGGAGGAGA | ASSLRDGGQLY | 0.794345 |
| GSTAGNGGNGGNATNGTIAGQP | ASSLRDGGQLY | 0.8185154 |
| GTGGDGGNGGNGGTGG | ASSLRDGGQLY | 0.40917242 |
| GVRRPAGIPADAPV | ASSLRDGGQLY | 0.8512139 |
| GYRQVTDAHLLTLARR | ASSLRDGGQLY | 0.7121272 |
| IAKMGQDGHDRG | ASSLRDGGQLY | 0.7053548 |
| KAEPSIEIQYT | ASSLRDGGQLY | 0.8376788 |
| KPVRIRMAADEDTVAVT | ASSLRDGGQLY | 0.84166235 |
| MPPRIAGMRLLVIKPEP | ASSLRDGGQLY | 0.71217895 |
| MKFYAVALFRSRGVPPTR | ASSLRDGGQLY | 0.7220989 |
| NAGGFGILGAVAHSP | ASSLRDGGQLY | 0.71997344 |
| NEGREITLAMAEM | ASSLRDGGQLY | 0.8382005 |
| NELVEWSHPLGHRWRWE | ASSLRDGGQLY | 0.85330486 |
| NLGAGGTGGNGGNPG | ASSLRDGGQLY | 0.7583521 |
| NTEWPKWETVMAADPVD | ASSLRDGGQLY | 0.84722996 |
| PASAAPAETQP | ASSLRDGGQLY | 0.7950758 |
| QQSSDGKNGVRAAEITGE | ASSLRDGGQLY | 0.38313973 |
| RAPDAVTTSWWKE | ASSLRDGGQLY | 0.77432966 |
| SGFGNVGAHTSGWFNQ | ASSLRDGGQLY | 0.8331585 |
| SGVAGAVGQAASVGGLKV | ASSLRDGGQLY | 0.8376197 |
| SNGGAGTGSAGGNGGT | ASSLRDGGQLY | 0.8596152 |
| TGGAGGEGGAAG | ASSLRDGGQLY | 0.80135566 |
| TGGDGGAAGAGGAPGHG | ASSLRDGGQLY | 0.72162294 |
| TMPPLLAAIANAGRHGV | ASSLRDGGQLY | 0.85112226 |
| TSSTAGNSGGA | ASSLRDGGQLY | 0.8599409 |
| VPATVPVGILC | ASSLRDGGQLY | 0.84673154 |
| VQRPDSGGVVLTGRIS | ASSLRDGGQLY | 0.74293476 |
| WTPNPVPATKGVPAGLVQAT | ASSLRDGGQLY | 0.6839037 |
| YALKRALAKFSDRFDVVII | ASSLRDGGQLY | 0.8055252 |
| YLIGSVIV | ASSLRDGGQLY | 0.8420202 |
| YGQLVAVID | ASSLRDGGQLY | 0.6150545 |
| VTVGGMTLPA | ASSLRDGGQLY | 0.7914145 |
| VLALPAV | ASSLRDGGQLY | 0.82608616 |
| VIPPTLNYET | ASSLRDGGQLY | 0.47213215 |

| | | |
|---|---|---|
| VGMGGMHPSSG | ASSLRDGGQLY | 0.82846045 |
| VGLAGGAATA | ASSLRDGGQLY | 0.84416056 |
| VGGAGGAGGSGP | ASSLRDGGQLY | 0.7727215 |
| VFAAFMV | ASSLRDGGQLY | 0.73780465 |
| TLYDKLR | ASSLRDGGQLY | 0.84281355 |
| TLAVFTVL | ASSLRDGGQLY | 0.864509 |
| TGSVQPARV | ASSLRDGGQLY | 0.8530299 |
| TGGGAGSGGT | ASSLRDGGQLY | 0.43222624 |
| SHLETSL | ASSLRDGGQLY | 0.23820229 |
| SFTTGMT | ASSLRDGGQLY | 0.7037351 |
| QVEVAKRMGVRQPTVS | ASSLRDGGQLY | 0.8225106 |
| QGSGGPALAM | ASSLRDGGQLY | 0.7636577 |
| MLAPSNP | ASSLRDGGQLY | 0.7530901 |
| LVNNAAI | ASSLRDGGQLY | 0.81455743 |
| LTGEINTLA | ASSLRDGGQLY | 0.84719616 |
| LSIFGPV | ASSLRDGGQLY | 0.7048811 |
| LLALLANL | ASSLRDGGQLY | 0.4135016 |
| LISNASDALDKLR | ASSLRDGGQLY | 0.8147199 |
| LINEAVP | ASSLRDGGQLY | 0.8381994 |
| LGYAFSGL | ASSLRDGGQLY | 0.70833284 |
| LDFVAVAL | ASSLRDGGQLY | 0.7876435 |
| LAQPGAE | ASSLRDGGQLY | 0.69406176 |
| ISNLGAVTGF | ASSLRDGGQLY | 0.33299872 |
| HQEDPGRI | ASSLRDGGQLY | 0.8068867 |
| GTPGEPTP | ASSLRDGGQLY | 0.6532873 |
| GFFNSGS | ASSLRDGGQLY | 0.8485639 |
| GAGGGGLTAGAN | ASSLRDGGQLY | 0.67582387 |
| FVDALVNM | ASSLRDGGQLY | 0.81824106 |
| FLTVAPEL | ASSLRDGGQLY | 0.81799734 |
| FETLTAL | ASSLRDGGQLY | 0.818913 |
| DVINPIEL | ASSLRDGGQLY | 0.7626328 |
| APVSGAPV | ASSLRDGGQLY | 0.6777587 |
| ALVGGLAW | ASSLRDGGQLY | 0.6075201 |
| ALAAAAATAI | ASSLRDGGQLY | 0.33644795 |
| AFGLGLLL | ASSLRDGGQLY | 0.8331022 |
| AFFAAASL | ASSLRDGGQLY | 0.14639464 |
| AAAALPAFSPP | ASSLRDGGQLY | 0.84723425 |
| VVLAPAL | ASSLRDGGQLY | 0.84531254 |
| TIGHVDHGKTTL | ASSLRDGGQLY | 0.6846803 |
| LISNASDALDK | ASSLRDGGQLY | 0.8118073 |
| LISNASDAL | ASSLRDGGQLY | 0.67628825 |
| ISNASDALDK | ASSLRDGGQLY | 0.57451385 |
| NQAETLVYQTE | ASSLRDGGQLY | 0.84101576 |
| TGGTGGTAGAG | ASSLRDGGQLY | 0.8269149 |
| TGSTAGGAGGA | ASSLRDGGQLY | 0.6897991 |
| DPLAVPVAAGPV | ASSLRDGGQLY | 0.79936385 |
| GGTGGTPGNGGN | ASSLRDGGQLY | 0.8061506 |
| PAGLAGVDPATL | ASSLRDGGQLY | 0.6946534 |
| GNNLVVEAPHAGASV | ASSLRDGGQLY | 0.7801813 |
| KVTRIMSASTLEQPA | ASSLRDGGQLY | 0.84812784 |
| LGFALPIALPRRTVV | ASSLRDGGQLY | 0.7987714 |

| | | |
|---|---|---|
| TRLGNLTARKLVVLV | ASSLRDGGQLY | 0.7171912 |
| WSQQRRLIYFPSAGP | ASSLRDGGQLY | 0.8305881 |
| GPNGNPDPMAAAVDIR | ASSLRDGGQLY | 0.59411675 |
| TQRCVSVADLRAAASR | ASSLRDGGQLY | 0.8238704 |
| QFVLDVAEHPVEVLTETV | ASSLRDGGQLY | 0.74999106 |
| GGAGGTGGAGGNGGTQAGDG | ASSLRDGGQLY | 0.7822988 |
| PLLATLRAIAVDVDAKPAQV | ASSLRDGGQLY | 0.8327511 |
| APKFRTGGYMIDFWGVGYQVAK | ASSLRDGGQLY | 0.5098767 |
| GAGGAGGAGGAAGGPGGTGGTG | ASSLRDGGQLY | 0.66958183 |
| IGALGLLATDVAPAGSVAFGVA | ASSLRDGGQLY | 0.7613949 |
| AGGAGGIGGNGGAGGAGGVLMG | ASSLRDGGQLY | 0.82809424 |
| ADELIGLL | ASSLRDGGQLY | 0.8434978 |
| ADRALKA | ASSLRDGGQLY | 0.7205119 |
| AGAVGQVGGAGGA | ASSLRDGGQLY | 0.7988402 |
| AGGGGGLGGAP | ASSLRDGGQLY | 0.7571998 |
| AGGLLYGNGGAG | ASSLRDGGQLY | 0.7264031 |
| AGLESIGL | ASSLRDGGQLY | 0.7878504 |
| AGLKTGII | ASSLRDGGQLY | 0.7331201 |
| ASLAALLL | ASSLRDGGQLY | 0.64143956 |
| ASVINISSI | ASSLRDGGQLY | 0.81217957 |
| ATPPGPA | ASSLRDGGQLY | 0.718167 |
| ATVASALV | ASSLRDGGQLY | 0.8034427 |
| ATVASLIL | ASSLRDGGQLY | 0.85427076 |
| AVNGPVLIRH | ASSLRDGGQLY | 0.7255267 |
| DGAAGAVGI | ASSLRDGGQLY | 0.8129692 |
| DPLLHGI | ASSLRDGGQLY | 0.8086194 |
| ELGAALGAL | ASSLRDGGQLY | 0.8069916 |
| ELGRTQAV | ASSLRDGGQLY | 0.7923976 |
| FVAGGYLS | ASSLRDGGQLY | 0.7070565 |
| GASGGAGGAGGW | ASSLRDGGQLY | 0.8494059 |
| GGAGAGGDGGAGGI | ASSLRDGGQLY | 0.83918536 |
| GTGHNGSQPGMGGNG | ASSLRDGGQLY | 0.83476377 |
| GYAAASASAS | ASSLRDGGQLY | 0.816802 |
| HQVDVAAVQQG | ASSLRDGGQLY | 0.8185633 |
| IGAAIAHT | ASSLRDGGQLY | 0.83257663 |
| IGMIEVAQL | ASSLRDGGQLY | 0.7388021 |
| IRLAPEHTSAMD | ASSLRDGGQLY | 0.46387208 |
| IYNVSVL | ASSLRDGGQLY | 0.78745496 |
| KQGFLGM | ASSLRDGGQLY | 0.7924527 |
| LESVITQL | ASSLRDGGQLY | 0.69222796 |
| LGLGALVVV | ASSLRDGGQLY | 0.31098616 |
| LIGNGGNGGSGGI | ASSLRDGGQLY | 0.8500106 |
| LNAINAPTL | ASSLRDGGQLY | 0.8625737 |
| LQLLNTGV | ASSLRDGGQLY | 0.6762531 |
| LQTSIEF | ASSLRDGGQLY | 0.8571212 |
| MSFAVSEL | ASSLRDGGQLY | 0.49989 |
| NTGALGL | ASSLRDGGQLY | 0.835927 |
| PPGAGGAGGAGTT | ASSLRDGGQLY | 0.8502301 |
| QLVGLGI | ASSLRDGGQLY | 0.68495417 |
| RFLEQVHQL | ASSLRDGGQLY | 0.83940065 |
| RGVGYVLR | ASSLRDGGQLY | 0.66951036 |

| | | |
|---|---|---|
| SATGVFAGVI | ASSLRDGGQLY | 0.7152692 |
| SLPAITLP | ASSLRDGGQLY | 0.8536171 |
| STALAFTV | ASSLRDGGQLY | 0.76984715 |
| SVLTVGLGK | ASSLRDGGQLY | 0.7867929 |
| TALAGAGSGI | ASSLRDGGQLY | 0.8025226 |
| TGLPFTGI | ASSLRDGGQLY | 0.66441196 |
| VAEHVTV | ASSLRDGGQLY | 0.79698294 |
| VDIVLNSL | ASSLRDGGQLY | 0.8429371 |
| VLTGGGAVL | ASSLRDGGQLY | 0.5970699 |
| VNVGLAYGI | ASSLRDGGQLY | 0.8569169 |
| VSSAAVGAV | ASSLRDGGQLY | 0.67819905 |
| VTSVYEAGV | ASSLRDGGQLY | 0.86158556 |
| WVALDRGA | ASSLRDGGQLY | 0.82664883 |
| YLVGSVVI | ASSLRDGGQLY | 0.8060255 |
| GPPGTGKTLL | ASSLRDGGQLY | 0.70650446 |
| GVGMTKF | ASSLRDGGQLY | 0.8512295 |
| IGIGGSDL | ASSLRDGGQLY | 0.79661 |
| IGTLLAEL | ASSLRDGGQLY | 0.7973586 |
| IYTSGTT | ASSLRDGGQLY | 0.84443814 |
| LSFPAGV | ASSLRDGGQLY | 0.6263404 |
| LYTSGTT | ASSLRDGGQLY | 0.8370376 |
| VGAFPFGF | ASSLRDGGQLY | 0.7588723 |
| VGAGTVEF | ASSLRDGGQLY | 0.7921688 |
| VGKSTLLL | ASSLRDGGQLY | 0.6651851 |
| AALAHAGGGIV | ASSLRDGGQLY | 0.4904976 |
| AALVAVSLL | ASSLRDGGQLY | 0.7735815 |
| AAVSRAVAAL | ASSLRDGGQLY | 0.5724929 |
| AGGSSGIGGF | ASSLRDGGQLY | 0.7815639 |
| AGLTVANANG | ASSLRDGGQLY | 0.82568103 |
| AGLVGAVL | ASSLRDGGQLY | 0.6457838 |
| ALSQAFTV | ASSLRDGGQLY | 0.8364993 |
| ALTFSVSL | ASSLRDGGQLY | 0.8370292 |
| ALTGAALVAL | ASSLRDGGQLY | 0.4924228 |
| ASFSIGAGVL | ASSLRDGGQLY | 0.71832377 |
| DGSVYKI | ASSLRDGGQLY | 0.74189806 |
| EQLAEIV | ASSLRDGGQLY | 0.7444285 |
| FAVSELTVM | ASSLRDGGQLY | 0.4341442 |
| GAAPLAAGL | ASSLRDGGQLY | 0.8113208 |
| GADGGDGGSGGAGG | ASSLRDGGQLY | 0.65628344 |
| GFGLDTGGAGG | ASSLRDGGQLY | 0.20331906 |
| GLASGGLL | ASSLRDGGQLY | 0.78166187 |
| GRIAGAAVLL | ASSLRDGGQLY | 0.7320476 |
| GSAGSLNTSF | ASSLRDGGQLY | 0.6997584 |
| GTGQIGI | ASSLRDGGQLY | 0.8152604 |
| GTLIALIL | ASSLRDGGQLY | 0.5928773 |
| IAATMGFVVG | ASSLRDGGQLY | 0.6118286 |
| IFAQALL | ASSLRDGGQLY | 0.79696864 |
| IGAAVALGF | ASSLRDGGQLY | 0.83989584 |
| IGLTPLAL | ASSLRDGGQLY | 0.53895015 |
| ILGSNTSTL | ASSLRDGGQLY | 0.7597395 |
| IVFSLTVL | ASSLRDGGQLY | 0.83987916 |

| | | |
|---|---|---|
| LASKATGF | ASSLRDGGQLY | 0.7737532 |
| LASVALIL | ASSLRDGGQLY | 0.84005576 |
| LFAVGLTL | ASSLRDGGQLY | 0.84587336 |
| LFNGGIGTYI | ASSLRDGGQLY | 0.6873175 |
| LTLGAGII | ASSLRDGGQLY | 0.8296542 |
| LVAAALGDI | ASSLRDGGQLY | 0.84175014 |
| NTAFVQLGI | ASSLRDGGQLY | 0.72491664 |
| QLFSGAEL | ASSLRDGGQLY | 0.7570127 |
| SAATFITL | ASSLRDGGQLY | 0.59938484 |
| SNYGLANLGV | ASSLRDGGQLY | 0.7262254 |
| SYGGAYAVM | ASSLRDGGQLY | 0.37033877 |
| TTGVGGLTLG | ASSLRDGGQLY | 0.762066 |
| TTQTIYI | ASSLRDGGQLY | 0.8352889 |
| VAAGAGLGAVI | ASSLRDGGQLY | 0.7101457 |
| VAAVVTLL | ASSLRDGGQLY | 0.5303146 |
| VGFLGGLI | ASSLRDGGQLY | 0.5546448 |
| VGVIPILL | ASSLRDGGQLY | 0.7465546 |
| VIFASSLI | ASSLRDGGQLY | 0.66060245 |
| VKGGDGGAAAG | ASSLRDGGQLY | 0.41261995 |
| VTKTFTNI | ASSLRDGGQLY | 0.72096527 |
| VVAAAILL | ASSLRDGGQLY | 0.8330301 |
| YAALDVEL | ASSLRDGGQLY | 0.8289877 |
| LASVALLL | ASSLRDGGQLY | 0.4694692 |
| LDAVHRV | ASSLRDGGQLY | 0.84653896 |
| QGLSSSIF | ASSLRDGGQLY | 0.79314244 |
| RGTFANIRL | ASSLRDGGQLY | 0.83876264 |
| RQLWWGHRI | ASSLRDGGQLY | 0.7600362 |
| YGPPGTGKTLL | ASSLRDGGQLY | 0.61701983 |
| AALVGAPANVA | ASSLRDGGQLY | 0.8179142 |
| AATFRRTE | ASSLRDGGQLY | 0.6654105 |
| AGALGGGI | ASSLRDGGQLY | 0.67504144 |
| AGGAGGAGGIGAG | ASSLRDGGQLY | 0.8316277 |
| AGSGALGL | ASSLRDGGQLY | 0.7071332 |
| AQIADQI | ASSLRDGGQLY | 0.6293509 |
| ASAVAVTYL | ASSLRDGGQLY | 0.6594295 |
| ASTVLGGTGG | ASSLRDGGQLY | 0.8440186 |
| AVGGEPAK | ASSLRDGGQLY | 0.5709011 |
| CEQGTFGGTC | ASSLRDGGQLY | 0.56904626 |
| DEGKAPAG | ASSLRDGGQLY | 0.614481 |
| DTVVFPAL | ASSLRDGGQLY | 0.7237907 |
| FEGITMAG | ASSLRDGGQLY | 0.83648676 |
| FPLVVQD | ASSLRDGGQLY | 0.80241966 |
| GAFTTPQ | ASSLRDGGQLY | 0.69381344 |
| GFGGFGTAGGVGGA | ASSLRDGGQLY | 0.78182006 |
| GFLAQFL | ASSLRDGGQLY | 0.806808 |
| GGAGGAGGNAGVA | ASSLRDGGQLY | 0.7726532 |
| GGVGTAGAGGAGGAGG | ASSLRDGGQLY | 0.7818592 |
| GNGGGGDAGNAGSGGN | ASSLRDGGQLY | 0.61376786 |
| GTGGAGGAGNDAGS | ASSLRDGGQLY | 0.841885 |
| GTTNTGWFNS | ASSLRDGGQLY | 0.8146155 |
| IFMGPLFTL | ASSLRDGGQLY | 0.82889646 |

| | | |
|---|---|---|
| ILAHAIEAI | ASSLRDGGQLY | 0.8299885 |
| IRSAIREAL | ASSLRDGGQLY | 0.76876783 |
| ITFAGGG | ASSLRDGGQLY | 0.7774667 |
| ITGMSGHQEV | ASSLRDGGQLY | 0.4007818 |
| IVAGLACTL | ASSLRDGGQLY | 0.7107576 |
| IVGGGLAGA | ASSLRDGGQLY | 0.74809813 |
| IVTGVLNI | ASSLRDGGQLY | 0.77101237 |
| LAQQGCI | ASSLRDGGQLY | 0.80314875 |
| LCLHVEN | ASSLRDGGQLY | 0.81538284 |
| LDAISDGS | ASSLRDGGQLY | 0.64451814 |
| LGGFAAML | ASSLRDGGQLY | 0.8203229 |
| LGGKNPMI | ASSLRDGGQLY | 0.83468294 |
| LGPKGVAAIL | ASSLRDGGQLY | 0.48155093 |
| LMVGVSI | ASSLRDGGQLY | 0.71713954 |
| LVGAAFVL | ASSLRDGGQLY | 0.85716367 |
| LVLAVAPL | ASSLRDGGQLY | 0.7648617 |
| MAAIAVIDAA | ASSLRDGGQLY | 0.70082617 |
| NNGINAELQ | ASSLRDGGQLY | 0.7292681 |
| RGEPLVLVHG | ASSLRDGGQLY | 0.83588636 |
| RGGAGGAAGLI | ASSLRDGGQLY | 0.694805 |
| RSYRMHPAV | ASSLRDGGQLY | 0.8466003 |
| SAAHLLAGQLA | ASSLRDGGQLY | 0.29994488 |
| SAAVGSY | ASSLRDGGQLY | 0.72277504 |
| SPHAVLHRI | ASSLRDGGQLY | 0.64668137 |
| SQIAKVL | ASSLRDGGQLY | 0.32952935 |
| TALGHLVP | ASSLRDGGQLY | 0.8193698 |
| TAPTTSGAA | ASSLRDGGQLY | 0.72115797 |
| TDGDDGNGGSPGT | ASSLRDGGQLY | 0.81547236 |
| TDGLGEN | ASSLRDGGQLY | 0.7544608 |
| TGGPPLPGQA | ASSLRDGGQLY | 0.8349351 |
| TIPAITI | ASSLRDGGQLY | 0.75707024 |
| TIPGFTI | ASSLRDGGQLY | 0.77196753 |
| TSIASGALA | ASSLRDGGQLY | 0.67781216 |
| TVTLPGAAI | ASSLRDGGQLY | 0.8094383 |
| VASGLSAALP | ASSLRDGGQLY | 0.83751106 |
| VGAGVIGI | ASSLRDGGQLY | 0.83533204 |
| VGPKRFGV | ASSLRDGGQLY | 0.43472698 |
| VNVAFPD | ASSLRDGGQLY | 0.71452713 |
| VSMTNSGVS | ASSLRDGGQLY | 0.75667906 |
| VTDVLGDLSV | ASSLRDGGQLY | 0.40797174 |
| VVVVGAGFAGL | ASSLRDGGQLY | 0.30382058 |
| VYTVGTVV | ASSLRDGGQLY | 0.6730024 |
| WTTFSGP | ASSLRDGGQLY | 0.6581751 |
| YDELVEAF | ASSLRDGGQLY | 0.815976 |
| YGTGGAGGNGG | ASSLRDGGQLY | 0.81381434 |
| YTNVMAV | ASSLRDGGQLY | 0.71513814 |
| FLEQVHQL | ASSLRDGGQLY | 0.80283505 |
| GLSLLAVAV | ASSLRDGGQLY | 0.738042 |
| LEQVHQL | ASSLRDGGQLY | 0.1334947 |
| LNAAGAI | ASSLRDGGQLY | 0.70072854 |
| PAVHASL | ASSLRDGGQLY | 0.5047464 |

| | | |
|---|---|---|
| TELEDDA | ASSLRDGGQLY | 0.84922767 |
| TPPGPPN | ASSLRDGGQLY | 0.64063096 |
| TPPPPPPP | ASSLRDGGQLY | 0.6353841 |
| TQSPALI | ASSPGTVSGNTLY | 0.7438735 |
| EFAAAFPQ | ASSPGTVSGNTLY | 0.7488941 |
| LLSGNLDVL | ASSPGTVSGNTLY | 0.74490005 |
| GPANLGNYNI | ASSPGTVSGNTLY | 0.60255134 |
| TLGPTLP | ASSPGTVSGNTLY | 0.25103715 |
| VYAYSSCCK | ASSPGTVSGNTLY | 0.82095945 |
| APAPAPAPNI | ASSPGTVSGNTLY | 0.56287026 |
| PGGPVPIP | ASSPGTVSGNTLY | 0.59468585 |
| FLTVALEP | ASSPGTVSGNTLY | 0.46410057 |
| SQYFGANL | ASSPGTVSGNTLY | 0.35217568 |
| GGVKPAA | ASSPGTVSGNTLY | 0.4611676 |
| GGTGGAGGAGGW | ASSPGTVSGNTLY | 0.832237 |
| GVGGVAGLGGAGG | ASSPGTVSGNTLY | 0.6910491 |
| VLLVLATTI | ASSPGTVSGNTLY | 0.70281535 |
| IGIFAAML | ASSPGTVSGNTLY | 0.84910357 |
| AVGDLID | ASSPGTVSGNTLY | 0.71401846 |
| NISPLPL | ASSPGTVSGNTLY | 0.70763457 |
| VTAVFVAI | ASSPGTVSGNTLY | 0.65652406 |
| SGDDNDAAL | ASSPGTVSGNTLY | 0.5941402 |
| AVAALAHS | ASSPGTVSGNTLY | 0.61711794 |
| GGVTVMATP | ASSPGTVSGNTLY | 0.63146484 |
| AGLAANVRD | ASSPGTVSGNTLY | 0.8507638 |
| TNRPDLIDPALLRPG | ASSPGTVSGNTLY | 0.74923575 |
| LANLGVTAL | ASSPGTVSGNTLY | 0.60224485 |
| TGERIEL | ASSPGTVSGNTLY | 0.78985107 |
| NGEELAA | ASSPGTVSGNTLY | 0.84262675 |
| VLSLVWG | ASSPGTVSGNTLY | 0.8609606 |
| IGYALWL | ASSPGTVSGNTLY | 0.6050156 |
| LSLLLAPL | ASSPGTVSGNTLY | 0.6949359 |
| TLDEALK | ASSPGTVSGNTLY | 0.7318162 |
| TDGAGTAEGG | ASSPGTVSGNTLY | 0.71999186 |
| NAPGGAGGNGG | ASSPGTVSGNTLY | 0.703006 |
| VSVFMGMGGNF | ASSPGTVSGNTLY | 0.5570062 |
| LASYLAAL | ASSPGTVSGNTLY | 0.8291272 |
| LVGGTPAL | ASSPGTVSGNTLY | 0.85424966 |
| VRNADVL | ASSPGTVSGNTLY | 0.7186559 |
| TCSSPFLVSAM | ASSPGTVSGNTLY | 0.62378466 |
| VHITDAE | ASSPGTVSGNTLY | 0.7118848 |
| EYAAPSF | ASSPGTVSGNTLY | 0.59602845 |
| VVDAFHGGL | ASSPGTVSGNTLY | 0.64018154 |
| ASVVDPQ | ASSPGTVSGNTLY | 0.74651587 |
| SYNEFGGLS | ASSPGTVSGNTLY | 0.45808625 |
| DPDGSVY | ASSPGTVSGNTLY | 0.7292299 |
| MVSTIHATE | ASSPGTVSGNTLY | 0.8498664 |
| LAAGVADPV | ASSPGTVSGNTLY | 0.6687371 |
| TKEQIAA | ASSPGTVSGNTLY | 0.7968318 |
| SLASPAAL | ASSPGTVSGNTLY | 0.8312154 |
| IGLVFYGV | ASSPGTVSGNTLY | 0.79936457 |

| | | |
|---|---|---|
| FLLDASNL | ASSPGTVSGNTLY | 0.83575296 |
| YQVGNFP | ASSPGTVSGNTLY | 0.8005982 |
| ESLLSQST | ASSPGTVSGNTLY | 0.7899661 |
| LFTVGFAAVG | ASSPGTVSGNTLY | 0.7580966 |
| ATPPTPT | ASSPGTVSGNTLY | 0.78317404 |
| TDLATLG | ASSPGTVSGNTLY | 0.73669505 |
| SAGGTGR | ASSPGTVSGNTLY | 0.72992676 |
| GFSNSGS | ASSPGTVSGNTLY | 0.7050537 |
| GGNGGSGGTSV | ASSPGTVSGNTLY | 0.81101906 |
| GTSVATGGAGN | ASSPGTVSGNTLY | 0.7501478 |
| NGGNGGSGGTS | ASSPGTVSGNTLY | 0.8291381 |
| AVVKFPH | ASSPGTVSGNTLY | 0.43474323 |
| TPPPPPP | ASSPGTVSGNTLY | 0.5584965 |
| AAPAAAAAPA | ASSPGTVSGNTLY | 0.8181609 |
| IALNEDL | ASSPGTVSGNTLY | 0.20260793 |
| IVNVSSM | ASSPGTVSGNTLY | 0.3411635 |
| TGIPPAPR | ASSPGTVSGNTLY | 0.76945496 |
| FDLGGGTF | ASSPGTVSGNTLY | 0.82873034 |
| FDLGGGTFDVS | ASSPGTVSGNTLY | 0.2053264 |
| NEPTAAA | ASSPGTVSGNTLY | 0.61600566 |
| NEPTAAALA | ASSPGTVSGNTLY | 0.51784515 |
| GSIINISSI | ASSPGTVSGNTLY | 0.82148874 |
| VAPGNAGTA | ASSPGTVSGNTLY | 0.7412211 |
| TGGGASLE | ASSPGTVSGNTLY | 0.43899408 |
| VAVYNHYKRI | ASSPGTVSGNTLY | 0.8186672 |
| TDPAPATT | ASSPGTVSGNTLY | 0.5043121 |
| TNRPDILDPALLRPG | ASSPGTVSGNTLY | 0.620862 |
| LTTTTTS | ASSPGTVSGNTLY | 0.83459723 |
| PTGSGKTL | ASSPGTVSGNTLY | 0.8356933 |
| TGSGKTL | ASSPGTVSGNTLY | 0.59908724 |
| TIGHVDHGK | ASSPGTVSGNTLY | 0.8129417 |
| TIGHVDHGKTT | ASSPGTVSGNTLY | 0.6577937 |
| GHVDHGKTT | ASSPGTVSGNTLY | 0.86031556 |
| AQFEHTLL | ASSPGTVSGNTLY | 0.7079201 |
| IVNQDEL | ASSPGTVSGNTLY | 0.48395902 |
| TNGSQFF | ASSPGTVSGNTLY | 0.81755525 |
| TGKGIVE | ASSPGTVSGNTLY | 0.806723 |
| LISNASDA | ASSPGTVSGNTLY | 0.7641978 |
| AELRTLGELTKIAKAHGAQ | ASSPGTVSGNTLY | 0.8035085 |
| APNAVAAAEPSW | ASSPGTVSGNTLY | 0.7009186 |
| APQVAGAATAVL | ASSPGTVSGNTLY | 0.7052211 |
| ARRDPAAAAAILPTDA | ASSPGTVSGNTLY | 0.83023965 |
| DAAARDVLAEAGLAEYF | ASSPGTVSGNTLY | 0.8085382 |
| DDQIVLSAKANLAKVEQ | ASSPGTVSGNTLY | 0.57984143 |
| DELLPDTGEDSDEDGH | ASSPGTVSGNTLY | 0.8400736 |
| ELPEAPANRATVPQD | ASSPGTVSGNTLY | 0.7283685 |
| FGQSKIERVTGIG | ASSPGTVSGNTLY | 0.64681995 |
| GAGGTGSTAGGAGGAGGAGGLY | ASSPGTVSGNTLY | 0.85158753 |
| GHGGTGAAGGNGGTGGAGGSGI | ASSPGTVSGNTLY | 0.7565248 |
| GNGGNGGAGGEGGPGGAGGAGA | ASSPGTVSGNTLY | 0.8118606 |
| GSTAGNGGNGGNATNGTIAGQP | ASSPGTVSGNTLY | 0.8146923 |

| | | |
|---|---|---|
| GTGGDGGNGGNGGTGG | ASSPGTVSGNTLY | 0.81935143 |
| GVRRPAGIPADAPV | ASSPGTVSGNTLY | 0.5225629 |
| GYRQVTDAHLLTLARR | ASSPGTVSGNTLY | 0.79565203 |
| IAKMGQDGHDRG | ASSPGTVSGNTLY | 0.7330684 |
| KAEPSIEIQYT | ASSPGTVSGNTLY | 0.823565 |
| KPVRIRMAADEDTVAVT | ASSPGTVSGNTLY | 0.7543516 |
| MPPRIAGMRLLVIKPEP | ASSPGTVSGNTLY | 0.6628574 |
| MKFYAVALFRSRGVPPTR | ASSPGTVSGNTLY | 0.80616546 |
| NAGGFGILGAVAHSP | ASSPGTVSGNTLY | 0.83854616 |
| NEGREITLAMAEM | ASSPGTVSGNTLY | 0.7839129 |
| NELVEWSHPLGHRWRWE | ASSPGTVSGNTLY | 0.76264554 |
| NLGAGGTGGNGGNPG | ASSPGTVSGNTLY | 0.84503365 |
| NTEWPKWETVMAADPVD | ASSPGTVSGNTLY | 0.63647676 |
| PASAAPAETQP | ASSPGTVSGNTLY | 0.73484683 |
| QQSSDGKNGVRAAEITGE | ASSPGTVSGNTLY | 0.8176581 |
| RAPDAVTTSWWKE | ASSPGTVSGNTLY | 0.83092 |
| SGFGNVGAHTSGWFNQ | ASSPGTVSGNTLY | 0.75272954 |
| SGVAGAVGQAASVGGLKV | ASSPGTVSGNTLY | 0.8338139 |
| SNGGAGTGSAGGNGGT | ASSPGTVSGNTLY | 0.644777 |
| TGGAGGEGGAAG | ASSPGTVSGNTLY | 0.79492027 |
| TGGDGGAAGAGGAPGHG | ASSPGTVSGNTLY | 0.6147467 |
| TMPPLLAAIANAGRHGV | ASSPGTVSGNTLY | 0.78073543 |
| TSSTAGNSGGA | ASSPGTVSGNTLY | 0.56021273 |
| VPATVPVGILC | ASSPGTVSGNTLY | 0.7901118 |
| VQRPDSGGVVLTGRIS | ASSPGTVSGNTLY | 0.80879724 |
| WTPNPVPATKGVPAGLVQAT | ASSPGTVSGNTLY | 0.812559 |
| YALKRALAKFSDRFDVVII | ASSPGTVSGNTLY | 0.32158783 |
| YLIGSVIV | ASSPGTVSGNTLY | 0.80339205 |
| YGQLVAVID | ASSPGTVSGNTLY | 0.8159272 |
| VTVGGMTLPA | ASSPGTVSGNTLY | 0.8573412 |
| VLALPAV | ASSPGTVSGNTLY | 0.734185 |
| VIPPTLNYET | ASSPGTVSGNTLY | 0.81514955 |
| VGMGGMHPSSG | ASSPGTVSGNTLY | 0.6724829 |
| VGLAGGAATA | ASSPGTVSGNTLY | 0.749359 |
| VGGAGGAGGSGP | ASSPGTVSGNTLY | 0.5055969 |
| VFAAFMV | ASSPGTVSGNTLY | 0.66541773 |
| TLYDKLR | ASSPGTVSGNTLY | 0.6611875 |
| TLAVFTVL | ASSPGTVSGNTLY | 0.60417545 |
| TGSVQPARV | ASSPGTVSGNTLY | 0.7966678 |
| TGGGAGSGGT | ASSPGTVSGNTLY | 0.81585073 |
| SHLETSL | ASSPGTVSGNTLY | 0.82173693 |
| SFTTGMT | ASSPGTVSGNTLY | 0.8272518 |
| QVEVAKRMGVRQPTVS | ASSPGTVSGNTLY | 0.8381778 |
| QGSGGPALAM | ASSPGTVSGNTLY | 0.836886 |
| MLAPSNP | ASSPGTVSGNTLY | 0.66165 |
| LVNNAAI | ASSPGTVSGNTLY | 0.7393842 |
| LTGEINTLA | ASSPGTVSGNTLY | 0.51368487 |
| LSIFGPV | ASSPGTVSGNTLY | 0.72391427 |
| LLALLANL | ASSPGTVSGNTLY | 0.6675869 |
| LISNASDALDKLR | ASSPGTVSGNTLY | 0.77792156 |
| LINEAVP | ASSPGTVSGNTLY | 0.17771547 |

| | | |
|---|---|---|
| LGYAFSGL | ASSPGTVSGNTLY | 0.7315172 |
| LDFVAVAL | ASSPGTVSGNTLY | 0.84938884 |
| LAQPGAE | ASSPGTVSGNTLY | 0.6319678 |
| ISNLGAVTGF | ASSPGTVSGNTLY | 0.78279084 |
| HQEDPGRI | ASSPGTVSGNTLY | 0.77359474 |
| GTPGEPTP | ASSPGTVSGNTLY | 0.6736388 |
| GFFNSGS | ASSPGTVSGNTLY | 0.80103827 |
| GAGGGGLTAGAN | ASSPGTVSGNTLY | 0.63432896 |
| FVDALVNM | ASSPGTVSGNTLY | 0.809382 |
| FLTVAPEL | ASSPGTVSGNTLY | 0.8466152 |
| FETLTAL | ASSPGTVSGNTLY | 0.7731647 |
| DVINPIEL | ASSPGTVSGNTLY | 0.71048117 |
| APVSGAPV | ASSPGTVSGNTLY | 0.7660469 |
| ALVGGLAW | ASSPGTVSGNTLY | 0.47281963 |
| ALAAAAATAI | ASSPGTVSGNTLY | 0.74647576 |
| AFGLGLLL | ASSPGTVSGNTLY | 0.18399693 |
| AFFAAASL | ASSPGTVSGNTLY | 0.73984003 |
| AAAALPAFSPP | ASSPGTVSGNTLY | 0.6169867 |
| VVLAPAL | ASSPGTVSGNTLY | 0.77547145 |
| TIGHVDHGKTTL | ASSPGTVSGNTLY | 0.75308555 |
| LISNASDALDK | ASSPGTVSGNTLY | 0.8003504 |
| LISNASDAL | ASSPGTVSGNTLY | 0.5902528 |
| ISNASDALDK | ASSPGTVSGNTLY | 0.7424686 |
| NQAETLVYQTE | ASSPGTVSGNTLY | 0.66222775 |
| TGGTGGTAGAG | ASSPGTVSGNTLY | 0.5224644 |
| TGSTAGGAGGA | ASSPGTVSGNTLY | 0.815493 |
| DPLAVPVAAGPV | ASSPGTVSGNTLY | 0.74716496 |
| GGTGGTPGNGGN | ASSPGTVSGNTLY | 0.7053136 |
| PAGLAGVDPATL | ASSPGTVSGNTLY | 0.83364344 |
| GNNLVVEAPHAGASV | ASSPGTVSGNTLY | 0.77325976 |
| KVTRIMSASTLEQPA | ASSPGTVSGNTLY | 0.5411322 |
| LGFALPIALPRRTVV | ASSPGTVSGNTLY | 0.6830785 |
| TRLGNLTARKLVVLV | ASSPGTVSGNTLY | 0.7461102 |
| WSQQRRLIYFPSAGP | ASSPGTVSGNTLY | 0.70130825 |
| GPNGNPDPMAAAVDIR | ASSPGTVSGNTLY | 0.19489284 |
| TQRCVSVADLRAAASR | ASSPGTVSGNTLY | 0.8048432 |
| QFVLDVAEHPVEVLTETV | ASSPGTVSGNTLY | 0.6520403 |
| GGAGGTGGAGGNGGTQAGDG | ASSPGTVSGNTLY | 0.5977773 |
| PLLATLRAIAVDVDAKPAQV | ASSPGTVSGNTLY | 0.68559694 |
| APKFRTGGYMIDFWGVGYQVAK | ASSPGTVSGNTLY | 0.8161695 |
| GAGGAGGAGGAAGGPGGTGGTG | ASSPGTVSGNTLY | 0.6658581 |
| IGALGLLATDVAPAGSVAFGVA | ASSPGTVSGNTLY | 0.7319534 |
| AGGAGGIGGNGGAGGAGGVLMG | ASSPGTVSGNTLY | 0.79062843 |
| ADELIGLL | ASSPGTVSGNTLY | 0.7889053 |
| ADRALKA | ASSPGTVSGNTLY | 0.83719385 |
| AGAVGQVGGAGGA | ASSPGTVSGNTLY | 0.781502 |
| AGGGGGLGGAP | ASSPGTVSGNTLY | 0.7529359 |
| AGGLLYGNGGAG | ASSPGTVSGNTLY | 0.8115711 |
| AGLESIGL | ASSPGTVSGNTLY | 0.7712353 |
| AGLKTGII | ASSPGTVSGNTLY | 0.509993 |
| ASLAALLL | ASSPGTVSGNTLY | 0.4301691 |

| | | |
|---|---|---|
| ASVINISSI | ASSPGTVSGNTLY | 0.7530331 |
| ATPPGPA | ASSPGTVSGNTLY | 0.4248441 |
| ATVASALV | ASSPGTVSGNTLY | 0.7656373 |
| ATVASLIL | ASSPGTVSGNTLY | 0.84813255 |
| AVNGPVLIRH | ASSPGTVSGNTLY | 0.7490749 |
| DGAAGAVGI | ASSPGTVSGNTLY | 0.51657295 |
| DPLLHGI | ASSPGTVSGNTLY | 0.70644855 |
| ELGAALGAL | ASSPGTVSGNTLY | 0.69455665 |
| ELGRTQAV | ASSPGTVSGNTLY | 0.79861665 |
| FVAGGYLS | ASSPGTVSGNTLY | 0.8108755 |
| GASGGAGGAGGW | ASSPGTVSGNTLY | 0.8420216 |
| GGAGAGGDGGAGGI | ASSPGTVSGNTLY | 0.8154089 |
| GTGHNGSQPGMGGNG | ASSPGTVSGNTLY | 0.6582139 |
| GYAAASASAS | ASSPGTVSGNTLY | 0.806852 |
| HQVDVAAVQQG | ASSPGTVSGNTLY | 0.727205 |
| IGAAIAHT | ASSPGTVSGNTLY | 0.75891274 |
| IGMIEVAQL | ASSPGTVSGNTLY | 0.7686907 |
| IRLAPEHTSAMD | ASSPGTVSGNTLY | 0.78041714 |
| IYNVSVL | ASSPGTVSGNTLY | 0.6954823 |
| KQGFLGM | ASSPGTVSGNTLY | 0.722896 |
| LESVITQL | ASSPGTVSGNTLY | 0.5187392 |
| LGLGALVVV | ASSPGTVSGNTLY | 0.80483603 |
| LIGNGGNGGSGGI | ASSPGTVSGNTLY | 0.7752255 |
| LNAINAPTL | ASSPGTVSGNTLY | 0.49669376 |
| LQLLNTGV | ASSPGTVSGNTLY | 0.70368916 |
| LQTSIEF | ASSPGTVSGNTLY | 0.8537429 |
| MSFAVSEL | ASSPGTVSGNTLY | 0.8383724 |
| NTGALGL | ASSPGTVSGNTLY | 0.6192497 |
| PPGAGGAGGAGTT | ASSPGTVSGNTLY | 0.6287504 |
| QLVGLGI | ASSPGTVSGNTLY | 0.7565678 |
| RFLEQVHQL | ASSPGTVSGNTLY | 0.4573562 |
| RGVGYVLR | ASSPGTVSGNTLY | 0.8049667 |
| SATGVFAGVI | ASSPGTVSGNTLY | 0.7124607 |
| SLPAITLP | ASSPGTVSGNTLY | 0.7060392 |
| STALAFTV | ASSPGTVSGNTLY | 0.67951876 |
| SVLTVGLGK | ASSPGTVSGNTLY | 0.77747524 |
| TALAGAGSGI | ASSPGTVSGNTLY | 0.68689996 |
| TGLPFTGI | ASSPGTVSGNTLY | 0.8128041 |
| VAEHVTV | ASSPGTVSGNTLY | 0.2480452 |
| VDIVLNSL | ASSPGTVSGNTLY | 0.8207507 |
| VLTGGGAVL | ASSPGTVSGNTLY | 0.568076 |
| VNVGLAYGI | ASSPGTVSGNTLY | 0.65183944 |
| VSSAAVGAV | ASSPGTVSGNTLY | 0.36872727 |
| VTSVYEAGV | ASSPGTVSGNTLY | 0.78710467 |
| WVALDRGA | ASSPGTVSGNTLY | 0.8138584 |
| YLVGSVVI | ASSPGTVSGNTLY | 0.828463 |
| GPPGTGKTLL | ASSPGTVSGNTLY | 0.6557655 |
| GVGMTKF | ASSPGTVSGNTLY | 0.81203103 |
| IGIGGSDL | ASSPGTVSGNTLY | 0.84794766 |
| IGTLLAEL | ASSPGTVSGNTLY | 0.68535864 |
| IYTSGTT | ASSPGTVSGNTLY | 0.4200206 |

| | | |
|---|---|---|
| LSFPAGV | ASSPGTVSGNTLY | 0.7845104 |
| LYTSGTT | ASSPGTVSGNTLY | 0.7255043 |
| VGAFPFGF | ASSPGTVSGNTLY | 0.82058454 |
| VGAGTVEF | ASSPGTVSGNTLY | 0.79701436 |
| VGKSTLLL | ASSPGTVSGNTLY | 0.5051988 |
| AALAHAGGGIV | ASSPGTVSGNTLY | 0.8167335 |
| AALVAVSLL | ASSPGTVSGNTLY | 0.82997346 |
| AAVSRAVAAL | ASSPGTVSGNTLY | 0.57837784 |
| AGGSSGIGGF | ASSPGTVSGNTLY | 0.6875345 |
| AGLTVANANG | ASSPGTVSGNTLY | 0.7365637 |
| AGLVGAVL | ASSPGTVSGNTLY | 0.4278372 |
| ALSQAFTV | ASSPGTVSGNTLY | 0.7990175 |
| ALTFSVSL | ASSPGTVSGNTLY | 0.6532148 |
| ALTGAALVAL | ASSPGTVSGNTLY | 0.6464237 |
| ASFSIGAGVL | ASSPGTVSGNTLY | 0.75524735 |
| DGSVYKI | ASSPGTVSGNTLY | 0.65393794 |
| EQLAEIV | ASSPGTVSGNTLY | 0.68096966 |
| FAVSELTVM | ASSPGTVSGNTLY | 0.69660544 |
| GAAPLAAGL | ASSPGTVSGNTLY | 0.7775551 |
| GADGGDGGSGGAGG | ASSPGTVSGNTLY | 0.7937474 |
| GFGLDTGGAGG | ASSPGTVSGNTLY | 0.7431639 |
| GLASGGLL | ASSPGTVSGNTLY | 0.794204 |
| GRIAGAAVLL | ASSPGTVSGNTLY | 0.83865225 |
| GSAGSLNTSF | ASSPGTVSGNTLY | 0.48465472 |
| GTGQIGI | ASSPGTVSGNTLY | 0.6267463 |
| GTLIALIL | ASSPGTVSGNTLY | 0.7881784 |
| IAATMGFVVG | ASSPGTVSGNTLY | 0.51702344 |
| IFAQALL | ASSPGTVSGNTLY | 0.80868876 |
| IGAAVALGF | ASSPGTVSGNTLY | 0.7560159 |
| IGLTPLAL | ASSPGTVSGNTLY | 0.785148 |
| ILGSNTSTL | ASSPGTVSGNTLY | 0.74616927 |
| IVFSLTVL | ASSPGTVSGNTLY | 0.85569465 |
| LASKATGF | ASSPGTVSGNTLY | 0.8112712 |
| LASVALIL | ASSPGTVSGNTLY | 0.7723782 |
| LFAVGLTL | ASSPGTVSGNTLY | 0.75740457 |
| LFNGGIGTYI | ASSPGTVSGNTLY | 0.6132184 |
| LTLGAGII | ASSPGTVSGNTLY | 0.81919396 |
| LVAAALGDI | ASSPGTVSGNTLY | 0.80479586 |
| NTAFVQLGI | ASSPGTVSGNTLY | 0.5474654 |
| QLFSGAEL | ASSPGTVSGNTLY | 0.7094121 |
| SAATFITL | ASSPGTVSGNTLY | 0.8420001 |
| SNYGLANLGV | ASSPGTVSGNTLY | 0.83982205 |
| SYGGAYAVM | ASSPGTVSGNTLY | 0.47308648 |
| TTGVGGLTLG | ASSPGTVSGNTLY | 0.68341494 |
| TTQTIYI | ASSPGTVSGNTLY | 0.47048017 |
| VAAGAGLGAVI | ASSPGTVSGNTLY | 0.68614554 |
| VAAVVTLL | ASSPGTVSGNTLY | 0.54406786 |
| VGFLGGLI | ASSPGTVSGNTLY | 0.7910626 |
| VGVIPILL | ASSPGTVSGNTLY | 0.6851876 |
| VIFASSLI | ASSPGTVSGNTLY | 0.78197515 |
| VKGGDGGAAAG | ASSPGTVSGNTLY | 0.7687787 |

| | | |
|---|---|---|
| VTKTFTNI | ASSPGTVSGNTLY | 0.8386384 |
| VVAAAILL | ASSPGTVSGNTLY | 0.64544994 |
| YAALDVEL | ASSPGTVSGNTLY | 0.7052918 |
| LASVALLL | ASSPGTVSGNTLY | 0.45235825 |
| LDAVHRV | ASSPGTVSGNTLY | 0.5371915 |
| QGLSSSIF | ASSPGTVSGNTLY | 0.6621422 |
| RGTFANIRL | ASSPGTVSGNTLY | 0.6922002 |
| RQLWWGHRI | ASSPGTVSGNTLY | 0.3118718 |
| YGPPGTGKTLL | ASSPGTVSGNTLY | 0.8440985 |
| AALVGAPANVA | ASSPGTVSGNTLY | 0.7403231 |
| AATFRRTE | ASSPGTVSGNTLY | 0.8128286 |
| AGALGGGI | ASSPGTVSGNTLY | 0.6720322 |
| AGGAGGAGGIGAG | ASSPGTVSGNTLY | 0.6796978 |
| AGSGALGL | ASSPGTVSGNTLY | 0.61991274 |
| AQIADQI | ASSPGTVSGNTLY | 0.78945273 |
| ASAVAVTYL | ASSPGTVSGNTLY | 0.582633 |
| ASTVLGGTGG | ASSPGTVSGNTLY | 0.49005914 |
| AVGGEPAK | ASSPGTVSGNTLY | 0.7084717 |
| CEQGTFGGTC | ASSPGTVSGNTLY | 0.6437153 |
| DEGKAPAG | ASSPGTVSGNTLY | 0.8459562 |
| DTVVFPAL | ASSPGTVSGNTLY | 0.8212737 |
| FEGITMAG | ASSPGTVSGNTLY | 0.76955384 |
| FPLVVQD | ASSPGTVSGNTLY | 0.8407966 |
| GAFTTPQ | ASSPGTVSGNTLY | 0.2639187 |
| GFGGFGTAGGVGGA | ASSPGTVSGNTLY | 0.69640064 |
| GFLAQFL | ASSPGTVSGNTLY | 0.8477392 |
| GGAGGAGGNAGVA | ASSPGTVSGNTLY | 0.77862096 |
| GGVGTAGAGGAGGAGG | ASSPGTVSGNTLY | 0.72652423 |
| GNGGGGDAGNAGSGGN | ASSPGTVSGNTLY | 0.83396906 |
| GTGGAGGAGNDAGS | ASSPGTVSGNTLY | 0.5939032 |
| GTTNTGWFNS | ASSPGTVSGNTLY | 0.6982656 |
| IFMGPLFTL | ASSPGTVSGNTLY | 0.7355751 |
| ILAHAIEAI | ASSPGTVSGNTLY | 0.82058007 |
| IRSAIREAL | ASSPGTVSGNTLY | 0.7844049 |
| ITFAGGG | ASSPGTVSGNTLY | 0.5806544 |
| ITGMSGHQEV | ASSPGTVSGNTLY | 0.8072597 |
| IVAGLACTL | ASSPGTVSGNTLY | 0.85627544 |
| IVGGGLAGA | ASSPGTVSGNTLY | 0.6821352 |
| IVTGVLNI | ASSPGTVSGNTLY | 0.8222383 |
| LAQQGCI | ASSPGTVSGNTLY | 0.7860217 |
| LCLHVEN | ASSPGTVSGNTLY | 0.74095 |
| LDAISDGS | ASSPGTVSGNTLY | 0.60041094 |
| LGGFAAML | ASSPGTVSGNTLY | 0.6201223 |
| LGGKNPMI | ASSPGTVSGNTLY | 0.7409964 |
| LGPKGVAAIL | ASSPGTVSGNTLY | 0.54214853 |
| LMVGVSI | ASSPGTVSGNTLY | 0.7494471 |
| LVGAAFVL | ASSPGTVSGNTLY | 0.72946703 |
| LVLAVAPL | ASSPGTVSGNTLY | 0.8619449 |
| MAAIAVIDAA | ASSPGTVSGNTLY | 0.79337204 |
| NNGINAELQ | ASSPGTVSGNTLY | 0.7943431 |
| RGEPLVLVHG | ASSPGTVSGNTLY | 0.5535958 |

| | | |
|---|---|---|
| RGGAGGAAGLI | ASSPGTVSGNTLY | 0.7876401 |
| RSYRMHPAV | ASSPGTVSGNTLY | 0.7835009 |
| SAAHLLAGQLA | ASSPGTVSGNTLY | 0.5878739 |
| SAAVGSY | ASSPGTVSGNTLY | 0.84114057 |
| SPHAVLHRI | ASSPGTVSGNTLY | 0.6257827 |
| SQIAKVL | ASSPGTVSGNTLY | 0.7117263 |
| TALGHLVP | ASSPGTVSGNTLY | 0.8186332 |
| TAPTTSGAA | ASSPGTVSGNTLY | 0.81938785 |
| TDGDDGNGGSPGT | ASSPGTVSGNTLY | 0.70540226 |
| TDGLGEN | ASSPGTVSGNTLY | 0.73663515 |
| TGGPPLPGQA | ASSPGTVSGNTLY | 0.5737522 |
| TIPAITI | ASSPGTVSGNTLY | 0.81982815 |
| TIPGFTI | ASSPGTVSGNTLY | 0.82812726 |
| TSIASGALA | ASSPGTVSGNTLY | 0.7103101 |
| TVTLPGAAI | ASSPGTVSGNTLY | 0.85144174 |
| VASGLSAALP | ASSPGTVSGNTLY | 0.81081307 |
| VGAGVIGI | ASSPGTVSGNTLY | 0.45354787 |
| VGPKRFGV | ASSPGTVSGNTLY | 0.7562091 |
| VNVAFPD | ASSPGTVSGNTLY | 0.83568454 |
| VSMTNSGVS | ASSPGTVSGNTLY | 0.82548493 |
| VTDVLGDLSV | ASSPGTVSGNTLY | 0.8430217 |
| VVVVGAGFAGL | ASSPGTVSGNTLY | 0.7827903 |
| VYTVGTVV | ASSPGTVSGNTLY | 0.68401444 |
| WTTFSGP | ASSPGTVSGNTLY | 0.8408034 |
| YDELVEAF | ASSPGTVSGNTLY | 0.85896754 |
| YGTGGAGGNGG | ASSPGTVSGNTLY | 0.8188939 |
| YTNVMAV | ASSPGTVSGNTLY | 0.6837015 |
| FLEQVHQL | ASSPGTVSGNTLY | 0.6555424 |
| GLSLLAVAV | ASSPGTVSGNTLY | 0.77512884 |
| LEQVHQL | ASSPGTVSGNTLY | 0.8071884 |
| LNAAGAI | ASSPGTVSGNTLY | 0.5818322 |
| PAVHASL | ASSPGTVSGNTLY | 0.75557023 |
| TELEDDA | ASSPGTVSGNTLY | 0.76209134 |
| TPPGPPN | ASSPGTVSGNTLY | 0.42202127 |
| TPPPPPPP | ASSPGTVSGNTLY | 0.80984867 |
| TQSPALI | ASSRTGGYEQY | 0.83299553 |
| EFAAAFPQ | ASSRTGGYEQY | 0.7837728 |
| LLSGNLDVL | ASSRTGGYEQY | 0.63449633 |
| GPANLGNYNI | ASSRTGGYEQY | 0.8283453 |
| TLGPTLP | ASSRTGGYEQY | 0.8556151 |
| VYAYSSCCK | ASSRTGGYEQY | 0.84555036 |
| APAPAPAPNI | ASSRTGGYEQY | 0.4451116 |
| PGGPVPIP | ASSRTGGYEQY | 0.27718502 |
| FLTVALEP | ASSRTGGYEQY | 0.6837736 |
| SQYFGANL | ASSRTGGYEQY | 0.7852764 |
| GGVKPAA | ASSRTGGYEQY | 0.6900915 |
| GGTGGAGGAGGW | ASSRTGGYEQY | 0.62081885 |
| GVGGVAGLGGAGG | ASSRTGGYEQY | 0.7568449 |
| VLLVLATTI | ASSRTGGYEQY | 0.8308184 |
| IGIFAAML | ASSRTGGYEQY | 0.6093787 |
| AVGDLID | ASSRTGGYEQY | 0.5350759 |

| | | |
|---|---|---|
| NISPLPL | ASSRTGGYEQY | 0.81500375 |
| VTAVFVAI | ASSRTGGYEQY | 0.8183547 |
| SGDDNDAAL | ASSRTGGYEQY | 0.607337 |
| AVAALAHS | ASSRTGGYEQY | 0.6790164 |
| GGVTVMATP | ASSRTGGYEQY | 0.71358144 |
| AGLAANVRD | ASSRTGGYEQY | 0.671093 |
| TNRPDLIDPALLRPG | ASSRTGGYEQY | 0.76777387 |
| LANLGVTAL | ASSRTGGYEQY | 0.5384897 |
| TGERIEL | ASSRTGGYEQY | 0.8370652 |
| NGEELAA | ASSRTGGYEQY | 0.5435647 |
| VLSLVWG | ASSRTGGYEQY | 0.80882955 |
| IGYALWL | ASSRTGGYEQY | 0.7692686 |
| LSLLLAPL | ASSRTGGYEQY | 0.801544 |
| TLDEALK | ASSRTGGYEQY | 0.66789275 |
| TDGAGTAEGG | ASSRTGGYEQY | 0.65796006 |
| NAPGGAGGNGG | ASSRTGGYEQY | 0.5665941 |
| VSVFMGMGGNF | ASSRTGGYEQY | 0.39911464 |
| LASYLAAL | ASSRTGGYEQY | 0.82797736 |
| LVGGTPAL | ASSRTGGYEQY | 0.33910817 |
| VRNADVL | ASSRTGGYEQY | 0.83103573 |
| TCSSPFLVSAM | ASSRTGGYEQY | 0.7953718 |
| VHITDAE | ASSRTGGYEQY | 0.64791584 |
| EYAAPSF | ASSRTGGYEQY | 0.80098665 |
| VVDAFHGGL | ASSRTGGYEQY | 0.47851652 |
| ASVVDPQ | ASSRTGGYEQY | 0.7085165 |
| SYNEFGGLS | ASSRTGGYEQY | 0.7858019 |
| DPDGSVY | ASSRTGGYEQY | 0.81768894 |
| MVSTIHATE | ASSRTGGYEQY | 0.6337987 |
| LAAGVADPV | ASSRTGGYEQY | 0.80181205 |
| TKEQIAA | ASSRTGGYEQY | 0.79028106 |
| SLASPAAL | ASSRTGGYEQY | 0.6719922 |
| IGLVFYGV | ASSRTGGYEQY | 0.7781568 |
| FLLDASNL | ASSRTGGYEQY | 0.8200928 |
| YQVGNFP | ASSRTGGYEQY | 0.79613227 |
| ESLLSQST | ASSRTGGYEQY | 0.5834001 |
| LFTVGFAAVG | ASSRTGGYEQY | 0.789711 |
| ATPPTPT | ASSRTGGYEQY | 0.608032 |
| TDLATLG | ASSRTGGYEQY | 0.7764511 |
| SAGGTGR | ASSRTGGYEQY | 0.7644173 |
| GFSNSGS | ASSRTGGYEQY | 0.6979934 |
| GGNGGSGGTSV | ASSRTGGYEQY | 0.81911486 |
| GTSVATGGAGN | ASSRTGGYEQY | 0.6808146 |
| NGGNGGSGGTS | ASSRTGGYEQY | 0.6085267 |
| AVVKFPH | ASSRTGGYEQY | 0.6935214 |
| TPPPPPP | ASSRTGGYEQY | 0.7557005 |
| AAPAAAAAPA | ASSRTGGYEQY | 0.81504685 |
| IALNEDL | ASSRTGGYEQY | 0.5561097 |
| IVNVSSM | ASSRTGGYEQY | 0.77330804 |
| TGIPPAPR | ASSRTGGYEQY | 0.7109244 |
| FDLGGGTF | ASSRTGGYEQY | 0.5926801 |
| FDLGGGTFDVS | ASSRTGGYEQY | 0.7806654 |

| | | |
|---|---|---|
| NEPTAAA | ASSRTGGYEQY | 0.7395602 |
| NEPTAAALA | ASSRTGGYEQY | 0.56538534 |
| GSIINISSI | ASSRTGGYEQY | 0.8100635 |
| VAPGNAGTA | ASSRTGGYEQY | 0.62139165 |
| TGGGASLE | ASSRTGGYEQY | 0.7869069 |
| VAVYNHYKRI | ASSRTGGYEQY | 0.8447015 |
| TDPAPATT | ASSRTGGYEQY | 0.5576542 |
| TNRPDILDPALLRPG | ASSRTGGYEQY | 0.8110186 |
| LTTTTTS | ASSRTGGYEQY | 0.7859684 |
| PTGSGKTL | ASSRTGGYEQY | 0.7722962 |
| TGSGKTL | ASSRTGGYEQY | 0.7127372 |
| TIGHVDHGK | ASSRTGGYEQY | 0.67622715 |
| TIGHVDHGKTT | ASSRTGGYEQY | 0.8436623 |
| GHVDHGKTT | ASSRTGGYEQY | 0.8068891 |
| AQFEHTLL | ASSRTGGYEQY | 0.8153198 |
| IVNQDEL | ASSRTGGYEQY | 0.80473435 |
| TNGSQFF | ASSRTGGYEQY | 0.80870265 |
| TGKGIVE | ASSRTGGYEQY | 0.8110404 |
| LISNASDA | ASSRTGGYEQY | 0.61748636 |
| AELRTLGELTKIAKAHGAQ | ASSRTGGYEQY | 0.45540464 |
| APNAVAAAEPSW | ASSRTGGYEQY | 0.6934135 |
| APQVAGAATAVL | ASSRTGGYEQY | 0.7147981 |
| ARRDPAAAAAILPTDA | ASSRTGGYEQY | 0.7349885 |
| DAAARDVLAEAGLAEYF | ASSRTGGYEQY | 0.4573019 |
| DDQIVLSAKANLAKVEQ | ASSRTGGYEQY | 0.8238013 |
| DELLPDTGEDSDEDGH | ASSRTGGYEQY | 0.86176956 |
| ELPEAPANRATVPQD | ASSRTGGYEQY | 0.6258242 |
| FGQSKIERVTGIG | ASSRTGGYEQY | 0.8517406 |
| GAGGTGSTAGGAGGAGGAGGLY | ASSRTGGYEQY | 0.5528752 |
| GHGGTGAAGGNGGTGGAGGSGI | ASSRTGGYEQY | 0.78852206 |
| GNGGNGGAGGEGGPGGAGGAGA | ASSRTGGYEQY | 0.8327222 |
| GSTAGNGGNGGNATNGTIAGQP | ASSRTGGYEQY | 0.5654294 |
| GTGGDGGNGGNGGTGG | ASSRTGGYEQY | 0.7964398 |
| GVRRPAGIPADAPV | ASSRTGGYEQY | 0.6911515 |
| GYRQVTDAHLLTLARR | ASSRTGGYEQY | 0.70376384 |
| IAKMGQDGHDRG | ASSRTGGYEQY | 0.8302781 |
| KAEPSIEIQYT | ASSRTGGYEQY | 0.6804888 |
| KPVRIRMAADEDTVAVT | ASSRTGGYEQY | 0.7981149 |
| MPPRIAGMRLLVIKPEP | ASSRTGGYEQY | 0.7215765 |
| MKFYAVALFRSRGVPPTR | ASSRTGGYEQY | 0.61144626 |
| NAGGFGILGAVAHSP | ASSRTGGYEQY | 0.80126053 |
| NEGREITLAMAEM | ASSRTGGYEQY | 0.8280009 |
| NELVEWSHPLGHRWRWE | ASSRTGGYEQY | 0.69034284 |
| NLGAGGTGGNGGNPG | ASSRTGGYEQY | 0.854131 |
| NTEWPKWETVMAADPVD | ASSRTGGYEQY | 0.62404096 |
| PASAAPAETQP | ASSRTGGYEQY | 0.85735655 |
| QQSSDGKNGVRAAEITGE | ASSRTGGYEQY | 0.820796 |
| RAPDAVTTSWWKE | ASSRTGGYEQY | 0.7313223 |
| SGFGNVGAHTSGWFNQ | ASSRTGGYEQY | 0.706055 |
| SGVAGAVGQAASVGGLKV | ASSRTGGYEQY | 0.814728 |
| SNGGAGTGSAGGNGGT | ASSRTGGYEQY | 0.7049113 |

| | | |
|---|---|---|
| TGGAGGEGGAAG | ASSRTGGYEQY | 0.7842593 |
| TGGDGGAAGAGGAPGHG | ASSRTGGYEQY | 0.8471075 |
| TMPPLLAAIANAGRHGV | ASSRTGGYEQY | 0.35011584 |
| TSSTAGNSGGA | ASSRTGGYEQY | 0.78949594 |
| VPATVPVGILC | ASSRTGGYEQY | 0.7269254 |
| VQRPDSGGVVLTGRIS | ASSRTGGYEQY | 0.7013169 |
| WTPNPVPATKGVPAGLVQAT | ASSRTGGYEQY | 0.5703528 |
| YALKRALAKFSDRFDVVII | ASSRTGGYEQY | 0.34411445 |
| YLIGSVIV | ASSRTGGYEQY | 0.63767004 |
| YGQLVAVID | ASSRTGGYEQY | 0.5062138 |
| VTVGGMTLPA | ASSRTGGYEQY | 0.76101285 |
| VLALPAV | ASSRTGGYEQY | 0.74588084 |
| VIPPTLNYET | ASSRTGGYEQY | 0.5253466 |
| VGMGGMHPSSG | ASSRTGGYEQY | 0.7751887 |
| VGLAGGAATA | ASSRTGGYEQY | 0.717882 |
| VGGAGGAGGSGP | ASSRTGGYEQY | 0.3725028 |
| VFAAFMV | ASSRTGGYEQY | 0.6566733 |
| TLYDKLR | ASSRTGGYEQY | 0.63837075 |
| TLAVFTVL | ASSRTGGYEQY | 0.643432 |
| TGSVQPARV | ASSRTGGYEQY | 0.36933845 |
| TGGGAGSGGT | ASSRTGGYEQY | 0.73533595 |
| SHLETSL | ASSRTGGYEQY | 0.6490902 |
| SFTTGMT | ASSRTGGYEQY | 0.21565714 |
| QVEVAKRMGVRQPTVS | ASSRTGGYEQY | 0.73757964 |
| QGSGGPALAM | ASSRTGGYEQY | 0.6101176 |
| MLAPSNP | ASSRTGGYEQY | 0.56559515 |
| LVNNAAI | ASSRTGGYEQY | 0.77174187 |
| LTGEINTLA | ASSRTGGYEQY | 0.5094825 |
| LSIFGPV | ASSRTGGYEQY | 0.5107777 |
| LLALLANL | ASSRTGGYEQY | 0.7638155 |
| LISNASDALDKLR | ASSRTGGYEQY | 0.78151166 |
| LINEAVP | ASSRTGGYEQY | 0.43243358 |
| LGYAFSGL | ASSRTGGYEQY | 0.70897543 |
| LDFVAVAL | ASSRTGGYEQY | 0.78322655 |
| LAQPGAE | ASSRTGGYEQY | 0.7183677 |
| ISNLGAVTGF | ASSRTGGYEQY | 0.7851627 |
| HQEDPGRI | ASSRTGGYEQY | 0.7887194 |
| GTPGEPTP | ASSRTGGYEQY | 0.5186459 |
| GFFNSGS | ASSRTGGYEQY | 0.77337885 |
| GAGGGGLTAGAN | ASSRTGGYEQY | 0.7912496 |
| FVDALVNM | ASSRTGGYEQY | 0.5877613 |
| FLTVAPEL | ASSRTGGYEQY | 0.696797 |
| FETLTAL | ASSRTGGYEQY | 0.3829485 |
| DVINPIEL | ASSRTGGYEQY | 0.64171994 |
| APVSGAPV | ASSRTGGYEQY | 0.3060604 |
| ALVGGLAW | ASSRTGGYEQY | 0.7474278 |
| ALAAAAATAI | ASSRTGGYEQY | 0.781116 |
| AFGLGLLL | ASSRTGGYEQY | 0.68551445 |
| AFFAAASL | ASSRTGGYEQY | 0.41468093 |
| AAAALPAFSPP | ASSRTGGYEQY | 0.4342361 |
| VVLAPAL | ASSRTGGYEQY | 0.707691 |

| | | |
|---|---|---|
| TIGHVDHGKTTL | ASSRTGGYEQY | 0.4064026 |
| LISNASDALDK | ASSRTGGYEQY | 0.31290245 |
| LISNASDAL | ASSRTGGYEQY | 0.66342217 |
| ISNASDALDK | ASSRTGGYEQY | 0.76892316 |
| NQAETLVYQTE | ASSRTGGYEQY | 0.77241474 |
| TGGTGGTAGAG | ASSRTGGYEQY | 0.46634758 |
| TGSTAGGAGGA | ASSRTGGYEQY | 0.7888993 |
| DPLAVPVAAGPV | ASSRTGGYEQY | 0.74792075 |
| GGTGGTPGNGGN | ASSRTGGYEQY | 0.7849394 |
| PAGLAGVDPATL | ASSRTGGYEQY | 0.684957 |
| GNNLVVEAPHAGASV | ASSRTGGYEQY | 0.43092886 |
| KVTRIMSASTLEQPA | ASSRTGGYEQY | 0.7689 |
| LGFALPIALPRRTVV | ASSRTGGYEQY | 0.67081547 |
| TRLGNLTARKLVVLV | ASSRTGGYEQY | 0.72538966 |
| WSQQRRLIYFPSAGP | ASSRTGGYEQY | 0.7603473 |
| GPNGNPDPMAAAVDIR | ASSRTGGYEQY | 0.6685241 |
| TQRCVSVADLRAAASR | ASSRTGGYEQY | 0.56382656 |
| QFVLDVAEHPVEVLTETV | ASSRTGGYEQY | 0.7108786 |
| GGAGGTGGAGGNGGTQAGDG | ASSRTGGYEQY | 0.7788508 |
| PLLATLRAIAVDVDAKPAQV | ASSRTGGYEQY | 0.5409436 |
| APKFRTGGYMIDFWGVGYQVAK | ASSRTGGYEQY | 0.40328056 |
| GAGGAGGAGGAAGGPGGTGGTG | ASSRTGGYEQY | 0.5796627 |
| IGALGLLATDVAPAGSVAFGVA | ASSRTGGYEQY | 0.6824002 |
| AGGAGGIGGNGGAGGAGGVLMG | ASSRTGGYEQY | 0.78196883 |
| ADELIGLL | ASSRTGGYEQY | 0.7023392 |
| ADRALKA | ASSRTGGYEQY | 0.67801285 |
| AGAVGQVGGAGGA | ASSRTGGYEQY | 0.75428754 |
| AGGGGGLGGAP | ASSRTGGYEQY | 0.74623966 |
| AGGLLYGNGGAG | ASSRTGGYEQY | 0.71090347 |
| AGLESIGL | ASSRTGGYEQY | 0.6801163 |
| AGLKTGII | ASSRTGGYEQY | 0.6068431 |
| ASLAALLL | ASSRTGGYEQY | 0.7739817 |
| ASVINISSI | ASSRTGGYEQY | 0.7624823 |
| ATPPGPA | ASSRTGGYEQY | 0.74114776 |
| ATVASALV | ASSRTGGYEQY | 0.7688345 |
| ATVASLIL | ASSRTGGYEQY | 0.7085111 |
| AVNGPVLIRH | ASSRTGGYEQY | 0.70587325 |
| DGAAGAVGI | ASSRTGGYEQY | 0.36809182 |
| DPLLHGI | ASSRTGGYEQY | 0.56321776 |
| ELGAALGAL | ASSRTGGYEQY | 0.70014715 |
| ELGRTQAV | ASSRTGGYEQY | 0.7353144 |
| FVAGGYLS | ASSRTGGYEQY | 0.7418901 |
| GASGGAGGAGGW | ASSRTGGYEQY | 0.7061698 |
| GGAGAGGDGGAGGI | ASSRTGGYEQY | 0.60044676 |
| GTGHNGSQPGMGGNG | ASSRTGGYEQY | 0.75949335 |
| GYAAASASAS | ASSRTGGYEQY | 0.7715049 |
| HQVDVAAVQQG | ASSRTGGYEQY | 0.43968755 |
| IGAAIAHT | ASSRTGGYEQY | 0.66820574 |
| IGMIEVAQL | ASSRTGGYEQY | 0.78664017 |
| IRLAPEHTSAMD | ASSRTGGYEQY | 0.73770666 |
| IYNVSVL | ASSRTGGYEQY | 0.6593132 |

| | | |
|---|---|---|
| KQGFLGM | ASSRTGGYEQY | 0.617911 |
| LESVITQL | ASSRTGGYEQY | 0.7801126 |
| LGLGALVVV | ASSRTGGYEQY | 0.60971236 |
| LIGNGGNGGSGGI | ASSRTGGYEQY | 0.7858021 |
| LNAINAPTL | ASSRTGGYEQY | 0.257275 |
| LQLLNTGV | ASSRTGGYEQY | 0.5776341 |
| LQTSIEF | ASSRTGGYEQY | 0.5821694 |
| MSFAVSEL | ASSRTGGYEQY | 0.22161093 |
| NTGALGL | ASSRTGGYEQY | 0.77384746 |
| PPGAGGAGGAGTT | ASSRTGGYEQY | 0.44130653 |
| QLVGLGI | ASSRTGGYEQY | 0.72917116 |
| RFLEQVHQL | ASSRTGGYEQY | 0.68847674 |
| RGVGYVLR | ASSRTGGYEQY | 0.7719052 |
| SATGVFAGVI | ASSRTGGYEQY | 0.74204797 |
| SLPAITLP | ASSRTGGYEQY | 0.5609999 |
| STALAFTV | ASSRTGGYEQY | 0.47215462 |
| SVLTVGLGK | ASSRTGGYEQY | 0.7453104 |
| TALAGAGSGI | ASSRTGGYEQY | 0.769791 |
| TGLPFTGI | ASSRTGGYEQY | 0.77012956 |
| VAEHVTV | ASSRTGGYEQY | 0.317407 |
| VDIVLNSL | ASSRTGGYEQY | 0.7308036 |
| VLTGGGAVL | ASSRTGGYEQY | 0.6706153 |
| VNVGLAYGI | ASSRTGGYEQY | 0.37731606 |
| VSSAAVGAV | ASSRTGGYEQY | 0.22639501 |
| VTSVYEAGV | ASSRTGGYEQY | 0.6185296 |
| WVALDRGA | ASSRTGGYEQY | 0.68773806 |
| YLVGSVVI | ASSRTGGYEQY | 0.76227826 |
| GPPGTGKTLL | ASSRTGGYEQY | 0.78067285 |
| GVGMTKF | ASSRTGGYEQY | 0.60668445 |
| IGIGGSDL | ASSRTGGYEQY | 0.763295 |
| IGTLLAEL | ASSRTGGYEQY | 0.7068867 |
| IYTSGTT | ASSRTGGYEQY | 0.16831793 |
| LSFPAGV | ASSRTGGYEQY | 0.566661 |
| LYTSGTT | ASSRTGGYEQY | 0.37691954 |
| VGAFPFGF | ASSRTGGYEQY | 0.7629535 |
| VGAGTVEF | ASSRTGGYEQY | 0.46406502 |
| VGKSTLLL | ASSRTGGYEQY | 0.643564 |
| AALAHAGGGIV | ASSRTGGYEQY | 0.6843755 |
| AALVAVSLL | ASSRTGGYEQY | 0.74625975 |
| AAVSRAVAAL | ASSRTGGYEQY | 0.7199482 |
| AGGSSGIGGF | ASSRTGGYEQY | 0.50784886 |
| AGLTVANANG | ASSRTGGYEQY | 0.34729934 |
| AGLVGAVL | ASSRTGGYEQY | 0.75090384 |
| ALSQAFTV | ASSRTGGYEQY | 0.60794884 |
| ALTFSVSL | ASSRTGGYEQY | 0.44931558 |
| ALTGAALVAL | ASSRTGGYEQY | 0.38250208 |
| ASFSIGAGVL | ASSRTGGYEQY | 0.7605137 |
| DGSVYKI | ASSRTGGYEQY | 0.29198304 |
| EQLAEIV | ASSRTGGYEQY | 0.7725007 |
| FAVSELTVM | ASSRTGGYEQY | 0.6160188 |
| GAAPLAAGL | ASSRTGGYEQY | 0.67772853 |

| | | |
|---|---|---|
| GADGGDGGSGGAGG | ASSRTGGYEQY | 0.7953377 |
| GFGLDTGGAGG | ASSRTGGYEQY | 0.736407 |
| GLASGGLL | ASSRTGGYEQY | 0.69192106 |
| GRIAGAAVLL | ASSRTGGYEQY | 0.5434743 |
| GSAGSLNTSF | ASSRTGGYEQY | 0.4578163 |
| GTGQIGI | ASSRTGGYEQY | 0.57050806 |
| GTLIALIL | ASSRTGGYEQY | 0.530404 |
| IAATMGFVVG | ASSRTGGYEQY | 0.7825372 |
| IFAQALL | ASSRTGGYEQY | 0.64088345 |
| IGAAVALGF | ASSRTGGYEQY | 0.3642628 |
| IGLTPLAL | ASSRTGGYEQY | 0.68297195 |
| ILGSNTSTL | ASSRTGGYEQY | 0.78205264 |
| IVFSLTVL | ASSRTGGYEQY | 0.7974769 |
| LASKATGF | ASSRTGGYEQY | 0.52206826 |
| LASVALIL | ASSRTGGYEQY | 0.5475824 |
| LFAVGLTL | ASSRTGGYEQY | 0.60747266 |
| LFNGGIGTYI | ASSRTGGYEQY | 0.68476987 |
| LTLGAGII | ASSRTGGYEQY | 0.4694828 |
| LVAAALGDI | ASSRTGGYEQY | 0.43658078 |
| NTAFVQLGI | ASSRTGGYEQY | 0.7640096 |
| QLFSGAEL | ASSRTGGYEQY | 0.78236574 |
| SAATFITL | ASSRTGGYEQY | 0.6422765 |
| SNYGLANLGV | ASSRTGGYEQY | 0.51678634 |
| SYGGAYAVM | ASSRTGGYEQY | 0.6672174 |
| TTGVGGLTLG | ASSRTGGYEQY | 0.54861414 |
| TTQTIYI | ASSRTGGYEQY | 0.5859715 |
| VAAGAGLGAVI | ASSRTGGYEQY | 0.63812876 |
| VAAVVTLL | ASSRTGGYEQY | 0.3564198 |
| VGFLGGLI | ASSRTGGYEQY | 0.6609411 |
| VGVIPILL | ASSRTGGYEQY | 0.8001251 |
| VIFASSLI | ASSRTGGYEQY | 0.67358065 |
| VKGGDGGAAAG | ASSRTGGYEQY | 0.73742104 |
| VTKTFTNI | ASSRTGGYEQY | 0.78898 |
| VVAAAILL | ASSRTGGYEQY | 0.7548393 |
| YAALDVEL | ASSRTGGYEQY | 0.782705 |
| LASVALLL | ASSRTGGYEQY | 0.7580752 |
| LDAVHRV | ASSRTGGYEQY | 0.6267176 |
| QGLSSSIF | ASSRTGGYEQY | 0.70436865 |
| RGTFANIRL | ASSRTGGYEQY | 0.7668246 |
| RQLWWGHRI | ASSRTGGYEQY | 0.6607591 |
| YGPPGTGKTLL | ASSRTGGYEQY | 0.7006423 |
| AALVGAPANVA | ASSRTGGYEQY | 0.63146865 |
| AATFRRTE | ASSRTGGYEQY | 0.75475466 |
| AGALGGGI | ASSRTGGYEQY | 0.7139348 |
| AGGAGGAGGIGAG | ASSRTGGYEQY | 0.7695731 |
| AGSGALGL | ASSRTGGYEQY | 0.24058688 |
| AQIADQI | ASSRTGGYEQY | 0.45020628 |
| ASAVAVTYL | ASSRTGGYEQY | 0.774379 |
| ASTVLGGTGG | ASSRTGGYEQY | 0.19925074 |
| AVGGEPAK | ASSRTGGYEQY | 0.26249266 |
| CEQGTFGGTC | ASSRTGGYEQY | 0.7059791 |

| | | |
|---|---|---|
| DEGKAPAG | ASSRTGGYEQY | 0.78404367 |
| DTVVFPAL | ASSRTGGYEQY | 0.22101492 |
| FEGITMAG | ASSRTGGYEQY | 0.5476475 |
| FPLVVQD | ASSRTGGYEQY | 0.40700588 |
| GAFTTPQ | ASSRTGGYEQY | 0.73852026 |
| GFGGFGTAGGVGGA | ASSRTGGYEQY | 0.6373332 |
| GFLAQFL | ASSRTGGYEQY | 0.3263747 |
| GGAGGAGGNAGVA | ASSRTGGYEQY | 0.7363813 |
| GGVGTAGAGGAGGAGG | ASSRTGGYEQY | 0.45450753 |
| GNGGGGDAGNAGSGGN | ASSRTGGYEQY | 0.6860361 |
| GTGGAGGAGNDAGS | ASSRTGGYEQY | 0.775572 |
| GTTNTGWFNS | ASSRTGGYEQY | 0.78636533 |
| IFMGPLFTL | ASSRTGGYEQY | 0.44106922 |
| ILAHAIEAI | ASSRTGGYEQY | 0.7537284 |
| IRSAIREAL | ASSRTGGYEQY | 0.5683601 |
| ITFAGGG | ASSRTGGYEQY | 0.80350775 |
| ITGMSGHQEV | ASSRTGGYEQY | 0.6561694 |
| IVAGLACTL | ASSRTGGYEQY | 0.31614628 |
| IVGGGLAGA | ASSRTGGYEQY | 0.7591319 |
| IVTGVLNI | ASSRTGGYEQY | 0.7348993 |
| LAQQGCI | ASSRTGGYEQY | 0.58409715 |
| LCLHVEN | ASSRTGGYEQY | 0.7552397 |
| LDAISDGS | ASSRTGGYEQY | 0.5889251 |
| LGGFAAML | ASSRTGGYEQY | 0.64163435 |
| LGGKNPMI | ASSRTGGYEQY | 0.76566017 |
| LGPKGVAAIL | ASSRTGGYEQY | 0.7462883 |
| LMVGVSI | ASSRTGGYEQY | 0.49820402 |
| LVGAAFVL | ASSRTGGYEQY | 0.7729214 |
| LVLAVAPL | ASSRTGGYEQY | 0.68505394 |
| MAAIAVIDAA | ASSRTGGYEQY | 0.45222545 |
| NNGINAELQ | ASSRTGGYEQY | 0.797308 |
| RGEPLVLVHG | ASSRTGGYEQY | 0.6407381 |
| RGGAGGAAGLI | ASSRTGGYEQY | 0.76844037 |
| RSYRMHPAV | ASSRTGGYEQY | 0.73044395 |
| SAAHLLAGQLA | ASSRTGGYEQY | 0.7677673 |
| SAAVGSY | ASSRTGGYEQY | 0.44035292 |
| SPHAVLHRI | ASSRTGGYEQY | 0.6470734 |
| SQIAKVL | ASSRTGGYEQY | 0.7102596 |
| TALGHLVP | ASSRTGGYEQY | 0.7533809 |
| TAPTTSGAA | ASSRTGGYEQY | 0.6335642 |
| TDGDDGNGGSPGT | ASSRTGGYEQY | 0.74027646 |
| TDGLGEN | ASSRTGGYEQY | 0.739146 |
| TGGPPLPGQA | ASSRTGGYEQY | 0.7844083 |
| TIPAITI | ASSRTGGYEQY | 0.67269444 |
| TIPGFTI | ASSRTGGYEQY | 0.66998196 |
| TSIASGALA | ASSRTGGYEQY | 0.7906778 |
| TVTLPGAAI | ASSRTGGYEQY | 0.3976457 |
| VASGLSAALP | ASSRTGGYEQY | 0.68678355 |
| VGAGVIGI | ASSRTGGYEQY | 0.76173127 |
| VGPKRFGV | ASSRTGGYEQY | 0.7729211 |
| VNVAFPD | ASSRTGGYEQY | 0.64075696 |

| | | |
|---|---|---|
| VSMTNSGVS | ASSRTGGYEQY | 0.77815026 |
| VTDVLGDLSV | ASSRTGGYEQY | 0.6736015 |
| VVVVGAGFAGL | ASSRTGGYEQY | 0.7456447 |
| VYTVGTVV | ASSRTGGYEQY | 0.68778473 |
| WTTFSGP | ASSRTGGYEQY | 0.6770536 |
| YDELVEAF | ASSRTGGYEQY | 0.5539432 |
| YGTGGAGGNGG | ASSRTGGYEQY | 0.7435336 |
| YTNVMAV | ASSRTGGYEQY | 0.7730165 |
| FLEQVHQL | ASSRTGGYEQY | 0.7508279 |
| GLSLLAVAV | ASSRTGGYEQY | 0.261984 |
| LEQVHQL | ASSRTGGYEQY | 0.75791997 |
| LNAAGAI | ASSRTGGYEQY | 0.753763 |
| PAVHASL | ASSRTGGYEQY | 0.7905843 |
| TELEDDA | ASSRTGGYEQY | 0.70305 |
| TPPGPPN | ASSRTGGYEQY | 0.74485564 |
| TPPPPPPP | ASSRTGGYEQY | 0.65626156 |
| TQSPALI | ASSFGPADSDYT | 0.7037381 |
| EFAAAFPQ | ASSFGPADSDYT | 0.41159207 |
| LLSGNLDVL | ASSFGPADSDYT | 0.5507808 |
| GPANLGNYNI | ASSFGPADSDYT | 0.6692093 |
| TLGPTLP | ASSFGPADSDYT | 0.57209164 |
| VYAYSSCCK | ASSFGPADSDYT | 0.7760575 |
| APAPAPAPNI | ASSFGPADSDYT | 0.78087866 |
| PGGPVPIP | ASSFGPADSDYT | 0.7742375 |
| FLTVALEP | ASSFGPADSDYT | 0.78024745 |
| SQYFGANL | ASSFGPADSDYT | 0.7866376 |
| GGVKPAA | ASSFGPADSDYT | 0.74660194 |
| GGTGGAGGAGGW | ASSFGPADSDYT | 0.6258496 |
| GVGGVAGLGGAGG | ASSFGPADSDYT | 0.65750504 |
| VLLVLATTI | ASSFGPADSDYT | 0.3743415 |
| IGIFAAML | ASSFGPADSDYT | 0.5913813 |
| AVGDLID | ASSFGPADSDYT | 0.6203056 |
| NISPLPL | ASSFGPADSDYT | 0.7118584 |
| VTAVFVAI | ASSFGPADSDYT | 0.18275853 |
| SGDDNDAAL | ASSFGPADSDYT | 0.73145497 |
| AVAALAHS | ASSFGPADSDYT | 0.787487 |
| GGVTVMATP | ASSFGPADSDYT | 0.6884771 |
| AGLAANVRD | ASSFGPADSDYT | 0.73075986 |
| TNRPDLIDPALLRPG | ASSFGPADSDYT | 0.7575769 |
| LANLGVTAL | ASSFGPADSDYT | 0.59525275 |
| TGERIEL | ASSFGPADSDYT | 0.7607707 |
| NGEELAA | ASSFGPADSDYT | 0.5904 |
| VLSLVWG | ASSFGPADSDYT | 0.7661443 |
| IGYALWL | ASSFGPADSDYT | 0.78940547 |
| LSLLLAPL | ASSFGPADSDYT | 0.7325797 |
| TLDEALK | ASSFGPADSDYT | 0.6156307 |
| TDGAGTAEGG | ASSFGPADSDYT | 0.694315 |
| NAPGGAGGNGG | ASSFGPADSDYT | 0.3486821 |
| VSVFMGMGGNF | ASSFGPADSDYT | 0.663923 |
| LASYLAAL | ASSFGPADSDYT | 0.44856435 |
| LVGGTPAL | ASSFGPADSDYT | 0.7125472 |

| | | |
|---|---|---|
| VRNADVL | ASSFGPADSDYT | 0.51246274 |
| TCSSPFLVSAM | ASSFGPADSDYT | 0.74022096 |
| VHITDAE | ASSFGPADSDYT | 0.71154785 |
| EYAAPSF | ASSFGPADSDYT | 0.7327159 |
| VVDAFHGGL | ASSFGPADSDYT | 0.53451645 |
| ASVVDPQ | ASSFGPADSDYT | 0.73506916 |
| SYNEFGGLS | ASSFGPADSDYT | 0.74483323 |
| DPDGSVY | ASSFGPADSDYT | 0.38596332 |
| MVSTIHATE | ASSFGPADSDYT | 0.77741885 |
| LAAGVADPV | ASSFGPADSDYT | 0.63363427 |
| TKEQIAA | ASSFGPADSDYT | 0.6985777 |
| SLASPAAL | ASSFGPADSDYT | 0.76795983 |
| IGLVFYGV | ASSFGPADSDYT | 0.6775514 |
| FLLDASNL | ASSFGPADSDYT | 0.43031257 |
| YQVGNFP | ASSFGPADSDYT | 0.6695027 |
| ESLLSQST | ASSFGPADSDYT | 0.6945094 |
| LFTVGFAAVG | ASSFGPADSDYT | 0.66353846 |
| ATPPTPT | ASSFGPADSDYT | 0.21716093 |
| TDLATLG | ASSFGPADSDYT | 0.7408704 |
| SAGGTGR | ASSFGPADSDYT | 0.4555124 |
| GFSNSGS | ASSFGPADSDYT | 0.63880897 |
| GGNGGSGGTSV | ASSFGPADSDYT | 0.58704793 |
| GTSVATGGAGN | ASSFGPADSDYT | 0.77168846 |
| NGGNGGSGGTS | ASSFGPADSDYT | 0.702693 |
| AVVKFPH | ASSFGPADSDYT | 0.5678034 |
| TPPPPPP | ASSFGPADSDYT | 0.74321413 |
| AAPAAAAAPA | ASSFGPADSDYT | 0.71590436 |
| IALNEDL | ASSFGPADSDYT | 0.7719109 |
| IVNVSSM | ASSFGPADSDYT | 0.6703619 |
| TGIPPAPR | ASSFGPADSDYT | 0.7045754 |
| FDLGGGTF | ASSFGPADSDYT | 0.65997434 |
| FDLGGGTFDVS | ASSFGPADSDYT | 0.6086569 |
| NEPTAAA | ASSFGPADSDYT | 0.4717179 |
| NEPTAAALA | ASSFGPADSDYT | 0.55694985 |
| GSIINISSI | ASSFGPADSDYT | 0.66162 |
| VAPGNAGTA | ASSFGPADSDYT | 0.39201874 |
| TGGGASLE | ASSFGPADSDYT | 0.7245065 |
| VAVYNHYKRI | ASSFGPADSDYT | 0.76234883 |
| TDPAPATT | ASSFGPADSDYT | 0.6908766 |
| TNRPDILDPALLRPG | ASSFGPADSDYT | 0.3077863 |
| LTTTTTS | ASSFGPADSDYT | 0.63550144 |
| PTGSGKTL | ASSFGPADSDYT | 0.6063962 |
| TGSGKTL | ASSFGPADSDYT | 0.7622217 |
| TIGHVDHGK | ASSFGPADSDYT | 0.7430173 |
| TIGHVDHGKTT | ASSFGPADSDYT | 0.77838564 |
| GHVDHGKTT | ASSFGPADSDYT | 0.76946723 |
| AQFEHTLL | ASSFGPADSDYT | 0.62793845 |
| IVNQDEL | ASSFGPADSDYT | 0.7640531 |
| TNGSQFF | ASSFGPADSDYT | 0.6110897 |
| TGKGIVE | ASSFGPADSDYT | 0.7310798 |
| LISNASDA | ASSFGPADSDYT | 0.69383234 |

| | | |
|---|---|---|
| AELRTLGELTKIAKAHGAQ | ASSFGPADSDYT | 0.73679394 |
| APNAVAAAEPSW | ASSFGPADSDYT | 0.5290147 |
| APQVAGAATAVL | ASSFGPADSDYT | 0.61934525 |
| ARRDPAAAAAILPTDA | ASSFGPADSDYT | 0.40104815 |
| DAAARDVLAEAGLAEYF | ASSFGPADSDYT | 0.7296242 |
| DDQIVLSAKANLAKVEQ | ASSFGPADSDYT | 0.49828523 |
| DELLPDTGEDSDEDGH | ASSFGPADSDYT | 0.39427283 |
| ELPEAPANRATVPQD | ASSFGPADSDYT | 0.64313453 |
| FGQSKIERVTGIG | ASSFGPADSDYT | 0.79437035 |
| GAGGTGSTAGGAGGAGGAGGLY | ASSFGPADSDYT | 0.79071 |
| GHGGTGAAGGNGGTGGAGGSGI | ASSFGPADSDYT | 0.5861996 |
| GNGGNGGAGGEGGPGGAGGAGA | ASSFGPADSDYT | 0.5499413 |
| GSTAGNGGNGGNATNGTIAGQP | ASSFGPADSDYT | 0.74850357 |
| GTGGDGGNGGNGGTGG | ASSFGPADSDYT | 0.48423532 |
| GVRRPAGIPADAPV | ASSFGPADSDYT | 0.71394426 |
| GYRQVTDAHLLTLARR | ASSFGPADSDYT | 0.7120581 |
| IAKMGQDGHDRG | ASSFGPADSDYT | 0.705656 |
| KAEPSIEIQYT | ASSFGPADSDYT | 0.48521227 |
| KPVRIRMAADEDTVAVT | ASSFGPADSDYT | 0.7506077 |
| MPPRIAGMRLLVIKPEP | ASSFGPADSDYT | 0.65382326 |
| MKFYAVALFRSRGVPPTR | ASSFGPADSDYT | 0.70791066 |
| NAGGFGILGAVAHSP | ASSFGPADSDYT | 0.3095829 |
| NEGREITLAMAEM | ASSFGPADSDYT | 0.7613286 |
| NELVEWSHPLGHRWRWE | ASSFGPADSDYT | 0.5308739 |
| NLGAGGTGGNGGNPG | ASSFGPADSDYT | 0.5807774 |
| NTEWPKWETVMAADPVD | ASSFGPADSDYT | 0.25397086 |
| PASAAPAETQP | ASSFGPADSDYT | 0.6542326 |
| QQSSDGKNGVRAAEITGE | ASSFGPADSDYT | 0.76456565 |
| RAPDAVTTSWWKE | ASSFGPADSDYT | 0.7845231 |
| SGFGNVGAHTSGWFNQ | ASSFGPADSDYT | 0.6685252 |
| SGVAGAVGQAASVGGLKV | ASSFGPADSDYT | 0.728136 |
| SNGGAGTGSAGGNGGT | ASSFGPADSDYT | 0.79369783 |
| TGGAGGEGGAAG | ASSFGPADSDYT | 0.6727094 |
| TGGDGGAAGAGGAPGHG | ASSFGPADSDYT | 0.3136936 |
| TMPPLLAAIANAGRHGV | ASSFGPADSDYT | 0.67652476 |
| TSSTAGNSGGA | ASSFGPADSDYT | 0.6914072 |
| VPATVPVGILC | ASSFGPADSDYT | 0.77707183 |
| VQRPDSGGVVLTGRIS | ASSFGPADSDYT | 0.7436689 |
| WTPNPVPATKGVPAGLVQAT | ASSFGPADSDYT | 0.40505382 |
| YALKRALAKFSDRFDVVII | ASSFGPADSDYT | 0.68723774 |
| YLIGSVIV | ASSFGPADSDYT | 0.7719506 |
| YGQLVAVID | ASSFGPADSDYT | 0.46397322 |
| VTVGGMTLPA | ASSFGPADSDYT | 0.5406133 |
| VLALPAV | ASSFGPADSDYT | 0.49259502 |
| VIPPTLNYET | ASSFGPADSDYT | 0.3905772 |
| VGMGGMHPSSG | ASSFGPADSDYT | 0.7094839 |
| VGLAGGAATA | ASSFGPADSDYT | 0.6479003 |
| VGGAGGAGGSGP | ASSFGPADSDYT | 0.5851928 |
| VFAAFMV | ASSFGPADSDYT | 0.6471181 |
| TLYDKLR | ASSFGPADSDYT | 0.5298786 |
| TLAVFTVL | ASSFGPADSDYT | 0.54629225 |

| | | |
|---|---|---|
| TGSVQPARV | ASSFGPADSDYT | 0.6053836 |
| TGGGAGSGGT | ASSFGPADSDYT | 0.7034459 |
| SHLETSL | ASSFGPADSDYT | 0.76453686 |
| SFTTGMT | ASSFGPADSDYT | 0.683166 |
| QVEVAKRMGVRQPTVS | ASSFGPADSDYT | 0.6692012 |
| QGSGGPALAM | ASSFGPADSDYT | 0.7939085 |
| MLAPSNP | ASSFGPADSDYT | 0.40801638 |
| LVNNAAI | ASSFGPADSDYT | 0.5090145 |
| LTGEINTLA | ASSFGPADSDYT | 0.73903024 |
| LSIFGPV | ASSFGPADSDYT | 0.58964413 |
| LLALLANL | ASSFGPADSDYT | 0.77456653 |
| LISNASDALDKLR | ASSFGPADSDYT | 0.7203666 |
| LINEAVP | ASSFGPADSDYT | 0.7292006 |
| LGYAFSGL | ASSFGPADSDYT | 0.7321054 |
| LDFVAVAL | ASSFGPADSDYT | 0.79300046 |
| LAQPGAE | ASSFGPADSDYT | 0.7472074 |
| ISNLGAVTGF | ASSFGPADSDYT | 0.56715846 |
| HQEDPGRI | ASSFGPADSDYT | 0.7346228 |
| GTPGEPTP | ASSFGPADSDYT | 0.32586095 |
| GFFNSGS | ASSFGPADSDYT | 0.74335456 |
| GAGGGGLTAGAN | ASSFGPADSDYT | 0.74312997 |
| FVDALVNM | ASSFGPADSDYT | 0.56427723 |
| FLTVAPEL | ASSFGPADSDYT | 0.64644235 |
| FETLTAL | ASSFGPADSDYT | 0.7929539 |
| DVINPIEL | ASSFGPADSDYT | 0.7733065 |
| APVSGAPV | ASSFGPADSDYT | 0.5447551 |
| ALVGGLAW | ASSFGPADSDYT | 0.5804157 |
| ALAAAAATAI | ASSFGPADSDYT | 0.4683341 |
| AFGLGLLL | ASSFGPADSDYT | 0.62306637 |
| AFFAAASL | ASSFGPADSDYT | 0.29094976 |
| AAAALPAFSPP | ASSFGPADSDYT | 0.75322354 |
| VVLAPAL | ASSFGPADSDYT | 0.6451454 |
| TIGHVDHGKTTL | ASSFGPADSDYT | 0.75236046 |
| LISNASDALDK | ASSFGPADSDYT | 0.73027486 |
| LISNASDAL | ASSFGPADSDYT | 0.78037214 |
| ISNASDALDK | ASSFGPADSDYT | 0.5678133 |
| NQAETLVYQTE | ASSFGPADSDYT | 0.6312503 |
| TGGTGGTAGAG | ASSFGPADSDYT | 0.27350596 |
| TGSTAGGAGGA | ASSFGPADSDYT | 0.5144153 |
| DPLAVPVAAGPV | ASSFGPADSDYT | 0.5678048 |
| GGTGGTPGNGGN | ASSFGPADSDYT | 0.6466098 |
| PAGLAGVDPATL | ASSFGPADSDYT | 0.38004974 |
| GNNLVVEAPHAGASV | ASSFGPADSDYT | 0.7852562 |
| KVTRIMSASTLEQPA | ASSFGPADSDYT | 0.6464521 |
| LGFALPIALPRRTVV | ASSFGPADSDYT | 0.7509664 |
| TRLGNLTARKLVVLV | ASSFGPADSDYT | 0.5724203 |
| WSQQRRLIYFPSAGP | ASSFGPADSDYT | 0.45621407 |
| GPNGNPDPMAAAVDIR | ASSFGPADSDYT | 0.6014024 |
| TQRCVSVADLRAAASR | ASSFGPADSDYT | 0.7583513 |
| QFVLDVAEHPVEVLTETV | ASSFGPADSDYT | 0.4615185 |
| GGAGGTGGAGGNGGTQAGDG | ASSFGPADSDYT | 0.33293143 |

| | | |
|---|---|---|
| PLLATLRAIAVDVDAKPAQV | ASSFGPADSDYT | 0.5570035 |
| APKFRTGGYMIDFWGVGYQVAK | ASSFGPADSDYT | 0.6382995 |
| GAGGAGGAGGAAGGPGGTGGTG | ASSFGPADSDYT | 0.78811544 |
| IGALGLLATDVAPAGSVAFGVA | ASSFGPADSDYT | 0.7820829 |
| AGGAGGIGGNGGAGGAGGVLMG | ASSFGPADSDYT | 0.70917845 |
| ADELIGLL | ASSFGPADSDYT | 0.7763419 |
| ADRALKA | ASSFGPADSDYT | 0.19367988 |
| AGAVGQVGGAGGA | ASSFGPADSDYT | 0.5913801 |
| AGGGGGLGGAP | ASSFGPADSDYT | 0.78349364 |
| AGGLLYGNGGAG | ASSFGPADSDYT | 0.7520594 |
| AGLESIGL | ASSFGPADSDYT | 0.690956 |
| AGLKTGII | ASSFGPADSDYT | 0.78388906 |
| ASLAALLL | ASSFGPADSDYT | 0.55477774 |
| ASVINISSI | ASSFGPADSDYT | 0.66193664 |
| ATPPGPA | ASSFGPADSDYT | 0.6891973 |
| ATVASALV | ASSFGPADSDYT | 0.7419419 |
| ATVASLIL | ASSFGPADSDYT | 0.74960184 |
| AVNGPVLIRH | ASSFGPADSDYT | 0.3784778 |
| DGAAGAVGI | ASSFGPADSDYT | 0.7247615 |
| DPLLHGI | ASSFGPADSDYT | 0.8014518 |
| ELGAALGAL | ASSFGPADSDYT | 0.6467941 |
| ELGRTQAV | ASSFGPADSDYT | 0.7754427 |
| FVAGGYLS | ASSFGPADSDYT | 0.68740267 |
| GASGGAGGAGGW | ASSFGPADSDYT | 0.7172682 |
| GGAGAGGDGGAGGI | ASSFGPADSDYT | 0.5427466 |
| GTGHNGSQPGMGGNG | ASSFGPADSDYT | 0.53256595 |
| GYAAASASAS | ASSFGPADSDYT | 0.73359275 |
| HQVDVAAVQQG | ASSFGPADSDYT | 0.5137122 |
| IGAAIAHT | ASSFGPADSDYT | 0.667238 |
| IGMIEVAQL | ASSFGPADSDYT | 0.634884 |
| IRLAPEHTSAMD | ASSFGPADSDYT | 0.8061646 |
| IYNVSVL | ASSFGPADSDYT | 0.7381991 |
| KQGFLGM | ASSFGPADSDYT | 0.74251664 |
| LESVITQL | ASSFGPADSDYT | 0.539544 |
| LGLGALVVV | ASSFGPADSDYT | 0.7165483 |
| LIGNGGNGGSGGI | ASSFGPADSDYT | 0.75272363 |
| LNAINAPTL | ASSFGPADSDYT | 0.43536258 |
| LQLLNTGV | ASSFGPADSDYT | 0.7817838 |
| LQTSIEF | ASSFGPADSDYT | 0.5059497 |
| MSFAVSEL | ASSFGPADSDYT | 0.7152365 |
| NTGALGL | ASSFGPADSDYT | 0.68640625 |
| PPGAGGAGGAGTT | ASSFGPADSDYT | 0.7623805 |
| QLVGLGI | ASSFGPADSDYT | 0.6707418 |
| RFLEQVHQL | ASSFGPADSDYT | 0.76027834 |
| RGVGYVLR | ASSFGPADSDYT | 0.46827352 |
| SATGVFAGVI | ASSFGPADSDYT | 0.7503355 |
| SLPAITLP | ASSFGPADSDYT | 0.7812469 |
| STALAFTV | ASSFGPADSDYT | 0.6670161 |
| SVLTVGLGK | ASSFGPADSDYT | 0.78751624 |
| TALAGAGSGI | ASSFGPADSDYT | 0.69990444 |
| TGLPFTGI | ASSFGPADSDYT | 0.25345844 |

| | | |
|---|---|---|
| VAEHVTV | ASSFGPADSDYT | 0.6734096 |
| VDIVLNSL | ASSFGPADSDYT | 0.7780434 |
| VLTGGGAVL | ASSFGPADSDYT | 0.78824854 |
| VNVGLAYGI | ASSFGPADSDYT | 0.76979876 |
| VSSAAVGAV | ASSFGPADSDYT | 0.7261725 |
| VTSVYEAGV | ASSFGPADSDYT | 0.6775099 |
| WVALDRGA | ASSFGPADSDYT | 0.78753126 |
| YLVGSVVI | ASSFGPADSDYT | 0.80523825 |
| GPPGTGKTLL | ASSFGPADSDYT | 0.7340621 |
| GVGMTKF | ASSFGPADSDYT | 0.5111973 |
| IGIGGSDL | ASSFGPADSDYT | 0.45438945 |
| IGTLLAEL | ASSFGPADSDYT | 0.74647963 |
| IYTSGTT | ASSFGPADSDYT | 0.7748985 |
| LSFPAGV | ASSFGPADSDYT | 0.3960877 |
| LYTSGTT | ASSFGPADSDYT | 0.69080853 |
| VGAFPFGF | ASSFGPADSDYT | 0.7165024 |
| VGAGTVEF | ASSFGPADSDYT | 0.2588631 |
| VGKSTLLL | ASSFGPADSDYT | 0.74318373 |
| AALAHAGGGIV | ASSFGPADSDYT | 0.7856662 |
| AALVAVSLL | ASSFGPADSDYT | 0.704914 |
| AAVSRAVAAL | ASSFGPADSDYT | 0.44732928 |
| AGGSSGIGGF | ASSFGPADSDYT | 0.78361166 |
| AGLTVANANG | ASSFGPADSDYT | 0.797153 |
| AGLVGAVL | ASSFGPADSDYT | 0.7803719 |
| ALSQAFTV | ASSFGPADSDYT | 0.33057573 |
| ALTFSVSL | ASSFGPADSDYT | 0.21394266 |
| ALTGAALVAL | ASSFGPADSDYT | 0.67901266 |
| ASFSIGAGVL | ASSFGPADSDYT | 0.7508918 |
| DGSVYKI | ASSFGPADSDYT | 0.5835556 |
| EQLAEIV | ASSFGPADSDYT | 0.46419203 |
| FAVSELTVM | ASSFGPADSDYT | 0.7302122 |
| GAAPLAAGL | ASSFGPADSDYT | 0.7753244 |
| GADGGDGGSGGAGG | ASSFGPADSDYT | 0.3464578 |
| GFGLDTGGAGG | ASSFGPADSDYT | 0.4307246 |
| GLASGGLL | ASSFGPADSDYT | 0.7541722 |
| GRIAGAAVLL | ASSFGPADSDYT | 0.7586469 |
| GSAGSLNTSF | ASSFGPADSDYT | 0.5392171 |
| GTGQIGI | ASSFGPADSDYT | 0.64894986 |
| GTLIALIL | ASSFGPADSDYT | 0.72885615 |
| IAATMGFVVG | ASSFGPADSDYT | 0.71543735 |
| IFAQALL | ASSFGPADSDYT | 0.7309867 |
| IGAAVALGF | ASSFGPADSDYT | 0.39884728 |
| IGLTPLAL | ASSFGPADSDYT | 0.7830264 |
| ILGSNTSTL | ASSFGPADSDYT | 0.43428308 |
| IVFSLTVL | ASSFGPADSDYT | 0.7686698 |
| LASKATGF | ASSFGPADSDYT | 0.6985743 |
| LASVALIL | ASSFGPADSDYT | 0.71618116 |
| LFAVGLTL | ASSFGPADSDYT | 0.62802 |
| LFNGGIGTYI | ASSFGPADSDYT | 0.49434078 |
| LTLGAGII | ASSFGPADSDYT | 0.44752294 |
| LVAAALGDI | ASSFGPADSDYT | 0.50244963 |

| | | |
|---|---|---|
| NTAFVQLGI | ASSFGPADSDYT | 0.78061163 |
| QLFSGAEL | ASSFGPADSDYT | 0.68729347 |
| SAATFITL | ASSFGPADSDYT | 0.7560296 |
| SNYGLANLGV | ASSFGPADSDYT | 0.7337203 |
| SYGGAYAVM | ASSFGPADSDYT | 0.51401 |
| TTGVGGLTLG | ASSFGPADSDYT | 0.74615455 |
| TTQTIYI | ASSFGPADSDYT | 0.36685118 |
| VAAGAGLGAVI | ASSFGPADSDYT | 0.6735733 |
| VAAVVTLL | ASSFGPADSDYT | 0.65396166 |
| VGFLGGLI | ASSFGPADSDYT | 0.7033001 |
| VGVIPILL | ASSFGPADSDYT | 0.5775638 |
| VIFASSLI | ASSFGPADSDYT | 0.73409975 |
| VKGGDGGAAAG | ASSFGPADSDYT | 0.7444565 |
| VTKTFTNI | ASSFGPADSDYT | 0.5393318 |
| VVAAAILL | ASSFGPADSDYT | 0.67711985 |
| YAALDVEL | ASSFGPADSDYT | 0.77676374 |
| LASVALLL | ASSFGPADSDYT | 0.720969 |
| LDAVHRV | ASSFGPADSDYT | 0.45339683 |
| QGLSSSIF | ASSFGPADSDYT | 0.7249571 |
| RGTFANIRL | ASSFGPADSDYT | 0.5416054 |
| RQLWWGHRI | ASSFGPADSDYT | 0.7263507 |
| YGPPGTGKTLL | ASSFGPADSDYT | 0.7156111 |
| AALVGAPANVA | ASSFGPADSDYT | 0.5918845 |
| AATFRRTE | ASSFGPADSDYT | 0.75731 |
| AGALGGGI | ASSFGPADSDYT | 0.36876112 |
| AGGAGGAGGIGAG | ASSFGPADSDYT | 0.6387447 |
| AGSGALGL | ASSFGPADSDYT | 0.5981603 |
| AQIADQI | ASSFGPADSDYT | 0.7121373 |
| ASAVAVTYL | ASSFGPADSDYT | 0.7400241 |
| ASTVLGGTGG | ASSFGPADSDYT | 0.45856255 |
| AVGGEPAK | ASSFGPADSDYT | 0.72554266 |
| CEQGTFGGTC | ASSFGPADSDYT | 0.63430715 |
| DEGKAPAG | ASSFGPADSDYT | 0.4958622 |
| DTVVFPAL | ASSFGPADSDYT | 0.64427173 |
| FEGITMAG | ASSFGPADSDYT | 0.6100855 |
| FPLVVQD | ASSFGPADSDYT | 0.40088123 |
| GAFTTPQ | ASSFGPADSDYT | 0.73918104 |
| GFGGFGTAGGVGGA | ASSFGPADSDYT | 0.5309212 |
| GFLAQFL | ASSFGPADSDYT | 0.70932305 |
| GGAGGAGGNAGVA | ASSFGPADSDYT | 0.79331076 |
| GGVGTAGAGGAGGAGG | ASSFGPADSDYT | 0.36939776 |
| GNGGGGDAGNAGSGGN | ASSFGPADSDYT | 0.7663568 |
| GTGGAGGAGNDAGS | ASSFGPADSDYT | 0.71068597 |
| GTTNTGWFNS | ASSFGPADSDYT | 0.72114563 |
| IFMGPLFTL | ASSFGPADSDYT | 0.63605046 |
| ILAHAIEAI | ASSFGPADSDYT | 0.57220054 |
| IRSAIREAL | ASSFGPADSDYT | 0.7782227 |
| ITFAGGG | ASSFGPADSDYT | 0.74966383 |
| ITGMSGHQEV | ASSFGPADSDYT | 0.6922851 |
| IVAGLACTL | ASSFGPADSDYT | 0.7086115 |
| IVGGGLAGA | ASSFGPADSDYT | 0.75436616 |

| | | |
|---|---|---|
| IVTGVLNI | ASSFGPADSDYT | 0.76409954 |
| LAQQGCI | ASSFGPADSDYT | 0.60879815 |
| LCLHVEN | ASSFGPADSDYT | 0.33895087 |
| LDAISDGS | ASSFGPADSDYT | 0.64709944 |
| LGGFAAML | ASSFGPADSDYT | 0.66375554 |
| LGGKNPMI | ASSFGPADSDYT | 0.5334933 |
| LGPKGVAAIL | ASSFGPADSDYT | 0.32036394 |
| LMVGVSI | ASSFGPADSDYT | 0.7565401 |
| LVGAAFVL | ASSFGPADSDYT | 0.80362654 |
| LVLAVAPL | ASSFGPADSDYT | 0.5554515 |
| MAAIAVIDAA | ASSFGPADSDYT | 0.7906249 |
| NNGINAELQ | ASSFGPADSDYT | 0.36354363 |
| RGEPLVLVHG | ASSFGPADSDYT | 0.7563899 |
| RGGAGGAAGLI | ASSFGPADSDYT | 0.77521074 |
| RSYRMHPAV | ASSFGPADSDYT | 0.5028751 |
| SAAHLLAGQLA | ASSFGPADSDYT | 0.6850158 |
| SAAVGSY | ASSFGPADSDYT | 0.65292025 |
| SPHAVLHRI | ASSFGPADSDYT | 0.5654628 |
| SQIAKVL | ASSFGPADSDYT | 0.7763213 |
| TALGHLVP | ASSFGPADSDYT | 0.5205877 |
| TAPTTSGAA | ASSFGPADSDYT | 0.70453036 |
| TDGDDGNGGSPGT | ASSFGPADSDYT | 0.525051 |
| TDGLGEN | ASSFGPADSDYT | 0.4958758 |
| TGGPPLPGQA | ASSFGPADSDYT | 0.7120626 |
| TIPAITI | ASSFGPADSDYT | 0.73787946 |
| TIPGFTI | ASSFGPADSDYT | 0.650528 |
| TSIASGALA | ASSFGPADSDYT | 0.8031008 |
| TVTLPGAAI | ASSFGPADSDYT | 0.59416115 |
| VASGLSAALP | ASSFGPADSDYT | 0.7954674 |
| VGAGVIGI | ASSFGPADSDYT | 0.77334595 |
| VGPKRFGV | ASSFGPADSDYT | 0.71408117 |
| VNVAFPD | ASSFGPADSDYT | 0.6284213 |
| VSMTNSGVS | ASSFGPADSDYT | 0.77648944 |
| VTDVLGDLSV | ASSFGPADSDYT | 0.57892185 |
| VVVVGAGFAGL | ASSFGPADSDYT | 0.7250278 |
| VYTVGTVV | ASSFGPADSDYT | 0.76864296 |
| WTTFSGP | ASSFGPADSDYT | 0.4344292 |
| YDELVEAF | ASSFGPADSDYT | 0.82478344 |
| YGTGGAGGNGG | ASSFGPADSDYT | 0.78497314 |
| YTNVMAV | ASSFGPADSDYT | 0.695345 |
| FLEQVHQL | ASSFGPADSDYT | 0.64664924 |
| GLSLLAVAV | ASSFGPADSDYT | 0.32030892 |
| LEQVHQL | ASSFGPADSDYT | 0.6574271 |
| LNAAGAI | ASSFGPADSDYT | 0.53403497 |
| PAVHASL | ASSFGPADSDYT | 0.8025501 |
| TELEDDA | ASSFGPADSDYT | 0.79895335 |
| TPPGPPN | ASSFGPADSDYT | 0.5944842 |
| TPPPPPPP | ASSFGPADSDYT | 0.799507 |
| TQSPALI | ASSRLGVSQNTLY | 0.67518103 |
| EFAAAFPQ | ASSRLGVSQNTLY | 0.4372201 |
| LLSGNLDVL | ASSRLGVSQNTLY | 0.7298155 |

| | | |
|---|---|---|
| GPANLGNYNI | ASSRLGVSQNTLY | 0.6314053 |
| TLGPTLP | ASSRLGVSQNTLY | 0.72122616 |
| VYAYSSCCK | ASSRLGVSQNTLY | 0.3626951 |
| APAPAPAPNI | ASSRLGVSQNTLY | 0.7855317 |
| PGGPVPIP | ASSRLGVSQNTLY | 0.67799896 |
| FLTVALEP | ASSRLGVSQNTLY | 0.2140466 |
| SQYFGANL | ASSRLGVSQNTLY | 0.7952469 |
| GGVKPAA | ASSRLGVSQNTLY | 0.6387987 |
| GGTGGAGGAGGW | ASSRLGVSQNTLY | 0.6162298 |
| GVGGVAGLGGAGG | ASSRLGVSQNTLY | 0.82369083 |
| VLLVLATTI | ASSRLGVSQNTLY | 0.577093 |
| IGIFAAML | ASSRLGVSQNTLY | 0.4641586 |
| AVGDLID | ASSRLGVSQNTLY | 0.8060702 |
| NISPLPL | ASSRLGVSQNTLY | 0.82148147 |
| VTAVFVAI | ASSRLGVSQNTLY | 0.38695645 |
| SGDDNDAAL | ASSRLGVSQNTLY | 0.7685338 |
| AVAALAHS | ASSRLGVSQNTLY | 0.81135637 |
| GGVTVMATP | ASSRLGVSQNTLY | 0.77000284 |
| AGLAANVRD | ASSRLGVSQNTLY | 0.82261145 |
| TNRPDLIDPALLRPG | ASSRLGVSQNTLY | 0.82673407 |
| LANLGVTAL | ASSRLGVSQNTLY | 0.53682876 |
| TGERIEL | ASSRLGVSQNTLY | 0.8229647 |
| NGEELAA | ASSRLGVSQNTLY | 0.82970685 |
| VLSLVWG | ASSRLGVSQNTLY | 0.58469963 |
| IGYALWL | ASSRLGVSQNTLY | 0.7342926 |
| LSLLLAPL | ASSRLGVSQNTLY | 0.48073098 |
| TLDEALK | ASSRLGVSQNTLY | 0.6515645 |
| TDGAGTAEGG | ASSRLGVSQNTLY | 0.3526631 |
| NAPGGAGGNGG | ASSRLGVSQNTLY | 0.80100256 |
| VSVFMGMGGNF | ASSRLGVSQNTLY | 0.8078157 |
| LASYLAAL | ASSRLGVSQNTLY | 0.74823487 |
| LVGGTPAL | ASSRLGVSQNTLY | 0.5224127 |
| VRNADVL | ASSRLGVSQNTLY | 0.4289245 |
| TCSSPFLVSAM | ASSRLGVSQNTLY | 0.7715281 |
| VHITDAE | ASSRLGVSQNTLY | 0.4496957 |
| EYAAPSF | ASSRLGVSQNTLY | 0.3531819 |
| VVDAFHGGL | ASSRLGVSQNTLY | 0.6999346 |
| ASVVDPQ | ASSRLGVSQNTLY | 0.7873006 |
| SYNEFGGLS | ASSRLGVSQNTLY | 0.79443073 |
| DPDGSVY | ASSRLGVSQNTLY | 0.55623996 |
| MVSTIHATE | ASSRLGVSQNTLY | 0.836918 |
| LAAGVADPV | ASSRLGVSQNTLY | 0.7978147 |
| TKEQIAA | ASSRLGVSQNTLY | 0.8073544 |
| SLASPAAL | ASSRLGVSQNTLY | 0.6427143 |
| IGLVFYGV | ASSRLGVSQNTLY | 0.45859912 |
| FLLDASNL | ASSRLGVSQNTLY | 0.8191266 |
| YQVGNFP | ASSRLGVSQNTLY | 0.7499122 |
| ESLLSQST | ASSRLGVSQNTLY | 0.7555931 |
| LFTVGFAAVG | ASSRLGVSQNTLY | 0.8113812 |
| ATPPTPT | ASSRLGVSQNTLY | 0.7096664 |
| TDLATLG | ASSRLGVSQNTLY | 0.612581 |

| | | |
|---|---|---|
| SAGGTGR | ASSRLGVSQNTLY | 0.73135716 |
| GFSNSGS | ASSRLGVSQNTLY | 0.8335819 |
| GGNGGSGGTSV | ASSRLGVSQNTLY | 0.5919225 |
| GTSVATGGAGN | ASSRLGVSQNTLY | 0.38236907 |
| NGGNGGSGGTS | ASSRLGVSQNTLY | 0.6253586 |
| AVVKFPH | ASSRLGVSQNTLY | 0.70555365 |
| TPPPPPP | ASSRLGVSQNTLY | 0.824267 |
| AAPAAAAAPA | ASSRLGVSQNTLY | 0.7416027 |
| IALNEDL | ASSRLGVSQNTLY | 0.66026485 |
| IVNVSSM | ASSRLGVSQNTLY | 0.7903069 |
| TGIPPAPR | ASSRLGVSQNTLY | 0.7422123 |
| FDLGGGTF | ASSRLGVSQNTLY | 0.7345829 |
| FDLGGGTFDVS | ASSRLGVSQNTLY | 0.72258675 |
| NEPTAAA | ASSRLGVSQNTLY | 0.688068 |
| NEPTAAALA | ASSRLGVSQNTLY | 0.82886213 |
| GSIINISSI | ASSRLGVSQNTLY | 0.80892754 |
| VAPGNAGTA | ASSRLGVSQNTLY | 0.7757625 |
| TGGGASLE | ASSRLGVSQNTLY | 0.80960953 |
| VAVYNHYKRI | ASSRLGVSQNTLY | 0.70184076 |
| TDPAPATT | ASSRLGVSQNTLY | 0.7386801 |
| TNRPDILDPALLRPG | ASSRLGVSQNTLY | 0.44771641 |
| LTTTTTS | ASSRLGVSQNTLY | 0.5942972 |
| PTGSGKTL | ASSRLGVSQNTLY | 0.7431811 |
| TGSGKTL | ASSRLGVSQNTLY | 0.7936263 |
| TIGHVDHGK | ASSRLGVSQNTLY | 0.79842603 |
| TIGHVDHGKTT | ASSRLGVSQNTLY | 0.7045232 |
| GHVDHGKTT | ASSRLGVSQNTLY | 0.61896086 |
| AQFEHTLL | ASSRLGVSQNTLY | 0.77896476 |
| IVNQDEL | ASSRLGVSQNTLY | 0.81181085 |
| TNGSQFF | ASSRLGVSQNTLY | 0.51833034 |
| TGKGIVE | ASSRLGVSQNTLY | 0.69268453 |
| LISNASDA | ASSRLGVSQNTLY | 0.83847797 |
| AELRTLGELTKIAKAHGAQ | ASSRLGVSQNTLY | 0.78492403 |
| APNAVAAAEPSW | ASSRLGVSQNTLY | 0.71491575 |
| APQVAGAATAVL | ASSRLGVSQNTLY | 0.56071514 |
| ARRDPAAAAAILPTDA | ASSRLGVSQNTLY | 0.8171865 |
| DAAARDVLAEAGLAEYF | ASSRLGVSQNTLY | 0.66331106 |
| DDQIVLSAKANLAKVEQ | ASSRLGVSQNTLY | 0.8346303 |
| DELLPDTGEDSDEDGH | ASSRLGVSQNTLY | 0.26399237 |
| ELPEAPANRATVPQD | ASSRLGVSQNTLY | 0.6459609 |
| FGQSKIERVTGIG | ASSRLGVSQNTLY | 0.618973 |
| GAGGTGSTAGGAGGAGGAGGLY | ASSRLGVSQNTLY | 0.20945047 |
| GHGGTGAAGGNGGTGGAGGSGI | ASSRLGVSQNTLY | 0.8192954 |
| GNGGNGGAGGEGGPGGAGGAGA | ASSRLGVSQNTLY | 0.43280184 |
| GSTAGNGGNGGNATNGTIAGQP | ASSRLGVSQNTLY | 0.701911 |
| GTGGDGGNGGNGGTGG | ASSRLGVSQNTLY | 0.69858384 |
| GVRRPAGIPADAPV | ASSRLGVSQNTLY | 0.8198557 |
| GYRQVTDAHLLTLARR | ASSRLGVSQNTLY | 0.7867309 |
| IAKMGQDGHDRG | ASSRLGVSQNTLY | 0.536979 |
| KAEPSIEIQYT | ASSRLGVSQNTLY | 0.4633808 |
| KPVRIRMAADEDTVAVT | ASSRLGVSQNTLY | 0.7965812 |

| | | |
|---|---|---|
| MPPRIAGMRLLVIKPEP | ASSRLGVSQNTLY | 0.80409455 |
| MKFYAVALFRSRGVPPTR | ASSRLGVSQNTLY | 0.80864614 |
| NAGGFGILGAVAHSP | ASSRLGVSQNTLY | 0.2711735 |
| NEGREITLAMAEM | ASSRLGVSQNTLY | 0.7904749 |
| NELVEWSHPLGHRWRWE | ASSRLGVSQNTLY | 0.70736283 |
| NLGAGGTGGNGGNPG | ASSRLGVSQNTLY | 0.45399904 |
| NTEWPKWETVMAADPVD | ASSRLGVSQNTLY | 0.25761324 |
| PASAAPAETQP | ASSRLGVSQNTLY | 0.7090507 |
| QQSSDGKNGVRAAEITGE | ASSRLGVSQNTLY | 0.7431339 |
| RAPDAVTTSWWKE | ASSRLGVSQNTLY | 0.8040471 |
| SGFGNVGAHTSGWFNQ | ASSRLGVSQNTLY | 0.81210446 |
| SGVAGAVGQAASVGGLKV | ASSRLGVSQNTLY | 0.7053951 |
| SNGGAGTGSAGGNGGT | ASSRLGVSQNTLY | 0.7911961 |
| TGGAGGEGGAAG | ASSRLGVSQNTLY | 0.71740603 |
| TGGDGGAAGAGGAPGHG | ASSRLGVSQNTLY | 0.13290557 |
| TMPPLLAAIANAGRHGV | ASSRLGVSQNTLY | 0.5520716 |
| TSSTAGNSGGA | ASSRLGVSQNTLY | 0.41619128 |
| VPATVPVGILC | ASSRLGVSQNTLY | 0.7987571 |
| VQRPDSGGVVLTGRIS | ASSRLGVSQNTLY | 0.5617704 |
| WTPNPVPATKGVPAGLVQAT | ASSRLGVSQNTLY | 0.6981626 |
| YALKRALAKFSDRFDVVII | ASSRLGVSQNTLY | 0.7366382 |
| YLIGSVIV | ASSRLGVSQNTLY | 0.7795411 |
| YGQLVAVID | ASSRLGVSQNTLY | 0.74127173 |
| VTVGGMTLPA | ASSRLGVSQNTLY | 0.5470239 |
| VLALPAV | ASSRLGVSQNTLY | 0.45097208 |
| VIPPTLNYET | ASSRLGVSQNTLY | 0.7872426 |
| VGMGGMHPSSG | ASSRLGVSQNTLY | 0.611109 |
| VGLAGGAATA | ASSRLGVSQNTLY | 0.40812647 |
| VGGAGGAGGSGP | ASSRLGVSQNTLY | 0.48121655 |
| VFAAFMV | ASSRLGVSQNTLY | 0.82449317 |
| TLYDKLR | ASSRLGVSQNTLY | 0.2571801 |
| TLAVFTVL | ASSRLGVSQNTLY | 0.8063569 |
| TGSVQPARV | ASSRLGVSQNTLY | 0.63519967 |
| TGGGAGSGGT | ASSRLGVSQNTLY | 0.7373937 |
| SHLETSL | ASSRLGVSQNTLY | 0.839603 |
| SFTTGMT | ASSRLGVSQNTLY | 0.7626183 |
| QVEVAKRMGVRQPTVS | ASSRLGVSQNTLY | 0.7630094 |
| QGSGGPALAM | ASSRLGVSQNTLY | 0.48936167 |
| MLAPSNP | ASSRLGVSQNTLY | 0.5938466 |
| LVNNAAI | ASSRLGVSQNTLY | 0.58447665 |
| LTGEINTLA | ASSRLGVSQNTLY | 0.50385475 |
| LSIFGPV | ASSRLGVSQNTLY | 0.8137839 |
| LLALLANL | ASSRLGVSQNTLY | 0.70312005 |
| LISNASDALDKLR | ASSRLGVSQNTLY | 0.37196818 |
| LINEAVP | ASSRLGVSQNTLY | 0.70301485 |
| LGYAFSGL | ASSRLGVSQNTLY | 0.8327217 |
| LDFVAVAL | ASSRLGVSQNTLY | 0.8436172 |
| LAQPGAE | ASSRLGVSQNTLY | 0.57824326 |
| ISNLGAVTGF | ASSRLGVSQNTLY | 0.44304347 |
| HQEDPGRI | ASSRLGVSQNTLY | 0.52555925 |
| GTPGEPTP | ASSRLGVSQNTLY | 0.7406467 |

| | | |
|---|---|---|
| GFFNSGS | ASSRLGVSQNTLY | 0.44286942 |
| GAGGGGLTAGAN | ASSRLGVSQNTLY | 0.42675507 |
| FVDALVNM | ASSRLGVSQNTLY | 0.8156675 |
| FLTVAPEL | ASSRLGVSQNTLY | 0.8231314 |
| FETLTAL | ASSRLGVSQNTLY | 0.6609034 |
| DVINPIEL | ASSRLGVSQNTLY | 0.40799177 |
| APVSGAPV | ASSRLGVSQNTLY | 0.7112392 |
| ALVGGLAW | ASSRLGVSQNTLY | 0.5878949 |
| ALAAAAATAI | ASSRLGVSQNTLY | 0.6629578 |
| AFGLGLLL | ASSRLGVSQNTLY | 0.5176568 |
| AFFAAASL | ASSRLGVSQNTLY | 0.43519285 |
| AAAALPAFSPP | ASSRLGVSQNTLY | 0.69095874 |
| VVLAPAL | ASSRLGVSQNTLY | 0.840201 |
| TIGHVDHGKTTL | ASSRLGVSQNTLY | 0.70340073 |
| LISNASDALDK | ASSRLGVSQNTLY | 0.76416636 |
| LISNASDAL | ASSRLGVSQNTLY | 0.8387781 |
| ISNASDALDK | ASSRLGVSQNTLY | 0.8103994 |
| NQAETLVYQTE | ASSRLGVSQNTLY | 0.8315727 |
| TGGTGGTAGAG | ASSRLGVSQNTLY | 0.7950672 |
| TGSTAGGAGGA | ASSRLGVSQNTLY | 0.6834427 |
| DPLAVPVAAGPV | ASSRLGVSQNTLY | 0.73101014 |
| GGTGGTPGNGGN | ASSRLGVSQNTLY | 0.82662994 |
| PAGLAGVDPATL | ASSRLGVSQNTLY | 0.6878548 |
| GNNLVVEAPHAGASV | ASSRLGVSQNTLY | 0.768756 |
| KVTRIMSASTLEQPA | ASSRLGVSQNTLY | 0.6790581 |
| LGFALPIALPRRTVV | ASSRLGVSQNTLY | 0.79479194 |
| TRLGNLTARKLVVLV | ASSRLGVSQNTLY | 0.7175082 |
| WSQQRRLIYFPSAGP | ASSRLGVSQNTLY | 0.823524 |
| GPNGNPDPMAAAVDIR | ASSRLGVSQNTLY | 0.22853631 |
| TQRCVSVADLRAAASR | ASSRLGVSQNTLY | 0.44659442 |
| QFVLDVAEHPVEVLTETV | ASSRLGVSQNTLY | 0.82426965 |
| GGAGGTGGAGGNGGTQAGDG | ASSRLGVSQNTLY | 0.1612408 |
| PLLATLRAIAVDVDAKPAQV | ASSRLGVSQNTLY | 0.22964847 |
| APKFRTGGYMIDFWGVGYQVAK | ASSRLGVSQNTLY | 0.76780546 |
| GAGGAGGAGGAAGGPGGTGGTG | ASSRLGVSQNTLY | 0.8241776 |
| IGALGLLATDVAPAGSVAFGVA | ASSRLGVSQNTLY | 0.18550238 |
| AGGAGGIGGNGGAGGAGGVLMG | ASSRLGVSQNTLY | 0.5348592 |
| ADELIGLL | ASSRLGVSQNTLY | 0.37250355 |
| ADRALKA | ASSRLGVSQNTLY | 0.74591243 |
| AGAVGQVGGAGGA | ASSRLGVSQNTLY | 0.69577205 |
| AGGGGGLGGAP | ASSRLGVSQNTLY | 0.31761003 |
| AGGLLYGNGGAG | ASSRLGVSQNTLY | 0.7885274 |
| AGLESIGL | ASSRLGVSQNTLY | 0.47221625 |
| AGLKTGII | ASSRLGVSQNTLY | 0.7734306 |
| ASLAALLL | ASSRLGVSQNTLY | 0.8208094 |
| ASVINISSI | ASSRLGVSQNTLY | 0.8313167 |
| ATPPGPA | ASSRLGVSQNTLY | 0.5262296 |
| ATVASALV | ASSRLGVSQNTLY | 0.7852893 |
| ATVASLIL | ASSRLGVSQNTLY | 0.6587248 |
| AVNGPVLIRH | ASSRLGVSQNTLY | 0.8477633 |
| DGAAGAVGI | ASSRLGVSQNTLY | 0.69251287 |

| | | |
|---|---|---|
| DPLLHGI | ASSRLGVSQNTLY | 0.3448584 |
| ELGAALGAL | ASSRLGVSQNTLY | 0.807359 |
| ELGRTQAV | ASSRLGVSQNTLY | 0.8074517 |
| FVAGGYLS | ASSRLGVSQNTLY | 0.59894013 |
| GASGGAGGAGGW | ASSRLGVSQNTLY | 0.80961156 |
| GGAGAGGDGGAGGI | ASSRLGVSQNTLY | 0.6628028 |
| GTGHNGSQPGMGGNG | ASSRLGVSQNTLY | 0.70341563 |
| GYAAASASAS | ASSRLGVSQNTLY | 0.8128409 |
| HQVDVAAVQQG | ASSRLGVSQNTLY | 0.79817164 |
| IGAAIAHT | ASSRLGVSQNTLY | 0.4835434 |
| IGMIEVAQL | ASSRLGVSQNTLY | 0.8029145 |
| IRLAPEHTSAMD | ASSRLGVSQNTLY | 0.72912514 |
| IYNVSVL | ASSRLGVSQNTLY | 0.46793014 |
| KQGFLGM | ASSRLGVSQNTLY | 0.8404876 |
| LESVITQL | ASSRLGVSQNTLY | 0.6551504 |
| LGLGALVVV | ASSRLGVSQNTLY | 0.8191945 |
| LIGNGGNGGSGGI | ASSRLGVSQNTLY | 0.77796245 |
| LNAINAPTL | ASSRLGVSQNTLY | 0.80364174 |
| LQLLNTGV | ASSRLGVSQNTLY | 0.501572 |
| LQTSIEF | ASSRLGVSQNTLY | 0.7140688 |
| MSFAVSEL | ASSRLGVSQNTLY | 0.7250378 |
| NTGALGL | ASSRLGVSQNTLY | 0.7825434 |
| PPGAGGAGGAGTT | ASSRLGVSQNTLY | 0.64921117 |
| QLVGLGI | ASSRLGVSQNTLY | 0.7812999 |
| RFLEQVHQL | ASSRLGVSQNTLY | 0.78926545 |
| RGVGYVLR | ASSRLGVSQNTLY | 0.832027 |
| SATGVFAGVI | ASSRLGVSQNTLY | 0.7439522 |
| SLPAITLP | ASSRLGVSQNTLY | 0.6957385 |
| STALAFTV | ASSRLGVSQNTLY | 0.8397293 |
| SVLTVGLGK | ASSRLGVSQNTLY | 0.53158 |
| TALAGAGSGI | ASSRLGVSQNTLY | 0.7115568 |
| TGLPFTGI | ASSRLGVSQNTLY | 0.8031447 |
| VAEHVTV | ASSRLGVSQNTLY | 0.808204 |
| VDIVLNSL | ASSRLGVSQNTLY | 0.699862 |
| VLTGGGAVL | ASSRLGVSQNTLY | 0.81912625 |
| VNVGLAYGI | ASSRLGVSQNTLY | 0.61877275 |
| VSSAAVGAV | ASSRLGVSQNTLY | 0.77123564 |
| VTSVYEAGV | ASSRLGVSQNTLY | 0.730999 |
| WVALDRGA | ASSRLGVSQNTLY | 0.6753114 |
| YLVGSVVI | ASSRLGVSQNTLY | 0.594412 |
| GPPGTGKTLL | ASSRLGVSQNTLY | 0.78295416 |
| GVGMTKF | ASSRLGVSQNTLY | 0.8196995 |
| IGIGGSDL | ASSRLGVSQNTLY | 0.7665989 |
| IGTLLAEL | ASSRLGVSQNTLY | 0.23834151 |
| IYTSGTT | ASSRLGVSQNTLY | 0.79656446 |
| LSFPAGV | ASSRLGVSQNTLY | 0.6869744 |
| LYTSGTT | ASSRLGVSQNTLY | 0.8434645 |
| VGAFPFGF | ASSRLGVSQNTLY | 0.7260759 |
| VGAGTVEF | ASSRLGVSQNTLY | 0.7230369 |
| VGKSTLLL | ASSRLGVSQNTLY | 0.6183977 |
| AALAHAGGGIV | ASSRLGVSQNTLY | 0.704579 |

| | | |
|---|---|---|
| AALVAVSLL | ASSRLGVSQNTLY | 0.43428043 |
| AAVSRAVAAL | ASSRLGVSQNTLY | 0.61794406 |
| AGGSSGIGGF | ASSRLGVSQNTLY | 0.71191025 |
| AGLTVANANG | ASSRLGVSQNTLY | 0.670529 |
| AGLVGAVL | ASSRLGVSQNTLY | 0.8107266 |
| ALSQAFTV | ASSRLGVSQNTLY | 0.82776827 |
| ALTFSVSL | ASSRLGVSQNTLY | 0.8085948 |
| ALTGAALVAL | ASSRLGVSQNTLY | 0.8216766 |
| ASFSIGAGVL | ASSRLGVSQNTLY | 0.84138477 |
| DGSVYKI | ASSRLGVSQNTLY | 0.7931669 |
| EQLAEIV | ASSRLGVSQNTLY | 0.6078615 |
| FAVSELTVM | ASSRLGVSQNTLY | 0.6889949 |
| GAAPLAAGL | ASSRLGVSQNTLY | 0.46720746 |
| GADGGDGGSGGAGG | ASSRLGVSQNTLY | 0.630627 |
| GFGLDTGGAGG | ASSRLGVSQNTLY | 0.6620716 |
| GLASGGLL | ASSRLGVSQNTLY | 0.74972093 |
| GRIAGAAVLL | ASSRLGVSQNTLY | 0.1572773 |
| GSAGSLNTSF | ASSRLGVSQNTLY | 0.778351 |
| GTGQIGI | ASSRLGVSQNTLY | 0.8303908 |
| GTLIALIL | ASSRLGVSQNTLY | 0.7069869 |
| IAATMGFVVG | ASSRLGVSQNTLY | 0.74725866 |
| IFAQALL | ASSRLGVSQNTLY | 0.81275684 |
| IGAAVALGF | ASSRLGVSQNTLY | 0.6909801 |
| IGLTPLAL | ASSRLGVSQNTLY | 0.79221046 |
| ILGSNTSTL | ASSRLGVSQNTLY | 0.65443635 |
| IVFSLTVL | ASSRLGVSQNTLY | 0.79820275 |
| LASKATGF | ASSRLGVSQNTLY | 0.831554 |
| LASVALIL | ASSRLGVSQNTLY | 0.77002084 |
| LFAVGLTL | ASSRLGVSQNTLY | 0.612424 |
| LFNGGIGTYI | ASSRLGVSQNTLY | 0.6576637 |
| LTLGAGII | ASSRLGVSQNTLY | 0.39072126 |
| LVAAALGDI | ASSRLGVSQNTLY | 0.6804422 |
| NTAFVQLGI | ASSRLGVSQNTLY | 0.6756494 |
| QLFSGAEL | ASSRLGVSQNTLY | 0.75632787 |
| SAATFITL | ASSRLGVSQNTLY | 0.5417939 |
| SNYGLANLGV | ASSRLGVSQNTLY | 0.75923526 |
| SYGGAYAVM | ASSRLGVSQNTLY | 0.73214746 |
| TTGVGGLTLG | ASSRLGVSQNTLY | 0.7728394 |
| TTQTIYI | ASSRLGVSQNTLY | 0.5517463 |
| VAAGAGLGAVI | ASSRLGVSQNTLY | 0.7909941 |
| VAAVVTLL | ASSRLGVSQNTLY | 0.7430627 |
| VGFLGGLI | ASSRLGVSQNTLY | 0.34502125 |
| VGVIPILL | ASSRLGVSQNTLY | 0.8225906 |
| VIFASSLI | ASSRLGVSQNTLY | 0.6899409 |
| VKGGDGGAAAG | ASSRLGVSQNTLY | 0.7575762 |
| VTKTFTNI | ASSRLGVSQNTLY | 0.7898605 |
| VVAAAILL | ASSRLGVSQNTLY | 0.7272158 |
| YAALDVEL | ASSRLGVSQNTLY | 0.4007079 |
| LASVALLL | ASSRLGVSQNTLY | 0.707639 |
| LDAVHRV | ASSRLGVSQNTLY | 0.74800885 |
| QGLSSSIF | ASSRLGVSQNTLY | 0.7313541 |

| | | |
|---|---|---|
| RGTFANIRL | ASSRLGVSQNTLY | 0.21923 |
| RQLWWGHRI | ASSRLGVSQNTLY | 0.78535056 |
| YGPPGTGKTLL | ASSRLGVSQNTLY | 0.4538881 |
| AALVGAPANVA | ASSRLGVSQNTLY | 0.6518423 |
| AATFRRTE | ASSRLGVSQNTLY | 0.707152 |
| AGALGGGI | ASSRLGVSQNTLY | 0.8199079 |
| AGGAGGAGGIGAG | ASSRLGVSQNTLY | 0.7469208 |
| AGSGALGL | ASSRLGVSQNTLY | 0.42676198 |
| AQIADQI | ASSRLGVSQNTLY | 0.79926264 |
| ASAVAVTYL | ASSRLGVSQNTLY | 0.7238438 |
| ASTVLGGTGG | ASSRLGVSQNTLY | 0.8016585 |
| AVGGEPAK | ASSRLGVSQNTLY | 0.6673571 |
| CEQGTFGGTC | ASSRLGVSQNTLY | 0.7162509 |
| DEGKAPAG | ASSRLGVSQNTLY | 0.6390984 |
| DTVVFPAL | ASSRLGVSQNTLY | 0.5488597 |
| FEGITMAG | ASSRLGVSQNTLY | 0.5872756 |
| FPLVVQD | ASSRLGVSQNTLY | 0.4954324 |
| GAFTTPQ | ASSRLGVSQNTLY | 0.5804881 |
| GFGGFGTAGGVGGA | ASSRLGVSQNTLY | 0.4010304 |
| GFLAQFL | ASSRLGVSQNTLY | 0.74567825 |
| GGAGGAGGNAGVA | ASSRLGVSQNTLY | 0.8094504 |
| GGVGTAGAGGAGGAGG | ASSRLGVSQNTLY | 0.68649065 |
| GNGGGGDAGNAGSGGN | ASSRLGVSQNTLY | 0.34731966 |
| GTGGAGGAGNDAGS | ASSRLGVSQNTLY | 0.7023064 |
| GTTNTGWFNS | ASSRLGVSQNTLY | 0.73243237 |
| IFMGPLFTL | ASSRLGVSQNTLY | 0.8153672 |
| ILAHAIEAI | ASSRLGVSQNTLY | 0.7712743 |
| IRSAIREAL | ASSRLGVSQNTLY | 0.8246843 |
| ITFAGGG | ASSRLGVSQNTLY | 0.79179233 |
| ITGMSGHQEV | ASSRLGVSQNTLY | 0.62235296 |
| IVAGLACTL | ASSRLGVSQNTLY | 0.8217752 |
| IVGGGLAGA | ASSRLGVSQNTLY | 0.68559766 |
| IVTGVLNI | ASSRLGVSQNTLY | 0.77592486 |
| LAQQGCI | ASSRLGVSQNTLY | 0.753451 |
| LCLHVEN | ASSRLGVSQNTLY | 0.7845393 |
| LDAISDGS | ASSRLGVSQNTLY | 0.54922557 |
| LGGFAAML | ASSRLGVSQNTLY | 0.65978 |
| LGGKNPMI | ASSRLGVSQNTLY | 0.353666 |
| LGPKGVAAIL | ASSRLGVSQNTLY | 0.7827722 |
| LMVGVSI | ASSRLGVSQNTLY | 0.44385743 |
| LVGAAFVL | ASSRLGVSQNTLY | 0.4787258 |
| LVLAVAPL | ASSRLGVSQNTLY | 0.645491 |
| MAAIAVIDAA | ASSRLGVSQNTLY | 0.8390316 |
| NNGINAELQ | ASSRLGVSQNTLY | 0.8201349 |
| RGEPLVLVHG | ASSRLGVSQNTLY | 0.6524401 |
| RGGAGGAAGLI | ASSRLGVSQNTLY | 0.5534696 |
| RSYRMHPAV | ASSRLGVSQNTLY | 0.79528457 |
| SAAHLLAGQLA | ASSRLGVSQNTLY | 0.5439792 |
| SAAVGSY | ASSRLGVSQNTLY | 0.78582835 |
| SPHAVLHRI | ASSRLGVSQNTLY | 0.75568616 |
| SQIAKVL | ASSRLGVSQNTLY | 0.73143935 |

| | | |
|---|---|---|
| TALGHLVP | ASSRLGVSQNTLY | 0.42634192 |
| TAPTTSGAA | ASSRLGVSQNTLY | 0.80219364 |
| TDGDDGNGGSPGT | ASSRLGVSQNTLY | 0.6600712 |
| TDGLGEN | ASSRLGVSQNTLY | 0.74030983 |
| TGGPPLPGQA | ASSRLGVSQNTLY | 0.33766928 |
| TIPAITI | ASSRLGVSQNTLY | 0.7936865 |
| TIPGFTI | ASSRLGVSQNTLY | 0.57856494 |
| TSIASGALA | ASSRLGVSQNTLY | 0.5963518 |
| TVTLPGAAI | ASSRLGVSQNTLY | 0.24597934 |
| VASGLSAALP | ASSRLGVSQNTLY | 0.6731038 |
| VGAGVIGI | ASSRLGVSQNTLY | 0.8078352 |
| VGPKRFGV | ASSRLGVSQNTLY | 0.8330927 |
| VNVAFPD | ASSRLGVSQNTLY | 0.7493607 |
| VSMTNSGVS | ASSRLGVSQNTLY | 0.75965613 |
| VTDVLGDLSV | ASSRLGVSQNTLY | 0.82662016 |
| VVVVGAGFAGL | ASSRLGVSQNTLY | 0.7350372 |
| VYTVGTVV | ASSRLGVSQNTLY | 0.2925417 |
| WTTFSGP | ASSRLGVSQNTLY | 0.6929691 |
| YDELVEAF | ASSRLGVSQNTLY | 0.72317034 |
| YGTGGAGGNGG | ASSRLGVSQNTLY | 0.82101965 |
| YTNVMAV | ASSRLGVSQNTLY | 0.7677923 |
| FLEQVHQL | ASSRLGVSQNTLY | 0.53599155 |
| GLSLLAVAV | ASSRLGVSQNTLY | 0.64486456 |
| LEQVHQL | ASSRLGVSQNTLY | 0.7861924 |
| LNAAGAI | ASSRLGVSQNTLY | 0.42913365 |
| PAVHASL | ASSRLGVSQNTLY | 0.6217369 |
| TELEDDA | ASSRLGVSQNTLY | 0.45565742 |
| TPPGPPN | ASSRLGVSQNTLY | 0.5317126 |
| TPPPPPPP | ASSRLGVSQNTLY | 0.72126913 |
| TQSPALI | ASSRGQGGDEQY | 0.71152186 |
| EFAAAFPQ | ASSRGQGGDEQY | 0.5214919 |
| LLSGNLDVL | ASSRGQGGDEQY | 0.69678605 |
| GPANLGNYNI | ASSRGQGGDEQY | 0.4986784 |
| TLGPTLP | ASSRGQGGDEQY | 0.5622232 |
| VYAYSSCCK | ASSRGQGGDEQY | 0.63252234 |
| APAPAPAPNI | ASSRGQGGDEQY | 0.70852095 |
| PGGPVPIP | ASSRGQGGDEQY | 0.81656635 |
| FLTVALEP | ASSRGQGGDEQY | 0.6739577 |
| SQYFGANL | ASSRGQGGDEQY | 0.69263244 |
| GGVKPAA | ASSRGQGGDEQY | 0.8420813 |
| GGTGGAGGAGGW | ASSRGQGGDEQY | 0.34937543 |
| GVGGVAGLGGAGG | ASSRGQGGDEQY | 0.49202475 |
| VLLVLATTI | ASSRGQGGDEQY | 0.7732661 |
| IGIFAAML | ASSRGQGGDEQY | 0.6658512 |
| AVGDLID | ASSRGQGGDEQY | 0.80227315 |
| NISPLPL | ASSRGQGGDEQY | 0.73225343 |
| VTAVFVAI | ASSRGQGGDEQY | 0.78654486 |
| SGDDNDAAL | ASSRGQGGDEQY | 0.73460543 |
| AVAALAHS | ASSRGQGGDEQY | 0.8403652 |
| GGVTVMATP | ASSRGQGGDEQY | 0.79797673 |
| AGLAANVRD | ASSRGQGGDEQY | 0.6267097 |

| | | |
|---|---|---|
| TNRPDLIDPALLRPG | ASSRGQGGDEQY | 0.79766464 |
| LANLGVTAL | ASSRGQGGDEQY | 0.31353092 |
| TGERIEL | ASSRGQGGDEQY | 0.7784817 |
| NGEELAA | ASSRGQGGDEQY | 0.77925754 |
| VLSLVWG | ASSRGQGGDEQY | 0.63079596 |
| IGYALWL | ASSRGQGGDEQY | 0.7014082 |
| LSLLLAPL | ASSRGQGGDEQY | 0.84025615 |
| TLDEALK | ASSRGQGGDEQY | 0.823632 |
| TDGAGTAEGG | ASSRGQGGDEQY | 0.49112016 |
| NAPGGAGGNGG | ASSRGQGGDEQY | 0.6497098 |
| VSVFMGMGGNF | ASSRGQGGDEQY | 0.50739396 |
| LASYLAAL | ASSRGQGGDEQY | 0.6095215 |
| LVGGTPAL | ASSRGQGGDEQY | 0.29183957 |
| VRNADVL | ASSRGQGGDEQY | 0.8055273 |
| TCSSPFLVSAM | ASSRGQGGDEQY | 0.6863604 |
| VHITDAE | ASSRGQGGDEQY | 0.787096 |
| EYAAPSF | ASSRGQGGDEQY | 0.77522177 |
| VVDAFHGGL | ASSRGQGGDEQY | 0.82808745 |
| ASVVDPQ | ASSRGQGGDEQY | 0.63306546 |
| SYNEFGGLS | ASSRGQGGDEQY | 0.64807785 |
| DPDGSVY | ASSRGQGGDEQY | 0.28695512 |
| MVSTIHATE | ASSRGQGGDEQY | 0.4203521 |
| LAAGVADPV | ASSRGQGGDEQY | 0.58207595 |
| TKEQIAA | ASSRGQGGDEQY | 0.60144424 |
| SLASPAAL | ASSRGQGGDEQY | 0.3335812 |
| IGLVFYGV | ASSRGQGGDEQY | 0.8234675 |
| FLLDASNL | ASSRGQGGDEQY | 0.67771006 |
| YQVGNFP | ASSRGQGGDEQY | 0.8071959 |
| ESLLSQST | ASSRGQGGDEQY | 0.53903794 |
| LFTVGFAAVG | ASSRGQGGDEQY | 0.51428235 |
| ATPPTPT | ASSRGQGGDEQY | 0.6600008 |
| TDLATLG | ASSRGQGGDEQY | 0.7886091 |
| SAGGTGR | ASSRGQGGDEQY | 0.5222773 |
| GFSNSGS | ASSRGQGGDEQY | 0.27049416 |
| GGNGGSGGTSV | ASSRGQGGDEQY | 0.48020136 |
| GTSVATGGAGN | ASSRGQGGDEQY | 0.61381614 |
| NGGNGGSGGTS | ASSRGQGGDEQY | 0.8384175 |
| AVVKFPH | ASSRGQGGDEQY | 0.8196889 |
| TPPPPPP | ASSRGQGGDEQY | 0.757975 |
| AAPAAAAAPA | ASSRGQGGDEQY | 0.8337745 |
| IALNEDL | ASSRGQGGDEQY | 0.17183027 |
| IVNVSSM | ASSRGQGGDEQY | 0.5748128 |
| TGIPPAPR | ASSRGQGGDEQY | 0.80159706 |
| FDLGGGTF | ASSRGQGGDEQY | 0.7797061 |
| FDLGGGTFDVS | ASSRGQGGDEQY | 0.72882354 |
| NEPTAAA | ASSRGQGGDEQY | 0.83033335 |
| NEPTAAALA | ASSRGQGGDEQY | 0.52173615 |
| GSIINISSI | ASSRGQGGDEQY | 0.68247277 |
| VAPGNAGTA | ASSRGQGGDEQY | 0.7339598 |
| TGGGASLE | ASSRGQGGDEQY | 0.80314755 |
| VAVYNHYKRI | ASSRGQGGDEQY | 0.7941398 |

| | | |
|---|---|---|
| TDPAPATT | ASSRGQGGDEQY | 0.38866782 |
| TNRPDILDPALLRPG | ASSRGQGGDEQY | 0.7213174 |
| LTTTTTS | ASSRGQGGDEQY | 0.8520472 |
| PTGSGKTL | ASSRGQGGDEQY | 0.6793364 |
| TGSGKTL | ASSRGQGGDEQY | 0.8039176 |
| TIGHVDHGK | ASSRGQGGDEQY | 0.7158576 |
| TIGHVDHGKTT | ASSRGQGGDEQY | 0.70229125 |
| GHVDHGKTT | ASSRGQGGDEQY | 0.6195029 |
| AQFEHTLL | ASSRGQGGDEQY | 0.6378315 |
| IVNQDEL | ASSRGQGGDEQY | 0.78263277 |
| TNGSQFF | ASSRGQGGDEQY | 0.54854035 |
| TGKGIVE | ASSRGQGGDEQY | 0.68975616 |
| LISNASDA | ASSRGQGGDEQY | 0.6865718 |
| AELRTLGELTKIAKAHGAQ | ASSRGQGGDEQY | 0.852041 |
| APNAVAAAEPSW | ASSRGQGGDEQY | 0.7760032 |
| APQVAGAATAVL | ASSRGQGGDEQY | 0.7820019 |
| ARRDPAAAAAILPTDA | ASSRGQGGDEQY | 0.5983336 |
| DAAARDVLAEAGLAEYF | ASSRGQGGDEQY | 0.7785729 |
| DDQIVLSAKANLAKVEQ | ASSRGQGGDEQY | 0.80773354 |
| DELLPDTGEDSDEDGH | ASSRGQGGDEQY | 0.43538612 |
| ELPEAPANRATVPQD | ASSRGQGGDEQY | 0.8219211 |
| FGQSKIERVTGIG | ASSRGQGGDEQY | 0.54786134 |
| GAGGTGSTAGGAGGAGGAGGLY | ASSRGQGGDEQY | 0.749988 |
| GHGGTGAAGGNGGTGGAGGSGI | ASSRGQGGDEQY | 0.66517484 |
| GNGGNGGAGGEGGPGGAGGAGA | ASSRGQGGDEQY | 0.804187 |
| GSTAGNGGNGGNATNGTIAGQP | ASSRGQGGDEQY | 0.75511956 |
| GTGGDGGNGGNGGTGG | ASSRGQGGDEQY | 0.8043839 |
| GVRRPAGIPADAPV | ASSRGQGGDEQY | 0.5361904 |
| GYRQVTDAHLLTLARR | ASSRGQGGDEQY | 0.8016542 |
| IAKMGQDGHDRG | ASSRGQGGDEQY | 0.82551026 |
| KAEPSIEIQYT | ASSRGQGGDEQY | 0.74160516 |
| KPVRIRMAADEDTVAVT | ASSRGQGGDEQY | 0.8419687 |
| MPPRIAGMRLLVIKPEP | ASSRGQGGDEQY | 0.66528803 |
| MKFYAVALFRSRGVPPTR | ASSRGQGGDEQY | 0.24957266 |
| NAGGFGILGAVAHSP | ASSRGQGGDEQY | 0.68950737 |
| NEGREITLAMAEM | ASSRGQGGDEQY | 0.82394207 |
| NELVEWSHPLGHRWRWE | ASSRGQGGDEQY | 0.83779323 |
| NLGAGGTGGNGGNPG | ASSRGQGGDEQY | 0.76786774 |
| NTEWPKWETVMAADPVD | ASSRGQGGDEQY | 0.74964035 |
| PASAAPAETQP | ASSRGQGGDEQY | 0.72222155 |
| QQSSDGKNGVRAAEITGE | ASSRGQGGDEQY | 0.8324895 |
| RAPDAVTTSWWKE | ASSRGQGGDEQY | 0.8428065 |
| SGFGNVGAHTSGWFNQ | ASSRGQGGDEQY | 0.7654437 |
| SGVAGAVGQAASVGGLKV | ASSRGQGGDEQY | 0.53959876 |
| SNGGAGTGSAGGNGGT | ASSRGQGGDEQY | 0.42112637 |
| TGGAGGEGGAAG | ASSRGQGGDEQY | 0.7854564 |
| TGGDGGAAGAGGAPGHG | ASSRGQGGDEQY | 0.8220721 |
| TMPPLLAAIANAGRHGV | ASSRGQGGDEQY | 0.42869 |
| TSSTAGNSGGA | ASSRGQGGDEQY | 0.7465733 |
| VPATVPVGILC | ASSRGQGGDEQY | 0.7587233 |
| VQRPDSGGVVLTGRIS | ASSRGQGGDEQY | 0.25578877 |

| | | |
|---|---|---|
| WTPNPVPATKGVPAGLVQAT | ASSRGQGGDEQY | 0.7652335 |
| YALKRALAKFSDRFDVVII | ASSRGQGGDEQY | 0.8343469 |
| YLIGSVIV | ASSRGQGGDEQY | 0.7583827 |
| YGQLVAVID | ASSRGQGGDEQY | 0.4133746 |
| VTVGGMTLPA | ASSRGQGGDEQY | 0.8275211 |
| VLALPAV | ASSRGQGGDEQY | 0.8464523 |
| VIPPTLNYET | ASSRGQGGDEQY | 0.8375875 |
| VGMGGMHPSSG | ASSRGQGGDEQY | 0.28037673 |
| VGLAGGAATA | ASSRGQGGDEQY | 0.19747554 |
| VGGAGGAGGSGP | ASSRGQGGDEQY | 0.7581743 |
| VFAAFMV | ASSRGQGGDEQY | 0.772173 |
| TLYDKLR | ASSRGQGGDEQY | 0.5377794 |
| TLAVFTVL | ASSRGQGGDEQY | 0.48648947 |
| TGSVQPARV | ASSRGQGGDEQY | 0.7860704 |
| TGGGAGSGGT | ASSRGQGGDEQY | 0.82030225 |
| SHLETSL | ASSRGQGGDEQY | 0.44172013 |
| SFTTGMT | ASSRGQGGDEQY | 0.37969056 |
| QVEVAKRMGVRQPTVS | ASSRGQGGDEQY | 0.8100604 |
| QGSGGPALAM | ASSRGQGGDEQY | 0.79707384 |
| MLAPSNP | ASSRGQGGDEQY | 0.591179 |
| LVNNAAI | ASSRGQGGDEQY | 0.72206956 |
| LTGEINTLA | ASSRGQGGDEQY | 0.7762307 |
| LSIFGPV | ASSRGQGGDEQY | 0.76161385 |
| LLALLANL | ASSRGQGGDEQY | 0.7805889 |
| LISNASDALDKLR | ASSRGQGGDEQY | 0.39500818 |
| LINEAVP | ASSRGQGGDEQY | 0.82572633 |
| LGYAFSGL | ASSRGQGGDEQY | 0.5182569 |
| LDFVAVAL | ASSRGQGGDEQY | 0.81274915 |
| LAQPGAE | ASSRGQGGDEQY | 0.73660743 |
| ISNLGAVTGF | ASSRGQGGDEQY | 0.7646599 |
| HQEDPGRI | ASSRGQGGDEQY | 0.72632194 |
| GTPGEPTP | ASSRGQGGDEQY | 0.45484447 |
| GFFNSGS | ASSRGQGGDEQY | 0.51209617 |
| GAGGGGLTAGAN | ASSRGQGGDEQY | 0.5041536 |
| FVDALVNM | ASSRGQGGDEQY | 0.82387245 |
| FLTVAPEL | ASSRGQGGDEQY | 0.7838068 |
| FETLTAL | ASSRGQGGDEQY | 0.811802 |
| DVINPIEL | ASSRGQGGDEQY | 0.77190983 |
| APVSGAPV | ASSRGQGGDEQY | 0.5651392 |
| ALVGGLAW | ASSRGQGGDEQY | 0.80424416 |
| ALAAAAATAI | ASSRGQGGDEQY | 0.3767367 |
| AFGLGLLL | ASSRGQGGDEQY | 0.7418755 |
| AFFAAASL | ASSRGQGGDEQY | 0.690907 |
| AAAALPAFSPP | ASSRGQGGDEQY | 0.74076796 |
| VVLAPAL | ASSRGQGGDEQY | 0.6086214 |
| TIGHVDHGKTTL | ASSRGQGGDEQY | 0.7936976 |
| LISNASDALDK | ASSRGQGGDEQY | 0.7832804 |
| LISNASDAL | ASSRGQGGDEQY | 0.42003292 |
| ISNASDALDK | ASSRGQGGDEQY | 0.6844436 |
| NQAETLVYQTE | ASSRGQGGDEQY | 0.8132341 |
| TGGTGGTAGAG | ASSRGQGGDEQY | 0.73345965 |

| | | |
|---|---|---|
| TGSTAGGAGGA | ASSRGQGGDEQY | 0.42147017 |
| DPLAVPVAAGPV | ASSRGQGGDEQY | 0.7532385 |
| GGTGGTPGNGGN | ASSRGQGGDEQY | 0.62213457 |
| PAGLAGVDPATL | ASSRGQGGDEQY | 0.7797539 |
| GNNLVVEAPHAGASV | ASSRGQGGDEQY | 0.7608814 |
| KVTRIMSASTLEQPA | ASSRGQGGDEQY | 0.52323925 |
| LGFALPIALPRRTVV | ASSRGQGGDEQY | 0.8020829 |
| TRLGNLTARKLVVLV | ASSRGQGGDEQY | 0.313206 |
| WSQQRRLIYFPSAGP | ASSRGQGGDEQY | 0.652315 |
| GPNGNPDPMAAAVDIR | ASSRGQGGDEQY | 0.67245793 |
| TQRCVSVADLRAAASR | ASSRGQGGDEQY | 0.7383592 |
| QFVLDVAEHPVEVLTETV | ASSRGQGGDEQY | 0.7839564 |
| GGAGGTGGAGGNGGTQAGDG | ASSRGQGGDEQY | 0.49817967 |
| PLLATLRAIAVDVDAKPAQV | ASSRGQGGDEQY | 0.749463 |
| APKFRTGGYMIDFWGVGYQVAK | ASSRGQGGDEQY | 0.6391158 |
| GAGGAGGAGGAAGGPGGTGGTG | ASSRGQGGDEQY | 0.50839627 |
| IGALGLLATDVAPAGSVAFGVA | ASSRGQGGDEQY | 0.71166277 |
| AGGAGGIGGNGGAGGAGGVLMG | ASSRGQGGDEQY | 0.6162325 |
| ADELIGLL | ASSRGQGGDEQY | 0.40118796 |
| ADRALKA | ASSRGQGGDEQY | 0.79385257 |
| AGAVGQVGGAGGA | ASSRGQGGDEQY | 0.4969117 |
| AGGGGGLGGAP | ASSRGQGGDEQY | 0.7095871 |
| AGGLLYGNGGAG | ASSRGQGGDEQY | 0.8245573 |
| AGLESIGL | ASSRGQGGDEQY | 0.37426353 |
| AGLKTGII | ASSRGQGGDEQY | 0.8146622 |
| ASLAALLL | ASSRGQGGDEQY | 0.752679 |
| ASVINISSI | ASSRGQGGDEQY | 0.7653105 |
| ATPPGPA | ASSRGQGGDEQY | 0.67763174 |
| ATVASALV | ASSRGQGGDEQY | 0.5476754 |
| ATVASLIL | ASSRGQGGDEQY | 0.8240407 |
| AVNGPVLIRH | ASSRGQGGDEQY | 0.78127575 |
| DGAAGAVGI | ASSRGQGGDEQY | 0.62395346 |
| DPLLHGI | ASSRGQGGDEQY | 0.75953215 |
| ELGAALGAL | ASSRGQGGDEQY | 0.8019552 |
| ELGRTQAV | ASSRGQGGDEQY | 0.8161773 |
| FVAGGYLS | ASSRGQGGDEQY | 0.64223045 |
| GASGGAGGAGGW | ASSRGQGGDEQY | 0.35195178 |
| GGAGAGGDGGAGGI | ASSRGQGGDEQY | 0.68024695 |
| GTGHNGSQPGMGGNG | ASSRGQGGDEQY | 0.719558 |
| GYAAASASAS | ASSRGQGGDEQY | 0.59412616 |
| HQVDVAAVQQG | ASSRGQGGDEQY | 0.2864107 |
| IGAAIAHT | ASSRGQGGDEQY | 0.79068744 |
| IGMIEVAQL | ASSRGQGGDEQY | 0.84883773 |
| IRLAPEHTSAMD | ASSRGQGGDEQY | 0.5396467 |
| IYNVSVL | ASSRGQGGDEQY | 0.8448279 |
| KQGFLGM | ASSRGQGGDEQY | 0.39772403 |
| LESVITQL | ASSRGQGGDEQY | 0.7825895 |
| LGLGALVVV | ASSRGQGGDEQY | 0.7926614 |
| LIGNGGNGGSGGI | ASSRGQGGDEQY | 0.5423685 |
| LNAINAPTL | ASSRGQGGDEQY | 0.6790261 |
| LQLLNTGV | ASSRGQGGDEQY | 0.70742714 |

| | | |
|---|---|---|
| LQTSIEF | ASSRGQGGDEQY | 0.6173538 |
| MSFAVSEL | ASSRGQGGDEQY | 0.8143993 |
| NTGALGL | ASSRGQGGDEQY | 0.48973173 |
| PPGAGGAGGAGTT | ASSRGQGGDEQY | 0.7357092 |
| QLVGLGI | ASSRGQGGDEQY | 0.58447695 |
| RFLEQVHQL | ASSRGQGGDEQY | 0.50636786 |
| RGVGYVLR | ASSRGQGGDEQY | 0.79025865 |
| SATGVFAGVI | ASSRGQGGDEQY | 0.73978263 |
| SLPAITLP | ASSRGQGGDEQY | 0.73466504 |
| STALAFTV | ASSRGQGGDEQY | 0.8383223 |
| SVLTVGLGK | ASSRGQGGDEQY | 0.6385992 |
| TALAGAGSGI | ASSRGQGGDEQY | 0.83784497 |
| TGLPFTGI | ASSRGQGGDEQY | 0.825024 |
| VAEHVTV | ASSRGQGGDEQY | 0.7706089 |
| VDIVLNSL | ASSRGQGGDEQY | 0.6654756 |
| VLTGGGAVL | ASSRGQGGDEQY | 0.8116635 |
| VNVGLAYGI | ASSRGQGGDEQY | 0.52627707 |
| VSSAAVGAV | ASSRGQGGDEQY | 0.75113684 |
| VTSVYEAGV | ASSRGQGGDEQY | 0.82078266 |
| WVALDRGA | ASSRGQGGDEQY | 0.79530156 |
| YLVGSVVI | ASSRGQGGDEQY | 0.8947445 |
| GPPGTGKTLL | ASSRGQGGDEQY | 0.87455386 |
| GVGMTKF | ASSRGQGGDEQY | 0.86620164 |
| IGIGGSDL | ASSRGQGGDEQY | 0.8249837 |
| IGTLLAEL | ASSRGQGGDEQY | 0.69687796 |
| IYTSGTT | ASSRGQGGDEQY | 0.86568516 |
| LSFPAGV | ASSRGQGGDEQY | 0.77907914 |
| LYTSGTT | ASSRGQGGDEQY | 0.87983763 |
| VGAFPFGF | ASSRGQGGDEQY | 0.8911078 |
| VGAGTVEF | ASSRGQGGDEQY | 0.8229654 |
| VGKSTLLL | ASSRGQGGDEQY | 0.89243984 |
| AALAHAGGGIV | ASSRGQGGDEQY | 0.8947482 |
| AALVAVSLL | ASSRGQGGDEQY | 0.73749715 |
| AAVSRAVAAL | ASSRGQGGDEQY | 0.85269195 |
| AGGSSGIGGF | ASSRGQGGDEQY | 0.8070059 |
| AGLTVANANG | ASSRGQGGDEQY | 0.8507206 |
| AGLVGAVL | ASSRGQGGDEQY | 0.5030356 |
| ALSQAFTV | ASSRGQGGDEQY | 0.8847329 |
| ALTFSVSL | ASSRGQGGDEQY | 0.8145052 |
| ALTGAALVAL | ASSRGQGGDEQY | 0.3475209 |
| ASFSIGAGVL | ASSRGQGGDEQY | 0.8804454 |
| DGSVYKI | ASSRGQGGDEQY | 0.8535493 |
| EQLAEIV | ASSRGQGGDEQY | 0.85491467 |
| FAVSELTVM | ASSRGQGGDEQY | 0.8907176 |
| GAAPLAAGL | ASSRGQGGDEQY | 0.8015356 |
| GADGGDGGSGGAGG | ASSRGQGGDEQY | 0.80185044 |
| GFGLDTGGAGG | ASSRGQGGDEQY | 0.8831284 |
| GLASGGLL | ASSRGQGGDEQY | 0.8956573 |
| GRIAGAAVLL | ASSRGQGGDEQY | 0.77412 |
| GSAGSLNTSF | ASSRGQGGDEQY | 0.86883736 |
| GTGQIGI | ASSRGQGGDEQY | 0.8956413 |

| | | |
|---|---|---|
| GTLIALIL | ASSRGQGGDEQY | 0.8823425 |
| IAATMGFVVG | ASSRGQGGDEQY | 0.89685607 |
| IFAQALL | ASSRGQGGDEQY | 0.8999938 |
| IGAAVALGF | ASSRGQGGDEQY | 0.8191929 |
| IGLTPLAL | ASSRGQGGDEQY | 0.89041376 |
| ILGSNTSTL | ASSRGQGGDEQY | 0.8957189 |
| IVFSLTVL | ASSRGQGGDEQY | 0.84266937 |
| LASKATGF | ASSRGQGGDEQY | 0.85986066 |
| LASVALIL | ASSRGQGGDEQY | 0.8184364 |
| LFAVGLTL | ASSRGQGGDEQY | 0.8673768 |
| LFNGGIGTYI | ASSRGQGGDEQY | 0.6409439 |
| LTLGAGII | ASSRGQGGDEQY | 0.8704041 |
| LVAAALGDI | ASSRGQGGDEQY | 0.8961263 |
| NTAFVQLGI | ASSRGQGGDEQY | 0.85493547 |
| QLFSGAEL | ASSRGQGGDEQY | 0.76595044 |
| SAATFITL | ASSRGQGGDEQY | 0.7303121 |
| SNYGLANLGV | ASSRGQGGDEQY | 0.8669884 |
| SYGGAYAVM | ASSRGQGGDEQY | 0.8056674 |
| TTGVGGLTLG | ASSRGQGGDEQY | 0.6162255 |
| TTQTIYI | ASSRGQGGDEQY | 0.8358449 |
| VAAGAGLGAVI | ASSRGQGGDEQY | 0.8955702 |
| VAAVVTLL | ASSRGQGGDEQY | 0.89553535 |
| VGFLGGLI | ASSRGQGGDEQY | 0.72884583 |
| VGVIPILL | ASSRGQGGDEQY | 0.8972456 |
| VIFASSLI | ASSRGQGGDEQY | 0.8767952 |
| VKGGDGGAAAG | ASSRGQGGDEQY | 0.8930573 |
| VTKTFTNI | ASSRGQGGDEQY | 0.8695326 |
| VVAAAILL | ASSRGQGGDEQY | 0.7938944 |
| YAALDVEL | ASSRGQGGDEQY | 0.8829696 |
| LASVALLL | ASSRGQGGDEQY | 0.8465198 |
| LDAVHRV | ASSRGQGGDEQY | 0.8406707 |
| QGLSSSIF | ASSRGQGGDEQY | 0.8906703 |
| RGTFANIRL | ASSRGQGGDEQY | 0.86013556 |
| RQLWWGHRI | ASSRGQGGDEQY | 0.8114398 |
| YGPPGTGKTLL | ASSRGQGGDEQY | 0.8290804 |
| AALVGAPANVA | ASSRGQGGDEQY | 0.8958092 |
| AATFRRTE | ASSRGQGGDEQY | 0.7730986 |
| AGALGGGI | ASSRGQGGDEQY | 0.7412582 |
| AGGAGGAGGIGAG | ASSRGQGGDEQY | 0.8175316 |
| AGSGALGL | ASSRGQGGDEQY | 0.86737084 |
| AQIADQI | ASSRGQGGDEQY | 0.89434934 |
| ASAVAVTYL | ASSRGQGGDEQY | 0.8846415 |
| ASTVLGGTGG | ASSRGQGGDEQY | 0.83235586 |
| AVGGEPAK | ASSRGQGGDEQY | 0.8810166 |
| CEQGTFGGTC | ASSRGQGGDEQY | 0.8813368 |
| DEGKAPAG | ASSRGQGGDEQY | 0.8666947 |
| DTVVFPAL | ASSRGQGGDEQY | 0.88770807 |
| FEGITMAG | ASSRGQGGDEQY | 0.80536497 |
| FPLVVQD | ASSRGQGGDEQY | 0.8934525 |
| GAFTTPQ | ASSRGQGGDEQY | 0.87956303 |
| GFGGFGTAGGVGGA | ASSRGQGGDEQY | 0.890219 |

| | | |
|---|---|---|
| GFLAQFL | ASSRGQGGDEQY | 0.89460593 |
| GGAGGAGGNAGVA | ASSRGQGGDEQY | 0.87383366 |
| GGVGTAGAGGAGGAGG | ASSRGQGGDEQY | 0.8635322 |
| GNGGGGDAGNAGSGGN | ASSRGQGGDEQY | 0.6302522 |
| GTGGAGGAGNDAGS | ASSRGQGGDEQY | 0.8484468 |
| GTTNTGWFNS | ASSRGQGGDEQY | 0.8726315 |
| IFMGPLFTL | ASSRGQGGDEQY | 0.86987126 |
| ILAHAIEAI | ASSRGQGGDEQY | 0.88619554 |
| IRSAIREAL | ASSRGQGGDEQY | 0.8887192 |
| ITFAGGG | ASSRGQGGDEQY | 0.82831556 |
| ITGMSGHQEV | ASSRGQGGDEQY | 0.89589334 |
| IVAGLACTL | ASSRGQGGDEQY | 0.8921076 |
| IVGGGLAGA | ASSRGQGGDEQY | 0.7169663 |
| IVTGVLNI | ASSRGQGGDEQY | 0.8468305 |
| LAQQGCI | ASSRGQGGDEQY | 0.90007496 |
| LCLHVEN | ASSRGQGGDEQY | 0.8611474 |
| LDAISDGS | ASSRGQGGDEQY | 0.8476254 |
| LGGFAAML | ASSRGQGGDEQY | 0.8445689 |
| LGGKNPMI | ASSRGQGGDEQY | 0.89284533 |
| LGPKGVAAIL | ASSRGQGGDEQY | 0.83647 |
| LMVGVSI | ASSRGQGGDEQY | 0.8963125 |
| LVGAAFVL | ASSRGQGGDEQY | 0.40781927 |
| LVLAVAPL | ASSRGQGGDEQY | 0.8441872 |
| MAAIAVIDAA | ASSRGQGGDEQY | 0.81957054 |
| NNGINAELQ | ASSRGQGGDEQY | 0.44832647 |
| RGEPLVLVHG | ASSRGQGGDEQY | 0.88993144 |
| RGGAGGAAGLI | ASSRGQGGDEQY | 0.8401634 |
| RSYRMHPAV | ASSRGQGGDEQY | 0.884207 |
| SAAHLLAGQLA | ASSRGQGGDEQY | 0.85838896 |
| SAAVGSY | ASSRGQGGDEQY | 0.89029753 |
| SPHAVLHRI | ASSRGQGGDEQY | 0.8565129 |
| SQIAKVL | ASSRGQGGDEQY | 0.87035096 |
| TALGHLVP | ASSRGQGGDEQY | 0.8149946 |
| TAPTTSGAA | ASSRGQGGDEQY | 0.88271904 |
| TDGDDGNGGSPGT | ASSRGQGGDEQY | 0.8969334 |
| TDGLGEN | ASSRGQGGDEQY | 0.88983107 |
| TGGPPLPGQA | ASSRGQGGDEQY | 0.59217924 |
| TIPAITI | ASSRGQGGDEQY | 0.8489193 |
| TIPGFTI | ASSRGQGGDEQY | 0.85928 |
| TSIASGALA | ASSRGQGGDEQY | 0.52266115 |
| TVTLPGAAI | ASSRGQGGDEQY | 0.53793526 |
| VASGLSAALP | ASSRGQGGDEQY | 0.81997013 |
| VGAGVIGI | ASSRGQGGDEQY | 0.8408215 |
| VGPKRFGV | ASSRGQGGDEQY | 0.8873487 |
| VNVAFPD | ASSRGQGGDEQY | 0.8878354 |
| VSMTNSGVS | ASSRGQGGDEQY | 0.82863325 |
| VTDVLGDLSV | ASSRGQGGDEQY | 0.8837122 |
| VVVVGAGFAGL | ASSRGQGGDEQY | 0.86929524 |
| VYTVGTVV | ASSRGQGGDEQY | 0.1065416 |
| WTTFSGP | ASSRGQGGDEQY | 0.8330221 |
| YDELVEAF | ASSRGQGGDEQY | 0.74902844 |

| | | |
|---|---|---|
| YGTGGAGGNGG | ASSRGQGGDEQY | 0.8965788 |
| YTNVMAV | ASSRGQGGDEQY | 0.81736475 |
| FLEQVHQL | ASSRGQGGDEQY | 0.82272184 |
| GLSLLAVAV | ASSRGQGGDEQY | 0.85807085 |
| LEQVHQL | ASSRGQGGDEQY | 0.86035395 |
| LNAAGAI | ASSRGQGGDEQY | 0.8606539 |
| PAVHASL | ASSRGQGGDEQY | 0.79415 |
| TELEDDA | ASSRGQGGDEQY | 0.5227536 |
| TPPGPPN | ASSRGQGGDEQY | 0.88748103 |
| TPPPPPPP | ASSRGQGGDEQY | 0.78226566 |
| TQSPALI | ASSLVQGANTEVF | 0.79029083 |
| EFAAAFPQ | ASSLVQGANTEVF | 0.68850756 |
| LLSGNLDVL | ASSLVQGANTEVF | 0.85742074 |
| GPANLGNYNI | ASSLVQGANTEVF | 0.59532344 |
| TLGPTLP | ASSLVQGANTEVF | 0.89393616 |
| VYAYSSCCK | ASSLVQGANTEVF | 0.8239755 |
| APAPAPAPNI | ASSLVQGANTEVF | 0.8454664 |
| PGGPVPIP | ASSLVQGANTEVF | 0.9000287 |
| FLTVALEP | ASSLVQGANTEVF | 0.8535465 |
| SQYFGANL | ASSLVQGANTEVF | 0.8402641 |
| GGVKPAA | ASSLVQGANTEVF | 0.8147297 |
| GGTGGAGGAGGW | ASSLVQGANTEVF | 0.7837535 |
| GVGGVAGLGGAGG | ASSLVQGANTEVF | 0.8196988 |
| VLLVLATTI | ASSLVQGANTEVF | 0.82316697 |
| IGIFAAML | ASSLVQGANTEVF | 0.8978925 |
| AVGDLID | ASSLVQGANTEVF | 0.84620315 |
| NISPLPL | ASSLVQGANTEVF | 0.7237183 |
| VTAVFVAI | ASSLVQGANTEVF | 0.87379 |
| SGDDNDAAL | ASSLVQGANTEVF | 0.89517117 |
| AVAALAHS | ASSLVQGANTEVF | 0.8991277 |
| GGVTVMATP | ASSLVQGANTEVF | 0.8202733 |
| AGLAANVRD | ASSLVQGANTEVF | 0.8381838 |
| TNRPDLIDPALLRPG | ASSLVQGANTEVF | 0.8339901 |
| LANLGVTAL | ASSLVQGANTEVF | 0.84981436 |
| TGERIEL | ASSLVQGANTEVF | 0.829519 |
| NGEELAA | ASSLVQGANTEVF | 0.75794613 |
| VLSLVWG | ASSLVQGANTEVF | 0.88907075 |
| IGYALWL | ASSLVQGANTEVF | 0.8981738 |
| LSLLLAPL | ASSLVQGANTEVF | 0.85095257 |
| TLDEALK | ASSLVQGANTEVF | 0.57856226 |
| TDGAGTAEGG | ASSLVQGANTEVF | 0.84278834 |
| NAPGGAGGNGG | ASSLVQGANTEVF | 0.79249 |
| VSVFMGMGGNF | ASSLVQGANTEVF | 0.8226655 |
| LASYLAAL | ASSLVQGANTEVF | 0.86369383 |
| LVGGTPAL | ASSLVQGANTEVF | 0.69517374 |
| VRNADVL | ASSLVQGANTEVF | 0.8582653 |
| TCSSPFLVSAM | ASSLVQGANTEVF | 0.90131867 |
| VHITDAE | ASSLVQGANTEVF | 0.83299357 |
| EYAAPSF | ASSLVQGANTEVF | 0.87621444 |
| VVDAFHGGL | ASSLVQGANTEVF | 0.8949636 |
| ASVVDPQ | ASSLVQGANTEVF | 0.87997794 |

| | | |
|---|---|---|
| SYNEFGGLS | ASSLVQGANTEVF | 0.8953957 |
| DPDGSVY | ASSLVQGANTEVF | 0.88277423 |
| MVSTIHATE | ASSLVQGANTEVF | 0.8683559 |
| LAAGVADPV | ASSLVQGANTEVF | 0.8631835 |
| TKEQIAA | ASSLVQGANTEVF | 0.8886442 |
| SLASPAAL | ASSLVQGANTEVF | 0.85399765 |
| IGLVFYGV | ASSLVQGANTEVF | 0.85675323 |
| FLLDASNL | ASSLVQGANTEVF | 0.8287521 |
| YQVGNFP | ASSLVQGANTEVF | 0.88964915 |
| ESLLSQST | ASSLVQGANTEVF | 0.8617101 |
| LFTVGFAAVG | ASSLVQGANTEVF | 0.8880179 |
| ATPPTPT | ASSLVQGANTEVF | 0.49881336 |
| TDLATLG | ASSLVQGANTEVF | 0.77452046 |
| SAGGTGR | ASSLVQGANTEVF | 0.88483846 |
| GFSNSGS | ASSLVQGANTEVF | 0.20734733 |
| GGNGGSGGTSV | ASSLVQGANTEVF | 0.4951026 |
| GTSVATGGAGN | ASSLVQGANTEVF | 0.8714168 |
| NGGNGGSGGTS | ASSLVQGANTEVF | 0.8907455 |
| AVVKFPH | ASSLVQGANTEVF | 0.2903092 |
| TPPPPPP | ASSLVQGANTEVF | 0.7995301 |
| AAPAAAAAPA | ASSLVQGANTEVF | 0.6923121 |
| IALNEDL | ASSLVQGANTEVF | 0.8867916 |
| IVNVSSM | ASSLVQGANTEVF | 0.86557406 |
| TGIPPAPR | ASSLVQGANTEVF | 0.58243436 |
| FDLGGGTF | ASSLVQGANTEVF | 0.88695896 |
| FDLGGGTFDVS | ASSLVQGANTEVF | 0.58466244 |
| NEPTAAA | ASSLVQGANTEVF | 0.82967937 |
| NEPTAAALA | ASSLVQGANTEVF | 0.8895583 |
| GSIINISSI | ASSLVQGANTEVF | 0.89397675 |
| VAPGNAGTA | ASSLVQGANTEVF | 0.79364073 |
| TGGGASLE | ASSLVQGANTEVF | 0.88660085 |
| VAVYNHYKRI | ASSLVQGANTEVF | 0.8233763 |
| TDPAPATT | ASSLVQGANTEVF | 0.9023054 |
| TNRPDILDPALLRPG | ASSLVQGANTEVF | 0.84686655 |
| LTTTTTS | ASSLVQGANTEVF | 0.6758511 |
| PTGSGKTL | ASSLVQGANTEVF | 0.8897192 |
| TGSGKTL | ASSLVQGANTEVF | 0.87888896 |
| TIGHVDHGK | ASSLVQGANTEVF | 0.87890816 |
| TIGHVDHGKTT | ASSLVQGANTEVF | 0.8801429 |
| GHVDHGKTT | ASSLVQGANTEVF | 0.82348824 |
| AQFEHTLL | ASSLVQGANTEVF | 0.84336114 |
| IVNQDEL | ASSLVQGANTEVF | 0.890787 |
| TNGSQFF | ASSLVQGANTEVF | 0.88487697 |
| TGKGIVE | ASSLVQGANTEVF | 0.79841596 |
| LISNASDA | ASSLVQGANTEVF | 0.89680016 |
| AELRTLGELTKIAKAHGAQ | ASSLVQGANTEVF | 0.85599554 |
| APNAVAAAEPSW | ASSLVQGANTEVF | 0.82240915 |
| APQVAGAATAVL | ASSLVQGANTEVF | 0.9001659 |
| ARRDPAAAAAILPTDA | ASSLVQGANTEVF | 0.8727052 |
| DAAARDVLAEAGLAEYF | ASSLVQGANTEVF | 0.88598275 |
| DDQIVLSAKANLAKVEQ | ASSLVQGANTEVF | 0.8854037 |

| | | |
|---|---|---|
| DELLPDTGEDSDEDGH | ASSLVQGANTEVF | 0.88820666 |
| ELPEAPANRATVPQD | ASSLVQGANTEVF | 0.771988 |
| FGQSKIERVTGIG | ASSLVQGANTEVF | 0.87777114 |
| GAGGTGSTAGGAGGAGGAGGLY | ASSLVQGANTEVF | 0.87252605 |
| GHGGTGAAGGNGGTGGAGGSGI | ASSLVQGANTEVF | 0.88934404 |
| GNGGNGGAGGEGGPGGAGGAGA | ASSLVQGANTEVF | 0.8625345 |
| GSTAGNGGNGGNATNGTIAGQP | ASSLVQGANTEVF | 0.8385155 |
| GTGGDGGNGGNGGTGG | ASSLVQGANTEVF | 0.8812063 |
| GVRRPAGIPADAPV | ASSLVQGANTEVF | 0.89647555 |
| GYRQVTDAHLLTLARR | ASSLVQGANTEVF | 0.8672906 |
| IAKMGQDGHDRG | ASSLVQGANTEVF | 0.880532 |
| KAEPSIEIQYT | ASSLVQGANTEVF | 0.89894974 |
| KPVRIRMAADEDTVAVT | ASSLVQGANTEVF | 0.82576406 |
| MPPRIAGMRLLVIKPEP | ASSLVQGANTEVF | 0.8574505 |
| MKFYAVALFRSRGVPPTR | ASSLVQGANTEVF | 0.8903479 |
| NAGGFGILGAVAHSP | ASSLVQGANTEVF | 0.89684886 |
| NEGREITLAMAEM | ASSLVQGANTEVF | 0.8523002 |
| NELVEWSHPLGHRWRWE | ASSLVQGANTEVF | 0.8937405 |
| NLGAGGTGGNGGNPG | ASSLVQGANTEVF | 0.8406086 |
| NTEWPKWETVMAADPVD | ASSLVQGANTEVF | 0.881361 |
| PASAAPAETQP | ASSLVQGANTEVF | 0.81667006 |
| QQSSDGKNGVRAAEITGE | ASSLVQGANTEVF | 0.8637165 |
| RAPDAVTTSWWKE | ASSLVQGANTEVF | 0.80463403 |
| SGFGNVGAHTSGWFNQ | ASSLVQGANTEVF | 0.87629247 |
| SGVAGAVGQAASVGGLKV | ASSLVQGANTEVF | 0.88797045 |
| SNGGAGTGSAGGNGGT | ASSLVQGANTEVF | 0.88399947 |
| TGGAGGEGGAAG | ASSLVQGANTEVF | 0.4655713 |
| TGGDGGAAGAGGAPGHG | ASSLVQGANTEVF | 0.8891704 |
| TMPPLLAAIANAGRHGV | ASSLVQGANTEVF | 0.88389707 |
| TSSTAGNSGGA | ASSLVQGANTEVF | 0.897018 |
| VPATVPVGILC | ASSLVQGANTEVF | 0.8618572 |
| VQRPDSGGVVLTGRIS | ASSLVQGANTEVF | 0.88165355 |
| WTPNPVPATKGVPAGLVQAT | ASSLVQGANTEVF | 0.83542854 |
| YALKRALAKFSDRFDVVII | ASSLVQGANTEVF | 0.8653812 |
| YLIGSVIV | ASSLVQGANTEVF | 0.7522043 |
| YGQLVAVID | ASSLVQGANTEVF | 0.85079455 |
| VTVGGMTLPA | ASSLVQGANTEVF | 0.8444376 |
| VLALPAV | ASSLVQGANTEVF | 0.8168834 |
| VIPPTLNYET | ASSLVQGANTEVF | 0.88088983 |
| VGMGGMHPSSG | ASSLVQGANTEVF | 0.89441895 |
| VGLAGGAATA | ASSLVQGANTEVF | 0.89175034 |
| VGGAGGAGGSGP | ASSLVQGANTEVF | 0.8908252 |
| VFAAFMV | ASSLVQGANTEVF | 0.89451104 |
| TLYDKLR | ASSLVQGANTEVF | 0.8949879 |
| TLAVFTVL | ASSLVQGANTEVF | 0.8325932 |
| TGSVQPARV | ASSLVQGANTEVF | 0.8668771 |
| TGGGAGSGGT | ASSLVQGANTEVF | 0.7384096 |
| SHLETSL | ASSLVQGANTEVF | 0.8446372 |
| SFTTGMT | ASSLVQGANTEVF | 0.8335604 |
| QVEVAKRMGVRQPTVS | ASSLVQGANTEVF | 0.8750529 |
| QGSGGPALAM | ASSLVQGANTEVF | 0.16442281 |

| | | |
|---|---|---|
| MLAPSNP | ASSLVQGANTEVF | 0.85263014 |
| LVNNAAI | ASSLVQGANTEVF | 0.8988759 |
| LTGEINTLA | ASSLVQGANTEVF | 0.81232536 |
| LSIFGPV | ASSLVQGANTEVF | 0.88255215 |
| LLALLANL | ASSLVQGANTEVF | 0.8747914 |
| LISNASDALDKLR | ASSLVQGANTEVF | 0.8192152 |
| LINEAVP | ASSLVQGANTEVF | 0.88337183 |
| LGYAFSGL | ASSLVQGANTEVF | 0.8158734 |
| LDFVAVAL | ASSLVQGANTEVF | 0.8855689 |
| LAQPGAE | ASSLVQGANTEVF | 0.89843774 |
| ISNLGAVTGF | ASSLVQGANTEVF | 0.87195265 |
| HQEDPGRI | ASSLVQGANTEVF | 0.87035835 |
| GTPGEPTP | ASSLVQGANTEVF | 0.86566234 |
| GFFNSGS | ASSLVQGANTEVF | 0.6627292 |
| GAGGGGLTAGAN | ASSLVQGANTEVF | 0.87540215 |
| FVDALVNM | ASSLVQGANTEVF | 0.73363525 |
| FLTVAPEL | ASSLVQGANTEVF | 0.85882556 |
| FETLTAL | ASSLVQGANTEVF | 0.7896622 |
| DVINPIEL | ASSLVQGANTEVF | 0.8775124 |
| APVSGAPV | ASSLVQGANTEVF | 0.8655653 |
| ALVGGLAW | ASSLVQGANTEVF | 0.88078535 |
| ALAAAAATAI | ASSLVQGANTEVF | 0.7785959 |
| AFGLGLLL | ASSLVQGANTEVF | 0.86109674 |
| AFFAAASL | ASSLVQGANTEVF | 0.8347589 |
| AAAALPAFSPP | ASSLVQGANTEVF | 0.72578907 |
| VVLAPAL | ASSLVQGANTEVF | 0.8901123 |
| TIGHVDHGKTTL | ASSLVQGANTEVF | 0.8658209 |
| LISNASDALDK | ASSLVQGANTEVF | 0.84192157 |
| LISNASDAL | ASSLVQGANTEVF | 0.8944255 |
| ISNASDALDK | ASSLVQGANTEVF | 0.8801321 |
| NQAETLVYQTE | ASSLVQGANTEVF | 0.5784064 |
| TGGTGGTAGAG | ASSLVQGANTEVF | 0.8464511 |
| TGSTAGGAGGA | ASSLVQGANTEVF | 0.86098313 |
| DPLAVPVAAGPV | ASSLVQGANTEVF | 0.8600651 |
| GGTGGTPGNGGN | ASSLVQGANTEVF | 0.2358769 |
| PAGLAGVDPATL | ASSLVQGANTEVF | 0.8821851 |
| GNNLVVEAPHAGASV | ASSLVQGANTEVF | 0.8165719 |
| KVTRIMSASTLEQPA | ASSLVQGANTEVF | 0.7822034 |
| LGFALPIALPRRTVV | ASSLVQGANTEVF | 0.8278736 |
| TRLGNLTARKLVVLV | ASSLVQGANTEVF | 0.8907965 |
| WSQQRRLIYFPSAGP | ASSLVQGANTEVF | 0.8483181 |
| GPNGNPDPMAAAVDIR | ASSLVQGANTEVF | 0.8648199 |
| TQRCVSVADLRAAASR | ASSLVQGANTEVF | 0.8880847 |
| QFVLDVAEHPVEVLTETV | ASSLVQGANTEVF | 0.8822367 |
| GGAGGTGGAGGNGGTQAGDG | ASSLVQGANTEVF | 0.89495707 |
| PLLATLRAIAVDVDAKPAQV | ASSLVQGANTEVF | 0.88420975 |
| APKFRTGGYMIDFWGVGYQVAK | ASSLVQGANTEVF | 0.8685957 |
| GAGGAGGAGGAAGGPGGTGGTG | ASSLVQGANTEVF | 0.8857131 |
| IGALGLLATDVAPAGSVAFGVA | ASSLVQGANTEVF | 0.8504304 |
| AGGAGGIGGNGGAGGAGGVLMG | ASSLVQGANTEVF | 0.7418008 |
| ADELIGLL | ASSLVQGANTEVF | 0.6925467 |

| | | |
|---|---|---|
| ADRALKA | ASSLVQGANTEVF | 0.81802535 |
| AGAVGQVGGAGGA | ASSLVQGANTEVF | 0.67709994 |
| AGGGGGLGGAP | ASSLVQGANTEVF | 0.8739053 |
| AGGLLYGNGGAG | ASSLVQGANTEVF | 0.8971552 |
| AGLESIGL | ASSLVQGANTEVF | 0.8541588 |
| AGLKTGII | ASSLVQGANTEVF | 0.74671423 |
| ASLAALLL | ASSLVQGANTEVF | 0.84763503 |
| ASVINISSI | ASSLVQGANTEVF | 0.846001 |
| ATPPGPA | ASSLVQGANTEVF | 0.8863841 |
| ATVASALV | ASSLVQGANTEVF | 0.88228 |
| ATVASLIL | ASSLVQGANTEVF | 0.896546 |
| AVNGPVLIRH | ASSLVQGANTEVF | 0.8900157 |
| DGAAGAVGI | ASSLVQGANTEVF | 0.8302093 |
| DPLLHGI | ASSLVQGANTEVF | 0.88734543 |
| ELGAALGAL | ASSLVQGANTEVF | 0.82900345 |
| ELGRTQAV | ASSLVQGANTEVF | 0.8668381 |
| FVAGGYLS | ASSLVQGANTEVF | 0.8691201 |
| GASGGAGGAGGW | ASSLVQGANTEVF | 0.8733479 |
| GGAGAGGDGGAGGI | ASSLVQGANTEVF | 0.8610724 |
| GTGHNGSQPGMGGNG | ASSLVQGANTEVF | 0.8567632 |
| GYAAASASAS | ASSLVQGANTEVF | 0.55165875 |
| HQVDVAAVQQG | ASSLVQGANTEVF | 0.8805007 |
| IGAAIAHT | ASSLVQGANTEVF | 0.7023176 |
| IGMIEVAQL | ASSLVQGANTEVF | 0.66354465 |
| IRLAPEHTSAMD | ASSLVQGANTEVF | 0.8209669 |
| IYNVSVL | ASSLVQGANTEVF | 0.89866483 |
| KQGFLGM | ASSLVQGANTEVF | 0.8986554 |
| LESVITQL | ASSLVQGANTEVF | 0.81221175 |
| LGLGALVVV | ASSLVQGANTEVF | 0.798922 |
| LIGNGGNGGSGGI | ASSLVQGANTEVF | 0.86354387 |
| LNAINAPTL | ASSLVQGANTEVF | 0.7124458 |
| LQLLNTGV | ASSLVQGANTEVF | 0.877669 |
| LQTSIEF | ASSLVQGANTEVF | 0.85013336 |
| MSFAVSEL | ASSLVQGANTEVF | 0.8461576 |
| NTGALGL | ASSLVQGANTEVF | 0.80548316 |
| PPGAGGAGGAGTT | ASSLVQGANTEVF | 0.8764075 |
| QLVGLGI | ASSLVQGANTEVF | 0.829623 |
| RFLEQVHQL | ASSLVQGANTEVF | 0.8892756 |
| RGVGYVLR | ASSLVQGANTEVF | 0.5511465 |
| SATGVFAGVI | ASSLVQGANTEVF | 0.8844031 |
| SLPAITLP | ASSLVQGANTEVF | 0.79515195 |
| STALAFTV | ASSLVQGANTEVF | 0.81485957 |
| SVLTVGLGK | ASSLVQGANTEVF | 0.4369558 |
| TALAGAGSGI | ASSLVQGANTEVF | 0.8666396 |
| TGLPFTGI | ASSLVQGANTEVF | 0.88236725 |
| VAEHVTV | ASSLVQGANTEVF | 0.89233005 |
| VDIVLNSL | ASSLVQGANTEVF | 0.8303045 |
| VLTGGGAVL | ASSLVQGANTEVF | 0.88538057 |
| VNVGLAYGI | ASSLVQGANTEVF | 0.89350677 |
| VSSAAVGAV | ASSLVQGANTEVF | 0.83996165 |
| VTSVYEAGV | ASSLVQGANTEVF | 0.58839726 |

| | | |
|---|---|---|
| WVALDRGA | ASSLVQGANTEVF | 0.8771169 |
| YLVGSVVI | ASSLVQGANTEVF | 0.8377162 |
| GPPGTGKTLL | ASSLVQGANTEVF | 0.89074993 |
| GVGMTKF | ASSLVQGANTEVF | 0.88159424 |
| IGIGGSDL | ASSLVQGANTEVF | 0.75398016 |
| IGTLLAEL | ASSLVQGANTEVF | 0.88606155 |
| IYTSGTT | ASSLVQGANTEVF | 0.8853572 |
| LSFPAGV | ASSLVQGANTEVF | 0.76877075 |
| LYTSGTT | ASSLVQGANTEVF | 0.82539904 |
| VGAFPFGF | ASSLVQGANTEVF | 0.8288317 |
| VGAGTVEF | ASSLVQGANTEVF | 0.6798529 |
| VGKSTLLL | ASSLVQGANTEVF | 0.88141525 |
| AALAHAGGGIV | ASSLVQGANTEVF | 0.82146573 |
| AALVAVSLL | ASSLVQGANTEVF | 0.84596884 |
| AAVSRAVAAL | ASSLVQGANTEVF | 0.8565208 |
| AGGSSGIGGF | ASSLVQGANTEVF | 0.83937687 |
| AGLTVANANG | ASSLVQGANTEVF | 0.8214609 |
| AGLVGAVL | ASSLVQGANTEVF | 0.843577 |
| ALSQAFTV | ASSLVQGANTEVF | 0.87157 |
| ALTFSVSL | ASSLVQGANTEVF | 0.88134813 |
| ALTGAALVAL | ASSLVQGANTEVF | 0.8501445 |
| ASFSIGAGVL | ASSLVQGANTEVF | 0.8705224 |
| DGSVYKI | ASSLVQGANTEVF | 0.8975019 |
| EQLAEIV | ASSLVQGANTEVF | 0.74012476 |
| FAVSELTVM | ASSLVQGANTEVF | 0.7845713 |
| GAAPLAAGL | ASSLVQGANTEVF | 0.88285846 |
| GADGGDGGSGGAGG | ASSLVQGANTEVF | 0.76405525 |
| GFGLDTGGAGG | ASSLVQGANTEVF | 0.88930374 |
| GLASGGLL | ASSLVQGANTEVF | 0.86158967 |
| GRIAGAAVLL | ASSLVQGANTEVF | 0.8744519 |
| GSAGSLNTSF | ASSLVQGANTEVF | 0.86088204 |
| GTGQIGI | ASSLVQGANTEVF | 0.89984506 |
| GTLIALIL | ASSLVQGANTEVF | 0.88809097 |
| IAATMGFVVG | ASSLVQGANTEVF | 0.8514888 |
| IFAQALL | ASSLVQGANTEVF | 0.87590796 |
| IGAAVALGF | ASSLVQGANTEVF | 0.7525054 |
| IGLTPLAL | ASSLVQGANTEVF | 0.88752395 |
| ILGSNTSTL | ASSLVQGANTEVF | 0.88690895 |
| IVFSLTVL | ASSLVQGANTEVF | 0.787515 |
| LASKATGF | ASSLVQGANTEVF | 0.8432234 |
| LASVALIL | ASSLVQGANTEVF | 0.89808595 |
| LFAVGLTL | ASSLVQGANTEVF | 0.8953395 |
| LFNGGIGTYI | ASSLVQGANTEVF | 0.6908377 |
| LTLGAGII | ASSLVQGANTEVF | 0.834553 |
| LVAAALGDI | ASSLVQGANTEVF | 0.72619534 |
| NTAFVQLGI | ASSLVQGANTEVF | 0.8212459 |
| QLFSGAEL | ASSLVQGANTEVF | 0.6540384 |
| SAATFITL | ASSLVQGANTEVF | 0.88295376 |
| SNYGLANLGV | ASSLVQGANTEVF | 0.8446634 |
| SYGGAYAVM | ASSLVQGANTEVF | 0.873983 |
| TTGVGGLTLG | ASSLVQGANTEVF | 0.8745018 |

| | | |
|---|---|---|
| TTQTIYI | ASSLVQGANTEVF | 0.8959194 |
| VAAGAGLGAVI | ASSLVQGANTEVF | 0.82399505 |
| VAAVVTLL | ASSLVQGANTEVF | 0.84719294 |
| VGFLGGLI | ASSLVQGANTEVF | 0.6730982 |
| VGVIPILL | ASSLVQGANTEVF | 0.75709426 |
| VIFASSLI | ASSLVQGANTEVF | 0.795101 |
| VKGGDGGAAAG | ASSLVQGANTEVF | 0.84067374 |
| VTKTFTNI | ASSLVQGANTEVF | 0.6354009 |
| VVAAAILL | ASSLVQGANTEVF | 0.89591855 |
| YAALDVEL | ASSLVQGANTEVF | 0.8623589 |
| LASVALLL | ASSLVQGANTEVF | 0.88598245 |
| LDAVHRV | ASSLVQGANTEVF | 0.8312669 |
| QGLSSSIF | ASSLVQGANTEVF | 0.8299228 |
| RGTFANIRL | ASSLVQGANTEVF | 0.8280988 |
| RQLWWGHRI | ASSLVQGANTEVF | 0.885841 |
| YGPPGTGKTLL | ASSLVQGANTEVF | 0.79004216 |
| AALVGAPANVA | ASSLVQGANTEVF | 0.3595694 |
| AATFRRTE | ASSLVQGANTEVF | 0.8186652 |
| AGALGGGI | ASSLVQGANTEVF | 0.70431006 |
| AGGAGGAGGIGAG | ASSLVQGANTEVF | 0.8957374 |
| AGSGALGL | ASSLVQGANTEVF | 0.89091504 |
| AQIADQI | ASSLVQGANTEVF | 0.8687304 |
| ASAVAVTYL | ASSLVQGANTEVF | 0.89083874 |
| ASTVLGGTGG | ASSLVQGANTEVF | 0.22901271 |
| AVGGEPAK | ASSLVQGANTEVF | 0.83594203 |
| CEQGTFGGTC | ASSLVQGANTEVF | 0.8964772 |
| DEGKAPAG | ASSLVQGANTEVF | 0.880546 |
| DTVVFPAL | ASSLVQGANTEVF | 0.85615957 |
| FEGITMAG | ASSLVQGANTEVF | 0.8940817 |
| FPLVVQD | ASSLVQGANTEVF | 0.799737 |
| GAFTTPQ | ASSLVQGANTEVF | 0.86192304 |
| GFGGFGTAGGVGGA | ASSLVQGANTEVF | 0.8751893 |
| GFLAQFL | ASSLVQGANTEVF | 0.8859625 |
| GGAGGAGGNAGVA | ASSLVQGANTEVF | 0.8764509 |
| GGVGTAGAGGAGGAGG | ASSLVQGANTEVF | 0.72246456 |
| GNGGGGDAGNAGSGGN | ASSLVQGANTEVF | 0.8767444 |
| GTGGAGGAGNDAGS | ASSLVQGANTEVF | 0.9005884 |
| GTTNTGWFNS | ASSLVQGANTEVF | 0.849564 |
| IFMGPLFTL | ASSLVQGANTEVF | 0.885957 |
| ILAHAIEAI | ASSLVQGANTEVF | 0.8794438 |
| IRSAIREAL | ASSLVQGANTEVF | 0.8422515 |
| ITFAGGG | ASSLVQGANTEVF | 0.8063663 |
| ITGMSGHQEV | ASSLVQGANTEVF | 0.8178471 |
| IVAGLACTL | ASSLVQGANTEVF | 0.85525304 |
| IVGGGLAGA | ASSLVQGANTEVF | 0.77942944 |
| IVTGVLNI | ASSLVQGANTEVF | 0.86351883 |
| LAQQGCI | ASSLVQGANTEVF | 0.8436879 |
| LCLHVEN | ASSLVQGANTEVF | 0.9034084 |
| LDAISDGS | ASSLVQGANTEVF | 0.87917936 |
| LGGFAAML | ASSLVQGANTEVF | 0.8841667 |
| LGGKNPMI | ASSLVQGANTEVF | 0.7963684 |

| | | |
|---|---|---|
| LGPKGVAAIL | ASSLVQGANTEVF | 0.87503976 |
| LMVGVSI | ASSLVQGANTEVF | 0.8719269 |
| LVGAAFVL | ASSLVQGANTEVF | 0.58169734 |
| LVLAVAPL | ASSLVQGANTEVF | 0.89424425 |
| MAAIAVIDAA | ASSLVQGANTEVF | 0.7911582 |
| NNGINAELQ | ASSLVQGANTEVF | 0.8508898 |
| RGEPLVLVHG | ASSLVQGANTEVF | 0.8793333 |
| RGGAGGAAGLI | ASSLVQGANTEVF | 0.88331795 |
| RSYRMHPAV | ASSLVQGANTEVF | 0.8526816 |
| SAAHLLAGQLA | ASSLVQGANTEVF | 0.856358 |
| SAAVGSY | ASSLVQGANTEVF | 0.8099959 |
| SPHAVLHRI | ASSLVQGANTEVF | 0.8862032 |
| SQIAKVL | ASSLVQGANTEVF | 0.8949123 |
| TALGHLVP | ASSLVQGANTEVF | 0.84504884 |
| TAPTTSGAA | ASSLVQGANTEVF | 0.8976009 |
| TDGDDGNGGSPGT | ASSLVQGANTEVF | 0.8834621 |
| TDGLGEN | ASSLVQGANTEVF | 0.57794136 |
| TGGPPLPGQA | ASSLVQGANTEVF | 0.8649862 |
| TIPAITI | ASSLVQGANTEVF | 0.8934795 |
| TIPGFTI | ASSLVQGANTEVF | 0.8903613 |
| TSIASGALA | ASSLVQGANTEVF | 0.8949205 |
| TVTLPGAAI | ASSLVQGANTEVF | 0.8880404 |
| VASGLSAALP | ASSLVQGANTEVF | 0.82725465 |
| VGAGVIGI | ASSLVQGANTEVF | 0.8945186 |
| VGPKRFGV | ASSLVQGANTEVF | 0.9029576 |
| VNVAFPD | ASSLVQGANTEVF | 0.88919455 |
| VSMTNSGVS | ASSLVQGANTEVF | 0.817934 |
| VTDVLGDLSV | ASSLVQGANTEVF | 0.8086562 |
| VVVVGAGFAGL | ASSLVQGANTEVF | 0.87826574 |
| VYTVGTVV | ASSLVQGANTEVF | 0.8843173 |
| WTTFSGP | ASSLVQGANTEVF | 0.7579186 |
| YDELVEAF | ASSLVQGANTEVF | 0.8727133 |
| YGTGGAGGNGG | ASSLVQGANTEVF | 0.8705338 |
| YTNVMAV | ASSLVQGANTEVF | 0.6521865 |
| FLEQVHQL | ASSLVQGANTEVF | 0.87253606 |
| GLSLLAVAV | ASSLVQGANTEVF | 0.89515615 |
| LEQVHQL | ASSLVQGANTEVF | 0.86665463 |
| LNAAGAI | ASSLVQGANTEVF | 0.7821416 |
| PAVHASL | ASSLVQGANTEVF | 0.8942809 |
| TELEDDA | ASSLVQGANTEVF | 0.9013448 |
| TPPGPPN | ASSLVQGANTEVF | 0.89632124 |
| TPPPPPPP | ASSLVQGANTEVF | 0.63183904 |
| TQSPALI | ASGGTGGARAEQF | 0.3197829 |
| EFAAAFPQ | ASGGTGGARAEQF | 0.8523474 |
| LLSGNLDVL | ASGGTGGARAEQF | 0.87579894 |
| GPANLGNYNI | ASGGTGGARAEQF | 0.85051215 |
| TLGPTLP | ASGGTGGARAEQF | 0.8271066 |
| VYAYSSCCK | ASGGTGGARAEQF | 0.8748398 |
| APAPAPAPNI | ASGGTGGARAEQF | 0.89117485 |
| PGGPVPIP | ASGGTGGARAEQF | 0.81905115 |
| FLTVALEP | ASGGTGGARAEQF | 0.76978415 |

| | | |
|---|---|---|
| SQYFGANL | ASGGTGGARAEQF | 0.8844153 |
| GGVKPAA | ASGGTGGARAEQF | 0.88947284 |
| GGTGGAGGAGGW | ASGGTGGARAEQF | 0.77605766 |
| GVGGVAGLGGAGG | ASGGTGGARAEQF | 0.85081244 |
| VLLVLATTI | ASGGTGGARAEQF | 0.8571671 |
| IGIFAAML | ASGGTGGARAEQF | 0.8377757 |
| AVGDLID | ASGGTGGARAEQF | 0.86100954 |
| NISPLPL | ASGGTGGARAEQF | 0.72777665 |
| VTAVFVAI | ASGGTGGARAEQF | 0.8909149 |
| SGDDNDAAL | ASGGTGGARAEQF | 0.71868086 |
| AVAALAHS | ASGGTGGARAEQF | 0.88949835 |
| GGVTVMATP | ASGGTGGARAEQF | 0.8771862 |
| AGLAANVRD | ASGGTGGARAEQF | 0.8807384 |
| TNRPDLIDPALLRPG | ASGGTGGARAEQF | 0.84674495 |
| LANLGVTAL | ASGGTGGARAEQF | 0.8092183 |
| TGERIEL | ASGGTGGARAEQF | 0.751438 |
| NGEELAA | ASGGTGGARAEQF | 0.6993672 |
| VLSLVWG | ASGGTGGARAEQF | 0.8953641 |
| IGYALWL | ASGGTGGARAEQF | 0.8397647 |
| LSLLLAPL | ASGGTGGARAEQF | 0.8910661 |
| TLDEALK | ASGGTGGARAEQF | 0.88313174 |
| TDGAGTAEGG | ASGGTGGARAEQF | 0.81299347 |
| NAPGGAGGNGG | ASGGTGGARAEQF | 0.8810843 |
| VSVFMGMGGNF | ASGGTGGARAEQF | 0.80275935 |
| LASYLAAL | ASGGTGGARAEQF | 0.83689773 |
| LVGGTPAL | ASGGTGGARAEQF | 0.8630418 |
| VRNADVL | ASGGTGGARAEQF | 0.881757 |
| TCSSPFLVSAM | ASGGTGGARAEQF | 0.81442326 |
| VHITDAE | ASGGTGGARAEQF | 0.87861085 |
| EYAAPSF | ASGGTGGARAEQF | 0.8732409 |
| VVDAFHGGL | ASGGTGGARAEQF | 0.77868545 |
| ASVVDPQ | ASGGTGGARAEQF | 0.8641031 |
| SYNEFGGLS | ASGGTGGARAEQF | 0.8903602 |
| DPDGSVY | ASGGTGGARAEQF | 0.8754754 |
| MVSTIHATE | ASGGTGGARAEQF | 0.76922864 |
| LAAGVADPV | ASGGTGGARAEQF | 0.88333 |
| TKEQIAA | ASGGTGGARAEQF | 0.8105923 |
| SLASPAAL | ASGGTGGARAEQF | 0.8721236 |
| IGLVFYGV | ASGGTGGARAEQF | 0.8713985 |
| FLLDASNL | ASGGTGGARAEQF | 0.8537303 |
| YQVGNFP | ASGGTGGARAEQF | 0.89207566 |
| ESLLSQST | ASGGTGGARAEQF | 0.61272544 |
| LFTVGFAAVG | ASGGTGGARAEQF | 0.82637763 |
| ATPPTPT | ASGGTGGARAEQF | 0.8471007 |
| TDLATLG | ASGGTGGARAEQF | 0.86800987 |
| SAGGTGR | ASGGTGGARAEQF | 0.8885987 |
| GFSNSGS | ASGGTGGARAEQF | 0.7796407 |
| GGNGGSGGTSV | ASGGTGGARAEQF | 0.86592734 |
| GTSVATGGAGN | ASGGTGGARAEQF | 0.845335 |
| NGGNGGSGGTS | ASGGTGGARAEQF | 0.79547614 |
| AVVKFPH | ASGGTGGARAEQF | 0.8577597 |

| | | |
|---|---|---|
| TPPPPPP | ASGGTGGARAEQF | 0.841362 |
| AAPAAAAAPA | ASGGTGGARAEQF | 0.7766316 |
| IALNEDL | ASGGTGGARAEQF | 0.88357556 |
| IVNVSSM | ASGGTGGARAEQF | 0.83838046 |
| TGIPPAPR | ASGGTGGARAEQF | 0.8757026 |
| FDLGGGTF | ASGGTGGARAEQF | 0.8919821 |
| FDLGGGTFDVS | ASGGTGGARAEQF | 0.8018191 |
| NEPTAAA | ASGGTGGARAEQF | 0.8882929 |
| NEPTAAALA | ASGGTGGARAEQF | 0.87063193 |
| GSIINISSI | ASGGTGGARAEQF | 0.87949646 |
| VAPGNAGTA | ASGGTGGARAEQF | 0.8437536 |
| TGGGASLE | ASGGTGGARAEQF | 0.7875105 |
| VAVYNHYKRI | ASGGTGGARAEQF | 0.8938588 |
| TDPAPATT | ASGGTGGARAEQF | 0.8888536 |
| TNRPDILDPALLRPG | ASGGTGGARAEQF | 0.89138097 |
| LTTTTTS | ASGGTGGARAEQF | 0.88375676 |
| PTGSGKTL | ASGGTGGARAEQF | 0.8798382 |
| TGSGKTL | ASGGTGGARAEQF | 0.88953805 |
| TIGHVDHGK | ASGGTGGARAEQF | 0.82118857 |
| TIGHVDHGKTT | ASGGTGGARAEQF | 0.6613138 |
| GHVDHGKTT | ASGGTGGARAEQF | 0.85132384 |
| AQFEHTLL | ASGGTGGARAEQF | 0.8218207 |
| IVNQDEL | ASGGTGGARAEQF | 0.834992 |
| TNGSQFF | ASGGTGGARAEQF | 0.6234088 |
| TGKGIVE | ASGGTGGARAEQF | 0.88827866 |
| LISNASDA | ASGGTGGARAEQF | 0.9026808 |
| AELRTLGELTKIAKAHGAQ | ASGGTGGARAEQF | 0.83618563 |
| APNAVAAAEPSW | ASGGTGGARAEQF | 0.8992051 |
| APQVAGAATAVL | ASGGTGGARAEQF | 0.6983777 |
| ARRDPAAAAAILPTDA | ASGGTGGARAEQF | 0.88987446 |
| DAAARDVLAEAGLAEYF | ASGGTGGARAEQF | 0.8918917 |
| DDQIVLSAKANLAKVEQ | ASGGTGGARAEQF | 0.78376216 |
| DELLPDTGEDSDEDGH | ASGGTGGARAEQF | 0.88231725 |
| ELPEAPANRATVPQD | ASGGTGGARAEQF | 0.81373525 |
| FGQSKIERVTGIG | ASGGTGGARAEQF | 0.8411087 |
| GAGGTGSTAGGAGGAGGAGGLY | ASGGTGGARAEQF | 0.89608926 |
| GHGGTGAAGGNGGTGGAGGSGI | ASGGTGGARAEQF | 0.83755517 |
| GNGGNGGAGGEGGPGGAGGAGA | ASGGTGGARAEQF | 0.8828465 |
| GSTAGNGGNGGNATNGTIAGQP | ASGGTGGARAEQF | 0.8571171 |
| GTGGDGGNGGNGGTGG | ASGGTGGARAEQF | 0.80058 |
| GVRRPAGIPADAPV | ASGGTGGARAEQF | 0.8794515 |
| GYRQVTDAHLLTLARR | ASGGTGGARAEQF | 0.89244914 |
| IAKMGQDGHDRG | ASGGTGGARAEQF | 0.8247209 |
| KAEPSIEIQYT | ASGGTGGARAEQF | 0.9019309 |
| KPVRIRMAADEDTVAVT | ASGGTGGARAEQF | 0.8038054 |
| MPPRIAGMRLLVIKPEP | ASGGTGGARAEQF | 0.8980757 |
| MKFYAVALFRSRGVPPTR | ASGGTGGARAEQF | 0.88790184 |
| NAGGFGILGAVAHSP | ASGGTGGARAEQF | 0.862936 |
| NEGREITLAMAEM | ASGGTGGARAEQF | 0.84238315 |
| NELVEWSHPLGHRWRWE | ASGGTGGARAEQF | 0.89351916 |
| NLGAGGTGGNGGNPG | ASGGTGGARAEQF | 0.8548244 |

| | | |
|---|---|---|
| NTEWPKWETVMAADPVD | ASGGTGGARAEQF | 0.8875829 |
| PASAAPAETQP | ASGGTGGARAEQF | 0.8957583 |
| QQSSDGKNGVRAAEITGE | ASGGTGGARAEQF | 0.5644964 |
| RAPDAVTTSWWKE | ASGGTGGARAEQF | 0.8742664 |
| SGFGNVGAHTSGWFNQ | ASGGTGGARAEQF | 0.79328334 |
| SGVAGAVGQAASVGGLKV | ASGGTGGARAEQF | 0.74080145 |
| SNGGAGTGSAGGNGGT | ASGGTGGARAEQF | 0.6295599 |
| TGGAGGEGGAAG | ASGGTGGARAEQF | 0.42783147 |
| TGGDGGAAGAGGAPGHG | ASGGTGGARAEQF | 0.69963264 |
| TMPPLLAAIANAGRHGV | ASGGTGGARAEQF | 0.5477302 |
| TSSTAGNSGGA | ASGGTGGARAEQF | 0.8452096 |
| VPATVPVGILC | ASGGTGGARAEQF | 0.85605454 |
| VQRPDSGGVVLTGRIS | ASGGTGGARAEQF | 0.6090294 |
| WTPNPVPATKGVPAGLVQAT | ASGGTGGARAEQF | 0.8795165 |
| YALKRALAKFSDRFDVVII | ASGGTGGARAEQF | 0.83196354 |
| YLIGSVIV | ASGGTGGARAEQF | 0.46483582 |
| YGQLVAVID | ASGGTGGARAEQF | 0.71975887 |
| VTVGGMTLPA | ASGGTGGARAEQF | 0.6192298 |
| VLALPAV | ASGGTGGARAEQF | 0.70054317 |
| VIPPTLNYET | ASGGTGGARAEQF | 0.3403683 |
| VGMGGMHPSSG | ASGGTGGARAEQF | 0.82602966 |
| VGLAGGAATA | ASGGTGGARAEQF | 0.62718636 |
| VGGAGGAGGSGP | ASGGTGGARAEQF | 0.18172568 |
| VFAAFMV | ASGGTGGARAEQF | 0.7985879 |
| TLYDKLR | ASGGTGGARAEQF | 0.7434653 |
| TLAVFTVL | ASGGTGGARAEQF | 0.71111596 |
| TGSVQPARV | ASGGTGGARAEQF | 0.8590058 |
| TGGGAGSGGT | ASGGTGGARAEQF | 0.49911252 |
| SHLETSL | ASGGTGGARAEQF | 0.60420567 |
| SFTTGMT | ASGGTGGARAEQF | 0.8512076 |
| QVEVAKRMGVRQPTVS | ASGGTGGARAEQF | 0.8757812 |
| QGSGGPALAM | ASGGTGGARAEQF | 0.52272046 |
| MLAPSNP | ASGGTGGARAEQF | 0.7812512 |
| LVNNAAI | ASGGTGGARAEQF | 0.8597872 |
| LTGEINTLA | ASGGTGGARAEQF | 0.7985678 |
| LSIFGPV | ASGGTGGARAEQF | 0.8594537 |
| LLALLANL | ASGGTGGARAEQF | 0.87415934 |
| LISNASDALDKLR | ASGGTGGARAEQF | 0.64004195 |
| LINEAVP | ASGGTGGARAEQF | 0.86942536 |
| LGYAFSGL | ASGGTGGARAEQF | 0.88749325 |
| LDFVAVAL | ASGGTGGARAEQF | 0.70533204 |
| LAQPGAE | ASGGTGGARAEQF | 0.7571783 |
| ISNLGAVTGF | ASGGTGGARAEQF | 0.5529806 |
| HQEDPGRI | ASGGTGGARAEQF | 0.7301464 |
| GTPGEPTP | ASGGTGGARAEQF | 0.37087935 |
| GFFNSGS | ASGGTGGARAEQF | 0.7971654 |
| GAGGGGLTAGAN | ASGGTGGARAEQF | 0.8594332 |
| FVDALVNM | ASGGTGGARAEQF | 0.7101909 |
| FLTVAPEL | ASGGTGGARAEQF | 0.5348307 |
| FETLTAL | ASGGTGGARAEQF | 0.47866964 |
| DVINPIEL | ASGGTGGARAEQF | 0.7744863 |

| | | |
|---|---|---|
| APVSGAPV | ASGGTGGARAEQF | 0.5678902 |
| ALVGGLAW | ASGGTGGARAEQF | 0.2989713 |
| ALAAAAATAI | ASGGTGGARAEQF | 0.7375875 |
| AFGLGLLL | ASGGTGGARAEQF | 0.8621943 |
| AFFAAASL | ASGGTGGARAEQF | 0.8402942 |
| AAAALPAFSPP | ASGGTGGARAEQF | 0.47724286 |
| VVLAPAL | ASGGTGGARAEQF | 0.8891765 |
| TIGHVDHGKTTL | ASGGTGGARAEQF | 0.82855636 |
| LISNASDALDK | ASGGTGGARAEQF | 0.80174744 |
| LISNASDAL | ASGGTGGARAEQF | 0.661263 |
| ISNASDALDK | ASGGTGGARAEQF | 0.5576134 |
| NQAETLVYQTE | ASGGTGGARAEQF | 0.86195886 |
| TGGTGGTAGAG | ASGGTGGARAEQF | 0.7277396 |
| TGSTAGGAGGA | ASGGTGGARAEQF | 0.6950176 |
| DPLAVPVAAGPV | ASGGTGGARAEQF | 0.85190165 |
| GGTGGTPGNGGN | ASGGTGGARAEQF | 0.74866843 |
| PAGLAGVDPATL | ASGGTGGARAEQF | 0.6087861 |
| GNNLVVEAPHAGASV | ASGGTGGARAEQF | 0.65979636 |
| KVTRIMSASTLEQPA | ASGGTGGARAEQF | 0.84950155 |
| LGFALPIALPRRTVV | ASGGTGGARAEQF | 0.6074811 |
| TRLGNLTARKLVVLV | ASGGTGGARAEQF | 0.49260277 |
| WSQQRRLIYFPSAGP | ASGGTGGARAEQF | 0.58721805 |
| GPNGNPDPMAAAVDIR | ASGGTGGARAEQF | 0.74315554 |
| TQRCVSVADLRAAASR | ASGGTGGARAEQF | 0.87150955 |
| QFVLDVAEHPVEVLTETV | ASGGTGGARAEQF | 0.78622 |
| GGAGGTGGAGGNGGTQAGDG | ASGGTGGARAEQF | 0.6174604 |
| PLLATLRAIAVDVDAKPAQV | ASGGTGGARAEQF | 0.8209934 |
| APKFRTGGYMIDFWGVGYQVAK | ASGGTGGARAEQF | 0.78169894 |
| GAGGAGGAGGAAGGPGGTGGTG | ASGGTGGARAEQF | 0.74681616 |
| IGALGLLATDVAPAGSVAFGVA | ASGGTGGARAEQF | 0.8286189 |
| AGGAGGIGGNGGAGGAGGVLMG | ASGGTGGARAEQF | 0.62926567 |
| ADELIGLL | ASGGTGGARAEQF | 0.84435034 |
| ADRALKA | ASGGTGGARAEQF | 0.8056072 |
| AGAVGQVGGAGGA | ASGGTGGARAEQF | 0.8210543 |
| AGGGGGLGGAP | ASGGTGGARAEQF | 0.8593615 |
| AGGLLYGNGGAG | ASGGTGGARAEQF | 0.799688 |
| AGLESIGL | ASGGTGGARAEQF | 0.7312317 |
| AGLKTGII | ASGGTGGARAEQF | 0.34835702 |
| ASLAALLL | ASGGTGGARAEQF | 0.6678926 |
| ASVINISSI | ASGGTGGARAEQF | 0.8138024 |
| ATPPGPA | ASGGTGGARAEQF | 0.78480595 |
| ATVASALV | ASGGTGGARAEQF | 0.8549604 |
| ATVASLIL | ASGGTGGARAEQF | 0.78968596 |
| AVNGPVLIRH | ASGGTGGARAEQF | 0.6498234 |
| DGAAGAVGI | ASGGTGGARAEQF | 0.8485295 |
| DPLLHGI | ASGGTGGARAEQF | 0.8546351 |
| ELGAALGAL | ASGGTGGARAEQF | 0.40363657 |
| ELGRTQAV | ASGGTGGARAEQF | 0.7369672 |
| FVAGGYLS | ASGGTGGARAEQF | 0.8985678 |
| GASGGAGGAGGW | ASGGTGGARAEQF | 0.7819125 |
| GGAGAGGDGGAGGI | ASGGTGGARAEQF | 0.73358786 |

| | | |
|---|---|---|
| GTGHNGSQPGMGGNG | ASGGTGGARAEQF | 0.62943906 |
| GYAAASASAS | ASGGTGGARAEQF | 0.87360907 |
| HQVDVAAVQQG | ASGGTGGARAEQF | 0.67231286 |
| IGAAIAHT | ASGGTGGARAEQF | 0.893492 |
| IGMIEVAQL | ASGGTGGARAEQF | 0.19839358 |
| IRLAPEHTSAMD | ASGGTGGARAEQF | 0.5658388 |
| IYNVSVL | ASGGTGGARAEQF | 0.5158286 |
| KQGFLGM | ASGGTGGARAEQF | 0.2676957 |
| LESVITQL | ASGGTGGARAEQF | 0.8609686 |
| LGLGALVVV | ASGGTGGARAEQF | 0.58844364 |
| LIGNGGNGGSGGI | ASGGTGGARAEQF | 0.8213002 |
| LNAINAPTL | ASGGTGGARAEQF | 0.7522726 |
| LQLLNTGV | ASGGTGGARAEQF | 0.86681163 |
| LQTSIEF | ASGGTGGARAEQF | 0.7545181 |
| MSFAVSEL | ASGGTGGARAEQF | 0.7158233 |
| NTGALGL | ASGGTGGARAEQF | 0.55025566 |
| PPGAGGAGGAGTT | ASGGTGGARAEQF | 0.8457108 |
| QLVGLGI | ASGGTGGARAEQF | 0.8706537 |
| RFLEQVHQL | ASGGTGGARAEQF | 0.7561724 |
| RGVGYVLR | ASGGTGGARAEQF | 0.43121326 |
| SATGVFAGVI | ASGGTGGARAEQF | 0.7523439 |
| SLPAITLP | ASGGTGGARAEQF | 0.7736321 |
| STALAFTV | ASGGTGGARAEQF | 0.2847654 |
| SVLTVGLGK | ASGGTGGARAEQF | 0.2841791 |
| TALAGAGSGI | ASGGTGGARAEQF | 0.64740145 |
| TGLPFTGI | ASGGTGGARAEQF | 0.7202585 |
| VAEHVTV | ASGGTGGARAEQF | 0.86057544 |
| VDIVLNSL | ASGGTGGARAEQF | 0.85195047 |
| VLTGGGAVL | ASGGTGGARAEQF | 0.6781926 |
| VNVGLAYGI | ASGGTGGARAEQF | 0.8134234 |
| VSSAAVGAV | ASGGTGGARAEQF | 0.79271513 |
| VTSVYEAGV | ASGGTGGARAEQF | 0.093459584 |
| WVALDRGA | ASGGTGGARAEQF | 0.6211786 |
| YLVGSVVI | ASGGTGGARAEQF | 0.469144 |
| GPPGTGKTLL | ASGGTGGARAEQF | 0.8712369 |
| GVGMTKF | ASGGTGGARAEQF | 0.69186985 |
| IGIGGSDL | ASGGTGGARAEQF | 0.68103623 |
| IGTLLAEL | ASGGTGGARAEQF | 0.736001 |
| IYTSGTT | ASGGTGGARAEQF | 0.7745514 |
| LSFPAGV | ASGGTGGARAEQF | 0.74566495 |
| LYTSGTT | ASGGTGGARAEQF | 0.49446273 |
| VGAFPFGF | ASGGTGGARAEQF | 0.27202833 |
| VGAGTVEF | ASGGTGGARAEQF | 0.85220057 |
| VGKSTLLL | ASGGTGGARAEQF | 0.4750818 |
| AALAHAGGGIV | ASGGTGGARAEQF | 0.61005783 |
| AALVAVSLL | ASGGTGGARAEQF | 0.3355958 |
| AAVSRAVAAL | ASGGTGGARAEQF | 0.78431153 |
| AGGSSGIGGF | ASGGTGGARAEQF | 0.36094996 |
| AGLTVANANG | ASGGTGGARAEQF | 0.8215864 |
| AGLVGAVL | ASGGTGGARAEQF | 0.6774106 |
| ALSQAFTV | ASGGTGGARAEQF | 0.7616534 |

| | | |
|---|---|---|
| ALTFSVSL | ASGGTGGARAEQF | 0.89297855 |
| ALTGAALVAL | ASGGTGGARAEQF | 0.75641763 |
| ASFSIGAGVL | ASGGTGGARAEQF | 0.7117983 |
| DGSVYKI | ASGGTGGARAEQF | 0.6538198 |
| EQLAEIV | ASGGTGGARAEQF | 0.55115277 |
| FAVSELTVM | ASGGTGGARAEQF | 0.59231186 |
| GAAPLAAGL | ASGGTGGARAEQF | 0.6453714 |
| GADGGDGGSGGAGG | ASGGTGGARAEQF | 0.8831198 |
| GFGLDTGGAGG | ASGGTGGARAEQF | 0.6369449 |
| GLASGGLL | ASGGTGGARAEQF | 0.46013218 |
| GRIAGAAVLL | ASGGTGGARAEQF | 0.6556061 |
| GSAGSLNTSF | ASGGTGGARAEQF | 0.8829874 |
| GTGQIGI | ASGGTGGARAEQF | 0.88560545 |
| GTLIALIL | ASGGTGGARAEQF | 0.662629 |
| IAATMGFVVG | ASGGTGGARAEQF | 0.59053445 |
| IFAQALL | ASGGTGGARAEQF | 0.601087 |
| IGAAVALGF | ASGGTGGARAEQF | 0.74724054 |
| IGLTPLAL | ASGGTGGARAEQF | 0.57236063 |
| ILGSNTSTL | ASGGTGGARAEQF | 0.46679696 |
| IVFSLTVL | ASGGTGGARAEQF | 0.8089106 |
| LASKATGF | ASGGTGGARAEQF | 0.8852217 |
| LASVALIL | ASGGTGGARAEQF | 0.68473303 |
| LFAVGLTL | ASGGTGGARAEQF | 0.34932265 |
| LFNGGIGTYI | ASGGTGGARAEQF | 0.7390373 |
| LTLGAGII | ASGGTGGARAEQF | 0.54823756 |
| LVAAALGDI | ASGGTGGARAEQF | 0.62472117 |
| NTAFVQLGI | ASGGTGGARAEQF | 0.68300736 |
| QLFSGAEL | ASGGTGGARAEQF | 0.4099351 |
| SAATFITL | ASGGTGGARAEQF | 0.751655 |
| SNYGLANLGV | ASGGTGGARAEQF | 0.8844366 |
| SYGGAYAVM | ASGGTGGARAEQF | 0.7006105 |
| TTGVGGLTLG | ASGGTGGARAEQF | 0.80250466 |
| TTQTIYI | ASGGTGGARAEQF | 0.8717125 |
| VAAGAGLGAVI | ASGGTGGARAEQF | 0.8316603 |
| VAAVVTLL | ASGGTGGARAEQF | 0.8684099 |
| VGFLGGLI | ASGGTGGARAEQF | 0.839808 |
| VGVIPILL | ASGGTGGARAEQF | 0.76832235 |
| VIFASSLI | ASGGTGGARAEQF | 0.75323534 |
| VKGGDGGAAAG | ASGGTGGARAEQF | 0.8628758 |
| VTKTFTNI | ASGGTGGARAEQF | 0.75608015 |
| VVAAAILL | ASGGTGGARAEQF | 0.72471815 |
| YAALDVEL | ASGGTGGARAEQF | 0.63834745 |
| LASVALLL | ASGGTGGARAEQF | 0.860931 |
| LDAVHRV | ASGGTGGARAEQF | 0.6818157 |
| QGLSSSIF | ASGGTGGARAEQF | 0.8547615 |
| RGTFANIRL | ASGGTGGARAEQF | 0.3259852 |
| RQLWWGHRI | ASGGTGGARAEQF | 0.47544506 |
| YGPPGTGKTLL | ASGGTGGARAEQF | 0.84273857 |
| AALVGAPANVA | ASGGTGGARAEQF | 0.11064379 |
| AATFRRTE | ASGGTGGARAEQF | 0.23317698 |
| AGALGGGI | ASGGTGGARAEQF | 0.73475623 |

| | | |
|---|---|---|
| AGGAGGAGGIGAG | ASGGTGGARAEQF | 0.8527795 |
| AGSGALGL | ASGGTGGARAEQF | 0.1420763 |
| AQIADQI | ASGGTGGARAEQF | 0.58155215 |
| ASAVAVTYL | ASGGTGGARAEQF | 0.37624374 |
| ASTVLGGTGG | ASGGTGGARAEQF | 0.7756982 |
| AVGGEPAK | ASGGTGGARAEQF | 0.7689488 |
| CEQGTFGGTC | ASGGTGGARAEQF | 0.3359639 |
| DEGKAPAG | ASGGTGGARAEQF | 0.8170562 |
| DTVVFPAL | ASGGTGGARAEQF | 0.324552 |
| FEGITMAG | ASGGTGGARAEQF | 0.65064526 |
| FPLVVQD | ASGGTGGARAEQF | 0.86137307 |
| GAFTTPQ | ASGGTGGARAEQF | 0.8779694 |
| GFGGFGTAGGVGGA | ASGGTGGARAEQF | 0.62709016 |
| GFLAQFL | ASGGTGGARAEQF | 0.8339677 |
| GGAGGAGGNAGVA | ASGGTGGARAEQF | 0.6059277 |
| GGVGTAGAGGAGGAGG | ASGGTGGARAEQF | 0.89641386 |
| GNGGGGDAGNAGSGGN | ASGGTGGARAEQF | 0.7220553 |
| GTGGAGGAGNDAGS | ASGGTGGARAEQF | 0.38024563 |
| GTTNTGWFNS | ASGGTGGARAEQF | 0.8360266 |
| IFMGPLFTL | ASGGTGGARAEQF | 0.814986 |
| ILAHAIEAI | ASGGTGGARAEQF | 0.78233075 |
| IRSAIREAL | ASGGTGGARAEQF | 0.8241297 |
| ITFAGGG | ASGGTGGARAEQF | 0.5507959 |
| ITGMSGHQEV | ASGGTGGARAEQF | 0.7043965 |
| IVAGLACTL | ASGGTGGARAEQF | 0.85521674 |
| IVGGGLAGA | ASGGTGGARAEQF | 0.85752594 |
| IVTGVLNI | ASGGTGGARAEQF | 0.5214983 |
| LAQQGCI | ASGGTGGARAEQF | 0.8453294 |
| LCLHVEN | ASGGTGGARAEQF | 0.7377285 |
| LDAISDGS | ASGGTGGARAEQF | 0.62367797 |
| LGGFAAML | ASGGTGGARAEQF | 0.8862348 |
| LGGKNPMI | ASGGTGGARAEQF | 0.6682791 |
| LGPKGVAAIL | ASGGTGGARAEQF | 0.843059 |
| LMVGVSI | ASGGTGGARAEQF | 0.8288376 |
| LVGAAFVL | ASGGTGGARAEQF | 0.8434928 |
| LVLAVAPL | ASGGTGGARAEQF | 0.5371281 |
| MAAIAVIDAA | ASGGTGGARAEQF | 0.7989135 |
| NNGINAELQ | ASGGTGGARAEQF | 0.76705384 |
| RGEPLVLVHG | ASGGTGGARAEQF | 0.83583 |
| RGGAGGAAGLI | ASGGTGGARAEQF | 0.71738505 |
| RSYRMHPAV | ASGGTGGARAEQF | 0.72257674 |
| SAAHLLAGQLA | ASGGTGGARAEQF | 0.8121144 |
| SAAVGSY | ASGGTGGARAEQF | 0.87770706 |
| SPHAVLHRI | ASGGTGGARAEQF | 0.78834707 |
| SQIAKVL | ASGGTGGARAEQF | 0.8076265 |
| TALGHLVP | ASGGTGGARAEQF | 0.88023764 |
| TAPTTSGAA | ASGGTGGARAEQF | 0.60423195 |
| TDGDDGNGGSPGT | ASGGTGGARAEQF | 0.72224104 |
| TDGLGEN | ASGGTGGARAEQF | 0.8394406 |
| TGGPPLPGQA | ASGGTGGARAEQF | 0.8681465 |
| TIPAITI | ASGGTGGARAEQF | 0.65726966 |

| | | |
|---|---|---|
| TIPGFTI | ASGGTGGARAEQF | 0.863656 |
| TSIASGALA | ASGGTGGARAEQF | 0.65336955 |
| TVTLPGAAI | ASGGTGGARAEQF | 0.8237158 |
| VASGLSAALP | ASGGTGGARAEQF | 0.681831 |
| VGAGVIGI | ASGGTGGARAEQF | 0.7807183 |
| VGPKRFGV | ASGGTGGARAEQF | 0.62936175 |
| VNVAFPD | ASGGTGGARAEQF | 0.8418165 |
| VSMTNSGVS | ASGGTGGARAEQF | 0.8438647 |
| VTDVLGDLSV | ASGGTGGARAEQF | 0.8462287 |
| VVVVGAGFAGL | ASGGTGGARAEQF | 0.21484528 |
| VYTVGTVV | ASGGTGGARAEQF | 0.8381187 |
| WTTFSGP | ASGGTGGARAEQF | 0.8190098 |
| YDELVEAF | ASGGTGGARAEQF | 0.8909499 |
| YGTGGAGGNGG | ASGGTGGARAEQF | 0.6953391 |
| YTNVMAV | ASGGTGGARAEQF | 0.7822741 |
| FLEQVHQL | ASGGTGGARAEQF | 0.6978071 |
| GLSLLAVAV | ASGGTGGARAEQF | 0.6789695 |
| LEQVHQL | ASGGTGGARAEQF | 0.4214148 |
| LNAAGAI | ASGGTGGARAEQF | 0.6945101 |
| PAVHASL | ASGGTGGARAEQF | 0.6892796 |
| TELEDDA | ASGGTGGARAEQF | 0.62265646 |
| TPPGPPN | ASGGTGGARAEQF | 0.84220815 |
| TPPPPPPP | ASGGTGGARAEQF | 0.865857 |
| TQSPALI | ASSQDEGKNTEVF | 0.86889803 |
| EFAAAFPQ | ASSQDEGKNTEVF | 0.85943913 |
| LLSGNLDVL | ASSQDEGKNTEVF | 0.8791011 |
| GPANLGNYNI | ASSQDEGKNTEVF | 0.81098574 |
| TLGPTLP | ASSQDEGKNTEVF | 0.6495387 |
| VYAYSSCCK | ASSQDEGKNTEVF | 0.7416673 |
| APAPAPAPNI | ASSQDEGKNTEVF | 0.4819923 |
| PGGPVPIP | ASSQDEGKNTEVF | 0.73787665 |
| FLTVALEP | ASSQDEGKNTEVF | 0.6710577 |
| SQYFGANL | ASSQDEGKNTEVF | 0.798722 |
| GGVKPAA | ASSQDEGKNTEVF | 0.120671816 |
| GGTGGAGGAGGW | ASSQDEGKNTEVF | 0.78125596 |
| GVGGVAGLGGAGG | ASSQDEGKNTEVF | 0.89215994 |
| VLLVLATTI | ASSQDEGKNTEVF | 0.62942904 |
| IGIFAAML | ASSQDEGKNTEVF | 0.78441215 |
| AVGDLID | ASSQDEGKNTEVF | 0.8308883 |
| NISPLPL | ASSQDEGKNTEVF | 0.64294255 |
| VTAVFVAI | ASSQDEGKNTEVF | 0.82794756 |
| SGDDNDAAL | ASSQDEGKNTEVF | 0.56532764 |
| AVAALAHS | ASSQDEGKNTEVF | 0.79816574 |
| GGVTVMATP | ASSQDEGKNTEVF | 0.88568753 |
| AGLAANVRD | ASSQDEGKNTEVF | 0.78824437 |
| TNRPDLIDPALLRPG | ASSQDEGKNTEVF | 0.7444674 |
| LANLGVTAL | ASSQDEGKNTEVF | 0.69665724 |
| TGERIEL | ASSQDEGKNTEVF | 0.41369668 |
| NGEELAA | ASSQDEGKNTEVF | 0.78801066 |
| VLSLVWG | ASSQDEGKNTEVF | 0.51593906 |
| IGYALWL | ASSQDEGKNTEVF | 0.7438961 |

| | | |
|---|---|---|
| LSLLLAPL | ASSQDEGKNTEVF | 0.5746341 |
| TLDEALK | ASSQDEGKNTEVF | 0.8166505 |
| TDGAGTAEGG | ASSQDEGKNTEVF | 0.80221 |
| NAPGGAGGNGG | ASSQDEGKNTEVF | 0.7978888 |
| VSVFMGMGGNF | ASSQDEGKNTEVF | 0.5785242 |
| LASYLAAL | ASSQDEGKNTEVF | 0.76070416 |
| LVGGTPAL | ASSQDEGKNTEVF | 0.6976532 |
| VRNADVL | ASSQDEGKNTEVF | 0.50601923 |
| TCSSPFLVSAM | ASSQDEGKNTEVF | 0.8519937 |
| VHITDAE | ASSQDEGKNTEVF | 0.6915037 |
| EYAAPSF | ASSQDEGKNTEVF | 0.7499187 |
| VVDAFHGGL | ASSQDEGKNTEVF | 0.8631579 |
| ASVVDPQ | ASSQDEGKNTEVF | 0.7999751 |
| SYNEFGGLS | ASSQDEGKNTEVF | 0.33634588 |
| DPDGSVY | ASSQDEGKNTEVF | 0.71336806 |
| MVSTIHATE | ASSQDEGKNTEVF | 0.76451176 |
| LAAGVADPV | ASSQDEGKNTEVF | 0.70843875 |
| TKEQIAA | ASSQDEGKNTEVF | 0.15808882 |
| SLASPAAL | ASSQDEGKNTEVF | 0.8354523 |
| IGLVFYGV | ASSQDEGKNTEVF | 0.6238409 |
| FLLDASNL | ASSQDEGKNTEVF | 0.5872258 |
| YQVGNFP | ASSQDEGKNTEVF | 0.6672043 |
| ESLLSQST | ASSQDEGKNTEVF | 0.84628373 |
| LFTVGFAAVG | ASSQDEGKNTEVF | 0.66319084 |
| ATPPTPT | ASSQDEGKNTEVF | 0.665496 |
| TDLATLG | ASSQDEGKNTEVF | 0.8345318 |
| SAGGTGR | ASSQDEGKNTEVF | 0.83712006 |
| GFSNSGS | ASSQDEGKNTEVF | 0.8736822 |
| GGNGGSGGTSV | ASSQDEGKNTEVF | 0.8026064 |
| GTSVATGGAGN | ASSQDEGKNTEVF | 0.69658947 |
| NGGNGGSGGTS | ASSQDEGKNTEVF | 0.7927938 |
| AVVKFPH | ASSQDEGKNTEVF | 0.6049497 |
| TPPPPPP | ASSQDEGKNTEVF | 0.53339565 |
| AAPAAAAAPA | ASSQDEGKNTEVF | 0.31076792 |
| IALNEDL | ASSQDEGKNTEVF | 0.6533685 |
| IVNVSSM | ASSQDEGKNTEVF | 0.39299828 |
| TGIPPAPR | ASSQDEGKNTEVF | 0.79303586 |
| FDLGGGTF | ASSQDEGKNTEVF | 0.83040226 |
| FDLGGGTFDVS | ASSQDEGKNTEVF | 0.69079524 |
| NEPTAAA | ASSQDEGKNTEVF | 0.47836 |
| NEPTAAALA | ASSQDEGKNTEVF | 0.7157202 |
| GSIINISSI | ASSQDEGKNTEVF | 0.7014955 |
| VAPGNAGTA | ASSQDEGKNTEVF | 0.8361194 |
| TGGGASLE | ASSQDEGKNTEVF | 0.79610246 |
| VAVYNHYKRI | ASSQDEGKNTEVF | 0.8851708 |
| TDPAPATT | ASSQDEGKNTEVF | 0.860373 |
| TNRPDILDPALLRPG | ASSQDEGKNTEVF | 0.68415254 |
| LTTTTTS | ASSQDEGKNTEVF | 0.8342335 |
| PTGSGKTL | ASSQDEGKNTEVF | 0.6470124 |
| TGSGKTL | ASSQDEGKNTEVF | 0.78667367 |
| TIGHVDHGK | ASSQDEGKNTEVF | 0.76333255 |

| | | |
|---|---|---|
| TIGHVDHGKTT | ASSQDEGKNTEVF | 0.7595264 |
| GHVDHGKTT | ASSQDEGKNTEVF | 0.7164241 |
| AQFEHTLL | ASSQDEGKNTEVF | 0.7377378 |
| IVNQDEL | ASSQDEGKNTEVF | 0.28590965 |
| TNGSQFF | ASSQDEGKNTEVF | 0.79795253 |
| TGKGIVE | ASSQDEGKNTEVF | 0.5449333 |
| LISNASDA | ASSQDEGKNTEVF | 0.30948538 |
| AELRTLGELTKIAKAHGAQ | ASSQDEGKNTEVF | 0.6898403 |
| APNAVAAAEPSW | ASSQDEGKNTEVF | 0.8898171 |
| APQVAGAATAVL | ASSQDEGKNTEVF | 0.8599692 |
| ARRDPAAAAAILPTDA | ASSQDEGKNTEVF | 0.6225238 |
| DAAARDVLAEAGLAEYF | ASSQDEGKNTEVF | 0.5652024 |
| DDQIVLSAKANLAKVEQ | ASSQDEGKNTEVF | 0.7988877 |
| DELLPDTGEDSDEDGH | ASSQDEGKNTEVF | 0.38215962 |
| ELPEAPANRATVPQD | ASSQDEGKNTEVF | 0.76762885 |
| FGQSKIERVTGIG | ASSQDEGKNTEVF | 0.74310696 |
| GAGGTGSTAGGAGGAGGAGGLY | ASSQDEGKNTEVF | 0.76038885 |
| GHGGTGAAGGNGGTGGAGGSGI | ASSQDEGKNTEVF | 0.5265537 |
| GNGGNGGAGGEGGPGGAGGAGA | ASSQDEGKNTEVF | 0.8051044 |
| GSTAGNGGNGGNATNGTIAGQP | ASSQDEGKNTEVF | 0.72276306 |
| GTGGDGGNGGNGGTGG | ASSQDEGKNTEVF | 0.7487533 |
| GVRRPAGIPADAPV | ASSQDEGKNTEVF | 0.21076998 |
| GYRQVTDAHLLTLARR | ASSQDEGKNTEVF | 0.8531506 |
| IAKMGQDGHDRG | ASSQDEGKNTEVF | 0.62388396 |
| KAEPSIEIQYT | ASSQDEGKNTEVF | 0.5464213 |
| KPVRIRMAADEDTVAVT | ASSQDEGKNTEVF | 0.26030955 |
| MPPRIAGMRLLVIKPEP | ASSQDEGKNTEVF | 0.74894977 |
| MKFYAVALFRSRGVPPTR | ASSQDEGKNTEVF | 0.82261014 |
| NAGGFGILGAVAHSP | ASSQDEGKNTEVF | 0.85419834 |
| NEGREITLAMAEM | ASSQDEGKNTEVF | 0.6485262 |
| NELVEWSHPLGHRWRWE | ASSQDEGKNTEVF | 0.84695005 |
| NLGAGGTGGNGGNPG | ASSQDEGKNTEVF | 0.84855795 |
| NTEWPKWETVMAADPVD | ASSQDEGKNTEVF | 0.7343143 |
| PASAAPAETQP | ASSQDEGKNTEVF | 0.38436717 |
| QQSSDGKNGVRAAEITGE | ASSQDEGKNTEVF | 0.769902 |
| RAPDAVTTSWWKE | ASSQDEGKNTEVF | 0.7199316 |
| SGFGNVGAHTSGWFNQ | ASSQDEGKNTEVF | 0.8539227 |
| SGVAGAVGQAASVGGLKV | ASSQDEGKNTEVF | 0.78156835 |
| SNGGAGTGSAGGNGGT | ASSQDEGKNTEVF | 0.5039859 |
| TGGAGGEGGAAG | ASSQDEGKNTEVF | 0.8019216 |
| TGGDGGAAGAGGAPGHG | ASSQDEGKNTEVF | 0.8331078 |
| TMPPLLAAIANAGRHGV | ASSQDEGKNTEVF | 0.50362146 |
| TSSTAGNSGGA | ASSQDEGKNTEVF | 0.6347275 |
| VPATVPVGILC | ASSQDEGKNTEVF | 0.65721655 |
| VQRPDSGGVVLTGRIS | ASSQDEGKNTEVF | 0.394539 |
| WTPNPVPATKGVPAGLVQAT | ASSQDEGKNTEVF | 0.8344391 |
| YALKRALAKFSDRFDVVII | ASSQDEGKNTEVF | 0.69361097 |
| YLIGSVIV | ASSQDEGKNTEVF | 0.6428894 |
| YGQLVAVID | ASSQDEGKNTEVF | 0.7420511 |
| VTVGGMTLPA | ASSQDEGKNTEVF | 0.66522133 |
| VLALPAV | ASSQDEGKNTEVF | 0.6129899 |

| | | |
|---|---|---|
| VIPPTLNYET | ASSQDEGKNTEVF | 0.7106746 |
| VGMGGMHPSSG | ASSQDEGKNTEVF | 0.80691075 |
| VGLAGGAATA | ASSQDEGKNTEVF | 0.8119992 |
| VGGAGGAGGSGP | ASSQDEGKNTEVF | 0.7315991 |
| VFAAFMV | ASSQDEGKNTEVF | 0.68748355 |
| TLYDKLR | ASSQDEGKNTEVF | 0.88578063 |
| TLAVFTVL | ASSQDEGKNTEVF | 0.45626736 |
| TGSVQPARV | ASSQDEGKNTEVF | 0.62380695 |
| TGGGAGSGGT | ASSQDEGKNTEVF | 0.79041123 |
| SHLETSL | ASSQDEGKNTEVF | 0.4905421 |
| SFTTGMT | ASSQDEGKNTEVF | 0.82939464 |
| QVEVAKRMGVRQPTVS | ASSQDEGKNTEVF | 0.7967806 |
| QGSGGPALAM | ASSQDEGKNTEVF | 0.80343294 |
| MLAPSNP | ASSQDEGKNTEVF | 0.7346502 |
| LVNNAAI | ASSQDEGKNTEVF | 0.89641577 |
| LTGEINTLA | ASSQDEGKNTEVF | 0.8362913 |
| LSIFGPV | ASSQDEGKNTEVF | 0.71722555 |
| LLALLANL | ASSQDEGKNTEVF | 0.84499586 |
| LISNASDALDKLR | ASSQDEGKNTEVF | 0.47558925 |
| LINEAVP | ASSQDEGKNTEVF | 0.8495673 |
| LGYAFSGL | ASSQDEGKNTEVF | 0.7818816 |
| LDFVAVAL | ASSQDEGKNTEVF | 0.5425215 |
| LAQPGAE | ASSQDEGKNTEVF | 0.7198814 |
| ISNLGAVTGF | ASSQDEGKNTEVF | 0.87208045 |
| HQEDPGRI | ASSQDEGKNTEVF | 0.857628 |
| GTPGEPTP | ASSQDEGKNTEVF | 0.37112665 |
| GFFNSGS | ASSQDEGKNTEVF | 0.6564796 |
| GAGGGGLTAGAN | ASSQDEGKNTEVF | 0.39076167 |
| FVDALVNM | ASSQDEGKNTEVF | 0.5494163 |
| FLTVAPEL | ASSQDEGKNTEVF | 0.48972043 |
| FETLTAL | ASSQDEGKNTEVF | 0.8111949 |
| DVINPIEL | ASSQDEGKNTEVF | 0.7094597 |
| APVSGAPV | ASSQDEGKNTEVF | 0.8134676 |
| ALVGGLAW | ASSQDEGKNTEVF | 0.7804066 |
| ALAAAAATAI | ASSQDEGKNTEVF | 0.8715597 |
| AFGLGLLL | ASSQDEGKNTEVF | 0.61402446 |
| AFFAAASL | ASSQDEGKNTEVF | 0.7734555 |
| AAAALPAFSPP | ASSQDEGKNTEVF | 0.3423161 |
| VVLAPAL | ASSQDEGKNTEVF | 0.5110545 |
| TIGHVDHGKTTL | ASSQDEGKNTEVF | 0.5680149 |
| LISNASDALDK | ASSQDEGKNTEVF | 0.7395595 |
| LISNASDAL | ASSQDEGKNTEVF | 0.2589966 |
| ISNASDALDK | ASSQDEGKNTEVF | 0.86649024 |
| NQAETLVYQTE | ASSQDEGKNTEVF | 0.7339228 |
| TGGTGGTAGAG | ASSQDEGKNTEVF | 0.8205215 |
| TGSTAGGAGGA | ASSQDEGKNTEVF | 0.6865959 |
| DPLAVPVAAGPV | ASSQDEGKNTEVF | 0.6298996 |
| GGTGGTPGNGGN | ASSQDEGKNTEVF | 0.6532035 |
| PAGLAGVDPATL | ASSQDEGKNTEVF | 0.830568 |
| GNNLVVEAPHAGASV | ASSQDEGKNTEVF | 0.62993395 |
| KVTRIMSASTLEQPA | ASSQDEGKNTEVF | 0.23947698 |

| | | |
|---|---|---|
| LGFALPIALPRRTVV | ASSQDEGKNTEVF | 0.54023594 |
| TRLGNLTARKLVVLV | ASSQDEGKNTEVF | 0.48683274 |
| WSQQRRLIYFPSAGP | ASSQDEGKNTEVF | 0.87531143 |
| GPNGNPDPMAAAVDIR | ASSQDEGKNTEVF | 0.85258126 |
| TQRCVSVADLRAAASR | ASSQDEGKNTEVF | 0.7828783 |
| QFVLDVAEHPVEVLTETV | ASSQDEGKNTEVF | 0.87050855 |
| GGAGGTGGAGGNGGTQAGDG | ASSQDEGKNTEVF | 0.16772662 |
| PLLATLRAIAVDVDAKPAQV | ASSQDEGKNTEVF | 0.71401227 |
| APKFRTGGYMIDFWGVGYQVAK | ASSQDEGKNTEVF | 0.8711762 |
| GAGGAGGAGGAAGGPGGTGGTG | ASSQDEGKNTEVF | 0.79730654 |
| IGALGLLATDVAPAGSVAFGVA | ASSQDEGKNTEVF | 0.75516474 |
| AGGAGGIGGNGGAGGAGGVLMG | ASSQDEGKNTEVF | 0.8636194 |
| ADELIGLL | ASSQDEGKNTEVF | 0.6348797 |
| ADRALKA | ASSQDEGKNTEVF | 0.716982 |
| AGAVGQVGGAGGA | ASSQDEGKNTEVF | 0.7642925 |
| AGGGGGLGGAP | ASSQDEGKNTEVF | 0.82817554 |
| AGGLLYGNGGAG | ASSQDEGKNTEVF | 0.7547498 |
| AGLESIGL | ASSQDEGKNTEVF | 0.4993058 |
| AGLKTGII | ASSQDEGKNTEVF | 0.7478255 |
| ASLAALLL | ASSQDEGKNTEVF | 0.9090051 |
| ASVINISSI | ASSQDEGKNTEVF | 0.7158968 |
| ATPPGPA | ASSQDEGKNTEVF | 0.831445 |
| ATVASALV | ASSQDEGKNTEVF | 0.79826486 |
| ATVASLIL | ASSQDEGKNTEVF | 0.72793114 |
| AVNGPVLIRH | ASSQDEGKNTEVF | 0.583359 |
| DGAAGAVGI | ASSQDEGKNTEVF | 0.62088263 |
| DPLLHGI | ASSQDEGKNTEVF | 0.78598666 |
| ELGAALGAL | ASSQDEGKNTEVF | 0.5213616 |
| ELGRTQAV | ASSQDEGKNTEVF | 0.72444963 |
| FVAGGYLS | ASSQDEGKNTEVF | 0.64838576 |
| GASGGAGGAGGW | ASSQDEGKNTEVF | 0.9023905 |
| GGAGAGGDGGAGGI | ASSQDEGKNTEVF | 0.8198473 |
| GTGHNGSQPGMGGNG | ASSQDEGKNTEVF | 0.8131968 |
| GYAAASASAS | ASSQDEGKNTEVF | 0.5660436 |
| HQVDVAAVQQG | ASSQDEGKNTEVF | 0.7853937 |
| IGAAIAHT | ASSQDEGKNTEVF | 0.8117787 |
| IGMIEVAQL | ASSQDEGKNTEVF | 0.35391334 |
| IRLAPEHTSAMD | ASSQDEGKNTEVF | 0.8718227 |
| IYNVSVL | ASSQDEGKNTEVF | 0.5680132 |
| KQGFLGM | ASSQDEGKNTEVF | 0.7159314 |
| LESVITQL | ASSQDEGKNTEVF | 0.80285335 |
| LGLGALVVV | ASSQDEGKNTEVF | 0.8671466 |
| LIGNGGNGGSGGI | ASSQDEGKNTEVF | 0.69077307 |
| LNAINAPTL | ASSQDEGKNTEVF | 0.7330629 |
| LQLLNTGV | ASSQDEGKNTEVF | 0.5944309 |
| LQTSIEF | ASSQDEGKNTEVF | 0.80884004 |
| MSFAVSEL | ASSQDEGKNTEVF | 0.86334383 |
| NTGALGL | ASSQDEGKNTEVF | 0.7203765 |
| PPGAGGAGGAGTT | ASSQDEGKNTEVF | 0.89181054 |
| QLVGLGI | ASSQDEGKNTEVF | 0.7497558 |
| RFLEQVHQL | ASSQDEGKNTEVF | 0.3585521 |

| | | |
|---|---|---|
| RGVGYVLR | ASSQDEGKNTEVF | 0.71552783 |
| SATGVFAGVI | ASSQDEGKNTEVF | 0.8741084 |
| SLPAITLP | ASSQDEGKNTEVF | 0.86707735 |
| STALAFTV | ASSQDEGKNTEVF | 0.8737032 |
| SVLTVGLGK | ASSQDEGKNTEVF | 0.8300619 |
| TALAGAGSGI | ASSQDEGKNTEVF | 0.6767595 |
| TGLPFTGI | ASSQDEGKNTEVF | 0.8677163 |
| VAEHVTV | ASSQDEGKNTEVF | 0.8775233 |
| VDIVLNSL | ASSQDEGKNTEVF | 0.8064735 |
| VLTGGGAVL | ASSQDEGKNTEVF | 0.5102615 |
| VNVGLAYGI | ASSQDEGKNTEVF | 0.4673832 |
| VSSAAVGAV | ASSQDEGKNTEVF | 0.8037161 |
| VTSVYEAGV | ASSQDEGKNTEVF | 0.84885 |
| WVALDRGA | ASSQDEGKNTEVF | 0.41433394 |
| YLVGSVVI | ASSQDEGKNTEVF | 0.79442984 |
| GPPGTGKTLL | ASSQDEGKNTEVF | 0.7857058 |
| GVGMTKF | ASSQDEGKNTEVF | 0.39700496 |
| IGIGGSDL | ASSQDEGKNTEVF | 0.7935766 |
| IGTLLAEL | ASSQDEGKNTEVF | 0.882051 |
| IYTSGTT | ASSQDEGKNTEVF | 0.75054073 |
| LSFPAGV | ASSQDEGKNTEVF | 0.54384065 |
| LYTSGTT | ASSQDEGKNTEVF | 0.870617 |
| VGAFPFGF | ASSQDEGKNTEVF | 0.8986925 |
| VGAGTVEF | ASSQDEGKNTEVF | 0.87182355 |
| VGKSTLLL | ASSQDEGKNTEVF | 0.39024213 |
| AALAHAGGGIV | ASSQDEGKNTEVF | 0.16995445 |
| AALVAVSLL | ASSQDEGKNTEVF | 0.7250187 |
| AAVSRAVAAL | ASSQDEGKNTEVF | 0.824908 |
| AGGSSGIGGF | ASSQDEGKNTEVF | 0.71638834 |
| AGLTVANANG | ASSQDEGKNTEVF | 0.62846625 |
| AGLVGAVL | ASSQDEGKNTEVF | 0.8186996 |
| ALSQAFTV | ASSQDEGKNTEVF | 0.8538555 |
| ALTFSVSL | ASSQDEGKNTEVF | 0.5975352 |
| ALTGAALVAL | ASSQDEGKNTEVF | 0.3742437 |
| ASFSIGAGVL | ASSQDEGKNTEVF | 0.8452995 |
| DGSVYKI | ASSQDEGKNTEVF | 0.80195785 |
| EQLAEIV | ASSQDEGKNTEVF | 0.60962087 |
| FAVSELTVM | ASSQDEGKNTEVF | 0.71374017 |
| GAAPLAAGL | ASSQDEGKNTEVF | 0.75763285 |
| GADGGDGGSGGAGG | ASSQDEGKNTEVF | 0.6387602 |
| GFGLDTGGAGG | ASSQDEGKNTEVF | 0.7896936 |
| GLASGGLL | ASSQDEGKNTEVF | 0.44600427 |
| GRIAGAAVLL | ASSQDEGKNTEVF | 0.8812151 |
| GSAGSLNTSF | ASSQDEGKNTEVF | 0.46723375 |
| GTGQIGI | ASSQDEGKNTEVF | 0.8256124 |
| GTLIALIL | ASSQDEGKNTEVF | 0.8110454 |
| IAATMGFVVG | ASSQDEGKNTEVF | 0.7974099 |
| IFAQALL | ASSQDEGKNTEVF | 0.7193662 |
| IGAAVALGF | ASSQDEGKNTEVF | 0.5952929 |
| IGLTPLAL | ASSQDEGKNTEVF | 0.52071005 |
| ILGSNTSTL | ASSQDEGKNTEVF | 0.28980416 |

| | | |
|---|---|---|
| IVFSLTVL | ASSQDEGKNTEVF | 0.8752264 |
| LASKATGF | ASSQDEGKNTEVF | 0.72045195 |
| LASVALIL | ASSQDEGKNTEVF | 0.8572268 |
| LFAVGLTL | ASSQDEGKNTEVF | 0.82576287 |
| LFNGGIGTYI | ASSQDEGKNTEVF | 0.6305438 |
| LTLGAGII | ASSQDEGKNTEVF | 0.7960919 |
| LVAAALGDI | ASSQDEGKNTEVF | 0.5207366 |
| NTAFVQLGI | ASSQDEGKNTEVF | 0.64862657 |
| QLFSGAEL | ASSQDEGKNTEVF | 0.7481368 |
| SAATFITL | ASSQDEGKNTEVF | 0.8029992 |
| SNYGLANLGV | ASSQDEGKNTEVF | 0.6115329 |
| SYGGAYAVM | ASSQDEGKNTEVF | 0.7744481 |
| TTGVGGLTLG | ASSQDEGKNTEVF | 0.7877296 |
| TTQTIYI | ASSQDEGKNTEVF | 0.50334233 |
| VAAGAGLGAVI | ASSQDEGKNTEVF | 0.75707936 |
| VAAVVTLL | ASSQDEGKNTEVF | 0.85424984 |
| VGFLGGLI | ASSQDEGKNTEVF | 0.7558413 |
| VGVIPILL | ASSQDEGKNTEVF | 0.5353836 |
| VIFASSLI | ASSQDEGKNTEVF | 0.8414984 |
| VKGGDGGAAAG | ASSQDEGKNTEVF | 0.5818498 |
| VTKTFTNI | ASSQDEGKNTEVF | 0.79818094 |
| VVAAAILL | ASSQDEGKNTEVF | 0.78275275 |
| YAALDVEL | ASSQDEGKNTEVF | 0.66058815 |
| LASVALLL | ASSQDEGKNTEVF | 0.84982896 |
| LDAVHRV | ASSQDEGKNTEVF | 0.38715684 |
| QGLSSSIF | ASSQDEGKNTEVF | 0.6178262 |
| RGTFANIRL | ASSQDEGKNTEVF | 0.66545206 |
| RQLWWGHRI | ASSQDEGKNTEVF | 0.79046655 |
| YGPPGTGKTLL | ASSQDEGKNTEVF | 0.82831573 |
| AALVGAPANVA | ASSQDEGKNTEVF | 0.56588906 |
| AATFRRTE | ASSQDEGKNTEVF | 0.7389026 |
| AGALGGGI | ASSQDEGKNTEVF | 0.71092147 |
| AGGAGGAGGIGAG | ASSQDEGKNTEVF | 0.5702747 |
| AGSGALGL | ASSQDEGKNTEVF | 0.6491309 |
| AQIADQI | ASSQDEGKNTEVF | 0.6751927 |
| ASAVAVTYL | ASSQDEGKNTEVF | 0.4471506 |
| ASTVLGGTGG | ASSQDEGKNTEVF | 0.8241781 |
| AVGGEPAK | ASSQDEGKNTEVF | 0.6475018 |
| CEQGTFGGTC | ASSQDEGKNTEVF | 0.8190284 |
| DEGKAPAG | ASSQDEGKNTEVF | 0.87265694 |
| DTVVFPAL | ASSQDEGKNTEVF | 0.5081308 |
| FEGITMAG | ASSQDEGKNTEVF | 0.81194085 |
| FPLVVQD | ASSQDEGKNTEVF | 0.7851193 |
| GAFTTPQ | ASSQDEGKNTEVF | 0.8250457 |
| GFGGFGTAGGVGGA | ASSQDEGKNTEVF | 0.74399185 |
| GFLAQFL | ASSQDEGKNTEVF | 0.65458864 |
| GGAGGAGGNAGVA | ASSQDEGKNTEVF | 0.8767674 |
| GGVGTAGAGGAGGAGG | ASSQDEGKNTEVF | 0.8355014 |
| GNGGGGDAGNAGSGGN | ASSQDEGKNTEVF | 0.7858963 |
| GTGGAGGAGNDAGS | ASSQDEGKNTEVF | 0.8372849 |
| GTTNTGWFNS | ASSQDEGKNTEVF | 0.8237561 |

| | | |
|---|---|---|
| IFMGPLFTL | ASSQDEGKNTEVF | 0.82799804 |
| ILAHAIEAI | ASSQDEGKNTEVF | 0.6135359 |
| IRSAIREAL | ASSQDEGKNTEVF | 0.32216004 |
| ITFAGGG | ASSQDEGKNTEVF | 0.73626375 |
| ITGMSGHQEV | ASSQDEGKNTEVF | 0.7138828 |
| IVAGLACTL | ASSQDEGKNTEVF | 0.593868 |
| IVGGGLAGA | ASSQDEGKNTEVF | 0.26217124 |
| IVTGVLNI | ASSQDEGKNTEVF | 0.8558619 |
| LAQQGCI | ASSQDEGKNTEVF | 0.89706373 |
| LCLHVEN | ASSQDEGKNTEVF | 0.6468353 |
| LDAISDGS | ASSQDEGKNTEVF | 0.8898791 |
| LGGFAAML | ASSQDEGKNTEVF | 0.4206136 |
| LGGKNPMI | ASSQDEGKNTEVF | 0.83360386 |
| LGPKGVAAIL | ASSQDEGKNTEVF | 0.8597131 |
| LMVGVSI | ASSQDEGKNTEVF | 0.511209 |
| LVGAAFVL | ASSQDEGKNTEVF | 0.8152777 |
| LVLAVAPL | ASSQDEGKNTEVF | 0.6422254 |
| MAAIAVIDAA | ASSQDEGKNTEVF | 0.6939314 |
| NNGINAELQ | ASSQDEGKNTEVF | 0.8698238 |
| RGEPLVLVHG | ASSQDEGKNTEVF | 0.69974387 |
| RGGAGGAAGLI | ASSQDEGKNTEVF | 0.8218675 |
| RSYRMHPAV | ASSQDEGKNTEVF | 0.69267476 |
| SAAHLLAGQLA | ASSQDEGKNTEVF | 0.56142473 |
| SAAVGSY | ASSQDEGKNTEVF | 0.8012382 |
| SPHAVLHRI | ASSQDEGKNTEVF | 0.84287214 |
| SQIAKVL | ASSQDEGKNTEVF | 0.6262979 |
| TALGHLVP | ASSQDEGKNTEVF | 0.8595182 |
| TAPTTSGAA | ASSQDEGKNTEVF | 0.6612355 |
| TDGDDGNGGSPGT | ASSQDEGKNTEVF | 0.89718884 |
| TDGLGEN | ASSQDEGKNTEVF | 0.8516217 |
| TGGPPLPGQA | ASSQDEGKNTEVF | 0.7618196 |
| TIPAITI | ASSQDEGKNTEVF | 0.63957965 |
| TIPGFTI | ASSQDEGKNTEVF | 0.86082816 |
| TSIASGALA | ASSQDEGKNTEVF | 0.6888845 |
| TVTLPGAAI | ASSQDEGKNTEVF | 0.8344873 |
| VASGLSAALP | ASSQDEGKNTEVF | 0.86639905 |
| VGAGVIGI | ASSQDEGKNTEVF | 0.647474 |
| VGPKRFGV | ASSQDEGKNTEVF | 0.8556705 |
| VNVAFPD | ASSQDEGKNTEVF | 0.7522467 |
| VSMTNSGVS | ASSQDEGKNTEVF | 0.7834172 |
| VTDVLGDLSV | ASSQDEGKNTEVF | 0.67458206 |
| VVVVGAGFAGL | ASSQDEGKNTEVF | 0.4681065 |
| VYTVGTVV | ASSQDEGKNTEVF | 0.7419284 |
| WTTFSGP | ASSQDEGKNTEVF | 0.58399457 |
| YDELVEAF | ASSQDEGKNTEVF | 0.7333573 |
| YGTGGAGGNGG | ASSQDEGKNTEVF | 0.82999486 |
| YTNVMAV | ASSQDEGKNTEVF | 0.66819805 |
| FLEQVHQL | ASSQDEGKNTEVF | 0.85424846 |
| GLSLLAVAV | ASSQDEGKNTEVF | 0.8491318 |
| LEQVHQL | ASSQDEGKNTEVF | 0.46287677 |
| LNAAGAI | ASSQDEGKNTEVF | 0.71115303 |

| | | |
|---|---|---|
| PAVHASL | ASSQDEGKNTEVF | 0.68803537 |
| TELEDDA | ASSQDEGKNTEVF | 0.721457 |
| TPPGPPN | ASSQDEGKNTEVF | 0.39718702 |
| TPPPPPPP | ASSQDEGKNTEVF | 0.8151804 |
| TQSPALI | ASSPQGSAETLY | 0.6060606 |
| EFAAAFPQ | ASSPQGSAETLY | 0.15904835 |
| LLSGNLDVL | ASSPQGSAETLY | 0.7742529 |
| GPANLGNYNI | ASSPQGSAETLY | 0.7508323 |
| TLGPTLP | ASSPQGSAETLY | 0.695804 |
| VYAYSSCCK | ASSPQGSAETLY | 0.82037044 |
| APAPAPAPNI | ASSPQGSAETLY | 0.34755346 |
| PGGPVPIP | ASSPQGSAETLY | 0.6614783 |
| FLTVALEP | ASSPQGSAETLY | 0.8173174 |
| SQYFGANL | ASSPQGSAETLY | 0.84571004 |
| GGVKPAA | ASSPQGSAETLY | 0.5952419 |
| GGTGGAGGAGGW | ASSPQGSAETLY | 0.76157427 |
| GVGGVAGLGGAGG | ASSPQGSAETLY | 0.8501034 |
| VLLVLATTI | ASSPQGSAETLY | 0.7809213 |
| IGIFAAML | ASSPQGSAETLY | 0.8435661 |
| AVGDLID | ASSPQGSAETLY | 0.85655606 |
| NISPLPL | ASSPQGSAETLY | 0.6400341 |
| VTAVFVAI | ASSPQGSAETLY | 0.8344258 |
| SGDDNDAAL | ASSPQGSAETLY | 0.8574313 |
| AVAALAHS | ASSPQGSAETLY | 0.74607366 |
| GGVTVMATP | ASSPQGSAETLY | 0.76694095 |
| AGLAANVRD | ASSPQGSAETLY | 0.6465956 |
| TNRPDLIDPALLRPG | ASSPQGSAETLY | 0.7041287 |
| LANLGVTAL | ASSPQGSAETLY | 0.2699299 |
| TGERIEL | ASSPQGSAETLY | 0.76072073 |
| NGEELAA | ASSPQGSAETLY | 0.8412608 |
| VLSLVWG | ASSPQGSAETLY | 0.6878377 |
| IGYALWL | ASSPQGSAETLY | 0.54705364 |
| LSLLLAPL | ASSPQGSAETLY | 0.5531713 |
| TLDEALK | ASSPQGSAETLY | 0.71308553 |
| TDGAGTAEGG | ASSPQGSAETLY | 0.6947921 |
| NAPGGAGGNGG | ASSPQGSAETLY | 0.3234006 |
| VSVFMGMGGNF | ASSPQGSAETLY | 0.6168035 |
| LASYLAAL | ASSPQGSAETLY | 0.84533465 |
| LVGGTPAL | ASSPQGSAETLY | 0.8399533 |
| VRNADVL | ASSPQGSAETLY | 0.39578512 |
| TCSSPFLVSAM | ASSPQGSAETLY | 0.8508046 |
| VHITDAE | ASSPQGSAETLY | 0.79508686 |
| EYAAPSF | ASSPQGSAETLY | 0.841336 |
| VVDAFHGGL | ASSPQGSAETLY | 0.76941615 |
| ASVVDPQ | ASSPQGSAETLY | 0.63889027 |
| SYNEFGGLS | ASSPQGSAETLY | 0.8028065 |
| DPDGSVY | ASSPQGSAETLY | 0.6559044 |
| MVSTIHATE | ASSPQGSAETLY | 0.610708 |
| LAAGVADPV | ASSPQGSAETLY | 0.8414163 |
| TKEQIAA | ASSPQGSAETLY | 0.699257 |
| SLASPAAL | ASSPQGSAETLY | 0.6378646 |

| | | |
|---|---|---|
| IGLVFYGV | ASSPQGSAETLY | 0.5629826 |
| FLLDASNL | ASSPQGSAETLY | 0.8484923 |
| YQVGNFP | ASSPQGSAETLY | 0.53904146 |
| ESLLSQST | ASSPQGSAETLY | 0.55306536 |
| LFTVGFAAVG | ASSPQGSAETLY | 0.53113997 |
| ATPPTPT | ASSPQGSAETLY | 0.7345043 |
| TDLATLG | ASSPQGSAETLY | 0.84089345 |
| SAGGTGR | ASSPQGSAETLY | 0.7976937 |
| GFSNSGS | ASSPQGSAETLY | 0.64436895 |
| GGNGGSGGTSV | ASSPQGSAETLY | 0.78143036 |
| GTSVATGGAGN | ASSPQGSAETLY | 0.8091888 |
| NGGNGGSGGTS | ASSPQGSAETLY | 0.7735863 |
| AVVKFPH | ASSPQGSAETLY | 0.82707787 |
| TPPPPPP | ASSPQGSAETLY | 0.5876045 |
| AAPAAAAAPA | ASSPQGSAETLY | 0.8453554 |
| IALNEDL | ASSPQGSAETLY | 0.8078194 |
| IVNVSSM | ASSPQGSAETLY | 0.8309468 |
| TGIPPAPR | ASSPQGSAETLY | 0.83361614 |
| FDLGGGTF | ASSPQGSAETLY | 0.7885094 |
| FDLGGGTFDVS | ASSPQGSAETLY | 0.76200706 |
| NEPTAAA | ASSPQGSAETLY | 0.3334936 |
| NEPTAAALA | ASSPQGSAETLY | 0.6653291 |
| GSIINISSI | ASSPQGSAETLY | 0.774869 |
| VAPGNAGTA | ASSPQGSAETLY | 0.7649898 |
| TGGGASLE | ASSPQGSAETLY | 0.8011001 |
| VAVYNHYKRI | ASSPQGSAETLY | 0.8156823 |
| TDPAPATT | ASSPQGSAETLY | 0.6389788 |
| TNRPDILDPALLRPG | ASSPQGSAETLY | 0.8416685 |
| LTTTTTS | ASSPQGSAETLY | 0.8294549 |
| PTGSGKTL | ASSPQGSAETLY | 0.44116378 |
| TGSGKTL | ASSPQGSAETLY | 0.69039005 |
| TIGHVDHGK | ASSPQGSAETLY | 0.8663871 |
| TIGHVDHGKTT | ASSPQGSAETLY | 0.7395822 |
| GHVDHGKTT | ASSPQGSAETLY | 0.7218895 |
| AQFEHTLL | ASSPQGSAETLY | 0.70004034 |
| IVNQDEL | ASSPQGSAETLY | 0.8368636 |
| TNGSQFF | ASSPQGSAETLY | 0.5919213 |
| TGKGIVE | ASSPQGSAETLY | 0.86736923 |
| LISNASDA | ASSPQGSAETLY | 0.22487922 |
| AELRTLGELTKIAKAHGAQ | ASSPQGSAETLY | 0.6659516 |
| APNAVAAAEPSW | ASSPQGSAETLY | 0.61222064 |
| APQVAGAATAVL | ASSPQGSAETLY | 0.34281412 |
| ARRDPAAAAAILPTDA | ASSPQGSAETLY | 0.84124935 |
| DAAARDVLAEAGLAEYF | ASSPQGSAETLY | 0.6814362 |
| DDQIVLSAKANLAKVEQ | ASSPQGSAETLY | 0.81631416 |
| DELLPDTGEDSDEDGH | ASSPQGSAETLY | 0.7366131 |
| ELPEAPANRATVPQD | ASSPQGSAETLY | 0.8502424 |
| FGQSKIERVTGIG | ASSPQGSAETLY | 0.72963864 |
| GAGGTGSTAGGAGGAGGAGGLY | ASSPQGSAETLY | 0.75843847 |
| GHGGTGAAGGNGGTGGAGGSGI | ASSPQGSAETLY | 0.6275035 |
| GNGGNGGAGGEGGPGGAGGAGA | ASSPQGSAETLY | 0.8286307 |

| | | |
|---|---|---|
| GSTAGNGGNGGNATNGTIAGQP | ASSPQGSAETLY | 0.8512249 |
| GTGGDGGNGGNGGTGG | ASSPQGSAETLY | 0.8192969 |
| GVRRPAGIPADAPV | ASSPQGSAETLY | 0.53327763 |
| GYRQVTDAHLLTLARR | ASSPQGSAETLY | 0.646032 |
| IAKMGQDGHDRG | ASSPQGSAETLY | 0.7811942 |
| KAEPSIEIQYT | ASSPQGSAETLY | 0.30909517 |
| KPVRIRMAADEDTVAVT | ASSPQGSAETLY | 0.239867 |
| MPPRIAGMRLLVIKPEP | ASSPQGSAETLY | 0.6614357 |
| MKFYAVALFRSRGVPPTR | ASSPQGSAETLY | 0.586257 |
| NAGGFGILGAVAHSP | ASSPQGSAETLY | 0.83703065 |
| NEGREITLAMAEM | ASSPQGSAETLY | 0.8194643 |
| NELVEWSHPLGHRWRWE | ASSPQGSAETLY | 0.70422643 |
| NLGAGGTGGNGGNPG | ASSPQGSAETLY | 0.80245686 |
| NTEWPKWETVMAADPVD | ASSPQGSAETLY | 0.75692815 |
| PASAAPAETQP | ASSPQGSAETLY | 0.11618955 |
| QQSSDGKNGVRAAEITGE | ASSPQGSAETLY | 0.6849756 |
| RAPDAVTTSWWKE | ASSPQGSAETLY | 0.50597507 |
| SGFGNVGAHTSGWFNQ | ASSPQGSAETLY | 0.85844684 |
| SGVAGAVGQAASVGGLKV | ASSPQGSAETLY | 0.59359396 |
| SNGGAGTGSAGGNGGT | ASSPQGSAETLY | 0.603709 |
| TGGAGGEGGAAG | ASSPQGSAETLY | 0.73675156 |
| TGGDGGAAGAGGAPGHG | ASSPQGSAETLY | 0.7049979 |
| TMPPLLAAIANAGRHGV | ASSPQGSAETLY | 0.74106157 |
| TSSTAGNSGGA | ASSPQGSAETLY | 0.5807657 |
| VPATVPVGILC | ASSPQGSAETLY | 0.17639516 |
| VQRPDSGGVVLTGRIS | ASSPQGSAETLY | 0.84027636 |
| WTPNPVPATKGVPAGLVQAT | ASSPQGSAETLY | 0.4190003 |
| YALKRALAKFSDRFDVVII | ASSPQGSAETLY | 0.6269281 |
| YLIGSVIV | ASSPQGSAETLY | 0.31758195 |
| YGQLVAVID | ASSPQGSAETLY | 0.20332086 |
| VTVGGMTLPA | ASSPQGSAETLY | 0.47184336 |
| VLALPAV | ASSPQGSAETLY | 0.8275662 |
| VIPPTLNYET | ASSPQGSAETLY | 0.67345953 |
| VGMGGMHPSSG | ASSPQGSAETLY | 0.6907942 |
| VGLAGGAATA | ASSPQGSAETLY | 0.85882497 |
| VGGAGGAGGSGP | ASSPQGSAETLY | 0.67316335 |
| VFAAFMV | ASSPQGSAETLY | 0.6882011 |
| TLYDKLR | ASSPQGSAETLY | 0.68772686 |
| TLAVFTVL | ASSPQGSAETLY | 0.57721245 |
| TGSVQPARV | ASSPQGSAETLY | 0.62312317 |
| TGGGAGSGGT | ASSPQGSAETLY | 0.62633663 |
| SHLETSL | ASSPQGSAETLY | 0.8672031 |
| SFTTGMT | ASSPQGSAETLY | 0.6826292 |
| QVEVAKRMGVRQPTVS | ASSPQGSAETLY | 0.5417124 |
| QGSGGPALAM | ASSPQGSAETLY | 0.74525625 |
| MLAPSNP | ASSPQGSAETLY | 0.8536267 |
| LVNNAAI | ASSPQGSAETLY | 0.8648181 |
| LTGEINTLA | ASSPQGSAETLY | 0.6066611 |
| LSIFGPV | ASSPQGSAETLY | 0.73257196 |
| LLALLANL | ASSPQGSAETLY | 0.7293223 |
| LISNASDALDKLR | ASSPQGSAETLY | 0.7201694 |

| | | |
|---|---|---|
| LINEAVP | ASSPQGSAETLY | 0.69189537 |
| LGYAFSGL | ASSPQGSAETLY | 0.5385572 |
| LDFVAVAL | ASSPQGSAETLY | 0.82290375 |
| LAQPGAE | ASSPQGSAETLY | 0.8532652 |
| ISNLGAVTGF | ASSPQGSAETLY | 0.7009798 |
| HQEDPGRI | ASSPQGSAETLY | 0.54508865 |
| GTPGEPTP | ASSPQGSAETLY | 0.72154033 |
| GFFNSGS | ASSPQGSAETLY | 0.58115304 |
| GAGGGGLTAGAN | ASSPQGSAETLY | 0.654207 |
| FVDALVNM | ASSPQGSAETLY | 0.7626937 |
| FLTVAPEL | ASSPQGSAETLY | 0.42338127 |
| FETLTAL | ASSPQGSAETLY | 0.7169542 |
| DVINPIEL | ASSPQGSAETLY | 0.86616063 |
| APVSGAPV | ASSPQGSAETLY | 0.6210785 |
| ALVGGLAW | ASSPQGSAETLY | 0.76099116 |
| ALAAAAATAI | ASSPQGSAETLY | 0.83185256 |
| AFGLGLLL | ASSPQGSAETLY | 0.7891362 |
| AFFAAASL | ASSPQGSAETLY | 0.83970016 |
| AAAALPAFSPP | ASSPQGSAETLY | 0.7965499 |
| VVLAPAL | ASSPQGSAETLY | 0.77928627 |
| TIGHVDHGKTTL | ASSPQGSAETLY | 0.76576656 |
| LISNASDALDK | ASSPQGSAETLY | 0.72913015 |
| LISNASDAL | ASSPQGSAETLY | 0.7411784 |
| ISNASDALDK | ASSPQGSAETLY | 0.71501887 |
| NQAETLVYQTE | ASSPQGSAETLY | 0.6669299 |
| TGGTGGTAGAG | ASSPQGSAETLY | 0.8432814 |
| TGSTAGGAGGA | ASSPQGSAETLY | 0.76918393 |
| DPLAVPVAAGPV | ASSPQGSAETLY | 0.79753447 |
| GGTGGTPGNGGN | ASSPQGSAETLY | 0.50104547 |
| PAGLAGVDPATL | ASSPQGSAETLY | 0.56072044 |
| GNNLVVEAPHAGASV | ASSPQGSAETLY | 0.8062975 |
| KVTRIMSASTLEQPA | ASSPQGSAETLY | 0.1756965 |
| LGFALPIALPRRTVV | ASSPQGSAETLY | 0.25991753 |
| TRLGNLTARKLVVLV | ASSPQGSAETLY | 0.76438963 |
| WSQQRRLIYFPSAGP | ASSPQGSAETLY | 0.84520364 |
| GPNGNPDPMAAAVDIR | ASSPQGSAETLY | 0.18057115 |
| TQRCVSVADLRAAASR | ASSPQGSAETLY | 0.63512164 |
| QFVLDVAEHPVEVLTETV | ASSPQGSAETLY | 0.4646144 |
| GGAGGTGGAGGNGGTQAGDG | ASSPQGSAETLY | 0.8141707 |
| PLLATLRAIAVDVDAKPAQV | ASSPQGSAETLY | 0.7694528 |
| APKFRTGGYMIDFWGVGYQVAK | ASSPQGSAETLY | 0.3808075 |
| GAGGAGGAGGAAGGPGGTGGTG | ASSPQGSAETLY | 0.8096493 |
| IGALGLLATDVAPAGSVAFGVA | ASSPQGSAETLY | 0.4018963 |
| AGGAGGIGGNGGAGGAGGVLMG | ASSPQGSAETLY | 0.41637993 |
| ADELIGLL | ASSPQGSAETLY | 0.8442714 |
| ADRALKA | ASSPQGSAETLY | 0.8294668 |
| AGAVGQVGGAGGA | ASSPQGSAETLY | 0.57052326 |
| AGGGGGLGGAP | ASSPQGSAETLY | 0.8280883 |
| AGGLLYGNGGAG | ASSPQGSAETLY | 0.62217724 |
| AGLESIGL | ASSPQGSAETLY | 0.86480284 |
| AGLKTGII | ASSPQGSAETLY | 0.70564616 |

| | | |
|---|---|---|
| ASLAALLL | ASSPQGSAETLY | 0.4367473 |
| ASVINISSI | ASSPQGSAETLY | 0.8124585 |
| ATPPGPA | ASSPQGSAETLY | 0.77136123 |
| ATVASALV | ASSPQGSAETLY | 0.7858913 |
| ATVASLIL | ASSPQGSAETLY | 0.7911814 |
| AVNGPVLIRH | ASSPQGSAETLY | 0.6238365 |
| DGAAGAVGI | ASSPQGSAETLY | 0.6921153 |
| DPLLHGI | ASSPQGSAETLY | 0.8328058 |
| ELGAALGAL | ASSPQGSAETLY | 0.83436775 |
| ELGRTQAV | ASSPQGSAETLY | 0.5434726 |
| FVAGGYLS | ASSPQGSAETLY | 0.8360764 |
| GASGGAGGAGGW | ASSPQGSAETLY | 0.71475124 |
| GGAGAGGDGGAGGI | ASSPQGSAETLY | 0.6710901 |
| GTGHNGSQPGMGGNG | ASSPQGSAETLY | 0.8674052 |
| GYAAASASAS | ASSPQGSAETLY | 0.7411211 |
| HQVDVAAVQQG | ASSPQGSAETLY | 0.8129501 |
| IGAAIAHT | ASSPQGSAETLY | 0.8309558 |
| IGMIEVAQL | ASSPQGSAETLY | 0.7792956 |
| IRLAPEHTSAMD | ASSPQGSAETLY | 0.45967376 |
| IYNVSVL | ASSPQGSAETLY | 0.7616948 |
| KQGFLGM | ASSPQGSAETLY | 0.73873097 |
| LESVITQL | ASSPQGSAETLY | 0.8244734 |
| LGLGALVVV | ASSPQGSAETLY | 0.74423635 |
| LIGNGGNGGSGGI | ASSPQGSAETLY | 0.4692785 |
| LNAINAPTL | ASSPQGSAETLY | 0.78122103 |
| LQLLNTGV | ASSPQGSAETLY | 0.8471893 |
| LQTSIEF | ASSPQGSAETLY | 0.7935208 |
| MSFAVSEL | ASSPQGSAETLY | 0.7973706 |
| NTGALGL | ASSPQGSAETLY | 0.8533582 |
| PPGAGGAGGAGTT | ASSPQGSAETLY | 0.64533985 |
| QLVGLGI | ASSPQGSAETLY | 0.72949064 |
| RFLEQVHQL | ASSPQGSAETLY | 0.8361374 |
| RGVGYVLR | ASSPQGSAETLY | 0.8391109 |
| SATGVFAGVI | ASSPQGSAETLY | 0.7135019 |
| SLPAITLP | ASSPQGSAETLY | 0.84279853 |
| STALAFTV | ASSPQGSAETLY | 0.5634134 |
| SVLTVGLGK | ASSPQGSAETLY | 0.7978234 |
| TALAGAGSGI | ASSPQGSAETLY | 0.57509077 |
| TGLPFTGI | ASSPQGSAETLY | 0.7970154 |
| VAEHVTV | ASSPQGSAETLY | 0.54911345 |
| VDIVLNSL | ASSPQGSAETLY | 0.80692536 |
| VLTGGGAVL | ASSPQGSAETLY | 0.81162477 |
| VNVGLAYGI | ASSPQGSAETLY | 0.8294312 |
| VSSAAVGAV | ASSPQGSAETLY | 0.25666213 |
| VTSVYEAGV | ASSPQGSAETLY | 0.8153187 |
| WVALDRGA | ASSPQGSAETLY | 0.80343795 |
| YLVGSVVI | ASSPQGSAETLY | 0.86187667 |
| GPPGTGKTLL | ASSPQGSAETLY | 0.7328827 |
| GVGMTKF | ASSPQGSAETLY | 0.80780816 |
| IGIGGSDL | ASSPQGSAETLY | 0.6935984 |
| IGTLLAEL | ASSPQGSAETLY | 0.71321297 |

| | | |
|---|---|---|
| IYTSGTT | ASSPQGSAETLY | 0.53777254 |
| LSFPAGV | ASSPQGSAETLY | 0.6644804 |
| LYTSGTT | ASSPQGSAETLY | 0.65480137 |
| VGAFPFGF | ASSPQGSAETLY | 0.58853245 |
| VGAGTVEF | ASSPQGSAETLY | 0.7901833 |
| VGKSTLLL | ASSPQGSAETLY | 0.8350334 |
| AALAHAGGGIV | ASSPQGSAETLY | 0.83542776 |
| AALVAVSLL | ASSPQGSAETLY | 0.8406037 |
| AAVSRAVAAL | ASSPQGSAETLY | 0.8359729 |
| AGGSSGIGGF | ASSPQGSAETLY | 0.8334713 |
| AGLTVANANG | ASSPQGSAETLY | 0.65449786 |
| AGLVGAVL | ASSPQGSAETLY | 0.706425 |
| ALSQAFTV | ASSPQGSAETLY | 0.5312771 |
| ALTFSVSL | ASSPQGSAETLY | 0.71801794 |
| ALTGAALVAL | ASSPQGSAETLY | 0.6628547 |
| ASFSIGAGVL | ASSPQGSAETLY | 0.77981865 |
| DGSVYKI | ASSPQGSAETLY | 0.14237522 |
| EQLAEIV | ASSPQGSAETLY | 0.6920173 |
| FAVSELTVM | ASSPQGSAETLY | 0.8625821 |
| GAAPLAAGL | ASSPQGSAETLY | 0.5274153 |
| GADGGDGGSGGAGG | ASSPQGSAETLY | 0.79096556 |
| GFGLDTGGAGG | ASSPQGSAETLY | 0.7256863 |
| GLASGGLL | ASSPQGSAETLY | 0.64552915 |
| GRIAGAAVLL | ASSPQGSAETLY | 0.817404 |
| GSAGSLNTSF | ASSPQGSAETLY | 0.59649813 |
| GTGQIGI | ASSPQGSAETLY | 0.79518104 |
| GTLIALIL | ASSPQGSAETLY | 0.8679228 |
| IAATMGFVVG | ASSPQGSAETLY | 0.76509035 |
| IFAQALL | ASSPQGSAETLY | 0.7519372 |
| IGAAVALGF | ASSPQGSAETLY | 0.76763964 |
| IGLTPLAL | ASSPQGSAETLY | 0.43953836 |
| ILGSNTSTL | ASSPQGSAETLY | 0.77489483 |
| IVFSLTVL | ASSPQGSAETLY | 0.11955619 |
| LASKATGF | ASSPQGSAETLY | 0.7197001 |
| LASVALIL | ASSPQGSAETLY | 0.57221717 |
| LFAVGLTL | ASSPQGSAETLY | 0.78440857 |
| LFNGGIGTYI | ASSPQGSAETLY | 0.76204264 |
| LTLGAGII | ASSPQGSAETLY | 0.82136875 |
| LVAAALGDI | ASSPQGSAETLY | 0.57941264 |
| NTAFVQLGI | ASSPQGSAETLY | 0.67627084 |
| QLFSGAEL | ASSPQGSAETLY | 0.5585077 |
| SAATFITL | ASSPQGSAETLY | 0.54168767 |
| SNYGLANLGV | ASSPQGSAETLY | 0.8042524 |
| SYGGAYAVM | ASSPQGSAETLY | 0.75093704 |
| TTGVGGLTLG | ASSPQGSAETLY | 0.6811803 |
| TTQTIYI | ASSPQGSAETLY | 0.8480246 |
| VAAGAGLGAVI | ASSPQGSAETLY | 0.78979945 |
| VAAVVTLL | ASSPQGSAETLY | 0.4921146 |
| VGFLGGLI | ASSPQGSAETLY | 0.6714034 |
| VGVIPILL | ASSPQGSAETLY | 0.7347298 |
| VIFASSLI | ASSPQGSAETLY | 0.70436 |

| | | |
|---|---|---|
| VKGGDGGAAAG | ASSPQGSAETLY | 0.13290483 |
| VTKTFTNI | ASSPQGSAETLY | 0.78144896 |
| VVAAAILL | ASSPQGSAETLY | 0.6754143 |
| YAALDVEL | ASSPQGSAETLY | 0.56880873 |
| LASVALLL | ASSPQGSAETLY | 0.66268826 |
| LDAVHRV | ASSPQGSAETLY | 0.81114763 |
| QGLSSSIF | ASSPQGSAETLY | 0.41739583 |
| RGTFANIRL | ASSPQGSAETLY | 0.7176359 |
| RQLWWGHRI | ASSPQGSAETLY | 0.8036953 |
| YGPPGTGKTLL | ASSPQGSAETLY | 0.8327792 |
| AALVGAPANVA | ASSPQGSAETLY | 0.8501998 |
| AATFRRTE | ASSPQGSAETLY | 0.7948963 |
| AGALGGGI | ASSPQGSAETLY | 0.7362476 |
| AGGAGGAGGIGAG | ASSPQGSAETLY | 0.80401087 |
| AGSGALGL | ASSPQGSAETLY | 0.78389156 |
| AQIADQI | ASSPQGSAETLY | 0.346817 |
| ASAVAVTYL | ASSPQGSAETLY | 0.28760353 |
| ASTVLGGTGG | ASSPQGSAETLY | 0.7441388 |
| AVGGEPAK | ASSPQGSAETLY | 0.3291949 |
| CEQGTFGGTC | ASSPQGSAETLY | 0.77210075 |
| DEGKAPAG | ASSPQGSAETLY | 0.847026 |
| DTVVFPAL | ASSPQGSAETLY | 0.7281994 |
| FEGITMAG | ASSPQGSAETLY | 0.54139405 |
| FPLVVQD | ASSPQGSAETLY | 0.7166705 |
| GAFTTPQ | ASSPQGSAETLY | 0.63994473 |
| GFGGFGTAGGVGGA | ASSPQGSAETLY | 0.798693 |
| GFLAQFL | ASSPQGSAETLY | 0.80262417 |
| GGAGGAGGNAGVA | ASSPQGSAETLY | 0.83286244 |
| GGVGTAGAGGAGGAGG | ASSPQGSAETLY | 0.82988274 |
| GNGGGGDAGNAGSGGN | ASSPQGSAETLY | 0.65959287 |
| GTGGAGGAGNDAGS | ASSPQGSAETLY | 0.8047358 |
| GTTNTGWFNS | ASSPQGSAETLY | 0.68802905 |
| IFMGPLFTL | ASSPQGSAETLY | 0.7176246 |
| ILAHAIEAI | ASSPQGSAETLY | 0.7435726 |
| IRSAIREAL | ASSPQGSAETLY | 0.7643859 |
| ITFAGGG | ASSPQGSAETLY | 0.7313327 |
| ITGMSGHQEV | ASSPQGSAETLY | 0.69589686 |
| IVAGLACTL | ASSPQGSAETLY | 0.46204415 |
| IVGGGLAGA | ASSPQGSAETLY | 0.7700251 |
| IVTGVLNI | ASSPQGSAETLY | 0.7866173 |
| LAQQGCI | ASSPQGSAETLY | 0.3625549 |
| LCLHVEN | ASSPQGSAETLY | 0.6874951 |
| LDAISDGS | ASSPQGSAETLY | 0.8602474 |
| LGGFAAML | ASSPQGSAETLY | 0.8492428 |
| LGGKNPMI | ASSPQGSAETLY | 0.57556325 |
| LGPKGVAAIL | ASSPQGSAETLY | 0.623812 |
| LMVGVSI | ASSPQGSAETLY | 0.7170774 |
| LVGAAFVL | ASSPQGSAETLY | 0.3630048 |
| LVLAVAPL | ASSPQGSAETLY | 0.7843374 |
| MAAIAVIDAA | ASSPQGSAETLY | 0.7028333 |
| NNGINAELQ | ASSPQGSAETLY | 0.70408165 |

| | | |
|---|---|---|
| RGEPLVLVHG | ASSPQGSAETLY | 0.6770156 |
| RGGAGGAAGLI | ASSPQGSAETLY | 0.7614374 |
| RSYRMHPAV | ASSPQGSAETLY | 0.6579596 |
| SAAHLLAGQLA | ASSPQGSAETLY | 0.80632854 |
| SAAVGSY | ASSPQGSAETLY | 0.17371725 |
| SPHAVLHRI | ASSPQGSAETLY | 0.8134253 |
| SQIAKVL | ASSPQGSAETLY | 0.5804155 |
| TALGHLVP | ASSPQGSAETLY | 0.635961 |
| TAPTTSGAA | ASSPQGSAETLY | 0.3784426 |
| TDGDDGNGGSPGT | ASSPQGSAETLY | 0.76916265 |
| TDGLGEN | ASSPQGSAETLY | 0.8065603 |
| TGGPPLPGQA | ASSPQGSAETLY | 0.8279656 |
| TIPAITI | ASSPQGSAETLY | 0.4831961 |
| TIPGFTI | ASSPQGSAETLY | 0.83148754 |
| TSIASGALA | ASSPQGSAETLY | 0.86210656 |
| TVTLPGAAI | ASSPQGSAETLY | 0.6740501 |
| VASGLSAALP | ASSPQGSAETLY | 0.39911982 |
| VGAGVIGI | ASSPQGSAETLY | 0.7947462 |
| VGPKRFGV | ASSPQGSAETLY | 0.6582629 |
| VNVAFPD | ASSPQGSAETLY | 0.81307304 |
| VSMTNSGVS | ASSPQGSAETLY | 0.76926225 |
| VTDVLGDLSV | ASSPQGSAETLY | 0.48429227 |
| VVVVGAGFAGL | ASSPQGSAETLY | 0.822181 |
| VYTVGTVV | ASSPQGSAETLY | 0.83093387 |
| WTTFSGP | ASSPQGSAETLY | 0.6018468 |
| YDELVEAF | ASSPQGSAETLY | 0.69429517 |
| YGTGGAGGNGG | ASSPQGSAETLY | 0.7024889 |
| YTNVMAV | ASSPQGSAETLY | 0.26014507 |
| FLEQVHQL | ASSPQGSAETLY | 0.8277149 |
| GLSLLAVAV | ASSPQGSAETLY | 0.6288436 |
| LEQVHQL | ASSPQGSAETLY | 0.67578757 |
| LNAAGAI | ASSPQGSAETLY | 0.75173616 |
| PAVHASL | ASSPQGSAETLY | 0.6848905 |
| TELEDDA | ASSPQGSAETLY | 0.70074224 |
| TPPGPPN | ASSPQGSAETLY | 0.7110708 |
| TPPPPPPP | ASSPQGSAETLY | 0.77935934 |
| TQSPALI | ASVGGNTLY | 0.7715021 |
| EFAAAFPQ | ASVGGNTLY | 0.7336823 |
| LLSGNLDVL | ASVGGNTLY | 0.74562925 |
| GPANLGNYNI | ASVGGNTLY | 0.8517521 |
| TLGPTLP | ASVGGNTLY | 0.47569633 |
| VYAYSSCCK | ASVGGNTLY | 0.673408 |
| APAPAPAPNI | ASVGGNTLY | 0.7819413 |
| PGGPVPIP | ASVGGNTLY | 0.4285746 |
| FLTVALEP | ASVGGNTLY | 0.7888473 |
| SQYFGANL | ASVGGNTLY | 0.764976 |
| GGVKPAA | ASVGGNTLY | 0.77828383 |
| GGTGGAGGAGGW | ASVGGNTLY | 0.69464666 |
| GVGGVAGLGGAGG | ASVGGNTLY | 0.8715284 |
| VLLVLATTI | ASVGGNTLY | 0.8189219 |
| IGIFAAML | ASVGGNTLY | 0.74461234 |

| | | |
|---|---|---|
| AVGDLID | ASVGGNTLY | 0.714396 |
| NISPLPL | ASVGGNTLY | 0.6106437 |
| VTAVFVAI | ASVGGNTLY | 0.833707 |
| SGDDNDAAL | ASVGGNTLY | 0.79096055 |
| AVAALAHS | ASVGGNTLY | 0.53928196 |
| GGVTVMATP | ASVGGNTLY | 0.70769966 |
| AGLAANVRD | ASVGGNTLY | 0.8566098 |
| TNRPDLIDPALLRPG | ASVGGNTLY | 0.8309432 |
| LANLGVTAL | ASVGGNTLY | 0.35698494 |
| TGERIEL | ASVGGNTLY | 0.67698836 |
| NGEELAA | ASVGGNTLY | 0.41020137 |
| VLSLVWG | ASVGGNTLY | 0.68358505 |
| IGYALWL | ASVGGNTLY | 0.60480225 |
| LSLLLAPL | ASVGGNTLY | 0.7892377 |
| TLDEALK | ASVGGNTLY | 0.6942848 |
| TDGAGTAEGG | ASVGGNTLY | 0.74514025 |
| NAPGGAGGNGG | ASVGGNTLY | 0.7657064 |
| VSVFMGMGGNF | ASVGGNTLY | 0.8553562 |
| LASYLAAL | ASVGGNTLY | 0.64813834 |
| LVGGTPAL | ASVGGNTLY | 0.737287 |
| VRNADVL | ASVGGNTLY | 0.4027 |
| TCSSPFLVSAM | ASVGGNTLY | 0.48187584 |
| VHITDAE | ASVGGNTLY | 0.63540167 |
| EYAAPSF | ASVGGNTLY | 0.7250493 |
| VVDAFHGGL | ASVGGNTLY | 0.24648629 |
| ASVVDPQ | ASVGGNTLY | 0.8491054 |
| SYNEFGGLS | ASVGGNTLY | 0.75498444 |
| DPDGSVY | ASVGGNTLY | 0.8210876 |
| MVSTIHATE | ASVGGNTLY | 0.7061884 |
| LAAGVADPV | ASVGGNTLY | 0.7319232 |
| TKEQIAA | ASVGGNTLY | 0.6353912 |
| SLASPAAL | ASVGGNTLY | 0.7947633 |
| IGLVFYGV | ASVGGNTLY | 0.5561632 |
| FLLDASNL | ASVGGNTLY | 0.43843278 |
| YQVGNFP | ASVGGNTLY | 0.71896434 |
| ESLLSQST | ASVGGNTLY | 0.55713516 |
| LFTVGFAAVG | ASVGGNTLY | 0.83974254 |
| ATPPTPT | ASVGGNTLY | 0.8261116 |
| TDLATLG | ASVGGNTLY | 0.77116483 |
| SAGGTGR | ASVGGNTLY | 0.8331223 |
| GFSNSGS | ASVGGNTLY | 0.27889818 |
| GGNGGSGGTSV | ASVGGNTLY | 0.6908247 |
| GTSVATGGAGN | ASVGGNTLY | 0.85692644 |
| NGGNGGSGGTS | ASVGGNTLY | 0.77149093 |
| AVVKFPH | ASVGGNTLY | 0.7263575 |
| TPPPPPP | ASVGGNTLY | 0.8400977 |
| AAPAAAAAPA | ASVGGNTLY | 0.63266546 |
| IALNEDL | ASVGGNTLY | 0.7102661 |
| IVNVSSM | ASVGGNTLY | 0.7307712 |
| TGIPPAPR | ASVGGNTLY | 0.8087278 |
| FDLGGGTF | ASVGGNTLY | 0.7701736 |

| | | |
|---|---|---|
| FDLGGGTFDVS | ASVGGNTLY | 0.6080882 |
| NEPTAAA | ASVGGNTLY | 0.7978174 |
| NEPTAAALA | ASVGGNTLY | 0.8784284 |
| GSIINISSI | ASVGGNTLY | 0.6858972 |
| VAPGNAGTA | ASVGGNTLY | 0.8176197 |
| TGGGASLE | ASVGGNTLY | 0.799182 |
| VAVYNHYKRI | ASVGGNTLY | 0.7065998 |
| TDPAPATT | ASVGGNTLY | 0.59042937 |
| TNRPDILDPALLRPG | ASVGGNTLY | 0.6240455 |
| LTTTTTS | ASVGGNTLY | 0.68860465 |
| PTGSGKTL | ASVGGNTLY | 0.54479456 |
| TGSGKTL | ASVGGNTLY | 0.7454071 |
| TIGHVDHGK | ASVGGNTLY | 0.70351565 |
| TIGHVDHGKTT | ASVGGNTLY | 0.87870634 |
| GHVDHGKTT | ASVGGNTLY | 0.79220474 |
| AQFEHTLL | ASVGGNTLY | 0.7887646 |
| IVNQDEL | ASVGGNTLY | 0.5628298 |
| TNGSQFF | ASVGGNTLY | 0.79296213 |
| TGKGIVE | ASVGGNTLY | 0.7704936 |
| LISNASDA | ASVGGNTLY | 0.5150895 |
| AELRTLGELTKIAKAHGAQ | ASVGGNTLY | 0.85745484 |
| APNAVAAAEPSW | ASVGGNTLY | 0.64556277 |
| APQVAGAATAVL | ASVGGNTLY | 0.6673945 |
| ARRDPAAAAAILPTDA | ASVGGNTLY | 0.8278591 |
| DAAARDVLAEAGLAEYF | ASVGGNTLY | 0.81402135 |
| DDQIVLSAKANLAKVEQ | ASVGGNTLY | 0.69545054 |
| DELLPDTGEDSDEDGH | ASVGGNTLY | 0.67485344 |
| ELPEAPANRATVPQD | ASVGGNTLY | 0.59264135 |
| FGQSKIERVTGIG | ASVGGNTLY | 0.8191747 |
| GAGGTGSTAGGAGGAGGAGGLY | ASVGGNTLY | 0.8307199 |
| GHGGTGAAGGNGGTGGAGGSGI | ASVGGNTLY | 0.63534164 |
| GNGGNGGAGGEGGPGGAGGAGA | ASVGGNTLY | 0.84352267 |
| GSTAGNGGNGGNATNGTIAGQP | ASVGGNTLY | 0.79491085 |
| GTGGDGGNGGNGGTGG | ASVGGNTLY | 0.4952252 |
| GVRRPAGIPADAPV | ASVGGNTLY | 0.7304083 |
| GYRQVTDAHLLTLARR | ASVGGNTLY | 0.8415113 |
| IAKMGQDGHDRG | ASVGGNTLY | 0.824741 |
| KAEPSIEIQYT | ASVGGNTLY | 0.86165833 |
| KPVRIRMAADEDTVAVT | ASVGGNTLY | 0.8009928 |
| MPPRIAGMRLLVIKPEP | ASVGGNTLY | 0.6484523 |
| MKFYAVALFRSRGVPPTR | ASVGGNTLY | 0.84960103 |
| NAGGFGILGAVAHSP | ASVGGNTLY | 0.86770964 |
| NEGREITLAMAEM | ASVGGNTLY | 0.8216872 |
| NELVEWSHPLGHRWRWE | ASVGGNTLY | 0.68756694 |
| NLGAGGTGGNGGNPG | ASVGGNTLY | 0.64141876 |
| NTEWPKWETVMAADPVD | ASVGGNTLY | 0.7726737 |
| PASAAPAETQP | ASVGGNTLY | 0.8032601 |
| QQSSDGKNGVRAAEITGE | ASVGGNTLY | 0.5680569 |
| RAPDAVTTSWWKE | ASVGGNTLY | 0.75806355 |
| SGFGNVGAHTSGWFNQ | ASVGGNTLY | 0.7666904 |
| SGVAGAVGQAASVGGLKV | ASVGGNTLY | 0.48153287 |

| | | |
|---|---|---|
| SNGGAGTGSAGGNGGT | ASVGGNTLY | 0.80113375 |
| TGGAGGEGGAAG | ASVGGNTLY | 0.8438253 |
| TGGDGGAAGAGGAPGHG | ASVGGNTLY | 0.7333376 |
| TMPPLLAAIANAGRHGV | ASVGGNTLY | 0.6410339 |
| TSSTAGNSGGA | ASVGGNTLY | 0.83666086 |
| VPATVPVGILC | ASVGGNTLY | 0.8761066 |
| VQRPDSGGVVLTGRIS | ASVGGNTLY | 0.8437577 |
| WTPNPVPATKGVPAGLVQAT | ASVGGNTLY | 0.5030097 |
| YALKRALAKFSDRFDVVII | ASVGGNTLY | 0.27918518 |
| YLIGSVIV | ASVGGNTLY | 0.61520797 |
| YGQLVAVID | ASVGGNTLY | 0.7630601 |
| VTVGGMTLPA | ASVGGNTLY | 0.712932 |
| VLALPAV | ASVGGNTLY | 0.62099934 |
| VIPPTLNYET | ASVGGNTLY | 0.7477821 |
| VGMGGMHPSSG | ASVGGNTLY | 0.8375132 |
| VGLAGGAATA | ASVGGNTLY | 0.6103305 |
| VGGAGGAGGSGP | ASVGGNTLY | 0.49968004 |
| VFAAFMV | ASVGGNTLY | 0.7888681 |
| TLYDKLR | ASVGGNTLY | 0.8042166 |
| TLAVFTVL | ASVGGNTLY | 0.6042347 |
| TGSVQPARV | ASVGGNTLY | 0.67714465 |
| TGGGAGSGGT | ASVGGNTLY | 0.66534126 |
| SHLETSL | ASVGGNTLY | 0.39775681 |
| SFTTGMT | ASVGGNTLY | 0.76178634 |
| QVEVAKRMGVRQPTVS | ASVGGNTLY | 0.5659864 |
| QGSGGPALAM | ASVGGNTLY | 0.85576713 |
| MLAPSNP | ASVGGNTLY | 0.57692456 |
| LVNNAAI | ASVGGNTLY | 0.8087564 |
| LTGEINTLA | ASVGGNTLY | 0.77970964 |
| LSIFGPV | ASVGGNTLY | 0.79550153 |
| LLALLANL | ASVGGNTLY | 0.68212575 |
| LISNASDALDKLR | ASVGGNTLY | 0.6497964 |
| LINEAVP | ASVGGNTLY | 0.5115153 |
| LGYAFSGL | ASVGGNTLY | 0.29845503 |
| LDFVAVAL | ASVGGNTLY | 0.8413167 |
| LAQPGAE | ASVGGNTLY | 0.17770603 |
| ISNLGAVTGF | ASVGGNTLY | 0.833142 |
| HQEDPGRI | ASVGGNTLY | 0.8224308 |
| GTPGEPTP | ASVGGNTLY | 0.63630486 |
| GFFNSGS | ASVGGNTLY | 0.7931578 |
| GAGGGGLTAGAN | ASVGGNTLY | 0.48594663 |
| FVDALVNM | ASVGGNTLY | 0.5546524 |
| FLTVAPEL | ASVGGNTLY | 0.8113267 |
| FETLTAL | ASVGGNTLY | 0.805959 |
| DVINPIEL | ASVGGNTLY | 0.63413227 |
| APVSGAPV | ASVGGNTLY | 0.77697897 |
| ALVGGLAW | ASVGGNTLY | 0.7688355 |
| ALAAAAATAI | ASVGGNTLY | 0.6519289 |
| AFGLGLLL | ASVGGNTLY | 0.7585544 |
| AFFAAASL | ASVGGNTLY | 0.83226997 |
| AAAALPAFSPP | ASVGGNTLY | 0.789768 |

| | | |
|---|---|---|
| VVLAPAL | ASVGGNTLY | 0.57577884 |
| TIGHVDHGKTTL | ASVGGNTLY | 0.8145669 |
| LISNASDALDK | ASVGGNTLY | 0.5425081 |
| LISNASDAL | ASVGGNTLY | 0.76817936 |
| ISNASDALDK | ASVGGNTLY | 0.75087494 |
| NQAETLVYQTE | ASVGGNTLY | 0.72522247 |
| TGGTGGTAGAG | ASVGGNTLY | 0.81473285 |
| TGSTAGGAGGA | ASVGGNTLY | 0.61461973 |
| DPLAVPVAAGPV | ASVGGNTLY | 0.5815481 |
| GGTGGTPGNGGN | ASVGGNTLY | 0.66657525 |
| PAGLAGVDPATL | ASVGGNTLY | 0.77993524 |
| GNNLVVEAPHAGASV | ASVGGNTLY | 0.8203408 |
| KVTRIMSASTLEQPA | ASVGGNTLY | 0.5750968 |
| LGFALPIALPRRTVV | ASVGGNTLY | 0.7525125 |
| TRLGNLTARKLVVLV | ASVGGNTLY | 0.7156571 |
| WSQQRRLIYFPSAGP | ASVGGNTLY | 0.5960472 |
| GPNGNPDPMAAAVDIR | ASVGGNTLY | 0.7475466 |
| TQRCVSVADLRAAASR | ASVGGNTLY | 0.72507256 |
| QFVLDVAEHPVEVLTETV | ASVGGNTLY | 0.58545715 |
| GGAGGTGGAGGNGGTQAGDG | ASVGGNTLY | 0.802605 |
| PLLATLRAIAVDVDAKPAQV | ASVGGNTLY | 0.60233015 |
| APKFRTGGYMIDFWGVGYQVAK | ASVGGNTLY | 0.81803006 |
| GAGGAGGAGGAAGGPGGTGGTG | ASVGGNTLY | 0.855356 |
| IGALGLLATDVAPAGSVAFGVA | ASVGGNTLY | 0.66072285 |
| AGGAGGIGGNGGAGGAGGVLMG | ASVGGNTLY | 0.8122414 |
| ADELIGLL | ASVGGNTLY | 0.7802069 |
| ADRALKA | ASVGGNTLY | 0.78476334 |
| AGAVGQVGGAGGA | ASVGGNTLY | 0.72620064 |
| AGGGGGLGGAP | ASVGGNTLY | 0.6523706 |
| AGGLLYGNGGAG | ASVGGNTLY | 0.8433596 |
| AGLESIGL | ASVGGNTLY | 0.8253547 |
| AGLKTGII | ASVGGNTLY | 0.82862246 |
| ASLAALLL | ASVGGNTLY | 0.78757405 |
| ASVINISSI | ASVGGNTLY | 0.77402365 |
| ATPPGPA | ASVGGNTLY | 0.80175704 |
| ATVASALV | ASVGGNTLY | 0.61273396 |
| ATVASLIL | ASVGGNTLY | 0.44393897 |
| AVNGPVLIRH | ASVGGNTLY | 0.69958407 |
| DGAAGAVGI | ASVGGNTLY | 0.72523683 |
| DPLLHGI | ASVGGNTLY | 0.7077477 |
| ELGAALGAL | ASVGGNTLY | 0.43611312 |
| ELGRTQAV | ASVGGNTLY | 0.8342004 |
| FVAGGYLS | ASVGGNTLY | 0.8733668 |
| GASGGAGGAGGW | ASVGGNTLY | 0.67774326 |
| GGAGAGGDGGAGGI | ASVGGNTLY | 0.8545897 |
| GTGHNGSQPGMGGNG | ASVGGNTLY | 0.4956534 |
| GYAAASASAS | ASVGGNTLY | 0.81744176 |
| HQVDVAAVQQG | ASVGGNTLY | 0.84159976 |
| IGAAIAHT | ASVGGNTLY | 0.5739246 |
| IGMIEVAQL | ASVGGNTLY | 0.81246114 |
| IRLAPEHTSAMD | ASVGGNTLY | 0.61465275 |

| | | |
|---|---|---|
| IYNVSVL | ASVGGNTLY | 0.6964253 |
| KQGFLGM | ASVGGNTLY | 0.84117556 |
| LESVITQL | ASVGGNTLY | 0.7141655 |
| LGLGALVVV | ASVGGNTLY | 0.7989457 |
| LIGNGGNGGSGGI | ASVGGNTLY | 0.70928633 |
| LNAINAPTL | ASVGGNTLY | 0.6036358 |
| LQLLNTGV | ASVGGNTLY | 0.7839319 |
| LQTSIEF | ASVGGNTLY | 0.84429455 |
| MSFAVSEL | ASVGGNTLY | 0.5964277 |
| NTGALGL | ASVGGNTLY | 0.86353534 |
| PPGAGGAGGAGTT | ASVGGNTLY | 0.65217507 |
| QLVGLGI | ASVGGNTLY | 0.8715043 |
| RFLEQVHQL | ASVGGNTLY | 0.8094466 |
| RGVGYVLR | ASVGGNTLY | 0.71971405 |
| SATGVFAGVI | ASVGGNTLY | 0.69187284 |
| SLPAITLP | ASVGGNTLY | 0.8380375 |
| STALAFTV | ASVGGNTLY | 0.6809554 |
| SVLTVGLGK | ASVGGNTLY | 0.81957424 |
| TALAGAGSGI | ASVGGNTLY | 0.85331476 |
| TGLPFTGI | ASVGGNTLY | 0.25911498 |
| VAEHVTV | ASVGGNTLY | 0.83861345 |
| VDIVLNSL | ASVGGNTLY | 0.80240655 |
| VLTGGGAVL | ASVGGNTLY | 0.7978545 |
| VNVGLAYGI | ASVGGNTLY | 0.5666907 |
| VSSAAVGAV | ASVGGNTLY | 0.25044248 |
| VTSVYEAGV | ASVGGNTLY | 0.5852948 |
| WVALDRGA | ASVGGNTLY | 0.4629397 |
| YLVGSVVI | ASVGGNTLY | 0.8171845 |
| GPPGTGKTLL | ASVGGNTLY | 0.7904992 |
| GVGMTKF | ASVGGNTLY | 0.5032598 |
| IGIGGSDL | ASVGGNTLY | 0.7704431 |
| IGTLLAEL | ASVGGNTLY | 0.8310623 |
| IYTSGTT | ASVGGNTLY | 0.2660836 |
| LSFPAGV | ASVGGNTLY | 0.6634445 |
| LYTSGTT | ASVGGNTLY | 0.54022104 |
| VGAFPFGF | ASVGGNTLY | 0.5750878 |
| VGAGTVEF | ASVGGNTLY | 0.1838143 |
| VGKSTLLL | ASVGGNTLY | 0.7709577 |
| AALAHAGGGIV | ASVGGNTLY | 0.60431933 |
| AALVAVSLL | ASVGGNTLY | 0.14339352 |
| AAVSRAVAAL | ASVGGNTLY | 0.7688153 |
| AGGSSGIGGF | ASVGGNTLY | 0.45385996 |
| AGLTVANANG | ASVGGNTLY | 0.50571305 |
| AGLVGAVL | ASVGGNTLY | 0.78784335 |
| ALSQAFTV | ASVGGNTLY | 0.5174291 |
| ALTFSVSL | ASVGGNTLY | 0.46479052 |
| ALTGAALVAL | ASVGGNTLY | 0.7822342 |
| ASFSIGAGVL | ASVGGNTLY | 0.8112465 |
| DGSVYKI | ASVGGNTLY | 0.38081563 |
| EQLAEIV | ASVGGNTLY | 0.73780704 |
| FAVSELTVM | ASVGGNTLY | 0.8526311 |

| | | |
|---|---|---|
| GAAPLAAGL | ASVGGNTLY | 0.7126925 |
| GADGGDGGSGGAGG | ASVGGNTLY | 0.84595096 |
| GFGLDTGGAGG | ASVGGNTLY | 0.8549179 |
| GLASGGLL | ASVGGNTLY | 0.6127704 |
| GRIAGAAVLL | ASVGGNTLY | 0.8263806 |
| GSAGSLNTSF | ASVGGNTLY | 0.8418265 |
| GTGQIGI | ASVGGNTLY | 0.5554962 |
| GTLIALIL | ASVGGNTLY | 0.63512355 |
| IAATMGFVVG | ASVGGNTLY | 0.27771223 |
| IFAQALL | ASVGGNTLY | 0.7067686 |
| IGAAVALGF | ASVGGNTLY | 0.38424394 |
| IGLTPLAL | ASVGGNTLY | 0.7627945 |
| ILGSNTSTL | ASVGGNTLY | 0.8376217 |
| IVFSLTVL | ASVGGNTLY | 0.6672199 |
| LASKATGF | ASVGGNTLY | 0.31487107 |
| LASVALIL | ASVGGNTLY | 0.2885511 |
| LFAVGLTL | ASVGGNTLY | 0.7478789 |
| LFNGGIGTYI | ASVGGNTLY | 0.31037015 |
| LTLGAGII | ASVGGNTLY | 0.25675622 |
| LVAAALGDI | ASVGGNTLY | 0.51798004 |
| NTAFVQLGI | ASVGGNTLY | 0.8155769 |
| QLFSGAEL | ASVGGNTLY | 0.82239175 |
| SAATFITL | ASVGGNTLY | 0.49556813 |
| SNYGLANLGV | ASVGGNTLY | 0.85471284 |
| SYGGAYAVM | ASVGGNTLY | 0.81696624 |
| TTGVGGLTLG | ASVGGNTLY | 0.8646446 |
| TTQTIYI | ASVGGNTLY | 0.67868423 |
| VAAGAGLGAVI | ASVGGNTLY | 0.28262877 |
| VAAVVTLL | ASVGGNTLY | 0.82806975 |
| VGFLGGLI | ASVGGNTLY | 0.6139042 |
| VGVIPILL | ASVGGNTLY | 0.7423513 |
| VIFASSLI | ASVGGNTLY | 0.79945225 |
| VKGGDGGAAAG | ASVGGNTLY | 0.62824774 |
| VTKTFTNI | ASVGGNTLY | 0.4213899 |
| VVAAAILL | ASVGGNTLY | 0.75858027 |
| YAALDVEL | ASVGGNTLY | 0.8574718 |
| LASVALLL | ASVGGNTLY | 0.46720505 |
| LDAVHRV | ASVGGNTLY | 0.33320498 |
| QGLSSSIF | ASVGGNTLY | 0.6183617 |
| RGTFANIRL | ASVGGNTLY | 0.643683 |
| RQLWWGHRI | ASVGGNTLY | 0.8152699 |
| YGPPGTGKTLL | ASVGGNTLY | 0.7104227 |
| AALVGAPANVA | ASVGGNTLY | 0.64047754 |
| AATFRRTE | ASVGGNTLY | 0.7570197 |
| AGALGGGI | ASVGGNTLY | 0.7628177 |
| AGGAGGAGGIGAG | ASVGGNTLY | 0.7034749 |
| AGSGALGL | ASVGGNTLY | 0.7945497 |
| AQIADQI | ASVGGNTLY | 0.53289545 |
| ASAVAVTYL | ASVGGNTLY | 0.84246355 |
| ASTVLGGTGG | ASVGGNTLY | 0.80276334 |
| AVGGEPAK | ASVGGNTLY | 0.7916559 |

| | | |
|---|---|---|
| CEQGTFGGTC | ASVGGNTLY | 0.8416046 |
| DEGKAPAG | ASVGGNTLY | 0.60576576 |
| DTVVFPAL | ASVGGNTLY | 0.7393503 |
| FEGITMAG | ASVGGNTLY | 0.3721001 |
| FPLVVQD | ASVGGNTLY | 0.57053804 |
| GAFTTPQ | ASVGGNTLY | 0.712349 |
| GFGGFGTAGGVGGA | ASVGGNTLY | 0.77290165 |
| GFLAQFL | ASVGGNTLY | 0.8136054 |
| GGAGGAGGNAGVA | ASVGGNTLY | 0.7885235 |
| GGVGTAGAGGAGGAGG | ASVGGNTLY | 0.5590033 |
| GNGGGGDAGNAGSGGN | ASVGGNTLY | 0.85557246 |
| GTGGAGGAGNDAGS | ASVGGNTLY | 0.84531456 |
| GTTNTGWFNS | ASVGGNTLY | 0.38658524 |
| IFMGPLFTL | ASVGGNTLY | 0.6098167 |
| ILAHAIEAI | ASVGGNTLY | 0.84318537 |
| IRSAIREAL | ASVGGNTLY | 0.7812318 |
| ITFAGGG | ASVGGNTLY | 0.60531807 |
| ITGMSGHQEV | ASVGGNTLY | 0.7014141 |
| IVAGLACTL | ASVGGNTLY | 0.8431717 |
| IVGGGLAGA | ASVGGNTLY | 0.66920555 |
| IVTGVLNI | ASVGGNTLY | 0.8558416 |
| LAQQGCI | ASVGGNTLY | 0.23571174 |
| LCLHVEN | ASVGGNTLY | 0.70452476 |
| LDAISDGS | ASVGGNTLY | 0.5932267 |
| LGGFAAML | ASVGGNTLY | 0.14174055 |
| LGGKNPMI | ASVGGNTLY | 0.8352998 |
| LGPKGVAAIL | ASVGGNTLY | 0.55073446 |
| LMVGVSI | ASVGGNTLY | 0.76481384 |
| LVGAAFVL | ASVGGNTLY | 0.5103927 |
| LVLAVAPL | ASVGGNTLY | 0.8109638 |
| MAAIAVIDAA | ASVGGNTLY | 0.7653534 |
| NNGINAELQ | ASVGGNTLY | 0.586897 |
| RGEPLVLVHG | ASVGGNTLY | 0.3434004 |
| RGGAGGAAGLI | ASVGGNTLY | 0.7831096 |
| RSYRMHPAV | ASVGGNTLY | 0.8530655 |
| SAAHLLAGQLA | ASVGGNTLY | 0.8394923 |
| SAAVGSY | ASVGGNTLY | 0.22320008 |
| SPHAVLHRI | ASVGGNTLY | 0.8056916 |
| SQIAKVL | ASVGGNTLY | 0.6855358 |
| TALGHLVP | ASVGGNTLY | 0.19002452 |
| TAPTTSGAA | ASVGGNTLY | 0.16385457 |
| TDGDDGNGGSPGT | ASVGGNTLY | 0.6105852 |
| TDGLGEN | ASVGGNTLY | 0.62546504 |
| TGGPPLPGQA | ASVGGNTLY | 0.7616972 |
| TIPAITI | ASVGGNTLY | 0.7775203 |
| TIPGFTI | ASVGGNTLY | 0.5202881 |
| TSIASGALA | ASVGGNTLY | 0.79006076 |
| TVTLPGAAI | ASVGGNTLY | 0.6739385 |
| VASGLSAALP | ASVGGNTLY | 0.094839446 |
| VGAGVIGI | ASVGGNTLY | 0.4638791 |
| VGPKRFGV | ASVGGNTLY | 0.23609202 |

| | | |
|---|---|---|
| VNVAFPD | ASVGGNTLY | 0.8501873 |
| VSMTNSGVS | ASVGGNTLY | 0.37756062 |
| VTDVLGDLSV | ASVGGNTLY | 0.6339592 |
| VVVVGAGFAGL | ASVGGNTLY | 0.6077975 |
| VYTVGTVV | ASVGGNTLY | 0.805902 |
| WTTFSGP | ASVGGNTLY | 0.73147124 |
| YDELVEAF | ASVGGNTLY | 0.45012164 |
| YGTGGAGGNGG | ASVGGNTLY | 0.24236709 |
| YTNVMAV | ASVGGNTLY | 0.7668932 |
| FLEQVHQL | ASVGGNTLY | 0.5203086 |
| GLSLLAVAV | ASVGGNTLY | 0.42654908 |
| LEQVHQL | ASVGGNTLY | 0.4252004 |
| LNAAGAI | ASVGGNTLY | 0.43543816 |
| PAVHASL | ASVGGNTLY | 0.26302087 |
| TELEDDA | ASVGGNTLY | 0.81622565 |
| TPPGPPN | ASVGGNTLY | 0.44698146 |
| TPPPPPPP | ASVGGNTLY | 0.57157326 |
| TQSPALI | ASSPRGWSSYEQY | 0.8612801 |
| EFAAAFPQ | ASSPRGWSSYEQY | 0.7720401 |
| LLSGNLDVL | ASSPRGWSSYEQY | 0.63432634 |
| GPANLGNYNI | ASSPRGWSSYEQY | 0.47842643 |
| TLGPTLP | ASSPRGWSSYEQY | 0.4087296 |
| VYAYSSCCK | ASSPRGWSSYEQY | 0.5246871 |
| APAPAPAPNI | ASSPRGWSSYEQY | 0.4807022 |
| PGGPVPIP | ASSPRGWSSYEQY | 0.8435806 |
| FLTVALEP | ASSPRGWSSYEQY | 0.7546639 |
| SQYFGANL | ASSPRGWSSYEQY | 0.32389405 |
| GGVKPAA | ASSPRGWSSYEQY | 0.82742494 |
| GGTGGAGGAGGW | ASSPRGWSSYEQY | 0.8490486 |
| GVGGVAGLGGAGG | ASSPRGWSSYEQY | 0.8701639 |
| VLLVLATTI | ASSPRGWSSYEQY | 0.36028185 |
| IGIFAAML | ASSPRGWSSYEQY | 0.64200383 |
| AVGDLID | ASSPRGWSSYEQY | 0.69610417 |
| NISPLPL | ASSPRGWSSYEQY | 0.6459983 |
| VTAVFVAI | ASSPRGWSSYEQY | 0.396159 |
| SGDDNDAAL | ASSPRGWSSYEQY | 0.3790834 |
| AVAALAHS | ASSPRGWSSYEQY | 0.83600277 |
| GGVTVMATP | ASSPRGWSSYEQY | 0.86640084 |
| AGLAANVRD | ASSPRGWSSYEQY | 0.6200131 |
| TNRPDLIDPALLRPG | ASSPRGWSSYEQY | 0.23546784 |
| LANLGVTAL | ASSPRGWSSYEQY | 0.6031926 |
| TGERIEL | ASSPRGWSSYEQY | 0.45077395 |
| NGEELAA | ASSPRGWSSYEQY | 0.4387337 |
| VLSLVWG | ASSPRGWSSYEQY | 0.5840621 |
| IGYALWL | ASSPRGWSSYEQY | 0.2743718 |
| LSLLLAPL | ASSPRGWSSYEQY | 0.64653593 |
| TLDEALK | ASSPRGWSSYEQY | 0.86163485 |
| TDGAGTAEGG | ASSPRGWSSYEQY | 0.72836846 |
| NAPGGAGGNGG | ASSPRGWSSYEQY | 0.79848593 |
| VSVFMGMGGNF | ASSPRGWSSYEQY | 0.839663 |
| LASYLAAL | ASSPRGWSSYEQY | 0.79259664 |

| | | |
|---|---|---|
| LVGGTPAL | ASSPRGWSSYEQY | 0.8424803 |
| VRNADVL | ASSPRGWSSYEQY | 0.82289696 |
| TCSSPFLVSAM | ASSPRGWSSYEQY | 0.63450897 |
| VHITDAE | ASSPRGWSSYEQY | 0.6657744 |
| EYAAPSF | ASSPRGWSSYEQY | 0.8193227 |
| VVDAFHGGL | ASSPRGWSSYEQY | 0.55554664 |
| ASVVDPQ | ASSPRGWSSYEQY | 0.7110938 |
| SYNEFGGLS | ASSPRGWSSYEQY | 0.7272965 |
| DPDGSVY | ASSPRGWSSYEQY | 0.8079755 |
| MVSTIHATE | ASSPRGWSSYEQY | 0.6333724 |
| LAAGVADPV | ASSPRGWSSYEQY | 0.8514092 |
| TKEQIAA | ASSPRGWSSYEQY | 0.1575638 |
| SLASPAAL | ASSPRGWSSYEQY | 0.37282807 |
| IGLVFYGV | ASSPRGWSSYEQY | 0.8427933 |
| FLLDASNL | ASSPRGWSSYEQY | 0.12521529 |
| YQVGNFP | ASSPRGWSSYEQY | 0.24805552 |
| ESLLSQST | ASSPRGWSSYEQY | 0.73335934 |
| LFTVGFAAVG | ASSPRGWSSYEQY | 0.84834146 |
| ATPPTPT | ASSPRGWSSYEQY | 0.12817214 |
| TDLATLG | ASSPRGWSSYEQY | 0.5455408 |
| SAGGTGR | ASSPRGWSSYEQY | 0.253784 |
| GFSNSGS | ASSPRGWSSYEQY | 0.78019506 |
| GGNGGSGGTSV | ASSPRGWSSYEQY | 0.60517716 |
| GTSVATGGAGN | ASSPRGWSSYEQY | 0.21930948 |
| NGGNGGSGGTS | ASSPRGWSSYEQY | 0.7762034 |
| AVVKFPH | ASSPRGWSSYEQY | 0.3583725 |
| TPPPPPP | ASSPRGWSSYEQY | 0.59545475 |
| AAPAAAAAPA | ASSPRGWSSYEQY | 0.81423634 |
| IALNEDL | ASSPRGWSSYEQY | 0.85772336 |
| IVNVSSM | ASSPRGWSSYEQY | 0.27351284 |
| TGIPPAPR | ASSPRGWSSYEQY | 0.7589979 |
| FDLGGGTF | ASSPRGWSSYEQY | 0.6139216 |
| FDLGGGTFDVS | ASSPRGWSSYEQY | 0.8796817 |
| NEPTAAA | ASSPRGWSSYEQY | 0.60186553 |
| NEPTAAALA | ASSPRGWSSYEQY | 0.26097262 |
| GSIINISSI | ASSPRGWSSYEQY | 0.818633 |
| VAPGNAGTA | ASSPRGWSSYEQY | 0.82340527 |
| TGGGASLE | ASSPRGWSSYEQY | 0.5927969 |
| VAVYNHYKRI | ASSPRGWSSYEQY | 0.81944096 |
| TDPAPATT | ASSPRGWSSYEQY | 0.6901766 |
| TNRPDILDPALLRPG | ASSPRGWSSYEQY | 0.611605 |
| LTTTTTS | ASSPRGWSSYEQY | 0.7808401 |
| PTGSGKTL | ASSPRGWSSYEQY | 0.79190916 |
| TGSGKTL | ASSPRGWSSYEQY | 0.41444147 |
| TIGHVDHGK | ASSPRGWSSYEQY | 0.82638794 |
| TIGHVDHGKTT | ASSPRGWSSYEQY | 0.6780157 |
| GHVDHGKTT | ASSPRGWSSYEQY | 0.3150907 |
| AQFEHTLL | ASSPRGWSSYEQY | 0.87857115 |
| IVNQDEL | ASSPRGWSSYEQY | 0.6131613 |
| TNGSQFF | ASSPRGWSSYEQY | 0.80403644 |
| TGKGIVE | ASSPRGWSSYEQY | 0.8178297 |

| | | |
|---|---|---|
| LISNASDA | ASSPRGWSSYEQY | 0.8517767 |
| AELRTLGELTKIAKAHGAQ | ASSPRGWSSYEQY | 0.4170546 |
| APNAVAAAEPSW | ASSPRGWSSYEQY | 0.78519547 |
| APQVAGAATAVL | ASSPRGWSSYEQY | 0.72872543 |
| ARRDPAAAAAILPTDA | ASSPRGWSSYEQY | 0.78654593 |
| DAAARDVLAEAGLAEYF | ASSPRGWSSYEQY | 0.7359353 |
| DDQIVLSAKANLAKVEQ | ASSPRGWSSYEQY | 0.74117494 |
| DELLPDTGEDSDEDGH | ASSPRGWSSYEQY | 0.8010955 |
| ELPEAPANRATVPQD | ASSPRGWSSYEQY | 0.83778465 |
| FGQSKIERVTGIG | ASSPRGWSSYEQY | 0.678777 |
| GAGGTGSTAGGAGGAGGAGGLY | ASSPRGWSSYEQY | 0.73404163 |
| GHGGTGAAGGNGGTGGAGGSGI | ASSPRGWSSYEQY | 0.8547021 |
| GNGGNGGAGGEGGPGGAGGAGA | ASSPRGWSSYEQY | 0.33476764 |
| GSTAGNGGNGGNATNGTIAGQP | ASSPRGWSSYEQY | 0.6773005 |
| GTGGDGGNGGNGGTGG | ASSPRGWSSYEQY | 0.8254329 |
| GVRRPAGIPADAPV | ASSPRGWSSYEQY | 0.8376969 |
| GYRQVTDAHLLTLARR | ASSPRGWSSYEQY | 0.6911583 |
| IAKMGQDGHDRG | ASSPRGWSSYEQY | 0.8589642 |
| KAEPSIEIQYT | ASSPRGWSSYEQY | 0.5181807 |
| KPVRIRMAADEDTVAVT | ASSPRGWSSYEQY | 0.749364 |
| MPPRIAGMRLLVIKPEP | ASSPRGWSSYEQY | 0.7417302 |
| MKFYAVALFRSRGVPPTR | ASSPRGWSSYEQY | 0.53574854 |
| NAGGFGILGAVAHSP | ASSPRGWSSYEQY | 0.40895084 |
| NEGREITLAMAEM | ASSPRGWSSYEQY | 0.7288476 |
| NELVEWSHPLGHRWRWE | ASSPRGWSSYEQY | 0.80817926 |
| NLGAGGTGGNGGNPG | ASSPRGWSSYEQY | 0.7344599 |
| NTEWPKWETVMAADPVD | ASSPRGWSSYEQY | 0.21074417 |
| PASAAPAETQP | ASSPRGWSSYEQY | 0.7307316 |
| QQSSDGKNGVRAAEITGE | ASSPRGWSSYEQY | 0.75125873 |
| RAPDAVTTSWWKE | ASSPRGWSSYEQY | 0.8640169 |
| SGFGNVGAHTSGWFNQ | ASSPRGWSSYEQY | 0.6472454 |
| SGVAGAVGQAASVGGLKV | ASSPRGWSSYEQY | 0.8385179 |
| SNGGAGTGSAGGNGGT | ASSPRGWSSYEQY | 0.6094536 |
| TGGAGGEGGAAG | ASSPRGWSSYEQY | 0.72569656 |
| TGGDGGAAGAGGAPGHG | ASSPRGWSSYEQY | 0.29285738 |
| TMPPLLAAIANAGRHGV | ASSPRGWSSYEQY | 0.44920903 |
| TSSTAGNSGGA | ASSPRGWSSYEQY | 0.6219551 |
| VPATVPVGILC | ASSPRGWSSYEQY | 0.4833171 |
| VQRPDSGGVVLTGRIS | ASSPRGWSSYEQY | 0.82354236 |
| WTPNPVPATKGVPAGLVQAT | ASSPRGWSSYEQY | 0.83757424 |
| YALKRALAKFSDRFDVVII | ASSPRGWSSYEQY | 0.79404026 |
| YLIGSVIV | ASSPRGWSSYEQY | 0.8341395 |
| YGQLVAVID | ASSPRGWSSYEQY | 0.8581294 |
| VTVGGMTLPA | ASSPRGWSSYEQY | 0.8137654 |
| VLALPAV | ASSPRGWSSYEQY | 0.65359586 |
| VIPPTLNYET | ASSPRGWSSYEQY | 0.607142 |
| VGMGGMHPSSG | ASSPRGWSSYEQY | 0.2616824 |
| VGLAGGAATA | ASSPRGWSSYEQY | 0.4698993 |
| VGGAGGAGGSGP | ASSPRGWSSYEQY | 0.6221764 |
| VFAAFMV | ASSPRGWSSYEQY | 0.587585 |
| TLYDKLR | ASSPRGWSSYEQY | 0.10570892 |

| | | |
|---|---|---|
| TLAVFTVL | ASSPRGWSSYEQY | 0.78265333 |
| TGSVQPARV | ASSPRGWSSYEQY | 0.84513855 |
| TGGGAGSGGT | ASSPRGWSSYEQY | 0.6575091 |
| SHLETSL | ASSPRGWSSYEQY | 0.7916932 |
| SFTTGMT | ASSPRGWSSYEQY | 0.7735987 |
| QVEVAKRMGVRQPTVS | ASSPRGWSSYEQY | 0.56860644 |
| QGSGGPALAM | ASSPRGWSSYEQY | 0.7780324 |
| MLAPSNP | ASSPRGWSSYEQY | 0.5321607 |
| LVNNAAI | ASSPRGWSSYEQY | 0.81685925 |
| LTGEINTLA | ASSPRGWSSYEQY | 0.8624145 |
| LSIFGPV | ASSPRGWSSYEQY | 0.7793114 |
| LLALLANL | ASSPRGWSSYEQY | 0.7241956 |
| LISNASDALDKLR | ASSPRGWSSYEQY | 0.758796 |
| LINEAVP | ASSPRGWSSYEQY | 0.43642008 |
| LGYAFSGL | ASSPRGWSSYEQY | 0.7422242 |
| LDFVAVAL | ASSPRGWSSYEQY | 0.13052124 |
| LAQPGAE | ASSPRGWSSYEQY | 0.7423639 |
| ISNLGAVTGF | ASSPRGWSSYEQY | 0.5122498 |
| HQEDPGRI | ASSPRGWSSYEQY | 0.7380357 |
| GTPGEPTP | ASSPRGWSSYEQY | 0.6719438 |
| GFFNSGS | ASSPRGWSSYEQY | 0.76828396 |
| GAGGGGLTAGAN | ASSPRGWSSYEQY | 0.46522754 |
| FVDALVNM | ASSPRGWSSYEQY | 0.7858782 |
| FLTVAPEL | ASSPRGWSSYEQY | 0.76424134 |
| FETLTAL | ASSPRGWSSYEQY | 0.31113568 |
| DVINPIEL | ASSPRGWSSYEQY | 0.842182 |
| APVSGAPV | ASSPRGWSSYEQY | 0.67103165 |
| ALVGGLAW | ASSPRGWSSYEQY | 0.66634953 |
| ALAAAAATAI | ASSPRGWSSYEQY | 0.8361679 |
| AFGLGLLL | ASSPRGWSSYEQY | 0.671612 |
| AFFAAASL | ASSPRGWSSYEQY | 0.38239 |
| AAAALPAFSPP | ASSPRGWSSYEQY | 0.5945166 |
| VVLAPAL | ASSPRGWSSYEQY | 0.73969615 |
| TIGHVDHGKTTL | ASSPRGWSSYEQY | 0.67727494 |
| LISNASDALDK | ASSPRGWSSYEQY | 0.12510876 |
| LISNASDAL | ASSPRGWSSYEQY | 0.7762766 |
| ISNASDALDK | ASSPRGWSSYEQY | 0.31514728 |
| NQAETLVYQTE | ASSPRGWSSYEQY | 0.56599385 |
| TGGTGGTAGAG | ASSPRGWSSYEQY | 0.58900356 |
| TGSTAGGAGGA | ASSPRGWSSYEQY | 0.80158514 |
| DPLAVPVAAGPV | ASSPRGWSSYEQY | 0.6709349 |
| GGTGGTPGNGGN | ASSPRGWSSYEQY | 0.47897163 |
| PAGLAGVDPATL | ASSPRGWSSYEQY | 0.7864579 |
| GNNLVVEAPHAGASV | ASSPRGWSSYEQY | 0.6640625 |
| KVTRIMSASTLEQPA | ASSPRGWSSYEQY | 0.8006545 |
| LGFALPIALPRRTVV | ASSPRGWSSYEQY | 0.72068304 |
| TRLGNLTARKLVVLV | ASSPRGWSSYEQY | 0.7305882 |
| WSQQRRLIYFPSAGP | ASSPRGWSSYEQY | 0.7621193 |
| GPNGNPDPMAAAVDIR | ASSPRGWSSYEQY | 0.4736107 |
| TQRCVSVADLRAAASR | ASSPRGWSSYEQY | 0.41752243 |
| QFVLDVAEHPVEVLTETV | ASSPRGWSSYEQY | 0.51810485 |

| | | |
|---|---|---|
| GGAGGTGGAGGNGGTQAGDG | ASSPRGWSSYEQY | 0.5022744 |
| PLLATLRAIAVDVDAKPAQV | ASSPRGWSSYEQY | 0.3739874 |
| APKFRTGGYMIDFWGVGYQVAK | ASSPRGWSSYEQY | 0.7820164 |
| GAGGAGGAGGAAGGPGGTGGTG | ASSPRGWSSYEQY | 0.84068364 |
| IGALGLLATDVAPAGSVAFGVA | ASSPRGWSSYEQY | 0.73659396 |
| AGGAGGIGGNGGAGGAGGVLMG | ASSPRGWSSYEQY | 0.23235 |
| ADELIGLL | ASSPRGWSSYEQY | 0.61531866 |
| ADRALKA | ASSPRGWSSYEQY | 0.5927783 |
| AGAVGQVGGAGGA | ASSPRGWSSYEQY | 0.7678545 |
| AGGGGGLGGAP | ASSPRGWSSYEQY | 0.8181145 |
| AGGLLYGNGGAG | ASSPRGWSSYEQY | 0.85184044 |
| AGLESIGL | ASSPRGWSSYEQY | 0.79677916 |
| AGLKTGII | ASSPRGWSSYEQY | 0.595616 |
| ASLAALLL | ASSPRGWSSYEQY | 0.8156364 |
| ASVINISSI | ASSPRGWSSYEQY | 0.59838283 |
| ATPPGPA | ASSPRGWSSYEQY | 0.7404871 |
| ATVASALV | ASSPRGWSSYEQY | 0.7228011 |
| ATVASLIL | ASSPRGWSSYEQY | 0.7804272 |
| AVNGPVLIRH | ASSPRGWSSYEQY | 0.50656337 |
| DGAAGAVGI | ASSPRGWSSYEQY | 0.60726887 |
| DPLLHGI | ASSPRGWSSYEQY | 0.30203074 |
| ELGAALGAL | ASSPRGWSSYEQY | 0.78298855 |
| ELGRTQAV | ASSPRGWSSYEQY | 0.39971843 |
| FVAGGYLS | ASSPRGWSSYEQY | 0.3447246 |
| GASGGAGGAGGW | ASSPRGWSSYEQY | 0.6144246 |
| GGAGAGGDGGAGGI | ASSPRGWSSYEQY | 0.861794 |
| GTGHNGSQPGMGGNG | ASSPRGWSSYEQY | 0.8374115 |
| GYAAASASAS | ASSPRGWSSYEQY | 0.5646558 |
| HQVDVAAVQQG | ASSPRGWSSYEQY | 0.5402583 |
| IGAAIAHT | ASSPRGWSSYEQY | 0.8055266 |
| IGMIEVAQL | ASSPRGWSSYEQY | 0.5679074 |
| IRLAPEHTSAMD | ASSPRGWSSYEQY | 0.8217175 |
| IYNVSVL | ASSPRGWSSYEQY | 0.67636275 |
| KQGFLGM | ASSPRGWSSYEQY | 0.6269046 |
| LESVITQL | ASSPRGWSSYEQY | 0.62125385 |
| LGLGALVVV | ASSPRGWSSYEQY | 0.8111611 |
| LIGNGGNGGSGGI | ASSPRGWSSYEQY | 0.57258254 |
| LNAINAPTL | ASSPRGWSSYEQY | 0.7475221 |
| LQLLNTGV | ASSPRGWSSYEQY | 0.17158264 |
| LQTSIEF | ASSPRGWSSYEQY | 0.79518676 |
| MSFAVSEL | ASSPRGWSSYEQY | 0.3984602 |
| NTGALGL | ASSPRGWSSYEQY | 0.3967904 |
| PPGAGGAGGAGTT | ASSPRGWSSYEQY | 0.18424699 |
| QLVGLGI | ASSPRGWSSYEQY | 0.6275446 |
| RFLEQVHQL | ASSPRGWSSYEQY | 0.80959177 |
| RGVGYVLR | ASSPRGWSSYEQY | 0.850157 |
| SATGVFAGVI | ASSPRGWSSYEQY | 0.72736585 |
| SLPAITLP | ASSPRGWSSYEQY | 0.79817206 |
| STALAFTV | ASSPRGWSSYEQY | 0.85092574 |
| SVLTVGLGK | ASSPRGWSSYEQY | 0.5391247 |
| TALAGAGSGI | ASSPRGWSSYEQY | 0.19338329 |

| | | |
|---|---|---|
| TGLPFTGI | ASSPRGWSSYEQY | 0.6969228 |
| VAEHVTV | ASSPRGWSSYEQY | 0.6873067 |
| VDIVLNSL | ASSPRGWSSYEQY | 0.8375499 |
| VLTGGGAVL | ASSPRGWSSYEQY | 0.8223947 |
| VNVGLAYGI | ASSPRGWSSYEQY | 0.25971633 |
| VSSAAVGAV | ASSPRGWSSYEQY | 0.8111042 |
| VTSVYEAGV | ASSPRGWSSYEQY | 0.7618439 |
| WVALDRGA | ASSPRGWSSYEQY | 0.3259835 |
| YLVGSVVI | ASSPRGWSSYEQY | 0.5298875 |
| GPPGTGKTLL | ASSPRGWSSYEQY | 0.45808613 |
| GVGMTKF | ASSPRGWSSYEQY | 0.3163507 |
| IGIGGSDL | ASSPRGWSSYEQY | 0.72864395 |
| IGTLLAEL | ASSPRGWSSYEQY | 0.6325095 |
| IYTSGTT | ASSPRGWSSYEQY | 0.5809785 |
| LSFPAGV | ASSPRGWSSYEQY | 0.62747014 |
| LYTSGTT | ASSPRGWSSYEQY | 0.5216694 |
| VGAFPFGF | ASSPRGWSSYEQY | 0.50725114 |
| VGAGTVEF | ASSPRGWSSYEQY | 0.5707807 |
| VGKSTLLL | ASSPRGWSSYEQY | 0.78804153 |
| AALAHAGGGIV | ASSPRGWSSYEQY | 0.8275367 |
| AALVAVSLL | ASSPRGWSSYEQY | 0.6608274 |
| AAVSRAVAAL | ASSPRGWSSYEQY | 0.7301127 |
| AGGSSGIGGF | ASSPRGWSSYEQY | 0.8371948 |
| AGLTVANANG | ASSPRGWSSYEQY | 0.23735568 |
| AGLVGAVL | ASSPRGWSSYEQY | 0.3779205 |
| ALSQAFTV | ASSPRGWSSYEQY | 0.7686011 |
| ALTFSVSL | ASSPRGWSSYEQY | 0.64231884 |
| ALTGAALVAL | ASSPRGWSSYEQY | 0.8317766 |
| ASFSIGAGVL | ASSPRGWSSYEQY | 0.7103014 |
| DGSVYKI | ASSPRGWSSYEQY | 0.6948332 |
| EQLAEIV | ASSPRGWSSYEQY | 0.6655922 |
| FAVSELTVM | ASSPRGWSSYEQY | 0.8768661 |
| GAAPLAAGL | ASSPRGWSSYEQY | 0.7370405 |
| GADGGDGGSGGAGG | ASSPRGWSSYEQY | 0.61967176 |
| GFGLDTGGAGG | ASSPRGWSSYEQY | 0.7886738 |
| GLASGGLL | ASSPRGWSSYEQY | 0.3076864 |
| GRIAGAAVLL | ASSPRGWSSYEQY | 0.829265 |
| GSAGSLNTSF | ASSPRGWSSYEQY | 0.8317292 |
| GTGQIGI | ASSPRGWSSYEQY | 0.4186718 |
| GTLIALIL | ASSPRGWSSYEQY | 0.6205671 |
| IAATMGFVVG | ASSPRGWSSYEQY | 0.8628942 |
| IFAQALL | ASSPRGWSSYEQY | 0.8151371 |
| IGAAVALGF | ASSPRGWSSYEQY | 0.5383398 |
| IGLTPLAL | ASSPRGWSSYEQY | 0.42644027 |
| ILGSNTSTL | ASSPRGWSSYEQY | 0.36860764 |
| IVFSLTVL | ASSPRGWSSYEQY | 0.56841534 |
| LASKATGF | ASSPRGWSSYEQY | 0.2055847 |
| LASVALIL | ASSPRGWSSYEQY | 0.76873434 |
| LFAVGLTL | ASSPRGWSSYEQY | 0.5717555 |
| LFNGGIGTYI | ASSPRGWSSYEQY | 0.82983536 |
| LTLGAGII | ASSPRGWSSYEQY | 0.7638804 |

| | | |
|---|---|---|
| LVAAALGDI | ASSPRGWSSYEQY | 0.84247506 |
| NTAFVQLGI | ASSPRGWSSYEQY | 0.608119 |
| QLFSGAEL | ASSPRGWSSYEQY | 0.60041595 |
| SAATFITL | ASSPRGWSSYEQY | 0.24435577 |
| SNYGLANLGV | ASSPRGWSSYEQY | 0.3402575 |
| SYGGAYAVM | ASSPRGWSSYEQY | 0.50983536 |
| TTGVGGLTLG | ASSPRGWSSYEQY | 0.672943 |
| TTQTIYI | ASSPRGWSSYEQY | 0.18353269 |
| VAAGAGLGAVI | ASSPRGWSSYEQY | 0.85629493 |
| VAAVVTLL | ASSPRGWSSYEQY | 0.51461315 |
| VGFLGGLI | ASSPRGWSSYEQY | 0.7897291 |
| VGVIPILL | ASSPRGWSSYEQY | 0.5303509 |
| VIFASSLI | ASSPRGWSSYEQY | 0.3291892 |
| VKGGDGGAAAG | ASSPRGWSSYEQY | 0.60886514 |
| VTKTFTNI | ASSPRGWSSYEQY | 0.7902423 |
| VVAAAILL | ASSPRGWSSYEQY | 0.33808208 |
| YAALDVEL | ASSPRGWSSYEQY | 0.20479093 |
| LASVALLL | ASSPRGWSSYEQY | 0.3478533 |
| LDAVHRV | ASSPRGWSSYEQY | 0.5533998 |
| QGLSSSIF | ASSPRGWSSYEQY | 0.869063 |
| RGTFANIRL | ASSPRGWSSYEQY | 0.83991146 |
| RQLWWGHRI | ASSPRGWSSYEQY | 0.7091259 |
| YGPPGTGKTLL | ASSPRGWSSYEQY | 0.86479175 |
| AALVGAPANVA | ASSPRGWSSYEQY | 0.12813291 |
| AATFRRTE | ASSPRGWSSYEQY | 0.48778328 |
| AGALGGGI | ASSPRGWSSYEQY | 0.8453552 |
| AGGAGGAGGIGAG | ASSPRGWSSYEQY | 0.78677696 |
| AGSGALGL | ASSPRGWSSYEQY | 0.61138177 |
| AQIADQI | ASSPRGWSSYEQY | 0.82507145 |
| ASAVAVTYL | ASSPRGWSSYEQY | 0.50213206 |
| ASTVLGGTGG | ASSPRGWSSYEQY | 0.67437506 |
| AVGGEPAK | ASSPRGWSSYEQY | 0.64425987 |
| CEQGTFGGTC | ASSPRGWSSYEQY | 0.80263865 |
| DEGKAPAG | ASSPRGWSSYEQY | 0.7547661 |
| DTVVFPAL | ASSPRGWSSYEQY | 0.24129143 |
| FEGITMAG | ASSPRGWSSYEQY | 0.7988498 |
| FPLVVQD | ASSPRGWSSYEQY | 0.875378 |
| GAFTTPQ | ASSPRGWSSYEQY | 0.58182836 |
| GFGGFGTAGGVGGA | ASSPRGWSSYEQY | 0.81526804 |
| GFLAQFL | ASSPRGWSSYEQY | 0.68606913 |
| GGAGGAGGNAGVA | ASSPRGWSSYEQY | 0.6362818 |
| GGVGTAGAGGAGGAGG | ASSPRGWSSYEQY | 0.5012541 |
| GNGGGGDAGNAGSGGN | ASSPRGWSSYEQY | 0.30218703 |
| GTGGAGGAGNDAGS | ASSPRGWSSYEQY | 0.79029644 |
| GTTNTGWFNS | ASSPRGWSSYEQY | 0.3505445 |
| IFMGPLFTL | ASSPRGWSSYEQY | 0.74240434 |
| ILAHAIEAI | ASSPRGWSSYEQY | 0.62738633 |
| IRSAIREAL | ASSPRGWSSYEQY | 0.8814783 |
| ITFAGGG | ASSPRGWSSYEQY | 0.7833609 |
| ITGMSGHQEV | ASSPRGWSSYEQY | 0.7268308 |
| IVAGLACTL | ASSPRGWSSYEQY | 0.5257699 |

| | | |
|---|---|---|
| IVGGGLAGA | ASSPRGWSSYEQY | 0.7726693 |
| IVTGVLNI | ASSPRGWSSYEQY | 0.78218734 |
| LAQQGCI | ASSPRGWSSYEQY | 0.29338 |
| LCLHVEN | ASSPRGWSSYEQY | 0.82422554 |
| LDAISDGS | ASSPRGWSSYEQY | 0.34768677 |
| LGGFAAML | ASSPRGWSSYEQY | 0.73713803 |
| LGGKNPMI | ASSPRGWSSYEQY | 0.68359727 |
| LGPKGVAAIL | ASSPRGWSSYEQY | 0.83252937 |
| LMVGVSI | ASSPRGWSSYEQY | 0.71568066 |
| LVGAAFVL | ASSPRGWSSYEQY | 0.7660396 |
| LVLAVAPL | ASSPRGWSSYEQY | 0.44172648 |
| MAAIAVIDAA | ASSPRGWSSYEQY | 0.7996491 |
| NNGINAELQ | ASSPRGWSSYEQY | 0.8281597 |
| RGEPLVLVHG | ASSPRGWSSYEQY | 0.6528733 |
| RGGAGGAAGLI | ASSPRGWSSYEQY | 0.86513215 |
| RSYRMHPAV | ASSPRGWSSYEQY | 0.74187696 |
| SAAHLLAGQLA | ASSPRGWSSYEQY | 0.18669869 |
| SAAVGSY | ASSPRGWSSYEQY | 0.62805206 |
| SPHAVLHRI | ASSPRGWSSYEQY | 0.83274394 |
| SQIAKVL | ASSPRGWSSYEQY | 0.8570311 |
| TALGHLVP | ASSPRGWSSYEQY | 0.8351003 |
| TAPTTSGAA | ASSPRGWSSYEQY | 0.81674653 |
| TDGDDGNGGSPGT | ASSPRGWSSYEQY | 0.64520836 |
| TDGLGEN | ASSPRGWSSYEQY | 0.8509872 |
| TGGPPLPGQA | ASSPRGWSSYEQY | 0.8861862 |
| TIPAITI | ASSPRGWSSYEQY | 0.7788568 |
| TIPGFTI | ASSPRGWSSYEQY | 0.37236804 |
| TSIASGALA | ASSPRGWSSYEQY | 0.33193028 |
| TVTLPGAAI | ASSPRGWSSYEQY | 0.8057521 |
| VASGLSAALP | ASSPRGWSSYEQY | 0.82790065 |
| VGAGVIGI | ASSPRGWSSYEQY | 0.40216362 |
| VGPKRFGV | ASSPRGWSSYEQY | 0.6049015 |
| VNVAFPD | ASSPRGWSSYEQY | 0.6985178 |
| VSMTNSGVS | ASSPRGWSSYEQY | 0.15917197 |
| VTDVLGDLSV | ASSPRGWSSYEQY | 0.79003155 |
| VVVVGAGFAGL | ASSPRGWSSYEQY | 0.8665184 |
| VYTVGTVV | ASSPRGWSSYEQY | 0.77696514 |
| WTTFSGP | ASSPRGWSSYEQY | 0.31369823 |
| YDELVEAF | ASSPRGWSSYEQY | 0.8322304 |
| YGTGGAGGNGG | ASSPRGWSSYEQY | 0.8491043 |
| YTNVMAV | ASSPRGWSSYEQY | 0.85887504 |
| FLEQVHQL | ASSPRGWSSYEQY | 0.25962096 |
| GLSLLAVAV | ASSPRGWSSYEQY | 0.13641651 |
| LEQVHQL | ASSPRGWSSYEQY | 0.7338897 |
| LNAAGAI | ASSPRGWSSYEQY | 0.79552925 |
| PAVHASL | ASSPRGWSSYEQY | 0.5945493 |
| TELEDDA | ASSPRGWSSYEQY | 0.4114508 |
| TPPGPPN | ASSPRGWSSYEQY | 0.7332759 |
| TPPPPPPP | ASSPRGWSSYEQY | 0.81956995 |
| TQSPALI | ASRSGTGGTQY | 0.31465578 |
| EFAAAFPQ | ASRSGTGGTQY | 0.32226261 |

| | | |
|---|---|---|
| LLSGNLDVL | ASRSGTGGTQY | 0.8062182 |
| GPANLGNYNI | ASRSGTGGTQY | 0.8370774 |
| TLGPTLP | ASRSGTGGTQY | 0.3476559 |
| VYAYSSCCK | ASRSGTGGTQY | 0.6430181 |
| APAPAPAPNI | ASRSGTGGTQY | 0.7616815 |
| PGGPVPIP | ASRSGTGGTQY | 0.7350798 |
| FLTVALEP | ASRSGTGGTQY | 0.717063 |
| SQYFGANL | ASRSGTGGTQY | 0.28223544 |
| GGVKPAA | ASRSGTGGTQY | 0.7890121 |
| GGTGGAGGAGGW | ASRSGTGGTQY | 0.31213373 |
| GVGGVAGLGGAGG | ASRSGTGGTQY | 0.83581686 |
| VLLVLATTI | ASRSGTGGTQY | 0.7014811 |
| IGIFAAML | ASRSGTGGTQY | 0.76335114 |
| AVGDLID | ASRSGTGGTQY | 0.6395082 |
| NISPLPL | ASRSGTGGTQY | 0.2736066 |
| VTAVFVAI | ASRSGTGGTQY | 0.40934426 |
| SGDDNDAAL | ASRSGTGGTQY | 0.45617306 |
| AVAALAHS | ASRSGTGGTQY | 0.84083444 |
| GGVTVMATP | ASRSGTGGTQY | 0.27458695 |
| AGLAANVRD | ASRSGTGGTQY | 0.80084836 |
| TNRPDLIDPALLRPG | ASRSGTGGTQY | 0.74220616 |
| LANLGVTAL | ASRSGTGGTQY | 0.32627457 |
| TGERIEL | ASRSGTGGTQY | 0.7789912 |
| NGEELAA | ASRSGTGGTQY | 0.24823816 |
| VLSLVWG | ASRSGTGGTQY | 0.7368833 |
| IGYALWL | ASRSGTGGTQY | 0.7434368 |
| LSLLLAPL | ASRSGTGGTQY | 0.74751866 |
| TLDEALK | ASRSGTGGTQY | 0.5126134 |
| TDGAGTAEGG | ASRSGTGGTQY | 0.80036145 |
| NAPGGAGGNGG | ASRSGTGGTQY | 0.77418363 |
| VSVFMGMGGNF | ASRSGTGGTQY | 0.40374136 |
| LASYLAAL | ASRSGTGGTQY | 0.7859085 |
| LVGGTPAL | ASRSGTGGTQY | 0.80167353 |
| VRNADVL | ASRSGTGGTQY | 0.78657657 |
| TCSSPFLVSAM | ASRSGTGGTQY | 0.22748101 |
| VHITDAE | ASRSGTGGTQY | 0.6816722 |
| EYAAPSF | ASRSGTGGTQY | 0.4924214 |
| VVDAFHGGL | ASRSGTGGTQY | 0.7397895 |
| ASVVDPQ | ASRSGTGGTQY | 0.67201257 |
| SYNEFGGLS | ASRSGTGGTQY | 0.5897365 |
| DPDGSVY | ASRSGTGGTQY | 0.7609966 |
| MVSTIHATE | ASRSGTGGTQY | 0.31468326 |
| LAAGVADPV | ASRSGTGGTQY | 0.5460249 |
| TKEQIAA | ASRSGTGGTQY | 0.5736169 |
| SLASPAAL | ASRSGTGGTQY | 0.7397911 |
| IGLVFYGV | ASRSGTGGTQY | 0.7582029 |
| FLLDASNL | ASRSGTGGTQY | 0.3815369 |
| YQVGNFP | ASRSGTGGTQY | 0.7362558 |
| ESLLSQST | ASRSGTGGTQY | 0.6882744 |
| LFTVGFAAVG | ASRSGTGGTQY | 0.49600014 |
| ATPPTPT | ASRSGTGGTQY | 0.76429504 |

| | | |
|---|---|---|
| TDLATLG | ASRSGTGGTQY | 0.54719126 |
| SAGGTGR | ASRSGTGGTQY | 0.22769976 |
| GFSNSGS | ASRSGTGGTQY | 0.8076619 |
| GGNGGSGGTSV | ASRSGTGGTQY | 0.4127294 |
| GTSVATGGAGN | ASRSGTGGTQY | 0.7088555 |
| NGGNGGSGGTS | ASRSGTGGTQY | 0.87265635 |
| AVVKFPH | ASRSGTGGTQY | 0.308801 |
| TPPPPPP | ASRSGTGGTQY | 0.81358945 |
| AAPAAAAAPA | ASRSGTGGTQY | 0.63381803 |
| IALNEDL | ASRSGTGGTQY | 0.7235409 |
| IVNVSSM | ASRSGTGGTQY | 0.6735996 |
| TGIPPAPR | ASRSGTGGTQY | 0.44661802 |
| FDLGGGTF | ASRSGTGGTQY | 0.8372403 |
| FDLGGGTFDVS | ASRSGTGGTQY | 0.7251005 |
| NEPTAAA | ASRSGTGGTQY | 0.7995564 |
| NEPTAAALA | ASRSGTGGTQY | 0.74210024 |
| GSIINISSI | ASRSGTGGTQY | 0.8034861 |
| VAPGNAGTA | ASRSGTGGTQY | 0.7914379 |
| TGGGASLE | ASRSGTGGTQY | 0.42067337 |
| VAVYNHYKRI | ASRSGTGGTQY | 0.25239044 |
| TDPAPATT | ASRSGTGGTQY | 0.6421318 |
| TNRPDILDPALLRPG | ASRSGTGGTQY | 0.62234235 |
| LTTTTTS | ASRSGTGGTQY | 0.43626732 |
| PTGSGKTL | ASRSGTGGTQY | 0.22985864 |
| TGSGKTL | ASRSGTGGTQY | 0.7958906 |
| TIGHVDHGK | ASRSGTGGTQY | 0.8781042 |
| TIGHVDHGKTT | ASRSGTGGTQY | 0.5038357 |
| GHVDHGKTT | ASRSGTGGTQY | 0.86904675 |
| AQFEHTLL | ASRSGTGGTQY | 0.39522916 |
| IVNQDEL | ASRSGTGGTQY | 0.77336323 |
| TNGSQFF | ASRSGTGGTQY | 0.81772566 |
| TGKGIVE | ASRSGTGGTQY | 0.32401517 |
| LISNASDA | ASRSGTGGTQY | 0.7331939 |
| AELRTLGELTKIAKAHGAQ | ASRSGTGGTQY | 0.4520651 |
| APNAVAAAEPSW | ASRSGTGGTQY | 0.40099153 |
| APQVAGAATAVL | ASRSGTGGTQY | 0.8292276 |
| ARRDPAAAAAILPTDA | ASRSGTGGTQY | 0.5760567 |
| DAAARDVLAEAGLAEYF | ASRSGTGGTQY | 0.78767586 |
| DDQIVLSAKANLAKVEQ | ASRSGTGGTQY | 0.49085343 |
| DELLPDTGEDSDEDGH | ASRSGTGGTQY | 0.46595252 |
| ELPEAPANRATVPQD | ASRSGTGGTQY | 0.7350627 |
| FGQSKIERVTGIG | ASRSGTGGTQY | 0.81609213 |
| GAGGTGSTAGGAGGAGGAGGLY | ASRSGTGGTQY | 0.69743955 |
| GHGGTGAAGGNGGTGGAGGSGI | ASRSGTGGTQY | 0.883 |
| GNGGNGGAGGEGGPGGAGGAGA | ASRSGTGGTQY | 0.46518987 |
| GSTAGNGGNGGNATNGTIAGQP | ASRSGTGGTQY | 0.86751646 |
| GTGGDGGNGGNGGTGG | ASRSGTGGTQY | 0.8578014 |
| GVRRPAGIPADAPV | ASRSGTGGTQY | 0.73813164 |
| GYRQVTDAHLLTLARR | ASRSGTGGTQY | 0.6041571 |
| IAKMGQDGHDRG | ASRSGTGGTQY | 0.81300884 |
| KAEPSIEIQYT | ASRSGTGGTQY | 0.57610834 |

| | | |
|---|---|---|
| KPVRIRMAADEDTVAVT | ASRSGTGGTQY | 0.78288394 |
| MPPRIAGMRLLVIKPEP | ASRSGTGGTQY | 0.8439757 |
| MKFYAVALFRSRGVPPTR | ASRSGTGGTQY | 0.61403465 |
| NAGGFGILGAVAHSP | ASRSGTGGTQY | 0.89470506 |
| NEGREITLAMAEM | ASRSGTGGTQY | 0.8636966 |
| NELVEWSHPLGHRWRWE | ASRSGTGGTQY | 0.8189857 |
| NLGAGGTGGNGGNPG | ASRSGTGGTQY | 0.70913965 |
| NTEWPKWETVMAADPVD | ASRSGTGGTQY | 0.45682642 |
| PASAAPAETQP | ASRSGTGGTQY | 0.78996414 |
| QQSSDGKNGVRAAEITGE | ASRSGTGGTQY | 0.6237961 |
| RAPDAVTTSWWKE | ASRSGTGGTQY | 0.8660067 |
| SGFGNVGAHTSGWFNQ | ASRSGTGGTQY | 0.87162566 |
| SGVAGAVGQAASVGGLKV | ASRSGTGGTQY | 0.693051 |
| SNGGAGTGSAGGNGGT | ASRSGTGGTQY | 0.88686216 |
| TGGAGGEGGAAG | ASRSGTGGTQY | 0.8832972 |
| TGGDGGAAGAGGAPGHG | ASRSGTGGTQY | 0.45016348 |
| TMPPLLAAIANAGRHGV | ASRSGTGGTQY | 0.8007605 |
| TSSTAGNSGGA | ASRSGTGGTQY | 0.7553692 |
| VPATVPVGILC | ASRSGTGGTQY | 0.790995 |
| VQRPDSGGVVLTGRIS | ASRSGTGGTQY | 0.34659597 |
| WTPNPVPATKGVPAGLVQAT | ASRSGTGGTQY | 0.8713608 |
| YALKRALAKFSDRFDVVII | ASRSGTGGTQY | 0.7908164 |
| YLIGSVIV | ASRSGTGGTQY | 0.14573848 |
| YGQLVAVID | ASRSGTGGTQY | 0.8383012 |
| VTVGGMTLPA | ASRSGTGGTQY | 0.7777074 |
| VLALPAV | ASRSGTGGTQY | 0.7415866 |
| VIPPTLNYET | ASRSGTGGTQY | 0.885551 |
| VGMGGMHPSSG | ASRSGTGGTQY | 0.6191585 |
| VGLAGGAATA | ASRSGTGGTQY | 0.6355605 |
| VGGAGGAGGSGP | ASRSGTGGTQY | 0.8725959 |
| VFAAFMV | ASRSGTGGTQY | 0.88989925 |
| TLYDKLR | ASRSGTGGTQY | 0.5086448 |
| TLAVFTVL | ASRSGTGGTQY | 0.8609531 |
| TGSVQPARV | ASRSGTGGTQY | 0.8904709 |
| TGGGAGSGGT | ASRSGTGGTQY | 0.8599979 |
| SHLETSL | ASRSGTGGTQY | 0.8873204 |
| SFTTGMT | ASRSGTGGTQY | 0.89464325 |
| QVEVAKRMGVRQPTVS | ASRSGTGGTQY | 0.7286599 |
| QGSGGPALAM | ASRSGTGGTQY | 0.8838425 |
| MLAPSNP | ASRSGTGGTQY | 0.8927891 |
| LVNNAAI | ASRSGTGGTQY | 0.7262075 |
| LTGEINTLA | ASRSGTGGTQY | 0.81774116 |
| LSIFGPV | ASRSGTGGTQY | 0.65177876 |
| LLALLANL | ASRSGTGGTQY | 0.81066 |
| LISNASDALDKLR | ASRSGTGGTQY | 0.32921034 |
| LINEAVP | ASRSGTGGTQY | 0.85666674 |
| LGYAFSGL | ASRSGTGGTQY | 0.8884865 |
| LDFVAVAL | ASRSGTGGTQY | 0.79748833 |
| LAQPGAE | ASRSGTGGTQY | 0.67642486 |
| ISNLGAVTGF | ASRSGTGGTQY | 0.6088739 |
| HQEDPGRI | ASRSGTGGTQY | 0.84610933 |

| | | |
|---|---|---|
| GTPGEPTP | ASRSGTGGTQY | 0.7225716 |
| GFFNSGS | ASRSGTGGTQY | 0.3458364 |
| GAGGGGLTAGAN | ASRSGTGGTQY | 0.7943978 |
| FVDALVNM | ASRSGTGGTQY | 0.87760925 |
| FLTVAPEL | ASRSGTGGTQY | 0.8780255 |
| FETLTAL | ASRSGTGGTQY | 0.5047096 |
| DVINPIEL | ASRSGTGGTQY | 0.89548063 |
| APVSGAPV | ASRSGTGGTQY | 0.8756461 |
| ALVGGLAW | ASRSGTGGTQY | 0.89527595 |
| ALAAAAATAI | ASRSGTGGTQY | 0.83017564 |
| AFGLGLLL | ASRSGTGGTQY | 0.6737075 |
| AFFAAASL | ASRSGTGGTQY | 0.88055456 |
| AAAALPAFSPP | ASRSGTGGTQY | 0.8086406 |
| VVLAPAL | ASRSGTGGTQY | 0.70831466 |
| TIGHVDHGKTTL | ASRSGTGGTQY | 0.8790635 |
| LISNASDALDK | ASRSGTGGTQY | 0.80422115 |
| LISNASDAL | ASRSGTGGTQY | 0.7030109 |
| ISNASDALDK | ASRSGTGGTQY | 0.7947876 |
| NQAETLVYQTE | ASRSGTGGTQY | 0.8974484 |
| TGGTGGTAGAG | ASRSGTGGTQY | 0.6882274 |
| TGSTAGGAGGA | ASRSGTGGTQY | 0.57099724 |
| DPLAVPVAAGPV | ASRSGTGGTQY | 0.7103698 |
| GGTGGTPGNGGN | ASRSGTGGTQY | 0.82833457 |
| PAGLAGVDPATL | ASRSGTGGTQY | 0.8833438 |
| GNNLVVEAPHAGASV | ASRSGTGGTQY | 0.8485242 |
| KVTRIMSASTLEQPA | ASRSGTGGTQY | 0.79464567 |
| LGFALPIALPRRTVV | ASRSGTGGTQY | 0.8579091 |
| TRLGNLTARKLVVLV | ASRSGTGGTQY | 0.8610174 |
| WSQQRRLIYFPSAGP | ASRSGTGGTQY | 0.8344242 |
| GPNGNPDPMAAAVDIR | ASRSGTGGTQY | 0.8818652 |
| TQRCVSVADLRAAASR | ASRSGTGGTQY | 0.7368474 |
| QFVLDVAEHPVEVLTETV | ASRSGTGGTQY | 0.8946471 |
| GGAGGTGGAGGNGGTQAGDG | ASRSGTGGTQY | 0.8669987 |
| PLLATLRAIAVDVDAKPAQV | ASRSGTGGTQY | 0.8733707 |
| APKFRTGGYMIDFWGVGYQVAK | ASRSGTGGTQY | 0.8835082 |
| GAGGAGGAGGAAGGPGGTGGTG | ASRSGTGGTQY | 0.8459388 |
| IGALGLLATDVAPAGSVAFGVA | ASRSGTGGTQY | 0.8550446 |
| AGGAGGIGGNGGAGGAGGVLMG | ASRSGTGGTQY | 0.3608295 |
| ADELIGLL | ASRSGTGGTQY | 0.78608686 |
| ADRALKA | ASRSGTGGTQY | 0.8584791 |
| AGAVGQVGGAGGA | ASRSGTGGTQY | 0.8588026 |
| AGGGGGLGGAP | ASRSGTGGTQY | 0.8713037 |
| AGGLLYGNGGAG | ASRSGTGGTQY | 0.846617 |
| AGLESIGL | ASRSGTGGTQY | 0.74311364 |
| AGLKTGII | ASRSGTGGTQY | 0.8896151 |
| ASLAALLL | ASRSGTGGTQY | 0.8865336 |
| ASVINISSI | ASRSGTGGTQY | 0.514557 |
| ATPPGPA | ASRSGTGGTQY | 0.80130994 |
| ATVASALV | ASRSGTGGTQY | 0.8997072 |
| ATVASLIL | ASRSGTGGTQY | 0.85802203 |
| AVNGPVLIRH | ASRSGTGGTQY | 0.78457904 |

| | | |
|---|---|---|
| DGAAGAVGI | ASRSGTGGTQY | 0.7491382 |
| DPLLHGI | ASRSGTGGTQY | 0.88919705 |
| ELGAALGAL | ASRSGTGGTQY | 0.7592055 |
| ELGRTQAV | ASRSGTGGTQY | 0.89709973 |
| FVAGGYLS | ASRSGTGGTQY | 0.204617 |
| GASGGAGGAGGW | ASRSGTGGTQY | 0.76803255 |
| GGAGAGGDGGAGGI | ASRSGTGGTQY | 0.7573802 |
| GTGHNGSQPGMGGNG | ASRSGTGGTQY | 0.28982142 |
| GYAAASASAS | ASRSGTGGTQY | 0.8771352 |
| HQVDVAAVQQG | ASRSGTGGTQY | 0.7082716 |
| IGAAIAHT | ASRSGTGGTQY | 0.8689965 |
| IGMIEVAQL | ASRSGTGGTQY | 0.8305951 |
| IRLAPEHTSAMD | ASRSGTGGTQY | 0.88008153 |
| IYNVSVL | ASRSGTGGTQY | 0.8505941 |
| KQGFLGM | ASRSGTGGTQY | 0.8169751 |
| LESVITQL | ASRSGTGGTQY | 0.6069002 |
| LGLGALVVV | ASRSGTGGTQY | 0.86818194 |
| LIGNGGNGGSGGI | ASRSGTGGTQY | 0.89110744 |
| LNAINAPTL | ASRSGTGGTQY | 0.88157535 |
| LQLLNTGV | ASRSGTGGTQY | 0.34553242 |
| LQTSIEF | ASRSGTGGTQY | 0.832238 |
| MSFAVSEL | ASRSGTGGTQY | 0.80350745 |
| NTGALGL | ASRSGTGGTQY | 0.2278386 |
| PPGAGGAGGAGTT | ASRSGTGGTQY | 0.23462984 |
| QLVGLGI | ASRSGTGGTQY | 0.8146802 |
| RFLEQVHQL | ASRSGTGGTQY | 0.80196106 |
| RGVGYVLR | ASRSGTGGTQY | 0.8810366 |
| SATGVFAGVI | ASRSGTGGTQY | 0.8716148 |
| SLPAITLP | ASRSGTGGTQY | 0.78878546 |
| STALAFTV | ASRSGTGGTQY | 0.8782423 |
| SVLTVGLGK | ASRSGTGGTQY | 0.8336669 |
| TALAGAGSGI | ASRSGTGGTQY | 0.08761262 |
| TGLPFTGI | ASRSGTGGTQY | 0.76714027 |
| VAEHVTV | ASRSGTGGTQY | 0.44155997 |
| VDIVLNSL | ASRSGTGGTQY | 0.8856863 |
| VLTGGGAVL | ASRSGTGGTQY | 0.711407 |
| VNVGLAYGI | ASRSGTGGTQY | 0.74492776 |
| VSSAAVGAV | ASRSGTGGTQY | 0.81568074 |
| VTSVYEAGV | ASRSGTGGTQY | 0.83969337 |
| WVALDRGA | ASRSGTGGTQY | 0.83208305 |
| YLVGSVVI | ASRSGTGGTQY | 0.69870555 |
| GPPGTGKTLL | ASRSGTGGTQY | 0.18936981 |
| GVGMTKF | ASRSGTGGTQY | 0.867668 |
| IGIGGSDL | ASRSGTGGTQY | 0.45730788 |
| IGTLLAEL | ASRSGTGGTQY | 0.5587699 |
| IYTSGTT | ASRSGTGGTQY | 0.3581283 |
| LSFPAGV | ASRSGTGGTQY | 0.45469016 |
| LYTSGTT | ASRSGTGGTQY | 0.38257682 |
| VGAFPFGF | ASRSGTGGTQY | 0.8788796 |
| VGAGTVEF | ASRSGTGGTQY | 0.8084527 |
| VGKSTLLL | ASRSGTGGTQY | 0.7999342 |

| | | |
|---|---|---|
| AALAHAGGGIV | ASRSGTGGTQY | 0.8978525 |
| AALVAVSLL | ASRSGTGGTQY | 0.82111794 |
| AAVSRAVAAL | ASRSGTGGTQY | 0.8072708 |
| AGGSSGIGGF | ASRSGTGGTQY | 0.64006686 |
| AGLTVANANG | ASRSGTGGTQY | 0.63993007 |
| AGLVGAVL | ASRSGTGGTQY | 0.6889708 |
| ALSQAFTV | ASRSGTGGTQY | 0.6630079 |
| ALTFSVSL | ASRSGTGGTQY | 0.8982622 |
| ALTGAALVAL | ASRSGTGGTQY | 0.764159 |
| ASFSIGAGVL | ASRSGTGGTQY | 0.4722839 |
| DGSVYKI | ASRSGTGGTQY | 0.8238164 |
| EQLAEIV | ASRSGTGGTQY | 0.89633787 |
| FAVSELTVM | ASRSGTGGTQY | 0.9050545 |
| GAAPLAAGL | ASRSGTGGTQY | 0.67274356 |
| GADGGDGGSGGAGG | ASRSGTGGTQY | 0.78141725 |
| GFGLDTGGAGG | ASRSGTGGTQY | 0.77508146 |
| GLASGGLL | ASRSGTGGTQY | 0.8200593 |
| GRIAGAAVLL | ASRSGTGGTQY | 0.7763916 |
| GSAGSLNTSF | ASRSGTGGTQY | 0.47307953 |
| GTGQIGI | ASRSGTGGTQY | 0.8875933 |
| GTLIALIL | ASRSGTGGTQY | 0.8963373 |
| IAATMGFVVG | ASRSGTGGTQY | 0.78850466 |
| IFAQALL | ASRSGTGGTQY | 0.48535466 |
| IGAAVALGF | ASRSGTGGTQY | 0.7981636 |
| IGLTPLAL | ASRSGTGGTQY | 0.69658697 |
| ILGSNTSTL | ASRSGTGGTQY | 0.7544821 |
| IVFSLTVL | ASRSGTGGTQY | 0.78869987 |
| LASKATGF | ASRSGTGGTQY | 0.53304446 |
| LASVALIL | ASRSGTGGTQY | 0.8108326 |
| LFAVGLTL | ASRSGTGGTQY | 0.9025154 |
| LFNGGIGTYI | ASRSGTGGTQY | 0.7806345 |
| LTLGAGII | ASRSGTGGTQY | 0.86736995 |
| LVAAALGDI | ASRSGTGGTQY | 0.89121944 |
| NTAFVQLGI | ASRSGTGGTQY | 0.8751991 |
| QLFSGAEL | ASRSGTGGTQY | 0.89448273 |
| SAATFITL | ASRSGTGGTQY | 0.8742931 |
| SNYGLANLGV | ASRSGTGGTQY | 0.78863645 |
| SYGGAYAVM | ASRSGTGGTQY | 0.84465337 |
| TTGVGGLTLG | ASRSGTGGTQY | 0.86435705 |
| TTQTIYI | ASRSGTGGTQY | 0.807652 |
| VAAGAGLGAVI | ASRSGTGGTQY | 0.8135692 |
| VAAVVTLL | ASRSGTGGTQY | 0.7574079 |
| VGFLGGLI | ASRSGTGGTQY | 0.877838 |
| VGVIPILL | ASRSGTGGTQY | 0.84044373 |
| VIFASSLI | ASRSGTGGTQY | 0.88507706 |
| VKGGDGGAAAG | ASRSGTGGTQY | 0.34809816 |
| VTKTFTNI | ASRSGTGGTQY | 0.5408778 |
| VVAAAILL | ASRSGTGGTQY | 0.8820857 |
| YAALDVEL | ASRSGTGGTQY | 0.14545482 |
| LASVALLL | ASRSGTGGTQY | 0.21632268 |
| LDAVHRV | ASRSGTGGTQY | 0.8594801 |

| | | |
|---|---|---|
| QGLSSSIF | ASRSGTGGTQY | 0.8943726 |
| RGTFANIRL | ASRSGTGGTQY | 0.14534591 |
| RQLWWGHRI | ASRSGTGGTQY | 0.5615684 |
| YGPPGTGKTLL | ASRSGTGGTQY | 0.56947815 |
| AALVGAPANVA | ASRSGTGGTQY | 0.8621823 |
| AATFRRTE | ASRSGTGGTQY | 0.8108776 |
| AGALGGGI | ASRSGTGGTQY | 0.39896083 |
| AGGAGGAGGIGAG | ASRSGTGGTQY | 0.86767095 |
| AGSGALGL | ASRSGTGGTQY | 0.43854785 |
| AQIADQI | ASRSGTGGTQY | 0.7093401 |
| ASAVAVTYL | ASRSGTGGTQY | 0.8788576 |
| ASTVLGGTGG | ASRSGTGGTQY | 0.89012265 |
| AVGGEPAK | ASRSGTGGTQY | 0.62615913 |
| CEQGTFGGTC | ASRSGTGGTQY | 0.87480503 |
| DEGKAPAG | ASRSGTGGTQY | 0.7768315 |
| DTVVFPAL | ASRSGTGGTQY | 0.90457535 |
| FEGITMAG | ASRSGTGGTQY | 0.75603384 |
| FPLVVQD | ASRSGTGGTQY | 0.47521436 |
| GAFTTPQ | ASRSGTGGTQY | 0.87942994 |
| GFGGFGTAGGVGGA | ASRSGTGGTQY | 0.8783143 |
| GFLAQFL | ASRSGTGGTQY | 0.795419 |
| GGAGGAGGNAGVA | ASRSGTGGTQY | 0.87828016 |
| GGVGTAGAGGAGGAGG | ASRSGTGGTQY | 0.75253534 |
| GNGGGGDAGNAGSGGN | ASRSGTGGTQY | 0.7968535 |
| GTGGAGGAGNDAGS | ASRSGTGGTQY | 0.87954926 |
| GTTNTGWFNS | ASRSGTGGTQY | 0.8829107 |
| IFMGPLFTL | ASRSGTGGTQY | 0.5668514 |
| ILAHAIEAI | ASRSGTGGTQY | 0.8837715 |
| IRSAIREAL | ASRSGTGGTQY | 0.80797595 |
| ITFAGGG | ASRSGTGGTQY | 0.6450863 |
| ITGMSGHQEV | ASRSGTGGTQY | 0.9026606 |
| IVAGLACTL | ASRSGTGGTQY | 0.8281135 |
| IVGGGLAGA | ASRSGTGGTQY | 0.88356537 |
| IVTGVLNI | ASRSGTGGTQY | 0.873049 |
| LAQQGCI | ASRSGTGGTQY | 0.8855995 |
| LCLHVEN | ASRSGTGGTQY | 0.5256257 |
| LDAISDGS | ASRSGTGGTQY | 0.8561323 |
| LGGFAAML | ASRSGTGGTQY | 0.84265953 |
| LGGKNPMI | ASRSGTGGTQY | 0.8707812 |
| LGPKGVAAIL | ASRSGTGGTQY | 0.8217006 |
| LMVGVSI | ASRSGTGGTQY | 0.6559981 |
| LVGAAFVL | ASRSGTGGTQY | 0.8641994 |
| LVLAVAPL | ASRSGTGGTQY | 0.89355487 |
| MAAIAVIDAA | ASRSGTGGTQY | 0.8514409 |
| NNGINAELQ | ASRSGTGGTQY | 0.84861207 |
| RGEPLVLVHG | ASRSGTGGTQY | 0.8980477 |
| RGGAGGAAGLI | ASRSGTGGTQY | 0.66678596 |
| RSYRMHPAV | ASRSGTGGTQY | 0.7816333 |
| SAAHLLAGQLA | ASRSGTGGTQY | 0.883054 |
| SAAVGSY | ASRSGTGGTQY | 0.88605154 |
| SPHAVLHRI | ASRSGTGGTQY | 0.78806686 |

| | | |
|---|---|---|
| SQIAKVL | ASRSGTGGTQY | 0.8953805 |
| TALGHLVP | ASRSGTGGTQY | 0.7373965 |
| TAPTTSGAA | ASRSGTGGTQY | 0.8646966 |
| TDGDDGNGGSPGT | ASRSGTGGTQY | 0.779783 |
| TDGLGEN | ASRSGTGGTQY | 0.84343195 |
| TGGPPLPGQA | ASRSGTGGTQY | 0.70085394 |
| TIPAITI | ASRSGTGGTQY | 0.86219174 |
| TIPGFTI | ASRSGTGGTQY | 0.8831296 |
| TSIASGALA | ASRSGTGGTQY | 0.87346125 |
| TVTLPGAAI | ASRSGTGGTQY | 0.19387421 |
| VASGLSAALP | ASRSGTGGTQY | 0.87592953 |
| VGAGVIGI | ASRSGTGGTQY | 0.8563309 |
| VGPKRFGV | ASRSGTGGTQY | 0.90200365 |
| VNVAFPD | ASRSGTGGTQY | 0.831697 |
| VSMTNSGVS | ASRSGTGGTQY | 0.86161584 |
| VTDVLGDLSV | ASRSGTGGTQY | 0.7422023 |
| VVVVGAGFAGL | ASRSGTGGTQY | 0.8101468 |
| VYTVGTVV | ASRSGTGGTQY | 0.61609745 |
| WTTFSGP | ASRSGTGGTQY | 0.7751327 |
| YDELVEAF | ASRSGTGGTQY | 0.7942301 |
| YGTGGAGGNGG | ASRSGTGGTQY | 0.7169923 |
| YTNVMAV | ASRSGTGGTQY | 0.8721286 |
| FLEQVHQL | ASRSGTGGTQY | 0.88932455 |
| GLSLLAVAV | ASRSGTGGTQY | 0.8853946 |
| LEQVHQL | ASRSGTGGTQY | 0.88908565 |
| LNAAGAI | ASRSGTGGTQY | 0.8976753 |
| PAVHASL | ASRSGTGGTQY | 0.88154286 |
| TELEDDA | ASRSGTGGTQY | 0.7537416 |
| TPPGPPN | ASRSGTGGTQY | 0.8151578 |
| TPPPPPPP | ASRSGTGGTQY | 0.5515933 |
| TQSPALI | ASSWGGENEQY | 0.79407096 |
| EFAAAFPQ | ASSWGGENEQY | 0.76575756 |
| LLSGNLDVL | ASSWGGENEQY | 0.83942944 |
| GPANLGNYNI | ASSWGGENEQY | 0.10998602 |
| TLGPTLP | ASSWGGENEQY | 0.84382737 |
| VYAYSSCCK | ASSWGGENEQY | 0.89814687 |
| APAPAPAPNI | ASSWGGENEQY | 0.76559794 |
| PGGPVPIP | ASSWGGENEQY | 0.8564416 |
| FLTVALEP | ASSWGGENEQY | 0.86348957 |
| SQYFGANL | ASSWGGENEQY | 0.7829093 |
| GGVKPAA | ASSWGGENEQY | 0.8754312 |
| GGTGGAGGAGGW | ASSWGGENEQY | 0.69335294 |
| GVGGVAGLGGAGG | ASSWGGENEQY | 0.8687405 |
| VLLVLATTI | ASSWGGENEQY | 0.89939445 |
| IGIFAAML | ASSWGGENEQY | 0.85231423 |
| AVGDLID | ASSWGGENEQY | 0.8458571 |
| NISPLPL | ASSWGGENEQY | 0.814212 |
| VTAVFVAI | ASSWGGENEQY | 0.37556756 |
| SGDDNDAAL | ASSWGGENEQY | 0.85385615 |
| AVAALAHS | ASSWGGENEQY | 0.13167076 |
| GGVTVMATP | ASSWGGENEQY | 0.8314718 |

| | | |
|---|---|---|
| AGLAANVRD | ASSWGGENEQY | 0.6752742 |
| TNRPDLIDPALLRPG | ASSWGGENEQY | 0.8352814 |
| LANLGVTAL | ASSWGGENEQY | 0.82499063 |
| TGERIEL | ASSWGGENEQY | 0.8718604 |
| NGEELAA | ASSWGGENEQY | 0.63277155 |
| VLSLVWG | ASSWGGENEQY | 0.83023393 |
| IGYALWL | ASSWGGENEQY | 0.77005804 |
| LSLLLAPL | ASSWGGENEQY | 0.50662637 |
| TLDEALK | ASSWGGENEQY | 0.8841408 |
| TDGAGTAEGG | ASSWGGENEQY | 0.8236023 |
| NAPGGAGGNGG | ASSWGGENEQY | 0.8321252 |
| VSVFMGMGGNF | ASSWGGENEQY | 0.88390434 |
| LASYLAAL | ASSWGGENEQY | 0.84194887 |
| LVGGTPAL | ASSWGGENEQY | 0.41546395 |
| VRNADVL | ASSWGGENEQY | 0.82542396 |
| TCSSPFLVSAM | ASSWGGENEQY | 0.8276919 |
| VHITDAE | ASSWGGENEQY | 0.8043594 |
| EYAAPSF | ASSWGGENEQY | 0.12642646 |
| VVDAFHGGL | ASSWGGENEQY | 0.86901796 |
| ASVVDPQ | ASSWGGENEQY | 0.6383753 |
| SYNEFGGLS | ASSWGGENEQY | 0.7383063 |
| DPDGSVY | ASSWGGENEQY | 0.7702588 |
| MVSTIHATE | ASSWGGENEQY | 0.88082194 |
| LAAGVADPV | ASSWGGENEQY | 0.7945653 |
| TKEQIAA | ASSWGGENEQY | 0.7199436 |
| SLASPAAL | ASSWGGENEQY | 0.88072056 |
| IGLVFYGV | ASSWGGENEQY | 0.84523004 |
| FLLDASNL | ASSWGGENEQY | 0.8918501 |
| YQVGNFP | ASSWGGENEQY | 0.83726543 |
| ESLLSQST | ASSWGGENEQY | 0.8411555 |
| LFTVGFAAVG | ASSWGGENEQY | 0.83847946 |
| ATPPTPT | ASSWGGENEQY | 0.8186699 |
| TDLATLG | ASSWGGENEQY | 0.49512604 |
| SAGGTGR | ASSWGGENEQY | 0.39744717 |
| GFSNSGS | ASSWGGENEQY | 0.822258 |
| GGNGGSGGTSV | ASSWGGENEQY | 0.37039912 |
| GTSVATGGAGN | ASSWGGENEQY | 0.8253045 |
| NGGNGGSGGTS | ASSWGGENEQY | 0.88436353 |
| AVVKFPH | ASSWGGENEQY | 0.83821934 |
| TPPPPPP | ASSWGGENEQY | 0.53591037 |
| AAPAAAAAPA | ASSWGGENEQY | 0.7997633 |
| IALNEDL | ASSWGGENEQY | 0.77931476 |
| IVNVSSM | ASSWGGENEQY | 0.8739705 |
| TGIPPAPR | ASSWGGENEQY | 0.87616783 |
| FDLGGGTF | ASSWGGENEQY | 0.89017916 |
| FDLGGGTFDVS | ASSWGGENEQY | 0.87458825 |
| NEPTAAA | ASSWGGENEQY | 0.7484586 |
| NEPTAAALA | ASSWGGENEQY | 0.87950236 |
| GSIINISSI | ASSWGGENEQY | 0.78808117 |
| VAPGNAGTA | ASSWGGENEQY | 0.83525646 |
| TGGGASLE | ASSWGGENEQY | 0.8349166 |

| | | |
|---|---|---|
| VAVYNHYKRI | ASSWGGENEQY | 0.86212826 |
| TDPAPATT | ASSWGGENEQY | 0.7832964 |
| TNRPDILDPALLRPG | ASSWGGENEQY | 0.7806059 |
| LTTTTTS | ASSWGGENEQY | 0.4079814 |
| PTGSGKTL | ASSWGGENEQY | 0.8687276 |
| TGSGKTL | ASSWGGENEQY | 0.84385407 |
| TIGHVDHGK | ASSWGGENEQY | 0.5258107 |
| TIGHVDHGKTT | ASSWGGENEQY | 0.75010014 |
| GHVDHGKTT | ASSWGGENEQY | 0.89964473 |
| AQFEHTLL | ASSWGGENEQY | 0.8931359 |
| IVNQDEL | ASSWGGENEQY | 0.70534664 |
| TNGSQFF | ASSWGGENEQY | 0.66479945 |
| TGKGIVE | ASSWGGENEQY | 0.8509605 |
| LISNASDA | ASSWGGENEQY | 0.46420258 |
| AELRTLGELTKIAKAHGAQ | ASSWGGENEQY | 0.8571346 |
| APNAVAAAEPSW | ASSWGGENEQY | 0.8203976 |
| APQVAGAATAVL | ASSWGGENEQY | 0.8330507 |
| ARRDPAAAAAILPTDA | ASSWGGENEQY | 0.7376311 |
| DAAARDVLAEAGLAEYF | ASSWGGENEQY | 0.8605827 |
| DDQIVLSAKANLAKVEQ | ASSWGGENEQY | 0.7798066 |
| DELLPDTGEDSDEDGH | ASSWGGENEQY | 0.8684951 |
| ELPEAPANRATVPQD | ASSWGGENEQY | 0.15956186 |
| FGQSKIERVTGIG | ASSWGGENEQY | 0.8656156 |
| GAGGTGSTAGGAGGAGGAGGLY | ASSWGGENEQY | 0.68380183 |
| GHGGTGAAGGNGGTGGAGGSGI | ASSWGGENEQY | 0.72686785 |
| GNGGNGGAGGEGGPGGAGGAGA | ASSWGGENEQY | 0.37574196 |
| GSTAGNGGNGGNATNGTIAGQP | ASSWGGENEQY | 0.8007709 |
| GTGGDGGNGGNGGTGG | ASSWGGENEQY | 0.8805599 |
| GVRRPAGIPADAPV | ASSWGGENEQY | 0.88952124 |
| GYRQVTDAHLLTLARR | ASSWGGENEQY | 0.7486026 |
| IAKMGQDGHDRG | ASSWGGENEQY | 0.87907577 |
| KAEPSIEIQYT | ASSWGGENEQY | 0.90060174 |
| KPVRIRMAADEDTVAVT | ASSWGGENEQY | 0.8036754 |
| MPPRIAGMRLLVIKPEP | ASSWGGENEQY | 0.31038508 |
| MKFYAVALFRSRGVPPTR | ASSWGGENEQY | 0.8302009 |
| NAGGFGILGAVAHSP | ASSWGGENEQY | 0.81245697 |
| NEGREITLAMAEM | ASSWGGENEQY | 0.8800225 |
| NELVEWSHPLGHRWRWE | ASSWGGENEQY | 0.87552196 |
| NLGAGGTGGNGGNPG | ASSWGGENEQY | 0.59069884 |
| NTEWPKWETVMAADPVD | ASSWGGENEQY | 0.8500049 |
| PASAAPAETQP | ASSWGGENEQY | 0.88099223 |
| QQSSDGKNGVRAAEITGE | ASSWGGENEQY | 0.69752866 |
| RAPDAVTTSWWKE | ASSWGGENEQY | 0.75237286 |
| SGFGNVGAHTSGWFNQ | ASSWGGENEQY | 0.6732447 |
| SGVAGAVGQAASVGGLKV | ASSWGGENEQY | 0.3626458 |
| SNGGAGTGSAGGNGGT | ASSWGGENEQY | 0.85878724 |
| TGGAGGEGGAAG | ASSWGGENEQY | 0.7838976 |
| TGGDGGAAGAGGAPGHG | ASSWGGENEQY | 0.6746577 |
| TMPPLLAAIANAGRHGV | ASSWGGENEQY | 0.821126 |
| TSSTAGNSGGA | ASSWGGENEQY | 0.6817008 |
| VPATVPVGILC | ASSWGGENEQY | 0.66708684 |

| | | |
|---|---|---|
| VQRPDSGGVVLTGRIS | ASSWGGENEQY | 0.7465848 |
| WTPNPVPATKGVPAGLVQAT | ASSWGGENEQY | 0.8233 |
| YALKRALAKFSDRFDVVII | ASSWGGENEQY | 0.87247086 |
| YLIGSVIV | ASSWGGENEQY | 0.8232079 |
| YGQLVAVID | ASSWGGENEQY | 0.8346497 |
| VTVGGMTLPA | ASSWGGENEQY | 0.8953126 |
| VLALPAV | ASSWGGENEQY | 0.5052788 |
| VIPPTLNYET | ASSWGGENEQY | 0.6298063 |
| VGMGGMHPSSG | ASSWGGENEQY | 0.8703151 |
| VGLAGGAATA | ASSWGGENEQY | 0.59974366 |
| VGGAGGAGGSGP | ASSWGGENEQY | 0.87766385 |
| VFAAFMV | ASSWGGENEQY | 0.8252318 |
| TLYDKLR | ASSWGGENEQY | 0.8562646 |
| TLAVFTVL | ASSWGGENEQY | 0.832564 |
| TGSVQPARV | ASSWGGENEQY | 0.90278476 |
| TGGGAGSGGT | ASSWGGENEQY | 0.8741542 |
| SHLETSL | ASSWGGENEQY | 0.7898541 |
| SFTTGMT | ASSWGGENEQY | 0.85866725 |
| QVEVAKRMGVRQPTVS | ASSWGGENEQY | 0.5344137 |
| QGSGGPALAM | ASSWGGENEQY | 0.88335305 |
| MLAPSNP | ASSWGGENEQY | 0.8659371 |
| LVNNAAI | ASSWGGENEQY | 0.6876304 |
| LTGEINTLA | ASSWGGENEQY | 0.7994236 |
| LSIFGPV | ASSWGGENEQY | 0.9000323 |
| LLALLANL | ASSWGGENEQY | 0.8904553 |
| LISNASDALDKLR | ASSWGGENEQY | 0.4495657 |
| LINEAVP | ASSWGGENEQY | 0.75192267 |
| LGYAFSGL | ASSWGGENEQY | 0.4735947 |
| LDFVAVAL | ASSWGGENEQY | 0.8058901 |
| LAQPGAE | ASSWGGENEQY | 0.4101863 |
| ISNLGAVTGF | ASSWGGENEQY | 0.87368524 |
| HQEDPGRI | ASSWGGENEQY | 0.76031923 |
| GTPGEPTP | ASSWGGENEQY | 0.8613458 |
| GFFNSGS | ASSWGGENEQY | 0.8305601 |
| GAGGGGLTAGAN | ASSWGGENEQY | 0.8867402 |
| FVDALVNM | ASSWGGENEQY | 0.7552049 |
| FLTVAPEL | ASSWGGENEQY | 0.7333628 |
| FETLTAL | ASSWGGENEQY | 0.4181891 |
| DVINPIEL | ASSWGGENEQY | 0.50389004 |
| APVSGAPV | ASSWGGENEQY | 0.7630331 |
| ALVGGLAW | ASSWGGENEQY | 0.7474761 |
| ALAAAAATAI | ASSWGGENEQY | 0.23946017 |
| AFGLGLLL | ASSWGGENEQY | 0.895002 |
| AFFAAASL | ASSWGGENEQY | 0.8311227 |
| AAAALPAFSPP | ASSWGGENEQY | 0.86984146 |
| VVLAPAL | ASSWGGENEQY | 0.65501004 |
| TIGHVDHGKTTL | ASSWGGENEQY | 0.74149835 |
| LISNASDALDK | ASSWGGENEQY | 0.7414983 |
| LISNASDAL | ASSWGGENEQY | 0.8659943 |
| ISNASDALDK | ASSWGGENEQY | 0.6674812 |
| NQAETLVYQTE | ASSWGGENEQY | 0.29798707 |

| | | |
|---|---|---|
| TGGTGGTAGAG | ASSWGGENEQY | 0.7521336 |
| TGSTAGGAGGA | ASSWGGENEQY | 0.56483877 |
| DPLAVPVAAGPV | ASSWGGENEQY | 0.8995148 |
| GGTGGTPGNGGN | ASSWGGENEQY | 0.8857419 |
| PAGLAGVDPATL | ASSWGGENEQY | 0.8494307 |
| GNNLVVEAPHAGASV | ASSWGGENEQY | 0.8925961 |
| KVTRIMSASTLEQPA | ASSWGGENEQY | 0.19641612 |
| LGFALPIALPRRTVV | ASSWGGENEQY | 0.6933762 |
| TRLGNLTARKLVVLV | ASSWGGENEQY | 0.89128417 |
| WSQQRRLIYFPSAGP | ASSWGGENEQY | 0.86969256 |
| GPNGNPDPMAAAVDIR | ASSWGGENEQY | 0.79586786 |
| TQRCVSVADLRAAASR | ASSWGGENEQY | 0.8895025 |
| QFVLDVAEHPVEVLTETV | ASSWGGENEQY | 0.6613747 |
| GGAGGTGGAGGNGGTQAGDG | ASSWGGENEQY | 0.7898264 |
| PLLATLRAIAVDVDAKPAQV | ASSWGGENEQY | 0.82160807 |
| APKFRTGGYMIDFWGVGYQVAK | ASSWGGENEQY | 0.8695523 |
| GAGGAGGAGGAAGGPGGTGGTG | ASSWGGENEQY | 0.864218 |
| IGALGLLATDVAPAGSVAFGVA | ASSWGGENEQY | 0.5916102 |
| AGGAGGIGGNGGAGGAGGVLMG | ASSWGGENEQY | 0.8471413 |
| ADELIGLL | ASSWGGENEQY | 0.905218 |
| ADRALKA | ASSWGGENEQY | 0.7778809 |
| AGAVGQVGGAGGA | ASSWGGENEQY | 0.88353884 |
| AGGGGGLGGAP | ASSWGGENEQY | 0.8543977 |
| AGGLLYGNGGAG | ASSWGGENEQY | 0.83059716 |
| AGLESIGL | ASSWGGENEQY | 0.6681845 |
| AGLKTGII | ASSWGGENEQY | 0.74558526 |
| ASLAALLL | ASSWGGENEQY | 0.8277512 |
| ASVINISSI | ASSWGGENEQY | 0.6785425 |
| ATPPGPA | ASSWGGENEQY | 0.8118321 |
| ATVASALV | ASSWGGENEQY | 0.7706597 |
| ATVASLIL | ASSWGGENEQY | 0.9083786 |
| AVNGPVLIRH | ASSWGGENEQY | 0.8710711 |
| DGAAGAVGI | ASSWGGENEQY | 0.8660846 |
| DPLLHGI | ASSWGGENEQY | 0.6280993 |
| ELGAALGAL | ASSWGGENEQY | 0.8668957 |
| ELGRTQAV | ASSWGGENEQY | 0.86166745 |
| FVAGGYLS | ASSWGGENEQY | 0.4476037 |
| GASGGAGGAGGW | ASSWGGENEQY | 0.89493597 |
| GGAGAGGDGGAGGI | ASSWGGENEQY | 0.6447574 |
| GTGHNGSQPGMGGNG | ASSWGGENEQY | 0.8172538 |
| GYAAASASAS | ASSWGGENEQY | 0.85983133 |
| HQVDVAAVQQG | ASSWGGENEQY | 0.8773408 |
| IGAAIAHT | ASSWGGENEQY | 0.80299103 |
| IGMIEVAQL | ASSWGGENEQY | 0.82820785 |
| IRLAPEHTSAMD | ASSWGGENEQY | 0.61335635 |
| IYNVSVL | ASSWGGENEQY | 0.8832886 |
| KQGFLGM | ASSWGGENEQY | 0.8855834 |
| LESVITQL | ASSWGGENEQY | 0.8360036 |
| LGLGALVVV | ASSWGGENEQY | 0.8977354 |
| LIGNGGNGGSGGI | ASSWGGENEQY | 0.838248 |
| LNAINAPTL | ASSWGGENEQY | 0.34965247 |

| | | |
|---|---|---|
| LQLLNTGV | ASSWGGENEQY | 0.8160994 |
| LQTSIEF | ASSWGGENEQY | 0.8888257 |
| MSFAVSEL | ASSWGGENEQY | 0.892823 |
| NTGALGL | ASSWGGENEQY | 0.88722086 |
| PPGAGGAGGAGTT | ASSWGGENEQY | 0.86643547 |
| QLVGLGI | ASSWGGENEQY | 0.7941022 |
| RFLEQVHQL | ASSWGGENEQY | 0.8943375 |
| RGVGYVLR | ASSWGGENEQY | 0.9053849 |
| SATGVFAGVI | ASSWGGENEQY | 0.8690127 |
| SLPAITLP | ASSWGGENEQY | 0.73818743 |
| STALAFTV | ASSWGGENEQY | 0.654876 |
| SVLTVGLGK | ASSWGGENEQY | 0.8616863 |
| TALAGAGSGI | ASSWGGENEQY | 0.88405395 |
| TGLPFTGI | ASSWGGENEQY | 0.59520537 |
| VAEHVTV | ASSWGGENEQY | 0.8453789 |
| VDIVLNSL | ASSWGGENEQY | 0.8471501 |
| VLTGGGAVL | ASSWGGENEQY | 0.3719672 |
| VNVGLAYGI | ASSWGGENEQY | 0.86367464 |
| VSSAAVGAV | ASSWGGENEQY | 0.89523363 |
| VTSVYEAGV | ASSWGGENEQY | 0.83550704 |
| WVALDRGA | ASSWGGENEQY | 0.7250794 |
| YLVGSVVI | ASSWGGENEQY | 0.89201194 |
| GPPGTGKTLL | ASSWGGENEQY | 0.90538764 |
| GVGMTKF | ASSWGGENEQY | 0.8930703 |
| IGIGGSDL | ASSWGGENEQY | 0.32382756 |
| IGTLLAEL | ASSWGGENEQY | 0.24905717 |
| IYTSGTT | ASSWGGENEQY | 0.7918931 |
| LSFPAGV | ASSWGGENEQY | 0.8602905 |
| LYTSGTT | ASSWGGENEQY | 0.75230134 |
| VGAFPFGF | ASSWGGENEQY | 0.68832076 |
| VGAGTVEF | ASSWGGENEQY | 0.85735357 |
| VGKSTLLL | ASSWGGENEQY | 0.88555944 |
| AALAHAGGGIV | ASSWGGENEQY | 0.60987973 |
| AALVAVSLL | ASSWGGENEQY | 0.5656644 |
| AAVSRAVAAL | ASSWGGENEQY | 0.86968553 |
| AGGSSGIGGF | ASSWGGENEQY | 0.87008345 |
| AGLTVANANG | ASSWGGENEQY | 0.6865268 |
| AGLVGAVL | ASSWGGENEQY | 0.8130702 |
| ALSQAFTV | ASSWGGENEQY | 0.78851426 |
| ALTFSVSL | ASSWGGENEQY | 0.7802224 |
| ALTGAALVAL | ASSWGGENEQY | 0.8492379 |
| ASFSIGAGVL | ASSWGGENEQY | 0.5224699 |
| DGSVYKI | ASSWGGENEQY | 0.8910818 |
| EQLAEIV | ASSWGGENEQY | 0.5820507 |
| FAVSELTVM | ASSWGGENEQY | 0.8883939 |
| GAAPLAAGL | ASSWGGENEQY | 0.8380316 |
| GADGGDGGSGGAGG | ASSWGGENEQY | 0.85300756 |
| GFGLDTGGAGG | ASSWGGENEQY | 0.7887385 |
| GLASGGLL | ASSWGGENEQY | 0.69178534 |
| GRIAGAAVLL | ASSWGGENEQY | 0.6451233 |
| GSAGSLNTSF | ASSWGGENEQY | 0.4565342 |

| | | |
|---|---|---|
| GTGQIGI | ASSWGGENEQY | 0.8894235 |
| GTLIALIL | ASSWGGENEQY | 0.30253226 |
| IAATMGFVVG | ASSWGGENEQY | 0.8840126 |
| IFAQALL | ASSWGGENEQY | 0.8596021 |
| IGAAVALGF | ASSWGGENEQY | 0.6334327 |
| IGLTPLAL | ASSWGGENEQY | 0.8720096 |
| ILGSNTSTL | ASSWGGENEQY | 0.61021066 |
| IVFSLTVL | ASSWGGENEQY | 0.77172625 |
| LASKATGF | ASSWGGENEQY | 0.8765561 |
| LASVALIL | ASSWGGENEQY | 0.8600855 |
| LFAVGLTL | ASSWGGENEQY | 0.68534005 |
| LFNGGIGTYI | ASSWGGENEQY | 0.86024404 |
| LTLGAGII | ASSWGGENEQY | 0.8608599 |
| LVAAALGDI | ASSWGGENEQY | 0.7491485 |
| NTAFVQLGI | ASSWGGENEQY | 0.8163124 |
| QLFSGAEL | ASSWGGENEQY | 0.87692213 |
| SAATFITL | ASSWGGENEQY | 0.8485353 |
| SNYGLANLGV | ASSWGGENEQY | 0.6409267 |
| SYGGAYAVM | ASSWGGENEQY | 0.8676053 |
| TTGVGGLTLG | ASSWGGENEQY | 0.71468186 |
| TTQTIYI | ASSWGGENEQY | 0.8523239 |
| VAAGAGLGAVI | ASSWGGENEQY | 0.83307976 |
| VAAVVTLL | ASSWGGENEQY | 0.78083473 |
| VGFLGGLI | ASSWGGENEQY | 0.8841274 |
| VGVIPILL | ASSWGGENEQY | 0.5236252 |
| VIFASSLI | ASSWGGENEQY | 0.75141686 |
| VKGGDGGAAAG | ASSWGGENEQY | 0.76775724 |
| VTKTFTNI | ASSWGGENEQY | 0.79078484 |
| VVAAAILL | ASSWGGENEQY | 0.87021905 |
| YAALDVEL | ASSWGGENEQY | 0.63397795 |
| LASVALLL | ASSWGGENEQY | 0.8467032 |
| LDAVHRV | ASSWGGENEQY | 0.7708172 |
| QGLSSSIF | ASSWGGENEQY | 0.58211976 |
| RGTFANIRL | ASSWGGENEQY | 0.79322237 |
| RQLWWGHRI | ASSWGGENEQY | 0.8040007 |
| YGPPGTGKTLL | ASSWGGENEQY | 0.63191617 |
| AALVGAPANVA | ASSWGGENEQY | 0.868512 |
| AATFRRTE | ASSWGGENEQY | 0.66518545 |
| AGALGGGI | ASSWGGENEQY | 0.8464066 |
| AGGAGGAGGIGAG | ASSWGGENEQY | 0.9002295 |
| AGSGALGL | ASSWGGENEQY | 0.71898305 |
| AQIADQI | ASSWGGENEQY | 0.8873478 |
| ASAVAVTYL | ASSWGGENEQY | 0.85855764 |
| ASTVLGGTGG | ASSWGGENEQY | 0.8676417 |
| AVGGEPAK | ASSWGGENEQY | 0.7609317 |
| CEQGTFGGTC | ASSWGGENEQY | 0.7175409 |
| DEGKAPAG | ASSWGGENEQY | 0.8920937 |
| DTVVFPAL | ASSWGGENEQY | 0.86252755 |
| FEGITMAG | ASSWGGENEQY | 0.8793788 |
| FPLVVQD | ASSWGGENEQY | 0.8652714 |
| GAFTTPQ | ASSWGGENEQY | 0.864985 |

| | | |
|---|---|---|
| GFGGFGTAGGVGGA | ASSWGGENEQY | 0.8740989 |
| GFLAQFL | ASSWGGENEQY | 0.7304421 |
| GGAGGAGGNAGVA | ASSWGGENEQY | 0.5080613 |
| GGVGTAGAGGAGGAGG | ASSWGGENEQY | 0.7440374 |
| GNGGGGDAGNAGSGGN | ASSWGGENEQY | 0.80356944 |
| GTGGAGGAGNDAGS | ASSWGGENEQY | 0.74493897 |
| GTTNTGWFNS | ASSWGGENEQY | 0.35561755 |
| IFMGPLFTL | ASSWGGENEQY | 0.8768513 |
| ILAHAIEAI | ASSWGGENEQY | 0.9044751 |
| IRSAIREAL | ASSWGGENEQY | 0.67226005 |
| ITFAGGG | ASSWGGENEQY | 0.9011611 |
| ITGMSGHQEV | ASSWGGENEQY | 0.6076687 |
| IVAGLACTL | ASSWGGENEQY | 0.87087876 |
| IVGGGLAGA | ASSWGGENEQY | 0.8841333 |
| IVTGVLNI | ASSWGGENEQY | 0.6439444 |
| LAQQGCI | ASSWGGENEQY | 0.84125507 |
| LCLHVEN | ASSWGGENEQY | 0.7298866 |
| LDAISDGS | ASSWGGENEQY | 0.79587543 |
| LGGFAAML | ASSWGGENEQY | 0.88853633 |
| LGGKNPMI | ASSWGGENEQY | 0.6805115 |
| LGPKGVAAIL | ASSWGGENEQY | 0.8549517 |
| LMVGVSI | ASSWGGENEQY | 0.7225803 |
| LVGAAFVL | ASSWGGENEQY | 0.6311096 |
| LVLAVAPL | ASSWGGENEQY | 0.84242785 |
| MAAIAVIDAA | ASSWGGENEQY | 0.88698673 |
| NNGINAELQ | ASSWGGENEQY | 0.7916434 |
| RGEPLVLVHG | ASSWGGENEQY | 0.90360194 |
| RGGAGGAAGLI | ASSWGGENEQY | 0.6990002 |
| RSYRMHPAV | ASSWGGENEQY | 0.8976366 |
| SAAHLLAGQLA | ASSWGGENEQY | 0.8862402 |
| SAAVGSY | ASSWGGENEQY | 0.83271503 |
| SPHAVLHRI | ASSWGGENEQY | 0.7688191 |
| SQIAKVL | ASSWGGENEQY | 0.8797546 |
| TALGHLVP | ASSWGGENEQY | 0.7191203 |
| TAPTTSGAA | ASSWGGENEQY | 0.8757385 |
| TDGDDGNGGSPGT | ASSWGGENEQY | 0.8881507 |
| TDGLGEN | ASSWGGENEQY | 0.6810694 |
| TGGPPLPGQA | ASSWGGENEQY | 0.8349583 |
| TIPAITI | ASSWGGENEQY | 0.79535544 |
| TIPGFTI | ASSWGGENEQY | 0.7366165 |
| TSIASGALA | ASSWGGENEQY | 0.72453654 |
| TVTLPGAAI | ASSWGGENEQY | 0.43830192 |
| VASGLSAALP | ASSWGGENEQY | 0.7539682 |
| VGAGVIGI | ASSWGGENEQY | 0.6659866 |
| VGPKRFGV | ASSWGGENEQY | 0.7889143 |
| VNVAFPD | ASSWGGENEQY | 0.805757 |
| VSMTNSGVS | ASSWGGENEQY | 0.73101056 |
| VTDVLGDLSV | ASSWGGENEQY | 0.84393543 |
| VVVVGAGFAGL | ASSWGGENEQY | 0.84299076 |
| VYTVGTVV | ASSWGGENEQY | 0.569074 |
| WTTFSGP | ASSWGGENEQY | 0.72377354 |

| | | |
|---|---|---|
| YDELVEAF | ASSWGGENEQY | 0.6886816 |
| YGTGGAGGNGG | ASSWGGENEQY | 0.74891055 |
| YTNVMAV | ASSWGGENEQY | 0.37727618 |
| FLEQVHQL | ASSWGGENEQY | 0.8239135 |
| GLSLLAVAV | ASSWGGENEQY | 0.6741428 |
| LEQVHQL | ASSWGGENEQY | 0.19335946 |
| LNAAGAI | ASSWGGENEQY | 0.76738405 |
| PAVHASL | ASSWGGENEQY | 0.76283765 |
| TELEDDA | ASSWGGENEQY | 0.76640415 |
| TPPGPPN | ASSWGGENEQY | 0.8317587 |
| TPPPPPPP | ASSWGGENEQY | 0.46864873 |
| TQSPALI | AFRGGDTQY | 0.6600158 |
| EFAAAFPQ | AFRGGDTQY | 0.82159793 |
| LLSGNLDVL | AFRGGDTQY | 0.8411475 |
| GPANLGNYNI | AFRGGDTQY | 0.57144207 |
| TLGPTLP | AFRGGDTQY | 0.80827034 |
| VYAYSSCCK | AFRGGDTQY | 0.83327985 |
| APAPAPAPNI | AFRGGDTQY | 0.8040466 |
| PGGPVPIP | AFRGGDTQY | 0.8188554 |
| FLTVALEP | AFRGGDTQY | 0.83962965 |
| SQYFGANL | AFRGGDTQY | 0.6454824 |
| GGVKPAA | AFRGGDTQY | 0.83642423 |
| GGTGGAGGAGGW | AFRGGDTQY | 0.84627473 |
| GVGGVAGLGGAGG | AFRGGDTQY | 0.7321559 |
| VLLVLATTI | AFRGGDTQY | 0.7706294 |
| IGIFAAML | AFRGGDTQY | 0.6780713 |
| AVGDLID | AFRGGDTQY | 0.6911189 |
| NISPLPL | AFRGGDTQY | 0.3912669 |
| VTAVFVAI | AFRGGDTQY | 0.80373013 |
| SGDDNDAAL | AFRGGDTQY | 0.825796 |
| AVAALAHS | AFRGGDTQY | 0.735326 |
| GGVTVMATP | AFRGGDTQY | 0.61590564 |
| AGLAANVRD | AFRGGDTQY | 0.6697805 |
| TNRPDLIDPALLRPG | AFRGGDTQY | 0.72650653 |
| LANLGVTAL | AFRGGDTQY | 0.74646056 |
| TGERIEL | AFRGGDTQY | 0.33509642 |
| NGEELAA | AFRGGDTQY | 0.6619471 |
| VLSLVWG | AFRGGDTQY | 0.82861686 |
| IGYALWL | AFRGGDTQY | 0.83237696 |
| LSLLLAPL | AFRGGDTQY | 0.42594647 |
| TLDEALK | AFRGGDTQY | 0.8430959 |
| TDGAGTAEGG | AFRGGDTQY | 0.8032851 |
| NAPGGAGGNGG | AFRGGDTQY | 0.8407645 |
| VSVFMGMGGNF | AFRGGDTQY | 0.77184707 |
| LASYLAAL | AFRGGDTQY | 0.6750568 |
| LVGGTPAL | AFRGGDTQY | 0.7995327 |
| VRNADVL | AFRGGDTQY | 0.7276912 |
| TCSSPFLVSAM | AFRGGDTQY | 0.51789284 |
| VHITDAE | AFRGGDTQY | 0.8206078 |
| EYAAPSF | AFRGGDTQY | 0.7550298 |
| VVDAFHGGL | AFRGGDTQY | 0.6759399 |

| | | |
|---|---|---|
| ASVVDPQ | AFRGGDTQY | 0.6198988 |
| SYNEFGGLS | AFRGGDTQY | 0.8297109 |
| DPDGSVY | AFRGGDTQY | 0.5608005 |
| MVSTIHATE | AFRGGDTQY | 0.690647 |
| LAAGVADPV | AFRGGDTQY | 0.58846253 |
| TKEQIAA | AFRGGDTQY | 0.78678244 |
| SLASPAAL | AFRGGDTQY | 0.8300642 |
| IGLVFYGV | AFRGGDTQY | 0.79402846 |
| FLLDASNL | AFRGGDTQY | 0.67057514 |
| YQVGNFP | AFRGGDTQY | 0.80213785 |
| ESLLSQST | AFRGGDTQY | 0.7937707 |
| LFTVGFAAVG | AFRGGDTQY | 0.7763431 |
| ATPPTPT | AFRGGDTQY | 0.8213317 |
| TDLATLG | AFRGGDTQY | 0.6547789 |
| SAGGTGR | AFRGGDTQY | 0.82752526 |
| GFSNSGS | AFRGGDTQY | 0.78536624 |
| GGNGGSGGTSV | AFRGGDTQY | 0.82453334 |
| GTSVATGGAGN | AFRGGDTQY | 0.81740516 |
| NGGNGGSGGTS | AFRGGDTQY | 0.789221 |
| AVVKFPH | AFRGGDTQY | 0.75969297 |
| TPPPPPP | AFRGGDTQY | 0.34208483 |
| AAPAAAAAPA | AFRGGDTQY | 0.7144711 |
| IALNEDL | AFRGGDTQY | 0.8042251 |
| IVNVSSM | AFRGGDTQY | 0.78463674 |
| TGIPPAPR | AFRGGDTQY | 0.81768584 |
| FDLGGGTF | AFRGGDTQY | 0.784389 |
| FDLGGGTFDVS | AFRGGDTQY | 0.67786026 |
| NEPTAAA | AFRGGDTQY | 0.83022666 |
| NEPTAAALA | AFRGGDTQY | 0.81891173 |
| GSIINISSI | AFRGGDTQY | 0.5063661 |
| VAPGNAGTA | AFRGGDTQY | 0.71024036 |
| TGGGASLE | AFRGGDTQY | 0.85108757 |
| VAVYNHYKRI | AFRGGDTQY | 0.7631626 |
| TDPAPATT | AFRGGDTQY | 0.75274754 |
| TNRPDILDPALLRPG | AFRGGDTQY | 0.65421945 |
| LTTTTTS | AFRGGDTQY | 0.8441499 |
| PTGSGKTL | AFRGGDTQY | 0.59794277 |
| TGSGKTL | AFRGGDTQY | 0.8420578 |
| TIGHVDHGK | AFRGGDTQY | 0.24609213 |
| TIGHVDHGKTT | AFRGGDTQY | 0.6661743 |
| GHVDHGKTT | AFRGGDTQY | 0.5961114 |
| AQFEHTLL | AFRGGDTQY | 0.5066948 |
| IVNQDEL | AFRGGDTQY | 0.8253746 |
| TNGSQFF | AFRGGDTQY | 0.70300394 |
| TGKGIVE | AFRGGDTQY | 0.8196047 |
| LISNASDA | AFRGGDTQY | 0.7716474 |
| AELRTLGELTKIAKAHGAQ | AFRGGDTQY | 0.8159342 |
| APNAVAAAEPSW | AFRGGDTQY | 0.7861075 |
| APQVAGAATAVL | AFRGGDTQY | 0.7906136 |
| ARRDPAAAAAILPTDA | AFRGGDTQY | 0.6405405 |
| DAAARDVLAEAGLAEYF | AFRGGDTQY | 0.8172819 |

| | | |
|---|---|---|
| DDQIVLSAKANLAKVEQ | AFRGGDTQY | 0.8355094 |
| DELLPDTGEDSDEDGH | AFRGGDTQY | 0.7680105 |
| ELPEAPANRATVPQD | AFRGGDTQY | 0.5130707 |
| FGQSKIERVTGIG | AFRGGDTQY | 0.596947 |
| GAGGTGSTAGGAGGAGGAGGLY | AFRGGDTQY | 0.78645283 |
| GHGGTGAAGGNGGTGGAGGSGI | AFRGGDTQY | 0.34120288 |
| GNGGNGGAGGEGGPGGAGGAGA | AFRGGDTQY | 0.2725961 |
| GSTAGNGGNGGNATNGTIAGQP | AFRGGDTQY | 0.7319628 |
| GTGGDGGNGGNGGTGG | AFRGGDTQY | 0.62572545 |
| GVRRPAGIPADAPV | AFRGGDTQY | 0.8418926 |
| GYRQVTDAHLLTLARR | AFRGGDTQY | 0.8180713 |
| IAKMGQDGHDRG | AFRGGDTQY | 0.76491094 |
| KAEPSIEIQYT | AFRGGDTQY | 0.819296 |
| KPVRIRMAADEDTVAVT | AFRGGDTQY | 0.75587785 |
| MPPRIAGMRLLVIKPEP | AFRGGDTQY | 0.15774697 |
| MKFYAVALFRSRGVPPTR | AFRGGDTQY | 0.70181537 |
| NAGGFGILGAVAHSP | AFRGGDTQY | 0.59203124 |
| NEGREITLAMAEM | AFRGGDTQY | 0.8401644 |
| NELVEWSHPLGHRWRWE | AFRGGDTQY | 0.68488955 |
| NLGAGGTGGNGGNPG | AFRGGDTQY | 0.6204098 |
| NTEWPKWETVMAADPVD | AFRGGDTQY | 0.7743497 |
| PASAAPAETQP | AFRGGDTQY | 0.7432085 |
| QQSSDGKNGVRAAEITGE | AFRGGDTQY | 0.7379129 |
| RAPDAVTTSWWKE | AFRGGDTQY | 0.5938586 |
| SGFGNVGAHTSGWFNQ | AFRGGDTQY | 0.20961651 |
| SGVAGAVGQAASVGGLKV | AFRGGDTQY | 0.8184206 |
| SNGGAGTGSAGGNGGT | AFRGGDTQY | 0.43688872 |
| TGGAGGEGGAAG | AFRGGDTQY | 0.6278686 |
| TGGDGGAAGAGGAPGHG | AFRGGDTQY | 0.2897772 |
| TMPPLLAAIANAGRHGV | AFRGGDTQY | 0.46425438 |
| TSSTAGNSGGA | AFRGGDTQY | 0.45252016 |
| VPATVPVGILC | AFRGGDTQY | 0.80815643 |
| VQRPDSGGVVLTGRIS | AFRGGDTQY | 0.74342 |
| WTPNPVPATKGVPAGLVQAT | AFRGGDTQY | 0.6915014 |
| YALKRALAKFSDRFDVVII | AFRGGDTQY | 0.8441547 |
| YLIGSVIV | AFRGGDTQY | 0.75989187 |
| YGQLVAVID | AFRGGDTQY | 0.71151507 |
| VTVGGMTLPA | AFRGGDTQY | 0.6942371 |
| VLALPAV | AFRGGDTQY | 0.657441 |
| VIPPTLNYET | AFRGGDTQY | 0.67251784 |
| VGMGGMHPSSG | AFRGGDTQY | 0.6219261 |
| VGLAGGAATA | AFRGGDTQY | 0.8497574 |
| VGGAGGAGGSGP | AFRGGDTQY | 0.67196286 |
| VFAAFMV | AFRGGDTQY | 0.54862213 |
| TLYDKLR | AFRGGDTQY | 0.70104474 |
| TLAVFTVL | AFRGGDTQY | 0.84641206 |
| TGSVQPARV | AFRGGDTQY | 0.8580861 |
| TGGGAGSGGT | AFRGGDTQY | 0.68298435 |
| SHLETSL | AFRGGDTQY | 0.7226678 |
| SFTTGMT | AFRGGDTQY | 0.6914531 |
| QVEVAKRMGVRQPTVS | AFRGGDTQY | 0.76854026 |

| | | |
|---|---|---|
| QGSGGPALAM | AFRGGDTQY | 0.6861991 |
| MLAPSNP | AFRGGDTQY | 0.5606051 |
| LVNNAAI | AFRGGDTQY | 0.80786455 |
| LTGEINTLA | AFRGGDTQY | 0.85104924 |
| LSIFGPV | AFRGGDTQY | 0.71144307 |
| LLALLANL | AFRGGDTQY | 0.49002957 |
| LISNASDALDKLR | AFRGGDTQY | 0.7655235 |
| LINEAVP | AFRGGDTQY | 0.64445406 |
| LGYAFSGL | AFRGGDTQY | 0.75036174 |
| LDFVAVAL | AFRGGDTQY | 0.78663415 |
| LAQPGAE | AFRGGDTQY | 0.46076494 |
| ISNLGAVTGF | AFRGGDTQY | 0.6829934 |
| HQEDPGRI | AFRGGDTQY | 0.8483672 |
| GTPGEPTP | AFRGGDTQY | 0.7177708 |
| GFFNSGS | AFRGGDTQY | 0.781545 |
| GAGGGGLTAGAN | AFRGGDTQY | 0.8365849 |
| FVDALVNM | AFRGGDTQY | 0.81490004 |
| FLTVAPEL | AFRGGDTQY | 0.84202814 |
| FETLTAL | AFRGGDTQY | 0.8113228 |
| DVINPIEL | AFRGGDTQY | 0.7185029 |
| APVSGAPV | AFRGGDTQY | 0.7430806 |
| ALVGGLAW | AFRGGDTQY | 0.7919494 |
| ALAAAAATAI | AFRGGDTQY | 0.7822162 |
| AFGLGLLL | AFRGGDTQY | 0.7407796 |
| AFFAAASL | AFRGGDTQY | 0.6011255 |
| AAAALPAFSPP | AFRGGDTQY | 0.8423524 |
| VVLAPAL | AFRGGDTQY | 0.75463253 |
| TIGHVDHGKTTL | AFRGGDTQY | 0.8231154 |
| LISNASDALDK | AFRGGDTQY | 0.6135751 |
| LISNASDAL | AFRGGDTQY | 0.6039168 |
| ISNASDALDK | AFRGGDTQY | 0.8233662 |
| NQAETLVYQTE | AFRGGDTQY | 0.20263207 |
| TGGTGGTAGAG | AFRGGDTQY | 0.26643994 |
| TGSTAGGAGGA | AFRGGDTQY | 0.7693182 |
| DPLAVPVAAGPV | AFRGGDTQY | 0.8449723 |
| GGTGGTPGNGGN | AFRGGDTQY | 0.21398295 |
| PAGLAGVDPATL | AFRGGDTQY | 0.6224258 |
| GNNLVVEAPHAGASV | AFRGGDTQY | 0.5178234 |
| KVTRIMSASTLEQPA | AFRGGDTQY | 0.7907398 |
| LGFALPIALPRRTVV | AFRGGDTQY | 0.76899624 |
| TRLGNLTARKLVVLV | AFRGGDTQY | 0.51282305 |
| WSQQRRLIYFPSAGP | AFRGGDTQY | 0.8147465 |
| GPNGNPDPMAAAVDIR | AFRGGDTQY | 0.41379404 |
| TQRCVSVADLRAAASR | AFRGGDTQY | 0.5415346 |
| QFVLDVAEHPVEVLTETV | AFRGGDTQY | 0.8123129 |
| GGAGGTGGAGGNGGTQAGDG | AFRGGDTQY | 0.83198524 |
| PLLATLRAIAVDVDAKPAQV | AFRGGDTQY | 0.6217549 |
| APKFRTGGYMIDFWGVGYQVAK | AFRGGDTQY | 0.8249573 |
| GAGGAGGAGGAAGGPGGTGGTG | AFRGGDTQY | 0.67525774 |
| IGALGLLATDVAPAGSVAFGVA | AFRGGDTQY | 0.85655093 |
| AGGAGGIGGNGGAGGAGGVLMG | AFRGGDTQY | 0.73870075 |

| | | |
|---|---|---|
| ADELIGLL | AFRGGDTQY | 0.49469882 |
| ADRALKA | AFRGGDTQY | 0.8155761 |
| AGAVGQVGGAGGA | AFRGGDTQY | 0.8068746 |
| AGGGGGLGGAP | AFRGGDTQY | 0.78320223 |
| AGGLLYGNGGAG | AFRGGDTQY | 0.81466776 |
| AGLESIGL | AFRGGDTQY | 0.65834516 |
| AGLKTGII | AFRGGDTQY | 0.75629663 |
| ASLAALLL | AFRGGDTQY | 0.84060836 |
| ASVINISSI | AFRGGDTQY | 0.8236041 |
| ATPPGPA | AFRGGDTQY | 0.6212469 |
| ATVASALV | AFRGGDTQY | 0.80623615 |
| ATVASLIL | AFRGGDTQY | 0.73208654 |
| AVNGPVLIRH | AFRGGDTQY | 0.68533736 |
| DGAAGAVGI | AFRGGDTQY | 0.8394755 |
| DPLLHGI | AFRGGDTQY | 0.7716826 |
| ELGAALGAL | AFRGGDTQY | 0.824633 |
| ELGRTQAV | AFRGGDTQY | 0.82109845 |
| FVAGGYLS | AFRGGDTQY | 0.83614516 |
| GASGGAGGAGGW | AFRGGDTQY | 0.56616575 |
| GGAGAGGDGGAGGI | AFRGGDTQY | 0.8121018 |
| GTGHNGSQPGMGGNG | AFRGGDTQY | 0.76410264 |
| GYAAASASAS | AFRGGDTQY | 0.81450194 |
| HQVDVAAVQQG | AFRGGDTQY | 0.7541692 |
| IGAAIAHT | AFRGGDTQY | 0.5499455 |
| IGMIEVAQL | AFRGGDTQY | 0.7868488 |
| IRLAPEHTSAMD | AFRGGDTQY | 0.83372384 |
| IYNVSVL | AFRGGDTQY | 0.7594395 |
| KQGFLGM | AFRGGDTQY | 0.77234113 |
| LESVITQL | AFRGGDTQY | 0.84715766 |
| LGLGALVVV | AFRGGDTQY | 0.71599865 |
| LIGNGGNGGSGGI | AFRGGDTQY | 0.7297616 |
| LNAINAPTL | AFRGGDTQY | 0.8077625 |
| LQLLNTGV | AFRGGDTQY | 0.82870317 |
| LQTSIEF | AFRGGDTQY | 0.7213925 |
| MSFAVSEL | AFRGGDTQY | 0.84298044 |
| NTGALGL | AFRGGDTQY | 0.6233655 |
| PPGAGGAGGAGTT | AFRGGDTQY | 0.8214317 |
| QLVGLGI | AFRGGDTQY | 0.6851241 |
| RFLEQVHQL | AFRGGDTQY | 0.8064208 |
| RGVGYVLR | AFRGGDTQY | 0.6780134 |
| SATGVFAGVI | AFRGGDTQY | 0.81751245 |
| SLPAITLP | AFRGGDTQY | 0.8299623 |
| STALAFTV | AFRGGDTQY | 0.8254796 |
| SVLTVGLGK | AFRGGDTQY | 0.24942808 |
| TALAGAGSGI | AFRGGDTQY | 0.8313813 |
| TGLPFTGI | AFRGGDTQY | 0.82463866 |
| VAEHVTV | AFRGGDTQY | 0.85248667 |
| VDIVLNSL | AFRGGDTQY | 0.7462056 |
| VLTGGGAVL | AFRGGDTQY | 0.7893649 |
| VNVGLAYGI | AFRGGDTQY | 0.74276453 |
| VSSAAVGAV | AFRGGDTQY | 0.72367024 |

| | | |
|---|---|---|
| VTSVYEAGV | AFRGGDTQY | 0.6263212 |
| WVALDRGA | AFRGGDTQY | 0.76064044 |
| YLVGSVVI | AFRGGDTQY | 0.7131233 |
| GPPGTGKTLL | AFRGGDTQY | 0.70291513 |
| GVGMTKF | AFRGGDTQY | 0.81207573 |
| IGIGGSDL | AFRGGDTQY | 0.82463175 |
| IGTLLAEL | AFRGGDTQY | 0.8202678 |
| IYTSGTT | AFRGGDTQY | 0.8393663 |
| LSFPAGV | AFRGGDTQY | 0.84388113 |
| LYTSGTT | AFRGGDTQY | 0.82896125 |
| VGAFPFGF | AFRGGDTQY | 0.6807379 |
| VGAGTVEF | AFRGGDTQY | 0.7594705 |
| VGKSTLLL | AFRGGDTQY | 0.6105517 |
| AALAHAGGGIV | AFRGGDTQY | 0.72118765 |
| AALVAVSLL | AFRGGDTQY | 0.7049052 |
| AAVSRAVAAL | AFRGGDTQY | 0.79623157 |
| AGGSSGIGGF | AFRGGDTQY | 0.17903276 |
| AGLTVANANG | AFRGGDTQY | 0.7282295 |
| AGLVGAVL | AFRGGDTQY | 0.84757054 |
| ALSQAFTV | AFRGGDTQY | 0.6128451 |
| ALTFSVSL | AFRGGDTQY | 0.799019 |
| ALTGAALVAL | AFRGGDTQY | 0.76500934 |
| ASFSIGAGVL | AFRGGDTQY | 0.64921284 |
| DGSVYKI | AFRGGDTQY | 0.8280296 |
| EQLAEIV | AFRGGDTQY | 0.59140486 |
| FAVSELTVM | AFRGGDTQY | 0.79192275 |
| GAAPLAAGL | AFRGGDTQY | 0.8419428 |
| GADGGDGGSGGAGG | AFRGGDTQY | 0.76240623 |
| GFGLDTGGAGG | AFRGGDTQY | 0.7734386 |
| GLASGGLL | AFRGGDTQY | 0.71937317 |
| GRIAGAAVLL | AFRGGDTQY | 0.47324443 |
| GSAGSLNTSF | AFRGGDTQY | 0.79477674 |
| GTGQIGI | AFRGGDTQY | 0.31586248 |
| GTLIALIL | AFRGGDTQY | 0.75095224 |
| IAATMGFVVG | AFRGGDTQY | 0.6275467 |
| IFAQALL | AFRGGDTQY | 0.78978145 |
| IGAAVALGF | AFRGGDTQY | 0.7625881 |
| IGLTPLAL | AFRGGDTQY | 0.7897602 |
| ILGSNTSTL | AFRGGDTQY | 0.64784884 |
| IVFSLTVL | AFRGGDTQY | 0.69916534 |
| LASKATGF | AFRGGDTQY | 0.5733021 |
| LASVALIL | AFRGGDTQY | 0.55234164 |
| LFAVGLTL | AFRGGDTQY | 0.82418865 |
| LFNGGIGTYI | AFRGGDTQY | 0.71891826 |
| LTLGAGII | AFRGGDTQY | 0.6911855 |
| LVAAALGDI | AFRGGDTQY | 0.8324739 |
| NTAFVQLGI | AFRGGDTQY | 0.77214885 |
| QLFSGAEL | AFRGGDTQY | 0.47551173 |
| SAATFITL | AFRGGDTQY | 0.77177995 |
| SNYGLANLGV | AFRGGDTQY | 0.75617397 |
| SYGGAYAVM | AFRGGDTQY | 0.7588835 |

| | | |
|---|---|---|
| TTGVGGLTLG | AFRGGDTQY | 0.19792172 |
| TTQTIYI | AFRGGDTQY | 0.78533477 |
| VAAGAGLGAVI | AFRGGDTQY | 0.68114823 |
| VAAVVTLL | AFRGGDTQY | 0.6387317 |
| VGFLGGLI | AFRGGDTQY | 0.6651011 |
| VGVIPILL | AFRGGDTQY | 0.8215743 |
| VIFASSLI | AFRGGDTQY | 0.6719968 |
| VKGGDGGAAAG | AFRGGDTQY | 0.78140897 |
| VTKTFTNI | AFRGGDTQY | 0.8177734 |
| VVAAAILL | AFRGGDTQY | 0.815259 |
| YAALDVEL | AFRGGDTQY | 0.84363306 |
| LASVALLL | AFRGGDTQY | 0.8143929 |
| LDAVHRV | AFRGGDTQY | 0.69973683 |
| QGLSSSIF | AFRGGDTQY | 0.7862678 |
| RGTFANIRL | AFRGGDTQY | 0.7573119 |
| RQLWWGHRI | AFRGGDTQY | 0.33533695 |
| YGPPGTGKTLL | AFRGGDTQY | 0.28004014 |
| AALVGAPANVA | AFRGGDTQY | 0.806606 |
| AATFRRTE | AFRGGDTQY | 0.31877264 |
| AGALGGGI | AFRGGDTQY | 0.7807324 |
| AGGAGGAGGIGAG | AFRGGDTQY | 0.8269799 |
| AGSGALGL | AFRGGDTQY | 0.75827575 |
| AQIADQI | AFRGGDTQY | 0.5942062 |
| ASAVAVTYL | AFRGGDTQY | 0.70594317 |
| ASTVLGGTGG | AFRGGDTQY | 0.7337446 |
| AVGGEPAK | AFRGGDTQY | 0.82862586 |
| CEQGTFGGTC | AFRGGDTQY | 0.81443363 |
| DEGKAPAG | AFRGGDTQY | 0.8410138 |
| DTVVFPAL | AFRGGDTQY | 0.828112 |
| FEGITMAG | AFRGGDTQY | 0.6954889 |
| FPLVVQD | AFRGGDTQY | 0.814583 |
| GAFTTPQ | AFRGGDTQY | 0.68724006 |
| GFGGFGTAGGVGGA | AFRGGDTQY | 0.74465394 |
| GFLAQFL | AFRGGDTQY | 0.7177344 |
| GGAGGAGGNAGVA | AFRGGDTQY | 0.7718857 |
| GGVGTAGAGGAGGAGG | AFRGGDTQY | 0.7673534 |
| GNGGGGDAGNAGSGGN | AFRGGDTQY | 0.65890074 |
| GTGGAGGAGNDAGS | AFRGGDTQY | 0.46868712 |
| GTTNTGWFNS | AFRGGDTQY | 0.79935765 |
| IFMGPLFTL | AFRGGDTQY | 0.81465816 |
| ILAHAIEAI | AFRGGDTQY | 0.3615603 |
| IRSAIREAL | AFRGGDTQY | 0.64162076 |
| ITFAGGG | AFRGGDTQY | 0.85576093 |
| ITGMSGHQEV | AFRGGDTQY | 0.8442278 |
| IVAGLACTL | AFRGGDTQY | 0.6521013 |
| IVGGGLAGA | AFRGGDTQY | 0.6703046 |
| IVTGVLNI | AFRGGDTQY | 0.7325252 |
| LAQQGCI | AFRGGDTQY | 0.376605 |
| LCLHVEN | AFRGGDTQY | 0.77734375 |
| LDAISDGS | AFRGGDTQY | 0.73821914 |
| LGGFAAML | AFRGGDTQY | 0.7415577 |

| | | |
|---|---|---|
| LGGKNPMI | AFRGGDTQY | 0.6130855 |
| LGPKGVAAIL | AFRGGDTQY | 0.74957716 |
| LMVGVSI | AFRGGDTQY | 0.68138504 |
| LVGAAFVL | AFRGGDTQY | 0.81411695 |
| LVLAVAPL | AFRGGDTQY | 0.20406683 |
| MAAIAVIDAA | AFRGGDTQY | 0.78108215 |
| NNGINAELQ | AFRGGDTQY | 0.63528776 |
| RGEPLVLVHG | AFRGGDTQY | 0.6743009 |
| RGGAGGAAGLI | AFRGGDTQY | 0.47529027 |
| RSYRMHPAV | AFRGGDTQY | 0.7493441 |
| SAAHLLAGQLA | AFRGGDTQY | 0.8346366 |
| SAAVGSY | AFRGGDTQY | 0.8185761 |
| SPHAVLHRI | AFRGGDTQY | 0.4381664 |
| SQIAKVL | AFRGGDTQY | 0.8295444 |
| TALGHLVP | AFRGGDTQY | 0.85524964 |
| TAPTTSGAA | AFRGGDTQY | 0.6792954 |
| TDGDDGNGGSPGT | AFRGGDTQY | 0.46599662 |
| TDGLGEN | AFRGGDTQY | 0.8045678 |
| TGGPPLPGQA | AFRGGDTQY | 0.72701436 |
| TIPAITI | AFRGGDTQY | 0.7985827 |
| TIPGFTI | AFRGGDTQY | 0.80822444 |
| TSIASGALA | AFRGGDTQY | 0.6011318 |
| TVTLPGAAI | AFRGGDTQY | 0.8256241 |
| VASGLSAALP | AFRGGDTQY | 0.8281338 |
| VGAGVIGI | AFRGGDTQY | 0.6507534 |
| VGPKRFGV | AFRGGDTQY | 0.6595519 |
| VNVAFPD | AFRGGDTQY | 0.65307856 |
| VSMTNSGVS | AFRGGDTQY | 0.33151746 |
| VTDVLGDLSV | AFRGGDTQY | 0.81214064 |
| VVVVGAGFAGL | AFRGGDTQY | 0.64895 |
| VYTVGTVV | AFRGGDTQY | 0.67900205 |
| WTTFSGP | AFRGGDTQY | 0.79193515 |
| YDELVEAF | AFRGGDTQY | 0.6746268 |
| YGTGGAGGNGG | AFRGGDTQY | 0.6941048 |
| YTNVMAV | AFRGGDTQY | 0.71679866 |
| FLEQVHQL | AFRGGDTQY | 0.7797146 |
| GLSLLAVAV | AFRGGDTQY | 0.77907723 |
| LEQVHQL | AFRGGDTQY | 0.76803124 |
| LNAAGAI | AFRGGDTQY | 0.7599841 |
| PAVHASL | AFRGGDTQY | 0.84215134 |
| TELEDDA | AFRGGDTQY | 0.5655287 |
| TPPGPPN | AFRGGDTQY | 0.65950537 |
| TPPPPPPP | AFRGGDTQY | 0.7938038 |
| TQSPALI | ASSQPGHSGNTLY | 0.44749543 |
| EFAAAFPQ | ASSQPGHSGNTLY | 0.8128439 |
| LLSGNLDVL | ASSQPGHSGNTLY | 0.7458184 |
| GPANLGNYNI | ASSQPGHSGNTLY | 0.7930829 |
| TLGPTLP | ASSQPGHSGNTLY | 0.7407932 |
| VYAYSSCCK | ASSQPGHSGNTLY | 0.8526093 |
| APAPAPAPNI | ASSQPGHSGNTLY | 0.8283604 |
| PGGPVPIP | ASSQPGHSGNTLY | 0.73986506 |

| | | |
|---|---|---|
| FLTVALEP | ASSQPGHSGNTLY | 0.8150063 |
| SQYFGANL | ASSQPGHSGNTLY | 0.63414663 |
| GGVKPAA | ASSQPGHSGNTLY | 0.8300082 |
| GGTGGAGGAGGW | ASSQPGHSGNTLY | 0.8038715 |
| GVGGVAGLGGAGG | ASSQPGHSGNTLY | 0.62078476 |
| VLLVLATTI | ASSQPGHSGNTLY | 0.7615044 |
| IGIFAAML | ASSQPGHSGNTLY | 0.8349199 |
| AVGDLID | ASSQPGHSGNTLY | 0.83157265 |
| NISPLPL | ASSQPGHSGNTLY | 0.37160566 |
| VTAVFVAI | ASSQPGHSGNTLY | 0.7290974 |
| SGDDNDAAL | ASSQPGHSGNTLY | 0.36879945 |
| AVAALAHS | ASSQPGHSGNTLY | 0.65426 |
| GGVTVMATP | ASSQPGHSGNTLY | 0.62056804 |
| AGLAANVRD | ASSQPGHSGNTLY | 0.81223804 |
| TNRPDLIDPALLRPG | ASSQPGHSGNTLY | 0.72493684 |
| LANLGVTAL | ASSQPGHSGNTLY | 0.7867333 |
| TGERIEL | ASSQPGHSGNTLY | 0.7874691 |
| NGEELAA | ASSQPGHSGNTLY | 0.83790493 |
| VLSLVWG | ASSQPGHSGNTLY | 0.68660295 |
| IGYALWL | ASSQPGHSGNTLY | 0.74593616 |
| LSLLLAPL | ASSQPGHSGNTLY | 0.4763354 |
| TLDEALK | ASSQPGHSGNTLY | 0.49877435 |
| TDGAGTAEGG | ASSQPGHSGNTLY | 0.65053093 |
| NAPGGAGGNGG | ASSQPGHSGNTLY | 0.7236742 |
| VSVFMGMGGNF | ASSQPGHSGNTLY | 0.31435624 |
| LASYLAAL | ASSQPGHSGNTLY | 0.837795 |
| LVGGTPAL | ASSQPGHSGNTLY | 0.7960242 |
| VRNADVL | ASSQPGHSGNTLY | 0.8149233 |
| TCSSPFLVSAM | ASSQPGHSGNTLY | 0.6619984 |
| VHITDAE | ASSQPGHSGNTLY | 0.73686713 |
| EYAAPSF | ASSQPGHSGNTLY | 0.7042111 |
| VVDAFHGGL | ASSQPGHSGNTLY | 0.8096017 |
| ASVVDPQ | ASSQPGHSGNTLY | 0.67582774 |
| SYNEFGGLS | ASSQPGHSGNTLY | 0.45630443 |
| DPDGSVY | ASSQPGHSGNTLY | 0.68395334 |
| MVSTIHATE | ASSQPGHSGNTLY | 0.53140706 |
| LAAGVADPV | ASSQPGHSGNTLY | 0.8426305 |
| TKEQIAA | ASSQPGHSGNTLY | 0.82836646 |
| SLASPAAL | ASSQPGHSGNTLY | 0.80727315 |
| IGLVFYGV | ASSQPGHSGNTLY | 0.824241 |
| FLLDASNL | ASSQPGHSGNTLY | 0.37487382 |
| YQVGNFP | ASSQPGHSGNTLY | 0.7047789 |
| ESLLSQST | ASSQPGHSGNTLY | 0.84501815 |
| LFTVGFAAVG | ASSQPGHSGNTLY | 0.7979831 |
| ATPPTPT | ASSQPGHSGNTLY | 0.7380222 |
| TDLATLG | ASSQPGHSGNTLY | 0.8378383 |
| SAGGTGR | ASSQPGHSGNTLY | 0.6513649 |
| GFSNSGS | ASSQPGHSGNTLY | 0.7559942 |
| GGNGGSGGTSV | ASSQPGHSGNTLY | 0.77406967 |
| GTSVATGGAGN | ASSQPGHSGNTLY | 0.77794576 |
| NGGNGGSGGTS | ASSQPGHSGNTLY | 0.7833281 |

| | | |
|---|---|---|
| AVVKFPH | ASSQPGHSGNTLY | 0.62114114 |
| TPPPPPP | ASSQPGHSGNTLY | 0.76734763 |
| AAPAAAAAPA | ASSQPGHSGNTLY | 0.8561723 |
| IALNEDL | ASSQPGHSGNTLY | 0.74650073 |
| IVNVSSM | ASSQPGHSGNTLY | 0.8315971 |
| TGIPPAPR | ASSQPGHSGNTLY | 0.80104077 |
| FDLGGGTF | ASSQPGHSGNTLY | 0.8139081 |
| FDLGGGTFDVS | ASSQPGHSGNTLY | 0.6767245 |
| NEPTAAA | ASSQPGHSGNTLY | 0.7151732 |
| NEPTAAALA | ASSQPGHSGNTLY | 0.6919701 |
| GSIINISSI | ASSQPGHSGNTLY | 0.63083947 |
| VAPGNAGTA | ASSQPGHSGNTLY | 0.7211563 |
| TGGGASLE | ASSQPGHSGNTLY | 0.6835823 |
| VAVYNHYKRI | ASSQPGHSGNTLY | 0.8624693 |
| TDPAPATT | ASSQPGHSGNTLY | 0.803717 |
| TNRPDILDPALLRPG | ASSQPGHSGNTLY | 0.80220103 |
| LTTTTTS | ASSQPGHSGNTLY | 0.6717695 |
| PTGSGKTL | ASSQPGHSGNTLY | 0.7704248 |
| TGSGKTL | ASSQPGHSGNTLY | 0.78586805 |
| TIGHVDHGK | ASSQPGHSGNTLY | 0.4999414 |
| TIGHVDHGKTT | ASSQPGHSGNTLY | 0.84181017 |
| GHVDHGKTT | ASSQPGHSGNTLY | 0.6723496 |
| AQFEHTLL | ASSQPGHSGNTLY | 0.6660149 |
| IVNQDEL | ASSQPGHSGNTLY | 0.81597733 |
| TNGSQFF | ASSQPGHSGNTLY | 0.83436805 |
| TGKGIVE | ASSQPGHSGNTLY | 0.70976686 |
| LISNASDA | ASSQPGHSGNTLY | 0.6510328 |
| AELRTLGELTKIAKAHGAQ | ASSQPGHSGNTLY | 0.6890808 |
| APNAVAAAEPSW | ASSQPGHSGNTLY | 0.82994324 |
| APQVAGAATAVL | ASSQPGHSGNTLY | 0.8383171 |
| ARRDPAAAAAILPTDA | ASSQPGHSGNTLY | 0.7639776 |
| DAAARDVLAEAGLAEYF | ASSQPGHSGNTLY | 0.84552777 |
| DDQIVLSAKANLAKVEQ | ASSQPGHSGNTLY | 0.77131695 |
| DELLPDTGEDSDEDGH | ASSQPGHSGNTLY | 0.5307101 |
| ELPEAPANRATVPQD | ASSQPGHSGNTLY | 0.7721168 |
| FGQSKIERVTGIG | ASSQPGHSGNTLY | 0.81804645 |
| GAGGTGSTAGGAGGAGGAGGLY | ASSQPGHSGNTLY | 0.82600784 |
| GHGGTGAAGGNGGTGGAGGSGI | ASSQPGHSGNTLY | 0.8533574 |
| GNGGNGGAGGEGGPGGAGGAGA | ASSQPGHSGNTLY | 0.8015889 |
| GSTAGNGGNGGNATNGTIAGQP | ASSQPGHSGNTLY | 0.6916841 |
| GTGGDGGNGGNGGTGG | ASSQPGHSGNTLY | 0.8413497 |
| GVRRPAGIPADAPV | ASSQPGHSGNTLY | 0.8525404 |
| GYRQVTDAHLLTLARR | ASSQPGHSGNTLY | 0.82035923 |
| IAKMGQDGHDRG | ASSQPGHSGNTLY | 0.72900283 |
| KAEPSIEIQYT | ASSQPGHSGNTLY | 0.66224104 |
| KPVRIRMAADEDTVAVT | ASSQPGHSGNTLY | 0.75069875 |
| MPPRIAGMRLLVIKPEP | ASSQPGHSGNTLY | 0.8231305 |
| MKFYAVALFRSRGVPPTR | ASSQPGHSGNTLY | 0.58621657 |
| NAGGFGILGAVAHSP | ASSQPGHSGNTLY | 0.76944447 |
| NEGREITLAMAEM | ASSQPGHSGNTLY | 0.80230165 |
| NELVEWSHPLGHRWRWE | ASSQPGHSGNTLY | 0.54018795 |

| | | |
|---|---|---|
| NLGAGGTGGNGGNPG | ASSQPGHSGNTLY | 0.8008769 |
| NTEWPKWETVMAADPVD | ASSQPGHSGNTLY | 0.84157175 |
| PASAAPAETQP | ASSQPGHSGNTLY | 0.69719124 |
| QQSSDGKNGVRAAEITGE | ASSQPGHSGNTLY | 0.7214086 |
| RAPDAVTTSWWKE | ASSQPGHSGNTLY | 0.833257 |
| SGFGNVGAHTSGWFNQ | ASSQPGHSGNTLY | 0.8630438 |
| SGVAGAVGQAASVGGLKV | ASSQPGHSGNTLY | 0.8360684 |
| SNGGAGTGSAGGNGGT | ASSQPGHSGNTLY | 0.4904587 |
| TGGAGGEGGAAG | ASSQPGHSGNTLY | 0.32900983 |
| TGGDGGAAGAGGAPGHG | ASSQPGHSGNTLY | 0.6153731 |
| TMPPLLAAIANAGRHGV | ASSQPGHSGNTLY | 0.8079065 |
| TSSTAGNSGGA | ASSQPGHSGNTLY | 0.7010962 |
| VPATVPVGILC | ASSQPGHSGNTLY | 0.7247188 |
| VQRPDSGGVVLTGRIS | ASSQPGHSGNTLY | 0.8081802 |
| WTPNPVPATKGVPAGLVQAT | ASSQPGHSGNTLY | 0.8332927 |
| YALKRALAKFSDRFDVVII | ASSQPGHSGNTLY | 0.6941668 |
| YLIGSVIV | ASSQPGHSGNTLY | 0.5682468 |
| YGQLVAVID | ASSQPGHSGNTLY | 0.7913872 |
| VTVGGMTLPA | ASSQPGHSGNTLY | 0.8092567 |
| VLALPAV | ASSQPGHSGNTLY | 0.6442247 |
| VIPPTLNYET | ASSQPGHSGNTLY | 0.76498675 |
| VGMGGMHPSSG | ASSQPGHSGNTLY | 0.6150528 |
| VGLAGGAATA | ASSQPGHSGNTLY | 0.523774 |
| VGGAGGAGGSGP | ASSQPGHSGNTLY | 0.7751547 |
| VFAAFMV | ASSQPGHSGNTLY | 0.66917586 |
| TLYDKLR | ASSQPGHSGNTLY | 0.8347722 |
| TLAVFTVL | ASSQPGHSGNTLY | 0.68741965 |
| TGSVQPARV | ASSQPGHSGNTLY | 0.8131726 |
| TGGGAGSGGT | ASSQPGHSGNTLY | 0.7979933 |
| SHLETSL | ASSQPGHSGNTLY | 0.7949432 |
| SFTTGMT | ASSQPGHSGNTLY | 0.73436385 |
| QVEVAKRMGVRQPTVS | ASSQPGHSGNTLY | 0.67961687 |
| QGSGGPALAM | ASSQPGHSGNTLY | 0.58053267 |
| MLAPSNP | ASSQPGHSGNTLY | 0.33498275 |
| LVNNAAI | ASSQPGHSGNTLY | 0.8285573 |
| LTGEINTLA | ASSQPGHSGNTLY | 0.5874179 |
| LSIFGPV | ASSQPGHSGNTLY | 0.8360326 |
| LLALLANL | ASSQPGHSGNTLY | 0.8161608 |
| LISNASDALDKLR | ASSQPGHSGNTLY | 0.637904 |
| LINEAVP | ASSQPGHSGNTLY | 0.79253983 |
| LGYAFSGL | ASSQPGHSGNTLY | 0.68631375 |
| LDFVAVAL | ASSQPGHSGNTLY | 0.5219435 |
| LAQPGAE | ASSQPGHSGNTLY | 0.8360202 |
| ISNLGAVTGF | ASSQPGHSGNTLY | 0.79315096 |
| HQEDPGRI | ASSQPGHSGNTLY | 0.6536498 |
| GTPGEPTP | ASSQPGHSGNTLY | 0.77927864 |
| GFFNSGS | ASSQPGHSGNTLY | 0.769915 |
| GAGGGGLTAGAN | ASSQPGHSGNTLY | 0.66940355 |
| FVDALVNM | ASSQPGHSGNTLY | 0.7117704 |
| FLTVAPEL | ASSQPGHSGNTLY | 0.8269503 |
| FETLTAL | ASSQPGHSGNTLY | 0.7538045 |

| | | |
|---|---|---|
| DVINPIEL | ASSQPGHSGNTLY | 0.7109844 |
| APVSGAPV | ASSQPGHSGNTLY | 0.7975079 |
| ALVGGLAW | ASSQPGHSGNTLY | 0.596868 |
| ALAAAAATAI | ASSQPGHSGNTLY | 0.80863535 |
| AFGLGLLL | ASSQPGHSGNTLY | 0.73497015 |
| AFFAAASL | ASSQPGHSGNTLY | 0.7290524 |
| AAAALPAFSPP | ASSQPGHSGNTLY | 0.8006478 |
| VVLAPAL | ASSQPGHSGNTLY | 0.58884114 |
| TIGHVDHGKTTL | ASSQPGHSGNTLY | 0.6480235 |
| LISNASDALDK | ASSQPGHSGNTLY | 0.7108996 |
| LISNASDAL | ASSQPGHSGNTLY | 0.7897067 |
| ISNASDALDK | ASSQPGHSGNTLY | 0.82293576 |
| NQAETLVYQTE | ASSQPGHSGNTLY | 0.65003884 |
| TGGTGGTAGAG | ASSQPGHSGNTLY | 0.77172387 |
| TGSTAGGAGGA | ASSQPGHSGNTLY | 0.7081097 |
| DPLAVPVAAGPV | ASSQPGHSGNTLY | 0.6407769 |
| GGTGGTPGNGGN | ASSQPGHSGNTLY | 0.7225475 |
| PAGLAGVDPATL | ASSQPGHSGNTLY | 0.7135101 |
| GNNLVVEAPHAGASV | ASSQPGHSGNTLY | 0.71461546 |
| KVTRIMSASTLEQPA | ASSQPGHSGNTLY | 0.77998585 |
| LGFALPIALPRRTVV | ASSQPGHSGNTLY | 0.6648519 |
| TRLGNLTARKLVVLV | ASSQPGHSGNTLY | 0.7694959 |
| WSQQRRLIYFPSAGP | ASSQPGHSGNTLY | 0.81338763 |
| GPNGNPDPMAAAVDIR | ASSQPGHSGNTLY | 0.7303833 |
| TQRCVSVADLRAAASR | ASSQPGHSGNTLY | 0.8090186 |
| QFVLDVAEHPVEVLTETV | ASSQPGHSGNTLY | 0.7974782 |
| GGAGGTGGAGGNGGTQAGDG | ASSQPGHSGNTLY | 0.80173844 |
| PLLATLRAIAVDVDAKPAQV | ASSQPGHSGNTLY | 0.7234297 |
| APKFRTGGYMIDFWGVGYQVAK | ASSQPGHSGNTLY | 0.6992618 |
| GAGGAGGAGGAAGGPGGTGGTG | ASSQPGHSGNTLY | 0.83740175 |
| IGALGLLATDVAPAGSVAFGVA | ASSQPGHSGNTLY | 0.80787677 |
| AGGAGGIGGNGGAGGAGGVLMG | ASSQPGHSGNTLY | 0.8240044 |
| ADELIGLL | ASSQPGHSGNTLY | 0.7994044 |
| ADRALKA | ASSQPGHSGNTLY | 0.78877646 |
| AGAVGQVGGAGGA | ASSQPGHSGNTLY | 0.81279725 |
| AGGGGGLGGAP | ASSQPGHSGNTLY | 0.72208756 |
| AGGLLYGNGGAG | ASSQPGHSGNTLY | 0.49608067 |
| AGLESIGL | ASSQPGHSGNTLY | 0.72845596 |
| AGLKTGII | ASSQPGHSGNTLY | 0.7748519 |
| ASLAALLL | ASSQPGHSGNTLY | 0.707001 |
| ASVINISSI | ASSQPGHSGNTLY | 0.45726815 |
| ATPPGPA | ASSQPGHSGNTLY | 0.8331726 |
| ATVASALV | ASSQPGHSGNTLY | 0.8609202 |
| ATVASLIL | ASSQPGHSGNTLY | 0.663134 |
| AVNGPVLIRH | ASSQPGHSGNTLY | 0.8545857 |
| DGAAGAVGI | ASSQPGHSGNTLY | 0.49325112 |
| DPLLHGI | ASSQPGHSGNTLY | 0.80219024 |
| ELGAALGAL | ASSQPGHSGNTLY | 0.8346803 |
| ELGRTQAV | ASSQPGHSGNTLY | 0.6092082 |
| FVAGGYLS | ASSQPGHSGNTLY | 0.8201581 |
| GASGGAGGAGGW | ASSQPGHSGNTLY | 0.6732491 |

| | | |
|---|---|---|
| GGAGAGGDGGAGGI | ASSQPGHSGNTLY | 0.7405596 |
| GTGHNGSQPGMGGNG | ASSQPGHSGNTLY | 0.8425683 |
| GYAAASASAS | ASSQPGHSGNTLY | 0.71450955 |
| HQVDVAAVQQG | ASSQPGHSGNTLY | 0.76238835 |
| IGAAIAHT | ASSQPGHSGNTLY | 0.76359886 |
| IGMIEVAQL | ASSQPGHSGNTLY | 0.6603184 |
| IRLAPEHTSAMD | ASSQPGHSGNTLY | 0.75474626 |
| IYNVSVL | ASSQPGHSGNTLY | 0.84034973 |
| KQGFLGM | ASSQPGHSGNTLY | 0.56840473 |
| LESVITQL | ASSQPGHSGNTLY | 0.8503799 |
| LGLGALVVV | ASSQPGHSGNTLY | 0.65391725 |
| LIGNGGNGGSGGI | ASSQPGHSGNTLY | 0.85622597 |
| LNAINAPTL | ASSQPGHSGNTLY | 0.80164015 |
| LQLLNTGV | ASSQPGHSGNTLY | 0.7770035 |
| LQTSIEF | ASSQPGHSGNTLY | 0.6913601 |
| MSFAVSEL | ASSQPGHSGNTLY | 0.8392233 |
| NTGALGL | ASSQPGHSGNTLY | 0.7401166 |
| PPGAGGAGGAGTT | ASSQPGHSGNTLY | 0.80226815 |
| QLVGLGI | ASSQPGHSGNTLY | 0.83592486 |
| RFLEQVHQL | ASSQPGHSGNTLY | 0.53735536 |
| RGVGYVLR | ASSQPGHSGNTLY | 0.7802 |
| SATGVFAGVI | ASSQPGHSGNTLY | 0.7266867 |
| SLPAITLP | ASSQPGHSGNTLY | 0.7316991 |
| STALAFTV | ASSQPGHSGNTLY | 0.6180726 |
| SVLTVGLGK | ASSQPGHSGNTLY | 0.37869304 |
| TALAGAGSGI | ASSQPGHSGNTLY | 0.6908947 |
| TGLPFTGI | ASSQPGHSGNTLY | 0.5163714 |
| VAEHVTV | ASSQPGHSGNTLY | 0.7537948 |
| VDIVLNSL | ASSQPGHSGNTLY | 0.74337554 |
| VLTGGGAVL | ASSQPGHSGNTLY | 0.56994635 |
| VNVGLAYGI | ASSQPGHSGNTLY | 0.7805098 |
| VSSAAVGAV | ASSQPGHSGNTLY | 0.78504163 |
| VTSVYEAGV | ASSQPGHSGNTLY | 0.4146312 |
| WVALDRGA | ASSQPGHSGNTLY | 0.6737561 |
| YLVGSVVI | ASSQPGHSGNTLY | 0.56703985 |
| GPPGTGKTLL | ASSQPGHSGNTLY | 0.670413 |
| GVGMTKF | ASSQPGHSGNTLY | 0.2952187 |
| IGIGGSDL | ASSQPGHSGNTLY | 0.76030296 |
| IGTLLAEL | ASSQPGHSGNTLY | 0.578922 |
| IYTSGTT | ASSQPGHSGNTLY | 0.19104135 |
| LSFPAGV | ASSQPGHSGNTLY | 0.6879903 |
| LYTSGTT | ASSQPGHSGNTLY | 0.64837974 |
| VGAFPFGF | ASSQPGHSGNTLY | 0.65991116 |
| VGAGTVEF | ASSQPGHSGNTLY | 0.7703644 |
| VGKSTLLL | ASSQPGHSGNTLY | 0.35626984 |
| AALAHAGGGIV | ASSQPGHSGNTLY | 0.58499366 |
| AALVAVSLL | ASSQPGHSGNTLY | 0.75111914 |
| AAVSRAVAAL | ASSQPGHSGNTLY | 0.7754168 |
| AGGSSGIGGF | ASSQPGHSGNTLY | 0.53272706 |
| AGLTVANANG | ASSQPGHSGNTLY | 0.7289337 |
| AGLVGAVL | ASSQPGHSGNTLY | 0.7836634 |

| | | |
|---|---|---|
| ALSQAFTV | ASSQPGHSGNTLY | 0.7207866 |
| ALTFSVSL | ASSQPGHSGNTLY | 0.7694137 |
| ALTGAALVAL | ASSQPGHSGNTLY | 0.7736639 |
| ASFSIGAGVL | ASSQPGHSGNTLY | 0.5839089 |
| DGSVYKI | ASSQPGHSGNTLY | 0.77212846 |
| EQLAEIV | ASSQPGHSGNTLY | 0.7879853 |
| FAVSELTVM | ASSQPGHSGNTLY | 0.6228157 |
| GAAPLAAGL | ASSQPGHSGNTLY | 0.7184313 |
| GADGGDGGSGGAGG | ASSQPGHSGNTLY | 0.48868185 |
| GFGLDTGGAGG | ASSQPGHSGNTLY | 0.6958991 |
| GLASGGLL | ASSQPGHSGNTLY | 0.28519365 |
| GRIAGAAVLL | ASSQPGHSGNTLY | 0.72947776 |
| GSAGSLNTSF | ASSQPGHSGNTLY | 0.7741556 |
| GTGQIGI | ASSQPGHSGNTLY | 0.68466985 |
| GTLIALIL | ASSQPGHSGNTLY | 0.47365445 |
| IAATMGFVVG | ASSQPGHSGNTLY | 0.44694373 |
| IFAQALL | ASSQPGHSGNTLY | 0.64300597 |
| IGAAVALGF | ASSQPGHSGNTLY | 0.62696725 |
| IGLTPLAL | ASSQPGHSGNTLY | 0.28952166 |
| ILGSNTSTL | ASSQPGHSGNTLY | 0.49663675 |
| IVFSLTVL | ASSQPGHSGNTLY | 0.77575946 |
| LASKATGF | ASSQPGHSGNTLY | 0.7777596 |
| LASVALIL | ASSQPGHSGNTLY | 0.48900762 |
| LFAVGLTL | ASSQPGHSGNTLY | 0.7823106 |
| LFNGGIGTYI | ASSQPGHSGNTLY | 0.7432493 |
| LTLGAGII | ASSQPGHSGNTLY | 0.7877078 |
| LVAAALGDI | ASSQPGHSGNTLY | 0.6899799 |
| NTAFVQLGI | ASSQPGHSGNTLY | 0.48085195 |
| QLFSGAEL | ASSQPGHSGNTLY | 0.7672967 |
| SAATFITL | ASSQPGHSGNTLY | 0.676414 |
| SNYGLANLGV | ASSQPGHSGNTLY | 0.5527116 |
| SYGGAYAVM | ASSQPGHSGNTLY | 0.766008 |
| TTGVGGLTLG | ASSQPGHSGNTLY | 0.6867916 |
| TTQTIYI | ASSQPGHSGNTLY | 0.57777727 |
| VAAGAGLGAVI | ASSQPGHSGNTLY | 0.66990566 |
| VAAVVTLL | ASSQPGHSGNTLY | 0.75163126 |
| VGFLGGLI | ASSQPGHSGNTLY | 0.54442596 |
| VGVIPILL | ASSQPGHSGNTLY | 0.41051492 |
| VIFASSLI | ASSQPGHSGNTLY | 0.5876285 |
| VKGGDGGAAAG | ASSQPGHSGNTLY | 0.71887887 |
| VTKTFTNI | ASSQPGHSGNTLY | 0.77101123 |
| VVAAAILL | ASSQPGHSGNTLY | 0.7213316 |
| YAALDVEL | ASSQPGHSGNTLY | 0.602382 |
| LASVALLL | ASSQPGHSGNTLY | 0.75246763 |
| LDAVHRV | ASSQPGHSGNTLY | 0.7393309 |
| QGLSSSIF | ASSQPGHSGNTLY | 0.7176394 |
| RGTFANIRL | ASSQPGHSGNTLY | 0.72212243 |
| RQLWWGHRI | ASSQPGHSGNTLY | 0.61637944 |
| YGPPGTGKTLL | ASSQPGHSGNTLY | 0.74772817 |
| AALVGAPANVA | ASSQPGHSGNTLY | 0.75443166 |
| AATFRRTE | ASSQPGHSGNTLY | 0.7458662 |

| | | |
|---|---|---|
| AGALGGGI | ASSQPGHSGNTLY | 0.75482523 |
| AGGAGGAGGIGAG | ASSQPGHSGNTLY | 0.67397845 |
| AGSGALGL | ASSQPGHSGNTLY | 0.7277769 |
| AQIADQI | ASSQPGHSGNTLY | 0.32172835 |
| ASAVAVTYL | ASSQPGHSGNTLY | 0.59892905 |
| ASTVLGGTGG | ASSQPGHSGNTLY | 0.71442366 |
| AVGGEPAK | ASSQPGHSGNTLY | 0.73238295 |
| CEQGTFGGTC | ASSQPGHSGNTLY | 0.74391997 |
| DEGKAPAG | ASSQPGHSGNTLY | 0.7368696 |
| DTVVFPAL | ASSQPGHSGNTLY | 0.5976615 |
| FEGITMAG | ASSQPGHSGNTLY | 0.74848396 |
| FPLVVQD | ASSQPGHSGNTLY | 0.7784562 |
| GAFTTPQ | ASSQPGHSGNTLY | 0.38754177 |
| GFGGFGTAGGVGGA | ASSQPGHSGNTLY | 0.5779669 |
| GFLAQFL | ASSQPGHSGNTLY | 0.7872051 |
| GGAGGAGGNAGVA | ASSQPGHSGNTLY | 0.7250977 |
| GGVGTAGAGGAGGAGG | ASSQPGHSGNTLY | 0.6748638 |
| GNGGGGDAGNAGSGGN | ASSQPGHSGNTLY | 0.6672702 |
| GTGGAGGAGNDAGS | ASSQPGHSGNTLY | 0.78288245 |
| GTTNTGWFNS | ASSQPGHSGNTLY | 0.5817721 |
| IFMGPLFTL | ASSQPGHSGNTLY | 0.7811266 |
| ILAHAIEAI | ASSQPGHSGNTLY | 0.2801456 |
| IRSAIREAL | ASSQPGHSGNTLY | 0.6170512 |
| ITFAGGG | ASSQPGHSGNTLY | 0.5039717 |
| ITGMSGHQEV | ASSQPGHSGNTLY | 0.24280323 |
| IVAGLACTL | ASSQPGHSGNTLY | 0.77000535 |
| IVGGGLAGA | ASSQPGHSGNTLY | 0.67756915 |
| IVTGVLNI | ASSQPGHSGNTLY | 0.75846434 |
| LAQQGCI | ASSQPGHSGNTLY | 0.70007837 |
| LCLHVEN | ASSQPGHSGNTLY | 0.75910485 |
| LDAISDGS | ASSQPGHSGNTLY | 0.723959 |
| LGGFAAML | ASSQPGHSGNTLY | 0.70181453 |
| LGGKNPMI | ASSQPGHSGNTLY | 0.45367718 |
| LGPKGVAAIL | ASSQPGHSGNTLY | 0.75731623 |
| LMVGVSI | ASSQPGHSGNTLY | 0.7613164 |
| LVGAAFVL | ASSQPGHSGNTLY | 0.7098334 |
| LVLAVAPL | ASSQPGHSGNTLY | 0.40449202 |
| MAAIAVIDAA | ASSQPGHSGNTLY | 0.6229008 |
| NNGINAELQ | ASSQPGHSGNTLY | 0.70165825 |
| RGEPLVLVHG | ASSQPGHSGNTLY | 0.24521881 |
| RGGAGGAAGLI | ASSQPGHSGNTLY | 0.22018902 |
| RSYRMHPAV | ASSQPGHSGNTLY | 0.59438205 |
| SAAHLLAGQLA | ASSQPGHSGNTLY | 0.5152437 |
| SAAVGSY | ASSQPGHSGNTLY | 0.76597226 |
| SPHAVLHRI | ASSQPGHSGNTLY | 0.753803 |
| SQIAKVL | ASSQPGHSGNTLY | 0.665014 |
| TALGHLVP | ASSQPGHSGNTLY | 0.7523571 |
| TAPTTSGAA | ASSQPGHSGNTLY | 0.74206513 |
| TDGDDGNGGSPGT | ASSQPGHSGNTLY | 0.1724837 |
| TDGLGEN | ASSQPGHSGNTLY | 0.6370052 |
| TGGPPLPGQA | ASSQPGHSGNTLY | 0.4390471 |

| | | |
|---|---|---|
| TIPAITI | ASSQPGHSGNTLY | 0.77626276 |
| TIPGFTI | ASSQPGHSGNTLY | 0.45821333 |
| TSIASGALA | ASSQPGHSGNTLY | 0.5976821 |
| TVTLPGAAI | ASSQPGHSGNTLY | 0.69704235 |
| VASGLSAALP | ASSQPGHSGNTLY | 0.7254381 |
| VGAGVIGI | ASSQPGHSGNTLY | 0.71352726 |
| VGPKRFGV | ASSQPGHSGNTLY | 0.44571474 |
| VNVAFPD | ASSQPGHSGNTLY | 0.2132959 |
| VSMTNSGVS | ASSQPGHSGNTLY | 0.763304 |
| VTDVLGDLSV | ASSQPGHSGNTLY | 0.36395448 |
| VVVVGAGFAGL | ASSQPGHSGNTLY | 0.52394927 |
| VYTVGTVV | ASSQPGHSGNTLY | 0.31839022 |
| WTTFSGP | ASSQPGHSGNTLY | 0.3285374 |
| YDELVEAF | ASSQPGHSGNTLY | 0.40268922 |
| YGTGGAGGNGG | ASSQPGHSGNTLY | 0.75302947 |
| YTNVMAV | ASSQPGHSGNTLY | 0.65129554 |
| FLEQVHQL | ASSQPGHSGNTLY | 0.6715056 |
| GLSLLAVAV | ASSQPGHSGNTLY | 0.79270184 |
| LEQVHQL | ASSQPGHSGNTLY | 0.7276166 |
| LNAAGAI | ASSQPGHSGNTLY | 0.6734078 |
| PAVHASL | ASSQPGHSGNTLY | 0.6172314 |
| TELEDDA | ASSQPGHSGNTLY | 0.5170187 |
| TPPGPPN | ASSQPGHSGNTLY | 0.55700254 |
| TPPPPPPP | ASSQPGHSGNTLY | 0.62052214 |
| TQSPALI | ASSQEGIGGHYEQY | 0.7897482 |
| EFAAAFPQ | ASSQEGIGGHYEQY | 0.5797466 |
| LLSGNLDVL | ASSQEGIGGHYEQY | 0.473068 |
| GPANLGNYNI | ASSQEGIGGHYEQY | 0.68928534 |
| TLGPTLP | ASSQEGIGGHYEQY | 0.7898247 |
| VYAYSSCCK | ASSQEGIGGHYEQY | 0.79694396 |
| APAPAPAPNI | ASSQEGIGGHYEQY | 0.508941 |
| PGGPVPIP | ASSQEGIGGHYEQY | 0.68211126 |
| FLTVALEP | ASSQEGIGGHYEQY | 0.6656371 |
| SQYFGANL | ASSQEGIGGHYEQY | 0.70249975 |
| GGVKPAA | ASSQEGIGGHYEQY | 0.53436565 |
| GGTGGAGGAGGW | ASSQEGIGGHYEQY | 0.43493444 |
| GVGGVAGLGGAGG | ASSQEGIGGHYEQY | 0.73181784 |
| VLLVLATTI | ASSQEGIGGHYEQY | 0.7940539 |
| IGIFAAML | ASSQEGIGGHYEQY | 0.6410253 |
| AVGDLID | ASSQEGIGGHYEQY | 0.46204907 |
| NISPLPL | ASSQEGIGGHYEQY | 0.67429596 |
| VTAVFVAI | ASSQEGIGGHYEQY | 0.56328833 |
| SGDDNDAAL | ASSQEGIGGHYEQY | 0.5895251 |
| AVAALAHS | ASSQEGIGGHYEQY | 0.70624334 |
| GGVTVMATP | ASSQEGIGGHYEQY | 0.35559878 |
| AGLAANVRD | ASSQEGIGGHYEQY | 0.6777384 |
| TNRPDLIDPALLRPG | ASSQEGIGGHYEQY | 0.796242 |
| LANLGVTAL | ASSQEGIGGHYEQY | 0.6957623 |
| TGERIEL | ASSQEGIGGHYEQY | 0.7555091 |
| NGEELAA | ASSQEGIGGHYEQY | 0.7786509 |
| VLSLVWG | ASSQEGIGGHYEQY | 0.7490173 |

| | | |
|---|---|---|
| IGYALWL | ASSQEGIGGHYEQY | 0.77947795 |
| LSLLLAPL | ASSQEGIGGHYEQY | 0.74169964 |
| TLDEALK | ASSQEGIGGHYEQY | 0.6415241 |
| TDGAGTAEGG | ASSQEGIGGHYEQY | 0.6909374 |
| NAPGGAGGNGG | ASSQEGIGGHYEQY | 0.73496187 |
| VSVFMGMGGNF | ASSQEGIGGHYEQY | 0.7028341 |
| LASYLAAL | ASSQEGIGGHYEQY | 0.70060647 |
| LVGGTPAL | ASSQEGIGGHYEQY | 0.6128368 |
| VRNADVL | ASSQEGIGGHYEQY | 0.7775234 |
| TCSSPFLVSAM | ASSQEGIGGHYEQY | 0.6537778 |
| VHITDAE | ASSQEGIGGHYEQY | 0.76883745 |
| EYAAPSF | ASSQEGIGGHYEQY | 0.3512006 |
| VVDAFHGGL | ASSQEGIGGHYEQY | 0.49296933 |
| ASVVDPQ | ASSQEGIGGHYEQY | 0.76502055 |
| SYNEFGGLS | ASSQEGIGGHYEQY | 0.19011566 |
| DPDGSVY | ASSQEGIGGHYEQY | 0.22961168 |
| MVSTIHATE | ASSQEGIGGHYEQY | 0.6962764 |
| LAAGVADPV | ASSQEGIGGHYEQY | 0.78475213 |
| TKEQIAA | ASSQEGIGGHYEQY | 0.18907265 |
| SLASPAAL | ASSQEGIGGHYEQY | 0.51118326 |
| IGLVFYGV | ASSQEGIGGHYEQY | 0.38821414 |
| FLLDASNL | ASSQEGIGGHYEQY | 0.69718707 |
| YQVGNFP | ASSQEGIGGHYEQY | 0.7051411 |
| ESLLSQST | ASSQEGIGGHYEQY | 0.26362902 |
| LFTVGFAAVG | ASSQEGIGGHYEQY | 0.75018054 |
| ATPPTPT | ASSQEGIGGHYEQY | 0.37839043 |
| TDLATLG | ASSQEGIGGHYEQY | 0.41912663 |
| SAGGTGR | ASSQEGIGGHYEQY | 0.75108135 |
| GFSNSGS | ASSQEGIGGHYEQY | 0.77049255 |
| GGNGGSGGTSV | ASSQEGIGGHYEQY | 0.36870775 |
| GTSVATGGAGN | ASSQEGIGGHYEQY | 0.7509599 |
| NGGNGGSGGTS | ASSQEGIGGHYEQY | 0.6199771 |
| AVVKFPH | ASSQEGIGGHYEQY | 0.8004408 |
| TPPPPPP | ASSQEGIGGHYEQY | 0.6759244 |
| AAPAAAAAPA | ASSQEGIGGHYEQY | 0.32538977 |
| IALNEDL | ASSQEGIGGHYEQY | 0.7724032 |
| IVNVSSM | ASSQEGIGGHYEQY | 0.7391593 |
| TGIPPAPR | ASSQEGIGGHYEQY | 0.6637862 |
| FDLGGGTF | ASSQEGIGGHYEQY | 0.75238216 |
| FDLGGGTFDVS | ASSQEGIGGHYEQY | 0.49656507 |
| NEPTAAA | ASSQEGIGGHYEQY | 0.68819404 |
| NEPTAAALA | ASSQEGIGGHYEQY | 0.77508616 |
| GSIINISSI | ASSQEGIGGHYEQY | 0.74583834 |
| VAPGNAGTA | ASSQEGIGGHYEQY | 0.41301444 |
| TGGGASLE | ASSQEGIGGHYEQY | 0.7449294 |
| VAVYNHYKRI | ASSQEGIGGHYEQY | 0.71139747 |
| TDPAPATT | ASSQEGIGGHYEQY | 0.5789118 |
| TNRPDILDPALLRPG | ASSQEGIGGHYEQY | 0.792647 |
| LTTTTTS | ASSQEGIGGHYEQY | 0.62477785 |
| PTGSGKTL | ASSQEGIGGHYEQY | 0.76606 |
| TGSGKTL | ASSQEGIGGHYEQY | 0.76545936 |

| | | |
|---|---|---|
| TIGHVDHGK | ASSQEGIGGHYEQY | 0.7842474 |
| TIGHVDHGKTT | ASSQEGIGGHYEQY | 0.34471413 |
| GHVDHGKTT | ASSQEGIGGHYEQY | 0.74004424 |
| AQFEHTLL | ASSQEGIGGHYEQY | 0.71479976 |
| IVNQDEL | ASSQEGIGGHYEQY | 0.7504529 |
| TNGSQFF | ASSQEGIGGHYEQY | 0.6752528 |
| TGKGIVE | ASSQEGIGGHYEQY | 0.5703402 |
| LISNASDA | ASSQEGIGGHYEQY | 0.7460034 |
| AELRTLGELTKIAKAHGAQ | ASSQEGIGGHYEQY | 0.78237885 |
| APNAVAAAEPSW | ASSQEGIGGHYEQY | 0.6562593 |
| APQVAGAATAVL | ASSQEGIGGHYEQY | 0.706522 |
| ARRDPAAAAAILPTDA | ASSQEGIGGHYEQY | 0.7856117 |
| DAAARDVLAEAGLAEYF | ASSQEGIGGHYEQY | 0.5351893 |
| DDQIVLSAKANLAKVEQ | ASSQEGIGGHYEQY | 0.70212084 |
| DELLPDTGEDSDEDGH | ASSQEGIGGHYEQY | 0.7488409 |
| ELPEAPANRATVPQD | ASSQEGIGGHYEQY | 0.7619231 |
| FGQSKIERVTGIG | ASSQEGIGGHYEQY | 0.64961344 |
| GAGGTGSTAGGAGGAGGAGGLY | ASSQEGIGGHYEQY | 0.76948583 |
| GHGGTGAAGGNGGTGGAGGSGI | ASSQEGIGGHYEQY | 0.52816737 |
| GNGGNGGAGGEGGPGGAGGAGA | ASSQEGIGGHYEQY | 0.7525774 |
| GSTAGNGGNGGNATNGTIAGQP | ASSQEGIGGHYEQY | 0.65661085 |
| GTGGDGGNGGNGGTGG | ASSQEGIGGHYEQY | 0.7337615 |
| GVRRPAGIPADAPV | ASSQEGIGGHYEQY | 0.4933053 |
| GYRQVTDAHLLTLARR | ASSQEGIGGHYEQY | 0.74847144 |
| IAKMGQDGHDRG | ASSQEGIGGHYEQY | 0.769367 |
| KAEPSIEIQYT | ASSQEGIGGHYEQY | 0.76660216 |
| KPVRIRMAADEDTVAVT | ASSQEGIGGHYEQY | 0.26434588 |
| MPPRIAGMRLLVIKPEP | ASSQEGIGGHYEQY | 0.755044 |
| MKFYAVALFRSRGVPPTR | ASSQEGIGGHYEQY | 0.75549936 |
| NAGGFGILGAVAHSP | ASSQEGIGGHYEQY | 0.7951929 |
| NEGREITLAMAEM | ASSQEGIGGHYEQY | 0.6747746 |
| NELVEWSHPLGHRWRWE | ASSQEGIGGHYEQY | 0.7575364 |
| NLGAGGTGGNGGNPG | ASSQEGIGGHYEQY | 0.6667792 |
| NTEWPKWETVMAADPVD | ASSQEGIGGHYEQY | 0.63054717 |
| PASAAPAETQP | ASSQEGIGGHYEQY | 0.5202802 |
| QQSSDGKNGVRAAEITGE | ASSQEGIGGHYEQY | 0.63109875 |
| RAPDAVTTSWWKE | ASSQEGIGGHYEQY | 0.63051903 |
| SGFGNVGAHTSGWFNQ | ASSQEGIGGHYEQY | 0.5972314 |
| SGVAGAVGQAASVGGLKV | ASSQEGIGGHYEQY | 0.7612608 |
| SNGGAGTGSAGGNGGT | ASSQEGIGGHYEQY | 0.7536627 |
| TGGAGGEGGAAG | ASSQEGIGGHYEQY | 0.77265984 |
| TGGDGGAAGAGGAPGHG | ASSQEGIGGHYEQY | 0.78240967 |
| TMPPLLAAIANAGRHGV | ASSQEGIGGHYEQY | 0.7798408 |
| TSSTAGNSGGA | ASSQEGIGGHYEQY | 0.7419865 |
| VPATVPVGILC | ASSQEGIGGHYEQY | 0.62242687 |
| VQRPDSGGVVLTGRIS | ASSQEGIGGHYEQY | 0.66418374 |
| WTPNPVPATKGVPAGLVQAT | ASSQEGIGGHYEQY | 0.4510752 |
| YALKRALAKFSDRFDVVII | ASSQEGIGGHYEQY | 0.6324538 |
| YLIGSVIV | ASSQEGIGGHYEQY | 0.6271012 |
| YGQLVAVID | ASSQEGIGGHYEQY | 0.72358465 |
| VTVGGMTLPA | ASSQEGIGGHYEQY | 0.18121256 |

| | | |
|---|---|---|
| VLALPAV | ASSQEGIGGHYEQY | 0.71660185 |
| VIPPTLNYET | ASSQEGIGGHYEQY | 0.78980863 |
| VGMGGMHPSSG | ASSQEGIGGHYEQY | 0.6274954 |
| VGLAGGAATA | ASSQEGIGGHYEQY | 0.7586856 |
| VGGAGGAGGSGP | ASSQEGIGGHYEQY | 0.71042854 |
| VFAAFMV | ASSQEGIGGHYEQY | 0.5668955 |
| TLYDKLR | ASSQEGIGGHYEQY | 0.7668641 |
| TLAVFTVL | ASSQEGIGGHYEQY | 0.54839706 |
| TGSVQPARV | ASSQEGIGGHYEQY | 0.7582679 |
| TGGGAGSGGT | ASSQEGIGGHYEQY | 0.7925166 |
| SHLETSL | ASSQEGIGGHYEQY | 0.7506093 |
| SFTTGMT | ASSQEGIGGHYEQY | 0.6741828 |
| QVEVAKRMGVRQPTVS | ASSQEGIGGHYEQY | 0.70586723 |
| QGSGGPALAM | ASSQEGIGGHYEQY | 0.34774914 |
| MLAPSNP | ASSQEGIGGHYEQY | 0.6872065 |
| LVNNAAI | ASSQEGIGGHYEQY | 0.18618566 |
| LTGEINTLA | ASSQEGIGGHYEQY | 0.7106291 |
| LSIFGPV | ASSQEGIGGHYEQY | 0.5089512 |
| LLALLANL | ASSQEGIGGHYEQY | 0.73463386 |
| LISNASDALDKLR | ASSQEGIGGHYEQY | 0.73338675 |
| LINEAVP | ASSQEGIGGHYEQY | 0.70933187 |
| LGYAFSGL | ASSQEGIGGHYEQY | 0.6077646 |
| LDFVAVAL | ASSQEGIGGHYEQY | 0.6542671 |
| LAQPGAE | ASSQEGIGGHYEQY | 0.66571844 |
| ISNLGAVTGF | ASSQEGIGGHYEQY | 0.47150254 |
| HQEDPGRI | ASSQEGIGGHYEQY | 0.7512129 |
| GTPGEPTP | ASSQEGIGGHYEQY | 0.61273503 |
| GFFNSGS | ASSQEGIGGHYEQY | 0.65454715 |
| GAGGGGLTAGAN | ASSQEGIGGHYEQY | 0.7809725 |
| FVDALVNM | ASSQEGIGGHYEQY | 0.7024993 |
| FLTVAPEL | ASSQEGIGGHYEQY | 0.29697672 |
| FETLTAL | ASSQEGIGGHYEQY | 0.70236826 |
| DVINPIEL | ASSQEGIGGHYEQY | 0.7116554 |
| APVSGAPV | ASSQEGIGGHYEQY | 0.68553895 |
| ALVGGLAW | ASSQEGIGGHYEQY | 0.19007349 |
| ALAAAAATAI | ASSQEGIGGHYEQY | 0.7128307 |
| AFGLGLLL | ASSQEGIGGHYEQY | 0.60460234 |
| AFFAAASL | ASSQEGIGGHYEQY | 0.54021573 |
| AAAALPAFSPP | ASSQEGIGGHYEQY | 0.58571166 |
| VVLAPAL | ASSQEGIGGHYEQY | 0.7688948 |
| TIGHVDHGKTTL | ASSQEGIGGHYEQY | 0.39368996 |
| LISNASDALDK | ASSQEGIGGHYEQY | 0.6850936 |
| LISNASDAL | ASSQEGIGGHYEQY | 0.7340692 |
| ISNASDALDK | ASSQEGIGGHYEQY | 0.75612533 |
| NQAETLVYQTE | ASSQEGIGGHYEQY | 0.77988046 |
| TGGTGGTAGAG | ASSQEGIGGHYEQY | 0.7250843 |
| TGSTAGGAGGA | ASSQEGIGGHYEQY | 0.64630306 |
| DPLAVPVAAGPV | ASSQEGIGGHYEQY | 0.74866444 |
| GGTGGTPGNGGN | ASSQEGIGGHYEQY | 0.63386303 |
| PAGLAGVDPATL | ASSQEGIGGHYEQY | 0.28757474 |
| GNNLVVEAPHAGASV | ASSQEGIGGHYEQY | 0.29773542 |

| | | |
|---|---|---|
| KVTRIMSASTLEQPA | ASSQEGIGGHYEQY | 0.7152724 |
| LGFALPIALPRRTVV | ASSQEGIGGHYEQY | 0.30225247 |
| TRLGNLTARKLVVLV | ASSQEGIGGHYEQY | 0.7357161 |
| WSQQRRLIYFPSAGP | ASSQEGIGGHYEQY | 0.75602394 |
| GPNGNPDPMAAAVDIR | ASSQEGIGGHYEQY | 0.5795426 |
| TQRCVSVADLRAAASR | ASSQEGIGGHYEQY | 0.41068056 |
| QFVLDVAEHPVEVLTETV | ASSQEGIGGHYEQY | 0.658061 |
| GGAGGTGGAGGNGGTQAGDG | ASSQEGIGGHYEQY | 0.45988578 |
| PLLATLRAIAVDVDAKPAQV | ASSQEGIGGHYEQY | 0.76360404 |
| APKFRTGGYMIDFWGVGYQVAK | ASSQEGIGGHYEQY | 0.75174284 |
| GAGGAGGAGGAAGGPGGTGGTG | ASSQEGIGGHYEQY | 0.7808198 |
| IGALGLLATDVAPAGSVAFGVA | ASSQEGIGGHYEQY | 0.76207316 |
| AGGAGGIGGNGGAGGAGGVLMG | ASSQEGIGGHYEQY | 0.68028235 |
| ADELIGLL | ASSQEGIGGHYEQY | 0.7524627 |
| ADRALKA | ASSQEGIGGHYEQY | 0.6415185 |
| AGAVGQVGGAGGA | ASSQEGIGGHYEQY | 0.69079113 |
| AGGGGGLGGAP | ASSQEGIGGHYEQY | 0.67068803 |
| AGGLLYGNGGAG | ASSQEGIGGHYEQY | 0.6764407 |
| AGLESIGL | ASSQEGIGGHYEQY | 0.5922647 |
| AGLKTGII | ASSQEGIGGHYEQY | 0.6293386 |
| ASLAALLL | ASSQEGIGGHYEQY | 0.36052197 |
| ASVINISSI | ASSQEGIGGHYEQY | 0.68568146 |
| ATPPGPA | ASSQEGIGGHYEQY | 0.7187259 |
| ATVASALV | ASSQEGIGGHYEQY | 0.30067942 |
| ATVASLIL | ASSQEGIGGHYEQY | 0.60922074 |
| AVNGPVLIRH | ASSQEGIGGHYEQY | 0.7853147 |
| DGAAGAVGI | ASSQEGIGGHYEQY | 0.7867224 |
| DPLLHGI | ASSQEGIGGHYEQY | 0.5947809 |
| ELGAALGAL | ASSQEGIGGHYEQY | 0.5451808 |
| ELGRTQAV | ASSQEGIGGHYEQY | 0.7150395 |
| FVAGGYLS | ASSQEGIGGHYEQY | 0.37345642 |
| GASGGAGGAGGW | ASSQEGIGGHYEQY | 0.6892191 |
| GGAGAGGDGGAGGI | ASSQEGIGGHYEQY | 0.69308734 |
| GTGHNGSQPGMGGNG | ASSQEGIGGHYEQY | 0.684225 |
| GYAAASASAS | ASSQEGIGGHYEQY | 0.5781611 |
| HQVDVAAVQQG | ASSQEGIGGHYEQY | 0.72874534 |
| IGAAIAHT | ASSQEGIGGHYEQY | 0.67569816 |
| IGMIEVAQL | ASSQEGIGGHYEQY | 0.7032745 |
| IRLAPEHTSAMD | ASSQEGIGGHYEQY | 0.19155541 |
| IYNVSVL | ASSQEGIGGHYEQY | 0.6509833 |
| KQGFLGM | ASSQEGIGGHYEQY | 0.55129623 |
| LESVITQL | ASSQEGIGGHYEQY | 0.49887735 |
| LGLGALVVV | ASSQEGIGGHYEQY | 0.25379154 |
| LIGNGGNGGSGGI | ASSQEGIGGHYEQY | 0.6883067 |
| LNAINAPTL | ASSQEGIGGHYEQY | 0.7755046 |
| LQLLNTGV | ASSQEGIGGHYEQY | 0.76773757 |
| LQTSIEF | ASSQEGIGGHYEQY | 0.49267206 |
| MSFAVSEL | ASSQEGIGGHYEQY | 0.7583293 |
| NTGALGL | ASSQEGIGGHYEQY | 0.78264636 |
| PPGAGGAGGAGTT | ASSQEGIGGHYEQY | 0.65666044 |
| QLVGLGI | ASSQEGIGGHYEQY | 0.25613368 |

| | | |
|---|---|---|
| RFLEQVHQL | ASSQEGIGGHYEQY | 0.74628633 |
| RGVGYVLR | ASSQEGIGGHYEQY | 0.6884311 |
| SATGVFAGVI | ASSQEGIGGHYEQY | 0.7382356 |
| SLPAITLP | ASSQEGIGGHYEQY | 0.76823175 |
| STALAFTV | ASSQEGIGGHYEQY | 0.478449 |
| SVLTVGLGK | ASSQEGIGGHYEQY | 0.7700652 |
| TALAGAGSGI | ASSQEGIGGHYEQY | 0.7774074 |
| TGLPFTGI | ASSQEGIGGHYEQY | 0.5106896 |
| VAEHVTV | ASSQEGIGGHYEQY | 0.5879378 |
| VDIVLNSL | ASSQEGIGGHYEQY | 0.58347553 |
| VLTGGGAVL | ASSQEGIGGHYEQY | 0.2944385 |
| VNVGLAYGI | ASSQEGIGGHYEQY | 0.7591055 |
| VSSAAVGAV | ASSQEGIGGHYEQY | 0.6474836 |
| VTSVYEAGV | ASSQEGIGGHYEQY | 0.6570399 |
| WVALDRGA | ASSQEGIGGHYEQY | 0.7085719 |
| YLVGSVVI | ASSQEGIGGHYEQY | 0.6135145 |
| GPPGTGKTLL | ASSQEGIGGHYEQY | 0.6053437 |
| GVGMTKF | ASSQEGIGGHYEQY | 0.6706135 |
| IGIGGSDL | ASSQEGIGGHYEQY | 0.7003981 |
| IGTLLAEL | ASSQEGIGGHYEQY | 0.7376649 |
| IYTSGTT | ASSQEGIGGHYEQY | 0.6845267 |
| LSFPAGV | ASSQEGIGGHYEQY | 0.71462107 |
| LYTSGTT | ASSQEGIGGHYEQY | 0.78603417 |
| VGAFPFGF | ASSQEGIGGHYEQY | 0.39560097 |
| VGAGTVEF | ASSQEGIGGHYEQY | 0.56452334 |
| VGKSTLLL | ASSQEGIGGHYEQY | 0.73396814 |
| AALAHAGGGIV | ASSQEGIGGHYEQY | 0.4362839 |
| AALVAVSLL | ASSQEGIGGHYEQY | 0.7588849 |
| AAVSRAVAAL | ASSQEGIGGHYEQY | 0.7361834 |
| AGGSSGIGGF | ASSQEGIGGHYEQY | 0.7326318 |
| AGLTVANANG | ASSQEGIGGHYEQY | 0.6451643 |
| AGLVGAVL | ASSQEGIGGHYEQY | 0.7911182 |
| ALSQAFTV | ASSQEGIGGHYEQY | 0.76660466 |
| ALTFSVSL | ASSQEGIGGHYEQY | 0.68583566 |
| ALTGAALVAL | ASSQEGIGGHYEQY | 0.72062266 |
| ASFSIGAGVL | ASSQEGIGGHYEQY | 0.49214697 |
| DGSVYKI | ASSQEGIGGHYEQY | 0.7580936 |
| EQLAEIV | ASSQEGIGGHYEQY | 0.75429374 |
| FAVSELTVM | ASSQEGIGGHYEQY | 0.5205469 |
| GAAPLAAGL | ASSQEGIGGHYEQY | 0.6857059 |
| GADGGDGGSGGAGG | ASSQEGIGGHYEQY | 0.7731948 |
| GFGLDTGGAGG | ASSQEGIGGHYEQY | 0.77195334 |
| GLASGGLL | ASSQEGIGGHYEQY | 0.33612972 |
| GRIAGAAVLL | ASSQEGIGGHYEQY | 0.6391594 |
| GSAGSLNTSF | ASSQEGIGGHYEQY | 0.32506287 |
| GTGQIGI | ASSQEGIGGHYEQY | 0.5221498 |
| GTLIALIL | ASSQEGIGGHYEQY | 0.44948375 |
| IAATMGFVVG | ASSQEGIGGHYEQY | 0.74208844 |
| IFAQALL | ASSQEGIGGHYEQY | 0.6716376 |
| IGAAVALGF | ASSQEGIGGHYEQY | 0.7498449 |
| IGLTPLAL | ASSQEGIGGHYEQY | 0.7267079 |

| | | |
|---|---|---|
| ILGSNTSTL | ASSQEGIGGHYEQY | 0.7824197 |
| IVFSLTVL | ASSQEGIGGHYEQY | 0.59707856 |
| LASKATGF | ASSQEGIGGHYEQY | 0.7082773 |
| LASVALIL | ASSQEGIGGHYEQY | 0.31840396 |
| LFAVGLTL | ASSQEGIGGHYEQY | 0.42428386 |
| LFNGGIGTYI | ASSQEGIGGHYEQY | 0.48460293 |
| LTLGAGII | ASSQEGIGGHYEQY | 0.6777415 |
| LVAAALGDI | ASSQEGIGGHYEQY | 0.22941849 |
| NTAFVQLGI | ASSQEGIGGHYEQY | 0.7812751 |
| QLFSGAEL | ASSQEGIGGHYEQY | 0.715761 |
| SAATFITL | ASSQEGIGGHYEQY | 0.758139 |
| SNYGLANLGV | ASSQEGIGGHYEQY | 0.59404904 |
| SYGGAYAVM | ASSQEGIGGHYEQY | 0.57993656 |
| TTGVGGLTLG | ASSQEGIGGHYEQY | 0.5838767 |
| TTQTIYI | ASSQEGIGGHYEQY | 0.75714445 |
| VAAGAGLGAVI | ASSQEGIGGHYEQY | 0.4159513 |
| VAAVVTLL | ASSQEGIGGHYEQY | 0.3651495 |
| VGFLGGLI | ASSQEGIGGHYEQY | 0.5654682 |
| VGVIPILL | ASSQEGIGGHYEQY | 0.46467385 |
| VIFASSLI | ASSQEGIGGHYEQY | 0.7914308 |
| VKGGDGGAAAG | ASSQEGIGGHYEQY | 0.77592415 |
| VTKTFTNI | ASSQEGIGGHYEQY | 0.72702324 |
| VVAAAILL | ASSQEGIGGHYEQY | 0.77169 |
| YAALDVEL | ASSQEGIGGHYEQY | 0.21357404 |
| LASVALLL | ASSQEGIGGHYEQY | 0.5584724 |
| LDAVHRV | ASSQEGIGGHYEQY | 0.7880943 |
| QGLSSSIF | ASSQEGIGGHYEQY | 0.7384641 |
| RGTFANIRL | ASSQEGIGGHYEQY | 0.6903188 |
| RQLWWGHRI | ASSQEGIGGHYEQY | 0.7760123 |
| YGPPGTGKTLL | ASSQEGIGGHYEQY | 0.6516218 |
| AALVGAPANVA | ASSQEGIGGHYEQY | 0.6901604 |
| AATFRRTE | ASSQEGIGGHYEQY | 0.7001256 |
| AGALGGGI | ASSQEGIGGHYEQY | 0.7260097 |
| AGGAGGAGGIGAG | ASSQEGIGGHYEQY | 0.6972792 |
| AGSGALGL | ASSQEGIGGHYEQY | 0.4359213 |
| AQIADQI | ASSQEGIGGHYEQY | 0.74856555 |
| ASAVAVTYL | ASSQEGIGGHYEQY | 0.7982421 |
| ASTVLGGTGG | ASSQEGIGGHYEQY | 0.6883651 |
| AVGGEPAK | ASSQEGIGGHYEQY | 0.75794435 |
| CEQGTFGGTC | ASSQEGIGGHYEQY | 0.74373114 |
| DEGKAPAG | ASSQEGIGGHYEQY | 0.75087255 |
| DTVVFPAL | ASSQEGIGGHYEQY | 0.6095079 |
| FEGITMAG | ASSQEGIGGHYEQY | 0.5493201 |
| FPLVVQD | ASSQEGIGGHYEQY | 0.67947495 |
| GAFTTPQ | ASSQEGIGGHYEQY | 0.48793787 |
| GFGGFGTAGGVGGA | ASSQEGIGGHYEQY | 0.70831656 |
| GFLAQFL | ASSQEGIGGHYEQY | 0.6572391 |
| GGAGGAGGNAGVA | ASSQEGIGGHYEQY | 0.80286336 |
| GGVGTAGAGGAGGAGG | ASSQEGIGGHYEQY | 0.75762314 |
| GNGGGGDAGNAGSGGN | ASSQEGIGGHYEQY | 0.72393847 |
| GTGGAGGAGNDAGS | ASSQEGIGGHYEQY | 0.52498364 |

| | | |
|---|---|---|
| GTTNTGWFNS | ASSQEGIGGHYEQY | 0.69006264 |
| IFMGPLFTL | ASSQEGIGGHYEQY | 0.72969556 |
| ILAHAIEAI | ASSQEGIGGHYEQY | 0.38225782 |
| IRSAIREAL | ASSQEGIGGHYEQY | 0.7858084 |
| ITFAGGG | ASSQEGIGGHYEQY | 0.53727484 |
| ITGMSGHQEV | ASSQEGIGGHYEQY | 0.6496177 |
| IVAGLACTL | ASSQEGIGGHYEQY | 0.7413543 |
| IVGGGLAGA | ASSQEGIGGHYEQY | 0.77285844 |
| IVTGVLNI | ASSQEGIGGHYEQY | 0.6335734 |
| LAQQGCI | ASSQEGIGGHYEQY | 0.6597754 |
| LCLHVEN | ASSQEGIGGHYEQY | 0.5551121 |
| LDAISDGS | ASSQEGIGGHYEQY | 0.7565967 |
| LGGFAAML | ASSQEGIGGHYEQY | 0.7734345 |
| LGGKNPMI | ASSQEGIGGHYEQY | 0.6801151 |
| LGPKGVAAIL | ASSQEGIGGHYEQY | 0.7805496 |
| LMVGVSI | ASSQEGIGGHYEQY | 0.6776136 |
| LVGAAFVL | ASSQEGIGGHYEQY | 0.31149203 |
| LVLAVAPL | ASSQEGIGGHYEQY | 0.6860899 |
| MAAIAVIDAA | ASSQEGIGGHYEQY | 0.74964535 |
| NNGINAELQ | ASSQEGIGGHYEQY | 0.77504504 |
| RGEPLVLVHG | ASSQEGIGGHYEQY | 0.7888675 |
| RGGAGGAAGLI | ASSQEGIGGHYEQY | 0.7137819 |
| RSYRMHPAV | ASSQEGIGGHYEQY | 0.65734136 |
| SAAHLLAGQLA | ASSQEGIGGHYEQY | 0.7842659 |
| SAAVGSY | ASSQEGIGGHYEQY | 0.8008187 |
| SPHAVLHRI | ASSQEGIGGHYEQY | 0.7578503 |
| SQIAKVL | ASSQEGIGGHYEQY | 0.5595399 |
| TALGHLVP | ASSQEGIGGHYEQY | 0.52173495 |
| TAPTTSGAA | ASSQEGIGGHYEQY | 0.74801743 |
| TDGDDGNGGSPGT | ASSQEGIGGHYEQY | 0.7536448 |
| TDGLGEN | ASSQEGIGGHYEQY | 0.50097644 |
| TGGPPLPGQA | ASSQEGIGGHYEQY | 0.71129876 |
| TIPAITI | ASSQEGIGGHYEQY | 0.7443898 |
| TIPGFTI | ASSQEGIGGHYEQY | 0.37230897 |
| TSIASGALA | ASSQEGIGGHYEQY | 0.75659394 |
| TVTLPGAAI | ASSQEGIGGHYEQY | 0.7703825 |
| VASGLSAALP | ASSQEGIGGHYEQY | 0.68944615 |
| VGAGVIGI | ASSQEGIGGHYEQY | 0.57396966 |
| VGPKRFGV | ASSQEGIGGHYEQY | 0.77719665 |
| VNVAFPD | ASSQEGIGGHYEQY | 0.7986884 |
| VSMTNSGVS | ASSQEGIGGHYEQY | 0.78220975 |
| VTDVLGDLSV | ASSQEGIGGHYEQY | 0.34997702 |
| VVVVGAGFAGL | ASSQEGIGGHYEQY | 0.23647362 |
| VYTVGTVV | ASSQEGIGGHYEQY | 0.61137784 |
| WTTFSGP | ASSQEGIGGHYEQY | 0.75551164 |
| YDELVEAF | ASSQEGIGGHYEQY | 0.6966715 |
| YGTGGAGGNGG | ASSQEGIGGHYEQY | 0.59017587 |
| YTNVMAV | ASSQEGIGGHYEQY | 0.72402924 |
| FLEQVHQL | ASSQEGIGGHYEQY | 0.7785733 |
| GLSLLAVAV | ASSQEGIGGHYEQY | 0.5381901 |
| LEQVHQL | ASSQEGIGGHYEQY | 0.35170424 |

| | | |
|---|---|---|
| LNAAGAI | ASSQEGIGGHYEQY | 0.75769144 |
| PAVHASL | ASSQEGIGGHYEQY | 0.7556644 |
| TELEDDA | ASSQEGIGGHYEQY | 0.5442569 |
| TPPGPPN | ASSQEGIGGHYEQY | 0.6662419 |
| TPPPPPPP | ASSQEGIGGHYEQY | 0.6623206 |
| TQSPALI | TCSAYDNYEQY | 0.5519662 |
| EFAAAFPQ | TCSAYDNYEQY | 0.7129961 |
| LLSGNLDVL | TCSAYDNYEQY | 0.4203807 |
| GPANLGNYNI | TCSAYDNYEQY | 0.7656522 |
| TLGPTLP | TCSAYDNYEQY | 0.4274047 |
| VYAYSSCCK | TCSAYDNYEQY | 0.73267424 |
| APAPAPAPNI | TCSAYDNYEQY | 0.72950923 |
| PGGPVPIP | TCSAYDNYEQY | 0.7252486 |
| FLTVALEP | TCSAYDNYEQY | 0.659516 |
| SQYFGANL | TCSAYDNYEQY | 0.5587327 |
| GGVKPAA | TCSAYDNYEQY | 0.40688896 |
| GGTGGAGGAGGW | TCSAYDNYEQY | 0.28313032 |
| GVGGVAGLGGAGG | TCSAYDNYEQY | 0.7651267 |
| VLLVLATTI | TCSAYDNYEQY | 0.2437831 |
| IGIFAAML | TCSAYDNYEQY | 0.76195943 |
| AVGDLID | TCSAYDNYEQY | 0.75244623 |
| NISPLPL | TCSAYDNYEQY | 0.4840397 |
| VTAVFVAI | TCSAYDNYEQY | 0.7187333 |
| SGDDNDAAL | TCSAYDNYEQY | 0.4166243 |
| AVAALAHS | TCSAYDNYEQY | 0.58694506 |
| GGVTVMATP | TCSAYDNYEQY | 0.755497 |
| AGLAANVRD | TCSAYDNYEQY | 0.7445564 |
| TNRPDLIDPALLRPG | TCSAYDNYEQY | 0.55034024 |
| LANLGVTAL | TCSAYDNYEQY | 0.71129274 |
| TGERIEL | TCSAYDNYEQY | 0.74353284 |
| NGEELAA | TCSAYDNYEQY | 0.59636986 |
| VLSLVWG | TCSAYDNYEQY | 0.6973516 |
| IGYALWL | TCSAYDNYEQY | 0.7745291 |
| LSLLLAPL | TCSAYDNYEQY | 0.71462774 |
| TLDEALK | TCSAYDNYEQY | 0.45724368 |
| TDGAGTAEGG | TCSAYDNYEQY | 0.7608868 |
| NAPGGAGGNGG | TCSAYDNYEQY | 0.5680415 |
| VSVFMGMGGNF | TCSAYDNYEQY | 0.7355026 |
| LASYLAAL | TCSAYDNYEQY | 0.6577765 |
| LVGGTPAL | TCSAYDNYEQY | 0.69388866 |
| VRNADVL | TCSAYDNYEQY | 0.7551926 |
| TCSSPFLVSAM | TCSAYDNYEQY | 0.41758418 |
| VHITDAE | TCSAYDNYEQY | 0.5756061 |
| EYAAPSF | TCSAYDNYEQY | 0.6234086 |
| VVDAFHGGL | TCSAYDNYEQY | 0.718331 |
| ASVVDPQ | TCSAYDNYEQY | 0.7621599 |
| SYNEFGGLS | TCSAYDNYEQY | 0.56576943 |
| DPDGSVY | TCSAYDNYEQY | 0.720355 |
| MVSTIHATE | TCSAYDNYEQY | 0.68433505 |
| LAAGVADPV | TCSAYDNYEQY | 0.5947559 |
| TKEQIAA | TCSAYDNYEQY | 0.6676299 |

| | | |
|---|---|---|
| SLASPAAL | TCSAYDNYEQY | 0.6238843 |
| IGLVFYGV | TCSAYDNYEQY | 0.47228152 |
| FLLDASNL | TCSAYDNYEQY | 0.7453171 |
| YQVGNFP | TCSAYDNYEQY | 0.55415475 |
| ESLLSQST | TCSAYDNYEQY | 0.6979649 |
| LFTVGFAAVG | TCSAYDNYEQY | 0.79139805 |
| ATPPTPT | TCSAYDNYEQY | 0.48784193 |
| TDLATLG | TCSAYDNYEQY | 0.7221476 |
| SAGGTGR | TCSAYDNYEQY | 0.7206321 |
| GFSNSGS | TCSAYDNYEQY | 0.72598577 |
| GGNGGSGGTSV | TCSAYDNYEQY | 0.6823678 |
| GTSVATGGAGN | TCSAYDNYEQY | 0.5614516 |
| NGGNGGSGGTS | TCSAYDNYEQY | 0.7700473 |
| AVVKFPH | TCSAYDNYEQY | 0.7527024 |
| TPPPPPP | TCSAYDNYEQY | 0.7502934 |
| AAPAAAAAPA | TCSAYDNYEQY | 0.73539126 |
| IALNEDL | TCSAYDNYEQY | 0.7504436 |
| IVNVSSM | TCSAYDNYEQY | 0.75515497 |
| TGIPPAPR | TCSAYDNYEQY | 0.6201465 |
| FDLGGGTF | TCSAYDNYEQY | 0.35960454 |
| FDLGGGTFDVS | TCSAYDNYEQY | 0.67547077 |
| NEPTAAA | TCSAYDNYEQY | 0.6741643 |
| NEPTAAALA | TCSAYDNYEQY | 0.60024726 |
| GSIINISSI | TCSAYDNYEQY | 0.28555238 |
| VAPGNAGTA | TCSAYDNYEQY | 0.772176 |
| TGGGASLE | TCSAYDNYEQY | 0.79567134 |
| VAVYNHYKRI | TCSAYDNYEQY | 0.5694382 |
| TDPAPATT | TCSAYDNYEQY | 0.7919147 |
| TNRPDILDPALLRPG | TCSAYDNYEQY | 0.4003635 |
| LTTTTTS | TCSAYDNYEQY | 0.746585 |
| PTGSGKTL | TCSAYDNYEQY | 0.77446467 |
| TGSGKTL | TCSAYDNYEQY | 0.47399193 |
| TIGHVDHGK | TCSAYDNYEQY | 0.7472404 |
| TIGHVDHGKTT | TCSAYDNYEQY | 0.52787423 |
| GHVDHGKTT | TCSAYDNYEQY | 0.5105858 |
| AQFEHTLL | TCSAYDNYEQY | 0.7806654 |
| IVNQDEL | TCSAYDNYEQY | 0.66188097 |
| TNGSQFF | TCSAYDNYEQY | 0.7340777 |
| TGKGIVE | TCSAYDNYEQY | 0.651636 |
| LISNASDA | TCSAYDNYEQY | 0.5219909 |
| AELRTLGELTKIAKAHGAQ | TCSAYDNYEQY | 0.7039741 |
| APNAVAAAEPSW | TCSAYDNYEQY | 0.77709323 |
| APQVAGAATAVL | TCSAYDNYEQY | 0.48801565 |
| ARRDPAAAAAILPTDA | TCSAYDNYEQY | 0.79937065 |
| DAAARDVLAEAGLAEYF | TCSAYDNYEQY | 0.5925547 |
| DDQIVLSAKANLAKVEQ | TCSAYDNYEQY | 0.7939755 |
| DELLPDTGEDSDEDGH | TCSAYDNYEQY | 0.7607217 |
| ELPEAPANRATVPQD | TCSAYDNYEQY | 0.70286083 |
| FGQSKIERVTGIG | TCSAYDNYEQY | 0.62345916 |
| GAGGTGSTAGGAGGAGGAGGLY | TCSAYDNYEQY | 0.7743169 |
| GHGGTGAAGGNGGTGGAGGSGI | TCSAYDNYEQY | 0.67093146 |

| | | |
|---|---|---|
| GNGGNGGAGGEGGPGGAGGAGA | TCSAYDNYEQY | 0.71915394 |
| GSTAGNGGNGGNATNGTIAGQP | TCSAYDNYEQY | 0.76431406 |
| GTGGDGGNGGNGGTGG | TCSAYDNYEQY | 0.4946361 |
| GVRRPAGIPADAPV | TCSAYDNYEQY | 0.6858706 |
| GYRQVTDAHLLTLARR | TCSAYDNYEQY | 0.65981746 |
| IAKMGQDGHDRG | TCSAYDNYEQY | 0.6373166 |
| KAEPSIEIQYT | TCSAYDNYEQY | 0.5733354 |
| KPVRIRMAADEDTVAVT | TCSAYDNYEQY | 0.47346705 |
| MPPRIAGMRLLVIKPEP | TCSAYDNYEQY | 0.60449547 |
| MKFYAVALFRSRGVPPTR | TCSAYDNYEQY | 0.53073144 |
| NAGGFGILGAVAHSP | TCSAYDNYEQY | 0.67416906 |
| NEGREITLAMAEM | TCSAYDNYEQY | 0.67001575 |
| NELVEWSHPLGHRWRWE | TCSAYDNYEQY | 0.5815714 |
| NLGAGGTGGNGGNPG | TCSAYDNYEQY | 0.67770237 |
| NTEWPKWETVMAADPVD | TCSAYDNYEQY | 0.65454566 |
| PASAAPAETQP | TCSAYDNYEQY | 0.49680084 |
| QQSSDGKNGVRAAEITGE | TCSAYDNYEQY | 0.6150589 |
| RAPDAVTTSWWKE | TCSAYDNYEQY | 0.6488975 |
| SGFGNVGAHTSGWFNQ | TCSAYDNYEQY | 0.63026524 |
| SGVAGAVGQAASVGGLKV | TCSAYDNYEQY | 0.47412786 |
| SNGGAGTGSAGGNGGT | TCSAYDNYEQY | 0.6667068 |
| TGGAGGEGGAAG | TCSAYDNYEQY | 0.60641396 |
| TGGDGGAAGAGGAPGHG | TCSAYDNYEQY | 0.33215454 |
| TMPPLLAAIANAGRHGV | TCSAYDNYEQY | 0.6599529 |
| TSSTAGNSGGA | TCSAYDNYEQY | 0.62038755 |
| VPATVPVGILC | TCSAYDNYEQY | 0.610679 |
| VQRPDSGGVVLTGRIS | TCSAYDNYEQY | 0.6814433 |
| WTPNPVPATKGVPAGLVQAT | TCSAYDNYEQY | 0.618168 |
| YALKRALAKFSDRFDVVII | TCSAYDNYEQY | 0.5480537 |
| YLIGSVIV | TCSAYDNYEQY | 0.6692443 |
| YGQLVAVID | TCSAYDNYEQY | 0.6744319 |
| VTVGGMTLPA | TCSAYDNYEQY | 0.48930025 |
| VLALPAV | TCSAYDNYEQY | 0.6524442 |
| VIPPTLNYET | TCSAYDNYEQY | 0.67598706 |
| VGMGGMHPSSG | TCSAYDNYEQY | 0.6487162 |
| VGLAGGAATA | TCSAYDNYEQY | 0.6860171 |
| VGGAGGAGGSGP | TCSAYDNYEQY | 0.68702507 |
| VFAAFMV | TCSAYDNYEQY | 0.57450676 |
| TLYDKLR | TCSAYDNYEQY | 0.6806935 |
| TLAVFTVL | TCSAYDNYEQY | 0.68720984 |
| TGSVQPARV | TCSAYDNYEQY | 0.5873232 |
| TGGGAGSGGT | TCSAYDNYEQY | 0.6425996 |
| SHLETSL | TCSAYDNYEQY | 0.4920956 |
| SFTTGMT | TCSAYDNYEQY | 0.62940323 |
| QVEVAKRMGVRQPTVS | TCSAYDNYEQY | 0.5013843 |
| QGSGGPALAM | TCSAYDNYEQY | 0.6628339 |
| MLAPSNP | TCSAYDNYEQY | 0.6850598 |
| LVNNAAI | TCSAYDNYEQY | 0.6362939 |
| LTGEINTLA | TCSAYDNYEQY | 0.5263876 |
| LSIFGPV | TCSAYDNYEQY | 0.51267266 |
| LLALLANL | TCSAYDNYEQY | 0.6568033 |

| | | |
|---|---|---|
| LISNASDALDKLR | TCSAYDNYEQY | 0.51145554 |
| LINEAVP | TCSAYDNYEQY | 0.5101652 |
| LGYAFSGL | TCSAYDNYEQY | 0.63246703 |
| LDFVAVAL | TCSAYDNYEQY | 0.6683066 |
| LAQPGAE | TCSAYDNYEQY | 0.67446804 |
| ISNLGAVTGF | TCSAYDNYEQY | 0.5367369 |
| HQEDPGRI | TCSAYDNYEQY | 0.68827385 |
| GTPGEPTP | TCSAYDNYEQY | 0.6611427 |
| GFFNSGS | TCSAYDNYEQY | 0.6819025 |
| GAGGGGLTAGAN | TCSAYDNYEQY | 0.6363485 |
| FVDALVNM | TCSAYDNYEQY | 0.5290059 |
| FLTVAPEL | TCSAYDNYEQY | 0.6775422 |
| FETLTAL | TCSAYDNYEQY | 0.61646044 |
| DVINPIEL | TCSAYDNYEQY | 0.63690805 |
| APVSGAPV | TCSAYDNYEQY | 0.67559046 |
| ALVGGLAW | TCSAYDNYEQY | 0.6486273 |
| ALAAAAATAI | TCSAYDNYEQY | 0.5607142 |
| AFGLGLLL | TCSAYDNYEQY | 0.6208039 |
| AFFAAASL | TCSAYDNYEQY | 0.68447906 |
| AAAALPAFSPP | TCSAYDNYEQY | 0.5527119 |
| VVLAPAL | TCSAYDNYEQY | 0.5092031 |
| TIGHVDHGKTTL | TCSAYDNYEQY | 0.60611725 |
| LISNASDALDK | TCSAYDNYEQY | 0.62656057 |
| LISNASDAL | TCSAYDNYEQY | 0.68056554 |
| ISNASDALDK | TCSAYDNYEQY | 0.6650796 |
| NQAETLVYQTE | TCSAYDNYEQY | 0.6110165 |
| TGGTGGTAGAG | TCSAYDNYEQY | 0.6638495 |
| TGSTAGGAGGA | TCSAYDNYEQY | 0.6637523 |
| DPLAVPVAAGPV | TCSAYDNYEQY | 0.64557076 |
| GGTGGTPGNGGN | TCSAYDNYEQY | 0.6490781 |
| PAGLAGVDPATL | TCSAYDNYEQY | 0.6040979 |
| GNNLVVEAPHAGASV | TCSAYDNYEQY | 0.6795848 |
| KVTRIMSASTLEQPA | TCSAYDNYEQY | 0.6754664 |
| LGFALPIALPRRTVV | TCSAYDNYEQY | 0.67064595 |
| TRLGNLTARKLVVLV | TCSAYDNYEQY | 0.6790026 |
| WSQQRRLIYFPSAGP | TCSAYDNYEQY | 0.64431506 |
| GPNGNPDPMAAAVDIR | TCSAYDNYEQY | 0.64819527 |
| TQRCVSVADLRAAASR | TCSAYDNYEQY | 0.4110549 |
| QFVLDVAEHPVEVLTETV | TCSAYDNYEQY | 0.598949 |
| GGAGGTGGAGGNGGTQAGDG | TCSAYDNYEQY | 0.64109695 |
| PLLATLRAIAVDVDAKPAQV | TCSAYDNYEQY | 0.65997195 |
| APKFRTGGYMIDFWGVGYQVAK | TCSAYDNYEQY | 0.6690337 |
| GAGGAGGAGGAAGGPGGTGGTG | TCSAYDNYEQY | 0.65422523 |
| IGALGLLATDVAPAGSVAFGVA | TCSAYDNYEQY | 0.5794061 |
| AGGAGGIGGNGGAGGAGGVLMG | TCSAYDNYEQY | 0.6655908 |
| ADELIGLL | TCSAYDNYEQY | 0.6788159 |
| ADRALKA | TCSAYDNYEQY | 0.5118554 |
| AGAVGQVGGAGGA | TCSAYDNYEQY | 0.6148522 |
| AGGGGGLGGAP | TCSAYDNYEQY | 0.6926693 |
| AGGLLYGNGGAG | TCSAYDNYEQY | 0.6525395 |
| AGLESIGL | TCSAYDNYEQY | 0.6290695 |

| | | |
|---|---|---|
| AGLKTGII | TCSAYDNYEQY | 0.5973893 |
| ASLAALLL | TCSAYDNYEQY | 0.68076956 |
| ASVINISSI | TCSAYDNYEQY | 0.62179184 |
| ATPPGPA | TCSAYDNYEQY | 0.68584365 |
| ATVASALV | TCSAYDNYEQY | 0.37251505 |
| ATVASLIL | TCSAYDNYEQY | 0.62303007 |
| AVNGPVLIRH | TCSAYDNYEQY | 0.5352231 |
| DGAAGAVGI | TCSAYDNYEQY | 0.34104094 |
| DPLLHGI | TCSAYDNYEQY | 0.6760767 |
| ELGAALGAL | TCSAYDNYEQY | 0.5173849 |
| ELGRTQAV | TCSAYDNYEQY | 0.6481395 |
| FVAGGYLS | TCSAYDNYEQY | 0.6402216 |
| GASGGAGGAGGW | TCSAYDNYEQY | 0.67835885 |
| GGAGAGGDGGAGGI | TCSAYDNYEQY | 0.6577692 |
| GTGHNGSQPGMGGNG | TCSAYDNYEQY | 0.607092 |
| GYAAASASAS | TCSAYDNYEQY | 0.57098186 |
| HQVDVAAVQQG | TCSAYDNYEQY | 0.66344976 |
| IGAAIAHT | TCSAYDNYEQY | 0.67241746 |
| IGMIEVAQL | TCSAYDNYEQY | 0.68117833 |
| IRLAPEHTSAMD | TCSAYDNYEQY | 0.3902303 |
| IYNVSVL | TCSAYDNYEQY | 0.6530459 |
| KQGFLGM | TCSAYDNYEQY | 0.5992019 |
| LESVITQL | TCSAYDNYEQY | 0.5538305 |
| LGLGALVVV | TCSAYDNYEQY | 0.3887537 |
| LIGNGGNGGSGGI | TCSAYDNYEQY | 0.6267499 |
| LNAINAPTL | TCSAYDNYEQY | 0.63893425 |
| LQLLNTGV | TCSAYDNYEQY | 0.6693757 |
| LQTSIEF | TCSAYDNYEQY | 0.6737313 |
| MSFAVSEL | TCSAYDNYEQY | 0.6288595 |
| NTGALGL | TCSAYDNYEQY | 0.6658833 |
| PPGAGGAGGAGTT | TCSAYDNYEQY | 0.618513 |
| QLVGLGI | TCSAYDNYEQY | 0.27968055 |
| RFLEQVHQL | TCSAYDNYEQY | 0.5862863 |
| RGVGYVLR | TCSAYDNYEQY | 0.49508798 |
| SATGVFAGVI | TCSAYDNYEQY | 0.6804397 |
| SLPAITLP | TCSAYDNYEQY | 0.5543885 |
| STALAFTV | TCSAYDNYEQY | 0.6154553 |
| SVLTVGLGK | TCSAYDNYEQY | 0.64464027 |
| TALAGAGSGI | TCSAYDNYEQY | 0.65455127 |
| TGLPFTGI | TCSAYDNYEQY | 0.6390923 |
| VAEHVTV | TCSAYDNYEQY | 0.5500081 |
| VDIVLNSL | TCSAYDNYEQY | 0.51918906 |
| VLTGGGAVL | TCSAYDNYEQY | 0.66500777 |
| VNVGLAYGI | TCSAYDNYEQY | 0.62437475 |
| VSSAAVGAV | TCSAYDNYEQY | 0.49155876 |
| VTSVYEAGV | TCSAYDNYEQY | 0.50818145 |
| WVALDRGA | TCSAYDNYEQY | 0.6742945 |
| YLVGSVVI | TCSAYDNYEQY | 0.4532402 |
| GPPGTGKTLL | TCSAYDNYEQY | 0.68076 |
| GVGMTKF | TCSAYDNYEQY | 0.6041422 |
| IGIGGSDL | TCSAYDNYEQY | 0.6243603 |

| | | |
|---|---|---|
| IGTLLAEL | TCSAYDNYEQY | 0.6919355 |
| IYTSGTT | TCSAYDNYEQY | 0.638945 |
| LSFPAGV | TCSAYDNYEQY | 0.6411711 |
| LYTSGTT | TCSAYDNYEQY | 0.5520396 |
| VGAFPFGF | TCSAYDNYEQY | 0.5815898 |
| VGAGTVEF | TCSAYDNYEQY | 0.5776402 |
| VGKSTLLL | TCSAYDNYEQY | 0.58147395 |
| AALAHAGGGIV | TCSAYDNYEQY | 0.6856574 |
| AALVAVSLL | TCSAYDNYEQY | 0.6390686 |
| AAVSRAVAAL | TCSAYDNYEQY | 0.49849188 |
| AGGSSGIGGF | TCSAYDNYEQY | 0.6523029 |
| AGLTVANANG | TCSAYDNYEQY | 0.6888083 |
| AGLVGAVL | TCSAYDNYEQY | 0.6908485 |
| ALSQAFTV | TCSAYDNYEQY | 0.549979 |
| ALTFSVSL | TCSAYDNYEQY | 0.5785887 |
| ALTGAALVAL | TCSAYDNYEQY | 0.569085 |
| ASFSIGAGVL | TCSAYDNYEQY | 0.62232554 |
| DGSVYKI | TCSAYDNYEQY | 0.5349059 |
| EQLAEIV | TCSAYDNYEQY | 0.5269865 |
| FAVSELTVM | TCSAYDNYEQY | 0.67271453 |
| GAAPLAAGL | TCSAYDNYEQY | 0.6857348 |
| GADGGDGGSGGAGG | TCSAYDNYEQY | 0.61895347 |
| GFGLDTGGAGG | TCSAYDNYEQY | 0.5375805 |
| GLASGGLL | TCSAYDNYEQY | 0.6265217 |
| GRIAGAAVLL | TCSAYDNYEQY | 0.5691746 |
| GSAGSLNTSF | TCSAYDNYEQY | 0.5925488 |
| GTGQIGI | TCSAYDNYEQY | 0.56231964 |
| GTLIALIL | TCSAYDNYEQY | 0.48675188 |
| IAATMGFVVG | TCSAYDNYEQY | 0.645552 |
| IFAQALL | TCSAYDNYEQY | 0.69050026 |
| IGAAVALGF | TCSAYDNYEQY | 0.61556554 |
| IGLTPLAL | TCSAYDNYEQY | 0.6625712 |
| ILGSNTSTL | TCSAYDNYEQY | 0.6902782 |
| IVFSLTVL | TCSAYDNYEQY | 0.6739408 |
| LASKATGF | TCSAYDNYEQY | 0.6840856 |
| LASVALIL | TCSAYDNYEQY | 0.6673322 |
| LFAVGLTL | TCSAYDNYEQY | 0.63708055 |
| LFNGGIGTYI | TCSAYDNYEQY | 0.6314249 |
| LTLGAGII | TCSAYDNYEQY | 0.68199694 |
| LVAAALGDI | TCSAYDNYEQY | 0.6260227 |
| NTAFVQLGI | TCSAYDNYEQY | 0.6519424 |
| QLFSGAEL | TCSAYDNYEQY | 0.6111571 |
| SAATFITL | TCSAYDNYEQY | 0.6689626 |
| SNYGLANLGV | TCSAYDNYEQY | 0.6195233 |
| SYGGAYAVM | TCSAYDNYEQY | 0.6832297 |
| TTGVGGLTLG | TCSAYDNYEQY | 0.36108917 |
| TTQTIYI | TCSAYDNYEQY | 0.5341543 |
| VAAGAGLGAVI | TCSAYDNYEQY | 0.6803671 |
| VAAVVTLL | TCSAYDNYEQY | 0.2910666 |
| VGFLGGLI | TCSAYDNYEQY | 0.36540538 |
| VGVIPILL | TCSAYDNYEQY | 0.6438454 |

| | | |
|---|---|---|
| VIFASSLI | TCSAYDNYEQY | 0.67885566 |
| VKGGDGGAAAG | TCSAYDNYEQY | 0.34205073 |
| VTKTFTNI | TCSAYDNYEQY | 0.5805922 |
| VVAAAILL | TCSAYDNYEQY | 0.47572598 |
| YAALDVEL | TCSAYDNYEQY | 0.67578214 |
| LASVALLL | TCSAYDNYEQY | 0.61784685 |
| LDAVHRV | TCSAYDNYEQY | 0.46949458 |
| QGLSSSIF | TCSAYDNYEQY | 0.67085516 |
| RGTFANIRL | TCSAYDNYEQY | 0.5533793 |
| RQLWWGHRI | TCSAYDNYEQY | 0.65798736 |
| YGPPGTGKTLL | TCSAYDNYEQY | 0.6786519 |
| AALVGAPANVA | TCSAYDNYEQY | 0.6867157 |
| AATFRRTE | TCSAYDNYEQY | 0.51909614 |
| AGALGGGI | TCSAYDNYEQY | 0.6641424 |
| AGGAGGAGGIGAG | TCSAYDNYEQY | 0.5986296 |
| AGSGALGL | TCSAYDNYEQY | 0.69539905 |
| AQIADQI | TCSAYDNYEQY | 0.6083728 |
| ASAVAVTYL | TCSAYDNYEQY | 0.48669362 |
| ASTVLGGTGG | TCSAYDNYEQY | 0.6794176 |
| AVGGEPAK | TCSAYDNYEQY | 0.6714902 |
| CEQGTFGGTC | TCSAYDNYEQY | 0.57231617 |
| DEGKAPAG | TCSAYDNYEQY | 0.6691554 |
| DTVVFPAL | TCSAYDNYEQY | 0.6248362 |
| FEGITMAG | TCSAYDNYEQY | 0.6313758 |
| FPLVVQD | TCSAYDNYEQY | 0.6793902 |
| GAFTTPQ | TCSAYDNYEQY | 0.66964793 |
| GFGGFGTAGGVGGA | TCSAYDNYEQY | 0.58786595 |
| GFLAQFL | TCSAYDNYEQY | 0.6771413 |
| GGAGGAGGNAGVA | TCSAYDNYEQY | 0.63213164 |
| GGVGTAGAGGAGGAGG | TCSAYDNYEQY | 0.5150558 |
| GNGGGGDAGNAGSGGN | TCSAYDNYEQY | 0.690698 |
| GTGGAGGAGNDAGS | TCSAYDNYEQY | 0.6311696 |
| GTTNTGWFNS | TCSAYDNYEQY | 0.6773385 |
| IFMGPLFTL | TCSAYDNYEQY | 0.66725373 |
| ILAHAIEAI | TCSAYDNYEQY | 0.66698897 |
| IRSAIREAL | TCSAYDNYEQY | 0.5726303 |
| ITFAGGG | TCSAYDNYEQY | 0.61687636 |
| ITGMSGHQEV | TCSAYDNYEQY | 0.6394297 |
| IVAGLACTL | TCSAYDNYEQY | 0.6659374 |
| IVGGGLAGA | TCSAYDNYEQY | 0.6301385 |
| IVTGVLNI | TCSAYDNYEQY | 0.6594452 |
| LAQQGCI | TCSAYDNYEQY | 0.66961 |
| LCLHVEN | TCSAYDNYEQY | 0.6890744 |
| LDAISDGS | TCSAYDNYEQY | 0.6190858 |
| LGGFAAML | TCSAYDNYEQY | 0.6360873 |
| LGGKNPMI | TCSAYDNYEQY | 0.68866396 |
| LGPKGVAAIL | TCSAYDNYEQY | 0.56495273 |
| LMVGVSI | TCSAYDNYEQY | 0.6303208 |
| LVGAAFVL | TCSAYDNYEQY | 0.66198146 |
| LVLAVAPL | TCSAYDNYEQY | 0.6736722 |
| MAAIAVIDAA | TCSAYDNYEQY | 0.6356958 |

| | | |
|---|---|---|
| NNGINAELQ | TCSAYDNYEQY | 0.6822386 |
| RGEPLVLVHG | TCSAYDNYEQY | 0.60854995 |
| RGGAGGAAGLI | TCSAYDNYEQY | 0.6631149 |
| RSYRMHPAV | TCSAYDNYEQY | 0.63291276 |
| SAAHLLAGQLA | TCSAYDNYEQY | 0.6354993 |
| SAAVGSY | TCSAYDNYEQY | 0.5866914 |
| SPHAVLHRI | TCSAYDNYEQY | 0.6557704 |
| SQIAKVL | TCSAYDNYEQY | 0.6792867 |
| TALGHLVP | TCSAYDNYEQY | 0.66504407 |
| TAPTTSGAA | TCSAYDNYEQY | 0.3674308 |
| TDGDDGNGGSPGT | TCSAYDNYEQY | 0.67431295 |
| TDGLGEN | TCSAYDNYEQY | 0.66094244 |
| TGGPPLPGQA | TCSAYDNYEQY | 0.69138205 |
| TIPAITI | TCSAYDNYEQY | 0.6193887 |
| TIPGFTI | TCSAYDNYEQY | 0.6461787 |
| TSIASGALA | TCSAYDNYEQY | 0.589907 |
| TVTLPGAAI | TCSAYDNYEQY | 0.6474645 |
| VASGLSAALP | TCSAYDNYEQY | 0.48007512 |
| VGAGVIGI | TCSAYDNYEQY | 0.58438265 |
| VGPKRFGV | TCSAYDNYEQY | 0.62971187 |
| VNVAFPD | TCSAYDNYEQY | 0.58962935 |
| VSMTNSGVS | TCSAYDNYEQY | 0.67246306 |
| VTDVLGDLSV | TCSAYDNYEQY | 0.67987126 |
| VVVVGAGFAGL | TCSAYDNYEQY | 0.67634225 |
| VYTVGTVV | TCSAYDNYEQY | 0.68212146 |
| WTTFSGP | TCSAYDNYEQY | 0.6871182 |
| YDELVEAF | TCSAYDNYEQY | 0.68047035 |
| YGTGGAGGNGG | TCSAYDNYEQY | 0.58517003 |
| YTNVMAV | TCSAYDNYEQY | 0.6410971 |
| FLEQVHQL | TCSAYDNYEQY | 0.48834366 |
| GLSLLAVAV | TCSAYDNYEQY | 0.62475663 |
| LEQVHQL | TCSAYDNYEQY | 0.6132704 |
| LNAAGAI | TCSAYDNYEQY | 0.6571568 |
| PAVHASL | TCSAYDNYEQY | 0.30054176 |
| TELEDDA | TCSAYDNYEQY | 0.6498991 |
| TPPGPPN | TCSAYDNYEQY | 0.68955237 |
| TPPPPPPP | TCSAYDNYEQY | 0.6122062 |
| TQSPALI | ASSRRDRGFSQNTLY | 0.6566113 |
| EFAAAFPQ | ASSRRDRGFSQNTLY | 0.6735286 |
| LLSGNLDVL | ASSRRDRGFSQNTLY | 0.60107964 |
| GPANLGNYNI | ASSRRDRGFSQNTLY | 0.6682384 |
| TLGPTLP | ASSRRDRGFSQNTLY | 0.60139817 |
| VYAYSSCCK | ASSRRDRGFSQNTLY | 0.6768659 |
| APAPAPAPNI | ASSRRDRGFSQNTLY | 0.6871765 |
| PGGPVPIP | ASSRRDRGFSQNTLY | 0.66153204 |
| FLTVALEP | ASSRRDRGFSQNTLY | 0.59543884 |
| SQYFGANL | ASSRRDRGFSQNTLY | 0.6283642 |
| GGVKPAA | ASSRRDRGFSQNTLY | 0.49936604 |
| GGTGGAGGAGGW | ASSRRDRGFSQNTLY | 0.6215446 |
| GVGGVAGLGGAGG | ASSRRDRGFSQNTLY | 0.62617916 |
| VLLVLATTI | ASSRRDRGFSQNTLY | 0.6452296 |

| | | |
|---|---|---|
| IGIFAAML | ASSRRDRGFSQNTLY | 0.5548538 |
| AVGDLID | ASSRRDRGFSQNTLY | 0.64568186 |
| NISPLPL | ASSRRDRGFSQNTLY | 0.6401404 |
| VTAVFVAI | ASSRRDRGFSQNTLY | 0.6344342 |
| SGDDNDAAL | ASSRRDRGFSQNTLY | 0.5637563 |
| AVAALAHS | ASSRRDRGFSQNTLY | 0.6670457 |
| GGVTVMATP | ASSRRDRGFSQNTLY | 0.63881266 |
| AGLAANVRD | ASSRRDRGFSQNTLY | 0.4651745 |
| TNRPDLIDPALLRPG | ASSRRDRGFSQNTLY | 0.6793575 |
| LANLGVTAL | ASSRRDRGFSQNTLY | 0.61932844 |
| TGERIEL | ASSRRDRGFSQNTLY | 0.64228934 |
| NGEELAA | ASSRRDRGFSQNTLY | 0.673398 |
| VLSLVWG | ASSRRDRGFSQNTLY | 0.6397399 |
| IGYALWL | ASSRRDRGFSQNTLY | 0.55651516 |
| LSLLLAPL | ASSRRDRGFSQNTLY | 0.6232979 |
| TLDEALK | ASSRRDRGFSQNTLY | 0.64438474 |
| TDGAGTAEGG | ASSRRDRGFSQNTLY | 0.6153429 |
| NAPGGAGGNGG | ASSRRDRGFSQNTLY | 0.3913356 |
| VSVFMGMGGNF | ASSRRDRGFSQNTLY | 0.6726803 |
| LASYLAAL | ASSRRDRGFSQNTLY | 0.51035225 |
| LVGGTPAL | ASSRRDRGFSQNTLY | 0.59885 |
| VRNADVL | ASSRRDRGFSQNTLY | 0.6184643 |
| TCSSPFLVSAM | ASSRRDRGFSQNTLY | 0.6781955 |
| VHITDAE | ASSRRDRGFSQNTLY | 0.63648266 |
| EYAAPSF | ASSRRDRGFSQNTLY | 0.57696474 |
| VVDAFHGGL | ASSRRDRGFSQNTLY | 0.6649548 |
| ASVVDPQ | ASSRRDRGFSQNTLY | 0.64271474 |
| SYNEFGGLS | ASSRRDRGFSQNTLY | 0.6754112 |
| DPDGSVY | ASSRRDRGFSQNTLY | 0.6379714 |
| MVSTIHATE | ASSRRDRGFSQNTLY | 0.6096016 |
| LAAGVADPV | ASSRRDRGFSQNTLY | 0.6320326 |
| TKEQIAA | ASSRRDRGFSQNTLY | 0.59150743 |
| SLASPAAL | ASSRRDRGFSQNTLY | 0.58259904 |
| IGLVFYGV | ASSRRDRGFSQNTLY | 0.53720874 |
| FLLDASNL | ASSRRDRGFSQNTLY | 0.62410116 |
| YQVGNFP | ASSRRDRGFSQNTLY | 0.51097524 |
| ESLLSQST | ASSRRDRGFSQNTLY | 0.63099587 |
| LFTVGFAAVG | ASSRRDRGFSQNTLY | 0.68485236 |
| ATPPTPT | ASSRRDRGFSQNTLY | 0.64030766 |
| TDLATLG | ASSRRDRGFSQNTLY | 0.42917353 |
| SAGGTGR | ASSRRDRGFSQNTLY | 0.5964034 |
| GFSNSGS | ASSRRDRGFSQNTLY | 0.64813423 |
| GGNGGSGGTSV | ASSRRDRGFSQNTLY | 0.67681396 |
| GTSVATGGAGN | ASSRRDRGFSQNTLY | 0.6719961 |
| NGGNGGSGGTS | ASSRRDRGFSQNTLY | 0.6856708 |
| AVVKFPH | ASSRRDRGFSQNTLY | 0.6665952 |
| TPPPPPP | ASSRRDRGFSQNTLY | 0.5882038 |
| AAPAAAAAPA | ASSRRDRGFSQNTLY | 0.6753342 |
| IALNEDL | ASSRRDRGFSQNTLY | 0.62844956 |
| IVNVSSM | ASSRRDRGFSQNTLY | 0.6574834 |
| TGIPPAPR | ASSRRDRGFSQNTLY | 0.6568222 |

| | | |
|---|---|---|
| FDLGGGTF | ASSRRDRGFSQNTLY | 0.6574832 |
| FDLGGGTFDVS | ASSRRDRGFSQNTLY | 0.5865978 |
| NEPTAAA | ASSRRDRGFSQNTLY | 0.615054 |
| NEPTAAALA | ASSRRDRGFSQNTLY | 0.4631737 |
| GSIINISSI | ASSRRDRGFSQNTLY | 0.6609313 |
| VAPGNAGTA | ASSRRDRGFSQNTLY | 0.5876032 |
| TGGGASLE | ASSRRDRGFSQNTLY | 0.55218637 |
| VAVYNHYKRI | ASSRRDRGFSQNTLY | 0.60230947 |
| TDPAPATT | ASSRRDRGFSQNTLY | 0.6844175 |
| TNRPDILDPALLRPG | ASSRRDRGFSQNTLY | 0.6848837 |
| LTTTTTS | ASSRRDRGFSQNTLY | 0.5850064 |
| PTGSGKTL | ASSRRDRGFSQNTLY | 0.5661926 |
| TGSGKTL | ASSRRDRGFSQNTLY | 0.65781355 |
| TIGHVDHGK | ASSRRDRGFSQNTLY | 0.51438344 |
| TIGHVDHGKTT | ASSRRDRGFSQNTLY | 0.66860247 |
| GHVDHGKTT | ASSRRDRGFSQNTLY | 0.63899636 |
| AQFEHTLL | ASSRRDRGFSQNTLY | 0.6421137 |
| IVNQDEL | ASSRRDRGFSQNTLY | 0.52805114 |
| TNGSQFF | ASSRRDRGFSQNTLY | 0.6647278 |
| TGKGIVE | ASSRRDRGFSQNTLY | 0.61035264 |
| LISNASDA | ASSRRDRGFSQNTLY | 0.6602812 |
| AELRTLGELTKIAKAHGAQ | ASSRRDRGFSQNTLY | 0.47601414 |
| APNAVAAAEPSW | ASSRRDRGFSQNTLY | 0.6725627 |
| APQVAGAATAVL | ASSRRDRGFSQNTLY | 0.553317 |
| ARRDPAAAAAILPTDA | ASSRRDRGFSQNTLY | 0.5677444 |
| DAAARDVLAEAGLAEYF | ASSRRDRGFSQNTLY | 0.35620952 |
| DDQIVLSAKANLAKVEQ | ASSRRDRGFSQNTLY | 0.6111801 |
| DELLPDTGEDSDEDGH | ASSRRDRGFSQNTLY | 0.67427135 |
| ELPEAPANRATVPQD | ASSRRDRGFSQNTLY | 0.68244565 |
| FGQSKIERVTGIG | ASSRRDRGFSQNTLY | 0.6403425 |
| GAGGTGSTAGGAGGAGGAGGLY | ASSRRDRGFSQNTLY | 0.667204 |
| GHGGTGAAGGNGGTGGAGGSGI | ASSRRDRGFSQNTLY | 0.68479234 |
| GNGGNGGAGGEGGPGGAGGAGA | ASSRRDRGFSQNTLY | 0.6290282 |
| GSTAGNGGNGGNATNGTIAGQP | ASSRRDRGFSQNTLY | 0.3849321 |
| GTGGDGGNGGNGGTGG | ASSRRDRGFSQNTLY | 0.6374666 |
| GVRRPAGIPADAPV | ASSRRDRGFSQNTLY | 0.622625 |
| GYRQVTDAHLLTLARR | ASSRRDRGFSQNTLY | 0.68357027 |
| IAKMGQDGHDRG | ASSRRDRGFSQNTLY | 0.65180004 |
| KAEPSIEIQYT | ASSRRDRGFSQNTLY | 0.5235034 |
| KPVRIRMAADEDTVAVT | ASSRRDRGFSQNTLY | 0.620651 |
| MPPRIAGMRLLVIKPEP | ASSRRDRGFSQNTLY | 0.6741581 |
| MKFYAVALFRSRGVPPTR | ASSRRDRGFSQNTLY | 0.52093226 |
| NAGGFGILGAVAHSP | ASSRRDRGFSQNTLY | 0.55632395 |
| NEGREITLAMAEM | ASSRRDRGFSQNTLY | 0.5311519 |
| NELVEWSHPLGHRWRWE | ASSRRDRGFSQNTLY | 0.557665 |
| NLGAGGTGGNGGNPG | ASSRRDRGFSQNTLY | 0.6498823 |
| NTEWPKWETVMAADPVD | ASSRRDRGFSQNTLY | 0.6205327 |
| PASAAPAETQP | ASSRRDRGFSQNTLY | 0.59017223 |
| QQSSDGKNGVRAAEITGE | ASSRRDRGFSQNTLY | 0.6380731 |
| RAPDAVTTSWWKE | ASSRRDRGFSQNTLY | 0.5754044 |
| SGFGNVGAHTSGWFNQ | ASSRRDRGFSQNTLY | 0.59397733 |

| | | |
|---|---|---|
| SGVAGAVGQAASVGGLKV | ASSRRDRGFSQNTLY | 0.62521446 |
| SNGGAGTGSAGGNGGT | ASSRRDRGFSQNTLY | 0.6680291 |
| TGGAGGEGGAAG | ASSRRDRGFSQNTLY | 0.67249274 |
| TGGDGGAAGAGGAPGHG | ASSRRDRGFSQNTLY | 0.63553536 |
| TMPPLLAAIANAGRHGV | ASSRRDRGFSQNTLY | 0.6335391 |
| TSSTAGNSGGA | ASSRRDRGFSQNTLY | 0.68988687 |
| VPATVPVGILC | ASSRRDRGFSQNTLY | 0.5109074 |
| VQRPDSGGVVLTGRIS | ASSRRDRGFSQNTLY | 0.54780877 |
| WTPNPVPATKGVPAGLVQAT | ASSRRDRGFSQNTLY | 0.6677377 |
| YALKRALAKFSDRFDVVII | ASSRRDRGFSQNTLY | 0.60278076 |
| YLIGSVIV | ASSRRDRGFSQNTLY | 0.6780696 |
| YGQLVAVID | ASSRRDRGFSQNTLY | 0.637732 |
| VTVGGMTLPA | ASSRRDRGFSQNTLY | 0.6665971 |
| VLALPAV | ASSRRDRGFSQNTLY | 0.64606714 |
| VIPPTLNYET | ASSRRDRGFSQNTLY | 0.69091433 |
| VGMGGMHPSSG | ASSRRDRGFSQNTLY | 0.6756245 |
| VGLAGGAATA | ASSRRDRGFSQNTLY | 0.6037112 |
| VGGAGGAGGSGP | ASSRRDRGFSQNTLY | 0.67255235 |
| VFAAFMV | ASSRRDRGFSQNTLY | 0.40739194 |
| TLYDKLR | ASSRRDRGFSQNTLY | 0.67269886 |
| TLAVFTVL | ASSRRDRGFSQNTLY | 0.6649705 |
| TGSVQPARV | ASSRRDRGFSQNTLY | 0.57590234 |
| TGGGAGSGGT | ASSRRDRGFSQNTLY | 0.6301118 |
| SHLETSL | ASSRRDRGFSQNTLY | 0.68550307 |
| SFTTGMT | ASSRRDRGFSQNTLY | 0.6812586 |
| QVEVAKRMGVRQPTVS | ASSRRDRGFSQNTLY | 0.5189317 |
| QGSGGPALAM | ASSRRDRGFSQNTLY | 0.60134375 |
| MLAPSNP | ASSRRDRGFSQNTLY | 0.49274844 |
| LVNNAAI | ASSRRDRGFSQNTLY | 0.503608 |
| LTGEINTLA | ASSRRDRGFSQNTLY | 0.39937627 |
| LSIFGPV | ASSRRDRGFSQNTLY | 0.67080677 |
| LLALLANL | ASSRRDRGFSQNTLY | 0.6217618 |
| LISNASDALDKLR | ASSRRDRGFSQNTLY | 0.6592436 |
| LINEAVP | ASSRRDRGFSQNTLY | 0.6509896 |
| LGYAFSGL | ASSRRDRGFSQNTLY | 0.68272406 |
| LDFVAVAL | ASSRRDRGFSQNTLY | 0.5905602 |
| LAQPGAE | ASSRRDRGFSQNTLY | 0.6056944 |
| ISNLGAVTGF | ASSRRDRGFSQNTLY | 0.4131176 |
| HQEDPGRI | ASSRRDRGFSQNTLY | 0.5527499 |
| GTPGEPTP | ASSRRDRGFSQNTLY | 0.579478 |
| GFFNSGS | ASSRRDRGFSQNTLY | 0.61408615 |
| GAGGGGLTAGAN | ASSRRDRGFSQNTLY | 0.504472 |
| FVDALVNM | ASSRRDRGFSQNTLY | 0.6887618 |
| FLTVAPEL | ASSRRDRGFSQNTLY | 0.62428665 |
| FETLTAL | ASSRRDRGFSQNTLY | 0.6720324 |
| DVINPIEL | ASSRRDRGFSQNTLY | 0.5701343 |
| APVSGAPV | ASSRRDRGFSQNTLY | 0.5598068 |
| ALVGGLAW | ASSRRDRGFSQNTLY | 0.6028404 |
| ALAAAAATAI | ASSRRDRGFSQNTLY | 0.65698093 |
| AFGLGLLL | ASSRRDRGFSQNTLY | 0.54015803 |
| AFFAAASL | ASSRRDRGFSQNTLY | 0.44124296 |

| | | |
|---|---|---|
| AAAALPAFSPP | ASSRRDRGFSQNTLY | 0.5407995 |
| VVLAPAL | ASSRRDRGFSQNTLY | 0.6212537 |
| TIGHVDHGKTTL | ASSRRDRGFSQNTLY | 0.6886252 |
| LISNASDALDK | ASSRRDRGFSQNTLY | 0.68430185 |
| LISNASDAL | ASSRRDRGFSQNTLY | 0.6524645 |
| ISNASDALDK | ASSRRDRGFSQNTLY | 0.6875398 |
| NQAETLVYQTE | ASSRRDRGFSQNTLY | 0.3053363 |
| TGGTGGTAGAG | ASSRRDRGFSQNTLY | 0.5920003 |
| TGSTAGGAGGA | ASSRRDRGFSQNTLY | 0.6792792 |
| DPLAVPVAAGPV | ASSRRDRGFSQNTLY | 0.6656052 |
| GGTGGTPGNGGN | ASSRRDRGFSQNTLY | 0.64036644 |
| PAGLAGVDPATL | ASSRRDRGFSQNTLY | 0.6865289 |
| GNNLVVEAPHAGASV | ASSRRDRGFSQNTLY | 0.54088116 |
| KVTRIMSASTLEQPA | ASSRRDRGFSQNTLY | 0.59449303 |
| LGFALPIALPRRTVV | ASSRRDRGFSQNTLY | 0.64856184 |
| TRLGNLTARKLVVLV | ASSRRDRGFSQNTLY | 0.6752661 |
| WSQQRRLIYFPSAGP | ASSRRDRGFSQNTLY | 0.6711822 |
| GPNGNPDPMAAAVDIR | ASSRRDRGFSQNTLY | 0.46239686 |
| TQRCVSVADLRAAASR | ASSRRDRGFSQNTLY | 0.64783883 |
| QFVLDVAEHPVEVLTETV | ASSRRDRGFSQNTLY | 0.6970463 |
| GGAGGTGGAGGNGGTQAGDG | ASSRRDRGFSQNTLY | 0.6071682 |
| PLLATLRAIAVDVDAKPAQV | ASSRRDRGFSQNTLY | 0.6824864 |
| APKFRTGGYMIDFWGVGYQVAK | ASSRRDRGFSQNTLY | 0.6352303 |
| GAGGAGGAGGAAGGPGGTGGTG | ASSRRDRGFSQNTLY | 0.6293831 |
| IGALGLLATDVAPAGSVAFGVA | ASSRRDRGFSQNTLY | 0.5881454 |
| AGGAGGIGGNGGAGGAGGVLMG | ASSRRDRGFSQNTLY | 0.5666215 |
| ADELIGLL | ASSRRDRGFSQNTLY | 0.6577604 |
| ADRALKA | ASSRRDRGFSQNTLY | 0.5740188 |
| AGAVGQVGGAGGA | ASSRRDRGFSQNTLY | 0.625953 |
| AGGGGGLGGAP | ASSRRDRGFSQNTLY | 0.6364065 |
| AGGLLYGNGGAG | ASSRRDRGFSQNTLY | 0.6949861 |
| AGLESIGL | ASSRRDRGFSQNTLY | 0.656871 |
| AGLKTGII | ASSRRDRGFSQNTLY | 0.6715 |
| ASLAALLL | ASSRRDRGFSQNTLY | 0.53169805 |
| ASVINISSI | ASSRRDRGFSQNTLY | 0.6524991 |
| ATPPGPA | ASSRRDRGFSQNTLY | 0.66954565 |
| ATVASALV | ASSRRDRGFSQNTLY | 0.5096731 |
| ATVASLIL | ASSRRDRGFSQNTLY | 0.6818624 |
| AVNGPVLIRH | ASSRRDRGFSQNTLY | 0.55261385 |
| DGAAGAVGI | ASSRRDRGFSQNTLY | 0.6460724 |
| DPLLHGI | ASSRRDRGFSQNTLY | 0.6526748 |
| ELGAALGAL | ASSRRDRGFSQNTLY | 0.6727402 |
| ELGRTQAV | ASSRRDRGFSQNTLY | 0.6481799 |
| FVAGGYLS | ASSRRDRGFSQNTLY | 0.66433954 |
| GASGGAGGAGGW | ASSRRDRGFSQNTLY | 0.5524314 |
| GGAGAGGDGGAGGI | ASSRRDRGFSQNTLY | 0.674651 |
| GTGHNGSQPGMGGNG | ASSRRDRGFSQNTLY | 0.6803312 |
| GYAAASASAS | ASSRRDRGFSQNTLY | 0.6512741 |
| HQVDVAAVQQG | ASSRRDRGFSQNTLY | 0.69175214 |
| IGAAIAHT | ASSRRDRGFSQNTLY | 0.6595979 |
| IGMIEVAQL | ASSRRDRGFSQNTLY | 0.36233076 |

| | | |
|---|---|---|
| IRLAPEHTSAMD | ASSRRDRGFSQNTLY | 0.63038 |
| IYNVSVL | ASSRRDRGFSQNTLY | 0.6804784 |
| KQGFLGM | ASSRRDRGFSQNTLY | 0.68606836 |
| LESVITQL | ASSRRDRGFSQNTLY | 0.68444955 |
| LGLGALVVV | ASSRRDRGFSQNTLY | 0.6548103 |
| LIGNGGNGGSGGI | ASSRRDRGFSQNTLY | 0.6116165 |
| LNAINAPTL | ASSRRDRGFSQNTLY | 0.6877365 |
| LQLLNTGV | ASSRRDRGFSQNTLY | 0.6953909 |
| LQTSIEF | ASSRRDRGFSQNTLY | 0.6591122 |
| MSFAVSEL | ASSRRDRGFSQNTLY | 0.56467164 |
| NTGALGL | ASSRRDRGFSQNTLY | 0.5295594 |
| PPGAGGAGGAGTT | ASSRRDRGFSQNTLY | 0.6670984 |
| QLVGLGI | ASSRRDRGFSQNTLY | 0.67947125 |
| RFLEQVHQL | ASSRRDRGFSQNTLY | 0.5011714 |
| RGVGYVLR | ASSRRDRGFSQNTLY | 0.6459384 |
| SATGVFAGVI | ASSRRDRGFSQNTLY | 0.6514201 |
| SLPAITLP | ASSRRDRGFSQNTLY | 0.37324324 |
| STALAFTV | ASSRRDRGFSQNTLY | 0.6670444 |
| SVLTVGLGK | ASSRRDRGFSQNTLY | 0.6852723 |
| TALAGAGSGI | ASSRRDRGFSQNTLY | 0.6643637 |
| TGLPFTGI | ASSRRDRGFSQNTLY | 0.51784676 |
| VAEHVTV | ASSRRDRGFSQNTLY | 0.6824763 |
| VDIVLNSL | ASSRRDRGFSQNTLY | 0.6928377 |
| VLTGGGAVL | ASSRRDRGFSQNTLY | 0.69134605 |
| VNVGLAYGI | ASSRRDRGFSQNTLY | 0.41213724 |
| VSSAAVGAV | ASSRRDRGFSQNTLY | 0.32867986 |
| VTSVYEAGV | ASSRRDRGFSQNTLY | 0.63745004 |
| WVALDRGA | ASSRRDRGFSQNTLY | 0.661042 |
| YLVGSVVI | ASSRRDRGFSQNTLY | 0.59408724 |
| GPPGTGKTLL | ASSRRDRGFSQNTLY | 0.553179 |
| GVGMTKF | ASSRRDRGFSQNTLY | 0.6637542 |
| IGIGGSDL | ASSRRDRGFSQNTLY | 0.6807533 |
| IGTLLAEL | ASSRRDRGFSQNTLY | 0.518249 |
| IYTSGTT | ASSRRDRGFSQNTLY | 0.45978343 |
| LSFPAGV | ASSRRDRGFSQNTLY | 0.6777912 |
| LYTSGTT | ASSRRDRGFSQNTLY | 0.6719588 |
| VGAFPFGF | ASSRRDRGFSQNTLY | 0.56176484 |
| VGAGTVEF | ASSRRDRGFSQNTLY | 0.6156823 |
| VGKSTLLL | ASSRRDRGFSQNTLY | 0.6449337 |
| AALAHAGGGIV | ASSRRDRGFSQNTLY | 0.6610724 |
| AALVAVSLL | ASSRRDRGFSQNTLY | 0.6547697 |
| AAVSRAVAAL | ASSRRDRGFSQNTLY | 0.477863 |
| AGGSSGIGGF | ASSRRDRGFSQNTLY | 0.6834855 |
| AGLTVANANG | ASSRRDRGFSQNTLY | 0.56262136 |
| AGLVGAVL | ASSRRDRGFSQNTLY | 0.66809213 |
| ALSQAFTV | ASSRRDRGFSQNTLY | 0.6376516 |
| ALTFSVSL | ASSRRDRGFSQNTLY | 0.6609635 |
| ALTGAALVAL | ASSRRDRGFSQNTLY | 0.6268679 |
| ASFSIGAGVL | ASSRRDRGFSQNTLY | 0.5472929 |
| DGSVYKI | ASSRRDRGFSQNTLY | 0.5463556 |
| EQLAEIV | ASSRRDRGFSQNTLY | 0.518416 |

| | | |
|---|---|---|
| FAVSELTVM | ASSRRDRGFSQNTLY | 0.6784111 |
| GAAPLAAGL | ASSRRDRGFSQNTLY | 0.66675293 |
| GADGGDGGSGGAGG | ASSRRDRGFSQNTLY | 0.68124115 |
| GFGLDTGGAGG | ASSRRDRGFSQNTLY | 0.6572629 |
| GLASGGLL | ASSRRDRGFSQNTLY | 0.57098097 |
| GRIAGAAVLL | ASSRRDRGFSQNTLY | 0.66720116 |
| GSAGSLNTSF | ASSRRDRGFSQNTLY | 0.5092039 |
| GTGQIGI | ASSRRDRGFSQNTLY | 0.64100343 |
| GTLIALIL | ASSRRDRGFSQNTLY | 0.6245753 |
| IAATMGFVVG | ASSRRDRGFSQNTLY | 0.6565769 |
| IFAQALL | ASSRRDRGFSQNTLY | 0.59331477 |
| IGAAVALGF | ASSRRDRGFSQNTLY | 0.67291117 |
| IGLTPLAL | ASSRRDRGFSQNTLY | 0.66269886 |
| ILGSNTSTL | ASSRRDRGFSQNTLY | 0.5512133 |
| IVFSLTVL | ASSRRDRGFSQNTLY | 0.64339435 |
| LASKATGF | ASSRRDRGFSQNTLY | 0.67975795 |
| LASVALIL | ASSRRDRGFSQNTLY | 0.65530986 |
| LFAVGLTL | ASSRRDRGFSQNTLY | 0.50718856 |
| LFNGGIGTYI | ASSRRDRGFSQNTLY | 0.659153 |
| LTLGAGII | ASSRRDRGFSQNTLY | 0.57150054 |
| LVAAALGDI | ASSRRDRGFSQNTLY | 0.6582713 |
| NTAFVQLGI | ASSRRDRGFSQNTLY | 0.6555506 |
| QLFSGAEL | ASSRRDRGFSQNTLY | 0.5805922 |
| SAATFITL | ASSRRDRGFSQNTLY | 0.6728499 |
| SNYGLANLGV | ASSRRDRGFSQNTLY | 0.45719618 |
| SYGGAYAVM | ASSRRDRGFSQNTLY | 0.608196 |
| TTGVGGLTLG | ASSRRDRGFSQNTLY | 0.61139446 |
| TTQTIYI | ASSRRDRGFSQNTLY | 0.64129305 |
| VAAGAGLGAVI | ASSRRDRGFSQNTLY | 0.66873085 |
| VAAVVTLL | ASSRRDRGFSQNTLY | 0.5225659 |
| VGFLGGLI | ASSRRDRGFSQNTLY | 0.6504692 |
| VGVIPILL | ASSRRDRGFSQNTLY | 0.5861662 |
| VIFASSLI | ASSRRDRGFSQNTLY | 0.5136769 |
| VKGGDGGAAAG | ASSRRDRGFSQNTLY | 0.6494934 |
| VTKTFTNI | ASSRRDRGFSQNTLY | 0.62260354 |
| VVAAAILL | ASSRRDRGFSQNTLY | 0.49011204 |
| YAALDVEL | ASSRRDRGFSQNTLY | 0.67168677 |
| LASVALLL | ASSRRDRGFSQNTLY | 0.57626784 |
| LDAVHRV | ASSRRDRGFSQNTLY | 0.6315857 |
| QGLSSSIF | ASSRRDRGFSQNTLY | 0.69084466 |
| RGTFANIRL | ASSRRDRGFSQNTLY | 0.4557931 |
| RQLWWGHRI | ASSRRDRGFSQNTLY | 0.66871107 |
| YGPPGTGKTLL | ASSRRDRGFSQNTLY | 0.6464361 |
| AALVGAPANVA | ASSRRDRGFSQNTLY | 0.65335435 |
| AATFRRTE | ASSRRDRGFSQNTLY | 0.62987745 |
| AGALGGGI | ASSRRDRGFSQNTLY | 0.5743081 |
| AGGAGGAGGIGAG | ASSRRDRGFSQNTLY | 0.6862581 |
| AGSGALGL | ASSRRDRGFSQNTLY | 0.66394436 |
| AQIADQI | ASSRRDRGFSQNTLY | 0.6356453 |
| ASAVAVTYL | ASSRRDRGFSQNTLY | 0.662259 |
| ASTVLGGTGG | ASSRRDRGFSQNTLY | 0.6655438 |

| | | |
|---|---|---|
| AVGGEPAK | ASSRRDRGFSQNTLY | 0.6767625 |
| CEQGTFGGTC | ASSRRDRGFSQNTLY | 0.59904194 |
| DEGKAPAG | ASSRRDRGFSQNTLY | 0.4377941 |
| DTVVFPAL | ASSRRDRGFSQNTLY | 0.61841464 |
| FEGITMAG | ASSRRDRGFSQNTLY | 0.6302445 |
| FPLVVQD | ASSRRDRGFSQNTLY | 0.5951915 |
| GAFTTPQ | ASSRRDRGFSQNTLY | 0.41390955 |
| GFGGFGTAGGVGGA | ASSRRDRGFSQNTLY | 0.6686927 |
| GFLAQFL | ASSRRDRGFSQNTLY | 0.69401747 |
| GGAGGAGGNAGVA | ASSRRDRGFSQNTLY | 0.5777109 |
| GGVGTAGAGGAGGAGG | ASSRRDRGFSQNTLY | 0.6952597 |
| GNGGGGDAGNAGSGGN | ASSRRDRGFSQNTLY | 0.49286878 |
| GTGGAGGAGNDAGS | ASSRRDRGFSQNTLY | 0.6584326 |
| GTTNTGWFNS | ASSRRDRGFSQNTLY | 0.67469 |
| IFMGPLFTL | ASSRRDRGFSQNTLY | 0.5382005 |
| ILAHAIEAI | ASSRRDRGFSQNTLY | 0.6487577 |
| IRSAIREAL | ASSRRDRGFSQNTLY | 0.63789606 |
| ITFAGGG | ASSRRDRGFSQNTLY | 0.5960197 |
| ITGMSGHQEV | ASSRRDRGFSQNTLY | 0.67998075 |
| IVAGLACTL | ASSRRDRGFSQNTLY | 0.5445187 |
| IVGGGLAGA | ASSRRDRGFSQNTLY | 0.65917706 |
| IVTGVLNI | ASSRRDRGFSQNTLY | 0.5957447 |
| LAQQGCI | ASSRRDRGFSQNTLY | 0.53467727 |
| LCLHVEN | ASSRRDRGFSQNTLY | 0.6675729 |
| LDAISDGS | ASSRRDRGFSQNTLY | 0.6667478 |
| LGGFAAML | ASSRRDRGFSQNTLY | 0.6394478 |
| LGGKNPMI | ASSRRDRGFSQNTLY | 0.69519854 |
| LGPKGVAAIL | ASSRRDRGFSQNTLY | 0.5881763 |
| LMVGVSI | ASSRRDRGFSQNTLY | 0.6944683 |
| LVGAAFVL | ASSRRDRGFSQNTLY | 0.6796845 |
| LVLAVAPL | ASSRRDRGFSQNTLY | 0.6466534 |
| MAAIAVIDAA | ASSRRDRGFSQNTLY | 0.61833256 |
| NNGINAELQ | ASSRRDRGFSQNTLY | 0.6754791 |
| RGEPLVLVHG | ASSRRDRGFSQNTLY | 0.5479132 |
| RGGAGGAAGLI | ASSRRDRGFSQNTLY | 0.6533172 |
| RSYRMHPAV | ASSRRDRGFSQNTLY | 0.68776596 |
| SAAHLLAGQLA | ASSRRDRGFSQNTLY | 0.69781476 |
| SAAVGSY | ASSRRDRGFSQNTLY | 0.7432734 |
| SPHAVLHRI | ASSRRDRGFSQNTLY | 0.7327211 |
| SQIAKVL | ASSRRDRGFSQNTLY | 0.712995 |
| TALGHLVP | ASSRRDRGFSQNTLY | 0.69398904 |
| TAPTTSGAA | ASSRRDRGFSQNTLY | 0.55995226 |
| TDGDDGNGGSPGT | ASSRRDRGFSQNTLY | 0.71503353 |
| TDGLGEN | ASSRRDRGFSQNTLY | 0.67645407 |
| TGGPPLPGQA | ASSRRDRGFSQNTLY | 0.7223481 |
| TIPAITI | ASSRRDRGFSQNTLY | 0.73600316 |
| TIPGFTI | ASSRRDRGFSQNTLY | 0.6763263 |
| TSIASGALA | ASSRRDRGFSQNTLY | 0.74007523 |
| TVTLPGAAI | ASSRRDRGFSQNTLY | 0.725755 |
| VASGLSAALP | ASSRRDRGFSQNTLY | 0.6404679 |
| VGAGVIGI | ASSRRDRGFSQNTLY | 0.70867884 |

| | | |
|---|---|---|
| VGPKRFGV | ASSRRDRGFSQNTLY | 0.70828146 |
| VNVAFPD | ASSRRDRGFSQNTLY | 0.7212174 |
| VSMTNSGVS | ASSRRDRGFSQNTLY | 0.6245942 |
| VTDVLGDLSV | ASSRRDRGFSQNTLY | 0.73433304 |
| VVVVGAGFAGL | ASSRRDRGFSQNTLY | 0.69398075 |
| VYTVGTVV | ASSRRDRGFSQNTLY | 0.36089826 |
| WTTFSGP | ASSRRDRGFSQNTLY | 0.7269298 |
| YDELVEAF | ASSRRDRGFSQNTLY | 0.7068628 |
| YGTGGAGGNGG | ASSRRDRGFSQNTLY | 0.7210058 |
| YTNVMAV | ASSRRDRGFSQNTLY | 0.7397797 |
| FLEQVHQL | ASSRRDRGFSQNTLY | 0.5915757 |
| GLSLLAVAV | ASSRRDRGFSQNTLY | 0.68393326 |
| LEQVHQL | ASSRRDRGFSQNTLY | 0.73084986 |
| LNAAGAI | ASSRRDRGFSQNTLY | 0.7394099 |
| PAVHASL | ASSRRDRGFSQNTLY | 0.6567062 |
| TELEDDA | ASSRRDRGFSQNTLY | 0.72845125 |
| TPPGPPN | ASSRRDRGFSQNTLY | 0.7349634 |
| TPPPPPPP | ASSRRDRGFSQNTLY | 0.73395956 |
| TQSPALI | ASSRGTDSAETLY | 0.74237394 |
| EFAAAFPQ | ASSRGTDSAETLY | 0.7444344 |
| LLSGNLDVL | ASSRGTDSAETLY | 0.67027843 |
| GPANLGNYNI | ASSRGTDSAETLY | 0.7386034 |
| TLGPTLP | ASSRGTDSAETLY | 0.74669075 |
| VYAYSSCCK | ASSRGTDSAETLY | 0.7024993 |
| APAPAPAPNI | ASSRGTDSAETLY | 0.71303195 |
| PGGPVPIP | ASSRGTDSAETLY | 0.69648254 |
| FLTVALEP | ASSRGTDSAETLY | 0.7021125 |
| SQYFGANL | ASSRGTDSAETLY | 0.5226899 |
| GGVKPAA | ASSRGTDSAETLY | 0.72288465 |
| GGTGGAGGAGGW | ASSRGTDSAETLY | 0.74240756 |
| GVGGVAGLGGAGG | ASSRGTDSAETLY | 0.69878936 |
| VLLVLATTI | ASSRGTDSAETLY | 0.68027186 |
| IGIFAAML | ASSRGTDSAETLY | 0.66271234 |
| AVGDLID | ASSRGTDSAETLY | 0.72177756 |
| NISPLPL | ASSRGTDSAETLY | 0.70273095 |
| VTAVFVAI | ASSRGTDSAETLY | 0.46719998 |
| SGDDNDAAL | ASSRGTDSAETLY | 0.7053619 |
| AVAALAHS | ASSRGTDSAETLY | 0.73466325 |
| GGVTVMATP | ASSRGTDSAETLY | 0.735229 |
| AGLAANVRD | ASSRGTDSAETLY | 0.5310484 |
| TNRPDLIDPALLRPG | ASSRGTDSAETLY | 0.7432039 |
| LANLGVTAL | ASSRGTDSAETLY | 0.7252603 |
| TGERIEL | ASSRGTDSAETLY | 0.74538624 |
| NGEELAA | ASSRGTDSAETLY | 0.7188666 |
| VLSLVWG | ASSRGTDSAETLY | 0.6982156 |
| IGYALWL | ASSRGTDSAETLY | 0.7275412 |
| LSLLLAPL | ASSRGTDSAETLY | 0.693354 |
| TLDEALK | ASSRGTDSAETLY | 0.66754854 |
| TDGAGTAEGG | ASSRGTDSAETLY | 0.7373066 |
| NAPGGAGGNGG | ASSRGTDSAETLY | 0.72065085 |
| VSVFMGMGGNF | ASSRGTDSAETLY | 0.6622833 |

| | | |
|---|---|---|
| LASYLAAL | ASSRGTDSAETLY | 0.67779875 |
| LVGGTPAL | ASSRGTDSAETLY | 0.74849033 |
| VRNADVL | ASSRGTDSAETLY | 0.64428043 |
| TCSSPFLVSAM | ASSRGTDSAETLY | 0.7078965 |
| VHITDAE | ASSRGTDSAETLY | 0.60067934 |
| EYAAPSF | ASSRGTDSAETLY | 0.70441437 |
| VVDAFHGGL | ASSRGTDSAETLY | 0.7370353 |
| ASVVDPQ | ASSRGTDSAETLY | 0.7339785 |
| SYNEFGGLS | ASSRGTDSAETLY | 0.691934 |
| DPDGSVY | ASSRGTDSAETLY | 0.7284354 |
| MVSTIHATE | ASSRGTDSAETLY | 0.73112106 |
| LAAGVADPV | ASSRGTDSAETLY | 0.72010946 |
| TKEQIAA | ASSRGTDSAETLY | 0.7329388 |
| SLASPAAL | ASSRGTDSAETLY | 0.67598933 |
| IGLVFYGV | ASSRGTDSAETLY | 0.7454703 |
| FLLDASNL | ASSRGTDSAETLY | 0.72944975 |
| YQVGNFP | ASSRGTDSAETLY | 0.7382304 |
| ESLLSQST | ASSRGTDSAETLY | 0.7369361 |
| LFTVGFAAVG | ASSRGTDSAETLY | 0.7240478 |
| ATPPTPT | ASSRGTDSAETLY | 0.7240026 |
| TDLATLG | ASSRGTDSAETLY | 0.524642 |
| SAGGTGR | ASSRGTDSAETLY | 0.6519332 |
| GFSNSGS | ASSRGTDSAETLY | 0.7248989 |
| GGNGGSGGTSV | ASSRGTDSAETLY | 0.7248529 |
| GTSVATGGAGN | ASSRGTDSAETLY | 0.7321184 |
| NGGNGGSGGTS | ASSRGTDSAETLY | 0.7278757 |
| AVVKFPH | ASSRGTDSAETLY | 0.69190204 |
| TPPPPPP | ASSRGTDSAETLY | 0.73226887 |
| AAPAAAAAPA | ASSRGTDSAETLY | 0.73193455 |
| IALNEDL | ASSRGTDSAETLY | 0.60732615 |
| IVNVSSM | ASSRGTDSAETLY | 0.7094184 |
| TGIPPAPR | ASSRGTDSAETLY | 0.7476679 |
| FDLGGGTF | ASSRGTDSAETLY | 0.7145181 |
| FDLGGGTFDVS | ASSRGTDSAETLY | 0.70293665 |
| NEPTAAA | ASSRGTDSAETLY | 0.69680774 |
| NEPTAAALA | ASSRGTDSAETLY | 0.74098074 |
| GSIINISSI | ASSRGTDSAETLY | 0.6834698 |
| VAPGNAGTA | ASSRGTDSAETLY | 0.7462785 |
| TGGGASLE | ASSRGTDSAETLY | 0.38424206 |
| VAVYNHYKRI | ASSRGTDSAETLY | 0.7087461 |
| TDPAPATT | ASSRGTDSAETLY | 0.7023901 |
| TNRPDILDPALLRPG | ASSRGTDSAETLY | 0.6440511 |
| LTTTTTS | ASSRGTDSAETLY | 0.7405746 |
| PTGSGKTL | ASSRGTDSAETLY | 0.69212127 |
| TGSGKTL | ASSRGTDSAETLY | 0.72750735 |
| TIGHVDHGK | ASSRGTDSAETLY | 0.72304535 |
| TIGHVDHGKTT | ASSRGTDSAETLY | 0.7402471 |
| GHVDHGKTT | ASSRGTDSAETLY | 0.7159846 |
| AQFEHTLL | ASSRGTDSAETLY | 0.7104993 |
| IVNQDEL | ASSRGTDSAETLY | 0.6881918 |
| TNGSQFF | ASSRGTDSAETLY | 0.73551464 |

| | | |
|---|---|---|
| TGKGIVE | ASSRGTDSAETLY | 0.73708713 |
| LISNASDA | ASSRGTDSAETLY | 0.74009824 |
| AELRTLGELTKIAKAHGAQ | ASSRGTDSAETLY | 0.5966607 |
| APNAVAAAEPSW | ASSRGTDSAETLY | 0.7105032 |
| APQVAGAATAVL | ASSRGTDSAETLY | 0.7186755 |
| ARRDPAAAAAILPTDA | ASSRGTDSAETLY | 0.41545203 |
| DAAARDVLAEAGLAEYF | ASSRGTDSAETLY | 0.41840327 |
| DDQIVLSAKANLAKVEQ | ASSRGTDSAETLY | 0.72082186 |
| DELLPDTGEDSDEDGH | ASSRGTDSAETLY | 0.6948553 |
| ELPEAPANRATVPQD | ASSRGTDSAETLY | 0.73642254 |
| FGQSKIERVTGIG | ASSRGTDSAETLY | 0.7306553 |
| GAGGTGSTAGGAGGAGGAGGLY | ASSRGTDSAETLY | 0.7064148 |
| GHGGTGAAGGNGGTGGAGGSGI | ASSRGTDSAETLY | 0.73119974 |
| GNGGNGGAGGEGGPGGAGGAGA | ASSRGTDSAETLY | 0.70874095 |
| GSTAGNGGNGGNATNGTIAGQP | ASSRGTDSAETLY | 0.24491848 |
| GTGGDGGNGGNGGTGG | ASSRGTDSAETLY | 0.6793616 |
| GVRRPAGIPADAPV | ASSRGTDSAETLY | 0.64021003 |
| GYRQVTDAHLLTLARR | ASSRGTDSAETLY | 0.7399055 |
| IAKMGQDGHDRG | ASSRGTDSAETLY | 0.7089865 |
| KAEPSIEIQYT | ASSRGTDSAETLY | 0.6713062 |
| KPVRIRMAADEDTVAVT | ASSRGTDSAETLY | 0.70981455 |
| MPPRIAGMRLLVIKPEP | ASSRGTDSAETLY | 0.7088336 |
| MKFYAVALFRSRGVPPTR | ASSRGTDSAETLY | 0.708394 |
| NAGGFGILGAVAHSP | ASSRGTDSAETLY | 0.67244124 |
| NEGREITLAMAEM | ASSRGTDSAETLY | 0.35045338 |
| NELVEWSHPLGHRWRWE | ASSRGTDSAETLY | 0.733042 |
| NLGAGGTGGNGGNPG | ASSRGTDSAETLY | 0.5472854 |
| NTEWPKWETVMAADPVD | ASSRGTDSAETLY | 0.6826676 |
| PASAAPAETQP | ASSRGTDSAETLY | 0.47161314 |
| QQSSDGKNGVRAAEITGE | ASSRGTDSAETLY | 0.68464875 |
| RAPDAVTTSWWKE | ASSRGTDSAETLY | 0.5386622 |
| SGFGNVGAHTSGWFNQ | ASSRGTDSAETLY | 0.74000335 |
| SGVAGAVGQAASVGGLKV | ASSRGTDSAETLY | 0.6900506 |
| SNGGAGTGSAGGNGGT | ASSRGTDSAETLY | 0.6878203 |
| TGGAGGEGGAAG | ASSRGTDSAETLY | 0.7446438 |
| TGGDGGAAGAGGAPGHG | ASSRGTDSAETLY | 0.68520683 |
| TMPPLLAAIANAGRHGV | ASSRGTDSAETLY | 0.69829726 |
| TSSTAGNSGGA | ASSRGTDSAETLY | 0.6943537 |
| VPATVPVGILC | ASSRGTDSAETLY | 0.6907805 |
| VQRPDSGGVVLTGRIS | ASSRGTDSAETLY | 0.71467066 |
| WTPNPVPATKGVPAGLVQAT | ASSRGTDSAETLY | 0.689057 |
| YALKRALAKFSDRFDVVII | ASSRGTDSAETLY | 0.7420889 |
| YLIGSVIV | ASSRGTDSAETLY | 0.71545064 |
| YGQLVAVID | ASSRGTDSAETLY | 0.6787502 |
| VTVGGMTLPA | ASSRGTDSAETLY | 0.72338116 |
| VLALPAV | ASSRGTDSAETLY | 0.74263096 |
| VIPPTLNYET | ASSRGTDSAETLY | 0.75060534 |
| VGMGGMHPSSG | ASSRGTDSAETLY | 0.688784 |
| VGLAGGAATA | ASSRGTDSAETLY | 0.6979605 |
| VGGAGGAGGSGP | ASSRGTDSAETLY | 0.6938101 |
| VFAAFMV | ASSRGTDSAETLY | 0.69879365 |

| | | |
|---|---|---|
| TLYDKLR | ASSRGTDSAETLY | 0.6955131 |
| TLAVFTVL | ASSRGTDSAETLY | 0.6141895 |
| TGSVQPARV | ASSRGTDSAETLY | 0.74192405 |
| TGGGAGSGGT | ASSRGTDSAETLY | 0.7415204 |
| SHLETSL | ASSRGTDSAETLY | 0.7142409 |
| SFTTGMT | ASSRGTDSAETLY | 0.69135547 |
| QVEVAKRMGVRQPTVS | ASSRGTDSAETLY | 0.70126045 |
| QGSGGPALAM | ASSRGTDSAETLY | 0.66768086 |
| MLAPSNP | ASSRGTDSAETLY | 0.69416225 |
| LVNNAAI | ASSRGTDSAETLY | 0.7108073 |
| LTGEINTLA | ASSRGTDSAETLY | 0.62248886 |
| LSIFGPV | ASSRGTDSAETLY | 0.7078135 |
| LLALLANL | ASSRGTDSAETLY | 0.7494115 |
| LISNASDALDKLR | ASSRGTDSAETLY | 0.68884563 |
| LINEAVP | ASSRGTDSAETLY | 0.72451496 |
| LGYAFSGL | ASSRGTDSAETLY | 0.74452996 |
| LDFVAVAL | ASSRGTDSAETLY | 0.73824835 |
| LAQPGAE | ASSRGTDSAETLY | 0.74442303 |
| ISNLGAVTGF | ASSRGTDSAETLY | 0.7251568 |
| HQEDPGRI | ASSRGTDSAETLY | 0.7207139 |
| GTPGEPTP | ASSRGTDSAETLY | 0.714852 |
| GFFNSGS | ASSRGTDSAETLY | 0.7232956 |
| GAGGGGLTAGAN | ASSRGTDSAETLY | 0.70685256 |
| FVDALVNM | ASSRGTDSAETLY | 0.708121 |
| FLTVAPEL | ASSRGTDSAETLY | 0.68115187 |
| FETLTAL | ASSRGTDSAETLY | 0.73516756 |
| DVINPIEL | ASSRGTDSAETLY | 0.7210742 |
| APVSGAPV | ASSRGTDSAETLY | 0.73642945 |
| ALVGGLAW | ASSRGTDSAETLY | 0.6128376 |
| ALAAAAATAI | ASSRGTDSAETLY | 0.681334 |
| AFGLGLLL | ASSRGTDSAETLY | 0.73718524 |
| AFFAAASL | ASSRGTDSAETLY | 0.2641069 |
| AAAALPAFSPP | ASSRGTDSAETLY | 0.41111517 |
| VVLAPAL | ASSRGTDSAETLY | 0.7282117 |
| TIGHVDHGKTTL | ASSRGTDSAETLY | 0.74131763 |
| LISNASDALDK | ASSRGTDSAETLY | 0.31742686 |
| LISNASDAL | ASSRGTDSAETLY | 0.62875605 |
| ISNASDALDK | ASSRGTDSAETLY | 0.6150435 |
| NQAETLVYQTE | ASSRGTDSAETLY | 0.7298287 |
| TGGTGGTAGAG | ASSRGTDSAETLY | 0.71536374 |
| TGSTAGGAGGA | ASSRGTDSAETLY | 0.5381477 |
| DPLAVPVAAGPV | ASSRGTDSAETLY | 0.73558795 |
| GGTGGTPGNGGN | ASSRGTDSAETLY | 0.62677914 |
| PAGLAGVDPATL | ASSRGTDSAETLY | 0.59694946 |
| GNNLVVEAPHAGASV | ASSRGTDSAETLY | 0.7411556 |
| KVTRIMSASTLEQPA | ASSRGTDSAETLY | 0.7394413 |
| LGFALPIALPRRTVV | ASSRGTDSAETLY | 0.68803036 |
| TRLGNLTARKLVVLV | ASSRGTDSAETLY | 0.7304598 |
| WSQQRRLIYFPSAGP | ASSRGTDSAETLY | 0.687256 |
| GPNGNPDPMAAAVDIR | ASSRGTDSAETLY | 0.7512213 |
| TQRCVSVADLRAAASR | ASSRGTDSAETLY | 0.7004242 |

| | | |
|---|---|---|
| QFVLDVAEHPVEVLTETV | ASSRGTDSAETLY | 0.635846 |
| GGAGGTGGAGGNGGTQAGDG | ASSRGTDSAETLY | 0.73870087 |
| PLLATLRAIAVDVDAKPAQV | ASSRGTDSAETLY | 0.7285757 |
| APKFRTGGYMIDFWGVGYQVAK | ASSRGTDSAETLY | 0.721789 |
| GAGGAGGAGGAAGGPGGTGGTG | ASSRGTDSAETLY | 0.7362092 |
| IGALGLLATDVAPAGSVAFGVA | ASSRGTDSAETLY | 0.70605826 |
| AGGAGGIGGNGGAGGAGGVLMG | ASSRGTDSAETLY | 0.7001621 |
| ADELIGLL | ASSRGTDSAETLY | 0.73540926 |
| ADRALKA | ASSRGTDSAETLY | 0.73650986 |
| AGAVGQVGGAGGA | ASSRGTDSAETLY | 0.61512667 |
| AGGGGGLGGAP | ASSRGTDSAETLY | 0.741956 |
| AGGLLYGNGGAG | ASSRGTDSAETLY | 0.6971221 |
| AGLESIGL | ASSRGTDSAETLY | 0.6773755 |
| AGLKTGII | ASSRGTDSAETLY | 0.7492974 |
| ASLAALLL | ASSRGTDSAETLY | 0.72647 |
| ASVINISSI | ASSRGTDSAETLY | 0.73855984 |
| ATPPGPA | ASSRGTDSAETLY | 0.7402809 |
| ATVASALV | ASSRGTDSAETLY | 0.728919 |
| ATVASLIL | ASSRGTDSAETLY | 0.63148075 |
| AVNGPVLIRH | ASSRGTDSAETLY | 0.7168834 |
| DGAAGAVGI | ASSRGTDSAETLY | 0.7052889 |
| DPLLHGI | ASSRGTDSAETLY | 0.7326639 |
| ELGAALGAL | ASSRGTDSAETLY | 0.72083884 |
| ELGRTQAV | ASSRGTDSAETLY | 0.6786109 |
| FVAGGYLS | ASSRGTDSAETLY | 0.73318666 |
| GASGGAGGAGGW | ASSRGTDSAETLY | 0.7433631 |
| GGAGAGGDGGAGGI | ASSRGTDSAETLY | 0.7284659 |
| GTGHNGSQPGMGGNG | ASSRGTDSAETLY | 0.7182432 |
| GYAAASASAS | ASSRGTDSAETLY | 0.74598765 |
| HQVDVAAVQQG | ASSRGTDSAETLY | 0.70465314 |
| IGAAIAHT | ASSRGTDSAETLY | 0.6969903 |
| IGMIEVAQL | ASSRGTDSAETLY | 0.7393132 |
| IRLAPEHTSAMD | ASSRGTDSAETLY | 0.73405325 |
| IYNVSVL | ASSRGTDSAETLY | 0.71640265 |
| KQGFLGM | ASSRGTDSAETLY | 0.7420132 |
| LESVITQL | ASSRGTDSAETLY | 0.6674671 |
| LGLGALVVV | ASSRGTDSAETLY | 0.7223393 |
| LIGNGGNGGSGGI | ASSRGTDSAETLY | 0.67791855 |
| LNAINAPTL | ASSRGTDSAETLY | 0.7183347 |
| LQLLNTGV | ASSRGTDSAETLY | 0.6862343 |
| LQTSIEF | ASSRGTDSAETLY | 0.73021424 |
| MSFAVSEL | ASSRGTDSAETLY | 0.74038815 |
| NTGALGL | ASSRGTDSAETLY | 0.7306679 |
| PPGAGGAGGAGTT | ASSRGTDSAETLY | 0.37542886 |
| QLVGLGI | ASSRGTDSAETLY | 0.737753 |
| RFLEQVHQL | ASSRGTDSAETLY | 0.71995413 |
| RGVGYVLR | ASSRGTDSAETLY | 0.7479651 |
| SATGVFAGVI | ASSRGTDSAETLY | 0.7146764 |
| SLPAITLP | ASSRGTDSAETLY | 0.7218496 |
| STALAFTV | ASSRGTDSAETLY | 0.7023239 |
| SVLTVGLGK | ASSRGTDSAETLY | 0.7242794 |

| | | |
|---|---|---|
| TALAGAGSGI | ASSRGTDSAETLY | 0.6432346 |
| TGLPFTGI | ASSRGTDSAETLY | 0.6932425 |
| VAEHVTV | ASSRGTDSAETLY | 0.67695427 |
| VDIVLNSL | ASSRGTDSAETLY | 0.6763791 |
| VLTGGGAVL | ASSRGTDSAETLY | 0.7316954 |
| VNVGLAYGI | ASSRGTDSAETLY | 0.7403906 |
| VSSAAVGAV | ASSRGTDSAETLY | 0.73591626 |
| VTSVYEAGV | ASSRGTDSAETLY | 0.7353027 |
| WVALDRGA | ASSRGTDSAETLY | 0.7417826 |
| YLVGSVVI | ASSRGTDSAETLY | 0.7432806 |
| GPPGTGKTLL | ASSRGTDSAETLY | 0.7076392 |
| GVGMTKF | ASSRGTDSAETLY | 0.69250107 |
| IGIGGSDL | ASSRGTDSAETLY | 0.6779847 |
| IGTLLAEL | ASSRGTDSAETLY | 0.700052 |
| IYTSGTT | ASSRGTDSAETLY | 0.7058518 |
| LSFPAGV | ASSRGTDSAETLY | 0.72017646 |
| LYTSGTT | ASSRGTDSAETLY | 0.24635702 |
| VGAFPFGF | ASSRGTDSAETLY | 0.7082243 |
| VGAGTVEF | ASSRGTDSAETLY | 0.74339664 |
| VGKSTLLL | ASSRGTDSAETLY | 0.6674347 |
| AALAHAGGGIV | ASSRGTDSAETLY | 0.72409964 |
| AALVAVSLL | ASSRGTDSAETLY | 0.71379316 |
| AAVSRAVAAL | ASSRGTDSAETLY | 0.6995498 |
| AGGSSGIGGF | ASSRGTDSAETLY | 0.72553694 |
| AGLTVANANG | ASSRGTDSAETLY | 0.67548347 |
| AGLVGAVL | ASSRGTDSAETLY | 0.7329513 |
| ALSQAFTV | ASSRGTDSAETLY | 0.7488575 |
| ALTFSVSL | ASSRGTDSAETLY | 0.7243032 |
| ALTGAALVAL | ASSRGTDSAETLY | 0.7026227 |
| ASFSIGAGVL | ASSRGTDSAETLY | 0.71174026 |
| DGSVYKI | ASSRGTDSAETLY | 0.5514826 |
| EQLAEIV | ASSRGTDSAETLY | 0.7104346 |
| FAVSELTVM | ASSRGTDSAETLY | 0.5281116 |
| GAAPLAAGL | ASSRGTDSAETLY | 0.70653474 |
| GADGGDGGSGGAGG | ASSRGTDSAETLY | 0.7067242 |
| GFGLDTGGAGG | ASSRGTDSAETLY | 0.72195625 |
| GLASGGLL | ASSRGTDSAETLY | 0.70605254 |
| GRIAGAAVLL | ASSRGTDSAETLY | 0.73016953 |
| GSAGSLNTSF | ASSRGTDSAETLY | 0.6669346 |
| GTGQIGI | ASSRGTDSAETLY | 0.71797127 |
| GTLIALIL | ASSRGTDSAETLY | 0.69724226 |
| IAATMGFVVG | ASSRGTDSAETLY | 0.6449852 |
| IFAQALL | ASSRGTDSAETLY | 0.74134076 |
| IGAAVALGF | ASSRGTDSAETLY | 0.7286724 |
| IGLTPLAL | ASSRGTDSAETLY | 0.709321 |
| ILGSNTSTL | ASSRGTDSAETLY | 0.7377259 |
| IVFSLTVL | ASSRGTDSAETLY | 0.7273679 |
| LASKATGF | ASSRGTDSAETLY | 0.64506173 |
| LASVALIL | ASSRGTDSAETLY | 0.70441085 |
| LFAVGLTL | ASSRGTDSAETLY | 0.7090024 |
| LFNGGIGTYI | ASSRGTDSAETLY | 0.71427286 |

| | | |
|---|---|---|
| LTLGAGII | ASSRGTDSAETLY | 0.36066246 |
| LVAAALGDI | ASSRGTDSAETLY | 0.7348207 |
| NTAFVQLGI | ASSRGTDSAETLY | 0.6756948 |
| QLFSGAEL | ASSRGTDSAETLY | 0.6621148 |
| SAATFITL | ASSRGTDSAETLY | 0.7131239 |
| SNYGLANLGV | ASSRGTDSAETLY | 0.73510826 |
| SYGGAYAVM | ASSRGTDSAETLY | 0.6774396 |
| TTGVGGLTLG | ASSRGTDSAETLY | 0.7172651 |
| TTQTIYI | ASSRGTDSAETLY | 0.7373102 |
| VAAGAGLGAVI | ASSRGTDSAETLY | 0.7268567 |
| VAAVVTLL | ASSRGTDSAETLY | 0.73583084 |
| VGFLGGLI | ASSRGTDSAETLY | 0.7291943 |
| VGVIPILL | ASSRGTDSAETLY | 0.7173121 |
| VIFASSLI | ASSRGTDSAETLY | 0.7213205 |
| VKGGDGGAAAG | ASSRGTDSAETLY | 0.70930576 |
| VTKTFTNI | ASSRGTDSAETLY | 0.6174067 |
| VVAAAILL | ASSRGTDSAETLY | 0.6216891 |
| YAALDVEL | ASSRGTDSAETLY | 0.7198033 |
| LASVALLL | ASSRGTDSAETLY | 0.51257515 |
| LDAVHRV | ASSRGTDSAETLY | 0.7177068 |
| QGLSSSIF | ASSRGTDSAETLY | 0.7405616 |
| RGTFANIRL | ASSRGTDSAETLY | 0.7172662 |
| RQLWWGHRI | ASSRGTDSAETLY | 0.6522006 |
| YGPPGTGKTLL | ASSRGTDSAETLY | 0.7069399 |
| AALVGAPANVA | ASSRGTDSAETLY | 0.6908501 |
| AATFRRTE | ASSRGTDSAETLY | 0.73794657 |
| AGALGGGI | ASSRGTDSAETLY | 0.74117696 |
| AGGAGGAGGIGAG | ASSRGTDSAETLY | 0.74123603 |
| AGSGALGL | ASSRGTDSAETLY | 0.7367667 |
| AQIADQI | ASSRGTDSAETLY | 0.68522686 |
| ASAVAVTYL | ASSRGTDSAETLY | 0.73777306 |
| ASTVLGGTGG | ASSRGTDSAETLY | 0.7027706 |
| AVGGEPAK | ASSRGTDSAETLY | 0.70860577 |
| CEQGTFGGTC | ASSRGTDSAETLY | 0.7236433 |
| DEGKAPAG | ASSRGTDSAETLY | 0.7283945 |
| DTVVFPAL | ASSRGTDSAETLY | 0.71994567 |
| FEGITMAG | ASSRGTDSAETLY | 0.70811737 |
| FPLVVQD | ASSRGTDSAETLY | 0.6266103 |
| GAFTTPQ | ASSRGTDSAETLY | 0.7318765 |
| GFGGFGTAGGVGGA | ASSRGTDSAETLY | 0.7025064 |
| GFLAQFL | ASSRGTDSAETLY | 0.5608578 |
| GGAGGAGGNAGVA | ASSRGTDSAETLY | 0.69111276 |
| GGVGTAGAGGAGGAGG | ASSRGTDSAETLY | 0.74474573 |
| GNGGGGDAGNAGSGGN | ASSRGTDSAETLY | 0.7354845 |
| GTGGAGGAGNDAGS | ASSRGTDSAETLY | 0.6561104 |
| GTTNTGWFNS | ASSRGTDSAETLY | 0.69025093 |
| IFMGPLFTL | ASSRGTDSAETLY | 0.70675814 |
| ILAHAIEAI | ASSRGTDSAETLY | 0.52385074 |
| IRSAIREAL | ASSRGTDSAETLY | 0.73610616 |
| ITFAGGG | ASSRGTDSAETLY | 0.6977525 |
| ITGMSGHQEV | ASSRGTDSAETLY | 0.7058959 |

| | | |
|---|---|---|
| IVAGLACTL | ASSRGTDSAETLY | 0.6704823 |
| IVGGGLAGA | ASSRGTDSAETLY | 0.7252393 |
| IVTGVLNI | ASSRGTDSAETLY | 0.69724184 |
| LAQQGCI | ASSRGTDSAETLY | 0.73839164 |
| LCLHVEN | ASSRGTDSAETLY | 0.30767134 |
| LDAISDGS | ASSRGTDSAETLY | 0.7354675 |
| LGGFAAML | ASSRGTDSAETLY | 0.6537568 |
| LGGKNPMI | ASSRGTDSAETLY | 0.708578 |
| LGPKGVAAIL | ASSRGTDSAETLY | 0.64334935 |
| LMVGVSI | ASSRGTDSAETLY | 0.71476066 |
| LVGAAFVL | ASSRGTDSAETLY | 0.7418735 |
| LVLAVAPL | ASSRGTDSAETLY | 0.7397394 |
| MAAIAVIDAA | ASSRGTDSAETLY | 0.68621695 |
| NNGINAELQ | ASSRGTDSAETLY | 0.73153275 |
| RGEPLVLVHG | ASSRGTDSAETLY | 0.74530387 |
| RGGAGGAAGLI | ASSRGTDSAETLY | 0.69611573 |
| RSYRMHPAV | ASSRGTDSAETLY | 0.6227753 |
| SAAHLLAGQLA | ASSRGTDSAETLY | 0.72673863 |
| SAAVGSY | ASSRGTDSAETLY | 0.69905734 |
| SPHAVLHRI | ASSRGTDSAETLY | 0.73734534 |
| SQIAKVL | ASSRGTDSAETLY | 0.72337365 |
| TALGHLVP | ASSRGTDSAETLY | 0.6407693 |
| TAPTTSGAA | ASSRGTDSAETLY | 0.724015 |
| TDGDDGNGGSPGT | ASSRGTDSAETLY | 0.7349404 |
| TDGLGEN | ASSRGTDSAETLY | 0.6697377 |
| TGGPPLPGQA | ASSRGTDSAETLY | 0.70764965 |
| TIPAITI | ASSRGTDSAETLY | 0.6852417 |
| TIPGFTI | ASSRGTDSAETLY | 0.59377325 |
| TSIASGALA | ASSRGTDSAETLY | 0.7294632 |
| TVTLPGAAI | ASSRGTDSAETLY | 0.6804505 |
| VASGLSAALP | ASSRGTDSAETLY | 0.68117166 |
| VGAGVIGI | ASSRGTDSAETLY | 0.71767855 |
| VGPKRFGV | ASSRGTDSAETLY | 0.70097977 |
| VNVAFPD | ASSRGTDSAETLY | 0.6864288 |
| VSMTNSGVS | ASSRGTDSAETLY | 0.70353186 |
| VTDVLGDLSV | ASSRGTDSAETLY | 0.7265811 |
| VVVVGAGFAGL | ASSRGTDSAETLY | 0.7334032 |
| VYTVGTVV | ASSRGTDSAETLY | 0.7070215 |
| WTTFSGP | ASSRGTDSAETLY | 0.7060069 |
| YDELVEAF | ASSRGTDSAETLY | 0.7436235 |
| YGTGGAGGNGG | ASSRGTDSAETLY | 0.6108955 |
| YTNVMAV | ASSRGTDSAETLY | 0.64426994 |
| FLEQVHQL | ASSRGTDSAETLY | 0.7321732 |
| GLSLLAVAV | ASSRGTDSAETLY | 0.6452961 |
| LEQVHQL | ASSRGTDSAETLY | 0.73637986 |
| LNAAGAI | ASSRGTDSAETLY | 0.7064899 |
| PAVHASL | ASSRGTDSAETLY | 0.7241131 |
| TELEDDA | ASSRGTDSAETLY | 0.70479786 |
| TPPGPPN | ASSRGTDSAETLY | 0.7455661 |
| TPPPPPPP | ASSRGTDSAETLY | 0.73926455 |
| TQSPALI | ASSQDRGANTQY | 0.7163138 |

| | | |
|---|---|---|
| EFAAAFPQ | ASSQDRGANTQY | 0.7285874 |
| LLSGNLDVL | ASSQDRGANTQY | 0.6354389 |
| GPANLGNYNI | ASSQDRGANTQY | 0.73145604 |
| TLGPTLP | ASSQDRGANTQY | 0.73034346 |
| VYAYSSCCK | ASSQDRGANTQY | 0.65812695 |
| APAPAPAPNI | ASSQDRGANTQY | 0.7006955 |
| PGGPVPIP | ASSQDRGANTQY | 0.7469715 |
| FLTVALEP | ASSQDRGANTQY | 0.7426323 |
| SQYFGANL | ASSQDRGANTQY | 0.61400384 |
| GGVKPAA | ASSQDRGANTQY | 0.69656295 |
| GGTGGAGGAGGW | ASSQDRGANTQY | 0.47138542 |
| GVGGVAGLGGAGG | ASSQDRGANTQY | 0.6995406 |
| VLLVLATTI | ASSQDRGANTQY | 0.62389505 |
| IGIFAAML | ASSQDRGANTQY | 0.7328659 |
| AVGDLID | ASSQDRGANTQY | 0.69394225 |
| NISPLPL | ASSQDRGANTQY | 0.7150674 |
| VTAVFVAI | ASSQDRGANTQY | 0.72015107 |
| SGDDNDAAL | ASSQDRGANTQY | 0.74538255 |
| AVAALAHS | ASSQDRGANTQY | 0.69268906 |
| GGVTVMATP | ASSQDRGANTQY | 0.70483077 |
| AGLAANVRD | ASSQDRGANTQY | 0.63949907 |
| TNRPDLIDPALLRPG | ASSQDRGANTQY | 0.6012743 |
| LANLGVTAL | ASSQDRGANTQY | 0.7154136 |
| TGERIEL | ASSQDRGANTQY | 0.70846236 |
| NGEELAA | ASSQDRGANTQY | 0.44404802 |
| VLSLVWG | ASSQDRGANTQY | 0.7419894 |
| IGYALWL | ASSQDRGANTQY | 0.71838844 |
| LSLLLAPL | ASSQDRGANTQY | 0.7354853 |
| TLDEALK | ASSQDRGANTQY | 0.6814268 |
| TDGAGTAEGG | ASSQDRGANTQY | 0.7161293 |
| NAPGGAGGNGG | ASSQDRGANTQY | 0.7020467 |
| VSVFMGMGGNF | ASSQDRGANTQY | 0.7269707 |
| LASYLAAL | ASSQDRGANTQY | 0.6965257 |
| LVGGTPAL | ASSQDRGANTQY | 0.6242814 |
| VRNADVL | ASSQDRGANTQY | 0.70450556 |
| TCSSPFLVSAM | ASSQDRGANTQY | 0.66666305 |
| VHITDAE | ASSQDRGANTQY | 0.7468136 |
| EYAAPSF | ASSQDRGANTQY | 0.74084884 |
| VVDAFHGGL | ASSQDRGANTQY | 0.72420144 |
| ASVVDPQ | ASSQDRGANTQY | 0.74162126 |
| SYNEFGGLS | ASSQDRGANTQY | 0.40902936 |
| DPDGSVY | ASSQDRGANTQY | 0.68073595 |
| MVSTIHATE | ASSQDRGANTQY | 0.73663044 |
| LAAGVADPV | ASSQDRGANTQY | 0.73541194 |
| TKEQIAA | ASSQDRGANTQY | 0.70256007 |
| SLASPAAL | ASSQDRGANTQY | 0.7449188 |
| IGLVFYGV | ASSQDRGANTQY | 0.67326283 |
| FLLDASNL | ASSQDRGANTQY | 0.7080667 |
| YQVGNFP | ASSQDRGANTQY | 0.7320756 |
| ESLLSQST | ASSQDRGANTQY | 0.7372743 |
| LFTVGFAAVG | ASSQDRGANTQY | 0.7377858 |

| | | |
|---|---|---|
| ATPPTPT | ASSQDRGANTQY | 0.6972126 |
| TDLATLG | ASSQDRGANTQY | 0.7199881 |
| SAGGTGR | ASSQDRGANTQY | 0.7515216 |
| GFSNSGS | ASSQDRGANTQY | 0.70465636 |
| GGNGGSGGTSV | ASSQDRGANTQY | 0.73920834 |
| GTSVATGGAGN | ASSQDRGANTQY | 0.7254977 |
| NGGNGGSGGTS | ASSQDRGANTQY | 0.70530444 |
| AVVKFPH | ASSQDRGANTQY | 0.66649204 |
| TPPPPPP | ASSQDRGANTQY | 0.6878023 |
| AAPAAAAAPA | ASSQDRGANTQY | 0.712989 |
| IALNEDL | ASSQDRGANTQY | 0.6451615 |
| IVNVSSM | ASSQDRGANTQY | 0.71627176 |
| TGIPPAPR | ASSQDRGANTQY | 0.69649684 |
| FDLGGGTF | ASSQDRGANTQY | 0.75277734 |
| FDLGGGTFDVS | ASSQDRGANTQY | 0.72801983 |
| NEPTAAA | ASSQDRGANTQY | 0.7365949 |
| NEPTAAALA | ASSQDRGANTQY | 0.6687982 |
| GSIINISSI | ASSQDRGANTQY | 0.7322434 |
| VAPGNAGTA | ASSQDRGANTQY | 0.7274257 |
| TGGGASLE | ASSQDRGANTQY | 0.65027314 |
| VAVYNHYKRI | ASSQDRGANTQY | 0.737453 |
| TDPAPATT | ASSQDRGANTQY | 0.660192 |
| TNRPDILDPALLRPG | ASSQDRGANTQY | 0.7021116 |
| LTTTTTS | ASSQDRGANTQY | 0.7249216 |
| PTGSGKTL | ASSQDRGANTQY | 0.73240113 |
| TGSGKTL | ASSQDRGANTQY | 0.7111278 |
| TIGHVDHGK | ASSQDRGANTQY | 0.7068888 |
| TIGHVDHGKTT | ASSQDRGANTQY | 0.67725 |
| GHVDHGKTT | ASSQDRGANTQY | 0.73710275 |
| AQFEHTLL | ASSQDRGANTQY | 0.7417418 |
| IVNQDEL | ASSQDRGANTQY | 0.7050705 |
| TNGSQFF | ASSQDRGANTQY | 0.7468121 |
| TGKGIVE | ASSQDRGANTQY | 0.7252109 |
| LISNASDA | ASSQDRGANTQY | 0.57235265 |
| AELRTLGELTKIAKAHGAQ | ASSQDRGANTQY | 0.7158075 |
| APNAVAAAEPSW | ASSQDRGANTQY | 0.7368763 |
| APQVAGAATAVL | ASSQDRGANTQY | 0.74344146 |
| ARRDPAAAAAILPTDA | ASSQDRGANTQY | 0.74774766 |
| DAAARDVLAEAGLAEYF | ASSQDRGANTQY | 0.7270093 |
| DDQIVLSAKANLAKVEQ | ASSQDRGANTQY | 0.6916254 |
| DELLPDTGEDSDEDGH | ASSQDRGANTQY | 0.7412101 |
| ELPEAPANRATVPQD | ASSQDRGANTQY | 0.7488284 |
| FGQSKIERVTGIG | ASSQDRGANTQY | 0.7299329 |
| GAGGTGSTAGGAGGAGGAGGLY | ASSQDRGANTQY | 0.68660295 |
| GHGGTGAAGGNGGTGGAGGSGI | ASSQDRGANTQY | 0.6600486 |
| GNGGNGGAGGEGGPGGAGGAGA | ASSQDRGANTQY | 0.73094237 |
| GSTAGNGGNGGNATNGTIAGQP | ASSQDRGANTQY | 0.7426442 |
| GTGGDGGNGGNGGTGG | ASSQDRGANTQY | 0.67275834 |
| GVRRPAGIPADAPV | ASSQDRGANTQY | 0.717816 |
| GYRQVTDAHLLTLARR | ASSQDRGANTQY | 0.71623313 |
| IAKMGQDGHDRG | ASSQDRGANTQY | 0.60254574 |

| | | |
|---|---|---|
| KAEPSIEIQYT | ASSQDRGANTQY | 0.7258767 |
| KPVRIRMAADEDTVAVT | ASSQDRGANTQY | 0.7392955 |
| MPPRIAGMRLLVIKPEP | ASSQDRGANTQY | 0.72453153 |
| MKFYAVALFRSRGVPPTR | ASSQDRGANTQY | 0.67989707 |
| NAGGFGILGAVAHSP | ASSQDRGANTQY | 0.73615 |
| NEGREITLAMAEM | ASSQDRGANTQY | 0.7505831 |
| NELVEWSHPLGHRWRWE | ASSQDRGANTQY | 0.745314 |
| NLGAGGTGGNGGNPG | ASSQDRGANTQY | 0.5969057 |
| NTEWPKWETVMAADPVD | ASSQDRGANTQY | 0.5124434 |
| PASAAPAETQP | ASSQDRGANTQY | 0.67346483 |
| QQSSDGKNGVRAAEITGE | ASSQDRGANTQY | 0.7225805 |
| RAPDAVTTSWWKE | ASSQDRGANTQY | 0.70307946 |
| SGFGNVGAHTSGWFNQ | ASSQDRGANTQY | 0.6945756 |
| SGVAGAVGQAASVGGLKV | ASSQDRGANTQY | 0.7284109 |
| SNGGAGTGSAGGNGGT | ASSQDRGANTQY | 0.7367877 |
| TGGAGGEGGAAG | ASSQDRGANTQY | 0.68663776 |
| TGGDGGAAGAGGAPGHG | ASSQDRGANTQY | 0.6761637 |
| TMPPLLAAIANAGRHGV | ASSQDRGANTQY | 0.7371212 |
| TSSTAGNSGGA | ASSQDRGANTQY | 0.73558915 |
| VPATVPVGILC | ASSQDRGANTQY | 0.6655188 |
| VQRPDSGGVVLTGRIS | ASSQDRGANTQY | 0.71432406 |
| WTPNPVPATKGVPAGLVQAT | ASSQDRGANTQY | 0.6659356 |
| YALKRALAKFSDRFDVVII | ASSQDRGANTQY | 0.66724217 |
| YLIGSVIV | ASSQDRGANTQY | 0.7160938 |
| YGQLVAVID | ASSQDRGANTQY | 0.6426064 |
| VTVGGMTLPA | ASSQDRGANTQY | 0.74240935 |
| VLALPAV | ASSQDRGANTQY | 0.6674852 |
| VIPPTLNYET | ASSQDRGANTQY | 0.7365688 |
| VGMGGMHPSSG | ASSQDRGANTQY | 0.72401947 |
| VGLAGGAATA | ASSQDRGANTQY | 0.732991 |
| VGGAGGAGGSGP | ASSQDRGANTQY | 0.69999963 |
| VFAAFMV | ASSQDRGANTQY | 0.67484194 |
| TLYDKLR | ASSQDRGANTQY | 0.6641648 |
| TLAVFTVL | ASSQDRGANTQY | 0.57743454 |
| TGSVQPARV | ASSQDRGANTQY | 0.73798525 |
| TGGGAGSGGT | ASSQDRGANTQY | 0.694157 |
| SHLETSL | ASSQDRGANTQY | 0.74110067 |
| SFTTGMT | ASSQDRGANTQY | 0.7376988 |
| QVEVAKRMGVRQPTVS | ASSQDRGANTQY | 0.6843004 |
| QGSGGPALAM | ASSQDRGANTQY | 0.736909 |
| MLAPSNP | ASSQDRGANTQY | 0.6732352 |
| LVNNAAI | ASSQDRGANTQY | 0.67793727 |
| LTGEINTLA | ASSQDRGANTQY | 0.72573376 |
| LSIFGPV | ASSQDRGANTQY | 0.72915184 |
| LLALLANL | ASSQDRGANTQY | 0.67688465 |
| LISNASDALDKLR | ASSQDRGANTQY | 0.7339072 |
| LINEAVP | ASSQDRGANTQY | 0.7238457 |
| LGYAFSGL | ASSQDRGANTQY | 0.6755463 |
| LDFVAVAL | ASSQDRGANTQY | 0.71806014 |
| LAQPGAE | ASSQDRGANTQY | 0.7346687 |
| ISNLGAVTGF | ASSQDRGANTQY | 0.73186415 |

| | | |
|---|---|---|
| HQEDPGRI | ASSQDRGANTQY | 0.6930189 |
| GTPGEPTP | ASSQDRGANTQY | 0.7335713 |
| GFFNSGS | ASSQDRGANTQY | 0.6549674 |
| GAGGGGLTAGAN | ASSQDRGANTQY | 0.7178056 |
| FVDALVNM | ASSQDRGANTQY | 0.72787195 |
| FLTVAPEL | ASSQDRGANTQY | 0.7011931 |
| FETLTAL | ASSQDRGANTQY | 0.73802507 |
| DVINPIEL | ASSQDRGANTQY | 0.65901303 |
| APVSGAPV | ASSQDRGANTQY | 0.6864046 |
| ALVGGLAW | ASSQDRGANTQY | 0.6916319 |
| ALAAAAATAI | ASSQDRGANTQY | 0.72143745 |
| AFGLGLLL | ASSQDRGANTQY | 0.73355746 |
| AFFAAASL | ASSQDRGANTQY | 0.6483717 |
| AAAALPAFSPP | ASSQDRGANTQY | 0.7122677 |
| VVLAPAL | ASSQDRGANTQY | 0.70567185 |
| TIGHVDHGKTTL | ASSQDRGANTQY | 0.66496134 |
| LISNASDALDK | ASSQDRGANTQY | 0.72465265 |
| LISNASDAL | ASSQDRGANTQY | 0.71610385 |
| ISNASDALDK | ASSQDRGANTQY | 0.67392534 |
| NQAETLVYQTE | ASSQDRGANTQY | 0.7299942 |
| TGGTGGTAGAG | ASSQDRGANTQY | 0.70244455 |
| TGSTAGGAGGA | ASSQDRGANTQY | 0.72950184 |
| DPLAVPVAAGPV | ASSQDRGANTQY | 0.74560535 |
| GGTGGTPGNGGN | ASSQDRGANTQY | 0.69425046 |
| PAGLAGVDPATL | ASSQDRGANTQY | 0.74080086 |
| GNNLVVEAPHAGASV | ASSQDRGANTQY | 0.72572637 |
| KVTRIMSASTLEQPA | ASSQDRGANTQY | 0.72983795 |
| LGFALPIALPRRTVV | ASSQDRGANTQY | 0.71091264 |
| TRLGNLTARKLVVLV | ASSQDRGANTQY | 0.6894114 |
| WSQQRRLIYFPSAGP | ASSQDRGANTQY | 0.7402568 |
| GPNGNPDPMAAAVDIR | ASSQDRGANTQY | 0.73131 |
| TQRCVSVADLRAAASR | ASSQDRGANTQY | 0.71594834 |
| QFVLDVAEHPVEVLTETV | ASSQDRGANTQY | 0.72631216 |
| GGAGGTGGAGGNGGTQAGDG | ASSQDRGANTQY | 0.7269418 |
| PLLATLRAIAVDVDAKPAQV | ASSQDRGANTQY | 0.73137105 |
| APKFRTGGYMIDFWGVGYQVAK | ASSQDRGANTQY | 0.67543817 |
| GAGGAGGAGGAAGGPGGTGGTG | ASSQDRGANTQY | 0.6215982 |
| IGALGLLATDVAPAGSVAFGVA | ASSQDRGANTQY | 0.7096381 |
| AGGAGGIGGNGGAGGAGGVLMG | ASSQDRGANTQY | 0.66819847 |
| ADELIGLL | ASSQDRGANTQY | 0.718034 |
| ADRALKA | ASSQDRGANTQY | 0.49952412 |
| AGAVGQVGGAGGA | ASSQDRGANTQY | 0.7298831 |
| AGGGGGLGGAP | ASSQDRGANTQY | 0.74897015 |
| AGGLLYGNGGAG | ASSQDRGANTQY | 0.66459095 |
| AGLESIGL | ASSQDRGANTQY | 0.7480822 |
| AGLKTGII | ASSQDRGANTQY | 0.66213256 |
| ASLAALLL | ASSQDRGANTQY | 0.7328718 |
| ASVINISSI | ASSQDRGANTQY | 0.73591965 |
| ATPPGPA | ASSQDRGANTQY | 0.63288677 |
| ATVASALV | ASSQDRGANTQY | 0.71895874 |
| ATVASLIL | ASSQDRGANTQY | 0.7038987 |

| | | |
|---|---|---|
| AVNGPVLIRH | ASSQDRGANTQY | 0.71419966 |
| DGAAGAVGI | ASSQDRGANTQY | 0.7417362 |
| DPLLHGI | ASSQDRGANTQY | 0.6905635 |
| ELGAALGAL | ASSQDRGANTQY | 0.72310966 |
| ELGRTQAV | ASSQDRGANTQY | 0.72042954 |
| FVAGGYLS | ASSQDRGANTQY | 0.69589436 |
| GASGGAGGAGGW | ASSQDRGANTQY | 0.7346604 |
| GGAGAGGDGGAGGI | ASSQDRGANTQY | 0.7339587 |
| GTGHNGSQPGMGGNG | ASSQDRGANTQY | 0.6758857 |
| GYAAASASAS | ASSQDRGANTQY | 0.7486438 |
| HQVDVAAVQQG | ASSQDRGANTQY | 0.6809702 |
| IGAAIAHT | ASSQDRGANTQY | 0.74645513 |
| IGMIEVAQL | ASSQDRGANTQY | 0.73101735 |
| IRLAPEHTSAMD | ASSQDRGANTQY | 0.7097554 |
| IYNVSVL | ASSQDRGANTQY | 0.7140269 |
| KQGFLGM | ASSQDRGANTQY | 0.7352965 |
| LESVITQL | ASSQDRGANTQY | 0.7022532 |
| LGLGALVVV | ASSQDRGANTQY | 0.7280437 |
| LIGNGGNGGSGGI | ASSQDRGANTQY | 0.7448869 |
| LNAINAPTL | ASSQDRGANTQY | 0.3336879 |
| LQLLNTGV | ASSQDRGANTQY | 0.7537137 |
| LQTSIEF | ASSQDRGANTQY | 0.63533294 |
| MSFAVSEL | ASSQDRGANTQY | 0.6942987 |
| NTGALGL | ASSQDRGANTQY | 0.45356983 |
| PPGAGGAGGAGTT | ASSQDRGANTQY | 0.3266731 |
| QLVGLGI | ASSQDRGANTQY | 0.4888472 |
| RFLEQVHQL | ASSQDRGANTQY | 0.40259942 |
| RGVGYVLR | ASSQDRGANTQY | 0.6974379 |
| SATGVFAGVI | ASSQDRGANTQY | 0.7206725 |
| SLPAITLP | ASSQDRGANTQY | 0.41211665 |
| STALAFTV | ASSQDRGANTQY | 0.6986184 |
| SVLTVGLGK | ASSQDRGANTQY | 0.7635629 |
| TALAGAGSGI | ASSQDRGANTQY | 0.27551565 |
| TGLPFTGI | ASSQDRGANTQY | 0.5558806 |
| VAEHVTV | ASSQDRGANTQY | 0.38526338 |
| VDIVLNSL | ASSQDRGANTQY | 0.5271058 |
| VLTGGGAVL | ASSQDRGANTQY | 0.18047814 |
| VNVGLAYGI | ASSQDRGANTQY | 0.714535 |
| VSSAAVGAV | ASSQDRGANTQY | 0.36760494 |
| VTSVYEAGV | ASSQDRGANTQY | 0.17892383 |
| WVALDRGA | ASSQDRGANTQY | 0.58450687 |
| YLVGSVVI | ASSQDRGANTQY | 0.4990417 |
| GPPGTGKTLL | ASSQDRGANTQY | 0.5741275 |
| GVGMTKF | ASSQDRGANTQY | 0.7230593 |
| IGIGGSDL | ASSQDRGANTQY | 0.36399746 |
| IGTLLAEL | ASSQDRGANTQY | 0.37185034 |
| IYTSGTT | ASSQDRGANTQY | 0.6544243 |
| LSFPAGV | ASSQDRGANTQY | 0.74877346 |
| LYTSGTT | ASSQDRGANTQY | 0.4075075 |
| VGAFPFGF | ASSQDRGANTQY | 0.6206672 |
| VGAGTVEF | ASSQDRGANTQY | 0.7610744 |

| | | |
|---|---|---|
| VGKSTLLL | ASSQDRGANTQY | 0.5606773 |
| AALAHAGGGIV | ASSQDRGANTQY | 0.7672638 |
| AALVAVSLL | ASSQDRGANTQY | 0.78059185 |
| AAVSRAVAAL | ASSQDRGANTQY | 0.45324636 |
| AGGSSGIGGF | ASSQDRGANTQY | 0.7469373 |
| AGLTVANANG | ASSQDRGANTQY | 0.709237 |
| AGLVGAVL | ASSQDRGANTQY | 0.5181908 |
| ALSQAFTV | ASSQDRGANTQY | 0.58854616 |
| ALTFSVSL | ASSQDRGANTQY | 0.30886465 |
| ALTGAALVAL | ASSQDRGANTQY | 0.5263248 |
| ASFSIGAGVL | ASSQDRGANTQY | 0.30886522 |
| DGSVYKI | ASSQDRGANTQY | 0.6190035 |
| EQLAEIV | ASSQDRGANTQY | 0.744773 |
| FAVSELTVM | ASSQDRGANTQY | 0.53899574 |
| GAAPLAAGL | ASSQDRGANTQY | 0.28521234 |
| GADGGDGGSGGAGG | ASSQDRGANTQY | 0.26554754 |
| GFGLDTGGAGG | ASSQDRGANTQY | 0.60074323 |
| GLASGGLL | ASSQDRGANTQY | 0.42190403 |
| GRIAGAAVLL | ASSQDRGANTQY | 0.28396955 |
| GSAGSLNTSF | ASSQDRGANTQY | 0.47007114 |
| GTGQIGI | ASSQDRGANTQY | 0.7135219 |
| GTLIALIL | ASSQDRGANTQY | 0.73127556 |
| IAATMGFVVG | ASSQDRGANTQY | 0.36437157 |
| IFAQALL | ASSQDRGANTQY | 0.7728845 |
| IGAAVALGF | ASSQDRGANTQY | 0.6692366 |
| IGLTPLAL | ASSQDRGANTQY | 0.75361836 |
| ILGSNTSTL | ASSQDRGANTQY | 0.5400456 |
| IVFSLTVL | ASSQDRGANTQY | 0.32526684 |
| LASKATGF | ASSQDRGANTQY | 0.7169391 |
| LASVALIL | ASSQDRGANTQY | 0.5101725 |
| LFAVGLTL | ASSQDRGANTQY | 0.6331346 |
| LFNGGIGTYI | ASSQDRGANTQY | 0.6968312 |
| LTLGAGII | ASSQDRGANTQY | 0.5301678 |
| LVAAALGDI | ASSQDRGANTQY | 0.39997995 |
| NTAFVQLGI | ASSQDRGANTQY | 0.56187814 |
| QLFSGAEL | ASSQDRGANTQY | 0.72229886 |
| SAATFITL | ASSQDRGANTQY | 0.3313256 |
| SNYGLANLGV | ASSQDRGANTQY | 0.38057134 |
| SYGGAYAVM | ASSQDRGANTQY | 0.55999327 |
| TTGVGGLTLG | ASSQDRGANTQY | 0.55236024 |
| TTQTIYI | ASSQDRGANTQY | 0.7547002 |
| VAAGAGLGAVI | ASSQDRGANTQY | 0.6656005 |
| VAAVVTLL | ASSQDRGANTQY | 0.47833756 |
| VGFLGGLI | ASSQDRGANTQY | 0.69517773 |
| VGVIPILL | ASSQDRGANTQY | 0.70819163 |
| VIFASSLI | ASSQDRGANTQY | 0.59480876 |
| VKGGDGGAAAG | ASSQDRGANTQY | 0.58857095 |
| VTKTFTNI | ASSQDRGANTQY | 0.4286488 |
| VVAAAILL | ASSQDRGANTQY | 0.72279525 |
| YAALDVEL | ASSQDRGANTQY | 0.71087635 |
| LASVALLL | ASSQDRGANTQY | 0.7324908 |

| | | |
|---|---|---|
| LDAVHRV | ASSQDRGANTQY | 0.7394784 |
| QGLSSSIF | ASSQDRGANTQY | 0.65175223 |
| RGTFANIRL | ASSQDRGANTQY | 0.57860196 |
| RQLWWGHRI | ASSQDRGANTQY | 0.23951289 |
| YGPPGTGKTLL | ASSQDRGANTQY | 0.40732387 |
| AALVGAPANVA | ASSQDRGANTQY | 0.5554619 |
| AATFRRTE | ASSQDRGANTQY | 0.63090813 |
| AGALGGGI | ASSQDRGANTQY | 0.6909741 |
| AGGAGGAGGIGAG | ASSQDRGANTQY | 0.7375736 |
| AGSGALGL | ASSQDRGANTQY | 0.3692434 |
| AQIADQI | ASSQDRGANTQY | 0.703454 |
| ASAVAVTYL | ASSQDRGANTQY | 0.74706626 |
| ASTVLGGTGG | ASSQDRGANTQY | 0.29356962 |
| AVGGEPAK | ASSQDRGANTQY | 0.51861006 |
| CEQGTFGGTC | ASSQDRGANTQY | 0.77715695 |
| DEGKAPAG | ASSQDRGANTQY | 0.6072345 |
| DTVVFPAL | ASSQDRGANTQY | 0.44476545 |
| FEGITMAG | ASSQDRGANTQY | 0.57608265 |
| FPLVVQD | ASSQDRGANTQY | 0.74272156 |
| GAFTTPQ | ASSQDRGANTQY | 0.6495606 |
| GFGGFGTAGGVGGA | ASSQDRGANTQY | 0.7510907 |
| GFLAQFL | ASSQDRGANTQY | 0.22058037 |
| GGAGGAGGNAGVA | ASSQDRGANTQY | 0.5539756 |
| GGVGTAGAGGAGGAGG | ASSQDRGANTQY | 0.39187825 |
| GNGGGGDAGNAGSGGN | ASSQDRGANTQY | 0.19373405 |
| GTGGAGGAGNDAGS | ASSQDRGANTQY | 0.71530575 |
| GTTNTGWFNS | ASSQDRGANTQY | 0.4979809 |
| IFMGPLFTL | ASSQDRGANTQY | 0.68804926 |
| ILAHAIEAI | ASSQDRGANTQY | 0.45564735 |
| IRSAIREAL | ASSQDRGANTQY | 0.7456689 |
| ITFAGGG | ASSQDRGANTQY | 0.5977383 |
| ITGMSGHQEV | ASSQDRGANTQY | 0.5104572 |
| IVAGLACTL | ASSQDRGANTQY | 0.39915764 |
| IVGGGLAGA | ASSQDRGANTQY | 0.65699124 |
| IVTGVLNI | ASSQDRGANTQY | 0.71810746 |
| LAQQGCI | ASSQDRGANTQY | 0.7516619 |
| LCLHVEN | ASSQDRGANTQY | 0.29005334 |
| LDAISDGS | ASSQDRGANTQY | 0.62272674 |
| LGGFAAML | ASSQDRGANTQY | 0.55774367 |
| LGGKNPMI | ASSQDRGANTQY | 0.21589503 |
| LGPKGVAAIL | ASSQDRGANTQY | 0.21903914 |
| LMVGVSI | ASSQDRGANTQY | 0.41074613 |
| LVGAAFVL | ASSQDRGANTQY | 0.47194248 |
| LVLAVAPL | ASSQDRGANTQY | 0.6562264 |
| MAAIAVIDAA | ASSQDRGANTQY | 0.65793073 |
| NNGINAELQ | ASSQDRGANTQY | 0.4841904 |
| RGEPLVLVHG | ASSQDRGANTQY | 0.62616235 |
| RGGAGGAAGLI | ASSQDRGANTQY | 0.6316035 |
| RSYRMHPAV | ASSQDRGANTQY | 0.13653727 |
| SAAHLLAGQLA | ASSQDRGANTQY | 0.43547225 |
| SAAVGSY | ASSQDRGANTQY | 0.28111455 |

| | | |
|---|---|---|
| SPHAVLHRI | ASSQDRGANTQY | 0.78634983 |
| SQIAKVL | ASSQDRGANTQY | 0.3726269 |
| TALGHLVP | ASSQDRGANTQY | 0.38935745 |
| TAPTTSGAA | ASSQDRGANTQY | 0.5472992 |
| TDGDDGNGGSPGT | ASSQDRGANTQY | 0.64964485 |
| TDGLGEN | ASSQDRGANTQY | 0.5623753 |
| TGGPPLPGQA | ASSQDRGANTQY | 0.37964204 |
| TIPAITI | ASSQDRGANTQY | 0.21650335 |
| TIPGFTI | ASSQDRGANTQY | 0.6641811 |
| TSIASGALA | ASSQDRGANTQY | 0.38092393 |
| TVTLPGAAI | ASSQDRGANTQY | 0.4276802 |
| VASGLSAALP | ASSQDRGANTQY | 0.24966113 |
| VGAGVIGI | ASSQDRGANTQY | 0.5584444 |
| VGPKRFGV | ASSQDRGANTQY | 0.30575642 |
| VNVAFPD | ASSQDRGANTQY | 0.7436638 |
| VSMTNSGVS | ASSQDRGANTQY | 0.40568388 |
| VTDVLGDLSV | ASSQDRGANTQY | 0.49075893 |
| VVVVGAGFAGL | ASSQDRGANTQY | 0.7752034 |
| VYTVGTVV | ASSQDRGANTQY | 0.6337154 |
| WTTFSGP | ASSQDRGANTQY | 0.5025942 |
| YDELVEAF | ASSQDRGANTQY | 0.46527642 |
| YGTGGAGGNGG | ASSQDRGANTQY | 0.3247341 |
| YTNVMAV | ASSQDRGANTQY | 0.347022 |
| FLEQVHQL | ASSQDRGANTQY | 0.46798772 |
| GLSLLAVAV | ASSQDRGANTQY | 0.74956983 |
| LEQVHQL | ASSQDRGANTQY | 0.62172425 |
| LNAAGAI | ASSQDRGANTQY | 0.3227206 |
| PAVHASL | ASSQDRGANTQY | 0.72762775 |
| TELEDDA | ASSQDRGANTQY | 0.77197564 |
| TPPGPPN | ASSQDRGANTQY | 0.7693927 |
| TPPPPPPP | ASSQDRGANTQY | 0.29221502 |
| TQSPALI | ASGRLGALY | 0.4950819 |
| EFAAAFPQ | ASGRLGALY | 0.52650267 |
| LLSGNLDVL | ASGRLGALY | 0.52372 |
| GPANLGNYNI | ASGRLGALY | 0.3991975 |
| TLGPTLP | ASGRLGALY | 0.39872804 |
| VYAYSSCCK | ASGRLGALY | 0.6796007 |
| APAPAPAPNI | ASGRLGALY | 0.79061353 |
| PGGPVPIP | ASGRLGALY | 0.53303367 |
| FLTVALEP | ASGRLGALY | 0.20739096 |
| SQYFGANL | ASGRLGALY | 0.51217914 |
| GGVKPAA | ASGRLGALY | 0.3767124 |
| GGTGGAGGAGGW | ASGRLGALY | 0.35898522 |
| GVGGVAGLGGAGG | ASGRLGALY | 0.50982773 |
| VLLVLATTI | ASGRLGALY | 0.2728464 |
| IGIFAAML | ASGRLGALY | 0.58252114 |
| AVGDLID | ASGRLGALY | 0.78024817 |
| NISPLPL | ASGRLGALY | 0.4861377 |
| VTAVFVAI | ASGRLGALY | 0.7114368 |
| SGDDNDAAL | ASGRLGALY | 0.7535492 |
| AVAALAHS | ASGRLGALY | 0.6764996 |

| | | |
|---|---|---|
| GGVTVMATP | ASGRLGALY | 0.7538672 |
| AGLAANVRD | ASGRLGALY | 0.68551743 |
| TNRPDLIDPALLRPG | ASGRLGALY | 0.59309435 |
| LANLGVTAL | ASGRLGALY | 0.5613289 |
| TGERIEL | ASGRLGALY | 0.7135341 |
| NGEELAA | ASGRLGALY | 0.51983076 |
| VLSLVWG | ASGRLGALY | 0.5430007 |
| IGYALWL | ASGRLGALY | 0.51169395 |
| LSLLLAPL | ASGRLGALY | 0.7026757 |
| TLDEALK | ASGRLGALY | 0.47479796 |
| TDGAGTAEGG | ASGRLGALY | 0.7665642 |
| NAPGGAGGNGG | ASGRLGALY | 0.24926849 |
| VSVFMGMGGNF | ASGRLGALY | 0.36508316 |
| LASYLAAL | ASGRLGALY | 0.7449137 |
| LVGGTPAL | ASGRLGALY | 0.15726624 |
| VRNADVL | ASGRLGALY | 0.272908 |
| TCSSPFLVSAM | ASGRLGALY | 0.50897384 |
| VHITDAE | ASGRLGALY | 0.73707366 |
| EYAAPSF | ASGRLGALY | 0.16308486 |
| VVDAFHGGL | ASGRLGALY | 0.4379916 |
| ASVVDPQ | ASGRLGALY | 0.30108532 |
| SYNEFGGLS | ASGRLGALY | 0.7699318 |
| DPDGSVY | ASGRLGALY | 0.61188024 |
| MVSTIHATE | ASGRLGALY | 0.31945413 |
| LAAGVADPV | ASGRLGALY | 0.7417171 |
| TKEQIAA | ASGRLGALY | 0.2229987 |
| SLASPAAL | ASGRLGALY | 0.47730774 |
| IGLVFYGV | ASGRLGALY | 0.7122607 |
| FLLDASNL | ASGRLGALY | 0.751767 |
| YQVGNFP | ASGRLGALY | 0.28436202 |
| ESLLSQST | ASGRLGALY | 0.6955265 |
| LFTVGFAAVG | ASGRLGALY | 0.4228208 |
| ATPPTPT | ASGRLGALY | 0.7880378 |
| TDLATLG | ASGRLGALY | 0.47312862 |
| SAGGTGR | ASGRLGALY | 0.28106707 |
| GFSNSGS | ASGRLGALY | 0.7484335 |
| GGNGGSGGTSV | ASGRLGALY | 0.7275987 |
| GTSVATGGAGN | ASGRLGALY | 0.58872896 |
| NGGNGGSGGTS | ASGRLGALY | 0.68563837 |
| AVVKFPH | ASGRLGALY | 0.58952606 |
| TPPPPPP | ASGRLGALY | 0.5507043 |
| AAPAAAAAPA | ASGRLGALY | 0.71973336 |
| IALNEDL | ASGRLGALY | 0.6561868 |
| IVNVSSM | ASGRLGALY | 0.4257025 |
| TGIPPAPR | ASGRLGALY | 0.7576015 |
| FDLGGGTF | ASGRLGALY | 0.5871335 |
| FDLGGGTFDVS | ASGRLGALY | 0.35029572 |
| NEPTAAA | ASGRLGALY | 0.78229177 |
| NEPTAAALA | ASGRLGALY | 0.59287935 |
| GSIINISSI | ASGRLGALY | 0.6945854 |
| VAPGNAGTA | ASGRLGALY | 0.70251024 |

| | | |
|---|---|---|
| TGGGASLE | ASGRLGALY | 0.77630687 |
| VAVYNHYKRI | ASGRLGALY | 0.36749172 |
| TDPAPATT | ASGRLGALY | 0.54284763 |
| TNRPDILDPALLRPG | ASGRLGALY | 0.55344236 |
| LTTTTTS | ASGRLGALY | 0.71883076 |
| PTGSGKTL | ASGRLGALY | 0.6137243 |
| TGSGKTL | ASGRLGALY | 0.44636226 |
| TIGHVDHGK | ASGRLGALY | 0.6847635 |
| TIGHVDHGKTT | ASGRLGALY | 0.7495431 |
| GHVDHGKTT | ASGRLGALY | 0.52886295 |
| AQFEHTLL | ASGRLGALY | 0.5983968 |
| IVNQDEL | ASGRLGALY | 0.77315104 |
| TNGSQFF | ASGRLGALY | 0.35560948 |
| TGKGIVE | ASGRLGALY | 0.60693395 |
| LISNASDA | ASGRLGALY | 0.68911624 |
| AELRTLGELTKIAKAHGAQ | ASGRLGALY | 0.7160916 |
| APNAVAAAEPSW | ASGRLGALY | 0.5914206 |
| APQVAGAATAVL | ASGRLGALY | 0.7034311 |
| ARRDPAAAAAILPTDA | ASGRLGALY | 0.41256386 |
| DAAARDVLAEAGLAEYF | ASGRLGALY | 0.6342387 |
| DDQIVLSAKANLAKVEQ | ASGRLGALY | 0.4737571 |
| DELLPDTGEDSDEDGH | ASGRLGALY | 0.5137602 |
| ELPEAPANRATVPQD | ASGRLGALY | 0.3415868 |
| FGQSKIERVTGIG | ASGRLGALY | 0.56582034 |
| GAGGTGSTAGGAGGAGGAGGLY | ASGRLGALY | 0.6765884 |
| GHGGTGAAGGNGGTGGAGGSGI | ASGRLGALY | 0.6531838 |
| GNGGNGGAGGEGGPGGAGGAGA | ASGRLGALY | 0.20947215 |
| GSTAGNGGNGGNATNGTIAGQP | ASGRLGALY | 0.6594216 |
| GTGGDGGNGGNGGTGG | ASGRLGALY | 0.6367378 |
| GVRRPAGIPADAPV | ASGRLGALY | 0.7871609 |
| GYRQVTDAHLLTLARR | ASGRLGALY | 0.51920414 |
| IAKMGQDGHDRG | ASGRLGALY | 0.7241724 |
| KAEPSIEIQYT | ASGRLGALY | 0.35970426 |
| KPVRIRMAADEDTVAVT | ASGRLGALY | 0.59010273 |
| MPPRIAGMRLLVIKPEP | ASGRLGALY | 0.2547871 |
| MKFYAVALFRSRGVPPTR | ASGRLGALY | 0.4100234 |
| NAGGFGILGAVAHSP | ASGRLGALY | 0.47202942 |
| NEGREITLAMAEM | ASGRLGALY | 0.3997177 |
| NELVEWSHPLGHRWRWE | ASGRLGALY | 0.7087957 |
| NLGAGGTGGNGGNPG | ASGRLGALY | 0.6976652 |
| NTEWPKWETVMAADPVD | ASGRLGALY | 0.7385986 |
| PASAAPAETQP | ASGRLGALY | 0.7511481 |
| QQSSDGKNGVRAAEITGE | ASGRLGALY | 0.7613951 |
| RAPDAVTTSWWKE | ASGRLGALY | 0.76326144 |
| SGFGNVGAHTSGWFNQ | ASGRLGALY | 0.46797663 |
| SGVAGAVGQAASVGGLKV | ASGRLGALY | 0.47434217 |
| SNGGAGTGSAGGNGGT | ASGRLGALY | 0.26648688 |
| TGGAGGEGGAAG | ASGRLGALY | 0.5033665 |
| TGGDGGAAGAGGAPGHG | ASGRLGALY | 0.5019556 |
| TMPPLLAAIANAGRHGV | ASGRLGALY | 0.5993414 |
| TSSTAGNSGGA | ASGRLGALY | 0.17299652 |

| | | |
|---|---|---|
| VPATVPVGILC | ASGRLGALY | 0.6224667 |
| VQRPDSGGVVLTGRIS | ASGRLGALY | 0.7627115 |
| WTPNPVPATKGVPAGLVQAT | ASGRLGALY | 0.4623818 |
| YALKRALAKFSDRFDVVII | ASGRLGALY | 0.661916 |
| YLIGSVIV | ASGRLGALY | 0.6790645 |
| YGQLVAVID | ASGRLGALY | 0.43522906 |
| VTVGGMTLPA | ASGRLGALY | 0.6793781 |
| VLALPAV | ASGRLGALY | 0.48919296 |
| VIPPTLNYET | ASGRLGALY | 0.7459641 |
| VGMGGMHPSSG | ASGRLGALY | 0.76258713 |
| VGLAGGAATA | ASGRLGALY | 0.6611576 |
| VGGAGGAGGSGP | ASGRLGALY | 0.49040878 |
| VFAAFMV | ASGRLGALY | 0.66732335 |
| TLYDKLR | ASGRLGALY | 0.32100546 |
| TLAVFTVL | ASGRLGALY | 0.52025926 |
| TGSVQPARV | ASGRLGALY | 0.26992324 |
| TGGGAGSGGT | ASGRLGALY | 0.5781731 |
| SHLETSL | ASGRLGALY | 0.35791188 |
| SFTTGMT | ASGRLGALY | 0.64391994 |
| QVEVAKRMGVRQPTVS | ASGRLGALY | 0.59465796 |
| QGSGGPALAM | ASGRLGALY | 0.5768783 |
| MLAPSNP | ASGRLGALY | 0.32687113 |
| LVNNAAI | ASGRLGALY | 0.71466994 |
| LTGEINTLA | ASGRLGALY | 0.63190293 |
| LSIFGPV | ASGRLGALY | 0.3245666 |
| LLALLANL | ASGRLGALY | 0.6815272 |
| LISNASDALDKLR | ASGRLGALY | 0.50458556 |
| LINEAVP | ASGRLGALY | 0.52708256 |
| LGYAFSGL | ASGRLGALY | 0.73268616 |
| LDFVAVAL | ASGRLGALY | 0.6632595 |
| LAQPGAE | ASGRLGALY | 0.24956965 |
| ISNLGAVTGF | ASGRLGALY | 0.46854317 |
| HQEDPGRI | ASGRLGALY | 0.5934972 |
| GTPGEPTP | ASGRLGALY | 0.51131994 |
| GFFNSGS | ASGRLGALY | 0.18124743 |
| GAGGGGLTAGAN | ASGRLGALY | 0.666275 |
| FVDALVNM | ASGRLGALY | 0.36426717 |
| FLTVAPEL | ASGRLGALY | 0.41227686 |
| FETLTAL | ASGRLGALY | 0.5291039 |
| DVINPIEL | ASGRLGALY | 0.7370825 |
| APVSGAPV | ASGRLGALY | 0.42848247 |
| ALVGGLAW | ASGRLGALY | 0.5826495 |
| ALAAAAATAI | ASGRLGALY | 0.6248903 |
| AFGLGLLL | ASGRLGALY | 0.6401627 |
| AFFAAASL | ASGRLGALY | 0.7312236 |
| AAAALPAFSPP | ASGRLGALY | 0.5696855 |
| VVLAPAL | ASGRLGALY | 0.47590455 |
| TIGHVDHGKTTL | ASGRLGALY | 0.6763507 |
| LISNASDALDK | ASGRLGALY | 0.45096266 |
| LISNASDAL | ASGRLGALY | 0.40683568 |
| ISNASDALDK | ASGRLGALY | 0.32315978 |

| | | |
|---|---|---|
| NQAETLVYQTE | ASGRLGALY | 0.3407682 |
| TGGTGGTAGAG | ASGRLGALY | 0.2996627 |
| TGSTAGGAGGA | ASGRLGALY | 0.6293919 |
| DPLAVPVAAGPV | ASGRLGALY | 0.7885316 |
| GGTGGTPGNGGN | ASGRLGALY | 0.57695866 |
| PAGLAGVDPATL | ASGRLGALY | 0.26537237 |
| GNNLVVEAPHAGASV | ASGRLGALY | 0.4431947 |
| KVTRIMSASTLEQPA | ASGRLGALY | 0.50919515 |
| LGFALPIALPRRTVV | ASGRLGALY | 0.6875186 |
| TRLGNLTARKLVVLV | ASGRLGALY | 0.67984337 |
| WSQQRRLIYFPSAGP | ASGRLGALY | 0.76607037 |
| GPNGNPDPMAAAVDIR | ASGRLGALY | 0.68808556 |
| TQRCVSVADLRAAASR | ASGRLGALY | 0.5046491 |
| QFVLDVAEHPVEVLTETV | ASGRLGALY | 0.6759034 |
| GGAGGTGGAGGNGGTQAGDG | ASGRLGALY | 0.5289988 |
| PLLATLRAIAVDVDAKPAQV | ASGRLGALY | 0.5158175 |
| APKFRTGGYMIDFWGVGYQVAK | ASGRLGALY | 0.5744464 |
| GAGGAGGAGGAAGGPGGTGGTG | ASGRLGALY | 0.61877656 |
| IGALGLLATDVAPAGSVAFGVA | ASGRLGALY | 0.5081883 |
| AGGAGGIGGNGGAGGAGGVLMG | ASGRLGALY | 0.5062305 |
| ADELIGLL | ASGRLGALY | 0.24132316 |
| ADRALKA | ASGRLGALY | 0.63117427 |
| AGAVGQVGGAGGA | ASGRLGALY | 0.4013551 |
| AGGGGGLGGAP | ASGRLGALY | 0.37014103 |
| AGGLLYGNGGAG | ASGRLGALY | 0.48188394 |
| AGLESIGL | ASGRLGALY | 0.7693131 |
| AGLKTGII | ASGRLGALY | 0.7555336 |
| ASLAALLL | ASGRLGALY | 0.47209477 |
| ASVINISSI | ASGRLGALY | 0.43185815 |
| ATPPGPA | ASGRLGALY | 0.6210988 |
| ATVASALV | ASGRLGALY | 0.3354549 |
| ATVASLIL | ASGRLGALY | 0.6974168 |
| AVNGPVLIRH | ASGRLGALY | 0.57511234 |
| DGAAGAVGI | ASGRLGALY | 0.58071405 |
| DPLLHGI | ASGRLGALY | 0.47500739 |
| ELGAALGAL | ASGRLGALY | 0.67570674 |
| ELGRTQAV | ASGRLGALY | 0.48090065 |
| FVAGGYLS | ASGRLGALY | 0.67709416 |
| GASGGAGGAGGW | ASGRLGALY | 0.15581453 |
| GGAGAGGDGGAGGI | ASGRLGALY | 0.7391907 |
| GTGHNGSQPGMGGNG | ASGRLGALY | 0.32716048 |
| GYAAASASAS | ASGRLGALY | 0.34556305 |
| HQVDVAAVQQG | ASGRLGALY | 0.19780028 |
| IGAAIAHT | ASGRLGALY | 0.4849642 |
| IGMIEVAQL | ASGRLGALY | 0.64490914 |
| IRLAPEHTSAMD | ASGRLGALY | 0.7426617 |
| IYNVSVL | ASGRLGALY | 0.5129799 |
| KQGFLGM | ASGRLGALY | 0.65404737 |
| LESVITQL | ASGRLGALY | 0.6765929 |
| LGLGALVVV | ASGRLGALY | 0.43140298 |
| LIGNGGNGGSGGI | ASGRLGALY | 0.24940284 |

| | | |
|---|---|---|
| LNAINAPTL | ASGRLGALY | 0.69934094 |
| LQLLNTGV | ASGRLGALY | 0.46278924 |
| LQTSIEF | ASGRLGALY | 0.75585985 |
| MSFAVSEL | ASGRLGALY | 0.62999034 |
| NTGALGL | ASGRLGALY | 0.26477897 |
| PPGAGGAGGAGTT | ASGRLGALY | 0.77231324 |
| QLVGLGI | ASGRLGALY | 0.7061151 |
| RFLEQVHQL | ASGRLGALY | 0.3182727 |
| RGVGYVLR | ASGRLGALY | 0.3142437 |
| SATGVFAGVI | ASGRLGALY | 0.41076684 |
| SLPAITLP | ASGRLGALY | 0.23612322 |
| STALAFTV | ASGRLGALY | 0.6882793 |
| SVLTVGLGK | ASGRLGALY | 0.46299344 |
| TALAGAGSGI | ASGRLGALY | 0.557799 |
| TGLPFTGI | ASGRLGALY | 0.5851413 |
| VAEHVTV | ASGRLGALY | 0.5152183 |
| VDIVLNSL | ASGRLGALY | 0.49085575 |
| VLTGGGAVL | ASGRLGALY | 0.54327124 |
| VNVGLAYGI | ASGRLGALY | 0.62204325 |
| VSSAAVGAV | ASGRLGALY | 0.6738641 |
| VTSVYEAGV | ASGRLGALY | 0.54047054 |
| WVALDRGA | ASGRLGALY | 0.59657013 |
| YLVGSVVI | ASGRLGALY | 0.7535875 |
| GPPGTGKTLL | ASGRLGALY | 0.3436973 |
| GVGMTKF | ASGRLGALY | 0.47865194 |
| IGIGGSDL | ASGRLGALY | 0.6956688 |
| IGTLLAEL | ASGRLGALY | 0.50333893 |
| IYTSGTT | ASGRLGALY | 0.73652184 |
| LSFPAGV | ASGRLGALY | 0.63793015 |
| LYTSGTT | ASGRLGALY | 0.62652016 |
| VGAFPFGF | ASGRLGALY | 0.51367855 |
| VGAGTVEF | ASGRLGALY | 0.7690096 |
| VGKSTLLL | ASGRLGALY | 0.70699275 |
| AALAHAGGGIV | ASGRLGALY | 0.57433116 |
| AALVAVSLL | ASGRLGALY | 0.6411928 |
| AAVSRAVAAL | ASGRLGALY | 0.3088486 |
| AGGSSGIGGF | ASGRLGALY | 0.66616225 |
| AGLTVANANG | ASGRLGALY | 0.6916882 |
| AGLVGAVL | ASGRLGALY | 0.37295145 |
| ALSQAFTV | ASGRLGALY | 0.54460955 |
| ALTFSVSL | ASGRLGALY | 0.74412054 |
| ALTGAALVAL | ASGRLGALY | 0.71491027 |
| ASFSIGAGVL | ASGRLGALY | 0.4236857 |
| DGSVYKI | ASGRLGALY | 0.44085193 |
| EQLAEIV | ASGRLGALY | 0.29837602 |
| FAVSELTVM | ASGRLGALY | 0.3816561 |
| GAAPLAAGL | ASGRLGALY | 0.3169073 |
| GADGGDGGSGGAGG | ASGRLGALY | 0.62628704 |
| GFGLDTGGAGG | ASGRLGALY | 0.43561995 |
| GLASGGLL | ASGRLGALY | 0.7057047 |
| GRIAGAAVLL | ASGRLGALY | 0.6245277 |

| | | |
|---|---|---|
| GSAGSLNTSF | ASGRLGALY | 0.74397886 |
| GTGQIGI | ASGRLGALY | 0.37694976 |
| GTLIALIL | ASGRLGALY | 0.5202871 |
| IAATMGFVVG | ASGRLGALY | 0.2582965 |
| IFAQALL | ASGRLGALY | 0.35208696 |
| IGAAVALGF | ASGRLGALY | 0.34954962 |
| IGLTPLAL | ASGRLGALY | 0.5765951 |
| ILGSNTSTL | ASGRLGALY | 0.19424827 |
| IVFSLTVL | ASGRLGALY | 0.77783513 |
| LASKATGF | ASGRLGALY | 0.5332782 |
| LASVALIL | ASGRLGALY | 0.67361546 |
| LFAVGLTL | ASGRLGALY | 0.5109766 |
| LFNGGIGTYI | ASGRLGALY | 0.37348822 |
| LTLGAGII | ASGRLGALY | 0.4568665 |
| LVAAALGDI | ASGRLGALY | 0.70078254 |
| NTAFVQLGI | ASGRLGALY | 0.3101283 |
| QLFSGAEL | ASGRLGALY | 0.1997314 |
| SAATFITL | ASGRLGALY | 0.37924916 |
| SNYGLANLGV | ASGRLGALY | 0.25772297 |
| SYGGAYAVM | ASGRLGALY | 0.7827319 |
| TTGVGGLTLG | ASGRLGALY | 0.73895955 |
| TTQTIYI | ASGRLGALY | 0.6508464 |
| VAAGAGLGAVI | ASGRLGALY | 0.7797069 |
| VAAVVTLL | ASGRLGALY | 0.16935265 |
| VGFLGGLI | ASGRLGALY | 0.51452756 |
| VGVIPILL | ASGRLGALY | 0.76131374 |
| VIFASSLI | ASGRLGALY | 0.59543186 |
| VKGGDGGAAAG | ASGRLGALY | 0.49068105 |
| VTKTFTNI | ASGRLGALY | 0.73780143 |
| VVAAAILL | ASGRLGALY | 0.48099023 |
| YAALDVEL | ASGRLGALY | 0.51537323 |
| LASVALLL | ASGRLGALY | 0.57459223 |
| LDAVHRV | ASGRLGALY | 0.66521883 |
| QGLSSSIF | ASGRLGALY | 0.5929322 |
| RGTFANIRL | ASGRLGALY | 0.27580595 |
| RQLWWGHRI | ASGRLGALY | 0.7196548 |
| YGPPGTGKTLL | ASGRLGALY | 0.7825967 |
| AALVGAPANVA | ASGRLGALY | 0.5149123 |
| AATFRRTE | ASGRLGALY | 0.64261556 |
| AGALGGGI | ASGRLGALY | 0.6636389 |
| AGGAGGAGGIGAG | ASGRLGALY | 0.60445154 |
| AGSGALGL | ASGRLGALY | 0.3788656 |
| AQIADQI | ASGRLGALY | 0.31445813 |
| ASAVAVTYL | ASGRLGALY | 0.71860605 |
| ASTVLGGTGG | ASGRLGALY | 0.31204686 |
| AVGGEPAK | ASGRLGALY | 0.5936358 |
| CEQGTFGGTC | ASGRLGALY | 0.57418513 |
| DEGKAPAG | ASGRLGALY | 0.80428886 |
| DTVVFPAL | ASGRLGALY | 0.65137887 |
| FEGITMAG | ASGRLGALY | 0.6904336 |
| FPLVVQD | ASGRLGALY | 0.33331496 |

| | | |
|---|---|---|
| GAFTTPQ | ASGRLGALY | 0.5984485 |
| GFGGFGTAGGVGGA | ASGRLGALY | 0.6883192 |
| GFLAQFL | ASGRLGALY | 0.2750522 |
| GGAGGAGGNAGVA | ASGRLGALY | 0.73002446 |
| GGVGTAGAGGAGGAGG | ASGRLGALY | 0.37706405 |
| GNGGGGDAGNAGSGGN | ASGRLGALY | 0.6247845 |
| GTGGAGGAGNDAGS | ASGRLGALY | 0.67434776 |
| GTTNTGWFNS | ASGRLGALY | 0.7440898 |
| IFMGPLFTL | ASGRLGALY | 0.49891588 |
| ILAHAIEAI | ASGRLGALY | 0.6717303 |
| IRSAIREAL | ASGRLGALY | 0.36802825 |
| ITFAGGG | ASGRLGALY | 0.69307804 |
| ITGMSGHQEV | ASGRLGALY | 0.7414537 |
| IVAGLACTL | ASGRLGALY | 0.582111 |
| IVGGGLAGA | ASGRLGALY | 0.76820016 |
| IVTGVLNI | ASGRLGALY | 0.74279755 |
| LAQQGCI | ASGRLGALY | 0.25979486 |
| LCLHVEN | ASGRLGALY | 0.5426446 |
| LDAISDGS | ASGRLGALY | 0.7426537 |
| LGGFAAML | ASGRLGALY | 0.736372 |
| LGGKNPMI | ASGRLGALY | 0.7016908 |
| LGPKGVAAIL | ASGRLGALY | 0.62082714 |
| LMVGVSI | ASGRLGALY | 0.44971 |
| LVGAAFVL | ASGRLGALY | 0.76770985 |
| LVLAVAPL | ASGRLGALY | 0.8058187 |
| MAAIAVIDAA | ASGRLGALY | 0.7143369 |
| NNGINAELQ | ASGRLGALY | 0.37088263 |
| RGEPLVLVHG | ASGRLGALY | 0.4125548 |
| RGGAGGAAGLI | ASGRLGALY | 0.6820453 |
| RSYRMHPAV | ASGRLGALY | 0.73021555 |
| SAAHLLAGQLA | ASGRLGALY | 0.3367603 |
| SAAVGSY | ASGRLGALY | 0.5960938 |
| SPHAVLHRI | ASGRLGALY | 0.604249 |
| SQIAKVL | ASGRLGALY | 0.2239537 |
| TALGHLVP | ASGRLGALY | 0.63529676 |
| TAPTTSGAA | ASGRLGALY | 0.7271536 |
| TDGDDGNGGSPGT | ASGRLGALY | 0.6866915 |
| TDGLGEN | ASGRLGALY | 0.42242113 |
| TGGPPLPGQA | ASGRLGALY | 0.7280886 |
| TIPAITI | ASGRLGALY | 0.7493225 |
| TIPGFTI | ASGRLGALY | 0.797377 |
| TSIASGALA | ASGRLGALY | 0.29634064 |
| TVTLPGAAI | ASGRLGALY | 0.17308293 |
| VASGLSAALP | ASGRLGALY | 0.54956776 |
| VGAGVIGI | ASGRLGALY | 0.69509536 |
| VGPKRFGV | ASGRLGALY | 0.5398572 |
| VNVAFPD | ASGRLGALY | 0.44462258 |
| VSMTNSGVS | ASGRLGALY | 0.63286453 |
| VTDVLGDLSV | ASGRLGALY | 0.74602795 |
| VVVVGAGFAGL | ASGRLGALY | 0.35094088 |
| VYTVGTVV | ASGRLGALY | 0.28888336 |

| | | |
|---|---|---|
| WTTFSGP | ASGRLGALY | 0.67214835 |
| YDELVEAF | ASGRLGALY | 0.7350961 |
| YGTGGAGGNGG | ASGRLGALY | 0.36841327 |
| YTNVMAV | ASGRLGALY | 0.44320595 |
| FLEQVHQL | ASGRLGALY | 0.5680718 |
| GLSLLAVAV | ASGRLGALY | 0.5921393 |
| LEQVHQL | ASGRLGALY | 0.631356 |
| LNAAGAI | ASGRLGALY | 0.312949 |
| PAVHASL | ASGRLGALY | 0.6907064 |
| TELEDDA | ASGRLGALY | 0.274687 |
| TPPGPPN | ASGRLGALY | 0.6455326 |
| TPPPPPPP | ASGRLGALY | 0.5938705 |
| TQSPALI | ASSMTGENTLY | 0.6504328 |
| EFAAAFPQ | ASSMTGENTLY | 0.45338595 |
| LLSGNLDVL | ASSMTGENTLY | 0.32127717 |
| GPANLGNYNI | ASSMTGENTLY | 0.35630974 |
| TLGPTLP | ASSMTGENTLY | 0.27177873 |
| VYAYSSCCK | ASSMTGENTLY | 0.7159358 |
| APAPAPAPNI | ASSMTGENTLY | 0.44901752 |
| PGGPVPIP | ASSMTGENTLY | 0.7256137 |
| FLTVALEP | ASSMTGENTLY | 0.63875747 |
| SQYFGANL | ASSMTGENTLY | 0.3714899 |
| GGVKPAA | ASSMTGENTLY | 0.64234614 |
| GGTGGAGGAGGW | ASSMTGENTLY | 0.301 |
| GVGGVAGLGGAGG | ASSMTGENTLY | 0.55251527 |
| VLLVLATTI | ASSMTGENTLY | 0.5589366 |
| IGIFAAML | ASSMTGENTLY | 0.738938 |
| AVGDLID | ASSMTGENTLY | 0.33397138 |
| NISPLPL | ASSMTGENTLY | 0.660105 |
| VTAVFVAI | ASSMTGENTLY | 0.6369269 |
| SGDDNDAAL | ASSMTGENTLY | 0.33931533 |
| AVAALAHS | ASSMTGENTLY | 0.70537555 |
| GGVTVMATP | ASSMTGENTLY | 0.7227524 |
| AGLAANVRD | ASSMTGENTLY | 0.6508248 |
| TNRPDLIDPALLRPG | ASSMTGENTLY | 0.2760989 |
| LANLGVTAL | ASSMTGENTLY | 0.6708036 |
| TGERIEL | ASSMTGENTLY | 0.36946404 |
| NGEELAA | ASSMTGENTLY | 0.64176744 |
| VLSLVWG | ASSMTGENTLY | 0.5424294 |
| IGYALWL | ASSMTGENTLY | 0.4768917 |
| LSLLLAPL | ASSMTGENTLY | 0.7401267 |
| TLDEALK | ASSMTGENTLY | 0.30869085 |
| TDGAGTAEGG | ASSMTGENTLY | 0.36821634 |
| NAPGGAGGNGG | ASSMTGENTLY | 0.49154112 |
| VSVFMGMGGNF | ASSMTGENTLY | 0.58916324 |
| LASYLAAL | ASSMTGENTLY | 0.7213266 |
| LVGGTPAL | ASSMTGENTLY | 0.36055836 |
| VRNADVL | ASSMTGENTLY | 0.58164793 |
| TCSSPFLVSAM | ASSMTGENTLY | 0.5387175 |
| VHITDAE | ASSMTGENTLY | 0.38023105 |
| EYAAPSF | ASSMTGENTLY | 0.64853114 |

| | | |
|---|---|---|
| VVDAFHGGL | ASSMTGENTLY | 0.48272043 |
| ASVVDPQ | ASSMTGENTLY | 0.28372544 |
| SYNEFGGLS | ASSMTGENTLY | 0.67403215 |
| DPDGSVY | ASSMTGENTLY | 0.46042588 |
| MVSTIHATE | ASSMTGENTLY | 0.6668178 |
| LAAGVADPV | ASSMTGENTLY | 0.76572114 |
| TKEQIAA | ASSMTGENTLY | 0.29759997 |
| SLASPAAL | ASSMTGENTLY | 0.64551336 |
| IGLVFYGV | ASSMTGENTLY | 0.63096416 |
| FLLDASNL | ASSMTGENTLY | 0.6031203 |
| YQVGNFP | ASSMTGENTLY | 0.5738428 |
| ESLLSQST | ASSMTGENTLY | 0.4080948 |
| LFTVGFAAVG | ASSMTGENTLY | 0.76946706 |
| ATPPTPT | ASSMTGENTLY | 0.7150912 |
| TDLATLG | ASSMTGENTLY | 0.65254295 |
| SAGGTGR | ASSMTGENTLY | 0.70340884 |
| GFSNSGS | ASSMTGENTLY | 0.6452167 |
| GGNGGSGGTSV | ASSMTGENTLY | 0.72415704 |
| GTSVATGGAGN | ASSMTGENTLY | 0.4267407 |
| NGGNGGSGGTS | ASSMTGENTLY | 0.27377182 |
| AVVKFPH | ASSMTGENTLY | 0.5308935 |
| TPPPPPP | ASSMTGENTLY | 0.48357517 |
| AAPAAAAAPA | ASSMTGENTLY | 0.41663012 |
| IALNEDL | ASSMTGENTLY | 0.30100882 |
| IVNVSSM | ASSMTGENTLY | 0.723904 |
| TGIPPAPR | ASSMTGENTLY | 0.7906277 |
| FDLGGGTF | ASSMTGENTLY | 0.42647693 |
| FDLGGGTFDVS | ASSMTGENTLY | 0.7995813 |
| NEPTAAA | ASSMTGENTLY | 0.32232058 |
| NEPTAAALA | ASSMTGENTLY | 0.7120254 |
| GSIINISSI | ASSMTGENTLY | 0.7235054 |
| VAPGNAGTA | ASSMTGENTLY | 0.3462736 |
| TGGGASLE | ASSMTGENTLY | 0.66096914 |
| VAVYNHYKRI | ASSMTGENTLY | 0.3450818 |
| TDPAPATT | ASSMTGENTLY | 0.44449744 |
| TNRPDILDPALLRPG | ASSMTGENTLY | 0.7405668 |
| LTTTTTS | ASSMTGENTLY | 0.5521048 |
| PTGSGKTL | ASSMTGENTLY | 0.66872114 |
| TGSGKTL | ASSMTGENTLY | 0.5948667 |
| TIGHVDHGK | ASSMTGENTLY | 0.40813124 |
| TIGHVDHGKTT | ASSMTGENTLY | 0.6573273 |
| GHVDHGKTT | ASSMTGENTLY | 0.6715907 |
| AQFEHTLL | ASSMTGENTLY | 0.57060444 |
| IVNQDEL | ASSMTGENTLY | 0.80115795 |
| TNGSQFF | ASSMTGENTLY | 0.30757663 |
| TGKGIVE | ASSMTGENTLY | 0.74577653 |
| LISNASDA | ASSMTGENTLY | 0.725716 |
| AELRTLGELTKIAKAHGAQ | ASSMTGENTLY | 0.5963142 |
| APNAVAAAEPSW | ASSMTGENTLY | 0.45308298 |
| APQVAGAATAVL | ASSMTGENTLY | 0.71530247 |
| ARRDPAAAAAILPTDA | ASSMTGENTLY | 0.5138502 |

| | | |
|---|---|---|
| DAAARDVLAEAGLAEYF | ASSMTGENTLY | 0.61428666 |
| DDQIVLSAKANLAKVEQ | ASSMTGENTLY | 0.7734468 |
| DELLPDTGEDSDEDGH | ASSMTGENTLY | 0.2726742 |
| ELPEAPANRATVPQD | ASSMTGENTLY | 0.8409841 |
| FGQSKIERVTGIG | ASSMTGENTLY | 0.78568566 |
| GAGGTGSTAGGAGGAGGAGGLY | ASSMTGENTLY | 0.747577 |
| GHGGTGAAGGNGGTGGAGGSGI | ASSMTGENTLY | 0.4786299 |
| GNGGNGGAGGEGGPGGAGGAGA | ASSMTGENTLY | 0.24345592 |
| GSTAGNGGNGGNATNGTIAGQP | ASSMTGENTLY | 0.44681704 |
| GTGGDGGNGGNGGTGG | ASSMTGENTLY | 0.43066198 |
| GVRRPAGIPADAPV | ASSMTGENTLY | 0.83562875 |
| GYRQVTDAHLLTLARR | ASSMTGENTLY | 0.8118827 |
| IAKMGQDGHDRG | ASSMTGENTLY | 0.44216663 |
| KAEPSIEIQYT | ASSMTGENTLY | 0.799851 |
| KPVRIRMAADEDTVAVT | ASSMTGENTLY | 0.8254911 |
| MPPRIAGMRLLVIKPEP | ASSMTGENTLY | 0.2225241 |
| MKFYAVALFRSRGVPPTR | ASSMTGENTLY | 0.58170843 |
| NAGGFGILGAVAHSP | ASSMTGENTLY | 0.38893905 |
| NEGREITLAMAEM | ASSMTGENTLY | 0.570367 |
| NELVEWSHPLGHRWRWE | ASSMTGENTLY | 0.13792148 |
| NLGAGGTGGNGGNPG | ASSMTGENTLY | 0.80155075 |
| NTEWPKWETVMAADPVD | ASSMTGENTLY | 0.49719793 |
| PASAAPAETQP | ASSMTGENTLY | 0.15248743 |
| QQSSDGKNGVRAAEITGE | ASSMTGENTLY | 0.6978517 |
| RAPDAVTTSWWKE | ASSMTGENTLY | 0.6069591 |
| SGFGNVGAHTSGWFNQ | ASSMTGENTLY | 0.5736538 |
| SGVAGAVGQAASVGGLKV | ASSMTGENTLY | 0.83223444 |
| SNGGAGTGSAGGNGGT | ASSMTGENTLY | 0.5526159 |
| TGGAGGEGGAAG | ASSMTGENTLY | 0.39219236 |
| TGGDGGAAGAGGAPGHG | ASSMTGENTLY | 0.786055 |
| TMPPLLAAIANAGRHGV | ASSMTGENTLY | 0.8491379 |
| TSSTAGNSGGA | ASSMTGENTLY | 0.34538335 |
| VPATVPVGILC | ASSMTGENTLY | 0.7703609 |
| VQRPDSGGVVLTGRIS | ASSMTGENTLY | 0.85511637 |
| WTPNPVPATKGVPAGLVQAT | ASSMTGENTLY | 0.7387562 |
| YALKRALAKFSDRFDVVII | ASSMTGENTLY | 0.8401562 |
| YLIGSVIV | ASSMTGENTLY | 0.8512838 |
| YGQLVAVID | ASSMTGENTLY | 0.4727016 |
| VTVGGMTLPA | ASSMTGENTLY | 0.8335054 |
| VLALPAV | ASSMTGENTLY | 0.81577563 |
| VIPPTLNYET | ASSMTGENTLY | 0.508783 |
| VGMGGMHPSSG | ASSMTGENTLY | 0.689955 |
| VGLAGGAATA | ASSMTGENTLY | 0.28297 |
| VGGAGGAGGSGP | ASSMTGENTLY | 0.571344 |
| VFAAFMV | ASSMTGENTLY | 0.38819322 |
| TLYDKLR | ASSMTGENTLY | 0.79097754 |
| TLAVFTVL | ASSMTGENTLY | 0.8286968 |
| TGSVQPARV | ASSMTGENTLY | 0.6564012 |
| TGGGAGSGGT | ASSMTGENTLY | 0.3528148 |
| SHLETSL | ASSMTGENTLY | 0.2774273 |
| SFTTGMT | ASSMTGENTLY | 0.73528254 |

| | | |
|---|---|---|
| QVEVAKRMGVRQPTVS | ASSMTGENTLY | 0.44067693 |
| QGSGGPALAM | ASSMTGENTLY | 0.29593122 |
| MLAPSNP | ASSMTGENTLY | 0.53926885 |
| LVNNAAI | ASSMTGENTLY | 0.8118398 |
| LTGEINTLA | ASSMTGENTLY | 0.80913794 |
| LSIFGPV | ASSMTGENTLY | 0.43591157 |
| LLALLANL | ASSMTGENTLY | 0.8553592 |
| LISNASDALDKLR | ASSMTGENTLY | 0.82597625 |
| LINEAVP | ASSMTGENTLY | 0.85913277 |
| LGYAFSGL | ASSMTGENTLY | 0.6283971 |
| LDFVAVAL | ASSMTGENTLY | 0.2877677 |
| LAQPGAE | ASSMTGENTLY | 0.84129965 |
| ISNLGAVTGF | ASSMTGENTLY | 0.69316095 |
| HQEDPGRI | ASSMTGENTLY | 0.78080976 |
| GTPGEPTP | ASSMTGENTLY | 0.76500076 |
| GFFNSGS | ASSMTGENTLY | 0.55892485 |
| GAGGGGLTAGAN | ASSMTGENTLY | 0.56669724 |
| FVDALVNM | ASSMTGENTLY | 0.76039374 |
| FLTVAPEL | ASSMTGENTLY | 0.82641137 |
| FETLTAL | ASSMTGENTLY | 0.40851682 |
| DVINPIEL | ASSMTGENTLY | 0.32631135 |
| APVSGAPV | ASSMTGENTLY | 0.69026166 |
| ALVGGLAW | ASSMTGENTLY | 0.70859164 |
| ALAAAAATAI | ASSMTGENTLY | 0.8412522 |
| AFGLGLLL | ASSMTGENTLY | 0.6919597 |
| AFFAAASL | ASSMTGENTLY | 0.59803486 |
| AAAALPAFSPP | ASSMTGENTLY | 0.7837509 |
| VVLAPAL | ASSMTGENTLY | 0.78503525 |
| TIGHVDHGKTTL | ASSMTGENTLY | 0.72446257 |
| LISNASDALDK | ASSMTGENTLY | 0.7928229 |
| LISNASDAL | ASSMTGENTLY | 0.5464543 |
| ISNASDALDK | ASSMTGENTLY | 0.8205448 |
| NQAETLVYQTE | ASSMTGENTLY | 0.80468875 |
| TGGTGGTAGAG | ASSMTGENTLY | 0.76978713 |
| TGSTAGGAGGA | ASSMTGENTLY | 0.8341893 |
| DPLAVPVAAGPV | ASSMTGENTLY | 0.74952626 |
| GGTGGTPGNGGN | ASSMTGENTLY | 0.68517363 |
| PAGLAGVDPATL | ASSMTGENTLY | 0.31635624 |
| GNNLVVEAPHAGASV | ASSMTGENTLY | 0.5957595 |
| KVTRIMSASTLEQPA | ASSMTGENTLY | 0.69006294 |
| LGFALPIALPRRTVV | ASSMTGENTLY | 0.76660275 |
| TRLGNLTARKLVVLV | ASSMTGENTLY | 0.8091527 |
| WSQQRRLIYFPSAGP | ASSMTGENTLY | 0.75457335 |
| GPNGNPDPMAAAVDIR | ASSMTGENTLY | 0.49233225 |
| TQRCVSVADLRAAASR | ASSMTGENTLY | 0.84183013 |
| QFVLDVAEHPVEVLTETV | ASSMTGENTLY | 0.8342175 |
| GGAGGTGGAGGNGGTQAGDG | ASSMTGENTLY | 0.35041592 |
| PLLATLRAIAVDVDAKPAQV | ASSMTGENTLY | 0.64071745 |
| APKFRTGGYMIDFWGVGYQVAK | ASSMTGENTLY | 0.8528982 |
| GAGGAGGAGGAAGGPGGTGGTG | ASSMTGENTLY | 0.7722496 |
| IGALGLLATDVAPAGSVAFGVA | ASSMTGENTLY | 0.51972246 |

| | | |
|---|---|---|
| AGGAGGIGGNGGAGGAGGVLMG | ASSMTGENTLY | 0.60132086 |
| ADELIGLL | ASSMTGENTLY | 0.8551753 |
| ADRALKA | ASSMTGENTLY | 0.7960895 |
| AGAVGQVGGAGGA | ASSMTGENTLY | 0.84845036 |
| AGGGGGLGGAP | ASSMTGENTLY | 0.19125673 |
| AGGLLYGNGGAG | ASSMTGENTLY | 0.54091 |
| AGLESIGL | ASSMTGENTLY | 0.45139724 |
| AGLKTGII | ASSMTGENTLY | 0.16447413 |
| ASLAALLL | ASSMTGENTLY | 0.81437397 |
| ASVINISSI | ASSMTGENTLY | 0.3841651 |
| ATPPGPA | ASSMTGENTLY | 0.7676343 |
| ATVASALV | ASSMTGENTLY | 0.5005137 |
| ATVASLIL | ASSMTGENTLY | 0.8301813 |
| AVNGPVLIRH | ASSMTGENTLY | 0.76322263 |
| DGAAGAVGI | ASSMTGENTLY | 0.5513662 |
| DPLLHGI | ASSMTGENTLY | 0.33889747 |
| ELGAALGAL | ASSMTGENTLY | 0.75204 |
| ELGRTQAV | ASSMTGENTLY | 0.84696853 |
| FVAGGYLS | ASSMTGENTLY | 0.8182107 |
| GASGGAGGAGGW | ASSMTGENTLY | 0.22117439 |
| GGAGAGGDGGAGGI | ASSMTGENTLY | 0.79286695 |
| GTGHNGSQPGMGGNG | ASSMTGENTLY | 0.6583502 |
| GYAAASASAS | ASSMTGENTLY | 0.2185978 |
| HQVDVAAVQQG | ASSMTGENTLY | 0.2038588 |
| IGAAIAHT | ASSMTGENTLY | 0.48851967 |
| IGMIEVAQL | ASSMTGENTLY | 0.68241423 |
| IRLAPEHTSAMD | ASSMTGENTLY | 0.77380943 |
| IYNVSVL | ASSMTGENTLY | 0.7905363 |
| KQGFLGM | ASSMTGENTLY | 0.57122564 |
| LESVITQL | ASSMTGENTLY | 0.7876149 |
| LGLGALVVV | ASSMTGENTLY | 0.7091584 |
| LIGNGGNGGSGGI | ASSMTGENTLY | 0.11140008 |
| LNAINAPTL | ASSMTGENTLY | 0.46032205 |
| LQLLNTGV | ASSMTGENTLY | 0.23451245 |
| LQTSIEF | ASSMTGENTLY | 0.8416835 |
| MSFAVSEL | ASSMTGENTLY | 0.46228588 |
| NTGALGL | ASSMTGENTLY | 0.51385725 |
| PPGAGGAGGAGTT | ASSMTGENTLY | 0.6627997 |
| QLVGLGI | ASSMTGENTLY | 0.8059572 |
| RFLEQVHQL | ASSMTGENTLY | 0.700306 |
| RGVGYVLR | ASSMTGENTLY | 0.4192028 |
| SATGVFAGVI | ASSMTGENTLY | 0.258408 |
| SLPAITLP | ASSMTGENTLY | 0.74658245 |
| STALAFTV | ASSMTGENTLY | 0.45279336 |
| SVLTVGLGK | ASSMTGENTLY | 0.37337077 |
| TALAGAGSGI | ASSMTGENTLY | 0.20433857 |
| TGLPFTGI | ASSMTGENTLY | 0.78350616 |
| VAEHVTV | ASSMTGENTLY | 0.2080963 |
| VDIVLNSL | ASSMTGENTLY | 0.81608665 |
| VLTGGGAVL | ASSMTGENTLY | 0.39477992 |
| VNVGLAYGI | ASSMTGENTLY | 0.63677573 |

| | | |
|---|---|---|
| VSSAAVGAV | ASSMTGENTLY | 0.86030203 |
| VTSVYEAGV | ASSMTGENTLY | 0.7896744 |
| WVALDRGA | ASSMTGENTLY | 0.6272418 |
| YLVGSVVI | ASSMTGENTLY | 0.34456402 |
| GPPGTGKTLL | ASSMTGENTLY | 0.32927346 |
| GVGMTKF | ASSMTGENTLY | 0.38107407 |
| IGIGGSDL | ASSMTGENTLY | 0.42547855 |
| IGTLLAEL | ASSMTGENTLY | 0.8311362 |
| IYTSGTT | ASSMTGENTLY | 0.64605206 |
| LSFPAGV | ASSMTGENTLY | 0.21956706 |
| LYTSGTT | ASSMTGENTLY | 0.76902467 |
| VGAFPFGF | ASSMTGENTLY | 0.8468387 |
| VGAGTVEF | ASSMTGENTLY | 0.86626214 |
| VGKSTLLL | ASSMTGENTLY | 0.37838003 |
| AALAHAGGGIV | ASSMTGENTLY | 0.5498398 |
| AALVAVSLL | ASSMTGENTLY | 0.6160256 |
| AAVSRAVAAL | ASSMTGENTLY | 0.72959733 |
| AGGSSGIGGF | ASSMTGENTLY | 0.40252602 |
| AGLTVANANG | ASSMTGENTLY | 0.38756162 |
| AGLVGAVL | ASSMTGENTLY | 0.78897846 |
| ALSQAFTV | ASSMTGENTLY | 0.87370634 |
| ALTFSVSL | ASSMTGENTLY | 0.6152213 |
| ALTGAALVAL | ASSMTGENTLY | 0.173779 |
| ASFSIGAGVL | ASSMTGENTLY | 0.6322073 |
| DGSVYKI | ASSMTGENTLY | 0.4818439 |
| EQLAEIV | ASSMTGENTLY | 0.50935876 |
| FAVSELTVM | ASSMTGENTLY | 0.67297447 |
| GAAPLAAGL | ASSMTGENTLY | 0.27692026 |
| GADGGDGGSGGAGG | ASSMTGENTLY | 0.6302955 |
| GFGLDTGGAGG | ASSMTGENTLY | 0.86480296 |
| GLASGGLL | ASSMTGENTLY | 0.6292067 |
| GRIAGAAVLL | ASSMTGENTLY | 0.7105143 |
| GSAGSLNTSF | ASSMTGENTLY | 0.84676623 |
| GTGQIGI | ASSMTGENTLY | 0.8025491 |
| GTLIALIL | ASSMTGENTLY | 0.8532671 |
| IAATMGFVVG | ASSMTGENTLY | 0.8369343 |
| IFAQALL | ASSMTGENTLY | 0.5422564 |
| IGAAVALGF | ASSMTGENTLY | 0.67663777 |
| IGLTPLAL | ASSMTGENTLY | 0.83110094 |
| ILGSNTSTL | ASSMTGENTLY | 0.54734415 |
| IVFSLTVL | ASSMTGENTLY | 0.65627986 |
| LASKATGF | ASSMTGENTLY | 0.51459944 |
| LASVALIL | ASSMTGENTLY | 0.7971233 |
| LFAVGLTL | ASSMTGENTLY | 0.615534 |
| LFNGGIGTYI | ASSMTGENTLY | 0.8536991 |
| LTLGAGII | ASSMTGENTLY | 0.17219536 |
| LVAAALGDI | ASSMTGENTLY | 0.32789385 |
| NTAFVQLGI | ASSMTGENTLY | 0.84435385 |
| QLFSGAEL | ASSMTGENTLY | 0.1257579 |
| SAATFITL | ASSMTGENTLY | 0.21786849 |
| SNYGLANLGV | ASSMTGENTLY | 0.60670555 |

| | | |
|---|---|---|
| SYGGAYAVM | ASSMTGENTLY | 0.84882843 |
| TTGVGGLTLG | ASSMTGENTLY | 0.136374 |
| TTQTIYI | ASSMTGENTLY | 0.45435804 |
| VAAGAGLGAVI | ASSMTGENTLY | 0.23847055 |
| VAAVVTLL | ASSMTGENTLY | 0.8295553 |
| VGFLGGLI | ASSMTGENTLY | 0.6801091 |
| VGVIPILL | ASSMTGENTLY | 0.26640615 |
| VIFASSLI | ASSMTGENTLY | 0.81865156 |
| VKGGDGGAAAG | ASSMTGENTLY | 0.19731371 |
| VTKTFTNI | ASSMTGENTLY | 0.74328876 |
| VVAAAILL | ASSMTGENTLY | 0.80740535 |
| YAALDVEL | ASSMTGENTLY | 0.8652423 |
| LASVALLL | ASSMTGENTLY | 0.29073852 |
| LDAVHRV | ASSMTGENTLY | 0.8103777 |
| QGLSSSIF | ASSMTGENTLY | 0.5440631 |
| RGTFANIRL | ASSMTGENTLY | 0.8769526 |
| RQLWWGHRI | ASSMTGENTLY | 0.5392198 |
| YGPPGTGKTLL | ASSMTGENTLY | 0.2554012 |
| AALVGAPANVA | ASSMTGENTLY | 0.8237104 |
| AATFRRTE | ASSMTGENTLY | 0.8056546 |
| AGALGGGI | ASSMTGENTLY | 0.6704655 |
| AGGAGGAGGIGAG | ASSMTGENTLY | 0.8399665 |
| AGSGALGL | ASSMTGENTLY | 0.67766654 |
| AQIADQI | ASSMTGENTLY | 0.6286138 |
| ASAVAVTYL | ASSMTGENTLY | 0.79483116 |
| ASTVLGGTGG | ASSMTGENTLY | 0.78743374 |
| AVGGEPAK | ASSMTGENTLY | 0.41470945 |
| CEQGTFGGTC | ASSMTGENTLY | 0.8391607 |
| DEGKAPAG | ASSMTGENTLY | 0.6783714 |
| DTVVFPAL | ASSMTGENTLY | 0.29621917 |
| FEGITMAG | ASSMTGENTLY | 0.8672679 |
| FPLVVQD | ASSMTGENTLY | 0.6674441 |
| GAFTTPQ | ASSMTGENTLY | 0.8034183 |
| GFGGFGTAGGVGGA | ASSMTGENTLY | 0.762744 |
| GFLAQFL | ASSMTGENTLY | 0.86915565 |
| GGAGGAGGNAGVA | ASSMTGENTLY | 0.37759337 |
| GGVGTAGAGGAGGAGG | ASSMTGENTLY | 0.6738373 |
| GNGGGGDAGNAGSGGN | ASSMTGENTLY | 0.7187412 |
| GTGGAGGAGNDAGS | ASSMTGENTLY | 0.8051577 |
| GTTNTGWFNS | ASSMTGENTLY | 0.68661034 |
| IFMGPLFTL | ASSMTGENTLY | 0.6569214 |
| ILAHAIEAI | ASSMTGENTLY | 0.7589668 |
| IRSAIREAL | ASSMTGENTLY | 0.82408863 |
| ITFAGGG | ASSMTGENTLY | 0.62201333 |
| ITGMSGHQEV | ASSMTGENTLY | 0.7423874 |
| IVAGLACTL | ASSMTGENTLY | 0.86126006 |
| IVGGGLAGA | ASSMTGENTLY | 0.2908249 |
| IVTGVLNI | ASSMTGENTLY | 0.65932107 |
| LAQQGCI | ASSMTGENTLY | 0.7659025 |
| LCLHVEN | ASSMTGENTLY | 0.8331711 |
| LDAISDGS | ASSMTGENTLY | 0.6400815 |

| | | |
|---|---|---|
| LGGFAAML | ASSMTGENTLY | 0.8224975 |
| LGGKNPMI | ASSMTGENTLY | 0.63616294 |
| LGPKGVAAIL | ASSMTGENTLY | 0.7352797 |
| LMVGVSI | ASSMTGENTLY | 0.73077816 |
| LVGAAFVL | ASSMTGENTLY | 0.61073065 |
| LVLAVAPL | ASSMTGENTLY | 0.35015452 |
| MAAIAVIDAA | ASSMTGENTLY | 0.7230418 |
| NNGINAELQ | ASSMTGENTLY | 0.81501794 |
| RGEPLVLVHG | ASSMTGENTLY | 0.7447681 |
| RGGAGGAAGLI | ASSMTGENTLY | 0.15506741 |
| RSYRMHPAV | ASSMTGENTLY | 0.81000024 |
| SAAHLLAGQLA | ASSMTGENTLY | 0.72776383 |
| SAAVGSY | ASSMTGENTLY | 0.8732358 |
| SPHAVLHRI | ASSMTGENTLY | 0.65193546 |
| SQIAKVL | ASSMTGENTLY | 0.8164312 |
| TALGHLVP | ASSMTGENTLY | 0.44087923 |
| TAPTTSGAA | ASSMTGENTLY | 0.72429943 |
| TDGDDGNGGSPGT | ASSMTGENTLY | 0.3083219 |
| TDGLGEN | ASSMTGENTLY | 0.45009622 |
| TGGPPLPGQA | ASSMTGENTLY | 0.6891432 |
| TIPAITI | ASSMTGENTLY | 0.5070418 |
| TIPGFTI | ASSMTGENTLY | 0.8345562 |
| TSIASGALA | ASSMTGENTLY | 0.8362393 |
| TVTLPGAAI | ASSMTGENTLY | 0.8336887 |
| VASGLSAALP | ASSMTGENTLY | 0.8324853 |
| VGAGVIGI | ASSMTGENTLY | 0.85874414 |
| VGPKRFGV | ASSMTGENTLY | 0.82519335 |
| VNVAFPD | ASSMTGENTLY | 0.503441 |
| VSMTNSGVS | ASSMTGENTLY | 0.5730618 |
| VTDVLGDLSV | ASSMTGENTLY | 0.30594206 |
| VVVVGAGFAGL | ASSMTGENTLY | 0.56723106 |
| VYTVGTVV | ASSMTGENTLY | 0.56676173 |
| WTTFSGP | ASSMTGENTLY | 0.71548116 |
| YDELVEAF | ASSMTGENTLY | 0.126605 |
| YGTGGAGGNGG | ASSMTGENTLY | 0.8178882 |
| YTNVMAV | ASSMTGENTLY | 0.83589804 |
| FLEQVHQL | ASSMTGENTLY | 0.70000434 |
| GLSLLAVAV | ASSMTGENTLY | 0.73619807 |
| LEQVHQL | ASSMTGENTLY | 0.8075113 |
| LNAAGAI | ASSMTGENTLY | 0.5598823 |
| PAVHASL | ASSMTGENTLY | 0.76800716 |
| TELEDDA | ASSMTGENTLY | 0.5333204 |
| TPPGPPN | ASSMTGENTLY | 0.82293904 |
| TPPPPPPP | ASSMTGENTLY | 0.84493256 |
| TQSPALI | ASSDGGGGEQY | 0.7292669 |
| EFAAAFPQ | ASSDGGGGEQY | 0.7212311 |
| LLSGNLDVL | ASSDGGGGEQY | 0.73577595 |
| GPANLGNYNI | ASSDGGGGEQY | 0.25935552 |
| TLGPTLP | ASSDGGGGEQY | 0.7404543 |
| VYAYSSCCK | ASSDGGGGEQY | 0.31712806 |
| APAPAPAPNI | ASSDGGGGEQY | 0.70111823 |

| | | |
|---|---|---|
| PGGPVPIP | ASSDGGGGEQY | 0.3951794 |
| FLTVALEP | ASSDGGGGEQY | 0.75140464 |
| SQYFGANL | ASSDGGGGEQY | 0.6863076 |
| GGVKPAA | ASSDGGGGEQY | 0.6978556 |
| GGTGGAGGAGGW | ASSDGGGGEQY | 0.3413752 |
| GVGGVAGLGGAGG | ASSDGGGGEQY | 0.8431294 |
| VLLVLATTI | ASSDGGGGEQY | 0.7829691 |
| IGIFAAML | ASSDGGGGEQY | 0.2943235 |
| AVGDLID | ASSDGGGGEQY | 0.8177308 |
| NISPLPL | ASSDGGGGEQY | 0.50589347 |
| VTAVFVAI | ASSDGGGGEQY | 0.72815883 |
| SGDDNDAAL | ASSDGGGGEQY | 0.81736416 |
| AVAALAHS | ASSDGGGGEQY | 0.72387576 |
| GGVTVMATP | ASSDGGGGEQY | 0.22316387 |
| AGLAANVRD | ASSDGGGGEQY | 0.6456666 |
| TNRPDLIDPALLRPG | ASSDGGGGEQY | 0.6590363 |
| LANLGVTAL | ASSDGGGGEQY | 0.6221117 |
| TGERIEL | ASSDGGGGEQY | 0.15778421 |
| NGEELAA | ASSDGGGGEQY | 0.73590237 |
| VLSLVWG | ASSDGGGGEQY | 0.2907921 |
| IGYALWL | ASSDGGGGEQY | 0.5329016 |
| LSLLLAPL | ASSDGGGGEQY | 0.60852146 |
| TLDEALK | ASSDGGGGEQY | 0.82667804 |
| TDGAGTAEGG | ASSDGGGGEQY | 0.82403713 |
| NAPGGAGGNGG | ASSDGGGGEQY | 0.5390352 |
| VSVFMGMGGNF | ASSDGGGGEQY | 0.7890597 |
| LASYLAAL | ASSDGGGGEQY | 0.65429205 |
| LVGGTPAL | ASSDGGGGEQY | 0.8432843 |
| VRNADVL | ASSDGGGGEQY | 0.6233121 |
| TCSSPFLVSAM | ASSDGGGGEQY | 0.6221669 |
| VHITDAE | ASSDGGGGEQY | 0.69556797 |
| EYAAPSF | ASSDGGGGEQY | 0.45139214 |
| VVDAFHGGL | ASSDGGGGEQY | 0.52065635 |
| ASVVDPQ | ASSDGGGGEQY | 0.4787929 |
| SYNEFGGLS | ASSDGGGGEQY | 0.4563091 |
| DPDGSVY | ASSDGGGGEQY | 0.2663491 |
| MVSTIHATE | ASSDGGGGEQY | 0.68470097 |
| LAAGVADPV | ASSDGGGGEQY | 0.8370559 |
| TKEQIAA | ASSDGGGGEQY | 0.7465686 |
| SLASPAAL | ASSDGGGGEQY | 0.21029592 |
| IGLVFYGV | ASSDGGGGEQY | 0.5401883 |
| FLLDASNL | ASSDGGGGEQY | 0.6176816 |
| YQVGNFP | ASSDGGGGEQY | 0.8025086 |
| ESLLSQST | ASSDGGGGEQY | 0.82547206 |
| LFTVGFAAVG | ASSDGGGGEQY | 0.86152416 |
| ATPPTPT | ASSDGGGGEQY | 0.7981342 |
| TDLATLG | ASSDGGGGEQY | 0.5458579 |
| SAGGTGR | ASSDGGGGEQY | 0.7973129 |
| GFSNSGS | ASSDGGGGEQY | 0.6225795 |
| GGNGGSGGTSV | ASSDGGGGEQY | 0.6039468 |
| GTSVATGGAGN | ASSDGGGGEQY | 0.60173595 |

| | | |
|---|---|---|
| NGGNGGSGGTS | ASSDGGGGEQY | 0.78880167 |
| AVVKFPH | ASSDGGGGEQY | 0.5026725 |
| TPPPPPP | ASSDGGGGEQY | 0.4426314 |
| AAPAAAAAPA | ASSDGGGGEQY | 0.19368811 |
| IALNEDL | ASSDGGGGEQY | 0.69982684 |
| IVNVSSM | ASSDGGGGEQY | 0.49779752 |
| TGIPPAPR | ASSDGGGGEQY | 0.47716942 |
| FDLGGGTF | ASSDGGGGEQY | 0.42097923 |
| FDLGGGTFDVS | ASSDGGGGEQY | 0.8600918 |
| NEPTAAA | ASSDGGGGEQY | 0.8506677 |
| NEPTAAALA | ASSDGGGGEQY | 0.58561873 |
| GSIINISSI | ASSDGGGGEQY | 0.43281078 |
| VAPGNAGTA | ASSDGGGGEQY | 0.7990622 |
| TGGGASLE | ASSDGGGGEQY | 0.43471962 |
| VAVYNHYKRI | ASSDGGGGEQY | 0.7797462 |
| TDPAPATT | ASSDGGGGEQY | 0.75274205 |
| TNRPDILDPALLRPG | ASSDGGGGEQY | 0.73981845 |
| LTTTTTS | ASSDGGGGEQY | 0.5304018 |
| PTGSGKTL | ASSDGGGGEQY | 0.79062617 |
| TGSGKTL | ASSDGGGGEQY | 0.5967475 |
| TIGHVDHGK | ASSDGGGGEQY | 0.7745573 |
| TIGHVDHGKTT | ASSDGGGGEQY | 0.15418577 |
| GHVDHGKTT | ASSDGGGGEQY | 0.81489503 |
| AQFEHTLL | ASSDGGGGEQY | 0.38357985 |
| IVNQDEL | ASSDGGGGEQY | 0.38633996 |
| TNGSQFF | ASSDGGGGEQY | 0.17775492 |
| TGKGIVE | ASSDGGGGEQY | 0.5148005 |
| LISNASDA | ASSDGGGGEQY | 0.8149216 |
| AELRTLGELTKIAKAHGAQ | ASSDGGGGEQY | 0.84050536 |
| APNAVAAAEPSW | ASSDGGGGEQY | 0.66789585 |
| APQVAGAATAVL | ASSDGGGGEQY | 0.79296255 |
| ARRDPAAAAAILPTDA | ASSDGGGGEQY | 0.86042726 |
| DAAARDVLAEAGLAEYF | ASSDGGGGEQY | 0.5935801 |
| DDQIVLSAKANLAKVEQ | ASSDGGGGEQY | 0.18546018 |
| DELLPDTGEDSDEDGH | ASSDGGGGEQY | 0.76398027 |
| ELPEAPANRATVPQD | ASSDGGGGEQY | 0.62937224 |
| FGQSKIERVTGIG | ASSDGGGGEQY | 0.85236824 |
| GAGGTGSTAGGAGGAGGAGGLY | ASSDGGGGEQY | 0.78237504 |
| GHGGTGAAGGNGGTGGAGGSGI | ASSDGGGGEQY | 0.31488144 |
| GNGGNGGAGGEGGPGGAGGAGA | ASSDGGGGEQY | 0.8401121 |
| GSTAGNGGNGGNATNGTIAGQP | ASSDGGGGEQY | 0.78485954 |
| GTGGDGGNGGNGGTGG | ASSDGGGGEQY | 0.36090326 |
| GVRRPAGIPADAPV | ASSDGGGGEQY | 0.31205845 |
| GYRQVTDAHLLTLARR | ASSDGGGGEQY | 0.3224224 |
| IAKMGQDGHDRG | ASSDGGGGEQY | 0.31959152 |
| KAEPSIEIQYT | ASSDGGGGEQY | 0.7312889 |
| KPVRIRMAADEDTVAVT | ASSDGGGGEQY | 0.56425524 |
| MPPRIAGMRLLVIKPEP | ASSDGGGGEQY | 0.591811 |
| MKFYAVALFRSRGVPPTR | ASSDGGGGEQY | 0.6530865 |
| NAGGFGILGAVAHSP | ASSDGGGGEQY | 0.465048 |
| NEGREITLAMAEM | ASSDGGGGEQY | 0.47518557 |

| | | |
|---|---|---|
| NELVEWSHPLGHRWRWE | ASSDGGGGEQY | 0.56193835 |
| NLGAGGTGGNGGNPG | ASSDGGGGEQY | 0.6744994 |
| NTEWPKWETVMAADPVD | ASSDGGGGEQY | 0.8081965 |
| PASAAPAETQP | ASSDGGGGEQY | 0.6707715 |
| QQSSDGKNGVRAAEITGE | ASSDGGGGEQY | 0.64607203 |
| RAPDAVTTSWWKE | ASSDGGGGEQY | 0.84198475 |
| SGFGNVGAHTSGWFNQ | ASSDGGGGEQY | 0.31978694 |
| SGVAGAVGQAASVGGLKV | ASSDGGGGEQY | 0.4817886 |
| SNGGAGTGSAGGNGGT | ASSDGGGGEQY | 0.80527973 |
| TGGAGGEGGAAG | ASSDGGGGEQY | 0.62412524 |
| TGGDGGAAGAGGAPGHG | ASSDGGGGEQY | 0.8417495 |
| TMPPLLAAIANAGRHGV | ASSDGGGGEQY | 0.6877171 |
| TSSTAGNSGGA | ASSDGGGGEQY | 0.75073177 |
| VPATVPVGILC | ASSDGGGGEQY | 0.6981621 |
| VQRPDSGGVVLTGRIS | ASSDGGGGEQY | 0.86371654 |
| WTPNPVPATKGVPAGLVQAT | ASSDGGGGEQY | 0.780076 |
| YALKRALAKFSDRFDVVII | ASSDGGGGEQY | 0.50419915 |
| YLIGSVIV | ASSDGGGGEQY | 0.84595585 |
| YGQLVAVID | ASSDGGGGEQY | 0.2350116 |
| VTVGGMTLPA | ASSDGGGGEQY | 0.7879525 |
| VLALPAV | ASSDGGGGEQY | 0.68431056 |
| VIPPTLNYET | ASSDGGGGEQY | 0.4898659 |
| VGMGGMHPSSG | ASSDGGGGEQY | 0.6300068 |
| VGLAGGAATA | ASSDGGGGEQY | 0.8416548 |
| VGGAGGAGGSGP | ASSDGGGGEQY | 0.82584226 |
| VFAAFMV | ASSDGGGGEQY | 0.48856002 |
| TLYDKLR | ASSDGGGGEQY | 0.5359335 |
| TLAVFTVL | ASSDGGGGEQY | 0.3052162 |
| TGSVQPARV | ASSDGGGGEQY | 0.49539736 |
| TGGGAGSGGT | ASSDGGGGEQY | 0.20843016 |
| SHLETSL | ASSDGGGGEQY | 0.7550774 |
| SFTTGMT | ASSDGGGGEQY | 0.49060783 |
| QVEVAKRMGVRQPTVS | ASSDGGGGEQY | 0.8381998 |
| QGSGGPALAM | ASSDGGGGEQY | 0.7050623 |
| MLAPSNP | ASSDGGGGEQY | 0.8232175 |
| LVNNAAI | ASSDGGGGEQY | 0.44183683 |
| LTGEINTLA | ASSDGGGGEQY | 0.4597138 |
| LSIFGPV | ASSDGGGGEQY | 0.19877197 |
| LLALLANL | ASSDGGGGEQY | 0.518208 |
| LISNASDALDKLR | ASSDGGGGEQY | 0.39746356 |
| LINEAVP | ASSDGGGGEQY | 0.60149956 |
| LGYAFSGL | ASSDGGGGEQY | 0.21235827 |
| LDFVAVAL | ASSDGGGGEQY | 0.8601298 |
| LAQPGAE | ASSDGGGGEQY | 0.67782235 |
| ISNLGAVTGF | ASSDGGGGEQY | 0.76116556 |
| HQEDPGRI | ASSDGGGGEQY | 0.4212147 |
| GTPGEPTP | ASSDGGGGEQY | 0.35165316 |
| GFFNSGS | ASSDGGGGEQY | 0.5744668 |
| GAGGGGLTAGAN | ASSDGGGGEQY | 0.8010366 |
| FVDALVNM | ASSDGGGGEQY | 0.40391076 |
| FLTVAPEL | ASSDGGGGEQY | 0.15032157 |

| | | |
|---|---|---|
| FETLTAL | ASSDGGGGEQY | 0.36289626 |
| DVINPIEL | ASSDGGGGEQY | 0.2500044 |
| APVSGAPV | ASSDGGGGEQY | 0.87017965 |
| ALVGGLAW | ASSDGGGGEQY | 0.8302504 |
| ALAAAAATAI | ASSDGGGGEQY | 0.7347271 |
| AFGLGLLL | ASSDGGGGEQY | 0.8571855 |
| AFFAAASL | ASSDGGGGEQY | 0.12790442 |
| AAAALPAFSPP | ASSDGGGGEQY | 0.55925155 |
| VVLAPAL | ASSDGGGGEQY | 0.85430473 |
| TIGHVDHGKTTL | ASSDGGGGEQY | 0.7658341 |
| LISNASDALDK | ASSDGGGGEQY | 0.55038655 |
| LISNASDAL | ASSDGGGGEQY | 0.83299804 |
| ISNASDALDK | ASSDGGGGEQY | 0.45770085 |
| NQAETLVYQTE | ASSDGGGGEQY | 0.60333914 |
| TGGTGGTAGAG | ASSDGGGGEQY | 0.6931304 |
| TGSTAGGAGGA | ASSDGGGGEQY | 0.71978426 |
| DPLAVPVAAGPV | ASSDGGGGEQY | 0.7484325 |
| GGTGGTPGNGGN | ASSDGGGGEQY | 0.20928507 |
| PAGLAGVDPATL | ASSDGGGGEQY | 0.7712509 |
| GNNLVVEAPHAGASV | ASSDGGGGEQY | 0.8668976 |
| KVTRIMSASTLEQPA | ASSDGGGGEQY | 0.62878853 |
| LGFALPIALPRRTVV | ASSDGGGGEQY | 0.79957324 |
| TRLGNLTARKLVVLV | ASSDGGGGEQY | 0.7583695 |
| WSQQRRLIYFPSAGP | ASSDGGGGEQY | 0.7841094 |
| GPNGNPDPMAAAVDIR | ASSDGGGGEQY | 0.441468 |
| TQRCVSVADLRAAASR | ASSDGGGGEQY | 0.44662604 |
| QFVLDVAEHPVEVLTETV | ASSDGGGGEQY | 0.8381503 |
| GGAGGTGGAGGNGGTQAGDG | ASSDGGGGEQY | 0.49068376 |
| PLLATLRAIAVDVDAKPAQV | ASSDGGGGEQY | 0.5884508 |
| APKFRTGGYMIDFWGVGYQVAK | ASSDGGGGEQY | 0.64433026 |
| GAGGAGGAGGAAGGPGGTGGTG | ASSDGGGGEQY | 0.87700355 |
| IGALGLLATDVAPAGSVAFGVA | ASSDGGGGEQY | 0.7501222 |
| AGGAGGIGGNGGAGGAGGVLMG | ASSDGGGGEQY | 0.81464523 |
| ADELIGLL | ASSDGGGGEQY | 0.45886838 |
| ADRALKA | ASSDGGGGEQY | 0.7136874 |
| AGAVGQVGGAGGA | ASSDGGGGEQY | 0.8147038 |
| AGGGGGLGGAP | ASSDGGGGEQY | 0.22037704 |
| AGGLLYGNGGAG | ASSDGGGGEQY | 0.8261841 |
| AGLESIGL | ASSDGGGGEQY | 0.4165161 |
| AGLKTGII | ASSDGGGGEQY | 0.7624661 |
| ASLAALLL | ASSDGGGGEQY | 0.7188579 |
| ASVINISSI | ASSDGGGGEQY | 0.8484786 |
| ATPPGPA | ASSDGGGGEQY | 0.64219415 |
| ATVASALV | ASSDGGGGEQY | 0.83291185 |
| ATVASLIL | ASSDGGGGEQY | 0.36891568 |
| AVNGPVLIRH | ASSDGGGGEQY | 0.79193777 |
| DGAAGAVGI | ASSDGGGGEQY | 0.8360809 |
| DPLLHGI | ASSDGGGGEQY | 0.79659706 |
| ELGAALGAL | ASSDGGGGEQY | 0.86681414 |
| ELGRTQAV | ASSDGGGGEQY | 0.78540206 |
| FVAGGYLS | ASSDGGGGEQY | 0.18158183 |

| | | |
|---|---|---|
| GASGGAGGAGGW | ASSDGGGGEQY | 0.60462725 |
| GGAGAGGDGGAGGI | ASSDGGGGEQY | 0.8321987 |
| GTGHNGSQPGMGGNG | ASSDGGGGEQY | 0.8538096 |
| GYAAASASAS | ASSDGGGGEQY | 0.7948606 |
| HQVDVAAVQQG | ASSDGGGGEQY | 0.7734816 |
| IGAAIAHT | ASSDGGGGEQY | 0.63527954 |
| IGMIEVAQL | ASSDGGGGEQY | 0.8511872 |
| IRLAPEHTSAMD | ASSDGGGGEQY | 0.8802333 |
| IYNVSVL | ASSDGGGGEQY | 0.81457484 |
| KQGFLGM | ASSDGGGGEQY | 0.37117612 |
| LESVITQL | ASSDGGGGEQY | 0.35522228 |
| LGLGALVVV | ASSDGGGGEQY | 0.75878507 |
| LIGNGGNGGSGGI | ASSDGGGGEQY | 0.8509902 |
| LNAINAPTL | ASSDGGGGEQY | 0.26978466 |
| LQLLNTGV | ASSDGGGGEQY | 0.7440127 |
| LQTSIEF | ASSDGGGGEQY | 0.70397866 |
| MSFAVSEL | ASSDGGGGEQY | 0.15461755 |
| NTGALGL | ASSDGGGGEQY | 0.7489798 |
| PPGAGGAGGAGTT | ASSDGGGGEQY | 0.8495356 |
| QLVGLGI | ASSDGGGGEQY | 0.7401819 |
| RFLEQVHQL | ASSDGGGGEQY | 0.41565818 |
| RGVGYVLR | ASSDGGGGEQY | 0.8322912 |
| SATGVFAGVI | ASSDGGGGEQY | 0.85482025 |
| SLPAITLP | ASSDGGGGEQY | 0.8653182 |
| STALAFTV | ASSDGGGGEQY | 0.20887037 |
| SVLTVGLGK | ASSDGGGGEQY | 0.1348231 |
| TALAGAGSGI | ASSDGGGGEQY | 0.7060996 |
| TGLPFTGI | ASSDGGGGEQY | 0.8269007 |
| VAEHVTV | ASSDGGGGEQY | 0.51961714 |
| VDIVLNSL | ASSDGGGGEQY | 0.46872157 |
| VLTGGGAVL | ASSDGGGGEQY | 0.7858325 |
| VNVGLAYGI | ASSDGGGGEQY | 0.8379152 |
| VSSAAVGAV | ASSDGGGGEQY | 0.26654524 |
| VTSVYEAGV | ASSDGGGGEQY | 0.29101792 |
| WVALDRGA | ASSDGGGGEQY | 0.7535245 |
| YLVGSVVI | ASSDGGGGEQY | 0.7916663 |
| GPPGTGKTLL | ASSDGGGGEQY | 0.41036105 |
| GVGMTKF | ASSDGGGGEQY | 0.6470411 |
| IGIGGSDL | ASSDGGGGEQY | 0.7343353 |
| IGTLLAEL | ASSDGGGGEQY | 0.7722292 |
| IYTSGTT | ASSDGGGGEQY | 0.7679031 |
| LSFPAGV | ASSDGGGGEQY | 0.32159954 |
| LYTSGTT | ASSDGGGGEQY | 0.82002157 |
| VGAFPFGF | ASSDGGGGEQY | 0.26232082 |
| VGAGTVEF | ASSDGGGGEQY | 0.8028121 |
| VGKSTLLL | ASSDGGGGEQY | 0.6825346 |
| AALAHAGGGIV | ASSDGGGGEQY | 0.6836854 |
| AALVAVSLL | ASSDGGGGEQY | 0.5809344 |
| AAVSRAVAAL | ASSDGGGGEQY | 0.40029734 |
| AGGSSGIGGF | ASSDGGGGEQY | 0.38304144 |
| AGLTVANANG | ASSDGGGGEQY | 0.3683966 |

| | | |
|---|---|---|
| AGLVGAVL | ASSDGGGGEQY | 0.8398224 |
| ALSQAFTV | ASSDGGGGEQY | 0.81557846 |
| ALTFSVSL | ASSDGGGGEQY | 0.78918886 |
| ALTGAALVAL | ASSDGGGGEQY | 0.70745176 |
| ASFSIGAGVL | ASSDGGGGEQY | 0.33914995 |
| DGSVYKI | ASSDGGGGEQY | 0.7473471 |
| EQLAEIV | ASSDGGGGEQY | 0.35136306 |
| FAVSELTVM | ASSDGGGGEQY | 0.6472359 |
| GAAPLAAGL | ASSDGGGGEQY | 0.7279366 |
| GADGGDGGSGGAGG | ASSDGGGGEQY | 0.8162629 |
| GFGLDTGGAGG | ASSDGGGGEQY | 0.32340464 |
| GLASGGLL | ASSDGGGGEQY | 0.73355556 |
| GRIAGAAVLL | ASSDGGGGEQY | 0.7351567 |
| GSAGSLNTSF | ASSDGGGGEQY | 0.35029006 |
| GTGQIGI | ASSDGGGGEQY | 0.74134064 |
| GTLIALIL | ASSDGGGGEQY | 0.8068377 |
| IAATMGFVVG | ASSDGGGGEQY | 0.71290165 |
| IFAQALL | ASSDGGGGEQY | 0.27482957 |
| IGAAVALGF | ASSDGGGGEQY | 0.7777678 |
| IGLTPLAL | ASSDGGGGEQY | 0.53479433 |
| ILGSNTSTL | ASSDGGGGEQY | 0.7516344 |
| IVFSLTVL | ASSDGGGGEQY | 0.6218289 |
| LASKATGF | ASSDGGGGEQY | 0.47054023 |
| LASVALIL | ASSDGGGGEQY | 0.8064594 |
| LFAVGLTL | ASSDGGGGEQY | 0.30865163 |
| LFNGGIGTYI | ASSDGGGGEQY | 0.5502544 |
| LTLGAGII | ASSDGGGGEQY | 0.59699845 |
| LVAAALGDI | ASSDGGGGEQY | 0.629653 |
| NTAFVQLGI | ASSDGGGGEQY | 0.8176124 |
| QLFSGAEL | ASSDGGGGEQY | 0.38103956 |
| SAATFITL | ASSDGGGGEQY | 0.70394766 |
| SNYGLANLGV | ASSDGGGGEQY | 0.5559646 |
| SYGGAYAVM | ASSDGGGGEQY | 0.37920403 |
| TTGVGGLTLG | ASSDGGGGEQY | 0.66689694 |
| TTQTIYI | ASSDGGGGEQY | 0.5378095 |
| VAAGAGLGAVI | ASSDGGGGEQY | 0.3592419 |
| VAAVVTLL | ASSDGGGGEQY | 0.6799711 |
| VGFLGGLI | ASSDGGGGEQY | 0.37486902 |
| VGVIPILL | ASSDGGGGEQY | 0.7531966 |
| VIFASSLI | ASSDGGGGEQY | 0.8659613 |
| VKGGDGGAAAG | ASSDGGGGEQY | 0.35640368 |
| VTKTFTNI | ASSDGGGGEQY | 0.78500897 |
| VVAAAILL | ASSDGGGGEQY | 0.78077155 |
| YAALDVEL | ASSDGGGGEQY | 0.7136959 |
| LASVALLL | ASSDGGGGEQY | 0.5990078 |
| LDAVHRV | ASSDGGGGEQY | 0.38100463 |
| QGLSSSIF | ASSDGGGGEQY | 0.84740686 |
| RGTFANIRL | ASSDGGGGEQY | 0.7971759 |
| RQLWWGHRI | ASSDGGGGEQY | 0.7875048 |
| YGPPGTGKTLL | ASSDGGGGEQY | 0.7799253 |
| AALVGAPANVA | ASSDGGGGEQY | 0.7613803 |

| | | |
|---|---|---|
| AATFRRTE | ASSDGGGGEQY | 0.8302502 |
| AGALGGGI | ASSDGGGGEQY | 0.4775871 |
| AGGAGGAGGIGAG | ASSDGGGGEQY | 0.262811 |
| AGSGALGL | ASSDGGGGEQY | 0.50582814 |
| AQIADQI | ASSDGGGGEQY | 0.51021266 |
| ASAVAVTYL | ASSDGGGGEQY | 0.36497912 |
| ASTVLGGTGG | ASSDGGGGEQY | 0.24682987 |
| AVGGEPAK | ASSDGGGGEQY | 0.8114892 |
| CEQGTFGGTC | ASSDGGGGEQY | 0.8711135 |
| DEGKAPAG | ASSDGGGGEQY | 0.5592115 |
| DTVVFPAL | ASSDGGGGEQY | 0.8800995 |
| FEGITMAG | ASSDGGGGEQY | 0.34655166 |
| FPLVVQD | ASSDGGGGEQY | 0.8125802 |
| GAFTTPQ | ASSDGGGGEQY | 0.8089411 |
| GFGGFGTAGGVGGA | ASSDGGGGEQY | 0.5269846 |
| GFLAQFL | ASSDGGGGEQY | 0.7249555 |
| GGAGGAGGNAGVA | ASSDGGGGEQY | 0.34565207 |
| GGVGTAGAGGAGGAGG | ASSDGGGGEQY | 0.5642089 |
| GNGGGGDAGNAGSGGN | ASSDGGGGEQY | 0.82216465 |
| GTGGAGGAGNDAGS | ASSDGGGGEQY | 0.5344344 |
| GTTNTGWFNS | ASSDGGGGEQY | 0.7030306 |
| IFMGPLFTL | ASSDGGGGEQY | 0.59604317 |
| ILAHAIEAI | ASSDGGGGEQY | 0.42082614 |
| IRSAIREAL | ASSDGGGGEQY | 0.6570158 |
| ITFAGGG | ASSDGGGGEQY | 0.7280785 |
| ITGMSGHQEV | ASSDGGGGEQY | 0.72650015 |
| IVAGLACTL | ASSDGGGGEQY | 0.8796692 |
| IVGGGLAGA | ASSDGGGGEQY | 0.27739057 |
| IVTGVLNI | ASSDGGGGEQY | 0.83598614 |
| LAQQGCI | ASSDGGGGEQY | 0.84688926 |
| LCLHVEN | ASSDGGGGEQY | 0.7427456 |
| LDAISDGS | ASSDGGGGEQY | 0.49431717 |
| LGGFAAML | ASSDGGGGEQY | 0.8192032 |
| LGGKNPMI | ASSDGGGGEQY | 0.5481626 |
| LGPKGVAAIL | ASSDGGGGEQY | 0.79228723 |
| LMVGVSI | ASSDGGGGEQY | 0.8423192 |
| LVGAAFVL | ASSDGGGGEQY | 0.15503289 |
| LVLAVAPL | ASSDGGGGEQY | 0.76468605 |
| MAAIAVIDAA | ASSDGGGGEQY | 0.3687826 |
| NNGINAELQ | ASSDGGGGEQY | 0.36550102 |
| RGEPLVLVHG | ASSDGGGGEQY | 0.1985579 |
| RGGAGGAAGLI | ASSDGGGGEQY | 0.13709727 |
| RSYRMHPAV | ASSDGGGGEQY | 0.25893247 |
| SAAHLLAGQLA | ASSDGGGGEQY | 0.16712025 |
| SAAVGSY | ASSDGGGGEQY | 0.64823925 |
| SPHAVLHRI | ASSDGGGGEQY | 0.45584482 |
| SQIAKVL | ASSDGGGGEQY | 0.20733885 |
| TALGHLVP | ASSDGGGGEQY | 0.51171255 |
| TAPTTSGAA | ASSDGGGGEQY | 0.6580361 |
| TDGDDGNGGSPGT | ASSDGGGGEQY | 0.12983374 |
| TDGLGEN | ASSDGGGGEQY | 0.26817128 |

| | | |
|---|---|---|
| TGGPPLPGQA | ASSDGGGGEQY | 0.21811655 |
| TIPAITI | ASSDGGGGEQY | 0.22260404 |
| TIPGFTI | ASSDGGGGEQY | 0.10229129 |
| TSIASGALA | ASSDGGGGEQY | 0.39200854 |
| TVTLPGAAI | ASSDGGGGEQY | 0.19413999 |
| VASGLSAALP | ASSDGGGGEQY | 0.07730671 |
| VGAGVIGI | ASSDGGGGEQY | 0.32361454 |
| VGPKRFGV | ASSDGGGGEQY | 0.25095773 |
| VNVAFPD | ASSDGGGGEQY | 0.20242818 |
| VSMTNSGVS | ASSDGGGGEQY | 0.57221663 |
| VTDVLGDLSV | ASSDGGGGEQY | 0.12927333 |
| VVVVGAGFAGL | ASSDGGGGEQY | 0.17324524 |
| VYTVGTVV | ASSDGGGGEQY | 0.47792494 |
| WTTFSGP | ASSDGGGGEQY | 0.6622659 |
| YDELVEAF | ASSDGGGGEQY | 0.15131715 |
| YGTGGAGGNGG | ASSDGGGGEQY | 0.36836553 |
| YTNVMAV | ASSDGGGGEQY | 0.7298114 |
| FLEQVHQL | ASSDGGGGEQY | 0.31573752 |
| GLSLLAVAV | ASSDGGGGEQY | 0.5828675 |
| LEQVHQL | ASSDGGGGEQY | 0.64504373 |
| LNAAGAI | ASSDGGGGEQY | 0.1905838 |
| PAVHASL | ASSDGGGGEQY | 0.58478445 |
| TELEDDA | ASSDGGGGEQY | 0.5838269 |
| TPPGPPN | ASSDGGGGEQY | 0.28095895 |
| TPPPPPPP | ASSDGGGGEQY | 0.2778628 |
| TQSPALI | ASSQDQLGDSYEQY | 0.16699117 |
| EFAAAFPQ | ASSQDQLGDSYEQY | 0.26212645 |
| LLSGNLDVL | ASSQDQLGDSYEQY | 0.10214543 |
| GPANLGNYNI | ASSQDQLGDSYEQY | 0.50658095 |
| TLGPTLP | ASSQDQLGDSYEQY | 0.55494386 |
| VYAYSSCCK | ASSQDQLGDSYEQY | 0.23137477 |
| APAPAPAPNI | ASSQDQLGDSYEQY | 0.12485312 |
| PGGPVPIP | ASSQDQLGDSYEQY | 0.117706284 |
| FLTVALEP | ASSQDQLGDSYEQY | 0.2561559 |
| SQYFGANL | ASSQDQLGDSYEQY | 0.17263319 |
| GGVKPAA | ASSQDQLGDSYEQY | 0.10979657 |
| GGTGGAGGAGGW | ASSQDQLGDSYEQY | 0.2665679 |
| GVGGVAGLGGAGG | ASSQDQLGDSYEQY | 0.47528383 |
| VLLVLATTI | ASSQDQLGDSYEQY | 0.51095474 |
| IGIFAAML | ASSQDQLGDSYEQY | 0.13344854 |
| AVGDLID | ASSQDQLGDSYEQY | 0.7850446 |
| NISPLPL | ASSQDQLGDSYEQY | 0.550015 |
| VTAVFVAI | ASSQDQLGDSYEQY | 0.6503775 |
| SGDDNDAAL | ASSQDQLGDSYEQY | 0.2997455 |
| AVAALAHS | ASSQDQLGDSYEQY | 0.18035842 |
| GGVTVMATP | ASSQDQLGDSYEQY | 0.7420789 |
| AGLAANVRD | ASSQDQLGDSYEQY | 0.25129497 |
| TNRPDLIDPALLRPG | ASSQDQLGDSYEQY | 0.23358192 |
| LANLGVTAL | ASSQDQLGDSYEQY | 0.48600048 |
| TGERIEL | ASSQDQLGDSYEQY | 0.23879561 |
| NGEELAA | ASSQDQLGDSYEQY | 0.19195364 |

| | | |
|---|---|---|
| VLSLVWG | ASSQDQLGDSYEQY | 0.44731638 |
| IGYALWL | ASSQDQLGDSYEQY | 0.5322908 |
| LSLLLAPL | ASSQDQLGDSYEQY | 0.13797276 |
| TLDEALK | ASSQDQLGDSYEQY | 0.17135927 |
| TDGAGTAEGG | ASSQDQLGDSYEQY | 0.15655863 |
| NAPGGAGGNGG | ASSQDQLGDSYEQY | 0.28402382 |
| VSVFMGMGGNF | ASSQDQLGDSYEQY | 0.6327411 |
| LASYLAAL | ASSQDQLGDSYEQY | 0.34238914 |
| LVGGTPAL | ASSQDQLGDSYEQY | 0.2887655 |
| VRNADVL | ASSQDQLGDSYEQY | 0.47738117 |
| TCSSPFLVSAM | ASSQDQLGDSYEQY | 0.47549853 |
| VHITDAE | ASSQDQLGDSYEQY | 0.32696742 |
| EYAAPSF | ASSQDQLGDSYEQY | 0.5871637 |
| VVDAFHGGL | ASSQDQLGDSYEQY | 0.20852497 |
| ASVVDPQ | ASSQDQLGDSYEQY | 0.5557578 |
| SYNEFGGLS | ASSQDQLGDSYEQY | 0.49307632 |
| DPDGSVY | ASSQDQLGDSYEQY | 0.45489693 |
| MVSTIHATE | ASSQDQLGDSYEQY | 0.58692664 |
| LAAGVADPV | ASSQDQLGDSYEQY | 0.47292995 |
| TKEQIAA | ASSQDQLGDSYEQY | 0.32462364 |
| SLASPAAL | ASSQDQLGDSYEQY | 0.10351728 |
| IGLVFYGV | ASSQDQLGDSYEQY | 0.279219 |
| FLLDASNL | ASSQDQLGDSYEQY | 0.39588225 |
| YQVGNFP | ASSQDQLGDSYEQY | 0.42736065 |
| ESLLSQST | ASSQDQLGDSYEQY | 0.45878077 |
| LFTVGFAAVG | ASSQDQLGDSYEQY | 0.4009521 |
| ATPPTPT | ASSQDQLGDSYEQY | 0.18115085 |
| TDLATLG | ASSQDQLGDSYEQY | 0.6419381 |
| SAGGTGR | ASSQDQLGDSYEQY | 0.6568071 |
| GFSNSGS | ASSQDQLGDSYEQY | 0.12804513 |
| GGNGGSGGTSV | ASSQDQLGDSYEQY | 0.27564403 |
| GTSVATGGAGN | ASSQDQLGDSYEQY | 0.7793609 |
| NGGNGGSGGTS | ASSQDQLGDSYEQY | 0.5141096 |
| AVVKFPH | ASSQDQLGDSYEQY | 0.22226708 |
| TPPPPPP | ASSQDQLGDSYEQY | 0.23052135 |
| AAPAAAAAPA | ASSQDQLGDSYEQY | 0.7741358 |
| IALNEDL | ASSQDQLGDSYEQY | 0.16491754 |
| IVNVSSM | ASSQDQLGDSYEQY | 0.6963224 |
| TGIPPAPR | ASSQDQLGDSYEQY | 0.0931481 |
| FDLGGGTF | ASSQDQLGDSYEQY | 0.22765768 |
| FDLGGGTFDVS | ASSQDQLGDSYEQY | 0.1772506 |
| NEPTAAA | ASSQDQLGDSYEQY | 0.09716611 |
| NEPTAAALA | ASSQDQLGDSYEQY | 0.4474038 |
| GSIINISSI | ASSQDQLGDSYEQY | 0.19799043 |
| VAPGNAGTA | ASSQDQLGDSYEQY | 0.46412605 |
| TGGGASLE | ASSQDQLGDSYEQY | 0.2578233 |
| VAVYNHYKRI | ASSQDQLGDSYEQY | 0.52644026 |
| TDPAPATT | ASSQDQLGDSYEQY | 0.49796486 |
| TNRPDILDPALLRPG | ASSQDQLGDSYEQY | 0.26719102 |
| LTTTTTS | ASSQDQLGDSYEQY | 0.19209656 |
| PTGSGKTL | ASSQDQLGDSYEQY | 0.33613366 |

| | | |
|---|---|---|
| TGSGKTL | ASSQDQLGDSYEQY | 0.64007413 |
| TIGHVDHGK | ASSQDQLGDSYEQY | 0.43733847 |
| TIGHVDHGKTT | ASSQDQLGDSYEQY | 0.120608285 |
| GHVDHGKTT | ASSQDQLGDSYEQY | 0.25883347 |
| AQFEHTLL | ASSQDQLGDSYEQY | 0.2404574 |
| IVNQDEL | ASSQDQLGDSYEQY | 0.1043732 |
| TNGSQFF | ASSQDQLGDSYEQY | 0.081122614 |
| TGKGIVE | ASSQDQLGDSYEQY | 0.16767797 |
| LISNASDA | ASSQDQLGDSYEQY | 0.25195324 |
| AELRTLGELTKIAKAHGAQ | ASSQDQLGDSYEQY | 0.50134885 |
| APNAVAAAEPSW | ASSQDQLGDSYEQY | 0.5420915 |
| APQVAGAATAVL | ASSQDQLGDSYEQY | 0.20549619 |
| ARRDPAAAAAILPTDA | ASSQDQLGDSYEQY | 0.5710905 |
| DAAARDVLAEAGLAEYF | ASSQDQLGDSYEQY | 0.37215343 |
| DDQIVLSAKANLAKVEQ | ASSQDQLGDSYEQY | 0.070466295 |
| DELLPDTGEDSDEDGH | ASSQDQLGDSYEQY | 0.23228803 |
| ELPEAPANRATVPQD | ASSQDQLGDSYEQY | 0.12772955 |
| FGQSKIERVTGIG | ASSQDQLGDSYEQY | 0.5446167 |
| GAGGTGSTAGGAGGAGGAGGLY | ASSQDQLGDSYEQY | 0.1631207 |
| GHGGTGAAGGNGGTGGAGGSGI | ASSQDQLGDSYEQY | 0.20977509 |
| GNGGNGGAGGEGGPGGAGGAGA | ASSQDQLGDSYEQY | 0.25551394 |
| GSTAGNGGNGGNATNGTIAGQP | ASSQDQLGDSYEQY | 0.6511657 |
| GTGGDGGNGGNGGTGG | ASSQDQLGDSYEQY | 0.33603653 |
| GVRRPAGIPADAPV | ASSQDQLGDSYEQY | 0.18232861 |
| GYRQVTDAHLLTLARR | ASSQDQLGDSYEQY | 0.080867015 |
| IAKMGQDGHDRG | ASSQDQLGDSYEQY | 0.38107046 |
| KAEPSIEIQYT | ASSQDQLGDSYEQY | 0.16720591 |
| KPVRIRMAADEDTVAVT | ASSQDQLGDSYEQY | 0.16884546 |
| MPPRIAGMRLLVIKPEP | ASSQDQLGDSYEQY | 0.11307796 |
| MKFYAVALFRSRGVPPTR | ASSQDQLGDSYEQY | 0.09501997 |
| NAGGFGILGAVAHSP | ASSQDQLGDSYEQY | 0.13025421 |
| NEGREITLAMAEM | ASSQDQLGDSYEQY | 0.4990044 |
| NELVEWSHPLGHRWRWE | ASSQDQLGDSYEQY | 0.26014906 |
| NLGAGGTGGNGGNPG | ASSQDQLGDSYEQY | 0.23521546 |
| NTEWPKWETVMAADPVD | ASSQDQLGDSYEQY | 0.74199176 |
| PASAAPAETQP | ASSQDQLGDSYEQY | 0.62491745 |
| QQSSDGKNGVRAAEITGE | ASSQDQLGDSYEQY | 0.25581524 |
| RAPDAVTTSWWKE | ASSQDQLGDSYEQY | 0.21365298 |
| SGFGNVGAHTSGWFNQ | ASSQDQLGDSYEQY | 0.15356487 |
| SGVAGAVGQAASVGGLKV | ASSQDQLGDSYEQY | 0.16430904 |
| SNGGAGTGSAGGNGGT | ASSQDQLGDSYEQY | 0.1744338 |
| TGGAGGEGGAAG | ASSQDQLGDSYEQY | 0.70842904 |
| TGGDGGAAGAGGAPGHG | ASSQDQLGDSYEQY | 0.21475115 |
| TMPPLLAAIANAGRHGV | ASSQDQLGDSYEQY | 0.16340071 |
| TSSTAGNSGGA | ASSQDQLGDSYEQY | 0.3334544 |
| VPATVPVGILC | ASSQDQLGDSYEQY | 0.6839569 |
| VQRPDSGGVVLTGRIS | ASSQDQLGDSYEQY | 0.796556 |
| WTPNPVPATKGVPAGLVQAT | ASSQDQLGDSYEQY | 0.15463749 |
| YALKRALAKFSDRFDVVII | ASSQDQLGDSYEQY | 0.24680826 |
| YLIGSVIV | ASSQDQLGDSYEQY | 0.25973004 |
| YGQLVAVID | ASSQDQLGDSYEQY | 0.29535496 |

| | | |
|---|---|---|
| VTVGGMTLPA | ASSQDQLGDSYEQY | 0.22356929 |
| VLALPAV | ASSQDQLGDSYEQY | 0.16459633 |
| VIPPTLNYET | ASSQDQLGDSYEQY | 0.5042426 |
| VGMGGMHPSSG | ASSQDQLGDSYEQY | 0.79626095 |
| VGLAGGAATA | ASSQDQLGDSYEQY | 0.24084231 |
| VGGAGGAGGSGP | ASSQDQLGDSYEQY | 0.124963395 |
| VFAAFMV | ASSQDQLGDSYEQY | 0.24237722 |
| TLYDKLR | ASSQDQLGDSYEQY | 0.160175 |
| TLAVFTVL | ASSQDQLGDSYEQY | 0.16901892 |
| TGSVQPARV | ASSQDQLGDSYEQY | 0.32785213 |
| TGGGAGSGGT | ASSQDQLGDSYEQY | 0.12401405 |
| SHLETSL | ASSQDQLGDSYEQY | 0.24583179 |
| SFTTGMT | ASSQDQLGDSYEQY | 0.7930869 |
| QVEVAKRMGVRQPTVS | ASSQDQLGDSYEQY | 0.2354807 |
| QGSGGPALAM | ASSQDQLGDSYEQY | 0.5520872 |
| MLAPSNP | ASSQDQLGDSYEQY | 0.6816708 |
| LVNNAAI | ASSQDQLGDSYEQY | 0.43768927 |
| LTGEINTLA | ASSQDQLGDSYEQY | 0.71544003 |
| LSIFGPV | ASSQDQLGDSYEQY | 0.6431831 |
| LLALLANL | ASSQDQLGDSYEQY | 0.28175044 |
| LISNASDALDKLR | ASSQDQLGDSYEQY | 0.34539926 |
| LINEAVP | ASSQDQLGDSYEQY | 0.44338584 |
| LGYAFSGL | ASSQDQLGDSYEQY | 0.24263564 |
| LDFVAVAL | ASSQDQLGDSYEQY | 0.28699178 |
| LAQPGAE | ASSQDQLGDSYEQY | 0.23454815 |
| ISNLGAVTGF | ASSQDQLGDSYEQY | 0.53029263 |
| HQEDPGRI | ASSQDQLGDSYEQY | 0.39488488 |
| GTPGEPTP | ASSQDQLGDSYEQY | 0.558945 |
| GFFNSGS | ASSQDQLGDSYEQY | 0.122943744 |
| GAGGGGLTAGAN | ASSQDQLGDSYEQY | 0.1500177 |
| FVDALVNM | ASSQDQLGDSYEQY | 0.67808545 |
| FLTVAPEL | ASSQDQLGDSYEQY | 0.08353322 |
| FETLTAL | ASSQDQLGDSYEQY | 0.09347693 |
| DVINPIEL | ASSQDQLGDSYEQY | 0.32595012 |
| APVSGAPV | ASSQDQLGDSYEQY | 0.76821524 |
| ALVGGLAW | ASSQDQLGDSYEQY | 0.08196065 |
| ALAAAAATAI | ASSQDQLGDSYEQY | 0.19617425 |
| AFGLGLLL | ASSQDQLGDSYEQY | 0.14061253 |
| AFFAAASL | ASSQDQLGDSYEQY | 0.46171278 |
| AAAALPAFSPP | ASSQDQLGDSYEQY | 0.26616532 |
| VVLAPAL | ASSQDQLGDSYEQY | 0.13084726 |
| TIGHVDHGKTTL | ASSQDQLGDSYEQY | 0.4346074 |
| LISNASDALDK | ASSQDQLGDSYEQY | 0.11074804 |
| LISNASDAL | ASSQDQLGDSYEQY | 0.1446698 |
| ISNASDALDK | ASSQDQLGDSYEQY | 0.5130939 |
| NQAETLVYQTE | ASSQDQLGDSYEQY | 0.74508166 |
| TGGTGGTAGAG | ASSQDQLGDSYEQY | 0.15589254 |
| TGSTAGGAGGA | ASSQDQLGDSYEQY | 0.51077914 |
| DPLAVPVAAGPV | ASSQDQLGDSYEQY | 0.18935183 |
| GGTGGTPGNGGN | ASSQDQLGDSYEQY | 0.82501704 |
| PAGLAGVDPATL | ASSQDQLGDSYEQY | 0.21251535 |

| | | |
|---|---|---|
| GNNLVVEAPHAGASV | ASSQDQLGDSYEQY | 0.12386038 |
| KVTRIMSASTLEQPA | ASSQDQLGDSYEQY | 0.5395268 |
| LGFALPIALPRRTVV | ASSQDQLGDSYEQY | 0.6074501 |
| TRLGNLTARKLVVLV | ASSQDQLGDSYEQY | 0.2838901 |
| WSQQRRLIYFPSAGP | ASSQDQLGDSYEQY | 0.5955734 |
| GPNGNPDPMAAAVDIR | ASSQDQLGDSYEQY | 0.23135187 |
| TQRCVSVADLRAAASR | ASSQDQLGDSYEQY | 0.22304553 |
| QFVLDVAEHPVEVLTETV | ASSQDQLGDSYEQY | 0.58029664 |
| GGAGGTGGAGGNGGTQAGDG | ASSQDQLGDSYEQY | 0.5856881 |
| PLLATLRAIAVDVDAKPAQV | ASSQDQLGDSYEQY | 0.16717497 |
| APKFRTGGYMIDFWGVGYQVAK | ASSQDQLGDSYEQY | 0.4969859 |
| GAGGAGGAGGAAGGPGGTGGTG | ASSQDQLGDSYEQY | 0.2957766 |
| IGALGLLATDVAPAGSVAFGVA | ASSQDQLGDSYEQY | 0.17945908 |
| AGGAGGIGGNGGAGGAGGVLMG | ASSQDQLGDSYEQY | 0.7619225 |
| ADELIGLL | ASSQDQLGDSYEQY | 0.32212266 |
| ADRALKA | ASSQDQLGDSYEQY | 0.53497505 |
| AGAVGQVGGAGGA | ASSQDQLGDSYEQY | 0.32921582 |
| AGGGGGLGGAP | ASSQDQLGDSYEQY | 0.71670246 |
| AGGLLYGNGGAG | ASSQDQLGDSYEQY | 0.15656762 |
| AGLESIGL | ASSQDQLGDSYEQY | 0.45552287 |
| AGLKTGII | ASSQDQLGDSYEQY | 0.33846486 |
| ASLAALLL | ASSQDQLGDSYEQY | 0.53809357 |
| ASVINISSI | ASSQDQLGDSYEQY | 0.2349681 |
| ATPPGPA | ASSQDQLGDSYEQY | 0.15935192 |
| ATVASALV | ASSQDQLGDSYEQY | 0.3458595 |
| ATVASLIL | ASSQDQLGDSYEQY | 0.61570215 |
| AVNGPVLIRH | ASSQDQLGDSYEQY | 0.30667126 |
| DGAAGAVGI | ASSQDQLGDSYEQY | 0.4684086 |
| DPLLHGI | ASSQDQLGDSYEQY | 0.7656073 |
| ELGAALGAL | ASSQDQLGDSYEQY | 0.16012506 |
| ELGRTQAV | ASSQDQLGDSYEQY | 0.2753529 |
| FVAGGYLS | ASSQDQLGDSYEQY | 0.48261195 |
| GASGGAGGAGGW | ASSQDQLGDSYEQY | 0.6301853 |
| GGAGAGGDGGAGGI | ASSQDQLGDSYEQY | 0.26499903 |
| GTGHNGSQPGMGGNG | ASSQDQLGDSYEQY | 0.70499647 |
| GYAAASASAS | ASSQDQLGDSYEQY | 0.2521472 |
| HQVDVAAVQQG | ASSQDQLGDSYEQY | 0.4609556 |
| IGAAIAHT | ASSQDQLGDSYEQY | 0.18343492 |
| IGMIEVAQL | ASSQDQLGDSYEQY | 0.2748776 |
| IRLAPEHTSAMD | ASSQDQLGDSYEQY | 0.1400091 |
| IYNVSVL | ASSQDQLGDSYEQY | 0.3636796 |
| KQGFLGM | ASSQDQLGDSYEQY | 0.61495876 |
| LESVITQL | ASSQDQLGDSYEQY | 0.4489169 |
| LGLGALVVV | ASSQDQLGDSYEQY | 0.09482029 |
| LIGNGGNGGSGGI | ASSQDQLGDSYEQY | 0.5259296 |
| LNAINAPTL | ASSQDQLGDSYEQY | 0.48950627 |
| LQLLNTGV | ASSQDQLGDSYEQY | 0.8177841 |
| LQTSIEF | ASSQDQLGDSYEQY | 0.32618594 |
| MSFAVSEL | ASSQDQLGDSYEQY | 0.4902121 |
| NTGALGL | ASSQDQLGDSYEQY | 0.20869839 |
| PPGAGGAGGAGTT | ASSQDQLGDSYEQY | 0.28730124 |

| | | |
|---|---|---|
| QLVGLGI | ASSQDQLGDSYEQY | 0.12806784 |
| RFLEQVHQL | ASSQDQLGDSYEQY | 0.22943808 |
| RGVGYVLR | ASSQDQLGDSYEQY | 0.3019809 |
| SATGVFAGVI | ASSQDQLGDSYEQY | 0.15701821 |
| SLPAITLP | ASSQDQLGDSYEQY | 0.71252435 |
| STALAFTV | ASSQDQLGDSYEQY | 0.6958035 |
| SVLTVGLGK | ASSQDQLGDSYEQY | 0.6388648 |
| TALAGAGSGI | ASSQDQLGDSYEQY | 0.7204495 |
| TGLPFTGI | ASSQDQLGDSYEQY | 0.74763596 |
| VAEHVTV | ASSQDQLGDSYEQY | 0.46670985 |
| VDIVLNSL | ASSQDQLGDSYEQY | 0.19505495 |
| VLTGGGAVL | ASSQDQLGDSYEQY | 0.278112 |
| VNVGLAYGI | ASSQDQLGDSYEQY | 0.12437017 |
| VSSAAVGAV | ASSQDQLGDSYEQY | 0.24196416 |
| VTSVYEAGV | ASSQDQLGDSYEQY | 0.2134246 |
| WVALDRGA | ASSQDQLGDSYEQY | 0.33624882 |
| YLVGSVVI | ASSQDQLGDSYEQY | 0.07760181 |
| GPPGTGKTLL | ASSQDQLGDSYEQY | 0.5627515 |
| GVGMTKF | ASSQDQLGDSYEQY | 0.7030604 |
| IGIGGSDL | ASSQDQLGDSYEQY | 0.2140722 |
| IGTLLAEL | ASSQDQLGDSYEQY | 0.3819139 |
| IYTSGTT | ASSQDQLGDSYEQY | 0.59831 |
| LSFPAGV | ASSQDQLGDSYEQY | 0.1829676 |
| LYTSGTT | ASSQDQLGDSYEQY | 0.59519625 |
| VGAFPFGF | ASSQDQLGDSYEQY | 0.16315833 |
| VGAGTVEF | ASSQDQLGDSYEQY | 0.47156274 |
| VGKSTLLL | ASSQDQLGDSYEQY | 0.6368603 |
| AALAHAGGGIV | ASSQDQLGDSYEQY | 0.35055828 |
| AALVAVSLL | ASSQDQLGDSYEQY | 0.3590464 |
| AAVSRAVAAL | ASSQDQLGDSYEQY | 0.2962057 |
| AGGSSGIGGF | ASSQDQLGDSYEQY | 0.14037848 |
| AGLTVANANG | ASSQDQLGDSYEQY | 0.40079293 |
| AGLVGAVL | ASSQDQLGDSYEQY | 0.07881212 |
| ALSQAFTV | ASSQDQLGDSYEQY | 0.39681327 |
| ALTFSVSL | ASSQDQLGDSYEQY | 0.1567795 |
| ALTGAALVAL | ASSQDQLGDSYEQY | 0.32674462 |
| ASFSIGAGVL | ASSQDQLGDSYEQY | 0.3142342 |
| DGSVYKI | ASSQDQLGDSYEQY | 0.45512208 |
| EQLAEIV | ASSQDQLGDSYEQY | 0.14693023 |
| FAVSELTVM | ASSQDQLGDSYEQY | 0.28211668 |
| GAAPLAAGL | ASSQDQLGDSYEQY | 0.20899145 |
| GADGGDGGSGGAGG | ASSQDQLGDSYEQY | 0.13442668 |
| GFGLDTGGAGG | ASSQDQLGDSYEQY | 0.6519301 |
| GLASGGLL | ASSQDQLGDSYEQY | 0.2508122 |
| GRIAGAAVLL | ASSQDQLGDSYEQY | 0.33031034 |
| GSAGSLNTSF | ASSQDQLGDSYEQY | 0.48153472 |
| GTGQIGI | ASSQDQLGDSYEQY | 0.3087557 |
| GTLIALIL | ASSQDQLGDSYEQY | 0.13898921 |
| IAATMGFVVG | ASSQDQLGDSYEQY | 0.31720525 |
| IFAQALL | ASSQDQLGDSYEQY | 0.28213638 |
| IGAAVALGF | ASSQDQLGDSYEQY | 0.20923428 |

| | | |
|---|---|---|
| IGLTPLAL | ASSQDQLGDSYEQY | 0.07741589 |
| ILGSNTSTL | ASSQDQLGDSYEQY | 0.47500432 |
| IVFSLTVL | ASSQDQLGDSYEQY | 0.1825438 |
| LASKATGF | ASSQDQLGDSYEQY | 0.25593713 |
| LASVALIL | ASSQDQLGDSYEQY | 0.22175491 |
| LFAVGLTL | ASSQDQLGDSYEQY | 0.5973784 |
| LFNGGIGTYI | ASSQDQLGDSYEQY | 0.19230673 |
| LTLGAGII | ASSQDQLGDSYEQY | 0.21125335 |
| LVAAALGDI | ASSQDQLGDSYEQY | 0.5659348 |
| NTAFVQLGI | ASSQDQLGDSYEQY | 0.3867731 |
| QLFSGAEL | ASSQDQLGDSYEQY | 0.64450324 |
| SAATFITL | ASSQDQLGDSYEQY | 0.37166128 |
| SNYGLANLGV | ASSQDQLGDSYEQY | 0.27474594 |
| SYGGAYAVM | ASSQDQLGDSYEQY | 0.3774563 |
| TTGVGGLTLG | ASSQDQLGDSYEQY | 0.31728545 |
| TTQTIYI | ASSQDQLGDSYEQY | 0.09535572 |
| VAAGAGLGAVI | ASSQDQLGDSYEQY | 0.10153659 |
| VAAVVTLL | ASSQDQLGDSYEQY | 0.20759724 |
| VGFLGGLI | ASSQDQLGDSYEQY | 0.09847861 |
| VGVIPILL | ASSQDQLGDSYEQY | 0.33070868 |
| VIFASSLI | ASSQDQLGDSYEQY | 0.5518272 |
| VKGGDGGAAAG | ASSQDQLGDSYEQY | 0.40220678 |
| VTKTFTNI | ASSQDQLGDSYEQY | 0.1329898 |
| VVAAAILL | ASSQDQLGDSYEQY | 0.24093142 |
| YAALDVEL | ASSQDQLGDSYEQY | 0.23241076 |
| LASVALLL | ASSQDQLGDSYEQY | 0.45847374 |
| LDAVHRV | ASSQDQLGDSYEQY | 0.31707877 |
| QGLSSSIF | ASSQDQLGDSYEQY | 0.78395665 |
| RGTFANIRL | ASSQDQLGDSYEQY | 0.58460593 |
| RQLWWGHRI | ASSQDQLGDSYEQY | 0.2035883 |
| YGPPGTGKTLL | ASSQDQLGDSYEQY | 0.54182386 |
| AALVGAPANVA | ASSQDQLGDSYEQY | 0.22146586 |
| AATFRRTE | ASSQDQLGDSYEQY | 0.31083417 |
| AGALGGGI | ASSQDQLGDSYEQY | 0.27527577 |
| AGGAGGAGGIGAG | ASSQDQLGDSYEQY | 0.39693397 |
| AGSGALGL | ASSQDQLGDSYEQY | 0.2844297 |
| AQIADQI | ASSQDQLGDSYEQY | 0.21468055 |
| ASAVAVTYL | ASSQDQLGDSYEQY | 0.12923232 |
| ASTVLGGTGG | ASSQDQLGDSYEQY | 0.31976798 |
| AVGGEPAK | ASSQDQLGDSYEQY | 0.20441024 |
| CEQGTFGGTC | ASSQDQLGDSYEQY | 0.108216904 |
| DEGKAPAG | ASSQDQLGDSYEQY | 0.20319283 |
| DTVVFPAL | ASSQDQLGDSYEQY | 0.8177439 |
| FEGITMAG | ASSQDQLGDSYEQY | 0.7747501 |
| FPLVVQD | ASSQDQLGDSYEQY | 0.17780747 |
| GAFTTPQ | ASSQDQLGDSYEQY | 0.17447981 |
| GFGGFGTAGGVGGA | ASSQDQLGDSYEQY | 0.6838688 |
| GFLAQFL | ASSQDQLGDSYEQY | 0.13586533 |
| GGAGGAGGNAGVA | ASSQDQLGDSYEQY | 0.35280925 |
| GGVGTAGAGGAGGAGG | ASSQDQLGDSYEQY | 0.39066404 |
| GNGGGGDAGNAGSGGN | ASSQDQLGDSYEQY | 0.35440743 |

| | | |
|---|---|---|
| GTGGAGGAGNDAGS | ASSQDQLGDSYEQY | 0.19308293 |
| GTTNTGWFNS | ASSQDQLGDSYEQY | 0.43405792 |
| IFMGPLFTL | ASSQDQLGDSYEQY | 0.2404319 |
| ILAHAIEAI | ASSQDQLGDSYEQY | 0.4550804 |
| IRSAIREAL | ASSQDQLGDSYEQY | 0.0809478 |
| ITFAGGG | ASSQDQLGDSYEQY | 0.50758016 |
| ITGMSGHQEV | ASSQDQLGDSYEQY | 0.14893314 |
| IVAGLACTL | ASSQDQLGDSYEQY | 0.2130268 |
| IVGGGLAGA | ASSQDQLGDSYEQY | 0.106314816 |
| IVTGVLNI | ASSQDQLGDSYEQY | 0.273189 |
| LAQQGCI | ASSQDQLGDSYEQY | 0.5929798 |
| LCLHVEN | ASSQDQLGDSYEQY | 0.705011 |
| LDAISDGS | ASSQDQLGDSYEQY | 0.13771652 |
| LGGFAAML | ASSQDQLGDSYEQY | 0.58294624 |
| LGGKNPMI | ASSQDQLGDSYEQY | 0.7606483 |
| LGPKGVAAIL | ASSQDQLGDSYEQY | 0.21334225 |
| LMVGVSI | ASSQDQLGDSYEQY | 0.12551416 |
| LVGAAFVL | ASSQDQLGDSYEQY | 0.37077284 |
| LVLAVAPL | ASSQDQLGDSYEQY | 0.3508743 |
| MAAIAVIDAA | ASSQDQLGDSYEQY | 0.57869 |
| NNGINAELQ | ASSQDQLGDSYEQY | 0.6266478 |
| RGEPLVLVHG | ASSQDQLGDSYEQY | 0.11315921 |
| RGGAGGAAGLI | ASSQDQLGDSYEQY | 0.41855332 |
| RSYRMHPAV | ASSQDQLGDSYEQY | 0.61838573 |
| SAAHLLAGQLA | ASSQDQLGDSYEQY | 0.16025327 |
| SAAVGSY | ASSQDQLGDSYEQY | 0.20061368 |
| SPHAVLHRI | ASSQDQLGDSYEQY | 0.21053942 |
| SQIAKVL | ASSQDQLGDSYEQY | 0.096298374 |
| TALGHLVP | ASSQDQLGDSYEQY | 0.42740566 |
| TAPTTSGAA | ASSQDQLGDSYEQY | 0.20832919 |
| TDGDDGNGGSPGT | ASSQDQLGDSYEQY | 0.21588463 |
| TDGLGEN | ASSQDQLGDSYEQY | 0.24335843 |
| TGGPPLPGQA | ASSQDQLGDSYEQY | 0.19338071 |
| TIPAITI | ASSQDQLGDSYEQY | 0.2028594 |
| TIPGFTI | ASSQDQLGDSYEQY | 0.21642396 |
| TSIASGALA | ASSQDQLGDSYEQY | 0.33745518 |
| TVTLPGAAI | ASSQDQLGDSYEQY | 0.5280427 |
| VASGLSAALP | ASSQDQLGDSYEQY | 0.36899644 |
| VGAGVIGI | ASSQDQLGDSYEQY | 0.43203723 |
| VGPKRFGV | ASSQDQLGDSYEQY | 0.6790204 |
| VNVAFPD | ASSQDQLGDSYEQY | 0.13503803 |
| VSMTNSGVS | ASSQDQLGDSYEQY | 0.17857282 |
| VTDVLGDLSV | ASSQDQLGDSYEQY | 0.5383389 |
| VVVVGAGFAGL | ASSQDQLGDSYEQY | 0.13881764 |
| VYTVGTVV | ASSQDQLGDSYEQY | 0.6270651 |
| WTTFSGP | ASSQDQLGDSYEQY | 0.361839 |
| YDELVEAF | ASSQDQLGDSYEQY | 0.39255172 |
| YGTGGAGGNGG | ASSQDQLGDSYEQY | 0.46966723 |
| YTNVMAV | ASSQDQLGDSYEQY | 0.83758736 |
| FLEQVHQL | ASSQDQLGDSYEQY | 0.41415012 |
| GLSLLAVAV | ASSQDQLGDSYEQY | 0.23687927 |

| | | |
|---|---|---|
| LEQVHQL | ASSQDQLGDSYEQY | 0.38438845 |
| LNAAGAI | ASSQDQLGDSYEQY | 0.18396565 |
| PAVHASL | ASSQDQLGDSYEQY | 0.5693363 |
| TELEDDA | ASSQDQLGDSYEQY | 0.45462957 |
| TPPGPPN | ASSQDQLGDSYEQY | 0.15086877 |
| TPPPPPPP | ASSQDQLGDSYEQY | 0.23317564 |
| TQSPALI | ASSLRGVNYAEQF | 0.65610576 |
| EFAAAFPQ | ASSLRGVNYAEQF | 0.6158814 |
| LLSGNLDVL | ASSLRGVNYAEQF | 0.11940921 |
| GPANLGNYNI | ASSLRGVNYAEQF | 0.18587044 |
| TLGPTLP | ASSLRGVNYAEQF | 0.14867777 |
| VYAYSSCCK | ASSLRGVNYAEQF | 0.27761176 |
| APAPAPAPNI | ASSLRGVNYAEQF | 0.16094364 |
| PGGPVPIP | ASSLRGVNYAEQF | 0.4985269 |
| FLTVALEP | ASSLRGVNYAEQF | 0.21631637 |
| SQYFGANL | ASSLRGVNYAEQF | 0.64370406 |
| GGVKPAA | ASSLRGVNYAEQF | 0.3519693 |
| GGTGGAGGAGGW | ASSLRGVNYAEQF | 0.4987948 |
| GVGGVAGLGGAGG | ASSLRGVNYAEQF | 0.19440867 |
| VLLVLATTI | ASSLRGVNYAEQF | 0.23137045 |
| IGIFAAML | ASSLRGVNYAEQF | 0.119317695 |
| AVGDLID | ASSLRGVNYAEQF | 0.13170426 |
| NISPLPL | ASSLRGVNYAEQF | 0.16764264 |
| VTAVFVAI | ASSLRGVNYAEQF | 0.24590702 |
| SGDDNDAAL | ASSLRGVNYAEQF | 0.09258906 |
| AVAALAHS | ASSLRGVNYAEQF | 0.74272025 |
| GGVTVMATP | ASSLRGVNYAEQF | 0.26586744 |
| AGLAANVRD | ASSLRGVNYAEQF | 0.38919342 |
| TNRPDLIDPALLRPG | ASSLRGVNYAEQF | 0.22799361 |
| LANLGVTAL | ASSLRGVNYAEQF | 0.20478594 |
| TGERIEL | ASSLRGVNYAEQF | 0.20324522 |
| NGEELAA | ASSLRGVNYAEQF | 0.55222535 |
| VLSLVWG | ASSLRGVNYAEQF | 0.15339778 |
| IGYALWL | ASSLRGVNYAEQF | 0.106120005 |
| LSLLLAPL | ASSLRGVNYAEQF | 0.21818952 |
| TLDEALK | ASSLRGVNYAEQF | 0.15051788 |
| TDGAGTAEGG | ASSLRGVNYAEQF | 0.76520205 |
| NAPGGAGGNGG | ASSLRGVNYAEQF | 0.5842383 |
| VSVFMGMGGNF | ASSLRGVNYAEQF | 0.33378038 |
| LASYLAAL | ASSLRGVNYAEQF | 0.55310535 |
| LVGGTPAL | ASSLRGVNYAEQF | 0.0914914 |
| VRNADVL | ASSLRGVNYAEQF | 0.25773776 |
| TCSSPFLVSAM | ASSLRGVNYAEQF | 0.73292655 |
| VHITDAE | ASSLRGVNYAEQF | 0.35591364 |
| EYAAPSF | ASSLRGVNYAEQF | 0.2470982 |
| VVDAFHGGL | ASSLRGVNYAEQF | 0.5689326 |
| ASVVDPQ | ASSLRGVNYAEQF | 0.17574535 |
| SYNEFGGLS | ASSLRGVNYAEQF | 0.20903993 |
| DPDGSVY | ASSLRGVNYAEQF | 0.27574503 |
| MVSTIHATE | ASSLRGVNYAEQF | 0.37452126 |
| LAAGVADPV | ASSLRGVNYAEQF | 0.2618323 |

| | | |
|---|---|---|
| TKEQIAA | ASSLRGVNYAEQF | 0.14671631 |
| SLASPAAL | ASSLRGVNYAEQF | 0.39727795 |
| IGLVFYGV | ASSLRGVNYAEQF | 0.84563595 |
| FLLDASNL | ASSLRGVNYAEQF | 0.20542261 |
| YQVGNFP | ASSLRGVNYAEQF | 0.58691585 |
| ESLLSQST | ASSLRGVNYAEQF | 0.37048674 |
| LFTVGFAAVG | ASSLRGVNYAEQF | 0.29054648 |
| ATPPTPT | ASSLRGVNYAEQF | 0.16234459 |
| TDLATLG | ASSLRGVNYAEQF | 0.1466223 |
| SAGGTGR | ASSLRGVNYAEQF | 0.27325746 |
| GFSNSGS | ASSLRGVNYAEQF | 0.15862069 |
| GGNGGSGGTSV | ASSLRGVNYAEQF | 0.28568393 |
| GTSVATGGAGN | ASSLRGVNYAEQF | 0.25949055 |
| NGGNGGSGGTS | ASSLRGVNYAEQF | 0.85745054 |
| AVVKFPH | ASSLRGVNYAEQF | 0.43568486 |
| TPPPPPP | ASSLRGVNYAEQF | 0.4573987 |
| AAPAAAAAPA | ASSLRGVNYAEQF | 0.14579026 |
| IALNEDL | ASSLRGVNYAEQF | 0.31055933 |
| IVNVSSM | ASSLRGVNYAEQF | 0.47483823 |
| TGIPPAPR | ASSLRGVNYAEQF | 0.13562463 |
| FDLGGGTF | ASSLRGVNYAEQF | 0.71029836 |
| FDLGGGTFDVS | ASSLRGVNYAEQF | 0.16219804 |
| NEPTAAA | ASSLRGVNYAEQF | 0.39838028 |
| NEPTAAALA | ASSLRGVNYAEQF | 0.440841 |
| GSIINISSI | ASSLRGVNYAEQF | 0.7290045 |
| VAPGNAGTA | ASSLRGVNYAEQF | 0.24128714 |
| TGGGASLE | ASSLRGVNYAEQF | 0.316077 |
| VAVYNHYKRI | ASSLRGVNYAEQF | 0.14264226 |
| TDPAPATT | ASSLRGVNYAEQF | 0.5790154 |
| TNRPDILDPALLRPG | ASSLRGVNYAEQF | 0.6071213 |
| LTTTTTS | ASSLRGVNYAEQF | 0.19021007 |
| PTGSGKTL | ASSLRGVNYAEQF | 0.77374005 |
| TGSGKTL | ASSLRGVNYAEQF | 0.37663424 |
| TIGHVDHGK | ASSLRGVNYAEQF | 0.14674126 |
| TIGHVDHGKTT | ASSLRGVNYAEQF | 0.256306 |
| GHVDHGKTT | ASSLRGVNYAEQF | 0.67768514 |
| AQFEHTLL | ASSLRGVNYAEQF | 0.744913 |
| IVNQDEL | ASSLRGVNYAEQF | 0.4634654 |
| TNGSQFF | ASSLRGVNYAEQF | 0.42048824 |
| TGKGIVE | ASSLRGVNYAEQF | 0.21832442 |
| LISNASDA | ASSLRGVNYAEQF | 0.70704 |
| AELRTLGELTKIAKAHGAQ | ASSLRGVNYAEQF | 0.79221416 |
| APNAVAAAEPSW | ASSLRGVNYAEQF | 0.5390387 |
| APQVAGAATAVL | ASSLRGVNYAEQF | 0.20572914 |
| ARRDPAAAAAILPTDA | ASSLRGVNYAEQF | 0.21699 |
| DAAARDVLAEAGLAEYF | ASSLRGVNYAEQF | 0.31022546 |
| DDQIVLSAKANLAKVEQ | ASSLRGVNYAEQF | 0.6317222 |
| DELLPDTGEDSDEDGH | ASSLRGVNYAEQF | 0.15021795 |
| ELPEAPANRATVPQD | ASSLRGVNYAEQF | 0.32026324 |
| FGQSKIERVTGIG | ASSLRGVNYAEQF | 0.31474996 |
| GAGGTGSTAGGAGGAGGAGGLY | ASSLRGVNYAEQF | 0.11563509 |

| | | |
|---|---|---|
| GHGGTGAAGGNGGTGGAGGSGI | ASSLRGVNYAEQF | 0.49506658 |
| GNGGNGGAGGEGGPGGAGGAGA | ASSLRGVNYAEQF | 0.8065566 |
| GSTAGNGGNGGNATNGTIAGQP | ASSLRGVNYAEQF | 0.2959299 |
| GTGGDGGNGGNGGTGG | ASSLRGVNYAEQF | 0.17040382 |
| GVRRPAGIPADAPV | ASSLRGVNYAEQF | 0.70415294 |
| GYRQVTDAHLLTLARR | ASSLRGVNYAEQF | 0.83252394 |
| IAKMGQDGHDRG | ASSLRGVNYAEQF | 0.51366276 |
| KAEPSIEIQYT | ASSLRGVNYAEQF | 0.12702955 |
| KPVRIRMAADEDTVAVT | ASSLRGVNYAEQF | 0.08819128 |
| MPPRIAGMRLLVIKPEP | ASSLRGVNYAEQF | 0.26432902 |
| MKFYAVALFRSRGVPPTR | ASSLRGVNYAEQF | 0.62412477 |
| NAGGFGILGAVAHSP | ASSLRGVNYAEQF | 0.22484253 |
| NEGREITLAMAEM | ASSLRGVNYAEQF | 0.1606612 |
| NELVEWSHPLGHRWRWE | ASSLRGVNYAEQF | 0.4354148 |
| NLGAGGTGGNGGNPG | ASSLRGVNYAEQF | 0.6139473 |
| NTEWPKWETVMAADPVD | ASSLRGVNYAEQF | 0.15495549 |
| PASAAPAETQP | ASSLRGVNYAEQF | 0.15127794 |
| QQSSDGKNGVRAAEITGE | ASSLRGVNYAEQF | 0.4359711 |
| RAPDAVTTSWWKE | ASSLRGVNYAEQF | 0.54854727 |
| SGFGNVGAHTSGWFNQ | ASSLRGVNYAEQF | 0.14168504 |
| SGVAGAVGQAASVGGLKV | ASSLRGVNYAEQF | 0.20158303 |
| SNGGAGTGSAGGNGGT | ASSLRGVNYAEQF | 0.27911353 |
| TGGAGGEGGAAG | ASSLRGVNYAEQF | 0.2151939 |
| TGGDGGAAGAGGAPGHG | ASSLRGVNYAEQF | 0.43324682 |
| TMPPLLAAIANAGRHGV | ASSLRGVNYAEQF | 0.15225539 |
| TSSTAGNSGGA | ASSLRGVNYAEQF | 0.71351516 |
| VPATVPVGILC | ASSLRGVNYAEQF | 0.13174802 |
| VQRPDSGGVVLTGRIS | ASSLRGVNYAEQF | 0.57759625 |
| WTPNPVPATKGVPAGLVQAT | ASSLRGVNYAEQF | 0.3389467 |
| YALKRALAKFSDRFDVVII | ASSLRGVNYAEQF | 0.35260472 |
| YLIGSVIV | ASSLRGVNYAEQF | 0.2281709 |
| YGQLVAVID | ASSLRGVNYAEQF | 0.15544571 |
| VTVGGMTLPA | ASSLRGVNYAEQF | 0.15774137 |
| VLALPAV | ASSLRGVNYAEQF | 0.102138594 |
| VIPPTLNYET | ASSLRGVNYAEQF | 0.67770267 |
| VGMGGMHPSSG | ASSLRGVNYAEQF | 0.0860944 |
| VGLAGGAATA | ASSLRGVNYAEQF | 0.5733119 |
| VGGAGGAGGSGP | ASSLRGVNYAEQF | 0.28109995 |
| VFAAFMV | ASSLRGVNYAEQF | 0.1681846 |
| TLYDKLR | ASSLRGVNYAEQF | 0.40000552 |
| TLAVFTVL | ASSLRGVNYAEQF | 0.12831494 |
| TGSVQPARV | ASSLRGVNYAEQF | 0.15172465 |
| TGGGAGSGGT | ASSLRGVNYAEQF | 0.4900439 |
| SHLETSL | ASSLRGVNYAEQF | 0.41906726 |
| SFTTGMT | ASSLRGVNYAEQF | 0.18565516 |
| QVEVAKRMGVRQPTVS | ASSLRGVNYAEQF | 0.34401572 |
| QGSGGPALAM | ASSLRGVNYAEQF | 0.40690714 |
| MLAPSNP | ASSLRGVNYAEQF | 0.1904883 |
| LVNNAAI | ASSLRGVNYAEQF | 0.26835892 |
| LTGEINTLA | ASSLRGVNYAEQF | 0.61858994 |
| LSIFGPV | ASSLRGVNYAEQF | 0.3217605 |

| | | |
|---|---|---|
| LLALLANL | ASSLRGVNYAEQF | 0.13516511 |
| LISNASDALDKLR | ASSLRGVNYAEQF | 0.46825466 |
| LINEAVP | ASSLRGVNYAEQF | 0.15823387 |
| LGYAFSGL | ASSLRGVNYAEQF | 0.41538012 |
| LDFVAVAL | ASSLRGVNYAEQF | 0.32010692 |
| LAQPGAE | ASSLRGVNYAEQF | 0.23844692 |
| ISNLGAVTGF | ASSLRGVNYAEQF | 0.5038485 |
| HQEDPGRI | ASSLRGVNYAEQF | 0.18913072 |
| GTPGEPTP | ASSLRGVNYAEQF | 0.19365236 |
| GFFNSGS | ASSLRGVNYAEQF | 0.209218 |
| GAGGGGLTAGAN | ASSLRGVNYAEQF | 0.2855928 |
| FVDALVNM | ASSLRGVNYAEQF | 0.5174889 |
| FLTVAPEL | ASSLRGVNYAEQF | 0.14373253 |
| FETLTAL | ASSLRGVNYAEQF | 0.36605245 |
| DVINPIEL | ASSLRGVNYAEQF | 0.19283612 |
| APVSGAPV | ASSLRGVNYAEQF | 0.14765583 |
| ALVGGLAW | ASSLRGVNYAEQF | 0.28220892 |
| ALAAAAATAI | ASSLRGVNYAEQF | 0.22195028 |
| AFGLGLLL | ASSLRGVNYAEQF | 0.16490722 |
| AFFAAASL | ASSLRGVNYAEQF | 0.4012074 |
| AAAALPAFSPP | ASSLRGVNYAEQF | 0.19095829 |
| VVLAPAL | ASSLRGVNYAEQF | 0.36546558 |
| TIGHVDHGKTTL | ASSLRGVNYAEQF | 0.7670143 |
| LISNASDALDK | ASSLRGVNYAEQF | 0.19049492 |
| LISNASDAL | ASSLRGVNYAEQF | 0.49858803 |
| ISNASDALDK | ASSLRGVNYAEQF | 0.46217716 |
| NQAETLVYQTE | ASSLRGVNYAEQF | 0.4570057 |
| TGGTGGTAGAG | ASSLRGVNYAEQF | 0.23209076 |
| TGSTAGGAGGA | ASSLRGVNYAEQF | 0.18231302 |
| DPLAVPVAAGPV | ASSLRGVNYAEQF | 0.7286863 |
| GGTGGTPGNGGN | ASSLRGVNYAEQF | 0.453534 |
| PAGLAGVDPATL | ASSLRGVNYAEQF | 0.54398894 |
| GNNLVVEAPHAGASV | ASSLRGVNYAEQF | 0.47611308 |
| KVTRIMSASTLEQPA | ASSLRGVNYAEQF | 0.55671316 |
| LGFALPIALPRRTVV | ASSLRGVNYAEQF | 0.5254941 |
| TRLGNLTARKLVVLV | ASSLRGVNYAEQF | 0.18857956 |
| WSQQRRLIYFPSAGP | ASSLRGVNYAEQF | 0.11912249 |
| GPNGNPDPMAAAVDIR | ASSLRGVNYAEQF | 0.2213921 |
| TQRCVSVADLRAAASR | ASSLRGVNYAEQF | 0.24213845 |
| QFVLDVAEHPVEVLTETV | ASSLRGVNYAEQF | 0.20369779 |
| GGAGGTGGAGGNGGTQAGDG | ASSLRGVNYAEQF | 0.14006546 |
| PLLATLRAIAVDVDAKPAQV | ASSLRGVNYAEQF | 0.5979628 |
| APKFRTGGYMIDFWGVGYQVAK | ASSLRGVNYAEQF | 0.8494903 |
| GAGGAGGAGGAAGGPGGTGGTG | ASSLRGVNYAEQF | 0.20849366 |
| IGALGLLATDVAPAGSVAFGVA | ASSLRGVNYAEQF | 0.6730893 |
| AGGAGGIGGNGGAGGAGGVLMG | ASSLRGVNYAEQF | 0.12678696 |
| ADELIGLL | ASSLRGVNYAEQF | 0.46498942 |
| ADRALKA | ASSLRGVNYAEQF | 0.62786615 |
| AGAVGQVGGAGGA | ASSLRGVNYAEQF | 0.15425763 |
| AGGGGGLGGAP | ASSLRGVNYAEQF | 0.3727255 |
| AGGLLYGNGGAG | ASSLRGVNYAEQF | 0.18559471 |

| | | |
|---|---|---|
| AGLESIGL | ASSLRGVNYAEQF | 0.21657556 |
| AGLKTGII | ASSLRGVNYAEQF | 0.6323624 |
| ASLAALLL | ASSLRGVNYAEQF | 0.22231066 |
| ASVINISSI | ASSLRGVNYAEQF | 0.34462097 |
| ATPPGPA | ASSLRGVNYAEQF | 0.20270228 |
| ATVASALV | ASSLRGVNYAEQF | 0.16567951 |
| ATVASLIL | ASSLRGVNYAEQF | 0.3210045 |
| AVNGPVLIRH | ASSLRGVNYAEQF | 0.60046107 |
| DGAAGAVGI | ASSLRGVNYAEQF | 0.18696715 |
| DPLLHGI | ASSLRGVNYAEQF | 0.7666985 |
| ELGAALGAL | ASSLRGVNYAEQF | 0.17981178 |
| ELGRTQAV | ASSLRGVNYAEQF | 0.8114337 |
| FVAGGYLS | ASSLRGVNYAEQF | 0.77706534 |
| GASGGAGGAGGW | ASSLRGVNYAEQF | 0.36293542 |
| GGAGAGGDGGAGGI | ASSLRGVNYAEQF | 0.23199111 |
| GTGHNGSQPGMGGNG | ASSLRGVNYAEQF | 0.6140207 |
| GYAAASASAS | ASSLRGVNYAEQF | 0.20448774 |
| HQVDVAAVQQG | ASSLRGVNYAEQF | 0.55497646 |
| IGAAIAHT | ASSLRGVNYAEQF | 0.5289352 |
| IGMIEVAQL | ASSLRGVNYAEQF | 0.48637858 |
| IRLAPEHTSAMD | ASSLRGVNYAEQF | 0.83584476 |
| IYNVSVL | ASSLRGVNYAEQF | 0.80426085 |
| KQGFLGM | ASSLRGVNYAEQF | 0.8242667 |
| LESVITQL | ASSLRGVNYAEQF | 0.71342576 |
| LGLGALVVV | ASSLRGVNYAEQF | 0.35423124 |
| LIGNGGNGGSGGI | ASSLRGVNYAEQF | 0.7154957 |
| LNAINAPTL | ASSLRGVNYAEQF | 0.6268343 |
| LQLLNTGV | ASSLRGVNYAEQF | 0.86769974 |
| LQTSIEF | ASSLRGVNYAEQF | 0.8596451 |
| MSFAVSEL | ASSLRGVNYAEQF | 0.6390338 |
| NTGALGL | ASSLRGVNYAEQF | 0.8267751 |
| PPGAGGAGGAGTT | ASSLRGVNYAEQF | 0.84137446 |
| QLVGLGI | ASSLRGVNYAEQF | 0.46490213 |
| RFLEQVHQL | ASSLRGVNYAEQF | 0.7768369 |
| RGVGYVLR | ASSLRGVNYAEQF | 0.6976352 |
| SATGVFAGVI | ASSLRGVNYAEQF | 0.76530856 |
| SLPAITLP | ASSLRGVNYAEQF | 0.32827157 |
| STALAFTV | ASSLRGVNYAEQF | 0.8292913 |
| SVLTVGLGK | ASSLRGVNYAEQF | 0.7315584 |
| TALAGAGSGI | ASSLRGVNYAEQF | 0.295448 |
| TGLPFTGI | ASSLRGVNYAEQF | 0.847842 |
| VAEHVTV | ASSLRGVNYAEQF | 0.64187944 |
| VDIVLNSL | ASSLRGVNYAEQF | 0.72121733 |
| VLTGGGAVL | ASSLRGVNYAEQF | 0.8441438 |
| VNVGLAYGI | ASSLRGVNYAEQF | 0.45105636 |
| VSSAAVGAV | ASSLRGVNYAEQF | 0.5918599 |
| VTSVYEAGV | ASSLRGVNYAEQF | 0.81681603 |
| WVALDRGA | ASSLRGVNYAEQF | 0.861282 |
| YLVGSVVI | ASSLRGVNYAEQF | 0.48677287 |
| GPPGTGKTLL | ASSLRGVNYAEQF | 0.8039119 |
| GVGMTKF | ASSLRGVNYAEQF | 0.8777642 |

| | | |
|---|---|---|
| IGIGGSDL | ASSLRGVNYAEQF | 0.81003624 |
| IGTLLAEL | ASSLRGVNYAEQF | 0.87388587 |
| IYTSGTT | ASSLRGVNYAEQF | 0.8821989 |
| LSFPAGV | ASSLRGVNYAEQF | 0.64917314 |
| LYTSGTT | ASSLRGVNYAEQF | 0.8663711 |
| VGAFPFGF | ASSLRGVNYAEQF | 0.8703114 |
| VGAGTVEF | ASSLRGVNYAEQF | 0.65890396 |
| VGKSTLLL | ASSLRGVNYAEQF | 0.755813 |
| AALAHAGGGIV | ASSLRGVNYAEQF | 0.5821836 |
| AALVAVSLL | ASSLRGVNYAEQF | 0.7952849 |
| AAVSRAVAAL | ASSLRGVNYAEQF | 0.43172562 |
| AGGSSGIGGF | ASSLRGVNYAEQF | 0.80453515 |
| AGLTVANANG | ASSLRGVNYAEQF | 0.8589635 |
| AGLVGAVL | ASSLRGVNYAEQF | 0.75073016 |
| ALSQAFTV | ASSLRGVNYAEQF | 0.51404876 |
| ALTFSVSL | ASSLRGVNYAEQF | 0.4692229 |
| ALTGAALVAL | ASSLRGVNYAEQF | 0.7944665 |
| ASFSIGAGVL | ASSLRGVNYAEQF | 0.4809184 |
| DGSVYKI | ASSLRGVNYAEQF | 0.4598605 |
| EQLAEIV | ASSLRGVNYAEQF | 0.4564972 |
| FAVSELTVM | ASSLRGVNYAEQF | 0.8224543 |
| GAAPLAAGL | ASSLRGVNYAEQF | 0.8442844 |
| GADGGDGGSGGAGG | ASSLRGVNYAEQF | 0.6204739 |
| GFGLDTGGAGG | ASSLRGVNYAEQF | 0.8613359 |
| GLASGGLL | ASSLRGVNYAEQF | 0.8590944 |
| GRIAGAAVLL | ASSLRGVNYAEQF | 0.8542502 |
| GSAGSLNTSF | ASSLRGVNYAEQF | 0.66217613 |
| GTGQIGI | ASSLRGVNYAEQF | 0.42587066 |
| GTLIALIL | ASSLRGVNYAEQF | 0.86899555 |
| IAATMGFVVG | ASSLRGVNYAEQF | 0.73972476 |
| IFAQALL | ASSLRGVNYAEQF | 0.83371174 |
| IGAAVALGF | ASSLRGVNYAEQF | 0.853695 |
| IGLTPLAL | ASSLRGVNYAEQF | 0.7181225 |
| ILGSNTSTL | ASSLRGVNYAEQF | 0.59970593 |
| IVFSLTVL | ASSLRGVNYAEQF | 0.7843055 |
| LASKATGF | ASSLRGVNYAEQF | 0.8963002 |
| LASVALIL | ASSLRGVNYAEQF | 0.5805599 |
| LFAVGLTL | ASSLRGVNYAEQF | 0.3723479 |
| LFNGGIGTYI | ASSLRGVNYAEQF | 0.68227124 |
| LTLGAGII | ASSLRGVNYAEQF | 0.68458587 |
| LVAAALGDI | ASSLRGVNYAEQF | 0.86719596 |
| NTAFVQLGI | ASSLRGVNYAEQF | 0.773543 |
| QLFSGAEL | ASSLRGVNYAEQF | 0.72201693 |
| SAATFITL | ASSLRGVNYAEQF | 0.8218721 |
| SNYGLANLGV | ASSLRGVNYAEQF | 0.80492115 |
| SYGGAYAVM | ASSLRGVNYAEQF | 0.77225745 |
| TTGVGGLTLG | ASSLRGVNYAEQF | 0.85957205 |
| TTQTIYI | ASSLRGVNYAEQF | 0.66600055 |
| VAAGAGLGAVI | ASSLRGVNYAEQF | 0.8859052 |
| VAAVVTLL | ASSLRGVNYAEQF | 0.8526472 |
| VGFLGGLI | ASSLRGVNYAEQF | 0.83188295 |

| | | |
|---|---|---|
| VGVIPILL | ASSLRGVNYAEQF | 0.86689764 |
| VIFASSLI | ASSLRGVNYAEQF | 0.7086549 |
| VKGGDGGAAAG | ASSLRGVNYAEQF | 0.79774255 |
| VTKTFTNI | ASSLRGVNYAEQF | 0.5355444 |
| VVAAAILL | ASSLRGVNYAEQF | 0.61694777 |
| YAALDVEL | ASSLRGVNYAEQF | 0.8214147 |
| LASVALLL | ASSLRGVNYAEQF | 0.8196355 |
| LDAVHRV | ASSLRGVNYAEQF | 0.8685637 |
| QGLSSSIF | ASSLRGVNYAEQF | 0.81818056 |
| RGTFANIRL | ASSLRGVNYAEQF | 0.7001926 |
| RQLWWGHRI | ASSLRGVNYAEQF | 0.88504523 |
| YGPPGTGKTLL | ASSLRGVNYAEQF | 0.8619673 |
| AALVGAPANVA | ASSLRGVNYAEQF | 0.6076621 |
| AATFRRTE | ASSLRGVNYAEQF | 0.76962835 |
| AGALGGGI | ASSLRGVNYAEQF | 0.86065173 |
| AGGAGGAGGIGAG | ASSLRGVNYAEQF | 0.8401638 |
| AGSGALGL | ASSLRGVNYAEQF | 0.7338664 |
| AQIADQI | ASSLRGVNYAEQF | 0.78155136 |
| ASAVAVTYL | ASSLRGVNYAEQF | 0.878428 |
| ASTVLGGTGG | ASSLRGVNYAEQF | 0.802497 |
| AVGGEPAK | ASSLRGVNYAEQF | 0.8852803 |
| CEQGTFGGTC | ASSLRGVNYAEQF | 0.36728832 |
| DEGKAPAG | ASSLRGVNYAEQF | 0.75748986 |
| DTVVFPAL | ASSLRGVNYAEQF | 0.6551031 |
| FEGITMAG | ASSLRGVNYAEQF | 0.18360595 |
| FPLVVQD | ASSLRGVNYAEQF | 0.85876703 |
| GAFTTPQ | ASSLRGVNYAEQF | 0.66879153 |
| GFGGFGTAGGVGGA | ASSLRGVNYAEQF | 0.8086145 |
| GFLAQFL | ASSLRGVNYAEQF | 0.7252829 |
| GGAGGAGGNAGVA | ASSLRGVNYAEQF | 0.855913 |
| GGVGTAGAGGAGGAGG | ASSLRGVNYAEQF | 0.7973956 |
| GNGGGGDAGNAGSGGN | ASSLRGVNYAEQF | 0.74857664 |
| GTGGAGGAGNDAGS | ASSLRGVNYAEQF | 0.47258264 |
| GTTNTGWFNS | ASSLRGVNYAEQF | 0.8096909 |
| IFMGPLFTL | ASSLRGVNYAEQF | 0.89068455 |
| ILAHAIEAI | ASSLRGVNYAEQF | 0.8765628 |
| IRSAIREAL | ASSLRGVNYAEQF | 0.28272775 |
| ITFAGGG | ASSLRGVNYAEQF | 0.83980083 |
| ITGMSGHQEV | ASSLRGVNYAEQF | 0.73477685 |
| IVAGLACTL | ASSLRGVNYAEQF | 0.40973783 |
| IVGGGLAGA | ASSLRGVNYAEQF | 0.44540694 |
| IVTGVLNI | ASSLRGVNYAEQF | 0.69820845 |
| LAQQGCI | ASSLRGVNYAEQF | 0.6943565 |
| LCLHVEN | ASSLRGVNYAEQF | 0.8225813 |
| LDAISDGS | ASSLRGVNYAEQF | 0.83208394 |
| LGGFAAML | ASSLRGVNYAEQF | 0.71888614 |
| LGGKNPMI | ASSLRGVNYAEQF | 0.8303234 |
| LGPKGVAAIL | ASSLRGVNYAEQF | 0.7467198 |
| LMVGVSI | ASSLRGVNYAEQF | 0.07941529 |
| LVGAAFVL | ASSLRGVNYAEQF | 0.632967 |
| LVLAVAPL | ASSLRGVNYAEQF | 0.5284736 |

| | | |
|---|---|---|
| MAAIAVIDAA | ASSLRGVNYAEQF | 0.8650943 |
| NNGINAELQ | ASSLRGVNYAEQF | 0.6545704 |
| RGEPLVLVHG | ASSLRGVNYAEQF | 0.7599288 |
| RGGAGGAAGLI | ASSLRGVNYAEQF | 0.7629925 |
| RSYRMHPAV | ASSLRGVNYAEQF | 0.82435286 |
| SAAHLLAGQLA | ASSLRGVNYAEQF | 0.7725123 |
| SAAVGSY | ASSLRGVNYAEQF | 0.6258781 |
| SPHAVLHRI | ASSLRGVNYAEQF | 0.37804613 |
| SQIAKVL | ASSLRGVNYAEQF | 0.80775845 |
| TALGHLVP | ASSLRGVNYAEQF | 0.6959479 |
| TAPTTSGAA | ASSLRGVNYAEQF | 0.51541877 |
| TDGDDGNGGSPGT | ASSLRGVNYAEQF | 0.65396816 |
| TDGLGEN | ASSLRGVNYAEQF | 0.83324325 |
| TGGPPLPGQA | ASSLRGVNYAEQF | 0.28216892 |
| TIPAITI | ASSLRGVNYAEQF | 0.86799794 |
| TIPGFTI | ASSLRGVNYAEQF | 0.45554745 |
| TSIASGALA | ASSLRGVNYAEQF | 0.7120818 |
| TVTLPGAAI | ASSLRGVNYAEQF | 0.8630148 |
| VASGLSAALP | ASSLRGVNYAEQF | 0.798392 |
| VGAGVIGI | ASSLRGVNYAEQF | 0.7432629 |
| VGPKRFGV | ASSLRGVNYAEQF | 0.6229 |
| VNVAFPD | ASSLRGVNYAEQF | 0.66891354 |
| VSMTNSGVS | ASSLRGVNYAEQF | 0.6892793 |
| VTDVLGDLSV | ASSLRGVNYAEQF | 0.59991825 |
| VVVVGAGFAGL | ASSLRGVNYAEQF | 0.8626515 |
| VYTVGTVV | ASSLRGVNYAEQF | 0.827009 |
| WTTFSGP | ASSLRGVNYAEQF | 0.5185954 |
| YDELVEAF | ASSLRGVNYAEQF | 0.8264729 |
| YGTGGAGGNGG | ASSLRGVNYAEQF | 0.8818929 |
| YTNVMAV | ASSLRGVNYAEQF | 0.88937557 |
| FLEQVHQL | ASSLRGVNYAEQF | 0.65123177 |
| GLSLLAVAV | ASSLRGVNYAEQF | 0.5953058 |
| LEQVHQL | ASSLRGVNYAEQF | 0.68674195 |
| LNAAGAI | ASSLRGVNYAEQF | 0.73021173 |
| PAVHASL | ASSLRGVNYAEQF | 0.48837844 |
| TELEDDA | ASSLRGVNYAEQF | 0.572904 |
| TPPGPPN | ASSLRGVNYAEQF | 0.8808541 |
| TPPPPPPP | ASSLRGVNYAEQF | 0.8946857 |
| TQSPALI | ASSATGGLNQDTQY | 0.7487884 |
| EFAAAFPQ | ASSATGGLNQDTQY | 0.3636728 |
| LLSGNLDVL | ASSATGGLNQDTQY | 0.7232992 |
| GPANLGNYNI | ASSATGGLNQDTQY | 0.61838937 |
| TLGPTLP | ASSATGGLNQDTQY | 0.6581155 |
| VYAYSSCCK | ASSATGGLNQDTQY | 0.7147726 |
| APAPAPAPNI | ASSATGGLNQDTQY | 0.47163516 |
| PGGPVPIP | ASSATGGLNQDTQY | 0.7264447 |
| FLTVALEP | ASSATGGLNQDTQY | 0.86825514 |
| SQYFGANL | ASSATGGLNQDTQY | 0.76073605 |
| GGVKPAA | ASSATGGLNQDTQY | 0.78833807 |
| GGTGGAGGAGGW | ASSATGGLNQDTQY | 0.87603664 |
| GVGGVAGLGGAGG | ASSATGGLNQDTQY | 0.84479 |

| | | |
|---|---|---|
| VLLVLATTI | ASSATGGLNQDTQY | 0.8744637 |
| IGIFAAML | ASSATGGLNQDTQY | 0.85882163 |
| AVGDLID | ASSATGGLNQDTQY | 0.76587117 |
| NISPLPL | ASSATGGLNQDTQY | 0.7497917 |
| VTAVFVAI | ASSATGGLNQDTQY | 0.8607379 |
| SGDDNDAAL | ASSATGGLNQDTQY | 0.7150014 |
| AVAALAHS | ASSATGGLNQDTQY | 0.81376386 |
| GGVTVMATP | ASSATGGLNQDTQY | 0.7771538 |
| AGLAANVRD | ASSATGGLNQDTQY | 0.8440647 |
| TNRPDLIDPALLRPG | ASSATGGLNQDTQY | 0.7218034 |
| LANLGVTAL | ASSATGGLNQDTQY | 0.8877673 |
| TGERIEL | ASSATGGLNQDTQY | 0.19860317 |
| NGEELAA | ASSATGGLNQDTQY | 0.54917157 |
| VLSLVWG | ASSATGGLNQDTQY | 0.8614596 |
| IGYALWL | ASSATGGLNQDTQY | 0.1599206 |
| LSLLLAPL | ASSATGGLNQDTQY | 0.44614756 |
| TLDEALK | ASSATGGLNQDTQY | 0.8306703 |
| TDGAGTAEGG | ASSATGGLNQDTQY | 0.8513955 |
| NAPGGAGGNGG | ASSATGGLNQDTQY | 0.15482435 |
| VSVFMGMGGNF | ASSATGGLNQDTQY | 0.6788608 |
| LASYLAAL | ASSATGGLNQDTQY | 0.33331102 |
| LVGGTPAL | ASSATGGLNQDTQY | 0.7699885 |
| VRNADVL | ASSATGGLNQDTQY | 0.74876165 |
| TCSSPFLVSAM | ASSATGGLNQDTQY | 0.39201972 |
| VHITDAE | ASSATGGLNQDTQY | 0.8493004 |
| EYAAPSF | ASSATGGLNQDTQY | 0.58910966 |
| VVDAFHGGL | ASSATGGLNQDTQY | 0.76486593 |
| ASVVDPQ | ASSATGGLNQDTQY | 0.86521673 |
| SYNEFGGLS | ASSATGGLNQDTQY | 0.89042544 |
| DPDGSVY | ASSATGGLNQDTQY | 0.5777804 |
| MVSTIHATE | ASSATGGLNQDTQY | 0.82007897 |
| LAAGVADPV | ASSATGGLNQDTQY | 0.7008797 |
| TKEQIAA | ASSATGGLNQDTQY | 0.8954508 |
| SLASPAAL | ASSATGGLNQDTQY | 0.6870903 |
| IGLVFYGV | ASSATGGLNQDTQY | 0.35039985 |
| FLLDASNL | ASSATGGLNQDTQY | 0.8438964 |
| YQVGNFP | ASSATGGLNQDTQY | 0.81722313 |
| ESLLSQST | ASSATGGLNQDTQY | 0.7456627 |
| LFTVGFAAVG | ASSATGGLNQDTQY | 0.85832417 |
| ATPPTPT | ASSATGGLNQDTQY | 0.65985465 |
| TDLATLG | ASSATGGLNQDTQY | 0.7361618 |
| SAGGTGR | ASSATGGLNQDTQY | 0.82196724 |
| GFSNSGS | ASSATGGLNQDTQY | 0.8431962 |
| GGNGGSGGTSV | ASSATGGLNQDTQY | 0.582878 |
| GTSVATGGAGN | ASSATGGLNQDTQY | 0.8833599 |
| NGGNGGSGGTS | ASSATGGLNQDTQY | 0.77717876 |
| AVVKFPH | ASSATGGLNQDTQY | 0.51559687 |
| TPPPPPP | ASSATGGLNQDTQY | 0.8862329 |
| AAPAAAAAPA | ASSATGGLNQDTQY | 0.7820486 |
| IALNEDL | ASSATGGLNQDTQY | 0.85429156 |
| IVNVSSM | ASSATGGLNQDTQY | 0.8435717 |

| | | |
|---|---|---|
| TGIPPAPR | ASSATGGLNQDTQY | 0.8816539 |
| FDLGGGTF | ASSATGGLNQDTQY | 0.6036094 |
| FDLGGGTFDVS | ASSATGGLNQDTQY | 0.76418203 |
| NEPTAAA | ASSATGGLNQDTQY | 0.7676614 |
| NEPTAAALA | ASSATGGLNQDTQY | 0.8349538 |
| GSIINISSI | ASSATGGLNQDTQY | 0.66574246 |
| VAPGNAGTA | ASSATGGLNQDTQY | 0.784927 |
| TGGGASLE | ASSATGGLNQDTQY | 0.85358834 |
| VAVYNHYKRI | ASSATGGLNQDTQY | 0.86624384 |
| TDPAPATT | ASSATGGLNQDTQY | 0.80500305 |
| TNRPDILDPALLRPG | ASSATGGLNQDTQY | 0.82336235 |
| LTTTTTS | ASSATGGLNQDTQY | 0.8866854 |
| PTGSGKTL | ASSATGGLNQDTQY | 0.6567844 |
| TGSGKTL | ASSATGGLNQDTQY | 0.77348834 |
| TIGHVDHGK | ASSATGGLNQDTQY | 0.85072255 |
| TIGHVDHGKTT | ASSATGGLNQDTQY | 0.8795473 |
| GHVDHGKTT | ASSATGGLNQDTQY | 0.76893973 |
| AQFEHTLL | ASSATGGLNQDTQY | 0.8800989 |
| IVNQDEL | ASSATGGLNQDTQY | 0.71101767 |
| TNGSQFF | ASSATGGLNQDTQY | 0.8280119 |
| TGKGIVE | ASSATGGLNQDTQY | 0.6488458 |
| LISNASDA | ASSATGGLNQDTQY | 0.64847267 |
| AELRTLGELTKIAKAHGAQ | ASSATGGLNQDTQY | 0.59388804 |
| APNAVAAAEPSW | ASSATGGLNQDTQY | 0.8003251 |
| APQVAGAATAVL | ASSATGGLNQDTQY | 0.85241085 |
| ARRDPAAAAAILPTDA | ASSATGGLNQDTQY | 0.78541887 |
| DAAARDVLAEAGLAEYF | ASSATGGLNQDTQY | 0.35815507 |
| DDQIVLSAKANLAKVEQ | ASSATGGLNQDTQY | 0.8158209 |
| DELLPDTGEDSDEDGH | ASSATGGLNQDTQY | 0.75457215 |
| ELPEAPANRATVPQD | ASSATGGLNQDTQY | 0.89545643 |
| FGQSKIERVTGIG | ASSATGGLNQDTQY | 0.7500189 |
| GAGGTGSTAGGAGGAGGAGGLY | ASSATGGLNQDTQY | 0.8353738 |
| GHGGTGAAGGNGGTGGAGGSGI | ASSATGGLNQDTQY | 0.7399386 |
| GNGGNGGAGGEGGPGGAGGAGA | ASSATGGLNQDTQY | 0.8062835 |
| GSTAGNGGNGGNATNGTIAGQP | ASSATGGLNQDTQY | 0.43705028 |
| GTGGDGGNGGNGGTGG | ASSATGGLNQDTQY | 0.59197754 |
| GVRRPAGIPADAPV | ASSATGGLNQDTQY | 0.74464065 |
| GYRQVTDAHLLTLARR | ASSATGGLNQDTQY | 0.66948545 |
| IAKMGQDGHDRG | ASSATGGLNQDTQY | 0.8491061 |
| KAEPSIEIQYT | ASSATGGLNQDTQY | 0.88167167 |
| KPVRIRMAADEDTVAVT | ASSATGGLNQDTQY | 0.824239 |
| MPPRIAGMRLLVIKPEP | ASSATGGLNQDTQY | 0.8580674 |
| MKFYAVALFRSRGVPPTR | ASSATGGLNQDTQY | 0.8709451 |
| NAGGFGILGAVAHSP | ASSATGGLNQDTQY | 0.8702687 |
| NEGREITLAMAEM | ASSATGGLNQDTQY | 0.72545683 |
| NELVEWSHPLGHRWRWE | ASSATGGLNQDTQY | 0.68461466 |
| NLGAGGTGGNGGNPG | ASSATGGLNQDTQY | 0.49522018 |
| NTEWPKWETVMAADPVD | ASSATGGLNQDTQY | 0.6782688 |
| PASAAPAETQP | ASSATGGLNQDTQY | 0.7113112 |
| QQSSDGKNGVRAAEITGE | ASSATGGLNQDTQY | 0.7583614 |
| RAPDAVTTSWWKE | ASSATGGLNQDTQY | 0.1228597 |

| | | |
|---|---|---|
| SGFGNVGAHTSGWFNQ | ASSATGGLNQDTQY | 0.8254454 |
| SGVAGAVGQAASVGGLKV | ASSATGGLNQDTQY | 0.8664812 |
| SNGGAGTGSAGGNGGT | ASSATGGLNQDTQY | 0.7284048 |
| TGGAGGEGGAAG | ASSATGGLNQDTQY | 0.819717 |
| TGGDGGAAGAGGAPGHG | ASSATGGLNQDTQY | 0.81112224 |
| TMPPLLAAIANAGRHGV | ASSATGGLNQDTQY | 0.6993768 |
| TSSTAGNSGGA | ASSATGGLNQDTQY | 0.8139611 |
| VPATVPVGILC | ASSATGGLNQDTQY | 0.69722056 |
| VQRPDSGGVVLTGRIS | ASSATGGLNQDTQY | 0.8618258 |
| WTPNPVPATKGVPAGLVQAT | ASSATGGLNQDTQY | 0.86758935 |
| YALKRALAKFSDRFDVVII | ASSATGGLNQDTQY | 0.81825316 |
| YLIGSVIV | ASSATGGLNQDTQY | 0.80249095 |
| YGQLVAVID | ASSATGGLNQDTQY | 0.8119346 |
| VTVGGMTLPA | ASSATGGLNQDTQY | 0.6228418 |
| VLALPAV | ASSATGGLNQDTQY | 0.80756235 |
| VIPPTLNYET | ASSATGGLNQDTQY | 0.53329265 |
| VGMGGMHPSSG | ASSATGGLNQDTQY | 0.7910502 |
| VGLAGGAATA | ASSATGGLNQDTQY | 0.6894555 |
| VGGAGGAGGSGP | ASSATGGLNQDTQY | 0.80761063 |
| VFAAFMV | ASSATGGLNQDTQY | 0.7449516 |
| TLYDKLR | ASSATGGLNQDTQY | 0.839668 |
| TLAVFTVL | ASSATGGLNQDTQY | 0.65205026 |
| TGSVQPARV | ASSATGGLNQDTQY | 0.8407066 |
| TGGGAGSGGT | ASSATGGLNQDTQY | 0.80158406 |
| SHLETSL | ASSATGGLNQDTQY | 0.3956323 |
| SFTTGMT | ASSATGGLNQDTQY | 0.8380904 |
| QVEVAKRMGVRQPTVS | ASSATGGLNQDTQY | 0.795339 |
| QGSGGPALAM | ASSATGGLNQDTQY | 0.72666556 |
| MLAPSNP | ASSATGGLNQDTQY | 0.83319867 |
| LVNNAAI | ASSATGGLNQDTQY | 0.795613 |
| LTGEINTLA | ASSATGGLNQDTQY | 0.588598 |
| LSIFGPV | ASSATGGLNQDTQY | 0.6529393 |
| LLALLANL | ASSATGGLNQDTQY | 0.81453145 |
| LISNASDALDKLR | ASSATGGLNQDTQY | 0.75914574 |
| LINEAVP | ASSATGGLNQDTQY | 0.171385 |
| LGYAFSGL | ASSATGGLNQDTQY | 0.8232061 |
| LDFVAVAL | ASSATGGLNQDTQY | 0.5031803 |
| LAQPGAE | ASSATGGLNQDTQY | 0.6277523 |
| ISNLGAVTGF | ASSATGGLNQDTQY | 0.793471 |
| HQEDPGRI | ASSATGGLNQDTQY | 0.85425633 |
| GTPGEPTP | ASSATGGLNQDTQY | 0.7272723 |
| GFFNSGS | ASSATGGLNQDTQY | 0.58030117 |
| GAGGGGLTAGAN | ASSATGGLNQDTQY | 0.8673307 |
| FVDALVNM | ASSATGGLNQDTQY | 0.78253853 |
| FLTVAPEL | ASSATGGLNQDTQY | 0.83056706 |
| FETLTAL | ASSATGGLNQDTQY | 0.7908418 |
| DVINPIEL | ASSATGGLNQDTQY | 0.74471587 |
| APVSGAPV | ASSATGGLNQDTQY | 0.81189513 |
| ALVGGLAW | ASSATGGLNQDTQY | 0.57924616 |
| ALAAAAATAI | ASSATGGLNQDTQY | 0.7430811 |
| AFGLGLLL | ASSATGGLNQDTQY | 0.6933931 |

| | | |
|---|---|---|
| AFFAAASL | ASSATGGLNQDTQY | 0.6131983 |
| AAAALPAFSPP | ASSATGGLNQDTQY | 0.70532346 |
| VVLAPAL | ASSATGGLNQDTQY | 0.8029996 |
| TIGHVDHGKTTL | ASSATGGLNQDTQY | 0.88763803 |
| LISNASDALDK | ASSATGGLNQDTQY | 0.63720787 |
| LISNASDAL | ASSATGGLNQDTQY | 0.42290747 |
| ISNASDALDK | ASSATGGLNQDTQY | 0.76093787 |
| NQAETLVYQTE | ASSATGGLNQDTQY | 0.7389548 |
| TGGTGGTAGAG | ASSATGGLNQDTQY | 0.85313267 |
| TGSTAGGAGGA | ASSATGGLNQDTQY | 0.8036376 |
| DPLAVPVAAGPV | ASSATGGLNQDTQY | 0.8837565 |
| GGTGGTPGNGGN | ASSATGGLNQDTQY | 0.8272029 |
| PAGLAGVDPATL | ASSATGGLNQDTQY | 0.68196774 |
| GNNLVVEAPHAGASV | ASSATGGLNQDTQY | 0.8699323 |
| KVTRIMSASTLEQPA | ASSATGGLNQDTQY | 0.7216362 |
| LGFALPIALPRRTVV | ASSATGGLNQDTQY | 0.71656597 |
| TRLGNLTARKLVVLV | ASSATGGLNQDTQY | 0.8129535 |
| WSQQRRLIYFPSAGP | ASSATGGLNQDTQY | 0.8113667 |
| GPNGNPDPMAAAVDIR | ASSATGGLNQDTQY | 0.7447777 |
| TQRCVSVADLRAAASR | ASSATGGLNQDTQY | 0.7539166 |
| QFVLDVAEHPVEVLTETV | ASSATGGLNQDTQY | 0.42156395 |
| GGAGGTGGAGGNGGTQAGDG | ASSATGGLNQDTQY | 0.863189 |
| PLLATLRAIAVDVDAKPAQV | ASSATGGLNQDTQY | 0.5081854 |
| APKFRTGGYMIDFWGVGYQVAK | ASSATGGLNQDTQY | 0.53836614 |
| GAGGAGGAGGAAGGPGGTGGTG | ASSATGGLNQDTQY | 0.69111073 |
| IGALGLLATDVAPAGSVAFGVA | ASSATGGLNQDTQY | 0.8840466 |
| AGGAGGIGGNGGAGGAGGVLMG | ASSATGGLNQDTQY | 0.8612388 |
| ADELIGLL | ASSATGGLNQDTQY | 0.6917565 |
| ADRALKA | ASSATGGLNQDTQY | 0.6607761 |
| AGAVGQVGGAGGA | ASSATGGLNQDTQY | 0.8409097 |
| AGGGGGLGGAP | ASSATGGLNQDTQY | 0.67588544 |
| AGGLLYGNGGAG | ASSATGGLNQDTQY | 0.8595335 |
| AGLESIGL | ASSATGGLNQDTQY | 0.7447952 |
| AGLKTGII | ASSATGGLNQDTQY | 0.6931491 |
| ASLAALLL | ASSATGGLNQDTQY | 0.58462477 |
| ASVINISSI | ASSATGGLNQDTQY | 0.84129876 |
| ATPPGPA | ASSATGGLNQDTQY | 0.6935808 |
| ATVASALV | ASSATGGLNQDTQY | 0.8563511 |
| ATVASLIL | ASSATGGLNQDTQY | 0.3557709 |
| AVNGPVLIRH | ASSATGGLNQDTQY | 0.8408162 |
| DGAAGAVGI | ASSATGGLNQDTQY | 0.58336866 |
| DPLLHGI | ASSATGGLNQDTQY | 0.61577743 |
| ELGAALGAL | ASSATGGLNQDTQY | 0.22437151 |
| ELGRTQAV | ASSATGGLNQDTQY | 0.70813096 |
| FVAGGYLS | ASSATGGLNQDTQY | 0.8207418 |
| GASGGAGGAGGW | ASSATGGLNQDTQY | 0.8759807 |
| GGAGAGGDGGAGGI | ASSATGGLNQDTQY | 0.8253099 |
| GTGHNGSQPGMGGNG | ASSATGGLNQDTQY | 0.8340621 |
| GYAAASASAS | ASSATGGLNQDTQY | 0.85540515 |
| HQVDVAAVQQG | ASSATGGLNQDTQY | 0.7212151 |
| IGAAIAHT | ASSATGGLNQDTQY | 0.30528468 |

| | | |
|---|---|---|
| IGMIEVAQL | ASSATGGLNQDTQY | 0.7864963 |
| IRLAPEHTSAMD | ASSATGGLNQDTQY | 0.7859739 |
| IYNVSVL | ASSATGGLNQDTQY | 0.8336872 |
| KQGFLGM | ASSATGGLNQDTQY | 0.8471581 |
| LESVITQL | ASSATGGLNQDTQY | 0.5636656 |
| LGLGALVVV | ASSATGGLNQDTQY | 0.8366858 |
| LIGNGGNGGSGGI | ASSATGGLNQDTQY | 0.7424977 |
| LNAINAPTL | ASSATGGLNQDTQY | 0.45392254 |
| LQLLNTGV | ASSATGGLNQDTQY | 0.7270671 |
| LQTSIEF | ASSATGGLNQDTQY | 0.5256821 |
| MSFAVSEL | ASSATGGLNQDTQY | 0.34956974 |
| NTGALGL | ASSATGGLNQDTQY | 0.8199408 |
| PPGAGGAGGAGTT | ASSATGGLNQDTQY | 0.72245836 |
| QLVGLGI | ASSATGGLNQDTQY | 0.6883194 |
| RFLEQVHQL | ASSATGGLNQDTQY | 0.728415 |
| RGVGYVLR | ASSATGGLNQDTQY | 0.64064854 |
| SATGVFAGVI | ASSATGGLNQDTQY | 0.67501324 |
| SLPAITLP | ASSATGGLNQDTQY | 0.6664778 |
| STALAFTV | ASSATGGLNQDTQY | 0.7734481 |
| SVLTVGLGK | ASSATGGLNQDTQY | 0.8593651 |
| TALAGAGSGI | ASSATGGLNQDTQY | 0.6905708 |
| TGLPFTGI | ASSATGGLNQDTQY | 0.7601762 |
| VAEHVTV | ASSATGGLNQDTQY | 0.86948323 |
| VDIVLNSL | ASSATGGLNQDTQY | 0.3504977 |
| VLTGGGAVL | ASSATGGLNQDTQY | 0.55088603 |
| VNVGLAYGI | ASSATGGLNQDTQY | 0.8226906 |
| VSSAAVGAV | ASSATGGLNQDTQY | 0.7221842 |
| VTSVYEAGV | ASSATGGLNQDTQY | 0.8571336 |
| WVALDRGA | ASSATGGLNQDTQY | 0.76630926 |
| YLVGSVVI | ASSATGGLNQDTQY | 0.8086624 |
| GPPGTGKTLL | ASSATGGLNQDTQY | 0.78187275 |
| GVGMTKF | ASSATGGLNQDTQY | 0.88949835 |
| IGIGGSDL | ASSATGGLNQDTQY | 0.81428456 |
| IGTLLAEL | ASSATGGLNQDTQY | 0.71193635 |
| IYTSGTT | ASSATGGLNQDTQY | 0.8444621 |
| LSFPAGV | ASSATGGLNQDTQY | 0.42002743 |
| LYTSGTT | ASSATGGLNQDTQY | 0.8344964 |
| VGAFPFGF | ASSATGGLNQDTQY | 0.84651995 |
| VGAGTVEF | ASSATGGLNQDTQY | 0.60841715 |
| VGKSTLLL | ASSATGGLNQDTQY | 0.72571623 |
| AALAHAGGGIV | ASSATGGLNQDTQY | 0.8919935 |
| AALVAVSLL | ASSATGGLNQDTQY | 0.869851 |
| AAVSRAVAAL | ASSATGGLNQDTQY | 0.7223201 |
| AGGSSGIGGF | ASSATGGLNQDTQY | 0.62626404 |
| AGLTVANANG | ASSATGGLNQDTQY | 0.5529134 |
| AGLVGAVL | ASSATGGLNQDTQY | 0.64027 |
| ALSQAFTV | ASSATGGLNQDTQY | 0.32338518 |
| ALTFSVSL | ASSATGGLNQDTQY | 0.8205972 |
| ALTGAALVAL | ASSATGGLNQDTQY | 0.73573613 |
| ASFSIGAGVL | ASSATGGLNQDTQY | 0.8600808 |
| DGSVYKI | ASSATGGLNQDTQY | 0.82430077 |

| | | |
|---|---|---|
| EQLAEIV | ASSATGGLNQDTQY | 0.86980593 |
| FAVSELTVM | ASSATGGLNQDTQY | 0.7313317 |
| GAAPLAAGL | ASSATGGLNQDTQY | 0.7023386 |
| GADGGDGGSGGAGG | ASSATGGLNQDTQY | 0.34115303 |
| GFGLDTGGAGG | ASSATGGLNQDTQY | 0.35882568 |
| GLASGGLL | ASSATGGLNQDTQY | 0.6668668 |
| GRIAGAAVLL | ASSATGGLNQDTQY | 0.6704677 |
| GSAGSLNTSF | ASSATGGLNQDTQY | 0.2885967 |
| GTGQIGI | ASSATGGLNQDTQY | 0.85944027 |
| GTLIALIL | ASSATGGLNQDTQY | 0.6955581 |
| IAATMGFVVG | ASSATGGLNQDTQY | 0.84359574 |
| IFAQALL | ASSATGGLNQDTQY | 0.696375 |
| IGAAVALGF | ASSATGGLNQDTQY | 0.55570316 |
| IGLTPLAL | ASSATGGLNQDTQY | 0.724094 |
| ILGSNTSTL | ASSATGGLNQDTQY | 0.8357709 |
| IVFSLTVL | ASSATGGLNQDTQY | 0.5427507 |
| LASKATGF | ASSATGGLNQDTQY | 0.34462684 |
| LASVALIL | ASSATGGLNQDTQY | 0.5788511 |
| LFAVGLTL | ASSATGGLNQDTQY | 0.75504994 |
| LFNGGIGTYI | ASSATGGLNQDTQY | 0.89393735 |
| LTLGAGII | ASSATGGLNQDTQY | 0.8605269 |
| LVAAALGDI | ASSATGGLNQDTQY | 0.7965785 |
| NTAFVQLGI | ASSATGGLNQDTQY | 0.90285033 |
| QLFSGAEL | ASSATGGLNQDTQY | 0.142394 |
| SAATFITL | ASSATGGLNQDTQY | 0.70727074 |
| SNYGLANLGV | ASSATGGLNQDTQY | 0.86678165 |
| SYGGAYAVM | ASSATGGLNQDTQY | 0.8216924 |
| TTGVGGLTLG | ASSATGGLNQDTQY | 0.74770135 |
| TTQTIYI | ASSATGGLNQDTQY | 0.8709642 |
| VAAGAGLGAVI | ASSATGGLNQDTQY | 0.594784 |
| VAAVVTLL | ASSATGGLNQDTQY | 0.74722034 |
| VGFLGGLI | ASSATGGLNQDTQY | 0.77388954 |
| VGVIPILL | ASSATGGLNQDTQY | 0.85938776 |
| VIFASSLI | ASSATGGLNQDTQY | 0.7234554 |
| VKGGDGGAAAG | ASSATGGLNQDTQY | 0.45962268 |
| VTKTFTNI | ASSATGGLNQDTQY | 0.83029294 |
| VVAAAILL | ASSATGGLNQDTQY | 0.8968432 |
| YAALDVEL | ASSATGGLNQDTQY | 0.7057209 |
| LASVALLL | ASSATGGLNQDTQY | 0.858563 |
| LDAVHRV | ASSATGGLNQDTQY | 0.7846241 |
| QGLSSSIF | ASSATGGLNQDTQY | 0.6605597 |
| RGTFANIRL | ASSATGGLNQDTQY | 0.6760706 |
| RQLWWGHRI | ASSATGGLNQDTQY | 0.5510879 |
| YGPPGTGKTLL | ASSATGGLNQDTQY | 0.8364879 |
| AALVGAPANVA | ASSATGGLNQDTQY | 0.5035194 |
| AATFRRTE | ASSATGGLNQDTQY | 0.7580838 |
| AGALGGGI | ASSATGGLNQDTQY | 0.75836384 |
| AGGAGGAGGIGAG | ASSATGGLNQDTQY | 0.9016479 |
| AGSGALGL | ASSATGGLNQDTQY | 0.835642 |
| AQIADQI | ASSATGGLNQDTQY | 0.82100976 |
| ASAVAVTYL | ASSATGGLNQDTQY | 0.65616715 |

| | | |
|---|---|---|
| ASTVLGGTGG | ASSATGGLNQDTQY | 0.83665746 |
| AVGGEPAK | ASSATGGLNQDTQY | 0.8175372 |
| CEQGTFGGTC | ASSATGGLNQDTQY | 0.47269017 |
| DEGKAPAG | ASSATGGLNQDTQY | 0.86193573 |
| DTVVFPAL | ASSATGGLNQDTQY | 0.5320285 |
| FEGITMAG | ASSATGGLNQDTQY | 0.74213654 |
| FPLVVQD | ASSATGGLNQDTQY | 0.75041926 |
| GAFTTPQ | ASSATGGLNQDTQY | 0.86606944 |
| GFGGFGTAGGVGGA | ASSATGGLNQDTQY | 0.7905868 |
| GFLAQFL | ASSATGGLNQDTQY | 0.8400667 |
| GGAGGAGGNAGVA | ASSATGGLNQDTQY | 0.6028291 |
| GGVGTAGAGGAGGAGG | ASSATGGLNQDTQY | 0.8411944 |
| GNGGGGDAGNAGSGGN | ASSATGGLNQDTQY | 0.86948043 |
| GTGGAGGAGNDAGS | ASSATGGLNQDTQY | 0.7761209 |
| GTTNTGWFNS | ASSATGGLNQDTQY | 0.899244 |
| IFMGPLFTL | ASSATGGLNQDTQY | 0.70153975 |
| ILAHAIEAI | ASSATGGLNQDTQY | 0.28068295 |
| IRSAIREAL | ASSATGGLNQDTQY | 0.7534703 |
| ITFAGGG | ASSATGGLNQDTQY | 0.86113906 |
| ITGMSGHQEV | ASSATGGLNQDTQY | 0.8779113 |
| IVAGLACTL | ASSATGGLNQDTQY | 0.8314834 |
| IVGGGLAGA | ASSATGGLNQDTQY | 0.85005987 |
| IVTGVLNI | ASSATGGLNQDTQY | 0.7059653 |
| LAQQGCI | ASSATGGLNQDTQY | 0.8721833 |
| LCLHVEN | ASSATGGLNQDTQY | 0.89644915 |
| LDAISDGS | ASSATGGLNQDTQY | 0.8418357 |
| LGGFAAML | ASSATGGLNQDTQY | 0.5029938 |
| LGGKNPMI | ASSATGGLNQDTQY | 0.43098342 |
| LGPKGVAAIL | ASSATGGLNQDTQY | 0.846478 |
| LMVGVSI | ASSATGGLNQDTQY | 0.85279536 |
| LVGAAFVL | ASSATGGLNQDTQY | 0.5169455 |
| LVLAVAPL | ASSATGGLNQDTQY | 0.7308855 |
| MAAIAVIDAA | ASSATGGLNQDTQY | 0.75372595 |
| NNGINAELQ | ASSATGGLNQDTQY | 0.2592642 |
| RGEPLVLVHG | ASSATGGLNQDTQY | 0.7688359 |
| RGGAGGAAGLI | ASSATGGLNQDTQY | 0.8835938 |
| RSYRMHPAV | ASSATGGLNQDTQY | 0.8387315 |
| SAAHLLAGQLA | ASSATGGLNQDTQY | 0.43569094 |
| SAAVGSY | ASSATGGLNQDTQY | 0.86728954 |
| SPHAVLHRI | ASSATGGLNQDTQY | 0.8750076 |
| SQIAKVL | ASSATGGLNQDTQY | 0.8851305 |
| TALGHLVP | ASSATGGLNQDTQY | 0.3695132 |
| TAPTTSGAA | ASSATGGLNQDTQY | 0.17352046 |
| TDGDDGNGGSPGT | ASSATGGLNQDTQY | 0.79669815 |
| TDGLGEN | ASSATGGLNQDTQY | 0.83951795 |
| TGGPPLPGQA | ASSATGGLNQDTQY | 0.64125556 |
| TIPAITI | ASSATGGLNQDTQY | 0.5753609 |
| TIPGFTI | ASSATGGLNQDTQY | 0.7106688 |
| TSIASGALA | ASSATGGLNQDTQY | 0.8527639 |
| TVTLPGAAI | ASSATGGLNQDTQY | 0.5847609 |
| VASGLSAALP | ASSATGGLNQDTQY | 0.5466422 |

| | | |
|---|---|---|
| VGAGVIGI | ASSATGGLNQDTQY | 0.7979296 |
| VGPKRFGV | ASSATGGLNQDTQY | 0.8711652 |
| VNVAFPD | ASSATGGLNQDTQY | 0.60419077 |
| VSMTNSGVS | ASSATGGLNQDTQY | 0.7567444 |
| VTDVLGDLSV | ASSATGGLNQDTQY | 0.8012694 |
| VVVVGAGFAGL | ASSATGGLNQDTQY | 0.7473053 |
| VYTVGTVV | ASSATGGLNQDTQY | 0.7810348 |
| WTTFSGP | ASSATGGLNQDTQY | 0.4234973 |
| YDELVEAF | ASSATGGLNQDTQY | 0.82926965 |
| YGTGGAGGNGG | ASSATGGLNQDTQY | 0.55306566 |
| YTNVMAV | ASSATGGLNQDTQY | 0.8643142 |
| FLEQVHQL | ASSATGGLNQDTQY | 0.8110417 |
| GLSLLAVAV | ASSATGGLNQDTQY | 0.8365782 |
| LEQVHQL | ASSATGGLNQDTQY | 0.7451754 |
| LNAAGAI | ASSATGGLNQDTQY | 0.44453925 |
| PAVHASL | ASSATGGLNQDTQY | 0.5953734 |
| TELEDDA | ASSATGGLNQDTQY | 0.59771854 |
| TPPGPPN | ASSATGGLNQDTQY | 0.8745229 |
| TPPPPPPP | ASSATGGLNQDTQY | 0.7372676 |
| TQSPALI | ASSQDPGTKNTLY | 0.86525786 |
| EFAAAFPQ | ASSQDPGTKNTLY | 0.808465 |
| LLSGNLDVL | ASSQDPGTKNTLY | 0.5527005 |
| GPANLGNYNI | ASSQDPGTKNTLY | 0.8497448 |
| TLGPTLP | ASSQDPGTKNTLY | 0.4393952 |
| VYAYSSCCK | ASSQDPGTKNTLY | 0.8454209 |
| APAPAPAPNI | ASSQDPGTKNTLY | 0.5678757 |
| PGGPVPIP | ASSQDPGTKNTLY | 0.83561444 |
| FLTVALEP | ASSQDPGTKNTLY | 0.7299727 |
| SQYFGANL | ASSQDPGTKNTLY | 0.84168255 |
| GGVKPAA | ASSQDPGTKNTLY | 0.8196584 |
| GGTGGAGGAGGW | ASSQDPGTKNTLY | 0.48317862 |
| GVGGVAGLGGAGG | ASSQDPGTKNTLY | 0.82930803 |
| VLLVLATTI | ASSQDPGTKNTLY | 0.8355381 |
| IGIFAAML | ASSQDPGTKNTLY | 0.8466022 |
| AVGDLID | ASSQDPGTKNTLY | 0.45263618 |
| NISPLPL | ASSQDPGTKNTLY | 0.77570426 |
| VTAVFVAI | ASSQDPGTKNTLY | 0.6536515 |
| SGDDNDAAL | ASSQDPGTKNTLY | 0.8102838 |
| AVAALAHS | ASSQDPGTKNTLY | 0.7910751 |
| GGVTVMATP | ASSQDPGTKNTLY | 0.636842 |
| AGLAANVRD | ASSQDPGTKNTLY | 0.78391737 |
| TNRPDLIDPALLRPG | ASSQDPGTKNTLY | 0.3165043 |
| LANLGVTAL | ASSQDPGTKNTLY | 0.6987943 |
| TGERIEL | ASSQDPGTKNTLY | 0.7011642 |
| NGEELAA | ASSQDPGTKNTLY | 0.79759336 |
| VLSLVWG | ASSQDPGTKNTLY | 0.8293381 |
| IGYALWL | ASSQDPGTKNTLY | 0.47246674 |
| LSLLLAPL | ASSQDPGTKNTLY | 0.7912951 |
| TLDEALK | ASSQDPGTKNTLY | 0.7242551 |
| TDGAGTAEGG | ASSQDPGTKNTLY | 0.6019056 |
| NAPGGAGGNGG | ASSQDPGTKNTLY | 0.8246408 |

| | | |
|---|---|---|
| VSVFMGMGGNF | ASSQDPGTKNTLY | 0.63925946 |
| LASYLAAL | ASSQDPGTKNTLY | 0.3420087 |
| LVGGTPAL | ASSQDPGTKNTLY | 0.851979 |
| VRNADVL | ASSQDPGTKNTLY | 0.55672246 |
| TCSSPFLVSAM | ASSQDPGTKNTLY | 0.7829766 |
| VHITDAE | ASSQDPGTKNTLY | 0.6840068 |
| EYAAPSF | ASSQDPGTKNTLY | 0.58220434 |
| VVDAFHGGL | ASSQDPGTKNTLY | 0.8616278 |
| ASVVDPQ | ASSQDPGTKNTLY | 0.7293278 |
| SYNEFGGLS | ASSQDPGTKNTLY | 0.826197 |
| DPDGSVY | ASSQDPGTKNTLY | 0.72570115 |
| MVSTIHATE | ASSQDPGTKNTLY | 0.6149148 |
| LAAGVADPV | ASSQDPGTKNTLY | 0.86051655 |
| TKEQIAA | ASSQDPGTKNTLY | 0.8204054 |
| SLASPAAL | ASSQDPGTKNTLY | 0.81326365 |
| IGLVFYGV | ASSQDPGTKNTLY | 0.7834468 |
| FLLDASNL | ASSQDPGTKNTLY | 0.8477825 |
| YQVGNFP | ASSQDPGTKNTLY | 0.85393614 |
| ESLLSQST | ASSQDPGTKNTLY | 0.6320266 |
| LFTVGFAAVG | ASSQDPGTKNTLY | 0.29730856 |
| ATPPTPT | ASSQDPGTKNTLY | 0.73610234 |
| TDLATLG | ASSQDPGTKNTLY | 0.6868302 |
| SAGGTGR | ASSQDPGTKNTLY | 0.6297984 |
| GFSNSGS | ASSQDPGTKNTLY | 0.3310167 |
| GGNGGSGGTSV | ASSQDPGTKNTLY | 0.8188051 |
| GTSVATGGAGN | ASSQDPGTKNTLY | 0.89931905 |
| NGGNGGSGGTS | ASSQDPGTKNTLY | 0.6186218 |
| AVVKFPH | ASSQDPGTKNTLY | 0.8974496 |
| TPPPPPP | ASSQDPGTKNTLY | 0.53643477 |
| AAPAAAAAPA | ASSQDPGTKNTLY | 0.83739316 |
| IALNEDL | ASSQDPGTKNTLY | 0.8483605 |
| IVNVSSM | ASSQDPGTKNTLY | 0.52792764 |
| TGIPPAPR | ASSQDPGTKNTLY | 0.7505315 |
| FDLGGGTF | ASSQDPGTKNTLY | 0.6914878 |
| FDLGGGTFDVS | ASSQDPGTKNTLY | 0.68899083 |
| NEPTAAA | ASSQDPGTKNTLY | 0.8644769 |
| NEPTAAALA | ASSQDPGTKNTLY | 0.5933897 |
| GSIINISSI | ASSQDPGTKNTLY | 0.82823783 |
| VAPGNAGTA | ASSQDPGTKNTLY | 0.7355448 |
| TGGGASLE | ASSQDPGTKNTLY | 0.63174164 |
| VAVYNHYKRI | ASSQDPGTKNTLY | 0.83734834 |
| TDPAPATT | ASSQDPGTKNTLY | 0.85910475 |
| TNRPDILDPALLRPG | ASSQDPGTKNTLY | 0.8045227 |
| LTTTTTS | ASSQDPGTKNTLY | 0.89247966 |
| PTGSGKTL | ASSQDPGTKNTLY | 0.7201551 |
| TGSGKTL | ASSQDPGTKNTLY | 0.89109224 |
| TIGHVDHGK | ASSQDPGTKNTLY | 0.88434327 |
| TIGHVDHGKTT | ASSQDPGTKNTLY | 0.79171073 |
| GHVDHGKTT | ASSQDPGTKNTLY | 0.7554928 |
| AQFEHTLL | ASSQDPGTKNTLY | 0.8510891 |
| IVNQDEL | ASSQDPGTKNTLY | 0.69115317 |

| | | |
|---|---|---|
| TNGSQFF | ASSQDPGTKNTLY | 0.85837483 |
| TGKGIVE | ASSQDPGTKNTLY | 0.87051684 |
| LISNASDA | ASSQDPGTKNTLY | 0.71571827 |
| AELRTLGELTKIAKAHGAQ | ASSQDPGTKNTLY | 0.82881415 |
| APNAVAAAEPSW | ASSQDPGTKNTLY | 0.7974454 |
| APQVAGAATAVL | ASSQDPGTKNTLY | 0.75655377 |
| ARRDPAAAAAILPTDA | ASSQDPGTKNTLY | 0.7071826 |
| DAAARDVLAEAGLAEYF | ASSQDPGTKNTLY | 0.31751907 |
| DDQIVLSAKANLAKVEQ | ASSQDPGTKNTLY | 0.7814274 |
| DELLPDTGEDSDEDGH | ASSQDPGTKNTLY | 0.64482945 |
| ELPEAPANRATVPQD | ASSQDPGTKNTLY | 0.80518794 |
| FGQSKIERVTGIG | ASSQDPGTKNTLY | 0.77547455 |
| GAGGTGSTAGGAGGAGGAGGLY | ASSQDPGTKNTLY | 0.6859789 |
| GHGGTGAAGGNGGTGGAGGSGI | ASSQDPGTKNTLY | 0.83169186 |
| GNGGNGGAGGEGGPGGAGGAGA | ASSQDPGTKNTLY | 0.8316368 |
| GSTAGNGGNGGNATNGTIAGQP | ASSQDPGTKNTLY | 0.5553847 |
| GTGGDGGNGGNGGTGG | ASSQDPGTKNTLY | 0.76127636 |
| GVRRPAGIPADAPV | ASSQDPGTKNTLY | 0.7209367 |
| GYRQVTDAHLLTLARR | ASSQDPGTKNTLY | 0.7518789 |
| IAKMGQDGHDRG | ASSQDPGTKNTLY | 0.39231092 |
| KAEPSIEIQYT | ASSQDPGTKNTLY | 0.8151839 |
| KPVRIRMAADEDTVAVT | ASSQDPGTKNTLY | 0.720916 |
| MPPRIAGMRLLVIKPEP | ASSQDPGTKNTLY | 0.16579863 |
| MKFYAVALFRSRGVPPTR | ASSQDPGTKNTLY | 0.75575167 |
| NAGGFGILGAVAHSP | ASSQDPGTKNTLY | 0.7610916 |
| NEGREITLAMAEM | ASSQDPGTKNTLY | 0.74821186 |
| NELVEWSHPLGHRWRWE | ASSQDPGTKNTLY | 0.8256705 |
| NLGAGGTGGNGGNPG | ASSQDPGTKNTLY | 0.22626686 |
| NTEWPKWETVMAADPVD | ASSQDPGTKNTLY | 0.6134807 |
| PASAAPAETQP | ASSQDPGTKNTLY | 0.82299984 |
| QQSSDGKNGVRAAEITGE | ASSQDPGTKNTLY | 0.82735074 |
| RAPDAVTTSWWKE | ASSQDPGTKNTLY | 0.48861915 |
| SGFGNVGAHTSGWFNQ | ASSQDPGTKNTLY | 0.81240344 |
| SGVAGAVGQAASVGGLKV | ASSQDPGTKNTLY | 0.8263723 |
| SNGGAGTGSAGGNGGT | ASSQDPGTKNTLY | 0.7920654 |
| TGGAGGEGGAAG | ASSQDPGTKNTLY | 0.81922215 |
| TGGDGGAAGAGGAPGHG | ASSQDPGTKNTLY | 0.8252741 |
| TMPPLLAAIANAGRHGV | ASSQDPGTKNTLY | 0.61437815 |
| TSSTAGNSGGA | ASSQDPGTKNTLY | 0.8229719 |
| VPATVPVGILC | ASSQDPGTKNTLY | 0.83217835 |
| VQRPDSGGVVLTGRIS | ASSQDPGTKNTLY | 0.66256994 |
| WTPNPVPATKGVPAGLVQAT | ASSQDPGTKNTLY | 0.79811764 |
| YALKRALAKFSDRFDVVII | ASSQDPGTKNTLY | 0.660882 |
| YLIGSVIV | ASSQDPGTKNTLY | 0.68321365 |
| YGQLVAVID | ASSQDPGTKNTLY | 0.20566785 |
| VTVGGMTLPA | ASSQDPGTKNTLY | 0.81378275 |
| VLALPAV | ASSQDPGTKNTLY | 0.80700874 |
| VIPPTLNYET | ASSQDPGTKNTLY | 0.7762497 |
| VGMGGMHPSSG | ASSQDPGTKNTLY | 0.66360205 |
| VGLAGGAATA | ASSQDPGTKNTLY | 0.6711076 |
| VGGAGGAGGSGP | ASSQDPGTKNTLY | 0.71702003 |

| | | |
|---|---|---|
| VFAAFMV | ASSQDPGTKNTLY | 0.74955845 |
| TLYDKLR | ASSQDPGTKNTLY | 0.2562781 |
| TLAVFTVL | ASSQDPGTKNTLY | 0.65813637 |
| TGSVQPARV | ASSQDPGTKNTLY | 0.8240225 |
| TGGGAGSGGT | ASSQDPGTKNTLY | 0.8243396 |
| SHLETSL | ASSQDPGTKNTLY | 0.43482754 |
| SFTTGMT | ASSQDPGTKNTLY | 0.8294096 |
| QVEVAKRMGVRQPTVS | ASSQDPGTKNTLY | 0.80824125 |
| QGSGGPALAM | ASSQDPGTKNTLY | 0.82896215 |
| MLAPSNP | ASSQDPGTKNTLY | 0.7494131 |
| LVNNAAI | ASSQDPGTKNTLY | 0.71038234 |
| LTGEINTLA | ASSQDPGTKNTLY | 0.8157146 |
| LSIFGPV | ASSQDPGTKNTLY | 0.7741269 |
| LLALLANL | ASSQDPGTKNTLY | 0.37845132 |
| LISNASDALDKLR | ASSQDPGTKNTLY | 0.81683344 |
| LINEAVP | ASSQDPGTKNTLY | 0.72699463 |
| LGYAFSGL | ASSQDPGTKNTLY | 0.71932304 |
| LDFVAVAL | ASSQDPGTKNTLY | 0.75206447 |
| LAQPGAE | ASSQDPGTKNTLY | 0.80103236 |
| ISNLGAVTGF | ASSQDPGTKNTLY | 0.65837085 |
| HQEDPGRI | ASSQDPGTKNTLY | 0.40722954 |
| GTPGEPTP | ASSQDPGTKNTLY | 0.57776594 |
| GFFNSGS | ASSQDPGTKNTLY | 0.8028839 |
| GAGGGGLTAGAN | ASSQDPGTKNTLY | 0.8239371 |
| FVDALVNM | ASSQDPGTKNTLY | 0.7819572 |
| FLTVAPEL | ASSQDPGTKNTLY | 0.6203016 |
| FETLTAL | ASSQDPGTKNTLY | 0.81907016 |
| DVINPIEL | ASSQDPGTKNTLY | 0.787531 |
| APVSGAPV | ASSQDPGTKNTLY | 0.77088624 |
| ALVGGLAW | ASSQDPGTKNTLY | 0.8158256 |
| ALAAAAATAI | ASSQDPGTKNTLY | 0.7464218 |
| AFGLGLLL | ASSQDPGTKNTLY | 0.8042476 |
| AFFAAASL | ASSQDPGTKNTLY | 0.79880387 |
| AAAALPAFSPP | ASSQDPGTKNTLY | 0.80837595 |
| VVLAPAL | ASSQDPGTKNTLY | 0.814072 |
| TIGHVDHGKTTL | ASSQDPGTKNTLY | 0.72047216 |
| LISNASDALDK | ASSQDPGTKNTLY | 0.73898554 |
| LISNASDAL | ASSQDPGTKNTLY | 0.25505167 |
| ISNASDALDK | ASSQDPGTKNTLY | 0.7232585 |
| NQAETLVYQTE | ASSQDPGTKNTLY | 0.80908996 |
| TGGTGGTAGAG | ASSQDPGTKNTLY | 0.7992995 |
| TGSTAGGAGGA | ASSQDPGTKNTLY | 0.8161721 |
| DPLAVPVAAGPV | ASSQDPGTKNTLY | 0.75954247 |
| GGTGGTPGNGGN | ASSQDPGTKNTLY | 0.6475832 |
| PAGLAGVDPATL | ASSQDPGTKNTLY | 0.8201792 |
| GNNLVVEAPHAGASV | ASSQDPGTKNTLY | 0.8205431 |
| KVTRIMSASTLEQPA | ASSQDPGTKNTLY | 0.496621 |
| LGFALPIALPRRTVV | ASSQDPGTKNTLY | 0.73042655 |
| TRLGNLTARKLVVLV | ASSQDPGTKNTLY | 0.8345009 |
| WSQQRRLIYFPSAGP | ASSQDPGTKNTLY | 0.80140114 |
| GPNGNPDPMAAAVDIR | ASSQDPGTKNTLY | 0.7731838 |

| | | |
|---|---|---|
| TQRCVSVADLRAAASR | ASSQDPGTKNTLY | 0.663187 |
| QFVLDVAEHPVEVLTETV | ASSQDPGTKNTLY | 0.8274424 |
| GGAGGTGGAGGNGGTQAGDG | ASSQDPGTKNTLY | 0.62940085 |
| PLLATLRAIAVDVDAKPAQV | ASSQDPGTKNTLY | 0.83221304 |
| APKFRTGGYMIDFWGVGYQVAK | ASSQDPGTKNTLY | 0.2153113 |
| GAGGAGGAGGAAGGPGGTGGTG | ASSQDPGTKNTLY | 0.66163266 |
| IGALGLLATDVAPAGSVAFGVA | ASSQDPGTKNTLY | 0.52612096 |
| AGGAGGIGGNGGAGGAGGVLMG | ASSQDPGTKNTLY | 0.38620967 |
| ADELIGLL | ASSQDPGTKNTLY | 0.81934595 |
| ADRALKA | ASSQDPGTKNTLY | 0.7370273 |
| AGAVGQVGGAGGA | ASSQDPGTKNTLY | 0.8234951 |
| AGGGGGLGGAP | ASSQDPGTKNTLY | 0.7981281 |
| AGGLLYGNGGAG | ASSQDPGTKNTLY | 0.8130451 |
| AGLESIGL | ASSQDPGTKNTLY | 0.79857314 |
| AGLKTGII | ASSQDPGTKNTLY | 0.772853 |
| ASLAALLL | ASSQDPGTKNTLY | 0.48490882 |
| ASVINISSI | ASSQDPGTKNTLY | 0.8152183 |
| ATPPGPA | ASSQDPGTKNTLY | 0.82109094 |
| ATVASALV | ASSQDPGTKNTLY | 0.7654278 |
| ATVASLIL | ASSQDPGTKNTLY | 0.40009803 |
| AVNGPVLIRH | ASSQDPGTKNTLY | 0.73317647 |
| DGAAGAVGI | ASSQDPGTKNTLY | 0.79180205 |
| DPLLHGI | ASSQDPGTKNTLY | 0.1820322 |
| ELGAALGAL | ASSQDPGTKNTLY | 0.2113521 |
| ELGRTQAV | ASSQDPGTKNTLY | 0.66978526 |
| FVAGGYLS | ASSQDPGTKNTLY | 0.65031993 |
| GASGGAGGAGGW | ASSQDPGTKNTLY | 0.82924205 |
| GGAGAGGDGGAGGI | ASSQDPGTKNTLY | 0.82373935 |
| GTGHNGSQPGMGGNG | ASSQDPGTKNTLY | 0.7852079 |
| GYAAASASAS | ASSQDPGTKNTLY | 0.80739415 |
| HQVDVAAVQQG | ASSQDPGTKNTLY | 0.79780793 |
| IGAAIAHT | ASSQDPGTKNTLY | 0.15509048 |
| IGMIEVAQL | ASSQDPGTKNTLY | 0.6910201 |
| IRLAPEHTSAMD | ASSQDPGTKNTLY | 0.6123845 |
| IYNVSVL | ASSQDPGTKNTLY | 0.8239851 |
| KQGFLGM | ASSQDPGTKNTLY | 0.6776303 |
| LESVITQL | ASSQDPGTKNTLY | 0.7200764 |
| LGLGALVVV | ASSQDPGTKNTLY | 0.80215245 |
| LIGNGGNGGSGGI | ASSQDPGTKNTLY | 0.7943361 |
| LNAINAPTL | ASSQDPGTKNTLY | 0.7838658 |
| LQLLNTGV | ASSQDPGTKNTLY | 0.5456409 |
| LQTSIEF | ASSQDPGTKNTLY | 0.16752441 |
| MSFAVSEL | ASSQDPGTKNTLY | 0.8174384 |
| NTGALGL | ASSQDPGTKNTLY | 0.25142983 |
| PPGAGGAGGAGTT | ASSQDPGTKNTLY | 0.5213164 |
| QLVGLGI | ASSQDPGTKNTLY | 0.22377652 |
| RFLEQVHQL | ASSQDPGTKNTLY | 0.16765717 |
| RGVGYVLR | ASSQDPGTKNTLY | 0.4152674 |
| SATGVFAGVI | ASSQDPGTKNTLY | 0.80470395 |
| SLPAITLP | ASSQDPGTKNTLY | 0.7681494 |
| STALAFTV | ASSQDPGTKNTLY | 0.7624146 |

| | | |
|---|---|---|
| SVLTVGLGK | ASSQDPGTKNTLY | 0.8304999 |
| TALAGAGSGI | ASSQDPGTKNTLY | 0.8059753 |
| TGLPFTGI | ASSQDPGTKNTLY | 0.75977457 |
| VAEHVTV | ASSQDPGTKNTLY | 0.61394423 |
| VDIVLNSL | ASSQDPGTKNTLY | 0.61530286 |
| VLTGGGAVL | ASSQDPGTKNTLY | 0.53366125 |
| VNVGLAYGI | ASSQDPGTKNTLY | 0.5545068 |
| VSSAAVGAV | ASSQDPGTKNTLY | 0.83323836 |
| VTSVYEAGV | ASSQDPGTKNTLY | 0.6634033 |
| WVALDRGA | ASSQDPGTKNTLY | 0.5427625 |
| YLVGSVVI | ASSQDPGTKNTLY | 0.7509026 |
| GPPGTGKTLL | ASSQDPGTKNTLY | 0.83032835 |
| GVGMTKF | ASSQDPGTKNTLY | 0.833292 |
| IGIGGSDL | ASSQDPGTKNTLY | 0.67776644 |
| IGTLLAEL | ASSQDPGTKNTLY | 0.76350164 |
| IYTSGTT | ASSQDPGTKNTLY | 0.6998875 |
| LSFPAGV | ASSQDPGTKNTLY | 0.8016335 |
| LYTSGTT | ASSQDPGTKNTLY | 0.6755312 |
| VGAFPFGF | ASSQDPGTKNTLY | 0.38536072 |
| VGAGTVEF | ASSQDPGTKNTLY | 0.78853106 |
| VGKSTLLL | ASSQDPGTKNTLY | 0.8325362 |
| AALAHAGGGIV | ASSQDPGTKNTLY | 0.6321095 |
| AALVAVSLL | ASSQDPGTKNTLY | 0.6419283 |
| AAVSRAVAAL | ASSQDPGTKNTLY | 0.79028946 |
| AGGSSGIGGF | ASSQDPGTKNTLY | 0.6685709 |
| AGLTVANANG | ASSQDPGTKNTLY | 0.77026856 |
| AGLVGAVL | ASSQDPGTKNTLY | 0.7647522 |
| ALSQAFTV | ASSQDPGTKNTLY | 0.54958963 |
| ALTFSVSL | ASSQDPGTKNTLY | 0.70299196 |
| ALTGAALVAL | ASSQDPGTKNTLY | 0.8322594 |
| ASFSIGAGVL | ASSQDPGTKNTLY | 0.7816336 |
| DGSVYKI | ASSQDPGTKNTLY | 0.7977039 |
| EQLAEIV | ASSQDPGTKNTLY | 0.8305181 |
| FAVSELTVM | ASSQDPGTKNTLY | 0.81733775 |
| GAAPLAAGL | ASSQDPGTKNTLY | 0.8310078 |
| GADGGDGGSGGAGG | ASSQDPGTKNTLY | 0.81543726 |
| GFGLDTGGAGG | ASSQDPGTKNTLY | 0.6687245 |
| GLASGGLL | ASSQDPGTKNTLY | 0.746452 |
| GRIAGAAVLL | ASSQDPGTKNTLY | 0.79910296 |
| GSAGSLNTSF | ASSQDPGTKNTLY | 0.80394363 |
| GTGQIGI | ASSQDPGTKNTLY | 0.7839722 |
| GTLIALIL | ASSQDPGTKNTLY | 0.64577883 |
| IAATMGFVVG | ASSQDPGTKNTLY | 0.83150995 |
| IFAQALL | ASSQDPGTKNTLY | 0.7483219 |
| IGAAVALGF | ASSQDPGTKNTLY | 0.81231487 |
| IGLTPLAL | ASSQDPGTKNTLY | 0.50753176 |
| ILGSNTSTL | ASSQDPGTKNTLY | 0.47974962 |
| IVFSLTVL | ASSQDPGTKNTLY | 0.82071435 |
| LASKATGF | ASSQDPGTKNTLY | 0.15987308 |
| LASVALIL | ASSQDPGTKNTLY | 0.20847788 |
| LFAVGLTL | ASSQDPGTKNTLY | 0.7690904 |

| | | |
|---|---|---|
| LFNGGIGTYI | ASSQDPGTKNTLY | 0.8272006 |
| LTLGAGII | ASSQDPGTKNTLY | 0.15955074 |
| LVAAALGDI | ASSQDPGTKNTLY | 0.5510906 |
| NTAFVQLGI | ASSQDPGTKNTLY | 0.42069414 |
| QLFSGAEL | ASSQDPGTKNTLY | 0.7792189 |
| SAATFITL | ASSQDPGTKNTLY | 0.77692056 |
| SNYGLANLGV | ASSQDPGTKNTLY | 0.37870836 |
| SYGGAYAVM | ASSQDPGTKNTLY | 0.8089082 |
| TTGVGGLTLG | ASSQDPGTKNTLY | 0.52566385 |
| TTQTIYI | ASSQDPGTKNTLY | 0.28107756 |
| VAAGAGLGAVI | ASSQDPGTKNTLY | 0.80490714 |
| VAAVVTLL | ASSQDPGTKNTLY | 0.8243489 |
| VGFLGGLI | ASSQDPGTKNTLY | 0.58412945 |
| VGVIPILL | ASSQDPGTKNTLY | 0.81771344 |
| VIFASSLI | ASSQDPGTKNTLY | 0.68503904 |
| VKGGDGGAAAG | ASSQDPGTKNTLY | 0.83424205 |
| VTKTFTNI | ASSQDPGTKNTLY | 0.7443102 |
| VVAAAILL | ASSQDPGTKNTLY | 0.39918804 |
| YAALDVEL | ASSQDPGTKNTLY | 0.820812 |
| LASVALLL | ASSQDPGTKNTLY | 0.80010116 |
| LDAVHRV | ASSQDPGTKNTLY | 0.7184639 |
| QGLSSSIF | ASSQDPGTKNTLY | 0.8194829 |
| RGTFANIRL | ASSQDPGTKNTLY | 0.55006725 |
| RQLWWGHRI | ASSQDPGTKNTLY | 0.7925983 |
| YGPPGTGKTLL | ASSQDPGTKNTLY | 0.8314626 |
| AALVGAPANVA | ASSQDPGTKNTLY | 0.82151854 |
| AATFRRTE | ASSQDPGTKNTLY | 0.48322594 |
| AGALGGGI | ASSQDPGTKNTLY | 0.7916856 |
| AGGAGGAGGIGAG | ASSQDPGTKNTLY | 0.792415 |
| AGSGALGL | ASSQDPGTKNTLY | 0.7212944 |
| AQIADQI | ASSQDPGTKNTLY | 0.8282686 |
| ASAVAVTYL | ASSQDPGTKNTLY | 0.7207084 |
| ASTVLGGTGG | ASSQDPGTKNTLY | 0.8162266 |
| AVGGEPAK | ASSQDPGTKNTLY | 0.80425465 |
| CEQGTFGGTC | ASSQDPGTKNTLY | 0.8215492 |
| DEGKAPAG | ASSQDPGTKNTLY | 0.35568792 |
| DTVVFPAL | ASSQDPGTKNTLY | 0.8098227 |
| FEGITMAG | ASSQDPGTKNTLY | 0.7812277 |
| FPLVVQD | ASSQDPGTKNTLY | 0.8143897 |
| GAFTTPQ | ASSQDPGTKNTLY | 0.7969394 |
| GFGGFGTAGGVGGA | ASSQDPGTKNTLY | 0.32771754 |
| GFLAQFL | ASSQDPGTKNTLY | 0.7811231 |
| GGAGGAGGNAGVA | ASSQDPGTKNTLY | 0.8243992 |
| GGVGTAGAGGAGGAGG | ASSQDPGTKNTLY | 0.7298504 |
| GNGGGGDAGNAGSGGN | ASSQDPGTKNTLY | 0.7938275 |
| GTGGAGGAGNDAGS | ASSQDPGTKNTLY | 0.8331768 |
| GTTNTGWFNS | ASSQDPGTKNTLY | 0.67899203 |
| IFMGPLFTL | ASSQDPGTKNTLY | 0.7620517 |
| ILAHAIEAI | ASSQDPGTKNTLY | 0.7863896 |
| IRSAIREAL | ASSQDPGTKNTLY | 0.82181674 |
| ITFAGGG | ASSQDPGTKNTLY | 0.73350626 |

| | | |
|---|---|---|
| ITGMSGHQEV | ASSQDPGTKNTLY | 0.8277675 |
| IVAGLACTL | ASSQDPGTKNTLY | 0.5771949 |
| IVGGGLAGA | ASSQDPGTKNTLY | 0.8204351 |
| IVTGVLNI | ASSQDPGTKNTLY | 0.6926557 |
| LAQQGCI | ASSQDPGTKNTLY | 0.8192218 |
| LCLHVEN | ASSQDPGTKNTLY | 0.69423616 |
| LDAISDGS | ASSQDPGTKNTLY | 0.82710266 |
| LGGFAAML | ASSQDPGTKNTLY | 0.8235751 |
| LGGKNPMI | ASSQDPGTKNTLY | 0.826103 |
| LGPKGVAAIL | ASSQDPGTKNTLY | 0.21551692 |
| LMVGVSI | ASSQDPGTKNTLY | 0.82535493 |
| LVGAAFVL | ASSQDPGTKNTLY | 0.825605 |
| LVLAVAPL | ASSQDPGTKNTLY | 0.8358164 |
| MAAIAVIDAA | ASSQDPGTKNTLY | 0.7282899 |
| NNGINAELQ | ASSQDPGTKNTLY | 0.7983744 |
| RGEPLVLVHG | ASSQDPGTKNTLY | 0.74082553 |
| RGGAGGAAGLI | ASSQDPGTKNTLY | 0.7220887 |
| RSYRMHPAV | ASSQDPGTKNTLY | 0.6832347 |
| SAAHLLAGQLA | ASSQDPGTKNTLY | 0.76104194 |
| SAAVGSY | ASSQDPGTKNTLY | 0.7666875 |
| SPHAVLHRI | ASSQDPGTKNTLY | 0.75087374 |
| SQIAKVL | ASSQDPGTKNTLY | 0.8184583 |
| TALGHLVP | ASSQDPGTKNTLY | 0.8230954 |
| TAPTTSGAA | ASSQDPGTKNTLY | 0.81469065 |
| TDGDDGNGGSPGT | ASSQDPGTKNTLY | 0.830228 |
| TDGLGEN | ASSQDPGTKNTLY | 0.8235997 |
| TGGPPLPGQA | ASSQDPGTKNTLY | 0.80575144 |
| TIPAITI | ASSQDPGTKNTLY | 0.7222165 |
| TIPGFTI | ASSQDPGTKNTLY | 0.76313573 |
| TSIASGALA | ASSQDPGTKNTLY | 0.68063414 |
| TVTLPGAAI | ASSQDPGTKNTLY | 0.7201488 |
| VASGLSAALP | ASSQDPGTKNTLY | 0.69558346 |
| VGAGVIGI | ASSQDPGTKNTLY | 0.805676 |
| VGPKRFGV | ASSQDPGTKNTLY | 0.15104342 |
| VNVAFPD | ASSQDPGTKNTLY | 0.7890464 |
| VSMTNSGVS | ASSQDPGTKNTLY | 0.8326746 |
| VTDVLGDLSV | ASSQDPGTKNTLY | 0.69178575 |
| VVVVGAGFAGL | ASSQDPGTKNTLY | 0.8000812 |
| VYTVGTVV | ASSQDPGTKNTLY | 0.7740344 |
| WTTFSGP | ASSQDPGTKNTLY | 0.6780926 |
| YDELVEAF | ASSQDPGTKNTLY | 0.8247739 |
| YGTGGAGGNGG | ASSQDPGTKNTLY | 0.52687687 |
| YTNVMAV | ASSQDPGTKNTLY | 0.8017608 |
| FLEQVHQL | ASSQDPGTKNTLY | 0.82985544 |
| GLSLLAVAV | ASSQDPGTKNTLY | 0.78600466 |
| LEQVHQL | ASSQDPGTKNTLY | 0.78630984 |
| LNAAGAI | ASSQDPGTKNTLY | 0.7362131 |
| PAVHASL | ASSQDPGTKNTLY | 0.32234466 |
| TELEDDA | ASSQDPGTKNTLY | 0.79925966 |
| TPPGPPN | ASSQDPGTKNTLY | 0.14938325 |
| TPPPPPPP | ASSQDPGTKNTLY | 0.79279006 |

| | | |
|---|---|---|
| TQSPALI | ASSSPTKNTLY | 0.40224636 |
| EFAAAFPQ | ASSSPTKNTLY | 0.80980664 |
| LLSGNLDVL | ASSSPTKNTLY | 0.8015712 |
| GPANLGNYNI | ASSSPTKNTLY | 0.78421235 |
| TLGPTLP | ASSSPTKNTLY | 0.7438191 |
| VYAYSSCCK | ASSSPTKNTLY | 0.7367486 |
| APAPAPAPNI | ASSSPTKNTLY | 0.55365086 |
| PGGPVPIP | ASSSPTKNTLY | 0.43052036 |
| FLTVALEP | ASSSPTKNTLY | 0.8219643 |
| SQYFGANL | ASSSPTKNTLY | 0.637305 |
| GGVKPAA | ASSSPTKNTLY | 0.7318926 |
| GGTGGAGGAGGW | ASSSPTKNTLY | 0.82278585 |
| GVGGVAGLGGAGG | ASSSPTKNTLY | 0.75960934 |
| VLLVLATTI | ASSSPTKNTLY | 0.35305125 |
| IGIFAAML | ASSSPTKNTLY | 0.79741776 |
| AVGDLID | ASSSPTKNTLY | 0.78312105 |
| NISPLPL | ASSSPTKNTLY | 0.7316748 |
| VTAVFVAI | ASSSPTKNTLY | 0.15294144 |
| SGDDNDAAL | ASSSPTKNTLY | 0.7566121 |
| AVAALAHS | ASSSPTKNTLY | 0.71995556 |
| GGVTVMATP | ASSSPTKNTLY | 0.71450746 |
| AGLAANVRD | ASSSPTKNTLY | 0.6931691 |
| TNRPDLIDPALLRPG | ASSSPTKNTLY | 0.82643074 |
| LANLGVTAL | ASSSPTKNTLY | 0.23648186 |
| TGERIEL | ASSSPTKNTLY | 0.7898581 |
| NGEELAA | ASSSPTKNTLY | 0.8171431 |
| VLSLVWG | ASSSPTKNTLY | 0.79471153 |
| IGYALWL | ASSSPTKNTLY | 0.83252823 |
| LSLLLAPL | ASSSPTKNTLY | 0.77401155 |
| TLDEALK | ASSSPTKNTLY | 0.65332764 |
| TDGAGTAEGG | ASSSPTKNTLY | 0.77902687 |
| NAPGGAGGNGG | ASSSPTKNTLY | 0.758237 |
| VSVFMGMGGNF | ASSSPTKNTLY | 0.25559354 |
| LASYLAAL | ASSSPTKNTLY | 0.2175292 |
| LVGGTPAL | ASSSPTKNTLY | 0.80623007 |
| VRNADVL | ASSSPTKNTLY | 0.2214738 |
| TCSSPFLVSAM | ASSSPTKNTLY | 0.7856862 |
| VHITDAE | ASSSPTKNTLY | 0.8113226 |
| EYAAPSF | ASSSPTKNTLY | 0.70896864 |
| VVDAFHGGL | ASSSPTKNTLY | 0.4970564 |
| ASVVDPQ | ASSSPTKNTLY | 0.7242284 |
| SYNEFGGLS | ASSSPTKNTLY | 0.71035194 |
| DPDGSVY | ASSSPTKNTLY | 0.81973636 |
| MVSTIHATE | ASSSPTKNTLY | 0.7669132 |
| LAAGVADPV | ASSSPTKNTLY | 0.83043367 |
| TKEQIAA | ASSSPTKNTLY | 0.8277521 |
| SLASPAAL | ASSSPTKNTLY | 0.7823663 |
| IGLVFYGV | ASSSPTKNTLY | 0.81624466 |
| FLLDASNL | ASSSPTKNTLY | 0.7276317 |
| YQVGNFP | ASSSPTKNTLY | 0.78250897 |
| ESLLSQST | ASSSPTKNTLY | 0.71684766 |

| | | |
|---|---|---|
| LFTVGFAAVG | ASSSPTKNTLY | 0.7659396 |
| ATPPTPT | ASSSPTKNTLY | 0.73068064 |
| TDLATLG | ASSSPTKNTLY | 0.64538854 |
| SAGGTGR | ASSSPTKNTLY | 0.40479904 |
| GFSNSGS | ASSSPTKNTLY | 0.7654803 |
| GGNGGSGGTSV | ASSSPTKNTLY | 0.7811482 |
| GTSVATGGAGN | ASSSPTKNTLY | 0.26707578 |
| NGGNGGSGGTS | ASSSPTKNTLY | 0.665399 |
| AVVKFPH | ASSSPTKNTLY | 0.8328946 |
| TPPPPPP | ASSSPTKNTLY | 0.83065724 |
| AAPAAAAAPA | ASSSPTKNTLY | 0.7178236 |
| IALNEDL | ASSSPTKNTLY | 0.5475842 |
| IVNVSSM | ASSSPTKNTLY | 0.78984785 |
| TGIPPAPR | ASSSPTKNTLY | 0.26830566 |
| FDLGGGTF | ASSSPTKNTLY | 0.76290274 |
| FDLGGGTFDVS | ASSSPTKNTLY | 0.79287237 |
| NEPTAAA | ASSSPTKNTLY | 0.7850853 |
| NEPTAAALA | ASSSPTKNTLY | 0.5840689 |
| GSIINISSI | ASSSPTKNTLY | 0.7896805 |
| VAPGNAGTA | ASSSPTKNTLY | 0.74071264 |
| TGGGASLE | ASSSPTKNTLY | 0.7770941 |
| VAVYNHYKRI | ASSSPTKNTLY | 0.16207647 |
| TDPAPATT | ASSSPTKNTLY | 0.76052576 |
| TNRPDILDPALLRPG | ASSSPTKNTLY | 0.7007057 |
| LTTTTTS | ASSSPTKNTLY | 0.563812 |
| PTGSGKTL | ASSSPTKNTLY | 0.32425487 |
| TGSGKTL | ASSSPTKNTLY | 0.7910601 |
| TIGHVDHGK | ASSSPTKNTLY | 0.82753813 |
| TIGHVDHGKTT | ASSSPTKNTLY | 0.8201661 |
| GHVDHGKTT | ASSSPTKNTLY | 0.44459742 |
| AQFEHTLL | ASSSPTKNTLY | 0.82738674 |
| IVNQDEL | ASSSPTKNTLY | 0.8302989 |
| TNGSQFF | ASSSPTKNTLY | 0.7286302 |
| TGKGIVE | ASSSPTKNTLY | 0.2538982 |
| LISNASDA | ASSSPTKNTLY | 0.8061831 |
| AELRTLGELTKIAKAHGAQ | ASSSPTKNTLY | 0.75715613 |
| APNAVAAAEPSW | ASSSPTKNTLY | 0.8086378 |
| APQVAGAATAVL | ASSSPTKNTLY | 0.80752265 |
| ARRDPAAAAAILPTDA | ASSSPTKNTLY | 0.671973 |
| DAAARDVLAEAGLAEYF | ASSSPTKNTLY | 0.8160349 |
| DDQIVLSAKANLAKVEQ | ASSSPTKNTLY | 0.8101798 |
| DELLPDTGEDSDEDGH | ASSSPTKNTLY | 0.63852286 |
| ELPEAPANRATVPQD | ASSSPTKNTLY | 0.70942986 |
| FGQSKIERVTGIG | ASSSPTKNTLY | 0.6928973 |
| GAGGTGSTAGGAGGAGGAGGLY | ASSSPTKNTLY | 0.31905788 |
| GHGGTGAAGGNGGTGGAGGSGI | ASSSPTKNTLY | 0.7812239 |
| GNGGNGGAGGEGGPGGAGGAGA | ASSSPTKNTLY | 0.7402003 |
| GSTAGNGGNGGNATNGTIAGQP | ASSSPTKNTLY | 0.6610036 |
| GTGGDGGNGGNGGTGG | ASSSPTKNTLY | 0.8070478 |
| GVRRPAGIPADAPV | ASSSPTKNTLY | 0.5796713 |
| GYRQVTDAHLLTLARR | ASSSPTKNTLY | 0.66224885 |

| | | |
|---|---|---|
| IAKMGQDGHDRG | ASSSPTKNTLY | 0.69056904 |
| KAEPSIEIQYT | ASSSPTKNTLY | 0.78796023 |
| KPVRIRMAADEDTVAVT | ASSSPTKNTLY | 0.7963139 |
| MPPRIAGMRLLVIKPEP | ASSSPTKNTLY | 0.799091 |
| MKFYAVALFRSRGVPPTR | ASSSPTKNTLY | 0.79858124 |
| NAGGFGILGAVAHSP | ASSSPTKNTLY | 0.8326302 |
| NEGREITLAMAEM | ASSSPTKNTLY | 0.48821872 |
| NELVEWSHPLGHRWRWE | ASSSPTKNTLY | 0.6267526 |
| NLGAGGTGGNGGNPG | ASSSPTKNTLY | 0.7993773 |
| NTEWPKWETVMAADPVD | ASSSPTKNTLY | 0.35668442 |
| PASAAPAETQP | ASSSPTKNTLY | 0.8126643 |
| QQSSDGKNGVRAAEITGE | ASSSPTKNTLY | 0.7954369 |
| RAPDAVTTSWWKE | ASSSPTKNTLY | 0.80405086 |
| SGFGNVGAHTSGWFNQ | ASSSPTKNTLY | 0.7683692 |
| SGVAGAVGQAASVGGLKV | ASSSPTKNTLY | 0.83493996 |
| SNGGAGTGSAGGNGGT | ASSSPTKNTLY | 0.8251815 |
| TGGAGGEGGAAG | ASSSPTKNTLY | 0.7737975 |
| TGGDGGAAGAGGAPGHG | ASSSPTKNTLY | 0.7971199 |
| TMPPLLAAIANAGRHGV | ASSSPTKNTLY | 0.5782942 |
| TSSTAGNSGGA | ASSSPTKNTLY | 0.8250317 |
| VPATVPVGILC | ASSSPTKNTLY | 0.80693406 |
| VQRPDSGGVVLTGRIS | ASSSPTKNTLY | 0.70846844 |
| WTPNPVPATKGVPAGLVQAT | ASSSPTKNTLY | 0.79253423 |
| YALKRALAKFSDRFDVVII | ASSSPTKNTLY | 0.8198028 |
| YLIGSVIV | ASSSPTKNTLY | 0.82926697 |
| YGQLVAVID | ASSSPTKNTLY | 0.22878926 |
| VTVGGMTLPA | ASSSPTKNTLY | 0.7791456 |
| VLALPAV | ASSSPTKNTLY | 0.22987825 |
| VIPPTLNYET | ASSSPTKNTLY | 0.5853759 |
| VGMGGMHPSSG | ASSSPTKNTLY | 0.54200816 |
| VGLAGGAATA | ASSSPTKNTLY | 0.7978689 |
| VGGAGGAGGSGP | ASSSPTKNTLY | 0.7687217 |
| VFAAFMV | ASSSPTKNTLY | 0.8069217 |
| TLYDKLR | ASSSPTKNTLY | 0.7942658 |
| TLAVFTVL | ASSSPTKNTLY | 0.8285992 |
| TGSVQPARV | ASSSPTKNTLY | 0.68759656 |
| TGGGAGSGGT | ASSSPTKNTLY | 0.74960816 |
| SHLETSL | ASSSPTKNTLY | 0.3562275 |
| SFTTGMT | ASSSPTKNTLY | 0.23380913 |
| QVEVAKRMGVRQPTVS | ASSSPTKNTLY | 0.6215061 |
| QGSGGPALAM | ASSSPTKNTLY | 0.72948 |
| MLAPSNP | ASSSPTKNTLY | 0.1823809 |
| LVNNAAI | ASSSPTKNTLY | 0.82740164 |
| LTGEINTLA | ASSSPTKNTLY | 0.8075131 |
| LSIFGPV | ASSSPTKNTLY | 0.81768036 |
| LLALLANL | ASSSPTKNTLY | 0.5409119 |
| LISNASDALDKLR | ASSSPTKNTLY | 0.7639876 |
| LINEAVP | ASSSPTKNTLY | 0.6920278 |
| LGYAFSGL | ASSSPTKNTLY | 0.8192451 |
| LDFVAVAL | ASSSPTKNTLY | 0.6577735 |
| LAQPGAE | ASSSPTKNTLY | 0.44616705 |

| | | |
|---|---|---|
| ISNLGAVTGF | ASSSPTKNTLY | 0.71914536 |
| HQEDPGRI | ASSSPTKNTLY | 0.6573226 |
| GTPGEPTP | ASSSPTKNTLY | 0.8242597 |
| GFFNSGS | ASSSPTKNTLY | 0.8208729 |
| GAGGGGLTAGAN | ASSSPTKNTLY | 0.812783 |
| FVDALVNM | ASSSPTKNTLY | 0.80948573 |
| FLTVAPEL | ASSSPTKNTLY | 0.2150023 |
| FETLTAL | ASSSPTKNTLY | 0.6400901 |
| DVINPIEL | ASSSPTKNTLY | 0.8334703 |
| APVSGAPV | ASSSPTKNTLY | 0.771236 |
| ALVGGLAW | ASSSPTKNTLY | 0.7777102 |
| ALAAAAATAI | ASSSPTKNTLY | 0.82575583 |
| AFGLGLLL | ASSSPTKNTLY | 0.72461545 |
| AFFAAASL | ASSSPTKNTLY | 0.7285235 |
| AAAALPAFSPP | ASSSPTKNTLY | 0.7689644 |
| VVLAPAL | ASSSPTKNTLY | 0.76007 |
| TIGHVDHGKTTL | ASSSPTKNTLY | 0.7716634 |
| LISNASDALDK | ASSSPTKNTLY | 0.60961497 |
| LISNASDAL | ASSSPTKNTLY | 0.7681842 |
| ISNASDALDK | ASSSPTKNTLY | 0.8353048 |
| NQAETLVYQTE | ASSSPTKNTLY | 0.7990905 |
| TGGTGGTAGAG | ASSSPTKNTLY | 0.8201052 |
| TGSTAGGAGGA | ASSSPTKNTLY | 0.8080981 |
| DPLAVPVAAGPV | ASSSPTKNTLY | 0.8117088 |
| GGTGGTPGNGGN | ASSSPTKNTLY | 0.74132663 |
| PAGLAGVDPATL | ASSSPTKNTLY | 0.71682525 |
| GNNLVVEAPHAGASV | ASSSPTKNTLY | 0.72676957 |
| KVTRIMSASTLEQPA | ASSSPTKNTLY | 0.6787361 |
| LGFALPIALPRRTVV | ASSSPTKNTLY | 0.7569289 |
| TRLGNLTARKLVVLV | ASSSPTKNTLY | 0.6993221 |
| WSQQRRLIYFPSAGP | ASSSPTKNTLY | 0.83704436 |
| GPNGNPDPMAAAVDIR | ASSSPTKNTLY | 0.8167754 |
| TQRCVSVADLRAAASR | ASSSPTKNTLY | 0.7937869 |
| QFVLDVAEHPVEVLTETV | ASSSPTKNTLY | 0.666482 |
| GGAGGTGGAGGNGGTQAGDG | ASSSPTKNTLY | 0.76314354 |
| PLLATLRAIAVDVDAKPAQV | ASSSPTKNTLY | 0.8006276 |
| APKFRTGGYMIDFWGVGYQVAK | ASSSPTKNTLY | 0.44744706 |
| GAGGAGGAGGAAGGPGGTGGTG | ASSSPTKNTLY | 0.82843935 |
| IGALGLLATDVAPAGSVAFGVA | ASSSPTKNTLY | 0.68237054 |
| AGGAGGIGGNGGAGGAGGVLMG | ASSSPTKNTLY | 0.697491 |
| ADELIGLL | ASSSPTKNTLY | 0.8168216 |
| ADRALKA | ASSSPTKNTLY | 0.8276843 |
| AGAVGQVGGAGGA | ASSSPTKNTLY | 0.6711727 |
| AGGGGGLGGAP | ASSSPTKNTLY | 0.6938498 |
| AGGLLYGNGGAG | ASSSPTKNTLY | 0.70263505 |
| AGLESIGL | ASSSPTKNTLY | 0.7994873 |
| AGLKTGII | ASSSPTKNTLY | 0.829063 |
| ASLAALLL | ASSSPTKNTLY | 0.7599377 |
| ASVINISSI | ASSSPTKNTLY | 0.8266041 |
| ATPPGPA | ASSSPTKNTLY | 0.7788658 |
| ATVASALV | ASSSPTKNTLY | 0.39507824 |

| | | |
|---|---|---|
| ATVASLIL | ASSSPTKNTLY | 0.7181352 |
| AVNGPVLIRH | ASSSPTKNTLY | 0.81300265 |
| DGAAGAVGI | ASSSPTKNTLY | 0.821403 |
| DPLLHGI | ASSSPTKNTLY | 0.8317087 |
| ELGAALGAL | ASSSPTKNTLY | 0.81046927 |
| ELGRTQAV | ASSSPTKNTLY | 0.7118273 |
| FVAGGYLS | ASSSPTKNTLY | 0.82960916 |
| GASGGAGGAGGW | ASSSPTKNTLY | 0.83140665 |
| GGAGAGGDGGAGGI | ASSSPTKNTLY | 0.8211308 |
| GTGHNGSQPGMGGNG | ASSSPTKNTLY | 0.6693473 |
| GYAAASASAS | ASSSPTKNTLY | 0.52256995 |
| HQVDVAAVQQG | ASSSPTKNTLY | 0.7771046 |
| IGAAIAHT | ASSSPTKNTLY | 0.8198025 |
| IGMIEVAQL | ASSSPTKNTLY | 0.5541344 |
| IRLAPEHTSAMD | ASSSPTKNTLY | 0.7841064 |
| IYNVSVL | ASSSPTKNTLY | 0.81051576 |
| KQGFLGM | ASSSPTKNTLY | 0.5144944 |
| LESVITQL | ASSSPTKNTLY | 0.7981335 |
| LGLGALVVV | ASSSPTKNTLY | 0.82909364 |
| LIGNGGNGGSGGI | ASSSPTKNTLY | 0.7325828 |
| LNAINAPTL | ASSSPTKNTLY | 0.74292004 |
| LQLLNTGV | ASSSPTKNTLY | 0.82464266 |
| LQTSIEF | ASSSPTKNTLY | 0.8369563 |
| MSFAVSEL | ASSSPTKNTLY | 0.8266878 |
| NTGALGL | ASSSPTKNTLY | 0.35251126 |
| PPGAGGAGGAGTT | ASSSPTKNTLY | 0.27795267 |
| QLVGLGI | ASSSPTKNTLY | 0.67789114 |
| RFLEQVHQL | ASSSPTKNTLY | 0.80297303 |
| RGVGYVLR | ASSSPTKNTLY | 0.6938058 |
| SATGVFAGVI | ASSSPTKNTLY | 0.6480549 |
| SLPAITLP | ASSSPTKNTLY | 0.815019 |
| STALAFTV | ASSSPTKNTLY | 0.82739186 |
| SVLTVGLGK | ASSSPTKNTLY | 0.6497859 |
| TALAGAGSGI | ASSSPTKNTLY | 0.34379333 |
| TGLPFTGI | ASSSPTKNTLY | 0.81591415 |
| VAEHVTV | ASSSPTKNTLY | 0.81407225 |
| VDIVLNSL | ASSSPTKNTLY | 0.71217024 |
| VLTGGGAVL | ASSSPTKNTLY | 0.7899742 |
| VNVGLAYGI | ASSSPTKNTLY | 0.5754751 |
| VSSAAVGAV | ASSSPTKNTLY | 0.27158067 |
| VTSVYEAGV | ASSSPTKNTLY | 0.79681206 |
| WVALDRGA | ASSSPTKNTLY | 0.6113779 |
| YLVGSVVI | ASSSPTKNTLY | 0.8295422 |
| GPPGTGKTLL | ASSSPTKNTLY | 0.7017197 |
| GVGMTKF | ASSSPTKNTLY | 0.79061365 |
| IGIGGSDL | ASSSPTKNTLY | 0.8132229 |
| IGTLLAEL | ASSSPTKNTLY | 0.7704077 |
| IYTSGTT | ASSSPTKNTLY | 0.7652852 |
| LSFPAGV | ASSSPTKNTLY | 0.6869085 |
| LYTSGTT | ASSSPTKNTLY | 0.55859584 |
| VGAFPFGF | ASSSPTKNTLY | 0.23151234 |

| | | |
|---|---|---|
| VGAGTVEF | ASSSPTKNTLY | 0.82586163 |
| VGKSTLLL | ASSSPTKNTLY | 0.16869345 |
| AALAHAGGGIV | ASSSPTKNTLY | 0.8242383 |
| AALVAVSLL | ASSSPTKNTLY | 0.81717855 |
| AAVSRAVAAL | ASSSPTKNTLY | 0.6444681 |
| AGGSSGIGGF | ASSSPTKNTLY | 0.789451 |
| AGLTVANANG | ASSSPTKNTLY | 0.6309556 |
| AGLVGAVL | ASSSPTKNTLY | 0.4296805 |
| ALSQAFTV | ASSSPTKNTLY | 0.8139903 |
| ALTFSVSL | ASSSPTKNTLY | 0.80416304 |
| ALTGAALVAL | ASSSPTKNTLY | 0.7115803 |
| ASFSIGAGVL | ASSSPTKNTLY | 0.77038324 |
| DGSVYKI | ASSSPTKNTLY | 0.79194874 |
| EQLAEIV | ASSSPTKNTLY | 0.68690574 |
| FAVSELTVM | ASSSPTKNTLY | 0.7281065 |
| GAAPLAAGL | ASSSPTKNTLY | 0.8289274 |
| GADGGDGGSGGAGG | ASSSPTKNTLY | 0.7526041 |
| GFGLDTGGAGG | ASSSPTKNTLY | 0.7256479 |
| GLASGGLL | ASSSPTKNTLY | 0.80217266 |
| GRIAGAAVLL | ASSSPTKNTLY | 0.5891045 |
| GSAGSLNTSF | ASSSPTKNTLY | 0.8149542 |
| GTGQIGI | ASSSPTKNTLY | 0.7302551 |
| GTLIALIL | ASSSPTKNTLY | 0.7420683 |
| IAATMGFVVG | ASSSPTKNTLY | 0.7962748 |
| IFAQALL | ASSSPTKNTLY | 0.51295716 |
| IGAAVALGF | ASSSPTKNTLY | 0.6574404 |
| IGLTPLAL | ASSSPTKNTLY | 0.6718149 |
| ILGSNTSTL | ASSSPTKNTLY | 0.7843373 |
| IVFSLTVL | ASSSPTKNTLY | 0.8226709 |
| LASKATGF | ASSSPTKNTLY | 0.70721775 |
| LASVALIL | ASSSPTKNTLY | 0.7748434 |
| LFAVGLTL | ASSSPTKNTLY | 0.6697178 |
| LFNGGIGTYI | ASSSPTKNTLY | 0.6059668 |
| LTLGAGII | ASSSPTKNTLY | 0.7294165 |
| LVAAALGDI | ASSSPTKNTLY | 0.72724557 |
| NTAFVQLGI | ASSSPTKNTLY | 0.6151622 |
| QLFSGAEL | ASSSPTKNTLY | 0.7798202 |
| SAATFITL | ASSSPTKNTLY | 0.59184027 |
| SNYGLANLGV | ASSSPTKNTLY | 0.7047521 |
| SYGGAYAVM | ASSSPTKNTLY | 0.83095217 |
| TTGVGGLTLG | ASSSPTKNTLY | 0.7629967 |
| TTQTIYI | ASSSPTKNTLY | 0.78773165 |
| VAAGAGLGAVI | ASSSPTKNTLY | 0.7977879 |
| VAAVVTLL | ASSSPTKNTLY | 0.81048745 |
| VGFLGGLI | ASSSPTKNTLY | 0.72446036 |
| VGVIPILL | ASSSPTKNTLY | 0.72017807 |
| VIFASSLI | ASSSPTKNTLY | 0.82941604 |
| VKGGDGGAAAG | ASSSPTKNTLY | 0.804942 |
| VTKTFTNI | ASSSPTKNTLY | 0.82586277 |
| VVAAAILL | ASSSPTKNTLY | 0.8104341 |
| YAALDVEL | ASSSPTKNTLY | 0.81560785 |

| | | |
|---|---|---|
| LASVALLL | ASSSPTKNTLY | 0.81254023 |
| LDAVHRV | ASSSPTKNTLY | 0.7610812 |
| QGLSSSIF | ASSSPTKNTLY | 0.3755238 |
| RGTFANIRL | ASSSPTKNTLY | 0.72579575 |
| RQLWWGHRI | ASSSPTKNTLY | 0.7704392 |
| YGPPGTGKTLL | ASSSPTKNTLY | 0.7158592 |
| AALVGAPANVA | ASSSPTKNTLY | 0.2821772 |
| AATFRRTE | ASSSPTKNTLY | 0.82992643 |
| AGALGGGI | ASSSPTKNTLY | 0.8357911 |
| AGGAGGAGGIGAG | ASSSPTKNTLY | 0.5134059 |
| AGSGALGL | ASSSPTKNTLY | 0.82857555 |
| AQIADQI | ASSSPTKNTLY | 0.53388155 |
| ASAVAVTYL | ASSSPTKNTLY | 0.79396784 |
| ASTVLGGTGG | ASSSPTKNTLY | 0.8298539 |
| AVGGEPAK | ASSSPTKNTLY | 0.6387656 |
| CEQGTFGGTC | ASSSPTKNTLY | 0.8152066 |
| DEGKAPAG | ASSSPTKNTLY | 0.65429264 |
| DTVVFPAL | ASSSPTKNTLY | 0.72804546 |
| FEGITMAG | ASSSPTKNTLY | 0.8311001 |
| FPLVVQD | ASSSPTKNTLY | 0.6877558 |
| GAFTTPQ | ASSSPTKNTLY | 0.76002806 |
| GFGGFGTAGGVGGA | ASSSPTKNTLY | 0.73444045 |
| GFLAQFL | ASSSPTKNTLY | 0.5580562 |
| GGAGGAGGNAGVA | ASSSPTKNTLY | 0.75847316 |
| GGVGTAGAGGAGGAGG | ASSSPTKNTLY | 0.8270134 |
| GNGGGGDAGNAGSGGN | ASSSPTKNTLY | 0.46078542 |
| GTGGAGGAGNDAGS | ASSSPTKNTLY | 0.82976514 |
| GTTNTGWFNS | ASSSPTKNTLY | 0.65728676 |
| IFMGPLFTL | ASSSPTKNTLY | 0.8368398 |
| ILAHAIEAI | ASSSPTKNTLY | 0.81493044 |
| IRSAIREAL | ASSSPTKNTLY | 0.79971206 |
| ITFAGGG | ASSSPTKNTLY | 0.66583276 |
| ITGMSGHQEV | ASSSPTKNTLY | 0.8313339 |
| IVAGLACTL | ASSSPTKNTLY | 0.7820823 |
| IVGGGLAGA | ASSSPTKNTLY | 0.81204563 |
| IVTGVLNI | ASSSPTKNTLY | 0.8183714 |
| LAQQGCI | ASSSPTKNTLY | 0.81266105 |
| LCLHVEN | ASSSPTKNTLY | 0.8567991 |
| LDAISDGS | ASSSPTKNTLY | 0.836169 |
| LGGFAAML | ASSSPTKNTLY | 0.8298842 |
| LGGKNPMI | ASSSPTKNTLY | 0.8120611 |
| LGPKGVAAIL | ASSSPTKNTLY | 0.63074106 |
| LMVGVSI | ASSSPTKNTLY | 0.8301016 |
| LVGAAFVL | ASSSPTKNTLY | 0.7794392 |
| LVLAVAPL | ASSSPTKNTLY | 0.8205364 |
| MAAIAVIDAA | ASSSPTKNTLY | 0.84729517 |
| NNGINAELQ | ASSSPTKNTLY | 0.80551887 |
| RGEPLVLVHG | ASSSPTKNTLY | 0.8578548 |
| RGGAGGAAGLI | ASSSPTKNTLY | 0.8511952 |
| RSYRMHPAV | ASSSPTKNTLY | 0.7379471 |
| SAAHLLAGQLA | ASSSPTKNTLY | 0.8267948 |

| | | |
|---|---|---|
| SAAVGSY | ASSSPTKNTLY | 0.8155144 |
| SPHAVLHRI | ASSSPTKNTLY | 0.82927257 |
| SQIAKVL | ASSSPTKNTLY | 0.6528549 |
| TALGHLVP | ASSSPTKNTLY | 0.84850425 |
| TAPTTSGAA | ASSSPTKNTLY | 0.80164844 |
| TDGDDGNGGSPGT | ASSSPTKNTLY | 0.2005373 |
| TDGLGEN | ASSSPTKNTLY | 0.8369312 |
| TGGPPLPGQA | ASSSPTKNTLY | 0.8336758 |
| TIPAITI | ASSSPTKNTLY | 0.8231776 |
| TIPGFTI | ASSSPTKNTLY | 0.8489331 |
| TSIASGALA | ASSSPTKNTLY | 0.24480146 |
| TVTLPGAAI | ASSSPTKNTLY | 0.7955649 |
| VASGLSAALP | ASSSPTKNTLY | 0.85173786 |
| VGAGVIGI | ASSSPTKNTLY | 0.85364574 |
| VGPKRFGV | ASSSPTKNTLY | 0.7562398 |
| VNVAFPD | ASSSPTKNTLY | 0.8376336 |
| VSMTNSGVS | ASSSPTKNTLY | 0.8509834 |
| VTDVLGDLSV | ASSSPTKNTLY | 0.83837163 |
| VVVVGAGFAGL | ASSSPTKNTLY | 0.8518142 |
| VYTVGTVV | ASSSPTKNTLY | 0.85503733 |
| WTTFSGP | ASSSPTKNTLY | 0.74719703 |
| YDELVEAF | ASSSPTKNTLY | 0.8537042 |
| YGTGGAGGNGG | ASSSPTKNTLY | 0.85783213 |
| YTNVMAV | ASSSPTKNTLY | 0.820387 |
| FLEQVHQL | ASSSPTKNTLY | 0.8362125 |
| GLSLLAVAV | ASSSPTKNTLY | 0.7911155 |
| LEQVHQL | ASSSPTKNTLY | 0.80985224 |
| LNAAGAI | ASSSPTKNTLY | 0.35907626 |
| PAVHASL | ASSSPTKNTLY | 0.8361494 |
| TELEDDA | ASSSPTKNTLY | 0.85078704 |
| TPPGPPN | ASSSPTKNTLY | 0.8139242 |
| TPPPPPPP | ASSSPTKNTLY | 0.7835493 |
| TQSPALI | ASSPGTGESQNTLY | 0.77501 |
| EFAAAFPQ | ASSPGTGESQNTLY | 0.80395406 |
| LLSGNLDVL | ASSPGTGESQNTLY | 0.81061584 |
| GPANLGNYNI | ASSPGTGESQNTLY | 0.3994416 |
| TLGPTLP | ASSPGTGESQNTLY | 0.80024135 |
| VYAYSSCCK | ASSPGTGESQNTLY | 0.85358775 |
| APAPAPAPNI | ASSPGTGESQNTLY | 0.8511282 |
| PGGPVPIP | ASSPGTGESQNTLY | 0.68713504 |
| FLTVALEP | ASSPGTGESQNTLY | 0.8552809 |
| SQYFGANL | ASSPGTGESQNTLY | 0.84229106 |
| GGVKPAA | ASSPGTGESQNTLY | 0.8518678 |
| GGTGGAGGAGGW | ASSPGTGESQNTLY | 0.8205475 |
| GVGGVAGLGGAGG | ASSPGTGESQNTLY | 0.7980697 |
| VLLVLATTI | ASSPGTGESQNTLY | 0.8424227 |
| IGIFAAML | ASSPGTGESQNTLY | 0.81065166 |
| AVGDLID | ASSPGTGESQNTLY | 0.72848296 |
| NISPLPL | ASSPGTGESQNTLY | 0.8530922 |
| VTAVFVAI | ASSPGTGESQNTLY | 0.80891085 |
| SGDDNDAAL | ASSPGTGESQNTLY | 0.79677546 |

| | | |
|---|---|---|
| AVAALAHS | ASSPGTGESQNTLY | 0.7745445 |
| GGVTVMATP | ASSPGTGESQNTLY | 0.85153437 |
| AGLAANVRD | ASSPGTGESQNTLY | 0.7770798 |
| TNRPDLIDPALLRPG | ASSPGTGESQNTLY | 0.6724194 |
| LANLGVTAL | ASSPGTGESQNTLY | 0.72996837 |
| TGERIEL | ASSPGTGESQNTLY | 0.8301692 |
| NGEELAA | ASSPGTGESQNTLY | 0.85258406 |
| VLSLVWG | ASSPGTGESQNTLY | 0.8419993 |
| IGYALWL | ASSPGTGESQNTLY | 0.7572919 |
| LSLLLAPL | ASSPGTGESQNTLY | 0.84286 |
| TLDEALK | ASSPGTGESQNTLY | 0.8368087 |
| TDGAGTAEGG | ASSPGTGESQNTLY | 0.8344527 |
| NAPGGAGGNGG | ASSPGTGESQNTLY | 0.8407055 |
| VSVFMGMGGNF | ASSPGTGESQNTLY | 0.78865784 |
| LASYLAAL | ASSPGTGESQNTLY | 0.8525001 |
| LVGGTPAL | ASSPGTGESQNTLY | 0.84091866 |
| VRNADVL | ASSPGTGESQNTLY | 0.8493352 |
| TCSSPFLVSAM | ASSPGTGESQNTLY | 0.85136145 |
| VHITDAE | ASSPGTGESQNTLY | 0.82223725 |
| EYAAPSF | ASSPGTGESQNTLY | 0.82957697 |
| VVDAFHGGL | ASSPGTGESQNTLY | 0.542905 |
| ASVVDPQ | ASSPGTGESQNTLY | 0.7396257 |
| SYNEFGGLS | ASSPGTGESQNTLY | 0.84112626 |
| DPDGSVY | ASSPGTGESQNTLY | 0.8366301 |
| MVSTIHATE | ASSPGTGESQNTLY | 0.8481204 |
| LAAGVADPV | ASSPGTGESQNTLY | 0.8376816 |
| TKEQIAA | ASSPGTGESQNTLY | 0.7913027 |
| SLASPAAL | ASSPGTGESQNTLY | 0.84680676 |
| IGLVFYGV | ASSPGTGESQNTLY | 0.85036236 |
| FLLDASNL | ASSPGTGESQNTLY | 0.71417224 |
| YQVGNFP | ASSPGTGESQNTLY | 0.8139754 |
| ESLLSQST | ASSPGTGESQNTLY | 0.86045855 |
| LFTVGFAAVG | ASSPGTGESQNTLY | 0.82533026 |
| ATPPTPT | ASSPGTGESQNTLY | 0.82547164 |
| TDLATLG | ASSPGTGESQNTLY | 0.7988553 |
| SAGGTGR | ASSPGTGESQNTLY | 0.84779835 |
| GFSNSGS | ASSPGTGESQNTLY | 0.7654674 |
| GGNGGSGGTSV | ASSPGTGESQNTLY | 0.86018777 |
| GTSVATGGAGN | ASSPGTGESQNTLY | 0.35082138 |
| NGGNGGSGGTS | ASSPGTGESQNTLY | 0.78834015 |
| AVVKFPH | ASSPGTGESQNTLY | 0.74279547 |
| TPPPPPP | ASSPGTGESQNTLY | 0.5812228 |
| AAPAAAAAPA | ASSPGTGESQNTLY | 0.85357106 |
| IALNEDL | ASSPGTGESQNTLY | 0.80568993 |
| IVNVSSM | ASSPGTGESQNTLY | 0.8473369 |
| TGIPPAPR | ASSPGTGESQNTLY | 0.8376059 |
| FDLGGGTF | ASSPGTGESQNTLY | 0.8549628 |
| FDLGGGTFDVS | ASSPGTGESQNTLY | 0.82594883 |
| NEPTAAA | ASSPGTGESQNTLY | 0.8221479 |
| NEPTAAALA | ASSPGTGESQNTLY | 0.7233101 |
| GSIINISSI | ASSPGTGESQNTLY | 0.8505525 |

| | | |
|---|---|---|
| VAPGNAGTA | ASSPGTGESQNTLY | 0.85220027 |
| TGGGASLE | ASSPGTGESQNTLY | 0.8361641 |
| VAVYNHYKRI | ASSPGTGESQNTLY | 0.71422875 |
| TDPAPATT | ASSPGTGESQNTLY | 0.80169433 |
| TNRPDILDPALLRPG | ASSPGTGESQNTLY | 0.8387865 |
| LTTTTTS | ASSPGTGESQNTLY | 0.21461117 |
| PTGSGKTL | ASSPGTGESQNTLY | 0.35977268 |
| TGSGKTL | ASSPGTGESQNTLY | 0.79878014 |
| TIGHVDHGK | ASSPGTGESQNTLY | 0.7716534 |
| TIGHVDHGKTT | ASSPGTGESQNTLY | 0.85514104 |
| GHVDHGKTT | ASSPGTGESQNTLY | 0.8504882 |
| AQFEHTLL | ASSPGTGESQNTLY | 0.8212858 |
| IVNQDEL | ASSPGTGESQNTLY | 0.83874655 |
| TNGSQFF | ASSPGTGESQNTLY | 0.82985103 |
| TGKGIVE | ASSPGTGESQNTLY | 0.1427642 |
| LISNASDA | ASSPGTGESQNTLY | 0.7890581 |
| AELRTLGELTKIAKAHGAQ | ASSPGTGESQNTLY | 0.75773084 |
| APNAVAAAEPSW | ASSPGTGESQNTLY | 0.8557135 |
| APQVAGAATAVL | ASSPGTGESQNTLY | 0.8026954 |
| ARRDPAAAAAILPTDA | ASSPGTGESQNTLY | 0.789312 |
| DAAARDVLAEAGLAEYF | ASSPGTGESQNTLY | 0.82736343 |
| DDQIVLSAKANLAKVEQ | ASSPGTGESQNTLY | 0.8181448 |
| DELLPDTGEDSDEDGH | ASSPGTGESQNTLY | 0.8246012 |
| ELPEAPANRATVPQD | ASSPGTGESQNTLY | 0.7522317 |
| FGQSKIERVTGIG | ASSPGTGESQNTLY | 0.19810352 |
| GAGGTGSTAGGAGGAGGAGGLY | ASSPGTGESQNTLY | 0.85266954 |
| GHGGTGAAGGNGGTGGAGGSGI | ASSPGTGESQNTLY | 0.45670006 |
| GNGGNGGAGGEGGPGGAGGAGA | ASSPGTGESQNTLY | 0.78773755 |
| GSTAGNGGNGGNATNGTIAGQP | ASSPGTGESQNTLY | 0.43444356 |
| GTGGDGGNGGNGGTGG | ASSPGTGESQNTLY | 0.17909269 |
| GVRRPAGIPADAPV | ASSPGTGESQNTLY | 0.62115526 |
| GYRQVTDAHLLTLARR | ASSPGTGESQNTLY | 0.8469001 |
| IAKMGQDGHDRG | ASSPGTGESQNTLY | 0.8165499 |
| KAEPSIEIQYT | ASSPGTGESQNTLY | 0.81582713 |
| KPVRIRMAADEDTVAVT | ASSPGTGESQNTLY | 0.85880363 |
| MPPRIAGMRLLVIKPEP | ASSPGTGESQNTLY | 0.8190174 |
| MKFYAVALFRSRGVPPTR | ASSPGTGESQNTLY | 0.8119043 |
| NAGGFGILGAVAHSP | ASSPGTGESQNTLY | 0.80068463 |
| NEGREITLAMAEM | ASSPGTGESQNTLY | 0.79377127 |
| NELVEWSHPLGHRWRWE | ASSPGTGESQNTLY | 0.7807287 |
| NLGAGGTGGNGGNPG | ASSPGTGESQNTLY | 0.78260255 |
| NTEWPKWETVMAADPVD | ASSPGTGESQNTLY | 0.85874045 |
| PASAAPAETQP | ASSPGTGESQNTLY | 0.79836726 |
| QQSSDGKNGVRAAEITGE | ASSPGTGESQNTLY | 0.7296597 |
| RAPDAVTTSWWKE | ASSPGTGESQNTLY | 0.8090943 |
| SGFGNVGAHTSGWFNQ | ASSPGTGESQNTLY | 0.85714746 |
| SGVAGAVGQAASVGGLKV | ASSPGTGESQNTLY | 0.85817915 |
| SNGGAGTGSAGGNGGT | ASSPGTGESQNTLY | 0.79428697 |
| TGGAGGEGGAAG | ASSPGTGESQNTLY | 0.7968872 |
| TGGDGGAAGAGGAPGHG | ASSPGTGESQNTLY | 0.8104478 |
| TMPPLLAAIANAGRHGV | ASSPGTGESQNTLY | 0.8271284 |

| | | |
|---|---|---|
| TSSTAGNSGGA | ASSPGTGESQNTLY | 0.7942289 |
| VPATVPVGILC | ASSPGTGESQNTLY | 0.669893 |
| VQRPDSGGVVLTGRIS | ASSPGTGESQNTLY | 0.8445977 |
| WTPNPVPATKGVPAGLVQAT | ASSPGTGESQNTLY | 0.8536241 |
| YALKRALAKFSDRFDVVII | ASSPGTGESQNTLY | 0.79816335 |
| YLIGSVIV | ASSPGTGESQNTLY | 0.78194386 |
| YGQLVAVID | ASSPGTGESQNTLY | 0.8235598 |
| VTVGGMTLPA | ASSPGTGESQNTLY | 0.7855085 |
| VLALPAV | ASSPGTGESQNTLY | 0.8061211 |
| VIPPTLNYET | ASSPGTGESQNTLY | 0.8095428 |
| VGMGGMHPSSG | ASSPGTGESQNTLY | 0.7317625 |
| VGLAGGAATA | ASSPGTGESQNTLY | 0.819774 |
| VGGAGGAGGSGP | ASSPGTGESQNTLY | 0.8582926 |
| VFAAFMV | ASSPGTGESQNTLY | 0.809736 |
| TLYDKLR | ASSPGTGESQNTLY | 0.8362664 |
| TLAVFTVL | ASSPGTGESQNTLY | 0.8544107 |
| TGSVQPARV | ASSPGTGESQNTLY | 0.8444241 |
| TGGGAGSGGT | ASSPGTGESQNTLY | 0.8546963 |
| SHLETSL | ASSPGTGESQNTLY | 0.8427094 |
| SFTTGMT | ASSPGTGESQNTLY | 0.8297102 |
| QVEVAKRMGVRQPTVS | ASSPGTGESQNTLY | 0.8315627 |
| QGSGGPALAM | ASSPGTGESQNTLY | 0.8335482 |
| MLAPSNP | ASSPGTGESQNTLY | 0.8300319 |
| LVNNAAI | ASSPGTGESQNTLY | 0.8205121 |
| LTGEINTLA | ASSPGTGESQNTLY | 0.7937165 |
| LSIFGPV | ASSPGTGESQNTLY | 0.8561035 |
| LLALLANL | ASSPGTGESQNTLY | 0.8285596 |
| LISNASDALDKLR | ASSPGTGESQNTLY | 0.84330976 |
| LINEAVP | ASSPGTGESQNTLY | 0.7088149 |
| LGYAFSGL | ASSPGTGESQNTLY | 0.7350081 |
| LDFVAVAL | ASSPGTGESQNTLY | 0.84602994 |
| LAQPGAE | ASSPGTGESQNTLY | 0.15315093 |
| ISNLGAVTGF | ASSPGTGESQNTLY | 0.31435314 |
| HQEDPGRI | ASSPGTGESQNTLY | 0.83521986 |
| GTPGEPTP | ASSPGTGESQNTLY | 0.8515736 |
| GFFNSGS | ASSPGTGESQNTLY | 0.15469697 |
| GAGGGGLTAGAN | ASSPGTGESQNTLY | 0.7565915 |
| FVDALVNM | ASSPGTGESQNTLY | 0.68682945 |
| FLTVAPEL | ASSPGTGESQNTLY | 0.8338011 |
| FETLTAL | ASSPGTGESQNTLY | 0.83605504 |
| DVINPIEL | ASSPGTGESQNTLY | 0.6034013 |
| APVSGAPV | ASSPGTGESQNTLY | 0.8462815 |
| ALVGGLAW | ASSPGTGESQNTLY | 0.71165246 |
| ALAAAAATAI | ASSPGTGESQNTLY | 0.44060886 |
| AFGLGLLL | ASSPGTGESQNTLY | 0.85240144 |
| AFFAAASL | ASSPGTGESQNTLY | 0.8501425 |
| AAAALPAFSPP | ASSPGTGESQNTLY | 0.78392774 |
| VVLAPAL | ASSPGTGESQNTLY | 0.8489115 |
| TIGHVDHGKTTL | ASSPGTGESQNTLY | 0.7824179 |
| LISNASDALDK | ASSPGTGESQNTLY | 0.8581679 |
| LISNASDAL | ASSPGTGESQNTLY | 0.82135737 |

| | | |
|---|---|---|
| ISNASDALDK | ASSPGTGESQNTLY | 0.65548795 |
| NQAETLVYQTE | ASSPGTGESQNTLY | 0.84885395 |
| TGGTGGTAGAG | ASSPGTGESQNTLY | 0.8375765 |
| TGSTAGGAGGA | ASSPGTGESQNTLY | 0.83391905 |
| DPLAVPVAAGPV | ASSPGTGESQNTLY | 0.84462273 |
| GGTGGTPGNGGN | ASSPGTGESQNTLY | 0.7209037 |
| PAGLAGVDPATL | ASSPGTGESQNTLY | 0.8234211 |
| GNNLVVEAPHAGASV | ASSPGTGESQNTLY | 0.85441136 |
| KVTRIMSASTLEQPA | ASSPGTGESQNTLY | 0.8535422 |
| LGFALPIALPRRTVV | ASSPGTGESQNTLY | 0.64407146 |
| TRLGNLTARKLVVLV | ASSPGTGESQNTLY | 0.84852624 |
| WSQQRRLIYFPSAGP | ASSPGTGESQNTLY | 0.8190465 |
| GPNGNPDPMAAAVDIR | ASSPGTGESQNTLY | 0.814082 |
| TQRCVSVADLRAAASR | ASSPGTGESQNTLY | 0.8577013 |
| QFVLDVAEHPVEVLTETV | ASSPGTGESQNTLY | 0.8089123 |
| GGAGGTGGAGGNGGTQAGDG | ASSPGTGESQNTLY | 0.8473878 |
| PLLATLRAIAVDVDAKPAQV | ASSPGTGESQNTLY | 0.84976995 |
| APKFRTGGYMIDFWGVGYQVAK | ASSPGTGESQNTLY | 0.8251188 |
| GAGGAGGAGGAAGGPGGTGGTG | ASSPGTGESQNTLY | 0.60521346 |
| IGALGLLATDVAPAGSVAFGVA | ASSPGTGESQNTLY | 0.8203969 |
| AGGAGGIGGNGGAGGAGGVLMG | ASSPGTGESQNTLY | 0.8262585 |
| ADELIGLL | ASSPGTGESQNTLY | 0.8496975 |
| ADRALKA | ASSPGTGESQNTLY | 0.8246122 |
| AGAVGQVGGAGGA | ASSPGTGESQNTLY | 0.606684 |
| AGGGGGLGGAP | ASSPGTGESQNTLY | 0.84025866 |
| AGGLLYGNGGAG | ASSPGTGESQNTLY | 0.8542396 |
| AGLESIGL | ASSPGTGESQNTLY | 0.83843684 |
| AGLKTGII | ASSPGTGESQNTLY | 0.8355677 |
| ASLAALLL | ASSPGTGESQNTLY | 0.8562258 |
| ASVINISSI | ASSPGTGESQNTLY | 0.8076285 |
| ATPPGPA | ASSPGTGESQNTLY | 0.8166598 |
| ATVASALV | ASSPGTGESQNTLY | 0.8506993 |
| ATVASLIL | ASSPGTGESQNTLY | 0.8510224 |
| AVNGPVLIRH | ASSPGTGESQNTLY | 0.8162204 |
| DGAAGAVGI | ASSPGTGESQNTLY | 0.85202765 |
| DPLLHGI | ASSPGTGESQNTLY | 0.730786 |
| ELGAALGAL | ASSPGTGESQNTLY | 0.8440243 |
| ELGRTQAV | ASSPGTGESQNTLY | 0.7760353 |
| FVAGGYLS | ASSPGTGESQNTLY | 0.8460996 |
| GASGGAGGAGGW | ASSPGTGESQNTLY | 0.7728472 |
| GGAGAGGDGGAGGI | ASSPGTGESQNTLY | 0.8512809 |
| GTGHNGSQPGMGGNG | ASSPGTGESQNTLY | 0.84805155 |
| GYAAASASAS | ASSPGTGESQNTLY | 0.8532171 |
| HQVDVAAVQQG | ASSPGTGESQNTLY | 0.35263443 |
| IGAAIAHT | ASSPGTGESQNTLY | 0.84936166 |
| IGMIEVAQL | ASSPGTGESQNTLY | 0.8467242 |
| IRLAPEHTSAMD | ASSPGTGESQNTLY | 0.85851806 |
| IYNVSVL | ASSPGTGESQNTLY | 0.8199275 |
| KQGFLGM | ASSPGTGESQNTLY | 0.83931243 |
| LESVITQL | ASSPGTGESQNTLY | 0.80780494 |
| LGLGALVVV | ASSPGTGESQNTLY | 0.8055087 |

| | | |
|---|---|---|
| LIGNGGNGGSGGI | ASSPGTGESQNTLY | 0.7644311 |
| LNAINAPTL | ASSPGTGESQNTLY | 0.79736114 |
| LQLLNTGV | ASSPGTGESQNTLY | 0.7944249 |
| LQTSIEF | ASSPGTGESQNTLY | 0.79659224 |
| MSFAVSEL | ASSPGTGESQNTLY | 0.84217274 |
| NTGALGL | ASSPGTGESQNTLY | 0.84997463 |
| PPGAGGAGGAGTT | ASSPGTGESQNTLY | 0.8509289 |
| QLVGLGI | ASSPGTGESQNTLY | 0.8541511 |
| RFLEQVHQL | ASSPGTGESQNTLY | 0.8512692 |
| RGVGYVLR | ASSPGTGESQNTLY | 0.8484349 |
| SATGVFAGVI | ASSPGTGESQNTLY | 0.7912154 |
| SLPAITLP | ASSPGTGESQNTLY | 0.8150782 |
| STALAFTV | ASSPGTGESQNTLY | 0.78288823 |
| SVLTVGLGK | ASSPGTGESQNTLY | 0.82325804 |
| TALAGAGSGI | ASSPGTGESQNTLY | 0.797259 |
| TGLPFTGI | ASSPGTGESQNTLY | 0.84031016 |
| VAEHVTV | ASSPGTGESQNTLY | 0.1348743 |
| VDIVLNSL | ASSPGTGESQNTLY | 0.8147459 |
| VLTGGGAVL | ASSPGTGESQNTLY | 0.85826373 |
| VNVGLAYGI | ASSPGTGESQNTLY | 0.7573768 |
| VSSAAVGAV | ASSPGTGESQNTLY | 0.8401026 |
| VTSVYEAGV | ASSPGTGESQNTLY | 0.8233993 |
| WVALDRGA | ASSPGTGESQNTLY | 0.8028401 |
| YLVGSVVI | ASSPGTGESQNTLY | 0.8480822 |
| GPPGTGKTLL | ASSPGTGESQNTLY | 0.75916415 |
| GVGMTKF | ASSPGTGESQNTLY | 0.8380471 |
| IGIGGSDL | ASSPGTGESQNTLY | 0.85776114 |
| IGTLLAEL | ASSPGTGESQNTLY | 0.8394181 |
| IYTSGTT | ASSPGTGESQNTLY | 0.82095766 |
| LSFPAGV | ASSPGTGESQNTLY | 0.82082224 |
| LYTSGTT | ASSPGTGESQNTLY | 0.5805709 |
| VGAFPFGF | ASSPGTGESQNTLY | 0.83491236 |
| VGAGTVEF | ASSPGTGESQNTLY | 0.13113727 |
| VGKSTLLL | ASSPGTGESQNTLY | 0.8259379 |
| AALAHAGGGIV | ASSPGTGESQNTLY | 0.7361564 |
| AALVAVSLL | ASSPGTGESQNTLY | 0.8405144 |
| AAVSRAVAAL | ASSPGTGESQNTLY | 0.83555114 |
| AGGSSGIGGF | ASSPGTGESQNTLY | 0.8433628 |
| AGLTVANANG | ASSPGTGESQNTLY | 0.7934053 |
| AGLVGAVL | ASSPGTGESQNTLY | 0.80352885 |
| ALSQAFTV | ASSPGTGESQNTLY | 0.75293726 |
| ALTFSVSL | ASSPGTGESQNTLY | 0.7222855 |
| ALTGAALVAL | ASSPGTGESQNTLY | 0.8474226 |
| ASFSIGAGVL | ASSPGTGESQNTLY | 0.81567854 |
| DGSVYKI | ASSPGTGESQNTLY | 0.8047686 |
| EQLAEIV | ASSPGTGESQNTLY | 0.85498565 |
| FAVSELTVM | ASSPGTGESQNTLY | 0.83941567 |
| GAAPLAAGL | ASSPGTGESQNTLY | 0.64913166 |
| GADGGDGGSGGAGG | ASSPGTGESQNTLY | 0.81994927 |
| GFGLDTGGAGG | ASSPGTGESQNTLY | 0.8317863 |
| GLASGGLL | ASSPGTGESQNTLY | 0.81901014 |

| | | |
|---|---|---|
| GRIAGAAVLL | ASSPGTGESQNTLY | 0.13686463 |
| GSAGSLNTSF | ASSPGTGESQNTLY | 0.83294433 |
| GTGQIGI | ASSPGTGESQNTLY | 0.81397605 |
| GTLIALIL | ASSPGTGESQNTLY | 0.7835535 |
| IAATMGFVVG | ASSPGTGESQNTLY | 0.8044071 |
| IFAQALL | ASSPGTGESQNTLY | 0.84793556 |
| IGAAVALGF | ASSPGTGESQNTLY | 0.28971064 |
| IGLTPLAL | ASSPGTGESQNTLY | 0.80780065 |
| ILGSNTSTL | ASSPGTGESQNTLY | 0.84342504 |
| IVFSLTVL | ASSPGTGESQNTLY | 0.84803367 |
| LASKATGF | ASSPGTGESQNTLY | 0.8552556 |
| LASVALIL | ASSPGTGESQNTLY | 0.8345696 |
| LFAVGLTL | ASSPGTGESQNTLY | 0.8052671 |
| LFNGGIGTYI | ASSPGTGESQNTLY | 0.84036297 |
| LTLGAGII | ASSPGTGESQNTLY | 0.8220391 |
| LVAAALGDI | ASSPGTGESQNTLY | 0.58634675 |
| NTAFVQLGI | ASSPGTGESQNTLY | 0.46266758 |
| QLFSGAEL | ASSPGTGESQNTLY | 0.83181787 |
| SAATFITL | ASSPGTGESQNTLY | 0.49676058 |
| SNYGLANLGV | ASSPGTGESQNTLY | 0.83670807 |
| SYGGAYAVM | ASSPGTGESQNTLY | 0.8513653 |
| TTGVGGLTLG | ASSPGTGESQNTLY | 0.7995733 |
| TTQTIYI | ASSPGTGESQNTLY | 0.72255 |
| VAAGAGLGAVI | ASSPGTGESQNTLY | 0.8231505 |
| VAAVVTLL | ASSPGTGESQNTLY | 0.8052458 |
| VGFLGGLI | ASSPGTGESQNTLY | 0.8452047 |
| VGVIPILL | ASSPGTGESQNTLY | 0.8393481 |
| VIFASSLI | ASSPGTGESQNTLY | 0.8547078 |
| VKGGDGGAAAG | ASSPGTGESQNTLY | 0.8530499 |
| VTKTFTNI | ASSPGTGESQNTLY | 0.8116703 |
| VVAAAILL | ASSPGTGESQNTLY | 0.84372425 |
| YAALDVEL | ASSPGTGESQNTLY | 0.8088012 |
| LASVALLL | ASSPGTGESQNTLY | 0.82193494 |
| LDAVHRV | ASSPGTGESQNTLY | 0.8270012 |
| QGLSSSIF | ASSPGTGESQNTLY | 0.82636166 |
| RGTFANIRL | ASSPGTGESQNTLY | 0.81019115 |
| RQLWWGHRI | ASSPGTGESQNTLY | 0.81828153 |
| YGPPGTGKTLL | ASSPGTGESQNTLY | 0.6815802 |
| AALVGAPANVA | ASSPGTGESQNTLY | 0.83376837 |
| AATFRRTE | ASSPGTGESQNTLY | 0.81840897 |
| AGALGGGI | ASSPGTGESQNTLY | 0.47590703 |
| AGGAGGAGGIGAG | ASSPGTGESQNTLY | 0.80629015 |
| AGSGALGL | ASSPGTGESQNTLY | 0.85599816 |
| AQIADQI | ASSPGTGESQNTLY | 0.85281014 |
| ASAVAVTYL | ASSPGTGESQNTLY | 0.7782679 |
| ASTVLGGTGG | ASSPGTGESQNTLY | 0.74567604 |
| AVGGEPAK | ASSPGTGESQNTLY | 0.82180464 |
| CEQGTFGGTC | ASSPGTGESQNTLY | 0.55187094 |
| DEGKAPAG | ASSPGTGESQNTLY | 0.83448493 |
| DTVVFPAL | ASSPGTGESQNTLY | 0.8235107 |
| FEGITMAG | ASSPGTGESQNTLY | 0.8210358 |

| | | |
|---|---|---|
| FPLVVQD | ASSPGTGESQNTLY | 0.7820553 |
| GAFTTPQ | ASSPGTGESQNTLY | 0.8361831 |
| GFGGFGTAGGVGGA | ASSPGTGESQNTLY | 0.803546 |
| GFLAQFL | ASSPGTGESQNTLY | 0.8320908 |
| GGAGGAGGNAGVA | ASSPGTGESQNTLY | 0.16167897 |
| GGVGTAGAGGAGGAGG | ASSPGTGESQNTLY | 0.8374814 |
| GNGGGGDAGNAGSGGN | ASSPGTGESQNTLY | 0.78521687 |
| GTGGAGGAGNDAGS | ASSPGTGESQNTLY | 0.7716857 |
| GTTNTGWFNS | ASSPGTGESQNTLY | 0.50034606 |
| IFMGPLFTL | ASSPGTGESQNTLY | 0.83688116 |
| ILAHAIEAI | ASSPGTGESQNTLY | 0.8495285 |
| IRSAIREAL | ASSPGTGESQNTLY | 0.84940004 |
| ITFAGGG | ASSPGTGESQNTLY | 0.7239151 |
| ITGMSGHQEV | ASSPGTGESQNTLY | 0.85258025 |
| IVAGLACTL | ASSPGTGESQNTLY | 0.85459244 |
| IVGGGLAGA | ASSPGTGESQNTLY | 0.8114703 |
| IVTGVLNI | ASSPGTGESQNTLY | 0.48007724 |
| LAQQGCI | ASSPGTGESQNTLY | 0.8391042 |
| LCLHVEN | ASSPGTGESQNTLY | 0.8007728 |
| LDAISDGS | ASSPGTGESQNTLY | 0.84281814 |
| LGGFAAML | ASSPGTGESQNTLY | 0.82429576 |
| LGGKNPMI | ASSPGTGESQNTLY | 0.77192724 |
| LGPKGVAAIL | ASSPGTGESQNTLY | 0.8440911 |
| LMVGVSI | ASSPGTGESQNTLY | 0.85046065 |
| LVGAAFVL | ASSPGTGESQNTLY | 0.78024405 |
| LVLAVAPL | ASSPGTGESQNTLY | 0.80879176 |
| MAAIAVIDAA | ASSPGTGESQNTLY | 0.8122517 |
| NNGINAELQ | ASSPGTGESQNTLY | 0.52626663 |
| RGEPLVLVHG | ASSPGTGESQNTLY | 0.8466362 |
| RGGAGGAAGLI | ASSPGTGESQNTLY | 0.800865 |
| RSYRMHPAV | ASSPGTGESQNTLY | 0.791417 |
| SAAHLLAGQLA | ASSPGTGESQNTLY | 0.83582294 |
| SAAVGSY | ASSPGTGESQNTLY | 0.80079 |
| SPHAVLHRI | ASSPGTGESQNTLY | 0.79472435 |
| SQIAKVL | ASSPGTGESQNTLY | 0.8162814 |
| TALGHLVP | ASSPGTGESQNTLY | 0.8357411 |
| TAPTTSGAA | ASSPGTGESQNTLY | 0.83701086 |
| TDGDDGNGGSPGT | ASSPGTGESQNTLY | 0.82815623 |
| TDGLGEN | ASSPGTGESQNTLY | 0.8214375 |
| TGGPPLPGQA | ASSPGTGESQNTLY | 0.85712326 |
| TIPAITI | ASSPGTGESQNTLY | 0.6751069 |
| TIPGFTI | ASSPGTGESQNTLY | 0.7878103 |
| TSIASGALA | ASSPGTGESQNTLY | 0.83444357 |
| TVTLPGAAI | ASSPGTGESQNTLY | 0.61748874 |
| VASGLSAALP | ASSPGTGESQNTLY | 0.84025955 |
| VGAGVIGI | ASSPGTGESQNTLY | 0.833815 |
| VGPKRFGV | ASSPGTGESQNTLY | 0.83803475 |
| VNVAFPD | ASSPGTGESQNTLY | 0.8182844 |
| VSMTNSGVS | ASSPGTGESQNTLY | 0.8595191 |
| VTDVLGDLSV | ASSPGTGESQNTLY | 0.8499535 |
| VVVVGAGFAGL | ASSPGTGESQNTLY | 0.83358353 |

| | | |
|---|---|---|
| VYTVGTVV | ASSPGTGESQNTLY | 0.8211045 |
| WTTFSGP | ASSPGTGESQNTLY | 0.7016466 |
| YDELVEAF | ASSPGTGESQNTLY | 0.8516822 |
| YGTGGAGGNGG | ASSPGTGESQNTLY | 0.83955014 |
| YTNVMAV | ASSPGTGESQNTLY | 0.77149415 |
| FLEQVHQL | ASSPGTGESQNTLY | 0.8249483 |
| GLSLLAVAV | ASSPGTGESQNTLY | 0.8558011 |
| LEQVHQL | ASSPGTGESQNTLY | 0.85289603 |
| LNAAGAI | ASSPGTGESQNTLY | 0.45017737 |
| PAVHASL | ASSPGTGESQNTLY | 0.8164387 |
| TELEDDA | ASSPGTGESQNTLY | 0.5210447 |
| TPPGPPN | ASSPGTGESQNTLY | 0.7742738 |
| TPPPPPPP | ASSPGTGESQNTLY | 0.7757818 |
| TQSPALI | ASGDAGQVYNNQAPL | 0.8400632 |
| EFAAAFPQ | ASGDAGQVYNNQAPL | 0.8185568 |
| LLSGNLDVL | ASGDAGQVYNNQAPL | 0.82366276 |
| GPANLGNYNI | ASGDAGQVYNNQAPL | 0.83474565 |
| TLGPTLP | ASGDAGQVYNNQAPL | 0.85680425 |
| VYAYSSCCK | ASGDAGQVYNNQAPL | 0.8050139 |
| APAPAPAPNI | ASGDAGQVYNNQAPL | 0.8312954 |
| PGGPVPIP | ASGDAGQVYNNQAPL | 0.5988333 |
| FLTVALEP | ASGDAGQVYNNQAPL | 0.46539682 |
| SQYFGANL | ASGDAGQVYNNQAPL | 0.79246926 |
| GGVKPAA | ASGDAGQVYNNQAPL | 0.8155942 |
| GGTGGAGGAGGW | ASGDAGQVYNNQAPL | 0.21312696 |
| GVGGVAGLGGAGG | ASGDAGQVYNNQAPL | 0.8519933 |
| VLLVLATTI | ASGDAGQVYNNQAPL | 0.83536124 |
| IGIFAAML | ASGDAGQVYNNQAPL | 0.8475858 |
| AVGDLID | ASGDAGQVYNNQAPL | 0.79130507 |
| NISPLPL | ASGDAGQVYNNQAPL | 0.82484686 |
| VTAVFVAI | ASGDAGQVYNNQAPL | 0.79611987 |
| SGDDNDAAL | ASGDAGQVYNNQAPL | 0.8439984 |
| AVAALAHS | ASGDAGQVYNNQAPL | 0.7901939 |
| GGVTVMATP | ASGDAGQVYNNQAPL | 0.69921947 |
| AGLAANVRD | ASGDAGQVYNNQAPL | 0.805938 |
| TNRPDLIDPALLRPG | ASGDAGQVYNNQAPL | 0.7709205 |
| LANLGVTAL | ASGDAGQVYNNQAPL | 0.85241544 |
| TGERIEL | ASGDAGQVYNNQAPL | 0.85009396 |
| NGEELAA | ASGDAGQVYNNQAPL | 0.84006035 |
| VLSLVWG | ASGDAGQVYNNQAPL | 0.8494054 |
| IGYALWL | ASGDAGQVYNNQAPL | 0.24963987 |
| LSLLLAPL | ASGDAGQVYNNQAPL | 0.7865639 |
| TLDEALK | ASGDAGQVYNNQAPL | 0.8563541 |
| TDGAGTAEGG | ASGDAGQVYNNQAPL | 0.83393073 |
| NAPGGAGGNGG | ASGDAGQVYNNQAPL | 0.82603025 |
| VSVFMGMGGNF | ASGDAGQVYNNQAPL | 0.8535027 |
| LASYLAAL | ASGDAGQVYNNQAPL | 0.7998193 |
| LVGGTPAL | ASGDAGQVYNNQAPL | 0.818919 |
| VRNADVL | ASGDAGQVYNNQAPL | 0.8269176 |
| TCSSPFLVSAM | ASGDAGQVYNNQAPL | 0.8437396 |
| VHITDAE | ASGDAGQVYNNQAPL | 0.83072114 |

| | | |
|---|---|---|
| EYAAPSF | ASGDAGQVYNNQAPL | 0.79946923 |
| VVDAFHGGL | ASGDAGQVYNNQAPL | 0.8347263 |
| ASVVDPQ | ASGDAGQVYNNQAPL | 0.8617167 |
| SYNEFGGLS | ASGDAGQVYNNQAPL | 0.82199824 |
| DPDGSVY | ASGDAGQVYNNQAPL | 0.84773934 |
| MVSTIHATE | ASGDAGQVYNNQAPL | 0.8425796 |
| LAAGVADPV | ASGDAGQVYNNQAPL | 0.81425756 |
| TKEQIAA | ASGDAGQVYNNQAPL | 0.7955267 |
| SLASPAAL | ASGDAGQVYNNQAPL | 0.80287045 |
| IGLVFYGV | ASGDAGQVYNNQAPL | 0.8101214 |
| FLLDASNL | ASGDAGQVYNNQAPL | 0.7543754 |
| YQVGNFP | ASGDAGQVYNNQAPL | 0.82575727 |
| ESLLSQST | ASGDAGQVYNNQAPL | 0.81145513 |
| LFTVGFAAVG | ASGDAGQVYNNQAPL | 0.8611982 |
| ATPPTPT | ASGDAGQVYNNQAPL | 0.84178966 |
| TDLATLG | ASGDAGQVYNNQAPL | 0.8397635 |
| SAGGTGR | ASGDAGQVYNNQAPL | 0.775049 |
| GFSNSGS | ASGDAGQVYNNQAPL | 0.8403618 |
| GGNGGSGGTSV | ASGDAGQVYNNQAPL | 0.83819234 |
| GTSVATGGAGN | ASGDAGQVYNNQAPL | 0.732646 |
| NGGNGGSGGTS | ASGDAGQVYNNQAPL | 0.85618675 |
| AVVKFPH | ASGDAGQVYNNQAPL | 0.7931293 |
| TPPPPPP | ASGDAGQVYNNQAPL | 0.7944281 |
| AAPAAAAAPA | ASGDAGQVYNNQAPL | 0.8479376 |
| IALNEDL | ASGDAGQVYNNQAPL | 0.8495821 |
| IVNVSSM | ASGDAGQVYNNQAPL | 0.80969405 |
| TGIPPAPR | ASGDAGQVYNNQAPL | 0.78910345 |
| FDLGGGTF | ASGDAGQVYNNQAPL | 0.79499555 |
| FDLGGGTFDVS | ASGDAGQVYNNQAPL | 0.84596443 |
| NEPTAAA | ASGDAGQVYNNQAPL | 0.8524229 |
| NEPTAAALA | ASGDAGQVYNNQAPL | 0.76425403 |
| GSIINISSI | ASGDAGQVYNNQAPL | 0.85463125 |
| VAPGNAGTA | ASGDAGQVYNNQAPL | 0.8270867 |
| TGGGASLE | ASGDAGQVYNNQAPL | 0.6444782 |
| VAVYNHYKRI | ASGDAGQVYNNQAPL | 0.821278 |
| TDPAPATT | ASGDAGQVYNNQAPL | 0.8506749 |
| TNRPDILDPALLRPG | ASGDAGQVYNNQAPL | 0.8486785 |
| LTTTTTS | ASGDAGQVYNNQAPL | 0.8567381 |
| PTGSGKTL | ASGDAGQVYNNQAPL | 0.83943135 |
| TGSGKTL | ASGDAGQVYNNQAPL | 0.8077799 |
| TIGHVDHGK | ASGDAGQVYNNQAPL | 0.85495555 |
| TIGHVDHGKTT | ASGDAGQVYNNQAPL | 0.8559905 |
| GHVDHGKTT | ASGDAGQVYNNQAPL | 0.8464053 |
| AQFEHTLL | ASGDAGQVYNNQAPL | 0.79399943 |
| IVNQDEL | ASGDAGQVYNNQAPL | 0.73982286 |
| TNGSQFF | ASGDAGQVYNNQAPL | 0.8397659 |
| TGKGIVE | ASGDAGQVYNNQAPL | 0.84524244 |
| LISNASDA | ASGDAGQVYNNQAPL | 0.7422211 |
| AELRTLGELTKIAKAHGAQ | ASGDAGQVYNNQAPL | 0.8323471 |
| APNAVAAAEPSW | ASGDAGQVYNNQAPL | 0.8365144 |
| APQVAGAATAVL | ASGDAGQVYNNQAPL | 0.74476004 |

| | | |
|---|---|---|
| ARRDPAAAAAILPTDA | ASGDAGQVYNNQAPL | 0.83603245 |
| DAAARDVLAEAGLAEYF | ASGDAGQVYNNQAPL | 0.8533913 |
| DDQIVLSAKANLAKVEQ | ASGDAGQVYNNQAPL | 0.82185686 |
| DELLPDTGEDSDEDGH | ASGDAGQVYNNQAPL | 0.8027729 |
| ELPEAPANRATVPQD | ASGDAGQVYNNQAPL | 0.8534349 |
| FGQSKIERVTGIG | ASGDAGQVYNNQAPL | 0.8617029 |
| GAGGTGSTAGGAGGAGGAGGLY | ASGDAGQVYNNQAPL | 0.85481155 |
| GHGGTGAAGGNGGTGGAGGSGI | ASGDAGQVYNNQAPL | 0.667906 |
| GNGGNGGAGGEGGPGGAGGAGA | ASGDAGQVYNNQAPL | 0.48553073 |
| GSTAGNGGNGGNATNGTIAGQP | ASGDAGQVYNNQAPL | 0.78457886 |
| GTGGDGGNGGNGGTGG | ASGDAGQVYNNQAPL | 0.83125305 |
| GVRRPAGIPADAPV | ASGDAGQVYNNQAPL | 0.8167958 |
| GYRQVTDAHLLTLARR | ASGDAGQVYNNQAPL | 0.78813076 |
| IAKMGQDGHDRG | ASGDAGQVYNNQAPL | 0.83846456 |
| KAEPSIEIQYT | ASGDAGQVYNNQAPL | 0.85301185 |
| KPVRIRMAADEDTVAVT | ASGDAGQVYNNQAPL | 0.8051721 |
| MPPRIAGMRLLVIKPEP | ASGDAGQVYNNQAPL | 0.6008241 |
| MKFYAVALFRSRGVPPTR | ASGDAGQVYNNQAPL | 0.84434205 |
| NAGGFGILGAVAHSP | ASGDAGQVYNNQAPL | 0.8434536 |
| NEGREITLAMAEM | ASGDAGQVYNNQAPL | 0.7925396 |
| NELVEWSHPLGHRWRWE | ASGDAGQVYNNQAPL | 0.8192252 |
| NLGAGGTGGNGGNPG | ASGDAGQVYNNQAPL | 0.79058206 |
| NTEWPKWETVMAADPVD | ASGDAGQVYNNQAPL | 0.3452363 |
| PASAAPAETQP | ASGDAGQVYNNQAPL | 0.8331578 |
| QQSSDGKNGVRAAEITGE | ASGDAGQVYNNQAPL | 0.7578446 |
| RAPDAVTTSWWKE | ASGDAGQVYNNQAPL | 0.85805607 |
| SGFGNVGAHTSGWFNQ | ASGDAGQVYNNQAPL | 0.78851575 |
| SGVAGAVGQAASVGGLKV | ASGDAGQVYNNQAPL | 0.83998173 |
| SNGGAGTGSAGGNGGT | ASGDAGQVYNNQAPL | 0.8446014 |
| TGGAGGEGGAAG | ASGDAGQVYNNQAPL | 0.84248185 |
| TGGDGGAAGAGGAPGHG | ASGDAGQVYNNQAPL | 0.8191073 |
| TMPPLLAAIANAGRHGV | ASGDAGQVYNNQAPL | 0.8050524 |
| TSSTAGNSGGA | ASGDAGQVYNNQAPL | 0.7562768 |
| VPATVPVGILC | ASGDAGQVYNNQAPL | 0.5137611 |
| VQRPDSGGVVLTGRIS | ASGDAGQVYNNQAPL | 0.8535712 |
| WTPNPVPATKGVPAGLVQAT | ASGDAGQVYNNQAPL | 0.19381283 |
| YALKRALAKFSDRFDVVII | ASGDAGQVYNNQAPL | 0.85316604 |
| YLIGSVIV | ASGDAGQVYNNQAPL | 0.8504721 |
| YGQLVAVID | ASGDAGQVYNNQAPL | 0.79679275 |
| VTVGGMTLPA | ASGDAGQVYNNQAPL | 0.8430344 |
| VLALPAV | ASGDAGQVYNNQAPL | 0.7230551 |
| VIPPTLNYET | ASGDAGQVYNNQAPL | 0.7322966 |
| VGMGGMHPSSG | ASGDAGQVYNNQAPL | 0.84176743 |
| VGLAGGAATA | ASGDAGQVYNNQAPL | 0.84317774 |
| VGGAGGAGGSGP | ASGDAGQVYNNQAPL | 0.7974739 |
| VFAAFMV | ASGDAGQVYNNQAPL | 0.83336127 |
| TLYDKLR | ASGDAGQVYNNQAPL | 0.8377321 |
| TLAVFTVL | ASGDAGQVYNNQAPL | 0.78591675 |
| TGSVQPARV | ASGDAGQVYNNQAPL | 0.83350617 |
| TGGGAGSGGT | ASGDAGQVYNNQAPL | 0.8531056 |
| SHLETSL | ASGDAGQVYNNQAPL | 0.8339535 |

| | | |
|---|---|---|
| SFTTGMT | ASGDAGQVYNNQAPL | 0.80571353 |
| QVEVAKRMGVRQPTVS | ASGDAGQVYNNQAPL | 0.8476353 |
| QGSGGPALAM | ASGDAGQVYNNQAPL | 0.7635517 |
| MLAPSNP | ASGDAGQVYNNQAPL | 0.8389696 |
| LVNNAAI | ASGDAGQVYNNQAPL | 0.8293382 |
| LTGEINTLA | ASGDAGQVYNNQAPL | 0.8237721 |
| LSIFGPV | ASGDAGQVYNNQAPL | 0.84465235 |
| LLALLANL | ASGDAGQVYNNQAPL | 0.6090553 |
| LISNASDALDKLR | ASGDAGQVYNNQAPL | 0.7815531 |
| LINEAVP | ASGDAGQVYNNQAPL | 0.79942954 |
| LGYAFSGL | ASGDAGQVYNNQAPL | 0.8380083 |
| LDFVAVAL | ASGDAGQVYNNQAPL | 0.84881926 |
| LAQPGAE | ASGDAGQVYNNQAPL | 0.7852887 |
| ISNLGAVTGF | ASGDAGQVYNNQAPL | 0.82982934 |
| HQEDPGRI | ASGDAGQVYNNQAPL | 0.8128276 |
| GTPGEPTP | ASGDAGQVYNNQAPL | 0.7746538 |
| GFFNSGS | ASGDAGQVYNNQAPL | 0.8172908 |
| GAGGGGLTAGAN | ASGDAGQVYNNQAPL | 0.8189759 |
| FVDALVNM | ASGDAGQVYNNQAPL | 0.76078886 |
| FLTVAPEL | ASGDAGQVYNNQAPL | 0.8428607 |
| FETLTAL | ASGDAGQVYNNQAPL | 0.75911343 |
| DVINPIEL | ASGDAGQVYNNQAPL | 0.8412647 |
| APVSGAPV | ASGDAGQVYNNQAPL | 0.8515842 |
| ALVGGLAW | ASGDAGQVYNNQAPL | 0.79742694 |
| ALAAAAATAI | ASGDAGQVYNNQAPL | 0.8445945 |
| AFGLGLLL | ASGDAGQVYNNQAPL | 0.83722985 |
| AFFAAASL | ASGDAGQVYNNQAPL | 0.8432341 |
| AAAALPAFSPP | ASGDAGQVYNNQAPL | 0.822139 |
| VVLAPAL | ASGDAGQVYNNQAPL | 0.810815 |
| TIGHVDHGKTTL | ASGDAGQVYNNQAPL | 0.85591537 |
| LISNASDALDK | ASGDAGQVYNNQAPL | 0.84767246 |
| LISNASDAL | ASGDAGQVYNNQAPL | 0.8400202 |
| ISNASDALDK | ASGDAGQVYNNQAPL | 0.84116215 |
| NQAETLVYQTE | ASGDAGQVYNNQAPL | 0.8406462 |
| TGGTGGTAGAG | ASGDAGQVYNNQAPL | 0.8419974 |
| TGSTAGGAGGA | ASGDAGQVYNNQAPL | 0.7898164 |
| DPLAVPVAAGPV | ASGDAGQVYNNQAPL | 0.6463951 |
| GGTGGTPGNGGN | ASGDAGQVYNNQAPL | 0.817962 |
| PAGLAGVDPATL | ASGDAGQVYNNQAPL | 0.8231499 |
| GNNLVVEAPHAGASV | ASGDAGQVYNNQAPL | 0.8187167 |
| KVTRIMSASTLEQPA | ASGDAGQVYNNQAPL | 0.4450248 |
| LGFALPIALPRRTVV | ASGDAGQVYNNQAPL | 0.85403776 |
| TRLGNLTARKLVVLV | ASGDAGQVYNNQAPL | 0.85959136 |
| WSQQRRLIYFPSAGP | ASGDAGQVYNNQAPL | 0.7887949 |
| GPNGNPDPMAAAVDIR | ASGDAGQVYNNQAPL | 0.8562432 |
| TQRCVSVADLRAAASR | ASGDAGQVYNNQAPL | 0.730091 |
| QFVLDVAEHPVEVLTETV | ASGDAGQVYNNQAPL | 0.8446797 |
| GGAGGTGGAGGNGGTQAGDG | ASGDAGQVYNNQAPL | 0.8536675 |
| PLLATLRAIAVDVDAKPAQV | ASGDAGQVYNNQAPL | 0.771887 |
| APKFRTGGYMIDFWGVGYQVAK | ASGDAGQVYNNQAPL | 0.84705925 |
| GAGGAGGAGGAAGGPGGTGGTG | ASGDAGQVYNNQAPL | 0.7976574 |

| | | |
|---|---|---|
| IGALGLLATDVAPAGSVAFGVA | ASGDAGQVYNNQAPL | 0.8239694 |
| AGGAGGIGGNGGAGGAGGVLMG | ASGDAGQVYNNQAPL | 0.855069 |
| ADELIGLL | ASGDAGQVYNNQAPL | 0.81307864 |
| ADRALKA | ASGDAGQVYNNQAPL | 0.8405368 |
| AGAVGQVGGAGGA | ASGDAGQVYNNQAPL | 0.8184805 |
| AGGGGGLGGAP | ASGDAGQVYNNQAPL | 0.76576746 |
| AGGLLYGNGGAG | ASGDAGQVYNNQAPL | 0.84094894 |
| AGLESIGL | ASGDAGQVYNNQAPL | 0.85334635 |
| AGLKTGII | ASGDAGQVYNNQAPL | 0.7445535 |
| ASLAALLL | ASGDAGQVYNNQAPL | 0.85463035 |
| ASVINISSI | ASGDAGQVYNNQAPL | 0.80108744 |
| ATPPGPA | ASGDAGQVYNNQAPL | 0.8623185 |
| ATVASALV | ASGDAGQVYNNQAPL | 0.8421921 |
| ATVASLIL | ASGDAGQVYNNQAPL | 0.82549256 |
| AVNGPVLIRH | ASGDAGQVYNNQAPL | 0.80347574 |
| DGAAGAVGI | ASGDAGQVYNNQAPL | 0.8541744 |
| DPLLHGI | ASGDAGQVYNNQAPL | 0.81856424 |
| ELGAALGAL | ASGDAGQVYNNQAPL | 0.84319943 |
| ELGRTQAV | ASGDAGQVYNNQAPL | 0.8555027 |
| FVAGGYLS | ASGDAGQVYNNQAPL | 0.54380476 |
| GASGGAGGAGGW | ASGDAGQVYNNQAPL | 0.87894183 |
| GGAGAGGDGGAGGI | ASGDAGQVYNNQAPL | 0.8417732 |
| GTGHNGSQPGMGGNG | ASGDAGQVYNNQAPL | 0.8451315 |
| GYAAASASAS | ASGDAGQVYNNQAPL | 0.7168528 |
| HQVDVAAVQQG | ASGDAGQVYNNQAPL | 0.39631093 |
| IGAAIAHT | ASGDAGQVYNNQAPL | 0.77265644 |
| IGMIEVAQL | ASGDAGQVYNNQAPL | 0.62332726 |
| IRLAPEHTSAMD | ASGDAGQVYNNQAPL | 0.8592647 |
| IYNVSVL | ASGDAGQVYNNQAPL | 0.85867184 |
| KQGFLGM | ASGDAGQVYNNQAPL | 0.7385775 |
| LESVITQL | ASGDAGQVYNNQAPL | 0.8671373 |
| LGLGALVVV | ASGDAGQVYNNQAPL | 0.87471145 |
| LIGNGGNGGSGGI | ASGDAGQVYNNQAPL | 0.39209455 |
| LNAINAPTL | ASGDAGQVYNNQAPL | 0.80841935 |
| LQLLNTGV | ASGDAGQVYNNQAPL | 0.67078114 |
| LQTSIEF | ASGDAGQVYNNQAPL | 0.77719104 |
| MSFAVSEL | ASGDAGQVYNNQAPL | 0.31936336 |
| NTGALGL | ASGDAGQVYNNQAPL | 0.8524965 |
| PPGAGGAGGAGTT | ASGDAGQVYNNQAPL | 0.7717249 |
| QLVGLGI | ASGDAGQVYNNQAPL | 0.14984204 |
| RFLEQVHQL | ASGDAGQVYNNQAPL | 0.83045566 |
| RGVGYVLR | ASGDAGQVYNNQAPL | 0.7913899 |
| SATGVFAGVI | ASGDAGQVYNNQAPL | 0.7237987 |
| SLPAITLP | ASGDAGQVYNNQAPL | 0.8639415 |
| STALAFTV | ASGDAGQVYNNQAPL | 0.76507527 |
| SVLTVGLGK | ASGDAGQVYNNQAPL | 0.6396141 |
| TALAGAGSGI | ASGDAGQVYNNQAPL | 0.85422647 |
| TGLPFTGI | ASGDAGQVYNNQAPL | 0.8691532 |
| VAEHVTV | ASGDAGQVYNNQAPL | 0.515937 |
| VDIVLNSL | ASGDAGQVYNNQAPL | 0.84167635 |
| VLTGGGAVL | ASGDAGQVYNNQAPL | 0.8809502 |

| | | |
|---|---|---|
| VNVGLAYGI | ASGDAGQVYNNQAPL | 0.8172747 |
| VSSAAVGAV | ASGDAGQVYNNQAPL | 0.87396806 |
| VTSVYEAGV | ASGDAGQVYNNQAPL | 0.87958944 |
| WVALDRGA | ASGDAGQVYNNQAPL | 0.7652838 |
| YLVGSVVI | ASGDAGQVYNNQAPL | 0.86543345 |
| GPPGTGKTLL | ASGDAGQVYNNQAPL | 0.87066793 |
| GVGMTKF | ASGDAGQVYNNQAPL | 0.77187884 |
| IGIGGSDL | ASGDAGQVYNNQAPL | 0.81487453 |
| IGTLLAEL | ASGDAGQVYNNQAPL | 0.5208326 |
| IYTSGTT | ASGDAGQVYNNQAPL | 0.8140373 |
| LSFPAGV | ASGDAGQVYNNQAPL | 0.34656686 |
| LYTSGTT | ASGDAGQVYNNQAPL | 0.84458256 |
| VGAFPFGF | ASGDAGQVYNNQAPL | 0.8684527 |
| VGAGTVEF | ASGDAGQVYNNQAPL | 0.78866655 |
| VGKSTLLL | ASGDAGQVYNNQAPL | 0.5913648 |
| AALAHAGGGIV | ASGDAGQVYNNQAPL | 0.6148694 |
| AALVAVSLL | ASGDAGQVYNNQAPL | 0.8501485 |
| AAVSRAVAAL | ASGDAGQVYNNQAPL | 0.6918584 |
| AGGSSGIGGF | ASGDAGQVYNNQAPL | 0.3694747 |
| AGLTVANANG | ASGDAGQVYNNQAPL | 0.77500355 |
| AGLVGAVL | ASGDAGQVYNNQAPL | 0.86639416 |
| ALSQAFTV | ASGDAGQVYNNQAPL | 0.8690908 |
| ALTFSVSL | ASGDAGQVYNNQAPL | 0.6294211 |
| ALTGAALVAL | ASGDAGQVYNNQAPL | 0.87669027 |
| ASFSIGAGVL | ASGDAGQVYNNQAPL | 0.8661071 |
| DGSVYKI | ASGDAGQVYNNQAPL | 0.8798447 |
| EQLAEIV | ASGDAGQVYNNQAPL | 0.8122297 |
| FAVSELTVM | ASGDAGQVYNNQAPL | 0.6252111 |
| GAAPLAAGL | ASGDAGQVYNNQAPL | 0.8694269 |
| GADGGDGGSGGAGG | ASGDAGQVYNNQAPL | 0.80638194 |
| GFGLDTGGAGG | ASGDAGQVYNNQAPL | 0.7599298 |
| GLASGGLL | ASGDAGQVYNNQAPL | 0.86915547 |
| GRIAGAAVLL | ASGDAGQVYNNQAPL | 0.75602293 |
| GSAGSLNTSF | ASGDAGQVYNNQAPL | 0.704651 |
| GTGQIGI | ASGDAGQVYNNQAPL | 0.8213438 |
| GTLIALIL | ASGDAGQVYNNQAPL | 0.8808286 |
| IAATMGFVVG | ASGDAGQVYNNQAPL | 0.6933903 |
| IFAQALL | ASGDAGQVYNNQAPL | 0.5211785 |
| IGAAVALGF | ASGDAGQVYNNQAPL | 0.7562576 |
| IGLTPLAL | ASGDAGQVYNNQAPL | 0.8202859 |
| ILGSNTSTL | ASGDAGQVYNNQAPL | 0.8693198 |
| IVFSLTVL | ASGDAGQVYNNQAPL | 0.83556634 |
| LASKATGF | ASGDAGQVYNNQAPL | 0.80800223 |
| LASVALIL | ASGDAGQVYNNQAPL | 0.8478438 |
| LFAVGLTL | ASGDAGQVYNNQAPL | 0.8599925 |
| LFNGGIGTYI | ASGDAGQVYNNQAPL | 0.8319248 |
| LTLGAGII | ASGDAGQVYNNQAPL | 0.86772454 |
| LVAAALGDI | ASGDAGQVYNNQAPL | 0.7438617 |
| NTAFVQLGI | ASGDAGQVYNNQAPL | 0.8781181 |
| QLFSGAEL | ASGDAGQVYNNQAPL | 0.8675775 |
| SAATFITL | ASGDAGQVYNNQAPL | 0.861956 |

| | | |
|---|---|---|
| SNYGLANLGV | ASGDAGQVYNNQAPL | 0.87254035 |
| SYGGAYAVM | ASGDAGQVYNNQAPL | 0.851467 |
| TTGVGGLTLG | ASGDAGQVYNNQAPL | 0.84387654 |
| TTQTIYI | ASGDAGQVYNNQAPL | 0.45459554 |
| VAAGAGLGAVI | ASGDAGQVYNNQAPL | 0.73676467 |
| VAAVVTLL | ASGDAGQVYNNQAPL | 0.84343284 |
| VGFLGGLI | ASGDAGQVYNNQAPL | 0.84499454 |
| VGVIPILL | ASGDAGQVYNNQAPL | 0.8596225 |
| VIFASSLI | ASGDAGQVYNNQAPL | 0.8386719 |
| VKGGDGGAAAG | ASGDAGQVYNNQAPL | 0.75010073 |
| VTKTFTNI | ASGDAGQVYNNQAPL | 0.87945163 |
| VVAAAILL | ASGDAGQVYNNQAPL | 0.87201303 |
| YAALDVEL | ASGDAGQVYNNQAPL | 0.47171223 |
| LASVALLL | ASGDAGQVYNNQAPL | 0.8084603 |
| LDAVHRV | ASGDAGQVYNNQAPL | 0.88116574 |
| QGLSSSIF | ASGDAGQVYNNQAPL | 0.8469887 |
| RGTFANIRL | ASGDAGQVYNNQAPL | 0.7797555 |
| RQLWWGHRI | ASGDAGQVYNNQAPL | 0.78135735 |
| YGPPGTGKTLL | ASGDAGQVYNNQAPL | 0.873312 |
| AALVGAPANVA | ASGDAGQVYNNQAPL | 0.7802325 |
| AATFRRTE | ASGDAGQVYNNQAPL | 0.88057905 |
| AGALGGGI | ASGDAGQVYNNQAPL | 0.28835103 |
| AGGAGGAGGIGAG | ASGDAGQVYNNQAPL | 0.8003039 |
| AGSGALGL | ASGDAGQVYNNQAPL | 0.75619954 |
| AQIADQI | ASGDAGQVYNNQAPL | 0.22868297 |
| ASAVAVTYL | ASGDAGQVYNNQAPL | 0.85139894 |
| ASTVLGGTGG | ASGDAGQVYNNQAPL | 0.70383334 |
| AVGGEPAK | ASGDAGQVYNNQAPL | 0.8638376 |
| CEQGTFGGTC | ASGDAGQVYNNQAPL | 0.7499546 |
| DEGKAPAG | ASGDAGQVYNNQAPL | 0.8669684 |
| DTVVFPAL | ASGDAGQVYNNQAPL | 0.83927035 |
| FEGITMAG | ASGDAGQVYNNQAPL | 0.83424574 |
| FPLVVQD | ASGDAGQVYNNQAPL | 0.5988635 |
| GAFTTPQ | ASGDAGQVYNNQAPL | 0.8455026 |
| GFGGFGTAGGVGGA | ASGDAGQVYNNQAPL | 0.880082 |
| GFLAQFL | ASGDAGQVYNNQAPL | 0.86832196 |
| GGAGGAGGNAGVA | ASGDAGQVYNNQAPL | 0.3385256 |
| GGVGTAGAGGAGGAGG | ASGDAGQVYNNQAPL | 0.833275 |
| GNGGGGDAGNAGSGGN | ASGDAGQVYNNQAPL | 0.77203023 |
| GTGGAGGAGNDAGS | ASGDAGQVYNNQAPL | 0.22886996 |
| GTTNTGWFNS | ASGDAGQVYNNQAPL | 0.22659433 |
| IFMGPLFTL | ASGDAGQVYNNQAPL | 0.80691296 |
| ILAHAIEAI | ASGDAGQVYNNQAPL | 0.8137933 |
| IRSAIREAL | ASGDAGQVYNNQAPL | 0.86151826 |
| ITFAGGG | ASGDAGQVYNNQAPL | 0.8606616 |
| ITGMSGHQEV | ASGDAGQVYNNQAPL | 0.78868735 |
| IVAGLACTL | ASGDAGQVYNNQAPL | 0.8643681 |
| IVGGGLAGA | ASGDAGQVYNNQAPL | 0.8417527 |
| IVTGVLNI | ASGDAGQVYNNQAPL | 0.10227999 |
| LAQQGCI | ASGDAGQVYNNQAPL | 0.77819943 |
| LCLHVEN | ASGDAGQVYNNQAPL | 0.35571903 |

| | | |
|---|---|---|
| LDAISDGS | ASGDAGQVYNNQAPL | 0.8767159 |
| LGGFAAML | ASGDAGQVYNNQAPL | 0.688897 |
| LGGKNPMI | ASGDAGQVYNNQAPL | 0.7343035 |
| LGPKGVAAIL | ASGDAGQVYNNQAPL | 0.8215977 |
| LMVGVSI | ASGDAGQVYNNQAPL | 0.84436584 |
| LVGAAFVL | ASGDAGQVYNNQAPL | 0.8302711 |
| LVLAVAPL | ASGDAGQVYNNQAPL | 0.735474 |
| MAAIAVIDAA | ASGDAGQVYNNQAPL | 0.22508591 |
| NNGINAELQ | ASGDAGQVYNNQAPL | 0.85258377 |
| RGEPLVLVHG | ASGDAGQVYNNQAPL | 0.622913 |
| RGGAGGAAGLI | ASGDAGQVYNNQAPL | 0.5244995 |
| RSYRMHPAV | ASGDAGQVYNNQAPL | 0.5042648 |
| SAAHLLAGQLA | ASGDAGQVYNNQAPL | 0.56972605 |
| SAAVGSY | ASGDAGQVYNNQAPL | 0.4661954 |
| SPHAVLHRI | ASGDAGQVYNNQAPL | 0.863596 |
| SQIAKVL | ASGDAGQVYNNQAPL | 0.77308285 |
| TALGHLVP | ASGDAGQVYNNQAPL | 0.78733855 |
| TAPTTSGAA | ASGDAGQVYNNQAPL | 0.8762983 |
| TDGDDGNGGSPGT | ASGDAGQVYNNQAPL | 0.8390055 |
| TDGLGEN | ASGDAGQVYNNQAPL | 0.81067634 |
| TGGPPLPGQA | ASGDAGQVYNNQAPL | 0.7049811 |
| TIPAITI | ASGDAGQVYNNQAPL | 0.597227 |
| TIPGFTI | ASGDAGQVYNNQAPL | 0.6443982 |
| TSIASGALA | ASGDAGQVYNNQAPL | 0.6699785 |
| TVTLPGAAI | ASGDAGQVYNNQAPL | 0.88183284 |
| VASGLSAALP | ASGDAGQVYNNQAPL | 0.8005941 |
| VGAGVIGI | ASGDAGQVYNNQAPL | 0.47857016 |
| VGPKRFGV | ASGDAGQVYNNQAPL | 0.85522556 |
| VNVAFPD | ASGDAGQVYNNQAPL | 0.8786392 |
| VSMTNSGVS | ASGDAGQVYNNQAPL | 0.88500977 |
| VTDVLGDLSV | ASGDAGQVYNNQAPL | 0.64116824 |
| VVVVGAGFAGL | ASGDAGQVYNNQAPL | 0.7779216 |
| VYTVGTVV | ASGDAGQVYNNQAPL | 0.7786865 |
| WTTFSGP | ASGDAGQVYNNQAPL | 0.80642664 |
| YDELVEAF | ASGDAGQVYNNQAPL | 0.7402796 |
| YGTGGAGGNGG | ASGDAGQVYNNQAPL | 0.5103537 |
| YTNVMAV | ASGDAGQVYNNQAPL | 0.87278515 |
| FLEQVHQL | ASGDAGQVYNNQAPL | 0.8872299 |
| GLSLLAVAV | ASGDAGQVYNNQAPL | 0.7902658 |
| LEQVHQL | ASGDAGQVYNNQAPL | 0.3425473 |
| LNAAGAI | ASGDAGQVYNNQAPL | 0.8127253 |
| PAVHASL | ASGDAGQVYNNQAPL | 0.7120285 |
| TELEDDA | ASGDAGQVYNNQAPL | 0.72823644 |
| TPPGPPN | ASGDAGQVYNNQAPL | 0.77848756 |
| TPPPPPPP | ASGDAGQVYNNQAPL | 0.42876565 |
| TQSPALI | ASSDANTEVF | 0.79565847 |
| EFAAAFPQ | ASSDANTEVF | 0.8826058 |
| LLSGNLDVL | ASSDANTEVF | 0.79610974 |
| GPANLGNYNI | ASSDANTEVF | 0.85298765 |
| TLGPTLP | ASSDANTEVF | 0.87025213 |
| VYAYSSCCK | ASSDANTEVF | 0.8533592 |

| | | |
|---|---|---|
| APAPAPAPNI | ASSDANTEVF | 0.87414354 |
| PGGPVPIP | ASSDANTEVF | 0.8674319 |
| FLTVALEP | ASSDANTEVF | 0.80491555 |
| SQYFGANL | ASSDANTEVF | 0.836041 |
| GGVKPAA | ASSDANTEVF | 0.85059094 |
| GGTGGAGGAGGW | ASSDANTEVF | 0.80977505 |
| GVGGVAGLGGAGG | ASSDANTEVF | 0.8284023 |
| VLLVLATTI | ASSDANTEVF | 0.78068286 |
| IGIFAAML | ASSDANTEVF | 0.8608907 |
| AVGDLID | ASSDANTEVF | 0.8382588 |
| NISPLPL | ASSDANTEVF | 0.8773273 |
| VTAVFVAI | ASSDANTEVF | 0.30628246 |
| SGDDNDAAL | ASSDANTEVF | 0.55571425 |
| AVAALAHS | ASSDANTEVF | 0.86868924 |
| GGVTVMATP | ASSDANTEVF | 0.18120408 |
| AGLAANVRD | ASSDANTEVF | 0.27026367 |
| TNRPDLIDPALLRPG | ASSDANTEVF | 0.8350923 |
| LANLGVTAL | ASSDANTEVF | 0.879238 |
| TGERIEL | ASSDANTEVF | 0.1605619 |
| NGEELAA | ASSDANTEVF | 0.67360234 |
| VLSLVWG | ASSDANTEVF | 0.4977394 |
| IGYALWL | ASSDANTEVF | 0.8649795 |
| LSLLLAPL | ASSDANTEVF | 0.78484 |
| TLDEALK | ASSDANTEVF | 0.45273528 |
| TDGAGTAEGG | ASSDANTEVF | 0.8568099 |
| NAPGGAGGNGG | ASSDANTEVF | 0.45149407 |
| VSVFMGMGGNF | ASSDANTEVF | 0.76903284 |
| LASYLAAL | ASSDANTEVF | 0.8615155 |
| LVGGTPAL | ASSDANTEVF | 0.8770666 |
| VRNADVL | ASSDANTEVF | 0.54919195 |
| TCSSPFLVSAM | ASSDANTEVF | 0.8555986 |
| VHITDAE | ASSDANTEVF | 0.8000185 |
| EYAAPSF | ASSDANTEVF | 0.8856085 |
| VVDAFHGGL | ASSDANTEVF | 0.75941694 |
| ASVVDPQ | ASSDANTEVF | 0.43797332 |
| SYNEFGGLS | ASSDANTEVF | 0.8621344 |
| DPDGSVY | ASSDANTEVF | 0.86108935 |
| MVSTIHATE | ASSDANTEVF | 0.77066 |
| LAAGVADPV | ASSDANTEVF | 0.86388826 |
| TKEQIAA | ASSDANTEVF | 0.8177581 |
| SLASPAAL | ASSDANTEVF | 0.79388714 |
| IGLVFYGV | ASSDANTEVF | 0.8647332 |
| FLLDASNL | ASSDANTEVF | 0.8673391 |
| YQVGNFP | ASSDANTEVF | 0.58882076 |
| ESLLSQST | ASSDANTEVF | 0.865033 |
| LFTVGFAAVG | ASSDANTEVF | 0.82402194 |
| ATPPTPT | ASSDANTEVF | 0.67134917 |
| TDLATLG | ASSDANTEVF | 0.88275445 |
| SAGGTGR | ASSDANTEVF | 0.8043932 |
| GFSNSGS | ASSDANTEVF | 0.8653921 |
| GGNGGSGGTSV | ASSDANTEVF | 0.8482241 |

| | | |
|---|---|---|
| GTSVATGGAGN | ASSDANTEVF | 0.87933433 |
| NGGNGGSGGTS | ASSDANTEVF | 0.4611361 |
| AVVKFPH | ASSDANTEVF | 0.8447531 |
| TPPPPPP | ASSDANTEVF | 0.8305347 |
| AAPAAAAAPA | ASSDANTEVF | 0.86245066 |
| IALNEDL | ASSDANTEVF | 0.80334973 |
| IVNVSSM | ASSDANTEVF | 0.68890965 |
| TGIPPAPR | ASSDANTEVF | 0.85213244 |
| FDLGGGTF | ASSDANTEVF | 0.8732039 |
| FDLGGGTFDVS | ASSDANTEVF | 0.83452433 |
| NEPTAAA | ASSDANTEVF | 0.85333765 |
| NEPTAAALA | ASSDANTEVF | 0.87997204 |
| GSIINISSI | ASSDANTEVF | 0.57291496 |
| VAPGNAGTA | ASSDANTEVF | 0.8034178 |
| TGGGASLE | ASSDANTEVF | 0.8645377 |
| VAVYNHYKRI | ASSDANTEVF | 0.8754525 |
| TDPAPATT | ASSDANTEVF | 0.809603 |
| TNRPDILDPALLRPG | ASSDANTEVF | 0.8785275 |
| LTTTTTS | ASSDANTEVF | 0.73623514 |
| PTGSGKTL | ASSDANTEVF | 0.85906154 |
| TGSGKTL | ASSDANTEVF | 0.7716801 |
| TIGHVDHGK | ASSDANTEVF | 0.829085 |
| TIGHVDHGKTT | ASSDANTEVF | 0.6781785 |
| GHVDHGKTT | ASSDANTEVF | 0.8440602 |
| AQFEHTLL | ASSDANTEVF | 0.8631029 |
| IVNQDEL | ASSDANTEVF | 0.85885674 |
| TNGSQFF | ASSDANTEVF | 0.23111157 |
| TGKGIVE | ASSDANTEVF | 0.8527411 |
| LISNASDA | ASSDANTEVF | 0.8655916 |
| AELRTLGELTKIAKAHGAQ | ASSDANTEVF | 0.8828776 |
| APNAVAAAEPSW | ASSDANTEVF | 0.82441044 |
| APQVAGAATAVL | ASSDANTEVF | 0.85594785 |
| ARRDPAAAAAILPTDA | ASSDANTEVF | 0.7267692 |
| DAAARDVLAEAGLAEYF | ASSDANTEVF | 0.82408106 |
| DDQIVLSAKANLAKVEQ | ASSDANTEVF | 0.65897727 |
| DELLPDTGEDSDEDGH | ASSDANTEVF | 0.736788 |
| ELPEAPANRATVPQD | ASSDANTEVF | 0.7904568 |
| FGQSKIERVTGIG | ASSDANTEVF | 0.6972176 |
| GAGGTGSTAGGAGGAGGAGGLY | ASSDANTEVF | 0.8644672 |
| GHGGTGAAGGNGGTGGAGGSGI | ASSDANTEVF | 0.87159216 |
| GNGGNGGAGGEGGPGGAGGAGA | ASSDANTEVF | 0.8638696 |
| GSTAGNGGNGGNATNGTIAGQP | ASSDANTEVF | 0.8737268 |
| GTGGDGGNGGNGGTGG | ASSDANTEVF | 0.8769678 |
| GVRRPAGIPADAPV | ASSDANTEVF | 0.8666538 |
| GYRQVTDAHLLTLARR | ASSDANTEVF | 0.73868597 |
| IAKMGQDGHDRG | ASSDANTEVF | 0.7767946 |
| KAEPSIEIQYT | ASSDANTEVF | 0.5207532 |
| KPVRIRMAADEDTVAVT | ASSDANTEVF | 0.7789734 |
| MPPRIAGMRLLVIKPEP | ASSDANTEVF | 0.7437639 |
| MKFYAVALFRSRGVPPTR | ASSDANTEVF | 0.8318826 |
| NAGGFGILGAVAHSP | ASSDANTEVF | 0.12355399 |

| | | |
|---|---|---|
| NEGREITLAMAEM | ASSDANTEVF | 0.841231 |
| NELVEWSHPLGHRWRWE | ASSDANTEVF | 0.88081026 |
| NLGAGGTGGNGGNPG | ASSDANTEVF | 0.7981545 |
| NTEWPKWETVMAADPVD | ASSDANTEVF | 0.8518614 |
| PASAAPAETQP | ASSDANTEVF | 0.85022545 |
| QQSSDGKNGVRAAEITGE | ASSDANTEVF | 0.7825819 |
| RAPDAVTTSWWKE | ASSDANTEVF | 0.86165667 |
| SGFGNVGAHTSGWFNQ | ASSDANTEVF | 0.706625 |
| SGVAGAVGQAASVGGLKV | ASSDANTEVF | 0.85990787 |
| SNGGAGTGSAGGNGGT | ASSDANTEVF | 0.8773767 |
| TGGAGGEGGAAG | ASSDANTEVF | 0.83490884 |
| TGGDGGAAGAGGAPGHG | ASSDANTEVF | 0.838531 |
| TMPPLLAAIANAGRHGV | ASSDANTEVF | 0.81453484 |
| TSSTAGNSGGA | ASSDANTEVF | 0.34869707 |
| VPATVPVGILC | ASSDANTEVF | 0.8463881 |
| VQRPDSGGVVLTGRIS | ASSDANTEVF | 0.1276801 |
| WTPNPVPATKGVPAGLVQAT | ASSDANTEVF | 0.8264065 |
| YALKRALAKFSDRFDVVII | ASSDANTEVF | 0.5966376 |
| YLIGSVIV | ASSDANTEVF | 0.8284722 |
| YGQLVAVID | ASSDANTEVF | 0.8189424 |
| VTVGGMTLPA | ASSDANTEVF | 0.86266965 |
| VLALPAV | ASSDANTEVF | 0.66670996 |
| VIPPTLNYET | ASSDANTEVF | 0.8324475 |
| VGMGGMHPSSG | ASSDANTEVF | 0.7929133 |
| VGLAGGAATA | ASSDANTEVF | 0.5063689 |
| VGGAGGAGGSGP | ASSDANTEVF | 0.8644166 |
| VFAAFMV | ASSDANTEVF | 0.7967584 |
| TLYDKLR | ASSDANTEVF | 0.820199 |
| TLAVFTVL | ASSDANTEVF | 0.8657075 |
| TGSVQPARV | ASSDANTEVF | 0.8061918 |
| TGGGAGSGGT | ASSDANTEVF | 0.34459713 |
| SHLETSL | ASSDANTEVF | 0.80299276 |
| SFTTGMT | ASSDANTEVF | 0.82063663 |
| QVEVAKRMGVRQPTVS | ASSDANTEVF | 0.78145266 |
| QGSGGPALAM | ASSDANTEVF | 0.13497852 |
| MLAPSNP | ASSDANTEVF | 0.85768765 |
| LVNNAAI | ASSDANTEVF | 0.6658911 |
| LTGEINTLA | ASSDANTEVF | 0.74325216 |
| LSIFGPV | ASSDANTEVF | 0.7463608 |
| LLALLANL | ASSDANTEVF | 0.8650793 |
| LISNASDALDKLR | ASSDANTEVF | 0.8360234 |
| LINEAVP | ASSDANTEVF | 0.7459021 |
| LGYAFSGL | ASSDANTEVF | 0.85749125 |
| LDFVAVAL | ASSDANTEVF | 0.82579947 |
| LAQPGAE | ASSDANTEVF | 0.8765589 |
| ISNLGAVTGF | ASSDANTEVF | 0.83651394 |
| HQEDPGRI | ASSDANTEVF | 0.83668786 |
| GTPGEPTP | ASSDANTEVF | 0.85468084 |
| GFFNSGS | ASSDANTEVF | 0.80999494 |
| GAGGGGLTAGAN | ASSDANTEVF | 0.5108777 |
| FVDALVNM | ASSDANTEVF | 0.5035217 |

| | | |
|---|---|---|
| FLTVAPEL | ASSDANTEVF | 0.80100983 |
| FETLTAL | ASSDANTEVF | 0.437406 |
| DVINPIEL | ASSDANTEVF | 0.83281296 |
| APVSGAPV | ASSDANTEVF | 0.8719081 |
| ALVGGLAW | ASSDANTEVF | 0.8444313 |
| ALAAAAATAI | ASSDANTEVF | 0.4504308 |
| AFGLGLLL | ASSDANTEVF | 0.7907816 |
| AFFAAASL | ASSDANTEVF | 0.7188071 |
| AAAALPAFSPP | ASSDANTEVF | 0.8532106 |
| VVLAPAL | ASSDANTEVF | 0.8594864 |
| TIGHVDHGKTTL | ASSDANTEVF | 0.87935483 |
| LISNASDALDK | ASSDANTEVF | 0.866595 |
| LISNASDAL | ASSDANTEVF | 0.75852966 |
| ISNASDALDK | ASSDANTEVF | 0.86658907 |
| NQAETLVYQTE | ASSDANTEVF | 0.78234494 |
| TGGTGGTAGAG | ASSDANTEVF | 0.8311279 |
| TGSTAGGAGGA | ASSDANTEVF | 0.813861 |
| DPLAVPVAAGPV | ASSDANTEVF | 0.8536962 |
| GGTGGTPGNGGN | ASSDANTEVF | 0.8143513 |
| PAGLAGVDPATL | ASSDANTEVF | 0.763193 |
| GNNLVVEAPHAGASV | ASSDANTEVF | 0.49297947 |
| KVTRIMSASTLEQPA | ASSDANTEVF | 0.8597375 |
| LGFALPIALPRRTVV | ASSDANTEVF | 0.794986 |
| TRLGNLTARKLVVLV | ASSDANTEVF | 0.61928415 |
| WSQQRRLIYFPSAGP | ASSDANTEVF | 0.7908999 |
| GPNGNPDPMAAAVDIR | ASSDANTEVF | 0.87958056 |
| TQRCVSVADLRAAASR | ASSDANTEVF | 0.8779628 |
| QFVLDVAEHPVEVLTETV | ASSDANTEVF | 0.7408226 |
| GGAGGTGGAGGNGGTQAGDG | ASSDANTEVF | 0.68713516 |
| PLLATLRAIAVDVDAKPAQV | ASSDANTEVF | 0.84947973 |
| APKFRTGGYMIDFWGVGYQVAK | ASSDANTEVF | 0.590046 |
| GAGGAGGAGGAAGGPGGTGGTG | ASSDANTEVF | 0.85419077 |
| IGALGLLATDVAPAGSVAFGVA | ASSDANTEVF | 0.81803954 |
| AGGAGGIGGNGGAGGAGGVLMG | ASSDANTEVF | 0.8206772 |
| ADELIGLL | ASSDANTEVF | 0.758272 |
| ADRALKA | ASSDANTEVF | 0.8512307 |
| AGAVGQVGGAGGA | ASSDANTEVF | 0.79420686 |
| AGGGGGLGGAP | ASSDANTEVF | 0.8569516 |
| AGGLLYGNGGAG | ASSDANTEVF | 0.18943469 |
| AGLESIGL | ASSDANTEVF | 0.85843635 |
| AGLKTGII | ASSDANTEVF | 0.69177103 |
| ASLAALLL | ASSDANTEVF | 0.74159515 |
| ASVINISSI | ASSDANTEVF | 0.26338944 |
| ATPPGPA | ASSDANTEVF | 0.8199659 |
| ATVASALV | ASSDANTEVF | 0.8604582 |
| ATVASLIL | ASSDANTEVF | 0.8745489 |
| AVNGPVLIRH | ASSDANTEVF | 0.7713618 |
| DGAAGAVGI | ASSDANTEVF | 0.8625157 |
| DPLLHGI | ASSDANTEVF | 0.8834427 |
| ELGAALGAL | ASSDANTEVF | 0.765562 |
| ELGRTQAV | ASSDANTEVF | 0.2412253 |

| | | |
|---|---|---|
| FVAGGYLS | ASSDANTEVF | 0.819628 |
| GASGGAGGAGGW | ASSDANTEVF | 0.8178694 |
| GGAGAGGDGGAGGI | ASSDANTEVF | 0.8686364 |
| GTGHNGSQPGMGGNG | ASSDANTEVF | 0.86857677 |
| GYAAASASAS | ASSDANTEVF | 0.5773517 |
| HQVDVAAVQQG | ASSDANTEVF | 0.848944 |
| IGAAIAHT | ASSDANTEVF | 0.86093724 |
| IGMIEVAQL | ASSDANTEVF | 0.7055548 |
| IRLAPEHTSAMD | ASSDANTEVF | 0.6945847 |
| IYNVSVL | ASSDANTEVF | 0.7212194 |
| KQGFLGM | ASSDANTEVF | 0.33323073 |
| LESVITQL | ASSDANTEVF | 0.8443664 |
| LGLGALVVV | ASSDANTEVF | 0.77630866 |
| LIGNGGNGGSGGI | ASSDANTEVF | 0.6862546 |
| LNAINAPTL | ASSDANTEVF | 0.80312115 |
| LQLLNTGV | ASSDANTEVF | 0.67625386 |
| LQTSIEF | ASSDANTEVF | 0.68297184 |
| MSFAVSEL | ASSDANTEVF | 0.7292712 |
| NTGALGL | ASSDANTEVF | 0.8195438 |
| PPGAGGAGGAGTT | ASSDANTEVF | 0.86124194 |
| QLVGLGI | ASSDANTEVF | 0.81707495 |
| RFLEQVHQL | ASSDANTEVF | 0.84559155 |
| RGVGYVLR | ASSDANTEVF | 0.87439775 |
| SATGVFAGVI | ASSDANTEVF | 0.53782165 |
| SLPAITLP | ASSDANTEVF | 0.7261104 |
| STALAFTV | ASSDANTEVF | 0.86076915 |
| SVLTVGLGK | ASSDANTEVF | 0.6667532 |
| TALAGAGSGI | ASSDANTEVF | 0.8648489 |
| TGLPFTGI | ASSDANTEVF | 0.84041923 |
| VAEHVTV | ASSDANTEVF | 0.85123146 |
| VDIVLNSL | ASSDANTEVF | 0.8300557 |
| VLTGGGAVL | ASSDANTEVF | 0.8836714 |
| VNVGLAYGI | ASSDANTEVF | 0.84159786 |
| VSSAAVGAV | ASSDANTEVF | 0.78800833 |
| VTSVYEAGV | ASSDANTEVF | 0.8310406 |
| WVALDRGA | ASSDANTEVF | 0.61901104 |
| YLVGSVVI | ASSDANTEVF | 0.86952496 |
| GPPGTGKTLL | ASSDANTEVF | 0.8619867 |
| GVGMTKF | ASSDANTEVF | 0.67114073 |
| IGIGGSDL | ASSDANTEVF | 0.80067503 |
| IGTLLAEL | ASSDANTEVF | 0.88106716 |
| IYTSGTT | ASSDANTEVF | 0.87126803 |
| LSFPAGV | ASSDANTEVF | 0.5095537 |
| LYTSGTT | ASSDANTEVF | 0.72790045 |
| VGAFPFGF | ASSDANTEVF | 0.60875076 |
| VGAGTVEF | ASSDANTEVF | 0.8151035 |
| VGKSTLLL | ASSDANTEVF | 0.4095549 |
| AALAHAGGGIV | ASSDANTEVF | 0.85808045 |
| AALVAVSLL | ASSDANTEVF | 0.7595897 |
| AAVSRAVAAL | ASSDANTEVF | 0.86223996 |
| AGGSSGIGGF | ASSDANTEVF | 0.8352642 |

| | | |
|---|---|---|
| AGLTVANANG | ASSDANTEVF | 0.8711506 |
| AGLVGAVL | ASSDANTEVF | 0.7676581 |
| ALSQAFTV | ASSDANTEVF | 0.75204486 |
| ALTFSVSL | ASSDANTEVF | 0.40101695 |
| ALTGAALVAL | ASSDANTEVF | 0.45651913 |
| ASFSIGAGVL | ASSDANTEVF | 0.72337043 |
| DGSVYKI | ASSDANTEVF | 0.7762034 |
| EQLAEIV | ASSDANTEVF | 0.26909012 |
| FAVSELTVM | ASSDANTEVF | 0.8804531 |
| GAAPLAAGL | ASSDANTEVF | 0.7977362 |
| GADGGDGGSGGAGG | ASSDANTEVF | 0.85587215 |
| GFGLDTGGAGG | ASSDANTEVF | 0.6608341 |
| GLASGGLL | ASSDANTEVF | 0.6607137 |
| GRIAGAAVLL | ASSDANTEVF | 0.72705203 |
| GSAGSLNTSF | ASSDANTEVF | 0.85989124 |
| GTGQIGI | ASSDANTEVF | 0.6338805 |
| GTLIALIL | ASSDANTEVF | 0.273332 |
| IAATMGFVVG | ASSDANTEVF | 0.7481638 |
| IFAQALL | ASSDANTEVF | 0.57389563 |
| IGAAVALGF | ASSDANTEVF | 0.8816732 |
| IGLTPLAL | ASSDANTEVF | 0.86867785 |
| ILGSNTSTL | ASSDANTEVF | 0.82830095 |
| IVFSLTVL | ASSDANTEVF | 0.8824791 |
| LASKATGF | ASSDANTEVF | 0.16921112 |
| LASVALIL | ASSDANTEVF | 0.6821935 |
| LFAVGLTL | ASSDANTEVF | 0.8809618 |
| LFNGGIGTYI | ASSDANTEVF | 0.8431473 |
| LTLGAGII | ASSDANTEVF | 0.79769063 |
| LVAAALGDI | ASSDANTEVF | 0.8677128 |
| NTAFVQLGI | ASSDANTEVF | 0.6929215 |
| QLFSGAEL | ASSDANTEVF | 0.78227115 |
| SAATFITL | ASSDANTEVF | 0.78270257 |
| SNYGLANLGV | ASSDANTEVF | 0.85808957 |
| SYGGAYAVM | ASSDANTEVF | 0.8308835 |
| TTGVGGLTLG | ASSDANTEVF | 0.5198324 |
| TTQTIYI | ASSDANTEVF | 0.8380581 |
| VAAGAGLGAVI | ASSDANTEVF | 0.8833952 |
| VAAVVTLL | ASSDANTEVF | 0.74928397 |
| VGFLGGLI | ASSDANTEVF | 0.8651698 |
| VGVIPILL | ASSDANTEVF | 0.8310852 |
| VIFASSLI | ASSDANTEVF | 0.8359999 |
| VKGGDGGAAAG | ASSDANTEVF | 0.70929027 |
| VTKTFTNI | ASSDANTEVF | 0.68394625 |
| VVAAAILL | ASSDANTEVF | 0.8313377 |
| YAALDVEL | ASSDANTEVF | 0.69971085 |
| LASVALLL | ASSDANTEVF | 0.8122395 |
| LDAVHRV | ASSDANTEVF | 0.7732451 |
| QGLSSSIF | ASSDANTEVF | 0.890068 |
| RGTFANIRL | ASSDANTEVF | 0.8562398 |
| RQLWWGHRI | ASSDANTEVF | 0.8415904 |
| YGPPGTGKTLL | ASSDANTEVF | 0.6467686 |

| | | |
|---|---|---|
| AALVGAPANVA | ASSDANTEVF | 0.8511688 |
| AATFRRTE | ASSDANTEVF | 0.85089326 |
| AGALGGGI | ASSDANTEVF | 0.43823105 |
| AGGAGGAGGIGAG | ASSDANTEVF | 0.88031626 |
| AGSGALGL | ASSDANTEVF | 0.62177783 |
| AQIADQI | ASSDANTEVF | 0.8322388 |
| ASAVAVTYL | ASSDANTEVF | 0.8618902 |
| ASTVLGGTGG | ASSDANTEVF | 0.870218 |
| AVGGEPAK | ASSDANTEVF | 0.7973174 |
| CEQGTFGGTC | ASSDANTEVF | 0.8366982 |
| DEGKAPAG | ASSDANTEVF | 0.59024906 |
| DTVVFPAL | ASSDANTEVF | 0.8705234 |
| FEGITMAG | ASSDANTEVF | 0.8669396 |
| FPLVVQD | ASSDANTEVF | 0.8281911 |
| GAFTTPQ | ASSDANTEVF | 0.882528 |
| GFGGFGTAGGVGGA | ASSDANTEVF | 0.84845006 |
| GFLAQFL | ASSDANTEVF | 0.35431543 |
| GGAGGAGGNAGVA | ASSDANTEVF | 0.7880221 |
| GGVGTAGAGGAGGAGG | ASSDANTEVF | 0.8735254 |
| GNGGGGDAGNAGSGGN | ASSDANTEVF | 0.87567437 |
| GTGGAGGAGNDAGS | ASSDANTEVF | 0.8615023 |
| GTTNTGWFNS | ASSDANTEVF | 0.84290624 |
| IFMGPLFTL | ASSDANTEVF | 0.79770577 |
| ILAHAIEAI | ASSDANTEVF | 0.87842685 |
| IRSAIREAL | ASSDANTEVF | 0.8899203 |
| ITFAGGG | ASSDANTEVF | 0.86015195 |
| ITGMSGHQEV | ASSDANTEVF | 0.7878814 |
| IVAGLACTL | ASSDANTEVF | 0.76813495 |
| IVGGGLAGA | ASSDANTEVF | 0.8442262 |
| IVTGVLNI | ASSDANTEVF | 0.86566985 |
| LAQQGCI | ASSDANTEVF | 0.5967678 |
| LCLHVEN | ASSDANTEVF | 0.8331442 |
| LDAISDGS | ASSDANTEVF | 0.8399743 |
| LGGFAAML | ASSDANTEVF | 0.32423866 |
| LGGKNPMI | ASSDANTEVF | 0.85620093 |
| LGPKGVAAIL | ASSDANTEVF | 0.87469995 |
| LMVGVSI | ASSDANTEVF | 0.8302614 |
| LVGAAFVL | ASSDANTEVF | 0.68786657 |
| LVLAVAPL | ASSDANTEVF | 0.8733462 |
| MAAIAVIDAA | ASSDANTEVF | 0.88427484 |
| NNGINAELQ | ASSDANTEVF | 0.87603164 |
| RGEPLVLVHG | ASSDANTEVF | 0.29967928 |
| RGGAGGAAGLI | ASSDANTEVF | 0.24703139 |
| RSYRMHPAV | ASSDANTEVF | 0.80378604 |
| SAAHLLAGQLA | ASSDANTEVF | 0.8534578 |
| SAAVGSY | ASSDANTEVF | 0.7522639 |
| SPHAVLHRI | ASSDANTEVF | 0.67097116 |
| SQIAKVL | ASSDANTEVF | 0.82615817 |
| TALGHLVP | ASSDANTEVF | 0.8706141 |
| TAPTTSGAA | ASSDANTEVF | 0.54251933 |
| TDGDDGNGGSPGT | ASSDANTEVF | 0.6189133 |

| | | |
|---|---|---|
| TDGLGEN | ASSDANTEVF | 0.84863776 |
| TGGPPLPGQA | ASSDANTEVF | 0.8666343 |
| TIPAITI | ASSDANTEVF | 0.6723648 |
| TIPGFTI | ASSDANTEVF | 0.78415406 |
| TSIASGALA | ASSDANTEVF | 0.80486864 |
| TVTLPGAAI | ASSDANTEVF | 0.8424992 |
| VASGLSAALP | ASSDANTEVF | 0.8403224 |
| VGAGVIGI | ASSDANTEVF | 0.62140125 |
| VGPKRFGV | ASSDANTEVF | 0.86935323 |
| VNVAFPD | ASSDANTEVF | 0.5138502 |
| VSMTNSGVS | ASSDANTEVF | 0.8720306 |
| VTDVLGDLSV | ASSDANTEVF | 0.8389058 |
| VVVVGAGFAGL | ASSDANTEVF | 0.83219755 |
| VYTVGTVV | ASSDANTEVF | 0.784517 |
| WTTFSGP | ASSDANTEVF | 0.6928222 |
| YDELVEAF | ASSDANTEVF | 0.6588107 |
| YGTGGAGGNGG | ASSDANTEVF | 0.6090715 |
| YTNVMAV | ASSDANTEVF | 0.87368685 |
| FLEQVHQL | ASSDANTEVF | 0.39637798 |
| GLSLLAVAV | ASSDANTEVF | 0.8683616 |
| LEQVHQL | ASSDANTEVF | 0.8124349 |
| LNAAGAI | ASSDANTEVF | 0.5907054 |
| PAVHASL | ASSDANTEVF | 0.856512 |
| TELEDDA | ASSDANTEVF | 0.5747499 |
| TPPGPPN | ASSDANTEVF | 0.8003419 |
| TPPPPPPP | ASSDANTEVF | 0.86019576 |
| TQSPALI | AWSLNTEVF | 0.8532374 |
| EFAAAFPQ | AWSLNTEVF | 0.670779 |
| LLSGNLDVL | AWSLNTEVF | 0.8501991 |
| GPANLGNYNI | AWSLNTEVF | 0.8429526 |
| TLGPTLP | AWSLNTEVF | 0.7242905 |
| VYAYSSCCK | AWSLNTEVF | 0.8275453 |
| APAPAPAPNI | AWSLNTEVF | 0.8660399 |
| PGGPVPIP | AWSLNTEVF | 0.833933 |
| FLTVALEP | AWSLNTEVF | 0.4817841 |
| SQYFGANL | AWSLNTEVF | 0.8322902 |
| GGVKPAA | AWSLNTEVF | 0.7273021 |
| GGTGGAGGAGGW | AWSLNTEVF | 0.8425671 |
| GVGGVAGLGGAGG | AWSLNTEVF | 0.82148194 |
| VLLVLATTI | AWSLNTEVF | 0.8000679 |
| IGIFAAML | AWSLNTEVF | 0.84900653 |
| AVGDLID | AWSLNTEVF | 0.5560786 |
| NISPLPL | AWSLNTEVF | 0.7291312 |
| VTAVFVAI | AWSLNTEVF | 0.7801873 |
| SGDDNDAAL | AWSLNTEVF | 0.7838669 |
| AVAALAHS | AWSLNTEVF | 0.85233 |
| GGVTVMATP | AWSLNTEVF | 0.61924356 |
| AGLAANVRD | AWSLNTEVF | 0.8329175 |
| TNRPDLIDPALLRPG | AWSLNTEVF | 0.7427094 |
| LANLGVTAL | AWSLNTEVF | 0.619759 |
| TGERIEL | AWSLNTEVF | 0.8228141 |

| | | |
|---|---|---|
| NGEELAA | AWSLNTEVF | 0.7758101 |
| VLSLVWG | AWSLNTEVF | 0.57879376 |
| IGYALWL | AWSLNTEVF | 0.85796434 |
| LSLLLAPL | AWSLNTEVF | 0.556818 |
| TLDEALK | AWSLNTEVF | 0.8337106 |
| TDGAGTAEGG | AWSLNTEVF | 0.87704426 |
| NAPGGAGGNGG | AWSLNTEVF | 0.70648694 |
| VSVFMGMGGNF | AWSLNTEVF | 0.8672534 |
| LASYLAAL | AWSLNTEVF | 0.8432611 |
| LVGGTPAL | AWSLNTEVF | 0.8512671 |
| VRNADVL | AWSLNTEVF | 0.7627439 |
| TCSSPFLVSAM | AWSLNTEVF | 0.70174193 |
| VHITDAE | AWSLNTEVF | 0.87253004 |
| EYAAPSF | AWSLNTEVF | 0.8449205 |
| VVDAFHGGL | AWSLNTEVF | 0.86667484 |
| ASVVDPQ | AWSLNTEVF | 0.85686725 |
| SYNEFGGLS | AWSLNTEVF | 0.85729474 |
| DPDGSVY | AWSLNTEVF | 0.86315846 |
| MVSTIHATE | AWSLNTEVF | 0.74992126 |
| LAAGVADPV | AWSLNTEVF | 0.48729914 |
| TKEQIAA | AWSLNTEVF | 0.75203836 |
| SLASPAAL | AWSLNTEVF | 0.789842 |
| IGLVFYGV | AWSLNTEVF | 0.6876297 |
| FLLDASNL | AWSLNTEVF | 0.50102305 |
| YQVGNFP | AWSLNTEVF | 0.8665651 |
| ESLLSQST | AWSLNTEVF | 0.88553596 |
| LFTVGFAAVG | AWSLNTEVF | 0.6817193 |
| ATPPTPT | AWSLNTEVF | 0.88436323 |
| TDLATLG | AWSLNTEVF | 0.56187624 |
| SAGGTGR | AWSLNTEVF | 0.85235953 |
| GFSNSGS | AWSLNTEVF | 0.8725643 |
| GGNGGSGGTSV | AWSLNTEVF | 0.6428057 |
| GTSVATGGAGN | AWSLNTEVF | 0.8429414 |
| NGGNGGSGGTS | AWSLNTEVF | 0.5829002 |
| AVVKFPH | AWSLNTEVF | 0.73511255 |
| TPPPPPP | AWSLNTEVF | 0.8719437 |
| AAPAAAAAPA | AWSLNTEVF | 0.7245563 |
| IALNEDL | AWSLNTEVF | 0.8515041 |
| IVNVSSM | AWSLNTEVF | 0.7289967 |
| TGIPPAPR | AWSLNTEVF | 0.632205 |
| FDLGGGTF | AWSLNTEVF | 0.8300482 |
| FDLGGGTFDVS | AWSLNTEVF | 0.87649417 |
| NEPTAAA | AWSLNTEVF | 0.817997 |
| NEPTAAALA | AWSLNTEVF | 0.88838345 |
| GSIINISSI | AWSLNTEVF | 0.6744442 |
| VAPGNAGTA | AWSLNTEVF | 0.87946963 |
| TGGGASLE | AWSLNTEVF | 0.87357986 |
| VAVYNHYKRI | AWSLNTEVF | 0.82764417 |
| TDPAPATT | AWSLNTEVF | 0.777128 |
| TNRPDILDPALLRPG | AWSLNTEVF | 0.86182964 |
| LTTTTTS | AWSLNTEVF | 0.76386034 |

| | | |
|---|---|---|
| PTGSGKTL | AWSLNTEVF | 0.8620055 |
| TGSGKTL | AWSLNTEVF | 0.876504 |
| TIGHVDHGK | AWSLNTEVF | 0.39937294 |
| TIGHVDHGKTT | AWSLNTEVF | 0.85198045 |
| GHVDHGKTT | AWSLNTEVF | 0.7701472 |
| AQFEHTLL | AWSLNTEVF | 0.7164396 |
| IVNQDEL | AWSLNTEVF | 0.5728552 |
| TNGSQFF | AWSLNTEVF | 0.22262728 |
| TGKGIVE | AWSLNTEVF | 0.66097975 |
| LISNASDA | AWSLNTEVF | 0.45119223 |
| AELRTLGELTKIAKAHGAQ | AWSLNTEVF | 0.848764 |
| APNAVAAAEPSW | AWSLNTEVF | 0.79020166 |
| APQVAGAATAVL | AWSLNTEVF | 0.5017209 |
| ARRDPAAAAAILPTDA | AWSLNTEVF | 0.8426025 |
| DAAARDVLAEAGLAEYF | AWSLNTEVF | 0.7738564 |
| DDQIVLSAKANLAKVEQ | AWSLNTEVF | 0.34319764 |
| DELLPDTGEDSDEDGH | AWSLNTEVF | 0.6425663 |
| ELPEAPANRATVPQD | AWSLNTEVF | 0.5020328 |
| FGQSKIERVTGIG | AWSLNTEVF | 0.6921015 |
| GAGGTGSTAGGAGGAGGAGGLY | AWSLNTEVF | 0.2961207 |
| GHGGTGAAGGNGGTGGAGGSGI | AWSLNTEVF | 0.7913761 |
| GNGGNGGAGGEGGPGGAGGAGA | AWSLNTEVF | 0.6165348 |
| GSTAGNGGNGGNATNGTIAGQP | AWSLNTEVF | 0.14192122 |
| GTGGDGGNGGNGGTGG | AWSLNTEVF | 0.7721219 |
| GVRRPAGIPADAPV | AWSLNTEVF | 0.5534713 |
| GYRQVTDAHLLTLARR | AWSLNTEVF | 0.52181804 |
| IAKMGQDGHDRG | AWSLNTEVF | 0.82140636 |
| KAEPSIEIQYT | AWSLNTEVF | 0.4849956 |
| KPVRIRMAADEDTVAVT | AWSLNTEVF | 0.31178722 |
| MPPRIAGMRLLVIKPEP | AWSLNTEVF | 0.8109447 |
| MKFYAVALFRSRGVPPTR | AWSLNTEVF | 0.7896106 |
| NAGGFGILGAVAHSP | AWSLNTEVF | 0.25077042 |
| NEGREITLAMAEM | AWSLNTEVF | 0.7415452 |
| NELVEWSHPLGHRWRWE | AWSLNTEVF | 0.83702767 |
| NLGAGGTGGNGGNPG | AWSLNTEVF | 0.67937607 |
| NTEWPKWETVMAADPVD | AWSLNTEVF | 0.82304806 |
| PASAAPAETQP | AWSLNTEVF | 0.83398426 |
| QQSSDGKNGVRAAEITGE | AWSLNTEVF | 0.64833194 |
| RAPDAVTTSWWKE | AWSLNTEVF | 0.8304668 |
| SGFGNVGAHTSGWFNQ | AWSLNTEVF | 0.86682546 |
| SGVAGAVGQAASVGGLKV | AWSLNTEVF | 0.5553671 |
| SNGGAGTGSAGGNGGT | AWSLNTEVF | 0.68565625 |
| TGGAGGEGGAAG | AWSLNTEVF | 0.32155526 |
| TGGDGGAAGAGGAPGHG | AWSLNTEVF | 0.7008345 |
| TMPPLLAAIANAGRHGV | AWSLNTEVF | 0.2647508 |
| TSSTAGNSGGA | AWSLNTEVF | 0.7948427 |
| VPATVPVGILC | AWSLNTEVF | 0.8088895 |
| VQRPDSGGVVLTGRIS | AWSLNTEVF | 0.6512145 |
| WTPNPVPATKGVPAGLVQAT | AWSLNTEVF | 0.41197178 |
| YALKRALAKFSDRFDVVII | AWSLNTEVF | 0.33709145 |
| YLIGSVIV | AWSLNTEVF | 0.72257984 |

| | | |
|---|---|---|
| YGQLVAVID | AWSLNTEVF | 0.3431129 |
| VTVGGMTLPA | AWSLNTEVF | 0.24307765 |
| VLALPAV | AWSLNTEVF | 0.6787063 |
| VIPPTLNYET | AWSLNTEVF | 0.80805624 |
| VGMGGMHPSSG | AWSLNTEVF | 0.78669286 |
| VGLAGGAATA | AWSLNTEVF | 0.51877785 |
| VGGAGGAGGSGP | AWSLNTEVF | 0.86591554 |
| VFAAFMV | AWSLNTEVF | 0.8367955 |
| TLYDKLR | AWSLNTEVF | 0.8391758 |
| TLAVFTVL | AWSLNTEVF | 0.4816986 |
| TGSVQPARV | AWSLNTEVF | 0.38632888 |
| TGGGAGSGGT | AWSLNTEVF | 0.8632376 |
| SHLETSL | AWSLNTEVF | 0.6751636 |
| SFTTGMT | AWSLNTEVF | 0.69117767 |
| QVEVAKRMGVRQPTVS | AWSLNTEVF | 0.8176687 |
| QGSGGPALAM | AWSLNTEVF | 0.50083065 |
| MLAPSNP | AWSLNTEVF | 0.4747342 |
| LVNNAAI | AWSLNTEVF | 0.76861554 |
| LTGEINTLA | AWSLNTEVF | 0.84646165 |
| LSIFGPV | AWSLNTEVF | 0.5326656 |
| LLALLANL | AWSLNTEVF | 0.31159896 |
| LISNASDALDKLR | AWSLNTEVF | 0.54844093 |
| LINEAVP | AWSLNTEVF | 0.61692023 |
| LGYAFSGL | AWSLNTEVF | 0.8078556 |
| LDFVAVAL | AWSLNTEVF | 0.7149888 |
| LAQPGAE | AWSLNTEVF | 0.6116695 |
| ISNLGAVTGF | AWSLNTEVF | 0.75343823 |
| HQEDPGRI | AWSLNTEVF | 0.75210094 |
| GTPGEPTP | AWSLNTEVF | 0.7249526 |
| GFFNSGS | AWSLNTEVF | 0.67771363 |
| GAGGGGLTAGAN | AWSLNTEVF | 0.63659614 |
| FVDALVNM | AWSLNTEVF | 0.8385675 |
| FLTVAPEL | AWSLNTEVF | 0.79205424 |
| FETLTAL | AWSLNTEVF | 0.7657607 |
| DVINPIEL | AWSLNTEVF | 0.8235034 |
| APVSGAPV | AWSLNTEVF | 0.60783947 |
| ALVGGLAW | AWSLNTEVF | 0.74022704 |
| ALAAAAATAI | AWSLNTEVF | 0.29662657 |
| AFGLGLLL | AWSLNTEVF | 0.51285875 |
| AFFAAASL | AWSLNTEVF | 0.7228058 |
| AAAALPAFSPP | AWSLNTEVF | 0.7681555 |
| VVLAPAL | AWSLNTEVF | 0.8245926 |
| TIGHVDHGKTTL | AWSLNTEVF | 0.6638308 |
| LISNASDALDK | AWSLNTEVF | 0.58258486 |
| LISNASDAL | AWSLNTEVF | 0.7838059 |
| ISNASDALDK | AWSLNTEVF | 0.8448839 |
| NQAETLVYQTE | AWSLNTEVF | 0.37203288 |
| TGGTGGTAGAG | AWSLNTEVF | 0.6210376 |
| TGSTAGGAGGA | AWSLNTEVF | 0.88114524 |
| DPLAVPVAAGPV | AWSLNTEVF | 0.806299 |
| GGTGGTPGNGGN | AWSLNTEVF | 0.64093566 |

| | | |
|---|---|---|
| PAGLAGVDPATL | AWSLNTEVF | 0.6069492 |
| GNNLVVEAPHAGASV | AWSLNTEVF | 0.8689721 |
| KVTRIMSASTLEQPA | AWSLNTEVF | 0.67139935 |
| LGFALPIALPRRTVV | AWSLNTEVF | 0.87137985 |
| TRLGNLTARKLVVLV | AWSLNTEVF | 0.19282973 |
| WSQQRRLIYFPSAGP | AWSLNTEVF | 0.602391 |
| GPNGNPDPMAAAVDIR | AWSLNTEVF | 0.46962857 |
| TQRCVSVADLRAAASR | AWSLNTEVF | 0.1513103 |
| QFVLDVAEHPVEVLTETV | AWSLNTEVF | 0.8093112 |
| GGAGGTGGAGGNGGTQAGDG | AWSLNTEVF | 0.5670464 |
| PLLATLRAIAVDVDAKPAQV | AWSLNTEVF | 0.7810373 |
| APKFRTGGYMIDFWGVGYQVAK | AWSLNTEVF | 0.6584835 |
| GAGGAGGAGGAAGGPGGTGGTG | AWSLNTEVF | 0.82243794 |
| IGALGLLATDVAPAGSVAFGVA | AWSLNTEVF | 0.7999371 |
| AGGAGGIGGNGGAGGAGGVLMG | AWSLNTEVF | 0.65478384 |
| ADELIGLL | AWSLNTEVF | 0.43205014 |
| ADRALKA | AWSLNTEVF | 0.76762044 |
| AGAVGQVGGAGGA | AWSLNTEVF | 0.82965124 |
| AGGGGGLGGAP | AWSLNTEVF | 0.7691945 |
| AGGLLYGNGGAG | AWSLNTEVF | 0.17671525 |
| AGLESIGL | AWSLNTEVF | 0.79968786 |
| AGLKTGII | AWSLNTEVF | 0.62040985 |
| ASLAALLL | AWSLNTEVF | 0.2259527 |
| ASVINISSI | AWSLNTEVF | 0.18192133 |
| ATPPGPA | AWSLNTEVF | 0.54763114 |
| ATVASALV | AWSLNTEVF | 0.6843158 |
| ATVASLIL | AWSLNTEVF | 0.8158711 |
| AVNGPVLIRH | AWSLNTEVF | 0.80678165 |
| DGAAGAVGI | AWSLNTEVF | 0.6216737 |
| DPLLHGI | AWSLNTEVF | 0.79258704 |
| ELGAALGAL | AWSLNTEVF | 0.6972375 |
| ELGRTQAV | AWSLNTEVF | 0.09277915 |
| FVAGGYLS | AWSLNTEVF | 0.50437 |
| GASGGAGGAGGW | AWSLNTEVF | 0.30544925 |
| GGAGAGGDGGAGGI | AWSLNTEVF | 0.8356619 |
| GTGHNGSQPGMGGNG | AWSLNTEVF | 0.46407276 |
| GYAAASASAS | AWSLNTEVF | 0.6808672 |
| HQVDVAAVQQG | AWSLNTEVF | 0.67996585 |
| IGAAIAHT | AWSLNTEVF | 0.8170831 |
| IGMIEVAQL | AWSLNTEVF | 0.71684366 |
| IRLAPEHTSAMD | AWSLNTEVF | 0.39586663 |
| IYNVSVL | AWSLNTEVF | 0.20940304 |
| KQGFLGM | AWSLNTEVF | 0.7630192 |
| LESVITQL | AWSLNTEVF | 0.46490115 |
| LGLGALVVV | AWSLNTEVF | 0.2790052 |
| LIGNGGNGGSGGI | AWSLNTEVF | 0.5050659 |
| LNAINAPTL | AWSLNTEVF | 0.51069254 |
| LQLLNTGV | AWSLNTEVF | 0.23346542 |
| LQTSIEF | AWSLNTEVF | 0.7841418 |
| MSFAVSEL | AWSLNTEVF | 0.4792114 |
| NTGALGL | AWSLNTEVF | 0.6857759 |

| | | |
|---|---|---|
| PPGAGGAGGAGTT | AWSLNTEVF | 0.8685714 |
| QLVGLGI | AWSLNTEVF | 0.8078599 |
| RFLEQVHQL | AWSLNTEVF | 0.70894885 |
| RGVGYVLR | AWSLNTEVF | 0.43737593 |
| SATGVFAGVI | AWSLNTEVF | 0.5089686 |
| SLPAITLP | AWSLNTEVF | 0.43919015 |
| STALAFTV | AWSLNTEVF | 0.391517 |
| SVLTVGLGK | AWSLNTEVF | 0.8364578 |
| TALAGAGSGI | AWSLNTEVF | 0.6985718 |
| TGLPFTGI | AWSLNTEVF | 0.4083438 |
| VAEHVTV | AWSLNTEVF | 0.7602515 |
| VDIVLNSL | AWSLNTEVF | 0.8632941 |
| VLTGGGAVL | AWSLNTEVF | 0.87846124 |
| VNVGLAYGI | AWSLNTEVF | 0.512954 |
| VSSAAVGAV | AWSLNTEVF | 0.46198905 |
| VTSVYEAGV | AWSLNTEVF | 0.5260401 |
| WVALDRGA | AWSLNTEVF | 0.73295903 |
| YLVGSVVI | AWSLNTEVF | 0.41377103 |
| GPPGTGKTLL | AWSLNTEVF | 0.30347684 |
| GVGMTKF | AWSLNTEVF | 0.8084097 |
| IGIGGSDL | AWSLNTEVF | 0.8886591 |
| IGTLLAEL | AWSLNTEVF | 0.61576676 |
| IYTSGTT | AWSLNTEVF | 0.36921465 |
| LSFPAGV | AWSLNTEVF | 0.67378134 |
| LYTSGTT | AWSLNTEVF | 0.44605318 |
| VGAFPFGF | AWSLNTEVF | 0.5276196 |
| VGAGTVEF | AWSLNTEVF | 0.4936633 |
| VGKSTLLL | AWSLNTEVF | 0.31511828 |
| AALAHAGGGIV | AWSLNTEVF | 0.5895032 |
| AALVAVSLL | AWSLNTEVF | 0.85263526 |
| AAVSRAVAAL | AWSLNTEVF | 0.73234 |
| AGGSSGIGGF | AWSLNTEVF | 0.76625854 |
| AGLTVANANG | AWSLNTEVF | 0.85993856 |
| AGLVGAVL | AWSLNTEVF | 0.8193282 |
| ALSQAFTV | AWSLNTEVF | 0.8495177 |
| ALTFSVSL | AWSLNTEVF | 0.83881193 |
| ALTGAALVAL | AWSLNTEVF | 0.6663188 |
| ASFSIGAGVL | AWSLNTEVF | 0.7272973 |
| DGSVYKI | AWSLNTEVF | 0.8454251 |
| EQLAEIV | AWSLNTEVF | 0.6533819 |
| FAVSELTVM | AWSLNTEVF | 0.73135537 |
| GAAPLAAGL | AWSLNTEVF | 0.6700371 |
| GADGGDGGSGGAGG | AWSLNTEVF | 0.8284751 |
| GFGLDTGGAGG | AWSLNTEVF | 0.5727396 |
| GLASGGLL | AWSLNTEVF | 0.8697241 |
| GRIAGAAVLL | AWSLNTEVF | 0.17538106 |
| GSAGSLNTSF | AWSLNTEVF | 0.289246 |
| GTGQIGI | AWSLNTEVF | 0.8524293 |
| GTLIALIL | AWSLNTEVF | 0.10763241 |
| IAATMGFVVG | AWSLNTEVF | 0.18064079 |
| IFAQALL | AWSLNTEVF | 0.69538075 |

| | | |
|---|---|---|
| IGAAVALGF | AWSLNTEVF | 0.8521666 |
| IGLTPLAL | AWSLNTEVF | 0.11565353 |
| ILGSNTSTL | AWSLNTEVF | 0.3795342 |
| IVFSLTVL | AWSLNTEVF | 0.24812475 |
| LASKATGF | AWSLNTEVF | 0.7278475 |
| LASVALIL | AWSLNTEVF | 0.6135135 |
| LFAVGLTL | AWSLNTEVF | 0.21887481 |
| LFNGGIGTYI | AWSLNTEVF | 0.777104 |
| LTLGAGII | AWSLNTEVF | 0.4049296 |
| LVAAALGDI | AWSLNTEVF | 0.68988097 |
| NTAFVQLGI | AWSLNTEVF | 0.81062907 |
| QLFSGAEL | AWSLNTEVF | 0.871942 |
| SAATFITL | AWSLNTEVF | 0.38589123 |
| SNYGLANLGV | AWSLNTEVF | 0.74390244 |
| SYGGAYAVM | AWSLNTEVF | 0.6006131 |
| TTGVGGLTLG | AWSLNTEVF | 0.8867509 |
| TTQTIYI | AWSLNTEVF | 0.6196986 |
| VAAGAGLGAVI | AWSLNTEVF | 0.2695048 |
| VAAVVTLL | AWSLNTEVF | 0.808526 |
| VGFLGGLI | AWSLNTEVF | 0.83446103 |
| VGVIPILL | AWSLNTEVF | 0.5290604 |
| VIFASSLI | AWSLNTEVF | 0.84003645 |
| VKGGDGGAAAG | AWSLNTEVF | 0.58296436 |
| VTKTFTNI | AWSLNTEVF | 0.6573045 |
| VVAAAILL | AWSLNTEVF | 0.8427375 |
| YAALDVEL | AWSLNTEVF | 0.8077142 |
| LASVALLL | AWSLNTEVF | 0.32474425 |
| LDAVHRV | AWSLNTEVF | 0.83143723 |
| QGLSSSIF | AWSLNTEVF | 0.7163576 |
| RGTFANIRL | AWSLNTEVF | 0.4030236 |
| RQLWWGHRI | AWSLNTEVF | 0.8696756 |
| YGPPGTGKTLL | AWSLNTEVF | 0.55135155 |
| AALVGAPANVA | AWSLNTEVF | 0.82436204 |
| AATFRRTE | AWSLNTEVF | 0.7750327 |
| AGALGGGI | AWSLNTEVF | 0.8587483 |
| AGGAGGAGGIGAG | AWSLNTEVF | 0.29071563 |
| AGSGALGL | AWSLNTEVF | 0.77497584 |
| AQIADQI | AWSLNTEVF | 0.65525234 |
| ASAVAVTYL | AWSLNTEVF | 0.774631 |
| ASTVLGGTGG | AWSLNTEVF | 0.66171575 |
| AVGGEPAK | AWSLNTEVF | 0.7009139 |
| CEQGTFGGTC | AWSLNTEVF | 0.8120259 |
| DEGKAPAG | AWSLNTEVF | 0.83147216 |
| DTVVFPAL | AWSLNTEVF | 0.69536555 |
| FEGITMAG | AWSLNTEVF | 0.6002924 |
| FPLVVQD | AWSLNTEVF | 0.86465174 |
| GAFTTPQ | AWSLNTEVF | 0.38305026 |
| GFGGFGTAGGVGGA | AWSLNTEVF | 0.66086626 |
| GFLAQFL | AWSLNTEVF | 0.7805093 |
| GGAGGAGGNAGVA | AWSLNTEVF | 0.82585263 |
| GGVGTAGAGGAGGAGG | AWSLNTEVF | 0.6947646 |

| | | |
|---|---|---|
| GNGGGGDAGNAGSGGN | AWSLNTEVF | 0.8587844 |
| GTGGAGGAGNDAGS | AWSLNTEVF | 0.44521284 |
| GTTNTGWFNS | AWSLNTEVF | 0.8184571 |
| IFMGPLFTL | AWSLNTEVF | 0.7373864 |
| ILAHAIEAI | AWSLNTEVF | 0.6119286 |
| IRSAIREAL | AWSLNTEVF | 0.44672233 |
| ITFAGGG | AWSLNTEVF | 0.81096554 |
| ITGMSGHQEV | AWSLNTEVF | 0.8260767 |
| IVAGLACTL | AWSLNTEVF | 0.816357 |
| IVGGGLAGA | AWSLNTEVF | 0.22324243 |
| IVTGVLNI | AWSLNTEVF | 0.79244965 |
| LAQQGCI | AWSLNTEVF | 0.8282472 |
| LCLHVEN | AWSLNTEVF | 0.8834245 |
| LDAISDGS | AWSLNTEVF | 0.6476067 |
| LGGFAAML | AWSLNTEVF | 0.7555568 |
| LGGKNPMI | AWSLNTEVF | 0.5607271 |
| LGPKGVAAIL | AWSLNTEVF | 0.707193 |
| LMVGVSI | AWSLNTEVF | 0.30841845 |
| LVGAAFVL | AWSLNTEVF | 0.51743007 |
| LVLAVAPL | AWSLNTEVF | 0.6125546 |
| MAAIAVIDAA | AWSLNTEVF | 0.5676258 |
| NNGINAELQ | AWSLNTEVF | 0.84408826 |
| RGEPLVLVHG | AWSLNTEVF | 0.81851715 |
| RGGAGGAAGLI | AWSLNTEVF | 0.80570495 |
| RSYRMHPAV | AWSLNTEVF | 0.83458126 |
| SAAHLLAGQLA | AWSLNTEVF | 0.8719607 |
| SAAVGSY | AWSLNTEVF | 0.75358164 |
| SPHAVLHRI | AWSLNTEVF | 0.58159316 |
| SQIAKVL | AWSLNTEVF | 0.63682234 |
| TALGHLVP | AWSLNTEVF | 0.3695257 |
| TAPTTSGAA | AWSLNTEVF | 0.54285073 |
| TDGDDGNGGSPGT | AWSLNTEVF | 0.6063028 |
| TDGLGEN | AWSLNTEVF | 0.65428543 |
| TGGPPLPGQA | AWSLNTEVF | 0.099184215 |
| TIPAITI | AWSLNTEVF | 0.81824213 |
| TIPGFTI | AWSLNTEVF | 0.85697925 |
| TSIASGALA | AWSLNTEVF | 0.6623273 |
| TVTLPGAAI | AWSLNTEVF | 0.74895287 |
| VASGLSAALP | AWSLNTEVF | 0.82181543 |
| VGAGVIGI | AWSLNTEVF | 0.60735327 |
| VGPKRFGV | AWSLNTEVF | 0.7630136 |
| VNVAFPD | AWSLNTEVF | 0.4772925 |
| VSMTNSGVS | AWSLNTEVF | 0.78875995 |
| VTDVLGDLSV | AWSLNTEVF | 0.8474002 |
| VVVVGAGFAGL | AWSLNTEVF | 0.7701621 |
| VYTVGTVV | AWSLNTEVF | 0.53395444 |
| WTTFSGP | AWSLNTEVF | 0.65710175 |
| YDELVEAF | AWSLNTEVF | 0.24149597 |
| YGTGGAGGNGG | AWSLNTEVF | 0.6051499 |
| YTNVMAV | AWSLNTEVF | 0.13715424 |
| FLEQVHQL | AWSLNTEVF | 0.7573005 |

| | | |
|---|---|---|
| GLSLLAVAV | AWSLNTEVF | 0.46254736 |
| LEQVHQL | AWSLNTEVF | 0.7274315 |
| LNAAGAI | AWSLNTEVF | 0.7223598 |
| PAVHASL | AWSLNTEVF | 0.73209834 |
| TELEDDA | AWSLNTEVF | 0.51080495 |
| TPPGPPN | AWSLNTEVF | 0.7812397 |
| TPPPPPPP | AWSLNTEVF | 0.7690356 |
| TQSPALI | ASSRGQGAGNTLY | 0.23880115 |
| EFAAAFPQ | ASSRGQGAGNTLY | 0.85387474 |
| LLSGNLDVL | ASSRGQGAGNTLY | 0.5842434 |
| GPANLGNYNI | ASSRGQGAGNTLY | 0.7579059 |
| TLGPTLP | ASSRGQGAGNTLY | 0.83274364 |
| VYAYSSCCK | ASSRGQGAGNTLY | 0.63760066 |
| APAPAPAPNI | ASSRGQGAGNTLY | 0.33642378 |
| PGGPVPIP | ASSRGQGAGNTLY | 0.6732469 |
| FLTVALEP | ASSRGQGAGNTLY | 0.72925746 |
| SQYFGANL | ASSRGQGAGNTLY | 0.5906738 |
| GGVKPAA | ASSRGQGAGNTLY | 0.14116241 |
| GGTGGAGGAGGW | ASSRGQGAGNTLY | 0.8165208 |
| GVGGVAGLGGAGG | ASSRGQGAGNTLY | 0.39928398 |
| VLLVLATTI | ASSRGQGAGNTLY | 0.62541485 |
| IGIFAAML | ASSRGQGAGNTLY | 0.6391472 |
| AVGDLID | ASSRGQGAGNTLY | 0.8065454 |
| NISPLPL | ASSRGQGAGNTLY | 0.67500937 |
| VTAVFVAI | ASSRGQGAGNTLY | 0.41778046 |
| SGDDNDAAL | ASSRGQGAGNTLY | 0.7747469 |
| AVAALAHS | ASSRGQGAGNTLY | 0.60888517 |
| GGVTVMATP | ASSRGQGAGNTLY | 0.81232405 |
| AGLAANVRD | ASSRGQGAGNTLY | 0.6756147 |
| TNRPDLIDPALLRPG | ASSRGQGAGNTLY | 0.6613963 |
| LANLGVTAL | ASSRGQGAGNTLY | 0.76299775 |
| TGERIEL | ASSRGQGAGNTLY | 0.48118466 |
| NGEELAA | ASSRGQGAGNTLY | 0.4690777 |
| VLSLVWG | ASSRGQGAGNTLY | 0.39958054 |
| IGYALWL | ASSRGQGAGNTLY | 0.63075733 |
| LSLLLAPL | ASSRGQGAGNTLY | 0.3228379 |
| TLDEALK | ASSRGQGAGNTLY | 0.74487215 |
| TDGAGTAEGG | ASSRGQGAGNTLY | 0.80621564 |
| NAPGGAGGNGG | ASSRGQGAGNTLY | 0.5699477 |
| VSVFMGMGGNF | ASSRGQGAGNTLY | 0.309317 |
| LASYLAAL | ASSRGQGAGNTLY | 0.6015624 |
| LVGGTPAL | ASSRGQGAGNTLY | 0.59642816 |
| VRNADVL | ASSRGQGAGNTLY | 0.7923087 |
| TCSSPFLVSAM | ASSRGQGAGNTLY | 0.7722169 |
| VHITDAE | ASSRGQGAGNTLY | 0.8815607 |
| EYAAPSF | ASSRGQGAGNTLY | 0.82976174 |
| VVDAFHGGL | ASSRGQGAGNTLY | 0.6349093 |
| ASVVDPQ | ASSRGQGAGNTLY | 0.8283242 |
| SYNEFGGLS | ASSRGQGAGNTLY | 0.6444238 |
| DPDGSVY | ASSRGQGAGNTLY | 0.7919624 |
| MVSTIHATE | ASSRGQGAGNTLY | 0.7490321 |

| | | |
|---|---|---|
| LAAGVADPV | ASSRGQGAGNTLY | 0.7723828 |
| TKEQIAA | ASSRGQGAGNTLY | 0.41484392 |
| SLASPAAL | ASSRGQGAGNTLY | 0.6355648 |
| IGLVFYGV | ASSRGQGAGNTLY | 0.29785663 |
| FLLDASNL | ASSRGQGAGNTLY | 0.8112423 |
| YQVGNFP | ASSRGQGAGNTLY | 0.4931524 |
| ESLLSQST | ASSRGQGAGNTLY | 0.30179372 |
| LFTVGFAAVG | ASSRGQGAGNTLY | 0.65037125 |
| ATPPTPT | ASSRGQGAGNTLY | 0.8642658 |
| TDLATLG | ASSRGQGAGNTLY | 0.8187618 |
| SAGGTGR | ASSRGQGAGNTLY | 0.57117605 |
| GFSNSGS | ASSRGQGAGNTLY | 0.4706096 |
| GGNGGSGGTSV | ASSRGQGAGNTLY | 0.82382023 |
| GTSVATGGAGN | ASSRGQGAGNTLY | 0.44408423 |
| NGGNGGSGGTS | ASSRGQGAGNTLY | 0.79480207 |
| AVVKFPH | ASSRGQGAGNTLY | 0.7379581 |
| TPPPPPP | ASSRGQGAGNTLY | 0.7251924 |
| AAPAAAAAPA | ASSRGQGAGNTLY | 0.44284308 |
| IALNEDL | ASSRGQGAGNTLY | 0.7904435 |
| IVNVSSM | ASSRGQGAGNTLY | 0.6646569 |
| TGIPPAPR | ASSRGQGAGNTLY | 0.69261634 |
| FDLGGGTF | ASSRGQGAGNTLY | 0.14727671 |
| FDLGGGTFDVS | ASSRGQGAGNTLY | 0.7777184 |
| NEPTAAA | ASSRGQGAGNTLY | 0.51085085 |
| NEPTAAALA | ASSRGQGAGNTLY | 0.4633805 |
| GSIINISSI | ASSRGQGAGNTLY | 0.18122607 |
| VAPGNAGTA | ASSRGQGAGNTLY | 0.726269 |
| TGGGASLE | ASSRGQGAGNTLY | 0.8138747 |
| VAVYNHYKRI | ASSRGQGAGNTLY | 0.8436126 |
| TDPAPATT | ASSRGQGAGNTLY | 0.7449965 |
| TNRPDILDPALLRPG | ASSRGQGAGNTLY | 0.7879169 |
| LTTTTTS | ASSRGQGAGNTLY | 0.8513813 |
| PTGSGKTL | ASSRGQGAGNTLY | 0.66910845 |
| TGSGKTL | ASSRGQGAGNTLY | 0.19765556 |
| TIGHVDHGK | ASSRGQGAGNTLY | 0.67871886 |
| TIGHVDHGKTT | ASSRGQGAGNTLY | 0.80183464 |
| GHVDHGKTT | ASSRGQGAGNTLY | 0.8187783 |
| AQFEHTLL | ASSRGQGAGNTLY | 0.81468606 |
| IVNQDEL | ASSRGQGAGNTLY | 0.4259983 |
| TNGSQFF | ASSRGQGAGNTLY | 0.71814847 |
| TGKGIVE | ASSRGQGAGNTLY | 0.78174394 |
| LISNASDA | ASSRGQGAGNTLY | 0.30850655 |
| AELRTLGELTKIAKAHGAQ | ASSRGQGAGNTLY | 0.57058924 |
| APNAVAAAEPSW | ASSRGQGAGNTLY | 0.61309385 |
| APQVAGAATAVL | ASSRGQGAGNTLY | 0.34814376 |
| ARRDPAAAAAILPTDA | ASSRGQGAGNTLY | 0.7006945 |
| DAAARDVLAEAGLAEYF | ASSRGQGAGNTLY | 0.676324 |
| DDQIVLSAKANLAKVEQ | ASSRGQGAGNTLY | 0.34422562 |
| DELLPDTGEDSDEDGH | ASSRGQGAGNTLY | 0.70849246 |
| ELPEAPANRATVPQD | ASSRGQGAGNTLY | 0.35827136 |
| FGQSKIERVTGIG | ASSRGQGAGNTLY | 0.36886764 |

| | | |
|---|---|---|
| GAGGTGSTAGGAGGAGGAGGLY | ASSRGQGAGNTLY | 0.4575305 |
| GHGGTGAAGGNGGTGGAGGSGI | ASSRGQGAGNTLY | 0.70320773 |
| GNGGNGGAGGEGGPGGAGGAGA | ASSRGQGAGNTLY | 0.81636345 |
| GSTAGNGGNGGNATNGTIAGQP | ASSRGQGAGNTLY | 0.72341615 |
| GTGGDGGNGGNGGTGG | ASSRGQGAGNTLY | 0.7744082 |
| GVRRPAGIPADAPV | ASSRGQGAGNTLY | 0.85885423 |
| GYRQVTDAHLLTLARR | ASSRGQGAGNTLY | 0.20819001 |
| IAKMGQDGHDRG | ASSRGQGAGNTLY | 0.39755642 |
| KAEPSIEIQYT | ASSRGQGAGNTLY | 0.735811 |
| KPVRIRMAADEDTVAVT | ASSRGQGAGNTLY | 0.5644254 |
| MPPRIAGMRLLVIKPEP | ASSRGQGAGNTLY | 0.8205058 |
| MKFYAVALFRSRGVPPTR | ASSRGQGAGNTLY | 0.7606636 |
| NAGGFGILGAVAHSP | ASSRGQGAGNTLY | 0.7443542 |
| NEGREITLAMAEM | ASSRGQGAGNTLY | 0.7566582 |
| NELVEWSHPLGHRWRWE | ASSRGQGAGNTLY | 0.87634915 |
| NLGAGGTGGNGGNPG | ASSRGQGAGNTLY | 0.73542285 |
| NTEWPKWETVMAADPVD | ASSRGQGAGNTLY | 0.7071525 |
| PASAAPAETQP | ASSRGQGAGNTLY | 0.8404428 |
| QQSSDGKNGVRAAEITGE | ASSRGQGAGNTLY | 0.34146982 |
| RAPDAVTTSWWKE | ASSRGQGAGNTLY | 0.80890375 |
| SGFGNVGAHTSGWFNQ | ASSRGQGAGNTLY | 0.79744583 |
| SGVAGAVGQAASVGGLKV | ASSRGQGAGNTLY | 0.46859437 |
| SNGGAGTGSAGGNGGT | ASSRGQGAGNTLY | 0.67714095 |
| TGGAGGEGGAAG | ASSRGQGAGNTLY | 0.8530591 |
| TGGDGGAAGAGGAPGHG | ASSRGQGAGNTLY | 0.85063744 |
| TMPPLLAAIANAGRHGV | ASSRGQGAGNTLY | 0.38372108 |
| TSSTAGNSGGA | ASSRGQGAGNTLY | 0.60496384 |
| VPATVPVGILC | ASSRGQGAGNTLY | 0.33843613 |
| VQRPDSGGVVLTGRIS | ASSRGQGAGNTLY | 0.5527961 |
| WTPNPVPATKGVPAGLVQAT | ASSRGQGAGNTLY | 0.26776886 |
| YALKRALAKFSDRFDVVII | ASSRGQGAGNTLY | 0.766876 |
| YLIGSVIV | ASSRGQGAGNTLY | 0.6047501 |
| YGQLVAVID | ASSRGQGAGNTLY | 0.8486848 |
| VTVGGMTLPA | ASSRGQGAGNTLY | 0.7493427 |
| VLALPAV | ASSRGQGAGNTLY | 0.82328403 |
| VIPPTLNYET | ASSRGQGAGNTLY | 0.5713723 |
| VGMGGMHPSSG | ASSRGQGAGNTLY | 0.63440454 |
| VGLAGGAATA | ASSRGQGAGNTLY | 0.24278693 |
| VGGAGGAGGSGP | ASSRGQGAGNTLY | 0.2896348 |
| VFAAFMV | ASSRGQGAGNTLY | 0.47672033 |
| TLYDKLR | ASSRGQGAGNTLY | 0.5010854 |
| TLAVFTVL | ASSRGQGAGNTLY | 0.17840354 |
| TGSVQPARV | ASSRGQGAGNTLY | 0.8664882 |
| TGGGAGSGGT | ASSRGQGAGNTLY | 0.5642994 |
| SHLETSL | ASSRGQGAGNTLY | 0.7824656 |
| SFTTGMT | ASSRGQGAGNTLY | 0.46141517 |
| QVEVAKRMGVRQPTVS | ASSRGQGAGNTLY | 0.3468104 |
| QGSGGPALAM | ASSRGQGAGNTLY | 0.6015778 |
| MLAPSNP | ASSRGQGAGNTLY | 0.7559141 |
| LVNNAAI | ASSRGQGAGNTLY | 0.41611177 |
| LTGEINTLA | ASSRGQGAGNTLY | 0.24438292 |

| | | |
|---|---|---|
| LSIFGPV | ASSRGQGAGNTLY | 0.35818785 |
| LLALLANL | ASSRGQGAGNTLY | 0.6118683 |
| LISNASDALDKLR | ASSRGQGAGNTLY | 0.87942606 |
| LINEAVP | ASSRGQGAGNTLY | 0.8416311 |
| LGYAFSGL | ASSRGQGAGNTLY | 0.72578895 |
| LDFVAVAL | ASSRGQGAGNTLY | 0.8762047 |
| LAQPGAE | ASSRGQGAGNTLY | 0.12236427 |
| ISNLGAVTGF | ASSRGQGAGNTLY | 0.5565004 |
| HQEDPGRI | ASSRGQGAGNTLY | 0.8377573 |
| GTPGEPTP | ASSRGQGAGNTLY | 0.74876434 |
| GFFNSGS | ASSRGQGAGNTLY | 0.65652514 |
| GAGGGGLTAGAN | ASSRGQGAGNTLY | 0.82808745 |
| FVDALVNM | ASSRGQGAGNTLY | 0.51903903 |
| FLTVAPEL | ASSRGQGAGNTLY | 0.54303396 |
| FETLTAL | ASSRGQGAGNTLY | 0.5986015 |
| DVINPIEL | ASSRGQGAGNTLY | 0.81923133 |
| APVSGAPV | ASSRGQGAGNTLY | 0.7137518 |
| ALVGGLAW | ASSRGQGAGNTLY | 0.33320373 |
| ALAAAAATAI | ASSRGQGAGNTLY | 0.7219691 |
| AFGLGLLL | ASSRGQGAGNTLY | 0.8862298 |
| AFFAAASL | ASSRGQGAGNTLY | 0.58083975 |
| AAAALPAFSPP | ASSRGQGAGNTLY | 0.8276961 |
| VVLAPAL | ASSRGQGAGNTLY | 0.6873816 |
| TIGHVDHGKTTL | ASSRGQGAGNTLY | 0.7369431 |
| LISNASDALDK | ASSRGQGAGNTLY | 0.5621705 |
| LISNASDAL | ASSRGQGAGNTLY | 0.45480922 |
| ISNASDALDK | ASSRGQGAGNTLY | 0.7981614 |
| NQAETLVYQTE | ASSRGQGAGNTLY | 0.3569854 |
| TGGTGGTAGAG | ASSRGQGAGNTLY | 0.65055436 |
| TGSTAGGAGGA | ASSRGQGAGNTLY | 0.6092003 |
| DPLAVPVAAGPV | ASSRGQGAGNTLY | 0.8880483 |
| GGTGGTPGNGGN | ASSRGQGAGNTLY | 0.7996371 |
| PAGLAGVDPATL | ASSRGQGAGNTLY | 0.7438965 |
| GNNLVVEAPHAGASV | ASSRGQGAGNTLY | 0.51069844 |
| KVTRIMSASTLEQPA | ASSRGQGAGNTLY | 0.7471832 |
| LGFALPIALPRRTVV | ASSRGQGAGNTLY | 0.8077122 |
| TRLGNLTARKLVVLV | ASSRGQGAGNTLY | 0.34456563 |
| WSQQRRLIYFPSAGP | ASSRGQGAGNTLY | 0.8381649 |
| GPNGNPDPMAAAVDIR | ASSRGQGAGNTLY | 0.3471825 |
| TQRCVSVADLRAAASR | ASSRGQGAGNTLY | 0.7599809 |
| QFVLDVAEHPVEVLTETV | ASSRGQGAGNTLY | 0.6846335 |
| GGAGGTGGAGGNGGTQAGDG | ASSRGQGAGNTLY | 0.8694346 |
| PLLATLRAIAVDVDAKPAQV | ASSRGQGAGNTLY | 0.6993607 |
| APKFRTGGYMIDFWGVGYQVAK | ASSRGQGAGNTLY | 0.7941085 |
| GAGGAGGAGGAAGGPGGTGGTG | ASSRGQGAGNTLY | 0.44598272 |
| IGALGLLATDVAPAGSVAFGVA | ASSRGQGAGNTLY | 0.7851132 |
| AGGAGGIGGNGGAGGAGGVLMG | ASSRGQGAGNTLY | 0.83684146 |
| ADELIGLL | ASSRGQGAGNTLY | 0.750968 |
| ADRALKA | ASSRGQGAGNTLY | 0.88451374 |
| AGAVGQVGGAGGA | ASSRGQGAGNTLY | 0.6181321 |
| AGGGGGLGGAP | ASSRGQGAGNTLY | 0.20883334 |

| | | |
|---|---|---|
| AGGLLYGNGGAG | ASSRGQGAGNTLY | 0.61717147 |
| AGLESIGL | ASSRGQGAGNTLY | 0.83897036 |
| AGLKTGII | ASSRGQGAGNTLY | 0.86001474 |
| ASLAALLL | ASSRGQGAGNTLY | 0.779109 |
| ASVINISSI | ASSRGQGAGNTLY | 0.7587102 |
| ATPPGPA | ASSRGQGAGNTLY | 0.674773 |
| ATVASALV | ASSRGQGAGNTLY | 0.8576628 |
| ATVASLIL | ASSRGQGAGNTLY | 0.87636393 |
| AVNGPVLIRH | ASSRGQGAGNTLY | 0.7309361 |
| DGAAGAVGI | ASSRGQGAGNTLY | 0.35751998 |
| DPLLHGI | ASSRGQGAGNTLY | 0.34422702 |
| ELGAALGAL | ASSRGQGAGNTLY | 0.79391 |
| ELGRTQAV | ASSRGQGAGNTLY | 0.844303 |
| FVAGGYLS | ASSRGQGAGNTLY | 0.32207724 |
| GASGGAGGAGGW | ASSRGQGAGNTLY | 0.6522905 |
| GGAGAGGDGGAGGI | ASSRGQGAGNTLY | 0.7096878 |
| GTGHNGSQPGMGGNG | ASSRGQGAGNTLY | 0.19503763 |
| GYAAASASAS | ASSRGQGAGNTLY | 0.7999538 |
| HQVDVAAVQQG | ASSRGQGAGNTLY | 0.8623265 |
| IGAAIAHT | ASSRGQGAGNTLY | 0.7594838 |
| IGMIEVAQL | ASSRGQGAGNTLY | 0.30624187 |
| IRLAPEHTSAMD | ASSRGQGAGNTLY | 0.84323156 |
| IYNVSVL | ASSRGQGAGNTLY | 0.866811 |
| KQGFLGM | ASSRGQGAGNTLY | 0.85581315 |
| LESVITQL | ASSRGQGAGNTLY | 0.19373015 |
| LGLGALVVV | ASSRGQGAGNTLY | 0.14522131 |
| LIGNGGNGGSGGI | ASSRGQGAGNTLY | 0.6807929 |
| LNAINAPTL | ASSRGQGAGNTLY | 0.83209914 |
| LQLLNTGV | ASSRGQGAGNTLY | 0.496257 |
| LQTSIEF | ASSRGQGAGNTLY | 0.3730952 |
| MSFAVSEL | ASSRGQGAGNTLY | 0.7720601 |
| NTGALGL | ASSRGQGAGNTLY | 0.8433941 |
| PPGAGGAGGAGTT | ASSRGQGAGNTLY | 0.40225038 |
| QLVGLGI | ASSRGQGAGNTLY | 0.27009258 |
| RFLEQVHQL | ASSRGQGAGNTLY | 0.8438789 |
| RGVGYVLR | ASSRGQGAGNTLY | 0.8096314 |
| SATGVFAGVI | ASSRGQGAGNTLY | 0.4857257 |
| SLPAITLP | ASSRGQGAGNTLY | 0.67108977 |
| STALAFTV | ASSRGQGAGNTLY | 0.7500606 |
| SVLTVGLGK | ASSRGQGAGNTLY | 0.78862894 |
| TALAGAGSGI | ASSRGQGAGNTLY | 0.75488627 |
| TGLPFTGI | ASSRGQGAGNTLY | 0.2868962 |
| VAEHVTV | ASSRGQGAGNTLY | 0.85681504 |
| VDIVLNSL | ASSRGQGAGNTLY | 0.3669359 |
| VLTGGGAVL | ASSRGQGAGNTLY | 0.80632854 |
| VNVGLAYGI | ASSRGQGAGNTLY | 0.71189535 |
| VSSAAVGAV | ASSRGQGAGNTLY | 0.7621587 |
| VTSVYEAGV | ASSRGQGAGNTLY | 0.65775526 |
| WVALDRGA | ASSRGQGAGNTLY | 0.34045947 |
| YLVGSVVI | ASSRGQGAGNTLY | 0.42601186 |
| GPPGTGKTLL | ASSRGQGAGNTLY | 0.2981561 |

| | | |
|---|---|---|
| GVGMTKF | ASSRGQGAGNTLY | 0.84304196 |
| IGIGGSDL | ASSRGQGAGNTLY | 0.38375148 |
| IGTLLAEL | ASSRGQGAGNTLY | 0.80888695 |
| IYTSGTT | ASSRGQGAGNTLY | 0.67172146 |
| LSFPAGV | ASSRGQGAGNTLY | 0.4622247 |
| LYTSGTT | ASSRGQGAGNTLY | 0.7888251 |
| VGAFPFGF | ASSRGQGAGNTLY | 0.23189366 |
| VGAGTVEF | ASSRGQGAGNTLY | 0.7537855 |
| VGKSTLLL | ASSRGQGAGNTLY | 0.72079664 |
| AALAHAGGGIV | ASSRGQGAGNTLY | 0.7705464 |
| AALVAVSLL | ASSRGQGAGNTLY | 0.5005398 |
| AAVSRAVAAL | ASSRGQGAGNTLY | 0.78380966 |
| AGGSSGIGGF | ASSRGQGAGNTLY | 0.7846964 |
| AGLTVANANG | ASSRGQGAGNTLY | 0.36684096 |
| AGLVGAVL | ASSRGQGAGNTLY | 0.7612872 |
| ALSQAFTV | ASSRGQGAGNTLY | 0.8520644 |
| ALTFSVSL | ASSRGQGAGNTLY | 0.7331883 |
| ALTGAALVAL | ASSRGQGAGNTLY | 0.32185346 |
| ASFSIGAGVL | ASSRGQGAGNTLY | 0.70041704 |
| DGSVYKI | ASSRGQGAGNTLY | 0.599541 |
| EQLAEIV | ASSRGQGAGNTLY | 0.78590465 |
| FAVSELTVM | ASSRGQGAGNTLY | 0.76373345 |
| GAAPLAAGL | ASSRGQGAGNTLY | 0.4715887 |
| GADGGDGGSGGAGG | ASSRGQGAGNTLY | 0.7511304 |
| GFGLDTGGAGG | ASSRGQGAGNTLY | 0.27056587 |
| GLASGGLL | ASSRGQGAGNTLY | 0.498473 |
| GRIAGAAVLL | ASSRGQGAGNTLY | 0.6050506 |
| GSAGSLNTSF | ASSRGQGAGNTLY | 0.67589974 |
| GTGQIGI | ASSRGQGAGNTLY | 0.77171624 |
| GTLIALIL | ASSRGQGAGNTLY | 0.43234617 |
| IAATMGFVVG | ASSRGQGAGNTLY | 0.72869503 |
| IFAQALL | ASSRGQGAGNTLY | 0.57430905 |
| IGAAVALGF | ASSRGQGAGNTLY | 0.4190818 |
| IGLTPLAL | ASSRGQGAGNTLY | 0.70187104 |
| ILGSNTSTL | ASSRGQGAGNTLY | 0.5609784 |
| IVFSLTVL | ASSRGQGAGNTLY | 0.23468181 |
| LASKATGF | ASSRGQGAGNTLY | 0.8118131 |
| LASVALIL | ASSRGQGAGNTLY | 0.3296426 |
| LFAVGLTL | ASSRGQGAGNTLY | 0.63987136 |
| LFNGGIGTYI | ASSRGQGAGNTLY | 0.8743879 |
| LTLGAGII | ASSRGQGAGNTLY | 0.23913783 |
| LVAAALGDI | ASSRGQGAGNTLY | 0.78179204 |
| NTAFVQLGI | ASSRGQGAGNTLY | 0.71632016 |
| QLFSGAEL | ASSRGQGAGNTLY | 0.75096893 |
| SAATFITL | ASSRGQGAGNTLY | 0.5309025 |
| SNYGLANLGV | ASSRGQGAGNTLY | 0.5306629 |
| SYGGAYAVM | ASSRGQGAGNTLY | 0.84727454 |
| TTGVGGLTLG | ASSRGQGAGNTLY | 0.684281 |
| TTQTIYI | ASSRGQGAGNTLY | 0.6689478 |
| VAAGAGLGAVI | ASSRGQGAGNTLY | 0.7927748 |
| VAAVVTLL | ASSRGQGAGNTLY | 0.8491017 |

| | | |
|---|---|---|
| VGFLGGLI | ASSRGQGAGNTLY | 0.7924531 |
| VGVIPILL | ASSRGQGAGNTLY | 0.49795434 |
| VIFASSLI | ASSRGQGAGNTLY | 0.21250004 |
| VKGGDGGAAAG | ASSRGQGAGNTLY | 0.6554352 |
| VTKTFTNI | ASSRGQGAGNTLY | 0.6836271 |
| VVAAAILL | ASSRGQGAGNTLY | 0.5128341 |
| YAALDVEL | ASSRGQGAGNTLY | 0.19019699 |
| LASVALLL | ASSRGQGAGNTLY | 0.8290534 |
| LDAVHRV | ASSRGQGAGNTLY | 0.8825582 |
| QGLSSSIF | ASSRGQGAGNTLY | 0.31100994 |
| RGTFANIRL | ASSRGQGAGNTLY | 0.8784012 |
| RQLWWGHRI | ASSRGQGAGNTLY | 0.37117893 |
| YGPPGTGKTLL | ASSRGQGAGNTLY | 0.7052634 |
| AALVGAPANVA | ASSRGQGAGNTLY | 0.83538866 |
| AATFRRTE | ASSRGQGAGNTLY | 0.32173085 |
| AGALGGGI | ASSRGQGAGNTLY | 0.6578827 |
| AGGAGGAGGIGAG | ASSRGQGAGNTLY | 0.56566405 |
| AGSGALGL | ASSRGQGAGNTLY | 0.451977 |
| AQIADQI | ASSRGQGAGNTLY | 0.8433029 |
| ASAVAVTYL | ASSRGQGAGNTLY | 0.40579695 |
| ASTVLGGTGG | ASSRGQGAGNTLY | 0.7755376 |
| AVGGEPAK | ASSRGQGAGNTLY | 0.4935997 |
| CEQGTFGGTC | ASSRGQGAGNTLY | 0.46191996 |
| DEGKAPAG | ASSRGQGAGNTLY | 0.7780832 |
| DTVVFPAL | ASSRGQGAGNTLY | 0.80843264 |
| FEGITMAG | ASSRGQGAGNTLY | 0.67396283 |
| FPLVVQD | ASSRGQGAGNTLY | 0.8719473 |
| GAFTTPQ | ASSRGQGAGNTLY | 0.5360028 |
| GFGGFGTAGGVGGA | ASSRGQGAGNTLY | 0.88161874 |
| GFLAQFL | ASSRGQGAGNTLY | 0.8576821 |
| GGAGGAGGNAGVA | ASSRGQGAGNTLY | 0.7614405 |
| GGVGTAGAGGAGGAGG | ASSRGQGAGNTLY | 0.617433 |
| GNGGGGDAGNAGSGGN | ASSRGQGAGNTLY | 0.81734216 |
| GTGGAGGAGNDAGS | ASSRGQGAGNTLY | 0.51663357 |
| GTTNTGWFNS | ASSRGQGAGNTLY | 0.64903814 |
| IFMGPLFTL | ASSRGQGAGNTLY | 0.82817024 |
| ILAHAIEAI | ASSRGQGAGNTLY | 0.63686335 |
| IRSAIREAL | ASSRGQGAGNTLY | 0.804999 |
| ITFAGGG | ASSRGQGAGNTLY | 0.77513355 |
| ITGMSGHQEV | ASSRGQGAGNTLY | 0.7526446 |
| IVAGLACTL | ASSRGQGAGNTLY | 0.71508735 |
| IVGGGLAGA | ASSRGQGAGNTLY | 0.45259923 |
| IVTGVLNI | ASSRGQGAGNTLY | 0.6908138 |
| LAQQGCI | ASSRGQGAGNTLY | 0.6614491 |
| LCLHVEN | ASSRGQGAGNTLY | 0.7743367 |
| LDAISDGS | ASSRGQGAGNTLY | 0.7968398 |
| LGGFAAML | ASSRGQGAGNTLY | 0.70022947 |
| LGGKNPMI | ASSRGQGAGNTLY | 0.80211437 |
| LGPKGVAAIL | ASSRGQGAGNTLY | 0.78144383 |
| LMVGVSI | ASSRGQGAGNTLY | 0.47847337 |
| LVGAAFVL | ASSRGQGAGNTLY | 0.7484803 |

| | | |
|---|---|---|
| LVLAVAPL | ASSRGQGAGNTLY | 0.6546999 |
| MAAIAVIDAA | ASSRGQGAGNTLY | 0.7478788 |
| NNGINAELQ | ASSRGQGAGNTLY | 0.40010536 |
| RGEPLVLVHG | ASSRGQGAGNTLY | 0.79241997 |
| RGGAGGAAGLI | ASSRGQGAGNTLY | 0.71005106 |
| RSYRMHPAV | ASSRGQGAGNTLY | 0.2977148 |
| SAAHLLAGQLA | ASSRGQGAGNTLY | 0.77657855 |
| SAAVGSY | ASSRGQGAGNTLY | 0.7575325 |
| SPHAVLHRI | ASSRGQGAGNTLY | 0.7318113 |
| SQIAKVL | ASSRGQGAGNTLY | 0.7942849 |
| TALGHLVP | ASSRGQGAGNTLY | 0.6862313 |
| TAPTTSGAA | ASSRGQGAGNTLY | 0.6341522 |
| TDGDDGNGGSPGT | ASSRGQGAGNTLY | 0.78987056 |
| TDGLGEN | ASSRGQGAGNTLY | 0.806461 |
| TGGPPLPGQA | ASSRGQGAGNTLY | 0.5954534 |
| TIPAITI | ASSRGQGAGNTLY | 0.774035 |
| TIPGFTI | ASSRGQGAGNTLY | 0.8025298 |
| TSIASGALA | ASSRGQGAGNTLY | 0.77766335 |
| TVTLPGAAI | ASSRGQGAGNTLY | 0.8043142 |
| VASGLSAALP | ASSRGQGAGNTLY | 0.81074476 |
| VGAGVIGI | ASSRGQGAGNTLY | 0.68425965 |
| VGPKRFGV | ASSRGQGAGNTLY | 0.80173594 |
| VNVAFPD | ASSRGQGAGNTLY | 0.80228 |
| VSMTNSGVS | ASSRGQGAGNTLY | 0.7278993 |
| VTDVLGDLSV | ASSRGQGAGNTLY | 0.7697376 |
| VVVVGAGFAGL | ASSRGQGAGNTLY | 0.5737947 |
| VYTVGTVV | ASSRGQGAGNTLY | 0.7458008 |
| WTTFSGP | ASSRGQGAGNTLY | 0.45834482 |
| YDELVEAF | ASSRGQGAGNTLY | 0.77467644 |
| YGTGGAGGNGG | ASSRGQGAGNTLY | 0.80345833 |
| YTNVMAV | ASSRGQGAGNTLY | 0.74292177 |
| FLEQVHQL | ASSRGQGAGNTLY | 0.6459116 |
| GLSLLAVAV | ASSRGQGAGNTLY | 0.6738537 |
| LEQVHQL | ASSRGQGAGNTLY | 0.76293397 |
| LNAAGAI | ASSRGQGAGNTLY | 0.71223795 |
| PAVHASL | ASSRGQGAGNTLY | 0.5255171 |
| TELEDDA | ASSRGQGAGNTLY | 0.7067516 |
| TPPGPPN | ASSRGQGAGNTLY | 0.7937893 |
| TPPPPPPP | ASSRGQGAGNTLY | 0.79596484 |
| TQSPALI | TCSPGGGTGQLY | 0.6261941 |
| EFAAAFPQ | TCSPGGGTGQLY | 0.8045621 |
| LLSGNLDVL | TCSPGGGTGQLY | 0.7858061 |
| GPANLGNYNI | TCSPGGGTGQLY | 0.80218667 |
| TLGPTLP | TCSPGGGTGQLY | 0.74202657 |
| VYAYSSCCK | TCSPGGGTGQLY | 0.63261676 |
| APAPAPAPNI | TCSPGGGTGQLY | 0.7860706 |
| PGGPVPIP | TCSPGGGTGQLY | 0.746482 |
| FLTVALEP | TCSPGGGTGQLY | 0.7686217 |
| SQYFGANL | TCSPGGGTGQLY | 0.79576707 |
| GGVKPAA | TCSPGGGTGQLY | 0.725561 |
| GGTGGAGGAGGW | TCSPGGGTGQLY | 0.7035419 |

| | | |
|---|---|---|
| GVGGVAGLGGAGG | TCSPGGGTGQLY | 0.71473604 |
| VLLVLATTI | TCSPGGGTGQLY | 0.81034505 |
| IGIFAAML | TCSPGGGTGQLY | 0.65636617 |
| AVGDLID | TCSPGGGTGQLY | 0.63751507 |
| NISPLPL | TCSPGGGTGQLY | 0.72714293 |
| VTAVFVAI | TCSPGGGTGQLY | 0.7628106 |
| SGDDNDAAL | TCSPGGGTGQLY | 0.8012048 |
| AVAALAHS | TCSPGGGTGQLY | 0.7834489 |
| GGVTVMATP | TCSPGGGTGQLY | 0.739915 |
| AGLAANVRD | TCSPGGGTGQLY | 0.78163695 |
| TNRPDLIDPALLRPG | TCSPGGGTGQLY | 0.78978354 |
| LANLGVTAL | TCSPGGGTGQLY | 0.7718641 |
| TGERIEL | TCSPGGGTGQLY | 0.7797513 |
| NGEELAA | TCSPGGGTGQLY | 0.7042179 |
| VLSLVWG | TCSPGGGTGQLY | 0.8073579 |
| IGYALWL | TCSPGGGTGQLY | 0.7863001 |
| LSLLLAPL | TCSPGGGTGQLY | 0.7968284 |
| TLDEALK | TCSPGGGTGQLY | 0.80207384 |
| TDGAGTAEGG | TCSPGGGTGQLY | 0.7815195 |
| NAPGGAGGNGG | TCSPGGGTGQLY | 0.77719563 |
| VSVFMGMGGNF | TCSPGGGTGQLY | 0.5570941 |
| LASYLAAL | TCSPGGGTGQLY | 0.6037792 |
| LVGGTPAL | TCSPGGGTGQLY | 0.7781745 |
| VRNADVL | TCSPGGGTGQLY | 0.7736286 |
| TCSSPFLVSAM | TCSPGGGTGQLY | 0.790925 |
| VHITDAE | TCSPGGGTGQLY | 0.77370346 |
| EYAAPSF | TCSPGGGTGQLY | 0.71320593 |
| VVDAFHGGL | TCSPGGGTGQLY | 0.79699016 |
| ASVVDPQ | TCSPGGGTGQLY | 0.79955065 |
| SYNEFGGLS | TCSPGGGTGQLY | 0.52610874 |
| DPDGSVY | TCSPGGGTGQLY | 0.72949004 |
| MVSTIHATE | TCSPGGGTGQLY | 0.8084735 |
| LAAGVADPV | TCSPGGGTGQLY | 0.76772785 |
| TKEQIAA | TCSPGGGTGQLY | 0.7401372 |
| SLASPAAL | TCSPGGGTGQLY | 0.7082267 |
| IGLVFYGV | TCSPGGGTGQLY | 0.79852414 |
| FLLDASNL | TCSPGGGTGQLY | 0.76097745 |
| YQVGNFP | TCSPGGGTGQLY | 0.81150794 |
| ESLLSQST | TCSPGGGTGQLY | 0.34773552 |
| LFTVGFAAVG | TCSPGGGTGQLY | 0.7309339 |
| ATPPTPT | TCSPGGGTGQLY | 0.70324314 |
| TDLATLG | TCSPGGGTGQLY | 0.41676325 |
| SAGGTGR | TCSPGGGTGQLY | 0.79884773 |
| GFSNSGS | TCSPGGGTGQLY | 0.5954081 |
| GGNGGSGGTSV | TCSPGGGTGQLY | 0.7833344 |
| GTSVATGGAGN | TCSPGGGTGQLY | 0.7379873 |
| NGGNGGSGGTS | TCSPGGGTGQLY | 0.8037733 |
| AVVKFPH | TCSPGGGTGQLY | 0.75837547 |
| TPPPPPP | TCSPGGGTGQLY | 0.7479888 |
| AAPAAAAAPA | TCSPGGGTGQLY | 0.6636716 |
| IALNEDL | TCSPGGGTGQLY | 0.79160976 |

| | | |
|---|---|---|
| IVNVSSM | TCSPGGGTGQLY | 0.80295265 |
| TGIPPAPR | TCSPGGGTGQLY | 0.8017808 |
| FDLGGGTF | TCSPGGGTGQLY | 0.4867472 |
| FDLGGGTFDVS | TCSPGGGTGQLY | 0.7795479 |
| NEPTAAA | TCSPGGGTGQLY | 0.7667182 |
| NEPTAAALA | TCSPGGGTGQLY | 0.37674052 |
| GSIINISSI | TCSPGGGTGQLY | 0.4681672 |
| VAPGNAGTA | TCSPGGGTGQLY | 0.7369762 |
| TGGGASLE | TCSPGGGTGQLY | 0.716524 |
| VAVYNHYKRI | TCSPGGGTGQLY | 0.79316616 |
| TDPAPATT | TCSPGGGTGQLY | 0.7878555 |
| TNRPDILDPALLRPG | TCSPGGGTGQLY | 0.745136 |
| LTTTTTS | TCSPGGGTGQLY | 0.7933477 |
| PTGSGKTL | TCSPGGGTGQLY | 0.77587867 |
| TGSGKTL | TCSPGGGTGQLY | 0.17906418 |
| TIGHVDHGK | TCSPGGGTGQLY | 0.6647047 |
| TIGHVDHGKTT | TCSPGGGTGQLY | 0.47404683 |
| GHVDHGKTT | TCSPGGGTGQLY | 0.80628026 |
| AQFEHTLL | TCSPGGGTGQLY | 0.676708 |
| IVNQDEL | TCSPGGGTGQLY | 0.67566067 |
| TNGSQFF | TCSPGGGTGQLY | 0.7561185 |
| TGKGIVE | TCSPGGGTGQLY | 0.75435406 |
| LISNASDA | TCSPGGGTGQLY | 0.75417143 |
| AELRTLGELTKIAKAHGAQ | TCSPGGGTGQLY | 0.6828495 |
| APNAVAAAEPSW | TCSPGGGTGQLY | 0.35494363 |
| APQVAGAATAVL | TCSPGGGTGQLY | 0.79381895 |
| ARRDPAAAAAILPTDA | TCSPGGGTGQLY | 0.57850766 |
| DAAARDVLAEAGLAEYF | TCSPGGGTGQLY | 0.6360506 |
| DDQIVLSAKANLAKVEQ | TCSPGGGTGQLY | 0.43366915 |
| DELLPDTGEDSDEDGH | TCSPGGGTGQLY | 0.76091933 |
| ELPEAPANRATVPQD | TCSPGGGTGQLY | 0.38687778 |
| FGQSKIERVTGIG | TCSPGGGTGQLY | 0.79771006 |
| GAGGTGSTAGGAGGAGGAGGLY | TCSPGGGTGQLY | 0.7096158 |
| GHGGTGAAGGNGGTGGAGGSGI | TCSPGGGTGQLY | 0.74049926 |
| GNGGNGGAGGEGGPGGAGGAGA | TCSPGGGTGQLY | 0.8080136 |
| GSTAGNGGNGGNATNGTIAGQP | TCSPGGGTGQLY | 0.741578 |
| GTGGDGGNGGNGGTGG | TCSPGGGTGQLY | 0.74661267 |
| GVRRPAGIPADAPV | TCSPGGGTGQLY | 0.69304365 |
| GYRQVTDAHLLTLARR | TCSPGGGTGQLY | 0.64089996 |
| IAKMGQDGHDRG | TCSPGGGTGQLY | 0.6789037 |
| KAEPSIEIQYT | TCSPGGGTGQLY | 0.67557824 |
| KPVRIRMAADEDTVAVT | TCSPGGGTGQLY | 0.8096402 |
| MPPRIAGMRLLVIKPEP | TCSPGGGTGQLY | 0.7511765 |
| MKFYAVALFRSRGVPPTR | TCSPGGGTGQLY | 0.38626346 |
| NAGGFGILGAVAHSP | TCSPGGGTGQLY | 0.75531006 |
| NEGREITLAMAEM | TCSPGGGTGQLY | 0.80948114 |
| NELVEWSHPLGHRWRWE | TCSPGGGTGQLY | 0.8104589 |
| NLGAGGTGGNGGNPG | TCSPGGGTGQLY | 0.6802255 |
| NTEWPKWETVMAADPVD | TCSPGGGTGQLY | 0.68429875 |
| PASAAPAETQP | TCSPGGGTGQLY | 0.70967007 |
| QQSSDGKNGVRAAEITGE | TCSPGGGTGQLY | 0.74892795 |

| | | |
|---|---|---|
| RAPDAVTTSWWKE | TCSPGGGTGQLY | 0.69055426 |
| SGFGNVGAHTSGWFNQ | TCSPGGGTGQLY | 0.5712041 |
| SGVAGAVGQAASVGGLKV | TCSPGGGTGQLY | 0.80166435 |
| SNGGAGTGSAGGNGGT | TCSPGGGTGQLY | 0.80447173 |
| TGGAGGEGGAAG | TCSPGGGTGQLY | 0.7435165 |
| TGGDGGAAGAGGAPGHG | TCSPGGGTGQLY | 0.38886982 |
| TMPPLLAAIANAGRHGV | TCSPGGGTGQLY | 0.7576133 |
| TSSTAGNSGGA | TCSPGGGTGQLY | 0.6796006 |
| VPATVPVGILC | TCSPGGGTGQLY | 0.72759616 |
| VQRPDSGGVVLTGRIS | TCSPGGGTGQLY | 0.726369 |
| WTPNPVPATKGVPAGLVQAT | TCSPGGGTGQLY | 0.48353797 |
| YALKRALAKFSDRFDVVII | TCSPGGGTGQLY | 0.750749 |
| YLIGSVIV | TCSPGGGTGQLY | 0.8123052 |
| YGQLVAVID | TCSPGGGTGQLY | 0.70788944 |
| VTVGGMTLPA | TCSPGGGTGQLY | 0.7626302 |
| VLALPAV | TCSPGGGTGQLY | 0.8046806 |
| VIPPTLNYET | TCSPGGGTGQLY | 0.7915414 |
| VGMGGMHPSSG | TCSPGGGTGQLY | 0.8035041 |
| VGLAGGAATA | TCSPGGGTGQLY | 0.7872591 |
| VGGAGGAGGSGP | TCSPGGGTGQLY | 0.7632866 |
| VFAAFMV | TCSPGGGTGQLY | 0.7769079 |
| TLYDKLR | TCSPGGGTGQLY | 0.774446 |
| TLAVFTVL | TCSPGGGTGQLY | 0.7642374 |
| TGSVQPARV | TCSPGGGTGQLY | 0.7516167 |
| TGGGAGSGGT | TCSPGGGTGQLY | 0.7220614 |
| SHLETSL | TCSPGGGTGQLY | 0.7890303 |
| SFTTGMT | TCSPGGGTGQLY | 0.75921834 |
| QVEVAKRMGVRQPTVS | TCSPGGGTGQLY | 0.797951 |
| QGSGGPALAM | TCSPGGGTGQLY | 0.47586158 |
| MLAPSNP | TCSPGGGTGQLY | 0.6068221 |
| LVNNAAI | TCSPGGGTGQLY | 0.7954012 |
| LTGEINTLA | TCSPGGGTGQLY | 0.23247409 |
| LSIFGPV | TCSPGGGTGQLY | 0.44073194 |
| LLALLANL | TCSPGGGTGQLY | 0.76827466 |
| LISNASDALDKLR | TCSPGGGTGQLY | 0.8030891 |
| LINEAVP | TCSPGGGTGQLY | 0.24990562 |
| LGYAFSGL | TCSPGGGTGQLY | 0.6132902 |
| LDFVAVAL | TCSPGGGTGQLY | 0.5824988 |
| LAQPGAE | TCSPGGGTGQLY | 0.7810738 |
| ISNLGAVTGF | TCSPGGGTGQLY | 0.75174886 |
| HQEDPGRI | TCSPGGGTGQLY | 0.44600254 |
| GTPGEPTP | TCSPGGGTGQLY | 0.7933872 |
| GFFNSGS | TCSPGGGTGQLY | 0.42159885 |
| GAGGGGLTAGAN | TCSPGGGTGQLY | 0.7303812 |
| FVDALVNM | TCSPGGGTGQLY | 0.7989212 |
| FLTVAPEL | TCSPGGGTGQLY | 0.80105186 |
| FETLTAL | TCSPGGGTGQLY | 0.5780185 |
| DVINPIEL | TCSPGGGTGQLY | 0.7928992 |
| APVSGAPV | TCSPGGGTGQLY | 0.723521 |
| ALVGGLAW | TCSPGGGTGQLY | 0.81257945 |
| ALAAAAATAI | TCSPGGGTGQLY | 0.7387837 |

| | | |
|---|---|---|
| AFGLGLLL | TCSPGGGTGQLY | 0.51762044 |
| AFFAAASL | TCSPGGGTGQLY | 0.7934383 |
| AAAALPAFSPP | TCSPGGGTGQLY | 0.7764506 |
| VVLAPAL | TCSPGGGTGQLY | 0.7583643 |
| TIGHVDHGKTTL | TCSPGGGTGQLY | 0.78289235 |
| LISNASDALDK | TCSPGGGTGQLY | 0.7091993 |
| LISNASDAL | TCSPGGGTGQLY | 0.7461239 |
| ISNASDALDK | TCSPGGGTGQLY | 0.7949504 |
| NQAETLVYQTE | TCSPGGGTGQLY | 0.79882216 |
| TGGTGGTAGAG | TCSPGGGTGQLY | 0.6175797 |
| TGSTAGGAGGA | TCSPGGGTGQLY | 0.7993933 |
| DPLAVPVAAGPV | TCSPGGGTGQLY | 0.75075877 |
| GGTGGTPGNGGN | TCSPGGGTGQLY | 0.66288096 |
| PAGLAGVDPATL | TCSPGGGTGQLY | 0.8097488 |
| GNNLVVEAPHAGASV | TCSPGGGTGQLY | 0.7317499 |
| KVTRIMSASTLEQPA | TCSPGGGTGQLY | 0.79663146 |
| LGFALPIALPRRTVV | TCSPGGGTGQLY | 0.7949778 |
| TRLGNLTARKLVVLV | TCSPGGGTGQLY | 0.786481 |
| WSQQRRLIYFPSAGP | TCSPGGGTGQLY | 0.5038253 |
| GPNGNPDPMAAAVDIR | TCSPGGGTGQLY | 0.72710055 |
| TQRCVSVADLRAAASR | TCSPGGGTGQLY | 0.7620126 |
| QFVLDVAEHPVEVLTETV | TCSPGGGTGQLY | 0.7948377 |
| GGAGGTGGAGGNGGTQAGDG | TCSPGGGTGQLY | 0.7295866 |
| PLLATLRAIAVDVDAKPAQV | TCSPGGGTGQLY | 0.70711184 |
| APKFRTGGYMIDFWGVGYQVAK | TCSPGGGTGQLY | 0.79108036 |
| GAGGAGGAGGAAGGPGGTGGTG | TCSPGGGTGQLY | 0.80470693 |
| IGALGLLATDVAPAGSVAFGVA | TCSPGGGTGQLY | 0.77951884 |
| AGGAGGIGGNGGAGGAGGVLMG | TCSPGGGTGQLY | 0.7608632 |
| ADELIGLL | TCSPGGGTGQLY | 0.8051609 |
| ADRALKA | TCSPGGGTGQLY | 0.6418686 |
| AGAVGQVGGAGGA | TCSPGGGTGQLY | 0.75013065 |
| AGGGGGLGGAP | TCSPGGGTGQLY | 0.79742885 |
| AGGLLYGNGGAG | TCSPGGGTGQLY | 0.79894334 |
| AGLESIGL | TCSPGGGTGQLY | 0.75359917 |
| AGLKTGII | TCSPGGGTGQLY | 0.8060713 |
| ASLAALLL | TCSPGGGTGQLY | 0.68076634 |
| ASVINISSI | TCSPGGGTGQLY | 0.7831389 |
| ATPPGPA | TCSPGGGTGQLY | 0.6917926 |
| ATVASALV | TCSPGGGTGQLY | 0.7664371 |
| ATVASLIL | TCSPGGGTGQLY | 0.68523663 |
| AVNGPVLIRH | TCSPGGGTGQLY | 0.78248495 |
| DGAAGAVGI | TCSPGGGTGQLY | 0.7924279 |
| DPLLHGI | TCSPGGGTGQLY | 0.7877213 |
| ELGAALGAL | TCSPGGGTGQLY | 0.31709638 |
| ELGRTQAV | TCSPGGGTGQLY | 0.7913093 |
| FVAGGYLS | TCSPGGGTGQLY | 0.7757567 |
| GASGGAGGAGGW | TCSPGGGTGQLY | 0.8086684 |
| GGAGAGGDGGAGGI | TCSPGGGTGQLY | 0.7707841 |
| GTGHNGSQPGMGGNG | TCSPGGGTGQLY | 0.77577287 |
| GYAAASASAS | TCSPGGGTGQLY | 0.7205858 |
| HQVDVAAVQQG | TCSPGGGTGQLY | 0.76009655 |

| | | |
|---|---|---|
| IGAAIAHT | TCSPGGGTGQLY | 0.63614506 |
| IGMIEVAQL | TCSPGGGTGQLY | 0.6731987 |
| IRLAPEHTSAMD | TCSPGGGTGQLY | 0.72687876 |
| IYNVSVL | TCSPGGGTGQLY | 0.69815135 |
| KQGFLGM | TCSPGGGTGQLY | 0.7897613 |
| LESVITQL | TCSPGGGTGQLY | 0.80302066 |
| LGLGALVVV | TCSPGGGTGQLY | 0.79485273 |
| LIGNGGNGGSGGI | TCSPGGGTGQLY | 0.80076087 |
| LNAINAPTL | TCSPGGGTGQLY | 0.8024399 |
| LQLLNTGV | TCSPGGGTGQLY | 0.8008497 |
| LQTSIEF | TCSPGGGTGQLY | 0.7132089 |
| MSFAVSEL | TCSPGGGTGQLY | 0.7069763 |
| NTGALGL | TCSPGGGTGQLY | 0.5879811 |
| PPGAGGAGGAGTT | TCSPGGGTGQLY | 0.7324071 |
| QLVGLGI | TCSPGGGTGQLY | 0.7199602 |
| RFLEQVHQL | TCSPGGGTGQLY | 0.7752504 |
| RGVGYVLR | TCSPGGGTGQLY | 0.20418265 |
| SATGVFAGVI | TCSPGGGTGQLY | 0.7474413 |
| SLPAITLP | TCSPGGGTGQLY | 0.8087807 |
| STALAFTV | TCSPGGGTGQLY | 0.7302464 |
| SVLTVGLGK | TCSPGGGTGQLY | 0.7716502 |
| TALAGAGSGI | TCSPGGGTGQLY | 0.7668158 |
| TGLPFTGI | TCSPGGGTGQLY | 0.7321019 |
| VAEHVTV | TCSPGGGTGQLY | 0.7885221 |
| VDIVLNSL | TCSPGGGTGQLY | 0.70669913 |
| VLTGGGAVL | TCSPGGGTGQLY | 0.7952684 |
| VNVGLAYGI | TCSPGGGTGQLY | 0.8073897 |
| VSSAAVGAV | TCSPGGGTGQLY | 0.7742973 |
| VTSVYEAGV | TCSPGGGTGQLY | 0.749666 |
| WVALDRGA | TCSPGGGTGQLY | 0.7542157 |
| YLVGSVVI | TCSPGGGTGQLY | 0.40446007 |
| GPPGTGKTLL | TCSPGGGTGQLY | 0.76939225 |
| GVGMTKF | TCSPGGGTGQLY | 0.62748015 |
| IGIGGSDL | TCSPGGGTGQLY | 0.746088 |
| IGTLLAEL | TCSPGGGTGQLY | 0.6751444 |
| IYTSGTT | TCSPGGGTGQLY | 0.77763355 |
| LSFPAGV | TCSPGGGTGQLY | 0.7658862 |
| LYTSGTT | TCSPGGGTGQLY | 0.78720695 |
| VGAFPFGF | TCSPGGGTGQLY | 0.6372483 |
| VGAGTVEF | TCSPGGGTGQLY | 0.78574693 |
| VGKSTLLL | TCSPGGGTGQLY | 0.76839066 |
| AALAHAGGGIV | TCSPGGGTGQLY | 0.5819323 |
| AALVAVSLL | TCSPGGGTGQLY | 0.7829398 |
| AAVSRAVAAL | TCSPGGGTGQLY | 0.7587313 |
| AGGSSGIGGF | TCSPGGGTGQLY | 0.73107874 |
| AGLTVANANG | TCSPGGGTGQLY | 0.7973234 |
| AGLVGAVL | TCSPGGGTGQLY | 0.768973 |
| ALSQAFTV | TCSPGGGTGQLY | 0.38321725 |
| ALTFSVSL | TCSPGGGTGQLY | 0.73111784 |
| ALTGAALVAL | TCSPGGGTGQLY | 0.755435 |
| ASFSIGAGVL | TCSPGGGTGQLY | 0.7581965 |

| | | |
|---|---|---|
| DGSVYKI | TCSPGGGTGQLY | 0.24463081 |
| EQLAEIV | TCSPGGGTGQLY | 0.7881153 |
| FAVSELTVM | TCSPGGGTGQLY | 0.6604704 |
| GAAPLAAGL | TCSPGGGTGQLY | 0.6526065 |
| GADGGDGGSGGAGG | TCSPGGGTGQLY | 0.7178725 |
| GFGLDTGGAGG | TCSPGGGTGQLY | 0.788621 |
| GLASGGLL | TCSPGGGTGQLY | 0.7577081 |
| GRIAGAAVLL | TCSPGGGTGQLY | 0.7028276 |
| GSAGSLNTSF | TCSPGGGTGQLY | 0.7843144 |
| GTGQIGI | TCSPGGGTGQLY | 0.79085946 |
| GTLIALIL | TCSPGGGTGQLY | 0.80520093 |
| IAATMGFVVG | TCSPGGGTGQLY | 0.7630332 |
| IFAQALL | TCSPGGGTGQLY | 0.76570845 |
| IGAAVALGF | TCSPGGGTGQLY | 0.77320814 |
| IGLTPLAL | TCSPGGGTGQLY | 0.69111526 |
| ILGSNTSTL | TCSPGGGTGQLY | 0.6920544 |
| IVFSLTVL | TCSPGGGTGQLY | 0.6287388 |
| LASKATGF | TCSPGGGTGQLY | 0.7463236 |
| LASVALIL | TCSPGGGTGQLY | 0.63731575 |
| LFAVGLTL | TCSPGGGTGQLY | 0.7668756 |
| LFNGGIGTYI | TCSPGGGTGQLY | 0.80254203 |
| LTLGAGII | TCSPGGGTGQLY | 0.73692834 |
| LVAAALGDI | TCSPGGGTGQLY | 0.51836455 |
| NTAFVQLGI | TCSPGGGTGQLY | 0.74174255 |
| QLFSGAEL | TCSPGGGTGQLY | 0.64844716 |
| SAATFITL | TCSPGGGTGQLY | 0.79367495 |
| SNYGLANLGV | TCSPGGGTGQLY | 0.78853333 |
| SYGGAYAVM | TCSPGGGTGQLY | 0.80476815 |
| TTGVGGLTLG | TCSPGGGTGQLY | 0.799955 |
| TTQTIYI | TCSPGGGTGQLY | 0.7226349 |
| VAAGAGLGAVI | TCSPGGGTGQLY | 0.7919061 |
| VAAVVTLL | TCSPGGGTGQLY | 0.721919 |
| VGFLGGLI | TCSPGGGTGQLY | 0.71621346 |
| VGVIPILL | TCSPGGGTGQLY | 0.7632057 |
| VIFASSLI | TCSPGGGTGQLY | 0.773587 |
| VKGGDGGAAAG | TCSPGGGTGQLY | 0.7464868 |
| VTKTFTNI | TCSPGGGTGQLY | 0.73864985 |
| VVAAAILL | TCSPGGGTGQLY | 0.43742126 |
| YAALDVEL | TCSPGGGTGQLY | 0.7899606 |
| LASVALLL | TCSPGGGTGQLY | 0.7718265 |
| LDAVHRV | TCSPGGGTGQLY | 0.67457616 |
| QGLSSSIF | TCSPGGGTGQLY | 0.73276883 |
| RGTFANIRL | TCSPGGGTGQLY | 0.8067359 |
| RQLWWGHRI | TCSPGGGTGQLY | 0.8032057 |
| YGPPGTGKTLL | TCSPGGGTGQLY | 0.7111506 |
| AALVGAPANVA | TCSPGGGTGQLY | 0.7036029 |
| AATFRRTE | TCSPGGGTGQLY | 0.76178306 |
| AGALGGGI | TCSPGGGTGQLY | 0.5980377 |
| AGGAGGAGGIGAG | TCSPGGGTGQLY | 0.7869754 |
| AGSGALGL | TCSPGGGTGQLY | 0.7504704 |
| AQIADQI | TCSPGGGTGQLY | 0.7518981 |

| | | |
|---|---|---|
| ASAVAVTYL | TCSPGGGTGQLY | 0.6881319 |
| ASTVLGGTGG | TCSPGGGTGQLY | 0.7790841 |
| AVGGEPAK | TCSPGGGTGQLY | 0.73106354 |
| CEQGTFGGTC | TCSPGGGTGQLY | 0.7859467 |
| DEGKAPAG | TCSPGGGTGQLY | 0.24039605 |
| DTVVFPAL | TCSPGGGTGQLY | 0.785619 |
| FEGITMAG | TCSPGGGTGQLY | 0.674837 |
| FPLVVQD | TCSPGGGTGQLY | 0.69069064 |
| GAFTTPQ | TCSPGGGTGQLY | 0.42234284 |
| GFGGFGTAGGVGGA | TCSPGGGTGQLY | 0.7064435 |
| GFLAQFL | TCSPGGGTGQLY | 0.78380805 |
| GGAGGAGGNAGVA | TCSPGGGTGQLY | 0.7979449 |
| GGVGTAGAGGAGGAGG | TCSPGGGTGQLY | 0.7543533 |
| GNGGGGDAGNAGSGGN | TCSPGGGTGQLY | 0.78893685 |
| GTGGAGGAGNDAGS | TCSPGGGTGQLY | 0.8082578 |
| GTTNTGWFNS | TCSPGGGTGQLY | 0.73774004 |
| IFMGPLFTL | TCSPGGGTGQLY | 0.3702556 |
| ILAHAIEAI | TCSPGGGTGQLY | 0.75974977 |
| IRSAIREAL | TCSPGGGTGQLY | 0.719002 |
| ITFAGGG | TCSPGGGTGQLY | 0.79287434 |
| ITGMSGHQEV | TCSPGGGTGQLY | 0.78060716 |
| IVAGLACTL | TCSPGGGTGQLY | 0.6191534 |
| IVGGGLAGA | TCSPGGGTGQLY | 0.7810768 |
| IVTGVLNI | TCSPGGGTGQLY | 0.7924183 |
| LAQQGCI | TCSPGGGTGQLY | 0.69413054 |
| LCLHVEN | TCSPGGGTGQLY | 0.61276555 |
| LDAISDGS | TCSPGGGTGQLY | 0.5835603 |
| LGGFAAML | TCSPGGGTGQLY | 0.4424492 |
| LGGKNPMI | TCSPGGGTGQLY | 0.78999954 |
| LGPKGVAAIL | TCSPGGGTGQLY | 0.7175338 |
| LMVGVSI | TCSPGGGTGQLY | 0.6692425 |
| LVGAAFVL | TCSPGGGTGQLY | 0.7602656 |
| LVLAVAPL | TCSPGGGTGQLY | 0.69673795 |
| MAAIAVIDAA | TCSPGGGTGQLY | 0.6832812 |
| NNGINAELQ | TCSPGGGTGQLY | 0.7155645 |
| RGEPLVLVHG | TCSPGGGTGQLY | 0.7586477 |
| RGGAGGAAGLI | TCSPGGGTGQLY | 0.78679264 |
| RSYRMHPAV | TCSPGGGTGQLY | 0.71885926 |
| SAAHLLAGQLA | TCSPGGGTGQLY | 0.68139243 |
| SAAVGSY | TCSPGGGTGQLY | 0.80563974 |
| SPHAVLHRI | TCSPGGGTGQLY | 0.56037486 |
| SQIAKVL | TCSPGGGTGQLY | 0.70806414 |
| TALGHLVP | TCSPGGGTGQLY | 0.78102934 |
| TAPTTSGAA | TCSPGGGTGQLY | 0.6980696 |
| TDGDDGNGGSPGT | TCSPGGGTGQLY | 0.7927627 |
| TDGLGEN | TCSPGGGTGQLY | 0.7720735 |
| TGGPPLPGQA | TCSPGGGTGQLY | 0.7769457 |
| TIPAITI | TCSPGGGTGQLY | 0.7463474 |
| TIPGFTI | TCSPGGGTGQLY | 0.809945 |
| TSIASGALA | TCSPGGGTGQLY | 0.7883438 |
| TVTLPGAAI | TCSPGGGTGQLY | 0.7017009 |

| | | |
|---|---|---|
| VASGLSAALP | TCSPGGGTGQLY | 0.7707639 |
| VGAGVIGI | TCSPGGGTGQLY | 0.50936556 |
| VGPKRFGV | TCSPGGGTGQLY | 0.7973499 |
| VNVAFPD | TCSPGGGTGQLY | 0.7823081 |
| VSMTNSGVS | TCSPGGGTGQLY | 0.66003144 |
| VTDVLGDLSV | TCSPGGGTGQLY | 0.75109816 |
| VVVVGAGFAGL | TCSPGGGTGQLY | 0.81271815 |
| VYTVGTVV | TCSPGGGTGQLY | 0.79980075 |
| WTTFSGP | TCSPGGGTGQLY | 0.67432785 |
| YDELVEAF | TCSPGGGTGQLY | 0.7135081 |
| YGTGGAGGNGG | TCSPGGGTGQLY | 0.6201004 |
| YTNVMAV | TCSPGGGTGQLY | 0.73045695 |
| FLEQVHQL | TCSPGGGTGQLY | 0.50239 |
| GLSLLAVAV | TCSPGGGTGQLY | 0.7905057 |
| LEQVHQL | TCSPGGGTGQLY | 0.72571385 |
| LNAAGAI | TCSPGGGTGQLY | 0.76967776 |
| PAVHASL | TCSPGGGTGQLY | 0.77026093 |
| TELEDDA | TCSPGGGTGQLY | 0.8055221 |
| TPPGPPN | TCSPGGGTGQLY | 0.70965827 |
| TPPPPPPP | TCSPGGGTGQLY | 0.74465317 |
| TQSPALI | ASSQEGTGDSDYT | 0.49269533 |
| EFAAAFPQ | ASSQEGTGDSDYT | 0.69426 |
| LLSGNLDVL | ASSQEGTGDSDYT | 0.726593 |
| GPANLGNYNI | ASSQEGTGDSDYT | 0.7116968 |
| TLGPTLP | ASSQEGTGDSDYT | 0.3473202 |
| VYAYSSCCK | ASSQEGTGDSDYT | 0.8033482 |
| APAPAPAPNI | ASSQEGTGDSDYT | 0.76253796 |
| PGGPVPIP | ASSQEGTGDSDYT | 0.7892289 |
| FLTVALEP | ASSQEGTGDSDYT | 0.68784523 |
| SQYFGANL | ASSQEGTGDSDYT | 0.6799909 |
| GGVKPAA | ASSQEGTGDSDYT | 0.71086675 |
| GGTGGAGGAGGW | ASSQEGTGDSDYT | 0.7894489 |
| GVGGVAGLGGAGG | ASSQEGTGDSDYT | 0.65711933 |
| VLLVLATTI | ASSQEGTGDSDYT | 0.34081453 |
| IGIFAAML | ASSQEGTGDSDYT | 0.6058674 |
| AVGDLID | ASSQEGTGDSDYT | 0.524729 |
| NISPLPL | ASSQEGTGDSDYT | 0.80611277 |
| VTAVFVAI | ASSQEGTGDSDYT | 0.8001373 |
| SGDDNDAAL | ASSQEGTGDSDYT | 0.77354014 |
| AVAALAHS | ASSQEGTGDSDYT | 0.80611 |
| GGVTVMATP | ASSQEGTGDSDYT | 0.25793022 |
| AGLAANVRD | ASSQEGTGDSDYT | 0.6643319 |
| TNRPDLIDPALLRPG | ASSQEGTGDSDYT | 0.80146873 |
| LANLGVTAL | ASSQEGTGDSDYT | 0.7857449 |
| TGERIEL | ASSQEGTGDSDYT | 0.74532413 |
| NGEELAA | ASSQEGTGDSDYT | 0.804909 |
| VLSLVWG | ASSQEGTGDSDYT | 0.69290876 |
| IGYALWL | ASSQEGTGDSDYT | 0.7499829 |
| LSLLLAPL | ASSQEGTGDSDYT | 0.7533939 |
| TLDEALK | ASSQEGTGDSDYT | 0.79794663 |
| TDGAGTAEGG | ASSQEGTGDSDYT | 0.77050257 |

| | | |
|---|---|---|
| NAPGGAGGNGG | ASSQEGTGDSDYT | 0.6216868 |
| VSVFMGMGGNF | ASSQEGTGDSDYT | 0.76635885 |
| LASYLAAL | ASSQEGTGDSDYT | 0.81364834 |
| LVGGTPAL | ASSQEGTGDSDYT | 0.7215407 |
| VRNADVL | ASSQEGTGDSDYT | 0.7904466 |
| TCSSPFLVSAM | ASSQEGTGDSDYT | 0.77062464 |
| VHITDAE | ASSQEGTGDSDYT | 0.7637066 |
| EYAAPSF | ASSQEGTGDSDYT | 0.684908 |
| VVDAFHGGL | ASSQEGTGDSDYT | 0.71087337 |
| ASVVDPQ | ASSQEGTGDSDYT | 0.79100037 |
| SYNEFGGLS | ASSQEGTGDSDYT | 0.66467375 |
| DPDGSVY | ASSQEGTGDSDYT | 0.74365306 |
| MVSTIHATE | ASSQEGTGDSDYT | 0.7063756 |
| LAAGVADPV | ASSQEGTGDSDYT | 0.8176466 |
| TKEQIAA | ASSQEGTGDSDYT | 0.7839447 |
| SLASPAAL | ASSQEGTGDSDYT | 0.7821468 |
| IGLVFYGV | ASSQEGTGDSDYT | 0.6819699 |
| FLLDASNL | ASSQEGTGDSDYT | 0.7892343 |
| YQVGNFP | ASSQEGTGDSDYT | 0.7848748 |
| ESLLSQST | ASSQEGTGDSDYT | 0.45005804 |
| LFTVGFAAVG | ASSQEGTGDSDYT | 0.8070441 |
| ATPPTPT | ASSQEGTGDSDYT | 0.67499936 |
| TDLATLG | ASSQEGTGDSDYT | 0.7481013 |
| SAGGTGR | ASSQEGTGDSDYT | 0.7867552 |
| GFSNSGS | ASSQEGTGDSDYT | 0.79257905 |
| GGNGGSGGTSV | ASSQEGTGDSDYT | 0.7572843 |
| GTSVATGGAGN | ASSQEGTGDSDYT | 0.78426874 |
| NGGNGGSGGTS | ASSQEGTGDSDYT | 0.6416562 |
| AVVKFPH | ASSQEGTGDSDYT | 0.7981508 |
| TPPPPPP | ASSQEGTGDSDYT | 0.79567516 |
| AAPAAAAAPA | ASSQEGTGDSDYT | 0.75028473 |
| IALNEDL | ASSQEGTGDSDYT | 0.8102269 |
| IVNVSSM | ASSQEGTGDSDYT | 0.7532912 |
| TGIPPAPR | ASSQEGTGDSDYT | 0.42038655 |
| FDLGGGTF | ASSQEGTGDSDYT | 0.7577343 |
| FDLGGGTFDVS | ASSQEGTGDSDYT | 0.7975402 |
| NEPTAAA | ASSQEGTGDSDYT | 0.79891664 |
| NEPTAAALA | ASSQEGTGDSDYT | 0.8051521 |
| GSIINISSI | ASSQEGTGDSDYT | 0.7642107 |
| VAPGNAGTA | ASSQEGTGDSDYT | 0.74594915 |
| TGGGASLE | ASSQEGTGDSDYT | 0.8066741 |
| VAVYNHYKRI | ASSQEGTGDSDYT | 0.81324565 |
| TDPAPATT | ASSQEGTGDSDYT | 0.7799988 |
| TNRPDILDPALLRPG | ASSQEGTGDSDYT | 0.711608 |
| LTTTTTS | ASSQEGTGDSDYT | 0.68415415 |
| PTGSGKTL | ASSQEGTGDSDYT | 0.7861142 |
| TGSGKTL | ASSQEGTGDSDYT | 0.79682994 |
| TIGHVDHGK | ASSQEGTGDSDYT | 0.623947 |
| TIGHVDHGKTT | ASSQEGTGDSDYT | 0.7754445 |
| GHVDHGKTT | ASSQEGTGDSDYT | 0.7746395 |
| AQFEHTLL | ASSQEGTGDSDYT | 0.4219228 |

| | | |
|---|---|---|
| IVNQDEL | ASSQEGTGDSDYT | 0.77515125 |
| TNGSQFF | ASSQEGTGDSDYT | 0.8008309 |
| TGKGIVE | ASSQEGTGDSDYT | 0.7742687 |
| LISNASDA | ASSQEGTGDSDYT | 0.6833524 |
| AELRTLGELTKIAKAHGAQ | ASSQEGTGDSDYT | 0.8039589 |
| APNAVAAAEPSW | ASSQEGTGDSDYT | 0.8125176 |
| APQVAGAATAVL | ASSQEGTGDSDYT | 0.8087255 |
| ARRDPAAAAAILPTDA | ASSQEGTGDSDYT | 0.42180985 |
| DAAARDVLAEAGLAEYF | ASSQEGTGDSDYT | 0.2992051 |
| DDQIVLSAKANLAKVEQ | ASSQEGTGDSDYT | 0.76340485 |
| DELLPDTGEDSDEDGH | ASSQEGTGDSDYT | 0.77461946 |
| ELPEAPANRATVPQD | ASSQEGTGDSDYT | 0.7186915 |
| FGQSKIERVTGIG | ASSQEGTGDSDYT | 0.6956812 |
| GAGGTGSTAGGAGGAGGAGGLY | ASSQEGTGDSDYT | 0.7640883 |
| GHGGTGAAGGNGGTGGAGGSGI | ASSQEGTGDSDYT | 0.7988064 |
| GNGGNGGAGGEGGPGGAGGAGA | ASSQEGTGDSDYT | 0.6227084 |
| GSTAGNGGNGGNATNGTIAGQP | ASSQEGTGDSDYT | 0.58872914 |
| GTGGDGGNGGNGGTGG | ASSQEGTGDSDYT | 0.75790846 |
| GVRRPAGIPADAPV | ASSQEGTGDSDYT | 0.78703964 |
| GYRQVTDAHLLTLARR | ASSQEGTGDSDYT | 0.6890774 |
| IAKMGQDGHDRG | ASSQEGTGDSDYT | 0.7413759 |
| KAEPSIEIQYT | ASSQEGTGDSDYT | 0.76327085 |
| KPVRIRMAADEDTVAVT | ASSQEGTGDSDYT | 0.7601518 |
| MPPRIAGMRLLVIKPEP | ASSQEGTGDSDYT | 0.772314 |
| MKFYAVALFRSRGVPPTR | ASSQEGTGDSDYT | 0.6509644 |
| NAGGFGILGAVAHSP | ASSQEGTGDSDYT | 0.80121267 |
| NEGREITLAMAEM | ASSQEGTGDSDYT | 0.61861324 |
| NELVEWSHPLGHRWRWE | ASSQEGTGDSDYT | 0.7995322 |
| NLGAGGTGGNGGNPG | ASSQEGTGDSDYT | 0.77345157 |
| NTEWPKWETVMAADPVD | ASSQEGTGDSDYT | 0.7811116 |
| PASAAPAETQP | ASSQEGTGDSDYT | 0.73700804 |
| QQSSDGKNGVRAAEITGE | ASSQEGTGDSDYT | 0.6736847 |
| RAPDAVTTSWWKE | ASSQEGTGDSDYT | 0.5947273 |
| SGFGNVGAHTSGWFNQ | ASSQEGTGDSDYT | 0.629878 |
| SGVAGAVGQAASVGGLKV | ASSQEGTGDSDYT | 0.80208737 |
| SNGGAGTGSAGGNGGT | ASSQEGTGDSDYT | 0.7303071 |
| TGGAGGEGGAAG | ASSQEGTGDSDYT | 0.80111337 |
| TGGDGGAAGAGGAPGHG | ASSQEGTGDSDYT | 0.78905416 |
| TMPPLLAAIANAGRHGV | ASSQEGTGDSDYT | 0.67191297 |
| TSSTAGNSGGA | ASSQEGTGDSDYT | 0.78975666 |
| VPATVPVGILC | ASSQEGTGDSDYT | 0.633863 |
| VQRPDSGGVVLTGRIS | ASSQEGTGDSDYT | 0.7703441 |
| WTPNPVPATKGVPAGLVQAT | ASSQEGTGDSDYT | 0.7838662 |
| YALKRALAKFSDRFDVVII | ASSQEGTGDSDYT | 0.7841095 |
| YLIGSVIV | ASSQEGTGDSDYT | 0.6673846 |
| YGQLVAVID | ASSQEGTGDSDYT | 0.7837926 |
| VTVGGMTLPA | ASSQEGTGDSDYT | 0.7820896 |
| VLALPAV | ASSQEGTGDSDYT | 0.651862 |
| VIPPTLNYET | ASSQEGTGDSDYT | 0.76693594 |
| VGMGGMHPSSG | ASSQEGTGDSDYT | 0.79863286 |
| VGLAGGAATA | ASSQEGTGDSDYT | 0.7861765 |

| | | |
|---|---|---|
| VGGAGGAGGSGP | ASSQEGTGDSDYT | 0.6108668 |
| VFAAFMV | ASSQEGTGDSDYT | 0.7821016 |
| TLYDKLR | ASSQEGTGDSDYT | 0.6850047 |
| TLAVFTVL | ASSQEGTGDSDYT | 0.77311695 |
| TGSVQPARV | ASSQEGTGDSDYT | 0.7674968 |
| TGGGAGSGGT | ASSQEGTGDSDYT | 0.734094 |
| SHLETSL | ASSQEGTGDSDYT | 0.7912923 |
| SFTTGMT | ASSQEGTGDSDYT | 0.44127768 |
| QVEVAKRMGVRQPTVS | ASSQEGTGDSDYT | 0.6930878 |
| QGSGGPALAM | ASSQEGTGDSDYT | 0.74491155 |
| MLAPSNP | ASSQEGTGDSDYT | 0.7687647 |
| LVNNAAI | ASSQEGTGDSDYT | 0.7856332 |
| LTGEINTLA | ASSQEGTGDSDYT | 0.6714673 |
| LSIFGPV | ASSQEGTGDSDYT | 0.7706969 |
| LLALLANL | ASSQEGTGDSDYT | 0.72965574 |
| LISNASDALDKLR | ASSQEGTGDSDYT | 0.6606248 |
| LINEAVP | ASSQEGTGDSDYT | 0.76573825 |
| LGYAFSGL | ASSQEGTGDSDYT | 0.728143 |
| LDFVAVAL | ASSQEGTGDSDYT | 0.6296715 |
| LAQPGAE | ASSQEGTGDSDYT | 0.79377687 |
| ISNLGAVTGF | ASSQEGTGDSDYT | 0.635731 |
| HQEDPGRI | ASSQEGTGDSDYT | 0.78775275 |
| GTPGEPTP | ASSQEGTGDSDYT | 0.8068819 |
| GFFNSGS | ASSQEGTGDSDYT | 0.63609385 |
| GAGGGGLTAGAN | ASSQEGTGDSDYT | 0.79864395 |
| FVDALVNM | ASSQEGTGDSDYT | 0.77305436 |
| FLTVAPEL | ASSQEGTGDSDYT | 0.78511864 |
| FETLTAL | ASSQEGTGDSDYT | 0.72908497 |
| DVINPIEL | ASSQEGTGDSDYT | 0.6695074 |
| APVSGAPV | ASSQEGTGDSDYT | 0.79347146 |
| ALVGGLAW | ASSQEGTGDSDYT | 0.78821397 |
| ALAAAAATAI | ASSQEGTGDSDYT | 0.7559222 |
| AFGLGLLL | ASSQEGTGDSDYT | 0.7788299 |
| AFFAAASL | ASSQEGTGDSDYT | 0.7742243 |
| AAAALPAFSPP | ASSQEGTGDSDYT | 0.78809977 |
| VVLAPAL | ASSQEGTGDSDYT | 0.71830606 |
| TIGHVDHGKTTL | ASSQEGTGDSDYT | 0.5730705 |
| LISNASDALDK | ASSQEGTGDSDYT | 0.7359127 |
| LISNASDAL | ASSQEGTGDSDYT | 0.75267494 |
| ISNASDALDK | ASSQEGTGDSDYT | 0.6417303 |
| NQAETLVYQTE | ASSQEGTGDSDYT | 0.5074136 |
| TGGTGGTAGAG | ASSQEGTGDSDYT | 0.79275537 |
| TGSTAGGAGGA | ASSQEGTGDSDYT | 0.8140168 |
| DPLAVPVAAGPV | ASSQEGTGDSDYT | 0.6608064 |
| GGTGGTPGNGGN | ASSQEGTGDSDYT | 0.8091396 |
| PAGLAGVDPATL | ASSQEGTGDSDYT | 0.5769124 |
| GNNLVVEAPHAGASV | ASSQEGTGDSDYT | 0.7952297 |
| KVTRIMSASTLEQPA | ASSQEGTGDSDYT | 0.798476 |
| LGFALPIALPRRTVV | ASSQEGTGDSDYT | 0.65207195 |
| TRLGNLTARKLVVLV | ASSQEGTGDSDYT | 0.7787357 |
| WSQQRRLIYFPSAGP | ASSQEGTGDSDYT | 0.6712239 |

| | | |
|---|---|---|
| GPNGNPDPMAAAVDIR | ASSQEGTGDSDYT | 0.7008927 |
| TQRCVSVADLRAAASR | ASSQEGTGDSDYT | 0.80402 |
| QFVLDVAEHPVEVLTETV | ASSQEGTGDSDYT | 0.6879647 |
| GGAGGTGGAGGNGGTQAGDG | ASSQEGTGDSDYT | 0.7832626 |
| PLLATLRAIAVDVDAKPAQV | ASSQEGTGDSDYT | 0.73292536 |
| APKFRTGGYMIDFWGVGYQVAK | ASSQEGTGDSDYT | 0.6633963 |
| GAGGAGGAGGAAGGPGGTGGTG | ASSQEGTGDSDYT | 0.7812268 |
| IGALGLLATDVAPAGSVAFGVA | ASSQEGTGDSDYT | 0.7953459 |
| AGGAGGIGGNGGAGGAGGVLMG | ASSQEGTGDSDYT | 0.76900053 |
| ADELIGLL | ASSQEGTGDSDYT | 0.8115965 |
| ADRALKA | ASSQEGTGDSDYT | 0.7020278 |
| AGAVGQVGGAGGA | ASSQEGTGDSDYT | 0.8031658 |
| AGGGGGLGGAP | ASSQEGTGDSDYT | 0.78904045 |
| AGGLLYGNGGAG | ASSQEGTGDSDYT | 0.75776654 |
| AGLESIGL | ASSQEGTGDSDYT | 0.73841995 |
| AGLKTGII | ASSQEGTGDSDYT | 0.7936976 |
| ASLAALLL | ASSQEGTGDSDYT | 0.7099891 |
| ASVINISSI | ASSQEGTGDSDYT | 0.7829608 |
| ATPPGPA | ASSQEGTGDSDYT | 0.8085908 |
| ATVASALV | ASSQEGTGDSDYT | 0.57398057 |
| ATVASLIL | ASSQEGTGDSDYT | 0.84994483 |
| AVNGPVLIRH | ASSQEGTGDSDYT | 0.8120809 |
| DGAAGAVGI | ASSQEGTGDSDYT | 0.7368092 |
| DPLLHGI | ASSQEGTGDSDYT | 0.6640893 |
| ELGAALGAL | ASSQEGTGDSDYT | 0.2975135 |
| ELGRTQAV | ASSQEGTGDSDYT | 0.72416246 |
| FVAGGYLS | ASSQEGTGDSDYT | 0.57906437 |
| GASGGAGGAGGW | ASSQEGTGDSDYT | 0.82477987 |
| GGAGAGGDGGAGGI | ASSQEGTGDSDYT | 0.82339144 |
| GTGHNGSQPGMGGNG | ASSQEGTGDSDYT | 0.68530416 |
| GYAAASASAS | ASSQEGTGDSDYT | 0.8527729 |
| HQVDVAAVQQG | ASSQEGTGDSDYT | 0.8620894 |
| IGAAIAHT | ASSQEGTGDSDYT | 0.3899308 |
| IGMIEVAQL | ASSQEGTGDSDYT | 0.7151923 |
| IRLAPEHTSAMD | ASSQEGTGDSDYT | 0.60170144 |
| IYNVSVL | ASSQEGTGDSDYT | 0.74151826 |
| KQGFLGM | ASSQEGTGDSDYT | 0.2466156 |
| LESVITQL | ASSQEGTGDSDYT | 0.83772945 |
| LGLGALVVV | ASSQEGTGDSDYT | 0.6586331 |
| LIGNGGNGGSGGI | ASSQEGTGDSDYT | 0.12345658 |
| LNAINAPTL | ASSQEGTGDSDYT | 0.74255633 |
| LQLLNTGV | ASSQEGTGDSDYT | 0.7115215 |
| LQTSIEF | ASSQEGTGDSDYT | 0.7228562 |
| MSFAVSEL | ASSQEGTGDSDYT | 0.84777427 |
| NTGALGL | ASSQEGTGDSDYT | 0.48917162 |
| PPGAGGAGGAGTT | ASSQEGTGDSDYT | 0.57067484 |
| QLVGLGI | ASSQEGTGDSDYT | 0.797081 |
| RFLEQVHQL | ASSQEGTGDSDYT | 0.86803377 |
| RGVGYVLR | ASSQEGTGDSDYT | 0.47835404 |
| SATGVFAGVI | ASSQEGTGDSDYT | 0.7951123 |
| SLPAITLP | ASSQEGTGDSDYT | 0.8670221 |

| | | |
|---|---|---|
| STALAFTV | ASSQEGTGDSDYT | 0.8024766 |
| SVLTVGLGK | ASSQEGTGDSDYT | 0.8687235 |
| TALAGAGSGI | ASSQEGTGDSDYT | 0.8828923 |
| TGLPFTGI | ASSQEGTGDSDYT | 0.54413736 |
| VAEHVTV | ASSQEGTGDSDYT | 0.84551275 |
| VDIVLNSL | ASSQEGTGDSDYT | 0.8678135 |
| VLTGGGAVL | ASSQEGTGDSDYT | 0.71869516 |
| VNVGLAYGI | ASSQEGTGDSDYT | 0.7475215 |
| VSSAAVGAV | ASSQEGTGDSDYT | 0.6043922 |
| VTSVYEAGV | ASSQEGTGDSDYT | 0.67293847 |
| WVALDRGA | ASSQEGTGDSDYT | 0.2646371 |
| YLVGSVVI | ASSQEGTGDSDYT | 0.7734523 |
| GPPGTGKTLL | ASSQEGTGDSDYT | 0.8290832 |
| GVGMTKF | ASSQEGTGDSDYT | 0.6846847 |
| IGIGGSDL | ASSQEGTGDSDYT | 0.5115705 |
| IGTLLAEL | ASSQEGTGDSDYT | 0.5259955 |
| IYTSGTT | ASSQEGTGDSDYT | 0.6809207 |
| LSFPAGV | ASSQEGTGDSDYT | 0.72534907 |
| LYTSGTT | ASSQEGTGDSDYT | 0.27309138 |
| VGAFPFGF | ASSQEGTGDSDYT | 0.6682982 |
| VGAGTVEF | ASSQEGTGDSDYT | 0.8412442 |
| VGKSTLLL | ASSQEGTGDSDYT | 0.85789406 |
| AALAHAGGGIV | ASSQEGTGDSDYT | 0.27508608 |
| AALVAVSLL | ASSQEGTGDSDYT | 0.858356 |
| AAVSRAVAAL | ASSQEGTGDSDYT | 0.81892157 |
| AGGSSGIGGF | ASSQEGTGDSDYT | 0.8690188 |
| AGLTVANANG | ASSQEGTGDSDYT | 0.7496528 |
| AGLVGAVL | ASSQEGTGDSDYT | 0.58563715 |
| ALSQAFTV | ASSQEGTGDSDYT | 0.8172417 |
| ALTFSVSL | ASSQEGTGDSDYT | 0.6902716 |
| ALTGAALVAL | ASSQEGTGDSDYT | 0.5312582 |
| ASFSIGAGVL | ASSQEGTGDSDYT | 0.842862 |
| DGSVYKI | ASSQEGTGDSDYT | 0.67737997 |
| EQLAEIV | ASSQEGTGDSDYT | 0.6155169 |
| FAVSELTVM | ASSQEGTGDSDYT | 0.62258387 |
| GAAPLAAGL | ASSQEGTGDSDYT | 0.8654122 |
| GADGGDGGSGGAGG | ASSQEGTGDSDYT | 0.42679223 |
| GFGLDTGGAGG | ASSQEGTGDSDYT | 0.63494027 |
| GLASGGLL | ASSQEGTGDSDYT | 0.44474426 |
| GRIAGAAVLL | ASSQEGTGDSDYT | 0.7588763 |
| GSAGSLNTSF | ASSQEGTGDSDYT | 0.84891045 |
| GTGQIGI | ASSQEGTGDSDYT | 0.8026006 |
| GTLIALIL | ASSQEGTGDSDYT | 0.65305483 |
| IAATMGFVVG | ASSQEGTGDSDYT | 0.8323157 |
| IFAQALL | ASSQEGTGDSDYT | 0.81802976 |
| IGAAVALGF | ASSQEGTGDSDYT | 0.7470462 |
| IGLTPLAL | ASSQEGTGDSDYT | 0.83784807 |
| ILGSNTSTL | ASSQEGTGDSDYT | 0.61622065 |
| IVFSLTVL | ASSQEGTGDSDYT | 0.85741144 |
| LASKATGF | ASSQEGTGDSDYT | 0.7862679 |
| LASVALIL | ASSQEGTGDSDYT | 0.86381423 |

| | | |
|---|---|---|
| LFAVGLTL | ASSQEGTGDSDYT | 0.8478632 |
| LFNGGIGTYI | ASSQEGTGDSDYT | 0.7948212 |
| LTLGAGII | ASSQEGTGDSDYT | 0.7391304 |
| LVAAALGDI | ASSQEGTGDSDYT | 0.24205123 |
| NTAFVQLGI | ASSQEGTGDSDYT | 0.6371677 |
| QLFSGAEL | ASSQEGTGDSDYT | 0.8100233 |
| SAATFITL | ASSQEGTGDSDYT | 0.7554689 |
| SNYGLANLGV | ASSQEGTGDSDYT | 0.82327616 |
| SYGGAYAVM | ASSQEGTGDSDYT | 0.83571184 |
| TTGVGGLTLG | ASSQEGTGDSDYT | 0.5970167 |
| TTQTIYI | ASSQEGTGDSDYT | 0.85966456 |
| VAAGAGLGAVI | ASSQEGTGDSDYT | 0.83816886 |
| VAAVVTLL | ASSQEGTGDSDYT | 0.35376275 |
| VGFLGGLI | ASSQEGTGDSDYT | 0.6799873 |
| VGVIPILL | ASSQEGTGDSDYT | 0.88031465 |
| VIFASSLI | ASSQEGTGDSDYT | 0.7761649 |
| VKGGDGGAAAG | ASSQEGTGDSDYT | 0.7249086 |
| VTKTFTNI | ASSQEGTGDSDYT | 0.6967378 |
| VVAAAILL | ASSQEGTGDSDYT | 0.86689436 |
| YAALDVEL | ASSQEGTGDSDYT | 0.6327115 |
| LASVALLL | ASSQEGTGDSDYT | 0.8700279 |
| LDAVHRV | ASSQEGTGDSDYT | 0.16581725 |
| QGLSSSIF | ASSQEGTGDSDYT | 0.7476193 |
| RGTFANIRL | ASSQEGTGDSDYT | 0.57058764 |
| RQLWWGHRI | ASSQEGTGDSDYT | 0.38241807 |
| YGPPGTGKTLL | ASSQEGTGDSDYT | 0.84405136 |
| AALVGAPANVA | ASSQEGTGDSDYT | 0.6442706 |
| AATFRRTE | ASSQEGTGDSDYT | 0.84169126 |
| AGALGGGI | ASSQEGTGDSDYT | 0.7046424 |
| AGGAGGAGGIGAG | ASSQEGTGDSDYT | 0.8281156 |
| AGSGALGL | ASSQEGTGDSDYT | 0.7429124 |
| AQIADQI | ASSQEGTGDSDYT | 0.7654069 |
| ASAVAVTYL | ASSQEGTGDSDYT | 0.56266737 |
| ASTVLGGTGG | ASSQEGTGDSDYT | 0.82947004 |
| AVGGEPAK | ASSQEGTGDSDYT | 0.8601615 |
| CEQGTFGGTC | ASSQEGTGDSDYT | 0.8312193 |
| DEGKAPAG | ASSQEGTGDSDYT | 0.35924178 |
| DTVVFPAL | ASSQEGTGDSDYT | 0.65852284 |
| FEGITMAG | ASSQEGTGDSDYT | 0.77396464 |
| FPLVVQD | ASSQEGTGDSDYT | 0.20253617 |
| GAFTTPQ | ASSQEGTGDSDYT | 0.20574398 |
| GFGGFGTAGGVGGA | ASSQEGTGDSDYT | 0.63602793 |
| GFLAQFL | ASSQEGTGDSDYT | 0.60650086 |
| GGAGGAGGNAGVA | ASSQEGTGDSDYT | 0.846558 |
| GGVGTAGAGGAGGAGG | ASSQEGTGDSDYT | 0.8175422 |
| GNGGGGDAGNAGSGGN | ASSQEGTGDSDYT | 0.7445394 |
| GTGGAGGAGNDAGS | ASSQEGTGDSDYT | 0.8151158 |
| GTTNTGWFNS | ASSQEGTGDSDYT | 0.7533468 |
| IFMGPLFTL | ASSQEGTGDSDYT | 0.10606858 |
| ILAHAIEAI | ASSQEGTGDSDYT | 0.6425803 |
| IRSAIREAL | ASSQEGTGDSDYT | 0.4236397 |

| | | |
|---|---|---|
| ITFAGGG | ASSQEGTGDSDYT | 0.87603414 |
| ITGMSGHQEV | ASSQEGTGDSDYT | 0.56678843 |
| IVAGLACTL | ASSQEGTGDSDYT | 0.5590029 |
| IVGGGLAGA | ASSQEGTGDSDYT | 0.7562511 |
| IVTGVLNI | ASSQEGTGDSDYT | 0.7330907 |
| LAQQGCI | ASSQEGTGDSDYT | 0.73646104 |
| LCLHVEN | ASSQEGTGDSDYT | 0.5384692 |
| LDAISDGS | ASSQEGTGDSDYT | 0.15225004 |
| LGGFAAML | ASSQEGTGDSDYT | 0.82927376 |
| LGGKNPMI | ASSQEGTGDSDYT | 0.42875683 |
| LGPKGVAAIL | ASSQEGTGDSDYT | 0.5434779 |
| LMVGVSI | ASSQEGTGDSDYT | 0.22827132 |
| LVGAAFVL | ASSQEGTGDSDYT | 0.41097867 |
| LVLAVAPL | ASSQEGTGDSDYT | 0.39804137 |
| MAAIAVIDAA | ASSQEGTGDSDYT | 0.86709124 |
| NNGINAELQ | ASSQEGTGDSDYT | 0.6910751 |
| RGEPLVLVHG | ASSQEGTGDSDYT | 0.6338807 |
| RGGAGGAAGLI | ASSQEGTGDSDYT | 0.8382288 |
| RSYRMHPAV | ASSQEGTGDSDYT | 0.7549586 |
| SAAHLLAGQLA | ASSQEGTGDSDYT | 0.67675734 |
| SAAVGSY | ASSQEGTGDSDYT | 0.6336149 |
| SPHAVLHRI | ASSQEGTGDSDYT | 0.5909864 |
| SQIAKVL | ASSQEGTGDSDYT | 0.54326856 |
| TALGHLVP | ASSQEGTGDSDYT | 0.5174625 |
| TAPTTSGAA | ASSQEGTGDSDYT | 0.8658761 |
| TDGDDGNGGSPGT | ASSQEGTGDSDYT | 0.7250528 |
| TDGLGEN | ASSQEGTGDSDYT | 0.4202327 |
| TGGPPLPGQA | ASSQEGTGDSDYT | 0.83499455 |
| TIPAITI | ASSQEGTGDSDYT | 0.8646692 |
| TIPGFTI | ASSQEGTGDSDYT | 0.8780363 |
| TSIASGALA | ASSQEGTGDSDYT | 0.55281144 |
| TVTLPGAAI | ASSQEGTGDSDYT | 0.6943611 |
| VASGLSAALP | ASSQEGTGDSDYT | 0.6646116 |
| VGAGVIGI | ASSQEGTGDSDYT | 0.734121 |
| VGPKRFGV | ASSQEGTGDSDYT | 0.6733618 |
| VNVAFPD | ASSQEGTGDSDYT | 0.43100384 |
| VSMTNSGVS | ASSQEGTGDSDYT | 0.8431709 |
| VTDVLGDLSV | ASSQEGTGDSDYT | 0.8873702 |
| VVVVGAGFAGL | ASSQEGTGDSDYT | 0.6072036 |
| VYTVGTVV | ASSQEGTGDSDYT | 0.41967303 |
| WTTFSGP | ASSQEGTGDSDYT | 0.7274961 |
| YDELVEAF | ASSQEGTGDSDYT | 0.5391704 |
| YGTGGAGGNGG | ASSQEGTGDSDYT | 0.67176163 |
| YTNVMAV | ASSQEGTGDSDYT | 0.79132926 |
| FLEQVHQL | ASSQEGTGDSDYT | 0.3614588 |
| GLSLLAVAV | ASSQEGTGDSDYT | 0.61271423 |
| LEQVHQL | ASSQEGTGDSDYT | 0.875837 |
| LNAAGAI | ASSQEGTGDSDYT | 0.6854261 |
| PAVHASL | ASSQEGTGDSDYT | 0.7750846 |
| TELEDDA | ASSQEGTGDSDYT | 0.839756 |
| TPPGPPN | ASSQEGTGDSDYT | 0.80414724 |

| | | |
|---|---|---|
| TPPPPPPP | ASSQEGTGDSDYT | 0.87420374 |
| TQSPALI | ASSLDWGGNYAEQF | 0.8457868 |
| EFAAAFPQ | ASSLDWGGNYAEQF | 0.7110113 |
| LLSGNLDVL | ASSLDWGGNYAEQF | 0.7259164 |
| GPANLGNYNI | ASSLDWGGNYAEQF | 0.7402848 |
| TLGPTLP | ASSLDWGGNYAEQF | 0.7634383 |
| VYAYSSCCK | ASSLDWGGNYAEQF | 0.7276803 |
| APAPAPAPNI | ASSLDWGGNYAEQF | 0.6775832 |
| PGGPVPIP | ASSLDWGGNYAEQF | 0.8563309 |
| FLTVALEP | ASSLDWGGNYAEQF | 0.7757883 |
| SQYFGANL | ASSLDWGGNYAEQF | 0.83779156 |
| GGVKPAA | ASSLDWGGNYAEQF | 0.5058108 |
| GGTGGAGGAGGW | ASSLDWGGNYAEQF | 0.49529657 |
| GVGGVAGLGGAGG | ASSLDWGGNYAEQF | 0.8336024 |
| VLLVLATTI | ASSLDWGGNYAEQF | 0.13214572 |
| IGIFAAML | ASSLDWGGNYAEQF | 0.2710564 |
| AVGDLID | ASSLDWGGNYAEQF | 0.7978791 |
| NISPLPL | ASSLDWGGNYAEQF | 0.8578328 |
| VTAVFVAI | ASSLDWGGNYAEQF | 0.17917848 |
| SGDDNDAAL | ASSLDWGGNYAEQF | 0.5088655 |
| AVAALAHS | ASSLDWGGNYAEQF | 0.48848793 |
| GGVTVMATP | ASSLDWGGNYAEQF | 0.8518115 |
| AGLAANVRD | ASSLDWGGNYAEQF | 0.7660824 |
| TNRPDLIDPALLRPG | ASSLDWGGNYAEQF | 0.34936658 |
| LANLGVTAL | ASSLDWGGNYAEQF | 0.83017975 |
| TGERIEL | ASSLDWGGNYAEQF | 0.3018667 |
| NGEELAA | ASSLDWGGNYAEQF | 0.42918396 |
| VLSLVWG | ASSLDWGGNYAEQF | 0.8360677 |
| IGYALWL | ASSLDWGGNYAEQF | 0.8650317 |
| LSLLLAPL | ASSLDWGGNYAEQF | 0.4828952 |
| TLDEALK | ASSLDWGGNYAEQF | 0.84871185 |
| TDGAGTAEGG | ASSLDWGGNYAEQF | 0.62156785 |
| NAPGGAGGNGG | ASSLDWGGNYAEQF | 0.88676625 |
| VSVFMGMGGNF | ASSLDWGGNYAEQF | 0.6971345 |
| LASYLAAL | ASSLDWGGNYAEQF | 0.4199193 |
| LVGGTPAL | ASSLDWGGNYAEQF | 0.8242531 |
| VRNADVL | ASSLDWGGNYAEQF | 0.82903874 |
| TCSSPFLVSAM | ASSLDWGGNYAEQF | 0.78059167 |
| VHITDAE | ASSLDWGGNYAEQF | 0.8313541 |
| EYAAPSF | ASSLDWGGNYAEQF | 0.74056745 |
| VVDAFHGGL | ASSLDWGGNYAEQF | 0.70585805 |
| ASVVDPQ | ASSLDWGGNYAEQF | 0.8606661 |
| SYNEFGGLS | ASSLDWGGNYAEQF | 0.8446684 |
| DPDGSVY | ASSLDWGGNYAEQF | 0.46975428 |
| MVSTIHATE | ASSLDWGGNYAEQF | 0.8124448 |
| LAAGVADPV | ASSLDWGGNYAEQF | 0.71782625 |
| TKEQIAA | ASSLDWGGNYAEQF | 0.61468565 |
| SLASPAAL | ASSLDWGGNYAEQF | 0.8728596 |
| IGLVFYGV | ASSLDWGGNYAEQF | 0.7993808 |
| FLLDASNL | ASSLDWGGNYAEQF | 0.832549 |
| YQVGNFP | ASSLDWGGNYAEQF | 0.82836336 |

| | | |
|---|---|---|
| ESLLSQST | ASSLDWGGNYAEQF | 0.8712917 |
| LFTVGFAAVG | ASSLDWGGNYAEQF | 0.37917843 |
| ATPPTPT | ASSLDWGGNYAEQF | 0.80585194 |
| TDLATLG | ASSLDWGGNYAEQF | 0.7372584 |
| SAGGTGR | ASSLDWGGNYAEQF | 0.8435594 |
| GFSNSGS | ASSLDWGGNYAEQF | 0.8183304 |
| GGNGGSGGTSV | ASSLDWGGNYAEQF | 0.44838747 |
| GTSVATGGAGN | ASSLDWGGNYAEQF | 0.7884507 |
| NGGNGGSGGTS | ASSLDWGGNYAEQF | 0.84526366 |
| AVVKFPH | ASSLDWGGNYAEQF | 0.75668526 |
| TPPPPPP | ASSLDWGGNYAEQF | 0.8037287 |
| AAPAAAAAPA | ASSLDWGGNYAEQF | 0.87786925 |
| IALNEDL | ASSLDWGGNYAEQF | 0.60402083 |
| IVNVSSM | ASSLDWGGNYAEQF | 0.704947 |
| TGIPPAPR | ASSLDWGGNYAEQF | 0.8544878 |
| FDLGGGTF | ASSLDWGGNYAEQF | 0.8523375 |
| FDLGGGTFDVS | ASSLDWGGNYAEQF | 0.7659942 |
| NEPTAAA | ASSLDWGGNYAEQF | 0.85916513 |
| NEPTAAALA | ASSLDWGGNYAEQF | 0.5206387 |
| GSIINISSI | ASSLDWGGNYAEQF | 0.81770134 |
| VAPGNAGTA | ASSLDWGGNYAEQF | 0.57252765 |
| TGGGASLE | ASSLDWGGNYAEQF | 0.81021327 |
| VAVYNHYKRI | ASSLDWGGNYAEQF | 0.5711018 |
| TDPAPATT | ASSLDWGGNYAEQF | 0.8086075 |
| TNRPDILDPALLRPG | ASSLDWGGNYAEQF | 0.81641936 |
| LTTTTTS | ASSLDWGGNYAEQF | 0.82393384 |
| PTGSGKTL | ASSLDWGGNYAEQF | 0.20584893 |
| TGSGKTL | ASSLDWGGNYAEQF | 0.8283903 |
| TIGHVDHGK | ASSLDWGGNYAEQF | 0.84264815 |
| TIGHVDHGKTT | ASSLDWGGNYAEQF | 0.88925964 |
| GHVDHGKTT | ASSLDWGGNYAEQF | 0.7506597 |
| AQFEHTLL | ASSLDWGGNYAEQF | 0.8009115 |
| IVNQDEL | ASSLDWGGNYAEQF | 0.68800414 |
| TNGSQFF | ASSLDWGGNYAEQF | 0.7495406 |
| TGKGIVE | ASSLDWGGNYAEQF | 0.5378513 |
| LISNASDA | ASSLDWGGNYAEQF | 0.72860694 |
| AELRTLGELTKIAKAHGAQ | ASSLDWGGNYAEQF | 0.6603462 |
| APNAVAAAEPSW | ASSLDWGGNYAEQF | 0.6403034 |
| APQVAGAATAVL | ASSLDWGGNYAEQF | 0.8310609 |
| ARRDPAAAAAILPTDA | ASSLDWGGNYAEQF | 0.8548144 |
| DAAARDVLAEAGLAEYF | ASSLDWGGNYAEQF | 0.8253494 |
| DDQIVLSAKANLAKVEQ | ASSLDWGGNYAEQF | 0.84298563 |
| DELLPDTGEDSDEDGH | ASSLDWGGNYAEQF | 0.8539631 |
| ELPEAPANRATVPQD | ASSLDWGGNYAEQF | 0.86606467 |
| FGQSKIERVTGIG | ASSLDWGGNYAEQF | 0.64973974 |
| GAGGTGSTAGGAGGAGGAGGLY | ASSLDWGGNYAEQF | 0.7178377 |
| GHGGTGAAGGNGGTGGAGGSGI | ASSLDWGGNYAEQF | 0.52508616 |
| GNGGNGGAGGEGGPGGAGGAGA | ASSLDWGGNYAEQF | 0.6301723 |
| GSTAGNGGNGGNATNGTIAGQP | ASSLDWGGNYAEQF | 0.65892637 |
| GTGGDGGNGGNGGTGG | ASSLDWGGNYAEQF | 0.7808386 |
| GVRRPAGIPADAPV | ASSLDWGGNYAEQF | 0.13716508 |

| | | |
|---|---|---|
| GYRQVTDAHLLTLARR | ASSLDWGGNYAEQF | 0.7020476 |
| IAKMGQDGHDRG | ASSLDWGGNYAEQF | 0.8655566 |
| KAEPSIEIQYT | ASSLDWGGNYAEQF | 0.5764724 |
| KPVRIRMAADEDTVAVT | ASSLDWGGNYAEQF | 0.8085896 |
| MPPRIAGMRLLVIKPEP | ASSLDWGGNYAEQF | 0.72532284 |
| MKFYAVALFRSRGVPPTR | ASSLDWGGNYAEQF | 0.62895286 |
| NAGGFGILGAVAHSP | ASSLDWGGNYAEQF | 0.82354665 |
| NEGREITLAMAEM | ASSLDWGGNYAEQF | 0.5166315 |
| NELVEWSHPLGHRWRWE | ASSLDWGGNYAEQF | 0.84046644 |
| NLGAGGTGGNGGNPG | ASSLDWGGNYAEQF | 0.8757716 |
| NTEWPKWETVMAADPVD | ASSLDWGGNYAEQF | 0.7261024 |
| PASAAPAETQP | ASSLDWGGNYAEQF | 0.7746346 |
| QQSSDGKNGVRAAEITGE | ASSLDWGGNYAEQF | 0.74422574 |
| RAPDAVTTSWWKE | ASSLDWGGNYAEQF | 0.34517023 |
| SGFGNVGAHTSGWFNQ | ASSLDWGGNYAEQF | 0.7824874 |
| SGVAGAVGQAASVGGLKV | ASSLDWGGNYAEQF | 0.23248431 |
| SNGGAGTGSAGGNGGT | ASSLDWGGNYAEQF | 0.7341323 |
| TGGAGGEGGAAG | ASSLDWGGNYAEQF | 0.4860497 |
| TGGDGGAAGAGGAPGHG | ASSLDWGGNYAEQF | 0.7950703 |
| TMPPLLAAIANAGRHGV | ASSLDWGGNYAEQF | 0.7365918 |
| TSSTAGNSGGA | ASSLDWGGNYAEQF | 0.8064467 |
| VPATVPVGILC | ASSLDWGGNYAEQF | 0.5656216 |
| VQRPDSGGVVLTGRIS | ASSLDWGGNYAEQF | 0.71230817 |
| WTPNPVPATKGVPAGLVQAT | ASSLDWGGNYAEQF | 0.5358912 |
| YALKRALAKFSDRFDVVII | ASSLDWGGNYAEQF | 0.39364633 |
| YLIGSVIV | ASSLDWGGNYAEQF | 0.82360965 |
| YGQLVAVID | ASSLDWGGNYAEQF | 0.7718161 |
| VTVGGMTLPA | ASSLDWGGNYAEQF | 0.66341066 |
| VLALPAV | ASSLDWGGNYAEQF | 0.8255153 |
| VIPPTLNYET | ASSLDWGGNYAEQF | 0.8015054 |
| VGMGGMHPSSG | ASSLDWGGNYAEQF | 0.40224344 |
| VGLAGGAATA | ASSLDWGGNYAEQF | 0.70829344 |
| VGGAGGAGGSGP | ASSLDWGGNYAEQF | 0.7387372 |
| VFAAFMV | ASSLDWGGNYAEQF | 0.72980356 |
| TLYDKLR | ASSLDWGGNYAEQF | 0.14047496 |
| TLAVFTVL | ASSLDWGGNYAEQF | 0.77923334 |
| TGSVQPARV | ASSLDWGGNYAEQF | 0.59846574 |
| TGGGAGSGGT | ASSLDWGGNYAEQF | 0.5547688 |
| SHLETSL | ASSLDWGGNYAEQF | 0.61461776 |
| SFTTGMT | ASSLDWGGNYAEQF | 0.8406262 |
| QVEVAKRMGVRQPTVS | ASSLDWGGNYAEQF | 0.6844023 |
| QGSGGPALAM | ASSLDWGGNYAEQF | 0.8143605 |
| MLAPSNP | ASSLDWGGNYAEQF | 0.8371184 |
| LVNNAAI | ASSLDWGGNYAEQF | 0.8155614 |
| LTGEINTLA | ASSLDWGGNYAEQF | 0.8702313 |
| LSIFGPV | ASSLDWGGNYAEQF | 0.8179109 |
| LLALLANL | ASSLDWGGNYAEQF | 0.70782626 |
| LISNASDALDKLR | ASSLDWGGNYAEQF | 0.7915733 |
| LINEAVP | ASSLDWGGNYAEQF | 0.7715479 |
| LGYAFSGL | ASSLDWGGNYAEQF | 0.26249582 |
| LDFVAVAL | ASSLDWGGNYAEQF | 0.2764898 |

| | | |
|---|---|---|
| LAQPGAE | ASSLDWGGNYAEQF | 0.77239144 |
| ISNLGAVTGF | ASSLDWGGNYAEQF | 0.24293907 |
| HQEDPGRI | ASSLDWGGNYAEQF | 0.7894982 |
| GTPGEPTP | ASSLDWGGNYAEQF | 0.8793176 |
| GFFNSGS | ASSLDWGGNYAEQF | 0.7887911 |
| GAGGGGLTAGAN | ASSLDWGGNYAEQF | 0.4499467 |
| FVDALVNM | ASSLDWGGNYAEQF | 0.67682815 |
| FLTVAPEL | ASSLDWGGNYAEQF | 0.69540477 |
| FETLTAL | ASSLDWGGNYAEQF | 0.8500291 |
| DVINPIEL | ASSLDWGGNYAEQF | 0.8357539 |
| APVSGAPV | ASSLDWGGNYAEQF | 0.8644953 |
| ALVGGLAW | ASSLDWGGNYAEQF | 0.8368156 |
| ALAAAAATAI | ASSLDWGGNYAEQF | 0.6572495 |
| AFGLGLLL | ASSLDWGGNYAEQF | 0.8321613 |
| AFFAAASL | ASSLDWGGNYAEQF | 0.66414046 |
| AAAALPAFSPP | ASSLDWGGNYAEQF | 0.7253238 |
| VVLAPAL | ASSLDWGGNYAEQF | 0.7240819 |
| TIGHVDHGKTTL | ASSLDWGGNYAEQF | 0.81252754 |
| LISNASDALDK | ASSLDWGGNYAEQF | 0.7827167 |
| LISNASDAL | ASSLDWGGNYAEQF | 0.63895535 |
| ISNASDALDK | ASSLDWGGNYAEQF | 0.39718404 |
| NQAETLVYQTE | ASSLDWGGNYAEQF | 0.82583165 |
| TGGTGGTAGAG | ASSLDWGGNYAEQF | 0.8077911 |
| TGSTAGGAGGA | ASSLDWGGNYAEQF | 0.33853808 |
| DPLAVPVAAGPV | ASSLDWGGNYAEQF | 0.6181296 |
| GGTGGTPGNGGN | ASSLDWGGNYAEQF | 0.8782379 |
| PAGLAGVDPATL | ASSLDWGGNYAEQF | 0.86275536 |
| GNNLVVEAPHAGASV | ASSLDWGGNYAEQF | 0.5157511 |
| KVTRIMSASTLEQPA | ASSLDWGGNYAEQF | 0.54110223 |
| LGFALPIALPRRTVV | ASSLDWGGNYAEQF | 0.73160636 |
| TRLGNLTARKLVVLV | ASSLDWGGNYAEQF | 0.28205842 |
| WSQQRRLIYFPSAGP | ASSLDWGGNYAEQF | 0.82126856 |
| GPNGNPDPMAAAVDIR | ASSLDWGGNYAEQF | 0.7233247 |
| TQRCVSVADLRAAASR | ASSLDWGGNYAEQF | 0.6969163 |
| QFVLDVAEHPVEVLTETV | ASSLDWGGNYAEQF | 0.55844057 |
| GGAGGTGGAGGNGGTQAGDG | ASSLDWGGNYAEQF | 0.7510344 |
| PLLATLRAIAVDVDAKPAQV | ASSLDWGGNYAEQF | 0.62394 |
| APKFRTGGYMIDFWGVGYQVAK | ASSLDWGGNYAEQF | 0.8521143 |
| GAGGAGGAGGAAGGPGGTGGTG | ASSLDWGGNYAEQF | 0.13324872 |
| IGALGLLATDVAPAGSVAFGVA | ASSLDWGGNYAEQF | 0.787493 |
| AGGAGGIGGNGGAGGAGGVLMG | ASSLDWGGNYAEQF | 0.55418533 |
| ADELIGLL | ASSLDWGGNYAEQF | 0.66455424 |
| ADRALKA | ASSLDWGGNYAEQF | 0.41965353 |
| AGAVGQVGGAGGA | ASSLDWGGNYAEQF | 0.69513285 |
| AGGGGGLGGAP | ASSLDWGGNYAEQF | 0.83231676 |
| AGGLLYGNGGAG | ASSLDWGGNYAEQF | 0.8444501 |
| AGLESIGL | ASSLDWGGNYAEQF | 0.44449407 |
| AGLKTGII | ASSLDWGGNYAEQF | 0.82460743 |
| ASLAALLL | ASSLDWGGNYAEQF | 0.8791928 |
| ASVINISSI | ASSLDWGGNYAEQF | 0.6400249 |
| ATPPGPA | ASSLDWGGNYAEQF | 0.3166133 |

| | | |
|---|---|---|
| ATVASALV | ASSLDWGGNYAEQF | 0.816306 |
| ATVASLIL | ASSLDWGGNYAEQF | 0.72822636 |
| AVNGPVLIRH | ASSLDWGGNYAEQF | 0.83643144 |
| DGAAGAVGI | ASSLDWGGNYAEQF | 0.81500447 |
| DPLLHGI | ASSLDWGGNYAEQF | 0.48613292 |
| ELGAALGAL | ASSLDWGGNYAEQF | 0.8501789 |
| ELGRTQAV | ASSLDWGGNYAEQF | 0.8360766 |
| FVAGGYLS | ASSLDWGGNYAEQF | 0.56740725 |
| GASGGAGGAGGW | ASSLDWGGNYAEQF | 0.5785643 |
| GGAGAGGDGGAGGI | ASSLDWGGNYAEQF | 0.5616702 |
| GTGHNGSQPGMGGNG | ASSLDWGGNYAEQF | 0.3414184 |
| GYAAASASAS | ASSLDWGGNYAEQF | 0.79623556 |
| HQVDVAAVQQG | ASSLDWGGNYAEQF | 0.5503857 |
| IGAAIAHT | ASSLDWGGNYAEQF | 0.63598746 |
| IGMIEVAQL | ASSLDWGGNYAEQF | 0.7579173 |
| IRLAPEHTSAMD | ASSLDWGGNYAEQF | 0.56419396 |
| IYNVSVL | ASSLDWGGNYAEQF | 0.66429186 |
| KQGFLGM | ASSLDWGGNYAEQF | 0.6532989 |
| LESVITQL | ASSLDWGGNYAEQF | 0.7791757 |
| LGLGALVVV | ASSLDWGGNYAEQF | 0.7656361 |
| LIGNGGNGGSGGI | ASSLDWGGNYAEQF | 0.7402079 |
| LNAINAPTL | ASSLDWGGNYAEQF | 0.76642233 |
| LQLLNTGV | ASSLDWGGNYAEQF | 0.8598487 |
| LQTSIEF | ASSLDWGGNYAEQF | 0.44146836 |
| MSFAVSEL | ASSLDWGGNYAEQF | 0.56313276 |
| NTGALGL | ASSLDWGGNYAEQF | 0.82303494 |
| PPGAGGAGGAGTT | ASSLDWGGNYAEQF | 0.36440665 |
| QLVGLGI | ASSLDWGGNYAEQF | 0.82202625 |
| RFLEQVHQL | ASSLDWGGNYAEQF | 0.7298755 |
| RGVGYVLR | ASSLDWGGNYAEQF | 0.77978724 |
| SATGVFAGVI | ASSLDWGGNYAEQF | 0.7449715 |
| SLPAITLP | ASSLDWGGNYAEQF | 0.87980247 |
| STALAFTV | ASSLDWGGNYAEQF | 0.854172 |
| SVLTVGLGK | ASSLDWGGNYAEQF | 0.6608311 |
| TALAGAGSGI | ASSLDWGGNYAEQF | 0.8278265 |
| TGLPFTGI | ASSLDWGGNYAEQF | 0.54470146 |
| VAEHVTV | ASSLDWGGNYAEQF | 0.82290304 |
| VDIVLNSL | ASSLDWGGNYAEQF | 0.84771216 |
| VLTGGGAVL | ASSLDWGGNYAEQF | 0.5289583 |
| VNVGLAYGI | ASSLDWGGNYAEQF | 0.7096374 |
| VSSAAVGAV | ASSLDWGGNYAEQF | 0.8711588 |
| VTSVYEAGV | ASSLDWGGNYAEQF | 0.84839493 |
| WVALDRGA | ASSLDWGGNYAEQF | 0.26181242 |
| YLVGSVVI | ASSLDWGGNYAEQF | 0.669715 |
| GPPGTGKTLL | ASSLDWGGNYAEQF | 0.2840368 |
| GVGMTKF | ASSLDWGGNYAEQF | 0.70500684 |
| IGIGGSDL | ASSLDWGGNYAEQF | 0.49181747 |
| IGTLLAEL | ASSLDWGGNYAEQF | 0.79263705 |
| IYTSGTT | ASSLDWGGNYAEQF | 0.7207342 |
| LSFPAGV | ASSLDWGGNYAEQF | 0.8240511 |
| LYTSGTT | ASSLDWGGNYAEQF | 0.77847373 |

| | | |
|---|---|---|
| VGAFPFGF | ASSLDWGGNYAEQF | 0.8636887 |
| VGAGTVEF | ASSLDWGGNYAEQF | 0.67494196 |
| VGKSTLLL | ASSLDWGGNYAEQF | 0.70570874 |
| AALAHAGGGIV | ASSLDWGGNYAEQF | 0.41382447 |
| AALVAVSLL | ASSLDWGGNYAEQF | 0.37679613 |
| AAVSRAVAAL | ASSLDWGGNYAEQF | 0.6050007 |
| AGGSSGIGGF | ASSLDWGGNYAEQF | 0.64906955 |
| AGLTVANANG | ASSLDWGGNYAEQF | 0.21599463 |
| AGLVGAVL | ASSLDWGGNYAEQF | 0.85778415 |
| ALSQAFTV | ASSLDWGGNYAEQF | 0.81297195 |
| ALTFSVSL | ASSLDWGGNYAEQF | 0.82951665 |
| ALTGAALVAL | ASSLDWGGNYAEQF | 0.62450886 |
| ASFSIGAGVL | ASSLDWGGNYAEQF | 0.737955 |
| DGSVYKI | ASSLDWGGNYAEQF | 0.6493809 |
| EQLAEIV | ASSLDWGGNYAEQF | 0.83629906 |
| FAVSELTVM | ASSLDWGGNYAEQF | 0.56874424 |
| GAAPLAAGL | ASSLDWGGNYAEQF | 0.30992284 |
| GADGGDGGSGGAGG | ASSLDWGGNYAEQF | 0.6516807 |
| GFGLDTGGAGG | ASSLDWGGNYAEQF | 0.38563144 |
| GLASGGLL | ASSLDWGGNYAEQF | 0.85211897 |
| GRIAGAAVLL | ASSLDWGGNYAEQF | 0.8211143 |
| GSAGSLNTSF | ASSLDWGGNYAEQF | 0.8016646 |
| GTGQIGI | ASSLDWGGNYAEQF | 0.85303056 |
| GTLIALIL | ASSLDWGGNYAEQF | 0.23754005 |
| IAATMGFVVG | ASSLDWGGNYAEQF | 0.6232996 |
| IFAQALL | ASSLDWGGNYAEQF | 0.8707867 |
| IGAAVALGF | ASSLDWGGNYAEQF | 0.7728909 |
| IGLTPLAL | ASSLDWGGNYAEQF | 0.7360269 |
| ILGSNTSTL | ASSLDWGGNYAEQF | 0.8716402 |
| IVFSLTVL | ASSLDWGGNYAEQF | 0.5935457 |
| LASKATGF | ASSLDWGGNYAEQF | 0.7410377 |
| LASVALIL | ASSLDWGGNYAEQF | 0.77649224 |
| LFAVGLTL | ASSLDWGGNYAEQF | 0.7907746 |
| LFNGGIGTYI | ASSLDWGGNYAEQF | 0.8361406 |
| LTLGAGII | ASSLDWGGNYAEQF | 0.55042887 |
| LVAAALGDI | ASSLDWGGNYAEQF | 0.7961439 |
| NTAFVQLGI | ASSLDWGGNYAEQF | 0.87623316 |
| QLFSGAEL | ASSLDWGGNYAEQF | 0.731078 |
| SAATFITL | ASSLDWGGNYAEQF | 0.83546484 |
| SNYGLANLGV | ASSLDWGGNYAEQF | 0.811182 |
| SYGGAYAVM | ASSLDWGGNYAEQF | 0.7980267 |
| TTGVGGLTLG | ASSLDWGGNYAEQF | 0.62391686 |
| TTQTIYI | ASSLDWGGNYAEQF | 0.6376848 |
| VAAGAGLGAVI | ASSLDWGGNYAEQF | 0.6981745 |
| VAAVVTLL | ASSLDWGGNYAEQF | 0.5617023 |
| VGFLGGLI | ASSLDWGGNYAEQF | 0.7907971 |
| VGVIPILL | ASSLDWGGNYAEQF | 0.63232136 |
| VIFASSLI | ASSLDWGGNYAEQF | 0.8929986 |
| VKGGDGGAAAG | ASSLDWGGNYAEQF | 0.8116987 |
| VTKTFTNI | ASSLDWGGNYAEQF | 0.8069701 |
| VVAAAILL | ASSLDWGGNYAEQF | 0.52899003 |

| | | |
|---|---|---|
| YAALDVEL | ASSLDWGGNYAEQF | 0.7917851 |
| LASVALLL | ASSLDWGGNYAEQF | 0.7615407 |
| LDAVHRV | ASSLDWGGNYAEQF | 0.41277853 |
| QGLSSSIF | ASSLDWGGNYAEQF | 0.8514231 |
| RGTFANIRL | ASSLDWGGNYAEQF | 0.64008176 |
| RQLWWGHRI | ASSLDWGGNYAEQF | 0.67240024 |
| YGPPGTGKTLL | ASSLDWGGNYAEQF | 0.8466977 |
| AALVGAPANVA | ASSLDWGGNYAEQF | 0.85055816 |
| AATFRRTE | ASSLDWGGNYAEQF | 0.6999999 |
| AGALGGGI | ASSLDWGGNYAEQF | 0.6365243 |
| AGGAGGAGGIGAG | ASSLDWGGNYAEQF | 0.5596033 |
| AGSGALGL | ASSLDWGGNYAEQF | 0.83798325 |
| AQIADQI | ASSLDWGGNYAEQF | 0.860188 |
| ASAVAVTYL | ASSLDWGGNYAEQF | 0.7778388 |
| ASTVLGGTGG | ASSLDWGGNYAEQF | 0.87195265 |
| AVGGEPAK | ASSLDWGGNYAEQF | 0.84868616 |
| CEQGTFGGTC | ASSLDWGGNYAEQF | 0.45637065 |
| DEGKAPAG | ASSLDWGGNYAEQF | 0.73569185 |
| DTVVFPAL | ASSLDWGGNYAEQF | 0.8311771 |
| FEGITMAG | ASSLDWGGNYAEQF | 0.82585 |
| FPLVVQD | ASSLDWGGNYAEQF | 0.8790882 |
| GAFTTPQ | ASSLDWGGNYAEQF | 0.83796096 |
| GFGGFGTAGGVGGA | ASSLDWGGNYAEQF | 0.6169089 |
| GFLAQFL | ASSLDWGGNYAEQF | 0.85384023 |
| GGAGGAGGNAGVA | ASSLDWGGNYAEQF | 0.88477945 |
| GGVGTAGAGGAGGAGG | ASSLDWGGNYAEQF | 0.84896153 |
| GNGGGGDAGNAGSGGN | ASSLDWGGNYAEQF | 0.712108 |
| GTGGAGGAGNDAGS | ASSLDWGGNYAEQF | 0.67233455 |
| GTTNTGWFNS | ASSLDWGGNYAEQF | 0.7353312 |
| IFMGPLFTL | ASSLDWGGNYAEQF | 0.84426826 |
| ILAHAIEAI | ASSLDWGGNYAEQF | 0.62699187 |
| IRSAIREAL | ASSLDWGGNYAEQF | 0.7652958 |
| ITFAGGG | ASSLDWGGNYAEQF | 0.77504253 |
| ITGMSGHQEV | ASSLDWGGNYAEQF | 0.42201263 |
| IVAGLACTL | ASSLDWGGNYAEQF | 0.78888375 |
| IVGGGLAGA | ASSLDWGGNYAEQF | 0.8534957 |
| IVTGVLNI | ASSLDWGGNYAEQF | 0.68950033 |
| LAQQGCI | ASSLDWGGNYAEQF | 0.6593236 |
| LCLHVEN | ASSLDWGGNYAEQF | 0.83720326 |
| LDAISDGS | ASSLDWGGNYAEQF | 0.8878326 |
| LGGFAAML | ASSLDWGGNYAEQF | 0.8655444 |
| LGGKNPMI | ASSLDWGGNYAEQF | 0.4018194 |
| LGPKGVAAIL | ASSLDWGGNYAEQF | 0.20632724 |
| LMVGVSI | ASSLDWGGNYAEQF | 0.5820738 |
| LVGAAFVL | ASSLDWGGNYAEQF | 0.8295822 |
| LVLAVAPL | ASSLDWGGNYAEQF | 0.630675 |
| MAAIAVIDAA | ASSLDWGGNYAEQF | 0.6695727 |
| NNGINAELQ | ASSLDWGGNYAEQF | 0.8107493 |
| RGEPLVLVHG | ASSLDWGGNYAEQF | 0.84941554 |
| RGGAGGAAGLI | ASSLDWGGNYAEQF | 0.584162 |
| RSYRMHPAV | ASSLDWGGNYAEQF | 0.48367077 |

| | | |
|---|---|---|
| SAAHLLAGQLA | ASSLDWGGNYAEQF | 0.81029123 |
| SAAVGSY | ASSLDWGGNYAEQF | 0.8512225 |
| SPHAVLHRI | ASSLDWGGNYAEQF | 0.5097797 |
| SQIAKVL | ASSLDWGGNYAEQF | 0.718198 |
| TALGHLVP | ASSLDWGGNYAEQF | 0.50474155 |
| TAPTTSGAA | ASSLDWGGNYAEQF | 0.54000247 |
| TDGDDGNGGSPGT | ASSLDWGGNYAEQF | 0.7262103 |
| TDGLGEN | ASSLDWGGNYAEQF | 0.5813478 |
| TGGPPLPGQA | ASSLDWGGNYAEQF | 0.84909576 |
| TIPAITI | ASSLDWGGNYAEQF | 0.56089497 |
| TIPGFTI | ASSLDWGGNYAEQF | 0.84428054 |
| TSIASGALA | ASSLDWGGNYAEQF | 0.792516 |
| TVTLPGAAI | ASSLDWGGNYAEQF | 0.7853299 |
| VASGLSAALP | ASSLDWGGNYAEQF | 0.7237097 |
| VGAGVIGI | ASSLDWGGNYAEQF | 0.63068616 |
| VGPKRFGV | ASSLDWGGNYAEQF | 0.43404397 |
| VNVAFPD | ASSLDWGGNYAEQF | 0.27877104 |
| VSMTNSGVS | ASSLDWGGNYAEQF | 0.85630095 |
| VTDVLGDLSV | ASSLDWGGNYAEQF | 0.49028924 |
| VVVVGAGFAGL | ASSLDWGGNYAEQF | 0.8621597 |
| VYTVGTVV | ASSLDWGGNYAEQF | 0.8355639 |
| WTTFSGP | ASSLDWGGNYAEQF | 0.5398551 |
| YDELVEAF | ASSLDWGGNYAEQF | 0.81807554 |
| YGTGGAGGNGG | ASSLDWGGNYAEQF | 0.5901456 |
| YTNVMAV | ASSLDWGGNYAEQF | 0.43479076 |
| FLEQVHQL | ASSLDWGGNYAEQF | 0.84548104 |
| GLSLLAVAV | ASSLDWGGNYAEQF | 0.8176726 |
| LEQVHQL | ASSLDWGGNYAEQF | 0.6246551 |
| LNAAGAI | ASSLDWGGNYAEQF | 0.820667 |
| PAVHASL | ASSLDWGGNYAEQF | 0.732903 |
| TELEDDA | ASSLDWGGNYAEQF | 0.55841863 |
| TPPGPPN | ASSLDWGGNYAEQF | 0.7070934 |
| TPPPPPPP | ASSLDWGGNYAEQF | 0.84714746 |
| TQSPALI | ASSPDWGVSYEQY | 0.7761099 |
| EFAAAFPQ | ASSPDWGVSYEQY | 0.64703023 |
| LLSGNLDVL | ASSPDWGVSYEQY | 0.8031446 |
| GPANLGNYNI | ASSPDWGVSYEQY | 0.524503 |
| TLGPTLP | ASSPDWGVSYEQY | 0.77300286 |
| VYAYSSCCK | ASSPDWGVSYEQY | 0.73773956 |
| APAPAPAPNI | ASSPDWGVSYEQY | 0.71619815 |
| PGGPVPIP | ASSPDWGVSYEQY | 0.8308697 |
| FLTVALEP | ASSPDWGVSYEQY | 0.57865345 |
| SQYFGANL | ASSPDWGVSYEQY | 0.55067307 |
| GGVKPAA | ASSPDWGVSYEQY | 0.6926189 |
| GGTGGAGGAGGW | ASSPDWGVSYEQY | 0.76374114 |
| GVGGVAGLGGAGG | ASSPDWGVSYEQY | 0.83942723 |
| VLLVLATTI | ASSPDWGVSYEQY | 0.549561 |
| IGIFAAML | ASSPDWGVSYEQY | 0.753856 |
| AVGDLID | ASSPDWGVSYEQY | 0.65016425 |
| NISPLPL | ASSPDWGVSYEQY | 0.5736177 |
| VTAVFVAI | ASSPDWGVSYEQY | 0.7803019 |

| | | |
|---|---|---|
| SGDDNDAAL | ASSPDWGVSYEQY | 0.6965611 |
| AVAALAHS | ASSPDWGVSYEQY | 0.6661527 |
| GGVTVMATP | ASSPDWGVSYEQY | 0.7848385 |
| AGLAANVRD | ASSPDWGVSYEQY | 0.6223779 |
| TNRPDLIDPALLRPG | ASSPDWGVSYEQY | 0.7850823 |
| LANLGVTAL | ASSPDWGVSYEQY | 0.8382597 |
| TGERIEL | ASSPDWGVSYEQY | 0.7205126 |
| NGEELAA | ASSPDWGVSYEQY | 0.83714247 |
| VLSLVWG | ASSPDWGVSYEQY | 0.7920519 |
| IGYALWL | ASSPDWGVSYEQY | 0.8120146 |
| LSLLLAPL | ASSPDWGVSYEQY | 0.6573749 |
| TLDEALK | ASSPDWGVSYEQY | 0.5708164 |
| TDGAGTAEGG | ASSPDWGVSYEQY | 0.85204667 |
| NAPGGAGGNGG | ASSPDWGVSYEQY | 0.8205383 |
| VSVFMGMGGNF | ASSPDWGVSYEQY | 0.8383485 |
| LASYLAAL | ASSPDWGVSYEQY | 0.8269515 |
| LVGGTPAL | ASSPDWGVSYEQY | 0.8170781 |
| VRNADVL | ASSPDWGVSYEQY | 0.8310893 |
| TCSSPFLVSAM | ASSPDWGVSYEQY | 0.6349983 |
| VHITDAE | ASSPDWGVSYEQY | 0.4535672 |
| EYAAPSF | ASSPDWGVSYEQY | 0.67678773 |
| VVDAFHGGL | ASSPDWGVSYEQY | 0.6677693 |
| ASVVDPQ | ASSPDWGVSYEQY | 0.66477066 |
| SYNEFGGLS | ASSPDWGVSYEQY | 0.4075903 |
| DPDGSVY | ASSPDWGVSYEQY | 0.8560287 |
| MVSTIHATE | ASSPDWGVSYEQY | 0.8883876 |
| LAAGVADPV | ASSPDWGVSYEQY | 0.6480055 |
| TKEQIAA | ASSPDWGVSYEQY | 0.88143414 |
| SLASPAAL | ASSPDWGVSYEQY | 0.42614502 |
| IGLVFYGV | ASSPDWGVSYEQY | 0.8189404 |
| FLLDASNL | ASSPDWGVSYEQY | 0.86380076 |
| YQVGNFP | ASSPDWGVSYEQY | 0.59904957 |
| ESLLSQST | ASSPDWGVSYEQY | 0.81833917 |
| LFTVGFAAVG | ASSPDWGVSYEQY | 0.59217536 |
| ATPPTPT | ASSPDWGVSYEQY | 0.69840366 |
| TDLATLG | ASSPDWGVSYEQY | 0.8640722 |
| SAGGTGR | ASSPDWGVSYEQY | 0.6649505 |
| GFSNSGS | ASSPDWGVSYEQY | 0.7525085 |
| GGNGGSGGTSV | ASSPDWGVSYEQY | 0.74457604 |
| GTSVATGGAGN | ASSPDWGVSYEQY | 0.5944121 |
| NGGNGGSGGTS | ASSPDWGVSYEQY | 0.7638767 |
| AVVKFPH | ASSPDWGVSYEQY | 0.86117744 |
| TPPPPPP | ASSPDWGVSYEQY | 0.54342043 |
| AAPAAAAAPA | ASSPDWGVSYEQY | 0.882136 |
| IALNEDL | ASSPDWGVSYEQY | 0.5910752 |
| IVNVSSM | ASSPDWGVSYEQY | 0.8813534 |
| TGIPPAPR | ASSPDWGVSYEQY | 0.8383251 |
| FDLGGGTF | ASSPDWGVSYEQY | 0.73220754 |
| FDLGGGTFDVS | ASSPDWGVSYEQY | 0.6799271 |
| NEPTAAA | ASSPDWGVSYEQY | 0.8604304 |
| NEPTAAALA | ASSPDWGVSYEQY | 0.7501677 |

| | | |
|---|---|---|
| GSIINISSI | ASSPDWGVSYEQY | 0.82835233 |
| VAPGNAGTA | ASSPDWGVSYEQY | 0.873575 |
| TGGGASLE | ASSPDWGVSYEQY | 0.55141693 |
| VAVYNHYKRI | ASSPDWGVSYEQY | 0.85168964 |
| TDPAPATT | ASSPDWGVSYEQY | 0.81158507 |
| TNRPDILDPALLRPG | ASSPDWGVSYEQY | 0.83476216 |
| LTTTTTS | ASSPDWGVSYEQY | 0.74569964 |
| PTGSGKTL | ASSPDWGVSYEQY | 0.30759078 |
| TGSGKTL | ASSPDWGVSYEQY | 0.78529704 |
| TIGHVDHGK | ASSPDWGVSYEQY | 0.6106635 |
| TIGHVDHGKTT | ASSPDWGVSYEQY | 0.8421376 |
| GHVDHGKTT | ASSPDWGVSYEQY | 0.86203694 |
| AQFEHTLL | ASSPDWGVSYEQY | 0.6922612 |
| IVNQDEL | ASSPDWGVSYEQY | 0.82424325 |
| TNGSQFF | ASSPDWGVSYEQY | 0.8339219 |
| TGKGIVE | ASSPDWGVSYEQY | 0.56911236 |
| LISNASDA | ASSPDWGVSYEQY | 0.783911 |
| AELRTLGELTKIAKAHGAQ | ASSPDWGVSYEQY | 0.6305288 |
| APNAVAAAEPSW | ASSPDWGVSYEQY | 0.79418325 |
| APQVAGAATAVL | ASSPDWGVSYEQY | 0.36047244 |
| ARRDPAAAAAILPTDA | ASSPDWGVSYEQY | 0.83713967 |
| DAAARDVLAEAGLAEYF | ASSPDWGVSYEQY | 0.67401296 |
| DDQIVLSAKANLAKVEQ | ASSPDWGVSYEQY | 0.17091937 |
| DELLPDTGEDSDEDGH | ASSPDWGVSYEQY | 0.8443492 |
| ELPEAPANRATVPQD | ASSPDWGVSYEQY | 0.7502984 |
| FGQSKIERVTGIG | ASSPDWGVSYEQY | 0.73873067 |
| GAGGTGSTAGGAGGAGGAGGLY | ASSPDWGVSYEQY | 0.8413356 |
| GHGGTGAAGGNGGTGGAGGSGI | ASSPDWGVSYEQY | 0.40833703 |
| GNGGNGGAGGEGGPGGAGGAGA | ASSPDWGVSYEQY | 0.52648675 |
| GSTAGNGGNGGNATNGTIAGQP | ASSPDWGVSYEQY | 0.8271101 |
| GTGGDGGNGGNGGTGG | ASSPDWGVSYEQY | 0.8667354 |
| GVRRPAGIPADAPV | ASSPDWGVSYEQY | 0.4450163 |
| GYRQVTDAHLLTLARR | ASSPDWGVSYEQY | 0.81962687 |
| IAKMGQDGHDRG | ASSPDWGVSYEQY | 0.8706999 |
| KAEPSIEIQYT | ASSPDWGVSYEQY | 0.81387997 |
| KPVRIRMAADEDTVAVT | ASSPDWGVSYEQY | 0.8764394 |
| MPPRIAGMRLLVIKPEP | ASSPDWGVSYEQY | 0.88362163 |
| MKFYAVALFRSRGVPPTR | ASSPDWGVSYEQY | 0.7261081 |
| NAGGFGILGAVAHSP | ASSPDWGVSYEQY | 0.8665236 |
| NEGREITLAMAEM | ASSPDWGVSYEQY | 0.8630301 |
| NELVEWSHPLGHRWRWE | ASSPDWGVSYEQY | 0.7261057 |
| NLGAGGTGGNGGNPG | ASSPDWGVSYEQY | 0.79729855 |
| NTEWPKWETVMAADPVD | ASSPDWGVSYEQY | 0.5953843 |
| PASAAPAETQP | ASSPDWGVSYEQY | 0.7759439 |
| QQSSDGKNGVRAAEITGE | ASSPDWGVSYEQY | 0.3771426 |
| RAPDAVTTSWWKE | ASSPDWGVSYEQY | 0.8087306 |
| SGFGNVGAHTSGWFNQ | ASSPDWGVSYEQY | 0.8741919 |
| SGVAGAVGQAASVGGLKV | ASSPDWGVSYEQY | 0.7688831 |
| SNGGAGTGSAGGNGGT | ASSPDWGVSYEQY | 0.52087647 |
| TGGAGGEGGAAG | ASSPDWGVSYEQY | 0.5329467 |
| TGGDGGAAGAGGAPGHG | ASSPDWGVSYEQY | 0.82131314 |

| | | |
|---|---|---|
| TMPPLLAAIANAGRHGV | ASSPDWGVSYEQY | 0.50297594 |
| TSSTAGNSGGA | ASSPDWGVSYEQY | 0.3744207 |
| VPATVPVGILC | ASSPDWGVSYEQY | 0.5976297 |
| VQRPDSGGVVLTGRIS | ASSPDWGVSYEQY | 0.8405032 |
| WTPNPVPATKGVPAGLVQAT | ASSPDWGVSYEQY | 0.86014175 |
| YALKRALAKFSDRFDVVII | ASSPDWGVSYEQY | 0.5396812 |
| YLIGSVIV | ASSPDWGVSYEQY | 0.86859745 |
| YGQLVAVID | ASSPDWGVSYEQY | 0.8435959 |
| VTVGGMTLPA | ASSPDWGVSYEQY | 0.8685696 |
| VLALPAV | ASSPDWGVSYEQY | 0.7750908 |
| VIPPTLNYET | ASSPDWGVSYEQY | 0.55474174 |
| VGMGGMHPSSG | ASSPDWGVSYEQY | 0.85967106 |
| VGLAGGAATA | ASSPDWGVSYEQY | 0.7361717 |
| VGGAGGAGGSGP | ASSPDWGVSYEQY | 0.7817209 |
| VFAAFMV | ASSPDWGVSYEQY | 0.85313094 |
| TLYDKLR | ASSPDWGVSYEQY | 0.5460048 |
| TLAVFTVL | ASSPDWGVSYEQY | 0.6318645 |
| TGSVQPARV | ASSPDWGVSYEQY | 0.77385116 |
| TGGGAGSGGT | ASSPDWGVSYEQY | 0.89023817 |
| SHLETSL | ASSPDWGVSYEQY | 0.6237142 |
| SFTTGMT | ASSPDWGVSYEQY | 0.46722656 |
| QVEVAKRMGVRQPTVS | ASSPDWGVSYEQY | 0.60054153 |
| QGSGGPALAM | ASSPDWGVSYEQY | 0.7049176 |
| MLAPSNP | ASSPDWGVSYEQY | 0.8699144 |
| LVNNAAI | ASSPDWGVSYEQY | 0.8321722 |
| LTGEINTLA | ASSPDWGVSYEQY | 0.77877414 |
| LSIFGPV | ASSPDWGVSYEQY | 0.82971776 |
| LLALLANL | ASSPDWGVSYEQY | 0.8249229 |
| LISNASDALDKLR | ASSPDWGVSYEQY | 0.80472094 |
| LINEAVP | ASSPDWGVSYEQY | 0.8063368 |
| LGYAFSGL | ASSPDWGVSYEQY | 0.68775475 |
| LDFVAVAL | ASSPDWGVSYEQY | 0.88540393 |
| LAQPGAE | ASSPDWGVSYEQY | 0.85250825 |
| ISNLGAVTGF | ASSPDWGVSYEQY | 0.8598059 |
| HQEDPGRI | ASSPDWGVSYEQY | 0.87112594 |
| GTPGEPTP | ASSPDWGVSYEQY | 0.70369315 |
| GFFNSGS | ASSPDWGVSYEQY | 0.8232292 |
| GAGGGGLTAGAN | ASSPDWGVSYEQY | 0.51115775 |
| FVDALVNM | ASSPDWGVSYEQY | 0.554374 |
| FLTVAPEL | ASSPDWGVSYEQY | 0.7937617 |
| FETLTAL | ASSPDWGVSYEQY | 0.81327766 |
| DVINPIEL | ASSPDWGVSYEQY | 0.8528473 |
| APVSGAPV | ASSPDWGVSYEQY | 0.8390006 |
| ALVGGLAW | ASSPDWGVSYEQY | 0.73093337 |
| ALAAAAATAI | ASSPDWGVSYEQY | 0.863789 |
| AFGLGLLL | ASSPDWGVSYEQY | 0.86517185 |
| AFFAAASL | ASSPDWGVSYEQY | 0.65007424 |
| AAAALPAFSPP | ASSPDWGVSYEQY | 0.7674922 |
| VVLAPAL | ASSPDWGVSYEQY | 0.87317675 |
| TIGHVDHGKTTL | ASSPDWGVSYEQY | 0.82547027 |
| LISNASDALDK | ASSPDWGVSYEQY | 0.75527024 |

| | | |
|---|---|---|
| LISNASDAL | ASSPDWGVSYEQY | 0.7945876 |
| ISNASDALDK | ASSPDWGVSYEQY | 0.86503196 |
| NQAETLVYQTE | ASSPDWGVSYEQY | 0.7577183 |
| TGGTGGTAGAG | ASSPDWGVSYEQY | 0.87806594 |
| TGSTAGGAGGA | ASSPDWGVSYEQY | 0.30647188 |
| DPLAVPVAAGPV | ASSPDWGVSYEQY | 0.8139378 |
| GGTGGTPGNGGN | ASSPDWGVSYEQY | 0.74370563 |
| PAGLAGVDPATL | ASSPDWGVSYEQY | 0.1840088 |
| GNNLVVEAPHAGASV | ASSPDWGVSYEQY | 0.8707485 |
| KVTRIMSASTLEQPA | ASSPDWGVSYEQY | 0.7072911 |
| LGFALPIALPRRTVV | ASSPDWGVSYEQY | 0.8158437 |
| TRLGNLTARKLVVLV | ASSPDWGVSYEQY | 0.7075217 |
| WSQQRRLIYFPSAGP | ASSPDWGVSYEQY | 0.86871064 |
| GPNGNPDPMAAAVDIR | ASSPDWGVSYEQY | 0.82129025 |
| TQRCVSVADLRAAASR | ASSPDWGVSYEQY | 0.7059345 |
| QFVLDVAEHPVEVLTETV | ASSPDWGVSYEQY | 0.6261867 |
| GGAGGTGGAGGNGGTQAGDG | ASSPDWGVSYEQY | 0.8097439 |
| PLLATLRAIAVDVDAKPAQV | ASSPDWGVSYEQY | 0.8753942 |
| APKFRTGGYMIDFWGVGYQVAK | ASSPDWGVSYEQY | 0.88457644 |
| GAGGAGGAGGAAGGPGGTGGTG | ASSPDWGVSYEQY | 0.2457636 |
| IGALGLLATDVAPAGSVAFGVA | ASSPDWGVSYEQY | 0.8336697 |
| AGGAGGIGGNGGAGGAGGVLMG | ASSPDWGVSYEQY | 0.78763145 |
| ADELIGLL | ASSPDWGVSYEQY | 0.18111576 |
| ADRALKA | ASSPDWGVSYEQY | 0.2598937 |
| AGAVGQVGGAGGA | ASSPDWGVSYEQY | 0.666834 |
| AGGGGGLGGAP | ASSPDWGVSYEQY | 0.7482674 |
| AGGLLYGNGGAG | ASSPDWGVSYEQY | 0.8228207 |
| AGLESIGL | ASSPDWGVSYEQY | 0.83868605 |
| AGLKTGII | ASSPDWGVSYEQY | 0.7466583 |
| ASLAALLL | ASSPDWGVSYEQY | 0.83896935 |
| ASVINISSI | ASSPDWGVSYEQY | 0.7210603 |
| ATPPGPA | ASSPDWGVSYEQY | 0.08664916 |
| ATVASALV | ASSPDWGVSYEQY | 0.64566475 |
| ATVASLIL | ASSPDWGVSYEQY | 0.574975 |
| AVNGPVLIRH | ASSPDWGVSYEQY | 0.86805093 |
| DGAAGAVGI | ASSPDWGVSYEQY | 0.60792536 |
| DPLLHGI | ASSPDWGVSYEQY | 0.7570939 |
| ELGAALGAL | ASSPDWGVSYEQY | 0.79861295 |
| ELGRTQAV | ASSPDWGVSYEQY | 0.82041216 |
| FVAGGYLS | ASSPDWGVSYEQY | 0.79174095 |
| GASGGAGGAGGW | ASSPDWGVSYEQY | 0.68478787 |
| GGAGAGGDGGAGGI | ASSPDWGVSYEQY | 0.21269591 |
| GTGHNGSQPGMGGNG | ASSPDWGVSYEQY | 0.82746756 |
| GYAAASASAS | ASSPDWGVSYEQY | 0.59184146 |
| HQVDVAAVQQG | ASSPDWGVSYEQY | 0.49553367 |
| IGAAIAHT | ASSPDWGVSYEQY | 0.6303332 |
| IGMIEVAQL | ASSPDWGVSYEQY | 0.78352827 |
| IRLAPEHTSAMD | ASSPDWGVSYEQY | 0.277245 |
| IYNVSVL | ASSPDWGVSYEQY | 0.8716886 |
| KQGFLGM | ASSPDWGVSYEQY | 0.51956433 |
| LESVITQL | ASSPDWGVSYEQY | 0.7217942 |

| | | |
|---|---|---|
| LGLGALVVV | ASSPDWGVSYEQY | 0.87715375 |
| LIGNGGNGGSGGI | ASSPDWGVSYEQY | 0.77780974 |
| LNAINAPTL | ASSPDWGVSYEQY | 0.7729669 |
| LQLLNTGV | ASSPDWGVSYEQY | 0.5817028 |
| LQTSIEF | ASSPDWGVSYEQY | 0.7018201 |
| MSFAVSEL | ASSPDWGVSYEQY | 0.641997 |
| NTGALGL | ASSPDWGVSYEQY | 0.59450686 |
| PPGAGGAGGAGTT | ASSPDWGVSYEQY | 0.86080694 |
| QLVGLGI | ASSPDWGVSYEQY | 0.84503317 |
| RFLEQVHQL | ASSPDWGVSYEQY | 0.63873434 |
| RGVGYVLR | ASSPDWGVSYEQY | 0.87361515 |
| SATGVFAGVI | ASSPDWGVSYEQY | 0.872716 |
| SLPAITLP | ASSPDWGVSYEQY | 0.88029027 |
| STALAFTV | ASSPDWGVSYEQY | 0.6699553 |
| SVLTVGLGK | ASSPDWGVSYEQY | 0.63330126 |
| TALAGAGSGI | ASSPDWGVSYEQY | 0.7636151 |
| TGLPFTGI | ASSPDWGVSYEQY | 0.7417842 |
| VAEHVTV | ASSPDWGVSYEQY | 0.51308167 |
| VDIVLNSL | ASSPDWGVSYEQY | 0.6104137 |
| VLTGGGAVL | ASSPDWGVSYEQY | 0.8801012 |
| VNVGLAYGI | ASSPDWGVSYEQY | 0.88097024 |
| VSSAAVGAV | ASSPDWGVSYEQY | 0.7286443 |
| VTSVYEAGV | ASSPDWGVSYEQY | 0.32947975 |
| WVALDRGA | ASSPDWGVSYEQY | 0.7510178 |
| YLVGSVVI | ASSPDWGVSYEQY | 0.6567497 |
| GPPGTGKTLL | ASSPDWGVSYEQY | 0.72858185 |
| GVGMTKF | ASSPDWGVSYEQY | 0.7291181 |
| IGIGGSDL | ASSPDWGVSYEQY | 0.55531347 |
| IGTLLAEL | ASSPDWGVSYEQY | 0.78419614 |
| IYTSGTT | ASSPDWGVSYEQY | 0.8721839 |
| LSFPAGV | ASSPDWGVSYEQY | 0.7624761 |
| LYTSGTT | ASSPDWGVSYEQY | 0.81355476 |
| VGAFPFGF | ASSPDWGVSYEQY | 0.87500983 |
| VGAGTVEF | ASSPDWGVSYEQY | 0.8394431 |
| VGKSTLLL | ASSPDWGVSYEQY | 0.87290645 |
| AALAHAGGGIV | ASSPDWGVSYEQY | 0.8491554 |
| AALVAVSLL | ASSPDWGVSYEQY | 0.7747644 |
| AAVSRAVAAL | ASSPDWGVSYEQY | 0.781973 |
| AGGSSGIGGF | ASSPDWGVSYEQY | 0.8466878 |
| AGLTVANANG | ASSPDWGVSYEQY | 0.7754785 |
| AGLVGAVL | ASSPDWGVSYEQY | 0.81241184 |
| ALSQAFTV | ASSPDWGVSYEQY | 0.8245798 |
| ALTFSVSL | ASSPDWGVSYEQY | 0.84503484 |
| ALTGAALVAL | ASSPDWGVSYEQY | 0.76649857 |
| ASFSIGAGVL | ASSPDWGVSYEQY | 0.8744479 |
| DGSVYKI | ASSPDWGVSYEQY | 0.18936417 |
| EQLAEIV | ASSPDWGVSYEQY | 0.47006172 |
| FAVSELTVM | ASSPDWGVSYEQY | 0.8592646 |
| GAAPLAAGL | ASSPDWGVSYEQY | 0.1260654 |
| GADGGDGGSGGAGG | ASSPDWGVSYEQY | 0.43138987 |
| GFGLDTGGAGG | ASSPDWGVSYEQY | 0.843477 |

| | | |
|---|---|---|
| GLASGGLL | ASSPDWGVSYEQY | 0.8501791 |
| GRIAGAAVLL | ASSPDWGVSYEQY | 0.12852955 |
| GSAGSLNTSF | ASSPDWGVSYEQY | 0.6210057 |
| GTGQIGI | ASSPDWGVSYEQY | 0.35896182 |
| GTLIALIL | ASSPDWGVSYEQY | 0.82881796 |
| IAATMGFVVG | ASSPDWGVSYEQY | 0.77071846 |
| IFAQALL | ASSPDWGVSYEQY | 0.33410817 |
| IGAAVALGF | ASSPDWGVSYEQY | 0.85841703 |
| IGLTPLAL | ASSPDWGVSYEQY | 0.6132387 |
| ILGSNTSTL | ASSPDWGVSYEQY | 0.7288729 |
| IVFSLTVL | ASSPDWGVSYEQY | 0.87114227 |
| LASKATGF | ASSPDWGVSYEQY | 0.8795738 |
| LASVALIL | ASSPDWGVSYEQY | 0.56885797 |
| LFAVGLTL | ASSPDWGVSYEQY | 0.8413638 |
| LFNGGIGTYI | ASSPDWGVSYEQY | 0.74443513 |
| LTLGAGII | ASSPDWGVSYEQY | 0.88760054 |
| LVAAALGDI | ASSPDWGVSYEQY | 0.74300927 |
| NTAFVQLGI | ASSPDWGVSYEQY | 0.3946138 |
| QLFSGAEL | ASSPDWGVSYEQY | 0.85986876 |
| SAATFITL | ASSPDWGVSYEQY | 0.8447169 |
| SNYGLANLGV | ASSPDWGVSYEQY | 0.7569594 |
| SYGGAYAVM | ASSPDWGVSYEQY | 0.8474889 |
| TTGVGGLTLG | ASSPDWGVSYEQY | 0.7685447 |
| TTQTIYI | ASSPDWGVSYEQY | 0.7799715 |
| VAAGAGLGAVI | ASSPDWGVSYEQY | 0.8520225 |
| VAAVVTLL | ASSPDWGVSYEQY | 0.8387052 |
| VGFLGGLI | ASSPDWGVSYEQY | 0.58099616 |
| VGVIPILL | ASSPDWGVSYEQY | 0.8863539 |
| VIFASSLI | ASSPDWGVSYEQY | 0.75356203 |
| VKGGDGGAAAG | ASSPDWGVSYEQY | 0.5371553 |
| VTKTFTNI | ASSPDWGVSYEQY | 0.8891795 |
| VVAAAILL | ASSPDWGVSYEQY | 0.814911 |
| YAALDVEL | ASSPDWGVSYEQY | 0.8594296 |
| LASVALLL | ASSPDWGVSYEQY | 0.8562175 |
| LDAVHRV | ASSPDWGVSYEQY | 0.86254275 |
| QGLSSSIF | ASSPDWGVSYEQY | 0.47669536 |
| RGTFANIRL | ASSPDWGVSYEQY | 0.7742921 |
| RQLWWGHRI | ASSPDWGVSYEQY | 0.77292615 |
| YGPPGTGKTLL | ASSPDWGVSYEQY | 0.8527181 |
| AALVGAPANVA | ASSPDWGVSYEQY | 0.7709813 |
| AATFRRTE | ASSPDWGVSYEQY | 0.6515806 |
| AGALGGGI | ASSPDWGVSYEQY | 0.8554054 |
| AGGAGGAGGIGAG | ASSPDWGVSYEQY | 0.8746538 |
| AGSGALGL | ASSPDWGVSYEQY | 0.8160466 |
| AQIADQI | ASSPDWGVSYEQY | 0.80642664 |
| ASAVAVTYL | ASSPDWGVSYEQY | 0.8857455 |
| ASTVLGGTGG | ASSPDWGVSYEQY | 0.64376426 |
| AVGGEPAK | ASSPDWGVSYEQY | 0.7887323 |
| CEQGTFGGTC | ASSPDWGVSYEQY | 0.86944544 |
| DEGKAPAG | ASSPDWGVSYEQY | 0.86790013 |
| DTVVFPAL | ASSPDWGVSYEQY | 0.821728 |

| | | |
|---|---|---|
| FEGITMAG | ASSPDWGVSYEQY | 0.8722073 |
| FPLVVQD | ASSPDWGVSYEQY | 0.6428117 |
| GAFTTPQ | ASSPDWGVSYEQY | 0.8090799 |
| GFGGFGTAGGVGGA | ASSPDWGVSYEQY | 0.68846816 |
| GFLAQFL | ASSPDWGVSYEQY | 0.68791723 |
| GGAGGAGGNAGVA | ASSPDWGVSYEQY | 0.47877795 |
| GGVGTAGAGGAGGAGG | ASSPDWGVSYEQY | 0.80426264 |
| GNGGGGDAGNAGSGGN | ASSPDWGVSYEQY | 0.8497549 |
| GTGGAGGAGNDAGS | ASSPDWGVSYEQY | 0.7998855 |
| GTTNTGWFNS | ASSPDWGVSYEQY | 0.2831259 |
| IFMGPLFTL | ASSPDWGVSYEQY | 0.82922924 |
| ILAHAIEAI | ASSPDWGVSYEQY | 0.81706405 |
| IRSAIREAL | ASSPDWGVSYEQY | 0.88231343 |
| ITFAGGG | ASSPDWGVSYEQY | 0.78208566 |
| ITGMSGHQEV | ASSPDWGVSYEQY | 0.8540864 |
| IVAGLACTL | ASSPDWGVSYEQY | 0.66401386 |
| IVGGGLAGA | ASSPDWGVSYEQY | 0.82633567 |
| IVTGVLNI | ASSPDWGVSYEQY | 0.5487857 |
| LAQQGCI | ASSPDWGVSYEQY | 0.50730145 |
| LCLHVEN | ASSPDWGVSYEQY | 0.7649182 |
| LDAISDGS | ASSPDWGVSYEQY | 0.69278055 |
| LGGFAAML | ASSPDWGVSYEQY | 0.83303726 |
| LGGKNPMI | ASSPDWGVSYEQY | 0.86430776 |
| LGPKGVAAIL | ASSPDWGVSYEQY | 0.8139378 |
| LMVGVSI | ASSPDWGVSYEQY | 0.8516336 |
| LVGAAFVL | ASSPDWGVSYEQY | 0.8751825 |
| LVLAVAPL | ASSPDWGVSYEQY | 0.8765106 |
| MAAIAVIDAA | ASSPDWGVSYEQY | 0.7393958 |
| NNGINAELQ | ASSPDWGVSYEQY | 0.67282957 |
| RGEPLVLVHG | ASSPDWGVSYEQY | 0.4302767 |
| RGGAGGAAGLI | ASSPDWGVSYEQY | 0.74606377 |
| RSYRMHPAV | ASSPDWGVSYEQY | 0.7233174 |
| SAAHLLAGQLA | ASSPDWGVSYEQY | 0.8144082 |
| SAAVGSY | ASSPDWGVSYEQY | 0.12149944 |
| SPHAVLHRI | ASSPDWGVSYEQY | 0.79117244 |
| SQIAKVL | ASSPDWGVSYEQY | 0.8594833 |
| TALGHLVP | ASSPDWGVSYEQY | 0.59855986 |
| TAPTTSGAA | ASSPDWGVSYEQY | 0.83041257 |
| TDGDDGNGGSPGT | ASSPDWGVSYEQY | 0.8254984 |
| TDGLGEN | ASSPDWGVSYEQY | 0.73157144 |
| TGGPPLPGQA | ASSPDWGVSYEQY | 0.8212849 |
| TIPAITI | ASSPDWGVSYEQY | 0.7195913 |
| TIPGFTI | ASSPDWGVSYEQY | 0.8634393 |
| TSIASGALA | ASSPDWGVSYEQY | 0.8720706 |
| TVTLPGAAI | ASSPDWGVSYEQY | 0.8350556 |
| VASGLSAALP | ASSPDWGVSYEQY | 0.7654947 |
| VGAGVIGI | ASSPDWGVSYEQY | 0.8301306 |
| VGPKRFGV | ASSPDWGVSYEQY | 0.43149137 |
| VNVAFPD | ASSPDWGVSYEQY | 0.79732645 |
| VSMTNSGVS | ASSPDWGVSYEQY | 0.31151623 |
| VTDVLGDLSV | ASSPDWGVSYEQY | 0.8116703 |

| | | |
|---|---|---|
| VVVVGAGFAGL | ASSPDWGVSYEQY | 0.6945904 |
| VYTVGTVV | ASSPDWGVSYEQY | 0.8073787 |
| WTTFSGP | ASSPDWGVSYEQY | 0.74004847 |
| YDELVEAF | ASSPDWGVSYEQY | 0.8489586 |
| YGTGGAGGNGG | ASSPDWGVSYEQY | 0.68192756 |
| YTNVMAV | ASSPDWGVSYEQY | 0.8333397 |
| FLEQVHQL | ASSPDWGVSYEQY | 0.7561652 |
| GLSLLAVAV | ASSPDWGVSYEQY | 0.3761525 |
| LEQVHQL | ASSPDWGVSYEQY | 0.85938424 |
| LNAAGAI | ASSPDWGVSYEQY | 0.8351638 |
| PAVHASL | ASSPDWGVSYEQY | 0.7421392 |
| TELEDDA | ASSPDWGVSYEQY | 0.85735756 |
| TPPGPPN | ASSPDWGVSYEQY | 0.82380253 |
| TPPPPPPP | ASSPDWGVSYEQY | 0.48935077 |
| TQSPALI | AWRPGVGENYAEQF | 0.6366141 |
| EFAAAFPQ | AWRPGVGENYAEQF | 0.80454385 |
| LLSGNLDVL | AWRPGVGENYAEQF | 0.7846614 |
| GPANLGNYNI | AWRPGVGENYAEQF | 0.14099042 |
| TLGPTLP | AWRPGVGENYAEQF | 0.8345977 |
| VYAYSSCCK | AWRPGVGENYAEQF | 0.516057 |
| APAPAPAPNI | AWRPGVGENYAEQF | 0.68705976 |
| PGGPVPIP | AWRPGVGENYAEQF | 0.78146994 |
| FLTVALEP | AWRPGVGENYAEQF | 0.85864425 |
| SQYFGANL | AWRPGVGENYAEQF | 0.6147752 |
| GGVKPAA | AWRPGVGENYAEQF | 0.681068 |
| GGTGGAGGAGGW | AWRPGVGENYAEQF | 0.8479496 |
| GVGGVAGLGGAGG | AWRPGVGENYAEQF | 0.76706517 |
| VLLVLATTI | AWRPGVGENYAEQF | 0.8284321 |
| IGIFAAML | AWRPGVGENYAEQF | 0.73000205 |
| AVGDLID | AWRPGVGENYAEQF | 0.81071746 |
| NISPLPL | AWRPGVGENYAEQF | 0.81145763 |
| VTAVFVAI | AWRPGVGENYAEQF | 0.601635 |
| SGDDNDAAL | AWRPGVGENYAEQF | 0.6714306 |
| AVAALAHS | AWRPGVGENYAEQF | 0.637216 |
| GGVTVMATP | AWRPGVGENYAEQF | 0.6341589 |
| AGLAANVRD | AWRPGVGENYAEQF | 0.5741032 |
| TNRPDLIDPALLRPG | AWRPGVGENYAEQF | 0.82996136 |
| LANLGVTAL | AWRPGVGENYAEQF | 0.8893672 |
| TGERIEL | AWRPGVGENYAEQF | 0.741408 |
| NGEELAA | AWRPGVGENYAEQF | 0.43046808 |
| VLSLVWG | AWRPGVGENYAEQF | 0.76231575 |
| IGYALWL | AWRPGVGENYAEQF | 0.7261181 |
| LSLLLAPL | AWRPGVGENYAEQF | 0.83808285 |
| TLDEALK | AWRPGVGENYAEQF | 0.84471524 |
| TDGAGTAEGG | AWRPGVGENYAEQF | 0.8686514 |
| NAPGGAGGNGG | AWRPGVGENYAEQF | 0.83383566 |
| VSVFMGMGGNF | AWRPGVGENYAEQF | 0.6923498 |
| LASYLAAL | AWRPGVGENYAEQF | 0.8688207 |
| LVGGTPAL | AWRPGVGENYAEQF | 0.78068674 |
| VRNADVL | AWRPGVGENYAEQF | 0.7227008 |
| TCSSPFLVSAM | AWRPGVGENYAEQF | 0.8210076 |

| | | |
|---|---|---|
| VHITDAE | AWRPGVGENYAEQF | 0.83996046 |
| EYAAPSF | AWRPGVGENYAEQF | 0.7076757 |
| VVDAFHGGL | AWRPGVGENYAEQF | 0.7755765 |
| ASVVDPQ | AWRPGVGENYAEQF | 0.5075736 |
| SYNEFGGLS | AWRPGVGENYAEQF | 0.86003226 |
| DPDGSVY | AWRPGVGENYAEQF | 0.27365527 |
| MVSTIHATE | AWRPGVGENYAEQF | 0.56764907 |
| LAAGVADPV | AWRPGVGENYAEQF | 0.7403737 |
| TKEQIAA | AWRPGVGENYAEQF | 0.87300396 |
| SLASPAAL | AWRPGVGENYAEQF | 0.8608654 |
| IGLVFYGV | AWRPGVGENYAEQF | 0.7090587 |
| FLLDASNL | AWRPGVGENYAEQF | 0.6873656 |
| YQVGNFP | AWRPGVGENYAEQF | 0.8264627 |
| ESLLSQST | AWRPGVGENYAEQF | 0.62226146 |
| LFTVGFAAVG | AWRPGVGENYAEQF | 0.86962134 |
| ATPPTPT | AWRPGVGENYAEQF | 0.7644379 |
| TDLATLG | AWRPGVGENYAEQF | 0.6977523 |
| SAGGTGR | AWRPGVGENYAEQF | 0.7099763 |
| GFSNSGS | AWRPGVGENYAEQF | 0.83599436 |
| GGNGGSGGTSV | AWRPGVGENYAEQF | 0.6883929 |
| GTSVATGGAGN | AWRPGVGENYAEQF | 0.8558252 |
| NGGNGGSGGTS | AWRPGVGENYAEQF | 0.17207313 |
| AVVKFPH | AWRPGVGENYAEQF | 0.84369195 |
| TPPPPPP | AWRPGVGENYAEQF | 0.5699078 |
| AAPAAAAAPA | AWRPGVGENYAEQF | 0.65643585 |
| IALNEDL | AWRPGVGENYAEQF | 0.28555307 |
| IVNVSSM | AWRPGVGENYAEQF | 0.78804076 |
| TGIPPAPR | AWRPGVGENYAEQF | 0.826193 |
| FDLGGGTF | AWRPGVGENYAEQF | 0.8716147 |
| FDLGGGTFDVS | AWRPGVGENYAEQF | 0.7907464 |
| NEPTAAA | AWRPGVGENYAEQF | 0.8298491 |
| NEPTAAALA | AWRPGVGENYAEQF | 0.85603976 |
| GSIINISSI | AWRPGVGENYAEQF | 0.7398136 |
| VAPGNAGTA | AWRPGVGENYAEQF | 0.24395846 |
| TGGGASLE | AWRPGVGENYAEQF | 0.8184673 |
| VAVYNHYKRI | AWRPGVGENYAEQF | 0.7459638 |
| TDPAPATT | AWRPGVGENYAEQF | 0.8613357 |
| TNRPDILDPALLRPG | AWRPGVGENYAEQF | 0.82419413 |
| LTTTTTS | AWRPGVGENYAEQF | 0.5918368 |
| PTGSGKTL | AWRPGVGENYAEQF | 0.81182796 |
| TGSGKTL | AWRPGVGENYAEQF | 0.80118906 |
| TIGHVDHGK | AWRPGVGENYAEQF | 0.548375 |
| TIGHVDHGKTT | AWRPGVGENYAEQF | 0.7195308 |
| GHVDHGKTT | AWRPGVGENYAEQF | 0.6402657 |
| AQFEHTLL | AWRPGVGENYAEQF | 0.42435712 |
| IVNQDEL | AWRPGVGENYAEQF | 0.8078507 |
| TNGSQFF | AWRPGVGENYAEQF | 0.7339611 |
| TGKGIVE | AWRPGVGENYAEQF | 0.65546525 |
| LISNASDA | AWRPGVGENYAEQF | 0.76939034 |
| AELRTLGELTKIAKAHGAQ | AWRPGVGENYAEQF | 0.6492486 |
| APNAVAAAEPSW | AWRPGVGENYAEQF | 0.64280504 |

| | | |
|---|---|---|
| APQVAGAATAVL | AWRPGVGENYAEQF | 0.6100134 |
| ARRDPAAAAAILPTDA | AWRPGVGENYAEQF | 0.79118234 |
| DAAARDVLAEAGLAEYF | AWRPGVGENYAEQF | 0.85449624 |
| DDQIVLSAKANLAKVEQ | AWRPGVGENYAEQF | 0.68317974 |
| DELLPDTGEDSDEDGH | AWRPGVGENYAEQF | 0.8182562 |
| ELPEAPANRATVPQD | AWRPGVGENYAEQF | 0.8625719 |
| FGQSKIERVTGIG | AWRPGVGENYAEQF | 0.31755123 |
| GAGGTGSTAGGAGGAGGAGGLY | AWRPGVGENYAEQF | 0.5684089 |
| GHGGTGAAGGNGGTGGAGGSGI | AWRPGVGENYAEQF | 0.83835787 |
| GNGGNGGAGGEGGPGGAGGAGA | AWRPGVGENYAEQF | 0.6299045 |
| GSTAGNGGNGGNATNGTIAGQP | AWRPGVGENYAEQF | 0.86183304 |
| GTGGDGGNGGNGGTGG | AWRPGVGENYAEQF | 0.7422844 |
| GVRRPAGIPADAPV | AWRPGVGENYAEQF | 0.82834625 |
| GYRQVTDAHLLTLARR | AWRPGVGENYAEQF | 0.7365278 |
| IAKMGQDGHDRG | AWRPGVGENYAEQF | 0.8824119 |
| KAEPSIEIQYT | AWRPGVGENYAEQF | 0.84430504 |
| KPVRIRMAADEDTVAVT | AWRPGVGENYAEQF | 0.77437043 |
| MPPRIAGMRLLVIKPEP | AWRPGVGENYAEQF | 0.83009857 |
| MKFYAVALFRSRGVPPTR | AWRPGVGENYAEQF | 0.45356542 |
| NAGGFGILGAVAHSP | AWRPGVGENYAEQF | 0.8374642 |
| NEGREITLAMAEM | AWRPGVGENYAEQF | 0.8596176 |
| NELVEWSHPLGHRWRWE | AWRPGVGENYAEQF | 0.61520946 |
| NLGAGGTGGNGGNPG | AWRPGVGENYAEQF | 0.7745929 |
| NTEWPKWETVMAADPVD | AWRPGVGENYAEQF | 0.88521993 |
| PASAAPAETQP | AWRPGVGENYAEQF | 0.87122643 |
| QQSSDGKNGVRAAEITGE | AWRPGVGENYAEQF | 0.6588509 |
| RAPDAVTTSWWKE | AWRPGVGENYAEQF | 0.7200791 |
| SGFGNVGAHTSGWFNQ | AWRPGVGENYAEQF | 0.46351033 |
| SGVAGAVGQAASVGGLKV | AWRPGVGENYAEQF | 0.715788 |
| SNGGAGTGSAGGNGGT | AWRPGVGENYAEQF | 0.292776 |
| TGGAGGEGGAAG | AWRPGVGENYAEQF | 0.8474488 |
| TGGDGGAAGAGGAPGHG | AWRPGVGENYAEQF | 0.7634139 |
| TMPPLLAAIANAGRHGV | AWRPGVGENYAEQF | 0.8434719 |
| TSSTAGNSGGA | AWRPGVGENYAEQF | 0.8372638 |
| VPATVPVGILC | AWRPGVGENYAEQF | 0.8727661 |
| VQRPDSGGVVLTGRIS | AWRPGVGENYAEQF | 0.7717788 |
| WTPNPVPATKGVPAGLVQAT | AWRPGVGENYAEQF | 0.6805413 |
| YALKRALAKFSDRFDVVII | AWRPGVGENYAEQF | 0.5398009 |
| YLIGSVIV | AWRPGVGENYAEQF | 0.27688548 |
| YGQLVAVID | AWRPGVGENYAEQF | 0.6769282 |
| VTVGGMTLPA | AWRPGVGENYAEQF | 0.6278854 |
| VLALPAV | AWRPGVGENYAEQF | 0.19494207 |
| VIPPTLNYET | AWRPGVGENYAEQF | 0.87018764 |
| VGMGGMHPSSG | AWRPGVGENYAEQF | 0.76216155 |
| VGLAGGAATA | AWRPGVGENYAEQF | 0.84662724 |
| VGGAGGAGGSGP | AWRPGVGENYAEQF | 0.7409152 |
| VFAAFMV | AWRPGVGENYAEQF | 0.48935962 |
| TLYDKLR | AWRPGVGENYAEQF | 0.71169806 |
| TLAVFTVL | AWRPGVGENYAEQF | 0.83868426 |
| TGSVQPARV | AWRPGVGENYAEQF | 0.5226337 |
| TGGGAGSGGT | AWRPGVGENYAEQF | 0.46378648 |

| | | |
|---|---|---|
| SHLETSL | AWRPGVGENYAEQF | 0.60281575 |
| SFTTGMT | AWRPGVGENYAEQF | 0.7478671 |
| QVEVAKRMGVRQPTVS | AWRPGVGENYAEQF | 0.8842091 |
| QGSGGPALAM | AWRPGVGENYAEQF | 0.8645495 |
| MLAPSNP | AWRPGVGENYAEQF | 0.81698245 |
| LVNNAAI | AWRPGVGENYAEQF | 0.8909541 |
| LTGEINTLA | AWRPGVGENYAEQF | 0.15630965 |
| LSIFGPV | AWRPGVGENYAEQF | 0.7411277 |
| LLALLANL | AWRPGVGENYAEQF | 0.86351347 |
| LISNASDALDKLR | AWRPGVGENYAEQF | 0.83793575 |
| LINEAVP | AWRPGVGENYAEQF | 0.7603858 |
| LGYAFSGL | AWRPGVGENYAEQF | 0.87272173 |
| LDFVAVAL | AWRPGVGENYAEQF | 0.58727956 |
| LAQPGAE | AWRPGVGENYAEQF | 0.78167665 |
| ISNLGAVTGF | AWRPGVGENYAEQF | 0.7850324 |
| HQEDPGRI | AWRPGVGENYAEQF | 0.86567867 |
| GTPGEPTP | AWRPGVGENYAEQF | 0.8167953 |
| GFFNSGS | AWRPGVGENYAEQF | 0.42470545 |
| GAGGGGLTAGAN | AWRPGVGENYAEQF | 0.85186136 |
| FVDALVNM | AWRPGVGENYAEQF | 0.8842609 |
| FLTVAPEL | AWRPGVGENYAEQF | 0.7280331 |
| FETLTAL | AWRPGVGENYAEQF | 0.8306075 |
| DVINPIEL | AWRPGVGENYAEQF | 0.81697917 |
| APVSGAPV | AWRPGVGENYAEQF | 0.5484164 |
| ALVGGLAW | AWRPGVGENYAEQF | 0.7191614 |
| ALAAAAATAI | AWRPGVGENYAEQF | 0.6100855 |
| AFGLGLLL | AWRPGVGENYAEQF | 0.8272388 |
| AFFAAASL | AWRPGVGENYAEQF | 0.55789906 |
| AAAALPAFSPP | AWRPGVGENYAEQF | 0.80098724 |
| VVLAPAL | AWRPGVGENYAEQF | 0.80577624 |
| TIGHVDHGKTTL | AWRPGVGENYAEQF | 0.89108306 |
| LISNASDALDK | AWRPGVGENYAEQF | 0.83920175 |
| LISNASDAL | AWRPGVGENYAEQF | 0.84082884 |
| ISNASDALDK | AWRPGVGENYAEQF | 0.61406964 |
| NQAETLVYQTE | AWRPGVGENYAEQF | 0.8464893 |
| TGGTGGTAGAG | AWRPGVGENYAEQF | 0.8163804 |
| TGSTAGGAGGA | AWRPGVGENYAEQF | 0.5233587 |
| DPLAVPVAAGPV | AWRPGVGENYAEQF | 0.86036897 |
| GGTGGTPGNGGN | AWRPGVGENYAEQF | 0.59502625 |
| PAGLAGVDPATL | AWRPGVGENYAEQF | 0.8051244 |
| GNNLVVEAPHAGASV | AWRPGVGENYAEQF | 0.7567492 |
| KVTRIMSASTLEQPA | AWRPGVGENYAEQF | 0.85136074 |
| LGFALPIALPRRTVV | AWRPGVGENYAEQF | 0.8184892 |
| TRLGNLTARKLVVLV | AWRPGVGENYAEQF | 0.79480433 |
| WSQQRRLIYFPSAGP | AWRPGVGENYAEQF | 0.5634214 |
| GPNGNPDPMAAAVDIR | AWRPGVGENYAEQF | 0.86219144 |
| TQRCVSVADLRAAASR | AWRPGVGENYAEQF | 0.8694288 |
| QFVLDVAEHPVEVLTETV | AWRPGVGENYAEQF | 0.76317656 |
| GGAGGTGGAGGNGGTQAGDG | AWRPGVGENYAEQF | 0.8848438 |
| PLLATLRAIAVDVDAKPAQV | AWRPGVGENYAEQF | 0.78833044 |
| APKFRTGGYMIDFWGVGYQVAK | AWRPGVGENYAEQF | 0.2714481 |

| | | |
|---|---|---|
| GAGGAGGAGGAAGGPGGTGGTG | AWRPGVGENYAEQF | 0.76865053 |
| IGALGLLATDVAPAGSVAFGVA | AWRPGVGENYAEQF | 0.86254835 |
| AGGAGGIGGNGGAGGAGGVLMG | AWRPGVGENYAEQF | 0.8561833 |
| ADELIGLL | AWRPGVGENYAEQF | 0.8425992 |
| ADRALKA | AWRPGVGENYAEQF | 0.8495045 |
| AGAVGQVGGAGGA | AWRPGVGENYAEQF | 0.7613044 |
| AGGGGGLGGAP | AWRPGVGENYAEQF | 0.8719446 |
| AGGLLYGNGGAG | AWRPGVGENYAEQF | 0.8947792 |
| AGLESIGL | AWRPGVGENYAEQF | 0.86069816 |
| AGLKTGII | AWRPGVGENYAEQF | 0.72167635 |
| ASLAALLL | AWRPGVGENYAEQF | 0.5954106 |
| ASVINISSI | AWRPGVGENYAEQF | 0.85156286 |
| ATPPGPA | AWRPGVGENYAEQF | 0.8543452 |
| ATVASALV | AWRPGVGENYAEQF | 0.6391928 |
| ATVASLIL | AWRPGVGENYAEQF | 0.77016866 |
| AVNGPVLIRH | AWRPGVGENYAEQF | 0.73249304 |
| DGAAGAVGI | AWRPGVGENYAEQF | 0.2386562 |
| DPLLHGI | AWRPGVGENYAEQF | 0.8246899 |
| ELGAALGAL | AWRPGVGENYAEQF | 0.86835754 |
| ELGRTQAV | AWRPGVGENYAEQF | 0.8611909 |
| FVAGGYLS | AWRPGVGENYAEQF | 0.38848072 |
| GASGGAGGAGGW | AWRPGVGENYAEQF | 0.86432135 |
| GGAGAGGDGGAGGI | AWRPGVGENYAEQF | 0.86297846 |
| GTGHNGSQPGMGGNG | AWRPGVGENYAEQF | 0.8849569 |
| GYAAASASAS | AWRPGVGENYAEQF | 0.3791649 |
| HQVDVAAVQQG | AWRPGVGENYAEQF | 0.19443911 |
| IGAAIAHT | AWRPGVGENYAEQF | 0.78299516 |
| IGMIEVAQL | AWRPGVGENYAEQF | 0.8197038 |
| IRLAPEHTSAMD | AWRPGVGENYAEQF | 0.63658947 |
| IYNVSVL | AWRPGVGENYAEQF | 0.63402915 |
| KQGFLGM | AWRPGVGENYAEQF | 0.7057419 |
| LESVITQL | AWRPGVGENYAEQF | 0.8616892 |
| LGLGALVVV | AWRPGVGENYAEQF | 0.57378244 |
| LIGNGGNGGSGGI | AWRPGVGENYAEQF | 0.59317195 |
| LNAINAPTL | AWRPGVGENYAEQF | 0.79751384 |
| LQLLNTGV | AWRPGVGENYAEQF | 0.87690735 |
| LQTSIEF | AWRPGVGENYAEQF | 0.64718425 |
| MSFAVSEL | AWRPGVGENYAEQF | 0.7690415 |
| NTGALGL | AWRPGVGENYAEQF | 0.76103604 |
| PPGAGGAGGAGTT | AWRPGVGENYAEQF | 0.72734773 |
| QLVGLGI | AWRPGVGENYAEQF | 0.7825955 |
| RFLEQVHQL | AWRPGVGENYAEQF | 0.51463974 |
| RGVGYVLR | AWRPGVGENYAEQF | 0.84424704 |
| SATGVFAGVI | AWRPGVGENYAEQF | 0.62768936 |
| SLPAITLP | AWRPGVGENYAEQF | 0.8575622 |
| STALAFTV | AWRPGVGENYAEQF | 0.8023989 |
| SVLTVGLGK | AWRPGVGENYAEQF | 0.84780324 |
| TALAGAGSGI | AWRPGVGENYAEQF | 0.75417393 |
| TGLPFTGI | AWRPGVGENYAEQF | 0.6160581 |
| VAEHVTV | AWRPGVGENYAEQF | 0.56431174 |
| VDIVLNSL | AWRPGVGENYAEQF | 0.5839969 |

| | | |
|---|---|---|
| VLTGGGAVL | AWRPGVGENYAEQF | 0.86448246 |
| VNVGLAYGI | AWRPGVGENYAEQF | 0.500422 |
| VSSAAVGAV | AWRPGVGENYAEQF | 0.8674075 |
| VTSVYEAGV | AWRPGVGENYAEQF | 0.82924235 |
| WVALDRGA | AWRPGVGENYAEQF | 0.6358489 |
| YLVGSVVI | AWRPGVGENYAEQF | 0.85884583 |
| GPPGTGKTLL | AWRPGVGENYAEQF | 0.3735006 |
| GVGMTKF | AWRPGVGENYAEQF | 0.8211001 |
| IGIGGSDL | AWRPGVGENYAEQF | 0.7358366 |
| IGTLLAEL | AWRPGVGENYAEQF | 0.8533893 |
| IYTSGTT | AWRPGVGENYAEQF | 0.7781141 |
| LSFPAGV | AWRPGVGENYAEQF | 0.85168314 |
| LYTSGTT | AWRPGVGENYAEQF | 0.83748275 |
| VGAFPFGF | AWRPGVGENYAEQF | 0.42434612 |
| VGAGTVEF | AWRPGVGENYAEQF | 0.8498368 |
| VGKSTLLL | AWRPGVGENYAEQF | 0.8490202 |
| AALAHAGGGIV | AWRPGVGENYAEQF | 0.8589648 |
| AALVAVSLL | AWRPGVGENYAEQF | 0.40231532 |
| AAVSRAVAAL | AWRPGVGENYAEQF | 0.7977506 |
| AGGSSGIGGF | AWRPGVGENYAEQF | 0.69603825 |
| AGLTVANANG | AWRPGVGENYAEQF | 0.82305866 |
| AGLVGAVL | AWRPGVGENYAEQF | 0.80305845 |
| ALSQAFTV | AWRPGVGENYAEQF | 0.7177555 |
| ALTFSVSL | AWRPGVGENYAEQF | 0.8466749 |
| ALTGAALVAL | AWRPGVGENYAEQF | 0.31926057 |
| ASFSIGAGVL | AWRPGVGENYAEQF | 0.7194445 |
| DGSVYKI | AWRPGVGENYAEQF | 0.7656603 |
| EQLAEIV | AWRPGVGENYAEQF | 0.78667027 |
| FAVSELTVM | AWRPGVGENYAEQF | 0.8537794 |
| GAAPLAAGL | AWRPGVGENYAEQF | 0.55970865 |
| GADGGDGGSGGAGG | AWRPGVGENYAEQF | 0.817306 |
| GFGLDTGGAGG | AWRPGVGENYAEQF | 0.7253637 |
| GLASGGLL | AWRPGVGENYAEQF | 0.67031395 |
| GRIAGAAVLL | AWRPGVGENYAEQF | 0.84974223 |
| GSAGSLNTSF | AWRPGVGENYAEQF | 0.6882031 |
| GTGQIGI | AWRPGVGENYAEQF | 0.40045643 |
| GTLIALIL | AWRPGVGENYAEQF | 0.8558169 |
| IAATMGFVVG | AWRPGVGENYAEQF | 0.52334833 |
| IFAQALL | AWRPGVGENYAEQF | 0.7968358 |
| IGAAVALGF | AWRPGVGENYAEQF | 0.84201866 |
| IGLTPLAL | AWRPGVGENYAEQF | 0.41058418 |
| ILGSNTSTL | AWRPGVGENYAEQF | 0.86294806 |
| IVFSLTVL | AWRPGVGENYAEQF | 0.7984276 |
| LASKATGF | AWRPGVGENYAEQF | 0.8039093 |
| LASVALIL | AWRPGVGENYAEQF | 0.609045 |
| LFAVGLTL | AWRPGVGENYAEQF | 0.5801488 |
| LFNGGIGTYI | AWRPGVGENYAEQF | 0.8669801 |
| LTLGAGII | AWRPGVGENYAEQF | 0.8308355 |
| LVAAALGDI | AWRPGVGENYAEQF | 0.7893288 |
| NTAFVQLGI | AWRPGVGENYAEQF | 0.8071121 |
| QLFSGAEL | AWRPGVGENYAEQF | 0.8431359 |

| | | |
|---|---|---|
| SAATFITL | AWRPGVGENYAEQF | 0.86339116 |
| SNYGLANLGV | AWRPGVGENYAEQF | 0.68918 |
| SYGGAYAVM | AWRPGVGENYAEQF | 0.45505854 |
| TTGVGGLTLG | AWRPGVGENYAEQF | 0.7365694 |
| TTQTIYI | AWRPGVGENYAEQF | 0.62059164 |
| VAAGAGLGAVI | AWRPGVGENYAEQF | 0.7173898 |
| VAAVVTLL | AWRPGVGENYAEQF | 0.45440543 |
| VGFLGGLI | AWRPGVGENYAEQF | 0.83537847 |
| VGVIPILL | AWRPGVGENYAEQF | 0.8918717 |
| VIFASSLI | AWRPGVGENYAEQF | 0.5852776 |
| VKGGDGGAAAG | AWRPGVGENYAEQF | 0.8865994 |
| VTKTFTNI | AWRPGVGENYAEQF | 0.5329861 |
| VVAAAILL | AWRPGVGENYAEQF | 0.84220576 |
| YAALDVEL | AWRPGVGENYAEQF | 0.846277 |
| LASVALLL | AWRPGVGENYAEQF | 0.56688094 |
| LDAVHRV | AWRPGVGENYAEQF | 0.77590483 |
| QGLSSSIF | AWRPGVGENYAEQF | 0.5787121 |
| RGTFANIRL | AWRPGVGENYAEQF | 0.64257133 |
| RQLWWGHRI | AWRPGVGENYAEQF | 0.8661871 |
| YGPPGTGKTLL | AWRPGVGENYAEQF | 0.48651585 |
| AALVGAPANVA | AWRPGVGENYAEQF | 0.81172466 |
| AATFRRTE | AWRPGVGENYAEQF | 0.7463753 |
| AGALGGGI | AWRPGVGENYAEQF | 0.63986444 |
| AGGAGGAGGIGAG | AWRPGVGENYAEQF | 0.85054266 |
| AGSGALGL | AWRPGVGENYAEQF | 0.8411342 |
| AQIADQI | AWRPGVGENYAEQF | 0.78086555 |
| ASAVAVTYL | AWRPGVGENYAEQF | 0.8928479 |
| ASTVLGGTGG | AWRPGVGENYAEQF | 0.6892822 |
| AVGGEPAK | AWRPGVGENYAEQF | 0.8815087 |
| CEQGTFGGTC | AWRPGVGENYAEQF | 0.87830323 |
| DEGKAPAG | AWRPGVGENYAEQF | 0.8037926 |
| DTVVFPAL | AWRPGVGENYAEQF | 0.7636038 |
| FEGITMAG | AWRPGVGENYAEQF | 0.8558295 |
| FPLVVQD | AWRPGVGENYAEQF | 0.7583624 |
| GAFTTPQ | AWRPGVGENYAEQF | 0.80595875 |
| GFGGFGTAGGVGGA | AWRPGVGENYAEQF | 0.8807806 |
| GFLAQFL | AWRPGVGENYAEQF | 0.75907457 |
| GGAGGAGGNAGVA | AWRPGVGENYAEQF | 0.8462911 |
| GGVGTAGAGGAGGAGG | AWRPGVGENYAEQF | 0.82944983 |
| GNGGGGDAGNAGSGGN | AWRPGVGENYAEQF | 0.74513364 |
| GTGGAGGAGNDAGS | AWRPGVGENYAEQF | 0.7604548 |
| GTTNTGWFNS | AWRPGVGENYAEQF | 0.34868366 |
| IFMGPLFTL | AWRPGVGENYAEQF | 0.7807344 |
| ILAHAIEAI | AWRPGVGENYAEQF | 0.69637066 |
| IRSAIREAL | AWRPGVGENYAEQF | 0.8049314 |
| ITFAGGG | AWRPGVGENYAEQF | 0.8230478 |
| ITGMSGHQEV | AWRPGVGENYAEQF | 0.7215762 |
| IVAGLACTL | AWRPGVGENYAEQF | 0.8474284 |
| IVGGGLAGA | AWRPGVGENYAEQF | 0.82124937 |
| IVTGVLNI | AWRPGVGENYAEQF | 0.66080326 |
| LAQQGCI | AWRPGVGENYAEQF | 0.7992555 |

| | | |
|---|---|---|
| LCLHVEN | AWRPGVGENYAEQF | 0.7457241 |
| LDAISDGS | AWRPGVGENYAEQF | 0.7763028 |
| LGGFAAML | AWRPGVGENYAEQF | 0.5502503 |
| LGGKNPMI | AWRPGVGENYAEQF | 0.83376 |
| LGPKGVAAIL | AWRPGVGENYAEQF | 0.7651956 |
| LMVGVSI | AWRPGVGENYAEQF | 0.17157328 |
| LVGAAFVL | AWRPGVGENYAEQF | 0.8071883 |
| LVLAVAPL | AWRPGVGENYAEQF | 0.79534036 |
| MAAIAVIDAA | AWRPGVGENYAEQF | 0.76703686 |
| NNGINAELQ | AWRPGVGENYAEQF | 0.8420979 |
| RGEPLVLVHG | AWRPGVGENYAEQF | 0.20432192 |
| RGGAGGAAGLI | AWRPGVGENYAEQF | 0.6949904 |
| RSYRMHPAV | AWRPGVGENYAEQF | 0.8432504 |
| SAAHLLAGQLA | AWRPGVGENYAEQF | 0.84616196 |
| SAAVGSY | AWRPGVGENYAEQF | 0.572654 |
| SPHAVLHRI | AWRPGVGENYAEQF | 0.8322263 |
| SQIAKVL | AWRPGVGENYAEQF | 0.8352708 |
| TALGHLVP | AWRPGVGENYAEQF | 0.8277533 |
| TAPTTSGAA | AWRPGVGENYAEQF | 0.8264454 |
| TDGDDGNGGSPGT | AWRPGVGENYAEQF | 0.8368855 |
| TDGLGEN | AWRPGVGENYAEQF | 0.70843834 |
| TGGPPLPGQA | AWRPGVGENYAEQF | 0.8436176 |
| TIPAITI | AWRPGVGENYAEQF | 0.8476082 |
| TIPGFTI | AWRPGVGENYAEQF | 0.72520065 |
| TSIASGALA | AWRPGVGENYAEQF | 0.8150303 |
| TVTLPGAAI | AWRPGVGENYAEQF | 0.68765116 |
| VASGLSAALP | AWRPGVGENYAEQF | 0.73861206 |
| VGAGVIGI | AWRPGVGENYAEQF | 0.2452931 |
| VGPKRFGV | AWRPGVGENYAEQF | 0.832893 |
| VNVAFPD | AWRPGVGENYAEQF | 0.8345635 |
| VSMTNSGVS | AWRPGVGENYAEQF | 0.79753816 |
| VTDVLGDLSV | AWRPGVGENYAEQF | 0.7407278 |
| VVVVGAGFAGL | AWRPGVGENYAEQF | 0.7549727 |
| VYTVGTVV | AWRPGVGENYAEQF | 0.7723621 |
| WTTFSGP | AWRPGVGENYAEQF | 0.759144 |
| YDELVEAF | AWRPGVGENYAEQF | 0.24682562 |
| YGTGGAGGNGG | AWRPGVGENYAEQF | 0.7650422 |
| YTNVMAV | AWRPGVGENYAEQF | 0.8425589 |
| FLEQVHQL | AWRPGVGENYAEQF | 0.8352063 |
| GLSLLAVAV | AWRPGVGENYAEQF | 0.50502825 |
| LEQVHQL | AWRPGVGENYAEQF | 0.84687984 |
| LNAAGAI | AWRPGVGENYAEQF | 0.8312048 |
| PAVHASL | AWRPGVGENYAEQF | 0.84179175 |
| TELEDDA | AWRPGVGENYAEQF | 0.67063147 |
| TPPGPPN | AWRPGVGENYAEQF | 0.74713063 |
| TPPPPPPP | AWRPGVGENYAEQF | 0.83126277 |
| TQSPALI | AWSLWQGGNTEVF | 0.8068502 |
| EFAAAFPQ | AWSLWQGGNTEVF | 0.5069581 |
| LLSGNLDVL | AWSLWQGGNTEVF | 0.8341076 |
| GPANLGNYNI | AWSLWQGGNTEVF | 0.7779659 |
| TLGPTLP | AWSLWQGGNTEVF | 0.7691726 |

| | | |
|---|---|---|
| VYAYSSCCK | AWSLWQGGNTEVF | 0.7611517 |
| APAPAPAPNI | AWSLWQGGNTEVF | 0.83387434 |
| PGGPVPIP | AWSLWQGGNTEVF | 0.7259459 |
| FLTVALEP | AWSLWQGGNTEVF | 0.62549734 |
| SQYFGANL | AWSLWQGGNTEVF | 0.4919201 |
| GGVKPAA | AWSLWQGGNTEVF | 0.81544787 |
| GGTGGAGGAGGW | AWSLWQGGNTEVF | 0.84217584 |
| GVGGVAGLGGAGG | AWSLWQGGNTEVF | 0.7908894 |
| VLLVLATTI | AWSLWQGGNTEVF | 0.7330171 |
| IGIFAAML | AWSLWQGGNTEVF | 0.8329226 |
| AVGDLID | AWSLWQGGNTEVF | 0.77424365 |
| NISPLPL | AWSLWQGGNTEVF | 0.7950381 |
| VTAVFVAI | AWSLWQGGNTEVF | 0.8248589 |
| SGDDNDAAL | AWSLWQGGNTEVF | 0.7655399 |
| AVAALAHS | AWSLWQGGNTEVF | 0.83567166 |
| GGVTVMATP | AWSLWQGGNTEVF | 0.79901284 |
| AGLAANVRD | AWSLWQGGNTEVF | 0.81167495 |
| TNRPDLIDPALLRPG | AWSLWQGGNTEVF | 0.82941085 |
| LANLGVTAL | AWSLWQGGNTEVF | 0.8035788 |
| TGERIEL | AWSLWQGGNTEVF | 0.7892875 |
| NGEELAA | AWSLWQGGNTEVF | 0.34753266 |
| VLSLVWG | AWSLWQGGNTEVF | 0.6369301 |
| IGYALWL | AWSLWQGGNTEVF | 0.83245265 |
| LSLLLAPL | AWSLWQGGNTEVF | 0.82223684 |
| TLDEALK | AWSLWQGGNTEVF | 0.83200675 |
| TDGAGTAEGG | AWSLWQGGNTEVF | 0.74783635 |
| NAPGGAGGNGG | AWSLWQGGNTEVF | 0.76399374 |
| VSVFMGMGGNF | AWSLWQGGNTEVF | 0.81477976 |
| LASYLAAL | AWSLWQGGNTEVF | 0.83559436 |
| LVGGTPAL | AWSLWQGGNTEVF | 0.5488417 |
| VRNADVL | AWSLWQGGNTEVF | 0.78612256 |
| TCSSPFLVSAM | AWSLWQGGNTEVF | 0.8507043 |
| VHITDAE | AWSLWQGGNTEVF | 0.8202803 |
| EYAAPSF | AWSLWQGGNTEVF | 0.8051119 |
| VVDAFHGGL | AWSLWQGGNTEVF | 0.62823254 |
| ASVVDPQ | AWSLWQGGNTEVF | 0.83765733 |
| SYNEFGGLS | AWSLWQGGNTEVF | 0.64781487 |
| DPDGSVY | AWSLWQGGNTEVF | 0.8502265 |
| MVSTIHATE | AWSLWQGGNTEVF | 0.20904522 |
| LAAGVADPV | AWSLWQGGNTEVF | 0.6229571 |
| TKEQIAA | AWSLWQGGNTEVF | 0.5763388 |
| SLASPAAL | AWSLWQGGNTEVF | 0.5503043 |
| IGLVFYGV | AWSLWQGGNTEVF | 0.8405502 |
| FLLDASNL | AWSLWQGGNTEVF | 0.6611916 |
| YQVGNFP | AWSLWQGGNTEVF | 0.82492906 |
| ESLLSQST | AWSLWQGGNTEVF | 0.8209828 |
| LFTVGFAAVG | AWSLWQGGNTEVF | 0.8370255 |
| ATPPTPT | AWSLWQGGNTEVF | 0.8208164 |
| TDLATLG | AWSLWQGGNTEVF | 0.7939224 |
| SAGGTGR | AWSLWQGGNTEVF | 0.54995716 |
| GFSNSGS | AWSLWQGGNTEVF | 0.8371181 |

| | | |
|---|---|---|
| GGNGGSGGTSV | AWSLWQGGNTEVF | 0.82936704 |
| GTSVATGGAGN | AWSLWQGGNTEVF | 0.77726376 |
| NGGNGGSGGTS | AWSLWQGGNTEVF | 0.53584373 |
| AVVKFPH | AWSLWQGGNTEVF | 0.77777714 |
| TPPPPPP | AWSLWQGGNTEVF | 0.82170576 |
| AAPAAAAAPA | AWSLWQGGNTEVF | 0.20440768 |
| IALNEDL | AWSLWQGGNTEVF | 0.21937749 |
| IVNVSSM | AWSLWQGGNTEVF | 0.7967559 |
| TGIPPAPR | AWSLWQGGNTEVF | 0.69776785 |
| FDLGGGTF | AWSLWQGGNTEVF | 0.8465331 |
| FDLGGGTFDVS | AWSLWQGGNTEVF | 0.84169376 |
| NEPTAAA | AWSLWQGGNTEVF | 0.8024317 |
| NEPTAAALA | AWSLWQGGNTEVF | 0.83085716 |
| GSIINISSI | AWSLWQGGNTEVF | 0.81829286 |
| VAPGNAGTA | AWSLWQGGNTEVF | 0.1439903 |
| TGGGASLE | AWSLWQGGNTEVF | 0.66038066 |
| VAVYNHYKRI | AWSLWQGGNTEVF | 0.6562283 |
| TDPAPATT | AWSLWQGGNTEVF | 0.8303203 |
| TNRPDILDPALLRPG | AWSLWQGGNTEVF | 0.7740343 |
| LTTTTTS | AWSLWQGGNTEVF | 0.7615942 |
| PTGSGKTL | AWSLWQGGNTEVF | 0.8154353 |
| TGSGKTL | AWSLWQGGNTEVF | 0.8129495 |
| TIGHVDHGK | AWSLWQGGNTEVF | 0.790652 |
| TIGHVDHGKTT | AWSLWQGGNTEVF | 0.6515442 |
| GHVDHGKTT | AWSLWQGGNTEVF | 0.1646816 |
| AQFEHTLL | AWSLWQGGNTEVF | 0.8373112 |
| IVNQDEL | AWSLWQGGNTEVF | 0.2707522 |
| TNGSQFF | AWSLWQGGNTEVF | 0.6755992 |
| TGKGIVE | AWSLWQGGNTEVF | 0.23214963 |
| LISNASDA | AWSLWQGGNTEVF | 0.17421183 |
| AELRTLGELTKIAKAHGAQ | AWSLWQGGNTEVF | 0.3286595 |
| APNAVAAAEPSW | AWSLWQGGNTEVF | 0.80737853 |
| APQVAGAATAVL | AWSLWQGGNTEVF | 0.7895254 |
| ARRDPAAAAAILPTDA | AWSLWQGGNTEVF | 0.7952268 |
| DAAARDVLAEAGLAEYF | AWSLWQGGNTEVF | 0.8492653 |
| DDQIVLSAKANLAKVEQ | AWSLWQGGNTEVF | 0.81386054 |
| DELLPDTGEDSDEDGH | AWSLWQGGNTEVF | 0.7938323 |
| ELPEAPANRATVPQD | AWSLWQGGNTEVF | 0.6809323 |
| FGQSKIERVTGIG | AWSLWQGGNTEVF | 0.71292293 |
| GAGGTGSTAGGAGGAGGAGGLY | AWSLWQGGNTEVF | 0.6754887 |
| GHGGTGAAGGNGGTGGAGGSGI | AWSLWQGGNTEVF | 0.62045985 |
| GNGGNGGAGGEGGPGGAGGAGA | AWSLWQGGNTEVF | 0.84563047 |
| GSTAGNGGNGGNATNGTIAGQP | AWSLWQGGNTEVF | 0.6248368 |
| GTGGDGGNGGNGGTGG | AWSLWQGGNTEVF | 0.4822612 |
| GVRRPAGIPADAPV | AWSLWQGGNTEVF | 0.62836957 |
| GYRQVTDAHLLTLARR | AWSLWQGGNTEVF | 0.8435904 |
| IAKMGQDGHDRG | AWSLWQGGNTEVF | 0.85020435 |
| KAEPSIEIQYT | AWSLWQGGNTEVF | 0.7946552 |
| KPVRIRMAADEDTVAVT | AWSLWQGGNTEVF | 0.6334627 |
| MPPRIAGMRLLVIKPEP | AWSLWQGGNTEVF | 0.7231341 |
| MKFYAVALFRSRGVPPTR | AWSLWQGGNTEVF | 0.8243584 |

| | | |
|---|---|---|
| NAGGFGILGAVAHSP | AWSLWQGGNTEVF | 0.704131 |
| NEGREITLAMAEM | AWSLWQGGNTEVF | 0.41023892 |
| NELVEWSHPLGHRWRWE | AWSLWQGGNTEVF | 0.8243541 |
| NLGAGGTGGNGGNPG | AWSLWQGGNTEVF | 0.84259635 |
| NTEWPKWETVMAADPVD | AWSLWQGGNTEVF | 0.69715 |
| PASAAPAETQP | AWSLWQGGNTEVF | 0.65361357 |
| QQSSDGKNGVRAAEITGE | AWSLWQGGNTEVF | 0.8127813 |
| RAPDAVTTSWWKE | AWSLWQGGNTEVF | 0.74973696 |
| SGFGNVGAHTSGWFNQ | AWSLWQGGNTEVF | 0.8031759 |
| SGVAGAVGQAASVGGLKV | AWSLWQGGNTEVF | 0.80000246 |
| SNGGAGTGSAGGNGGT | AWSLWQGGNTEVF | 0.56171817 |
| TGGAGGEGGAAG | AWSLWQGGNTEVF | 0.7475404 |
| TGGDGGAAGAGGAPGHG | AWSLWQGGNTEVF | 0.8497053 |
| TMPPLLAAIANAGRHGV | AWSLWQGGNTEVF | 0.79708624 |
| TSSTAGNSGGA | AWSLWQGGNTEVF | 0.81241065 |
| VPATVPVGILC | AWSLWQGGNTEVF | 0.84597456 |
| VQRPDSGGVVLTGRIS | AWSLWQGGNTEVF | 0.8359928 |
| WTPNPVPATKGVPAGLVQAT | AWSLWQGGNTEVF | 0.84687495 |
| YALKRALAKFSDRFDVVII | AWSLWQGGNTEVF | 0.82990795 |
| YLIGSVIV | AWSLWQGGNTEVF | 0.6844041 |
| YGQLVAVID | AWSLWQGGNTEVF | 0.7996454 |
| VTVGGMTLPA | AWSLWQGGNTEVF | 0.81294274 |
| VLALPAV | AWSLWQGGNTEVF | 0.8233834 |
| VIPPTLNYET | AWSLWQGGNTEVF | 0.7967057 |
| VGMGGMHPSSG | AWSLWQGGNTEVF | 0.65056664 |
| VGLAGGAATA | AWSLWQGGNTEVF | 0.8444801 |
| VGGAGGAGGSGP | AWSLWQGGNTEVF | 0.7718745 |
| VFAAFMV | AWSLWQGGNTEVF | 0.82445467 |
| TLYDKLR | AWSLWQGGNTEVF | 0.5463302 |
| TLAVFTVL | AWSLWQGGNTEVF | 0.52770936 |
| TGSVQPARV | AWSLWQGGNTEVF | 0.837291 |
| TGGGAGSGGT | AWSLWQGGNTEVF | 0.15368408 |
| SHLETSL | AWSLWQGGNTEVF | 0.18126923 |
| SFTTGMT | AWSLWQGGNTEVF | 0.8127293 |
| QVEVAKRMGVRQPTVS | AWSLWQGGNTEVF | 0.8433944 |
| QGSGGPALAM | AWSLWQGGNTEVF | 0.15987237 |
| MLAPSNP | AWSLWQGGNTEVF | 0.6127422 |
| LVNNAAI | AWSLWQGGNTEVF | 0.539521 |
| LTGEINTLA | AWSLWQGGNTEVF | 0.73715806 |
| LSIFGPV | AWSLWQGGNTEVF | 0.81067693 |
| LLALLANL | AWSLWQGGNTEVF | 0.431654 |
| LISNASDALDKLR | AWSLWQGGNTEVF | 0.81617725 |
| LINEAVP | AWSLWQGGNTEVF | 0.5573077 |
| LGYAFSGL | AWSLWQGGNTEVF | 0.30726174 |
| LDFVAVAL | AWSLWQGGNTEVF | 0.8341901 |
| LAQPGAE | AWSLWQGGNTEVF | 0.8403018 |
| ISNLGAVTGF | AWSLWQGGNTEVF | 0.7155438 |
| HQEDPGRI | AWSLWQGGNTEVF | 0.83747214 |
| GTPGEPTP | AWSLWQGGNTEVF | 0.71146935 |
| GFFNSGS | AWSLWQGGNTEVF | 0.8503013 |
| GAGGGGLTAGAN | AWSLWQGGNTEVF | 0.79284495 |

| | | |
|---|---|---|
| FVDALVNM | AWSLWQGGNTEVF | 0.41671115 |
| FLTVAPEL | AWSLWQGGNTEVF | 0.8371549 |
| FETLTAL | AWSLWQGGNTEVF | 0.8140078 |
| DVINPIEL | AWSLWQGGNTEVF | 0.79609156 |
| APVSGAPV | AWSLWQGGNTEVF | 0.83757913 |
| ALVGGLAW | AWSLWQGGNTEVF | 0.51016885 |
| ALAAAAATAI | AWSLWQGGNTEVF | 0.8077209 |
| AFGLGLLL | AWSLWQGGNTEVF | 0.8453264 |
| AFFAAASL | AWSLWQGGNTEVF | 0.8430748 |
| AAAALPAFSPP | AWSLWQGGNTEVF | 0.52243626 |
| VVLAPAL | AWSLWQGGNTEVF | 0.8302052 |
| TIGHVDHGKTTL | AWSLWQGGNTEVF | 0.80652285 |
| LISNASDALDK | AWSLWQGGNTEVF | 0.7609 |
| LISNASDAL | AWSLWQGGNTEVF | 0.84666765 |
| ISNASDALDK | AWSLWQGGNTEVF | 0.7261628 |
| NQAETLVYQTE | AWSLWQGGNTEVF | 0.83589005 |
| TGGTGGTAGAG | AWSLWQGGNTEVF | 0.8247309 |
| TGSTAGGAGGA | AWSLWQGGNTEVF | 0.8117267 |
| DPLAVPVAAGPV | AWSLWQGGNTEVF | 0.49476382 |
| GGTGGTPGNGGN | AWSLWQGGNTEVF | 0.8152962 |
| PAGLAGVDPATL | AWSLWQGGNTEVF | 0.80984753 |
| GNNLVVEAPHAGASV | AWSLWQGGNTEVF | 0.8258387 |
| KVTRIMSASTLEQPA | AWSLWQGGNTEVF | 0.7799097 |
| LGFALPIALPRRTVV | AWSLWQGGNTEVF | 0.46761972 |
| TRLGNLTARKLVVLV | AWSLWQGGNTEVF | 0.8123736 |
| WSQQRRLIYFPSAGP | AWSLWQGGNTEVF | 0.84103024 |
| GPNGNPDPMAAAVDIR | AWSLWQGGNTEVF | 0.79669094 |
| TQRCVSVADLRAAASR | AWSLWQGGNTEVF | 0.77189916 |
| QFVLDVAEHPVEVLTETV | AWSLWQGGNTEVF | 0.8479636 |
| GGAGGTGGAGGNGGTQAGDG | AWSLWQGGNTEVF | 0.7259501 |
| PLLATLRAIAVDVDAKPAQV | AWSLWQGGNTEVF | 0.7726227 |
| APKFRTGGYMIDFWGVGYQVAK | AWSLWQGGNTEVF | 0.8302283 |
| GAGGAGGAGGAAGGPGGTGGTG | AWSLWQGGNTEVF | 0.8313899 |
| IGALGLLATDVAPAGSVAFGVA | AWSLWQGGNTEVF | 0.72074056 |
| AGGAGGIGGNGGAGGAGGVLMG | AWSLWQGGNTEVF | 0.84431034 |
| ADELIGLL | AWSLWQGGNTEVF | 0.67528975 |
| ADRALKA | AWSLWQGGNTEVF | 0.8415659 |
| AGAVGQVGGAGGA | AWSLWQGGNTEVF | 0.7656318 |
| AGGGGGLGGAP | AWSLWQGGNTEVF | 0.8266622 |
| AGGLLYGNGGAG | AWSLWQGGNTEVF | 0.77652466 |
| AGLESIGL | AWSLWQGGNTEVF | 0.8436656 |
| AGLKTGII | AWSLWQGGNTEVF | 0.8424279 |
| ASLAALLL | AWSLWQGGNTEVF | 0.8421385 |
| ASVINISSI | AWSLWQGGNTEVF | 0.1934383 |
| ATPPGPA | AWSLWQGGNTEVF | 0.84139407 |
| ATVASALV | AWSLWQGGNTEVF | 0.83520496 |
| ATVASLIL | AWSLWQGGNTEVF | 0.8508726 |
| AVNGPVLIRH | AWSLWQGGNTEVF | 0.7888692 |
| DGAAGAVGI | AWSLWQGGNTEVF | 0.8086116 |
| DPLLHGI | AWSLWQGGNTEVF | 0.8084588 |
| ELGAALGAL | AWSLWQGGNTEVF | 0.7216495 |

| | | |
|---|---|---|
| ELGRTQAV | AWSLWQGGNTEVF | 0.7025632 |
| FVAGGYLS | AWSLWQGGNTEVF | 0.790888 |
| GASGGAGGAGGW | AWSLWQGGNTEVF | 0.7969928 |
| GGAGAGGDGGAGGI | AWSLWQGGNTEVF | 0.7850007 |
| GTGHNGSQPGMGGNG | AWSLWQGGNTEVF | 0.8356776 |
| GYAAASASAS | AWSLWQGGNTEVF | 0.840171 |
| HQVDVAAVQQG | AWSLWQGGNTEVF | 0.8407798 |
| IGAAIAHT | AWSLWQGGNTEVF | 0.84260714 |
| IGMIEVAQL | AWSLWQGGNTEVF | 0.84209466 |
| IRLAPEHTSAMD | AWSLWQGGNTEVF | 0.8007918 |
| IYNVSVL | AWSLWQGGNTEVF | 0.73743427 |
| KQGFLGM | AWSLWQGGNTEVF | 0.76613235 |
| LESVITQL | AWSLWQGGNTEVF | 0.7341581 |
| LGLGALVVV | AWSLWQGGNTEVF | 0.7773497 |
| LIGNGGNGGSGGI | AWSLWQGGNTEVF | 0.73633754 |
| LNAINAPTL | AWSLWQGGNTEVF | 0.8248461 |
| LQLLNTGV | AWSLWQGGNTEVF | 0.14159638 |
| LQTSIEF | AWSLWQGGNTEVF | 0.8128575 |
| MSFAVSEL | AWSLWQGGNTEVF | 0.8462831 |
| NTGALGL | AWSLWQGGNTEVF | 0.74965477 |
| PPGAGGAGGAGTT | AWSLWQGGNTEVF | 0.8037957 |
| QLVGLGI | AWSLWQGGNTEVF | 0.79212743 |
| RFLEQVHQL | AWSLWQGGNTEVF | 0.7438853 |
| RGVGYVLR | AWSLWQGGNTEVF | 0.83656275 |
| SATGVFAGVI | AWSLWQGGNTEVF | 0.57726705 |
| SLPAITLP | AWSLWQGGNTEVF | 0.8031156 |
| STALAFTV | AWSLWQGGNTEVF | 0.84672797 |
| SVLTVGLGK | AWSLWQGGNTEVF | 0.8146037 |
| TALAGAGSGI | AWSLWQGGNTEVF | 0.7879504 |
| TGLPFTGI | AWSLWQGGNTEVF | 0.73750865 |
| VAEHVTV | AWSLWQGGNTEVF | 0.34042308 |
| VDIVLNSL | AWSLWQGGNTEVF | 0.8131183 |
| VLTGGGAVL | AWSLWQGGNTEVF | 0.14013733 |
| VNVGLAYGI | AWSLWQGGNTEVF | 0.8088644 |
| VSSAAVGAV | AWSLWQGGNTEVF | 0.6016128 |
| VTSVYEAGV | AWSLWQGGNTEVF | 0.82818294 |
| WVALDRGA | AWSLWQGGNTEVF | 0.8214563 |
| YLVGSVVI | AWSLWQGGNTEVF | 0.82306445 |
| GPPGTGKTLL | AWSLWQGGNTEVF | 0.75788504 |
| GVGMTKF | AWSLWQGGNTEVF | 0.76491743 |
| IGIGGSDL | AWSLWQGGNTEVF | 0.6818911 |
| IGTLLAEL | AWSLWQGGNTEVF | 0.55255365 |
| IYTSGTT | AWSLWQGGNTEVF | 0.83909845 |
| LSFPAGV | AWSLWQGGNTEVF | 0.7285527 |
| LYTSGTT | AWSLWQGGNTEVF | 0.76642156 |
| VGAFPFGF | AWSLWQGGNTEVF | 0.84267414 |
| VGAGTVEF | AWSLWQGGNTEVF | 0.8065456 |
| VGKSTLLL | AWSLWQGGNTEVF | 0.38016903 |
| AALAHAGGGIV | AWSLWQGGNTEVF | 0.8170526 |
| AALVAVSLL | AWSLWQGGNTEVF | 0.80780876 |
| AAVSRAVAAL | AWSLWQGGNTEVF | 0.7976774 |

| | | |
|---|---|---|
| AGGSSGIGGF | AWSLWQGGNTEVF | 0.13922562 |
| AGLTVANANG | AWSLWQGGNTEVF | 0.81259847 |
| AGLVGAVL | AWSLWQGGNTEVF | 0.758169 |
| ALSQAFTV | AWSLWQGGNTEVF | 0.74476695 |
| ALTFSVSL | AWSLWQGGNTEVF | 0.71438074 |
| ALTGAALVAL | AWSLWQGGNTEVF | 0.8387841 |
| ASFSIGAGVL | AWSLWQGGNTEVF | 0.33769596 |
| DGSVYKI | AWSLWQGGNTEVF | 0.78775775 |
| EQLAEIV | AWSLWQGGNTEVF | 0.8364286 |
| FAVSELTVM | AWSLWQGGNTEVF | 0.83528435 |
| GAAPLAAGL | AWSLWQGGNTEVF | 0.84473985 |
| GADGGDGGSGGAGG | AWSLWQGGNTEVF | 0.79969 |
| GFGLDTGGAGG | AWSLWQGGNTEVF | 0.73219013 |
| GLASGGLL | AWSLWQGGNTEVF | 0.7877587 |
| GRIAGAAVLL | AWSLWQGGNTEVF | 0.7165962 |
| GSAGSLNTSF | AWSLWQGGNTEVF | 0.31974357 |
| GTGQIGI | AWSLWQGGNTEVF | 0.25315887 |
| GTLIALIL | AWSLWQGGNTEVF | 0.8270034 |
| IAATMGFVVG | AWSLWQGGNTEVF | 0.22838551 |
| IFAQALL | AWSLWQGGNTEVF | 0.81021184 |
| IGAAVALGF | AWSLWQGGNTEVF | 0.81233764 |
| IGLTPLAL | AWSLWQGGNTEVF | 0.7451197 |
| ILGSNTSTL | AWSLWQGGNTEVF | 0.58207667 |
| IVFSLTVL | AWSLWQGGNTEVF | 0.7956776 |
| LASKATGF | AWSLWQGGNTEVF | 0.78069735 |
| LASVALIL | AWSLWQGGNTEVF | 0.8361991 |
| LFAVGLTL | AWSLWQGGNTEVF | 0.80933696 |
| LFNGGIGTYI | AWSLWQGGNTEVF | 0.8468547 |
| LTLGAGII | AWSLWQGGNTEVF | 0.8433754 |
| LVAAALGDI | AWSLWQGGNTEVF | 0.7939198 |
| NTAFVQLGI | AWSLWQGGNTEVF | 0.83453274 |
| QLFSGAEL | AWSLWQGGNTEVF | 0.7421572 |
| SAATFITL | AWSLWQGGNTEVF | 0.7762894 |
| SNYGLANLGV | AWSLWQGGNTEVF | 0.76557565 |
| SYGGAYAVM | AWSLWQGGNTEVF | 0.8027877 |
| TTGVGGLTLG | AWSLWQGGNTEVF | 0.7380627 |
| TTQTIYI | AWSLWQGGNTEVF | 0.73643696 |
| VAAGAGLGAVI | AWSLWQGGNTEVF | 0.38557398 |
| VAAVVTLL | AWSLWQGGNTEVF | 0.8105363 |
| VGFLGGLI | AWSLWQGGNTEVF | 0.8151373 |
| VGVIPILL | AWSLWQGGNTEVF | 0.27633813 |
| VIFASSLI | AWSLWQGGNTEVF | 0.72746974 |
| VKGGDGGAAAG | AWSLWQGGNTEVF | 0.84986174 |
| VTKTFTNI | AWSLWQGGNTEVF | 0.8421829 |
| VVAAAILL | AWSLWQGGNTEVF | 0.7616619 |
| YAALDVEL | AWSLWQGGNTEVF | 0.63902783 |
| LASVALLL | AWSLWQGGNTEVF | 0.8159319 |
| LDAVHRV | AWSLWQGGNTEVF | 0.32581058 |
| QGLSSSIF | AWSLWQGGNTEVF | 0.7797024 |
| RGTFANIRL | AWSLWQGGNTEVF | 0.8160105 |
| RQLWWGHRI | AWSLWQGGNTEVF | 0.8124998 |

| | | |
|---|---|---|
| YGPPGTGKTLL | AWSLWQGGNTEVF | 0.6294069 |
| AALVGAPANVA | AWSLWQGGNTEVF | 0.8132765 |
| AATFRRTE | AWSLWQGGNTEVF | 0.78307265 |
| AGALGGGI | AWSLWQGGNTEVF | 0.78376174 |
| AGGAGGAGGIGAG | AWSLWQGGNTEVF | 0.16226915 |
| AGSGALGL | AWSLWQGGNTEVF | 0.792351 |
| AQIADQI | AWSLWQGGNTEVF | 0.7638817 |
| ASAVAVTYL | AWSLWQGGNTEVF | 0.6712953 |
| ASTVLGGTGG | AWSLWQGGNTEVF | 0.47507387 |
| AVGGEPAK | AWSLWQGGNTEVF | 0.7830828 |
| CEQGTFGGTC | AWSLWQGGNTEVF | 0.8433461 |
| DEGKAPAG | AWSLWQGGNTEVF | 0.8385601 |
| DTVVFPAL | AWSLWQGGNTEVF | 0.5618413 |
| FEGITMAG | AWSLWQGGNTEVF | 0.83794355 |
| FPLVVQD | AWSLWQGGNTEVF | 0.8500499 |
| GAFTTPQ | AWSLWQGGNTEVF | 0.7812576 |
| GFGGFGTAGGVGGA | AWSLWQGGNTEVF | 0.39218372 |
| GFLAQFL | AWSLWQGGNTEVF | 0.81371987 |
| GGAGGAGGNAGVA | AWSLWQGGNTEVF | 0.7756666 |
| GGVGTAGAGGAGGAGG | AWSLWQGGNTEVF | 0.80940354 |
| GNGGGGDAGNAGSGGN | AWSLWQGGNTEVF | 0.8232131 |
| GTGGAGGAGNDAGS | AWSLWQGGNTEVF | 0.7315535 |
| GTTNTGWFNS | AWSLWQGGNTEVF | 0.8169075 |
| IFMGPLFTL | AWSLWQGGNTEVF | 0.8355205 |
| ILAHAIEAI | AWSLWQGGNTEVF | 0.7422439 |
| IRSAIREAL | AWSLWQGGNTEVF | 0.73147637 |
| ITFAGGG | AWSLWQGGNTEVF | 0.6367737 |
| ITGMSGHQEV | AWSLWQGGNTEVF | 0.3573422 |
| IVAGLACTL | AWSLWQGGNTEVF | 0.8278688 |
| IVGGGLAGA | AWSLWQGGNTEVF | 0.77031887 |
| IVTGVLNI | AWSLWQGGNTEVF | 0.6765537 |
| LAQQGCI | AWSLWQGGNTEVF | 0.8190789 |
| LCLHVEN | AWSLWQGGNTEVF | 0.6570543 |
| LDAISDGS | AWSLWQGGNTEVF | 0.6801032 |
| LGGFAAML | AWSLWQGGNTEVF | 0.7456602 |
| LGGKNPMI | AWSLWQGGNTEVF | 0.7951844 |
| LGPKGVAAIL | AWSLWQGGNTEVF | 0.8237848 |
| LMVGVSI | AWSLWQGGNTEVF | 0.8039721 |
| LVGAAFVL | AWSLWQGGNTEVF | 0.73451364 |
| LVLAVAPL | AWSLWQGGNTEVF | 0.8478027 |
| MAAIAVIDAA | AWSLWQGGNTEVF | 0.5908991 |
| NNGINAELQ | AWSLWQGGNTEVF | 0.6289106 |
| RGEPLVLVHG | AWSLWQGGNTEVF | 0.82018596 |
| RGGAGGAAGLI | AWSLWQGGNTEVF | 0.39262095 |
| RSYRMHPAV | AWSLWQGGNTEVF | 0.83102334 |
| SAAHLLAGQLA | AWSLWQGGNTEVF | 0.81698537 |
| SAAVGSY | AWSLWQGGNTEVF | 0.820599 |
| SPHAVLHRI | AWSLWQGGNTEVF | 0.7876363 |
| SQIAKVL | AWSLWQGGNTEVF | 0.8495987 |
| TALGHLVP | AWSLWQGGNTEVF | 0.8392681 |
| TAPTTSGAA | AWSLWQGGNTEVF | 0.75839984 |

| | | |
|---|---|---|
| TDGDDGNGGSPGT | AWSLWQGGNTEVF | 0.8060923 |
| TDGLGEN | AWSLWQGGNTEVF | 0.51766694 |
| TGGPPLPGQA | AWSLWQGGNTEVF | 0.8321764 |
| TIPAITI | AWSLWQGGNTEVF | 0.807045 |
| TIPGFTI | AWSLWQGGNTEVF | 0.73407936 |
| TSIASGALA | AWSLWQGGNTEVF | 0.8109284 |
| TVTLPGAAI | AWSLWQGGNTEVF | 0.8427294 |
| VASGLSAALP | AWSLWQGGNTEVF | 0.84429955 |
| VGAGVIGI | AWSLWQGGNTEVF | 0.26131353 |
| VGPKRFGV | AWSLWQGGNTEVF | 0.80037355 |
| VNVAFPD | AWSLWQGGNTEVF | 0.29805416 |
| VSMTNSGVS | AWSLWQGGNTEVF | 0.7145599 |
| VTDVLGDLSV | AWSLWQGGNTEVF | 0.6081047 |
| VVVVGAGFAGL | AWSLWQGGNTEVF | 0.8280983 |
| VYTVGTVV | AWSLWQGGNTEVF | 0.79369473 |
| WTTFSGP | AWSLWQGGNTEVF | 0.8115722 |
| YDELVEAF | AWSLWQGGNTEVF | 0.8138433 |
| YGTGGAGGNGG | AWSLWQGGNTEVF | 0.8447028 |
| YTNVMAV | AWSLWQGGNTEVF | 0.74530244 |
| FLEQVHQL | AWSLWQGGNTEVF | 0.79849696 |
| GLSLLAVAV | AWSLWQGGNTEVF | 0.4580791 |
| LEQVHQL | AWSLWQGGNTEVF | 0.33237237 |
| LNAAGAI | AWSLWQGGNTEVF | 0.71512973 |
| PAVHASL | AWSLWQGGNTEVF | 0.7323599 |
| TELEDDA | AWSLWQGGNTEVF | 0.20294847 |
| TPPGPPN | AWSLWQGGNTEVF | 0.8311816 |
| TPPPPPPP | AWSLWQGGNTEVF | 0.8252166 |
| TQSPALI | ASRDWGLYAEQF | 0.8332293 |
| EFAAAFPQ | ASRDWGLYAEQF | 0.7056391 |
| LLSGNLDVL | ASRDWGLYAEQF | 0.7582681 |
| GPANLGNYNI | ASRDWGLYAEQF | 0.74132836 |
| TLGPTLP | ASRDWGLYAEQF | 0.8343706 |
| VYAYSSCCK | ASRDWGLYAEQF | 0.781299 |
| APAPAPAPNI | ASRDWGLYAEQF | 0.46802765 |
| PGGPVPIP | ASRDWGLYAEQF | 0.6657909 |
| FLTVALEP | ASRDWGLYAEQF | 0.70830834 |
| SQYFGANL | ASRDWGLYAEQF | 0.8429976 |
| GGVKPAA | ASRDWGLYAEQF | 0.83691853 |
| GGTGGAGGAGGW | ASRDWGLYAEQF | 0.82965344 |
| GVGGVAGLGGAGG | ASRDWGLYAEQF | 0.8248677 |
| VLLVLATTI | ASRDWGLYAEQF | 0.25373662 |
| IGIFAAML | ASRDWGLYAEQF | 0.7446454 |
| AVGDLID | ASRDWGLYAEQF | 0.8433632 |
| NISPLPL | ASRDWGLYAEQF | 0.8175042 |
| VTAVFVAI | ASRDWGLYAEQF | 0.8001945 |
| SGDDNDAAL | ASRDWGLYAEQF | 0.84370327 |
| AVAALAHS | ASRDWGLYAEQF | 0.7720754 |
| GGVTVMATP | ASRDWGLYAEQF | 0.78869927 |
| AGLAANVRD | ASRDWGLYAEQF | 0.7958893 |
| TNRPDLIDPALLRPG | ASRDWGLYAEQF | 0.8048858 |
| LANLGVTAL | ASRDWGLYAEQF | 0.79853946 |

| | | |
|---|---|---|
| TGERIEL | ASRDWGLYAEQF | 0.70315313 |
| NGEELAA | ASRDWGLYAEQF | 0.78505033 |
| VLSLVWG | ASRDWGLYAEQF | 0.8516165 |
| IGYALWL | ASRDWGLYAEQF | 0.81309897 |
| LSLLLAPL | ASRDWGLYAEQF | 0.84113336 |
| TLDEALK | ASRDWGLYAEQF | 0.8251598 |
| TDGAGTAEGG | ASRDWGLYAEQF | 0.8199541 |
| NAPGGAGGNGG | ASRDWGLYAEQF | 0.76208425 |
| VSVFMGMGGNF | ASRDWGLYAEQF | 0.7970729 |
| LASYLAAL | ASRDWGLYAEQF | 0.7658217 |
| LVGGTPAL | ASRDWGLYAEQF | 0.7572582 |
| VRNADVL | ASRDWGLYAEQF | 0.75998896 |
| TCSSPFLVSAM | ASRDWGLYAEQF | 0.6968272 |
| VHITDAE | ASRDWGLYAEQF | 0.8528047 |
| EYAAPSF | ASRDWGLYAEQF | 0.8280041 |
| VVDAFHGGL | ASRDWGLYAEQF | 0.8187642 |
| ASVVDPQ | ASRDWGLYAEQF | 0.75060534 |
| SYNEFGGLS | ASRDWGLYAEQF | 0.80966145 |
| DPDGSVY | ASRDWGLYAEQF | 0.8249925 |
| MVSTIHATE | ASRDWGLYAEQF | 0.44230986 |
| LAAGVADPV | ASRDWGLYAEQF | 0.8446483 |
| TKEQIAA | ASRDWGLYAEQF | 0.7319283 |
| SLASPAAL | ASRDWGLYAEQF | 0.71100426 |
| IGLVFYGV | ASRDWGLYAEQF | 0.7908198 |
| FLLDASNL | ASRDWGLYAEQF | 0.8404678 |
| YQVGNFP | ASRDWGLYAEQF | 0.7415013 |
| ESLLSQST | ASRDWGLYAEQF | 0.74656796 |
| LFTVGFAAVG | ASRDWGLYAEQF | 0.7400613 |
| ATPPTPT | ASRDWGLYAEQF | 0.8245941 |
| TDLATLG | ASRDWGLYAEQF | 0.84390116 |
| SAGGTGR | ASRDWGLYAEQF | 0.7347282 |
| GFSNSGS | ASRDWGLYAEQF | 0.8471697 |
| GGNGGSGGTSV | ASRDWGLYAEQF | 0.6807775 |
| GTSVATGGAGN | ASRDWGLYAEQF | 0.4427957 |
| NGGNGGSGGTS | ASRDWGLYAEQF | 0.78356 |
| AVVKFPH | ASRDWGLYAEQF | 0.8355825 |
| TPPPPPP | ASRDWGLYAEQF | 0.8405971 |
| AAPAAAAAPA | ASRDWGLYAEQF | 0.8474947 |
| IALNEDL | ASRDWGLYAEQF | 0.8114461 |
| IVNVSSM | ASRDWGLYAEQF | 0.79400253 |
| TGIPPAPR | ASRDWGLYAEQF | 0.84370565 |
| FDLGGGTF | ASRDWGLYAEQF | 0.8410431 |
| FDLGGGTFDVS | ASRDWGLYAEQF | 0.83027637 |
| NEPTAAA | ASRDWGLYAEQF | 0.7177645 |
| NEPTAAALA | ASRDWGLYAEQF | 0.5100497 |
| GSIINISSI | ASRDWGLYAEQF | 0.79587805 |
| VAPGNAGTA | ASRDWGLYAEQF | 0.84085023 |
| TGGGASLE | ASRDWGLYAEQF | 0.54146695 |
| VAVYNHYKRI | ASRDWGLYAEQF | 0.81024903 |
| TDPAPATT | ASRDWGLYAEQF | 0.8234017 |
| TNRPDILDPALLRPG | ASRDWGLYAEQF | 0.61934465 |

| | | |
|---|---|---|
| LTTTTTS | ASRDWGLYAEQF | 0.8176539 |
| PTGSGKTL | ASRDWGLYAEQF | 0.8458491 |
| TGSGKTL | ASRDWGLYAEQF | 0.7258439 |
| TIGHVDHGK | ASRDWGLYAEQF | 0.7572454 |
| TIGHVDHGKTT | ASRDWGLYAEQF | 0.8425879 |
| GHVDHGKTT | ASRDWGLYAEQF | 0.8531567 |
| AQFEHTLL | ASRDWGLYAEQF | 0.837034 |
| IVNQDEL | ASRDWGLYAEQF | 0.40418696 |
| TNGSQFF | ASRDWGLYAEQF | 0.2914886 |
| TGKGIVE | ASRDWGLYAEQF | 0.7422913 |
| LISNASDA | ASRDWGLYAEQF | 0.82362133 |
| AELRTLGELTKIAKAHGAQ | ASRDWGLYAEQF | 0.72596335 |
| APNAVAAAEPSW | ASRDWGLYAEQF | 0.7268181 |
| APQVAGAATAVL | ASRDWGLYAEQF | 0.8334573 |
| ARRDPAAAAAILPTDA | ASRDWGLYAEQF | 0.84017575 |
| DAAARDVLAEAGLAEYF | ASRDWGLYAEQF | 0.73730135 |
| DDQIVLSAKANLAKVEQ | ASRDWGLYAEQF | 0.45172805 |
| DELLPDTGEDSDEDGH | ASRDWGLYAEQF | 0.82112724 |
| ELPEAPANRATVPQD | ASRDWGLYAEQF | 0.81449014 |
| FGQSKIERVTGIG | ASRDWGLYAEQF | 0.7621597 |
| GAGGTGSTAGGAGGAGGAGGLY | ASRDWGLYAEQF | 0.8184193 |
| GHGGTGAAGGNGGTGGAGGSGI | ASRDWGLYAEQF | 0.7339144 |
| GNGGNGGAGGEGGPGGAGGAGA | ASRDWGLYAEQF | 0.23468645 |
| GSTAGNGGNGGNATNGTIAGQP | ASRDWGLYAEQF | 0.8246475 |
| GTGGDGGNGGNGGTGG | ASRDWGLYAEQF | 0.6706314 |
| GVRRPAGIPADAPV | ASRDWGLYAEQF | 0.84895694 |
| GYRQVTDAHLLTLARR | ASRDWGLYAEQF | 0.7449398 |
| IAKMGQDGHDRG | ASRDWGLYAEQF | 0.8359885 |
| KAEPSIEIQYT | ASRDWGLYAEQF | 0.830108 |
| KPVRIRMAADEDTVAVT | ASRDWGLYAEQF | 0.81485474 |
| MPPRIAGMRLLVIKPEP | ASRDWGLYAEQF | 0.8008068 |
| MKFYAVALFRSRGVPPTR | ASRDWGLYAEQF | 0.73763883 |
| NAGGFGILGAVAHSP | ASRDWGLYAEQF | 0.6737697 |
| NEGREITLAMAEM | ASRDWGLYAEQF | 0.29655412 |
| NELVEWSHPLGHRWRWE | ASRDWGLYAEQF | 0.84022975 |
| NLGAGGTGGNGGNPG | ASRDWGLYAEQF | 0.1645752 |
| NTEWPKWETVMAADPVD | ASRDWGLYAEQF | 0.84000653 |
| PASAAPAETQP | ASRDWGLYAEQF | 0.8340869 |
| QQSSDGKNGVRAAEITGE | ASRDWGLYAEQF | 0.7223693 |
| RAPDAVTTSWWKE | ASRDWGLYAEQF | 0.81820977 |
| SGFGNVGAHTSGWFNQ | ASRDWGLYAEQF | 0.72294796 |
| SGVAGAVGQAASVGGLKV | ASRDWGLYAEQF | 0.5764531 |
| SNGGAGTGSAGGNGGT | ASRDWGLYAEQF | 0.8374356 |
| TGGAGGEGGAAG | ASRDWGLYAEQF | 0.8081327 |
| TGGDGGAAGAGGAPGHG | ASRDWGLYAEQF | 0.75871325 |
| TMPPLLAAIANAGRHGV | ASRDWGLYAEQF | 0.7813743 |
| TSSTAGNSGGA | ASRDWGLYAEQF | 0.8212792 |
| VPATVPVGILC | ASRDWGLYAEQF | 0.6946691 |
| VQRPDSGGVVLTGRIS | ASRDWGLYAEQF | 0.7313504 |
| WTPNPVPATKGVPAGLVQAT | ASRDWGLYAEQF | 0.84455884 |
| YALKRALAKFSDRFDVVII | ASRDWGLYAEQF | 0.7676492 |

| | | |
|---|---|---|
| YLIGSVIV | ASRDWGLYAEQF | 0.787314 |
| YGQLVAVID | ASRDWGLYAEQF | 0.83180803 |
| VTVGGMTLPA | ASRDWGLYAEQF | 0.6861494 |
| VLALPAV | ASRDWGLYAEQF | 0.8330079 |
| VIPPTLNYET | ASRDWGLYAEQF | 0.77885437 |
| VGMGGMHPSSG | ASRDWGLYAEQF | 0.76890343 |
| VGLAGGAATA | ASRDWGLYAEQF | 0.8322516 |
| VGGAGGAGGSGP | ASRDWGLYAEQF | 0.40018827 |
| VFAAFMV | ASRDWGLYAEQF | 0.7444944 |
| TLYDKLR | ASRDWGLYAEQF | 0.76937044 |
| TLAVFTVL | ASRDWGLYAEQF | 0.8196672 |
| TGSVQPARV | ASRDWGLYAEQF | 0.8371943 |
| TGGGAGSGGT | ASRDWGLYAEQF | 0.75749147 |
| SHLETSL | ASRDWGLYAEQF | 0.8110771 |
| SFTTGMT | ASRDWGLYAEQF | 0.726398 |
| QVEVAKRMGVRQPTVS | ASRDWGLYAEQF | 0.6808926 |
| QGSGGPALAM | ASRDWGLYAEQF | 0.65391374 |
| MLAPSNP | ASRDWGLYAEQF | 0.7332197 |
| LVNNAAI | ASRDWGLYAEQF | 0.7500544 |
| LTGEINTLA | ASRDWGLYAEQF | 0.81219447 |
| LSIFGPV | ASRDWGLYAEQF | 0.68843544 |
| LLALLANL | ASRDWGLYAEQF | 0.8062744 |
| LISNASDALDKLR | ASRDWGLYAEQF | 0.84378195 |
| LINEAVP | ASRDWGLYAEQF | 0.7811781 |
| LGYAFSGL | ASRDWGLYAEQF | 0.82865787 |
| LDFVAVAL | ASRDWGLYAEQF | 0.8272202 |
| LAQPGAE | ASRDWGLYAEQF | 0.8338352 |
| ISNLGAVTGF | ASRDWGLYAEQF | 0.7418932 |
| HQEDPGRI | ASRDWGLYAEQF | 0.7753932 |
| GTPGEPTP | ASRDWGLYAEQF | 0.843817 |
| GFFNSGS | ASRDWGLYAEQF | 0.8320124 |
| GAGGGGLTAGAN | ASRDWGLYAEQF | 0.7879939 |
| FVDALVNM | ASRDWGLYAEQF | 0.8208791 |
| FLTVAPEL | ASRDWGLYAEQF | 0.82967055 |
| FETLTAL | ASRDWGLYAEQF | 0.82632464 |
| DVINPIEL | ASRDWGLYAEQF | 0.79098547 |
| APVSGAPV | ASRDWGLYAEQF | 0.49946427 |
| ALVGGLAW | ASRDWGLYAEQF | 0.77803904 |
| ALAAAAATAI | ASRDWGLYAEQF | 0.82043 |
| AFGLGLLL | ASRDWGLYAEQF | 0.6955479 |
| AFFAAASL | ASRDWGLYAEQF | 0.2703061 |
| AAAALPAFSPP | ASRDWGLYAEQF | 0.8403394 |
| VVLAPAL | ASRDWGLYAEQF | 0.8516097 |
| TIGHVDHGKTTL | ASRDWGLYAEQF | 0.657596 |
| LISNASDALDK | ASRDWGLYAEQF | 0.8436945 |
| LISNASDAL | ASRDWGLYAEQF | 0.5324122 |
| ISNASDALDK | ASRDWGLYAEQF | 0.82962245 |
| NQAETLVYQTE | ASRDWGLYAEQF | 0.8391607 |
| TGGTGGTAGAG | ASRDWGLYAEQF | 0.7077172 |
| TGSTAGGAGGA | ASRDWGLYAEQF | 0.81318545 |
| DPLAVPVAAGPV | ASRDWGLYAEQF | 0.7630415 |

| | | |
|---|---|---|
| GGTGGTPGNGGN | ASRDWGLYAEQF | 0.7894879 |
| PAGLAGVDPATL | ASRDWGLYAEQF | 0.8461714 |
| GNNLVVEAPHAGASV | ASRDWGLYAEQF | 0.7383733 |
| KVTRIMSASTLEQPA | ASRDWGLYAEQF | 0.79375255 |
| LGFALPIALPRRTVV | ASRDWGLYAEQF | 0.7333107 |
| TRLGNLTARKLVVLV | ASRDWGLYAEQF | 0.62407905 |
| WSQQRRLIYFPSAGP | ASRDWGLYAEQF | 0.7896973 |
| GPNGNPDPMAAAVDIR | ASRDWGLYAEQF | 0.8313732 |
| TQRCVSVADLRAAASR | ASRDWGLYAEQF | 0.54440665 |
| QFVLDVAEHPVEVLTETV | ASRDWGLYAEQF | 0.83828974 |
| GGAGGTGGAGGNGGTQAGDG | ASRDWGLYAEQF | 0.77526873 |
| PLLATLRAIAVDVDAKPAQV | ASRDWGLYAEQF | 0.8515977 |
| APKFRTGGYMIDFWGVGYQVAK | ASRDWGLYAEQF | 0.83471143 |
| GAGGAGGAGGAAGGPGGTGGTG | ASRDWGLYAEQF | 0.8205288 |
| IGALGLLATDVAPAGSVAFGVA | ASRDWGLYAEQF | 0.72246677 |
| AGGAGGIGGNGGAGGAGGVLMG | ASRDWGLYAEQF | 0.8461243 |
| ADELIGLL | ASRDWGLYAEQF | 0.79919034 |
| ADRALKA | ASRDWGLYAEQF | 0.82187885 |
| AGAVGQVGGAGGA | ASRDWGLYAEQF | 0.8312806 |
| AGGGGGLGGAP | ASRDWGLYAEQF | 0.6840606 |
| AGGLLYGNGGAG | ASRDWGLYAEQF | 0.7235301 |
| AGLESIGL | ASRDWGLYAEQF | 0.7070246 |
| AGLKTGII | ASRDWGLYAEQF | 0.7091564 |
| ASLAALLL | ASRDWGLYAEQF | 0.68579674 |
| ASVINISSI | ASRDWGLYAEQF | 0.5856825 |
| ATPPGPA | ASRDWGLYAEQF | 0.7111212 |
| ATVASALV | ASRDWGLYAEQF | 0.651581 |
| ATVASLIL | ASRDWGLYAEQF | 0.7004913 |
| AVNGPVLIRH | ASRDWGLYAEQF | 0.7223147 |
| DGAAGAVGI | ASRDWGLYAEQF | 0.6917572 |
| DPLLHGI | ASRDWGLYAEQF | 0.72880334 |
| ELGAALGAL | ASRDWGLYAEQF | 0.7297518 |
| ELGRTQAV | ASRDWGLYAEQF | 0.60703397 |
| FVAGGYLS | ASRDWGLYAEQF | 0.6800314 |
| GASGGAGGAGGW | ASRDWGLYAEQF | 0.6893112 |
| GGAGAGGDGGAGGI | ASRDWGLYAEQF | 0.7020347 |
| GTGHNGSQPGMGGNG | ASRDWGLYAEQF | 0.56375796 |
| GYAAASASAS | ASRDWGLYAEQF | 0.72293484 |
| HQVDVAAVQQG | ASRDWGLYAEQF | 0.674214 |
| IGAAIAHT | ASRDWGLYAEQF | 0.34232014 |
| IGMIEVAQL | ASRDWGLYAEQF | 0.71270466 |
| IRLAPEHTSAMD | ASRDWGLYAEQF | 0.7021785 |
| IYNVSVL | ASRDWGLYAEQF | 0.69014704 |
| KQGFLGM | ASRDWGLYAEQF | 0.72176695 |
| LESVITQL | ASRDWGLYAEQF | 0.5021056 |
| LGLGALVVV | ASRDWGLYAEQF | 0.6780869 |
| LIGNGGNGGSGGI | ASRDWGLYAEQF | 0.71844155 |
| LNAINAPTL | ASRDWGLYAEQF | 0.72753215 |
| LQLLNTGV | ASRDWGLYAEQF | 0.6448604 |
| LQTSIEF | ASRDWGLYAEQF | 0.7087252 |
| MSFAVSEL | ASRDWGLYAEQF | 0.7248858 |

| | | |
|---|---|---|
| NTGALGL | ASRDWGLYAEQF | 0.7127175 |
| PPGAGGAGGAGTT | ASRDWGLYAEQF | 0.7269947 |
| QLVGLGI | ASRDWGLYAEQF | 0.7304414 |
| RFLEQVHQL | ASRDWGLYAEQF | 0.6860719 |
| RGVGYVLR | ASRDWGLYAEQF | 0.7241453 |
| SATGVFAGVI | ASRDWGLYAEQF | 0.73160005 |
| SLPAITLP | ASRDWGLYAEQF | 0.6938261 |
| STALAFTV | ASRDWGLYAEQF | 0.7079279 |
| SVLTVGLGK | ASRDWGLYAEQF | 0.6928619 |
| TALAGAGSGI | ASRDWGLYAEQF | 0.6972252 |
| TGLPFTGI | ASRDWGLYAEQF | 0.45102525 |
| VAEHVTV | ASRDWGLYAEQF | 0.7047933 |
| VDIVLNSL | ASRDWGLYAEQF | 0.7240444 |
| VLTGGGAVL | ASRDWGLYAEQF | 0.6817248 |
| VNVGLAYGI | ASRDWGLYAEQF | 0.641955 |
| VSSAAVGAV | ASRDWGLYAEQF | 0.6476073 |
| VTSVYEAGV | ASRDWGLYAEQF | 0.6877692 |
| WVALDRGA | ASRDWGLYAEQF | 0.7030262 |
| YLVGSVVI | ASRDWGLYAEQF | 0.5322971 |
| GPPGTGKTLL | ASRDWGLYAEQF | 0.6720001 |
| GVGMTKF | ASRDWGLYAEQF | 0.7238457 |
| IGIGGSDL | ASRDWGLYAEQF | 0.7247725 |
| IGTLLAEL | ASRDWGLYAEQF | 0.5268657 |
| IYTSGTT | ASRDWGLYAEQF | 0.7273057 |
| LSFPAGV | ASRDWGLYAEQF | 0.718486 |
| LYTSGTT | ASRDWGLYAEQF | 0.7272142 |
| VGAFPFGF | ASRDWGLYAEQF | 0.7123793 |
| VGAGTVEF | ASRDWGLYAEQF | 0.657452 |
| VGKSTLLL | ASRDWGLYAEQF | 0.71540654 |
| AALAHAGGGIV | ASRDWGLYAEQF | 0.6718508 |
| AALVAVSLL | ASRDWGLYAEQF | 0.64118046 |
| AAVSRAVAAL | ASRDWGLYAEQF | 0.72608274 |
| AGGSSGIGGF | ASRDWGLYAEQF | 0.6929499 |
| AGLTVANANG | ASRDWGLYAEQF | 0.65638435 |
| AGLVGAVL | ASRDWGLYAEQF | 0.6415954 |
| ALSQAFTV | ASRDWGLYAEQF | 0.72674936 |
| ALTFSVSL | ASRDWGLYAEQF | 0.61540264 |
| ALTGAALVAL | ASRDWGLYAEQF | 0.65752596 |
| ASFSIGAGVL | ASRDWGLYAEQF | 0.6359307 |
| DGSVYKI | ASRDWGLYAEQF | 0.69978285 |
| EQLAEIV | ASRDWGLYAEQF | 0.72205806 |
| FAVSELTVM | ASRDWGLYAEQF | 0.716434 |
| GAAPLAAGL | ASRDWGLYAEQF | 0.665006 |
| GADGGDGGSGGAGG | ASRDWGLYAEQF | 0.7139846 |
| GFGLDTGGAGG | ASRDWGLYAEQF | 0.7146257 |
| GLASGGLL | ASRDWGLYAEQF | 0.7124737 |
| GRIAGAAVLL | ASRDWGLYAEQF | 0.7236867 |
| GSAGSLNTSF | ASRDWGLYAEQF | 0.63662946 |
| GTGQIGI | ASRDWGLYAEQF | 0.72509426 |
| GTLIALIL | ASRDWGLYAEQF | 0.71618205 |
| IAATMGFVVG | ASRDWGLYAEQF | 0.7282763 |

| | | |
|---|---|---|
| IFAQALL | ASRDWGLYAEQF | 0.72376 |
| IGAAVALGF | ASRDWGLYAEQF | 0.70048696 |
| IGLTPLAL | ASRDWGLYAEQF | 0.7085427 |
| ILGSNTSTL | ASRDWGLYAEQF | 0.50456744 |
| IVFSLTVL | ASRDWGLYAEQF | 0.69978464 |
| LASKATGF | ASRDWGLYAEQF | 0.71648675 |
| LASVALIL | ASRDWGLYAEQF | 0.7023787 |
| LFAVGLTL | ASRDWGLYAEQF | 0.72230643 |
| LFNGGIGTYI | ASRDWGLYAEQF | 0.7239959 |
| LTLGAGII | ASRDWGLYAEQF | 0.6698941 |
| LVAAALGDI | ASRDWGLYAEQF | 0.72825396 |
| NTAFVQLGI | ASRDWGLYAEQF | 0.7185205 |
| QLFSGAEL | ASRDWGLYAEQF | 0.59063405 |
| SAATFITL | ASRDWGLYAEQF | 0.68770504 |
| SNYGLANLGV | ASRDWGLYAEQF | 0.7286991 |
| SYGGAYAVM | ASRDWGLYAEQF | 0.7052141 |
| TTGVGGLTLG | ASRDWGLYAEQF | 0.7005682 |
| TTQTIYI | ASRDWGLYAEQF | 0.6962081 |
| VAAGAGLGAVI | ASRDWGLYAEQF | 0.72963285 |
| VAAVVTLL | ASRDWGLYAEQF | 0.66324615 |
| VGFLGGLI | ASRDWGLYAEQF | 0.7305605 |
| VGVIPILL | ASRDWGLYAEQF | 0.4172083 |
| VIFASSLI | ASRDWGLYAEQF | 0.6901777 |
| VKGGDGGAAAG | ASRDWGLYAEQF | 0.6506064 |
| VTKTFTNI | ASRDWGLYAEQF | 0.58244497 |
| VVAAAILL | ASRDWGLYAEQF | 0.7257571 |
| YAALDVEL | ASRDWGLYAEQF | 0.70452183 |
| LASVALLL | ASRDWGLYAEQF | 0.72279835 |
| LDAVHRV | ASRDWGLYAEQF | 0.7096466 |
| QGLSSSIF | ASRDWGLYAEQF | 0.7231158 |
| RGTFANIRL | ASRDWGLYAEQF | 0.6969915 |
| RQLWWGHRI | ASRDWGLYAEQF | 0.7106619 |
| YGPPGTGKTLL | ASRDWGLYAEQF | 0.6495888 |
| AALVGAPANVA | ASRDWGLYAEQF | 0.7251055 |
| AATFRRTE | ASRDWGLYAEQF | 0.73112345 |
| AGALGGGI | ASRDWGLYAEQF | 0.7171028 |
| AGGAGGAGGIGAG | ASRDWGLYAEQF | 0.62780356 |
| AGSGALGL | ASRDWGLYAEQF | 0.66185516 |
| AQIADQI | ASRDWGLYAEQF | 0.70858026 |
| ASAVAVTYL | ASRDWGLYAEQF | 0.38280937 |
| ASTVLGGTGG | ASRDWGLYAEQF | 0.45938608 |
| AVGGEPAK | ASRDWGLYAEQF | 0.66624343 |
| CEQGTFGGTC | ASRDWGLYAEQF | 0.6412342 |
| DEGKAPAG | ASRDWGLYAEQF | 0.7280147 |
| DTVVFPAL | ASRDWGLYAEQF | 0.71853757 |
| FEGITMAG | ASRDWGLYAEQF | 0.6952377 |
| FPLVVQD | ASRDWGLYAEQF | 0.71951115 |
| GAFTTPQ | ASRDWGLYAEQF | 0.6988811 |
| GFGGFGTAGGVGGA | ASRDWGLYAEQF | 0.26870582 |
| GFLAQFL | ASRDWGLYAEQF | 0.68779045 |
| GGAGGAGGNAGVA | ASRDWGLYAEQF | 0.6394088 |

| | | |
|---|---|---|
| GGVGTAGAGGAGGAGG | ASRDWGLYAEQF | 0.7315963 |
| GNGGGGDAGNAGSGGN | ASRDWGLYAEQF | 0.6664806 |
| GTGGAGGAGNDAGS | ASRDWGLYAEQF | 0.6411556 |
| GTTNTGWFNS | ASRDWGLYAEQF | 0.6986153 |
| IFMGPLFTL | ASRDWGLYAEQF | 0.694373 |
| ILAHAIEAI | ASRDWGLYAEQF | 0.69641227 |
| IRSAIREAL | ASRDWGLYAEQF | 0.619286 |
| ITFAGGG | ASRDWGLYAEQF | 0.3324652 |
| ITGMSGHQEV | ASRDWGLYAEQF | 0.7241285 |
| IVAGLACTL | ASRDWGLYAEQF | 0.5372282 |
| IVGGGLAGA | ASRDWGLYAEQF | 0.6692526 |
| IVTGVLNI | ASRDWGLYAEQF | 0.5191338 |
| LAQQGCI | ASRDWGLYAEQF | 0.51657325 |
| LCLHVEN | ASRDWGLYAEQF | 0.63537514 |
| LDAISDGS | ASRDWGLYAEQF | 0.7288481 |
| LGGFAAML | ASRDWGLYAEQF | 0.6748424 |
| LGGKNPMI | ASRDWGLYAEQF | 0.6652712 |
| LGPKGVAAIL | ASRDWGLYAEQF | 0.7280375 |
| LMVGVSI | ASRDWGLYAEQF | 0.69258076 |
| LVGAAFVL | ASRDWGLYAEQF | 0.6770594 |
| LVLAVAPL | ASRDWGLYAEQF | 0.67222667 |
| MAAIAVIDAA | ASRDWGLYAEQF | 0.6677183 |
| NNGINAELQ | ASRDWGLYAEQF | 0.67491144 |
| RGEPLVLVHG | ASRDWGLYAEQF | 0.67033756 |
| RGGAGGAAGLI | ASRDWGLYAEQF | 0.732071 |
| RSYRMHPAV | ASRDWGLYAEQF | 0.69772387 |
| SAAHLLAGQLA | ASRDWGLYAEQF | 0.6567749 |
| SAAVGSY | ASRDWGLYAEQF | 0.7052158 |
| SPHAVLHRI | ASRDWGLYAEQF | 0.73056227 |
| SQIAKVL | ASRDWGLYAEQF | 0.7324469 |
| TALGHLVP | ASRDWGLYAEQF | 0.65047383 |
| TAPTTSGAA | ASRDWGLYAEQF | 0.7032618 |
| TDGDDGNGGSPGT | ASRDWGLYAEQF | 0.6947715 |
| TDGLGEN | ASRDWGLYAEQF | 0.69591916 |
| TGGPPLPGQA | ASRDWGLYAEQF | 0.69076306 |
| TIPAITI | ASRDWGLYAEQF | 0.6118018 |
| TIPGFTI | ASRDWGLYAEQF | 0.7187137 |
| TSIASGALA | ASRDWGLYAEQF | 0.7324417 |
| TVTLPGAAI | ASRDWGLYAEQF | 0.6951646 |
| VASGLSAALP | ASRDWGLYAEQF | 0.65228415 |
| VGAGVIGI | ASRDWGLYAEQF | 0.6930114 |
| VGPKRFGV | ASRDWGLYAEQF | 0.6461634 |
| VNVAFPD | ASRDWGLYAEQF | 0.6749903 |
| VSMTNSGVS | ASRDWGLYAEQF | 0.7112961 |
| VTDVLGDLSV | ASRDWGLYAEQF | 0.5831474 |
| VVVVGAGFAGL | ASRDWGLYAEQF | 0.677567 |
| VYTVGTVV | ASRDWGLYAEQF | 0.7315617 |
| WTTFSGP | ASRDWGLYAEQF | 0.6808744 |
| YDELVEAF | ASRDWGLYAEQF | 0.70415235 |
| YGTGGAGGNGG | ASRDWGLYAEQF | 0.7238289 |
| YTNVMAV | ASRDWGLYAEQF | 0.7139571 |

| | | |
|---|---|---|
| FLEQVHQL | ASRDWGLYAEQF | 0.7277244 |
| GLSLLAVAV | ASRDWGLYAEQF | 0.7213807 |
| LEQVHQL | ASRDWGLYAEQF | 0.7103174 |
| LNAAGAI | ASRDWGLYAEQF | 0.70468915 |
| PAVHASL | ASRDWGLYAEQF | 0.69763637 |
| TELEDDA | ASRDWGLYAEQF | 0.70505124 |
| TPPGPPN | ASRDWGLYAEQF | 0.6913758 |
| TPPPPPPP | ASRDWGLYAEQF | 0.68231636 |
| TQSPALI | ASSDAQTSNERLF | 0.7285033 |
| EFAAAFPQ | ASSDAQTSNERLF | 0.7024976 |
| LLSGNLDVL | ASSDAQTSNERLF | 0.72207934 |
| GPANLGNYNI | ASSDAQTSNERLF | 0.67722094 |
| TLGPTLP | ASSDAQTSNERLF | 0.65717906 |
| VYAYSSCCK | ASSDAQTSNERLF | 0.7194383 |
| APAPAPAPNI | ASSDAQTSNERLF | 0.30680272 |
| PGGPVPIP | ASSDAQTSNERLF | 0.43513215 |
| FLTVALEP | ASSDAQTSNERLF | 0.7060149 |
| SQYFGANL | ASSDAQTSNERLF | 0.7271414 |
| GGVKPAA | ASSDAQTSNERLF | 0.30510646 |
| GGTGGAGGAGGW | ASSDAQTSNERLF | 0.66549134 |
| GVGGVAGLGGAGG | ASSDAQTSNERLF | 0.6165467 |
| VLLVLATTI | ASSDAQTSNERLF | 0.71367776 |
| IGIFAAML | ASSDAQTSNERLF | 0.70177925 |
| AVGDLID | ASSDAQTSNERLF | 0.5729016 |
| NISPLPL | ASSDAQTSNERLF | 0.71903807 |
| VTAVFVAI | ASSDAQTSNERLF | 0.5880637 |
| SGDDNDAAL | ASSDAQTSNERLF | 0.5026188 |
| AVAALAHS | ASSDAQTSNERLF | 0.72635174 |
| GGVTVMATP | ASSDAQTSNERLF | 0.72667825 |
| AGLAANVRD | ASSDAQTSNERLF | 0.6631695 |
| TNRPDLIDPALLRPG | ASSDAQTSNERLF | 0.7240354 |
| LANLGVTAL | ASSDAQTSNERLF | 0.67307705 |
| TGERIEL | ASSDAQTSNERLF | 0.7316801 |
| NGEELAA | ASSDAQTSNERLF | 0.6946716 |
| VLSLVWG | ASSDAQTSNERLF | 0.6026734 |
| IGYALWL | ASSDAQTSNERLF | 0.7184275 |
| LSLLLAPL | ASSDAQTSNERLF | 0.7102771 |
| TLDEALK | ASSDAQTSNERLF | 0.7217499 |
| TDGAGTAEGG | ASSDAQTSNERLF | 0.71734905 |
| NAPGGAGGNGG | ASSDAQTSNERLF | 0.6652663 |
| VSVFMGMGGNF | ASSDAQTSNERLF | 0.67847204 |
| LASYLAAL | ASSDAQTSNERLF | 0.72643924 |
| LVGGTPAL | ASSDAQTSNERLF | 0.7242046 |
| VRNADVL | ASSDAQTSNERLF | 0.619944 |
| TCSSPFLVSAM | ASSDAQTSNERLF | 0.7245481 |
| VHITDAE | ASSDAQTSNERLF | 0.68752694 |
| EYAAPSF | ASSDAQTSNERLF | 0.68037516 |
| VVDAFHGGL | ASSDAQTSNERLF | 0.72953904 |
| ASVVDPQ | ASSDAQTSNERLF | 0.71421486 |
| SYNEFGGLS | ASSDAQTSNERLF | 0.7218648 |
| DPDGSVY | ASSDAQTSNERLF | 0.72482324 |

| | | |
|---|---|---|
| MVSTIHATE | ASSDAQTSNERLF | 0.7259284 |
| LAAGVADPV | ASSDAQTSNERLF | 0.63247275 |
| TKEQIAA | ASSDAQTSNERLF | 0.7153125 |
| SLASPAAL | ASSDAQTSNERLF | 0.7055076 |
| IGLVFYGV | ASSDAQTSNERLF | 0.7198329 |
| FLLDASNL | ASSDAQTSNERLF | 0.70678294 |
| YQVGNFP | ASSDAQTSNERLF | 0.5983902 |
| ESLLSQST | ASSDAQTSNERLF | 0.71523005 |
| LFTVGFAAVG | ASSDAQTSNERLF | 0.7282506 |
| ATPPTPT | ASSDAQTSNERLF | 0.7087012 |
| TDLATLG | ASSDAQTSNERLF | 0.71528506 |
| SAGGTGR | ASSDAQTSNERLF | 0.72906005 |
| GFSNSGS | ASSDAQTSNERLF | 0.69071287 |
| GGNGGSGGTSV | ASSDAQTSNERLF | 0.69468904 |
| GTSVATGGAGN | ASSDAQTSNERLF | 0.725213 |
| NGGNGGSGGTS | ASSDAQTSNERLF | 0.72756845 |
| AVVKFPH | ASSDAQTSNERLF | 0.70080245 |
| TPPPPPP | ASSDAQTSNERLF | 0.7273108 |
| AAPAAAAAPA | ASSDAQTSNERLF | 0.6607298 |
| IALNEDL | ASSDAQTSNERLF | 0.7174493 |
| IVNVSSM | ASSDAQTSNERLF | 0.6510022 |
| TGIPPAPR | ASSDAQTSNERLF | 0.71864796 |
| FDLGGGTF | ASSDAQTSNERLF | 0.62691975 |
| FDLGGGTFDVS | ASSDAQTSNERLF | 0.71913755 |
| NEPTAAA | ASSDAQTSNERLF | 0.7215588 |
| NEPTAAALA | ASSDAQTSNERLF | 0.72063786 |
| GSIINISSI | ASSDAQTSNERLF | 0.4825262 |
| VAPGNAGTA | ASSDAQTSNERLF | 0.71996844 |
| TGGGASLE | ASSDAQTSNERLF | 0.71890795 |
| VAVYNHYKRI | ASSDAQTSNERLF | 0.73305047 |
| TDPAPATT | ASSDAQTSNERLF | 0.7027323 |
| TNRPDILDPALLRPG | ASSDAQTSNERLF | 0.7181456 |
| LTTTTTS | ASSDAQTSNERLF | 0.689266 |
| PTGSGKTL | ASSDAQTSNERLF | 0.6931807 |
| TGSGKTL | ASSDAQTSNERLF | 0.64022565 |
| TIGHVDHGK | ASSDAQTSNERLF | 0.6921353 |
| TIGHVDHGKTT | ASSDAQTSNERLF | 0.6635625 |
| GHVDHGKTT | ASSDAQTSNERLF | 0.6408297 |
| AQFEHTLL | ASSDAQTSNERLF | 0.7215391 |
| IVNQDEL | ASSDAQTSNERLF | 0.7267104 |
| TNGSQFF | ASSDAQTSNERLF | 0.7216095 |
| TGKGIVE | ASSDAQTSNERLF | 0.72483754 |
| LISNASDA | ASSDAQTSNERLF | 0.7255241 |
| AELRTLGELTKIAKAHGAQ | ASSDAQTSNERLF | 0.72693837 |
| APNAVAAAEPSW | ASSDAQTSNERLF | 0.6760362 |
| APQVAGAATAVL | ASSDAQTSNERLF | 0.70255566 |
| ARRDPAAAAAILPTDA | ASSDAQTSNERLF | 0.6438718 |
| DAAARDVLAEAGLAEYF | ASSDAQTSNERLF | 0.6873356 |
| DDQIVLSAKANLAKVEQ | ASSDAQTSNERLF | 0.6775011 |
| DELLPDTGEDSDEDGH | ASSDAQTSNERLF | 0.7087029 |
| ELPEAPANRATVPQD | ASSDAQTSNERLF | 0.28130233 |

| | | |
|---|---|---|
| FGQSKIERVTGIG | ASSDAQTSNERLF | 0.6879171 |
| GAGGTGSTAGGAGGAGGAGGLY | ASSDAQTSNERLF | 0.73142505 |
| GHGGTGAAGGNGGTGGAGGSGI | ASSDAQTSNERLF | 0.6203538 |
| GNGGNGGAGGEGGPGGAGGAGA | ASSDAQTSNERLF | 0.7183708 |
| GSTAGNGGNGGNATNGTIAGQP | ASSDAQTSNERLF | 0.68809265 |
| GTGGDGGNGGNGGTGG | ASSDAQTSNERLF | 0.6722349 |
| GVRRPAGIPADAPV | ASSDAQTSNERLF | 0.7225819 |
| GYRQVTDAHLLTLARR | ASSDAQTSNERLF | 0.6642639 |
| IAKMGQDGHDRG | ASSDAQTSNERLF | 0.71631706 |
| KAEPSIEIQYT | ASSDAQTSNERLF | 0.7305016 |
| KPVRIRMAADEDTVAVT | ASSDAQTSNERLF | 0.70577264 |
| MPPRIAGMRLLVIKPEP | ASSDAQTSNERLF | 0.71567833 |
| MKFYAVALFRSRGVPPTR | ASSDAQTSNERLF | 0.7065248 |
| NAGGFGILGAVAHSP | ASSDAQTSNERLF | 0.601467 |
| NEGREITLAMAEM | ASSDAQTSNERLF | 0.716079 |
| NELVEWSHPLGHRWRWE | ASSDAQTSNERLF | 0.323362 |
| NLGAGGTGGNGGNPG | ASSDAQTSNERLF | 0.6891429 |
| NTEWPKWETVMAADPVD | ASSDAQTSNERLF | 0.6664963 |
| PASAAPAETQP | ASSDAQTSNERLF | 0.70372206 |
| QQSSDGKNGVRAAEITGE | ASSDAQTSNERLF | 0.69607073 |
| RAPDAVTTSWWKE | ASSDAQTSNERLF | 0.7163819 |
| SGFGNVGAHTSGWFNQ | ASSDAQTSNERLF | 0.6610799 |
| SGVAGAVGQAASVGGLKV | ASSDAQTSNERLF | 0.6684736 |
| SNGGAGTGSAGGNGGT | ASSDAQTSNERLF | 0.6369014 |
| TGGAGGEGGAAG | ASSDAQTSNERLF | 0.64025754 |
| TGGDGGAAGAGGAPGHG | ASSDAQTSNERLF | 0.7200278 |
| TMPPLLAAIANAGRHGV | ASSDAQTSNERLF | 0.7069964 |
| TSSTAGNSGGA | ASSDAQTSNERLF | 0.6737708 |
| VPATVPVGILC | ASSDAQTSNERLF | 0.7258573 |
| VQRPDSGGVVLTGRIS | ASSDAQTSNERLF | 0.71046734 |
| WTPNPVPATKGVPAGLVQAT | ASSDAQTSNERLF | 0.61350125 |
| YALKRALAKFSDRFDVVII | ASSDAQTSNERLF | 0.6813806 |
| YLIGSVIV | ASSDAQTSNERLF | 0.70219004 |
| YGQLVAVID | ASSDAQTSNERLF | 0.6979633 |
| VTVGGMTLPA | ASSDAQTSNERLF | 0.29756963 |
| VLALPAV | ASSDAQTSNERLF | 0.70576227 |
| VIPPTLNYET | ASSDAQTSNERLF | 0.683554 |
| VGMGGMHPSSG | ASSDAQTSNERLF | 0.6243179 |
| VGLAGGAATA | ASSDAQTSNERLF | 0.67150116 |
| VGGAGGAGGSGP | ASSDAQTSNERLF | 0.715911 |
| VFAAFMV | ASSDAQTSNERLF | 0.5844755 |
| TLYDKLR | ASSDAQTSNERLF | 0.7034967 |
| TLAVFTVL | ASSDAQTSNERLF | 0.72153014 |
| TGSVQPARV | ASSDAQTSNERLF | 0.7239476 |
| TGGGAGSGGT | ASSDAQTSNERLF | 0.7296468 |
| SHLETSL | ASSDAQTSNERLF | 0.71366966 |
| SFTTGMT | ASSDAQTSNERLF | 0.69546705 |
| QVEVAKRMGVRQPTVS | ASSDAQTSNERLF | 0.72158366 |
| QGSGGPALAM | ASSDAQTSNERLF | 0.7169776 |
| MLAPSNP | ASSDAQTSNERLF | 0.5378292 |
| LVNNAAI | ASSDAQTSNERLF | 0.46833992 |

| | | |
|---|---|---|
| LTGEINTLA | ASSDAQTSNERLF | 0.7087521 |
| LSIFGPV | ASSDAQTSNERLF | 0.5124167 |
| LLALLANL | ASSDAQTSNERLF | 0.70821524 |
| LISNASDALDKLR | ASSDAQTSNERLF | 0.7306467 |
| LINEAVP | ASSDAQTSNERLF | 0.69520915 |
| LGYAFSGL | ASSDAQTSNERLF | 0.65850633 |
| LDFVAVAL | ASSDAQTSNERLF | 0.69121826 |
| LAQPGAE | ASSDAQTSNERLF | 0.6825944 |
| ISNLGAVTGF | ASSDAQTSNERLF | 0.7218932 |
| HQEDPGRI | ASSDAQTSNERLF | 0.72372335 |
| GTPGEPTP | ASSDAQTSNERLF | 0.7275202 |
| GFFNSGS | ASSDAQTSNERLF | 0.7198686 |
| GAGGGGLTAGAN | ASSDAQTSNERLF | 0.6699164 |
| FVDALVNM | ASSDAQTSNERLF | 0.71575356 |
| FLTVAPEL | ASSDAQTSNERLF | 0.6799934 |
| FETLTAL | ASSDAQTSNERLF | 0.7010347 |
| DVINPIEL | ASSDAQTSNERLF | 0.7085201 |
| APVSGAPV | ASSDAQTSNERLF | 0.70961434 |
| ALVGGLAW | ASSDAQTSNERLF | 0.6993406 |
| ALAAAAATAI | ASSDAQTSNERLF | 0.6982343 |
| AFGLGLLL | ASSDAQTSNERLF | 0.6378336 |
| AFFAAASL | ASSDAQTSNERLF | 0.7217513 |
| AAAALPAFSPP | ASSDAQTSNERLF | 0.6917353 |
| VVLAPAL | ASSDAQTSNERLF | 0.5483134 |
| TIGHVDHGKTTL | ASSDAQTSNERLF | 0.6837036 |
| LISNASDALDK | ASSDAQTSNERLF | 0.7300469 |
| LISNASDAL | ASSDAQTSNERLF | 0.72862625 |
| ISNASDALDK | ASSDAQTSNERLF | 0.63295174 |
| NQAETLVYQTE | ASSDAQTSNERLF | 0.67113435 |
| TGGTGGTAGAG | ASSDAQTSNERLF | 0.6959457 |
| TGSTAGGAGGA | ASSDAQTSNERLF | 0.5029315 |
| DPLAVPVAAGPV | ASSDAQTSNERLF | 0.71548736 |
| GGTGGTPGNGGN | ASSDAQTSNERLF | 0.6887386 |
| PAGLAGVDPATL | ASSDAQTSNERLF | 0.6743887 |
| GNNLVVEAPHAGASV | ASSDAQTSNERLF | 0.6839552 |
| KVTRIMSASTLEQPA | ASSDAQTSNERLF | 0.6946362 |
| LGFALPIALPRRTVV | ASSDAQTSNERLF | 0.6553648 |
| TRLGNLTARKLVVLV | ASSDAQTSNERLF | 0.72397757 |
| WSQQRRLIYFPSAGP | ASSDAQTSNERLF | 0.31254208 |
| GPNGNPDPMAAAVDIR | ASSDAQTSNERLF | 0.71349263 |
| TQRCVSVADLRAAASR | ASSDAQTSNERLF | 0.6428908 |
| QFVLDVAEHPVEVLTETV | ASSDAQTSNERLF | 0.65531373 |
| GGAGGTGGAGGNGGTQAGDG | ASSDAQTSNERLF | 0.59622735 |
| PLLATLRAIAVDVDAKPAQV | ASSDAQTSNERLF | 0.7079312 |
| APKFRTGGYMIDFWGVGYQVAK | ASSDAQTSNERLF | 0.72313625 |
| GAGGAGGAGGAAGGPGGTGGTG | ASSDAQTSNERLF | 0.72062397 |
| IGALGLLATDVAPAGSVAFGVA | ASSDAQTSNERLF | 0.60359997 |
| AGGAGGIGGNGGAGGAGGVLMG | ASSDAQTSNERLF | 0.7250691 |
| ADELIGLL | ASSDAQTSNERLF | 0.7303841 |
| ADRALKA | ASSDAQTSNERLF | 0.6768954 |
| AGAVGQVGGAGGA | ASSDAQTSNERLF | 0.59420854 |

| | | |
|---|---|---|
| AGGGGGLGGAP | ASSDAQTSNERLF | 0.7173621 |
| AGGLLYGNGGAG | ASSDAQTSNERLF | 0.6991794 |
| AGLESIGL | ASSDAQTSNERLF | 0.7187995 |
| AGLKTGII | ASSDAQTSNERLF | 0.7124849 |
| ASLAALLL | ASSDAQTSNERLF | 0.610706 |
| ASVINISSI | ASSDAQTSNERLF | 0.7283802 |
| ATPPGPA | ASSDAQTSNERLF | 0.72279173 |
| ATVASALV | ASSDAQTSNERLF | 0.65733755 |
| ATVASLIL | ASSDAQTSNERLF | 0.69911677 |
| AVNGPVLIRH | ASSDAQTSNERLF | 0.68862706 |
| DGAAGAVGI | ASSDAQTSNERLF | 0.50616455 |
| DPLLHGI | ASSDAQTSNERLF | 0.72446376 |
| ELGAALGAL | ASSDAQTSNERLF | 0.6497607 |
| ELGRTQAV | ASSDAQTSNERLF | 0.6785477 |
| FVAGGYLS | ASSDAQTSNERLF | 0.7065283 |
| GASGGAGGAGGW | ASSDAQTSNERLF | 0.68831265 |
| GGAGAGGDGGAGGI | ASSDAQTSNERLF | 0.68855655 |
| GTGHNGSQPGMGGNG | ASSDAQTSNERLF | 0.6892052 |
| GYAAASASAS | ASSDAQTSNERLF | 0.7060212 |
| HQVDVAAVQQG | ASSDAQTSNERLF | 0.70007694 |
| IGAAIAHT | ASSDAQTSNERLF | 0.69525826 |
| IGMIEVAQL | ASSDAQTSNERLF | 0.70771456 |
| IRLAPEHTSAMD | ASSDAQTSNERLF | 0.7278946 |
| IYNVSVL | ASSDAQTSNERLF | 0.602391 |
| KQGFLGM | ASSDAQTSNERLF | 0.663509 |
| LESVITQL | ASSDAQTSNERLF | 0.7053518 |
| LGLGALVVV | ASSDAQTSNERLF | 0.5617236 |
| LIGNGGNGGSGGI | ASSDAQTSNERLF | 0.7141799 |
| LNAINAPTL | ASSDAQTSNERLF | 0.6955707 |
| LQLLNTGV | ASSDAQTSNERLF | 0.7082964 |
| LQTSIEF | ASSDAQTSNERLF | 0.6926065 |
| MSFAVSEL | ASSDAQTSNERLF | 0.73219717 |
| NTGALGL | ASSDAQTSNERLF | 0.72364545 |
| PPGAGGAGGAGTT | ASSDAQTSNERLF | 0.71172 |
| QLVGLGI | ASSDAQTSNERLF | 0.71003497 |
| RFLEQVHQL | ASSDAQTSNERLF | 0.6805974 |
| RGVGYVLR | ASSDAQTSNERLF | 0.7260211 |
| SATGVFAGVI | ASSDAQTSNERLF | 0.72048104 |
| SLPAITLP | ASSDAQTSNERLF | 0.6166649 |
| STALAFTV | ASSDAQTSNERLF | 0.68431973 |
| SVLTVGLGK | ASSDAQTSNERLF | 0.72905916 |
| TALAGAGSGI | ASSDAQTSNERLF | 0.72277266 |
| TGLPFTGI | ASSDAQTSNERLF | 0.5074125 |
| VAEHVTV | ASSDAQTSNERLF | 0.6750289 |
| VDIVLNSL | ASSDAQTSNERLF | 0.5081941 |
| VLTGGGAVL | ASSDAQTSNERLF | 0.6751418 |
| VNVGLAYGI | ASSDAQTSNERLF | 0.66981 |
| VSSAAVGAV | ASSDAQTSNERLF | 0.71463835 |
| VTSVYEAGV | ASSDAQTSNERLF | 0.69087154 |
| WVALDRGA | ASSDAQTSNERLF | 0.7121 |
| YLVGSVVI | ASSDAQTSNERLF | 0.7088013 |

| | | |
|---|---|---|
| GPPGTGKTLL | ASSDAQTSNERLF | 0.72944367 |
| GVGMTKF | ASSDAQTSNERLF | 0.67852336 |
| IGIGGSDL | ASSDAQTSNERLF | 0.6937804 |
| IGTLLAEL | ASSDAQTSNERLF | 0.61043525 |
| IYTSGTT | ASSDAQTSNERLF | 0.5384048 |
| LSFPAGV | ASSDAQTSNERLF | 0.6824531 |
| LYTSGTT | ASSDAQTSNERLF | 0.69170856 |
| VGAFPFGF | ASSDAQTSNERLF | 0.40471107 |
| VGAGTVEF | ASSDAQTSNERLF | 0.7229009 |
| VGKSTLLL | ASSDAQTSNERLF | 0.7142521 |
| AALAHAGGGIV | ASSDAQTSNERLF | 0.72089493 |
| AALVAVSLL | ASSDAQTSNERLF | 0.6920843 |
| AAVSRAVAAL | ASSDAQTSNERLF | 0.71093583 |
| AGGSSGIGGF | ASSDAQTSNERLF | 0.6688287 |
| AGLTVANANG | ASSDAQTSNERLF | 0.7142518 |
| AGLVGAVL | ASSDAQTSNERLF | 0.65485275 |
| ALSQAFTV | ASSDAQTSNERLF | 0.633959 |
| ALTFSVSL | ASSDAQTSNERLF | 0.69263387 |
| ALTGAALVAL | ASSDAQTSNERLF | 0.63909924 |
| ASFSIGAGVL | ASSDAQTSNERLF | 0.72652584 |
| DGSVYKI | ASSDAQTSNERLF | 0.72339153 |
| EQLAEIV | ASSDAQTSNERLF | 0.70869356 |
| FAVSELTVM | ASSDAQTSNERLF | 0.7237463 |
| GAAPLAAGL | ASSDAQTSNERLF | 0.5094024 |
| GADGGDGGSGGAGG | ASSDAQTSNERLF | 0.684902 |
| GFGLDTGGAGG | ASSDAQTSNERLF | 0.729553 |
| GLASGGLL | ASSDAQTSNERLF | 0.7109521 |
| GRIAGAAVLL | ASSDAQTSNERLF | 0.69509387 |
| GSAGSLNTSF | ASSDAQTSNERLF | 0.7287654 |
| GTGQIGI | ASSDAQTSNERLF | 0.66135174 |
| GTLIALIL | ASSDAQTSNERLF | 0.69900894 |
| IAATMGFVVG | ASSDAQTSNERLF | 0.7006109 |
| IFAQALL | ASSDAQTSNERLF | 0.72196805 |
| IGAAVALGF | ASSDAQTSNERLF | 0.7136413 |
| IGLTPLAL | ASSDAQTSNERLF | 0.6809469 |
| ILGSNTSTL | ASSDAQTSNERLF | 0.71461433 |
| IVFSLTVL | ASSDAQTSNERLF | 0.73183197 |
| LASKATGF | ASSDAQTSNERLF | 0.67989016 |
| LASVALIL | ASSDAQTSNERLF | 0.72280407 |
| LFAVGLTL | ASSDAQTSNERLF | 0.71753275 |
| LFNGGIGTYI | ASSDAQTSNERLF | 0.69739914 |
| LTLGAGII | ASSDAQTSNERLF | 0.6593889 |
| LVAAALGDI | ASSDAQTSNERLF | 0.67794335 |
| NTAFVQLGI | ASSDAQTSNERLF | 0.6787805 |
| QLFSGAEL | ASSDAQTSNERLF | 0.6349346 |
| SAATFITL | ASSDAQTSNERLF | 0.70332736 |
| SNYGLANLGV | ASSDAQTSNERLF | 0.6881363 |
| SYGGAYAVM | ASSDAQTSNERLF | 0.7336446 |
| TTGVGGLTLG | ASSDAQTSNERLF | 0.7168268 |
| TTQTIYI | ASSDAQTSNERLF | 0.709695 |
| VAAGAGLGAVI | ASSDAQTSNERLF | 0.6416906 |

| | | |
|---|---|---|
| VAAVVTLL | ASSDAQTSNERLF | 0.7124024 |
| VGFLGGLI | ASSDAQTSNERLF | 0.71174407 |
| VGVIPILL | ASSDAQTSNERLF | 0.6508241 |
| VIFASSLI | ASSDAQTSNERLF | 0.7292291 |
| VKGGDGGAAAG | ASSDAQTSNERLF | 0.6729187 |
| VTKTFTNI | ASSDAQTSNERLF | 0.6564672 |
| VVAAAILL | ASSDAQTSNERLF | 0.71885276 |
| YAALDVEL | ASSDAQTSNERLF | 0.72856075 |
| LASVALLL | ASSDAQTSNERLF | 0.69413066 |
| LDAVHRV | ASSDAQTSNERLF | 0.6609747 |
| QGLSSSIF | ASSDAQTSNERLF | 0.65596133 |
| RGTFANIRL | ASSDAQTSNERLF | 0.723036 |
| RQLWWGHRI | ASSDAQTSNERLF | 0.7240094 |
| YGPPGTGKTLL | ASSDAQTSNERLF | 0.69603604 |
| AALVGAPANVA | ASSDAQTSNERLF | 0.72874224 |
| AATFRRTE | ASSDAQTSNERLF | 0.716592 |
| AGALGGGI | ASSDAQTSNERLF | 0.6079646 |
| AGGAGGAGGIGAG | ASSDAQTSNERLF | 0.70528173 |
| AGSGALGL | ASSDAQTSNERLF | 0.72199744 |
| AQIADQI | ASSDAQTSNERLF | 0.71719885 |
| ASAVAVTYL | ASSDAQTSNERLF | 0.7333451 |
| ASTVLGGTGG | ASSDAQTSNERLF | 0.71872205 |
| AVGGEPAK | ASSDAQTSNERLF | 0.6750628 |
| CEQGTFGGTC | ASSDAQTSNERLF | 0.72630286 |
| DEGKAPAG | ASSDAQTSNERLF | 0.7318333 |
| DTVVFPAL | ASSDAQTSNERLF | 0.72431576 |
| FEGITMAG | ASSDAQTSNERLF | 0.68929285 |
| FPLVVQD | ASSDAQTSNERLF | 0.66322595 |
| GAFTTPQ | ASSDAQTSNERLF | 0.71310204 |
| GFGGFGTAGGVGGA | ASSDAQTSNERLF | 0.7156862 |
| GFLAQFL | ASSDAQTSNERLF | 0.67554176 |
| GGAGGAGGNAGVA | ASSDAQTSNERLF | 0.7040098 |
| GGVGTAGAGGAGGAGG | ASSDAQTSNERLF | 0.70784175 |
| GNGGGGDAGNAGSGGN | ASSDAQTSNERLF | 0.6426217 |
| GTGGAGGAGNDAGS | ASSDAQTSNERLF | 0.71549493 |
| GTTNTGWFNS | ASSDAQTSNERLF | 0.72685254 |
| IFMGPLFTL | ASSDAQTSNERLF | 0.70058286 |
| ILAHAIEAI | ASSDAQTSNERLF | 0.67668974 |
| IRSAIREAL | ASSDAQTSNERLF | 0.72416294 |
| ITFAGGG | ASSDAQTSNERLF | 0.73414564 |
| ITGMSGHQEV | ASSDAQTSNERLF | 0.72814196 |
| IVAGLACTL | ASSDAQTSNERLF | 0.6083828 |
| IVGGGLAGA | ASSDAQTSNERLF | 0.48838824 |
| IVTGVLNI | ASSDAQTSNERLF | 0.6401826 |
| LAQQGCI | ASSDAQTSNERLF | 0.7188467 |
| LCLHVEN | ASSDAQTSNERLF | 0.6924753 |
| LDAISDGS | ASSDAQTSNERLF | 0.67959 |
| LGGFAAML | ASSDAQTSNERLF | 0.7090114 |
| LGGKNPMI | ASSDAQTSNERLF | 0.72407365 |
| LGPKGVAAIL | ASSDAQTSNERLF | 0.6892762 |
| LMVGVSI | ASSDAQTSNERLF | 0.5870883 |

| | | |
|---|---|---|
| LVGAAFVL | ASSDAQTSNERLF | 0.7164787 |
| LVLAVAPL | ASSDAQTSNERLF | 0.7194994 |
| MAAIAVIDAA | ASSDAQTSNERLF | 0.64244324 |
| NNGINAELQ | ASSDAQTSNERLF | 0.68157864 |
| RGEPLVLVHG | ASSDAQTSNERLF | 0.63624793 |
| RGGAGGAAGLI | ASSDAQTSNERLF | 0.5439891 |
| RSYRMHPAV | ASSDAQTSNERLF | 0.7023114 |
| SAAHLLAGQLA | ASSDAQTSNERLF | 0.6463403 |
| SAAVGSY | ASSDAQTSNERLF | 0.723431 |
| SPHAVLHRI | ASSDAQTSNERLF | 0.6629189 |
| SQIAKVL | ASSDAQTSNERLF | 0.7226307 |
| TALGHLVP | ASSDAQTSNERLF | 0.7156476 |
| TAPTTSGAA | ASSDAQTSNERLF | 0.7204867 |
| TDGDDGNGGSPGT | ASSDAQTSNERLF | 0.6884237 |
| TDGLGEN | ASSDAQTSNERLF | 0.6755001 |
| TGGPPLPGQA | ASSDAQTSNERLF | 0.6385173 |
| TIPAITI | ASSDAQTSNERLF | 0.48671263 |
| TIPGFTI | ASSDAQTSNERLF | 0.7279393 |
| TSIASGALA | ASSDAQTSNERLF | 0.49633217 |
| TVTLPGAAI | ASSDAQTSNERLF | 0.7290616 |
| VASGLSAALP | ASSDAQTSNERLF | 0.72452813 |
| VGAGVIGI | ASSDAQTSNERLF | 0.6518622 |
| VGPKRFGV | ASSDAQTSNERLF | 0.71260273 |
| VNVAFPD | ASSDAQTSNERLF | 0.6341969 |
| VSMTNSGVS | ASSDAQTSNERLF | 0.6352803 |
| VTDVLGDLSV | ASSDAQTSNERLF | 0.72201097 |
| VVVVGAGFAGL | ASSDAQTSNERLF | 0.7160659 |
| VYTVGTVV | ASSDAQTSNERLF | 0.67317736 |
| WTTFSGP | ASSDAQTSNERLF | 0.7164555 |
| YDELVEAF | ASSDAQTSNERLF | 0.7051315 |
| YGTGGAGGNGG | ASSDAQTSNERLF | 0.679689 |
| YTNVMAV | ASSDAQTSNERLF | 0.7071359 |
| FLEQVHQL | ASSDAQTSNERLF | 0.7225198 |
| GLSLLAVAV | ASSDAQTSNERLF | 0.71857715 |
| LEQVHQL | ASSDAQTSNERLF | 0.67230827 |
| LNAAGAI | ASSDAQTSNERLF | 0.71919614 |
| PAVHASL | ASSDAQTSNERLF | 0.63008076 |
| TELEDDA | ASSDAQTSNERLF | 0.7077675 |
| TPPGPPN | ASSDAQTSNERLF | 0.7004095 |
| TPPPPPPP | ASSDAQTSNERLF | 0.70194274 |
| TQSPALI | ASSEQNSGNTLY | 0.7150886 |
| EFAAAFPQ | ASSEQNSGNTLY | 0.6508244 |
| LLSGNLDVL | ASSEQNSGNTLY | 0.65840924 |
| GPANLGNYNI | ASSEQNSGNTLY | 0.69926095 |
| TLGPTLP | ASSEQNSGNTLY | 0.7085202 |
| VYAYSSCCK | ASSEQNSGNTLY | 0.7207256 |
| APAPAPAPNI | ASSEQNSGNTLY | 0.6468437 |
| PGGPVPIP | ASSEQNSGNTLY | 0.70468605 |
| FLTVALEP | ASSEQNSGNTLY | 0.69114256 |
| SQYFGANL | ASSEQNSGNTLY | 0.6611371 |
| GGVKPAA | ASSEQNSGNTLY | 0.7127678 |

| | | |
|---|---|---|
| GGTGGAGGAGGW | ASSEQNSGNTLY | 0.704289 |
| GVGGVAGLGGAGG | ASSEQNSGNTLY | 0.6760877 |
| VLLVLATTI | ASSEQNSGNTLY | 0.7198956 |
| IGIFAAML | ASSEQNSGNTLY | 0.6601568 |
| AVGDLID | ASSEQNSGNTLY | 0.7172758 |
| NISPLPL | ASSEQNSGNTLY | 0.7276202 |
| VTAVFVAI | ASSEQNSGNTLY | 0.6977817 |
| SGDDNDAAL | ASSEQNSGNTLY | 0.71983397 |
| AVAALAHS | ASSEQNSGNTLY | 0.7096889 |
| GGVTVMATP | ASSEQNSGNTLY | 0.7141403 |
| AGLAANVRD | ASSEQNSGNTLY | 0.69448227 |
| TNRPDLIDPALLRPG | ASSEQNSGNTLY | 0.6790463 |
| LANLGVTAL | ASSEQNSGNTLY | 0.723203 |
| TGERIEL | ASSEQNSGNTLY | 0.71768516 |
| NGEELAA | ASSEQNSGNTLY | 0.72691905 |
| VLSLVWG | ASSEQNSGNTLY | 0.7116685 |
| IGYALWL | ASSEQNSGNTLY | 0.72202104 |
| LSLLLAPL | ASSEQNSGNTLY | 0.71555007 |
| TLDEALK | ASSEQNSGNTLY | 0.6623815 |
| TDGAGTAEGG | ASSEQNSGNTLY | 0.6070519 |
| NAPGGAGGNGG | ASSEQNSGNTLY | 0.6829123 |
| VSVFMGMGGNF | ASSEQNSGNTLY | 0.68954647 |
| LASYLAAL | ASSEQNSGNTLY | 0.7078399 |
| LVGGTPAL | ASSEQNSGNTLY | 0.5428539 |
| VRNADVL | ASSEQNSGNTLY | 0.7227553 |
| TCSSPFLVSAM | ASSEQNSGNTLY | 0.7332655 |
| VHITDAE | ASSEQNSGNTLY | 0.67454624 |
| EYAAPSF | ASSEQNSGNTLY | 0.7296988 |
| VVDAFHGGL | ASSEQNSGNTLY | 0.638224 |
| ASVVDPQ | ASSEQNSGNTLY | 0.7192057 |
| SYNEFGGLS | ASSEQNSGNTLY | 0.72603905 |
| DPDGSVY | ASSEQNSGNTLY | 0.634464 |
| MVSTIHATE | ASSEQNSGNTLY | 0.71905744 |
| LAAGVADPV | ASSEQNSGNTLY | 0.693307 |
| TKEQIAA | ASSEQNSGNTLY | 0.698781 |
| SLASPAAL | ASSEQNSGNTLY | 0.7307692 |
| IGLVFYGV | ASSEQNSGNTLY | 0.69283426 |
| FLLDASNL | ASSEQNSGNTLY | 0.7175127 |
| YQVGNFP | ASSEQNSGNTLY | 0.69890535 |
| ESLLSQST | ASSEQNSGNTLY | 0.66782904 |
| LFTVGFAAVG | ASSEQNSGNTLY | 0.71347463 |
| ATPPTPT | ASSEQNSGNTLY | 0.7300736 |
| TDLATLG | ASSEQNSGNTLY | 0.6423306 |
| SAGGTGR | ASSEQNSGNTLY | 0.7313309 |
| GFSNSGS | ASSEQNSGNTLY | 0.68562955 |
| GGNGGSGGTSV | ASSEQNSGNTLY | 0.73155886 |
| GTSVATGGAGN | ASSEQNSGNTLY | 0.7177947 |
| NGGNGGSGGTS | ASSEQNSGNTLY | 0.6915074 |
| AVVKFPH | ASSEQNSGNTLY | 0.6920605 |
| TPPPPPP | ASSEQNSGNTLY | 0.72716933 |
| AAPAAAAAPA | ASSEQNSGNTLY | 0.6859419 |

| | | |
|---|---|---|
| IALNEDL | ASSEQNSGNTLY | 0.7237163 |
| IVNVSSM | ASSEQNSGNTLY | 0.7295074 |
| TGIPPAPR | ASSEQNSGNTLY | 0.52148664 |
| FDLGGGTF | ASSEQNSGNTLY | 0.78301054 |
| FDLGGGTFDVS | ASSEQNSGNTLY | 0.7370761 |
| NEPTAAA | ASSEQNSGNTLY | 0.6801692 |
| NEPTAAALA | ASSEQNSGNTLY | 0.59736514 |
| GSIINISSI | ASSEQNSGNTLY | 0.42004836 |
| VAPGNAGTA | ASSEQNSGNTLY | 0.6296079 |
| TGGGASLE | ASSEQNSGNTLY | 0.53994465 |
| VAVYNHYKRI | ASSEQNSGNTLY | 0.7734089 |
| TDPAPATT | ASSEQNSGNTLY | 0.7615559 |
| TNRPDILDPALLRPG | ASSEQNSGNTLY | 0.5654849 |
| LTTTTTS | ASSEQNSGNTLY | 0.7848168 |
| PTGSGKTL | ASSEQNSGNTLY | 0.7584373 |
| TGSGKTL | ASSEQNSGNTLY | 0.4504337 |
| TIGHVDHGK | ASSEQNSGNTLY | 0.68919426 |
| TIGHVDHGKTT | ASSEQNSGNTLY | 0.5783236 |
| GHVDHGKTT | ASSEQNSGNTLY | 0.6411408 |
| AQFEHTLL | ASSEQNSGNTLY | 0.38734794 |
| IVNQDEL | ASSEQNSGNTLY | 0.7481755 |
| TNGSQFF | ASSEQNSGNTLY | 0.62745684 |
| TGKGIVE | ASSEQNSGNTLY | 0.25968248 |
| LISNASDA | ASSEQNSGNTLY | 0.72356987 |
| AELRTLGELTKIAKAHGAQ | ASSEQNSGNTLY | 0.7097353 |
| APNAVAAAEPSW | ASSEQNSGNTLY | 0.68083394 |
| APQVAGAATAVL | ASSEQNSGNTLY | 0.7707591 |
| ARRDPAAAAAILPTDA | ASSEQNSGNTLY | 0.53294015 |
| DAAARDVLAEAGLAEYF | ASSEQNSGNTLY | 0.600497 |
| DDQIVLSAKANLAKVEQ | ASSEQNSGNTLY | 0.7719326 |
| DELLPDTGEDSDEDGH | ASSEQNSGNTLY | 0.7781664 |
| ELPEAPANRATVPQD | ASSEQNSGNTLY | 0.5164174 |
| FGQSKIERVTGIG | ASSEQNSGNTLY | 0.7368698 |
| GAGGTGSTAGGAGGAGGAGGLY | ASSEQNSGNTLY | 0.7641002 |
| GHGGTGAAGGNGGTGGAGGSGI | ASSEQNSGNTLY | 0.7233354 |
| GNGGNGGAGGEGGPGGAGGAGA | ASSEQNSGNTLY | 0.77038825 |
| GSTAGNGGNGGNATNGTIAGQP | ASSEQNSGNTLY | 0.7769195 |
| GTGGDGGNGGNGGTGG | ASSEQNSGNTLY | 0.638237 |
| GVRRPAGIPADAPV | ASSEQNSGNTLY | 0.7784085 |
| GYRQVTDAHLLTLARR | ASSEQNSGNTLY | 0.78810924 |
| IAKMGQDGHDRG | ASSEQNSGNTLY | 0.65082866 |
| KAEPSIEIQYT | ASSEQNSGNTLY | 0.69427323 |
| KPVRIRMAADEDTVAVT | ASSEQNSGNTLY | 0.47242337 |
| MPPRIAGMRLLVIKPEP | ASSEQNSGNTLY | 0.68582106 |
| MKFYAVALFRSRGVPPTR | ASSEQNSGNTLY | 0.44344988 |
| NAGGFGILGAVAHSP | ASSEQNSGNTLY | 0.7514435 |
| NEGREITLAMAEM | ASSEQNSGNTLY | 0.76272696 |
| NELVEWSHPLGHRWRWE | ASSEQNSGNTLY | 0.67174876 |
| NLGAGGTGGNGGNPG | ASSEQNSGNTLY | 0.52285564 |
| NTEWPKWETVMAADPVD | ASSEQNSGNTLY | 0.4296279 |
| PASAAPAETQP | ASSEQNSGNTLY | 0.7037594 |

| | | |
|---|---|---|
| QQSSDGKNGVRAAEITGE | ASSEQNSGNTLY | 0.5253862 |
| RAPDAVTTSWWKE | ASSEQNSGNTLY | 0.33191592 |
| SGFGNVGAHTSGWFNQ | ASSEQNSGNTLY | 0.6925584 |
| SGVAGAVGQAASVGGLKV | ASSEQNSGNTLY | 0.76907444 |
| SNGGAGTGSAGGNGGT | ASSEQNSGNTLY | 0.7537403 |
| TGGAGGEGGAAG | ASSEQNSGNTLY | 0.5261203 |
| TGGDGGAAGAGGAPGHG | ASSEQNSGNTLY | 0.78736186 |
| TMPPLLAAIANAGRHGV | ASSEQNSGNTLY | 0.7637949 |
| TSSTAGNSGGA | ASSEQNSGNTLY | 0.7408538 |
| VPATVPVGILC | ASSEQNSGNTLY | 0.6253989 |
| VQRPDSGGVVLTGRIS | ASSEQNSGNTLY | 0.5010414 |
| WTPNPVPATKGVPAGLVQAT | ASSEQNSGNTLY | 0.77809024 |
| YALKRALAKFSDRFDVVII | ASSEQNSGNTLY | 0.7045933 |
| YLIGSVIV | ASSEQNSGNTLY | 0.6621572 |
| YGQLVAVID | ASSEQNSGNTLY | 0.76273763 |
| VTVGGMTLPA | ASSEQNSGNTLY | 0.70115626 |
| VLALPAV | ASSEQNSGNTLY | 0.61883944 |
| VIPPTLNYET | ASSEQNSGNTLY | 0.7013491 |
| VGMGGMHPSSG | ASSEQNSGNTLY | 0.73804355 |
| VGLAGGAATA | ASSEQNSGNTLY | 0.6259169 |
| VGGAGGAGGSGP | ASSEQNSGNTLY | 0.42470884 |
| VFAAFMV | ASSEQNSGNTLY | 0.61738944 |
| TLYDKLR | ASSEQNSGNTLY | 0.7050463 |
| TLAVFTVL | ASSEQNSGNTLY | 0.77542233 |
| TGSVQPARV | ASSEQNSGNTLY | 0.70760864 |
| TGGGAGSGGT | ASSEQNSGNTLY | 0.6097621 |
| SHLETSL | ASSEQNSGNTLY | 0.75728 |
| SFTTGMT | ASSEQNSGNTLY | 0.71206254 |
| QVEVAKRMGVRQPTVS | ASSEQNSGNTLY | 0.69225895 |
| QGSGGPALAM | ASSEQNSGNTLY | 0.7260798 |
| MLAPSNP | ASSEQNSGNTLY | 0.653972 |
| LVNNAAI | ASSEQNSGNTLY | 0.7454717 |
| LTGEINTLA | ASSEQNSGNTLY | 0.72741526 |
| LSIFGPV | ASSEQNSGNTLY | 0.7348876 |
| LLALLANL | ASSEQNSGNTLY | 0.769576 |
| LISNASDALDKLR | ASSEQNSGNTLY | 0.7391634 |
| LINEAVP | ASSEQNSGNTLY | 0.67535114 |
| LGYAFSGL | ASSEQNSGNTLY | 0.3266023 |
| LDFVAVAL | ASSEQNSGNTLY | 0.67929804 |
| LAQPGAE | ASSEQNSGNTLY | 0.74943864 |
| ISNLGAVTGF | ASSEQNSGNTLY | 0.7354994 |
| HQEDPGRI | ASSEQNSGNTLY | 0.772736 |
| GTPGEPTP | ASSEQNSGNTLY | 0.71150243 |
| GFFNSGS | ASSEQNSGNTLY | 0.63599825 |
| GAGGGGLTAGAN | ASSEQNSGNTLY | 0.7588526 |
| FVDALVNM | ASSEQNSGNTLY | 0.7664605 |
| FLTVAPEL | ASSEQNSGNTLY | 0.39727122 |
| FETLTAL | ASSEQNSGNTLY | 0.68077517 |
| DVINPIEL | ASSEQNSGNTLY | 0.79321826 |
| APVSGAPV | ASSEQNSGNTLY | 0.7407904 |
| ALVGGLAW | ASSEQNSGNTLY | 0.6782473 |

| | | |
|---|---|---|
| ALAAAAATAI | ASSEQNSGNTLY | 0.59730744 |
| AFGLGLLL | ASSEQNSGNTLY | 0.7848469 |
| AFFAAASL | ASSEQNSGNTLY | 0.6319433 |
| AAAALPAFSPP | ASSEQNSGNTLY | 0.7884227 |
| VVLAPAL | ASSEQNSGNTLY | 0.27030358 |
| TIGHVDHGKTTL | ASSEQNSGNTLY | 0.5322689 |
| LISNASDALDK | ASSEQNSGNTLY | 0.4288044 |
| LISNASDAL | ASSEQNSGNTLY | 0.2756459 |
| ISNASDALDK | ASSEQNSGNTLY | 0.7664646 |
| NQAETLVYQTE | ASSEQNSGNTLY | 0.55892277 |
| TGGTGGTAGAG | ASSEQNSGNTLY | 0.7530439 |
| TGSTAGGAGGA | ASSEQNSGNTLY | 0.7134114 |
| DPLAVPVAAGPV | ASSEQNSGNTLY | 0.7750015 |
| GGTGGTPGNGGN | ASSEQNSGNTLY | 0.7140812 |
| PAGLAGVDPATL | ASSEQNSGNTLY | 0.70604193 |
| GNNLVVEAPHAGASV | ASSEQNSGNTLY | 0.48813057 |
| KVTRIMSASTLEQPA | ASSEQNSGNTLY | 0.7558472 |
| LGFALPIALPRRTVV | ASSEQNSGNTLY | 0.77506435 |
| TRLGNLTARKLVVLV | ASSEQNSGNTLY | 0.643878 |
| WSQQRRLIYFPSAGP | ASSEQNSGNTLY | 0.4359749 |
| GPNGNPDPMAAAVDIR | ASSEQNSGNTLY | 0.71599346 |
| TQRCVSVADLRAAASR | ASSEQNSGNTLY | 0.6875425 |
| QFVLDVAEHPVEVLTETV | ASSEQNSGNTLY | 0.3069675 |
| GGAGGTGGAGGNGGTQAGDG | ASSEQNSGNTLY | 0.29401505 |
| PLLATLRAIAVDVDAKPAQV | ASSEQNSGNTLY | 0.6038233 |
| APKFRTGGYMIDFWGVGYQVAK | ASSEQNSGNTLY | 0.6945355 |
| GAGGAGGAGGAAGGPGGTGGTG | ASSEQNSGNTLY | 0.77531946 |
| IGALGLLATDVAPAGSVAFGVA | ASSEQNSGNTLY | 0.7716583 |
| AGGAGGIGGNGGAGGAGGVLMG | ASSEQNSGNTLY | 0.6373346 |
| ADELIGLL | ASSEQNSGNTLY | 0.74346685 |
| ADRALKA | ASSEQNSGNTLY | 0.7445357 |
| AGAVGQVGGAGGA | ASSEQNSGNTLY | 0.20068018 |
| AGGGGGLGGAP | ASSEQNSGNTLY | 0.5958686 |
| AGGLLYGNGGAG | ASSEQNSGNTLY | 0.4729732 |
| AGLESIGL | ASSEQNSGNTLY | 0.781512 |
| AGLKTGII | ASSEQNSGNTLY | 0.66038704 |
| ASLAALLL | ASSEQNSGNTLY | 0.67501056 |
| ASVINISSI | ASSEQNSGNTLY | 0.6833961 |
| ATPPGPA | ASSEQNSGNTLY | 0.7469133 |
| ATVASALV | ASSEQNSGNTLY | 0.71271443 |
| ATVASLIL | ASSEQNSGNTLY | 0.5372555 |
| AVNGPVLIRH | ASSEQNSGNTLY | 0.31827104 |
| DGAAGAVGI | ASSEQNSGNTLY | 0.75350094 |
| DPLLHGI | ASSEQNSGNTLY | 0.53610843 |
| ELGAALGAL | ASSEQNSGNTLY | 0.58285046 |
| ELGRTQAV | ASSEQNSGNTLY | 0.3644556 |
| FVAGGYLS | ASSEQNSGNTLY | 0.73232764 |
| GASGGAGGAGGW | ASSEQNSGNTLY | 0.37388617 |
| GGAGAGGDGGAGGI | ASSEQNSGNTLY | 0.7391214 |
| GTGHNGSQPGMGGNG | ASSEQNSGNTLY | 0.65884197 |
| GYAAASASAS | ASSEQNSGNTLY | 0.7206181 |

| | | |
|---|---|---|
| HQVDVAAVQQG | ASSEQNSGNTLY | 0.78992355 |
| IGAAIAHT | ASSEQNSGNTLY | 0.74437404 |
| IGMIEVAQL | ASSEQNSGNTLY | 0.66354954 |
| IRLAPEHTSAMD | ASSEQNSGNTLY | 0.60877675 |
| IYNVSVL | ASSEQNSGNTLY | 0.5151541 |
| KQGFLGM | ASSEQNSGNTLY | 0.5432125 |
| LESVITQL | ASSEQNSGNTLY | 0.6174034 |
| LGLGALVVV | ASSEQNSGNTLY | 0.7853209 |
| LIGNGGNGGSGGI | ASSEQNSGNTLY | 0.574609 |
| LNAINAPTL | ASSEQNSGNTLY | 0.43061936 |
| LQLLNTGV | ASSEQNSGNTLY | 0.58082175 |
| LQTSIEF | ASSEQNSGNTLY | 0.7847144 |
| MSFAVSEL | ASSEQNSGNTLY | 0.78442585 |
| NTGALGL | ASSEQNSGNTLY | 0.61783826 |
| PPGAGGAGGAGTT | ASSEQNSGNTLY | 0.52029175 |
| QLVGLGI | ASSEQNSGNTLY | 0.55345386 |
| RFLEQVHQL | ASSEQNSGNTLY | 0.7205512 |
| RGVGYVLR | ASSEQNSGNTLY | 0.49576712 |
| SATGVFAGVI | ASSEQNSGNTLY | 0.4523223 |
| SLPAITLP | ASSEQNSGNTLY | 0.71200204 |
| STALAFTV | ASSEQNSGNTLY | 0.78970236 |
| SVLTVGLGK | ASSEQNSGNTLY | 0.63936186 |
| TALAGAGSGI | ASSEQNSGNTLY | 0.34689367 |
| TGLPFTGI | ASSEQNSGNTLY | 0.7048598 |
| VAEHVTV | ASSEQNSGNTLY | 0.57768506 |
| VDIVLNSL | ASSEQNSGNTLY | 0.6004553 |
| VLTGGGAVL | ASSEQNSGNTLY | 0.6458121 |
| VNVGLAYGI | ASSEQNSGNTLY | 0.4069398 |
| VSSAAVGAV | ASSEQNSGNTLY | 0.707631 |
| VTSVYEAGV | ASSEQNSGNTLY | 0.7824343 |
| WVALDRGA | ASSEQNSGNTLY | 0.70292723 |
| YLVGSVVI | ASSEQNSGNTLY | 0.7362391 |
| GPPGTGKTLL | ASSEQNSGNTLY | 0.7803359 |
| GVGMTKF | ASSEQNSGNTLY | 0.7543426 |
| IGIGGSDL | ASSEQNSGNTLY | 0.7789225 |
| IGTLLAEL | ASSEQNSGNTLY | 0.77345014 |
| IYTSGTT | ASSEQNSGNTLY | 0.68437696 |
| LSFPAGV | ASSEQNSGNTLY | 0.6653737 |
| LYTSGTT | ASSEQNSGNTLY | 0.77672017 |
| VGAFPFGF | ASSEQNSGNTLY | 0.71570486 |
| VGAGTVEF | ASSEQNSGNTLY | 0.6863892 |
| VGKSTLLL | ASSEQNSGNTLY | 0.60081786 |
| AALAHAGGGIV | ASSEQNSGNTLY | 0.7774923 |
| AALVAVSLL | ASSEQNSGNTLY | 0.57646656 |
| AAVSRAVAAL | ASSEQNSGNTLY | 0.77114797 |
| AGGSSGIGGF | ASSEQNSGNTLY | 0.30686364 |
| AGLTVANANG | ASSEQNSGNTLY | 0.46753848 |
| AGLVGAVL | ASSEQNSGNTLY | 0.7688166 |
| ALSQAFTV | ASSEQNSGNTLY | 0.1959776 |
| ALTFSVSL | ASSEQNSGNTLY | 0.26955014 |
| ALTGAALVAL | ASSEQNSGNTLY | 0.6534892 |

| | | |
|---|---|---|
| ASFSIGAGVL | ASSEQNSGNTLY | 0.7708807 |
| DGSVYKI | ASSEQNSGNTLY | 0.21207711 |
| EQLAEIV | ASSEQNSGNTLY | 0.56357265 |
| FAVSELTVM | ASSEQNSGNTLY | 0.36499426 |
| GAAPLAAGL | ASSEQNSGNTLY | 0.70012915 |
| GADGGDGGSGGAGG | ASSEQNSGNTLY | 0.71755785 |
| GFGLDTGGAGG | ASSEQNSGNTLY | 0.36133444 |
| GLASGGLL | ASSEQNSGNTLY | 0.745262 |
| GRIAGAAVLL | ASSEQNSGNTLY | 0.34128484 |
| GSAGSLNTSF | ASSEQNSGNTLY | 0.6529826 |
| GTGQIGI | ASSEQNSGNTLY | 0.76487774 |
| GTLIALIL | ASSEQNSGNTLY | 0.7831521 |
| IAATMGFVVG | ASSEQNSGNTLY | 0.5679002 |
| IFAQALL | ASSEQNSGNTLY | 0.7551677 |
| IGAAVALGF | ASSEQNSGNTLY | 0.6054031 |
| IGLTPLAL | ASSEQNSGNTLY | 0.79058886 |
| ILGSNTSTL | ASSEQNSGNTLY | 0.6649116 |
| IVFSLTVL | ASSEQNSGNTLY | 0.3788047 |
| LASKATGF | ASSEQNSGNTLY | 0.7631041 |
| LASVALIL | ASSEQNSGNTLY | 0.7298286 |
| LFAVGLTL | ASSEQNSGNTLY | 0.6770437 |
| LFNGGIGTYI | ASSEQNSGNTLY | 0.76240873 |
| LTLGAGII | ASSEQNSGNTLY | 0.49199837 |
| LVAAALGDI | ASSEQNSGNTLY | 0.6903719 |
| NTAFVQLGI | ASSEQNSGNTLY | 0.77639735 |
| QLFSGAEL | ASSEQNSGNTLY | 0.7726843 |
| SAATFITL | ASSEQNSGNTLY | 0.4658025 |
| SNYGLANLGV | ASSEQNSGNTLY | 0.7408526 |
| SYGGAYAVM | ASSEQNSGNTLY | 0.70256305 |
| TTGVGGLTLG | ASSEQNSGNTLY | 0.55796576 |
| TTQTIYI | ASSEQNSGNTLY | 0.7835326 |
| VAAGAGLGAVI | ASSEQNSGNTLY | 0.6165211 |
| VAAVVTLL | ASSEQNSGNTLY | 0.7537079 |
| VGFLGGLI | ASSEQNSGNTLY | 0.732095 |
| VGVIPILL | ASSEQNSGNTLY | 0.7709439 |
| VIFASSLI | ASSEQNSGNTLY | 0.51176083 |
| VKGGDGGAAAG | ASSEQNSGNTLY | 0.72893035 |
| VTKTFTNI | ASSEQNSGNTLY | 0.7090111 |
| VVAAAILL | ASSEQNSGNTLY | 0.7618437 |
| YAALDVEL | ASSEQNSGNTLY | 0.67219865 |
| LASVALLL | ASSEQNSGNTLY | 0.7004557 |
| LDAVHRV | ASSEQNSGNTLY | 0.73073983 |
| QGLSSSIF | ASSEQNSGNTLY | 0.77595943 |
| RGTFANIRL | ASSEQNSGNTLY | 0.6844723 |
| RQLWWGHRI | ASSEQNSGNTLY | 0.73512447 |
| YGPPGTGKTLL | ASSEQNSGNTLY | 0.7838971 |
| AALVGAPANVA | ASSEQNSGNTLY | 0.5328125 |
| AATFRRTE | ASSEQNSGNTLY | 0.6583319 |
| AGALGGGI | ASSEQNSGNTLY | 0.74228466 |
| AGGAGGAGGIGAG | ASSEQNSGNTLY | 0.7744731 |
| AGSGALGL | ASSEQNSGNTLY | 0.6176033 |

| | | |
|---|---|---|
| AQIADQI | ASSEQNSGNTLY | 0.77757037 |
| ASAVAVTYL | ASSEQNSGNTLY | 0.6704984 |
| ASTVLGGTGG | ASSEQNSGNTLY | 0.7742678 |
| AVGGEPAK | ASSEQNSGNTLY | 0.7166928 |
| CEQGTFGGTC | ASSEQNSGNTLY | 0.7270535 |
| DEGKAPAG | ASSEQNSGNTLY | 0.62492025 |
| DTVVFPAL | ASSEQNSGNTLY | 0.7735448 |
| FEGITMAG | ASSEQNSGNTLY | 0.7653167 |
| FPLVVQD | ASSEQNSGNTLY | 0.7748443 |
| GAFTTPQ | ASSEQNSGNTLY | 0.26227358 |
| GFGGFGTAGGVGGA | ASSEQNSGNTLY | 0.76478636 |
| GFLAQFL | ASSEQNSGNTLY | 0.7733151 |
| GGAGGAGGNAGVA | ASSEQNSGNTLY | 0.7889231 |
| GGVGTAGAGGAGGAGG | ASSEQNSGNTLY | 0.6188843 |
| GNGGGGDAGNAGSGGN | ASSEQNSGNTLY | 0.7116264 |
| GTGGAGGAGNDAGS | ASSEQNSGNTLY | 0.67771995 |
| GTTNTGWFNS | ASSEQNSGNTLY | 0.6347624 |
| IFMGPLFTL | ASSEQNSGNTLY | 0.45727915 |
| ILAHAIEAI | ASSEQNSGNTLY | 0.64524055 |
| IRSAIREAL | ASSEQNSGNTLY | 0.6951152 |
| ITFAGGG | ASSEQNSGNTLY | 0.6271941 |
| ITGMSGHQEV | ASSEQNSGNTLY | 0.7702433 |
| IVAGLACTL | ASSEQNSGNTLY | 0.77445495 |
| IVGGGLAGA | ASSEQNSGNTLY | 0.77716255 |
| IVTGVLNI | ASSEQNSGNTLY | 0.77567863 |
| LAQQGCI | ASSEQNSGNTLY | 0.78015184 |
| LCLHVEN | ASSEQNSGNTLY | 0.728616 |
| LDAISDGS | ASSEQNSGNTLY | 0.6038598 |
| LGGFAAML | ASSEQNSGNTLY | 0.7178689 |
| LGGKNPMI | ASSEQNSGNTLY | 0.47745812 |
| LGPKGVAAIL | ASSEQNSGNTLY | 0.7041869 |
| LMVGVSI | ASSEQNSGNTLY | 0.64416254 |
| LVGAAFVL | ASSEQNSGNTLY | 0.7360019 |
| LVLAVAPL | ASSEQNSGNTLY | 0.20180185 |
| MAAIAVIDAA | ASSEQNSGNTLY | 0.74897635 |
| NNGINAELQ | ASSEQNSGNTLY | 0.78814006 |
| RGEPLVLVHG | ASSEQNSGNTLY | 0.6672367 |
| RGGAGGAAGLI | ASSEQNSGNTLY | 0.7275273 |
| RSYRMHPAV | ASSEQNSGNTLY | 0.7605329 |
| SAAHLLAGQLA | ASSEQNSGNTLY | 0.6096107 |
| SAAVGSY | ASSEQNSGNTLY | 0.7526994 |
| SPHAVLHRI | ASSEQNSGNTLY | 0.55725455 |
| SQIAKVL | ASSEQNSGNTLY | 0.7339212 |
| TALGHLVP | ASSEQNSGNTLY | 0.7806569 |
| TAPTTSGAA | ASSEQNSGNTLY | 0.7298105 |
| TDGDDGNGGSPGT | ASSEQNSGNTLY | 0.65287924 |
| TDGLGEN | ASSEQNSGNTLY | 0.6493702 |
| TGGPPLPGQA | ASSEQNSGNTLY | 0.41052812 |
| TIPAITI | ASSEQNSGNTLY | 0.72297305 |
| TIPGFTI | ASSEQNSGNTLY | 0.4935649 |
| TSIASGALA | ASSEQNSGNTLY | 0.7137457 |

| | | |
|---|---|---|
| TVTLPGAAI | ASSEQNSGNTLY | 0.5138811 |
| VASGLSAALP | ASSEQNSGNTLY | 0.75107706 |
| VGAGVIGI | ASSEQNSGNTLY | 0.74219394 |
| VGPKRFGV | ASSEQNSGNTLY | 0.69843024 |
| VNVAFPD | ASSEQNSGNTLY | 0.5939199 |
| VSMTNSGVS | ASSEQNSGNTLY | 0.7234122 |
| VTDVLGDLSV | ASSEQNSGNTLY | 0.68039083 |
| VVVVGAGFAGL | ASSEQNSGNTLY | 0.47332963 |
| VYTVGTVV | ASSEQNSGNTLY | 0.7649874 |
| WTTFSGP | ASSEQNSGNTLY | 0.57784545 |
| YDELVEAF | ASSEQNSGNTLY | 0.6951999 |
| YGTGGAGGNGG | ASSEQNSGNTLY | 0.77287984 |
| YTNVMAV | ASSEQNSGNTLY | 0.7211406 |
| FLEQVHQL | ASSEQNSGNTLY | 0.3041427 |
| GLSLLAVAV | ASSEQNSGNTLY | 0.707102 |
| LEQVHQL | ASSEQNSGNTLY | 0.7197869 |
| LNAAGAI | ASSEQNSGNTLY | 0.6536735 |
| PAVHASL | ASSEQNSGNTLY | 0.22188312 |
| TELEDDA | ASSEQNSGNTLY | 0.7557163 |
| TPPGPPN | ASSEQNSGNTLY | 0.5577946 |
| TPPPPPPP | ASSEQNSGNTLY | 0.5962758 |
| TQSPALI | ASRSGQYAEQF | 0.62276185 |
| EFAAAFPQ | ASRSGQYAEQF | 0.7674179 |
| LLSGNLDVL | ASRSGQYAEQF | 0.6159853 |
| GPANLGNYNI | ASRSGQYAEQF | 0.61744815 |
| TLGPTLP | ASRSGQYAEQF | 0.75964165 |
| VYAYSSCCK | ASRSGQYAEQF | 0.7525002 |
| APAPAPAPNI | ASRSGQYAEQF | 0.780271 |
| PGGPVPIP | ASRSGQYAEQF | 0.72998714 |
| FLTVALEP | ASRSGQYAEQF | 0.59002805 |
| SQYFGANL | ASRSGQYAEQF | 0.7217345 |
| GGVKPAA | ASRSGQYAEQF | 0.52357686 |
| GGTGGAGGAGGW | ASRSGQYAEQF | 0.5386021 |
| GVGGVAGLGGAGG | ASRSGQYAEQF | 0.33974984 |
| VLLVLATTI | ASRSGQYAEQF | 0.61316204 |
| IGIFAAML | ASRSGQYAEQF | 0.40365717 |
| AVGDLID | ASRSGQYAEQF | 0.7308198 |
| NISPLPL | ASRSGQYAEQF | 0.7442088 |
| VTAVFVAI | ASRSGQYAEQF | 0.66115 |
| SGDDNDAAL | ASRSGQYAEQF | 0.41558293 |
| AVAALAHS | ASRSGQYAEQF | 0.659951 |
| GGVTVMATP | ASRSGQYAEQF | 0.67389333 |
| AGLAANVRD | ASRSGQYAEQF | 0.7607698 |
| TNRPDLIDPALLRPG | ASRSGQYAEQF | 0.7129952 |
| LANLGVTAL | ASRSGQYAEQF | 0.7908281 |
| TGERIEL | ASRSGQYAEQF | 0.7721483 |
| NGEELAA | ASRSGQYAEQF | 0.66630495 |
| VLSLVWG | ASRSGQYAEQF | 0.7532606 |
| IGYALWL | ASRSGQYAEQF | 0.6095372 |
| LSLLLAPL | ASRSGQYAEQF | 0.7226462 |
| TLDEALK | ASRSGQYAEQF | 0.6829529 |

| | | |
|---|---|---|
| TDGAGTAEGG | ASRSGQYAEQF | 0.67247987 |
| NAPGGAGGNGG | ASRSGQYAEQF | 0.66904056 |
| VSVFMGMGGNF | ASRSGQYAEQF | 0.6784667 |
| LASYLAAL | ASRSGQYAEQF | 0.29424948 |
| LVGGTPAL | ASRSGQYAEQF | 0.71114725 |
| VRNADVL | ASRSGQYAEQF | 0.5093168 |
| TCSSPFLVSAM | ASRSGQYAEQF | 0.31254885 |
| VHITDAE | ASRSGQYAEQF | 0.6598539 |
| EYAAPSF | ASRSGQYAEQF | 0.7889972 |
| VVDAFHGGL | ASRSGQYAEQF | 0.77122283 |
| ASVVDPQ | ASRSGQYAEQF | 0.6572231 |
| SYNEFGGLS | ASRSGQYAEQF | 0.5374366 |
| DPDGSVY | ASRSGQYAEQF | 0.7547725 |
| MVSTIHATE | ASRSGQYAEQF | 0.41415763 |
| LAAGVADPV | ASRSGQYAEQF | 0.69578487 |
| TKEQIAA | ASRSGQYAEQF | 0.729711 |
| SLASPAAL | ASRSGQYAEQF | 0.72042 |
| IGLVFYGV | ASRSGQYAEQF | 0.4738756 |
| FLLDASNL | ASRSGQYAEQF | 0.7469027 |
| YQVGNFP | ASRSGQYAEQF | 0.6973489 |
| ESLLSQST | ASRSGQYAEQF | 0.6623938 |
| LFTVGFAAVG | ASRSGQYAEQF | 0.304102 |
| ATPPTPT | ASRSGQYAEQF | 0.76204586 |
| TDLATLG | ASRSGQYAEQF | 0.6296812 |
| SAGGTGR | ASRSGQYAEQF | 0.45305097 |
| GFSNSGS | ASRSGQYAEQF | 0.26371133 |
| GGNGGSGGTSV | ASRSGQYAEQF | 0.6683768 |
| GTSVATGGAGN | ASRSGQYAEQF | 0.7593075 |
| NGGNGGSGGTS | ASRSGQYAEQF | 0.7705053 |
| AVVKFPH | ASRSGQYAEQF | 0.6079956 |
| TPPPPPP | ASRSGQYAEQF | 0.77563155 |
| AAPAAAAAPA | ASRSGQYAEQF | 0.7645976 |
| IALNEDL | ASRSGQYAEQF | 0.69726515 |
| IVNVSSM | ASRSGQYAEQF | 0.34699845 |
| TGIPPAPR | ASRSGQYAEQF | 0.7121738 |
| FDLGGGTF | ASRSGQYAEQF | 0.70721436 |
| FDLGGGTFDVS | ASRSGQYAEQF | 0.76642334 |
| NEPTAAA | ASRSGQYAEQF | 0.7453501 |
| NEPTAAALA | ASRSGQYAEQF | 0.525356 |
| GSIINISSI | ASRSGQYAEQF | 0.7262654 |
| VAPGNAGTA | ASRSGQYAEQF | 0.75730765 |
| TGGGASLE | ASRSGQYAEQF | 0.5211253 |
| VAVYNHYKRI | ASRSGQYAEQF | 0.5110426 |
| TDPAPATT | ASRSGQYAEQF | 0.59213245 |
| TNRPDILDPALLRPG | ASRSGQYAEQF | 0.43064672 |
| LTTTTTS | ASRSGQYAEQF | 0.7574378 |
| PTGSGKTL | ASRSGQYAEQF | 0.68755686 |
| TGSGKTL | ASRSGQYAEQF | 0.6484642 |
| TIGHVDHGK | ASRSGQYAEQF | 0.68787277 |
| TIGHVDHGKTT | ASRSGQYAEQF | 0.62624246 |
| GHVDHGKTT | ASRSGQYAEQF | 0.55959404 |

| | | |
|---|---|---|
| AQFEHTLL | ASRSGQYAEQF | 0.6653631 |
| IVNQDEL | ASRSGQYAEQF | 0.74696606 |
| TNGSQFF | ASRSGQYAEQF | 0.7477375 |
| TGKGIVE | ASRSGQYAEQF | 0.71294206 |
| LISNASDA | ASRSGQYAEQF | 0.64128625 |
| AELRTLGELTKIAKAHGAQ | ASRSGQYAEQF | 0.78566456 |
| APNAVAAAEPSW | ASRSGQYAEQF | 0.47208762 |
| APQVAGAATAVL | ASRSGQYAEQF | 0.6193186 |
| ARRDPAAAAAILPTDA | ASRSGQYAEQF | 0.7394552 |
| DAAARDVLAEAGLAEYF | ASRSGQYAEQF | 0.48776534 |
| DDQIVLSAKANLAKVEQ | ASRSGQYAEQF | 0.76018375 |
| DELLPDTGEDSDEDGH | ASRSGQYAEQF | 0.74538565 |
| ELPEAPANRATVPQD | ASRSGQYAEQF | 0.74138784 |
| FGQSKIERVTGIG | ASRSGQYAEQF | 0.7035702 |
| GAGGTGSTAGGAGGAGGAGGLY | ASRSGQYAEQF | 0.79150045 |
| GHGGTGAAGGNGGTGGAGGSGI | ASRSGQYAEQF | 0.75659966 |
| GNGGNGGAGGEGGPGGAGGAGA | ASRSGQYAEQF | 0.6662191 |
| GSTAGNGGNGGNATNGTIAGQP | ASRSGQYAEQF | 0.7703672 |
| GTGGDGGNGGNGGTGG | ASRSGQYAEQF | 0.40704483 |
| GVRRPAGIPADAPV | ASRSGQYAEQF | 0.77290636 |
| GYRQVTDAHLLTLARR | ASRSGQYAEQF | 0.7277123 |
| IAKMGQDGHDRG | ASRSGQYAEQF | 0.5447167 |
| KAEPSIEIQYT | ASRSGQYAEQF | 0.6990874 |
| KPVRIRMAADEDTVAVT | ASRSGQYAEQF | 0.76639736 |
| MPPRIAGMRLLVIKPEP | ASRSGQYAEQF | 0.77660584 |
| MKFYAVALFRSRGVPPTR | ASRSGQYAEQF | 0.38487107 |
| NAGGFGILGAVAHSP | ASRSGQYAEQF | 0.64634037 |
| NEGREITLAMAEM | ASRSGQYAEQF | 0.35708466 |
| NELVEWSHPLGHRWRWE | ASRSGQYAEQF | 0.41177586 |
| NLGAGGTGGNGGNPG | ASRSGQYAEQF | 0.41404456 |
| NTEWPKWETVMAADPVD | ASRSGQYAEQF | 0.73801124 |
| PASAAPAETQP | ASRSGQYAEQF | 0.66843635 |
| QQSSDGKNGVRAAEITGE | ASRSGQYAEQF | 0.75889146 |
| RAPDAVTTSWWKE | ASRSGQYAEQF | 0.7218848 |
| SGFGNVGAHTSGWFNQ | ASRSGQYAEQF | 0.77154213 |
| SGVAGAVGQAASVGGLKV | ASRSGQYAEQF | 0.57919174 |
| SNGGAGTGSAGGNGGT | ASRSGQYAEQF | 0.7208779 |
| TGGAGGEGGAAG | ASRSGQYAEQF | 0.3356127 |
| TGGDGGAAGAGGAPGHG | ASRSGQYAEQF | 0.47627264 |
| TMPPLLAAIANAGRHGV | ASRSGQYAEQF | 0.48386675 |
| TSSTAGNSGGA | ASRSGQYAEQF | 0.70677567 |
| VPATVPVGILC | ASRSGQYAEQF | 0.3215508 |
| VQRPDSGGVVLTGRIS | ASRSGQYAEQF | 0.7715807 |
| WTPNPVPATKGVPAGLVQAT | ASRSGQYAEQF | 0.69394994 |
| YALKRALAKFSDRFDVVII | ASRSGQYAEQF | 0.7341182 |
| YLIGSVIV | ASRSGQYAEQF | 0.63384354 |
| YGQLVAVID | ASRSGQYAEQF | 0.58989924 |
| VTVGGMTLPA | ASRSGQYAEQF | 0.6234181 |
| VLALPAV | ASRSGQYAEQF | 0.75959086 |
| VIPPTLNYET | ASRSGQYAEQF | 0.61091775 |
| VGMGGMHPSSG | ASRSGQYAEQF | 0.2520656 |

| | | |
|---|---|---|
| VGLAGGAATA | ASRSGQYAEQF | 0.4704365 |
| VGGAGGAGGSGP | ASRSGQYAEQF | 0.47668517 |
| VFAAFMV | ASRSGQYAEQF | 0.7797712 |
| TLYDKLR | ASRSGQYAEQF | 0.7664057 |
| TLAVFTVL | ASRSGQYAEQF | 0.7309419 |
| TGSVQPARV | ASRSGQYAEQF | 0.77290004 |
| TGGGAGSGGT | ASRSGQYAEQF | 0.22113957 |
| SHLETSL | ASRSGQYAEQF | 0.64845276 |
| SFTTGMT | ASRSGQYAEQF | 0.78094256 |
| QVEVAKRMGVRQPTVS | ASRSGQYAEQF | 0.71964383 |
| QGSGGPALAM | ASRSGQYAEQF | 0.6991848 |
| MLAPSNP | ASRSGQYAEQF | 0.7706134 |
| LVNNAAI | ASRSGQYAEQF | 0.63612235 |
| LTGEINTLA | ASRSGQYAEQF | 0.66394424 |
| LSIFGPV | ASRSGQYAEQF | 0.716879 |
| LLALLANL | ASRSGQYAEQF | 0.7311959 |
| LISNASDALDKLR | ASRSGQYAEQF | 0.6695406 |
| LINEAVP | ASRSGQYAEQF | 0.45099846 |
| LGYAFSGL | ASRSGQYAEQF | 0.68694746 |
| LDFVAVAL | ASRSGQYAEQF | 0.7955196 |
| LAQPGAE | ASRSGQYAEQF | 0.6964766 |
| ISNLGAVTGF | ASRSGQYAEQF | 0.7516 |
| HQEDPGRI | ASRSGQYAEQF | 0.7253599 |
| GTPGEPTP | ASRSGQYAEQF | 0.7155117 |
| GFFNSGS | ASRSGQYAEQF | 0.58033776 |
| GAGGGGLTAGAN | ASRSGQYAEQF | 0.5847173 |
| FVDALVNM | ASRSGQYAEQF | 0.7386782 |
| FLTVAPEL | ASRSGQYAEQF | 0.5772989 |
| FETLTAL | ASRSGQYAEQF | 0.67786765 |
| DVINPIEL | ASRSGQYAEQF | 0.616238 |
| APVSGAPV | ASRSGQYAEQF | 0.7929311 |
| ALVGGLAW | ASRSGQYAEQF | 0.7429119 |
| ALAAAAATAI | ASRSGQYAEQF | 0.73103416 |
| AFGLGLLL | ASRSGQYAEQF | 0.5683669 |
| AFFAAASL | ASRSGQYAEQF | 0.6828468 |
| AAAALPAFSPP | ASRSGQYAEQF | 0.7533065 |
| VVLAPAL | ASRSGQYAEQF | 0.334714 |
| TIGHVDHGKTTL | ASRSGQYAEQF | 0.7756164 |
| LISNASDALDK | ASRSGQYAEQF | 0.54010546 |
| LISNASDAL | ASRSGQYAEQF | 0.696122 |
| ISNASDALDK | ASRSGQYAEQF | 0.749822 |
| NQAETLVYQTE | ASRSGQYAEQF | 0.7844878 |
| TGGTGGTAGAG | ASRSGQYAEQF | 0.6317742 |
| TGSTAGGAGGA | ASRSGQYAEQF | 0.6859446 |
| DPLAVPVAAGPV | ASRSGQYAEQF | 0.57335246 |
| GGTGGTPGNGGN | ASRSGQYAEQF | 0.7274892 |
| PAGLAGVDPATL | ASRSGQYAEQF | 0.7753421 |
| GNNLVVEAPHAGASV | ASRSGQYAEQF | 0.7040059 |
| KVTRIMSASTLEQPA | ASRSGQYAEQF | 0.7894684 |
| LGFALPIALPRRTVV | ASRSGQYAEQF | 0.6938809 |
| TRLGNLTARKLVVLV | ASRSGQYAEQF | 0.31596723 |

| | | |
|---|---|---|
| WSQQRRLIYFPSAGP | ASRSGQYAEQF | 0.6388707 |
| GPNGNPDPMAAAVDIR | ASRSGQYAEQF | 0.77582 |
| TQRCVSVADLRAAASR | ASRSGQYAEQF | 0.77762216 |
| QFVLDVAEHPVEVLTETV | ASRSGQYAEQF | 0.7808173 |
| GGAGGTGGAGGNGGTQAGDG | ASRSGQYAEQF | 0.7628838 |
| PLLATLRAIAVDVDAKPAQV | ASRSGQYAEQF | 0.6487435 |
| APKFRTGGYMIDFWGVGYQVAK | ASRSGQYAEQF | 0.7760948 |
| GAGGAGGAGGAAGGPGGTGGTG | ASRSGQYAEQF | 0.7824542 |
| IGALGLLATDVAPAGSVAFGVA | ASRSGQYAEQF | 0.72749615 |
| AGGAGGIGGNGGAGGAGGVLMG | ASRSGQYAEQF | 0.4958121 |
| ADELIGLL | ASRSGQYAEQF | 0.4592433 |
| ADRALKA | ASRSGQYAEQF | 0.7330958 |
| AGAVGQVGGAGGA | ASRSGQYAEQF | 0.7672391 |
| AGGGGGLGGAP | ASRSGQYAEQF | 0.38848564 |
| AGGLLYGNGGAG | ASRSGQYAEQF | 0.74050903 |
| AGLESIGL | ASRSGQYAEQF | 0.732134 |
| AGLKTGII | ASRSGQYAEQF | 0.37264404 |
| ASLAALLL | ASRSGQYAEQF | 0.75424427 |
| ASVINISSI | ASRSGQYAEQF | 0.7855872 |
| ATPPGPA | ASRSGQYAEQF | 0.663188 |
| ATVASALV | ASRSGQYAEQF | 0.5714972 |
| ATVASLIL | ASRSGQYAEQF | 0.7760444 |
| AVNGPVLIRH | ASRSGQYAEQF | 0.7915821 |
| DGAAGAVGI | ASRSGQYAEQF | 0.7758841 |
| DPLLHGI | ASRSGQYAEQF | 0.37998348 |
| ELGAALGAL | ASRSGQYAEQF | 0.23220924 |
| ELGRTQAV | ASRSGQYAEQF | 0.70683175 |
| FVAGGYLS | ASRSGQYAEQF | 0.7636408 |
| GASGGAGGAGGW | ASRSGQYAEQF | 0.67748296 |
| GGAGAGGDGGAGGI | ASRSGQYAEQF | 0.5987141 |
| GTGHNGSQPGMGGNG | ASRSGQYAEQF | 0.75606287 |
| GYAAASASAS | ASRSGQYAEQF | 0.7688054 |
| HQVDVAAVQQG | ASRSGQYAEQF | 0.5465647 |
| IGAAIAHT | ASRSGQYAEQF | 0.35014087 |
| IGMIEVAQL | ASRSGQYAEQF | 0.76426303 |
| IRLAPEHTSAMD | ASRSGQYAEQF | 0.7356036 |
| IYNVSVL | ASRSGQYAEQF | 0.6208708 |
| KQGFLGM | ASRSGQYAEQF | 0.6861491 |
| LESVITQL | ASRSGQYAEQF | 0.7169775 |
| LGLGALVVV | ASRSGQYAEQF | 0.624153 |
| LIGNGGNGGSGGI | ASRSGQYAEQF | 0.7428205 |
| LNAINAPTL | ASRSGQYAEQF | 0.4569614 |
| LQLLNTGV | ASRSGQYAEQF | 0.7852322 |
| LQTSIEF | ASRSGQYAEQF | 0.44664487 |
| MSFAVSEL | ASRSGQYAEQF | 0.73834443 |
| NTGALGL | ASRSGQYAEQF | 0.74716425 |
| PPGAGGAGGAGTT | ASRSGQYAEQF | 0.72147083 |
| QLVGLGI | ASRSGQYAEQF | 0.6838264 |
| RFLEQVHQL | ASRSGQYAEQF | 0.60583335 |
| RGVGYVLR | ASRSGQYAEQF | 0.5131419 |
| SATGVFAGVI | ASRSGQYAEQF | 0.33856118 |

| | | |
|---|---|---|
| SLPAITLP | ASRSGQYAEQF | 0.78094095 |
| STALAFTV | ASRSGQYAEQF | 0.7117417 |
| SVLTVGLGK | ASRSGQYAEQF | 0.7730118 |
| TALAGAGSGI | ASRSGQYAEQF | 0.7342018 |
| TGLPFTGI | ASRSGQYAEQF | 0.5896205 |
| VAEHVTV | ASRSGQYAEQF | 0.70939535 |
| VDIVLNSL | ASRSGQYAEQF | 0.5303719 |
| VLTGGGAVL | ASRSGQYAEQF | 0.6096975 |
| VNVGLAYGI | ASRSGQYAEQF | 0.69804096 |
| VSSAAVGAV | ASRSGQYAEQF | 0.7416917 |
| VTSVYEAGV | ASRSGQYAEQF | 0.5836921 |
| WVALDRGA | ASRSGQYAEQF | 0.69267154 |
| YLVGSVVI | ASRSGQYAEQF | 0.7293551 |
| GPPGTGKTLL | ASRSGQYAEQF | 0.4623301 |
| GVGMTKF | ASRSGQYAEQF | 0.6927671 |
| IGIGGSDL | ASRSGQYAEQF | 0.77335775 |
| IGTLLAEL | ASRSGQYAEQF | 0.6706927 |
| IYTSGTT | ASRSGQYAEQF | 0.52350134 |
| LSFPAGV | ASRSGQYAEQF | 0.7574055 |
| LYTSGTT | ASRSGQYAEQF | 0.5932235 |
| VGAFPFGF | ASRSGQYAEQF | 0.74615896 |
| VGAGTVEF | ASRSGQYAEQF | 0.7138262 |
| VGKSTLLL | ASRSGQYAEQF | 0.60485554 |
| AALAHAGGGIV | ASRSGQYAEQF | 0.7530388 |
| AALVAVSLL | ASRSGQYAEQF | 0.3586874 |
| AAVSRAVAAL | ASRSGQYAEQF | 0.5694605 |
| AGGSSGIGGF | ASRSGQYAEQF | 0.640069 |
| AGLTVANANG | ASRSGQYAEQF | 0.7143998 |
| AGLVGAVL | ASRSGQYAEQF | 0.7542542 |
| ALSQAFTV | ASRSGQYAEQF | 0.59400356 |
| ALTFSVSL | ASRSGQYAEQF | 0.7002354 |
| ALTGAALVAL | ASRSGQYAEQF | 0.656999 |
| ASFSIGAGVL | ASRSGQYAEQF | 0.5376798 |
| DGSVYKI | ASRSGQYAEQF | 0.5905025 |
| EQLAEIV | ASRSGQYAEQF | 0.57848513 |
| FAVSELTVM | ASRSGQYAEQF | 0.4533456 |
| GAAPLAAGL | ASRSGQYAEQF | 0.7443527 |
| GADGGDGGSGGAGG | ASRSGQYAEQF | 0.58030033 |
| GFGLDTGGAGG | ASRSGQYAEQF | 0.72534037 |
| GLASGGLL | ASRSGQYAEQF | 0.78305966 |
| GRIAGAAVLL | ASRSGQYAEQF | 0.5042265 |
| GSAGSLNTSF | ASRSGQYAEQF | 0.71698874 |
| GTGQIGI | ASRSGQYAEQF | 0.7297666 |
| GTLIALIL | ASRSGQYAEQF | 0.7594422 |
| IAATMGFVVG | ASRSGQYAEQF | 0.695516 |
| IFAQALL | ASRSGQYAEQF | 0.6305731 |
| IGAAVALGF | ASRSGQYAEQF | 0.7827381 |
| IGLTPLAL | ASRSGQYAEQF | 0.74925417 |
| ILGSNTSTL | ASRSGQYAEQF | 0.69014204 |
| IVFSLTVL | ASRSGQYAEQF | 0.761637 |
| LASKATGF | ASRSGQYAEQF | 0.76261485 |

| | | |
|---|---|---|
| LASVALIL | ASRSGQYAEQF | 0.75064665 |
| LFAVGLTL | ASRSGQYAEQF | 0.6418794 |
| LFNGGIGTYI | ASRSGQYAEQF | 0.33212695 |
| LTLGAGII | ASRSGQYAEQF | 0.68547857 |
| LVAAALGDI | ASRSGQYAEQF | 0.6945412 |
| NTAFVQLGI | ASRSGQYAEQF | 0.5236651 |
| QLFSGAEL | ASRSGQYAEQF | 0.28341857 |
| SAATFITL | ASRSGQYAEQF | 0.77781236 |
| SNYGLANLGV | ASRSGQYAEQF | 0.7911659 |
| SYGGAYAVM | ASRSGQYAEQF | 0.6283841 |
| TTGVGGLTLG | ASRSGQYAEQF | 0.7868006 |
| TTQTIYI | ASRSGQYAEQF | 0.41919848 |
| VAAGAGLGAVI | ASRSGQYAEQF | 0.75426584 |
| VAAVVTLL | ASRSGQYAEQF | 0.77672017 |
| VGFLGGLI | ASRSGQYAEQF | 0.53656 |
| VGVIPILL | ASRSGQYAEQF | 0.7535623 |
| VIFASSLI | ASRSGQYAEQF | 0.5954606 |
| VKGGDGGAAAG | ASRSGQYAEQF | 0.63911843 |
| VTKTFTNI | ASRSGQYAEQF | 0.7789003 |
| VVAAAILL | ASRSGQYAEQF | 0.6711001 |
| YAALDVEL | ASRSGQYAEQF | 0.7455224 |
| LASVALLL | ASRSGQYAEQF | 0.65711284 |
| LDAVHRV | ASRSGQYAEQF | 0.522459 |
| QGLSSSIF | ASRSGQYAEQF | 0.7189671 |
| RGTFANIRL | ASRSGQYAEQF | 0.76226366 |
| RQLWWGHRI | ASRSGQYAEQF | 0.59285545 |
| YGPPGTGKTLL | ASRSGQYAEQF | 0.7718656 |
| AALVGAPANVA | ASRSGQYAEQF | 0.61113423 |
| AATFRRTE | ASRSGQYAEQF | 0.79333293 |
| AGALGGGI | ASRSGQYAEQF | 0.77392244 |
| AGGAGGAGGIGAG | ASRSGQYAEQF | 0.732491 |
| AGSGALGL | ASRSGQYAEQF | 0.586355 |
| AQIADQI | ASRSGQYAEQF | 0.7781557 |
| ASAVAVTYL | ASRSGQYAEQF | 0.66889346 |
| ASTVLGGTGG | ASRSGQYAEQF | 0.7429395 |
| AVGGEPAK | ASRSGQYAEQF | 0.769683 |
| CEQGTFGGTC | ASRSGQYAEQF | 0.51363057 |
| DEGKAPAG | ASRSGQYAEQF | 0.6732446 |
| DTVVFPAL | ASRSGQYAEQF | 0.6460186 |
| FEGITMAG | ASRSGQYAEQF | 0.62434286 |
| FPLVVQD | ASRSGQYAEQF | 0.60741246 |
| GAFTTPQ | ASRSGQYAEQF | 0.4467324 |
| GFGGFGTAGGVGGA | ASRSGQYAEQF | 0.6086036 |
| GFLAQFL | ASRSGQYAEQF | 0.5804111 |
| GGAGGAGGNAGVA | ASRSGQYAEQF | 0.6757563 |
| GGVGTAGAGGAGGAGG | ASRSGQYAEQF | 0.651384 |
| GNGGGGDAGNAGSGGN | ASRSGQYAEQF | 0.5926412 |
| GTGGAGGAGNDAGS | ASRSGQYAEQF | 0.67634165 |
| GTTNTGWFNS | ASRSGQYAEQF | 0.6651273 |
| IFMGPLFTL | ASRSGQYAEQF | 0.48186874 |
| ILAHAIEAI | ASRSGQYAEQF | 0.6253319 |

| | | |
|---|---|---|
| IRSAIREAL | ASRSGQYAEQF | 0.55867213 |
| ITFAGGG | ASRSGQYAEQF | 0.6161099 |
| ITGMSGHQEV | ASRSGQYAEQF | 0.41693884 |
| IVAGLACTL | ASRSGQYAEQF | 0.65186733 |
| IVGGGLAGA | ASRSGQYAEQF | 0.61331415 |
| IVTGVLNI | ASRSGQYAEQF | 0.3151603 |
| LAQQGCI | ASRSGQYAEQF | 0.6383207 |
| LCLHVEN | ASRSGQYAEQF | 0.63512695 |
| LDAISDGS | ASRSGQYAEQF | 0.63130677 |
| LGGFAAML | ASRSGQYAEQF | 0.6648884 |
| LGGKNPMI | ASRSGQYAEQF | 0.50570434 |
| LGPKGVAAIL | ASRSGQYAEQF | 0.5963769 |
| LMVGVSI | ASRSGQYAEQF | 0.665303 |
| LVGAAFVL | ASRSGQYAEQF | 0.67346644 |
| LVLAVAPL | ASRSGQYAEQF | 0.54964995 |
| MAAIAVIDAA | ASRSGQYAEQF | 0.64647114 |
| NNGINAELQ | ASRSGQYAEQF | 0.6706241 |
| RGEPLVLVHG | ASRSGQYAEQF | 0.64048636 |
| RGGAGGAAGLI | ASRSGQYAEQF | 0.663087 |
| RSYRMHPAV | ASRSGQYAEQF | 0.6691525 |
| SAAHLLAGQLA | ASRSGQYAEQF | 0.49473935 |
| SAAVGSY | ASRSGQYAEQF | 0.67599523 |
| SPHAVLHRI | ASRSGQYAEQF | 0.6786255 |
| SQIAKVL | ASRSGQYAEQF | 0.61551076 |
| TALGHLVP | ASRSGQYAEQF | 0.6211132 |
| TAPTTSGAA | ASRSGQYAEQF | 0.50547576 |
| TDGDDGNGGSPGT | ASRSGQYAEQF | 0.60948133 |
| TDGLGEN | ASRSGQYAEQF | 0.4598299 |
| TGGPPLPGQA | ASRSGQYAEQF | 0.6502653 |
| TIPAITI | ASRSGQYAEQF | 0.6710613 |
| TIPGFTI | ASRSGQYAEQF | 0.61856675 |
| TSIASGALA | ASRSGQYAEQF | 0.5029732 |
| TVTLPGAAI | ASRSGQYAEQF | 0.52333343 |
| VASGLSAALP | ASRSGQYAEQF | 0.5945076 |
| VGAGVIGI | ASRSGQYAEQF | 0.5588654 |
| VGPKRFGV | ASRSGQYAEQF | 0.3892126 |
| VNVAFPD | ASRSGQYAEQF | 0.52072525 |
| VSMTNSGVS | ASRSGQYAEQF | 0.67033476 |
| VTDVLGDLSV | ASRSGQYAEQF | 0.6692873 |
| VVVVGAGFAGL | ASRSGQYAEQF | 0.54252815 |
| VYTVGTVV | ASRSGQYAEQF | 0.67730975 |
| WTTFSGP | ASRSGQYAEQF | 0.6650554 |
| YDELVEAF | ASRSGQYAEQF | 0.6724402 |
| YGTGGAGGNGG | ASRSGQYAEQF | 0.6213963 |
| YTNVMAV | ASRSGQYAEQF | 0.5330334 |
| FLEQVHQL | ASRSGQYAEQF | 0.67300177 |
| GLSLLAVAV | ASRSGQYAEQF | 0.62015563 |
| LEQVHQL | ASRSGQYAEQF | 0.59834415 |
| LNAAGAI | ASRSGQYAEQF | 0.66263866 |
| PAVHASL | ASRSGQYAEQF | 0.6339126 |
| TELEDDA | ASRSGQYAEQF | 0.58719623 |

| | | |
|---|---|---|
| TPPGPPN | ASRSGQYAEQF | 0.62672496 |
| TPPPPPPP | ASRSGQYAEQF | 0.6457685 |
| TQSPALI | ASGGGTGGSAETLY | 0.5473935 |
| EFAAAFPQ | ASGGGTGGSAETLY | 0.5013311 |
| LLSGNLDVL | ASGGGTGGSAETLY | 0.580449 |
| GPANLGNYNI | ASGGGTGGSAETLY | 0.64109606 |
| TLGPTLP | ASGGGTGGSAETLY | 0.6724818 |
| VYAYSSCCK | ASGGGTGGSAETLY | 0.6311156 |
| APAPAPAPNI | ASGGGTGGSAETLY | 0.5648311 |
| PGGPVPIP | ASGGGTGGSAETLY | 0.66037863 |
| FLTVALEP | ASGGGTGGSAETLY | 0.65138286 |
| SQYFGANL | ASGGGTGGSAETLY | 0.6355422 |
| GGVKPAA | ASGGGTGGSAETLY | 0.6105344 |
| GGTGGAGGAGGW | ASGGGTGGSAETLY | 0.59760886 |
| GVGGVAGLGGAGG | ASGGGTGGSAETLY | 0.64960706 |
| VLLVLATTI | ASGGGTGGSAETLY | 0.6519196 |
| IGIFAAML | ASGGGTGGSAETLY | 0.64527786 |
| AVGDLID | ASGGGTGGSAETLY | 0.65700454 |
| NISPLPL | ASGGGTGGSAETLY | 0.6383043 |
| VTAVFVAI | ASGGGTGGSAETLY | 0.62396014 |
| SGDDNDAAL | ASGGGTGGSAETLY | 0.44808742 |
| AVAALAHS | ASGGGTGGSAETLY | 0.5444231 |
| GGVTVMATP | ASGGGTGGSAETLY | 0.6478625 |
| AGLAANVRD | ASGGGTGGSAETLY | 0.6497519 |
| TNRPDLIDPALLRPG | ASGGGTGGSAETLY | 0.6652261 |
| LANLGVTAL | ASGGGTGGSAETLY | 0.61389893 |
| TGERIEL | ASGGGTGGSAETLY | 0.5817113 |
| NGEELAA | ASGGGTGGSAETLY | 0.6352916 |
| VLSLVWG | ASGGGTGGSAETLY | 0.67301655 |
| IGYALWL | ASGGGTGGSAETLY | 0.46339124 |
| LSLLLAPL | ASGGGTGGSAETLY | 0.59197426 |
| TLDEALK | ASGGGTGGSAETLY | 0.6819202 |
| TDGAGTAEGG | ASGGGTGGSAETLY | 0.6518243 |
| NAPGGAGGNGG | ASGGGTGGSAETLY | 0.6276324 |
| VSVFMGMGGNF | ASGGGTGGSAETLY | 0.5830145 |
| LASYLAAL | ASGGGTGGSAETLY | 0.6827679 |
| LVGGTPAL | ASGGGTGGSAETLY | 0.57051516 |
| VRNADVL | ASGGGTGGSAETLY | 0.67748725 |
| TCSSPFLVSAM | ASGGGTGGSAETLY | 0.37144825 |
| VHITDAE | ASGGGTGGSAETLY | 0.5498822 |
| EYAAPSF | ASGGGTGGSAETLY | 0.43187487 |
| VVDAFHGGL | ASGGGTGGSAETLY | 0.36944532 |
| ASVVDPQ | ASGGGTGGSAETLY | 0.664031 |
| SYNEFGGLS | ASGGGTGGSAETLY | 0.5756364 |
| DPDGSVY | ASGGGTGGSAETLY | 0.6613938 |
| MVSTIHATE | ASGGGTGGSAETLY | 0.63089 |
| LAAGVADPV | ASGGGTGGSAETLY | 0.67016995 |
| TKEQIAA | ASGGGTGGSAETLY | 0.64125633 |
| SLASPAAL | ASGGGTGGSAETLY | 0.62944156 |
| IGLVFYGV | ASGGGTGGSAETLY | 0.519198 |
| FLLDASNL | ASGGGTGGSAETLY | 0.66058916 |

| | | |
|---|---|---|
| YQVGNFP | ASGGGTGGSAETLY | 0.65606004 |
| ESLLSQST | ASGGGTGGSAETLY | 0.60508925 |
| LFTVGFAAVG | ASGGGTGGSAETLY | 0.48752576 |
| ATPPTPT | ASGGGTGGSAETLY | 0.6138886 |
| TDLATLG | ASGGGTGGSAETLY | 0.63267195 |
| SAGGTGR | ASGGGTGGSAETLY | 0.39547387 |
| GFSNSGS | ASGGGTGGSAETLY | 0.35019842 |
| GGNGGSGGTSV | ASGGGTGGSAETLY | 0.5803924 |
| GTSVATGGAGN | ASGGGTGGSAETLY | 0.55050135 |
| NGGNGGSGGTS | ASGGGTGGSAETLY | 0.67297554 |
| AVVKFPH | ASGGGTGGSAETLY | 0.66455835 |
| TPPPPPP | ASGGGTGGSAETLY | 0.6212146 |
| AAPAAAAAPA | ASGGGTGGSAETLY | 0.655663 |
| IALNEDL | ASGGGTGGSAETLY | 0.64649206 |
| IVNVSSM | ASGGGTGGSAETLY | 0.29327992 |
| TGIPPAPR | ASGGGTGGSAETLY | 0.54954314 |
| FDLGGGTF | ASGGGTGGSAETLY | 0.48272336 |
| FDLGGGTFDVS | ASGGGTGGSAETLY | 0.6681651 |
| NEPTAAA | ASGGGTGGSAETLY | 0.5442036 |
| NEPTAAALA | ASGGGTGGSAETLY | 0.5776037 |
| GSIINISSI | ASGGGTGGSAETLY | 0.6428267 |
| VAPGNAGTA | ASGGGTGGSAETLY | 0.6543597 |
| TGGGASLE | ASGGGTGGSAETLY | 0.63214695 |
| VAVYNHYKRI | ASGGGTGGSAETLY | 0.5247712 |
| TDPAPATT | ASGGGTGGSAETLY | 0.370456 |
| TNRPDILDPALLRPG | ASGGGTGGSAETLY | 0.66387904 |
| LTTTTTS | ASGGGTGGSAETLY | 0.5110267 |
| PTGSGKTL | ASGGGTGGSAETLY | 0.5526271 |
| TGSGKTL | ASGGGTGGSAETLY | 0.4214614 |
| TIGHVDHGK | ASGGGTGGSAETLY | 0.58985084 |
| TIGHVDHGKTT | ASGGGTGGSAETLY | 0.45011747 |
| GHVDHGKTT | ASGGGTGGSAETLY | 0.64526385 |
| AQFEHTLL | ASGGGTGGSAETLY | 0.60371435 |
| IVNQDEL | ASGGGTGGSAETLY | 0.6234501 |
| TNGSQFF | ASGGGTGGSAETLY | 0.6806777 |
| TGKGIVE | ASGGGTGGSAETLY | 0.65611017 |
| LISNASDA | ASGGGTGGSAETLY | 0.61484945 |
| AELRTLGELTKIAKAHGAQ | ASGGGTGGSAETLY | 0.6139651 |
| APNAVAAAEPSW | ASGGGTGGSAETLY | 0.54000014 |
| APQVAGAATAVL | ASGGGTGGSAETLY | 0.5873903 |
| ARRDPAAAAAILPTDA | ASGGGTGGSAETLY | 0.5924019 |
| DAAARDVLAEAGLAEYF | ASGGGTGGSAETLY | 0.6782092 |
| DDQIVLSAKANLAKVEQ | ASGGGTGGSAETLY | 0.5505336 |
| DELLPDTGEDSDEDGH | ASGGGTGGSAETLY | 0.49694967 |
| ELPEAPANRATVPQD | ASGGGTGGSAETLY | 0.57555026 |
| FGQSKIERVTGIG | ASGGGTGGSAETLY | 0.6772344 |
| GAGGTGSTAGGAGGAGGAGGLY | ASGGGTGGSAETLY | 0.6816324 |
| GHGGTGAAGGNGGTGGAGGSGI | ASGGGTGGSAETLY | 0.5543054 |
| GNGGNGGAGGEGGPGGAGGAGA | ASGGGTGGSAETLY | 0.5390322 |
| GSTAGNGGNGGNATNGTIAGQP | ASGGGTGGSAETLY | 0.5800367 |
| GTGGDGGNGGNGGTGG | ASGGGTGGSAETLY | 0.6280673 |

| | | |
|---|---|---|
| GVRRPAGIPADAPV | ASGGGTGGSAETLY | 0.52561486 |
| GYRQVTDAHLLTLARR | ASGGGTGGSAETLY | 0.52433324 |
| IAKMGQDGHDRG | ASGGGTGGSAETLY | 0.63435715 |
| KAEPSIEIQYT | ASGGGTGGSAETLY | 0.6850953 |
| KPVRIRMAADEDTVAVT | ASGGGTGGSAETLY | 0.5829036 |
| MPPRIAGMRLLVIKPEP | ASGGGTGGSAETLY | 0.45597282 |
| MKFYAVALFRSRGVPPTR | ASGGGTGGSAETLY | 0.64366925 |
| NAGGFGILGAVAHSP | ASGGGTGGSAETLY | 0.5736752 |
| NEGREITLAMAEM | ASGGGTGGSAETLY | 0.60657907 |
| NELVEWSHPLGHRWRWE | ASGGGTGGSAETLY | 0.6151792 |
| NLGAGGTGGNGGNPG | ASGGGTGGSAETLY | 0.42788735 |
| NTEWPKWETVMAADPVD | ASGGGTGGSAETLY | 0.62622535 |
| PASAAPAETQP | ASGGGTGGSAETLY | 0.67829376 |
| QQSSDGKNGVRAAEITGE | ASGGGTGGSAETLY | 0.6388209 |
| RAPDAVTTSWWKE | ASGGGTGGSAETLY | 0.65268993 |
| SGFGNVGAHTSGWFNQ | ASGGGTGGSAETLY | 0.6751608 |
| SGVAGAVGQAASVGGLKV | ASGGGTGGSAETLY | 0.66020155 |
| SNGGAGTGSAGGNGGT | ASGGGTGGSAETLY | 0.67073846 |
| TGGAGGEGGAAG | ASGGGTGGSAETLY | 0.6712252 |
| TGGDGGAAGAGGAPGHG | ASGGGTGGSAETLY | 0.6079162 |
| TMPPLLAAIANAGRHGV | ASGGGTGGSAETLY | 0.6091161 |
| TSSTAGNSGGA | ASGGGTGGSAETLY | 0.66488093 |
| VPATVPVGILC | ASGGGTGGSAETLY | 0.6472611 |
| VQRPDSGGVVLTGRIS | ASGGGTGGSAETLY | 0.6378882 |
| WTPNPVPATKGVPAGLVQAT | ASGGGTGGSAETLY | 0.5631991 |
| YALKRALAKFSDRFDVVII | ASGGGTGGSAETLY | 0.6731508 |
| YLIGSVIV | ASGGGTGGSAETLY | 0.5955673 |
| YGQLVAVID | ASGGGTGGSAETLY | 0.6759378 |
| VTVGGMTLPA | ASGGGTGGSAETLY | 0.40784392 |
| VLALPAV | ASGGGTGGSAETLY | 0.5665999 |
| VIPPTLNYET | ASGGGTGGSAETLY | 0.66440845 |
| VGMGGMHPSSG | ASGGGTGGSAETLY | 0.3076429 |
| VGLAGGAATA | ASGGGTGGSAETLY | 0.35575864 |
| VGGAGGAGGSGP | ASGGGTGGSAETLY | 0.6147249 |
| VFAAFMV | ASGGGTGGSAETLY | 0.669631 |
| TLYDKLR | ASGGGTGGSAETLY | 0.3077852 |
| TLAVFTVL | ASGGGTGGSAETLY | 0.5705962 |
| TGSVQPARV | ASGGGTGGSAETLY | 0.45114857 |
| TGGGAGSGGT | ASGGGTGGSAETLY | 0.6170291 |
| SHLETSL | ASGGGTGGSAETLY | 0.6360427 |
| SFTTGMT | ASGGGTGGSAETLY | 0.40885368 |
| QVEVAKRMGVRQPTVS | ASGGGTGGSAETLY | 0.6531849 |
| QGSGGPALAM | ASGGGTGGSAETLY | 0.43707618 |
| MLAPSNP | ASGGGTGGSAETLY | 0.5742636 |
| LVNNAAI | ASGGGTGGSAETLY | 0.66606617 |
| LTGEINTLA | ASGGGTGGSAETLY | 0.6787306 |
| LSIFGPV | ASGGGTGGSAETLY | 0.47098407 |
| LLALLANL | ASGGGTGGSAETLY | 0.66256785 |
| LISNASDALDKLR | ASGGGTGGSAETLY | 0.52852625 |
| LINEAVP | ASGGGTGGSAETLY | 0.68486655 |
| LGYAFSGL | ASGGGTGGSAETLY | 0.6225934 |

| | | |
|---|---|---|
| LDFVAVAL | ASGGGTGGSAETLY | 0.40532258 |
| LAQPGAE | ASGGGTGGSAETLY | 0.66656494 |
| ISNLGAVTGF | ASGGGTGGSAETLY | 0.63943905 |
| HQEDPGRI | ASGGGTGGSAETLY | 0.59500957 |
| GTPGEPTP | ASGGGTGGSAETLY | 0.6678654 |
| GFFNSGS | ASGGGTGGSAETLY | 0.49497074 |
| GAGGGGLTAGAN | ASGGGTGGSAETLY | 0.6414007 |
| FVDALVNM | ASGGGTGGSAETLY | 0.67201936 |
| FLTVAPEL | ASGGGTGGSAETLY | 0.6628866 |
| FETLTAL | ASGGGTGGSAETLY | 0.49212942 |
| DVINPIEL | ASGGGTGGSAETLY | 0.6490183 |
| APVSGAPV | ASGGGTGGSAETLY | 0.63505435 |
| ALVGGLAW | ASGGGTGGSAETLY | 0.5485826 |
| ALAAAAATAI | ASGGGTGGSAETLY | 0.6746056 |
| AFGLGLLL | ASGGGTGGSAETLY | 0.5731803 |
| AFFAAASL | ASGGGTGGSAETLY | 0.6662514 |
| AAAALPAFSPP | ASGGGTGGSAETLY | 0.65572083 |
| VVLAPAL | ASGGGTGGSAETLY | 0.67908245 |
| TIGHVDHGKTTL | ASGGGTGGSAETLY | 0.45362943 |
| LISNASDALDK | ASGGGTGGSAETLY | 0.62329936 |
| LISNASDAL | ASGGGTGGSAETLY | 0.63421583 |
| ISNASDALDK | ASGGGTGGSAETLY | 0.665246 |
| NQAETLVYQTE | ASGGGTGGSAETLY | 0.57839453 |
| TGGTGGTAGAG | ASGGGTGGSAETLY | 0.60259026 |
| TGSTAGGAGGA | ASGGGTGGSAETLY | 0.6428021 |
| DPLAVPVAAGPV | ASGGGTGGSAETLY | 0.67362154 |
| GGTGGTPGNGGN | ASGGGTGGSAETLY | 0.60292804 |
| PAGLAGVDPATL | ASGGGTGGSAETLY | 0.60664564 |
| GNNLVVEAPHAGASV | ASGGGTGGSAETLY | 0.67606485 |
| KVTRIMSASTLEQPA | ASGGGTGGSAETLY | 0.56570655 |
| LGFALPIALPRRTVV | ASGGGTGGSAETLY | 0.62590253 |
| TRLGNLTARKLVVLV | ASGGGTGGSAETLY | 0.6500451 |
| WSQQRRLIYFPSAGP | ASGGGTGGSAETLY | 0.6667114 |
| GPNGNPDPMAAAVDIR | ASGGGTGGSAETLY | 0.59021753 |
| TQRCVSVADLRAAASR | ASGGGTGGSAETLY | 0.670472 |
| QFVLDVAEHPVEVLTETV | ASGGGTGGSAETLY | 0.51040876 |
| GGAGGTGGAGGNGGTQAGDG | ASGGGTGGSAETLY | 0.67002386 |
| PLLATLRAIAVDVDAKPAQV | ASGGGTGGSAETLY | 0.55925727 |
| APKFRTGGYMIDFWGVGYQVAK | ASGGGTGGSAETLY | 0.64825714 |
| GAGGAGGAGGAAGGPGGTGGTG | ASGGGTGGSAETLY | 0.56801283 |
| IGALGLLATDVAPAGSVAFGVA | ASGGGTGGSAETLY | 0.6655724 |
| AGGAGGIGGNGGAGGAGGVLMG | ASGGGTGGSAETLY | 0.6679059 |
| ADELIGLL | ASGGGTGGSAETLY | 0.67205584 |
| ADRALKA | ASGGGTGGSAETLY | 0.37709484 |
| AGAVGQVGGAGGA | ASGGGTGGSAETLY | 0.66406727 |
| AGGGGGLGGAP | ASGGGTGGSAETLY | 0.6742188 |
| AGGLLYGNGGAG | ASGGGTGGSAETLY | 0.6812018 |
| AGLESIGL | ASGGGTGGSAETLY | 0.6044902 |
| AGLKTGII | ASGGGTGGSAETLY | 0.64292276 |
| ASLAALLL | ASGGGTGGSAETLY | 0.6229969 |
| ASVINISSI | ASGGGTGGSAETLY | 0.60937476 |

| | | |
|---|---|---|
| ATPPGPA | ASGGGTGGSAETLY | 0.51425374 |
| ATVASALV | ASGGGTGGSAETLY | 0.5975237 |
| ATVASLIL | ASGGGTGGSAETLY | 0.6044705 |
| AVNGPVLIRH | ASGGGTGGSAETLY | 0.5994083 |
| DGAAGAVGI | ASGGGTGGSAETLY | 0.6692226 |
| DPLLHGI | ASGGGTGGSAETLY | 0.66483366 |
| ELGAALGAL | ASGGGTGGSAETLY | 0.66619396 |
| ELGRTQAV | ASGGGTGGSAETLY | 0.6773499 |
| FVAGGYLS | ASGGGTGGSAETLY | 0.6754148 |
| GASGGAGGAGGW | ASGGGTGGSAETLY | 0.6482559 |
| GGAGAGGDGGAGGI | ASGGGTGGSAETLY | 0.6056378 |
| GTGHNGSQPGMGGNG | ASGGGTGGSAETLY | 0.619959 |
| GYAAASASAS | ASGGGTGGSAETLY | 0.52912885 |
| HQVDVAAVQQG | ASGGGTGGSAETLY | 0.6066145 |
| IGAAIAHT | ASGGGTGGSAETLY | 0.61107457 |
| IGMIEVAQL | ASGGGTGGSAETLY | 0.6469898 |
| IRLAPEHTSAMD | ASGGGTGGSAETLY | 0.30784822 |
| IYNVSVL | ASGGGTGGSAETLY | 0.64511025 |
| KQGFLGM | ASGGGTGGSAETLY | 0.67896104 |
| LESVITQL | ASGGGTGGSAETLY | 0.612285 |
| LGLGALVVV | ASGGGTGGSAETLY | 0.65429294 |
| LIGNGGNGGSGGI | ASGGGTGGSAETLY | 0.6441033 |
| LNAINAPTL | ASGGGTGGSAETLY | 0.5547171 |
| LQLLNTGV | ASGGGTGGSAETLY | 0.6690817 |
| LQTSIEF | ASGGGTGGSAETLY | 0.5451923 |
| MSFAVSEL | ASGGGTGGSAETLY | 0.6499797 |
| NTGALGL | ASGGGTGGSAETLY | 0.6756115 |
| PPGAGGAGGAGTT | ASGGGTGGSAETLY | 0.6526679 |
| QLVGLGI | ASGGGTGGSAETLY | 0.54692274 |
| RFLEQVHQL | ASGGGTGGSAETLY | 0.603635 |
| RGVGYVLR | ASGGGTGGSAETLY | 0.44002634 |
| SATGVFAGVI | ASGGGTGGSAETLY | 0.6056282 |
| SLPAITLP | ASGGGTGGSAETLY | 0.50545794 |
| STALAFTV | ASGGGTGGSAETLY | 0.63671327 |
| SVLTVGLGK | ASGGGTGGSAETLY | 0.5230286 |
| TALAGAGSGI | ASGGGTGGSAETLY | 0.6586187 |
| TGLPFTGI | ASGGGTGGSAETLY | 0.647467 |
| VAEHVTV | ASGGGTGGSAETLY | 0.63534206 |
| VDIVLNSL | ASGGGTGGSAETLY | 0.58162796 |
| VLTGGGAVL | ASGGGTGGSAETLY | 0.6351955 |
| VNVGLAYGI | ASGGGTGGSAETLY | 0.60979044 |
| VSSAAVGAV | ASGGGTGGSAETLY | 0.51797354 |
| VTSVYEAGV | ASGGGTGGSAETLY | 0.66358423 |
| WVALDRGA | ASGGGTGGSAETLY | 0.5578023 |
| YLVGSVVI | ASGGGTGGSAETLY | 0.5977803 |
| GPPGTGKTLL | ASGGGTGGSAETLY | 0.6724092 |
| GVGMTKF | ASGGGTGGSAETLY | 0.64386415 |
| IGIGGSDL | ASGGGTGGSAETLY | 0.436728 |
| IGTLLAEL | ASGGGTGGSAETLY | 0.6370582 |
| IYTSGTT | ASGGGTGGSAETLY | 0.6424128 |
| LSFPAGV | ASGGGTGGSAETLY | 0.62757015 |

| | | |
|---|---|---|
| LYTSGTT | ASGGGTGGSAETLY | 0.32090265 |
| VGAFPFGF | ASGGGTGGSAETLY | 0.63500047 |
| VGAGTVEF | ASGGGTGGSAETLY | 0.54680634 |
| VGKSTLLL | ASGGGTGGSAETLY | 0.57153594 |
| AALAHAGGGIV | ASGGGTGGSAETLY | 0.5841186 |
| AALVAVSLL | ASGGGTGGSAETLY | 0.6691494 |
| AAVSRAVAAL | ASGGGTGGSAETLY | 0.61041254 |
| AGGSSGIGGF | ASGGGTGGSAETLY | 0.6196441 |
| AGLTVANANG | ASGGGTGGSAETLY | 0.6546929 |
| AGLVGAVL | ASGGGTGGSAETLY | 0.6543921 |
| ALSQAFTV | ASGGGTGGSAETLY | 0.674143 |
| ALTFSVSL | ASGGGTGGSAETLY | 0.6593393 |
| ALTGAALVAL | ASGGGTGGSAETLY | 0.5414152 |
| ASFSIGAGVL | ASGGGTGGSAETLY | 0.62757766 |
| DGSVYKI | ASGGGTGGSAETLY | 0.56351376 |
| EQLAEIV | ASGGGTGGSAETLY | 0.43661293 |
| FAVSELTVM | ASGGGTGGSAETLY | 0.42651895 |
| GAAPLAAGL | ASGGGTGGSAETLY | 0.644479 |
| GADGGDGGSGGAGG | ASGGGTGGSAETLY | 0.45063654 |
| GFGLDTGGAGG | ASGGGTGGSAETLY | 0.6517876 |
| GLASGGLL | ASGGGTGGSAETLY | 0.65203863 |
| GRIAGAAVLL | ASGGGTGGSAETLY | 0.55079305 |
| GSAGSLNTSF | ASGGGTGGSAETLY | 0.42274976 |
| GTGQIGI | ASGGGTGGSAETLY | 0.5896448 |
| GTLIALIL | ASGGGTGGSAETLY | 0.5423342 |
| IAATMGFVVG | ASGGGTGGSAETLY | 0.66283447 |
| IFAQALL | ASGGGTGGSAETLY | 0.63312745 |
| IGAAVALGF | ASGGGTGGSAETLY | 0.6826198 |
| IGLTPLAL | ASGGGTGGSAETLY | 0.66976297 |
| ILGSNTSTL | ASGGGTGGSAETLY | 0.6166948 |
| IVFSLTVL | ASGGGTGGSAETLY | 0.660763 |
| LASKATGF | ASGGGTGGSAETLY | 0.56639576 |
| LASVALIL | ASGGGTGGSAETLY | 0.60176367 |
| LFAVGLTL | ASGGGTGGSAETLY | 0.58226967 |
| LFNGGIGTYI | ASGGGTGGSAETLY | 0.6104654 |
| LTLGAGII | ASGGGTGGSAETLY | 0.60324 |
| LVAAALGDI | ASGGGTGGSAETLY | 0.5688137 |
| NTAFVQLGI | ASGGGTGGSAETLY | 0.43890712 |
| QLFSGAEL | ASGGGTGGSAETLY | 0.65860665 |
| SAATFITL | ASGGGTGGSAETLY | 0.64394605 |
| SNYGLANLGV | ASGGGTGGSAETLY | 0.38798785 |
| SYGGAYAVM | ASGGGTGGSAETLY | 0.5855362 |
| TTGVGGLTLG | ASGGGTGGSAETLY | 0.6801369 |
| TTQTIYI | ASGGGTGGSAETLY | 0.675763 |
| VAAGAGLGAVI | ASGGGTGGSAETLY | 0.5880258 |
| VAAVVTLL | ASGGGTGGSAETLY | 0.5837027 |
| VGFLGGLI | ASGGGTGGSAETLY | 0.6510951 |
| VGVIPILL | ASGGGTGGSAETLY | 0.51249534 |
| VIFASSLI | ASGGGTGGSAETLY | 0.6132189 |
| VKGGDGGAAAG | ASGGGTGGSAETLY | 0.6303118 |
| VTKTFTNI | ASGGGTGGSAETLY | 0.6309755 |

| | | |
|---|---|---|
| VVAAAILL | ASGGGTGGSAETLY | 0.5061601 |
| YAALDVEL | ASGGGTGGSAETLY | 0.6497922 |
| LASVALLL | ASGGGTGGSAETLY | 0.62644863 |
| LDAVHRV | ASGGGTGGSAETLY | 0.6168777 |
| QGLSSSIF | ASGGGTGGSAETLY | 0.33309767 |
| RGTFANIRL | ASGGGTGGSAETLY | 0.646054 |
| RQLWWGHRI | ASGGGTGGSAETLY | 0.56960595 |
| YGPPGTGKTLL | ASGGGTGGSAETLY | 0.51332784 |
| AALVGAPANVA | ASGGGTGGSAETLY | 0.356563 |
| AATFRRTE | ASGGGTGGSAETLY | 0.62001204 |
| AGALGGGI | ASGGGTGGSAETLY | 0.6724721 |
| AGGAGGAGGIGAG | ASGGGTGGSAETLY | 0.66962266 |
| AGSGALGL | ASGGGTGGSAETLY | 0.532775 |
| AQIADQI | ASGGGTGGSAETLY | 0.66963995 |
| ASAVAVTYL | ASGGGTGGSAETLY | 0.67883193 |
| ASTVLGGTGG | ASGGGTGGSAETLY | 0.59478664 |
| AVGGEPAK | ASGGGTGGSAETLY | 0.38181207 |
| CEQGTFGGTC | ASGGGTGGSAETLY | 0.646977 |
| DEGKAPAG | ASGGGTGGSAETLY | 0.63884735 |
| DTVVFPAL | ASGGGTGGSAETLY | 0.6520464 |
| FEGITMAG | ASGGGTGGSAETLY | 0.66893774 |
| FPLVVQD | ASGGGTGGSAETLY | 0.54931813 |
| GAFTTPQ | ASGGGTGGSAETLY | 0.6313308 |
| GFGGFGTAGGVGGA | ASGGGTGGSAETLY | 0.66957355 |
| GFLAQFL | ASGGGTGGSAETLY | 0.5275741 |
| GGAGGAGGNAGVA | ASGGGTGGSAETLY | 0.5007218 |
| GGVGTAGAGGAGGAGG | ASGGGTGGSAETLY | 0.56491524 |
| GNGGGGDAGNAGSGGN | ASGGGTGGSAETLY | 0.4382811 |
| GTGGAGGAGNDAGS | ASGGGTGGSAETLY | 0.6582126 |
| GTTNTGWFNS | ASGGGTGGSAETLY | 0.6027899 |
| IFMGPLFTL | ASGGGTGGSAETLY | 0.6121054 |
| ILAHAIEAI | ASGGGTGGSAETLY | 0.6335617 |
| IRSAIREAL | ASGGGTGGSAETLY | 0.5974431 |
| ITFAGGG | ASGGGTGGSAETLY | 0.6005517 |
| ITGMSGHQEV | ASGGGTGGSAETLY | 0.62211883 |
| IVAGLACTL | ASGGGTGGSAETLY | 0.65351176 |
| IVGGGLAGA | ASGGGTGGSAETLY | 0.6546235 |
| IVTGVLNI | ASGGGTGGSAETLY | 0.61786306 |
| LAQQGCI | ASGGGTGGSAETLY | 0.6164243 |
| LCLHVEN | ASGGGTGGSAETLY | 0.67694205 |
| LDAISDGS | ASGGGTGGSAETLY | 0.5119414 |
| LGGFAAML | ASGGGTGGSAETLY | 0.58063054 |
| LGGKNPMI | ASGGGTGGSAETLY | 0.6523903 |
| LGPKGVAAIL | ASGGGTGGSAETLY | 0.527384 |
| LMVGVSI | ASGGGTGGSAETLY | 0.6573779 |
| LVGAAFVL | ASGGGTGGSAETLY | 0.6503137 |
| LVLAVAPL | ASGGGTGGSAETLY | 0.65379333 |
| MAAIAVIDAA | ASGGGTGGSAETLY | 0.6283511 |
| NNGINAELQ | ASGGGTGGSAETLY | 0.68099535 |
| RGEPLVLVHG | ASGGGTGGSAETLY | 0.6600827 |
| RGGAGGAAGLI | ASGGGTGGSAETLY | 0.58725417 |

| | | |
|---|---|---|
| RSYRMHPAV | ASGGGTGGSAETLY | 0.6696459 |
| SAAHLLAGQLA | ASGGGTGGSAETLY | 0.46273303 |
| SAAVGSY | ASGGGTGGSAETLY | 0.6683827 |
| SPHAVLHRI | ASGGGTGGSAETLY | 0.6643963 |
| SQIAKVL | ASGGGTGGSAETLY | 0.5573468 |
| TALGHLVP | ASGGGTGGSAETLY | 0.6429223 |
| TAPTTSGAA | ASGGGTGGSAETLY | 0.6681893 |
| TDGDDGNGGSPGT | ASGGGTGGSAETLY | 0.67332244 |
| TDGLGEN | ASGGGTGGSAETLY | 0.46837002 |
| TGGPPLPGQA | ASGGGTGGSAETLY | 0.5966384 |
| TIPAITI | ASGGGTGGSAETLY | 0.44947475 |
| TIPGFTI | ASGGGTGGSAETLY | 0.44177654 |
| TSIASGALA | ASGGGTGGSAETLY | 0.464957 |
| TVTLPGAAI | ASGGGTGGSAETLY | 0.6539581 |
| VASGLSAALP | ASGGGTGGSAETLY | 0.6264612 |
| VGAGVIGI | ASGGGTGGSAETLY | 0.66805303 |
| VGPKRFGV | ASGGGTGGSAETLY | 0.6512425 |
| VNVAFPD | ASGGGTGGSAETLY | 0.6720911 |
| VSMTNSGVS | ASGGGTGGSAETLY | 0.58412594 |
| VTDVLGDLSV | ASGGGTGGSAETLY | 0.642592 |
| VVVVGAGFAGL | ASGGGTGGSAETLY | 0.39410678 |
| VYTVGTVV | ASGGGTGGSAETLY | 0.55204165 |
| WTTFSGP | ASGGGTGGSAETLY | 0.47390908 |
| YDELVEAF | ASGGGTGGSAETLY | 0.6337153 |
| YGTGGAGGNGG | ASGGGTGGSAETLY | 0.3747804 |
| YTNVMAV | ASGGGTGGSAETLY | 0.6756289 |
| FLEQVHQL | ASGGGTGGSAETLY | 0.62743306 |
| GLSLLAVAV | ASGGGTGGSAETLY | 0.6608186 |
| LEQVHQL | ASGGGTGGSAETLY | 0.57689124 |
| LNAAGAI | ASGGGTGGSAETLY | 0.56921226 |
| PAVHASL | ASGGGTGGSAETLY | 0.60860544 |
| TELEDDA | ASGGGTGGSAETLY | 0.6630439 |
| TPPGPPN | ASGGGTGGSAETLY | 0.52524626 |
| TPPPPPPP | ASGGGTGGSAETLY | 0.39466682 |
| TQSPALI | ASRTVYAEQF | 0.57834023 |
| EFAAAFPQ | ASRTVYAEQF | 0.48515826 |
| LLSGNLDVL | ASRTVYAEQF | 0.676311 |
| GPANLGNYNI | ASRTVYAEQF | 0.67204 |
| TLGPTLP | ASRTVYAEQF | 0.6508626 |
| VYAYSSCCK | ASRTVYAEQF | 0.67739135 |
| APAPAPAPNI | ASRTVYAEQF | 0.32282665 |
| PGGPVPIP | ASRTVYAEQF | 0.57753 |
| FLTVALEP | ASRTVYAEQF | 0.675835 |
| SQYFGANL | ASRTVYAEQF | 0.64626026 |
| GGVKPAA | ASRTVYAEQF | 0.6388808 |
| GGTGGAGGAGGW | ASRTVYAEQF | 0.6695089 |
| GVGGVAGLGGAGG | ASRTVYAEQF | 0.6030709 |
| VLLVLATTI | ASRTVYAEQF | 0.630481 |
| IGIFAAML | ASRTVYAEQF | 0.6376067 |
| AVGDLID | ASRTVYAEQF | 0.6298261 |
| NISPLPL | ASRTVYAEQF | 0.59914416 |

| | | |
|---|---|---|
| VTAVFVAI | ASRTVYAEQF | 0.49899337 |
| SGDDNDAAL | ASRTVYAEQF | 0.63248956 |
| AVAALAHS | ASRTVYAEQF | 0.6853486 |
| GGVTVMATP | ASRTVYAEQF | 0.6348482 |
| AGLAANVRD | ASRTVYAEQF | 0.6652582 |
| TNRPDLIDPALLRPG | ASRTVYAEQF | 0.6483983 |
| LANLGVTAL | ASRTVYAEQF | 0.658028 |
| TGERIEL | ASRTVYAEQF | 0.59433174 |
| NGEELAA | ASRTVYAEQF | 0.5622742 |
| VLSLVWG | ASRTVYAEQF | 0.636341 |
| IGYALWL | ASRTVYAEQF | 0.5227063 |
| LSLLLAPL | ASRTVYAEQF | 0.62182665 |
| TLDEALK | ASRTVYAEQF | 0.5780572 |
| TDGAGTAEGG | ASRTVYAEQF | 0.686201 |
| NAPGGAGGNGG | ASRTVYAEQF | 0.6591141 |
| VSVFMGMGGNF | ASRTVYAEQF | 0.6515503 |
| LASYLAAL | ASRTVYAEQF | 0.58126146 |
| LVGGTPAL | ASRTVYAEQF | 0.6127553 |
| VRNADVL | ASRTVYAEQF | 0.64870316 |
| TCSSPFLVSAM | ASRTVYAEQF | 0.4457999 |
| VHITDAE | ASRTVYAEQF | 0.6721724 |
| EYAAPSF | ASRTVYAEQF | 0.5680633 |
| VVDAFHGGL | ASRTVYAEQF | 0.5863962 |
| ASVVDPQ | ASRTVYAEQF | 0.661101 |
| SYNEFGGLS | ASRTVYAEQF | 0.679911 |
| DPDGSVY | ASRTVYAEQF | 0.60144955 |
| MVSTIHATE | ASRTVYAEQF | 0.62487966 |
| LAAGVADPV | ASRTVYAEQF | 0.59400654 |
| TKEQIAA | ASRTVYAEQF | 0.6544746 |
| SLASPAAL | ASRTVYAEQF | 0.6725656 |
| IGLVFYGV | ASRTVYAEQF | 0.65007925 |
| FLLDASNL | ASRTVYAEQF | 0.68298125 |
| YQVGNFP | ASRTVYAEQF | 0.60221446 |
| ESLLSQST | ASRTVYAEQF | 0.40690988 |
| LFTVGFAAVG | ASRTVYAEQF | 0.60898185 |
| ATPPTPT | ASRTVYAEQF | 0.6644261 |
| TDLATLG | ASRTVYAEQF | 0.6722304 |
| SAGGTGR | ASRTVYAEQF | 0.67963827 |
| GFSNSGS | ASRTVYAEQF | 0.63007444 |
| GGNGGSGGTSV | ASRTVYAEQF | 0.5938484 |
| GTSVATGGAGN | ASRTVYAEQF | 0.67622817 |
| NGGNGGSGGTS | ASRTVYAEQF | 0.67875654 |
| AVVKFPH | ASRTVYAEQF | 0.6566931 |
| TPPPPPP | ASRTVYAEQF | 0.56388235 |
| AAPAAAAAPA | ASRTVYAEQF | 0.5173283 |
| IALNEDL | ASRTVYAEQF | 0.6441464 |
| IVNVSSM | ASRTVYAEQF | 0.65839326 |
| TGIPPAPR | ASRTVYAEQF | 0.50133634 |
| FDLGGGTF | ASRTVYAEQF | 0.6430172 |
| FDLGGGTFDVS | ASRTVYAEQF | 0.65292716 |
| NEPTAAA | ASRTVYAEQF | 0.4429315 |

| | | |
|---|---|---|
| NEPTAAALA | ASRTVYAEQF | 0.63904375 |
| GSIINISSI | ASRTVYAEQF | 0.67196345 |
| VAPGNAGTA | ASRTVYAEQF | 0.5771515 |
| TGGGASLE | ASRTVYAEQF | 0.55133486 |
| VAVYNHYKRI | ASRTVYAEQF | 0.6722656 |
| TDPAPATT | ASRTVYAEQF | 0.6844826 |
| TNRPDILDPALLRPG | ASRTVYAEQF | 0.66843 |
| LTTTTTS | ASRTVYAEQF | 0.46733832 |
| PTGSGKTL | ASRTVYAEQF | 0.3478857 |
| TGSGKTL | ASRTVYAEQF | 0.6186819 |
| TIGHVDHGK | ASRTVYAEQF | 0.67048895 |
| TIGHVDHGKTT | ASRTVYAEQF | 0.62812304 |
| GHVDHGKTT | ASRTVYAEQF | 0.5819411 |
| AQFEHTLL | ASRTVYAEQF | 0.64519393 |
| IVNQDEL | ASRTVYAEQF | 0.6698884 |
| TNGSQFF | ASRTVYAEQF | 0.5368168 |
| TGKGIVE | ASRTVYAEQF | 0.4578849 |
| LISNASDA | ASRTVYAEQF | 0.64752173 |
| AELRTLGELTKIAKAHGAQ | ASRTVYAEQF | 0.6640806 |
| APNAVAAAEPSW | ASRTVYAEQF | 0.5725508 |
| APQVAGAATAVL | ASRTVYAEQF | 0.6269126 |
| ARRDPAAAAAILPTDA | ASRTVYAEQF | 0.6285087 |
| DAAARDVLAEAGLAEYF | ASRTVYAEQF | 0.57759523 |
| DDQIVLSAKANLAKVEQ | ASRTVYAEQF | 0.6435911 |
| DELLPDTGEDSDEDGH | ASRTVYAEQF | 0.5442455 |
| ELPEAPANRATVPQD | ASRTVYAEQF | 0.6737604 |
| FGQSKIERVTGIG | ASRTVYAEQF | 0.52723205 |
| GAGGTGSTAGGAGGAGGAGGLY | ASRTVYAEQF | 0.6325096 |
| GHGGTGAAGGNGGTGGAGGSGI | ASRTVYAEQF | 0.65591526 |
| GNGGNGGAGGEGGPGGAGGAGA | ASRTVYAEQF | 0.64276695 |
| GSTAGNGGNGGNATNGTIAGQP | ASRTVYAEQF | 0.62280256 |
| GTGGDGGNGGNGGTGG | ASRTVYAEQF | 0.5889762 |
| GVRRPAGIPADAPV | ASRTVYAEQF | 0.5010488 |
| GYRQVTDAHLLTLARR | ASRTVYAEQF | 0.46086505 |
| IAKMGQDGHDRG | ASRTVYAEQF | 0.669687 |
| KAEPSIEIQYT | ASRTVYAEQF | 0.5825153 |
| KPVRIRMAADEDTVAVT | ASRTVYAEQF | 0.67195547 |
| MPPRIAGMRLLVIKPEP | ASRTVYAEQF | 0.65488756 |
| MKFYAVALFRSRGVPPTR | ASRTVYAEQF | 0.5811213 |
| NAGGFGILGAVAHSP | ASRTVYAEQF | 0.644987 |
| NEGREITLAMAEM | ASRTVYAEQF | 0.51343656 |
| NELVEWSHPLGHRWRWE | ASRTVYAEQF | 0.56064576 |
| NLGAGGTGGNGGNPG | ASRTVYAEQF | 0.6526748 |
| NTEWPKWETVMAADPVD | ASRTVYAEQF | 0.64336246 |
| PASAAPAETQP | ASRTVYAEQF | 0.57611424 |
| QQSSDGKNGVRAAEITGE | ASRTVYAEQF | 0.6301989 |
| RAPDAVTTSWWKE | ASRTVYAEQF | 0.64818645 |
| SGFGNVGAHTSGWFNQ | ASRTVYAEQF | 0.50090885 |
| SGVAGAVGQAASVGGLKV | ASRTVYAEQF | 0.59634566 |
| SNGGAGTGSAGGNGGT | ASRTVYAEQF | 0.66967666 |
| TGGAGGEGGAAG | ASRTVYAEQF | 0.62561136 |

| | | |
|---|---|---|
| TGGDGGAAGAGGAPGHG | ASRTVYAEQF | 0.52903575 |
| TMPPLLAAIANAGRHGV | ASRTVYAEQF | 0.654952 |
| TSSTAGNSGGA | ASRTVYAEQF | 0.5472446 |
| VPATVPVGILC | ASRTVYAEQF | 0.6550115 |
| VQRPDSGGVVLTGRIS | ASRTVYAEQF | 0.62980497 |
| WTPNPVPATKGVPAGLVQAT | ASRTVYAEQF | 0.6115712 |
| YALKRALAKFSDRFDVVII | ASRTVYAEQF | 0.64927757 |
| YLIGSVIV | ASRTVYAEQF | 0.45395938 |
| YGQLVAVID | ASRTVYAEQF | 0.5462779 |
| VTVGGMTLPA | ASRTVYAEQF | 0.58569455 |
| VLALPAV | ASRTVYAEQF | 0.648015 |
| VIPPTLNYET | ASRTVYAEQF | 0.65969336 |
| VGMGGMHPSSG | ASRTVYAEQF | 0.5622654 |
| VGLAGGAATA | ASRTVYAEQF | 0.6366478 |
| VGGAGGAGGSGP | ASRTVYAEQF | 0.62301123 |
| VFAAFMV | ASRTVYAEQF | 0.5686409 |
| TLYDKLR | ASRTVYAEQF | 0.57556224 |
| TLAVFTVL | ASRTVYAEQF | 0.5704122 |
| TGSVQPARV | ASRTVYAEQF | 0.46971622 |
| TGGGAGSGGT | ASRTVYAEQF | 0.6453321 |
| SHLETSL | ASRTVYAEQF | 0.55491173 |
| SFTTGMT | ASRTVYAEQF | 0.6243572 |
| QVEVAKRMGVRQPTVS | ASRTVYAEQF | 0.6477899 |
| QGSGGPALAM | ASRTVYAEQF | 0.5118674 |
| MLAPSNP | ASRTVYAEQF | 0.6324087 |
| LVNNAAI | ASRTVYAEQF | 0.6416026 |
| LTGEINTLA | ASRTVYAEQF | 0.6558126 |
| LSIFGPV | ASRTVYAEQF | 0.6328567 |
| LLALLANL | ASRTVYAEQF | 0.58132297 |
| LISNASDALDKLR | ASRTVYAEQF | 0.66771233 |
| LINEAVP | ASRTVYAEQF | 0.6595952 |
| LGYAFSGL | ASRTVYAEQF | 0.6378303 |
| LDFVAVAL | ASRTVYAEQF | 0.6439925 |
| LAQPGAE | ASRTVYAEQF | 0.6658938 |
| ISNLGAVTGF | ASRTVYAEQF | 0.65714717 |
| HQEDPGRI | ASRTVYAEQF | 0.6099989 |
| GTPGEPTP | ASRTVYAEQF | 0.4150223 |
| GFFNSGS | ASRTVYAEQF | 0.6349761 |
| GAGGGGLTAGAN | ASRTVYAEQF | 0.6343156 |
| FVDALVNM | ASRTVYAEQF | 0.57099664 |
| FLTVAPEL | ASRTVYAEQF | 0.4007257 |
| FETLTAL | ASRTVYAEQF | 0.6710592 |
| DVINPIEL | ASRTVYAEQF | 0.68505335 |
| APVSGAPV | ASRTVYAEQF | 0.6018144 |
| ALVGGLAW | ASRTVYAEQF | 0.68069524 |
| ALAAAAATAI | ASRTVYAEQF | 0.42337912 |
| AFGLGLLL | ASRTVYAEQF | 0.6593565 |
| AFFAAASL | ASRTVYAEQF | 0.67313004 |
| AAAALPAFSPP | ASRTVYAEQF | 0.56567925 |
| VVLAPAL | ASRTVYAEQF | 0.6615963 |
| TIGHVDHGKTTL | ASRTVYAEQF | 0.51721185 |

| | | |
|---|---|---|
| LISNASDALDK | ASRTVYAEQF | 0.5735293 |
| LISNASDAL | ASRTVYAEQF | 0.6742146 |
| ISNASDALDK | ASRTVYAEQF | 0.61505085 |
| NQAETLVYQTE | ASRTVYAEQF | 0.62925655 |
| TGGTGGTAGAG | ASRTVYAEQF | 0.6212347 |
| TGSTAGGAGGA | ASRTVYAEQF | 0.58338237 |
| DPLAVPVAAGPV | ASRTVYAEQF | 0.6271971 |
| GGTGGTPGNGGN | ASRTVYAEQF | 0.67051435 |
| PAGLAGVDPATL | ASRTVYAEQF | 0.5672577 |
| GNNLVVEAPHAGASV | ASRTVYAEQF | 0.677437 |
| KVTRIMSASTLEQPA | ASRTVYAEQF | 0.566196 |
| LGFALPIALPRRTVV | ASRTVYAEQF | 0.68452823 |
| TRLGNLTARKLVVLV | ASRTVYAEQF | 0.66684854 |
| WSQQRRLIYFPSAGP | ASRTVYAEQF | 0.6390127 |
| GPNGNPDPMAAAVDIR | ASRTVYAEQF | 0.57050717 |
| TQRCVSVADLRAAASR | ASRTVYAEQF | 0.6706685 |
| QFVLDVAEHPVEVLTETV | ASRTVYAEQF | 0.6266763 |
| GGAGGTGGAGGNGGTQAGDG | ASRTVYAEQF | 0.6184112 |
| PLLATLRAIAVDVDAKPAQV | ASRTVYAEQF | 0.6613129 |
| APKFRTGGYMIDFWGVGYQVAK | ASRTVYAEQF | 0.46528816 |
| GAGGAGGAGGAAGGPGGTGGTG | ASRTVYAEQF | 0.6407016 |
| IGALGLLATDVAPAGSVAFGVA | ASRTVYAEQF | 0.6043311 |
| AGGAGGIGGNGGAGGAGGVLMG | ASRTVYAEQF | 0.6184449 |
| ADELIGLL | ASRTVYAEQF | 0.54785466 |
| ADRALKA | ASRTVYAEQF | 0.44373733 |
| AGAVGQVGGAGGA | ASRTVYAEQF | 0.601354 |
| AGGGGGLGGAP | ASRTVYAEQF | 0.5189913 |
| AGGLLYGNGGAG | ASRTVYAEQF | 0.62771374 |
| AGLESIGL | ASRTVYAEQF | 0.6311232 |
| AGLKTGII | ASRTVYAEQF | 0.54210585 |
| ASLAALLL | ASRTVYAEQF | 0.63900185 |
| ASVINISSI | ASRTVYAEQF | 0.6225464 |
| ATPPGPA | ASRTVYAEQF | 0.44262967 |
| ATVASALV | ASRTVYAEQF | 0.57226706 |
| ATVASLIL | ASRTVYAEQF | 0.57991886 |
| AVNGPVLIRH | ASRTVYAEQF | 0.5902163 |
| DGAAGAVGI | ASRTVYAEQF | 0.40776348 |
| DPLLHGI | ASRTVYAEQF | 0.6231859 |
| ELGAALGAL | ASRTVYAEQF | 0.53951293 |
| ELGRTQAV | ASRTVYAEQF | 0.3692435 |
| FVAGGYLS | ASRTVYAEQF | 0.6204805 |
| GASGGAGGAGGW | ASRTVYAEQF | 0.5665858 |
| GGAGAGGDGGAGGI | ASRTVYAEQF | 0.55467355 |
| GTGHNGSQPGMGGNG | ASRTVYAEQF | 0.63878024 |
| GYAAASASAS | ASRTVYAEQF | 0.5848848 |
| HQVDVAAVQQG | ASRTVYAEQF | 0.5271713 |
| IGAAIAHT | ASRTVYAEQF | 0.6197846 |
| IGMIEVAQL | ASRTVYAEQF | 0.64824 |
| IRLAPEHTSAMD | ASRTVYAEQF | 0.4848144 |
| IYNVSVL | ASRTVYAEQF | 0.59747756 |
| KQGFLGM | ASRTVYAEQF | 0.64343476 |

| | | |
|---|---|---|
| LESVITQL | ASRTVYAEQF | 0.6220685 |
| LGLGALVVV | ASRTVYAEQF | 0.65229523 |
| LIGNGGNGGSGGI | ASRTVYAEQF | 0.65397716 |
| LNAINAPTL | ASRTVYAEQF | 0.5183109 |
| LQLLNTGV | ASRTVYAEQF | 0.6397572 |
| LQTSIEF | ASRTVYAEQF | 0.64446723 |
| MSFAVSEL | ASRTVYAEQF | 0.5690532 |
| NTGALGL | ASRTVYAEQF | 0.60844046 |
| PPGAGGAGGAGTT | ASRTVYAEQF | 0.49049997 |
| QLVGLGI | ASRTVYAEQF | 0.5798698 |
| RFLEQVHQL | ASRTVYAEQF | 0.46354473 |
| RGVGYVLR | ASRTVYAEQF | 0.61041284 |
| SATGVFAGVI | ASRTVYAEQF | 0.64555794 |
| SLPAITLP | ASRTVYAEQF | 0.59693867 |
| STALAFTV | ASRTVYAEQF | 0.47464603 |
| SVLTVGLGK | ASRTVYAEQF | 0.4916701 |
| TALAGAGSGI | ASRTVYAEQF | 0.59784555 |
| TGLPFTGI | ASRTVYAEQF | 0.5080579 |
| VAEHVTV | ASRTVYAEQF | 0.5058119 |
| VDIVLNSL | ASRTVYAEQF | 0.5551318 |
| VLTGGGAVL | ASRTVYAEQF | 0.64260024 |
| VNVGLAYGI | ASRTVYAEQF | 0.6425728 |
| VSSAAVGAV | ASRTVYAEQF | 0.5283308 |
| VTSVYEAGV | ASRTVYAEQF | 0.6434569 |
| WVALDRGA | ASRTVYAEQF | 0.60996187 |
| YLVGSVVI | ASRTVYAEQF | 0.6452225 |
| GPPGTGKTLL | ASRTVYAEQF | 0.61501765 |
| GVGMTKF | ASRTVYAEQF | 0.49081796 |
| IGIGGSDL | ASRTVYAEQF | 0.62894 |
| IGTLLAEL | ASRTVYAEQF | 0.58697623 |
| IYTSGTT | ASRTVYAEQF | 0.6343782 |
| LSFPAGV | ASRTVYAEQF | 0.6395651 |
| LYTSGTT | ASRTVYAEQF | 0.5948816 |
| VGAFPFGF | ASRTVYAEQF | 0.54225755 |
| VGAGTVEF | ASRTVYAEQF | 0.5739855 |
| VGKSTLLL | ASRTVYAEQF | 0.6447096 |
| AALAHAGGGIV | ASRTVYAEQF | 0.5101748 |
| AALVAVSLL | ASRTVYAEQF | 0.46921277 |
| AAVSRAVAAL | ASRTVYAEQF | 0.58410585 |
| AGGSSGIGGF | ASRTVYAEQF | 0.6062492 |
| AGLTVANANG | ASRTVYAEQF | 0.64670146 |
| AGLVGAVL | ASRTVYAEQF | 0.6212491 |
| ALSQAFTV | ASRTVYAEQF | 0.56866264 |
| ALTFSVSL | ASRTVYAEQF | 0.6275043 |
| ALTGAALVAL | ASRTVYAEQF | 0.63484573 |
| ASFSIGAGVL | ASRTVYAEQF | 0.61476576 |
| DGSVYKI | ASRTVYAEQF | 0.58224547 |
| EQLAEIV | ASRTVYAEQF | 0.54617757 |
| FAVSELTVM | ASRTVYAEQF | 0.6405075 |
| GAAPLAAGL | ASRTVYAEQF | 0.6373825 |
| GADGGDGGSGGAGG | ASRTVYAEQF | 0.6330136 |

| | | |
|---|---|---|
| GFGLDTGGAGG | ASRTVYAEQF | 0.6366326 |
| GLASGGLL | ASRTVYAEQF | 0.60029626 |
| GRIAGAAVLL | ASRTVYAEQF | 0.61217886 |
| GSAGSLNTSF | ASRTVYAEQF | 0.5089144 |
| GTGQIGI | ASRTVYAEQF | 0.5439128 |
| GTLIALIL | ASRTVYAEQF | 0.58145046 |
| IAATMGFVVG | ASRTVYAEQF | 0.60660595 |
| IFAQALL | ASRTVYAEQF | 0.62549776 |
| IGAAVALGF | ASRTVYAEQF | 0.61028004 |
| IGLTPLAL | ASRTVYAEQF | 0.5358248 |
| ILGSNTSTL | ASRTVYAEQF | 0.6333894 |
| IVFSLTVL | ASRTVYAEQF | 0.63531065 |
| LASKATGF | ASRTVYAEQF | 0.49009794 |
| LASVALIL | ASRTVYAEQF | 0.5815414 |
| LFAVGLTL | ASRTVYAEQF | 0.64382327 |
| LFNGGIGTYI | ASRTVYAEQF | 0.6117905 |
| LTLGAGII | ASRTVYAEQF | 0.59263813 |
| LVAAALGDI | ASRTVYAEQF | 0.5913181 |
| NTAFVQLGI | ASRTVYAEQF | 0.63820726 |
| QLFSGAEL | ASRTVYAEQF | 0.6097684 |
| SAATFITL | ASRTVYAEQF | 0.642617 |
| SNYGLANLGV | ASRTVYAEQF | 0.40400574 |
| SYGGAYAVM | ASRTVYAEQF | 0.5744395 |
| TTGVGGLTLG | ASRTVYAEQF | 0.5544731 |
| TTQTIYI | ASRTVYAEQF | 0.3700558 |
| VAAGAGLGAVI | ASRTVYAEQF | 0.6447673 |
| VAAVVTLL | ASRTVYAEQF | 0.5379755 |
| VGFLGGLI | ASRTVYAEQF | 0.6227644 |
| VGVIPILL | ASRTVYAEQF | 0.57966876 |
| VIFASSLI | ASRTVYAEQF | 0.64163035 |
| VKGGDGGAAAG | ASRTVYAEQF | 0.60820806 |
| VTKTFTNI | ASRTVYAEQF | 0.5801551 |
| VVAAAILL | ASRTVYAEQF | 0.5488287 |
| YAALDVEL | ASRTVYAEQF | 0.63460505 |
| LASVALLL | ASRTVYAEQF | 0.640839 |
| LDAVHRV | ASRTVYAEQF | 0.6418943 |
| QGLSSSIF | ASRTVYAEQF | 0.4264121 |
| RGTFANIRL | ASRTVYAEQF | 0.6223526 |
| RQLWWGHRI | ASRTVYAEQF | 0.60946673 |
| YGPPGTGKTLL | ASRTVYAEQF | 0.52361214 |
| AALVGAPANVA | ASRTVYAEQF | 0.4440867 |
| AATFRRTE | ASRTVYAEQF | 0.53992236 |
| AGALGGGI | ASRTVYAEQF | 0.56737363 |
| AGGAGGAGGIGAG | ASRTVYAEQF | 0.6291061 |
| AGSGALGL | ASRTVYAEQF | 0.63536626 |
| AQIADQI | ASRTVYAEQF | 0.5820716 |
| ASAVAVTYL | ASRTVYAEQF | 0.61962175 |
| ASTVLGGTGG | ASRTVYAEQF | 0.6059767 |
| AVGGEPAK | ASRTVYAEQF | 0.3106333 |
| CEQGTFGGTC | ASRTVYAEQF | 0.54007196 |
| DEGKAPAG | ASRTVYAEQF | 0.44579986 |

| | | |
|---|---|---|
| DTVVFPAL | ASRTVYAEQF | 0.64230335 |
| FEGITMAG | ASRTVYAEQF | 0.50038373 |
| FPLVVQD | ASRTVYAEQF | 0.55063874 |
| GAFTTPQ | ASRTVYAEQF | 0.6024855 |
| GFGGFGTAGGVGGA | ASRTVYAEQF | 0.59528875 |
| GFLAQFL | ASRTVYAEQF | 0.6007576 |
| GGAGGAGGNAGVA | ASRTVYAEQF | 0.51797426 |
| GGVGTAGAGGAGGAGG | ASRTVYAEQF | 0.50707525 |
| GNGGGGDAGNAGSGGN | ASRTVYAEQF | 0.6387318 |
| GTGGAGGAGNDAGS | ASRTVYAEQF | 0.5834236 |
| GTTNTGWFNS | ASRTVYAEQF | 0.49834824 |
| IFMGPLFTL | ASRTVYAEQF | 0.53031576 |
| ILAHAIEAI | ASRTVYAEQF | 0.6339501 |
| IRSAIREAL | ASRTVYAEQF | 0.41443333 |
| ITFAGGG | ASRTVYAEQF | 0.6466004 |
| ITGMSGHQEV | ASRTVYAEQF | 0.5495363 |
| IVAGLACTL | ASRTVYAEQF | 0.58485276 |
| IVGGGLAGA | ASRTVYAEQF | 0.6440321 |
| IVTGVLNI | ASRTVYAEQF | 0.5947646 |
| LAQQGCI | ASRTVYAEQF | 0.59871864 |
| LCLHVEN | ASRTVYAEQF | 0.5382444 |
| LDAISDGS | ASRTVYAEQF | 0.5019736 |
| LGGFAAML | ASRTVYAEQF | 0.5514552 |
| LGGKNPMI | ASRTVYAEQF | 0.522843 |
| LGPKGVAAIL | ASRTVYAEQF | 0.6476813 |
| LMVGVSI | ASRTVYAEQF | 0.5968686 |
| LVGAAFVL | ASRTVYAEQF | 0.46214345 |
| LVLAVAPL | ASRTVYAEQF | 0.61117053 |
| MAAIAVIDAA | ASRTVYAEQF | 0.64481235 |
| NNGINAELQ | ASRTVYAEQF | 0.6499758 |
| RGEPLVLVHG | ASRTVYAEQF | 0.5126449 |
| RGGAGGAAGLI | ASRTVYAEQF | 0.5470733 |
| RSYRMHPAV | ASRTVYAEQF | 0.57392144 |
| SAAHLLAGQLA | ASRTVYAEQF | 0.5896493 |
| SAAVGSY | ASRTVYAEQF | 0.53524876 |
| SPHAVLHRI | ASRTVYAEQF | 0.49900872 |
| SQIAKVL | ASRTVYAEQF | 0.63517386 |
| TALGHLVP | ASRTVYAEQF | 0.6496994 |
| TAPTTSGAA | ASRTVYAEQF | 0.58677727 |
| TDGDDGNGGSPGT | ASRTVYAEQF | 0.4939918 |
| TDGLGEN | ASRTVYAEQF | 0.59235996 |
| TGGPPLPGQA | ASRTVYAEQF | 0.51335406 |
| TIPAITI | ASRTVYAEQF | 0.5576849 |
| TIPGFTI | ASRTVYAEQF | 0.57541597 |
| TSIASGALA | ASRTVYAEQF | 0.452298 |
| TVTLPGAAI | ASRTVYAEQF | 0.58602023 |
| VASGLSAALP | ASRTVYAEQF | 0.6514393 |
| VGAGVIGI | ASRTVYAEQF | 0.5591408 |
| VGPKRFGV | ASRTVYAEQF | 0.6140336 |
| VNVAFPD | ASRTVYAEQF | 0.64438933 |
| VSMTNSGVS | ASRTVYAEQF | 0.62612087 |

| | | |
|---|---|---|
| VTDVLGDLSV | ASRTVYAEQF | 0.6432046 |
| VVVVGAGFAGL | ASRTVYAEQF | 0.61840045 |
| VYTVGTVV | ASRTVYAEQF | 0.60991997 |
| WTTFSGP | ASRTVYAEQF | 0.6036489 |
| YDELVEAF | ASRTVYAEQF | 0.6272507 |
| YGTGGAGGNGG | ASRTVYAEQF | 0.59731805 |
| YTNVMAV | ASRTVYAEQF | 0.61066425 |
| FLEQVHQL | ASRTVYAEQF | 0.58408165 |
| GLSLLAVAV | ASRTVYAEQF | 0.6245878 |
| LEQVHQL | ASRTVYAEQF | 0.60009027 |
| LNAAGAI | ASRTVYAEQF | 0.6387397 |
| PAVHASL | ASRTVYAEQF | 0.3922034 |
| TELEDDA | ASRTVYAEQF | 0.5167021 |
| TPPGPPN | ASRTVYAEQF | 0.63437253 |
| TPPPPPPP | ASRTVYAEQF | 0.35456842 |
| TQSPALI | ASREGAENTEVF | 0.4618283 |
| EFAAAFPQ | ASREGAENTEVF | 0.5925941 |
| LLSGNLDVL | ASREGAENTEVF | 0.6401093 |
| GPANLGNYNI | ASREGAENTEVF | 0.3614688 |
| TLGPTLP | ASREGAENTEVF | 0.5417118 |
| VYAYSSCCK | ASREGAENTEVF | 0.48043633 |
| APAPAPAPNI | ASREGAENTEVF | 0.6332667 |
| PGGPVPIP | ASREGAENTEVF | 0.57703 |
| FLTVALEP | ASREGAENTEVF | 0.4303179 |
| SQYFGANL | ASREGAENTEVF | 0.6306248 |
| GGVKPAA | ASREGAENTEVF | 0.4937892 |
| GGTGGAGGAGGW | ASREGAENTEVF | 0.6143169 |
| GVGGVAGLGGAGG | ASREGAENTEVF | 0.6420822 |
| VLLVLATTI | ASREGAENTEVF | 0.64412916 |
| IGIFAAML | ASREGAENTEVF | 0.48301452 |
| AVGDLID | ASREGAENTEVF | 0.62831926 |
| NISPLPL | ASREGAENTEVF | 0.54050833 |
| VTAVFVAI | ASREGAENTEVF | 0.65471244 |
| SGDDNDAAL | ASREGAENTEVF | 0.58442163 |
| AVAALAHS | ASREGAENTEVF | 0.4556371 |
| GGVTVMATP | ASREGAENTEVF | 0.6269207 |
| AGLAANVRD | ASREGAENTEVF | 0.6212883 |
| TNRPDLIDPALLRPG | ASREGAENTEVF | 0.57310283 |
| LANLGVTAL | ASREGAENTEVF | 0.6136168 |
| TGERIEL | ASREGAENTEVF | 0.57823867 |
| NGEELAA | ASREGAENTEVF | 0.5752173 |
| VLSLVWG | ASREGAENTEVF | 0.6287871 |
| IGYALWL | ASREGAENTEVF | 0.6133237 |
| LSLLLAPL | ASREGAENTEVF | 0.54390466 |
| TLDEALK | ASREGAENTEVF | 0.6421465 |
| TDGAGTAEGG | ASREGAENTEVF | 0.60287607 |
| NAPGGAGGNGG | ASREGAENTEVF | 0.51796216 |
| VSVFMGMGGNF | ASREGAENTEVF | 0.6509776 |
| LASYLAAL | ASREGAENTEVF | 0.5893013 |
| LVGGTPAL | ASREGAENTEVF | 0.63653755 |
| VRNADVL | ASREGAENTEVF | 0.6311464 |

| | | |
|---|---|---|
| TCSSPFLVSAM | ASREGAENTEVF | 0.6419667 |
| VHITDAE | ASREGAENTEVF | 0.4776571 |
| EYAAPSF | ASREGAENTEVF | 0.6004531 |
| VVDAFHGGL | ASREGAENTEVF | 0.60848475 |
| ASVVDPQ | ASREGAENTEVF | 0.62969255 |
| SYNEFGGLS | ASREGAENTEVF | 0.58849823 |
| DPDGSVY | ASREGAENTEVF | 0.6235839 |
| MVSTIHATE | ASREGAENTEVF | 0.62818563 |
| LAAGVADPV | ASREGAENTEVF | 0.6473816 |
| TKEQIAA | ASREGAENTEVF | 0.5964508 |
| SLASPAAL | ASREGAENTEVF | 0.57304144 |
| IGLVFYGV | ASREGAENTEVF | 0.6475977 |
| FLLDASNL | ASREGAENTEVF | 0.5053778 |
| YQVGNFP | ASREGAENTEVF | 0.6045913 |
| ESLLSQST | ASREGAENTEVF | 0.63254815 |
| LFTVGFAAVG | ASREGAENTEVF | 0.6430795 |
| ATPPTPT | ASREGAENTEVF | 0.58973414 |
| TDLATLG | ASREGAENTEVF | 0.63600576 |
| SAGGTGR | ASREGAENTEVF | 0.5493071 |
| GFSNSGS | ASREGAENTEVF | 0.6135149 |
| GGNGGSGGTSV | ASREGAENTEVF | 0.5427077 |
| GTSVATGGAGN | ASREGAENTEVF | 0.59914577 |
| NGGNGGSGGTS | ASREGAENTEVF | 0.5327596 |
| AVVKFPH | ASREGAENTEVF | 0.61046225 |
| TPPPPPP | ASREGAENTEVF | 0.63406855 |
| AAPAAAAAPA | ASREGAENTEVF | 0.62016743 |
| IALNEDL | ASREGAENTEVF | 0.46107578 |
| IVNVSSM | ASREGAENTEVF | 0.62965167 |
| TGIPPAPR | ASREGAENTEVF | 0.622501 |
| FDLGGGTF | ASREGAENTEVF | 0.6467862 |
| FDLGGGTFDVS | ASREGAENTEVF | 0.6085431 |
| NEPTAAA | ASREGAENTEVF | 0.6277839 |
| NEPTAAALA | ASREGAENTEVF | 0.5761081 |
| GSIINISSI | ASREGAENTEVF | 0.6170049 |
| VAPGNAGTA | ASREGAENTEVF | 0.48102397 |
| TGGGASLE | ASREGAENTEVF | 0.5554601 |
| VAVYNHYKRI | ASREGAENTEVF | 0.5819201 |
| TDPAPATT | ASREGAENTEVF | 0.54210806 |
| TNRPDILDPALLRPG | ASREGAENTEVF | 0.63120973 |
| LTTTTTS | ASREGAENTEVF | 0.6375621 |
| PTGSGKTL | ASREGAENTEVF | 0.634834 |
| TGSGKTL | ASREGAENTEVF | 0.63802886 |
| TIGHVDHGK | ASREGAENTEVF | 0.6416776 |
| TIGHVDHGKTT | ASREGAENTEVF | 0.63984746 |
| GHVDHGKTT | ASREGAENTEVF | 0.5575503 |
| AQFEHTLL | ASREGAENTEVF | 0.5782686 |
| IVNQDEL | ASREGAENTEVF | 0.47967017 |
| TNGSQFF | ASREGAENTEVF | 0.55510753 |
| TGKGIVE | ASREGAENTEVF | 0.5586478 |
| LISNASDA | ASREGAENTEVF | 0.61598146 |
| AELRTLGELTKIAKAHGAQ | ASREGAENTEVF | 0.3405591 |

| | | |
|---|---|---|
| APNAVAAAEPSW | ASREGAENTEVF | 0.58583915 |
| APQVAGAATAVL | ASREGAENTEVF | 0.64985764 |
| ARRDPAAAAAILPTDA | ASREGAENTEVF | 0.5907322 |
| DAAARDVLAEAGLAEYF | ASREGAENTEVF | 0.62403953 |
| DDQIVLSAKANLAKVEQ | ASREGAENTEVF | 0.62304974 |
| DELLPDTGEDSDEDGH | ASREGAENTEVF | 0.5494163 |
| ELPEAPANRATVPQD | ASREGAENTEVF | 0.6285762 |
| FGQSKIERVTGIG | ASREGAENTEVF | 0.56276464 |
| GAGGTGSTAGGAGGAGGAGGLY | ASREGAENTEVF | 0.64250726 |
| GHGGTGAAGGNGGTGGAGGSGI | ASREGAENTEVF | 0.64947426 |
| GNGGNGGAGGEGGPGGAGGAGA | ASREGAENTEVF | 0.6134516 |
| GSTAGNGGNGGNATNGTIAGQP | ASREGAENTEVF | 0.53295 |
| GTGGDGGNGGNGGTGG | ASREGAENTEVF | 0.6138706 |
| GVRRPAGIPADAPV | ASREGAENTEVF | 0.46394858 |
| GYRQVTDAHLLTLARR | ASREGAENTEVF | 0.5693382 |
| IAKMGQDGHDRG | ASREGAENTEVF | 0.576377 |
| KAEPSIEIQYT | ASREGAENTEVF | 0.58880603 |
| KPVRIRMAADEDTVAVT | ASREGAENTEVF | 0.5250524 |
| MPPRIAGMRLLVIKPEP | ASREGAENTEVF | 0.6223516 |
| MKFYAVALFRSRGVPPTR | ASREGAENTEVF | 0.61217815 |
| NAGGFGILGAVAHSP | ASREGAENTEVF | 0.61430657 |
| NEGREITLAMAEM | ASREGAENTEVF | 0.52106166 |
| NELVEWSHPLGHRWRWE | ASREGAENTEVF | 0.63290757 |
| NLGAGGTGGNGGNPG | ASREGAENTEVF | 0.6161729 |
| NTEWPKWETVMAADPVD | ASREGAENTEVF | 0.4569642 |
| PASAAPAETQP | ASREGAENTEVF | 0.632721 |
| QQSSDGKNGVRAAEITGE | ASREGAENTEVF | 0.57514274 |
| RAPDAVTTSWWKE | ASREGAENTEVF | 0.57204986 |
| SGFGNVGAHTSGWFNQ | ASREGAENTEVF | 0.63640046 |
| SGVAGAVGQAASVGGLKV | ASREGAENTEVF | 0.59852034 |
| SNGGAGTGSAGGNGGT | ASREGAENTEVF | 0.4831455 |
| TGGAGGEGGAAG | ASREGAENTEVF | 0.5573279 |
| TGGDGGAAGAGGAPGHG | ASREGAENTEVF | 0.5947431 |
| TMPPLLAAIANAGRHGV | ASREGAENTEVF | 0.59317136 |
| TSSTAGNSGGA | ASREGAENTEVF | 0.3924801 |
| VPATVPVGILC | ASREGAENTEVF | 0.61377746 |
| VQRPDSGGVVLTGRIS | ASREGAENTEVF | 0.51982594 |
| WTPNPVPATKGVPAGLVQAT | ASREGAENTEVF | 0.533165 |
| YALKRALAKFSDRFDVVII | ASREGAENTEVF | 0.56338614 |
| YLIGSVIV | ASREGAENTEVF | 0.63768184 |
| YGQLVAVID | ASREGAENTEVF | 0.62364453 |
| VTVGGMTLPA | ASREGAENTEVF | 0.5753342 |
| VLALPAV | ASREGAENTEVF | 0.61172605 |
| VIPPTLNYET | ASREGAENTEVF | 0.61311656 |
| VGMGGMHPSSG | ASREGAENTEVF | 0.6437374 |
| VGLAGGAATA | ASREGAENTEVF | 0.605734 |
| VGGAGGAGGSGP | ASREGAENTEVF | 0.60189694 |
| VFAAFMV | ASREGAENTEVF | 0.6121487 |
| TLYDKLR | ASREGAENTEVF | 0.5975399 |
| TLAVFTVL | ASREGAENTEVF | 0.55362093 |
| TGSVQPARV | ASREGAENTEVF | 0.54930997 |

| | | |
|---|---|---|
| TGGGAGSGGT | ASREGAENTEVF | 0.58330905 |
| SHLETSL | ASREGAENTEVF | 0.5326253 |
| SFTTGMT | ASREGAENTEVF | 0.6091701 |
| QVEVAKRMGVRQPTVS | ASREGAENTEVF | 0.64017946 |
| QGSGGPALAM | ASREGAENTEVF | 0.5649167 |
| MLAPSNP | ASREGAENTEVF | 0.42972875 |
| LVNNAAI | ASREGAENTEVF | 0.55293053 |
| LTGEINTLA | ASREGAENTEVF | 0.54219335 |
| LSIFGPV | ASREGAENTEVF | 0.6378911 |
| LLALLANL | ASREGAENTEVF | 0.61981535 |
| LISNASDALDKLR | ASREGAENTEVF | 0.6431044 |
| LINEAVP | ASREGAENTEVF | 0.6339365 |
| LGYAFSGL | ASREGAENTEVF | 0.5669775 |
| LDFVAVAL | ASREGAENTEVF | 0.6331501 |
| LAQPGAE | ASREGAENTEVF | 0.57182014 |
| ISNLGAVTGF | ASREGAENTEVF | 0.6167105 |
| HQEDPGRI | ASREGAENTEVF | 0.60823953 |
| GTPGEPTP | ASREGAENTEVF | 0.602646 |
| GFFNSGS | ASREGAENTEVF | 0.5220467 |
| GAGGGGLTAGAN | ASREGAENTEVF | 0.5914279 |
| FVDALVNM | ASREGAENTEVF | 0.48793268 |
| FLTVAPEL | ASREGAENTEVF | 0.61378694 |
| FETLTAL | ASREGAENTEVF | 0.5728233 |
| DVINPIEL | ASREGAENTEVF | 0.5482456 |
| APVSGAPV | ASREGAENTEVF | 0.5731791 |
| ALVGGLAW | ASREGAENTEVF | 0.6488714 |
| ALAAAAATAI | ASREGAENTEVF | 0.647918 |
| AFGLGLLL | ASREGAENTEVF | 0.5493289 |
| AFFAAASL | ASREGAENTEVF | 0.5412639 |
| AAAALPAFSPP | ASREGAENTEVF | 0.59983957 |
| VVLAPAL | ASREGAENTEVF | 0.5303296 |
| TIGHVDHGKTTL | ASREGAENTEVF | 0.61965954 |
| LISNASDALDK | ASREGAENTEVF | 0.5942018 |
| LISNASDAL | ASREGAENTEVF | 0.5870467 |
| ISNASDALDK | ASREGAENTEVF | 0.5267075 |
| NQAETLVYQTE | ASREGAENTEVF | 0.61922824 |
| TGGTGGTAGAG | ASREGAENTEVF | 0.5667926 |
| TGSTAGGAGGA | ASREGAENTEVF | 0.6193118 |
| DPLAVPVAAGPV | ASREGAENTEVF | 0.44020367 |
| GGTGGTPGNGGN | ASREGAENTEVF | 0.6373725 |
| PAGLAGVDPATL | ASREGAENTEVF | 0.5082808 |
| GNNLVVEAPHAGASV | ASREGAENTEVF | 0.5400333 |
| KVTRIMSASTLEQPA | ASREGAENTEVF | 0.37791002 |
| LGFALPIALPRRTVV | ASREGAENTEVF | 0.61193806 |
| TRLGNLTARKLVVLV | ASREGAENTEVF | 0.62626547 |
| WSQQRRLIYFPSAGP | ASREGAENTEVF | 0.6388965 |
| GPNGNPDPMAAAVDIR | ASREGAENTEVF | 0.61316776 |
| TQRCVSVADLRAAASR | ASREGAENTEVF | 0.613001 |
| QFVLDVAEHPVEVLTETV | ASREGAENTEVF | 0.64782846 |
| GGAGGTGGAGGNGGTQAGDG | ASREGAENTEVF | 0.5836564 |
| PLLATLRAIAVDVDAKPAQV | ASREGAENTEVF | 0.42873836 |

| | | |
|---|---|---|
| APKFRTGGYMIDFWGVGYQVAK | ASREGAENTEVF | 0.6059445 |
| GAGGAGGAGGAAGGPGGTGGTG | ASREGAENTEVF | 0.5766169 |
| IGALGLLATDVAPAGSVAFGVA | ASREGAENTEVF | 0.63942915 |
| AGGAGGIGGNGGAGGAGGVLMG | ASREGAENTEVF | 0.62024873 |
| ADELIGLL | ASREGAENTEVF | 0.47371447 |
| ADRALKA | ASREGAENTEVF | 0.6026846 |
| AGAVGQVGGAGGA | ASREGAENTEVF | 0.6333253 |
| AGGGGGLGGAP | ASREGAENTEVF | 0.48234007 |
| AGGLLYGNGGAG | ASREGAENTEVF | 0.54429686 |
| AGLESIGL | ASREGAENTEVF | 0.56378716 |
| AGLKTGII | ASREGAENTEVF | 0.53032523 |
| ASLAALLL | ASREGAENTEVF | 0.59647655 |
| ASVINISSI | ASREGAENTEVF | 0.55499417 |
| ATPPGPA | ASREGAENTEVF | 0.5551329 |
| ATVASALV | ASREGAENTEVF | 0.5901454 |
| ATVASLIL | ASREGAENTEVF | 0.510647 |
| AVNGPVLIRH | ASREGAENTEVF | 0.56630296 |
| DGAAGAVGI | ASREGAENTEVF | 0.5776085 |
| DPLLHGI | ASREGAENTEVF | 0.6120998 |
| ELGAALGAL | ASREGAENTEVF | 0.6256687 |
| ELGRTQAV | ASREGAENTEVF | 0.5683087 |
| FVAGGYLS | ASREGAENTEVF | 0.6111212 |
| GASGGAGGAGGW | ASREGAENTEVF | 0.64931506 |
| GGAGAGGDGGAGGI | ASREGAENTEVF | 0.47747302 |
| GTGHNGSQPGMGGNG | ASREGAENTEVF | 0.5407325 |
| GYAAASASAS | ASREGAENTEVF | 0.6170341 |
| HQVDVAAVQQG | ASREGAENTEVF | 0.57507294 |
| IGAAIAHT | ASREGAENTEVF | 0.6362293 |
| IGMIEVAQL | ASREGAENTEVF | 0.61141515 |
| IRLAPEHTSAMD | ASREGAENTEVF | 0.6210998 |
| IYNVSVL | ASREGAENTEVF | 0.5917394 |
| KQGFLGM | ASREGAENTEVF | 0.6471125 |
| LESVITQL | ASREGAENTEVF | 0.63029814 |
| LGLGALVVV | ASREGAENTEVF | 0.58548355 |
| LIGNGGNGGSGGI | ASREGAENTEVF | 0.6090201 |
| LNAINAPTL | ASREGAENTEVF | 0.46727413 |
| LQLLNTGV | ASREGAENTEVF | 0.62312216 |
| LQTSIEF | ASREGAENTEVF | 0.6269189 |
| MSFAVSEL | ASREGAENTEVF | 0.52741235 |
| NTGALGL | ASREGAENTEVF | 0.576114 |
| PPGAGGAGGAGTT | ASREGAENTEVF | 0.65189433 |
| QLVGLGI | ASREGAENTEVF | 0.63637 |
| RFLEQVHQL | ASREGAENTEVF | 0.56655586 |
| RGVGYVLR | ASREGAENTEVF | 0.5566488 |
| SATGVFAGVI | ASREGAENTEVF | 0.54776204 |
| SLPAITLP | ASREGAENTEVF | 0.55644095 |
| STALAFTV | ASREGAENTEVF | 0.43764898 |
| SVLTVGLGK | ASREGAENTEVF | 0.6266087 |
| TALAGAGSGI | ASREGAENTEVF | 0.5872187 |
| TGLPFTGI | ASREGAENTEVF | 0.62440187 |
| VAEHVTV | ASREGAENTEVF | 0.6128498 |

| | | |
|---|---|---|
| VDIVLNSL | ASREGAENTEVF | 0.64792037 |
| VLTGGGAVL | ASREGAENTEVF | 0.5495403 |
| VNVGLAYGI | ASREGAENTEVF | 0.5812305 |
| VSSAAVGAV | ASREGAENTEVF | 0.4197447 |
| VTSVYEAGV | ASREGAENTEVF | 0.6051145 |
| WVALDRGA | ASREGAENTEVF | 0.52573895 |
| YLVGSVVI | ASREGAENTEVF | 0.583302 |
| GPPGTGKTLL | ASREGAENTEVF | 0.48276672 |
| GVGMTKF | ASREGAENTEVF | 0.6458211 |
| IGIGGSDL | ASREGAENTEVF | 0.5908926 |
| IGTLLAEL | ASREGAENTEVF | 0.6385402 |
| IYTSGTT | ASREGAENTEVF | 0.52411926 |
| LSFPAGV | ASREGAENTEVF | 0.5172516 |
| LYTSGTT | ASREGAENTEVF | 0.5592598 |
| VGAFPFGF | ASREGAENTEVF | 0.6285502 |
| VGAGTVEF | ASREGAENTEVF | 0.4764972 |
| VGKSTLLL | ASREGAENTEVF | 0.43059176 |
| AALAHAGGGIV | ASREGAENTEVF | 0.557163 |
| AALVAVSLL | ASREGAENTEVF | 0.5542794 |
| AAVSRAVAAL | ASREGAENTEVF | 0.64356625 |
| AGGSSGIGGF | ASREGAENTEVF | 0.6417607 |
| AGLTVANANG | ASREGAENTEVF | 0.60369116 |
| AGLVGAVL | ASREGAENTEVF | 0.63899547 |
| ALSQAFTV | ASREGAENTEVF | 0.33902138 |
| ALTFSVSL | ASREGAENTEVF | 0.5327802 |
| ALTGAALVAL | ASREGAENTEVF | 0.64308035 |
| ASFSIGAGVL | ASREGAENTEVF | 0.62045634 |
| DGSVYKI | ASREGAENTEVF | 0.60283995 |
| EQLAEIV | ASREGAENTEVF | 0.6495115 |
| FAVSELTVM | ASREGAENTEVF | 0.52470356 |
| GAAPLAAGL | ASREGAENTEVF | 0.59382135 |
| GADGGDGGSGGAGG | ASREGAENTEVF | 0.6080265 |
| GFGLDTGGAGG | ASREGAENTEVF | 0.63380146 |
| GLASGGLL | ASREGAENTEVF | 0.6285088 |
| GRIAGAAVLL | ASREGAENTEVF | 0.45999664 |
| GSAGSLNTSF | ASREGAENTEVF | 0.62077296 |
| GTGQIGI | ASREGAENTEVF | 0.6511321 |
| GTLIALIL | ASREGAENTEVF | 0.56828165 |
| IAATMGFVVG | ASREGAENTEVF | 0.63779235 |
| IFAQALL | ASREGAENTEVF | 0.60969096 |
| IGAAVALGF | ASREGAENTEVF | 0.6184554 |
| IGLTPLAL | ASREGAENTEVF | 0.54206604 |
| ILGSNTSTL | ASREGAENTEVF | 0.5271333 |
| IVFSLTVL | ASREGAENTEVF | 0.6301807 |
| LASKATGF | ASREGAENTEVF | 0.514927 |
| LASVALIL | ASREGAENTEVF | 0.60154057 |
| LFAVGLTL | ASREGAENTEVF | 0.58669275 |
| LFNGGIGTYI | ASREGAENTEVF | 0.65839005 |
| LTLGAGII | ASREGAENTEVF | 0.621757 |
| LVAAALGDI | ASREGAENTEVF | 0.6294311 |
| NTAFVQLGI | ASREGAENTEVF | 0.540367 |

| | | |
|---|---|---|
| QLFSGAEL | ASREGAENTEVF | 0.60995317 |
| SAATFITL | ASREGAENTEVF | 0.6203563 |
| SNYGLANLGV | ASREGAENTEVF | 0.50680417 |
| SYGGAYAVM | ASREGAENTEVF | 0.6440195 |
| TTGVGGLTLG | ASREGAENTEVF | 0.5347459 |
| TTQTIYI | ASREGAENTEVF | 0.58287144 |
| VAAGAGLGAVI | ASREGAENTEVF | 0.6140376 |
| VAAVVTLL | ASREGAENTEVF | 0.62959087 |
| VGFLGGLI | ASREGAENTEVF | 0.60473835 |
| VGVIPILL | ASREGAENTEVF | 0.64103967 |
| VIFASSLI | ASREGAENTEVF | 0.5296992 |
| VKGGDGGAAAG | ASREGAENTEVF | 0.6280203 |
| VTKTFTNI | ASREGAENTEVF | 0.64031583 |
| VVAAAILL | ASREGAENTEVF | 0.59325725 |
| YAALDVEL | ASREGAENTEVF | 0.6463033 |
| LASVALLL | ASREGAENTEVF | 0.6219223 |
| LDAVHRV | ASREGAENTEVF | 0.40527475 |
| QGLSSSIF | ASREGAENTEVF | 0.61248565 |
| RGTFANIRL | ASREGAENTEVF | 0.6431383 |
| RQLWWGHRI | ASREGAENTEVF | 0.6397794 |
| YGPPGTGKTLL | ASREGAENTEVF | 0.64247537 |
| AALVGAPANVA | ASREGAENTEVF | 0.6011241 |
| AATFRRTE | ASREGAENTEVF | 0.5767812 |
| AGALGGGI | ASREGAENTEVF | 0.6463577 |
| AGGAGGAGGIGAG | ASREGAENTEVF | 0.65954876 |
| AGSGALGL | ASREGAENTEVF | 0.6256667 |
| AQIADQI | ASREGAENTEVF | 0.5387759 |
| ASAVAVTYL | ASREGAENTEVF | 0.5119624 |
| ASTVLGGTGG | ASREGAENTEVF | 0.62754023 |
| AVGGEPAK | ASREGAENTEVF | 0.634585 |
| CEQGTFGGTC | ASREGAENTEVF | 0.50894773 |
| DEGKAPAG | ASREGAENTEVF | 0.6102905 |
| DTVVFPAL | ASREGAENTEVF | 0.61840224 |
| FEGITMAG | ASREGAENTEVF | 0.40120354 |
| FPLVVQD | ASREGAENTEVF | 0.60571754 |
| GAFTTPQ | ASREGAENTEVF | 0.6374124 |
| GFGGFGTAGGVGGA | ASREGAENTEVF | 0.6193454 |
| GFLAQFL | ASREGAENTEVF | 0.48770326 |
| GGAGGAGGNAGVA | ASREGAENTEVF | 0.641502 |
| GGVGTAGAGGAGGAGG | ASREGAENTEVF | 0.6508227 |
| GNGGGDAGNAGSGGN | ASREGAENTEVF | 0.6479935 |
| GTGGAGGAGNDAGS | ASREGAENTEVF | 0.4267112 |
| GTTNTGWFNS | ASREGAENTEVF | 0.36441186 |
| IFMGPLFTL | ASREGAENTEVF | 0.61739445 |
| ILAHAIEAI | ASREGAENTEVF | 0.6186998 |
| IRSAIREAL | ASREGAENTEVF | 0.5447112 |
| ITFAGGG | ASREGAENTEVF | 0.50407267 |
| ITGMSGHQEV | ASREGAENTEVF | 0.6037706 |
| IVAGLACTL | ASREGAENTEVF | 0.6399033 |
| IVGGGLAGA | ASREGAENTEVF | 0.4773972 |
| IVTGVLNI | ASREGAENTEVF | 0.4879926 |

| | | |
|---|---|---|
| LAQQGCI | ASREGAENTEVF | 0.63222826 |
| LCLHVEN | ASREGAENTEVF | 0.6328089 |
| LDAISDGS | ASREGAENTEVF | 0.5186015 |
| LGGFAAML | ASREGAENTEVF | 0.57358825 |
| LGGKNPMI | ASREGAENTEVF | 0.62216294 |
| LGPKGVAAIL | ASREGAENTEVF | 0.6261743 |
| LMVGVSI | ASREGAENTEVF | 0.60729504 |
| LVGAAFVL | ASREGAENTEVF | 0.48428375 |
| LVLAVAPL | ASREGAENTEVF | 0.6296233 |
| MAAIAVIDAA | ASREGAENTEVF | 0.49339145 |
| NNGINAELQ | ASREGAENTEVF | 0.628775 |
| RGEPLVLVHG | ASREGAENTEVF | 0.6029187 |
| RGGAGGAAGLI | ASREGAENTEVF | 0.6153182 |
| RSYRMHPAV | ASREGAENTEVF | 0.58302903 |
| SAAHLLAGQLA | ASREGAENTEVF | 0.52159166 |
| SAAVGSY | ASREGAENTEVF | 0.48072135 |
| SPHAVLHRI | ASREGAENTEVF | 0.5237076 |
| SQIAKVL | ASREGAENTEVF | 0.64292604 |
| TALGHLVP | ASREGAENTEVF | 0.6090287 |
| TAPTTSGAA | ASREGAENTEVF | 0.6281722 |
| TDGDDGNGGSPGT | ASREGAENTEVF | 0.6316012 |
| TDGLGEN | ASREGAENTEVF | 0.53163344 |
| TGGPPLPGQA | ASREGAENTEVF | 0.624494 |
| TIPAITI | ASREGAENTEVF | 0.46965176 |
| TIPGFTI | ASREGAENTEVF | 0.6187212 |
| TSIASGALA | ASREGAENTEVF | 0.5869435 |
| TVTLPGAAI | ASREGAENTEVF | 0.62118536 |
| VASGLSAALP | ASREGAENTEVF | 0.5525894 |
| VGAGVIGI | ASREGAENTEVF | 0.63144416 |
| VGPKRFGV | ASREGAENTEVF | 0.6203853 |
| VNVAFPD | ASREGAENTEVF | 0.53170836 |
| VSMTNSGVS | ASREGAENTEVF | 0.60502726 |
| VTDVLGDLSV | ASREGAENTEVF | 0.6317848 |
| VVVVGAGFAGL | ASREGAENTEVF | 0.6283672 |
| VYTVGTVV | ASREGAENTEVF | 0.48955554 |
| WTTFSGP | ASREGAENTEVF | 0.61568946 |
| YDELVEAF | ASREGAENTEVF | 0.5386746 |
| YGTGGAGGNGG | ASREGAENTEVF | 0.61042714 |
| YTNVMAV | ASREGAENTEVF | 0.6147555 |
| FLEQVHQL | ASREGAENTEVF | 0.5762969 |
| GLSLLAVAV | ASREGAENTEVF | 0.6275527 |
| LEQVHQL | ASREGAENTEVF | 0.4767058 |
| LNAAGAI | ASREGAENTEVF | 0.5534042 |
| PAVHASL | ASREGAENTEVF | 0.5802969 |
| TELEDDA | ASREGAENTEVF | 0.61534745 |
| TPPGPPN | ASREGAENTEVF | 0.6257824 |
| TPPPPPPP | ASREGAENTEVF | 0.5028176 |
| TQSPALI | ASSLRDSYNNQAPL | 0.6091291 |
| EFAAAFPQ | ASSLRDSYNNQAPL | 0.57874984 |
| LLSGNLDVL | ASSLRDSYNNQAPL | 0.51384246 |
| GPANLGNYNI | ASSLRDSYNNQAPL | 0.592774 |

| | | |
|---|---|---|
| TLGPTLP | ASSLRDSYNNQAPL | 0.58170104 |
| VYAYSSCCK | ASSLRDSYNNQAPL | 0.46058044 |
| APAPAPAPNI | ASSLRDSYNNQAPL | 0.62954223 |
| PGGPVPIP | ASSLRDSYNNQAPL | 0.5378597 |
| FLTVALEP | ASSLRDSYNNQAPL | 0.6009857 |
| SQYFGANL | ASSLRDSYNNQAPL | 0.64148885 |
| GGVKPAA | ASSLRDSYNNQAPL | 0.43633828 |
| GGTGGAGGAGGW | ASSLRDSYNNQAPL | 0.63296074 |
| GVGGVAGLGGAGG | ASSLRDSYNNQAPL | 0.60551405 |
| VLLVLATTI | ASSLRDSYNNQAPL | 0.5935111 |
| IGIFAAML | ASSLRDSYNNQAPL | 0.59185845 |
| AVGDLID | ASSLRDSYNNQAPL | 0.5410348 |
| NISPLPL | ASSLRDSYNNQAPL | 0.6370847 |
| VTAVFVAI | ASSLRDSYNNQAPL | 0.63035953 |
| SGDDNDAAL | ASSLRDSYNNQAPL | 0.61734563 |
| AVAALAHS | ASSLRDSYNNQAPL | 0.6147727 |
| GGVTVMATP | ASSLRDSYNNQAPL | 0.63089395 |
| AGLAANVRD | ASSLRDSYNNQAPL | 0.6364577 |
| TNRPDLIDPALLRPG | ASSLRDSYNNQAPL | 0.556928 |
| LANLGVTAL | ASSLRDSYNNQAPL | 0.44046062 |
| TGERIEL | ASSLRDSYNNQAPL | 0.58553195 |
| NGEELAA | ASSLRDSYNNQAPL | 0.5897309 |
| VLSLVWG | ASSLRDSYNNQAPL | 0.56727326 |
| IGYALWL | ASSLRDSYNNQAPL | 0.45333153 |
| LSLLLAPL | ASSLRDSYNNQAPL | 0.6243694 |
| TLDEALK | ASSLRDSYNNQAPL | 0.6562224 |
| TDGAGTAEGG | ASSLRDSYNNQAPL | 0.53742814 |
| NAPGGAGGNGG | ASSLRDSYNNQAPL | 0.6479919 |
| VSVFMGMGGNF | ASSLRDSYNNQAPL | 0.48367214 |
| LASYLAAL | ASSLRDSYNNQAPL | 0.6318182 |
| LVGGTPAL | ASSLRDSYNNQAPL | 0.6424332 |
| VRNADVL | ASSLRDSYNNQAPL | 0.505641 |
| TCSSPFLVSAM | ASSLRDSYNNQAPL | 0.607666 |
| VHITDAE | ASSLRDSYNNQAPL | 0.57317585 |
| EYAAPSF | ASSLRDSYNNQAPL | 0.55177677 |
| VVDAFHGGL | ASSLRDSYNNQAPL | 0.6409817 |
| ASVVDPQ | ASSLRDSYNNQAPL | 0.5504746 |
| SYNEFGGLS | ASSLRDSYNNQAPL | 0.61945534 |
| DPDGSVY | ASSLRDSYNNQAPL | 0.5566031 |
| MVSTIHATE | ASSLRDSYNNQAPL | 0.5389515 |
| LAAGVADPV | ASSLRDSYNNQAPL | 0.62387294 |
| TKEQIAA | ASSLRDSYNNQAPL | 0.6276871 |
| SLASPAAL | ASSLRDSYNNQAPL | 0.60339177 |
| IGLVFYGV | ASSLRDSYNNQAPL | 0.6583912 |
| FLLDASNL | ASSLRDSYNNQAPL | 0.5423988 |
| YQVGNFP | ASSLRDSYNNQAPL | 0.6380891 |
| ESLLSQST | ASSLRDSYNNQAPL | 0.62670016 |
| LFTVGFAAVG | ASSLRDSYNNQAPL | 0.5954101 |
| ATPPTPT | ASSLRDSYNNQAPL | 0.58652663 |
| TDLATLG | ASSLRDSYNNQAPL | 0.63916636 |
| SAGGTGR | ASSLRDSYNNQAPL | 0.5730428 |

| | | |
|---|---|---|
| GFSNSGS | ASSLRDSYNNQAPL | 0.605839 |
| GGNGGSGGTSV | ASSLRDSYNNQAPL | 0.6440493 |
| GTSVATGGAGN | ASSLRDSYNNQAPL | 0.53406924 |
| NGGNGGSGGTS | ASSLRDSYNNQAPL | 0.8257605 |
| AVVKFPH | ASSLRDSYNNQAPL | 0.7256027 |
| TPPPPPP | ASSLRDSYNNQAPL | 0.7844044 |
| AAPAAAAAPA | ASSLRDSYNNQAPL | 0.60354686 |
| IALNEDL | ASSLRDSYNNQAPL | 0.326238 |
| IVNVSSM | ASSLRDSYNNQAPL | 0.7021127 |
| TGIPPAPR | ASSLRDSYNNQAPL | 0.49495584 |
| FDLGGGTF | ASSLRDSYNNQAPL | 0.75004077 |
| FDLGGGTFDVS | ASSLRDSYNNQAPL | 0.8059077 |
| NEPTAAA | ASSLRDSYNNQAPL | 0.584805 |
| NEPTAAALA | ASSLRDSYNNQAPL | 0.8187642 |
| GSIINISSI | ASSLRDSYNNQAPL | 0.8187672 |
| VAPGNAGTA | ASSLRDSYNNQAPL | 0.44074315 |
| TGGGASLE | ASSLRDSYNNQAPL | 0.69703186 |
| VAVYNHYKRI | ASSLRDSYNNQAPL | 0.48246008 |
| TDPAPATT | ASSLRDSYNNQAPL | 0.7008766 |
| TNRPDILDPALLRPG | ASSLRDSYNNQAPL | 0.18317133 |
| LTTTTTS | ASSLRDSYNNQAPL | 0.7901752 |
| PTGSGKTL | ASSLRDSYNNQAPL | 0.5639391 |
| TGSGKTL | ASSLRDSYNNQAPL | 0.14921296 |
| TIGHVDHGK | ASSLRDSYNNQAPL | 0.72917104 |
| TIGHVDHGKTT | ASSLRDSYNNQAPL | 0.6304566 |
| GHVDHGKTT | ASSLRDSYNNQAPL | 0.6740735 |
| AQFEHTLL | ASSLRDSYNNQAPL | 0.81159323 |
| IVNQDEL | ASSLRDSYNNQAPL | 0.39885032 |
| TNGSQFF | ASSLRDSYNNQAPL | 0.39550662 |
| TGKGIVE | ASSLRDSYNNQAPL | 0.7852247 |
| LISNASDA | ASSLRDSYNNQAPL | 0.84222627 |
| AELRTLGELTKIAKAHGAQ | ASSLRDSYNNQAPL | 0.4004416 |
| APNAVAAAEPSW | ASSLRDSYNNQAPL | 0.74310756 |
| APQVAGAATAVL | ASSLRDSYNNQAPL | 0.85161364 |
| ARRDPAAAAAILPTDA | ASSLRDSYNNQAPL | 0.6800541 |
| DAAARDVLAEAGLAEYF | ASSLRDSYNNQAPL | 0.8319043 |
| DDQIVLSAKANLAKVEQ | ASSLRDSYNNQAPL | 0.83798015 |
| DELLPDTGEDSDEDGH | ASSLRDSYNNQAPL | 0.60084367 |
| ELPEAPANRATVPQD | ASSLRDSYNNQAPL | 0.82898694 |
| FGQSKIERVTGIG | ASSLRDSYNNQAPL | 0.82165575 |
| GAGGTGSTAGGAGGAGGAGGLY | ASSLRDSYNNQAPL | 0.6910057 |
| GHGGTGAAGGNGGTGGAGGSGI | ASSLRDSYNNQAPL | 0.72049475 |
| GNGGNGGAGGEGGPGGAGGAGA | ASSLRDSYNNQAPL | 0.53111774 |
| GSTAGNGGNGGNATNGTIAGQP | ASSLRDSYNNQAPL | 0.70895183 |
| GTGGDGGNGGNGGTGG | ASSLRDSYNNQAPL | 0.31847346 |
| GVRRPAGIPADAPV | ASSLRDSYNNQAPL | 0.72218657 |
| GYRQVTDAHLLTLARR | ASSLRDSYNNQAPL | 0.8401977 |
| IAKMGQDGHDRG | ASSLRDSYNNQAPL | 0.6729743 |
| KAEPSIEIQYT | ASSLRDSYNNQAPL | 0.45462358 |
| KPVRIRMAADEDTVAVT | ASSLRDSYNNQAPL | 0.4443881 |
| MPPRIAGMRLLVIKPEP | ASSLRDSYNNQAPL | 0.70436954 |

| | | |
|---|---|---|
| MKFYAVALFRSRGVPPTR | ASSLRDSYNNQAPL | 0.6250285 |
| NAGGFGILGAVAHSP | ASSLRDSYNNQAPL | 0.32620847 |
| NEGREITLAMAEM | ASSLRDSYNNQAPL | 0.62729895 |
| NELVEWSHPLGHRWRWE | ASSLRDSYNNQAPL | 0.8152993 |
| NLGAGGTGGNGGNPG | ASSLRDSYNNQAPL | 0.819968 |
| NTEWPKWETVMAADPVD | ASSLRDSYNNQAPL | 0.37334442 |
| PASAAPAETQP | ASSLRDSYNNQAPL | 0.8526106 |
| QQSSDGKNGVRAAEITGE | ASSLRDSYNNQAPL | 0.74842393 |
| RAPDAVTTSWWKE | ASSLRDSYNNQAPL | 0.8240701 |
| SGFGNVGAHTSGWFNQ | ASSLRDSYNNQAPL | 0.70472085 |
| SGVAGAVGQAASVGGLKV | ASSLRDSYNNQAPL | 0.54689634 |
| SNGGAGTGSAGGNGGT | ASSLRDSYNNQAPL | 0.80963147 |
| TGGAGGEGGAAG | ASSLRDSYNNQAPL | 0.64549315 |
| TGGDGGAAGAGGAPGHG | ASSLRDSYNNQAPL | 0.5723774 |
| TMPPLLAAIANAGRHGV | ASSLRDSYNNQAPL | 0.8144408 |
| TSSTAGNSGGA | ASSLRDSYNNQAPL | 0.5238215 |
| VPATVPVGILC | ASSLRDSYNNQAPL | 0.5185423 |
| VQRPDSGGVVLTGRIS | ASSLRDSYNNQAPL | 0.6397344 |
| WTPNPVPATKGVPAGLVQAT | ASSLRDSYNNQAPL | 0.82083285 |
| YALKRALAKFSDRFDVVII | ASSLRDSYNNQAPL | 0.5290331 |
| YLIGSVIV | ASSLRDSYNNQAPL | 0.44624314 |
| YGQLVAVID | ASSLRDSYNNQAPL | 0.6105187 |
| VTVGGMTLPA | ASSLRDSYNNQAPL | 0.6906295 |
| VLALPAV | ASSLRDSYNNQAPL | 0.8408065 |
| VIPPTLNYET | ASSLRDSYNNQAPL | 0.7694807 |
| VGMGGMHPSSG | ASSLRDSYNNQAPL | 0.69415474 |
| VGLAGGAATA | ASSLRDSYNNQAPL | 0.7833078 |
| VGGAGGAGGSGP | ASSLRDSYNNQAPL | 0.7790636 |
| VFAAFMV | ASSLRDSYNNQAPL | 0.7207699 |
| TLYDKLR | ASSLRDSYNNQAPL | 0.7376873 |
| TLAVFTVL | ASSLRDSYNNQAPL | 0.54763997 |
| TGSVQPARV | ASSLRDSYNNQAPL | 0.81741226 |
| TGGGAGSGGT | ASSLRDSYNNQAPL | 0.7755699 |
| SHLETSL | ASSLRDSYNNQAPL | 0.81965685 |
| SFTTGMT | ASSLRDSYNNQAPL | 0.8121909 |
| QVEVAKRMGVRQPTVS | ASSLRDSYNNQAPL | 0.70251817 |
| QGSGGPALAM | ASSLRDSYNNQAPL | 0.7632648 |
| MLAPSNP | ASSLRDSYNNQAPL | 0.28026035 |
| LVNNAAI | ASSLRDSYNNQAPL | 0.6075374 |
| LTGEINTLA | ASSLRDSYNNQAPL | 0.7100924 |
| LSIFGPV | ASSLRDSYNNQAPL | 0.75439155 |
| LLALLANL | ASSLRDSYNNQAPL | 0.7607082 |
| LISNASDALDKLR | ASSLRDSYNNQAPL | 0.8072444 |
| LINEAVP | ASSLRDSYNNQAPL | 0.6014931 |
| LGYAFSGL | ASSLRDSYNNQAPL | 0.7850979 |
| LDFVAVAL | ASSLRDSYNNQAPL | 0.8468554 |
| LAQPGAE | ASSLRDSYNNQAPL | 0.383664 |
| ISNLGAVTGF | ASSLRDSYNNQAPL | 0.67417985 |
| HQEDPGRI | ASSLRDSYNNQAPL | 0.8493718 |
| GTPGEPTP | ASSLRDSYNNQAPL | 0.7408718 |
| GFFNSGS | ASSLRDSYNNQAPL | 0.6463653 |

| | | |
|---|---|---|
| GAGGGGLTAGAN | ASSLRDSYNNQAPL | 0.7252258 |
| FVDALVNM | ASSLRDSYNNQAPL | 0.8089142 |
| FLTVAPEL | ASSLRDSYNNQAPL | 0.6202538 |
| FETLTAL | ASSLRDSYNNQAPL | 0.83430976 |
| DVINPIEL | ASSLRDSYNNQAPL | 0.21060228 |
| APVSGAPV | ASSLRDSYNNQAPL | 0.71230567 |
| ALVGGLAW | ASSLRDSYNNQAPL | 0.5939386 |
| ALAAAAATAI | ASSLRDSYNNQAPL | 0.24845941 |
| AFGLGLLL | ASSLRDSYNNQAPL | 0.77997273 |
| AFFAAASL | ASSLRDSYNNQAPL | 0.67491734 |
| AAAALPAFSPP | ASSLRDSYNNQAPL | 0.82135993 |
| VVLAPAL | ASSLRDSYNNQAPL | 0.642897 |
| TIGHVDHGKTTL | ASSLRDSYNNQAPL | 0.8213869 |
| LISNASDALDK | ASSLRDSYNNQAPL | 0.7063849 |
| LISNASDAL | ASSLRDSYNNQAPL | 0.7069492 |
| ISNASDALDK | ASSLRDSYNNQAPL | 0.6146573 |
| NQAETLVYQTE | ASSLRDSYNNQAPL | 0.71502745 |
| TGGTGGTAGAG | ASSLRDSYNNQAPL | 0.80135596 |
| TGSTAGGAGGA | ASSLRDSYNNQAPL | 0.81589365 |
| DPLAVPVAAGPV | ASSLRDSYNNQAPL | 0.28618717 |
| GGTGGTPGNGGN | ASSLRDSYNNQAPL | 0.6642182 |
| PAGLAGVDPATL | ASSLRDSYNNQAPL | 0.6880169 |
| GNNLVVEAPHAGASV | ASSLRDSYNNQAPL | 0.20187701 |
| KVTRIMSASTLEQPA | ASSLRDSYNNQAPL | 0.19913481 |
| LGFALPIALPRRTVV | ASSLRDSYNNQAPL | 0.51361245 |
| TRLGNLTARKLVVLV | ASSLRDSYNNQAPL | 0.6105501 |
| WSQQRRLIYFPSAGP | ASSLRDSYNNQAPL | 0.7865927 |
| GPNGNPDPMAAAVDIR | ASSLRDSYNNQAPL | 0.781554 |
| TQRCVSVADLRAAASR | ASSLRDSYNNQAPL | 0.6209704 |
| QFVLDVAEHPVEVLTETV | ASSLRDSYNNQAPL | 0.7769569 |
| GGAGGTGGAGGNGGTQAGDG | ASSLRDSYNNQAPL | 0.6972306 |
| PLLATLRAIAVDVDAKPAQV | ASSLRDSYNNQAPL | 0.09966852 |
| APKFRTGGYMIDFWGVGYQVAK | ASSLRDSYNNQAPL | 0.6151618 |
| GAGGAGGAGGAAGGPGGTGGTG | ASSLRDSYNNQAPL | 0.46299452 |
| IGALGLLATDVAPAGSVAFGVA | ASSLRDSYNNQAPL | 0.8308205 |
| AGGAGGIGGNGGAGGAGGVLMG | ASSLRDSYNNQAPL | 0.51897216 |
| ADELIGLL | ASSLRDSYNNQAPL | 0.55540603 |
| ADRALKA | ASSLRDSYNNQAPL | 0.7079507 |
| AGAVGQVGGAGGA | ASSLRDSYNNQAPL | 0.7157594 |
| AGGGGGLGGAP | ASSLRDSYNNQAPL | 0.7118275 |
| AGGLLYGNGGAG | ASSLRDSYNNQAPL | 0.4915855 |
| AGLESIGL | ASSLRDSYNNQAPL | 0.16136704 |
| AGLKTGII | ASSLRDSYNNQAPL | 0.7827507 |
| ASLAALLL | ASSLRDSYNNQAPL | 0.33640587 |
| ASVINISSI | ASSLRDSYNNQAPL | 0.5867702 |
| ATPPGPA | ASSLRDSYNNQAPL | 0.36579537 |
| ATVASALV | ASSLRDSYNNQAPL | 0.18123701 |
| ATVASLIL | ASSLRDSYNNQAPL | 0.34084105 |
| AVNGPVLIRH | ASSLRDSYNNQAPL | 0.8086939 |
| DGAAGAVGI | ASSLRDSYNNQAPL | 0.6824827 |
| DPLLHGI | ASSLRDSYNNQAPL | 0.64950585 |

| | | |
|---|---|---|
| ELGAALGAL | ASSLRDSYNNQAPL | 0.85364044 |
| ELGRTQAV | ASSLRDSYNNQAPL | 0.6877239 |
| FVAGGYLS | ASSLRDSYNNQAPL | 0.6823648 |
| GASGGAGGAGGW | ASSLRDSYNNQAPL | 0.5881235 |
| GGAGAGGDGGAGGI | ASSLRDSYNNQAPL | 0.5098722 |
| GTGHNGSQPGMGGNG | ASSLRDSYNNQAPL | 0.46870947 |
| GYAAASASAS | ASSLRDSYNNQAPL | 0.5547369 |
| HQVDVAAVQQG | ASSLRDSYNNQAPL | 0.83752453 |
| IGAAIAHT | ASSLRDSYNNQAPL | 0.7279198 |
| IGMIEVAQL | ASSLRDSYNNQAPL | 0.4524195 |
| IRLAPEHTSAMD | ASSLRDSYNNQAPL | 0.78931534 |
| IYNVSVL | ASSLRDSYNNQAPL | 0.8424908 |
| KQGFLGM | ASSLRDSYNNQAPL | 0.84666896 |
| LESVITQL | ASSLRDSYNNQAPL | 0.58582175 |
| LGLGALVVV | ASSLRDSYNNQAPL | 0.6606405 |
| LIGNGGNGGSGGI | ASSLRDSYNNQAPL | 0.7044325 |
| LNAINAPTL | ASSLRDSYNNQAPL | 0.66774595 |
| LQLLNTGV | ASSLRDSYNNQAPL | 0.6266565 |
| LQTSIEF | ASSLRDSYNNQAPL | 0.4287705 |
| MSFAVSEL | ASSLRDSYNNQAPL | 0.811569 |
| NTGALGL | ASSLRDSYNNQAPL | 0.85276747 |
| PPGAGGAGGAGTT | ASSLRDSYNNQAPL | 0.66722 |
| QLVGLGI | ASSLRDSYNNQAPL | 0.25908512 |
| RFLEQVHQL | ASSLRDSYNNQAPL | 0.627627 |
| RGVGYVLR | ASSLRDSYNNQAPL | 0.4976048 |
| SATGVFAGVI | ASSLRDSYNNQAPL | 0.58655804 |
| SLPAITLP | ASSLRDSYNNQAPL | 0.6912455 |
| STALAFTV | ASSLRDSYNNQAPL | 0.42034698 |
| SVLTVGLGK | ASSLRDSYNNQAPL | 0.6989373 |
| TALAGAGSGI | ASSLRDSYNNQAPL | 0.85213006 |
| TGLPFTGI | ASSLRDSYNNQAPL | 0.6227537 |
| VAEHVTV | ASSLRDSYNNQAPL | 0.77974385 |
| VDIVLNSL | ASSLRDSYNNQAPL | 0.8315881 |
| VLTGGGAVL | ASSLRDSYNNQAPL | 0.80130076 |
| VNVGLAYGI | ASSLRDSYNNQAPL | 0.83701205 |
| VSSAAVGAV | ASSLRDSYNNQAPL | 0.7634127 |
| VTSVYEAGV | ASSLRDSYNNQAPL | 0.71114224 |
| WVALDRGA | ASSLRDSYNNQAPL | 0.7508812 |
| YLVGSVVI | ASSLRDSYNNQAPL | 0.78076494 |
| GPPGTGKTLL | ASSLRDSYNNQAPL | 0.68108505 |
| GVGMTKF | ASSLRDSYNNQAPL | 0.691571 |
| IGIGGSDL | ASSLRDSYNNQAPL | 0.63855743 |
| IGTLLAEL | ASSLRDSYNNQAPL | 0.8158082 |
| IYTSGTT | ASSLRDSYNNQAPL | 0.6729844 |
| LSFPAGV | ASSLRDSYNNQAPL | 0.81101537 |
| LYTSGTT | ASSLRDSYNNQAPL | 0.32483333 |
| VGAFPFGF | ASSLRDSYNNQAPL | 0.40985292 |
| VGAGTVEF | ASSLRDSYNNQAPL | 0.8124788 |
| VGKSTLLL | ASSLRDSYNNQAPL | 0.14590861 |
| AALAHAGGGIV | ASSLRDSYNNQAPL | 0.27592647 |
| AALVAVSLL | ASSLRDSYNNQAPL | 0.7432116 |

| | | |
|---|---|---|
| AAVSRAVAAL | ASSLRDSYNNQAPL | 0.8129214 |
| AGGSSGIGGF | ASSLRDSYNNQAPL | 0.1370562 |
| AGLTVANANG | ASSLRDSYNNQAPL | 0.48173323 |
| AGLVGAVL | ASSLRDSYNNQAPL | 0.49153537 |
| ALSQAFTV | ASSLRDSYNNQAPL | 0.77454954 |
| ALTFSVSL | ASSLRDSYNNQAPL | 0.67867863 |
| ALTGAALVAL | ASSLRDSYNNQAPL | 0.29925162 |
| ASFSIGAGVL | ASSLRDSYNNQAPL | 0.81550837 |
| DGSVYKI | ASSLRDSYNNQAPL | 0.32553926 |
| EQLAEIV | ASSLRDSYNNQAPL | 0.53185564 |
| FAVSELTVM | ASSLRDSYNNQAPL | 0.8078103 |
| GAAPLAAGL | ASSLRDSYNNQAPL | 0.8323325 |
| GADGGDGGSGGAGG | ASSLRDSYNNQAPL | 0.5037679 |
| GFGLDTGGAGG | ASSLRDSYNNQAPL | 0.8067502 |
| GLASGGLL | ASSLRDSYNNQAPL | 0.63892436 |
| GRIAGAAVLL | ASSLRDSYNNQAPL | 0.86254066 |
| GSAGSLNTSF | ASSLRDSYNNQAPL | 0.6499543 |
| GTGQIGI | ASSLRDSYNNQAPL | 0.36249962 |
| GTLIALIL | ASSLRDSYNNQAPL | 0.8118045 |
| IAATMGFVVG | ASSLRDSYNNQAPL | 0.79995406 |
| IFAQALL | ASSLRDSYNNQAPL | 0.7798703 |
| IGAAVALGF | ASSLRDSYNNQAPL | 0.7874577 |
| IGLTPLAL | ASSLRDSYNNQAPL | 0.67164373 |
| ILGSNTSTL | ASSLRDSYNNQAPL | 0.69066465 |
| IVFSLTVL | ASSLRDSYNNQAPL | 0.8072785 |
| LASKATGF | ASSLRDSYNNQAPL | 0.7610889 |
| LASVALIL | ASSLRDSYNNQAPL | 0.39088166 |
| LFAVGLTL | ASSLRDSYNNQAPL | 0.8319894 |
| LFNGGIGTYI | ASSLRDSYNNQAPL | 0.66584414 |
| LTLGAGII | ASSLRDSYNNQAPL | 0.5758023 |
| LVAAALGDI | ASSLRDSYNNQAPL | 0.8422502 |
| NTAFVQLGI | ASSLRDSYNNQAPL | 0.74201244 |
| QLFSGAEL | ASSLRDSYNNQAPL | 0.8112223 |
| SAATFITL | ASSLRDSYNNQAPL | 0.7842534 |
| SNYGLANLGV | ASSLRDSYNNQAPL | 0.8120743 |
| SYGGAYAVM | ASSLRDSYNNQAPL | 0.33956826 |
| TTGVGGLTLG | ASSLRDSYNNQAPL | 0.73389035 |
| TTQTIYI | ASSLRDSYNNQAPL | 0.7277678 |
| VAAGAGLGAVI | ASSLRDSYNNQAPL | 0.80852675 |
| VAAVVTLL | ASSLRDSYNNQAPL | 0.6618135 |
| VGFLGGLI | ASSLRDSYNNQAPL | 0.454925 |
| VGVIPILL | ASSLRDSYNNQAPL | 0.788203 |
| VIFASSLI | ASSLRDSYNNQAPL | 0.8386247 |
| VKGGDGGAAAG | ASSLRDSYNNQAPL | 0.7440355 |
| VTKTFTNI | ASSLRDSYNNQAPL | 0.71167827 |
| VVAAAILL | ASSLRDSYNNQAPL | 0.85437995 |
| YAALDVEL | ASSLRDSYNNQAPL | 0.53950125 |
| LASVALLL | ASSLRDSYNNQAPL | 0.70670164 |
| LDAVHRV | ASSLRDSYNNQAPL | 0.81487274 |
| QGLSSSIF | ASSLRDSYNNQAPL | 0.8092787 |
| RGTFANIRL | ASSLRDSYNNQAPL | 0.7563225 |

| | | |
|---|---|---|
| RQLWWGHRI | ASSLRDSYNNQAPL | 0.8083745 |
| YGPPGTGKTLL | ASSLRDSYNNQAPL | 0.50754315 |
| AALVGAPANVA | ASSLRDSYNNQAPL | 0.7749612 |
| AATFRRTE | ASSLRDSYNNQAPL | 0.5949317 |
| AGALGGGI | ASSLRDSYNNQAPL | 0.70264566 |
| AGGAGGAGGIGAG | ASSLRDSYNNQAPL | 0.353334 |
| AGSGALGL | ASSLRDSYNNQAPL | 0.7353785 |
| AQIADQI | ASSLRDSYNNQAPL | 0.79686105 |
| ASAVAVTYL | ASSLRDSYNNQAPL | 0.7881948 |
| ASTVLGGTGG | ASSLRDSYNNQAPL | 0.21188417 |
| AVGGEPAK | ASSLRDSYNNQAPL | 0.7815007 |
| CEQGTFGGTC | ASSLRDSYNNQAPL | 0.79013693 |
| DEGKAPAG | ASSLRDSYNNQAPL | 0.854988 |
| DTVVFPAL | ASSLRDSYNNQAPL | 0.7113329 |
| FEGITMAG | ASSLRDSYNNQAPL | 0.8177894 |
| FPLVVQD | ASSLRDSYNNQAPL | 0.5318977 |
| GAFTTPQ | ASSLRDSYNNQAPL | 0.72696805 |
| GFGGFGTAGGVGGA | ASSLRDSYNNQAPL | 0.47229117 |
| GFLAQFL | ASSLRDSYNNQAPL | 0.5997368 |
| GGAGGAGGNAGVA | ASSLRDSYNNQAPL | 0.6143724 |
| GGVGTAGAGGAGGAGG | ASSLRDSYNNQAPL | 0.5618341 |
| GNGGGGDAGNAGSGGN | ASSLRDSYNNQAPL | 0.7740121 |
| GTGGAGGAGNDAGS | ASSLRDSYNNQAPL | 0.7906965 |
| GTTNTGWFNS | ASSLRDSYNNQAPL | 0.8168228 |
| IFMGPLFTL | ASSLRDSYNNQAPL | 0.8275884 |
| ILAHAIEAI | ASSLRDSYNNQAPL | 0.842155 |
| IRSAIREAL | ASSLRDSYNNQAPL | 0.83369803 |
| ITFAGGG | ASSLRDSYNNQAPL | 0.5933894 |
| ITGMSGHQEV | ASSLRDSYNNQAPL | 0.6393192 |
| IVAGLACTL | ASSLRDSYNNQAPL | 0.35658866 |
| IVGGGLAGA | ASSLRDSYNNQAPL | 0.6578258 |
| IVTGVLNI | ASSLRDSYNNQAPL | 0.6341392 |
| LAQQGCI | ASSLRDSYNNQAPL | 0.73953176 |
| LCLHVEN | ASSLRDSYNNQAPL | 0.121030614 |
| LDAISDGS | ASSLRDSYNNQAPL | 0.7276275 |
| LGGFAAML | ASSLRDSYNNQAPL | 0.83781266 |
| LGGKNPMI | ASSLRDSYNNQAPL | 0.46933818 |
| LGPKGVAAIL | ASSLRDSYNNQAPL | 0.7786946 |
| LMVGVSI | ASSLRDSYNNQAPL | 0.7448983 |
| LVGAAFVL | ASSLRDSYNNQAPL | 0.6005988 |
| LVLAVAPL | ASSLRDSYNNQAPL | 0.8004775 |
| MAAIAVIDAA | ASSLRDSYNNQAPL | 0.61605173 |
| NNGINAELQ | ASSLRDSYNNQAPL | 0.8220822 |
| RGEPLVLVHG | ASSLRDSYNNQAPL | 0.8359214 |
| RGGAGGAAGLI | ASSLRDSYNNQAPL | 0.77482176 |
| RSYRMHPAV | ASSLRDSYNNQAPL | 0.63589597 |
| SAAHLLAGQLA | ASSLRDSYNNQAPL | 0.7464763 |
| SAAVGSY | ASSLRDSYNNQAPL | 0.28789213 |
| SPHAVLHRI | ASSLRDSYNNQAPL | 0.68076193 |
| SQIAKVL | ASSLRDSYNNQAPL | 0.12836403 |
| TALGHLVP | ASSLRDSYNNQAPL | 0.72400916 |

| | | |
|---|---|---|
| TAPTTSGAA | ASSLRDSYNNQAPL | 0.56857723 |
| TDGDDGNGGSPGT | ASSLRDSYNNQAPL | 0.78181803 |
| TDGLGEN | ASSLRDSYNNQAPL | 0.6986192 |
| TGGPPLPGQA | ASSLRDSYNNQAPL | 0.7646312 |
| TIPAITI | ASSLRDSYNNQAPL | 0.49458462 |
| TIPGFTI | ASSLRDSYNNQAPL | 0.7262949 |
| TSIASGALA | ASSLRDSYNNQAPL | 0.64661336 |
| TVTLPGAAI | ASSLRDSYNNQAPL | 0.4152551 |
| VASGLSAALP | ASSLRDSYNNQAPL | 0.7771327 |
| VGAGVIGI | ASSLRDSYNNQAPL | 0.7423185 |
| VGPKRFGV | ASSLRDSYNNQAPL | 0.6580453 |
| VNVAFPD | ASSLRDSYNNQAPL | 0.83009106 |
| VSMTNSGVS | ASSLRDSYNNQAPL | 0.7858349 |
| VTDVLGDLSV | ASSLRDSYNNQAPL | 0.3141563 |
| VVVVGAGFAGL | ASSLRDSYNNQAPL | 0.6271843 |
| VYTVGTVV | ASSLRDSYNNQAPL | 0.7486272 |
| WTTFSGP | ASSLRDSYNNQAPL | 0.6570357 |
| YDELVEAF | ASSLRDSYNNQAPL | 0.12448655 |
| YGTGGAGGNGG | ASSLRDSYNNQAPL | 0.77427924 |
| YTNVMAV | ASSLRDSYNNQAPL | 0.5927401 |
| FLEQVHQL | ASSLRDSYNNQAPL | 0.49806505 |
| GLSLLAVAV | ASSLRDSYNNQAPL | 0.61613584 |
| LEQVHQL | ASSLRDSYNNQAPL | 0.82632565 |
| LNAAGAI | ASSLRDSYNNQAPL | 0.49591404 |
| PAVHASL | ASSLRDSYNNQAPL | 0.67106813 |
| TELEDDA | ASSLRDSYNNQAPL | 0.7698333 |
| TPPGPPN | ASSLRDSYNNQAPL | 0.7363385 |
| TPPPPPPP | ASSLRDSYNNQAPL | 0.825032 |
| TQSPALI | ASSLSGNTLY | 0.73195434 |
| EFAAAFPQ | ASSLSGNTLY | 0.72857773 |
| LLSGNLDVL | ASSLSGNTLY | 0.78920007 |
| GPANLGNYNI | ASSLSGNTLY | 0.67860603 |
| TLGPTLP | ASSLSGNTLY | 0.43161538 |
| VYAYSSCCK | ASSLSGNTLY | 0.37440354 |
| APAPAPAPNI | ASSLSGNTLY | 0.60122657 |
| PGGPVPIP | ASSLSGNTLY | 0.36650887 |
| FLTVALEP | ASSLSGNTLY | 0.75349987 |
| SQYFGANL | ASSLSGNTLY | 0.83580697 |
| GGVKPAA | ASSLSGNTLY | 0.63144815 |
| GGTGGAGGAGGW | ASSLSGNTLY | 0.4096392 |
| GVGGVAGLGGAGG | ASSLSGNTLY | 0.67635137 |
| VLLVLATTI | ASSLSGNTLY | 0.5088292 |
| IGIFAAML | ASSLSGNTLY | 0.7999792 |
| AVGDLID | ASSLSGNTLY | 0.7820139 |
| NISPLPL | ASSLSGNTLY | 0.8283836 |
| VTAVFVAI | ASSLSGNTLY | 0.79588723 |
| SGDDNDAAL | ASSLSGNTLY | 0.5896089 |
| AVAALAHS | ASSLSGNTLY | 0.8237088 |
| GGVTVMATP | ASSLSGNTLY | 0.67733884 |
| AGLAANVRD | ASSLSGNTLY | 0.6633955 |
| TNRPDLIDPALLRPG | ASSLSGNTLY | 0.7308807 |

| | | |
|---|---|---|
| LANLGVTAL | ASSLSGNTLY | 0.71714735 |
| TGERIEL | ASSLSGNTLY | 0.56766164 |
| NGEELAA | ASSLSGNTLY | 0.70770645 |
| VLSLVWG | ASSLSGNTLY | 0.34398183 |
| IGYALWL | ASSLSGNTLY | 0.73084277 |
| LSLLLAPL | ASSLSGNTLY | 0.48507124 |
| TLDEALK | ASSLSGNTLY | 0.4198039 |
| TDGAGTAEGG | ASSLSGNTLY | 0.645153 |
| NAPGGAGGNGG | ASSLSGNTLY | 0.85169375 |
| VSVFMGMGGNF | ASSLSGNTLY | 0.8496342 |
| LASYLAAL | ASSLSGNTLY | 0.58928 |
| LVGGTPAL | ASSLSGNTLY | 0.5798645 |
| VRNADVL | ASSLSGNTLY | 0.7302791 |
| TCSSPFLVSAM | ASSLSGNTLY | 0.40530586 |
| VHITDAE | ASSLSGNTLY | 0.7909858 |
| EYAAPSF | ASSLSGNTLY | 0.66208184 |
| VVDAFHGGL | ASSLSGNTLY | 0.6659219 |
| ASVVDPQ | ASSLSGNTLY | 0.6613343 |
| SYNEFGGLS | ASSLSGNTLY | 0.77015847 |
| DPDGSVY | ASSLSGNTLY | 0.6112065 |
| MVSTIHATE | ASSLSGNTLY | 0.79285955 |
| LAAGVADPV | ASSLSGNTLY | 0.13532452 |
| TKEQIAA | ASSLSGNTLY | 0.7740824 |
| SLASPAAL | ASSLSGNTLY | 0.48610628 |
| IGLVFYGV | ASSLSGNTLY | 0.5251804 |
| FLLDASNL | ASSLSGNTLY | 0.25156912 |
| YQVGNFP | ASSLSGNTLY | 0.7211022 |
| ESLLSQST | ASSLSGNTLY | 0.77216834 |
| LFTVGFAAVG | ASSLSGNTLY | 0.8295083 |
| ATPPTPT | ASSLSGNTLY | 0.60321045 |
| TDLATLG | ASSLSGNTLY | 0.7809702 |
| SAGGTGR | ASSLSGNTLY | 0.82348275 |
| GFSNSGS | ASSLSGNTLY | 0.6371834 |
| GGNGGSGGTSV | ASSLSGNTLY | 0.27090237 |
| GTSVATGGAGN | ASSLSGNTLY | 0.77454686 |
| NGGNGGSGGTS | ASSLSGNTLY | 0.5916727 |
| AVVKFPH | ASSLSGNTLY | 0.8108587 |
| TPPPPPP | ASSLSGNTLY | 0.7513927 |
| AAPAAAAAPA | ASSLSGNTLY | 0.4040997 |
| IALNEDL | ASSLSGNTLY | 0.8278903 |
| IVNVSSM | ASSLSGNTLY | 0.83190316 |
| TGIPPAPR | ASSLSGNTLY | 0.4856683 |
| FDLGGGTF | ASSLSGNTLY | 0.5464372 |
| FDLGGGTFDVS | ASSLSGNTLY | 0.60446244 |
| NEPTAAA | ASSLSGNTLY | 0.2635863 |
| NEPTAAALA | ASSLSGNTLY | 0.74810535 |
| GSIINISSI | ASSLSGNTLY | 0.6318928 |
| VAPGNAGTA | ASSLSGNTLY | 0.53077257 |
| TGGGASLE | ASSLSGNTLY | 0.7314187 |
| VAVYNHYKRI | ASSLSGNTLY | 0.5521986 |
| TDPAPATT | ASSLSGNTLY | 0.57876563 |

| | | |
|---|---|---|
| TNRPDILDPALLRPG | ASSLSGNTLY | 0.5716536 |
| LTTTTTS | ASSLSGNTLY | 0.5779435 |
| PTGSGKTL | ASSLSGNTLY | 0.77318996 |
| TGSGKTL | ASSLSGNTLY | 0.61864364 |
| TIGHVDHGK | ASSLSGNTLY | 0.746336 |
| TIGHVDHGKTT | ASSLSGNTLY | 0.8415352 |
| GHVDHGKTT | ASSLSGNTLY | 0.31397328 |
| AQFEHTLL | ASSLSGNTLY | 0.5129911 |
| IVNQDEL | ASSLSGNTLY | 0.77842593 |
| TNGSQFF | ASSLSGNTLY | 0.5171976 |
| TGKGIVE | ASSLSGNTLY | 0.79964507 |
| LISNASDA | ASSLSGNTLY | 0.6666687 |
| AELRTLGELTKIAKAHGAQ | ASSLSGNTLY | 0.74402076 |
| APNAVAAAEPSW | ASSLSGNTLY | 0.64870775 |
| APQVAGAATAVL | ASSLSGNTLY | 0.84870064 |
| ARRDPAAAAAILPTDA | ASSLSGNTLY | 0.8155541 |
| DAAARDVLAEAGLAEYF | ASSLSGNTLY | 0.68996894 |
| DDQIVLSAKANLAKVEQ | ASSLSGNTLY | 0.6875584 |
| DELLPDTGEDSDEDGH | ASSLSGNTLY | 0.47356644 |
| ELPEAPANRATVPQD | ASSLSGNTLY | 0.77924377 |
| FGQSKIERVTGIG | ASSLSGNTLY | 0.8099896 |
| GAGGTGSTAGGAGGAGGAGGLY | ASSLSGNTLY | 0.46813864 |
| GHGGTGAAGGNGGTGGAGGSGI | ASSLSGNTLY | 0.6883939 |
| GNGGNGGAGGEGGPGGAGGAGA | ASSLSGNTLY | 0.8306718 |
| GSTAGNGGNGGNATNGTIAGQP | ASSLSGNTLY | 0.8371262 |
| GTGGDGGNGGNGGTGG | ASSLSGNTLY | 0.4172266 |
| GVRRPAGIPADAPV | ASSLSGNTLY | 0.6529652 |
| GYRQVTDAHLLTLARR | ASSLSGNTLY | 0.42423975 |
| IAKMGQDGHDRG | ASSLSGNTLY | 0.6392407 |
| KAEPSIEIQYT | ASSLSGNTLY | 0.41774166 |
| KPVRIRMAADEDTVAVT | ASSLSGNTLY | 0.773339 |
| MPPRIAGMRLLVIKPEP | ASSLSGNTLY | 0.6421454 |
| MKFYAVALFRSRGVPPTR | ASSLSGNTLY | 0.8009641 |
| NAGGFGILGAVAHSP | ASSLSGNTLY | 0.7557844 |
| NEGREITLAMAEM | ASSLSGNTLY | 0.81662226 |
| NELVEWSHPLGHRWRWE | ASSLSGNTLY | 0.6362505 |
| NLGAGGTGGNGGNPG | ASSLSGNTLY | 0.6640223 |
| NTEWPKWETVMAADPVD | ASSLSGNTLY | 0.46898484 |
| PASAAPAETQP | ASSLSGNTLY | 0.30248678 |
| QQSSDGKNGVRAAEITGE | ASSLSGNTLY | 0.49059498 |
| RAPDAVTTSWWKE | ASSLSGNTLY | 0.5886509 |
| SGFGNVGAHTSGWFNQ | ASSLSGNTLY | 0.18410408 |
| SGVAGAVGQAASVGGLKV | ASSLSGNTLY | 0.8378675 |
| SNGGAGTGSAGGNGGT | ASSLSGNTLY | 0.7447613 |
| TGGAGGEGGAAG | ASSLSGNTLY | 0.74952614 |
| TGGDGGAAGAGGAPGHG | ASSLSGNTLY | 0.55145925 |
| TMPPLLAAIANAGRHGV | ASSLSGNTLY | 0.5087653 |
| TSSTAGNSGGA | ASSLSGNTLY | 0.5770159 |
| VPATVPVGILC | ASSLSGNTLY | 0.8026494 |
| VQRPDSGGVVLTGRIS | ASSLSGNTLY | 0.45477894 |
| WTPNPVPATKGVPAGLVQAT | ASSLSGNTLY | 0.23427106 |

| | | |
|---|---|---|
| YALKRALAKFSDRFDVVII | ASSLSGNTLY | 0.57045853 |
| YLIGSVIV | ASSLSGNTLY | 0.32985878 |
| YGQLVAVID | ASSLSGNTLY | 0.8383305 |
| VTVGGMTLPA | ASSLSGNTLY | 0.82021 |
| VLALPAV | ASSLSGNTLY | 0.7826015 |
| VIPPTLNYET | ASSLSGNTLY | 0.8384972 |
| VGMGGMHPSSG | ASSLSGNTLY | 0.19273254 |
| VGLAGGAATA | ASSLSGNTLY | 0.5613452 |
| VGGAGGAGGSGP | ASSLSGNTLY | 0.84322214 |
| VFAAFMV | ASSLSGNTLY | 0.7726915 |
| TLYDKLR | ASSLSGNTLY | 0.68039644 |
| TLAVFTVL | ASSLSGNTLY | 0.82957965 |
| TGSVQPARV | ASSLSGNTLY | 0.5364597 |
| TGGGAGSGGT | ASSLSGNTLY | 0.66172594 |
| SHLETSL | ASSLSGNTLY | 0.7173587 |
| SFTTGMT | ASSLSGNTLY | 0.7954968 |
| QVEVAKRMGVRQPTVS | ASSLSGNTLY | 0.6911081 |
| QGSGGPALAM | ASSLSGNTLY | 0.37783322 |
| MLAPSNP | ASSLSGNTLY | 0.8059598 |
| LVNNAAI | ASSLSGNTLY | 0.84262204 |
| LTGEINTLA | ASSLSGNTLY | 0.67744017 |
| LSIFGPV | ASSLSGNTLY | 0.75914955 |
| LLALLANL | ASSLSGNTLY | 0.7798867 |
| LISNASDALDKLR | ASSLSGNTLY | 0.7037815 |
| LINEAVP | ASSLSGNTLY | 0.55851364 |
| LGYAFSGL | ASSLSGNTLY | 0.4944747 |
| LDFVAVAL | ASSLSGNTLY | 0.71352136 |
| LAQPGAE | ASSLSGNTLY | 0.43693253 |
| ISNLGAVTGF | ASSLSGNTLY | 0.7509788 |
| HQEDPGRI | ASSLSGNTLY | 0.6482771 |
| GTPGEPTP | ASSLSGNTLY | 0.86693925 |
| GFFNSGS | ASSLSGNTLY | 0.79282343 |
| GAGGGGLTAGAN | ASSLSGNTLY | 0.77259445 |
| FVDALVNM | ASSLSGNTLY | 0.43607903 |
| FLTVAPEL | ASSLSGNTLY | 0.74099755 |
| FETLTAL | ASSLSGNTLY | 0.74976957 |
| DVINPIEL | ASSLSGNTLY | 0.36244375 |
| APVSGAPV | ASSLSGNTLY | 0.8352275 |
| ALVGGLAW | ASSLSGNTLY | 0.49188566 |
| ALAAAAATAI | ASSLSGNTLY | 0.6899358 |
| AFGLGLLL | ASSLSGNTLY | 0.77093226 |
| AFFAAASL | ASSLSGNTLY | 0.8207314 |
| AAAALPAFSPP | ASSLSGNTLY | 0.67259526 |
| VVLAPAL | ASSLSGNTLY | 0.66166484 |
| TIGHVDHGKTTL | ASSLSGNTLY | 0.48787647 |
| LISNASDALDK | ASSLSGNTLY | 0.80309784 |
| LISNASDAL | ASSLSGNTLY | 0.83423376 |
| ISNASDALDK | ASSLSGNTLY | 0.54774237 |
| NQAETLVYQTE | ASSLSGNTLY | 0.84184825 |
| TGGTGGTAGAG | ASSLSGNTLY | 0.7156522 |
| TGSTAGGAGGA | ASSLSGNTLY | 0.35164303 |

| | | |
|---|---|---|
| DPLAVPVAAGPV | ASSLSGNTLY | 0.757936 |
| GGTGGTPGNGGN | ASSLSGNTLY | 0.8269786 |
| PAGLAGVDPATL | ASSLSGNTLY | 0.8097515 |
| GNNLVVEAPHAGASV | ASSLSGNTLY | 0.72346604 |
| KVTRIMSASTLEQPA | ASSLSGNTLY | 0.707594 |
| LGFALPIALPRRTVV | ASSLSGNTLY | 0.64300174 |
| TRLGNLTARKLVVLV | ASSLSGNTLY | 0.8448634 |
| WSQQRRLIYFPSAGP | ASSLSGNTLY | 0.862558 |
| GPNGNPDPMAAAVDIR | ASSLSGNTLY | 0.82173204 |
| TQRCVSVADLRAAASR | ASSLSGNTLY | 0.6538319 |
| QFVLDVAEHPVEVLTETV | ASSLSGNTLY | 0.5884219 |
| GGAGGTGGAGGNGGTQAGDG | ASSLSGNTLY | 0.7876659 |
| PLLATLRAIAVDVDAKPAQV | ASSLSGNTLY | 0.8052782 |
| APKFRTGGYMIDFWGVGYQVAK | ASSLSGNTLY | 0.5487926 |
| GAGGAGGAGGAAGGPGGTGGTG | ASSLSGNTLY | 0.7337822 |
| IGALGLLATDVAPAGSVAFGVA | ASSLSGNTLY | 0.7124566 |
| AGGAGGIGGNGGAGGAGGVLMG | ASSLSGNTLY | 0.33522183 |
| ADELIGLL | ASSLSGNTLY | 0.7574011 |
| ADRALKA | ASSLSGNTLY | 0.8134927 |
| AGAVGQVGGAGGA | ASSLSGNTLY | 0.7574544 |
| AGGGGGLGGAP | ASSLSGNTLY | 0.5019592 |
| AGGLLYGNGGAG | ASSLSGNTLY | 0.82892436 |
| AGLESIGL | ASSLSGNTLY | 0.8476914 |
| AGLKTGII | ASSLSGNTLY | 0.84952253 |
| ASLAALLL | ASSLSGNTLY | 0.29556742 |
| ASVINISSI | ASSLSGNTLY | 0.20027246 |
| ATPPGPA | ASSLSGNTLY | 0.66110706 |
| ATVASALV | ASSLSGNTLY | 0.7710637 |
| ATVASLIL | ASSLSGNTLY | 0.63506734 |
| AVNGPVLIRH | ASSLSGNTLY | 0.59345984 |
| DGAAGAVGI | ASSLSGNTLY | 0.7311806 |
| DPLLHGI | ASSLSGNTLY | 0.82199913 |
| ELGAALGAL | ASSLSGNTLY | 0.5292028 |
| ELGRTQAV | ASSLSGNTLY | 0.38928652 |
| FVAGGYLS | ASSLSGNTLY | 0.75184673 |
| GASGGAGGAGGW | ASSLSGNTLY | 0.82956326 |
| GGAGAGGDGGAGGI | ASSLSGNTLY | 0.5401851 |
| GTGHNGSQPGMGGNG | ASSLSGNTLY | 0.68529665 |
| GYAAASASAS | ASSLSGNTLY | 0.6516767 |
| HQVDVAAVQQG | ASSLSGNTLY | 0.55353403 |
| IGAAIAHT | ASSLSGNTLY | 0.69768953 |
| IGMIEVAQL | ASSLSGNTLY | 0.42802602 |
| IRLAPEHTSAMD | ASSLSGNTLY | 0.7968459 |
| IYNVSVL | ASSLSGNTLY | 0.41684183 |
| KQGFLGM | ASSLSGNTLY | 0.7933896 |
| LESVITQL | ASSLSGNTLY | 0.7447983 |
| LGLGALVVV | ASSLSGNTLY | 0.7810991 |
| LIGNGGNGGSGGI | ASSLSGNTLY | 0.6270813 |
| LNAINAPTL | ASSLSGNTLY | 0.48994523 |
| LQLLNTGV | ASSLSGNTLY | 0.46023238 |
| LQTSIEF | ASSLSGNTLY | 0.3403033 |

| | | |
|---|---|---|
| MSFAVSEL | ASSLSGNTLY | 0.80247176 |
| NTGALGL | ASSLSGNTLY | 0.18163033 |
| PPGAGGAGGAGTT | ASSLSGNTLY | 0.8206503 |
| QLVGLGI | ASSLSGNTLY | 0.7145877 |
| RFLEQVHQL | ASSLSGNTLY | 0.56852794 |
| RGVGYVLR | ASSLSGNTLY | 0.78640175 |
| SATGVFAGVI | ASSLSGNTLY | 0.34184247 |
| SLPAITLP | ASSLSGNTLY | 0.6566457 |
| STALAFTV | ASSLSGNTLY | 0.7471958 |
| SVLTVGLGK | ASSLSGNTLY | 0.81351423 |
| TALAGAGSGI | ASSLSGNTLY | 0.5246079 |
| TGLPFTGI | ASSLSGNTLY | 0.8045038 |
| VAEHVTV | ASSLSGNTLY | 0.7845291 |
| VDIVLNSL | ASSLSGNTLY | 0.57649374 |
| VLTGGGAVL | ASSLSGNTLY | 0.75431156 |
| VNVGLAYGI | ASSLSGNTLY | 0.8083031 |
| VSSAAVGAV | ASSLSGNTLY | 0.7574756 |
| VTSVYEAGV | ASSLSGNTLY | 0.4606893 |
| WVALDRGA | ASSLSGNTLY | 0.7490988 |
| YLVGSVVI | ASSLSGNTLY | 0.57541955 |
| GPPGTGKTLL | ASSLSGNTLY | 0.75013435 |
| GVGMTKF | ASSLSGNTLY | 0.7174721 |
| IGIGGSDL | ASSLSGNTLY | 0.6945901 |
| IGTLLAEL | ASSLSGNTLY | 0.8353697 |
| IYTSGTT | ASSLSGNTLY | 0.3727382 |
| LSFPAGV | ASSLSGNTLY | 0.59448934 |
| LYTSGTT | ASSLSGNTLY | 0.660841 |
| VGAFPFGF | ASSLSGNTLY | 0.7184669 |
| VGAGTVEF | ASSLSGNTLY | 0.80947435 |
| VGKSTLLL | ASSLSGNTLY | 0.47653022 |
| AALAHAGGGIV | ASSLSGNTLY | 0.7657776 |
| AALVAVSLL | ASSLSGNTLY | 0.72456765 |
| AAVSRAVAAL | ASSLSGNTLY | 0.52164567 |
| AGGSSGIGGF | ASSLSGNTLY | 0.780624 |
| AGLTVANANG | ASSLSGNTLY | 0.6289195 |
| AGLVGAVL | ASSLSGNTLY | 0.50489604 |
| ALSQAFTV | ASSLSGNTLY | 0.8024267 |
| ALTFSVSL | ASSLSGNTLY | 0.5822548 |
| ALTGAALVAL | ASSLSGNTLY | 0.77341676 |
| ASFSIGAGVL | ASSLSGNTLY | 0.8086208 |
| DGSVYKI | ASSLSGNTLY | 0.4808546 |
| EQLAEIV | ASSLSGNTLY | 0.79555 |
| FAVSELTVM | ASSLSGNTLY | 0.7611434 |
| GAAPLAAGL | ASSLSGNTLY | 0.7371472 |
| GADGGDGGSGGAGG | ASSLSGNTLY | 0.54314524 |
| GFGLDTGGAGG | ASSLSGNTLY | 0.5714494 |
| GLASGGLL | ASSLSGNTLY | 0.8308616 |
| GRIAGAAVLL | ASSLSGNTLY | 0.7848743 |
| GSAGSLNTSF | ASSLSGNTLY | 0.7720057 |
| GTGQIGI | ASSLSGNTLY | 0.7825223 |
| GTLIALIL | ASSLSGNTLY | 0.7641344 |

| | | |
|---|---|---|
| IAATMGFVVG | ASSLSGNTLY | 0.8206525 |
| IFAQALL | ASSLSGNTLY | 0.55119884 |
| IGAAVALGF | ASSLSGNTLY | 0.4391336 |
| IGLTPLAL | ASSLSGNTLY | 0.63460827 |
| ILGSNTSTL | ASSLSGNTLY | 0.5591552 |
| IVFSLTVL | ASSLSGNTLY | 0.6087526 |
| LASKATGF | ASSLSGNTLY | 0.33855897 |
| LASVALIL | ASSLSGNTLY | 0.8093101 |
| LFAVGLTL | ASSLSGNTLY | 0.8635634 |
| LFNGGIGTYI | ASSLSGNTLY | 0.4564334 |
| LTLGAGII | ASSLSGNTLY | 0.85533035 |
| LVAAALGDI | ASSLSGNTLY | 0.44272852 |
| NTAFVQLGI | ASSLSGNTLY | 0.7741801 |
| QLFSGAEL | ASSLSGNTLY | 0.8110739 |
| SAATFITL | ASSLSGNTLY | 0.4665502 |
| SNYGLANLGV | ASSLSGNTLY | 0.72658384 |
| SYGGAYAVM | ASSLSGNTLY | 0.5206742 |
| TTGVGGLTLG | ASSLSGNTLY | 0.54858327 |
| TTQTIYI | ASSLSGNTLY | 0.818724 |
| VAAGAGLGAVI | ASSLSGNTLY | 0.5420917 |
| VAAVVTLL | ASSLSGNTLY | 0.7652991 |
| VGFLGGLI | ASSLSGNTLY | 0.70324314 |
| VGVIPILL | ASSLSGNTLY | 0.5229925 |
| VIFASSLI | ASSLSGNTLY | 0.7527753 |
| VKGGDGGAAAG | ASSLSGNTLY | 0.8083192 |
| VTKTFTNI | ASSLSGNTLY | 0.6347903 |
| VVAAAILL | ASSLSGNTLY | 0.8633813 |
| YAALDVEL | ASSLSGNTLY | 0.4800044 |
| LASVALLL | ASSLSGNTLY | 0.8239881 |
| LDAVHRV | ASSLSGNTLY | 0.8325943 |
| QGLSSSIF | ASSLSGNTLY | 0.7137841 |
| RGTFANIRL | ASSLSGNTLY | 0.6281369 |
| RQLWWGHRI | ASSLSGNTLY | 0.8096638 |
| YGPPGTGKTLL | ASSLSGNTLY | 0.7414787 |
| AALVGAPANVA | ASSLSGNTLY | 0.72606695 |
| AATFRRTE | ASSLSGNTLY | 0.8349423 |
| AGALGGGI | ASSLSGNTLY | 0.58078027 |
| AGGAGGAGGIGAG | ASSLSGNTLY | 0.6946945 |
| AGSGALGL | ASSLSGNTLY | 0.67226124 |
| AQIADQI | ASSLSGNTLY | 0.7001153 |
| ASAVAVTYL | ASSLSGNTLY | 0.5274226 |
| ASTVLGGTGG | ASSLSGNTLY | 0.3612073 |
| AVGGEPAK | ASSLSGNTLY | 0.67856514 |
| CEQGTFGGTC | ASSLSGNTLY | 0.5010048 |
| DEGKAPAG | ASSLSGNTLY | 0.71760094 |
| DTVVFPAL | ASSLSGNTLY | 0.6955277 |
| FEGITMAG | ASSLSGNTLY | 0.5230336 |
| FPLVVQD | ASSLSGNTLY | 0.6927837 |
| GAFTTPQ | ASSLSGNTLY | 0.7099538 |
| GFGGFGTAGGVGGA | ASSLSGNTLY | 0.47647735 |
| GFLAQFL | ASSLSGNTLY | 0.62209165 |

| | | |
|---|---|---|
| GGAGGAGGNAGVA | ASSLSGNTLY | 0.5251278 |
| GGVGTAGAGGAGGAGG | ASSLSGNTLY | 0.6507681 |
| GNGGGGDAGNAGSGGN | ASSLSGNTLY | 0.3704332 |
| GTGGAGGAGNDAGS | ASSLSGNTLY | 0.67071104 |
| GTTNTGWFNS | ASSLSGNTLY | 0.6090718 |
| IFMGPLFTL | ASSLSGNTLY | 0.2820998 |
| ILAHAIEAI | ASSLSGNTLY | 0.68518436 |
| IRSAIREAL | ASSLSGNTLY | 0.5648875 |
| ITFAGGG | ASSLSGNTLY | 0.6592301 |
| ITGMSGHQEV | ASSLSGNTLY | 0.7085291 |
| IVAGLACTL | ASSLSGNTLY | 0.6264258 |
| IVGGGLAGA | ASSLSGNTLY | 0.5057326 |
| IVTGVLNI | ASSLSGNTLY | 0.6775063 |
| LAQQGCI | ASSLSGNTLY | 0.71000916 |
| LCLHVEN | ASSLSGNTLY | 0.42802477 |
| LDAISDGS | ASSLSGNTLY | 0.6874062 |
| LGGFAAML | ASSLSGNTLY | 0.7144607 |
| LGGKNPMI | ASSLSGNTLY | 0.6050751 |
| LGPKGVAAIL | ASSLSGNTLY | 0.71870816 |
| LMVGVSI | ASSLSGNTLY | 0.7208673 |
| LVGAAFVL | ASSLSGNTLY | 0.6205445 |
| LVLAVAPL | ASSLSGNTLY | 0.70221055 |
| MAAIAVIDAA | ASSLSGNTLY | 0.7088301 |
| NNGINAELQ | ASSLSGNTLY | 0.6111008 |
| RGEPLVLVHG | ASSLSGNTLY | 0.6390137 |
| RGGAGGAAGLI | ASSLSGNTLY | 0.6068804 |
| RSYRMHPAV | ASSLSGNTLY | 0.68146527 |
| SAAHLLAGQLA | ASSLSGNTLY | 0.45057052 |
| SAAVGSY | ASSLSGNTLY | 0.6498102 |
| SPHAVLHRI | ASSLSGNTLY | 0.703917 |
| SQIAKVL | ASSLSGNTLY | 0.605338 |
| TALGHLVP | ASSLSGNTLY | 0.4663653 |
| TAPTTSGAA | ASSLSGNTLY | 0.4781295 |
| TDGDDGNGGSPGT | ASSLSGNTLY | 0.63159347 |
| TDGLGEN | ASSLSGNTLY | 0.6198733 |
| TGGPPLPGQA | ASSLSGNTLY | 0.36010876 |
| TIPAITI | ASSLSGNTLY | 0.60326815 |
| TIPGFTI | ASSLSGNTLY | 0.68597215 |
| TSIASGALA | ASSLSGNTLY | 0.699144 |
| TVTLPGAAI | ASSLSGNTLY | 0.45577568 |
| VASGLSAALP | ASSLSGNTLY | 0.71365595 |
| VGAGVIGI | ASSLSGNTLY | 0.71071106 |
| VGPKRFGV | ASSLSGNTLY | 0.7144991 |
| VNVAFPD | ASSLSGNTLY | 0.6426864 |
| VSMTNSGVS | ASSLSGNTLY | 0.5792977 |
| VTDVLGDLSV | ASSLSGNTLY | 0.70355487 |
| VVVVGAGFAGL | ASSLSGNTLY | 0.6014123 |
| VYTVGTVV | ASSLSGNTLY | 0.61024535 |
| WTTFSGP | ASSLSGNTLY | 0.69829535 |
| YDELVEAF | ASSLSGNTLY | 0.53658116 |
| YGTGGAGGNGG | ASSLSGNTLY | 0.4925834 |

| | | |
|---|---|---|
| YTNVMAV | ASSLSGNTLY | 0.6737378 |
| FLEQVHQL | ASSLSGNTLY | 0.71180403 |
| GLSLLAVAV | ASSLSGNTLY | 0.50977814 |
| LEQVHQL | ASSLSGNTLY | 0.40676558 |
| LNAAGAI | ASSLSGNTLY | 0.6306515 |
| PAVHASL | ASSLSGNTLY | 0.6408031 |
| TELEDDA | ASSLSGNTLY | 0.7113694 |
| TPPGPPN | ASSLSGNTLY | 0.67034596 |
| TPPPPPPP | ASSLSGNTLY | 0.64938694 |
| TQSPALI | ASARDNNQAPL | 0.7056292 |
| EFAAAFPQ | ASARDNNQAPL | 0.6710964 |
| LLSGNLDVL | ASARDNNQAPL | 0.65111923 |
| GPANLGNYNI | ASARDNNQAPL | 0.7116233 |
| TLGPTLP | ASARDNNQAPL | 0.5195528 |
| VYAYSSCCK | ASARDNNQAPL | 0.7082827 |
| APAPAPAPNI | ASARDNNQAPL | 0.67753506 |
| PGGPVPIP | ASARDNNQAPL | 0.71020347 |
| FLTVALEP | ASARDNNQAPL | 0.71487355 |
| SQYFGANL | ASARDNNQAPL | 0.65333176 |
| GGVKPAA | ASARDNNQAPL | 0.6711383 |
| GGTGGAGGAGGW | ASARDNNQAPL | 0.39036012 |
| GVGGVAGLGGAGG | ASARDNNQAPL | 0.61825 |
| VLLVLATTI | ASARDNNQAPL | 0.6968522 |
| IGIFAAML | ASARDNNQAPL | 0.6580955 |
| AVGDLID | ASARDNNQAPL | 0.70993847 |
| NISPLPL | ASARDNNQAPL | 0.70613164 |
| VTAVFVAI | ASARDNNQAPL | 0.57278425 |
| SGDDNDAAL | ASARDNNQAPL | 0.7203827 |
| AVAALAHS | ASARDNNQAPL | 0.7113205 |
| GGVTVMATP | ASARDNNQAPL | 0.42599455 |
| AGLAANVRD | ASARDNNQAPL | 0.6423324 |
| TNRPDLIDPALLRPG | ASARDNNQAPL | 0.7121402 |
| LANLGVTAL | ASARDNNQAPL | 0.7024097 |
| TGERIEL | ASARDNNQAPL | 0.5828427 |
| NGEELAA | ASARDNNQAPL | 0.68562937 |
| VLSLVWG | ASARDNNQAPL | 0.7183119 |
| IGYALWL | ASARDNNQAPL | 0.6312601 |
| LSLLLAPL | ASARDNNQAPL | 0.7119762 |
| TLDEALK | ASARDNNQAPL | 0.34912178 |
| TDGAGTAEGG | ASARDNNQAPL | 0.66557693 |
| NAPGGAGGNGG | ASARDNNQAPL | 0.5731368 |
| VSVFMGMGGNF | ASARDNNQAPL | 0.32500494 |
| LASYLAAL | ASARDNNQAPL | 0.6671284 |
| LVGGTPAL | ASARDNNQAPL | 0.6598404 |
| VRNADVL | ASARDNNQAPL | 0.71497226 |
| TCSSPFLVSAM | ASARDNNQAPL | 0.61735845 |
| VHITDAE | ASARDNNQAPL | 0.70300376 |
| EYAAPSF | ASARDNNQAPL | 0.6661711 |
| VVDAFHGGL | ASARDNNQAPL | 0.6917047 |
| ASVVDPQ | ASARDNNQAPL | 0.57435703 |
| SYNEFGGLS | ASARDNNQAPL | 0.63310415 |

| | | |
|---|---|---|
| DPDGSVY | ASARDNNQAPL | 0.72028506 |
| MVSTIHATE | ASARDNNQAPL | 0.70682406 |
| LAAGVADPV | ASARDNNQAPL | 0.34277645 |
| TKEQIAA | ASARDNNQAPL | 0.68168485 |
| SLASPAAL | ASARDNNQAPL | 0.5823047 |
| IGLVFYGV | ASARDNNQAPL | 0.36958146 |
| FLLDASNL | ASARDNNQAPL | 0.37572464 |
| YQVGNFP | ASARDNNQAPL | 0.4883746 |
| ESLLSQST | ASARDNNQAPL | 0.6024685 |
| LFTVGFAAVG | ASARDNNQAPL | 0.6695424 |
| ATPPTPT | ASARDNNQAPL | 0.68110657 |
| TDLATLG | ASARDNNQAPL | 0.5598278 |
| SAGGTGR | ASARDNNQAPL | 0.6856112 |
| GFSNSGS | ASARDNNQAPL | 0.63040227 |
| GGNGGSGGTSV | ASARDNNQAPL | 0.25459158 |
| GTSVATGGAGN | ASARDNNQAPL | 0.5704566 |
| NGGNGGSGGTS | ASARDNNQAPL | 0.49788085 |
| AVVKFPH | ASARDNNQAPL | 0.7188049 |
| TPPPPPP | ASARDNNQAPL | 0.5622827 |
| AAPAAAAAPA | ASARDNNQAPL | 0.5455644 |
| IALNEDL | ASARDNNQAPL | 0.61541903 |
| IVNVSSM | ASARDNNQAPL | 0.6860832 |
| TGIPPAPR | ASARDNNQAPL | 0.64906895 |
| FDLGGGTF | ASARDNNQAPL | 0.5376174 |
| FDLGGGTFDVS | ASARDNNQAPL | 0.3812684 |
| NEPTAAA | ASARDNNQAPL | 0.6653919 |
| NEPTAAALA | ASARDNNQAPL | 0.572278 |
| GSIINISSI | ASARDNNQAPL | 0.5933993 |
| VAPGNAGTA | ASARDNNQAPL | 0.5725342 |
| TGGGASLE | ASARDNNQAPL | 0.6023549 |
| VAVYNHYKRI | ASARDNNQAPL | 0.4020748 |
| TDPAPATT | ASARDNNQAPL | 0.7149471 |
| TNRPDILDPALLRPG | ASARDNNQAPL | 0.64294255 |
| LTTTTTS | ASARDNNQAPL | 0.57137847 |
| PTGSGKTL | ASARDNNQAPL | 0.7081003 |
| TGSGKTL | ASARDNNQAPL | 0.684242 |
| TIGHVDHGK | ASARDNNQAPL | 0.59863555 |
| TIGHVDHGKTT | ASARDNNQAPL | 0.5556809 |
| GHVDHGKTT | ASARDNNQAPL | 0.47654295 |
| AQFEHTLL | ASARDNNQAPL | 0.517505 |
| IVNQDEL | ASARDNNQAPL | 0.5239849 |
| TNGSQFF | ASARDNNQAPL | 0.71395344 |
| TGKGIVE | ASARDNNQAPL | 0.6829332 |
| LISNASDA | ASARDNNQAPL | 0.5718392 |
| AELRTLGELTKIAKAHGAQ | ASARDNNQAPL | 0.69593644 |
| APNAVAAAEPSW | ASARDNNQAPL | 0.70760894 |
| APQVAGAATAVL | ASARDNNQAPL | 0.71407276 |
| ARRDPAAAAAILPTDA | ASARDNNQAPL | 0.59949076 |
| DAAARDVLAEAGLAEYF | ASARDNNQAPL | 0.6534582 |
| DDQIVLSAKANLAKVEQ | ASARDNNQAPL | 0.68257797 |
| DELLPDTGEDSDEDGH | ASARDNNQAPL | 0.60121465 |

| | | |
|---|---|---|
| ELPEAPANRATVPQD | ASARDNNQAPL | 0.6569612 |
| FGQSKIERVTGIG | ASARDNNQAPL | 0.47476536 |
| GAGGTGSTAGGAGGAGGAGGLY | ASARDNNQAPL | 0.70510674 |
| GHGGTGAAGGNGGTGGAGGSGI | ASARDNNQAPL | 0.7266841 |
| GNGGNGGAGGEGGPGGAGGAGA | ASARDNNQAPL | 0.621428 |
| GSTAGNGGNGGNATNGTIAGQP | ASARDNNQAPL | 0.44318962 |
| GTGGDGGNGGNGGTGG | ASARDNNQAPL | 0.5652306 |
| GVRRPAGIPADAPV | ASARDNNQAPL | 0.49781817 |
| GYRQVTDAHLLTLARR | ASARDNNQAPL | 0.53523535 |
| IAKMGQDGHDRG | ASARDNNQAPL | 0.6698093 |
| KAEPSIEIQYT | ASARDNNQAPL | 0.45308244 |
| KPVRIRMAADEDTVAVT | ASARDNNQAPL | 0.61995316 |
| MPPRIAGMRLLVIKPEP | ASARDNNQAPL | 0.70973945 |
| MKFYAVALFRSRGVPPTR | ASARDNNQAPL | 0.63912344 |
| NAGGFGILGAVAHSP | ASARDNNQAPL | 0.70301414 |
| NEGREITLAMAEM | ASARDNNQAPL | 0.6980281 |
| NELVEWSHPLGHRWRWE | ASARDNNQAPL | 0.68171453 |
| NLGAGGTGGNGGNPG | ASARDNNQAPL | 0.71568453 |
| NTEWPKWETVMAADPVD | ASARDNNQAPL | 0.71794826 |
| PASAAPAETQP | ASARDNNQAPL | 0.6514397 |
| QQSSDGKNGVRAAEITGE | ASARDNNQAPL | 0.6521209 |
| RAPDAVTTSWWKE | ASARDNNQAPL | 0.6931503 |
| SGFGNVGAHTSGWFNQ | ASARDNNQAPL | 0.6219895 |
| SGVAGAVGQAASVGGLKV | ASARDNNQAPL | 0.6075524 |
| SNGGAGTGSAGGNGGT | ASARDNNQAPL | 0.62078935 |
| TGGAGGEGGAAG | ASARDNNQAPL | 0.6866693 |
| TGGDGGAAGAGGAPGHG | ASARDNNQAPL | 0.62208915 |
| TMPPLLAAIANAGRHGV | ASARDNNQAPL | 0.72061515 |
| TSSTAGNSGGA | ASARDNNQAPL | 0.4020458 |
| VPATVPVGILC | ASARDNNQAPL | 0.42113888 |
| VQRPDSGGVVLTGRIS | ASARDNNQAPL | 0.70432633 |
| WTPNPVPATKGVPAGLVQAT | ASARDNNQAPL | 0.284424 |
| YALKRALAKFSDRFDVVII | ASARDNNQAPL | 0.45731348 |
| YLIGSVIV | ASARDNNQAPL | 0.6757415 |
| YGQLVAVID | ASARDNNQAPL | 0.7017303 |
| VTVGGMTLPA | ASARDNNQAPL | 0.28851357 |
| VLALPAV | ASARDNNQAPL | 0.5551473 |
| VIPPTLNYET | ASARDNNQAPL | 0.49820152 |
| VGMGGMHPSSG | ASARDNNQAPL | 0.7089548 |
| VGLAGGAATA | ASARDNNQAPL | 0.61494416 |
| VGGAGGAGGSGP | ASARDNNQAPL | 0.3285388 |
| VFAAFMV | ASARDNNQAPL | 0.7125927 |
| TLYDKLR | ASARDNNQAPL | 0.46415734 |
| TLAVFTVL | ASARDNNQAPL | 0.6330249 |
| TGSVQPARV | ASARDNNQAPL | 0.7041597 |
| TGGGAGSGGT | ASARDNNQAPL | 0.7234509 |
| SHLETSL | ASARDNNQAPL | 0.56181645 |
| SFTTGMT | ASARDNNQAPL | 0.6983597 |
| QVEVAKRMGVRQPTVS | ASARDNNQAPL | 0.6124572 |
| QGSGGPALAM | ASARDNNQAPL | 0.718925 |
| MLAPSNP | ASARDNNQAPL | 0.57168245 |

| | | |
|---|---|---|
| LVNNAAI | ASARDNNQAPL | 0.40954906 |
| LTGEINTLA | ASARDNNQAPL | 0.708969 |
| LSIFGPV | ASARDNNQAPL | 0.709643 |
| LLALLANL | ASARDNNQAPL | 0.68097353 |
| LISNASDALDKLR | ASARDNNQAPL | 0.70580304 |
| LINEAVP | ASARDNNQAPL | 0.65106416 |
| LGYAFSGL | ASARDNNQAPL | 0.6167997 |
| LDFVAVAL | ASARDNNQAPL | 0.6953728 |
| LAQPGAE | ASARDNNQAPL | 0.70455056 |
| ISNLGAVTGF | ASARDNNQAPL | 0.4756211 |
| HQEDPGRI | ASARDNNQAPL | 0.7066765 |
| GTPGEPTP | ASARDNNQAPL | 0.61467326 |
| GFFNSGS | ASARDNNQAPL | 0.646399 |
| GAGGGGLTAGAN | ASARDNNQAPL | 0.70800877 |
| FVDALVNM | ASARDNNQAPL | 0.6903007 |
| FLTVAPEL | ASARDNNQAPL | 0.689282 |
| FETLTAL | ASARDNNQAPL | 0.6607877 |
| DVINPIEL | ASARDNNQAPL | 0.7267562 |
| APVSGAPV | ASARDNNQAPL | 0.37927842 |
| ALVGGLAW | ASARDNNQAPL | 0.70090437 |
| ALAAAAATAI | ASARDNNQAPL | 0.66174304 |
| AFGLGLLL | ASARDNNQAPL | 0.7139231 |
| AFFAAASL | ASARDNNQAPL | 0.692873 |
| AAAALPAFSPP | ASARDNNQAPL | 0.61833537 |
| VVLAPAL | ASARDNNQAPL | 0.6921588 |
| TIGHVDHGKTTL | ASARDNNQAPL | 0.7021489 |
| LISNASDALDK | ASARDNNQAPL | 0.66302145 |
| LISNASDAL | ASARDNNQAPL | 0.70367557 |
| ISNASDALDK | ASARDNNQAPL | 0.72111875 |
| NQAETLVYQTE | ASARDNNQAPL | 0.6135757 |
| TGGTGGTAGAG | ASARDNNQAPL | 0.6424628 |
| TGSTAGGAGGA | ASARDNNQAPL | 0.7018001 |
| DPLAVPVAAGPV | ASARDNNQAPL | 0.71948457 |
| GGTGGTPGNGGN | ASARDNNQAPL | 0.67452765 |
| PAGLAGVDPATL | ASARDNNQAPL | 0.71360993 |
| GNNLVVEAPHAGASV | ASARDNNQAPL | 0.57253677 |
| KVTRIMSASTLEQPA | ASARDNNQAPL | 0.6821474 |
| LGFALPIALPRRTVV | ASARDNNQAPL | 0.6447811 |
| TRLGNLTARKLVVLV | ASARDNNQAPL | 0.65786004 |
| WSQQRRLIYFPSAGP | ASARDNNQAPL | 0.41074437 |
| GPNGNPDPMAAAVDIR | ASARDNNQAPL | 0.6646559 |
| TQRCVSVADLRAAASR | ASARDNNQAPL | 0.68812585 |
| QFVLDVAEHPVEVLTETV | ASARDNNQAPL | 0.6831103 |
| GGAGGTGGAGGNGGTQAGDG | ASARDNNQAPL | 0.43556428 |
| PLLATLRAIAVDVDAKPAQV | ASARDNNQAPL | 0.6586464 |
| APKFRTGGYMIDFWGVGYQVAK | ASARDNNQAPL | 0.6895611 |
| GAGGAGGAGGAAGGPGGTGGTG | ASARDNNQAPL | 0.72367907 |
| IGALGLLATDVAPAGSVAFGVA | ASARDNNQAPL | 0.6298352 |
| AGGAGGIGGNGGAGGAGGVLMG | ASARDNNQAPL | 0.7117915 |
| ADELIGLL | ASARDNNQAPL | 0.5835861 |
| ADRALKA | ASARDNNQAPL | 0.6534407 |

| | | |
|---|---|---|
| AGAVGQVGGAGGA | ASARDNNQAPL | 0.46264273 |
| AGGGGGLGGAP | ASARDNNQAPL | 0.6669003 |
| AGGLLYGNGGAG | ASARDNNQAPL | 0.5956415 |
| AGLESIGL | ASARDNNQAPL | 0.52151227 |
| AGLKTGII | ASARDNNQAPL | 0.6953769 |
| ASLAALLL | ASARDNNQAPL | 0.71674544 |
| ASVINISSI | ASARDNNQAPL | 0.6866503 |
| ATPPGPA | ASARDNNQAPL | 0.70328844 |
| ATVASALV | ASARDNNQAPL | 0.7055546 |
| ATVASLIL | ASARDNNQAPL | 0.7197413 |
| AVNGPVLIRH | ASARDNNQAPL | 0.5320848 |
| DGAAGAVGI | ASARDNNQAPL | 0.6328226 |
| DPLLHGI | ASARDNNQAPL | 0.39724502 |
| ELGAALGAL | ASARDNNQAPL | 0.59615624 |
| ELGRTQAV | ASARDNNQAPL | 0.55442286 |
| FVAGGYLS | ASARDNNQAPL | 0.6292031 |
| GASGGAGGAGGW | ASARDNNQAPL | 0.27021748 |
| GGAGAGGDGGAGGI | ASARDNNQAPL | 0.68296194 |
| GTGHNGSQPGMGGNG | ASARDNNQAPL | 0.7147676 |
| GYAAASASAS | ASARDNNQAPL | 0.6083727 |
| HQVDVAAVQQG | ASARDNNQAPL | 0.6924385 |
| IGAAIAHT | ASARDNNQAPL | 0.6721666 |
| IGMIEVAQL | ASARDNNQAPL | 0.5564551 |
| IRLAPEHTSAMD | ASARDNNQAPL | 0.6840692 |
| IYNVSVL | ASARDNNQAPL | 0.54316336 |
| KQGFLGM | ASARDNNQAPL | 0.714352 |
| LESVITQL | ASARDNNQAPL | 0.70455885 |
| LGLGALVVV | ASARDNNQAPL | 0.677078 |
| LIGNGGNGGSGGI | ASARDNNQAPL | 0.6861329 |
| LNAINAPTL | ASARDNNQAPL | 0.6900301 |
| LQLLNTGV | ASARDNNQAPL | 0.34393072 |
| LQTSIEF | ASARDNNQAPL | 0.6931063 |
| MSFAVSEL | ASARDNNQAPL | 0.58895123 |
| NTGALGL | ASARDNNQAPL | 0.6555153 |
| PPGAGGAGGAGTT | ASARDNNQAPL | 0.49647665 |
| QLVGLGI | ASARDNNQAPL | 0.68487626 |
| RFLEQVHQL | ASARDNNQAPL | 0.6157391 |
| RGVGYVLR | ASARDNNQAPL | 0.6889596 |
| SATGVFAGVI | ASARDNNQAPL | 0.5402155 |
| SLPAITLP | ASARDNNQAPL | 0.6668606 |
| STALAFTV | ASARDNNQAPL | 0.65479815 |
| SVLTVGLGK | ASARDNNQAPL | 0.43938404 |
| TALAGAGSGI | ASARDNNQAPL | 0.7074913 |
| TGLPFTGI | ASARDNNQAPL | 0.65726614 |
| VAEHVTV | ASARDNNQAPL | 0.59478056 |
| VDIVLNSL | ASARDNNQAPL | 0.69320965 |
| VLTGGGAVL | ASARDNNQAPL | 0.6870061 |
| VNVGLAYGI | ASARDNNQAPL | 0.35755467 |
| VSSAAVGAV | ASARDNNQAPL | 0.6332969 |
| VTSVYEAGV | ASARDNNQAPL | 0.6648004 |
| WVALDRGA | ASARDNNQAPL | 0.5836812 |

| | | |
|---|---|---|
| YLVGSVVI | ASARDNNQAPL | 0.3365927 |
| GPPGTGKTLL | ASARDNNQAPL | 0.68988633 |
| GVGMTKF | ASARDNNQAPL | 0.6401545 |
| IGIGGSDL | ASARDNNQAPL | 0.5103916 |
| IGTLLAEL | ASARDNNQAPL | 0.6026061 |
| IYTSGTT | ASARDNNQAPL | 0.71484387 |
| LSFPAGV | ASARDNNQAPL | 0.6638142 |
| LYTSGTT | ASARDNNQAPL | 0.69101846 |
| VGAFPFGF | ASARDNNQAPL | 0.7055248 |
| VGAGTVEF | ASARDNNQAPL | 0.6279596 |
| VGKSTLLL | ASARDNNQAPL | 0.7002903 |
| AALAHAGGGIV | ASARDNNQAPL | 0.6493229 |
| AALVAVSLL | ASARDNNQAPL | 0.64914453 |
| AAVSRAVAAL | ASARDNNQAPL | 0.700029 |
| AGGSSGIGGF | ASARDNNQAPL | 0.6392653 |
| AGLTVANANG | ASARDNNQAPL | 0.522336 |
| AGLVGAVL | ASARDNNQAPL | 0.4585616 |
| ALSQAFTV | ASARDNNQAPL | 0.6223811 |
| ALTFSVSL | ASARDNNQAPL | 0.50969946 |
| ALTGAALVAL | ASARDNNQAPL | 0.65763146 |
| ASFSIGAGVL | ASARDNNQAPL | 0.7213173 |
| DGSVYKI | ASARDNNQAPL | 0.6648573 |
| EQLAEIV | ASARDNNQAPL | 0.4966542 |
| FAVSELTVM | ASARDNNQAPL | 0.60952187 |
| GAAPLAAGL | ASARDNNQAPL | 0.47819752 |
| GADGGDGGSGGAGG | ASARDNNQAPL | 0.70537734 |
| GFGLDTGGAGG | ASARDNNQAPL | 0.69560087 |
| GLASGGLL | ASARDNNQAPL | 0.72135234 |
| GRIAGAAVLL | ASARDNNQAPL | 0.6812134 |
| GSAGSLNTSF | ASARDNNQAPL | 0.55837286 |
| GTGQIGI | ASARDNNQAPL | 0.7095523 |
| GTLIALIL | ASARDNNQAPL | 0.60833275 |
| IAATMGFVVG | ASARDNNQAPL | 0.6901779 |
| IFAQALL | ASARDNNQAPL | 0.67043895 |
| IGAAVALGF | ASARDNNQAPL | 0.6763314 |
| IGLTPLAL | ASARDNNQAPL | 0.6738962 |
| ILGSNTSTL | ASARDNNQAPL | 0.6885176 |
| IVFSLTVL | ASARDNNQAPL | 0.40133983 |
| LASKATGF | ASARDNNQAPL | 0.7045102 |
| LASVALIL | ASARDNNQAPL | 0.54198885 |
| LFAVGLTL | ASARDNNQAPL | 0.49223495 |
| LFNGGIGTYI | ASARDNNQAPL | 0.6013192 |
| LTLGAGII | ASARDNNQAPL | 0.7141193 |
| LVAAALGDI | ASARDNNQAPL | 0.71052265 |
| NTAFVQLGI | ASARDNNQAPL | 0.57679665 |
| QLFSGAEL | ASARDNNQAPL | 0.50425583 |
| SAATFITL | ASARDNNQAPL | 0.6851057 |
| SNYGLANLGV | ASARDNNQAPL | 0.48985738 |
| SYGGAYAVM | ASARDNNQAPL | 0.70259786 |
| TTGVGGLTLG | ASARDNNQAPL | 0.6217431 |
| TTQTIYI | ASARDNNQAPL | 0.62195575 |

| | | |
|---|---|---|
| VAAGAGLGAVI | ASARDNNQAPL | 0.64239097 |
| VAAVVTLL | ASARDNNQAPL | 0.68157923 |
| VGFLGGLI | ASARDNNQAPL | 0.59663165 |
| VGVIPILL | ASARDNNQAPL | 0.7126004 |
| VIFASSLI | ASARDNNQAPL | 0.34846318 |
| VKGGDGGAAAG | ASARDNNQAPL | 0.68695235 |
| VTKTFTNI | ASARDNNQAPL | 0.4612087 |
| VVAAAILL | ASARDNNQAPL | 0.4928814 |
| YAALDVEL | ASARDNNQAPL | 0.3625424 |
| LASVALLL | ASARDNNQAPL | 0.68972504 |
| LDAVHRV | ASARDNNQAPL | 0.6923505 |
| QGLSSSIF | ASARDNNQAPL | 0.7024976 |
| RGTFANIRL | ASARDNNQAPL | 0.65792835 |
| RQLWWGHRI | ASARDNNQAPL | 0.70223606 |
| YGPPGTGKTLL | ASARDNNQAPL | 0.71084785 |
| AALVGAPANVA | ASARDNNQAPL | 0.536882 |
| AATFRRTE | ASARDNNQAPL | 0.31984568 |
| AGALGGGI | ASARDNNQAPL | 0.68348444 |
| AGGAGGAGGIGAG | ASARDNNQAPL | 0.6914342 |
| AGSGALGL | ASARDNNQAPL | 0.7083356 |
| AQIADQI | ASARDNNQAPL | 0.69757026 |
| ASAVAVTYL | ASARDNNQAPL | 0.43925864 |
| ASTVLGGTGG | ASARDNNQAPL | 0.70140344 |
| AVGGEPAK | ASARDNNQAPL | 0.6929415 |
| CEQGTFGGTC | ASARDNNQAPL | 0.4686878 |
| DEGKAPAG | ASARDNNQAPL | 0.615798 |
| DTVVFPAL | ASARDNNQAPL | 0.6349119 |
| FEGITMAG | ASARDNNQAPL | 0.51952183 |
| FPLVVQD | ASARDNNQAPL | 0.652099 |
| GAFTTPQ | ASARDNNQAPL | 0.5631354 |
| GFGGFGTAGGVGGA | ASARDNNQAPL | 0.576696 |
| GFLAQFL | ASARDNNQAPL | 0.61568815 |
| GGAGGAGGNAGVA | ASARDNNQAPL | 0.51499605 |
| GGVGTAGAGGAGGAGG | ASARDNNQAPL | 0.5713882 |
| GNGGGGDAGNAGSGGN | ASARDNNQAPL | 0.5647131 |
| GTGGAGGAGNDAGS | ASARDNNQAPL | 0.6429456 |
| GTTNTGWFNS | ASARDNNQAPL | 0.67978084 |
| IFMGPLFTL | ASARDNNQAPL | 0.63678694 |
| ILAHAIEAI | ASARDNNQAPL | 0.70263404 |
| IRSAIREAL | ASARDNNQAPL | 0.6980407 |
| ITFAGGG | ASARDNNQAPL | 0.4122542 |
| ITGMSGHQEV | ASARDNNQAPL | 0.52994883 |
| IVAGLACTL | ASARDNNQAPL | 0.703605 |
| IVGGGLAGA | ASARDNNQAPL | 0.51600456 |
| IVTGVLNI | ASARDNNQAPL | 0.70923865 |
| LAQQGCI | ASARDNNQAPL | 0.5964652 |
| LCLHVEN | ASARDNNQAPL | 0.6350999 |
| LDAISDGS | ASARDNNQAPL | 0.6790122 |
| LGGFAAML | ASARDNNQAPL | 0.71931726 |
| LGGKNPMI | ASARDNNQAPL | 0.7039249 |
| LGPKGVAAIL | ASARDNNQAPL | 0.65693253 |

| | | |
|---|---|---|
| LMVGVSI | ASARDNNQAPL | 0.7056601 |
| LVGAAFVL | ASARDNNQAPL | 0.55871105 |
| LVLAVAPL | ASARDNNQAPL | 0.7042774 |
| MAAIAVIDAA | ASARDNNQAPL | 0.70873964 |
| NNGINAELQ | ASARDNNQAPL | 0.44521612 |
| RGEPLVLVHG | ASARDNNQAPL | 0.61719 |
| RGGAGGAAGLI | ASARDNNQAPL | 0.7124666 |
| RSYRMHPAV | ASARDNNQAPL | 0.715753 |
| SAAHLLAGQLA | ASARDNNQAPL | 0.50633764 |
| SAAVGSY | ASARDNNQAPL | 0.60459524 |
| SPHAVLHRI | ASARDNNQAPL | 0.48977298 |
| SQIAKVL | ASARDNNQAPL | 0.590914 |
| TALGHLVP | ASARDNNQAPL | 0.46919125 |
| TAPTTSGAA | ASARDNNQAPL | 0.68523365 |
| TDGDDGNGGSPGT | ASARDNNQAPL | 0.59851587 |
| TDGLGEN | ASARDNNQAPL | 0.7175896 |
| TGGPPLPGQA | ASARDNNQAPL | 0.650408 |
| TIPAITI | ASARDNNQAPL | 0.6812235 |
| TIPGFTI | ASARDNNQAPL | 0.60403275 |
| TSIASGALA | ASARDNNQAPL | 0.5944007 |
| TVTLPGAAI | ASARDNNQAPL | 0.5273256 |
| VASGLSAALP | ASARDNNQAPL | 0.49734318 |
| VGAGVIGI | ASARDNNQAPL | 0.56477416 |
| VGPKRFGV | ASARDNNQAPL | 0.5704906 |
| VNVAFPD | ASARDNNQAPL | 0.32932916 |
| VSMTNSGVS | ASARDNNQAPL | 0.70783055 |
| VTDVLGDLSV | ASARDNNQAPL | 0.67947286 |
| VVVVGAGFAGL | ASARDNNQAPL | 0.6545346 |
| VYTVGTVV | ASARDNNQAPL | 0.51147145 |
| WTTFSGP | ASARDNNQAPL | 0.64289963 |
| YDELVEAF | ASARDNNQAPL | 0.5585271 |
| YGTGGAGGNGG | ASARDNNQAPL | 0.7121147 |
| YTNVMAV | ASARDNNQAPL | 0.47647107 |
| FLEQVHQL | ASARDNNQAPL | 0.37165043 |
| GLSLLAVAV | ASARDNNQAPL | 0.6224835 |
| LEQVHQL | ASARDNNQAPL | 0.4767039 |
| LNAAGAI | ASARDNNQAPL | 0.71490395 |
| PAVHASL | ASARDNNQAPL | 0.6970142 |
| TELEDDA | ASARDNNQAPL | 0.66927433 |
| TPPGPPN | ASARDNNQAPL | 0.7218926 |
| TPPPPPPP | ASARDNNQAPL | 0.28948742 |
| TQSPALI | ASSSRWGVEQY | 0.53546745 |
| EFAAAFPQ | ASSSRWGVEQY | 0.71274436 |
| LLSGNLDVL | ASSSRWGVEQY | 0.6698348 |
| GPANLGNYNI | ASSSRWGVEQY | 0.61470324 |
| TLGPTLP | ASSSRWGVEQY | 0.7055987 |
| VYAYSSCCK | ASSSRWGVEQY | 0.53181624 |
| APAPAPAPNI | ASSSRWGVEQY | 0.6087905 |
| PGGPVPIP | ASSSRWGVEQY | 0.68370074 |
| FLTVALEP | ASSSRWGVEQY | 0.70024 |
| SQYFGANL | ASSSRWGVEQY | 0.69003886 |

| | | |
|---|---|---|
| GGVKPAA | ASSSRWGVEQY | 0.50003797 |
| GGTGGAGGAGGW | ASSSRWGVEQY | 0.7050582 |
| GVGGVAGLGGAGG | ASSSRWGVEQY | 0.70983404 |
| VLLVLATTI | ASSSRWGVEQY | 0.6346774 |
| IGIFAAML | ASSSRWGVEQY | 0.68477887 |
| AVGDLID | ASSSRWGVEQY | 0.6803229 |
| NISPLPL | ASSSRWGVEQY | 0.6761154 |
| VTAVFVAI | ASSSRWGVEQY | 0.52279353 |
| SGDDNDAAL | ASSSRWGVEQY | 0.4333681 |
| AVAALAHS | ASSSRWGVEQY | 0.6668405 |
| GGVTVMATP | ASSSRWGVEQY | 0.48161256 |
| AGLAANVRD | ASSSRWGVEQY | 0.69251835 |
| TNRPDLIDPALLRPG | ASSSRWGVEQY | 0.617437 |
| LANLGVTAL | ASSSRWGVEQY | 0.7192769 |
| TGERIEL | ASSSRWGVEQY | 0.6864784 |
| NGEELAA | ASSSRWGVEQY | 0.67672586 |
| VLSLVWG | ASSSRWGVEQY | 0.47105289 |
| IGYALWL | ASSSRWGVEQY | 0.68113124 |
| LSLLLAPL | ASSSRWGVEQY | 0.66347605 |
| TLDEALK | ASSSRWGVEQY | 0.4022534 |
| TDGAGTAEGG | ASSSRWGVEQY | 0.69835496 |
| NAPGGAGGNGG | ASSSRWGVEQY | 0.434393 |
| VSVFMGMGGNF | ASSSRWGVEQY | 0.64483464 |
| LASYLAAL | ASSSRWGVEQY | 0.6816848 |
| LVGGTPAL | ASSSRWGVEQY | 0.72155523 |
| VRNADVL | ASSSRWGVEQY | 0.6087141 |
| TCSSPFLVSAM | ASSSRWGVEQY | 0.6504697 |
| VHITDAE | ASSSRWGVEQY | 0.46600422 |
| EYAAPSF | ASSSRWGVEQY | 0.6995379 |
| VVDAFHGGL | ASSSRWGVEQY | 0.7063063 |
| ASVVDPQ | ASSSRWGVEQY | 0.70063335 |
| SYNEFGGLS | ASSSRWGVEQY | 0.72190785 |
| DPDGSVY | ASSSRWGVEQY | 0.69351375 |
| MVSTIHATE | ASSSRWGVEQY | 0.43157393 |
| LAAGVADPV | ASSSRWGVEQY | 0.6908924 |
| TKEQIAA | ASSSRWGVEQY | 0.70771706 |
| SLASPAAL | ASSSRWGVEQY | 0.69921017 |
| IGLVFYGV | ASSSRWGVEQY | 0.6926559 |
| FLLDASNL | ASSSRWGVEQY | 0.71539813 |
| YQVGNFP | ASSSRWGVEQY | 0.61923 |
| ESLLSQST | ASSSRWGVEQY | 0.70350647 |
| LFTVGFAAVG | ASSSRWGVEQY | 0.72295666 |
| ATPPTPT | ASSSRWGVEQY | 0.70394397 |
| TDLATLG | ASSSRWGVEQY | 0.6341036 |
| SAGGTGR | ASSSRWGVEQY | 0.58061457 |
| GFSNSGS | ASSSRWGVEQY | 0.68900293 |
| GGNGGSGGTSV | ASSSRWGVEQY | 0.69363284 |
| GTSVATGGAGN | ASSSRWGVEQY | 0.6022489 |
| NGGNGGSGGTS | ASSSRWGVEQY | 0.6232425 |
| AVVKFPH | ASSSRWGVEQY | 0.6275334 |
| TPPPPPP | ASSSRWGVEQY | 0.4898459 |

| | | |
|---|---|---|
| AAPAAAAAPA | ASSSRWGVEQY | 0.6931449 |
| IALNEDL | ASSSRWGVEQY | 0.7078999 |
| IVNVSSM | ASSSRWGVEQY | 0.686444 |
| TGIPPAPR | ASSSRWGVEQY | 0.56995606 |
| FDLGGGTF | ASSSRWGVEQY | 0.7007649 |
| FDLGGGTFDVS | ASSSRWGVEQY | 0.70671356 |
| NEPTAAA | ASSSRWGVEQY | 0.7161569 |
| NEPTAAALA | ASSSRWGVEQY | 0.35084808 |
| GSIINISSI | ASSSRWGVEQY | 0.2948076 |
| VAPGNAGTA | ASSSRWGVEQY | 0.6660309 |
| TGGGASLE | ASSSRWGVEQY | 0.7031269 |
| VAVYNHYKRI | ASSSRWGVEQY | 0.57908106 |
| TDPAPATT | ASSSRWGVEQY | 0.61836886 |
| TNRPDILDPALLRPG | ASSSRWGVEQY | 0.6863091 |
| LTTTTTS | ASSSRWGVEQY | 0.6878574 |
| PTGSGKTL | ASSSRWGVEQY | 0.60206795 |
| TGSGKTL | ASSSRWGVEQY | 0.45686316 |
| TIGHVDHGK | ASSSRWGVEQY | 0.6973825 |
| TIGHVDHGKTT | ASSSRWGVEQY | 0.7149584 |
| GHVDHGKTT | ASSSRWGVEQY | 0.5388584 |
| AQFEHTLL | ASSSRWGVEQY | 0.6267851 |
| IVNQDEL | ASSSRWGVEQY | 0.6249982 |
| TNGSQFF | ASSSRWGVEQY | 0.63359416 |
| TGKGIVE | ASSSRWGVEQY | 0.6245588 |
| LISNASDA | ASSSRWGVEQY | 0.43316305 |
| AELRTLGELTKIAKAHGAQ | ASSSRWGVEQY | 0.6916846 |
| APNAVAAAEPSW | ASSSRWGVEQY | 0.40208223 |
| APQVAGAATAVL | ASSSRWGVEQY | 0.6960921 |
| ARRDPAAAAAILPTDA | ASSSRWGVEQY | 0.699799 |
| DAAARDVLAEAGLAEYF | ASSSRWGVEQY | 0.6869998 |
| DDQIVLSAKANLAKVEQ | ASSSRWGVEQY | 0.585088 |
| DELLPDTGEDSDEDGH | ASSSRWGVEQY | 0.5525186 |
| ELPEAPANRATVPQD | ASSSRWGVEQY | 0.44993937 |
| FGQSKIERVTGIG | ASSSRWGVEQY | 0.451532 |
| GAGGTGSTAGGAGGAGGAGGLY | ASSSRWGVEQY | 0.71630925 |
| GHGGTGAAGGNGGTGGAGGSGI | ASSSRWGVEQY | 0.63023186 |
| GNGGNGGAGGEGGPGGAGGAGA | ASSSRWGVEQY | 0.714633 |
| GSTAGNGGNGGNATNGTIAGQP | ASSSRWGVEQY | 0.6347101 |
| GTGGDGGNGGNGGTGG | ASSSRWGVEQY | 0.55223656 |
| GVRRPAGIPADAPV | ASSSRWGVEQY | 0.68850446 |
| GYRQVTDAHLLTLARR | ASSSRWGVEQY | 0.5182873 |
| IAKMGQDGHDRG | ASSSRWGVEQY | 0.6677557 |
| KAEPSIEIQYT | ASSSRWGVEQY | 0.6851553 |
| KPVRIRMAADEDTVAVT | ASSSRWGVEQY | 0.7128147 |
| MPPRIAGMRLLVIKPEP | ASSSRWGVEQY | 0.5548171 |
| MKFYAVALFRSRGVPPTR | ASSSRWGVEQY | 0.69961655 |
| NAGGFGILGAVAHSP | ASSSRWGVEQY | 0.6799716 |
| NEGREITLAMAEM | ASSSRWGVEQY | 0.5804862 |
| NELVEWSHPLGHRWRWE | ASSSRWGVEQY | 0.6859381 |
| NLGAGGTGGNGGNPG | ASSSRWGVEQY | 0.6978643 |
| NTEWPKWETVMAADPVD | ASSSRWGVEQY | 0.6928343 |

| | | |
|---|---|---|
| PASAAPAETQP | ASSSRWGVEQY | 0.48384285 |
| QQSSDGKNGVRAAEITGE | ASSSRWGVEQY | 0.6474306 |
| RAPDAVTTSWWKE | ASSSRWGVEQY | 0.5638671 |
| SGFGNVGAHTSGWFNQ | ASSSRWGVEQY | 0.66354287 |
| SGVAGAVGQAASVGGLKV | ASSSRWGVEQY | 0.6550313 |
| SNGGAGTGSAGGNGGT | ASSSRWGVEQY | 0.6488356 |
| TGGAGGEGGAAG | ASSSRWGVEQY | 0.70132315 |
| TGGDGGAAGAGGAPGHG | ASSSRWGVEQY | 0.5504353 |
| TMPPLLAAIANAGRHGV | ASSSRWGVEQY | 0.5664582 |
| TSSTAGNSGGA | ASSSRWGVEQY | 0.6354246 |
| VPATVPVGILC | ASSSRWGVEQY | 0.62829375 |
| VQRPDSGGVVLTGRIS | ASSSRWGVEQY | 0.6967163 |
| WTPNPVPATKGVPAGLVQAT | ASSSRWGVEQY | 0.45328408 |
| YALKRALAKFSDRFDVVII | ASSSRWGVEQY | 0.6742857 |
| YLIGSVIV | ASSSRWGVEQY | 0.62148094 |
| YGQLVAVID | ASSSRWGVEQY | 0.48344374 |
| VTVGGMTLPA | ASSSRWGVEQY | 0.6841285 |
| VLALPAV | ASSSRWGVEQY | 0.63260466 |
| VIPPTLNYET | ASSSRWGVEQY | 0.56967485 |
| VGMGGMHPSSG | ASSSRWGVEQY | 0.6974367 |
| VGLAGGAATA | ASSSRWGVEQY | 0.5844412 |
| VGGAGGAGGSGP | ASSSRWGVEQY | 0.6601763 |
| VFAAFMV | ASSSRWGVEQY | 0.6826087 |
| TLYDKLR | ASSSRWGVEQY | 0.6316149 |
| TLAVFTVL | ASSSRWGVEQY | 0.69406474 |
| TGSVQPARV | ASSSRWGVEQY | 0.66024494 |
| TGGGAGSGGT | ASSSRWGVEQY | 0.69952834 |
| SHLETSL | ASSSRWGVEQY | 0.5311787 |
| SFTTGMT | ASSSRWGVEQY | 0.54923296 |
| QVEVAKRMGVRQPTVS | ASSSRWGVEQY | 0.70062727 |
| QGSGGPALAM | ASSSRWGVEQY | 0.66990405 |
| MLAPSNP | ASSSRWGVEQY | 0.70387197 |
| LVNNAAI | ASSSRWGVEQY | 0.70238435 |
| LTGEINTLA | ASSSRWGVEQY | 0.71252275 |
| LSIFGPV | ASSSRWGVEQY | 0.7152244 |
| LLALLANL | ASSSRWGVEQY | 0.5542144 |
| LISNASDALDKLR | ASSSRWGVEQY | 0.48112732 |
| LINEAVP | ASSSRWGVEQY | 0.6019316 |
| LGYAFSGL | ASSSRWGVEQY | 0.56371206 |
| LDFVAVAL | ASSSRWGVEQY | 0.6389824 |
| LAQPGAE | ASSSRWGVEQY | 0.3951603 |
| ISNLGAVTGF | ASSSRWGVEQY | 0.6988117 |
| HQEDPGRI | ASSSRWGVEQY | 0.7198592 |
| GTPGEPTP | ASSSRWGVEQY | 0.526425 |
| GFFNSGS | ASSSRWGVEQY | 0.72054 |
| GAGGGGLTAGAN | ASSSRWGVEQY | 0.4644042 |
| FVDALVNM | ASSSRWGVEQY | 0.6741928 |
| FLTVAPEL | ASSSRWGVEQY | 0.7040022 |
| FETLTAL | ASSSRWGVEQY | 0.45530346 |
| DVINPIEL | ASSSRWGVEQY | 0.6624106 |
| APVSGAPV | ASSSRWGVEQY | 0.52307 |

| | | |
|---|---|---|
| ALVGGLAW | ASSSRWGVEQY | 0.5794339 |
| ALAAAAATAI | ASSSRWGVEQY | 0.7107265 |
| AFGLGLLL | ASSSRWGVEQY | 0.5404327 |
| AFFAAASL | ASSSRWGVEQY | 0.69226646 |
| AAAALPAFSPP | ASSSRWGVEQY | 0.675897 |
| VVLAPAL | ASSSRWGVEQY | 0.47818738 |
| TIGHVDHGKTTL | ASSSRWGVEQY | 0.6910533 |
| LISNASDALDK | ASSSRWGVEQY | 0.7106341 |
| LISNASDAL | ASSSRWGVEQY | 0.6514499 |
| ISNASDALDK | ASSSRWGVEQY | 0.7222815 |
| NQAETLVYQTE | ASSSRWGVEQY | 0.53713655 |
| TGGTGGTAGAG | ASSSRWGVEQY | 0.7080256 |
| TGSTAGGAGGA | ASSSRWGVEQY | 0.7102046 |
| DPLAVPVAAGPV | ASSSRWGVEQY | 0.6387094 |
| GGTGGTPGNGGN | ASSSRWGVEQY | 0.61760795 |
| PAGLAGVDPATL | ASSSRWGVEQY | 0.6908448 |
| GNNLVVEAPHAGASV | ASSSRWGVEQY | 0.6941362 |
| KVTRIMSASTLEQPA | ASSSRWGVEQY | 0.7077719 |
| LGFALPIALPRRTVV | ASSSRWGVEQY | 0.7163644 |
| TRLGNLTARKLVVLV | ASSSRWGVEQY | 0.2641224 |
| WSQQRRLIYFPSAGP | ASSSRWGVEQY | 0.82128495 |
| GPNGNPDPMAAAVDIR | ASSSRWGVEQY | 0.6977471 |
| TQRCVSVADLRAAASR | ASSSRWGVEQY | 0.6961807 |
| QFVLDVAEHPVEVLTETV | ASSSRWGVEQY | 0.4118408 |
| GGAGGTGGAGGNGGTQAGDG | ASSSRWGVEQY | 0.2122176 |
| PLLATLRAIAVDVDAKPAQV | ASSSRWGVEQY | 0.5432918 |
| APKFRTGGYMIDFWGVGYQVAK | ASSSRWGVEQY | 0.3474549 |
| GAGGAGGAGGAAGGPGGTGGTG | ASSSRWGVEQY | 0.79902714 |
| IGALGLLATDVAPAGSVAFGVA | ASSSRWGVEQY | 0.72437394 |
| AGGAGGIGGNGGAGGAGGVLMG | ASSSRWGVEQY | 0.4856338 |
| ADELIGLL | ASSSRWGVEQY | 0.7607626 |
| ADRALKA | ASSSRWGVEQY | 0.7892549 |
| AGAVGQVGGAGGA | ASSSRWGVEQY | 0.2227647 |
| AGGGGGLGGAP | ASSSRWGVEQY | 0.4916383 |
| AGGLLYGNGGAG | ASSSRWGVEQY | 0.33584967 |
| AGLESIGL | ASSSRWGVEQY | 0.50091493 |
| AGLKTGII | ASSSRWGVEQY | 0.15445726 |
| ASLAALLL | ASSSRWGVEQY | 0.74543244 |
| ASVINISSI | ASSSRWGVEQY | 0.4625803 |
| ATPPGPA | ASSSRWGVEQY | 0.1333037 |
| ATVASALV | ASSSRWGVEQY | 0.5438899 |
| ATVASLIL | ASSSRWGVEQY | 0.47890836 |
| AVNGPVLIRH | ASSSRWGVEQY | 0.4201969 |
| DGAAGAVGI | ASSSRWGVEQY | 0.76372576 |
| DPLLHGI | ASSSRWGVEQY | 0.56289554 |
| ELGAALGAL | ASSSRWGVEQY | 0.30011627 |
| ELGRTQAV | ASSSRWGVEQY | 0.7178823 |
| FVAGGYLS | ASSSRWGVEQY | 0.8144691 |
| GASGGAGGAGGW | ASSSRWGVEQY | 0.28167635 |
| GGAGAGGDGGAGGI | ASSSRWGVEQY | 0.7013239 |
| GTGHNGSQPGMGGNG | ASSSRWGVEQY | 0.8406063 |

| | | |
|---|---|---|
| GYAAASASAS | ASSSRWGVEQY | 0.57967716 |
| HQVDVAAVQQG | ASSSRWGVEQY | 0.7894256 |
| IGAAIAHT | ASSSRWGVEQY | 0.8175974 |
| IGMIEVAQL | ASSSRWGVEQY | 0.41463703 |
| IRLAPEHTSAMD | ASSSRWGVEQY | 0.794308 |
| IYNVSVL | ASSSRWGVEQY | 0.79717016 |
| KQGFLGM | ASSSRWGVEQY | 0.52088356 |
| LESVITQL | ASSSRWGVEQY | 0.5903711 |
| LGLGALVVV | ASSSRWGVEQY | 0.32561934 |
| LIGNGGNGGSGGI | ASSSRWGVEQY | 0.56173694 |
| LNAINAPTL | ASSSRWGVEQY | 0.22732297 |
| LQLLNTGV | ASSSRWGVEQY | 0.70651394 |
| LQTSIEF | ASSSRWGVEQY | 0.7933134 |
| MSFAVSEL | ASSSRWGVEQY | 0.5700941 |
| NTGALGL | ASSSRWGVEQY | 0.24193247 |
| PPGAGGAGGAGTT | ASSSRWGVEQY | 0.26847073 |
| QLVGLGI | ASSSRWGVEQY | 0.658207 |
| RFLEQVHQL | ASSSRWGVEQY | 0.42189774 |
| RGVGYVLR | ASSSRWGVEQY | 0.20944461 |
| SATGVFAGVI | ASSSRWGVEQY | 0.59437513 |
| SLPAITLP | ASSSRWGVEQY | 0.77294827 |
| STALAFTV | ASSSRWGVEQY | 0.7879008 |
| SVLTVGLGK | ASSSRWGVEQY | 0.294054 |
| TALAGAGSGI | ASSSRWGVEQY | 0.83209884 |
| TGLPFTGI | ASSSRWGVEQY | 0.79603755 |
| VAEHVTV | ASSSRWGVEQY | 0.83003616 |
| VDIVLNSL | ASSSRWGVEQY | 0.5457413 |
| VLTGGGAVL | ASSSRWGVEQY | 0.31314516 |
| VNVGLAYGI | ASSSRWGVEQY | 0.81198615 |
| VSSAAVGAV | ASSSRWGVEQY | 0.562057 |
| VTSVYEAGV | ASSSRWGVEQY | 0.5027 |
| WVALDRGA | ASSSRWGVEQY | 0.76555824 |
| YLVGSVVI | ASSSRWGVEQY | 0.41436622 |
| GPPGTGKTLL | ASSSRWGVEQY | 0.41777408 |
| GVGMTKF | ASSSRWGVEQY | 0.72616625 |
| IGIGGSDL | ASSSRWGVEQY | 0.79402006 |
| IGTLLAEL | ASSSRWGVEQY | 0.3044043 |
| IYTSGTT | ASSSRWGVEQY | 0.38466758 |
| LSFPAGV | ASSSRWGVEQY | 0.41706127 |
| LYTSGTT | ASSSRWGVEQY | 0.5774287 |
| VGAFPFGF | ASSSRWGVEQY | 0.8139899 |
| VGAGTVEF | ASSSRWGVEQY | 0.63835186 |
| VGKSTLLL | ASSSRWGVEQY | 0.5831 |
| AALAHAGGGIV | ASSSRWGVEQY | 0.7371571 |
| AALVAVSLL | ASSSRWGVEQY | 0.73685396 |
| AAVSRAVAAL | ASSSRWGVEQY | 0.6103928 |
| AGGSSGIGGF | ASSSRWGVEQY | 0.7556237 |
| AGLTVANANG | ASSSRWGVEQY | 0.42655542 |
| AGLVGAVL | ASSSRWGVEQY | 0.7807075 |
| ALSQAFTV | ASSSRWGVEQY | 0.73961365 |
| ALTFSVSL | ASSSRWGVEQY | 0.7666213 |

| | | |
|---|---|---|
| ALTGAALVAL | ASSSRWGVEQY | 0.7923356 |
| ASFSIGAGVL | ASSSRWGVEQY | 0.74266565 |
| DGSVYKI | ASSSRWGVEQY | 0.62137437 |
| EQLAEIV | ASSSRWGVEQY | 0.21720937 |
| FAVSELTVM | ASSSRWGVEQY | 0.48104012 |
| GAAPLAAGL | ASSSRWGVEQY | 0.6847811 |
| GADGGDGGSGGAGG | ASSSRWGVEQY | 0.6580385 |
| GFGLDTGGAGG | ASSSRWGVEQY | 0.7645165 |
| GLASGGLL | ASSSRWGVEQY | 0.74737597 |
| GRIAGAAVLL | ASSSRWGVEQY | 0.39275044 |
| GSAGSLNTSF | ASSSRWGVEQY | 0.8078064 |
| GTGQIGI | ASSSRWGVEQY | 0.8001303 |
| GTLIALIL | ASSSRWGVEQY | 0.25621897 |
| IAATMGFVVG | ASSSRWGVEQY | 0.5899362 |
| IFAQALL | ASSSRWGVEQY | 0.8353481 |
| IGAAVALGF | ASSSRWGVEQY | 0.76942825 |
| IGLTPLAL | ASSSRWGVEQY | 0.47776178 |
| ILGSNTSTL | ASSSRWGVEQY | 0.56175375 |
| IVFSLTVL | ASSSRWGVEQY | 0.83199084 |
| LASKATGF | ASSSRWGVEQY | 0.51098037 |
| LASVALIL | ASSSRWGVEQY | 0.8089216 |
| LFAVGLTL | ASSSRWGVEQY | 0.17542139 |
| LFNGGIGTYI | ASSSRWGVEQY | 0.55023503 |
| LTLGAGII | ASSSRWGVEQY | 0.39066237 |
| LVAAALGDI | ASSSRWGVEQY | 0.16326095 |
| NTAFVQLGI | ASSSRWGVEQY | 0.6960137 |
| QLFSGAEL | ASSSRWGVEQY | 0.39899707 |
| SAATFITL | ASSSRWGVEQY | 0.7237625 |
| SNYGLANLGV | ASSSRWGVEQY | 0.45917726 |
| SYGGAYAVM | ASSSRWGVEQY | 0.7764331 |
| TTGVGGLTLG | ASSSRWGVEQY | 0.73849785 |
| TTQTIYI | ASSSRWGVEQY | 0.56050557 |
| VAAGAGLGAVI | ASSSRWGVEQY | 0.3587711 |
| VAAVVTLL | ASSSRWGVEQY | 0.63631636 |
| VGFLGGLI | ASSSRWGVEQY | 0.7943069 |
| VGVIPILL | ASSSRWGVEQY | 0.73833036 |
| VIFASSLI | ASSSRWGVEQY | 0.2116144 |
| VKGGDGGAAAG | ASSSRWGVEQY | 0.5440175 |
| VTKTFTNI | ASSSRWGVEQY | 0.56350684 |
| VVAAAILL | ASSSRWGVEQY | 0.17010674 |
| YAALDVEL | ASSSRWGVEQY | 0.15134692 |
| LASVALLL | ASSSRWGVEQY | 0.39804864 |
| LDAVHRV | ASSSRWGVEQY | 0.6150044 |
| QGLSSSIF | ASSSRWGVEQY | 0.7354047 |
| RGTFANIRL | ASSSRWGVEQY | 0.7493398 |
| RQLWWGHRI | ASSSRWGVEQY | 0.52743757 |
| YGPPGTGKTLL | ASSSRWGVEQY | 0.75450736 |
| AALVGAPANVA | ASSSRWGVEQY | 0.67496693 |
| AATFRRTE | ASSSRWGVEQY | 0.11847061 |
| AGALGGGI | ASSSRWGVEQY | 0.42622638 |
| AGGAGGAGGIGAG | ASSSRWGVEQY | 0.22439402 |

| | | |
|---|---|---|
| AGSGALGL | ASSSRWGVEQY | 0.79732597 |
| AQIADQI | ASSSRWGVEQY | 0.30837482 |
| ASAVAVTYL | ASSSRWGVEQY | 0.370114 |
| ASTVLGGTGG | ASSSRWGVEQY | 0.663699 |
| AVGGEPAK | ASSSRWGVEQY | 0.7699855 |
| CEQGTFGGTC | ASSSRWGVEQY | 0.6410361 |
| DEGKAPAG | ASSSRWGVEQY | 0.37088853 |
| DTVVFPAL | ASSSRWGVEQY | 0.17097083 |
| FEGITMAG | ASSSRWGVEQY | 0.715203 |
| FPLVVQD | ASSSRWGVEQY | 0.3955016 |
| GAFTTPQ | ASSSRWGVEQY | 0.30604097 |
| GFGGFGTAGGVGGA | ASSSRWGVEQY | 0.21339741 |
| GFLAQFL | ASSSRWGVEQY | 0.22876404 |
| GGAGGAGGNAGVA | ASSSRWGVEQY | 0.24995251 |
| GGVGTAGAGGAGGAGG | ASSSRWGVEQY | 0.766417 |
| GNGGGGDAGNAGSGGN | ASSSRWGVEQY | 0.57447624 |
| GTGGAGGAGNDAGS | ASSSRWGVEQY | 0.4512391 |
| GTTNTGWFNS | ASSSRWGVEQY | 0.8102566 |
| IFMGPLFTL | ASSSRWGVEQY | 0.77150065 |
| ILAHAIEAI | ASSSRWGVEQY | 0.59316814 |
| IRSAIREAL | ASSSRWGVEQY | 0.3539234 |
| ITFAGGG | ASSSRWGVEQY | 0.31882784 |
| ITGMSGHQEV | ASSSRWGVEQY | 0.2989849 |
| IVAGLACTL | ASSSRWGVEQY | 0.33175856 |
| IVGGGLAGA | ASSSRWGVEQY | 0.79933643 |
| IVTGVLNI | ASSSRWGVEQY | 0.5348997 |
| LAQQGCI | ASSSRWGVEQY | 0.26216614 |
| LCLHVEN | ASSSRWGVEQY | 0.76113296 |
| LDAISDGS | ASSSRWGVEQY | 0.81716967 |
| LGGFAAML | ASSSRWGVEQY | 0.8415065 |
| LGGKNPMI | ASSSRWGVEQY | 0.31301612 |
| LGPKGVAAIL | ASSSRWGVEQY | 0.4920804 |
| LMVGVSI | ASSSRWGVEQY | 0.53552854 |
| LVGAAFVL | ASSSRWGVEQY | 0.602295 |
| LVLAVAPL | ASSSRWGVEQY | 0.4163097 |
| MAAIAVIDAA | ASSSRWGVEQY | 0.2844425 |
| NNGINAELQ | ASSSRWGVEQY | 0.77936685 |
| RGEPLVLVHG | ASSSRWGVEQY | 0.85199463 |
| RGGAGGAAGLI | ASSSRWGVEQY | 0.5094744 |
| RSYRMHPAV | ASSSRWGVEQY | 0.1891788 |
| SAAHLLAGQLA | ASSSRWGVEQY | 0.55827254 |
| SAAVGSY | ASSSRWGVEQY | 0.38439995 |
| SPHAVLHRI | ASSSRWGVEQY | 0.45212087 |
| SQIAKVL | ASSSRWGVEQY | 0.6412095 |
| TALGHLVP | ASSSRWGVEQY | 0.25426844 |
| TAPTTSGAA | ASSSRWGVEQY | 0.50724167 |
| TDGDDGNGGSPGT | ASSSRWGVEQY | 0.84394944 |
| TDGLGEN | ASSSRWGVEQY | 0.6115557 |
| TGGPPLPGQA | ASSSRWGVEQY | 0.7421053 |
| TIPAITI | ASSSRWGVEQY | 0.7904203 |
| TIPGFTI | ASSSRWGVEQY | 0.7240486 |

| | | |
|---|---|---|
| TSIASGALA | ASSSRWGVEQY | 0.8085295 |
| TVTLPGAAI | ASSSRWGVEQY | 0.80583364 |
| VASGLSAALP | ASSSRWGVEQY | 0.52822614 |
| VGAGVIGI | ASSSRWGVEQY | 0.6286026 |
| VGPKRFGV | ASSSRWGVEQY | 0.7439239 |
| VNVAFPD | ASSSRWGVEQY | 0.59186447 |
| VSMTNSGVS | ASSSRWGVEQY | 0.6317966 |
| VTDVLGDLSV | ASSSRWGVEQY | 0.4756407 |
| VVVVGAGFAGL | ASSSRWGVEQY | 0.7731254 |
| VYTVGTVV | ASSSRWGVEQY | 0.6314445 |
| WTTFSGP | ASSSRWGVEQY | 0.8211724 |
| YDELVEAF | ASSSRWGVEQY | 0.22004685 |
| YGTGGAGGNGG | ASSSRWGVEQY | 0.2673388 |
| YTNVMAV | ASSSRWGVEQY | 0.7933717 |
| FLEQVHQL | ASSSRWGVEQY | 0.16215871 |
| GLSLLAVAV | ASSSRWGVEQY | 0.19294924 |
| LEQVHQL | ASSSRWGVEQY | 0.64994603 |
| LNAAGAI | ASSSRWGVEQY | 0.8288752 |
| PAVHASL | ASSSRWGVEQY | 0.14887443 |
| TELEDDA | ASSSRWGVEQY | 0.3516385 |
| TPPGPPN | ASSSRWGVEQY | 0.26305193 |
| TPPPPPPP | ASSSRWGVEQY | 0.77475715 |
| TQSPALI | ASSSRLGENYAEQF | 0.61659575 |
| EFAAAFPQ | ASSSRLGENYAEQF | 0.25368512 |
| LLSGNLDVL | ASSSRLGENYAEQF | 0.7400139 |
| GPANLGNYNI | ASSSRLGENYAEQF | 0.20771085 |
| TLGPTLP | ASSSRLGENYAEQF | 0.47435933 |
| VYAYSSCCK | ASSSRLGENYAEQF | 0.7571584 |
| APAPAPAPNI | ASSSRLGENYAEQF | 0.8305926 |
| PGGPVPIP | ASSSRLGENYAEQF | 0.2789139 |
| FLTVALEP | ASSSRLGENYAEQF | 0.77488446 |
| SQYFGANL | ASSSRLGENYAEQF | 0.49385145 |
| GGVKPAA | ASSSRLGENYAEQF | 0.84879017 |
| GGTGGAGGAGGW | ASSSRLGENYAEQF | 0.42395315 |
| GVGGVAGLGGAGG | ASSSRLGENYAEQF | 0.22879678 |
| VLLVLATTI | ASSSRLGENYAEQF | 0.76032704 |
| IGIFAAML | ASSSRLGENYAEQF | 0.78545505 |
| AVGDLID | ASSSRLGENYAEQF | 0.6091023 |
| NISPLPL | ASSSRLGENYAEQF | 0.79303604 |
| VTAVFVAI | ASSSRLGENYAEQF | 0.60196114 |
| SGDDNDAAL | ASSSRLGENYAEQF | 0.5918757 |
| AVAALAHS | ASSSRLGENYAEQF | 0.76385474 |
| GGVTVMATP | ASSSRLGENYAEQF | 0.7586976 |
| AGLAANVRD | ASSSRLGENYAEQF | 0.3054547 |
| TNRPDLIDPALLRPG | ASSSRLGENYAEQF | 0.792442 |
| LANLGVTAL | ASSSRLGENYAEQF | 0.6290796 |
| TGERIEL | ASSSRLGENYAEQF | 0.3684963 |
| NGEELAA | ASSSRLGENYAEQF | 0.82769936 |
| VLSLVWG | ASSSRLGENYAEQF | 0.64312696 |
| IGYALWL | ASSSRLGENYAEQF | 0.76036656 |
| LSLLLAPL | ASSSRLGENYAEQF | 0.67409694 |

| | | |
|---|---|---|
| TLDEALK | ASSSRLGENYAEQF | 0.8445367 |
| TDGAGTAEGG | ASSSRLGENYAEQF | 0.23520115 |
| NAPGGAGGNGG | ASSSRLGENYAEQF | 0.71948224 |
| VSVFMGMGGNF | ASSSRLGENYAEQF | 0.61430764 |
| LASYLAAL | ASSSRLGENYAEQF | 0.77684325 |
| LVGGTPAL | ASSSRLGENYAEQF | 0.65726745 |
| VRNADVL | ASSSRLGENYAEQF | 0.4043442 |
| TCSSPFLVSAM | ASSSRLGENYAEQF | 0.6976821 |
| VHITDAE | ASSSRLGENYAEQF | 0.7907277 |
| EYAAPSF | ASSSRLGENYAEQF | 0.6004003 |
| VVDAFHGGL | ASSSRLGENYAEQF | 0.7102928 |
| ASVVDPQ | ASSSRLGENYAEQF | 0.8306572 |
| SYNEFGGLS | ASSSRLGENYAEQF | 0.30261585 |
| DPDGSVY | ASSSRLGENYAEQF | 0.6291704 |
| MVSTIHATE | ASSSRLGENYAEQF | 0.7463188 |
| LAAGVADPV | ASSSRLGENYAEQF | 0.79030955 |
| TKEQIAA | ASSSRLGENYAEQF | 0.5659368 |
| SLASPAAL | ASSSRLGENYAEQF | 0.7908405 |
| IGLVFYGV | ASSSRLGENYAEQF | 0.43719563 |
| FLLDASNL | ASSSRLGENYAEQF | 0.6997072 |
| YQVGNFP | ASSSRLGENYAEQF | 0.5352666 |
| ESLLSQST | ASSSRLGENYAEQF | 0.6050395 |
| LFTVGFAAVG | ASSSRLGENYAEQF | 0.28163454 |
| ATPPTPT | ASSSRLGENYAEQF | 0.65664434 |
| TDLATLG | ASSSRLGENYAEQF | 0.7592693 |
| SAGGTGR | ASSSRLGENYAEQF | 0.73223305 |
| GFSNSGS | ASSSRLGENYAEQF | 0.16594902 |
| GGNGGSGGTSV | ASSSRLGENYAEQF | 0.748233 |
| GTSVATGGAGN | ASSSRLGENYAEQF | 0.723233 |
| NGGNGGSGGTS | ASSSRLGENYAEQF | 0.8436558 |
| AVVKFPH | ASSSRLGENYAEQF | 0.61473393 |
| TPPPPPP | ASSSRLGENYAEQF | 0.7767693 |
| AAPAAAAAPA | ASSSRLGENYAEQF | 0.37203065 |
| IALNEDL | ASSSRLGENYAEQF | 0.6447121 |
| IVNVSSM | ASSSRLGENYAEQF | 0.3325286 |
| TGIPPAPR | ASSSRLGENYAEQF | 0.497412 |
| FDLGGGTF | ASSSRLGENYAEQF | 0.58194184 |
| FDLGGGTFDVS | ASSSRLGENYAEQF | 0.37727174 |
| NEPTAAA | ASSSRLGENYAEQF | 0.805616 |
| NEPTAAALA | ASSSRLGENYAEQF | 0.78818 |
| GSIINISSI | ASSSRLGENYAEQF | 0.8011381 |
| VAPGNAGTA | ASSSRLGENYAEQF | 0.8063229 |
| TGGGASLE | ASSSRLGENYAEQF | 0.82141423 |
| VAVYNHYKRI | ASSSRLGENYAEQF | 0.7590188 |
| TDPAPATT | ASSSRLGENYAEQF | 0.44469905 |
| TNRPDILDPALLRPG | ASSSRLGENYAEQF | 0.50375074 |
| LTTTTTS | ASSSRLGENYAEQF | 0.23377961 |
| PTGSGKTL | ASSSRLGENYAEQF | 0.4581625 |
| TGSGKTL | ASSSRLGENYAEQF | 0.47606546 |
| TIGHVDHGK | ASSSRLGENYAEQF | 0.6526432 |
| TIGHVDHGKTT | ASSSRLGENYAEQF | 0.14028803 |

| | | |
|---|---|---|
| GHVDHGKTT | ASSSRLGENYAEQF | 0.7405682 |
| AQFEHTLL | ASSSRLGENYAEQF | 0.8042229 |
| IVNQDEL | ASSSRLGENYAEQF | 0.56405306 |
| TNGSQFF | ASSSRLGENYAEQF | 0.7249253 |
| TGKGIVE | ASSSRLGENYAEQF | 0.73085725 |
| LISNASDA | ASSSRLGENYAEQF | 0.4250161 |
| AELRTLGELTKIAKAHGAQ | ASSSRLGENYAEQF | 0.7660756 |
| APNAVAAAEPSW | ASSSRLGENYAEQF | 0.41096476 |
| APQVAGAATAVL | ASSSRLGENYAEQF | 0.76159143 |
| ARRDPAAAAAILPTDA | ASSSRLGENYAEQF | 0.8070923 |
| DAAARDVLAEAGLAEYF | ASSSRLGENYAEQF | 0.6500816 |
| DDQIVLSAKANLAKVEQ | ASSSRLGENYAEQF | 0.6730991 |
| DELLPDTGEDSDEDGH | ASSSRLGENYAEQF | 0.6608921 |
| ELPEAPANRATVPQD | ASSSRLGENYAEQF | 0.2107777 |
| FGQSKIERVTGIG | ASSSRLGENYAEQF | 0.6811582 |
| GAGGTGSTAGGAGGAGGAGGLY | ASSSRLGENYAEQF | 0.15748295 |
| GHGGTGAAGGNGGTGGAGGSGI | ASSSRLGENYAEQF | 0.6617404 |
| GNGGNGGAGGEGGPGGAGGAGA | ASSSRLGENYAEQF | 0.3145023 |
| GSTAGNGGNGGNATNGTIAGQP | ASSSRLGENYAEQF | 0.6635984 |
| GTGGDGGNGGNGGTGG | ASSSRLGENYAEQF | 0.58642375 |
| GVRRPAGIPADAPV | ASSSRLGENYAEQF | 0.72588277 |
| GYRQVTDAHLLTLARR | ASSSRLGENYAEQF | 0.32231325 |
| IAKMGQDGHDRG | ASSSRLGENYAEQF | 0.66543686 |
| KAEPSIEIQYT | ASSSRLGENYAEQF | 0.5245415 |
| KPVRIRMAADEDTVAVT | ASSSRLGENYAEQF | 0.23818885 |
| MPPRIAGMRLLVIKPEP | ASSSRLGENYAEQF | 0.7756175 |
| MKFYAVALFRSRGVPPTR | ASSSRLGENYAEQF | 0.560436 |
| NAGGFGILGAVAHSP | ASSSRLGENYAEQF | 0.6816437 |
| NEGREITLAMAEM | ASSSRLGENYAEQF | 0.7642403 |
| NELVEWSHPLGHRWRWE | ASSSRLGENYAEQF | 0.6555586 |
| NLGAGGTGGNGGNPG | ASSSRLGENYAEQF | 0.2091387 |
| NTEWPKWETVMAADPVD | ASSSRLGENYAEQF | 0.6045972 |
| PASAAPAETQP | ASSSRLGENYAEQF | 0.6049072 |
| QQSSDGKNGVRAAEITGE | ASSSRLGENYAEQF | 0.4828897 |
| RAPDAVTTSWWKE | ASSSRLGENYAEQF | 0.14056386 |
| SGFGNVGAHTSGWFNQ | ASSSRLGENYAEQF | 0.75206494 |
| SGVAGAVGQAASVGGLKV | ASSSRLGENYAEQF | 0.3750593 |
| SNGGAGTGSAGGNGGT | ASSSRLGENYAEQF | 0.5028488 |
| TGGAGGEGGAAG | ASSSRLGENYAEQF | 0.46415707 |
| TGGDGGAAGAGGAPGHG | ASSSRLGENYAEQF | 0.7986027 |
| TMPPLLAAIANAGRHGV | ASSSRLGENYAEQF | 0.73615444 |
| TSSTAGNSGGA | ASSSRLGENYAEQF | 0.4770982 |
| VPATVPVGILC | ASSSRLGENYAEQF | 0.7620852 |
| VQRPDSGGVVLTGRIS | ASSSRLGENYAEQF | 0.6763818 |
| WTPNPVPATKGVPAGLVQAT | ASSSRLGENYAEQF | 0.80932844 |
| YALKRALAKFSDRFDVVII | ASSSRLGENYAEQF | 0.67146033 |
| YLIGSVIV | ASSSRLGENYAEQF | 0.5760186 |
| YGQLVAVID | ASSSRLGENYAEQF | 0.6222736 |
| VTVGGMTLPA | ASSSRLGENYAEQF | 0.57706594 |
| VLALPAV | ASSSRLGENYAEQF | 0.19887628 |
| VIPPTLNYET | ASSSRLGENYAEQF | 0.21949948 |

| | | |
|---|---|---|
| VGMGGMHPSSG | ASSSRLGENYAEQF | 0.55726194 |
| VGLAGGAATA | ASSSRLGENYAEQF | 0.22011754 |
| VGGAGGAGGSGP | ASSSRLGENYAEQF | 0.63250536 |
| VFAAFMV | ASSSRLGENYAEQF | 0.7981583 |
| TLYDKLR | ASSSRLGENYAEQF | 0.72533023 |
| TLAVFTVL | ASSSRLGENYAEQF | 0.23083654 |
| TGSVQPARV | ASSSRLGENYAEQF | 0.43782955 |
| TGGGAGSGGT | ASSSRLGENYAEQF | 0.43464863 |
| SHLETSL | ASSSRLGENYAEQF | 0.72828805 |
| SFTTGMT | ASSSRLGENYAEQF | 0.7314596 |
| QVEVAKRMGVRQPTVS | ASSSRLGENYAEQF | 0.8338069 |
| QGSGGPALAM | ASSSRLGENYAEQF | 0.78375596 |
| MLAPSNP | ASSSRLGENYAEQF | 0.4507426 |
| LVNNAAI | ASSSRLGENYAEQF | 0.78075826 |
| LTGEINTLA | ASSSRLGENYAEQF | 0.52469635 |
| LSIFGPV | ASSSRLGENYAEQF | 0.603257 |
| LLALLANL | ASSSRLGENYAEQF | 0.5478562 |
| LISNASDALDKLR | ASSSRLGENYAEQF | 0.739511 |
| LINEAVP | ASSSRLGENYAEQF | 0.56275654 |
| LGYAFSGL | ASSSRLGENYAEQF | 0.40452737 |
| LDFVAVAL | ASSSRLGENYAEQF | 0.24186338 |
| LAQPGAE | ASSSRLGENYAEQF | 0.6605207 |
| ISNLGAVTGF | ASSSRLGENYAEQF | 0.6277331 |
| HQEDPGRI | ASSSRLGENYAEQF | 0.25767744 |
| GTPGEPTP | ASSSRLGENYAEQF | 0.4123094 |
| GFFNSGS | ASSSRLGENYAEQF | 0.83854556 |
| GAGGGGLTAGAN | ASSSRLGENYAEQF | 0.8273805 |
| FVDALVNM | ASSSRLGENYAEQF | 0.4350251 |
| FLTVAPEL | ASSSRLGENYAEQF | 0.36052966 |
| FETLTAL | ASSSRLGENYAEQF | 0.7466978 |
| DVINPIEL | ASSSRLGENYAEQF | 0.32930464 |
| APVSGAPV | ASSSRLGENYAEQF | 0.6851641 |
| ALVGGLAW | ASSSRLGENYAEQF | 0.68308043 |
| ALAAAAATAI | ASSSRLGENYAEQF | 0.6748004 |
| AFGLGLLL | ASSSRLGENYAEQF | 0.44832367 |
| AFFAAASL | ASSSRLGENYAEQF | 0.68458134 |
| AAAALPAFSPP | ASSSRLGENYAEQF | 0.51787484 |
| VVLAPAL | ASSSRLGENYAEQF | 0.7467178 |
| TIGHVDHGKTTL | ASSSRLGENYAEQF | 0.13918953 |
| LISNASDALDK | ASSSRLGENYAEQF | 0.7553734 |
| LISNASDAL | ASSSRLGENYAEQF | 0.30220187 |
| ISNASDALDK | ASSSRLGENYAEQF | 0.42428607 |
| NQAETLVYQTE | ASSSRLGENYAEQF | 0.19455037 |
| TGGTGGTAGAG | ASSSRLGENYAEQF | 0.58516514 |
| TGSTAGGAGGA | ASSSRLGENYAEQF | 0.76644427 |
| DPLAVPVAAGPV | ASSSRLGENYAEQF | 0.79898524 |
| GGTGGTPGNGGN | ASSSRLGENYAEQF | 0.42648 |
| PAGLAGVDPATL | ASSSRLGENYAEQF | 0.7409133 |
| GNNLVVEAPHAGASV | ASSSRLGENYAEQF | 0.83192086 |
| KVTRIMSASTLEQPA | ASSSRLGENYAEQF | 0.43093896 |
| LGFALPIALPRRTVV | ASSSRLGENYAEQF | 0.17869705 |

| | | |
|---|---|---|
| TRLGNLTARKLVVLV | ASSSRLGENYAEQF | 0.7152118 |
| WSQQRRLIYFPSAGP | ASSSRLGENYAEQF | 0.65628827 |
| GPNGNPDPMAAAVDIR | ASSSRLGENYAEQF | 0.79890656 |
| TQRCVSVADLRAAASR | ASSSRLGENYAEQF | 0.79980505 |
| QFVLDVAEHPVEVLTETV | ASSSRLGENYAEQF | 0.2697204 |
| GGAGGTGGAGGNGGTQAGDG | ASSSRLGENYAEQF | 0.7923588 |
| PLLATLRAIAVDVDAKPAQV | ASSSRLGENYAEQF | 0.7848904 |
| APKFRTGGYMIDFWGVGYQVAK | ASSSRLGENYAEQF | 0.31769207 |
| GAGGAGGAGGAAGGPGGTGGTG | ASSSRLGENYAEQF | 0.3409853 |
| IGALGLLATDVAPAGSVAFGVA | ASSSRLGENYAEQF | 0.38101548 |
| AGGAGGIGGNGGAGGAGGVLMG | ASSSRLGENYAEQF | 0.22151412 |
| ADELIGLL | ASSSRLGENYAEQF | 0.6910877 |
| ADRALKA | ASSSRLGENYAEQF | 0.4435609 |
| AGAVGQVGGAGGA | ASSSRLGENYAEQF | 0.41331798 |
| AGGGGGLGGAP | ASSSRLGENYAEQF | 0.62448305 |
| AGGLLYGNGGAG | ASSSRLGENYAEQF | 0.35147977 |
| AGLESIGL | ASSSRLGENYAEQF | 0.45571905 |
| AGLKTGII | ASSSRLGENYAEQF | 0.39790648 |
| ASLAALLL | ASSSRLGENYAEQF | 0.5594187 |
| ASVINISSI | ASSSRLGENYAEQF | 0.7273662 |
| ATPPGPA | ASSSRLGENYAEQF | 0.6343855 |
| ATVASALV | ASSSRLGENYAEQF | 0.7672601 |
| ATVASLIL | ASSSRLGENYAEQF | 0.8044035 |
| AVNGPVLIRH | ASSSRLGENYAEQF | 0.27842152 |
| DGAAGAVGI | ASSSRLGENYAEQF | 0.4172737 |
| DPLLHGI | ASSSRLGENYAEQF | 0.7543441 |
| ELGAALGAL | ASSSRLGENYAEQF | 0.35251266 |
| ELGRTQAV | ASSSRLGENYAEQF | 0.78633654 |
| FVAGGYLS | ASSSRLGENYAEQF | 0.5732405 |
| GASGGAGGAGGW | ASSSRLGENYAEQF | 0.70022225 |
| GGAGAGGDGGAGGI | ASSSRLGENYAEQF | 0.6716402 |
| GTGHNGSQPGMGGNG | ASSSRLGENYAEQF | 0.83618736 |
| GYAAASASAS | ASSSRLGENYAEQF | 0.72646815 |
| HQVDVAAVQQG | ASSSRLGENYAEQF | 0.48498172 |
| IGAAIAHT | ASSSRLGENYAEQF | 0.72090465 |
| IGMIEVAQL | ASSSRLGENYAEQF | 0.31824905 |
| IRLAPEHTSAMD | ASSSRLGENYAEQF | 0.745032 |
| IYNVSVL | ASSSRLGENYAEQF | 0.74809813 |
| KQGFLGM | ASSSRLGENYAEQF | 0.37694812 |
| LESVITQL | ASSSRLGENYAEQF | 0.60128677 |
| LGLGALVVV | ASSSRLGENYAEQF | 0.81330025 |
| LIGNGGNGGSGGI | ASSSRLGENYAEQF | 0.7637116 |
| LNAINAPTL | ASSSRLGENYAEQF | 0.23688939 |
| LQLLNTGV | ASSSRLGENYAEQF | 0.46709877 |
| LQTSIEF | ASSSRLGENYAEQF | 0.28177193 |
| MSFAVSEL | ASSSRLGENYAEQF | 0.507763 |
| NTGALGL | ASSSRLGENYAEQF | 0.25312182 |
| PPGAGGAGGAGTT | ASSSRLGENYAEQF | 0.74196726 |
| QLVGLGI | ASSSRLGENYAEQF | 0.46123186 |
| RFLEQVHQL | ASSSRLGENYAEQF | 0.8166359 |
| RGVGYVLR | ASSSRLGENYAEQF | 0.67340875 |

| | | |
|---|---|---|
| SATGVFAGVI | ASSSRLGENYAEQF | 0.78860366 |
| SLPAITLP | ASSSRLGENYAEQF | 0.4410686 |
| STALAFTV | ASSSRLGENYAEQF | 0.40848276 |
| SVLTVGLGK | ASSSRLGENYAEQF | 0.23459579 |
| TALAGAGSGI | ASSSRLGENYAEQF | 0.2611336 |
| TGLPFTGI | ASSSRLGENYAEQF | 0.34456474 |
| VAEHVTV | ASSSRLGENYAEQF | 0.52553976 |
| VDIVLNSL | ASSSRLGENYAEQF | 0.17627285 |
| VLTGGGAVL | ASSSRLGENYAEQF | 0.83279324 |
| VNVGLAYGI | ASSSRLGENYAEQF | 0.62265295 |
| VSSAAVGAV | ASSSRLGENYAEQF | 0.71350497 |
| VTSVYEAGV | ASSSRLGENYAEQF | 0.42353314 |
| WVALDRGA | ASSSRLGENYAEQF | 0.3664239 |
| YLVGSVVI | ASSSRLGENYAEQF | 0.45582658 |
| GPPGTGKTLL | ASSSRLGENYAEQF | 0.77927893 |
| GVGMTKF | ASSSRLGENYAEQF | 0.29269898 |
| IGIGGSDL | ASSSRLGENYAEQF | 0.16800626 |
| IGTLLAEL | ASSSRLGENYAEQF | 0.45347774 |
| IYTSGTT | ASSSRLGENYAEQF | 0.22498792 |
| LSFPAGV | ASSSRLGENYAEQF | 0.83109045 |
| LYTSGTT | ASSSRLGENYAEQF | 0.7828618 |
| VGAFPFGF | ASSSRLGENYAEQF | 0.67534006 |
| VGAGTVEF | ASSSRLGENYAEQF | 0.82112473 |
| VGKSTLLL | ASSSRLGENYAEQF | 0.15253134 |
| AALAHAGGGIV | ASSSRLGENYAEQF | 0.47495916 |
| AALVAVSLL | ASSSRLGENYAEQF | 0.8357103 |
| AAVSRAVAAL | ASSSRLGENYAEQF | 0.6544157 |
| AGGSSGIGGF | ASSSRLGENYAEQF | 0.5265384 |
| AGLTVANANG | ASSSRLGENYAEQF | 0.79450524 |
| AGLVGAVL | ASSSRLGENYAEQF | 0.37793785 |
| ALSQAFTV | ASSSRLGENYAEQF | 0.49366456 |
| ALTFSVSL | ASSSRLGENYAEQF | 0.4928583 |
| ALTGAALVAL | ASSSRLGENYAEQF | 0.64126027 |
| ASFSIGAGVL | ASSSRLGENYAEQF | 0.6421387 |
| DGSVYKI | ASSSRLGENYAEQF | 0.26884547 |
| EQLAEIV | ASSSRLGENYAEQF | 0.74430525 |
| FAVSELTVM | ASSSRLGENYAEQF | 0.83379555 |
| GAAPLAAGL | ASSSRLGENYAEQF | 0.50768423 |
| GADGGDGGSGGAGG | ASSSRLGENYAEQF | 0.7944508 |
| GFGLDTGGAGG | ASSSRLGENYAEQF | 0.7137008 |
| GLASGGLL | ASSSRLGENYAEQF | 0.7870937 |
| GRIAGAAVLL | ASSSRLGENYAEQF | 0.40744823 |
| GSAGSLNTSF | ASSSRLGENYAEQF | 0.3321057 |
| GTGQIGI | ASSSRLGENYAEQF | 0.6493089 |
| GTLIALIL | ASSSRLGENYAEQF | 0.36617744 |
| IAATMGFVVG | ASSSRLGENYAEQF | 0.60715544 |
| IFAQALL | ASSSRLGENYAEQF | 0.49894553 |
| IGAAVALGF | ASSSRLGENYAEQF | 0.85325885 |
| IGLTPLAL | ASSSRLGENYAEQF | 0.7551429 |
| ILGSNTSTL | ASSSRLGENYAEQF | 0.72637403 |
| IVFSLTVL | ASSSRLGENYAEQF | 0.31790742 |

| | | |
|---|---|---|
| LASKATGF | ASSSRLGENYAEQF | 0.661119 |
| LASVALIL | ASSSRLGENYAEQF | 0.7417918 |
| LFAVGLTL | ASSSRLGENYAEQF | 0.23997603 |
| LFNGGIGTYI | ASSSRLGENYAEQF | 0.8153865 |
| LTLGAGII | ASSSRLGENYAEQF | 0.3712715 |
| LVAAALGDI | ASSSRLGENYAEQF | 0.70248014 |
| NTAFVQLGI | ASSSRLGENYAEQF | 0.7095409 |
| QLFSGAEL | ASSSRLGENYAEQF | 0.8298528 |
| SAATFITL | ASSSRLGENYAEQF | 0.5004252 |
| SNYGLANLGV | ASSSRLGENYAEQF | 0.61642396 |
| SYGGAYAVM | ASSSRLGENYAEQF | 0.310241 |
| TTGVGGLTLG | ASSSRLGENYAEQF | 0.77804315 |
| TTQTIYI | ASSSRLGENYAEQF | 0.7973603 |
| VAAGAGLGAVI | ASSSRLGENYAEQF | 0.7464869 |
| VAAVVTLL | ASSSRLGENYAEQF | 0.83465046 |
| VGFLGGLI | ASSSRLGENYAEQF | 0.74203146 |
| VGVIPILL | ASSSRLGENYAEQF | 0.22955742 |
| VIFASSLI | ASSSRLGENYAEQF | 0.56725836 |
| VKGGDGGAAAG | ASSSRLGENYAEQF | 0.81394005 |
| VTKTFTNI | ASSSRLGENYAEQF | 0.79553807 |
| VVAAAILL | ASSSRLGENYAEQF | 0.7101948 |
| YAALDVEL | ASSSRLGENYAEQF | 0.7233976 |
| LASVALLL | ASSSRLGENYAEQF | 0.483947 |
| LDAVHRV | ASSSRLGENYAEQF | 0.8123005 |
| QGLSSSIF | ASSSRLGENYAEQF | 0.85509604 |
| RGTFANIRL | ASSSRLGENYAEQF | 0.7901047 |
| RQLWWGHRI | ASSSRLGENYAEQF | 0.5716473 |
| YGPPGTGKTLL | ASSSRLGENYAEQF | 0.4659807 |
| AALVGAPANVA | ASSSRLGENYAEQF | 0.6705334 |
| AATFRRTE | ASSSRLGENYAEQF | 0.8011564 |
| AGALGGGI | ASSSRLGENYAEQF | 0.3208468 |
| AGGAGGAGGIGAG | ASSSRLGENYAEQF | 0.66595864 |
| AGSGALGL | ASSSRLGENYAEQF | 0.59356505 |
| AQIADQI | ASSSRLGENYAEQF | 0.1970227 |
| ASAVAVTYL | ASSSRLGENYAEQF | 0.72888047 |
| ASTVLGGTGG | ASSSRLGENYAEQF | 0.8125612 |
| AVGGEPAK | ASSSRLGENYAEQF | 0.6353862 |
| CEQGTFGGTC | ASSSRLGENYAEQF | 0.36738682 |
| DEGKAPAG | ASSSRLGENYAEQF | 0.79974854 |
| DTVVFPAL | ASSSRLGENYAEQF | 0.8368875 |
| FEGITMAG | ASSSRLGENYAEQF | 0.7952533 |
| FPLVVQD | ASSSRLGENYAEQF | 0.20962058 |
| GAFTTPQ | ASSSRLGENYAEQF | 0.1506096 |
| GFGGFGTAGGVGGA | ASSSRLGENYAEQF | 0.55090934 |
| GFLAQFL | ASSSRLGENYAEQF | 0.81977564 |
| GGAGGAGGNAGVA | ASSSRLGENYAEQF | 0.4540686 |
| GGVGTAGAGGAGGAGG | ASSSRLGENYAEQF | 0.3705246 |
| GNGGGGDAGNAGSGGN | ASSSRLGENYAEQF | 0.7221253 |
| GTGGAGGAGNDAGS | ASSSRLGENYAEQF | 0.7758045 |
| GTTNTGWFNS | ASSSRLGENYAEQF | 0.2821494 |
| IFMGPLFTL | ASSSRLGENYAEQF | 0.2969972 |

| | | |
|---|---|---|
| ILAHAIEAI | ASSSRLGENYAEQF | 0.691039 |
| IRSAIREAL | ASSSRLGENYAEQF | 0.80297303 |
| ITFAGGG | ASSSRLGENYAEQF | 0.3423162 |
| ITGMSGHQEV | ASSSRLGENYAEQF | 0.5154022 |
| IVAGLACTL | ASSSRLGENYAEQF | 0.5128148 |
| IVGGGLAGA | ASSSRLGENYAEQF | 0.71783197 |
| IVTGVLNI | ASSSRLGENYAEQF | 0.6807197 |
| LAQQGCI | ASSSRLGENYAEQF | 0.3520573 |
| LCLHVEN | ASSSRLGENYAEQF | 0.8010345 |
| LDAISDGS | ASSSRLGENYAEQF | 0.2654562 |
| LGGFAAML | ASSSRLGENYAEQF | 0.7877037 |
| LGGKNPMI | ASSSRLGENYAEQF | 0.6169143 |
| LGPKGVAAIL | ASSSRLGENYAEQF | 0.6445124 |
| LMVGVSI | ASSSRLGENYAEQF | 0.49752575 |
| LVGAAFVL | ASSSRLGENYAEQF | 0.3981471 |
| LVLAVAPL | ASSSRLGENYAEQF | 0.2951602 |
| MAAIAVIDAA | ASSSRLGENYAEQF | 0.2418727 |
| NNGINAELQ | ASSSRLGENYAEQF | 0.7944056 |
| RGEPLVLVHG | ASSSRLGENYAEQF | 0.26476797 |
| RGGAGGAAGLI | ASSSRLGENYAEQF | 0.76926905 |
| RSYRMHPAV | ASSSRLGENYAEQF | 0.59879434 |
| SAAHLLAGQLA | ASSSRLGENYAEQF | 0.3085017 |
| SAAVGSY | ASSSRLGENYAEQF | 0.72502893 |
| SPHAVLHRI | ASSSRLGENYAEQF | 0.25906378 |
| SQIAKVL | ASSSRLGENYAEQF | 0.41947225 |
| TALGHLVP | ASSSRLGENYAEQF | 0.78662896 |
| TAPTTSGAA | ASSSRLGENYAEQF | 0.7326942 |
| TDGDDGNGGSPGT | ASSSRLGENYAEQF | 0.33231908 |
| TDGLGEN | ASSSRLGENYAEQF | 0.6892035 |
| TGGPPLPGQA | ASSSRLGENYAEQF | 0.67687786 |
| TIPAITI | ASSSRLGENYAEQF | 0.3347168 |
| TIPGFTI | ASSSRLGENYAEQF | 0.62895644 |
| TSIASGALA | ASSSRLGENYAEQF | 0.7730355 |
| TVTLPGAAI | ASSSRLGENYAEQF | 0.6456241 |
| VASGLSAALP | ASSSRLGENYAEQF | 0.26161063 |
| VGAGVIGI | ASSSRLGENYAEQF | 0.7223582 |
| VGPKRFGV | ASSSRLGENYAEQF | 0.41233546 |
| VNVAFPD | ASSSRLGENYAEQF | 0.7191427 |
| VSMTNSGVS | ASSSRLGENYAEQF | 0.59049 |
| VTDVLGDLSV | ASSSRLGENYAEQF | 0.4950161 |
| VVVVGAGFAGL | ASSSRLGENYAEQF | 0.77499306 |
| VYTVGTVV | ASSSRLGENYAEQF | 0.3337622 |
| WTTFSGP | ASSSRLGENYAEQF | 0.4207716 |
| YDELVEAF | ASSSRLGENYAEQF | 0.55724835 |
| YGTGGAGGNGG | ASSSRLGENYAEQF | 0.5101237 |
| YTNVMAV | ASSSRLGENYAEQF | 0.764331 |
| FLEQVHQL | ASSSRLGENYAEQF | 0.2925178 |
| GLSLLAVAV | ASSSRLGENYAEQF | 0.6736026 |
| LEQVHQL | ASSSRLGENYAEQF | 0.46908432 |
| LNAAGAI | ASSSRLGENYAEQF | 0.29743055 |
| PAVHASL | ASSSRLGENYAEQF | 0.60506046 |

| | | |
|---|---|---|
| TELEDDA | ASSSRLGENYAEQF | 0.4816947 |
| TPPGPPN | ASSSRLGENYAEQF | 0.30008075 |
| TPPPPPPP | ASSSRLGENYAEQF | 0.7041762 |
| TQSPALI | ASSDAGQPNYAEQF | 0.31230056 |
| EFAAAFPQ | ASSDAGQPNYAEQF | 0.68342257 |
| LLSGNLDVL | ASSDAGQPNYAEQF | 0.7932428 |
| GPANLGNYNI | ASSDAGQPNYAEQF | 0.48234513 |
| TLGPTLP | ASSDAGQPNYAEQF | 0.7605541 |
| VYAYSSCCK | ASSDAGQPNYAEQF | 0.7507359 |
| APAPAPAPNI | ASSDAGQPNYAEQF | 0.70403683 |
| PGGPVPIP | ASSDAGQPNYAEQF | 0.40391245 |
| FLTVALEP | ASSDAGQPNYAEQF | 0.36810154 |
| SQYFGANL | ASSDAGQPNYAEQF | 0.8148824 |
| GGVKPAA | ASSDAGQPNYAEQF | 0.7598013 |
| GGTGGAGGAGGW | ASSDAGQPNYAEQF | 0.7403069 |
| GVGGVAGLGGAGG | ASSDAGQPNYAEQF | 0.78101486 |
| VLLVLATTI | ASSDAGQPNYAEQF | 0.7653826 |
| IGIFAAML | ASSDAGQPNYAEQF | 0.77338517 |
| AVGDLID | ASSDAGQPNYAEQF | 0.49085462 |
| NISPLPL | ASSDAGQPNYAEQF | 0.26717204 |
| VTAVFVAI | ASSDAGQPNYAEQF | 0.41614845 |
| SGDDNDAAL | ASSDAGQPNYAEQF | 0.45800295 |
| AVAALAHS | ASSDAGQPNYAEQF | 0.41029054 |
| GGVTVMATP | ASSDAGQPNYAEQF | 0.27090916 |
| AGLAANVRD | ASSDAGQPNYAEQF | 0.7792955 |
| TNRPDLIDPALLRPG | ASSDAGQPNYAEQF | 0.85091126 |
| LANLGVTAL | ASSDAGQPNYAEQF | 0.35400724 |
| TGERIEL | ASSDAGQPNYAEQF | 0.8351612 |
| NGEELAA | ASSDAGQPNYAEQF | 0.27028182 |
| VLSLVWG | ASSDAGQPNYAEQF | 0.76573086 |
| IGYALWL | ASSDAGQPNYAEQF | 0.78182936 |
| LSLLLAPL | ASSDAGQPNYAEQF | 0.37251866 |
| TLDEALK | ASSDAGQPNYAEQF | 0.63855445 |
| TDGAGTAEGG | ASSDAGQPNYAEQF | 0.2933702 |
| NAPGGAGGNGG | ASSDAGQPNYAEQF | 0.4801945 |
| VSVFMGMGGNF | ASSDAGQPNYAEQF | 0.7814789 |
| LASYLAAL | ASSDAGQPNYAEQF | 0.42610782 |
| LVGGTPAL | ASSDAGQPNYAEQF | 0.6420196 |
| VRNADVL | ASSDAGQPNYAEQF | 0.4740415 |
| TCSSPFLVSAM | ASSDAGQPNYAEQF | 0.37261248 |
| VHITDAE | ASSDAGQPNYAEQF | 0.6085992 |
| EYAAPSF | ASSDAGQPNYAEQF | 0.75321853 |
| VVDAFHGGL | ASSDAGQPNYAEQF | 0.50321656 |
| ASVVDPQ | ASSDAGQPNYAEQF | 0.85217357 |
| SYNEFGGLS | ASSDAGQPNYAEQF | 0.2664999 |
| DPDGSVY | ASSDAGQPNYAEQF | 0.82660794 |
| MVSTIHATE | ASSDAGQPNYAEQF | 0.8180895 |
| LAAGVADPV | ASSDAGQPNYAEQF | 0.6634064 |
| TKEQIAA | ASSDAGQPNYAEQF | 0.4965641 |
| SLASPAAL | ASSDAGQPNYAEQF | 0.78538954 |
| IGLVFYGV | ASSDAGQPNYAEQF | 0.53984845 |

| | | |
|---|---|---|
| FLLDASNL | ASSDAGQPNYAEQF | 0.74396086 |
| YQVGNFP | ASSDAGQPNYAEQF | 0.78587675 |
| ESLLSQST | ASSDAGQPNYAEQF | 0.52153 |
| LFTVGFAAVG | ASSDAGQPNYAEQF | 0.70436317 |
| ATPPTPT | ASSDAGQPNYAEQF | 0.6628617 |
| TDLATLG | ASSDAGQPNYAEQF | 0.65223855 |
| SAGGTGR | ASSDAGQPNYAEQF | 0.5977716 |
| GFSNSGS | ASSDAGQPNYAEQF | 0.34698498 |
| GGNGGSGGTSV | ASSDAGQPNYAEQF | 0.615165 |
| GTSVATGGAGN | ASSDAGQPNYAEQF | 0.54569894 |
| NGGNGGSGGTS | ASSDAGQPNYAEQF | 0.6860366 |
| AVVKFPH | ASSDAGQPNYAEQF | 0.66324043 |
| TPPPPPP | ASSDAGQPNYAEQF | 0.5829394 |
| AAPAAAAAPA | ASSDAGQPNYAEQF | 0.7117859 |
| IALNEDL | ASSDAGQPNYAEQF | 0.7124304 |
| IVNVSSM | ASSDAGQPNYAEQF | 0.42341924 |
| TGIPPAPR | ASSDAGQPNYAEQF | 0.644466 |
| FDLGGGTF | ASSDAGQPNYAEQF | 0.56126356 |
| FDLGGGTFDVS | ASSDAGQPNYAEQF | 0.623703 |
| NEPTAAA | ASSDAGQPNYAEQF | 0.36465508 |
| NEPTAAALA | ASSDAGQPNYAEQF | 0.6866428 |
| GSIINISSI | ASSDAGQPNYAEQF | 0.60263526 |
| VAPGNAGTA | ASSDAGQPNYAEQF | 0.27125156 |
| TGGGASLE | ASSDAGQPNYAEQF | 0.66184676 |
| VAVYNHYKRI | ASSDAGQPNYAEQF | 0.6681076 |
| TDPAPATT | ASSDAGQPNYAEQF | 0.6073786 |
| TNRPDILDPALLRPG | ASSDAGQPNYAEQF | 0.6991615 |
| LTTTTTS | ASSDAGQPNYAEQF | 0.382475 |
| PTGSGKTL | ASSDAGQPNYAEQF | 0.53788835 |
| TGSGKTL | ASSDAGQPNYAEQF | 0.6969292 |
| TIGHVDHGK | ASSDAGQPNYAEQF | 0.70451283 |
| TIGHVDHGKTT | ASSDAGQPNYAEQF | 0.45733622 |
| GHVDHGKTT | ASSDAGQPNYAEQF | 0.6759937 |
| AQFEHTLL | ASSDAGQPNYAEQF | 0.70435333 |
| IVNQDEL | ASSDAGQPNYAEQF | 0.6512083 |
| TNGSQFF | ASSDAGQPNYAEQF | 0.68537325 |
| TGKGIVE | ASSDAGQPNYAEQF | 0.7012644 |
| LISNASDA | ASSDAGQPNYAEQF | 0.5886717 |
| AELRTLGELTKIAKAHGAQ | ASSDAGQPNYAEQF | 0.7002444 |
| APNAVAAAEPSW | ASSDAGQPNYAEQF | 0.7093983 |
| APQVAGAATAVL | ASSDAGQPNYAEQF | 0.60555464 |
| ARRDPAAAAAILPTDA | ASSDAGQPNYAEQF | 0.66670686 |
| DAAARDVLAEAGLAEYF | ASSDAGQPNYAEQF | 0.4718397 |
| DDQIVLSAKANLAKVEQ | ASSDAGQPNYAEQF | 0.62906873 |
| DELLPDTGEDSDEDGH | ASSDAGQPNYAEQF | 0.3380384 |
| ELPEAPANRATVPQD | ASSDAGQPNYAEQF | 0.6810018 |
| FGQSKIERVTGIG | ASSDAGQPNYAEQF | 0.6893041 |
| GAGGTGSTAGGAGGAGGAGGLY | ASSDAGQPNYAEQF | 0.6212328 |
| GHGGTGAAGGNGGTGGAGGSGI | ASSDAGQPNYAEQF | 0.5052308 |
| GNGGNGGAGGEGGPGGAGGAGA | ASSDAGQPNYAEQF | 0.4778601 |
| GSTAGNGGNGGNATNGTIAGQP | ASSDAGQPNYAEQF | 0.6057776 |

| | | |
|---|---|---|
| GTGGDGGNGGNGGTGG | ASSDAGQPNYAEQF | 0.6363251 |
| GVRRPAGIPADAPV | ASSDAGQPNYAEQF | 0.3182028 |
| GYRQVTDAHLLTLARR | ASSDAGQPNYAEQF | 0.61071897 |
| IAKMGQDGHDRG | ASSDAGQPNYAEQF | 0.7047769 |
| KAEPSIEIQYT | ASSDAGQPNYAEQF | 0.69531035 |
| KPVRIRMAADEDTVAVT | ASSDAGQPNYAEQF | 0.4276459 |
| MPPRIAGMRLLVIKPEP | ASSDAGQPNYAEQF | 0.70874643 |
| MKFYAVALFRSRGVPPTR | ASSDAGQPNYAEQF | 0.70066744 |
| NAGGFGILGAVAHSP | ASSDAGQPNYAEQF | 0.6830212 |
| NEGREITLAMAEM | ASSDAGQPNYAEQF | 0.58048654 |
| NELVEWSHPLGHRWRWE | ASSDAGQPNYAEQF | 0.47323078 |
| NLGAGGTGGNGGNPG | ASSDAGQPNYAEQF | 0.6966499 |
| NTEWPKWETVMAADPVD | ASSDAGQPNYAEQF | 0.6408967 |
| PASAAPAETQP | ASSDAGQPNYAEQF | 0.47402382 |
| QQSSDGKNGVRAAEITGE | ASSDAGQPNYAEQF | 0.70299065 |
| RAPDAVTTSWWKE | ASSDAGQPNYAEQF | 0.614312 |
| SGFGNVGAHTSGWFNQ | ASSDAGQPNYAEQF | 0.61949086 |
| SGVAGAVGQAASVGGLKV | ASSDAGQPNYAEQF | 0.6386831 |
| SNGGAGTGSAGGNGGT | ASSDAGQPNYAEQF | 0.6779282 |
| TGGAGGEGGAAG | ASSDAGQPNYAEQF | 0.58541 |
| TGGDGGAAGAGGAPGHG | ASSDAGQPNYAEQF | 0.41723534 |
| TMPPLLAAIANAGRHGV | ASSDAGQPNYAEQF | 0.53364164 |
| TSSTAGNSGGA | ASSDAGQPNYAEQF | 0.65707785 |
| VPATVPVGILC | ASSDAGQPNYAEQF | 0.69959486 |
| VQRPDSGGVVLTGRIS | ASSDAGQPNYAEQF | 0.62651986 |
| WTPNPVPATKGVPAGLVQAT | ASSDAGQPNYAEQF | 0.59569514 |
| YALKRALAKFSDRFDVVII | ASSDAGQPNYAEQF | 0.6862606 |
| YLIGSVIV | ASSDAGQPNYAEQF | 0.6341888 |
| YGQLVAVID | ASSDAGQPNYAEQF | 0.6546676 |
| VTVGGMTLPA | ASSDAGQPNYAEQF | 0.7006905 |
| VLALPAV | ASSDAGQPNYAEQF | 0.60148835 |
| VIPPTLNYET | ASSDAGQPNYAEQF | 0.68633956 |
| VGMGGMHPSSG | ASSDAGQPNYAEQF | 0.6763197 |
| VGLAGGAATA | ASSDAGQPNYAEQF | 0.6815145 |
| VGGAGGAGGSGP | ASSDAGQPNYAEQF | 0.6942231 |
| VFAAFMV | ASSDAGQPNYAEQF | 0.6306404 |
| TLYDKLR | ASSDAGQPNYAEQF | 0.646117 |
| TLAVFTVL | ASSDAGQPNYAEQF | 0.30062595 |
| TGSVQPARV | ASSDAGQPNYAEQF | 0.6184306 |
| TGGGAGSGGT | ASSDAGQPNYAEQF | 0.6902642 |
| SHLETSL | ASSDAGQPNYAEQF | 0.6750395 |
| SFTTGMT | ASSDAGQPNYAEQF | 0.69420546 |
| QVEVAKRMGVRQPTVS | ASSDAGQPNYAEQF | 0.64375806 |
| QGSGGPALAM | ASSDAGQPNYAEQF | 0.58812296 |
| MLAPSNP | ASSDAGQPNYAEQF | 0.7031678 |
| LVNNAAI | ASSDAGQPNYAEQF | 0.6975076 |
| LTGEINTLA | ASSDAGQPNYAEQF | 0.40709516 |
| LSIFGPV | ASSDAGQPNYAEQF | 0.63100815 |
| LLALLANL | ASSDAGQPNYAEQF | 0.71171963 |
| LISNASDALDKLR | ASSDAGQPNYAEQF | 0.6756034 |
| LINEAVP | ASSDAGQPNYAEQF | 0.6480204 |

| | | |
|---|---|---|
| LGYAFSGL | ASSDAGQPNYAEQF | 0.53406376 |
| LDFVAVAL | ASSDAGQPNYAEQF | 0.71100336 |
| LAQPGAE | ASSDAGQPNYAEQF | 0.5765693 |
| ISNLGAVTGF | ASSDAGQPNYAEQF | 0.7122906 |
| HQEDPGRI | ASSDAGQPNYAEQF | 0.28705293 |
| GTPGEPTP | ASSDAGQPNYAEQF | 0.49817443 |
| GFFNSGS | ASSDAGQPNYAEQF | 0.38639346 |
| GAGGGGLTAGAN | ASSDAGQPNYAEQF | 0.31914794 |
| FVDALVNM | ASSDAGQPNYAEQF | 0.6877544 |
| FLTVAPEL | ASSDAGQPNYAEQF | 0.58157027 |
| FETLTAL | ASSDAGQPNYAEQF | 0.7016163 |
| DVINPIEL | ASSDAGQPNYAEQF | 0.66342795 |
| APVSGAPV | ASSDAGQPNYAEQF | 0.69424695 |
| ALVGGLAW | ASSDAGQPNYAEQF | 0.6597947 |
| ALAAAAATAI | ASSDAGQPNYAEQF | 0.68502015 |
| AFGLGLLL | ASSDAGQPNYAEQF | 0.4073802 |
| AFFAAASL | ASSDAGQPNYAEQF | 0.69406617 |
| AAAALPAFSPP | ASSDAGQPNYAEQF | 0.7090547 |
| VVLAPAL | ASSDAGQPNYAEQF | 0.61307704 |
| TIGHVDHGKTTL | ASSDAGQPNYAEQF | 0.4877894 |
| LISNASDALDK | ASSDAGQPNYAEQF | 0.6335418 |
| LISNASDAL | ASSDAGQPNYAEQF | 0.6645125 |
| ISNASDALDK | ASSDAGQPNYAEQF | 0.2812177 |
| NQAETLVYQTE | ASSDAGQPNYAEQF | 0.2878893 |
| TGGTGGTAGAG | ASSDAGQPNYAEQF | 0.60198 |
| TGSTAGGAGGA | ASSDAGQPNYAEQF | 0.53455645 |
| DPLAVPVAAGPV | ASSDAGQPNYAEQF | 0.7082766 |
| GGTGGTPGNGGN | ASSDAGQPNYAEQF | 0.6964581 |
| PAGLAGVDPATL | ASSDAGQPNYAEQF | 0.6519865 |
| GNNLVVEAPHAGASV | ASSDAGQPNYAEQF | 0.67919403 |
| KVTRIMSASTLEQPA | ASSDAGQPNYAEQF | 0.6835133 |
| LGFALPIALPRRTVV | ASSDAGQPNYAEQF | 0.2584307 |
| TRLGNLTARKLVVLV | ASSDAGQPNYAEQF | 0.5566823 |
| WSQQRRLIYFPSAGP | ASSDAGQPNYAEQF | 0.4814415 |
| GPNGNPDPMAAAVDIR | ASSDAGQPNYAEQF | 0.7102853 |
| TQRCVSVADLRAAASR | ASSDAGQPNYAEQF | 0.5733207 |
| QFVLDVAEHPVEVLTETV | ASSDAGQPNYAEQF | 0.60836935 |
| GGAGGTGGAGGNGGTQAGDG | ASSDAGQPNYAEQF | 0.66679275 |
| PLLATLRAIAVDVDAKPAQV | ASSDAGQPNYAEQF | 0.67871475 |
| APKFRTGGYMIDFWGVGYQVAK | ASSDAGQPNYAEQF | 0.65124583 |
| GAGGAGGAGGAAGGPGGTGGTG | ASSDAGQPNYAEQF | 0.4663593 |
| IGALGLLATDVAPAGSVAFGVA | ASSDAGQPNYAEQF | 0.2624474 |
| AGGAGGIGGNGGAGGAGGVLMG | ASSDAGQPNYAEQF | 0.69811624 |
| ADELIGLL | ASSDAGQPNYAEQF | 0.35144922 |
| ADRALKA | ASSDAGQPNYAEQF | 0.52282643 |
| AGAVGQVGGAGGA | ASSDAGQPNYAEQF | 0.33688933 |
| AGGGGGLGGAP | ASSDAGQPNYAEQF | 0.27536315 |
| AGGLLYGNGGAG | ASSDAGQPNYAEQF | 0.3822167 |
| AGLESIGL | ASSDAGQPNYAEQF | 0.68002415 |
| AGLKTGII | ASSDAGQPNYAEQF | 0.64038515 |
| ASLAALLL | ASSDAGQPNYAEQF | 0.64335376 |

| | | |
|---|---|---|
| ASVINISSI | ASSDAGQPNYAEQF | 0.7120571 |
| ATPPGPA | ASSDAGQPNYAEQF | 0.67974377 |
| ATVASALV | ASSDAGQPNYAEQF | 0.64418447 |
| ATVASLIL | ASSDAGQPNYAEQF | 0.5826507 |
| AVNGPVLIRH | ASSDAGQPNYAEQF | 0.55626124 |
| DGAAGAVGI | ASSDAGQPNYAEQF | 0.46339944 |
| DPLLHGI | ASSDAGQPNYAEQF | 0.53169644 |
| ELGAALGAL | ASSDAGQPNYAEQF | 0.7143367 |
| ELGRTQAV | ASSDAGQPNYAEQF | 0.49561828 |
| FVAGGYLS | ASSDAGQPNYAEQF | 0.42134246 |
| GASGGAGGAGGW | ASSDAGQPNYAEQF | 0.53258026 |
| GGAGAGGDGGAGGI | ASSDAGQPNYAEQF | 0.7099323 |
| GTGHNGSQPGMGGNG | ASSDAGQPNYAEQF | 0.7092035 |
| GYAAASASAS | ASSDAGQPNYAEQF | 0.5839027 |
| HQVDVAAVQQG | ASSDAGQPNYAEQF | 0.55321467 |
| IGAAIAHT | ASSDAGQPNYAEQF | 0.5691153 |
| IGMIEVAQL | ASSDAGQPNYAEQF | 0.67074454 |
| IRLAPEHTSAMD | ASSDAGQPNYAEQF | 0.50569916 |
| IYNVSVL | ASSDAGQPNYAEQF | 0.40068173 |
| KQGFLGM | ASSDAGQPNYAEQF | 0.6670345 |
| LESVITQL | ASSDAGQPNYAEQF | 0.71599436 |
| LGLGALVVV | ASSDAGQPNYAEQF | 0.5654321 |
| LIGNGGNGGSGGI | ASSDAGQPNYAEQF | 0.39866364 |
| LNAINAPTL | ASSDAGQPNYAEQF | 0.66920006 |
| LQLLNTGV | ASSDAGQPNYAEQF | 0.56813973 |
| LQTSIEF | ASSDAGQPNYAEQF | 0.6221452 |
| MSFAVSEL | ASSDAGQPNYAEQF | 0.62783736 |
| NTGALGL | ASSDAGQPNYAEQF | 0.44869208 |
| PPGAGGAGGAGTT | ASSDAGQPNYAEQF | 0.6030679 |
| QLVGLGI | ASSDAGQPNYAEQF | 0.708192 |
| RFLEQVHQL | ASSDAGQPNYAEQF | 0.64021146 |
| RGVGYVLR | ASSDAGQPNYAEQF | 0.6789602 |
| SATGVFAGVI | ASSDAGQPNYAEQF | 0.7038535 |
| SLPAITLP | ASSDAGQPNYAEQF | 0.6915473 |
| STALAFTV | ASSDAGQPNYAEQF | 0.70415694 |
| SVLTVGLGK | ASSDAGQPNYAEQF | 0.701496 |
| TALAGAGSGI | ASSDAGQPNYAEQF | 0.6282451 |
| TGLPFTGI | ASSDAGQPNYAEQF | 0.6265038 |
| VAEHVTV | ASSDAGQPNYAEQF | 0.6724639 |
| VDIVLNSL | ASSDAGQPNYAEQF | 0.68670166 |
| VLTGGGAVL | ASSDAGQPNYAEQF | 0.64245677 |
| VNVGLAYGI | ASSDAGQPNYAEQF | 0.52994853 |
| VSSAAVGAV | ASSDAGQPNYAEQF | 0.705779 |
| VTSVYEAGV | ASSDAGQPNYAEQF | 0.5801129 |
| WVALDRGA | ASSDAGQPNYAEQF | 0.6964785 |
| YLVGSVVI | ASSDAGQPNYAEQF | 0.48974943 |
| GPPGTGKTLL | ASSDAGQPNYAEQF | 0.49099836 |
| GVGMTKF | ASSDAGQPNYAEQF | 0.69456965 |
| IGIGGSDL | ASSDAGQPNYAEQF | 0.25669748 |
| IGTLLAEL | ASSDAGQPNYAEQF | 0.29289123 |
| IYTSGTT | ASSDAGQPNYAEQF | 0.62910116 |

| | | |
|---|---|---|
| LSFPAGV | ASSDAGQPNYAEQF | 0.70409214 |
| LYTSGTT | ASSDAGQPNYAEQF | 0.2642996 |
| VGAFPFGF | ASSDAGQPNYAEQF | 0.51737314 |
| VGAGTVEF | ASSDAGQPNYAEQF | 0.41675496 |
| VGKSTLLL | ASSDAGQPNYAEQF | 0.62563014 |
| AALAHAGGGIV | ASSDAGQPNYAEQF | 0.6407592 |
| AALVAVSLL | ASSDAGQPNYAEQF | 0.38864204 |
| AAVSRAVAAL | ASSDAGQPNYAEQF | 0.68038344 |
| AGGSSGIGGF | ASSDAGQPNYAEQF | 0.3954341 |
| AGLTVANANG | ASSDAGQPNYAEQF | 0.43868575 |
| AGLVGAVL | ASSDAGQPNYAEQF | 0.67876744 |
| ALSQAFTV | ASSDAGQPNYAEQF | 0.7039504 |
| ALTFSVSL | ASSDAGQPNYAEQF | 0.51753 |
| ALTGAALVAL | ASSDAGQPNYAEQF | 0.6832347 |
| ASFSIGAGVL | ASSDAGQPNYAEQF | 0.558648 |
| DGSVYKI | ASSDAGQPNYAEQF | 0.70924544 |
| EQLAEIV | ASSDAGQPNYAEQF | 0.6170659 |
| FAVSELTVM | ASSDAGQPNYAEQF | 0.34968215 |
| GAAPLAAGL | ASSDAGQPNYAEQF | 0.68759084 |
| GADGGDGGSGGAGG | ASSDAGQPNYAEQF | 0.6665943 |
| GFGLDTGGAGG | ASSDAGQPNYAEQF | 0.62934864 |
| GLASGGLL | ASSDAGQPNYAEQF | 0.69628096 |
| GRIAGAAVLL | ASSDAGQPNYAEQF | 0.44821626 |
| GSAGSLNTSF | ASSDAGQPNYAEQF | 0.65347564 |
| GTGQIGI | ASSDAGQPNYAEQF | 0.70669675 |
| GTLIALIL | ASSDAGQPNYAEQF | 0.7066726 |
| IAATMGFVVG | ASSDAGQPNYAEQF | 0.42448786 |
| IFAQALL | ASSDAGQPNYAEQF | 0.65603566 |
| IGAAVALGF | ASSDAGQPNYAEQF | 0.6628225 |
| IGLTPLAL | ASSDAGQPNYAEQF | 0.5747 |
| ILGSNTSTL | ASSDAGQPNYAEQF | 0.70257425 |
| IVFSLTVL | ASSDAGQPNYAEQF | 0.6011944 |
| LASKATGF | ASSDAGQPNYAEQF | 0.6903231 |
| LASVALIL | ASSDAGQPNYAEQF | 0.6687131 |
| LFAVGLTL | ASSDAGQPNYAEQF | 0.7090218 |
| LFNGGIGTYI | ASSDAGQPNYAEQF | 0.3958197 |
| LTLGAGII | ASSDAGQPNYAEQF | 0.67214507 |
| LVAAALGDI | ASSDAGQPNYAEQF | 0.6560463 |
| NTAFVQLGI | ASSDAGQPNYAEQF | 0.6932703 |
| QLFSGAEL | ASSDAGQPNYAEQF | 0.60785323 |
| SAATFITL | ASSDAGQPNYAEQF | 0.4238202 |
| SNYGLANLGV | ASSDAGQPNYAEQF | 0.66063845 |
| SYGGAYAVM | ASSDAGQPNYAEQF | 0.70115316 |
| TTGVGGLTLG | ASSDAGQPNYAEQF | 0.64625114 |
| TTQTIYI | ASSDAGQPNYAEQF | 0.6825137 |
| VAAGAGLGAVI | ASSDAGQPNYAEQF | 0.7093959 |
| VAAVVTLL | ASSDAGQPNYAEQF | 0.49478862 |
| VGFLGGLI | ASSDAGQPNYAEQF | 0.6480199 |
| VGVIPILL | ASSDAGQPNYAEQF | 0.6830621 |
| VIFASSLI | ASSDAGQPNYAEQF | 0.7017754 |
| VKGGDGGAAAG | ASSDAGQPNYAEQF | 0.57821524 |

| | | |
|---|---|---|
| VTKTFTNI | ASSDAGQPNYAEQF | 0.7090218 |
| VVAAAILL | ASSDAGQPNYAEQF | 0.54868853 |
| YAALDVEL | ASSDAGQPNYAEQF | 0.7023004 |
| LASVALLL | ASSDAGQPNYAEQF | 0.59266955 |
| LDAVHRV | ASSDAGQPNYAEQF | 0.6875597 |
| QGLSSSIF | ASSDAGQPNYAEQF | 0.57017815 |
| RGTFANIRL | ASSDAGQPNYAEQF | 0.7027027 |
| RQLWWGHRI | ASSDAGQPNYAEQF | 0.6938252 |
| YGPPGTGKTLL | ASSDAGQPNYAEQF | 0.70769477 |
| AALVGAPANVA | ASSDAGQPNYAEQF | 0.28272268 |
| AATFRRTE | ASSDAGQPNYAEQF | 0.6947386 |
| AGALGGGI | ASSDAGQPNYAEQF | 0.7066027 |
| AGGAGGAGGIGAG | ASSDAGQPNYAEQF | 0.71521825 |
| AGSGALGL | ASSDAGQPNYAEQF | 0.60160136 |
| AQIADQI | ASSDAGQPNYAEQF | 0.6771649 |
| ASAVAVTYL | ASSDAGQPNYAEQF | 0.62555873 |
| ASTVLGGTGG | ASSDAGQPNYAEQF | 0.54760236 |
| AVGGEPAK | ASSDAGQPNYAEQF | 0.5694382 |
| CEQGTFGGTC | ASSDAGQPNYAEQF | 0.56557816 |
| DEGKAPAG | ASSDAGQPNYAEQF | 0.64153963 |
| DTVVFPAL | ASSDAGQPNYAEQF | 0.59089756 |
| FEGITMAG | ASSDAGQPNYAEQF | 0.6974411 |
| FPLVVQD | ASSDAGQPNYAEQF | 0.70169294 |
| GAFTTPQ | ASSDAGQPNYAEQF | 0.6957768 |
| GFGGFGTAGGVGGA | ASSDAGQPNYAEQF | 0.70375943 |
| GFLAQFL | ASSDAGQPNYAEQF | 0.7018744 |
| GGAGGAGGNAGVA | ASSDAGQPNYAEQF | 0.6698839 |
| GGVGTAGAGGAGGAGG | ASSDAGQPNYAEQF | 0.56061906 |
| GNGGGGDAGNAGSGGN | ASSDAGQPNYAEQF | 0.64785886 |
| GTGGAGGAGNDAGS | ASSDAGQPNYAEQF | 0.5344113 |
| GTTNTGWFNS | ASSDAGQPNYAEQF | 0.6325941 |
| IFMGPLFTL | ASSDAGQPNYAEQF | 0.5831681 |
| ILAHAIEAI | ASSDAGQPNYAEQF | 0.67697996 |
| IRSAIREAL | ASSDAGQPNYAEQF | 0.2559356 |
| ITFAGGG | ASSDAGQPNYAEQF | 0.6773238 |
| ITGMSGHQEV | ASSDAGQPNYAEQF | 0.71193534 |
| IVAGLACTL | ASSDAGQPNYAEQF | 0.57265687 |
| IVGGGLAGA | ASSDAGQPNYAEQF | 0.6686357 |
| IVTGVLNI | ASSDAGQPNYAEQF | 0.653944 |
| LAQQGCI | ASSDAGQPNYAEQF | 0.59022033 |
| LCLHVEN | ASSDAGQPNYAEQF | 0.6945398 |
| LDAISDGS | ASSDAGQPNYAEQF | 0.5131404 |
| LGGFAAML | ASSDAGQPNYAEQF | 0.6578831 |
| LGGKNPMI | ASSDAGQPNYAEQF | 0.7033719 |
| LGPKGVAAIL | ASSDAGQPNYAEQF | 0.6627359 |
| LMVGVSI | ASSDAGQPNYAEQF | 0.6721983 |
| LVGAAFVL | ASSDAGQPNYAEQF | 0.5857064 |
| LVLAVAPL | ASSDAGQPNYAEQF | 0.3455019 |
| MAAIAVIDAA | ASSDAGQPNYAEQF | 0.6882138 |
| NNGINAELQ | ASSDAGQPNYAEQF | 0.256617 |
| RGEPLVLVHG | ASSDAGQPNYAEQF | 0.65679586 |

| | | |
|---|---|---|
| RGGAGGAAGLI | ASSDAGQPNYAEQF | 0.48216078 |
| RSYRMHPAV | ASSDAGQPNYAEQF | 0.68332875 |
| SAAHLLAGQLA | ASSDAGQPNYAEQF | 0.6745914 |
| SAAVGSY | ASSDAGQPNYAEQF | 0.67265487 |
| SPHAVLHRI | ASSDAGQPNYAEQF | 0.60451794 |
| SQIAKVL | ASSDAGQPNYAEQF | 0.63873756 |
| TALGHLVP | ASSDAGQPNYAEQF | 0.5786245 |
| TAPTTSGAA | ASSDAGQPNYAEQF | 0.42066818 |
| TDGDDGNGGSPGT | ASSDAGQPNYAEQF | 0.682003 |
| TDGLGEN | ASSDAGQPNYAEQF | 0.5508034 |
| TGGPPLPGQA | ASSDAGQPNYAEQF | 0.6190276 |
| TIPAITI | ASSDAGQPNYAEQF | 0.7037843 |
| TIPGFTI | ASSDAGQPNYAEQF | 0.63376915 |
| TSIASGALA | ASSDAGQPNYAEQF | 0.35166496 |
| TVTLPGAAI | ASSDAGQPNYAEQF | 0.647037 |
| VASGLSAALP | ASSDAGQPNYAEQF | 0.66838336 |
| VGAGVIGI | ASSDAGQPNYAEQF | 0.6135622 |
| VGPKRFGV | ASSDAGQPNYAEQF | 0.25744772 |
| VNVAFPD | ASSDAGQPNYAEQF | 0.67492604 |
| VSMTNSGVS | ASSDAGQPNYAEQF | 0.59803784 |
| VTDVLGDLSV | ASSDAGQPNYAEQF | 0.56876117 |
| VVVVGAGFAGL | ASSDAGQPNYAEQF | 0.54481554 |
| VYTVGTVV | ASSDAGQPNYAEQF | 0.69518805 |
| WTTFSGP | ASSDAGQPNYAEQF | 0.38902524 |
| YDELVEAF | ASSDAGQPNYAEQF | 0.6363802 |
| YGTGGAGGNGG | ASSDAGQPNYAEQF | 0.6938448 |
| YTNVMAV | ASSDAGQPNYAEQF | 0.7050973 |
| FLEQVHQL | ASSDAGQPNYAEQF | 0.7101182 |
| GLSLLAVAV | ASSDAGQPNYAEQF | 0.6557464 |
| LEQVHQL | ASSDAGQPNYAEQF | 0.49742725 |
| LNAAGAI | ASSDAGQPNYAEQF | 0.6928717 |
| PAVHASL | ASSDAGQPNYAEQF | 0.58518296 |
| TELEDDA | ASSDAGQPNYAEQF | 0.39384687 |
| TPPGPPN | ASSDAGQPNYAEQF | 0.34202713 |
| TPPPPPPP | ASSDAGQPNYAEQF | 0.64795625 |
| TQSPALI | ASSAGGAQNTLY | 0.37207693 |
| EFAAAFPQ | ASSAGGAQNTLY | 0.66380703 |
| LLSGNLDVL | ASSAGGAQNTLY | 0.6862747 |
| GPANLGNYNI | ASSAGGAQNTLY | 0.5903717 |
| TLGPTLP | ASSAGGAQNTLY | 0.39132276 |
| VYAYSSCCK | ASSAGGAQNTLY | 0.627665 |
| APAPAPAPNI | ASSAGGAQNTLY | 0.60788566 |
| PGGPVPIP | ASSAGGAQNTLY | 0.6858823 |
| FLTVALEP | ASSAGGAQNTLY | 0.64373815 |
| SQYFGANL | ASSAGGAQNTLY | 0.7146542 |
| GGVKPAA | ASSAGGAQNTLY | 0.70088315 |
| GGTGGAGGAGGW | ASSAGGAQNTLY | 0.63242626 |
| GVGGVAGLGGAGG | ASSAGGAQNTLY | 0.687632 |
| VLLVLATTI | ASSAGGAQNTLY | 0.59438634 |
| IGIFAAML | ASSAGGAQNTLY | 0.62255794 |
| AVGDLID | ASSAGGAQNTLY | 0.62990904 |

| | | |
|---|---|---|
| NISPLPL | ASSAGGAQNTLY | 0.6388837 |
| VTAVFVAI | ASSAGGAQNTLY | 0.62975776 |
| SGDDNDAAL | ASSAGGAQNTLY | 0.6153294 |
| AVAALAHS | ASSAGGAQNTLY | 0.3745258 |
| GGVTVMATP | ASSAGGAQNTLY | 0.69541067 |
| AGLAANVRD | ASSAGGAQNTLY | 0.6610719 |
| TNRPDLIDPALLRPG | ASSAGGAQNTLY | 0.34941432 |
| LANLGVTAL | ASSAGGAQNTLY | 0.6518666 |
| TGERIEL | ASSAGGAQNTLY | 0.7101115 |
| NGEELAA | ASSAGGAQNTLY | 0.7043177 |
| VLSLVWG | ASSAGGAQNTLY | 0.5487845 |
| IGYALWL | ASSAGGAQNTLY | 0.52376294 |
| LSLLLAPL | ASSAGGAQNTLY | 0.67226297 |
| TLDEALK | ASSAGGAQNTLY | 0.3479883 |
| TDGAGTAEGG | ASSAGGAQNTLY | 0.64137876 |
| NAPGGAGGNGG | ASSAGGAQNTLY | 0.6699922 |
| VSVFMGMGGNF | ASSAGGAQNTLY | 0.65282464 |
| LASYLAAL | ASSAGGAQNTLY | 0.5190157 |
| LVGGTPAL | ASSAGGAQNTLY | 0.66017413 |
| VRNADVL | ASSAGGAQNTLY | 0.5926874 |
| TCSSPFLVSAM | ASSAGGAQNTLY | 0.66233635 |
| VHITDAE | ASSAGGAQNTLY | 0.26357266 |
| EYAAPSF | ASSAGGAQNTLY | 0.62205243 |
| VVDAFHGGL | ASSAGGAQNTLY | 0.60918134 |
| ASVVDPQ | ASSAGGAQNTLY | 0.44208774 |
| SYNEFGGLS | ASSAGGAQNTLY | 0.31671637 |
| DPDGSVY | ASSAGGAQNTLY | 0.63472855 |
| MVSTIHATE | ASSAGGAQNTLY | 0.6817791 |
| LAAGVADPV | ASSAGGAQNTLY | 0.6930838 |
| TKEQIAA | ASSAGGAQNTLY | 0.5322688 |
| SLASPAAL | ASSAGGAQNTLY | 0.70446265 |
| IGLVFYGV | ASSAGGAQNTLY | 0.6952157 |
| FLLDASNL | ASSAGGAQNTLY | 0.63398206 |
| YQVGNFP | ASSAGGAQNTLY | 0.359874 |
| ESLLSQST | ASSAGGAQNTLY | 0.6751109 |
| LFTVGFAAVG | ASSAGGAQNTLY | 0.6584791 |
| ATPPTPT | ASSAGGAQNTLY | 0.67835987 |
| TDLATLG | ASSAGGAQNTLY | 0.68641967 |
| SAGGTGR | ASSAGGAQNTLY | 0.5527954 |
| GFSNSGS | ASSAGGAQNTLY | 0.69447726 |
| GGNGGSGGTSV | ASSAGGAQNTLY | 0.7033996 |
| GTSVATGGAGN | ASSAGGAQNTLY | 0.5894954 |
| NGGNGGSGGTS | ASSAGGAQNTLY | 0.48974335 |
| AVVKFPH | ASSAGGAQNTLY | 0.49921674 |
| TPPPPPP | ASSAGGAQNTLY | 0.29671842 |
| AAPAAAAAPA | ASSAGGAQNTLY | 0.70437974 |
| IALNEDL | ASSAGGAQNTLY | 0.63110197 |
| IVNVSSM | ASSAGGAQNTLY | 0.53938967 |
| TGIPPAPR | ASSAGGAQNTLY | 0.67976296 |
| FDLGGGTF | ASSAGGAQNTLY | 0.55769455 |
| FDLGGGTFDVS | ASSAGGAQNTLY | 0.59782565 |

| | | |
|---|---|---|
| NEPTAAA | ASSAGGAQNTLY | 0.5784998 |
| NEPTAAALA | ASSAGGAQNTLY | 0.6072346 |
| GSIINISSI | ASSAGGAQNTLY | 0.67024356 |
| VAPGNAGTA | ASSAGGAQNTLY | 0.638713 |
| TGGGASLE | ASSAGGAQNTLY | 0.6348491 |
| VAVYNHYKRI | ASSAGGAQNTLY | 0.70823336 |
| TDPAPATT | ASSAGGAQNTLY | 0.4100594 |
| TNRPDILDPALLRPG | ASSAGGAQNTLY | 0.595749 |
| LTTTTTS | ASSAGGAQNTLY | 0.6650114 |
| PTGSGKTL | ASSAGGAQNTLY | 0.41860384 |
| TGSGKTL | ASSAGGAQNTLY | 0.6843474 |
| TIGHVDHGK | ASSAGGAQNTLY | 0.679664 |
| TIGHVDHGKTT | ASSAGGAQNTLY | 0.6822144 |
| GHVDHGKTT | ASSAGGAQNTLY | 0.6266305 |
| AQFEHTLL | ASSAGGAQNTLY | 0.71489096 |
| IVNQDEL | ASSAGGAQNTLY | 0.691703 |
| TNGSQFF | ASSAGGAQNTLY | 0.60851103 |
| TGKGIVE | ASSAGGAQNTLY | 0.6740196 |
| LISNASDA | ASSAGGAQNTLY | 0.45792896 |
| AELRTLGELTKIAKAHGAQ | ASSAGGAQNTLY | 0.7037302 |
| APNAVAAAEPSW | ASSAGGAQNTLY | 0.68460083 |
| APQVAGAATAVL | ASSAGGAQNTLY | 0.49414158 |
| ARRDPAAAAAILPTDA | ASSAGGAQNTLY | 0.6584875 |
| DAAARDVLAEAGLAEYF | ASSAGGAQNTLY | 0.6963949 |
| DDQIVLSAKANLAKVEQ | ASSAGGAQNTLY | 0.7009189 |
| DELLPDTGEDSDEDGH | ASSAGGAQNTLY | 0.35141715 |
| ELPEAPANRATVPQD | ASSAGGAQNTLY | 0.6418793 |
| FGQSKIERVTGIG | ASSAGGAQNTLY | 0.31053397 |
| GAGGTGSTAGGAGGAGGAGGLY | ASSAGGAQNTLY | 0.47575235 |
| GHGGTGAAGGNGGTGGAGGSGI | ASSAGGAQNTLY | 0.49582455 |
| GNGGNGGAGGEGGPGGAGGAGA | ASSAGGAQNTLY | 0.6768085 |
| GSTAGNGGNGGNATNGTIAGQP | ASSAGGAQNTLY | 0.63634896 |
| GTGGDGGNGGNGGTGG | ASSAGGAQNTLY | 0.6890298 |
| GVRRPAGIPADAPV | ASSAGGAQNTLY | 0.6637396 |
| GYRQVTDAHLLTLARR | ASSAGGAQNTLY | 0.7056155 |
| IAKMGQDGHDRG | ASSAGGAQNTLY | 0.5902746 |
| KAEPSIEIQYT | ASSAGGAQNTLY | 0.66956115 |
| KPVRIRMAADEDTVAVT | ASSAGGAQNTLY | 0.36205015 |
| MPPRIAGMRLLVIKPEP | ASSAGGAQNTLY | 0.32249528 |
| MKFYAVALFRSRGVPPTR | ASSAGGAQNTLY | 0.48305386 |
| NAGGFGILGAVAHSP | ASSAGGAQNTLY | 0.64915764 |
| NEGREITLAMAEM | ASSAGGAQNTLY | 0.2763569 |
| NELVEWSHPLGHRWRWE | ASSAGGAQNTLY | 0.68777937 |
| NLGAGGTGGNGGNPG | ASSAGGAQNTLY | 0.66364217 |
| NTEWPKWETVMAADPVD | ASSAGGAQNTLY | 0.6886945 |
| PASAAPAETQP | ASSAGGAQNTLY | 0.58566284 |
| QQSSDGKNGVRAAEITGE | ASSAGGAQNTLY | 0.6138254 |
| RAPDAVTTSWWKE | ASSAGGAQNTLY | 0.57275325 |
| SGFGNVGAHTSGWFNQ | ASSAGGAQNTLY | 0.6886398 |
| SGVAGAVGQAASVGGLKV | ASSAGGAQNTLY | 0.56611407 |
| SNGGAGTGSAGGNGGT | ASSAGGAQNTLY | 0.35258603 |

| | | |
|---|---|---|
| TGGAGGEGGAAG | ASSAGGAQNTLY | 0.48755714 |
| TGGDGGAAGAGGAPGHG | ASSAGGAQNTLY | 0.50578105 |
| TMPPLLAAIANAGRHGV | ASSAGGAQNTLY | 0.7008559 |
| TSSTAGNSGGA | ASSAGGAQNTLY | 0.6948763 |
| VPATVPVGILC | ASSAGGAQNTLY | 0.6818509 |
| VQRPDSGGVVLTGRIS | ASSAGGAQNTLY | 0.69820076 |
| WTPNPVPATKGVPAGLVQAT | ASSAGGAQNTLY | 0.28463575 |
| YALKRALAKFSDRFDVVII | ASSAGGAQNTLY | 0.55072236 |
| YLIGSVIV | ASSAGGAQNTLY | 0.7144971 |
| YGQLVAVID | ASSAGGAQNTLY | 0.63311714 |
| VTVGGMTLPA | ASSAGGAQNTLY | 0.6545816 |
| VLALPAV | ASSAGGAQNTLY | 0.69963336 |
| VIPPTLNYET | ASSAGGAQNTLY | 0.62923455 |
| VGMGGMHPSSG | ASSAGGAQNTLY | 0.62511504 |
| VGLAGGAATA | ASSAGGAQNTLY | 0.61483717 |
| VGGAGGAGGSGP | ASSAGGAQNTLY | 0.6832642 |
| VFAAFMV | ASSAGGAQNTLY | 0.5916824 |
| TLYDKLR | ASSAGGAQNTLY | 0.5532238 |
| TLAVFTVL | ASSAGGAQNTLY | 0.65053326 |
| TGSVQPARV | ASSAGGAQNTLY | 0.7141553 |
| TGGGAGSGGT | ASSAGGAQNTLY | 0.641049 |
| SHLETSL | ASSAGGAQNTLY | 0.68748236 |
| SFTTGMT | ASSAGGAQNTLY | 0.6626208 |
| QVEVAKRMGVRQPTVS | ASSAGGAQNTLY | 0.6834129 |
| QGSGGPALAM | ASSAGGAQNTLY | 0.5796689 |
| MLAPSNP | ASSAGGAQNTLY | 0.6201484 |
| LVNNAAI | ASSAGGAQNTLY | 0.64917946 |
| LTGEINTLA | ASSAGGAQNTLY | 0.5962645 |
| LSIFGPV | ASSAGGAQNTLY | 0.61798704 |
| LLALLANL | ASSAGGAQNTLY | 0.5855156 |
| LISNASDALDKLR | ASSAGGAQNTLY | 0.7155205 |
| LINEAVP | ASSAGGAQNTLY | 0.68453515 |
| LGYAFSGL | ASSAGGAQNTLY | 0.63127923 |
| LDFVAVAL | ASSAGGAQNTLY | 0.56900024 |
| LAQPGAE | ASSAGGAQNTLY | 0.63258064 |
| ISNLGAVTGF | ASSAGGAQNTLY | 0.6818863 |
| HQEDPGRI | ASSAGGAQNTLY | 0.39448047 |
| GTPGEPTP | ASSAGGAQNTLY | 0.7095245 |
| GFFNSGS | ASSAGGAQNTLY | 0.5915876 |
| GAGGGGLTAGAN | ASSAGGAQNTLY | 0.54455286 |
| FVDALVNM | ASSAGGAQNTLY | 0.69374126 |
| FLTVAPEL | ASSAGGAQNTLY | 0.7127664 |
| FETLTAL | ASSAGGAQNTLY | 0.57410705 |
| DVINPIEL | ASSAGGAQNTLY | 0.60242414 |
| APVSGAPV | ASSAGGAQNTLY | 0.5679978 |
| ALVGGLAW | ASSAGGAQNTLY | 0.68222165 |
| ALAAAAATAI | ASSAGGAQNTLY | 0.7015317 |
| AFGLGLLL | ASSAGGAQNTLY | 0.6444706 |
| AFFAAASL | ASSAGGAQNTLY | 0.7120012 |
| AAAALPAFSPP | ASSAGGAQNTLY | 0.5837681 |
| VVLAPAL | ASSAGGAQNTLY | 0.36218202 |

| | | |
|---|---|---|
| TIGHVDHGKTTL | ASSAGGAQNTLY | 0.6303823 |
| LISNASDALDK | ASSAGGAQNTLY | 0.6843312 |
| LISNASDAL | ASSAGGAQNTLY | 0.69392 |
| ISNASDALDK | ASSAGGAQNTLY | 0.7099656 |
| NQAETLVYQTE | ASSAGGAQNTLY | 0.67145556 |
| TGGTGGTAGAG | ASSAGGAQNTLY | 0.5848478 |
| TGSTAGGAGGA | ASSAGGAQNTLY | 0.70417845 |
| DPLAVPVAAGPV | ASSAGGAQNTLY | 0.70615566 |
| GGTGGTPGNGGN | ASSAGGAQNTLY | 0.68593967 |
| PAGLAGVDPATL | ASSAGGAQNTLY | 0.5715591 |
| GNNLVVEAPHAGASV | ASSAGGAQNTLY | 0.45471132 |
| KVTRIMSASTLEQPA | ASSAGGAQNTLY | 0.6433617 |
| LGFALPIALPRRTVV | ASSAGGAQNTLY | 0.6846212 |
| TRLGNLTARKLVVLV | ASSAGGAQNTLY | 0.4421869 |
| WSQQRRLIYFPSAGP | ASSAGGAQNTLY | 0.6715516 |
| GPNGNPDPMAAAVDIR | ASSAGGAQNTLY | 0.67352533 |
| TQRCVSVADLRAAASR | ASSAGGAQNTLY | 0.43053654 |
| QFVLDVAEHPVEVLTETV | ASSAGGAQNTLY | 0.65582895 |
| GGAGGTGGAGGNGGTQAGDG | ASSAGGAQNTLY | 0.7072983 |
| PLLATLRAIAVDVDAKPAQV | ASSAGGAQNTLY | 0.5657943 |
| APKFRTGGYMIDFWGVGYQVAK | ASSAGGAQNTLY | 0.5832038 |
| GAGGAGGAGGAAGGPGGTGGTG | ASSAGGAQNTLY | 0.699227 |
| IGALGLLATDVAPAGSVAFGVA | ASSAGGAQNTLY | 0.7161315 |
| AGGAGGIGGNGGAGGAGGVLMG | ASSAGGAQNTLY | 0.69789803 |
| ADELIGLL | ASSAGGAQNTLY | 0.408935 |
| ADRALKA | ASSAGGAQNTLY | 0.31466135 |
| AGAVGQVGGAGGA | ASSAGGAQNTLY | 0.5651792 |
| AGGGGGLGGAP | ASSAGGAQNTLY | 0.69621044 |
| AGGLLYGNGGAG | ASSAGGAQNTLY | 0.61639416 |
| AGLESIGL | ASSAGGAQNTLY | 0.5287007 |
| AGLKTGII | ASSAGGAQNTLY | 0.673506 |
| ASLAALLL | ASSAGGAQNTLY | 0.6988131 |
| ASVINISSI | ASSAGGAQNTLY | 0.50284165 |
| ATPPGPA | ASSAGGAQNTLY | 0.32786125 |
| ATVASALV | ASSAGGAQNTLY | 0.6655522 |
| ATVASLIL | ASSAGGAQNTLY | 0.679745 |
| AVNGPVLIRH | ASSAGGAQNTLY | 0.6146761 |
| DGAAGAVGI | ASSAGGAQNTLY | 0.64611834 |
| DPLLHGI | ASSAGGAQNTLY | 0.52981097 |
| ELGAALGAL | ASSAGGAQNTLY | 0.45401698 |
| ELGRTQAV | ASSAGGAQNTLY | 0.68128693 |
| FVAGGYLS | ASSAGGAQNTLY | 0.52013993 |
| GASGGAGGAGGW | ASSAGGAQNTLY | 0.7039734 |
| GGAGAGGDGGAGGI | ASSAGGAQNTLY | 0.5270411 |
| GTGHNGSQPGMGGNG | ASSAGGAQNTLY | 0.6802082 |
| GYAAASASAS | ASSAGGAQNTLY | 0.69161224 |
| HQVDVAAVQQG | ASSAGGAQNTLY | 0.68313473 |
| IGAAIAHT | ASSAGGAQNTLY | 0.62790346 |
| IGMIEVAQL | ASSAGGAQNTLY | 0.61086476 |
| IRLAPEHTSAMD | ASSAGGAQNTLY | 0.50127363 |
| IYNVSVL | ASSAGGAQNTLY | 0.36225608 |

| | | |
|---|---|---|
| KQGFLGM | ASSAGGAQNTLY | 0.7050508 |
| LESVITQL | ASSAGGAQNTLY | 0.2593652 |
| LGLGALVVV | ASSAGGAQNTLY | 0.70829546 |
| LIGNGGNGGSGGI | ASSAGGAQNTLY | 0.67706156 |
| LNAINAPTL | ASSAGGAQNTLY | 0.51030636 |
| LQLLNTGV | ASSAGGAQNTLY | 0.6675278 |
| LQTSIEF | ASSAGGAQNTLY | 0.48529756 |
| MSFAVSEL | ASSAGGAQNTLY | 0.54317796 |
| NTGALGL | ASSAGGAQNTLY | 0.683858 |
| PPGAGGAGGAGTT | ASSAGGAQNTLY | 0.67539907 |
| QLVGLGI | ASSAGGAQNTLY | 0.58982706 |
| RFLEQVHQL | ASSAGGAQNTLY | 0.63684916 |
| RGVGYVLR | ASSAGGAQNTLY | 0.6715821 |
| SATGVFAGVI | ASSAGGAQNTLY | 0.57787055 |
| SLPAITLP | ASSAGGAQNTLY | 0.59742224 |
| STALAFTV | ASSAGGAQNTLY | 0.7002034 |
| SVLTVGLGK | ASSAGGAQNTLY | 0.61001635 |
| TALAGAGSGI | ASSAGGAQNTLY | 0.57260054 |
| TGLPFTGI | ASSAGGAQNTLY | 0.67467713 |
| VAEHVTV | ASSAGGAQNTLY | 0.5555117 |
| VDIVLNSL | ASSAGGAQNTLY | 0.6889077 |
| VLTGGGAVL | ASSAGGAQNTLY | 0.6371497 |
| VNVGLAYGI | ASSAGGAQNTLY | 0.65606445 |
| VSSAAVGAV | ASSAGGAQNTLY | 0.6663071 |
| VTSVYEAGV | ASSAGGAQNTLY | 0.35612324 |
| WVALDRGA | ASSAGGAQNTLY | 0.4899215 |
| YLVGSVVI | ASSAGGAQNTLY | 0.60774153 |
| GPPGTGKTLL | ASSAGGAQNTLY | 0.6649991 |
| GVGMTKF | ASSAGGAQNTLY | 0.68887234 |
| IGIGGSDL | ASSAGGAQNTLY | 0.5832412 |
| IGTLLAEL | ASSAGGAQNTLY | 0.66447645 |
| IYTSGTT | ASSAGGAQNTLY | 0.61854994 |
| LSFPAGV | ASSAGGAQNTLY | 0.5393284 |
| LYTSGTT | ASSAGGAQNTLY | 0.5616079 |
| VGAFPFGF | ASSAGGAQNTLY | 0.5169466 |
| VGAGTVEF | ASSAGGAQNTLY | 0.4930935 |
| VGKSTLLL | ASSAGGAQNTLY | 0.6965463 |
| AALAHAGGGIV | ASSAGGAQNTLY | 0.47755128 |
| AALVAVSLL | ASSAGGAQNTLY | 0.64941204 |
| AAVSRAVAAL | ASSAGGAQNTLY | 0.71264744 |
| AGGSSGIGGF | ASSAGGAQNTLY | 0.6435894 |
| AGLTVANANG | ASSAGGAQNTLY | 0.6549444 |
| AGLVGAVL | ASSAGGAQNTLY | 0.65057313 |
| ALSQAFTV | ASSAGGAQNTLY | 0.68863964 |
| ALTFSVSL | ASSAGGAQNTLY | 0.62281394 |
| ALTGAALVAL | ASSAGGAQNTLY | 0.6385626 |
| ASFSIGAGVL | ASSAGGAQNTLY | 0.70127153 |
| DGSVYKI | ASSAGGAQNTLY | 0.68499 |
| EQLAEIV | ASSAGGAQNTLY | 0.6988151 |
| FAVSELTVM | ASSAGGAQNTLY | 0.6789489 |
| GAAPLAAGL | ASSAGGAQNTLY | 0.6904915 |

| | | |
|---|---|---|
| GADGGDGGSGGAGG | ASSAGGAQNTLY | 0.68036497 |
| GFGLDTGGAGG | ASSAGGAQNTLY | 0.64902055 |
| GLASGGLL | ASSAGGAQNTLY | 0.37997785 |
| GRIAGAAVLL | ASSAGGAQNTLY | 0.59875226 |
| GSAGSLNTSF | ASSAGGAQNTLY | 0.67723525 |
| GTGQIGI | ASSAGGAQNTLY | 0.53619415 |
| GTLIALIL | ASSAGGAQNTLY | 0.32820562 |
| IAATMGFVVG | ASSAGGAQNTLY | 0.7080624 |
| IFAQALL | ASSAGGAQNTLY | 0.71386683 |
| IGAAVALGF | ASSAGGAQNTLY | 0.5167915 |
| IGLTPLAL | ASSAGGAQNTLY | 0.7100933 |
| ILGSNTSTL | ASSAGGAQNTLY | 0.4133825 |
| IVFSLTVL | ASSAGGAQNTLY | 0.6728409 |
| LASKATGF | ASSAGGAQNTLY | 0.70638293 |
| LASVALIL | ASSAGGAQNTLY | 0.5607027 |
| LFAVGLTL | ASSAGGAQNTLY | 0.6909808 |
| LFNGGIGTYI | ASSAGGAQNTLY | 0.56972474 |
| LTLGAGII | ASSAGGAQNTLY | 0.602939 |
| LVAAALGDI | ASSAGGAQNTLY | 0.7082356 |
| NTAFVQLGI | ASSAGGAQNTLY | 0.64429307 |
| QLFSGAEL | ASSAGGAQNTLY | 0.6819931 |
| SAATFITL | ASSAGGAQNTLY | 0.6077018 |
| SNYGLANLGV | ASSAGGAQNTLY | 0.53155255 |
| SYGGAYAVM | ASSAGGAQNTLY | 0.661554 |
| TTGVGGLTLG | ASSAGGAQNTLY | 0.7116032 |
| TTQTIYI | ASSAGGAQNTLY | 0.5392888 |
| VAAGAGLGAVI | ASSAGGAQNTLY | 0.70258385 |
| VAAVVTLL | ASSAGGAQNTLY | 0.61767566 |
| VGFLGGLI | ASSAGGAQNTLY | 0.7160299 |
| VGVIPILL | ASSAGGAQNTLY | 0.6930475 |
| VIFASSLI | ASSAGGAQNTLY | 0.66988546 |
| VKGGDGGAAAG | ASSAGGAQNTLY | 0.5844064 |
| VTKTFTNI | ASSAGGAQNTLY | 0.7036998 |
| VVAAAILL | ASSAGGAQNTLY | 0.657063 |
| YAALDVEL | ASSAGGAQNTLY | 0.69536257 |
| LASVALLL | ASSAGGAQNTLY | 0.70144594 |
| LDAVHRV | ASSAGGAQNTLY | 0.7336551 |
| QGLSSSIF | ASSAGGAQNTLY | 0.8950654 |
| RGTFANIRL | ASSAGGAQNTLY | 0.8278922 |
| RQLWWGHRI | ASSAGGAQNTLY | 0.8448079 |
| YGPPGTGKTLL | ASSAGGAQNTLY | 0.72778547 |
| AALVGAPANVA | ASSAGGAQNTLY | 0.591117 |
| AATFRRTE | ASSAGGAQNTLY | 0.82892334 |
| AGALGGGI | ASSAGGAQNTLY | 0.6430838 |
| AGGAGGAGGIGAG | ASSAGGAQNTLY | 0.82233346 |
| AGSGALGL | ASSAGGAQNTLY | 0.8784296 |
| AQIADQI | ASSAGGAQNTLY | 0.74514437 |
| ASAVAVTYL | ASSAGGAQNTLY | 0.8698467 |
| ASTVLGGTGG | ASSAGGAQNTLY | 0.87698776 |
| AVGGEPAK | ASSAGGAQNTLY | 0.6714298 |
| CEQGTFGGTC | ASSAGGAQNTLY | 0.7989208 |

| | | |
|---|---|---|
| DEGKAPAG | ASSAGGAQNTLY | 0.7742827 |
| DTVVFPAL | ASSAGGAQNTLY | 0.8126632 |
| FEGITMAG | ASSAGGAQNTLY | 0.6300793 |
| FPLVVQD | ASSAGGAQNTLY | 0.8635484 |
| GAFTTPQ | ASSAGGAQNTLY | 0.6916611 |
| GFGGFGTAGGVGGA | ASSAGGAQNTLY | 0.19284746 |
| GFLAQFL | ASSAGGAQNTLY | 0.84883714 |
| GGAGGAGGNAGVA | ASSAGGAQNTLY | 0.7892295 |
| GGVGTAGAGGAGGAGG | ASSAGGAQNTLY | 0.7760024 |
| GNGGGGDAGNAGSGGN | ASSAGGAQNTLY | 0.85852075 |
| GTGGAGGAGNDAGS | ASSAGGAQNTLY | 0.639109 |
| GTTNTGWFNS | ASSAGGAQNTLY | 0.7564651 |
| IFMGPLFTL | ASSAGGAQNTLY | 0.8454801 |
| ILAHAIEAI | ASSAGGAQNTLY | 0.8691592 |
| IRSAIREAL | ASSAGGAQNTLY | 0.6798377 |
| ITFAGGG | ASSAGGAQNTLY | 0.808742 |
| ITGMSGHQEV | ASSAGGAQNTLY | 0.8853559 |
| IVAGLACTL | ASSAGGAQNTLY | 0.83501416 |
| IVGGGLAGA | ASSAGGAQNTLY | 0.8881765 |
| IVTGVLNI | ASSAGGAQNTLY | 0.8949418 |
| LAQQGCI | ASSAGGAQNTLY | 0.77945375 |
| LCLHVEN | ASSAGGAQNTLY | 0.8709055 |
| LDAISDGS | ASSAGGAQNTLY | 0.8943666 |
| LGGFAAML | ASSAGGAQNTLY | 0.8291286 |
| LGGKNPMI | ASSAGGAQNTLY | 0.7994609 |
| LGPKGVAAIL | ASSAGGAQNTLY | 0.67054904 |
| LMVGVSI | ASSAGGAQNTLY | 0.80323017 |
| LVGAAFVL | ASSAGGAQNTLY | 0.39713115 |
| LVLAVAPL | ASSAGGAQNTLY | 0.80020076 |
| MAAIAVIDAA | ASSAGGAQNTLY | 0.8917396 |
| NNGINAELQ | ASSAGGAQNTLY | 0.7388856 |
| RGEPLVLVHG | ASSAGGAQNTLY | 0.6439167 |
| RGGAGGAAGLI | ASSAGGAQNTLY | 0.5887722 |
| RSYRMHPAV | ASSAGGAQNTLY | 0.843534 |
| SAAHLLAGQLA | ASSAGGAQNTLY | 0.7027209 |
| SAAVGSY | ASSAGGAQNTLY | 0.552924 |
| SPHAVLHRI | ASSAGGAQNTLY | 0.8110245 |
| SQIAKVL | ASSAGGAQNTLY | 0.87029564 |
| TALGHLVP | ASSAGGAQNTLY | 0.87805146 |
| TAPTTSGAA | ASSAGGAQNTLY | 0.532464 |
| TDGDDGNGGSPGT | ASSAGGAQNTLY | 0.89038473 |
| TDGLGEN | ASSAGGAQNTLY | 0.84135103 |
| TGGPPLPGQA | ASSAGGAQNTLY | 0.889517 |
| TIPAITI | ASSAGGAQNTLY | 0.8386331 |
| TIPGFTI | ASSAGGAQNTLY | 0.7461314 |
| TSIASGALA | ASSAGGAQNTLY | 0.84791887 |
| TVTLPGAAI | ASSAGGAQNTLY | 0.7025315 |
| VASGLSAALP | ASSAGGAQNTLY | 0.7388649 |
| VGAGVIGI | ASSAGGAQNTLY | 0.87010777 |
| VGPKRFGV | ASSAGGAQNTLY | 0.7849709 |
| VNVAFPD | ASSAGGAQNTLY | 0.6912577 |

| | | |
|---|---|---|
| VSMTNSGVS | ASSAGGAQNTLY | 0.68523526 |
| VTDVLGDLSV | ASSAGGAQNTLY | 0.8901062 |
| VVVVGAGFAGL | ASSAGGAQNTLY | 0.63957006 |
| VYTVGTVV | ASSAGGAQNTLY | 0.66883796 |
| WTTFSGP | ASSAGGAQNTLY | 0.6818066 |
| YDELVEAF | ASSAGGAQNTLY | 0.78873086 |
| YGTGGAGGNGG | ASSAGGAQNTLY | 0.86857224 |
| YTNVMAV | ASSAGGAQNTLY | 0.875154 |
| FLEQVHQL | ASSAGGAQNTLY | 0.81114435 |
| GLSLLAVAV | ASSAGGAQNTLY | 0.8213688 |
| LEQVHQL | ASSAGGAQNTLY | 0.872316 |
| LNAAGAI | ASSAGGAQNTLY | 0.83926296 |
| PAVHASL | ASSAGGAQNTLY | 0.8363241 |
| TELEDDA | ASSAGGAQNTLY | 0.6608902 |
| TPPGPPN | ASSAGGAQNTLY | 0.8904215 |
| TPPPPPPP | ASSAGGAQNTLY | 0.84733295 |
| TQSPALI | ASGRDWGGEDTQY | 0.87730694 |
| EFAAAFPQ | ASGRDWGGEDTQY | 0.884101 |
| LLSGNLDVL | ASGRDWGGEDTQY | 0.87211156 |
| GPANLGNYNI | ASGRDWGGEDTQY | 0.8279069 |
| TLGPTLP | ASGRDWGGEDTQY | 0.36590034 |
| VYAYSSCCK | ASGRDWGGEDTQY | 0.7898205 |
| APAPAPAPNI | ASGRDWGGEDTQY | 0.7996403 |
| PGGPVPIP | ASGRDWGGEDTQY | 0.826303 |
| FLTVALEP | ASGRDWGGEDTQY | 0.8394381 |
| SQYFGANL | ASGRDWGGEDTQY | 0.8870964 |
| GGVKPAA | ASGRDWGGEDTQY | 0.71533656 |
| GGTGGAGGAGGW | ASGRDWGGEDTQY | 0.8756752 |
| GVGGVAGLGGAGG | ASGRDWGGEDTQY | 0.882362 |
| VLLVLATTI | ASGRDWGGEDTQY | 0.5541272 |
| IGIFAAML | ASGRDWGGEDTQY | 0.7604717 |
| AVGDLID | ASGRDWGGEDTQY | 0.9039218 |
| NISPLPL | ASGRDWGGEDTQY | 0.7911378 |
| VTAVFVAI | ASGRDWGGEDTQY | 0.7613883 |
| SGDDNDAAL | ASGRDWGGEDTQY | 0.77909255 |
| AVAALAHS | ASGRDWGGEDTQY | 0.8563558 |
| GGVTVMATP | ASGRDWGGEDTQY | 0.7380373 |
| AGLAANVRD | ASGRDWGGEDTQY | 0.89905584 |
| TNRPDLIDPALLRPG | ASGRDWGGEDTQY | 0.32708955 |
| LANLGVTAL | ASGRDWGGEDTQY | 0.81651354 |
| TGERIEL | ASGRDWGGEDTQY | 0.71994257 |
| NGEELAA | ASGRDWGGEDTQY | 0.358862 |
| VLSLVWG | ASGRDWGGEDTQY | 0.8564731 |
| IGYALWL | ASGRDWGGEDTQY | 0.81237936 |
| LSLLLAPL | ASGRDWGGEDTQY | 0.84987485 |
| TLDEALK | ASGRDWGGEDTQY | 0.82605964 |
| TDGAGTAEGG | ASGRDWGGEDTQY | 0.8688086 |
| NAPGGAGGNGG | ASGRDWGGEDTQY | 0.8116126 |
| VSVFMGMGGNF | ASGRDWGGEDTQY | 0.8124552 |
| LASYLAAL | ASGRDWGGEDTQY | 0.7987553 |
| LVGGTPAL | ASGRDWGGEDTQY | 0.8308245 |

| | | |
|---|---|---|
| VRNADVL | ASGRDWGGEDTQY | 0.88698494 |
| TCSSPFLVSAM | ASGRDWGGEDTQY | 0.89235145 |
| VHITDAE | ASGRDWGGEDTQY | 0.56371176 |
| EYAAPSF | ASGRDWGGEDTQY | 0.7879542 |
| VVDAFHGGL | ASGRDWGGEDTQY | 0.7757492 |
| ASVVDPQ | ASGRDWGGEDTQY | 0.3944486 |
| SYNEFGGLS | ASGRDWGGEDTQY | 0.3103541 |
| DPDGSVY | ASGRDWGGEDTQY | 0.7411929 |
| MVSTIHATE | ASGRDWGGEDTQY | 0.76912665 |
| LAAGVADPV | ASGRDWGGEDTQY | 0.8487062 |
| TKEQIAA | ASGRDWGGEDTQY | 0.8620822 |
| SLASPAAL | ASGRDWGGEDTQY | 0.737352 |
| IGLVFYGV | ASGRDWGGEDTQY | 0.85638654 |
| FLLDASNL | ASGRDWGGEDTQY | 0.7980501 |
| YQVGNFP | ASGRDWGGEDTQY | 0.09139439 |
| ESLLSQST | ASGRDWGGEDTQY | 0.80482274 |
| LFTVGFAAVG | ASGRDWGGEDTQY | 0.6612663 |
| ATPPTPT | ASGRDWGGEDTQY | 0.897617 |
| TDLATLG | ASGRDWGGEDTQY | 0.7144972 |
| SAGGTGR | ASGRDWGGEDTQY | 0.7063339 |
| GFSNSGS | ASGRDWGGEDTQY | 0.77942586 |
| GGNGGSGGTSV | ASGRDWGGEDTQY | 0.78665274 |
| GTSVATGGAGN | ASGRDWGGEDTQY | 0.7901746 |
| NGGNGGSGGTS | ASGRDWGGEDTQY | 0.6897728 |
| AVVKFPH | ASGRDWGGEDTQY | 0.2302955 |
| TPPPPPP | ASGRDWGGEDTQY | 0.8548029 |
| AAPAAAAAPA | ASGRDWGGEDTQY | 0.698534 |
| IALNEDL | ASGRDWGGEDTQY | 0.74024564 |
| IVNVSSM | ASGRDWGGEDTQY | 0.583086 |
| TGIPPAPR | ASGRDWGGEDTQY | 0.37653208 |
| FDLGGGTF | ASGRDWGGEDTQY | 0.65592 |
| FDLGGGTFDVS | ASGRDWGGEDTQY | 0.8929974 |
| NEPTAAA | ASGRDWGGEDTQY | 0.71119666 |
| NEPTAAALA | ASGRDWGGEDTQY | 0.73284805 |
| GSIINISSI | ASGRDWGGEDTQY | 0.89924634 |
| VAPGNAGTA | ASGRDWGGEDTQY | 0.7083703 |
| TGGGASLE | ASGRDWGGEDTQY | 0.75358224 |
| VAVYNHYKRI | ASGRDWGGEDTQY | 0.8055284 |
| TDPAPATT | ASGRDWGGEDTQY | 0.7078553 |
| TNRPDILDPALLRPG | ASGRDWGGEDTQY | 0.75084454 |
| LTTTTTS | ASGRDWGGEDTQY | 0.7585929 |
| PTGSGKTL | ASGRDWGGEDTQY | 0.8885106 |
| TGSGKTL | ASGRDWGGEDTQY | 0.8224197 |
| TIGHVDHGK | ASGRDWGGEDTQY | 0.75514054 |
| TIGHVDHGKTT | ASGRDWGGEDTQY | 0.87938637 |
| GHVDHGKTT | ASGRDWGGEDTQY | 0.8848444 |
| AQFEHTLL | ASGRDWGGEDTQY | 0.89706534 |
| IVNQDEL | ASGRDWGGEDTQY | 0.7173978 |
| TNGSQFF | ASGRDWGGEDTQY | 0.82103026 |
| TGKGIVE | ASGRDWGGEDTQY | 0.81087756 |
| LISNASDA | ASGRDWGGEDTQY | 0.7376169 |

| | | |
|---|---|---|
| AELRTLGELTKIAKAHGAQ | ASGRDWGGEDTQY | 0.817563 |
| APNAVAAAEPSW | ASGRDWGGEDTQY | 0.68129206 |
| APQVAGAATAVL | ASGRDWGGEDTQY | 0.87962997 |
| ARRDPAAAAAILPTDA | ASGRDWGGEDTQY | 0.88295645 |
| DAAARDVLAEAGLAEYF | ASGRDWGGEDTQY | 0.8276001 |
| DDQIVLSAKANLAKVEQ | ASGRDWGGEDTQY | 0.69752795 |
| DELLPDTGEDSDEDGH | ASGRDWGGEDTQY | 0.73935616 |
| ELPEAPANRATVPQD | ASGRDWGGEDTQY | 0.65867555 |
| FGQSKIERVTGIG | ASGRDWGGEDTQY | 0.68834597 |
| GAGGTGSTAGGAGGAGGAGGLY | ASGRDWGGEDTQY | 0.81757194 |
| GHGGTGAAGGNGGTGGAGGSGI | ASGRDWGGEDTQY | 0.54465413 |
| GNGGNGGAGGEGGPGGAGGAGA | ASGRDWGGEDTQY | 0.79921436 |
| GSTAGNGGNGGNATNGTIAGQP | ASGRDWGGEDTQY | 0.9016187 |
| GTGGDGGNGGNGGTGG | ASGRDWGGEDTQY | 0.684034 |
| GVRRPAGIPADAPV | ASGRDWGGEDTQY | 0.8599472 |
| GYRQVTDAHLLTLARR | ASGRDWGGEDTQY | 0.8856306 |
| IAKMGQDGHDRG | ASGRDWGGEDTQY | 0.8527477 |
| KAEPSIEIQYT | ASGRDWGGEDTQY | 0.8742751 |
| KPVRIRMAADEDTVAVT | ASGRDWGGEDTQY | 0.8409251 |
| MPPRIAGMRLLVIKPEP | ASGRDWGGEDTQY | 0.8554835 |
| MKFYAVALFRSRGVPPTR | ASGRDWGGEDTQY | 0.82786584 |
| NAGGFGILGAVAHSP | ASGRDWGGEDTQY | 0.8276299 |
| NEGREITLAMAEM | ASGRDWGGEDTQY | 0.7588278 |
| NELVEWSHPLGHRWRWE | ASGRDWGGEDTQY | 0.7741388 |
| NLGAGGTGGNGGNPG | ASGRDWGGEDTQY | 0.76598054 |
| NTEWPKWETVMAADPVD | ASGRDWGGEDTQY | 0.8551873 |
| PASAAPAETQP | ASGRDWGGEDTQY | 0.81737894 |
| QQSSDGKNGVRAAEITGE | ASGRDWGGEDTQY | 0.855742 |
| RAPDAVTTSWWKE | ASGRDWGGEDTQY | 0.49418658 |
| SGFGNVGAHTSGWFNQ | ASGRDWGGEDTQY | 0.67246133 |
| SGVAGAVGQAASVGGLKV | ASGRDWGGEDTQY | 0.86433774 |
| SNGGAGTGSAGGNGGT | ASGRDWGGEDTQY | 0.18417999 |
| TGGAGGEGGAAG | ASGRDWGGEDTQY | 0.39812484 |
| TGGDGGAAGAGGAPGHG | ASGRDWGGEDTQY | 0.8409034 |
| TMPPLLAAIANAGRHGV | ASGRDWGGEDTQY | 0.876647 |
| TSSTAGNSGGA | ASGRDWGGEDTQY | 0.21702863 |
| VPATVPVGILC | ASGRDWGGEDTQY | 0.7568429 |
| VQRPDSGGVVLTGRIS | ASGRDWGGEDTQY | 0.5688981 |
| WTPNPVPATKGVPAGLVQAT | ASGRDWGGEDTQY | 0.8984158 |
| YALKRALAKFSDRFDVVII | ASGRDWGGEDTQY | 0.7967119 |
| YLIGSVIV | ASGRDWGGEDTQY | 0.5002688 |
| YGQLVAVID | ASGRDWGGEDTQY | 0.8543805 |
| VTVGGMTLPA | ASGRDWGGEDTQY | 0.64046943 |
| VLALPAV | ASGRDWGGEDTQY | 0.64596003 |
| VIPPTLNYET | ASGRDWGGEDTQY | 0.8723756 |
| VGMGGMHPSSG | ASGRDWGGEDTQY | 0.8743902 |
| VGLAGGAATA | ASGRDWGGEDTQY | 0.6901046 |
| VGGAGGAGGSGP | ASGRDWGGEDTQY | 0.84595895 |
| VFAAFMV | ASGRDWGGEDTQY | 0.7295432 |
| TLYDKLR | ASGRDWGGEDTQY | 0.89828944 |
| TLAVFTVL | ASGRDWGGEDTQY | 0.7616925 |

| | | |
|---|---|---|
| TGSVQPARV | ASGRDWGGEDTQY | 0.59400165 |
| TGGGAGSGGT | ASGRDWGGEDTQY | 0.8744174 |
| SHLETSL | ASGRDWGGEDTQY | 0.8567935 |
| SFTTGMT | ASGRDWGGEDTQY | 0.8460159 |
| QVEVAKRMGVRQPTVS | ASGRDWGGEDTQY | 0.8439772 |
| QGSGGPALAM | ASGRDWGGEDTQY | 0.817111 |
| MLAPSNP | ASGRDWGGEDTQY | 0.74817145 |
| LVNNAAI | ASGRDWGGEDTQY | 0.8614749 |
| LTGEINTLA | ASGRDWGGEDTQY | 0.85393703 |
| LSIFGPV | ASGRDWGGEDTQY | 0.668843 |
| LLALLANL | ASGRDWGGEDTQY | 0.8914257 |
| LISNASDALDKLR | ASGRDWGGEDTQY | 0.77822447 |
| LINEAVP | ASGRDWGGEDTQY | 0.73028296 |
| LGYAFSGL | ASGRDWGGEDTQY | 0.8956411 |
| LDFVAVAL | ASGRDWGGEDTQY | 0.84411085 |
| LAQPGAE | ASGRDWGGEDTQY | 0.8562043 |
| ISNLGAVTGF | ASGRDWGGEDTQY | 0.85100234 |
| HQEDPGRI | ASGRDWGGEDTQY | 0.8354993 |
| GTPGEPTP | ASGRDWGGEDTQY | 0.68830127 |
| GFFNSGS | ASGRDWGGEDTQY | 0.84335756 |
| GAGGGGLTAGAN | ASGRDWGGEDTQY | 0.8205576 |
| FVDALVNM | ASGRDWGGEDTQY | 0.86687815 |
| FLTVAPEL | ASGRDWGGEDTQY | 0.83014727 |
| FETLTAL | ASGRDWGGEDTQY | 0.6625727 |
| DVINPIEL | ASGRDWGGEDTQY | 0.8483597 |
| APVSGAPV | ASGRDWGGEDTQY | 0.8903161 |
| ALVGGLAW | ASGRDWGGEDTQY | 0.85572505 |
| ALAAAAATAI | ASGRDWGGEDTQY | 0.8343828 |
| AFGLGLLL | ASGRDWGGEDTQY | 0.88074476 |
| AFFAAASL | ASGRDWGGEDTQY | 0.7161737 |
| AAAALPAFSPP | ASGRDWGGEDTQY | 0.8040153 |
| VVLAPAL | ASGRDWGGEDTQY | 0.8884759 |
| TIGHVDHGKTTL | ASGRDWGGEDTQY | 0.8744961 |
| LISNASDALDK | ASGRDWGGEDTQY | 0.8242328 |
| LISNASDAL | ASGRDWGGEDTQY | 0.8827332 |
| ISNASDALDK | ASGRDWGGEDTQY | 0.7607248 |
| NQAETLVYQTE | ASGRDWGGEDTQY | 0.842597 |
| TGGTGGTAGAG | ASGRDWGGEDTQY | 0.7070245 |
| TGSTAGGAGGA | ASGRDWGGEDTQY | 0.815699 |
| DPLAVPVAAGPV | ASGRDWGGEDTQY | 0.63777244 |
| GGTGGTPGNGGN | ASGRDWGGEDTQY | 0.8447928 |
| PAGLAGVDPATL | ASGRDWGGEDTQY | 0.8643685 |
| GNNLVVEAPHAGASV | ASGRDWGGEDTQY | 0.8484393 |
| KVTRIMSASTLEQPA | ASGRDWGGEDTQY | 0.40224302 |
| LGFALPIALPRRTVV | ASGRDWGGEDTQY | 0.84765995 |
| TRLGNLTARKLVVLV | ASGRDWGGEDTQY | 0.8458072 |
| WSQQRRLIYFPSAGP | ASGRDWGGEDTQY | 0.8870535 |
| GPNGNPDPMAAAVDIR | ASGRDWGGEDTQY | 0.8035829 |
| TQRCVSVADLRAAASR | ASGRDWGGEDTQY | 0.8603833 |
| QFVLDVAEHPVEVLTETV | ASGRDWGGEDTQY | 0.79203475 |
| GGAGGTGGAGGNGGTQAGDG | ASGRDWGGEDTQY | 0.83093995 |

| | | |
|---|---|---|
| PLLATLRAIAVDVDAKPAQV | ASGRDWGGEDTQY | 0.6241512 |
| APKFRTGGYMIDFWGVGYQVAK | ASGRDWGGEDTQY | 0.7306209 |
| GAGGAGGAGGAAGGPGGTGGTG | ASGRDWGGEDTQY | 0.7193078 |
| IGALGLLATDVAPAGSVAFGVA | ASGRDWGGEDTQY | 0.66135323 |
| AGGAGGIGGNGGAGGAGGVLMG | ASGRDWGGEDTQY | 0.83447427 |
| ADELIGLL | ASGRDWGGEDTQY | 0.8625889 |
| ADRALKA | ASGRDWGGEDTQY | 0.8788377 |
| AGAVGQVGGAGGA | ASGRDWGGEDTQY | 0.8710139 |
| AGGGGGLGGAP | ASGRDWGGEDTQY | 0.8798398 |
| AGGLLYGNGGAG | ASGRDWGGEDTQY | 0.8852416 |
| AGLESIGL | ASGRDWGGEDTQY | 0.70805144 |
| AGLKTGII | ASGRDWGGEDTQY | 0.7933162 |
| ASLAALLL | ASGRDWGGEDTQY | 0.6285023 |
| ASVINISSI | ASGRDWGGEDTQY | 0.77880496 |
| ATPPGPA | ASGRDWGGEDTQY | 0.7572488 |
| ATVASALV | ASGRDWGGEDTQY | 0.812682 |
| ATVASLIL | ASGRDWGGEDTQY | 0.16562141 |
| AVNGPVLIRH | ASGRDWGGEDTQY | 0.76729107 |
| DGAAGAVGI | ASGRDWGGEDTQY | 0.8901955 |
| DPLLHGI | ASGRDWGGEDTQY | 0.61656654 |
| ELGAALGAL | ASGRDWGGEDTQY | 0.8452852 |
| ELGRTQAV | ASGRDWGGEDTQY | 0.81970704 |
| FVAGGYLS | ASGRDWGGEDTQY | 0.75029767 |
| GASGGAGGAGGW | ASGRDWGGEDTQY | 0.8504752 |
| GGAGAGGDGGAGGI | ASGRDWGGEDTQY | 0.72687864 |
| GTGHNGSQPGMGGNG | ASGRDWGGEDTQY | 0.8564951 |
| GYAAASASAS | ASGRDWGGEDTQY | 0.8862656 |
| HQVDVAAVQQG | ASGRDWGGEDTQY | 0.8389474 |
| IGAAIAHT | ASGRDWGGEDTQY | 0.81749034 |
| IGMIEVAQL | ASGRDWGGEDTQY | 0.8191443 |
| IRLAPEHTSAMD | ASGRDWGGEDTQY | 0.55883116 |
| IYNVSVL | ASGRDWGGEDTQY | 0.82128084 |
| KQGFLGM | ASGRDWGGEDTQY | 0.1698603 |
| LESVITQL | ASGRDWGGEDTQY | 0.7825012 |
| LGLGALVVV | ASGRDWGGEDTQY | 0.66324323 |
| LIGNGGNGGSGGI | ASGRDWGGEDTQY | 0.82124066 |
| LNAINAPTL | ASGRDWGGEDTQY | 0.8040359 |
| LQLLNTGV | ASGRDWGGEDTQY | 0.8674284 |
| LQTSIEF | ASGRDWGGEDTQY | 0.66597843 |
| MSFAVSEL | ASGRDWGGEDTQY | 0.7975701 |
| NTGALGL | ASGRDWGGEDTQY | 0.7426769 |
| PPGAGGAGGAGTT | ASGRDWGGEDTQY | 0.6734407 |
| QLVGLGI | ASGRDWGGEDTQY | 0.8809221 |
| RFLEQVHQL | ASGRDWGGEDTQY | 0.84726286 |
| RGVGYVLR | ASGRDWGGEDTQY | 0.7713375 |
| SATGVFAGVI | ASGRDWGGEDTQY | 0.8774568 |
| SLPAITLP | ASGRDWGGEDTQY | 0.8393905 |
| STALAFTV | ASGRDWGGEDTQY | 0.6300664 |
| SVLTVGLGK | ASGRDWGGEDTQY | 0.7407584 |
| TALAGAGSGI | ASGRDWGGEDTQY | 0.80752397 |
| TGLPFTGI | ASGRDWGGEDTQY | 0.7433368 |

| | | |
|---|---|---|
| VAEHVTV | ASGRDWGGEDTQY | 0.17884158 |
| VDIVLNSL | ASGRDWGGEDTQY | 0.8776231 |
| VLTGGGAVL | ASGRDWGGEDTQY | 0.74011284 |
| VNVGLAYGI | ASGRDWGGEDTQY | 0.652665 |
| VSSAAVGAV | ASGRDWGGEDTQY | 0.7362335 |
| VTSVYEAGV | ASGRDWGGEDTQY | 0.8470895 |
| WVALDRGA | ASGRDWGGEDTQY | 0.643353 |
| YLVGSVVI | ASGRDWGGEDTQY | 0.7641579 |
| GPPGTGKTLL | ASGRDWGGEDTQY | 0.83881575 |
| GVGMTKF | ASGRDWGGEDTQY | 0.8458899 |
| IGIGGSDL | ASGRDWGGEDTQY | 0.8724477 |
| IGTLLAEL | ASGRDWGGEDTQY | 0.86391973 |
| IYTSGTT | ASGRDWGGEDTQY | 0.810904 |
| LSFPAGV | ASGRDWGGEDTQY | 0.8680903 |
| LYTSGTT | ASGRDWGGEDTQY | 0.8600558 |
| VGAFPFGF | ASGRDWGGEDTQY | 0.5872467 |
| VGAGTVEF | ASGRDWGGEDTQY | 0.5105985 |
| VGKSTLLL | ASGRDWGGEDTQY | 0.7974991 |
| AALAHAGGGIV | ASGRDWGGEDTQY | 0.5265839 |
| AALVAVSLL | ASGRDWGGEDTQY | 0.8222792 |
| AAVSRAVAAL | ASGRDWGGEDTQY | 0.90246236 |
| AGGSSGIGGF | ASGRDWGGEDTQY | 0.82829773 |
| AGLTVANANG | ASGRDWGGEDTQY | 0.61604977 |
| AGLVGAVL | ASGRDWGGEDTQY | 0.7910284 |
| ALSQAFTV | ASGRDWGGEDTQY | 0.7859483 |
| ALTFSVSL | ASGRDWGGEDTQY | 0.82440305 |
| ALTGAALVAL | ASGRDWGGEDTQY | 0.8687965 |
| ASFSIGAGVL | ASGRDWGGEDTQY | 0.88128656 |
| DGSVYKI | ASGRDWGGEDTQY | 0.8441159 |
| EQLAEIV | ASGRDWGGEDTQY | 0.72063524 |
| FAVSELTVM | ASGRDWGGEDTQY | 0.8451878 |
| GAAPLAAGL | ASGRDWGGEDTQY | 0.76134837 |
| GADGGDGGSGGAGG | ASGRDWGGEDTQY | 0.83167267 |
| GFGLDTGGAGG | ASGRDWGGEDTQY | 0.83619344 |
| GLASGGLL | ASGRDWGGEDTQY | 0.8705232 |
| GRIAGAAVLL | ASGRDWGGEDTQY | 0.82558703 |
| GSAGSLNTSF | ASGRDWGGEDTQY | 0.79070497 |
| GTGQIGI | ASGRDWGGEDTQY | 0.59317875 |
| GTLIALIL | ASGRDWGGEDTQY | 0.86100554 |
| IAATMGFVVG | ASGRDWGGEDTQY | 0.75524235 |
| IFAQALL | ASGRDWGGEDTQY | 0.59345996 |
| IGAAVALGF | ASGRDWGGEDTQY | 0.74111104 |
| IGLTPLAL | ASGRDWGGEDTQY | 0.8860675 |
| ILGSNTSTL | ASGRDWGGEDTQY | 0.88624626 |
| IVFSLTVL | ASGRDWGGEDTQY | 0.6663336 |
| LASKATGF | ASGRDWGGEDTQY | 0.7228997 |
| LASVALIL | ASGRDWGGEDTQY | 0.8011875 |
| LFAVGLTL | ASGRDWGGEDTQY | 0.51660323 |
| LFNGGIGTYI | ASGRDWGGEDTQY | 0.8721585 |
| LTLGAGII | ASGRDWGGEDTQY | 0.7458041 |
| LVAAALGDI | ASGRDWGGEDTQY | 0.78167117 |

| | | |
|---|---|---|
| NTAFVQLGI | ASGRDWGGEDTQY | 0.771628 |
| QLFSGAEL | ASGRDWGGEDTQY | 0.8452194 |
| SAATFITL | ASGRDWGGEDTQY | 0.7724551 |
| SNYGLANLGV | ASGRDWGGEDTQY | 0.87670666 |
| SYGGAYAVM | ASGRDWGGEDTQY | 0.24336834 |
| TTGVGGLTLG | ASGRDWGGEDTQY | 0.8694384 |
| TTQTIYI | ASGRDWGGEDTQY | 0.640386 |
| VAAGAGLGAVI | ASGRDWGGEDTQY | 0.7417732 |
| VAAVVTLL | ASGRDWGGEDTQY | 0.5298739 |
| VGFLGGLI | ASGRDWGGEDTQY | 0.8431567 |
| VGVIPILL | ASGRDWGGEDTQY | 0.8588508 |
| VIFASSLI | ASGRDWGGEDTQY | 0.8819522 |
| VKGGDGGAAAG | ASGRDWGGEDTQY | 0.6954136 |
| VTKTFTNI | ASGRDWGGEDTQY | 0.85928977 |
| VVAAAILL | ASGRDWGGEDTQY | 0.88670516 |
| YAALDVEL | ASGRDWGGEDTQY | 0.7268913 |
| LASVALLL | ASGRDWGGEDTQY | 0.5192963 |
| LDAVHRV | ASGRDWGGEDTQY | 0.8299681 |
| QGLSSSIF | ASGRDWGGEDTQY | 0.77398324 |
| RGTFANIRL | ASGRDWGGEDTQY | 0.87812936 |
| RQLWWGHRI | ASGRDWGGEDTQY | 0.8384133 |
| YGPPGTGKTLL | ASGRDWGGEDTQY | 0.5963999 |
| AALVGAPANVA | ASGRDWGGEDTQY | 0.8628217 |
| AATFRRTE | ASGRDWGGEDTQY | 0.8831836 |
| AGALGGGI | ASGRDWGGEDTQY | 0.6783934 |
| AGGAGGAGGIGAG | ASGRDWGGEDTQY | 0.7862312 |
| AGSGALGL | ASGRDWGGEDTQY | 0.8201145 |
| AQIADQI | ASGRDWGGEDTQY | 0.48098138 |
| ASAVAVTYL | ASGRDWGGEDTQY | 0.86635196 |
| ASTVLGGTGG | ASGRDWGGEDTQY | 0.71421695 |
| AVGGEPAK | ASGRDWGGEDTQY | 0.7583647 |
| CEQGTFGGTC | ASGRDWGGEDTQY | 0.8228635 |
| DEGKAPAG | ASGRDWGGEDTQY | 0.7955922 |
| DTVVFPAL | ASGRDWGGEDTQY | 0.7495166 |
| FEGITMAG | ASGRDWGGEDTQY | 0.7819352 |
| FPLVVQD | ASGRDWGGEDTQY | 0.8502786 |
| GAFTTPQ | ASGRDWGGEDTQY | 0.852022 |
| GFGGFGTAGGVGGA | ASGRDWGGEDTQY | 0.8350117 |
| GFLAQFL | ASGRDWGGEDTQY | 0.8429244 |
| GGAGGAGGNAGVA | ASGRDWGGEDTQY | 0.8758511 |
| GGVGTAGAGGAGGAGG | ASGRDWGGEDTQY | 0.6343574 |
| GNGGGGDAGNAGSGGN | ASGRDWGGEDTQY | 0.7100824 |
| GTGGAGGAGNDAGS | ASGRDWGGEDTQY | 0.86269426 |
| GTTNTGWFNS | ASGRDWGGEDTQY | 0.62058645 |
| IFMGPLFTL | ASGRDWGGEDTQY | 0.87350327 |
| ILAHAIEAI | ASGRDWGGEDTQY | 0.8064115 |
| IRSAIREAL | ASGRDWGGEDTQY | 0.8142809 |
| ITFAGGG | ASGRDWGGEDTQY | 0.77140343 |
| ITGMSGHQEV | ASGRDWGGEDTQY | 0.90071404 |
| IVAGLACTL | ASGRDWGGEDTQY | 0.8498088 |
| IVGGGLAGA | ASGRDWGGEDTQY | 0.839803 |

| | | |
|---|---|---|
| IVTGVLNI | ASGRDWGGEDTQY | 0.81668425 |
| LAQQGCI | ASGRDWGGEDTQY | 0.7552514 |
| LCLHVEN | ASGRDWGGEDTQY | 0.8735375 |
| LDAISDGS | ASGRDWGGEDTQY | 0.8573462 |
| LGGFAAML | ASGRDWGGEDTQY | 0.5892091 |
| LGGKNPMI | ASGRDWGGEDTQY | 0.74982953 |
| LGPKGVAAIL | ASGRDWGGEDTQY | 0.8920858 |
| LMVGVSI | ASGRDWGGEDTQY | 0.87408555 |
| LVGAAFVL | ASGRDWGGEDTQY | 0.4674558 |
| LVLAVAPL | ASGRDWGGEDTQY | 0.76409477 |
| MAAIAVIDAA | ASGRDWGGEDTQY | 0.6013125 |
| NNGINAELQ | ASGRDWGGEDTQY | 0.7723058 |
| RGEPLVLVHG | ASGRDWGGEDTQY | 0.6736555 |
| RGGAGGAAGLI | ASGRDWGGEDTQY | 0.8339635 |
| RSYRMHPAV | ASGRDWGGEDTQY | 0.758396 |
| SAAHLLAGQLA | ASGRDWGGEDTQY | 0.81143856 |
| SAAVGSY | ASGRDWGGEDTQY | 0.8219087 |
| SPHAVLHRI | ASGRDWGGEDTQY | 0.87543297 |
| SQIAKVL | ASGRDWGGEDTQY | 0.7260163 |
| TALGHLVP | ASGRDWGGEDTQY | 0.78829527 |
| TAPTTSGAA | ASGRDWGGEDTQY | 0.65454614 |
| TDGDDGNGGSPGT | ASGRDWGGEDTQY | 0.4626591 |
| TDGLGEN | ASGRDWGGEDTQY | 0.7490866 |
| TGGPPLPGQA | ASGRDWGGEDTQY | 0.7637738 |
| TIPAITI | ASGRDWGGEDTQY | 0.36117348 |
| TIPGFTI | ASGRDWGGEDTQY | 0.8973694 |
| TSIASGALA | ASGRDWGGEDTQY | 0.79664075 |
| TVTLPGAAI | ASGRDWGGEDTQY | 0.84198296 |
| VASGLSAALP | ASGRDWGGEDTQY | 0.8151573 |
| VGAGVIGI | ASGRDWGGEDTQY | 0.7796719 |
| VGPKRFGV | ASGRDWGGEDTQY | 0.716247 |
| VNVAFPD | ASGRDWGGEDTQY | 0.83458006 |
| VSMTNSGVS | ASGRDWGGEDTQY | 0.6882941 |
| VTDVLGDLSV | ASGRDWGGEDTQY | 0.6643927 |
| VVVVGAGFAGL | ASGRDWGGEDTQY | 0.82940155 |
| VYTVGTVV | ASGRDWGGEDTQY | 0.72722715 |
| WTTFSGP | ASGRDWGGEDTQY | 0.88565135 |
| YDELVEAF | ASGRDWGGEDTQY | 0.8716709 |
| YGTGGAGGNGG | ASGRDWGGEDTQY | 0.8297224 |
| YTNVMAV | ASGRDWGGEDTQY | 0.8835268 |
| FLEQVHQL | ASGRDWGGEDTQY | 0.31373885 |
| GLSLLAVAV | ASGRDWGGEDTQY | 0.80995536 |
| LEQVHQL | ASGRDWGGEDTQY | 0.8769592 |
| LNAAGAI | ASGRDWGGEDTQY | 0.8460161 |
| PAVHASL | ASGRDWGGEDTQY | 0.786718 |
| TELEDDA | ASGRDWGGEDTQY | 0.87194514 |
| TPPGPPN | ASGRDWGGEDTQY | 0.7058865 |
| TPPPPPPP | ASGRDWGGEDTQY | 0.75964355 |
| TQSPALI | ASSTRTGSAETLY | 0.81585 |
| EFAAAFPQ | ASSTRTGSAETLY | 0.87370384 |
| LLSGNLDVL | ASSTRTGSAETLY | 0.86566484 |

| | | |
|---|---|---|
| GPANLGNYNI | ASSTRTGSAETLY | 0.7137125 |
| TLGPTLP | ASSTRTGSAETLY | 0.85362166 |
| VYAYSSCCK | ASSTRTGSAETLY | 0.9017525 |
| APAPAPAPNI | ASSTRTGSAETLY | 0.7505146 |
| PGGPVPIP | ASSTRTGSAETLY | 0.8699739 |
| FLTVALEP | ASSTRTGSAETLY | 0.8178377 |
| SQYFGANL | ASSTRTGSAETLY | 0.6514362 |
| GGVKPAA | ASSTRTGSAETLY | 0.6822161 |
| GGTGGAGGAGGW | ASSTRTGSAETLY | 0.6887975 |
| GVGGVAGLGGAGG | ASSTRTGSAETLY | 0.7906274 |
| VLLVLATTI | ASSTRTGSAETLY | 0.6486373 |
| IGIFAAML | ASSTRTGSAETLY | 0.8294157 |
| AVGDLID | ASSTRTGSAETLY | 0.80368876 |
| NISPLPL | ASSTRTGSAETLY | 0.90347075 |
| VTAVFVAI | ASSTRTGSAETLY | 0.83848035 |
| SGDDNDAAL | ASSTRTGSAETLY | 0.8465929 |
| AVAALAHS | ASSTRTGSAETLY | 0.62422216 |
| GGVTVMATP | ASSTRTGSAETLY | 0.8489727 |
| AGLAANVRD | ASSTRTGSAETLY | 0.826199 |
| TNRPDLIDPALLRPG | ASSTRTGSAETLY | 0.61326444 |
| LANLGVTAL | ASSTRTGSAETLY | 0.88556314 |
| TGERIEL | ASSTRTGSAETLY | 0.66675484 |
| NGEELAA | ASSTRTGSAETLY | 0.80722386 |
| VLSLVWG | ASSTRTGSAETLY | 0.87234676 |
| IGYALWL | ASSTRTGSAETLY | 0.8497871 |
| LSLLLAPL | ASSTRTGSAETLY | 0.782917 |
| TLDEALK | ASSTRTGSAETLY | 0.78224206 |
| TDGAGTAEGG | ASSTRTGSAETLY | 0.71207225 |
| NAPGGAGGNGG | ASSTRTGSAETLY | 0.86109424 |
| VSVFMGMGGNF | ASSTRTGSAETLY | 0.86863554 |
| LASYLAAL | ASSTRTGSAETLY | 0.7244823 |
| LVGGTPAL | ASSTRTGSAETLY | 0.88130677 |
| VRNADVL | ASSTRTGSAETLY | 0.875105 |
| TCSSPFLVSAM | ASSTRTGSAETLY | 0.56960636 |
| VHITDAE | ASSTRTGSAETLY | 0.7927345 |
| EYAAPSF | ASSTRTGSAETLY | 0.87364537 |
| VVDAFHGGL | ASSTRTGSAETLY | 0.8747719 |
| ASVVDPQ | ASSTRTGSAETLY | 0.89574623 |
| SYNEFGGLS | ASSTRTGSAETLY | 0.85940033 |
| DPDGSVY | ASSTRTGSAETLY | 0.71616095 |
| MVSTIHATE | ASSTRTGSAETLY | 0.8857105 |
| LAAGVADPV | ASSTRTGSAETLY | 0.90879405 |
| TKEQIAA | ASSTRTGSAETLY | 0.8540065 |
| SLASPAAL | ASSTRTGSAETLY | 0.75668335 |
| IGLVFYGV | ASSTRTGSAETLY | 0.7652465 |
| FLLDASNL | ASSTRTGSAETLY | 0.85356605 |
| YQVGNFP | ASSTRTGSAETLY | 0.85982335 |
| ESLLSQST | ASSTRTGSAETLY | 0.69998324 |
| LFTVGFAAVG | ASSTRTGSAETLY | 0.795678 |
| ATPPTPT | ASSTRTGSAETLY | 0.80942464 |
| TDLATLG | ASSTRTGSAETLY | 0.5162059 |

| | | |
|---|---|---|
| SAGGTGR | ASSTRTGSAETLY | 0.8765059 |
| GFSNSGS | ASSTRTGSAETLY | 0.8846025 |
| GGNGGSGGTSV | ASSTRTGSAETLY | 0.84378207 |
| GTSVATGGAGN | ASSTRTGSAETLY | 0.6915976 |
| NGGNGGSGGTS | ASSTRTGSAETLY | 0.8774065 |
| AVVKFPH | ASSTRTGSAETLY | 0.89053655 |
| TPPPPPP | ASSTRTGSAETLY | 0.8882576 |
| AAPAAAAAPA | ASSTRTGSAETLY | 0.5913591 |
| IALNEDL | ASSTRTGSAETLY | 0.3369619 |
| IVNVSSM | ASSTRTGSAETLY | 0.7267833 |
| TGIPPAPR | ASSTRTGSAETLY | 0.8252816 |
| FDLGGGTF | ASSTRTGSAETLY | 0.80463827 |
| FDLGGGTFDVS | ASSTRTGSAETLY | 0.74708915 |
| NEPTAAA | ASSTRTGSAETLY | 0.8089269 |
| NEPTAAALA | ASSTRTGSAETLY | 0.87231183 |
| GSIINISSI | ASSTRTGSAETLY | 0.6969703 |
| VAPGNAGTA | ASSTRTGSAETLY | 0.6059395 |
| TGGGASLE | ASSTRTGSAETLY | 0.87228453 |
| VAVYNHYKRI | ASSTRTGSAETLY | 0.86748695 |
| TDPAPATT | ASSTRTGSAETLY | 0.6989819 |
| TNRPDILDPALLRPG | ASSTRTGSAETLY | 0.7403559 |
| LTTTTTS | ASSTRTGSAETLY | 0.7477949 |
| PTGSGKTL | ASSTRTGSAETLY | 0.75956184 |
| TGSGKTL | ASSTRTGSAETLY | 0.80270827 |
| TIGHVDHGK | ASSTRTGSAETLY | 0.64412457 |
| TIGHVDHGKTT | ASSTRTGSAETLY | 0.8964565 |
| GHVDHGKTT | ASSTRTGSAETLY | 0.6486601 |
| AQFEHTLL | ASSTRTGSAETLY | 0.8718302 |
| IVNQDEL | ASSTRTGSAETLY | 0.8185611 |
| TNGSQFF | ASSTRTGSAETLY | 0.86268425 |
| TGKGIVE | ASSTRTGSAETLY | 0.738952 |
| LISNASDA | ASSTRTGSAETLY | 0.7005449 |
| AELRTLGELTKIAKAHGAQ | ASSTRTGSAETLY | 0.65663624 |
| APNAVAAAEPSW | ASSTRTGSAETLY | 0.50252634 |
| APQVAGAATAVL | ASSTRTGSAETLY | 0.8710666 |
| ARRDPAAAAAILPTDA | ASSTRTGSAETLY | 0.24321511 |
| DAAARDVLAEAGLAEYF | ASSTRTGSAETLY | 0.8742541 |
| DDQIVLSAKANLAKVEQ | ASSTRTGSAETLY | 0.82811916 |
| DELLPDTGEDSDEDGH | ASSTRTGSAETLY | 0.75970066 |
| ELPEAPANRATVPQD | ASSTRTGSAETLY | 0.8618754 |
| FGQSKIERVTGIG | ASSTRTGSAETLY | 0.5951538 |
| GAGGTGSTAGGAGGAGGAGGLY | ASSTRTGSAETLY | 0.7160505 |
| GHGGTGAAGGNGGTGGAGGSGI | ASSTRTGSAETLY | 0.8283137 |
| GNGGNGGAGGEGGPGGAGGAGA | ASSTRTGSAETLY | 0.8483108 |
| GSTAGNGGNGGNATNGTIAGQP | ASSTRTGSAETLY | 0.72884065 |
| GTGGDGGNGGNGGTGG | ASSTRTGSAETLY | 0.86125875 |
| GVRRPAGIPADAPV | ASSTRTGSAETLY | 0.8370775 |
| GYRQVTDAHLLTLARR | ASSTRTGSAETLY | 0.7417823 |
| IAKMGQDGHDRG | ASSTRTGSAETLY | 0.83962667 |
| KAEPSIEIQYT | ASSTRTGSAETLY | 0.8727504 |
| KPVRIRMAADEDTVAVT | ASSTRTGSAETLY | 0.849726 |

| | | |
|---|---|---|
| MPPRIAGMRLLVIKPEP | ASSTRTGSAETLY | 0.65413666 |
| MKFYAVALFRSRGVPPTR | ASSTRTGSAETLY | 0.85840255 |
| NAGGFGILGAVAHSP | ASSTRTGSAETLY | 0.6596366 |
| NEGREITLAMAEM | ASSTRTGSAETLY | 0.8287776 |
| NELVEWSHPLGHRWRWE | ASSTRTGSAETLY | 0.8396033 |
| NLGAGGTGGNGGNPG | ASSTRTGSAETLY | 0.81219643 |
| NTEWPKWETVMAADPVD | ASSTRTGSAETLY | 0.8818308 |
| PASAAPAETQP | ASSTRTGSAETLY | 0.73630136 |
| QQSSDGKNGVRAAEITGE | ASSTRTGSAETLY | 0.71825564 |
| RAPDAVTTSWWKE | ASSTRTGSAETLY | 0.7677132 |
| SGFGNVGAHTSGWFNQ | ASSTRTGSAETLY | 0.8262627 |
| SGVAGAVGQAASVGGLKV | ASSTRTGSAETLY | 0.8566235 |
| SNGGAGTGSAGGNGGT | ASSTRTGSAETLY | 0.6291348 |
| TGGAGGEGGAAG | ASSTRTGSAETLY | 0.81892866 |
| TGGDGGAAGAGGAPGHG | ASSTRTGSAETLY | 0.77267194 |
| TMPPLLAAIANAGRHGV | ASSTRTGSAETLY | 0.63647103 |
| TSSTAGNSGGA | ASSTRTGSAETLY | 0.8525027 |
| VPATVPVGILC | ASSTRTGSAETLY | 0.81519157 |
| VQRPDSGGVVLTGRIS | ASSTRTGSAETLY | 0.7094214 |
| WTPNPVPATKGVPAGLVQAT | ASSTRTGSAETLY | 0.86555195 |
| YALKRALAKFSDRFDVVII | ASSTRTGSAETLY | 0.7652187 |
| YLIGSVIV | ASSTRTGSAETLY | 0.8699683 |
| YGQLVAVID | ASSTRTGSAETLY | 0.90440154 |
| VTVGGMTLPA | ASSTRTGSAETLY | 0.6801056 |
| VLALPAV | ASSTRTGSAETLY | 0.87305343 |
| VIPPTLNYET | ASSTRTGSAETLY | 0.8291254 |
| VGMGGMHPSSG | ASSTRTGSAETLY | 0.8163314 |
| VGLAGGAATA | ASSTRTGSAETLY | 0.767088 |
| VGGAGGAGGSGP | ASSTRTGSAETLY | 0.75298995 |
| VFAAFMV | ASSTRTGSAETLY | 0.88848215 |
| TLYDKLR | ASSTRTGSAETLY | 0.86452854 |
| TLAVFTVL | ASSTRTGSAETLY | 0.8605865 |
| TGSVQPARV | ASSTRTGSAETLY | 0.864805 |
| TGGGAGSGGT | ASSTRTGSAETLY | 0.8404683 |
| SHLETSL | ASSTRTGSAETLY | 0.85808754 |
| SFTTGMT | ASSTRTGSAETLY | 0.6687838 |
| QVEVAKRMGVRQPTVS | ASSTRTGSAETLY | 0.56016445 |
| QGSGGPALAM | ASSTRTGSAETLY | 0.77754253 |
| MLAPSNP | ASSTRTGSAETLY | 0.73986554 |
| LVNNAAI | ASSTRTGSAETLY | 0.7996994 |
| LTGEINTLA | ASSTRTGSAETLY | 0.592757 |
| LSIFGPV | ASSTRTGSAETLY | 0.85297567 |
| LLALLANL | ASSTRTGSAETLY | 0.89964974 |
| LISNASDALDKLR | ASSTRTGSAETLY | 0.7510586 |
| LINEAVP | ASSTRTGSAETLY | 0.9040302 |
| LGYAFSGL | ASSTRTGSAETLY | 0.60904837 |
| LDFVAVAL | ASSTRTGSAETLY | 0.868897 |
| LAQPGAE | ASSTRTGSAETLY | 0.87478113 |
| ISNLGAVTGF | ASSTRTGSAETLY | 0.63976705 |
| HQEDPGRI | ASSTRTGSAETLY | 0.83978176 |
| GTPGEPTP | ASSTRTGSAETLY | 0.7605227 |

| | | |
|---|---|---|
| GFFNSGS | ASSTRTGSAETLY | 0.76054686 |
| GAGGGGLTAGAN | ASSTRTGSAETLY | 0.88161343 |
| FVDALVNM | ASSTRTGSAETLY | 0.7785495 |
| FLTVAPEL | ASSTRTGSAETLY | 0.8654494 |
| FETLTAL | ASSTRTGSAETLY | 0.77353543 |
| DVINPIEL | ASSTRTGSAETLY | 0.704781 |
| APVSGAPV | ASSTRTGSAETLY | 0.8468907 |
| ALVGGLAW | ASSTRTGSAETLY | 0.87840486 |
| ALAAAAATAI | ASSTRTGSAETLY | 0.7185725 |
| AFGLGLLL | ASSTRTGSAETLY | 0.90734816 |
| AFFAAASL | ASSTRTGSAETLY | 0.77597487 |
| AAAALPAFSPP | ASSTRTGSAETLY | 0.90735567 |
| VVLAPAL | ASSTRTGSAETLY | 0.8807295 |
| TIGHVDHGKTTL | ASSTRTGSAETLY | 0.7820902 |
| LISNASDALDK | ASSTRTGSAETLY | 0.8076376 |
| LISNASDAL | ASSTRTGSAETLY | 0.8502831 |
| ISNASDALDK | ASSTRTGSAETLY | 0.7326057 |
| NQAETLVYQTE | ASSTRTGSAETLY | 0.85169256 |
| TGGTGGTAGAG | ASSTRTGSAETLY | 0.8955691 |
| TGSTAGGAGGA | ASSTRTGSAETLY | 0.78041226 |
| DPLAVPVAAGPV | ASSTRTGSAETLY | 0.9097462 |
| GGTGGTPGNGGN | ASSTRTGSAETLY | 0.87720954 |
| PAGLAGVDPATL | ASSTRTGSAETLY | 0.8804499 |
| GNNLVVEAPHAGASV | ASSTRTGSAETLY | 0.78400296 |
| KVTRIMSASTLEQPA | ASSTRTGSAETLY | 0.434457 |
| LGFALPIALPRRTVV | ASSTRTGSAETLY | 0.87295306 |
| TRLGNLTARKLVVLV | ASSTRTGSAETLY | 0.7055973 |
| WSQQRRLIYFPSAGP | ASSTRTGSAETLY | 0.88892233 |
| GPNGNPDPMAAAVDIR | ASSTRTGSAETLY | 0.8926324 |
| TQRCVSVADLRAAASR | ASSTRTGSAETLY | 0.80079174 |
| QFVLDVAEHPVEVLTETV | ASSTRTGSAETLY | 0.9051259 |
| GGAGGTGGAGGNGGTQAGDG | ASSTRTGSAETLY | 0.90770674 |
| PLLATLRAIAVDVDAKPAQV | ASSTRTGSAETLY | 0.55423546 |
| APKFRTGGYMIDFWGVGYQVAK | ASSTRTGSAETLY | 0.845981 |
| GAGGAGGAGGAAGGPGGTGGTG | ASSTRTGSAETLY | 0.814268 |
| IGALGLLATDVAPAGSVAFGVA | ASSTRTGSAETLY | 0.84177345 |
| AGGAGGIGGNGGAGGAGGVLMG | ASSTRTGSAETLY | 0.5031108 |
| ADELIGLL | ASSTRTGSAETLY | 0.89699835 |
| ADRALKA | ASSTRTGSAETLY | 0.82987 |
| AGAVGQVGGAGGA | ASSTRTGSAETLY | 0.14232026 |
| AGGGGGLGGAP | ASSTRTGSAETLY | 0.86351407 |
| AGGLLYGNGGAG | ASSTRTGSAETLY | 0.842947 |
| AGLESIGL | ASSTRTGSAETLY | 0.80024934 |
| AGLKTGII | ASSTRTGSAETLY | 0.8983325 |
| ASLAALLL | ASSTRTGSAETLY | 0.5942224 |
| ASVINISSI | ASSTRTGSAETLY | 0.766135 |
| ATPPGPA | ASSTRTGSAETLY | 0.88958585 |
| ATVASALV | ASSTRTGSAETLY | 0.90590614 |
| ATVASLIL | ASSTRTGSAETLY | 0.66169417 |
| AVNGPVLIRH | ASSTRTGSAETLY | 0.878361 |
| DGAAGAVGI | ASSTRTGSAETLY | 0.91165435 |

| | | |
|---|---|---|
| DPLLHGI | ASSTRTGSAETLY | 0.8825948 |
| ELGAALGAL | ASSTRTGSAETLY | 0.9069608 |
| ELGRTQAV | ASSTRTGSAETLY | 0.9127842 |
| FVAGGYLS | ASSTRTGSAETLY | 0.8057579 |
| GASGGAGGAGGW | ASSTRTGSAETLY | 0.89903927 |
| GGAGAGGDGGAGGI | ASSTRTGSAETLY | 0.90921307 |
| GTGHNGSQPGMGGNG | ASSTRTGSAETLY | 0.83629787 |
| GYAAASASAS | ASSTRTGSAETLY | 0.86655045 |
| HQVDVAAVQQG | ASSTRTGSAETLY | 0.80303234 |
| IGAAIAHT | ASSTRTGSAETLY | 0.84393716 |
| IGMIEVAQL | ASSTRTGSAETLY | 0.3490531 |
| IRLAPEHTSAMD | ASSTRTGSAETLY | 0.8728214 |
| IYNVSVL | ASSTRTGSAETLY | 0.9018141 |
| KQGFLGM | ASSTRTGSAETLY | 0.83348083 |
| LESVITQL | ASSTRTGSAETLY | 0.74180377 |
| LGLGALVVV | ASSTRTGSAETLY | 0.7553363 |
| LIGNGGNGGSGGI | ASSTRTGSAETLY | 0.8654689 |
| LNAINAPTL | ASSTRTGSAETLY | 0.850247 |
| LQLLNTGV | ASSTRTGSAETLY | 0.43284804 |
| LQTSIEF | ASSTRTGSAETLY | 0.77400076 |
| MSFAVSEL | ASSTRTGSAETLY | 0.90344596 |
| NTGALGL | ASSTRTGSAETLY | 0.9076149 |
| PPGAGGAGGAGTT | ASSTRTGSAETLY | 0.49531978 |
| QLVGLGI | ASSTRTGSAETLY | 0.9029453 |
| RFLEQVHQL | ASSTRTGSAETLY | 0.9022595 |
| RGVGYVLR | ASSTRTGSAETLY | 0.9128189 |
| SATGVFAGVI | ASSTRTGSAETLY | 0.8754577 |
| SLPAITLP | ASSTRTGSAETLY | 0.7731981 |
| STALAFTV | ASSTRTGSAETLY | 0.8962535 |
| SVLTVGLGK | ASSTRTGSAETLY | 0.8331499 |
| TALAGAGSGI | ASSTRTGSAETLY | 0.73692214 |
| TGLPFTGI | ASSTRTGSAETLY | 0.90285397 |
| VAEHVTV | ASSTRTGSAETLY | 0.8314667 |
| VDIVLNSL | ASSTRTGSAETLY | 0.78352034 |
| VLTGGGAVL | ASSTRTGSAETLY | 0.83005077 |
| VNVGLAYGI | ASSTRTGSAETLY | 0.911028 |
| VSSAAVGAV | ASSTRTGSAETLY | 0.7352384 |
| VTSVYEAGV | ASSTRTGSAETLY | 0.67112005 |
| WVALDRGA | ASSTRTGSAETLY | 0.7578792 |
| YLVGSVVI | ASSTRTGSAETLY | 0.8721876 |
| GPPGTGKTLL | ASSTRTGSAETLY | 0.90051126 |
| GVGMTKF | ASSTRTGSAETLY | 0.8813648 |
| IGIGGSDL | ASSTRTGSAETLY | 0.83813936 |
| IGTLLAEL | ASSTRTGSAETLY | 0.8854908 |
| IYTSGTT | ASSTRTGSAETLY | 0.8968538 |
| LSFPAGV | ASSTRTGSAETLY | 0.86790264 |
| LYTSGTT | ASSTRTGSAETLY | 0.8971745 |
| VGAFPFGF | ASSTRTGSAETLY | 0.77609265 |
| VGAGTVEF | ASSTRTGSAETLY | 0.9075427 |
| VGKSTLLL | ASSTRTGSAETLY | 0.8939017 |
| AALAHAGGGIV | ASSTRTGSAETLY | 0.904305 |

| | | |
|---|---|---|
| AALVAVSLL | ASSTRTGSAETLY | 0.90102696 |
| AAVSRAVAAL | ASSTRTGSAETLY | 0.8873406 |
| AGGSSGIGGF | ASSTRTGSAETLY | 0.88203764 |
| AGLTVANANG | ASSTRTGSAETLY | 0.34518868 |
| AGLVGAVL | ASSTRTGSAETLY | 0.8329145 |
| ALSQAFTV | ASSTRTGSAETLY | 0.87751627 |
| ALTFSVSL | ASSTRTGSAETLY | 0.8748723 |
| ALTGAALVAL | ASSTRTGSAETLY | 0.88673824 |
| ASFSIGAGVL | ASSTRTGSAETLY | 0.89585906 |
| DGSVYKI | ASSTRTGSAETLY | 0.81416523 |
| EQLAEIV | ASSTRTGSAETLY | 0.9085554 |
| FAVSELTVM | ASSTRTGSAETLY | 0.89747775 |
| GAAPLAAGL | ASSTRTGSAETLY | 0.5382569 |
| GADGGDGGSGGAGG | ASSTRTGSAETLY | 0.8348784 |
| GFGLDTGGAGG | ASSTRTGSAETLY | 0.9104496 |
| GLASGGLL | ASSTRTGSAETLY | 0.8781902 |
| GRIAGAAVLL | ASSTRTGSAETLY | 0.84883994 |
| GSAGSLNTSF | ASSTRTGSAETLY | 0.8420859 |
| GTGQIGI | ASSTRTGSAETLY | 0.91300803 |
| GTLIALIL | ASSTRTGSAETLY | 0.8111228 |
| IAATMGFVVG | ASSTRTGSAETLY | 0.9135806 |
| IFAQALL | ASSTRTGSAETLY | 0.20034228 |
| IGAAVALGF | ASSTRTGSAETLY | 0.8402852 |
| IGLTPLAL | ASSTRTGSAETLY | 0.81966615 |
| ILGSNTSTL | ASSTRTGSAETLY | 0.40822485 |
| IVFSLTVL | ASSTRTGSAETLY | 0.8984318 |
| LASKATGF | ASSTRTGSAETLY | 0.83771944 |
| LASVALIL | ASSTRTGSAETLY | 0.9003973 |
| LFAVGLTL | ASSTRTGSAETLY | 0.8545994 |
| LFNGGIGTYI | ASSTRTGSAETLY | 0.89279056 |
| LTLGAGII | ASSTRTGSAETLY | 0.86410564 |
| LVAAALGDI | ASSTRTGSAETLY | 0.87641 |
| NTAFVQLGI | ASSTRTGSAETLY | 0.7072964 |
| QLFSGAEL | ASSTRTGSAETLY | 0.8966108 |
| SAATFITL | ASSTRTGSAETLY | 0.9076507 |
| SNYGLANLGV | ASSTRTGSAETLY | 0.8988111 |
| SYGGAYAVM | ASSTRTGSAETLY | 0.60042596 |
| TTGVGGLTLG | ASSTRTGSAETLY | 0.82812965 |
| TTQTIYI | ASSTRTGSAETLY | 0.85761976 |
| VAAGAGLGAVI | ASSTRTGSAETLY | 0.23351982 |
| VAAVVTLL | ASSTRTGSAETLY | 0.2759987 |
| VGFLGGLI | ASSTRTGSAETLY | 0.8315312 |
| VGVIPILL | ASSTRTGSAETLY | 0.779449 |
| VIFASSLI | ASSTRTGSAETLY | 0.9028387 |
| VKGGDGGAAAG | ASSTRTGSAETLY | 0.8952748 |
| VTKTFTNI | ASSTRTGSAETLY | 0.85140246 |
| VVAAAILL | ASSTRTGSAETLY | 0.8968482 |
| YAALDVEL | ASSTRTGSAETLY | 0.87686145 |
| LASVALLL | ASSTRTGSAETLY | 0.08290954 |
| LDAVHRV | ASSTRTGSAETLY | 0.84292054 |
| QGLSSSIF | ASSTRTGSAETLY | 0.61062515 |

| | | |
|---|---|---|
| RGTFANIRL | ASSTRTGSAETLY | 0.91260695 |
| RQLWWGHRI | ASSTRTGSAETLY | 0.7154942 |
| YGPPGTGKTLL | ASSTRTGSAETLY | 0.78229207 |
| AALVGAPANVA | ASSTRTGSAETLY | 0.86322933 |
| AATFRRTE | ASSTRTGSAETLY | 0.8701464 |
| AGALGGGI | ASSTRTGSAETLY | 0.8685649 |
| AGGAGGAGGIGAG | ASSTRTGSAETLY | 0.7723576 |
| AGSGALGL | ASSTRTGSAETLY | 0.1454152 |
| AQIADQI | ASSTRTGSAETLY | 0.90077305 |
| ASAVAVTYL | ASSTRTGSAETLY | 0.61433876 |
| ASTVLGGTGG | ASSTRTGSAETLY | 0.664009 |
| AVGGEPAK | ASSTRTGSAETLY | 0.5307647 |
| CEQGTFGGTC | ASSTRTGSAETLY | 0.35949963 |
| DEGKAPAG | ASSTRTGSAETLY | 0.6055124 |
| DTVVFPAL | ASSTRTGSAETLY | 0.9046091 |
| FEGITMAG | ASSTRTGSAETLY | 0.8173397 |
| FPLVVQD | ASSTRTGSAETLY | 0.8201983 |
| GAFTTPQ | ASSTRTGSAETLY | 0.9052772 |
| GFGGFGTAGGVGGA | ASSTRTGSAETLY | 0.8703648 |
| GFLAQFL | ASSTRTGSAETLY | 0.83119816 |
| GGAGGAGGNAGVA | ASSTRTGSAETLY | 0.79682946 |
| GGVGTAGAGGAGGAGG | ASSTRTGSAETLY | 0.73867166 |
| GNGGGGDAGNAGSGGN | ASSTRTGSAETLY | 0.7453477 |
| GTGGAGGAGNDAGS | ASSTRTGSAETLY | 0.7239296 |
| GTTNTGWFNS | ASSTRTGSAETLY | 0.911939 |
| IFMGPLFTL | ASSTRTGSAETLY | 0.83993125 |
| ILAHAIEAI | ASSTRTGSAETLY | 0.6473106 |
| IRSAIREAL | ASSTRTGSAETLY | 0.88538826 |
| ITFAGGG | ASSTRTGSAETLY | 0.91092837 |
| ITGMSGHQEV | ASSTRTGSAETLY | 0.91611224 |
| IVAGLACTL | ASSTRTGSAETLY | 0.7479956 |
| IVGGGLAGA | ASSTRTGSAETLY | 0.8574653 |
| IVTGVLNI | ASSTRTGSAETLY | 0.8433419 |
| LAQQGCI | ASSTRTGSAETLY | 0.8513982 |
| LCLHVEN | ASSTRTGSAETLY | 0.83786166 |
| LDAISDGS | ASSTRTGSAETLY | 0.47560495 |
| LGGFAAML | ASSTRTGSAETLY | 0.9040669 |
| LGGKNPMI | ASSTRTGSAETLY | 0.92010283 |
| LGPKGVAAIL | ASSTRTGSAETLY | 0.8174536 |
| LMVGVSI | ASSTRTGSAETLY | 0.7093438 |
| LVGAAFVL | ASSTRTGSAETLY | 0.854392 |
| LVLAVAPL | ASSTRTGSAETLY | 0.7786411 |
| MAAIAVIDAA | ASSTRTGSAETLY | 0.8193929 |
| NNGINAELQ | ASSTRTGSAETLY | 0.87835175 |
| RGEPLVLVHG | ASSTRTGSAETLY | 0.533064 |
| RGGAGGAAGLI | ASSTRTGSAETLY | 0.807691 |
| RSYRMHPAV | ASSTRTGSAETLY | 0.91596854 |
| SAAHLLAGQLA | ASSTRTGSAETLY | 0.83310556 |
| SAAVGSY | ASSTRTGSAETLY | 0.8660374 |
| SPHAVLHRI | ASSTRTGSAETLY | 0.89810437 |
| SQIAKVL | ASSTRTGSAETLY | 0.87949324 |

| | | |
|---|---|---|
| TALGHLVP | ASSTRTGSAETLY | 0.9055741 |
| TAPTTSGAA | ASSTRTGSAETLY | 0.90436924 |
| TDGDDGNGGSPGT | ASSTRTGSAETLY | 0.84256244 |
| TDGLGEN | ASSTRTGSAETLY | 0.8717985 |
| TGGPPLPGQA | ASSTRTGSAETLY | 0.8465051 |
| TIPAITI | ASSTRTGSAETLY | 0.87103796 |
| TIPGFTI | ASSTRTGSAETLY | 0.8585375 |
| TSIASGALA | ASSTRTGSAETLY | 0.8157963 |
| TVTLPGAAI | ASSTRTGSAETLY | 0.90131825 |
| VASGLSAALP | ASSTRTGSAETLY | 0.88154435 |
| VGAGVIGI | ASSTRTGSAETLY | 0.9042666 |
| VGPKRFGV | ASSTRTGSAETLY | 0.70222914 |
| VNVAFPD | ASSTRTGSAETLY | 0.6785208 |
| VSMTNSGVS | ASSTRTGSAETLY | 0.89731944 |
| VTDVLGDLSV | ASSTRTGSAETLY | 0.18177344 |
| VVVVGAGFAGL | ASSTRTGSAETLY | 0.3371784 |
| VYTVGTVV | ASSTRTGSAETLY | 0.8787446 |
| WTTFSGP | ASSTRTGSAETLY | 0.9115849 |
| YDELVEAF | ASSTRTGSAETLY | 0.16516264 |
| YGTGGAGGNGG | ASSTRTGSAETLY | 0.72620535 |
| YTNVMAV | ASSTRTGSAETLY | 0.6913132 |
| FLEQVHQL | ASSTRTGSAETLY | 0.903555 |
| GLSLLAVAV | ASSTRTGSAETLY | 0.8590468 |
| LEQVHQL | ASSTRTGSAETLY | 0.5196838 |
| LNAAGAI | ASSTRTGSAETLY | 0.8904232 |
| PAVHASL | ASSTRTGSAETLY | 0.64948446 |
| TELEDDA | ASSTRTGSAETLY | 0.6362087 |
| TPPGPPN | ASSTRTGSAETLY | 0.8960368 |
| TPPPPPPP | ASSTRTGSAETLY | 0.9099277 |
| TQSPALI | ASSRTGTNTGQLY | 0.7326566 |
| EFAAAFPQ | ASSRTGTNTGQLY | 0.8996359 |
| LLSGNLDVL | ASSRTGTNTGQLY | 0.8367734 |
| GPANLGNYNI | ASSRTGTNTGQLY | 0.91504025 |
| TLGPTLP | ASSRTGTNTGQLY | 0.81538516 |
| VYAYSSCCK | ASSRTGTNTGQLY | 0.5327984 |
| APAPAPAPNI | ASSRTGTNTGQLY | 0.8933827 |
| PGGPVPIP | ASSRTGTNTGQLY | 0.8757359 |
| FLTVALEP | ASSRTGTNTGQLY | 0.86956275 |
| SQYFGANL | ASSRTGTNTGQLY | 0.89828765 |
| GGVKPAA | ASSRTGTNTGQLY | 0.84178823 |
| GGTGGAGGAGGW | ASSRTGTNTGQLY | 0.8316753 |
| GVGGVAGLGGAGG | ASSRTGTNTGQLY | 0.90165615 |
| VLLVLATTI | ASSRTGTNTGQLY | 0.89519083 |
| IGIFAAML | ASSRTGTNTGQLY | 0.60786164 |
| AVGDLID | ASSRTGTNTGQLY | 0.8915691 |
| NISPLPL | ASSRTGTNTGQLY | 0.86066556 |
| VTAVFVAI | ASSRTGTNTGQLY | 0.82018304 |
| SGDDNDAAL | ASSRTGTNTGQLY | 0.9146105 |
| AVAALAHS | ASSRTGTNTGQLY | 0.88047886 |
| GGVTVMATP | ASSRTGTNTGQLY | 0.89762723 |
| AGLAANVRD | ASSRTGTNTGQLY | 0.8946911 |

| | | |
|---|---|---|
| TNRPDLIDPALLRPG | ASSRTGTNTGQLY | 0.9155966 |
| LANLGVTAL | ASSRTGTNTGQLY | 0.45935762 |
| TGERIEL | ASSRTGTNTGQLY | 0.8807095 |
| NGEELAA | ASSRTGTNTGQLY | 0.8668647 |
| VLSLVWG | ASSRTGTNTGQLY | 0.89430046 |
| IGYALWL | ASSRTGTNTGQLY | 0.87395287 |
| LSLLLAPL | ASSRTGTNTGQLY | 0.64491415 |
| TLDEALK | ASSRTGTNTGQLY | 0.8791107 |
| TDGAGTAEGG | ASSRTGTNTGQLY | 0.90556335 |
| NAPGGAGGNGG | ASSRTGTNTGQLY | 0.87424636 |
| VSVFMGMGGNF | ASSRTGTNTGQLY | 0.8839437 |
| LASYLAAL | ASSRTGTNTGQLY | 0.9112965 |
| LVGGTPAL | ASSRTGTNTGQLY | 0.78562534 |
| VRNADVL | ASSRTGTNTGQLY | 0.84578186 |
| TCSSPFLVSAM | ASSRTGTNTGQLY | 0.9025357 |
| VHITDAE | ASSRTGTNTGQLY | 0.90128446 |
| EYAAPSF | ASSRTGTNTGQLY | 0.84482896 |
| VVDAFHGGL | ASSRTGTNTGQLY | 0.908524 |
| ASVVDPQ | ASSRTGTNTGQLY | 0.7823733 |
| SYNEFGGLS | ASSRTGTNTGQLY | 0.8890972 |
| DPDGSVY | ASSRTGTNTGQLY | 0.7701074 |
| MVSTIHATE | ASSRTGTNTGQLY | 0.8852863 |
| LAAGVADPV | ASSRTGTNTGQLY | 0.732939 |
| TKEQIAA | ASSRTGTNTGQLY | 0.88701886 |
| SLASPAAL | ASSRTGTNTGQLY | 0.8973106 |
| IGLVFYGV | ASSRTGTNTGQLY | 0.8923725 |
| FLLDASNL | ASSRTGTNTGQLY | 0.32805195 |
| YQVGNFP | ASSRTGTNTGQLY | 0.89207196 |
| ESLLSQST | ASSRTGTNTGQLY | 0.901662 |
| LFTVGFAAVG | ASSRTGTNTGQLY | 0.9169958 |
| ATPPTPT | ASSRTGTNTGQLY | 0.87308407 |
| TDLATLG | ASSRTGTNTGQLY | 0.8904896 |
| SAGGTGR | ASSRTGTNTGQLY | 0.8412172 |
| GFSNSGS | ASSRTGTNTGQLY | 0.8501508 |
| GGNGGSGGTSV | ASSRTGTNTGQLY | 0.7727034 |
| GTSVATGGAGN | ASSRTGTNTGQLY | 0.84338045 |
| NGGNGGSGGTS | ASSRTGTNTGQLY | 0.8260901 |
| AVVKFPH | ASSRTGTNTGQLY | 0.76175 |
| TPPPPPP | ASSRTGTNTGQLY | 0.89959717 |
| AAPAAAAAPA | ASSRTGTNTGQLY | 0.9019933 |
| IALNEDL | ASSRTGTNTGQLY | 0.8968195 |
| IVNVSSM | ASSRTGTNTGQLY | 0.9045775 |
| TGIPPAPR | ASSRTGTNTGQLY | 0.90465254 |
| FDLGGGTF | ASSRTGTNTGQLY | 0.9050064 |
| FDLGGGTFDVS | ASSRTGTNTGQLY | 0.80053043 |
| NEPTAAA | ASSRTGTNTGQLY | 0.8518119 |
| NEPTAAALA | ASSRTGTNTGQLY | 0.7094816 |
| GSIINISSI | ASSRTGTNTGQLY | 0.8262205 |
| VAPGNAGTA | ASSRTGTNTGQLY | 0.8016218 |
| TGGGASLE | ASSRTGTNTGQLY | 0.8749899 |
| VAVYNHYKRI | ASSRTGTNTGQLY | 0.1223999 |

| | | |
|---|---|---|
| TDPAPATT | ASSRTGTNTGQLY | 0.8630197 |
| TNRPDILDPALLRPG | ASSRTGTNTGQLY | 0.9089557 |
| LTTTTTS | ASSRTGTNTGQLY | 0.78097475 |
| PTGSGKTL | ASSRTGTNTGQLY | 0.8954079 |
| TGSGKTL | ASSRTGTNTGQLY | 0.8472324 |
| TIGHVDHGK | ASSRTGTNTGQLY | 0.8139551 |
| TIGHVDHGKTT | ASSRTGTNTGQLY | 0.89687407 |
| GHVDHGKTT | ASSRTGTNTGQLY | 0.7371244 |
| AQFEHTLL | ASSRTGTNTGQLY | 0.89413047 |
| IVNQDEL | ASSRTGTNTGQLY | 0.91282624 |
| TNGSQFF | ASSRTGTNTGQLY | 0.86943537 |
| TGKGIVE | ASSRTGTNTGQLY | 0.87105864 |
| LISNASDA | ASSRTGTNTGQLY | 0.86374164 |
| AELRTLGELTKIAKAHGAQ | ASSRTGTNTGQLY | 0.426767 |
| APNAVAAAEPSW | ASSRTGTNTGQLY | 0.8751239 |
| APQVAGAATAVL | ASSRTGTNTGQLY | 0.103016704 |
| ARRDPAAAAAILPTDA | ASSRTGTNTGQLY | 0.8555814 |
| DAAARDVLAEAGLAEYF | ASSRTGTNTGQLY | 0.70101285 |
| DDQIVLSAKANLAKVEQ | ASSRTGTNTGQLY | 0.8768908 |
| DELLPDTGEDSDEDGH | ASSRTGTNTGQLY | 0.8580839 |
| ELPEAPANRATVPQD | ASSRTGTNTGQLY | 0.8970033 |
| FGQSKIERVTGIG | ASSRTGTNTGQLY | 0.76992977 |
| GAGGTGSTAGGAGGAGGAGGLY | ASSRTGTNTGQLY | 0.8377348 |
| GHGGTGAAGGNGGTGGAGGSGI | ASSRTGTNTGQLY | 0.79102945 |
| GNGGNGGAGGEGGPGGAGGAGA | ASSRTGTNTGQLY | 0.6248595 |
| GSTAGNGGNGGNATNGTIAGQP | ASSRTGTNTGQLY | 0.89472437 |
| GTGGDGGNGGNGGTGG | ASSRTGTNTGQLY | 0.86272496 |
| GVRRPAGIPADAPV | ASSRTGTNTGQLY | 0.83007795 |
| GYRQVTDAHLLTLARR | ASSRTGTNTGQLY | 0.90051407 |
| IAKMGQDGHDRG | ASSRTGTNTGQLY | 0.86558867 |
| KAEPSIEIQYT | ASSRTGTNTGQLY | 0.57382435 |
| KPVRIRMAADEDTVAVT | ASSRTGTNTGQLY | 0.8435426 |
| MPPRIAGMRLLVIKPEP | ASSRTGTNTGQLY | 0.8602526 |
| MKFYAVALFRSRGVPPTR | ASSRTGTNTGQLY | 0.8358476 |
| NAGGFGILGAVAHSP | ASSRTGTNTGQLY | 0.103699595 |
| NEGREITLAMAEM | ASSRTGTNTGQLY | 0.8762859 |
| NELVEWSHPLGHRWRWE | ASSRTGTNTGQLY | 0.82548255 |
| NLGAGGTGGNGGNPG | ASSRTGTNTGQLY | 0.7558703 |
| NTEWPKWETVMAADPVD | ASSRTGTNTGQLY | 0.7738405 |
| PASAAPAETQP | ASSRTGTNTGQLY | 0.89501834 |
| QQSSDGKNGVRAAEITGE | ASSRTGTNTGQLY | 0.7469631 |
| RAPDAVTTSWWKE | ASSRTGTNTGQLY | 0.8823532 |
| SGFGNVGAHTSGWFNQ | ASSRTGTNTGQLY | 0.8947121 |
| SGVAGAVGQAASVGGLKV | ASSRTGTNTGQLY | 0.8971462 |
| SNGGAGTGSAGGNGGT | ASSRTGTNTGQLY | 0.91000354 |
| TGGAGGEGGAAG | ASSRTGTNTGQLY | 0.8804729 |
| TGGDGGAAGAGGAPGHG | ASSRTGTNTGQLY | 0.8702054 |
| TMPPLLAAIANAGRHGV | ASSRTGTNTGQLY | 0.88922113 |
| TSSTAGNSGGA | ASSRTGTNTGQLY | 0.8879619 |
| VPATVPVGILC | ASSRTGTNTGQLY | 0.39823353 |
| VQRPDSGGVVLTGRIS | ASSRTGTNTGQLY | 0.45432818 |

| | | |
|---|---|---|
| WTPNPVPATKGVPAGLVQAT | ASSRTGTNTGQLY | 0.87147665 |
| YALKRALAKFSDRFDVVII | ASSRTGTNTGQLY | 0.39882976 |
| YLIGSVIV | ASSRTGTNTGQLY | 0.8728474 |
| YGQLVAVID | ASSRTGTNTGQLY | 0.9111923 |
| VTVGGMTLPA | ASSRTGTNTGQLY | 0.8750075 |
| VLALPAV | ASSRTGTNTGQLY | 0.71930814 |
| VIPPTLNYET | ASSRTGTNTGQLY | 0.84817564 |
| VGMGGMHPSSG | ASSRTGTNTGQLY | 0.8134308 |
| VGLAGGAATA | ASSRTGTNTGQLY | 0.89428747 |
| VGGAGGAGGSGP | ASSRTGTNTGQLY | 0.9052607 |
| VFAAFMV | ASSRTGTNTGQLY | 0.9123846 |
| TLYDKLR | ASSRTGTNTGQLY | 0.89992225 |
| TLAVFTVL | ASSRTGTNTGQLY | 0.8213882 |
| TGSVQPARV | ASSRTGTNTGQLY | 0.89284503 |
| TGGGAGSGGT | ASSRTGTNTGQLY | 0.8276074 |
| SHLETSL | ASSRTGTNTGQLY | 0.85706854 |
| SFTTGMT | ASSRTGTNTGQLY | 0.8568003 |
| QVEVAKRMGVRQPTVS | ASSRTGTNTGQLY | 0.8838369 |
| QGSGGPALAM | ASSRTGTNTGQLY | 0.8627311 |
| MLAPSNP | ASSRTGTNTGQLY | 0.8184583 |
| LVNNAAI | ASSRTGTNTGQLY | 0.6084438 |
| LTGEINTLA | ASSRTGTNTGQLY | 0.9044634 |
| LSIFGPV | ASSRTGTNTGQLY | 0.8843787 |
| LLALLANL | ASSRTGTNTGQLY | 0.65981126 |
| LISNASDALDKLR | ASSRTGTNTGQLY | 0.839622 |
| LINEAVP | ASSRTGTNTGQLY | 0.9095958 |
| LGYAFSGL | ASSRTGTNTGQLY | 0.91095006 |
| LDFVAVAL | ASSRTGTNTGQLY | 0.7710024 |
| LAQPGAE | ASSRTGTNTGQLY | 0.7541046 |
| ISNLGAVTGF | ASSRTGTNTGQLY | 0.87140524 |
| HQEDPGRI | ASSRTGTNTGQLY | 0.41244102 |
| GTPGEPTP | ASSRTGTNTGQLY | 0.8858453 |
| GFFNSGS | ASSRTGTNTGQLY | 0.8461453 |
| GAGGGGLTAGAN | ASSRTGTNTGQLY | 0.8366382 |
| FVDALVNM | ASSRTGTNTGQLY | 0.7918345 |
| FLTVAPEL | ASSRTGTNTGQLY | 0.8680501 |
| FETLTAL | ASSRTGTNTGQLY | 0.81408405 |
| DVINPIEL | ASSRTGTNTGQLY | 0.90185535 |
| APVSGAPV | ASSRTGTNTGQLY | 0.1645312 |
| ALVGGLAW | ASSRTGTNTGQLY | 0.87747496 |
| ALAAAAATAI | ASSRTGTNTGQLY | 0.7735472 |
| AFGLGLLL | ASSRTGTNTGQLY | 0.83090985 |
| AFFAAASL | ASSRTGTNTGQLY | 0.46651417 |
| AAAALPAFSPP | ASSRTGTNTGQLY | 0.87217695 |
| VVLAPAL | ASSRTGTNTGQLY | 0.8983154 |
| TIGHVDHGKTTL | ASSRTGTNTGQLY | 0.9020157 |
| LISNASDALDK | ASSRTGTNTGQLY | 0.69071543 |
| LISNASDAL | ASSRTGTNTGQLY | 0.8935118 |
| ISNASDALDK | ASSRTGTNTGQLY | 0.9169063 |
| NQAETLVYQTE | ASSRTGTNTGQLY | 0.8140686 |
| TGGTGGTAGAG | ASSRTGTNTGQLY | 0.3373262 |

| | | |
|---|---|---|
| TGSTAGGAGGA | ASSRTGTNTGQLY | 0.8850604 |
| DPLAVPVAAGPV | ASSRTGTNTGQLY | 0.85856074 |
| GGTGGTPGNGGN | ASSRTGTNTGQLY | 0.8950047 |
| PAGLAGVDPATL | ASSRTGTNTGQLY | 0.89378905 |
| GNNLVVEAPHAGASV | ASSRTGTNTGQLY | 0.68549013 |
| KVTRIMSASTLEQPA | ASSRTGTNTGQLY | 0.9037972 |
| LGFALPIALPRRTVV | ASSRTGTNTGQLY | 0.89801747 |
| TRLGNLTARKLVVLV | ASSRTGTNTGQLY | 0.80230165 |
| WSQQRRLIYFPSAGP | ASSRTGTNTGQLY | 0.82748365 |
| GPNGNPDPMAAAVDIR | ASSRTGTNTGQLY | 0.8023201 |
| TQRCVSVADLRAAASR | ASSRTGTNTGQLY | 0.3431723 |
| QFVLDVAEHPVEVLTETV | ASSRTGTNTGQLY | 0.8946315 |
| GGAGGTGGAGGNGGTQAGDG | ASSRTGTNTGQLY | 0.79094994 |
| PLLATLRAIAVDVDAKPAQV | ASSRTGTNTGQLY | 0.7948661 |
| APKFRTGGYMIDFWGVGYQVAK | ASSRTGTNTGQLY | 0.8650263 |
| GAGGAGGAGGAAGGPGGTGGTG | ASSRTGTNTGQLY | 0.76263505 |
| IGALGLLATDVAPAGSVAFGVA | ASSRTGTNTGQLY | 0.7945734 |
| AGGAGGIGGNGGAGGAGGVLMG | ASSRTGTNTGQLY | 0.80008996 |
| ADELIGLL | ASSRTGTNTGQLY | 0.86664903 |
| ADRALKA | ASSRTGTNTGQLY | 0.8793084 |
| AGAVGQVGGAGGA | ASSRTGTNTGQLY | 0.86877435 |
| AGGGGGLGGAP | ASSRTGTNTGQLY | 0.88313836 |
| AGGLLYGNGGAG | ASSRTGTNTGQLY | 0.90580285 |
| AGLESIGL | ASSRTGTNTGQLY | 0.6252457 |
| AGLKTGII | ASSRTGTNTGQLY | 0.7850759 |
| ASLAALLL | ASSRTGTNTGQLY | 0.892704 |
| ASVINISSI | ASSRTGTNTGQLY | 0.58909476 |
| ATPPGPA | ASSRTGTNTGQLY | 0.8922807 |
| ATVASALV | ASSRTGTNTGQLY | 0.87361276 |
| ATVASLIL | ASSRTGTNTGQLY | 0.88084424 |
| AVNGPVLIRH | ASSRTGTNTGQLY | 0.8767061 |
| DGAAGAVGI | ASSRTGTNTGQLY | 0.91238153 |
| DPLLHGI | ASSRTGTNTGQLY | 0.8960141 |
| ELGAALGAL | ASSRTGTNTGQLY | 0.858801 |
| ELGRTQAV | ASSRTGTNTGQLY | 0.86989695 |
| FVAGGYLS | ASSRTGTNTGQLY | 0.7822082 |
| GASGGAGGAGGW | ASSRTGTNTGQLY | 0.8966494 |
| GGAGAGGDGGAGGI | ASSRTGTNTGQLY | 0.8981365 |
| GTGHNGSQPGMGGNG | ASSRTGTNTGQLY | 0.6948793 |
| GYAAASASAS | ASSRTGTNTGQLY | 0.8362185 |
| HQVDVAAVQQG | ASSRTGTNTGQLY | 0.9122317 |
| IGAAIAHT | ASSRTGTNTGQLY | 0.8976402 |
| IGMIEVAQL | ASSRTGTNTGQLY | 0.4134831 |
| IRLAPEHTSAMD | ASSRTGTNTGQLY | 0.8204634 |
| IYNVSVL | ASSRTGTNTGQLY | 0.5835624 |
| KQGFLGM | ASSRTGTNTGQLY | 0.8706814 |
| LESVITQL | ASSRTGTNTGQLY | 0.7005966 |
| LGLGALVVV | ASSRTGTNTGQLY | 0.8875895 |
| LIGNGGNGGSGGI | ASSRTGTNTGQLY | 0.8387302 |
| LNAINAPTL | ASSRTGTNTGQLY | 0.8957671 |
| LQLLNTGV | ASSRTGTNTGQLY | 0.8690387 |

| | | |
|---|---|---|
| LQTSIEF | ASSRTGTNTGQLY | 0.9089458 |
| MSFAVSEL | ASSRTGTNTGQLY | 0.8333339 |
| NTGALGL | ASSRTGTNTGQLY | 0.84803903 |
| PPGAGGAGGAGTT | ASSRTGTNTGQLY | 0.5700667 |
| QLVGLGI | ASSRTGTNTGQLY | 0.5499519 |
| RFLEQVHQL | ASSRTGTNTGQLY | 0.7997267 |
| RGVGYVLR | ASSRTGTNTGQLY | 0.8338541 |
| SATGVFAGVI | ASSRTGTNTGQLY | 0.29903582 |
| SLPAITLP | ASSRTGTNTGQLY | 0.908675 |
| STALAFTV | ASSRTGTNTGQLY | 0.87545496 |
| SVLTVGLGK | ASSRTGTNTGQLY | 0.89969015 |
| TALAGAGSGI | ASSRTGTNTGQLY | 0.8211126 |
| TGLPFTGI | ASSRTGTNTGQLY | 0.8434078 |
| VAEHVTV | ASSRTGTNTGQLY | 0.7776584 |
| VDIVLNSL | ASSRTGTNTGQLY | 0.89379877 |
| VLTGGGAVL | ASSRTGTNTGQLY | 0.689558 |
| VNVGLAYGI | ASSRTGTNTGQLY | 0.5989966 |
| VSSAAVGAV | ASSRTGTNTGQLY | 0.85775316 |
| VTSVYEAGV | ASSRTGTNTGQLY | 0.7411492 |
| WVALDRGA | ASSRTGTNTGQLY | 0.9077594 |
| YLVGSVVI | ASSRTGTNTGQLY | 0.90046227 |
| GPPGTGKTLL | ASSRTGTNTGQLY | 0.8761393 |
| GVGMTKF | ASSRTGTNTGQLY | 0.907202 |
| IGIGGSDL | ASSRTGTNTGQLY | 0.31524673 |
| IGTLLAEL | ASSRTGTNTGQLY | 0.80190694 |
| IYTSGTT | ASSRTGTNTGQLY | 0.9134227 |
| LSFPAGV | ASSRTGTNTGQLY | 0.88548124 |
| LYTSGTT | ASSRTGTNTGQLY | 0.8499085 |
| VGAFPFGF | ASSRTGTNTGQLY | 0.9060011 |
| VGAGTVEF | ASSRTGTNTGQLY | 0.78347087 |
| VGKSTLLL | ASSRTGTNTGQLY | 0.8481694 |
| AALAHAGGGIV | ASSRTGTNTGQLY | 0.8509708 |
| AALVAVSLL | ASSRTGTNTGQLY | 0.8830741 |
| AAVSRAVAAL | ASSRTGTNTGQLY | 0.8861346 |
| AGGSSGIGGF | ASSRTGTNTGQLY | 0.74308205 |
| AGLTVANANG | ASSRTGTNTGQLY | 0.8878145 |
| AGLVGAVL | ASSRTGTNTGQLY | 0.9136971 |
| ALSQAFTV | ASSRTGTNTGQLY | 0.8358409 |
| ALTFSVSL | ASSRTGTNTGQLY | 0.90129757 |
| ALTGAALVAL | ASSRTGTNTGQLY | 0.8829428 |
| ASFSIGAGVL | ASSRTGTNTGQLY | 0.8847493 |
| DGSVYKI | ASSRTGTNTGQLY | 0.7682607 |
| EQLAEIV | ASSRTGTNTGQLY | 0.8045365 |
| FAVSELTVM | ASSRTGTNTGQLY | 0.84742975 |
| GAAPLAAGL | ASSRTGTNTGQLY | 0.77042013 |
| GADGGDGGSGGAGG | ASSRTGTNTGQLY | 0.8701272 |
| GFGLDTGGAGG | ASSRTGTNTGQLY | 0.80951625 |
| GLASGGLL | ASSRTGTNTGQLY | 0.9198288 |
| GRIAGAAVLL | ASSRTGTNTGQLY | 0.89122194 |
| GSAGSLNTSF | ASSRTGTNTGQLY | 0.87606454 |
| GTGQIGI | ASSRTGTNTGQLY | 0.71439075 |

| | | |
|---|---|---|
| GTLIALIL | ASSRTGTNTGQLY | 0.88569456 |
| IAATMGFVVG | ASSRTGTNTGQLY | 0.8784951 |
| IFAQALL | ASSRTGTNTGQLY | 0.5727465 |
| IGAAVALGF | ASSRTGTNTGQLY | 0.9098119 |
| IGLTPLAL | ASSRTGTNTGQLY | 0.7812916 |
| ILGSNTSTL | ASSRTGTNTGQLY | 0.8412733 |
| IVFSLTVL | ASSRTGTNTGQLY | 0.90240407 |
| LASKATGF | ASSRTGTNTGQLY | 0.90930367 |
| LASVALIL | ASSRTGTNTGQLY | 0.8248807 |
| LFAVGLTL | ASSRTGTNTGQLY | 0.8393758 |
| LFNGGIGTYI | ASSRTGTNTGQLY | 0.70708555 |
| LTLGAGII | ASSRTGTNTGQLY | 0.9024906 |
| LVAAALGDI | ASSRTGTNTGQLY | 0.9028859 |
| NTAFVQLGI | ASSRTGTNTGQLY | 0.87436616 |
| QLFSGAEL | ASSRTGTNTGQLY | 0.9104459 |
| SAATFITL | ASSRTGTNTGQLY | 0.8956119 |
| SNYGLANLGV | ASSRTGTNTGQLY | 0.60698277 |
| SYGGAYAVM | ASSRTGTNTGQLY | 0.86269605 |
| TTGVGGLTLG | ASSRTGTNTGQLY | 0.8950243 |
| TTQTIYI | ASSRTGTNTGQLY | 0.89970624 |
| VAAGAGLGAVI | ASSRTGTNTGQLY | 0.91404694 |
| VAAVVTLL | ASSRTGTNTGQLY | 0.8856209 |
| VGFLGGLI | ASSRTGTNTGQLY | 0.81854117 |
| VGVIPILL | ASSRTGTNTGQLY | 0.90837914 |
| VIFASSLI | ASSRTGTNTGQLY | 0.9185391 |
| VKGGDGGAAAG | ASSRTGTNTGQLY | 0.9002619 |
| VTKTFTNI | ASSRTGTNTGQLY | 0.8576351 |
| VVAAAILL | ASSRTGTNTGQLY | 0.8450552 |
| YAALDVEL | ASSRTGTNTGQLY | 0.86759716 |
| LASVALLL | ASSRTGTNTGQLY | 0.8933489 |
| LDAVHRV | ASSRTGTNTGQLY | 0.7759606 |
| QGLSSSIF | ASSRTGTNTGQLY | 0.86592746 |
| RGTFANIRL | ASSRTGTNTGQLY | 0.8852481 |
| RQLWWGHRI | ASSRTGTNTGQLY | 0.66542643 |
| YGPPGTGKTLL | ASSRTGTNTGQLY | 0.8889076 |
| AALVGAPANVA | ASSRTGTNTGQLY | 0.9013367 |
| AATFRRTE | ASSRTGTNTGQLY | 0.83482945 |
| AGALGGGI | ASSRTGTNTGQLY | 0.8312831 |
| AGGAGGAGGIGAG | ASSRTGTNTGQLY | 0.9019387 |
| AGSGALGL | ASSRTGTNTGQLY | 0.9173895 |
| AQIADQI | ASSRTGTNTGQLY | 0.90569264 |
| ASAVAVTYL | ASSRTGTNTGQLY | 0.5028201 |
| ASTVLGGTGG | ASSRTGTNTGQLY | 0.46869522 |
| AVGGEPAK | ASSRTGTNTGQLY | 0.78089345 |
| CEQGTFGGTC | ASSRTGTNTGQLY | 0.89599234 |
| DEGKAPAG | ASSRTGTNTGQLY | 0.8087173 |
| DTVVFPAL | ASSRTGTNTGQLY | 0.78484046 |
| FEGITMAG | ASSRTGTNTGQLY | 0.8647423 |
| FPLVVQD | ASSRTGTNTGQLY | 0.9037874 |
| GAFTTPQ | ASSRTGTNTGQLY | 0.7892922 |
| GFGGFGTAGGVGGA | ASSRTGTNTGQLY | 0.6953179 |

| | | |
|---|---|---|
| GFLAQFL | ASSRTGTNTGQLY | 0.88292223 |
| GGAGGAGGNAGVA | ASSRTGTNTGQLY | 0.89994335 |
| GGVGTAGAGGAGGAGG | ASSRTGTNTGQLY | 0.76127756 |
| GNGGGGDAGNAGSGGN | ASSRTGTNTGQLY | 0.83682644 |
| GTGGAGGAGNDAGS | ASSRTGTNTGQLY | 0.7668812 |
| GTTNTGWFNS | ASSRTGTNTGQLY | 0.75297606 |
| IFMGPLFTL | ASSRTGTNTGQLY | 0.8717264 |
| ILAHAIEAI | ASSRTGTNTGQLY | 0.7327163 |
| IRSAIREAL | ASSRTGTNTGQLY | 0.9025748 |
| ITFAGGG | ASSRTGTNTGQLY | 0.74077356 |
| ITGMSGHQEV | ASSRTGTNTGQLY | 0.90214574 |
| IVAGLACTL | ASSRTGTNTGQLY | 0.8815345 |
| IVGGGLAGA | ASSRTGTNTGQLY | 0.8828889 |
| IVTGVLNI | ASSRTGTNTGQLY | 0.8365556 |
| LAQQGCI | ASSRTGTNTGQLY | 0.81123513 |
| LCLHVEN | ASSRTGTNTGQLY | 0.6578204 |
| LDAISDGS | ASSRTGTNTGQLY | 0.5741905 |
| LGGFAAML | ASSRTGTNTGQLY | 0.90134436 |
| LGGKNPMI | ASSRTGTNTGQLY | 0.21334778 |
| LGPKGVAAIL | ASSRTGTNTGQLY | 0.9006551 |
| LMVGVSI | ASSRTGTNTGQLY | 0.89056826 |
| LVGAAFVL | ASSRTGTNTGQLY | 0.69668484 |
| LVLAVAPL | ASSRTGTNTGQLY | 0.889122 |
| MAAIAVIDAA | ASSRTGTNTGQLY | 0.7311922 |
| NNGINAELQ | ASSRTGTNTGQLY | 0.7680263 |
| RGEPLVLVHG | ASSRTGTNTGQLY | 0.9018203 |
| RGGAGGAAGLI | ASSRTGTNTGQLY | 0.88807684 |
| RSYRMHPAV | ASSRTGTNTGQLY | 0.79770875 |
| SAAHLLAGQLA | ASSRTGTNTGQLY | 0.88530356 |
| SAAVGSY | ASSRTGTNTGQLY | 0.8709226 |
| SPHAVLHRI | ASSRTGTNTGQLY | 0.82778215 |
| SQIAKVL | ASSRTGTNTGQLY | 0.8383411 |
| TALGHLVP | ASSRTGTNTGQLY | 0.899348 |
| TAPTTSGAA | ASSRTGTNTGQLY | 0.875329 |
| TDGDDGNGGSPGT | ASSRTGTNTGQLY | 0.7712685 |
| TDGLGEN | ASSRTGTNTGQLY | 0.88763773 |
| TGGPPLPGQA | ASSRTGTNTGQLY | 0.74777657 |
| TIPAITI | ASSRTGTNTGQLY | 0.8777444 |
| TIPGFTI | ASSRTGTNTGQLY | 0.8483742 |
| TSIASGALA | ASSRTGTNTGQLY | 0.8682444 |
| TVTLPGAAI | ASSRTGTNTGQLY | 0.88744813 |
| VASGLSAALP | ASSRTGTNTGQLY | 0.7426808 |
| VGAGVIGI | ASSRTGTNTGQLY | 0.7909297 |
| VGPKRFGV | ASSRTGTNTGQLY | 0.8408859 |
| VNVAFPD | ASSRTGTNTGQLY | 0.8601254 |
| VSMTNSGVS | ASSRTGTNTGQLY | 0.89450604 |
| VTDVLGDLSV | ASSRTGTNTGQLY | 0.75221276 |
| VVVVGAGFAGL | ASSRTGTNTGQLY | 0.87745863 |
| VYTVGTVV | ASSRTGTNTGQLY | 0.826851 |
| WTTFSGP | ASSRTGTNTGQLY | 0.7480631 |
| YDELVEAF | ASSRTGTNTGQLY | 0.85706943 |

| | | |
|---|---|---|
| YGTGGAGGNGG | ASSRTGTNTGQLY | 0.8553108 |
| YTNVMAV | ASSRTGTNTGQLY | 0.8117795 |
| FLEQVHQL | ASSRTGTNTGQLY | 0.88624024 |
| GLSLLAVAV | ASSRTGTNTGQLY | 0.6960795 |
| LEQVHQL | ASSRTGTNTGQLY | 0.8875618 |
| LNAAGAI | ASSRTGTNTGQLY | 0.9103574 |
| PAVHASL | ASSRTGTNTGQLY | 0.850947 |
| TELEDDA | ASSRTGTNTGQLY | 0.8990423 |
| TPPGPPN | ASSRTGTNTGQLY | 0.87730396 |
| TPPPPPPP | ASSRTGTNTGQLY | 0.8757471 |
| TQSPALI | ASSPTGNNSPLY | 0.82275575 |
| EFAAAFPQ | ASSPTGNNSPLY | 0.81807595 |
| LLSGNLDVL | ASSPTGNNSPLY | 0.9010614 |
| GPANLGNYNI | ASSPTGNNSPLY | 0.8927655 |
| TLGPTLP | ASSPTGNNSPLY | 0.89741635 |
| VYAYSSCCK | ASSPTGNNSPLY | 0.88298357 |
| APAPAPAPNI | ASSPTGNNSPLY | 0.897874 |
| PGGPVPIP | ASSPTGNNSPLY | 0.8938569 |
| FLTVALEP | ASSPTGNNSPLY | 0.8105939 |
| SQYFGANL | ASSPTGNNSPLY | 0.62606406 |
| GGVKPAA | ASSPTGNNSPLY | 0.82019365 |
| GGTGGAGGAGGW | ASSPTGNNSPLY | 0.8329529 |
| GVGGVAGLGGAGG | ASSPTGNNSPLY | 0.8442522 |
| VLLVLATTI | ASSPTGNNSPLY | 0.5904116 |
| IGIFAAML | ASSPTGNNSPLY | 0.9027941 |
| AVGDLID | ASSPTGNNSPLY | 0.91634285 |
| NISPLPL | ASSPTGNNSPLY | 0.79616654 |
| VTAVFVAI | ASSPTGNNSPLY | 0.91198814 |
| SGDDNDAAL | ASSPTGNNSPLY | 0.6688253 |
| AVAALAHS | ASSPTGNNSPLY | 0.89313936 |
| GGVTVMATP | ASSPTGNNSPLY | 0.9073702 |
| AGLAANVRD | ASSPTGNNSPLY | 0.7581103 |
| TNRPDLIDPALLRPG | ASSPTGNNSPLY | 0.89000106 |
| LANLGVTAL | ASSPTGNNSPLY | 0.7732663 |
| TGERIEL | ASSPTGNNSPLY | 0.8216398 |
| NGEELAA | ASSPTGNNSPLY | 0.90507436 |
| VLSLVWG | ASSPTGNNSPLY | 0.8367605 |
| IGYALWL | ASSPTGNNSPLY | 0.8738191 |
| LSLLLAPL | ASSPTGNNSPLY | 0.8115045 |
| TLDEALK | ASSPTGNNSPLY | 0.7113384 |
| TDGAGTAEGG | ASSPTGNNSPLY | 0.85970396 |
| NAPGGAGGNGG | ASSPTGNNSPLY | 0.9096246 |
| VSVFMGMGGNF | ASSPTGNNSPLY | 0.8061018 |
| LASYLAAL | ASSPTGNNSPLY | 0.9183843 |
| LVGGTPAL | ASSPTGNNSPLY | 0.80950207 |
| VRNADVL | ASSPTGNNSPLY | 0.9133121 |
| TCSSPFLVSAM | ASSPTGNNSPLY | 0.897378 |
| VHITDAE | ASSPTGNNSPLY | 0.8574096 |
| EYAAPSF | ASSPTGNNSPLY | 0.8364453 |
| VVDAFHGGL | ASSPTGNNSPLY | 0.90334964 |
| ASVVDPQ | ASSPTGNNSPLY | 0.85662675 |

| | | |
|---|---|---|
| SYNEFGGLS | ASSPTGNNSPLY | 0.89267695 |
| DPDGSVY | ASSPTGNNSPLY | 0.9108314 |
| MVSTIHATE | ASSPTGNNSPLY | 0.168906 |
| LAAGVADPV | ASSPTGNNSPLY | 0.7687813 |
| TKEQIAA | ASSPTGNNSPLY | 0.65727335 |
| SLASPAAL | ASSPTGNNSPLY | 0.46947384 |
| IGLVFYGV | ASSPTGNNSPLY | 0.19938016 |
| FLLDASNL | ASSPTGNNSPLY | 0.12969832 |
| YQVGNFP | ASSPTGNNSPLY | 0.27248368 |
| ESLLSQST | ASSPTGNNSPLY | 0.15896104 |
| LFTVGFAAVG | ASSPTGNNSPLY | 0.68238986 |
| ATPPTPT | ASSPTGNNSPLY | 0.4309215 |
| TDLATLG | ASSPTGNNSPLY | 0.22038811 |
| SAGGTGR | ASSPTGNNSPLY | 0.72676814 |
| GFSNSGS | ASSPTGNNSPLY | 0.72615665 |
| GGNGGSGGTSV | ASSPTGNNSPLY | 0.14076081 |
| GTSVATGGAGN | ASSPTGNNSPLY | 0.32466915 |
| NGGNGGSGGTS | ASSPTGNNSPLY | 0.19749425 |
| AVVKFPH | ASSPTGNNSPLY | 0.27894342 |
| TPPPPPP | ASSPTGNNSPLY | 0.12143322 |
| AAPAAAAAPA | ASSPTGNNSPLY | 0.53948975 |
| IALNEDL | ASSPTGNNSPLY | 0.2822709 |
| IVNVSSM | ASSPTGNNSPLY | 0.11630885 |
| TGIPPAPR | ASSPTGNNSPLY | 0.34472752 |
| FDLGGGTF | ASSPTGNNSPLY | 0.28361204 |
| FDLGGGTFDVS | ASSPTGNNSPLY | 0.21907206 |
| NEPTAAA | ASSPTGNNSPLY | 0.72189033 |
| NEPTAAALA | ASSPTGNNSPLY | 0.2238811 |
| GSIINISSI | ASSPTGNNSPLY | 0.16635905 |
| VAPGNAGTA | ASSPTGNNSPLY | 0.64101505 |
| TGGGASLE | ASSPTGNNSPLY | 0.73571086 |
| VAVYNHYKRI | ASSPTGNNSPLY | 0.15323107 |
| TDPAPATT | ASSPTGNNSPLY | 0.6397114 |
| TNRPDILDPALLRPG | ASSPTGNNSPLY | 0.7927179 |
| LTTTTTS | ASSPTGNNSPLY | 0.39224887 |
| PTGSGKTL | ASSPTGNNSPLY | 0.70209783 |
| TGSGKTL | ASSPTGNNSPLY | 0.76491624 |
| TIGHVDHGK | ASSPTGNNSPLY | 0.21639645 |
| TIGHVDHGKTT | ASSPTGNNSPLY | 0.6479106 |
| GHVDHGKTT | ASSPTGNNSPLY | 0.7839593 |
| AQFEHTLL | ASSPTGNNSPLY | 0.25343564 |
| IVNQDEL | ASSPTGNNSPLY | 0.3656842 |
| TNGSQFF | ASSPTGNNSPLY | 0.17327726 |
| TGKGIVE | ASSPTGNNSPLY | 0.30252677 |
| LISNASDA | ASSPTGNNSPLY | 0.13220456 |
| AELRTLGELTKIAKAHGAQ | ASSPTGNNSPLY | 0.63522905 |
| APNAVAAAEPSW | ASSPTGNNSPLY | 0.5962173 |
| APQVAGAATAVL | ASSPTGNNSPLY | 0.34694886 |
| ARRDPAAAAAILPTDA | ASSPTGNNSPLY | 0.17490935 |
| DAAARDVLAEAGLAEYF | ASSPTGNNSPLY | 0.1693986 |
| DDQIVLSAKANLAKVEQ | ASSPTGNNSPLY | 0.5279395 |

| | | |
|---|---|---|
| DELLPDTGEDSDEDGH | ASSPTGNNSPLY | 0.35201168 |
| ELPEAPANRATVPQD | ASSPTGNNSPLY | 0.14580336 |
| FGQSKIERVTGIG | ASSPTGNNSPLY | 0.2655732 |
| GAGGTGSTAGGAGGAGGAGGLY | ASSPTGNNSPLY | 0.6164895 |
| GHGGTGAAGGNGGTGGAGGSGI | ASSPTGNNSPLY | 0.6473589 |
| GNGGNGGAGGEGGPGGAGGAGA | ASSPTGNNSPLY | 0.14562613 |
| GSTAGNGGNGGNATNGTIAGQP | ASSPTGNNSPLY | 0.7761314 |
| GTGGDGGNGGNGGTGG | ASSPTGNNSPLY | 0.7745521 |
| GVRRPAGIPADAPV | ASSPTGNNSPLY | 0.8062702 |
| GYRQVTDAHLLTLARR | ASSPTGNNSPLY | 0.3259604 |
| IAKMGQDGHDRG | ASSPTGNNSPLY | 0.16881649 |
| KAEPSIEIQYT | ASSPTGNNSPLY | 0.7646606 |
| KPVRIRMAADEDTVAVT | ASSPTGNNSPLY | 0.4618465 |
| MPPRIAGMRLLVIKPEP | ASSPTGNNSPLY | 0.16965732 |
| MKFYAVALFRSRGVPPTR | ASSPTGNNSPLY | 0.59073037 |
| NAGGFGILGAVAHSP | ASSPTGNNSPLY | 0.22370863 |
| NEGREITLAMAEM | ASSPTGNNSPLY | 0.20872593 |
| NELVEWSHPLGHRWRWE | ASSPTGNNSPLY | 0.6490569 |
| NLGAGGTGGNGGNPG | ASSPTGNNSPLY | 0.65893275 |
| NTEWPKWETVMAADPVD | ASSPTGNNSPLY | 0.18844603 |
| PASAAPAETQP | ASSPTGNNSPLY | 0.1555146 |
| QQSSDGKNGVRAAEITGE | ASSPTGNNSPLY | 0.23426446 |
| RAPDAVTTSWWKE | ASSPTGNNSPLY | 0.5193962 |
| SGFGNVGAHTSGWFNQ | ASSPTGNNSPLY | 0.6624509 |
| SGVAGAVGQAASVGGLKV | ASSPTGNNSPLY | 0.38673198 |
| SNGGAGTGSAGGNGGT | ASSPTGNNSPLY | 0.38846302 |
| TGGAGGEGGAAG | ASSPTGNNSPLY | 0.70274615 |
| TGGDGGAAGAGGAPGHG | ASSPTGNNSPLY | 0.53580546 |
| TMPPLLAAIANAGRHGV | ASSPTGNNSPLY | 0.34595257 |
| TSSTAGNSGGA | ASSPTGNNSPLY | 0.6879908 |
| VPATVPVGILC | ASSPTGNNSPLY | 0.2262303 |
| VQRPDSGGVVLTGRIS | ASSPTGNNSPLY | 0.6403576 |
| WTPNPVPATKGVPAGLVQAT | ASSPTGNNSPLY | 0.56633055 |
| YALKRALAKFSDRFDVVII | ASSPTGNNSPLY | 0.61633176 |
| YLIGSVIV | ASSPTGNNSPLY | 0.7176362 |
| YGQLVAVID | ASSPTGNNSPLY | 0.42285442 |
| VTVGGMTLPA | ASSPTGNNSPLY | 0.44606444 |
| VLALPAV | ASSPTGNNSPLY | 0.12862268 |
| VIPPTLNYET | ASSPTGNNSPLY | 0.28068346 |
| VGMGGMHPSSG | ASSPTGNNSPLY | 0.62229264 |
| VGLAGGAATA | ASSPTGNNSPLY | 0.5522251 |
| VGGAGGAGGSGP | ASSPTGNNSPLY | 0.6590314 |
| VFAAFMV | ASSPTGNNSPLY | 0.50718546 |
| TLYDKLR | ASSPTGNNSPLY | 0.23585325 |
| TLAVFTVL | ASSPTGNNSPLY | 0.74227417 |
| TGSVQPARV | ASSPTGNNSPLY | 0.767261 |
| TGGGAGSGGT | ASSPTGNNSPLY | 0.14249173 |
| SHLETSL | ASSPTGNNSPLY | 0.24597605 |
| SFTTGMT | ASSPTGNNSPLY | 0.78874546 |
| QVEVAKRMGVRQPTVS | ASSPTGNNSPLY | 0.68876505 |
| QGSGGPALAM | ASSPTGNNSPLY | 0.3091379 |

| | | |
|---|---|---|
| MLAPSNP | ASSPTGNNSPLY | 0.3277266 |
| LVNNAAI | ASSPTGNNSPLY | 0.75610226 |
| LTGEINTLA | ASSPTGNNSPLY | 0.29773086 |
| LSIFGPV | ASSPTGNNSPLY | 0.7851163 |
| LLALLANL | ASSPTGNNSPLY | 0.12276887 |
| LISNASDALDKLR | ASSPTGNNSPLY | 0.3441686 |
| LINEAVP | ASSPTGNNSPLY | 0.22375174 |
| LGYAFSGL | ASSPTGNNSPLY | 0.1272394 |
| LDFVAVAL | ASSPTGNNSPLY | 0.48088133 |
| LAQPGAE | ASSPTGNNSPLY | 0.23056403 |
| ISNLGAVTGF | ASSPTGNNSPLY | 0.6958296 |
| HQEDPGRI | ASSPTGNNSPLY | 0.3270589 |
| GTPGEPTP | ASSPTGNNSPLY | 0.51291937 |
| GFFNSGS | ASSPTGNNSPLY | 0.62616926 |
| GAGGGGLTAGAN | ASSPTGNNSPLY | 0.40250307 |
| FVDALVNM | ASSPTGNNSPLY | 0.17264777 |
| FLTVAPEL | ASSPTGNNSPLY | 0.4827279 |
| FETLTAL | ASSPTGNNSPLY | 0.73275316 |
| DVINPIEL | ASSPTGNNSPLY | 0.57783437 |
| APVSGAPV | ASSPTGNNSPLY | 0.13748375 |
| ALVGGLAW | ASSPTGNNSPLY | 0.5650519 |
| ALAAAAATAI | ASSPTGNNSPLY | 0.28714025 |
| AFGLGLLL | ASSPTGNNSPLY | 0.116836786 |
| AFFAAASL | ASSPTGNNSPLY | 0.12389187 |
| AAAALPAFSPP | ASSPTGNNSPLY | 0.31055653 |
| VVLAPAL | ASSPTGNNSPLY | 0.2746582 |
| TIGHVDHGKTTL | ASSPTGNNSPLY | 0.7022571 |
| LISNASDALDK | ASSPTGNNSPLY | 0.66804284 |
| LISNASDAL | ASSPTGNNSPLY | 0.32092634 |
| ISNASDALDK | ASSPTGNNSPLY | 0.69958806 |
| NQAETLVYQTE | ASSPTGNNSPLY | 0.50525326 |
| TGGTGGTAGAG | ASSPTGNNSPLY | 0.11252569 |
| TGSTAGGAGGA | ASSPTGNNSPLY | 0.26541492 |
| DPLAVPVAAGPV | ASSPTGNNSPLY | 0.14609556 |
| GGTGGTPGNGGN | ASSPTGNNSPLY | 0.7132468 |
| PAGLAGVDPATL | ASSPTGNNSPLY | 0.15823105 |
| GNNLVVEAPHAGASV | ASSPTGNNSPLY | 0.23011021 |
| KVTRIMSASTLEQPA | ASSPTGNNSPLY | 0.40902516 |
| LGFALPIALPRRTVV | ASSPTGNNSPLY | 0.62418866 |
| TRLGNLTARKLVVLV | ASSPTGNNSPLY | 0.49664852 |
| WSQQRRLIYFPSAGP | ASSPTGNNSPLY | 0.19982558 |
| GPNGNPDPMAAAVDIR | ASSPTGNNSPLY | 0.12138362 |
| TQRCVSVADLRAAASR | ASSPTGNNSPLY | 0.5040445 |
| QFVLDVAEHPVEVLTETV | ASSPTGNNSPLY | 0.13593629 |
| GGAGGTGGAGGNGGTQAGDG | ASSPTGNNSPLY | 0.16272864 |
| PLLATLRAIAVDVDAKPAQV | ASSPTGNNSPLY | 0.1311778 |
| APKFRTGGYMIDFWGVGYQVAK | ASSPTGNNSPLY | 0.112853274 |
| GAGGAGGAGGAAGGPGGTGGTG | ASSPTGNNSPLY | 0.14175881 |
| IGALGLLATDVAPAGSVAFGVA | ASSPTGNNSPLY | 0.65381014 |
| AGGAGGIGGNGGAGGAGGVLMG | ASSPTGNNSPLY | 0.35950094 |
| ADELIGLL | ASSPTGNNSPLY | 0.3404854 |

| | | |
|---|---|---|
| ADRALKA | ASSPTGNNSPLY | 0.76744473 |
| AGAVGQVGGAGGA | ASSPTGNNSPLY | 0.71051496 |
| AGGGGGLGGAP | ASSPTGNNSPLY | 0.36493456 |
| AGGLLYGNGGAG | ASSPTGNNSPLY | 0.18946402 |
| AGLESIGL | ASSPTGNNSPLY | 0.15921725 |
| AGLKTGII | ASSPTGNNSPLY | 0.16248 |
| ASLAALLL | ASSPTGNNSPLY | 0.1674363 |
| ASVINISSI | ASSPTGNNSPLY | 0.76909477 |
| ATPPGPA | ASSPTGNNSPLY | 0.34649277 |
| ATVASALV | ASSPTGNNSPLY | 0.14206459 |
| ATVASLIL | ASSPTGNNSPLY | 0.60797846 |
| AVNGPVLIRH | ASSPTGNNSPLY | 0.7741393 |
| DGAAGAVGI | ASSPTGNNSPLY | 0.8304589 |
| DPLLHGI | ASSPTGNNSPLY | 0.17818934 |
| ELGAALGAL | ASSPTGNNSPLY | 0.36133593 |
| ELGRTQAV | ASSPTGNNSPLY | 0.31019914 |
| FVAGGYLS | ASSPTGNNSPLY | 0.44911647 |
| GASGGAGGAGGW | ASSPTGNNSPLY | 0.29373908 |
| GGAGAGGDGGAGGI | ASSPTGNNSPLY | 0.14811161 |
| GTGHNGSQPGMGGNG | ASSPTGNNSPLY | 0.6396257 |
| GYAAASASAS | ASSPTGNNSPLY | 0.83759606 |
| HQVDVAAVQQG | ASSPTGNNSPLY | 0.22746895 |
| IGAAIAHT | ASSPTGNNSPLY | 0.13422796 |
| IGMIEVAQL | ASSPTGNNSPLY | 0.4044593 |
| IRLAPEHTSAMD | ASSPTGNNSPLY | 0.20691386 |
| IYNVSVL | ASSPTGNNSPLY | 0.27207643 |
| KQGFLGM | ASSPTGNNSPLY | 0.3695068 |
| LESVITQL | ASSPTGNNSPLY | 0.13700679 |
| LGLGALVVV | ASSPTGNNSPLY | 0.26046017 |
| LIGNGGNGGSGGI | ASSPTGNNSPLY | 0.80230665 |
| LNAINAPTL | ASSPTGNNSPLY | 0.37167853 |
| LQLLNTGV | ASSPTGNNSPLY | 0.6565102 |
| LQTSIEF | ASSPTGNNSPLY | 0.7526344 |
| MSFAVSEL | ASSPTGNNSPLY | 0.64154184 |
| NTGALGL | ASSPTGNNSPLY | 0.7900971 |
| PPGAGGAGGAGTT | ASSPTGNNSPLY | 0.78538704 |
| QLVGLGI | ASSPTGNNSPLY | 0.2511156 |
| RFLEQVHQL | ASSPTGNNSPLY | 0.41350895 |
| RGVGYVLR | ASSPTGNNSPLY | 0.65026575 |
| SATGVFAGVI | ASSPTGNNSPLY | 0.42180854 |
| SLPAITLP | ASSPTGNNSPLY | 0.41258755 |
| STALAFTV | ASSPTGNNSPLY | 0.25685963 |
| SVLTVGLGK | ASSPTGNNSPLY | 0.73484415 |
| TALAGAGSGI | ASSPTGNNSPLY | 0.39227802 |
| TGLPFTGI | ASSPTGNNSPLY | 0.7524897 |
| VAEHVTV | ASSPTGNNSPLY | 0.16286111 |
| VDIVLNSL | ASSPTGNNSPLY | 0.15569183 |
| VLTGGGAVL | ASSPTGNNSPLY | 0.75683284 |
| VNVGLAYGI | ASSPTGNNSPLY | 0.117715396 |
| VSSAAVGAV | ASSPTGNNSPLY | 0.14366469 |
| VTSVYEAGV | ASSPTGNNSPLY | 0.4633175 |

| | | |
|---|---|---|
| WVALDRGA | ASSPTGNNSPLY | 0.79369354 |
| YLVGSVVI | ASSPTGNNSPLY | 0.11529413 |
| GPPGTGKTLL | ASSPTGNNSPLY | 0.16712417 |
| GVGMTKF | ASSPTGNNSPLY | 0.1577232 |
| IGIGGSDL | ASSPTGNNSPLY | 0.663935 |
| IGTLLAEL | ASSPTGNNSPLY | 0.31204158 |
| IYTSGTT | ASSPTGNNSPLY | 0.13254777 |
| LSFPAGV | ASSPTGNNSPLY | 0.5771297 |
| LYTSGTT | ASSPTGNNSPLY | 0.13352735 |
| VGAFPFGF | ASSPTGNNSPLY | 0.14548104 |
| VGAGTVEF | ASSPTGNNSPLY | 0.4929191 |
| VGKSTLLL | ASSPTGNNSPLY | 0.7972002 |
| AALAHAGGGIV | ASSPTGNNSPLY | 0.15409622 |
| AALVAVSLL | ASSPTGNNSPLY | 0.5621463 |
| AAVSRAVAAL | ASSPTGNNSPLY | 0.386217 |
| AGGSSGIGGF | ASSPTGNNSPLY | 0.8341463 |
| AGLTVANANG | ASSPTGNNSPLY | 0.24018942 |
| AGLVGAVL | ASSPTGNNSPLY | 0.14509091 |
| ALSQAFTV | ASSPTGNNSPLY | 0.6652576 |
| ALTFSVSL | ASSPTGNNSPLY | 0.6901481 |
| ALTGAALVAL | ASSPTGNNSPLY | 0.28613135 |
| ASFSIGAGVL | ASSPTGNNSPLY | 0.7653458 |
| DGSVYKI | ASSPTGNNSPLY | 0.3757575 |
| EQLAEIV | ASSPTGNNSPLY | 0.37089813 |
| FAVSELTVM | ASSPTGNNSPLY | 0.70815134 |
| GAAPLAAGL | ASSPTGNNSPLY | 0.66566575 |
| GADGGDGGSGGAGG | ASSPTGNNSPLY | 0.14525987 |
| GFGLDTGGAGG | ASSPTGNNSPLY | 0.5031347 |
| GLASGGLL | ASSPTGNNSPLY | 0.42498383 |
| GRIAGAAVLL | ASSPTGNNSPLY | 0.21439253 |
| GSAGSLNTSF | ASSPTGNNSPLY | 0.756754 |
| GTGQIGI | ASSPTGNNSPLY | 0.41352913 |
| GTLIALIL | ASSPTGNNSPLY | 0.68332297 |
| IAATMGFVVG | ASSPTGNNSPLY | 0.42904782 |
| IFAQALL | ASSPTGNNSPLY | 0.78335804 |
| IGAAVALGF | ASSPTGNNSPLY | 0.1369881 |
| IGLTPLAL | ASSPTGNNSPLY | 0.6810317 |
| ILGSNTSTL | ASSPTGNNSPLY | 0.45506075 |
| IVFSLTVL | ASSPTGNNSPLY | 0.66679263 |
| LASKATGF | ASSPTGNNSPLY | 0.6005504 |
| LASVALIL | ASSPTGNNSPLY | 0.1866596 |
| LFAVGLTL | ASSPTGNNSPLY | 0.47648865 |
| LFNGGIGTYI | ASSPTGNNSPLY | 0.704257 |
| LTLGAGII | ASSPTGNNSPLY | 0.3209971 |
| LVAAALGDI | ASSPTGNNSPLY | 0.5626416 |
| NTAFVQLGI | ASSPTGNNSPLY | 0.8059258 |
| QLFSGAEL | ASSPTGNNSPLY | 0.1834539 |
| SAATFITL | ASSPTGNNSPLY | 0.29844528 |
| SNYGLANLGV | ASSPTGNNSPLY | 0.57972616 |
| SYGGAYAVM | ASSPTGNNSPLY | 0.7357134 |
| TTGVGGLTLG | ASSPTGNNSPLY | 0.33498847 |

| | | |
|---|---|---|
| TTQTIYI | ASSPTGNNSPLY | 0.76187825 |
| VAAGAGLGAVI | ASSPTGNNSPLY | 0.19843726 |
| VAAVVTLL | ASSPTGNNSPLY | 0.6655253 |
| VGFLGGLI | ASSPTGNNSPLY | 0.4466043 |
| VGVIPILL | ASSPTGNNSPLY | 0.5374023 |
| VIFASSLI | ASSPTGNNSPLY | 0.15457818 |
| VKGGDGGAAAG | ASSPTGNNSPLY | 0.6612632 |
| VTKTFTNI | ASSPTGNNSPLY | 0.7258111 |
| VVAAAILL | ASSPTGNNSPLY | 0.6884047 |
| YAALDVEL | ASSPTGNNSPLY | 0.12051454 |
| LASVALLL | ASSPTGNNSPLY | 0.6107569 |
| LDAVHRV | ASSPTGNNSPLY | 0.70689833 |
| QGLSSSIF | ASSPTGNNSPLY | 0.82704055 |
| RGTFANIRL | ASSPTGNNSPLY | 0.43383312 |
| RQLWWGHRI | ASSPTGNNSPLY | 0.567847 |
| YGPPGTGKTLL | ASSPTGNNSPLY | 0.24040717 |
| AALVGAPANVA | ASSPTGNNSPLY | 0.31469458 |
| AATFRRTE | ASSPTGNNSPLY | 0.21663022 |
| AGALGGGI | ASSPTGNNSPLY | 0.378657 |
| AGGAGGAGGIGAG | ASSPTGNNSPLY | 0.39112267 |
| AGSGALGL | ASSPTGNNSPLY | 0.22689162 |
| AQIADQI | ASSPTGNNSPLY | 0.7149386 |
| ASAVAVTYL | ASSPTGNNSPLY | 0.7576032 |
| ASTVLGGTGG | ASSPTGNNSPLY | 0.6857226 |
| AVGGEPAK | ASSPTGNNSPLY | 0.77727544 |
| CEQGTFGGTC | ASSPTGNNSPLY | 0.7925024 |
| DEGKAPAG | ASSPTGNNSPLY | 0.6058407 |
| DTVVFPAL | ASSPTGNNSPLY | 0.19589548 |
| FEGITMAG | ASSPTGNNSPLY | 0.29333287 |
| FPLVVQD | ASSPTGNNSPLY | 0.1529957 |
| GAFTTPQ | ASSPTGNNSPLY | 0.24827258 |
| GFGGFGTAGGVGGA | ASSPTGNNSPLY | 0.21305412 |
| GFLAQFL | ASSPTGNNSPLY | 0.46344775 |
| GGAGGAGGNAGVA | ASSPTGNNSPLY | 0.11382358 |
| GGVGTAGAGGAGGAGG | ASSPTGNNSPLY | 0.71988636 |
| GNGGGGDAGNAGSGGN | ASSPTGNNSPLY | 0.7685562 |
| GTGGAGGAGNDAGS | ASSPTGNNSPLY | 0.35246846 |
| GTTNTGWFNS | ASSPTGNNSPLY | 0.564774 |
| IFMGPLFTL | ASSPTGNNSPLY | 0.5569972 |
| ILAHAIEAI | ASSPTGNNSPLY | 0.2566046 |
| IRSAIREAL | ASSPTGNNSPLY | 0.7182641 |
| ITFAGGG | ASSPTGNNSPLY | 0.19078442 |
| ITGMSGHQEV | ASSPTGNNSPLY | 0.63467956 |
| IVAGLACTL | ASSPTGNNSPLY | 0.7153867 |
| IVGGGLAGA | ASSPTGNNSPLY | 0.43721238 |
| IVTGVLNI | ASSPTGNNSPLY | 0.59727716 |
| LAQQGCI | ASSPTGNNSPLY | 0.39684796 |
| LCLHVEN | ASSPTGNNSPLY | 0.13214575 |
| LDAISDGS | ASSPTGNNSPLY | 0.6143173 |
| LGGFAAML | ASSPTGNNSPLY | 0.1121183 |
| LGGKNPMI | ASSPTGNNSPLY | 0.51448095 |

| | | |
|---|---|---|
| LGPKGVAAIL | ASSPTGNNSPLY | 0.16007335 |
| LMVGVSI | ASSPTGNNSPLY | 0.5255873 |
| LVGAAFVL | ASSPTGNNSPLY | 0.35746026 |
| LVLAVAPL | ASSPTGNNSPLY | 0.5168369 |
| MAAIAVIDAA | ASSPTGNNSPLY | 0.22005698 |
| NNGINAELQ | ASSPTGNNSPLY | 0.45376647 |
| RGEPLVLVHG | ASSPTGNNSPLY | 0.27011782 |
| RGGAGGAAGLI | ASSPTGNNSPLY | 0.14331616 |
| RSYRMHPAV | ASSPTGNNSPLY | 0.72686356 |
| SAAHLLAGQLA | ASSPTGNNSPLY | 0.27202043 |
| SAAVGSY | ASSPTGNNSPLY | 0.43621475 |
| SPHAVLHRI | ASSPTGNNSPLY | 0.61244416 |
| SQIAKVL | ASSPTGNNSPLY | 0.30910695 |
| TALGHLVP | ASSPTGNNSPLY | 0.1288544 |
| TAPTTSGAA | ASSPTGNNSPLY | 0.59654045 |
| TDGDDGNGGSPGT | ASSPTGNNSPLY | 0.42318314 |
| TDGLGEN | ASSPTGNNSPLY | 0.28689364 |
| TGGPPLPGQA | ASSPTGNNSPLY | 0.11272031 |
| TIPAITI | ASSPTGNNSPLY | 0.4970122 |
| TIPGFTI | ASSPTGNNSPLY | 0.21817246 |
| TSIASGALA | ASSPTGNNSPLY | 0.24375625 |
| TVTLPGAAI | ASSPTGNNSPLY | 0.19143566 |
| VASGLSAALP | ASSPTGNNSPLY | 0.7369442 |
| VGAGVIGI | ASSPTGNNSPLY | 0.2413486 |
| VGPKRFGV | ASSPTGNNSPLY | 0.36272508 |
| VNVAFPD | ASSPTGNNSPLY | 0.668476 |
| VSMTNSGVS | ASSPTGNNSPLY | 0.41375107 |
| VTDVLGDLSV | ASSPTGNNSPLY | 0.7925688 |
| VVVVGAGFAGL | ASSPTGNNSPLY | 0.33502975 |
| VYTVGTVV | ASSPTGNNSPLY | 0.3036626 |
| WTTFSGP | ASSPTGNNSPLY | 0.4213915 |
| YDELVEAF | ASSPTGNNSPLY | 0.35278013 |
| YGTGGAGGNGG | ASSPTGNNSPLY | 0.13817011 |
| YTNVMAV | ASSPTGNNSPLY | 0.13231423 |
| FLEQVHQL | ASSPTGNNSPLY | 0.5032733 |
| GLSLLAVAV | ASSPTGNNSPLY | 0.13439219 |
| LEQVHQL | ASSPTGNNSPLY | 0.41037714 |
| LNAAGAI | ASSPTGNNSPLY | 0.7010778 |
| PAVHASL | ASSPTGNNSPLY | 0.5023002 |
| TELEDDA | ASSPTGNNSPLY | 0.15287887 |
| TPPGPPN | ASSPTGNNSPLY | 0.28148487 |
| TPPPPPPP | ASSPTGNNSPLY | 0.1870369 |
| TQSPALI | ASSPRTFSNERLF | 0.6493367 |
| EFAAAFPQ | ASSPRTFSNERLF | 0.6283628 |
| LLSGNLDVL | ASSPRTFSNERLF | 0.81932604 |
| GPANLGNYNI | ASSPRTFSNERLF | 0.67090863 |
| TLGPTLP | ASSPRTFSNERLF | 0.26311085 |
| VYAYSSCCK | ASSPRTFSNERLF | 0.67362 |
| APAPAPAPNI | ASSPRTFSNERLF | 0.30415857 |
| PGGPVPIP | ASSPRTFSNERLF | 0.40837437 |
| FLTVALEP | ASSPRTFSNERLF | 0.30619997 |

| | | |
|---|---|---|
| SQYFGANL | ASSPRTFSNERLF | 0.54092216 |
| GGVKPAA | ASSPRTFSNERLF | 0.36110353 |
| GGTGGAGGAGGW | ASSPRTFSNERLF | 0.251122 |
| GVGGVAGLGGAGG | ASSPRTFSNERLF | 0.13820966 |
| VLLVLATTI | ASSPRTFSNERLF | 0.61251605 |
| IGIFAAML | ASSPRTFSNERLF | 0.49790472 |
| AVGDLID | ASSPRTFSNERLF | 0.20070878 |
| NISPLPL | ASSPRTFSNERLF | 0.30150008 |
| VTAVFVAI | ASSPRTFSNERLF | 0.807403 |
| SGDDNDAAL | ASSPRTFSNERLF | 0.80176306 |
| AVAALAHS | ASSPRTFSNERLF | 0.25497967 |
| GGVTVMATP | ASSPRTFSNERLF | 0.1708082 |
| AGLAANVRD | ASSPRTFSNERLF | 0.70307577 |
| TNRPDLIDPALLRPG | ASSPRTFSNERLF | 0.13475706 |
| LANLGVTAL | ASSPRTFSNERLF | 0.526013 |
| TGERIEL | ASSPRTFSNERLF | 0.47829825 |
| NGEELAA | ASSPRTFSNERLF | 0.41412753 |
| VLSLVWG | ASSPRTFSNERLF | 0.24373558 |
| IGYALWL | ASSPRTFSNERLF | 0.53590083 |
| LSLLLAPL | ASSPRTFSNERLF | 0.2825322 |
| TLDEALK | ASSPRTFSNERLF | 0.6007425 |
| TDGAGTAEGG | ASSPRTFSNERLF | 0.113412276 |
| NAPGGAGGNGG | ASSPRTFSNERLF | 0.3173049 |
| VSVFMGMGGNF | ASSPRTFSNERLF | 0.2222403 |
| LASYLAAL | ASSPRTFSNERLF | 0.20527986 |
| LVGGTPAL | ASSPRTFSNERLF | 0.14104405 |
| VRNADVL | ASSPRTFSNERLF | 0.28428176 |
| TCSSPFLVSAM | ASSPRTFSNERLF | 0.77306587 |
| VHITDAE | ASSPRTFSNERLF | 0.73067987 |
| EYAAPSF | ASSPRTFSNERLF | 0.2439025 |
| VVDAFHGGL | ASSPRTFSNERLF | 0.7204293 |
| ASVVDPQ | ASSPRTFSNERLF | 0.82272804 |
| SYNEFGGLS | ASSPRTFSNERLF | 0.2810402 |
| DPDGSVY | ASSPRTFSNERLF | 0.12642929 |
| MVSTIHATE | ASSPRTFSNERLF | 0.41911095 |
| LAAGVADPV | ASSPRTFSNERLF | 0.18856466 |
| TKEQIAA | ASSPRTFSNERLF | 0.66933143 |
| SLASPAAL | ASSPRTFSNERLF | 0.7898731 |
| IGLVFYGV | ASSPRTFSNERLF | 0.15088467 |
| FLLDASNL | ASSPRTFSNERLF | 0.63104093 |
| YQVGNFP | ASSPRTFSNERLF | 0.741871 |
| ESLLSQST | ASSPRTFSNERLF | 0.22304061 |
| LFTVGFAAVG | ASSPRTFSNERLF | 0.1769765 |
| ATPPTPT | ASSPRTFSNERLF | 0.16714326 |
| TDLATLG | ASSPRTFSNERLF | 0.14493859 |
| SAGGTGR | ASSPRTFSNERLF | 0.40883225 |
| GFSNSGS | ASSPRTFSNERLF | 0.2841326 |
| GGNGGSGGTSV | ASSPRTFSNERLF | 0.19464403 |
| GTSVATGGAGN | ASSPRTFSNERLF | 0.3858838 |
| NGGNGGSGGTS | ASSPRTFSNERLF | 0.18057448 |
| AVVKFPH | ASSPRTFSNERLF | 0.1994713 |

| | | |
|---|---|---|
| TPPPPPP | ASSPRTFSNERLF | 0.20480427 |
| AAPAAAAAPA | ASSPRTFSNERLF | 0.25044042 |
| IALNEDL | ASSPRTFSNERLF | 0.62207884 |
| IVNVSSM | ASSPRTFSNERLF | 0.36890072 |
| TGIPPAPR | ASSPRTFSNERLF | 0.582653 |
| FDLGGGTF | ASSPRTFSNERLF | 0.7666266 |
| FDLGGGTFDVS | ASSPRTFSNERLF | 0.13448252 |
| NEPTAAA | ASSPRTFSNERLF | 0.18909723 |
| NEPTAAALA | ASSPRTFSNERLF | 0.62995756 |
| GSIINISSI | ASSPRTFSNERLF | 0.14656927 |
| VAPGNAGTA | ASSPRTFSNERLF | 0.73704225 |
| TGGGASLE | ASSPRTFSNERLF | 0.41833705 |
| VAVYNHYKRI | ASSPRTFSNERLF | 0.55336624 |
| TDPAPATT | ASSPRTFSNERLF | 0.44629997 |
| TNRPDILDPALLRPG | ASSPRTFSNERLF | 0.82155114 |
| LTTTTTS | ASSPRTFSNERLF | 0.5955988 |
| PTGSGKTL | ASSPRTFSNERLF | 0.2517225 |
| TGSGKTL | ASSPRTFSNERLF | 0.222687 |
| TIGHVDHGK | ASSPRTFSNERLF | 0.16626936 |
| TIGHVDHGKTT | ASSPRTFSNERLF | 0.7020589 |
| GHVDHGKTT | ASSPRTFSNERLF | 0.65837 |
| AQFEHTLL | ASSPRTFSNERLF | 0.17484057 |
| IVNQDEL | ASSPRTFSNERLF | 0.3847108 |
| TNGSQFF | ASSPRTFSNERLF | 0.7297443 |
| TGKGIVE | ASSPRTFSNERLF | 0.7316406 |
| LISNASDA | ASSPRTFSNERLF | 0.13448288 |
| AELRTLGELTKIAKAHGAQ | ASSPRTFSNERLF | 0.23233363 |
| APNAVAAAEPSW | ASSPRTFSNERLF | 0.14525682 |
| APQVAGAATAVL | ASSPRTFSNERLF | 0.36270136 |
| ARRDPAAAAAILPTDA | ASSPRTFSNERLF | 0.15075691 |
| DAAARDVLAEAGLAEYF | ASSPRTFSNERLF | 0.6430491 |
| DDQIVLSAKANLAKVEQ | ASSPRTFSNERLF | 0.28904867 |
| DELLPDTGEDSDEDGH | ASSPRTFSNERLF | 0.7738433 |
| ELPEAPANRATVPQD | ASSPRTFSNERLF | 0.42189157 |
| FGQSKIERVTGIG | ASSPRTFSNERLF | 0.6386685 |
| GAGGTGSTAGGAGGAGGAGGLY | ASSPRTFSNERLF | 0.24888141 |
| GHGGTGAAGGNGGTGGAGGSGI | ASSPRTFSNERLF | 0.2389479 |
| GNGGNGGAGGEGGPGGAGGAGA | ASSPRTFSNERLF | 0.15124233 |
| GSTAGNGGNGGNATNGTIAGQP | ASSPRTFSNERLF | 0.13910697 |
| GTGGDGGNGGNGGTGG | ASSPRTFSNERLF | 0.20879243 |
| GVRRPAGIPADAPV | ASSPRTFSNERLF | 0.22755703 |
| GYRQVTDAHLLTLARR | ASSPRTFSNERLF | 0.117561854 |
| IAKMGQDGHDRG | ASSPRTFSNERLF | 0.77921075 |
| KAEPSIEIQYT | ASSPRTFSNERLF | 0.50886667 |
| KPVRIRMAADEDTVAVT | ASSPRTFSNERLF | 0.5358262 |
| MPPRIAGMRLLVIKPEP | ASSPRTFSNERLF | 0.2024327 |
| MKFYAVALFRSRGVPPTR | ASSPRTFSNERLF | 0.2681624 |
| NAGGFGILGAVAHSP | ASSPRTFSNERLF | 0.18425333 |
| NEGREITLAMAEM | ASSPRTFSNERLF | 0.7429468 |
| NELVEWSHPLGHRWRWE | ASSPRTFSNERLF | 0.1517792 |
| NLGAGGTGGNGGNPG | ASSPRTFSNERLF | 0.12788145 |

| | | |
|---|---|---|
| NTEWPKWETVMAADPVD | ASSPRTFSNERLF | 0.23636043 |
| PASAAPAETQP | ASSPRTFSNERLF | 0.14259803 |
| QQSSDGKNGVRAAEITGE | ASSPRTFSNERLF | 0.81181586 |
| RAPDAVTTSWWKE | ASSPRTFSNERLF | 0.7158903 |
| SGFGNVGAHTSGWFNQ | ASSPRTFSNERLF | 0.5410428 |
| SGVAGAVGQAASVGGLKV | ASSPRTFSNERLF | 0.76252913 |
| SNGGAGTGSAGGNGGT | ASSPRTFSNERLF | 0.123470835 |
| TGGAGGEGGAAG | ASSPRTFSNERLF | 0.19483155 |
| TGGDGGAAGAGGAPGHG | ASSPRTFSNERLF | 0.8042604 |
| TMPPLLAAIANAGRHGV | ASSPRTFSNERLF | 0.4708956 |
| TSSTAGNSGGA | ASSPRTFSNERLF | 0.33883715 |
| VPATVPVGILC | ASSPRTFSNERLF | 0.6870254 |
| VQRPDSGGVVLTGRIS | ASSPRTFSNERLF | 0.20796666 |
| WTPNPVPATKGVPAGLVQAT | ASSPRTFSNERLF | 0.30172986 |
| YALKRALAKFSDRFDVVII | ASSPRTFSNERLF | 0.2797413 |
| YLIGSVIV | ASSPRTFSNERLF | 0.42427915 |
| YGQLVAVID | ASSPRTFSNERLF | 0.5059703 |
| VTVGGMTLPA | ASSPRTFSNERLF | 0.1534743 |
| VLALPAV | ASSPRTFSNERLF | 0.5128622 |
| VIPPTLNYET | ASSPRTFSNERLF | 0.8228413 |
| VGMGGMHPSSG | ASSPRTFSNERLF | 0.39768308 |
| VGLAGGAATA | ASSPRTFSNERLF | 0.68227255 |
| VGGAGGAGGSGP | ASSPRTFSNERLF | 0.4516877 |
| VFAAFMV | ASSPRTFSNERLF | 0.76814985 |
| TLYDKLR | ASSPRTFSNERLF | 0.20879416 |
| TLAVFTVL | ASSPRTFSNERLF | 0.1857758 |
| TGSVQPARV | ASSPRTFSNERLF | 0.43781236 |
| TGGGAGSGGT | ASSPRTFSNERLF | 0.22918567 |
| SHLETSL | ASSPRTFSNERLF | 0.46396267 |
| SFTTGMT | ASSPRTFSNERLF | 0.23462607 |
| QVEVAKRMGVRQPTVS | ASSPRTFSNERLF | 0.8439457 |
| QGSGGPALAM | ASSPRTFSNERLF | 0.6853138 |
| MLAPSNP | ASSPRTFSNERLF | 0.39283484 |
| LVNNAAI | ASSPRTFSNERLF | 0.16134131 |
| LTGEINTLA | ASSPRTFSNERLF | 0.4433449 |
| LSIFGPV | ASSPRTFSNERLF | 0.573001 |
| LLALLANL | ASSPRTFSNERLF | 0.13673428 |
| LISNASDALDKLR | ASSPRTFSNERLF | 0.7720045 |
| LINEAVP | ASSPRTFSNERLF | 0.1896703 |
| LGYAFSGL | ASSPRTFSNERLF | 0.4289967 |
| LDFVAVAL | ASSPRTFSNERLF | 0.57988465 |
| LAQPGAE | ASSPRTFSNERLF | 0.8065301 |
| ISNLGAVTGF | ASSPRTFSNERLF | 0.2847513 |
| HQEDPGRI | ASSPRTFSNERLF | 0.30494204 |
| GTPGEPTP | ASSPRTFSNERLF | 0.15316778 |
| GFFNSGS | ASSPRTFSNERLF | 0.7056314 |
| GAGGGGLTAGAN | ASSPRTFSNERLF | 0.7474791 |
| FVDALVNM | ASSPRTFSNERLF | 0.31906903 |
| FLTVAPEL | ASSPRTFSNERLF | 0.803668 |
| FETLTAL | ASSPRTFSNERLF | 0.54512906 |
| DVINPIEL | ASSPRTFSNERLF | 0.1452576 |

| | | |
|---|---|---|
| APVSGAPV | ASSPRTFSNERLF | 0.3314889 |
| ALVGGLAW | ASSPRTFSNERLF | 0.6140001 |
| ALAAAAATAI | ASSPRTFSNERLF | 0.7904255 |
| AFGLGLLL | ASSPRTFSNERLF | 0.67738974 |
| AFFAAASL | ASSPRTFSNERLF | 0.71438414 |
| AAAALPAFSPP | ASSPRTFSNERLF | 0.2942077 |
| VVLAPAL | ASSPRTFSNERLF | 0.7806809 |
| TIGHVDHGKTTL | ASSPRTFSNERLF | 0.83416426 |
| LISNASDALDK | ASSPRTFSNERLF | 0.5706984 |
| LISNASDAL | ASSPRTFSNERLF | 0.36346877 |
| ISNASDALDK | ASSPRTFSNERLF | 0.26942343 |
| NQAETLVYQTE | ASSPRTFSNERLF | 0.4401875 |
| TGGTGGTAGAG | ASSPRTFSNERLF | 0.71764636 |
| TGSTAGGAGGA | ASSPRTFSNERLF | 0.21081315 |
| DPLAVPVAAGPV | ASSPRTFSNERLF | 0.51528394 |
| GGTGGTPGNGGN | ASSPRTFSNERLF | 0.5422442 |
| PAGLAGVDPATL | ASSPRTFSNERLF | 0.1399773 |
| GNNLVVEAPHAGASV | ASSPRTFSNERLF | 0.67900497 |
| KVTRIMSASTLEQPA | ASSPRTFSNERLF | 0.79238224 |
| LGFALPIALPRRTVV | ASSPRTFSNERLF | 0.3584416 |
| TRLGNLTARKLVVLV | ASSPRTFSNERLF | 0.2677305 |
| WSQQRRLIYFPSAGP | ASSPRTFSNERLF | 0.76392746 |
| GPNGNPDPMAAAVDIR | ASSPRTFSNERLF | 0.8312467 |
| TQRCVSVADLRAAASR | ASSPRTFSNERLF | 0.73806787 |
| QFVLDVAEHPVEVLTETV | ASSPRTFSNERLF | 0.13374119 |
| GGAGGTGGAGGNGGTQAGDG | ASSPRTFSNERLF | 0.12405031 |
| PLLATLRAIAVDVDAKPAQV | ASSPRTFSNERLF | 0.3409325 |
| APKFRTGGYMIDFWGVGYQVAK | ASSPRTFSNERLF | 0.66694987 |
| GAGGAGGAGGAAGGPGGTGGTG | ASSPRTFSNERLF | 0.21113876 |
| IGALGLLATDVAPAGSVAFGVA | ASSPRTFSNERLF | 0.20273757 |
| AGGAGGIGGNGGAGGAGGVLMG | ASSPRTFSNERLF | 0.7182633 |
| ADELIGLL | ASSPRTFSNERLF | 0.6934496 |
| ADRALKA | ASSPRTFSNERLF | 0.18097492 |
| AGAVGQVGGAGGA | ASSPRTFSNERLF | 0.15118596 |
| AGGGGGLGGAP | ASSPRTFSNERLF | 0.6127383 |
| AGGLLYGNGGAG | ASSPRTFSNERLF | 0.69246954 |
| AGLESIGL | ASSPRTFSNERLF | 0.21167889 |
| AGLKTGII | ASSPRTFSNERLF | 0.41389337 |
| ASLAALLL | ASSPRTFSNERLF | 0.2530723 |
| ASVINISSI | ASSPRTFSNERLF | 0.5476827 |
| ATPPGPA | ASSPRTFSNERLF | 0.49468863 |
| ATVASALV | ASSPRTFSNERLF | 0.1699295 |
| ATVASLIL | ASSPRTFSNERLF | 0.75290674 |
| AVNGPVLIRH | ASSPRTFSNERLF | 0.16005234 |
| DGAAGAVGI | ASSPRTFSNERLF | 0.68992716 |
| DPLLHGI | ASSPRTFSNERLF | 0.5252217 |
| ELGAALGAL | ASSPRTFSNERLF | 0.41343996 |
| ELGRTQAV | ASSPRTFSNERLF | 0.35992706 |
| FVAGGYLS | ASSPRTFSNERLF | 0.23754835 |
| GASGGAGGAGGW | ASSPRTFSNERLF | 0.15021695 |
| GGAGAGGDGGAGGI | ASSPRTFSNERLF | 0.13380757 |

| | | |
|---|---|---|
| GTGHNGSQPGMGGNG | ASSPRTFSNERLF | 0.7541687 |
| GYAAASASAS | ASSPRTFSNERLF | 0.1127255 |
| HQVDVAAVQQG | ASSPRTFSNERLF | 0.6901554 |
| IGAAIAHT | ASSPRTFSNERLF | 0.3900919 |
| IGMIEVAQL | ASSPRTFSNERLF | 0.15441921 |
| IRLAPEHTSAMD | ASSPRTFSNERLF | 0.47659382 |
| IYNVSVL | ASSPRTFSNERLF | 0.15452453 |
| KQGFLGM | ASSPRTFSNERLF | 0.26750383 |
| LESVITQL | ASSPRTFSNERLF | 0.79160637 |
| LGLGALVVV | ASSPRTFSNERLF | 0.54662097 |
| LIGNGGNGGSGGI | ASSPRTFSNERLF | 0.19551043 |
| LNAINAPTL | ASSPRTFSNERLF | 0.49851328 |
| LQLLNTGV | ASSPRTFSNERLF | 0.51476276 |
| LQTSIEF | ASSPRTFSNERLF | 0.20482458 |
| MSFAVSEL | ASSPRTFSNERLF | 0.3399119 |
| NTGALGL | ASSPRTFSNERLF | 0.7000873 |
| PPGAGGAGGAGTT | ASSPRTFSNERLF | 0.41726625 |
| QLVGLGI | ASSPRTFSNERLF | 0.16213965 |
| RFLEQVHQL | ASSPRTFSNERLF | 0.5490202 |
| RGVGYVLR | ASSPRTFSNERLF | 0.21140476 |
| SATGVFAGVI | ASSPRTFSNERLF | 0.57396746 |
| SLPAITLP | ASSPRTFSNERLF | 0.26727882 |
| STALAFTV | ASSPRTFSNERLF | 0.3383336 |
| SVLTVGLGK | ASSPRTFSNERLF | 0.57878524 |
| TALAGAGSGI | ASSPRTFSNERLF | 0.17995575 |
| TGLPFTGI | ASSPRTFSNERLF | 0.21111792 |
| VAEHVTV | ASSPRTFSNERLF | 0.35321733 |
| VDIVLNSL | ASSPRTFSNERLF | 0.30190048 |
| VLTGGGAVL | ASSPRTFSNERLF | 0.5460272 |
| VNVGLAYGI | ASSPRTFSNERLF | 0.18785527 |
| VSSAAVGAV | ASSPRTFSNERLF | 0.46019414 |
| VTSVYEAGV | ASSPRTFSNERLF | 0.24298835 |
| WVALDRGA | ASSPRTFSNERLF | 0.17714404 |
| YLVGSVVI | ASSPRTFSNERLF | 0.3643784 |
| GPPGTGKTLL | ASSPRTFSNERLF | 0.29670727 |
| GVGMTKF | ASSPRTFSNERLF | 0.20018414 |
| IGIGGSDL | ASSPRTFSNERLF | 0.4924599 |
| IGTLLAEL | ASSPRTFSNERLF | 0.16174932 |
| IYTSGTT | ASSPRTFSNERLF | 0.39042756 |
| LSFPAGV | ASSPRTFSNERLF | 0.7104442 |
| LYTSGTT | ASSPRTFSNERLF | 0.32233518 |
| VGAFPFGF | ASSPRTFSNERLF | 0.6041439 |
| VGAGTVEF | ASSPRTFSNERLF | 0.4662555 |
| VGKSTLLL | ASSPRTFSNERLF | 0.634358 |
| AALAHAGGGIV | ASSPRTFSNERLF | 0.21807769 |
| AALVAVSLL | ASSPRTFSNERLF | 0.19393218 |
| AAVSRAVAAL | ASSPRTFSNERLF | 0.64916885 |
| AGGSSGIGGF | ASSPRTFSNERLF | 0.503976 |
| AGLTVANANG | ASSPRTFSNERLF | 0.713655 |
| AGLVGAVL | ASSPRTFSNERLF | 0.6443304 |
| ALSQAFTV | ASSPRTFSNERLF | 0.6536807 |

| | | |
|---|---|---|
| ALTFSVSL | ASSPRTFSNERLF | 0.64729446 |
| ALTGAALVAL | ASSPRTFSNERLF | 0.34766972 |
| ASFSIGAGVL | ASSPRTFSNERLF | 0.15444411 |
| DGSVYKI | ASSPRTFSNERLF | 0.21407545 |
| EQLAEIV | ASSPRTFSNERLF | 0.24600697 |
| FAVSELTVM | ASSPRTFSNERLF | 0.20000121 |
| GAAPLAAGL | ASSPRTFSNERLF | 0.14717709 |
| GADGGDGGSGGAGG | ASSPRTFSNERLF | 0.7313751 |
| GFGLDTGGAGG | ASSPRTFSNERLF | 0.82277817 |
| GLASGGLL | ASSPRTFSNERLF | 0.16545539 |
| GRIAGAAVLL | ASSPRTFSNERLF | 0.8164158 |
| GSAGSLNTSF | ASSPRTFSNERLF | 0.17144881 |
| GTGQIGI | ASSPRTFSNERLF | 0.51753277 |
| GTLIALIL | ASSPRTFSNERLF | 0.76219344 |
| IAATMGFVVG | ASSPRTFSNERLF | 0.17474626 |
| IFAQALL | ASSPRTFSNERLF | 0.42915493 |
| IGAAVALGF | ASSPRTFSNERLF | 0.18628204 |
| IGLTPLAL | ASSPRTFSNERLF | 0.18585792 |
| ILGSNTSTL | ASSPRTFSNERLF | 0.7706326 |
| IVFSLTVL | ASSPRTFSNERLF | 0.233316 |
| LASKATGF | ASSPRTFSNERLF | 0.41025972 |
| LASVALIL | ASSPRTFSNERLF | 0.23088682 |
| LFAVGLTL | ASSPRTFSNERLF | 0.15645626 |
| LFNGGIGTYI | ASSPRTFSNERLF | 0.332878 |
| LTLGAGII | ASSPRTFSNERLF | 0.7320341 |
| LVAAALGDI | ASSPRTFSNERLF | 0.3835692 |
| NTAFVQLGI | ASSPRTFSNERLF | 0.828653 |
| QLFSGAEL | ASSPRTFSNERLF | 0.18413754 |
| SAATFITL | ASSPRTFSNERLF | 0.80644035 |
| SNYGLANLGV | ASSPRTFSNERLF | 0.7770755 |
| SYGGAYAVM | ASSPRTFSNERLF | 0.542454 |
| TTGVGGLTLG | ASSPRTFSNERLF | 0.271846 |
| TTQTIYI | ASSPRTFSNERLF | 0.7478285 |
| VAAGAGLGAVI | ASSPRTFSNERLF | 0.42186144 |
| VAAVVTLL | ASSPRTFSNERLF | 0.6767713 |
| VGFLGGLI | ASSPRTFSNERLF | 0.69456625 |
| VGVIPILL | ASSPRTFSNERLF | 0.3830326 |
| VIFASSLI | ASSPRTFSNERLF | 0.80017984 |
| VKGGDGGAAAG | ASSPRTFSNERLF | 0.6837195 |
| VTKTFTNI | ASSPRTFSNERLF | 0.6733057 |
| VVAAAILL | ASSPRTFSNERLF | 0.48053277 |
| YAALDVEL | ASSPRTFSNERLF | 0.37493974 |
| LASVALLL | ASSPRTFSNERLF | 0.5404039 |
| LDAVHRV | ASSPRTFSNERLF | 0.41294822 |
| QGLSSSIF | ASSPRTFSNERLF | 0.78588766 |
| RGTFANIRL | ASSPRTFSNERLF | 0.7458428 |
| RQLWWGHRI | ASSPRTFSNERLF | 0.49126878 |
| YGPPGTGKTLL | ASSPRTFSNERLF | 0.76609504 |
| AALVGAPANVA | ASSPRTFSNERLF | 0.77543855 |
| AATFRRTE | ASSPRTFSNERLF | 0.31918555 |
| AGALGGGI | ASSPRTFSNERLF | 0.5367477 |

| | | |
|---|---|---|
| AGGAGGAGGIGAG | ASSPRTFSNERLF | 0.4809442 |
| AGSGALGL | ASSPRTFSNERLF | 0.534192 |
| AQIADQI | ASSPRTFSNERLF | 0.18883221 |
| ASAVAVTYL | ASSPRTFSNERLF | 0.7161793 |
| ASTVLGGTGG | ASSPRTFSNERLF | 0.45359787 |
| AVGGEPAK | ASSPRTFSNERLF | 0.18534927 |
| CEQGTFGGTC | ASSPRTFSNERLF | 0.6363727 |
| DEGKAPAG | ASSPRTFSNERLF | 0.5430398 |
| DTVVFPAL | ASSPRTFSNERLF | 0.54272324 |
| FEGITMAG | ASSPRTFSNERLF | 0.76752377 |
| FPLVVQD | ASSPRTFSNERLF | 0.5590559 |
| GAFTTPQ | ASSPRTFSNERLF | 0.44195846 |
| GFGGFGTAGGVGGA | ASSPRTFSNERLF | 0.70370054 |
| GFLAQFL | ASSPRTFSNERLF | 0.7882067 |
| GGAGGAGGNAGVA | ASSPRTFSNERLF | 0.415249 |
| GGVGTAGAGGAGGAGG | ASSPRTFSNERLF | 0.6260971 |
| GNGGGGDAGNAGSGGN | ASSPRTFSNERLF | 0.80047095 |
| GTGGAGGAGNDAGS | ASSPRTFSNERLF | 0.67862177 |
| GTTNTGWFNS | ASSPRTFSNERLF | 0.78958416 |
| IFMGPLFTL | ASSPRTFSNERLF | 0.8094315 |
| ILAHAIEAI | ASSPRTFSNERLF | 0.48217428 |
| IRSAIREAL | ASSPRTFSNERLF | 0.7725871 |
| ITFAGGG | ASSPRTFSNERLF | 0.79732615 |
| ITGMSGHQEV | ASSPRTFSNERLF | 0.6026602 |
| IVAGLACTL | ASSPRTFSNERLF | 0.5685622 |
| IVGGGLAGA | ASSPRTFSNERLF | 0.4016749 |
| IVTGVLNI | ASSPRTFSNERLF | 0.6022792 |
| LAQQGCI | ASSPRTFSNERLF | 0.3816598 |
| LCLHVEN | ASSPRTFSNERLF | 0.66064763 |
| LDAISDGS | ASSPRTFSNERLF | 0.7982259 |
| LGGFAAML | ASSPRTFSNERLF | 0.54250187 |
| LGGKNPMI | ASSPRTFSNERLF | 0.32459232 |
| LGPKGVAAIL | ASSPRTFSNERLF | 0.29389092 |
| LMVGVSI | ASSPRTFSNERLF | 0.67930675 |
| LVGAAFVL | ASSPRTFSNERLF | 0.4053728 |
| LVLAVAPL | ASSPRTFSNERLF | 0.36209908 |
| MAAIAVIDAA | ASSPRTFSNERLF | 0.5263097 |
| NNGINAELQ | ASSPRTFSNERLF | 0.7757546 |
| RGEPLVLVHG | ASSPRTFSNERLF | 0.77788717 |
| RGGAGGAAGLI | ASSPRTFSNERLF | 0.3556382 |
| RSYRMHPAV | ASSPRTFSNERLF | 0.8083982 |
| SAAHLLAGQLA | ASSPRTFSNERLF | 0.7358383 |
| SAAVGSY | ASSPRTFSNERLF | 0.79202235 |
| SPHAVLHRI | ASSPRTFSNERLF | 0.64416957 |
| SQIAKVL | ASSPRTFSNERLF | 0.37453756 |
| TALGHLVP | ASSPRTFSNERLF | 0.78907746 |
| TAPTTSGAA | ASSPRTFSNERLF | 0.5503133 |
| TDGDDGNGGSPGT | ASSPRTFSNERLF | 0.6114428 |
| TDGLGEN | ASSPRTFSNERLF | 0.7271908 |
| TGGPPLPGQA | ASSPRTFSNERLF | 0.63736236 |
| TIPAITI | ASSPRTFSNERLF | 0.4357947 |

| | | |
|---|---|---|
| TIPGFTI | ASSPRTFSNERLF | 0.5812731 |
| TSIASGALA | ASSPRTFSNERLF | 0.7720402 |
| TVTLPGAAI | ASSPRTFSNERLF | 0.3457866 |
| VASGLSAALP | ASSPRTFSNERLF | 0.4594443 |
| VGAGVIGI | ASSPRTFSNERLF | 0.5688231 |
| VGPKRFGV | ASSPRTFSNERLF | 0.6179428 |
| VNVAFPD | ASSPRTFSNERLF | 0.78287673 |
| VSMTNSGVS | ASSPRTFSNERLF | 0.6939844 |
| VTDVLGDLSV | ASSPRTFSNERLF | 0.5130786 |
| VVVVGAGFAGL | ASSPRTFSNERLF | 0.68491435 |
| VYTVGTVV | ASSPRTFSNERLF | 0.72550994 |
| WTTFSGP | ASSPRTFSNERLF | 0.6175018 |
| YDELVEAF | ASSPRTFSNERLF | 0.68221605 |
| YGTGGAGGNGG | ASSPRTFSNERLF | 0.44684237 |
| YTNVMAV | ASSPRTFSNERLF | 0.7541307 |
| FLEQVHQL | ASSPRTFSNERLF | 0.7172613 |
| GLSLLAVAV | ASSPRTFSNERLF | 0.72924066 |
| LEQVHQL | ASSPRTFSNERLF | 0.7801956 |
| LNAAGAI | ASSPRTFSNERLF | 0.71967375 |
| PAVHASL | ASSPRTFSNERLF | 0.6371201 |
| TELEDDA | ASSPRTFSNERLF | 0.2769771 |
| TPPGPPN | ASSPRTFSNERLF | 0.56110865 |
| TPPPPPPP | ASSPRTFSNERLF | 0.59539163 |
| TQSPALI | ASRFDRQAPL | 0.66299856 |
| EFAAAFPQ | ASRFDRQAPL | 0.71027756 |
| LLSGNLDVL | ASRFDRQAPL | 0.7509398 |
| GPANLGNYNI | ASRFDRQAPL | 0.4873029 |
| TLGPTLP | ASRFDRQAPL | 0.77427614 |
| VYAYSSCCK | ASRFDRQAPL | 0.7911849 |
| APAPAPAPNI | ASRFDRQAPL | 0.30187798 |
| PGGPVPIP | ASRFDRQAPL | 0.5341881 |
| FLTVALEP | ASRFDRQAPL | 0.82028556 |
| SQYFGANL | ASRFDRQAPL | 0.6863924 |
| GGVKPAA | ASRFDRQAPL | 0.5511045 |
| GGTGGAGGAGGW | ASRFDRQAPL | 0.5626327 |
| GVGGVAGLGGAGG | ASRFDRQAPL | 0.82120556 |
| VLLVLATTI | ASRFDRQAPL | 0.59826183 |
| IGIFAAML | ASRFDRQAPL | 0.8043842 |
| AVGDLID | ASRFDRQAPL | 0.22252476 |
| NISPLPL | ASRFDRQAPL | 0.5617777 |
| VTAVFVAI | ASRFDRQAPL | 0.47975555 |
| SGDDNDAAL | ASRFDRQAPL | 0.19565286 |
| AVAALAHS | ASRFDRQAPL | 0.7504085 |
| GGVTVMATP | ASRFDRQAPL | 0.5238221 |
| AGLAANVRD | ASRFDRQAPL | 0.7217126 |
| TNRPDLIDPALLRPG | ASRFDRQAPL | 0.558917 |
| LANLGVTAL | ASRFDRQAPL | 0.7393931 |
| TGERIEL | ASRFDRQAPL | 0.57287216 |
| NGEELAA | ASRFDRQAPL | 0.6060788 |
| VLSLVWG | ASRFDRQAPL | 0.4625723 |
| IGYALWL | ASRFDRQAPL | 0.71981883 |

| | | |
|---|---|---|
| LSLLLAPL | ASRFDRQAPL | 0.7747973 |
| TLDEALK | ASRFDRQAPL | 0.7553366 |
| TDGAGTAEGG | ASRFDRQAPL | 0.2846559 |
| NAPGGAGGNGG | ASRFDRQAPL | 0.62646127 |
| VSVFMGMGGNF | ASRFDRQAPL | 0.5940475 |
| LASYLAAL | ASRFDRQAPL | 0.3217244 |
| LVGGTPAL | ASRFDRQAPL | 0.25868154 |
| VRNADVL | ASRFDRQAPL | 0.5210118 |
| TCSSPFLVSAM | ASRFDRQAPL | 0.55617213 |
| VHITDAE | ASRFDRQAPL | 0.7275346 |
| EYAAPSF | ASRFDRQAPL | 0.71160966 |
| VVDAFHGGL | ASRFDRQAPL | 0.4896495 |
| ASVVDPQ | ASRFDRQAPL | 0.7209445 |
| SYNEFGGLS | ASRFDRQAPL | 0.6233956 |
| DPDGSVY | ASRFDRQAPL | 0.11411525 |
| MVSTIHATE | ASRFDRQAPL | 0.5493947 |
| LAAGVADPV | ASRFDRQAPL | 0.31643614 |
| TKEQIAA | ASRFDRQAPL | 0.802786 |
| SLASPAAL | ASRFDRQAPL | 0.40481362 |
| IGLVFYGV | ASRFDRQAPL | 0.45462275 |
| FLLDASNL | ASRFDRQAPL | 0.552957 |
| YQVGNFP | ASRFDRQAPL | 0.69871986 |
| ESLLSQST | ASRFDRQAPL | 0.63052243 |
| LFTVGFAAVG | ASRFDRQAPL | 0.412033 |
| ATPPTPT | ASRFDRQAPL | 0.27009898 |
| TDLATLG | ASRFDRQAPL | 0.7252602 |
| SAGGTGR | ASRFDRQAPL | 0.5298531 |
| GFSNSGS | ASRFDRQAPL | 0.4325834 |
| GGNGGSGGTSV | ASRFDRQAPL | 0.2835705 |
| GTSVATGGAGN | ASRFDRQAPL | 0.70683444 |
| NGGNGGSGGTS | ASRFDRQAPL | 0.32811666 |
| AVVKFPH | ASRFDRQAPL | 0.7522496 |
| TPPPPPP | ASRFDRQAPL | 0.4441477 |
| AAPAAAAAPA | ASRFDRQAPL | 0.54433 |
| IALNEDL | ASRFDRQAPL | 0.82966626 |
| IVNVSSM | ASRFDRQAPL | 0.65731037 |
| TGIPPAPR | ASRFDRQAPL | 0.53579104 |
| FDLGGGTF | ASRFDRQAPL | 0.51322985 |
| FDLGGGTFDVS | ASRFDRQAPL | 0.38871062 |
| NEPTAAA | ASRFDRQAPL | 0.43095493 |
| NEPTAAALA | ASRFDRQAPL | 0.48394144 |
| GSIINISSI | ASRFDRQAPL | 0.7949806 |
| VAPGNAGTA | ASRFDRQAPL | 0.5818659 |
| TGGGASLE | ASRFDRQAPL | 0.36534163 |
| VAVYNHYKRI | ASRFDRQAPL | 0.71430236 |
| TDPAPATT | ASRFDRQAPL | 0.8078394 |
| TNRPDILDPALLRPG | ASRFDRQAPL | 0.82784814 |
| LTTTTTS | ASRFDRQAPL | 0.39783525 |
| PTGSGKTL | ASRFDRQAPL | 0.5951834 |
| TGSGKTL | ASRFDRQAPL | 0.53445154 |
| TIGHVDHGK | ASRFDRQAPL | 0.5469492 |

| | | |
|---|---|---|
| TIGHVDHGKTT | ASRFDRQAPL | 0.5047393 |
| GHVDHGKTT | ASRFDRQAPL | 0.42999482 |
| AQFEHTLL | ASRFDRQAPL | 0.7285124 |
| IVNQDEL | ASRFDRQAPL | 0.85493004 |
| TNGSQFF | ASRFDRQAPL | 0.5543668 |
| TGKGIVE | ASRFDRQAPL | 0.25787055 |
| LISNASDA | ASRFDRQAPL | 0.52105653 |
| AELRTLGELTKIAKAHGAQ | ASRFDRQAPL | 0.38409144 |
| APNAVAAAEPSW | ASRFDRQAPL | 0.39472386 |
| APQVAGAATAVL | ASRFDRQAPL | 0.6173799 |
| ARRDPAAAAAILPTDA | ASRFDRQAPL | 0.27447426 |
| DAAARDVLAEAGLAEYF | ASRFDRQAPL | 0.6009342 |
| DDQIVLSAKANLAKVEQ | ASRFDRQAPL | 0.8276629 |
| DELLPDTGEDSDEDGH | ASRFDRQAPL | 0.5349301 |
| ELPEAPANRATVPQD | ASRFDRQAPL | 0.70320106 |
| FGQSKIERVTGIG | ASRFDRQAPL | 0.7883669 |
| GAGGTGSTAGGAGGAGGAGGLY | ASRFDRQAPL | 0.6941742 |
| GHGGTGAAGGNGGTGGAGGSGI | ASRFDRQAPL | 0.79286826 |
| GNGGNGGAGGEGGPGGAGGAGA | ASRFDRQAPL | 0.76055264 |
| GSTAGNGGNGGNATNGTIAGQP | ASRFDRQAPL | 0.6106968 |
| GTGGDGGNGGNGGTGG | ASRFDRQAPL | 0.6099739 |
| GVRRPAGIPADAPV | ASRFDRQAPL | 0.780836 |
| GYRQVTDAHLLTLARR | ASRFDRQAPL | 0.5279561 |
| IAKMGQDGHDRG | ASRFDRQAPL | 0.58933944 |
| KAEPSIEIQYT | ASRFDRQAPL | 0.5018239 |
| KPVRIRMAADEDTVAVT | ASRFDRQAPL | 0.7624823 |
| MPPRIAGMRLLVIKPEP | ASRFDRQAPL | 0.60372746 |
| MKFYAVALFRSRGVPPTR | ASRFDRQAPL | 0.8183618 |
| NAGGFGILGAVAHSP | ASRFDRQAPL | 0.29995245 |
| NEGREITLAMAEM | ASRFDRQAPL | 0.40592778 |
| NELVEWSHPLGHRWRWE | ASRFDRQAPL | 0.772305 |
| NLGAGGTGGNGGNPG | ASRFDRQAPL | 0.16170962 |
| NTEWPKWETVMAADPVD | ASRFDRQAPL | 0.28043124 |
| PASAAPAETQP | ASRFDRQAPL | 0.6001628 |
| QQSSDGKNGVRAAEITGE | ASRFDRQAPL | 0.7830194 |
| RAPDAVTTSWWKE | ASRFDRQAPL | 0.17863756 |
| SGFGNVGAHTSGWFNQ | ASRFDRQAPL | 0.48490578 |
| SGVAGAVGQAASVGGLKV | ASRFDRQAPL | 0.3583308 |
| SNGGAGTGSAGGNGGT | ASRFDRQAPL | 0.77434516 |
| TGGAGGEGGAAG | ASRFDRQAPL | 0.6221425 |
| TGGDGGAAGAGGAPGHG | ASRFDRQAPL | 0.3206859 |
| TMPPLLAAIANAGRHGV | ASRFDRQAPL | 0.7194061 |
| TSSTAGNSGGA | ASRFDRQAPL | 0.24784318 |
| VPATVPVGILC | ASRFDRQAPL | 0.6460003 |
| VQRPDSGGVVLTGRIS | ASRFDRQAPL | 0.7196463 |
| WTPNPVPATKGVPAGLVQAT | ASRFDRQAPL | 0.81761986 |
| YALKRALAKFSDRFDVVII | ASRFDRQAPL | 0.36682415 |
| YLIGSVIV | ASRFDRQAPL | 0.72198325 |
| YGQLVAVID | ASRFDRQAPL | 0.4984988 |
| VTVGGMTLPA | ASRFDRQAPL | 0.85502625 |
| VLALPAV | ASRFDRQAPL | 0.5408065 |

| | | |
|---|---|---|
| VIPPTLNYET | ASRFDRQAPL | 0.33766863 |
| VGMGGMHPSSG | ASRFDRQAPL | 0.76365775 |
| VGLAGGAATA | ASRFDRQAPL | 0.700609 |
| VGGAGGAGGSGP | ASRFDRQAPL | 0.65101147 |
| VFAAFMV | ASRFDRQAPL | 0.75454515 |
| TLYDKLR | ASRFDRQAPL | 0.6052147 |
| TLAVFTVL | ASRFDRQAPL | 0.510496 |
| TGSVQPARV | ASRFDRQAPL | 0.7553817 |
| TGGGAGSGGT | ASRFDRQAPL | 0.6712437 |
| SHLETSL | ASRFDRQAPL | 0.48648077 |
| SFTTGMT | ASRFDRQAPL | 0.7680322 |
| QVEVAKRMGVRQPTVS | ASRFDRQAPL | 0.60100424 |
| QGSGGPALAM | ASRFDRQAPL | 0.42761356 |
| MLAPSNP | ASRFDRQAPL | 0.8307277 |
| LVNNAAI | ASRFDRQAPL | 0.6345532 |
| LTGEINTLA | ASRFDRQAPL | 0.733937 |
| LSIFGPV | ASRFDRQAPL | 0.72013706 |
| LLALLANL | ASRFDRQAPL | 0.82516325 |
| LISNASDALDKLR | ASRFDRQAPL | 0.4587339 |
| LINEAVP | ASRFDRQAPL | 0.6719035 |
| LGYAFSGL | ASRFDRQAPL | 0.6205222 |
| LDFVAVAL | ASRFDRQAPL | 0.7351801 |
| LAQPGAE | ASRFDRQAPL | 0.61059326 |
| ISNLGAVTGF | ASRFDRQAPL | 0.6446979 |
| HQEDPGRI | ASRFDRQAPL | 0.7014475 |
| GTPGEPTP | ASRFDRQAPL | 0.8076853 |
| GFFNSGS | ASRFDRQAPL | 0.56079453 |
| GAGGGGLTAGAN | ASRFDRQAPL | 0.6511558 |
| FVDALVNM | ASRFDRQAPL | 0.8155029 |
| FLTVAPEL | ASRFDRQAPL | 0.4492757 |
| FETLTAL | ASRFDRQAPL | 0.58386487 |
| DVINPIEL | ASRFDRQAPL | 0.7258016 |
| APVSGAPV | ASRFDRQAPL | 0.7631039 |
| ALVGGLAW | ASRFDRQAPL | 0.5703586 |
| ALAAAAATAI | ASRFDRQAPL | 0.7804424 |
| AFGLGLLL | ASRFDRQAPL | 0.4929821 |
| AFFAAASL | ASRFDRQAPL | 0.6972366 |
| AAAALPAFSPP | ASRFDRQAPL | 0.55481017 |
| VVLAPAL | ASRFDRQAPL | 0.6037428 |
| TIGHVDHGKTTL | ASRFDRQAPL | 0.4052038 |
| LISNASDALDK | ASRFDRQAPL | 0.64600307 |
| LISNASDAL | ASRFDRQAPL | 0.7525522 |
| ISNASDALDK | ASRFDRQAPL | 0.7164154 |
| NQAETLVYQTE | ASRFDRQAPL | 0.2286819 |
| TGGTGGTAGAG | ASRFDRQAPL | 0.7203661 |
| TGSTAGGAGGA | ASRFDRQAPL | 0.72604054 |
| DPLAVPVAAGPV | ASRFDRQAPL | 0.8427006 |
| GGTGGTPGNGGN | ASRFDRQAPL | 0.5785279 |
| PAGLAGVDPATL | ASRFDRQAPL | 0.71842694 |
| GNNLVVEAPHAGASV | ASRFDRQAPL | 0.51564324 |
| KVTRIMSASTLEQPA | ASRFDRQAPL | 0.59085166 |

| | | |
|---|---|---|
| LGFALPIALPRRTVV | ASRFDRQAPL | 0.29402745 |
| TRLGNLTARKLVVLV | ASRFDRQAPL | 0.5005691 |
| WSQQRRLIYFPSAGP | ASRFDRQAPL | 0.513597 |
| GPNGNPDPMAAAVDIR | ASRFDRQAPL | 0.39943618 |
| TQRCVSVADLRAAASR | ASRFDRQAPL | 0.7519033 |
| QFVLDVAEHPVEVLTETV | ASRFDRQAPL | 0.73884284 |
| GGAGGTGGAGGNGGTQAGDG | ASRFDRQAPL | 0.7804782 |
| PLLATLRAIAVDVDAKPAQV | ASRFDRQAPL | 0.79134274 |
| APKFRTGGYMIDFWGVGYQVAK | ASRFDRQAPL | 0.8087497 |
| GAGGAGGAGGAAGGPGGTGGTG | ASRFDRQAPL | 0.75193906 |
| IGALGLLATDVAPAGSVAFGVA | ASRFDRQAPL | 0.4687675 |
| AGGAGGIGGNGGAGGAGGVLMG | ASRFDRQAPL | 0.62010723 |
| ADELIGLL | ASRFDRQAPL | 0.30296496 |
| ADRALKA | ASRFDRQAPL | 0.55541646 |
| AGAVGQVGGAGGA | ASRFDRQAPL | 0.5133346 |
| AGGGGGLGGAP | ASRFDRQAPL | 0.63801885 |
| AGGLLYGNGGAG | ASRFDRQAPL | 0.17392628 |
| AGLESIGL | ASRFDRQAPL | 0.68732417 |
| AGLKTGII | ASRFDRQAPL | 0.8060533 |
| ASLAALLL | ASRFDRQAPL | 0.5314802 |
| ASVINISSI | ASRFDRQAPL | 0.7260245 |
| ATPPGPA | ASRFDRQAPL | 0.70744324 |
| ATVASALV | ASRFDRQAPL | 0.43022066 |
| ATVASLIL | ASRFDRQAPL | 0.732225 |
| AVNGPVLIRH | ASRFDRQAPL | 0.4611256 |
| DGAAGAVGI | ASRFDRQAPL | 0.72690684 |
| DPLLHGI | ASRFDRQAPL | 0.8151192 |
| ELGAALGAL | ASRFDRQAPL | 0.6629334 |
| ELGRTQAV | ASRFDRQAPL | 0.5763902 |
| FVAGGYLS | ASRFDRQAPL | 0.6386912 |
| GASGGAGGAGGW | ASRFDRQAPL | 0.40553606 |
| GGAGAGGDGGAGGI | ASRFDRQAPL | 0.5950457 |
| GTGHNGSQPGMGGNG | ASRFDRQAPL | 0.41199133 |
| GYAAASASAS | ASRFDRQAPL | 0.61675894 |
| HQVDVAAVQQG | ASRFDRQAPL | 0.4161261 |
| IGAAIAHT | ASRFDRQAPL | 0.6568832 |
| IGMIEVAQL | ASRFDRQAPL | 0.6202651 |
| IRLAPEHTSAMD | ASRFDRQAPL | 0.6210151 |
| IYNVSVL | ASRFDRQAPL | 0.36053675 |
| KQGFLGM | ASRFDRQAPL | 0.7000483 |
| LESVITQL | ASRFDRQAPL | 0.5898626 |
| LGLGALVVV | ASRFDRQAPL | 0.35723543 |
| LIGNGGNGGSGGI | ASRFDRQAPL | 0.70718575 |
| LNAINAPTL | ASRFDRQAPL | 0.5811682 |
| LQLLNTGV | ASRFDRQAPL | 0.5871179 |
| LQTSIEF | ASRFDRQAPL | 0.78307843 |
| MSFAVSEL | ASRFDRQAPL | 0.64926267 |
| NTGALGL | ASRFDRQAPL | 0.28442568 |
| PPGAGGAGGAGTT | ASRFDRQAPL | 0.5488297 |
| QLVGLGI | ASRFDRQAPL | 0.6090263 |
| RFLEQVHQL | ASRFDRQAPL | 0.54589766 |

| | | |
|---|---|---|
| RGVGYVLR | ASRFDRQAPL | 0.19248904 |
| SATGVFAGVI | ASRFDRQAPL | 0.7023598 |
| SLPAITLP | ASRFDRQAPL | 0.42958322 |
| STALAFTV | ASRFDRQAPL | 0.37994578 |
| SVLTVGLGK | ASRFDRQAPL | 0.46305582 |
| TALAGAGSGI | ASRFDRQAPL | 0.7491056 |
| TGLPFTGI | ASRFDRQAPL | 0.65172946 |
| VAEHVTV | ASRFDRQAPL | 0.6277113 |
| VDIVLNSL | ASRFDRQAPL | 0.68510973 |
| VLTGGGAVL | ASRFDRQAPL | 0.68550694 |
| VNVGLAYGI | ASRFDRQAPL | 0.7886432 |
| VSSAAVGAV | ASRFDRQAPL | 0.71001863 |
| VTSVYEAGV | ASRFDRQAPL | 0.62825227 |
| WVALDRGA | ASRFDRQAPL | 0.7000608 |
| YLVGSVVI | ASRFDRQAPL | 0.60218155 |
| GPPGTGKTLL | ASRFDRQAPL | 0.38075095 |
| GVGMTKF | ASRFDRQAPL | 0.33912304 |
| IGIGGSDL | ASRFDRQAPL | 0.45384315 |
| IGTLLAEL | ASRFDRQAPL | 0.3094067 |
| IYTSGTT | ASRFDRQAPL | 0.6620246 |
| LSFPAGV | ASRFDRQAPL | 0.8064186 |
| LYTSGTT | ASRFDRQAPL | 0.63294697 |
| VGAFPFGF | ASRFDRQAPL | 0.34617904 |
| VGAGTVEF | ASRFDRQAPL | 0.5107324 |
| VGKSTLLL | ASRFDRQAPL | 0.6221312 |
| AALAHAGGGIV | ASRFDRQAPL | 0.7403556 |
| AALVAVSLL | ASRFDRQAPL | 0.7611787 |
| AAVSRAVAAL | ASRFDRQAPL | 0.8392282 |
| AGGSSGIGGF | ASRFDRQAPL | 0.7423874 |
| AGLTVANANG | ASRFDRQAPL | 0.4753324 |
| AGLVGAVL | ASRFDRQAPL | 0.70775384 |
| ALSQAFTV | ASRFDRQAPL | 0.52098054 |
| ALTFSVSL | ASRFDRQAPL | 0.6111806 |
| ALTGAALVAL | ASRFDRQAPL | 0.58881533 |
| ASFSIGAGVL | ASRFDRQAPL | 0.63220507 |
| DGSVYKI | ASRFDRQAPL | 0.5510442 |
| EQLAEIV | ASRFDRQAPL | 0.5068664 |
| FAVSELTVM | ASRFDRQAPL | 0.29850203 |
| GAAPLAAGL | ASRFDRQAPL | 0.7113803 |
| GADGGDGGSGGAGG | ASRFDRQAPL | 0.46684518 |
| GFGLDTGGAGG | ASRFDRQAPL | 0.4038177 |
| GLASGGLL | ASRFDRQAPL | 0.5028249 |
| GRIAGAAVLL | ASRFDRQAPL | 0.821943 |
| GSAGSLNTSF | ASRFDRQAPL | 0.81560373 |
| GTGQIGI | ASRFDRQAPL | 0.45179623 |
| GTLIALIL | ASRFDRQAPL | 0.4563741 |
| IAATMGFVVG | ASRFDRQAPL | 0.7041794 |
| IFAQALL | ASRFDRQAPL | 0.3525792 |
| IGAAVALGF | ASRFDRQAPL | 0.6974084 |
| IGLTPLAL | ASRFDRQAPL | 0.59982115 |
| ILGSNTSTL | ASRFDRQAPL | 0.5559734 |

| | | |
|---|---|---|
| IVFSLTVL | ASRFDRQAPL | 0.4717811 |
| LASKATGF | ASRFDRQAPL | 0.69281596 |
| LASVALIL | ASRFDRQAPL | 0.48963076 |
| LFAVGLTL | ASRFDRQAPL | 0.71564037 |
| LFNGGIGTYI | ASRFDRQAPL | 0.20119727 |
| LTLGAGII | ASRFDRQAPL | 0.72327995 |
| LVAAALGDI | ASRFDRQAPL | 0.3743495 |
| NTAFVQLGI | ASRFDRQAPL | 0.48840824 |
| QLFSGAEL | ASRFDRQAPL | 0.25599557 |
| SAATFITL | ASRFDRQAPL | 0.5419209 |
| SNYGLANLGV | ASRFDRQAPL | 0.7400862 |
| SYGGAYAVM | ASRFDRQAPL | 0.78041136 |
| TTGVGGLTLG | ASRFDRQAPL | 0.5485323 |
| TTQTIYI | ASRFDRQAPL | 0.7113915 |
| VAAGAGLGAVI | ASRFDRQAPL | 0.7826482 |
| VAAVVTLL | ASRFDRQAPL | 0.52223307 |
| VGFLGGLI | ASRFDRQAPL | 0.29319683 |
| VGVIPILL | ASRFDRQAPL | 0.7055457 |
| VIFASSLI | ASRFDRQAPL | 0.536922 |
| VKGGDGGAAAG | ASRFDRQAPL | 0.7845337 |
| VTKTFTNI | ASRFDRQAPL | 0.7278272 |
| VVAAAILL | ASRFDRQAPL | 0.28063044 |
| YAALDVEL | ASRFDRQAPL | 0.7671277 |
| LASVALLL | ASRFDRQAPL | 0.75193644 |
| LDAVHRV | ASRFDRQAPL | 0.3575112 |
| QGLSSSIF | ASRFDRQAPL | 0.40660152 |
| RGTFANIRL | ASRFDRQAPL | 0.4622169 |
| RQLWWGHRI | ASRFDRQAPL | 0.30501255 |
| YGPPGTGKTLL | ASRFDRQAPL | 0.68277514 |
| AALVGAPANVA | ASRFDRQAPL | 0.49483714 |
| AATFRRTE | ASRFDRQAPL | 0.5715947 |
| AGALGGGI | ASRFDRQAPL | 0.54959583 |
| AGGAGGAGGIGAG | ASRFDRQAPL | 0.5121787 |
| AGSGALGL | ASRFDRQAPL | 0.52437073 |
| AQIADQI | ASRFDRQAPL | 0.56815624 |
| ASAVAVTYL | ASRFDRQAPL | 0.64664954 |
| ASTVLGGTGG | ASRFDRQAPL | 0.72336316 |
| AVGGEPAK | ASRFDRQAPL | 0.5472786 |
| CEQGTFGGTC | ASRFDRQAPL | 0.62793595 |
| DEGKAPAG | ASRFDRQAPL | 0.799605 |
| DTVVFPAL | ASRFDRQAPL | 0.37601638 |
| FEGITMAG | ASRFDRQAPL | 0.41661954 |
| FPLVVQD | ASRFDRQAPL | 0.71432316 |
| GAFTTPQ | ASRFDRQAPL | 0.43840235 |
| GFGGFGTAGGVGGA | ASRFDRQAPL | 0.765121 |
| GFLAQFL | ASRFDRQAPL | 0.6332673 |
| GGAGGAGGNAGVA | ASRFDRQAPL | 0.66861254 |
| GGVGTAGAGGAGGAGG | ASRFDRQAPL | 0.6485209 |
| GNGGGGDAGNAGSGGN | ASRFDRQAPL | 0.82003415 |
| GTGGAGGAGNDAGS | ASRFDRQAPL | 0.73551387 |
| GTTNTGWFNS | ASRFDRQAPL | 0.5337218 |

| | | |
|---|---|---|
| IFMGPLFTL | ASRFDRQAPL | 0.75513965 |
| ILAHAIEAI | ASRFDRQAPL | 0.39032012 |
| IRSAIREAL | ASRFDRQAPL | 0.70859677 |
| ITFAGGG | ASRFDRQAPL | 0.71665096 |
| ITGMSGHQEV | ASRFDRQAPL | 0.3694013 |
| IVAGLACTL | ASRFDRQAPL | 0.5263245 |
| IVGGGLAGA | ASRFDRQAPL | 0.78434145 |
| IVTGVLNI | ASRFDRQAPL | 0.7527311 |
| LAQQGCI | ASRFDRQAPL | 0.35826322 |
| LCLHVEN | ASRFDRQAPL | 0.4404703 |
| LDAISDGS | ASRFDRQAPL | 0.39993343 |
| LGGFAAML | ASRFDRQAPL | 0.5189441 |
| LGGKNPMI | ASRFDRQAPL | 0.36322635 |
| LGPKGVAAIL | ASRFDRQAPL | 0.71944404 |
| LMVGVSI | ASRFDRQAPL | 0.5247826 |
| LVGAAFVL | ASRFDRQAPL | 0.7475531 |
| LVLAVAPL | ASRFDRQAPL | 0.6489197 |
| MAAIAVIDAA | ASRFDRQAPL | 0.77432334 |
| NNGINAELQ | ASRFDRQAPL | 0.4668107 |
| RGEPLVLVHG | ASRFDRQAPL | 0.58207774 |
| RGGAGGAAGLI | ASRFDRQAPL | 0.31009123 |
| RSYRMHPAV | ASRFDRQAPL | 0.48766172 |
| SAAHLLAGQLA | ASRFDRQAPL | 0.44255844 |
| SAAVGSY | ASRFDRQAPL | 0.5738212 |
| SPHAVLHRI | ASRFDRQAPL | 0.32458213 |
| SQIAKVL | ASRFDRQAPL | 0.8163694 |
| TALGHLVP | ASRFDRQAPL | 0.56959295 |
| TAPTTSGAA | ASRFDRQAPL | 0.72484 |
| TDGDDGNGGSPGT | ASRFDRQAPL | 0.520061 |
| TDGLGEN | ASRFDRQAPL | 0.46946406 |
| TGGPPLPGQA | ASRFDRQAPL | 0.46871123 |
| TIPAITI | ASRFDRQAPL | 0.72650295 |
| TIPGFTI | ASRFDRQAPL | 0.36296827 |
| TSIASGALA | ASRFDRQAPL | 0.21056178 |
| TVTLPGAAI | ASRFDRQAPL | 0.45216316 |
| VASGLSAALP | ASRFDRQAPL | 0.29543063 |
| VGAGVIGI | ASRFDRQAPL | 0.8184275 |
| VGPKRFGV | ASRFDRQAPL | 0.77883315 |
| VNVAFPD | ASRFDRQAPL | 0.63422525 |
| VSMTNSGVS | ASRFDRQAPL | 0.8137748 |
| VTDVLGDLSV | ASRFDRQAPL | 0.18727882 |
| VVVVGAGFAGL | ASRFDRQAPL | 0.57686573 |
| VYTVGTVV | ASRFDRQAPL | 0.8032426 |
| WTTFSGP | ASRFDRQAPL | 0.6905377 |
| YDELVEAF | ASRFDRQAPL | 0.58774763 |
| YGTGGAGGNGG | ASRFDRQAPL | 0.7729457 |
| YTNVMAV | ASRFDRQAPL | 0.43490076 |
| FLEQVHQL | ASRFDRQAPL | 0.5699452 |
| GLSLLAVAV | ASRFDRQAPL | 0.62337667 |
| LEQVHQL | ASRFDRQAPL | 0.69569546 |
| LNAAGAI | ASRFDRQAPL | 0.68881476 |

| | | |
|---|---|---|
| PAVHASL | ASRFDRQAPL | 0.32594478 |
| TELEDDA | ASRFDRQAPL | 0.71240467 |
| TPPGPPN | ASRFDRQAPL | 0.83233845 |
| TPPPPPPP | ASRFDRQAPL | 0.4995442 |
| TQSPALI | ASGPDRNTEVF | 0.7321425 |
| EFAAAFPQ | ASGPDRNTEVF | 0.6743548 |
| LLSGNLDVL | ASGPDRNTEVF | 0.6610441 |
| GPANLGNYNI | ASGPDRNTEVF | 0.4134512 |
| TLGPTLP | ASGPDRNTEVF | 0.3633139 |
| VYAYSSCCK | ASGPDRNTEVF | 0.6853225 |
| APAPAPAPNI | ASGPDRNTEVF | 0.36317128 |
| PGGPVPIP | ASGPDRNTEVF | 0.6031539 |
| FLTVALEP | ASGPDRNTEVF | 0.54597104 |
| SQYFGANL | ASGPDRNTEVF | 0.85316885 |
| GGVKPAA | ASGPDRNTEVF | 0.6853493 |
| GGTGGAGGAGGW | ASGPDRNTEVF | 0.71353066 |
| GVGGVAGLGGAGG | ASGPDRNTEVF | 0.37109005 |
| VLLVLATTI | ASGPDRNTEVF | 0.61172193 |
| IGIFAAML | ASGPDRNTEVF | 0.6984893 |
| AVGDLID | ASGPDRNTEVF | 0.33952036 |
| NISPLPL | ASGPDRNTEVF | 0.78652924 |
| VTAVFVAI | ASGPDRNTEVF | 0.4278072 |
| SGDDNDAAL | ASGPDRNTEVF | 0.62232697 |
| AVAALAHS | ASGPDRNTEVF | 0.7164378 |
| GGVTVMATP | ASGPDRNTEVF | 0.7915441 |
| AGLAANVRD | ASGPDRNTEVF | 0.5476111 |
| TNRPDLIDPALLRPG | ASGPDRNTEVF | 0.6704965 |
| LANLGVTAL | ASGPDRNTEVF | 0.3901399 |
| TGERIEL | ASGPDRNTEVF | 0.75589097 |
| NGEELAA | ASGPDRNTEVF | 0.773356 |
| VLSLVWG | ASGPDRNTEVF | 0.6704027 |
| IGYALWL | ASGPDRNTEVF | 0.83399075 |
| LSLLLAPL | ASGPDRNTEVF | 0.75530696 |
| TLDEALK | ASGPDRNTEVF | 0.3193202 |
| TDGAGTAEGG | ASGPDRNTEVF | 0.60452616 |
| NAPGGAGGNGG | ASGPDRNTEVF | 0.75646424 |
| VSVFMGMGGNF | ASGPDRNTEVF | 0.79841554 |
| LASYLAAL | ASGPDRNTEVF | 0.7751278 |
| LVGGTPAL | ASGPDRNTEVF | 0.72735155 |
| VRNADVL | ASGPDRNTEVF | 0.4598037 |
| TCSSPFLVSAM | ASGPDRNTEVF | 0.80450994 |
| VHITDAE | ASGPDRNTEVF | 0.8582088 |
| EYAAPSF | ASGPDRNTEVF | 0.7290337 |
| VVDAFHGGL | ASGPDRNTEVF | 0.45387593 |
| ASVVDPQ | ASGPDRNTEVF | 0.5119155 |
| SYNEFGGLS | ASGPDRNTEVF | 0.6823039 |
| DPDGSVY | ASGPDRNTEVF | 0.7435064 |
| MVSTIHATE | ASGPDRNTEVF | 0.37526095 |
| LAAGVADPV | ASGPDRNTEVF | 0.6517224 |
| TKEQIAA | ASGPDRNTEVF | 0.6737908 |
| SLASPAAL | ASGPDRNTEVF | 0.26015678 |

| | | |
|---|---|---|
| IGLVFYGV | ASGPDRNTEVF | 0.69206154 |
| FLLDASNL | ASGPDRNTEVF | 0.78490275 |
| YQVGNFP | ASGPDRNTEVF | 0.6540807 |
| ESLLSQST | ASGPDRNTEVF | 0.4343942 |
| LFTVGFAAVG | ASGPDRNTEVF | 0.7872739 |
| ATPPTPT | ASGPDRNTEVF | 0.83285743 |
| TDLATLG | ASGPDRNTEVF | 0.8183916 |
| SAGGTGR | ASGPDRNTEVF | 0.29488963 |
| GFSNSGS | ASGPDRNTEVF | 0.17418662 |
| GGNGGSGGTSV | ASGPDRNTEVF | 0.5640227 |
| GTSVATGGAGN | ASGPDRNTEVF | 0.7491362 |
| NGGNGGSGGTS | ASGPDRNTEVF | 0.57094437 |
| AVVKFPH | ASGPDRNTEVF | 0.46025342 |
| TPPPPPP | ASGPDRNTEVF | 0.69344985 |
| AAPAAAAAPA | ASGPDRNTEVF | 0.77518654 |
| IALNEDL | ASGPDRNTEVF | 0.4417563 |
| IVNVSSM | ASGPDRNTEVF | 0.37528127 |
| TGIPPAPR | ASGPDRNTEVF | 0.7246108 |
| FDLGGGTF | ASGPDRNTEVF | 0.74005365 |
| FDLGGGTFDVS | ASGPDRNTEVF | 0.35101777 |
| NEPTAAA | ASGPDRNTEVF | 0.4933588 |
| NEPTAAALA | ASGPDRNTEVF | 0.64772844 |
| GSIINISSI | ASGPDRNTEVF | 0.68043584 |
| VAPGNAGTA | ASGPDRNTEVF | 0.6479124 |
| TGGGASLE | ASGPDRNTEVF | 0.3551278 |
| VAVYNHYKRI | ASGPDRNTEVF | 0.74832624 |
| TDPAPATT | ASGPDRNTEVF | 0.31213236 |
| TNRPDILDPALLRPG | ASGPDRNTEVF | 0.71638036 |
| LTTTTTS | ASGPDRNTEVF | 0.612068 |
| PTGSGKTL | ASGPDRNTEVF | 0.653405 |
| TGSGKTL | ASGPDRNTEVF | 0.5391996 |
| TIGHVDHGK | ASGPDRNTEVF | 0.40592128 |
| TIGHVDHGKTT | ASGPDRNTEVF | 0.39361626 |
| GHVDHGKTT | ASGPDRNTEVF | 0.29156286 |
| AQFEHTLL | ASGPDRNTEVF | 0.7448687 |
| IVNQDEL | ASGPDRNTEVF | 0.6781869 |
| TNGSQFF | ASGPDRNTEVF | 0.7221795 |
| TGKGIVE | ASGPDRNTEVF | 0.67555285 |
| LISNASDA | ASGPDRNTEVF | 0.4095168 |
| AELRTLGELTKIAKAHGAQ | ASGPDRNTEVF | 0.650235 |
| APNAVAAAEPSW | ASGPDRNTEVF | 0.38949 |
| APQVAGAATAVL | ASGPDRNTEVF | 0.5953629 |
| ARRDPAAAAAILPTDA | ASGPDRNTEVF | 0.65567684 |
| DAAARDVLAEAGLAEYF | ASGPDRNTEVF | 0.71349853 |
| DDQIVLSAKANLAKVEQ | ASGPDRNTEVF | 0.4116422 |
| DELLPDTGEDSDEDGH | ASGPDRNTEVF | 0.6863817 |
| ELPEAPANRATVPQD | ASGPDRNTEVF | 0.6728997 |
| FGQSKIERVTGIG | ASGPDRNTEVF | 0.4499301 |
| GAGGTGSTAGGAGGAGGAGGLY | ASGPDRNTEVF | 0.65066075 |
| GHGGTGAAGGNGGTGGAGGSGI | ASGPDRNTEVF | 0.77275825 |
| GNGGNGGAGGEGGPGGAGGAGA | ASGPDRNTEVF | 0.6661832 |

| | | |
|---|---|---|
| GSTAGNGGNGGNATNGTIAGQP | ASGPDRNTEVF | 0.32049662 |
| GTGGDGGNGGNGGTGG | ASGPDRNTEVF | 0.7275927 |
| GVRRPAGIPADAPV | ASGPDRNTEVF | 0.41261622 |
| GYRQVTDAHLLTLARR | ASGPDRNTEVF | 0.6544596 |
| IAKMGQDGHDRG | ASGPDRNTEVF | 0.59083885 |
| KAEPSIEIQYT | ASGPDRNTEVF | 0.5753509 |
| KPVRIRMAADEDTVAVT | ASGPDRNTEVF | 0.74797046 |
| MPPRIAGMRLLVIKPEP | ASGPDRNTEVF | 0.4360617 |
| MKFYAVALFRSRGVPPTR | ASGPDRNTEVF | 0.4448989 |
| NAGGFGILGAVAHSP | ASGPDRNTEVF | 0.5426209 |
| NEGREITLAMAEM | ASGPDRNTEVF | 0.65054095 |
| NELVEWSHPLGHRWRWE | ASGPDRNTEVF | 0.7505406 |
| NLGAGGTGGNGGNPG | ASGPDRNTEVF | 0.3718617 |
| NTEWPKWETVMAADPVD | ASGPDRNTEVF | 0.63770694 |
| PASAAPAETQP | ASGPDRNTEVF | 0.5408992 |
| QQSSDGKNGVRAAEITGE | ASGPDRNTEVF | 0.41496912 |
| RAPDAVTTSWWKE | ASGPDRNTEVF | 0.6077268 |
| SGFGNVGAHTSGWFNQ | ASGPDRNTEVF | 0.5662582 |
| SGVAGAVGQAASVGGLKV | ASGPDRNTEVF | 0.39231747 |
| SNGGAGTGSAGGNGGT | ASGPDRNTEVF | 0.6847453 |
| TGGAGGEGGAAG | ASGPDRNTEVF | 0.50948936 |
| TGGDGGAAGAGGAPGHG | ASGPDRNTEVF | 0.697756 |
| TMPPLLAAIANAGRHGV | ASGPDRNTEVF | 0.78071976 |
| TSSTAGNSGGA | ASGPDRNTEVF | 0.3808283 |
| VPATVPVGILC | ASGPDRNTEVF | 0.6973043 |
| VQRPDSGGVVLTGRIS | ASGPDRNTEVF | 0.6621809 |
| WTPNPVPATKGVPAGLVQAT | ASGPDRNTEVF | 0.6469722 |
| YALKRALAKFSDRFDVVII | ASGPDRNTEVF | 0.58983445 |
| YLIGSVIV | ASGPDRNTEVF | 0.4241273 |
| YGQLVAVID | ASGPDRNTEVF | 0.7916799 |
| VTVGGMTLPA | ASGPDRNTEVF | 0.7380087 |
| VLALPAV | ASGPDRNTEVF | 0.7254659 |
| VIPPTLNYET | ASGPDRNTEVF | 0.7019205 |
| VGMGGMHPSSG | ASGPDRNTEVF | 0.7147633 |
| VGLAGGAATA | ASGPDRNTEVF | 0.7339845 |
| VGGAGGAGGSGP | ASGPDRNTEVF | 0.44037032 |
| VFAAFMV | ASGPDRNTEVF | 0.27674076 |
| TLYDKLR | ASGPDRNTEVF | 0.5759518 |
| TLAVFTVL | ASGPDRNTEVF | 0.5834553 |
| TGSVQPARV | ASGPDRNTEVF | 0.4715222 |
| TGGGAGSGGT | ASGPDRNTEVF | 0.3312156 |
| SHLETSL | ASGPDRNTEVF | 0.7583312 |
| SFTTGMT | ASGPDRNTEVF | 0.8369339 |
| QVEVAKRMGVRQPTVS | ASGPDRNTEVF | 0.51629686 |
| QGSGGPALAM | ASGPDRNTEVF | 0.8517939 |
| MLAPSNP | ASGPDRNTEVF | 0.36278322 |
| LVNNAAI | ASGPDRNTEVF | 0.73980945 |
| LTGEINTLA | ASGPDRNTEVF | 0.7785707 |
| LSIFGPV | ASGPDRNTEVF | 0.3675792 |
| LLALLANL | ASGPDRNTEVF | 0.7246879 |
| LISNASDALDKLR | ASGPDRNTEVF | 0.4527839 |

| | | |
|---|---|---|
| LINEAVP | ASGPDRNTEVF | 0.51082706 |
| LGYAFSGL | ASGPDRNTEVF | 0.76808536 |
| LDFVAVAL | ASGPDRNTEVF | 0.58582896 |
| LAQPGAE | ASGPDRNTEVF | 0.6884315 |
| ISNLGAVTGF | ASGPDRNTEVF | 0.5612977 |
| HQEDPGRI | ASGPDRNTEVF | 0.42215574 |
| GTPGEPTP | ASGPDRNTEVF | 0.6320262 |
| GFFNSGS | ASGPDRNTEVF | 0.74779093 |
| GAGGGGLTAGAN | ASGPDRNTEVF | 0.5850122 |
| FVDALVNM | ASGPDRNTEVF | 0.85538936 |
| FLTVAPEL | ASGPDRNTEVF | 0.3934965 |
| FETLTAL | ASGPDRNTEVF | 0.8025304 |
| DVINPIEL | ASGPDRNTEVF | 0.7830425 |
| APVSGAPV | ASGPDRNTEVF | 0.6106165 |
| ALVGGLAW | ASGPDRNTEVF | 0.5278262 |
| ALAAAAATAI | ASGPDRNTEVF | 0.7785428 |
| AFGLGLLL | ASGPDRNTEVF | 0.5279624 |
| AFFAAASL | ASGPDRNTEVF | 0.6798129 |
| AAAALPAFSPP | ASGPDRNTEVF | 0.79296625 |
| VVLAPAL | ASGPDRNTEVF | 0.6275874 |
| TIGHVDHGKTTL | ASGPDRNTEVF | 0.7646345 |
| LISNASDALDK | ASGPDRNTEVF | 0.72870666 |
| LISNASDAL | ASGPDRNTEVF | 0.7236154 |
| ISNASDALDK | ASGPDRNTEVF | 0.6931727 |
| NQAETLVYQTE | ASGPDRNTEVF | 0.57130516 |
| TGGTGGTAGAG | ASGPDRNTEVF | 0.68343455 |
| TGSTAGGAGGA | ASGPDRNTEVF | 0.6179694 |
| DPLAVPVAAGPV | ASGPDRNTEVF | 0.7476067 |
| GGTGGTPGNGGN | ASGPDRNTEVF | 0.7589535 |
| PAGLAGVDPATL | ASGPDRNTEVF | 0.6854366 |
| GNNLVVEAPHAGASV | ASGPDRNTEVF | 0.772226 |
| KVTRIMSASTLEQPA | ASGPDRNTEVF | 0.7433338 |
| LGFALPIALPRRTVV | ASGPDRNTEVF | 0.5261141 |
| TRLGNLTARKLVVLV | ASGPDRNTEVF | 0.6940748 |
| WSQQRRLIYFPSAGP | ASGPDRNTEVF | 0.6592509 |
| GPNGNPDPMAAAVDIR | ASGPDRNTEVF | 0.71623135 |
| TQRCVSVADLRAAASR | ASGPDRNTEVF | 0.39720863 |
| QFVLDVAEHPVEVLTETV | ASGPDRNTEVF | 0.7554506 |
| GGAGGTGGAGGNGGTQAGDG | ASGPDRNTEVF | 0.6248814 |
| PLLATLRAIAVDVDAKPAQV | ASGPDRNTEVF | 0.28533962 |
| APKFRTGGYMIDFWGVGYQVAK | ASGPDRNTEVF | 0.7407957 |
| GAGGAGGAGGAAGGPGGTGGTG | ASGPDRNTEVF | 0.7122371 |
| IGALGLLATDVAPAGSVAFGVA | ASGPDRNTEVF | 0.69536316 |
| AGGAGGIGGNGGAGGAGGVLMG | ASGPDRNTEVF | 0.7567656 |
| ADELIGLL | ASGPDRNTEVF | 0.5580727 |
| ADRALKA | ASGPDRNTEVF | 0.63175243 |
| AGAVGQVGGAGGA | ASGPDRNTEVF | 0.7463755 |
| AGGGGGLGGAP | ASGPDRNTEVF | 0.7761378 |
| AGGLLYGNGGAG | ASGPDRNTEVF | 0.61573815 |
| AGLESIGL | ASGPDRNTEVF | 0.7245666 |
| AGLKTGII | ASGPDRNTEVF | 0.77109885 |

| | | |
|---|---|---|
| ASLAALLL | ASGPDRNTEVF | 0.7259338 |
| ASVINISSI | ASGPDRNTEVF | 0.7693517 |
| ATPPGPA | ASGPDRNTEVF | 0.78002566 |
| ATVASALV | ASGPDRNTEVF | 0.62966675 |
| ATVASLIL | ASGPDRNTEVF | 0.77149165 |
| AVNGPVLIRH | ASGPDRNTEVF | 0.77273387 |
| DGAAGAVGI | ASGPDRNTEVF | 0.7131419 |
| DPLLHGI | ASGPDRNTEVF | 0.7177244 |
| ELGAALGAL | ASGPDRNTEVF | 0.6371224 |
| ELGRTQAV | ASGPDRNTEVF | 0.68951833 |
| FVAGGYLS | ASGPDRNTEVF | 0.5134225 |
| GASGGAGGAGGW | ASGPDRNTEVF | 0.714234 |
| GGAGAGGDGGAGGI | ASGPDRNTEVF | 0.7740014 |
| GTGHNGSQPGMGGNG | ASGPDRNTEVF | 0.68144834 |
| GYAAASASAS | ASGPDRNTEVF | 0.5641908 |
| HQVDVAAVQQG | ASGPDRNTEVF | 0.61381406 |
| IGAAIAHT | ASGPDRNTEVF | 0.7089119 |
| IGMIEVAQL | ASGPDRNTEVF | 0.66803056 |
| IRLAPEHTSAMD | ASGPDRNTEVF | 0.5055958 |
| IYNVSVL | ASGPDRNTEVF | 0.643311 |
| KQGFLGM | ASGPDRNTEVF | 0.7610519 |
| LESVITQL | ASGPDRNTEVF | 0.7612455 |
| LGLGALVVV | ASGPDRNTEVF | 0.52689034 |
| LIGNGGNGGSGGI | ASGPDRNTEVF | 0.7800147 |
| LNAINAPTL | ASGPDRNTEVF | 0.7358935 |
| LQLLNTGV | ASGPDRNTEVF | 0.7714918 |
| LQTSIEF | ASGPDRNTEVF | 0.72268707 |
| MSFAVSEL | ASGPDRNTEVF | 0.6445217 |
| NTGALGL | ASGPDRNTEVF | 0.75327456 |
| PPGAGGAGGAGTT | ASGPDRNTEVF | 0.6638057 |
| QLVGLGI | ASGPDRNTEVF | 0.6590377 |
| RFLEQVHQL | ASGPDRNTEVF | 0.76121366 |
| RGVGYVLR | ASGPDRNTEVF | 0.6981962 |
| SATGVFAGVI | ASGPDRNTEVF | 0.61670893 |
| SLPAITLP | ASGPDRNTEVF | 0.6131166 |
| STALAFTV | ASGPDRNTEVF | 0.77774054 |
| SVLTVGLGK | ASGPDRNTEVF | 0.5576358 |
| TALAGAGSGI | ASGPDRNTEVF | 0.68889266 |
| TGLPFTGI | ASGPDRNTEVF | 0.6216712 |
| VAEHVTV | ASGPDRNTEVF | 0.70685524 |
| VDIVLNSL | ASGPDRNTEVF | 0.7723888 |
| VLTGGGAVL | ASGPDRNTEVF | 0.7468405 |
| VNVGLAYGI | ASGPDRNTEVF | 0.67101353 |
| VSSAAVGAV | ASGPDRNTEVF | 0.7387043 |
| VTSVYEAGV | ASGPDRNTEVF | 0.7497438 |
| WVALDRGA | ASGPDRNTEVF | 0.7246871 |
| YLVGSVVI | ASGPDRNTEVF | 0.7452031 |
| GPPGTGKTLL | ASGPDRNTEVF | 0.5901247 |
| GVGMTKF | ASGPDRNTEVF | 0.7714417 |
| IGIGGSDL | ASGPDRNTEVF | 0.75244486 |
| IGTLLAEL | ASGPDRNTEVF | 0.7668984 |

| | | |
|---|---|---|
| IYTSGTT | ASGPDRNTEVF | 0.76442397 |
| LSFPAGV | ASGPDRNTEVF | 0.7396188 |
| LYTSGTT | ASGPDRNTEVF | 0.73132503 |
| VGAFPFGF | ASGPDRNTEVF | 0.50462437 |
| VGAGTVEF | ASGPDRNTEVF | 0.5827259 |
| VGKSTLLL | ASGPDRNTEVF | 0.7363889 |
| AALAHAGGGIV | ASGPDRNTEVF | 0.7125183 |
| AALVAVSLL | ASGPDRNTEVF | 0.7604302 |
| AAVSRAVAAL | ASGPDRNTEVF | 0.7412841 |
| AGGSSGIGGF | ASGPDRNTEVF | 0.6345091 |
| AGLTVANANG | ASGPDRNTEVF | 0.7619896 |
| AGLVGAVL | ASGPDRNTEVF | 0.76804996 |
| ALSQAFTV | ASGPDRNTEVF | 0.51651853 |
| ALTFSVSL | ASGPDRNTEVF | 0.6683538 |
| ALTGAALVAL | ASGPDRNTEVF | 0.78273094 |
| ASFSIGAGVL | ASGPDRNTEVF | 0.70776963 |
| DGSVYKI | ASGPDRNTEVF | 0.70416594 |
| EQLAEIV | ASGPDRNTEVF | 0.686906 |
| FAVSELTVM | ASGPDRNTEVF | 0.773429 |
| GAAPLAAGL | ASGPDRNTEVF | 0.6562041 |
| GADGGDGGSGGAGG | ASGPDRNTEVF | 0.78557205 |
| GFGLDTGGAGG | ASGPDRNTEVF | 0.36137557 |
| GLASGGLL | ASGPDRNTEVF | 0.7077016 |
| GRIAGAAVLL | ASGPDRNTEVF | 0.6573663 |
| GSAGSLNTSF | ASGPDRNTEVF | 0.5033606 |
| GTGQIGI | ASGPDRNTEVF | 0.7574988 |
| GTLIALIL | ASGPDRNTEVF | 0.6598207 |
| IAATMGFVVG | ASGPDRNTEVF | 0.74283457 |
| IFAQALL | ASGPDRNTEVF | 0.694695 |
| IGAAVALGF | ASGPDRNTEVF | 0.7628625 |
| IGLTPLAL | ASGPDRNTEVF | 0.6969789 |
| ILGSNTSTL | ASGPDRNTEVF | 0.7125721 |
| IVFSLTVL | ASGPDRNTEVF | 0.6864462 |
| LASKATGF | ASGPDRNTEVF | 0.75250244 |
| LASVALIL | ASGPDRNTEVF | 0.77213067 |
| LFAVGLTL | ASGPDRNTEVF | 0.7699131 |
| LFNGGIGTYI | ASGPDRNTEVF | 0.52743953 |
| LTLGAGII | ASGPDRNTEVF | 0.69097626 |
| LVAAALGDI | ASGPDRNTEVF | 0.7174115 |
| NTAFVQLGI | ASGPDRNTEVF | 0.47590968 |
| QLFSGAEL | ASGPDRNTEVF | 0.44248623 |
| SAATFITL | ASGPDRNTEVF | 0.6636473 |
| SNYGLANLGV | ASGPDRNTEVF | 0.64920926 |
| SYGGAYAVM | ASGPDRNTEVF | 0.7625003 |
| TTGVGGLTLG | ASGPDRNTEVF | 0.7477411 |
| TTQTIYI | ASGPDRNTEVF | 0.69759274 |
| VAAGAGLGAVI | ASGPDRNTEVF | 0.7425209 |
| VAAVVTLL | ASGPDRNTEVF | 0.7088646 |
| VGFLGGLI | ASGPDRNTEVF | 0.19917768 |
| VGVIPILL | ASGPDRNTEVF | 0.6286506 |
| VIFASSLI | ASGPDRNTEVF | 0.54785794 |

| | | |
|---|---|---|
| VKGGDGGAAAG | ASGPDRNTEVF | 0.77408993 |
| VTKTFTNI | ASGPDRNTEVF | 0.6273336 |
| VVAAAILL | ASGPDRNTEVF | 0.5900315 |
| YAALDVEL | ASGPDRNTEVF | 0.7145113 |
| LASVALLL | ASGPDRNTEVF | 0.6938919 |
| LDAVHRV | ASGPDRNTEVF | 0.68443775 |
| QGLSSSIF | ASGPDRNTEVF | 0.596106 |
| RGTFANIRL | ASGPDRNTEVF | 0.39929846 |
| RQLWWGHRI | ASGPDRNTEVF | 0.75677794 |
| YGPPGTGKTLL | ASGPDRNTEVF | 0.5472606 |
| AALVGAPANVA | ASGPDRNTEVF | 0.63527775 |
| AATFRRTE | ASGPDRNTEVF | 0.41878742 |
| AGALGGGI | ASGPDRNTEVF | 0.63450694 |
| AGGAGGAGGIGAG | ASGPDRNTEVF | 0.47796738 |
| AGSGALGL | ASGPDRNTEVF | 0.7626191 |
| AQIADQI | ASGPDRNTEVF | 0.6661871 |
| ASAVAVTYL | ASGPDRNTEVF | 0.6693667 |
| ASTVLGGTGG | ASGPDRNTEVF | 0.77799535 |
| AVGGEPAK | ASGPDRNTEVF | 0.6663455 |
| CEQGTFGGTC | ASGPDRNTEVF | 0.6828191 |
| DEGKAPAG | ASGPDRNTEVF | 0.7041967 |
| DTVVFPAL | ASGPDRNTEVF | 0.63466424 |
| FEGITMAG | ASGPDRNTEVF | 0.66101146 |
| FPLVVQD | ASGPDRNTEVF | 0.659112 |
| GAFTTPQ | ASGPDRNTEVF | 0.77763593 |
| GFGGFGTAGGVGGA | ASGPDRNTEVF | 0.72490215 |
| GFLAQFL | ASGPDRNTEVF | 0.5337455 |
| GGAGGAGGNAGVA | ASGPDRNTEVF | 0.738775 |
| GGVGTAGAGGAGGAGG | ASGPDRNTEVF | 0.78074586 |
| GNGGGGDAGNAGSGGN | ASGPDRNTEVF | 0.7851659 |
| GTGGAGGAGNDAGS | ASGPDRNTEVF | 0.6379756 |
| GTTNTGWFNS | ASGPDRNTEVF | 0.66682494 |
| IFMGPLFTL | ASGPDRNTEVF | 0.67178047 |
| ILAHAIEAI | ASGPDRNTEVF | 0.6711507 |
| IRSAIREAL | ASGPDRNTEVF | 0.6567894 |
| ITFAGGG | ASGPDRNTEVF | 0.6211649 |
| ITGMSGHQEV | ASGPDRNTEVF | 0.7599969 |
| IVAGLACTL | ASGPDRNTEVF | 0.7794596 |
| IVGGGLAGA | ASGPDRNTEVF | 0.69912755 |
| IVTGVLNI | ASGPDRNTEVF | 0.43187484 |
| LAQQGCI | ASGPDRNTEVF | 0.70561635 |
| LCLHVEN | ASGPDRNTEVF | 0.5938469 |
| LDAISDGS | ASGPDRNTEVF | 0.6693727 |
| LGGFAAML | ASGPDRNTEVF | 0.696507 |
| LGGKNPMI | ASGPDRNTEVF | 0.4725398 |
| LGPKGVAAIL | ASGPDRNTEVF | 0.6989465 |
| LMVGVSI | ASGPDRNTEVF | 0.78144276 |
| LVGAAFVL | ASGPDRNTEVF | 0.62968755 |
| LVLAVAPL | ASGPDRNTEVF | 0.73581004 |
| MAAIAVIDAA | ASGPDRNTEVF | 0.77294147 |
| NNGINAELQ | ASGPDRNTEVF | 0.7570599 |

| | | |
|---|---|---|
| RGEPLVLVHG | ASGPDRNTEVF | 0.7713183 |
| RGGAGGAAGLI | ASGPDRNTEVF | 0.7430521 |
| RSYRMHPAV | ASGPDRNTEVF | 0.7431044 |
| SAAHLLAGQLA | ASGPDRNTEVF | 0.7214912 |
| SAAVGSY | ASGPDRNTEVF | 0.75341344 |
| SPHAVLHRI | ASGPDRNTEVF | 0.7154531 |
| SQIAKVL | ASGPDRNTEVF | 0.7124238 |
| TALGHLVP | ASGPDRNTEVF | 0.6697212 |
| TAPTTSGAA | ASGPDRNTEVF | 0.7556768 |
| TDGDDGNGGSPGT | ASGPDRNTEVF | 0.7007862 |
| TDGLGEN | ASGPDRNTEVF | 0.77147603 |
| TGGPPLPGQA | ASGPDRNTEVF | 0.55728185 |
| TIPAITI | ASGPDRNTEVF | 0.6310711 |
| TIPGFTI | ASGPDRNTEVF | 0.7588762 |
| TSIASGALA | ASGPDRNTEVF | 0.28284404 |
| TVTLPGAAI | ASGPDRNTEVF | 0.43151167 |
| VASGLSAALP | ASGPDRNTEVF | 0.72088 |
| VGAGVIGI | ASGPDRNTEVF | 0.77162766 |
| VGPKRFGV | ASGPDRNTEVF | 0.2888776 |
| VNVAFPD | ASGPDRNTEVF | 0.6213955 |
| VSMTNSGVS | ASGPDRNTEVF | 0.5875933 |
| VTDVLGDLSV | ASGPDRNTEVF | 0.7519053 |
| VVVVGAGFAGL | ASGPDRNTEVF | 0.7010639 |
| VYTVGTVV | ASGPDRNTEVF | 0.5141256 |
| WTTFSGP | ASGPDRNTEVF | 0.75932205 |
| YDELVEAF | ASGPDRNTEVF | 0.4336415 |
| YGTGGAGGNGG | ASGPDRNTEVF | 0.6618583 |
| YTNVMAV | ASGPDRNTEVF | 0.76419926 |
| FLEQVHQL | ASGPDRNTEVF | 0.77180135 |
| GLSLLAVAV | ASGPDRNTEVF | 0.6215248 |
| LEQVHQL | ASGPDRNTEVF | 0.75615263 |
| LNAAGAI | ASGPDRNTEVF | 0.6088108 |
| PAVHASL | ASGPDRNTEVF | 0.7887306 |
| TELEDDA | ASGPDRNTEVF | 0.7072695 |
| TPPGPPN | ASGPDRNTEVF | 0.58987033 |
| TPPPPPPP | ASGPDRNTEVF | 0.7501307 |
| TQSPALI | ASSLGGGAETLY | 0.7560468 |
| EFAAAFPQ | ASSLGGGAETLY | 0.7311204 |
| LLSGNLDVL | ASSLGGGAETLY | 0.7373044 |
| GPANLGNYNI | ASSLGGGAETLY | 0.70202297 |
| TLGPTLP | ASSLGGGAETLY | 0.694592 |
| VYAYSSCCK | ASSLGGGAETLY | 0.7574201 |
| APAPAPAPNI | ASSLGGGAETLY | 0.7589861 |
| PGGPVPIP | ASSLGGGAETLY | 0.64757675 |
| FLTVALEP | ASSLGGGAETLY | 0.7713524 |
| SQYFGANL | ASSLGGGAETLY | 0.691583 |
| GGVKPAA | ASSLGGGAETLY | 0.6624438 |
| GGTGGAGGAGGW | ASSLGGGAETLY | 0.7740258 |
| GVGGVAGLGGAGG | ASSLGGGAETLY | 0.7156391 |
| VLLVLATTI | ASSLGGGAETLY | 0.7646079 |
| IGIFAAML | ASSLGGGAETLY | 0.75838566 |

| | | |
|---|---|---|
| AVGDLID | ASSLGGGAETLY | 0.7675201 |
| NISPLPL | ASSLGGGAETLY | 0.5930634 |
| VTAVFVAI | ASSLGGGAETLY | 0.7015084 |
| SGDDNDAAL | ASSLGGGAETLY | 0.7183672 |
| AVAALAHS | ASSLGGGAETLY | 0.75877136 |
| GGVTVMATP | ASSLGGGAETLY | 0.70807874 |
| AGLAANVRD | ASSLGGGAETLY | 0.62907743 |
| TNRPDLIDPALLRPG | ASSLGGGAETLY | 0.75442374 |
| LANLGVTAL | ASSLGGGAETLY | 0.7733254 |
| TGERIEL | ASSLGGGAETLY | 0.72993636 |
| NGEELAA | ASSLGGGAETLY | 0.71725005 |
| VLSLVWG | ASSLGGGAETLY | 0.777452 |
| IGYALWL | ASSLGGGAETLY | 0.6884388 |
| LSLLLAPL | ASSLGGGAETLY | 0.70357215 |
| TLDEALK | ASSLGGGAETLY | 0.75569165 |
| TDGAGTAEGG | ASSLGGGAETLY | 0.7656825 |
| NAPGGAGGNGG | ASSLGGGAETLY | 0.71434885 |
| VSVFMGMGGNF | ASSLGGGAETLY | 0.76877296 |
| LASYLAAL | ASSLGGGAETLY | 0.5984955 |
| LVGGTPAL | ASSLGGGAETLY | 0.7427079 |
| VRNADVL | ASSLGGGAETLY | 0.5953108 |
| TCSSPFLVSAM | ASSLGGGAETLY | 0.7166058 |
| VHITDAE | ASSLGGGAETLY | 0.6039641 |
| EYAAPSF | ASSLGGGAETLY | 0.74175864 |
| VVDAFHGGL | ASSLGGGAETLY | 0.7497548 |
| ASVVDPQ | ASSLGGGAETLY | 0.753307 |
| SYNEFGGLS | ASSLGGGAETLY | 0.34912103 |
| DPDGSVY | ASSLGGGAETLY | 0.7476217 |
| MVSTIHATE | ASSLGGGAETLY | 0.74373174 |
| LAAGVADPV | ASSLGGGAETLY | 0.78150713 |
| TKEQIAA | ASSLGGGAETLY | 0.71148765 |
| SLASPAAL | ASSLGGGAETLY | 0.73933434 |
| IGLVFYGV | ASSLGGGAETLY | 0.67668426 |
| FLLDASNL | ASSLGGGAETLY | 0.7162682 |
| YQVGNFP | ASSLGGGAETLY | 0.53642416 |
| ESLLSQST | ASSLGGGAETLY | 0.6778258 |
| LFTVGFAAVG | ASSLGGGAETLY | 0.6294342 |
| ATPPTPT | ASSLGGGAETLY | 0.6243702 |
| TDLATLG | ASSLGGGAETLY | 0.74726474 |
| SAGGTGR | ASSLGGGAETLY | 0.7705948 |
| GFSNSGS | ASSLGGGAETLY | 0.754825 |
| GGNGGSGGTSV | ASSLGGGAETLY | 0.767258 |
| GTSVATGGAGN | ASSLGGGAETLY | 0.7773255 |
| NGGNGGSGGTS | ASSLGGGAETLY | 0.77356243 |
| AVVKFPH | ASSLGGGAETLY | 0.6532525 |
| TPPPPPP | ASSLGGGAETLY | 0.68281066 |
| AAPAAAAAPA | ASSLGGGAETLY | 0.56413484 |
| IALNEDL | ASSLGGGAETLY | 0.684967 |
| IVNVSSM | ASSLGGGAETLY | 0.6574702 |
| TGIPPAPR | ASSLGGGAETLY | 0.73629105 |
| FDLGGGTF | ASSLGGGAETLY | 0.27818057 |

| | | |
|---|---|---|
| FDLGGGTFDVS | ASSLGGGAETLY | 0.6788696 |
| NEPTAAA | ASSLGGGAETLY | 0.77946055 |
| NEPTAAALA | ASSLGGGAETLY | 0.6100977 |
| GSIINISSI | ASSLGGGAETLY | 0.7311078 |
| VAPGNAGTA | ASSLGGGAETLY | 0.7268293 |
| TGGGASLE | ASSLGGGAETLY | 0.6439235 |
| VAVYNHYKRI | ASSLGGGAETLY | 0.7531467 |
| TDPAPATT | ASSLGGGAETLY | 0.6543082 |
| TNRPDILDPALLRPG | ASSLGGGAETLY | 0.753021 |
| LTTTTTS | ASSLGGGAETLY | 0.7768066 |
| PTGSGKTL | ASSLGGGAETLY | 0.7326989 |
| TGSGKTL | ASSLGGGAETLY | 0.68967825 |
| TIGHVDHGK | ASSLGGGAETLY | 0.71983826 |
| TIGHVDHGKTT | ASSLGGGAETLY | 0.5365933 |
| GHVDHGKTT | ASSLGGGAETLY | 0.72540885 |
| AQFEHTLL | ASSLGGGAETLY | 0.5667076 |
| IVNQDEL | ASSLGGGAETLY | 0.6933961 |
| TNGSQFF | ASSLGGGAETLY | 0.6741873 |
| TGKGIVE | ASSLGGGAETLY | 0.7412176 |
| LISNASDA | ASSLGGGAETLY | 0.7106712 |
| AELRTLGELTKIAKAHGAQ | ASSLGGGAETLY | 0.73664474 |
| APNAVAAAEPSW | ASSLGGGAETLY | 0.58990353 |
| APQVAGAATAVL | ASSLGGGAETLY | 0.7027391 |
| ARRDPAAAAAILPTDA | ASSLGGGAETLY | 0.6448988 |
| DAAARDVLAEAGLAEYF | ASSLGGGAETLY | 0.5901546 |
| DDQIVLSAKANLAKVEQ | ASSLGGGAETLY | 0.74449366 |
| DELLPDTGEDSDEDGH | ASSLGGGAETLY | 0.71975106 |
| ELPEAPANRATVPQD | ASSLGGGAETLY | 0.65832096 |
| FGQSKIERVTGIG | ASSLGGGAETLY | 0.76191986 |
| GAGGTGSTAGGAGGAGGAGGLY | ASSLGGGAETLY | 0.73356795 |
| GHGGTGAAGGNGGTGGAGGSGI | ASSLGGGAETLY | 0.50761557 |
| GNGGNGGAGGEGGPGGAGGAGA | ASSLGGGAETLY | 0.66750836 |
| GSTAGNGGNGGNATNGTIAGQP | ASSLGGGAETLY | 0.71924114 |
| GTGGDGGNGGNGGTGG | ASSLGGGAETLY | 0.7229901 |
| GVRRPAGIPADAPV | ASSLGGGAETLY | 0.27639166 |
| GYRQVTDAHLLTLARR | ASSLGGGAETLY | 0.74204135 |
| IAKMGQDGHDRG | ASSLGGGAETLY | 0.66395056 |
| KAEPSIEIQYT | ASSLGGGAETLY | 0.57085735 |
| KPVRIRMAADEDTVAVT | ASSLGGGAETLY | 0.65046483 |
| MPPRIAGMRLLVIKPEP | ASSLGGGAETLY | 0.7535393 |
| MKFYAVALFRSRGVPPTR | ASSLGGGAETLY | 0.6742257 |
| NAGGFGILGAVAHSP | ASSLGGGAETLY | 0.68602186 |
| NEGREITLAMAEM | ASSLGGGAETLY | 0.7416386 |
| NELVEWSHPLGHRWRWE | ASSLGGGAETLY | 0.748312 |
| NLGAGGTGGNGGNPG | ASSLGGGAETLY | 0.77060735 |
| NTEWPKWETVMAADPVD | ASSLGGGAETLY | 0.7398994 |
| PASAAPAETQP | ASSLGGGAETLY | 0.7085579 |
| QQSSDGKNGVRAAEITGE | ASSLGGGAETLY | 0.73562473 |
| RAPDAVTTSWWKE | ASSLGGGAETLY | 0.6812881 |
| SGFGNVGAHTSGWFNQ | ASSLGGGAETLY | 0.55948246 |
| SGVAGAVGQAASVGGLKV | ASSLGGGAETLY | 0.4835794 |

| | | |
|---|---|---|
| SNGGAGTGSAGGNGGT | ASSLGGGAETLY | 0.71244895 |
| TGGAGGEGGAAG | ASSLGGGAETLY | 0.5358995 |
| TGGDGGAAGAGGAPGHG | ASSLGGGAETLY | 0.7338217 |
| TMPPLLAAIANAGRHGV | ASSLGGGAETLY | 0.77196753 |
| TSSTAGNSGGA | ASSLGGGAETLY | 0.6969741 |
| VPATVPVGILC | ASSLGGGAETLY | 0.5629535 |
| VQRPDSGGVVLTGRIS | ASSLGGGAETLY | 0.66938186 |
| WTPNPVPATKGVPAGLVQAT | ASSLGGGAETLY | 0.6410673 |
| YALKRALAKFSDRFDVVII | ASSLGGGAETLY | 0.7562654 |
| YLIGSVIV | ASSLGGGAETLY | 0.7587105 |
| YGQLVAVID | ASSLGGGAETLY | 0.7794653 |
| VTVGGMTLPA | ASSLGGGAETLY | 0.75851595 |
| VLALPAV | ASSLGGGAETLY | 0.64453214 |
| VIPPTLNYET | ASSLGGGAETLY | 0.75791204 |
| VGMGGMHPSSG | ASSLGGGAETLY | 0.6722791 |
| VGLAGGAATA | ASSLGGGAETLY | 0.6802895 |
| VGGAGGAGGSGP | ASSLGGGAETLY | 0.7280098 |
| VFAAFMV | ASSLGGGAETLY | 0.728402 |
| TLYDKLR | ASSLGGGAETLY | 0.7279104 |
| TLAVFTVL | ASSLGGGAETLY | 0.7126712 |
| TGSVQPARV | ASSLGGGAETLY | 0.54683167 |
| TGGGAGSGGT | ASSLGGGAETLY | 0.7525135 |
| SHLETSL | ASSLGGGAETLY | 0.70948154 |
| SFTTGMT | ASSLGGGAETLY | 0.5397381 |
| QVEVAKRMGVRQPTVS | ASSLGGGAETLY | 0.67949444 |
| QGSGGPALAM | ASSLGGGAETLY | 0.78010726 |
| MLAPSNP | ASSLGGGAETLY | 0.77181125 |
| LVNNAAI | ASSLGGGAETLY | 0.61054903 |
| LTGEINTLA | ASSLGGGAETLY | 0.6492038 |
| LSIFGPV | ASSLGGGAETLY | 0.7002858 |
| LLALLANL | ASSLGGGAETLY | 0.53244656 |
| LISNASDALDKLR | ASSLGGGAETLY | 0.7558968 |
| LINEAVP | ASSLGGGAETLY | 0.66541505 |
| LGYAFSGL | ASSLGGGAETLY | 0.6701636 |
| LDFVAVAL | ASSLGGGAETLY | 0.6539196 |
| LAQPGAE | ASSLGGGAETLY | 0.725936 |
| ISNLGAVTGF | ASSLGGGAETLY | 0.65398484 |
| HQEDPGRI | ASSLGGGAETLY | 0.7559929 |
| GTPGEPTP | ASSLGGGAETLY | 0.28867632 |
| GFFNSGS | ASSLGGGAETLY | 0.74482644 |
| GAGGGGLTAGAN | ASSLGGGAETLY | 0.6037727 |
| FVDALVNM | ASSLGGGAETLY | 0.65136045 |
| FLTVAPEL | ASSLGGGAETLY | 0.46736446 |
| FETLTAL | ASSLGGGAETLY | 0.6949738 |
| DVINPIEL | ASSLGGGAETLY | 0.747044 |
| APVSGAPV | ASSLGGGAETLY | 0.76417863 |
| ALVGGLAW | ASSLGGGAETLY | 0.65340346 |
| ALAAAAATAI | ASSLGGGAETLY | 0.7463275 |
| AFGLGLLL | ASSLGGGAETLY | 0.77793705 |
| AFFAAASL | ASSLGGGAETLY | 0.66598845 |
| AAAALPAFSPP | ASSLGGGAETLY | 0.4922889 |

| | | |
|---|---|---|
| VVLAPAL | ASSLGGGAETLY | 0.7208421 |
| TIGHVDHGKTTL | ASSLGGGAETLY | 0.6493539 |
| LISNASDALDK | ASSLGGGAETLY | 0.7573379 |
| LISNASDAL | ASSLGGGAETLY | 0.7472632 |
| ISNASDALDK | ASSLGGGAETLY | 0.5290013 |
| NQAETLVYQTE | ASSLGGGAETLY | 0.74237514 |
| TGGTGGTAGAG | ASSLGGGAETLY | 0.7645707 |
| TGSTAGGAGGA | ASSLGGGAETLY | 0.62566894 |
| DPLAVPVAAGPV | ASSLGGGAETLY | 0.6114169 |
| GGTGGTPGNGGN | ASSLGGGAETLY | 0.6252019 |
| PAGLAGVDPATL | ASSLGGGAETLY | 0.40808365 |
| GNNLVVEAPHAGASV | ASSLGGGAETLY | 0.7438213 |
| KVTRIMSASTLEQPA | ASSLGGGAETLY | 0.6355558 |
| LGFALPIALPRRTVV | ASSLGGGAETLY | 0.66114056 |
| TRLGNLTARKLVVLV | ASSLGGGAETLY | 0.71517974 |
| WSQQRRLIYFPSAGP | ASSLGGGAETLY | 0.68003887 |
| GPNGNPDPMAAAVDIR | ASSLGGGAETLY | 0.6794983 |
| TQRCVSVADLRAAASR | ASSLGGGAETLY | 0.69178057 |
| QFVLDVAEHPVEVLTETV | ASSLGGGAETLY | 0.70794237 |
| GGAGGTGGAGGNGGTQAGDG | ASSLGGGAETLY | 0.73523533 |
| PLLATLRAIAVDVDAKPAQV | ASSLGGGAETLY | 0.6463165 |
| APKFRTGGYMIDFWGVGYQVAK | ASSLGGGAETLY | 0.67725563 |
| GAGGAGGAGGAAGGPGGTGGTG | ASSLGGGAETLY | 0.7732007 |
| IGALGLLATDVAPAGSVAFGVA | ASSLGGGAETLY | 0.5749118 |
| AGGAGGIGGNGGAGGAGGVLMG | ASSLGGGAETLY | 0.6428282 |
| ADELIGLL | ASSLGGGAETLY | 0.7316582 |
| ADRALKA | ASSLGGGAETLY | 0.57911444 |
| AGAVGQVGGAGGA | ASSLGGGAETLY | 0.7536736 |
| AGGGGGLGGAP | ASSLGGGAETLY | 0.7077766 |
| AGGLLYGNGGAG | ASSLGGGAETLY | 0.7422561 |
| AGLESIGL | ASSLGGGAETLY | 0.68355453 |
| AGLKTGII | ASSLGGGAETLY | 0.780835 |
| ASLAALLL | ASSLGGGAETLY | 0.74645066 |
| ASVINISSI | ASSLGGGAETLY | 0.69401515 |
| ATPPGPA | ASSLGGGAETLY | 0.75177246 |
| ATVASALV | ASSLGGGAETLY | 0.60527575 |
| ATVASLIL | ASSLGGGAETLY | 0.75460285 |
| AVNGPVLIRH | ASSLGGGAETLY | 0.7443673 |
| DGAAGAVGI | ASSLGGGAETLY | 0.59077287 |
| DPLLHGI | ASSLGGGAETLY | 0.6980789 |
| ELGAALGAL | ASSLGGGAETLY | 0.77807415 |
| ELGRTQAV | ASSLGGGAETLY | 0.7686342 |
| FVAGGYLS | ASSLGGGAETLY | 0.57572055 |
| GASGGAGGAGGW | ASSLGGGAETLY | 0.63483155 |
| GGAGAGGDGGAGGI | ASSLGGGAETLY | 0.5516331 |
| GTGHNGSQPGMGGNG | ASSLGGGAETLY | 0.6857049 |
| GYAAASASAS | ASSLGGGAETLY | 0.62112635 |
| HQVDVAAVQQG | ASSLGGGAETLY | 0.7444043 |
| IGAAIAHT | ASSLGGGAETLY | 0.68769616 |
| IGMIEVAQL | ASSLGGGAETLY | 0.7383389 |
| IRLAPEHTSAMD | ASSLGGGAETLY | 0.7382862 |

| | | |
|---|---|---|
| IYNVSVL | ASSLGGGAETLY | 0.7720122 |
| KQGFLGM | ASSLGGGAETLY | 0.67835426 |
| LESVITQL | ASSLGGGAETLY | 0.699199 |
| LGLGALVVV | ASSLGGGAETLY | 0.5455198 |
| LIGNGGNGGSGGI | ASSLGGGAETLY | 0.64855725 |
| LNAINAPTL | ASSLGGGAETLY | 0.6872948 |
| LQLLNTGV | ASSLGGGAETLY | 0.6737553 |
| LQTSIEF | ASSLGGGAETLY | 0.4145661 |
| MSFAVSEL | ASSLGGGAETLY | 0.7744984 |
| NTGALGL | ASSLGGGAETLY | 0.71616524 |
| PPGAGGAGGAGTT | ASSLGGGAETLY | 0.75183 |
| QLVGLGI | ASSLGGGAETLY | 0.6596897 |
| RFLEQVHQL | ASSLGGGAETLY | 0.6876433 |
| RGVGYVLR | ASSLGGGAETLY | 0.65392673 |
| SATGVFAGVI | ASSLGGGAETLY | 0.7529907 |
| SLPAITLP | ASSLGGGAETLY | 0.608627 |
| STALAFTV | ASSLGGGAETLY | 0.45446062 |
| SVLTVGLGK | ASSLGGGAETLY | 0.6518979 |
| TALAGAGSGI | ASSLGGGAETLY | 0.48432478 |
| TGLPFTGI | ASSLGGGAETLY | 0.78052014 |
| VAEHVTV | ASSLGGGAETLY | 0.7673328 |
| VDIVLNSL | ASSLGGGAETLY | 0.73073214 |
| VLTGGGAVL | ASSLGGGAETLY | 0.7819047 |
| VNVGLAYGI | ASSLGGGAETLY | 0.3370291 |
| VSSAAVGAV | ASSLGGGAETLY | 0.64158 |
| VTSVYEAGV | ASSLGGGAETLY | 0.7697729 |
| WVALDRGA | ASSLGGGAETLY | 0.7294701 |
| YLVGSVVI | ASSLGGGAETLY | 0.7080561 |
| GPPGTGKTLL | ASSLGGGAETLY | 0.77180266 |
| GVGMTKF | ASSLGGGAETLY | 0.61956024 |
| IGIGGSDL | ASSLGGGAETLY | 0.72118187 |
| IGTLLAEL | ASSLGGGAETLY | 0.6998073 |
| IYTSGTT | ASSLGGGAETLY | 0.76685876 |
| LSFPAGV | ASSLGGGAETLY | 0.73604715 |
| LYTSGTT | ASSLGGGAETLY | 0.6307709 |
| VGAFPFGF | ASSLGGGAETLY | 0.7354157 |
| VGAGTVEF | ASSLGGGAETLY | 0.7897835 |
| VGKSTLLL | ASSLGGGAETLY | 0.6773239 |
| AALAHAGGGIV | ASSLGGGAETLY | 0.7562962 |
| AALVAVSLL | ASSLGGGAETLY | 0.7358105 |
| AAVSRAVAAL | ASSLGGGAETLY | 0.67402756 |
| AGGSSGIGGF | ASSLGGGAETLY | 0.6413637 |
| AGLTVANANG | ASSLGGGAETLY | 0.6314653 |
| AGLVGAVL | ASSLGGGAETLY | 0.70099306 |
| ALSQAFTV | ASSLGGGAETLY | 0.5576743 |
| ALTFSVSL | ASSLGGGAETLY | 0.70983344 |
| ALTGAALVAL | ASSLGGGAETLY | 0.6703346 |
| ASFSIGAGVL | ASSLGGGAETLY | 0.7940755 |
| DGSVYKI | ASSLGGGAETLY | 0.7469972 |
| EQLAEIV | ASSLGGGAETLY | 0.7382082 |
| FAVSELTVM | ASSLGGGAETLY | 0.58658457 |

| | | |
|---|---|---|
| GAAPLAAGL | ASSLGGGAETLY | 0.73837745 |
| GADGGDGGSGGAGG | ASSLGGGAETLY | 0.7238155 |
| GFGLDTGGAGG | ASSLGGGAETLY | 0.560532 |
| GLASGGLL | ASSLGGGAETLY | 0.7727418 |
| GRIAGAAVLL | ASSLGGGAETLY | 0.6444533 |
| GSAGSLNTSF | ASSLGGGAETLY | 0.65960425 |
| GTGQIGI | ASSLGGGAETLY | 0.745222 |
| GTLIALIL | ASSLGGGAETLY | 0.76662153 |
| IAATMGFVVG | ASSLGGGAETLY | 0.7164923 |
| IFAQALL | ASSLGGGAETLY | 0.68831694 |
| IGAAVALGF | ASSLGGGAETLY | 0.6188334 |
| IGLTPLAL | ASSLGGGAETLY | 0.76213133 |
| ILGSNTSTL | ASSLGGGAETLY | 0.7701589 |
| IVFSLTVL | ASSLGGGAETLY | 0.70147437 |
| LASKATGF | ASSLGGGAETLY | 0.78654784 |
| LASVALIL | ASSLGGGAETLY | 0.73132306 |
| LFAVGLTL | ASSLGGGAETLY | 0.5623966 |
| LFNGGIGTYI | ASSLGGGAETLY | 0.7374357 |
| LTLGAGII | ASSLGGGAETLY | 0.7612002 |
| LVAAALGDI | ASSLGGGAETLY | 0.7618534 |
| NTAFVQLGI | ASSLGGGAETLY | 0.7789639 |
| QLFSGAEL | ASSLGGGAETLY | 0.72332686 |
| SAATFITL | ASSLGGGAETLY | 0.66315377 |
| SNYGLANLGV | ASSLGGGAETLY | 0.77271396 |
| SYGGAYAVM | ASSLGGGAETLY | 0.7861453 |
| TTGVGGLTLG | ASSLGGGAETLY | 0.74743557 |
| TTQTIYI | ASSLGGGAETLY | 0.6522731 |
| VAAGAGLGAVI | ASSLGGGAETLY | 0.6334677 |
| VAAVVTLL | ASSLGGGAETLY | 0.74237144 |
| VGFLGGLI | ASSLGGGAETLY | 0.7519029 |
| VGVIPILL | ASSLGGGAETLY | 0.63065284 |
| VIFASSLI | ASSLGGGAETLY | 0.7297436 |
| VKGGDGGAAAG | ASSLGGGAETLY | 0.72971857 |
| VTKTFTNI | ASSLGGGAETLY | 0.5224965 |
| VVAAAILL | ASSLGGGAETLY | 0.7228557 |
| YAALDVEL | ASSLGGGAETLY | 0.76278156 |
| LASVALLL | ASSLGGGAETLY | 0.73870397 |
| LDAVHRV | ASSLGGGAETLY | 0.6436661 |
| QGLSSSIF | ASSLGGGAETLY | 0.7679685 |
| RGTFANIRL | ASSLGGGAETLY | 0.78910434 |
| RQLWWGHRI | ASSLGGGAETLY | 0.7821176 |
| YGPPGTGKTLL | ASSLGGGAETLY | 0.49528646 |
| AALVGAPANVA | ASSLGGGAETLY | 0.3468215 |
| AATFRRTE | ASSLGGGAETLY | 0.6715151 |
| AGALGGGI | ASSLGGGAETLY | 0.737443 |
| AGGAGGAGGIGAG | ASSLGGGAETLY | 0.6827888 |
| AGSGALGL | ASSLGGGAETLY | 0.6524616 |
| AQIADQI | ASSLGGGAETLY | 0.7190754 |
| ASAVAVTYL | ASSLGGGAETLY | 0.7596965 |
| ASTVLGGTGG | ASSLGGGAETLY | 0.6708054 |
| AVGGEPAK | ASSLGGGAETLY | 0.58728206 |

| | | |
|---|---|---|
| CEQGTFGGTC | ASSLGGGAETLY | 0.7235211 |
| DEGKAPAG | ASSLGGGAETLY | 0.7561896 |
| DTVVFPAL | ASSLGGGAETLY | 0.61653775 |
| FEGITMAG | ASSLGGGAETLY | 0.68820894 |
| FPLVVQD | ASSLGGGAETLY | 0.6931083 |
| GAFTTPQ | ASSLGGGAETLY | 0.6666418 |
| GFGGFGTAGGVGGA | ASSLGGGAETLY | 0.6978911 |
| GFLAQFL | ASSLGGGAETLY | 0.618745 |
| GGAGGAGGNAGVA | ASSLGGGAETLY | 0.76801926 |
| GGVGTAGAGGAGGAGG | ASSLGGGAETLY | 0.6149281 |
| GNGGGGDAGNAGSGGN | ASSLGGGAETLY | 0.7480298 |
| GTGGAGGAGNDAGS | ASSLGGGAETLY | 0.73394114 |
| GTTNTGWFNS | ASSLGGGAETLY | 0.74461234 |
| IFMGPLFTL | ASSLGGGAETLY | 0.6938676 |
| ILAHAIEAI | ASSLGGGAETLY | 0.6344339 |
| IRSAIREAL | ASSLGGGAETLY | 0.59090185 |
| ITFAGGG | ASSLGGGAETLY | 0.47965688 |
| ITGMSGHQEV | ASSLGGGAETLY | 0.76673895 |
| IVAGLACTL | ASSLGGGAETLY | 0.6068843 |
| IVGGGLAGA | ASSLGGGAETLY | 0.7740817 |
| IVTGVLNI | ASSLGGGAETLY | 0.7481778 |
| LAQQGCI | ASSLGGGAETLY | 0.65045816 |
| LCLHVEN | ASSLGGGAETLY | 0.74564534 |
| LDAISDGS | ASSLGGGAETLY | 0.59438413 |
| LGGFAAML | ASSLGGGAETLY | 0.68496305 |
| LGGKNPMI | ASSLGGGAETLY | 0.72841656 |
| LGPKGVAAIL | ASSLGGGAETLY | 0.7414251 |
| LMVGVSI | ASSLGGGAETLY | 0.64377713 |
| LVGAAFVL | ASSLGGGAETLY | 0.7533477 |
| LVLAVAPL | ASSLGGGAETLY | 0.735199 |
| MAAIAVIDAA | ASSLGGGAETLY | 0.64743066 |
| NNGINAELQ | ASSLGGGAETLY | 0.7316105 |
| RGEPLVLVHG | ASSLGGGAETLY | 0.7606434 |
| RGGAGGAAGLI | ASSLGGGAETLY | 0.75141203 |
| RSYRMHPAV | ASSLGGGAETLY | 0.62642336 |
| SAAHLLAGQLA | ASSLGGGAETLY | 0.73196524 |
| SAAVGSY | ASSLGGGAETLY | 0.59943235 |
| SPHAVLHRI | ASSLGGGAETLY | 0.7188436 |
| SQIAKVL | ASSLGGGAETLY | 0.71790546 |
| TALGHLVP | ASSLGGGAETLY | 0.6974073 |
| TAPTTSGAA | ASSLGGGAETLY | 0.75675964 |
| TDGDDGNGGSPGT | ASSLGGGAETLY | 0.50176096 |
| TDGLGEN | ASSLGGGAETLY | 0.6237299 |
| TGGPPLPGQA | ASSLGGGAETLY | 0.6741146 |
| TIPAITI | ASSLGGGAETLY | 0.7364444 |
| TIPGFTI | ASSLGGGAETLY | 0.7437906 |
| TSIASGALA | ASSLGGGAETLY | 0.5785677 |
| TVTLPGAAI | ASSLGGGAETLY | 0.7186836 |
| VASGLSAALP | ASSLGGGAETLY | 0.7087134 |
| VGAGVIGI | ASSLGGGAETLY | 0.64264375 |
| VGPKRFGV | ASSLGGGAETLY | 0.73620874 |

| | | |
|---|---|---|
| VNVAFPD | ASSLGGGAETLY | 0.70295227 |
| VSMTNSGVS | ASSLGGGAETLY | 0.6047576 |
| VTDVLGDLSV | ASSLGGGAETLY | 0.7575582 |
| VVVVGAGFAGL | ASSLGGGAETLY | 0.66448605 |
| VYTVGTVV | ASSLGGGAETLY | 0.7469285 |
| WTTFSGP | ASSLGGGAETLY | 0.6999035 |
| YDELVEAF | ASSLGGGAETLY | 0.630773 |
| YGTGGAGGNGG | ASSLGGGAETLY | 0.7501102 |
| YTNVMAV | ASSLGGGAETLY | 0.71672523 |
| FLEQVHQL | ASSLGGGAETLY | 0.7454914 |
| GLSLLAVAV | ASSLGGGAETLY | 0.6979232 |
| LEQVHQL | ASSLGGGAETLY | 0.64499915 |
| LNAAGAI | ASSLGGGAETLY | 0.7549875 |
| PAVHASL | ASSLGGGAETLY | 0.7547745 |
| TELEDDA | ASSLGGGAETLY | 0.7233826 |
| TPPGPPN | ASSLGGGAETLY | 0.73962235 |
| TPPPPPPP | ASSLGGGAETLY | 0.72028595 |
| TQSPALI | ASSLSGARYEQY | 0.7519185 |
| EFAAAFPQ | ASSLSGARYEQY | 0.6320015 |
| LLSGNLDVL | ASSLSGARYEQY | 0.5475699 |
| GPANLGNYNI | ASSLSGARYEQY | 0.6681285 |
| TLGPTLP | ASSLSGARYEQY | 0.6757581 |
| VYAYSSCCK | ASSLSGARYEQY | 0.6778762 |
| APAPAPAPNI | ASSLSGARYEQY | 0.5297981 |
| PGGPVPIP | ASSLSGARYEQY | 0.75513065 |
| FLTVALEP | ASSLSGARYEQY | 0.7903152 |
| SQYFGANL | ASSLSGARYEQY | 0.6444698 |
| GGVKPAA | ASSLSGARYEQY | 0.78927344 |
| GGTGGAGGAGGW | ASSLSGARYEQY | 0.5719906 |
| GVGGVAGLGGAGG | ASSLSGARYEQY | 0.75135726 |
| VLLVLATTI | ASSLSGARYEQY | 0.76382166 |
| IGIFAAML | ASSLSGARYEQY | 0.59615284 |
| AVGDLID | ASSLSGARYEQY | 0.7315284 |
| NISPLPL | ASSLSGARYEQY | 0.6561391 |
| VTAVFVAI | ASSLSGARYEQY | 0.6823111 |
| SGDDNDAAL | ASSLSGARYEQY | 0.76982546 |
| AVAALAHS | ASSLSGARYEQY | 0.673002 |
| GGVTVMATP | ASSLSGARYEQY | 0.7468668 |
| AGLAANVRD | ASSLSGARYEQY | 0.7007655 |
| TNRPDLIDPALLRPG | ASSLSGARYEQY | 0.65429014 |
| LANLGVTAL | ASSLSGARYEQY | 0.7543866 |
| TGERIEL | ASSLSGARYEQY | 0.7614561 |
| NGEELAA | ASSLSGARYEQY | 0.6674075 |
| VLSLVWG | ASSLSGARYEQY | 0.7831646 |
| IGYALWL | ASSLSGARYEQY | 0.6550675 |
| LSLLLAPL | ASSLSGARYEQY | 0.78078336 |
| TLDEALK | ASSLSGARYEQY | 0.75644076 |
| TDGAGTAEGG | ASSLSGARYEQY | 0.6984476 |
| NAPGGAGGNGG | ASSLSGARYEQY | 0.6977413 |
| VSVFMGMGGNF | ASSLSGARYEQY | 0.7593978 |
| LASYLAAL | ASSLSGARYEQY | 0.6889788 |

| | | |
|---|---|---|
| LVGGTPAL | ASSLSGARYEQY | 0.7393451 |
| VRNADVL | ASSLSGARYEQY | 0.77831614 |
| TCSSPFLVSAM | ASSLSGARYEQY | 0.51373374 |
| VHITDAE | ASSLSGARYEQY | 0.66173303 |
| EYAAPSF | ASSLSGARYEQY | 0.63191855 |
| VVDAFHGGL | ASSLSGARYEQY | 0.6126375 |
| ASVVDPQ | ASSLSGARYEQY | 0.5475327 |
| SYNEFGGLS | ASSLSGARYEQY | 0.43295902 |
| DPDGSVY | ASSLSGARYEQY | 0.5977391 |
| MVSTIHATE | ASSLSGARYEQY | 0.5396949 |
| LAAGVADPV | ASSLSGARYEQY | 0.63976496 |
| TKEQIAA | ASSLSGARYEQY | 0.64123034 |
| SLASPAAL | ASSLSGARYEQY | 0.54400474 |
| IGLVFYGV | ASSLSGARYEQY | 0.6623778 |
| FLLDASNL | ASSLSGARYEQY | 0.66504014 |
| YQVGNFP | ASSLSGARYEQY | 0.4506644 |
| ESLLSQST | ASSLSGARYEQY | 0.58496475 |
| LFTVGFAAVG | ASSLSGARYEQY | 0.57404375 |
| ATPPTPT | ASSLSGARYEQY | 0.57743716 |
| TDLATLG | ASSLSGARYEQY | 0.3734316 |
| SAGGTGR | ASSLSGARYEQY | 0.6496731 |
| GFSNSGS | ASSLSGARYEQY | 0.5527763 |
| GGNGGSGGTSV | ASSLSGARYEQY | 0.3154363 |
| GTSVATGGAGN | ASSLSGARYEQY | 0.59411687 |
| NGGNGGSGGTS | ASSLSGARYEQY | 0.599205 |
| AVVKFPH | ASSLSGARYEQY | 0.6043891 |
| TPPPPPP | ASSLSGARYEQY | 0.6553469 |
| AAPAAAAAPA | ASSLSGARYEQY | 0.40139088 |
| IALNEDL | ASSLSGARYEQY | 0.54986405 |
| IVNVSSM | ASSLSGARYEQY | 0.6514878 |
| TGIPPAPR | ASSLSGARYEQY | 0.6623924 |
| FDLGGGTF | ASSLSGARYEQY | 0.5273424 |
| FDLGGGTFDVS | ASSLSGARYEQY | 0.63482124 |
| NEPTAAA | ASSLSGARYEQY | 0.66301864 |
| NEPTAAALA | ASSLSGARYEQY | 0.63770574 |
| GSIINISSI | ASSLSGARYEQY | 0.64564943 |
| VAPGNAGTA | ASSLSGARYEQY | 0.654193 |
| TGGGASLE | ASSLSGARYEQY | 0.5137601 |
| VAVYNHYKRI | ASSLSGARYEQY | 0.65752137 |
| TDPAPATT | ASSLSGARYEQY | 0.6628685 |
| TNRPDILDPALLRPG | ASSLSGARYEQY | 0.55828714 |
| LTTTTTS | ASSLSGARYEQY | 0.62423027 |
| PTGSGKTL | ASSLSGARYEQY | 0.5237461 |
| TGSGKTL | ASSLSGARYEQY | 0.5667131 |
| TIGHVDHGK | ASSLSGARYEQY | 0.3662301 |
| TIGHVDHGKTT | ASSLSGARYEQY | 0.6342165 |
| GHVDHGKTT | ASSLSGARYEQY | 0.6549473 |
| AQFEHTLL | ASSLSGARYEQY | 0.60388607 |
| IVNQDEL | ASSLSGARYEQY | 0.52835864 |
| TNGSQFF | ASSLSGARYEQY | 0.46917656 |
| TGKGIVE | ASSLSGARYEQY | 0.59264773 |

| | | |
|---|---|---|
| LISNASDA | ASSLSGARYEQY | 0.5800249 |
| AELRTLGELTKIAKAHGAQ | ASSLSGARYEQY | 0.34876895 |
| APNAVAAAEPSW | ASSLSGARYEQY | 0.57003033 |
| APQVAGAATAVL | ASSLSGARYEQY | 0.66287845 |
| ARRDPAAAAAILPTDA | ASSLSGARYEQY | 0.6558059 |
| DAAARDVLAEAGLAEYF | ASSLSGARYEQY | 0.43165055 |
| DDQIVLSAKANLAKVEQ | ASSLSGARYEQY | 0.6636083 |
| DELLPDTGEDSDEDGH | ASSLSGARYEQY | 0.6316558 |
| ELPEAPANRATVPQD | ASSLSGARYEQY | 0.65383166 |
| FGQSKIERVTGIG | ASSLSGARYEQY | 0.58059996 |
| GAGGTGSTAGGAGGAGGAGGLY | ASSLSGARYEQY | 0.5303576 |
| GHGGTGAAGGNGGTGGAGGSGI | ASSLSGARYEQY | 0.6447203 |
| GNGGNGGAGGEGGPGGAGGAGA | ASSLSGARYEQY | 0.6010648 |
| GSTAGNGGNGGNATNGTIAGQP | ASSLSGARYEQY | 0.47497806 |
| GTGGDGGNGGNGGTGG | ASSLSGARYEQY | 0.6497179 |
| GVRRPAGIPADAPV | ASSLSGARYEQY | 0.6178099 |
| GYRQVTDAHLLTLARR | ASSLSGARYEQY | 0.56419516 |
| IAKMGQDGHDRG | ASSLSGARYEQY | 0.5721589 |
| KAEPSIEIQYT | ASSLSGARYEQY | 0.6324969 |
| KPVRIRMAADEDTVAVT | ASSLSGARYEQY | 0.5312427 |
| MPPRIAGMRLLVIKPEP | ASSLSGARYEQY | 0.53299254 |
| MKFYAVALFRSRGVPPTR | ASSLSGARYEQY | 0.45192268 |
| NAGGFGILGAVAHSP | ASSLSGARYEQY | 0.6231331 |
| NEGREITLAMAEM | ASSLSGARYEQY | 0.66032577 |
| NELVEWSHPLGHRWRWE | ASSLSGARYEQY | 0.60628784 |
| NLGAGGTGGNGGNPG | ASSLSGARYEQY | 0.5101341 |
| NTEWPKWETVMAADPVD | ASSLSGARYEQY | 0.6420433 |
| PASAAPAETQP | ASSLSGARYEQY | 0.62798893 |
| QQSSDGKNGVRAAEITGE | ASSLSGARYEQY | 0.61106306 |
| RAPDAVTTSWWKE | ASSLSGARYEQY | 0.63411015 |
| SGFGNVGAHTSGWFNQ | ASSLSGARYEQY | 0.5788624 |
| SGVAGAVGQAASVGGLKV | ASSLSGARYEQY | 0.6372335 |
| SNGGAGTGSAGGNGGT | ASSLSGARYEQY | 0.6294753 |
| TGGAGGEGGAAG | ASSLSGARYEQY | 0.6389102 |
| TGGDGGAAGAGGAPGHG | ASSLSGARYEQY | 0.64104444 |
| TMPPLLAAIANAGRHGV | ASSLSGARYEQY | 0.62944436 |
| TSSTAGNSGGA | ASSLSGARYEQY | 0.57794464 |
| VPATVPVGILC | ASSLSGARYEQY | 0.3644167 |
| VQRPDSGGVVLTGRIS | ASSLSGARYEQY | 0.57512105 |
| WTPNPVPATKGVPAGLVQAT | ASSLSGARYEQY | 0.63192505 |
| YALKRALAKFSDRFDVVII | ASSLSGARYEQY | 0.630403 |
| YLIGSVIV | ASSLSGARYEQY | 0.6468839 |
| YGQLVAVID | ASSLSGARYEQY | 0.64039767 |
| VTVGGMTLPA | ASSLSGARYEQY | 0.545859 |
| VLALPAV | ASSLSGARYEQY | 0.6483454 |
| VIPPTLNYET | ASSLSGARYEQY | 0.65307426 |
| VGMGGMHPSSG | ASSLSGARYEQY | 0.43520823 |
| VGLAGGAATA | ASSLSGARYEQY | 0.5784508 |
| VGGAGGAGGSGP | ASSLSGARYEQY | 0.6657146 |
| VFAAFMV | ASSLSGARYEQY | 0.62193036 |
| TLYDKLR | ASSLSGARYEQY | 0.5756658 |

| | | |
|---|---|---|
| TLAVFTVL | ASSLSGARYEQY | 0.54283154 |
| TGSVQPARV | ASSLSGARYEQY | 0.6610898 |
| TGGGAGSGGT | ASSLSGARYEQY | 0.47855037 |
| SHLETSL | ASSLSGARYEQY | 0.6620163 |
| SFTTGMT | ASSLSGARYEQY | 0.3292349 |
| QVEVAKRMGVRQPTVS | ASSLSGARYEQY | 0.4676427 |
| QGSGGPALAM | ASSLSGARYEQY | 0.47001588 |
| MLAPSNP | ASSLSGARYEQY | 0.4014867 |
| LVNNAAI | ASSLSGARYEQY | 0.6566248 |
| LTGEINTLA | ASSLSGARYEQY | 0.549669 |
| LSIFGPV | ASSLSGARYEQY | 0.6490661 |
| LLALLANL | ASSLSGARYEQY | 0.62232125 |
| LISNASDALDKLR | ASSLSGARYEQY | 0.65364236 |
| LINEAVP | ASSLSGARYEQY | 0.62385386 |
| LGYAFSGL | ASSLSGARYEQY | 0.57767713 |
| LDFVAVAL | ASSLSGARYEQY | 0.46242067 |
| LAQPGAE | ASSLSGARYEQY | 0.65012896 |
| ISNLGAVTGF | ASSLSGARYEQY | 0.6576566 |
| HQEDPGRI | ASSLSGARYEQY | 0.5876859 |
| GTPGEPTP | ASSLSGARYEQY | 0.47990882 |
| GFFNSGS | ASSLSGARYEQY | 0.4953068 |
| GAGGGGLTAGAN | ASSLSGARYEQY | 0.63127923 |
| FVDALVNM | ASSLSGARYEQY | 0.36851734 |
| FLTVAPEL | ASSLSGARYEQY | 0.3309559 |
| FETLTAL | ASSLSGARYEQY | 0.5529463 |
| DVINPIEL | ASSLSGARYEQY | 0.5802187 |
| APVSGAPV | ASSLSGARYEQY | 0.6578788 |
| ALVGGLAW | ASSLSGARYEQY | 0.65392715 |
| ALAAAAATAI | ASSLSGARYEQY | 0.6004815 |
| AFGLGLLL | ASSLSGARYEQY | 0.6398208 |
| AFFAAASL | ASSLSGARYEQY | 0.635876 |
| AAAALPAFSPP | ASSLSGARYEQY | 0.30643618 |
| VVLAPAL | ASSLSGARYEQY | 0.53941166 |
| TIGHVDHGKTTL | ASSLSGARYEQY | 0.48119667 |
| LISNASDALDK | ASSLSGARYEQY | 0.6563417 |
| LISNASDAL | ASSLSGARYEQY | 0.5255593 |
| ISNASDALDK | ASSLSGARYEQY | 0.558781 |
| NQAETLVYQTE | ASSLSGARYEQY | 0.61750335 |
| TGGTGGTAGAG | ASSLSGARYEQY | 0.6152792 |
| TGSTAGGAGGA | ASSLSGARYEQY | 0.6065489 |
| DPLAVPVAAGPV | ASSLSGARYEQY | 0.45954317 |
| GGTGGTPGNGGN | ASSLSGARYEQY | 0.3201208 |
| PAGLAGVDPATL | ASSLSGARYEQY | 0.6554866 |
| GNNLVVEAPHAGASV | ASSLSGARYEQY | 0.42075697 |
| KVTRIMSASTLEQPA | ASSLSGARYEQY | 0.53157586 |
| LGFALPIALPRRTVV | ASSLSGARYEQY | 0.3387584 |
| TRLGNLTARKLVVLV | ASSLSGARYEQY | 0.5398909 |
| WSQQRRLIYFPSAGP | ASSLSGARYEQY | 0.4332111 |
| GPNGNPDPMAAAVDIR | ASSLSGARYEQY | 0.64028895 |
| TQRCVSVADLRAAASR | ASSLSGARYEQY | 0.59205407 |
| QFVLDVAEHPVEVLTETV | ASSLSGARYEQY | 0.60035473 |

| | | |
|---|---|---|
| GGAGGTGGAGGNGGTQAGDG | ASSLSGARYEQY | 0.66771644 |
| PLLATLRAIAVDVDAKPAQV | ASSLSGARYEQY | 0.62044877 |
| APKFRTGGYMIDFWGVGYQVAK | ASSLSGARYEQY | 0.59033614 |
| GAGGAGGAGGAAGGPGGTGGTG | ASSLSGARYEQY | 0.52988464 |
| IGALGLLATDVAPAGSVAFGVA | ASSLSGARYEQY | 0.49131268 |
| AGGAGGIGGNGGAGGAGGVLMG | ASSLSGARYEQY | 0.5411049 |
| ADELIGLL | ASSLSGARYEQY | 0.54675907 |
| ADRALKA | ASSLSGARYEQY | 0.66666377 |
| AGAVGQVGGAGGA | ASSLSGARYEQY | 0.49176198 |
| AGGGGGLGGAP | ASSLSGARYEQY | 0.4658145 |
| AGGLLYGNGGAG | ASSLSGARYEQY | 0.5473568 |
| AGLESIGL | ASSLSGARYEQY | 0.6624291 |
| AGLKTGII | ASSLSGARYEQY | 0.6680933 |
| ASLAALLL | ASSLSGARYEQY | 0.51091707 |
| ASVINISSI | ASSLSGARYEQY | 0.54291326 |
| ATPPGPA | ASSLSGARYEQY | 0.5640489 |
| ATVASALV | ASSLSGARYEQY | 0.62224895 |
| ATVASLIL | ASSLSGARYEQY | 0.49915504 |
| AVNGPVLIRH | ASSLSGARYEQY | 0.4565782 |
| DGAAGAVGI | ASSLSGARYEQY | 0.61680853 |
| DPLLHGI | ASSLSGARYEQY | 0.66478086 |
| ELGAALGAL | ASSLSGARYEQY | 0.552729 |
| ELGRTQAV | ASSLSGARYEQY | 0.5045372 |
| FVAGGYLS | ASSLSGARYEQY | 0.6064185 |
| GASGGAGGAGGW | ASSLSGARYEQY | 0.5367649 |
| GGAGAGGDGGAGGI | ASSLSGARYEQY | 0.5874463 |
| GTGHNGSQPGMGGNG | ASSLSGARYEQY | 0.6320317 |
| GYAAASASAS | ASSLSGARYEQY | 0.44591254 |
| HQVDVAAVQQG | ASSLSGARYEQY | 0.5988356 |
| IGAAIAHT | ASSLSGARYEQY | 0.66954815 |
| IGMIEVAQL | ASSLSGARYEQY | 0.5821229 |
| IRLAPEHTSAMD | ASSLSGARYEQY | 0.6296071 |
| IYNVSVL | ASSLSGARYEQY | 0.65729505 |
| KQGFLGM | ASSLSGARYEQY | 0.6444748 |
| LESVITQL | ASSLSGARYEQY | 0.6600457 |
| LGLGALVVV | ASSLSGARYEQY | 0.6333638 |
| LIGNGGNGGSGGI | ASSLSGARYEQY | 0.57310605 |
| LNAINAPTL | ASSLSGARYEQY | 0.5928018 |
| LQLLNTGV | ASSLSGARYEQY | 0.6396458 |
| LQTSIEF | ASSLSGARYEQY | 0.61665857 |
| MSFAVSEL | ASSLSGARYEQY | 0.5875857 |
| NTGALGL | ASSLSGARYEQY | 0.5012757 |
| PPGAGGAGGAGTT | ASSLSGARYEQY | 0.6617799 |
| QLVGLGI | ASSLSGARYEQY | 0.58229876 |
| RFLEQVHQL | ASSLSGARYEQY | 0.64050305 |
| RGVGYVLR | ASSLSGARYEQY | 0.43649784 |
| SATGVFAGVI | ASSLSGARYEQY | 0.50314724 |
| SLPAITLP | ASSLSGARYEQY | 0.64967465 |
| STALAFTV | ASSLSGARYEQY | 0.30983284 |
| SVLTVGLGK | ASSLSGARYEQY | 0.32284048 |
| TALAGAGSGI | ASSLSGARYEQY | 0.5796268 |

| | | |
|---|---|---|
| TGLPFTGI | ASSLSGARYEQY | 0.658702 |
| VAEHVTV | ASSLSGARYEQY | 0.3140877 |
| VDIVLNSL | ASSLSGARYEQY | 0.5116614 |
| VLTGGGAVL | ASSLSGARYEQY | 0.42812517 |
| VNVGLAYGI | ASSLSGARYEQY | 0.63243026 |
| VSSAAVGAV | ASSLSGARYEQY | 0.6387167 |
| VTSVYEAGV | ASSLSGARYEQY | 0.40413702 |
| WVALDRGA | ASSLSGARYEQY | 0.63797474 |
| YLVGSVVI | ASSLSGARYEQY | 0.44182593 |
| GPPGTGKTLL | ASSLSGARYEQY | 0.47596344 |
| GVGMTKF | ASSLSGARYEQY | 0.64893925 |
| IGIGGSDL | ASSLSGARYEQY | 0.6521536 |
| IGTLLAEL | ASSLSGARYEQY | 0.4816206 |
| IYTSGTT | ASSLSGARYEQY | 0.6506698 |
| LSFPAGV | ASSLSGARYEQY | 0.53277594 |
| LYTSGTT | ASSLSGARYEQY | 0.66850346 |
| VGAFPFGF | ASSLSGARYEQY | 0.57347023 |
| VGAGTVEF | ASSLSGARYEQY | 0.39666274 |
| VGKSTLLL | ASSLSGARYEQY | 0.6536599 |
| AALAHAGGGIV | ASSLSGARYEQY | 0.629281 |
| AALVAVSLL | ASSLSGARYEQY | 0.62681574 |
| AAVSRAVAAL | ASSLSGARYEQY | 0.6427406 |
| AGGSSGIGGF | ASSLSGARYEQY | 0.50711334 |
| AGLTVANANG | ASSLSGARYEQY | 0.6066778 |
| AGLVGAVL | ASSLSGARYEQY | 0.6544883 |
| ALSQAFTV | ASSLSGARYEQY | 0.6456721 |
| ALTFSVSL | ASSLSGARYEQY | 0.48521358 |
| ALTGAALVAL | ASSLSGARYEQY | 0.6483599 |
| ASFSIGAGVL | ASSLSGARYEQY | 0.6194935 |
| DGSVYKI | ASSLSGARYEQY | 0.52056676 |
| EQLAEIV | ASSLSGARYEQY | 0.66623 |
| FAVSELTVM | ASSLSGARYEQY | 0.59547424 |
| GAAPLAAGL | ASSLSGARYEQY | 0.6480305 |
| GADGGDGGSGGAGG | ASSLSGARYEQY | 0.64627194 |
| GFGLDTGGAGG | ASSLSGARYEQY | 0.6565654 |
| GLASGGLL | ASSLSGARYEQY | 0.46100628 |
| GRIAGAAVLL | ASSLSGARYEQY | 0.61885357 |
| GSAGSLNTSF | ASSLSGARYEQY | 0.6168793 |
| GTGQIGI | ASSLSGARYEQY | 0.6428577 |
| GTLIALIL | ASSLSGARYEQY | 0.58871984 |
| IAATMGFVVG | ASSLSGARYEQY | 0.5228028 |
| IFAQALL | ASSLSGARYEQY | 0.61464965 |
| IGAAVALGF | ASSLSGARYEQY | 0.6610514 |
| IGLTPLAL | ASSLSGARYEQY | 0.5774814 |
| ILGSNTSTL | ASSLSGARYEQY | 0.6295634 |
| IVFSLTVL | ASSLSGARYEQY | 0.6643424 |
| LASKATGF | ASSLSGARYEQY | 0.4865427 |
| LASVALIL | ASSLSGARYEQY | 0.60936517 |
| LFAVGLTL | ASSLSGARYEQY | 0.6354319 |
| LFNGGIGTYI | ASSLSGARYEQY | 0.6575919 |
| LTLGAGII | ASSLSGARYEQY | 0.5586604 |

| | | |
|---|---|---|
| LVAAALGDI | ASSLSGARYEQY | 0.64933366 |
| NTAFVQLGI | ASSLSGARYEQY | 0.55546623 |
| QLFSGAEL | ASSLSGARYEQY | 0.64034337 |
| SAATFITL | ASSLSGARYEQY | 0.5755231 |
| SNYGLANLGV | ASSLSGARYEQY | 0.638063 |
| SYGGAYAVM | ASSLSGARYEQY | 0.5683002 |
| TTGVGGLTLG | ASSLSGARYEQY | 0.6459727 |
| TTQTIYI | ASSLSGARYEQY | 0.6540061 |
| VAAGAGLGAVI | ASSLSGARYEQY | 0.6545382 |
| VAAVVTLL | ASSLSGARYEQY | 0.32596904 |
| VGFLGGLI | ASSLSGARYEQY | 0.65583646 |
| VGVIPILL | ASSLSGARYEQY | 0.64723086 |
| VIFASSLI | ASSLSGARYEQY | 0.66639346 |
| VKGGDGGAAAG | ASSLSGARYEQY | 0.5827538 |
| VTKTFTNI | ASSLSGARYEQY | 0.6383846 |
| VVAAAILL | ASSLSGARYEQY | 0.58855915 |
| YAALDVEL | ASSLSGARYEQY | 0.57701343 |
| LASVALLL | ASSLSGARYEQY | 0.5127536 |
| LDAVHRV | ASSLSGARYEQY | 0.5843746 |
| QGLSSSIF | ASSLSGARYEQY | 0.5975645 |
| RGTFANIRL | ASSLSGARYEQY | 0.56052476 |
| RQLWWGHRI | ASSLSGARYEQY | 0.6456993 |
| YGPPGTGKTLL | ASSLSGARYEQY | 0.65236336 |
| AALVGAPANVA | ASSLSGARYEQY | 0.66085947 |
| AATFRRTE | ASSLSGARYEQY | 0.6572179 |
| AGALGGGI | ASSLSGARYEQY | 0.6611043 |
| AGGAGGAGGIGAG | ASSLSGARYEQY | 0.6329186 |
| AGSGALGL | ASSLSGARYEQY | 0.58438194 |
| AQIADQI | ASSLSGARYEQY | 0.6021713 |
| ASAVAVTYL | ASSLSGARYEQY | 0.5121287 |
| ASTVLGGTGG | ASSLSGARYEQY | 0.5854062 |
| AVGGEPAK | ASSLSGARYEQY | 0.59655076 |
| CEQGTFGGTC | ASSLSGARYEQY | 0.63335234 |
| DEGKAPAG | ASSLSGARYEQY | 0.3071954 |
| DTVVFPAL | ASSLSGARYEQY | 0.6214727 |
| FEGITMAG | ASSLSGARYEQY | 0.6658366 |
| FPLVVQD | ASSLSGARYEQY | 0.5429788 |
| GAFTTPQ | ASSLSGARYEQY | 0.6312283 |
| GFGGFGTAGGVGGA | ASSLSGARYEQY | 0.6221994 |
| GFLAQFL | ASSLSGARYEQY | 0.55055845 |
| GGAGGAGGNAGVA | ASSLSGARYEQY | 0.64886487 |
| GGVGTAGAGGAGGAGG | ASSLSGARYEQY | 0.51073873 |
| GNGGGGDAGNAGSGGN | ASSLSGARYEQY | 0.62981665 |
| GTGGAGGAGNDAGS | ASSLSGARYEQY | 0.6650202 |
| GTTNTGWFNS | ASSLSGARYEQY | 0.62696326 |
| IFMGPLFTL | ASSLSGARYEQY | 0.61708117 |
| ILAHAIEAI | ASSLSGARYEQY | 0.59855795 |
| IRSAIREAL | ASSLSGARYEQY | 0.42842507 |
| ITFAGGG | ASSLSGARYEQY | 0.63209075 |
| ITGMSGHQEV | ASSLSGARYEQY | 0.35601324 |
| IVAGLACTL | ASSLSGARYEQY | 0.60423493 |

| | | |
|---|---|---|
| IVGGGLAGA | ASSLSGARYEQY | 0.4849242 |
| IVTGVLNI | ASSLSGARYEQY | 0.636634 |
| LAQQGCI | ASSLSGARYEQY | 0.6385212 |
| LCLHVEN | ASSLSGARYEQY | 0.61367714 |
| LDAISDGS | ASSLSGARYEQY | 0.51333094 |
| LGGFAAML | ASSLSGARYEQY | 0.5614575 |
| LGGKNPMI | ASSLSGARYEQY | 0.47742242 |
| LGPKGVAAIL | ASSLSGARYEQY | 0.490714 |
| LMVGVSI | ASSLSGARYEQY | 0.6471835 |
| LVGAAFVL | ASSLSGARYEQY | 0.5352805 |
| LVLAVAPL | ASSLSGARYEQY | 0.61722636 |
| MAAIAVIDAA | ASSLSGARYEQY | 0.6626649 |
| NNGINAELQ | ASSLSGARYEQY | 0.63819516 |
| RGEPLVLVHG | ASSLSGARYEQY | 0.4351822 |
| RGGAGGAAGLI | ASSLSGARYEQY | 0.61619127 |
| RSYRMHPAV | ASSLSGARYEQY | 0.597959 |
| SAAHLLAGQLA | ASSLSGARYEQY | 0.58855915 |
| SAAVGSY | ASSLSGARYEQY | 0.31613687 |
| SPHAVLHRI | ASSLSGARYEQY | 0.6174495 |
| SQIAKVL | ASSLSGARYEQY | 0.52292556 |
| TALGHLVP | ASSLSGARYEQY | 0.5602248 |
| TAPTTSGAA | ASSLSGARYEQY | 0.550542 |
| TDGDDGNGGSPGT | ASSLSGARYEQY | 0.6532139 |
| TDGLGEN | ASSLSGARYEQY | 0.5231478 |
| TGGPPLPGQA | ASSLSGARYEQY | 0.62562287 |
| TIPAITI | ASSLSGARYEQY | 0.64664996 |
| TIPGFTI | ASSLSGARYEQY | 0.6550085 |
| TSIASGALA | ASSLSGARYEQY | 0.6627052 |
| TVTLPGAAI | ASSLSGARYEQY | 0.6337681 |
| VASGLSAALP | ASSLSGARYEQY | 0.5587292 |
| VGAGVIGI | ASSLSGARYEQY | 0.624067 |
| VGPKRFGV | ASSLSGARYEQY | 0.56374216 |
| VNVAFPD | ASSLSGARYEQY | 0.359308 |
| VSMTNSGVS | ASSLSGARYEQY | 0.33183774 |
| VTDVLGDLSV | ASSLSGARYEQY | 0.61165035 |
| VVVVGAGFAGL | ASSLSGARYEQY | 0.34407312 |
| VYTVGTVV | ASSLSGARYEQY | 0.61300516 |
| WTTFSGP | ASSLSGARYEQY | 0.6448552 |
| YDELVEAF | ASSLSGARYEQY | 0.585281 |
| YGTGGAGGNGG | ASSLSGARYEQY | 0.43303502 |
| YTNVMAV | ASSLSGARYEQY | 0.56967276 |
| FLEQVHQL | ASSLSGARYEQY | 0.58756596 |
| GLSLLAVAV | ASSLSGARYEQY | 0.64934844 |
| LEQVHQL | ASSLSGARYEQY | 0.63606715 |
| LNAAGAI | ASSLSGARYEQY | 0.6602535 |
| PAVHASL | ASSLSGARYEQY | 0.6557317 |
| TELEDDA | ASSLSGARYEQY | 0.6038066 |
| TPPGPPN | ASSLSGARYEQY | 0.64046323 |
| TPPPPPPP | ASSLSGARYEQY | 0.5701223 |
| TQSPALI | ASSPGQKNTLY | 0.5917058 |
| EFAAAFPQ | ASSPGQKNTLY | 0.5608988 |

| | | |
|---|---|---|
| LLSGNLDVL | ASSPGQKNTLY | 0.60994303 |
| GPANLGNYNI | ASSPGQKNTLY | 0.58640707 |
| TLGPTLP | ASSPGQKNTLY | 0.5210648 |
| VYAYSSCCK | ASSPGQKNTLY | 0.38116395 |
| APAPAPAPNI | ASSPGQKNTLY | 0.578215 |
| PGGPVPIP | ASSPGQKNTLY | 0.62380624 |
| FLTVALEP | ASSPGQKNTLY | 0.35596564 |
| SQYFGANL | ASSPGQKNTLY | 0.5376111 |
| GGVKPAA | ASSPGQKNTLY | 0.66704214 |
| GGTGGAGGAGGW | ASSPGQKNTLY | 0.6618844 |
| GVGGVAGLGGAGG | ASSPGQKNTLY | 0.5418303 |
| VLLVLATTI | ASSPGQKNTLY | 0.5156628 |
| IGIFAAML | ASSPGQKNTLY | 0.6162953 |
| AVGDLID | ASSPGQKNTLY | 0.37273335 |
| NISPLPL | ASSPGQKNTLY | 0.59138244 |
| VTAVFVAI | ASSPGQKNTLY | 0.61967784 |
| SGDDNDAAL | ASSPGQKNTLY | 0.6285988 |
| AVAALAHS | ASSPGQKNTLY | 0.48826602 |
| GGVTVMATP | ASSPGQKNTLY | 0.6217524 |
| AGLAANVRD | ASSPGQKNTLY | 0.6129502 |
| TNRPDLIDPALLRPG | ASSPGQKNTLY | 0.619162 |
| LANLGVTAL | ASSPGQKNTLY | 0.32504657 |
| TGERIEL | ASSPGQKNTLY | 0.64625585 |
| NGEELAA | ASSPGQKNTLY | 0.54440904 |
| VLSLVWG | ASSPGQKNTLY | 0.504293 |
| IGYALWL | ASSPGQKNTLY | 0.36960602 |
| LSLLLAPL | ASSPGQKNTLY | 0.61100245 |
| TLDEALK | ASSPGQKNTLY | 0.6533432 |
| TDGAGTAEGG | ASSPGQKNTLY | 0.65086 |
| NAPGGAGGNGG | ASSPGQKNTLY | 0.3936129 |
| VSVFMGMGGNF | ASSPGQKNTLY | 0.65505713 |
| LASYLAAL | ASSPGQKNTLY | 0.6638302 |
| LVGGTPAL | ASSPGQKNTLY | 0.5917168 |
| VRNADVL | ASSPGQKNTLY | 0.39541218 |
| TCSSPFLVSAM | ASSPGQKNTLY | 0.65092534 |
| VHITDAE | ASSPGQKNTLY | 0.57157755 |
| EYAAPSF | ASSPGQKNTLY | 0.64828074 |
| VVDAFHGGL | ASSPGQKNTLY | 0.6383122 |
| ASVVDPQ | ASSPGQKNTLY | 0.50248337 |
| SYNEFGGLS | ASSPGQKNTLY | 0.66226923 |
| DPDGSVY | ASSPGQKNTLY | 0.6537838 |
| MVSTIHATE | ASSPGQKNTLY | 0.50979906 |
| LAAGVADPV | ASSPGQKNTLY | 0.52297497 |
| TKEQIAA | ASSPGQKNTLY | 0.53380525 |
| SLASPAAL | ASSPGQKNTLY | 0.36643392 |
| IGLVFYGV | ASSPGQKNTLY | 0.6574702 |
| FLLDASNL | ASSPGQKNTLY | 0.5804072 |
| YQVGNFP | ASSPGQKNTLY | 0.5865527 |
| ESLLSQST | ASSPGQKNTLY | 0.6277919 |
| LFTVGFAAVG | ASSPGQKNTLY | 0.5692027 |
| ATPPTPT | ASSPGQKNTLY | 0.5596647 |

| | | |
|---|---|---|
| TDLATLG | ASSPGQKNTLY | 0.5965182 |
| SAGGTGR | ASSPGQKNTLY | 0.638901 |
| GFSNSGS | ASSPGQKNTLY | 0.62954974 |
| GGNGGSGGTSV | ASSPGQKNTLY | 0.61883456 |
| GTSVATGGAGN | ASSPGQKNTLY | 0.61447775 |
| NGGNGGSGGTS | ASSPGQKNTLY | 0.6641055 |
| AVVKFPH | ASSPGQKNTLY | 0.4833557 |
| TPPPPPP | ASSPGQKNTLY | 0.56027865 |
| AAPAAAAAPA | ASSPGQKNTLY | 0.6382777 |
| IALNEDL | ASSPGQKNTLY | 0.39566374 |
| IVNVSSM | ASSPGQKNTLY | 0.64192206 |
| TGIPPAPR | ASSPGQKNTLY | 0.6309686 |
| FDLGGGTF | ASSPGQKNTLY | 0.6333878 |
| FDLGGGTFDVS | ASSPGQKNTLY | 0.5754779 |
| NEPTAAA | ASSPGQKNTLY | 0.66707385 |
| NEPTAAALA | ASSPGQKNTLY | 0.6556577 |
| GSIINISSI | ASSPGQKNTLY | 0.59475654 |
| VAPGNAGTA | ASSPGQKNTLY | 0.64307344 |
| TGGGASLE | ASSPGQKNTLY | 0.46293327 |
| VAVYNHYKRI | ASSPGQKNTLY | 0.6506572 |
| TDPAPATT | ASSPGQKNTLY | 0.6133696 |
| TNRPDILDPALLRPG | ASSPGQKNTLY | 0.53950983 |
| LTTTTTS | ASSPGQKNTLY | 0.6079433 |
| PTGSGKTL | ASSPGQKNTLY | 0.65031874 |
| TGSGKTL | ASSPGQKNTLY | 0.6526839 |
| TIGHVDHGK | ASSPGQKNTLY | 0.35464734 |
| TIGHVDHGKTT | ASSPGQKNTLY | 0.5923161 |
| GHVDHGKTT | ASSPGQKNTLY | 0.36234358 |
| AQFEHTLL | ASSPGQKNTLY | 0.47111857 |
| IVNQDEL | ASSPGQKNTLY | 0.45449454 |
| TNGSQFF | ASSPGQKNTLY | 0.6366852 |
| TGKGIVE | ASSPGQKNTLY | 0.5683091 |
| LISNASDA | ASSPGQKNTLY | 0.63176656 |
| AELRTLGELTKIAKAHGAQ | ASSPGQKNTLY | 0.61812556 |
| APNAVAAAEPSW | ASSPGQKNTLY | 0.66266835 |
| APQVAGAATAVL | ASSPGQKNTLY | 0.48991054 |
| ARRDPAAAAAILPTDA | ASSPGQKNTLY | 0.6089616 |
| DAAARDVLAEAGLAEYF | ASSPGQKNTLY | 0.38174605 |
| DDQIVLSAKANLAKVEQ | ASSPGQKNTLY | 0.47050238 |
| DELLPDTGEDSDEDGH | ASSPGQKNTLY | 0.48437157 |
| ELPEAPANRATVPQD | ASSPGQKNTLY | 0.6216791 |
| FGQSKIERVTGIG | ASSPGQKNTLY | 0.35052857 |
| GAGGTGSTAGGAGGAGGAGGLY | ASSPGQKNTLY | 0.65929234 |
| GHGGTGAAGGNGGTGGAGGSGI | ASSPGQKNTLY | 0.62357986 |
| GNGGNGGAGGEGGPGGAGGAGA | ASSPGQKNTLY | 0.6484351 |
| GSTAGNGGNGGNATNGTIAGQP | ASSPGQKNTLY | 0.58592975 |
| GTGGDGGNGGNGGTGG | ASSPGQKNTLY | 0.5876061 |
| GVRRPAGIPADAPV | ASSPGQKNTLY | 0.5859022 |
| GYRQVTDAHLLTLARR | ASSPGQKNTLY | 0.64399666 |
| IAKMGQDGHDRG | ASSPGQKNTLY | 0.5277915 |
| KAEPSIEIQYT | ASSPGQKNTLY | 0.3874714 |

| | | |
|---|---|---|
| KPVRIRMAADEDTVAVT | ASSPGQKNTLY | 0.54335934 |
| MPPRIAGMRLLVIKPEP | ASSPGQKNTLY | 0.5042989 |
| MKFYAVALFRSRGVPPTR | ASSPGQKNTLY | 0.6619384 |
| NAGGFGILGAVAHSP | ASSPGQKNTLY | 0.65438306 |
| NEGREITLAMAEM | ASSPGQKNTLY | 0.633087 |
| NELVEWSHPLGHRWRWE | ASSPGQKNTLY | 0.63486266 |
| NLGAGGTGGNGGNPG | ASSPGQKNTLY | 0.334434 |
| NTEWPKWETVMAADPVD | ASSPGQKNTLY | 0.5894508 |
| PASAAPAETQP | ASSPGQKNTLY | 0.66763234 |
| QQSSDGKNGVRAAEITGE | ASSPGQKNTLY | 0.63513476 |
| RAPDAVTTSWWKE | ASSPGQKNTLY | 0.5845669 |
| SGFGNVGAHTSGWFNQ | ASSPGQKNTLY | 0.6592805 |
| SGVAGAVGQAASVGGLKV | ASSPGQKNTLY | 0.5971901 |
| SNGGAGTGSAGGNGGT | ASSPGQKNTLY | 0.5817319 |
| TGGAGGEGGAAG | ASSPGQKNTLY | 0.6340048 |
| TGGDGGAAGAGGAPGHG | ASSPGQKNTLY | 0.59935796 |
| TMPPLLAAIANAGRHGV | ASSPGQKNTLY | 0.5969818 |
| TSSTAGNSGGA | ASSPGQKNTLY | 0.46601248 |
| VPATVPVGILC | ASSPGQKNTLY | 0.6206558 |
| VQRPDSGGVVLTGRIS | ASSPGQKNTLY | 0.67042714 |
| WTPNPVPATKGVPAGLVQAT | ASSPGQKNTLY | 0.6090035 |
| YALKRALAKFSDRFDVVII | ASSPGQKNTLY | 0.6558467 |
| YLIGSVIV | ASSPGQKNTLY | 0.6448939 |
| YGQLVAVID | ASSPGQKNTLY | 0.63435423 |
| VTVGGMTLPA | ASSPGQKNTLY | 0.5138258 |
| VLALPAV | ASSPGQKNTLY | 0.5713945 |
| VIPPTLNYET | ASSPGQKNTLY | 0.55909824 |
| VGMGGMHPSSG | ASSPGQKNTLY | 0.5412422 |
| VGLAGGAATA | ASSPGQKNTLY | 0.5627849 |
| VGGAGGAGGSGP | ASSPGQKNTLY | 0.56323063 |
| VFAAFMV | ASSPGQKNTLY | 0.67163134 |
| TLYDKLR | ASSPGQKNTLY | 0.63673353 |
| TLAVFTVL | ASSPGQKNTLY | 0.64708936 |
| TGSVQPARV | ASSPGQKNTLY | 0.54918396 |
| TGGGAGSGGT | ASSPGQKNTLY | 0.59079725 |
| SHLETSL | ASSPGQKNTLY | 0.63770324 |
| SFTTGMT | ASSPGQKNTLY | 0.4206792 |
| QVEVAKRMGVRQPTVS | ASSPGQKNTLY | 0.6594615 |
| QGSGGPALAM | ASSPGQKNTLY | 0.5493622 |
| MLAPSNP | ASSPGQKNTLY | 0.58675325 |
| LVNNAAI | ASSPGQKNTLY | 0.63838524 |
| LTGEINTLA | ASSPGQKNTLY | 0.65461683 |
| LSIFGPV | ASSPGQKNTLY | 0.54994816 |
| LLALLANL | ASSPGQKNTLY | 0.5338005 |
| LISNASDALDKLR | ASSPGQKNTLY | 0.56197 |
| LINEAVP | ASSPGQKNTLY | 0.64088523 |
| LGYAFSGL | ASSPGQKNTLY | 0.65758926 |
| LDFVAVAL | ASSPGQKNTLY | 0.5897838 |
| LAQPGAE | ASSPGQKNTLY | 0.6597451 |
| ISNLGAVTGF | ASSPGQKNTLY | 0.6331668 |
| HQEDPGRI | ASSPGQKNTLY | 0.38876745 |

| | | |
|---|---|---|
| GTPGEPTP | ASSPGQKNTLY | 0.57803416 |
| GFFNSGS | ASSPGQKNTLY | 0.6513611 |
| GAGGGGLTAGAN | ASSPGQKNTLY | 0.65530336 |
| FVDALVNM | ASSPGQKNTLY | 0.665742 |
| FLTVAPEL | ASSPGQKNTLY | 0.63585055 |
| FETLTAL | ASSPGQKNTLY | 0.5655265 |
| DVINPIEL | ASSPGQKNTLY | 0.660009 |
| APVSGAPV | ASSPGQKNTLY | 0.6626327 |
| ALVGGLAW | ASSPGQKNTLY | 0.6522447 |
| ALAAAAATAI | ASSPGQKNTLY | 0.52023065 |
| AFGLGLLL | ASSPGQKNTLY | 0.46948913 |
| AFFAAASL | ASSPGQKNTLY | 0.6198685 |
| AAAALPAFSPP | ASSPGQKNTLY | 0.65632 |
| VVLAPAL | ASSPGQKNTLY | 0.42707756 |
| TIGHVDHGKTTL | ASSPGQKNTLY | 0.6229479 |
| LISNASDALDK | ASSPGQKNTLY | 0.63335717 |
| LISNASDAL | ASSPGQKNTLY | 0.4131447 |
| ISNASDALDK | ASSPGQKNTLY | 0.6183547 |
| NQAETLVYQTE | ASSPGQKNTLY | 0.6602104 |
| TGGTGGTAGAG | ASSPGQKNTLY | 0.56964505 |
| TGSTAGGAGGA | ASSPGQKNTLY | 0.5692669 |
| DPLAVPVAAGPV | ASSPGQKNTLY | 0.6561589 |
| GGTGGTPGNGGN | ASSPGQKNTLY | 0.6713549 |
| PAGLAGVDPATL | ASSPGQKNTLY | 0.6562794 |
| GNNLVVEAPHAGASV | ASSPGQKNTLY | 0.44007096 |
| KVTRIMSASTLEQPA | ASSPGQKNTLY | 0.34387368 |
| LGFALPIALPRRTVV | ASSPGQKNTLY | 0.5472844 |
| TRLGNLTARKLVVLV | ASSPGQKNTLY | 0.6432022 |
| WSQQRRLIYFPSAGP | ASSPGQKNTLY | 0.6019219 |
| GPNGNPDPMAAAVDIR | ASSPGQKNTLY | 0.5648523 |
| TQRCVSVADLRAAASR | ASSPGQKNTLY | 0.64238155 |
| QFVLDVAEHPVEVLTETV | ASSPGQKNTLY | 0.6560578 |
| GGAGGTGGAGGNGGTQAGDG | ASSPGQKNTLY | 0.47340557 |
| PLLATLRAIAVDVDAKPAQV | ASSPGQKNTLY | 0.41745377 |
| APKFRTGGYMIDFWGVGYQVAK | ASSPGQKNTLY | 0.637232 |
| GAGGAGGAGGAAGGPGGTGGTG | ASSPGQKNTLY | 0.6273014 |
| IGALGLLATDVAPAGSVAFGVA | ASSPGQKNTLY | 0.54220426 |
| AGGAGGIGGNGGAGGAGGVLMG | ASSPGQKNTLY | 0.60729843 |
| ADELIGLL | ASSPGQKNTLY | 0.55681735 |
| ADRALKA | ASSPGQKNTLY | 0.45249045 |
| AGAVGQVGGAGGA | ASSPGQKNTLY | 0.6310277 |
| AGGGGGLGGAP | ASSPGQKNTLY | 0.48567438 |
| AGGLLYGNGGAG | ASSPGQKNTLY | 0.6570108 |
| AGLESIGL | ASSPGQKNTLY | 0.5102709 |
| AGLKTGII | ASSPGQKNTLY | 0.6299706 |
| ASLAALLL | ASSPGQKNTLY | 0.640938 |
| ASVINISSI | ASSPGQKNTLY | 0.62028295 |
| ATPPGPA | ASSPGQKNTLY | 0.57580227 |
| ATVASALV | ASSPGQKNTLY | 0.5500477 |
| ATVASLIL | ASSPGQKNTLY | 0.48430187 |
| AVNGPVLIRH | ASSPGQKNTLY | 0.3353879 |

| | | |
|---|---|---|
| DGAAGAVGI | ASSPGQKNTLY | 0.6589679 |
| DPLLHGI | ASSPGQKNTLY | 0.54086024 |
| ELGAALGAL | ASSPGQKNTLY | 0.653605 |
| ELGRTQAV | ASSPGQKNTLY | 0.6495889 |
| FVAGGYLS | ASSPGQKNTLY | 0.508179 |
| GASGGAGGAGGW | ASSPGQKNTLY | 0.6192949 |
| GGAGAGGDGGAGGI | ASSPGQKNTLY | 0.54293364 |
| GTGHNGSQPGMGGNG | ASSPGQKNTLY | 0.41272748 |
| GYAAASASAS | ASSPGQKNTLY | 0.637383 |
| HQVDVAAVQQG | ASSPGQKNTLY | 0.63099134 |
| IGAAIAHT | ASSPGQKNTLY | 0.49941772 |
| IGMIEVAQL | ASSPGQKNTLY | 0.6006057 |
| IRLAPEHTSAMD | ASSPGQKNTLY | 0.6221113 |
| IYNVSVL | ASSPGQKNTLY | 0.49493173 |
| KQGFLGM | ASSPGQKNTLY | 0.5841502 |
| LESVITQL | ASSPGQKNTLY | 0.65001035 |
| LGLGALVVV | ASSPGQKNTLY | 0.59374803 |
| LIGNGGNGGSGGI | ASSPGQKNTLY | 0.54262966 |
| LNAINAPTL | ASSPGQKNTLY | 0.6547969 |
| LQLLNTGV | ASSPGQKNTLY | 0.5134572 |
| LQTSIEF | ASSPGQKNTLY | 0.63491446 |
| MSFAVSEL | ASSPGQKNTLY | 0.59530705 |
| NTGALGL | ASSPGQKNTLY | 0.56892 |
| PPGAGGAGGAGTT | ASSPGQKNTLY | 0.65544605 |
| QLVGLGI | ASSPGQKNTLY | 0.4655202 |
| RFLEQVHQL | ASSPGQKNTLY | 0.5479362 |
| RGVGYVLR | ASSPGQKNTLY | 0.55486095 |
| SATGVFAGVI | ASSPGQKNTLY | 0.6279342 |
| SLPAITLP | ASSPGQKNTLY | 0.65459734 |
| STALAFTV | ASSPGQKNTLY | 0.54979247 |
| SVLTVGLGK | ASSPGQKNTLY | 0.6035981 |
| TALAGAGSGI | ASSPGQKNTLY | 0.56808054 |
| TGLPFTGI | ASSPGQKNTLY | 0.50033635 |
| VAEHVTV | ASSPGQKNTLY | 0.53804004 |
| VDIVLNSL | ASSPGQKNTLY | 0.55296314 |
| VLTGGGAVL | ASSPGQKNTLY | 0.5328676 |
| VNVGLAYGI | ASSPGQKNTLY | 0.61630577 |
| VSSAAVGAV | ASSPGQKNTLY | 0.56010693 |
| VTSVYEAGV | ASSPGQKNTLY | 0.63407946 |
| WVALDRGA | ASSPGQKNTLY | 0.66157347 |
| YLVGSVVI | ASSPGQKNTLY | 0.5887003 |
| GPPGTGKTLL | ASSPGQKNTLY | 0.6242169 |
| GVGMTKF | ASSPGQKNTLY | 0.6431918 |
| IGIGGSDL | ASSPGQKNTLY | 0.6348467 |
| IGTLLAEL | ASSPGQKNTLY | 0.6017842 |
| IYTSGTT | ASSPGQKNTLY | 0.55298495 |
| LSFPAGV | ASSPGQKNTLY | 0.66204023 |
| LYTSGTT | ASSPGQKNTLY | 0.6572156 |
| VGAFPFGF | ASSPGQKNTLY | 0.65558666 |
| VGAGTVEF | ASSPGQKNTLY | 0.6440085 |
| VGKSTLLL | ASSPGQKNTLY | 0.6297224 |

| | | |
|---|---|---|
| AALAHAGGGIV | ASSPGQKNTLY | 0.6461686 |
| AALVAVSLL | ASSPGQKNTLY | 0.58613217 |
| AAVSRAVAAL | ASSPGQKNTLY | 0.43393683 |
| AGGSSGIGGF | ASSPGQKNTLY | 0.6039558 |
| AGLTVANANG | ASSPGQKNTLY | 0.5674145 |
| AGLVGAVL | ASSPGQKNTLY | 0.56305575 |
| ALSQAFTV | ASSPGQKNTLY | 0.37463623 |
| ALTFSVSL | ASSPGQKNTLY | 0.65886503 |
| ALTGAALVAL | ASSPGQKNTLY | 0.6704134 |
| ASFSIGAGVL | ASSPGQKNTLY | 0.5445422 |
| DGSVYKI | ASSPGQKNTLY | 0.66475743 |
| EQLAEIV | ASSPGQKNTLY | 0.44803786 |
| FAVSELTVM | ASSPGQKNTLY | 0.65322983 |
| GAAPLAAGL | ASSPGQKNTLY | 0.65660965 |
| GADGGDGGSGGAGG | ASSPGQKNTLY | 0.52734995 |
| GFGLDTGGAGG | ASSPGQKNTLY | 0.6519562 |
| GLASGGLL | ASSPGQKNTLY | 0.5375907 |
| GRIAGAAVLL | ASSPGQKNTLY | 0.5878692 |
| GSAGSLNTSF | ASSPGQKNTLY | 0.6606591 |
| GTGQIGI | ASSPGQKNTLY | 0.6023783 |
| GTLIALIL | ASSPGQKNTLY | 0.61154157 |
| IAATMGFVVG | ASSPGQKNTLY | 0.5909818 |
| IFAQALL | ASSPGQKNTLY | 0.52940446 |
| IGAAVALGF | ASSPGQKNTLY | 0.600109 |
| IGLTPLAL | ASSPGQKNTLY | 0.654953 |
| ILGSNTSTL | ASSPGQKNTLY | 0.41698492 |
| IVFSLTVL | ASSPGQKNTLY | 0.6608453 |
| LASKATGF | ASSPGQKNTLY | 0.5318546 |
| LASVALIL | ASSPGQKNTLY | 0.6678123 |
| LFAVGLTL | ASSPGQKNTLY | 0.6414927 |
| LFNGGIGTYI | ASSPGQKNTLY | 0.625289 |
| LTLGAGII | ASSPGQKNTLY | 0.535743 |
| LVAAALGDI | ASSPGQKNTLY | 0.66058886 |
| NTAFVQLGI | ASSPGQKNTLY | 0.58435667 |
| QLFSGAEL | ASSPGQKNTLY | 0.6416619 |
| SAATFITL | ASSPGQKNTLY | 0.64730895 |
| SNYGLANLGV | ASSPGQKNTLY | 0.2818482 |
| SYGGAYAVM | ASSPGQKNTLY | 0.7362516 |
| TTGVGGLTLG | ASSPGQKNTLY | 0.61162615 |
| TTQTIYI | ASSPGQKNTLY | 0.5812929 |
| VAAGAGLGAVI | ASSPGQKNTLY | 0.36064112 |
| VAAVVTLL | ASSPGQKNTLY | 0.27055725 |
| VGFLGGLI | ASSPGQKNTLY | 0.40294898 |
| VGVIPILL | ASSPGQKNTLY | 0.31304055 |
| VIFASSLI | ASSPGQKNTLY | 0.7197226 |
| VKGGDGGAAAG | ASSPGQKNTLY | 0.6272198 |
| VTKTFTNI | ASSPGQKNTLY | 0.3541179 |
| VVAAAILL | ASSPGQKNTLY | 0.64437973 |
| YAALDVEL | ASSPGQKNTLY | 0.7182477 |
| LASVALLL | ASSPGQKNTLY | 0.23465994 |
| LDAVHRV | ASSPGQKNTLY | 0.5060603 |

| | | |
|---|---|---|
| QGLSSSIF | ASSPGQKNTLY | 0.29357928 |
| RGTFANIRL | ASSPGQKNTLY | 0.41591537 |
| RQLWWGHRI | ASSPGQKNTLY | 0.20042855 |
| YGPPGTGKTLL | ASSPGQKNTLY | 0.6277065 |
| AALVGAPANVA | ASSPGQKNTLY | 0.35669065 |
| AATFRRTE | ASSPGQKNTLY | 0.18361053 |
| AGALGGGI | ASSPGQKNTLY | 0.5289501 |
| AGGAGGAGGIGAG | ASSPGQKNTLY | 0.43591893 |
| AGSGALGL | ASSPGQKNTLY | 0.4502452 |
| AQIADQI | ASSPGQKNTLY | 0.6745648 |
| ASAVAVTYL | ASSPGQKNTLY | 0.31011096 |
| ASTVLGGTGG | ASSPGQKNTLY | 0.33686224 |
| AVGGEPAK | ASSPGQKNTLY | 0.61224633 |
| CEQGTFGGTC | ASSPGQKNTLY | 0.6906072 |
| DEGKAPAG | ASSPGQKNTLY | 0.3373358 |
| DTVVFPAL | ASSPGQKNTLY | 0.5623097 |
| FEGITMAG | ASSPGQKNTLY | 0.7384397 |
| FPLVVQD | ASSPGQKNTLY | 0.44781762 |
| GAFTTPQ | ASSPGQKNTLY | 0.71485585 |
| GFGGFGTAGGVGGA | ASSPGQKNTLY | 0.73603755 |
| GFLAQFL | ASSPGQKNTLY | 0.33786607 |
| GGAGGAGGNAGVA | ASSPGQKNTLY | 0.6900235 |
| GGVGTAGAGGAGGAGG | ASSPGQKNTLY | 0.67036015 |
| GNGGGGDAGNAGSGGN | ASSPGQKNTLY | 0.44674385 |
| GTGGAGGAGNDAGS | ASSPGQKNTLY | 0.45214438 |
| GTTNTGWFNS | ASSPGQKNTLY | 0.26585567 |
| IFMGPLFTL | ASSPGQKNTLY | 0.48357543 |
| ILAHAIEAI | ASSPGQKNTLY | 0.23022355 |
| IRSAIREAL | ASSPGQKNTLY | 0.6355839 |
| ITFAGGG | ASSPGQKNTLY | 0.71415854 |
| ITGMSGHQEV | ASSPGQKNTLY | 0.4267972 |
| IVAGLACTL | ASSPGQKNTLY | 0.25157902 |
| IVGGGLAGA | ASSPGQKNTLY | 0.2212422 |
| IVTGVLNI | ASSPGQKNTLY | 0.5089798 |
| LAQQGCI | ASSPGQKNTLY | 0.3118869 |
| LCLHVEN | ASSPGQKNTLY | 0.21384771 |
| LDAISDGS | ASSPGQKNTLY | 0.43809384 |
| LGGFAAML | ASSPGQKNTLY | 0.6422943 |
| LGGKNPMI | ASSPGQKNTLY | 0.66920114 |
| LGPKGVAAIL | ASSPGQKNTLY | 0.28912938 |
| LMVGVSI | ASSPGQKNTLY | 0.74145436 |
| LVGAAFVL | ASSPGQKNTLY | 0.6884739 |
| LVLAVAPL | ASSPGQKNTLY | 0.6743803 |
| MAAIAVIDAA | ASSPGQKNTLY | 0.46450192 |
| NNGINAELQ | ASSPGQKNTLY | 0.27910402 |
| RGEPLVLVHG | ASSPGQKNTLY | 0.7313285 |
| RGGAGGAAGLI | ASSPGQKNTLY | 0.4639051 |
| RSYRMHPAV | ASSPGQKNTLY | 0.46462515 |
| SAAHLLAGQLA | ASSPGQKNTLY | 0.623389 |
| SAAVGSY | ASSPGQKNTLY | 0.5072272 |
| SPHAVLHRI | ASSPGQKNTLY | 0.35033527 |

| | | |
|---|---|---|
| SQIAKVL | ASSPGQKNTLY | 0.6395216 |
| TALGHLVP | ASSPGQKNTLY | 0.6475582 |
| TAPTTSGAA | ASSPGQKNTLY | 0.29880556 |
| TDGDDGNGGSPGT | ASSPGQKNTLY | 0.3105539 |
| TDGLGEN | ASSPGQKNTLY | 0.4239673 |
| TGGPPLPGQA | ASSPGQKNTLY | 0.50590056 |
| TIPAITI | ASSPGQKNTLY | 0.69091415 |
| TIPGFTI | ASSPGQKNTLY | 0.5378139 |
| TSIASGALA | ASSPGQKNTLY | 0.46291095 |
| TVTLPGAAI | ASSPGQKNTLY | 0.6232306 |
| VASGLSAALP | ASSPGQKNTLY | 0.6419155 |
| VGAGVIGI | ASSPGQKNTLY | 0.52563107 |
| VGPKRFGV | ASSPGQKNTLY | 0.6739522 |
| VNVAFPD | ASSPGQKNTLY | 0.38460076 |
| VSMTNSGVS | ASSPGQKNTLY | 0.636068 |
| VTDVLGDLSV | ASSPGQKNTLY | 0.6422799 |
| VVVVGAGFAGL | ASSPGQKNTLY | 0.66052246 |
| VYTVGTVV | ASSPGQKNTLY | 0.7009611 |
| WTTFSGP | ASSPGQKNTLY | 0.6248538 |
| YDELVEAF | ASSPGQKNTLY | 0.49325585 |
| YGTGGAGGNGG | ASSPGQKNTLY | 0.21806243 |
| YTNVMAV | ASSPGQKNTLY | 0.40824583 |
| FLEQVHQL | ASSPGQKNTLY | 0.57461894 |
| GLSLLAVAV | ASSPGQKNTLY | 0.6111443 |
| LEQVHQL | ASSPGQKNTLY | 0.6694578 |
| LNAAGAI | ASSPGQKNTLY | 0.6506387 |
| PAVHASL | ASSPGQKNTLY | 0.36028123 |
| TELEDDA | ASSPGQKNTLY | 0.72046465 |
| TPPGPPN | ASSPGQKNTLY | 0.7065799 |
| TPPPPPPP | ASSPGQKNTLY | 0.24528533 |
| TQSPALI | ASGGWEQY | 0.42464107 |
| EFAAAFPQ | ASGGWEQY | 0.7533105 |
| LLSGNLDVL | ASGGWEQY | 0.65239716 |
| GPANLGNYNI | ASGGWEQY | 0.37609577 |
| TLGPTLP | ASGGWEQY | 0.45852017 |
| VYAYSSCCK | ASGGWEQY | 0.7570957 |
| APAPAPAPNI | ASGGWEQY | 0.3803581 |
| PGGPVPIP | ASGGWEQY | 0.72199184 |
| FLTVALEP | ASGGWEQY | 0.21094391 |
| SQYFGANL | ASGGWEQY | 0.40666533 |
| GGVKPAA | ASGGWEQY | 0.3541643 |
| GGTGGAGGAGGW | ASGGWEQY | 0.19989324 |
| GVGGVAGLGGAGG | ASGGWEQY | 0.6259307 |
| VLLVLATTI | ASGGWEQY | 0.34333947 |
| IGIFAAML | ASGGWEQY | 0.66492164 |
| AVGDLID | ASGGWEQY | 0.37590614 |
| NISPLPL | ASGGWEQY | 0.67810476 |
| VTAVFVAI | ASGGWEQY | 0.5213851 |
| SGDDNDAAL | ASGGWEQY | 0.4886408 |
| AVAALAHS | ASGGWEQY | 0.30843514 |
| GGVTVMATP | ASGGWEQY | 0.54240274 |

| | | |
|---|---|---|
| AGLAANVRD | ASGGWEQY | 0.7137102 |
| TNRPDLIDPALLRPG | ASGGWEQY | 0.6353317 |
| LANLGVTAL | ASGGWEQY | 0.24584995 |
| TGERIEL | ASGGWEQY | 0.4863625 |
| NGEELAA | ASGGWEQY | 0.3991406 |
| VLSLVWG | ASGGWEQY | 0.21846133 |
| IGYALWL | ASGGWEQY | 0.19767536 |
| LSLLLAPL | ASGGWEQY | 0.39667913 |
| TLDEALK | ASGGWEQY | 0.5272124 |
| TDGAGTAEGG | ASGGWEQY | 0.627198 |
| NAPGGAGGNGG | ASGGWEQY | 0.6173413 |
| VSVFMGMGGNF | ASGGWEQY | 0.3529239 |
| LASYLAAL | ASGGWEQY | 0.6419835 |
| LVGGTPAL | ASGGWEQY | 0.53289795 |
| VRNADVL | ASGGWEQY | 0.17385127 |
| TCSSPFLVSAM | ASGGWEQY | 0.3750672 |
| VHITDAE | ASGGWEQY | 0.23733622 |
| EYAAPSF | ASGGWEQY | 0.71060616 |
| VVDAFHGGL | ASGGWEQY | 0.30701816 |
| ASVVDPQ | ASGGWEQY | 0.37374297 |
| SYNEFGGLS | ASGGWEQY | 0.44509053 |
| DPDGSVY | ASGGWEQY | 0.6871276 |
| MVSTIHATE | ASGGWEQY | 0.5818863 |
| LAAGVADPV | ASGGWEQY | 0.3152272 |
| TKEQIAA | ASGGWEQY | 0.22056112 |
| SLASPAAL | ASGGWEQY | 0.53453636 |
| IGLVFYGV | ASGGWEQY | 0.34411317 |
| FLLDASNL | ASGGWEQY | 0.34031934 |
| YQVGNFP | ASGGWEQY | 0.25972918 |
| ESLLSQST | ASGGWEQY | 0.5854012 |
| LFTVGFAAVG | ASGGWEQY | 0.25312072 |
| ATPPTPT | ASGGWEQY | 0.6708019 |
| TDLATLG | ASGGWEQY | 0.3205734 |
| SAGGTGR | ASGGWEQY | 0.41567597 |
| GFSNSGS | ASGGWEQY | 0.749452 |
| GGNGGSGGTSV | ASGGWEQY | 0.6807602 |
| GTSVATGGAGN | ASGGWEQY | 0.45100015 |
| NGGNGGSGGTS | ASGGWEQY | 0.38745874 |
| AVVKFPH | ASGGWEQY | 0.2795774 |
| TPPPPPP | ASGGWEQY | 0.3243194 |
| AAPAAAAAPA | ASGGWEQY | 0.38961384 |
| IALNEDL | ASGGWEQY | 0.7157592 |
| IVNVSSM | ASGGWEQY | 0.47148097 |
| TGIPPAPR | ASGGWEQY | 0.2663716 |
| FDLGGGTF | ASGGWEQY | 0.5709779 |
| FDLGGGTFDVS | ASGGWEQY | 0.730423 |
| NEPTAAA | ASGGWEQY | 0.7163259 |
| NEPTAAALA | ASGGWEQY | 0.2694963 |
| GSIINISSI | ASGGWEQY | 0.44321898 |
| VAPGNAGTA | ASGGWEQY | 0.436509 |
| TGGGASLE | ASGGWEQY | 0.47927707 |

| | | |
|---|---|---|
| VAVYNHYKRI | ASGGWEQY | 0.36694166 |
| TDPAPATT | ASGGWEQY | 0.31904238 |
| TNRPDILDPALLRPG | ASGGWEQY | 0.6046164 |
| LTTTTTS | ASGGWEQY | 0.7746382 |
| PTGSGKTL | ASGGWEQY | 0.4113912 |
| TGSGKTL | ASGGWEQY | 0.20902307 |
| TIGHVDHGK | ASGGWEQY | 0.42404273 |
| TIGHVDHGKTT | ASGGWEQY | 0.3443113 |
| GHVDHGKTT | ASGGWEQY | 0.312117 |
| AQFEHTLL | ASGGWEQY | 0.45867902 |
| IVNQDEL | ASGGWEQY | 0.23365115 |
| TNGSQFF | ASGGWEQY | 0.5090498 |
| TGKGIVE | ASGGWEQY | 0.75476766 |
| LISNASDA | ASGGWEQY | 0.4255912 |
| AELRTLGELTKIAKAHGAQ | ASGGWEQY | 0.602016 |
| APNAVAAAEPSW | ASGGWEQY | 0.7098502 |
| APQVAGAATAVL | ASGGWEQY | 0.6083828 |
| ARRDPAAAAAILPTDA | ASGGWEQY | 0.7206565 |
| DAAARDVLAEAGLAEYF | ASGGWEQY | 0.715919 |
| DDQIVLSAKANLAKVEQ | ASGGWEQY | 0.49385223 |
| DELLPDTGEDSDEDGH | ASGGWEQY | 0.54382867 |
| ELPEAPANRATVPQD | ASGGWEQY | 0.637852 |
| FGQSKIERVTGIG | ASGGWEQY | 0.4102376 |
| GAGGTGSTAGGAGGAGGAGGLY | ASGGWEQY | 0.45609546 |
| GHGGTGAAGGNGGTGGAGGSGI | ASGGWEQY | 0.40267885 |
| GNGGNGGAGGEGGPGGAGGAGA | ASGGWEQY | 0.6777426 |
| GSTAGNGGNGGNATNGTIAGQP | ASGGWEQY | 0.52154016 |
| GTGGDGGNGGNGGTGG | ASGGWEQY | 0.73309207 |
| GVRRPAGIPADAPV | ASGGWEQY | 0.22588836 |
| GYRQVTDAHLLTLARR | ASGGWEQY | 0.3228014 |
| IAKMGQDGHDRG | ASGGWEQY | 0.7177235 |
| KAEPSIEIQYT | ASGGWEQY | 0.18479025 |
| KPVRIRMAADEDTVAVT | ASGGWEQY | 0.21188912 |
| MPPRIAGMRLLVIKPEP | ASGGWEQY | 0.41684973 |
| MKFYAVALFRSRGVPPTR | ASGGWEQY | 0.7264961 |
| NAGGFGILGAVAHSP | ASGGWEQY | 0.18435676 |
| NEGREITLAMAEM | ASGGWEQY | 0.37913406 |
| NELVEWSHPLGHRWRWE | ASGGWEQY | 0.25664034 |
| NLGAGGTGGNGGNPG | ASGGWEQY | 0.712062 |
| NTEWPKWETVMAADPVD | ASGGWEQY | 0.49099305 |
| PASAAPAETQP | ASGGWEQY | 0.25116396 |
| QQSSDGKNGVRAAEITGE | ASGGWEQY | 0.65142053 |
| RAPDAVTTSWWKE | ASGGWEQY | 0.21567962 |
| SGFGNVGAHTSGWFNQ | ASGGWEQY | 0.45043802 |
| SGVAGAVGQAASVGGLKV | ASGGWEQY | 0.64597243 |
| SNGGAGTGSAGGNGGT | ASGGWEQY | 0.7585064 |
| TGGAGGEGGAAG | ASGGWEQY | 0.25040698 |
| TGGDGGAAGAGGAPGHG | ASGGWEQY | 0.6342605 |
| TMPPLLAAIANAGRHGV | ASGGWEQY | 0.36017773 |
| TSSTAGNSGGA | ASGGWEQY | 0.7498835 |
| VPATVPVGILC | ASGGWEQY | 0.39290392 |

| | | |
|---|---|---|
| VQRPDSGGVVLTGRIS | ASGGWEQY | 0.22918475 |
| WTPNPVPATKGVPAGLVQAT | ASGGWEQY | 0.6974338 |
| YALKRALAKFSDRFDVVII | ASGGWEQY | 0.6463663 |
| YLIGSVIV | ASGGWEQY | 0.4883222 |
| YGQLVAVID | ASGGWEQY | 0.6958406 |
| VTVGGMTLPA | ASGGWEQY | 0.4659567 |
| VLALPAV | ASGGWEQY | 0.4519752 |
| VIPPTLNYET | ASGGWEQY | 0.66605973 |
| VGMGGMHPSSG | ASGGWEQY | 0.65657747 |
| VGLAGGAATA | ASGGWEQY | 0.35761505 |
| VGGAGGAGGSGP | ASGGWEQY | 0.67624295 |
| VFAAFMV | ASGGWEQY | 0.5051019 |
| TLYDKLR | ASGGWEQY | 0.2940813 |
| TLAVFTVL | ASGGWEQY | 0.755864 |
| TGSVQPARV | ASGGWEQY | 0.4863277 |
| TGGGAGSGGT | ASGGWEQY | 0.6610601 |
| SHLETSL | ASGGWEQY | 0.57939863 |
| SFTTGMT | ASGGWEQY | 0.7422576 |
| QVEVAKRMGVRQPTVS | ASGGWEQY | 0.30689207 |
| QGSGGPALAM | ASGGWEQY | 0.55642116 |
| MLAPSNP | ASGGWEQY | 0.4858806 |
| LVNNAAI | ASGGWEQY | 0.68139255 |
| LTGEINTLA | ASGGWEQY | 0.44958812 |
| LSIFGPV | ASGGWEQY | 0.43146527 |
| LLALLANL | ASGGWEQY | 0.59887874 |
| LISNASDALDKLR | ASGGWEQY | 0.7149124 |
| LINEAVP | ASGGWEQY | 0.448156 |
| LGYAFSGL | ASGGWEQY | 0.63799024 |
| LDFVAVAL | ASGGWEQY | 0.7442015 |
| LAQPGAE | ASGGWEQY | 0.2798701 |
| ISNLGAVTGF | ASGGWEQY | 0.4652363 |
| HQEDPGRI | ASGGWEQY | 0.63812304 |
| GTPGEPTP | ASGGWEQY | 0.7124065 |
| GFFNSGS | ASGGWEQY | 0.4839003 |
| GAGGGGLTAGAN | ASGGWEQY | 0.69694215 |
| FVDALVNM | ASGGWEQY | 0.41368243 |
| FLTVAPEL | ASGGWEQY | 0.5888535 |
| FETLTAL | ASGGWEQY | 0.30513373 |
| DVINPIEL | ASGGWEQY | 0.43662506 |
| APVSGAPV | ASGGWEQY | 0.33491915 |
| ALVGGLAW | ASGGWEQY | 0.50739896 |
| ALAAAAATAI | ASGGWEQY | 0.6652448 |
| AFGLGLLL | ASGGWEQY | 0.61342186 |
| AFFAAASL | ASGGWEQY | 0.20718977 |
| AAAALPAFSPP | ASGGWEQY | 0.6114732 |
| VVLAPAL | ASGGWEQY | 0.6470797 |
| TIGHVDHGKTTL | ASGGWEQY | 0.76698315 |
| LISNASDALDK | ASGGWEQY | 0.5138123 |
| LISNASDAL | ASGGWEQY | 0.6461921 |
| ISNASDALDK | ASGGWEQY | 0.34321812 |
| NQAETLVYQTE | ASGGWEQY | 0.47114938 |

| | | |
|---|---|---|
| TGGTGGTAGAG | ASGGWEQY | 0.2848787 |
| TGSTAGGAGGA | ASGGWEQY | 0.3499787 |
| DPLAVPVAAGPV | ASGGWEQY | 0.42594105 |
| GGTGGTPGNGGN | ASGGWEQY | 0.3146631 |
| PAGLAGVDPATL | ASGGWEQY | 0.7247907 |
| GNNLVVEAPHAGASV | ASGGWEQY | 0.712987 |
| KVTRIMSASTLEQPA | ASGGWEQY | 0.7031038 |
| LGFALPIALPRRTVV | ASGGWEQY | 0.7174201 |
| TRLGNLTARKLVVLV | ASGGWEQY | 0.7148325 |
| WSQQRRLIYFPSAGP | ASGGWEQY | 0.68011713 |
| GPNGNPDPMAAAVDIR | ASGGWEQY | 0.41319826 |
| TQRCVSVADLRAAASR | ASGGWEQY | 0.4649427 |
| QFVLDVAEHPVEVLTETV | ASGGWEQY | 0.23824644 |
| GGAGGTGGAGGNGGTQAGDG | ASGGWEQY | 0.41149664 |
| PLLATLRAIAVDVDAKPAQV | ASGGWEQY | 0.4391992 |
| APKFRTGGYMIDFWGVGYQVAK | ASGGWEQY | 0.5034391 |
| GAGGAGGAGGAAGGPGGTGGTG | ASGGWEQY | 0.18605913 |
| IGALGLLATDVAPAGSVAFGVA | ASGGWEQY | 0.64207166 |
| AGGAGGIGGNGGAGGAGGVLMG | ASGGWEQY | 0.7283682 |
| ADELIGLL | ASGGWEQY | 0.41511482 |
| ADRALKA | ASGGWEQY | 0.65585697 |
| AGAVGQVGGAGGA | ASGGWEQY | 0.6493087 |
| AGGGGGLGGAP | ASGGWEQY | 0.391621 |
| AGGLLYGNGGAG | ASGGWEQY | 0.6632322 |
| AGLESIGL | ASGGWEQY | 0.38357055 |
| AGLKTGII | ASGGWEQY | 0.6749967 |
| ASLAALLL | ASGGWEQY | 0.7330493 |
| ASVINISSI | ASGGWEQY | 0.5771803 |
| ATPPGPA | ASGGWEQY | 0.53651285 |
| ATVASALV | ASGGWEQY | 0.5535075 |
| ATVASLIL | ASGGWEQY | 0.27623367 |
| AVNGPVLIRH | ASGGWEQY | 0.55738246 |
| DGAAGAVGI | ASGGWEQY | 0.30479544 |
| DPLLHGI | ASGGWEQY | 0.54779744 |
| ELGAALGAL | ASGGWEQY | 0.28906283 |
| ELGRTQAV | ASGGWEQY | 0.5306243 |
| FVAGGYLS | ASGGWEQY | 0.49075228 |
| GASGGAGGAGGW | ASGGWEQY | 0.5450448 |
| GGAGAGGDGGAGGI | ASGGWEQY | 0.27419508 |
| GTGHNGSQPGMGGNG | ASGGWEQY | 0.52840424 |
| GYAAASASAS | ASGGWEQY | 0.49584058 |
| HQVDVAAVQQG | ASGGWEQY | 0.28149515 |
| IGAAIAHT | ASGGWEQY | 0.63751686 |
| IGMIEVAQL | ASGGWEQY | 0.39522156 |
| IRLAPEHTSAMD | ASGGWEQY | 0.5380671 |
| IYNVSVL | ASGGWEQY | 0.6816304 |
| KQGFLGM | ASGGWEQY | 0.53960925 |
| LESVITQL | ASGGWEQY | 0.22788785 |
| LGLGALVVV | ASGGWEQY | 0.46759915 |
| LIGNGGNGGSGGI | ASGGWEQY | 0.45555538 |
| LNAINAPTL | ASGGWEQY | 0.41054118 |

| | | |
|---|---|---|
| LQLLNTGV | ASGGWEQY | 0.18870853 |
| LQTSIEF | ASGGWEQY | 0.6222099 |
| MSFAVSEL | ASGGWEQY | 0.292906 |
| NTGALGL | ASGGWEQY | 0.34813294 |
| PPGAGGAGGAGTT | ASGGWEQY | 0.39239335 |
| QLVGLGI | ASGGWEQY | 0.66764164 |
| RFLEQVHQL | ASGGWEQY | 0.4758799 |
| RGVGYVLR | ASGGWEQY | 0.5392575 |
| SATGVFAGVI | ASGGWEQY | 0.63616693 |
| SLPAITLP | ASGGWEQY | 0.5183295 |
| STALAFTV | ASGGWEQY | 0.6956371 |
| SVLTVGLGK | ASGGWEQY | 0.57981104 |
| TALAGAGSGI | ASGGWEQY | 0.5300095 |
| TGLPFTGI | ASGGWEQY | 0.5893024 |
| VAEHVTV | ASGGWEQY | 0.39352113 |
| VDIVLNSL | ASGGWEQY | 0.23530987 |
| VLTGGGAVL | ASGGWEQY | 0.23670647 |
| VNVGLAYGI | ASGGWEQY | 0.24453081 |
| VSSAAVGAV | ASGGWEQY | 0.22758767 |
| VTSVYEAGV | ASGGWEQY | 0.5790354 |
| WVALDRGA | ASGGWEQY | 0.7185929 |
| YLVGSVVI | ASGGWEQY | 0.53444797 |
| GPPGTGKTLL | ASGGWEQY | 0.23651315 |
| GVGMTKF | ASGGWEQY | 0.41339052 |
| IGIGGSDL | ASGGWEQY | 0.4080745 |
| IGTLLAEL | ASGGWEQY | 0.64029855 |
| IYTSGTT | ASGGWEQY | 0.6369059 |
| LSFPAGV | ASGGWEQY | 0.7594838 |
| LYTSGTT | ASGGWEQY | 0.6686909 |
| VGAFPFGF | ASGGWEQY | 0.3988213 |
| VGAGTVEF | ASGGWEQY | 0.65211964 |
| VGKSTLLL | ASGGWEQY | 0.41547284 |
| AALAHAGGGIV | ASGGWEQY | 0.49815428 |
| AALVAVSLL | ASGGWEQY | 0.47335178 |
| AAVSRAVAAL | ASGGWEQY | 0.54137504 |
| AGGSSGIGGF | ASGGWEQY | 0.5162332 |
| AGLTVANANG | ASGGWEQY | 0.41940922 |
| AGLVGAVL | ASGGWEQY | 0.2278419 |
| ALSQAFTV | ASGGWEQY | 0.5780604 |
| ALTFSVSL | ASGGWEQY | 0.2595492 |
| ALTGAALVAL | ASGGWEQY | 0.23343062 |
| ASFSIGAGVL | ASGGWEQY | 0.41109747 |
| DGSVYKI | ASGGWEQY | 0.7363196 |
| EQLAEIV | ASGGWEQY | 0.7369367 |
| FAVSELTVM | ASGGWEQY | 0.34732163 |
| GAAPLAAGL | ASGGWEQY | 0.34322482 |
| GADGGDGGSGGAGG | ASGGWEQY | 0.69241846 |
| GFGLDTGGAGG | ASGGWEQY | 0.27369177 |
| GLASGGLL | ASGGWEQY | 0.6055972 |
| GRIAGAAVLL | ASGGWEQY | 0.55663514 |
| GSAGSLNTSF | ASGGWEQY | 0.54237556 |

| | | |
|---|---|---|
| GTGQIGI | ASGGWEQY | 0.3499301 |
| GTLIALIL | ASGGWEQY | 0.64135075 |
| IAATMGFVVG | ASGGWEQY | 0.42625502 |
| IFAQALL | ASGGWEQY | 0.6193063 |
| IGAAVALGF | ASGGWEQY | 0.18629262 |
| IGLTPLAL | ASGGWEQY | 0.655662 |
| ILGSNTSTL | ASGGWEQY | 0.29182655 |
| IVFSLTVL | ASGGWEQY | 0.3248394 |
| LASKATGF | ASGGWEQY | 0.20116729 |
| LASVALIL | ASGGWEQY | 0.42022955 |
| LFAVGLTL | ASGGWEQY | 0.5660366 |
| LFNGGIGTYI | ASGGWEQY | 0.7280419 |
| LTLGAGII | ASGGWEQY | 0.34663936 |
| LVAAALGDI | ASGGWEQY | 0.65897524 |
| NTAFVQLGI | ASGGWEQY | 0.62991107 |
| QLFSGAEL | ASGGWEQY | 0.37950864 |
| SAATFITL | ASGGWEQY | 0.23641762 |
| SNYGLANLGV | ASGGWEQY | 0.59994453 |
| SYGGAYAVM | ASGGWEQY | 0.49712875 |
| TTGVGGLTLG | ASGGWEQY | 0.72070205 |
| TTQTIYI | ASGGWEQY | 0.60810375 |
| VAAGAGLGAVI | ASGGWEQY | 0.23805828 |
| VAAVVTLL | ASGGWEQY | 0.67496705 |
| VGFLGGLI | ASGGWEQY | 0.64489436 |
| VGVIPILL | ASGGWEQY | 0.3064235 |
| VIFASSLI | ASGGWEQY | 0.29810873 |
| VKGGDGGAAAG | ASGGWEQY | 0.34187695 |
| VTKTFTNI | ASGGWEQY | 0.22341631 |
| VVAAAILL | ASGGWEQY | 0.5798613 |
| YAALDVEL | ASGGWEQY | 0.44146562 |
| LASVALLL | ASGGWEQY | 0.48124808 |
| LDAVHRV | ASGGWEQY | 0.46483856 |
| QGLSSSIF | ASGGWEQY | 0.41602686 |
| RGTFANIRL | ASGGWEQY | 0.37415278 |
| RQLWWGHRI | ASGGWEQY | 0.43173748 |
| YGPPGTGKTLL | ASGGWEQY | 0.5285078 |
| AALVGAPANVA | ASGGWEQY | 0.66187996 |
| AATFRRTE | ASGGWEQY | 0.45469067 |
| AGALGGGI | ASGGWEQY | 0.51631606 |
| AGGAGGAGGIGAG | ASGGWEQY | 0.7149895 |
| AGSGALGL | ASGGWEQY | 0.28835955 |
| AQIADQI | ASGGWEQY | 0.35154545 |
| ASAVAVTYL | ASGGWEQY | 0.64817774 |
| ASTVLGGTGG | ASGGWEQY | 0.30582047 |
| AVGGEPAK | ASGGWEQY | 0.6963178 |
| CEQGTFGGTC | ASGGWEQY | 0.53665644 |
| DEGKAPAG | ASGGWEQY | 0.5607412 |
| DTVVFPAL | ASGGWEQY | 0.54679555 |
| FEGITMAG | ASGGWEQY | 0.747067 |
| FPLVVQD | ASGGWEQY | 0.6140047 |
| GAFTTPQ | ASGGWEQY | 0.41219845 |

| | | |
|---|---|---|
| GFGGFGTAGGVGGA | ASGGWEQY | 0.67348695 |
| GFLAQFL | ASGGWEQY | 0.29423395 |
| GGAGGAGGNAGVA | ASGGWEQY | 0.6423073 |
| GGVGTAGAGGAGGAGG | ASGGWEQY | 0.6441761 |
| GNGGGGDAGNAGSGGN | ASGGWEQY | 0.2926044 |
| GTGGAGGAGNDAGS | ASGGWEQY | 0.45846117 |
| GTTNTGWFNS | ASGGWEQY | 0.703161 |
| IFMGPLFTL | ASGGWEQY | 0.6848271 |
| ILAHAIEAI | ASGGWEQY | 0.26985824 |
| IRSAIREAL | ASGGWEQY | 0.32906276 |
| ITFAGGG | ASGGWEQY | 0.30082124 |
| ITGMSGHQEV | ASGGWEQY | 0.43575096 |
| IVAGLACTL | ASGGWEQY | 0.25765955 |
| IVGGGLAGA | ASGGWEQY | 0.639665 |
| IVTGVLNI | ASGGWEQY | 0.35111785 |
| LAQQGCI | ASGGWEQY | 0.7230251 |
| LCLHVEN | ASGGWEQY | 0.549546 |
| LDAISDGS | ASGGWEQY | 0.6468665 |
| LGGFAAML | ASGGWEQY | 0.35367042 |
| LGGKNPMI | ASGGWEQY | 0.4221683 |
| LGPKGVAAIL | ASGGWEQY | 0.23310754 |
| LMVGVSI | ASGGWEQY | 0.3017138 |
| LVGAAFVL | ASGGWEQY | 0.27619708 |
| LVLAVAPL | ASGGWEQY | 0.52532154 |
| MAAIAVIDAA | ASGGWEQY | 0.2083279 |
| NNGINAELQ | ASGGWEQY | 0.7502382 |
| RGEPLVLVHG | ASGGWEQY | 0.4139751 |
| RGGAGGAAGLI | ASGGWEQY | 0.5476946 |
| RSYRMHPAV | ASGGWEQY | 0.38709247 |
| SAAHLLAGQLA | ASGGWEQY | 0.31627065 |
| SAAVGSY | ASGGWEQY | 0.382285 |
| SPHAVLHRI | ASGGWEQY | 0.66036844 |
| SQIAKVL | ASGGWEQY | 0.2761653 |
| TALGHLVP | ASGGWEQY | 0.19988415 |
| TAPTTSGAA | ASGGWEQY | 0.32123843 |
| TDGDDGNGGSPGT | ASGGWEQY | 0.23473375 |
| TDGLGEN | ASGGWEQY | 0.733714 |
| TGGPPLPGQA | ASGGWEQY | 0.6967466 |
| TIPAITI | ASGGWEQY | 0.54124737 |
| TIPGFTI | ASGGWEQY | 0.7369487 |
| TSIASGALA | ASGGWEQY | 0.18597463 |
| TVTLPGAAI | ASGGWEQY | 0.41566962 |
| VASGLSAALP | ASGGWEQY | 0.7306881 |
| VGAGVIGI | ASGGWEQY | 0.573682 |
| VGPKRFGV | ASGGWEQY | 0.40043172 |
| VNVAFPD | ASGGWEQY | 0.6671514 |
| VSMTNSGVS | ASGGWEQY | 0.35861364 |
| VTDVLGDLSV | ASGGWEQY | 0.4283775 |
| VVVVGAGFAGL | ASGGWEQY | 0.49328908 |
| VYTVGTVV | ASGGWEQY | 0.5841553 |
| WTTFSGP | ASGGWEQY | 0.43365476 |

| | | |
|---|---|---|
| YDELVEAF | ASGGWEQY | 0.24650313 |
| YGTGGAGGNGG | ASGGWEQY | 0.60877526 |
| YTNVMAV | ASGGWEQY | 0.7607338 |
| FLEQVHQL | ASGGWEQY | 0.39578593 |
| GLSLLAVAV | ASGGWEQY | 0.5438051 |
| LEQVHQL | ASGGWEQY | 0.57836616 |
| LNAAGAI | ASGGWEQY | 0.43881422 |
| PAVHASL | ASGGWEQY | 0.31299356 |
| TELEDDA | ASGGWEQY | 0.29638037 |
| TPPGPPN | ASGGWEQY | 0.5176101 |
| TPPPPPPP | ASGGWEQY | 0.27605414 |
| TQSPALI | ASSAWGGAYEQY | 0.46025547 |
| EFAAAFPQ | ASSAWGGAYEQY | 0.44893757 |
| LLSGNLDVL | ASSAWGGAYEQY | 0.76952225 |
| GPANLGNYNI | ASSAWGGAYEQY | 0.63919455 |
| TLGPTLP | ASSAWGGAYEQY | 0.5903684 |
| VYAYSSCCK | ASSAWGGAYEQY | 0.3013463 |
| APAPAPAPNI | ASSAWGGAYEQY | 0.48038357 |
| PGGPVPIP | ASSAWGGAYEQY | 0.64799345 |
| FLTVALEP | ASSAWGGAYEQY | 0.23343769 |
| SQYFGANL | ASSAWGGAYEQY | 0.707685 |
| GGVKPAA | ASSAWGGAYEQY | 0.3025645 |
| GGTGGAGGAGGW | ASSAWGGAYEQY | 0.60809803 |
| GVGGVAGLGGAGG | ASSAWGGAYEQY | 0.63096 |
| VLLVLATTI | ASSAWGGAYEQY | 0.73790574 |
| IGIFAAML | ASSAWGGAYEQY | 0.41660574 |
| AVGDLID | ASSAWGGAYEQY | 0.51750326 |
| NISPLPL | ASSAWGGAYEQY | 0.30634645 |
| VTAVFVAI | ASSAWGGAYEQY | 0.6749722 |
| SGDDNDAAL | ASSAWGGAYEQY | 0.7073555 |
| AVAALAHS | ASSAWGGAYEQY | 0.6141985 |
| GGVTVMATP | ASSAWGGAYEQY | 0.75931334 |
| AGLAANVRD | ASSAWGGAYEQY | 0.66657484 |
| TNRPDLIDPALLRPG | ASSAWGGAYEQY | 0.2414981 |
| LANLGVTAL | ASSAWGGAYEQY | 0.44597465 |
| TGERIEL | ASSAWGGAYEQY | 0.67072415 |
| NGEELAA | ASSAWGGAYEQY | 0.7210674 |
| VLSLVWG | ASSAWGGAYEQY | 0.66427577 |
| IGYALWL | ASSAWGGAYEQY | 0.6543816 |
| LSLLLAPL | ASSAWGGAYEQY | 0.381714 |
| TLDEALK | ASSAWGGAYEQY | 0.72325265 |
| TDGAGTAEGG | ASSAWGGAYEQY | 0.7536223 |
| NAPGGAGGNGG | ASSAWGGAYEQY | 0.62877095 |
| VSVFMGMGGNF | ASSAWGGAYEQY | 0.3670525 |
| LASYLAAL | ASSAWGGAYEQY | 0.42627543 |
| LVGGTPAL | ASSAWGGAYEQY | 0.5892048 |
| VRNADVL | ASSAWGGAYEQY | 0.70901656 |
| TCSSPFLVSAM | ASSAWGGAYEQY | 0.27100033 |
| VHITDAE | ASSAWGGAYEQY | 0.5111165 |
| EYAAPSF | ASSAWGGAYEQY | 0.5575615 |
| VVDAFHGGL | ASSAWGGAYEQY | 0.21983947 |

| | | |
|---|---|---|
| ASVVDPQ | ASSAWGGAYEQY | 0.4940586 |
| SYNEFGGLS | ASSAWGGAYEQY | 0.722857 |
| DPDGSVY | ASSAWGGAYEQY | 0.5661882 |
| MVSTIHATE | ASSAWGGAYEQY | 0.306709 |
| LAAGVADPV | ASSAWGGAYEQY | 0.7119535 |
| TKEQIAA | ASSAWGGAYEQY | 0.7299217 |
| SLASPAAL | ASSAWGGAYEQY | 0.70334333 |
| IGLVFYGV | ASSAWGGAYEQY | 0.26096457 |
| FLLDASNL | ASSAWGGAYEQY | 0.19111133 |
| YQVGNFP | ASSAWGGAYEQY | 0.48877013 |
| ESLLSQST | ASSAWGGAYEQY | 0.704619 |
| LFTVGFAAVG | ASSAWGGAYEQY | 0.48872086 |
| ATPPTPT | ASSAWGGAYEQY | 0.3558947 |
| TDLATLG | ASSAWGGAYEQY | 0.55367744 |
| SAGGTGR | ASSAWGGAYEQY | 0.688105 |
| GFSNSGS | ASSAWGGAYEQY | 0.27993318 |
| GGNGGSGGTSV | ASSAWGGAYEQY | 0.27727097 |
| GTSVATGGAGN | ASSAWGGAYEQY | 0.61840034 |
| NGGNGGSGGTS | ASSAWGGAYEQY | 0.6715392 |
| AVVKFPH | ASSAWGGAYEQY | 0.30723536 |
| TPPPPPP | ASSAWGGAYEQY | 0.38728178 |
| AAPAAAAAPA | ASSAWGGAYEQY | 0.47010654 |
| IALNEDL | ASSAWGGAYEQY | 0.3956259 |
| IVNVSSM | ASSAWGGAYEQY | 0.6023193 |
| TGIPPAPR | ASSAWGGAYEQY | 0.27305156 |
| FDLGGGTF | ASSAWGGAYEQY | 0.68751985 |
| FDLGGGTFDVS | ASSAWGGAYEQY | 0.2340596 |
| NEPTAAA | ASSAWGGAYEQY | 0.6126034 |
| NEPTAAALA | ASSAWGGAYEQY | 0.57456255 |
| GSIINISSI | ASSAWGGAYEQY | 0.55657184 |
| VAPGNAGTA | ASSAWGGAYEQY | 0.379248 |
| TGGGASLE | ASSAWGGAYEQY | 0.2849811 |
| VAVYNHYKRI | ASSAWGGAYEQY | 0.2799472 |
| TDPAPATT | ASSAWGGAYEQY | 0.24413805 |
| TNRPDILDPALLRPG | ASSAWGGAYEQY | 0.7125751 |
| LTTTTTS | ASSAWGGAYEQY | 0.4406325 |
| PTGSGKTL | ASSAWGGAYEQY | 0.68794787 |
| TGSGKTL | ASSAWGGAYEQY | 0.45443347 |
| TIGHVDHGK | ASSAWGGAYEQY | 0.30169645 |
| TIGHVDHGKTT | ASSAWGGAYEQY | 0.5950899 |
| GHVDHGKTT | ASSAWGGAYEQY | 0.2865115 |
| AQFEHTLL | ASSAWGGAYEQY | 0.34229594 |
| IVNQDEL | ASSAWGGAYEQY | 0.51720816 |
| TNGSQFF | ASSAWGGAYEQY | 0.6529137 |
| TGKGIVE | ASSAWGGAYEQY | 0.28764656 |
| LISNASDA | ASSAWGGAYEQY | 0.57805204 |
| AELRTLGELTKIAKAHGAQ | ASSAWGGAYEQY | 0.5715914 |
| APNAVAAAEPSW | ASSAWGGAYEQY | 0.28959802 |
| APQVAGAATAVL | ASSAWGGAYEQY | 0.5502431 |
| ARRDPAAAAAILPTDA | ASSAWGGAYEQY | 0.6896988 |
| DAAARDVLAEAGLAEYF | ASSAWGGAYEQY | 0.57311285 |

| | | |
|---|---|---|
| DDQIVLSAKANLAKVEQ | ASSAWGGAYEQY | 0.2339585 |
| DELLPDTGEDSDEDGH | ASSAWGGAYEQY | 0.6176204 |
| ELPEAPANRATVPQD | ASSAWGGAYEQY | 0.33852702 |
| FGQSKIERVTGIG | ASSAWGGAYEQY | 0.59276104 |
| GAGGTGSTAGGAGGAGGAGGLY | ASSAWGGAYEQY | 0.5028904 |
| GHGGTGAAGGNGGTGGAGGSGI | ASSAWGGAYEQY | 0.4335421 |
| GNGGNGGAGGEGGPGGAGGAGA | ASSAWGGAYEQY | 0.6836253 |
| GSTAGNGGNGGNATNGTIAGQP | ASSAWGGAYEQY | 0.27245137 |
| GTGGDGGNGGNGGTGG | ASSAWGGAYEQY | 0.36194158 |
| GVRRPAGIPADAPV | ASSAWGGAYEQY | 0.39935315 |
| GYRQVTDAHLLTLARR | ASSAWGGAYEQY | 0.48077887 |
| IAKMGQDGHDRG | ASSAWGGAYEQY | 0.6436031 |
| KAEPSIEIQYT | ASSAWGGAYEQY | 0.2867602 |
| KPVRIRMAADEDTVAVT | ASSAWGGAYEQY | 0.51598966 |
| MPPRIAGMRLLVIKPEP | ASSAWGGAYEQY | 0.40225345 |
| MKFYAVALFRSRGVPPTR | ASSAWGGAYEQY | 0.30437815 |
| NAGGFGILGAVAHSP | ASSAWGGAYEQY | 0.51406634 |
| NEGREITLAMAEM | ASSAWGGAYEQY | 0.36266598 |
| NELVEWSHPLGHRWRWE | ASSAWGGAYEQY | 0.2563896 |
| NLGAGGTGGNGGNPG | ASSAWGGAYEQY | 0.61124337 |
| NTEWPKWETVMAADPVD | ASSAWGGAYEQY | 0.4018482 |
| PASAAPAETQP | ASSAWGGAYEQY | 0.5704045 |
| QQSSDGKNGVRAAEITGE | ASSAWGGAYEQY | 0.720122 |
| RAPDAVTTSWWKE | ASSAWGGAYEQY | 0.28840193 |
| SGFGNVGAHTSGWFNQ | ASSAWGGAYEQY | 0.5595112 |
| SGVAGAVGQAASVGGLKV | ASSAWGGAYEQY | 0.5536875 |
| SNGGAGTGSAGGNGGT | ASSAWGGAYEQY | 0.61881924 |
| TGGAGGEGGAAG | ASSAWGGAYEQY | 0.4722841 |
| TGGDGGAAGAGGAPGHG | ASSAWGGAYEQY | 0.2980781 |
| TMPPLLAAIANAGRHGV | ASSAWGGAYEQY | 0.7132188 |
| TSSTAGNSGGA | ASSAWGGAYEQY | 0.6048053 |
| VPATVPVGILC | ASSAWGGAYEQY | 0.65862703 |
| VQRPDSGGVVLTGRIS | ASSAWGGAYEQY | 0.6810856 |
| WTPNPVPATKGVPAGLVQAT | ASSAWGGAYEQY | 0.64569426 |
| YALKRALAKFSDRFDVVII | ASSAWGGAYEQY | 0.6585917 |
| YLIGSVIV | ASSAWGGAYEQY | 0.38995117 |
| YGQLVAVID | ASSAWGGAYEQY | 0.2507839 |
| VTVGGMTLPA | ASSAWGGAYEQY | 0.4503622 |
| VLALPAV | ASSAWGGAYEQY | 0.4365667 |
| VIPPTLNYET | ASSAWGGAYEQY | 0.3241042 |
| VGMGGMHPSSG | ASSAWGGAYEQY | 0.26797524 |
| VGLAGGAATA | ASSAWGGAYEQY | 0.6957909 |
| VGGAGGAGGSGP | ASSAWGGAYEQY | 0.76408243 |
| VFAAFMV | ASSAWGGAYEQY | 0.41627365 |
| TLYDKLR | ASSAWGGAYEQY | 0.7263262 |
| TLAVFTVL | ASSAWGGAYEQY | 0.2413555 |
| TGSVQPARV | ASSAWGGAYEQY | 0.6526414 |
| TGGGAGSGGT | ASSAWGGAYEQY | 0.70252687 |
| SHLETSL | ASSAWGGAYEQY | 0.30317274 |
| SFTTGMT | ASSAWGGAYEQY | 0.5952346 |
| QVEVAKRMGVRQPTVS | ASSAWGGAYEQY | 0.26523834 |

| | | |
|---|---|---|
| QGSGGPALAM | ASSAWGGAYEQY | 0.3331839 |
| MLAPSNP | ASSAWGGAYEQY | 0.7011181 |
| LVNNAAI | ASSAWGGAYEQY | 0.46365422 |
| LTGEINTLA | ASSAWGGAYEQY | 0.5717667 |
| LSIFGPV | ASSAWGGAYEQY | 0.4333348 |
| LLALLANL | ASSAWGGAYEQY | 0.3307858 |
| LISNASDALDKLR | ASSAWGGAYEQY | 0.55918694 |
| LINEAVP | ASSAWGGAYEQY | 0.67810243 |
| LGYAFSGL | ASSAWGGAYEQY | 0.40074018 |
| LDFVAVAL | ASSAWGGAYEQY | 0.74803793 |
| LAQPGAE | ASSAWGGAYEQY | 0.28611168 |
| ISNLGAVTGF | ASSAWGGAYEQY | 0.7059458 |
| HQEDPGRI | ASSAWGGAYEQY | 0.74261075 |
| GTPGEPTP | ASSAWGGAYEQY | 0.556832 |
| GFFNSGS | ASSAWGGAYEQY | 0.43137574 |
| GAGGGGLTAGAN | ASSAWGGAYEQY | 0.6986511 |
| FVDALVNM | ASSAWGGAYEQY | 0.4416246 |
| FLTVAPEL | ASSAWGGAYEQY | 0.6443012 |
| FETLTAL | ASSAWGGAYEQY | 0.70846367 |
| DVINPIEL | ASSAWGGAYEQY | 0.5646131 |
| APVSGAPV | ASSAWGGAYEQY | 0.8795587 |
| ALVGGLAW | ASSAWGGAYEQY | 0.84383094 |
| ALAAAAATAI | ASSAWGGAYEQY | 0.81841207 |
| AFGLGLLL | ASSAWGGAYEQY | 0.6900881 |
| AFFAAASL | ASSAWGGAYEQY | 0.22196311 |
| AAAALPAFSPP | ASSAWGGAYEQY | 0.7877209 |
| VVLAPAL | ASSAWGGAYEQY | 0.5920801 |
| TIGHVDHGKTTL | ASSAWGGAYEQY | 0.8710145 |
| LISNASDALDK | ASSAWGGAYEQY | 0.87205565 |
| LISNASDAL | ASSAWGGAYEQY | 0.697863 |
| ISNASDALDK | ASSAWGGAYEQY | 0.862025 |
| NQAETLVYQTE | ASSAWGGAYEQY | 0.8676441 |
| TGGTGGTAGAG | ASSAWGGAYEQY | 0.5695606 |
| TGSTAGGAGGA | ASSAWGGAYEQY | 0.7848893 |
| DPLAVPVAAGPV | ASSAWGGAYEQY | 0.47731596 |
| GGTGGTPGNGGN | ASSAWGGAYEQY | 0.8058034 |
| PAGLAGVDPATL | ASSAWGGAYEQY | 0.32102227 |
| GNNLVVEAPHAGASV | ASSAWGGAYEQY | 0.8285798 |
| KVTRIMSASTLEQPA | ASSAWGGAYEQY | 0.7298331 |
| LGFALPIALPRRTVV | ASSAWGGAYEQY | 0.17240196 |
| TRLGNLTARKLVVLV | ASSAWGGAYEQY | 0.84732306 |
| WSQQRRLIYFPSAGP | ASSAWGGAYEQY | 0.8007575 |
| GPNGNPDPMAAAVDIR | ASSAWGGAYEQY | 0.7563375 |
| TQRCVSVADLRAAASR | ASSAWGGAYEQY | 0.8703201 |
| QFVLDVAEHPVEVLTETV | ASSAWGGAYEQY | 0.78715587 |
| GGAGGTGGAGGNGGTQAGDG | ASSAWGGAYEQY | 0.52654445 |
| PLLATLRAIAVDVDAKPAQV | ASSAWGGAYEQY | 0.82033026 |
| APKFRTGGYMIDFWGVGYQVAK | ASSAWGGAYEQY | 0.885494 |
| GAGGAGGAGGAAGGPGGTGGTG | ASSAWGGAYEQY | 0.4421248 |
| IGALGLLATDVAPAGSVAFGVA | ASSAWGGAYEQY | 0.83770496 |
| AGGAGGIGGNGGAGGAGGVLMG | ASSAWGGAYEQY | 0.8804756 |

| | | |
|---|---|---|
| ADELIGLL | ASSAWGGAYEQY | 0.80104893 |
| ADRALKA | ASSAWGGAYEQY | 0.89128155 |
| AGAVGQVGGAGGA | ASSAWGGAYEQY | 0.8935726 |
| AGGGGGLGGAP | ASSAWGGAYEQY | 0.65320605 |
| AGGLLYGNGGAG | ASSAWGGAYEQY | 0.88396376 |
| AGLESIGL | ASSAWGGAYEQY | 0.8577268 |
| AGLKTGII | ASSAWGGAYEQY | 0.77870464 |
| ASLAALLL | ASSAWGGAYEQY | 0.7904233 |
| ASVINISSI | ASSAWGGAYEQY | 0.49512076 |
| ATPPGPA | ASSAWGGAYEQY | 0.76474303 |
| ATVASALV | ASSAWGGAYEQY | 0.3740118 |
| ATVASLIL | ASSAWGGAYEQY | 0.83894706 |
| AVNGPVLIRH | ASSAWGGAYEQY | 0.85858303 |
| DGAAGAVGI | ASSAWGGAYEQY | 0.79173946 |
| DPLLHGI | ASSAWGGAYEQY | 0.58538 |
| ELGAALGAL | ASSAWGGAYEQY | 0.33038118 |
| ELGRTQAV | ASSAWGGAYEQY | 0.7995037 |
| FVAGGYLS | ASSAWGGAYEQY | 0.56389105 |
| GASGGAGGAGGW | ASSAWGGAYEQY | 0.31956244 |
| GGAGAGGDGGAGGI | ASSAWGGAYEQY | 0.7960784 |
| GTGHNGSQPGMGGNG | ASSAWGGAYEQY | 0.8476409 |
| GYAAASASAS | ASSAWGGAYEQY | 0.8644217 |
| HQVDVAAVQQG | ASSAWGGAYEQY | 0.6134913 |
| IGAAIAHT | ASSAWGGAYEQY | 0.8967531 |
| IGMIEVAQL | ASSAWGGAYEQY | 0.8635074 |
| IRLAPEHTSAMD | ASSAWGGAYEQY | 0.87066746 |
| IYNVSVL | ASSAWGGAYEQY | 0.76961064 |
| KQGFLGM | ASSAWGGAYEQY | 0.6057503 |
| LESVITQL | ASSAWGGAYEQY | 0.88395965 |
| LGLGALVVV | ASSAWGGAYEQY | 0.7716857 |
| LIGNGGNGGSGGI | ASSAWGGAYEQY | 0.7489717 |
| LNAINAPTL | ASSAWGGAYEQY | 0.8622475 |
| LQLLNTGV | ASSAWGGAYEQY | 0.67139375 |
| LQTSIEF | ASSAWGGAYEQY | 0.7193669 |
| MSFAVSEL | ASSAWGGAYEQY | 0.81047004 |
| NTGALGL | ASSAWGGAYEQY | 0.8808781 |
| PPGAGGAGGAGTT | ASSAWGGAYEQY | 0.5913096 |
| QLVGLGI | ASSAWGGAYEQY | 0.4661466 |
| RFLEQVHQL | ASSAWGGAYEQY | 0.6676539 |
| RGVGYVLR | ASSAWGGAYEQY | 0.79675967 |
| SATGVFAGVI | ASSAWGGAYEQY | 0.8846299 |
| SLPAITLP | ASSAWGGAYEQY | 0.8390957 |
| STALAFTV | ASSAWGGAYEQY | 0.7771568 |
| SVLTVGLGK | ASSAWGGAYEQY | 0.85272515 |
| TALAGAGSGI | ASSAWGGAYEQY | 0.85485935 |
| TGLPFTGI | ASSAWGGAYEQY | 0.8115614 |
| VAEHVTV | ASSAWGGAYEQY | 0.82455206 |
| VDIVLNSL | ASSAWGGAYEQY | 0.74278545 |
| VLTGGGAVL | ASSAWGGAYEQY | 0.8834182 |
| VNVGLAYGI | ASSAWGGAYEQY | 0.8595225 |
| VSSAAVGAV | ASSAWGGAYEQY | 0.8800542 |

| | | |
|---|---|---|
| VTSVYEAGV | ASSAWGGAYEQY | 0.88831675 |
| WVALDRGA | ASSAWGGAYEQY | 0.77817726 |
| YLVGSVVI | ASSAWGGAYEQY | 0.7893686 |
| GPPGTGKTLL | ASSAWGGAYEQY | 0.33348683 |
| GVGMTKF | ASSAWGGAYEQY | 0.71204257 |
| IGIGGSDL | ASSAWGGAYEQY | 0.82715946 |
| IGTLLAEL | ASSAWGGAYEQY | 0.8341257 |
| IYTSGTT | ASSAWGGAYEQY | 0.8650874 |
| LSFPAGV | ASSAWGGAYEQY | 0.83433306 |
| LYTSGTT | ASSAWGGAYEQY | 0.68798715 |
| VGAFPFGF | ASSAWGGAYEQY | 0.85792637 |
| VGAGTVEF | ASSAWGGAYEQY | 0.88213545 |
| VGKSTLLL | ASSAWGGAYEQY | 0.5537884 |
| AALAHAGGGIV | ASSAWGGAYEQY | 0.8157971 |
| AALVAVSLL | ASSAWGGAYEQY | 0.89753544 |
| AAVSRAVAAL | ASSAWGGAYEQY | 0.85529363 |
| AGGSSGIGGF | ASSAWGGAYEQY | 0.7198061 |
| AGLTVANANG | ASSAWGGAYEQY | 0.7401277 |
| AGLVGAVL | ASSAWGGAYEQY | 0.8899397 |
| ALSQAFTV | ASSAWGGAYEQY | 0.7811441 |
| ALTFSVSL | ASSAWGGAYEQY | 0.8758052 |
| ALTGAALVAL | ASSAWGGAYEQY | 0.26463228 |
| ASFSIGAGVL | ASSAWGGAYEQY | 0.8031366 |
| DGSVYKI | ASSAWGGAYEQY | 0.43841916 |
| EQLAEIV | ASSAWGGAYEQY | 0.2466912 |
| FAVSELTVM | ASSAWGGAYEQY | 0.86423004 |
| GAAPLAAGL | ASSAWGGAYEQY | 0.7143854 |
| GADGGDGGSGGAGG | ASSAWGGAYEQY | 0.8648471 |
| GFGLDTGGAGG | ASSAWGGAYEQY | 0.74108505 |
| GLASGGLL | ASSAWGGAYEQY | 0.88486195 |
| GRIAGAAVLL | ASSAWGGAYEQY | 0.82939696 |
| GSAGSLNTSF | ASSAWGGAYEQY | 0.79467595 |
| GTGQIGI | ASSAWGGAYEQY | 0.7231711 |
| GTLIALIL | ASSAWGGAYEQY | 0.8128255 |
| IAATMGFVVG | ASSAWGGAYEQY | 0.8710313 |
| IFAQALL | ASSAWGGAYEQY | 0.87287116 |
| IGAAVALGF | ASSAWGGAYEQY | 0.20756629 |
| IGLTPLAL | ASSAWGGAYEQY | 0.83857346 |
| ILGSNTSTL | ASSAWGGAYEQY | 0.7601313 |
| IVFSLTVL | ASSAWGGAYEQY | 0.2876824 |
| LASKATGF | ASSAWGGAYEQY | 0.22269279 |
| LASVALIL | ASSAWGGAYEQY | 0.64872754 |
| LFAVGLTL | ASSAWGGAYEQY | 0.7932075 |
| LFNGGIGTYI | ASSAWGGAYEQY | 0.8605103 |
| LTLGAGII | ASSAWGGAYEQY | 0.8521119 |
| LVAAALGDI | ASSAWGGAYEQY | 0.7734829 |
| NTAFVQLGI | ASSAWGGAYEQY | 0.84509635 |
| QLFSGAEL | ASSAWGGAYEQY | 0.7734567 |
| SAATFITL | ASSAWGGAYEQY | 0.106196694 |
| SNYGLANLGV | ASSAWGGAYEQY | 0.64558697 |
| SYGGAYAVM | ASSAWGGAYEQY | 0.5434328 |

| | | |
|---|---|---|
| TTGVGGLTLG | ASSAWGGAYEQY | 0.89104235 |
| TTQTIYI | ASSAWGGAYEQY | 0.7531225 |
| VAAGAGLGAVI | ASSAWGGAYEQY | 0.76383466 |
| VAAVVTLL | ASSAWGGAYEQY | 0.8146437 |
| VGFLGGLI | ASSAWGGAYEQY | 0.8530448 |
| VGVIPILL | ASSAWGGAYEQY | 0.8148171 |
| VIFASSLI | ASSAWGGAYEQY | 0.5843467 |
| VKGGDGGAAAG | ASSAWGGAYEQY | 0.22418319 |
| VTKTFTNI | ASSAWGGAYEQY | 0.80972165 |
| VVAAAILL | ASSAWGGAYEQY | 0.5381481 |
| YAALDVEL | ASSAWGGAYEQY | 0.4783787 |
| LASVALLL | ASSAWGGAYEQY | 0.6164335 |
| LDAVHRV | ASSAWGGAYEQY | 0.6578274 |
| QGLSSSIF | ASSAWGGAYEQY | 0.27266592 |
| RGTFANIRL | ASSAWGGAYEQY | 0.88277155 |
| RQLWWGHRI | ASSAWGGAYEQY | 0.69144267 |
| YGPPGTGKTLL | ASSAWGGAYEQY | 0.7718301 |
| AALVGAPANVA | ASSAWGGAYEQY | 0.88570154 |
| AATFRRTE | ASSAWGGAYEQY | 0.83741605 |
| AGALGGGI | ASSAWGGAYEQY | 0.78872585 |
| AGGAGGAGGIGAG | ASSAWGGAYEQY | 0.606369 |
| AGSGALGL | ASSAWGGAYEQY | 0.68330467 |
| AQIADQI | ASSAWGGAYEQY | 0.5436846 |
| ASAVAVTYL | ASSAWGGAYEQY | 0.52450967 |
| ASTVLGGTGG | ASSAWGGAYEQY | 0.88431394 |
| AVGGEPAK | ASSAWGGAYEQY | 0.8241579 |
| CEQGTFGGTC | ASSAWGGAYEQY | 0.5987899 |
| DEGKAPAG | ASSAWGGAYEQY | 0.8720534 |
| DTVVFPAL | ASSAWGGAYEQY | 0.8782405 |
| FEGITMAG | ASSAWGGAYEQY | 0.889964 |
| FPLVVQD | ASSAWGGAYEQY | 0.69602966 |
| GAFTTPQ | ASSAWGGAYEQY | 0.613222 |
| GFGGFGTAGGVGGA | ASSAWGGAYEQY | 0.74521446 |
| GFLAQFL | ASSAWGGAYEQY | 0.8051176 |
| GGAGGAGGNAGVA | ASSAWGGAYEQY | 0.73208153 |
| GGVGTAGAGGAGGAGG | ASSAWGGAYEQY | 0.55400753 |
| GNGGGGDAGNAGSGGN | ASSAWGGAYEQY | 0.8735727 |
| GTGGAGGAGNDAGS | ASSAWGGAYEQY | 0.9004339 |
| GTTNTGWFNS | ASSAWGGAYEQY | 0.7288023 |
| IFMGPLFTL | ASSAWGGAYEQY | 0.2239385 |
| ILAHAIEAI | ASSAWGGAYEQY | 0.7807995 |
| IRSAIREAL | ASSAWGGAYEQY | 0.57427096 |
| ITFAGGG | ASSAWGGAYEQY | 0.7218457 |
| ITGMSGHQEV | ASSAWGGAYEQY | 0.7150967 |
| IVAGLACTL | ASSAWGGAYEQY | 0.57366276 |
| IVGGGLAGA | ASSAWGGAYEQY | 0.7603262 |
| IVTGVLNI | ASSAWGGAYEQY | 0.88249415 |
| LAQQGCI | ASSAWGGAYEQY | 0.745065 |
| LCLHVEN | ASSAWGGAYEQY | 0.84670615 |
| LDAISDGS | ASSAWGGAYEQY | 0.8827819 |
| LGGFAAML | ASSAWGGAYEQY | 0.86144805 |

| | | |
|---|---|---|
| LGGKNPMI | ASSAWGGAYEQY | 0.8892925 |
| LGPKGVAAIL | ASSAWGGAYEQY | 0.8815828 |
| LMVGVSI | ASSAWGGAYEQY | 0.79689586 |
| LVGAAFVL | ASSAWGGAYEQY | 0.7499967 |
| LVLAVAPL | ASSAWGGAYEQY | 0.8706243 |
| MAAIAVIDAA | ASSAWGGAYEQY | 0.7888677 |
| NNGINAELQ | ASSAWGGAYEQY | 0.8015238 |
| RGEPLVLVHG | ASSAWGGAYEQY | 0.78374183 |
| RGGAGGAAGLI | ASSAWGGAYEQY | 0.86262643 |
| RSYRMHPAV | ASSAWGGAYEQY | 0.7761369 |
| SAAHLLAGQLA | ASSAWGGAYEQY | 0.8891019 |
| SAAVGSY | ASSAWGGAYEQY | 0.25563323 |
| SPHAVLHRI | ASSAWGGAYEQY | 0.5336409 |
| SQIAKVL | ASSAWGGAYEQY | 0.88236487 |
| TALGHLVP | ASSAWGGAYEQY | 0.10440553 |
| TAPTTSGAA | ASSAWGGAYEQY | 0.34252423 |
| TDGDDGNGGSPGT | ASSAWGGAYEQY | 0.8233973 |
| TDGLGEN | ASSAWGGAYEQY | 0.8697269 |
| TGGPPLPGQA | ASSAWGGAYEQY | 0.14052674 |
| TIPAITI | ASSAWGGAYEQY | 0.64325005 |
| TIPGFTI | ASSAWGGAYEQY | 0.38395327 |
| TSIASGALA | ASSAWGGAYEQY | 0.86970705 |
| TVTLPGAAI | ASSAWGGAYEQY | 0.76529026 |
| VASGLSAALP | ASSAWGGAYEQY | 0.41068202 |
| VGAGVIGI | ASSAWGGAYEQY | 0.8705011 |
| VGPKRFGV | ASSAWGGAYEQY | 0.39407402 |
| VNVAFPD | ASSAWGGAYEQY | 0.8118148 |
| VSMTNSGVS | ASSAWGGAYEQY | 0.8762874 |
| VTDVLGDLSV | ASSAWGGAYEQY | 0.8953769 |
| VVVVGAGFAGL | ASSAWGGAYEQY | 0.5954102 |
| VYTVGTVV | ASSAWGGAYEQY | 0.8669207 |
| WTTFSGP | ASSAWGGAYEQY | 0.715783 |
| YDELVEAF | ASSAWGGAYEQY | 0.8963525 |
| YGTGGAGGNGG | ASSAWGGAYEQY | 0.7048069 |
| YTNVMAV | ASSAWGGAYEQY | 0.4715852 |
| FLEQVHQL | ASSAWGGAYEQY | 0.86816025 |
| GLSLLAVAV | ASSAWGGAYEQY | 0.87271154 |
| LEQVHQL | ASSAWGGAYEQY | 0.67352426 |
| LNAAGAI | ASSAWGGAYEQY | 0.8744395 |
| PAVHASL | ASSAWGGAYEQY | 0.8133956 |
| TELEDDA | ASSAWGGAYEQY | 0.7940493 |
| TPPGPPN | ASSAWGGAYEQY | 0.883401 |
| TPPPPPPP | ASSAWGGAYEQY | 0.8664138 |
| TQSPALI | ASSFGRVEDTQY | 0.53246397 |
| EFAAAFPQ | ASSFGRVEDTQY | 0.8772087 |
| LLSGNLDVL | ASSFGRVEDTQY | 0.78685766 |
| GPANLGNYNI | ASSFGRVEDTQY | 0.5181377 |
| TLGPTLP | ASSFGRVEDTQY | 0.8938352 |
| VYAYSSCCK | ASSFGRVEDTQY | 0.82984024 |
| APAPAPAPNI | ASSFGRVEDTQY | 0.8650619 |
| PGGPVPIP | ASSFGRVEDTQY | 0.80831635 |

| | | |
|---|---|---|
| FLTVALEP | ASSFGRVEDTQY | 0.8902197 |
| SQYFGANL | ASSFGRVEDTQY | 0.39876288 |
| GGVKPAA | ASSFGRVEDTQY | 0.82521737 |
| GGTGGAGGAGGW | ASSFGRVEDTQY | 0.77960545 |
| GVGGVAGLGGAGG | ASSFGRVEDTQY | 0.8823595 |
| VLLVLATTI | ASSFGRVEDTQY | 0.8507368 |
| IGIFAAML | ASSFGRVEDTQY | 0.6382708 |
| AVGDLID | ASSFGRVEDTQY | 0.842281 |
| NISPLPL | ASSFGRVEDTQY | 0.87460494 |
| VTAVFVAI | ASSFGRVEDTQY | 0.8106574 |
| SGDDNDAAL | ASSFGRVEDTQY | 0.80286837 |
| AVAALAHS | ASSFGRVEDTQY | 0.8937868 |
| GGVTVMATP | ASSFGRVEDTQY | 0.59016454 |
| AGLAANVRD | ASSFGRVEDTQY | 0.78575486 |
| TNRPDLIDPALLRPG | ASSFGRVEDTQY | 0.8587346 |
| LANLGVTAL | ASSFGRVEDTQY | 0.87086535 |
| TGERIEL | ASSFGRVEDTQY | 0.832625 |
| NGEELAA | ASSFGRVEDTQY | 0.8833672 |
| VLSLVWG | ASSFGRVEDTQY | 0.63772494 |
| IGYALWL | ASSFGRVEDTQY | 0.86577344 |
| LSLLLAPL | ASSFGRVEDTQY | 0.7749387 |
| TLDEALK | ASSFGRVEDTQY | 0.7868919 |
| TDGAGTAEGG | ASSFGRVEDTQY | 0.5700915 |
| NAPGGAGGNGG | ASSFGRVEDTQY | 0.80375576 |
| VSVFMGMGGNF | ASSFGRVEDTQY | 0.8627829 |
| LASYLAAL | ASSFGRVEDTQY | 0.85158986 |
| LVGGTPAL | ASSFGRVEDTQY | 0.24474758 |
| VRNADVL | ASSFGRVEDTQY | 0.8655349 |
| TCSSPFLVSAM | ASSFGRVEDTQY | 0.86354876 |
| VHITDAE | ASSFGRVEDTQY | 0.8970326 |
| EYAAPSF | ASSFGRVEDTQY | 0.75227267 |
| VVDAFHGGL | ASSFGRVEDTQY | 0.85454273 |
| ASVVDPQ | ASSFGRVEDTQY | 0.6579072 |
| SYNEFGGLS | ASSFGRVEDTQY | 0.83463603 |
| DPDGSVY | ASSFGRVEDTQY | 0.48653188 |
| MVSTIHATE | ASSFGRVEDTQY | 0.5998349 |
| LAAGVADPV | ASSFGRVEDTQY | 0.79977715 |
| TKEQIAA | ASSFGRVEDTQY | 0.7371017 |
| SLASPAAL | ASSFGRVEDTQY | 0.86659026 |
| IGLVFYGV | ASSFGRVEDTQY | 0.87387055 |
| FLLDASNL | ASSFGRVEDTQY | 0.8546068 |
| YQVGNFP | ASSFGRVEDTQY | 0.8700214 |
| ESLLSQST | ASSFGRVEDTQY | 0.8892755 |
| LFTVGFAAVG | ASSFGRVEDTQY | 0.8893354 |
| ATPPTPT | ASSFGRVEDTQY | 0.6701007 |
| TDLATLG | ASSFGRVEDTQY | 0.79668653 |
| SAGGTGR | ASSFGRVEDTQY | 0.5592258 |
| GFSNSGS | ASSFGRVEDTQY | 0.7722163 |
| GGNGGSGGTSV | ASSFGRVEDTQY | 0.6837927 |
| GTSVATGGAGN | ASSFGRVEDTQY | 0.84013814 |
| NGGNGGSGGTS | ASSFGRVEDTQY | 0.105805606 |

| | | |
|---|---|---|
| AVVKFPH | ASSFGRVEDTQY | 0.83838177 |
| TPPPPPP | ASSFGRVEDTQY | 0.8872086 |
| AAPAAAAAPA | ASSFGRVEDTQY | 0.5937918 |
| IALNEDL | ASSFGRVEDTQY | 0.8289576 |
| IVNVSSM | ASSFGRVEDTQY | 0.8610182 |
| TGIPPAPR | ASSFGRVEDTQY | 0.71111727 |
| FDLGGGTF | ASSFGRVEDTQY | 0.83340704 |
| FDLGGGTFDVS | ASSFGRVEDTQY | 0.63639146 |
| NEPTAAA | ASSFGRVEDTQY | 0.872259 |
| NEPTAAALA | ASSFGRVEDTQY | 0.8774754 |
| GSIINISSI | ASSFGRVEDTQY | 0.808365 |
| VAPGNAGTA | ASSFGRVEDTQY | 0.69569206 |
| TGGGASLE | ASSFGRVEDTQY | 0.8066036 |
| VAVYNHYKRI | ASSFGRVEDTQY | 0.24089724 |
| TDPAPATT | ASSFGRVEDTQY | 0.77018684 |
| TNRPDILDPALLRPG | ASSFGRVEDTQY | 0.19995797 |
| LTTTTTS | ASSFGRVEDTQY | 0.823154 |
| PTGSGKTL | ASSFGRVEDTQY | 0.4169236 |
| TGSGKTL | ASSFGRVEDTQY | 0.8281605 |
| TIGHVDHGK | ASSFGRVEDTQY | 0.78158 |
| TIGHVDHGKTT | ASSFGRVEDTQY | 0.8227638 |
| GHVDHGKTT | ASSFGRVEDTQY | 0.4789381 |
| AQFEHTLL | ASSFGRVEDTQY | 0.8539471 |
| IVNQDEL | ASSFGRVEDTQY | 0.7767298 |
| TNGSQFF | ASSFGRVEDTQY | 0.30266404 |
| TGKGIVE | ASSFGRVEDTQY | 0.8811634 |
| LISNASDA | ASSFGRVEDTQY | 0.75308526 |
| AELRTLGELTKIAKAHGAQ | ASSFGRVEDTQY | 0.814538 |
| APNAVAAAEPSW | ASSFGRVEDTQY | 0.83769894 |
| APQVAGAATAVL | ASSFGRVEDTQY | 0.78923243 |
| ARRDPAAAAAILPTDA | ASSFGRVEDTQY | 0.28947207 |
| DAAARDVLAEAGLAEYF | ASSFGRVEDTQY | 0.74964225 |
| DDQIVLSAKANLAKVEQ | ASSFGRVEDTQY | 0.7672842 |
| DELLPDTGEDSDEDGH | ASSFGRVEDTQY | 0.68551326 |
| ELPEAPANRATVPQD | ASSFGRVEDTQY | 0.20314375 |
| FGQSKIERVTGIG | ASSFGRVEDTQY | 0.86766297 |
| GAGGTGSTAGGAGGAGGAGGLY | ASSFGRVEDTQY | 0.5093647 |
| GHGGTGAAGGNGGTGGAGGSGI | ASSFGRVEDTQY | 0.7100562 |
| GNGGNGGAGGEGGPGGAGGAGA | ASSFGRVEDTQY | 0.8068115 |
| GSTAGNGGNGGNATNGTIAGQP | ASSFGRVEDTQY | 0.8769096 |
| GTGGDGGNGGNGGTGG | ASSFGRVEDTQY | 0.7798499 |
| GVRRPAGIPADAPV | ASSFGRVEDTQY | 0.6711008 |
| GYRQVTDAHLLTLARR | ASSFGRVEDTQY | 0.8587718 |
| IAKMGQDGHDRG | ASSFGRVEDTQY | 0.74511725 |
| KAEPSIEIQYT | ASSFGRVEDTQY | 0.8815173 |
| KPVRIRMAADEDTVAVT | ASSFGRVEDTQY | 0.82320285 |
| MPPRIAGMRLLVIKPEP | ASSFGRVEDTQY | 0.6375625 |
| MKFYAVALFRSRGVPPTR | ASSFGRVEDTQY | 0.813796 |
| NAGGFGILGAVAHSP | ASSFGRVEDTQY | 0.7153598 |
| NEGREITLAMAEM | ASSFGRVEDTQY | 0.5867814 |
| NELVEWSHPLGHRWRWE | ASSFGRVEDTQY | 0.48053044 |

| | | |
|---|---|---|
| NLGAGGTGGNGGNPG | ASSFGRVEDTQY | 0.54582644 |
| NTEWPKWETVMAADPVD | ASSFGRVEDTQY | 0.38845485 |
| PASAAPAETQP | ASSFGRVEDTQY | 0.80830264 |
| QQSSDGKNGVRAAEITGE | ASSFGRVEDTQY | 0.89597785 |
| RAPDAVTTSWWKE | ASSFGRVEDTQY | 0.82504654 |
| SGFGNVGAHTSGWFNQ | ASSFGRVEDTQY | 0.3554578 |
| SGVAGAVGQAASVGGLKV | ASSFGRVEDTQY | 0.73251766 |
| SNGGAGTGSAGGNGGT | ASSFGRVEDTQY | 0.73066753 |
| TGGAGGEGGAAG | ASSFGRVEDTQY | 0.8560828 |
| TGGDGGAAGAGGAPGHG | ASSFGRVEDTQY | 0.81942123 |
| TMPPLLAAIANAGRHGV | ASSFGRVEDTQY | 0.8975347 |
| TSSTAGNSGGA | ASSFGRVEDTQY | 0.86073256 |
| VPATVPVGILC | ASSFGRVEDTQY | 0.7022298 |
| VQRPDSGGVVLTGRIS | ASSFGRVEDTQY | 0.8731289 |
| WTPNPVPATKGVPAGLVQAT | ASSFGRVEDTQY | 0.78772247 |
| YALKRALAKFSDRFDVVII | ASSFGRVEDTQY | 0.8008011 |
| YLIGSVIV | ASSFGRVEDTQY | 0.82265985 |
| YGQLVAVID | ASSFGRVEDTQY | 0.84619963 |
| VTVGGMTLPA | ASSFGRVEDTQY | 0.77631176 |
| VLALPAV | ASSFGRVEDTQY | 0.7378197 |
| VIPPTLNYET | ASSFGRVEDTQY | 0.27531606 |
| VGMGGMHPSSG | ASSFGRVEDTQY | 0.8561386 |
| VGLAGGAATA | ASSFGRVEDTQY | 0.31902066 |
| VGGAGGAGGSGP | ASSFGRVEDTQY | 0.54971087 |
| VFAAFMV | ASSFGRVEDTQY | 0.7635744 |
| TLYDKLR | ASSFGRVEDTQY | 0.89711916 |
| TLAVFTVL | ASSFGRVEDTQY | 0.87901497 |
| TGSVQPARV | ASSFGRVEDTQY | 0.6665153 |
| TGGGAGSGGT | ASSFGRVEDTQY | 0.532964 |
| SHLETSL | ASSFGRVEDTQY | 0.8501225 |
| SFTTGMT | ASSFGRVEDTQY | 0.61907065 |
| QVEVAKRMGVRQPTVS | ASSFGRVEDTQY | 0.8749229 |
| QGSGGPALAM | ASSFGRVEDTQY | 0.8215682 |
| MLAPSNP | ASSFGRVEDTQY | 0.78774375 |
| LVNNAAI | ASSFGRVEDTQY | 0.62617505 |
| LTGEINTLA | ASSFGRVEDTQY | 0.8483352 |
| LSIFGPV | ASSFGRVEDTQY | 0.6698421 |
| LLALLANL | ASSFGRVEDTQY | 0.8683253 |
| LISNASDALDKLR | ASSFGRVEDTQY | 0.14873973 |
| LINEAVP | ASSFGRVEDTQY | 0.8806786 |
| LGYAFSGL | ASSFGRVEDTQY | 0.6558578 |
| LDFVAVAL | ASSFGRVEDTQY | 0.5314092 |
| LAQPGAE | ASSFGRVEDTQY | 0.31106013 |
| ISNLGAVTGF | ASSFGRVEDTQY | 0.7735175 |
| HQEDPGRI | ASSFGRVEDTQY | 0.84013665 |
| GTPGEPTP | ASSFGRVEDTQY | 0.87972134 |
| GFFNSGS | ASSFGRVEDTQY | 0.7840867 |
| GAGGGGLTAGAN | ASSFGRVEDTQY | 0.8563914 |
| FVDALVNM | ASSFGRVEDTQY | 0.86858726 |
| FLTVAPEL | ASSFGRVEDTQY | 0.7654232 |
| FETLTAL | ASSFGRVEDTQY | 0.26330274 |

| | | |
|---|---|---|
| DVINPIEL | ASSFGRVEDTQY | 0.83806133 |
| APVSGAPV | ASSFGRVEDTQY | 0.8250983 |
| ALVGGLAW | ASSFGRVEDTQY | 0.88427556 |
| ALAAAAATAI | ASSFGRVEDTQY | 0.8132585 |
| AFGLGLLL | ASSFGRVEDTQY | 0.6159096 |
| AFFAAASL | ASSFGRVEDTQY | 0.7897256 |
| AAAALPAFSPP | ASSFGRVEDTQY | 0.87065506 |
| VVLAPAL | ASSFGRVEDTQY | 0.58118576 |
| TIGHVDHGKTTL | ASSFGRVEDTQY | 0.65897185 |
| LISNASDALDK | ASSFGRVEDTQY | 0.66319704 |
| LISNASDAL | ASSFGRVEDTQY | 0.648633 |
| ISNASDALDK | ASSFGRVEDTQY | 0.7305716 |
| NQAETLVYQTE | ASSFGRVEDTQY | 0.74632186 |
| TGGTGGTAGAG | ASSFGRVEDTQY | 0.6842423 |
| TGSTAGGAGGA | ASSFGRVEDTQY | 0.76406634 |
| DPLAVPVAAGPV | ASSFGRVEDTQY | 0.49846506 |
| GGTGGTPGNGGN | ASSFGRVEDTQY | 0.6885391 |
| PAGLAGVDPATL | ASSFGRVEDTQY | 0.6751317 |
| GNNLVVEAPHAGASV | ASSFGRVEDTQY | 0.7707222 |
| KVTRIMSASTLEQPA | ASSFGRVEDTQY | 0.8571944 |
| LGFALPIALPRRTVV | ASSFGRVEDTQY | 0.7653803 |
| TRLGNLTARKLVVLV | ASSFGRVEDTQY | 0.8367736 |
| WSQQRRLIYFPSAGP | ASSFGRVEDTQY | 0.8838933 |
| GPNGNPDPMAAAVDIR | ASSFGRVEDTQY | 0.31147102 |
| TQRCVSVADLRAAASR | ASSFGRVEDTQY | 0.5872368 |
| QFVLDVAEHPVEVLTETV | ASSFGRVEDTQY | 0.8650127 |
| GGAGGTGGAGGNGGTQAGDG | ASSFGRVEDTQY | 0.5549128 |
| PLLATLRAIAVDVDAKPAQV | ASSFGRVEDTQY | 0.8797924 |
| APKFRTGGYMIDFWGVGYQVAK | ASSFGRVEDTQY | 0.78457206 |
| GAGGAGGAGGAAGGPGGTGGTG | ASSFGRVEDTQY | 0.8487363 |
| IGALGLLATDVAPAGSVAFGVA | ASSFGRVEDTQY | 0.7895257 |
| AGGAGGIGGNGGAGGAGGVLMG | ASSFGRVEDTQY | 0.8949843 |
| ADELIGLL | ASSFGRVEDTQY | 0.8556141 |
| ADRALKA | ASSFGRVEDTQY | 0.7959431 |
| AGAVGQVGGAGGA | ASSFGRVEDTQY | 0.85833746 |
| AGGGGGLGGAP | ASSFGRVEDTQY | 0.47357672 |
| AGGLLYGNGGAG | ASSFGRVEDTQY | 0.86591065 |
| AGLESIGL | ASSFGRVEDTQY | 0.87064564 |
| AGLKTGII | ASSFGRVEDTQY | 0.6370255 |
| ASLAALLL | ASSFGRVEDTQY | 0.79270315 |
| ASVINISSI | ASSFGRVEDTQY | 0.88202965 |
| ATPPGPA | ASSFGRVEDTQY | 0.8885198 |
| ATVASALV | ASSFGRVEDTQY | 0.38121796 |
| ATVASLIL | ASSFGRVEDTQY | 0.779914 |
| AVNGPVLIRH | ASSFGRVEDTQY | 0.43778622 |
| DGAAGAVGI | ASSFGRVEDTQY | 0.47975528 |
| DPLLHGI | ASSFGRVEDTQY | 0.31167722 |
| ELGAALGAL | ASSFGRVEDTQY | 0.85278463 |
| ELGRTQAV | ASSFGRVEDTQY | 0.7214364 |
| FVAGGYLS | ASSFGRVEDTQY | 0.8743125 |
| GASGGAGGAGGW | ASSFGRVEDTQY | 0.8223145 |

| | | |
|---|---|---|
| GGAGAGGDGGAGGI | ASSFGRVEDTQY | 0.8700454 |
| GTGHNGSQPGMGGNG | ASSFGRVEDTQY | 0.6365385 |
| GYAAASASAS | ASSFGRVEDTQY | 0.7346459 |
| HQVDVAAVQQG | ASSFGRVEDTQY | 0.47371703 |
| IGAAIAHT | ASSFGRVEDTQY | 0.22239853 |
| IGMIEVAQL | ASSFGRVEDTQY | 0.62089103 |
| IRLAPEHTSAMD | ASSFGRVEDTQY | 0.72059524 |
| IYNVSVL | ASSFGRVEDTQY | 0.16021504 |
| KQGFLGM | ASSFGRVEDTQY | 0.89055014 |
| LESVITQL | ASSFGRVEDTQY | 0.78680724 |
| LGLGALVVV | ASSFGRVEDTQY | 0.8348727 |
| LIGNGGNGGSGGI | ASSFGRVEDTQY | 0.615192 |
| LNAINAPTL | ASSFGRVEDTQY | 0.60942537 |
| LQLLNTGV | ASSFGRVEDTQY | 0.7134977 |
| LQTSIEF | ASSFGRVEDTQY | 0.85432184 |
| MSFAVSEL | ASSFGRVEDTQY | 0.70643103 |
| NTGALGL | ASSFGRVEDTQY | 0.21403465 |
| PPGAGGAGGAGTT | ASSFGRVEDTQY | 0.63531643 |
| QLVGLGI | ASSFGRVEDTQY | 0.5315771 |
| RFLEQVHQL | ASSFGRVEDTQY | 0.88965 |
| RGVGYVLR | ASSFGRVEDTQY | 0.86951673 |
| SATGVFAGVI | ASSFGRVEDTQY | 0.84163845 |
| SLPAITLP | ASSFGRVEDTQY | 0.8925402 |
| STALAFTV | ASSFGRVEDTQY | 0.14377473 |
| SVLTVGLGK | ASSFGRVEDTQY | 0.6312144 |
| TALAGAGSGI | ASSFGRVEDTQY | 0.8898412 |
| TGLPFTGI | ASSFGRVEDTQY | 0.793352 |
| VAEHVTV | ASSFGRVEDTQY | 0.7429646 |
| VDIVLNSL | ASSFGRVEDTQY | 0.8856429 |
| VLTGGGAVL | ASSFGRVEDTQY | 0.6012113 |
| VNVGLAYGI | ASSFGRVEDTQY | 0.67104983 |
| VSSAAVGAV | ASSFGRVEDTQY | 0.80880845 |
| VTSVYEAGV | ASSFGRVEDTQY | 0.8635862 |
| WVALDRGA | ASSFGRVEDTQY | 0.85958564 |
| YLVGSVVI | ASSFGRVEDTQY | 0.524389 |
| GPPGTGKTLL | ASSFGRVEDTQY | 0.8455013 |
| GVGMTKF | ASSFGRVEDTQY | 0.8926277 |
| IGIGGSDL | ASSFGRVEDTQY | 0.7588066 |
| IGTLLAEL | ASSFGRVEDTQY | 0.8414415 |
| IYTSGTT | ASSFGRVEDTQY | 0.82910323 |
| LSFPAGV | ASSFGRVEDTQY | 0.58927447 |
| LYTSGTT | ASSFGRVEDTQY | 0.6978618 |
| VGAFPFGF | ASSFGRVEDTQY | 0.63018954 |
| VGAGTVEF | ASSFGRVEDTQY | 0.8447085 |
| VGKSTLLL | ASSFGRVEDTQY | 0.6897335 |
| AALAHAGGGIV | ASSFGRVEDTQY | 0.8273582 |
| AALVAVSLL | ASSFGRVEDTQY | 0.80788 |
| AAVSRAVAAL | ASSFGRVEDTQY | 0.89899325 |
| AGGSSGIGGF | ASSFGRVEDTQY | 0.8382709 |
| AGLTVANANG | ASSFGRVEDTQY | 0.8669628 |
| AGLVGAVL | ASSFGRVEDTQY | 0.6676948 |

| | | |
|---|---|---|
| ALSQAFTV | ASSFGRVEDTQY | 0.8275129 |
| ALTFSVSL | ASSFGRVEDTQY | 0.84037954 |
| ALTGAALVAL | ASSFGRVEDTQY | 0.29903007 |
| ASFSIGAGVL | ASSFGRVEDTQY | 0.8711331 |
| DGSVYKI | ASSFGRVEDTQY | 0.62872255 |
| EQLAEIV | ASSFGRVEDTQY | 0.82044196 |
| FAVSELTVM | ASSFGRVEDTQY | 0.861437 |
| GAAPLAAGL | ASSFGRVEDTQY | 0.8824098 |
| GADGGDGGSGGAGG | ASSFGRVEDTQY | 0.8229999 |
| GFGLDTGGAGG | ASSFGRVEDTQY | 0.8109671 |
| GLASGGLL | ASSFGRVEDTQY | 0.58892196 |
| GRIAGAAVLL | ASSFGRVEDTQY | 0.8706029 |
| GSAGSLNTSF | ASSFGRVEDTQY | 0.8803256 |
| GTGQIGI | ASSFGRVEDTQY | 0.82996756 |
| GTLIALIL | ASSFGRVEDTQY | 0.8962329 |
| IAATMGFVVG | ASSFGRVEDTQY | 0.8736198 |
| IFAQALL | ASSFGRVEDTQY | 0.2710622 |
| IGAAVALGF | ASSFGRVEDTQY | 0.75581545 |
| IGLTPLAL | ASSFGRVEDTQY | 0.88825035 |
| ILGSNTSTL | ASSFGRVEDTQY | 0.877465 |
| IVFSLTVL | ASSFGRVEDTQY | 0.7817656 |
| LASKATGF | ASSFGRVEDTQY | 0.8625513 |
| LASVALIL | ASSFGRVEDTQY | 0.67976886 |
| LFAVGLTL | ASSFGRVEDTQY | 0.88520944 |
| LFNGGIGTYI | ASSFGRVEDTQY | 0.89656734 |
| LTLGAGII | ASSFGRVEDTQY | 0.8681678 |
| LVAAALGDI | ASSFGRVEDTQY | 0.76899916 |
| NTAFVQLGI | ASSFGRVEDTQY | 0.64667773 |
| QLFSGAEL | ASSFGRVEDTQY | 0.8135926 |
| SAATFITL | ASSFGRVEDTQY | 0.885448 |
| SNYGLANLGV | ASSFGRVEDTQY | 0.57828945 |
| SYGGAYAVM | ASSFGRVEDTQY | 0.8387004 |
| TTGVGGLTLG | ASSFGRVEDTQY | 0.7819843 |
| TTQTIYI | ASSFGRVEDTQY | 0.22202867 |
| VAAGAGLGAVI | ASSFGRVEDTQY | 0.81439626 |
| VAAVVTLL | ASSFGRVEDTQY | 0.8868932 |
| VGFLGGLI | ASSFGRVEDTQY | 0.8431118 |
| VGVIPILL | ASSFGRVEDTQY | 0.568235 |
| VIFASSLI | ASSFGRVEDTQY | 0.87658787 |
| VKGGDGGAAAG | ASSFGRVEDTQY | 0.89152825 |
| VTKTFTNI | ASSFGRVEDTQY | 0.8903315 |
| VVAAAILL | ASSFGRVEDTQY | 0.3518961 |
| YAALDVEL | ASSFGRVEDTQY | 0.19529253 |
| LASVALLL | ASSFGRVEDTQY | 0.8229736 |
| LDAVHRV | ASSFGRVEDTQY | 0.85937726 |
| QGLSSSIF | ASSFGRVEDTQY | 0.61356956 |
| RGTFANIRL | ASSFGRVEDTQY | 0.59183407 |
| RQLWWGHRI | ASSFGRVEDTQY | 0.8460261 |
| YGPPGTGKTLL | ASSFGRVEDTQY | 0.87832123 |
| AALVGAPANVA | ASSFGRVEDTQY | 0.50036246 |
| AATFRRTE | ASSFGRVEDTQY | 0.323541 |

| | | |
|---|---|---|
| AGALGGGI | ASSFGRVEDTQY | 0.86752415 |
| AGGAGGAGGIGAG | ASSFGRVEDTQY | 0.8789759 |
| AGSGALGL | ASSFGRVEDTQY | 0.6880666 |
| AQIADQI | ASSFGRVEDTQY | 0.7715453 |
| ASAVAVTYL | ASSFGRVEDTQY | 0.78249633 |
| ASTVLGGTGG | ASSFGRVEDTQY | 0.83471817 |
| AVGGEPAK | ASSFGRVEDTQY | 0.8164145 |
| CEQGTFGGTC | ASSFGRVEDTQY | 0.5741447 |
| DEGKAPAG | ASSFGRVEDTQY | 0.869341 |
| DTVVFPAL | ASSFGRVEDTQY | 0.5636075 |
| FEGITMAG | ASSFGRVEDTQY | 0.8714436 |
| FPLVVQD | ASSFGRVEDTQY | 0.8162093 |
| GAFTTPQ | ASSFGRVEDTQY | 0.8466878 |
| GFGGFGTAGGVGGA | ASSFGRVEDTQY | 0.7811686 |
| GFLAQFL | ASSFGRVEDTQY | 0.64250034 |
| GGAGGAGGNAGVA | ASSFGRVEDTQY | 0.56540304 |
| GGVGTAGAGGAGGAGG | ASSFGRVEDTQY | 0.42999923 |
| GNGGGGDAGNAGSGGN | ASSFGRVEDTQY | 0.8781935 |
| GTGGAGGAGNDAGS | ASSFGRVEDTQY | 0.33150476 |
| GTTNTGWFNS | ASSFGRVEDTQY | 0.88330936 |
| IFMGPLFTL | ASSFGRVEDTQY | 0.7913694 |
| ILAHAIEAI | ASSFGRVEDTQY | 0.7096678 |
| IRSAIREAL | ASSFGRVEDTQY | 0.8606641 |
| ITFAGGG | ASSFGRVEDTQY | 0.41803586 |
| ITGMSGHQEV | ASSFGRVEDTQY | 0.7656288 |
| IVAGLACTL | ASSFGRVEDTQY | 0.8081271 |
| IVGGGLAGA | ASSFGRVEDTQY | 0.8685884 |
| IVTGVLNI | ASSFGRVEDTQY | 0.639977 |
| LAQQGCI | ASSFGRVEDTQY | 0.8646471 |
| LCLHVEN | ASSFGRVEDTQY | 0.8299056 |
| LDAISDGS | ASSFGRVEDTQY | 0.45908338 |
| LGGFAAML | ASSFGRVEDTQY | 0.8221086 |
| LGGKNPMI | ASSFGRVEDTQY | 0.8762396 |
| LGPKGVAAIL | ASSFGRVEDTQY | 0.8309599 |
| LMVGVSI | ASSFGRVEDTQY | 0.53793883 |
| LVGAAFVL | ASSFGRVEDTQY | 0.80864936 |
| LVLAVAPL | ASSFGRVEDTQY | 0.7390879 |
| MAAIAVIDAA | ASSFGRVEDTQY | 0.828678 |
| NNGINAELQ | ASSFGRVEDTQY | 0.84350264 |
| RGEPLVLVHG | ASSFGRVEDTQY | 0.69387037 |
| RGGAGGAAGLI | ASSFGRVEDTQY | 0.84161043 |
| RSYRMHPAV | ASSFGRVEDTQY | 0.47043633 |
| SAAHLLAGQLA | ASSFGRVEDTQY | 0.5880841 |
| SAAVGSY | ASSFGRVEDTQY | 0.80117536 |
| SPHAVLHRI | ASSFGRVEDTQY | 0.787876 |
| SQIAKVL | ASSFGRVEDTQY | 0.8744047 |
| TALGHLVP | ASSFGRVEDTQY | 0.61203825 |
| TAPTTSGAA | ASSFGRVEDTQY | 0.8135981 |
| TDGDDGNGGSPGT | ASSFGRVEDTQY | 0.5674511 |
| TDGLGEN | ASSFGRVEDTQY | 0.5249809 |
| TGGPPLPGQA | ASSFGRVEDTQY | 0.8468162 |

| | | |
|---|---|---|
| TIPAITI | ASSFGRVEDTQY | 0.75489557 |
| TIPGFTI | ASSFGRVEDTQY | 0.40814078 |
| TSIASGALA | ASSFGRVEDTQY | 0.8557826 |
| TVTLPGAAI | ASSFGRVEDTQY | 0.49003822 |
| VASGLSAALP | ASSFGRVEDTQY | 0.6914561 |
| VGAGVIGI | ASSFGRVEDTQY | 0.8901838 |
| VGPKRFGV | ASSFGRVEDTQY | 0.5827235 |
| VNVAFPD | ASSFGRVEDTQY | 0.86643016 |
| VSMTNSGVS | ASSFGRVEDTQY | 0.84845096 |
| VTDVLGDLSV | ASSFGRVEDTQY | 0.8498522 |
| VVVVGAGFAGL | ASSFGRVEDTQY | 0.71135956 |
| VYTVGTVV | ASSFGRVEDTQY | 0.5991628 |
| WTTFSGP | ASSFGRVEDTQY | 0.89579856 |
| YDELVEAF | ASSFGRVEDTQY | 0.8287013 |
| YGTGGAGGNGG | ASSFGRVEDTQY | 0.83456516 |
| YTNVMAV | ASSFGRVEDTQY | 0.85959196 |
| FLEQVHQL | ASSFGRVEDTQY | 0.8775923 |
| GLSLLAVAV | ASSFGRVEDTQY | 0.87271905 |
| LEQVHQL | ASSFGRVEDTQY | 0.73789966 |
| LNAAGAI | ASSFGRVEDTQY | 0.41840142 |
| PAVHASL | ASSFGRVEDTQY | 0.7494723 |
| TELEDDA | ASSFGRVEDTQY | 0.7590797 |
| TPPGPPN | ASSFGRVEDTQY | 0.75220376 |
| TPPPPPPP | ASSFGRVEDTQY | 0.38853484 |
| TQSPALI | ASSIAGTGGRSQNTLY | 0.8778345 |
| EFAAAFPQ | ASSIAGTGGRSQNTLY | 0.90265185 |
| LLSGNLDVL | ASSIAGTGGRSQNTLY | 0.6442385 |
| GPANLGNYNI | ASSIAGTGGRSQNTLY | 0.9004248 |
| TLGPTLP | ASSIAGTGGRSQNTLY | 0.5362562 |
| VYAYSSCCK | ASSIAGTGGRSQNTLY | 0.81255525 |
| APAPAPAPNI | ASSIAGTGGRSQNTLY | 0.88288015 |
| PGGPVPIP | ASSIAGTGGRSQNTLY | 0.6591671 |
| FLTVALEP | ASSIAGTGGRSQNTLY | 0.818153 |
| SQYFGANL | ASSIAGTGGRSQNTLY | 0.4909047 |
| GGVKPAA | ASSIAGTGGRSQNTLY | 0.6920629 |
| GGTGGAGGAGGW | ASSIAGTGGRSQNTLY | 0.8826603 |
| GVGGVAGLGGAGG | ASSIAGTGGRSQNTLY | 0.60864663 |
| VLLVLATTI | ASSIAGTGGRSQNTLY | 0.80750334 |
| IGIFAAML | ASSIAGTGGRSQNTLY | 0.7654527 |
| AVGDLID | ASSIAGTGGRSQNTLY | 0.63918066 |
| NISPLPL | ASSIAGTGGRSQNTLY | 0.8593128 |
| VTAVFVAI | ASSIAGTGGRSQNTLY | 0.86674917 |
| SGDDNDAAL | ASSIAGTGGRSQNTLY | 0.775612 |
| AVAALAHS | ASSIAGTGGRSQNTLY | 0.8968379 |
| GGVTVMATP | ASSIAGTGGRSQNTLY | 0.57235324 |
| AGLAANVRD | ASSIAGTGGRSQNTLY | 0.8905094 |
| TNRPDLIDPALLRPG | ASSIAGTGGRSQNTLY | 0.89075613 |
| LANLGVTAL | ASSIAGTGGRSQNTLY | 0.82445073 |
| TGERIEL | ASSIAGTGGRSQNTLY | 0.77421975 |
| NGEELAA | ASSIAGTGGRSQNTLY | 0.8716639 |
| VLSLVWG | ASSIAGTGGRSQNTLY | 0.76263046 |

| | | |
|---|---|---|
| IGYALWL | ASSIAGTGGRSQNTLY | 0.8150267 |
| LSLLLAPL | ASSIAGTGGRSQNTLY | 0.89289695 |
| TLDEALK | ASSIAGTGGRSQNTLY | 0.24266288 |
| TDGAGTAEGG | ASSIAGTGGRSQNTLY | 0.80135465 |
| NAPGGAGGNGG | ASSIAGTGGRSQNTLY | 0.75033945 |
| VSVFMGMGGNF | ASSIAGTGGRSQNTLY | 0.7168269 |
| LASYLAAL | ASSIAGTGGRSQNTLY | 0.40789583 |
| LVGGTPAL | ASSIAGTGGRSQNTLY | 0.22842419 |
| VRNADVL | ASSIAGTGGRSQNTLY | 0.5395503 |
| TCSSPFLVSAM | ASSIAGTGGRSQNTLY | 0.4211167 |
| VHITDAE | ASSIAGTGGRSQNTLY | 0.8533907 |
| EYAAPSF | ASSIAGTGGRSQNTLY | 0.7331519 |
| VVDAFHGGL | ASSIAGTGGRSQNTLY | 0.5182663 |
| ASVVDPQ | ASSIAGTGGRSQNTLY | 0.7786975 |
| SYNEFGGLS | ASSIAGTGGRSQNTLY | 0.7826088 |
| DPDGSVY | ASSIAGTGGRSQNTLY | 0.22736609 |
| MVSTIHATE | ASSIAGTGGRSQNTLY | 0.62211806 |
| LAAGVADPV | ASSIAGTGGRSQNTLY | 0.37586358 |
| TKEQIAA | ASSIAGTGGRSQNTLY | 0.52595973 |
| SLASPAAL | ASSIAGTGGRSQNTLY | 0.19364482 |
| IGLVFYGV | ASSIAGTGGRSQNTLY | 0.6926312 |
| FLLDASNL | ASSIAGTGGRSQNTLY | 0.6644902 |
| YQVGNFP | ASSIAGTGGRSQNTLY | 0.27025473 |
| ESLLSQST | ASSIAGTGGRSQNTLY | 0.68981916 |
| LFTVGFAAVG | ASSIAGTGGRSQNTLY | 0.49789974 |
| ATPPTPT | ASSIAGTGGRSQNTLY | 0.5447172 |
| TDLATLG | ASSIAGTGGRSQNTLY | 0.76357996 |
| SAGGTGR | ASSIAGTGGRSQNTLY | 0.44720095 |
| GFSNSGS | ASSIAGTGGRSQNTLY | 0.57254016 |
| GGNGGSGGTSV | ASSIAGTGGRSQNTLY | 0.77403045 |
| GTSVATGGAGN | ASSIAGTGGRSQNTLY | 0.80394405 |
| NGGNGGSGGTS | ASSIAGTGGRSQNTLY | 0.41680014 |
| AVVKFPH | ASSIAGTGGRSQNTLY | 0.69334 |
| TPPPPPP | ASSIAGTGGRSQNTLY | 0.8343516 |
| AAPAAAAAPA | ASSIAGTGGRSQNTLY | 0.62535006 |
| IALNEDL | ASSIAGTGGRSQNTLY | 0.79725075 |
| IVNVSSM | ASSIAGTGGRSQNTLY | 0.802226 |
| TGIPPAPR | ASSIAGTGGRSQNTLY | 0.39795244 |
| FDLGGGTF | ASSIAGTGGRSQNTLY | 0.79219174 |
| FDLGGGTFDVS | ASSIAGTGGRSQNTLY | 0.83344287 |
| NEPTAAA | ASSIAGTGGRSQNTLY | 0.5663703 |
| NEPTAAALA | ASSIAGTGGRSQNTLY | 0.6525767 |
| GSIINISSI | ASSIAGTGGRSQNTLY | 0.29082838 |
| VAPGNAGTA | ASSIAGTGGRSQNTLY | 0.5681335 |
| TGGGASLE | ASSIAGTGGRSQNTLY | 0.26580134 |
| VAVYNHYKRI | ASSIAGTGGRSQNTLY | 0.76389027 |
| TDPAPATT | ASSIAGTGGRSQNTLY | 0.79809743 |
| TNRPDILDPALLRPG | ASSIAGTGGRSQNTLY | 0.66195613 |
| LTTTTTS | ASSIAGTGGRSQNTLY | 0.26547387 |
| PTGSGKTL | ASSIAGTGGRSQNTLY | 0.27173683 |
| TGSGKTL | ASSIAGTGGRSQNTLY | 0.5449903 |

| | | |
|---|---|---|
| TIGHVDHGK | ASSIAGTGGRSQNTLY | 0.27295083 |
| TIGHVDHGKTT | ASSIAGTGGRSQNTLY | 0.24435353 |
| GHVDHGKTT | ASSIAGTGGRSQNTLY | 0.30324674 |
| AQFEHTLL | ASSIAGTGGRSQNTLY | 0.7532061 |
| IVNQDEL | ASSIAGTGGRSQNTLY | 0.7973483 |
| TNGSQFF | ASSIAGTGGRSQNTLY | 0.5477412 |
| TGKGIVE | ASSIAGTGGRSQNTLY | 0.7984552 |
| LISNASDA | ASSIAGTGGRSQNTLY | 0.84227204 |
| AELRTLGELTKIAKAHGAQ | ASSIAGTGGRSQNTLY | 0.7859385 |
| APNAVAAAEPSW | ASSIAGTGGRSQNTLY | 0.68864304 |
| APQVAGAATAVL | ASSIAGTGGRSQNTLY | 0.25239712 |
| ARRDPAAAAAILPTDA | ASSIAGTGGRSQNTLY | 0.850368 |
| DAAARDVLAEAGLAEYF | ASSIAGTGGRSQNTLY | 0.5212065 |
| DDQIVLSAKANLAKVEQ | ASSIAGTGGRSQNTLY | 0.78671336 |
| DELLPDTGEDSDEDGH | ASSIAGTGGRSQNTLY | 0.7174684 |
| ELPEAPANRATVPQD | ASSIAGTGGRSQNTLY | 0.65828973 |
| FGQSKIERVTGIG | ASSIAGTGGRSQNTLY | 0.49680763 |
| GAGGTGSTAGGAGGAGGAGGLY | ASSIAGTGGRSQNTLY | 0.7990109 |
| GHGGTGAAGGNGGTGGAGGSGI | ASSIAGTGGRSQNTLY | 0.7986176 |
| GNGGNGGAGGEGGPGGAGGAGA | ASSIAGTGGRSQNTLY | 0.4486111 |
| GSTAGNGGNGGNATNGTIAGQP | ASSIAGTGGRSQNTLY | 0.22531441 |
| GTGGDGGNGGNGGTGG | ASSIAGTGGRSQNTLY | 0.7056554 |
| GVRRPAGIPADAPV | ASSIAGTGGRSQNTLY | 0.6751043 |
| GYRQVTDAHLLTLARR | ASSIAGTGGRSQNTLY | 0.79301304 |
| IAKMGQDGHDRG | ASSIAGTGGRSQNTLY | 0.61984664 |
| KAEPSIEIQYT | ASSIAGTGGRSQNTLY | 0.58564615 |
| KPVRIRMAADEDTVAVT | ASSIAGTGGRSQNTLY | 0.7666391 |
| MPPRIAGMRLLVIKPEP | ASSIAGTGGRSQNTLY | 0.7257073 |
| MKFYAVALFRSRGVPPTR | ASSIAGTGGRSQNTLY | 0.65287626 |
| NAGGFGILGAVAHSP | ASSIAGTGGRSQNTLY | 0.7101056 |
| NEGREITLAMAEM | ASSIAGTGGRSQNTLY | 0.5662639 |
| NELVEWSHPLGHRWRWE | ASSIAGTGGRSQNTLY | 0.7929606 |
| NLGAGGTGGNGGNPG | ASSIAGTGGRSQNTLY | 0.7805437 |
| NTEWPKWETVMAADPVD | ASSIAGTGGRSQNTLY | 0.66156316 |
| PASAAPAETQP | ASSIAGTGGRSQNTLY | 0.790081 |
| QQSSDGKNGVRAAEITGE | ASSIAGTGGRSQNTLY | 0.60356325 |
| RAPDAVTTSWWKE | ASSIAGTGGRSQNTLY | 0.6567189 |
| SGFGNVGAHTSGWFNQ | ASSIAGTGGRSQNTLY | 0.32938552 |
| SGVAGAVGQAASVGGLKV | ASSIAGTGGRSQNTLY | 0.53340405 |
| SNGGAGTGSAGGNGGT | ASSIAGTGGRSQNTLY | 0.58977884 |
| TGGAGGEGGAAG | ASSIAGTGGRSQNTLY | 0.7167978 |
| TGGDGGAAGAGGAPGHG | ASSIAGTGGRSQNTLY | 0.7607719 |
| TMPPLLAAIANAGRHGV | ASSIAGTGGRSQNTLY | 0.6606668 |
| TSSTAGNSGGA | ASSIAGTGGRSQNTLY | 0.5717632 |
| VPATVPVGILC | ASSIAGTGGRSQNTLY | 0.79205626 |
| VQRPDSGGVVLTGRIS | ASSIAGTGGRSQNTLY | 0.8202785 |
| WTPNPVPATKGVPAGLVQAT | ASSIAGTGGRSQNTLY | 0.36350363 |
| YALKRALAKFSDRFDVVII | ASSIAGTGGRSQNTLY | 0.610046 |
| YLIGSVIV | ASSIAGTGGRSQNTLY | 0.79878056 |
| YGQLVAVID | ASSIAGTGGRSQNTLY | 0.8224294 |
| VTVGGMTLPA | ASSIAGTGGRSQNTLY | 0.5851972 |

| | | |
|---|---|---|
| VLALPAV | ASSIAGTGGRSQNTLY | 0.68195975 |
| VIPPTLNYET | ASSIAGTGGRSQNTLY | 0.8612174 |
| VGMGGMHPSSG | ASSIAGTGGRSQNTLY | 0.65855145 |
| VGLAGGAATA | ASSIAGTGGRSQNTLY | 0.8260665 |
| VGGAGGAGGSGP | ASSIAGTGGRSQNTLY | 0.2511604 |
| VFAAFMV | ASSIAGTGGRSQNTLY | 0.52298707 |
| TLYDKLR | ASSIAGTGGRSQNTLY | 0.41320044 |
| TLAVFTVL | ASSIAGTGGRSQNTLY | 0.14949693 |
| TGSVQPARV | ASSIAGTGGRSQNTLY | 0.7857319 |
| TGGGAGSGGT | ASSIAGTGGRSQNTLY | 0.39623117 |
| SHLETSL | ASSIAGTGGRSQNTLY | 0.7654412 |
| SFTTGMT | ASSIAGTGGRSQNTLY | 0.52633804 |
| QVEVAKRMGVRQPTVS | ASSIAGTGGRSQNTLY | 0.76408404 |
| QGSGGPALAM | ASSIAGTGGRSQNTLY | 0.73474675 |
| MLAPSNP | ASSIAGTGGRSQNTLY | 0.5426346 |
| LVNNAAI | ASSIAGTGGRSQNTLY | 0.27923787 |
| LTGEINTLA | ASSIAGTGGRSQNTLY | 0.7090992 |
| LSIFGPV | ASSIAGTGGRSQNTLY | 0.779853 |
| LLALLANL | ASSIAGTGGRSQNTLY | 0.7174176 |
| LISNASDALDKLR | ASSIAGTGGRSQNTLY | 0.23557806 |
| LINEAVP | ASSIAGTGGRSQNTLY | 0.7559991 |
| LGYAFSGL | ASSIAGTGGRSQNTLY | 0.58588904 |
| LDFVAVAL | ASSIAGTGGRSQNTLY | 0.36315364 |
| LAQPGAE | ASSIAGTGGRSQNTLY | 0.21298611 |
| ISNLGAVTGF | ASSIAGTGGRSQNTLY | 0.41060966 |
| HQEDPGRI | ASSIAGTGGRSQNTLY | 0.53398025 |
| GTPGEPTP | ASSIAGTGGRSQNTLY | 0.7567694 |
| GFFNSGS | ASSIAGTGGRSQNTLY | 0.80574477 |
| GAGGGGLTAGAN | ASSIAGTGGRSQNTLY | 0.5019051 |
| FVDALVNM | ASSIAGTGGRSQNTLY | 0.7978729 |
| FLTVAPEL | ASSIAGTGGRSQNTLY | 0.65267974 |
| FETLTAL | ASSIAGTGGRSQNTLY | 0.11747765 |
| DVINPIEL | ASSIAGTGGRSQNTLY | 0.5168554 |
| APVSGAPV | ASSIAGTGGRSQNTLY | 0.21220063 |
| ALVGGLAW | ASSIAGTGGRSQNTLY | 0.7836117 |
| ALAAAAATAI | ASSIAGTGGRSQNTLY | 0.37611765 |
| AFGLGLLL | ASSIAGTGGRSQNTLY | 0.67750585 |
| AFFAAASL | ASSIAGTGGRSQNTLY | 0.59234065 |
| AAAALPAFSPP | ASSIAGTGGRSQNTLY | 0.8326763 |
| VVLAPAL | ASSIAGTGGRSQNTLY | 0.7444606 |
| TIGHVDHGKTTL | ASSIAGTGGRSQNTLY | 0.38029212 |
| LISNASDALDK | ASSIAGTGGRSQNTLY | 0.19944844 |
| LISNASDAL | ASSIAGTGGRSQNTLY | 0.68486214 |
| ISNASDALDK | ASSIAGTGGRSQNTLY | 0.48880157 |
| NQAETLVYQTE | ASSIAGTGGRSQNTLY | 0.4320369 |
| TGGTGGTAGAG | ASSIAGTGGRSQNTLY | 0.3815378 |
| TGSTAGGAGGA | ASSIAGTGGRSQNTLY | 0.7315237 |
| DPLAVPVAAGPV | ASSIAGTGGRSQNTLY | 0.35944694 |
| GGTGGTPGNGGN | ASSIAGTGGRSQNTLY | 0.7383055 |
| PAGLAGVDPATL | ASSIAGTGGRSQNTLY | 0.298105 |
| GNNLVVEAPHAGASV | ASSIAGTGGRSQNTLY | 0.612018 |

| | | |
|---|---|---|
| KVTRIMSASTLEQPA | ASSIAGTGGRSQNTLY | 0.81438255 |
| LGFALPIALPRRTVV | ASSIAGTGGRSQNTLY | 0.82451904 |
| TRLGNLTARKLVVLV | ASSIAGTGGRSQNTLY | 0.50000215 |
| WSQQRRLIYFPSAGP | ASSIAGTGGRSQNTLY | 0.51980513 |
| GPNGNPDPMAAAVDIR | ASSIAGTGGRSQNTLY | 0.3691374 |
| TQRCVSVADLRAAASR | ASSIAGTGGRSQNTLY | 0.52487373 |
| QFVLDVAEHPVEVLTETV | ASSIAGTGGRSQNTLY | 0.5199318 |
| GGAGGTGGAGGNGGTQAGDG | ASSIAGTGGRSQNTLY | 0.8007171 |
| PLLATLRAIAVDVDAKPAQV | ASSIAGTGGRSQNTLY | 0.55052805 |
| APKFRTGGYMIDFWGVGYQVAK | ASSIAGTGGRSQNTLY | 0.27487883 |
| GAGGAGGAGGAAGGPGGTGGTG | ASSIAGTGGRSQNTLY | 0.7101376 |
| IGALGLLATDVAPAGSVAFGVA | ASSIAGTGGRSQNTLY | 0.82945013 |
| AGGAGGIGGNGGAGGAGGVLMG | ASSIAGTGGRSQNTLY | 0.8438442 |
| ADELIGLL | ASSIAGTGGRSQNTLY | 0.3202814 |
| ADRALKA | ASSIAGTGGRSQNTLY | 0.5612654 |
| AGAVGQVGGAGGA | ASSIAGTGGRSQNTLY | 0.610041 |
| AGGGGGLGGAP | ASSIAGTGGRSQNTLY | 0.68543684 |
| AGGLLYGNGGAG | ASSIAGTGGRSQNTLY | 0.36519557 |
| AGLESIGL | ASSIAGTGGRSQNTLY | 0.4343395 |
| AGLKTGII | ASSIAGTGGRSQNTLY | 0.7922356 |
| ASLAALLL | ASSIAGTGGRSQNTLY | 0.8671557 |
| ASVINISSI | ASSIAGTGGRSQNTLY | 0.5647508 |
| ATPPGPA | ASSIAGTGGRSQNTLY | 0.3826036 |
| ATVASALV | ASSIAGTGGRSQNTLY | 0.6326508 |
| ATVASLIL | ASSIAGTGGRSQNTLY | 0.4584833 |
| AVNGPVLIRH | ASSIAGTGGRSQNTLY | 0.49203146 |
| DGAAGAVGI | ASSIAGTGGRSQNTLY | 0.5790304 |
| DPLLHGI | ASSIAGTGGRSQNTLY | 0.26799423 |
| ELGAALGAL | ASSIAGTGGRSQNTLY | 0.6136626 |
| ELGRTQAV | ASSIAGTGGRSQNTLY | 0.7809601 |
| FVAGGYLS | ASSIAGTGGRSQNTLY | 0.69486374 |
| GASGGAGGAGGW | ASSIAGTGGRSQNTLY | 0.6715245 |
| GGAGAGGDGGAGGI | ASSIAGTGGRSQNTLY | 0.8105643 |
| GTGHNGSQPGMGGNG | ASSIAGTGGRSQNTLY | 0.72575307 |
| GYAAASASAS | ASSIAGTGGRSQNTLY | 0.82374895 |
| HQVDVAAVQQG | ASSIAGTGGRSQNTLY | 0.8408784 |
| IGAAIAHT | ASSIAGTGGRSQNTLY | 0.476676 |
| IGMIEVAQL | ASSIAGTGGRSQNTLY | 0.57996356 |
| IRLAPEHTSAMD | ASSIAGTGGRSQNTLY | 0.8086946 |
| IYNVSVL | ASSIAGTGGRSQNTLY | 0.60965455 |
| KQGFLGM | ASSIAGTGGRSQNTLY | 0.6697186 |
| LESVITQL | ASSIAGTGGRSQNTLY | 0.6483873 |
| LGLGALVVV | ASSIAGTGGRSQNTLY | 0.80198646 |
| LIGNGGNGGSGGI | ASSIAGTGGRSQNTLY | 0.6787622 |
| LNAINAPTL | ASSIAGTGGRSQNTLY | 0.848565 |
| LQLLNTGV | ASSIAGTGGRSQNTLY | 0.16701302 |
| LQTSIEF | ASSIAGTGGRSQNTLY | 0.43605152 |
| MSFAVSEL | ASSIAGTGGRSQNTLY | 0.8030429 |
| NTGALGL | ASSIAGTGGRSQNTLY | 0.14232773 |
| PPGAGGAGGAGTT | ASSIAGTGGRSQNTLY | 0.29247937 |
| QLVGLGI | ASSIAGTGGRSQNTLY | 0.7098975 |

| | | |
|---|---|---|
| RFLEQVHQL | ASSIAGTGGRSQNTLY | 0.80375147 |
| RGVGYVLR | ASSIAGTGGRSQNTLY | 0.15734212 |
| SATGVFAGVI | ASSIAGTGGRSQNTLY | 0.61529386 |
| SLPAITLP | ASSIAGTGGRSQNTLY | 0.20148383 |
| STALAFTV | ASSIAGTGGRSQNTLY | 0.52629036 |
| SVLTVGLGK | ASSIAGTGGRSQNTLY | 0.6127535 |
| TALAGAGSGI | ASSIAGTGGRSQNTLY | 0.22327477 |
| TGLPFTGI | ASSIAGTGGRSQNTLY | 0.72505397 |
| VAEHVTV | ASSIAGTGGRSQNTLY | 0.3309993 |
| VDIVLNSL | ASSIAGTGGRSQNTLY | 0.6435644 |
| VLTGGGAVL | ASSIAGTGGRSQNTLY | 0.77239585 |
| VNVGLAYGI | ASSIAGTGGRSQNTLY | 0.8556634 |
| VSSAAVGAV | ASSIAGTGGRSQNTLY | 0.29051685 |
| VTSVYEAGV | ASSIAGTGGRSQNTLY | 0.75413454 |
| WVALDRGA | ASSIAGTGGRSQNTLY | 0.5297359 |
| YLVGSVVI | ASSIAGTGGRSQNTLY | 0.85553813 |
| GPPGTGKTLL | ASSIAGTGGRSQNTLY | 0.46844393 |
| GVGMTKF | ASSIAGTGGRSQNTLY | 0.21889694 |
| IGIGGSDL | ASSIAGTGGRSQNTLY | 0.7758442 |
| IGTLLAEL | ASSIAGTGGRSQNTLY | 0.5980848 |
| IYTSGTT | ASSIAGTGGRSQNTLY | 0.55762815 |
| LSFPAGV | ASSIAGTGGRSQNTLY | 0.8345686 |
| LYTSGTT | ASSIAGTGGRSQNTLY | 0.38707614 |
| VGAFPFGF | ASSIAGTGGRSQNTLY | 0.62794507 |
| VGAGTVEF | ASSIAGTGGRSQNTLY | 0.7797345 |
| VGKSTLLL | ASSIAGTGGRSQNTLY | 0.7381573 |
| AALAHAGGGIV | ASSIAGTGGRSQNTLY | 0.46762532 |
| AALVAVSLL | ASSIAGTGGRSQNTLY | 0.7291143 |
| AAVSRAVAAL | ASSIAGTGGRSQNTLY | 0.7045431 |
| AGGSSGIGGF | ASSIAGTGGRSQNTLY | 0.33267632 |
| AGLTVANANG | ASSIAGTGGRSQNTLY | 0.8404794 |
| AGLVGAVL | ASSIAGTGGRSQNTLY | 0.4878074 |
| ALSQAFTV | ASSIAGTGGRSQNTLY | 0.78482443 |
| ALTFSVSL | ASSIAGTGGRSQNTLY | 0.7074559 |
| ALTGAALVAL | ASSIAGTGGRSQNTLY | 0.8603536 |
| ASFSIGAGVL | ASSIAGTGGRSQNTLY | 0.3673212 |
| DGSVYKI | ASSIAGTGGRSQNTLY | 0.75311 |
| EQLAEIV | ASSIAGTGGRSQNTLY | 0.7274939 |
| FAVSELTVM | ASSIAGTGGRSQNTLY | 0.75556195 |
| GAAPLAAGL | ASSIAGTGGRSQNTLY | 0.569713 |
| GADGGDGGSGGAGG | ASSIAGTGGRSQNTLY | 0.64281565 |
| GFGLDTGGAGG | ASSIAGTGGRSQNTLY | 0.6808474 |
| GLASGGLL | ASSIAGTGGRSQNTLY | 0.81091106 |
| GRIAGAAVLL | ASSIAGTGGRSQNTLY | 0.58927125 |
| GSAGSLNTSF | ASSIAGTGGRSQNTLY | 0.62483525 |
| GTGQIGI | ASSIAGTGGRSQNTLY | 0.8403481 |
| GTLIALIL | ASSIAGTGGRSQNTLY | 0.27710062 |
| IAATMGFVVG | ASSIAGTGGRSQNTLY | 0.6803781 |
| IFAQALL | ASSIAGTGGRSQNTLY | 0.7955673 |
| IGAAVALGF | ASSIAGTGGRSQNTLY | 0.7772566 |
| IGLTPLAL | ASSIAGTGGRSQNTLY | 0.5643561 |

| | | |
|---|---|---|
| ILGSNTSTL | ASSIAGTGGRSQNTLY | 0.8030726 |
| IVFSLTVL | ASSIAGTGGRSQNTLY | 0.5436939 |
| LASKATGF | ASSIAGTGGRSQNTLY | 0.70046544 |
| LASVALIL | ASSIAGTGGRSQNTLY | 0.52872527 |
| LFAVGLTL | ASSIAGTGGRSQNTLY | 0.61087316 |
| LFNGGIGTYI | ASSIAGTGGRSQNTLY | 0.3840773 |
| LTLGAGII | ASSIAGTGGRSQNTLY | 0.6971005 |
| LVAAALGDI | ASSIAGTGGRSQNTLY | 0.79879016 |
| NTAFVQLGI | ASSIAGTGGRSQNTLY | 0.7391872 |
| QLFSGAEL | ASSIAGTGGRSQNTLY | 0.25333217 |
| SAATFITL | ASSIAGTGGRSQNTLY | 0.77074265 |
| SNYGLANLGV | ASSIAGTGGRSQNTLY | 0.77622277 |
| SYGGAYAVM | ASSIAGTGGRSQNTLY | 0.86148083 |
| TTGVGGLTLG | ASSIAGTGGRSQNTLY | 0.55861044 |
| TTQTIYI | ASSIAGTGGRSQNTLY | 0.7879529 |
| VAAGAGLGAVI | ASSIAGTGGRSQNTLY | 0.6766576 |
| VAAVVTLL | ASSIAGTGGRSQNTLY | 0.66138095 |
| VGFLGGLI | ASSIAGTGGRSQNTLY | 0.31092072 |
| VGVIPILL | ASSIAGTGGRSQNTLY | 0.3471094 |
| VIFASSLI | ASSIAGTGGRSQNTLY | 0.68307203 |
| VKGGDGGAAAG | ASSIAGTGGRSQNTLY | 0.47860408 |
| VTKTFTNI | ASSIAGTGGRSQNTLY | 0.841092 |
| VVAAAILL | ASSIAGTGGRSQNTLY | 0.7950237 |
| YAALDVEL | ASSIAGTGGRSQNTLY | 0.73701894 |
| LASVALLL | ASSIAGTGGRSQNTLY | 0.824545 |
| LDAVHRV | ASSIAGTGGRSQNTLY | 0.8205063 |
| QGLSSSIF | ASSIAGTGGRSQNTLY | 0.67671365 |
| RGTFANIRL | ASSIAGTGGRSQNTLY | 0.54468584 |
| RQLWWGHRI | ASSIAGTGGRSQNTLY | 0.61713064 |
| YGPPGTGKTLL | ASSIAGTGGRSQNTLY | 0.28251982 |
| AALVGAPANVA | ASSIAGTGGRSQNTLY | 0.5217562 |
| AATFRRTE | ASSIAGTGGRSQNTLY | 0.53541565 |
| AGALGGGI | ASSIAGTGGRSQNTLY | 0.64573383 |
| AGGAGGAGGIGAG | ASSIAGTGGRSQNTLY | 0.13053644 |
| AGSGALGL | ASSIAGTGGRSQNTLY | 0.81803465 |
| AQIADQI | ASSIAGTGGRSQNTLY | 0.79865724 |
| ASAVAVTYL | ASSIAGTGGRSQNTLY | 0.60295266 |
| ASTVLGGTGG | ASSIAGTGGRSQNTLY | 0.78548175 |
| AVGGEPAK | ASSIAGTGGRSQNTLY | 0.7209091 |
| CEQGTFGGTC | ASSIAGTGGRSQNTLY | 0.47733584 |
| DEGKAPAG | ASSIAGTGGRSQNTLY | 0.7880671 |
| DTVVFPAL | ASSIAGTGGRSQNTLY | 0.51624095 |
| FEGITMAG | ASSIAGTGGRSQNTLY | 0.7753842 |
| FPLVVQD | ASSIAGTGGRSQNTLY | 0.80136657 |
| GAFTTPQ | ASSIAGTGGRSQNTLY | 0.72736406 |
| GFGGFGTAGGVGGA | ASSIAGTGGRSQNTLY | 0.56267333 |
| GFLAQFL | ASSIAGTGGRSQNTLY | 0.6860714 |
| GGAGGAGGNAGVA | ASSIAGTGGRSQNTLY | 0.2983814 |
| GGVGTAGAGGAGGAGG | ASSIAGTGGRSQNTLY | 0.58123124 |
| GNGGGGDAGNAGSGGN | ASSIAGTGGRSQNTLY | 0.21610327 |
| GTGGAGGAGNDAGS | ASSIAGTGGRSQNTLY | 0.7302222 |

| | | |
|---|---|---|
| GTTNTGWFNS | ASSIAGTGGRSQNTLY | 0.29989076 |
| IFMGPLFTL | ASSIAGTGGRSQNTLY | 0.747707 |
| ILAHAIEAI | ASSIAGTGGRSQNTLY | 0.688295 |
| IRSAIREAL | ASSIAGTGGRSQNTLY | 0.7099497 |
| ITFAGGG | ASSIAGTGGRSQNTLY | 0.4670422 |
| ITGMSGHQEV | ASSIAGTGGRSQNTLY | 0.7547465 |
| IVAGLACTL | ASSIAGTGGRSQNTLY | 0.7984612 |
| IVGGGLAGA | ASSIAGTGGRSQNTLY | 0.35425234 |
| IVTGVLNI | ASSIAGTGGRSQNTLY | 0.6727848 |
| LAQQGCI | ASSIAGTGGRSQNTLY | 0.6078606 |
| LCLHVEN | ASSIAGTGGRSQNTLY | 0.5613346 |
| LDAISDGS | ASSIAGTGGRSQNTLY | 0.77967656 |
| LGGFAAML | ASSIAGTGGRSQNTLY | 0.6286184 |
| LGGKNPMI | ASSIAGTGGRSQNTLY | 0.25775704 |
| LGPKGVAAIL | ASSIAGTGGRSQNTLY | 0.5945286 |
| LMVGVSI | ASSIAGTGGRSQNTLY | 0.7407043 |
| LVGAAFVL | ASSIAGTGGRSQNTLY | 0.4090494 |
| LVLAVAPL | ASSIAGTGGRSQNTLY | 0.14819355 |
| MAAIAVIDAA | ASSIAGTGGRSQNTLY | 0.5978749 |
| NNGINAELQ | ASSIAGTGGRSQNTLY | 0.32171658 |
| RGEPLVLVHG | ASSIAGTGGRSQNTLY | 0.4379939 |
| RGGAGGAAGLI | ASSIAGTGGRSQNTLY | 0.53864604 |
| RSYRMHPAV | ASSIAGTGGRSQNTLY | 0.79405004 |
| SAAHLLAGQLA | ASSIAGTGGRSQNTLY | 0.60580194 |
| SAAVGSY | ASSIAGTGGRSQNTLY | 0.6506327 |
| SPHAVLHRI | ASSIAGTGGRSQNTLY | 0.7426233 |
| SQIAKVL | ASSIAGTGGRSQNTLY | 0.63846165 |
| TALGHLVP | ASSIAGTGGRSQNTLY | 0.7825558 |
| TAPTTSGAA | ASSIAGTGGRSQNTLY | 0.7205986 |
| TDGDDGNGGSPGT | ASSIAGTGGRSQNTLY | 0.6353632 |
| TDGLGEN | ASSIAGTGGRSQNTLY | 0.6570706 |
| TGGPPLPGQA | ASSIAGTGGRSQNTLY | 0.49944192 |
| TIPAITI | ASSIAGTGGRSQNTLY | 0.529296 |
| TIPGFTI | ASSIAGTGGRSQNTLY | 0.55859447 |
| TSIASGALA | ASSIAGTGGRSQNTLY | 0.49881455 |
| TVTLPGAAI | ASSIAGTGGRSQNTLY | 0.5972056 |
| VASGLSAALP | ASSIAGTGGRSQNTLY | 0.6912555 |
| VGAGVIGI | ASSIAGTGGRSQNTLY | 0.7691351 |
| VGPKRFGV | ASSIAGTGGRSQNTLY | 0.5361981 |
| VNVAFPD | ASSIAGTGGRSQNTLY | 0.22979331 |
| VSMTNSGVS | ASSIAGTGGRSQNTLY | 0.55041444 |
| VTDVLGDLSV | ASSIAGTGGRSQNTLY | 0.47447157 |
| VVVVGAGFAGL | ASSIAGTGGRSQNTLY | 0.78182435 |
| VYTVGTVV | ASSIAGTGGRSQNTLY | 0.70068336 |
| WTTFSGP | ASSIAGTGGRSQNTLY | 0.86001265 |
| YDELVEAF | ASSIAGTGGRSQNTLY | 0.807671 |
| YGTGGAGGNGG | ASSIAGTGGRSQNTLY | 0.5886305 |
| YTNVMAV | ASSIAGTGGRSQNTLY | 0.7738792 |
| FLEQVHQL | ASSIAGTGGRSQNTLY | 0.49986008 |
| GLSLLAVAV | ASSIAGTGGRSQNTLY | 0.567227 |
| LEQVHQL | ASSIAGTGGRSQNTLY | 0.52375627 |

| | | |
|---|---|---|
| LNAAGAI | ASSIAGTGGRSQNTLY | 0.67909163 |
| PAVHASL | ASSIAGTGGRSQNTLY | 0.4411077 |
| TELEDDA | ASSIAGTGGRSQNTLY | 0.41615355 |
| TPPGPPN | ASSIAGTGGRSQNTLY | 0.25412095 |
| TPPPPPPP | ASSIAGTGGRSQNTLY | 0.789293 |
| TQSPALI | SSSRQGNTEVF | 0.32180133 |
| EFAAAFPQ | SSSRQGNTEVF | 0.35885537 |
| LLSGNLDVL | SSSRQGNTEVF | 0.5095372 |
| GPANLGNYNI | SSSRQGNTEVF | 0.84397566 |
| TLGPTLP | SSSRQGNTEVF | 0.8361957 |
| VYAYSSCCK | SSSRQGNTEVF | 0.54677576 |
| APAPAPAPNI | SSSRQGNTEVF | 0.4467304 |
| PGGPVPIP | SSSRQGNTEVF | 0.82497406 |
| FLTVALEP | SSSRQGNTEVF | 0.52118915 |
| SQYFGANL | SSSRQGNTEVF | 0.6604154 |
| GGVKPAA | SSSRQGNTEVF | 0.7041254 |
| GGTGGAGGAGGW | SSSRQGNTEVF | 0.54991615 |
| GVGGVAGLGGAGG | SSSRQGNTEVF | 0.45078158 |
| VLLVLATTI | SSSRQGNTEVF | 0.75785124 |
| IGIFAAML | SSSRQGNTEVF | 0.55767834 |
| AVGDLID | SSSRQGNTEVF | 0.60561764 |
| NISPLPL | SSSRQGNTEVF | 0.26811182 |
| VTAVFVAI | SSSRQGNTEVF | 0.7242313 |
| SGDDNDAAL | SSSRQGNTEVF | 0.44069445 |
| AVAALAHS | SSSRQGNTEVF | 0.36565167 |
| GGVTVMATP | SSSRQGNTEVF | 0.18387012 |
| AGLAANVRD | SSSRQGNTEVF | 0.55159676 |
| TNRPDLIDPALLRPG | SSSRQGNTEVF | 0.8040772 |
| LANLGVTAL | SSSRQGNTEVF | 0.82234704 |
| TGERIEL | SSSRQGNTEVF | 0.6903255 |
| NGEELAA | SSSRQGNTEVF | 0.7194761 |
| VLSLVWG | SSSRQGNTEVF | 0.82763565 |
| IGYALWL | SSSRQGNTEVF | 0.57481706 |
| LSLLLAPL | SSSRQGNTEVF | 0.18327835 |
| TLDEALK | SSSRQGNTEVF | 0.6617748 |
| TDGAGTAEGG | SSSRQGNTEVF | 0.7603522 |
| NAPGGAGGNGG | SSSRQGNTEVF | 0.72654366 |
| VSVFMGMGGNF | SSSRQGNTEVF | 0.80506486 |
| LASYLAAL | SSSRQGNTEVF | 0.31024322 |
| LVGGTPAL | SSSRQGNTEVF | 0.7412913 |
| VRNADVL | SSSRQGNTEVF | 0.641202 |
| TCSSPFLVSAM | SSSRQGNTEVF | 0.27700067 |
| VHITDAE | SSSRQGNTEVF | 0.4709916 |
| EYAAPSF | SSSRQGNTEVF | 0.39571744 |
| VVDAFHGGL | SSSRQGNTEVF | 0.23091376 |
| ASVVDPQ | SSSRQGNTEVF | 0.6838368 |
| SYNEFGGLS | SSSRQGNTEVF | 0.5944073 |
| DPDGSVY | SSSRQGNTEVF | 0.68004644 |
| MVSTIHATE | SSSRQGNTEVF | 0.5133149 |
| LAAGVADPV | SSSRQGNTEVF | 0.5106123 |
| TKEQIAA | SSSRQGNTEVF | 0.55593884 |

| | | |
|---|---|---|
| SLASPAAL | SSSRQGNTEVF | 0.6223435 |
| IGLVFYGV | SSSRQGNTEVF | 0.7285131 |
| FLLDASNL | SSSRQGNTEVF | 0.7738128 |
| YQVGNFP | SSSRQGNTEVF | 0.6701331 |
| ESLLSQST | SSSRQGNTEVF | 0.60989046 |
| LFTVGFAAVG | SSSRQGNTEVF | 0.8261521 |
| ATPPTPT | SSSRQGNTEVF | 0.33324385 |
| TDLATLG | SSSRQGNTEVF | 0.4018644 |
| SAGGTGR | SSSRQGNTEVF | 0.7341312 |
| GFSNSGS | SSSRQGNTEVF | 0.5718055 |
| GGNGGSGGTSV | SSSRQGNTEVF | 0.80037135 |
| GTSVATGGAGN | SSSRQGNTEVF | 0.6802452 |
| NGGNGGSGGTS | SSSRQGNTEVF | 0.7047647 |
| AVVKFPH | SSSRQGNTEVF | 0.7708769 |
| TPPPPPP | SSSRQGNTEVF | 0.83384323 |
| AAPAAAAAPA | SSSRQGNTEVF | 0.73922145 |
| IALNEDL | SSSRQGNTEVF | 0.4550114 |
| IVNVSSM | SSSRQGNTEVF | 0.8363928 |
| TGIPPAPR | SSSRQGNTEVF | 0.2821918 |
| FDLGGGTF | SSSRQGNTEVF | 0.75101256 |
| FDLGGGTFDVS | SSSRQGNTEVF | 0.7817115 |
| NEPTAAA | SSSRQGNTEVF | 0.36237139 |
| NEPTAAALA | SSSRQGNTEVF | 0.6278447 |
| GSIINISSI | SSSRQGNTEVF | 0.82129306 |
| VAPGNAGTA | SSSRQGNTEVF | 0.80286 |
| TGGGASLE | SSSRQGNTEVF | 0.47230095 |
| VAVYNHYKRI | SSSRQGNTEVF | 0.4006051 |
| TDPAPATT | SSSRQGNTEVF | 0.41440922 |
| TNRPDILDPALLRPG | SSSRQGNTEVF | 0.48763728 |
| LTTTTTS | SSSRQGNTEVF | 0.22231932 |
| PTGSGKTL | SSSRQGNTEVF | 0.7301785 |
| TGSGKTL | SSSRQGNTEVF | 0.6277107 |
| TIGHVDHGK | SSSRQGNTEVF | 0.8505231 |
| TIGHVDHGKTT | SSSRQGNTEVF | 0.6628628 |
| GHVDHGKTT | SSSRQGNTEVF | 0.78995717 |
| AQFEHTLL | SSSRQGNTEVF | 0.5929593 |
| IVNQDEL | SSSRQGNTEVF | 0.6445754 |
| TNGSQFF | SSSRQGNTEVF | 0.20093009 |
| TGKGIVE | SSSRQGNTEVF | 0.24337566 |
| LISNASDA | SSSRQGNTEVF | 0.3991915 |
| AELRTLGELTKIAKAHGAQ | SSSRQGNTEVF | 0.68952656 |
| APNAVAAAEPSW | SSSRQGNTEVF | 0.24691635 |
| APQVAGAATAVL | SSSRQGNTEVF | 0.81140864 |
| ARRDPAAAAAILPTDA | SSSRQGNTEVF | 0.57859826 |
| DAAARDVLAEAGLAEYF | SSSRQGNTEVF | 0.724139 |
| DDQIVLSAKANLAKVEQ | SSSRQGNTEVF | 0.46650934 |
| DELLPDTGEDSDEDGH | SSSRQGNTEVF | 0.28505263 |
| ELPEAPANRATVPQD | SSSRQGNTEVF | 0.5624001 |
| FGQSKIERVTGIG | SSSRQGNTEVF | 0.7959459 |
| GAGGTGSTAGGAGGAGGAGGLY | SSSRQGNTEVF | 0.33505177 |
| GHGGTGAAGGNGGTGGAGGSGI | SSSRQGNTEVF | 0.24602613 |

| | | |
|---|---|---|
| GNGGNGGAGGEGGPGGAGGAGA | SSSRQGNTEVF | 0.4155604 |
| GSTAGNGGNGGNATNGTIAGQP | SSSRQGNTEVF | 0.56359744 |
| GTGGDGGNGGNGGTGG | SSSRQGNTEVF | 0.8440163 |
| GVRRPAGIPADAPV | SSSRQGNTEVF | 0.79968655 |
| GYRQVTDAHLLTLARR | SSSRQGNTEVF | 0.6612798 |
| IAKMGQDGHDRG | SSSRQGNTEVF | 0.8459688 |
| KAEPSIEIQYT | SSSRQGNTEVF | 0.13816099 |
| KPVRIRMAADEDTVAVT | SSSRQGNTEVF | 0.502486 |
| MPPRIAGMRLLVIKPEP | SSSRQGNTEVF | 0.8289834 |
| MKFYAVALFRSRGVPPTR | SSSRQGNTEVF | 0.7067762 |
| NAGGFGILGAVAHSP | SSSRQGNTEVF | 0.61996514 |
| NEGREITLAMAEM | SSSRQGNTEVF | 0.80281925 |
| NELVEWSHPLGHRWRWE | SSSRQGNTEVF | 0.48310816 |
| NLGAGGTGGNGGNPG | SSSRQGNTEVF | 0.43267637 |
| NTEWPKWETVMAADPVD | SSSRQGNTEVF | 0.68772566 |
| PASAAPAETQP | SSSRQGNTEVF | 0.6106684 |
| QQSSDGKNGVRAAEITGE | SSSRQGNTEVF | 0.5253484 |
| RAPDAVTTSWWKE | SSSRQGNTEVF | 0.23474261 |
| SGFGNVGAHTSGWFNQ | SSSRQGNTEVF | 0.7328785 |
| SGVAGAVGQAASVGGLKV | SSSRQGNTEVF | 0.85524166 |
| SNGGAGTGSAGGNGGT | SSSRQGNTEVF | 0.59919727 |
| TGGAGGEGGAAG | SSSRQGNTEVF | 0.71379507 |
| TGGDGGAAGAGGAPGHG | SSSRQGNTEVF | 0.67741597 |
| TMPPLLAAIANAGRHGV | SSSRQGNTEVF | 0.7564999 |
| TSSTAGNSGGA | SSSRQGNTEVF | 0.49189296 |
| VPATVPVGILC | SSSRQGNTEVF | 0.36735207 |
| VQRPDSGGVVLTGRIS | SSSRQGNTEVF | 0.7906843 |
| WTPNPVPATKGVPAGLVQAT | SSSRQGNTEVF | 0.37170005 |
| YALKRALAKFSDRFDVVII | SSSRQGNTEVF | 0.67811275 |
| YLIGSVIV | SSSRQGNTEVF | 0.59742683 |
| YGQLVAVID | SSSRQGNTEVF | 0.8564194 |
| VTVGGMTLPA | SSSRQGNTEVF | 0.77778244 |
| VLALPAV | SSSRQGNTEVF | 0.68338907 |
| VIPPTLNYET | SSSRQGNTEVF | 0.5861188 |
| VGMGGMHPSSG | SSSRQGNTEVF | 0.6698447 |
| VGLAGGAATA | SSSRQGNTEVF | 0.7593292 |
| VGGAGGAGGSGP | SSSRQGNTEVF | 0.21842188 |
| VFAAFMV | SSSRQGNTEVF | 0.81350213 |
| TLYDKLR | SSSRQGNTEVF | 0.3691971 |
| TLAVFTVL | SSSRQGNTEVF | 0.56288683 |
| TGSVQPARV | SSSRQGNTEVF | 0.7118754 |
| TGGGAGSGGT | SSSRQGNTEVF | 0.8549385 |
| SHLETSL | SSSRQGNTEVF | 0.62820375 |
| SFTTGMT | SSSRQGNTEVF | 0.8052106 |
| QVEVAKRMGVRQPTVS | SSSRQGNTEVF | 0.44881958 |
| QGSGGPALAM | SSSRQGNTEVF | 0.76554364 |
| MLAPSNP | SSSRQGNTEVF | 0.81935775 |
| LVNNAAI | SSSRQGNTEVF | 0.7757793 |
| LTGEINTLA | SSSRQGNTEVF | 0.85338676 |
| LSIFGPV | SSSRQGNTEVF | 0.5807357 |
| LLALLANL | SSSRQGNTEVF | 0.20546693 |

| | | |
|---|---|---|
| LISNASDALDKLR | SSSRQGNTEVF | 0.5091554 |
| LINEAVP | SSSRQGNTEVF | 0.75603575 |
| LGYAFSGL | SSSRQGNTEVF | 0.8164173 |
| LDFVAVAL | SSSRQGNTEVF | 0.79586124 |
| LAQPGAE | SSSRQGNTEVF | 0.7693788 |
| ISNLGAVTGF | SSSRQGNTEVF | 0.56356055 |
| HQEDPGRI | SSSRQGNTEVF | 0.81595457 |
| GTPGEPTP | SSSRQGNTEVF | 0.8496052 |
| GFFNSGS | SSSRQGNTEVF | 0.7499943 |
| GAGGGGLTAGAN | SSSRQGNTEVF | 0.38176906 |
| FVDALVNM | SSSRQGNTEVF | 0.3660524 |
| FLTVAPEL | SSSRQGNTEVF | 0.7469629 |
| FETLTAL | SSSRQGNTEVF | 0.7726957 |
| DVINPIEL | SSSRQGNTEVF | 0.31312048 |
| APVSGAPV | SSSRQGNTEVF | 0.60177714 |
| ALVGGLAW | SSSRQGNTEVF | 0.70773304 |
| ALAAAAATAI | SSSRQGNTEVF | 0.18498388 |
| AFGLGLLL | SSSRQGNTEVF | 0.69665635 |
| AFFAAASL | SSSRQGNTEVF | 0.8245698 |
| AAAALPAFSPP | SSSRQGNTEVF | 0.615492 |
| VVLAPAL | SSSRQGNTEVF | 0.3006244 |
| TIGHVDHGKTTL | SSSRQGNTEVF | 0.8272872 |
| LISNASDALDK | SSSRQGNTEVF | 0.8400407 |
| LISNASDAL | SSSRQGNTEVF | 0.79792434 |
| ISNASDALDK | SSSRQGNTEVF | 0.30053052 |
| NQAETLVYQTE | SSSRQGNTEVF | 0.15571198 |
| TGGTGGTAGAG | SSSRQGNTEVF | 0.6590419 |
| TGSTAGGAGGA | SSSRQGNTEVF | 0.84368443 |
| DPLAVPVAAGPV | SSSRQGNTEVF | 0.6147199 |
| GGTGGTPGNGGN | SSSRQGNTEVF | 0.44902226 |
| PAGLAGVDPATL | SSSRQGNTEVF | 0.6795154 |
| GNNLVVEAPHAGASV | SSSRQGNTEVF | 0.78792727 |
| KVTRIMSASTLEQPA | SSSRQGNTEVF | 0.2878918 |
| LGFALPIALPRRTVV | SSSRQGNTEVF | 0.28153443 |
| TRLGNLTARKLVVLV | SSSRQGNTEVF | 0.67924744 |
| WSQQRRLIYFPSAGP | SSSRQGNTEVF | 0.8022326 |
| GPNGNPDPMAAAVDIR | SSSRQGNTEVF | 0.2934495 |
| TQRCVSVADLRAAASR | SSSRQGNTEVF | 0.57943034 |
| QFVLDVAEHPVEVLTETV | SSSRQGNTEVF | 0.7385913 |
| GGAGGTGGAGGNGGTQAGDG | SSSRQGNTEVF | 0.52312183 |
| PLLATLRAIAVDVDAKPAQV | SSSRQGNTEVF | 0.6895613 |
| APKFRTGGYMIDFWGVGYQVAK | SSSRQGNTEVF | 0.32775578 |
| GAGGAGGAGGAAGGPGGTGGTG | SSSRQGNTEVF | 0.74602014 |
| IGALGLLATDVAPAGSVAFGVA | SSSRQGNTEVF | 0.3297948 |
| AGGAGGIGGNGGAGGAGGVLMG | SSSRQGNTEVF | 0.7903814 |
| ADELIGLL | SSSRQGNTEVF | 0.5931723 |
| ADRALKA | SSSRQGNTEVF | 0.60219693 |
| AGAVGQVGGAGGA | SSSRQGNTEVF | 0.59114385 |
| AGGGGGLGGAP | SSSRQGNTEVF | 0.4139769 |
| AGGLLYGNGGAG | SSSRQGNTEVF | 0.37518376 |
| AGLESIGL | SSSRQGNTEVF | 0.44037855 |

| | | |
|---|---|---|
| AGLKTGII | SSSRQGNTEVF | 0.7859545 |
| ASLAALLL | SSSRQGNTEVF | 0.6632749 |
| ASVINISSI | SSSRQGNTEVF | 0.70783293 |
| ATPPGPA | SSSRQGNTEVF | 0.67809784 |
| ATVASALV | SSSRQGNTEVF | 0.29881462 |
| ATVASLIL | SSSRQGNTEVF | 0.73398197 |
| AVNGPVLIRH | SSSRQGNTEVF | 0.31252795 |
| DGAAGAVGI | SSSRQGNTEVF | 0.7584925 |
| DPLLHGI | SSSRQGNTEVF | 0.5544316 |
| ELGAALGAL | SSSRQGNTEVF | 0.6669691 |
| ELGRTQAV | SSSRQGNTEVF | 0.5473395 |
| FVAGGYLS | SSSRQGNTEVF | 0.66979396 |
| GASGGAGGAGGW | SSSRQGNTEVF | 0.7285734 |
| GGAGAGGDGGAGGI | SSSRQGNTEVF | 0.44258785 |
| GTGHNGSQPGMGGNG | SSSRQGNTEVF | 0.6312934 |
| GYAAASASAS | SSSRQGNTEVF | 0.7973609 |
| HQVDVAAVQQG | SSSRQGNTEVF | 0.7100744 |
| IGAAIAHT | SSSRQGNTEVF | 0.25815257 |
| IGMIEVAQL | SSSRQGNTEVF | 0.68148834 |
| IRLAPEHTSAMD | SSSRQGNTEVF | 0.43786246 |
| IYNVSVL | SSSRQGNTEVF | 0.7202457 |
| KQGFLGM | SSSRQGNTEVF | 0.57840466 |
| LESVITQL | SSSRQGNTEVF | 0.6263066 |
| LGLGALVVV | SSSRQGNTEVF | 0.5824131 |
| LIGNGGNGGSGGI | SSSRQGNTEVF | 0.34858716 |
| LNAINAPTL | SSSRQGNTEVF | 0.47317347 |
| LQLLNTGV | SSSRQGNTEVF | 0.42794365 |
| LQTSIEF | SSSRQGNTEVF | 0.73386824 |
| MSFAVSEL | SSSRQGNTEVF | 0.73225987 |
| NTGALGL | SSSRQGNTEVF | 0.3544163 |
| PPGAGGAGGAGTT | SSSRQGNTEVF | 0.7080256 |
| QLVGLGI | SSSRQGNTEVF | 0.4946462 |
| RFLEQVHQL | SSSRQGNTEVF | 0.31496897 |
| RGVGYVLR | SSSRQGNTEVF | 0.5851954 |
| SATGVFAGVI | SSSRQGNTEVF | 0.46588016 |
| SLPAITLP | SSSRQGNTEVF | 0.2753213 |
| STALAFTV | SSSRQGNTEVF | 0.62525004 |
| SVLTVGLGK | SSSRQGNTEVF | 0.4038812 |
| TALAGAGSGI | SSSRQGNTEVF | 0.65796983 |
| TGLPFTGI | SSSRQGNTEVF | 0.48536038 |
| VAEHVTV | SSSRQGNTEVF | 0.25534654 |
| VDIVLNSL | SSSRQGNTEVF | 0.76239914 |
| VLTGGGAVL | SSSRQGNTEVF | 0.5685801 |
| VNVGLAYGI | SSSRQGNTEVF | 0.63620114 |
| VSSAAVGAV | SSSRQGNTEVF | 0.68424803 |
| VTSVYEAGV | SSSRQGNTEVF | 0.45403805 |
| WVALDRGA | SSSRQGNTEVF | 0.7922609 |
| YLVGSVVI | SSSRQGNTEVF | 0.6970277 |
| GPPGTGKTLL | SSSRQGNTEVF | 0.70576644 |
| GVGMTKF | SSSRQGNTEVF | 0.5496652 |
| IGIGGSDL | SSSRQGNTEVF | 0.83396804 |

| | | |
|---|---|---|
| IGTLLAEL | SSSRQGNTEVF | 0.7580048 |
| IYTSGTT | SSSRQGNTEVF | 0.46730453 |
| LSFPAGV | SSSRQGNTEVF | 0.19196737 |
| LYTSGTT | SSSRQGNTEVF | 0.67414534 |
| VGAFPFGF | SSSRQGNTEVF | 0.66123724 |
| VGAGTVEF | SSSRQGNTEVF | 0.38022846 |
| VGKSTLLL | SSSRQGNTEVF | 0.22190855 |
| AALAHAGGGIV | SSSRQGNTEVF | 0.7921115 |
| AALVAVSLL | SSSRQGNTEVF | 0.8505987 |
| AAVSRAVAAL | SSSRQGNTEVF | 0.64210236 |
| AGGSSGIGGF | SSSRQGNTEVF | 0.83298767 |
| AGLTVANANG | SSSRQGNTEVF | 0.32759976 |
| AGLVGAVL | SSSRQGNTEVF | 0.6974923 |
| ALSQAFTV | SSSRQGNTEVF | 0.80718565 |
| ALTFSVSL | SSSRQGNTEVF | 0.4062345 |
| ALTGAALVAL | SSSRQGNTEVF | 0.6876999 |
| ASFSIGAGVL | SSSRQGNTEVF | 0.37271494 |
| DGSVYKI | SSSRQGNTEVF | 0.39599872 |
| EQLAEIV | SSSRQGNTEVF | 0.7738498 |
| FAVSELTVM | SSSRQGNTEVF | 0.61671495 |
| GAAPLAAGL | SSSRQGNTEVF | 0.6253475 |
| GADGGDGGSGGAGG | SSSRQGNTEVF | 0.5830105 |
| GFGLDTGGAGG | SSSRQGNTEVF | 0.35675424 |
| GLASGGLL | SSSRQGNTEVF | 0.5600445 |
| GRIAGAAVLL | SSSRQGNTEVF | 0.74821854 |
| GSAGSLNTSF | SSSRQGNTEVF | 0.7359593 |
| GTGQIGI | SSSRQGNTEVF | 0.84691495 |
| GTLIALIL | SSSRQGNTEVF | 0.40011916 |
| IAATMGFVVG | SSSRQGNTEVF | 0.8463805 |
| IFAQALL | SSSRQGNTEVF | 0.84620684 |
| IGAAVALGF | SSSRQGNTEVF | 0.72029686 |
| IGLTPLAL | SSSRQGNTEVF | 0.4838286 |
| ILGSNTSTL | SSSRQGNTEVF | 0.8298974 |
| IVFSLTVL | SSSRQGNTEVF | 0.669945 |
| LASKATGF | SSSRQGNTEVF | 0.66975474 |
| LASVALIL | SSSRQGNTEVF | 0.7113553 |
| LFAVGLTL | SSSRQGNTEVF | 0.44213843 |
| LFNGGIGTYI | SSSRQGNTEVF | 0.85322213 |
| LTLGAGII | SSSRQGNTEVF | 0.8239099 |
| LVAAALGDI | SSSRQGNTEVF | 0.74218756 |
| NTAFVQLGI | SSSRQGNTEVF | 0.65111566 |
| QLFSGAEL | SSSRQGNTEVF | 0.38479924 |
| SAATFITL | SSSRQGNTEVF | 0.6965732 |
| SNYGLANLGV | SSSRQGNTEVF | 0.60375035 |
| SYGGAYAVM | SSSRQGNTEVF | 0.8415027 |
| TTGVGGLTLG | SSSRQGNTEVF | 0.8371228 |
| TTQTIYI | SSSRQGNTEVF | 0.69763845 |
| VAAGAGLGAVI | SSSRQGNTEVF | 0.8384965 |
| VAAVVTLL | SSSRQGNTEVF | 0.838998 |
| VGFLGGLI | SSSRQGNTEVF | 0.40411466 |
| VGVIPILL | SSSRQGNTEVF | 0.75100476 |

| | | |
|---|---|---|
| VIFASSLI | SSSRQGNTEVF | 0.63065505 |
| VKGGDGGAAAG | SSSRQGNTEVF | 0.74528813 |
| VTKTFTNI | SSSRQGNTEVF | 0.26746783 |
| VVAAAILL | SSSRQGNTEVF | 0.8297915 |
| YAALDVEL | SSSRQGNTEVF | 0.72086453 |
| LASVALLL | SSSRQGNTEVF | 0.20106575 |
| LDAVHRV | SSSRQGNTEVF | 0.7926309 |
| QGLSSSIF | SSSRQGNTEVF | 0.7731717 |
| RGTFANIRL | SSSRQGNTEVF | 0.7899709 |
| RQLWWGHRI | SSSRQGNTEVF | 0.8475696 |
| YGPPGTGKTLL | SSSRQGNTEVF | 0.6556854 |
| AALVGAPANVA | SSSRQGNTEVF | 0.6552121 |
| AATFRRTE | SSSRQGNTEVF | 0.83137167 |
| AGALGGGI | SSSRQGNTEVF | 0.85314476 |
| AGGAGGAGGIGAG | SSSRQGNTEVF | 0.62372285 |
| AGSGALGL | SSSRQGNTEVF | 0.82347035 |
| AQIADQI | SSSRQGNTEVF | 0.8473872 |
| ASAVAVTYL | SSSRQGNTEVF | 0.805812 |
| ASTVLGGTGG | SSSRQGNTEVF | 0.85329145 |
| AVGGEPAK | SSSRQGNTEVF | 0.8563509 |
| CEQGTFGGTC | SSSRQGNTEVF | 0.5805014 |
| DEGKAPAG | SSSRQGNTEVF | 0.8483391 |
| DTVVFPAL | SSSRQGNTEVF | 0.856606 |
| FEGITMAG | SSSRQGNTEVF | 0.7502886 |
| FPLVVQD | SSSRQGNTEVF | 0.77261317 |
| GAFTTPQ | SSSRQGNTEVF | 0.41389325 |
| GFGGFGTAGGVGGA | SSSRQGNTEVF | 0.7056558 |
| GFLAQFL | SSSRQGNTEVF | 0.43166196 |
| GGAGGAGGNAGVA | SSSRQGNTEVF | 0.8208896 |
| GGVGTAGAGGAGGAGG | SSSRQGNTEVF | 0.8512678 |
| GNGGGGDAGNAGSGGN | SSSRQGNTEVF | 0.7978142 |
| GTGGAGGAGNDAGS | SSSRQGNTEVF | 0.5776137 |
| GTTNTGWFNS | SSSRQGNTEVF | 0.4324558 |
| IFMGPLFTL | SSSRQGNTEVF | 0.7542184 |
| ILAHAIEAI | SSSRQGNTEVF | 0.6944101 |
| IRSAIREAL | SSSRQGNTEVF | 0.2540601 |
| ITFAGGG | SSSRQGNTEVF | 0.6943017 |
| ITGMSGHQEV | SSSRQGNTEVF | 0.83536196 |
| IVAGLACTL | SSSRQGNTEVF | 0.83860624 |
| IVGGGLAGA | SSSRQGNTEVF | 0.59325504 |
| IVTGVLNI | SSSRQGNTEVF | 0.85768855 |
| LAQQGCI | SSSRQGNTEVF | 0.83084834 |
| LCLHVEN | SSSRQGNTEVF | 0.85280955 |
| LDAISDGS | SSSRQGNTEVF | 0.7713139 |
| LGGFAAML | SSSRQGNTEVF | 0.4567836 |
| LGGKNPMI | SSSRQGNTEVF | 0.8432367 |
| LGPKGVAAIL | SSSRQGNTEVF | 0.77703017 |
| LMVGVSI | SSSRQGNTEVF | 0.7338251 |
| LVGAAFVL | SSSRQGNTEVF | 0.8189994 |
| LVLAVAPL | SSSRQGNTEVF | 0.7560663 |
| MAAIAVIDAA | SSSRQGNTEVF | 0.7282332 |

| | | |
|---|---|---|
| NNGINAELQ | SSSRQGNTEVF | 0.7989385 |
| RGEPLVLVHG | SSSRQGNTEVF | 0.8266374 |
| RGGAGGAAGLI | SSSRQGNTEVF | 0.6492443 |
| RSYRMHPAV | SSSRQGNTEVF | 0.5360838 |
| SAAHLLAGQLA | SSSRQGNTEVF | 0.7201781 |
| SAAVGSY | SSSRQGNTEVF | 0.8105386 |
| SPHAVLHRI | SSSRQGNTEVF | 0.85055643 |
| SQIAKVL | SSSRQGNTEVF | 0.78341734 |
| TALGHLVP | SSSRQGNTEVF | 0.70688236 |
| TAPTTSGAA | SSSRQGNTEVF | 0.8373788 |
| TDGDDGNGGSPGT | SSSRQGNTEVF | 0.82429063 |
| TDGLGEN | SSSRQGNTEVF | 0.7588186 |
| TGGPPLPGQA | SSSRQGNTEVF | 0.7957736 |
| TIPAITI | SSSRQGNTEVF | 0.72609305 |
| TIPGFTI | SSSRQGNTEVF | 0.8295213 |
| TSIASGALA | SSSRQGNTEVF | 0.8152102 |
| TVTLPGAAI | SSSRQGNTEVF | 0.8431525 |
| VASGLSAALP | SSSRQGNTEVF | 0.8449721 |
| VGAGVIGI | SSSRQGNTEVF | 0.7944745 |
| VGPKRFGV | SSSRQGNTEVF | 0.75479233 |
| VNVAFPD | SSSRQGNTEVF | 0.36400408 |
| VSMTNSGVS | SSSRQGNTEVF | 0.692935 |
| VTDVLGDLSV | SSSRQGNTEVF | 0.79879373 |
| VVVVGAGFAGL | SSSRQGNTEVF | 0.8161002 |
| VYTVGTVV | SSSRQGNTEVF | 0.82392764 |
| WTTFSGP | SSSRQGNTEVF | 0.7939633 |
| YDELVEAF | SSSRQGNTEVF | 0.7162926 |
| YGTGGAGGNGG | SSSRQGNTEVF | 0.8167634 |
| YTNVMAV | SSSRQGNTEVF | 0.84530437 |
| FLEQVHQL | SSSRQGNTEVF | 0.5214343 |
| GLSLLAVAV | SSSRQGNTEVF | 0.7232172 |
| LEQVHQL | SSSRQGNTEVF | 0.8596581 |
| LNAAGAI | SSSRQGNTEVF | 0.81520855 |
| PAVHASL | SSSRQGNTEVF | 0.7433998 |
| TELEDDA | SSSRQGNTEVF | 0.72934747 |
| TPPGPPN | SSSRQGNTEVF | 0.85776234 |
| TPPPPPPP | SSSRQGNTEVF | 0.73176384 |
| TQSPALI | AWIPTGQDTQY | 0.8454921 |
| EFAAAFPQ | AWIPTGQDTQY | 0.34420764 |
| LLSGNLDVL | AWIPTGQDTQY | 0.7354399 |
| GPANLGNYNI | AWIPTGQDTQY | 0.36699688 |
| TLGPTLP | AWIPTGQDTQY | 0.29128 |
| VYAYSSCCK | AWIPTGQDTQY | 0.8432994 |
| APAPAPAPNI | AWIPTGQDTQY | 0.61398935 |
| PGGPVPIP | AWIPTGQDTQY | 0.83671117 |
| FLTVALEP | AWIPTGQDTQY | 0.72896063 |
| SQYFGANL | AWIPTGQDTQY | 0.84478 |
| GGVKPAA | AWIPTGQDTQY | 0.8067651 |
| GGTGGAGGAGGW | AWIPTGQDTQY | 0.77076447 |
| GVGGVAGLGGAGG | AWIPTGQDTQY | 0.5538519 |
| VLLVLATTI | AWIPTGQDTQY | 0.81520957 |

| | | |
|---|---|---|
| IGIFAAML | AWIPTGQDTQY | 0.82680035 |
| AVGDLID | AWIPTGQDTQY | 0.78388476 |
| NISPLPL | AWIPTGQDTQY | 0.43431783 |
| VTAVFVAI | AWIPTGQDTQY | 0.73187524 |
| SGDDNDAAL | AWIPTGQDTQY | 0.7583642 |
| AVAALAHS | AWIPTGQDTQY | 0.22368163 |
| GGVTVMATP | AWIPTGQDTQY | 0.17840384 |
| AGLAANVRD | AWIPTGQDTQY | 0.65158427 |
| TNRPDLIDPALLRPG | AWIPTGQDTQY | 0.7541401 |
| LANLGVTAL | AWIPTGQDTQY | 0.84038675 |
| TGERIEL | AWIPTGQDTQY | 0.8324477 |
| NGEELAA | AWIPTGQDTQY | 0.73235947 |
| VLSLVWG | AWIPTGQDTQY | 0.83009994 |
| IGYALWL | AWIPTGQDTQY | 0.8042765 |
| LSLLLAPL | AWIPTGQDTQY | 0.14076427 |
| TLDEALK | AWIPTGQDTQY | 0.6589994 |
| TDGAGTAEGG | AWIPTGQDTQY | 0.4475733 |
| NAPGGAGGNGG | AWIPTGQDTQY | 0.85100204 |
| VSVFMGMGGNF | AWIPTGQDTQY | 0.66094804 |
| LASYLAAL | AWIPTGQDTQY | 0.7235992 |
| LVGGTPAL | AWIPTGQDTQY | 0.7815454 |
| VRNADVL | AWIPTGQDTQY | 0.831295 |
| TCSSPFLVSAM | AWIPTGQDTQY | 0.8095189 |
| VHITDAE | AWIPTGQDTQY | 0.49644905 |
| EYAAPSF | AWIPTGQDTQY | 0.19992995 |
| VVDAFHGGL | AWIPTGQDTQY | 0.8150275 |
| ASVVDPQ | AWIPTGQDTQY | 0.4769482 |
| SYNEFGGLS | AWIPTGQDTQY | 0.603991 |
| DPDGSVY | AWIPTGQDTQY | 0.30177823 |
| MVSTIHATE | AWIPTGQDTQY | 0.6842514 |
| LAAGVADPV | AWIPTGQDTQY | 0.53486377 |
| TKEQIAA | AWIPTGQDTQY | 0.835544 |
| SLASPAAL | AWIPTGQDTQY | 0.72816515 |
| IGLVFYGV | AWIPTGQDTQY | 0.76800793 |
| FLLDASNL | AWIPTGQDTQY | 0.8580776 |
| YQVGNFP | AWIPTGQDTQY | 0.8254739 |
| ESLLSQST | AWIPTGQDTQY | 0.7413175 |
| LFTVGFAAVG | AWIPTGQDTQY | 0.6941669 |
| ATPPTPT | AWIPTGQDTQY | 0.61871237 |
| TDLATLG | AWIPTGQDTQY | 0.6538579 |
| SAGGTGR | AWIPTGQDTQY | 0.69754374 |
| GFSNSGS | AWIPTGQDTQY | 0.8475692 |
| GGNGGSGGTSV | AWIPTGQDTQY | 0.7189858 |
| GTSVATGGAGN | AWIPTGQDTQY | 0.3656851 |
| NGGNGGSGGTS | AWIPTGQDTQY | 0.7819867 |
| AVVKFPH | AWIPTGQDTQY | 0.85356873 |
| TPPPPPP | AWIPTGQDTQY | 0.86116123 |
| AAPAAAAAPA | AWIPTGQDTQY | 0.62319064 |
| IALNEDL | AWIPTGQDTQY | 0.6662931 |
| IVNVSSM | AWIPTGQDTQY | 0.67961836 |
| TGIPPAPR | AWIPTGQDTQY | 0.78956866 |

| | | |
|---|---|---|
| FDLGGGTF | AWIPTGQDTQY | 0.6743188 |
| FDLGGGTFDVS | AWIPTGQDTQY | 0.56595844 |
| NEPTAAA | AWIPTGQDTQY | 0.82141113 |
| NEPTAAALA | AWIPTGQDTQY | 0.86289006 |
| GSIINISSI | AWIPTGQDTQY | 0.7028901 |
| VAPGNAGTA | AWIPTGQDTQY | 0.42162424 |
| TGGGASLE | AWIPTGQDTQY | 0.75439703 |
| VAVYNHYKRI | AWIPTGQDTQY | 0.6862829 |
| TDPAPATT | AWIPTGQDTQY | 0.7233466 |
| TNRPDILDPALLRPG | AWIPTGQDTQY | 0.7602936 |
| LTTTTTS | AWIPTGQDTQY | 0.50840664 |
| PTGSGKTL | AWIPTGQDTQY | 0.78246504 |
| TGSGKTL | AWIPTGQDTQY | 0.85930264 |
| TIGHVDHGK | AWIPTGQDTQY | 0.76582634 |
| TIGHVDHGKTT | AWIPTGQDTQY | 0.8248776 |
| GHVDHGKTT | AWIPTGQDTQY | 0.8502005 |
| AQFEHTLL | AWIPTGQDTQY | 0.83511937 |
| IVNQDEL | AWIPTGQDTQY | 0.8589411 |
| TNGSQFF | AWIPTGQDTQY | 0.843091 |
| TGKGIVE | AWIPTGQDTQY | 0.74051714 |
| LISNASDA | AWIPTGQDTQY | 0.7137059 |
| AELRTLGELTKIAKAHGAQ | AWIPTGQDTQY | 0.84045297 |
| APNAVAAAEPSW | AWIPTGQDTQY | 0.7773726 |
| APQVAGAATAVL | AWIPTGQDTQY | 0.7641802 |
| ARRDPAAAAAILPTDA | AWIPTGQDTQY | 0.70755625 |
| DAAARDVLAEAGLAEYF | AWIPTGQDTQY | 0.8468036 |
| DDQIVLSAKANLAKVEQ | AWIPTGQDTQY | 0.74788946 |
| DELLPDTGEDSDEDGH | AWIPTGQDTQY | 0.84920025 |
| ELPEAPANRATVPQD | AWIPTGQDTQY | 0.34174803 |
| FGQSKIERVTGIG | AWIPTGQDTQY | 0.62824595 |
| GAGGTGSTAGGAGGAGGAGGLY | AWIPTGQDTQY | 0.8485259 |
| GHGGTGAAGGNGGTGGAGGSGI | AWIPTGQDTQY | 0.13715859 |
| GNGGNGGAGGEGGPGGAGGAGA | AWIPTGQDTQY | 0.24139643 |
| GSTAGNGGNGGNATNGTIAGQP | AWIPTGQDTQY | 0.76402086 |
| GTGGDGGNGGNGGTGG | AWIPTGQDTQY | 0.8516855 |
| GVRRPAGIPADAPV | AWIPTGQDTQY | 0.15060882 |
| GYRQVTDAHLLTLARR | AWIPTGQDTQY | 0.6222764 |
| IAKMGQDGHDRG | AWIPTGQDTQY | 0.51570773 |
| KAEPSIEIQYT | AWIPTGQDTQY | 0.80390894 |
| KPVRIRMAADEDTVAVT | AWIPTGQDTQY | 0.7772738 |
| MPPRIAGMRLLVIKPEP | AWIPTGQDTQY | 0.41783512 |
| MKFYAVALFRSRGVPPTR | AWIPTGQDTQY | 0.8411267 |
| NAGGFGILGAVAHSP | AWIPTGQDTQY | 0.31711364 |
| NEGREITLAMAEM | AWIPTGQDTQY | 0.68558174 |
| NELVEWSHPLGHRWRWE | AWIPTGQDTQY | 0.8360523 |
| NLGAGGTGGNGGNPG | AWIPTGQDTQY | 0.8556894 |
| NTEWPKWETVMAADPVD | AWIPTGQDTQY | 0.44021294 |
| PASAAPAETQP | AWIPTGQDTQY | 0.8457977 |
| QQSSDGKNGVRAAEITGE | AWIPTGQDTQY | 0.68613774 |
| RAPDAVTTSWWKE | AWIPTGQDTQY | 0.865703 |
| SGFGNVGAHTSGWFNQ | AWIPTGQDTQY | 0.71824795 |

| | | |
|---|---|---|
| SGVAGAVGQAASVGGLKV | AWIPTGQDTQY | 0.45703185 |
| SNGGAGTGSAGGNGGT | AWIPTGQDTQY | 0.84047866 |
| TGGAGGEGGAAG | AWIPTGQDTQY | 0.83908933 |
| TGGDGGAAGAGGAPGHG | AWIPTGQDTQY | 0.74233764 |
| TMPPLLAAIANAGRHGV | AWIPTGQDTQY | 0.8409194 |
| TSSTAGNSGGA | AWIPTGQDTQY | 0.6715145 |
| VPATVPVGILC | AWIPTGQDTQY | 0.78549355 |
| VQRPDSGGVVLTGRIS | AWIPTGQDTQY | 0.83827496 |
| WTPNPVPATKGVPAGLVQAT | AWIPTGQDTQY | 0.8270325 |
| YALKRALAKFSDRFDVVII | AWIPTGQDTQY | 0.53546816 |
| YLIGSVIV | AWIPTGQDTQY | 0.8256163 |
| YGQLVAVID | AWIPTGQDTQY | 0.7759881 |
| VTVGGMTLPA | AWIPTGQDTQY | 0.4739012 |
| VLALPAV | AWIPTGQDTQY | 0.860579 |
| VIPPTLNYET | AWIPTGQDTQY | 0.77806795 |
| VGMGGMHPSSG | AWIPTGQDTQY | 0.84001493 |
| VGLAGGAATA | AWIPTGQDTQY | 0.8225062 |
| VGGAGGAGGSGP | AWIPTGQDTQY | 0.85775006 |
| VFAAFMV | AWIPTGQDTQY | 0.5648506 |
| TLYDKLR | AWIPTGQDTQY | 0.812858 |
| TLAVFTVL | AWIPTGQDTQY | 0.79059696 |
| TGSVQPARV | AWIPTGQDTQY | 0.83959925 |
| TGGGAGSGGT | AWIPTGQDTQY | 0.8055887 |
| SHLETSL | AWIPTGQDTQY | 0.7206398 |
| SFTTGMT | AWIPTGQDTQY | 0.80069554 |
| QVEVAKRMGVRQPTVS | AWIPTGQDTQY | 0.84946376 |
| QGSGGPALAM | AWIPTGQDTQY | 0.72480536 |
| MLAPSNP | AWIPTGQDTQY | 0.7660704 |
| LVNNAAI | AWIPTGQDTQY | 0.86031806 |
| LTGEINTLA | AWIPTGQDTQY | 0.4747299 |
| LSIFGPV | AWIPTGQDTQY | 0.78495455 |
| LLALLANL | AWIPTGQDTQY | 0.8259723 |
| LISNASDALDKLR | AWIPTGQDTQY | 0.8335197 |
| LINEAVP | AWIPTGQDTQY | 0.76179785 |
| LGYAFSGL | AWIPTGQDTQY | 0.8392405 |
| LDFVAVAL | AWIPTGQDTQY | 0.6574017 |
| LAQPGAE | AWIPTGQDTQY | 0.8310326 |
| ISNLGAVTGF | AWIPTGQDTQY | 0.7523824 |
| HQEDPGRI | AWIPTGQDTQY | 0.80531347 |
| GTPGEPTP | AWIPTGQDTQY | 0.6189215 |
| GFFNSGS | AWIPTGQDTQY | 0.81259274 |
| GAGGGGLTAGAN | AWIPTGQDTQY | 0.8382555 |
| FVDALVNM | AWIPTGQDTQY | 0.8321711 |
| FLTVAPEL | AWIPTGQDTQY | 0.20433292 |
| FETLTAL | AWIPTGQDTQY | 0.84425306 |
| DVINPIEL | AWIPTGQDTQY | 0.8301408 |
| APVSGAPV | AWIPTGQDTQY | 0.86174875 |
| ALVGGLAW | AWIPTGQDTQY | 0.7327952 |
| ALAAAAATAI | AWIPTGQDTQY | 0.808259 |
| AFGLGLLL | AWIPTGQDTQY | 0.7148255 |
| AFFAAASL | AWIPTGQDTQY | 0.7913524 |

| | | |
|---|---|---|
| AAAALPAFSPP | AWIPTGQDTQY | 0.59703 |
| VVLAPAL | AWIPTGQDTQY | 0.6273256 |
| TIGHVDHGKTTL | AWIPTGQDTQY | 0.7813358 |
| LISNASDALDK | AWIPTGQDTQY | 0.72955614 |
| LISNASDAL | AWIPTGQDTQY | 0.8437846 |
| ISNASDALDK | AWIPTGQDTQY | 0.832808 |
| NQAETLVYQTE | AWIPTGQDTQY | 0.8375151 |
| TGGTGGTAGAG | AWIPTGQDTQY | 0.8496047 |
| TGSTAGGAGGA | AWIPTGQDTQY | 0.85846865 |
| DPLAVPVAAGPV | AWIPTGQDTQY | 0.8476635 |
| GGTGGTPGNGGN | AWIPTGQDTQY | 0.69595844 |
| PAGLAGVDPATL | AWIPTGQDTQY | 0.7764907 |
| GNNLVVEAPHAGASV | AWIPTGQDTQY | 0.56881523 |
| KVTRIMSASTLEQPA | AWIPTGQDTQY | 0.7392813 |
| LGFALPIALPRRTVV | AWIPTGQDTQY | 0.7436069 |
| TRLGNLTARKLVVLV | AWIPTGQDTQY | 0.8025315 |
| WSQQRRLIYFPSAGP | AWIPTGQDTQY | 0.14028259 |
| GPNGNPDPMAAAVDIR | AWIPTGQDTQY | 0.8200129 |
| TQRCVSVADLRAAASR | AWIPTGQDTQY | 0.8526837 |
| QFVLDVAEHPVEVLTETV | AWIPTGQDTQY | 0.72169316 |
| GGAGGTGGAGGNGGTQAGDG | AWIPTGQDTQY | 0.8215591 |
| PLLATLRAIAVDVDAKPAQV | AWIPTGQDTQY | 0.8306345 |
| APKFRTGGYMIDFWGVGYQVAK | AWIPTGQDTQY | 0.6783184 |
| GAGGAGGAGGAAGGPGGTGGTG | AWIPTGQDTQY | 0.83568937 |
| IGALGLLATDVAPAGSVAFGVA | AWIPTGQDTQY | 0.71423405 |
| AGGAGGIGGNGGAGGAGGVLMG | AWIPTGQDTQY | 0.84025383 |
| ADELIGLL | AWIPTGQDTQY | 0.85684025 |
| ADRALKA | AWIPTGQDTQY | 0.79605865 |
| AGAVGQVGGAGGA | AWIPTGQDTQY | 0.6662135 |
| AGGGGGLGGAP | AWIPTGQDTQY | 0.76910996 |
| AGGLLYGNGGAG | AWIPTGQDTQY | 0.3724392 |
| AGLESIGL | AWIPTGQDTQY | 0.727128 |
| AGLKTGII | AWIPTGQDTQY | 0.33508977 |
| ASLAALLL | AWIPTGQDTQY | 0.80368054 |
| ASVINISSI | AWIPTGQDTQY | 0.5558001 |
| ATPPGPA | AWIPTGQDTQY | 0.8238254 |
| ATVASALV | AWIPTGQDTQY | 0.7938937 |
| ATVASLIL | AWIPTGQDTQY | 0.7643547 |
| AVNGPVLIRH | AWIPTGQDTQY | 0.6272348 |
| DGAAGAVGI | AWIPTGQDTQY | 0.81857973 |
| DPLLHGI | AWIPTGQDTQY | 0.78639936 |
| ELGAALGAL | AWIPTGQDTQY | 0.5026672 |
| ELGRTQAV | AWIPTGQDTQY | 0.84100044 |
| FVAGGYLS | AWIPTGQDTQY | 0.69917834 |
| GASGGAGGAGGW | AWIPTGQDTQY | 0.7666366 |
| GGAGAGGDGGAGGI | AWIPTGQDTQY | 0.8411497 |
| GTGHNGSQPGMGGNG | AWIPTGQDTQY | 0.8102231 |
| GYAAASASAS | AWIPTGQDTQY | 0.43833816 |
| HQVDVAAVQQG | AWIPTGQDTQY | 0.78275985 |
| IGAAIAHT | AWIPTGQDTQY | 0.787871 |
| IGMIEVAQL | AWIPTGQDTQY | 0.74105513 |

| | | |
|---|---|---|
| IRLAPEHTSAMD | AWIPTGQDTQY | 0.18987815 |
| IYNVSVL | AWIPTGQDTQY | 0.808338 |
| KQGFLGM | AWIPTGQDTQY | 0.47892237 |
| LESVITQL | AWIPTGQDTQY | 0.61963546 |
| LGLGALVVV | AWIPTGQDTQY | 0.7230686 |
| LIGNGGNGGSGGI | AWIPTGQDTQY | 0.8489907 |
| LNAINAPTL | AWIPTGQDTQY | 0.76076853 |
| LQLLNTGV | AWIPTGQDTQY | 0.79206437 |
| LQTSIEF | AWIPTGQDTQY | 0.8145793 |
| MSFAVSEL | AWIPTGQDTQY | 0.79000247 |
| NTGALGL | AWIPTGQDTQY | 0.8503416 |
| PPGAGGAGGAGTT | AWIPTGQDTQY | 0.81133705 |
| QLVGLGI | AWIPTGQDTQY | 0.6040829 |
| RFLEQVHQL | AWIPTGQDTQY | 0.7959149 |
| RGVGYVLR | AWIPTGQDTQY | 0.6988723 |
| SATGVFAGVI | AWIPTGQDTQY | 0.4734033 |
| SLPAITLP | AWIPTGQDTQY | 0.53686047 |
| STALAFTV | AWIPTGQDTQY | 0.65869045 |
| SVLTVGLGK | AWIPTGQDTQY | 0.38636845 |
| TALAGAGSGI | AWIPTGQDTQY | 0.797142 |
| TGLPFTGI | AWIPTGQDTQY | 0.84235156 |
| VAEHVTV | AWIPTGQDTQY | 0.79079896 |
| VDIVLNSL | AWIPTGQDTQY | 0.30212995 |
| VLTGGGAVL | AWIPTGQDTQY | 0.67418325 |
| VNVGLAYGI | AWIPTGQDTQY | 0.7264061 |
| VSSAAVGAV | AWIPTGQDTQY | 0.83645105 |
| VTSVYEAGV | AWIPTGQDTQY | 0.8029178 |
| WVALDRGA | AWIPTGQDTQY | 0.8591704 |
| YLVGSVVI | AWIPTGQDTQY | 0.8402442 |
| GPPGTGKTLL | AWIPTGQDTQY | 0.744643 |
| GVGMTKF | AWIPTGQDTQY | 0.8393053 |
| IGIGGSDL | AWIPTGQDTQY | 0.72533906 |
| IGTLLAEL | AWIPTGQDTQY | 0.7516842 |
| IYTSGTT | AWIPTGQDTQY | 0.7279065 |
| LSFPAGV | AWIPTGQDTQY | 0.77533007 |
| LYTSGTT | AWIPTGQDTQY | 0.7022641 |
| VGAFPFGF | AWIPTGQDTQY | 0.66594934 |
| VGAGTVEF | AWIPTGQDTQY | 0.38029486 |
| VGKSTLLL | AWIPTGQDTQY | 0.77009535 |
| AALAHAGGGIV | AWIPTGQDTQY | 0.49638206 |
| AALVAVSLL | AWIPTGQDTQY | 0.37684333 |
| AAVSRAVAAL | AWIPTGQDTQY | 0.7235186 |
| AGGSSGIGGF | AWIPTGQDTQY | 0.8604685 |
| AGLTVANANG | AWIPTGQDTQY | 0.853935 |
| AGLVGAVL | AWIPTGQDTQY | 0.7025343 |
| ALSQAFTV | AWIPTGQDTQY | 0.61867326 |
| ALTFSVSL | AWIPTGQDTQY | 0.8179491 |
| ALTGAALVAL | AWIPTGQDTQY | 0.55091834 |
| ASFSIGAGVL | AWIPTGQDTQY | 0.8188459 |
| DGSVYKI | AWIPTGQDTQY | 0.8058708 |
| EQLAEIV | AWIPTGQDTQY | 0.7878382 |

| | | |
|---|---|---|
| FAVSELTVM | AWIPTGQDTQY | 0.58121854 |
| GAAPLAAGL | AWIPTGQDTQY | 0.8184384 |
| GADGGDGGSGGAGG | AWIPTGQDTQY | 0.7283987 |
| GFGLDTGGAGG | AWIPTGQDTQY | 0.81793594 |
| GLASGGLL | AWIPTGQDTQY | 0.17434792 |
| GRIAGAAVLL | AWIPTGQDTQY | 0.8271395 |
| GSAGSLNTSF | AWIPTGQDTQY | 0.6890102 |
| GTGQIGI | AWIPTGQDTQY | 0.563171 |
| GTLIALIL | AWIPTGQDTQY | 0.25336474 |
| IAATMGFVVG | AWIPTGQDTQY | 0.6814611 |
| IFAQALL | AWIPTGQDTQY | 0.8414477 |
| IGAAVALGF | AWIPTGQDTQY | 0.8464507 |
| IGLTPLAL | AWIPTGQDTQY | 0.61291134 |
| ILGSNTSTL | AWIPTGQDTQY | 0.8293191 |
| IVFSLTVL | AWIPTGQDTQY | 0.8554787 |
| LASKATGF | AWIPTGQDTQY | 0.75790226 |
| LASVALIL | AWIPTGQDTQY | 0.2865405 |
| LFAVGLTL | AWIPTGQDTQY | 0.8195446 |
| LFNGGIGTYI | AWIPTGQDTQY | 0.75903213 |
| LTLGAGII | AWIPTGQDTQY | 0.8415599 |
| LVAAALGDI | AWIPTGQDTQY | 0.8240758 |
| NTAFVQLGI | AWIPTGQDTQY | 0.58330214 |
| QLFSGAEL | AWIPTGQDTQY | 0.8324686 |
| SAATFITL | AWIPTGQDTQY | 0.84875405 |
| SNYGLANLGV | AWIPTGQDTQY | 0.57962894 |
| SYGGAYAVM | AWIPTGQDTQY | 0.5062116 |
| TTGVGGLTLG | AWIPTGQDTQY | 0.5513371 |
| TTQTIYI | AWIPTGQDTQY | 0.44430286 |
| VAAGAGLGAVI | AWIPTGQDTQY | 0.7945554 |
| VAAVVTLL | AWIPTGQDTQY | 0.75243306 |
| VGFLGGLI | AWIPTGQDTQY | 0.7687744 |
| VGVIPILL | AWIPTGQDTQY | 0.7862724 |
| VIFASSLI | AWIPTGQDTQY | 0.70635784 |
| VKGGDGGAAAG | AWIPTGQDTQY | 0.72441494 |
| VTKTFTNI | AWIPTGQDTQY | 0.7373597 |
| VVAAAILL | AWIPTGQDTQY | 0.79147357 |
| YAALDVEL | AWIPTGQDTQY | 0.8260149 |
| LASVALLL | AWIPTGQDTQY | 0.7762054 |
| LDAVHRV | AWIPTGQDTQY | 0.7635287 |
| QGLSSSIF | AWIPTGQDTQY | 0.8558959 |
| RGTFANIRL | AWIPTGQDTQY | 0.49419236 |
| RQLWWGHRI | AWIPTGQDTQY | 0.6391772 |
| YGPPGTGKTLL | AWIPTGQDTQY | 0.834599 |
| AALVGAPANVA | AWIPTGQDTQY | 0.55425835 |
| AATFRRTE | AWIPTGQDTQY | 0.8483207 |
| AGALGGGI | AWIPTGQDTQY | 0.78587806 |
| AGGAGGAGGIGAG | AWIPTGQDTQY | 0.8112546 |
| AGSGALGL | AWIPTGQDTQY | 0.7670307 |
| AQIADQI | AWIPTGQDTQY | 0.8572007 |
| ASAVAVTYL | AWIPTGQDTQY | 0.84503305 |
| ASTVLGGTGG | AWIPTGQDTQY | 0.682469 |

| | | |
|---|---|---|
| AVGGEPAK | AWIPTGQDTQY | 0.8473134 |
| CEQGTFGGTC | AWIPTGQDTQY | 0.39523095 |
| DEGKAPAG | AWIPTGQDTQY | 0.8218787 |
| DTVVFPAL | AWIPTGQDTQY | 0.8347218 |
| FEGITMAG | AWIPTGQDTQY | 0.6315565 |
| FPLVVQD | AWIPTGQDTQY | 0.78659934 |
| GAFTTPQ | AWIPTGQDTQY | 0.84727246 |
| GFGGFGTAGGVGGA | AWIPTGQDTQY | 0.8470844 |
| GFLAQFL | AWIPTGQDTQY | 0.3982587 |
| GGAGGAGGNAGVA | AWIPTGQDTQY | 0.71692383 |
| GGVGTAGAGGAGGAGG | AWIPTGQDTQY | 0.37888658 |
| GNGGGGDAGNAGSGGN | AWIPTGQDTQY | 0.4331382 |
| GTGGAGGAGNDAGS | AWIPTGQDTQY | 0.3234019 |
| GTTNTGWFNS | AWIPTGQDTQY | 0.8242573 |
| IFMGPLFTL | AWIPTGQDTQY | 0.74255097 |
| ILAHAIEAI | AWIPTGQDTQY | 0.8380718 |
| IRSAIREAL | AWIPTGQDTQY | 0.7922535 |
| ITFAGGG | AWIPTGQDTQY | 0.8465336 |
| ITGMSGHQEV | AWIPTGQDTQY | 0.68183064 |
| IVAGLACTL | AWIPTGQDTQY | 0.76228714 |
| IVGGGLAGA | AWIPTGQDTQY | 0.378879 |
| IVTGVLNI | AWIPTGQDTQY | 0.37561482 |
| LAQQGCI | AWIPTGQDTQY | 0.5133636 |
| LCLHVEN | AWIPTGQDTQY | 0.7711005 |
| LDAISDGS | AWIPTGQDTQY | 0.21689013 |
| LGGFAAML | AWIPTGQDTQY | 0.8520595 |
| LGGKNPMI | AWIPTGQDTQY | 0.80979246 |
| LGPKGVAAIL | AWIPTGQDTQY | 0.80929756 |
| LMVGVSI | AWIPTGQDTQY | 0.66101074 |
| LVGAAFVL | AWIPTGQDTQY | 0.6015427 |
| LVLAVAPL | AWIPTGQDTQY | 0.7293997 |
| MAAIAVIDAA | AWIPTGQDTQY | 0.8375554 |
| NNGINAELQ | AWIPTGQDTQY | 0.6199269 |
| RGEPLVLVHG | AWIPTGQDTQY | 0.23922297 |
| RGGAGGAAGLI | AWIPTGQDTQY | 0.5783189 |
| RSYRMHPAV | AWIPTGQDTQY | 0.47388473 |
| SAAHLLAGQLA | AWIPTGQDTQY | 0.857128 |
| SAAVGSY | AWIPTGQDTQY | 0.8400618 |
| SPHAVLHRI | AWIPTGQDTQY | 0.8191189 |
| SQIAKVL | AWIPTGQDTQY | 0.8473346 |
| TALGHLVP | AWIPTGQDTQY | 0.16195008 |
| TAPTTSGAA | AWIPTGQDTQY | 0.6438733 |
| TDGDDGNGGSPGT | AWIPTGQDTQY | 0.84991586 |
| TDGLGEN | AWIPTGQDTQY | 0.8171969 |
| TGGPPLPGQA | AWIPTGQDTQY | 0.7632983 |
| TIPAITI | AWIPTGQDTQY | 0.85275817 |
| TIPGFTI | AWIPTGQDTQY | 0.67846906 |
| TSIASGALA | AWIPTGQDTQY | 0.7402265 |
| TVTLPGAAI | AWIPTGQDTQY | 0.8194481 |
| VASGLSAALP | AWIPTGQDTQY | 0.7874038 |
| VGAGVIGI | AWIPTGQDTQY | 0.79401696 |

| | | |
|---|---|---|
| VGPKRFGV | AWIPTGQDTQY | 0.43770045 |
| VNVAFPD | AWIPTGQDTQY | 0.80513126 |
| VSMTNSGVS | AWIPTGQDTQY | 0.8625142 |
| VTDVLGDLSV | AWIPTGQDTQY | 0.78911674 |
| VVVVGAGFAGL | AWIPTGQDTQY | 0.818926 |
| VYTVGTVV | AWIPTGQDTQY | 0.8261472 |
| WTTFSGP | AWIPTGQDTQY | 0.81705076 |
| YDELVEAF | AWIPTGQDTQY | 0.669395 |
| YGTGGAGGNGG | AWIPTGQDTQY | 0.66465795 |
| YTNVMAV | AWIPTGQDTQY | 0.81130004 |
| FLEQVHQL | AWIPTGQDTQY | 0.7000791 |
| GLSLLAVAV | AWIPTGQDTQY | 0.7888367 |
| LEQVHQL | AWIPTGQDTQY | 0.7116326 |
| LNAAGAI | AWIPTGQDTQY | 0.8667654 |
| PAVHASL | AWIPTGQDTQY | 0.8246126 |
| TELEDDA | AWIPTGQDTQY | 0.8368011 |
| TPPGPPN | AWIPTGQDTQY | 0.6912987 |
| TPPPPPPP | AWIPTGQDTQY | 0.76357144 |
| TQSPALI | ASSLGFNQDTQY | 0.8277128 |
| EFAAAFPQ | ASSLGFNQDTQY | 0.40429598 |
| LLSGNLDVL | ASSLGFNQDTQY | 0.84441787 |
| GPANLGNYNI | ASSLGFNQDTQY | 0.6827611 |
| TLGPTLP | ASSLGFNQDTQY | 0.7726872 |
| VYAYSSCCK | ASSLGFNQDTQY | 0.82831323 |
| APAPAPAPNI | ASSLGFNQDTQY | 0.84984607 |
| PGGPVPIP | ASSLGFNQDTQY | 0.7791922 |
| FLTVALEP | ASSLGFNQDTQY | 0.7906157 |
| SQYFGANL | ASSLGFNQDTQY | 0.681289 |
| GGVKPAA | ASSLGFNQDTQY | 0.8353461 |
| GGTGGAGGAGGW | ASSLGFNQDTQY | 0.8549732 |
| GVGGVAGLGGAGG | ASSLGFNQDTQY | 0.8168409 |
| VLLVLATTI | ASSLGFNQDTQY | 0.85760903 |
| IGIFAAML | ASSLGFNQDTQY | 0.81745946 |
| AVGDLID | ASSLGFNQDTQY | 0.2492564 |
| NISPLPL | ASSLGFNQDTQY | 0.7473729 |
| VTAVFVAI | ASSLGFNQDTQY | 0.8434659 |
| SGDDNDAAL | ASSLGFNQDTQY | 0.8475212 |
| AVAALAHS | ASSLGFNQDTQY | 0.850769 |
| GGVTVMATP | ASSLGFNQDTQY | 0.8211404 |
| AGLAANVRD | ASSLGFNQDTQY | 0.66755354 |
| TNRPDLIDPALLRPG | ASSLGFNQDTQY | 0.85147035 |
| LANLGVTAL | ASSLGFNQDTQY | 0.8620871 |
| TGERIEL | ASSLGFNQDTQY | 0.83822954 |
| NGEELAA | ASSLGFNQDTQY | 0.6217449 |
| VLSLVWG | ASSLGFNQDTQY | 0.56071913 |
| IGYALWL | ASSLGFNQDTQY | 0.79901564 |
| LSLLLAPL | ASSLGFNQDTQY | 0.8467363 |
| TLDEALK | ASSLGFNQDTQY | 0.5668133 |
| TDGAGTAEGG | ASSLGFNQDTQY | 0.82152057 |
| NAPGGAGGNGG | ASSLGFNQDTQY | 0.8071592 |
| VSVFMGMGGNF | ASSLGFNQDTQY | 0.24425335 |

| | | |
|---|---|---|
| LASYLAAL | ASSLGFNQDTQY | 0.8223489 |
| LVGGTPAL | ASSLGFNQDTQY | 0.84857297 |
| VRNADVL | ASSLGFNQDTQY | 0.77424544 |
| TCSSPFLVSAM | ASSLGFNQDTQY | 0.6558197 |
| VHITDAE | ASSLGFNQDTQY | 0.84853894 |
| EYAAPSF | ASSLGFNQDTQY | 0.86073315 |
| VVDAFHGGL | ASSLGFNQDTQY | 0.8557861 |
| ASVVDPQ | ASSLGFNQDTQY | 0.47426337 |
| SYNEFGGLS | ASSLGFNQDTQY | 0.22552432 |
| DPDGSVY | ASSLGFNQDTQY | 0.7459048 |
| MVSTIHATE | ASSLGFNQDTQY | 0.83915496 |
| LAAGVADPV | ASSLGFNQDTQY | 0.7729168 |
| TKEQIAA | ASSLGFNQDTQY | 0.7318419 |
| SLASPAAL | ASSLGFNQDTQY | 0.8355943 |
| IGLVFYGV | ASSLGFNQDTQY | 0.8428745 |
| FLLDASNL | ASSLGFNQDTQY | 0.41872734 |
| YQVGNFP | ASSLGFNQDTQY | 0.3613867 |
| ESLLSQST | ASSLGFNQDTQY | 0.8179117 |
| LFTVGFAAVG | ASSLGFNQDTQY | 0.83807033 |
| ATPPTPT | ASSLGFNQDTQY | 0.59631467 |
| TDLATLG | ASSLGFNQDTQY | 0.78669643 |
| SAGGTGR | ASSLGFNQDTQY | 0.7695792 |
| GFSNSGS | ASSLGFNQDTQY | 0.6988578 |
| GGNGGSGGTSV | ASSLGFNQDTQY | 0.8030314 |
| GTSVATGGAGN | ASSLGFNQDTQY | 0.5539912 |
| NGGNGGSGGTS | ASSLGFNQDTQY | 0.84004086 |
| AVVKFPH | ASSLGFNQDTQY | 0.568738 |
| TPPPPPP | ASSLGFNQDTQY | 0.83022845 |
| AAPAAAAAPA | ASSLGFNQDTQY | 0.8062396 |
| IALNEDL | ASSLGFNQDTQY | 0.77291214 |
| IVNVSSM | ASSLGFNQDTQY | 0.7733501 |
| TGIPPAPR | ASSLGFNQDTQY | 0.70590955 |
| FDLGGGTF | ASSLGFNQDTQY | 0.5867659 |
| FDLGGGTFDVS | ASSLGFNQDTQY | 0.5107739 |
| NEPTAAA | ASSLGFNQDTQY | 0.83771825 |
| NEPTAAALA | ASSLGFNQDTQY | 0.7338907 |
| GSIINISSI | ASSLGFNQDTQY | 0.83887255 |
| VAPGNAGTA | ASSLGFNQDTQY | 0.82500476 |
| TGGGASLE | ASSLGFNQDTQY | 0.5875528 |
| VAVYNHYKRI | ASSLGFNQDTQY | 0.81064403 |
| TDPAPATT | ASSLGFNQDTQY | 0.55065584 |
| TNRPDILDPALLRPG | ASSLGFNQDTQY | 0.67670166 |
| LTTTTTS | ASSLGFNQDTQY | 0.8237834 |
| PTGSGKTL | ASSLGFNQDTQY | 0.8261535 |
| TGSGKTL | ASSLGFNQDTQY | 0.6053438 |
| TIGHVDHGK | ASSLGFNQDTQY | 0.8218181 |
| TIGHVDHGKTT | ASSLGFNQDTQY | 0.7993926 |
| GHVDHGKTT | ASSLGFNQDTQY | 0.4920626 |
| AQFEHTLL | ASSLGFNQDTQY | 0.74692726 |
| IVNQDEL | ASSLGFNQDTQY | 0.83977646 |
| TNGSQFF | ASSLGFNQDTQY | 0.78366077 |

| | | |
|---|---|---|
| TGKGIVE | ASSLGFNQDTQY | 0.56646687 |
| LISNASDA | ASSLGFNQDTQY | 0.8274808 |
| AELRTLGELTKIAKAHGAQ | ASSLGFNQDTQY | 0.70379305 |
| APNAVAAAEPSW | ASSLGFNQDTQY | 0.8121806 |
| APQVAGAATAVL | ASSLGFNQDTQY | 0.78890586 |
| ARRDPAAAAAILPTDA | ASSLGFNQDTQY | 0.72782123 |
| DAAARDVLAEAGLAEYF | ASSLGFNQDTQY | 0.8327675 |
| DDQIVLSAKANLAKVEQ | ASSLGFNQDTQY | 0.4820999 |
| DELLPDTGEDSDEDGH | ASSLGFNQDTQY | 0.55721164 |
| ELPEAPANRATVPQD | ASSLGFNQDTQY | 0.7329731 |
| FGQSKIERVTGIG | ASSLGFNQDTQY | 0.7840812 |
| GAGGTGSTAGGAGGAGGAGGLY | ASSLGFNQDTQY | 0.84103125 |
| GHGGTGAAGGNGGTGGAGGSGI | ASSLGFNQDTQY | 0.60218114 |
| GNGGNGGAGGEGGPGGAGGAGA | ASSLGFNQDTQY | 0.7871611 |
| GSTAGNGGNGGNATNGTIAGQP | ASSLGFNQDTQY | 0.7013557 |
| GTGGDGGNGGNGGTGG | ASSLGFNQDTQY | 0.5781194 |
| GVRRPAGIPADAPV | ASSLGFNQDTQY | 0.78128934 |
| GYRQVTDAHLLTLARR | ASSLGFNQDTQY | 0.68354833 |
| IAKMGQDGHDRG | ASSLGFNQDTQY | 0.56193244 |
| KAEPSIEIQYT | ASSLGFNQDTQY | 0.80370796 |
| KPVRIRMAADEDTVAVT | ASSLGFNQDTQY | 0.67949426 |
| MPPRIAGMRLLVIKPEP | ASSLGFNQDTQY | 0.76620054 |
| MKFYAVALFRSRGVPPTR | ASSLGFNQDTQY | 0.8561865 |
| NAGGFGILGAVAHSP | ASSLGFNQDTQY | 0.4668038 |
| NEGREITLAMAEM | ASSLGFNQDTQY | 0.8222563 |
| NELVEWSHPLGHRWRWE | ASSLGFNQDTQY | 0.81781256 |
| NLGAGGTGGNGGNPG | ASSLGFNQDTQY | 0.8175936 |
| NTEWPKWETVMAADPVD | ASSLGFNQDTQY | 0.76131344 |
| PASAAPAETQP | ASSLGFNQDTQY | 0.68501717 |
| QQSSDGKNGVRAAEITGE | ASSLGFNQDTQY | 0.854486 |
| RAPDAVTTSWWKE | ASSLGFNQDTQY | 0.8243761 |
| SGFGNVGAHTSGWFNQ | ASSLGFNQDTQY | 0.81829494 |
| SGVAGAVGQAASVGGLKV | ASSLGFNQDTQY | 0.8260695 |
| SNGGAGTGSAGGNGGT | ASSLGFNQDTQY | 0.8291376 |
| TGGAGGEGGAAG | ASSLGFNQDTQY | 0.8390963 |
| TGGDGGAAGAGGAPGHG | ASSLGFNQDTQY | 0.744554 |
| TMPPLLAAIANAGRHGV | ASSLGFNQDTQY | 0.49698296 |
| TSSTAGNSGGA | ASSLGFNQDTQY | 0.7617394 |
| VPATVPVGILC | ASSLGFNQDTQY | 0.73203707 |
| VQRPDSGGVVLTGRIS | ASSLGFNQDTQY | 0.68459296 |
| WTPNPVPATKGVPAGLVQAT | ASSLGFNQDTQY | 0.3767354 |
| YALKRALAKFSDRFDVVII | ASSLGFNQDTQY | 0.84267676 |
| YLIGSVIV | ASSLGFNQDTQY | 0.86400104 |
| YGQLVAVID | ASSLGFNQDTQY | 0.7374004 |
| VTVGGMTLPA | ASSLGFNQDTQY | 0.86433613 |
| VLALPAV | ASSLGFNQDTQY | 0.51577747 |
| VIPPTLNYET | ASSLGFNQDTQY | 0.8288275 |
| VGMGGMHPSSG | ASSLGFNQDTQY | 0.8473527 |
| VGLAGGAATA | ASSLGFNQDTQY | 0.6934994 |
| VGGAGGAGGSGP | ASSLGFNQDTQY | 0.8069236 |
| VFAAFMV | ASSLGFNQDTQY | 0.6484035 |

| | | |
|---|---|---|
| TLYDKLR | ASSLGFNQDTQY | 0.68610764 |
| TLAVFTVL | ASSLGFNQDTQY | 0.8435191 |
| TGSVQPARV | ASSLGFNQDTQY | 0.73365825 |
| TGGGAGSGGT | ASSLGFNQDTQY | 0.7882485 |
| SHLETSL | ASSLGFNQDTQY | 0.79418683 |
| SFTTGMT | ASSLGFNQDTQY | 0.6345959 |
| QVEVAKRMGVRQPTVS | ASSLGFNQDTQY | 0.7860307 |
| QGSGGPALAM | ASSLGFNQDTQY | 0.8407421 |
| MLAPSNP | ASSLGFNQDTQY | 0.6861696 |
| LVNNAAI | ASSLGFNQDTQY | 0.8612156 |
| LTGEINTLA | ASSLGFNQDTQY | 0.48926163 |
| LSIFGPV | ASSLGFNQDTQY | 0.85649717 |
| LLALLANL | ASSLGFNQDTQY | 0.84762114 |
| LISNASDALDKLR | ASSLGFNQDTQY | 0.8166631 |
| LINEAVP | ASSLGFNQDTQY | 0.7055301 |
| LGYAFSGL | ASSLGFNQDTQY | 0.85125065 |
| LDFVAVAL | ASSLGFNQDTQY | 0.77961516 |
| LAQPGAE | ASSLGFNQDTQY | 0.78652 |
| ISNLGAVTGF | ASSLGFNQDTQY | 0.8475727 |
| HQEDPGRI | ASSLGFNQDTQY | 0.50097525 |
| GTPGEPTP | ASSLGFNQDTQY | 0.7071389 |
| GFFNSGS | ASSLGFNQDTQY | 0.66905886 |
| GAGGGGLTAGAN | ASSLGFNQDTQY | 0.6353214 |
| FVDALVNM | ASSLGFNQDTQY | 0.59534097 |
| FLTVAPEL | ASSLGFNQDTQY | 0.38840613 |
| FETLTAL | ASSLGFNQDTQY | 0.6033925 |
| DVINPIEL | ASSLGFNQDTQY | 0.5377439 |
| APVSGAPV | ASSLGFNQDTQY | 0.7009881 |
| ALVGGLAW | ASSLGFNQDTQY | 0.6886076 |
| ALAAAAATAI | ASSLGFNQDTQY | 0.5697168 |
| AFGLGLLL | ASSLGFNQDTQY | 0.70178443 |
| AFFAAASL | ASSLGFNQDTQY | 0.6648506 |
| AAAALPAFSPP | ASSLGFNQDTQY | 0.44360024 |
| VVLAPAL | ASSLGFNQDTQY | 0.63280606 |
| TIGHVDHGKTTL | ASSLGFNQDTQY | 0.5697797 |
| LISNASDALDK | ASSLGFNQDTQY | 0.64496773 |
| LISNASDAL | ASSLGFNQDTQY | 0.42488754 |
| ISNASDALDK | ASSLGFNQDTQY | 0.6869929 |
| NQAETLVYQTE | ASSLGFNQDTQY | 0.6025973 |
| TGGTGGTAGAG | ASSLGFNQDTQY | 0.29001895 |
| TGSTAGGAGGA | ASSLGFNQDTQY | 0.67187876 |
| DPLAVPVAAGPV | ASSLGFNQDTQY | 0.6413516 |
| GGTGGTPGNGGN | ASSLGFNQDTQY | 0.6193977 |
| PAGLAGVDPATL | ASSLGFNQDTQY | 0.7041935 |
| GNNLVVEAPHAGASV | ASSLGFNQDTQY | 0.4194737 |
| KVTRIMSASTLEQPA | ASSLGFNQDTQY | 0.4777354 |
| LGFALPIALPRRTVV | ASSLGFNQDTQY | 0.6878072 |
| TRLGNLTARKLVVLV | ASSLGFNQDTQY | 0.6991254 |
| WSQQRRLIYFPSAGP | ASSLGFNQDTQY | 0.44412434 |
| GPNGNPDPMAAAVDIR | ASSLGFNQDTQY | 0.67600036 |
| TQRCVSVADLRAAASR | ASSLGFNQDTQY | 0.6959176 |

| | | |
|---|---|---|
| QFVLDVAEHPVEVLTETV | ASSLGFNQDTQY | 0.6621454 |
| GGAGGTGGAGGNGGTQAGDG | ASSLGFNQDTQY | 0.70289373 |
| PLLATLRAIAVDVDAKPAQV | ASSLGFNQDTQY | 0.70831954 |
| APKFRTGGYMIDFWGVGYQVAK | ASSLGFNQDTQY | 0.56350625 |
| GAGGAGGAGGAAGGPGGTGGTG | ASSLGFNQDTQY | 0.7053602 |
| IGALGLLATDVAPAGSVAFGVA | ASSLGFNQDTQY | 0.706323 |
| AGGAGGIGGNGGAGGAGGVLMG | ASSLGFNQDTQY | 0.57034326 |
| ADELIGLL | ASSLGFNQDTQY | 0.6533936 |
| ADRALKA | ASSLGFNQDTQY | 0.47130948 |
| AGAVGQVGGAGGA | ASSLGFNQDTQY | 0.6147789 |
| AGGGGGLGGAP | ASSLGFNQDTQY | 0.42960313 |
| AGGLLYGNGGAG | ASSLGFNQDTQY | 0.68647856 |
| AGLESIGL | ASSLGFNQDTQY | 0.7026949 |
| AGLKTGII | ASSLGFNQDTQY | 0.6392704 |
| ASLAALLL | ASSLGFNQDTQY | 0.49678206 |
| ASVINISSI | ASSLGFNQDTQY | 0.46038872 |
| ATPPGPA | ASSLGFNQDTQY | 0.64879024 |
| ATVASALV | ASSLGFNQDTQY | 0.506838 |
| ATVASLIL | ASSLGFNQDTQY | 0.3567018 |
| AVNGPVLIRH | ASSLGFNQDTQY | 0.63280416 |
| DGAAGAVGI | ASSLGFNQDTQY | 0.687891 |
| DPLLHGI | ASSLGFNQDTQY | 0.6833252 |
| ELGAALGAL | ASSLGFNQDTQY | 0.5539483 |
| ELGRTQAV | ASSLGFNQDTQY | 0.71367073 |
| FVAGGYLS | ASSLGFNQDTQY | 0.6979408 |
| GASGGAGGAGGW | ASSLGFNQDTQY | 0.70000577 |
| GGAGAGGDGGAGGI | ASSLGFNQDTQY | 0.5987913 |
| GTGHNGSQPGMGGNG | ASSLGFNQDTQY | 0.49312335 |
| GYAAASASAS | ASSLGFNQDTQY | 0.7070533 |
| HQVDVAAVQQG | ASSLGFNQDTQY | 0.6397233 |
| IGAAIAHT | ASSLGFNQDTQY | 0.6280959 |
| IGMIEVAQL | ASSLGFNQDTQY | 0.6964966 |
| IRLAPEHTSAMD | ASSLGFNQDTQY | 0.5976097 |
| IYNVSVL | ASSLGFNQDTQY | 0.59129965 |
| KQGFLGM | ASSLGFNQDTQY | 0.6607401 |
| LESVITQL | ASSLGFNQDTQY | 0.70226824 |
| LGLGALVVV | ASSLGFNQDTQY | 0.5681695 |
| LIGNGGNGGSGGI | ASSLGFNQDTQY | 0.48143315 |
| LNAINAPTL | ASSLGFNQDTQY | 0.5817207 |
| LQLLNTGV | ASSLGFNQDTQY | 0.64889157 |
| LQTSIEF | ASSLGFNQDTQY | 0.7002719 |
| MSFAVSEL | ASSLGFNQDTQY | 0.66764003 |
| NTGALGL | ASSLGFNQDTQY | 0.6181116 |
| PPGAGGAGGAGTT | ASSLGFNQDTQY | 0.6810907 |
| QLVGLGI | ASSLGFNQDTQY | 0.674397 |
| RFLEQVHQL | ASSLGFNQDTQY | 0.66003 |
| RGVGYVLR | ASSLGFNQDTQY | 0.66283965 |
| SATGVFAGVI | ASSLGFNQDTQY | 0.6023002 |
| SLPAITLP | ASSLGFNQDTQY | 0.70282245 |
| STALAFTV | ASSLGFNQDTQY | 0.6918509 |
| SVLTVGLGK | ASSLGFNQDTQY | 0.6886002 |

| | | |
|---|---|---|
| TALAGAGSGI | ASSLGFNQDTQY | 0.7017525 |
| TGLPFTGI | ASSLGFNQDTQY | 0.63483745 |
| VAEHVTV | ASSLGFNQDTQY | 0.6423317 |
| VDIVLNSL | ASSLGFNQDTQY | 0.4013796 |
| VLTGGGAVL | ASSLGFNQDTQY | 0.5875697 |
| VNVGLAYGI | ASSLGFNQDTQY | 0.6745018 |
| VSSAAVGAV | ASSLGFNQDTQY | 0.68015134 |
| VTSVYEAGV | ASSLGFNQDTQY | 0.6952679 |
| WVALDRGA | ASSLGFNQDTQY | 0.6321471 |
| YLVGSVVI | ASSLGFNQDTQY | 0.59479755 |
| GPPGTGKTLL | ASSLGFNQDTQY | 0.68540287 |
| GVGMTKF | ASSLGFNQDTQY | 0.69917107 |
| IGIGGSDL | ASSLGFNQDTQY | 0.49009076 |
| IGTLLAEL | ASSLGFNQDTQY | 0.6178758 |
| IYTSGTT | ASSLGFNQDTQY | 0.71662074 |
| LSFPAGV | ASSLGFNQDTQY | 0.67948306 |
| LYTSGTT | ASSLGFNQDTQY | 0.6326126 |
| VGAFPFGF | ASSLGFNQDTQY | 0.59539306 |
| VGAGTVEF | ASSLGFNQDTQY | 0.71277684 |
| VGKSTLLL | ASSLGFNQDTQY | 0.59443724 |
| AALAHAGGGIV | ASSLGFNQDTQY | 0.71186316 |
| AALVAVSLL | ASSLGFNQDTQY | 0.34939724 |
| AAVSRAVAAL | ASSLGFNQDTQY | 0.5824659 |
| AGGSSGIGGF | ASSLGFNQDTQY | 0.48958182 |
| AGLTVANANG | ASSLGFNQDTQY | 0.3186895 |
| AGLVGAVL | ASSLGFNQDTQY | 0.69800913 |
| ALSQAFTV | ASSLGFNQDTQY | 0.5393541 |
| ALTFSVSL | ASSLGFNQDTQY | 0.67047024 |
| ALTGAALVAL | ASSLGFNQDTQY | 0.62697506 |
| ASFSIGAGVL | ASSLGFNQDTQY | 0.7043191 |
| DGSVYKI | ASSLGFNQDTQY | 0.6760556 |
| EQLAEIV | ASSLGFNQDTQY | 0.6240437 |
| FAVSELTVM | ASSLGFNQDTQY | 0.5094111 |
| GAAPLAAGL | ASSLGFNQDTQY | 0.68189794 |
| GADGGDGGSGGAGG | ASSLGFNQDTQY | 0.69926053 |
| GFGLDTGGAGG | ASSLGFNQDTQY | 0.67262924 |
| GLASGGLL | ASSLGFNQDTQY | 0.35410964 |
| GRIAGAAVLL | ASSLGFNQDTQY | 0.66034555 |
| GSAGSLNTSF | ASSLGFNQDTQY | 0.6256793 |
| GTGQIGI | ASSLGFNQDTQY | 0.34289303 |
| GTLIALIL | ASSLGFNQDTQY | 0.3120108 |
| IAATMGFVVG | ASSLGFNQDTQY | 0.57193077 |
| IFAQALL | ASSLGFNQDTQY | 0.63948315 |
| IGAAVALGF | ASSLGFNQDTQY | 0.696581 |
| IGLTPLAL | ASSLGFNQDTQY | 0.68710953 |
| ILGSNTSTL | ASSLGFNQDTQY | 0.6262314 |
| IVFSLTVL | ASSLGFNQDTQY | 0.68568695 |
| LASKATGF | ASSLGFNQDTQY | 0.6544064 |
| LASVALIL | ASSLGFNQDTQY | 0.25475973 |
| LFAVGLTL | ASSLGFNQDTQY | 0.5479452 |
| LFNGGIGTYI | ASSLGFNQDTQY | 0.46168053 |

| | | |
|---|---|---|
| LTLGAGII | ASSLGFNQDTQY | 0.7069596 |
| LVAAALGDI | ASSLGFNQDTQY | 0.57450175 |
| NTAFVQLGI | ASSLGFNQDTQY | 0.6317123 |
| QLFSGAEL | ASSLGFNQDTQY | 0.6568382 |
| SAATFITL | ASSLGFNQDTQY | 0.68501174 |
| SNYGLANLGV | ASSLGFNQDTQY | 0.65336776 |
| SYGGAYAVM | ASSLGFNQDTQY | 0.4996471 |
| TTGVGGLTLG | ASSLGFNQDTQY | 0.3195012 |
| TTQTIYI | ASSLGFNQDTQY | 0.67739034 |
| VAAGAGLGAVI | ASSLGFNQDTQY | 0.46369246 |
| VAAVVTLL | ASSLGFNQDTQY | 0.46574977 |
| VGFLGGLI | ASSLGFNQDTQY | 0.45123142 |
| VGVIPILL | ASSLGFNQDTQY | 0.60358447 |
| VIFASSLI | ASSLGFNQDTQY | 0.36008546 |
| VKGGDGGAAAG | ASSLGFNQDTQY | 0.6898774 |
| VTKTFTNI | ASSLGFNQDTQY | 0.5788585 |
| VVAAAILL | ASSLGFNQDTQY | 0.6478946 |
| YAALDVEL | ASSLGFNQDTQY | 0.71489084 |
| LASVALLL | ASSLGFNQDTQY | 0.6806096 |
| LDAVHRV | ASSLGFNQDTQY | 0.64089054 |
| QGLSSSIF | ASSLGFNQDTQY | 0.52336776 |
| RGTFANIRL | ASSLGFNQDTQY | 0.5636644 |
| RQLWWGHRI | ASSLGFNQDTQY | 0.532588 |
| YGPPGTGKTLL | ASSLGFNQDTQY | 0.53995305 |
| AALVGAPANVA | ASSLGFNQDTQY | 0.704543 |
| AATFRRTE | ASSLGFNQDTQY | 0.6176576 |
| AGALGGGI | ASSLGFNQDTQY | 0.4614091 |
| AGGAGGAGGIGAG | ASSLGFNQDTQY | 0.63919044 |
| AGSGALGL | ASSLGFNQDTQY | 0.7138947 |
| AQIADQI | ASSLGFNQDTQY | 0.7149025 |
| ASAVAVTYL | ASSLGFNQDTQY | 0.57538223 |
| ASTVLGGTGG | ASSLGFNQDTQY | 0.5325834 |
| AVGGEPAK | ASSLGFNQDTQY | 0.5518981 |
| CEQGTFGGTC | ASSLGFNQDTQY | 0.66061056 |
| DEGKAPAG | ASSLGFNQDTQY | 0.52001166 |
| DTVVFPAL | ASSLGFNQDTQY | 0.49299088 |
| FEGITMAG | ASSLGFNQDTQY | 0.6865078 |
| FPLVVQD | ASSLGFNQDTQY | 0.71903443 |
| GAFTTPQ | ASSLGFNQDTQY | 0.6153223 |
| GFGGFGTAGGVGGA | ASSLGFNQDTQY | 0.41747656 |
| GFLAQFL | ASSLGFNQDTQY | 0.649645 |
| GGAGGAGGNAGVA | ASSLGFNQDTQY | 0.5391504 |
| GGVGTAGAGGAGGAGG | ASSLGFNQDTQY | 0.5915165 |
| GNGGGGDAGNAGSGGN | ASSLGFNQDTQY | 0.5599526 |
| GTGGAGGAGNDAGS | ASSLGFNQDTQY | 0.46057814 |
| GTTNTGWFNS | ASSLGFNQDTQY | 0.63554245 |
| IFMGPLFTL | ASSLGFNQDTQY | 0.7082548 |
| ILAHAIEAI | ASSLGFNQDTQY | 0.64983904 |
| IRSAIREAL | ASSLGFNQDTQY | 0.6705657 |
| ITFAGGG | ASSLGFNQDTQY | 0.7105204 |
| ITGMSGHQEV | ASSLGFNQDTQY | 0.6964817 |

| | | |
|---|---|---|
| IVAGLACTL | ASSLGFNQDTQY | 0.7101973 |
| IVGGGLAGA | ASSLGFNQDTQY | 0.7012745 |
| IVTGVLNI | ASSLGFNQDTQY | 0.64757085 |
| LAQQGCI | ASSLGFNQDTQY | 0.6382266 |
| LCLHVEN | ASSLGFNQDTQY | 0.7039784 |
| LDAISDGS | ASSLGFNQDTQY | 0.64728117 |
| LGGFAAML | ASSLGFNQDTQY | 0.6512526 |
| LGGKNPMI | ASSLGFNQDTQY | 0.6025607 |
| LGPKGVAAIL | ASSLGFNQDTQY | 0.70103216 |
| LMVGVSI | ASSLGFNQDTQY | 0.60662484 |
| LVGAAFVL | ASSLGFNQDTQY | 0.7123057 |
| LVLAVAPL | ASSLGFNQDTQY | 0.33586258 |
| MAAIAVIDAA | ASSLGFNQDTQY | 0.47520083 |
| NNGINAELQ | ASSLGFNQDTQY | 0.70494914 |
| RGEPLVLVHG | ASSLGFNQDTQY | 0.26166937 |
| RGGAGGAAGLI | ASSLGFNQDTQY | 0.308313 |
| RSYRMHPAV | ASSLGFNQDTQY | 0.6342629 |
| SAAHLLAGQLA | ASSLGFNQDTQY | 0.7053454 |
| SAAVGSY | ASSLGFNQDTQY | 0.26768947 |
| SPHAVLHRI | ASSLGFNQDTQY | 0.4813046 |
| SQIAKVL | ASSLGFNQDTQY | 0.3998587 |
| TALGHLVP | ASSLGFNQDTQY | 0.6549471 |
| TAPTTSGAA | ASSLGFNQDTQY | 0.63386214 |
| TDGDDGNGGSPGT | ASSLGFNQDTQY | 0.4241826 |
| TDGLGEN | ASSLGFNQDTQY | 0.6914789 |
| TGGPPLPGQA | ASSLGFNQDTQY | 0.40558964 |
| TIPAITI | ASSLGFNQDTQY | 0.64883447 |
| TIPGFTI | ASSLGFNQDTQY | 0.69546115 |
| TSIASGALA | ASSLGFNQDTQY | 0.7140869 |
| TVTLPGAAI | ASSLGFNQDTQY | 0.509023 |
| VASGLSAALP | ASSLGFNQDTQY | 0.68148345 |
| VGAGVIGI | ASSLGFNQDTQY | 0.5905708 |
| VGPKRFGV | ASSLGFNQDTQY | 0.7175716 |
| VNVAFPD | ASSLGFNQDTQY | 0.6174131 |
| VSMTNSGVS | ASSLGFNQDTQY | 0.42368394 |
| VTDVLGDLSV | ASSLGFNQDTQY | 0.6961149 |
| VVVVGAGFAGL | ASSLGFNQDTQY | 0.6979314 |
| VYTVGTVV | ASSLGFNQDTQY | 0.5584936 |
| WTTFSGP | ASSLGFNQDTQY | 0.70203644 |
| YDELVEAF | ASSLGFNQDTQY | 0.56983936 |
| YGTGGAGGNGG | ASSLGFNQDTQY | 0.6459476 |
| YTNVMAV | ASSLGFNQDTQY | 0.7031314 |
| FLEQVHQL | ASSLGFNQDTQY | 0.6984259 |
| GLSLLAVAV | ASSLGFNQDTQY | 0.48394746 |
| LEQVHQL | ASSLGFNQDTQY | 0.69248927 |
| LNAAGAI | ASSLGFNQDTQY | 0.6496303 |
| PAVHASL | ASSLGFNQDTQY | 0.490814 |
| TELEDDA | ASSLGFNQDTQY | 0.71280366 |
| TPPGPPN | ASSLGFNQDTQY | 0.60836565 |
| TPPPPPPP | ASSLGFNQDTQY | 0.6973243 |
| TQSPALI | ASSLRDWGDEQY | 0.68175924 |

| | | |
|---|---|---|
| EFAAAFPQ | ASSLRDWGDEQY | 0.7053608 |
| LLSGNLDVL | ASSLRDWGDEQY | 0.493204 |
| GPANLGNYNI | ASSLRDWGDEQY | 0.6515402 |
| TLGPTLP | ASSLRDWGDEQY | 0.6297576 |
| VYAYSSCCK | ASSLRDWGDEQY | 0.6903504 |
| APAPAPAPNI | ASSLRDWGDEQY | 0.6225092 |
| PGGPVPIP | ASSLRDWGDEQY | 0.5714029 |
| FLTVALEP | ASSLRDWGDEQY | 0.68453175 |
| SQYFGANL | ASSLRDWGDEQY | 0.70415115 |
| GGVKPAA | ASSLRDWGDEQY | 0.6481839 |
| GGTGGAGGAGGW | ASSLRDWGDEQY | 0.6534858 |
| GVGGVAGLGGAGG | ASSLRDWGDEQY | 0.71405625 |
| VLLVLATTI | ASSLRDWGDEQY | 0.50963193 |
| IGIFAAML | ASSLRDWGDEQY | 0.6387279 |
| AVGDLID | ASSLRDWGDEQY | 0.68382084 |
| NISPLPL | ASSLRDWGDEQY | 0.69829834 |
| VTAVFVAI | ASSLRDWGDEQY | 0.6365802 |
| SGDDNDAAL | ASSLRDWGDEQY | 0.7097583 |
| AVAALAHS | ASSLRDWGDEQY | 0.5736438 |
| GGVTVMATP | ASSLRDWGDEQY | 0.69339967 |
| AGLAANVRD | ASSLRDWGDEQY | 0.64130855 |
| TNRPDLIDPALLRPG | ASSLRDWGDEQY | 0.6167901 |
| LANLGVTAL | ASSLRDWGDEQY | 0.526014 |
| TGERIEL | ASSLRDWGDEQY | 0.68741816 |
| NGEELAA | ASSLRDWGDEQY | 0.6986438 |
| VLSLVWG | ASSLRDWGDEQY | 0.6924635 |
| IGYALWL | ASSLRDWGDEQY | 0.32037556 |
| LSLLLAPL | ASSLRDWGDEQY | 0.6931114 |
| TLDEALK | ASSLRDWGDEQY | 0.683776 |
| TDGAGTAEGG | ASSLRDWGDEQY | 0.71792173 |
| NAPGGAGGNGG | ASSLRDWGDEQY | 0.6242863 |
| VSVFMGMGGNF | ASSLRDWGDEQY | 0.65946126 |
| LASYLAAL | ASSLRDWGDEQY | 0.56582975 |
| LVGGTPAL | ASSLRDWGDEQY | 0.6468221 |
| VRNADVL | ASSLRDWGDEQY | 0.49567747 |
| TCSSPFLVSAM | ASSLRDWGDEQY | 0.54831135 |
| VHITDAE | ASSLRDWGDEQY | 0.64085615 |
| EYAAPSF | ASSLRDWGDEQY | 0.5946038 |
| VVDAFHGGL | ASSLRDWGDEQY | 0.69897866 |
| ASVVDPQ | ASSLRDWGDEQY | 0.70226026 |
| SYNEFGGLS | ASSLRDWGDEQY | 0.692989 |
| DPDGSVY | ASSLRDWGDEQY | 0.70430815 |
| MVSTIHATE | ASSLRDWGDEQY | 0.71260834 |
| LAAGVADPV | ASSLRDWGDEQY | 0.6904422 |
| TKEQIAA | ASSLRDWGDEQY | 0.61175287 |
| SLASPAAL | ASSLRDWGDEQY | 0.63140106 |
| IGLVFYGV | ASSLRDWGDEQY | 0.49496767 |
| FLLDASNL | ASSLRDWGDEQY | 0.63330543 |
| YQVGNFP | ASSLRDWGDEQY | 0.6078402 |
| ESLLSQST | ASSLRDWGDEQY | 0.6645293 |
| LFTVGFAAVG | ASSLRDWGDEQY | 0.25712532 |

| | | |
|---|---|---|
| ATPPTPT | ASSLRDWGDEQY | 0.68688726 |
| TDLATLG | ASSLRDWGDEQY | 0.71194947 |
| SAGGTGR | ASSLRDWGDEQY | 0.6155118 |
| GFSNSGS | ASSLRDWGDEQY | 0.66601264 |
| GGNGGSGGTSV | ASSLRDWGDEQY | 0.6936736 |
| GTSVATGGAGN | ASSLRDWGDEQY | 0.6029041 |
| NGGNGGSGGTS | ASSLRDWGDEQY | 0.6826667 |
| AVVKFPH | ASSLRDWGDEQY | 0.55669695 |
| TPPPPPP | ASSLRDWGDEQY | 0.6906589 |
| AAPAAAAAPA | ASSLRDWGDEQY | 0.7066884 |
| IALNEDL | ASSLRDWGDEQY | 0.6688671 |
| IVNVSSM | ASSLRDWGDEQY | 0.6039288 |
| TGIPPAPR | ASSLRDWGDEQY | 0.62982774 |
| FDLGGGTF | ASSLRDWGDEQY | 0.42282745 |
| FDLGGGTFDVS | ASSLRDWGDEQY | 0.6371757 |
| NEPTAAA | ASSLRDWGDEQY | 0.43048853 |
| NEPTAAALA | ASSLRDWGDEQY | 0.6570116 |
| GSIINISSI | ASSLRDWGDEQY | 0.53353906 |
| VAPGNAGTA | ASSLRDWGDEQY | 0.6676166 |
| TGGGASLE | ASSLRDWGDEQY | 0.6620695 |
| VAVYNHYKRI | ASSLRDWGDEQY | 0.65281403 |
| TDPAPATT | ASSLRDWGDEQY | 0.56028676 |
| TNRPDILDPALLRPG | ASSLRDWGDEQY | 0.6700885 |
| LTTTTTS | ASSLRDWGDEQY | 0.6562464 |
| PTGSGKTL | ASSLRDWGDEQY | 0.39824495 |
| TGSGKTL | ASSLRDWGDEQY | 0.7025935 |
| TIGHVDHGK | ASSLRDWGDEQY | 0.5901274 |
| TIGHVDHGKTT | ASSLRDWGDEQY | 0.655086 |
| GHVDHGKTT | ASSLRDWGDEQY | 0.6960833 |
| AQFEHTLL | ASSLRDWGDEQY | 0.6468805 |
| IVNQDEL | ASSLRDWGDEQY | 0.3989644 |
| TNGSQFF | ASSLRDWGDEQY | 0.63878703 |
| TGKGIVE | ASSLRDWGDEQY | 0.6560632 |
| LISNASDA | ASSLRDWGDEQY | 0.61728776 |
| AELRTLGELTKIAKAHGAQ | ASSLRDWGDEQY | 0.2833516 |
| APNAVAAAEPSW | ASSLRDWGDEQY | 0.6915085 |
| APQVAGAATAVL | ASSLRDWGDEQY | 0.48721015 |
| ARRDPAAAAAILPTDA | ASSLRDWGDEQY | 0.5918013 |
| DAAARDVLAEAGLAEYF | ASSLRDWGDEQY | 0.62792575 |
| DDQIVLSAKANLAKVEQ | ASSLRDWGDEQY | 0.69920945 |
| DELLPDTGEDSDEDGH | ASSLRDWGDEQY | 0.5159904 |
| ELPEAPANRATVPQD | ASSLRDWGDEQY | 0.521718 |
| FGQSKIERVTGIG | ASSLRDWGDEQY | 0.6838504 |
| GAGGTGSTAGGAGGAGGAGGLY | ASSLRDWGDEQY | 0.63604283 |
| GHGGTGAAGGNGGTGGAGGSGI | ASSLRDWGDEQY | 0.70106006 |
| GNGGNGGAGGEGGPGGAGGAGA | ASSLRDWGDEQY | 0.63030636 |
| GSTAGNGGNGGNATNGTIAGQP | ASSLRDWGDEQY | 0.61804116 |
| GTGGDGGNGGNGGTGG | ASSLRDWGDEQY | 0.6320875 |
| GVRRPAGIPADAPV | ASSLRDWGDEQY | 0.5293333 |
| GYRQVTDAHLLTLARR | ASSLRDWGDEQY | 0.48107925 |
| IAKMGQDGHDRG | ASSLRDWGDEQY | 0.42062002 |

| | | |
|---|---|---|
| KAEPSIEIQYT | ASSLRDWGDEQY | 0.57255197 |
| KPVRIRMAADEDTVAVT | ASSLRDWGDEQY | 0.41340327 |
| MPPRIAGMRLLVIKPEP | ASSLRDWGDEQY | 0.6614899 |
| MKFYAVALFRSRGVPPTR | ASSLRDWGDEQY | 0.69562614 |
| NAGGFGILGAVAHSP | ASSLRDWGDEQY | 0.6206184 |
| NEGREITLAMAEM | ASSLRDWGDEQY | 0.41169044 |
| NELVEWSHPLGHRWRWE | ASSLRDWGDEQY | 0.6025033 |
| NLGAGGTGGNGGNPG | ASSLRDWGDEQY | 0.63018405 |
| NTEWPKWETVMAADPVD | ASSLRDWGDEQY | 0.6952597 |
| PASAAPAETQP | ASSLRDWGDEQY | 0.69020414 |
| QQSSDGKNGVRAAEITGE | ASSLRDWGDEQY | 0.7187084 |
| RAPDAVTTSWWKE | ASSLRDWGDEQY | 0.6919095 |
| SGFGNVGAHTSGWFNQ | ASSLRDWGDEQY | 0.6136428 |
| SGVAGAVGQAASVGGLKV | ASSLRDWGDEQY | 0.6972211 |
| SNGGAGTGSAGGNGGT | ASSLRDWGDEQY | 0.61385316 |
| TGGAGGEGGAAG | ASSLRDWGDEQY | 0.6540674 |
| TGGDGGAAGAGGAPGHG | ASSLRDWGDEQY | 0.65772533 |
| TMPPLLAAIANAGRHGV | ASSLRDWGDEQY | 0.6712606 |
| TSSTAGNSGGA | ASSLRDWGDEQY | 0.5725566 |
| VPATVPVGILC | ASSLRDWGDEQY | 0.6164618 |
| VQRPDSGGVVLTGRIS | ASSLRDWGDEQY | 0.3824247 |
| WTPNPVPATKGVPAGLVQAT | ASSLRDWGDEQY | 0.68378764 |
| YALKRALAKFSDRFDVVII | ASSLRDWGDEQY | 0.42494133 |
| YLIGSVIV | ASSLRDWGDEQY | 0.41108066 |
| YGQLVAVID | ASSLRDWGDEQY | 0.62332535 |
| VTVGGMTLPA | ASSLRDWGDEQY | 0.71251833 |
| VLALPAV | ASSLRDWGDEQY | 0.7000765 |
| VIPPTLNYET | ASSLRDWGDEQY | 0.5834069 |
| VGMGGMHPSSG | ASSLRDWGDEQY | 0.535511 |
| VGLAGGAATA | ASSLRDWGDEQY | 0.6866944 |
| VGGAGGAGGSGP | ASSLRDWGDEQY | 0.5056514 |
| VFAAFMV | ASSLRDWGDEQY | 0.67979145 |
| TLYDKLR | ASSLRDWGDEQY | 0.66544807 |
| TLAVFTVL | ASSLRDWGDEQY | 0.6633291 |
| TGSVQPARV | ASSLRDWGDEQY | 0.49938846 |
| TGGGAGSGGT | ASSLRDWGDEQY | 0.68357337 |
| SHLETSL | ASSLRDWGDEQY | 0.6159733 |
| SFTTGMT | ASSLRDWGDEQY | 0.6634505 |
| QVEVAKRMGVRQPTVS | ASSLRDWGDEQY | 0.28202096 |
| QGSGGPALAM | ASSLRDWGDEQY | 0.68586445 |
| MLAPSNP | ASSLRDWGDEQY | 0.56900036 |
| LVNNAAI | ASSLRDWGDEQY | 0.5097523 |
| LTGEINTLA | ASSLRDWGDEQY | 0.3495155 |
| LSIFGPV | ASSLRDWGDEQY | 0.6326786 |
| LLALLANL | ASSLRDWGDEQY | 0.6831826 |
| LISNASDALDKLR | ASSLRDWGDEQY | 0.7032617 |
| LINEAVP | ASSLRDWGDEQY | 0.6181183 |
| LGYAFSGL | ASSLRDWGDEQY | 0.69169366 |
| LDFVAVAL | ASSLRDWGDEQY | 0.69464374 |
| LAQPGAE | ASSLRDWGDEQY | 0.6354867 |
| ISNLGAVTGF | ASSLRDWGDEQY | 0.35237563 |

| | | |
|---|---|---|
| HQEDPGRI | ASSLRDWGDEQY | 0.6565455 |
| GTPGEPTP | ASSLRDWGDEQY | 0.6619655 |
| GFFNSGS | ASSLRDWGDEQY | 0.69764876 |
| GAGGGGLTAGAN | ASSLRDWGDEQY | 0.66604495 |
| FVDALVNM | ASSLRDWGDEQY | 0.51937455 |
| FLTVAPEL | ASSLRDWGDEQY | 0.6565591 |
| FETLTAL | ASSLRDWGDEQY | 0.67953396 |
| DVINPIEL | ASSLRDWGDEQY | 0.47995088 |
| APVSGAPV | ASSLRDWGDEQY | 0.5421133 |
| ALVGGLAW | ASSLRDWGDEQY | 0.55513686 |
| ALAAAAATAI | ASSLRDWGDEQY | 0.41831392 |
| AFGLGLLL | ASSLRDWGDEQY | 0.65985817 |
| AFFAAASL | ASSLRDWGDEQY | 0.6341672 |
| AAAALPAFSPP | ASSLRDWGDEQY | 0.5511459 |
| VVLAPAL | ASSLRDWGDEQY | 0.65821224 |
| TIGHVDHGKTTL | ASSLRDWGDEQY | 0.5260153 |
| LISNASDALDK | ASSLRDWGDEQY | 0.5443668 |
| LISNASDAL | ASSLRDWGDEQY | 0.58222127 |
| ISNASDALDK | ASSLRDWGDEQY | 0.6536819 |
| NQAETLVYQTE | ASSLRDWGDEQY | 0.6924949 |
| TGGTGGTAGAG | ASSLRDWGDEQY | 0.646865 |
| TGSTAGGAGGA | ASSLRDWGDEQY | 0.6476936 |
| DPLAVPVAAGPV | ASSLRDWGDEQY | 0.7118383 |
| GGTGGTPGNGGN | ASSLRDWGDEQY | 0.38585836 |
| PAGLAGVDPATL | ASSLRDWGDEQY | 0.55318344 |
| GNNLVVEAPHAGASV | ASSLRDWGDEQY | 0.6813449 |
| KVTRIMSASTLEQPA | ASSLRDWGDEQY | 0.5206123 |
| LGFALPIALPRRTVV | ASSLRDWGDEQY | 0.6984745 |
| TRLGNLTARKLVVLV | ASSLRDWGDEQY | 0.6631415 |
| WSQQRRLIYFPSAGP | ASSLRDWGDEQY | 0.67955226 |
| GPNGNPDPMAAAVDIR | ASSLRDWGDEQY | 0.65447843 |
| TQRCVSVADLRAAASR | ASSLRDWGDEQY | 0.71402436 |
| QFVLDVAEHPVEVLTETV | ASSLRDWGDEQY | 0.68491685 |
| GGAGGTGGAGGNGGTQAGDG | ASSLRDWGDEQY | 0.6341243 |
| PLLATLRAIAVDVDAKPAQV | ASSLRDWGDEQY | 0.702688 |
| APKFRTGGYMIDFWGVGYQVAK | ASSLRDWGDEQY | 0.41118857 |
| GAGGAGGAGGAAGGPGGTGGTG | ASSLRDWGDEQY | 0.69181466 |
| IGALGLLATDVAPAGSVAFGVA | ASSLRDWGDEQY | 0.68217754 |
| AGGAGGIGGNGGAGGAGGVLMG | ASSLRDWGDEQY | 0.53485566 |
| ADELIGLL | ASSLRDWGDEQY | 0.6534358 |
| ADRALKA | ASSLRDWGDEQY | 0.7087159 |
| AGAVGQVGGAGGA | ASSLRDWGDEQY | 0.7054941 |
| AGGGGGLGGAP | ASSLRDWGDEQY | 0.46007362 |
| AGGLLYGNGGAG | ASSLRDWGDEQY | 0.61862236 |
| AGLESIGL | ASSLRDWGDEQY | 0.42066017 |
| AGLKTGII | ASSLRDWGDEQY | 0.4687722 |
| ASLAALLL | ASSLRDWGDEQY | 0.38527852 |
| ASVINISSI | ASSLRDWGDEQY | 0.6817535 |
| ATPPGPA | ASSLRDWGDEQY | 0.6164521 |
| ATVASALV | ASSLRDWGDEQY | 0.69715214 |
| ATVASLIL | ASSLRDWGDEQY | 0.6669133 |

| | | |
|---|---|---|
| AVNGPVLIRH | ASSLRDWGDEQY | 0.70218706 |
| DGAAGAVGI | ASSLRDWGDEQY | 0.57798374 |
| DPLLHGI | ASSLRDWGDEQY | 0.60817295 |
| ELGAALGAL | ASSLRDWGDEQY | 0.37066004 |
| ELGRTQAV | ASSLRDWGDEQY | 0.3868694 |
| FVAGGYLS | ASSLRDWGDEQY | 0.5345202 |
| GASGGAGGAGGW | ASSLRDWGDEQY | 0.59003735 |
| GGAGAGGDGGAGGI | ASSLRDWGDEQY | 0.3126643 |
| GTGHNGSQPGMGGNG | ASSLRDWGDEQY | 0.7091975 |
| GYAAASASAS | ASSLRDWGDEQY | 0.62869215 |
| HQVDVAAVQQG | ASSLRDWGDEQY | 0.67798954 |
| IGAAIAHT | ASSLRDWGDEQY | 0.52422285 |
| IGMIEVAQL | ASSLRDWGDEQY | 0.5330645 |
| IRLAPEHTSAMD | ASSLRDWGDEQY | 0.6098027 |
| IYNVSVL | ASSLRDWGDEQY | 0.67857707 |
| KQGFLGM | ASSLRDWGDEQY | 0.5634104 |
| LESVITQL | ASSLRDWGDEQY | 0.3440212 |
| LGLGALVVV | ASSLRDWGDEQY | 0.4909918 |
| LIGNGGNGGSGGI | ASSLRDWGDEQY | 0.53754693 |
| LNAINAPTL | ASSLRDWGDEQY | 0.71589434 |
| LQLLNTGV | ASSLRDWGDEQY | 0.7016172 |
| LQTSIEF | ASSLRDWGDEQY | 0.66568553 |
| MSFAVSEL | ASSLRDWGDEQY | 0.7131698 |
| NTGALGL | ASSLRDWGDEQY | 0.27995488 |
| PPGAGGAGGAGTT | ASSLRDWGDEQY | 0.5520958 |
| QLVGLGI | ASSLRDWGDEQY | 0.7003038 |
| RFLEQVHQL | ASSLRDWGDEQY | 0.67597663 |
| RGVGYVLR | ASSLRDWGDEQY | 0.6428113 |
| SATGVFAGVI | ASSLRDWGDEQY | 0.70181215 |
| SLPAITLP | ASSLRDWGDEQY | 0.5483871 |
| STALAFTV | ASSLRDWGDEQY | 0.6091187 |
| SVLTVGLGK | ASSLRDWGDEQY | 0.6471828 |
| TALAGAGSGI | ASSLRDWGDEQY | 0.68851155 |
| TGLPFTGI | ASSLRDWGDEQY | 0.6574081 |
| VAEHVTV | ASSLRDWGDEQY | 0.47519207 |
| VDIVLNSL | ASSLRDWGDEQY | 0.6545135 |
| VLTGGGAVL | ASSLRDWGDEQY | 0.72081405 |
| VNVGLAYGI | ASSLRDWGDEQY | 0.6228544 |
| VSSAAVGAV | ASSLRDWGDEQY | 0.6869316 |
| VTSVYEAGV | ASSLRDWGDEQY | 0.66404974 |
| WVALDRGA | ASSLRDWGDEQY | 0.61227334 |
| YLVGSVVI | ASSLRDWGDEQY | 0.57987785 |
| GPPGTGKTLL | ASSLRDWGDEQY | 0.5604832 |
| GVGMTKF | ASSLRDWGDEQY | 0.67397475 |
| IGIGGSDL | ASSLRDWGDEQY | 0.5532112 |
| IGTLLAEL | ASSLRDWGDEQY | 0.62254757 |
| IYTSGTT | ASSLRDWGDEQY | 0.63783026 |
| LSFPAGV | ASSLRDWGDEQY | 0.7201368 |
| LYTSGTT | ASSLRDWGDEQY | 0.68534803 |
| VGAFPFGF | ASSLRDWGDEQY | 0.6794605 |
| VGAGTVEF | ASSLRDWGDEQY | 0.56531584 |

| | | |
|---|---|---|
| VGKSTLLL | ASSLRDWGDEQY | 0.664691 |
| AALAHAGGGIV | ASSLRDWGDEQY | 0.68659586 |
| AALVAVSLL | ASSLRDWGDEQY | 0.42348522 |
| AAVSRAVAAL | ASSLRDWGDEQY | 0.7029548 |
| AGGSSGIGGF | ASSLRDWGDEQY | 0.5171379 |
| AGLTVANANG | ASSLRDWGDEQY | 0.64997464 |
| AGLVGAVL | ASSLRDWGDEQY | 0.65377903 |
| ALSQAFTV | ASSLRDWGDEQY | 0.7118104 |
| ALTFSVSL | ASSLRDWGDEQY | 0.6485837 |
| ALTGAALVAL | ASSLRDWGDEQY | 0.66123784 |
| ASFSIGAGVL | ASSLRDWGDEQY | 0.5440086 |
| DGSVYKI | ASSLRDWGDEQY | 0.69004536 |
| EQLAEIV | ASSLRDWGDEQY | 0.70428336 |
| FAVSELTVM | ASSLRDWGDEQY | 0.650638 |
| GAAPLAAGL | ASSLRDWGDEQY | 0.7205257 |
| GADGGDGGSGGAGG | ASSLRDWGDEQY | 0.63330007 |
| GFGLDTGGAGG | ASSLRDWGDEQY | 0.33263245 |
| GLASGGLL | ASSLRDWGDEQY | 0.6277633 |
| GRIAGAAVLL | ASSLRDWGDEQY | 0.7030343 |
| GSAGSLNTSF | ASSLRDWGDEQY | 0.7096169 |
| GTGQIGI | ASSLRDWGDEQY | 0.69760674 |
| GTLIALIL | ASSLRDWGDEQY | 0.66949505 |
| IAATMGFVVG | ASSLRDWGDEQY | 0.6205846 |
| IFAQALL | ASSLRDWGDEQY | 0.70968723 |
| IGAAVALGF | ASSLRDWGDEQY | 0.7168839 |
| IGLTPLAL | ASSLRDWGDEQY | 0.6731242 |
| ILGSNTSTL | ASSLRDWGDEQY | 0.5097196 |
| IVFSLTVL | ASSLRDWGDEQY | 0.4721238 |
| LASKATGF | ASSLRDWGDEQY | 0.6764034 |
| LASVALIL | ASSLRDWGDEQY | 0.702333 |
| LFAVGLTL | ASSLRDWGDEQY | 0.4455278 |
| LFNGGIGTYI | ASSLRDWGDEQY | 0.66782385 |
| LTLGAGII | ASSLRDWGDEQY | 0.6683415 |
| LVAAALGDI | ASSLRDWGDEQY | 0.34850875 |
| NTAFVQLGI | ASSLRDWGDEQY | 0.6874603 |
| QLFSGAEL | ASSLRDWGDEQY | 0.7097028 |
| SAATFITL | ASSLRDWGDEQY | 0.66430473 |
| SNYGLANLGV | ASSLRDWGDEQY | 0.48372138 |
| SYGGAYAVM | ASSLRDWGDEQY | 0.70536256 |
| TTGVGGLTLG | ASSLRDWGDEQY | 0.71385837 |
| TTQTIYI | ASSLRDWGDEQY | 0.71279585 |
| VAAGAGLGAVI | ASSLRDWGDEQY | 0.38009244 |
| VAAVVTLL | ASSLRDWGDEQY | 0.3071914 |
| VGFLGGLI | ASSLRDWGDEQY | 0.6484914 |
| VGVIPILL | ASSLRDWGDEQY | 0.6922768 |
| VIFASSLI | ASSLRDWGDEQY | 0.58177745 |
| VKGGDGGAAAG | ASSLRDWGDEQY | 0.5284107 |
| VTKTFTNI | ASSLRDWGDEQY | 0.678593 |
| VVAAAILL | ASSLRDWGDEQY | 0.70504296 |
| YAALDVEL | ASSLRDWGDEQY | 0.48721403 |
| LASVALLL | ASSLRDWGDEQY | 0.4251094 |

| | | |
|---|---|---|
| LDAVHRV | ASSLRDWGDEQY | 0.69027054 |
| QGLSSSIF | ASSLRDWGDEQY | 0.68674076 |
| RGTFANIRL | ASSLRDWGDEQY | 0.56951123 |
| RQLWWGHRI | ASSLRDWGDEQY | 0.6353323 |
| YGPPGTGKTLL | ASSLRDWGDEQY | 0.6530715 |
| AALVGAPANVA | ASSLRDWGDEQY | 0.61217546 |
| AATFRRTE | ASSLRDWGDEQY | 0.6701758 |
| AGALGGGI | ASSLRDWGDEQY | 0.46645686 |
| AGGAGGAGGIGAG | ASSLRDWGDEQY | 0.70387775 |
| AGSGALGL | ASSLRDWGDEQY | 0.4982733 |
| AQIADQI | ASSLRDWGDEQY | 0.6943002 |
| ASAVAVTYL | ASSLRDWGDEQY | 0.66807276 |
| ASTVLGGTGG | ASSLRDWGDEQY | 0.66214 |
| AVGGEPAK | ASSLRDWGDEQY | 0.6417805 |
| CEQGTFGGTC | ASSLRDWGDEQY | 0.5320147 |
| DEGKAPAG | ASSLRDWGDEQY | 0.5283492 |
| DTVVFPAL | ASSLRDWGDEQY | 0.41150028 |
| FEGITMAG | ASSLRDWGDEQY | 0.7049632 |
| FPLVVQD | ASSLRDWGDEQY | 0.5717772 |
| GAFTTPQ | ASSLRDWGDEQY | 0.7022544 |
| GFGGFGTAGGVGGA | ASSLRDWGDEQY | 0.6637994 |
| GFLAQFL | ASSLRDWGDEQY | 0.5626142 |
| GGAGGAGGNAGVA | ASSLRDWGDEQY | 0.68509525 |
| GGVGTAGAGGAGGAGG | ASSLRDWGDEQY | 0.4354456 |
| GNGGGGDAGNAGSGGN | ASSLRDWGDEQY | 0.6270827 |
| GTGGAGGAGNDAGS | ASSLRDWGDEQY | 0.65016735 |
| GTTNTGWFNS | ASSLRDWGDEQY | 0.6838229 |
| IFMGPLFTL | ASSLRDWGDEQY | 0.54497993 |
| ILAHAIEAI | ASSLRDWGDEQY | 0.6782156 |
| IRSAIREAL | ASSLRDWGDEQY | 0.67419744 |
| ITFAGGG | ASSLRDWGDEQY | 0.43174323 |
| ITGMSGHQEV | ASSLRDWGDEQY | 0.65852606 |
| IVAGLACTL | ASSLRDWGDEQY | 0.70193255 |
| IVGGGLAGA | ASSLRDWGDEQY | 0.66130686 |
| IVTGVLNI | ASSLRDWGDEQY | 0.48499513 |
| LAQQGCI | ASSLRDWGDEQY | 0.66379535 |
| LCLHVEN | ASSLRDWGDEQY | 0.5935102 |
| LDAISDGS | ASSLRDWGDEQY | 0.6797993 |
| LGGFAAML | ASSLRDWGDEQY | 0.6740223 |
| LGGKNPMI | ASSLRDWGDEQY | 0.52107275 |
| LGPKGVAAIL | ASSLRDWGDEQY | 0.6771715 |
| LMVGVSI | ASSLRDWGDEQY | 0.37376863 |
| LVGAAFVL | ASSLRDWGDEQY | 0.542567 |
| LVLAVAPL | ASSLRDWGDEQY | 0.60276324 |
| MAAIAVIDAA | ASSLRDWGDEQY | 0.6264416 |
| NNGINAELQ | ASSLRDWGDEQY | 0.6890548 |
| RGEPLVLVHG | ASSLRDWGDEQY | 0.5389892 |
| RGGAGGAAGLI | ASSLRDWGDEQY | 0.64840066 |
| RSYRMHPAV | ASSLRDWGDEQY | 0.604151 |
| SAAHLLAGQLA | ASSLRDWGDEQY | 0.5285877 |
| SAAVGSY | ASSLRDWGDEQY | 0.6397593 |

| | | |
|---|---|---|
| SPHAVLHRI | ASSLRDWGDEQY | 0.5986533 |
| SQIAKVL | ASSLRDWGDEQY | 0.4277565 |
| TALGHLVP | ASSLRDWGDEQY | 0.6874218 |
| TAPTTSGAA | ASSLRDWGDEQY | 0.49235594 |
| TDGDDGNGGSPGT | ASSLRDWGDEQY | 0.61309063 |
| TDGLGEN | ASSLRDWGDEQY | 0.7073544 |
| TGGPPLPGQA | ASSLRDWGDEQY | 0.43979195 |
| TIPAITI | ASSLRDWGDEQY | 0.68550956 |
| TIPGFTI | ASSLRDWGDEQY | 0.6577648 |
| TSIASGALA | ASSLRDWGDEQY | 0.6823987 |
| TVTLPGAAI | ASSLRDWGDEQY | 0.62176025 |
| VASGLSAALP | ASSLRDWGDEQY | 0.59097457 |
| VGAGVIGI | ASSLRDWGDEQY | 0.70981085 |
| VGPKRFGV | ASSLRDWGDEQY | 0.6629225 |
| VNVAFPD | ASSLRDWGDEQY | 0.59941584 |
| VSMTNSGVS | ASSLRDWGDEQY | 0.680585 |
| VTDVLGDLSV | ASSLRDWGDEQY | 0.6941403 |
| VVVVGAGFAGL | ASSLRDWGDEQY | 0.6938813 |
| VYTVGTVV | ASSLRDWGDEQY | 0.5991373 |
| WTTFSGP | ASSLRDWGDEQY | 0.37152207 |
| YDELVEAF | ASSLRDWGDEQY | 0.62479043 |
| YGTGGAGGNGG | ASSLRDWGDEQY | 0.61096376 |
| YTNVMAV | ASSLRDWGDEQY | 0.57226616 |
| FLEQVHQL | ASSLRDWGDEQY | 0.3533815 |
| GLSLLAVAV | ASSLRDWGDEQY | 0.6981719 |
| LEQVHQL | ASSLRDWGDEQY | 0.71894187 |
| LNAAGAI | ASSLRDWGDEQY | 0.5010439 |
| PAVHASL | ASSLRDWGDEQY | 0.7199167 |
| TELEDDA | ASSLRDWGDEQY | 0.46212107 |
| TPPGPPN | ASSLRDWGDEQY | 0.66984165 |
| TPPPPPPP | ASSLRDWGDEQY | 0.69874805 |
| TQSPALI | ASRPGTGGYEQY | 0.50101453 |
| EFAAAFPQ | ASRPGTGGYEQY | 0.6303141 |
| LLSGNLDVL | ASRPGTGGYEQY | 0.5873883 |
| GPANLGNYNI | ASRPGTGGYEQY | 0.58593744 |
| TLGPTLP | ASRPGTGGYEQY | 0.70193386 |
| VYAYSSCCK | ASRPGTGGYEQY | 0.53012663 |
| APAPAPAPNI | ASRPGTGGYEQY | 0.6736649 |
| PGGPVPIP | ASRPGTGGYEQY | 0.604144 |
| FLTVALEP | ASRPGTGGYEQY | 0.55925685 |
| SQYFGANL | ASRPGTGGYEQY | 0.68118525 |
| GGVKPAA | ASRPGTGGYEQY | 0.6813233 |
| GGTGGAGGAGGW | ASRPGTGGYEQY | 0.6048752 |
| GVGGVAGLGGAGG | ASRPGTGGYEQY | 0.71573424 |
| VLLVLATTI | ASRPGTGGYEQY | 0.54748756 |
| IGIFAAML | ASRPGTGGYEQY | 0.7155093 |
| AVGDLID | ASRPGTGGYEQY | 0.7049836 |
| NISPLPL | ASRPGTGGYEQY | 0.67313164 |
| VTAVFVAI | ASRPGTGGYEQY | 0.5975411 |
| SGDDNDAAL | ASRPGTGGYEQY | 0.7000693 |
| AVAALAHS | ASRPGTGGYEQY | 0.57861763 |

| | | |
|---|---|---|
| GGVTVMATP | ASRPGTGGYEQY | 0.6652622 |
| AGLAANVRD | ASRPGTGGYEQY | 0.70683753 |
| TNRPDLIDPALLRPG | ASRPGTGGYEQY | 0.20876008 |
| LANLGVTAL | ASRPGTGGYEQY | 0.84076446 |
| TGERIEL | ASRPGTGGYEQY | 0.7429769 |
| NGEELAA | ASRPGTGGYEQY | 0.58252233 |
| VLSLVWG | ASRPGTGGYEQY | 0.2993511 |
| IGYALWL | ASRPGTGGYEQY | 0.16699678 |
| LSLLLAPL | ASRPGTGGYEQY | 0.46396214 |
| TLDEALK | ASRPGTGGYEQY | 0.30467075 |
| TDGAGTAEGG | ASRPGTGGYEQY | 0.8051667 |
| NAPGGAGGNGG | ASRPGTGGYEQY | 0.7337793 |
| VSVFMGMGGNF | ASRPGTGGYEQY | 0.32043472 |
| LASYLAAL | ASRPGTGGYEQY | 0.78995466 |
| LVGGTPAL | ASRPGTGGYEQY | 0.75255144 |
| VRNADVL | ASRPGTGGYEQY | 0.19194111 |
| TCSSPFLVSAM | ASRPGTGGYEQY | 0.52074987 |
| VHITDAE | ASRPGTGGYEQY | 0.29167324 |
| EYAAPSF | ASRPGTGGYEQY | 0.44538286 |
| VVDAFHGGL | ASRPGTGGYEQY | 0.1905925 |
| ASVVDPQ | ASRPGTGGYEQY | 0.6619332 |
| SYNEFGGLS | ASRPGTGGYEQY | 0.42149845 |
| DPDGSVY | ASRPGTGGYEQY | 0.107451074 |
| MVSTIHATE | ASRPGTGGYEQY | 0.662629 |
| LAAGVADPV | ASRPGTGGYEQY | 0.3667021 |
| TKEQIAA | ASRPGTGGYEQY | 0.4116383 |
| SLASPAAL | ASRPGTGGYEQY | 0.72921014 |
| IGLVFYGV | ASRPGTGGYEQY | 0.33010083 |
| FLLDASNL | ASRPGTGGYEQY | 0.35028198 |
| YQVGNFP | ASRPGTGGYEQY | 0.75040853 |
| ESLLSQST | ASRPGTGGYEQY | 0.7738517 |
| LFTVGFAAVG | ASRPGTGGYEQY | 0.2598132 |
| ATPPTPT | ASRPGTGGYEQY | 0.60672235 |
| TDLATLG | ASRPGTGGYEQY | 0.8119165 |
| SAGGTGR | ASRPGTGGYEQY | 0.56563616 |
| GFSNSGS | ASRPGTGGYEQY | 0.7833308 |
| GGNGGSGGTSV | ASRPGTGGYEQY | 0.7953776 |
| GTSVATGGAGN | ASRPGTGGYEQY | 0.45987827 |
| NGGNGGSGGTS | ASRPGTGGYEQY | 0.77595043 |
| AVVKFPH | ASRPGTGGYEQY | 0.88314235 |
| TPPPPPP | ASRPGTGGYEQY | 0.4157998 |
| AAPAAAAAPA | ASRPGTGGYEQY | 0.52445513 |
| IALNEDL | ASRPGTGGYEQY | 0.19703901 |
| IVNVSSM | ASRPGTGGYEQY | 0.4419983 |
| TGIPPAPR | ASRPGTGGYEQY | 0.17318347 |
| FDLGGGTF | ASRPGTGGYEQY | 0.69950265 |
| FDLGGGTFDVS | ASRPGTGGYEQY | 0.7650869 |
| NEPTAAA | ASRPGTGGYEQY | 0.5283011 |
| NEPTAAALA | ASRPGTGGYEQY | 0.24854682 |
| GSIINISSI | ASRPGTGGYEQY | 0.19161075 |
| VAPGNAGTA | ASRPGTGGYEQY | 0.60338235 |

| | | |
|---|---|---|
| TGGGASLE | ASRPGTGGYEQY | 0.22818866 |
| VAVYNHYKRI | ASRPGTGGYEQY | 0.15088814 |
| TDPAPATT | ASRPGTGGYEQY | 0.4579045 |
| TNRPDILDPALLRPG | ASRPGTGGYEQY | 0.7250701 |
| LTTTTTS | ASRPGTGGYEQY | 0.7102543 |
| PTGSGKTL | ASRPGTGGYEQY | 0.3649757 |
| TGSGKTL | ASRPGTGGYEQY | 0.85393584 |
| TIGHVDHGK | ASRPGTGGYEQY | 0.74855363 |
| TIGHVDHGKTT | ASRPGTGGYEQY | 0.83359516 |
| GHVDHGKTT | ASRPGTGGYEQY | 0.48884118 |
| AQFEHTLL | ASRPGTGGYEQY | 0.22794943 |
| IVNQDEL | ASRPGTGGYEQY | 0.8301965 |
| TNGSQFF | ASRPGTGGYEQY | 0.4046229 |
| TGKGIVE | ASRPGTGGYEQY | 0.5546795 |
| LISNASDA | ASRPGTGGYEQY | 0.6300546 |
| AELRTLGELTKIAKAHGAQ | ASRPGTGGYEQY | 0.51808834 |
| APNAVAAAEPSW | ASRPGTGGYEQY | 0.34139934 |
| APQVAGAATAVL | ASRPGTGGYEQY | 0.6759445 |
| ARRDPAAAAAILPTDA | ASRPGTGGYEQY | 0.74618655 |
| DAAARDVLAEAGLAEYF | ASRPGTGGYEQY | 0.33728677 |
| DDQIVLSAKANLAKVEQ | ASRPGTGGYEQY | 0.237654 |
| DELLPDTGEDSDEDGH | ASRPGTGGYEQY | 0.36902905 |
| ELPEAPANRATVPQD | ASRPGTGGYEQY | 0.5315135 |
| FGQSKIERVTGIG | ASRPGTGGYEQY | 0.74723214 |
| GAGGTGSTAGGAGGAGGAGGLY | ASRPGTGGYEQY | 0.5702096 |
| GHGGTGAAGGNGGTGGAGGSGI | ASRPGTGGYEQY | 0.46096852 |
| GNGGNGGAGGEGGPGGAGGAGA | ASRPGTGGYEQY | 0.7062719 |
| GSTAGNGGNGGNATNGTIAGQP | ASRPGTGGYEQY | 0.6252688 |
| GTGGDGGNGGNGGTGG | ASRPGTGGYEQY | 0.5220931 |
| GVRRPAGIPADAPV | ASRPGTGGYEQY | 0.5478165 |
| GYRQVTDAHLLTLARR | ASRPGTGGYEQY | 0.465063 |
| IAKMGQDGHDRG | ASRPGTGGYEQY | 0.7399545 |
| KAEPSIEIQYT | ASRPGTGGYEQY | 0.70413595 |
| KPVRIRMAADEDTVAVT | ASRPGTGGYEQY | 0.6163378 |
| MPPRIAGMRLLVIKPEP | ASRPGTGGYEQY | 0.7662916 |
| MKFYAVALFRSRGVPPTR | ASRPGTGGYEQY | 0.49538445 |
| NAGGFGILGAVAHSP | ASRPGTGGYEQY | 0.5042471 |
| NEGREITLAMAEM | ASRPGTGGYEQY | 0.19095525 |
| NELVEWSHPLGHRWRWE | ASRPGTGGYEQY | 0.35537663 |
| NLGAGGTGGNGGNPG | ASRPGTGGYEQY | 0.5164432 |
| NTEWPKWETVMAADPVD | ASRPGTGGYEQY | 0.66408694 |
| PASAAPAETQP | ASRPGTGGYEQY | 0.6940892 |
| QQSSDGKNGVRAAEITGE | ASRPGTGGYEQY | 0.6047478 |
| RAPDAVTTSWWKE | ASRPGTGGYEQY | 0.39944112 |
| SGFGNVGAHTSGWFNQ | ASRPGTGGYEQY | 0.7011688 |
| SGVAGAVGQAASVGGLKV | ASRPGTGGYEQY | 0.80226386 |
| SNGGAGTGSAGGNGGT | ASRPGTGGYEQY | 0.22178137 |
| TGGAGGEGGAAG | ASRPGTGGYEQY | 0.5378093 |
| TGGDGGAAGAGGAPGHG | ASRPGTGGYEQY | 0.86981446 |
| TMPPLLAAIANAGRHGV | ASRPGTGGYEQY | 0.66574484 |
| TSSTAGNSGGA | ASRPGTGGYEQY | 0.39798146 |

| | | |
|---|---|---|
| VPATVPVGILC | ASRPGTGGYEQY | 0.49146324 |
| VQRPDSGGVVLTGRIS | ASRPGTGGYEQY | 0.86009777 |
| WTPNPVPATKGVPAGLVQAT | ASRPGTGGYEQY | 0.38348645 |
| YALKRALAKFSDRFDVVII | ASRPGTGGYEQY | 0.84507126 |
| YLIGSVIV | ASRPGTGGYEQY | 0.16727975 |
| YGQLVAVID | ASRPGTGGYEQY | 0.45191568 |
| VTVGGMTLPA | ASRPGTGGYEQY | 0.27030027 |
| VLALPAV | ASRPGTGGYEQY | 0.13162252 |
| VIPPTLNYET | ASRPGTGGYEQY | 0.75457716 |
| VGMGGMHPSSG | ASRPGTGGYEQY | 0.4121147 |
| VGLAGGAATA | ASRPGTGGYEQY | 0.75274754 |
| VGGAGGAGGSGP | ASRPGTGGYEQY | 0.4356917 |
| VFAAFMV | ASRPGTGGYEQY | 0.7645745 |
| TLYDKLR | ASRPGTGGYEQY | 0.7264382 |
| TLAVFTVL | ASRPGTGGYEQY | 0.47161537 |
| TGSVQPARV | ASRPGTGGYEQY | 0.27400777 |
| TGGGAGSGGT | ASRPGTGGYEQY | 0.6215849 |
| SHLETSL | ASRPGTGGYEQY | 0.7469826 |
| SFTTGMT | ASRPGTGGYEQY | 0.6559422 |
| QVEVAKRMGVRQPTVS | ASRPGTGGYEQY | 0.18709455 |
| QGSGGPALAM | ASRPGTGGYEQY | 0.6317315 |
| MLAPSNP | ASRPGTGGYEQY | 0.46055853 |
| LVNNAAI | ASRPGTGGYEQY | 0.18274549 |
| LTGEINTLA | ASRPGTGGYEQY | 0.10660826 |
| LSIFGPV | ASRPGTGGYEQY | 0.44042838 |
| LLALLANL | ASRPGTGGYEQY | 0.49493873 |
| LISNASDALDKLR | ASRPGTGGYEQY | 0.7396035 |
| LINEAVP | ASRPGTGGYEQY | 0.7647553 |
| LGYAFSGL | ASRPGTGGYEQY | 0.37364832 |
| LDFVAVAL | ASRPGTGGYEQY | 0.71612597 |
| LAQPGAE | ASRPGTGGYEQY | 0.59449565 |
| ISNLGAVTGF | ASRPGTGGYEQY | 0.08053239 |
| HQEDPGRI | ASRPGTGGYEQY | 0.33940288 |
| GTPGEPTP | ASRPGTGGYEQY | 0.18489599 |
| GFFNSGS | ASRPGTGGYEQY | 0.77773625 |
| GAGGGGLTAGAN | ASRPGTGGYEQY | 0.3802626 |
| FVDALVNM | ASRPGTGGYEQY | 0.5407467 |
| FLTVAPEL | ASRPGTGGYEQY | 0.46151567 |
| FETLTAL | ASRPGTGGYEQY | 0.7882568 |
| DVINPIEL | ASRPGTGGYEQY | 0.5720365 |
| APVSGAPV | ASRPGTGGYEQY | 0.23165041 |
| ALVGGLAW | ASRPGTGGYEQY | 0.11575686 |
| ALAAAAATAI | ASRPGTGGYEQY | 0.59794784 |
| AFGLGLLL | ASRPGTGGYEQY | 0.30307293 |
| AFFAAASL | ASRPGTGGYEQY | 0.31669658 |
| AAAALPAFSPP | ASRPGTGGYEQY | 0.24156055 |
| VVLAPAL | ASRPGTGGYEQY | 0.15110736 |
| TIGHVDHGKTTL | ASRPGTGGYEQY | 0.2524232 |
| LISNASDALDK | ASRPGTGGYEQY | 0.72834927 |
| LISNASDAL | ASRPGTGGYEQY | 0.26925457 |
| ISNASDALDK | ASRPGTGGYEQY | 0.4761075 |

| | | |
|---|---|---|
| NQAETLVYQTE | ASRPGTGGYEQY | 0.86000776 |
| TGGTGGTAGAG | ASRPGTGGYEQY | 0.75394523 |
| TGSTAGGAGGA | ASRPGTGGYEQY | 0.40404493 |
| DPLAVPVAAGPV | ASRPGTGGYEQY | 0.3691442 |
| GGTGGTPGNGGN | ASRPGTGGYEQY | 0.2796029 |
| PAGLAGVDPATL | ASRPGTGGYEQY | 0.33586633 |
| GNNLVVEAPHAGASV | ASRPGTGGYEQY | 0.36920235 |
| KVTRIMSASTLEQPA | ASRPGTGGYEQY | 0.7829497 |
| LGFALPIALPRRTVV | ASRPGTGGYEQY | 0.49905932 |
| TRLGNLTARKLVVLV | ASRPGTGGYEQY | 0.25470054 |
| WSQQRRLIYFPSAGP | ASRPGTGGYEQY | 0.6404849 |
| GPNGNPDPMAAAVDIR | ASRPGTGGYEQY | 0.8083097 |
| TQRCVSVADLRAAASR | ASRPGTGGYEQY | 0.86997145 |
| QFVLDVAEHPVEVLTETV | ASRPGTGGYEQY | 0.28314328 |
| GGAGGTGGAGGNGGTQAGDG | ASRPGTGGYEQY | 0.38537678 |
| PLLATLRAIAVDVDAKPAQV | ASRPGTGGYEQY | 0.48801786 |
| APKFRTGGYMIDFWGVGYQVAK | ASRPGTGGYEQY | 0.58890164 |
| GAGGAGGAGGAAGGPGGTGGTG | ASRPGTGGYEQY | 0.30182374 |
| IGALGLLATDVAPAGSVAFGVA | ASRPGTGGYEQY | 0.28899676 |
| AGGAGGIGGNGGAGGAGGVLMG | ASRPGTGGYEQY | 0.70101815 |
| ADELIGLL | ASRPGTGGYEQY | 0.8651324 |
| ADRALKA | ASRPGTGGYEQY | 0.4160626 |
| AGAVGQVGGAGGA | ASRPGTGGYEQY | 0.34119874 |
| AGGGGGLGGAP | ASRPGTGGYEQY | 0.44817787 |
| AGGLLYGNGGAG | ASRPGTGGYEQY | 0.2924733 |
| AGLESIGL | ASRPGTGGYEQY | 0.30025774 |
| AGLKTGII | ASRPGTGGYEQY | 0.42159408 |
| ASLAALLL | ASRPGTGGYEQY | 0.2063369 |
| ASVINISSI | ASRPGTGGYEQY | 0.4485332 |
| ATPPGPA | ASRPGTGGYEQY | 0.8415144 |
| ATVASALV | ASRPGTGGYEQY | 0.51749235 |
| ATVASLIL | ASRPGTGGYEQY | 0.7120348 |
| AVNGPVLIRH | ASRPGTGGYEQY | 0.8229538 |
| DGAAGAVGI | ASRPGTGGYEQY | 0.7005666 |
| DPLLHGI | ASRPGTGGYEQY | 0.81475556 |
| ELGAALGAL | ASRPGTGGYEQY | 0.77666384 |
| ELGRTQAV | ASRPGTGGYEQY | 0.43010402 |
| FVAGGYLS | ASRPGTGGYEQY | 0.4614095 |
| GASGGAGGAGGW | ASRPGTGGYEQY | 0.76488304 |
| GGAGAGGDGGAGGI | ASRPGTGGYEQY | 0.44304243 |
| GTGHNGSQPGMGGNG | ASRPGTGGYEQY | 0.51152384 |
| GYAAASASAS | ASRPGTGGYEQY | 0.4771598 |
| HQVDVAAVQQG | ASRPGTGGYEQY | 0.78682405 |
| IGAAIAHT | ASRPGTGGYEQY | 0.40676087 |
| IGMIEVAQL | ASRPGTGGYEQY | 0.80649155 |
| IRLAPEHTSAMD | ASRPGTGGYEQY | 0.14006062 |
| IYNVSVL | ASRPGTGGYEQY | 0.26178443 |
| KQGFLGM | ASRPGTGGYEQY | 0.80390537 |
| LESVITQL | ASRPGTGGYEQY | 0.09247534 |
| LGLGALVVV | ASRPGTGGYEQY | 0.12885839 |
| LIGNGGNGGSGGI | ASRPGTGGYEQY | 0.5514872 |

| | | |
|---|---|---|
| LNAINAPTL | ASRPGTGGYEQY | 0.8337169 |
| LQLLNTGV | ASRPGTGGYEQY | 0.10052668 |
| LQTSIEF | ASRPGTGGYEQY | 0.38291484 |
| MSFAVSEL | ASRPGTGGYEQY | 0.18042836 |
| NTGALGL | ASRPGTGGYEQY | 0.63879 |
| PPGAGGAGGAGTT | ASRPGTGGYEQY | 0.50161827 |
| QLVGLGI | ASRPGTGGYEQY | 0.15995252 |
| RFLEQVHQL | ASRPGTGGYEQY | 0.65715647 |
| RGVGYVLR | ASRPGTGGYEQY | 0.25352204 |
| SATGVFAGVI | ASRPGTGGYEQY | 0.3304006 |
| SLPAITLP | ASRPGTGGYEQY | 0.7689904 |
| STALAFTV | ASRPGTGGYEQY | 0.83750343 |
| SVLTVGLGK | ASRPGTGGYEQY | 0.2377109 |
| TALAGAGSGI | ASRPGTGGYEQY | 0.65275013 |
| TGLPFTGI | ASRPGTGGYEQY | 0.43049192 |
| VAEHVTV | ASRPGTGGYEQY | 0.88324594 |
| VDIVLNSL | ASRPGTGGYEQY | 0.35804445 |
| VLTGGGAVL | ASRPGTGGYEQY | 0.18298809 |
| VNVGLAYGI | ASRPGTGGYEQY | 0.7306663 |
| VSSAAVGAV | ASRPGTGGYEQY | 0.73944175 |
| VTSVYEAGV | ASRPGTGGYEQY | 0.46862888 |
| WVALDRGA | ASRPGTGGYEQY | 0.77303594 |
| YLVGSVVI | ASRPGTGGYEQY | 0.38777938 |
| GPPGTGKTLL | ASRPGTGGYEQY | 0.45180613 |
| GVGMTKF | ASRPGTGGYEQY | 0.75994575 |
| IGIGGSDL | ASRPGTGGYEQY | 0.70009184 |
| IGTLLAEL | ASRPGTGGYEQY | 0.28668234 |
| IYTSGTT | ASRPGTGGYEQY | 0.770288 |
| LSFPAGV | ASRPGTGGYEQY | 0.58021796 |
| LYTSGTT | ASRPGTGGYEQY | 0.21868014 |
| VGAFPFGF | ASRPGTGGYEQY | 0.87254524 |
| VGAGTVEF | ASRPGTGGYEQY | 0.38742515 |
| VGKSTLLL | ASRPGTGGYEQY | 0.71637636 |
| AALAHAGGGIV | ASRPGTGGYEQY | 0.63886225 |
| AALVAVSLL | ASRPGTGGYEQY | 0.78342915 |
| AAVSRAVAAL | ASRPGTGGYEQY | 0.2489737 |
| AGGSSGIGGF | ASRPGTGGYEQY | 0.75265586 |
| AGLTVANANG | ASRPGTGGYEQY | 0.6252719 |
| AGLVGAVL | ASRPGTGGYEQY | 0.6761862 |
| ALSQAFTV | ASRPGTGGYEQY | 0.54739296 |
| ALTFSVSL | ASRPGTGGYEQY | 0.599746 |
| ALTGAALVAL | ASRPGTGGYEQY | 0.6055978 |
| ASFSIGAGVL | ASRPGTGGYEQY | 0.78816533 |
| DGSVYKI | ASRPGTGGYEQY | 0.47907743 |
| EQLAEIV | ASRPGTGGYEQY | 0.4685248 |
| FAVSELTVM | ASRPGTGGYEQY | 0.8277596 |
| GAAPLAAGL | ASRPGTGGYEQY | 0.1959257 |
| GADGGDGGSGGAGG | ASRPGTGGYEQY | 0.57782483 |
| GFGLDTGGAGG | ASRPGTGGYEQY | 0.721061 |
| GLASGGLL | ASRPGTGGYEQY | 0.7146285 |
| GRIAGAAVLL | ASRPGTGGYEQY | 0.4961086 |

| | | |
|---|---|---|
| GSAGSLNTSF | ASRPGTGGYEQY | 0.807837 |
| GTGQIGI | ASRPGTGGYEQY | 0.3645774 |
| GTLIALIL | ASRPGTGGYEQY | 0.7529074 |
| IAATMGFVVG | ASRPGTGGYEQY | 0.6660284 |
| IFAQALL | ASRPGTGGYEQY | 0.49831384 |
| IGAAVALGF | ASRPGTGGYEQY | 0.27905488 |
| IGLTPLAL | ASRPGTGGYEQY | 0.73026276 |
| ILGSNTSTL | ASRPGTGGYEQY | 0.7521379 |
| IVFSLTVL | ASRPGTGGYEQY | 0.75682205 |
| LASKATGF | ASRPGTGGYEQY | 0.14319815 |
| LASVALIL | ASRPGTGGYEQY | 0.6955824 |
| LFAVGLTL | ASRPGTGGYEQY | 0.7696731 |
| LFNGGIGTYI | ASRPGTGGYEQY | 0.87000746 |
| LTLGAGII | ASRPGTGGYEQY | 0.39964175 |
| LVAAALGDI | ASRPGTGGYEQY | 0.73349833 |
| NTAFVQLGI | ASRPGTGGYEQY | 0.5568167 |
| QLFSGAEL | ASRPGTGGYEQY | 0.524239 |
| SAATFITL | ASRPGTGGYEQY | 0.1865614 |
| SNYGLANLGV | ASRPGTGGYEQY | 0.30791494 |
| SYGGAYAVM | ASRPGTGGYEQY | 0.5055318 |
| TTGVGGLTLG | ASRPGTGGYEQY | 0.3332981 |
| TTQTIYI | ASRPGTGGYEQY | 0.81559813 |
| VAAGAGLGAVI | ASRPGTGGYEQY | 0.7364756 |
| VAAVVTLL | ASRPGTGGYEQY | 0.75501496 |
| VGFLGGLI | ASRPGTGGYEQY | 0.7900882 |
| VGVIPILL | ASRPGTGGYEQY | 0.8493523 |
| VIFASSLI | ASRPGTGGYEQY | 0.6437483 |
| VKGGDGGAAAG | ASRPGTGGYEQY | 0.40883732 |
| VTKTFTNI | ASRPGTGGYEQY | 0.4653525 |
| VVAAAILL | ASRPGTGGYEQY | 0.23112786 |
| YAALDVEL | ASRPGTGGYEQY | 0.3584505 |
| LASVALLL | ASRPGTGGYEQY | 0.4216468 |
| LDAVHRV | ASRPGTGGYEQY | 0.47605932 |
| QGLSSSIF | ASRPGTGGYEQY | 0.090385266 |
| RGTFANIRL | ASRPGTGGYEQY | 0.77065206 |
| RQLWWGHRI | ASRPGTGGYEQY | 0.8057879 |
| YGPPGTGKTLL | ASRPGTGGYEQY | 0.44328904 |
| AALVGAPANVA | ASRPGTGGYEQY | 0.6495563 |
| AATFRRTE | ASRPGTGGYEQY | 0.74707144 |
| AGALGGGI | ASRPGTGGYEQY | 0.38106424 |
| AGGAGGAGGIGAG | ASRPGTGGYEQY | 0.6941683 |
| AGSGALGL | ASRPGTGGYEQY | 0.33535412 |
| AQIADQI | ASRPGTGGYEQY | 0.7208129 |
| ASAVAVTYL | ASRPGTGGYEQY | 0.8093712 |
| ASTVLGGTGG | ASRPGTGGYEQY | 0.6549565 |
| AVGGEPAK | ASRPGTGGYEQY | 0.3821547 |
| CEQGTFGGTC | ASRPGTGGYEQY | 0.5407481 |
| DEGKAPAG | ASRPGTGGYEQY | 0.18513754 |
| DTVVFPAL | ASRPGTGGYEQY | 0.43241966 |
| FEGITMAG | ASRPGTGGYEQY | 0.09258565 |
| FPLVVQD | ASRPGTGGYEQY | 0.61396325 |

| | | |
|---|---|---|
| GAFTTPQ | ASRPGTGGYEQY | 0.25956985 |
| GFGGFGTAGGVGGA | ASRPGTGGYEQY | 0.643737 |
| GFLAQFL | ASRPGTGGYEQY | 0.6023185 |
| GGAGGAGGNAGVA | ASRPGTGGYEQY | 0.5674567 |
| GGVGTAGAGGAGGAGG | ASRPGTGGYEQY | 0.29333013 |
| GNGGGGDAGNAGSGGN | ASRPGTGGYEQY | 0.5969192 |
| GTGGAGGAGNDAGS | ASRPGTGGYEQY | 0.6770996 |
| GTTNTGWFNS | ASRPGTGGYEQY | 0.24023521 |
| IFMGPLFTL | ASRPGTGGYEQY | 0.8452692 |
| ILAHAIEAI | ASRPGTGGYEQY | 0.4306513 |
| IRSAIREAL | ASRPGTGGYEQY | 0.5350676 |
| ITFAGGG | ASRPGTGGYEQY | 0.7794366 |
| ITGMSGHQEV | ASRPGTGGYEQY | 0.5049795 |
| IVAGLACTL | ASRPGTGGYEQY | 0.20701084 |
| IVGGGLAGA | ASRPGTGGYEQY | 0.45968482 |
| IVTGVLNI | ASRPGTGGYEQY | 0.60898185 |
| LAQQGCI | ASRPGTGGYEQY | 0.34689677 |
| LCLHVEN | ASRPGTGGYEQY | 0.100984484 |
| LDAISDGS | ASRPGTGGYEQY | 0.6951151 |
| LGGFAAML | ASRPGTGGYEQY | 0.22261539 |
| LGGKNPMI | ASRPGTGGYEQY | 0.40178382 |
| LGPKGVAAIL | ASRPGTGGYEQY | 0.4975017 |
| LMVGVSI | ASRPGTGGYEQY | 0.7412952 |
| LVGAAFVL | ASRPGTGGYEQY | 0.33186817 |
| LVLAVAPL | ASRPGTGGYEQY | 0.5202757 |
| MAAIAVIDAA | ASRPGTGGYEQY | 0.6270644 |
| NNGINAELQ | ASRPGTGGYEQY | 0.56098825 |
| RGEPLVLVHG | ASRPGTGGYEQY | 0.76413316 |
| RGGAGGAAGLI | ASRPGTGGYEQY | 0.6307073 |
| RSYRMHPAV | ASRPGTGGYEQY | 0.45519722 |
| SAAHLLAGQLA | ASRPGTGGYEQY | 0.656667 |
| SAAVGSY | ASRPGTGGYEQY | 0.37110198 |
| SPHAVLHRI | ASRPGTGGYEQY | 0.19252044 |
| SQIAKVL | ASRPGTGGYEQY | 0.22264236 |
| TALGHLVP | ASRPGTGGYEQY | 0.50229627 |
| TAPTTSGAA | ASRPGTGGYEQY | 0.18243857 |
| TDGDDGNGGSPGT | ASRPGTGGYEQY | 0.57013655 |
| TDGLGEN | ASRPGTGGYEQY | 0.7349811 |
| TGGPPLPGQA | ASRPGTGGYEQY | 0.53291094 |
| TIPAITI | ASRPGTGGYEQY | 0.16316392 |
| TIPGFTI | ASRPGTGGYEQY | 0.45966136 |
| TSIASGALA | ASRPGTGGYEQY | 0.3300357 |
| TVTLPGAAI | ASRPGTGGYEQY | 0.60609245 |
| VASGLSAALP | ASRPGTGGYEQY | 0.7140054 |
| VGAGVIGI | ASRPGTGGYEQY | 0.8584597 |
| VGPKRFGV | ASRPGTGGYEQY | 0.78636754 |
| VNVAFPD | ASRPGTGGYEQY | 0.47679225 |
| VSMTNSGVS | ASRPGTGGYEQY | 0.7371677 |
| VTDVLGDLSV | ASRPGTGGYEQY | 0.3685996 |
| VVVVGAGFAGL | ASRPGTGGYEQY | 0.72672397 |
| VYTVGTVV | ASRPGTGGYEQY | 0.49943903 |

| | | |
|---|---|---|
| WTTFSGP | ASRPGTGGYEQY | 0.633057 |
| YDELVEAF | ASRPGTGGYEQY | 0.31475008 |
| YGTGGAGGNGG | ASRPGTGGYEQY | 0.35392582 |
| YTNVMAV | ASRPGTGGYEQY | 0.18245447 |
| FLEQVHQL | ASRPGTGGYEQY | 0.65529704 |
| GLSLLAVAV | ASRPGTGGYEQY | 0.35264748 |
| LEQVHQL | ASRPGTGGYEQY | 0.16147937 |
| LNAAGAI | ASRPGTGGYEQY | 0.39279103 |
| PAVHASL | ASRPGTGGYEQY | 0.8652943 |
| TELEDDA | ASRPGTGGYEQY | 0.7873551 |
| TPPGPPN | ASRPGTGGYEQY | 0.395067 |
| TPPPPPPP | ASRPGTGGYEQY | 0.346166 |
| TQSPALI | ASSQDGNTLY | 0.7766822 |
| EFAAAFPQ | ASSQDGNTLY | 0.34066495 |
| LLSGNLDVL | ASSQDGNTLY | 0.69441175 |
| GPANLGNYNI | ASSQDGNTLY | 0.5660129 |
| TLGPTLP | ASSQDGNTLY | 0.5713995 |
| VYAYSSCCK | ASSQDGNTLY | 0.37838164 |
| APAPAPAPNI | ASSQDGNTLY | 0.71567833 |
| PGGPVPIP | ASSQDGNTLY | 0.50611794 |
| FLTVALEP | ASSQDGNTLY | 0.5172038 |
| SQYFGANL | ASSQDGNTLY | 0.11304317 |
| GGVKPAA | ASSQDGNTLY | 0.76627004 |
| GGTGGAGGAGGW | ASSQDGNTLY | 0.30056572 |
| GVGGVAGLGGAGG | ASSQDGNTLY | 0.24185842 |
| VLLVLATTI | ASSQDGNTLY | 0.13576494 |
| IGIFAAML | ASSQDGNTLY | 0.5136454 |
| AVGDLID | ASSQDGNTLY | 0.7864691 |
| NISPLPL | ASSQDGNTLY | 0.8051417 |
| VTAVFVAI | ASSQDGNTLY | 0.32683098 |
| SGDDNDAAL | ASSQDGNTLY | 0.718785 |
| AVAALAHS | ASSQDGNTLY | 0.8470073 |
| GGVTVMATP | ASSQDGNTLY | 0.43538937 |
| AGLAANVRD | ASSQDGNTLY | 0.15235513 |
| TNRPDLIDPALLRPG | ASSQDGNTLY | 0.5484702 |
| LANLGVTAL | ASSQDGNTLY | 0.6988282 |
| TGERIEL | ASSQDGNTLY | 0.72995996 |
| NGEELAA | ASSQDGNTLY | 0.78243876 |
| VLSLVWG | ASSQDGNTLY | 0.19896702 |
| IGYALWL | ASSQDGNTLY | 0.68532544 |
| LSLLLAPL | ASSQDGNTLY | 0.75074965 |
| TLDEALK | ASSQDGNTLY | 0.23359638 |
| TDGAGTAEGG | ASSQDGNTLY | 0.41015363 |
| NAPGGAGGNGG | ASSQDGNTLY | 0.39895523 |
| VSVFMGMGGNF | ASSQDGNTLY | 0.26642087 |
| LASYLAAL | ASSQDGNTLY | 0.6482299 |
| LVGGTPAL | ASSQDGNTLY | 0.47924778 |
| VRNADVL | ASSQDGNTLY | 0.41446 |
| TCSSPFLVSAM | ASSQDGNTLY | 0.4663167 |
| VHITDAE | ASSQDGNTLY | 0.37991735 |
| EYAAPSF | ASSQDGNTLY | 0.3506735 |

| | | |
|---|---|---|
| VVDAFHGGL | ASSQDGNTLY | 0.44948107 |
| ASVVDPQ | ASSQDGNTLY | 0.7106586 |
| SYNEFGGLS | ASSQDGNTLY | 0.74386513 |
| DPDGSVY | ASSQDGNTLY | 0.6663143 |
| MVSTIHATE | ASSQDGNTLY | 0.63610184 |
| LAAGVADPV | ASSQDGNTLY | 0.8296665 |
| TKEQIAA | ASSQDGNTLY | 0.2037426 |
| SLASPAAL | ASSQDGNTLY | 0.31580192 |
| IGLVFYGV | ASSQDGNTLY | 0.6589142 |
| FLLDASNL | ASSQDGNTLY | 0.328284 |
| YQVGNFP | ASSQDGNTLY | 0.7734859 |
| ESLLSQST | ASSQDGNTLY | 0.66060126 |
| LFTVGFAAVG | ASSQDGNTLY | 0.55763876 |
| ATPPTPT | ASSQDGNTLY | 0.5863511 |
| TDLATLG | ASSQDGNTLY | 0.8753592 |
| SAGGTGR | ASSQDGNTLY | 0.64298916 |
| GFSNSGS | ASSQDGNTLY | 0.47984183 |
| GGNGGSGGTSV | ASSQDGNTLY | 0.8186439 |
| GTSVATGGAGN | ASSQDGNTLY | 0.23452353 |
| NGGNGGSGGTS | ASSQDGNTLY | 0.7383506 |
| AVVKFPH | ASSQDGNTLY | 0.72604656 |
| TPPPPPP | ASSQDGNTLY | 0.22926793 |
| AAPAAAAAPA | ASSQDGNTLY | 0.46100253 |
| IALNEDL | ASSQDGNTLY | 0.7941986 |
| IVNVSSM | ASSQDGNTLY | 0.80115896 |
| TGIPPAPR | ASSQDGNTLY | 0.18934062 |
| FDLGGGTF | ASSQDGNTLY | 0.36684144 |
| FDLGGGTFDVS | ASSQDGNTLY | 0.20889804 |
| NEPTAAA | ASSQDGNTLY | 0.29934177 |
| NEPTAAALA | ASSQDGNTLY | 0.19499406 |
| GSIINISSI | ASSQDGNTLY | 0.5897821 |
| VAPGNAGTA | ASSQDGNTLY | 0.41333917 |
| TGGGASLE | ASSQDGNTLY | 0.78735316 |
| VAVYNHYKRI | ASSQDGNTLY | 0.5147536 |
| TDPAPATT | ASSQDGNTLY | 0.7450539 |
| TNRPDILDPALLRPG | ASSQDGNTLY | 0.3549654 |
| LTTTTTS | ASSQDGNTLY | 0.5617577 |
| PTGSGKTL | ASSQDGNTLY | 0.18737513 |
| TGSGKTL | ASSQDGNTLY | 0.23237005 |
| TIGHVDHGK | ASSQDGNTLY | 0.28573716 |
| TIGHVDHGKTT | ASSQDGNTLY | 0.54616606 |
| GHVDHGKTT | ASSQDGNTLY | 0.1359484 |
| AQFEHTLL | ASSQDGNTLY | 0.84066546 |
| IVNQDEL | ASSQDGNTLY | 0.40401134 |
| TNGSQFF | ASSQDGNTLY | 0.6067959 |
| TGKGIVE | ASSQDGNTLY | 0.42076337 |
| LISNASDA | ASSQDGNTLY | 0.25092217 |
| AELRTLGELTKIAKAHGAQ | ASSQDGNTLY | 0.40159717 |
| APNAVAAAEPSW | ASSQDGNTLY | 0.72355837 |
| APQVAGAATAVL | ASSQDGNTLY | 0.30199003 |
| ARRDPAAAAAILPTDA | ASSQDGNTLY | 0.17825413 |

| | | |
|---|---|---|
| DAAARDVLAEAGLAEYF | ASSQDGNTLY | 0.30075753 |
| DDQIVLSAKANLAKVEQ | ASSQDGNTLY | 0.4604466 |
| DELLPDTGEDSDEDGH | ASSQDGNTLY | 0.8467095 |
| ELPEAPANRATVPQD | ASSQDGNTLY | 0.7877196 |
| FGQSKIERVTGIG | ASSQDGNTLY | 0.5407761 |
| GAGGTGSTAGGAGGAGGAGGLY | ASSQDGNTLY | 0.815073 |
| GHGGTGAAGGNGGTGGAGGSGI | ASSQDGNTLY | 0.10303019 |
| GNGGNGGAGGEGGPGGAGGAGA | ASSQDGNTLY | 0.46561325 |
| GSTAGNGGNGGNATNGTIAGQP | ASSQDGNTLY | 0.8221116 |
| GTGGDGGNGGNGGTGG | ASSQDGNTLY | 0.57426023 |
| GVRRPAGIPADAPV | ASSQDGNTLY | 0.44474563 |
| GYRQVTDAHLLTLARR | ASSQDGNTLY | 0.7929839 |
| IAKMGQDGHDRG | ASSQDGNTLY | 0.41625953 |
| KAEPSIEIQYT | ASSQDGNTLY | 0.3663118 |
| KPVRIRMAADEDTVAVT | ASSQDGNTLY | 0.5638932 |
| MPPRIAGMRLLVIKPEP | ASSQDGNTLY | 0.63228613 |
| MKFYAVALFRSRGVPPTR | ASSQDGNTLY | 0.57980406 |
| NAGGFGILGAVAHSP | ASSQDGNTLY | 0.2010702 |
| NEGREITLAMAEM | ASSQDGNTLY | 0.61840045 |
| NELVEWSHPLGHRWRWE | ASSQDGNTLY | 0.88610786 |
| NLGAGGTGGNGGNPG | ASSQDGNTLY | 0.44198063 |
| NTEWPKWETVMAADPVD | ASSQDGNTLY | 0.7476984 |
| PASAAPAETQP | ASSQDGNTLY | 0.51881826 |
| QQSSDGKNGVRAAEITGE | ASSQDGNTLY | 0.6365045 |
| RAPDAVTTSWWKE | ASSQDGNTLY | 0.30663037 |
| SGFGNVGAHTSGWFNQ | ASSQDGNTLY | 0.25585872 |
| SGVAGAVGQAASVGGLKV | ASSQDGNTLY | 0.62057626 |
| SNGGAGTGSAGGNGGT | ASSQDGNTLY | 0.2406111 |
| TGGAGGEGGAAG | ASSQDGNTLY | 0.5913065 |
| TGGDGGAAGAGGAPGHG | ASSQDGNTLY | 0.47710747 |
| TMPPLLAAIANAGRHGV | ASSQDGNTLY | 0.8906513 |
| TSSTAGNSGGA | ASSQDGNTLY | 0.6946666 |
| VPATVPVGILC | ASSQDGNTLY | 0.60217965 |
| VQRPDSGGVVLTGRIS | ASSQDGNTLY | 0.38024783 |
| WTPNPVPATKGVPAGLVQAT | ASSQDGNTLY | 0.57026994 |
| YALKRALAKFSDRFDVVII | ASSQDGNTLY | 0.70726264 |
| YLIGSVIV | ASSQDGNTLY | 0.20054013 |
| YGQLVAVID | ASSQDGNTLY | 0.7612603 |
| VTVGGMTLPA | ASSQDGNTLY | 0.2512176 |
| VLALPAV | ASSQDGNTLY | 0.61532694 |
| VIPPTLNYET | ASSQDGNTLY | 0.62521 |
| VGMGGMHPSSG | ASSQDGNTLY | 0.83264726 |
| VGLAGGAATA | ASSQDGNTLY | 0.54188585 |
| VGGAGGAGGSGP | ASSQDGNTLY | 0.5980832 |
| VFAAFMV | ASSQDGNTLY | 0.32486755 |
| TLYDKLR | ASSQDGNTLY | 0.63220906 |
| TLAVFTVL | ASSQDGNTLY | 0.7882481 |
| TGSVQPARV | ASSQDGNTLY | 0.63669854 |
| TGGGAGSGGT | ASSQDGNTLY | 0.8425865 |
| SHLETSL | ASSQDGNTLY | 0.5378345 |
| SFTTGMT | ASSQDGNTLY | 0.1532157 |

| | | |
|---|---|---|
| QVEVAKRMGVRQPTVS | ASSQDGNTLY | 0.35076553 |
| QGSGGPALAM | ASSQDGNTLY | 0.81067765 |
| MLAPSNP | ASSQDGNTLY | 0.83860445 |
| LVNNAAI | ASSQDGNTLY | 0.846327 |
| LTGEINTLA | ASSQDGNTLY | 0.73093015 |
| LSIFGPV | ASSQDGNTLY | 0.46229395 |
| LLALLANL | ASSQDGNTLY | 0.8157973 |
| LISNASDALDKLR | ASSQDGNTLY | 0.8810917 |
| LINEAVP | ASSQDGNTLY | 0.62361926 |
| LGYAFSGL | ASSQDGNTLY | 0.22100589 |
| LDFVAVAL | ASSQDGNTLY | 0.21465145 |
| LAQPGAE | ASSQDGNTLY | 0.66783786 |
| ISNLGAVTGF | ASSQDGNTLY | 0.8128326 |
| HQEDPGRI | ASSQDGNTLY | 0.2336205 |
| GTPGEPTP | ASSQDGNTLY | 0.49373323 |
| GFFNSGS | ASSQDGNTLY | 0.5619228 |
| GAGGGGLTAGAN | ASSQDGNTLY | 0.13162214 |
| FVDALVNM | ASSQDGNTLY | 0.75943005 |
| FLTVAPEL | ASSQDGNTLY | 0.86229014 |
| FETLTAL | ASSQDGNTLY | 0.6171693 |
| DVINPIEL | ASSQDGNTLY | 0.22009274 |
| APVSGAPV | ASSQDGNTLY | 0.79189414 |
| ALVGGLAW | ASSQDGNTLY | 0.8607807 |
| ALAAAAATAI | ASSQDGNTLY | 0.805522 |
| AFGLGLLL | ASSQDGNTLY | 0.20756641 |
| AFFAAASL | ASSQDGNTLY | 0.11337805 |
| AAAALPAFSPP | ASSQDGNTLY | 0.5120449 |
| VVLAPAL | ASSQDGNTLY | 0.80181104 |
| TIGHVDHGKTTL | ASSQDGNTLY | 0.48605052 |
| LISNASDALDK | ASSQDGNTLY | 0.31323487 |
| LISNASDAL | ASSQDGNTLY | 0.7055322 |
| ISNASDALDK | ASSQDGNTLY | 0.78305626 |
| NQAETLVYQTE | ASSQDGNTLY | 0.19356924 |
| TGGTGGTAGAG | ASSQDGNTLY | 0.18957722 |
| TGSTAGGAGGA | ASSQDGNTLY | 0.82252425 |
| DPLAVPVAAGPV | ASSQDGNTLY | 0.7302453 |
| GGTGGTPGNGGN | ASSQDGNTLY | 0.25591317 |
| PAGLAGVDPATL | ASSQDGNTLY | 0.48891798 |
| GNNLVVEAPHAGASV | ASSQDGNTLY | 0.61104715 |
| KVTRIMSASTLEQPA | ASSQDGNTLY | 0.47821617 |
| LGFALPIALPRRTVV | ASSQDGNTLY | 0.61598474 |
| TRLGNLTARKLVVLV | ASSQDGNTLY | 0.18767092 |
| WSQQRRLIYFPSAGP | ASSQDGNTLY | 0.8387538 |
| GPNGNPDPMAAAVDIR | ASSQDGNTLY | 0.21447583 |
| TQRCVSVADLRAAASR | ASSQDGNTLY | 0.7495897 |
| QFVLDVAEHPVEVLTETV | ASSQDGNTLY | 0.551831 |
| GGAGGTGGAGGNGGTQAGDG | ASSQDGNTLY | 0.5455094 |
| PLLATLRAIAVDVDAKPAQV | ASSQDGNTLY | 0.45266524 |
| APKFRTGGYMIDFWGVGYQVAK | ASSQDGNTLY | 0.21727377 |
| GAGGAGGAGGAAGGPGGTGGTG | ASSQDGNTLY | 0.28584105 |
| IGALGLLATDVAPAGSVAFGVA | ASSQDGNTLY | 0.19260383 |

| | | |
|---|---|---|
| AGGAGGIGGNGGAGGAGGVLMG | ASSQDGNTLY | 0.7738771 |
| ADELIGLL | ASSQDGNTLY | 0.110933095 |
| ADRALKA | ASSQDGNTLY | 0.6793209 |
| AGAVGQVGGAGGA | ASSQDGNTLY | 0.53241384 |
| AGGGGGLGGAP | ASSQDGNTLY | 0.26944345 |
| AGGLLYGNGGAG | ASSQDGNTLY | 0.59201276 |
| AGLESIGL | ASSQDGNTLY | 0.19947414 |
| AGLKTGII | ASSQDGNTLY | 0.40167508 |
| ASLAALLL | ASSQDGNTLY | 0.6114212 |
| ASVINISSI | ASSQDGNTLY | 0.66259885 |
| ATPPGPA | ASSQDGNTLY | 0.33948392 |
| ATVASALV | ASSQDGNTLY | 0.65667593 |
| ATVASLIL | ASSQDGNTLY | 0.637584 |
| AVNGPVLIRH | ASSQDGNTLY | 0.26587677 |
| DGAAGAVGI | ASSQDGNTLY | 0.609588 |
| DPLLHGI | ASSQDGNTLY | 0.7850687 |
| ELGAALGAL | ASSQDGNTLY | 0.59646374 |
| ELGRTQAV | ASSQDGNTLY | 0.2043632 |
| FVAGGYLS | ASSQDGNTLY | 0.648556 |
| GASGGAGGAGGW | ASSQDGNTLY | 0.33823633 |
| GGAGAGGDGGAGGI | ASSQDGNTLY | 0.63294923 |
| GTGHNGSQPGMGGNG | ASSQDGNTLY | 0.5593157 |
| GYAAASASAS | ASSQDGNTLY | 0.43730068 |
| HQVDVAAVQQG | ASSQDGNTLY | 0.6740174 |
| IGAAIAHT | ASSQDGNTLY | 0.30105188 |
| IGMIEVAQL | ASSQDGNTLY | 0.336449 |
| IRLAPEHTSAMD | ASSQDGNTLY | 0.36612582 |
| IYNVSVL | ASSQDGNTLY | 0.6495136 |
| KQGFLGM | ASSQDGNTLY | 0.6509069 |
| LESVITQL | ASSQDGNTLY | 0.27123734 |
| LGLGALVVV | ASSQDGNTLY | 0.54898393 |
| LIGNGGNGGSGGI | ASSQDGNTLY | 0.38509566 |
| LNAINAPTL | ASSQDGNTLY | 0.24360901 |
| LQLLNTGV | ASSQDGNTLY | 0.5794188 |
| LQTSIEF | ASSQDGNTLY | 0.40275267 |
| MSFAVSEL | ASSQDGNTLY | 0.18495718 |
| NTGALGL | ASSQDGNTLY | 0.648077 |
| PPGAGGAGGAGTT | ASSQDGNTLY | 0.33294153 |
| QLVGLGI | ASSQDGNTLY | 0.5506037 |
| RFLEQVHQL | ASSQDGNTLY | 0.8872865 |
| RGVGYVLR | ASSQDGNTLY | 0.17855436 |
| SATGVFAGVI | ASSQDGNTLY | 0.680796 |
| SLPAITLP | ASSQDGNTLY | 0.57845324 |
| STALAFTV | ASSQDGNTLY | 0.559525 |
| SVLTVGLGK | ASSQDGNTLY | 0.5416614 |
| TALAGAGSGI | ASSQDGNTLY | 0.36178428 |
| TGLPFTGI | ASSQDGNTLY | 0.83919424 |
| VAEHVTV | ASSQDGNTLY | 0.5935542 |
| VDIVLNSL | ASSQDGNTLY | 0.6961297 |
| VLTGGGAVL | ASSQDGNTLY | 0.64040184 |
| VNVGLAYGI | ASSQDGNTLY | 0.80336016 |

| | | |
|---|---|---|
| VSSAAVGAV | ASSQDGNTLY | 0.6884655 |
| VTSVYEAGV | ASSQDGNTLY | 0.29433194 |
| WVALDRGA | ASSQDGNTLY | 0.16880041 |
| YLVGSVVI | ASSQDGNTLY | 0.5288433 |
| GPPGTGKTLL | ASSQDGNTLY | 0.56219894 |
| GVGMTKF | ASSQDGNTLY | 0.33391157 |
| IGIGGSDL | ASSQDGNTLY | 0.15091248 |
| IGTLLAEL | ASSQDGNTLY | 0.7851974 |
| IYTSGTT | ASSQDGNTLY | 0.8803689 |
| LSFPAGV | ASSQDGNTLY | 0.37320232 |
| LYTSGTT | ASSQDGNTLY | 0.8629743 |
| VGAFPFGF | ASSQDGNTLY | 0.23097086 |
| VGAGTVEF | ASSQDGNTLY | 0.6215776 |
| VGKSTLLL | ASSQDGNTLY | 0.81318367 |
| AALAHAGGGIV | ASSQDGNTLY | 0.2442967 |
| AALVAVSLL | ASSQDGNTLY | 0.58531487 |
| AAVSRAVAAL | ASSQDGNTLY | 0.31408066 |
| AGGSSGIGGF | ASSQDGNTLY | 0.3354495 |
| AGLTVANANG | ASSQDGNTLY | 0.77664214 |
| AGLVGAVL | ASSQDGNTLY | 0.48325258 |
| ALSQAFTV | ASSQDGNTLY | 0.62053 |
| ALTFSVSL | ASSQDGNTLY | 0.4204187 |
| ALTGAALVAL | ASSQDGNTLY | 0.30039614 |
| ASFSIGAGVL | ASSQDGNTLY | 0.5353601 |
| DGSVYKI | ASSQDGNTLY | 0.7442379 |
| EQLAEIV | ASSQDGNTLY | 0.33112592 |
| FAVSELTVM | ASSQDGNTLY | 0.8715761 |
| GAAPLAAGL | ASSQDGNTLY | 0.31033593 |
| GADGGDGGSGGAGG | ASSQDGNTLY | 0.88061535 |
| GFGLDTGGAGG | ASSQDGNTLY | 0.837906 |
| GLASGGLL | ASSQDGNTLY | 0.6342544 |
| GRIAGAAVLL | ASSQDGNTLY | 0.4050948 |
| GSAGSLNTSF | ASSQDGNTLY | 0.791173 |
| GTGQIGI | ASSQDGNTLY | 0.4250027 |
| GTLIALIL | ASSQDGNTLY | 0.5322281 |
| IAATMGFVVG | ASSQDGNTLY | 0.7349774 |
| IFAQALL | ASSQDGNTLY | 0.5340719 |
| IGAAVALGF | ASSQDGNTLY | 0.649225 |
| IGLTPLAL | ASSQDGNTLY | 0.6185471 |
| ILGSNTSTL | ASSQDGNTLY | 0.6287042 |
| IVFSLTVL | ASSQDGNTLY | 0.5824185 |
| LASKATGF | ASSQDGNTLY | 0.47487676 |
| LASVALIL | ASSQDGNTLY | 0.6039347 |
| LFAVGLTL | ASSQDGNTLY | 0.5504623 |
| LFNGGIGTYI | ASSQDGNTLY | 0.65184236 |
| LTLGAGII | ASSQDGNTLY | 0.6460182 |
| LVAAALGDI | ASSQDGNTLY | 0.5598823 |
| NTAFVQLGI | ASSQDGNTLY | 0.65274334 |
| QLFSGAEL | ASSQDGNTLY | 0.6469689 |
| SAATFITL | ASSQDGNTLY | 0.47414893 |
| SNYGLANLGV | ASSQDGNTLY | 0.59364104 |

| | | |
|---|---|---|
| SYGGAYAVM | ASSQDGNTLY | 0.5798686 |
| TTGVGGLTLG | ASSQDGNTLY | 0.60644084 |
| TTQTIYI | ASSQDGNTLY | 0.39968765 |
| VAAGAGLGAVI | ASSQDGNTLY | 0.64225054 |
| VAAVVTLL | ASSQDGNTLY | 0.5463816 |
| VGFLGGLI | ASSQDGNTLY | 0.34500083 |
| VGVIPILL | ASSQDGNTLY | 0.62230515 |
| VIFASSLI | ASSQDGNTLY | 0.6061799 |
| VKGGDGGAAAG | ASSQDGNTLY | 0.59269243 |
| VTKTFTNI | ASSQDGNTLY | 0.6415299 |
| VVAAAILL | ASSQDGNTLY | 0.5378556 |
| YAALDVEL | ASSQDGNTLY | 0.5250306 |
| LASVALLL | ASSQDGNTLY | 0.63493955 |
| LDAVHRV | ASSQDGNTLY | 0.651405 |
| QGLSSSIF | ASSQDGNTLY | 0.5236248 |
| RGTFANIRL | ASSQDGNTLY | 0.61602235 |
| RQLWWGHRI | ASSQDGNTLY | 0.65354955 |
| YGPPGTGKTLL | ASSQDGNTLY | 0.62181044 |
| AALVGAPANVA | ASSQDGNTLY | 0.65137994 |
| AATFRRTE | ASSQDGNTLY | 0.655932 |
| AGALGGGI | ASSQDGNTLY | 0.56993514 |
| AGGAGGAGGIGAG | ASSQDGNTLY | 0.65026176 |
| AGSGALGL | ASSQDGNTLY | 0.6545502 |
| AQIADQI | ASSQDGNTLY | 0.5880064 |
| ASAVAVTYL | ASSQDGNTLY | 0.62048966 |
| ASTVLGGTGG | ASSQDGNTLY | 0.49375445 |
| AVGGEPAK | ASSQDGNTLY | 0.5960927 |
| CEQGTFGGTC | ASSQDGNTLY | 0.47604764 |
| DEGKAPAG | ASSQDGNTLY | 0.61811525 |
| DTVVFPAL | ASSQDGNTLY | 0.6545889 |
| FEGITMAG | ASSQDGNTLY | 0.6009053 |
| FPLVVQD | ASSQDGNTLY | 0.49137488 |
| GAFTTPQ | ASSQDGNTLY | 0.5215818 |
| GFGGFGTAGGVGGA | ASSQDGNTLY | 0.58777523 |
| GFLAQFL | ASSQDGNTLY | 0.5512604 |
| GGAGGAGGNAGVA | ASSQDGNTLY | 0.42747957 |
| GGVGTAGAGGAGGAGG | ASSQDGNTLY | 0.5793408 |
| GNGGGGDAGNAGSGGN | ASSQDGNTLY | 0.6510099 |
| GTGGAGGAGNDAGS | ASSQDGNTLY | 0.64948875 |
| GTTNTGWFNS | ASSQDGNTLY | 0.5245039 |
| IFMGPLFTL | ASSQDGNTLY | 0.6561997 |
| ILAHAIEAI | ASSQDGNTLY | 0.63585496 |
| IRSAIREAL | ASSQDGNTLY | 0.6543781 |
| ITFAGGG | ASSQDGNTLY | 0.6072897 |
| ITGMSGHQEV | ASSQDGNTLY | 0.5053426 |
| IVAGLACTL | ASSQDGNTLY | 0.6448729 |
| IVGGGLAGA | ASSQDGNTLY | 0.5808513 |
| IVTGVLNI | ASSQDGNTLY | 0.6017233 |
| LAQQGCI | ASSQDGNTLY | 0.6455632 |
| LCLHVEN | ASSQDGNTLY | 0.59683126 |
| LDAISDGS | ASSQDGNTLY | 0.544834 |

| | | |
|---|---|---|
| LGGFAAML | ASSQDGNTLY | 0.58780515 |
| LGGKNPMI | ASSQDGNTLY | 0.64354 |
| LGPKGVAAIL | ASSQDGNTLY | 0.5367009 |
| LMVGVSI | ASSQDGNTLY | 0.5166882 |
| LVGAAFVL | ASSQDGNTLY | 0.5751507 |
| LVLAVAPL | ASSQDGNTLY | 0.6090871 |
| MAAIAVIDAA | ASSQDGNTLY | 0.6506887 |
| NNGINAELQ | ASSQDGNTLY | 0.63679844 |
| RGEPLVLVHG | ASSQDGNTLY | 0.5457284 |
| RGGAGGAAGLI | ASSQDGNTLY | 0.63206524 |
| RSYRMHPAV | ASSQDGNTLY | 0.6359271 |
| SAAHLLAGQLA | ASSQDGNTLY | 0.61490536 |
| SAAVGSY | ASSQDGNTLY | 0.6221681 |
| SPHAVLHRI | ASSQDGNTLY | 0.555483 |
| SQIAKVL | ASSQDGNTLY | 0.64043194 |
| TALGHLVP | ASSQDGNTLY | 0.63975114 |
| TAPTTSGAA | ASSQDGNTLY | 0.64097476 |
| TDGDDGNGGSPGT | ASSQDGNTLY | 0.6478392 |
| TDGLGEN | ASSQDGNTLY | 0.60942876 |
| TGGPPLPGQA | ASSQDGNTLY | 0.6132018 |
| TIPAITI | ASSQDGNTLY | 0.4758762 |
| TIPGFTI | ASSQDGNTLY | 0.57324636 |
| TSIASGALA | ASSQDGNTLY | 0.6147326 |
| TVTLPGAAI | ASSQDGNTLY | 0.616887 |
| VASGLSAALP | ASSQDGNTLY | 0.6434478 |
| VGAGVIGI | ASSQDGNTLY | 0.6396595 |
| VGPKRFGV | ASSQDGNTLY | 0.56016874 |
| VNVAFPD | ASSQDGNTLY | 0.64785236 |
| VSMTNSGVS | ASSQDGNTLY | 0.6534832 |
| VTDVLGDLSV | ASSQDGNTLY | 0.49813718 |
| VVVVGAGFAGL | ASSQDGNTLY | 0.57629395 |
| VYTVGTVV | ASSQDGNTLY | 0.6583798 |
| WTTFSGP | ASSQDGNTLY | 0.6213069 |
| YDELVEAF | ASSQDGNTLY | 0.59194815 |
| YGTGGAGGNGG | ASSQDGNTLY | 0.6044191 |
| YTNVMAV | ASSQDGNTLY | 0.6601928 |
| FLEQVHQL | ASSQDGNTLY | 0.58729756 |
| GLSLLAVAV | ASSQDGNTLY | 0.65867865 |
| LEQVHQL | ASSQDGNTLY | 0.41419938 |
| LNAAGAI | ASSQDGNTLY | 0.591672 |
| PAVHASL | ASSQDGNTLY | 0.5282104 |
| TELEDDA | ASSQDGNTLY | 0.36778072 |
| TPPGPPN | ASSQDGNTLY | 0.64755255 |
| TPPPPPPP | ASSQDGNTLY | 0.5982996 |
| TQSPALI | ASSLEGQSNERLF | 0.64159554 |
| EFAAAFPQ | ASSLEGQSNERLF | 0.6059029 |
| LLSGNLDVL | ASSLEGQSNERLF | 0.6536623 |
| GPANLGNYNI | ASSLEGQSNERLF | 0.61323524 |
| TLGPTLP | ASSLEGQSNERLF | 0.5982751 |
| VYAYSSCCK | ASSLEGQSNERLF | 0.5254241 |
| APAPAPAPNI | ASSLEGQSNERLF | 0.6414322 |

| | | |
|---|---|---|
| PGGPVPIP | ASSLEGQSNERLF | 0.6535028 |
| FLTVALEP | ASSLEGQSNERLF | 0.64282674 |
| SQYFGANL | ASSLEGQSNERLF | 0.45475492 |
| GGVKPAA | ASSLEGQSNERLF | 0.6035361 |
| GGTGGAGGAGGW | ASSLEGQSNERLF | 0.6225277 |
| GVGGVAGLGGAGG | ASSLEGQSNERLF | 0.39393207 |
| VLLVLATTI | ASSLEGQSNERLF | 0.38738492 |
| IGIFAAML | ASSLEGQSNERLF | 0.5384282 |
| AVGDLID | ASSLEGQSNERLF | 0.5707329 |
| NISPLPL | ASSLEGQSNERLF | 0.64101714 |
| VTAVFVAI | ASSLEGQSNERLF | 0.63053226 |
| SGDDNDAAL | ASSLEGQSNERLF | 0.59973025 |
| AVAALAHS | ASSLEGQSNERLF | 0.62242496 |
| GGVTVMATP | ASSLEGQSNERLF | 0.60945725 |
| AGLAANVRD | ASSLEGQSNERLF | 0.32178512 |
| TNRPDLIDPALLRPG | ASSLEGQSNERLF | 0.5433527 |
| LANLGVTAL | ASSLEGQSNERLF | 0.48749036 |
| TGERIEL | ASSLEGQSNERLF | 0.65740156 |
| NGEELAA | ASSLEGQSNERLF | 0.5455942 |
| VLSLVWG | ASSLEGQSNERLF | 0.5528336 |
| IGYALWL | ASSLEGQSNERLF | 0.6156687 |
| LSLLLAPL | ASSLEGQSNERLF | 0.61800337 |
| TLDEALK | ASSLEGQSNERLF | 0.60748535 |
| TDGAGTAEGG | ASSLEGQSNERLF | 0.52406466 |
| NAPGGAGGNGG | ASSLEGQSNERLF | 0.3893305 |
| VSVFMGMGGNF | ASSLEGQSNERLF | 0.64608127 |
| LASYLAAL | ASSLEGQSNERLF | 0.5006029 |
| LVGGTPAL | ASSLEGQSNERLF | 0.54365575 |
| VRNADVL | ASSLEGQSNERLF | 0.424849 |
| TCSSPFLVSAM | ASSLEGQSNERLF | 0.5892174 |
| VHITDAE | ASSLEGQSNERLF | 0.45792532 |
| EYAAPSF | ASSLEGQSNERLF | 0.6432336 |
| VVDAFHGGL | ASSLEGQSNERLF | 0.578035 |
| ASVVDPQ | ASSLEGQSNERLF | 0.5945384 |
| SYNEFGGLS | ASSLEGQSNERLF | 0.66079855 |
| DPDGSVY | ASSLEGQSNERLF | 0.61812055 |
| MVSTIHATE | ASSLEGQSNERLF | 0.59174734 |
| LAAGVADPV | ASSLEGQSNERLF | 0.57319224 |
| TKEQIAA | ASSLEGQSNERLF | 0.54445165 |
| SLASPAAL | ASSLEGQSNERLF | 0.524041 |
| IGLVFYGV | ASSLEGQSNERLF | 0.57524073 |
| FLLDASNL | ASSLEGQSNERLF | 0.655294 |
| YQVGNFP | ASSLEGQSNERLF | 0.60173684 |
| ESLLSQST | ASSLEGQSNERLF | 0.5128829 |
| LFTVGFAAVG | ASSLEGQSNERLF | 0.6049295 |
| ATPPTPT | ASSLEGQSNERLF | 0.65817595 |
| TDLATLG | ASSLEGQSNERLF | 0.65811175 |
| SAGGTGR | ASSLEGQSNERLF | 0.53811455 |
| GFSNSGS | ASSLEGQSNERLF | 0.56637406 |
| GGNGGSGGTSV | ASSLEGQSNERLF | 0.5837572 |
| GTSVATGGAGN | ASSLEGQSNERLF | 0.60399896 |

| | | |
|---|---|---|
| NGGNGGSGGTS | ASSLEGQSNERLF | 0.5286347 |
| AVVKFPH | ASSLEGQSNERLF | 0.52216345 |
| TPPPPPP | ASSLEGQSNERLF | 0.6289619 |
| AAPAAAAAPA | ASSLEGQSNERLF | 0.6652224 |
| IALNEDL | ASSLEGQSNERLF | 0.58449197 |
| IVNVSSM | ASSLEGQSNERLF | 0.46622783 |
| TGIPPAPR | ASSLEGQSNERLF | 0.6014498 |
| FDLGGGTF | ASSLEGQSNERLF | 0.5398405 |
| FDLGGGTFDVS | ASSLEGQSNERLF | 0.5583768 |
| NEPTAAA | ASSLEGQSNERLF | 0.5662168 |
| NEPTAAALA | ASSLEGQSNERLF | 0.47986394 |
| GSIINISSI | ASSLEGQSNERLF | 0.6229094 |
| VAPGNAGTA | ASSLEGQSNERLF | 0.66016483 |
| TGGGASLE | ASSLEGQSNERLF | 0.5935677 |
| VAVYNHYKRI | ASSLEGQSNERLF | 0.6339288 |
| TDPAPATT | ASSLEGQSNERLF | 0.6535122 |
| TNRPDILDPALLRPG | ASSLEGQSNERLF | 0.63609165 |
| LTTTTTS | ASSLEGQSNERLF | 0.6471444 |
| PTGSGKTL | ASSLEGQSNERLF | 0.63983905 |
| TGSGKTL | ASSLEGQSNERLF | 0.6266902 |
| TIGHVDHGK | ASSLEGQSNERLF | 0.6078083 |
| TIGHVDHGKTT | ASSLEGQSNERLF | 0.6417791 |
| GHVDHGKTT | ASSLEGQSNERLF | 0.60717344 |
| AQFEHTLL | ASSLEGQSNERLF | 0.6040854 |
| IVNQDEL | ASSLEGQSNERLF | 0.5760414 |
| TNGSQFF | ASSLEGQSNERLF | 0.6502737 |
| TGKGIVE | ASSLEGQSNERLF | 0.5619146 |
| LISNASDA | ASSLEGQSNERLF | 0.6558193 |
| AELRTLGELTKIAKAHGAQ | ASSLEGQSNERLF | 0.4123906 |
| APNAVAAAEPSW | ASSLEGQSNERLF | 0.49695048 |
| APQVAGAATAVL | ASSLEGQSNERLF | 0.63953435 |
| ARRDPAAAAAILPTDA | ASSLEGQSNERLF | 0.31639102 |
| DAAARDVLAEAGLAEYF | ASSLEGQSNERLF | 0.38040227 |
| DDQIVLSAKANLAKVEQ | ASSLEGQSNERLF | 0.59533733 |
| DELLPDTGEDSDEDGH | ASSLEGQSNERLF | 0.6524185 |
| ELPEAPANRATVPQD | ASSLEGQSNERLF | 0.32390448 |
| FGQSKIERVTGIG | ASSLEGQSNERLF | 0.52287483 |
| GAGGTGSTAGGAGGAGGAGGLY | ASSLEGQSNERLF | 0.47887647 |
| GHGGTGAAGGNGGTGGAGGSGI | ASSLEGQSNERLF | 0.6313689 |
| GNGGNGGAGGEGGPGGAGGAGA | ASSLEGQSNERLF | 0.61903256 |
| GSTAGNGGNGGNATNGTIAGQP | ASSLEGQSNERLF | 0.4447525 |
| GTGGDGGNGGNGGTGG | ASSLEGQSNERLF | 0.6428243 |
| GVRRPAGIPADAPV | ASSLEGQSNERLF | 0.43436894 |
| GYRQVTDAHLLTLARR | ASSLEGQSNERLF | 0.5812148 |
| IAKMGQDGHDRG | ASSLEGQSNERLF | 0.64644563 |
| KAEPSIEIQYT | ASSLEGQSNERLF | 0.6530659 |
| KPVRIRMAADEDTVAVT | ASSLEGQSNERLF | 0.5029429 |
| MPPRIAGMRLLVIKPEP | ASSLEGQSNERLF | 0.6380842 |
| MKFYAVALFRSRGVPPTR | ASSLEGQSNERLF | 0.56652856 |
| NAGGFGILGAVAHSP | ASSLEGQSNERLF | 0.6624185 |
| NEGREITLAMAEM | ASSLEGQSNERLF | 0.6009923 |

| | | |
|---|---|---|
| NELVEWSHPLGHRWRWE | ASSLEGQSNERLF | 0.48033494 |
| NLGAGGTGGNGGNPG | ASSLEGQSNERLF | 0.64734817 |
| NTEWPKWETVMAADPVD | ASSLEGQSNERLF | 0.6366073 |
| PASAAPAETQP | ASSLEGQSNERLF | 0.6077703 |
| QQSSDGKNGVRAAEITGE | ASSLEGQSNERLF | 0.6385031 |
| RAPDAVTTSWWKE | ASSLEGQSNERLF | 0.53683794 |
| SGFGNVGAHTSGWFNQ | ASSLEGQSNERLF | 0.60155267 |
| SGVAGAVGQAASVGGLKV | ASSLEGQSNERLF | 0.64842045 |
| SNGGAGTGSAGGNGGT | ASSLEGQSNERLF | 0.6371924 |
| TGGAGGEGGAAG | ASSLEGQSNERLF | 0.51240724 |
| TGGDGGAAGAGGAPGHG | ASSLEGQSNERLF | 0.6530516 |
| TMPPLLAAIANAGRHGV | ASSLEGQSNERLF | 0.6037384 |
| TSSTAGNSGGA | ASSLEGQSNERLF | 0.5619077 |
| VPATVPVGILC | ASSLEGQSNERLF | 0.65912575 |
| VQRPDSGGVVLTGRIS | ASSLEGQSNERLF | 0.58413047 |
| WTPNPVPATKGVPAGLVQAT | ASSLEGQSNERLF | 0.64175606 |
| YALKRALAKFSDRFDVVII | ASSLEGQSNERLF | 0.6495107 |
| YLIGSVIV | ASSLEGQSNERLF | 0.66005474 |
| YGQLVAVID | ASSLEGQSNERLF | 0.4887578 |
| VTVGGMTLPA | ASSLEGQSNERLF | 0.6218569 |
| VLALPAV | ASSLEGQSNERLF | 0.6059133 |
| VIPPTLNYET | ASSLEGQSNERLF | 0.6430049 |
| VGMGGMHPSSG | ASSLEGQSNERLF | 0.600139 |
| VGLAGGAATA | ASSLEGQSNERLF | 0.5775591 |
| VGGAGGAGGSGP | ASSLEGQSNERLF | 0.637808 |
| VFAAFMV | ASSLEGQSNERLF | 0.6539006 |
| TLYDKLR | ASSLEGQSNERLF | 0.6034839 |
| TLAVFTVL | ASSLEGQSNERLF | 0.62101114 |
| TGSVQPARV | ASSLEGQSNERLF | 0.65137416 |
| TGGGAGSGGT | ASSLEGQSNERLF | 0.5598797 |
| SHLETSL | ASSLEGQSNERLF | 0.61243 |
| SFTTGMT | ASSLEGQSNERLF | 0.64036286 |
| QVEVAKRMGVRQPTVS | ASSLEGQSNERLF | 0.6497461 |
| QGSGGPALAM | ASSLEGQSNERLF | 0.6006541 |
| MLAPSNP | ASSLEGQSNERLF | 0.64869034 |
| LVNNAAI | ASSLEGQSNERLF | 0.5375329 |
| LTGEINTLA | ASSLEGQSNERLF | 0.6257187 |
| LSIFGPV | ASSLEGQSNERLF | 0.5810888 |
| LLALLANL | ASSLEGQSNERLF | 0.61591566 |
| LISNASDALDKLR | ASSLEGQSNERLF | 0.5066881 |
| LINEAVP | ASSLEGQSNERLF | 0.6291303 |
| LGYAFSGL | ASSLEGQSNERLF | 0.6363512 |
| LDFVAVAL | ASSLEGQSNERLF | 0.6431345 |
| LAQPGAE | ASSLEGQSNERLF | 0.38246185 |
| ISNLGAVTGF | ASSLEGQSNERLF | 0.6345187 |
| HQEDPGRI | ASSLEGQSNERLF | 0.63954633 |
| GTPGEPTP | ASSLEGQSNERLF | 0.6588952 |
| GFFNSGS | ASSLEGQSNERLF | 0.5890494 |
| GAGGGGLTAGAN | ASSLEGQSNERLF | 0.63606304 |
| FVDALVNM | ASSLEGQSNERLF | 0.53476644 |
| FLTVAPEL | ASSLEGQSNERLF | 0.59926355 |

| | | |
|---|---|---|
| FETLTAL | ASSLEGQSNERLF | 0.4887974 |
| DVINPIEL | ASSLEGQSNERLF | 0.56466836 |
| APVSGAPV | ASSLEGQSNERLF | 0.5770734 |
| ALVGGLAW | ASSLEGQSNERLF | 0.55543286 |
| ALAAAAATAI | ASSLEGQSNERLF | 0.63746727 |
| AFGLGLLL | ASSLEGQSNERLF | 0.64505374 |
| AFFAAASL | ASSLEGQSNERLF | 0.64992106 |
| AAAALPAFSPP | ASSLEGQSNERLF | 0.64699554 |
| VVLAPAL | ASSLEGQSNERLF | 0.65154886 |
| TIGHVDHGKTTL | ASSLEGQSNERLF | 0.64732134 |
| LISNASDALDK | ASSLEGQSNERLF | 0.57794154 |
| LISNASDAL | ASSLEGQSNERLF | 0.60283846 |
| ISNASDALDK | ASSLEGQSNERLF | 0.476443 |
| NQAETLVYQTE | ASSLEGQSNERLF | 0.600837 |
| TGGTGGTAGAG | ASSLEGQSNERLF | 0.5770819 |
| TGSTAGGAGGA | ASSLEGQSNERLF | 0.6238738 |
| DPLAVPVAAGPV | ASSLEGQSNERLF | 0.321653 |
| GGTGGTPGNGGN | ASSLEGQSNERLF | 0.620084 |
| PAGLAGVDPATL | ASSLEGQSNERLF | 0.6567646 |
| GNNLVVEAPHAGASV | ASSLEGQSNERLF | 0.5618285 |
| KVTRIMSASTLEQPA | ASSLEGQSNERLF | 0.631027 |
| LGFALPIALPRRTVV | ASSLEGQSNERLF | 0.63087356 |
| TRLGNLTARKLVVLV | ASSLEGQSNERLF | 0.5539351 |
| WSQQRRLIYFPSAGP | ASSLEGQSNERLF | 0.63273865 |
| GPNGNPDPMAAAVDIR | ASSLEGQSNERLF | 0.56636775 |
| TQRCVSVADLRAAASR | ASSLEGQSNERLF | 0.64266634 |
| QFVLDVAEHPVEVLTETV | ASSLEGQSNERLF | 0.65886045 |
| GGAGGTGGAGGNGGTQAGDG | ASSLEGQSNERLF | 0.63136876 |
| PLLATLRAIAVDVDAKPAQV | ASSLEGQSNERLF | 0.59927887 |
| APKFRTGGYMIDFWGVGYQVAK | ASSLEGQSNERLF | 0.6255604 |
| GAGGAGGAGGAAGGPGGTGGTG | ASSLEGQSNERLF | 0.44383442 |
| IGALGLLATDVAPAGSVAFGVA | ASSLEGQSNERLF | 0.6175731 |
| AGGAGGIGGNGGAGGAGGVLMG | ASSLEGQSNERLF | 0.44749343 |
| ADELIGLL | ASSLEGQSNERLF | 0.6029476 |
| ADRALKA | ASSLEGQSNERLF | 0.5664694 |
| AGAVGQVGGAGGA | ASSLEGQSNERLF | 0.62910324 |
| AGGGGGLGGAP | ASSLEGQSNERLF | 0.61762935 |
| AGGLLYGNGGAG | ASSLEGQSNERLF | 0.60886645 |
| AGLESIGL | ASSLEGQSNERLF | 0.55424553 |
| AGLKTGII | ASSLEGQSNERLF | 0.6247853 |
| ASLAALLL | ASSLEGQSNERLF | 0.6039313 |
| ASVINISSI | ASSLEGQSNERLF | 0.5035348 |
| ATPPGPA | ASSLEGQSNERLF | 0.64097 |
| ATVASALV | ASSLEGQSNERLF | 0.58773535 |
| ATVASLIL | ASSLEGQSNERLF | 0.5927035 |
| AVNGPVLIRH | ASSLEGQSNERLF | 0.65340173 |
| DGAAGAVGI | ASSLEGQSNERLF | 0.6267345 |
| DPLLHGI | ASSLEGQSNERLF | 0.42474806 |
| ELGAALGAL | ASSLEGQSNERLF | 0.5908256 |
| ELGRTQAV | ASSLEGQSNERLF | 0.61482346 |
| FVAGGYLS | ASSLEGQSNERLF | 0.60301834 |

| | | |
|---|---|---|
| GASGGAGGAGGW | ASSLEGQSNERLF | 0.35933515 |
| GGAGAGGDGGAGGI | ASSLEGQSNERLF | 0.6276492 |
| GTGHNGSQPGMGGNG | ASSLEGQSNERLF | 0.56406033 |
| GYAAASASAS | ASSLEGQSNERLF | 0.54077363 |
| HQVDVAAVQQG | ASSLEGQSNERLF | 0.57646626 |
| IGAAIAHT | ASSLEGQSNERLF | 0.6408331 |
| IGMIEVAQL | ASSLEGQSNERLF | 0.5897576 |
| IRLAPEHTSAMD | ASSLEGQSNERLF | 0.60266703 |
| IYNVSVL | ASSLEGQSNERLF | 0.6221094 |
| KQGFLGM | ASSLEGQSNERLF | 0.637396 |
| LESVITQL | ASSLEGQSNERLF | 0.65303373 |
| LGLGALVVV | ASSLEGQSNERLF | 0.61103725 |
| LIGNGGNGGSGGI | ASSLEGQSNERLF | 0.59429175 |
| LNAINAPTL | ASSLEGQSNERLF | 0.62902915 |
| LQLLNTGV | ASSLEGQSNERLF | 0.5472383 |
| LQTSIEF | ASSLEGQSNERLF | 0.50739264 |
| MSFAVSEL | ASSLEGQSNERLF | 0.4452439 |
| NTGALGL | ASSLEGQSNERLF | 0.5940907 |
| PPGAGGAGGAGTT | ASSLEGQSNERLF | 0.46832848 |
| QLVGLGI | ASSLEGQSNERLF | 0.6257459 |
| RFLEQVHQL | ASSLEGQSNERLF | 0.64981073 |
| RGVGYVLR | ASSLEGQSNERLF | 0.59405565 |
| SATGVFAGVI | ASSLEGQSNERLF | 0.46987224 |
| SLPAITLP | ASSLEGQSNERLF | 0.57949376 |
| STALAFTV | ASSLEGQSNERLF | 0.56560946 |
| SVLTVGLGK | ASSLEGQSNERLF | 0.6378047 |
| TALAGAGSGI | ASSLEGQSNERLF | 0.64514947 |
| TGLPFTGI | ASSLEGQSNERLF | 0.6601518 |
| VAEHVTV | ASSLEGQSNERLF | 0.6439908 |
| VDIVLNSL | ASSLEGQSNERLF | 0.58232534 |
| VLTGGGAVL | ASSLEGQSNERLF | 0.63430226 |
| VNVGLAYGI | ASSLEGQSNERLF | 0.5935252 |
| VSSAAVGAV | ASSLEGQSNERLF | 0.6142743 |
| VTSVYEAGV | ASSLEGQSNERLF | 0.61553144 |
| WVALDRGA | ASSLEGQSNERLF | 0.60205853 |
| YLVGSVVI | ASSLEGQSNERLF | 0.58223176 |
| GPPGTGKTLL | ASSLEGQSNERLF | 0.6013924 |
| GVGMTKF | ASSLEGQSNERLF | 0.41942802 |
| IGIGGSDL | ASSLEGQSNERLF | 0.6296014 |
| IGTLLAEL | ASSLEGQSNERLF | 0.57515126 |
| IYTSGTT | ASSLEGQSNERLF | 0.43418413 |
| LSFPAGV | ASSLEGQSNERLF | 0.59990746 |
| LYTSGTT | ASSLEGQSNERLF | 0.65723985 |
| VGAFPFGF | ASSLEGQSNERLF | 0.6487843 |
| VGAGTVEF | ASSLEGQSNERLF | 0.57330173 |
| VGKSTLLL | ASSLEGQSNERLF | 0.5597323 |
| AALAHAGGGIV | ASSLEGQSNERLF | 0.6105221 |
| AALVAVSLL | ASSLEGQSNERLF | 0.5101712 |
| AAVSRAVAAL | ASSLEGQSNERLF | 0.6225954 |
| AGGSSGIGGF | ASSLEGQSNERLF | 0.5993705 |
| AGLTVANANG | ASSLEGQSNERLF | 0.60252106 |

| | | |
|---|---|---|
| AGLVGAVL | ASSLEGQSNERLF | 0.573045 |
| ALSQAFTV | ASSLEGQSNERLF | 0.6257734 |
| ALTFSVSL | ASSLEGQSNERLF | 0.57584304 |
| ALTGAALVAL | ASSLEGQSNERLF | 0.62495685 |
| ASFSIGAGVL | ASSLEGQSNERLF | 0.3332481 |
| DGSVYKI | ASSLEGQSNERLF | 0.6424925 |
| EQLAEIV | ASSLEGQSNERLF | 0.51990986 |
| FAVSELTVM | ASSLEGQSNERLF | 0.5125526 |
| GAAPLAAGL | ASSLEGQSNERLF | 0.3685806 |
| GADGGDGGSGGAGG | ASSLEGQSNERLF | 0.6152441 |
| GFGLDTGGAGG | ASSLEGQSNERLF | 0.63302475 |
| GLASGGLL | ASSLEGQSNERLF | 0.6382483 |
| GRIAGAAVLL | ASSLEGQSNERLF | 0.5604942 |
| GSAGSLNTSF | ASSLEGQSNERLF | 0.64117414 |
| GTGQIGI | ASSLEGQSNERLF | 0.64574665 |
| GTLIALIL | ASSLEGQSNERLF | 0.58377844 |
| IAATMGFVVG | ASSLEGQSNERLF | 0.3966686 |
| IFAQALL | ASSLEGQSNERLF | 0.6288202 |
| IGAAVALGF | ASSLEGQSNERLF | 0.60398066 |
| IGLTPLAL | ASSLEGQSNERLF | 0.6413579 |
| ILGSNTSTL | ASSLEGQSNERLF | 0.6202798 |
| IVFSLTVL | ASSLEGQSNERLF | 0.4985202 |
| LASKATGF | ASSLEGQSNERLF | 0.64533967 |
| LASVALIL | ASSLEGQSNERLF | 0.64520425 |
| LFAVGLTL | ASSLEGQSNERLF | 0.49262285 |
| LFNGGIGTYI | ASSLEGQSNERLF | 0.53434086 |
| LTLGAGII | ASSLEGQSNERLF | 0.559739 |
| LVAAALGDI | ASSLEGQSNERLF | 0.46615872 |
| NTAFVQLGI | ASSLEGQSNERLF | 0.6447244 |
| QLFSGAEL | ASSLEGQSNERLF | 0.5764984 |
| SAATFITL | ASSLEGQSNERLF | 0.58506644 |
| SNYGLANLGV | ASSLEGQSNERLF | 0.6173187 |
| SYGGAYAVM | ASSLEGQSNERLF | 0.58312464 |
| TTGVGGLTLG | ASSLEGQSNERLF | 0.5857721 |
| TTQTIYI | ASSLEGQSNERLF | 0.5932912 |
| VAAGAGLGAVI | ASSLEGQSNERLF | 0.6242053 |
| VAAVVTLL | ASSLEGQSNERLF | 0.62246495 |
| VGFLGGLI | ASSLEGQSNERLF | 0.6061419 |
| VGVIPILL | ASSLEGQSNERLF | 0.62042284 |
| VIFASSLI | ASSLEGQSNERLF | 0.6536559 |
| VKGGDGGAAAG | ASSLEGQSNERLF | 0.4622932 |
| VTKTFTNI | ASSLEGQSNERLF | 0.5868938 |
| VVAAAILL | ASSLEGQSNERLF | 0.62087476 |
| YAALDVEL | ASSLEGQSNERLF | 0.54419553 |
| LASVALLL | ASSLEGQSNERLF | 0.6386663 |
| LDAVHRV | ASSLEGQSNERLF | 0.615971 |
| QGLSSSIF | ASSLEGQSNERLF | 0.6262065 |
| RGTFANIRL | ASSLEGQSNERLF | 0.59763587 |
| RQLWWGHRI | ASSLEGQSNERLF | 0.6573446 |
| YGPPGTGKTLL | ASSLEGQSNERLF | 0.6390562 |
| AALVGAPANVA | ASSLEGQSNERLF | 0.61454856 |

| | | |
|---|---|---|
| AATFRRTE | ASSLEGQSNERLF | 0.64544237 |
| AGALGGGI | ASSLEGQSNERLF | 0.4647898 |
| AGGAGGAGGIGAG | ASSLEGQSNERLF | 0.6430341 |
| AGSGALGL | ASSLEGQSNERLF | 0.6294339 |
| AQIADQI | ASSLEGQSNERLF | 0.5330537 |
| ASAVAVTYL | ASSLEGQSNERLF | 0.60392797 |
| ASTVLGGTGG | ASSLEGQSNERLF | 0.65136534 |
| AVGGEPAK | ASSLEGQSNERLF | 0.6492171 |
| CEQGTFGGTC | ASSLEGQSNERLF | 0.53126884 |
| DEGKAPAG | ASSLEGQSNERLF | 0.5769299 |
| DTVVFPAL | ASSLEGQSNERLF | 0.44827974 |
| FEGITMAG | ASSLEGQSNERLF | 0.47816068 |
| FPLVVQD | ASSLEGQSNERLF | 0.47240406 |
| GAFTTPQ | ASSLEGQSNERLF | 0.6263466 |
| GFGGFGTAGGVGGA | ASSLEGQSNERLF | 0.58933663 |
| GFLAQFL | ASSLEGQSNERLF | 0.64323723 |
| GGAGGAGGNAGVA | ASSLEGQSNERLF | 0.62067485 |
| GGVGTAGAGGAGGAGG | ASSLEGQSNERLF | 0.6535952 |
| GNGGGDAGNAGSGGN | ASSLEGQSNERLF | 0.5670797 |
| GTGGAGGAGNDAGS | ASSLEGQSNERLF | 0.6002597 |
| GTTNTGWFNS | ASSLEGQSNERLF | 0.42688543 |
| IFMGPLFTL | ASSLEGQSNERLF | 0.5474293 |
| ILAHAIEAI | ASSLEGQSNERLF | 0.5328455 |
| IRSAIREAL | ASSLEGQSNERLF | 0.60082734 |
| ITFAGGG | ASSLEGQSNERLF | 0.36825517 |
| ITGMSGHQEV | ASSLEGQSNERLF | 0.6552413 |
| IVAGLACTL | ASSLEGQSNERLF | 0.59583044 |
| IVGGGLAGA | ASSLEGQSNERLF | 0.6419569 |
| IVTGVLNI | ASSLEGQSNERLF | 0.55943775 |
| LAQQGCI | ASSLEGQSNERLF | 0.55781066 |
| LCLHVEN | ASSLEGQSNERLF | 0.55588806 |
| LDAISDGS | ASSLEGQSNERLF | 0.63738114 |
| LGGFAAML | ASSLEGQSNERLF | 0.52389336 |
| LGGKNPMI | ASSLEGQSNERLF | 0.3918779 |
| LGPKGVAAIL | ASSLEGQSNERLF | 0.54085386 |
| LMVGVSI | ASSLEGQSNERLF | 0.48861188 |
| LVGAAFVL | ASSLEGQSNERLF | 0.6561053 |
| LVLAVAPL | ASSLEGQSNERLF | 0.65057385 |
| MAAIAVIDAA | ASSLEGQSNERLF | 0.62178 |
| NNGINAELQ | ASSLEGQSNERLF | 0.65464824 |
| RGEPLVLVHG | ASSLEGQSNERLF | 0.33092004 |
| RGGAGGAAGLI | ASSLEGQSNERLF | 0.56440437 |
| RSYRMHPAV | ASSLEGQSNERLF | 0.6564839 |
| SAAHLLAGQLA | ASSLEGQSNERLF | 0.62551695 |
| SAAVGSY | ASSLEGQSNERLF | 0.60601985 |
| SPHAVLHRI | ASSLEGQSNERLF | 0.6473219 |
| SQIAKVL | ASSLEGQSNERLF | 0.57400626 |
| TALGHLVP | ASSLEGQSNERLF | 0.608491 |
| TAPTTSGAA | ASSLEGQSNERLF | 0.59628505 |
| TDGDDGNGGSPGT | ASSLEGQSNERLF | 0.63993365 |
| TDGLGEN | ASSLEGQSNERLF | 0.62203085 |

| | | |
|---|---|---|
| TGGPPLPGQA | ASSLEGQSNERLF | 0.48751208 |
| TIPAITI | ASSLEGQSNERLF | 0.6361165 |
| TIPGFTI | ASSLEGQSNERLF | 0.6618398 |
| TSIASGALA | ASSLEGQSNERLF | 0.586017 |
| TVTLPGAAI | ASSLEGQSNERLF | 0.63467956 |
| VASGLSAALP | ASSLEGQSNERLF | 0.62687415 |
| VGAGVIGI | ASSLEGQSNERLF | 0.6175089 |
| VGPKRFGV | ASSLEGQSNERLF | 0.55970836 |
| VNVAFPD | ASSLEGQSNERLF | 0.52137685 |
| VSMTNSGVS | ASSLEGQSNERLF | 0.6293513 |
| VTDVLGDLSV | ASSLEGQSNERLF | 0.51910776 |
| VVVVGAGFAGL | ASSLEGQSNERLF | 0.6116489 |
| VYTVGTVV | ASSLEGQSNERLF | 0.5999138 |
| WTTFSGP | ASSLEGQSNERLF | 0.66563016 |
| YDELVEAF | ASSLEGQSNERLF | 0.63318825 |
| YGTGGAGGNGG | ASSLEGQSNERLF | 0.62574095 |
| YTNVMAV | ASSLEGQSNERLF | 0.52338284 |
| FLEQVHQL | ASSLEGQSNERLF | 0.61871517 |
| GLSLLAVAV | ASSLEGQSNERLF | 0.6298909 |
| LEQVHQL | ASSLEGQSNERLF | 0.45395565 |
| LNAAGAI | ASSLEGQSNERLF | 0.64691544 |
| PAVHASL | ASSLEGQSNERLF | 0.53661716 |
| TELEDDA | ASSLEGQSNERLF | 0.5975324 |
| TPPGPPN | ASSLEGQSNERLF | 0.6387788 |
| TPPPPPPP | ASSLEGQSNERLF | 0.6575534 |
| TQSPALI | ASSDREGNQAPL | 0.6053945 |
| EFAAAFPQ | ASSDREGNQAPL | 0.6164664 |
| LLSGNLDVL | ASSDREGNQAPL | 0.531562 |
| GPANLGNYNI | ASSDREGNQAPL | 0.6329671 |
| TLGPTLP | ASSDREGNQAPL | 0.64457786 |
| VYAYSSCCK | ASSDREGNQAPL | 0.6184202 |
| APAPAPAPNI | ASSDREGNQAPL | 0.65912527 |
| PGGPVPIP | ASSDREGNQAPL | 0.6156092 |
| FLTVALEP | ASSDREGNQAPL | 0.40834597 |
| SQYFGANL | ASSDREGNQAPL | 0.60991627 |
| GGVKPAA | ASSDREGNQAPL | 0.6523517 |
| GGTGGAGGAGGW | ASSDREGNQAPL | 0.64359033 |
| GVGGVAGLGGAGG | ASSDREGNQAPL | 0.65844524 |
| VLLVLATTI | ASSDREGNQAPL | 0.6286273 |
| IGIFAAML | ASSDREGNQAPL | 0.5828382 |
| AVGDLID | ASSDREGNQAPL | 0.65424716 |
| NISPLPL | ASSDREGNQAPL | 0.6638653 |
| VTAVFVAI | ASSDREGNQAPL | 0.6362107 |
| SGDDNDAAL | ASSDREGNQAPL | 0.5224087 |
| AVAALAHS | ASSDREGNQAPL | 0.493148 |
| GGVTVMATP | ASSDREGNQAPL | 0.6359373 |
| AGLAANVRD | ASSDREGNQAPL | 0.63789165 |
| TNRPDLIDPALLRPG | ASSDREGNQAPL | 0.49958953 |
| LANLGVTAL | ASSDREGNQAPL | 0.6198462 |
| TGERIEL | ASSDREGNQAPL | 0.62003136 |
| NGEELAA | ASSDREGNQAPL | 0.41587853 |

| | | |
|---|---|---|
| VLSLVWG | ASSDREGNQAPL | 0.62670445 |
| IGYALWL | ASSDREGNQAPL | 0.64585334 |
| LSLLLAPL | ASSDREGNQAPL | 0.63114387 |
| TLDEALK | ASSDREGNQAPL | 0.5522207 |
| TDGAGTAEGG | ASSDREGNQAPL | 0.6456742 |
| NAPGGAGGNGG | ASSDREGNQAPL | 0.6595574 |
| VSVFMGMGGNF | ASSDREGNQAPL | 0.6585334 |
| LASYLAAL | ASSDREGNQAPL | 0.45253134 |
| LVGGTPAL | ASSDREGNQAPL | 0.35485655 |
| VRNADVL | ASSDREGNQAPL | 0.59843147 |
| TCSSPFLVSAM | ASSDREGNQAPL | 0.6419721 |
| VHITDAE | ASSDREGNQAPL | 0.5978862 |
| EYAAPSF | ASSDREGNQAPL | 0.555124 |
| VVDAFHGGL | ASSDREGNQAPL | 0.6220275 |
| ASVVDPQ | ASSDREGNQAPL | 0.65046966 |
| SYNEFGGLS | ASSDREGNQAPL | 0.55841666 |
| DPDGSVY | ASSDREGNQAPL | 0.43577725 |
| MVSTIHATE | ASSDREGNQAPL | 0.63875973 |
| LAAGVADPV | ASSDREGNQAPL | 0.6354658 |
| TKEQIAA | ASSDREGNQAPL | 0.55871254 |
| SLASPAAL | ASSDREGNQAPL | 0.5843019 |
| IGLVFYGV | ASSDREGNQAPL | 0.6201187 |
| FLLDASNL | ASSDREGNQAPL | 0.6015022 |
| YQVGNFP | ASSDREGNQAPL | 0.6139013 |
| ESLLSQST | ASSDREGNQAPL | 0.5044303 |
| LFTVGFAAVG | ASSDREGNQAPL | 0.643224 |
| ATPPTPT | ASSDREGNQAPL | 0.5146473 |
| TDLATLG | ASSDREGNQAPL | 0.62432104 |
| SAGGTGR | ASSDREGNQAPL | 0.62059927 |
| GFSNSGS | ASSDREGNQAPL | 0.6284431 |
| GGNGGSGGTSV | ASSDREGNQAPL | 0.59137446 |
| GTSVATGGAGN | ASSDREGNQAPL | 0.55232424 |
| NGGNGGSGGTS | ASSDREGNQAPL | 0.5143787 |
| AVVKFPH | ASSDREGNQAPL | 0.4131974 |
| TPPPPPP | ASSDREGNQAPL | 0.6495028 |
| AAPAAAAAPA | ASSDREGNQAPL | 0.5921614 |
| IALNEDL | ASSDREGNQAPL | 0.64698523 |
| IVNVSSM | ASSDREGNQAPL | 0.6366484 |
| TGIPPAPR | ASSDREGNQAPL | 0.55117124 |
| FDLGGGTF | ASSDREGNQAPL | 0.6265628 |
| FDLGGGTFDVS | ASSDREGNQAPL | 0.44762024 |
| NEPTAAA | ASSDREGNQAPL | 0.60050815 |
| NEPTAAALA | ASSDREGNQAPL | 0.6188941 |
| GSIINISSI | ASSDREGNQAPL | 0.6421206 |
| VAPGNAGTA | ASSDREGNQAPL | 0.54335845 |
| TGGGASLE | ASSDREGNQAPL | 0.62633723 |
| VAVYNHYKRI | ASSDREGNQAPL | 0.6262722 |
| TDPAPATT | ASSDREGNQAPL | 0.5178663 |
| TNRPDILDPALLRPG | ASSDREGNQAPL | 0.63757265 |
| LTTTTTS | ASSDREGNQAPL | 0.649098 |
| PTGSGKTL | ASSDREGNQAPL | 0.6304083 |

| | | |
|---|---|---|
| TGSGKTL | ASSDREGNQAPL | 0.52406085 |
| TIGHVDHGK | ASSDREGNQAPL | 0.63803643 |
| TIGHVDHGKTT | ASSDREGNQAPL | 0.5522635 |
| GHVDHGKTT | ASSDREGNQAPL | 0.62272435 |
| AQFEHTLL | ASSDREGNQAPL | 0.6046496 |
| IVNQDEL | ASSDREGNQAPL | 0.577165 |
| TNGSQFF | ASSDREGNQAPL | 0.64426523 |
| TGKGIVE | ASSDREGNQAPL | 0.4501428 |
| LISNASDA | ASSDREGNQAPL | 0.544989 |
| AELRTLGELTKIAKAHGAQ | ASSDREGNQAPL | 0.5833598 |
| APNAVAAAEPSW | ASSDREGNQAPL | 0.62197506 |
| APQVAGAATAVL | ASSDREGNQAPL | 0.6385392 |
| ARRDPAAAAAILPTDA | ASSDREGNQAPL | 0.5461586 |
| DAAARDVLAEAGLAEYF | ASSDREGNQAPL | 0.61005926 |
| DDQIVLSAKANLAKVEQ | ASSDREGNQAPL | 0.61255705 |
| DELLPDTGEDSDEDGH | ASSDREGNQAPL | 0.56780374 |
| ELPEAPANRATVPQD | ASSDREGNQAPL | 0.62166023 |
| FGQSKIERVTGIG | ASSDREGNQAPL | 0.5995651 |
| GAGGTGSTAGGAGGAGGAGGLY | ASSDREGNQAPL | 0.44455382 |
| GHGGTGAAGGNGGTGGAGGSGI | ASSDREGNQAPL | 0.6366069 |
| GNGGNGGAGGEGGPGGAGGAGA | ASSDREGNQAPL | 0.54419816 |
| GSTAGNGGNGGNATNGTIAGQP | ASSDREGNQAPL | 0.63424826 |
| GTGGDGGNGGNGGTGG | ASSDREGNQAPL | 0.66174793 |
| GVRRPAGIPADAPV | ASSDREGNQAPL | 0.47519588 |
| GYRQVTDAHLLTLARR | ASSDREGNQAPL | 0.62334263 |
| IAKMGQDGHDRG | ASSDREGNQAPL | 0.61969864 |
| KAEPSIEIQYT | ASSDREGNQAPL | 0.62190366 |
| KPVRIRMAADEDTVAVT | ASSDREGNQAPL | 0.6021381 |
| MPPRIAGMRLLVIKPEP | ASSDREGNQAPL | 0.5745462 |
| MKFYAVALFRSRGVPPTR | ASSDREGNQAPL | 0.6539819 |
| NAGGFGILGAVAHSP | ASSDREGNQAPL | 0.6407573 |
| NEGREITLAMAEM | ASSDREGNQAPL | 0.6235329 |
| NELVEWSHPLGHRWRWE | ASSDREGNQAPL | 0.6319282 |
| NLGAGGTGGNGGNPG | ASSDREGNQAPL | 0.63697135 |
| NTEWPKWETVMAADPVD | ASSDREGNQAPL | 0.6406326 |
| PASAAPAETQP | ASSDREGNQAPL | 0.5601719 |
| QQSSDGKNGVRAAEITGE | ASSDREGNQAPL | 0.43574232 |
| RAPDAVTTSWWKE | ASSDREGNQAPL | 0.60144687 |
| SGFGNVGAHTSGWFNQ | ASSDREGNQAPL | 0.5867191 |
| SGVAGAVGQAASVGGLKV | ASSDREGNQAPL | 0.5871056 |
| SNGGAGTGSAGGNGGT | ASSDREGNQAPL | 0.39732054 |
| TGGAGGEGGAAG | ASSDREGNQAPL | 0.6492422 |
| TGGDGGAAGAGGAPGHG | ASSDREGNQAPL | 0.6617338 |
| TMPPLLAAIANAGRHGV | ASSDREGNQAPL | 0.5412577 |
| TSSTAGNSGGA | ASSDREGNQAPL | 0.6615125 |
| VPATVPVGILC | ASSDREGNQAPL | 0.50222003 |
| VQRPDSGGVVLTGRIS | ASSDREGNQAPL | 0.6448593 |
| WTPNPVPATKGVPAGLVQAT | ASSDREGNQAPL | 0.6516036 |
| YALKRALAKFSDRFDVVII | ASSDREGNQAPL | 0.5153318 |
| YLIGSVIV | ASSDREGNQAPL | 0.63913655 |
| YGQLVAVID | ASSDREGNQAPL | 0.5813153 |

| | | |
|---|---|---|
| VTVGGMTLPA | ASSDREGNQAPL | 0.56840557 |
| VLALPAV | ASSDREGNQAPL | 0.64893055 |
| VIPPTLNYET | ASSDREGNQAPL | 0.5835016 |
| VGMGGMHPSSG | ASSDREGNQAPL | 0.6404182 |
| VGLAGGAATA | ASSDREGNQAPL | 0.6034713 |
| VGGAGGAGGSGP | ASSDREGNQAPL | 0.542886 |
| VFAAFMV | ASSDREGNQAPL | 0.6326427 |
| TLYDKLR | ASSDREGNQAPL | 0.6407315 |
| TLAVFTVL | ASSDREGNQAPL | 0.57217145 |
| TGSVQPARV | ASSDREGNQAPL | 0.6626044 |
| TGGGAGSGGT | ASSDREGNQAPL | 0.5270612 |
| SHLETSL | ASSDREGNQAPL | 0.6588252 |
| SFTTGMT | ASSDREGNQAPL | 0.63615733 |
| QVEVAKRMGVRQPTVS | ASSDREGNQAPL | 0.60450065 |
| QGSGGPALAM | ASSDREGNQAPL | 0.5764044 |
| MLAPSNP | ASSDREGNQAPL | 0.644501 |
| LVNNAAI | ASSDREGNQAPL | 0.59389114 |
| LTGEINTLA | ASSDREGNQAPL | 0.62191904 |
| LSIFGPV | ASSDREGNQAPL | 0.65330845 |
| LLALLANL | ASSDREGNQAPL | 0.68964493 |
| LISNASDALDKLR | ASSDREGNQAPL | 0.7956955 |
| LINEAVP | ASSDREGNQAPL | 0.76102304 |
| LGYAFSGL | ASSDREGNQAPL | 0.7801763 |
| LDFVAVAL | ASSDREGNQAPL | 0.73008585 |
| LAQPGAE | ASSDREGNQAPL | 0.6190915 |
| ISNLGAVTGF | ASSDREGNQAPL | 0.7686541 |
| HQEDPGRI | ASSDREGNQAPL | 0.67721415 |
| GTPGEPTP | ASSDREGNQAPL | 0.79793155 |
| GFFNSGS | ASSDREGNQAPL | 0.79888284 |
| GAGGGGLTAGAN | ASSDREGNQAPL | 0.721843 |
| FVDALVNM | ASSDREGNQAPL | 0.79046226 |
| FLTVAPEL | ASSDREGNQAPL | 0.7983018 |
| FETLTAL | ASSDREGNQAPL | 0.6176726 |
| DVINPIEL | ASSDREGNQAPL | 0.7459852 |
| APVSGAPV | ASSDREGNQAPL | 0.7500837 |
| ALVGGLAW | ASSDREGNQAPL | 0.756145 |
| ALAAAAATAI | ASSDREGNQAPL | 0.5652396 |
| AFGLGLLL | ASSDREGNQAPL | 0.79120994 |
| AFFAAASL | ASSDREGNQAPL | 0.7180184 |
| AAAALPAFSPP | ASSDREGNQAPL | 0.3213625 |
| VVLAPAL | ASSDREGNQAPL | 0.77280354 |
| TIGHVDHGKTTL | ASSDREGNQAPL | 0.7488702 |
| LISNASDALDK | ASSDREGNQAPL | 0.74939454 |
| LISNASDAL | ASSDREGNQAPL | 0.79090655 |
| ISNASDALDK | ASSDREGNQAPL | 0.69416773 |
| NQAETLVYQTE | ASSDREGNQAPL | 0.7053038 |
| TGGTGGTAGAG | ASSDREGNQAPL | 0.77361226 |
| TGSTAGGAGGA | ASSDREGNQAPL | 0.7973237 |
| DPLAVPVAAGPV | ASSDREGNQAPL | 0.67012846 |
| GGTGGTPGNGGN | ASSDREGNQAPL | 0.7651692 |
| PAGLAGVDPATL | ASSDREGNQAPL | 0.80048645 |

| | | |
|---|---|---|
| GNNLVVEAPHAGASV | ASSDREGNQAPL | 0.76678765 |
| KVTRIMSASTLEQPA | ASSDREGNQAPL | 0.8053168 |
| LGFALPIALPRRTVV | ASSDREGNQAPL | 0.8083477 |
| TRLGNLTARKLVVLV | ASSDREGNQAPL | 0.73541045 |
| WSQQRRLIYFPSAGP | ASSDREGNQAPL | 0.79474235 |
| GPNGNPDPMAAAVDIR | ASSDREGNQAPL | 0.80381775 |
| TQRCVSVADLRAAASR | ASSDREGNQAPL | 0.7432359 |
| QFVLDVAEHPVEVLTETV | ASSDREGNQAPL | 0.75760305 |
| GGAGGTGGAGGNGGTQAGDG | ASSDREGNQAPL | 0.67014205 |
| PLLATLRAIAVDVDAKPAQV | ASSDREGNQAPL | 0.75645673 |
| APKFRTGGYMIDFWGVGYQVAK | ASSDREGNQAPL | 0.5805267 |
| GAGGAGGAGGAAGGPGGTGGTG | ASSDREGNQAPL | 0.7605485 |
| IGALGLLATDVAPAGSVAFGVA | ASSDREGNQAPL | 0.8036952 |
| AGGAGGIGGNGGAGGAGGVLMG | ASSDREGNQAPL | 0.74394166 |
| ADELIGLL | ASSDREGNQAPL | 0.62945527 |
| ADRALKA | ASSDREGNQAPL | 0.66308427 |
| AGAVGQVGGAGGA | ASSDREGNQAPL | 0.7683717 |
| AGGGGGLGGAP | ASSDREGNQAPL | 0.70997006 |
| AGGLLYGNGGAG | ASSDREGNQAPL | 0.59064245 |
| AGLESIGL | ASSDREGNQAPL | 0.714216 |
| AGLKTGII | ASSDREGNQAPL | 0.7958621 |
| ASLAALLL | ASSDREGNQAPL | 0.799542 |
| ASVINISSI | ASSDREGNQAPL | 0.6416671 |
| ATPPGPA | ASSDREGNQAPL | 0.80175567 |
| ATVASALV | ASSDREGNQAPL | 0.7922353 |
| ATVASLIL | ASSDREGNQAPL | 0.80445427 |
| AVNGPVLIRH | ASSDREGNQAPL | 0.778884 |
| DGAAGAVGI | ASSDREGNQAPL | 0.66843116 |
| DPLLHGI | ASSDREGNQAPL | 0.79648054 |
| ELGAALGAL | ASSDREGNQAPL | 0.7063266 |
| ELGRTQAV | ASSDREGNQAPL | 0.7584388 |
| FVAGGYLS | ASSDREGNQAPL | 0.791957 |
| GASGGAGGAGGW | ASSDREGNQAPL | 0.7436143 |
| GGAGAGGDGGAGGI | ASSDREGNQAPL | 0.7019773 |
| GTGHNGSQPGMGGNG | ASSDREGNQAPL | 0.7406838 |
| GYAAASASAS | ASSDREGNQAPL | 0.8064032 |
| HQVDVAAVQQG | ASSDREGNQAPL | 0.672215 |
| IGAAIAHT | ASSDREGNQAPL | 0.67287064 |
| IGMIEVAQL | ASSDREGNQAPL | 0.7290052 |
| IRLAPEHTSAMD | ASSDREGNQAPL | 0.7556052 |
| IYNVSVL | ASSDREGNQAPL | 0.7978796 |
| KQGFLGM | ASSDREGNQAPL | 0.7919774 |
| LESVITQL | ASSDREGNQAPL | 0.73375255 |
| LGLGALVVV | ASSDREGNQAPL | 0.77225935 |
| LIGNGGNGGSGGI | ASSDREGNQAPL | 0.7897499 |
| LNAINAPTL | ASSDREGNQAPL | 0.7718041 |
| LQLLNTGV | ASSDREGNQAPL | 0.7791258 |
| LQTSIEF | ASSDREGNQAPL | 0.68966055 |
| MSFAVSEL | ASSDREGNQAPL | 0.8022552 |
| NTGALGL | ASSDREGNQAPL | 0.7895983 |
| PPGAGGAGGAGTT | ASSDREGNQAPL | 0.7997192 |

| | | |
|---|---|---|
| QLVGLGI | ASSDREGNQAPL | 0.8006404 |
| RFLEQVHQL | ASSDREGNQAPL | 0.7642797 |
| RGVGYVLR | ASSDREGNQAPL | 0.77265346 |
| SATGVFAGVI | ASSDREGNQAPL | 0.5723815 |
| SLPAITLP | ASSDREGNQAPL | 0.7328144 |
| STALAFTV | ASSDREGNQAPL | 0.7535074 |
| SVLTVGLGK | ASSDREGNQAPL | 0.76279235 |
| TALAGAGSGI | ASSDREGNQAPL | 0.7790264 |
| TGLPFTGI | ASSDREGNQAPL | 0.79777443 |
| VAEHVTV | ASSDREGNQAPL | 0.71780264 |
| VDIVLNSL | ASSDREGNQAPL | 0.7976614 |
| VLTGGGAVL | ASSDREGNQAPL | 0.7961627 |
| VNVGLAYGI | ASSDREGNQAPL | 0.64361477 |
| VSSAAVGAV | ASSDREGNQAPL | 0.73726535 |
| VTSVYEAGV | ASSDREGNQAPL | 0.80383116 |
| WVALDRGA | ASSDREGNQAPL | 0.7752957 |
| YLVGSVVI | ASSDREGNQAPL | 0.73561466 |
| GPPGTGKTLL | ASSDREGNQAPL | 0.76159954 |
| GVGMTKF | ASSDREGNQAPL | 0.8076383 |
| IGIGGSDL | ASSDREGNQAPL | 0.75106287 |
| IGTLLAEL | ASSDREGNQAPL | 0.8022517 |
| IYTSGTT | ASSDREGNQAPL | 0.47852996 |
| LSFPAGV | ASSDREGNQAPL | 0.76284647 |
| LYTSGTT | ASSDREGNQAPL | 0.7398617 |
| VGAFPFGF | ASSDREGNQAPL | 0.36537087 |
| VGAGTVEF | ASSDREGNQAPL | 0.79464936 |
| VGKSTLLL | ASSDREGNQAPL | 0.7476021 |
| AALAHAGGGIV | ASSDREGNQAPL | 0.7845812 |
| AALVAVSLL | ASSDREGNQAPL | 0.7555013 |
| AAVSRAVAAL | ASSDREGNQAPL | 0.7985432 |
| AGGSSGIGGF | ASSDREGNQAPL | 0.75789464 |
| AGLTVANANG | ASSDREGNQAPL | 0.7695222 |
| AGLVGAVL | ASSDREGNQAPL | 0.71494144 |
| ALSQAFTV | ASSDREGNQAPL | 0.78418934 |
| ALTFSVSL | ASSDREGNQAPL | 0.79859483 |
| ALTGAALVAL | ASSDREGNQAPL | 0.80288297 |
| ASFSIGAGVL | ASSDREGNQAPL | 0.53688186 |
| DGSVYKI | ASSDREGNQAPL | 0.75155556 |
| EQLAEIV | ASSDREGNQAPL | 0.76115763 |
| FAVSELTVM | ASSDREGNQAPL | 0.44052497 |
| GAAPLAAGL | ASSDREGNQAPL | 0.46574336 |
| GADGGDGGSGGAGG | ASSDREGNQAPL | 0.71821964 |
| GFGLDTGGAGG | ASSDREGNQAPL | 0.7223365 |
| GLASGGLL | ASSDREGNQAPL | 0.7801924 |
| GRIAGAAVLL | ASSDREGNQAPL | 0.7834012 |
| GSAGSLNTSF | ASSDREGNQAPL | 0.739497 |
| GTGQIGI | ASSDREGNQAPL | 0.7803217 |
| GTLIALIL | ASSDREGNQAPL | 0.74925804 |
| IAATMGFVVG | ASSDREGNQAPL | 0.18358968 |
| IFAQALL | ASSDREGNQAPL | 0.7382648 |
| IGAAVALGF | ASSDREGNQAPL | 0.6160691 |

| | | |
|---|---|---|
| IGLTPLAL | ASSDREGNQAPL | 0.80993456 |
| ILGSNTSTL | ASSDREGNQAPL | 0.705694 |
| IVFSLTVL | ASSDREGNQAPL | 0.7127799 |
| LASKATGF | ASSDREGNQAPL | 0.75719863 |
| LASVALIL | ASSDREGNQAPL | 0.77351284 |
| LFAVGLTL | ASSDREGNQAPL | 0.7520942 |
| LFNGGIGTYI | ASSDREGNQAPL | 0.6893146 |
| LTLGAGII | ASSDREGNQAPL | 0.40799075 |
| LVAAALGDI | ASSDREGNQAPL | 0.7899729 |
| NTAFVQLGI | ASSDREGNQAPL | 0.7062191 |
| QLFSGAEL | ASSDREGNQAPL | 0.6786296 |
| SAATFITL | ASSDREGNQAPL | 0.6387832 |
| SNYGLANLGV | ASSDREGNQAPL | 0.7156671 |
| SYGGAYAVM | ASSDREGNQAPL | 0.6041044 |
| TTGVGGLTLG | ASSDREGNQAPL | 0.80013126 |
| TTQTIYI | ASSDREGNQAPL | 0.7021516 |
| VAAGAGLGAVI | ASSDREGNQAPL | 0.7302567 |
| VAAVVTLL | ASSDREGNQAPL | 0.8041859 |
| VGFLGGLI | ASSDREGNQAPL | 0.76141155 |
| VGVIPILL | ASSDREGNQAPL | 0.7343815 |
| VIFASSLI | ASSDREGNQAPL | 0.7122196 |
| VKGGDGGAAAG | ASSDREGNQAPL | 0.7021446 |
| VTKTFTNI | ASSDREGNQAPL | 0.7045971 |
| VVAAAILL | ASSDREGNQAPL | 0.72300816 |
| YAALDVEL | ASSDREGNQAPL | 0.80263525 |
| LASVALLL | ASSDREGNQAPL | 0.7770815 |
| LDAVHRV | ASSDREGNQAPL | 0.6659492 |
| QGLSSSIF | ASSDREGNQAPL | 0.79629064 |
| RGTFANIRL | ASSDREGNQAPL | 0.8037317 |
| RQLWWGHRI | ASSDREGNQAPL | 0.80762315 |
| YGPPGTGKTLL | ASSDREGNQAPL | 0.6878271 |
| AALVGAPANVA | ASSDREGNQAPL | 0.75969213 |
| AATFRRTE | ASSDREGNQAPL | 0.7521677 |
| AGALGGGI | ASSDREGNQAPL | 0.7377406 |
| AGGAGGAGGIGAG | ASSDREGNQAPL | 0.7267504 |
| AGSGALGL | ASSDREGNQAPL | 0.68376255 |
| AQIADQI | ASSDREGNQAPL | 0.7968048 |
| ASAVAVTYL | ASSDREGNQAPL | 0.8146541 |
| ASTVLGGTGG | ASSDREGNQAPL | 0.7558116 |
| AVGGEPAK | ASSDREGNQAPL | 0.667081 |
| CEQGTFGGTC | ASSDREGNQAPL | 0.7356119 |
| DEGKAPAG | ASSDREGNQAPL | 0.6938989 |
| DTVVFPAL | ASSDREGNQAPL | 0.7093955 |
| FEGITMAG | ASSDREGNQAPL | 0.7453122 |
| FPLVVQD | ASSDREGNQAPL | 0.6033773 |
| GAFTTPQ | ASSDREGNQAPL | 0.7576051 |
| GFGGFGTAGGVGGA | ASSDREGNQAPL | 0.8059437 |
| GFLAQFL | ASSDREGNQAPL | 0.73713887 |
| GGAGGAGGNAGVA | ASSDREGNQAPL | 0.77898914 |
| GGVGTAGAGGAGGAGG | ASSDREGNQAPL | 0.79757106 |
| GNGGGGDAGNAGSGGN | ASSDREGNQAPL | 0.7796327 |

| | | |
|---|---|---|
| GTGGAGGAGNDAGS | ASSDREGNQAPL | 0.79782146 |
| GTTNTGWFNS | ASSDREGNQAPL | 0.79719985 |
| IFMGPLFTL | ASSDREGNQAPL | 0.7776176 |
| ILAHAIEAI | ASSDREGNQAPL | 0.7631223 |
| IRSAIREAL | ASSDREGNQAPL | 0.7784275 |
| ITFAGGG | ASSDREGNQAPL | 0.74451894 |
| ITGMSGHQEV | ASSDREGNQAPL | 0.751976 |
| IVAGLACTL | ASSDREGNQAPL | 0.7348484 |
| IVGGGLAGA | ASSDREGNQAPL | 0.7851842 |
| IVTGVLNI | ASSDREGNQAPL | 0.7618481 |
| LAQQGCI | ASSDREGNQAPL | 0.8081031 |
| LCLHVEN | ASSDREGNQAPL | 0.4863513 |
| LDAISDGS | ASSDREGNQAPL | 0.6739983 |
| LGGFAAML | ASSDREGNQAPL | 0.7922338 |
| LGGKNPMI | ASSDREGNQAPL | 0.24226677 |
| LGPKGVAAIL | ASSDREGNQAPL | 0.52439106 |
| LMVGVSI | ASSDREGNQAPL | 0.7668724 |
| LVGAAFVL | ASSDREGNQAPL | 0.7951536 |
| LVLAVAPL | ASSDREGNQAPL | 0.24760908 |
| MAAIAVIDAA | ASSDREGNQAPL | 0.72137606 |
| NNGINAELQ | ASSDREGNQAPL | 0.63014853 |
| RGEPLVLVHG | ASSDREGNQAPL | 0.7966491 |
| RGGAGGAAGLI | ASSDREGNQAPL | 0.7540006 |
| RSYRMHPAV | ASSDREGNQAPL | 0.58219516 |
| SAAHLLAGQLA | ASSDREGNQAPL | 0.7949611 |
| SAAVGSY | ASSDREGNQAPL | 0.6508584 |
| SPHAVLHRI | ASSDREGNQAPL | 0.7363529 |
| SQIAKVL | ASSDREGNQAPL | 0.79628754 |
| TALGHLVP | ASSDREGNQAPL | 0.8071491 |
| TAPTTSGAA | ASSDREGNQAPL | 0.6679168 |
| TDGDDGNGGSPGT | ASSDREGNQAPL | 0.7883081 |
| TDGLGEN | ASSDREGNQAPL | 0.730104 |
| TGGPPLPGQA | ASSDREGNQAPL | 0.80875444 |
| TIPAITI | ASSDREGNQAPL | 0.74054027 |
| TIPGFTI | ASSDREGNQAPL | 0.6119628 |
| TSIASGALA | ASSDREGNQAPL | 0.7946228 |
| TVTLPGAAI | ASSDREGNQAPL | 0.7821622 |
| VASGLSAALP | ASSDREGNQAPL | 0.77615905 |
| VGAGVIGI | ASSDREGNQAPL | 0.78996193 |
| VGPKRFGV | ASSDREGNQAPL | 0.7534555 |
| VNVAFPD | ASSDREGNQAPL | 0.73848164 |
| VSMTNSGVS | ASSDREGNQAPL | 0.790697 |
| VTDVLGDLSV | ASSDREGNQAPL | 0.78183883 |
| VVVVGAGFAGL | ASSDREGNQAPL | 0.6767455 |
| VYTVGTVV | ASSDREGNQAPL | 0.8043233 |
| WTTFSGP | ASSDREGNQAPL | 0.74756396 |
| YDELVEAF | ASSDREGNQAPL | 0.71059966 |
| YGTGGAGGNGG | ASSDREGNQAPL | 0.8084593 |
| YTNVMAV | ASSDREGNQAPL | 0.7722877 |
| FLEQVHQL | ASSDREGNQAPL | 0.78408456 |
| GLSLLAVAV | ASSDREGNQAPL | 0.7962636 |

| | | |
|---|---|---|
| LEQVHQL | ASSDREGNQAPL | 0.806389 |
| LNAAGAI | ASSDREGNQAPL | 0.6657009 |
| PAVHASL | ASSDREGNQAPL | 0.7705923 |
| TELEDDA | ASSDREGNQAPL | 0.75285274 |
| TPPGPPN | ASSDREGNQAPL | 0.79369485 |
| TPPPPPPP | ASSDREGNQAPL | 0.76298463 |
| TQSPALI | ASTRGGTNERLF | 0.7274711 |
| EFAAAFPQ | ASTRGGTNERLF | 0.789536 |
| LLSGNLDVL | ASTRGGTNERLF | 0.79994184 |
| GPANLGNYNI | ASTRGGTNERLF | 0.7732972 |
| TLGPTLP | ASTRGGTNERLF | 0.7667681 |
| VYAYSSCCK | ASTRGGTNERLF | 0.80393785 |
| APAPAPAPNI | ASTRGGTNERLF | 0.70171154 |
| PGGPVPIP | ASTRGGTNERLF | 0.7581553 |
| FLTVALEP | ASTRGGTNERLF | 0.79275596 |
| SQYFGANL | ASTRGGTNERLF | 0.79442924 |
| GGVKPAA | ASTRGGTNERLF | 0.76995873 |
| GGTGGAGGAGGW | ASTRGGTNERLF | 0.79869473 |
| GVGGVAGLGGAGG | ASTRGGTNERLF | 0.7311668 |
| VLLVLATTI | ASTRGGTNERLF | 0.77073157 |
| IGIFAAML | ASTRGGTNERLF | 0.699018 |
| AVGDLID | ASTRGGTNERLF | 0.7596846 |
| NISPLPL | ASTRGGTNERLF | 0.6463045 |
| VTAVFVAI | ASTRGGTNERLF | 0.76816314 |
| SGDDNDAAL | ASTRGGTNERLF | 0.78742707 |
| AVAALAHS | ASTRGGTNERLF | 0.7794013 |
| GGVTVMATP | ASTRGGTNERLF | 0.44712543 |
| AGLAANVRD | ASTRGGTNERLF | 0.7834495 |
| TNRPDLIDPALLRPG | ASTRGGTNERLF | 0.78268194 |
| LANLGVTAL | ASTRGGTNERLF | 0.8103796 |
| TGERIEL | ASTRGGTNERLF | 0.7517972 |
| NGEELAA | ASTRGGTNERLF | 0.78825665 |
| VLSLVWG | ASTRGGTNERLF | 0.7193389 |
| IGYALWL | ASTRGGTNERLF | 0.77360636 |
| LSLLLAPL | ASTRGGTNERLF | 0.6270886 |
| TLDEALK | ASTRGGTNERLF | 0.6904439 |
| TDGAGTAEGG | ASTRGGTNERLF | 0.7137747 |
| NAPGGAGGNGG | ASTRGGTNERLF | 0.685133 |
| VSVFMGMGGNF | ASTRGGTNERLF | 0.7897638 |
| LASYLAAL | ASTRGGTNERLF | 0.78832495 |
| LVGGTPAL | ASTRGGTNERLF | 0.78765607 |
| VRNADVL | ASTRGGTNERLF | 0.79471976 |
| TCSSPFLVSAM | ASTRGGTNERLF | 0.79531205 |
| VHITDAE | ASTRGGTNERLF | 0.8057966 |
| EYAAPSF | ASTRGGTNERLF | 0.7208879 |
| VVDAFHGGL | ASTRGGTNERLF | 0.7534286 |
| ASVVDPQ | ASTRGGTNERLF | 0.59998685 |
| SYNEFGGLS | ASTRGGTNERLF | 0.7427449 |
| DPDGSVY | ASTRGGTNERLF | 0.7282113 |
| MVSTIHATE | ASTRGGTNERLF | 0.76785624 |
| LAAGVADPV | ASTRGGTNERLF | 0.21537718 |

| | | |
|---|---|---|
| TKEQIAA | ASTRGGTNERLF | 0.76347804 |
| SLASPAAL | ASTRGGTNERLF | 0.8034685 |
| IGLVFYGV | ASTRGGTNERLF | 0.69384336 |
| FLLDASNL | ASTRGGTNERLF | 0.78313696 |
| YQVGNFP | ASTRGGTNERLF | 0.7670742 |
| ESLLSQST | ASTRGGTNERLF | 0.71148807 |
| LFTVGFAAVG | ASTRGGTNERLF | 0.78096306 |
| ATPPTPT | ASTRGGTNERLF | 0.7204461 |
| TDLATLG | ASTRGGTNERLF | 0.79224885 |
| SAGGTGR | ASTRGGTNERLF | 0.80485904 |
| GFSNSGS | ASTRGGTNERLF | 0.7715703 |
| GGNGGSGGTSV | ASTRGGTNERLF | 0.7391273 |
| GTSVATGGAGN | ASTRGGTNERLF | 0.77589047 |
| NGGNGGSGGTS | ASTRGGTNERLF | 0.64798325 |
| AVVKFPH | ASTRGGTNERLF | 0.74926746 |
| TPPPPPP | ASTRGGTNERLF | 0.5715301 |
| AAPAAAAAPA | ASTRGGTNERLF | 0.7505995 |
| IALNEDL | ASTRGGTNERLF | 0.7226418 |
| IVNVSSM | ASTRGGTNERLF | 0.7677922 |
| TGIPPAPR | ASTRGGTNERLF | 0.7543248 |
| FDLGGGTF | ASTRGGTNERLF | 0.7803531 |
| FDLGGGTFDVS | ASTRGGTNERLF | 0.68261343 |
| NEPTAAA | ASTRGGTNERLF | 0.7766155 |
| NEPTAAALA | ASTRGGTNERLF | 0.7486132 |
| GSIINISSI | ASTRGGTNERLF | 0.6458397 |
| VAPGNAGTA | ASTRGGTNERLF | 0.788021 |
| TGGGASLE | ASTRGGTNERLF | 0.77108175 |
| VAVYNHYKRI | ASTRGGTNERLF | 0.7349235 |
| TDPAPATT | ASTRGGTNERLF | 0.79637456 |
| TNRPDILDPALLRPG | ASTRGGTNERLF | 0.77094066 |
| LTTTTTS | ASTRGGTNERLF | 0.64170754 |
| PTGSGKTL | ASTRGGTNERLF | 0.7248269 |
| TGSGKTL | ASTRGGTNERLF | 0.75637484 |
| TIGHVDHGK | ASTRGGTNERLF | 0.74573445 |
| TIGHVDHGKTT | ASTRGGTNERLF | 0.3402381 |
| GHVDHGKTT | ASTRGGTNERLF | 0.7798257 |
| AQFEHTLL | ASTRGGTNERLF | 0.71103466 |
| IVNQDEL | ASTRGGTNERLF | 0.67893636 |
| TNGSQFF | ASTRGGTNERLF | 0.7362714 |
| TGKGIVE | ASTRGGTNERLF | 0.7891982 |
| LISNASDA | ASTRGGTNERLF | 0.729267 |
| AELRTLGELTKIAKAHGAQ | ASTRGGTNERLF | 0.7652122 |
| APNAVAAAEPSW | ASTRGGTNERLF | 0.77297753 |
| APQVAGAATAVL | ASTRGGTNERLF | 0.779776 |
| ARRDPAAAAAILPTDA | ASTRGGTNERLF | 0.7959684 |
| DAAARDVLAEAGLAEYF | ASTRGGTNERLF | 0.77258 |
| DDQIVLSAKANLAKVEQ | ASTRGGTNERLF | 0.7748094 |
| DELLPDTGEDSDEDGH | ASTRGGTNERLF | 0.7884588 |
| ELPEAPANRATVPQD | ASTRGGTNERLF | 0.7531234 |
| FGQSKIERVTGIG | ASTRGGTNERLF | 0.6753917 |
| GAGGTGSTAGGAGGAGGAGGLY | ASTRGGTNERLF | 0.6487074 |

| | | |
|---|---|---|
| GHGGTGAAGGNGGTGGAGGSGI | ASTRGGTNERLF | 0.74656427 |
| GNGGNGGAGGEGGPGGAGGAGA | ASTRGGTNERLF | 0.6490488 |
| GSTAGNGGNGGNATNGTIAGQP | ASTRGGTNERLF | 0.7752247 |
| GTGGDGGNGGNGGTGG | ASTRGGTNERLF | 0.8094083 |
| GVRRPAGIPADAPV | ASTRGGTNERLF | 0.7634489 |
| GYRQVTDAHLLTLARR | ASTRGGTNERLF | 0.6281661 |
| IAKMGQDGHDRG | ASTRGGTNERLF | 0.7340104 |
| KAEPSIEIQYT | ASTRGGTNERLF | 0.74680114 |
| KPVRIRMAADEDTVAVT | ASTRGGTNERLF | 0.78307843 |
| MPPRIAGMRLLVIKPEP | ASTRGGTNERLF | 0.7996111 |
| MKFYAVALFRSRGVPPTR | ASTRGGTNERLF | 0.80991304 |
| NAGGFGILGAVAHSP | ASTRGGTNERLF | 0.78926 |
| NEGREITLAMAEM | ASTRGGTNERLF | 0.71310306 |
| NELVEWSHPLGHRWRWE | ASTRGGTNERLF | 0.78823733 |
| NLGAGGTGGNGGNPG | ASTRGGTNERLF | 0.7469084 |
| NTEWPKWETVMAADPVD | ASTRGGTNERLF | 0.76607037 |
| PASAAPAETQP | ASTRGGTNERLF | 0.77018523 |
| QQSSDGKNGVRAAEITGE | ASTRGGTNERLF | 0.77323824 |
| RAPDAVTTSWWKE | ASTRGGTNERLF | 0.73091114 |
| SGFGNVGAHTSGWFNQ | ASTRGGTNERLF | 0.74611866 |
| SGVAGAVGQAASVGGLKV | ASTRGGTNERLF | 0.6395892 |
| SNGGAGTGSAGGNGGT | ASTRGGTNERLF | 0.7992651 |
| TGGAGGEGGAAG | ASTRGGTNERLF | 0.6723271 |
| TGGDGGAAGAGGAPGHG | ASTRGGTNERLF | 0.657987 |
| TMPPLLAAIANAGRHGV | ASTRGGTNERLF | 0.729496 |
| TSSTAGNSGGA | ASTRGGTNERLF | 0.8007777 |
| VPATVPVGILC | ASTRGGTNERLF | 0.8008474 |
| VQRPDSGGVVLTGRIS | ASTRGGTNERLF | 0.7043067 |
| WTPNPVPATKGVPAGLVQAT | ASTRGGTNERLF | 0.71462536 |
| YALKRALAKFSDRFDVVII | ASTRGGTNERLF | 0.7674886 |
| YLIGSVIV | ASTRGGTNERLF | 0.64927316 |
| YGQLVAVID | ASTRGGTNERLF | 0.79305637 |
| VTVGGMTLPA | ASTRGGTNERLF | 0.74174833 |
| VLALPAV | ASTRGGTNERLF | 0.7327635 |
| VIPPTLNYET | ASTRGGTNERLF | 0.73753583 |
| VGMGGMHPSSG | ASTRGGTNERLF | 0.77589 |
| VGLAGGAATA | ASTRGGTNERLF | 0.7207725 |
| VGGAGGAGGSGP | ASTRGGTNERLF | 0.7955235 |
| VFAAFMV | ASTRGGTNERLF | 0.36618316 |
| TLYDKLR | ASTRGGTNERLF | 0.79244363 |
| TLAVFTVL | ASTRGGTNERLF | 0.67151564 |
| TGSVQPARV | ASTRGGTNERLF | 0.72693044 |
| TGGGAGSGGT | ASTRGGTNERLF | 0.47350436 |
| SHLETSL | ASTRGGTNERLF | 0.77493846 |
| SFTTGMT | ASTRGGTNERLF | 0.780504 |
| QVEVAKRMGVRQPTVS | ASTRGGTNERLF | 0.793381 |
| QGSGGPALAM | ASTRGGTNERLF | 0.73136777 |
| MLAPSNP | ASTRGGTNERLF | 0.7816868 |
| LVNNAAI | ASTRGGTNERLF | 0.7959768 |
| LTGEINTLA | ASTRGGTNERLF | 0.72406256 |
| LSIFGPV | ASTRGGTNERLF | 0.46021253 |

| | | |
|---|---|---|
| LLALLANL | ASTRGGTNERLF | 0.77594733 |
| LISNASDALDKLR | ASTRGGTNERLF | 0.764263 |
| LINEAVP | ASTRGGTNERLF | 0.7965294 |
| LGYAFSGL | ASTRGGTNERLF | 0.76755786 |
| LDFVAVAL | ASTRGGTNERLF | 0.63153005 |
| LAQPGAE | ASTRGGTNERLF | 0.796218 |
| ISNLGAVTGF | ASTRGGTNERLF | 0.7845084 |
| HQEDPGRI | ASTRGGTNERLF | 0.65265834 |
| GTPGEPTP | ASTRGGTNERLF | 0.7271684 |
| GFFNSGS | ASTRGGTNERLF | 0.7440536 |
| GAGGGGLTAGAN | ASTRGGTNERLF | 0.5334817 |
| FVDALVNM | ASTRGGTNERLF | 0.7865538 |
| FLTVAPEL | ASTRGGTNERLF | 0.7162315 |
| FETLTAL | ASTRGGTNERLF | 0.740861 |
| DVINPIEL | ASTRGGTNERLF | 0.76020086 |
| APVSGAPV | ASTRGGTNERLF | 0.73570925 |
| ALVGGLAW | ASTRGGTNERLF | 0.7343245 |
| ALAAAAATAI | ASTRGGTNERLF | 0.73190475 |
| AFGLGLLL | ASTRGGTNERLF | 0.77845985 |
| AFFAAASL | ASTRGGTNERLF | 0.7828331 |
| AAAALPAFSPP | ASTRGGTNERLF | 0.75341856 |
| VVLAPAL | ASTRGGTNERLF | 0.780199 |
| TIGHVDHGKTTL | ASTRGGTNERLF | 0.7966225 |
| LISNASDALDK | ASTRGGTNERLF | 0.6157691 |
| LISNASDAL | ASTRGGTNERLF | 0.7012012 |
| ISNASDALDK | ASTRGGTNERLF | 0.78501886 |
| NQAETLVYQTE | ASTRGGTNERLF | 0.70036393 |
| TGGTGGTAGAG | ASTRGGTNERLF | 0.7950313 |
| TGSTAGGAGGA | ASTRGGTNERLF | 0.75807893 |
| DPLAVPVAAGPV | ASTRGGTNERLF | 0.7727288 |
| GGTGGTPGNGGN | ASTRGGTNERLF | 0.7683859 |
| PAGLAGVDPATL | ASTRGGTNERLF | 0.8010684 |
| GNNLVVEAPHAGASV | ASTRGGTNERLF | 0.785421 |
| KVTRIMSASTLEQPA | ASTRGGTNERLF | 0.7688604 |
| LGFALPIALPRRTVV | ASTRGGTNERLF | 0.79010963 |
| TRLGNLTARKLVVLV | ASTRGGTNERLF | 0.6753818 |
| WSQQRRLIYFPSAGP | ASTRGGTNERLF | 0.7870985 |
| GPNGNPDPMAAAVDIR | ASTRGGTNERLF | 0.78982544 |
| TQRCVSVADLRAAASR | ASTRGGTNERLF | 0.6638582 |
| QFVLDVAEHPVEVLTETV | ASTRGGTNERLF | 0.7407473 |
| GGAGGTGGAGGNGGTQAGDG | ASTRGGTNERLF | 0.8043292 |
| PLLATLRAIAVDVDAKPAQV | ASTRGGTNERLF | 0.79779005 |
| APKFRTGGYMIDFWGVGYQVAK | ASTRGGTNERLF | 0.6899843 |
| GAGGAGGAGGAAGGPGGTGGTG | ASTRGGTNERLF | 0.72355103 |
| IGALGLLATDVAPAGSVAFGVA | ASTRGGTNERLF | 0.6342999 |
| AGGAGGIGGNGGAGGAGGVLMG | ASTRGGTNERLF | 0.7253292 |
| ADELIGLL | ASTRGGTNERLF | 0.60378706 |
| ADRALKA | ASTRGGTNERLF | 0.777928 |
| AGAVGQVGGAGGA | ASTRGGTNERLF | 0.72795516 |
| AGGGGGLGGAP | ASTRGGTNERLF | 0.78958964 |
| AGGLLYGNGGAG | ASTRGGTNERLF | 0.77393967 |

| | | |
|---|---|---|
| AGLESIGL | ASTRGGTNERLF | 0.80178833 |
| AGLKTGII | ASTRGGTNERLF | 0.72161204 |
| ASLAALLL | ASTRGGTNERLF | 0.7360867 |
| ASVINISSI | ASTRGGTNERLF | 0.5930091 |
| ATPPGPA | ASTRGGTNERLF | 0.6107198 |
| ATVASALV | ASTRGGTNERLF | 0.7286117 |
| ATVASLIL | ASTRGGTNERLF | 0.7402706 |
| AVNGPVLIRH | ASTRGGTNERLF | 0.41544765 |
| DGAAGAVGI | ASTRGGTNERLF | 0.80268574 |
| DPLLHGI | ASTRGGTNERLF | 0.7508854 |
| ELGAALGAL | ASTRGGTNERLF | 0.78916323 |
| ELGRTQAV | ASTRGGTNERLF | 0.7243528 |
| FVAGGYLS | ASTRGGTNERLF | 0.6947259 |
| GASGGAGGAGGW | ASTRGGTNERLF | 0.7166039 |
| GGAGAGGDGGAGGI | ASTRGGTNERLF | 0.7801717 |
| GTGHNGSQPGMGGNG | ASTRGGTNERLF | 0.6667999 |
| GYAAASASAS | ASTRGGTNERLF | 0.6119618 |
| HQVDVAAVQQG | ASTRGGTNERLF | 0.7164763 |
| IGAAIAHT | ASTRGGTNERLF | 0.7157364 |
| IGMIEVAQL | ASTRGGTNERLF | 0.80526304 |
| IRLAPEHTSAMD | ASTRGGTNERLF | 0.7949836 |
| IYNVSVL | ASTRGGTNERLF | 0.76014787 |
| KQGFLGM | ASTRGGTNERLF | 0.8070642 |
| LESVITQL | ASTRGGTNERLF | 0.28441793 |
| LGLGALVVV | ASTRGGTNERLF | 0.73442376 |
| LIGNGGNGGSGGI | ASTRGGTNERLF | 0.8009968 |
| LNAINAPTL | ASTRGGTNERLF | 0.78194475 |
| LQLLNTGV | ASTRGGTNERLF | 0.74648494 |
| LQTSIEF | ASTRGGTNERLF | 0.7991338 |
| MSFAVSEL | ASTRGGTNERLF | 0.68864685 |
| NTGALGL | ASTRGGTNERLF | 0.7496513 |
| PPGAGGAGGAGTT | ASTRGGTNERLF | 0.76139426 |
| QLVGLGI | ASTRGGTNERLF | 0.79195714 |
| RFLEQVHQL | ASTRGGTNERLF | 0.7753031 |
| RGVGYVLR | ASTRGGTNERLF | 0.62849283 |
| SATGVFAGVI | ASTRGGTNERLF | 0.78777134 |
| SLPAITLP | ASTRGGTNERLF | 0.80573463 |
| STALAFTV | ASTRGGTNERLF | 0.7172425 |
| SVLTVGLGK | ASTRGGTNERLF | 0.786957 |
| TALAGAGSGI | ASTRGGTNERLF | 0.77515966 |
| TGLPFTGI | ASTRGGTNERLF | 0.73672456 |
| VAEHVTV | ASTRGGTNERLF | 0.69986534 |
| VDIVLNSL | ASTRGGTNERLF | 0.6859573 |
| VLTGGGAVL | ASTRGGTNERLF | 0.7773824 |
| VNVGLAYGI | ASTRGGTNERLF | 0.64502823 |
| VSSAAVGAV | ASTRGGTNERLF | 0.7657272 |
| VTSVYEAGV | ASTRGGTNERLF | 0.75764734 |
| WVALDRGA | ASTRGGTNERLF | 0.8110423 |
| YLVGSVVI | ASTRGGTNERLF | 0.7753645 |
| GPPGTGKTLL | ASTRGGTNERLF | 0.78272116 |
| GVGMTKF | ASTRGGTNERLF | 0.67857146 |

| | | |
|---|---|---|
| IGIGGSDL | ASTRGGTNERLF | 0.7788433 |
| IGTLLAEL | ASTRGGTNERLF | 0.7726832 |
| IYTSGTT | ASTRGGTNERLF | 0.6622375 |
| LSFPAGV | ASTRGGTNERLF | 0.79642403 |
| LYTSGTT | ASTRGGTNERLF | 0.68413854 |
| VGAFPFGF | ASTRGGTNERLF | 0.7454921 |
| VGAGTVEF | ASTRGGTNERLF | 0.78815204 |
| VGKSTLLL | ASTRGGTNERLF | 0.80131257 |
| AALAHAGGGIV | ASTRGGTNERLF | 0.74992955 |
| AALVAVSLL | ASTRGGTNERLF | 0.7780177 |
| AAVSRAVAAL | ASTRGGTNERLF | 0.67250633 |
| AGGSSGIGGF | ASTRGGTNERLF | 0.79268086 |
| AGLTVANANG | ASTRGGTNERLF | 0.79290974 |
| AGLVGAVL | ASTRGGTNERLF | 0.76709735 |
| ALSQAFTV | ASTRGGTNERLF | 0.8093835 |
| ALTFSVSL | ASTRGGTNERLF | 0.7947806 |
| ALTGAALVAL | ASTRGGTNERLF | 0.5042927 |
| ASFSIGAGVL | ASTRGGTNERLF | 0.76617503 |
| DGSVYKI | ASTRGGTNERLF | 0.7954881 |
| EQLAEIV | ASTRGGTNERLF | 0.7930441 |
| FAVSELTVM | ASTRGGTNERLF | 0.8068781 |
| GAAPLAAGL | ASTRGGTNERLF | 0.7814169 |
| GADGGDGGSGGAGG | ASTRGGTNERLF | 0.71993876 |
| GFGLDTGGAGG | ASTRGGTNERLF | 0.8005053 |
| GLASGGLL | ASTRGGTNERLF | 0.8148663 |
| GRIAGAAVLL | ASTRGGTNERLF | 0.788241 |
| GSAGSLNTSF | ASTRGGTNERLF | 0.7113242 |
| GTGQIGI | ASTRGGTNERLF | 0.71136636 |
| GTLIALIL | ASTRGGTNERLF | 0.78153265 |
| IAATMGFVVG | ASTRGGTNERLF | 0.7888396 |
| IFAQALL | ASTRGGTNERLF | 0.68432224 |
| IGAAVALGF | ASTRGGTNERLF | 0.75392675 |
| IGLTPLAL | ASTRGGTNERLF | 0.7678299 |
| ILGSNTSTL | ASTRGGTNERLF | 0.5148052 |
| IVFSLTVL | ASTRGGTNERLF | 0.7806399 |
| LASKATGF | ASTRGGTNERLF | 0.79100037 |
| LASVALIL | ASTRGGTNERLF | 0.7847507 |
| LFAVGLTL | ASTRGGTNERLF | 0.69287884 |
| LFNGGIGTYI | ASTRGGTNERLF | 0.79201615 |
| LTLGAGII | ASTRGGTNERLF | 0.79999053 |
| LVAAALGDI | ASTRGGTNERLF | 0.8067397 |
| NTAFVQLGI | ASTRGGTNERLF | 0.5976021 |
| QLFSGAEL | ASTRGGTNERLF | 0.37662113 |
| SAATFITL | ASTRGGTNERLF | 0.73465943 |
| SNYGLANLGV | ASTRGGTNERLF | 0.7836941 |
| SYGGAYAVM | ASTRGGTNERLF | 0.7495018 |
| TTGVGGLTLG | ASTRGGTNERLF | 0.71144414 |
| TTQTIYI | ASTRGGTNERLF | 0.74995255 |
| VAAGAGLGAVI | ASTRGGTNERLF | 0.7963273 |
| VAAVVTLL | ASTRGGTNERLF | 0.6968725 |
| VGFLGGLI | ASTRGGTNERLF | 0.6758154 |

| | | |
|---|---|---|
| VGVIPILL | ASTRGGTNERLF | 0.7889435 |
| VIFASSLI | ASTRGGTNERLF | 0.79486215 |
| VKGGDGGAAAG | ASTRGGTNERLF | 0.6701559 |
| VTKTFTNI | ASTRGGTNERLF | 0.72781086 |
| VVAAAILL | ASTRGGTNERLF | 0.7517232 |
| YAALDVEL | ASTRGGTNERLF | 0.74933565 |
| LASVALLL | ASTRGGTNERLF | 0.76022756 |
| LDAVHRV | ASTRGGTNERLF | 0.6490134 |
| QGLSSSIF | ASTRGGTNERLF | 0.78795826 |
| RGTFANIRL | ASTRGGTNERLF | 0.6640116 |
| RQLWWGHRI | ASTRGGTNERLF | 0.7887557 |
| YGPPGTGKTLL | ASTRGGTNERLF | 0.7596886 |
| AALVGAPANVA | ASTRGGTNERLF | 0.7774019 |
| AATFRRTE | ASTRGGTNERLF | 0.7225523 |
| AGALGGGI | ASTRGGTNERLF | 0.6975082 |
| AGGAGGAGGIGAG | ASTRGGTNERLF | 0.67409474 |
| AGSGALGL | ASTRGGTNERLF | 0.5986202 |
| AQIADQI | ASTRGGTNERLF | 0.7905671 |
| ASAVAVTYL | ASTRGGTNERLF | 0.7217361 |
| ASTVLGGTGG | ASTRGGTNERLF | 0.78911734 |
| AVGGEPAK | ASTRGGTNERLF | 0.7811556 |
| CEQGTFGGTC | ASTRGGTNERLF | 0.6978544 |
| DEGKAPAG | ASTRGGTNERLF | 0.7826184 |
| DTVVFPAL | ASTRGGTNERLF | 0.62269455 |
| FEGITMAG | ASTRGGTNERLF | 0.7633816 |
| FPLVVQD | ASTRGGTNERLF | 0.7549494 |
| GAFTTPQ | ASTRGGTNERLF | 0.7935005 |
| GFGGFGTAGGVGGA | ASTRGGTNERLF | 0.70659685 |
| GFLAQFL | ASTRGGTNERLF | 0.78963614 |
| GGAGGAGGNAGVA | ASTRGGTNERLF | 0.77348566 |
| GGVGTAGAGGAGGAGG | ASTRGGTNERLF | 0.72190946 |
| GNGGGGDAGNAGSGGN | ASTRGGTNERLF | 0.7843528 |
| GTGGAGGAGNDAGS | ASTRGGTNERLF | 0.79190934 |
| GTTNTGWFNS | ASTRGGTNERLF | 0.78596115 |
| IFMGPLFTL | ASTRGGTNERLF | 0.6346973 |
| ILAHAIEAI | ASTRGGTNERLF | 0.7745315 |
| IRSAIREAL | ASTRGGTNERLF | 0.69254065 |
| ITFAGGG | ASTRGGTNERLF | 0.7675194 |
| ITGMSGHQEV | ASTRGGTNERLF | 0.76134 |
| IVAGLACTL | ASTRGGTNERLF | 0.74566 |
| IVGGGLAGA | ASTRGGTNERLF | 0.78961813 |
| IVTGVLNI | ASTRGGTNERLF | 0.62437415 |
| LAQQGCI | ASTRGGTNERLF | 0.70432353 |
| LCLHVEN | ASTRGGTNERLF | 0.73305565 |
| LDAISDGS | ASTRGGTNERLF | 0.7594056 |
| LGGFAAML | ASTRGGTNERLF | 0.7881851 |
| LGGKNPMI | ASTRGGTNERLF | 0.6653261 |
| LGPKGVAAIL | ASTRGGTNERLF | 0.76537645 |
| LMVGVSI | ASTRGGTNERLF | 0.7527121 |
| LVGAAFVL | ASTRGGTNERLF | 0.6956584 |
| LVLAVAPL | ASTRGGTNERLF | 0.78475565 |

| | | |
|---|---|---|
| MAAIAVIDAA | ASTRGGTNERLF | 0.7530841 |
| NNGINAELQ | ASTRGGTNERLF | 0.6522647 |
| RGEPLVLVHG | ASTRGGTNERLF | 0.78538144 |
| RGGAGGAAGLI | ASTRGGTNERLF | 0.70404506 |
| RSYRMHPAV | ASTRGGTNERLF | 0.7779025 |
| SAAHLLAGQLA | ASTRGGTNERLF | 0.8047011 |
| SAAVGSY | ASTRGGTNERLF | 0.62605685 |
| SPHAVLHRI | ASTRGGTNERLF | 0.7885041 |
| SQIAKVL | ASTRGGTNERLF | 0.7675164 |
| TALGHLVP | ASTRGGTNERLF | 0.76288766 |
| TAPTTSGAA | ASTRGGTNERLF | 0.74525523 |
| TDGDDGNGGSPGT | ASTRGGTNERLF | 0.7286062 |
| TDGLGEN | ASTRGGTNERLF | 0.8004257 |
| TGGPPLPGQA | ASTRGGTNERLF | 0.7843491 |
| TIPAITI | ASTRGGTNERLF | 0.7884004 |
| TIPGFTI | ASTRGGTNERLF | 0.7769456 |
| TSIASGALA | ASTRGGTNERLF | 0.7923963 |
| TVTLPGAAI | ASTRGGTNERLF | 0.7916583 |
| VASGLSAALP | ASTRGGTNERLF | 0.6984141 |
| VGAGVIGI | ASTRGGTNERLF | 0.5832414 |
| VGPKRFGV | ASTRGGTNERLF | 0.7375819 |
| VNVAFPD | ASTRGGTNERLF | 0.7084799 |
| VSMTNSGVS | ASTRGGTNERLF | 0.7468469 |
| VTDVLGDLSV | ASTRGGTNERLF | 0.56067383 |
| VVVVGAGFAGL | ASTRGGTNERLF | 0.7880275 |
| VYTVGTVV | ASTRGGTNERLF | 0.8100225 |
| WTTFSGP | ASTRGGTNERLF | 0.7185207 |
| YDELVEAF | ASTRGGTNERLF | 0.81056476 |
| YGTGGAGGNGG | ASTRGGTNERLF | 0.6553645 |
| YTNVMAV | ASTRGGTNERLF | 0.78472257 |
| FLEQVHQL | ASTRGGTNERLF | 0.7940935 |
| GLSLLAVAV | ASTRGGTNERLF | 0.6625226 |
| LEQVHQL | ASTRGGTNERLF | 0.7857757 |
| LNAAGAI | ASTRGGTNERLF | 0.721413 |
| PAVHASL | ASTRGGTNERLF | 0.7277364 |
| TELEDDA | ASTRGGTNERLF | 0.7944915 |
| TPPGPPN | ASTRGGTNERLF | 0.7349391 |
| TPPPPPPP | ASTRGGTNERLF | 0.7868992 |
| TQSPALI | ASSSDWGSAETLY | 0.7600991 |
| EFAAAFPQ | ASSSDWGSAETLY | 0.6989969 |
| LLSGNLDVL | ASSSDWGSAETLY | 0.780741 |
| GPANLGNYNI | ASSSDWGSAETLY | 0.79518473 |
| TLGPTLP | ASSSDWGSAETLY | 0.747214 |
| VYAYSSCCK | ASSSDWGSAETLY | 0.8144388 |
| APAPAPAPNI | ASSSDWGSAETLY | 0.6925434 |
| PGGPVPIP | ASSSDWGSAETLY | 0.80516213 |
| FLTVALEP | ASSSDWGSAETLY | 0.7955768 |
| SQYFGANL | ASSSDWGSAETLY | 0.7548642 |
| GGVKPAA | ASSSDWGSAETLY | 0.74288476 |
| GGTGGAGGAGGW | ASSSDWGSAETLY | 0.78923416 |
| GVGGVAGLGGAGG | ASSSDWGSAETLY | 0.74099076 |

| | | |
|---|---|---|
| VLLVLATTI | ASSSDWGSAETLY | 0.77176046 |
| IGIFAAML | ASSSDWGSAETLY | 0.8071773 |
| AVGDLID | ASSSDWGSAETLY | 0.10432029 |
| NISPLPL | ASSSDWGSAETLY | 0.14211671 |
| VTAVFVAI | ASSSDWGSAETLY | 0.102482624 |
| SGDDNDAAL | ASSSDWGSAETLY | 0.12229573 |
| AVAALAHS | ASSSDWGSAETLY | 0.114290155 |
| GGVTVMATP | ASSSDWGSAETLY | 0.10167343 |
| AGLAANVRD | ASSSDWGSAETLY | 0.11677794 |
| TNRPDLIDPALLRPG | ASSSDWGSAETLY | 0.10593949 |
| LANLGVTAL | ASSSDWGSAETLY | 0.10199519 |
| TGERIEL | ASSSDWGSAETLY | 0.11451698 |
| NGEELAA | ASSSDWGSAETLY | 0.11829357 |
| VLSLVWG | ASSSDWGSAETLY | 0.14492327 |
| IGYALWL | ASSSDWGSAETLY | 0.14253613 |
| LSLLLAPL | ASSSDWGSAETLY | 0.10274107 |
| TLDEALK | ASSSDWGSAETLY | 0.10673575 |
| TDGAGTAEGG | ASSSDWGSAETLY | 0.11555394 |
| NAPGGAGGNGG | ASSSDWGSAETLY | 0.109377176 |
| VSVFMGMGGNF | ASSSDWGSAETLY | 0.09733973 |
| LASYLAAL | ASSSDWGSAETLY | 0.116905905 |
| LVGGTPAL | ASSSDWGSAETLY | 0.09921346 |
| VRNADVL | ASSSDWGSAETLY | 0.096162915 |
| TCSSPFLVSAM | ASSSDWGSAETLY | 0.1162789 |
| VHITDAE | ASSSDWGSAETLY | 0.11227346 |
| EYAAPSF | ASSSDWGSAETLY | 0.10791895 |
| VVDAFHGGL | ASSSDWGSAETLY | 0.14950481 |
| ASVVDPQ | ASSSDWGSAETLY | 0.09633185 |
| SYNEFGGLS | ASSSDWGSAETLY | 0.10646603 |
| DPDGSVY | ASSSDWGSAETLY | 0.11433242 |
| MVSTIHATE | ASSSDWGSAETLY | 0.15958185 |
| LAAGVADPV | ASSSDWGSAETLY | 0.1069929 |
| TKEQIAA | ASSSDWGSAETLY | 0.1078259 |
| SLASPAAL | ASSSDWGSAETLY | 0.17281115 |
| IGLVFYGV | ASSSDWGSAETLY | 0.12227676 |
| FLLDASNL | ASSSDWGSAETLY | 0.120426886 |
| YQVGNFP | ASSSDWGSAETLY | 0.12547596 |
| ESLLSQST | ASSSDWGSAETLY | 0.100747846 |
| LFTVGFAAVG | ASSSDWGSAETLY | 0.126246 |
| ATPPTPT | ASSSDWGSAETLY | 0.11947063 |
| TDLATLG | ASSSDWGSAETLY | 0.12090551 |
| SAGGTGR | ASSSDWGSAETLY | 0.11527317 |
| GFSNSGS | ASSSDWGSAETLY | 0.107047915 |
| GGNGGSGGTSV | ASSSDWGSAETLY | 0.10507164 |
| GTSVATGGAGN | ASSSDWGSAETLY | 0.096811615 |
| NGGNGGSGGTS | ASSSDWGSAETLY | 0.11886942 |
| AVVKFPH | ASSSDWGSAETLY | 0.121626176 |
| TPPPPPP | ASSSDWGSAETLY | 0.11534855 |
| AAPAAAAAPA | ASSSDWGSAETLY | 0.09848794 |
| IALNEDL | ASSSDWGSAETLY | 0.1007073 |
| IVNVSSM | ASSSDWGSAETLY | 0.09967819 |

| | | |
|---|---|---|
| TGIPPAPR | ASSSDWGSAETLY | 0.104341835 |
| FDLGGGTF | ASSSDWGSAETLY | 0.097305894 |
| FDLGGGTFDVS | ASSSDWGSAETLY | 0.10536092 |
| NEPTAAA | ASSSDWGSAETLY | 0.16167176 |
| NEPTAAALA | ASSSDWGSAETLY | 0.16037878 |
| GSIINISSI | ASSSDWGSAETLY | 0.10724998 |
| VAPGNAGTA | ASSSDWGSAETLY | 0.14954893 |
| TGGGASLE | ASSSDWGSAETLY | 0.10506068 |
| VAVYNHYKRI | ASSSDWGSAETLY | 0.11136595 |
| TDPAPATT | ASSSDWGSAETLY | 0.11440435 |
| TNRPDILDPALLRPG | ASSSDWGSAETLY | 0.10630751 |
| LTTTTTS | ASSSDWGSAETLY | 0.22407278 |
| PTGSGKTL | ASSSDWGSAETLY | 0.11065984 |
| TGSGKTL | ASSSDWGSAETLY | 0.11654326 |
| TIGHVDHGK | ASSSDWGSAETLY | 0.14235316 |
| TIGHVDHGKTT | ASSSDWGSAETLY | 0.1145585 |
| GHVDHGKTT | ASSSDWGSAETLY | 0.10807401 |
| AQFEHTLL | ASSSDWGSAETLY | 0.11444907 |
| IVNQDEL | ASSSDWGSAETLY | 0.11769686 |
| TNGSQFF | ASSSDWGSAETLY | 0.10016873 |
| TGKGIVE | ASSSDWGSAETLY | 0.10551088 |
| LISNASDA | ASSSDWGSAETLY | 0.1033233 |
| AELRTLGELTKIAKAHGAQ | ASSSDWGSAETLY | 0.118841134 |
| APNAVAAAEPSW | ASSSDWGSAETLY | 0.17227885 |
| APQVAGAATAVL | ASSSDWGSAETLY | 0.11617965 |
| ARRDPAAAAAILPTDA | ASSSDWGSAETLY | 0.10286981 |
| DAAARDVLAEAGLAEYF | ASSSDWGSAETLY | 0.12650128 |
| DDQIVLSAKANLAKVEQ | ASSSDWGSAETLY | 0.12210416 |
| DELLPDTGEDSDEDGH | ASSSDWGSAETLY | 0.11975198 |
| ELPEAPANRATVPQD | ASSSDWGSAETLY | 0.13002762 |
| FGQSKIERVTGIG | ASSSDWGSAETLY | 0.104811385 |
| GAGGTGSTAGGAGGAGGAGGLY | ASSSDWGSAETLY | 0.12103796 |
| GHGGTGAAGGNGGTGGAGGSGI | ASSSDWGSAETLY | 0.14104486 |
| GNGGNGGAGGEGGPGGAGGAGA | ASSSDWGSAETLY | 0.12181977 |
| GSTAGNGGNGGNATNGTIAGQP | ASSSDWGSAETLY | 0.11446614 |
| GTGGDGGNGGNGGTGG | ASSSDWGSAETLY | 0.11427049 |
| GVRRPAGIPADAPV | ASSSDWGSAETLY | 0.112597294 |
| GYRQVTDAHLLTLARR | ASSSDWGSAETLY | 0.10210523 |
| IAKMGQDGHDRG | ASSSDWGSAETLY | 0.10705441 |
| KAEPSIEIQYT | ASSSDWGSAETLY | 0.1131838 |
| KPVRIRMAADEDTVAVT | ASSSDWGSAETLY | 0.111165784 |
| MPPRIAGMRLLVIKPEP | ASSSDWGSAETLY | 0.109416574 |
| MKFYAVALFRSRGVPPTR | ASSSDWGSAETLY | 0.12141346 |
| NAGGFGILGAVAHSP | ASSSDWGSAETLY | 0.104661494 |
| NEGREITLAMAEM | ASSSDWGSAETLY | 0.12522565 |
| NELVEWSHPLGHRWRWE | ASSSDWGSAETLY | 0.12851481 |
| NLGAGGTGGNGGNPG | ASSSDWGSAETLY | 0.10247266 |
| NTEWPKWETVMAADPVD | ASSSDWGSAETLY | 0.11427992 |
| PASAAPAETQP | ASSSDWGSAETLY | 0.14357837 |
| QQSSDGKNGVRAAEITGE | ASSSDWGSAETLY | 0.13088088 |
| RAPDAVTTSWWKE | ASSSDWGSAETLY | 0.1219345 |

| | | |
|---|---|---|
| SGFGNVGAHTSGWFNQ | ASSSDWGSAETLY | 0.10684245 |
| SGVAGAVGQAASVGGLKV | ASSSDWGSAETLY | 0.12865621 |
| SNGGAGTGSAGGNGGT | ASSSDWGSAETLY | 0.099416986 |
| TGGAGGEGGAAG | ASSSDWGSAETLY | 0.12957475 |
| TGGDGGAAGAGGAPGHG | ASSSDWGSAETLY | 0.10013379 |
| TMPPLLAAIANAGRHGV | ASSSDWGSAETLY | 0.104416594 |
| TSSTAGNSGGA | ASSSDWGSAETLY | 0.10280515 |
| VPATVPVGILC | ASSSDWGSAETLY | 0.0972846 |
| VQRPDSGGVVLTGRIS | ASSSDWGSAETLY | 0.14471176 |
| WTPNPVPATKGVPAGLVQAT | ASSSDWGSAETLY | 0.10283387 |
| YALKRALAKFSDRFDVVII | ASSSDWGSAETLY | 0.12263612 |
| YLIGSVIV | ASSSDWGSAETLY | 0.105021864 |
| YGQLVAVID | ASSSDWGSAETLY | 0.12843542 |
| VTVGGMTLPA | ASSSDWGSAETLY | 0.11394595 |
| VLALPAV | ASSSDWGSAETLY | 0.112260975 |
| VIPPTLNYET | ASSSDWGSAETLY | 0.107016854 |
| VGMGGMHPSSG | ASSSDWGSAETLY | 0.13670897 |
| VGLAGGAATA | ASSSDWGSAETLY | 0.12480698 |
| VGGAGGAGGSGP | ASSSDWGSAETLY | 0.115060024 |
| VFAAFMV | ASSSDWGSAETLY | 0.10160486 |
| TLYDKLR | ASSSDWGSAETLY | 0.09862743 |
| TLAVFTVL | ASSSDWGSAETLY | 0.12339777 |
| TGSVQPARV | ASSSDWGSAETLY | 0.098598085 |
| TGGGAGSGGT | ASSSDWGSAETLY | 0.095959514 |
| SHLETSL | ASSSDWGSAETLY | 0.0968509 |
| SFTTGMT | ASSSDWGSAETLY | 0.102773935 |
| QVEVAKRMGVRQPTVS | ASSSDWGSAETLY | 0.14488277 |
| QGSGGPALAM | ASSSDWGSAETLY | 0.14316729 |
| MLAPSNP | ASSSDWGSAETLY | 0.11348929 |
| LVNNAAI | ASSSDWGSAETLY | 0.11835373 |
| LTGEINTLA | ASSSDWGSAETLY | 0.119521625 |
| LSIFGPV | ASSSDWGSAETLY | 0.095266744 |
| LLALLANL | ASSSDWGSAETLY | 0.10732866 |
| LISNASDALDKLR | ASSSDWGSAETLY | 0.10290297 |
| LINEAVP | ASSSDWGSAETLY | 0.12303407 |
| LGYAFSGL | ASSSDWGSAETLY | 0.1014459 |
| LDFVAVAL | ASSSDWGSAETLY | 0.11327175 |
| LAQPGAE | ASSSDWGSAETLY | 0.12073183 |
| ISNLGAVTGF | ASSSDWGSAETLY | 0.11757126 |
| HQEDPGRI | ASSSDWGSAETLY | 0.11783225 |
| GTPGEPTP | ASSSDWGSAETLY | 0.100907706 |
| GFFNSGS | ASSSDWGSAETLY | 0.095587656 |
| GAGGGGLTAGAN | ASSSDWGSAETLY | 0.14303052 |
| FVDALVNM | ASSSDWGSAETLY | 0.100300886 |
| FLTVAPEL | ASSSDWGSAETLY | 0.11083121 |
| FETLTAL | ASSSDWGSAETLY | 0.09911198 |
| DVINPIEL | ASSSDWGSAETLY | 0.096802056 |
| APVSGAPV | ASSSDWGSAETLY | 0.10179927 |
| ALVGGLAW | ASSSDWGSAETLY | 0.11913994 |
| ALAAAAATAI | ASSSDWGSAETLY | 0.108636625 |
| AFGLGLLL | ASSSDWGSAETLY | 0.12188493 |

| | | |
|---|---|---|
| AFFAAASL | ASSSDWGSAETLY | 0.14254746 |
| AAAALPAFSPP | ASSSDWGSAETLY | 0.1824235 |
| VVLAPAL | ASSSDWGSAETLY | 0.11540986 |
| TIGHVDHGKTTL | ASSSDWGSAETLY | 0.11224791 |
| LISNASDALDK | ASSSDWGSAETLY | 0.106319785 |
| LISNASDAL | ASSSDWGSAETLY | 0.10606147 |
| ISNASDALDK | ASSSDWGSAETLY | 0.106901914 |
| NQAETLVYQTE | ASSSDWGSAETLY | 0.15466076 |
| TGGTGGTAGAG | ASSSDWGSAETLY | 0.10369822 |
| TGSTAGGAGGA | ASSSDWGSAETLY | 0.10312768 |
| DPLAVPVAAGPV | ASSSDWGSAETLY | 0.110600404 |
| GGTGGTPGNGGN | ASSSDWGSAETLY | 0.12587097 |
| PAGLAGVDPATL | ASSSDWGSAETLY | 0.13464469 |
| GNNLVVEAPHAGASV | ASSSDWGSAETLY | 0.10453985 |
| KVTRIMSASTLEQPA | ASSSDWGSAETLY | 0.108960636 |
| LGFALPIALPRRTVV | ASSSDWGSAETLY | 0.11059666 |
| TRLGNLTARKLVVLV | ASSSDWGSAETLY | 0.11912292 |
| WSQQRRLIYFPSAGP | ASSSDWGSAETLY | 0.10271123 |
| GPNGNPDPMAAAVDIR | ASSSDWGSAETLY | 0.105982095 |
| TQRCVSVADLRAAASR | ASSSDWGSAETLY | 0.116783984 |
| QFVLDVAEHPVEVLTETV | ASSSDWGSAETLY | 0.19646493 |
| GGAGGTGGAGGNGGTQAGDG | ASSSDWGSAETLY | 0.10986738 |
| PLLATLRAIAVDVDAKPAQV | ASSSDWGSAETLY | 0.10100177 |
| APKFRTGGYMIDFWGVGYQVAK | ASSSDWGSAETLY | 0.12014519 |
| GAGGAGGAGGAAGGPGGTGGTG | ASSSDWGSAETLY | 0.1014939 |
| IGALGLLATDVAPAGSVAFGVA | ASSSDWGSAETLY | 0.11052143 |
| AGGAGGIGGNGGAGGAGGVLMG | ASSSDWGSAETLY | 0.11809637 |
| ADELIGLL | ASSSDWGSAETLY | 0.10088132 |
| ADRALKA | ASSSDWGSAETLY | 0.107733175 |
| AGAVGQVGGAGGA | ASSSDWGSAETLY | 0.161389 |
| AGGGGGLGGAP | ASSSDWGSAETLY | 0.11535719 |
| AGGLLYGNGGAG | ASSSDWGSAETLY | 0.12202397 |
| AGLESIGL | ASSSDWGSAETLY | 0.13178013 |
| AGLKTGII | ASSSDWGSAETLY | 0.11822251 |
| ASLAALLL | ASSSDWGSAETLY | 0.16053985 |
| ASVINISSI | ASSSDWGSAETLY | 0.106942244 |
| ATPPGPA | ASSSDWGSAETLY | 0.11771229 |
| ATVASALV | ASSSDWGSAETLY | 0.10853016 |
| ATVASLIL | ASSSDWGSAETLY | 0.105652034 |
| AVNGPVLIRH | ASSSDWGSAETLY | 0.11938438 |
| DGAAGAVGI | ASSSDWGSAETLY | 0.13022143 |
| DPLLHGI | ASSSDWGSAETLY | 0.10561583 |
| ELGAALGAL | ASSSDWGSAETLY | 0.14760195 |
| ELGRTQAV | ASSSDWGSAETLY | 0.11759872 |
| FVAGGYLS | ASSSDWGSAETLY | 0.11707499 |
| GASGGAGGAGGW | ASSSDWGSAETLY | 0.1003297 |
| GGAGAGGDGGAGGI | ASSSDWGSAETLY | 0.10779783 |
| GTGHNGSQPGMGGNG | ASSSDWGSAETLY | 0.122191615 |
| GYAAASASAS | ASSSDWGSAETLY | 0.096825585 |
| HQVDVAAVQQG | ASSSDWGSAETLY | 0.0962187 |
| IGAAIAHT | ASSSDWGSAETLY | 0.1074163 |

| | | |
|---|---|---|
| IGMIEVAQL | ASSSDWGSAETLY | 0.1441993 |
| IRLAPEHTSAMD | ASSSDWGSAETLY | 0.095440775 |
| IYNVSVL | ASSSDWGSAETLY | 0.11309831 |
| KQGFLGM | ASSSDWGSAETLY | 0.10241643 |
| LESVITQL | ASSSDWGSAETLY | 0.11124663 |
| LGLGALVVV | ASSSDWGSAETLY | 0.11539838 |
| LIGNGGNGGSGGI | ASSSDWGSAETLY | 0.1035606 |
| LNAINAPTL | ASSSDWGSAETLY | 0.12166513 |
| LQLLNTGV | ASSSDWGSAETLY | 0.10038969 |
| LQTSIEF | ASSSDWGSAETLY | 0.09940184 |
| MSFAVSEL | ASSSDWGSAETLY | 0.1235606 |
| NTGALGL | ASSSDWGSAETLY | 0.1313399 |
| PPGAGGAGGAGTT | ASSSDWGSAETLY | 0.10237264 |
| QLVGLGI | ASSSDWGSAETLY | 0.12993191 |
| RFLEQVHQL | ASSSDWGSAETLY | 0.0998112 |
| RGVGYVLR | ASSSDWGSAETLY | 0.16042389 |
| SATGVFAGVI | ASSSDWGSAETLY | 0.12363147 |
| SLPAITLP | ASSSDWGSAETLY | 0.0980081 |
| STALAFTV | ASSSDWGSAETLY | 0.1254416 |
| SVLTVGLGK | ASSSDWGSAETLY | 0.11709043 |
| TALAGAGSGI | ASSSDWGSAETLY | 0.13025722 |
| TGLPFTGI | ASSSDWGSAETLY | 0.11374382 |
| VAEHVTV | ASSSDWGSAETLY | 0.103894316 |
| VDIVLNSL | ASSSDWGSAETLY | 0.11096773 |
| VLTGGGAVL | ASSSDWGSAETLY | 0.1399532 |
| VNVGLAYGI | ASSSDWGSAETLY | 0.12761138 |
| VSSAAVGAV | ASSSDWGSAETLY | 0.10308347 |
| VTSVYEAGV | ASSSDWGSAETLY | 0.118215695 |
| WVALDRGA | ASSSDWGSAETLY | 0.124921 |
| YLVGSVVI | ASSSDWGSAETLY | 0.11157592 |
| GPPGTGKTLL | ASSSDWGSAETLY | 0.13065793 |
| GVGMTKF | ASSSDWGSAETLY | 0.11392987 |
| IGIGGSDL | ASSSDWGSAETLY | 0.12384528 |
| IGTLLAEL | ASSSDWGSAETLY | 0.12170259 |
| IYTSGTT | ASSSDWGSAETLY | 0.112969644 |
| LSFPAGV | ASSSDWGSAETLY | 0.10552676 |
| LYTSGTT | ASSSDWGSAETLY | 0.10957717 |
| VGAFPFGF | ASSSDWGSAETLY | 0.11763049 |
| VGAGTVEF | ASSSDWGSAETLY | 0.12640399 |
| VGKSTLLL | ASSSDWGSAETLY | 0.10011319 |
| AALAHAGGGIV | ASSSDWGSAETLY | 0.09887581 |
| AALVAVSLL | ASSSDWGSAETLY | 0.110867016 |
| AAVSRAVAAL | ASSSDWGSAETLY | 0.14375632 |
| AGGSSGIGGF | ASSSDWGSAETLY | 0.10873893 |
| AGLTVANANG | ASSSDWGSAETLY | 0.11736277 |
| AGLVGAVL | ASSSDWGSAETLY | 0.22872853 |
| ALSQAFTV | ASSSDWGSAETLY | 0.10280541 |
| ALTFSVSL | ASSSDWGSAETLY | 0.1204766 |
| ALTGAALVAL | ASSSDWGSAETLY | 0.12214606 |
| ASFSIGAGVL | ASSSDWGSAETLY | 0.14635287 |
| DGSVYKI | ASSSDWGSAETLY | 0.11071683 |

| | | |
|---|---|---|
| EQLAEIV | ASSSDWGSAETLY | 0.124823056 |
| FAVSELTVM | ASSSDWGSAETLY | 0.106496505 |
| GAAPLAAGL | ASSSDWGSAETLY | 0.11876641 |
| GADGGDGGSGGAGG | ASSSDWGSAETLY | 0.096767955 |
| GFGLDTGGAGG | ASSSDWGSAETLY | 0.10921869 |
| GLASGGLL | ASSSDWGSAETLY | 0.09886807 |
| GRIAGAAVLL | ASSSDWGSAETLY | 0.11618307 |
| GSAGSLNTSF | ASSSDWGSAETLY | 0.13167879 |
| GTGQIGI | ASSSDWGSAETLY | 0.1296536 |
| GTLIALIL | ASSSDWGSAETLY | 0.09756405 |
| IAATMGFVVG | ASSSDWGSAETLY | 0.13955948 |
| IFAQALL | ASSSDWGSAETLY | 0.13837628 |
| IGAAVALGF | ASSSDWGSAETLY | 0.34338886 |
| IGLTPLAL | ASSSDWGSAETLY | 0.11023913 |
| ILGSNTSTL | ASSSDWGSAETLY | 0.13532893 |
| IVFSLTVL | ASSSDWGSAETLY | 0.114914864 |
| LASKATGF | ASSSDWGSAETLY | 0.11075365 |
| LASVALIL | ASSSDWGSAETLY | 0.09911669 |
| LFAVGLTL | ASSSDWGSAETLY | 0.10725746 |
| LFNGGIGTYI | ASSSDWGSAETLY | 0.11284965 |
| LTLGAGII | ASSSDWGSAETLY | 0.10616119 |
| LVAAALGDI | ASSSDWGSAETLY | 0.16532663 |
| NTAFVQLGI | ASSSDWGSAETLY | 0.14477304 |
| QLFSGAEL | ASSSDWGSAETLY | 0.14550194 |
| SAATFITL | ASSSDWGSAETLY | 0.15442146 |
| SNYGLANLGV | ASSSDWGSAETLY | 0.14160146 |
| SYGGAYAVM | ASSSDWGSAETLY | 0.11688323 |
| TTGVGGLTLG | ASSSDWGSAETLY | 0.10811187 |
| TTQTIYI | ASSSDWGSAETLY | 0.108884156 |
| VAAGAGLGAVI | ASSSDWGSAETLY | 0.10154892 |
| VAAVVTLL | ASSSDWGSAETLY | 0.107258074 |
| VGFLGGLI | ASSSDWGSAETLY | 0.107173115 |
| VGVIPILL | ASSSDWGSAETLY | 0.1279025 |
| VIFASSLI | ASSSDWGSAETLY | 0.09515306 |
| VKGGDGGAAAG | ASSSDWGSAETLY | 0.11134022 |
| VTKTFTNI | ASSSDWGSAETLY | 0.14924826 |
| VVAAAILL | ASSSDWGSAETLY | 0.099504896 |
| YAALDVEL | ASSSDWGSAETLY | 0.124778375 |
| LASVALLL | ASSSDWGSAETLY | 0.122185364 |
| LDAVHRV | ASSSDWGSAETLY | 0.099193364 |
| QGLSSSIF | ASSSDWGSAETLY | 0.15321948 |
| RGTFANIRL | ASSSDWGSAETLY | 0.101302534 |
| RQLWWGHRI | ASSSDWGSAETLY | 0.118909344 |
| YGPPGTGKTLL | ASSSDWGSAETLY | 0.146089 |
| AALVGAPANVA | ASSSDWGSAETLY | 0.115165286 |
| AATFRRTE | ASSSDWGSAETLY | 0.1053614 |
| AGALGGGI | ASSSDWGSAETLY | 0.11080466 |
| AGGAGGAGGIGAG | ASSSDWGSAETLY | 0.10707742 |
| AGSGALGL | ASSSDWGSAETLY | 0.108808406 |
| AQIADQI | ASSSDWGSAETLY | 0.09558075 |
| ASAVAVTYL | ASSSDWGSAETLY | 0.118546225 |

| | | |
|---|---|---|
| ASTVLGGTGG | ASSSDWGSAETLY | 0.10290007 |
| AVGGEPAK | ASSSDWGSAETLY | 0.13882886 |
| CEQGTFGGTC | ASSSDWGSAETLY | 0.12857981 |
| DEGKAPAG | ASSSDWGSAETLY | 0.11462019 |
| DTVVFPAL | ASSSDWGSAETLY | 0.10725774 |
| FEGITMAG | ASSSDWGSAETLY | 0.103624 |
| FPLVVQD | ASSSDWGSAETLY | 0.103303984 |
| GAFTTPQ | ASSSDWGSAETLY | 0.10211308 |
| GFGGFGTAGGVGGA | ASSSDWGSAETLY | 0.112116724 |
| GFLAQFL | ASSSDWGSAETLY | 0.1078679 |
| GGAGGAGGNAGVA | ASSSDWGSAETLY | 0.11147066 |
| GGVGTAGAGGAGGAGG | ASSSDWGSAETLY | 0.13366827 |
| GNGGGGDAGNAGSGGN | ASSSDWGSAETLY | 0.12081675 |
| GTGGAGGAGNDAGS | ASSSDWGSAETLY | 0.105222575 |
| GTTNTGWFNS | ASSSDWGSAETLY | 0.10858255 |
| IFMGPLFTL | ASSSDWGSAETLY | 0.11148682 |
| ILAHAIEAI | ASSSDWGSAETLY | 0.11233513 |
| IRSAIREAL | ASSSDWGSAETLY | 0.09520241 |
| ITFAGGG | ASSSDWGSAETLY | 0.113095276 |
| ITGMSGHQEV | ASSSDWGSAETLY | 0.1118491 |
| IVAGLACTL | ASSSDWGSAETLY | 0.10562516 |
| IVGGGLAGA | ASSSDWGSAETLY | 0.10598643 |
| IVTGVLNI | ASSSDWGSAETLY | 0.17234078 |
| LAQQGCI | ASSSDWGSAETLY | 0.09808984 |
| LCLHVEN | ASSSDWGSAETLY | 0.1110647 |
| LDAISDGS | ASSSDWGSAETLY | 0.13188489 |
| LGGFAAML | ASSSDWGSAETLY | 0.14514741 |
| LGGKNPMI | ASSSDWGSAETLY | 0.14024837 |
| LGPKGVAAIL | ASSSDWGSAETLY | 0.12037649 |
| LMVGVSI | ASSSDWGSAETLY | 0.10822146 |
| LVGAAFVL | ASSSDWGSAETLY | 0.123136684 |
| LVLAVAPL | ASSSDWGSAETLY | 0.11538622 |
| MAAIAVIDAA | ASSSDWGSAETLY | 0.09784959 |
| NNGINAELQ | ASSSDWGSAETLY | 0.09740477 |
| RGEPLVLVHG | ASSSDWGSAETLY | 0.106323116 |
| RGGAGGAAGLI | ASSSDWGSAETLY | 0.09811647 |
| RSYRMHPAV | ASSSDWGSAETLY | 0.13687082 |
| SAAHLLAGQLA | ASSSDWGSAETLY | 0.12345187 |
| SAAVGSY | ASSSDWGSAETLY | 0.10542074 |
| SPHAVLHRI | ASSSDWGSAETLY | 0.10017246 |
| SQIAKVL | ASSSDWGSAETLY | 0.105377994 |
| TALGHLVP | ASSSDWGSAETLY | 0.11826937 |
| TAPTTSGAA | ASSSDWGSAETLY | 0.1335869 |
| TDGDDGNGGSPGT | ASSSDWGSAETLY | 0.10233462 |
| TDGLGEN | ASSSDWGSAETLY | 0.33951885 |
| TGGPPLPGQA | ASSSDWGSAETLY | 0.15517008 |
| TIPAITI | ASSSDWGSAETLY | 0.109899215 |
| TIPGFTI | ASSSDWGSAETLY | 0.13982311 |
| TSIASGALA | ASSSDWGSAETLY | 0.10674451 |
| TVTLPGAAI | ASSSDWGSAETLY | 0.10416481 |
| VASGLSAALP | ASSSDWGSAETLY | 0.10571379 |

| | | |
|---|---|---|
| VGAGVIGI | ASSSDWGSAETLY | 0.104784906 |
| VGPKRFGV | ASSSDWGSAETLY | 0.10759042 |
| VNVAFPD | ASSSDWGSAETLY | 0.10616937 |
| VSMTNSGVS | ASSSDWGSAETLY | 0.10281126 |
| VTDVLGDLSV | ASSSDWGSAETLY | 0.104435004 |
| VVVVGAGFAGL | ASSSDWGSAETLY | 0.10138147 |
| VYTVGTVV | ASSSDWGSAETLY | 0.0971979 |
| WTTFSGP | ASSSDWGSAETLY | 0.10632126 |
| YDELVEAF | ASSSDWGSAETLY | 0.16953789 |
| YGTGGAGGNGG | ASSSDWGSAETLY | 0.2259842 |
| YTNVMAV | ASSSDWGSAETLY | 0.104309045 |
| FLEQVHQL | ASSSDWGSAETLY | 0.10753468 |
| GLSLLAVAV | ASSSDWGSAETLY | 0.16398177 |
| LEQVHQL | ASSSDWGSAETLY | 0.10449603 |
| LNAAGAI | ASSSDWGSAETLY | 0.109810285 |
| PAVHASL | ASSSDWGSAETLY | 0.119793296 |
| TELEDDA | ASSSDWGSAETLY | 0.11211063 |
| TPPGPPN | ASSSDWGSAETLY | 0.10155579 |
| TPPPPPPP | ASSSDWGSAETLY | 0.10452934 |
| TQSPALI | ASSLSGTEISNERLF | 0.104607224 |
| EFAAAFPQ | ASSLSGTEISNERLF | 0.12091903 |
| LLSGNLDVL | ASSLSGTEISNERLF | 0.0956864 |
| GPANLGNYNI | ASSLSGTEISNERLF | 0.11538363 |
| TLGPTLP | ASSLSGTEISNERLF | 0.106490575 |
| VYAYSSCCK | ASSLSGTEISNERLF | 0.105415456 |
| APAPAPAPNI | ASSLSGTEISNERLF | 0.097059436 |
| PGGPVPIP | ASSLSGTEISNERLF | 0.11505471 |
| FLTVALEP | ASSLSGTEISNERLF | 0.11330511 |
| SQYFGANL | ASSLSGTEISNERLF | 0.13451037 |
| GGVKPAA | ASSLSGTEISNERLF | 0.099972114 |
| GGTGGAGGAGGW | ASSLSGTEISNERLF | 0.11842395 |
| GVGGVAGLGGAGG | ASSLSGTEISNERLF | 0.12062018 |
| VLLVLATTI | ASSLSGTEISNERLF | 0.11399846 |
| IGIFAAML | ASSLSGTEISNERLF | 0.10127658 |
| AVGDLID | ASSLSGTEISNERLF | 0.12239081 |
| NISPLPL | ASSLSGTEISNERLF | 0.099397756 |
| VTAVFVAI | ASSLSGTEISNERLF | 0.11573679 |
| SGDDNDAAL | ASSLSGTEISNERLF | 0.12430644 |
| AVAALAHS | ASSLSGTEISNERLF | 0.10002941 |
| GGVTVMATP | ASSLSGTEISNERLF | 0.12524517 |
| AGLAANVRD | ASSLSGTEISNERLF | 0.13664588 |
| TNRPDLIDPALLRPG | ASSLSGTEISNERLF | 0.10273931 |
| LANLGVTAL | ASSLSGTEISNERLF | 0.105414756 |
| TGERIEL | ASSLSGTEISNERLF | 0.10869545 |
| NGEELAA | ASSLSGTEISNERLF | 0.095870115 |
| VLSLVWG | ASSLSGTEISNERLF | 0.12895116 |
| IGYALWL | ASSLSGTEISNERLF | 0.10577772 |
| LSLLLAPL | ASSLSGTEISNERLF | 0.113475904 |
| TLDEALK | ASSLSGTEISNERLF | 0.111597136 |
| TDGAGTAEGG | ASSLSGTEISNERLF | 0.10900797 |
| NAPGGAGGNGG | ASSLSGTEISNERLF | 0.11526083 |

| | | |
|---|---|---|
| VSVFMGMGGNF | ASSLSGTEISNERLF | 0.11810885 |
| LASYLAAL | ASSLSGTEISNERLF | 0.111206375 |
| LVGGTPAL | ASSLSGTEISNERLF | 0.10860535 |
| VRNADVL | ASSLSGTEISNERLF | 0.10579097 |
| TCSSPFLVSAM | ASSLSGTEISNERLF | 0.11052724 |
| VHITDAE | ASSLSGTEISNERLF | 0.17346115 |
| EYAAPSF | ASSLSGTEISNERLF | 0.09946 |
| VVDAFHGGL | ASSLSGTEISNERLF | 0.10533099 |
| ASVVDPQ | ASSLSGTEISNERLF | 0.1221556 |
| SYNEFGGLS | ASSLSGTEISNERLF | 0.10073862 |
| DPDGSVY | ASSLSGTEISNERLF | 0.11909104 |
| MVSTIHATE | ASSLSGTEISNERLF | 0.12636559 |
| LAAGVADPV | ASSLSGTEISNERLF | 0.12982905 |
| TKEQIAA | ASSLSGTEISNERLF | 0.10434546 |
| SLASPAAL | ASSLSGTEISNERLF | 0.14577185 |
| IGLVFYGV | ASSLSGTEISNERLF | 0.13995928 |
| FLLDASNL | ASSLSGTEISNERLF | 0.10530009 |
| YQVGNFP | ASSLSGTEISNERLF | 0.10060708 |
| ESLLSQST | ASSLSGTEISNERLF | 0.104711436 |
| LFTVGFAAVG | ASSLSGTEISNERLF | 0.1088135 |
| ATPPTPT | ASSLSGTEISNERLF | 0.12204887 |
| TDLATLG | ASSLSGTEISNERLF | 0.10574965 |
| SAGGTGR | ASSLSGTEISNERLF | 0.11142367 |
| GFSNSGS | ASSLSGTEISNERLF | 0.1217894 |
| GGNGGSGGTSV | ASSLSGTEISNERLF | 0.1316204 |
| GTSVATGGAGN | ASSLSGTEISNERLF | 0.10069213 |
| NGGNGGSGGTS | ASSLSGTEISNERLF | 0.10830259 |
| AVVKFPH | ASSLSGTEISNERLF | 0.1088154 |
| TPPPPPP | ASSLSGTEISNERLF | 0.10941458 |
| AAPAAAAAPA | ASSLSGTEISNERLF | 0.10678146 |
| IALNEDL | ASSLSGTEISNERLF | 0.12759049 |
| IVNVSSM | ASSLSGTEISNERLF | 0.11867048 |
| TGIPPAPR | ASSLSGTEISNERLF | 0.13203523 |
| FDLGGGTF | ASSLSGTEISNERLF | 0.14000145 |
| FDLGGGTFDVS | ASSLSGTEISNERLF | 0.16017437 |
| NEPTAAA | ASSLSGTEISNERLF | 0.10614745 |
| NEPTAAALA | ASSLSGTEISNERLF | 0.11672256 |
| GSIINISSI | ASSLSGTEISNERLF | 0.09877821 |
| VAPGNAGTA | ASSLSGTEISNERLF | 0.09770878 |
| TGGGASLE | ASSLSGTEISNERLF | 0.0992326 |
| VAVYNHYKRI | ASSLSGTEISNERLF | 0.112836234 |
| TDPAPATT | ASSLSGTEISNERLF | 0.09596246 |
| TNRPDILDPALLRPG | ASSLSGTEISNERLF | 0.12661587 |
| LTTTTTS | ASSLSGTEISNERLF | 0.11455425 |
| PTGSGKTL | ASSLSGTEISNERLF | 0.14480513 |
| TGSGKTL | ASSLSGTEISNERLF | 0.113919936 |
| TIGHVDHGK | ASSLSGTEISNERLF | 0.10557455 |
| TIGHVDHGKTT | ASSLSGTEISNERLF | 0.106680624 |
| GHVDHGKTT | ASSLSGTEISNERLF | 0.13868478 |
| AQFEHTLL | ASSLSGTEISNERLF | 0.10193872 |
| IVNQDEL | ASSLSGTEISNERLF | 0.09945681 |

| | | |
|---|---|---|
| TNGSQFF | ASSLSGTEISNERLF | 0.10484328 |
| TGKGIVE | ASSLSGTEISNERLF | 0.104117975 |
| LISNASDA | ASSLSGTEISNERLF | 0.13058329 |
| AELRTLGELTKIAKAHGAQ | ASSLSGTEISNERLF | 0.12119297 |
| APNAVAAAEPSW | ASSLSGTEISNERLF | 0.11515614 |
| APQVAGAATAVL | ASSLSGTEISNERLF | 0.11803207 |
| ARRDPAAAAAILPTDA | ASSLSGTEISNERLF | 0.09663666 |
| DAAARDVLAEAGLAEYF | ASSLSGTEISNERLF | 0.11087846 |
| DDQIVLSAKANLAKVEQ | ASSLSGTEISNERLF | 0.21605805 |
| DELLPDTGEDSDEDGH | ASSLSGTEISNERLF | 0.10995478 |
| ELPEAPANRATVPQD | ASSLSGTEISNERLF | 0.123662844 |
| FGQSKIERVTGIG | ASSLSGTEISNERLF | 0.13350643 |
| GAGGTGSTAGGAGGAGGAGGLY | ASSLSGTEISNERLF | 0.10931785 |
| GHGGTGAAGGNGGTGGAGGSGI | ASSLSGTEISNERLF | 0.11196417 |
| GNGGNGGAGGEGGPGGAGGAGA | ASSLSGTEISNERLF | 0.11376256 |
| GSTAGNGGNGGNATNGTIAGQP | ASSLSGTEISNERLF | 0.113694124 |
| GTGGDGGNGGNGGTGG | ASSLSGTEISNERLF | 0.100598715 |
| GVRRPAGIPADAPV | ASSLSGTEISNERLF | 0.10178493 |
| GYRQVTDAHLLTLARR | ASSLSGTEISNERLF | 0.12783873 |
| IAKMGQDGHDRG | ASSLSGTEISNERLF | 0.2545775 |
| KAEPSIEIQYT | ASSLSGTEISNERLF | 0.11591885 |
| KPVRIRMAADEDTVAVT | ASSLSGTEISNERLF | 0.13148029 |
| MPPRIAGMRLLVIKPEP | ASSLSGTEISNERLF | 0.12042951 |
| MKFYAVALFRSRGVPPTR | ASSLSGTEISNERLF | 0.10341485 |
| NAGGFGILGAVAHSP | ASSLSGTEISNERLF | 0.1102631 |
| NEGREITLAMAEM | ASSLSGTEISNERLF | 0.10357184 |
| NELVEWSHPLGHRWRWE | ASSLSGTEISNERLF | 0.1037115 |
| NLGAGGTGGNGGNPG | ASSLSGTEISNERLF | 0.10187494 |
| NTEWPKWETVMAADPVD | ASSLSGTEISNERLF | 0.10947322 |
| PASAAPAETQP | ASSLSGTEISNERLF | 0.11427898 |
| QQSSDGKNGVRAAEITGE | ASSLSGTEISNERLF | 0.26173282 |
| RAPDAVTTSWWKE | ASSLSGTEISNERLF | 0.11456691 |
| SGFGNVGAHTSGWFNQ | ASSLSGTEISNERLF | 0.12178907 |
| SGVAGAVGQAASVGGLKV | ASSLSGTEISNERLF | 0.10295857 |
| SNGGAGTGSAGGNGGT | ASSLSGTEISNERLF | 0.10405145 |
| TGGAGGEGGAAG | ASSLSGTEISNERLF | 0.10904452 |
| TGGDGGAAGAGGAPGHG | ASSLSGTEISNERLF | 0.1036526 |
| TMPPLLAAIANAGRHGV | ASSLSGTEISNERLF | 0.15514806 |
| TSSTAGNSGGA | ASSLSGTEISNERLF | 0.107931554 |
| VPATVPVGILC | ASSLSGTEISNERLF | 0.10732765 |
| VQRPDSGGVVLTGRIS | ASSLSGTEISNERLF | 0.12349849 |
| WTPNPVPATKGVPAGLVQAT | ASSLSGTEISNERLF | 0.19675492 |
| YALKRALAKFSDRFDVVII | ASSLSGTEISNERLF | 0.11008018 |
| YLIGSVIV | ASSLSGTEISNERLF | 0.12225933 |
| YGQLVAVID | ASSLSGTEISNERLF | 0.1034859 |
| VTVGGMTLPA | ASSLSGTEISNERLF | 0.12323302 |
| VLALPAV | ASSLSGTEISNERLF | 0.18641773 |
| VIPPTLNYET | ASSLSGTEISNERLF | 0.09692819 |
| VGMGGMHPSSG | ASSLSGTEISNERLF | 0.1817836 |
| VGLAGGAATA | ASSLSGTEISNERLF | 0.111576356 |
| VGGAGGAGGSGP | ASSLSGTEISNERLF | 0.10349181 |

| | | |
|---|---|---|
| VFAAFMV | ASSLSGTEISNERLF | 0.116548106 |
| TLYDKLR | ASSLSGTEISNERLF | 0.12434613 |
| TLAVFTVL | ASSLSGTEISNERLF | 0.12742162 |
| TGSVQPARV | ASSLSGTEISNERLF | 0.12007328 |
| TGGGAGSGGT | ASSLSGTEISNERLF | 0.10199167 |
| SHLETSL | ASSLSGTEISNERLF | 0.10290303 |
| SFTTGMT | ASSLSGTEISNERLF | 0.13719724 |
| QVEVAKRMGVRQPTVS | ASSLSGTEISNERLF | 0.1164069 |
| QGSGGPALAM | ASSLSGTEISNERLF | 0.13468565 |
| MLAPSNP | ASSLSGTEISNERLF | 0.105178654 |
| LVNNAAI | ASSLSGTEISNERLF | 0.111149736 |
| LTGEINTLA | ASSLSGTEISNERLF | 0.107195735 |
| LSIFGPV | ASSLSGTEISNERLF | 0.12437528 |
| LLALLANL | ASSLSGTEISNERLF | 0.10131766 |
| LISNASDALDKLR | ASSLSGTEISNERLF | 0.114443034 |
| LINEAVP | ASSLSGTEISNERLF | 0.1314348 |
| LGYAFSGL | ASSLSGTEISNERLF | 0.10056305 |
| LDFVAVAL | ASSLSGTEISNERLF | 0.10460566 |
| LAQPGAE | ASSLSGTEISNERLF | 0.15188518 |
| ISNLGAVTGF | ASSLSGTEISNERLF | 0.10687927 |
| HQEDPGRI | ASSLSGTEISNERLF | 0.10106423 |
| GTPGEPTP | ASSLSGTEISNERLF | 0.15232153 |
| GFFNSGS | ASSLSGTEISNERLF | 0.22499564 |
| GAGGGGLTAGAN | ASSLSGTEISNERLF | 0.11881358 |
| FVDALVNM | ASSLSGTEISNERLF | 0.10194335 |
| FLTVAPEL | ASSLSGTEISNERLF | 0.096879855 |
| FETLTAL | ASSLSGTEISNERLF | 0.11140286 |
| DVINPIEL | ASSLSGTEISNERLF | 0.12334862 |
| APVSGAPV | ASSLSGTEISNERLF | 0.110632755 |
| ALVGGLAW | ASSLSGTEISNERLF | 0.10179525 |
| ALAAAAATAI | ASSLSGTEISNERLF | 0.10929573 |
| AFGLGLLL | ASSLSGTEISNERLF | 0.13884111 |
| AFFAAASL | ASSLSGTEISNERLF | 0.1032215 |
| AAAALPAFSPP | ASSLSGTEISNERLF | 0.10060937 |
| VVLAPAL | ASSLSGTEISNERLF | 0.10611027 |
| TIGHVDHGKTTL | ASSLSGTEISNERLF | 0.13352007 |
| LISNASDALDK | ASSLSGTEISNERLF | 0.09892146 |
| LISNASDAL | ASSLSGTEISNERLF | 0.105485894 |
| ISNASDALDK | ASSLSGTEISNERLF | 0.1406165 |
| NQAETLVYQTE | ASSLSGTEISNERLF | 0.099697 |
| TGGTGGTAGAG | ASSLSGTEISNERLF | 0.11569897 |
| TGSTAGGAGGA | ASSLSGTEISNERLF | 0.10813814 |
| DPLAVPVAAGPV | ASSLSGTEISNERLF | 0.12519343 |
| GGTGGTPGNGGN | ASSLSGTEISNERLF | 0.106568635 |
| PAGLAGVDPATL | ASSLSGTEISNERLF | 0.11653075 |
| GNNLVVEAPHAGASV | ASSLSGTEISNERLF | 0.11799174 |
| KVTRIMSASTLEQPA | ASSLSGTEISNERLF | 0.118741676 |
| LGFALPIALPRRTVV | ASSLSGTEISNERLF | 0.11491388 |
| TRLGNLTARKLVVLV | ASSLSGTEISNERLF | 0.10360041 |
| WSQQRRLIYFPSAGP | ASSLSGTEISNERLF | 0.106413506 |
| GPNGNPDPMAAAVDIR | ASSLSGTEISNERLF | 0.09643685 |

| | | |
|---|---|---|
| TQRCVSVADLRAAASR | ASSLSGTEISNERLF | 0.14589158 |
| QFVLDVAEHPVEVLTETV | ASSLSGTEISNERLF | 0.095812686 |
| GGAGGTGGAGGNGGTQAGDG | ASSLSGTEISNERLF | 0.13305968 |
| PLLATLRAIAVDVDAKPAQV | ASSLSGTEISNERLF | 0.12896149 |
| APKFRTGGYMIDFWGVGYQVAK | ASSLSGTEISNERLF | 0.1071123 |
| GAGGAGGAGGAAGGPGGTGGTG | ASSLSGTEISNERLF | 0.11455608 |
| IGALGLLATDVAPAGSVAFGVA | ASSLSGTEISNERLF | 0.098694295 |
| AGGAGGIGGNGGAGGAGGVLMG | ASSLSGTEISNERLF | 0.10217573 |
| ADELIGLL | ASSLSGTEISNERLF | 0.108299226 |
| ADRALKA | ASSLSGTEISNERLF | 0.11825393 |
| AGAVGQVGGAGGA | ASSLSGTEISNERLF | 0.11021962 |
| AGGGGGLGGAP | ASSLSGTEISNERLF | 0.11737017 |
| AGGLLYGNGGAG | ASSLSGTEISNERLF | 0.12155277 |
| AGLESIGL | ASSLSGTEISNERLF | 0.10348381 |
| AGLKTGII | ASSLSGTEISNERLF | 0.10638045 |
| ASLAALLL | ASSLSGTEISNERLF | 0.14651854 |
| ASVINISSI | ASSLSGTEISNERLF | 0.11522812 |
| ATPPGPA | ASSLSGTEISNERLF | 0.101671025 |
| ATVASALV | ASSLSGTEISNERLF | 0.12964877 |
| ATVASLIL | ASSLSGTEISNERLF | 0.1032777 |
| AVNGPVLIRH | ASSLSGTEISNERLF | 0.12083451 |
| DGAAGAVGI | ASSLSGTEISNERLF | 0.11676516 |
| DPLLHGI | ASSLSGTEISNERLF | 0.114535145 |
| ELGAALGAL | ASSLSGTEISNERLF | 0.12882021 |
| ELGRTQAV | ASSLSGTEISNERLF | 0.101120174 |
| FVAGGYLS | ASSLSGTEISNERLF | 0.10225931 |
| GASGGAGGAGGW | ASSLSGTEISNERLF | 0.10776815 |
| GGAGAGGDGGAGGI | ASSLSGTEISNERLF | 0.13905413 |
| GTGHNGSQPGMGGNG | ASSLSGTEISNERLF | 0.13373648 |
| GYAAASASAS | ASSLSGTEISNERLF | 0.105554 |
| HQVDVAAVQQG | ASSLSGTEISNERLF | 0.11535824 |
| IGAAIAHT | ASSLSGTEISNERLF | 0.10711406 |
| IGMIEVAQL | ASSLSGTEISNERLF | 0.10806756 |
| IRLAPEHTSAMD | ASSLSGTEISNERLF | 0.11167851 |
| IYNVSVL | ASSLSGTEISNERLF | 0.10172628 |
| KQGFLGM | ASSLSGTEISNERLF | 0.105585955 |
| LESVITQL | ASSLSGTEISNERLF | 0.11386629 |
| LGLGALVVV | ASSLSGTEISNERLF | 0.10846385 |
| LIGNGGNGGSGGI | ASSLSGTEISNERLF | 0.12325244 |
| LNAINAPTL | ASSLSGTEISNERLF | 0.114937976 |
| LQLLNTGV | ASSLSGTEISNERLF | 0.1048425 |
| LQTSIEF | ASSLSGTEISNERLF | 0.11535863 |
| MSFAVSEL | ASSLSGTEISNERLF | 0.111352414 |
| NTGALGL | ASSLSGTEISNERLF | 0.117917985 |
| PPGAGGAGGAGTT | ASSLSGTEISNERLF | 0.11963914 |
| QLVGLGI | ASSLSGTEISNERLF | 0.10529509 |
| RFLEQVHQL | ASSLSGTEISNERLF | 0.13416074 |
| RGVGYVLR | ASSLSGTEISNERLF | 0.145731 |
| SATGVFAGVI | ASSLSGTEISNERLF | 0.11502901 |
| SLPAITLP | ASSLSGTEISNERLF | 0.11057487 |
| STALAFTV | ASSLSGTEISNERLF | 0.1191453 |

| | | |
|---|---|---|
| SVLTVGLGK | ASSLSGTEISNERLF | 0.13463676 |
| TALAGAGSGI | ASSLSGTEISNERLF | 0.10806662 |
| TGLPFTGI | ASSLSGTEISNERLF | 0.09882114 |
| VAEHVTV | ASSLSGTEISNERLF | 0.1110661 |
| VDIVLNSL | ASSLSGTEISNERLF | 0.10518109 |
| VLTGGGAVL | ASSLSGTEISNERLF | 0.10827012 |
| VNVGLAYGI | ASSLSGTEISNERLF | 0.09919957 |
| VSSAAVGAV | ASSLSGTEISNERLF | 0.14667942 |
| VTSVYEAGV | ASSLSGTEISNERLF | 0.2256872 |
| WVALDRGA | ASSLSGTEISNERLF | 0.10830539 |
| YLVGSVVI | ASSLSGTEISNERLF | 0.108880155 |
| GPPGTGKTLL | ASSLSGTEISNERLF | 0.098725975 |
| GVGMTKF | ASSLSGTEISNERLF | 0.12444614 |
| IGIGGSDL | ASSLSGTEISNERLF | 0.13351496 |
| IGTLLAEL | ASSLSGTEISNERLF | 0.10391183 |
| IYTSGTT | ASSLSGTEISNERLF | 0.124051645 |
| LSFPAGV | ASSLSGTEISNERLF | 0.10406807 |
| LYTSGTT | ASSLSGTEISNERLF | 0.109773904 |
| VGAFPFGF | ASSLSGTEISNERLF | 0.1330381 |
| VGAGTVEF | ASSLSGTEISNERLF | 0.11802265 |
| VGKSTLLL | ASSLSGTEISNERLF | 0.11042734 |
| AALAHAGGGIV | ASSLSGTEISNERLF | 0.107974894 |
| AALVAVSLL | ASSLSGTEISNERLF | 0.10549701 |
| AAVSRAVAAL | ASSLSGTEISNERLF | 0.113605246 |
| AGGSSGIGGF | ASSLSGTEISNERLF | 0.12808976 |
| AGLTVANANG | ASSLSGTEISNERLF | 0.10050128 |
| AGLVGAVL | ASSLSGTEISNERLF | 0.11806136 |
| ALSQAFTV | ASSLSGTEISNERLF | 0.10739495 |
| ALTFSVSL | ASSLSGTEISNERLF | 0.18920708 |
| ALTGAALVAL | ASSLSGTEISNERLF | 0.17778015 |
| ASFSIGAGVL | ASSLSGTEISNERLF | 0.119357154 |
| DGSVYKI | ASSLSGTEISNERLF | 0.10786014 |
| EQLAEIV | ASSLSGTEISNERLF | 0.18055137 |
| FAVSELTVM | ASSLSGTEISNERLF | 0.113797314 |
| GAAPLAAGL | ASSLSGTEISNERLF | 0.12306916 |
| GADGGDGGSGGAGG | ASSLSGTEISNERLF | 0.119326174 |
| GFGLDTGGAGG | ASSLSGTEISNERLF | 0.7448679 |
| GLASGGLL | ASSLSGTEISNERLF | 0.8029152 |
| GRIAGAAVLL | ASSLSGTEISNERLF | 0.7928967 |
| GSAGSLNTSF | ASSLSGTEISNERLF | 0.7883844 |
| GTGQIGI | ASSLSGTEISNERLF | 0.7680255 |
| GTLIALIL | ASSLSGTEISNERLF | 0.50537634 |
| IAATMGFVVG | ASSLSGTEISNERLF | 0.7723847 |
| IFAQALL | ASSLSGTEISNERLF | 0.7465353 |
| IGAAVALGF | ASSLSGTEISNERLF | 0.79443294 |
| IGLTPLAL | ASSLSGTEISNERLF | 0.7972321 |
| ILGSNTSTL | ASSLSGTEISNERLF | 0.7650202 |
| IVFSLTVL | ASSLSGTEISNERLF | 0.80224955 |
| LASKATGF | ASSLSGTEISNERLF | 0.8010067 |
| LASVALIL | ASSLSGTEISNERLF | 0.63376987 |
| LFAVGLTL | ASSLSGTEISNERLF | 0.7725657 |

| | | |
|---|---|---|
| LFNGGIGTYI | ASSLSGTEISNERLF | 0.75859094 |
| LTLGAGII | ASSLSGTEISNERLF | 0.7842949 |
| LVAAALGDI | ASSLSGTEISNERLF | 0.5703539 |
| NTAFVQLGI | ASSLSGTEISNERLF | 0.79943526 |
| QLFSGAEL | ASSLSGTEISNERLF | 0.7715329 |
| SAATFITL | ASSLSGTEISNERLF | 0.30572787 |
| SNYGLANLGV | ASSLSGTEISNERLF | 0.78442466 |
| SYGGAYAVM | ASSLSGTEISNERLF | 0.78376997 |
| TTGVGGLTLG | ASSLSGTEISNERLF | 0.77621716 |
| TTQTIYI | ASSLSGTEISNERLF | 0.80070114 |
| VAAGAGLGAVI | ASSLSGTEISNERLF | 0.7098297 |
| VAAVVTLL | ASSLSGTEISNERLF | 0.73691547 |
| VGFLGGLI | ASSLSGTEISNERLF | 0.7961588 |
| VGVIPILL | ASSLSGTEISNERLF | 0.8037075 |
| VIFASSLI | ASSLSGTEISNERLF | 0.7037191 |
| VKGGDGGAAAG | ASSLSGTEISNERLF | 0.7932566 |
| VTKTFTNI | ASSLSGTEISNERLF | 0.80249816 |
| VVAAAILL | ASSLSGTEISNERLF | 0.79360384 |
| YAALDVEL | ASSLSGTEISNERLF | 0.8004304 |
| LASVALLL | ASSLSGTEISNERLF | 0.803823 |
| LDAVHRV | ASSLSGTEISNERLF | 0.7399496 |
| QGLSSSIF | ASSLSGTEISNERLF | 0.8019481 |
| RGTFANIRL | ASSLSGTEISNERLF | 0.8017709 |
| RQLWWGHRI | ASSLSGTEISNERLF | 0.76907253 |
| YGPPGTGKTLL | ASSLSGTEISNERLF | 0.787406 |
| AALVGAPANVA | ASSLSGTEISNERLF | 0.7461645 |
| AATFRRTE | ASSLSGTEISNERLF | 0.7760343 |
| AGALGGGI | ASSLSGTEISNERLF | 0.5600897 |
| AGGAGGAGGIGAG | ASSLSGTEISNERLF | 0.7942658 |
| AGSGALGL | ASSLSGTEISNERLF | 0.8009968 |
| AQIADQI | ASSLSGTEISNERLF | 0.7789423 |
| ASAVAVTYL | ASSLSGTEISNERLF | 0.7367085 |
| ASTVLGGTGG | ASSLSGTEISNERLF | 0.74114835 |
| AVGGEPAK | ASSLSGTEISNERLF | 0.78549707 |
| CEQGTFGGTC | ASSLSGTEISNERLF | 0.77754855 |
| DEGKAPAG | ASSLSGTEISNERLF | 0.5865377 |
| DTVVFPAL | ASSLSGTEISNERLF | 0.75630414 |
| FEGITMAG | ASSLSGTEISNERLF | 0.80021393 |
| FPLVVQD | ASSLSGTEISNERLF | 0.8020328 |
| GAFTTPQ | ASSLSGTEISNERLF | 0.6396379 |
| GFGGFGTAGGVGGA | ASSLSGTEISNERLF | 0.80313045 |
| GFLAQFL | ASSLSGTEISNERLF | 0.7975885 |
| GGAGGAGGNAGVA | ASSLSGTEISNERLF | 0.80302596 |
| GGVGTAGAGGAGGAGG | ASSLSGTEISNERLF | 0.789431 |
| GNGGGGDAGNAGSGGN | ASSLSGTEISNERLF | 0.7407886 |
| GTGGAGGAGNDAGS | ASSLSGTEISNERLF | 0.8002895 |
| GTTNTGWFNS | ASSLSGTEISNERLF | 0.77419955 |
| IFMGPLFTL | ASSLSGTEISNERLF | 0.7471384 |
| ILAHAIEAI | ASSLSGTEISNERLF | 0.7994539 |
| IRSAIREAL | ASSLSGTEISNERLF | 0.7784849 |
| ITFAGGG | ASSLSGTEISNERLF | 0.7555301 |

| | | |
|---|---|---|
| ITGMSGHQEV | ASSLSGTEISNERLF | 0.76038396 |
| IVAGLACTL | ASSLSGTEISNERLF | 0.80361986 |
| IVGGGLAGA | ASSLSGTEISNERLF | 0.7244703 |
| IVTGVLNI | ASSLSGTEISNERLF | 0.73781246 |
| LAQQGCI | ASSLSGTEISNERLF | 0.75639236 |
| LCLHVEN | ASSLSGTEISNERLF | 0.7881605 |
| LDAISDGS | ASSLSGTEISNERLF | 0.8009795 |
| LGGFAAML | ASSLSGTEISNERLF | 0.7927564 |
| LGGKNPMI | ASSLSGTEISNERLF | 0.759642 |
| LGPKGVAAIL | ASSLSGTEISNERLF | 0.7929697 |
| LMVGVSI | ASSLSGTEISNERLF | 0.7967936 |
| LVGAAFVL | ASSLSGTEISNERLF | 0.79134965 |
| LVLAVAPL | ASSLSGTEISNERLF | 0.7965671 |
| MAAIAVIDAA | ASSLSGTEISNERLF | 0.75374234 |
| NNGINAELQ | ASSLSGTEISNERLF | 0.80197906 |
| RGEPLVLVHG | ASSLSGTEISNERLF | 0.7974938 |
| RGGAGGAAGLI | ASSLSGTEISNERLF | 0.7992028 |
| RSYRMHPAV | ASSLSGTEISNERLF | 0.79760015 |
| SAAHLLAGQLA | ASSLSGTEISNERLF | 0.78940374 |
| SAAVGSY | ASSLSGTEISNERLF | 0.7927168 |
| SPHAVLHRI | ASSLSGTEISNERLF | 0.59227765 |
| SQIAKVL | ASSLSGTEISNERLF | 0.7641151 |
| TALGHLVP | ASSLSGTEISNERLF | 0.790768 |
| TAPTTSGAA | ASSLSGTEISNERLF | 0.7905729 |
| TDGDDGNGGSPGT | ASSLSGTEISNERLF | 0.7957452 |
| TDGLGEN | ASSLSGTEISNERLF | 0.7845341 |
| TGGPPLPGQA | ASSLSGTEISNERLF | 0.7654287 |
| TIPAITI | ASSLSGTEISNERLF | 0.80083984 |
| TIPGFTI | ASSLSGTEISNERLF | 0.7989522 |
| TSIASGALA | ASSLSGTEISNERLF | 0.6519523 |
| TVTLPGAAI | ASSLSGTEISNERLF | 0.7728559 |
| VASGLSAALP | ASSLSGTEISNERLF | 0.80510604 |
| VGAGVIGI | ASSLSGTEISNERLF | 0.79005116 |
| VGPKRFGV | ASSLSGTEISNERLF | 0.78098357 |
| VNVAFPD | ASSLSGTEISNERLF | 0.77191097 |
| VSMTNSGVS | ASSLSGTEISNERLF | 0.80429995 |
| VTDVLGDLSV | ASSLSGTEISNERLF | 0.76672477 |
| VVVVGAGFAGL | ASSLSGTEISNERLF | 0.8043624 |
| VYTVGTVV | ASSLSGTEISNERLF | 0.38932174 |
| WTTFSGP | ASSLSGTEISNERLF | 0.7744007 |
| YDELVEAF | ASSLSGTEISNERLF | 0.7176627 |
| YGTGGAGGNGG | ASSLSGTEISNERLF | 0.6337304 |
| YTNVMAV | ASSLSGTEISNERLF | 0.8016379 |
| FLEQVHQL | ASSLSGTEISNERLF | 0.75096416 |
| GLSLLAVAV | ASSLSGTEISNERLF | 0.796875 |
| LEQVHQL | ASSLSGTEISNERLF | 0.7795896 |
| LNAAGAI | ASSLSGTEISNERLF | 0.80068356 |
| PAVHASL | ASSLSGTEISNERLF | 0.7860166 |
| TELEDDA | ASSLSGTEISNERLF | 0.7890482 |
| TPPGPPN | ASSLSGTEISNERLF | 0.72107244 |
| TPPPPPPP | ASSLSGTEISNERLF | 0.80133605 |

| | | |
|---|---|---|
| TQSPALI | ASSLRQGGSYEQY | 0.80164504 |
| EFAAAFPQ | ASSLRQGGSYEQY | 0.79564285 |
| LLSGNLDVL | ASSLRQGGSYEQY | 0.5927682 |
| GPANLGNYNI | ASSLRQGGSYEQY | 0.77601486 |
| TLGPTLP | ASSLRQGGSYEQY | 0.78893435 |
| VYAYSSCCK | ASSLRQGGSYEQY | 0.39276886 |
| APAPAPAPNI | ASSLRQGGSYEQY | 0.47610432 |
| PGGPVPIP | ASSLRQGGSYEQY | 0.7729455 |
| FLTVALEP | ASSLRQGGSYEQY | 0.7671144 |
| SQYFGANL | ASSLRQGGSYEQY | 0.8022295 |
| GGVKPAA | ASSLRQGGSYEQY | 0.79575515 |
| GGTGGAGGAGGW | ASSLRQGGSYEQY | 0.7842858 |
| GVGGVAGLGGAGG | ASSLRQGGSYEQY | 0.7981839 |
| VLLVLATTI | ASSLRQGGSYEQY | 0.7812929 |
| IGIFAAML | ASSLRQGGSYEQY | 0.19508545 |
| AVGDLID | ASSLRQGGSYEQY | 0.76854336 |
| NISPLPL | ASSLRQGGSYEQY | 0.65273035 |
| VTAVFVAI | ASSLRQGGSYEQY | 0.80178225 |
| SGDDNDAAL | ASSLRQGGSYEQY | 0.7487489 |
| AVAALAHS | ASSLRQGGSYEQY | 0.74838674 |
| GGVTVMATP | ASSLRQGGSYEQY | 0.7858141 |
| AGLAANVRD | ASSLRQGGSYEQY | 0.78584284 |
| TNRPDLIDPALLRPG | ASSLRQGGSYEQY | 0.78636456 |
| LANLGVTAL | ASSLRQGGSYEQY | 0.72967494 |
| TGERIEL | ASSLRQGGSYEQY | 0.3050522 |
| NGEELAA | ASSLRQGGSYEQY | 0.79879224 |
| VLSLVWG | ASSLRQGGSYEQY | 0.55693865 |
| IGYALWL | ASSLRQGGSYEQY | 0.71616507 |
| LSLLLAPL | ASSLRQGGSYEQY | 0.47537857 |
| TLDEALK | ASSLRQGGSYEQY | 0.7519312 |
| TDGAGTAEGG | ASSLRQGGSYEQY | 0.53294027 |
| NAPGGAGGNGG | ASSLRQGGSYEQY | 0.7978687 |
| VSVFMGMGGNF | ASSLRQGGSYEQY | 0.77300674 |
| LASYLAAL | ASSLRQGGSYEQY | 0.76810765 |
| LVGGTPAL | ASSLRQGGSYEQY | 0.802842 |
| VRNADVL | ASSLRQGGSYEQY | 0.7801462 |
| TCSSPFLVSAM | ASSLRQGGSYEQY | 0.7742342 |
| VHITDAE | ASSLRQGGSYEQY | 0.74796194 |
| EYAAPSF | ASSLRQGGSYEQY | 0.7502756 |
| VVDAFHGGL | ASSLRQGGSYEQY | 0.75124496 |
| ASVVDPQ | ASSLRQGGSYEQY | 0.7364651 |
| SYNEFGGLS | ASSLRQGGSYEQY | 0.80462843 |
| DPDGSVY | ASSLRQGGSYEQY | 0.7744844 |
| MVSTIHATE | ASSLRQGGSYEQY | 0.57844174 |
| LAAGVADPV | ASSLRQGGSYEQY | 0.78504366 |
| TKEQIAA | ASSLRQGGSYEQY | 0.80415547 |
| SLASPAAL | ASSLRQGGSYEQY | 0.8066913 |
| IGLVFYGV | ASSLRQGGSYEQY | 0.7488018 |
| FLLDASNL | ASSLRQGGSYEQY | 0.76905704 |
| YQVGNFP | ASSLRQGGSYEQY | 0.7748656 |
| ESLLSQST | ASSLRQGGSYEQY | 0.78396976 |

| | | |
|---|---|---|
| LFTVGFAAVG | ASSLRQGGSYEQY | 0.75861263 |
| ATPPTPT | ASSLRQGGSYEQY | 0.6652415 |
| TDLATLG | ASSLRQGGSYEQY | 0.79956025 |
| SAGGTGR | ASSLRQGGSYEQY | 0.80674416 |
| GFSNSGS | ASSLRQGGSYEQY | 0.7772212 |
| GGNGGSGGTSV | ASSLRQGGSYEQY | 0.6734795 |
| GTSVATGGAGN | ASSLRQGGSYEQY | 0.78370297 |
| NGGNGGSGGTS | ASSLRQGGSYEQY | 0.7501941 |
| AVVKFPH | ASSLRQGGSYEQY | 0.773505 |
| TPPPPPP | ASSLRQGGSYEQY | 0.78569555 |
| AAPAAAAAPA | ASSLRQGGSYEQY | 0.6526612 |
| IALNEDL | ASSLRQGGSYEQY | 0.77565724 |
| IVNVSSM | ASSLRQGGSYEQY | 0.80589855 |
| TGIPPAPR | ASSLRQGGSYEQY | 0.76752615 |
| FDLGGGTF | ASSLRQGGSYEQY | 0.78813785 |
| FDLGGGTFDVS | ASSLRQGGSYEQY | 0.8020904 |
| NEPTAAA | ASSLRQGGSYEQY | 0.7979982 |
| NEPTAAALA | ASSLRQGGSYEQY | 0.8039712 |
| GSIINISSI | ASSLRQGGSYEQY | 0.79761636 |
| VAPGNAGTA | ASSLRQGGSYEQY | 0.76903194 |
| TGGGASLE | ASSLRQGGSYEQY | 0.7887901 |
| VAVYNHYKRI | ASSLRQGGSYEQY | 0.78976095 |
| TDPAPATT | ASSLRQGGSYEQY | 0.7886742 |
| TNRPDILDPALLRPG | ASSLRQGGSYEQY | 0.78345156 |
| LTTTTTS | ASSLRQGGSYEQY | 0.7593528 |
| PTGSGKTL | ASSLRQGGSYEQY | 0.8006978 |
| TGSGKTL | ASSLRQGGSYEQY | 0.7926323 |
| TIGHVDHGK | ASSLRQGGSYEQY | 0.80241853 |
| TIGHVDHGKTT | ASSLRQGGSYEQY | 0.687953 |
| GHVDHGKTT | ASSLRQGGSYEQY | 0.7347704 |
| AQFEHTLL | ASSLRQGGSYEQY | 0.80053866 |
| IVNQDEL | ASSLRQGGSYEQY | 0.24959564 |
| TNGSQFF | ASSLRQGGSYEQY | 0.4307739 |
| TGKGIVE | ASSLRQGGSYEQY | 0.78924644 |
| LISNASDA | ASSLRQGGSYEQY | 0.8043721 |
| AELRTLGELTKIAKAHGAQ | ASSLRQGGSYEQY | 0.26030266 |
| APNAVAAAEPSW | ASSLRQGGSYEQY | 0.7037498 |
| APQVAGAATAVL | ASSLRQGGSYEQY | 0.71304893 |
| ARRDPAAAAAILPTDA | ASSLRQGGSYEQY | 0.788612 |
| DAAARDVLAEAGLAEYF | ASSLRQGGSYEQY | 0.7824881 |
| DDQIVLSAKANLAKVEQ | ASSLRQGGSYEQY | 0.5812447 |
| DELLPDTGEDSDEDGH | ASSLRQGGSYEQY | 0.7982483 |
| ELPEAPANRATVPQD | ASSLRQGGSYEQY | 0.62097526 |
| FGQSKIERVTGIG | ASSLRQGGSYEQY | 0.71407175 |
| GAGGTGSTAGGAGGAGGAGGLY | ASSLRQGGSYEQY | 0.8007171 |
| GHGGTGAAGGNGGTGGAGGSGI | ASSLRQGGSYEQY | 0.80340064 |
| GNGGNGGAGGEGGPGGAGGAGA | ASSLRQGGSYEQY | 0.71130705 |
| GSTAGNGGNGGNATNGTIAGQP | ASSLRQGGSYEQY | 0.80041885 |
| GTGGDGGNGGNGGTGG | ASSLRQGGSYEQY | 0.76163447 |
| GVRRPAGIPADAPV | ASSLRQGGSYEQY | 0.80669296 |
| GYRQVTDAHLLTLARR | ASSLRQGGSYEQY | 0.7753954 |

| | | |
|---|---|---|
| IAKMGQDGHDRG | ASSLRQGGSYEQY | 0.6225681 |
| KAEPSIEIQYT | ASSLRQGGSYEQY | 0.7985097 |
| KPVRIRMAADEDTVAVT | ASSLRQGGSYEQY | 0.79425985 |
| MPPRIAGMRLLVIKPEP | ASSLRQGGSYEQY | 0.7879953 |
| MKFYAVALFRSRGVPPTR | ASSLRQGGSYEQY | 0.7993159 |
| NAGGFGILGAVAHSP | ASSLRQGGSYEQY | 0.763247 |
| NEGREITLAMAEM | ASSLRQGGSYEQY | 0.7785286 |
| NELVEWSHPLGHRWRWE | ASSLRQGGSYEQY | 0.801811 |
| NLGAGGTGGNGGNPG | ASSLRQGGSYEQY | 0.79867786 |
| NTEWPKWETVMAADPVD | ASSLRQGGSYEQY | 0.6991135 |
| PASAAPAETQP | ASSLRQGGSYEQY | 0.79865 |
| QQSSDGKNGVRAAEITGE | ASSLRQGGSYEQY | 0.77772385 |
| RAPDAVTTSWWKE | ASSLRQGGSYEQY | 0.7429471 |
| SGFGNVGAHTSGWFNQ | ASSLRQGGSYEQY | 0.80561066 |
| SGVAGAVGQAASVGGLKV | ASSLRQGGSYEQY | 0.78206486 |
| SNGGAGTGSAGGNGGT | ASSLRQGGSYEQY | 0.80162555 |
| TGGAGGEGGAAG | ASSLRQGGSYEQY | 0.8011546 |
| TGGDGGAAGAGGAPGHG | ASSLRQGGSYEQY | 0.80256546 |
| TMPPLLAAIANAGRHGV | ASSLRQGGSYEQY | 0.64174604 |
| TSSTAGNSGGA | ASSLRQGGSYEQY | 0.7875705 |
| VPATVPVGILC | ASSLRQGGSYEQY | 0.7857646 |
| VQRPDSGGVVLTGRIS | ASSLRQGGSYEQY | 0.7959368 |
| WTPNPVPATKGVPAGLVQAT | ASSLRQGGSYEQY | 0.77502596 |
| YALKRALAKFSDRFDVVII | ASSLRQGGSYEQY | 0.69888866 |
| YLIGSVIV | ASSLRQGGSYEQY | 0.79464304 |
| YGQLVAVID | ASSLRQGGSYEQY | 0.80296445 |
| VTVGGMTLPA | ASSLRQGGSYEQY | 0.7846526 |
| VLALPAV | ASSLRQGGSYEQY | 0.7854721 |
| VIPPTLNYET | ASSLRQGGSYEQY | 0.8050642 |
| VGMGGMHPSSG | ASSLRQGGSYEQY | 0.7478893 |
| VGLAGGAATA | ASSLRQGGSYEQY | 0.77383757 |
| VGGAGGAGGSGP | ASSLRQGGSYEQY | 0.79850185 |
| VFAAFMV | ASSLRQGGSYEQY | 0.80038357 |
| TLYDKLR | ASSLRQGGSYEQY | 0.7804127 |
| TLAVFTVL | ASSLRQGGSYEQY | 0.8028617 |
| TGSVQPARV | ASSLRQGGSYEQY | 0.7511129 |
| TGGGAGSGGT | ASSLRQGGSYEQY | 0.79585195 |
| SHLETSL | ASSLRQGGSYEQY | 0.75808686 |
| SFTTGMT | ASSLRQGGSYEQY | 0.79261893 |
| QVEVAKRMGVRQPTVS | ASSLRQGGSYEQY | 0.7495891 |
| QGSGGPALAM | ASSLRQGGSYEQY | 0.79337925 |
| MLAPSNP | ASSLRQGGSYEQY | 0.80024534 |
| LVNNAAI | ASSLRQGGSYEQY | 0.7961454 |
| LTGEINTLA | ASSLRQGGSYEQY | 0.35837513 |
| LSIFGPV | ASSLRQGGSYEQY | 0.80063033 |
| LLALLANL | ASSLRQGGSYEQY | 0.7957956 |
| LISNASDALDKLR | ASSLRQGGSYEQY | 0.8070347 |
| LINEAVP | ASSLRQGGSYEQY | 0.78866804 |
| LGYAFSGL | ASSLRQGGSYEQY | 0.79609936 |
| LDFVAVAL | ASSLRQGGSYEQY | 0.7690362 |
| LAQPGAE | ASSLRQGGSYEQY | 0.7858149 |

| | | |
|---|---|---|
| ISNLGAVTGF | ASSLRQGGSYEQY | 0.7385576 |
| HQEDPGRI | ASSLRQGGSYEQY | 0.7656822 |
| GTPGEPTP | ASSLRQGGSYEQY | 0.7663562 |
| GFFNSGS | ASSLRQGGSYEQY | 0.7574605 |
| GAGGGGLTAGAN | ASSLRQGGSYEQY | 0.79937816 |
| FVDALVNM | ASSLRQGGSYEQY | 0.7997978 |
| FLTVAPEL | ASSLRQGGSYEQY | 0.7988229 |
| FETLTAL | ASSLRQGGSYEQY | 0.8023138 |
| DVINPIEL | ASSLRQGGSYEQY | 0.80397606 |
| APVSGAPV | ASSLRQGGSYEQY | 0.8012426 |
| ALVGGLAW | ASSLRQGGSYEQY | 0.7725208 |
| ALAAAAATAI | ASSLRQGGSYEQY | 0.77761817 |
| AFGLGLLL | ASSLRQGGSYEQY | 0.7214933 |
| AFFAAASL | ASSLRQGGSYEQY | 0.7739544 |
| AAAALPAFSPP | ASSLRQGGSYEQY | 0.76934934 |
| VVLAPAL | ASSLRQGGSYEQY | 0.79322994 |
| TIGHVDHGKTTL | ASSLRQGGSYEQY | 0.20987384 |
| LISNASDALDK | ASSLRQGGSYEQY | 0.7865819 |
| LISNASDAL | ASSLRQGGSYEQY | 0.8042519 |
| ISNASDALDK | ASSLRQGGSYEQY | 0.7550556 |
| NQAETLVYQTE | ASSLRQGGSYEQY | 0.7957276 |
| TGGTGGTAGAG | ASSLRQGGSYEQY | 0.78770137 |
| TGSTAGGAGGA | ASSLRQGGSYEQY | 0.7672576 |
| DPLAVPVAAGPV | ASSLRQGGSYEQY | 0.79911536 |
| GGTGGTPGNGGN | ASSLRQGGSYEQY | 0.74951184 |
| PAGLAGVDPATL | ASSLRQGGSYEQY | 0.79764473 |
| GNNLVVEAPHAGASV | ASSLRQGGSYEQY | 0.80538845 |
| KVTRIMSASTLEQPA | ASSLRQGGSYEQY | 0.78696847 |
| LGFALPIALPRRTVV | ASSLRQGGSYEQY | 0.78475726 |
| TRLGNLTARKLVVLV | ASSLRQGGSYEQY | 0.78542924 |
| WSQQRRLIYFPSAGP | ASSLRQGGSYEQY | 0.5695385 |
| GPNGNPDPMAAAVDIR | ASSLRQGGSYEQY | 0.7881218 |
| TQRCVSVADLRAAASR | ASSLRQGGSYEQY | 0.37383997 |
| QFVLDVAEHPVEVLTETV | ASSLRQGGSYEQY | 0.78340685 |
| GGAGGTGGAGGNGGTQAGDG | ASSLRQGGSYEQY | 0.7503012 |
| PLLATLRAIAVDVDAKPAQV | ASSLRQGGSYEQY | 0.78979725 |
| APKFRTGGYMIDFWGVGYQVAK | ASSLRQGGSYEQY | 0.78000236 |
| GAGGAGGAGGAAGGPGGTGGTG | ASSLRQGGSYEQY | 0.79300094 |
| IGALGLLATDVAPAGSVAFGVA | ASSLRQGGSYEQY | 0.72987586 |
| AGGAGGIGGNGGAGGAGGVLMG | ASSLRQGGSYEQY | 0.78330183 |
| ADELIGLL | ASSLRQGGSYEQY | 0.75981164 |
| ADRALKA | ASSLRQGGSYEQY | 0.6804594 |
| AGAVGQVGGAGGA | ASSLRQGGSYEQY | 0.79629725 |
| AGGGGGLGGAP | ASSLRQGGSYEQY | 0.77159107 |
| AGGLLYGNGGAG | ASSLRQGGSYEQY | 0.78060585 |
| AGLESIGL | ASSLRQGGSYEQY | 0.8012308 |
| AGLKTGII | ASSLRQGGSYEQY | 0.79220957 |
| ASLAALLL | ASSLRQGGSYEQY | 0.6143595 |
| ASVINISSI | ASSLRQGGSYEQY | 0.7801222 |
| ATPPGPA | ASSLRQGGSYEQY | 0.7848165 |
| ATVASALV | ASSLRQGGSYEQY | 0.78299075 |

| | | |
|---|---|---|
| ATVASLIL | ASSLRQGGSYEQY | 0.22335829 |
| AVNGPVLIRH | ASSLRQGGSYEQY | 0.7927023 |
| DGAAGAVGI | ASSLRQGGSYEQY | 0.73909867 |
| DPLLHGI | ASSLRQGGSYEQY | 0.7382984 |
| ELGAALGAL | ASSLRQGGSYEQY | 0.7627226 |
| ELGRTQAV | ASSLRQGGSYEQY | 0.79813784 |
| FVAGGYLS | ASSLRQGGSYEQY | 0.77139986 |
| GASGGAGGAGGW | ASSLRQGGSYEQY | 0.7827107 |
| GGAGAGGDGGAGGI | ASSLRQGGSYEQY | 0.7976818 |
| GTGHNGSQPGMGGNG | ASSLRQGGSYEQY | 0.79280573 |
| GYAAASASAS | ASSLRQGGSYEQY | 0.80365145 |
| HQVDVAAVQQG | ASSLRQGGSYEQY | 0.79295325 |
| IGAAIAHT | ASSLRQGGSYEQY | 0.7801137 |
| IGMIEVAQL | ASSLRQGGSYEQY | 0.7936008 |
| IRLAPEHTSAMD | ASSLRQGGSYEQY | 0.78327703 |
| IYNVSVL | ASSLRQGGSYEQY | 0.68231016 |
| KQGFLGM | ASSLRQGGSYEQY | 0.6341108 |
| LESVITQL | ASSLRQGGSYEQY | 0.7890814 |
| LGLGALVVV | ASSLRQGGSYEQY | 0.5764018 |
| LIGNGGNGGSGGI | ASSLRQGGSYEQY | 0.78976375 |
| LNAINAPTL | ASSLRQGGSYEQY | 0.80042547 |
| LQLLNTGV | ASSLRQGGSYEQY | 0.7861131 |
| LQTSIEF | ASSLRQGGSYEQY | 0.66553223 |
| MSFAVSEL | ASSLRQGGSYEQY | 0.77330756 |
| NTGALGL | ASSLRQGGSYEQY | 0.7614101 |
| PPGAGGAGGAGTT | ASSLRQGGSYEQY | 0.8004228 |
| QLVGLGI | ASSLRQGGSYEQY | 0.79883397 |
| RFLEQVHQL | ASSLRQGGSYEQY | 0.8042238 |
| RGVGYVLR | ASSLRQGGSYEQY | 0.7992271 |
| SATGVFAGVI | ASSLRQGGSYEQY | 0.7627418 |
| SLPAITLP | ASSLRQGGSYEQY | 0.7991909 |
| STALAFTV | ASSLRQGGSYEQY | 0.7779616 |
| SVLTVGLGK | ASSLRQGGSYEQY | 0.77190924 |
| TALAGAGSGI | ASSLRQGGSYEQY | 0.7794354 |
| TGLPFTGI | ASSLRQGGSYEQY | 0.79201186 |
| VAEHVTV | ASSLRQGGSYEQY | 0.76818454 |
| VDIVLNSL | ASSLRQGGSYEQY | 0.76559293 |
| VLTGGGAVL | ASSLRQGGSYEQY | 0.5827384 |
| VNVGLAYGI | ASSLRQGGSYEQY | 0.7948679 |
| VSSAAVGAV | ASSLRQGGSYEQY | 0.79464614 |
| VTSVYEAGV | ASSLRQGGSYEQY | 0.69700706 |
| WVALDRGA | ASSLRQGGSYEQY | 0.76757777 |
| YLVGSVVI | ASSLRQGGSYEQY | 0.80543655 |
| GPPGTGKTLL | ASSLRQGGSYEQY | 0.8044712 |
| GVGMTKF | ASSLRQGGSYEQY | 0.7530253 |
| IGIGGSDL | ASSLRQGGSYEQY | 0.75690854 |
| IGTLLAEL | ASSLRQGGSYEQY | 0.7881874 |
| IYTSGTT | ASSLRQGGSYEQY | 0.65297186 |
| LSFPAGV | ASSLRQGGSYEQY | 0.7943182 |
| LYTSGTT | ASSLRQGGSYEQY | 0.7825637 |
| VGAFPFGF | ASSLRQGGSYEQY | 0.77539325 |

| | | |
|---|---|---|
| VGAGTVEF | ASSLRQGGSYEQY | 0.7520615 |
| VGKSTLLL | ASSLRQGGSYEQY | 0.786512 |
| AALAHAGGGIV | ASSLRQGGSYEQY | 0.75859725 |
| AALVAVSLL | ASSLRQGGSYEQY | 0.79681873 |
| AAVSRAVAAL | ASSLRQGGSYEQY | 0.2774192 |
| AGGSSGIGGF | ASSLRQGGSYEQY | 0.7866978 |
| AGLTVANANG | ASSLRQGGSYEQY | 0.7409371 |
| AGLVGAVL | ASSLRQGGSYEQY | 0.7666433 |
| ALSQAFTV | ASSLRQGGSYEQY | 0.57063377 |
| ALTFSVSL | ASSLRQGGSYEQY | 0.77433866 |
| ALTGAALVAL | ASSLRQGGSYEQY | 0.79928815 |
| ASFSIGAGVL | ASSLRQGGSYEQY | 0.80099344 |
| DGSVYKI | ASSLRQGGSYEQY | 0.7441733 |
| EQLAEIV | ASSLRQGGSYEQY | 0.797866 |
| FAVSELTVM | ASSLRQGGSYEQY | 0.80590713 |
| GAAPLAAGL | ASSLRQGGSYEQY | 0.7749105 |
| GADGGDGGSGGAGG | ASSLRQGGSYEQY | 0.4871648 |
| GFGLDTGGAGG | ASSLRQGGSYEQY | 0.7924557 |
| GLASGGLL | ASSLRQGGSYEQY | 0.7731833 |
| GRIAGAAVLL | ASSLRQGGSYEQY | 0.79740465 |
| GSAGSLNTSF | ASSLRQGGSYEQY | 0.79419655 |
| GTGQIGI | ASSLRQGGSYEQY | 0.71939707 |
| GTLIALIL | ASSLRQGGSYEQY | 0.7970321 |
| IAATMGFVVG | ASSLRQGGSYEQY | 0.79884416 |
| IFAQALL | ASSLRQGGSYEQY | 0.75233877 |
| IGAAVALGF | ASSLRQGGSYEQY | 0.7508899 |
| IGLTPLAL | ASSLRQGGSYEQY | 0.70639133 |
| ILGSNTSTL | ASSLRQGGSYEQY | 0.4963444 |
| IVFSLTVL | ASSLRQGGSYEQY | 0.79302216 |
| LASKATGF | ASSLRQGGSYEQY | 0.76022243 |
| LASVALIL | ASSLRQGGSYEQY | 0.7518159 |
| LFAVGLTL | ASSLRQGGSYEQY | 0.7878123 |
| LFNGGIGTYI | ASSLRQGGSYEQY | 0.7420428 |
| LTLGAGII | ASSLRQGGSYEQY | 0.7606528 |
| LVAAALGDI | ASSLRQGGSYEQY | 0.7709761 |
| NTAFVQLGI | ASSLRQGGSYEQY | 0.77837896 |
| QLFSGAEL | ASSLRQGGSYEQY | 0.7930635 |
| SAATFITL | ASSLRQGGSYEQY | 0.77427495 |
| SNYGLANLGV | ASSLRQGGSYEQY | 0.77845216 |
| SYGGAYAVM | ASSLRQGGSYEQY | 0.80316186 |
| TTGVGGLTLG | ASSLRQGGSYEQY | 0.67007685 |
| TTQTIYI | ASSLRQGGSYEQY | 0.73025197 |
| VAAGAGLGAVI | ASSLRQGGSYEQY | 0.79667294 |
| VAAVVTLL | ASSLRQGGSYEQY | 0.7097314 |
| VGFLGGLI | ASSLRQGGSYEQY | 0.7962353 |
| VGVIPILL | ASSLRQGGSYEQY | 0.7800757 |
| VIFASSLI | ASSLRQGGSYEQY | 0.7950704 |
| VKGGDGGAAAG | ASSLRQGGSYEQY | 0.7865309 |
| VTKTFTNI | ASSLRQGGSYEQY | 0.8055645 |
| VVAAAILL | ASSLRQGGSYEQY | 0.80046105 |
| YAALDVEL | ASSLRQGGSYEQY | 0.76387537 |

| | | |
|---|---|---|
| LASVALLL | ASSLRQGGSYEQY | 0.7970519 |
| LDAVHRV | ASSLRQGGSYEQY | 0.66204315 |
| QGLSSSIF | ASSLRQGGSYEQY | 0.79970825 |
| RGTFANIRL | ASSLRQGGSYEQY | 0.7988622 |
| RQLWWGHRI | ASSLRQGGSYEQY | 0.73996186 |
| YGPPGTGKTLL | ASSLRQGGSYEQY | 0.78016114 |
| AALVGAPANVA | ASSLRQGGSYEQY | 0.8042177 |
| AATFRRTE | ASSLRQGGSYEQY | 0.8016088 |
| AGALGGGI | ASSLRQGGSYEQY | 0.6192975 |
| AGGAGGAGGIGAG | ASSLRQGGSYEQY | 0.76622427 |
| AGSGALGL | ASSLRQGGSYEQY | 0.6213043 |
| AQIADQI | ASSLRQGGSYEQY | 0.7683134 |
| ASAVAVTYL | ASSLRQGGSYEQY | 0.6278647 |
| ASTVLGGTGG | ASSLRQGGSYEQY | 0.7979471 |
| AVGGEPAK | ASSLRQGGSYEQY | 0.7716921 |
| CEQGTFGGTC | ASSLRQGGSYEQY | 0.7961973 |
| DEGKAPAG | ASSLRQGGSYEQY | 0.78985274 |
| DTVVFPAL | ASSLRQGGSYEQY | 0.804539 |
| FEGITMAG | ASSLRQGGSYEQY | 0.76883304 |
| FPLVVQD | ASSLRQGGSYEQY | 0.7734784 |
| GAFTTPQ | ASSLRQGGSYEQY | 0.6139991 |
| GFGGFGTAGGVGGA | ASSLRQGGSYEQY | 0.628584 |
| GFLAQFL | ASSLRQGGSYEQY | 0.74829435 |
| GGAGGAGGNAGVA | ASSLRQGGSYEQY | 0.7719274 |
| GGVGTAGAGGAGGAGG | ASSLRQGGSYEQY | 0.46379027 |
| GNGGGGDAGNAGSGGN | ASSLRQGGSYEQY | 0.80307233 |
| GTGGAGGAGNDAGS | ASSLRQGGSYEQY | 0.7924943 |
| GTTNTGWFNS | ASSLRQGGSYEQY | 0.7992387 |
| IFMGPLFTL | ASSLRQGGSYEQY | 0.7396387 |
| ILAHAIEAI | ASSLRQGGSYEQY | 0.7601346 |
| IRSAIREAL | ASSLRQGGSYEQY | 0.7697583 |
| ITFAGGG | ASSLRQGGSYEQY | 0.79647493 |
| ITGMSGHQEV | ASSLRQGGSYEQY | 0.7515855 |
| IVAGLACTL | ASSLRQGGSYEQY | 0.57224596 |
| IVGGGLAGA | ASSLRQGGSYEQY | 0.7360129 |
| IVTGVLNI | ASSLRQGGSYEQY | 0.69441736 |
| LAQQGCI | ASSLRQGGSYEQY | 0.8050978 |
| LCLHVEN | ASSLRQGGSYEQY | 0.8011324 |
| LDAISDGS | ASSLRQGGSYEQY | 0.7933922 |
| LGGFAAML | ASSLRQGGSYEQY | 0.80365807 |
| LGGKNPMI | ASSLRQGGSYEQY | 0.37209964 |
| LGPKGVAAIL | ASSLRQGGSYEQY | 0.7454938 |
| LMVGVSI | ASSLRQGGSYEQY | 0.8029078 |
| LVGAAFVL | ASSLRQGGSYEQY | 0.79523146 |
| LVLAVAPL | ASSLRQGGSYEQY | 0.77790725 |
| MAAIAVIDAA | ASSLRQGGSYEQY | 0.8043244 |
| NNGINAELQ | ASSLRQGGSYEQY | 0.74577665 |
| RGEPLVLVHG | ASSLRQGGSYEQY | 0.78250825 |
| RGGAGGAAGLI | ASSLRQGGSYEQY | 0.78353524 |
| RSYRMHPAV | ASSLRQGGSYEQY | 0.79441446 |
| SAAHLLAGQLA | ASSLRQGGSYEQY | 0.78971195 |

| | | |
|---|---|---|
| SAAVGSY | ASSLRQGGSYEQY | 0.7252735 |
| SPHAVLHRI | ASSLRQGGSYEQY | 0.7933265 |
| SQIAKVL | ASSLRQGGSYEQY | 0.80591285 |
| TALGHLVP | ASSLRQGGSYEQY | 0.77531135 |
| TAPTTSGAA | ASSLRQGGSYEQY | 0.80079854 |
| TDGDDGNGGSPGT | ASSLRQGGSYEQY | 0.79473007 |
| TDGLGEN | ASSLRQGGSYEQY | 0.7913281 |
| TGGPPLPGQA | ASSLRQGGSYEQY | 0.7542263 |
| TIPAITI | ASSLRQGGSYEQY | 0.76956487 |
| TIPGFTI | ASSLRQGGSYEQY | 0.78398246 |
| TSIASGALA | ASSLRQGGSYEQY | 0.7395524 |
| TVTLPGAAI | ASSLRQGGSYEQY | 0.7810969 |
| VASGLSAALP | ASSLRQGGSYEQY | 0.76628476 |
| VGAGVIGI | ASSLRQGGSYEQY | 0.80796105 |
| VGPKRFGV | ASSLRQGGSYEQY | 0.79768515 |
| VNVAFPD | ASSLRQGGSYEQY | 0.7951002 |
| VSMTNSGVS | ASSLRQGGSYEQY | 0.74733865 |
| VTDVLGDLSV | ASSLRQGGSYEQY | 0.789709 |
| VVVVGAGFAGL | ASSLRQGGSYEQY | 0.794929 |
| VYTVGTVV | ASSLRQGGSYEQY | 0.61672056 |
| WTTFSGP | ASSLRQGGSYEQY | 0.8039104 |
| YDELVEAF | ASSLRQGGSYEQY | 0.755012 |
| YGTGGAGGNGG | ASSLRQGGSYEQY | 0.771463 |
| YTNVMAV | ASSLRQGGSYEQY | 0.7967969 |
| FLEQVHQL | ASSLRQGGSYEQY | 0.80164576 |
| GLSLLAVAV | ASSLRQGGSYEQY | 0.78065395 |
| LEQVHQL | ASSLRQGGSYEQY | 0.78120965 |
| LNAAGAI | ASSLRQGGSYEQY | 0.7439063 |
| PAVHASL | ASSLRQGGSYEQY | 0.8021698 |
| TELEDDA | ASSLRQGGSYEQY | 0.8023823 |
| TPPGPPN | ASSLRQGGSYEQY | 0.7901225 |
| TPPPPPPP | ASSLRQGGSYEQY | 0.8054861 |
| TQSPALI | SSSPGQDTQY | 0.767755 |
| EFAAAFPQ | SSSPGQDTQY | 0.55814874 |
| LLSGNLDVL | SSSPGQDTQY | 0.78851116 |
| GPANLGNYNI | SSSPGQDTQY | 0.79873496 |
| TLGPTLP | SSSPGQDTQY | 0.80176944 |
| VYAYSSCCK | SSSPGQDTQY | 0.80433667 |
| APAPAPAPNI | SSSPGQDTQY | 0.789673 |
| PGGPVPIP | SSSPGQDTQY | 0.7730119 |
| FLTVALEP | SSSPGQDTQY | 0.80344474 |
| SQYFGANL | SSSPGQDTQY | 0.8075046 |
| GGVKPAA | SSSPGQDTQY | 0.79963326 |
| GGTGGAGGAGGW | SSSPGQDTQY | 0.7726294 |
| GVGGVAGLGGAGG | SSSPGQDTQY | 0.75458324 |
| VLLVLATTI | SSSPGQDTQY | 0.78901184 |
| IGIFAAML | SSSPGQDTQY | 0.8009441 |
| AVGDLID | SSSPGQDTQY | 0.737764 |
| NISPLPL | SSSPGQDTQY | 0.791332 |
| VTAVFVAI | SSSPGQDTQY | 0.79377794 |
| SGDDNDAAL | SSSPGQDTQY | 0.6096124 |

| | | |
|---|---|---|
| AVAALAHS | SSSPGQDTQY | 0.7896848 |
| GGVTVMATP | SSSPGQDTQY | 0.802639 |
| AGLAANVRD | SSSPGQDTQY | 0.78172016 |
| TNRPDLIDPALLRPG | SSSPGQDTQY | 0.7588862 |
| LANLGVTAL | SSSPGQDTQY | 0.8013252 |
| TGERIEL | SSSPGQDTQY | 0.8076396 |
| NGEELAA | SSSPGQDTQY | 0.8041864 |
| VLSLVWG | SSSPGQDTQY | 0.5389736 |
| IGYALWL | SSSPGQDTQY | 0.49078038 |
| LSLLLAPL | SSSPGQDTQY | 0.76367676 |
| TLDEALK | SSSPGQDTQY | 0.7965964 |
| TDGAGTAEGG | SSSPGQDTQY | 0.7618493 |
| NAPGGAGGNGG | SSSPGQDTQY | 0.75734484 |
| VSVFMGMGGNF | SSSPGQDTQY | 0.7926557 |
| LASYLAAL | SSSPGQDTQY | 0.80166113 |
| LVGGTPAL | SSSPGQDTQY | 0.74034786 |
| VRNADVL | SSSPGQDTQY | 0.67347074 |
| TCSSPFLVSAM | SSSPGQDTQY | 0.78814673 |
| VHITDAE | SSSPGQDTQY | 0.79917246 |
| EYAAPSF | SSSPGQDTQY | 0.72933644 |
| VVDAFHGGL | SSSPGQDTQY | 0.78124684 |
| ASVVDPQ | SSSPGQDTQY | 0.75622547 |
| SYNEFGGLS | SSSPGQDTQY | 0.7622248 |
| DPDGSVY | SSSPGQDTQY | 0.7888946 |
| MVSTIHATE | SSSPGQDTQY | 0.7413856 |
| LAAGVADPV | SSSPGQDTQY | 0.8003453 |
| TKEQIAA | SSSPGQDTQY | 0.7459451 |
| SLASPAAL | SSSPGQDTQY | 0.79986054 |
| IGLVFYGV | SSSPGQDTQY | 0.7890376 |
| FLLDASNL | SSSPGQDTQY | 0.7904036 |
| YQVGNFP | SSSPGQDTQY | 0.778008 |
| ESLLSQST | SSSPGQDTQY | 0.7637379 |
| LFTVGFAAVG | SSSPGQDTQY | 0.7093357 |
| ATPPTPT | SSSPGQDTQY | 0.6602993 |
| TDLATLG | SSSPGQDTQY | 0.8000406 |
| SAGGTGR | SSSPGQDTQY | 0.7129347 |
| GFSNSGS | SSSPGQDTQY | 0.801394 |
| GGNGGSGGTSV | SSSPGQDTQY | 0.7998218 |
| GTSVATGGAGN | SSSPGQDTQY | 0.72741306 |
| NGGNGGSGGTS | SSSPGQDTQY | 0.79592407 |
| AVVKFPH | SSSPGQDTQY | 0.73518926 |
| TPPPPPP | SSSPGQDTQY | 0.76683354 |
| AAPAAAAAPA | SSSPGQDTQY | 0.79821193 |
| IALNEDL | SSSPGQDTQY | 0.7944902 |
| IVNVSSM | SSSPGQDTQY | 0.74629474 |
| TGIPPAPR | SSSPGQDTQY | 0.7962797 |
| FDLGGGTF | SSSPGQDTQY | 0.79031104 |
| FDLGGGTFDVS | SSSPGQDTQY | 0.7544831 |
| NEPTAAA | SSSPGQDTQY | 0.7772735 |
| NEPTAAALA | SSSPGQDTQY | 0.80100954 |
| GSIINISSI | SSSPGQDTQY | 0.7928598 |

| | | |
|---|---|---|
| VAPGNAGTA | SSSPGQDTQY | 0.75352645 |
| TGGGASLE | SSSPGQDTQY | 0.79492015 |
| VAVYNHYKRI | SSSPGQDTQY | 0.74913 |
| TDPAPATT | SSSPGQDTQY | 0.7914697 |
| TNRPDILDPALLRPG | SSSPGQDTQY | 0.7851606 |
| LTTTTTS | SSSPGQDTQY | 0.7765454 |
| PTGSGKTL | SSSPGQDTQY | 0.7967325 |
| TGSGKTL | SSSPGQDTQY | 0.6358218 |
| TIGHVDHGK | SSSPGQDTQY | 0.754809 |
| TIGHVDHGKTT | SSSPGQDTQY | 0.7762324 |
| GHVDHGKTT | SSSPGQDTQY | 0.78335345 |
| AQFEHTLL | SSSPGQDTQY | 0.7989638 |
| IVNQDEL | SSSPGQDTQY | 0.7433839 |
| TNGSQFF | SSSPGQDTQY | 0.7889192 |
| TGKGIVE | SSSPGQDTQY | 0.77295375 |
| LISNASDA | SSSPGQDTQY | 0.7398126 |
| AELRTLGELTKIAKAHGAQ | SSSPGQDTQY | 0.78343475 |
| APNAVAAAEPSW | SSSPGQDTQY | 0.76873004 |
| APQVAGAATAVL | SSSPGQDTQY | 0.75173116 |
| ARRDPAAAAAILPTDA | SSSPGQDTQY | 0.7920451 |
| DAAARDVLAEAGLAEYF | SSSPGQDTQY | 0.728096 |
| DDQIVLSAKANLAKVEQ | SSSPGQDTQY | 0.79098797 |
| DELLPDTGEDSDEDGH | SSSPGQDTQY | 0.803622 |
| ELPEAPANRATVPQD | SSSPGQDTQY | 0.76572466 |
| FGQSKIERVTGIG | SSSPGQDTQY | 0.79953265 |
| GAGGTGSTAGGAGGAGGAGGLY | SSSPGQDTQY | 0.79400575 |
| GHGGTGAAGGNGGTGGAGGSGI | SSSPGQDTQY | 0.79180086 |
| GNGGNGGAGGEGGPGGAGGAGA | SSSPGQDTQY | 0.7775787 |
| GSTAGNGGNGGNATNGTIAGQP | SSSPGQDTQY | 0.75393593 |
| GTGGDGGNGGNGGTGG | SSSPGQDTQY | 0.8011343 |
| GVRRPAGIPADAPV | SSSPGQDTQY | 0.7950375 |
| GYRQVTDAHLLTLARR | SSSPGQDTQY | 0.79778636 |
| IAKMGQDGHDRG | SSSPGQDTQY | 0.79217875 |
| KAEPSIEIQYT | SSSPGQDTQY | 0.79475975 |
| KPVRIRMAADEDTVAVT | SSSPGQDTQY | 0.7974088 |
| MPPRIAGMRLLVIKPEP | SSSPGQDTQY | 0.762673 |
| MKFYAVALFRSRGVPPTR | SSSPGQDTQY | 0.69220495 |
| NAGGFGILGAVAHSP | SSSPGQDTQY | 0.7712741 |
| NEGREITLAMAEM | SSSPGQDTQY | 0.77702546 |
| NELVEWSHPLGHRWRWE | SSSPGQDTQY | 0.760523 |
| NLGAGGTGGNGGNPG | SSSPGQDTQY | 0.55901706 |
| NTEWPKWETVMAADPVD | SSSPGQDTQY | 0.7983905 |
| PASAAPAETQP | SSSPGQDTQY | 0.8074373 |
| QQSSDGKNGVRAAEITGE | SSSPGQDTQY | 0.7418928 |
| RAPDAVTTSWWKE | SSSPGQDTQY | 0.80614305 |
| SGFGNVGAHTSGWFNQ | SSSPGQDTQY | 0.6971878 |
| SGVAGAVGQAASVGGLKV | SSSPGQDTQY | 0.7973747 |
| SNGGAGTGSAGGNGGT | SSSPGQDTQY | 0.80097437 |
| TGGAGGEGGAAG | SSSPGQDTQY | 0.74150586 |
| TGGDGGAAGAGGAPGHG | SSSPGQDTQY | 0.7935659 |
| TMPPLLAAIANAGRHGV | SSSPGQDTQY | 0.7657354 |

| | | |
|---|---|---|
| TSSTAGNSGGA | SSSPGQDTQY | 0.7712021 |
| VPATVPVGILC | SSSPGQDTQY | 0.80363727 |
| VQRPDSGGVVLTGRIS | SSSPGQDTQY | 0.7617105 |
| WTPNPVPATKGVPAGLVQAT | SSSPGQDTQY | 0.7883499 |
| YALKRALAKFSDRFDVVII | SSSPGQDTQY | 0.77565587 |
| YLIGSVIV | SSSPGQDTQY | 0.7533814 |
| YGQLVAVID | SSSPGQDTQY | 0.7881212 |
| VTVGGMTLPA | SSSPGQDTQY | 0.80154335 |
| VLALPAV | SSSPGQDTQY | 0.7731985 |
| VIPPTLNYET | SSSPGQDTQY | 0.806978 |
| VGMGGMHPSSG | SSSPGQDTQY | 0.750706 |
| VGLAGGAATA | SSSPGQDTQY | 0.80382323 |
| VGGAGGAGGSGP | SSSPGQDTQY | 0.79960036 |
| VFAAFMV | SSSPGQDTQY | 0.78651494 |
| TLYDKLR | SSSPGQDTQY | 0.7758591 |
| TLAVFTVL | SSSPGQDTQY | 0.80147713 |
| TGSVQPARV | SSSPGQDTQY | 0.77683485 |
| TGGGAGSGGT | SSSPGQDTQY | 0.7932541 |
| SHLETSL | SSSPGQDTQY | 0.802441 |
| SFTTGMT | SSSPGQDTQY | 0.15984875 |
| QVEVAKRMGVRQPTVS | SSSPGQDTQY | 0.65119123 |
| QGSGGPALAM | SSSPGQDTQY | 0.3186649 |
| MLAPSNP | SSSPGQDTQY | 0.19858241 |
| LVNNAAI | SSSPGQDTQY | 0.20463856 |
| LTGEINTLA | SSSPGQDTQY | 0.1406419 |
| LSIFGPV | SSSPGQDTQY | 0.19253455 |
| LLALLANL | SSSPGQDTQY | 0.18102723 |
| LISNASDALDKLR | SSSPGQDTQY | 0.37997884 |
| LINEAVP | SSSPGQDTQY | 0.217082 |
| LGYAFSGL | SSSPGQDTQY | 0.20347416 |
| LDFVAVAL | SSSPGQDTQY | 0.6156424 |
| LAQPGAE | SSSPGQDTQY | 0.46852726 |
| ISNLGAVTGF | SSSPGQDTQY | 0.15050575 |
| HQEDPGRI | SSSPGQDTQY | 0.21535502 |
| GTPGEPTP | SSSPGQDTQY | 0.16942704 |
| GFFNSGS | SSSPGQDTQY | 0.18923397 |
| GAGGGGLTAGAN | SSSPGQDTQY | 0.13203178 |
| FVDALVNM | SSSPGQDTQY | 0.31503284 |
| FLTVAPEL | SSSPGQDTQY | 0.16327523 |
| FETLTAL | SSSPGQDTQY | 0.12875038 |
| DVINPIEL | SSSPGQDTQY | 0.18267946 |
| APVSGAPV | SSSPGQDTQY | 0.18329419 |
| ALVGGLAW | SSSPGQDTQY | 0.19562009 |
| ALAAAAATAI | SSSPGQDTQY | 0.5350983 |
| AFGLGLLL | SSSPGQDTQY | 0.13708426 |
| AFFAAASL | SSSPGQDTQY | 0.17121433 |
| AAAALPAFSPP | SSSPGQDTQY | 0.53114235 |
| VVLAPAL | SSSPGQDTQY | 0.66735893 |
| TIGHVDHGKTTL | SSSPGQDTQY | 0.15766479 |
| LISNASDALDK | SSSPGQDTQY | 0.34920982 |
| LISNASDAL | SSSPGQDTQY | 0.61877716 |

| | | |
|---|---|---|
| ISNASDALDK | SSSPGQDTQY | 0.30372256 |
| NQAETLVYQTE | SSSPGQDTQY | 0.2829749 |
| TGGTGGTAGAG | SSSPGQDTQY | 0.3353244 |
| TGSTAGGAGGA | SSSPGQDTQY | 0.15019101 |
| DPLAVPVAAGPV | SSSPGQDTQY | 0.48333284 |
| GGTGGTPGNGGN | SSSPGQDTQY | 0.6664804 |
| PAGLAGVDPATL | SSSPGQDTQY | 0.22079669 |
| GNNLVVEAPHAGASV | SSSPGQDTQY | 0.23399863 |
| KVTRIMSASTLEQPA | SSSPGQDTQY | 0.15890117 |
| LGFALPIALPRRTVV | SSSPGQDTQY | 0.15891775 |
| TRLGNLTARKLVVLV | SSSPGQDTQY | 0.134655 |
| WSQQRRLIYFPSAGP | SSSPGQDTQY | 0.46884456 |
| GPNGNPDPMAAAVDIR | SSSPGQDTQY | 0.40040404 |
| TQRCVSVADLRAAASR | SSSPGQDTQY | 0.23277393 |
| QFVLDVAEHPVEVLTETV | SSSPGQDTQY | 0.1474524 |
| GGAGGTGGAGGNGGTQAGDG | SSSPGQDTQY | 0.1708936 |
| PLLATLRAIAVDVDAKPAQV | SSSPGQDTQY | 0.18912494 |
| APKFRTGGYMIDFWGVGYQVAK | SSSPGQDTQY | 0.18880853 |
| GAGGAGGAGGAAGGPGGTGGTG | SSSPGQDTQY | 0.13786729 |
| IGALGLLATDVAPAGSVAFGVA | SSSPGQDTQY | 0.15388784 |
| AGGAGGIGGNGGAGGAGGVLMG | SSSPGQDTQY | 0.53654075 |
| ADELIGLL | SSSPGQDTQY | 0.5317799 |
| ADRALKA | SSSPGQDTQY | 0.14555785 |
| AGAVGQVGGAGGA | SSSPGQDTQY | 0.5851704 |
| AGGGGGLGGAP | SSSPGQDTQY | 0.36968598 |
| AGGLLYGNGGAG | SSSPGQDTQY | 0.59714437 |
| AGLESIGL | SSSPGQDTQY | 0.23391998 |
| AGLKTGII | SSSPGQDTQY | 0.16637273 |
| ASLAALLL | SSSPGQDTQY | 0.39507377 |
| ASVINISSI | SSSPGQDTQY | 0.26138592 |
| ATPPGPA | SSSPGQDTQY | 0.15551229 |
| ATVASALV | SSSPGQDTQY | 0.2813912 |
| ATVASLIL | SSSPGQDTQY | 0.23484707 |
| AVNGPVLIRH | SSSPGQDTQY | 0.19846246 |
| DGAAGAVGI | SSSPGQDTQY | 0.32772148 |
| DPLLHGI | SSSPGQDTQY | 0.28895897 |
| ELGAALGAL | SSSPGQDTQY | 0.15494187 |
| ELGRTQAV | SSSPGQDTQY | 0.16606037 |
| FVAGGYLS | SSSPGQDTQY | 0.15583625 |
| GASGGAGGAGGW | SSSPGQDTQY | 0.3852346 |
| GGAGAGGDGGAGGI | SSSPGQDTQY | 0.5761392 |
| GTGHNGSQPGMGGNG | SSSPGQDTQY | 0.23216441 |
| GYAAASASAS | SSSPGQDTQY | 0.18672332 |
| HQVDVAAVQQG | SSSPGQDTQY | 0.52473086 |
| IGAAIAHT | SSSPGQDTQY | 0.30252934 |
| IGMIEVAQL | SSSPGQDTQY | 0.22304548 |
| IRLAPEHTSAMD | SSSPGQDTQY | 0.26824003 |
| IYNVSVL | SSSPGQDTQY | 0.2167379 |
| KQGFLGM | SSSPGQDTQY | 0.29485223 |
| LESVITQL | SSSPGQDTQY | 0.26464847 |
| LGLGALVVV | SSSPGQDTQY | 0.25473475 |

| | | |
|---|---|---|
| LIGNGGNGGSGGI | SSSPGQDTQY | 0.27917016 |
| LNAINAPTL | SSSPGQDTQY | 0.29187468 |
| LQLLNTGV | SSSPGQDTQY | 0.22595716 |
| LQTSIEF | SSSPGQDTQY | 0.13872758 |
| MSFAVSEL | SSSPGQDTQY | 0.17013131 |
| NTGALGL | SSSPGQDTQY | 0.23365432 |
| PPGAGGAGGAGTT | SSSPGQDTQY | 0.33340758 |
| QLVGLGI | SSSPGQDTQY | 0.21805978 |
| RFLEQVHQL | SSSPGQDTQY | 0.2541628 |
| RGVGYVLR | SSSPGQDTQY | 0.17597735 |
| SATGVFAGVI | SSSPGQDTQY | 0.26639017 |
| SLPAITLP | SSSPGQDTQY | 0.5230409 |
| STALAFTV | SSSPGQDTQY | 0.14890131 |
| SVLTVGLGK | SSSPGQDTQY | 0.1808974 |
| TALAGAGSGI | SSSPGQDTQY | 0.6350227 |
| TGLPFTGI | SSSPGQDTQY | 0.3592462 |
| VAEHVTV | SSSPGQDTQY | 0.24822119 |
| VDIVLNSL | SSSPGQDTQY | 0.16963582 |
| VLTGGGAVL | SSSPGQDTQY | 0.63887453 |
| VNVGLAYGI | SSSPGQDTQY | 0.14516942 |
| VSSAAVGAV | SSSPGQDTQY | 0.5573546 |
| VTSVYEAGV | SSSPGQDTQY | 0.13416171 |
| WVALDRGA | SSSPGQDTQY | 0.18347967 |
| YLVGSVVI | SSSPGQDTQY | 0.17836191 |
| GPPGTGKTLL | SSSPGQDTQY | 0.13543692 |
| GVGMTKF | SSSPGQDTQY | 0.4382121 |
| IGIGGSDL | SSSPGQDTQY | 0.15976283 |
| IGTLLAEL | SSSPGQDTQY | 0.33006537 |
| IYTSGTT | SSSPGQDTQY | 0.20856702 |
| LSFPAGV | SSSPGQDTQY | 0.2611112 |
| LYTSGTT | SSSPGQDTQY | 0.2768926 |
| VGAFPFGF | SSSPGQDTQY | 0.17721625 |
| VGAGTVEF | SSSPGQDTQY | 0.15921041 |
| VGKSTLLL | SSSPGQDTQY | 0.40243644 |
| AALAHAGGGIV | SSSPGQDTQY | 0.26081342 |
| AALVAVSLL | SSSPGQDTQY | 0.23552202 |
| AAVSRAVAAL | SSSPGQDTQY | 0.1481956 |
| AGGSSGIGGF | SSSPGQDTQY | 0.14714363 |
| AGLTVANANG | SSSPGQDTQY | 0.35232812 |
| AGLVGAVL | SSSPGQDTQY | 0.13512602 |
| ALSQAFTV | SSSPGQDTQY | 0.1291725 |
| ALTFSVSL | SSSPGQDTQY | 0.19386053 |
| ALTGAALVAL | SSSPGQDTQY | 0.15247059 |
| ASFSIGAGVL | SSSPGQDTQY | 0.58799434 |
| DGSVYKI | SSSPGQDTQY | 0.5928179 |
| EQLAEIV | SSSPGQDTQY | 0.22540885 |
| FAVSELTVM | SSSPGQDTQY | 0.452058 |
| GAAPLAAGL | SSSPGQDTQY | 0.30372417 |
| GADGGDGGSGGAGG | SSSPGQDTQY | 0.12710781 |
| GFGLDTGGAGG | SSSPGQDTQY | 0.19046935 |
| GLASGGLL | SSSPGQDTQY | 0.150323 |

| | | |
|---|---|---|
| GRIAGAAVLL | SSSPGQDTQY | 0.2962987 |
| GSAGSLNTSF | SSSPGQDTQY | 0.14672692 |
| GTGQIGI | SSSPGQDTQY | 0.17371777 |
| GTLIALIL | SSSPGQDTQY | 0.27771884 |
| IAATMGFVVG | SSSPGQDTQY | 0.4170214 |
| IFAQALL | SSSPGQDTQY | 0.27071112 |
| IGAAVALGF | SSSPGQDTQY | 0.15932962 |
| IGLTPLAL | SSSPGQDTQY | 0.12881383 |
| ILGSNTSTL | SSSPGQDTQY | 0.40428293 |
| IVFSLTVL | SSSPGQDTQY | 0.14083676 |
| LASKATGF | SSSPGQDTQY | 0.17447431 |
| LASVALIL | SSSPGQDTQY | 0.13046661 |
| LFAVGLTL | SSSPGQDTQY | 0.13177994 |
| LFNGGIGTYI | SSSPGQDTQY | 0.1476337 |
| LTLGAGII | SSSPGQDTQY | 0.25123918 |
| LVAAALGDI | SSSPGQDTQY | 0.22463962 |
| NTAFVQLGI | SSSPGQDTQY | 0.22711244 |
| QLFSGAEL | SSSPGQDTQY | 0.6931168 |
| SAATFITL | SSSPGQDTQY | 0.6014197 |
| SNYGLANLGV | SSSPGQDTQY | 0.211293 |
| SYGGAYAVM | SSSPGQDTQY | 0.20093851 |
| TTGVGGLTLG | SSSPGQDTQY | 0.15833478 |
| TTQTIYI | SSSPGQDTQY | 0.1673661 |
| VAAGAGLGAVI | SSSPGQDTQY | 0.16688609 |
| VAAVVTLL | SSSPGQDTQY | 0.5579935 |
| VGFLGGLI | SSSPGQDTQY | 0.16899857 |
| VGVIPILL | SSSPGQDTQY | 0.14616331 |
| VIFASSLI | SSSPGQDTQY | 0.23916173 |
| VKGGDGGAAAG | SSSPGQDTQY | 0.4818893 |
| VTKTFTNI | SSSPGQDTQY | 0.7184648 |
| VVAAAILL | SSSPGQDTQY | 0.16088839 |
| YAALDVEL | SSSPGQDTQY | 0.22071937 |
| LASVALLL | SSSPGQDTQY | 0.17980188 |
| LDAVHRV | SSSPGQDTQY | 0.36730337 |
| QGLSSSIF | SSSPGQDTQY | 0.1601051 |
| RGTFANIRL | SSSPGQDTQY | 0.16075075 |
| RQLWWGHRI | SSSPGQDTQY | 0.29761308 |
| YGPPGTGKTLL | SSSPGQDTQY | 0.70965767 |
| AALVGAPANVA | SSSPGQDTQY | 0.17861743 |
| AATFRRTE | SSSPGQDTQY | 0.14374588 |
| AGALGGGI | SSSPGQDTQY | 0.29512787 |
| AGGAGGAGGIGAG | SSSPGQDTQY | 0.18076177 |
| AGSGALGL | SSSPGQDTQY | 0.235227 |
| AQIADQI | SSSPGQDTQY | 0.27926925 |
| ASAVAVTYL | SSSPGQDTQY | 0.14224775 |
| ASTVLGGTGG | SSSPGQDTQY | 0.1654102 |
| AVGGEPAK | SSSPGQDTQY | 0.7184919 |
| CEQGTFGGTC | SSSPGQDTQY | 0.31952536 |
| DEGKAPAG | SSSPGQDTQY | 0.2698759 |
| DTVVFPAL | SSSPGQDTQY | 0.5903673 |
| FEGITMAG | SSSPGQDTQY | 0.39117873 |

| | | |
|---|---|---|
| FPLVVQD | SSSPGQDTQY | 0.63387024 |
| GAFTTPQ | SSSPGQDTQY | 0.36179906 |
| GFGGFGTAGGVGGA | SSSPGQDTQY | 0.1769218 |
| GFLAQFL | SSSPGQDTQY | 0.22455156 |
| GGAGGAGGNAGVA | SSSPGQDTQY | 0.22590318 |
| GGVGTAGAGGAGGAGG | SSSPGQDTQY | 0.29774874 |
| GNGGGGDAGNAGSGGN | SSSPGQDTQY | 0.26390135 |
| GTGGAGGAGNDAGS | SSSPGQDTQY | 0.16189913 |
| GTTNTGWFNS | SSSPGQDTQY | 0.6641861 |
| IFMGPLFTL | SSSPGQDTQY | 0.26876587 |
| ILAHAIEAI | SSSPGQDTQY | 0.2552994 |
| IRSAIREAL | SSSPGQDTQY | 0.14614686 |
| ITFAGGG | SSSPGQDTQY | 0.16013147 |
| ITGMSGHQEV | SSSPGQDTQY | 0.44196895 |
| IVAGLACTL | SSSPGQDTQY | 0.13008454 |
| IVGGGLAGA | SSSPGQDTQY | 0.13205428 |
| IVTGVLNI | SSSPGQDTQY | 0.23669806 |
| LAQQGCI | SSSPGQDTQY | 0.61661375 |
| LCLHVEN | SSSPGQDTQY | 0.12941794 |
| LDAISDGS | SSSPGQDTQY | 0.16996415 |
| LGGFAAML | SSSPGQDTQY | 0.14184408 |
| LGGKNPMI | SSSPGQDTQY | 0.23545972 |
| LGPKGVAAIL | SSSPGQDTQY | 0.25695762 |
| LMVGVSI | SSSPGQDTQY | 0.14148648 |
| LVGAAFVL | SSSPGQDTQY | 0.26216656 |
| LVLAVAPL | SSSPGQDTQY | 0.13824734 |
| MAAIAVIDAA | SSSPGQDTQY | 0.13993882 |
| NNGINAELQ | SSSPGQDTQY | 0.24583182 |
| RGEPLVLVHG | SSSPGQDTQY | 0.42991483 |
| RGGAGGAAGLI | SSSPGQDTQY | 0.14326052 |
| RSYRMHPAV | SSSPGQDTQY | 0.4074164 |
| SAAHLLAGQLA | SSSPGQDTQY | 0.18072283 |
| SAAVGSY | SSSPGQDTQY | 0.72223854 |
| SPHAVLHRI | SSSPGQDTQY | 0.23136693 |
| SQIAKVL | SSSPGQDTQY | 0.14010163 |
| TALGHLVP | SSSPGQDTQY | 0.39178717 |
| TAPTTSGAA | SSSPGQDTQY | 0.23374847 |
| TDGDDGNGGSPGT | SSSPGQDTQY | 0.21957414 |
| TDGLGEN | SSSPGQDTQY | 0.45359164 |
| TGGPPLPGQA | SSSPGQDTQY | 0.18249637 |
| TIPAITI | SSSPGQDTQY | 0.30052602 |
| TIPGFTI | SSSPGQDTQY | 0.5648382 |
| TSIASGALA | SSSPGQDTQY | 0.28235537 |
| TVTLPGAAI | SSSPGQDTQY | 0.16176066 |
| VASGLSAALP | SSSPGQDTQY | 0.21952535 |
| VGAGVIGI | SSSPGQDTQY | 0.31142014 |
| VGPKRFGV | SSSPGQDTQY | 0.16822055 |
| VNVAFPD | SSSPGQDTQY | 0.5823258 |
| VSMTNSGVS | SSSPGQDTQY | 0.2093592 |
| VTDVLGDLSV | SSSPGQDTQY | 0.38848484 |
| VVVVGAGFAGL | SSSPGQDTQY | 0.31007177 |

| | | |
|---|---|---|
| VYTVGTVV | SSSPGQDTQY | 0.72019815 |
| WTTFSGP | SSSPGQDTQY | 0.14571881 |
| YDELVEAF | SSSPGQDTQY | 0.23434618 |
| YGTGGAGGNGG | SSSPGQDTQY | 0.29595748 |
| YTNVMAV | SSSPGQDTQY | 0.34055197 |
| FLEQVHQL | SSSPGQDTQY | 0.18280786 |
| GLSLLAVAV | SSSPGQDTQY | 0.15180236 |
| LEQVHQL | SSSPGQDTQY | 0.20366342 |
| LNAAGAI | SSSPGQDTQY | 0.5228529 |
| PAVHASL | SSSPGQDTQY | 0.18850733 |
| TELEDDA | SSSPGQDTQY | 0.20816757 |
| TPPGPPN | SSSPGQDTQY | 0.70772374 |
| TPPPPPPP | SSSPGQDTQY | 0.1589698 |
| TQSPALI | GAVLYNSPLY | 0.21956754 |
| EFAAAFPQ | GAVLYNSPLY | 0.28317192 |
| LLSGNLDVL | GAVLYNSPLY | 0.3223545 |
| GPANLGNYNI | GAVLYNSPLY | 0.19491656 |
| TLGPTLP | GAVLYNSPLY | 0.41592482 |
| VYAYSSCCK | GAVLYNSPLY | 0.1596704 |
| APAPAPAPNI | GAVLYNSPLY | 0.4690426 |
| PGGPVPIP | GAVLYNSPLY | 0.16508242 |
| FLTVALEP | GAVLYNSPLY | 0.34547883 |
| SQYFGANL | GAVLYNSPLY | 0.17320402 |
| GGVKPAA | GAVLYNSPLY | 0.5214822 |
| GGTGGAGGAGGW | GAVLYNSPLY | 0.5945034 |
| GVGGVAGLGGAGG | GAVLYNSPLY | 0.49404073 |
| VLLVLATTI | GAVLYNSPLY | 0.12934406 |
| IGIFAAML | GAVLYNSPLY | 0.57401145 |
| AVGDLID | GAVLYNSPLY | 0.54387796 |
| NISPLPL | GAVLYNSPLY | 0.7458018 |
| VTAVFVAI | GAVLYNSPLY | 0.23811245 |
| SGDDNDAAL | GAVLYNSPLY | 0.29627132 |
| AVAALAHS | GAVLYNSPLY | 0.3097629 |
| GGVTVMATP | GAVLYNSPLY | 0.19101535 |
| AGLAANVRD | GAVLYNSPLY | 0.1654844 |
| TNRPDLIDPALLRPG | GAVLYNSPLY | 0.23206311 |
| LANLGVTAL | GAVLYNSPLY | 0.27896795 |
| TGERIEL | GAVLYNSPLY | 0.21099994 |
| NGEELAA | GAVLYNSPLY | 0.53348225 |
| VLSLVWG | GAVLYNSPLY | 0.38970208 |
| IGYALWL | GAVLYNSPLY | 0.53341866 |
| LSLLLAPL | GAVLYNSPLY | 0.7017062 |
| TLDEALK | GAVLYNSPLY | 0.5569835 |
| TDGAGTAEGG | GAVLYNSPLY | 0.23771752 |
| NAPGGAGGNGG | GAVLYNSPLY | 0.18516187 |
| VSVFMGMGGNF | GAVLYNSPLY | 0.19252215 |
| LASYLAAL | GAVLYNSPLY | 0.15300517 |
| LVGGTPAL | GAVLYNSPLY | 0.17467761 |
| VRNADVL | GAVLYNSPLY | 0.197927 |
| TCSSPFLVSAM | GAVLYNSPLY | 0.2713024 |
| VHITDAE | GAVLYNSPLY | 0.12795337 |

| | | |
|---|---|---|
| EYAAPSF | GAVLYNSPLY | 0.31385988 |
| VVDAFHGGL | GAVLYNSPLY | 0.5203449 |
| ASVVDPQ | GAVLYNSPLY | 0.15844704 |
| SYNEFGGLS | GAVLYNSPLY | 0.32103264 |
| DPDGSVY | GAVLYNSPLY | 0.18722446 |
| MVSTIHATE | GAVLYNSPLY | 0.15618828 |
| LAAGVADPV | GAVLYNSPLY | 0.65184295 |
| TKEQIAA | GAVLYNSPLY | 0.15663227 |
| SLASPAAL | GAVLYNSPLY | 0.26150948 |
| IGLVFYGV | GAVLYNSPLY | 0.62753123 |
| FLLDASNL | GAVLYNSPLY | 0.2823144 |
| YQVGNFP | GAVLYNSPLY | 0.19638678 |
| ESLLSQST | GAVLYNSPLY | 0.18525344 |
| LFTVGFAAVG | GAVLYNSPLY | 0.1443772 |
| ATPPTPT | GAVLYNSPLY | 0.2346856 |
| TDLATLG | GAVLYNSPLY | 0.12710261 |
| SAGGTGR | GAVLYNSPLY | 0.34392 |
| GFSNSGS | GAVLYNSPLY | 0.1563011 |
| GGNGGSGGTSV | GAVLYNSPLY | 0.4178562 |
| GTSVATGGAGN | GAVLYNSPLY | 0.32373872 |
| NGGNGGSGGTS | GAVLYNSPLY | 0.22993304 |
| AVVKFPH | GAVLYNSPLY | 0.18712658 |
| TPPPPPP | GAVLYNSPLY | 0.17104664 |
| AAPAAAAAPA | GAVLYNSPLY | 0.16857141 |
| IALNEDL | GAVLYNSPLY | 0.15702863 |
| IVNVSSM | GAVLYNSPLY | 0.27099708 |
| TGIPPAPR | GAVLYNSPLY | 0.19653004 |
| FDLGGGTF | GAVLYNSPLY | 0.2063938 |
| FDLGGGTFDVS | GAVLYNSPLY | 0.48802888 |
| NEPTAAA | GAVLYNSPLY | 0.21809472 |
| NEPTAAALA | GAVLYNSPLY | 0.14378704 |
| GSIINISSI | GAVLYNSPLY | 0.44918787 |
| VAPGNAGTA | GAVLYNSPLY | 0.29150206 |
| TGGGASLE | GAVLYNSPLY | 0.24364835 |
| VAVYNHYKRI | GAVLYNSPLY | 0.12682036 |
| TDPAPATT | GAVLYNSPLY | 0.21178828 |
| TNRPDILDPALLRPG | GAVLYNSPLY | 0.1681291 |
| LTTTTTS | GAVLYNSPLY | 0.21118373 |
| PTGSGKTL | GAVLYNSPLY | 0.16412172 |
| TGSGKTL | GAVLYNSPLY | 0.61498284 |
| TIGHVDHGK | GAVLYNSPLY | 0.14217837 |
| TIGHVDHGKTT | GAVLYNSPLY | 0.37603673 |
| GHVDHGKTT | GAVLYNSPLY | 0.31507587 |
| AQFEHTLL | GAVLYNSPLY | 0.3644883 |
| IVNQDEL | GAVLYNSPLY | 0.6274785 |
| TNGSQFF | GAVLYNSPLY | 0.3432022 |
| TGKGIVE | GAVLYNSPLY | 0.21034703 |
| LISNASDA | GAVLYNSPLY | 0.19798462 |
| AELRTLGELTKIAKAHGAQ | GAVLYNSPLY | 0.2175551 |
| APNAVAAAEPSW | GAVLYNSPLY | 0.1326184 |
| APQVAGAATAVL | GAVLYNSPLY | 0.13344103 |

| | | |
|---|---|---|
| ARRDPAAAAAILPTDA | GAVLYNSPLY | 0.3267437 |
| DAAARDVLAEAGLAEYF | GAVLYNSPLY | 0.13254519 |
| DDQIVLSAKANLAKVEQ | GAVLYNSPLY | 0.32503873 |
| DELLPDTGEDSDEDGH | GAVLYNSPLY | 0.2747463 |
| ELPEAPANRATVPQD | GAVLYNSPLY | 0.23347491 |
| FGQSKIERVTGIG | GAVLYNSPLY | 0.14621364 |
| GAGGTGSTAGGAGGAGGAGGLY | GAVLYNSPLY | 0.19065383 |
| GHGGTGAAGGNGGTGGAGGSGI | GAVLYNSPLY | 0.20046861 |
| GNGGNGGAGGEGGPGGAGGAGA | GAVLYNSPLY | 0.40467754 |
| GSTAGNGGNGGNATNGTIAGQP | GAVLYNSPLY | 0.26995558 |
| GTGGDGGNGGNGGTGG | GAVLYNSPLY | 0.71665955 |
| GVRRPAGIPADAPV | GAVLYNSPLY | 0.6245415 |
| GYRQVTDAHLLTLARR | GAVLYNSPLY | 0.23668417 |
| IAKMGQDGHDRG | GAVLYNSPLY | 0.39207554 |
| KAEPSIEIQYT | GAVLYNSPLY | 0.18075684 |
| KPVRIRMAADEDTVAVT | GAVLYNSPLY | 0.15578151 |
| MPPRIAGMRLLVIKPEP | GAVLYNSPLY | 0.1713869 |
| MKFYAVALFRSRGVPPTR | GAVLYNSPLY | 0.2310851 |
| NAGGFGILGAVAHSP | GAVLYNSPLY | 0.17786096 |
| NEGREITLAMAEM | GAVLYNSPLY | 0.15406239 |
| NELVEWSHPLGHRWRWE | GAVLYNSPLY | 0.14768666 |
| NLGAGGTGGNGGNPG | GAVLYNSPLY | 0.19547659 |
| NTEWPKWETVMAADPVD | GAVLYNSPLY | 0.31603545 |
| PASAAPAETQP | GAVLYNSPLY | 0.13812171 |
| QQSSDGKNGVRAAEITGE | GAVLYNSPLY | 0.16036016 |
| RAPDAVTTSWWKE | GAVLYNSPLY | 0.7386551 |
| SGFGNVGAHTSGWFNQ | GAVLYNSPLY | 0.7476231 |
| SGVAGAVGQAASVGGLKV | GAVLYNSPLY | 0.19334342 |
| SNGGAGTGSAGGNGGT | GAVLYNSPLY | 0.18048434 |
| TGGAGGEGGAAG | GAVLYNSPLY | 0.3399825 |
| TGGDGGAAGAGGAPGHG | GAVLYNSPLY | 0.14236684 |
| TMPPLLAAIANAGRHGV | GAVLYNSPLY | 0.20564383 |
| TSSTAGNSGGA | GAVLYNSPLY | 0.38508314 |
| VPATVPVGILC | GAVLYNSPLY | 0.23667729 |
| VQRPDSGGVVLTGRIS | GAVLYNSPLY | 0.15789174 |
| WTPNPVPATKGVPAGLVQAT | GAVLYNSPLY | 0.23181602 |
| YALKRALAKFSDRFDVVII | GAVLYNSPLY | 0.21629031 |
| YLIGSVIV | GAVLYNSPLY | 0.25288188 |
| YGQLVAVID | GAVLYNSPLY | 0.1286044 |
| VTVGGMTLPA | GAVLYNSPLY | 0.19958702 |
| VLALPAV | GAVLYNSPLY | 0.16628508 |
| VIPPTLNYET | GAVLYNSPLY | 0.18039176 |
| VGMGGMHPSSG | GAVLYNSPLY | 0.14222333 |
| VGLAGGAATA | GAVLYNSPLY | 0.16725436 |
| VGGAGGAGGSGP | GAVLYNSPLY | 0.67197376 |
| VFAAFMV | GAVLYNSPLY | 0.50045407 |
| TLYDKLR | GAVLYNSPLY | 0.13465425 |
| TLAVFTVL | GAVLYNSPLY | 0.42918777 |
| TGSVQPARV | GAVLYNSPLY | 0.7155546 |
| TGGGAGSGGT | GAVLYNSPLY | 0.18979985 |
| SHLETSL | GAVLYNSPLY | 0.13792145 |

| | | |
|---|---|---|
| SFTTGMT | GAVLYNSPLY | 0.30090287 |
| QVEVAKRMGVRQPTVS | GAVLYNSPLY | 0.18427646 |
| QGSGGPALAM | GAVLYNSPLY | 0.31658265 |
| MLAPSNP | GAVLYNSPLY | 0.4749967 |
| LVNNAAI | GAVLYNSPLY | 0.14782144 |
| LTGEINTLA | GAVLYNSPLY | 0.35884959 |
| LSIFGPV | GAVLYNSPLY | 0.5768885 |
| LLALLANL | GAVLYNSPLY | 0.1599719 |
| LISNASDALDKLR | GAVLYNSPLY | 0.15236507 |
| LINEAVP | GAVLYNSPLY | 0.1515379 |
| LGYAFSGL | GAVLYNSPLY | 0.13148965 |
| LDFVAVAL | GAVLYNSPLY | 0.26655108 |
| LAQPGAE | GAVLYNSPLY | 0.18858811 |
| ISNLGAVTGF | GAVLYNSPLY | 0.19054808 |
| HQEDPGRI | GAVLYNSPLY | 0.22984767 |
| GTPGEPTP | GAVLYNSPLY | 0.17505416 |
| GFFNSGS | GAVLYNSPLY | 0.18686056 |
| GAGGGGLTAGAN | GAVLYNSPLY | 0.20776863 |
| FVDALVNM | GAVLYNSPLY | 0.22484091 |
| FLTVAPEL | GAVLYNSPLY | 0.300636 |
| FETLTAL | GAVLYNSPLY | 0.25094494 |
| DVINPIEL | GAVLYNSPLY | 0.17893502 |
| APVSGAPV | GAVLYNSPLY | 0.69504887 |
| ALVGGLAW | GAVLYNSPLY | 0.15443179 |
| ALAAAAATAI | GAVLYNSPLY | 0.15975468 |
| AFGLGLLL | GAVLYNSPLY | 0.35970646 |
| AFFAAASL | GAVLYNSPLY | 0.14293167 |
| AAAALPAFSPP | GAVLYNSPLY | 0.40482152 |
| VVLAPAL | GAVLYNSPLY | 0.28932124 |
| TIGHVDHGKTTL | GAVLYNSPLY | 0.27923906 |
| LISNASDALDK | GAVLYNSPLY | 0.23711497 |
| LISNASDAL | GAVLYNSPLY | 0.6529608 |
| ISNASDALDK | GAVLYNSPLY | 0.48198366 |
| NQAETLVYQTE | GAVLYNSPLY | 0.16612512 |
| TGGTGGTAGAG | GAVLYNSPLY | 0.22181892 |
| TGSTAGGAGGA | GAVLYNSPLY | 0.15196231 |
| DPLAVPVAAGPV | GAVLYNSPLY | 0.31441373 |
| GGTGGTPGNGGN | GAVLYNSPLY | 0.3056653 |
| PAGLAGVDPATL | GAVLYNSPLY | 0.17778899 |
| GNNLVVEAPHAGASV | GAVLYNSPLY | 0.30879766 |
| KVTRIMSASTLEQPA | GAVLYNSPLY | 0.34109125 |
| LGFALPIALPRRTVV | GAVLYNSPLY | 0.43104696 |
| TRLGNLTARKLVVLV | GAVLYNSPLY | 0.13884917 |
| WSQQRRLIYFPSAGP | GAVLYNSPLY | 0.20343709 |
| GPNGNPDPMAAAVDIR | GAVLYNSPLY | 0.14348488 |
| TQRCVSVADLRAAASR | GAVLYNSPLY | 0.20936267 |
| QFVLDVAEHPVEVLTETV | GAVLYNSPLY | 0.15451433 |
| GGAGGTGGAGGNGGTQAGDG | GAVLYNSPLY | 0.4241715 |
| PLLATLRAIAVDVDAKPAQV | GAVLYNSPLY | 0.23764515 |
| APKFRTGGYMIDFWGVGYQVAK | GAVLYNSPLY | 0.366692 |
| GAGGAGGAGGAAGGPGGTGGTG | GAVLYNSPLY | 0.33038488 |

| | | |
|---|---|---|
| IGALGLLATDVAPAGSVAFGVA | GAVLYNSPLY | 0.5204 |
| AGGAGGIGGNGGAGGAGGVLMG | GAVLYNSPLY | 0.18696894 |
| ADELIGLL | GAVLYNSPLY | 0.25802717 |
| ADRALKA | GAVLYNSPLY | 0.14218745 |
| AGAVGQVGGAGGA | GAVLYNSPLY | 0.14077765 |
| AGGGGGLGGAP | GAVLYNSPLY | 0.15322272 |
| AGGLLYGNGGAG | GAVLYNSPLY | 0.21243648 |
| AGLESIGL | GAVLYNSPLY | 0.13165127 |
| AGLKTGII | GAVLYNSPLY | 0.5777827 |
| ASLAALLL | GAVLYNSPLY | 0.32755142 |
| ASVINISSI | GAVLYNSPLY | 0.4260278 |
| ATPPGPA | GAVLYNSPLY | 0.1805459 |
| ATVASALV | GAVLYNSPLY | 0.15837444 |
| ATVASLIL | GAVLYNSPLY | 0.18680976 |
| AVNGPVLIRH | GAVLYNSPLY | 0.6058552 |
| DGAAGAVGI | GAVLYNSPLY | 0.14932822 |
| DPLLHGI | GAVLYNSPLY | 0.13934278 |
| ELGAALGAL | GAVLYNSPLY | 0.17157581 |
| ELGRTQAV | GAVLYNSPLY | 0.14750612 |
| FVAGGYLS | GAVLYNSPLY | 0.4666082 |
| GASGGAGGAGGW | GAVLYNSPLY | 0.48262995 |
| GGAGAGGDGGAGGI | GAVLYNSPLY | 0.36813435 |
| GTGHNGSQPGMGGNG | GAVLYNSPLY | 0.24169391 |
| GYAAASASAS | GAVLYNSPLY | 0.13290028 |
| HQVDVAAVQQG | GAVLYNSPLY | 0.1869269 |
| IGAAIAHT | GAVLYNSPLY | 0.70007944 |
| IGMIEVAQL | GAVLYNSPLY | 0.34158605 |
| IRLAPEHTSAMD | GAVLYNSPLY | 0.24862996 |
| IYNVSVL | GAVLYNSPLY | 0.44572425 |
| KQGFLGM | GAVLYNSPLY | 0.1916618 |
| LESVITQL | GAVLYNSPLY | 0.24032387 |
| LGLGALVVV | GAVLYNSPLY | 0.23591956 |
| LIGNGGNGGSGGI | GAVLYNSPLY | 0.1851768 |
| LNAINAPTL | GAVLYNSPLY | 0.19878227 |
| LQLLNTGV | GAVLYNSPLY | 0.14603876 |
| LQTSIEF | GAVLYNSPLY | 0.250628 |
| MSFAVSEL | GAVLYNSPLY | 0.7521491 |
| NTGALGL | GAVLYNSPLY | 0.35013247 |
| PPGAGGAGGAGTT | GAVLYNSPLY | 0.49647176 |
| QLVGLGI | GAVLYNSPLY | 0.2865587 |
| RFLEQVHQL | GAVLYNSPLY | 0.6928235 |
| RGVGYVLR | GAVLYNSPLY | 0.20516966 |
| SATGVFAGVI | GAVLYNSPLY | 0.1688557 |
| SLPAITLP | GAVLYNSPLY | 0.16839218 |
| STALAFTV | GAVLYNSPLY | 0.1701782 |
| SVLTVGLGK | GAVLYNSPLY | 0.21259926 |
| TALAGAGSGI | GAVLYNSPLY | 0.18709067 |
| TGLPFTGI | GAVLYNSPLY | 0.7806031 |
| VAEHVTV | GAVLYNSPLY | 0.35061127 |
| VDIVLNSL | GAVLYNSPLY | 0.2662471 |
| VLTGGGAVL | GAVLYNSPLY | 0.17330785 |

| | | |
|---|---|---|
| VNVGLAYGI | GAVLYNSPLY | 0.19566143 |
| VSSAAVGAV | GAVLYNSPLY | 0.2729705 |
| VTSVYEAGV | GAVLYNSPLY | 0.14924781 |
| WVALDRGA | GAVLYNSPLY | 0.60047704 |
| YLVGSVVI | GAVLYNSPLY | 0.17465931 |
| GPPGTGKTLL | GAVLYNSPLY | 0.20078631 |
| GVGMTKF | GAVLYNSPLY | 0.30286622 |
| IGIGGSDL | GAVLYNSPLY | 0.66773087 |
| IGTLLAEL | GAVLYNSPLY | 0.17423774 |
| IYTSGTT | GAVLYNSPLY | 0.18453409 |
| LSFPAGV | GAVLYNSPLY | 0.16415776 |
| LYTSGTT | GAVLYNSPLY | 0.39079678 |
| VGAFPFGF | GAVLYNSPLY | 0.6468059 |
| VGAGTVEF | GAVLYNSPLY | 0.2848534 |
| VGKSTLLL | GAVLYNSPLY | 0.4481933 |
| AALAHAGGGIV | GAVLYNSPLY | 0.22299328 |
| AALVAVSLL | GAVLYNSPLY | 0.14955018 |
| AAVSRAVAAL | GAVLYNSPLY | 0.20235758 |
| AGGSSGIGGF | GAVLYNSPLY | 0.24511114 |
| AGLTVANANG | GAVLYNSPLY | 0.62355727 |
| AGLVGAVL | GAVLYNSPLY | 0.5526526 |
| ALSQAFTV | GAVLYNSPLY | 0.1864704 |
| ALTFSVSL | GAVLYNSPLY | 0.18699013 |
| ALTGAALVAL | GAVLYNSPLY | 0.62760854 |
| ASFSIGAGVL | GAVLYNSPLY | 0.611435 |
| DGSVYKI | GAVLYNSPLY | 0.37398538 |
| EQLAEIV | GAVLYNSPLY | 0.19279565 |
| FAVSELTVM | GAVLYNSPLY | 0.19101755 |
| GAAPLAAGL | GAVLYNSPLY | 0.20701005 |
| GADGGDGGSGGAGG | GAVLYNSPLY | 0.5083046 |
| GFGLDTGGAGG | GAVLYNSPLY | 0.15643057 |
| GLASGGLL | GAVLYNSPLY | 0.21178766 |
| GRIAGAAVLL | GAVLYNSPLY | 0.43067306 |
| GSAGSLNTSF | GAVLYNSPLY | 0.14148684 |
| GTGQIGI | GAVLYNSPLY | 0.31298968 |
| GTLIALIL | GAVLYNSPLY | 0.55180025 |
| IAATMGFVVG | GAVLYNSPLY | 0.17270209 |
| IFAQALL | GAVLYNSPLY | 0.17759082 |
| IGAAVALGF | GAVLYNSPLY | 0.597413 |
| IGLTPLAL | GAVLYNSPLY | 0.78429264 |
| ILGSNTSTL | GAVLYNSPLY | 0.34557694 |
| IVFSLTVL | GAVLYNSPLY | 0.14649497 |
| LASKATGF | GAVLYNSPLY | 0.13310409 |
| LASVALIL | GAVLYNSPLY | 0.15851717 |
| LFAVGLTL | GAVLYNSPLY | 0.47256163 |
| LFNGGIGTYI | GAVLYNSPLY | 0.18443745 |
| LTLGAGII | GAVLYNSPLY | 0.18212132 |
| LVAAALGDI | GAVLYNSPLY | 0.35568032 |
| NTAFVQLGI | GAVLYNSPLY | 0.532236 |
| QLFSGAEL | GAVLYNSPLY | 0.16012102 |
| SAATFITL | GAVLYNSPLY | 0.15507254 |

| | | |
|---|---|---|
| SNYGLANLGV | GAVLYNSPLY | 0.18462794 |
| SYGGAYAVM | GAVLYNSPLY | 0.36825794 |
| TTGVGGLTLG | GAVLYNSPLY | 0.15378988 |
| TTQTIYI | GAVLYNSPLY | 0.30596387 |
| VAAGAGLGAVI | GAVLYNSPLY | 0.2117293 |
| VAAVVTLL | GAVLYNSPLY | 0.14647979 |
| VGFLGGLI | GAVLYNSPLY | 0.3201959 |
| VGVIPILL | GAVLYNSPLY | 0.17735197 |
| VIFASSLI | GAVLYNSPLY | 0.62162805 |
| VKGGDGGAAAG | GAVLYNSPLY | 0.16949801 |
| VTKTFTNI | GAVLYNSPLY | 0.28652504 |
| VVAAAILL | GAVLYNSPLY | 0.2509458 |
| YAALDVEL | GAVLYNSPLY | 0.23274826 |
| LASVALLL | GAVLYNSPLY | 0.24709147 |
| LDAVHRV | GAVLYNSPLY | 0.16315672 |
| QGLSSSIF | GAVLYNSPLY | 0.15242085 |
| RGTFANIRL | GAVLYNSPLY | 0.13274166 |
| RQLWWGHRI | GAVLYNSPLY | 0.360547 |
| YGPPGTGKTLL | GAVLYNSPLY | 0.12917437 |
| AALVGAPANVA | GAVLYNSPLY | 0.2836569 |
| AATFRRTE | GAVLYNSPLY | 0.38349292 |
| AGALGGGI | GAVLYNSPLY | 0.15329547 |
| AGGAGGAGGIGAG | GAVLYNSPLY | 0.23189075 |
| AGSGALGL | GAVLYNSPLY | 0.17477533 |
| AQIADQI | GAVLYNSPLY | 0.14724363 |
| ASAVAVTYL | GAVLYNSPLY | 0.5951549 |
| ASTVLGGTGG | GAVLYNSPLY | 0.2359996 |
| AVGGEPAK | GAVLYNSPLY | 0.17638578 |
| CEQGTFGGTC | GAVLYNSPLY | 0.21461418 |
| DEGKAPAG | GAVLYNSPLY | 0.33651453 |
| DTVVFPAL | GAVLYNSPLY | 0.17362379 |
| FEGITMAG | GAVLYNSPLY | 0.17503677 |
| FPLVVQD | GAVLYNSPLY | 0.5746961 |
| GAFTTPQ | GAVLYNSPLY | 0.1949977 |
| GFGGFGTAGGVGGA | GAVLYNSPLY | 0.16541126 |
| GFLAQFL | GAVLYNSPLY | 0.43521324 |
| GGAGGAGGNAGVA | GAVLYNSPLY | 0.15356956 |
| GGVGTAGAGGAGGAGG | GAVLYNSPLY | 0.3869357 |
| GNGGGGDAGNAGSGGN | GAVLYNSPLY | 0.17897204 |
| GTGGAGGAGNDAGS | GAVLYNSPLY | 0.22879237 |
| GTTNTGWFNS | GAVLYNSPLY | 0.33567888 |
| IFMGPLFTL | GAVLYNSPLY | 0.15978509 |
| ILAHAIEAI | GAVLYNSPLY | 0.17364015 |
| IRSAIREAL | GAVLYNSPLY | 0.1784527 |
| ITFAGGG | GAVLYNSPLY | 0.38388124 |
| ITGMSGHQEV | GAVLYNSPLY | 0.42774284 |
| IVAGLACTL | GAVLYNSPLY | 0.18268716 |
| IVGGGLAGA | GAVLYNSPLY | 0.30250287 |
| IVTGVLNI | GAVLYNSPLY | 0.19767849 |
| LAQQGCI | GAVLYNSPLY | 0.17352426 |
| LCLHVEN | GAVLYNSPLY | 0.19527654 |

| | | |
|---|---|---|
| LDAISDGS | GAVLYNSPLY | 0.17417115 |
| LGGFAAML | GAVLYNSPLY | 0.17391594 |
| LGGKNPMI | GAVLYNSPLY | 0.19809568 |
| LGPKGVAAIL | GAVLYNSPLY | 0.17214854 |
| LMVGVSI | GAVLYNSPLY | 0.25028548 |
| LVGAAFVL | GAVLYNSPLY | 0.528501 |
| LVLAVAPL | GAVLYNSPLY | 0.17635283 |
| MAAIAVIDAA | GAVLYNSPLY | 0.26610047 |
| NNGINAELQ | GAVLYNSPLY | 0.37841958 |
| RGEPLVLVHG | GAVLYNSPLY | 0.23839027 |
| RGGAGGAAGLI | GAVLYNSPLY | 0.20747308 |
| RSYRMHPAV | GAVLYNSPLY | 0.15850517 |
| SAAHLLAGQLA | GAVLYNSPLY | 0.43074295 |
| SAAVGSY | GAVLYNSPLY | 0.40545171 |
| SPHAVLHRI | GAVLYNSPLY | 0.3349107 |
| SQIAKVL | GAVLYNSPLY | 0.2567558 |
| TALGHLVP | GAVLYNSPLY | 0.24052301 |
| TAPTTSGAA | GAVLYNSPLY | 0.32964742 |
| TDGDDGNGGSPGT | GAVLYNSPLY | 0.23595592 |
| TDGLGEN | GAVLYNSPLY | 0.14054845 |
| TGGPPLPGQA | GAVLYNSPLY | 0.21082321 |
| TIPAITI | GAVLYNSPLY | 0.3077223 |
| TIPGFTI | GAVLYNSPLY | 0.1600196 |
| TSIASGALA | GAVLYNSPLY | 0.14420624 |
| TVTLPGAAI | GAVLYNSPLY | 0.5912936 |
| VASGLSAALP | GAVLYNSPLY | 0.76126 |
| VGAGVIGI | GAVLYNSPLY | 0.17271826 |
| VGPKRFGV | GAVLYNSPLY | 0.5253762 |
| VNVAFPD | GAVLYNSPLY | 0.14412092 |
| VSMTNSGVS | GAVLYNSPLY | 0.3329795 |
| VTDVLGDLSV | GAVLYNSPLY | 0.52294487 |
| VVVVGAGFAGL | GAVLYNSPLY | 0.17124778 |
| VYTVGTVV | GAVLYNSPLY | 0.30925685 |
| WTTFSGP | GAVLYNSPLY | 0.15593758 |
| YDELVEAF | GAVLYNSPLY | 0.17251611 |
| YGTGGAGGNGG | GAVLYNSPLY | 0.50565624 |
| YTNVMAV | GAVLYNSPLY | 0.20176354 |
| FLEQVHQL | GAVLYNSPLY | 0.17214552 |
| GLSLLAVAV | GAVLYNSPLY | 0.1925884 |
| LEQVHQL | GAVLYNSPLY | 0.16262534 |
| LNAAGAI | GAVLYNSPLY | 0.17968482 |
| PAVHASL | GAVLYNSPLY | 0.34827685 |
| TELEDDA | GAVLYNSPLY | 0.14786929 |
| TPPGPPN | GAVLYNSPLY | 0.6001462 |
| TPPPPPPP | GAVLYNSPLY | 0.15691644 |
| TQSPALI | ASSLSTGVNQDTQY | 0.7353591 |
| EFAAAFPQ | ASSLSTGVNQDTQY | 0.42039305 |
| LLSGNLDVL | ASSLSTGVNQDTQY | 0.3862584 |
| GPANLGNYNI | ASSLSTGVNQDTQY | 0.17145517 |
| TLGPTLP | ASSLSTGVNQDTQY | 0.6999704 |
| VYAYSSCCK | ASSLSTGVNQDTQY | 0.24226238 |

| | | |
|---|---|---|
| APAPAPAPNI | ASSLSTGVNQDTQY | 0.24816862 |
| PGGPVPIP | ASSLSTGVNQDTQY | 0.2689023 |
| FLTVALEP | ASSLSTGVNQDTQY | 0.11172806 |
| SQYFGANL | ASSLSTGVNQDTQY | 0.1451256 |
| GGVKPAA | ASSLSTGVNQDTQY | 0.10924599 |
| GGTGGAGGAGGW | ASSLSTGVNQDTQY | 0.124544896 |
| GVGGVAGLGGAGG | ASSLSTGVNQDTQY | 0.122196704 |
| VLLVLATTI | ASSLSTGVNQDTQY | 0.10760597 |
| IGIFAAML | ASSLSTGVNQDTQY | 0.11619247 |
| AVGDLID | ASSLSTGVNQDTQY | 0.115942344 |
| NISPLPL | ASSLSTGVNQDTQY | 0.10880755 |
| VTAVFVAI | ASSLSTGVNQDTQY | 0.118174806 |
| SGDDNDAAL | ASSLSTGVNQDTQY | 0.123391755 |
| AVAALAHS | ASSLSTGVNQDTQY | 0.13735747 |
| GGVTVMATP | ASSLSTGVNQDTQY | 0.14070776 |
| AGLAANVRD | ASSLSTGVNQDTQY | 0.11038168 |
| TNRPDLIDPALLRPG | ASSLSTGVNQDTQY | 0.1135817 |
| LANLGVTAL | ASSLSTGVNQDTQY | 0.1146931 |
| TGERIEL | ASSLSTGVNQDTQY | 0.11718211 |
| NGEELAA | ASSLSTGVNQDTQY | 0.10521192 |
| VLSLVWG | ASSLSTGVNQDTQY | 0.12153442 |
| IGYALWL | ASSLSTGVNQDTQY | 0.106181465 |
| LSLLLAPL | ASSLSTGVNQDTQY | 0.10412981 |
| TLDEALK | ASSLSTGVNQDTQY | 0.116130605 |
| TDGAGTAEGG | ASSLSTGVNQDTQY | 0.1199439 |
| NAPGGAGGNGG | ASSLSTGVNQDTQY | 0.11549947 |
| VSVFMGMGGNF | ASSLSTGVNQDTQY | 0.13317218 |
| LASYLAAL | ASSLSTGVNQDTQY | 0.10668308 |
| LVGGTPAL | ASSLSTGVNQDTQY | 0.11237119 |
| VRNADVL | ASSLSTGVNQDTQY | 0.11913765 |
| TCSSPFLVSAM | ASSLSTGVNQDTQY | 0.15469259 |
| VHITDAE | ASSLSTGVNQDTQY | 0.111796506 |
| EYAAPSF | ASSLSTGVNQDTQY | 0.11153697 |
| VVDAFHGGL | ASSLSTGVNQDTQY | 0.15627415 |
| ASVVDPQ | ASSLSTGVNQDTQY | 0.12365975 |
| SYNEFGGLS | ASSLSTGVNQDTQY | 0.117802136 |
| DPDGSVY | ASSLSTGVNQDTQY | 0.122495316 |
| MVSTIHATE | ASSLSTGVNQDTQY | 0.107266694 |
| LAAGVADPV | ASSLSTGVNQDTQY | 0.12799591 |
| TKEQIAA | ASSLSTGVNQDTQY | 0.11252293 |
| SLASPAAL | ASSLSTGVNQDTQY | 0.12876981 |
| IGLVFYGV | ASSLSTGVNQDTQY | 0.12283552 |
| FLLDASNL | ASSLSTGVNQDTQY | 0.11179328 |
| YQVGNFP | ASSLSTGVNQDTQY | 0.1098528 |
| ESLLSQST | ASSLSTGVNQDTQY | 0.10469533 |
| LFTVGFAAVG | ASSLSTGVNQDTQY | 0.11943798 |
| ATPPTPT | ASSLSTGVNQDTQY | 0.12152515 |
| TDLATLG | ASSLSTGVNQDTQY | 0.117638096 |
| SAGGTGR | ASSLSTGVNQDTQY | 0.10632886 |
| GFSNSGS | ASSLSTGVNQDTQY | 0.10962296 |
| GGNGGSGGTSV | ASSLSTGVNQDTQY | 0.10765164 |

| | | |
|---|---|---|
| GTSVATGGAGN | ASSLSTGVNQDTQY | 0.11041886 |
| NGGNGGSGGTS | ASSLSTGVNQDTQY | 0.105434224 |
| AVVKFPH | ASSLSTGVNQDTQY | 0.109746404 |
| TPPPPPP | ASSLSTGVNQDTQY | 0.15083098 |
| AAPAAAAAPA | ASSLSTGVNQDTQY | 0.14976262 |
| IALNEDL | ASSLSTGVNQDTQY | 0.12022529 |
| IVNVSSM | ASSLSTGVNQDTQY | 0.1332846 |
| TGIPPAPR | ASSLSTGVNQDTQY | 0.108778015 |
| FDLGGGTF | ASSLSTGVNQDTQY | 0.11772446 |
| FDLGGGTFDVS | ASSLSTGVNQDTQY | 0.118684165 |
| NEPTAAA | ASSLSTGVNQDTQY | 0.111961804 |
| NEPTAAALA | ASSLSTGVNQDTQY | 0.1428188 |
| GSIINISSI | ASSLSTGVNQDTQY | 0.11432941 |
| VAPGNAGTA | ASSLSTGVNQDTQY | 0.11020615 |
| TGGGASLE | ASSLSTGVNQDTQY | 0.13815098 |
| VAVYNHYKRI | ASSLSTGVNQDTQY | 0.11900333 |
| TDPAPATT | ASSLSTGVNQDTQY | 0.11570544 |
| TNRPDILDPALLRPG | ASSLSTGVNQDTQY | 0.109414704 |
| LTTTTTS | ASSLSTGVNQDTQY | 0.11657108 |
| PTGSGKTL | ASSLSTGVNQDTQY | 0.107092544 |
| TGSGKTL | ASSLSTGVNQDTQY | 0.114162594 |
| TIGHVDHGK | ASSLSTGVNQDTQY | 0.10814973 |
| TIGHVDHGKTT | ASSLSTGVNQDTQY | 0.11883293 |
| GHVDHGKTT | ASSLSTGVNQDTQY | 0.16402084 |
| AQFEHTLL | ASSLSTGVNQDTQY | 0.12391354 |
| IVNQDEL | ASSLSTGVNQDTQY | 0.10981072 |
| TNGSQFF | ASSLSTGVNQDTQY | 0.12268543 |
| TGKGIVE | ASSLSTGVNQDTQY | 0.1272866 |
| LISNASDA | ASSLSTGVNQDTQY | 0.12715581 |
| AELRTLGELTKIAKAHGAQ | ASSLSTGVNQDTQY | 0.12455369 |
| APNAVAAAEPSW | ASSLSTGVNQDTQY | 0.112705946 |
| APQVAGAATAVL | ASSLSTGVNQDTQY | 0.11850552 |
| ARRDPAAAAAILPTDA | ASSLSTGVNQDTQY | 0.13736764 |
| DAAARDVLAEAGLAEYF | ASSLSTGVNQDTQY | 0.12051182 |
| DDQIVLSAKANLAKVEQ | ASSLSTGVNQDTQY | 0.11475474 |
| DELLPDTGEDSDEDGH | ASSLSTGVNQDTQY | 0.12783939 |
| ELPEAPANRATVPQD | ASSLSTGVNQDTQY | 0.115821004 |
| FGQSKIERVTGIG | ASSLSTGVNQDTQY | 0.11101977 |
| GAGGTGSTAGGAGGAGGAGGLY | ASSLSTGVNQDTQY | 0.11043234 |
| GHGGTGAAGGNGGTGGAGGSGI | ASSLSTGVNQDTQY | 0.113366514 |
| GNGGNGGAGGEGGPGGAGGAGA | ASSLSTGVNQDTQY | 0.1140078 |
| GSTAGNGGNGGNATNGTIAGQP | ASSLSTGVNQDTQY | 0.110508665 |
| GTGGDGGNGGNGGTGG | ASSLSTGVNQDTQY | 0.12474998 |
| GVRRPAGIPADAPV | ASSLSTGVNQDTQY | 0.11109287 |
| GYRQVTDAHLLTLARR | ASSLSTGVNQDTQY | 0.11895531 |
| IAKMGQDGHDRG | ASSLSTGVNQDTQY | 0.12158753 |
| KAEPSIEIQYT | ASSLSTGVNQDTQY | 0.110454574 |
| KPVRIRMAADEDTVAVT | ASSLSTGVNQDTQY | 0.11956479 |
| MPPRIAGMRLLVIKPEP | ASSLSTGVNQDTQY | 0.13444456 |
| MKFYAVALFRSRGVPPTR | ASSLSTGVNQDTQY | 0.117724605 |
| NAGGFGILGAVAHSP | ASSLSTGVNQDTQY | 0.12092058 |

| | | |
|---|---|---|
| NEGREITLAMAEM | ASSLSTGVNQDTQY | 0.11118108 |
| NELVEWSHPLGHRWRWE | ASSLSTGVNQDTQY | 0.11794123 |
| NLGAGGTGGNGGNPG | ASSLSTGVNQDTQY | 0.1050255 |
| NTEWPKWETVMAADPVD | ASSLSTGVNQDTQY | 0.124864876 |
| PASAAPAETQP | ASSLSTGVNQDTQY | 0.10905813 |
| QQSSDGKNGVRAAEITGE | ASSLSTGVNQDTQY | 0.109908246 |
| RAPDAVTTSWWKE | ASSLSTGVNQDTQY | 0.1086761 |
| SGFGNVGAHTSGWFNQ | ASSLSTGVNQDTQY | 0.10541884 |
| SGVAGAVGQAASVGGLKV | ASSLSTGVNQDTQY | 0.13356687 |
| SNGGAGTGSAGGNGGT | ASSLSTGVNQDTQY | 0.10840548 |
| TGGAGGEGGAAG | ASSLSTGVNQDTQY | 0.120088756 |
| TGGDGGAAGAGGAPGHG | ASSLSTGVNQDTQY | 0.10935425 |
| TMPPLLAAIANAGRHGV | ASSLSTGVNQDTQY | 0.13100366 |
| TSSTAGNSGGA | ASSLSTGVNQDTQY | 0.11629997 |
| VPATVPVGILC | ASSLSTGVNQDTQY | 0.11657242 |
| VQRPDSGGVVLTGRIS | ASSLSTGVNQDTQY | 0.11242382 |
| WTPNPVPATKGVPAGLVQAT | ASSLSTGVNQDTQY | 0.13053356 |
| YALKRALAKFSDRFDVVII | ASSLSTGVNQDTQY | 0.11859387 |
| YLIGSVIV | ASSLSTGVNQDTQY | 0.116329856 |
| YGQLVAVID | ASSLSTGVNQDTQY | 0.10887681 |
| VTVGGMTLPA | ASSLSTGVNQDTQY | 0.10485104 |
| VLALPAV | ASSLSTGVNQDTQY | 0.1305647 |
| VIPPTLNYET | ASSLSTGVNQDTQY | 0.105449975 |
| VGMGGMHPSSG | ASSLSTGVNQDTQY | 0.103908025 |
| VGLAGGAATA | ASSLSTGVNQDTQY | 0.10495385 |
| VGGAGGAGGSGP | ASSLSTGVNQDTQY | 0.10827822 |
| VFAAFMV | ASSLSTGVNQDTQY | 0.13071878 |
| TLYDKLR | ASSLSTGVNQDTQY | 0.13616817 |
| TLAVFTVL | ASSLSTGVNQDTQY | 0.120177664 |
| TGSVQPARV | ASSLSTGVNQDTQY | 0.119097754 |
| TGGGAGSGGT | ASSLSTGVNQDTQY | 0.123740785 |
| SHLETSL | ASSLSTGVNQDTQY | 0.10384411 |
| SFTTGMT | ASSLSTGVNQDTQY | 0.11446372 |
| QVEVAKRMGVRQPTVS | ASSLSTGVNQDTQY | 0.10935981 |
| QGSGGPALAM | ASSLSTGVNQDTQY | 0.12337565 |
| MLAPSNP | ASSLSTGVNQDTQY | 0.10932706 |
| LVNNAAI | ASSLSTGVNQDTQY | 0.11169848 |
| LTGEINTLA | ASSLSTGVNQDTQY | 0.12828657 |
| LSIFGPV | ASSLSTGVNQDTQY | 0.11116106 |
| LLALLANL | ASSLSTGVNQDTQY | 0.11714453 |
| LISNASDALDKLR | ASSLSTGVNQDTQY | 0.10874 |
| LINEAVP | ASSLSTGVNQDTQY | 0.105386876 |
| LGYAFSGL | ASSLSTGVNQDTQY | 0.14191678 |
| LDFVAVAL | ASSLSTGVNQDTQY | 0.106194876 |
| LAQPGAE | ASSLSTGVNQDTQY | 0.116002515 |
| ISNLGAVTGF | ASSLSTGVNQDTQY | 0.10532523 |
| HQEDPGRI | ASSLSTGVNQDTQY | 0.10393122 |
| GTPGEPTP | ASSLSTGVNQDTQY | 0.1100404 |
| GFFNSGS | ASSLSTGVNQDTQY | 0.1196664 |
| GAGGGGLTAGAN | ASSLSTGVNQDTQY | 0.111750424 |
| FVDALVNM | ASSLSTGVNQDTQY | 0.118233405 |

| | | |
|---|---|---|
| FLTVAPEL | ASSLSTGVNQDTQY | 0.1275918 |
| FETLTAL | ASSLSTGVNQDTQY | 0.13217716 |
| DVINPIEL | ASSLSTGVNQDTQY | 0.118811995 |
| APVSGAPV | ASSLSTGVNQDTQY | 0.12021048 |
| ALVGGLAW | ASSLSTGVNQDTQY | 0.11332802 |
| ALAAAAATAI | ASSLSTGVNQDTQY | 0.11149419 |
| AFGLGLLL | ASSLSTGVNQDTQY | 0.11269562 |
| AFFAAASL | ASSLSTGVNQDTQY | 0.13943705 |
| AAAALPAFSPP | ASSLSTGVNQDTQY | 0.108487636 |
| VVLAPAL | ASSLSTGVNQDTQY | 0.10934824 |
| TIGHVDHGKTTL | ASSLSTGVNQDTQY | 0.115854 |
| LISNASDALDK | ASSLSTGVNQDTQY | 0.12324932 |
| LISNASDAL | ASSLSTGVNQDTQY | 0.12845501 |
| ISNASDALDK | ASSLSTGVNQDTQY | 0.1112447 |
| NQAETLVYQTE | ASSLSTGVNQDTQY | 0.114616044 |
| TGGTGGTAGAG | ASSLSTGVNQDTQY | 0.113114215 |
| TGSTAGGAGGA | ASSLSTGVNQDTQY | 0.12228331 |
| DPLAVPVAAGPV | ASSLSTGVNQDTQY | 0.1085367 |
| GGTGGTPGNGGN | ASSLSTGVNQDTQY | 0.115087256 |
| PAGLAGVDPATL | ASSLSTGVNQDTQY | 0.11692184 |
| GNNLVVEAPHAGASV | ASSLSTGVNQDTQY | 0.12819211 |
| KVTRIMSASTLEQPA | ASSLSTGVNQDTQY | 0.116247684 |
| LGFALPIALPRRTVV | ASSLSTGVNQDTQY | 0.10708666 |
| TRLGNLTARKLVVLV | ASSLSTGVNQDTQY | 0.124126315 |
| WSQQRRLIYFPSAGP | ASSLSTGVNQDTQY | 0.10899372 |
| GPNGNPDPMAAAVDIR | ASSLSTGVNQDTQY | 0.117524594 |
| TQRCVSVADLRAAASR | ASSLSTGVNQDTQY | 0.12110105 |
| QFVLDVAEHPVEVLTETV | ASSLSTGVNQDTQY | 0.109234214 |
| GGAGGTGGAGGNGGTQAGDG | ASSLSTGVNQDTQY | 0.11231203 |
| PLLATLRAIAVDVDAKPAQV | ASSLSTGVNQDTQY | 0.16087109 |
| APKFRTGGYMIDFWGVGYQVAK | ASSLSTGVNQDTQY | 0.11289203 |
| GAGGAGGAGGAAGGPGGTGGTG | ASSLSTGVNQDTQY | 0.11512146 |
| IGALGLLATDVAPAGSVAFGVA | ASSLSTGVNQDTQY | 0.12484588 |
| AGGAGGIGGNGGAGGAGGVLMG | ASSLSTGVNQDTQY | 0.11596947 |
| ADELIGLL | ASSLSTGVNQDTQY | 0.13804087 |
| ADRALKA | ASSLSTGVNQDTQY | 0.110418275 |
| AGAVGQVGGAGGA | ASSLSTGVNQDTQY | 0.12065316 |
| AGGGGGLGGAP | ASSLSTGVNQDTQY | 0.114039436 |
| AGGLLYGNGGAG | ASSLSTGVNQDTQY | 0.109865904 |
| AGLESIGL | ASSLSTGVNQDTQY | 0.1273326 |
| AGLKTGII | ASSLSTGVNQDTQY | 0.12686688 |
| ASLAALLL | ASSLSTGVNQDTQY | 0.11034118 |
| ASVINISSI | ASSLSTGVNQDTQY | 0.13503408 |
| ATPPGPA | ASSLSTGVNQDTQY | 0.124098785 |
| ATVASALV | ASSLSTGVNQDTQY | 0.11507945 |
| ATVASLIL | ASSLSTGVNQDTQY | 0.10819193 |
| AVNGPVLIRH | ASSLSTGVNQDTQY | 0.11362294 |
| DGAAGAVGI | ASSLSTGVNQDTQY | 0.118658334 |
| DPLLHGI | ASSLSTGVNQDTQY | 0.10772808 |
| ELGAALGAL | ASSLSTGVNQDTQY | 0.10453759 |
| ELGRTQAV | ASSLSTGVNQDTQY | 0.111459225 |

| | | |
|---|---|---|
| FVAGGYLS | ASSLSTGVNQDTQY | 0.13505135 |
| GASGGAGGAGGW | ASSLSTGVNQDTQY | 0.10408003 |
| GGAGAGGDGGAGGI | ASSLSTGVNQDTQY | 0.11788244 |
| GTGHNGSQPGMGGNG | ASSLSTGVNQDTQY | 0.10820383 |
| GYAAASASAS | ASSLSTGVNQDTQY | 0.116979785 |
| HQVDVAAVQQG | ASSLSTGVNQDTQY | 0.12572259 |
| IGAAIAHT | ASSLSTGVNQDTQY | 0.11103148 |
| IGMIEVAQL | ASSLSTGVNQDTQY | 0.12583788 |
| IRLAPEHTSAMD | ASSLSTGVNQDTQY | 0.1068536 |
| IYNVSVL | ASSLSTGVNQDTQY | 0.10727296 |
| KQGFLGM | ASSLSTGVNQDTQY | 0.12168576 |
| LESVITQL | ASSLSTGVNQDTQY | 0.11704182 |
| LGLGALVVV | ASSLSTGVNQDTQY | 0.11004393 |
| LIGNGGNGGSGGI | ASSLSTGVNQDTQY | 0.13099329 |
| LNAINAPTL | ASSLSTGVNQDTQY | 0.10709433 |
| LQLLNTGV | ASSLSTGVNQDTQY | 0.12391135 |
| LQTSIEF | ASSLSTGVNQDTQY | 0.12264693 |
| MSFAVSEL | ASSLSTGVNQDTQY | 0.10576395 |
| NTGALGL | ASSLSTGVNQDTQY | 0.12286728 |
| PPGAGGAGGAGTT | ASSLSTGVNQDTQY | 0.116770804 |
| QLVGLGI | ASSLSTGVNQDTQY | 0.13190827 |
| RFLEQVHQL | ASSLSTGVNQDTQY | 0.11203786 |
| RGVGYVLR | ASSLSTGVNQDTQY | 0.11150147 |
| SATGVFAGVI | ASSLSTGVNQDTQY | 0.116399325 |
| SLPAITLP | ASSLSTGVNQDTQY | 0.1380405 |
| STALAFTV | ASSLSTGVNQDTQY | 0.1263596 |
| SVLTVGLGK | ASSLSTGVNQDTQY | 0.11250758 |
| TALAGAGSGI | ASSLSTGVNQDTQY | 0.11936062 |
| TGLPFTGI | ASSLSTGVNQDTQY | 0.12531759 |
| VAEHVTV | ASSLSTGVNQDTQY | 0.11544834 |
| VDIVLNSL | ASSLSTGVNQDTQY | 0.12903441 |
| VLTGGGAVL | ASSLSTGVNQDTQY | 0.11649259 |
| VNVGLAYGI | ASSLSTGVNQDTQY | 0.12178669 |
| VSSAAVGAV | ASSLSTGVNQDTQY | 0.12295594 |
| VTSVYEAGV | ASSLSTGVNQDTQY | 0.10618239 |
| WVALDRGA | ASSLSTGVNQDTQY | 0.10978549 |
| YLVGSVVI | ASSLSTGVNQDTQY | 0.10973152 |
| GPPGTGKTLL | ASSLSTGVNQDTQY | 0.11727682 |
| GVGMTKF | ASSLSTGVNQDTQY | 0.12601206 |
| IGIGGSDL | ASSLSTGVNQDTQY | 0.107104674 |
| IGTLLAEL | ASSLSTGVNQDTQY | 0.10729275 |
| IYTSGTT | ASSLSTGVNQDTQY | 0.114173315 |
| LSFPAGV | ASSLSTGVNQDTQY | 0.13569704 |
| LYTSGTT | ASSLSTGVNQDTQY | 0.11370089 |
| VGAFPFGF | ASSLSTGVNQDTQY | 0.11872258 |
| VGAGTVEF | ASSLSTGVNQDTQY | 0.17821552 |
| VGKSTLLL | ASSLSTGVNQDTQY | 0.108916275 |
| AALAHAGGGIV | ASSLSTGVNQDTQY | 0.124753535 |
| AALVAVSLL | ASSLSTGVNQDTQY | 0.12075637 |
| AAVSRAVAAL | ASSLSTGVNQDTQY | 0.1269749 |
| AGGSSGIGGF | ASSLSTGVNQDTQY | 0.116493784 |

| | | |
|---|---|---|
| AGLTVANANG | ASSLSTGVNQDTQY | 0.120782845 |
| AGLVGAVL | ASSLSTGVNQDTQY | 0.11034999 |
| ALSQAFTV | ASSLSTGVNQDTQY | 0.120386906 |
| ALTFSVSL | ASSLSTGVNQDTQY | 0.10468681 |
| ALTGAALVAL | ASSLSTGVNQDTQY | 0.114345364 |
| ASFSIGAGVL | ASSLSTGVNQDTQY | 0.107852 |
| DGSVYKI | ASSLSTGVNQDTQY | 0.11772883 |
| EQLAEIV | ASSLSTGVNQDTQY | 0.12443125 |
| FAVSELTVM | ASSLSTGVNQDTQY | 0.12445323 |
| GAAPLAAGL | ASSLSTGVNQDTQY | 0.10417543 |
| GADGGDGGSGGAGG | ASSLSTGVNQDTQY | 0.14177927 |
| GFGLDTGGAGG | ASSLSTGVNQDTQY | 0.1310297 |
| GLASGGLL | ASSLSTGVNQDTQY | 0.16954151 |
| GRIAGAAVLL | ASSLSTGVNQDTQY | 0.117503375 |
| GSAGSLNTSF | ASSLSTGVNQDTQY | 0.13164604 |
| GTGQIGI | ASSLSTGVNQDTQY | 0.117978305 |
| GTLIALIL | ASSLSTGVNQDTQY | 0.11417478 |
| IAATMGFVVG | ASSLSTGVNQDTQY | 0.10732386 |
| IFAQALL | ASSLSTGVNQDTQY | 0.11070952 |
| IGAAVALGF | ASSLSTGVNQDTQY | 0.11442883 |
| IGLTPLAL | ASSLSTGVNQDTQY | 0.11246767 |
| ILGSNTSTL | ASSLSTGVNQDTQY | 0.1332988 |
| IVFSLTVL | ASSLSTGVNQDTQY | 0.12369156 |
| LASKATGF | ASSLSTGVNQDTQY | 0.14361157 |
| LASVALIL | ASSLSTGVNQDTQY | 0.13928904 |
| LFAVGLTL | ASSLSTGVNQDTQY | 0.12196556 |
| LFNGGIGTYI | ASSLSTGVNQDTQY | 0.1158022 |
| LTLGAGII | ASSLSTGVNQDTQY | 0.115088284 |
| LVAAALGDI | ASSLSTGVNQDTQY | 0.11130947 |
| NTAFVQLGI | ASSLSTGVNQDTQY | 0.10932833 |
| QLFSGAEL | ASSLSTGVNQDTQY | 0.114800915 |
| SAATFITL | ASSLSTGVNQDTQY | 0.11455436 |
| SNYGLANLGV | ASSLSTGVNQDTQY | 0.13327232 |
| SYGGAYAVM | ASSLSTGVNQDTQY | 0.103775546 |
| TTGVGGLTLG | ASSLSTGVNQDTQY | 0.109903425 |
| TTQTIYI | ASSLSTGVNQDTQY | 0.13102017 |
| VAAGAGLGAVI | ASSLSTGVNQDTQY | 0.10604203 |
| VAAVVTLL | ASSLSTGVNQDTQY | 0.123768315 |
| VGFLGGLI | ASSLSTGVNQDTQY | 0.11554078 |
| VGVIPILL | ASSLSTGVNQDTQY | 0.10692856 |
| VIFASSLI | ASSLSTGVNQDTQY | 0.1381711 |
| VKGGDGGAAAG | ASSLSTGVNQDTQY | 0.10885458 |
| VTKTFTNI | ASSLSTGVNQDTQY | 0.120100774 |
| VVAAAILL | ASSLSTGVNQDTQY | 0.12745443 |
| YAALDVEL | ASSLSTGVNQDTQY | 0.116604656 |
| LASVALLL | ASSLSTGVNQDTQY | 0.10982822 |
| LDAVHRV | ASSLSTGVNQDTQY | 0.11436299 |
| QGLSSSIF | ASSLSTGVNQDTQY | 0.110483944 |
| RGTFANIRL | ASSLSTGVNQDTQY | 0.11116831 |
| RQLWWGHRI | ASSLSTGVNQDTQY | 0.103637345 |
| YGPPGTGKTLL | ASSLSTGVNQDTQY | 0.11775782 |

| | | |
|---|---|---|
| AALVGAPANVA | ASSLSTGVNQDTQY | 0.109380506 |
| AATFRRTE | ASSLSTGVNQDTQY | 0.12898609 |
| AGALGGGI | ASSLSTGVNQDTQY | 0.12598057 |
| AGGAGGAGGIGAG | ASSLSTGVNQDTQY | 0.11808077 |
| AGSGALGL | ASSLSTGVNQDTQY | 0.111066714 |
| AQIADQI | ASSLSTGVNQDTQY | 0.10660773 |
| ASAVAVTYL | ASSLSTGVNQDTQY | 0.10655391 |
| ASTVLGGTGG | ASSLSTGVNQDTQY | 0.10941408 |
| AVGGEPAK | ASSLSTGVNQDTQY | 0.110449634 |
| CEQGTFGGTC | ASSLSTGVNQDTQY | 0.11167187 |
| DEGKAPAG | ASSLSTGVNQDTQY | 0.114985734 |
| DTVVFPAL | ASSLSTGVNQDTQY | 0.1290674 |
| FEGITMAG | ASSLSTGVNQDTQY | 0.12232662 |
| FPLVVQD | ASSLSTGVNQDTQY | 0.11015798 |
| GAFTTPQ | ASSLSTGVNQDTQY | 0.111926354 |
| GFGGFGTAGGVGGA | ASSLSTGVNQDTQY | 0.117337845 |
| GFLAQFL | ASSLSTGVNQDTQY | 0.1164132 |
| GGAGGAGGNAGVA | ASSLSTGVNQDTQY | 0.10364099 |
| GGVGTAGAGGAGGAGG | ASSLSTGVNQDTQY | 0.11533088 |
| GNGGGGDAGNAGSGGN | ASSLSTGVNQDTQY | 0.11589104 |
| GTGGAGGAGNDAGS | ASSLSTGVNQDTQY | 0.11061263 |
| GTTNTGWFNS | ASSLSTGVNQDTQY | 0.11428995 |
| IFMGPLFTL | ASSLSTGVNQDTQY | 0.14886801 |
| ILAHAIEAI | ASSLSTGVNQDTQY | 0.10595029 |
| IRSAIREAL | ASSLSTGVNQDTQY | 0.11375278 |
| ITFAGGG | ASSLSTGVNQDTQY | 0.12369517 |
| ITGMSGHQEV | ASSLSTGVNQDTQY | 0.15213658 |
| IVAGLACTL | ASSLSTGVNQDTQY | 0.13615717 |
| IVGGGLAGA | ASSLSTGVNQDTQY | 0.12994108 |
| IVTGVLNI | ASSLSTGVNQDTQY | 0.112100475 |
| LAQQGCI | ASSLSTGVNQDTQY | 0.12497977 |
| LCLHVEN | ASSLSTGVNQDTQY | 0.11976904 |
| LDAISDGS | ASSLSTGVNQDTQY | 0.10481235 |
| LGGFAAML | ASSLSTGVNQDTQY | 0.10427201 |
| LGGKNPMI | ASSLSTGVNQDTQY | 0.11657665 |
| LGPKGVAAIL | ASSLSTGVNQDTQY | 0.1052034 |
| LMVGVSI | ASSLSTGVNQDTQY | 0.13062103 |
| LVGAAFVL | ASSLSTGVNQDTQY | 0.12331062 |
| LVLAVAPL | ASSLSTGVNQDTQY | 0.11160629 |
| MAAIAVIDAA | ASSLSTGVNQDTQY | 0.10680309 |
| NNGINAELQ | ASSLSTGVNQDTQY | 0.11122713 |
| RGEPLVLVHG | ASSLSTGVNQDTQY | 0.12470519 |
| RGGAGGAAGLI | ASSLSTGVNQDTQY | 0.12454237 |
| RSYRMHPAV | ASSLSTGVNQDTQY | 0.10965987 |
| SAAHLLAGQLA | ASSLSTGVNQDTQY | 0.14048798 |
| SAAVGSY | ASSLSTGVNQDTQY | 0.15150869 |
| SPHAVLHRI | ASSLSTGVNQDTQY | 0.114617184 |
| SQIAKVL | ASSLSTGVNQDTQY | 0.13421863 |
| TALGHLVP | ASSLSTGVNQDTQY | 0.11337137 |
| TAPTTSGAA | ASSLSTGVNQDTQY | 0.106081635 |
| TDGDDGNGGSPGT | ASSLSTGVNQDTQY | 0.110472016 |

| | | |
|---|---|---|
| TDGLGEN | ASSLSTGVNQDTQY | 0.111370064 |
| TGGPPLPGQA | ASSLSTGVNQDTQY | 0.113972485 |
| TIPAITI | ASSLSTGVNQDTQY | 0.10954347 |
| TIPGFTI | ASSLSTGVNQDTQY | 0.110566385 |
| TSIASGALA | ASSLSTGVNQDTQY | 0.10946417 |
| TVTLPGAAI | ASSLSTGVNQDTQY | 0.115035266 |
| VASGLSAALP | ASSLSTGVNQDTQY | 0.10479845 |
| VGAGVIGI | ASSLSTGVNQDTQY | 0.1126018 |
| VGPKRFGV | ASSLSTGVNQDTQY | 0.16651759 |
| VNVAFPD | ASSLSTGVNQDTQY | 0.18164745 |
| VSMTNSGVS | ASSLSTGVNQDTQY | 0.11109287 |
| VTDVLGDLSV | ASSLSTGVNQDTQY | 0.11550435 |
| VVVVGAGFAGL | ASSLSTGVNQDTQY | 0.11906518 |
| VYTVGTVV | ASSLSTGVNQDTQY | 0.122184314 |
| WTTFSGP | ASSLSTGVNQDTQY | 0.113340065 |
| YDELVEAF | ASSLSTGVNQDTQY | 0.124537446 |
| YGTGGAGGNGG | ASSLSTGVNQDTQY | 0.11702151 |
| YTNVMAV | ASSLSTGVNQDTQY | 0.108157426 |
| FLEQVHQL | ASSLSTGVNQDTQY | 0.108766176 |
| GLSLLAVAV | ASSLSTGVNQDTQY | 0.11160948 |
| LEQVHQL | ASSLSTGVNQDTQY | 0.120121665 |
| LNAAGAI | ASSLSTGVNQDTQY | 0.104020186 |
| PAVHASL | ASSLSTGVNQDTQY | 0.11929774 |
| TELEDDA | ASSLSTGVNQDTQY | 0.11003259 |
| TPPGPPN | ASSLSTGVNQDTQY | 0.11637287 |
| TPPPPPPP | ASSLSTGVNQDTQY | 0.10534794 |
| TQSPALI | ASSDASSYNSPLY | 0.11522602 |
| EFAAAFPQ | ASSDASSYNSPLY | 0.114639916 |
| LLSGNLDVL | ASSDASSYNSPLY | 0.125218 |
| GPANLGNYNI | ASSDASSYNSPLY | 0.10480741 |
| TLGPTLP | ASSDASSYNSPLY | 0.11609784 |
| VYAYSSCCK | ASSDASSYNSPLY | 0.121428646 |
| APAPAPAPNI | ASSDASSYNSPLY | 0.115310565 |
| PGGPVPIP | ASSDASSYNSPLY | 0.10751767 |
| FLTVALEP | ASSDASSYNSPLY | 0.13072704 |
| SQYFGANL | ASSDASSYNSPLY | 0.10684897 |
| GGVKPAA | ASSDASSYNSPLY | 0.11836053 |
| GGTGGAGGAGGW | ASSDASSYNSPLY | 0.12634695 |
| GVGGVAGLGGAGG | ASSDASSYNSPLY | 0.10862004 |
| VLLVLATTI | ASSDASSYNSPLY | 0.12926203 |
| IGIFAAML | ASSDASSYNSPLY | 0.13172528 |
| AVGDLID | ASSDASSYNSPLY | 0.10996876 |
| NISPLPL | ASSDASSYNSPLY | 0.10913129 |
| VTAVFVAI | ASSDASSYNSPLY | 0.11175806 |
| SGDDNDAAL | ASSDASSYNSPLY | 0.10671432 |
| AVAALAHS | ASSDASSYNSPLY | 0.13423619 |
| GGVTVMATP | ASSDASSYNSPLY | 0.11057097 |
| AGLAANVRD | ASSDASSYNSPLY | 0.11660446 |
| TNRPDLIDPALLRPG | ASSDASSYNSPLY | 0.1181314 |
| LANLGVTAL | ASSDASSYNSPLY | 0.11478741 |
| TGERIEL | ASSDASSYNSPLY | 0.121992595 |

| | | |
|---|---|---|
| NGEELAA | ASSDASSYNSPLY | 0.12270154 |
| VLSLVWG | ASSDASSYNSPLY | 0.114266805 |
| IGYALWL | ASSDASSYNSPLY | 0.11125233 |
| LSLLLAPL | ASSDASSYNSPLY | 0.112449944 |
| TLDEALK | ASSDASSYNSPLY | 0.11253153 |
| TDGAGTAEGG | ASSDASSYNSPLY | 0.14426208 |
| NAPGGAGGNGG | ASSDASSYNSPLY | 0.10930072 |
| VSVFMGMGGNF | ASSDASSYNSPLY | 0.114756905 |
| LASYLAAL | ASSDASSYNSPLY | 0.12352762 |
| LVGGTPAL | ASSDASSYNSPLY | 0.10678655 |
| VRNADVL | ASSDASSYNSPLY | 0.11662028 |
| TCSSPFLVSAM | ASSDASSYNSPLY | 0.12569958 |
| VHITDAE | ASSDASSYNSPLY | 0.13237871 |
| EYAAPSF | ASSDASSYNSPLY | 0.11057883 |
| VVDAFHGGL | ASSDASSYNSPLY | 0.12179197 |
| ASVVDPQ | ASSDASSYNSPLY | 0.13397068 |
| SYNEFGGLS | ASSDASSYNSPLY | 0.109935656 |
| DPDGSVY | ASSDASSYNSPLY | 0.105825014 |
| MVSTIHATE | ASSDASSYNSPLY | 0.11014734 |
| LAAGVADPV | ASSDASSYNSPLY | 0.111700214 |
| TKEQIAA | ASSDASSYNSPLY | 0.12182466 |
| SLASPAAL | ASSDASSYNSPLY | 0.11209089 |
| IGLVFYGV | ASSDASSYNSPLY | 0.11714286 |
| FLLDASNL | ASSDASSYNSPLY | 0.11896645 |
| YQVGNFP | ASSDASSYNSPLY | 0.12049379 |
| ESLLSQST | ASSDASSYNSPLY | 0.105953164 |
| LFTVGFAAVG | ASSDASSYNSPLY | 0.11717028 |
| ATPPTPT | ASSDASSYNSPLY | 0.11823669 |
| TDLATLG | ASSDASSYNSPLY | 0.11631896 |
| SAGGTGR | ASSDASSYNSPLY | 0.113039196 |
| GFSNSGS | ASSDASSYNSPLY | 0.12248835 |
| GGNGGSGGTSV | ASSDASSYNSPLY | 0.118176535 |
| GTSVATGGAGN | ASSDASSYNSPLY | 0.11309782 |
| NGGNGGSGGTS | ASSDASSYNSPLY | 0.13730858 |
| AVVKFPH | ASSDASSYNSPLY | 0.15045501 |
| TPPPPPP | ASSDASSYNSPLY | 0.11203777 |
| AAPAAAAAPA | ASSDASSYNSPLY | 0.12445642 |
| IALNEDL | ASSDASSYNSPLY | 0.1059912 |
| IVNVSSM | ASSDASSYNSPLY | 0.105362274 |
| TGIPPAPR | ASSDASSYNSPLY | 0.105953224 |
| FDLGGGTF | ASSDASSYNSPLY | 0.120195314 |
| FDLGGGTFDVS | ASSDASSYNSPLY | 0.10420908 |
| NEPTAAA | ASSDASSYNSPLY | 0.12725401 |
| NEPTAAALA | ASSDASSYNSPLY | 0.11682024 |
| GSIINISSI | ASSDASSYNSPLY | 0.14048785 |
| VAPGNAGTA | ASSDASSYNSPLY | 0.120325975 |
| TGGGASLE | ASSDASSYNSPLY | 0.11002226 |
| VAVYNHYKRI | ASSDASSYNSPLY | 0.1151528 |
| TDPAPATT | ASSDASSYNSPLY | 0.13198885 |
| TNRPDILDPALLRPG | ASSDASSYNSPLY | 0.10995193 |
| LTTTTTS | ASSDASSYNSPLY | 0.10616919 |

| | | |
|---|---|---|
| PTGSGKTL | ASSDASSYNSPLY | 0.11100093 |
| TGSGKTL | ASSDASSYNSPLY | 0.10913456 |
| TIGHVDHGK | ASSDASSYNSPLY | 0.12133239 |
| TIGHVDHGKTT | ASSDASSYNSPLY | 0.121290535 |
| GHVDHGKTT | ASSDASSYNSPLY | 0.118962 |
| AQFEHTLL | ASSDASSYNSPLY | 0.11407726 |
| IVNQDEL | ASSDASSYNSPLY | 0.10478375 |
| TNGSQFF | ASSDASSYNSPLY | 0.12084039 |
| TGKGIVE | ASSDASSYNSPLY | 0.18033817 |
| LISNASDA | ASSDASSYNSPLY | 0.1154425 |
| AELRTLGELTKIAKAHGAQ | ASSDASSYNSPLY | 0.12209085 |
| APNAVAAAEPSW | ASSDASSYNSPLY | 0.13327166 |
| APQVAGAATAVL | ASSDASSYNSPLY | 0.11493564 |
| ARRDPAAAAAILPTDA | ASSDASSYNSPLY | 0.11654399 |
| DAAARDVLAEAGLAEYF | ASSDASSYNSPLY | 0.115089655 |
| DDQIVLSAKANLAKVEQ | ASSDASSYNSPLY | 0.11526394 |
| DELLPDTGEDSDEDGH | ASSDASSYNSPLY | 0.10710234 |
| ELPEAPANRATVPQD | ASSDASSYNSPLY | 0.10969325 |
| FGQSKIERVTGIG | ASSDASSYNSPLY | 0.12767512 |
| GAGGTGSTAGGAGGAGGAGGLY | ASSDASSYNSPLY | 0.13348356 |
| GHGGTGAAGGNGGTGGAGGSGI | ASSDASSYNSPLY | 0.11512953 |
| GNGGNGGAGGEGGPGGAGGAGA | ASSDASSYNSPLY | 0.13132416 |
| GSTAGNGGNGGNATNGTIAGQP | ASSDASSYNSPLY | 0.12151616 |
| GTGGDGGNGGNGGTGG | ASSDASSYNSPLY | 0.113943264 |
| GVRRPAGIPADAPV | ASSDASSYNSPLY | 0.116260454 |
| GYRQVTDAHLLTLARR | ASSDASSYNSPLY | 0.11163545 |
| IAKMGQDGHDRG | ASSDASSYNSPLY | 0.10648429 |
| KAEPSIEIQYT | ASSDASSYNSPLY | 0.10879142 |
| KPVRIRMAADEDTVAVT | ASSDASSYNSPLY | 0.11335312 |
| MPPRIAGMRLLVIKPEP | ASSDASSYNSPLY | 0.121909544 |
| MKFYAVALFRSRGVPPTR | ASSDASSYNSPLY | 0.14266972 |
| NAGGFGILGAVAHSP | ASSDASSYNSPLY | 0.11610884 |
| NEGREITLAMAEM | ASSDASSYNSPLY | 0.12768784 |
| NELVEWSHPLGHRWRWE | ASSDASSYNSPLY | 0.1116139 |
| NLGAGGTGGNGGNPG | ASSDASSYNSPLY | 0.10866742 |
| NTEWPKWETVMAADPVD | ASSDASSYNSPLY | 0.11638543 |
| PASAAPAETQP | ASSDASSYNSPLY | 0.111511186 |
| QQSSDGKNGVRAAEITGE | ASSDASSYNSPLY | 0.14721672 |
| RAPDAVTTSWWKE | ASSDASSYNSPLY | 0.11443273 |
| SGFGNVGAHTSGWFNQ | ASSDASSYNSPLY | 0.11384248 |
| SGVAGAVGQAASVGGLKV | ASSDASSYNSPLY | 0.12675829 |
| SNGGAGTGSAGGNGGT | ASSDASSYNSPLY | 0.1218828 |
| TGGAGGEGGAAG | ASSDASSYNSPLY | 0.114264295 |
| TGGDGGAAGAGGAPGHG | ASSDASSYNSPLY | 0.10997479 |
| TMPPLLAAIANAGRHGV | ASSDASSYNSPLY | 0.1124298 |
| TSSTAGNSGGA | ASSDASSYNSPLY | 0.12268525 |
| VPATVPVGILC | ASSDASSYNSPLY | 0.1479527 |
| VQRPDSGGVVLTGRIS | ASSDASSYNSPLY | 0.104533814 |
| WTPNPVPATKGVPAGLVQAT | ASSDASSYNSPLY | 0.1252089 |
| YALKRALAKFSDRFDVVII | ASSDASSYNSPLY | 0.11791329 |
| YLIGSVIV | ASSDASSYNSPLY | 0.11027088 |

| | | |
|---|---|---|
| YGQLVAVID | ASSDASSYNSPLY | 0.11909498 |
| VTVGGMTLPA | ASSDASSYNSPLY | 0.1236329 |
| VLALPAV | ASSDASSYNSPLY | 0.115028 |
| VIPPTLNYET | ASSDASSYNSPLY | 0.116615616 |
| VGMGGMHPSSG | ASSDASSYNSPLY | 0.10608645 |
| VGLAGGAATA | ASSDASSYNSPLY | 0.1097285 |
| VGGAGGAGGSGP | ASSDASSYNSPLY | 0.13162424 |
| VFAAFMV | ASSDASSYNSPLY | 0.12268629 |
| TLYDKLR | ASSDASSYNSPLY | 0.13437189 |
| TLAVFTVL | ASSDASSYNSPLY | 0.11324186 |
| TGSVQPARV | ASSDASSYNSPLY | 0.1187803 |
| TGGGAGSGGT | ASSDASSYNSPLY | 0.11125094 |
| SHLETSL | ASSDASSYNSPLY | 0.12452537 |
| SFTTGMT | ASSDASSYNSPLY | 0.10779172 |
| QVEVAKRMGVRQPTVS | ASSDASSYNSPLY | 0.11881391 |
| QGSGGPALAM | ASSDASSYNSPLY | 0.12634297 |
| MLAPSNP | ASSDASSYNSPLY | 0.10722437 |
| LVNNAAI | ASSDASSYNSPLY | 0.11113815 |
| LTGEINTLA | ASSDASSYNSPLY | 0.11906645 |
| LSIFGPV | ASSDASSYNSPLY | 0.11076747 |
| LLALLANL | ASSDASSYNSPLY | 0.10862714 |
| LISNASDALDKLR | ASSDASSYNSPLY | 0.13268542 |
| LINEAVP | ASSDASSYNSPLY | 0.15497856 |
| LGYAFSGL | ASSDASSYNSPLY | 0.1201879 |
| LDFVAVAL | ASSDASSYNSPLY | 0.10848491 |
| LAQPGAE | ASSDASSYNSPLY | 0.10509415 |
| ISNLGAVTGF | ASSDASSYNSPLY | 0.11178072 |
| HQEDPGRI | ASSDASSYNSPLY | 0.12466534 |
| GTPGEPTP | ASSDASSYNSPLY | 0.115108594 |
| GFFNSGS | ASSDASSYNSPLY | 0.10960175 |
| GAGGGGLTAGAN | ASSDASSYNSPLY | 0.110306546 |
| FVDALVNM | ASSDASSYNSPLY | 0.14262679 |
| FLTVAPEL | ASSDASSYNSPLY | 0.10937818 |
| FETLTAL | ASSDASSYNSPLY | 0.10832759 |
| DVINPIEL | ASSDASSYNSPLY | 0.10699838 |
| APVSGAPV | ASSDASSYNSPLY | 0.13432896 |
| ALVGGLAW | ASSDASSYNSPLY | 0.107594065 |
| ALAAAAATAI | ASSDASSYNSPLY | 0.11071332 |
| AFGLGLLL | ASSDASSYNSPLY | 0.12632997 |
| AFFAAASL | ASSDASSYNSPLY | 0.10626015 |
| AAAALPAFSPP | ASSDASSYNSPLY | 0.12220751 |
| VVLAPAL | ASSDASSYNSPLY | 0.11788104 |
| TIGHVDHGKTTL | ASSDASSYNSPLY | 0.12521641 |
| LISNASDALDK | ASSDASSYNSPLY | 0.1133375 |
| LISNASDAL | ASSDASSYNSPLY | 0.116926916 |
| ISNASDALDK | ASSDASSYNSPLY | 0.11582027 |
| NQAETLVYQTE | ASSDASSYNSPLY | 0.12321686 |
| TGGTGGTAGAG | ASSDASSYNSPLY | 0.116053976 |
| TGSTAGGAGGA | ASSDASSYNSPLY | 0.11145928 |
| DPLAVPVAAGPV | ASSDASSYNSPLY | 0.11123527 |
| GGTGGTPGNGGN | ASSDASSYNSPLY | 0.1041727 |

| | | |
|---|---|---|
| PAGLAGVDPATL | ASSDASSYNSPLY | 0.12271347 |
| GNNLVVEAPHAGASV | ASSDASSYNSPLY | 0.10388394 |
| KVTRIMSASTLEQPA | ASSDASSYNSPLY | 0.12808421 |
| LGFALPIALPRRTVV | ASSDASSYNSPLY | 0.1255985 |
| TRLGNLTARKLVVLV | ASSDASSYNSPLY | 0.11250096 |
| WSQQRRLIYFPSAGP | ASSDASSYNSPLY | 0.11629128 |
| GPNGNPDPMAAAVDIR | ASSDASSYNSPLY | 0.10709976 |
| TQRCVSVADLRAAASR | ASSDASSYNSPLY | 0.106117114 |
| QFVLDVAEHPVEVLTETV | ASSDASSYNSPLY | 0.114352085 |
| GGAGGTGGAGGNGGTQAGDG | ASSDASSYNSPLY | 0.12493261 |
| PLLATLRAIAVDVDAKPAQV | ASSDASSYNSPLY | 0.11358876 |
| APKFRTGGYMIDFWGVGYQVAK | ASSDASSYNSPLY | 0.1160539 |
| GAGGAGGAGGAAGGPGGTGGTG | ASSDASSYNSPLY | 0.11940818 |
| IGALGLLATDVAPAGSVAFGVA | ASSDASSYNSPLY | 0.10995887 |
| AGGAGGIGGNGGAGGAGGVLMG | ASSDASSYNSPLY | 0.11078813 |
| ADELIGLL | ASSDASSYNSPLY | 0.14441213 |
| ADRALKA | ASSDASSYNSPLY | 0.116138116 |
| AGAVGQVGGAGGA | ASSDASSYNSPLY | 0.10954096 |
| AGGGGGLGGAP | ASSDASSYNSPLY | 0.13479942 |
| AGGLLYGNGGAG | ASSDASSYNSPLY | 0.11003085 |
| AGLESIGL | ASSDASSYNSPLY | 0.12525705 |
| AGLKTGII | ASSDASSYNSPLY | 0.11887434 |
| ASLAALLL | ASSDASSYNSPLY | 0.11812727 |
| ASVINISSI | ASSDASSYNSPLY | 0.12953164 |
| ATPPGPA | ASSDASSYNSPLY | 0.10848713 |
| ATVASALV | ASSDASSYNSPLY | 0.10958557 |
| ATVASLIL | ASSDASSYNSPLY | 0.11534137 |
| AVNGPVLIRH | ASSDASSYNSPLY | 0.13896525 |
| DGAAGAVGI | ASSDASSYNSPLY | 0.14129403 |
| DPLLHGI | ASSDASSYNSPLY | 0.1147051 |
| ELGAALGAL | ASSDASSYNSPLY | 0.12084991 |
| ELGRTQAV | ASSDASSYNSPLY | 0.11242595 |
| FVAGGYLS | ASSDASSYNSPLY | 0.11309744 |
| GASGGAGGAGGW | ASSDASSYNSPLY | 0.11442259 |
| GGAGAGGDGGAGGI | ASSDASSYNSPLY | 0.108273715 |
| GTGHNGSQPGMGGNG | ASSDASSYNSPLY | 0.111630514 |
| GYAAASASAS | ASSDASSYNSPLY | 0.11763039 |
| HQVDVAAVQQG | ASSDASSYNSPLY | 0.11241238 |
| IGAAIAHT | ASSDASSYNSPLY | 0.12972374 |
| IGMIEVAQL | ASSDASSYNSPLY | 0.10777208 |
| IRLAPEHTSAMD | ASSDASSYNSPLY | 0.11290387 |
| IYNVSVL | ASSDASSYNSPLY | 0.11674846 |
| KQGFLGM | ASSDASSYNSPLY | 0.119950026 |
| LESVITQL | ASSDASSYNSPLY | 0.11402709 |
| LGLGALVVV | ASSDASSYNSPLY | 0.12141823 |
| LIGNGGNGGSGGI | ASSDASSYNSPLY | 0.11085101 |
| LNAINAPTL | ASSDASSYNSPLY | 0.13842374 |
| LQLLNTGV | ASSDASSYNSPLY | 0.1509548 |
| LQTSIEF | ASSDASSYNSPLY | 0.11540044 |
| MSFAVSEL | ASSDASSYNSPLY | 0.1132966 |
| NTGALGL | ASSDASSYNSPLY | 0.11304897 |

| | | |
|---|---|---|
| PPGAGGAGGAGTT | ASSDASSYNSPLY | 0.13324569 |
| QLVGLGI | ASSDASSYNSPLY | 0.114916936 |
| RFLEQVHQL | ASSDASSYNSPLY | 0.10643088 |
| RGVGYVLR | ASSDASSYNSPLY | 0.11944574 |
| SATGVFAGVI | ASSDASSYNSPLY | 0.11639507 |
| SLPAITLP | ASSDASSYNSPLY | 0.11116054 |
| STALAFTV | ASSDASSYNSPLY | 0.10708226 |
| SVLTVGLGK | ASSDASSYNSPLY | 0.13656262 |
| TALAGAGSGI | ASSDASSYNSPLY | 0.20496237 |
| TGLPFTGI | ASSDASSYNSPLY | 0.11467455 |
| VAEHVTV | ASSDASSYNSPLY | 0.10994176 |
| VDIVLNSL | ASSDASSYNSPLY | 0.10625804 |
| VLTGGGAVL | ASSDASSYNSPLY | 0.12734234 |
| VNVGLAYGI | ASSDASSYNSPLY | 0.12687407 |
| VSSAAVGAV | ASSDASSYNSPLY | 0.11220874 |
| VTSVYEAGV | ASSDASSYNSPLY | 0.12914625 |
| WVALDRGA | ASSDASSYNSPLY | 0.10897776 |
| YLVGSVVI | ASSDASSYNSPLY | 0.11829537 |
| GPPGTGKTLL | ASSDASSYNSPLY | 0.12743948 |
| GVGMTKF | ASSDASSYNSPLY | 0.12125026 |
| IGIGGSDL | ASSDASSYNSPLY | 0.111940406 |
| IGTLLAEL | ASSDASSYNSPLY | 0.11555165 |
| IYTSGTT | ASSDASSYNSPLY | 0.1146395 |
| LSFPAGV | ASSDASSYNSPLY | 0.1178992 |
| LYTSGTT | ASSDASSYNSPLY | 0.12339214 |
| VGAFPFGF | ASSDASSYNSPLY | 0.10576895 |
| VGAGTVEF | ASSDASSYNSPLY | 0.1220975 |
| VGKSTLLL | ASSDASSYNSPLY | 0.11118497 |
| AALAHAGGGIV | ASSDASSYNSPLY | 0.14753602 |
| AALVAVSLL | ASSDASSYNSPLY | 0.121790804 |
| AAVSRAVAAL | ASSDASSYNSPLY | 0.11923847 |
| AGGSSGIGGF | ASSDASSYNSPLY | 0.11357061 |
| AGLTVANANG | ASSDASSYNSPLY | 0.14151101 |
| AGLVGAVL | ASSDASSYNSPLY | 0.114826255 |
| ALSQAFTV | ASSDASSYNSPLY | 0.11821638 |
| ALTFSVSL | ASSDASSYNSPLY | 0.11988366 |
| ALTGAALVAL | ASSDASSYNSPLY | 0.10128031 |
| ASFSIGAGVL | ASSDASSYNSPLY | 0.14542902 |
| DGSVYKI | ASSDASSYNSPLY | 0.1859909 |
| EQLAEIV | ASSDASSYNSPLY | 0.23686746 |
| FAVSELTVM | ASSDASSYNSPLY | 0.11383019 |
| GAAPLAAGL | ASSDASSYNSPLY | 0.08980516 |
| GADGGDGGSGGAGG | ASSDASSYNSPLY | 0.15331393 |
| GFGLDTGGAGG | ASSDASSYNSPLY | 0.09049437 |
| GLASGGLL | ASSDASSYNSPLY | 0.0915859 |
| GRIAGAAVLL | ASSDASSYNSPLY | 0.16101807 |
| GSAGSLNTSF | ASSDASSYNSPLY | 0.12004916 |
| GTGQIGI | ASSDASSYNSPLY | 0.4094313 |
| GTLIALIL | ASSDASSYNSPLY | 0.26945078 |
| IAATMGFVVG | ASSDASSYNSPLY | 0.096232966 |
| IFAQALL | ASSDASSYNSPLY | 0.13215223 |

| | | |
|---|---|---|
| IGAAVALGF | ASSDASSYNSPLY | 0.1450005 |
| IGLTPLAL | ASSDASSYNSPLY | 0.14098813 |
| ILGSNTSTL | ASSDASSYNSPLY | 0.08144028 |
| IVFSLTVL | ASSDASSYNSPLY | 0.17216499 |
| LASKATGF | ASSDASSYNSPLY | 0.102399275 |
| LASVALIL | ASSDASSYNSPLY | 0.07855188 |
| LFAVGLTL | ASSDASSYNSPLY | 0.19921461 |
| LFNGGIGTYI | ASSDASSYNSPLY | 0.13917898 |
| LTLGAGII | ASSDASSYNSPLY | 0.10012123 |
| LVAAALGDI | ASSDASSYNSPLY | 0.33397388 |
| NTAFVQLGI | ASSDASSYNSPLY | 0.10048929 |
| QLFSGAEL | ASSDASSYNSPLY | 0.09662266 |
| SAATFITL | ASSDASSYNSPLY | 0.23826802 |
| SNYGLANLGV | ASSDASSYNSPLY | 0.31106955 |
| SYGGAYAVM | ASSDASSYNSPLY | 0.100064345 |
| TTGVGGLTLG | ASSDASSYNSPLY | 0.18693352 |
| TTQTIYI | ASSDASSYNSPLY | 0.5741066 |
| VAAGAGLGAVI | ASSDASSYNSPLY | 0.16173545 |
| VAAVVTLL | ASSDASSYNSPLY | 0.3114698 |
| VGFLGGLI | ASSDASSYNSPLY | 0.39279896 |
| VGVIPILL | ASSDASSYNSPLY | 0.12454251 |
| VIFASSLI | ASSDASSYNSPLY | 0.19579363 |
| VKGGDGGAAAG | ASSDASSYNSPLY | 0.36225754 |
| VTKTFTNI | ASSDASSYNSPLY | 0.14689551 |
| VVAAAILL | ASSDASSYNSPLY | 0.17463979 |
| YAALDVEL | ASSDASSYNSPLY | 0.118603475 |
| LASVALLL | ASSDASSYNSPLY | 0.17108162 |
| LDAVHRV | ASSDASSYNSPLY | 0.079416566 |
| QGLSSSIF | ASSDASSYNSPLY | 0.18110058 |
| RGTFANIRL | ASSDASSYNSPLY | 0.20717795 |
| RQLWWGHRI | ASSDASSYNSPLY | 0.17412762 |
| YGPPGTGKTLL | ASSDASSYNSPLY | 0.095537096 |
| AALVGAPANVA | ASSDASSYNSPLY | 0.106288806 |
| AATFRRTE | ASSDASSYNSPLY | 0.0866265 |
| AGALGGGI | ASSDASSYNSPLY | 0.13489518 |
| AGGAGGAGGIGAG | ASSDASSYNSPLY | 0.09557079 |
| AGSGALGL | ASSDASSYNSPLY | 0.08553707 |
| AQIADQI | ASSDASSYNSPLY | 0.26383358 |
| ASAVAVTYL | ASSDASSYNSPLY | 0.2435763 |
| ASTVLGGTGG | ASSDASSYNSPLY | 0.11372844 |
| AVGGEPAK | ASSDASSYNSPLY | 0.19002074 |
| CEQGTFGGTC | ASSDASSYNSPLY | 0.3367738 |
| DEGKAPAG | ASSDASSYNSPLY | 0.42597145 |
| DTVVFPAL | ASSDASSYNSPLY | 0.16339879 |
| FEGITMAG | ASSDASSYNSPLY | 0.08872245 |
| FPLVVQD | ASSDASSYNSPLY | 0.55381256 |
| GAFTTPQ | ASSDASSYNSPLY | 0.2155699 |
| GFGGFGTAGGVGGA | ASSDASSYNSPLY | 0.12640603 |
| GFLAQFL | ASSDASSYNSPLY | 0.3169625 |
| GGAGGAGGNAGVA | ASSDASSYNSPLY | 0.10303232 |
| GGVGTAGAGGAGGAGG | ASSDASSYNSPLY | 0.10600136 |

| | | |
|---|---|---|
| GNGGGGDAGNAGSGGN | ASSDASSYNSPLY | 0.37429404 |
| GTGGAGGAGNDAGS | ASSDASSYNSPLY | 0.29846364 |
| GTTNTGWFNS | ASSDASSYNSPLY | 0.10707841 |
| IFMGPLFTL | ASSDASSYNSPLY | 0.09424509 |
| ILAHAIEAI | ASSDASSYNSPLY | 0.12269139 |
| IRSAIREAL | ASSDASSYNSPLY | 0.22018342 |
| ITFAGGG | ASSDASSYNSPLY | 0.256441 |
| ITGMSGHQEV | ASSDASSYNSPLY | 0.16271307 |
| IVAGLACTL | ASSDASSYNSPLY | 0.13628912 |
| IVGGGLAGA | ASSDASSYNSPLY | 0.29663897 |
| IVTGVLNI | ASSDASSYNSPLY | 0.16175881 |
| LAQQGCI | ASSDASSYNSPLY | 0.16031912 |
| LCLHVEN | ASSDASSYNSPLY | 0.22430071 |
| LDAISDGS | ASSDASSYNSPLY | 0.08851382 |
| LGGFAAML | ASSDASSYNSPLY | 0.2814796 |
| LGGKNPMI | ASSDASSYNSPLY | 0.31638432 |
| LGPKGVAAIL | ASSDASSYNSPLY | 0.2416163 |
| LMVGVSI | ASSDASSYNSPLY | 0.3079634 |
| LVGAAFVL | ASSDASSYNSPLY | 0.1053768 |
| LVLAVAPL | ASSDASSYNSPLY | 0.23999159 |
| MAAIAVIDAA | ASSDASSYNSPLY | 0.101621374 |
| NNGINAELQ | ASSDASSYNSPLY | 0.15871474 |
| RGEPLVLVHG | ASSDASSYNSPLY | 0.23726057 |
| RGGAGGAAGLI | ASSDASSYNSPLY | 0.19728851 |
| RSYRMHPAV | ASSDASSYNSPLY | 0.28856784 |
| SAAHLLAGQLA | ASSDASSYNSPLY | 0.20742112 |
| SAAVGSY | ASSDASSYNSPLY | 0.13220352 |
| SPHAVLHRI | ASSDASSYNSPLY | 0.49701124 |
| SQIAKVL | ASSDASSYNSPLY | 0.5332885 |
| TALGHLVP | ASSDASSYNSPLY | 0.09685967 |
| TAPTTSGAA | ASSDASSYNSPLY | 0.107685946 |
| TDGDDGNGGSPGT | ASSDASSYNSPLY | 0.12816598 |
| TDGLGEN | ASSDASSYNSPLY | 0.2648521 |
| TGGPPLPGQA | ASSDASSYNSPLY | 0.22328547 |
| TIPAITI | ASSDASSYNSPLY | 0.1955238 |
| TIPGFTI | ASSDASSYNSPLY | 0.14474016 |
| TSIASGALA | ASSDASSYNSPLY | 0.098935805 |
| TVTLPGAAI | ASSDASSYNSPLY | 0.49267542 |
| VASGLSAALP | ASSDASSYNSPLY | 0.09156828 |
| VGAGVIGI | ASSDASSYNSPLY | 0.15767346 |
| VGPKRFGV | ASSDASSYNSPLY | 0.10396205 |
| VNVAFPD | ASSDASSYNSPLY | 0.07848696 |
| VSMTNSGVS | ASSDASSYNSPLY | 0.20339903 |
| VTDVLGDLSV | ASSDASSYNSPLY | 0.12538297 |
| VVVVGAGFAGL | ASSDASSYNSPLY | 0.31343237 |
| VYTVGTVV | ASSDASSYNSPLY | 0.14013901 |
| WTTFSGP | ASSDASSYNSPLY | 0.16291177 |
| YDELVEAF | ASSDASSYNSPLY | 0.22903052 |
| YGTGGAGGNGG | ASSDASSYNSPLY | 0.1579811 |
| YTNVMAV | ASSDASSYNSPLY | 0.1030272 |
| FLEQVHQL | ASSDASSYNSPLY | 0.18093039 |

| | | |
|---|---|---|
| GLSLLAVAV | ASSDASSYNSPLY | 0.47796917 |
| LEQVHQL | ASSDASSYNSPLY | 0.23242185 |
| LNAAGAI | ASSDASSYNSPLY | 0.08963119 |
| PAVHASL | ASSDASSYNSPLY | 0.13220541 |
| TELEDDA | ASSDASSYNSPLY | 0.13458486 |
| TPPGPPN | ASSDASSYNSPLY | 0.076070994 |
| TPPPPPPP | ASSDASSYNSPLY | 0.08317912 |
| TQSPALI | ASGDREVF | 0.07961937 |
| EFAAAFPQ | ASGDREVF | 0.08467852 |
| LLSGNLDVL | ASGDREVF | 0.3207749 |
| GPANLGNYNI | ASGDREVF | 0.2999192 |
| TLGPTLP | ASGDREVF | 0.18002139 |
| VYAYSSCCK | ASGDREVF | 0.33892375 |
| APAPAPAPNI | ASGDREVF | 0.13896942 |
| PGGPVPIP | ASGDREVF | 0.07402709 |
| FLTVALEP | ASGDREVF | 0.112762004 |
| SQYFGANL | ASGDREVF | 0.10762128 |
| GGVKPAA | ASGDREVF | 0.37231123 |
| GGTGGAGGAGGW | ASGDREVF | 0.08540362 |
| GVGGVAGLGGAGG | ASGDREVF | 0.13233374 |
| VLLVLATTI | ASGDREVF | 0.19716772 |
| IGIFAAML | ASGDREVF | 0.12782718 |
| AVGDLID | ASGDREVF | 0.26847148 |
| NISPLPL | ASGDREVF | 0.1043106 |
| VTAVFVAI | ASGDREVF | 0.092226595 |
| SGDDNDAAL | ASGDREVF | 0.2051373 |
| AVAALAHS | ASGDREVF | 0.079643264 |
| GGVTVMATP | ASGDREVF | 0.10608241 |
| AGLAANVRD | ASGDREVF | 0.095867634 |
| TNRPDLIDPALLRPG | ASGDREVF | 0.12775029 |
| LANLGVTAL | ASGDREVF | 0.08817896 |
| TGERIEL | ASGDREVF | 0.29812333 |
| NGEELAA | ASGDREVF | 0.18756592 |
| VLSLVWG | ASGDREVF | 0.1875808 |
| IGYALWL | ASGDREVF | 0.17799868 |
| LSLLLAPL | ASGDREVF | 0.1508449 |
| TLDEALK | ASGDREVF | 0.20811842 |
| TDGAGTAEGG | ASGDREVF | 0.1076638 |
| NAPGGAGGNGG | ASGDREVF | 0.11420473 |
| VSVFMGMGGNF | ASGDREVF | 0.09803717 |
| LASYLAAL | ASGDREVF | 0.10423555 |
| LVGGTPAL | ASGDREVF | 0.43735096 |
| VRNADVL | ASGDREVF | 0.15814716 |
| TCSSPFLVSAM | ASGDREVF | 0.0950271 |
| VHITDAE | ASGDREVF | 0.10194119 |
| EYAAPSF | ASGDREVF | 0.49192727 |
| VVDAFHGGL | ASGDREVF | 0.67117757 |
| ASVVDPQ | ASGDREVF | 0.112429194 |
| SYNEFGGLS | ASGDREVF | 0.15365046 |
| DPDGSVY | ASGDREVF | 0.17135064 |
| MVSTIHATE | ASGDREVF | 0.14880674 |

| | | |
|---|---|---|
| LAAGVADPV | ASGDREVF | 0.13503833 |
| TKEQIAA | ASGDREVF | 0.09836991 |
| SLASPAAL | ASGDREVF | 0.29187846 |
| IGLVFYGV | ASGDREVF | 0.77211 |
| FLLDASNL | ASGDREVF | 0.113574564 |
| YQVGNFP | ASGDREVF | 0.092705585 |
| ESLLSQST | ASGDREVF | 0.20728466 |
| LFTVGFAAVG | ASGDREVF | 0.09656226 |
| ATPPTPT | ASGDREVF | 0.1486666 |
| TDLATLG | ASGDREVF | 0.13551019 |
| SAGGTGR | ASGDREVF | 0.09031316 |
| GFSNSGS | ASGDREVF | 0.14026892 |
| GGNGGSGGTSV | ASGDREVF | 0.27314538 |
| GTSVATGGAGN | ASGDREVF | 0.11084812 |
| NGGNGGSGGTS | ASGDREVF | 0.39059317 |
| AVVKFPH | ASGDREVF | 0.4253421 |
| TPPPPPP | ASGDREVF | 0.2693001 |
| AAPAAAAAPA | ASGDREVF | 0.47665554 |
| IALNEDL | ASGDREVF | 0.37414223 |
| IVNVSSM | ASGDREVF | 0.13941927 |
| TGIPPAPR | ASGDREVF | 0.15059888 |
| FDLGGGTF | ASGDREVF | 0.32929605 |
| FDLGGGTFDVS | ASGDREVF | 0.2077119 |
| NEPTAAA | ASGDREVF | 0.24306166 |
| NEPTAAALA | ASGDREVF | 0.15953542 |
| GSIINISSI | ASGDREVF | 0.33826217 |
| VAPGNAGTA | ASGDREVF | 0.15504932 |
| TGGGASLE | ASGDREVF | 0.48398626 |
| VAVYNHYKRI | ASGDREVF | 0.10639848 |
| TDPAPATT | ASGDREVF | 0.1034904 |
| TNRPDILDPALLRPG | ASGDREVF | 0.3949827 |
| LTTTTTS | ASGDREVF | 0.09020866 |
| PTGSGKTL | ASGDREVF | 0.09827897 |
| TGSGKTL | ASGDREVF | 0.1590827 |
| TIGHVDHGK | ASGDREVF | 0.34213224 |
| TIGHVDHGKTT | ASGDREVF | 0.074293144 |
| GHVDHGKTT | ASGDREVF | 0.10523257 |
| AQFEHTLL | ASGDREVF | 0.11766151 |
| IVNQDEL | ASGDREVF | 0.108990975 |
| TNGSQFF | ASGDREVF | 0.13098653 |
| TGKGIVE | ASGDREVF | 0.09480816 |
| LISNASDA | ASGDREVF | 0.17899403 |
| AELRTLGELTKIAKAHGAQ | ASGDREVF | 0.0925484 |
| APNAVAAAEPSW | ASGDREVF | 0.085405886 |
| APQVAGAATAVL | ASGDREVF | 0.17035173 |
| ARRDPAAAAAILPTDA | ASGDREVF | 0.53689826 |
| DAAARDVLAEAGLAEYF | ASGDREVF | 0.11388749 |
| DDQIVLSAKANLAKVEQ | ASGDREVF | 0.20556268 |
| DELLPDTGEDSDEDGH | ASGDREVF | 0.12289675 |
| ELPEAPANRATVPQD | ASGDREVF | 0.76612234 |
| FGQSKIERVTGIG | ASGDREVF | 0.1406394 |

| | | |
|---|---|---|
| GAGGTGSTAGGAGGAGGAGGLY | ASGDREVF | 0.103552386 |
| GHGGTGAAGGNGGTGGAGGSGI | ASGDREVF | 0.22009623 |
| GNGGNGGAGGEGGPGGAGGAGA | ASGDREVF | 0.41319144 |
| GSTAGNGGNGGNATNGTIAGQP | ASGDREVF | 0.16678214 |
| GTGGDGGNGGNGGTGG | ASGDREVF | 0.37315798 |
| GVRRPAGIPADAPV | ASGDREVF | 0.09478058 |
| GYRQVTDAHLLTLARR | ASGDREVF | 0.16588756 |
| IAKMGQDGHDRG | ASGDREVF | 0.2390241 |
| KAEPSIEIQYT | ASGDREVF | 0.18830997 |
| KPVRIRMAADEDTVAVT | ASGDREVF | 0.08394378 |
| MPPRIAGMRLLVIKPEP | ASGDREVF | 0.17375258 |
| MKFYAVALFRSRGVPPTR | ASGDREVF | 0.12840587 |
| NAGGFGILGAVAHSP | ASGDREVF | 0.1348921 |
| NEGREITLAMAEM | ASGDREVF | 0.21118885 |
| NELVEWSHPLGHRWRWE | ASGDREVF | 0.16417503 |
| NLGAGGTGGNGGNPG | ASGDREVF | 0.3438277 |
| NTEWPKWETVMAADPVD | ASGDREVF | 0.1654415 |
| PASAAPAETQP | ASGDREVF | 0.13157468 |
| QQSSDGKNGVRAAEITGE | ASGDREVF | 0.09579925 |
| RAPDAVTTSWWKE | ASGDREVF | 0.26742572 |
| SGFGNVGAHTSGWFNQ | ASGDREVF | 0.20834255 |
| SGVAGAVGQAASVGGLKV | ASGDREVF | 0.1925373 |
| SNGGAGTGSAGGNGGT | ASGDREVF | 0.121601835 |
| TGGAGGEGGAAG | ASGDREVF | 0.10172866 |
| TGGDGGAAGAGGAPGHG | ASGDREVF | 0.24371406 |
| TMPPLLAAIANAGRHGV | ASGDREVF | 0.32449287 |
| TSSTAGNSGGA | ASGDREVF | 0.13333365 |
| VPATVPVGILC | ASGDREVF | 0.16742617 |
| VQRPDSGGVVLTGRIS | ASGDREVF | 0.5161135 |
| WTPNPVPATKGVPAGLVQAT | ASGDREVF | 0.119253054 |
| YALKRALAKFSDRFDVVII | ASGDREVF | 0.12687257 |
| YLIGSVIV | ASGDREVF | 0.226776 |
| YGQLVAVID | ASGDREVF | 0.41804886 |
| VTVGGMTLPA | ASGDREVF | 0.14187789 |
| VLALPAV | ASGDREVF | 0.4254042 |
| VIPPTLNYET | ASGDREVF | 0.08964909 |
| VGMGGMHPSSG | ASGDREVF | 0.2931774 |
| VGLAGGAATA | ASGDREVF | 0.14709339 |
| VGGAGGAGGSGP | ASGDREVF | 0.20788534 |
| VFAAFMV | ASGDREVF | 0.07927129 |
| TLYDKLR | ASGDREVF | 0.2379215 |
| TLAVFTVL | ASGDREVF | 0.354326 |
| TGSVQPARV | ASGDREVF | 0.35464856 |
| TGGGAGSGGT | ASGDREVF | 0.08249113 |
| SHLETSL | ASGDREVF | 0.14737172 |
| SFTTGMT | ASGDREVF | 0.36702955 |
| QVEVAKRMGVRQPTVS | ASGDREVF | 0.6852522 |
| QGSGGPALAM | ASGDREVF | 0.17108569 |
| MLAPSNP | ASGDREVF | 0.29422113 |
| LVNNAAI | ASGDREVF | 0.10636027 |
| LTGEINTLA | ASGDREVF | 0.11825619 |

| | | |
|---|---|---|
| LSIFGPV | ASGDREVF | 0.11431682 |
| LLALLANL | ASGDREVF | 0.1396628 |
| LISNASDALDKLR | ASGDREVF | 0.17758957 |
| LINEAVP | ASGDREVF | 0.11330288 |
| LGYAFSGL | ASGDREVF | 0.19311842 |
| LDFVAVAL | ASGDREVF | 0.45792925 |
| LAQPGAE | ASGDREVF | 0.17660889 |
| ISNLGAVTGF | ASGDREVF | 0.50588906 |
| HQEDPGRI | ASGDREVF | 0.7014776 |
| GTPGEPTP | ASGDREVF | 0.22090243 |
| GFFNSGS | ASGDREVF | 0.09796227 |
| GAGGGGLTAGAN | ASGDREVF | 0.16621472 |
| FVDALVNM | ASGDREVF | 0.09172929 |
| FLTVAPEL | ASGDREVF | 0.1447456 |
| FETLTAL | ASGDREVF | 0.0962216 |
| DVINPIEL | ASGDREVF | 0.22336519 |
| APVSGAPV | ASGDREVF | 0.074134886 |
| ALVGGLAW | ASGDREVF | 0.3589819 |
| ALAAAAATAI | ASGDREVF | 0.43220818 |
| AFGLGLLL | ASGDREVF | 0.08153157 |
| AFFAAASL | ASGDREVF | 0.28822684 |
| AAAALPAFSPP | ASGDREVF | 0.53071 |
| VVLAPAL | ASGDREVF | 0.09765443 |
| TIGHVDHGKTTL | ASGDREVF | 0.37947565 |
| LISNASDALDK | ASGDREVF | 0.108622216 |
| LISNASDAL | ASGDREVF | 0.28333977 |
| ISNASDALDK | ASGDREVF | 0.20028721 |
| NQAETLVYQTE | ASGDREVF | 0.21794814 |
| TGGTGGTAGAG | ASGDREVF | 0.18992202 |
| TGSTAGGAGGA | ASGDREVF | 0.21265662 |
| DPLAVPVAAGPV | ASGDREVF | 0.096512385 |
| GGTGGTPGNGGN | ASGDREVF | 0.2148726 |
| PAGLAGVDPATL | ASGDREVF | 0.08484553 |
| GNNLVVEAPHAGASV | ASGDREVF | 0.2348313 |
| KVTRIMSASTLEQPA | ASGDREVF | 0.10504554 |
| LGFALPIALPRRTVV | ASGDREVF | 0.24402277 |
| TRLGNLTARKLVVLV | ASGDREVF | 0.12690778 |
| WSQQRRLIYFPSAGP | ASGDREVF | 0.21265009 |
| GPNGNPDPMAAAVDIR | ASGDREVF | 0.14498916 |
| TQRCVSVADLRAAASR | ASGDREVF | 0.115754284 |
| QFVLDVAEHPVEVLTETV | ASGDREVF | 0.098348975 |
| GGAGGTGGAGGNGGTQAGDG | ASGDREVF | 0.09006407 |
| PLLATLRAIAVDVDAKPAQV | ASGDREVF | 0.33835125 |
| APKFRTGGYMIDFWGVGYQVAK | ASGDREVF | 0.13973217 |
| GAGGAGGAGGAAGGPGGTGGTG | ASGDREVF | 0.089134574 |
| IGALGLLATDVAPAGSVAFGVA | ASGDREVF | 0.21175101 |
| AGGAGGIGGNGGAGGAGGVLMG | ASGDREVF | 0.14379576 |
| ADELIGLL | ASGDREVF | 0.08888963 |
| ADRALKA | ASGDREVF | 0.20971406 |
| AGAVGQVGGAGGA | ASGDREVF | 0.21211469 |
| AGGGGGLGGAP | ASGDREVF | 0.13800333 |

| | | |
|---|---|---|
| AGGLLYGNGGAG | ASGDREVF | 0.07593874 |
| AGLESIGL | ASGDREVF | 0.099381045 |
| AGLKTGII | ASGDREVF | 0.13803896 |
| ASLAALLL | ASGDREVF | 0.08236239 |
| ASVINISSI | ASGDREVF | 0.0947482 |
| ATPPGPA | ASGDREVF | 0.33831745 |
| ATVASALV | ASGDREVF | 0.141691 |
| ATVASLIL | ASGDREVF | 0.107264854 |
| AVNGPVLIRH | ASGDREVF | 0.25115213 |
| DGAAGAVGI | ASGDREVF | 0.20115033 |
| DPLLHGI | ASGDREVF | 0.56726784 |
| ELGAALGAL | ASGDREVF | 0.10701296 |
| ELGRTQAV | ASGDREVF | 0.121235475 |
| FVAGGYLS | ASGDREVF | 0.20065084 |
| GASGGAGGAGGW | ASGDREVF | 0.18159713 |
| GGAGAGGDGGAGGI | ASGDREVF | 0.09508702 |
| GTGHNGSQPGMGGNG | ASGDREVF | 0.08337634 |
| GYAAASASAS | ASGDREVF | 0.17541102 |
| HQVDVAAVQQG | ASGDREVF | 0.11426688 |
| IGAAIAHT | ASGDREVF | 0.17084162 |
| IGMIEVAQL | ASGDREVF | 0.32844743 |
| IRLAPEHTSAMD | ASGDREVF | 0.09195231 |
| IYNVSVL | ASGDREVF | 0.1005655 |
| KQGFLGM | ASGDREVF | 0.13352318 |
| LESVITQL | ASGDREVF | 0.11450868 |
| LGLGALVVV | ASGDREVF | 0.28953552 |
| LIGNGGNGGSGGI | ASGDREVF | 0.107344344 |
| LNAINAPTL | ASGDREVF | 0.74656254 |
| LQLLNTGV | ASGDREVF | 0.17622286 |
| LQTSIEF | ASGDREVF | 0.10529766 |
| MSFAVSEL | ASGDREVF | 0.3043738 |
| NTGALGL | ASGDREVF | 0.16607338 |
| PPGAGGAGGAGTT | ASGDREVF | 0.15198219 |
| QLVGLGI | ASGDREVF | 0.17255358 |
| RFLEQVHQL | ASGDREVF | 0.088237286 |
| RGVGYVLR | ASGDREVF | 0.11622474 |
| SATGVFAGVI | ASGDREVF | 0.16945815 |
| SLPAITLP | ASGDREVF | 0.103269264 |
| STALAFTV | ASGDREVF | 0.162287 |
| SVLTVGLGK | ASGDREVF | 0.09011458 |
| TALAGAGSGI | ASGDREVF | 0.10083711 |
| TGLPFTGI | ASGDREVF | 0.22548747 |
| VAEHVTV | ASGDREVF | 0.2816651 |
| VDIVLNSL | ASGDREVF | 0.6490361 |
| VLTGGGAVL | ASGDREVF | 0.12344148 |
| VNVGLAYGI | ASGDREVF | 0.09494986 |
| VSSAAVGAV | ASGDREVF | 0.47728658 |
| VTSVYEAGV | ASGDREVF | 0.11147562 |
| WVALDRGA | ASGDREVF | 0.21177132 |
| YLVGSVVI | ASGDREVF | 0.12545164 |
| GPPGTGKTLL | ASGDREVF | 0.10008609 |

| | | |
|---|---|---|
| GVGMTKF | ASGDREVF | 0.13931873 |
| IGIGGSDL | ASGDREVF | 0.1934443 |
| IGTLLAEL | ASGDREVF | 0.082731284 |
| IYTSGTT | ASGDREVF | 0.2484991 |
| LSFPAGV | ASGDREVF | 0.073631875 |
| LYTSGTT | ASGDREVF | 0.096879266 |
| VGAFPFGF | ASGDREVF | 0.13439724 |
| VGAGTVEF | ASGDREVF | 0.10874797 |
| VGKSTLLL | ASGDREVF | 0.08861892 |
| AALAHAGGGIV | ASGDREVF | 0.15019406 |
| AALVAVSLL | ASGDREVF | 0.44099516 |
| AAVSRAVAAL | ASGDREVF | 0.38125086 |
| AGGSSGIGGF | ASGDREVF | 0.11339709 |
| AGLTVANANG | ASGDREVF | 0.29517108 |
| AGLVGAVL | ASGDREVF | 0.7161919 |
| ALSQAFTV | ASGDREVF | 0.19013304 |
| ALTFSVSL | ASGDREVF | 0.08714749 |
| ALTGAALVAL | ASGDREVF | 0.14426409 |
| ASFSIGAGVL | ASGDREVF | 0.07666006 |
| DGSVYKI | ASGDREVF | 0.09985918 |
| EQLAEIV | ASGDREVF | 0.6978849 |
| FAVSELTVM | ASGDREVF | 0.09090984 |
| GAAPLAAGL | ASGDREVF | 0.20567955 |
| GADGGDGGSGGAGG | ASGDREVF | 0.34994423 |
| GFGLDTGGAGG | ASGDREVF | 0.13674936 |
| GLASGGLL | ASGDREVF | 0.11360337 |
| GRIAGAAVLL | ASGDREVF | 0.10026658 |
| GSAGSLNTSF | ASGDREVF | 0.095168315 |
| GTGQIGI | ASGDREVF | 0.17606223 |
| GTLIALIL | ASGDREVF | 0.1411513 |
| IAATMGFVVG | ASGDREVF | 0.11137407 |
| IFAQALL | ASGDREVF | 0.15764755 |
| IGAAVALGF | ASGDREVF | 0.11067605 |
| IGLTPLAL | ASGDREVF | 0.12854436 |
| ILGSNTSTL | ASGDREVF | 0.11654669 |
| IVFSLTVL | ASGDREVF | 0.09630371 |
| LASKATGF | ASGDREVF | 0.22871605 |
| LASVALIL | ASGDREVF | 0.0871015 |
| LFAVGLTL | ASGDREVF | 0.1830095 |
| LFNGGIGTYI | ASGDREVF | 0.51561534 |
| LTLGAGII | ASGDREVF | 0.08053088 |
| LVAAALGDI | ASGDREVF | 0.108359076 |
| NTAFVQLGI | ASGDREVF | 0.12707448 |
| QLFSGAEL | ASGDREVF | 0.08714455 |
| SAATFITL | ASGDREVF | 0.39109126 |
| SNYGLANLGV | ASGDREVF | 0.13887668 |
| SYGGAYAVM | ASGDREVF | 0.25450686 |
| TTGVGGLTLG | ASGDREVF | 0.12754808 |
| TTQTIYI | ASGDREVF | 0.68393517 |
| VAAGAGLGAVI | ASGDREVF | 0.21275352 |
| VAAVVTLL | ASGDREVF | 0.1266738 |

| | | |
|---|---|---|
| VGFLGGLI | ASGDREVF | 0.0797251 |
| VGVIPILL | ASGDREVF | 0.09776829 |
| VIFASSLI | ASGDREVF | 0.27604938 |
| VKGGDGGAAAG | ASGDREVF | 0.33884034 |
| VTKTFTNI | ASGDREVF | 0.093714826 |
| VVAAAILL | ASGDREVF | 0.17490838 |
| YAALDVEL | ASGDREVF | 0.39612645 |
| LASVALLL | ASGDREVF | 0.32468262 |
| LDAVHRV | ASGDREVF | 0.093812585 |
| QGLSSSIF | ASGDREVF | 0.114915445 |
| RGTFANIRL | ASGDREVF | 0.15650071 |
| RQLWWGHRI | ASGDREVF | 0.16208166 |
| YGPPGTGKTLL | ASGDREVF | 0.09465084 |
| AALVGAPANVA | ASGDREVF | 0.38566327 |
| AATFRRTE | ASGDREVF | 0.21418032 |
| AGALGGGI | ASGDREVF | 0.6061272 |
| AGGAGGAGGIGAG | ASGDREVF | 0.17124318 |
| AGSGALGL | ASGDREVF | 0.2424547 |
| AQIADQI | ASGDREVF | 0.17370963 |
| ASAVAVTYL | ASGDREVF | 0.12120704 |
| ASTVLGGTGG | ASGDREVF | 0.10491436 |
| AVGGEPAK | ASGDREVF | 0.085900016 |
| CEQGTFGGTC | ASGDREVF | 0.10503989 |
| DEGKAPAG | ASGDREVF | 0.108998485 |
| DTVVFPAL | ASGDREVF | 0.07468664 |
| FEGITMAG | ASGDREVF | 0.12421076 |
| FPLVVQD | ASGDREVF | 0.17469838 |
| GAFTTPQ | ASGDREVF | 0.20335644 |
| GFGGFGTAGGVGGA | ASGDREVF | 0.12892456 |
| GFLAQFL | ASGDREVF | 0.14120346 |
| GGAGGAGGNAGVA | ASGDREVF | 0.090824775 |
| GGVGTAGAGGAGGAGG | ASGDREVF | 0.37590295 |
| GNGGGGDAGNAGSGGN | ASGDREVF | 0.085948594 |
| GTGGAGGAGNDAGS | ASGDREVF | 0.09457889 |
| GTTNTGWFNS | ASGDREVF | 0.1320666 |
| IFMGPLFTL | ASGDREVF | 0.09481865 |
| ILAHAIEAI | ASGDREVF | 0.70424235 |
| IRSAIREAL | ASGDREVF | 0.32685354 |
| ITFAGGG | ASGDREVF | 0.21145049 |
| ITGMSGHQEV | ASGDREVF | 0.47583148 |
| IVAGLACTL | ASGDREVF | 0.08398802 |
| IVGGGLAGA | ASGDREVF | 0.10534584 |
| IVTGVLNI | ASGDREVF | 0.54171485 |
| LAQQGCI | ASGDREVF | 0.12512271 |
| LCLHVEN | ASGDREVF | 0.23989484 |
| LDAISDGS | ASGDREVF | 0.25069702 |
| LGGFAAML | ASGDREVF | 0.1022408 |
| LGGKNPMI | ASGDREVF | 0.13988398 |
| LGPKGVAAIL | ASGDREVF | 0.10605712 |
| LMVGVSI | ASGDREVF | 0.24882388 |
| LVGAAFVL | ASGDREVF | 0.16005656 |

| | | |
|---|---|---|
| LVLAVAPL | ASGDREVF | 0.10026419 |
| MAAIAVIDAA | ASGDREVF | 0.290106 |
| NNGINAELQ | ASGDREVF | 0.78989065 |
| RGEPLVLVHG | ASGDREVF | 0.14811645 |
| RGGAGGAAGLI | ASGDREVF | 0.17403743 |
| RSYRMHPAV | ASGDREVF | 0.15735218 |
| SAAHLLAGQLA | ASGDREVF | 0.3236876 |
| SAAVGSY | ASGDREVF | 0.12683877 |
| SPHAVLHRI | ASGDREVF | 0.103928566 |
| SQIAKVL | ASGDREVF | 0.12144744 |
| TALGHLVP | ASGDREVF | 0.1203016 |
| TAPTTSGAA | ASGDREVF | 0.19797052 |
| TDGDDGNGGSPGT | ASGDREVF | 0.1346329 |
| TDGLGEN | ASGDREVF | 0.81627774 |
| TGGPPLPGQA | ASGDREVF | 0.3311125 |
| TIPAITI | ASGDREVF | 0.12103786 |
| TIPGFTI | ASGDREVF | 0.09043784 |
| TSIASGALA | ASGDREVF | 0.15941834 |
| TVTLPGAAI | ASGDREVF | 0.16171694 |
| VASGLSAALP | ASGDREVF | 0.102662444 |
| VGAGVIGI | ASGDREVF | 0.49394256 |
| VGPKRFGV | ASGDREVF | 0.122823186 |
| VNVAFPD | ASGDREVF | 0.09562653 |
| VSMTNSGVS | ASGDREVF | 0.17011358 |
| VTDVLGDLSV | ASGDREVF | 0.70600295 |
| VVVVGAGFAGL | ASGDREVF | 0.20866603 |
| VYTVGTVV | ASGDREVF | 0.12826884 |
| WTTFSGP | ASGDREVF | 0.09237242 |
| YDELVEAF | ASGDREVF | 0.43177602 |
| YGTGGAGGNGG | ASGDREVF | 0.4064354 |
| YTNVMAV | ASGDREVF | 0.11320318 |
| FLEQVHQL | ASGDREVF | 0.6344075 |
| GLSLLAVAV | ASGDREVF | 0.09668251 |
| LEQVHQL | ASGDREVF | 0.09527136 |
| LNAAGAI | ASGDREVF | 0.20011912 |
| PAVHASL | ASGDREVF | 0.16456133 |
| TELEDDA | ASGDREVF | 0.6122136 |
| TPPGPPN | ASGDREVF | 0.14364862 |
| TPPPPPPP | ASGDREVF | 0.36978942 |
| TQSPALI | ASSPPRGWNSPLY | 0.15017876 |
| EFAAAFPQ | ASSPPRGWNSPLY | 0.45886466 |
| LLSGNLDVL | ASSPPRGWNSPLY | 0.3152054 |
| GPANLGNYNI | ASSPPRGWNSPLY | 0.19166775 |
| TLGPTLP | ASSPPRGWNSPLY | 0.18411545 |
| VYAYSSCCK | ASSPPRGWNSPLY | 0.1274456 |
| APAPAPAPNI | ASSPPRGWNSPLY | 0.19532982 |
| PGGPVPIP | ASSPPRGWNSPLY | 0.101436265 |
| FLTVALEP | ASSPPRGWNSPLY | 0.12450191 |
| SQYFGANL | ASSPPRGWNSPLY | 0.28999007 |
| GGVKPAA | ASSPPRGWNSPLY | 0.23455314 |
| GGTGGAGGAGGW | ASSPPRGWNSPLY | 0.09877513 |

| | | |
|---|---|---|
| GVGGVAGLGGAGG | ASSPPRGWNSPLY | 0.2611822 |
| VLLVLATTI | ASSPPRGWNSPLY | 0.6016342 |
| IGIFAAML | ASSPPRGWNSPLY | 0.18941994 |
| AVGDLID | ASSPPRGWNSPLY | 0.112563536 |
| NISPLPL | ASSPPRGWNSPLY | 0.5190972 |
| VTAVFVAI | ASSPPRGWNSPLY | 0.7905155 |
| SGDDNDAAL | ASSPPRGWNSPLY | 0.32561564 |
| AVAALAHS | ASSPPRGWNSPLY | 0.09286759 |
| GGVTVMATP | ASSPPRGWNSPLY | 0.08833622 |
| AGLAANVRD | ASSPPRGWNSPLY | 0.19822468 |
| TNRPDLIDPALLRPG | ASSPPRGWNSPLY | 0.1289246 |
| LANLGVTAL | ASSPPRGWNSPLY | 0.11007287 |
| TGERIEL | ASSPPRGWNSPLY | 0.09141038 |
| NGEELAA | ASSPPRGWNSPLY | 0.27564305 |
| VLSLVWG | ASSPPRGWNSPLY | 0.23844594 |
| IGYALWL | ASSPPRGWNSPLY | 0.11839099 |
| LSLLLAPL | ASSPPRGWNSPLY | 0.08903817 |
| TLDEALK | ASSPPRGWNSPLY | 0.21655743 |
| TDGAGTAEGG | ASSPPRGWNSPLY | 0.2996061 |
| NAPGGAGGNGG | ASSPPRGWNSPLY | 0.113620095 |
| VSVFMGMGGNF | ASSPPRGWNSPLY | 0.13447526 |
| LASYLAAL | ASSPPRGWNSPLY | 0.19973963 |
| LVGGTPAL | ASSPPRGWNSPLY | 0.1237991 |
| VRNADVL | ASSPPRGWNSPLY | 0.14905907 |
| TCSSPFLVSAM | ASSPPRGWNSPLY | 0.10954047 |
| VHITDAE | ASSPPRGWNSPLY | 0.10375414 |
| EYAAPSF | ASSPPRGWNSPLY | 0.12458841 |
| VVDAFHGGL | ASSPPRGWNSPLY | 0.31421602 |
| ASVVDPQ | ASSPPRGWNSPLY | 0.20111638 |
| SYNEFGGLS | ASSPPRGWNSPLY | 0.21053141 |
| DPDGSVY | ASSPPRGWNSPLY | 0.22020894 |
| MVSTIHATE | ASSPPRGWNSPLY | 0.14018579 |
| LAAGVADPV | ASSPPRGWNSPLY | 0.094652735 |
| TKEQIAA | ASSPPRGWNSPLY | 0.08198699 |
| SLASPAAL | ASSPPRGWNSPLY | 0.45300403 |
| IGLVFYGV | ASSPPRGWNSPLY | 0.08365121 |
| FLLDASNL | ASSPPRGWNSPLY | 0.29458994 |
| YQVGNFP | ASSPPRGWNSPLY | 0.17293355 |
| ESLLSQST | ASSPPRGWNSPLY | 0.10377182 |
| LFTVGFAAVG | ASSPPRGWNSPLY | 0.15925902 |
| ATPPTPT | ASSPPRGWNSPLY | 0.077112585 |
| TDLATLG | ASSPPRGWNSPLY | 0.1386583 |
| SAGGTGR | ASSPPRGWNSPLY | 0.62130046 |
| GFSNSGS | ASSPPRGWNSPLY | 0.16866496 |
| GGNGGSGGTSV | ASSPPRGWNSPLY | 0.14876804 |
| GTSVATGGAGN | ASSPPRGWNSPLY | 0.21631965 |
| NGGNGGSGGTS | ASSPPRGWNSPLY | 0.2738693 |
| AVVKFPH | ASSPPRGWNSPLY | 0.13305408 |
| TPPPPPP | ASSPPRGWNSPLY | 0.18799442 |
| AAPAAAAAPA | ASSPPRGWNSPLY | 0.29463834 |
| IALNEDL | ASSPPRGWNSPLY | 0.22641543 |

| | | |
|---|---|---|
| IVNVSSM | ASSPPRGWNSPLY | 0.096593 |
| TGIPPAPR | ASSPPRGWNSPLY | 0.13925405 |
| FDLGGGTF | ASSPPRGWNSPLY | 0.112071484 |
| FDLGGGTFDVS | ASSPPRGWNSPLY | 0.20998393 |
| NEPTAAA | ASSPPRGWNSPLY | 0.12313084 |
| NEPTAAALA | ASSPPRGWNSPLY | 0.23269296 |
| GSIINISSI | ASSPPRGWNSPLY | 0.13632649 |
| VAPGNAGTA | ASSPPRGWNSPLY | 0.08266177 |
| TGGGASLE | ASSPPRGWNSPLY | 0.115449265 |
| VAVYNHYKRI | ASSPPRGWNSPLY | 0.21590081 |
| TDPAPATT | ASSPPRGWNSPLY | 0.14556924 |
| TNRPDILDPALLRPG | ASSPPRGWNSPLY | 0.16292994 |
| LTTTTTS | ASSPPRGWNSPLY | 0.105504975 |
| PTGSGKTL | ASSPPRGWNSPLY | 0.26302227 |
| TGSGKTL | ASSPPRGWNSPLY | 0.12628572 |
| TIGHVDHGK | ASSPPRGWNSPLY | 0.11864093 |
| TIGHVDHGKTT | ASSPPRGWNSPLY | 0.18296239 |
| GHVDHGKTT | ASSPPRGWNSPLY | 0.13475664 |
| AQFEHTLL | ASSPPRGWNSPLY | 0.12406321 |
| IVNQDEL | ASSPPRGWNSPLY | 0.30507636 |
| TNGSQFF | ASSPPRGWNSPLY | 0.09415325 |
| TGKGIVE | ASSPPRGWNSPLY | 0.18235976 |
| LISNASDA | ASSPPRGWNSPLY | 0.17679352 |
| AELRTLGELTKIAKAHGAQ | ASSPPRGWNSPLY | 0.123719305 |
| APNAVAAAEPSW | ASSPPRGWNSPLY | 0.2434604 |
| APQVAGAATAVL | ASSPPRGWNSPLY | 0.08867865 |
| ARRDPAAAAAILPTDA | ASSPPRGWNSPLY | 0.2489368 |
| DAAARDVLAEAGLAEYF | ASSPPRGWNSPLY | 0.11723408 |
| DDQIVLSAKANLAKVEQ | ASSPPRGWNSPLY | 0.1257757 |
| DELLPDTGEDSDEDGH | ASSPPRGWNSPLY | 0.11738159 |
| ELPEAPANRATVPQD | ASSPPRGWNSPLY | 0.20102692 |
| FGQSKIERVTGIG | ASSPPRGWNSPLY | 0.31510133 |
| GAGGTGSTAGGAGGAGGAGGLY | ASSPPRGWNSPLY | 0.093007624 |
| GHGGTGAAGGNGGTGGAGGSGI | ASSPPRGWNSPLY | 0.19536018 |
| GNGGNGGAGGEGGPGGAGGAGA | ASSPPRGWNSPLY | 0.27863398 |
| GSTAGNGGNGGNATNGTIAGQP | ASSPPRGWNSPLY | 0.15727761 |
| GTGGDGGNGGNGGTGG | ASSPPRGWNSPLY | 0.08663219 |
| GVRRPAGIPADAPV | ASSPPRGWNSPLY | 0.10996681 |
| GYRQVTDAHLLTLARR | ASSPPRGWNSPLY | 0.08607535 |
| IAKMGQDGHDRG | ASSPPRGWNSPLY | 0.13163759 |
| KAEPSIEIQYT | ASSPPRGWNSPLY | 0.10208601 |
| KPVRIRMAADEDTVAVT | ASSPPRGWNSPLY | 0.27359557 |
| MPPRIAGMRLLVIKPEP | ASSPPRGWNSPLY | 0.44430012 |
| MKFYAVALFRSRGVPPTR | ASSPPRGWNSPLY | 0.10936119 |
| NAGGFGILGAVAHSP | ASSPPRGWNSPLY | 0.4209203 |
| NEGREITLAMAEM | ASSPPRGWNSPLY | 0.10274736 |
| NELVEWSHPLGHRWRWE | ASSPPRGWNSPLY | 0.20473644 |
| NLGAGGTGGNGGNPG | ASSPPRGWNSPLY | 0.50296396 |
| NTEWPKWETVMAADPVD | ASSPPRGWNSPLY | 0.09996066 |
| PASAAPAETQP | ASSPPRGWNSPLY | 0.15475498 |
| QQSSDGKNGVRAAEITGE | ASSPPRGWNSPLY | 0.13694292 |

| | | |
|---|---|---|
| RAPDAVTTSWWKE | ASSPPRGWNSPLY | 0.10625491 |
| SGFGNVGAHTSGWFNQ | ASSPPRGWNSPLY | 0.44795293 |
| SGVAGAVGQAASVGGLKV | ASSPPRGWNSPLY | 0.12442848 |
| SNGGAGTGSAGGNGGT | ASSPPRGWNSPLY | 0.25574556 |
| TGGAGGEGGAAG | ASSPPRGWNSPLY | 0.10717932 |
| TGGDGGAAGAGGAPGHG | ASSPPRGWNSPLY | 0.0929807 |
| TMPPLLAAIANAGRHGV | ASSPPRGWNSPLY | 0.16275601 |
| TSSTAGNSGGA | ASSPPRGWNSPLY | 0.49233487 |
| VPATVPVGILC | ASSPPRGWNSPLY | 0.13678588 |
| VQRPDSGGVVLTGRIS | ASSPPRGWNSPLY | 0.3104363 |
| WTPNPVPATKGVPAGLVQAT | ASSPPRGWNSPLY | 0.14570051 |
| YALKRALAKFSDRFDVVII | ASSPPRGWNSPLY | 0.2224752 |
| YLIGSVIV | ASSPPRGWNSPLY | 0.58953536 |
| YGQLVAVID | ASSPPRGWNSPLY | 0.18560408 |
| VTVGGMTLPA | ASSPPRGWNSPLY | 0.13289942 |
| VLALPAV | ASSPPRGWNSPLY | 0.49327683 |
| VIPPTLNYET | ASSPPRGWNSPLY | 0.17365211 |
| VGMGGMHPSSG | ASSPPRGWNSPLY | 0.2930338 |
| VGLAGGAATA | ASSPPRGWNSPLY | 0.29854977 |
| VGGAGGAGGSGP | ASSPPRGWNSPLY | 0.14101535 |
| VFAAFMV | ASSPPRGWNSPLY | 0.50907665 |
| TLYDKLR | ASSPPRGWNSPLY | 0.29721147 |
| TLAVFTVL | ASSPPRGWNSPLY | 0.28746504 |
| TGSVQPARV | ASSPPRGWNSPLY | 0.1572558 |
| TGGGAGSGGT | ASSPPRGWNSPLY | 0.13213906 |
| SHLETSL | ASSPPRGWNSPLY | 0.20243752 |
| SFTTGMT | ASSPPRGWNSPLY | 0.13264376 |
| QVEVAKRMGVRQPTVS | ASSPPRGWNSPLY | 0.5082227 |
| QGSGGPALAM | ASSPPRGWNSPLY | 0.3046413 |
| MLAPSNP | ASSPPRGWNSPLY | 0.17606698 |
| LVNNAAI | ASSPPRGWNSPLY | 0.52573 |
| LTGEINTLA | ASSPPRGWNSPLY | 0.45921966 |
| LSIFGPV | ASSPPRGWNSPLY | 0.12650895 |
| LLALLANL | ASSPPRGWNSPLY | 0.17133103 |
| LISNASDALDKLR | ASSPPRGWNSPLY | 0.1778842 |
| LINEAVP | ASSPPRGWNSPLY | 0.19786096 |
| LGYAFSGL | ASSPPRGWNSPLY | 0.10930276 |
| LDFVAVAL | ASSPPRGWNSPLY | 0.3229077 |
| LAQPGAE | ASSPPRGWNSPLY | 0.14458346 |
| ISNLGAVTGF | ASSPPRGWNSPLY | 0.10677324 |
| HQEDPGRI | ASSPPRGWNSPLY | 0.23155782 |
| GTPGEPTP | ASSPPRGWNSPLY | 0.1836341 |
| GFFNSGS | ASSPPRGWNSPLY | 0.15587151 |
| GAGGGGLTAGAN | ASSPPRGWNSPLY | 0.51169485 |
| FVDALVNM | ASSPPRGWNSPLY | 0.1694916 |
| FLTVAPEL | ASSPPRGWNSPLY | 0.13421257 |
| FETLTAL | ASSPPRGWNSPLY | 0.35863948 |
| DVINPIEL | ASSPPRGWNSPLY | 0.58781636 |
| APVSGAPV | ASSPPRGWNSPLY | 0.13770542 |
| ALVGGLAW | ASSPPRGWNSPLY | 0.3172624 |
| ALAAAAATAI | ASSPPRGWNSPLY | 0.70069 |

| | | |
|---|---|---|
| AFGLGLLL | ASSPPRGWNSPLY | 0.29234537 |
| AFFAAASL | ASSPPRGWNSPLY | 0.5014999 |
| AAAALPAFSPP | ASSPPRGWNSPLY | 0.56074464 |
| VVLAPAL | ASSPPRGWNSPLY | 0.16820693 |
| TIGHVDHGKTTL | ASSPPRGWNSPLY | 0.4406822 |
| LISNASDALDK | ASSPPRGWNSPLY | 0.6074117 |
| LISNASDAL | ASSPPRGWNSPLY | 0.19640633 |
| ISNASDALDK | ASSPPRGWNSPLY | 0.23427334 |
| NQAETLVYQTE | ASSPPRGWNSPLY | 0.15346622 |
| TGGTGGTAGAG | ASSPPRGWNSPLY | 0.21997648 |
| TGSTAGGAGGA | ASSPPRGWNSPLY | 0.115329154 |
| DPLAVPVAAGPV | ASSPPRGWNSPLY | 0.41037065 |
| GGTGGTPGNGGN | ASSPPRGWNSPLY | 0.43291357 |
| PAGLAGVDPATL | ASSPPRGWNSPLY | 0.21708219 |
| GNNLVVEAPHAGASV | ASSPPRGWNSPLY | 0.13007917 |
| KVTRIMSASTLEQPA | ASSPPRGWNSPLY | 0.1295932 |
| LGFALPIALPRRTVV | ASSPPRGWNSPLY | 0.16880026 |
| TRLGNLTARKLVVLV | ASSPPRGWNSPLY | 0.16932726 |
| WSQQRRLIYFPSAGP | ASSPPRGWNSPLY | 0.13095449 |
| GPNGNPDPMAAAVDIR | ASSPPRGWNSPLY | 0.16818114 |
| TQRCVSVADLRAAASR | ASSPPRGWNSPLY | 0.4389059 |
| QFVLDVAEHPVEVLTETV | ASSPPRGWNSPLY | 0.42535734 |
| GGAGGTGGAGGNGGTQAGDG | ASSPPRGWNSPLY | 0.12698524 |
| PLLATLRAIAVDVDAKPAQV | ASSPPRGWNSPLY | 0.6047255 |
| APKFRTGGYMIDFWGVGYQVAK | ASSPPRGWNSPLY | 0.44057918 |
| GAGGAGGAGGAAGGPGGTGGTG | ASSPPRGWNSPLY | 0.5842698 |
| IGALGLLATDVAPAGSVAFGVA | ASSPPRGWNSPLY | 0.21759655 |
| AGGAGGIGGNGGAGGAGGVLMG | ASSPPRGWNSPLY | 0.1347065 |
| ADELIGLL | ASSPPRGWNSPLY | 0.6521222 |
| ADRALKA | ASSPPRGWNSPLY | 0.2505522 |
| AGAVGQVGGAGGA | ASSPPRGWNSPLY | 0.19701083 |
| AGGGGGLGGAP | ASSPPRGWNSPLY | 0.3774578 |
| AGGLLYGNGGAG | ASSPPRGWNSPLY | 0.16089489 |
| AGLESIGL | ASSPPRGWNSPLY | 0.14776292 |
| AGLKTGII | ASSPPRGWNSPLY | 0.3652639 |
| ASLAALLL | ASSPPRGWNSPLY | 0.37362063 |
| ASVINISSI | ASSPPRGWNSPLY | 0.12777068 |
| ATPPGPA | ASSPPRGWNSPLY | 0.14476168 |
| ATVASALV | ASSPPRGWNSPLY | 0.16120966 |
| ATVASLIL | ASSPPRGWNSPLY | 0.26524326 |
| AVNGPVLIRH | ASSPPRGWNSPLY | 0.50365996 |
| DGAAGAVGI | ASSPPRGWNSPLY | 0.2435084 |
| DPLLHGI | ASSPPRGWNSPLY | 0.21964821 |
| ELGAALGAL | ASSPPRGWNSPLY | 0.40775502 |
| ELGRTQAV | ASSPPRGWNSPLY | 0.31010753 |
| FVAGGYLS | ASSPPRGWNSPLY | 0.2227338 |
| GASGGAGGAGGW | ASSPPRGWNSPLY | 0.40732658 |
| GGAGAGGDGGAGGI | ASSPPRGWNSPLY | 0.14906844 |
| GTGHNGSQPGMGGNG | ASSPPRGWNSPLY | 0.36419702 |
| GYAAASASAS | ASSPPRGWNSPLY | 0.38182363 |
| HQVDVAAVQQG | ASSPPRGWNSPLY | 0.38260025 |

| | | |
|---|---|---|
| IGAAIAHT | ASSPPRGWNSPLY | 0.4737762 |
| IGMIEVAQL | ASSPPRGWNSPLY | 0.23361737 |
| IRLAPEHTSAMD | ASSPPRGWNSPLY | 0.25996053 |
| IYNVSVL | ASSPPRGWNSPLY | 0.118832424 |
| KQGFLGM | ASSPPRGWNSPLY | 0.28881836 |
| LESVITQL | ASSPPRGWNSPLY | 0.30625257 |
| LGLGALVVV | ASSPPRGWNSPLY | 0.2704243 |
| LIGNGGNGGSGGI | ASSPPRGWNSPLY | 0.42118746 |
| LNAINAPTL | ASSPPRGWNSPLY | 0.3442601 |
| LQLLNTGV | ASSPPRGWNSPLY | 0.15978502 |
| LQTSIEF | ASSPPRGWNSPLY | 0.5698024 |
| MSFAVSEL | ASSPPRGWNSPLY | 0.653803 |
| NTGALGL | ASSPPRGWNSPLY | 0.12756601 |
| PPGAGGAGGAGTT | ASSPPRGWNSPLY | 0.18545267 |
| QLVGLGI | ASSPPRGWNSPLY | 0.62934667 |
| RFLEQVHQL | ASSPPRGWNSPLY | 0.41992062 |
| RGVGYVLR | ASSPPRGWNSPLY | 0.22528589 |
| SATGVFAGVI | ASSPPRGWNSPLY | 0.23214276 |
| SLPAITLP | ASSPPRGWNSPLY | 0.3970612 |
| STALAFTV | ASSPPRGWNSPLY | 0.17790592 |
| SVLTVGLGK | ASSPPRGWNSPLY | 0.6016177 |
| TALAGAGSGI | ASSPPRGWNSPLY | 0.11102539 |
| TGLPFTGI | ASSPPRGWNSPLY | 0.214147 |
| VAEHVTV | ASSPPRGWNSPLY | 0.15676014 |
| VDIVLNSL | ASSPPRGWNSPLY | 0.108349495 |
| VLTGGGAVL | ASSPPRGWNSPLY | 0.3624066 |
| VNVGLAYGI | ASSPPRGWNSPLY | 0.16845451 |
| VSSAAVGAV | ASSPPRGWNSPLY | 0.37587106 |
| VTSVYEAGV | ASSPPRGWNSPLY | 0.21042511 |
| WVALDRGA | ASSPPRGWNSPLY | 0.32789832 |
| YLVGSVVI | ASSPPRGWNSPLY | 0.38513893 |
| GPPGTGKTLL | ASSPPRGWNSPLY | 0.21377875 |
| GVGMTKF | ASSPPRGWNSPLY | 0.15868306 |
| IGIGGSDL | ASSPPRGWNSPLY | 0.29171777 |
| IGTLLAEL | ASSPPRGWNSPLY | 0.5311966 |
| IYTSGTT | ASSPPRGWNSPLY | 0.3445804 |
| LSFPAGV | ASSPPRGWNSPLY | 0.11977884 |
| LYTSGTT | ASSPPRGWNSPLY | 0.200311 |
| VGAFPFGF | ASSPPRGWNSPLY | 0.20443912 |
| VGAGTVEF | ASSPPRGWNSPLY | 0.10941628 |
| VGKSTLLL | ASSPPRGWNSPLY | 0.11339916 |
| AALAHAGGGIV | ASSPPRGWNSPLY | 0.12609094 |
| AALVAVSLL | ASSPPRGWNSPLY | 0.18414815 |
| AAVSRAVAAL | ASSPPRGWNSPLY | 0.43674758 |
| AGGSSGIGGF | ASSPPRGWNSPLY | 0.42749742 |
| AGLTVANANG | ASSPPRGWNSPLY | 0.20977184 |
| AGLVGAVL | ASSPPRGWNSPLY | 0.47992843 |
| ALSQAFTV | ASSPPRGWNSPLY | 0.24432796 |
| ALTFSVSL | ASSPPRGWNSPLY | 0.09845124 |
| ALTGAALVAL | ASSPPRGWNSPLY | 0.18602805 |
| ASFSIGAGVL | ASSPPRGWNSPLY | 0.13042808 |

| | | |
|---|---|---|
| DGSVYKI | ASSPPRGWNSPLY | 0.4889021 |
| EQLAEIV | ASSPPRGWNSPLY | 0.13414085 |
| FAVSELTVM | ASSPPRGWNSPLY | 0.17095399 |
| GAAPLAAGL | ASSPPRGWNSPLY | 0.2333184 |
| GADGGDGGSGGAGG | ASSPPRGWNSPLY | 0.21094981 |
| GFGLDTGGAGG | ASSPPRGWNSPLY | 0.3209231 |
| GLASGGLL | ASSPPRGWNSPLY | 0.14544822 |
| GRIAGAAVLL | ASSPPRGWNSPLY | 0.11181933 |
| GSAGSLNTSF | ASSPPRGWNSPLY | 0.31197426 |
| GTGQIGI | ASSPPRGWNSPLY | 0.1209164 |
| GTLIALIL | ASSPPRGWNSPLY | 0.14762147 |
| IAATMGFVVG | ASSPPRGWNSPLY | 0.11891889 |
| IFAQALL | ASSPPRGWNSPLY | 0.3026113 |
| IGAAVALGF | ASSPPRGWNSPLY | 0.13131903 |
| IGLTPLAL | ASSPPRGWNSPLY | 0.46661276 |
| ILGSNTSTL | ASSPPRGWNSPLY | 0.19947776 |
| IVFSLTVL | ASSPPRGWNSPLY | 0.20463136 |
| LASKATGF | ASSPPRGWNSPLY | 0.5890032 |
| LASVALIL | ASSPPRGWNSPLY | 0.37441027 |
| LFAVGLTL | ASSPPRGWNSPLY | 0.23464833 |
| LFNGGIGTYI | ASSPPRGWNSPLY | 0.15781738 |
| LTLGAGII | ASSPPRGWNSPLY | 0.14550759 |
| LVAAALGDI | ASSPPRGWNSPLY | 0.13937442 |
| NTAFVQLGI | ASSPPRGWNSPLY | 0.14605439 |
| QLFSGAEL | ASSPPRGWNSPLY | 0.54824805 |
| SAATFITL | ASSPPRGWNSPLY | 0.23824756 |
| SNYGLANLGV | ASSPPRGWNSPLY | 0.13570756 |
| SYGGAYAVM | ASSPPRGWNSPLY | 0.25841582 |
| TTGVGGLTLG | ASSPPRGWNSPLY | 0.62768674 |
| TTQTIYI | ASSPPRGWNSPLY | 0.7560352 |
| VAAGAGLGAVI | ASSPPRGWNSPLY | 0.14319636 |
| VAAVVTLL | ASSPPRGWNSPLY | 0.2102801 |
| VGFLGGLI | ASSPPRGWNSPLY | 0.20426214 |
| VGVIPILL | ASSPPRGWNSPLY | 0.24201241 |
| VIFASSLI | ASSPPRGWNSPLY | 0.18340418 |
| VKGGDGGAAAG | ASSPPRGWNSPLY | 0.14217566 |
| VTKTFTNI | ASSPPRGWNSPLY | 0.34857818 |
| VVAAAILL | ASSPPRGWNSPLY | 0.8102349 |
| YAALDVEL | ASSPPRGWNSPLY | 0.168082 |
| LASVALLL | ASSPPRGWNSPLY | 0.12113995 |
| LDAVHRV | ASSPPRGWNSPLY | 0.22033009 |
| QGLSSSIF | ASSPPRGWNSPLY | 0.1306909 |
| RGTFANIRL | ASSPPRGWNSPLY | 0.17224762 |
| RQLWWGHRI | ASSPPRGWNSPLY | 0.24793012 |
| YGPPGTGKTLL | ASSPPRGWNSPLY | 0.12576889 |
| AALVGAPANVA | ASSPPRGWNSPLY | 0.19721246 |
| AATFRRTE | ASSPPRGWNSPLY | 0.71585554 |
| AGALGGGI | ASSPPRGWNSPLY | 0.17800042 |
| AGGAGGAGGIGAG | ASSPPRGWNSPLY | 0.40035295 |
| AGSGALGL | ASSPPRGWNSPLY | 0.5958108 |
| AQIADQI | ASSPPRGWNSPLY | 0.38924527 |

| | | |
|---|---|---|
| ASAVAVTYL | ASSPPRGWNSPLY | 0.6647866 |
| ASTVLGGTGG | ASSPPRGWNSPLY | 0.51856154 |
| AVGGEPAK | ASSPPRGWNSPLY | 0.20181242 |
| CEQGTFGGTC | ASSPPRGWNSPLY | 0.24280027 |
| DEGKAPAG | ASSPPRGWNSPLY | 0.4889931 |
| DTVVFPAL | ASSPPRGWNSPLY | 0.22260283 |
| FEGITMAG | ASSPPRGWNSPLY | 0.27574015 |
| FPLVVQD | ASSPPRGWNSPLY | 0.19808209 |
| GAFTTPQ | ASSPPRGWNSPLY | 0.50063604 |
| GFGGFGTAGGVGGA | ASSPPRGWNSPLY | 0.24788171 |
| GFLAQFL | ASSPPRGWNSPLY | 0.56861734 |
| GGAGGAGGNAGVA | ASSPPRGWNSPLY | 0.12927295 |
| GGVGTAGAGGAGGAGG | ASSPPRGWNSPLY | 0.14051925 |
| GNGGGGDAGNAGSGGN | ASSPPRGWNSPLY | 0.58912313 |
| GTGGAGGAGNDAGS | ASSPPRGWNSPLY | 0.10688882 |
| GTTNTGWFNS | ASSPPRGWNSPLY | 0.12537439 |
| IFMGPLFTL | ASSPPRGWNSPLY | 0.22612251 |
| ILAHAIEAI | ASSPPRGWNSPLY | 0.6627507 |
| IRSAIREAL | ASSPPRGWNSPLY | 0.10041404 |
| ITFAGGG | ASSPPRGWNSPLY | 0.1491061 |
| ITGMSGHQEV | ASSPPRGWNSPLY | 0.143236 |
| IVAGLACTL | ASSPPRGWNSPLY | 0.3187278 |
| IVGGGLAGA | ASSPPRGWNSPLY | 0.23914713 |
| IVTGVLNI | ASSPPRGWNSPLY | 0.1351078 |
| LAQQGCI | ASSPPRGWNSPLY | 0.3385821 |
| LCLHVEN | ASSPPRGWNSPLY | 0.11948063 |
| LDAISDGS | ASSPPRGWNSPLY | 0.14416498 |
| LGGFAAML | ASSPPRGWNSPLY | 0.33577013 |
| LGGKNPMI | ASSPPRGWNSPLY | 0.6552653 |
| LGPKGVAAIL | ASSPPRGWNSPLY | 0.1422759 |
| LMVGVSI | ASSPPRGWNSPLY | 0.39565426 |
| LVGAAFVL | ASSPPRGWNSPLY | 0.17550278 |
| LVLAVAPL | ASSPPRGWNSPLY | 0.81195766 |
| MAAIAVIDAA | ASSPPRGWNSPLY | 0.18014704 |
| NNGINAELQ | ASSPPRGWNSPLY | 0.13505284 |
| RGEPLVLVHG | ASSPPRGWNSPLY | 0.39243355 |
| RGGAGGAAGLI | ASSPPRGWNSPLY | 0.52742887 |
| RSYRMHPAV | ASSPPRGWNSPLY | 0.23060629 |
| SAAHLLAGQLA | ASSPPRGWNSPLY | 0.514723 |
| SAAVGSY | ASSPPRGWNSPLY | 0.18398164 |
| SPHAVLHRI | ASSPPRGWNSPLY | 0.20816223 |
| SQIAKVL | ASSPPRGWNSPLY | 0.46994513 |
| TALGHLVP | ASSPPRGWNSPLY | 0.33372596 |
| TAPTTSGAA | ASSPPRGWNSPLY | 0.13145424 |
| TDGDDGNGGSPGT | ASSPPRGWNSPLY | 0.37761712 |
| TDGLGEN | ASSPPRGWNSPLY | 0.22620702 |
| TGGPPLPGQA | ASSPPRGWNSPLY | 0.16237 |
| TIPAITI | ASSPPRGWNSPLY | 0.6353297 |
| TIPGFTI | ASSPPRGWNSPLY | 0.2762832 |
| TSIASGALA | ASSPPRGWNSPLY | 0.47290847 |
| TVTLPGAAI | ASSPPRGWNSPLY | 0.26712382 |

| | | |
|---|---|---|
| VASGLSAALP | ASSPPRGWNSPLY | 0.78739536 |
| VGAGVIGI | ASSPPRGWNSPLY | 0.14606242 |
| VGPKRFGV | ASSPPRGWNSPLY | 0.33930817 |
| VNVAFPD | ASSPPRGWNSPLY | 0.2635844 |
| VSMTNSGVS | ASSPPRGWNSPLY | 0.3651454 |
| VTDVLGDLSV | ASSPPRGWNSPLY | 0.260014 |
| VVVVGAGFAGL | ASSPPRGWNSPLY | 0.16912997 |
| VYTVGTVV | ASSPPRGWNSPLY | 0.31324548 |
| WTTFSGP | ASSPPRGWNSPLY | 0.5207894 |
| YDELVEAF | ASSPPRGWNSPLY | 0.1980949 |
| YGTGGAGGNGG | ASSPPRGWNSPLY | 0.29166463 |
| YTNVMAV | ASSPPRGWNSPLY | 0.7044962 |
| FLEQVHQL | ASSPPRGWNSPLY | 0.15983902 |
| GLSLLAVAV | ASSPPRGWNSPLY | 0.19522737 |
| LEQVHQL | ASSPPRGWNSPLY | 0.31763142 |
| LNAAGAI | ASSPPRGWNSPLY | 0.5210873 |
| PAVHASL | ASSPPRGWNSPLY | 0.21254495 |
| TELEDDA | ASSPPRGWNSPLY | 0.56951505 |
| TPPGPPN | ASSPPRGWNSPLY | 0.14714059 |
| TPPPPPPP | ASSPPRGWNSPLY | 0.37776533 |
| TQSPALI | ASSRDERGYAEQF | 0.25650543 |
| EFAAAFPQ | ASSRDERGYAEQF | 0.28989 |
| LLSGNLDVL | ASSRDERGYAEQF | 0.11463631 |
| GPANLGNYNI | ASSRDERGYAEQF | 0.32971358 |
| TLGPTLP | ASSRDERGYAEQF | 0.536813 |
| VYAYSSCCK | ASSRDERGYAEQF | 0.4324989 |
| APAPAPAPNI | ASSRDERGYAEQF | 0.10952556 |
| PGGPVPIP | ASSRDERGYAEQF | 0.38384628 |
| FLTVALEP | ASSRDERGYAEQF | 0.4490899 |
| SQYFGANL | ASSRDERGYAEQF | 0.7792869 |
| GGVKPAA | ASSRDERGYAEQF | 0.25135234 |
| GGTGGAGGAGGW | ASSRDERGYAEQF | 0.36419064 |
| GVGGVAGLGGAGG | ASSRDERGYAEQF | 0.15799688 |
| VLLVLATTI | ASSRDERGYAEQF | 0.2107526 |
| IGIFAAML | ASSRDERGYAEQF | 0.13851407 |
| AVGDLID | ASSRDERGYAEQF | 0.1921965 |
| NISPLPL | ASSRDERGYAEQF | 0.26889092 |
| VTAVFVAI | ASSRDERGYAEQF | 0.14988202 |
| SGDDNDAAL | ASSRDERGYAEQF | 0.41116095 |
| AVAALAHS | ASSRDERGYAEQF | 0.5820836 |
| GGVTVMATP | ASSRDERGYAEQF | 0.49956366 |
| AGLAANVRD | ASSRDERGYAEQF | 0.60603464 |
| TNRPDLIDPALLRPG | ASSRDERGYAEQF | 0.7315353 |
| LANLGVTAL | ASSRDERGYAEQF | 0.37688756 |
| TGERIEL | ASSRDERGYAEQF | 0.14408602 |
| NGEELAA | ASSRDERGYAEQF | 0.28926072 |
| VLSLVWG | ASSRDERGYAEQF | 0.12223798 |
| IGYALWL | ASSRDERGYAEQF | 0.19632404 |
| LSLLLAPL | ASSRDERGYAEQF | 0.15139188 |
| TLDEALK | ASSRDERGYAEQF | 0.2916304 |
| TDGAGTAEGG | ASSRDERGYAEQF | 0.09948341 |

| | | |
|---|---|---|
| NAPGGAGGNGG | ASSRDERGYAEQF | 0.4969257 |
| VSVFMGMGGNF | ASSRDERGYAEQF | 0.5696176 |
| LASYLAAL | ASSRDERGYAEQF | 0.14973056 |
| LVGGTPAL | ASSRDERGYAEQF | 0.37057132 |
| VRNADVL | ASSRDERGYAEQF | 0.5501516 |
| TCSSPFLVSAM | ASSRDERGYAEQF | 0.14629205 |
| VHITDAE | ASSRDERGYAEQF | 0.48552915 |
| EYAAPSF | ASSRDERGYAEQF | 0.14643295 |
| VVDAFHGGL | ASSRDERGYAEQF | 0.40015656 |
| ASVVDPQ | ASSRDERGYAEQF | 0.49751186 |
| SYNEFGGLS | ASSRDERGYAEQF | 0.27567023 |
| DPDGSVY | ASSRDERGYAEQF | 0.2702782 |
| MVSTIHATE | ASSRDERGYAEQF | 0.269807 |
| LAAGVADPV | ASSRDERGYAEQF | 0.13739091 |
| TKEQIAA | ASSRDERGYAEQF | 0.28522408 |
| SLASPAAL | ASSRDERGYAEQF | 0.13408753 |
| IGLVFYGV | ASSRDERGYAEQF | 0.31051838 |
| FLLDASNL | ASSRDERGYAEQF | 0.13902387 |
| YQVGNFP | ASSRDERGYAEQF | 0.28540498 |
| ESLLSQST | ASSRDERGYAEQF | 0.2090111 |
| LFTVGFAAVG | ASSRDERGYAEQF | 0.24753857 |
| ATPPTPT | ASSRDERGYAEQF | 0.15563181 |
| TDLATLG | ASSRDERGYAEQF | 0.2380724 |
| SAGGTGR | ASSRDERGYAEQF | 0.14611289 |
| GFSNSGS | ASSRDERGYAEQF | 0.12642102 |
| GGNGGSGGTSV | ASSRDERGYAEQF | 0.4847806 |
| GTSVATGGAGN | ASSRDERGYAEQF | 0.2024495 |
| NGGNGGSGGTS | ASSRDERGYAEQF | 0.23708233 |
| AVVKFPH | ASSRDERGYAEQF | 0.41080523 |
| TPPPPPP | ASSRDERGYAEQF | 0.23997715 |
| AAPAAAAAPA | ASSRDERGYAEQF | 0.12677771 |
| IALNEDL | ASSRDERGYAEQF | 0.27237314 |
| IVNVSSM | ASSRDERGYAEQF | 0.2470965 |
| TGIPPAPR | ASSRDERGYAEQF | 0.18525344 |
| FDLGGGTF | ASSRDERGYAEQF | 0.10304406 |
| FDLGGGTFDVS | ASSRDERGYAEQF | 0.24828902 |
| NEPTAAA | ASSRDERGYAEQF | 0.16363043 |
| NEPTAAALA | ASSRDERGYAEQF | 0.15472175 |
| GSIINISSI | ASSRDERGYAEQF | 0.1595689 |
| VAPGNAGTA | ASSRDERGYAEQF | 0.5284318 |
| TGGGASLE | ASSRDERGYAEQF | 0.5027289 |
| VAVYNHYKRI | ASSRDERGYAEQF | 0.18639797 |
| TDPAPATT | ASSRDERGYAEQF | 0.41315925 |
| TNRPDILDPALLRPG | ASSRDERGYAEQF | 0.29405966 |
| LTTTTTS | ASSRDERGYAEQF | 0.67184675 |
| PTGSGKTL | ASSRDERGYAEQF | 0.1881647 |
| TGSGKTL | ASSRDERGYAEQF | 0.18349117 |
| TIGHVDHGK | ASSRDERGYAEQF | 0.2593311 |
| TIGHVDHGKTT | ASSRDERGYAEQF | 0.25139722 |
| GHVDHGKTT | ASSRDERGYAEQF | 0.13432707 |
| AQFEHTLL | ASSRDERGYAEQF | 0.11345216 |

| | | |
|---|---|---|
| IVNQDEL | ASSRDERGYAEQF | 0.24205495 |
| TNGSQFF | ASSRDERGYAEQF | 0.13330719 |
| TGKGIVE | ASSRDERGYAEQF | 0.25085077 |
| LISNASDA | ASSRDERGYAEQF | 0.5141674 |
| AELRTLGELTKIAKAHGAQ | ASSRDERGYAEQF | 0.25044078 |
| APNAVAAAEPSW | ASSRDERGYAEQF | 0.13513377 |
| APQVAGAATAVL | ASSRDERGYAEQF | 0.16656293 |
| ARRDPAAAAAILPTDA | ASSRDERGYAEQF | 0.22463332 |
| DAAARDVLAEAGLAEYF | ASSRDERGYAEQF | 0.44005525 |
| DDQIVLSAKANLAKVEQ | ASSRDERGYAEQF | 0.23850581 |
| DELLPDTGEDSDEDGH | ASSRDERGYAEQF | 0.7822 |
| ELPEAPANRATVPQD | ASSRDERGYAEQF | 0.4208882 |
| FGQSKIERVTGIG | ASSRDERGYAEQF | 0.156582 |
| GAGGTGSTAGGAGGAGGAGGLY | ASSRDERGYAEQF | 0.40113506 |
| GHGGTGAAGGNGGTGGAGGSGI | ASSRDERGYAEQF | 0.22505912 |
| GNGGNGGAGGEGGPGGAGGAGA | ASSRDERGYAEQF | 0.26030582 |
| GSTAGNGGNGGNATNGTIAGQP | ASSRDERGYAEQF | 0.22499363 |
| GTGGDGGNGGNGGTGG | ASSRDERGYAEQF | 0.21087316 |
| GVRRPAGIPADAPV | ASSRDERGYAEQF | 0.18003377 |
| GYRQVTDAHLLTLARR | ASSRDERGYAEQF | 0.18337576 |
| IAKMGQDGHDRG | ASSRDERGYAEQF | 0.12900314 |
| KAEPSIEIQYT | ASSRDERGYAEQF | 0.21905467 |
| KPVRIRMAADEDTVAVT | ASSRDERGYAEQF | 0.14241606 |
| MPPRIAGMRLLVIKPEP | ASSRDERGYAEQF | 0.14017875 |
| MKFYAVALFRSRGVPPTR | ASSRDERGYAEQF | 0.20171365 |
| NAGGFGILGAVAHSP | ASSRDERGYAEQF | 0.7315559 |
| NEGREITLAMAEM | ASSRDERGYAEQF | 0.7271134 |
| NELVEWSHPLGHRWRWE | ASSRDERGYAEQF | 0.15328202 |
| NLGAGGTGGNGGNPG | ASSRDERGYAEQF | 0.14252119 |
| NTEWPKWETVMAADPVD | ASSRDERGYAEQF | 0.49653912 |
| PASAAPAETQP | ASSRDERGYAEQF | 0.12123566 |
| QQSSDGKNGVRAAEITGE | ASSRDERGYAEQF | 0.32250008 |
| RAPDAVTTSWWKE | ASSRDERGYAEQF | 0.27614242 |
| SGFGNVGAHTSGWFNQ | ASSRDERGYAEQF | 0.2471279 |
| SGVAGAVGQAASVGGLKV | ASSRDERGYAEQF | 0.17343539 |
| SNGGAGTGSAGGNGGT | ASSRDERGYAEQF | 0.25437906 |
| TGGAGGEGGAAG | ASSRDERGYAEQF | 0.13786612 |
| TGGDGGAAGAGGAPGHG | ASSRDERGYAEQF | 0.37530857 |
| TMPPLLAAIANAGRHGV | ASSRDERGYAEQF | 0.09851348 |
| TSSTAGNSGGA | ASSRDERGYAEQF | 0.27195913 |
| VPATVPVGILC | ASSRDERGYAEQF | 0.146222 |
| VQRPDSGGVVLTGRIS | ASSRDERGYAEQF | 0.16787174 |
| WTPNPVPATKGVPAGLVQAT | ASSRDERGYAEQF | 0.12305257 |
| YALKRALAKFSDRFDVVII | ASSRDERGYAEQF | 0.19217786 |
| YLIGSVIV | ASSRDERGYAEQF | 0.5399673 |
| YGQLVAVID | ASSRDERGYAEQF | 0.48115742 |
| VTVGGMTLPA | ASSRDERGYAEQF | 0.15619655 |
| VLALPAV | ASSRDERGYAEQF | 0.4605021 |
| VIPPTLNYET | ASSRDERGYAEQF | 0.73450464 |
| VGMGGMHPSSG | ASSRDERGYAEQF | 0.2243503 |
| VGLAGGAATA | ASSRDERGYAEQF | 0.12216184 |

| | | |
|---|---|---|
| VGGAGGAGGSGP | ASSRDERGYAEQF | 0.2826345 |
| VFAAFMV | ASSRDERGYAEQF | 0.12674035 |
| TLYDKLR | ASSRDERGYAEQF | 0.3949327 |
| TLAVFTVL | ASSRDERGYAEQF | 0.62098014 |
| TGSVQPARV | ASSRDERGYAEQF | 0.121316634 |
| TGGGAGSGGT | ASSRDERGYAEQF | 0.4097729 |
| SHLETSL | ASSRDERGYAEQF | 0.5029008 |
| SFTTGMT | ASSRDERGYAEQF | 0.14999615 |
| QVEVAKRMGVRQPTVS | ASSRDERGYAEQF | 0.14967029 |
| QGSGGPALAM | ASSRDERGYAEQF | 0.15042742 |
| MLAPSNP | ASSRDERGYAEQF | 0.123381086 |
| LVNNAAI | ASSRDERGYAEQF | 0.2631565 |
| LTGEINTLA | ASSRDERGYAEQF | 0.16977203 |
| LSIFGPV | ASSRDERGYAEQF | 0.16274852 |
| LLALLANL | ASSRDERGYAEQF | 0.21689674 |
| LISNASDALDKLR | ASSRDERGYAEQF | 0.15667789 |
| LINEAVP | ASSRDERGYAEQF | 0.17632213 |
| LGYAFSGL | ASSRDERGYAEQF | 0.16823761 |
| LDFVAVAL | ASSRDERGYAEQF | 0.18260749 |
| LAQPGAE | ASSRDERGYAEQF | 0.30257392 |
| ISNLGAVTGF | ASSRDERGYAEQF | 0.18209212 |
| HQEDPGRI | ASSRDERGYAEQF | 0.32577047 |
| GTPGEPTP | ASSRDERGYAEQF | 0.64539766 |
| GFFNSGS | ASSRDERGYAEQF | 0.11760078 |
| GAGGGGLTAGAN | ASSRDERGYAEQF | 0.15490735 |
| FVDALVNM | ASSRDERGYAEQF | 0.30878308 |
| FLTVAPEL | ASSRDERGYAEQF | 0.12219507 |
| FETLTAL | ASSRDERGYAEQF | 0.5557648 |
| DVINPIEL | ASSRDERGYAEQF | 0.20924984 |
| APVSGAPV | ASSRDERGYAEQF | 0.3388096 |
| ALVGGLAW | ASSRDERGYAEQF | 0.25975704 |
| ALAAAAATAI | ASSRDERGYAEQF | 0.7277579 |
| AFGLGLLL | ASSRDERGYAEQF | 0.35542858 |
| AFFAAASL | ASSRDERGYAEQF | 0.18461962 |
| AAAALPAFSPP | ASSRDERGYAEQF | 0.44313008 |
| VVLAPAL | ASSRDERGYAEQF | 0.14101529 |
| TIGHVDHGKTTL | ASSRDERGYAEQF | 0.40616393 |
| LISNASDALDK | ASSRDERGYAEQF | 0.35367996 |
| LISNASDAL | ASSRDERGYAEQF | 0.1474231 |
| ISNASDALDK | ASSRDERGYAEQF | 0.2138443 |
| NQAETLVYQTE | ASSRDERGYAEQF | 0.47459704 |
| TGGTGGTAGAG | ASSRDERGYAEQF | 0.47911128 |
| TGSTAGGAGGA | ASSRDERGYAEQF | 0.13083975 |
| DPLAVPVAAGPV | ASSRDERGYAEQF | 0.16275817 |
| GGTGGTPGNGGN | ASSRDERGYAEQF | 0.14422645 |
| PAGLAGVDPATL | ASSRDERGYAEQF | 0.20627043 |
| GNNLVVEAPHAGASV | ASSRDERGYAEQF | 0.13686791 |
| KVTRIMSASTLEQPA | ASSRDERGYAEQF | 0.45384133 |
| LGFALPIALPRRTVV | ASSRDERGYAEQF | 0.21654502 |
| TRLGNLTARKLVVLV | ASSRDERGYAEQF | 0.68649673 |
| WSQQRRLIYFPSAGP | ASSRDERGYAEQF | 0.2609825 |

| | | |
|---|---|---|
| GPNGNPDPMAAAVDIR | ASSRDERGYAEQF | 0.39817405 |
| TQRCVSVADLRAAASR | ASSRDERGYAEQF | 0.18080035 |
| QFVLDVAEHPVEVLTETV | ASSRDERGYAEQF | 0.1665812 |
| GGAGGTGGAGGNGGTQAGDG | ASSRDERGYAEQF | 0.13600235 |
| PLLATLRAIAVDVDAKPAQV | ASSRDERGYAEQF | 0.12515981 |
| APKFRTGGYMIDFWGVGYQVAK | ASSRDERGYAEQF | 0.1536181 |
| GAGGAGGAGGAAGGPGGTGGTG | ASSRDERGYAEQF | 0.16590944 |
| IGALGLLATDVAPAGSVAFGVA | ASSRDERGYAEQF | 0.10280067 |
| AGGAGGIGGNGGAGGAGGVLMG | ASSRDERGYAEQF | 0.6336197 |
| ADELIGLL | ASSRDERGYAEQF | 0.24849948 |
| ADRALKA | ASSRDERGYAEQF | 0.33538237 |
| AGAVGQVGGAGGA | ASSRDERGYAEQF | 0.17742142 |
| AGGGGGLGGAP | ASSRDERGYAEQF | 0.16846742 |
| AGGLLYGNGGAG | ASSRDERGYAEQF | 0.14097065 |
| AGLESIGL | ASSRDERGYAEQF | 0.43811017 |
| AGLKTGII | ASSRDERGYAEQF | 0.13149737 |
| ASLAALLL | ASSRDERGYAEQF | 0.11320938 |
| ASVINISSI | ASSRDERGYAEQF | 0.18353471 |
| ATPPGPA | ASSRDERGYAEQF | 0.1249048 |
| ATVASALV | ASSRDERGYAEQF | 0.721595 |
| ATVASLIL | ASSRDERGYAEQF | 0.47072852 |
| AVNGPVLIRH | ASSRDERGYAEQF | 0.31135738 |
| DGAAGAVGI | ASSRDERGYAEQF | 0.5056911 |
| DPLLHGI | ASSRDERGYAEQF | 0.11113003 |
| ELGAALGAL | ASSRDERGYAEQF | 0.1842168 |
| ELGRTQAV | ASSRDERGYAEQF | 0.66739273 |
| FVAGGYLS | ASSRDERGYAEQF | 0.23916844 |
| GASGGAGGAGGW | ASSRDERGYAEQF | 0.24208754 |
| GGAGAGGDGGAGGI | ASSRDERGYAEQF | 0.52359635 |
| GTGHNGSQPGMGGNG | ASSRDERGYAEQF | 0.14588334 |
| GYAAASASAS | ASSRDERGYAEQF | 0.18770301 |
| HQVDVAAVQQG | ASSRDERGYAEQF | 0.17843197 |
| IGAAIAHT | ASSRDERGYAEQF | 0.28332204 |
| IGMIEVAQL | ASSRDERGYAEQF | 0.26585594 |
| IRLAPEHTSAMD | ASSRDERGYAEQF | 0.1325325 |
| IYNVSVL | ASSRDERGYAEQF | 0.37460983 |
| KQGFLGM | ASSRDERGYAEQF | 0.7875976 |
| LESVITQL | ASSRDERGYAEQF | 0.20707023 |
| LGLGALVVV | ASSRDERGYAEQF | 0.51180285 |
| LIGNGGNGGSGGI | ASSRDERGYAEQF | 0.32610893 |
| LNAINAPTL | ASSRDERGYAEQF | 0.62453055 |
| LQLLNTGV | ASSRDERGYAEQF | 0.16129315 |
| LQTSIEF | ASSRDERGYAEQF | 0.14132202 |
| MSFAVSEL | ASSRDERGYAEQF | 0.24282451 |
| NTGALGL | ASSRDERGYAEQF | 0.16273418 |
| PPGAGGAGGAGTT | ASSRDERGYAEQF | 0.2500217 |
| QLVGLGI | ASSRDERGYAEQF | 0.19364165 |
| RFLEQVHQL | ASSRDERGYAEQF | 0.82085204 |
| RGVGYVLR | ASSRDERGYAEQF | 0.45176056 |
| SATGVFAGVI | ASSRDERGYAEQF | 0.2973349 |
| SLPAITLP | ASSRDERGYAEQF | 0.12663329 |

| | | |
|---|---|---|
| STALAFTV | ASSRDERGYAEQF | 0.21918203 |
| SVLTVGLGK | ASSRDERGYAEQF | 0.35335588 |
| TALAGAGSGI | ASSRDERGYAEQF | 0.13303116 |
| TGLPFTGI | ASSRDERGYAEQF | 0.6026925 |
| VAEHVTV | ASSRDERGYAEQF | 0.15597814 |
| VDIVLNSL | ASSRDERGYAEQF | 0.23335993 |
| VLTGGGAVL | ASSRDERGYAEQF | 0.31153542 |
| VNVGLAYGI | ASSRDERGYAEQF | 0.74499047 |
| VSSAAVGAV | ASSRDERGYAEQF | 0.2242771 |
| VTSVYEAGV | ASSRDERGYAEQF | 0.24207127 |
| WVALDRGA | ASSRDERGYAEQF | 0.13141182 |
| YLVGSVVI | ASSRDERGYAEQF | 0.48883456 |
| GPPGTGKTLL | ASSRDERGYAEQF | 0.55931604 |
| GVGMTKF | ASSRDERGYAEQF | 0.5490724 |
| IGIGGSDL | ASSRDERGYAEQF | 0.69340914 |
| IGTLLAEL | ASSRDERGYAEQF | 0.2635063 |
| IYTSGTT | ASSRDERGYAEQF | 0.13285042 |
| LSFPAGV | ASSRDERGYAEQF | 0.24046382 |
| LYTSGTT | ASSRDERGYAEQF | 0.47389728 |
| VGAFPFGF | ASSRDERGYAEQF | 0.66138595 |
| VGAGTVEF | ASSRDERGYAEQF | 0.3357614 |
| VGKSTLLL | ASSRDERGYAEQF | 0.40740135 |
| AALAHAGGGIV | ASSRDERGYAEQF | 0.17542286 |
| AALVAVSLL | ASSRDERGYAEQF | 0.6080874 |
| AAVSRAVAAL | ASSRDERGYAEQF | 0.7257265 |
| AGGSSGIGGF | ASSRDERGYAEQF | 0.41250998 |
| AGLTVANANG | ASSRDERGYAEQF | 0.20969312 |
| AGLVGAVL | ASSRDERGYAEQF | 0.19090398 |
| ALSQAFTV | ASSRDERGYAEQF | 0.27352515 |
| ALTFSVSL | ASSRDERGYAEQF | 0.4655464 |
| ALTGAALVAL | ASSRDERGYAEQF | 0.1546687 |
| ASFSIGAGVL | ASSRDERGYAEQF | 0.3516346 |
| DGSVYKI | ASSRDERGYAEQF | 0.27744988 |
| EQLAEIV | ASSRDERGYAEQF | 0.12142135 |
| FAVSELTVM | ASSRDERGYAEQF | 0.366328 |
| GAAPLAAGL | ASSRDERGYAEQF | 0.68864995 |
| GADGGDGGSGGAGG | ASSRDERGYAEQF | 0.27130795 |
| GFGLDTGGAGG | ASSRDERGYAEQF | 0.16265914 |
| GLASGGLL | ASSRDERGYAEQF | 0.65104043 |
| GRIAGAAVLL | ASSRDERGYAEQF | 0.78831226 |
| GSAGSLNTSF | ASSRDERGYAEQF | 0.5597706 |
| GTGQIGI | ASSRDERGYAEQF | 0.1268821 |
| GTLIALIL | ASSRDERGYAEQF | 0.11040084 |
| IAATMGFVVG | ASSRDERGYAEQF | 0.23283431 |
| IFAQALL | ASSRDERGYAEQF | 0.3550095 |
| IGAAVALGF | ASSRDERGYAEQF | 0.16036685 |
| IGLTPLAL | ASSRDERGYAEQF | 0.14071158 |
| ILGSNTSTL | ASSRDERGYAEQF | 0.42703402 |
| IVFSLTVL | ASSRDERGYAEQF | 0.45944715 |
| LASKATGF | ASSRDERGYAEQF | 0.15639012 |
| LASVALIL | ASSRDERGYAEQF | 0.13374504 |

| | | |
|---|---|---|
| LFAVGLTL | ASSRDERGYAEQF | 0.3531252 |
| LFNGGIGTYI | ASSRDERGYAEQF | 0.43560928 |
| LTLGAGII | ASSRDERGYAEQF | 0.1492597 |
| LVAAALGDI | ASSRDERGYAEQF | 0.18019384 |
| NTAFVQLGI | ASSRDERGYAEQF | 0.24790631 |
| QLFSGAEL | ASSRDERGYAEQF | 0.2755148 |
| SAATFITL | ASSRDERGYAEQF | 0.28168535 |
| SNYGLANLGV | ASSRDERGYAEQF | 0.14005177 |
| SYGGAYAVM | ASSRDERGYAEQF | 0.5104439 |
| TTGVGGLTLG | ASSRDERGYAEQF | 0.1429469 |
| TTQTIYI | ASSRDERGYAEQF | 0.43332836 |
| VAAGAGLGAVI | ASSRDERGYAEQF | 0.21367368 |
| VAAVVTLL | ASSRDERGYAEQF | 0.25958538 |
| VGFLGGLI | ASSRDERGYAEQF | 0.24059269 |
| VGVIPILL | ASSRDERGYAEQF | 0.17483419 |
| VIFASSLI | ASSRDERGYAEQF | 0.13732547 |
| VKGGDGGAAAG | ASSRDERGYAEQF | 0.10775442 |
| VTKTFTNI | ASSRDERGYAEQF | 0.5703725 |
| VVAAAILL | ASSRDERGYAEQF | 0.2554326 |
| YAALDVEL | ASSRDERGYAEQF | 0.37850356 |
| LASVALLL | ASSRDERGYAEQF | 0.27177295 |
| LDAVHRV | ASSRDERGYAEQF | 0.13873042 |
| QGLSSSIF | ASSRDERGYAEQF | 0.2529119 |
| RGTFANIRL | ASSRDERGYAEQF | 0.1136601 |
| RQLWWGHRI | ASSRDERGYAEQF | 0.19493626 |
| YGPPGTGKTLL | ASSRDERGYAEQF | 0.5048873 |
| AALVGAPANVA | ASSRDERGYAEQF | 0.30810177 |
| AATFRRTE | ASSRDERGYAEQF | 0.16135584 |
| AGALGGGI | ASSRDERGYAEQF | 0.30891347 |
| AGGAGGAGGIGAG | ASSRDERGYAEQF | 0.3141057 |
| AGSGALGL | ASSRDERGYAEQF | 0.16772695 |
| AQIADQI | ASSRDERGYAEQF | 0.2591591 |
| ASAVAVTYL | ASSRDERGYAEQF | 0.46582246 |
| ASTVLGGTGG | ASSRDERGYAEQF | 0.28202072 |
| AVGGEPAK | ASSRDERGYAEQF | 0.13518137 |
| CEQGTFGGTC | ASSRDERGYAEQF | 0.34647447 |
| DEGKAPAG | ASSRDERGYAEQF | 0.16369069 |
| DTVVFPAL | ASSRDERGYAEQF | 0.31985685 |
| FEGITMAG | ASSRDERGYAEQF | 0.19257452 |
| FPLVVQD | ASSRDERGYAEQF | 0.21401814 |
| GAFTTPQ | ASSRDERGYAEQF | 0.36663735 |
| GFGGFGTAGGVGGA | ASSRDERGYAEQF | 0.14937973 |
| GFLAQFL | ASSRDERGYAEQF | 0.15901127 |
| GGAGGAGGNAGVA | ASSRDERGYAEQF | 0.23666137 |
| GGVGTAGAGGAGGAGG | ASSRDERGYAEQF | 0.21169095 |
| GNGGGGDAGNAGSGGN | ASSRDERGYAEQF | 0.41476175 |
| GTGGAGGAGNDAGS | ASSRDERGYAEQF | 0.14117685 |
| GTTNTGWFNS | ASSRDERGYAEQF | 0.28510684 |
| IFMGPLFTL | ASSRDERGYAEQF | 0.17437646 |
| ILAHAIEAI | ASSRDERGYAEQF | 0.14333294 |
| IRSAIREAL | ASSRDERGYAEQF | 0.25457093 |

| | | |
|---|---|---|
| ITFAGGG | ASSRDERGYAEQF | 0.20509811 |
| ITGMSGHQEV | ASSRDERGYAEQF | 0.15551533 |
| IVAGLACTL | ASSRDERGYAEQF | 0.30630004 |
| IVGGGLAGA | ASSRDERGYAEQF | 0.15016674 |
| IVTGVLNI | ASSRDERGYAEQF | 0.27164084 |
| LAQQGCI | ASSRDERGYAEQF | 0.7307341 |
| LCLHVEN | ASSRDERGYAEQF | 0.16109385 |
| LDAISDGS | ASSRDERGYAEQF | 0.35056055 |
| LGGFAAML | ASSRDERGYAEQF | 0.22520082 |
| LGGKNPMI | ASSRDERGYAEQF | 0.33557338 |
| LGPKGVAAIL | ASSRDERGYAEQF | 0.16957757 |
| LMVGVSI | ASSRDERGYAEQF | 0.1607868 |
| LVGAAFVL | ASSRDERGYAEQF | 0.46074927 |
| LVLAVAPL | ASSRDERGYAEQF | 0.37239325 |
| MAAIAVIDAA | ASSRDERGYAEQF | 0.45561206 |
| NNGINAELQ | ASSRDERGYAEQF | 0.3101147 |
| RGEPLVLVHG | ASSRDERGYAEQF | 0.35004127 |
| RGGAGGAAGLI | ASSRDERGYAEQF | 0.42281413 |
| RSYRMHPAV | ASSRDERGYAEQF | 0.18275377 |
| SAAHLLAGQLA | ASSRDERGYAEQF | 0.118287496 |
| SAAVGSY | ASSRDERGYAEQF | 0.1626519 |
| SPHAVLHRI | ASSRDERGYAEQF | 0.13169247 |
| SQIAKVL | ASSRDERGYAEQF | 0.17298709 |
| TALGHLVP | ASSRDERGYAEQF | 0.13808997 |
| TAPTTSGAA | ASSRDERGYAEQF | 0.49137694 |
| TDGDDGNGGSPGT | ASSRDERGYAEQF | 0.7686703 |
| TDGLGEN | ASSRDERGYAEQF | 0.14923902 |
| TGGPPLPGQA | ASSRDERGYAEQF | 0.7262247 |
| TIPAITI | ASSRDERGYAEQF | 0.14086142 |
| TIPGFTI | ASSRDERGYAEQF | 0.37142807 |
| TSIASGALA | ASSRDERGYAEQF | 0.58485043 |
| TVTLPGAAI | ASSRDERGYAEQF | 0.13560215 |
| VASGLSAALP | ASSRDERGYAEQF | 0.25706747 |
| VGAGVIGI | ASSRDERGYAEQF | 0.18671212 |
| VGPKRFGV | ASSRDERGYAEQF | 0.17842695 |
| VNVAFPD | ASSRDERGYAEQF | 0.5412067 |
| VSMTNSGVS | ASSRDERGYAEQF | 0.17510699 |
| VTDVLGDLSV | ASSRDERGYAEQF | 0.2894898 |
| VVVVGAGFAGL | ASSRDERGYAEQF | 0.16673268 |
| VYTVGTVV | ASSRDERGYAEQF | 0.1371279 |
| WTTFSGP | ASSRDERGYAEQF | 0.22900067 |
| YDELVEAF | ASSRDERGYAEQF | 0.42145318 |
| YGTGGAGGNGG | ASSRDERGYAEQF | 0.18900728 |
| YTNVMAV | ASSRDERGYAEQF | 0.696443 |
| FLEQVHQL | ASSRDERGYAEQF | 0.15646017 |
| GLSLLAVAV | ASSRDERGYAEQF | 0.64692307 |
| LEQVHQL | ASSRDERGYAEQF | 0.677941 |
| LNAAGAI | ASSRDERGYAEQF | 0.26797068 |
| PAVHASL | ASSRDERGYAEQF | 0.19515795 |
| TELEDDA | ASSRDERGYAEQF | 0.630949 |
| TPPGPPN | ASSRDERGYAEQF | 0.20032117 |

| | | |
|---|---|---|
| TPPPPPPP | ASSRDERGYAEQF | 0.40534753 |
| TQSPALI | ASSFTGTNSEVF | 0.44690025 |
| EFAAAFPQ | ASSFTGTNSEVF | 0.15234269 |
| LLSGNLDVL | ASSFTGTNSEVF | 0.54093176 |
| GPANLGNYNI | ASSFTGTNSEVF | 0.29031414 |
| TLGPTLP | ASSFTGTNSEVF | 0.4315107 |
| VYAYSSCCK | ASSFTGTNSEVF | 0.181563 |
| APAPAPAPNI | ASSFTGTNSEVF | 0.14643057 |
| PGGPVPIP | ASSFTGTNSEVF | 0.2712583 |
| FLTVALEP | ASSFTGTNSEVF | 0.14528582 |
| SQYFGANL | ASSFTGTNSEVF | 0.76312286 |
| GGVKPAA | ASSFTGTNSEVF | 0.40318805 |
| GGTGGAGGAGGW | ASSFTGTNSEVF | 0.19869378 |
| GVGGVAGLGGAGG | ASSFTGTNSEVF | 0.40944964 |
| VLLVLATTI | ASSFTGTNSEVF | 0.5932962 |
| IGIFAAML | ASSFTGTNSEVF | 0.11781159 |
| AVGDLID | ASSFTGTNSEVF | 0.23980889 |
| NISPLPL | ASSFTGTNSEVF | 0.2529091 |
| VTAVFVAI | ASSFTGTNSEVF | 0.22767352 |
| SGDDNDAAL | ASSFTGTNSEVF | 0.102595195 |
| AVAALAHS | ASSFTGTNSEVF | 0.3634868 |
| GGVTVMATP | ASSFTGTNSEVF | 0.16383526 |
| AGLAANVRD | ASSFTGTNSEVF | 0.08482234 |
| TNRPDLIDPALLRPG | ASSFTGTNSEVF | 0.3334092 |
| LANLGVTAL | ASSFTGTNSEVF | 0.18618375 |
| TGERIEL | ASSFTGTNSEVF | 0.1866216 |
| NGEELAA | ASSFTGTNSEVF | 0.45225072 |
| VLSLVWG | ASSFTGTNSEVF | 0.22336733 |
| IGYALWL | ASSFTGTNSEVF | 0.16789365 |
| LSLLLAPL | ASSFTGTNSEVF | 0.36075616 |
| TLDEALK | ASSFTGTNSEVF | 0.48069495 |
| TDGAGTAEGG | ASSFTGTNSEVF | 0.15206008 |
| NAPGGAGGNGG | ASSFTGTNSEVF | 0.27335772 |
| VSVFMGMGGNF | ASSFTGTNSEVF | 0.6882609 |
| LASYLAAL | ASSFTGTNSEVF | 0.24508683 |
| LVGGTPAL | ASSFTGTNSEVF | 0.6032742 |
| VRNADVL | ASSFTGTNSEVF | 0.6298322 |
| TCSSPFLVSAM | ASSFTGTNSEVF | 0.20522803 |
| VHITDAE | ASSFTGTNSEVF | 0.48773104 |
| EYAAPSF | ASSFTGTNSEVF | 0.5762955 |
| VVDAFHGGL | ASSFTGTNSEVF | 0.2357316 |
| ASVVDPQ | ASSFTGTNSEVF | 0.23917666 |
| SYNEFGGLS | ASSFTGTNSEVF | 0.16299021 |
| DPDGSVY | ASSFTGTNSEVF | 0.31529334 |
| MVSTIHATE | ASSFTGTNSEVF | 0.14071561 |
| LAAGVADPV | ASSFTGTNSEVF | 0.38347203 |
| TKEQIAA | ASSFTGTNSEVF | 0.5475409 |
| SLASPAAL | ASSFTGTNSEVF | 0.22700386 |
| IGLVFYGV | ASSFTGTNSEVF | 0.10607798 |
| FLLDASNL | ASSFTGTNSEVF | 0.114398524 |
| YQVGNFP | ASSFTGTNSEVF | 0.30842376 |

| | | |
|---|---|---|
| ESLLSQST | ASSFTGTNSEVF | 0.16426697 |
| LFTVGFAAVG | ASSFTGTNSEVF | 0.1533058 |
| ATPPTPT | ASSFTGTNSEVF | 0.17111877 |
| TDLATLG | ASSFTGTNSEVF | 0.44539678 |
| SAGGTGR | ASSFTGTNSEVF | 0.50668734 |
| GFSNSGS | ASSFTGTNSEVF | 0.13494049 |
| GGNGGSGGTSV | ASSFTGTNSEVF | 0.65640104 |
| GTSVATGGAGN | ASSFTGTNSEVF | 0.7499234 |
| NGGNGGSGGTS | ASSFTGTNSEVF | 0.6374397 |
| AVVKFPH | ASSFTGTNSEVF | 0.30199647 |
| TPPPPPP | ASSFTGTNSEVF | 0.13940951 |
| AAPAAAAAPA | ASSFTGTNSEVF | 0.7073785 |
| IALNEDL | ASSFTGTNSEVF | 0.16915213 |
| IVNVSSM | ASSFTGTNSEVF | 0.3878287 |
| TGIPPAPR | ASSFTGTNSEVF | 0.48754317 |
| FDLGGGTF | ASSFTGTNSEVF | 0.2501309 |
| FDLGGGTFDVS | ASSFTGTNSEVF | 0.14926355 |
| NEPTAAA | ASSFTGTNSEVF | 0.5155066 |
| NEPTAAALA | ASSFTGTNSEVF | 0.6212518 |
| GSIINISSI | ASSFTGTNSEVF | 0.11894059 |
| VAPGNAGTA | ASSFTGTNSEVF | 0.15792976 |
| TGGGASLE | ASSFTGTNSEVF | 0.26659223 |
| VAVYNHYKRI | ASSFTGTNSEVF | 0.21272974 |
| TDPAPATT | ASSFTGTNSEVF | 0.5206089 |
| TNRPDILDPALLRPG | ASSFTGTNSEVF | 0.34765628 |
| LTTTTTS | ASSFTGTNSEVF | 0.23844412 |
| PTGSGKTL | ASSFTGTNSEVF | 0.38404432 |
| TGSGKTL | ASSFTGTNSEVF | 0.35653794 |
| TIGHVDHGK | ASSFTGTNSEVF | 0.2966311 |
| TIGHVDHGKTT | ASSFTGTNSEVF | 0.45710057 |
| GHVDHGKTT | ASSFTGTNSEVF | 0.13927676 |
| AQFEHTLL | ASSFTGTNSEVF | 0.6070475 |
| IVNQDEL | ASSFTGTNSEVF | 0.4744156 |
| TNGSQFF | ASSFTGTNSEVF | 0.46734452 |
| TGKGIVE | ASSFTGTNSEVF | 0.6046761 |
| LISNASDA | ASSFTGTNSEVF | 0.29803145 |
| AELRTLGELTKIAKAHGAQ | ASSFTGTNSEVF | 0.3365092 |
| APNAVAAAEPSW | ASSFTGTNSEVF | 0.12698637 |
| APQVAGAATAVL | ASSFTGTNSEVF | 0.21768057 |
| ARRDPAAAAAILPTDA | ASSFTGTNSEVF | 0.2792923 |
| DAAARDVLAEAGLAEYF | ASSFTGTNSEVF | 0.34015054 |
| DDQIVLSAKANLAKVEQ | ASSFTGTNSEVF | 0.510627 |
| DELLPDTGEDSDEDGH | ASSFTGTNSEVF | 0.5133233 |
| ELPEAPANRATVPQD | ASSFTGTNSEVF | 0.18247691 |
| FGQSKIERVTGIG | ASSFTGTNSEVF | 0.696429 |
| GAGGTGSTAGGAGGAGGAGGLY | ASSFTGTNSEVF | 0.62267333 |
| GHGGTGAAGGNGGTGGAGGSGI | ASSFTGTNSEVF | 0.12718551 |
| GNGGNGGAGGEGGPGGAGGAGA | ASSFTGTNSEVF | 0.1989786 |
| GSTAGNGGNGGNATNGTIAGQP | ASSFTGTNSEVF | 0.5359374 |
| GTGGDGGNGGNGGTGG | ASSFTGTNSEVF | 0.66807485 |
| GVRRPAGIPADAPV | ASSFTGTNSEVF | 0.2158218 |

| | | |
|---|---|---|
| GYRQVTDAHLLTLARR | ASSFTGTNSEVF | 0.31579903 |
| IAKMGQDGHDRG | ASSFTGTNSEVF | 0.7871134 |
| KAEPSIEIQYT | ASSFTGTNSEVF | 0.2765371 |
| KPVRIRMAADEDTVAVT | ASSFTGTNSEVF | 0.6273748 |
| MPPRIAGMRLLVIKPEP | ASSFTGTNSEVF | 0.1033116 |
| MKFYAVALFRSRGVPPTR | ASSFTGTNSEVF | 0.27665487 |
| NAGGFGILGAVAHSP | ASSFTGTNSEVF | 0.2548049 |
| NEGREITLAMAEM | ASSFTGTNSEVF | 0.087385036 |
| NELVEWSHPLGHRWRWE | ASSFTGTNSEVF | 0.42694837 |
| NLGAGGTGGNGGNPG | ASSFTGTNSEVF | 0.225179 |
| NTEWPKWETVMAADPVD | ASSFTGTNSEVF | 0.46916515 |
| PASAAPAETQP | ASSFTGTNSEVF | 0.18599007 |
| QQSSDGKNGVRAAEITGE | ASSFTGTNSEVF | 0.46708044 |
| RAPDAVTTSWWKE | ASSFTGTNSEVF | 0.3689416 |
| SGFGNVGAHTSGWFNQ | ASSFTGTNSEVF | 0.2777885 |
| SGVAGAVGQAASVGGLKV | ASSFTGTNSEVF | 0.18465987 |
| SNGGAGTGSAGGNGGT | ASSFTGTNSEVF | 0.29316008 |
| TGGAGGEGGAAG | ASSFTGTNSEVF | 0.6753036 |
| TGGDGGAAGAGGAPGHG | ASSFTGTNSEVF | 0.5878556 |
| TMPPLLAAIANAGRHGV | ASSFTGTNSEVF | 0.12032929 |
| TSSTAGNSGGA | ASSFTGTNSEVF | 0.4607747 |
| VPATVPVGILC | ASSFTGTNSEVF | 0.21868022 |
| VQRPDSGGVVLTGRIS | ASSFTGTNSEVF | 0.13648623 |
| WTPNPVPATKGVPAGLVQAT | ASSFTGTNSEVF | 0.10640717 |
| YALKRALAKFSDRFDVVII | ASSFTGTNSEVF | 0.17630154 |
| YLIGSVIV | ASSFTGTNSEVF | 0.22565493 |
| YGQLVAVID | ASSFTGTNSEVF | 0.34008026 |
| VTVGGMTLPA | ASSFTGTNSEVF | 0.45653397 |
| VLALPAV | ASSFTGTNSEVF | 0.18566239 |
| VIPPTLNYET | ASSFTGTNSEVF | 0.41500473 |
| VGMGGMHPSSG | ASSFTGTNSEVF | 0.22038999 |
| VGLAGGAATA | ASSFTGTNSEVF | 0.06747034 |
| VGGAGGAGGSGP | ASSFTGTNSEVF | 0.17229916 |
| VFAAFMV | ASSFTGTNSEVF | 0.11548089 |
| TLYDKLR | ASSFTGTNSEVF | 0.6440228 |
| TLAVFTVL | ASSFTGTNSEVF | 0.13277006 |
| TGSVQPARV | ASSFTGTNSEVF | 0.186757 |
| TGGGAGSGGT | ASSFTGTNSEVF | 0.21317154 |
| SHLETSL | ASSFTGTNSEVF | 0.647012 |
| SFTTGMT | ASSFTGTNSEVF | 0.3490464 |
| QVEVAKRMGVRQPTVS | ASSFTGTNSEVF | 0.1764795 |
| QGSGGPALAM | ASSFTGTNSEVF | 0.123024166 |
| MLAPSNP | ASSFTGTNSEVF | 0.32229453 |
| LVNNAAI | ASSFTGTNSEVF | 0.35124323 |
| LTGEINTLA | ASSFTGTNSEVF | 0.1775125 |
| LSIFGPV | ASSFTGTNSEVF | 0.16482797 |
| LLALLANL | ASSFTGTNSEVF | 0.7120027 |
| LISNASDALDKLR | ASSFTGTNSEVF | 0.13148457 |
| LINEAVP | ASSFTGTNSEVF | 0.5203198 |
| LGYAFSGL | ASSFTGTNSEVF | 0.17523064 |
| LDFVAVAL | ASSFTGTNSEVF | 0.18140215 |

| | | |
|---|---|---|
| LAQPGAE | ASSFTGTNSEVF | 0.68531907 |
| ISNLGAVTGF | ASSFTGTNSEVF | 0.5385779 |
| HQEDPGRI | ASSFTGTNSEVF | 0.22915667 |
| GTPGEPTP | ASSFTGTNSEVF | 0.1703971 |
| GFFNSGS | ASSFTGTNSEVF | 0.15554969 |
| GAGGGGLTAGAN | ASSFTGTNSEVF | 0.16195317 |
| FVDALVNM | ASSFTGTNSEVF | 0.17536072 |
| FLTVAPEL | ASSFTGTNSEVF | 0.62721175 |
| FETLTAL | ASSFTGTNSEVF | 0.30502957 |
| DVINPIEL | ASSFTGTNSEVF | 0.15575533 |
| APVSGAPV | ASSFTGTNSEVF | 0.5098636 |
| ALVGGLAW | ASSFTGTNSEVF | 0.6271099 |
| ALAAAAATAI | ASSFTGTNSEVF | 0.6804292 |
| AFGLGLLL | ASSFTGTNSEVF | 0.13939019 |
| AFFAAASL | ASSFTGTNSEVF | 0.25336987 |
| AAAALPAFSPP | ASSFTGTNSEVF | 0.30772653 |
| VVLAPAL | ASSFTGTNSEVF | 0.23685932 |
| TIGHVDHGKTTL | ASSFTGTNSEVF | 0.1916833 |
| LISNASDALDK | ASSFTGTNSEVF | 0.16911186 |
| LISNASDAL | ASSFTGTNSEVF | 0.5524467 |
| ISNASDALDK | ASSFTGTNSEVF | 0.84891653 |
| NQAETLVYQTE | ASSFTGTNSEVF | 0.2236211 |
| TGGTGGTAGAG | ASSFTGTNSEVF | 0.114179865 |
| TGSTAGGAGGA | ASSFTGTNSEVF | 0.19948977 |
| DPLAVPVAAGPV | ASSFTGTNSEVF | 0.13742362 |
| GGTGGTPGNGGN | ASSFTGTNSEVF | 0.14589502 |
| PAGLAGVDPATL | ASSFTGTNSEVF | 0.24981365 |
| GNNLVVEAPHAGASV | ASSFTGTNSEVF | 0.10818779 |
| KVTRIMSASTLEQPA | ASSFTGTNSEVF | 0.23848963 |
| LGFALPIALPRRTVV | ASSFTGTNSEVF | 0.69123846 |
| TRLGNLTARKLVVLV | ASSFTGTNSEVF | 0.22785927 |
| WSQQRRLIYFPSAGP | ASSFTGTNSEVF | 0.50472116 |
| GPNGNPDPMAAAVDIR | ASSFTGTNSEVF | 0.5686735 |
| TQRCVSVADLRAAASR | ASSFTGTNSEVF | 0.37296557 |
| QFVLDVAEHPVEVLTETV | ASSFTGTNSEVF | 0.5410366 |
| GGAGGTGGAGGNGGTQAGDG | ASSFTGTNSEVF | 0.71716195 |
| PLLATLRAIAVDVDAKPAQV | ASSFTGTNSEVF | 0.29712272 |
| APKFRTGGYMIDFWGVGYQVAK | ASSFTGTNSEVF | 0.28120756 |
| GAGGAGGAGGAAGGPGGTGGTG | ASSFTGTNSEVF | 0.6602415 |
| IGALGLLATDVAPAGSVAFGVA | ASSFTGTNSEVF | 0.19972678 |
| AGGAGGIGGNGGAGGAGGVLMG | ASSFTGTNSEVF | 0.27408046 |
| ADELIGLL | ASSFTGTNSEVF | 0.3047749 |
| ADRALKA | ASSFTGTNSEVF | 0.397529 |
| AGAVGQVGGAGGA | ASSFTGTNSEVF | 0.26571903 |
| AGGGGGLGGAP | ASSFTGTNSEVF | 0.73880875 |
| AGGLLYGNGGAG | ASSFTGTNSEVF | 0.109155916 |
| AGLESIGL | ASSFTGTNSEVF | 0.15105395 |
| AGLKTGII | ASSFTGTNSEVF | 0.6175038 |
| ASLAALLL | ASSFTGTNSEVF | 0.09089852 |
| ASVINISSI | ASSFTGTNSEVF | 0.13268656 |
| ATPPGPA | ASSFTGTNSEVF | 0.36750883 |

| | | |
|---|---|---|
| ATVASALV | ASSFTGTNSEVF | 0.63962865 |
| ATVASLIL | ASSFTGTNSEVF | 0.09477857 |
| AVNGPVLIRH | ASSFTGTNSEVF | 0.23545238 |
| DGAAGAVGI | ASSFTGTNSEVF | 0.13951555 |
| DPLLHGI | ASSFTGTNSEVF | 0.48522723 |
| ELGAALGAL | ASSFTGTNSEVF | 0.21916239 |
| ELGRTQAV | ASSFTGTNSEVF | 0.11626658 |
| FVAGGYLS | ASSFTGTNSEVF | 0.44083458 |
| GASGGAGGAGGW | ASSFTGTNSEVF | 0.13248758 |
| GGAGAGGDGGAGGI | ASSFTGTNSEVF | 0.39349604 |
| GTGHNGSQPGMGGNG | ASSFTGTNSEVF | 0.46351945 |
| GYAAASASAS | ASSFTGTNSEVF | 0.80806375 |
| HQVDVAAVQQG | ASSFTGTNSEVF | 0.13922231 |
| IGAAIAHT | ASSFTGTNSEVF | 0.39889115 |
| IGMIEVAQL | ASSFTGTNSEVF | 0.19322199 |
| IRLAPEHTSAMD | ASSFTGTNSEVF | 0.7828846 |
| IYNVSVL | ASSFTGTNSEVF | 0.2148254 |
| KQGFLGM | ASSFTGTNSEVF | 0.12223816 |
| LESVITQL | ASSFTGTNSEVF | 0.5337447 |
| LGLGALVVV | ASSFTGTNSEVF | 0.45280093 |
| LIGNGGNGGSGGI | ASSFTGTNSEVF | 0.3339212 |
| LNAINAPTL | ASSFTGTNSEVF | 0.6788441 |
| LQLLNTGV | ASSFTGTNSEVF | 0.268807 |
| LQTSIEF | ASSFTGTNSEVF | 0.1884374 |
| MSFAVSEL | ASSFTGTNSEVF | 0.42656946 |
| NTGALGL | ASSFTGTNSEVF | 0.37699234 |
| PPGAGGAGGAGTT | ASSFTGTNSEVF | 0.18453988 |
| QLVGLGI | ASSFTGTNSEVF | 0.5140276 |
| RFLEQVHQL | ASSFTGTNSEVF | 0.22723642 |
| RGVGYVLR | ASSFTGTNSEVF | 0.18264589 |
| SATGVFAGVI | ASSFTGTNSEVF | 0.6580118 |
| SLPAITLP | ASSFTGTNSEVF | 0.29333657 |
| STALAFTV | ASSFTGTNSEVF | 0.42447776 |
| SVLTVGLGK | ASSFTGTNSEVF | 0.34478447 |
| TALAGAGSGI | ASSFTGTNSEVF | 0.8288256 |
| TGLPFTGI | ASSFTGTNSEVF | 0.16720936 |
| VAEHVTV | ASSFTGTNSEVF | 0.36795607 |
| VDIVLNSL | ASSFTGTNSEVF | 0.2858152 |
| VLTGGGAVL | ASSFTGTNSEVF | 0.52917695 |
| VNVGLAYGI | ASSFTGTNSEVF | 0.26165557 |
| VSSAAVGAV | ASSFTGTNSEVF | 0.4720019 |
| VTSVYEAGV | ASSFTGTNSEVF | 0.4013217 |
| WVALDRGA | ASSFTGTNSEVF | 0.56382656 |
| YLVGSVVI | ASSFTGTNSEVF | 0.2882299 |
| GPPGTGKTLL | ASSFTGTNSEVF | 0.3540292 |
| GVGMTKF | ASSFTGTNSEVF | 0.6912408 |
| IGIGGSDL | ASSFTGTNSEVF | 0.17916952 |
| IGTLLAEL | ASSFTGTNSEVF | 0.24267653 |
| IYTSGTT | ASSFTGTNSEVF | 0.5064205 |
| LSFPAGV | ASSFTGTNSEVF | 0.6563707 |
| LYTSGTT | ASSFTGTNSEVF | 0.2799107 |

| | | |
|---|---|---|
| VGAFPFGF | ASSFTGTNSEVF | 0.60724753 |
| VGAGTVEF | ASSFTGTNSEVF | 0.21332039 |
| VGKSTLLL | ASSFTGTNSEVF | 0.30783403 |
| AALAHAGGGIV | ASSFTGTNSEVF | 0.35335332 |
| AALVAVSLL | ASSFTGTNSEVF | 0.21595293 |
| AAVSRAVAAL | ASSFTGTNSEVF | 0.120100915 |
| AGGSSGIGGF | ASSFTGTNSEVF | 0.24347615 |
| AGLTVANANG | ASSFTGTNSEVF | 0.46380687 |
| AGLVGAVL | ASSFTGTNSEVF | 0.32094795 |
| ALSQAFTV | ASSFTGTNSEVF | 0.12520793 |
| ALTFSVSL | ASSFTGTNSEVF | 0.32722685 |
| ALTGAALVAL | ASSFTGTNSEVF | 0.37811345 |
| ASFSIGAGVL | ASSFTGTNSEVF | 0.82919496 |
| DGSVYKI | ASSFTGTNSEVF | 0.24912018 |
| EQLAEIV | ASSFTGTNSEVF | 0.5849019 |
| FAVSELTVM | ASSFTGTNSEVF | 0.21092653 |
| GAAPLAAGL | ASSFTGTNSEVF | 0.30083868 |
| GADGGDGGSGGAGG | ASSFTGTNSEVF | 0.10415685 |
| GFGLDTGGAGG | ASSFTGTNSEVF | 0.21293357 |
| GLASGGLL | ASSFTGTNSEVF | 0.2127566 |
| GRIAGAAVLL | ASSFTGTNSEVF | 0.13925502 |
| GSAGSLNTSF | ASSFTGTNSEVF | 0.58460915 |
| GTGQIGI | ASSFTGTNSEVF | 0.6123804 |
| GTLIALIL | ASSFTGTNSEVF | 0.5268017 |
| IAATMGFVVG | ASSFTGTNSEVF | 0.62863266 |
| IFAQALL | ASSFTGTNSEVF | 0.68906116 |
| IGAAVALGF | ASSFTGTNSEVF | 0.5089087 |
| IGLTPLAL | ASSFTGTNSEVF | 0.17082009 |
| ILGSNTSTL | ASSFTGTNSEVF | 0.20359854 |
| IVFSLTVL | ASSFTGTNSEVF | 0.10469228 |
| LASKATGF | ASSFTGTNSEVF | 0.18224175 |
| LASVALIL | ASSFTGTNSEVF | 0.17947038 |
| LFAVGLTL | ASSFTGTNSEVF | 0.25523376 |
| LFNGGIGTYI | ASSFTGTNSEVF | 0.08680379 |
| LTLGAGII | ASSFTGTNSEVF | 0.59939325 |
| LVAAALGDI | ASSFTGTNSEVF | 0.63945097 |
| NTAFVQLGI | ASSFTGTNSEVF | 0.18379885 |
| QLFSGAEL | ASSFTGTNSEVF | 0.44516146 |
| SAATFITL | ASSFTGTNSEVF | 0.48461732 |
| SNYGLANLGV | ASSFTGTNSEVF | 0.15775986 |
| SYGGAYAVM | ASSFTGTNSEVF | 0.45414126 |
| TTGVGGLTLG | ASSFTGTNSEVF | 0.18791807 |
| TTQTIYI | ASSFTGTNSEVF | 0.5468463 |
| VAAGAGLGAVI | ASSFTGTNSEVF | 0.5181578 |
| VAAVVTLL | ASSFTGTNSEVF | 0.35699552 |
| VGFLGGLI | ASSFTGTNSEVF | 0.30803925 |
| VGVIPILL | ASSFTGTNSEVF | 0.3946195 |
| VIFASSLI | ASSFTGTNSEVF | 0.17032793 |
| VKGGDGGAAAG | ASSFTGTNSEVF | 0.34907496 |
| VTKTFTNI | ASSFTGTNSEVF | 0.23556688 |
| VVAAAILL | ASSFTGTNSEVF | 0.34082332 |

| | | |
|---|---|---|
| YAALDVEL | ASSFTGTNSEVF | 0.16590475 |
| LASVALLL | ASSFTGTNSEVF | 0.33316588 |
| LDAVHRV | ASSFTGTNSEVF | 0.23022799 |
| QGLSSSIF | ASSFTGTNSEVF | 0.38782647 |
| RGTFANIRL | ASSFTGTNSEVF | 0.13975666 |
| RQLWWGHRI | ASSFTGTNSEVF | 0.40251297 |
| YGPPGTGKTLL | ASSFTGTNSEVF | 0.38495964 |
| AALVGAPANVA | ASSFTGTNSEVF | 0.12679283 |
| AATFRRTE | ASSFTGTNSEVF | 0.54552555 |
| AGALGGGI | ASSFTGTNSEVF | 0.32496023 |
| AGGAGGAGGIGAG | ASSFTGTNSEVF | 0.24278742 |
| AGSGALGL | ASSFTGTNSEVF | 0.4787162 |
| AQIADQI | ASSFTGTNSEVF | 0.28187066 |
| ASAVAVTYL | ASSFTGTNSEVF | 0.14867844 |
| ASTVLGGTGG | ASSFTGTNSEVF | 0.17535727 |
| AVGGEPAK | ASSFTGTNSEVF | 0.36461318 |
| CEQGTFGGTC | ASSFTGTNSEVF | 0.20015812 |
| DEGKAPAG | ASSFTGTNSEVF | 0.09648097 |
| DTVVFPAL | ASSFTGTNSEVF | 0.3401278 |
| FEGITMAG | ASSFTGTNSEVF | 0.18510501 |
| FPLVVQD | ASSFTGTNSEVF | 0.14076042 |
| GAFTTPQ | ASSFTGTNSEVF | 0.18135518 |
| GFGGFGTAGGVGGA | ASSFTGTNSEVF | 0.5216995 |
| GFLAQFL | ASSFTGTNSEVF | 0.39390135 |
| GGAGGAGGNAGVA | ASSFTGTNSEVF | 0.28635257 |
| GGVGTAGAGGAGGAGG | ASSFTGTNSEVF | 0.45328692 |
| GNGGGGDAGNAGSGGN | ASSFTGTNSEVF | 0.3221267 |
| GTGGAGGAGNDAGS | ASSFTGTNSEVF | 0.45383814 |
| GTTNTGWFNS | ASSFTGTNSEVF | 0.31690136 |
| IFMGPLFTL | ASSFTGTNSEVF | 0.34531847 |
| ILAHAIEAI | ASSFTGTNSEVF | 0.4665228 |
| IRSAIREAL | ASSFTGTNSEVF | 0.2474996 |
| ITFAGGG | ASSFTGTNSEVF | 0.15904611 |
| ITGMSGHQEV | ASSFTGTNSEVF | 0.1541002 |
| IVAGLACTL | ASSFTGTNSEVF | 0.20975558 |
| IVGGGLAGA | ASSFTGTNSEVF | 0.1566269 |
| IVTGVLNI | ASSFTGTNSEVF | 0.3299631 |
| LAQQGCI | ASSFTGTNSEVF | 0.6261898 |
| LCLHVEN | ASSFTGTNSEVF | 0.2769267 |
| LDAISDGS | ASSFTGTNSEVF | 0.13544387 |
| LGGFAAML | ASSFTGTNSEVF | 0.22482853 |
| LGGKNPMI | ASSFTGTNSEVF | 0.21304733 |
| LGPKGVAAIL | ASSFTGTNSEVF | 0.4556582 |
| LMVGVSI | ASSFTGTNSEVF | 0.44973868 |
| LVGAAFVL | ASSFTGTNSEVF | 0.8047328 |
| LVLAVAPL | ASSFTGTNSEVF | 0.44301063 |
| MAAIAVIDAA | ASSFTGTNSEVF | 0.15462519 |
| NNGINAELQ | ASSFTGTNSEVF | 0.4474875 |
| RGEPLVLVHG | ASSFTGTNSEVF | 0.22454439 |
| RGGAGGAAGLI | ASSFTGTNSEVF | 0.37371963 |
| RSYRMHPAV | ASSFTGTNSEVF | 0.30460978 |

| | | |
|---|---|---|
| SAAHLLAGQLA | ASSFTGTNSEVF | 0.44513375 |
| SAAVGSY | ASSFTGTNSEVF | 0.20669857 |
| SPHAVLHRI | ASSFTGTNSEVF | 0.27213764 |
| SQIAKVL | ASSFTGTNSEVF | 0.14559126 |
| TALGHLVP | ASSFTGTNSEVF | 0.53205705 |
| TAPTTSGAA | ASSFTGTNSEVF | 0.20930408 |
| TDGDDGNGGSPGT | ASSFTGTNSEVF | 0.17523564 |
| TDGLGEN | ASSFTGTNSEVF | 0.20599702 |
| TGGPPLPGQA | ASSFTGTNSEVF | 0.6572904 |
| TIPAITI | ASSFTGTNSEVF | 0.7129028 |
| TIPGFTI | ASSFTGTNSEVF | 0.17434609 |
| TSIASGALA | ASSFTGTNSEVF | 0.16163974 |
| TVTLPGAAI | ASSFTGTNSEVF | 0.6316434 |
| VASGLSAALP | ASSFTGTNSEVF | 0.15288204 |
| VGAGVIGI | ASSFTGTNSEVF | 0.40013504 |
| VGPKRFGV | ASSFTGTNSEVF | 0.27146888 |
| VNVAFPD | ASSFTGTNSEVF | 0.18296246 |
| VSMTNSGVS | ASSFTGTNSEVF | 0.22669974 |
| VTDVLGDLSV | ASSFTGTNSEVF | 0.44019982 |
| VVVVGAGFAGL | ASSFTGTNSEVF | 0.16109437 |
| VYTVGTVV | ASSFTGTNSEVF | 0.45241475 |
| WTTFSGP | ASSFTGTNSEVF | 0.09633771 |
| YDELVEAF | ASSFTGTNSEVF | 0.37142476 |
| YGTGGAGGNGG | ASSFTGTNSEVF | 0.12622237 |
| YTNVMAV | ASSFTGTNSEVF | 0.16933915 |
| FLEQVHQL | ASSFTGTNSEVF | 0.10530523 |
| GLSLLAVAV | ASSFTGTNSEVF | 0.2852543 |
| LEQVHQL | ASSFTGTNSEVF | 0.42859414 |
| LNAAGAI | ASSFTGTNSEVF | 0.6419566 |
| PAVHASL | ASSFTGTNSEVF | 0.34835327 |
| TELEDDA | ASSFTGTNSEVF | 0.4104365 |
| TPPGPPN | ASSFTGTNSEVF | 0.64345044 |
| TPPPPPPP | ASSFTGTNSEVF | 0.1755884 |
| TQSPALI | ASSLDSGQNTEVF | 0.11976666 |
| EFAAAFPQ | ASSLDSGQNTEVF | 0.30905193 |
| LLSGNLDVL | ASSLDSGQNTEVF | 0.3986257 |
| GPANLGNYNI | ASSLDSGQNTEVF | 0.6060368 |
| TLGPTLP | ASSLDSGQNTEVF | 0.50492364 |
| VYAYSSCCK | ASSLDSGQNTEVF | 0.10121621 |
| APAPAPAPNI | ASSLDSGQNTEVF | 0.45406866 |
| PGGPVPIP | ASSLDSGQNTEVF | 0.4338513 |
| FLTVALEP | ASSLDSGQNTEVF | 0.13537075 |
| SQYFGANL | ASSLDSGQNTEVF | 0.20501232 |
| GGVKPAA | ASSLDSGQNTEVF | 0.19641392 |
| GGTGGAGGAGGW | ASSLDSGQNTEVF | 0.12571242 |
| GVGGVAGLGGAGG | ASSLDSGQNTEVF | 0.3859885 |
| VLLVLATTI | ASSLDSGQNTEVF | 0.16595784 |
| IGIFAAML | ASSLDSGQNTEVF | 0.2385052 |
| AVGDLID | ASSLDSGQNTEVF | 0.21115308 |
| NISPLPL | ASSLDSGQNTEVF | 0.20122637 |
| VTAVFVAI | ASSLDSGQNTEVF | 0.21605083 |

| | | |
|---|---|---|
| SGDDNDAAL | ASSLDSGQNTEVF | 0.21298516 |
| AVAALAHS | ASSLDSGQNTEVF | 0.3145197 |
| GGVTVMATP | ASSLDSGQNTEVF | 0.4878195 |
| AGLAANVRD | ASSLDSGQNTEVF | 0.20296033 |
| TNRPDLIDPALLRPG | ASSLDSGQNTEVF | 0.39364323 |
| LANLGVTAL | ASSLDSGQNTEVF | 0.5617621 |
| TGERIEL | ASSLDSGQNTEVF | 0.13394322 |
| NGEELAA | ASSLDSGQNTEVF | 0.15047325 |
| VLSLVWG | ASSLDSGQNTEVF | 0.37013596 |
| IGYALWL | ASSLDSGQNTEVF | 0.1910474 |
| LSLLLAPL | ASSLDSGQNTEVF | 0.5791459 |
| TLDEALK | ASSLDSGQNTEVF | 0.25433028 |
| TDGAGTAEGG | ASSLDSGQNTEVF | 0.2861229 |
| NAPGGAGGNGG | ASSLDSGQNTEVF | 0.32738492 |
| VSVFMGMGGNF | ASSLDSGQNTEVF | 0.75666285 |
| LASYLAAL | ASSLDSGQNTEVF | 0.397218 |
| LVGGTPAL | ASSLDSGQNTEVF | 0.23386486 |
| VRNADVL | ASSLDSGQNTEVF | 0.5134528 |
| TCSSPFLVSAM | ASSLDSGQNTEVF | 0.17008252 |
| VHITDAE | ASSLDSGQNTEVF | 0.4670685 |
| EYAAPSF | ASSLDSGQNTEVF | 0.5249543 |
| VVDAFHGGL | ASSLDSGQNTEVF | 0.12322098 |
| ASVVDPQ | ASSLDSGQNTEVF | 0.19041407 |
| SYNEFGGLS | ASSLDSGQNTEVF | 0.65212846 |
| DPDGSVY | ASSLDSGQNTEVF | 0.55013883 |
| MVSTIHATE | ASSLDSGQNTEVF | 0.18189326 |
| LAAGVADPV | ASSLDSGQNTEVF | 0.15503596 |
| TKEQIAA | ASSLDSGQNTEVF | 0.19646849 |
| SLASPAAL | ASSLDSGQNTEVF | 0.24637717 |
| IGLVFYGV | ASSLDSGQNTEVF | 0.1474009 |
| FLLDASNL | ASSLDSGQNTEVF | 0.4328723 |
| YQVGNFP | ASSLDSGQNTEVF | 0.22256085 |
| ESLLSQST | ASSLDSGQNTEVF | 0.71917903 |
| LFTVGFAAVG | ASSLDSGQNTEVF | 0.3055612 |
| ATPPTPT | ASSLDSGQNTEVF | 0.45242178 |
| TDLATLG | ASSLDSGQNTEVF | 0.21564318 |
| SAGGTGR | ASSLDSGQNTEVF | 0.19062892 |
| GFSNSGS | ASSLDSGQNTEVF | 0.13534759 |
| GGNGGSGGTSV | ASSLDSGQNTEVF | 0.16297135 |
| GTSVATGGAGN | ASSLDSGQNTEVF | 0.18446806 |
| NGGNGGSGGTS | ASSLDSGQNTEVF | 0.23085836 |
| AVVKFPH | ASSLDSGQNTEVF | 0.118473545 |
| TPPPPPP | ASSLDSGQNTEVF | 0.6101512 |
| AAPAAAAAPA | ASSLDSGQNTEVF | 0.22201556 |
| IALNEDL | ASSLDSGQNTEVF | 0.33605126 |
| IVNVSSM | ASSLDSGQNTEVF | 0.22842836 |
| TGIPPAPR | ASSLDSGQNTEVF | 0.18410935 |
| FDLGGGTF | ASSLDSGQNTEVF | 0.16058914 |
| FDLGGGTFDVS | ASSLDSGQNTEVF | 0.5014727 |
| NEPTAAA | ASSLDSGQNTEVF | 0.11852879 |
| NEPTAAALA | ASSLDSGQNTEVF | 0.120872095 |

| | | |
|---|---|---|
| GSIINISSI | ASSLDSGQNTEVF | 0.18867092 |
| VAPGNAGTA | ASSLDSGQNTEVF | 0.23081464 |
| TGGGASLE | ASSLDSGQNTEVF | 0.7532631 |
| VAVYNHYKRI | ASSLDSGQNTEVF | 0.5339279 |
| TDPAPATT | ASSLDSGQNTEVF | 0.28870404 |
| TNRPDILDPALLRPG | ASSLDSGQNTEVF | 0.7187841 |
| LTTTTTS | ASSLDSGQNTEVF | 0.08892721 |
| PTGSGKTL | ASSLDSGQNTEVF | 0.2292602 |
| TGSGKTL | ASSLDSGQNTEVF | 0.68417794 |
| TIGHVDHGK | ASSLDSGQNTEVF | 0.3309108 |
| TIGHVDHGKTT | ASSLDSGQNTEVF | 0.24038196 |
| GHVDHGKTT | ASSLDSGQNTEVF | 0.47334027 |
| AQFEHTLL | ASSLDSGQNTEVF | 0.1484133 |
| IVNQDEL | ASSLDSGQNTEVF | 0.20981906 |
| TNGSQFF | ASSLDSGQNTEVF | 0.23069613 |
| TGKGIVE | ASSLDSGQNTEVF | 0.44500944 |
| LISNASDA | ASSLDSGQNTEVF | 0.45710197 |
| AELRTLGELTKIAKAHGAQ | ASSLDSGQNTEVF | 0.13863225 |
| APNAVAAAEPSW | ASSLDSGQNTEVF | 0.49319458 |
| APQVAGAATAVL | ASSLDSGQNTEVF | 0.7736674 |
| ARRDPAAAAAILPTDA | ASSLDSGQNTEVF | 0.16163555 |
| DAAARDVLAEAGLAEYF | ASSLDSGQNTEVF | 0.54108363 |
| DDQIVLSAKANLAKVEQ | ASSLDSGQNTEVF | 0.30092558 |
| DELLPDTGEDSDEDGH | ASSLDSGQNTEVF | 0.38402075 |
| ELPEAPANRATVPQD | ASSLDSGQNTEVF | 0.16074024 |
| FGQSKIERVTGIG | ASSLDSGQNTEVF | 0.13085417 |
| GAGGTGSTAGGAGGAGGAGGLY | ASSLDSGQNTEVF | 0.4339369 |
| GHGGTGAAGGNGGTGGAGGSGI | ASSLDSGQNTEVF | 0.12499006 |
| GNGGNGGAGGEGGPGGAGGAGA | ASSLDSGQNTEVF | 0.32321328 |
| GSTAGNGGNGGNATNGTIAGQP | ASSLDSGQNTEVF | 0.254779 |
| GTGGDGGNGGNGGTGG | ASSLDSGQNTEVF | 0.8070388 |
| GVRRPAGIPADAPV | ASSLDSGQNTEVF | 0.38158938 |
| GYRQVTDAHLLTLARR | ASSLDSGQNTEVF | 0.3107707 |
| IAKMGQDGHDRG | ASSLDSGQNTEVF | 0.14206362 |
| KAEPSIEIQYT | ASSLDSGQNTEVF | 0.3618592 |
| KPVRIRMAADEDTVAVT | ASSLDSGQNTEVF | 0.45549104 |
| MPPRIAGMRLLVIKPEP | ASSLDSGQNTEVF | 0.1610274 |
| MKFYAVALFRSRGVPPTR | ASSLDSGQNTEVF | 0.52079123 |
| NAGGFGILGAVAHSP | ASSLDSGQNTEVF | 0.14935668 |
| NEGREITLAMAEM | ASSLDSGQNTEVF | 0.327307 |
| NELVEWSHPLGHRWRWE | ASSLDSGQNTEVF | 0.32292575 |
| NLGAGGTGGNGGNPG | ASSLDSGQNTEVF | 0.7742244 |
| NTEWPKWETVMAADPVD | ASSLDSGQNTEVF | 0.23659804 |
| PASAAPAETQP | ASSLDSGQNTEVF | 0.42791635 |
| QQSSDGKNGVRAAEITGE | ASSLDSGQNTEVF | 0.13323739 |
| RAPDAVTTSWWKE | ASSLDSGQNTEVF | 0.49539602 |
| SGFGNVGAHTSGWFNQ | ASSLDSGQNTEVF | 0.5435616 |
| SGVAGAVGQAASVGGLKV | ASSLDSGQNTEVF | 0.6356381 |
| SNGGAGTGSAGGNGGT | ASSLDSGQNTEVF | 0.7940132 |
| TGGAGGEGGAAG | ASSLDSGQNTEVF | 0.36910048 |
| TGGDGGAAGAGGAPGHG | ASSLDSGQNTEVF | 0.13846713 |

| | | |
|---|---|---|
| TMPPLLAAIANAGRHGV | ASSLDSGQNTEVF | 0.29082736 |
| TSSTAGNSGGA | ASSLDSGQNTEVF | 0.49346796 |
| VPATVPVGILC | ASSLDSGQNTEVF | 0.69542384 |
| VQRPDSGGVVLTGRIS | ASSLDSGQNTEVF | 0.43590605 |
| WTPNPVPATKGVPAGLVQAT | ASSLDSGQNTEVF | 0.4325122 |
| YALKRALAKFSDRFDVVII | ASSLDSGQNTEVF | 0.22890407 |
| YLIGSVIV | ASSLDSGQNTEVF | 0.61684346 |
| YGQLVAVID | ASSLDSGQNTEVF | 0.8386189 |
| VTVGGMTLPA | ASSLDSGQNTEVF | 0.4054572 |
| VLALPAV | ASSLDSGQNTEVF | 0.18707517 |
| VIPPTLNYET | ASSLDSGQNTEVF | 0.20634572 |
| VGMGGMHPSSG | ASSLDSGQNTEVF | 0.42683828 |
| VGLAGGAATA | ASSLDSGQNTEVF | 0.40621245 |
| VGGAGGAGGSGP | ASSLDSGQNTEVF | 0.17693762 |
| VFAAFMV | ASSLDSGQNTEVF | 0.21757644 |
| TLYDKLR | ASSLDSGQNTEVF | 0.26542938 |
| TLAVFTVL | ASSLDSGQNTEVF | 0.109138556 |
| TGSVQPARV | ASSLDSGQNTEVF | 0.37409964 |
| TGGGAGSGGT | ASSLDSGQNTEVF | 0.7220018 |
| SHLETSL | ASSLDSGQNTEVF | 0.36617744 |
| SFTTGMT | ASSLDSGQNTEVF | 0.13992815 |
| QVEVAKRMGVRQPTVS | ASSLDSGQNTEVF | 0.5842104 |
| QGSGGPALAM | ASSLDSGQNTEVF | 0.75020254 |
| MLAPSNP | ASSLDSGQNTEVF | 0.63800466 |
| LVNNAAI | ASSLDSGQNTEVF | 0.13659094 |
| LTGEINTLA | ASSLDSGQNTEVF | 0.08310559 |
| LSIFGPV | ASSLDSGQNTEVF | 0.32786232 |
| LLALLANL | ASSLDSGQNTEVF | 0.6652243 |
| LISNASDALDKLR | ASSLDSGQNTEVF | 0.208757 |
| LINEAVP | ASSLDSGQNTEVF | 0.15208013 |
| LGYAFSGL | ASSLDSGQNTEVF | 0.27414194 |
| LDFVAVAL | ASSLDSGQNTEVF | 0.46203542 |
| LAQPGAE | ASSLDSGQNTEVF | 0.1797764 |
| ISNLGAVTGF | ASSLDSGQNTEVF | 0.14111361 |
| HQEDPGRI | ASSLDSGQNTEVF | 0.38319045 |
| GTPGEPTP | ASSLDSGQNTEVF | 0.5505451 |
| GFFNSGS | ASSLDSGQNTEVF | 0.118618324 |
| GAGGGGLTAGAN | ASSLDSGQNTEVF | 0.16489276 |
| FVDALVNM | ASSLDSGQNTEVF | 0.34418088 |
| FLTVAPEL | ASSLDSGQNTEVF | 0.5471408 |
| FETLTAL | ASSLDSGQNTEVF | 0.31725854 |
| DVINPIEL | ASSLDSGQNTEVF | 0.13245946 |
| APVSGAPV | ASSLDSGQNTEVF | 0.38058382 |
| ALVGGLAW | ASSLDSGQNTEVF | 0.1352501 |
| ALAAAAATAI | ASSLDSGQNTEVF | 0.47844353 |
| AFGLGLLL | ASSLDSGQNTEVF | 0.30109894 |
| AFFAAASL | ASSLDSGQNTEVF | 0.3829638 |
| AAAALPAFSPP | ASSLDSGQNTEVF | 0.19140372 |
| VVLAPAL | ASSLDSGQNTEVF | 0.14149296 |
| TIGHVDHGKTTL | ASSLDSGQNTEVF | 0.13718137 |
| LISNASDALDK | ASSLDSGQNTEVF | 0.118895315 |

| | | |
|---|---|---|
| LISNASDAL | ASSLDSGQNTEVF | 0.60693073 |
| ISNASDALDK | ASSLDSGQNTEVF | 0.3749866 |
| NQAETLVYQTE | ASSLDSGQNTEVF | 0.49313638 |
| TGGTGGTAGAG | ASSLDSGQNTEVF | 0.24947476 |
| TGSTAGGAGGA | ASSLDSGQNTEVF | 0.15044543 |
| DPLAVPVAAGPV | ASSLDSGQNTEVF | 0.41305488 |
| GGTGGTPGNGGN | ASSLDSGQNTEVF | 0.12077303 |
| PAGLAGVDPATL | ASSLDSGQNTEVF | 0.3993435 |
| GNNLVVEAPHAGASV | ASSLDSGQNTEVF | 0.2711813 |
| KVTRIMSASTLEQPA | ASSLDSGQNTEVF | 0.38481277 |
| LGFALPIALPRRTVV | ASSLDSGQNTEVF | 0.20057991 |
| TRLGNLTARKLVVLV | ASSLDSGQNTEVF | 0.41497838 |
| WSQQRRLIYFPSAGP | ASSLDSGQNTEVF | 0.3906033 |
| GPNGNPDPMAAAVDIR | ASSLDSGQNTEVF | 0.16612491 |
| TQRCVSVADLRAAASR | ASSLDSGQNTEVF | 0.34431273 |
| QFVLDVAEHPVEVLTETV | ASSLDSGQNTEVF | 0.4716844 |
| GGAGGTGGAGGNGGTQAGDG | ASSLDSGQNTEVF | 0.43545324 |
| PLLATLRAIAVDVDAKPAQV | ASSLDSGQNTEVF | 0.11145633 |
| APKFRTGGYMIDFWGVGYQVAK | ASSLDSGQNTEVF | 0.35034275 |
| GAGGAGGAGGAAGGPGGTGGTG | ASSLDSGQNTEVF | 0.15113103 |
| IGALGLLATDVAPAGSVAFGVA | ASSLDSGQNTEVF | 0.29827356 |
| AGGAGGIGGNGGAGGAGGVLMG | ASSLDSGQNTEVF | 0.24957654 |
| ADELIGLL | ASSLDSGQNTEVF | 0.27647293 |
| ADRALKA | ASSLDSGQNTEVF | 0.43981978 |
| AGAVGQVGGAGGA | ASSLDSGQNTEVF | 0.14883362 |
| AGGGGGLGGAP | ASSLDSGQNTEVF | 0.18173194 |
| AGGLLYGNGGAG | ASSLDSGQNTEVF | 0.22824106 |
| AGLESIGL | ASSLDSGQNTEVF | 0.28586918 |
| AGLKTGII | ASSLDSGQNTEVF | 0.39418742 |
| ASLAALLL | ASSLDSGQNTEVF | 0.13125461 |
| ASVINISSI | ASSLDSGQNTEVF | 0.3549537 |
| ATPPGPA | ASSLDSGQNTEVF | 0.2328718 |
| ATVASALV | ASSLDSGQNTEVF | 0.14736696 |
| ATVASLIL | ASSLDSGQNTEVF | 0.3365789 |
| AVNGPVLIRH | ASSLDSGQNTEVF | 0.19127624 |
| DGAAGAVGI | ASSLDSGQNTEVF | 0.14454383 |
| DPLLHGI | ASSLDSGQNTEVF | 0.45429716 |
| ELGAALGAL | ASSLDSGQNTEVF | 0.19275755 |
| ELGRTQAV | ASSLDSGQNTEVF | 0.35831553 |
| FVAGGYLS | ASSLDSGQNTEVF | 0.74841213 |
| GASGGAGGAGGW | ASSLDSGQNTEVF | 0.15097792 |
| GGAGAGGDGGAGGI | ASSLDSGQNTEVF | 0.42849472 |
| GTGHNGSQPGMGGNG | ASSLDSGQNTEVF | 0.23470432 |
| GYAAASASAS | ASSLDSGQNTEVF | 0.30780035 |
| HQVDVAAVQQG | ASSLDSGQNTEVF | 0.22215028 |
| IGAAIAHT | ASSLDSGQNTEVF | 0.15590122 |
| IGMIEVAQL | ASSLDSGQNTEVF | 0.48106515 |
| IRLAPEHTSAMD | ASSLDSGQNTEVF | 0.36443785 |
| IYNVSVL | ASSLDSGQNTEVF | 0.42330825 |
| KQGFLGM | ASSLDSGQNTEVF | 0.30307877 |
| LESVITQL | ASSLDSGQNTEVF | 0.5802788 |

| | | |
|---|---|---|
| LGLGALVVV | ASSLDSGQNTEVF | 0.5073515 |
| LIGNGGNGGSGGI | ASSLDSGQNTEVF | 0.15752646 |
| LNAINAPTL | ASSLDSGQNTEVF | 0.113977805 |
| LQLLNTGV | ASSLDSGQNTEVF | 0.20257995 |
| LQTSIEF | ASSLDSGQNTEVF | 0.21041152 |
| MSFAVSEL | ASSLDSGQNTEVF | 0.20408563 |
| NTGALGL | ASSLDSGQNTEVF | 0.13538171 |
| PPGAGGAGGAGTT | ASSLDSGQNTEVF | 0.4614829 |
| QLVGLGI | ASSLDSGQNTEVF | 0.7542664 |
| RFLEQVHQL | ASSLDSGQNTEVF | 0.20339644 |
| RGVGYVLR | ASSLDSGQNTEVF | 0.79566026 |
| SATGVFAGVI | ASSLDSGQNTEVF | 0.1287676 |
| SLPAITLP | ASSLDSGQNTEVF | 0.38292813 |
| STALAFTV | ASSLDSGQNTEVF | 0.5170773 |
| SVLTVGLGK | ASSLDSGQNTEVF | 0.12458309 |
| TALAGAGSGI | ASSLDSGQNTEVF | 0.32504514 |
| TGLPFTGI | ASSLDSGQNTEVF | 0.19836548 |
| VAEHVTV | ASSLDSGQNTEVF | 0.17202665 |
| VDIVLNSL | ASSLDSGQNTEVF | 0.5355334 |
| VLTGGGAVL | ASSLDSGQNTEVF | 0.23296587 |
| VNVGLAYGI | ASSLDSGQNTEVF | 0.44551367 |
| VSSAAVGAV | ASSLDSGQNTEVF | 0.2128198 |
| VTSVYEAGV | ASSLDSGQNTEVF | 0.15818617 |
| WVALDRGA | ASSLDSGQNTEVF | 0.34743503 |
| YLVGSVVI | ASSLDSGQNTEVF | 0.50556767 |
| GPPGTGKTLL | ASSLDSGQNTEVF | 0.3384124 |
| GVGMTKF | ASSLDSGQNTEVF | 0.84394515 |
| IGIGGSDL | ASSLDSGQNTEVF | 0.177102 |
| IGTLLAEL | ASSLDSGQNTEVF | 0.7147385 |
| IYTSGTT | ASSLDSGQNTEVF | 0.75681686 |
| LSFPAGV | ASSLDSGQNTEVF | 0.27025723 |
| LYTSGTT | ASSLDSGQNTEVF | 0.24664421 |
| VGAFPFGF | ASSLDSGQNTEVF | 0.5217947 |
| VGAGTVEF | ASSLDSGQNTEVF | 0.25202918 |
| VGKSTLLL | ASSLDSGQNTEVF | 0.4085256 |
| AALAHAGGGIV | ASSLDSGQNTEVF | 0.58131456 |
| AALVAVSLL | ASSLDSGQNTEVF | 0.44484293 |
| AAVSRAVAAL | ASSLDSGQNTEVF | 0.8596519 |
| AGGSSGIGGF | ASSLDSGQNTEVF | 0.800259 |
| AGLTVANANG | ASSLDSGQNTEVF | 0.7663661 |
| AGLVGAVL | ASSLDSGQNTEVF | 0.65910494 |
| ALSQAFTV | ASSLDSGQNTEVF | 0.3830871 |
| ALTFSVSL | ASSLDSGQNTEVF | 0.68846756 |
| ALTGAALVAL | ASSLDSGQNTEVF | 0.53644687 |
| ASFSIGAGVL | ASSLDSGQNTEVF | 0.7790157 |
| DGSVYKI | ASSLDSGQNTEVF | 0.8416736 |
| EQLAEIV | ASSLDSGQNTEVF | 0.6136057 |
| FAVSELTVM | ASSLDSGQNTEVF | 0.81498355 |
| GAAPLAAGL | ASSLDSGQNTEVF | 0.7707145 |
| GADGGDGGSGGAGG | ASSLDSGQNTEVF | 0.34675676 |
| GFGLDTGGAGG | ASSLDSGQNTEVF | 0.7258636 |

| | | |
|---|---|---|
| GLASGGLL | ASSLDSGQNTEVF | 0.6926193 |
| GRIAGAAVLL | ASSLDSGQNTEVF | 0.7186123 |
| GSAGSLNTSF | ASSLDSGQNTEVF | 0.2786904 |
| GTGQIGI | ASSLDSGQNTEVF | 0.8194044 |
| GTLIALIL | ASSLDSGQNTEVF | 0.64999783 |
| IAATMGFVVG | ASSLDSGQNTEVF | 0.1531695 |
| IFAQALL | ASSLDSGQNTEVF | 0.78529584 |
| IGAAVALGF | ASSLDSGQNTEVF | 0.65735584 |
| IGLTPLAL | ASSLDSGQNTEVF | 0.6586815 |
| ILGSNTSTL | ASSLDSGQNTEVF | 0.83644724 |
| IVFSLTVL | ASSLDSGQNTEVF | 0.45310563 |
| LASKATGF | ASSLDSGQNTEVF | 0.502494 |
| LASVALIL | ASSLDSGQNTEVF | 0.80805683 |
| LFAVGLTL | ASSLDSGQNTEVF | 0.84895754 |
| LFNGGIGTYI | ASSLDSGQNTEVF | 0.4115669 |
| LTLGAGII | ASSLDSGQNTEVF | 0.7772262 |
| LVAAALGDI | ASSLDSGQNTEVF | 0.85222983 |
| NTAFVQLGI | ASSLDSGQNTEVF | 0.7669469 |
| QLFSGAEL | ASSLDSGQNTEVF | 0.86934984 |
| SAATFITL | ASSLDSGQNTEVF | 0.87613475 |
| SNYGLANLGV | ASSLDSGQNTEVF | 0.5780995 |
| SYGGAYAVM | ASSLDSGQNTEVF | 0.84456575 |
| TTGVGGLTLG | ASSLDSGQNTEVF | 0.8576517 |
| TTQTIYI | ASSLDSGQNTEVF | 0.64843506 |
| VAAGAGLGAVI | ASSLDSGQNTEVF | 0.7235671 |
| VAAVVTLL | ASSLDSGQNTEVF | 0.5233036 |
| VGFLGGLI | ASSLDSGQNTEVF | 0.68059266 |
| VGVIPILL | ASSLDSGQNTEVF | 0.35754794 |
| VIFASSLI | ASSLDSGQNTEVF | 0.77302563 |
| VKGGDGGAAAG | ASSLDSGQNTEVF | 0.8590056 |
| VTKTFTNI | ASSLDSGQNTEVF | 0.7036967 |
| VVAAAILL | ASSLDSGQNTEVF | 0.46365294 |
| YAALDVEL | ASSLDSGQNTEVF | 0.43942225 |
| LASVALLL | ASSLDSGQNTEVF | 0.77463526 |
| LDAVHRV | ASSLDSGQNTEVF | 0.58224773 |
| QGLSSSIF | ASSLDSGQNTEVF | 0.37130803 |
| RGTFANIRL | ASSLDSGQNTEVF | 0.65368706 |
| RQLWWGHRI | ASSLDSGQNTEVF | 0.81050944 |
| YGPPGTGKTLL | ASSLDSGQNTEVF | 0.82310456 |
| AALVGAPANVA | ASSLDSGQNTEVF | 0.45866397 |
| AATFRRTE | ASSLDSGQNTEVF | 0.8718287 |
| AGALGGGI | ASSLDSGQNTEVF | 0.8113872 |
| AGGAGGAGGIGAG | ASSLDSGQNTEVF | 0.865106 |
| AGSGALGL | ASSLDSGQNTEVF | 0.7492038 |
| AQIADQI | ASSLDSGQNTEVF | 0.5121448 |
| ASAVAVTYL | ASSLDSGQNTEVF | 0.81527567 |
| ASTVLGGTGG | ASSLDSGQNTEVF | 0.6608857 |
| AVGGEPAK | ASSLDSGQNTEVF | 0.7058668 |
| CEQGTFGGTC | ASSLDSGQNTEVF | 0.8238299 |
| DEGKAPAG | ASSLDSGQNTEVF | 0.66265684 |
| DTVVFPAL | ASSLDSGQNTEVF | 0.61797374 |

| | | |
|---|---|---|
| FEGITMAG | ASSLDSGQNTEVF | 0.6710378 |
| FPLVVQD | ASSLDSGQNTEVF | 0.88334954 |
| GAFTTPQ | ASSLDSGQNTEVF | 0.5569211 |
| GFGGFGTAGGVGGA | ASSLDSGQNTEVF | 0.46336 |
| GFLAQFL | ASSLDSGQNTEVF | 0.617886 |
| GGAGGAGGNAGVA | ASSLDSGQNTEVF | 0.7060783 |
| GGVGTAGAGGAGGAGG | ASSLDSGQNTEVF | 0.84661734 |
| GNGGGGDAGNAGSGGN | ASSLDSGQNTEVF | 0.7878671 |
| GTGGAGGAGNDAGS | ASSLDSGQNTEVF | 0.6952496 |
| GTTNTGWFNS | ASSLDSGQNTEVF | 0.78908205 |
| IFMGPLFTL | ASSLDSGQNTEVF | 0.82383907 |
| ILAHAIEAI | ASSLDSGQNTEVF | 0.7604462 |
| IRSAIREAL | ASSLDSGQNTEVF | 0.7726039 |
| ITFAGGG | ASSLDSGQNTEVF | 0.5856787 |
| ITGMSGHQEV | ASSLDSGQNTEVF | 0.8780464 |
| IVAGLACTL | ASSLDSGQNTEVF | 0.8298216 |
| IVGGGLAGA | ASSLDSGQNTEVF | 0.84809923 |
| IVTGVLNI | ASSLDSGQNTEVF | 0.8575675 |
| LAQQGCI | ASSLDSGQNTEVF | 0.74731064 |
| LCLHVEN | ASSLDSGQNTEVF | 0.78795576 |
| LDAISDGS | ASSLDSGQNTEVF | 0.42401147 |
| LGGFAAML | ASSLDSGQNTEVF | 0.5233614 |
| LGGKNPMI | ASSLDSGQNTEVF | 0.74540466 |
| LGPKGVAAIL | ASSLDSGQNTEVF | 0.79950774 |
| LMVGVSI | ASSLDSGQNTEVF | 0.8122337 |
| LVGAAFVL | ASSLDSGQNTEVF | 0.8042443 |
| LVLAVAPL | ASSLDSGQNTEVF | 0.5772147 |
| MAAIAVIDAA | ASSLDSGQNTEVF | 0.83878523 |
| NNGINAELQ | ASSLDSGQNTEVF | 0.8465231 |
| RGEPLVLVHG | ASSLDSGQNTEVF | 0.46539328 |
| RGGAGGAAGLI | ASSLDSGQNTEVF | 0.6691136 |
| RSYRMHPAV | ASSLDSGQNTEVF | 0.86566 |
| SAAHLLAGQLA | ASSLDSGQNTEVF | 0.7757335 |
| SAAVGSY | ASSLDSGQNTEVF | 0.6825044 |
| SPHAVLHRI | ASSLDSGQNTEVF | 0.6607708 |
| SQIAKVL | ASSLDSGQNTEVF | 0.8275442 |
| TALGHLVP | ASSLDSGQNTEVF | 0.6705881 |
| TAPTTSGAA | ASSLDSGQNTEVF | 0.8680759 |
| TDGDDGNGGSPGT | ASSLDSGQNTEVF | 0.24142519 |
| TDGLGEN | ASSLDSGQNTEVF | 0.7384889 |
| TGGPPLPGQA | ASSLDSGQNTEVF | 0.6491552 |
| TIPAITI | ASSLDSGQNTEVF | 0.2448218 |
| TIPGFTI | ASSLDSGQNTEVF | 0.84250283 |
| TSIASGALA | ASSLDSGQNTEVF | 0.5170322 |
| TVTLPGAAI | ASSLDSGQNTEVF | 0.7850082 |
| VASGLSAALP | ASSLDSGQNTEVF | 0.693086 |
| VGAGVIGI | ASSLDSGQNTEVF | 0.8494601 |
| VGPKRFGV | ASSLDSGQNTEVF | 0.76973027 |
| VNVAFPD | ASSLDSGQNTEVF | 0.56910586 |
| VSMTNSGVS | ASSLDSGQNTEVF | 0.5230446 |
| VTDVLGDLSV | ASSLDSGQNTEVF | 0.8188983 |

| | | |
|---|---|---|
| VVVVGAGFAGL | ASSLDSGQNTEVF | 0.8491111 |
| VYTVGTVV | ASSLDSGQNTEVF | 0.8702259 |
| WTTFSGP | ASSLDSGQNTEVF | 0.3043329 |
| YDELVEAF | ASSLDSGQNTEVF | 0.77327216 |
| YGTGGAGGNGG | ASSLDSGQNTEVF | 0.7386944 |
| YTNVMAV | ASSLDSGQNTEVF | 0.30871832 |
| FLEQVHQL | ASSLDSGQNTEVF | 0.22985007 |
| GLSLLAVAV | ASSLDSGQNTEVF | 0.694314 |
| LEQVHQL | ASSLDSGQNTEVF | 0.7135278 |
| LNAAGAI | ASSLDSGQNTEVF | 0.8076183 |
| PAVHASL | ASSLDSGQNTEVF | 0.81141925 |
| TELEDDA | ASSLDSGQNTEVF | 0.6935688 |
| TPPGPPN | ASSLDSGQNTEVF | 0.81507695 |
| TPPPPPPP | ASSLDSGQNTEVF | 0.7027139 |
| TQSPALI | ASSRGNNSGNTLY | 0.08721496 |
| EFAAAFPQ | ASSRGNNSGNTLY | 0.5940933 |
| LLSGNLDVL | ASSRGNNSGNTLY | 0.34669343 |
| GPANLGNYNI | ASSRGNNSGNTLY | 0.8629314 |
| TLGPTLP | ASSRGNNSGNTLY | 0.47649908 |
| VYAYSSCCK | ASSRGNNSGNTLY | 0.63354623 |
| APAPAPAPNI | ASSRGNNSGNTLY | 0.72264147 |
| PGGPVPIP | ASSRGNNSGNTLY | 0.74832094 |
| FLTVALEP | ASSRGNNSGNTLY | 0.7272069 |
| SQYFGANL | ASSRGNNSGNTLY | 0.6145309 |
| GGVKPAA | ASSRGNNSGNTLY | 0.21483822 |
| GGTGGAGGAGGW | ASSRGNNSGNTLY | 0.8203169 |
| GVGGVAGLGGAGG | ASSRGNNSGNTLY | 0.48032868 |
| VLLVLATTI | ASSRGNNSGNTLY | 0.43194366 |
| IGIFAAML | ASSRGNNSGNTLY | 0.41831905 |
| AVGDLID | ASSRGNNSGNTLY | 0.3967292 |
| NISPLPL | ASSRGNNSGNTLY | 0.33242637 |
| VTAVFVAI | ASSRGNNSGNTLY | 0.86004615 |
| SGDDNDAAL | ASSRGNNSGNTLY | 0.6682155 |
| AVAALAHS | ASSRGNNSGNTLY | 0.6922978 |
| GGVTVMATP | ASSRGNNSGNTLY | 0.87304926 |
| AGLAANVRD | ASSRGNNSGNTLY | 0.66970634 |
| TNRPDLIDPALLRPG | ASSRGNNSGNTLY | 0.70023584 |
| LANLGVTAL | ASSRGNNSGNTLY | 0.5341489 |
| TGERIEL | ASSRGNNSGNTLY | 0.53750545 |
| NGEELAA | ASSRGNNSGNTLY | 0.5894598 |
| VLSLVWG | ASSRGNNSGNTLY | 0.5329263 |
| IGYALWL | ASSRGNNSGNTLY | 0.8572462 |
| LSLLLAPL | ASSRGNNSGNTLY | 0.76751375 |
| TLDEALK | ASSRGNNSGNTLY | 0.4282969 |
| TDGAGTAEGG | ASSRGNNSGNTLY | 0.817199 |
| NAPGGAGGNGG | ASSRGNNSGNTLY | 0.86426556 |
| VSVFMGMGGNF | ASSRGNNSGNTLY | 0.8874117 |
| LASYLAAL | ASSRGNNSGNTLY | 0.44558012 |
| LVGGTPAL | ASSRGNNSGNTLY | 0.5888359 |
| VRNADVL | ASSRGNNSGNTLY | 0.66674674 |
| TCSSPFLVSAM | ASSRGNNSGNTLY | 0.6891686 |

| | | |
|---|---|---|
| VHITDAE | ASSRGNNSGNTLY | 0.6020013 |
| EYAAPSF | ASSRGNNSGNTLY | 0.50473285 |
| VVDAFHGGL | ASSRGNNSGNTLY | 0.86514395 |
| ASVVDPQ | ASSRGNNSGNTLY | 0.85508454 |
| SYNEFGGLS | ASSRGNNSGNTLY | 0.69898427 |
| DPDGSVY | ASSRGNNSGNTLY | 0.50593567 |
| MVSTIHATE | ASSRGNNSGNTLY | 0.686169 |
| LAAGVADPV | ASSRGNNSGNTLY | 0.5947883 |
| TKEQIAA | ASSRGNNSGNTLY | 0.6091049 |
| SLASPAAL | ASSRGNNSGNTLY | 0.6349512 |
| IGLVFYGV | ASSRGNNSGNTLY | 0.40271673 |
| FLLDASNL | ASSRGNNSGNTLY | 0.70996916 |
| YQVGNFP | ASSRGNNSGNTLY | 0.8753425 |
| ESLLSQST | ASSRGNNSGNTLY | 0.6185509 |
| LFTVGFAAVG | ASSRGNNSGNTLY | 0.8113595 |
| ATPPTPT | ASSRGNNSGNTLY | 0.8580195 |
| TDLATLG | ASSRGNNSGNTLY | 0.8251164 |
| SAGGTGR | ASSRGNNSGNTLY | 0.85848415 |
| GFSNSGS | ASSRGNNSGNTLY | 0.81205434 |
| GGNGGSGGTSV | ASSRGNNSGNTLY | 0.7217486 |
| GTSVATGGAGN | ASSRGNNSGNTLY | 0.77444243 |
| NGGNGGSGGTS | ASSRGNNSGNTLY | 0.7985705 |
| AVVKFPH | ASSRGNNSGNTLY | 0.6768315 |
| TPPPPPP | ASSRGNNSGNTLY | 0.72985095 |
| AAPAAAAAPA | ASSRGNNSGNTLY | 0.7265274 |
| IALNEDL | ASSRGNNSGNTLY | 0.7939841 |
| IVNVSSM | ASSRGNNSGNTLY | 0.768641 |
| TGIPPAPR | ASSRGNNSGNTLY | 0.8406638 |
| FDLGGGTF | ASSRGNNSGNTLY | 0.31021655 |
| FDLGGGTFDVS | ASSRGNNSGNTLY | 0.4716925 |
| NEPTAAA | ASSRGNNSGNTLY | 0.8431411 |
| NEPTAAALA | ASSRGNNSGNTLY | 0.15952164 |
| GSIINISSI | ASSRGNNSGNTLY | 0.2722424 |
| VAPGNAGTA | ASSRGNNSGNTLY | 0.8009255 |
| TGGGASLE | ASSRGNNSGNTLY | 0.859217 |
| VAVYNHYKRI | ASSRGNNSGNTLY | 0.16696304 |
| TDPAPATT | ASSRGNNSGNTLY | 0.5245566 |
| TNRPDILDPALLRPG | ASSRGNNSGNTLY | 0.43018192 |
| LTTTTTS | ASSRGNNSGNTLY | 0.839404 |
| PTGSGKTL | ASSRGNNSGNTLY | 0.69529176 |
| TGSGKTL | ASSRGNNSGNTLY | 0.32707307 |
| TIGHVDHGK | ASSRGNNSGNTLY | 0.8364272 |
| TIGHVDHGKTT | ASSRGNNSGNTLY | 0.4732802 |
| GHVDHGKTT | ASSRGNNSGNTLY | 0.6303398 |
| AQFEHTLL | ASSRGNNSGNTLY | 0.8571873 |
| IVNQDEL | ASSRGNNSGNTLY | 0.8550569 |
| TNGSQFF | ASSRGNNSGNTLY | 0.44666535 |
| TGKGIVE | ASSRGNNSGNTLY | 0.820864 |
| LISNASDA | ASSRGNNSGNTLY | 0.66099334 |
| AELRTLGELTKIAKAHGAQ | ASSRGNNSGNTLY | 0.8868011 |
| APNAVAAAEPSW | ASSRGNNSGNTLY | 0.66585654 |

| | | |
|---|---|---|
| APQVAGAATAVL | ASSRGNNSGNTLY | 0.41054618 |
| ARRDPAAAAAILPTDA | ASSRGNNSGNTLY | 0.8352556 |
| DAAARDVLAEAGLAEYF | ASSRGNNSGNTLY | 0.8373901 |
| DDQIVLSAKANLAKVEQ | ASSRGNNSGNTLY | 0.68051755 |
| DELLPDTGEDSDEDGH | ASSRGNNSGNTLY | 0.80994964 |
| ELPEAPANRATVPQD | ASSRGNNSGNTLY | 0.73550034 |
| FGQSKIERVTGIG | ASSRGNNSGNTLY | 0.683548 |
| GAGGTGSTAGGAGGAGGAGGLY | ASSRGNNSGNTLY | 0.8062073 |
| GHGGTGAAGGNGGTGGAGGSGI | ASSRGNNSGNTLY | 0.8139082 |
| GNGGNGGAGGEGGPGGAGGAGA | ASSRGNNSGNTLY | 0.5129069 |
| GSTAGNGGNGGNATNGTIAGQP | ASSRGNNSGNTLY | 0.8703283 |
| GTGGDGGNGGNGGTGG | ASSRGNNSGNTLY | 0.70845 |
| GVRRPAGIPADAPV | ASSRGNNSGNTLY | 0.48748922 |
| GYRQVTDAHLLTLARR | ASSRGNNSGNTLY | 0.8871908 |
| IAKMGQDGHDRG | ASSRGNNSGNTLY | 0.7583432 |
| KAEPSIEIQYT | ASSRGNNSGNTLY | 0.8384706 |
| KPVRIRMAADEDTVAVT | ASSRGNNSGNTLY | 0.8298292 |
| MPPRIAGMRLLVIKPEP | ASSRGNNSGNTLY | 0.8225395 |
| MKFYAVALFRSRGVPPTR | ASSRGNNSGNTLY | 0.42228192 |
| NAGGFGILGAVAHSP | ASSRGNNSGNTLY | 0.72816294 |
| NEGREITLAMAEM | ASSRGNNSGNTLY | 0.7152158 |
| NELVEWSHPLGHRWRWE | ASSRGNNSGNTLY | 0.8300023 |
| NLGAGGTGGNGGNPG | ASSRGNNSGNTLY | 0.74443257 |
| NTEWPKWETVMAADPVD | ASSRGNNSGNTLY | 0.64317286 |
| PASAAPAETQP | ASSRGNNSGNTLY | 0.8187468 |
| QQSSDGKNGVRAAEITGE | ASSRGNNSGNTLY | 0.8657843 |
| RAPDAVTTSWWKE | ASSRGNNSGNTLY | 0.80499417 |
| SGFGNVGAHTSGWFNQ | ASSRGNNSGNTLY | 0.73398507 |
| SGVAGAVGQAASVGGLKV | ASSRGNNSGNTLY | 0.8672075 |
| SNGGAGTGSAGGNGGT | ASSRGNNSGNTLY | 0.5183474 |
| TGGAGGEGGAAG | ASSRGNNSGNTLY | 0.72931325 |
| TGGDGGAAGAGGAPGHG | ASSRGNNSGNTLY | 0.85415834 |
| TMPPLLAAIANAGRHGV | ASSRGNNSGNTLY | 0.84357333 |
| TSSTAGNSGGA | ASSRGNNSGNTLY | 0.7587696 |
| VPATVPVGILC | ASSRGNNSGNTLY | 0.8529091 |
| VQRPDSGGVVLTGRIS | ASSRGNNSGNTLY | 0.56522906 |
| WTPNPVPATKGVPAGLVQAT | ASSRGNNSGNTLY | 0.73048985 |
| YALKRALAKFSDRFDVVII | ASSRGNNSGNTLY | 0.6165138 |
| YLIGSVIV | ASSRGNNSGNTLY | 0.72302544 |
| YGQLVAVID | ASSRGNNSGNTLY | 0.5109774 |
| VTVGGMTLPA | ASSRGNNSGNTLY | 0.75703454 |
| VLALPAV | ASSRGNNSGNTLY | 0.8315531 |
| VIPPTLNYET | ASSRGNNSGNTLY | 0.76118743 |
| VGMGGMHPSSG | ASSRGNNSGNTLY | 0.24950497 |
| VGLAGGAATA | ASSRGNNSGNTLY | 0.7978599 |
| VGGAGGAGGSGP | ASSRGNNSGNTLY | 0.69725806 |
| VFAAFMV | ASSRGNNSGNTLY | 0.86929107 |
| TLYDKLR | ASSRGNNSGNTLY | 0.7463451 |
| TLAVFTVL | ASSRGNNSGNTLY | 0.8039217 |
| TGSVQPARV | ASSRGNNSGNTLY | 0.57253116 |
| TGGGAGSGGT | ASSRGNNSGNTLY | 0.78629696 |

| | | |
|---|---|---|
| SHLETSL | ASSRGNNSGNTLY | 0.4270962 |
| SFTTGMT | ASSRGNNSGNTLY | 0.5643604 |
| QVEVAKRMGVRQPTVS | ASSRGNNSGNTLY | 0.6075578 |
| QGSGGPALAM | ASSRGNNSGNTLY | 0.57357264 |
| MLAPSNP | ASSRGNNSGNTLY | 0.7945529 |
| LVNNAAI | ASSRGNNSGNTLY | 0.85109705 |
| LTGEINTLA | ASSRGNNSGNTLY | 0.8287974 |
| LSIFGPV | ASSRGNNSGNTLY | 0.8484275 |
| LLALLANL | ASSRGNNSGNTLY | 0.8649925 |
| LISNASDALDKLR | ASSRGNNSGNTLY | 0.8653457 |
| LINEAVP | ASSRGNNSGNTLY | 0.65287495 |
| LGYAFSGL | ASSRGNNSGNTLY | 0.6462245 |
| LDFVAVAL | ASSRGNNSGNTLY | 0.45736945 |
| LAQPGAE | ASSRGNNSGNTLY | 0.6734154 |
| ISNLGAVTGF | ASSRGNNSGNTLY | 0.6816043 |
| HQEDPGRI | ASSRGNNSGNTLY | 0.7474624 |
| GTPGEPTP | ASSRGNNSGNTLY | 0.14328589 |
| GFFNSGS | ASSRGNNSGNTLY | 0.7289561 |
| GAGGGGLTAGAN | ASSRGNNSGNTLY | 0.8665887 |
| FVDALVNM | ASSRGNNSGNTLY | 0.6296259 |
| FLTVAPEL | ASSRGNNSGNTLY | 0.76937217 |
| FETLTAL | ASSRGNNSGNTLY | 0.7948515 |
| DVINPIEL | ASSRGNNSGNTLY | 0.69818306 |
| APVSGAPV | ASSRGNNSGNTLY | 0.8139417 |
| ALVGGLAW | ASSRGNNSGNTLY | 0.66299677 |
| ALAAAAATAI | ASSRGNNSGNTLY | 0.83778465 |
| AFGLGLLL | ASSRGNNSGNTLY | 0.877574 |
| AFFAAASL | ASSRGNNSGNTLY | 0.7757749 |
| AAAALPAFSPP | ASSRGNNSGNTLY | 0.6765936 |
| VVLAPAL | ASSRGNNSGNTLY | 0.7461896 |
| TIGHVDHGKTTL | ASSRGNNSGNTLY | 0.3922744 |
| LISNASDALDK | ASSRGNNSGNTLY | 0.7264529 |
| LISNASDAL | ASSRGNNSGNTLY | 0.16994387 |
| ISNASDALDK | ASSRGNNSGNTLY | 0.74358046 |
| NQAETLVYQTE | ASSRGNNSGNTLY | 0.5834073 |
| TGGTGGTAGAG | ASSRGNNSGNTLY | 0.74644697 |
| TGSTAGGAGGA | ASSRGNNSGNTLY | 0.7267631 |
| DPLAVPVAAGPV | ASSRGNNSGNTLY | 0.829426 |
| GGTGGTPGNGGN | ASSRGNNSGNTLY | 0.49738705 |
| PAGLAGVDPATL | ASSRGNNSGNTLY | 0.76976734 |
| GNNLVVEAPHAGASV | ASSRGNNSGNTLY | 0.6860415 |
| KVTRIMSASTLEQPA | ASSRGNNSGNTLY | 0.3759816 |
| LGFALPIALPRRTVV | ASSRGNNSGNTLY | 0.8495356 |
| TRLGNLTARKLVVLV | ASSRGNNSGNTLY | 0.7847613 |
| WSQQRRLIYFPSAGP | ASSRGNNSGNTLY | 0.7248752 |
| GPNGNPDPMAAAVDIR | ASSRGNNSGNTLY | 0.8179667 |
| TQRCVSVADLRAAASR | ASSRGNNSGNTLY | 0.7830297 |
| QFVLDVAEHPVEVLTETV | ASSRGNNSGNTLY | 0.53780997 |
| GGAGGTGGAGGNGGTQAGDG | ASSRGNNSGNTLY | 0.65911704 |
| PLLATLRAIAVDVDAKPAQV | ASSRGNNSGNTLY | 0.75152 |
| APKFRTGGYMIDFWGVGYQVAK | ASSRGNNSGNTLY | 0.69916606 |

| | | |
|---|---|---|
| GAGGAGGAGGAAGGPGGTGGTG | ASSRGNNSGNTLY | 0.14354347 |
| IGALGLLATDVAPAGSVAFGVA | ASSRGNNSGNTLY | 0.81195 |
| AGGAGGIGGNGGAGGAGGVLMG | ASSRGNNSGNTLY | 0.48256767 |
| ADELIGLL | ASSRGNNSGNTLY | 0.5952107 |
| ADRALKA | ASSRGNNSGNTLY | 0.7128078 |
| AGAVGQVGGAGGA | ASSRGNNSGNTLY | 0.82910657 |
| AGGGGGLGGAP | ASSRGNNSGNTLY | 0.6637616 |
| AGGLLYGNGGAG | ASSRGNNSGNTLY | 0.535241 |
| AGLESIGL | ASSRGNNSGNTLY | 0.8129848 |
| AGLKTGII | ASSRGNNSGNTLY | 0.74749124 |
| ASLAALLL | ASSRGNNSGNTLY | 0.8265589 |
| ASVINISSI | ASSRGNNSGNTLY | 0.71755165 |
| ATPPGPA | ASSRGNNSGNTLY | 0.79085886 |
| ATVASALV | ASSRGNNSGNTLY | 0.6885108 |
| ATVASLIL | ASSRGNNSGNTLY | 0.69100136 |
| AVNGPVLIRH | ASSRGNNSGNTLY | 0.48063785 |
| DGAAGAVGI | ASSRGNNSGNTLY | 0.4288794 |
| DPLLHGI | ASSRGNNSGNTLY | 0.69882333 |
| ELGAALGAL | ASSRGNNSGNTLY | 0.42309624 |
| ELGRTQAV | ASSRGNNSGNTLY | 0.7495305 |
| FVAGGYLS | ASSRGNNSGNTLY | 0.8732585 |
| GASGGAGGAGGW | ASSRGNNSGNTLY | 0.73423344 |
| GGAGAGGDGGAGGI | ASSRGNNSGNTLY | 0.40049317 |
| GTGHNGSQPGMGGNG | ASSRGNNSGNTLY | 0.70653886 |
| GYAAASASAS | ASSRGNNSGNTLY | 0.60595304 |
| HQVDVAAVQQG | ASSRGNNSGNTLY | 0.81115985 |
| IGAAIAHT | ASSRGNNSGNTLY | 0.81922776 |
| IGMIEVAQL | ASSRGNNSGNTLY | 0.8352798 |
| IRLAPEHTSAMD | ASSRGNNSGNTLY | 0.79102683 |
| IYNVSVL | ASSRGNNSGNTLY | 0.611047 |
| KQGFLGM | ASSRGNNSGNTLY | 0.83705497 |
| LESVITQL | ASSRGNNSGNTLY | 0.71314335 |
| LGLGALVVV | ASSRGNNSGNTLY | 0.75931156 |
| LIGNGGNGGSGGI | ASSRGNNSGNTLY | 0.7722677 |
| LNAINAPTL | ASSRGNNSGNTLY | 0.82021415 |
| LQLLNTGV | ASSRGNNSGNTLY | 0.58925456 |
| LQTSIEF | ASSRGNNSGNTLY | 0.66645336 |
| MSFAVSEL | ASSRGNNSGNTLY | 0.467983 |
| NTGALGL | ASSRGNNSGNTLY | 0.81867385 |
| PPGAGGAGGAGTT | ASSRGNNSGNTLY | 0.6993469 |
| QLVGLGI | ASSRGNNSGNTLY | 0.54273283 |
| RFLEQVHQL | ASSRGNNSGNTLY | 0.64224553 |
| RGVGYVLR | ASSRGNNSGNTLY | 0.8738143 |
| SATGVFAGVI | ASSRGNNSGNTLY | 0.85410047 |
| SLPAITLP | ASSRGNNSGNTLY | 0.6014155 |
| STALAFTV | ASSRGNNSGNTLY | 0.6009115 |
| SVLTVGLGK | ASSRGNNSGNTLY | 0.7677962 |
| TALAGAGSGI | ASSRGNNSGNTLY | 0.5108845 |
| TGLPFTGI | ASSRGNNSGNTLY | 0.84132797 |
| VAEHVTV | ASSRGNNSGNTLY | 0.709501 |
| VDIVLNSL | ASSRGNNSGNTLY | 0.67982733 |

| | | |
|---|---|---|
| VLTGGGAVL | ASSRGNNSGNTLY | 0.58644986 |
| VNVGLAYGI | ASSRGNNSGNTLY | 0.81754196 |
| VSSAAVGAV | ASSRGNNSGNTLY | 0.64352727 |
| VTSVYEAGV | ASSRGNNSGNTLY | 0.83593416 |
| WVALDRGA | ASSRGNNSGNTLY | 0.17217322 |
| YLVGSVVI | ASSRGNNSGNTLY | 0.840829 |
| GPPGTGKTLL | ASSRGNNSGNTLY | 0.51119405 |
| GVGMTKF | ASSRGNNSGNTLY | 0.59767884 |
| IGIGGSDL | ASSRGNNSGNTLY | 0.2840208 |
| IGTLLAEL | ASSRGNNSGNTLY | 0.72016776 |
| IYTSGTT | ASSRGNNSGNTLY | 0.8247183 |
| LSFPAGV | ASSRGNNSGNTLY | 0.8628423 |
| LYTSGTT | ASSRGNNSGNTLY | 0.6962253 |
| VGAFPFGF | ASSRGNNSGNTLY | 0.7949785 |
| VGAGTVEF | ASSRGNNSGNTLY | 0.87556535 |
| VGKSTLLL | ASSRGNNSGNTLY | 0.68347234 |
| AALAHAGGGIV | ASSRGNNSGNTLY | 0.30829608 |
| AALVAVSLL | ASSRGNNSGNTLY | 0.7538356 |
| AAVSRAVAAL | ASSRGNNSGNTLY | 0.68211484 |
| AGGSSGIGGF | ASSRGNNSGNTLY | 0.8558315 |
| AGLTVANANG | ASSRGNNSGNTLY | 0.78774107 |
| AGLVGAVL | ASSRGNNSGNTLY | 0.41946965 |
| ALSQAFTV | ASSRGNNSGNTLY | 0.7621144 |
| ALTFSVSL | ASSRGNNSGNTLY | 0.82240045 |
| ALTGAALVAL | ASSRGNNSGNTLY | 0.4634155 |
| ASFSIGAGVL | ASSRGNNSGNTLY | 0.6493119 |
| DGSVYKI | ASSRGNNSGNTLY | 0.55106795 |
| EQLAEIV | ASSRGNNSGNTLY | 0.26955405 |
| FAVSELTVM | ASSRGNNSGNTLY | 0.76698416 |
| GAAPLAAGL | ASSRGNNSGNTLY | 0.6749394 |
| GADGGDGGSGGAGG | ASSRGNNSGNTLY | 0.54002845 |
| GFGLDTGGAGG | ASSRGNNSGNTLY | 0.7266207 |
| GLASGGLL | ASSRGNNSGNTLY | 0.5535842 |
| GRIAGAAVLL | ASSRGNNSGNTLY | 0.6267153 |
| GSAGSLNTSF | ASSRGNNSGNTLY | 0.61581826 |
| GTGQIGI | ASSRGNNSGNTLY | 0.77312493 |
| GTLIALIL | ASSRGNNSGNTLY | 0.8382728 |
| IAATMGFVVG | ASSRGNNSGNTLY | 0.7324587 |
| IFAQALL | ASSRGNNSGNTLY | 0.74216104 |
| IGAAVALGF | ASSRGNNSGNTLY | 0.8554718 |
| IGLTPLAL | ASSRGNNSGNTLY | 0.38162664 |
| ILGSNTSTL | ASSRGNNSGNTLY | 0.5284908 |
| IVFSLTVL | ASSRGNNSGNTLY | 0.81681883 |
| LASKATGF | ASSRGNNSGNTLY | 0.5781174 |
| LASVALIL | ASSRGNNSGNTLY | 0.8302054 |
| LFAVGLTL | ASSRGNNSGNTLY | 0.72309613 |
| LFNGGIGTYI | ASSRGNNSGNTLY | 0.7800687 |
| LTLGAGII | ASSRGNNSGNTLY | 0.7064646 |
| LVAAALGDI | ASSRGNNSGNTLY | 0.8835905 |
| NTAFVQLGI | ASSRGNNSGNTLY | 0.8201448 |
| QLFSGAEL | ASSRGNNSGNTLY | 0.71146965 |

| | | |
|---|---|---|
| SAATFITL | ASSRGNNSGNTLY | 0.7591711 |
| SNYGLANLGV | ASSRGNNSGNTLY | 0.4353994 |
| SYGGAYAVM | ASSRGNNSGNTLY | 0.82171667 |
| TTGVGGLTLG | ASSRGNNSGNTLY | 0.8237004 |
| TTQTIYI | ASSRGNNSGNTLY | 0.5672745 |
| VAAGAGLGAVI | ASSRGNNSGNTLY | 0.6881465 |
| VAAVVTLL | ASSRGNNSGNTLY | 0.8832989 |
| VGFLGGLI | ASSRGNNSGNTLY | 0.84847766 |
| VGVIPILL | ASSRGNNSGNTLY | 0.52819526 |
| VIFASSLI | ASSRGNNSGNTLY | 0.5853654 |
| VKGGDGGAAAG | ASSRGNNSGNTLY | 0.47164196 |
| VTKTFTNI | ASSRGNNSGNTLY | 0.6935793 |
| VVAAAILL | ASSRGNNSGNTLY | 0.38158783 |
| YAALDVEL | ASSRGNNSGNTLY | 0.8225949 |
| LASVALLL | ASSRGNNSGNTLY | 0.64322144 |
| LDAVHRV | ASSRGNNSGNTLY | 0.78443635 |
| QGLSSSIF | ASSRGNNSGNTLY | 0.7751503 |
| RGTFANIRL | ASSRGNNSGNTLY | 0.8597492 |
| RQLWWGHRI | ASSRGNNSGNTLY | 0.6551548 |
| YGPPGTGKTLL | ASSRGNNSGNTLY | 0.6123824 |
| AALVGAPANVA | ASSRGNNSGNTLY | 0.36044598 |
| AATFRRTE | ASSRGNNSGNTLY | 0.4364817 |
| AGALGGGI | ASSRGNNSGNTLY | 0.6539221 |
| AGGAGGAGGIGAG | ASSRGNNSGNTLY | 0.61895394 |
| AGSGALGL | ASSRGNNSGNTLY | 0.24488804 |
| AQIADQI | ASSRGNNSGNTLY | 0.86338323 |
| ASAVAVTYL | ASSRGNNSGNTLY | 0.72613597 |
| ASTVLGGTGG | ASSRGNNSGNTLY | 0.8241388 |
| AVGGEPAK | ASSRGNNSGNTLY | 0.58216316 |
| CEQGTFGGTC | ASSRGNNSGNTLY | 0.57941175 |
| DEGKAPAG | ASSRGNNSGNTLY | 0.66308975 |
| DTVVFPAL | ASSRGNNSGNTLY | 0.7920115 |
| FEGITMAG | ASSRGNNSGNTLY | 0.40747747 |
| FPLVVQD | ASSRGNNSGNTLY | 0.4003424 |
| GAFTTPQ | ASSRGNNSGNTLY | 0.62756526 |
| GFGGFGTAGGVGGA | ASSRGNNSGNTLY | 0.6207063 |
| GFLAQFL | ASSRGNNSGNTLY | 0.87800217 |
| GGAGGAGGNAGVA | ASSRGNNSGNTLY | 0.8460831 |
| GGVGTAGAGGAGGAGG | ASSRGNNSGNTLY | 0.7659081 |
| GNGGGGDAGNAGSGGN | ASSRGNNSGNTLY | 0.87062776 |
| GTGGAGGAGNDAGS | ASSRGNNSGNTLY | 0.19841602 |
| GTTNTGWFNS | ASSRGNNSGNTLY | 0.5545493 |
| IFMGPLFTL | ASSRGNNSGNTLY | 0.83593 |
| ILAHAIEAI | ASSRGNNSGNTLY | 0.8105484 |
| IRSAIREAL | ASSRGNNSGNTLY | 0.69622403 |
| ITFAGGG | ASSRGNNSGNTLY | 0.85915774 |
| ITGMSGHQEV | ASSRGNNSGNTLY | 0.51385576 |
| IVAGLACTL | ASSRGNNSGNTLY | 0.65460676 |
| IVGGGLAGA | ASSRGNNSGNTLY | 0.72666025 |
| IVTGVLNI | ASSRGNNSGNTLY | 0.8370883 |
| LAQQGCI | ASSRGNNSGNTLY | 0.8132136 |

| | | |
|---|---|---|
| LCLHVEN | ASSRGNNSGNTLY | 0.41679367 |
| LDAISDGS | ASSRGNNSGNTLY | 0.7890692 |
| LGGFAAML | ASSRGNNSGNTLY | 0.88773096 |
| LGGKNPMI | ASSRGNNSGNTLY | 0.6546537 |
| LGPKGVAAIL | ASSRGNNSGNTLY | 0.8300299 |
| LMVGVSI | ASSRGNNSGNTLY | 0.773525 |
| LVGAAFVL | ASSRGNNSGNTLY | 0.7255465 |
| LVLAVAPL | ASSRGNNSGNTLY | 0.55993366 |
| MAAIAVIDAA | ASSRGNNSGNTLY | 0.59109765 |
| NNGINAELQ | ASSRGNNSGNTLY | 0.7632535 |
| RGEPLVLVHG | ASSRGNNSGNTLY | 0.39460605 |
| RGGAGGAAGLI | ASSRGNNSGNTLY | 0.7394208 |
| RSYRMHPAV | ASSRGNNSGNTLY | 0.6923504 |
| SAAHLLAGQLA | ASSRGNNSGNTLY | 0.89420617 |
| SAAVGSY | ASSRGNNSGNTLY | 0.8051547 |
| SPHAVLHRI | ASSRGNNSGNTLY | 0.8021623 |
| SQIAKVL | ASSRGNNSGNTLY | 0.5245409 |
| TALGHLVP | ASSRGNNSGNTLY | 0.8232757 |
| TAPTTSGAA | ASSRGNNSGNTLY | 0.80426544 |
| TDGDDGNGGSPGT | ASSRGNNSGNTLY | 0.55706733 |
| TDGLGEN | ASSRGNNSGNTLY | 0.85367185 |
| TGGPPLPGQA | ASSRGNNSGNTLY | 0.5910276 |
| TIPAITI | ASSRGNNSGNTLY | 0.74758697 |
| TIPGFTI | ASSRGNNSGNTLY | 0.77568126 |
| TSIASGALA | ASSRGNNSGNTLY | 0.8030076 |
| TVTLPGAAI | ASSRGNNSGNTLY | 0.7219167 |
| VASGLSAALP | ASSRGNNSGNTLY | 0.7776731 |
| VGAGVIGI | ASSRGNNSGNTLY | 0.502937 |
| VGPKRFGV | ASSRGNNSGNTLY | 0.8430858 |
| VNVAFPD | ASSRGNNSGNTLY | 0.8377634 |
| VSMTNSGVS | ASSRGNNSGNTLY | 0.6997057 |
| VTDVLGDLSV | ASSRGNNSGNTLY | 0.86879086 |
| VVVVGAGFAGL | ASSRGNNSGNTLY | 0.8017603 |
| VYTVGTVV | ASSRGNNSGNTLY | 0.3416523 |
| WTTFSGP | ASSRGNNSGNTLY | 0.7356217 |
| YDELVEAF | ASSRGNNSGNTLY | 0.8613628 |
| YGTGGAGGNGG | ASSRGNNSGNTLY | 0.86278296 |
| YTNVMAV | ASSRGNNSGNTLY | 0.8498228 |
| FLEQVHQL | ASSRGNNSGNTLY | 0.7736082 |
| GLSLLAVAV | ASSRGNNSGNTLY | 0.6844722 |
| LEQVHQL | ASSRGNNSGNTLY | 0.8625868 |
| LNAAGAI | ASSRGNNSGNTLY | 0.8955479 |
| PAVHASL | ASSRGNNSGNTLY | 0.81130207 |
| TELEDDA | ASSRGNNSGNTLY | 0.597316 |
| TPPGPPN | ASSRGNNSGNTLY | 0.547566 |
| TPPPPPPP | ASSRGNNSGNTLY | 0.80468494 |
| TQSPALI | ASSYRDENTGQLY | 0.8434636 |
| EFAAAFPQ | ASSYRDENTGQLY | 0.5321748 |
| LLSGNLDVL | ASSYRDENTGQLY | 0.7395433 |
| GPANLGNYNI | ASSYRDENTGQLY | 0.75896883 |
| TLGPTLP | ASSYRDENTGQLY | 0.2919268 |

| | | |
|---|---|---|
| VYAYSSCCK | ASSYRDENTGQLY | 0.7946263 |
| APAPAPAPNI | ASSYRDENTGQLY | 0.86539227 |
| PGGPVPIP | ASSYRDENTGQLY | 0.81258106 |
| FLTVALEP | ASSYRDENTGQLY | 0.49825215 |
| SQYFGANL | ASSYRDENTGQLY | 0.85113114 |
| GGVKPAA | ASSYRDENTGQLY | 0.87555003 |
| GGTGGAGGAGGW | ASSYRDENTGQLY | 0.871176 |
| GVGGVAGLGGAGG | ASSYRDENTGQLY | 0.341058 |
| VLLVLATTI | ASSYRDENTGQLY | 0.2139791 |
| IGIFAAML | ASSYRDENTGQLY | 0.6926758 |
| AVGDLID | ASSYRDENTGQLY | 0.7645857 |
| NISPLPL | ASSYRDENTGQLY | 0.57845724 |
| VTAVFVAI | ASSYRDENTGQLY | 0.5271764 |
| SGDDNDAAL | ASSYRDENTGQLY | 0.77555823 |
| AVAALAHS | ASSYRDENTGQLY | 0.8407694 |
| GGVTVMATP | ASSYRDENTGQLY | 0.419838 |
| AGLAANVRD | ASSYRDENTGQLY | 0.5105987 |
| TNRPDLIDPALLRPG | ASSYRDENTGQLY | 0.83264995 |
| LANLGVTAL | ASSYRDENTGQLY | 0.84177625 |
| TGERIEL | ASSYRDENTGQLY | 0.5296406 |
| NGEELAA | ASSYRDENTGQLY | 0.6845999 |
| VLSLVWG | ASSYRDENTGQLY | 0.7199744 |
| IGYALWL | ASSYRDENTGQLY | 0.725977 |
| LSLLLAPL | ASSYRDENTGQLY | 0.7758471 |
| TLDEALK | ASSYRDENTGQLY | 0.47433645 |
| TDGAGTAEGG | ASSYRDENTGQLY | 0.8485883 |
| NAPGGAGGNGG | ASSYRDENTGQLY | 0.50534695 |
| VSVFMGMGGNF | ASSYRDENTGQLY | 0.8365463 |
| LASYLAAL | ASSYRDENTGQLY | 0.7430837 |
| LVGGTPAL | ASSYRDENTGQLY | 0.7873213 |
| VRNADVL | ASSYRDENTGQLY | 0.66268045 |
| TCSSPFLVSAM | ASSYRDENTGQLY | 0.54191095 |
| VHITDAE | ASSYRDENTGQLY | 0.4915673 |
| EYAAPSF | ASSYRDENTGQLY | 0.44101796 |
| VVDAFHGGL | ASSYRDENTGQLY | 0.84560907 |
| ASVVDPQ | ASSYRDENTGQLY | 0.23218268 |
| SYNEFGGLS | ASSYRDENTGQLY | 0.84390795 |
| DPDGSVY | ASSYRDENTGQLY | 0.80594134 |
| MVSTIHATE | ASSYRDENTGQLY | 0.5457151 |
| LAAGVADPV | ASSYRDENTGQLY | 0.83391243 |
| TKEQIAA | ASSYRDENTGQLY | 0.36559972 |
| SLASPAAL | ASSYRDENTGQLY | 0.7298394 |
| IGLVFYGV | ASSYRDENTGQLY | 0.7874832 |
| FLLDASNL | ASSYRDENTGQLY | 0.8207282 |
| YQVGNFP | ASSYRDENTGQLY | 0.5669278 |
| ESLLSQST | ASSYRDENTGQLY | 0.8382741 |
| LFTVGFAAVG | ASSYRDENTGQLY | 0.7973042 |
| ATPPTPT | ASSYRDENTGQLY | 0.51548517 |
| TDLATLG | ASSYRDENTGQLY | 0.77297914 |
| SAGGTGR | ASSYRDENTGQLY | 0.8075182 |
| GFSNSGS | ASSYRDENTGQLY | 0.8095685 |

| | | |
|---|---|---|
| GGNGGSGGTSV | ASSYRDENTGQLY | 0.40577048 |
| GTSVATGGAGN | ASSYRDENTGQLY | 0.78201807 |
| NGGNGGSGGTS | ASSYRDENTGQLY | 0.55150425 |
| AVVKFPH | ASSYRDENTGQLY | 0.7636723 |
| TPPPPPP | ASSYRDENTGQLY | 0.75659716 |
| AAPAAAAAPA | ASSYRDENTGQLY | 0.56567425 |
| IALNEDL | ASSYRDENTGQLY | 0.83611876 |
| IVNVSSM | ASSYRDENTGQLY | 0.5137526 |
| TGIPPAPR | ASSYRDENTGQLY | 0.6123735 |
| FDLGGGTF | ASSYRDENTGQLY | 0.6213664 |
| FDLGGGTFDVS | ASSYRDENTGQLY | 0.69953054 |
| NEPTAAA | ASSYRDENTGQLY | 0.80389726 |
| NEPTAAALA | ASSYRDENTGQLY | 0.49099547 |
| GSIINISSI | ASSYRDENTGQLY | 0.76584435 |
| VAPGNAGTA | ASSYRDENTGQLY | 0.68617654 |
| TGGGASLE | ASSYRDENTGQLY | 0.46080393 |
| VAVYNHYKRI | ASSYRDENTGQLY | 0.79942715 |
| TDPAPATT | ASSYRDENTGQLY | 0.71631444 |
| TNRPDILDPALLRPG | ASSYRDENTGQLY | 0.4102642 |
| LTTTTTS | ASSYRDENTGQLY | 0.8215749 |
| PTGSGKTL | ASSYRDENTGQLY | 0.5553429 |
| TGSGKTL | ASSYRDENTGQLY | 0.7731296 |
| TIGHVDHGK | ASSYRDENTGQLY | 0.8718184 |
| TIGHVDHGKTT | ASSYRDENTGQLY | 0.41364002 |
| GHVDHGKTT | ASSYRDENTGQLY | 0.85228825 |
| AQFEHTLL | ASSYRDENTGQLY | 0.7820326 |
| IVNQDEL | ASSYRDENTGQLY | 0.7657326 |
| TNGSQFF | ASSYRDENTGQLY | 0.6762229 |
| TGKGIVE | ASSYRDENTGQLY | 0.5749883 |
| LISNASDA | ASSYRDENTGQLY | 0.85946393 |
| AELRTLGELTKIAKAHGAQ | ASSYRDENTGQLY | 0.7988909 |
| APNAVAAAEPSW | ASSYRDENTGQLY | 0.74898255 |
| APQVAGAATAVL | ASSYRDENTGQLY | 0.795158 |
| ARRDPAAAAAILPTDA | ASSYRDENTGQLY | 0.7958306 |
| DAAARDVLAEAGLAEYF | ASSYRDENTGQLY | 0.8288367 |
| DDQIVLSAKANLAKVEQ | ASSYRDENTGQLY | 0.5675707 |
| DELLPDTGEDSDEDGH | ASSYRDENTGQLY | 0.40189135 |
| ELPEAPANRATVPQD | ASSYRDENTGQLY | 0.6335838 |
| FGQSKIERVTGIG | ASSYRDENTGQLY | 0.6407765 |
| GAGGTGSTAGGAGGAGGAGGLY | ASSYRDENTGQLY | 0.6644645 |
| GHGGTGAAGGNGGTGGAGGSGI | ASSYRDENTGQLY | 0.42652145 |
| GNGGNGGAGGEGGPGGAGGAGA | ASSYRDENTGQLY | 0.7960155 |
| GSTAGNGGNGGNATNGTIAGQP | ASSYRDENTGQLY | 0.88850456 |
| GTGGDGGNGGNGGTGG | ASSYRDENTGQLY | 0.53748703 |
| GVRRPAGIPADAPV | ASSYRDENTGQLY | 0.88062066 |
| GYRQVTDAHLLTLARR | ASSYRDENTGQLY | 0.5031655 |
| IAKMGQDGHDRG | ASSYRDENTGQLY | 0.8003252 |
| KAEPSIEIQYT | ASSYRDENTGQLY | 0.82009506 |
| KPVRIRMAADEDTVAVT | ASSYRDENTGQLY | 0.5853126 |
| MPPRIAGMRLLVIKPEP | ASSYRDENTGQLY | 0.7106825 |
| MKFYAVALFRSRGVPPTR | ASSYRDENTGQLY | 0.6157412 |

| | | |
|---|---|---|
| NAGGFGILGAVAHSP | ASSYRDENTGQLY | 0.63255507 |
| NEGREITLAMAEM | ASSYRDENTGQLY | 0.8354948 |
| NELVEWSHPLGHRWRWE | ASSYRDENTGQLY | 0.5084131 |
| NLGAGGTGGNGGNPG | ASSYRDENTGQLY | 0.78446984 |
| NTEWPKWETVMAADPVD | ASSYRDENTGQLY | 0.66798127 |
| PASAAPAETQP | ASSYRDENTGQLY | 0.587659 |
| QQSSDGKNGVRAAEITGE | ASSYRDENTGQLY | 0.7970092 |
| RAPDAVTTSWWKE | ASSYRDENTGQLY | 0.8083302 |
| SGFGNVGAHTSGWFNQ | ASSYRDENTGQLY | 0.7389771 |
| SGVAGAVGQAASVGGLKV | ASSYRDENTGQLY | 0.8929008 |
| SNGGAGTGSAGGNGGT | ASSYRDENTGQLY | 0.5896965 |
| TGGAGGEGGAAG | ASSYRDENTGQLY | 0.8739831 |
| TGGDGGAAGAGGAPGHG | ASSYRDENTGQLY | 0.85069734 |
| TMPPLLAAIANAGRHGV | ASSYRDENTGQLY | 0.73948836 |
| TSSTAGNSGGA | ASSYRDENTGQLY | 0.6882724 |
| VPATVPVGILC | ASSYRDENTGQLY | 0.81027025 |
| VQRPDSGGVVLTGRIS | ASSYRDENTGQLY | 0.566042 |
| WTPNPVPATKGVPAGLVQAT | ASSYRDENTGQLY | 0.79451734 |
| YALKRALAKFSDRFDVVII | ASSYRDENTGQLY | 0.87411916 |
| YLIGSVIV | ASSYRDENTGQLY | 0.73894024 |
| YGQLVAVID | ASSYRDENTGQLY | 0.81810844 |
| VTVGGMTLPA | ASSYRDENTGQLY | 0.7987832 |
| VLALPAV | ASSYRDENTGQLY | 0.75404114 |
| VIPPTLNYET | ASSYRDENTGQLY | 0.7284534 |
| VGMGGMHPSSG | ASSYRDENTGQLY | 0.42763853 |
| VGLAGGAATA | ASSYRDENTGQLY | 0.7524706 |
| VGGAGGAGGSGP | ASSYRDENTGQLY | 0.66385305 |
| VFAAFMV | ASSYRDENTGQLY | 0.78361976 |
| TLYDKLR | ASSYRDENTGQLY | 0.7997941 |
| TLAVFTVL | ASSYRDENTGQLY | 0.7283339 |
| TGSVQPARV | ASSYRDENTGQLY | 0.81726086 |
| TGGGAGSGGT | ASSYRDENTGQLY | 0.7861899 |
| SHLETSL | ASSYRDENTGQLY | 0.6426084 |
| SFTTGMT | ASSYRDENTGQLY | 0.76784396 |
| QVEVAKRMGVRQPTVS | ASSYRDENTGQLY | 0.75478125 |
| QGSGGPALAM | ASSYRDENTGQLY | 0.78901297 |
| MLAPSNP | ASSYRDENTGQLY | 0.5737468 |
| LVNNAAI | ASSYRDENTGQLY | 0.8078025 |
| LTGEINTLA | ASSYRDENTGQLY | 0.7490593 |
| LSIFGPV | ASSYRDENTGQLY | 0.21229072 |
| LLALLANL | ASSYRDENTGQLY | 0.7963526 |
| LISNASDALDKLR | ASSYRDENTGQLY | 0.7761115 |
| LINEAVP | ASSYRDENTGQLY | 0.7601646 |
| LGYAFSGL | ASSYRDENTGQLY | 0.81458896 |
| LDFVAVAL | ASSYRDENTGQLY | 0.3324847 |
| LAQPGAE | ASSYRDENTGQLY | 0.6541014 |
| ISNLGAVTGF | ASSYRDENTGQLY | 0.8113076 |
| HQEDPGRI | ASSYRDENTGQLY | 0.8148644 |
| GTPGEPTP | ASSYRDENTGQLY | 0.6023185 |
| GFFNSGS | ASSYRDENTGQLY | 0.8011174 |
| GAGGGGLTAGAN | ASSYRDENTGQLY | 0.8057554 |

| | | |
|---|---|---|
| FVDALVNM | ASSYRDENTGQLY | 0.8024856 |
| FLTVAPEL | ASSYRDENTGQLY | 0.807485 |
| FETLTAL | ASSYRDENTGQLY | 0.81331396 |
| DVINPIEL | ASSYRDENTGQLY | 0.660427 |
| APVSGAPV | ASSYRDENTGQLY | 0.8144096 |
| ALVGGLAW | ASSYRDENTGQLY | 0.8215074 |
| ALAAAAATAI | ASSYRDENTGQLY | 0.6937339 |
| AFGLGLLL | ASSYRDENTGQLY | 0.7941718 |
| AFFAAASL | ASSYRDENTGQLY | 0.73938775 |
| AAAALPAFSPP | ASSYRDENTGQLY | 0.7362585 |
| VVLAPAL | ASSYRDENTGQLY | 0.32017177 |
| TIGHVDHGKTTL | ASSYRDENTGQLY | 0.7992092 |
| LISNASDALDK | ASSYRDENTGQLY | 0.81116056 |
| LISNASDAL | ASSYRDENTGQLY | 0.77358437 |
| ISNASDALDK | ASSYRDENTGQLY | 0.7270942 |
| NQAETLVYQTE | ASSYRDENTGQLY | 0.6800249 |
| TGGTGGTAGAG | ASSYRDENTGQLY | 0.76866496 |
| TGSTAGGAGGA | ASSYRDENTGQLY | 0.75926006 |
| DPLAVPVAAGPV | ASSYRDENTGQLY | 0.3353334 |
| GGTGGTPGNGGN | ASSYRDENTGQLY | 0.7357601 |
| PAGLAGVDPATL | ASSYRDENTGQLY | 0.81087005 |
| GNNLVVEAPHAGASV | ASSYRDENTGQLY | 0.8092765 |
| KVTRIMSASTLEQPA | ASSYRDENTGQLY | 0.49923328 |
| LGFALPIALPRRTVV | ASSYRDENTGQLY | 0.8194903 |
| TRLGNLTARKLVVLV | ASSYRDENTGQLY | 0.79880583 |
| WSQQRRLIYFPSAGP | ASSYRDENTGQLY | 0.8129988 |
| GPNGNPDPMAAAVDIR | ASSYRDENTGQLY | 0.6928711 |
| TQRCVSVADLRAAASR | ASSYRDENTGQLY | 0.7385105 |
| QFVLDVAEHPVEVLTETV | ASSYRDENTGQLY | 0.8046503 |
| GGAGGTGGAGGNGGTQAGDG | ASSYRDENTGQLY | 0.7720506 |
| PLLATLRAIAVDVDAKPAQV | ASSYRDENTGQLY | 0.5974716 |
| APKFRTGGYMIDFWGVGYQVAK | ASSYRDENTGQLY | 0.813889 |
| GAGGAGGAGGAAGGPGGTGGTG | ASSYRDENTGQLY | 0.76048195 |
| IGALGLLATDVAPAGSVAFGVA | ASSYRDENTGQLY | 0.72891116 |
| AGGAGGIGGNGGAGGAGGVLMG | ASSYRDENTGQLY | 0.73368174 |
| ADELIGLL | ASSYRDENTGQLY | 0.81641006 |
| ADRALKA | ASSYRDENTGQLY | 0.70741457 |
| AGAVGQVGGAGGA | ASSYRDENTGQLY | 0.62173367 |
| AGGGGGLGGAP | ASSYRDENTGQLY | 0.5501442 |
| AGGLLYGNGGAG | ASSYRDENTGQLY | 0.7730163 |
| AGLESIGL | ASSYRDENTGQLY | 0.81254745 |
| AGLKTGII | ASSYRDENTGQLY | 0.7826159 |
| ASLAALLL | ASSYRDENTGQLY | 0.728822 |
| ASVINISSI | ASSYRDENTGQLY | 0.80028015 |
| ATPPGPA | ASSYRDENTGQLY | 0.78387403 |
| ATVASALV | ASSYRDENTGQLY | 0.78928393 |
| ATVASLIL | ASSYRDENTGQLY | 0.7894935 |
| AVNGPVLIRH | ASSYRDENTGQLY | 0.7456889 |
| DGAAGAVGI | ASSYRDENTGQLY | 0.81350523 |
| DPLLHGI | ASSYRDENTGQLY | 0.80332804 |
| ELGAALGAL | ASSYRDENTGQLY | 0.80534935 |

| | | |
|---|---|---|
| ELGRTQAV | ASSYRDENTGQLY | 0.80371356 |
| FVAGGYLS | ASSYRDENTGQLY | 0.7582419 |
| GASGGAGGAGGW | ASSYRDENTGQLY | 0.7876555 |
| GGAGAGGDGGAGGI | ASSYRDENTGQLY | 0.32906252 |
| GTGHNGSQPGMGGNG | ASSYRDENTGQLY | 0.5319861 |
| GYAAASASAS | ASSYRDENTGQLY | 0.7944834 |
| HQVDVAAVQQG | ASSYRDENTGQLY | 0.7936969 |
| IGAAIAHT | ASSYRDENTGQLY | 0.8047071 |
| IGMIEVAQL | ASSYRDENTGQLY | 0.77081835 |
| IRLAPEHTSAMD | ASSYRDENTGQLY | 0.7340479 |
| IYNVSVL | ASSYRDENTGQLY | 0.7828667 |
| KQGFLGM | ASSYRDENTGQLY | 0.80601096 |
| LESVITQL | ASSYRDENTGQLY | 0.59437114 |
| LGLGALVVV | ASSYRDENTGQLY | 0.76377064 |
| LIGNGGNGGSGGI | ASSYRDENTGQLY | 0.82242715 |
| LNAINAPTL | ASSYRDENTGQLY | 0.7891408 |
| LQLLNTGV | ASSYRDENTGQLY | 0.7808565 |
| LQTSIEF | ASSYRDENTGQLY | 0.6947901 |
| MSFAVSEL | ASSYRDENTGQLY | 0.8071478 |
| NTGALGL | ASSYRDENTGQLY | 0.6927961 |
| PPGAGGAGGAGTT | ASSYRDENTGQLY | 0.8199681 |
| QLVGLGI | ASSYRDENTGQLY | 0.25771362 |
| RFLEQVHQL | ASSYRDENTGQLY | 0.7292489 |
| RGVGYVLR | ASSYRDENTGQLY | 0.59561706 |
| SATGVFAGVI | ASSYRDENTGQLY | 0.503746 |
| SLPAITLP | ASSYRDENTGQLY | 0.81466705 |
| STALAFTV | ASSYRDENTGQLY | 0.6832293 |
| SVLTVGLGK | ASSYRDENTGQLY | 0.7945496 |
| TALAGAGSGI | ASSYRDENTGQLY | 0.78308773 |
| TGLPFTGI | ASSYRDENTGQLY | 0.80995363 |
| VAEHVTV | ASSYRDENTGQLY | 0.79676867 |
| VDIVLNSL | ASSYRDENTGQLY | 0.74813193 |
| VLTGGGAVL | ASSYRDENTGQLY | 0.6315568 |
| VNVGLAYGI | ASSYRDENTGQLY | 0.809898 |
| VSSAAVGAV | ASSYRDENTGQLY | 0.79758 |
| VTSVYEAGV | ASSYRDENTGQLY | 0.7940173 |
| WVALDRGA | ASSYRDENTGQLY | 0.4943693 |
| YLVGSVVI | ASSYRDENTGQLY | 0.7583925 |
| GPPGTGKTLL | ASSYRDENTGQLY | 0.78356314 |
| GVGMTKF | ASSYRDENTGQLY | 0.27312985 |
| IGIGGSDL | ASSYRDENTGQLY | 0.31528825 |
| IGTLLAEL | ASSYRDENTGQLY | 0.7564628 |
| IYTSGTT | ASSYRDENTGQLY | 0.73912776 |
| LSFPAGV | ASSYRDENTGQLY | 0.81721795 |
| LYTSGTT | ASSYRDENTGQLY | 0.8129468 |
| VGAFPFGF | ASSYRDENTGQLY | 0.78721565 |
| VGAGTVEF | ASSYRDENTGQLY | 0.7990643 |
| VGKSTLLL | ASSYRDENTGQLY | 0.7862008 |
| AALAHAGGGIV | ASSYRDENTGQLY | 0.16674094 |
| AALVAVSLL | ASSYRDENTGQLY | 0.6536858 |
| AAVSRAVAAL | ASSYRDENTGQLY | 0.65834224 |

| | | |
|---|---|---|
| AGGSSGIGGF | ASSYRDENTGQLY | 0.80346954 |
| AGLTVANANG | ASSYRDENTGQLY | 0.72713315 |
| AGLVGAVL | ASSYRDENTGQLY | 0.7432208 |
| ALSQAFTV | ASSYRDENTGQLY | 0.7935812 |
| ALTFSVSL | ASSYRDENTGQLY | 0.7793715 |
| ALTGAALVAL | ASSYRDENTGQLY | 0.7791176 |
| ASFSIGAGVL | ASSYRDENTGQLY | 0.6656213 |
| DGSVYKI | ASSYRDENTGQLY | 0.2421848 |
| EQLAEIV | ASSYRDENTGQLY | 0.81156266 |
| FAVSELTVM | ASSYRDENTGQLY | 0.3722894 |
| GAAPLAAGL | ASSYRDENTGQLY | 0.64011467 |
| GADGGDGGSGGAGG | ASSYRDENTGQLY | 0.32258302 |
| GFGLDTGGAGG | ASSYRDENTGQLY | 0.41940486 |
| GLASGGLL | ASSYRDENTGQLY | 0.41311938 |
| GRIAGAAVLL | ASSYRDENTGQLY | 0.8051084 |
| GSAGSLNTSF | ASSYRDENTGQLY | 0.7691864 |
| GTGQIGI | ASSYRDENTGQLY | 0.76795554 |
| GTLIALIL | ASSYRDENTGQLY | 0.8215896 |
| IAATMGFVVG | ASSYRDENTGQLY | 0.77044606 |
| IFAQALL | ASSYRDENTGQLY | 0.78004503 |
| IGAAVALGF | ASSYRDENTGQLY | 0.6637448 |
| IGLTPLAL | ASSYRDENTGQLY | 0.72697747 |
| ILGSNTSTL | ASSYRDENTGQLY | 0.71840906 |
| IVFSLTVL | ASSYRDENTGQLY | 0.67419183 |
| LASKATGF | ASSYRDENTGQLY | 0.81797665 |
| LASVALIL | ASSYRDENTGQLY | 0.752813 |
| LFAVGLTL | ASSYRDENTGQLY | 0.5607816 |
| LFNGGIGTYI | ASSYRDENTGQLY | 0.7354964 |
| LTLGAGII | ASSYRDENTGQLY | 0.8183136 |
| LVAAALGDI | ASSYRDENTGQLY | 0.82269245 |
| NTAFVQLGI | ASSYRDENTGQLY | 0.7437169 |
| QLFSGAEL | ASSYRDENTGQLY | 0.6644652 |
| SAATFITL | ASSYRDENTGQLY | 0.6874 |
| SNYGLANLGV | ASSYRDENTGQLY | 0.786256 |
| SYGGAYAVM | ASSYRDENTGQLY | 0.68206054 |
| TTGVGGLTLG | ASSYRDENTGQLY | 0.50580525 |
| TTQTIYI | ASSYRDENTGQLY | 0.80764914 |
| VAAGAGLGAVI | ASSYRDENTGQLY | 0.81097394 |
| VAAVVTLL | ASSYRDENTGQLY | 0.7381158 |
| VGFLGGLI | ASSYRDENTGQLY | 0.70456135 |
| VGVIPILL | ASSYRDENTGQLY | 0.7806021 |
| VIFASSLI | ASSYRDENTGQLY | 0.712639 |
| VKGGDGGAAAG | ASSYRDENTGQLY | 0.7701558 |
| VTKTFTNI | ASSYRDENTGQLY | 0.7396945 |
| VVAAAILL | ASSYRDENTGQLY | 0.65290177 |
| YAALDVEL | ASSYRDENTGQLY | 0.76171964 |
| LASVALLL | ASSYRDENTGQLY | 0.82268643 |
| LDAVHRV | ASSYRDENTGQLY | 0.74632776 |
| QGLSSSIF | ASSYRDENTGQLY | 0.7917915 |
| RGTFANIRL | ASSYRDENTGQLY | 0.8190913 |
| RQLWWGHRI | ASSYRDENTGQLY | 0.81080675 |

| | | |
|---|---|---|
| YGPPGTGKTLL | ASSYRDENTGQLY | 0.81861424 |
| AALVGAPANVA | ASSYRDENTGQLY | 0.79572475 |
| AATFRRTE | ASSYRDENTGQLY | 0.7624663 |
| AGALGGGI | ASSYRDENTGQLY | 0.7648964 |
| AGGAGGAGGIGAG | ASSYRDENTGQLY | 0.7967217 |
| AGSGALGL | ASSYRDENTGQLY | 0.7905699 |
| AQIADQI | ASSYRDENTGQLY | 0.7834659 |
| ASAVAVTYL | ASSYRDENTGQLY | 0.71419114 |
| ASTVLGGTGG | ASSYRDENTGQLY | 0.81256175 |
| AVGGEPAK | ASSYRDENTGQLY | 0.76956093 |
| CEQGTFGGTC | ASSYRDENTGQLY | 0.80563617 |
| DEGKAPAG | ASSYRDENTGQLY | 0.5363352 |
| DTVVFPAL | ASSYRDENTGQLY | 0.63016856 |
| FEGITMAG | ASSYRDENTGQLY | 0.80960023 |
| FPLVVQD | ASSYRDENTGQLY | 0.18589768 |
| GAFTTPQ | ASSYRDENTGQLY | 0.24001688 |
| GFGGFGTAGGVGGA | ASSYRDENTGQLY | 0.79386175 |
| GFLAQFL | ASSYRDENTGQLY | 0.8154011 |
| GGAGGAGGNAGVA | ASSYRDENTGQLY | 0.20507385 |
| GGVGTAGAGGAGGAGG | ASSYRDENTGQLY | 0.59534836 |
| GNGGGGDAGNAGSGGN | ASSYRDENTGQLY | 0.614024 |
| GTGGAGGAGNDAGS | ASSYRDENTGQLY | 0.77107525 |
| GTTNTGWFNS | ASSYRDENTGQLY | 0.7787247 |
| IFMGPLFTL | ASSYRDENTGQLY | 0.41155833 |
| ILAHAIEAI | ASSYRDENTGQLY | 0.8039123 |
| IRSAIREAL | ASSYRDENTGQLY | 0.60846317 |
| ITFAGGG | ASSYRDENTGQLY | 0.453335 |
| ITGMSGHQEV | ASSYRDENTGQLY | 0.80846554 |
| IVAGLACTL | ASSYRDENTGQLY | 0.81283665 |
| IVGGGLAGA | ASSYRDENTGQLY | 0.71418655 |
| IVTGVLNI | ASSYRDENTGQLY | 0.80559695 |
| LAQQGCI | ASSYRDENTGQLY | 0.7173506 |
| LCLHVEN | ASSYRDENTGQLY | 0.82301724 |
| LDAISDGS | ASSYRDENTGQLY | 0.73102593 |
| LGGFAAML | ASSYRDENTGQLY | 0.5616533 |
| LGGKNPMI | ASSYRDENTGQLY | 0.8107475 |
| LGPKGVAAIL | ASSYRDENTGQLY | 0.80500185 |
| LMVGVSI | ASSYRDENTGQLY | 0.76291496 |
| LVGAAFVL | ASSYRDENTGQLY | 0.8034284 |
| LVLAVAPL | ASSYRDENTGQLY | 0.66343665 |
| MAAIAVIDAA | ASSYRDENTGQLY | 0.7797191 |
| NNGINAELQ | ASSYRDENTGQLY | 0.8140763 |
| RGEPLVLVHG | ASSYRDENTGQLY | 0.81299734 |
| RGGAGGAAGLI | ASSYRDENTGQLY | 0.58626145 |
| RSYRMHPAV | ASSYRDENTGQLY | 0.8102796 |
| SAAHLLAGQLA | ASSYRDENTGQLY | 0.783745 |
| SAAVGSY | ASSYRDENTGQLY | 0.7391772 |
| SPHAVLHRI | ASSYRDENTGQLY | 0.8204685 |
| SQIAKVL | ASSYRDENTGQLY | 0.7574271 |
| TALGHLVP | ASSYRDENTGQLY | 0.81286925 |
| TAPTTSGAA | ASSYRDENTGQLY | 0.8071263 |

| | | |
|---|---|---|
| TDGDDGNGGSPGT | ASSYRDENTGQLY | 0.78609335 |
| TDGLGEN | ASSYRDENTGQLY | 0.4376548 |
| TGGPPLPGQA | ASSYRDENTGQLY | 0.75161433 |
| TIPAITI | ASSYRDENTGQLY | 0.7803731 |
| TIPGFTI | ASSYRDENTGQLY | 0.79369414 |
| TSIASGALA | ASSYRDENTGQLY | 0.7777294 |
| TVTLPGAAI | ASSYRDENTGQLY | 0.64782596 |
| VASGLSAALP | ASSYRDENTGQLY | 0.80148804 |
| VGAGVIGI | ASSYRDENTGQLY | 0.8179927 |
| VGPKRFGV | ASSYRDENTGQLY | 0.78541577 |
| VNVAFPD | ASSYRDENTGQLY | 0.75181353 |
| VSMTNSGVS | ASSYRDENTGQLY | 0.8210126 |
| VTDVLGDLSV | ASSYRDENTGQLY | 0.7523978 |
| VVVVGAGFAGL | ASSYRDENTGQLY | 0.7703078 |
| VYTVGTVV | ASSYRDENTGQLY | 0.80191636 |
| WTTFSGP | ASSYRDENTGQLY | 0.80381244 |
| YDELVEAF | ASSYRDENTGQLY | 0.7580554 |
| YGTGGAGGNGG | ASSYRDENTGQLY | 0.81126076 |
| YTNVMAV | ASSYRDENTGQLY | 0.579813 |
| FLEQVHQL | ASSYRDENTGQLY | 0.8076439 |
| GLSLLAVAV | ASSYRDENTGQLY | 0.7389528 |
| LEQVHQL | ASSYRDENTGQLY | 0.7926917 |
| LNAAGAI | ASSYRDENTGQLY | 0.7345431 |
| PAVHASL | ASSYRDENTGQLY | 0.8118752 |
| TELEDDA | ASSYRDENTGQLY | 0.81333876 |
| TPPGPPN | ASSYRDENTGQLY | 0.8092937 |
| TPPPPPPP | ASSYRDENTGQLY | 0.26001838 |
| TQSPALI | ASSLGQGRSDTQY | 0.8124368 |
| EFAAAFPQ | ASSLGQGRSDTQY | 0.7890256 |
| LLSGNLDVL | ASSLGQGRSDTQY | 0.8216752 |
| GPANLGNYNI | ASSLGQGRSDTQY | 0.74685925 |
| TLGPTLP | ASSLGQGRSDTQY | 0.7707738 |
| VYAYSSCCK | ASSLGQGRSDTQY | 0.7702615 |
| APAPAPAPNI | ASSLGQGRSDTQY | 0.7565763 |
| PGGPVPIP | ASSLGQGRSDTQY | 0.6836419 |
| FLTVALEP | ASSLGQGRSDTQY | 0.75763583 |
| SQYFGANL | ASSLGQGRSDTQY | 0.75140315 |
| GGVKPAA | ASSLGQGRSDTQY | 0.74577165 |
| GGTGGAGGAGGW | ASSLGQGRSDTQY | 0.8065465 |
| GVGGVAGLGGAGG | ASSLGQGRSDTQY | 0.8142753 |
| VLLVLATTI | ASSLGQGRSDTQY | 0.81075716 |
| IGIFAAML | ASSLGQGRSDTQY | 0.8132663 |
| AVGDLID | ASSLGQGRSDTQY | 0.8172022 |
| NISPLPL | ASSLGQGRSDTQY | 0.8066836 |
| VTAVFVAI | ASSLGQGRSDTQY | 0.7427744 |
| SGDDNDAAL | ASSLGQGRSDTQY | 0.74273205 |
| AVAALAHS | ASSLGQGRSDTQY | 0.7185286 |
| GGVTVMATP | ASSLGQGRSDTQY | 0.75955236 |
| AGLAANVRD | ASSLGQGRSDTQY | 0.7374985 |
| TNRPDLIDPALLRPG | ASSLGQGRSDTQY | 0.79852575 |
| LANLGVTAL | ASSLGQGRSDTQY | 0.17472552 |

| | | |
|---|---|---|
| TGERIEL | ASSLGQGRSDTQY | 0.7793412 |
| NGEELAA | ASSLGQGRSDTQY | 0.81956786 |
| VLSLVWG | ASSLGQGRSDTQY | 0.7166672 |
| IGYALWL | ASSLGQGRSDTQY | 0.7929255 |
| LSLLLAPL | ASSLGQGRSDTQY | 0.78936046 |
| TLDEALK | ASSLGQGRSDTQY | 0.75805223 |
| TDGAGTAEGG | ASSLGQGRSDTQY | 0.8065918 |
| NAPGGAGGNGG | ASSLGQGRSDTQY | 0.6859124 |
| VSVFMGMGGNF | ASSLGQGRSDTQY | 0.79823256 |
| LASYLAAL | ASSLGQGRSDTQY | 0.82050955 |
| LVGGTPAL | ASSLGQGRSDTQY | 0.7970163 |
| VRNADVL | ASSLGQGRSDTQY | 0.75351465 |
| TCSSPFLVSAM | ASSLGQGRSDTQY | 0.7398045 |
| VHITDAE | ASSLGQGRSDTQY | 0.3952423 |
| EYAAPSF | ASSLGQGRSDTQY | 0.7710892 |
| VVDAFHGGL | ASSLGQGRSDTQY | 0.1918934 |
| ASVVDPQ | ASSLGQGRSDTQY | 0.7842113 |
| SYNEFGGLS | ASSLGQGRSDTQY | 0.64307433 |
| DPDGSVY | ASSLGQGRSDTQY | 0.79114985 |
| MVSTIHATE | ASSLGQGRSDTQY | 0.7920135 |
| LAAGVADPV | ASSLGQGRSDTQY | 0.7964704 |
| TKEQIAA | ASSLGQGRSDTQY | 0.7126534 |
| SLASPAAL | ASSLGQGRSDTQY | 0.7638607 |
| IGLVFYGV | ASSLGQGRSDTQY | 0.70317936 |
| FLLDASNL | ASSLGQGRSDTQY | 0.53438175 |
| YQVGNFP | ASSLGQGRSDTQY | 0.8125975 |
| ESLLSQST | ASSLGQGRSDTQY | 0.76959383 |
| LFTVGFAAVG | ASSLGQGRSDTQY | 0.77233994 |
| ATPPTPT | ASSLGQGRSDTQY | 0.8112302 |
| TDLATLG | ASSLGQGRSDTQY | 0.7901838 |
| SAGGTGR | ASSLGQGRSDTQY | 0.52606297 |
| GFSNSGS | ASSLGQGRSDTQY | 0.77523214 |
| GGNGGSGGTSV | ASSLGQGRSDTQY | 0.7802273 |
| GTSVATGGAGN | ASSLGQGRSDTQY | 0.7638148 |
| NGGNGGSGGTS | ASSLGQGRSDTQY | 0.18245305 |
| AVVKFPH | ASSLGQGRSDTQY | 0.7931015 |
| TPPPPPP | ASSLGQGRSDTQY | 0.7380638 |
| AAPAAAAAPA | ASSLGQGRSDTQY | 0.73317873 |
| IALNEDL | ASSLGQGRSDTQY | 0.75290585 |
| IVNVSSM | ASSLGQGRSDTQY | 0.81135833 |
| TGIPPAPR | ASSLGQGRSDTQY | 0.61001253 |
| FDLGGGTF | ASSLGQGRSDTQY | 0.72162664 |
| FDLGGGTFDVS | ASSLGQGRSDTQY | 0.81115746 |
| NEPTAAA | ASSLGQGRSDTQY | 0.8016025 |
| NEPTAAALA | ASSLGQGRSDTQY | 0.8137076 |
| GSIINISSI | ASSLGQGRSDTQY | 0.769578 |
| VAPGNAGTA | ASSLGQGRSDTQY | 0.71080506 |
| TGGGASLE | ASSLGQGRSDTQY | 0.7571702 |
| VAVYNHYKRI | ASSLGQGRSDTQY | 0.72908556 |
| TDPAPATT | ASSLGQGRSDTQY | 0.50563896 |
| TNRPDILDPALLRPG | ASSLGQGRSDTQY | 0.40868843 |

| | | |
|---|---|---|
| LTTTTTS | ASSLGQGRSDTQY | 0.79640895 |
| PTGSGKTL | ASSLGQGRSDTQY | 0.39413717 |
| TGSGKTL | ASSLGQGRSDTQY | 0.76857734 |
| TIGHVDHGK | ASSLGQGRSDTQY | 0.80663645 |
| TIGHVDHGKTT | ASSLGQGRSDTQY | 0.7411868 |
| GHVDHGKTT | ASSLGQGRSDTQY | 0.5858135 |
| AQFEHTLL | ASSLGQGRSDTQY | 0.7580372 |
| IVNQDEL | ASSLGQGRSDTQY | 0.69721055 |
| TNGSQFF | ASSLGQGRSDTQY | 0.8086565 |
| TGKGIVE | ASSLGQGRSDTQY | 0.7799581 |
| LISNASDA | ASSLGQGRSDTQY | 0.8163951 |
| AELRTLGELTKIAKAHGAQ | ASSLGQGRSDTQY | 0.8105597 |
| APNAVAAAEPSW | ASSLGQGRSDTQY | 0.76599634 |
| APQVAGAATAVL | ASSLGQGRSDTQY | 0.8117976 |
| ARRDPAAAAAILPTDA | ASSLGQGRSDTQY | 0.76103306 |
| DAAARDVLAEAGLAEYF | ASSLGQGRSDTQY | 0.75678766 |
| DDQIVLSAKANLAKVEQ | ASSLGQGRSDTQY | 0.78612936 |
| DELLPDTGEDSDEDGH | ASSLGQGRSDTQY | 0.79051346 |
| ELPEAPANRATVPQD | ASSLGQGRSDTQY | 0.69941723 |
| FGQSKIERVTGIG | ASSLGQGRSDTQY | 0.76209295 |
| GAGGTGSTAGGAGGAGGAGGLY | ASSLGQGRSDTQY | 0.57877934 |
| GHGGTGAAGGNGGTGGAGGSGI | ASSLGQGRSDTQY | 0.79628503 |
| GNGGNGGAGGEGGPGGAGGAGA | ASSLGQGRSDTQY | 0.79040366 |
| GSTAGNGGNGGNATNGTIAGQP | ASSLGQGRSDTQY | 0.40581244 |
| GTGGDGGNGGNGGTGG | ASSLGQGRSDTQY | 0.7341405 |
| GVRRPAGIPADAPV | ASSLGQGRSDTQY | 0.8205136 |
| GYRQVTDAHLLTLARR | ASSLGQGRSDTQY | 0.8118458 |
| IAKMGQDGHDRG | ASSLGQGRSDTQY | 0.7073809 |
| KAEPSIEIQYT | ASSLGQGRSDTQY | 0.6776802 |
| KPVRIRMAADEDTVAVT | ASSLGQGRSDTQY | 0.7834104 |
| MPPRIAGMRLLVIKPEP | ASSLGQGRSDTQY | 0.3990649 |
| MKFYAVALFRSRGVPPTR | ASSLGQGRSDTQY | 0.79411197 |
| NAGGFGILGAVAHSP | ASSLGQGRSDTQY | 0.7812065 |
| NEGREITLAMAEM | ASSLGQGRSDTQY | 0.78677964 |
| NELVEWSHPLGHRWRWE | ASSLGQGRSDTQY | 0.5922065 |
| NLGAGGTGGNGGNPG | ASSLGQGRSDTQY | 0.7924197 |
| NTEWPKWETVMAADPVD | ASSLGQGRSDTQY | 0.7509109 |
| PASAAPAETQP | ASSLGQGRSDTQY | 0.78561085 |
| QQSSDGKNGVRAAEITGE | ASSLGQGRSDTQY | 0.22073413 |
| RAPDAVTTSWWKE | ASSLGQGRSDTQY | 0.7858008 |
| SGFGNVGAHTSGWFNQ | ASSLGQGRSDTQY | 0.7305241 |
| SGVAGAVGQAASVGGLKV | ASSLGQGRSDTQY | 0.67458814 |
| SNGGAGTGSAGGNGGT | ASSLGQGRSDTQY | 0.3894956 |
| TGGAGGEGGAAG | ASSLGQGRSDTQY | 0.7687628 |
| TGGDGGAAGAGGAPGHG | ASSLGQGRSDTQY | 0.8139419 |
| TMPPLLAAIANAGRHGV | ASSLGQGRSDTQY | 0.81336087 |
| TSSTAGNSGGA | ASSLGQGRSDTQY | 0.68040925 |
| VPATVPVGILC | ASSLGQGRSDTQY | 0.8060836 |
| VQRPDSGGVVLTGRIS | ASSLGQGRSDTQY | 0.821259 |
| WTPNPVPATKGVPAGLVQAT | ASSLGQGRSDTQY | 0.77480245 |
| YALKRALAKFSDRFDVVII | ASSLGQGRSDTQY | 0.36152387 |

| | | |
|---|---|---|
| YLIGSVIV | ASSLGQGRSDTQY | 0.78454375 |
| YGQLVAVID | ASSLGQGRSDTQY | 0.748698 |
| VTVGGMTLPA | ASSLGQGRSDTQY | 0.801996 |
| VLALPAV | ASSLGQGRSDTQY | 0.7972763 |
| VIPPTLNYET | ASSLGQGRSDTQY | 0.7110659 |
| VGMGGMHPSSG | ASSLGQGRSDTQY | 0.7684475 |
| VGLAGGAATA | ASSLGQGRSDTQY | 0.8093407 |
| VGGAGGAGGSGP | ASSLGQGRSDTQY | 0.7119076 |
| VFAAFMV | ASSLGQGRSDTQY | 0.7345768 |
| TLYDKLR | ASSLGQGRSDTQY | 0.62631595 |
| TLAVFTVL | ASSLGQGRSDTQY | 0.41602814 |
| TGSVQPARV | ASSLGQGRSDTQY | 0.79452056 |
| TGGGAGSGGT | ASSLGQGRSDTQY | 0.75477046 |
| SHLETSL | ASSLGQGRSDTQY | 0.6493175 |
| SFTTGMT | ASSLGQGRSDTQY | 0.7970303 |
| QVEVAKRMGVRQPTVS | ASSLGQGRSDTQY | 0.68732 |
| QGSGGPALAM | ASSLGQGRSDTQY | 0.73191947 |
| MLAPSNP | ASSLGQGRSDTQY | 0.742792 |
| LVNNAAI | ASSLGQGRSDTQY | 0.77194035 |
| LTGEINTLA | ASSLGQGRSDTQY | 0.8005809 |
| LSIFGPV | ASSLGQGRSDTQY | 0.78095466 |
| LLALLANL | ASSLGQGRSDTQY | 0.76095474 |
| LISNASDALDKLR | ASSLGQGRSDTQY | 0.82046473 |
| LINEAVP | ASSLGQGRSDTQY | 0.5472332 |
| LGYAFSGL | ASSLGQGRSDTQY | 0.6386934 |
| LDFVAVAL | ASSLGQGRSDTQY | 0.79588246 |
| LAQPGAE | ASSLGQGRSDTQY | 0.56230885 |
| ISNLGAVTGF | ASSLGQGRSDTQY | 0.8038022 |
| HQEDPGRI | ASSLGQGRSDTQY | 0.78655267 |
| GTPGEPTP | ASSLGQGRSDTQY | 0.80263126 |
| GFFNSGS | ASSLGQGRSDTQY | 0.75746256 |
| GAGGGGLTAGAN | ASSLGQGRSDTQY | 0.82190776 |
| FVDALVNM | ASSLGQGRSDTQY | 0.81303394 |
| FLTVAPEL | ASSLGQGRSDTQY | 0.7541463 |
| FETLTAL | ASSLGQGRSDTQY | 0.7928006 |
| DVINPIEL | ASSLGQGRSDTQY | 0.46222174 |
| APVSGAPV | ASSLGQGRSDTQY | 0.802508 |
| ALVGGLAW | ASSLGQGRSDTQY | 0.7938215 |
| ALAAAAATAI | ASSLGQGRSDTQY | 0.727316 |
| AFGLGLLL | ASSLGQGRSDTQY | 0.7812948 |
| AFFAAASL | ASSLGQGRSDTQY | 0.8176124 |
| AAAALPAFSPP | ASSLGQGRSDTQY | 0.8166884 |
| VVLAPAL | ASSLGQGRSDTQY | 0.36809188 |
| TIGHVDHGKTTL | ASSLGQGRSDTQY | 0.77591264 |
| LISNASDALDK | ASSLGQGRSDTQY | 0.3153156 |
| LISNASDAL | ASSLGQGRSDTQY | 0.6654122 |
| ISNASDALDK | ASSLGQGRSDTQY | 0.6196549 |
| NQAETLVYQTE | ASSLGQGRSDTQY | 0.8016818 |
| TGGTGGTAGAG | ASSLGQGRSDTQY | 0.7685466 |
| TGSTAGGAGGA | ASSLGQGRSDTQY | 0.7860409 |
| DPLAVPVAAGPV | ASSLGQGRSDTQY | 0.78550637 |

| | | |
|---|---|---|
| GGTGGTPGNGGN | ASSLGQGRSDTQY | 0.81921744 |
| PAGLAGVDPATL | ASSLGQGRSDTQY | 0.7514989 |
| GNNLVVEAPHAGASV | ASSLGQGRSDTQY | 0.76262426 |
| KVTRIMSASTLEQPA | ASSLGQGRSDTQY | 0.5525367 |
| LGFALPIALPRRTVV | ASSLGQGRSDTQY | 0.46879792 |
| TRLGNLTARKLVVLV | ASSLGQGRSDTQY | 0.73584026 |
| WSQQRRLIYFPSAGP | ASSLGQGRSDTQY | 0.7230196 |
| GPNGNPDPMAAAVDIR | ASSLGQGRSDTQY | 0.2625529 |
| TQRCVSVADLRAAASR | ASSLGQGRSDTQY | 0.8126008 |
| QFVLDVAEHPVEVLTETV | ASSLGQGRSDTQY | 0.7949724 |
| GGAGGTGGAGGNGGTQAGDG | ASSLGQGRSDTQY | 0.8116429 |
| PLLATLRAIAVDVDAKPAQV | ASSLGQGRSDTQY | 0.70056254 |
| APKFRTGGYMIDFWGVGYQVAK | ASSLGQGRSDTQY | 0.7493639 |
| GAGGAGGAGGAAGGPGGTGGTG | ASSLGQGRSDTQY | 0.73885393 |
| IGALGLLATDVAPAGSVAFGVA | ASSLGQGRSDTQY | 0.79660594 |
| AGGAGGIGGNGGAGGAGGVLMG | ASSLGQGRSDTQY | 0.7112397 |
| ADELIGLL | ASSLGQGRSDTQY | 0.5664035 |
| ADRALKA | ASSLGQGRSDTQY | 0.6958806 |
| AGAVGQVGGAGGA | ASSLGQGRSDTQY | 0.7022272 |
| AGGGGGLGGAP | ASSLGQGRSDTQY | 0.81738937 |
| AGGLLYGNGGAG | ASSLGQGRSDTQY | 0.8132293 |
| AGLESIGL | ASSLGQGRSDTQY | 0.80076003 |
| AGLKTGII | ASSLGQGRSDTQY | 0.81352234 |
| ASLAALLL | ASSLGQGRSDTQY | 0.27468497 |
| ASVINISSI | ASSLGQGRSDTQY | 0.7155218 |
| ATPPGPA | ASSLGQGRSDTQY | 0.81199384 |
| ATVASALV | ASSLGQGRSDTQY | 0.78810287 |
| ATVASLIL | ASSLGQGRSDTQY | 0.77779037 |
| AVNGPVLIRH | ASSLGQGRSDTQY | 0.8172244 |
| DGAAGAVGI | ASSLGQGRSDTQY | 0.7370659 |
| DPLLHGI | ASSLGQGRSDTQY | 0.74287355 |
| ELGAALGAL | ASSLGQGRSDTQY | 0.7758131 |
| ELGRTQAV | ASSLGQGRSDTQY | 0.8047867 |
| FVAGGYLS | ASSLGQGRSDTQY | 0.7926147 |
| GASGGAGGAGGW | ASSLGQGRSDTQY | 0.6942125 |
| GGAGAGGDGGAGGI | ASSLGQGRSDTQY | 0.7696967 |
| GTGHNGSQPGMGGNG | ASSLGQGRSDTQY | 0.82488525 |
| GYAAASASAS | ASSLGQGRSDTQY | 0.77659684 |
| HQVDVAAVQQG | ASSLGQGRSDTQY | 0.8141506 |
| IGAAIAHT | ASSLGQGRSDTQY | 0.79689586 |
| IGMIEVAQL | ASSLGQGRSDTQY | 0.78613 |
| IRLAPEHTSAMD | ASSLGQGRSDTQY | 0.7217274 |
| IYNVSVL | ASSLGQGRSDTQY | 0.76385665 |
| KQGFLGM | ASSLGQGRSDTQY | 0.7595192 |
| LESVITQL | ASSLGQGRSDTQY | 0.69230103 |
| LGLGALVVV | ASSLGQGRSDTQY | 0.7411635 |
| LIGNGGNGGSGGI | ASSLGQGRSDTQY | 0.73398143 |
| LNAINAPTL | ASSLGQGRSDTQY | 0.8254716 |
| LQLLNTGV | ASSLGQGRSDTQY | 0.8049189 |
| LQTSIEF | ASSLGQGRSDTQY | 0.79859364 |
| MSFAVSEL | ASSLGQGRSDTQY | 0.7057334 |

| | | |
|---|---|---|
| NTGALGL | ASSLGQGRSDTQY | 0.7934449 |
| PPGAGGAGGAGTT | ASSLGQGRSDTQY | 0.8001252 |
| QLVGLGI | ASSLGQGRSDTQY | 0.6473806 |
| RFLEQVHQL | ASSLGQGRSDTQY | 0.816454 |
| RGVGYVLR | ASSLGQGRSDTQY | 0.7314824 |
| SATGVFAGVI | ASSLGQGRSDTQY | 0.75311184 |
| SLPAITLP | ASSLGQGRSDTQY | 0.76125205 |
| STALAFTV | ASSLGQGRSDTQY | 0.80495465 |
| SVLTVGLGK | ASSLGQGRSDTQY | 0.7606653 |
| TALAGAGSGI | ASSLGQGRSDTQY | 0.7523471 |
| TGLPFTGI | ASSLGQGRSDTQY | 0.7422075 |
| VAEHVTV | ASSLGQGRSDTQY | 0.8036521 |
| VDIVLNSL | ASSLGQGRSDTQY | 0.81714547 |
| VLTGGGAVL | ASSLGQGRSDTQY | 0.7525712 |
| VNVGLAYGI | ASSLGQGRSDTQY | 0.8183962 |
| VSSAAVGAV | ASSLGQGRSDTQY | 0.74927896 |
| VTSVYEAGV | ASSLGQGRSDTQY | 0.4597044 |
| WVALDRGA | ASSLGQGRSDTQY | 0.77780306 |
| YLVGSVVI | ASSLGQGRSDTQY | 0.81070316 |
| GPPGTGKTLL | ASSLGQGRSDTQY | 0.8133749 |
| GVGMTKF | ASSLGQGRSDTQY | 0.8184564 |
| IGIGGSDL | ASSLGQGRSDTQY | 0.7800876 |
| IGTLLAEL | ASSLGQGRSDTQY | 0.759506 |
| IYTSGTT | ASSLGQGRSDTQY | 0.8167608 |
| LSFPAGV | ASSLGQGRSDTQY | 0.8161071 |
| LYTSGTT | ASSLGQGRSDTQY | 0.80062973 |
| VGAFPFGF | ASSLGQGRSDTQY | 0.73297834 |
| VGAGTVEF | ASSLGQGRSDTQY | 0.5777658 |
| VGKSTLLL | ASSLGQGRSDTQY | 0.79609054 |
| AALAHAGGGIV | ASSLGQGRSDTQY | 0.8136342 |
| AALVAVSLL | ASSLGQGRSDTQY | 0.62241745 |
| AAVSRAVAAL | ASSLGQGRSDTQY | 0.7924682 |
| AGGSSGIGGF | ASSLGQGRSDTQY | 0.79214793 |
| AGLTVANANG | ASSLGQGRSDTQY | 0.6206557 |
| AGLVGAVL | ASSLGQGRSDTQY | 0.7861824 |
| ALSQAFTV | ASSLGQGRSDTQY | 0.8165417 |
| ALTFSVSL | ASSLGQGRSDTQY | 0.78280723 |
| ALTGAALVAL | ASSLGQGRSDTQY | 0.71041393 |
| ASFSIGAGVL | ASSLGQGRSDTQY | 0.81383246 |
| DGSVYKI | ASSLGQGRSDTQY | 0.82546765 |
| EQLAEIV | ASSLGQGRSDTQY | 0.815856 |
| FAVSELTVM | ASSLGQGRSDTQY | 0.44837433 |
| GAAPLAAGL | ASSLGQGRSDTQY | 0.3695117 |
| GADGGDGGSGGAGG | ASSLGQGRSDTQY | 0.70302796 |
| GFGLDTGGAGG | ASSLGQGRSDTQY | 0.78462195 |
| GLASGGLL | ASSLGQGRSDTQY | 0.71356237 |
| GRIAGAAVLL | ASSLGQGRSDTQY | 0.68678856 |
| GSAGSLNTSF | ASSLGQGRSDTQY | 0.8033092 |
| GTGQIGI | ASSLGQGRSDTQY | 0.81426054 |
| GTLIALIL | ASSLGQGRSDTQY | 0.7325175 |
| IAATMGFVVG | ASSLGQGRSDTQY | 0.4548838 |

| | | |
|---|---|---|
| IFAQALL | ASSLGQGRSDTQY | 0.8039205 |
| IGAAVALGF | ASSLGQGRSDTQY | 0.80314803 |
| IGLTPLAL | ASSLGQGRSDTQY | 0.7446086 |
| ILGSNTSTL | ASSLGQGRSDTQY | 0.7850214 |
| IVFSLTVL | ASSLGQGRSDTQY | 0.69540167 |
| LASKATGF | ASSLGQGRSDTQY | 0.48156434 |
| LASVALIL | ASSLGQGRSDTQY | 0.7954675 |
| LFAVGLTL | ASSLGQGRSDTQY | 0.6345945 |
| LFNGGIGTYI | ASSLGQGRSDTQY | 0.821046 |
| LTLGAGII | ASSLGQGRSDTQY | 0.7325051 |
| LVAAALGDI | ASSLGQGRSDTQY | 0.80769545 |
| NTAFVQLGI | ASSLGQGRSDTQY | 0.80013216 |
| QLFSGAEL | ASSLGQGRSDTQY | 0.80123544 |
| SAATFITL | ASSLGQGRSDTQY | 0.7719858 |
| SNYGLANLGV | ASSLGQGRSDTQY | 0.7294432 |
| SYGGAYAVM | ASSLGQGRSDTQY | 0.64092505 |
| TTGVGGLTLG | ASSLGQGRSDTQY | 0.42546386 |
| TTQTIYI | ASSLGQGRSDTQY | 0.811983 |
| VAAGAGLGAVI | ASSLGQGRSDTQY | 0.31743333 |
| VAAVVTLL | ASSLGQGRSDTQY | 0.8157675 |
| VGFLGGLI | ASSLGQGRSDTQY | 0.8103858 |
| VGVIPILL | ASSLGQGRSDTQY | 0.7488301 |
| VIFASSLI | ASSLGQGRSDTQY | 0.8029979 |
| VKGGDGGAAAG | ASSLGQGRSDTQY | 0.6466171 |
| VTKTFTNI | ASSLGQGRSDTQY | 0.6777289 |
| VVAAAILL | ASSLGQGRSDTQY | 0.8036734 |
| YAALDVEL | ASSLGQGRSDTQY | 0.7963238 |
| LASVALLL | ASSLGQGRSDTQY | 0.7307085 |
| LDAVHRV | ASSLGQGRSDTQY | 0.7960115 |
| QGLSSSIF | ASSLGQGRSDTQY | 0.79880273 |
| RGTFANIRL | ASSLGQGRSDTQY | 0.6549684 |
| RQLWWGHRI | ASSLGQGRSDTQY | 0.7637041 |
| YGPPGTGKTLL | ASSLGQGRSDTQY | 0.81401294 |
| AALVGAPANVA | ASSLGQGRSDTQY | 0.7936413 |
| AATFRRTE | ASSLGQGRSDTQY | 0.74324906 |
| AGALGGGI | ASSLGQGRSDTQY | 0.80533445 |
| AGGAGGAGGIGAG | ASSLGQGRSDTQY | 0.70469004 |
| AGSGALGL | ASSLGQGRSDTQY | 0.8017254 |
| AQIADQI | ASSLGQGRSDTQY | 0.7906177 |
| ASAVAVTYL | ASSLGQGRSDTQY | 0.73602664 |
| ASTVLGGTGG | ASSLGQGRSDTQY | 0.8089726 |
| AVGGEPAK | ASSLGQGRSDTQY | 0.48580256 |
| CEQGTFGGTC | ASSLGQGRSDTQY | 0.74396324 |
| DEGKAPAG | ASSLGQGRSDTQY | 0.7548349 |
| DTVVFPAL | ASSLGQGRSDTQY | 0.7897958 |
| FEGITMAG | ASSLGQGRSDTQY | 0.8068167 |
| FPLVVQD | ASSLGQGRSDTQY | 0.7079894 |
| GAFTTPQ | ASSLGQGRSDTQY | 0.7865926 |
| GFGGFGTAGGVGGA | ASSLGQGRSDTQY | 0.6874875 |
| GFLAQFL | ASSLGQGRSDTQY | 0.5985521 |
| GGAGGAGGNAGVA | ASSLGQGRSDTQY | 0.7613345 |

| | | |
|---|---|---|
| GGVGTAGAGGAGGAGG | ASSLGQGRSDTQY | 0.7449447 |
| GNGGGGDAGNAGSGGN | ASSLGQGRSDTQY | 0.68643445 |
| GTGGAGGAGNDAGS | ASSLGQGRSDTQY | 0.8037825 |
| GTTNTGWFNS | ASSLGQGRSDTQY | 0.68775517 |
| IFMGPLFTL | ASSLGQGRSDTQY | 0.7766925 |
| ILAHAIEAI | ASSLGQGRSDTQY | 0.81365204 |
| IRSAIREAL | ASSLGQGRSDTQY | 0.7069391 |
| ITFAGGG | ASSLGQGRSDTQY | 0.8058858 |
| ITGMSGHQEV | ASSLGQGRSDTQY | 0.7957222 |
| IVAGLACTL | ASSLGQGRSDTQY | 0.7990062 |
| IVGGGLAGA | ASSLGQGRSDTQY | 0.7527884 |
| IVTGVLNI | ASSLGQGRSDTQY | 0.75403845 |
| LAQQGCI | ASSLGQGRSDTQY | 0.8169697 |
| LCLHVEN | ASSLGQGRSDTQY | 0.80445254 |
| LDAISDGS | ASSLGQGRSDTQY | 0.7710936 |
| LGGFAAML | ASSLGQGRSDTQY | 0.7946472 |
| LGGKNPMI | ASSLGQGRSDTQY | 0.7977544 |
| LGPKGVAAIL | ASSLGQGRSDTQY | 0.80604184 |
| LMVGVSI | ASSLGQGRSDTQY | 0.75221145 |
| LVGAAFVL | ASSLGQGRSDTQY | 0.5581222 |
| LVLAVAPL | ASSLGQGRSDTQY | 0.7543904 |
| MAAIAVIDAA | ASSLGQGRSDTQY | 0.77589834 |
| NNGINAELQ | ASSLGQGRSDTQY | 0.7454076 |
| RGEPLVLVHG | ASSLGQGRSDTQY | 0.36530644 |
| RGGAGGAAGLI | ASSLGQGRSDTQY | 0.8083178 |
| RSYRMHPAV | ASSLGQGRSDTQY | 0.82365954 |
| SAAHLLAGQLA | ASSLGQGRSDTQY | 0.6151844 |
| SAAVGSY | ASSLGQGRSDTQY | 0.81823003 |
| SPHAVLHRI | ASSLGQGRSDTQY | 0.64015806 |
| SQIAKVL | ASSLGQGRSDTQY | 0.8000722 |
| TALGHLVP | ASSLGQGRSDTQY | 0.8098521 |
| TAPTTSGAA | ASSLGQGRSDTQY | 0.69560766 |
| TDGDDGNGGSPGT | ASSLGQGRSDTQY | 0.7738327 |
| TDGLGEN | ASSLGQGRSDTQY | 0.73539454 |
| TGGPPLPGQA | ASSLGQGRSDTQY | 0.76199764 |
| TIPAITI | ASSLGQGRSDTQY | 0.8155626 |
| TIPGFTI | ASSLGQGRSDTQY | 0.6937226 |
| TSIASGALA | ASSLGQGRSDTQY | 0.78570414 |
| TVTLPGAAI | ASSLGQGRSDTQY | 0.7504177 |
| VASGLSAALP | ASSLGQGRSDTQY | 0.66426414 |
| VGAGVIGI | ASSLGQGRSDTQY | 0.8018236 |
| VGPKRFGV | ASSLGQGRSDTQY | 0.8021593 |
| VNVAFPD | ASSLGQGRSDTQY | 0.6560289 |
| VSMTNSGVS | ASSLGQGRSDTQY | 0.81497467 |
| VTDVLGDLSV | ASSLGQGRSDTQY | 0.75546306 |
| VVVVGAGFAGL | ASSLGQGRSDTQY | 0.82468104 |
| VYTVGTVV | ASSLGQGRSDTQY | 0.8093933 |
| WTTFSGP | ASSLGQGRSDTQY | 0.7866599 |
| YDELVEAF | ASSLGQGRSDTQY | 0.7549659 |
| YGTGGAGGNGG | ASSLGQGRSDTQY | 0.81482923 |
| YTNVMAV | ASSLGQGRSDTQY | 0.7391881 |

| | | |
|---|---|---|
| FLEQVHQL | ASSLGQGRSDTQY | 0.7902187 |
| GLSLLAVAV | ASSLGQGRSDTQY | 0.81282747 |
| LEQVHQL | ASSLGQGRSDTQY | 0.64222515 |
| LNAAGAI | ASSLGQGRSDTQY | 0.86648065 |
| PAVHASL | ASSLGQGRSDTQY | 0.8081531 |
| TELEDDA | ASSLGQGRSDTQY | 0.79967797 |
| TPPGPPN | ASSLGQGRSDTQY | 0.72448564 |
| TPPPPPPP | ASSLGQGRSDTQY | 0.41543388 |
| TQSPALI | TCSAQAGGDSAETLY | 0.7197542 |
| EFAAAFPQ | TCSAQAGGDSAETLY | 0.66276455 |
| LLSGNLDVL | TCSAQAGGDSAETLY | 0.8121598 |
| GPANLGNYNI | TCSAQAGGDSAETLY | 0.84230757 |
| TLGPTLP | TCSAQAGGDSAETLY | 0.71031606 |
| VYAYSSCCK | TCSAQAGGDSAETLY | 0.8727723 |
| APAPAPAPNI | TCSAQAGGDSAETLY | 0.84146374 |
| PGGPVPIP | TCSAQAGGDSAETLY | 0.58228326 |
| FLTVALEP | TCSAQAGGDSAETLY | 0.7082013 |
| SQYFGANL | TCSAQAGGDSAETLY | 0.6379502 |
| GGVKPAA | TCSAQAGGDSAETLY | 0.79618067 |
| GGTGGAGGAGGW | TCSAQAGGDSAETLY | 0.30160022 |
| GVGGVAGLGGAGG | TCSAQAGGDSAETLY | 0.83084726 |
| VLLVLATTI | TCSAQAGGDSAETLY | 0.62623847 |
| IGIFAAML | TCSAQAGGDSAETLY | 0.15156929 |
| AVGDLID | TCSAQAGGDSAETLY | 0.8153599 |
| NISPLPL | TCSAQAGGDSAETLY | 0.7353232 |
| VTAVFVAI | TCSAQAGGDSAETLY | 0.79858613 |
| SGDDNDAAL | TCSAQAGGDSAETLY | 0.8608368 |
| AVAALAHS | TCSAQAGGDSAETLY | 0.51794755 |
| GGVTVMATP | TCSAQAGGDSAETLY | 0.65749246 |
| AGLAANVRD | TCSAQAGGDSAETLY | 0.84046084 |
| TNRPDLIDPALLRPG | TCSAQAGGDSAETLY | 0.87638456 |
| LANLGVTAL | TCSAQAGGDSAETLY | 0.6330849 |
| TGERIEL | TCSAQAGGDSAETLY | 0.79920834 |
| NGEELAA | TCSAQAGGDSAETLY | 0.8582707 |
| VLSLVWG | TCSAQAGGDSAETLY | 0.83164036 |
| IGYALWL | TCSAQAGGDSAETLY | 0.85395586 |
| LSLLLAPL | TCSAQAGGDSAETLY | 0.8646042 |
| TLDEALK | TCSAQAGGDSAETLY | 0.58813214 |
| TDGAGTAEGG | TCSAQAGGDSAETLY | 0.8645321 |
| NAPGGAGGNGG | TCSAQAGGDSAETLY | 0.8839884 |
| VSVFMGMGGNF | TCSAQAGGDSAETLY | 0.7656659 |
| LASYLAAL | TCSAQAGGDSAETLY | 0.7847127 |
| LVGGTPAL | TCSAQAGGDSAETLY | 0.6397008 |
| VRNADVL | TCSAQAGGDSAETLY | 0.7428808 |
| TCSSPFLVSAM | TCSAQAGGDSAETLY | 0.34858578 |
| VHITDAE | TCSAQAGGDSAETLY | 0.77091163 |
| EYAAPSF | TCSAQAGGDSAETLY | 0.86579907 |
| VVDAFHGGL | TCSAQAGGDSAETLY | 0.7427702 |
| ASVVDPQ | TCSAQAGGDSAETLY | 0.5978222 |
| SYNEFGGLS | TCSAQAGGDSAETLY | 0.52782375 |
| DPDGSVY | TCSAQAGGDSAETLY | 0.74561 |

| | | |
|---|---|---|
| MVSTIHATE | TCSAQAGGDSAETLY | 0.72531146 |
| LAAGVADPV | TCSAQAGGDSAETLY | 0.28893963 |
| TKEQIAA | TCSAQAGGDSAETLY | 0.74986523 |
| SLASPAAL | TCSAQAGGDSAETLY | 0.86930513 |
| IGLVFYGV | TCSAQAGGDSAETLY | 0.8637262 |
| FLLDASNL | TCSAQAGGDSAETLY | 0.5086928 |
| YQVGNFP | TCSAQAGGDSAETLY | 0.8852499 |
| ESLLSQST | TCSAQAGGDSAETLY | 0.7524198 |
| LFTVGFAAVG | TCSAQAGGDSAETLY | 0.83685315 |
| ATPPTPT | TCSAQAGGDSAETLY | 0.6903622 |
| TDLATLG | TCSAQAGGDSAETLY | 0.6761908 |
| SAGGTGR | TCSAQAGGDSAETLY | 0.83057606 |
| GFSNSGS | TCSAQAGGDSAETLY | 0.74314153 |
| GGNGGSGGTSV | TCSAQAGGDSAETLY | 0.6876165 |
| GTSVATGGAGN | TCSAQAGGDSAETLY | 0.8593856 |
| NGGNGGSGGTS | TCSAQAGGDSAETLY | 0.7834141 |
| AVVKFPH | TCSAQAGGDSAETLY | 0.64935416 |
| TPPPPPP | TCSAQAGGDSAETLY | 0.64112747 |
| AAPAAAAAPA | TCSAQAGGDSAETLY | 0.8319117 |
| IALNEDL | TCSAQAGGDSAETLY | 0.58980864 |
| IVNVSSM | TCSAQAGGDSAETLY | 0.64997995 |
| TGIPPAPR | TCSAQAGGDSAETLY | 0.5146133 |
| FDLGGGTF | TCSAQAGGDSAETLY | 0.75518835 |
| FDLGGGTFDVS | TCSAQAGGDSAETLY | 0.87151515 |
| NEPTAAA | TCSAQAGGDSAETLY | 0.7966814 |
| NEPTAAALA | TCSAQAGGDSAETLY | 0.66907847 |
| GSIINISSI | TCSAQAGGDSAETLY | 0.84676826 |
| VAPGNAGTA | TCSAQAGGDSAETLY | 0.8101689 |
| TGGGASLE | TCSAQAGGDSAETLY | 0.78322816 |
| VAVYNHYKRI | TCSAQAGGDSAETLY | 0.79673976 |
| TDPAPATT | TCSAQAGGDSAETLY | 0.6649499 |
| TNRPDILDPALLRPG | TCSAQAGGDSAETLY | 0.8271009 |
| LTTTTTS | TCSAQAGGDSAETLY | 0.8203478 |
| PTGSGKTL | TCSAQAGGDSAETLY | 0.84102684 |
| TGSGKTL | TCSAQAGGDSAETLY | 0.83285683 |
| TIGHVDHGK | TCSAQAGGDSAETLY | 0.7999283 |
| TIGHVDHGKTT | TCSAQAGGDSAETLY | 0.7746322 |
| GHVDHGKTT | TCSAQAGGDSAETLY | 0.39861533 |
| AQFEHTLL | TCSAQAGGDSAETLY | 0.51750576 |
| IVNQDEL | TCSAQAGGDSAETLY | 0.82161176 |
| TNGSQFF | TCSAQAGGDSAETLY | 0.77591455 |
| TGKGIVE | TCSAQAGGDSAETLY | 0.839614 |
| LISNASDA | TCSAQAGGDSAETLY | 0.80088806 |
| AELRTLGELTKIAKAHGAQ | TCSAQAGGDSAETLY | 0.68338245 |
| APNAVAAAEPSW | TCSAQAGGDSAETLY | 0.80693936 |
| APQVAGAATAVL | TCSAQAGGDSAETLY | 0.8582795 |
| ARRDPAAAAAILPTDA | TCSAQAGGDSAETLY | 0.48176712 |
| DAAARDVLAEAGLAEYF | TCSAQAGGDSAETLY | 0.7698196 |
| DDQIVLSAKANLAKVEQ | TCSAQAGGDSAETLY | 0.88806987 |
| DELLPDTGEDSDEDGH | TCSAQAGGDSAETLY | 0.78485805 |
| ELPEAPANRATVPQD | TCSAQAGGDSAETLY | 0.7652927 |

| | | |
|---|---|---|
| FGQSKIERVTGIG | TCSAQAGGDSAETLY | 0.7437509 |
| GAGGTGSTAGGAGGAGGAGGLY | TCSAQAGGDSAETLY | 0.84409285 |
| GHGGTGAAGGNGGTGGAGGSGI | TCSAQAGGDSAETLY | 0.60770357 |
| GNGGNGGAGGEGGPGGAGGAGA | TCSAQAGGDSAETLY | 0.87401223 |
| GSTAGNGGNGGNATNGTIAGQP | TCSAQAGGDSAETLY | 0.28163493 |
| GTGGDGGNGGNGGTGG | TCSAQAGGDSAETLY | 0.71344596 |
| GVRRPAGIPADAPV | TCSAQAGGDSAETLY | 0.5216684 |
| GYRQVTDAHLLTLARR | TCSAQAGGDSAETLY | 0.47198275 |
| IAKMGQDGHDRG | TCSAQAGGDSAETLY | 0.86264455 |
| KAEPSIEIQYT | TCSAQAGGDSAETLY | 0.6757501 |
| KPVRIRMAADEDTVAVT | TCSAQAGGDSAETLY | 0.8579316 |
| MPPRIAGMRLLVIKPEP | TCSAQAGGDSAETLY | 0.744889 |
| MKFYAVALFRSRGVPPTR | TCSAQAGGDSAETLY | 0.85704446 |
| NAGGFGILGAVAHSP | TCSAQAGGDSAETLY | 0.7693834 |
| NEGREITLAMAEM | TCSAQAGGDSAETLY | 0.7723907 |
| NELVEWSHPLGHRWRWE | TCSAQAGGDSAETLY | 0.69940686 |
| NLGAGGTGGNGGNPG | TCSAQAGGDSAETLY | 0.84883785 |
| NTEWPKWETVMAADPVD | TCSAQAGGDSAETLY | 0.8495079 |
| PASAAPAETQP | TCSAQAGGDSAETLY | 0.7962346 |
| QQSSDGKNGVRAAEITGE | TCSAQAGGDSAETLY | 0.44689712 |
| RAPDAVTTSWWKE | TCSAQAGGDSAETLY | 0.7110677 |
| SGFGNVGAHTSGWFNQ | TCSAQAGGDSAETLY | 0.7977929 |
| SGVAGAVGQAASVGGLKV | TCSAQAGGDSAETLY | 0.28472364 |
| SNGGAGTGSAGGNGGT | TCSAQAGGDSAETLY | 0.2404033 |
| TGGAGGEGGAAG | TCSAQAGGDSAETLY | 0.6129749 |
| TGGDGGAAGAGGAPGHG | TCSAQAGGDSAETLY | 0.6844741 |
| TMPPLLAAIANAGRHGV | TCSAQAGGDSAETLY | 0.8557067 |
| TSSTAGNSGGA | TCSAQAGGDSAETLY | 0.85149336 |
| VPATVPVGILC | TCSAQAGGDSAETLY | 0.74814975 |
| VQRPDSGGVVLTGRIS | TCSAQAGGDSAETLY | 0.8230512 |
| WTPNPVPATKGVPAGLVQAT | TCSAQAGGDSAETLY | 0.79631555 |
| YALKRALAKFSDRFDVVII | TCSAQAGGDSAETLY | 0.11766491 |
| YLIGSVIV | TCSAQAGGDSAETLY | 0.6117534 |
| YGQLVAVID | TCSAQAGGDSAETLY | 0.6087374 |
| VTVGGMTLPA | TCSAQAGGDSAETLY | 0.84806514 |
| VLALPAV | TCSAQAGGDSAETLY | 0.7000938 |
| VIPPTLNYET | TCSAQAGGDSAETLY | 0.65714103 |
| VGMGGMHPSSG | TCSAQAGGDSAETLY | 0.7876701 |
| VGLAGGAATA | TCSAQAGGDSAETLY | 0.7346798 |
| VGGAGGAGGSGP | TCSAQAGGDSAETLY | 0.7395171 |
| VFAAFMV | TCSAQAGGDSAETLY | 0.4941861 |
| TLYDKLR | TCSAQAGGDSAETLY | 0.27252382 |
| TLAVFTVL | TCSAQAGGDSAETLY | 0.8609805 |
| TGSVQPARV | TCSAQAGGDSAETLY | 0.51204693 |
| TGGGAGSGGT | TCSAQAGGDSAETLY | 0.6932345 |
| SHLETSL | TCSAQAGGDSAETLY | 0.31351537 |
| SFTTGMT | TCSAQAGGDSAETLY | 0.7247457 |
| QVEVAKRMGVRQPTVS | TCSAQAGGDSAETLY | 0.45368344 |
| QGSGGPALAM | TCSAQAGGDSAETLY | 0.8477993 |
| MLAPSNP | TCSAQAGGDSAETLY | 0.7390325 |
| LVNNAAI | TCSAQAGGDSAETLY | 0.7304765 |

| | | |
|---|---|---|
| LTGEINTLA | TCSAQAGGDSAETLY | 0.8837048 |
| LSIFGPV | TCSAQAGGDSAETLY | 0.72011477 |
| LLALLANL | TCSAQAGGDSAETLY | 0.7579758 |
| LISNASDALDKLR | TCSAQAGGDSAETLY | 0.72446835 |
| LINEAVP | TCSAQAGGDSAETLY | 0.65217465 |
| LGYAFSGL | TCSAQAGGDSAETLY | 0.7027523 |
| LDFVAVAL | TCSAQAGGDSAETLY | 0.71090424 |
| LAQPGAE | TCSAQAGGDSAETLY | 0.87400746 |
| ISNLGAVTGF | TCSAQAGGDSAETLY | 0.7596425 |
| HQEDPGRI | TCSAQAGGDSAETLY | 0.50404996 |
| GTPGEPTP | TCSAQAGGDSAETLY | 0.7815645 |
| GFFNSGS | TCSAQAGGDSAETLY | 0.8653378 |
| GAGGGGLTAGAN | TCSAQAGGDSAETLY | 0.8853017 |
| FVDALVNM | TCSAQAGGDSAETLY | 0.7093119 |
| FLTVAPEL | TCSAQAGGDSAETLY | 0.5912312 |
| FETLTAL | TCSAQAGGDSAETLY | 0.70378435 |
| DVINPIEL | TCSAQAGGDSAETLY | 0.7408583 |
| APVSGAPV | TCSAQAGGDSAETLY | 0.65583277 |
| ALVGGLAW | TCSAQAGGDSAETLY | 0.5905551 |
| ALAAAAATAI | TCSAQAGGDSAETLY | 0.81242347 |
| AFGLGLLL | TCSAQAGGDSAETLY | 0.8745779 |
| AFFAAASL | TCSAQAGGDSAETLY | 0.73223996 |
| AAAALPAFSPP | TCSAQAGGDSAETLY | 0.3937867 |
| VVLAPAL | TCSAQAGGDSAETLY | 0.74364 |
| TIGHVDHGKTTL | TCSAQAGGDSAETLY | 0.58195436 |
| LISNASDALDK | TCSAQAGGDSAETLY | 0.69343066 |
| LISNASDAL | TCSAQAGGDSAETLY | 0.79507685 |
| ISNASDALDK | TCSAQAGGDSAETLY | 0.5010071 |
| NQAETLVYQTE | TCSAQAGGDSAETLY | 0.7596973 |
| TGGTGGTAGAG | TCSAQAGGDSAETLY | 0.8885019 |
| TGSTAGGAGGA | TCSAQAGGDSAETLY | 0.70445156 |
| DPLAVPVAAGPV | TCSAQAGGDSAETLY | 0.845989 |
| GGTGGTPGNGGN | TCSAQAGGDSAETLY | 0.8687023 |
| PAGLAGVDPATL | TCSAQAGGDSAETLY | 0.83863676 |
| GNNLVVEAPHAGASV | TCSAQAGGDSAETLY | 0.8776673 |
| KVTRIMSASTLEQPA | TCSAQAGGDSAETLY | 0.793534 |
| LGFALPIALPRRTVV | TCSAQAGGDSAETLY | 0.761829 |
| TRLGNLTARKLVVLV | TCSAQAGGDSAETLY | 0.72248805 |
| WSQQRRLIYFPSAGP | TCSAQAGGDSAETLY | 0.81904453 |
| GPNGNPDPMAAAVDIR | TCSAQAGGDSAETLY | 0.7653127 |
| TQRCVSVADLRAAASR | TCSAQAGGDSAETLY | 0.7743742 |
| QFVLDVAEHPVEVLTETV | TCSAQAGGDSAETLY | 0.6936251 |
| GGAGGTGGAGGNGGTQAGDG | TCSAQAGGDSAETLY | 0.86236984 |
| PLLATLRAIAVDVDAKPAQV | TCSAQAGGDSAETLY | 0.70371056 |
| APKFRTGGYMIDFWGVGYQVAK | TCSAQAGGDSAETLY | 0.82647383 |
| GAGGAGGAGGAAGGPGGTGGTG | TCSAQAGGDSAETLY | 0.4363547 |
| IGALGLLATDVAPAGSVAFGVA | TCSAQAGGDSAETLY | 0.6542239 |
| AGGAGGIGGNGGAGGAGGVLMG | TCSAQAGGDSAETLY | 0.8472235 |
| ADELIGLL | TCSAQAGGDSAETLY | 0.12841915 |
| ADRALKA | TCSAQAGGDSAETLY | 0.2690633 |
| AGAVGQVGGAGGA | TCSAQAGGDSAETLY | 0.76818323 |

| | | |
|---|---|---|
| AGGGGGLGGAP | TCSAQAGGDSAETLY | 0.85292673 |
| AGGLLYGNGGAG | TCSAQAGGDSAETLY | 0.16256838 |
| AGLESIGL | TCSAQAGGDSAETLY | 0.60494936 |
| AGLKTGII | TCSAQAGGDSAETLY | 0.47589743 |
| ASLAALLL | TCSAQAGGDSAETLY | 0.77957904 |
| ASVINISSI | TCSAQAGGDSAETLY | 0.7893301 |
| ATPPGPA | TCSAQAGGDSAETLY | 0.32337445 |
| ATVASALV | TCSAQAGGDSAETLY | 0.8491433 |
| ATVASLIL | TCSAQAGGDSAETLY | 0.4304803 |
| AVNGPVLIRH | TCSAQAGGDSAETLY | 0.609434 |
| DGAAGAVGI | TCSAQAGGDSAETLY | 0.8584211 |
| DPLLHGI | TCSAQAGGDSAETLY | 0.8571254 |
| ELGAALGAL | TCSAQAGGDSAETLY | 0.6379385 |
| ELGRTQAV | TCSAQAGGDSAETLY | 0.83970976 |
| FVAGGYLS | TCSAQAGGDSAETLY | 0.6019968 |
| GASGGAGGAGGW | TCSAQAGGDSAETLY | 0.89324325 |
| GGAGAGGDGGAGGI | TCSAQAGGDSAETLY | 0.77001584 |
| GTGHNGSQPGMGGNG | TCSAQAGGDSAETLY | 0.44667506 |
| GYAAASASAS | TCSAQAGGDSAETLY | 0.8515303 |
| HQVDVAAVQQG | TCSAQAGGDSAETLY | 0.852445 |
| IGAAIAHT | TCSAQAGGDSAETLY | 0.81948066 |
| IGMIEVAQL | TCSAQAGGDSAETLY | 0.7910251 |
| IRLAPEHTSAMD | TCSAQAGGDSAETLY | 0.66988933 |
| IYNVSVL | TCSAQAGGDSAETLY | 0.73538435 |
| KQGFLGM | TCSAQAGGDSAETLY | 0.8633758 |
| LESVITQL | TCSAQAGGDSAETLY | 0.8517611 |
| LGLGALVVV | TCSAQAGGDSAETLY | 0.6117377 |
| LIGNGGNGGSGGI | TCSAQAGGDSAETLY | 0.8583062 |
| LNAINAPTL | TCSAQAGGDSAETLY | 0.7741015 |
| LQLLNTGV | TCSAQAGGDSAETLY | 0.65644413 |
| LQTSIEF | TCSAQAGGDSAETLY | 0.8800169 |
| MSFAVSEL | TCSAQAGGDSAETLY | 0.7164828 |
| NTGALGL | TCSAQAGGDSAETLY | 0.8473699 |
| PPGAGGAGGAGTT | TCSAQAGGDSAETLY | 0.84080875 |
| QLVGLGI | TCSAQAGGDSAETLY | 0.8148168 |
| RFLEQVHQL | TCSAQAGGDSAETLY | 0.5214124 |
| RGVGYVLR | TCSAQAGGDSAETLY | 0.7903589 |
| SATGVFAGVI | TCSAQAGGDSAETLY | 0.7878125 |
| SLPAITLP | TCSAQAGGDSAETLY | 0.8327557 |
| STALAFTV | TCSAQAGGDSAETLY | 0.7621298 |
| SVLTVGLGK | TCSAQAGGDSAETLY | 0.73879373 |
| TALAGAGSGI | TCSAQAGGDSAETLY | 0.82576966 |
| TGLPFTGI | TCSAQAGGDSAETLY | 0.8760229 |
| VAEHVTV | TCSAQAGGDSAETLY | 0.7616578 |
| VDIVLNSL | TCSAQAGGDSAETLY | 0.75921 |
| VLTGGGAVL | TCSAQAGGDSAETLY | 0.885925 |
| VNVGLAYGI | TCSAQAGGDSAETLY | 0.6705199 |
| VSSAAVGAV | TCSAQAGGDSAETLY | 0.79075015 |
| VTSVYEAGV | TCSAQAGGDSAETLY | 0.8489944 |
| WVALDRGA | TCSAQAGGDSAETLY | 0.8626943 |
| YLVGSVVI | TCSAQAGGDSAETLY | 0.7643507 |

| | | |
|---|---|---|
| GPPGTGKTLL | TCSAQAGGDSAETLY | 0.8565822 |
| GVGMTKF | TCSAQAGGDSAETLY | 0.5561034 |
| IGIGGSDL | TCSAQAGGDSAETLY | 0.84864146 |
| IGTLLAEL | TCSAQAGGDSAETLY | 0.6603055 |
| IYTSGTT | TCSAQAGGDSAETLY | 0.8173966 |
| LSFPAGV | TCSAQAGGDSAETLY | 0.65221494 |
| LYTSGTT | TCSAQAGGDSAETLY | 0.8498632 |
| VGAFPFGF | TCSAQAGGDSAETLY | 0.8441018 |
| VGAGTVEF | TCSAQAGGDSAETLY | 0.85392606 |
| VGKSTLLL | TCSAQAGGDSAETLY | 0.24675423 |
| AALAHAGGGIV | TCSAQAGGDSAETLY | 0.84578115 |
| AALVAVSLL | TCSAQAGGDSAETLY | 0.84590405 |
| AAVSRAVAAL | TCSAQAGGDSAETLY | 0.88364863 |
| AGGSSGIGGF | TCSAQAGGDSAETLY | 0.7215563 |
| AGLTVANANG | TCSAQAGGDSAETLY | 0.82400405 |
| AGLVGAVL | TCSAQAGGDSAETLY | 0.7762917 |
| ALSQAFTV | TCSAQAGGDSAETLY | 0.7648895 |
| ALTFSVSL | TCSAQAGGDSAETLY | 0.5211818 |
| ALTGAALVAL | TCSAQAGGDSAETLY | 0.751248 |
| ASFSIGAGVL | TCSAQAGGDSAETLY | 0.68687946 |
| DGSVYKI | TCSAQAGGDSAETLY | 0.68317914 |
| EQLAEIV | TCSAQAGGDSAETLY | 0.8218252 |
| FAVSELTVM | TCSAQAGGDSAETLY | 0.85716647 |
| GAAPLAAGL | TCSAQAGGDSAETLY | 0.8624308 |
| GADGGDGGSGGAGG | TCSAQAGGDSAETLY | 0.84519565 |
| GFGLDTGGAGG | TCSAQAGGDSAETLY | 0.8706387 |
| GLASGGLL | TCSAQAGGDSAETLY | 0.8316239 |
| GRIAGAAVLL | TCSAQAGGDSAETLY | 0.72753716 |
| GSAGSLNTSF | TCSAQAGGDSAETLY | 0.7076843 |
| GTGQIGI | TCSAQAGGDSAETLY | 0.593096 |
| GTLIALIL | TCSAQAGGDSAETLY | 0.7028539 |
| IAATMGFVVG | TCSAQAGGDSAETLY | 0.7312554 |
| IFAQALL | TCSAQAGGDSAETLY | 0.80162406 |
| IGAAVALGF | TCSAQAGGDSAETLY | 0.1275528 |
| IGLTPLAL | TCSAQAGGDSAETLY | 0.7213098 |
| ILGSNTSTL | TCSAQAGGDSAETLY | 0.88085073 |
| IVFSLTVL | TCSAQAGGDSAETLY | 0.67613864 |
| LASKATGF | TCSAQAGGDSAETLY | 0.80840063 |
| LASVALIL | TCSAQAGGDSAETLY | 0.7913142 |
| LFAVGLTL | TCSAQAGGDSAETLY | 0.63323003 |
| LFNGGIGTYI | TCSAQAGGDSAETLY | 0.824792 |
| LTLGAGII | TCSAQAGGDSAETLY | 0.6290732 |
| LVAAALGDI | TCSAQAGGDSAETLY | 0.84928703 |
| NTAFVQLGI | TCSAQAGGDSAETLY | 0.8820138 |
| QLFSGAEL | TCSAQAGGDSAETLY | 0.8078027 |
| SAATFITL | TCSAQAGGDSAETLY | 0.74221367 |
| SNYGLANLGV | TCSAQAGGDSAETLY | 0.7671532 |
| SYGGAYAVM | TCSAQAGGDSAETLY | 0.4373316 |
| TTGVGGLTLG | TCSAQAGGDSAETLY | 0.761493 |
| TTQTIYI | TCSAQAGGDSAETLY | 0.49227297 |
| VAAGAGLGAVI | TCSAQAGGDSAETLY | 0.751384 |

| | | |
|---|---|---|
| VAAVVTLL | TCSAQAGGDSAETLY | 0.63126683 |
| VGFLGGLI | TCSAQAGGDSAETLY | 0.83701277 |
| VGVIPILL | TCSAQAGGDSAETLY | 0.80739474 |
| VIFASSLI | TCSAQAGGDSAETLY | 0.7785311 |
| VKGGDGGAAAG | TCSAQAGGDSAETLY | 0.6501147 |
| VTKTFTNI | TCSAQAGGDSAETLY | 0.7389842 |
| VVAAAILL | TCSAQAGGDSAETLY | 0.6863539 |
| YAALDVEL | TCSAQAGGDSAETLY | 0.5608167 |
| LASVALLL | TCSAQAGGDSAETLY | 0.84623253 |
| LDAVHRV | TCSAQAGGDSAETLY | 0.75945467 |
| QGLSSSIF | TCSAQAGGDSAETLY | 0.73302364 |
| RGTFANIRL | TCSAQAGGDSAETLY | 0.86724555 |
| RQLWWGHRI | TCSAQAGGDSAETLY | 0.84179866 |
| YGPPGTGKTLL | TCSAQAGGDSAETLY | 0.40529257 |
| AALVGAPANVA | TCSAQAGGDSAETLY | 0.7233113 |
| AATFRRTE | TCSAQAGGDSAETLY | 0.76913416 |
| AGALGGGI | TCSAQAGGDSAETLY | 0.7794476 |
| AGGAGGAGGIGAG | TCSAQAGGDSAETLY | 0.16555816 |
| AGSGALGL | TCSAQAGGDSAETLY | 0.83314764 |
| AQIADQI | TCSAQAGGDSAETLY | 0.6571649 |
| ASAVAVTYL | TCSAQAGGDSAETLY | 0.5802896 |
| ASTVLGGTGG | TCSAQAGGDSAETLY | 0.7214186 |
| AVGGEPAK | TCSAQAGGDSAETLY | 0.86228263 |
| CEQGTFGGTC | TCSAQAGGDSAETLY | 0.68993163 |
| DEGKAPAG | TCSAQAGGDSAETLY | 0.8028381 |
| DTVVFPAL | TCSAQAGGDSAETLY | 0.84391665 |
| FEGITMAG | TCSAQAGGDSAETLY | 0.8271184 |
| FPLVVQD | TCSAQAGGDSAETLY | 0.8769387 |
| GAFTTPQ | TCSAQAGGDSAETLY | 0.84707105 |
| GFGGFGTAGGVGGA | TCSAQAGGDSAETLY | 0.6745991 |
| GFLAQFL | TCSAQAGGDSAETLY | 0.797181 |
| GGAGGAGGNAGVA | TCSAQAGGDSAETLY | 0.63745594 |
| GGVGTAGAGGAGGAGG | TCSAQAGGDSAETLY | 0.481218 |
| GNGGGGDAGNAGSGGN | TCSAQAGGDSAETLY | 0.36726126 |
| GTGGAGGAGNDAGS | TCSAQAGGDSAETLY | 0.7581748 |
| GTTNTGWFNS | TCSAQAGGDSAETLY | 0.35944223 |
| IFMGPLFTL | TCSAQAGGDSAETLY | 0.8187063 |
| ILAHAIEAI | TCSAQAGGDSAETLY | 0.8518543 |
| IRSAIREAL | TCSAQAGGDSAETLY | 0.684631 |
| ITFAGGG | TCSAQAGGDSAETLY | 0.5003922 |
| ITGMSGHQEV | TCSAQAGGDSAETLY | 0.7046294 |
| IVAGLACTL | TCSAQAGGDSAETLY | 0.6429963 |
| IVGGGLAGA | TCSAQAGGDSAETLY | 0.85825825 |
| IVTGVLNI | TCSAQAGGDSAETLY | 0.78989995 |
| LAQQGCI | TCSAQAGGDSAETLY | 0.8594346 |
| LCLHVEN | TCSAQAGGDSAETLY | 0.84521866 |
| LDAISDGS | TCSAQAGGDSAETLY | 0.7343396 |
| LGGFAAML | TCSAQAGGDSAETLY | 0.8522532 |
| LGGKNPMI | TCSAQAGGDSAETLY | 0.7007266 |
| LGPKGVAAIL | TCSAQAGGDSAETLY | 0.756749 |
| LMVGVSI | TCSAQAGGDSAETLY | 0.780442 |

| | | |
|---|---|---|
| LVGAAFVL | TCSAQAGGDSAETLY | 0.7885065 |
| LVLAVAPL | TCSAQAGGDSAETLY | 0.75163674 |
| MAAIAVIDAA | TCSAQAGGDSAETLY | 0.7665832 |
| NNGINAELQ | TCSAQAGGDSAETLY | 0.46473747 |
| RGEPLVLVHG | TCSAQAGGDSAETLY | 0.7695122 |
| RGGAGGAAGLI | TCSAQAGGDSAETLY | 0.7384472 |
| RSYRMHPAV | TCSAQAGGDSAETLY | 0.3067533 |
| SAAHLLAGQLA | TCSAQAGGDSAETLY | 0.69732153 |
| SAAVGSY | TCSAQAGGDSAETLY | 0.8883168 |
| SPHAVLHRI | TCSAQAGGDSAETLY | 0.86152375 |
| SQIAKVL | TCSAQAGGDSAETLY | 0.6274711 |
| TALGHLVP | TCSAQAGGDSAETLY | 0.6582582 |
| TAPTTSGAA | TCSAQAGGDSAETLY | 0.734795 |
| TDGDDGNGGSPGT | TCSAQAGGDSAETLY | 0.4822569 |
| TDGLGEN | TCSAQAGGDSAETLY | 0.8142048 |
| TGGPPLPGQA | TCSAQAGGDSAETLY | 0.7309307 |
| TIPAITI | TCSAQAGGDSAETLY | 0.768703 |
| TIPGFTI | TCSAQAGGDSAETLY | 0.58470416 |
| TSIASGALA | TCSAQAGGDSAETLY | 0.7865163 |
| TVTLPGAAI | TCSAQAGGDSAETLY | 0.74718547 |
| VASGLSAALP | TCSAQAGGDSAETLY | 0.78026307 |
| VGAGVIGI | TCSAQAGGDSAETLY | 0.23138312 |
| VGPKRFGV | TCSAQAGGDSAETLY | 0.82499063 |
| VNVAFPD | TCSAQAGGDSAETLY | 0.6546173 |
| VSMTNSGVS | TCSAQAGGDSAETLY | 0.5498253 |
| VTDVLGDLSV | TCSAQAGGDSAETLY | 0.3444587 |
| VVVVGAGFAGL | TCSAQAGGDSAETLY | 0.7448063 |
| VYTVGTVV | TCSAQAGGDSAETLY | 0.8455057 |
| WTTFSGP | TCSAQAGGDSAETLY | 0.8532504 |
| YDELVEAF | TCSAQAGGDSAETLY | 0.6186067 |
| YGTGGAGGNGG | TCSAQAGGDSAETLY | 0.83883107 |
| YTNVMAV | TCSAQAGGDSAETLY | 0.859723 |
| FLEQVHQL | TCSAQAGGDSAETLY | 0.7276153 |
| GLSLLAVAV | TCSAQAGGDSAETLY | 0.43514225 |
| LEQVHQL | TCSAQAGGDSAETLY | 0.83176124 |
| LNAAGAI | TCSAQAGGDSAETLY | 0.65711963 |
| PAVHASL | TCSAQAGGDSAETLY | 0.84765196 |
| TELEDDA | TCSAQAGGDSAETLY | 0.8308279 |
| TPPGPPN | TCSAQAGGDSAETLY | 0.58021975 |
| TPPPPPPP | TCSAQAGGDSAETLY | 0.8072793 |
| TQSPALI | ASSRGQTNTEVF | 0.868132 |
| EFAAAFPQ | ASSRGQTNTEVF | 0.60413563 |
| LLSGNLDVL | ASSRGQTNTEVF | 0.59954846 |
| GPANLGNYNI | ASSRGQTNTEVF | 0.645223 |
| TLGPTLP | ASSRGQTNTEVF | 0.49978393 |
| VYAYSSCCK | ASSRGQTNTEVF | 0.81666875 |
| APAPAPAPNI | ASSRGQTNTEVF | 0.6727356 |
| PGGPVPIP | ASSRGQTNTEVF | 0.69625497 |
| FLTVALEP | ASSRGQTNTEVF | 0.7857392 |
| SQYFGANL | ASSRGQTNTEVF | 0.6931877 |
| GGVKPAA | ASSRGQTNTEVF | 0.75546044 |

| | | |
|---|---|---|
| GGTGGAGGAGGW | ASSRGQTNTEVF | 0.77938145 |
| GVGGVAGLGGAGG | ASSRGQTNTEVF | 0.79304457 |
| VLLVLATTI | ASSRGQTNTEVF | 0.79476863 |
| IGIFAAML | ASSRGQTNTEVF | 0.7094434 |
| AVGDLID | ASSRGQTNTEVF | 0.7320783 |
| NISPLPL | ASSRGQTNTEVF | 0.88076186 |
| VTAVFVAI | ASSRGQTNTEVF | 0.5185021 |
| SGDDNDAAL | ASSRGQTNTEVF | 0.5787908 |
| AVAALAHS | ASSRGQTNTEVF | 0.8208212 |
| GGVTVMATP | ASSRGQTNTEVF | 0.5398278 |
| AGLAANVRD | ASSRGQTNTEVF | 0.8449111 |
| TNRPDLIDPALLRPG | ASSRGQTNTEVF | 0.79200053 |
| LANLGVTAL | ASSRGQTNTEVF | 0.8025991 |
| TGERIEL | ASSRGQTNTEVF | 0.65456474 |
| NGEELAA | ASSRGQTNTEVF | 0.8893266 |
| VLSLVWG | ASSRGQTNTEVF | 0.86410993 |
| IGYALWL | ASSRGQTNTEVF | 0.7105927 |
| LSLLLAPL | ASSRGQTNTEVF | 0.8299246 |
| TLDEALK | ASSRGQTNTEVF | 0.42932546 |
| TDGAGTAEGG | ASSRGQTNTEVF | 0.8304739 |
| NAPGGAGGNGG | ASSRGQTNTEVF | 0.8371252 |
| VSVFMGMGGNF | ASSRGQTNTEVF | 0.60638666 |
| LASYLAAL | ASSRGQTNTEVF | 0.73828423 |
| LVGGTPAL | ASSRGQTNTEVF | 0.8548527 |
| VRNADVL | ASSRGQTNTEVF | 0.8772919 |
| TCSSPFLVSAM | ASSRGQTNTEVF | 0.35623956 |
| VHITDAE | ASSRGQTNTEVF | 0.72804224 |
| EYAAPSF | ASSRGQTNTEVF | 0.34515154 |
| VVDAFHGGL | ASSRGQTNTEVF | 0.56883574 |
| ASVVDPQ | ASSRGQTNTEVF | 0.5039027 |
| SYNEFGGLS | ASSRGQTNTEVF | 0.79884446 |
| DPDGSVY | ASSRGQTNTEVF | 0.7616241 |
| MVSTIHATE | ASSRGQTNTEVF | 0.7990735 |
| LAAGVADPV | ASSRGQTNTEVF | 0.8218715 |
| TKEQIAA | ASSRGQTNTEVF | 0.87704873 |
| SLASPAAL | ASSRGQTNTEVF | 0.6751117 |
| IGLVFYGV | ASSRGQTNTEVF | 0.79780173 |
| FLLDASNL | ASSRGQTNTEVF | 0.5554131 |
| YQVGNFP | ASSRGQTNTEVF | 0.58591163 |
| ESLLSQST | ASSRGQTNTEVF | 0.5498854 |
| LFTVGFAAVG | ASSRGQTNTEVF | 0.75472844 |
| ATPPTPT | ASSRGQTNTEVF | 0.25273657 |
| TDLATLG | ASSRGQTNTEVF | 0.8666903 |
| SAGGTGR | ASSRGQTNTEVF | 0.817728 |
| GFSNSGS | ASSRGQTNTEVF | 0.85017216 |
| GGNGGSGGTSV | ASSRGQTNTEVF | 0.71069455 |
| GTSVATGGAGN | ASSRGQTNTEVF | 0.71590734 |
| NGGNGGSGGTS | ASSRGQTNTEVF | 0.7253649 |
| AVVKFPH | ASSRGQTNTEVF | 0.8464694 |
| TPPPPPP | ASSRGQTNTEVF | 0.66992664 |
| AAPAAAAPA | ASSRGQTNTEVF | 0.34684083 |

| | | |
|---|---|---|
| IALNEDL | ASSRGQTNTEVF | 0.54465216 |
| IVNVSSM | ASSRGQTNTEVF | 0.46257818 |
| TGIPPAPR | ASSRGQTNTEVF | 0.8651953 |
| FDLGGGTF | ASSRGQTNTEVF | 0.8563267 |
| FDLGGGTFDVS | ASSRGQTNTEVF | 0.8108277 |
| NEPTAAA | ASSRGQTNTEVF | 0.8576528 |
| NEPTAAALA | ASSRGQTNTEVF | 0.19750713 |
| GSIINISSI | ASSRGQTNTEVF | 0.6665108 |
| VAPGNAGTA | ASSRGQTNTEVF | 0.8746029 |
| TGGGASLE | ASSRGQTNTEVF | 0.8051293 |
| VAVYNHYKRI | ASSRGQTNTEVF | 0.784142 |
| TDPAPATT | ASSRGQTNTEVF | 0.87759626 |
| TNRPDILDPALLRPG | ASSRGQTNTEVF | 0.71506536 |
| LTTTTTS | ASSRGQTNTEVF | 0.78109366 |
| PTGSGKTL | ASSRGQTNTEVF | 0.8183142 |
| TGSGKTL | ASSRGQTNTEVF | 0.83295286 |
| TIGHVDHGK | ASSRGQTNTEVF | 0.78529036 |
| TIGHVDHGKTT | ASSRGQTNTEVF | 0.60664713 |
| GHVDHGKTT | ASSRGQTNTEVF | 0.8194066 |
| AQFEHTLL | ASSRGQTNTEVF | 0.88747764 |
| IVNQDEL | ASSRGQTNTEVF | 0.7777496 |
| TNGSQFF | ASSRGQTNTEVF | 0.85440207 |
| TGKGIVE | ASSRGQTNTEVF | 0.8251176 |
| LISNASDA | ASSRGQTNTEVF | 0.7635085 |
| AELRTLGELTKIAKAHGAQ | ASSRGQTNTEVF | 0.67446864 |
| APNAVAAAEPSW | ASSRGQTNTEVF | 0.6259488 |
| APQVAGAATAVL | ASSRGQTNTEVF | 0.73767936 |
| ARRDPAAAAAILPTDA | ASSRGQTNTEVF | 0.5661026 |
| DAAARDVLAEAGLAEYF | ASSRGQTNTEVF | 0.7985919 |
| DDQIVLSAKANLAKVEQ | ASSRGQTNTEVF | 0.69238037 |
| DELLPDTGEDSDEDGH | ASSRGQTNTEVF | 0.8967372 |
| ELPEAPANRATVPQD | ASSRGQTNTEVF | 0.82703036 |
| FGQSKIERVTGIG | ASSRGQTNTEVF | 0.8427366 |
| GAGGTGSTAGGAGGAGGAGGLY | ASSRGQTNTEVF | 0.6256074 |
| GHGGTGAAGGNGGTGGAGGSGI | ASSRGQTNTEVF | 0.7501923 |
| GNGGNGGAGGEGGPGGAGGAGA | ASSRGQTNTEVF | 0.7807867 |
| GSTAGNGGNGGNATNGTIAGQP | ASSRGQTNTEVF | 0.5287601 |
| GTGGDGGNGGNGGTGG | ASSRGQTNTEVF | 0.8626913 |
| GVRRPAGIPADAPV | ASSRGQTNTEVF | 0.65343225 |
| GYRQVTDAHLLTLARR | ASSRGQTNTEVF | 0.721476 |
| IAKMGQDGHDRG | ASSRGQTNTEVF | 0.8181459 |
| KAEPSIEIQYT | ASSRGQTNTEVF | 0.8377322 |
| KPVRIRMAADEDTVAVT | ASSRGQTNTEVF | 0.7610997 |
| MPPRIAGMRLLVIKPEP | ASSRGQTNTEVF | 0.70850474 |
| MKFYAVALFRSRGVPPTR | ASSRGQTNTEVF | 0.70539725 |
| NAGGFGILGAVAHSP | ASSRGQTNTEVF | 0.8126188 |
| NEGREITLAMAEM | ASSRGQTNTEVF | 0.875681 |
| NELVEWSHPLGHRWRWE | ASSRGQTNTEVF | 0.71901184 |
| NLGAGGTGGNGGNPG | ASSRGQTNTEVF | 0.86715645 |
| NTEWPKWETVMAADPVD | ASSRGQTNTEVF | 0.75228506 |
| PASAAPAETQP | ASSRGQTNTEVF | 0.39437237 |

| | | |
|---|---|---|
| QQSSDGKNGVRAAEITGE | ASSRGQTNTEVF | 0.7859286 |
| RAPDAVTTSWWKE | ASSRGQTNTEVF | 0.8596033 |
| SGFGNVGAHTSGWFNQ | ASSRGQTNTEVF | 0.8414701 |
| SGVAGAVGQAASVGGLKV | ASSRGQTNTEVF | 0.86889243 |
| SNGGAGTGSAGGNGGT | ASSRGQTNTEVF | 0.82193637 |
| TGGAGGEGGAAG | ASSRGQTNTEVF | 0.6848059 |
| TGGDGGAAGAGGAPGHG | ASSRGQTNTEVF | 0.86282104 |
| TMPPLLAAIANAGRHGV | ASSRGQTNTEVF | 0.8690183 |
| TSSTAGNSGGA | ASSRGQTNTEVF | 0.8600831 |
| VPATVPVGILC | ASSRGQTNTEVF | 0.66451985 |
| VQRPDSGGVVLTGRIS | ASSRGQTNTEVF | 0.550576 |
| WTPNPVPATKGVPAGLVQAT | ASSRGQTNTEVF | 0.81800884 |
| YALKRALAKFSDRFDVVII | ASSRGQTNTEVF | 0.8595705 |
| YLIGSVIV | ASSRGQTNTEVF | 0.6048412 |
| YGQLVAVID | ASSRGQTNTEVF | 0.8061054 |
| VTVGGMTLPA | ASSRGQTNTEVF | 0.77663505 |
| VLALPAV | ASSRGQTNTEVF | 0.43628186 |
| VIPPTLNYET | ASSRGQTNTEVF | 0.7848941 |
| VGMGGMHPSSG | ASSRGQTNTEVF | 0.8688174 |
| VGLAGGAATA | ASSRGQTNTEVF | 0.79229724 |
| VGGAGGAGGSGP | ASSRGQTNTEVF | 0.59024805 |
| VFAAFMV | ASSRGQTNTEVF | 0.85940677 |
| TLYDKLR | ASSRGQTNTEVF | 0.8913596 |
| TLAVFTVL | ASSRGQTNTEVF | 0.86967725 |
| TGSVQPARV | ASSRGQTNTEVF | 0.48114038 |
| TGGGAGSGGT | ASSRGQTNTEVF | 0.20085257 |
| SHLETSL | ASSRGQTNTEVF | 0.7272732 |
| SFTTGMT | ASSRGQTNTEVF | 0.806236 |
| QVEVAKRMGVRQPTVS | ASSRGQTNTEVF | 0.7590035 |
| QGSGGPALAM | ASSRGQTNTEVF | 0.7336583 |
| MLAPSNP | ASSRGQTNTEVF | 0.82594556 |
| LVNNAAI | ASSRGQTNTEVF | 0.85469854 |
| LTGEINTLA | ASSRGQTNTEVF | 0.6554852 |
| LSIFGPV | ASSRGQTNTEVF | 0.40828377 |
| LLALLANL | ASSRGQTNTEVF | 0.83531046 |
| LISNASDALDKLR | ASSRGQTNTEVF | 0.8550148 |
| LINEAVP | ASSRGQTNTEVF | 0.63984096 |
| LGYAFSGL | ASSRGQTNTEVF | 0.76193255 |
| LDFVAVAL | ASSRGQTNTEVF | 0.74574316 |
| LAQPGAE | ASSRGQTNTEVF | 0.6497294 |
| ISNLGAVTGF | ASSRGQTNTEVF | 0.7522253 |
| HQEDPGRI | ASSRGQTNTEVF | 0.51912767 |
| GTPGEPTP | ASSRGQTNTEVF | 0.8726287 |
| GFFNSGS | ASSRGQTNTEVF | 0.598233 |
| GAGGGGLTAGAN | ASSRGQTNTEVF | 0.8256502 |
| FVDALVNM | ASSRGQTNTEVF | 0.8325156 |
| FLTVAPEL | ASSRGQTNTEVF | 0.8212356 |
| FETLTAL | ASSRGQTNTEVF | 0.7549114 |
| DVINPIEL | ASSRGQTNTEVF | 0.6404054 |
| APVSGAPV | ASSRGQTNTEVF | 0.5648805 |
| ALVGGLAW | ASSRGQTNTEVF | 0.299272 |

| | | |
|---|---|---|
| ALAAAAATAI | ASSRGQTNTEVF | 0.8702791 |
| AFGLGLLL | ASSRGQTNTEVF | 0.7145173 |
| AFFAAASL | ASSRGQTNTEVF | 0.872662 |
| AAAALPAFSPP | ASSRGQTNTEVF | 0.85158825 |
| VVLAPAL | ASSRGQTNTEVF | 0.7149434 |
| TIGHVDHGKTTL | ASSRGQTNTEVF | 0.80396676 |
| LISNASDALDK | ASSRGQTNTEVF | 0.63123465 |
| LISNASDAL | ASSRGQTNTEVF | 0.6131362 |
| ISNASDALDK | ASSRGQTNTEVF | 0.80166507 |
| NQAETLVYQTE | ASSRGQTNTEVF | 0.83712554 |
| TGGTGGTAGAG | ASSRGQTNTEVF | 0.6634212 |
| TGSTAGGAGGA | ASSRGQTNTEVF | 0.824975 |
| DPLAVPVAAGPV | ASSRGQTNTEVF | 0.80516493 |
| GGTGGTPGNGGN | ASSRGQTNTEVF | 0.5163451 |
| PAGLAGVDPATL | ASSRGQTNTEVF | 0.7936981 |
| GNNLVVEAPHAGASV | ASSRGQTNTEVF | 0.85612893 |
| KVTRIMSASTLEQPA | ASSRGQTNTEVF | 0.8210441 |
| LGFALPIALPRRTVV | ASSRGQTNTEVF | 0.670985 |
| TRLGNLTARKLVVLV | ASSRGQTNTEVF | 0.845147 |
| WSQQRRLIYFPSAGP | ASSRGQTNTEVF | 0.63674855 |
| GPNGNPDPMAAAVDIR | ASSRGQTNTEVF | 0.7991169 |
| TQRCVSVADLRAAASR | ASSRGQTNTEVF | 0.8108138 |
| QFVLDVAEHPVEVLTETV | ASSRGQTNTEVF | 0.71981657 |
| GGAGGTGGAGGNGGTQAGDG | ASSRGQTNTEVF | 0.8691895 |
| PLLATLRAIAVDVDAKPAQV | ASSRGQTNTEVF | 0.3738143 |
| APKFRTGGYMIDFWGVGYQVAK | ASSRGQTNTEVF | 0.64573705 |
| GAGGAGGAGGAAGGPGGTGGTG | ASSRGQTNTEVF | 0.7462801 |
| IGALGLLATDVAPAGSVAFGVA | ASSRGQTNTEVF | 0.83514386 |
| AGGAGGIGGNGGAGGAGGVLMG | ASSRGQTNTEVF | 0.85282195 |
| ADELIGLL | ASSRGQTNTEVF | 0.6389705 |
| ADRALKA | ASSRGQTNTEVF | 0.76675284 |
| AGAVGQVGGAGGA | ASSRGQTNTEVF | 0.7294276 |
| AGGGGGLGGAP | ASSRGQTNTEVF | 0.64719075 |
| AGGLLYGNGGAG | ASSRGQTNTEVF | 0.78462744 |
| AGLESIGL | ASSRGQTNTEVF | 0.6685194 |
| AGLKTGII | ASSRGQTNTEVF | 0.56229675 |
| ASLAALLL | ASSRGQTNTEVF | 0.81931186 |
| ASVINISSI | ASSRGQTNTEVF | 0.7572473 |
| ATPPGPA | ASSRGQTNTEVF | 0.8133069 |
| ATVASALV | ASSRGQTNTEVF | 0.8557999 |
| ATVASLIL | ASSRGQTNTEVF | 0.6043962 |
| AVNGPVLIRH | ASSRGQTNTEVF | 0.8165722 |
| DGAAGAVGI | ASSRGQTNTEVF | 0.8173921 |
| DPLLHGI | ASSRGQTNTEVF | 0.8328236 |
| ELGAALGAL | ASSRGQTNTEVF | 0.7657205 |
| ELGRTQAV | ASSRGQTNTEVF | 0.63769865 |
| FVAGGYLS | ASSRGQTNTEVF | 0.86566526 |
| GASGGAGGAGGW | ASSRGQTNTEVF | 0.85106903 |
| GGAGAGGDGGAGGI | ASSRGQTNTEVF | 0.7855996 |
| GTGHNGSQPGMGGNG | ASSRGQTNTEVF | 0.8432114 |
| GYAAASASAS | ASSRGQTNTEVF | 0.80211323 |

| | | |
|---|---|---|
| HQVDVAAVQQG | ASSRGQTNTEVF | 0.8499435 |
| IGAAIAHT | ASSRGQTNTEVF | 0.6811104 |
| IGMIEVAQL | ASSRGQTNTEVF | 0.4846465 |
| IRLAPEHTSAMD | ASSRGQTNTEVF | 0.7798395 |
| IYNVSVL | ASSRGQTNTEVF | 0.7216141 |
| KQGFLGM | ASSRGQTNTEVF | 0.6626468 |
| LESVITQL | ASSRGQTNTEVF | 0.28162903 |
| LGLGALVVV | ASSRGQTNTEVF | 0.86107826 |
| LIGNGGNGGSGGI | ASSRGQTNTEVF | 0.8939252 |
| LNAINAPTL | ASSRGQTNTEVF | 0.6180979 |
| LQLLNTGV | ASSRGQTNTEVF | 0.8747101 |
| LQTSIEF | ASSRGQTNTEVF | 0.48035577 |
| MSFAVSEL | ASSRGQTNTEVF | 0.8410868 |
| NTGALGL | ASSRGQTNTEVF | 0.86829257 |
| PPGAGGAGGAGTT | ASSRGQTNTEVF | 0.57202065 |
| QLVGLGI | ASSRGQTNTEVF | 0.7976226 |
| RFLEQVHQL | ASSRGQTNTEVF | 0.6784041 |
| RGVGYVLR | ASSRGQTNTEVF | 0.70013857 |
| SATGVFAGVI | ASSRGQTNTEVF | 0.8771828 |
| SLPAITLP | ASSRGQTNTEVF | 0.7287333 |
| STALAFTV | ASSRGQTNTEVF | 0.78153527 |
| SVLTVGLGK | ASSRGQTNTEVF | 0.8066179 |
| TALAGAGSGI | ASSRGQTNTEVF | 0.6881988 |
| TGLPFTGI | ASSRGQTNTEVF | 0.82434934 |
| VAEHVTV | ASSRGQTNTEVF | 0.8387747 |
| VDIVLNSL | ASSRGQTNTEVF | 0.5573441 |
| VLTGGGAVL | ASSRGQTNTEVF | 0.8586836 |
| VNVGLAYGI | ASSRGQTNTEVF | 0.7108803 |
| VSSAAVGAV | ASSRGQTNTEVF | 0.887653 |
| VTSVYEAGV | ASSRGQTNTEVF | 0.84388554 |
| WVALDRGA | ASSRGQTNTEVF | 0.7628145 |
| YLVGSVVI | ASSRGQTNTEVF | 0.7347672 |
| GPPGTGKTLL | ASSRGQTNTEVF | 0.86403704 |
| GVGMTKF | ASSRGQTNTEVF | 0.7868885 |
| IGIGGSDL | ASSRGQTNTEVF | 0.8041585 |
| IGTLLAEL | ASSRGQTNTEVF | 0.8497232 |
| IYTSGTT | ASSRGQTNTEVF | 0.7118511 |
| LSFPAGV | ASSRGQTNTEVF | 0.9026923 |
| LYTSGTT | ASSRGQTNTEVF | 0.8603283 |
| VGAFPFGF | ASSRGQTNTEVF | 0.87697315 |
| VGAGTVEF | ASSRGQTNTEVF | 0.7923225 |
| VGKSTLLL | ASSRGQTNTEVF | 0.4698807 |
| AALAHAGGGIV | ASSRGQTNTEVF | 0.8110411 |
| AALVAVSLL | ASSRGQTNTEVF | 0.73637235 |
| AAVSRAVAAL | ASSRGQTNTEVF | 0.87194836 |
| AGGSSGIGGF | ASSRGQTNTEVF | 0.88742244 |
| AGLTVANANG | ASSRGQTNTEVF | 0.7908828 |
| AGLVGAVL | ASSRGQTNTEVF | 0.88783514 |
| ALSQAFTV | ASSRGQTNTEVF | 0.90117687 |
| ALTFSVSL | ASSRGQTNTEVF | 0.47055328 |
| ALTGAALVAL | ASSRGQTNTEVF | 0.83374226 |

| | | |
|---|---|---|
| ASFSIGAGVL | ASSRGQTNTEVF | 0.7119069 |
| DGSVYKI | ASSRGQTNTEVF | 0.819975 |
| EQLAEIV | ASSRGQTNTEVF | 0.29041335 |
| FAVSELTVM | ASSRGQTNTEVF | 0.88576365 |
| GAAPLAAGL | ASSRGQTNTEVF | 0.7641789 |
| GADGGDGGSGGAGG | ASSRGQTNTEVF | 0.14771055 |
| GFGLDTGGAGG | ASSRGQTNTEVF | 0.8518623 |
| GLASGGLL | ASSRGQTNTEVF | 0.82323384 |
| GRIAGAAVLL | ASSRGQTNTEVF | 0.81338835 |
| GSAGSLNTSF | ASSRGQTNTEVF | 0.87925434 |
| GTGQIGI | ASSRGQTNTEVF | 0.74919105 |
| GTLIALIL | ASSRGQTNTEVF | 0.7422478 |
| IAATMGFVVG | ASSRGQTNTEVF | 0.8611275 |
| IFAQALL | ASSRGQTNTEVF | 0.89835685 |
| IGAAVALGF | ASSRGQTNTEVF | 0.6927886 |
| IGLTPLAL | ASSRGQTNTEVF | 0.8524512 |
| ILGSNTSTL | ASSRGQTNTEVF | 0.90773714 |
| IVFSLTVL | ASSRGQTNTEVF | 0.863118 |
| LASKATGF | ASSRGQTNTEVF | 0.90018046 |
| LASVALIL | ASSRGQTNTEVF | 0.9110453 |
| LFAVGLTL | ASSRGQTNTEVF | 0.74711454 |
| LFNGGIGTYI | ASSRGQTNTEVF | 0.88422143 |
| LTLGAGII | ASSRGQTNTEVF | 0.88103455 |
| LVAAALGDI | ASSRGQTNTEVF | 0.8335823 |
| NTAFVQLGI | ASSRGQTNTEVF | 0.8409014 |
| QLFSGAEL | ASSRGQTNTEVF | 0.69114757 |
| SAATFITL | ASSRGQTNTEVF | 0.81251365 |
| SNYGLANLGV | ASSRGQTNTEVF | 0.45887312 |
| SYGGAYAVM | ASSRGQTNTEVF | 0.84208834 |
| TTGVGGLTLG | ASSRGQTNTEVF | 0.8917737 |
| TTQTIYI | ASSRGQTNTEVF | 0.8067709 |
| VAAGAGLGAVI | ASSRGQTNTEVF | 0.6309619 |
| VAAVVTLL | ASSRGQTNTEVF | 0.68881476 |
| VGFLGGLI | ASSRGQTNTEVF | 0.8597573 |
| VGVIPILL | ASSRGQTNTEVF | 0.79984915 |
| VIFASSLI | ASSRGQTNTEVF | 0.45007417 |
| VKGGDGGAAAG | ASSRGQTNTEVF | 0.7520834 |
| VTKTFTNI | ASSRGQTNTEVF | 0.8950204 |
| VVAAAILL | ASSRGQTNTEVF | 0.9013565 |
| YAALDVEL | ASSRGQTNTEVF | 0.6092405 |
| LASVALLL | ASSRGQTNTEVF | 0.88899577 |
| LDAVHRV | ASSRGQTNTEVF | 0.8895912 |
| QGLSSSIF | ASSRGQTNTEVF | 0.910075 |
| RGTFANIRL | ASSRGQTNTEVF | 0.8602529 |
| RQLWWGHRI | ASSRGQTNTEVF | 0.72552514 |
| YGPPGTGKTLL | ASSRGQTNTEVF | 0.87687063 |
| AALVGAPANVA | ASSRGQTNTEVF | 0.7906573 |
| AATFRRTE | ASSRGQTNTEVF | 0.7654186 |
| AGALGGGI | ASSRGQTNTEVF | 0.8940834 |
| AGGAGGAGGIGAG | ASSRGQTNTEVF | 0.8135514 |
| AGSGALGL | ASSRGQTNTEVF | 0.7585032 |

| | | |
|---|---|---|
| AQIADQI | ASSRGQTNTEVF | 0.78888255 |
| ASAVAVTYL | ASSRGQTNTEVF | 0.9126604 |
| ASTVLGGTGG | ASSRGQTNTEVF | 0.6598637 |
| AVGGEPAK | ASSRGQTNTEVF | 0.7236627 |
| CEQGTFGGTC | ASSRGQTNTEVF | 0.75265753 |
| DEGKAPAG | ASSRGQTNTEVF | 0.83627087 |
| DTVVFPAL | ASSRGQTNTEVF | 0.89275193 |
| FEGITMAG | ASSRGQTNTEVF | 0.87992644 |
| FPLVVQD | ASSRGQTNTEVF | 0.80666375 |
| GAFTTPQ | ASSRGQTNTEVF | 0.85818267 |
| GFGGFGTAGGVGGA | ASSRGQTNTEVF | 0.89112794 |
| GFLAQFL | ASSRGQTNTEVF | 0.85635877 |
| GGAGGAGGNAGVA | ASSRGQTNTEVF | 0.88905954 |
| GGVGTAGAGGAGGAGG | ASSRGQTNTEVF | 0.7282131 |
| GNGGGGDAGNAGSGGN | ASSRGQTNTEVF | 0.9066347 |
| GTGGAGGAGNDAGS | ASSRGQTNTEVF | 0.8853864 |
| GTTNTGWFNS | ASSRGQTNTEVF | 0.8944986 |
| IFMGPLFTL | ASSRGQTNTEVF | 0.8905546 |
| ILAHAIEAI | ASSRGQTNTEVF | 0.8759652 |
| IRSAIREAL | ASSRGQTNTEVF | 0.87028927 |
| ITFAGGG | ASSRGQTNTEVF | 0.504562 |
| ITGMSGHQEV | ASSRGQTNTEVF | 0.7495008 |
| IVAGLACTL | ASSRGQTNTEVF | 0.8468348 |
| IVGGGLAGA | ASSRGQTNTEVF | 0.85009515 |
| IVTGVLNI | ASSRGQTNTEVF | 0.865494 |
| LAQQGCI | ASSRGQTNTEVF | 0.8924359 |
| LCLHVEN | ASSRGQTNTEVF | 0.7726175 |
| LDAISDGS | ASSRGQTNTEVF | 0.9054097 |
| LGGFAAML | ASSRGQTNTEVF | 0.8857975 |
| LGGKNPMI | ASSRGQTNTEVF | 0.5624821 |
| LGPKGVAAIL | ASSRGQTNTEVF | 0.8105669 |
| LMVGVSI | ASSRGQTNTEVF | 0.90366995 |
| LVGAAFVL | ASSRGQTNTEVF | 0.85347444 |
| LVLAVAPL | ASSRGQTNTEVF | 0.7984798 |
| MAAIAVIDAA | ASSRGQTNTEVF | 0.8318259 |
| NNGINAELQ | ASSRGQTNTEVF | 0.90409964 |
| RGEPLVLVHG | ASSRGQTNTEVF | 0.77930725 |
| RGGAGGAAGLI | ASSRGQTNTEVF | 0.9090761 |
| RSYRMHPAV | ASSRGQTNTEVF | 0.2932948 |
| SAAHLLAGQLA | ASSRGQTNTEVF | 0.8459294 |
| SAAVGSY | ASSRGQTNTEVF | 0.8317336 |
| SPHAVLHRI | ASSRGQTNTEVF | 0.4760266 |
| SQIAKVL | ASSRGQTNTEVF | 0.8830778 |
| TALGHLVP | ASSRGQTNTEVF | 0.7928941 |
| TAPTTSGAA | ASSRGQTNTEVF | 0.88323784 |
| TDGDDGNGGSPGT | ASSRGQTNTEVF | 0.78058726 |
| TDGLGEN | ASSRGQTNTEVF | 0.89662427 |
| TGGPPLPGQA | ASSRGQTNTEVF | 0.8239579 |
| TIPAITI | ASSRGQTNTEVF | 0.8580221 |
| TIPGFTI | ASSRGQTNTEVF | 0.72613865 |
| TSIASGALA | ASSRGQTNTEVF | 0.8780188 |

| | | |
|---|---|---|
| TVTLPGAAI | ASSRGQTNTEVF | 0.9054431 |
| VASGLSAALP | ASSRGQTNTEVF | 0.90464306 |
| VGAGVIGI | ASSRGQTNTEVF | 0.5376071 |
| VGPKRFGV | ASSRGQTNTEVF | 0.8133751 |
| VNVAFPD | ASSRGQTNTEVF | 0.8502783 |
| VSMTNSGVS | ASSRGQTNTEVF | 0.32950738 |
| VTDVLGDLSV | ASSRGQTNTEVF | 0.26286098 |
| VVVVGAGFAGL | ASSRGQTNTEVF | 0.81245553 |
| VYTVGTVV | ASSRGQTNTEVF | 0.75840867 |
| WTTFSGP | ASSRGQTNTEVF | 0.88074535 |
| YDELVEAF | ASSRGQTNTEVF | 0.8690947 |
| YGTGGAGGNGG | ASSRGQTNTEVF | 0.8147531 |
| YTNVMAV | ASSRGQTNTEVF | 0.8763119 |
| FLEQVHQL | ASSRGQTNTEVF | 0.85841227 |
| GLSLLAVAV | ASSRGQTNTEVF | 0.090102136 |
| LEQVHQL | ASSRGQTNTEVF | 0.78046334 |
| LNAAGAI | ASSRGQTNTEVF | 0.45389366 |
| PAVHASL | ASSRGQTNTEVF | 0.90785456 |
| TELEDDA | ASSRGQTNTEVF | 0.7268176 |
| TPPGPPN | ASSRGQTNTEVF | 0.7079379 |
| TPPPPPPP | ASSRGQTNTEVF | 0.8382654 |
| TQSPALI | GARRRGGNTLY | 0.8374726 |
| EFAAAFPQ | GARRRGGNTLY | 0.8361099 |
| LLSGNLDVL | GARRRGGNTLY | 0.79255515 |
| GPANLGNYNI | GARRRGGNTLY | 0.21921323 |
| TLGPTLP | GARRRGGNTLY | 0.88668907 |
| VYAYSSCCK | GARRRGGNTLY | 0.61891603 |
| APAPAPAPNI | GARRRGGNTLY | 0.6976725 |
| PGGPVPIP | GARRRGGNTLY | 0.3760787 |
| FLTVALEP | GARRRGGNTLY | 0.46253228 |
| SQYFGANL | GARRRGGNTLY | 0.5401784 |
| GGVKPAA | GARRRGGNTLY | 0.89635533 |
| GGTGGAGGAGGW | GARRRGGNTLY | 0.7524842 |
| GVGGVAGLGGAGG | GARRRGGNTLY | 0.77974266 |
| VLLVLATTI | GARRRGGNTLY | 0.89256966 |
| IGIFAAML | GARRRGGNTLY | 0.8233148 |
| AVGDLID | GARRRGGNTLY | 0.7976784 |
| NISPLPL | GARRRGGNTLY | 0.80541563 |
| VTAVFVAI | GARRRGGNTLY | 0.675675 |
| SGDDNDAAL | GARRRGGNTLY | 0.7463299 |
| AVAALAHS | GARRRGGNTLY | 0.7515832 |
| GGVTVMATP | GARRRGGNTLY | 0.9082309 |
| AGLAANVRD | GARRRGGNTLY | 0.8385895 |
| TNRPDLIDPALLRPG | GARRRGGNTLY | 0.51348096 |
| LANLGVTAL | GARRRGGNTLY | 0.8850666 |
| TGERIEL | GARRRGGNTLY | 0.8993404 |
| NGEELAA | GARRRGGNTLY | 0.91126513 |
| VLSLVWG | GARRRGGNTLY | 0.6983572 |
| IGYALWL | GARRRGGNTLY | 0.8335867 |
| LSLLLAPL | GARRRGGNTLY | 0.82728684 |
| TLDEALK | GARRRGGNTLY | 0.8112111 |

| | | |
|---|---|---|
| TDGAGTAEGG | GARRRGGNTLY | 0.79909825 |
| NAPGGAGGNGG | GARRRGGNTLY | 0.6648269 |
| VSVFMGMGGNF | GARRRGGNTLY | 0.9020868 |
| LASYLAAL | GARRRGGNTLY | 0.91667855 |
| LVGGTPAL | GARRRGGNTLY | 0.82067215 |
| VRNADVL | GARRRGGNTLY | 0.39315873 |
| TCSSPFLVSAM | GARRRGGNTLY | 0.82244825 |
| VHITDAE | GARRRGGNTLY | 0.729657 |
| EYAAPSF | GARRRGGNTLY | 0.7741294 |
| VVDAFHGGL | GARRRGGNTLY | 0.84185547 |
| ASVVDPQ | GARRRGGNTLY | 0.4993496 |
| SYNEFGGLS | GARRRGGNTLY | 0.807637 |
| DPDGSVY | GARRRGGNTLY | 0.911126 |
| MVSTIHATE | GARRRGGNTLY | 0.7671421 |
| LAAGVADPV | GARRRGGNTLY | 0.84619087 |
| TKEQIAA | GARRRGGNTLY | 0.88589966 |
| SLASPAAL | GARRRGGNTLY | 0.8555544 |
| IGLVFYGV | GARRRGGNTLY | 0.8958188 |
| FLLDASNL | GARRRGGNTLY | 0.8883761 |
| YQVGNFP | GARRRGGNTLY | 0.8587147 |
| ESLLSQST | GARRRGGNTLY | 0.84959155 |
| LFTVGFAAVG | GARRRGGNTLY | 0.8235977 |
| ATPPTPT | GARRRGGNTLY | 0.83626693 |
| TDLATLG | GARRRGGNTLY | 0.83303297 |
| SAGGTGR | GARRRGGNTLY | 0.7900921 |
| GFSNSGS | GARRRGGNTLY | 0.87930846 |
| GGNGGSGGTSV | GARRRGGNTLY | 0.8637404 |
| GTSVATGGAGN | GARRRGGNTLY | 0.90029055 |
| NGGNGGSGGTS | GARRRGGNTLY | 0.58150065 |
| AVVKFPH | GARRRGGNTLY | 0.69532716 |
| TPPPPPP | GARRRGGNTLY | 0.88121825 |
| AAPAAAAAPA | GARRRGGNTLY | 0.24914008 |
| IALNEDL | GARRRGGNTLY | 0.34297645 |
| IVNVSSM | GARRRGGNTLY | 0.8636738 |
| TGIPPAPR | GARRRGGNTLY | 0.9055524 |
| FDLGGGTF | GARRRGGNTLY | 0.24296649 |
| FDLGGGTFDVS | GARRRGGNTLY | 0.7375089 |
| NEPTAAA | GARRRGGNTLY | 0.59561884 |
| NEPTAAALA | GARRRGGNTLY | 0.90237296 |
| GSIINISSI | GARRRGGNTLY | 0.8347132 |
| VAPGNAGTA | GARRRGGNTLY | 0.5195718 |
| TGGGASLE | GARRRGGNTLY | 0.885715 |
| VAVYNHYKRI | GARRRGGNTLY | 0.4272892 |
| TDPAPATT | GARRRGGNTLY | 0.70546746 |
| TNRPDILDPALLRPG | GARRRGGNTLY | 0.8866855 |
| LTTTTTS | GARRRGGNTLY | 0.8991089 |
| PTGSGKTL | GARRRGGNTLY | 0.6213062 |
| TGSGKTL | GARRRGGNTLY | 0.88677984 |
| TIGHVDHGK | GARRRGGNTLY | 0.79590523 |
| TIGHVDHGKTT | GARRRGGNTLY | 0.9134244 |
| GHVDHGKTT | GARRRGGNTLY | 0.79736406 |

| | | |
|---|---|---|
| AQFEHTLL | GARRRGGNTLY | 0.5340567 |
| IVNQDEL | GARRRGGNTLY | 0.8839281 |
| TNGSQFF | GARRRGGNTLY | 0.8621868 |
| TGKGIVE | GARRRGGNTLY | 0.8737318 |
| LISNASDA | GARRRGGNTLY | 0.8843781 |
| AELRTLGELTKIAKAHGAQ | GARRRGGNTLY | 0.83771646 |
| APNAVAAAEPSW | GARRRGGNTLY | 0.79123867 |
| APQVAGAATAVL | GARRRGGNTLY | 0.88380015 |
| ARRDPAAAAAILPTDA | GARRRGGNTLY | 0.8837429 |
| DAAARDVLAEAGLAEYF | GARRRGGNTLY | 0.646017 |
| DDQIVLSAKANLAKVEQ | GARRRGGNTLY | 0.8942629 |
| DELLPDTGEDSDEDGH | GARRRGGNTLY | 0.8355284 |
| ELPEAPANRATVPQD | GARRRGGNTLY | 0.76407987 |
| FGQSKIERVTGIG | GARRRGGNTLY | 0.9108361 |
| GAGGTGSTAGGAGGAGGAGGLY | GARRRGGNTLY | 0.85846806 |
| GHGGTGAAGGNGGTGGAGGSGI | GARRRGGNTLY | 0.87888265 |
| GNGGNGGAGGEGGPGGAGGAGA | GARRRGGNTLY | 0.89201057 |
| GSTAGNGGNGGNATNGTIAGQP | GARRRGGNTLY | 0.9091846 |
| GTGGDGGNGGNGGTGG | GARRRGGNTLY | 0.56415933 |
| GVRRPAGIPADAPV | GARRRGGNTLY | 0.851321 |
| GYRQVTDAHLLTLARR | GARRRGGNTLY | 0.8375205 |
| IAKMGQDGHDRG | GARRRGGNTLY | 0.8852786 |
| KAEPSIEIQYT | GARRRGGNTLY | 0.8460151 |
| KPVRIRMAADEDTVAVT | GARRRGGNTLY | 0.6813601 |
| MPPRIAGMRLLVIKPEP | GARRRGGNTLY | 0.8717718 |
| MKFYAVALFRSRGVPPTR | GARRRGGNTLY | 0.8941002 |
| NAGGFGILGAVAHSP | GARRRGGNTLY | 0.86444545 |
| NEGREITLAMAEM | GARRRGGNTLY | 0.8683366 |
| NELVEWSHPLGHRWRWE | GARRRGGNTLY | 0.9039062 |
| NLGAGGTGGNGGNPG | GARRRGGNTLY | 0.7308867 |
| NTEWPKWETVMAADPVD | GARRRGGNTLY | 0.8365444 |
| PASAAPAETQP | GARRRGGNTLY | 0.8962461 |
| QQSSDGKNGVRAAEITGE | GARRRGGNTLY | 0.8952165 |
| RAPDAVTTSWWKE | GARRRGGNTLY | 0.842093 |
| SGFGNVGAHTSGWFNQ | GARRRGGNTLY | 0.9043704 |
| SGVAGAVGQAASVGGLKV | GARRRGGNTLY | 0.74859107 |
| SNGGAGTGSAGGNGGT | GARRRGGNTLY | 0.8672831 |
| TGGAGGEGGAAG | GARRRGGNTLY | 0.7310495 |
| TGGDGGAAGAGGAPGHG | GARRRGGNTLY | 0.8400584 |
| TMPPLLAAIANAGRHGV | GARRRGGNTLY | 0.66569155 |
| TSSTAGNSGGA | GARRRGGNTLY | 0.850578 |
| VPATVPVGILC | GARRRGGNTLY | 0.87153506 |
| VQRPDSGGVVLTGRIS | GARRRGGNTLY | 0.8711313 |
| WTPNPVPATKGVPAGLVQAT | GARRRGGNTLY | 0.21660408 |
| YALKRALAKFSDRFDVVII | GARRRGGNTLY | 0.8706717 |
| YLIGSVIV | GARRRGGNTLY | 0.8853663 |
| YGQLVAVID | GARRRGGNTLY | 0.91095954 |
| VTVGGMTLPA | GARRRGGNTLY | 0.8483381 |
| VLALPAV | GARRRGGNTLY | 0.8872979 |
| VIPPTLNYET | GARRRGGNTLY | 0.77860016 |
| VGMGGMHPSSG | GARRRGGNTLY | 0.8471885 |

| | | |
|---|---|---|
| VGLAGGAATA | GARRRGGNTLY | 0.6957339 |
| VGGAGGAGGSGP | GARRRGGNTLY | 0.7987263 |
| VFAAFMV | GARRRGGNTLY | 0.77222645 |
| TLYDKLR | GARRRGGNTLY | 0.7118638 |
| TLAVFTVL | GARRRGGNTLY | 0.88246757 |
| TGSVQPARV | GARRRGGNTLY | 0.89053285 |
| TGGGAGSGGT | GARRRGGNTLY | 0.8797506 |
| SHLETSL | GARRRGGNTLY | 0.89232165 |
| SFTTGMT | GARRRGGNTLY | 0.89306545 |
| QVEVAKRMGVRQPTVS | GARRRGGNTLY | 0.9027872 |
| QGSGGPALAM | GARRRGGNTLY | 0.79667795 |
| MLAPSNP | GARRRGGNTLY | 0.7700022 |
| LVNNAAI | GARRRGGNTLY | 0.60543895 |
| LTGEINTLA | GARRRGGNTLY | 0.7851983 |
| LSIFGPV | GARRRGGNTLY | 0.79181695 |
| LLALLANL | GARRRGGNTLY | 0.84949815 |
| LISNASDALDKLR | GARRRGGNTLY | 0.14324653 |
| LINEAVP | GARRRGGNTLY | 0.82829964 |
| LGYAFSGL | GARRRGGNTLY | 0.9020044 |
| LDFVAVAL | GARRRGGNTLY | 0.77009726 |
| LAQPGAE | GARRRGGNTLY | 0.8754753 |
| ISNLGAVTGF | GARRRGGNTLY | 0.8197976 |
| HQEDPGRI | GARRRGGNTLY | 0.78123105 |
| GTPGEPTP | GARRRGGNTLY | 0.87789094 |
| GFFNSGS | GARRRGGNTLY | 0.7586302 |
| GAGGGGLTAGAN | GARRRGGNTLY | 0.8873693 |
| FVDALVNM | GARRRGGNTLY | 0.9045826 |
| FLTVAPEL | GARRRGGNTLY | 0.840829 |
| FETLTAL | GARRRGGNTLY | 0.8470925 |
| DVINPIEL | GARRRGGNTLY | 0.8665546 |
| APVSGAPV | GARRRGGNTLY | 0.4280126 |
| ALVGGLAW | GARRRGGNTLY | 0.85609806 |
| ALAAAAATAI | GARRRGGNTLY | 0.12043793 |
| AFGLGLLL | GARRRGGNTLY | 0.8180603 |
| AFFAAASL | GARRRGGNTLY | 0.72948617 |
| AAAALPAFSPP | GARRRGGNTLY | 0.8579523 |
| VVLAPAL | GARRRGGNTLY | 0.83166087 |
| TIGHVDHGKTTL | GARRRGGNTLY | 0.8908787 |
| LISNASDALDK | GARRRGGNTLY | 0.68519634 |
| LISNASDAL | GARRRGGNTLY | 0.8218604 |
| ISNASDALDK | GARRRGGNTLY | 0.75556016 |
| NQAETLVYQTE | GARRRGGNTLY | 0.6354005 |
| TGGTGGTAGAG | GARRRGGNTLY | 0.8763759 |
| TGSTAGGAGGA | GARRRGGNTLY | 0.8593594 |
| DPLAVPVAAGPV | GARRRGGNTLY | 0.7892393 |
| GGTGGTPGNGGN | GARRRGGNTLY | 0.8909385 |
| PAGLAGVDPATL | GARRRGGNTLY | 0.85714066 |
| GNNLVVEAPHAGASV | GARRRGGNTLY | 0.44762313 |
| KVTRIMSASTLEQPA | GARRRGGNTLY | 0.7884885 |
| LGFALPIALPRRTVV | GARRRGGNTLY | 0.836895 |
| TRLGNLTARKLVVLV | GARRRGGNTLY | 0.8259016 |

| | | |
|---|---|---|
| WSQQRRLIYFPSAGP | GARRRGGNTLY | 0.1276633 |
| GPNGNPDPMAAAVDIR | GARRRGGNTLY | 0.8634436 |
| TQRCVSVADLRAAASR | GARRRGGNTLY | 0.76646763 |
| QFVLDVAEHPVEVLTETV | GARRRGGNTLY | 0.67623925 |
| GGAGGTGGAGGNGGTQAGDG | GARRRGGNTLY | 0.7915671 |
| PLLATLRAIAVDVDAKPAQV | GARRRGGNTLY | 0.87804186 |
| APKFRTGGYMIDFWGVGYQVAK | GARRRGGNTLY | 0.82365453 |
| GAGGAGGAGGAAGGPGGTGGTG | GARRRGGNTLY | 0.8371633 |
| IGALGLLATDVAPAGSVAFGVA | GARRRGGNTLY | 0.8780726 |
| AGGAGGIGGNGGAGGAGGVLMG | GARRRGGNTLY | 0.88427186 |
| ADELIGLL | GARRRGGNTLY | 0.9007695 |
| ADRALKA | GARRRGGNTLY | 0.87023675 |
| AGAVGQVGGAGGA | GARRRGGNTLY | 0.86448 |
| AGGGGGLGGAP | GARRRGGNTLY | 0.8842075 |
| AGGLLYGNGGAG | GARRRGGNTLY | 0.8438108 |
| AGLESIGL | GARRRGGNTLY | 0.4771661 |
| AGLKTGII | GARRRGGNTLY | 0.49145395 |
| ASLAALLL | GARRRGGNTLY | 0.81137407 |
| ASVINISSI | GARRRGGNTLY | 0.4472042 |
| ATPPGPA | GARRRGGNTLY | 0.86078393 |
| ATVASALV | GARRRGGNTLY | 0.91001785 |
| ATVASLIL | GARRRGGNTLY | 0.8652506 |
| AVNGPVLIRH | GARRRGGNTLY | 0.57716966 |
| DGAAGAVGI | GARRRGGNTLY | 0.8173797 |
| DPLLHGI | GARRRGGNTLY | 0.73686796 |
| ELGAALGAL | GARRRGGNTLY | 0.8724289 |
| ELGRTQAV | GARRRGGNTLY | 0.90047 |
| FVAGGYLS | GARRRGGNTLY | 0.9016751 |
| GASGGAGGAGGW | GARRRGGNTLY | 0.8889453 |
| GGAGAGGDGGAGGI | GARRRGGNTLY | 0.7775879 |
| GTGHNGSQPGMGGNG | GARRRGGNTLY | 0.8814719 |
| GYAAASASAS | GARRRGGNTLY | 0.80306244 |
| HQVDVAAVQQG | GARRRGGNTLY | 0.7917174 |
| IGAAIAHT | GARRRGGNTLY | 0.83342826 |
| IGMIEVAQL | GARRRGGNTLY | 0.873368 |
| IRLAPEHTSAMD | GARRRGGNTLY | 0.86062586 |
| IYNVSVL | GARRRGGNTLY | 0.7954804 |
| KQGFLGM | GARRRGGNTLY | 0.50570446 |
| LESVITQL | GARRRGGNTLY | 0.89536893 |
| LGLGALVVV | GARRRGGNTLY | 0.8629738 |
| LIGNGGNGGSGGI | GARRRGGNTLY | 0.65123016 |
| LNAINAPTL | GARRRGGNTLY | 0.80316955 |
| LQLLNTGV | GARRRGGNTLY | 0.90317327 |
| LQTSIEF | GARRRGGNTLY | 0.9035127 |
| MSFAVSEL | GARRRGGNTLY | 0.7581997 |
| NTGALGL | GARRRGGNTLY | 0.7600696 |
| PPGAGGAGGAGTT | GARRRGGNTLY | 0.8394253 |
| QLVGLGI | GARRRGGNTLY | 0.62546784 |
| RFLEQVHQL | GARRRGGNTLY | 0.8831012 |
| RGVGYVLR | GARRRGGNTLY | 0.80634534 |
| SATGVFAGVI | GARRRGGNTLY | 0.7886149 |

| | | |
|---|---|---|
| SLPAITLP | GARRRGGNTLY | 0.80428934 |
| STALAFTV | GARRRGGNTLY | 0.8524467 |
| SVLTVGLGK | GARRRGGNTLY | 0.77407897 |
| TALAGAGSGI | GARRRGGNTLY | 0.89642966 |
| TGLPFTGI | GARRRGGNTLY | 0.2111787 |
| VAEHVTV | GARRRGGNTLY | 0.8755871 |
| VDIVLNSL | GARRRGGNTLY | 0.7020194 |
| VLTGGGAVL | GARRRGGNTLY | 0.81499994 |
| VNVGLAYGI | GARRRGGNTLY | 0.48019212 |
| VSSAAVGAV | GARRRGGNTLY | 0.825145 |
| VTSVYEAGV | GARRRGGNTLY | 0.86916876 |
| WVALDRGA | GARRRGGNTLY | 0.8943653 |
| YLVGSVVI | GARRRGGNTLY | 0.69392776 |
| GPPGTGKTLL | GARRRGGNTLY | 0.87632537 |
| GVGMTKF | GARRRGGNTLY | 0.91106856 |
| IGIGGSDL | GARRRGGNTLY | 0.75245064 |
| IGTLLAEL | GARRRGGNTLY | 0.3313885 |
| IYTSGTT | GARRRGGNTLY | 0.86353546 |
| LSFPAGV | GARRRGGNTLY | 0.80397165 |
| LYTSGTT | GARRRGGNTLY | 0.8835895 |
| VGAFPFGF | GARRRGGNTLY | 0.88540447 |
| VGAGTVEF | GARRRGGNTLY | 0.5929582 |
| VGKSTLLL | GARRRGGNTLY | 0.89876735 |
| AALAHAGGGIV | GARRRGGNTLY | 0.8823798 |
| AALVAVSLL | GARRRGGNTLY | 0.76016045 |
| AAVSRAVAAL | GARRRGGNTLY | 0.7174587 |
| AGGSSGIGGF | GARRRGGNTLY | 0.736127 |
| AGLTVANANG | GARRRGGNTLY | 0.43588907 |
| AGLVGAVL | GARRRGGNTLY | 0.89135027 |
| ALSQAFTV | GARRRGGNTLY | 0.74279296 |
| ALTFSVSL | GARRRGGNTLY | 0.7928593 |
| ALTGAALVAL | GARRRGGNTLY | 0.8334048 |
| ASFSIGAGVL | GARRRGGNTLY | 0.80176795 |
| DGSVYKI | GARRRGGNTLY | 0.7956712 |
| EQLAEIV | GARRRGGNTLY | 0.7936517 |
| FAVSELTVM | GARRRGGNTLY | 0.8357375 |
| GAAPLAAGL | GARRRGGNTLY | 0.86889744 |
| GADGGDGGSGGAGG | GARRRGGNTLY | 0.81599164 |
| GFGLDTGGAGG | GARRRGGNTLY | 0.8482766 |
| GLASGGLL | GARRRGGNTLY | 0.89095145 |
| GRIAGAAVLL | GARRRGGNTLY | 0.6359589 |
| GSAGSLNTSF | GARRRGGNTLY | 0.7704909 |
| GTGQIGI | GARRRGGNTLY | 0.8847118 |
| GTLIALIL | GARRRGGNTLY | 0.67041814 |
| IAATMGFVVG | GARRRGGNTLY | 0.8763491 |
| IFAQALL | GARRRGGNTLY | 0.8522435 |
| IGAAVALGF | GARRRGGNTLY | 0.85802644 |
| IGLTPLAL | GARRRGGNTLY | 0.83348477 |
| ILGSNTSTL | GARRRGGNTLY | 0.9077394 |
| IVFSLTVL | GARRRGGNTLY | 0.875743 |
| LASKATGF | GARRRGGNTLY | 0.8144908 |

| | | |
|---|---|---|
| LASVALIL | GARRRGGNTLY | 0.85233504 |
| LFAVGLTL | GARRRGGNTLY | 0.74105304 |
| LFNGGIGTYI | GARRRGGNTLY | 0.8861904 |
| LTLGAGII | GARRRGGNTLY | 0.8886563 |
| LVAAALGDI | GARRRGGNTLY | 0.66923654 |
| NTAFVQLGI | GARRRGGNTLY | 0.7996082 |
| QLFSGAEL | GARRRGGNTLY | 0.91102695 |
| SAATFITL | GARRRGGNTLY | 0.88804436 |
| SNYGLANLGV | GARRRGGNTLY | 0.5887915 |
| SYGGAYAVM | GARRRGGNTLY | 0.7658434 |
| TTGVGGLTLG | GARRRGGNTLY | 0.6548451 |
| TTQTIYI | GARRRGGNTLY | 0.86687434 |
| VAAGAGLGAVI | GARRRGGNTLY | 0.6028472 |
| VAAVVTLL | GARRRGGNTLY | 0.86983967 |
| VGFLGGLI | GARRRGGNTLY | 0.78419507 |
| VGVIPILL | GARRRGGNTLY | 0.8711967 |
| VIFASSLI | GARRRGGNTLY | 0.86000323 |
| VKGGDGGAAAG | GARRRGGNTLY | 0.9034171 |
| VTKTFTNI | GARRRGGNTLY | 0.8026752 |
| VVAAAILL | GARRRGGNTLY | 0.8076751 |
| YAALDVEL | GARRRGGNTLY | 0.52010953 |
| LASVALLL | GARRRGGNTLY | 0.58709085 |
| LDAVHRV | GARRRGGNTLY | 0.7763416 |
| QGLSSSIF | GARRRGGNTLY | 0.81171376 |
| RGTFANIRL | GARRRGGNTLY | 0.33535206 |
| RQLWWGHRI | GARRRGGNTLY | 0.9038904 |
| YGPPGTGKTLL | GARRRGGNTLY | 0.8384512 |
| AALVGAPANVA | GARRRGGNTLY | 0.88523376 |
| AATFRRTE | GARRRGGNTLY | 0.7814233 |
| AGALGGGI | GARRRGGNTLY | 0.78711736 |
| AGGAGGAGGIGAG | GARRRGGNTLY | 0.7812062 |
| AGSGALGL | GARRRGGNTLY | 0.8816784 |
| AQIADQI | GARRRGGNTLY | 0.6926482 |
| ASAVAVTYL | GARRRGGNTLY | 0.3315508 |
| ASTVLGGTGG | GARRRGGNTLY | 0.7790246 |
| AVGGEPAK | GARRRGGNTLY | 0.5215931 |
| CEQGTFGGTC | GARRRGGNTLY | 0.9081844 |
| DEGKAPAG | GARRRGGNTLY | 0.88653994 |
| DTVVFPAL | GARRRGGNTLY | 0.84473026 |
| FEGITMAG | GARRRGGNTLY | 0.90778613 |
| FPLVVQD | GARRRGGNTLY | 0.24106076 |
| GAFTTPQ | GARRRGGNTLY | 0.7647946 |
| GFGGFGTAGGVGGA | GARRRGGNTLY | 0.9078262 |
| GFLAQFL | GARRRGGNTLY | 0.8659004 |
| GGAGGAGGNAGVA | GARRRGGNTLY | 0.808924 |
| GGVGTAGAGGAGGAGG | GARRRGGNTLY | 0.8963785 |
| GNGGGGDAGNAGSGGN | GARRRGGNTLY | 0.74848706 |
| GTGGAGGAGNDAGS | GARRRGGNTLY | 0.8355124 |
| GTTNTGWFNS | GARRRGGNTLY | 0.83140576 |
| IFMGPLFTL | GARRRGGNTLY | 0.8839382 |
| ILAHAIEAI | GARRRGGNTLY | 0.877911 |

| | | |
|---|---|---|
| IRSAIREAL | GARRRGGNTLY | 0.6713496 |
| ITFAGGG | GARRRGGNTLY | 0.8811188 |
| ITGMSGHQEV | GARRRGGNTLY | 0.9059895 |
| IVAGLACTL | GARRRGGNTLY | 0.77348936 |
| IVGGGLAGA | GARRRGGNTLY | 0.88037026 |
| IVTGVLNI | GARRRGGNTLY | 0.85920334 |
| LAQQGCI | GARRRGGNTLY | 0.85191506 |
| LCLHVEN | GARRRGGNTLY | 0.72868073 |
| LDAISDGS | GARRRGGNTLY | 0.74317753 |
| LGGFAAML | GARRRGGNTLY | 0.825004 |
| LGGKNPMI | GARRRGGNTLY | 0.6886895 |
| LGPKGVAAIL | GARRRGGNTLY | 0.8416537 |
| LMVGVSI | GARRRGGNTLY | 0.79940224 |
| LVGAAFVL | GARRRGGNTLY | 0.9209676 |
| LVLAVAPL | GARRRGGNTLY | 0.8675588 |
| MAAIAVIDAA | GARRRGGNTLY | 0.86519754 |
| NNGINAELQ | GARRRGGNTLY | 0.7232383 |
| RGEPLVLVHG | GARRRGGNTLY | 0.8744732 |
| RGGAGGAAGLI | GARRRGGNTLY | 0.85638636 |
| RSYRMHPAV | GARRRGGNTLY | 0.47526437 |
| SAAHLLAGQLA | GARRRGGNTLY | 0.89999896 |
| SAAVGSY | GARRRGGNTLY | 0.7339203 |
| SPHAVLHRI | GARRRGGNTLY | 0.8292719 |
| SQIAKVL | GARRRGGNTLY | 0.8938113 |
| TALGHLVP | GARRRGGNTLY | 0.8866006 |
| TAPTTSGAA | GARRRGGNTLY | 0.81523556 |
| TDGDDGNGGSPGT | GARRRGGNTLY | 0.83124065 |
| TDGLGEN | GARRRGGNTLY | 0.7106029 |
| TGGPPLPGQA | GARRRGGNTLY | 0.89285433 |
| TIPAITI | GARRRGGNTLY | 0.88968027 |
| TIPGFTI | GARRRGGNTLY | 0.85072553 |
| TSIASGALA | GARRRGGNTLY | 0.90599984 |
| TVTLPGAAI | GARRRGGNTLY | 0.889295 |
| VASGLSAALP | GARRRGGNTLY | 0.5766581 |
| VGAGVIGI | GARRRGGNTLY | 0.8360287 |
| VGPKRFGV | GARRRGGNTLY | 0.88789225 |
| VNVAFPD | GARRRGGNTLY | 0.8891094 |
| VSMTNSGVS | GARRRGGNTLY | 0.9023928 |
| VTDVLGDLSV | GARRRGGNTLY | 0.870517 |
| VVVVGAGFAGL | GARRRGGNTLY | 0.7823193 |
| VYTVGTVV | GARRRGGNTLY | 0.89920545 |
| WTTFSGP | GARRRGGNTLY | 0.9224571 |
| YDELVEAF | GARRRGGNTLY | 0.8882446 |
| YGTGGAGGNGG | GARRRGGNTLY | 0.8318955 |
| YTNVMAV | GARRRGGNTLY | 0.842957 |
| FLEQVHQL | GARRRGGNTLY | 0.8575496 |
| GLSLLAVAV | GARRRGGNTLY | 0.88519096 |
| LEQVHQL | GARRRGGNTLY | 0.72911155 |
| LNAAGAI | GARRRGGNTLY | 0.8364186 |
| PAVHASL | GARRRGGNTLY | 0.85800195 |
| TELEDDA | GARRRGGNTLY | 0.46178484 |

| | | |
|---|---|---|
| TPPGPPN | GARRRGGNTLY | 0.86259365 |
| TPPPPPPP | GARRRGGNTLY | 0.88783556 |
| TQSPALI | ASSIDGGCTEVF | 0.8494371 |
| EFAAAFPQ | ASSIDGGCTEVF | 0.76847714 |
| LLSGNLDVL | ASSIDGGCTEVF | 0.88749975 |
| GPANLGNYNI | ASSIDGGCTEVF | 0.9103564 |
| TLGPTLP | ASSIDGGCTEVF | 0.90064406 |
| VYAYSSCCK | ASSIDGGCTEVF | 0.4680216 |
| APAPAPAPNI | ASSIDGGCTEVF | 0.31428528 |
| PGGPVPIP | ASSIDGGCTEVF | 0.8040798 |
| FLTVALEP | ASSIDGGCTEVF | 0.8708147 |
| SQYFGANL | ASSIDGGCTEVF | 0.80748117 |
| GGVKPAA | ASSIDGGCTEVF | 0.7753598 |
| GGTGGAGGAGGW | ASSIDGGCTEVF | 0.8274684 |
| GVGGVAGLGGAGG | ASSIDGGCTEVF | 0.8912184 |
| VLLVLATTI | ASSIDGGCTEVF | 0.71675587 |
| IGIFAAML | ASSIDGGCTEVF | 0.74347353 |
| AVGDLID | ASSIDGGCTEVF | 0.846601 |
| NISPLPL | ASSIDGGCTEVF | 0.89244473 |
| VTAVFVAI | ASSIDGGCTEVF | 0.6828043 |
| SGDDNDAAL | ASSIDGGCTEVF | 0.7944522 |
| AVAALAHS | ASSIDGGCTEVF | 0.79870665 |
| GGVTVMATP | ASSIDGGCTEVF | 0.8114053 |
| AGLAANVRD | ASSIDGGCTEVF | 0.8459184 |
| TNRPDLIDPALLRPG | ASSIDGGCTEVF | 0.751035 |
| LANLGVTAL | ASSIDGGCTEVF | 0.8818624 |
| TGERIEL | ASSIDGGCTEVF | 0.6356982 |
| NGEELAA | ASSIDGGCTEVF | 0.8976029 |
| VLSLVWG | ASSIDGGCTEVF | 0.8584126 |
| IGYALWL | ASSIDGGCTEVF | 0.87045515 |
| LSLLLAPL | ASSIDGGCTEVF | 0.78449845 |
| TLDEALK | ASSIDGGCTEVF | 0.72924745 |
| TDGAGTAEGG | ASSIDGGCTEVF | 0.6588041 |
| NAPGGAGGNGG | ASSIDGGCTEVF | 0.61771107 |
| VSVFMGMGGNF | ASSIDGGCTEVF | 0.8894756 |
| LASYLAAL | ASSIDGGCTEVF | 0.39578715 |
| LVGGTPAL | ASSIDGGCTEVF | 0.89131606 |
| VRNADVL | ASSIDGGCTEVF | 0.8689903 |
| TCSSPFLVSAM | ASSIDGGCTEVF | 0.6826068 |
| VHITDAE | ASSIDGGCTEVF | 0.8769014 |
| EYAAPSF | ASSIDGGCTEVF | 0.6904539 |
| VVDAFHGGL | ASSIDGGCTEVF | 0.7773061 |
| ASVVDPQ | ASSIDGGCTEVF | 0.8822148 |
| SYNEFGGLS | ASSIDGGCTEVF | 0.8831882 |
| DPDGSVY | ASSIDGGCTEVF | 0.7278983 |
| MVSTIHATE | ASSIDGGCTEVF | 0.8722799 |
| LAAGVADPV | ASSIDGGCTEVF | 0.849531 |
| TKEQIAA | ASSIDGGCTEVF | 0.7503866 |
| SLASPAAL | ASSIDGGCTEVF | 0.8531072 |
| IGLVFYGV | ASSIDGGCTEVF | 0.8888838 |
| FLLDASNL | ASSIDGGCTEVF | 0.87612814 |

| | | |
|---|---|---|
| YQVGNFP | ASSIDGGCTEVF | 0.6513177 |
| ESLLSQST | ASSIDGGCTEVF | 0.8606385 |
| LFTVGFAAVG | ASSIDGGCTEVF | 0.70756626 |
| ATPPTPT | ASSIDGGCTEVF | 0.8483768 |
| TDLATLG | ASSIDGGCTEVF | 0.8301556 |
| SAGGTGR | ASSIDGGCTEVF | 0.8336704 |
| GFSNSGS | ASSIDGGCTEVF | 0.8738478 |
| GGNGGSGGTSV | ASSIDGGCTEVF | 0.66890943 |
| GTSVATGGAGN | ASSIDGGCTEVF | 0.740998 |
| NGGNGGSGGTS | ASSIDGGCTEVF | 0.8172027 |
| AVVKFPH | ASSIDGGCTEVF | 0.8440348 |
| TPPPPPP | ASSIDGGCTEVF | 0.8804282 |
| AAPAAAAAPA | ASSIDGGCTEVF | 0.70363027 |
| IALNEDL | ASSIDGGCTEVF | 0.8532785 |
| IVNVSSM | ASSIDGGCTEVF | 0.8020209 |
| TGIPPAPR | ASSIDGGCTEVF | 0.7224909 |
| FDLGGGTF | ASSIDGGCTEVF | 0.8592415 |
| FDLGGGTFDVS | ASSIDGGCTEVF | 0.80373955 |
| NEPTAAA | ASSIDGGCTEVF | 0.71512187 |
| NEPTAAALA | ASSIDGGCTEVF | 0.8777485 |
| GSIINISSI | ASSIDGGCTEVF | 0.7003207 |
| VAPGNAGTA | ASSIDGGCTEVF | 0.88881946 |
| TGGGASLE | ASSIDGGCTEVF | 0.9081727 |
| VAVYNHYKRI | ASSIDGGCTEVF | 0.8081155 |
| TDPAPATT | ASSIDGGCTEVF | 0.8945386 |
| TNRPDILDPALLRPG | ASSIDGGCTEVF | 0.8517695 |
| LTTTTTS | ASSIDGGCTEVF | 0.84517384 |
| PTGSGKTL | ASSIDGGCTEVF | 0.8123142 |
| TGSGKTL | ASSIDGGCTEVF | 0.7346239 |
| TIGHVDHGK | ASSIDGGCTEVF | 0.89205223 |
| TIGHVDHGKTT | ASSIDGGCTEVF | 0.8799093 |
| GHVDHGKTT | ASSIDGGCTEVF | 0.88651776 |
| AQFEHTLL | ASSIDGGCTEVF | 0.8597674 |
| IVNQDEL | ASSIDGGCTEVF | 0.8598391 |
| TNGSQFF | ASSIDGGCTEVF | 0.882244 |
| TGKGIVE | ASSIDGGCTEVF | 0.75337076 |
| LISNASDA | ASSIDGGCTEVF | 0.63628614 |
| AELRTLGELTKIAKAHGAQ | ASSIDGGCTEVF | 0.79338706 |
| APNAVAAAEPSW | ASSIDGGCTEVF | 0.7661444 |
| APQVAGAATAVL | ASSIDGGCTEVF | 0.75689846 |
| ARRDPAAAAAILPTDA | ASSIDGGCTEVF | 0.6704602 |
| DAAARDVLAEAGLAEYF | ASSIDGGCTEVF | 0.8851806 |
| DDQIVLSAKANLAKVEQ | ASSIDGGCTEVF | 0.9147365 |
| DELLPDTGEDSDEDGH | ASSIDGGCTEVF | 0.7823043 |
| ELPEAPANRATVPQD | ASSIDGGCTEVF | 0.9104248 |
| FGQSKIERVTGIG | ASSIDGGCTEVF | 0.6278247 |
| GAGGTGSTAGGAGGAGGAGGLY | ASSIDGGCTEVF | 0.8840127 |
| GHGGTGAAGGNGGTGGAGGSGI | ASSIDGGCTEVF | 0.8938427 |
| GNGGNGGAGGEGGPGGAGGAGA | ASSIDGGCTEVF | 0.725049 |
| GSTAGNGGNGGNATNGTIAGQP | ASSIDGGCTEVF | 0.8764314 |
| GTGGDGGNGGNGGTGG | ASSIDGGCTEVF | 0.6835153 |

| | | |
|---|---|---|
| GVRRPAGIPADAPV | ASSIDGGCTEVF | 0.794428 |
| GYRQVTDAHLLTLARR | ASSIDGGCTEVF | 0.89593685 |
| IAKMGQDGHDRG | ASSIDGGCTEVF | 0.8150468 |
| KAEPSIEIQYT | ASSIDGGCTEVF | 0.8653215 |
| KPVRIRMAADEDTVAVT | ASSIDGGCTEVF | 0.8284733 |
| MPPRIAGMRLLVIKPEP | ASSIDGGCTEVF | 0.7536552 |
| MKFYAVALFRSRGVPPTR | ASSIDGGCTEVF | 0.85864455 |
| NAGGFGILGAVAHSP | ASSIDGGCTEVF | 0.9025855 |
| NEGREITLAMAEM | ASSIDGGCTEVF | 0.79679394 |
| NELVEWSHPLGHRWRWE | ASSIDGGCTEVF | 0.9210386 |
| NLGAGGTGGNGGNPG | ASSIDGGCTEVF | 0.7208061 |
| NTEWPKWETVMAADPVD | ASSIDGGCTEVF | 0.89651775 |
| PASAAPAETQP | ASSIDGGCTEVF | 0.88613546 |
| QQSSDGKNGVRAAEITGE | ASSIDGGCTEVF | 0.8289539 |
| RAPDAVTTSWWKE | ASSIDGGCTEVF | 0.8177413 |
| SGFGNVGAHTSGWFNQ | ASSIDGGCTEVF | 0.88035595 |
| SGVAGAVGQAASVGGLKV | ASSIDGGCTEVF | 0.8249123 |
| SNGGAGTGSAGGNGGT | ASSIDGGCTEVF | 0.8793731 |
| TGGAGGEGGAAG | ASSIDGGCTEVF | 0.9068012 |
| TGGDGGAAGAGGAPGHG | ASSIDGGCTEVF | 0.30277318 |
| TMPPLLAAIANAGRHGV | ASSIDGGCTEVF | 0.77242005 |
| TSSTAGNSGGA | ASSIDGGCTEVF | 0.5990208 |
| VPATVPVGILC | ASSIDGGCTEVF | 0.6018477 |
| VQRPDSGGVVLTGRIS | ASSIDGGCTEVF | 0.28892237 |
| WTPNPVPATKGVPAGLVQAT | ASSIDGGCTEVF | 0.19328664 |
| YALKRALAKFSDRFDVVII | ASSIDGGCTEVF | 0.5001668 |
| YLIGSVIV | ASSIDGGCTEVF | 0.25044966 |
| YGQLVAVID | ASSIDGGCTEVF | 0.8045915 |
| VTVGGMTLPA | ASSIDGGCTEVF | 0.6253733 |
| VLALPAV | ASSIDGGCTEVF | 0.30782816 |
| VIPPTLNYET | ASSIDGGCTEVF | 0.6804801 |
| VGMGGMHPSSG | ASSIDGGCTEVF | 0.7377452 |
| VGLAGGAATA | ASSIDGGCTEVF | 0.1834268 |
| VGGAGGAGGSGP | ASSIDGGCTEVF | 0.49969018 |
| VFAAFMV | ASSIDGGCTEVF | 0.25311854 |
| TLYDKLR | ASSIDGGCTEVF | 0.42787045 |
| TLAVFTVL | ASSIDGGCTEVF | 0.15474279 |
| TGSVQPARV | ASSIDGGCTEVF | 0.6183081 |
| TGGGAGSGGT | ASSIDGGCTEVF | 0.31211266 |
| SHLETSL | ASSIDGGCTEVF | 0.12751459 |
| SFTTGMT | ASSIDGGCTEVF | 0.54719454 |
| QVEVAKRMGVRQPTVS | ASSIDGGCTEVF | 0.4407335 |
| QGSGGPALAM | ASSIDGGCTEVF | 0.36706978 |
| MLAPSNP | ASSIDGGCTEVF | 0.6409935 |
| LVNNAAI | ASSIDGGCTEVF | 0.4735499 |
| LTGEINTLA | ASSIDGGCTEVF | 0.29415792 |
| LSIFGPV | ASSIDGGCTEVF | 0.6183117 |
| LLALLANL | ASSIDGGCTEVF | 0.68925256 |
| LISNASDALDKLR | ASSIDGGCTEVF | 0.27520385 |
| LINEAVP | ASSIDGGCTEVF | 0.50135076 |
| LGYAFSGL | ASSIDGGCTEVF | 0.7902888 |

| | | |
|---|---|---|
| LDFVAVAL | ASSIDGGCTEVF | 0.33793804 |
| LAQPGAE | ASSIDGGCTEVF | 0.78024924 |
| ISNLGAVTGF | ASSIDGGCTEVF | 0.8106799 |
| HQEDPGRI | ASSIDGGCTEVF | 0.3037219 |
| GTPGEPTP | ASSIDGGCTEVF | 0.6975745 |
| GFFNSGS | ASSIDGGCTEVF | 0.7458684 |
| GAGGGGLTAGAN | ASSIDGGCTEVF | 0.4308048 |
| FVDALVNM | ASSIDGGCTEVF | 0.48157352 |
| FLTVAPEL | ASSIDGGCTEVF | 0.23436272 |
| FETLTAL | ASSIDGGCTEVF | 0.46999794 |
| DVINPIEL | ASSIDGGCTEVF | 0.18833373 |
| APVSGAPV | ASSIDGGCTEVF | 0.5621557 |
| ALVGGLAW | ASSIDGGCTEVF | 0.6713946 |
| ALAAAAATAI | ASSIDGGCTEVF | 0.40408692 |
| AFGLGLLL | ASSIDGGCTEVF | 0.2021688 |
| AFFAAASL | ASSIDGGCTEVF | 0.2150201 |
| AAAALPAFSPP | ASSIDGGCTEVF | 0.56748843 |
| VVLAPAL | ASSIDGGCTEVF | 0.30678153 |
| TIGHVDHGKTTL | ASSIDGGCTEVF | 0.2433798 |
| LISNASDALDK | ASSIDGGCTEVF | 0.3258257 |
| LISNASDAL | ASSIDGGCTEVF | 0.57343996 |
| ISNASDALDK | ASSIDGGCTEVF | 0.63540274 |
| NQAETLVYQTE | ASSIDGGCTEVF | 0.28186405 |
| TGGTGGTAGAG | ASSIDGGCTEVF | 0.7891437 |
| TGSTAGGAGGA | ASSIDGGCTEVF | 0.79167557 |
| DPLAVPVAAGPV | ASSIDGGCTEVF | 0.77557254 |
| GGTGGTPGNGGN | ASSIDGGCTEVF | 0.49183178 |
| PAGLAGVDPATL | ASSIDGGCTEVF | 0.28678373 |
| GNNLVVEAPHAGASV | ASSIDGGCTEVF | 0.8035076 |
| KVTRIMSASTLEQPA | ASSIDGGCTEVF | 0.2666008 |
| LGFALPIALPRRTVV | ASSIDGGCTEVF | 0.5600363 |
| TRLGNLTARKLVVLV | ASSIDGGCTEVF | 0.6388308 |
| WSQQRRLIYFPSAGP | ASSIDGGCTEVF | 0.386764 |
| GPNGNPDPMAAAVDIR | ASSIDGGCTEVF | 0.25245768 |
| TQRCVSVADLRAAASR | ASSIDGGCTEVF | 0.56824744 |
| QFVLDVAEHPVEVLTETV | ASSIDGGCTEVF | 0.76834995 |
| GGAGGTGGAGGNGGTQAGDG | ASSIDGGCTEVF | 0.20912369 |
| PLLATLRAIAVDVDAKPAQV | ASSIDGGCTEVF | 0.2344316 |
| APKFRTGGYMIDFWGVGYQVAK | ASSIDGGCTEVF | 0.50113 |
| GAGGAGGAGGAAGGPGGTGGTG | ASSIDGGCTEVF | 0.49040398 |
| IGALGLLATDVAPAGSVAFGVA | ASSIDGGCTEVF | 0.72872597 |
| AGGAGGIGGNGGAGGAGGVLMG | ASSIDGGCTEVF | 0.6030727 |
| ADELIGLL | ASSIDGGCTEVF | 0.39740652 |
| ADRALKA | ASSIDGGCTEVF | 0.66901964 |
| AGAVGQVGGAGGA | ASSIDGGCTEVF | 0.6127796 |
| AGGGGGLGGAP | ASSIDGGCTEVF | 0.4667419 |
| AGGLLYGNGGAG | ASSIDGGCTEVF | 0.6486764 |
| AGLESIGL | ASSIDGGCTEVF | 0.2756374 |
| AGLKTGII | ASSIDGGCTEVF | 0.74157757 |
| ASLAALLL | ASSIDGGCTEVF | 0.70300627 |
| ASVINISSI | ASSIDGGCTEVF | 0.68893653 |

| | | |
|---|---|---|
| ATPPGPA | ASSIDGGCTEVF | 0.7680605 |
| ATVASALV | ASSIDGGCTEVF | 0.46504247 |
| ATVASLIL | ASSIDGGCTEVF | 0.48520035 |
| AVNGPVLIRH | ASSIDGGCTEVF | 0.16781835 |
| DGAAGAVGI | ASSIDGGCTEVF | 0.22167405 |
| DPLLHGI | ASSIDGGCTEVF | 0.5094307 |
| ELGAALGAL | ASSIDGGCTEVF | 0.52828985 |
| ELGRTQAV | ASSIDGGCTEVF | 0.72991014 |
| FVAGGYLS | ASSIDGGCTEVF | 0.6479856 |
| GASGGAGGAGGW | ASSIDGGCTEVF | 0.25010556 |
| GGAGAGGDGGAGGI | ASSIDGGCTEVF | 0.81927025 |
| GTGHNGSQPGMGGNG | ASSIDGGCTEVF | 0.77293515 |
| GYAAASASAS | ASSIDGGCTEVF | 0.17318359 |
| HQVDVAAVQQG | ASSIDGGCTEVF | 0.3482483 |
| IGAAIAHT | ASSIDGGCTEVF | 0.828769 |
| IGMIEVAQL | ASSIDGGCTEVF | 0.681453 |
| IRLAPEHTSAMD | ASSIDGGCTEVF | 0.34038952 |
| IYNVSVL | ASSIDGGCTEVF | 0.4588893 |
| KQGFLGM | ASSIDGGCTEVF | 0.8329791 |
| LESVITQL | ASSIDGGCTEVF | 0.5253815 |
| LGLGALVVV | ASSIDGGCTEVF | 0.80354017 |
| LIGNGGNGGSGGI | ASSIDGGCTEVF | 0.14449875 |
| LNAINAPTL | ASSIDGGCTEVF | 0.5098237 |
| LQLLNTGV | ASSIDGGCTEVF | 0.36998576 |
| LQTSIEF | ASSIDGGCTEVF | 0.14058416 |
| MSFAVSEL | ASSIDGGCTEVF | 0.56519294 |
| NTGALGL | ASSIDGGCTEVF | 0.44246793 |
| PPGAGGAGGAGTT | ASSIDGGCTEVF | 0.66348976 |
| QLVGLGI | ASSIDGGCTEVF | 0.34324986 |
| RFLEQVHQL | ASSIDGGCTEVF | 0.72569764 |
| RGVGYVLR | ASSIDGGCTEVF | 0.58414936 |
| SATGVFAGVI | ASSIDGGCTEVF | 0.4497932 |
| SLPAITLP | ASSIDGGCTEVF | 0.33975902 |
| STALAFTV | ASSIDGGCTEVF | 0.49124986 |
| SVLTVGLGK | ASSIDGGCTEVF | 0.8127244 |
| TALAGAGSGI | ASSIDGGCTEVF | 0.77791595 |
| TGLPFTGI | ASSIDGGCTEVF | 0.19911957 |
| VAEHVTV | ASSIDGGCTEVF | 0.71891505 |
| VDIVLNSL | ASSIDGGCTEVF | 0.37655362 |
| VLTGGGAVL | ASSIDGGCTEVF | 0.23134466 |
| VNVGLAYGI | ASSIDGGCTEVF | 0.18723652 |
| VSSAAVGAV | ASSIDGGCTEVF | 0.28652424 |
| VTSVYEAGV | ASSIDGGCTEVF | 0.38040096 |
| WVALDRGA | ASSIDGGCTEVF | 0.6496761 |
| YLVGSVVI | ASSIDGGCTEVF | 0.66596276 |
| GPPGTGKTLL | ASSIDGGCTEVF | 0.4020694 |
| GVGMTKF | ASSIDGGCTEVF | 0.5874126 |
| IGIGGSDL | ASSIDGGCTEVF | 0.5196815 |
| IGTLLAEL | ASSIDGGCTEVF | 0.08363564 |
| IYTSGTT | ASSIDGGCTEVF | 0.28372043 |
| LSFPAGV | ASSIDGGCTEVF | 0.1823232 |

| | | |
|---|---|---|
| LYTSGTT | ASSIDGGCTEVF | 0.8166522 |
| VGAFPFGF | ASSIDGGCTEVF | 0.23130731 |
| VGAGTVEF | ASSIDGGCTEVF | 0.26737067 |
| VGKSTLLL | ASSIDGGCTEVF | 0.4893942 |
| AALAHAGGGIV | ASSIDGGCTEVF | 0.69255185 |
| AALVAVSLL | ASSIDGGCTEVF | 0.49790347 |
| AAVSRAVAAL | ASSIDGGCTEVF | 0.28453216 |
| AGGSSGIGGF | ASSIDGGCTEVF | 0.17276534 |
| AGLTVANANG | ASSIDGGCTEVF | 0.52123946 |
| AGLVGAVL | ASSIDGGCTEVF | 0.31083643 |
| ALSQAFTV | ASSIDGGCTEVF | 0.2851792 |
| ALTFSVSL | ASSIDGGCTEVF | 0.30329248 |
| ALTGAALVAL | ASSIDGGCTEVF | 0.6379243 |
| ASFSIGAGVL | ASSIDGGCTEVF | 0.22681415 |
| DGSVYKI | ASSIDGGCTEVF | 0.73657 |
| EQLAEIV | ASSIDGGCTEVF | 0.28816745 |
| FAVSELTVM | ASSIDGGCTEVF | 0.37905568 |
| GAAPLAAGL | ASSIDGGCTEVF | 0.72234106 |
| GADGGDGGSGGAGG | ASSIDGGCTEVF | 0.6841953 |
| GFGLDTGGAGG | ASSIDGGCTEVF | 0.31617224 |
| GLASGGLL | ASSIDGGCTEVF | 0.34984177 |
| GRIAGAAVLL | ASSIDGGCTEVF | 0.22520426 |
| GSAGSLNTSF | ASSIDGGCTEVF | 0.24489021 |
| GTGQIGI | ASSIDGGCTEVF | 0.35046175 |
| GTLIALIL | ASSIDGGCTEVF | 0.81450856 |
| IAATMGFVVG | ASSIDGGCTEVF | 0.55927503 |
| IFAQALL | ASSIDGGCTEVF | 0.30819708 |
| IGAAVALGF | ASSIDGGCTEVF | 0.7344552 |
| IGLTPLAL | ASSIDGGCTEVF | 0.8175799 |
| ILGSNTSTL | ASSIDGGCTEVF | 0.81258583 |
| IVFSLTVL | ASSIDGGCTEVF | 0.20867448 |
| LASKATGF | ASSIDGGCTEVF | 0.3967328 |
| LASVALIL | ASSIDGGCTEVF | 0.42159727 |
| LFAVGLTL | ASSIDGGCTEVF | 0.3958323 |
| LFNGGIGTYI | ASSIDGGCTEVF | 0.4305599 |
| LTLGAGII | ASSIDGGCTEVF | 0.29032204 |
| LVAAALGDI | ASSIDGGCTEVF | 0.6949724 |
| NTAFVQLGI | ASSIDGGCTEVF | 0.8729286 |
| QLFSGAEL | ASSIDGGCTEVF | 0.38725707 |
| SAATFITL | ASSIDGGCTEVF | 0.15032312 |
| SNYGLANLGV | ASSIDGGCTEVF | 0.4674524 |
| SYGGAYAVM | ASSIDGGCTEVF | 0.22017643 |
| TTGVGGLTLG | ASSIDGGCTEVF | 0.270754 |
| TTQTIYI | ASSIDGGCTEVF | 0.32247534 |
| VAAGAGLGAVI | ASSIDGGCTEVF | 0.18010142 |
| VAAVVTLL | ASSIDGGCTEVF | 0.5071908 |
| VGFLGGLI | ASSIDGGCTEVF | 0.7609792 |
| VGVIPILL | ASSIDGGCTEVF | 0.25598162 |
| VIFASSLI | ASSIDGGCTEVF | 0.59325695 |
| VKGGDGGAAAG | ASSIDGGCTEVF | 0.7802947 |
| VTKTFTNI | ASSIDGGCTEVF | 0.60010576 |

| | | |
|---|---|---|
| VVAAAILL | ASSIDGGCTEVF | 0.7257886 |
| YAALDVEL | ASSIDGGCTEVF | 0.7977469 |
| LASVALLL | ASSIDGGCTEVF | 0.4895565 |
| LDAVHRV | ASSIDGGCTEVF | 0.5093176 |
| QGLSSSIF | ASSIDGGCTEVF | 0.73483694 |
| RGTFANIRL | ASSIDGGCTEVF | 0.5087856 |
| RQLWWGHRI | ASSIDGGCTEVF | 0.43342584 |
| YGPPGTGKTLL | ASSIDGGCTEVF | 0.39917636 |
| AALVGAPANVA | ASSIDGGCTEVF | 0.7332586 |
| AATFRRTE | ASSIDGGCTEVF | 0.4199186 |
| AGALGGGI | ASSIDGGCTEVF | 0.82242405 |
| AGGAGGAGGIGAG | ASSIDGGCTEVF | 0.1990177 |
| AGSGALGL | ASSIDGGCTEVF | 0.25373814 |
| AQIADQI | ASSIDGGCTEVF | 0.7297914 |
| ASAVAVTYL | ASSIDGGCTEVF | 0.12221051 |
| ASTVLGGTGG | ASSIDGGCTEVF | 0.2697727 |
| AVGGEPAK | ASSIDGGCTEVF | 0.5228985 |
| CEQGTFGGTC | ASSIDGGCTEVF | 0.75754786 |
| DEGKAPAG | ASSIDGGCTEVF | 0.13248377 |
| DTVVFPAL | ASSIDGGCTEVF | 0.36943674 |
| FEGITMAG | ASSIDGGCTEVF | 0.22096066 |
| FPLVVQD | ASSIDGGCTEVF | 0.7471795 |
| GAFTTPQ | ASSIDGGCTEVF | 0.40002522 |
| GFGGFGTAGGVGGA | ASSIDGGCTEVF | 0.16136794 |
| GFLAQFL | ASSIDGGCTEVF | 0.67128515 |
| GGAGGAGGNAGVA | ASSIDGGCTEVF | 0.17135449 |
| GGVGTAGAGGAGGAGG | ASSIDGGCTEVF | 0.5361814 |
| GNGGGGDAGNAGSGGN | ASSIDGGCTEVF | 0.66640955 |
| GTGGAGGAGNDAGS | ASSIDGGCTEVF | 0.8480302 |
| GTTNTGWFNS | ASSIDGGCTEVF | 0.23024414 |
| IFMGPLFTL | ASSIDGGCTEVF | 0.599321 |
| ILAHAIEAI | ASSIDGGCTEVF | 0.32643923 |
| IRSAIREAL | ASSIDGGCTEVF | 0.8404738 |
| ITFAGGG | ASSIDGGCTEVF | 0.32899392 |
| ITGMSGHQEV | ASSIDGGCTEVF | 0.24461208 |
| IVAGLACTL | ASSIDGGCTEVF | 0.74982643 |
| IVGGGLAGA | ASSIDGGCTEVF | 0.6728358 |
| IVTGVLNI | ASSIDGGCTEVF | 0.41679946 |
| LAQQGCI | ASSIDGGCTEVF | 0.76940864 |
| LCLHVEN | ASSIDGGCTEVF | 0.4388758 |
| LDAISDGS | ASSIDGGCTEVF | 0.44411924 |
| LGGFAAML | ASSIDGGCTEVF | 0.7101959 |
| LGGKNPMI | ASSIDGGCTEVF | 0.672904 |
| LGPKGVAAIL | ASSIDGGCTEVF | 0.30139166 |
| LMVGVSI | ASSIDGGCTEVF | 0.7195302 |
| LVGAAFVL | ASSIDGGCTEVF | 0.4637974 |
| LVLAVAPL | ASSIDGGCTEVF | 0.29627436 |
| MAAIAVIDAA | ASSIDGGCTEVF | 0.7541708 |
| NNGINAELQ | ASSIDGGCTEVF | 0.41953534 |
| RGEPLVLVHG | ASSIDGGCTEVF | 0.69265836 |
| RGGAGGAAGLI | ASSIDGGCTEVF | 0.5407817 |

| | | |
|---|---|---|
| RSYRMHPAV | ASSIDGGCTEVF | 0.85289097 |
| SAAHLLAGQLA | ASSIDGGCTEVF | 0.18185891 |
| SAAVGSY | ASSIDGGCTEVF | 0.66456723 |
| SPHAVLHRI | ASSIDGGCTEVF | 0.47450265 |
| SQIAKVL | ASSIDGGCTEVF | 0.71724427 |
| TALGHLVP | ASSIDGGCTEVF | 0.6299603 |
| TAPTTSGAA | ASSIDGGCTEVF | 0.36367413 |
| TDGDDGNGGSPGT | ASSIDGGCTEVF | 0.5555681 |
| TDGLGEN | ASSIDGGCTEVF | 0.7759161 |
| TGGPPLPGQA | ASSIDGGCTEVF | 0.5168198 |
| TIPAITI | ASSIDGGCTEVF | 0.6142285 |
| TIPGFTI | ASSIDGGCTEVF | 0.78562814 |
| TSIASGALA | ASSIDGGCTEVF | 0.3050189 |
| TVTLPGAAI | ASSIDGGCTEVF | 0.48013693 |
| VASGLSAALP | ASSIDGGCTEVF | 0.7310973 |
| VGAGVIGI | ASSIDGGCTEVF | 0.7688442 |
| VGPKRFGV | ASSIDGGCTEVF | 0.45383537 |
| VNVAFPD | ASSIDGGCTEVF | 0.77503735 |
| VSMTNSGVS | ASSIDGGCTEVF | 0.21665719 |
| VTDVLGDLSV | ASSIDGGCTEVF | 0.58566016 |
| VVVVGAGFAGL | ASSIDGGCTEVF | 0.43089157 |
| VYTVGTVV | ASSIDGGCTEVF | 0.39919284 |
| WTTFSGP | ASSIDGGCTEVF | 0.20945217 |
| YDELVEAF | ASSIDGGCTEVF | 0.46158895 |
| YGTGGAGGNGG | ASSIDGGCTEVF | 0.6477219 |
| YTNVMAV | ASSIDGGCTEVF | 0.6154962 |
| FLEQVHQL | ASSIDGGCTEVF | 0.20589292 |
| GLSLLAVAV | ASSIDGGCTEVF | 0.54242826 |
| LEQVHQL | ASSIDGGCTEVF | 0.6751511 |
| LNAAGAI | ASSIDGGCTEVF | 0.8481608 |
| PAVHASL | ASSIDGGCTEVF | 0.40033868 |
| TELEDDA | ASSIDGGCTEVF | 0.64590067 |
| TPPGPPN | ASSIDGGCTEVF | 0.2647188 |
| TPPPPPPP | ASSIDGGCTEVF | 0.43709272 |
| TQSPALI | ASSNWGYEQY | 0.1851059 |
| EFAAAFPQ | ASSNWGYEQY | 0.3218932 |
| LLSGNLDVL | ASSNWGYEQY | 0.3168542 |
| GPANLGNYNI | ASSNWGYEQY | 0.23824683 |
| TLGPTLP | ASSNWGYEQY | 0.768412 |
| VYAYSSCCK | ASSNWGYEQY | 0.7540666 |
| APAPAPAPNI | ASSNWGYEQY | 0.6022534 |
| PGGPVPIP | ASSNWGYEQY | 0.7959372 |
| FLTVALEP | ASSNWGYEQY | 0.80477655 |
| SQYFGANL | ASSNWGYEQY | 0.74638337 |
| GGVKPAA | ASSNWGYEQY | 0.29238766 |
| GGTGGAGGAGGW | ASSNWGYEQY | 0.2686433 |
| GVGGVAGLGGAGG | ASSNWGYEQY | 0.18438496 |
| VLLVLATTI | ASSNWGYEQY | 0.38102165 |
| IGIFAAML | ASSNWGYEQY | 0.29578844 |
| AVGDLID | ASSNWGYEQY | 0.5541066 |
| NISPLPL | ASSNWGYEQY | 0.1197236 |

| | | |
|---|---|---|
| VTAVFVAI | ASSNWGYEQY | 0.6494005 |
| SGDDNDAAL | ASSNWGYEQY | 0.80702305 |
| AVAALAHS | ASSNWGYEQY | 0.28119025 |
| GGVTVMATP | ASSNWGYEQY | 0.6255275 |
| AGLAANVRD | ASSNWGYEQY | 0.70562005 |
| TNRPDLIDPALLRPG | ASSNWGYEQY | 0.33314022 |
| LANLGVTAL | ASSNWGYEQY | 0.7154162 |
| TGERIEL | ASSNWGYEQY | 0.31477216 |
| NGEELAA | ASSNWGYEQY | 0.7140161 |
| VLSLVWG | ASSNWGYEQY | 0.6191314 |
| IGYALWL | ASSNWGYEQY | 0.5701904 |
| LSLLLAPL | ASSNWGYEQY | 0.553422 |
| TLDEALK | ASSNWGYEQY | 0.5218507 |
| TDGAGTAEGG | ASSNWGYEQY | 0.16393258 |
| NAPGGAGGNGG | ASSNWGYEQY | 0.59880817 |
| VSVFMGMGGNF | ASSNWGYEQY | 0.32121855 |
| LASYLAAL | ASSNWGYEQY | 0.42229405 |
| LVGGTPAL | ASSNWGYEQY | 0.2505457 |
| VRNADVL | ASSNWGYEQY | 0.6038835 |
| TCSSPFLVSAM | ASSNWGYEQY | 0.47661975 |
| VHITDAE | ASSNWGYEQY | 0.6335583 |
| EYAAPSF | ASSNWGYEQY | 0.2143793 |
| VVDAFHGGL | ASSNWGYEQY | 0.63931215 |
| ASVVDPQ | ASSNWGYEQY | 0.56723607 |
| SYNEFGGLS | ASSNWGYEQY | 0.23124649 |
| DPDGSVY | ASSNWGYEQY | 0.76847255 |
| MVSTIHATE | ASSNWGYEQY | 0.49796987 |
| LAAGVADPV | ASSNWGYEQY | 0.29599488 |
| TKEQIAA | ASSNWGYEQY | 0.5901061 |
| SLASPAAL | ASSNWGYEQY | 0.44897783 |
| IGLVFYGV | ASSNWGYEQY | 0.1715321 |
| FLLDASNL | ASSNWGYEQY | 0.28016356 |
| YQVGNFP | ASSNWGYEQY | 0.5334382 |
| ESLLSQST | ASSNWGYEQY | 0.2940223 |
| LFTVGFAAVG | ASSNWGYEQY | 0.19021718 |
| ATPPTPT | ASSNWGYEQY | 0.58592093 |
| TDLATLG | ASSNWGYEQY | 0.30047524 |
| SAGGTGR | ASSNWGYEQY | 0.21419868 |
| GFSNSGS | ASSNWGYEQY | 0.3492192 |
| GGNGGSGGTSV | ASSNWGYEQY | 0.730278 |
| GTSVATGGAGN | ASSNWGYEQY | 0.5001433 |
| NGGNGGSGGTS | ASSNWGYEQY | 0.4154942 |
| AVVKFPH | ASSNWGYEQY | 0.5766311 |
| TPPPPPP | ASSNWGYEQY | 0.5395138 |
| AAPAAAAAPA | ASSNWGYEQY | 0.73113346 |
| IALNEDL | ASSNWGYEQY | 0.48612133 |
| IVNVSSM | ASSNWGYEQY | 0.44575158 |
| TGIPPAPR | ASSNWGYEQY | 0.5910973 |
| FDLGGGTF | ASSNWGYEQY | 0.54660666 |
| FDLGGGTFDVS | ASSNWGYEQY | 0.36181715 |
| NEPTAAA | ASSNWGYEQY | 0.22011098 |

| | | |
|---|---|---|
| NEPTAAALA | ASSNWGYEQY | 0.39205214 |
| GSIINISSI | ASSNWGYEQY | 0.3342307 |
| VAPGNAGTA | ASSNWGYEQY | 0.43800318 |
| TGGGASLE | ASSNWGYEQY | 0.7992907 |
| VAVYNHYKRI | ASSNWGYEQY | 0.43929744 |
| TDPAPATT | ASSNWGYEQY | 0.22303253 |
| TNRPDILDPALLRPG | ASSNWGYEQY | 0.41028523 |
| LTTTTTS | ASSNWGYEQY | 0.17919306 |
| PTGSGKTL | ASSNWGYEQY | 0.5323526 |
| TGSGKTL | ASSNWGYEQY | 0.6015167 |
| TIGHVDHGK | ASSNWGYEQY | 0.8502197 |
| TIGHVDHGKTT | ASSNWGYEQY | 0.71159923 |
| GHVDHGKTT | ASSNWGYEQY | 0.34520143 |
| AQFEHTLL | ASSNWGYEQY | 0.652619 |
| IVNQDEL | ASSNWGYEQY | 0.32204226 |
| TNGSQFF | ASSNWGYEQY | 0.6494997 |
| TGKGIVE | ASSNWGYEQY | 0.47453195 |
| LISNASDA | ASSNWGYEQY | 0.66586584 |
| AELRTLGELTKIAKAHGAQ | ASSNWGYEQY | 0.46420503 |
| APNAVAAAEPSW | ASSNWGYEQY | 0.39076725 |
| APQVAGAATAVL | ASSNWGYEQY | 0.18302383 |
| ARRDPAAAAAILPTDA | ASSNWGYEQY | 0.8069596 |
| DAAARDVLAEAGLAEYF | ASSNWGYEQY | 0.2508613 |
| DDQIVLSAKANLAKVEQ | ASSNWGYEQY | 0.26884302 |
| DELLPDTGEDSDEDGH | ASSNWGYEQY | 0.39002505 |
| ELPEAPANRATVPQD | ASSNWGYEQY | 0.78302306 |
| FGQSKIERVTGIG | ASSNWGYEQY | 0.79989606 |
| GAGGTGSTAGGAGGAGGAGGLY | ASSNWGYEQY | 0.31343448 |
| GHGGTGAAGGNGGTGGAGGSGI | ASSNWGYEQY | 0.2715847 |
| GNGGNGGAGGEGGPGGAGGAGA | ASSNWGYEQY | 0.72461087 |
| GSTAGNGGNGGNATNGTIAGQP | ASSNWGYEQY | 0.23055202 |
| GTGGDGGNGGNGGTGG | ASSNWGYEQY | 0.6092185 |
| GVRRPAGIPADAPV | ASSNWGYEQY | 0.39105445 |
| GYRQVTDAHLLTLARR | ASSNWGYEQY | 0.42437696 |
| IAKMGQDGHDRG | ASSNWGYEQY | 0.39520827 |
| KAEPSIEIQYT | ASSNWGYEQY | 0.6844553 |
| KPVRIRMAADEDTVAVT | ASSNWGYEQY | 0.31876615 |
| MPPRIAGMRLLVIKPEP | ASSNWGYEQY | 0.60065365 |
| MKFYAVALFRSRGVPPTR | ASSNWGYEQY | 0.10612145 |
| NAGGFGILGAVAHSP | ASSNWGYEQY | 0.6737985 |
| NEGREITLAMAEM | ASSNWGYEQY | 0.21769477 |
| NELVEWSHPLGHRWRWE | ASSNWGYEQY | 0.27241793 |
| NLGAGGTGGNGGNPG | ASSNWGYEQY | 0.1734426 |
| NTEWPKWETVMAADPVD | ASSNWGYEQY | 0.5597001 |
| PASAAPAETQP | ASSNWGYEQY | 0.6856285 |
| QQSSDGKNGVRAAEITGE | ASSNWGYEQY | 0.7550385 |
| RAPDAVTTSWWKE | ASSNWGYEQY | 0.42035276 |
| SGFGNVGAHTSGWFNQ | ASSNWGYEQY | 0.69822353 |
| SGVAGAVGQAASVGGLKV | ASSNWGYEQY | 0.7215619 |
| SNGGAGTGSAGGNGGT | ASSNWGYEQY | 0.315445 |
| TGGAGGEGGAAG | ASSNWGYEQY | 0.1640968 |

| | | |
|---|---|---|
| TGGDGGAAGAGGAPGHG | ASSNWGYEQY | 0.4789104 |
| TMPPLLAAIANAGRHGV | ASSNWGYEQY | 0.7028532 |
| TSSTAGNSGGA | ASSNWGYEQY | 0.77748525 |
| VPATVPVGILC | ASSNWGYEQY | 0.6339654 |
| VQRPDSGGVVLTGRIS | ASSNWGYEQY | 0.17617875 |
| WTPNPVPATKGVPAGLVQAT | ASSNWGYEQY | 0.5296304 |
| YALKRALAKFSDRFDVVII | ASSNWGYEQY | 0.58896565 |
| YLIGSVIV | ASSNWGYEQY | 0.21909347 |
| YGQLVAVID | ASSNWGYEQY | 0.33541402 |
| VTVGGMTLPA | ASSNWGYEQY | 0.43671882 |
| VLALPAV | ASSNWGYEQY | 0.27949664 |
| VIPPTLNYET | ASSNWGYEQY | 0.57879 |
| VGMGGMHPSSG | ASSNWGYEQY | 0.33565402 |
| VGLAGGAATA | ASSNWGYEQY | 0.37400246 |
| VGGAGGAGGSGP | ASSNWGYEQY | 0.4592652 |
| VFAAFMV | ASSNWGYEQY | 0.37078598 |
| TLYDKLR | ASSNWGYEQY | 0.41072232 |
| TLAVFTVL | ASSNWGYEQY | 0.38922837 |
| TGSVQPARV | ASSNWGYEQY | 0.4415663 |
| TGGGAGSGGT | ASSNWGYEQY | 0.6452675 |
| SHLETSL | ASSNWGYEQY | 0.48200852 |
| SFTTGMT | ASSNWGYEQY | 0.6721848 |
| QVEVAKRMGVRQPTVS | ASSNWGYEQY | 0.77226996 |
| QGSGGPALAM | ASSNWGYEQY | 0.23514834 |
| MLAPSNP | ASSNWGYEQY | 0.36290208 |
| LVNNAAI | ASSNWGYEQY | 0.5847743 |
| LTGEINTLA | ASSNWGYEQY | 0.3310349 |
| LSIFGPV | ASSNWGYEQY | 0.6941576 |
| LLALLANL | ASSNWGYEQY | 0.527776 |
| LISNASDALDKLR | ASSNWGYEQY | 0.5823298 |
| LINEAVP | ASSNWGYEQY | 0.5328896 |
| LGYAFSGL | ASSNWGYEQY | 0.83760643 |
| LDFVAVAL | ASSNWGYEQY | 0.6368147 |
| LAQPGAE | ASSNWGYEQY | 0.5255507 |
| ISNLGAVTGF | ASSNWGYEQY | 0.791124 |
| HQEDPGRI | ASSNWGYEQY | 0.3709463 |
| GTPGEPTP | ASSNWGYEQY | 0.75249577 |
| GFFNSGS | ASSNWGYEQY | 0.72631055 |
| GAGGGGLTAGAN | ASSNWGYEQY | 0.18880211 |
| FVDALVNM | ASSNWGYEQY | 0.4463203 |
| FLTVAPEL | ASSNWGYEQY | 0.80365777 |
| FETLTAL | ASSNWGYEQY | 0.7546528 |
| DVINPIEL | ASSNWGYEQY | 0.28262654 |
| APVSGAPV | ASSNWGYEQY | 0.30379182 |
| ALVGGLAW | ASSNWGYEQY | 0.3044117 |
| ALAAAAATAI | ASSNWGYEQY | 0.40141386 |
| AFGLGLLL | ASSNWGYEQY | 0.27487925 |
| AFFAAASL | ASSNWGYEQY | 0.5256235 |
| AAAALPAFSPP | ASSNWGYEQY | 0.3534334 |
| VVLAPAL | ASSNWGYEQY | 0.7974757 |
| TIGHVDHGKTTL | ASSNWGYEQY | 0.4276288 |

| | | |
|---|---|---|
| LISNASDALDK | ASSNWGYEQY | 0.6654796 |
| LISNASDAL | ASSNWGYEQY | 0.3508872 |
| ISNASDALDK | ASSNWGYEQY | 0.4003152 |
| NQAETLVYQTE | ASSNWGYEQY | 0.20828524 |
| TGGTGGTAGAG | ASSNWGYEQY | 0.2386164 |
| TGSTAGGAGGA | ASSNWGYEQY | 0.32336336 |
| DPLAVPVAAGPV | ASSNWGYEQY | 0.54121095 |
| GGTGGTPGNGGN | ASSNWGYEQY | 0.14807574 |
| PAGLAGVDPATL | ASSNWGYEQY | 0.81863964 |
| GNNLVVEAPHAGASV | ASSNWGYEQY | 0.3814187 |
| KVTRIMSASTLEQPA | ASSNWGYEQY | 0.52060515 |
| LGFALPIALPRRTVV | ASSNWGYEQY | 0.339101 |
| TRLGNLTARKLVVLV | ASSNWGYEQY | 0.38666987 |
| WSQQRRLIYFPSAGP | ASSNWGYEQY | 0.2601687 |
| GPNGNPDPMAAAVDIR | ASSNWGYEQY | 0.7053055 |
| TQRCVSVADLRAAASR | ASSNWGYEQY | 0.17975554 |
| QFVLDVAEHPVEVLTETV | ASSNWGYEQY | 0.16551611 |
| GGAGGTGGAGGNGGTQAGDG | ASSNWGYEQY | 0.45244658 |
| PLLATLRAIAVDVDAKPAQV | ASSNWGYEQY | 0.20039956 |
| APKFRTGGYMIDFWGVGYQVAK | ASSNWGYEQY | 0.8125044 |
| GAGGAGGAGGAAGGPGGTGGTG | ASSNWGYEQY | 0.7140472 |
| IGALGLLATDVAPAGSVAFGVA | ASSNWGYEQY | 0.57124686 |
| AGGAGGIGGNGGAGGAGGVLMG | ASSNWGYEQY | 0.82899636 |
| ADELIGLL | ASSNWGYEQY | 0.122418895 |
| ADRALKA | ASSNWGYEQY | 0.36497194 |
| AGAVGQVGGAGGA | ASSNWGYEQY | 0.7874743 |
| AGGGGGLGGAP | ASSNWGYEQY | 0.3896901 |
| AGGLLYGNGGAG | ASSNWGYEQY | 0.3946832 |
| AGLESIGL | ASSNWGYEQY | 0.6769841 |
| AGLKTGII | ASSNWGYEQY | 0.27017605 |
| ASLAALLL | ASSNWGYEQY | 0.31903526 |
| ASVINISSI | ASSNWGYEQY | 0.38081518 |
| ATPPGPA | ASSNWGYEQY | 0.64503706 |
| ATVASALV | ASSNWGYEQY | 0.7168957 |
| ATVASLIL | ASSNWGYEQY | 0.24428445 |
| AVNGPVLIRH | ASSNWGYEQY | 0.5777806 |
| DGAAGAVGI | ASSNWGYEQY | 0.8472575 |
| DPLLHGI | ASSNWGYEQY | 0.37653247 |
| ELGAALGAL | ASSNWGYEQY | 0.53418374 |
| ELGRTQAV | ASSNWGYEQY | 0.46658003 |
| FVAGGYLS | ASSNWGYEQY | 0.4644984 |
| GASGGAGGAGGW | ASSNWGYEQY | 0.25566489 |
| GGAGAGGDGGAGGI | ASSNWGYEQY | 0.2280502 |
| GTGHNGSQPGMGGNG | ASSNWGYEQY | 0.6767589 |
| GYAAASASAS | ASSNWGYEQY | 0.16674511 |
| HQVDVAAVQQG | ASSNWGYEQY | 0.5421614 |
| IGAAIAHT | ASSNWGYEQY | 0.40213147 |
| IGMIEVAQL | ASSNWGYEQY | 0.85843146 |
| IRLAPEHTSAMD | ASSNWGYEQY | 0.67015874 |
| IYNVSVL | ASSNWGYEQY | 0.48195112 |
| KQGFLGM | ASSNWGYEQY | 0.23002501 |

| | | |
|---|---|---|
| LESVITQL | ASSNWGYEQY | 0.62416875 |
| LGLGALVVV | ASSNWGYEQY | 0.6355007 |
| LIGNGGNGGSGGI | ASSNWGYEQY | 0.19236192 |
| LNAINAPTL | ASSNWGYEQY | 0.76175565 |
| LQLLNTGV | ASSNWGYEQY | 0.26150897 |
| LQTSIEF | ASSNWGYEQY | 0.5305489 |
| MSFAVSEL | ASSNWGYEQY | 0.6363073 |
| NTGALGL | ASSNWGYEQY | 0.8272047 |
| PPGAGGAGGAGTT | ASSNWGYEQY | 0.382075 |
| QLVGLGI | ASSNWGYEQY | 0.6610985 |
| RFLEQVHQL | ASSNWGYEQY | 0.2612141 |
| RGVGYVLR | ASSNWGYEQY | 0.59723437 |
| SATGVFAGVI | ASSNWGYEQY | 0.72209907 |
| SLPAITLP | ASSNWGYEQY | 0.70932627 |
| STALAFTV | ASSNWGYEQY | 0.82417387 |
| SVLTVGLGK | ASSNWGYEQY | 0.72030485 |
| TALAGAGSGI | ASSNWGYEQY | 0.24129274 |
| TGLPFTGI | ASSNWGYEQY | 0.35425454 |
| VAEHVTV | ASSNWGYEQY | 0.7545814 |
| VDIVLNSL | ASSNWGYEQY | 0.7537116 |
| VLTGGGAVL | ASSNWGYEQY | 0.63237906 |
| VNVGLAYGI | ASSNWGYEQY | 0.7463964 |
| VSSAAVGAV | ASSNWGYEQY | 0.2787881 |
| VTSVYEAGV | ASSNWGYEQY | 0.7785092 |
| WVALDRGA | ASSNWGYEQY | 0.8654951 |
| YLVGSVVI | ASSNWGYEQY | 0.5533468 |
| GPPGTGKTLL | ASSNWGYEQY | 0.34132493 |
| GVGMTKF | ASSNWGYEQY | 0.3591231 |
| IGIGGSDL | ASSNWGYEQY | 0.6459751 |
| IGTLLAEL | ASSNWGYEQY | 0.67670214 |
| IYTSGTT | ASSNWGYEQY | 0.28275838 |
| LSFPAGV | ASSNWGYEQY | 0.49315986 |
| LYTSGTT | ASSNWGYEQY | 0.54760927 |
| VGAFPFGF | ASSNWGYEQY | 0.17036885 |
| VGAGTVEF | ASSNWGYEQY | 0.6692053 |
| VGKSTLLL | ASSNWGYEQY | 0.80048937 |
| AALAHAGGGIV | ASSNWGYEQY | 0.60986984 |
| AALVAVSLL | ASSNWGYEQY | 0.26877448 |
| AAVSRAVAAL | ASSNWGYEQY | 0.76498914 |
| AGGSSGIGGF | ASSNWGYEQY | 0.7912129 |
| AGLTVANANG | ASSNWGYEQY | 0.7913596 |
| AGLVGAVL | ASSNWGYEQY | 0.21089542 |
| ALSQAFTV | ASSNWGYEQY | 0.13066106 |
| ALTFSVSL | ASSNWGYEQY | 0.5761864 |
| ALTGAALVAL | ASSNWGYEQY | 0.76946926 |
| ASFSIGAGVL | ASSNWGYEQY | 0.4376886 |
| DGSVYKI | ASSNWGYEQY | 0.26784533 |
| EQLAEIV | ASSNWGYEQY | 0.6137638 |
| FAVSELTVM | ASSNWGYEQY | 0.6945587 |
| GAAPLAAGL | ASSNWGYEQY | 0.28467274 |
| GADGGDGGSGGAGG | ASSNWGYEQY | 0.23552415 |

| | | |
|---|---|---|
| GFGLDTGGAGG | ASSNWGYEQY | 0.7447911 |
| GLASGGLL | ASSNWGYEQY | 0.7376362 |
| GRIAGAAVLL | ASSNWGYEQY | 0.26570264 |
| GSAGSLNTSF | ASSNWGYEQY | 0.28426638 |
| GTGQIGI | ASSNWGYEQY | 0.42652315 |
| GTLIALIL | ASSNWGYEQY | 0.6437199 |
| IAATMGFVVG | ASSNWGYEQY | 0.50154126 |
| IFAQALL | ASSNWGYEQY | 0.256178 |
| IGAAVALGF | ASSNWGYEQY | 0.732803 |
| IGLTPLAL | ASSNWGYEQY | 0.22089893 |
| ILGSNTSTL | ASSNWGYEQY | 0.6118482 |
| IVFSLTVL | ASSNWGYEQY | 0.5587709 |
| LASKATGF | ASSNWGYEQY | 0.5408101 |
| LASVALIL | ASSNWGYEQY | 0.32706913 |
| LFAVGLTL | ASSNWGYEQY | 0.24804655 |
| LFNGGIGTYI | ASSNWGYEQY | 0.181769 |
| LTLGAGII | ASSNWGYEQY | 0.206512 |
| LVAAALGDI | ASSNWGYEQY | 0.7602464 |
| NTAFVQLGI | ASSNWGYEQY | 0.4962165 |
| QLFSGAEL | ASSNWGYEQY | 0.7465761 |
| SAATFITL | ASSNWGYEQY | 0.43306854 |
| SNYGLANLGV | ASSNWGYEQY | 0.28188196 |
| SYGGAYAVM | ASSNWGYEQY | 0.646471 |
| TTGVGGLTLG | ASSNWGYEQY | 0.19248238 |
| TTQTIYI | ASSNWGYEQY | 0.4945151 |
| VAAGAGLGAVI | ASSNWGYEQY | 0.4998783 |
| VAAVVTLL | ASSNWGYEQY | 0.63723606 |
| VGFLGGLI | ASSNWGYEQY | 0.27115303 |
| VGVIPILL | ASSNWGYEQY | 0.585878 |
| VIFASSLI | ASSNWGYEQY | 0.5369296 |
| VKGGDGGAAAG | ASSNWGYEQY | 0.28817645 |
| VTKTFTNI | ASSNWGYEQY | 0.58445764 |
| VVAAAILL | ASSNWGYEQY | 0.7303398 |
| YAALDVEL | ASSNWGYEQY | 0.58699757 |
| LASVALLL | ASSNWGYEQY | 0.17592287 |
| LDAVHRV | ASSNWGYEQY | 0.48836556 |
| QGLSSSIF | ASSNWGYEQY | 0.23226342 |
| RGTFANIRL | ASSNWGYEQY | 0.577152 |
| RQLWWGHRI | ASSNWGYEQY | 0.45827132 |
| YGPPGTGKTLL | ASSNWGYEQY | 0.44882834 |
| AALVGAPANVA | ASSNWGYEQY | 0.55124104 |
| AATFRRTE | ASSNWGYEQY | 0.35432056 |
| AGALGGGI | ASSNWGYEQY | 0.26120502 |
| AGGAGGAGGIGAG | ASSNWGYEQY | 0.3559728 |
| AGSGALGL | ASSNWGYEQY | 0.501255 |
| AQIADQI | ASSNWGYEQY | 0.54531443 |
| ASAVAVTYL | ASSNWGYEQY | 0.22675028 |
| ASTVLGGTGG | ASSNWGYEQY | 0.46475202 |
| AVGGEPAK | ASSNWGYEQY | 0.3441099 |
| CEQGTFGGTC | ASSNWGYEQY | 0.24924688 |
| DEGKAPAG | ASSNWGYEQY | 0.59409344 |

| | | |
|---|---|---|
| DTVVFPAL | ASSNWGYEQY | 0.51615083 |
| FEGITMAG | ASSNWGYEQY | 0.19653787 |
| FPLVVQD | ASSNWGYEQY | 0.66964614 |
| GAFTTPQ | ASSNWGYEQY | 0.30564627 |
| GFGGFGTAGGVGGA | ASSNWGYEQY | 0.518188 |
| GFLAQFL | ASSNWGYEQY | 0.79980475 |
| GGAGGAGGNAGVA | ASSNWGYEQY | 0.2509305 |
| GGVGTAGAGGAGGAGG | ASSNWGYEQY | 0.6006799 |
| GNGGGGDAGNAGSGGN | ASSNWGYEQY | 0.43075073 |
| GTGGAGGAGNDAGS | ASSNWGYEQY | 0.56073916 |
| GTTNTGWFNS | ASSNWGYEQY | 0.47284958 |
| IFMGPLFTL | ASSNWGYEQY | 0.26890618 |
| ILAHAIEAI | ASSNWGYEQY | 0.7451292 |
| IRSAIREAL | ASSNWGYEQY | 0.5699669 |
| ITFAGGG | ASSNWGYEQY | 0.6377307 |
| ITGMSGHQEV | ASSNWGYEQY | 0.680738 |
| IVAGLACTL | ASSNWGYEQY | 0.7853304 |
| IVGGGLAGA | ASSNWGYEQY | 0.6872227 |
| IVTGVLNI | ASSNWGYEQY | 0.29160526 |
| LAQQGCI | ASSNWGYEQY | 0.17730175 |
| LCLHVEN | ASSNWGYEQY | 0.39215332 |
| LDAISDGS | ASSNWGYEQY | 0.36567077 |
| LGGFAAML | ASSNWGYEQY | 0.38222522 |
| LGGKNPMI | ASSNWGYEQY | 0.24531603 |
| LGPKGVAAIL | ASSNWGYEQY | 0.69403493 |
| LMVGVSI | ASSNWGYEQY | 0.8351946 |
| LVGAAFVL | ASSNWGYEQY | 0.35412467 |
| LVLAVAPL | ASSNWGYEQY | 0.83958524 |
| MAAIAVIDAA | ASSNWGYEQY | 0.22555247 |
| NNGINAELQ | ASSNWGYEQY | 0.5621617 |
| RGEPLVLVHG | ASSNWGYEQY | 0.7379744 |
| RGGAGGAAGLI | ASSNWGYEQY | 0.2202293 |
| RSYRMHPAV | ASSNWGYEQY | 0.494187 |
| SAAHLLAGQLA | ASSNWGYEQY | 0.2050861 |
| SAAVGSY | ASSNWGYEQY | 0.2637462 |
| SPHAVLHRI | ASSNWGYEQY | 0.7567435 |
| SQIAKVL | ASSNWGYEQY | 0.37648517 |
| TALGHLVP | ASSNWGYEQY | 0.58213806 |
| TAPTTSGAA | ASSNWGYEQY | 0.39017767 |
| TDGDDGNGGSPGT | ASSNWGYEQY | 0.264792 |
| TDGLGEN | ASSNWGYEQY | 0.5499843 |
| TGGPPLPGQA | ASSNWGYEQY | 0.78612447 |
| TIPAITI | ASSNWGYEQY | 0.49856547 |
| TIPGFTI | ASSNWGYEQY | 0.8658055 |
| TSIASGALA | ASSNWGYEQY | 0.27955985 |
| TVTLPGAAI | ASSNWGYEQY | 0.7952806 |
| VASGLSAALP | ASSNWGYEQY | 0.8017163 |
| VGAGVIGI | ASSNWGYEQY | 0.4306823 |
| VGPKRFGV | ASSNWGYEQY | 0.3386584 |
| VNVAFPD | ASSNWGYEQY | 0.72165483 |
| VSMTNSGVS | ASSNWGYEQY | 0.4489189 |

| | | |
|---|---|---|
| VTDVLGDLSV | ASSNWGYEQY | 0.6308193 |
| VVVVGAGFAGL | ASSNWGYEQY | 0.79297554 |
| VYTVGTVV | ASSNWGYEQY | 0.72734296 |
| WTTFSGP | ASSNWGYEQY | 0.8794505 |
| YDELVEAF | ASSNWGYEQY | 0.85028946 |
| YGTGGAGGNGG | ASSNWGYEQY | 0.8024667 |
| YTNVMAV | ASSNWGYEQY | 0.7468742 |
| FLEQVHQL | ASSNWGYEQY | 0.3878003 |
| GLSLLAVAV | ASSNWGYEQY | 0.83453846 |
| LEQVHQL | ASSNWGYEQY | 0.6843362 |
| LNAAGAI | ASSNWGYEQY | 0.86864793 |
| PAVHASL | ASSNWGYEQY | 0.8527478 |
| TELEDDA | ASSNWGYEQY | 0.73449755 |
| TPPGPPN | ASSNWGYEQY | 0.87815094 |
| TPPPPPPP | ASSNWGYEQY | 0.8676123 |
| TQSPALI | ASSDGTGVSNERLF | 0.5313126 |
| EFAAAFPQ | ASSDGTGVSNERLF | 0.79434216 |
| LLSGNLDVL | ASSDGTGVSNERLF | 0.7972736 |
| GPANLGNYNI | ASSDGTGVSNERLF | 0.79001 |
| TLGPTLP | ASSDGTGVSNERLF | 0.48535064 |
| VYAYSSCCK | ASSDGTGVSNERLF | 0.86363757 |
| APAPAPAPNI | ASSDGTGVSNERLF | 0.7945551 |
| PGGPVPIP | ASSDGTGVSNERLF | 0.16443096 |
| FLTVALEP | ASSDGTGVSNERLF | 0.8290632 |
| SQYFGANL | ASSDGTGVSNERLF | 0.79120684 |
| GGVKPAA | ASSDGTGVSNERLF | 0.75745535 |
| GGTGGAGGAGGW | ASSDGTGVSNERLF | 0.87482405 |
| GVGGVAGLGGAGG | ASSDGTGVSNERLF | 0.3083005 |
| VLLVLATTI | ASSDGTGVSNERLF | 0.7135899 |
| IGIFAAML | ASSDGTGVSNERLF | 0.86995065 |
| AVGDLID | ASSDGTGVSNERLF | 0.87910986 |
| NISPLPL | ASSDGTGVSNERLF | 0.537176 |
| VTAVFVAI | ASSDGTGVSNERLF | 0.8588824 |
| SGDDNDAAL | ASSDGTGVSNERLF | 0.8731363 |
| AVAALAHS | ASSDGTGVSNERLF | 0.84787756 |
| GGVTVMATP | ASSDGTGVSNERLF | 0.8740144 |
| AGLAANVRD | ASSDGTGVSNERLF | 0.8794782 |
| TNRPDLIDPALLRPG | ASSDGTGVSNERLF | 0.6565836 |
| LANLGVTAL | ASSDGTGVSNERLF | 0.87345564 |
| TGERIEL | ASSDGTGVSNERLF | 0.88023853 |
| NGEELAA | ASSDGTGVSNERLF | 0.7486416 |
| VLSLVWG | ASSDGTGVSNERLF | 0.8313276 |
| IGYALWL | ASSDGTGVSNERLF | 0.74793106 |
| LSLLLAPL | ASSDGTGVSNERLF | 0.7781315 |
| TLDEALK | ASSDGTGVSNERLF | 0.3327021 |
| TDGAGTAEGG | ASSDGTGVSNERLF | 0.8537437 |
| NAPGGAGGNGG | ASSDGTGVSNERLF | 0.8697367 |
| VSVFMGMGGNF | ASSDGTGVSNERLF | 0.79130554 |
| LASYLAAL | ASSDGTGVSNERLF | 0.7305571 |
| LVGGTPAL | ASSDGTGVSNERLF | 0.73109984 |
| VRNADVL | ASSDGTGVSNERLF | 0.81602526 |

| | | |
|---|---|---|
| TCSSPFLVSAM | ASSDGTGVSNERLF | 0.76078594 |
| VHITDAE | ASSDGTGVSNERLF | 0.37450743 |
| EYAAPSF | ASSDGTGVSNERLF | 0.73901314 |
| VVDAFHGGL | ASSDGTGVSNERLF | 0.86926776 |
| ASVVDPQ | ASSDGTGVSNERLF | 0.8680826 |
| SYNEFGGLS | ASSDGTGVSNERLF | 0.3985374 |
| DPDGSVY | ASSDGTGVSNERLF | 0.879483 |
| MVSTIHATE | ASSDGTGVSNERLF | 0.8780051 |
| LAAGVADPV | ASSDGTGVSNERLF | 0.8818736 |
| TKEQIAA | ASSDGTGVSNERLF | 0.82636964 |
| SLASPAAL | ASSDGTGVSNERLF | 0.7651586 |
| IGLVFYGV | ASSDGTGVSNERLF | 0.86927617 |
| FLLDASNL | ASSDGTGVSNERLF | 0.7922753 |
| YQVGNFP | ASSDGTGVSNERLF | 0.582678 |
| ESLLSQST | ASSDGTGVSNERLF | 0.86217046 |
| LFTVGFAAVG | ASSDGTGVSNERLF | 0.7904235 |
| ATPPTPT | ASSDGTGVSNERLF | 0.74791545 |
| TDLATLG | ASSDGTGVSNERLF | 0.79839855 |
| SAGGTGR | ASSDGTGVSNERLF | 0.88197184 |
| GFSNSGS | ASSDGTGVSNERLF | 0.69075596 |
| GGNGGSGGTSV | ASSDGTGVSNERLF | 0.5466539 |
| GTSVATGGAGN | ASSDGTGVSNERLF | 0.65287435 |
| NGGNGGSGGTS | ASSDGTGVSNERLF | 0.8332145 |
| AVVKFPH | ASSDGTGVSNERLF | 0.8715362 |
| TPPPPPP | ASSDGTGVSNERLF | 0.84307647 |
| AAPAAAAAPA | ASSDGTGVSNERLF | 0.75948954 |
| IALNEDL | ASSDGTGVSNERLF | 0.8547572 |
| IVNVSSM | ASSDGTGVSNERLF | 0.83930004 |
| TGIPPAPR | ASSDGTGVSNERLF | 0.8308666 |
| FDLGGGTF | ASSDGTGVSNERLF | 0.86979586 |
| FDLGGGTFDVS | ASSDGTGVSNERLF | 0.7422122 |
| NEPTAAA | ASSDGTGVSNERLF | 0.88114697 |
| NEPTAAALA | ASSDGTGVSNERLF | 0.84898025 |
| GSIINISSI | ASSDGTGVSNERLF | 0.8673012 |
| VAPGNAGTA | ASSDGTGVSNERLF | 0.87262106 |
| TGGGASLE | ASSDGTGVSNERLF | 0.8312982 |
| VAVYNHYKRI | ASSDGTGVSNERLF | 0.83198214 |
| TDPAPATT | ASSDGTGVSNERLF | 0.3282511 |
| TNRPDILDPALLRPG | ASSDGTGVSNERLF | 0.7622668 |
| LTTTTTS | ASSDGTGVSNERLF | 0.84619015 |
| PTGSGKTL | ASSDGTGVSNERLF | 0.846017 |
| TGSGKTL | ASSDGTGVSNERLF | 0.8555005 |
| TIGHVDHGK | ASSDGTGVSNERLF | 0.8395178 |
| TIGHVDHGKTT | ASSDGTGVSNERLF | 0.74524176 |
| GHVDHGKTT | ASSDGTGVSNERLF | 0.8662445 |
| AQFEHTLL | ASSDGTGVSNERLF | 0.86623156 |
| IVNQDEL | ASSDGTGVSNERLF | 0.55153656 |
| TNGSQFF | ASSDGTGVSNERLF | 0.7720928 |
| TGKGIVE | ASSDGTGVSNERLF | 0.8847375 |
| LISNASDA | ASSDGTGVSNERLF | 0.8619362 |
| AELRTLGELTKIAKAHGAQ | ASSDGTGVSNERLF | 0.79673517 |

| | | |
|---|---|---|
| APNAVAAAEPSW | ASSDGTGVSNERLF | 0.6834577 |
| APQVAGAATAVL | ASSDGTGVSNERLF | 0.88818586 |
| ARRDPAAAAAILPTDA | ASSDGTGVSNERLF | 0.68591183 |
| DAAARDVLAEAGLAEYF | ASSDGTGVSNERLF | 0.88198864 |
| DDQIVLSAKANLAKVEQ | ASSDGTGVSNERLF | 0.2019731 |
| DELLPDTGEDSDEDGH | ASSDGTGVSNERLF | 0.7227267 |
| ELPEAPANRATVPQD | ASSDGTGVSNERLF | 0.68891716 |
| FGQSKIERVTGIG | ASSDGTGVSNERLF | 0.38391572 |
| GAGGTGSTAGGAGGAGGAGGLY | ASSDGTGVSNERLF | 0.86908984 |
| GHGGTGAAGGNGGTGGAGGSGI | ASSDGTGVSNERLF | 0.7423179 |
| GNGGNGGAGGEGGPGGAGGAGA | ASSDGTGVSNERLF | 0.85292727 |
| GSTAGNGGNGGNATNGTIAGQP | ASSDGTGVSNERLF | 0.83707523 |
| GTGGDGGNGGNGGTGG | ASSDGTGVSNERLF | 0.8671923 |
| GVRRPAGIPADAPV | ASSDGTGVSNERLF | 0.84371114 |
| GYRQVTDAHLLTLARR | ASSDGTGVSNERLF | 0.80841756 |
| IAKMGQDGHDRG | ASSDGTGVSNERLF | 0.55816543 |
| KAEPSIEIQYT | ASSDGTGVSNERLF | 0.86138785 |
| KPVRIRMAADEDTVAVT | ASSDGTGVSNERLF | 0.8727518 |
| MPPRIAGMRLLVIKPEP | ASSDGTGVSNERLF | 0.8633231 |
| MKFYAVALFRSRGVPPTR | ASSDGTGVSNERLF | 0.46350628 |
| NAGGFGILGAVAHSP | ASSDGTGVSNERLF | 0.78804505 |
| NEGREITLAMAEM | ASSDGTGVSNERLF | 0.8269138 |
| NELVEWSHPLGHRWRWE | ASSDGTGVSNERLF | 0.26159143 |
| NLGAGGTGGNGGNPG | ASSDGTGVSNERLF | 0.21913777 |
| NTEWPKWETVMAADPVD | ASSDGTGVSNERLF | 0.8134413 |
| PASAAPAETQP | ASSDGTGVSNERLF | 0.7397486 |
| QQSSDGKNGVRAAEITGE | ASSDGTGVSNERLF | 0.8736012 |
| RAPDAVTTSWWKE | ASSDGTGVSNERLF | 0.86698484 |
| SGFGNVGAHTSGWFNQ | ASSDGTGVSNERLF | 0.80209625 |
| SGVAGAVGQAASVGGLKV | ASSDGTGVSNERLF | 0.86349475 |
| SNGGAGTGSAGGNGGT | ASSDGTGVSNERLF | 0.81310767 |
| TGGAGGEGGAAG | ASSDGTGVSNERLF | 0.11899922 |
| TGGDGGAAGAGGAPGHG | ASSDGTGVSNERLF | 0.72928417 |
| TMPPLLAAIANAGRHGV | ASSDGTGVSNERLF | 0.56552327 |
| TSSTAGNSGGA | ASSDGTGVSNERLF | 0.8799659 |
| VPATVPVGILC | ASSDGTGVSNERLF | 0.68137264 |
| VQRPDSGGVVLTGRIS | ASSDGTGVSNERLF | 0.7355965 |
| WTPNPVPATKGVPAGLVQAT | ASSDGTGVSNERLF | 0.81173277 |
| YALKRALAKFSDRFDVVII | ASSDGTGVSNERLF | 0.8472168 |
| YLIGSVIV | ASSDGTGVSNERLF | 0.81337243 |
| YGQLVAVID | ASSDGTGVSNERLF | 0.68288875 |
| VTVGGMTLPA | ASSDGTGVSNERLF | 0.17860182 |
| VLALPAV | ASSDGTGVSNERLF | 0.85921156 |
| VIPPTLNYET | ASSDGTGVSNERLF | 0.43698427 |
| VGMGGMHPSSG | ASSDGTGVSNERLF | 0.6180136 |
| VGLAGGAATA | ASSDGTGVSNERLF | 0.31910053 |
| VGGAGGAGGSGP | ASSDGTGVSNERLF | 0.19482553 |
| VFAAFMV | ASSDGTGVSNERLF | 0.39669657 |
| TLYDKLR | ASSDGTGVSNERLF | 0.8647031 |
| TLAVFTVL | ASSDGTGVSNERLF | 0.81527615 |
| TGSVQPARV | ASSDGTGVSNERLF | 0.7882515 |

| | | |
|---|---|---|
| TGGGAGSGGT | ASSDGTGVSNERLF | 0.8844746 |
| SHLETSL | ASSDGTGVSNERLF | 0.83978593 |
| SFTTGMT | ASSDGTGVSNERLF | 0.7812572 |
| QVEVAKRMGVRQPTVS | ASSDGTGVSNERLF | 0.6524341 |
| QGSGGPALAM | ASSDGTGVSNERLF | 0.67724586 |
| MLAPSNP | ASSDGTGVSNERLF | 0.6994728 |
| LVNNAAI | ASSDGTGVSNERLF | 0.6265192 |
| LTGEINTLA | ASSDGTGVSNERLF | 0.88323325 |
| LSIFGPV | ASSDGTGVSNERLF | 0.72740716 |
| LLALLANL | ASSDGTGVSNERLF | 0.5865749 |
| LISNASDALDKLR | ASSDGTGVSNERLF | 0.808855 |
| LINEAVP | ASSDGTGVSNERLF | 0.88108784 |
| LGYAFSGL | ASSDGTGVSNERLF | 0.88951147 |
| LDFVAVAL | ASSDGTGVSNERLF | 0.7219304 |
| LAQPGAE | ASSDGTGVSNERLF | 0.78302693 |
| ISNLGAVTGF | ASSDGTGVSNERLF | 0.78071445 |
| HQEDPGRI | ASSDGTGVSNERLF | 0.82822824 |
| GTPGEPTP | ASSDGTGVSNERLF | 0.74838597 |
| GFFNSGS | ASSDGTGVSNERLF | 0.47619432 |
| GAGGGGLTAGAN | ASSDGTGVSNERLF | 0.872241 |
| FVDALVNM | ASSDGTGVSNERLF | 0.889969 |
| FLTVAPEL | ASSDGTGVSNERLF | 0.76239467 |
| FETLTAL | ASSDGTGVSNERLF | 0.74517405 |
| DVINPIEL | ASSDGTGVSNERLF | 0.8029914 |
| APVSGAPV | ASSDGTGVSNERLF | 0.7481232 |
| ALVGGLAW | ASSDGTGVSNERLF | 0.79471755 |
| ALAAAAATAI | ASSDGTGVSNERLF | 0.8378687 |
| AFGLGLLL | ASSDGTGVSNERLF | 0.5675152 |
| AFFAAASL | ASSDGTGVSNERLF | 0.7644106 |
| AAAALPAFSPP | ASSDGTGVSNERLF | 0.8881494 |
| VVLAPAL | ASSDGTGVSNERLF | 0.80341923 |
| TIGHVDHGKTTL | ASSDGTGVSNERLF | 0.84646744 |
| LISNASDALDK | ASSDGTGVSNERLF | 0.87678266 |
| LISNASDAL | ASSDGTGVSNERLF | 0.86144114 |
| ISNASDALDK | ASSDGTGVSNERLF | 0.88222516 |
| NQAETLVYQTE | ASSDGTGVSNERLF | 0.8800035 |
| TGGTGGTAGAG | ASSDGTGVSNERLF | 0.6958844 |
| TGSTAGGAGGA | ASSDGTGVSNERLF | 0.82473415 |
| DPLAVPVAAGPV | ASSDGTGVSNERLF | 0.8409872 |
| GGTGGTPGNGGN | ASSDGTGVSNERLF | 0.81716615 |
| PAGLAGVDPATL | ASSDGTGVSNERLF | 0.79724336 |
| GNNLVVEAPHAGASV | ASSDGTGVSNERLF | 0.715507 |
| KVTRIMSASTLEQPA | ASSDGTGVSNERLF | 0.8747225 |
| LGFALPIALPRRTVV | ASSDGTGVSNERLF | 0.83263755 |
| TRLGNLTARKLVVLV | ASSDGTGVSNERLF | 0.8733008 |
| WSQQRRLIYFPSAGP | ASSDGTGVSNERLF | 0.48058882 |
| GPNGNPDPMAAAVDIR | ASSDGTGVSNERLF | 0.5444739 |
| TQRCVSVADLRAAASR | ASSDGTGVSNERLF | 0.8690585 |
| QFVLDVAEHPVEVLTETV | ASSDGTGVSNERLF | 0.15989484 |
| GGAGGTGGAGGNGGTQAGDG | ASSDGTGVSNERLF | 0.21369931 |
| PLLATLRAIAVDVDAKPAQV | ASSDGTGVSNERLF | 0.8541402 |

| | | |
|---|---|---|
| APKFRTGGYMIDFWGVGYQVAK | ASSDGTGVSNERLF | 0.8797406 |
| GAGGAGGAGGAAGGPGGTGGTG | ASSDGTGVSNERLF | 0.18423094 |
| IGALGLLATDVAPAGSVAFGVA | ASSDGTGVSNERLF | 0.6536255 |
| AGGAGGIGGNGGAGGAGGVLMG | ASSDGTGVSNERLF | 0.55907893 |
| ADELIGLL | ASSDGTGVSNERLF | 0.8450399 |
| ADRALKA | ASSDGTGVSNERLF | 0.8201041 |
| AGAVGQVGGAGGA | ASSDGTGVSNERLF | 0.42104417 |
| AGGGGGLGGAP | ASSDGTGVSNERLF | 0.85545343 |
| AGGLLYGNGGAG | ASSDGTGVSNERLF | 0.5678394 |
| AGLESIGL | ASSDGTGVSNERLF | 0.4400736 |
| AGLKTGII | ASSDGTGVSNERLF | 0.8695591 |
| ASLAALLL | ASSDGTGVSNERLF | 0.88281214 |
| ASVINISSI | ASSDGTGVSNERLF | 0.67284364 |
| ATPPGPA | ASSDGTGVSNERLF | 0.8681961 |
| ATVASALV | ASSDGTGVSNERLF | 0.78148043 |
| ATVASLIL | ASSDGTGVSNERLF | 0.8889092 |
| AVNGPVLIRH | ASSDGTGVSNERLF | 0.7691829 |
| DGAAGAVGI | ASSDGTGVSNERLF | 0.47152078 |
| DPLLHGI | ASSDGTGVSNERLF | 0.87063164 |
| ELGAALGAL | ASSDGTGVSNERLF | 0.857884 |
| ELGRTQAV | ASSDGTGVSNERLF | 0.7845941 |
| FVAGGYLS | ASSDGTGVSNERLF | 0.87992287 |
| GASGGAGGAGGW | ASSDGTGVSNERLF | 0.70998 |
| GGAGAGGDGGAGGI | ASSDGTGVSNERLF | 0.80735475 |
| GTGHNGSQPGMGGNG | ASSDGTGVSNERLF | 0.8745455 |
| GYAAASASAS | ASSDGTGVSNERLF | 0.8688369 |
| HQVDVAAVQQG | ASSDGTGVSNERLF | 0.57794356 |
| IGAAIAHT | ASSDGTGVSNERLF | 0.862188 |
| IGMIEVAQL | ASSDGTGVSNERLF | 0.7912823 |
| IRLAPEHTSAMD | ASSDGTGVSNERLF | 0.7287793 |
| IYNVSVL | ASSDGTGVSNERLF | 0.8852295 |
| KQGFLGM | ASSDGTGVSNERLF | 0.8315668 |
| LESVITQL | ASSDGTGVSNERLF | 0.86775005 |
| LGLGALVVV | ASSDGTGVSNERLF | 0.8562256 |
| LIGNGGNGGSGGI | ASSDGTGVSNERLF | 0.8810733 |
| LNAINAPTL | ASSDGTGVSNERLF | 0.46373448 |
| LQLLNTGV | ASSDGTGVSNERLF | 0.85131776 |
| LQTSIEF | ASSDGTGVSNERLF | 0.82975745 |
| MSFAVSEL | ASSDGTGVSNERLF | 0.86559707 |
| NTGALGL | ASSDGTGVSNERLF | 0.83953166 |
| PPGAGGAGGAGTT | ASSDGTGVSNERLF | 0.5895913 |
| QLVGLGI | ASSDGTGVSNERLF | 0.83807796 |
| RFLEQVHQL | ASSDGTGVSNERLF | 0.87780356 |
| RGVGYVLR | ASSDGTGVSNERLF | 0.82289624 |
| SATGVFAGVI | ASSDGTGVSNERLF | 0.82444465 |
| SLPAITLP | ASSDGTGVSNERLF | 0.8850591 |
| STALAFTV | ASSDGTGVSNERLF | 0.7304406 |
| SVLTVGLGK | ASSDGTGVSNERLF | 0.7663434 |
| TALAGAGSGI | ASSDGTGVSNERLF | 0.8683127 |
| TGLPFTGI | ASSDGTGVSNERLF | 0.8689307 |
| VAEHVTV | ASSDGTGVSNERLF | 0.77696264 |

| | | |
|---|---|---|
| VDIVLNSL | ASSDGTGVSNERLF | 0.88307464 |
| VLTGGGAVL | ASSDGTGVSNERLF | 0.7266352 |
| VNVGLAYGI | ASSDGTGVSNERLF | 0.85827243 |
| VSSAAVGAV | ASSDGTGVSNERLF | 0.7854481 |
| VTSVYEAGV | ASSDGTGVSNERLF | 0.84999084 |
| WVALDRGA | ASSDGTGVSNERLF | 0.7027905 |
| YLVGSVVI | ASSDGTGVSNERLF | 0.8599524 |
| GPPGTGKTLL | ASSDGTGVSNERLF | 0.8744979 |
| GVGMTKF | ASSDGTGVSNERLF | 0.8646876 |
| IGIGGSDL | ASSDGTGVSNERLF | 0.21697979 |
| IGTLLAEL | ASSDGTGVSNERLF | 0.8719265 |
| IYTSGTT | ASSDGTGVSNERLF | 0.85415757 |
| LSFPAGV | ASSDGTGVSNERLF | 0.8922892 |
| LYTSGTT | ASSDGTGVSNERLF | 0.81872106 |
| VGAFPFGF | ASSDGTGVSNERLF | 0.85264766 |
| VGAGTVEF | ASSDGTGVSNERLF | 0.7944506 |
| VGKSTLLL | ASSDGTGVSNERLF | 0.79434997 |
| AALAHAGGGIV | ASSDGTGVSNERLF | 0.6635159 |
| AALVAVSLL | ASSDGTGVSNERLF | 0.8039721 |
| AAVSRAVAAL | ASSDGTGVSNERLF | 0.7914026 |
| AGGSSGIGGF | ASSDGTGVSNERLF | 0.7440207 |
| AGLTVANANG | ASSDGTGVSNERLF | 0.8651203 |
| AGLVGAVL | ASSDGTGVSNERLF | 0.8769981 |
| ALSQAFTV | ASSDGTGVSNERLF | 0.87252665 |
| ALTFSVSL | ASSDGTGVSNERLF | 0.8788073 |
| ALTGAALVAL | ASSDGTGVSNERLF | 0.879583 |
| ASFSIGAGVL | ASSDGTGVSNERLF | 0.86849815 |
| DGSVYKI | ASSDGTGVSNERLF | 0.7281545 |
| EQLAEIV | ASSDGTGVSNERLF | 0.7948211 |
| FAVSELTVM | ASSDGTGVSNERLF | 0.69003236 |
| GAAPLAAGL | ASSDGTGVSNERLF | 0.7995714 |
| GADGGDGGSGGAGG | ASSDGTGVSNERLF | 0.74449146 |
| GFGLDTGGAGG | ASSDGTGVSNERLF | 0.84340125 |
| GLASGGLL | ASSDGTGVSNERLF | 0.1361854 |
| GRIAGAAVLL | ASSDGTGVSNERLF | 0.84950036 |
| GSAGSLNTSF | ASSDGTGVSNERLF | 0.8817026 |
| GTGQIGI | ASSDGTGVSNERLF | 0.7145736 |
| GTLIALIL | ASSDGTGVSNERLF | 0.8437165 |
| IAATMGFVVG | ASSDGTGVSNERLF | 0.8255482 |
| IFAQALL | ASSDGTGVSNERLF | 0.753068 |
| IGAAVALGF | ASSDGTGVSNERLF | 0.8674543 |
| IGLTPLAL | ASSDGTGVSNERLF | 0.64981186 |
| ILGSNTSTL | ASSDGTGVSNERLF | 0.85055196 |
| IVFSLTVL | ASSDGTGVSNERLF | 0.8836979 |
| LASKATGF | ASSDGTGVSNERLF | 0.8427025 |
| LASVALIL | ASSDGTGVSNERLF | 0.8340458 |
| LFAVGLTL | ASSDGTGVSNERLF | 0.79584396 |
| LFNGGIGTYI | ASSDGTGVSNERLF | 0.37270275 |
| LTLGAGII | ASSDGTGVSNERLF | 0.8468219 |
| LVAAALGDI | ASSDGTGVSNERLF | 0.122460686 |
| NTAFVQLGI | ASSDGTGVSNERLF | 0.8332064 |

| | | |
|---|---|---|
| QLFSGAEL | ASSDGTGVSNERLF | 0.6155262 |
| SAATFITL | ASSDGTGVSNERLF | 0.8368447 |
| SNYGLANLGV | ASSDGTGVSNERLF | 0.81668776 |
| SYGGAYAVM | ASSDGTGVSNERLF | 0.85662544 |
| TTGVGGLTLG | ASSDGTGVSNERLF | 0.7072148 |
| TTQTIYI | ASSDGTGVSNERLF | 0.8006307 |
| VAAGAGLGAVI | ASSDGTGVSNERLF | 0.70840573 |
| VAAVVTLL | ASSDGTGVSNERLF | 0.5572628 |
| VGFLGGLI | ASSDGTGVSNERLF | 0.8750664 |
| VGVIPILL | ASSDGTGVSNERLF | 0.81507313 |
| VIFASSLI | ASSDGTGVSNERLF | 0.80440795 |
| VKGGDGGAAAG | ASSDGTGVSNERLF | 0.87166214 |
| VTKTFTNI | ASSDGTGVSNERLF | 0.8318852 |
| VVAAAILL | ASSDGTGVSNERLF | 0.51653874 |
| YAALDVEL | ASSDGTGVSNERLF | 0.8311779 |
| LASVALLL | ASSDGTGVSNERLF | 0.82591075 |
| LDAVHRV | ASSDGTGVSNERLF | 0.7912113 |
| QGLSSSIF | ASSDGTGVSNERLF | 0.124174304 |
| RGTFANIRL | ASSDGTGVSNERLF | 0.8457506 |
| RQLWWGHRI | ASSDGTGVSNERLF | 0.7151166 |
| YGPPGTGKTLL | ASSDGTGVSNERLF | 0.74545324 |
| AALVGAPANVA | ASSDGTGVSNERLF | 0.7172692 |
| AATFRRTE | ASSDGTGVSNERLF | 0.8680433 |
| AGALGGGI | ASSDGTGVSNERLF | 0.43279833 |
| AGGAGGAGGIGAG | ASSDGTGVSNERLF | 0.7858375 |
| AGSGALGL | ASSDGTGVSNERLF | 0.8742908 |
| AQIADQI | ASSDGTGVSNERLF | 0.85941017 |
| ASAVAVTYL | ASSDGTGVSNERLF | 0.87801516 |
| ASTVLGGTGG | ASSDGTGVSNERLF | 0.80335915 |
| AVGGEPAK | ASSDGTGVSNERLF | 0.80545145 |
| CEQGTFGGTC | ASSDGTGVSNERLF | 0.82690203 |
| DEGKAPAG | ASSDGTGVSNERLF | 0.8330581 |
| DTVVFPAL | ASSDGTGVSNERLF | 0.38836503 |
| FEGITMAG | ASSDGTGVSNERLF | 0.3029302 |
| FPLVVQD | ASSDGTGVSNERLF | 0.8462085 |
| GAFTTPQ | ASSDGTGVSNERLF | 0.2768221 |
| GFGGFGTAGGVGGA | ASSDGTGVSNERLF | 0.8184991 |
| GFLAQFL | ASSDGTGVSNERLF | 0.87528074 |
| GGAGGAGGNAGVA | ASSDGTGVSNERLF | 0.82746154 |
| GGVGTAGAGGAGGAGG | ASSDGTGVSNERLF | 0.5593893 |
| GNGGGGDAGNAGSGGN | ASSDGTGVSNERLF | 0.80104464 |
| GTGGAGGAGNDAGS | ASSDGTGVSNERLF | 0.7809691 |
| GTTNTGWFNS | ASSDGTGVSNERLF | 0.8673637 |
| IFMGPLFTL | ASSDGTGVSNERLF | 0.8631418 |
| ILAHAIEAI | ASSDGTGVSNERLF | 0.8875411 |
| IRSAIREAL | ASSDGTGVSNERLF | 0.8711216 |
| ITFAGGG | ASSDGTGVSNERLF | 0.7558582 |
| ITGMSGHQEV | ASSDGTGVSNERLF | 0.8618771 |
| IVAGLACTL | ASSDGTGVSNERLF | 0.77822006 |
| IVGGGLAGA | ASSDGTGVSNERLF | 0.82317007 |
| IVTGVLNI | ASSDGTGVSNERLF | 0.8026353 |

| | | |
|---|---|---|
| LAQQGCI | ASSDGTGVSNERLF | 0.85596275 |
| LCLHVEN | ASSDGTGVSNERLF | 0.75526536 |
| LDAISDGS | ASSDGTGVSNERLF | 0.7120179 |
| LGGFAAML | ASSDGTGVSNERLF | 0.4676506 |
| LGGKNPMI | ASSDGTGVSNERLF | 0.8645384 |
| LGPKGVAAIL | ASSDGTGVSNERLF | 0.8414331 |
| LMVGVSI | ASSDGTGVSNERLF | 0.48957142 |
| LVGAAFVL | ASSDGTGVSNERLF | 0.69006413 |
| LVLAVAPL | ASSDGTGVSNERLF | 0.8857969 |
| MAAIAVIDAA | ASSDGTGVSNERLF | 0.88229346 |
| NNGINAELQ | ASSDGTGVSNERLF | 0.7394556 |
| RGEPLVLVHG | ASSDGTGVSNERLF | 0.68591607 |
| RGGAGGAAGLI | ASSDGTGVSNERLF | 0.84572583 |
| RSYRMHPAV | ASSDGTGVSNERLF | 0.34585744 |
| SAAHLLAGQLA | ASSDGTGVSNERLF | 0.84106493 |
| SAAVGSY | ASSDGTGVSNERLF | 0.8214655 |
| SPHAVLHRI | ASSDGTGVSNERLF | 0.8069612 |
| SQIAKVL | ASSDGTGVSNERLF | 0.6660044 |
| TALGHLVP | ASSDGTGVSNERLF | 0.83261263 |
| TAPTTSGAA | ASSDGTGVSNERLF | 0.7576517 |
| TDGDDGNGGSPGT | ASSDGTGVSNERLF | 0.8644227 |
| TDGLGEN | ASSDGTGVSNERLF | 0.16963702 |
| TGGPPLPGQA | ASSDGTGVSNERLF | 0.8392326 |
| TIPAITI | ASSDGTGVSNERLF | 0.6905403 |
| TIPGFTI | ASSDGTGVSNERLF | 0.72948706 |
| TSIASGALA | ASSDGTGVSNERLF | 0.44123515 |
| TVTLPGAAI | ASSDGTGVSNERLF | 0.82344186 |
| VASGLSAALP | ASSDGTGVSNERLF | 0.8781159 |
| VGAGVIGI | ASSDGTGVSNERLF | 0.87311876 |
| VGPKRFGV | ASSDGTGVSNERLF | 0.62540734 |
| VNVAFPD | ASSDGTGVSNERLF | 0.87182236 |
| VSMTNSGVS | ASSDGTGVSNERLF | 0.8876648 |
| VTDVLGDLSV | ASSDGTGVSNERLF | 0.794277 |
| VVVVGAGFAGL | ASSDGTGVSNERLF | 0.31742555 |
| VYTVGTVV | ASSDGTGVSNERLF | 0.84218556 |
| WTTFSGP | ASSDGTGVSNERLF | 0.84490806 |
| YDELVEAF | ASSDGTGVSNERLF | 0.8643231 |
| YGTGGAGGNGG | ASSDGTGVSNERLF | 0.855368 |
| YTNVMAV | ASSDGTGVSNERLF | 0.648775 |
| FLEQVHQL | ASSDGTGVSNERLF | 0.8532382 |
| GLSLLAVAV | ASSDGTGVSNERLF | 0.87392807 |
| LEQVHQL | ASSDGTGVSNERLF | 0.71628845 |
| LNAAGAI | ASSDGTGVSNERLF | 0.78300816 |
| PAVHASL | ASSDGTGVSNERLF | 0.67851156 |
| TELEDDA | ASSDGTGVSNERLF | 0.4084672 |
| TPPGPPN | ASSDGTGVSNERLF | 0.8581524 |
| TPPPPPPP | ASSDGTGVSNERLF | 0.7585968 |
| TQSPALI | ASSQTGGNYAEQF | 0.7182105 |
| EFAAAFPQ | ASSQTGGNYAEQF | 0.8278582 |
| LLSGNLDVL | ASSQTGGNYAEQF | 0.6527443 |
| GPANLGNYNI | ASSQTGGNYAEQF | 0.70384884 |

| | | |
|---|---|---|
| TLGPTLP | ASSQTGGNYAEQF | 0.76246154 |
| VYAYSSCCK | ASSQTGGNYAEQF | 0.81202596 |
| APAPAPAPNI | ASSQTGGNYAEQF | 0.85178065 |
| PGGPVPIP | ASSQTGGNYAEQF | 0.8305218 |
| FLTVALEP | ASSQTGGNYAEQF | 0.83408356 |
| SQYFGANL | ASSQTGGNYAEQF | 0.8807925 |
| GGVKPAA | ASSQTGGNYAEQF | 0.58346546 |
| GGTGGAGGAGGW | ASSQTGGNYAEQF | 0.6353389 |
| GVGGVAGLGGAGG | ASSQTGGNYAEQF | 0.858046 |
| VLLVLATTI | ASSQTGGNYAEQF | 0.46243346 |
| IGIFAAML | ASSQTGGNYAEQF | 0.8642771 |
| AVGDLID | ASSQTGGNYAEQF | 0.80685186 |
| NISPLPL | ASSQTGGNYAEQF | 0.8428886 |
| VTAVFVAI | ASSQTGGNYAEQF | 0.84134257 |
| SGDDNDAAL | ASSQTGGNYAEQF | 0.88898486 |
| AVAALAHS | ASSQTGGNYAEQF | 0.87408936 |
| GGVTVMATP | ASSQTGGNYAEQF | 0.811175 |
| AGLAANVRD | ASSQTGGNYAEQF | 0.8508376 |
| TNRPDLIDPALLRPG | ASSQTGGNYAEQF | 0.5969602 |
| LANLGVTAL | ASSQTGGNYAEQF | 0.87148726 |
| TGERIEL | ASSQTGGNYAEQF | 0.848346 |
| NGEELAA | ASSQTGGNYAEQF | 0.7039796 |
| VLSLVWG | ASSQTGGNYAEQF | 0.8117429 |
| IGYALWL | ASSQTGGNYAEQF | 0.88164127 |
| LSLLLAPL | ASSQTGGNYAEQF | 0.87478113 |
| TLDEALK | ASSQTGGNYAEQF | 0.37164026 |
| TDGAGTAEGG | ASSQTGGNYAEQF | 0.78649986 |
| NAPGGAGGNGG | ASSQTGGNYAEQF | 0.35603637 |
| VSVFMGMGGNF | ASSQTGGNYAEQF | 0.74251974 |
| LASYLAAL | ASSQTGGNYAEQF | 0.52444774 |
| LVGGTPAL | ASSQTGGNYAEQF | 0.8629218 |
| VRNADVL | ASSQTGGNYAEQF | 0.7725573 |
| TCSSPFLVSAM | ASSQTGGNYAEQF | 0.86390936 |
| VHITDAE | ASSQTGGNYAEQF | 0.8312681 |
| EYAAPSF | ASSQTGGNYAEQF | 0.8763197 |
| VVDAFHGGL | ASSQTGGNYAEQF | 0.76170254 |
| ASVVDPQ | ASSQTGGNYAEQF | 0.75855803 |
| SYNEFGGLS | ASSQTGGNYAEQF | 0.40531057 |
| DPDGSVY | ASSQTGGNYAEQF | 0.39434528 |
| MVSTIHATE | ASSQTGGNYAEQF | 0.7809069 |
| LAAGVADPV | ASSQTGGNYAEQF | 0.7496366 |
| TKEQIAA | ASSQTGGNYAEQF | 0.26841068 |
| SLASPAAL | ASSQTGGNYAEQF | 0.8729917 |
| IGLVFYGV | ASSQTGGNYAEQF | 0.8522087 |
| FLLDASNL | ASSQTGGNYAEQF | 0.86038315 |
| YQVGNFP | ASSQTGGNYAEQF | 0.69784135 |
| ESLLSQST | ASSQTGGNYAEQF | 0.7812395 |
| LFTVGFAAVG | ASSQTGGNYAEQF | 0.7438984 |
| ATPPTPT | ASSQTGGNYAEQF | 0.8577101 |
| TDLATLG | ASSQTGGNYAEQF | 0.6787468 |
| SAGGTGR | ASSQTGGNYAEQF | 0.57889956 |

| | | |
|---|---|---|
| GFSNSGS | ASSQTGGNYAEQF | 0.78195024 |
| GGNGGSGGTSV | ASSQTGGNYAEQF | 0.7156427 |
| GTSVATGGAGN | ASSQTGGNYAEQF | 0.88287234 |
| NGGNGGSGGTS | ASSQTGGNYAEQF | 0.8722882 |
| AVVKFPH | ASSQTGGNYAEQF | 0.8523871 |
| TPPPPPP | ASSQTGGNYAEQF | 0.86956984 |
| AAPAAAAAPA | ASSQTGGNYAEQF | 0.27610067 |
| IALNEDL | ASSQTGGNYAEQF | 0.7231747 |
| IVNVSSM | ASSQTGGNYAEQF | 0.87844795 |
| TGIPPAPR | ASSQTGGNYAEQF | 0.8608856 |
| FDLGGGTF | ASSQTGGNYAEQF | 0.78541887 |
| FDLGGGTFDVS | ASSQTGGNYAEQF | 0.87752724 |
| NEPTAAA | ASSQTGGNYAEQF | 0.70944893 |
| NEPTAAALA | ASSQTGGNYAEQF | 0.76422215 |
| GSIINISSI | ASSQTGGNYAEQF | 0.8275901 |
| VAPGNAGTA | ASSQTGGNYAEQF | 0.84115124 |
| TGGGASLE | ASSQTGGNYAEQF | 0.8589586 |
| VAVYNHYKRI | ASSQTGGNYAEQF | 0.67592263 |
| TDPAPATT | ASSQTGGNYAEQF | 0.8395226 |
| TNRPDILDPALLRPG | ASSQTGGNYAEQF | 0.8894248 |
| LTTTTTS | ASSQTGGNYAEQF | 0.81013167 |
| PTGSGKTL | ASSQTGGNYAEQF | 0.8733251 |
| TGSGKTL | ASSQTGGNYAEQF | 0.8489329 |
| TIGHVDHGK | ASSQTGGNYAEQF | 0.84688294 |
| TIGHVDHGKTT | ASSQTGGNYAEQF | 0.71670204 |
| GHVDHGKTT | ASSQTGGNYAEQF | 0.77387923 |
| AQFEHTLL | ASSQTGGNYAEQF | 0.8043086 |
| IVNQDEL | ASSQTGGNYAEQF | 0.72257286 |
| TNGSQFF | ASSQTGGNYAEQF | 0.78475153 |
| TGKGIVE | ASSQTGGNYAEQF | 0.7609991 |
| LISNASDA | ASSQTGGNYAEQF | 0.8921533 |
| AELRTLGELTKIAKAHGAQ | ASSQTGGNYAEQF | 0.8496155 |
| APNAVAAAEPSW | ASSQTGGNYAEQF | 0.85013604 |
| APQVAGAATAVL | ASSQTGGNYAEQF | 0.7038971 |
| ARRDPAAAAAILPTDA | ASSQTGGNYAEQF | 0.8519163 |
| DAAARDVLAEAGLAEYF | ASSQTGGNYAEQF | 0.85056525 |
| DDQIVLSAKANLAKVEQ | ASSQTGGNYAEQF | 0.51279575 |
| DELLPDTGEDSDEDGH | ASSQTGGNYAEQF | 0.87829727 |
| ELPEAPANRATVPQD | ASSQTGGNYAEQF | 0.69246596 |
| FGQSKIERVTGIG | ASSQTGGNYAEQF | 0.781142 |
| GAGGTGSTAGGAGGAGGAGGLY | ASSQTGGNYAEQF | 0.83896923 |
| GHGGTGAAGGNGGTGGAGGSGI | ASSQTGGNYAEQF | 0.8844627 |
| GNGGNGGAGGEGGPGGAGGAGA | ASSQTGGNYAEQF | 0.7655879 |
| GSTAGNGGNGGNATNGTIAGQP | ASSQTGGNYAEQF | 0.7822162 |
| GTGGDGGNGGNGGTGG | ASSQTGGNYAEQF | 0.6538532 |
| GVRRPAGIPADAPV | ASSQTGGNYAEQF | 0.8733041 |
| GYRQVTDAHLLTLARR | ASSQTGGNYAEQF | 0.87565726 |
| IAKMGQDGHDRG | ASSQTGGNYAEQF | 0.8346485 |
| KAEPSIEIQYT | ASSQTGGNYAEQF | 0.8867507 |
| KPVRIRMAADEDTVAVT | ASSQTGGNYAEQF | 0.8407734 |
| MPPRIAGMRLLVIKPEP | ASSQTGGNYAEQF | 0.4297987 |

| | | |
|---|---|---|
| MKFYAVALFRSRGVPPTR | ASSQTGGNYAEQF | 0.8189399 |
| NAGGFGILGAVAHSP | ASSQTGGNYAEQF | 0.86739 |
| NEGREITLAMAEM | ASSQTGGNYAEQF | 0.87547815 |
| NELVEWSHPLGHRWRWE | ASSQTGGNYAEQF | 0.88021135 |
| NLGAGGTGGNGGNPG | ASSQTGGNYAEQF | 0.8518728 |
| NTEWPKWETVMAADPVD | ASSQTGGNYAEQF | 0.8020522 |
| PASAAPAETQP | ASSQTGGNYAEQF | 0.87891394 |
| QQSSDGKNGVRAAEITGE | ASSQTGGNYAEQF | 0.88919777 |
| RAPDAVTTSWWKE | ASSQTGGNYAEQF | 0.8694025 |
| SGFGNVGAHTSGWFNQ | ASSQTGGNYAEQF | 0.761793 |
| SGVAGAVGQAASVGGLKV | ASSQTGGNYAEQF | 0.639647 |
| SNGGAGTGSAGGNGGT | ASSQTGGNYAEQF | 0.8310784 |
| TGGAGGEGGAAG | ASSQTGGNYAEQF | 0.8711314 |
| TGGDGGAAGAGGAPGHG | ASSQTGGNYAEQF | 0.5832864 |
| TMPPLLAAIANAGRHGV | ASSQTGGNYAEQF | 0.820073 |
| TSSTAGNSGGA | ASSQTGGNYAEQF | 0.8434094 |
| VPATVPVGILC | ASSQTGGNYAEQF | 0.54056454 |
| VQRPDSGGVVLTGRIS | ASSQTGGNYAEQF | 0.85846555 |
| WTPNPVPATKGVPAGLVQAT | ASSQTGGNYAEQF | 0.88402426 |
| YALKRALAKFSDRFDVVII | ASSQTGGNYAEQF | 0.78681165 |
| YLIGSVIV | ASSQTGGNYAEQF | 0.77090275 |
| YGQLVAVID | ASSQTGGNYAEQF | 0.87520534 |
| VTVGGMTLPA | ASSQTGGNYAEQF | 0.8921008 |
| VLALPAV | ASSQTGGNYAEQF | 0.8797877 |
| VIPPTLNYET | ASSQTGGNYAEQF | 0.38005394 |
| VGMGGMHPSSG | ASSQTGGNYAEQF | 0.33173206 |
| VGLAGGAATA | ASSQTGGNYAEQF | 0.71396434 |
| VGGAGGAGGSGP | ASSQTGGNYAEQF | 0.86961514 |
| VFAAFMV | ASSQTGGNYAEQF | 0.748263 |
| TLYDKLR | ASSQTGGNYAEQF | 0.7296711 |
| TLAVFTVL | ASSQTGGNYAEQF | 0.8582823 |
| TGSVQPARV | ASSQTGGNYAEQF | 0.87296474 |
| TGGGAGSGGT | ASSQTGGNYAEQF | 0.67554593 |
| SHLETSL | ASSQTGGNYAEQF | 0.5480552 |
| SFTTGMT | ASSQTGGNYAEQF | 0.86091065 |
| QVEVAKRMGVRQPTVS | ASSQTGGNYAEQF | 0.8606351 |
| QGSGGPALAM | ASSQTGGNYAEQF | 0.7000635 |
| MLAPSNP | ASSQTGGNYAEQF | 0.82594943 |
| LVNNAAI | ASSQTGGNYAEQF | 0.7190739 |
| LTGEINTLA | ASSQTGGNYAEQF | 0.45209575 |
| LSIFGPV | ASSQTGGNYAEQF | 0.8371121 |
| LLALLANL | ASSQTGGNYAEQF | 0.6502777 |
| LISNASDALDKLR | ASSQTGGNYAEQF | 0.8796663 |
| LINEAVP | ASSQTGGNYAEQF | 0.7252352 |
| LGYAFSGL | ASSQTGGNYAEQF | 0.8721125 |
| LDFVAVAL | ASSQTGGNYAEQF | 0.8348093 |
| LAQPGAE | ASSQTGGNYAEQF | 0.8477895 |
| ISNLGAVTGF | ASSQTGGNYAEQF | 0.7879108 |
| HQEDPGRI | ASSQTGGNYAEQF | 0.7581213 |
| GTPGEPTP | ASSQTGGNYAEQF | 0.6454649 |
| GFFNSGS | ASSQTGGNYAEQF | 0.33704406 |

| | | |
|---|---|---|
| GAGGGGLTAGAN | ASSQTGGNYAEQF | 0.87383837 |
| FVDALVNM | ASSQTGGNYAEQF | 0.17358208 |
| FLTVAPEL | ASSQTGGNYAEQF | 0.8722795 |
| FETLTAL | ASSQTGGNYAEQF | 0.8608744 |
| DVINPIEL | ASSQTGGNYAEQF | 0.6480746 |
| APVSGAPV | ASSQTGGNYAEQF | 0.8559754 |
| ALVGGLAW | ASSQTGGNYAEQF | 0.70100564 |
| ALAAAAATAI | ASSQTGGNYAEQF | 0.6996969 |
| AFGLGLLL | ASSQTGGNYAEQF | 0.86752987 |
| AFFAAASL | ASSQTGGNYAEQF | 0.8419681 |
| AAAALPAFSPP | ASSQTGGNYAEQF | 0.7122544 |
| VVLAPAL | ASSQTGGNYAEQF | 0.8290302 |
| TIGHVDHGKTTL | ASSQTGGNYAEQF | 0.8471452 |
| LISNASDALDK | ASSQTGGNYAEQF | 0.7609525 |
| LISNASDAL | ASSQTGGNYAEQF | 0.7859124 |
| ISNASDALDK | ASSQTGGNYAEQF | 0.871727 |
| NQAETLVYQTE | ASSQTGGNYAEQF | 0.8189945 |
| TGGTGGTAGAG | ASSQTGGNYAEQF | 0.72569835 |
| TGSTAGGAGGA | ASSQTGGNYAEQF | 0.86504304 |
| DPLAVPVAAGPV | ASSQTGGNYAEQF | 0.6526573 |
| GGTGGTPGNGGN | ASSQTGGNYAEQF | 0.84949064 |
| PAGLAGVDPATL | ASSQTGGNYAEQF | 0.7839842 |
| GNNLVVEAPHAGASV | ASSQTGGNYAEQF | 0.762048 |
| KVTRIMSASTLEQPA | ASSQTGGNYAEQF | 0.86586714 |
| LGFALPIALPRRTVV | ASSQTGGNYAEQF | 0.6787237 |
| TRLGNLTARKLVVLV | ASSQTGGNYAEQF | 0.76536226 |
| WSQQRRLIYFPSAGP | ASSQTGGNYAEQF | 0.7567812 |
| GPNGNPDPMAAAVDIR | ASSQTGGNYAEQF | 0.81923217 |
| TQRCVSVADLRAAASR | ASSQTGGNYAEQF | 0.8684628 |
| QFVLDVAEHPVEVLTETV | ASSQTGGNYAEQF | 0.710265 |
| GGAGGTGGAGGNGGTQAGDG | ASSQTGGNYAEQF | 0.8419002 |
| PLLATLRAIAVDVDAKPAQV | ASSQTGGNYAEQF | 0.7246329 |
| APKFRTGGYMIDFWGVGYQVAK | ASSQTGGNYAEQF | 0.5853163 |
| GAGGAGGAGGAAGGPGGTGGTG | ASSQTGGNYAEQF | 0.7716787 |
| IGALGLLATDVAPAGSVAFGVA | ASSQTGGNYAEQF | 0.7991667 |
| AGGAGGIGGNGGAGGAGGVLMG | ASSQTGGNYAEQF | 0.76834786 |
| ADELIGLL | ASSQTGGNYAEQF | 0.8438978 |
| ADRALKA | ASSQTGGNYAEQF | 0.63580406 |
| AGAVGQVGGAGGA | ASSQTGGNYAEQF | 0.83203375 |
| AGGGGGLGGAP | ASSQTGGNYAEQF | 0.8834568 |
| AGGLLYGNGGAG | ASSQTGGNYAEQF | 0.8065936 |
| AGLESIGL | ASSQTGGNYAEQF | 0.8728162 |
| AGLKTGII | ASSQTGGNYAEQF | 0.8576293 |
| ASLAALLL | ASSQTGGNYAEQF | 0.8508152 |
| ASVINISSI | ASSQTGGNYAEQF | 0.7578399 |
| ATPPGPA | ASSQTGGNYAEQF | 0.730729 |
| ATVASALV | ASSQTGGNYAEQF | 0.87752676 |
| ATVASLIL | ASSQTGGNYAEQF | 0.8610748 |
| AVNGPVLIRH | ASSQTGGNYAEQF | 0.861702 |
| DGAAGAVGI | ASSQTGGNYAEQF | 0.84734905 |
| DPLLHGI | ASSQTGGNYAEQF | 0.8733939 |

| | | |
|---|---|---|
| ELGAALGAL | ASSQTGGNYAEQF | 0.86252666 |
| ELGRTQAV | ASSQTGGNYAEQF | 0.76558733 |
| FVAGGYLS | ASSQTGGNYAEQF | 0.49568158 |
| GASGGAGGAGGW | ASSQTGGNYAEQF | 0.7458752 |
| GGAGAGGDGGAGGI | ASSQTGGNYAEQF | 0.81338143 |
| GTGHNGSQPGMGGNG | ASSQTGGNYAEQF | 0.7999097 |
| GYAAASASAS | ASSQTGGNYAEQF | 0.3621704 |
| HQVDVAAVQQG | ASSQTGGNYAEQF | 0.86796695 |
| IGAAIAHT | ASSQTGGNYAEQF | 0.8901278 |
| IGMIEVAQL | ASSQTGGNYAEQF | 0.69910836 |
| IRLAPEHTSAMD | ASSQTGGNYAEQF | 0.887762 |
| IYNVSVL | ASSQTGGNYAEQF | 0.60237396 |
| KQGFLGM | ASSQTGGNYAEQF | 0.85442764 |
| LESVITQL | ASSQTGGNYAEQF | 0.86968374 |
| LGLGALVVV | ASSQTGGNYAEQF | 0.6870562 |
| LIGNGGNGGSGGI | ASSQTGGNYAEQF | 0.8399636 |
| LNAINAPTL | ASSQTGGNYAEQF | 0.73277926 |
| LQLLNTGV | ASSQTGGNYAEQF | 0.78798974 |
| LQTSIEF | ASSQTGGNYAEQF | 0.8782383 |
| MSFAVSEL | ASSQTGGNYAEQF | 0.7131797 |
| NTGALGL | ASSQTGGNYAEQF | 0.80637753 |
| PPGAGGAGGAGTT | ASSQTGGNYAEQF | 0.7411827 |
| QLVGLGI | ASSQTGGNYAEQF | 0.5883616 |
| RFLEQVHQL | ASSQTGGNYAEQF | 0.8111266 |
| RGVGYVLR | ASSQTGGNYAEQF | 0.86593324 |
| SATGVFAGVI | ASSQTGGNYAEQF | 0.7185877 |
| SLPAITLP | ASSQTGGNYAEQF | 0.88816625 |
| STALAFTV | ASSQTGGNYAEQF | 0.73287535 |
| SVLTVGLGK | ASSQTGGNYAEQF | 0.88769877 |
| TALAGAGSGI | ASSQTGGNYAEQF | 0.8657465 |
| TGLPFTGI | ASSQTGGNYAEQF | 0.8313859 |
| VAEHVTV | ASSQTGGNYAEQF | 0.7312233 |
| VDIVLNSL | ASSQTGGNYAEQF | 0.87687474 |
| VLTGGGAVL | ASSQTGGNYAEQF | 0.7612543 |
| VNVGLAYGI | ASSQTGGNYAEQF | 0.86053 |
| VSSAAVGAV | ASSQTGGNYAEQF | 0.8780375 |
| VTSVYEAGV | ASSQTGGNYAEQF | 0.33970574 |
| WVALDRGA | ASSQTGGNYAEQF | 0.85442126 |
| YLVGSVVI | ASSQTGGNYAEQF | 0.76670134 |
| GPPGTGKTLL | ASSQTGGNYAEQF | 0.7053298 |
| GVGMTKF | ASSQTGGNYAEQF | 0.48006204 |
| IGIGGSDL | ASSQTGGNYAEQF | 0.22358724 |
| IGTLLAEL | ASSQTGGNYAEQF | 0.65071535 |
| IYTSGTT | ASSQTGGNYAEQF | 0.42857164 |
| LSFPAGV | ASSQTGGNYAEQF | 0.8333003 |
| LYTSGTT | ASSQTGGNYAEQF | 0.7648393 |
| VGAFPFGF | ASSQTGGNYAEQF | 0.48114115 |
| VGAGTVEF | ASSQTGGNYAEQF | 0.83933365 |
| VGKSTLLL | ASSQTGGNYAEQF | 0.8367126 |
| AALAHAGGGIV | ASSQTGGNYAEQF | 0.2596156 |
| AALVAVSLL | ASSQTGGNYAEQF | 0.6346508 |

| | | |
|---|---|---|
| AAVSRAVAAL | ASSQTGGNYAEQF | 0.5275595 |
| AGGSSGIGGF | ASSQTGGNYAEQF | 0.5704427 |
| AGLTVANANG | ASSQTGGNYAEQF | 0.2581113 |
| AGLVGAVL | ASSQTGGNYAEQF | 0.76731706 |
| ALSQAFTV | ASSQTGGNYAEQF | 0.6052712 |
| ALTFSVSL | ASSQTGGNYAEQF | 0.13258159 |
| ALTGAALVAL | ASSQTGGNYAEQF | 0.64441246 |
| ASFSIGAGVL | ASSQTGGNYAEQF | 0.6494295 |
| DGSVYKI | ASSQTGGNYAEQF | 0.4885684 |
| EQLAEIV | ASSQTGGNYAEQF | 0.82404107 |
| FAVSELTVM | ASSQTGGNYAEQF | 0.4287409 |
| GAAPLAAGL | ASSQTGGNYAEQF | 0.4506551 |
| GADGGDGGSGGAGG | ASSQTGGNYAEQF | 0.8218937 |
| GFGLDTGGAGG | ASSQTGGNYAEQF | 0.8393856 |
| GLASGGLL | ASSQTGGNYAEQF | 0.32952625 |
| GRIAGAAVLL | ASSQTGGNYAEQF | 0.7779829 |
| GSAGSLNTSF | ASSQTGGNYAEQF | 0.853377 |
| GTGQIGI | ASSQTGGNYAEQF | 0.6939052 |
| GTLIALIL | ASSQTGGNYAEQF | 0.82866496 |
| IAATMGFVVG | ASSQTGGNYAEQF | 0.831038 |
| IFAQALL | ASSQTGGNYAEQF | 0.48727953 |
| IGAAVALGF | ASSQTGGNYAEQF | 0.830914 |
| IGLTPLAL | ASSQTGGNYAEQF | 0.8592633 |
| ILGSNTSTL | ASSQTGGNYAEQF | 0.56879497 |
| IVFSLTVL | ASSQTGGNYAEQF | 0.69405925 |
| LASKATGF | ASSQTGGNYAEQF | 0.3440297 |
| LASVALIL | ASSQTGGNYAEQF | 0.6325655 |
| LFAVGLTL | ASSQTGGNYAEQF | 0.2006214 |
| LFNGGIGTYI | ASSQTGGNYAEQF | 0.8031345 |
| LTLGAGII | ASSQTGGNYAEQF | 0.8046316 |
| LVAAALGDI | ASSQTGGNYAEQF | 0.64318776 |
| NTAFVQLGI | ASSQTGGNYAEQF | 0.3705798 |
| QLFSGAEL | ASSQTGGNYAEQF | 0.3781511 |
| SAATFITL | ASSQTGGNYAEQF | 0.7022163 |
| SNYGLANLGV | ASSQTGGNYAEQF | 0.46949428 |
| SYGGAYAVM | ASSQTGGNYAEQF | 0.20166568 |
| TTGVGGLTLG | ASSQTGGNYAEQF | 0.62821126 |
| TTQTIYI | ASSQTGGNYAEQF | 0.8164653 |
| VAAGAGLGAVI | ASSQTGGNYAEQF | 0.8179164 |
| VAAVVTLL | ASSQTGGNYAEQF | 0.32464504 |
| VGFLGGLI | ASSQTGGNYAEQF | 0.8571441 |
| VGVIPILL | ASSQTGGNYAEQF | 0.82267845 |
| VIFASSLI | ASSQTGGNYAEQF | 0.85423964 |
| VKGGDGGAAAG | ASSQTGGNYAEQF | 0.66963905 |
| VTKTFTNI | ASSQTGGNYAEQF | 0.40900502 |
| VVAAAILL | ASSQTGGNYAEQF | 0.8445505 |
| YAALDVEL | ASSQTGGNYAEQF | 0.6739522 |
| LASVALLL | ASSQTGGNYAEQF | 0.51048267 |
| LDAVHRV | ASSQTGGNYAEQF | 0.7894119 |
| QGLSSSIF | ASSQTGGNYAEQF | 0.5633266 |
| RGTFANIRL | ASSQTGGNYAEQF | 0.5335596 |

| | | |
|---|---|---|
| RQLWWGHRI | ASSQTGGNYAEQF | 0.7682724 |
| YGPPGTGKTLL | ASSQTGGNYAEQF | 0.7940123 |
| AALVGAPANVA | ASSQTGGNYAEQF | 0.4527653 |
| AATFRRTE | ASSQTGGNYAEQF | 0.30241624 |
| AGALGGGI | ASSQTGGNYAEQF | 0.38810676 |
| AGGAGGAGGIGAG | ASSQTGGNYAEQF | 0.6658442 |
| AGSGALGL | ASSQTGGNYAEQF | 0.8315584 |
| AQIADQI | ASSQTGGNYAEQF | 0.6929615 |
| ASAVAVTYL | ASSQTGGNYAEQF | 0.66029894 |
| ASTVLGGTGG | ASSQTGGNYAEQF | 0.7991302 |
| AVGGEPAK | ASSQTGGNYAEQF | 0.7889336 |
| CEQGTFGGTC | ASSQTGGNYAEQF | 0.6825021 |
| DEGKAPAG | ASSQTGGNYAEQF | 0.72742254 |
| DTVVFPAL | ASSQTGGNYAEQF | 0.6280818 |
| FEGITMAG | ASSQTGGNYAEQF | 0.80218416 |
| FPLVVQD | ASSQTGGNYAEQF | 0.78799486 |
| GAFTTPQ | ASSQTGGNYAEQF | 0.7732057 |
| GFGGFGTAGGVGGA | ASSQTGGNYAEQF | 0.8099935 |
| GFLAQFL | ASSQTGGNYAEQF | 0.7424507 |
| GGAGGAGGNAGVA | ASSQTGGNYAEQF | 0.6856901 |
| GGVGTAGAGGAGGAGG | ASSQTGGNYAEQF | 0.21451184 |
| GNGGGGDAGNAGSGGN | ASSQTGGNYAEQF | 0.5765345 |
| GTGGAGGAGNDAGS | ASSQTGGNYAEQF | 0.7173474 |
| GTTNTGWFNS | ASSQTGGNYAEQF | 0.7667171 |
| IFMGPLFTL | ASSQTGGNYAEQF | 0.79597336 |
| ILAHAIEAI | ASSQTGGNYAEQF | 0.71727216 |
| IRSAIREAL | ASSQTGGNYAEQF | 0.4723796 |
| ITFAGGG | ASSQTGGNYAEQF | 0.787555 |
| ITGMSGHQEV | ASSQTGGNYAEQF | 0.8436808 |
| IVAGLACTL | ASSQTGGNYAEQF | 0.29097074 |
| IVGGGLAGA | ASSQTGGNYAEQF | 0.5996921 |
| IVTGVLNI | ASSQTGGNYAEQF | 0.8619175 |
| LAQQGCI | ASSQTGGNYAEQF | 0.80120045 |
| LCLHVEN | ASSQTGGNYAEQF | 0.5737159 |
| LDAISDGS | ASSQTGGNYAEQF | 0.5755821 |
| LGGFAAML | ASSQTGGNYAEQF | 0.8569474 |
| LGGKNPMI | ASSQTGGNYAEQF | 0.5246947 |
| LGPKGVAAIL | ASSQTGGNYAEQF | 0.8500314 |
| LMVGVSI | ASSQTGGNYAEQF | 0.16647407 |
| LVGAAFVL | ASSQTGGNYAEQF | 0.5114825 |
| LVLAVAPL | ASSQTGGNYAEQF | 0.38120314 |
| MAAIAVIDAA | ASSQTGGNYAEQF | 0.17498226 |
| NNGINAELQ | ASSQTGGNYAEQF | 0.79921734 |
| RGEPLVLVHG | ASSQTGGNYAEQF | 0.51843596 |
| RGGAGGAAGLI | ASSQTGGNYAEQF | 0.8217552 |
| RSYRMHPAV | ASSQTGGNYAEQF | 0.70232433 |
| SAAHLLAGQLA | ASSQTGGNYAEQF | 0.81840694 |
| SAAVGSY | ASSQTGGNYAEQF | 0.8085401 |
| SPHAVLHRI | ASSQTGGNYAEQF | 0.6189002 |
| SQIAKVL | ASSQTGGNYAEQF | 0.3784887 |
| TALGHLVP | ASSQTGGNYAEQF | 0.7569032 |

| | | |
|---|---|---|
| TAPTTSGAA | ASSQTGGNYAEQF | 0.8052357 |
| TDGDDGNGGSPGT | ASSQTGGNYAEQF | 0.7655552 |
| TDGLGEN | ASSQTGGNYAEQF | 0.23642029 |
| TGGPPLPGQA | ASSQTGGNYAEQF | 0.6782082 |
| TIPAITI | ASSQTGGNYAEQF | 0.65416867 |
| TIPGFTI | ASSQTGGNYAEQF | 0.19997807 |
| TSIASGALA | ASSQTGGNYAEQF | 0.14799237 |
| TVTLPGAAI | ASSQTGGNYAEQF | 0.51326114 |
| VASGLSAALP | ASSQTGGNYAEQF | 0.64297616 |
| VGAGVIGI | ASSQTGGNYAEQF | 0.82361805 |
| VGPKRFGV | ASSQTGGNYAEQF | 0.8329444 |
| VNVAFPD | ASSQTGGNYAEQF | 0.61399007 |
| VSMTNSGVS | ASSQTGGNYAEQF | 0.8238643 |
| VTDVLGDLSV | ASSQTGGNYAEQF | 0.75132024 |
| VVVVGAGFAGL | ASSQTGGNYAEQF | 0.1176917 |
| VYTVGTVV | ASSQTGGNYAEQF | 0.63114697 |
| WTTFSGP | ASSQTGGNYAEQF | 0.25449887 |
| YDELVEAF | ASSQTGGNYAEQF | 0.81198144 |
| YGTGGAGGNGG | ASSQTGGNYAEQF | 0.370575 |
| YTNVMAV | ASSQTGGNYAEQF | 0.57921994 |
| FLEQVHQL | ASSQTGGNYAEQF | 0.7009679 |
| GLSLLAVAV | ASSQTGGNYAEQF | 0.8139669 |
| LEQVHQL | ASSQTGGNYAEQF | 0.7531915 |
| LNAAGAI | ASSQTGGNYAEQF | 0.4517095 |
| PAVHASL | ASSQTGGNYAEQF | 0.15859403 |
| TELEDDA | ASSQTGGNYAEQF | 0.7759553 |
| TPPGPPN | ASSQTGGNYAEQF | 0.34461147 |
| TPPPPPPP | ASSQTGGNYAEQF | 0.32323304 |
| TQSPALI | ASGDRLGGDEQY | 0.22622912 |
| EFAAAFPQ | ASGDRLGGDEQY | 0.16725503 |
| LLSGNLDVL | ASGDRLGGDEQY | 0.2930281 |
| GPANLGNYNI | ASGDRLGGDEQY | 0.78605056 |
| TLGPTLP | ASGDRLGGDEQY | 0.6312005 |
| VYAYSSCCK | ASGDRLGGDEQY | 0.6575632 |
| APAPAPAPNI | ASGDRLGGDEQY | 0.8546366 |
| PGGPVPIP | ASGDRLGGDEQY | 0.8110763 |
| FLTVALEP | ASGDRLGGDEQY | 0.66370094 |
| SQYFGANL | ASGDRLGGDEQY | 0.47901022 |
| GGVKPAA | ASGDRLGGDEQY | 0.34282345 |
| GGTGGAGGAGGW | ASGDRLGGDEQY | 0.3755871 |
| GVGGVAGLGGAGG | ASGDRLGGDEQY | 0.39806637 |
| VLLVLATTI | ASGDRLGGDEQY | 0.8479593 |
| IGIFAAML | ASGDRLGGDEQY | 0.5417047 |
| AVGDLID | ASGDRLGGDEQY | 0.3420899 |
| NISPLPL | ASGDRLGGDEQY | 0.7505592 |
| VTAVFVAI | ASGDRLGGDEQY | 0.8466284 |
| SGDDNDAAL | ASGDRLGGDEQY | 0.8663782 |
| AVAALAHS | ASGDRLGGDEQY | 0.3565517 |
| GGVTVMATP | ASGDRLGGDEQY | 0.5970503 |
| AGLAANVRD | ASGDRLGGDEQY | 0.58645296 |
| TNRPDLIDPALLRPG | ASGDRLGGDEQY | 0.7284104 |

| | | |
|---|---|---|
| LANLGVTAL | ASGDRLGGDEQY | 0.52629435 |
| TGERIEL | ASGDRLGGDEQY | 0.3009681 |
| NGEELAA | ASGDRLGGDEQY | 0.7912786 |
| VLSLVWG | ASGDRLGGDEQY | 0.86390007 |
| IGYALWL | ASGDRLGGDEQY | 0.53672045 |
| LSLLLAPL | ASGDRLGGDEQY | 0.44556966 |
| TLDEALK | ASGDRLGGDEQY | 0.6662774 |
| TDGAGTAEGG | ASGDRLGGDEQY | 0.49963143 |
| NAPGGAGGNGG | ASGDRLGGDEQY | 0.5605763 |
| VSVFMGMGGNF | ASGDRLGGDEQY | 0.69721544 |
| LASYLAAL | ASGDRLGGDEQY | 0.26793888 |
| LVGGTPAL | ASGDRLGGDEQY | 0.6406132 |
| VRNADVL | ASGDRLGGDEQY | 0.8590338 |
| TCSSPFLVSAM | ASGDRLGGDEQY | 0.7035736 |
| VHITDAE | ASGDRLGGDEQY | 0.79422873 |
| EYAAPSF | ASGDRLGGDEQY | 0.8461722 |
| VVDAFHGGL | ASGDRLGGDEQY | 0.7967085 |
| ASVVDPQ | ASGDRLGGDEQY | 0.8465421 |
| SYNEFGGLS | ASGDRLGGDEQY | 0.8321974 |
| DPDGSVY | ASGDRLGGDEQY | 0.5630269 |
| MVSTIHATE | ASGDRLGGDEQY | 0.71506906 |
| LAAGVADPV | ASGDRLGGDEQY | 0.7905958 |
| TKEQIAA | ASGDRLGGDEQY | 0.680819 |
| SLASPAAL | ASGDRLGGDEQY | 0.6686889 |
| IGLVFYGV | ASGDRLGGDEQY | 0.638985 |
| FLLDASNL | ASGDRLGGDEQY | 0.8225767 |
| YQVGNFP | ASGDRLGGDEQY | 0.72411764 |
| ESLLSQST | ASGDRLGGDEQY | 0.8278175 |
| LFTVGFAAVG | ASGDRLGGDEQY | 0.21300457 |
| ATPPTPT | ASGDRLGGDEQY | 0.32225817 |
| TDLATLG | ASGDRLGGDEQY | 0.83621186 |
| SAGGTGR | ASGDRLGGDEQY | 0.14010425 |
| GFSNSGS | ASGDRLGGDEQY | 0.17528039 |
| GGNGGSGGTSV | ASGDRLGGDEQY | 0.70323586 |
| GTSVATGGAGN | ASGDRLGGDEQY | 0.85184324 |
| NGGNGGSGGTS | ASGDRLGGDEQY | 0.1413243 |
| AVVKFPH | ASGDRLGGDEQY | 0.40455246 |
| TPPPPPP | ASGDRLGGDEQY | 0.26931807 |
| AAPAAAAAPA | ASGDRLGGDEQY | 0.79717517 |
| IALNEDL | ASGDRLGGDEQY | 0.6939891 |
| IVNVSSM | ASGDRLGGDEQY | 0.21366723 |
| TGIPPAPR | ASGDRLGGDEQY | 0.7689452 |
| FDLGGGTF | ASGDRLGGDEQY | 0.3262009 |
| FDLGGGTFDVS | ASGDRLGGDEQY | 0.39504924 |
| NEPTAAA | ASGDRLGGDEQY | 0.81870204 |
| NEPTAAALA | ASGDRLGGDEQY | 0.85230315 |
| GSIINISSI | ASGDRLGGDEQY | 0.29510313 |
| VAPGNAGTA | ASGDRLGGDEQY | 0.8158797 |
| TGGGASLE | ASGDRLGGDEQY | 0.5770077 |
| VAVYNHYKRI | ASGDRLGGDEQY | 0.8677092 |
| TDPAPATT | ASGDRLGGDEQY | 0.53927135 |

| | | |
|---|---|---|
| TNRPDILDPALLRPG | ASGDRLGGDEQY | 0.23242283 |
| LTTTTTS | ASGDRLGGDEQY | 0.80847013 |
| PTGSGKTL | ASGDRLGGDEQY | 0.8143474 |
| TGSGKTL | ASGDRLGGDEQY | 0.52103555 |
| TIGHVDHGK | ASGDRLGGDEQY | 0.82668674 |
| TIGHVDHGKTT | ASGDRLGGDEQY | 0.5748373 |
| GHVDHGKTT | ASGDRLGGDEQY | 0.6742182 |
| AQFEHTLL | ASGDRLGGDEQY | 0.842314 |
| IVNQDEL | ASGDRLGGDEQY | 0.7850684 |
| TNGSQFF | ASGDRLGGDEQY | 0.3064405 |
| TGKGIVE | ASGDRLGGDEQY | 0.7785257 |
| LISNASDA | ASGDRLGGDEQY | 0.71343005 |
| AELRTLGELTKIAKAHGAQ | ASGDRLGGDEQY | 0.4002394 |
| APNAVAAAEPSW | ASGDRLGGDEQY | 0.8537067 |
| APQVAGAATAVL | ASGDRLGGDEQY | 0.62496716 |
| ARRDPAAAAAILPTDA | ASGDRLGGDEQY | 0.82470655 |
| DAAARDVLAEAGLAEYF | ASGDRLGGDEQY | 0.7280631 |
| DDQIVLSAKANLAKVEQ | ASGDRLGGDEQY | 0.84682685 |
| DELLPDTGEDSDEDGH | ASGDRLGGDEQY | 0.23313981 |
| ELPEAPANRATVPQD | ASGDRLGGDEQY | 0.79700947 |
| FGQSKIERVTGIG | ASGDRLGGDEQY | 0.6960434 |
| GAGGTGSTAGGAGGAGGAGGLY | ASGDRLGGDEQY | 0.8138224 |
| GHGGTGAAGGNGGTGGAGGSGI | ASGDRLGGDEQY | 0.7369571 |
| GNGGNGGAGGEGGPGGAGGAGA | ASGDRLGGDEQY | 0.49867666 |
| GSTAGNGGNGGNATNGTIAGQP | ASGDRLGGDEQY | 0.71976906 |
| GTGGDGGNGGNGGTGG | ASGDRLGGDEQY | 0.8345859 |
| GVRRPAGIPADAPV | ASGDRLGGDEQY | 0.6248419 |
| GYRQVTDAHLLTLARR | ASGDRLGGDEQY | 0.71585715 |
| IAKMGQDGHDRG | ASGDRLGGDEQY | 0.8542955 |
| KAEPSIEIQYT | ASGDRLGGDEQY | 0.31721354 |
| KPVRIRMAADEDTVAVT | ASGDRLGGDEQY | 0.66697186 |
| MPPRIAGMRLLVIKPEP | ASGDRLGGDEQY | 0.7740095 |
| MKFYAVALFRSRGVPPTR | ASGDRLGGDEQY | 0.81507874 |
| NAGGFGILGAVAHSP | ASGDRLGGDEQY | 0.6096672 |
| NEGREITLAMAEM | ASGDRLGGDEQY | 0.8319926 |
| NELVEWSHPLGHRWRWE | ASGDRLGGDEQY | 0.5223268 |
| NLGAGGTGGNGGNPG | ASGDRLGGDEQY | 0.8156008 |
| NTEWPKWETVMAADPVD | ASGDRLGGDEQY | 0.684633 |
| PASAAPAETQP | ASGDRLGGDEQY | 0.76860064 |
| QQSSDGKNGVRAAEITGE | ASGDRLGGDEQY | 0.38551027 |
| RAPDAVTTSWWKE | ASGDRLGGDEQY | 0.80147564 |
| SGFGNVGAHTSGWFNQ | ASGDRLGGDEQY | 0.83540505 |
| SGVAGAVGQAASVGGLKV | ASGDRLGGDEQY | 0.8289052 |
| SNGGAGTGSAGGNGGT | ASGDRLGGDEQY | 0.16779277 |
| TGGAGGEGGAAG | ASGDRLGGDEQY | 0.82051384 |
| TGGDGGAAGAGGAPGHG | ASGDRLGGDEQY | 0.8319392 |
| TMPPLLAAIANAGRHGV | ASGDRLGGDEQY | 0.8630972 |
| TSSTAGNSGGA | ASGDRLGGDEQY | 0.6974735 |
| VPATVPVGILC | ASGDRLGGDEQY | 0.78119063 |
| VQRPDSGGVVLTGRIS | ASGDRLGGDEQY | 0.5676023 |
| WTPNPVPATKGVPAGLVQAT | ASGDRLGGDEQY | 0.64983165 |

| | | |
|---|---|---|
| YALKRALAKFSDRFDVVII | ASGDRLGGDEQY | 0.37982744 |
| YLIGSVIV | ASGDRLGGDEQY | 0.59738415 |
| YGQLVAVID | ASGDRLGGDEQY | 0.67701924 |
| VTVGGMTLPA | ASGDRLGGDEQY | 0.4989418 |
| VLALPAV | ASGDRLGGDEQY | 0.8375678 |
| VIPPTLNYET | ASGDRLGGDEQY | 0.8232249 |
| VGMGGMHPSSG | ASGDRLGGDEQY | 0.8304955 |
| VGLAGGAATA | ASGDRLGGDEQY | 0.8443053 |
| VGGAGGAGGSGP | ASGDRLGGDEQY | 0.8529867 |
| VFAAFMV | ASGDRLGGDEQY | 0.7585974 |
| TLYDKLR | ASGDRLGGDEQY | 0.58477175 |
| TLAVFTVL | ASGDRLGGDEQY | 0.652259 |
| TGSVQPARV | ASGDRLGGDEQY | 0.33704388 |
| TGGGAGSGGT | ASGDRLGGDEQY | 0.6207115 |
| SHLETSL | ASGDRLGGDEQY | 0.56762075 |
| SFTTGMT | ASGDRLGGDEQY | 0.7498457 |
| QVEVAKRMGVRQPTVS | ASGDRLGGDEQY | 0.12965326 |
| QGSGGPALAM | ASGDRLGGDEQY | 0.80396056 |
| MLAPSNP | ASGDRLGGDEQY | 0.84990954 |
| LVNNAAI | ASGDRLGGDEQY | 0.668967 |
| LTGEINTLA | ASGDRLGGDEQY | 0.758228 |
| LSIFGPV | ASGDRLGGDEQY | 0.79589057 |
| LLALLANL | ASGDRLGGDEQY | 0.566133 |
| LISNASDALDKLR | ASGDRLGGDEQY | 0.8142359 |
| LINEAVP | ASGDRLGGDEQY | 0.42259106 |
| LGYAFSGL | ASGDRLGGDEQY | 0.7873958 |
| LDFVAVAL | ASGDRLGGDEQY | 0.8490311 |
| LAQPGAE | ASGDRLGGDEQY | 0.7329126 |
| ISNLGAVTGF | ASGDRLGGDEQY | 0.63038194 |
| HQEDPGRI | ASGDRLGGDEQY | 0.6620896 |
| GTPGEPTP | ASGDRLGGDEQY | 0.21915245 |
| GFFNSGS | ASGDRLGGDEQY | 0.68310326 |
| GAGGGGLTAGAN | ASGDRLGGDEQY | 0.12191687 |
| FVDALVNM | ASGDRLGGDEQY | 0.75735056 |
| FLTVAPEL | ASGDRLGGDEQY | 0.3078462 |
| FETLTAL | ASGDRLGGDEQY | 0.71522415 |
| DVINPIEL | ASGDRLGGDEQY | 0.7300801 |
| APVSGAPV | ASGDRLGGDEQY | 0.77621603 |
| ALVGGLAW | ASGDRLGGDEQY | 0.42545 |
| ALAAAAATAI | ASGDRLGGDEQY | 0.70443106 |
| AFGLGLLL | ASGDRLGGDEQY | 0.52441555 |
| AFFAAASL | ASGDRLGGDEQY | 0.29294676 |
| AAAALPAFSPP | ASGDRLGGDEQY | 0.83624315 |
| VVLAPAL | ASGDRLGGDEQY | 0.4888811 |
| TIGHVDHGKTTL | ASGDRLGGDEQY | 0.75603163 |
| LISNASDALDK | ASGDRLGGDEQY | 0.8160064 |
| LISNASDAL | ASGDRLGGDEQY | 0.6714134 |
| ISNASDALDK | ASGDRLGGDEQY | 0.29493237 |
| NQAETLVYQTE | ASGDRLGGDEQY | 0.737437 |
| TGGTGGTAGAG | ASGDRLGGDEQY | 0.65495664 |
| TGSTAGGAGGA | ASGDRLGGDEQY | 0.5366794 |

| | | |
|---|---|---|
| DPLAVPVAAGPV | ASGDRLGGDEQY | 0.1311283 |
| GGTGGTPGNGGN | ASGDRLGGDEQY | 0.7660929 |
| PAGLAGVDPATL | ASGDRLGGDEQY | 0.41010106 |
| GNNLVVEAPHAGASV | ASGDRLGGDEQY | 0.66117567 |
| KVTRIMSASTLEQPA | ASGDRLGGDEQY | 0.54349035 |
| LGFALPIALPRRTVV | ASGDRLGGDEQY | 0.8259047 |
| TRLGNLTARKLVVLV | ASGDRLGGDEQY | 0.5524564 |
| WSQQRRLIYFPSAGP | ASGDRLGGDEQY | 0.5280541 |
| GPNGNPDPMAAAVDIR | ASGDRLGGDEQY | 0.78461385 |
| TQRCVSVADLRAAASR | ASGDRLGGDEQY | 0.7256261 |
| QFVLDVAEHPVEVLTETV | ASGDRLGGDEQY | 0.84558284 |
| GGAGGTGGAGGNGGTQAGDG | ASGDRLGGDEQY | 0.70724916 |
| PLLATLRAIAVDVDAKPAQV | ASGDRLGGDEQY | 0.6112448 |
| APKFRTGGYMIDFWGVGYQVAK | ASGDRLGGDEQY | 0.73453224 |
| GAGGAGGAGGAAGGPGGTGGTG | ASGDRLGGDEQY | 0.6960392 |
| IGALGLLATDVAPAGSVAFGVA | ASGDRLGGDEQY | 0.19493766 |
| AGGAGGIGGNGGAGGAGGVLMG | ASGDRLGGDEQY | 0.19942841 |
| ADELIGLL | ASGDRLGGDEQY | 0.7739654 |
| ADRALKA | ASGDRLGGDEQY | 0.18478331 |
| AGAVGQVGGAGGA | ASGDRLGGDEQY | 0.7172513 |
| AGGGGGLGGAP | ASGDRLGGDEQY | 0.8033771 |
| AGGLLYGNGGAG | ASGDRLGGDEQY | 0.7516331 |
| AGLESIGL | ASGDRLGGDEQY | 0.2632608 |
| AGLKTGII | ASGDRLGGDEQY | 0.564044 |
| ASLAALLL | ASGDRLGGDEQY | 0.48564604 |
| ASVINISSI | ASGDRLGGDEQY | 0.80020946 |
| ATPPGPA | ASGDRLGGDEQY | 0.76429653 |
| ATVASALV | ASGDRLGGDEQY | 0.8547989 |
| ATVASLIL | ASGDRLGGDEQY | 0.8343522 |
| AVNGPVLIRH | ASGDRLGGDEQY | 0.6244143 |
| DGAAGAVGI | ASGDRLGGDEQY | 0.8053951 |
| DPLLHGI | ASGDRLGGDEQY | 0.6171248 |
| ELGAALGAL | ASGDRLGGDEQY | 0.72817266 |
| ELGRTQAV | ASGDRLGGDEQY | 0.6182163 |
| FVAGGYLS | ASGDRLGGDEQY | 0.7817658 |
| GASGGAGGAGGW | ASGDRLGGDEQY | 0.5272994 |
| GGAGAGGDGGAGGI | ASGDRLGGDEQY | 0.51009476 |
| GTGHNGSQPGMGGNG | ASGDRLGGDEQY | 0.29869235 |
| GYAAASASAS | ASGDRLGGDEQY | 0.756005 |
| HQVDVAAVQQG | ASGDRLGGDEQY | 0.7507446 |
| IGAAIAHT | ASGDRLGGDEQY | 0.26466328 |
| IGMIEVAQL | ASGDRLGGDEQY | 0.5446625 |
| IRLAPEHTSAMD | ASGDRLGGDEQY | 0.8624662 |
| IYNVSVL | ASGDRLGGDEQY | 0.8508737 |
| KQGFLGM | ASGDRLGGDEQY | 0.5039566 |
| LESVITQL | ASGDRLGGDEQY | 0.44765475 |
| LGLGALVVV | ASGDRLGGDEQY | 0.81497645 |
| LIGNGGNGGSGGI | ASGDRLGGDEQY | 0.284043 |
| LNAINAPTL | ASGDRLGGDEQY | 0.7020453 |
| LQLLNTGV | ASGDRLGGDEQY | 0.76712245 |
| LQTSIEF | ASGDRLGGDEQY | 0.75944257 |

| | | |
|---|---|---|
| MSFAVSEL | ASGDRLGGDEQY | 0.47667477 |
| NTGALGL | ASGDRLGGDEQY | 0.7849481 |
| PPGAGGAGGAGTT | ASGDRLGGDEQY | 0.6968194 |
| QLVGLGI | ASGDRLGGDEQY | 0.7268486 |
| RFLEQVHQL | ASGDRLGGDEQY | 0.14691564 |
| RGVGYVLR | ASGDRLGGDEQY | 0.773215 |
| SATGVFAGVI | ASGDRLGGDEQY | 0.44952762 |
| SLPAITLP | ASGDRLGGDEQY | 0.47498238 |
| STALAFTV | ASGDRLGGDEQY | 0.210581 |
| SVLTVGLGK | ASGDRLGGDEQY | 0.68229544 |
| TALAGAGSGI | ASGDRLGGDEQY | 0.8497724 |
| TGLPFTGI | ASGDRLGGDEQY | 0.8425783 |
| VAEHVTV | ASGDRLGGDEQY | 0.39658204 |
| VDIVLNSL | ASGDRLGGDEQY | 0.82527775 |
| VLTGGGAVL | ASGDRLGGDEQY | 0.8621949 |
| VNVGLAYGI | ASGDRLGGDEQY | 0.6553276 |
| VSSAAVGAV | ASGDRLGGDEQY | 0.18245032 |
| VTSVYEAGV | ASGDRLGGDEQY | 0.7591522 |
| WVALDRGA | ASGDRLGGDEQY | 0.76523226 |
| YLVGSVVI | ASGDRLGGDEQY | 0.82821125 |
| GPPGTGKTLL | ASGDRLGGDEQY | 0.8368808 |
| GVGMTKF | ASGDRLGGDEQY | 0.3425069 |
| IGIGGSDL | ASGDRLGGDEQY | 0.78510517 |
| IGTLLAEL | ASGDRLGGDEQY | 0.81878734 |
| IYTSGTT | ASGDRLGGDEQY | 0.3974523 |
| LSFPAGV | ASGDRLGGDEQY | 0.48281658 |
| LYTSGTT | ASGDRLGGDEQY | 0.5355752 |
| VGAFPFGF | ASGDRLGGDEQY | 0.30751318 |
| VGAGTVEF | ASGDRLGGDEQY | 0.7384935 |
| VGKSTLLL | ASGDRLGGDEQY | 0.60397995 |
| AALAHAGGGIV | ASGDRLGGDEQY | 0.52866596 |
| AALVAVSLL | ASGDRLGGDEQY | 0.662294 |
| AAVSRAVAAL | ASGDRLGGDEQY | 0.4165184 |
| AGGSSGIGGF | ASGDRLGGDEQY | 0.46658564 |
| AGLTVANANG | ASGDRLGGDEQY | 0.522174 |
| AGLVGAVL | ASGDRLGGDEQY | 0.71574426 |
| ALSQAFTV | ASGDRLGGDEQY | 0.800306 |
| ALTFSVSL | ASGDRLGGDEQY | 0.7675807 |
| ALTGAALVAL | ASGDRLGGDEQY | 0.8028395 |
| ASFSIGAGVL | ASGDRLGGDEQY | 0.8512692 |
| DGSVYKI | ASGDRLGGDEQY | 0.3192206 |
| EQLAEIV | ASGDRLGGDEQY | 0.45684963 |
| FAVSELTVM | ASGDRLGGDEQY | 0.7922909 |
| GAAPLAAGL | ASGDRLGGDEQY | 0.31921875 |
| GADGGDGGSGGAGG | ASGDRLGGDEQY | 0.82235163 |
| GFGLDTGGAGG | ASGDRLGGDEQY | 0.74513495 |
| GLASGGLL | ASGDRLGGDEQY | 0.7792507 |
| GRIAGAAVLL | ASGDRLGGDEQY | 0.76029396 |
| GSAGSLNTSF | ASGDRLGGDEQY | 0.86499155 |
| GTGQIGI | ASGDRLGGDEQY | 0.7791596 |
| GTLIALIL | ASGDRLGGDEQY | 0.62554306 |

| | | |
|---|---|---|
| IAATMGFVVG | ASGDRLGGDEQY | 0.7913281 |
| IFAQALL | ASGDRLGGDEQY | 0.37849325 |
| IGAAVALGF | ASGDRLGGDEQY | 0.8226706 |
| IGLTPLAL | ASGDRLGGDEQY | 0.7847769 |
| ILGSNTSTL | ASGDRLGGDEQY | 0.4993934 |
| IVFSLTVL | ASGDRLGGDEQY | 0.6750049 |
| LASKATGF | ASGDRLGGDEQY | 0.8309531 |
| LASVALIL | ASGDRLGGDEQY | 0.84572035 |
| LFAVGLTL | ASGDRLGGDEQY | 0.21172231 |
| LFNGGIGTYI | ASGDRLGGDEQY | 0.55738217 |
| LTLGAGII | ASGDRLGGDEQY | 0.24759403 |
| LVAAALGDI | ASGDRLGGDEQY | 0.5856097 |
| NTAFVQLGI | ASGDRLGGDEQY | 0.2728429 |
| QLFSGAEL | ASGDRLGGDEQY | 0.79049814 |
| SAATFITL | ASGDRLGGDEQY | 0.56767464 |
| SNYGLANLGV | ASGDRLGGDEQY | 0.83128715 |
| SYGGAYAVM | ASGDRLGGDEQY | 0.7222107 |
| TTGVGGLTLG | ASGDRLGGDEQY | 0.81956077 |
| TTQTIYI | ASGDRLGGDEQY | 0.4692798 |
| VAAGAGLGAVI | ASGDRLGGDEQY | 0.602242 |
| VAAVVTLL | ASGDRLGGDEQY | 0.2275011 |
| VGFLGGLI | ASGDRLGGDEQY | 0.2764141 |
| VGVIPILL | ASGDRLGGDEQY | 0.41710642 |
| VIFASSLI | ASGDRLGGDEQY | 0.6181879 |
| VKGGDGGAAAG | ASGDRLGGDEQY | 0.19786744 |
| VTKTFTNI | ASGDRLGGDEQY | 0.8559391 |
| VVAAAILL | ASGDRLGGDEQY | 0.6867035 |
| YAALDVEL | ASGDRLGGDEQY | 0.78104657 |
| LASVALLL | ASGDRLGGDEQY | 0.5078692 |
| LDAVHRV | ASGDRLGGDEQY | 0.4518648 |
| QGLSSSIF | ASGDRLGGDEQY | 0.5457282 |
| RGTFANIRL | ASGDRLGGDEQY | 0.7952628 |
| RQLWWGHRI | ASGDRLGGDEQY | 0.3263377 |
| YGPPGTGKTLL | ASGDRLGGDEQY | 0.25034156 |
| AALVGAPANVA | ASGDRLGGDEQY | 0.6228443 |
| AATFRRTE | ASGDRLGGDEQY | 0.5039144 |
| AGALGGGI | ASGDRLGGDEQY | 0.8548543 |
| AGGAGGAGGIGAG | ASGDRLGGDEQY | 0.83264244 |
| AGSGALGL | ASGDRLGGDEQY | 0.7482033 |
| AQIADQI | ASGDRLGGDEQY | 0.82976997 |
| ASAVAVTYL | ASGDRLGGDEQY | 0.15618858 |
| ASTVLGGTGG | ASGDRLGGDEQY | 0.57109344 |
| AVGGEPAK | ASGDRLGGDEQY | 0.85560775 |
| CEQGTFGGTC | ASGDRLGGDEQY | 0.7533096 |
| DEGKAPAG | ASGDRLGGDEQY | 0.62054944 |
| DTVVFPAL | ASGDRLGGDEQY | 0.83450127 |
| FEGITMAG | ASGDRLGGDEQY | 0.5282395 |
| FPLVVQD | ASGDRLGGDEQY | 0.550954 |
| GAFTTPQ | ASGDRLGGDEQY | 0.63353956 |
| GFGGFGTAGGVGGA | ASGDRLGGDEQY | 0.7150296 |
| GFLAQFL | ASGDRLGGDEQY | 0.76696813 |

| | | |
|---|---|---|
| GGAGGAGGNAGVA | ASGDRLGGDEQY | 0.29109937 |
| GGVGTAGAGGAGGAGG | ASGDRLGGDEQY | 0.7205103 |
| GNGGGGDAGNAGSGGN | ASGDRLGGDEQY | 0.8660888 |
| GTGGAGGAGNDAGS | ASGDRLGGDEQY | 0.62409306 |
| GTTNTGWFNS | ASGDRLGGDEQY | 0.83699965 |
| IFMGPLFTL | ASGDRLGGDEQY | 0.7754647 |
| ILAHAIEAI | ASGDRLGGDEQY | 0.8246604 |
| IRSAIREAL | ASGDRLGGDEQY | 0.47354835 |
| ITFAGGG | ASGDRLGGDEQY | 0.45663768 |
| ITGMSGHQEV | ASGDRLGGDEQY | 0.6876932 |
| IVAGLACTL | ASGDRLGGDEQY | 0.46049395 |
| IVGGGLAGA | ASGDRLGGDEQY | 0.6784041 |
| IVTGVLNI | ASGDRLGGDEQY | 0.5845869 |
| LAQQGCI | ASGDRLGGDEQY | 0.86783516 |
| LCLHVEN | ASGDRLGGDEQY | 0.8080699 |
| LDAISDGS | ASGDRLGGDEQY | 0.775126 |
| LGGFAAML | ASGDRLGGDEQY | 0.40053692 |
| LGGKNPMI | ASGDRLGGDEQY | 0.7105752 |
| LGPKGVAAIL | ASGDRLGGDEQY | 0.7890883 |
| LMVGVSI | ASGDRLGGDEQY | 0.31170988 |
| LVGAAFVL | ASGDRLGGDEQY | 0.843018 |
| LVLAVAPL | ASGDRLGGDEQY | 0.43807042 |
| MAAIAVIDAA | ASGDRLGGDEQY | 0.7440096 |
| NNGINAELQ | ASGDRLGGDEQY | 0.8004036 |
| RGEPLVLVHG | ASGDRLGGDEQY | 0.84752584 |
| RGGAGGAAGLI | ASGDRLGGDEQY | 0.5688077 |
| RSYRMHPAV | ASGDRLGGDEQY | 0.68419886 |
| SAAHLLAGQLA | ASGDRLGGDEQY | 0.3776948 |
| SAAVGSY | ASGDRLGGDEQY | 0.8146024 |
| SPHAVLHRI | ASGDRLGGDEQY | 0.83234525 |
| SQIAKVL | ASGDRLGGDEQY | 0.7451103 |
| TALGHLVP | ASGDRLGGDEQY | 0.85290337 |
| TAPTTSGAA | ASGDRLGGDEQY | 0.76968575 |
| TDGDDGNGGSPGT | ASGDRLGGDEQY | 0.22923473 |
| TDGLGEN | ASGDRLGGDEQY | 0.5387943 |
| TGGPPLPGQA | ASGDRLGGDEQY | 0.8330023 |
| TIPAITI | ASGDRLGGDEQY | 0.85010797 |
| TIPGFTI | ASGDRLGGDEQY | 0.82734925 |
| TSIASGALA | ASGDRLGGDEQY | 0.76589584 |
| TVTLPGAAI | ASGDRLGGDEQY | 0.6478729 |
| VASGLSAALP | ASGDRLGGDEQY | 0.8521453 |
| VGAGVIGI | ASGDRLGGDEQY | 0.8668293 |
| VGPKRFGV | ASGDRLGGDEQY | 0.81693804 |
| VNVAFPD | ASGDRLGGDEQY | 0.5480644 |
| VSMTNSGVS | ASGDRLGGDEQY | 0.49029082 |
| VTDVLGDLSV | ASGDRLGGDEQY | 0.70736545 |
| VVVVGAGFAGL | ASGDRLGGDEQY | 0.8332629 |
| VYTVGTVV | ASGDRLGGDEQY | 0.34693775 |
| WTTFSGP | ASGDRLGGDEQY | 0.7390789 |
| YDELVEAF | ASGDRLGGDEQY | 0.7157347 |
| YGTGGAGGNGG | ASGDRLGGDEQY | 0.21540897 |

| | | |
|---|---|---|
| YTNVMAV | ASGDRLGGDEQY | 0.83317757 |
| FLEQVHQL | ASGDRLGGDEQY | 0.85499763 |
| GLSLLAVAV | ASGDRLGGDEQY | 0.6370869 |
| LEQVHQL | ASGDRLGGDEQY | 0.4569052 |
| LNAAGAI | ASGDRLGGDEQY | 0.842815 |
| PAVHASL | ASGDRLGGDEQY | 0.8647212 |
| TELEDDA | ASGDRLGGDEQY | 0.8171542 |
| TPPGPPN | ASGDRLGGDEQY | 0.23486486 |
| TPPPPPPP | ASGDRLGGDEQY | 0.17359112 |
| TQSPALI | TCSAENSGNTLY | 0.5212158 |
| EFAAAFPQ | TCSAENSGNTLY | 0.82960397 |
| LLSGNLDVL | TCSAENSGNTLY | 0.5318429 |
| GPANLGNYNI | TCSAENSGNTLY | 0.3710568 |
| TLGPTLP | TCSAENSGNTLY | 0.80430055 |
| VYAYSSCCK | TCSAENSGNTLY | 0.8383686 |
| APAPAPAPNI | TCSAENSGNTLY | 0.30987555 |
| PGGPVPIP | TCSAENSGNTLY | 0.29556638 |
| FLTVALEP | TCSAENSGNTLY | 0.8313025 |
| SQYFGANL | TCSAENSGNTLY | 0.8083379 |
| GGVKPAA | TCSAENSGNTLY | 0.4844511 |
| GGTGGAGGAGGW | TCSAENSGNTLY | 0.617851 |
| GVGGVAGLGGAGG | TCSAENSGNTLY | 0.5873245 |
| VLLVLATTI | TCSAENSGNTLY | 0.5904146 |
| IGIFAAML | TCSAENSGNTLY | 0.7829845 |
| AVGDLID | TCSAENSGNTLY | 0.37210965 |
| NISPLPL | TCSAENSGNTLY | 0.8537758 |
| VTAVFVAI | TCSAENSGNTLY | 0.34603107 |
| SGDDNDAAL | TCSAENSGNTLY | 0.7985586 |
| AVAALAHS | TCSAENSGNTLY | 0.70445514 |
| GGVTVMATP | TCSAENSGNTLY | 0.6928109 |
| AGLAANVRD | TCSAENSGNTLY | 0.5737629 |
| TNRPDLIDPALLRPG | TCSAENSGNTLY | 0.5565829 |
| LANLGVTAL | TCSAENSGNTLY | 0.31540054 |
| TGERIEL | TCSAENSGNTLY | 0.21591097 |
| NGEELAA | TCSAENSGNTLY | 0.8290931 |
| VLSLVWG | TCSAENSGNTLY | 0.16188364 |
| IGYALWL | TCSAENSGNTLY | 0.8122013 |
| LSLLLAPL | TCSAENSGNTLY | 0.7205929 |
| TLDEALK | TCSAENSGNTLY | 0.3877765 |
| TDGAGTAEGG | TCSAENSGNTLY | 0.75411767 |
| NAPGGAGGNGG | TCSAENSGNTLY | 0.28790492 |
| VSVFMGMGGNF | TCSAENSGNTLY | 0.40213406 |
| LASYLAAL | TCSAENSGNTLY | 0.82597166 |
| LVGGTPAL | TCSAENSGNTLY | 0.7603103 |
| VRNADVL | TCSAENSGNTLY | 0.4334742 |
| TCSSPFLVSAM | TCSAENSGNTLY | 0.70185363 |
| VHITDAE | TCSAENSGNTLY | 0.76544946 |
| EYAAPSF | TCSAENSGNTLY | 0.4889005 |
| VVDAFHGGL | TCSAENSGNTLY | 0.6093205 |
| ASVVDPQ | TCSAENSGNTLY | 0.84060085 |
| SYNEFGGLS | TCSAENSGNTLY | 0.6777758 |

| | | |
|---|---|---|
| DPDGSVY | TCSAENSGNTLY | 0.31596142 |
| MVSTIHATE | TCSAENSGNTLY | 0.8068259 |
| LAAGVADPV | TCSAENSGNTLY | 0.4664833 |
| TKEQIAA | TCSAENSGNTLY | 0.7798575 |
| SLASPAAL | TCSAENSGNTLY | 0.65364 |
| IGLVFYGV | TCSAENSGNTLY | 0.56854975 |
| FLLDASNL | TCSAENSGNTLY | 0.77684975 |
| YQVGNFP | TCSAENSGNTLY | 0.59044707 |
| ESLLSQST | TCSAENSGNTLY | 0.5136785 |
| LFTVGFAAVG | TCSAENSGNTLY | 0.54129237 |
| ATPPTPT | TCSAENSGNTLY | 0.64860344 |
| TDLATLG | TCSAENSGNTLY | 0.8125348 |
| SAGGTGR | TCSAENSGNTLY | 0.42260545 |
| GFSNSGS | TCSAENSGNTLY | 0.73604906 |
| GGNGGSGGTSV | TCSAENSGNTLY | 0.50570583 |
| GTSVATGGAGN | TCSAENSGNTLY | 0.3741138 |
| NGGNGGSGGTS | TCSAENSGNTLY | 0.59745145 |
| AVVKFPH | TCSAENSGNTLY | 0.61058736 |
| TPPPPPP | TCSAENSGNTLY | 0.3639542 |
| AAPAAAAAPA | TCSAENSGNTLY | 0.7542226 |
| IALNEDL | TCSAENSGNTLY | 0.30890715 |
| IVNVSSM | TCSAENSGNTLY | 0.6893359 |
| TGIPPAPR | TCSAENSGNTLY | 0.858431 |
| FDLGGGTF | TCSAENSGNTLY | 0.37398177 |
| FDLGGGTFDVS | TCSAENSGNTLY | 0.78072476 |
| NEPTAAA | TCSAENSGNTLY | 0.7973546 |
| NEPTAAALA | TCSAENSGNTLY | 0.767308 |
| GSIINISSI | TCSAENSGNTLY | 0.5619687 |
| VAPGNAGTA | TCSAENSGNTLY | 0.48625743 |
| TGGGASLE | TCSAENSGNTLY | 0.8414793 |
| VAVYNHYKRI | TCSAENSGNTLY | 0.7906647 |
| TDPAPATT | TCSAENSGNTLY | 0.8147368 |
| TNRPDILDPALLRPG | TCSAENSGNTLY | 0.8050114 |
| LTTTTTS | TCSAENSGNTLY | 0.82728314 |
| PTGSGKTL | TCSAENSGNTLY | 0.7989263 |
| TGSGKTL | TCSAENSGNTLY | 0.5815183 |
| TIGHVDHGK | TCSAENSGNTLY | 0.26118222 |
| TIGHVDHGKTT | TCSAENSGNTLY | 0.6051534 |
| GHVDHGKTT | TCSAENSGNTLY | 0.5992056 |
| AQFEHTLL | TCSAENSGNTLY | 0.5029998 |
| IVNQDEL | TCSAENSGNTLY | 0.25272626 |
| TNGSQFF | TCSAENSGNTLY | 0.843217 |
| TGKGIVE | TCSAENSGNTLY | 0.86658776 |
| LISNASDA | TCSAENSGNTLY | 0.5060153 |
| AELRTLGELTKIAKAHGAQ | TCSAENSGNTLY | 0.8574451 |
| APNAVAAAEPSW | TCSAENSGNTLY | 0.31603426 |
| APQVAGAATAVL | TCSAENSGNTLY | 0.7756294 |
| ARRDPAAAAAILPTDA | TCSAENSGNTLY | 0.8432295 |
| DAAARDVLAEAGLAEYF | TCSAENSGNTLY | 0.48386392 |
| DDQIVLSAKANLAKVEQ | TCSAENSGNTLY | 0.7672686 |
| DELLPDTGEDSDEDGH | TCSAENSGNTLY | 0.4072569 |

| | | |
|---|---|---|
| ELPEAPANRATVPQD | TCSAENSGNTLY | 0.48015553 |
| FGQSKIERVTGIG | TCSAENSGNTLY | 0.8348081 |
| GAGGTGSTAGGAGGAGGAGGLY | TCSAENSGNTLY | 0.51315546 |
| GHGGTGAAGGNGGTGGAGGSGI | TCSAENSGNTLY | 0.7121157 |
| GNGGNGGAGGEGGPGGAGGAGA | TCSAENSGNTLY | 0.49244007 |
| GSTAGNGGNGGNATNGTIAGQP | TCSAENSGNTLY | 0.3787648 |
| GTGGDGGNGGNGGTGG | TCSAENSGNTLY | 0.6565744 |
| GVRRPAGIPADAPV | TCSAENSGNTLY | 0.82295406 |
| GYRQVTDAHLLTLARR | TCSAENSGNTLY | 0.49919856 |
| IAKMGQDGHDRG | TCSAENSGNTLY | 0.8650433 |
| KAEPSIEIQYT | TCSAENSGNTLY | 0.39049032 |
| KPVRIRMAADEDTVAVT | TCSAENSGNTLY | 0.86340415 |
| MPPRIAGMRLLVIKPEP | TCSAENSGNTLY | 0.84302986 |
| MKFYAVALFRSRGVPPTR | TCSAENSGNTLY | 0.73501396 |
| NAGGFGILGAVAHSP | TCSAENSGNTLY | 0.54641616 |
| NEGREITLAMAEM | TCSAENSGNTLY | 0.8387912 |
| NELVEWSHPLGHRWRWE | TCSAENSGNTLY | 0.52888536 |
| NLGAGGTGGNGGNPG | TCSAENSGNTLY | 0.74464 |
| NTEWPKWETVMAADPVD | TCSAENSGNTLY | 0.8096744 |
| PASAAPAETQP | TCSAENSGNTLY | 0.44015902 |
| QQSSDGKNGVRAAEITGE | TCSAENSGNTLY | 0.86298466 |
| RAPDAVTTSWWKE | TCSAENSGNTLY | 0.8160825 |
| SGFGNVGAHTSGWFNQ | TCSAENSGNTLY | 0.7794807 |
| SGVAGAVGQAASVGGLKV | TCSAENSGNTLY | 0.6274877 |
| SNGGAGTGSAGGNGGT | TCSAENSGNTLY | 0.37124524 |
| TGGAGGEGGAAG | TCSAENSGNTLY | 0.6707577 |
| TGGDGGAAGAGGAPGHG | TCSAENSGNTLY | 0.5128626 |
| TMPPLLAAIANAGRHGV | TCSAENSGNTLY | 0.8465543 |
| TSSTAGNSGGA | TCSAENSGNTLY | 0.84096324 |
| VPATVPVGILC | TCSAENSGNTLY | 0.5937286 |
| VQRPDSGGVVLTGRIS | TCSAENSGNTLY | 0.80813426 |
| WTPNPVPATKGVPAGLVQAT | TCSAENSGNTLY | 0.81963426 |
| YALKRALAKFSDRFDVVII | TCSAENSGNTLY | 0.3828001 |
| YLIGSVIV | TCSAENSGNTLY | 0.76592815 |
| YGQLVAVID | TCSAENSGNTLY | 0.48563397 |
| VTVGGMTLPA | TCSAENSGNTLY | 0.6935023 |
| VLALPAV | TCSAENSGNTLY | 0.2705266 |
| VIPPTLNYET | TCSAENSGNTLY | 0.83515966 |
| VGMGGMHPSSG | TCSAENSGNTLY | 0.6941036 |
| VGLAGGAATA | TCSAENSGNTLY | 0.16548622 |
| VGGAGGAGGSGP | TCSAENSGNTLY | 0.79188794 |
| VFAAFMV | TCSAENSGNTLY | 0.63465303 |
| TLYDKLR | TCSAENSGNTLY | 0.6330394 |
| TLAVFTVL | TCSAENSGNTLY | 0.838403 |
| TGSVQPARV | TCSAENSGNTLY | 0.56685054 |
| TGGGAGSGGT | TCSAENSGNTLY | 0.488222 |
| SHLETSL | TCSAENSGNTLY | 0.8116854 |
| SFTTGMT | TCSAENSGNTLY | 0.83123875 |
| QVEVAKRMGVRQPTVS | TCSAENSGNTLY | 0.39309198 |
| QGSGGPALAM | TCSAENSGNTLY | 0.8058598 |
| MLAPSNP | TCSAENSGNTLY | 0.8554846 |

| | | |
|---|---|---|
| LVNNAAI | TCSAENSGNTLY | 0.74005735 |
| LTGEINTLA | TCSAENSGNTLY | 0.8702146 |
| LSIFGPV | TCSAENSGNTLY | 0.87835705 |
| LLALLANL | TCSAENSGNTLY | 0.69032985 |
| LISNASDALDKLR | TCSAENSGNTLY | 0.85396504 |
| LINEAVP | TCSAENSGNTLY | 0.8541543 |
| LGYAFSGL | TCSAENSGNTLY | 0.64419794 |
| LDFVAVAL | TCSAENSGNTLY | 0.71289027 |
| LAQPGAE | TCSAENSGNTLY | 0.42499435 |
| ISNLGAVTGF | TCSAENSGNTLY | 0.7539053 |
| HQEDPGRI | TCSAENSGNTLY | 0.44234017 |
| GTPGEPTP | TCSAENSGNTLY | 0.8011754 |
| GFFNSGS | TCSAENSGNTLY | 0.8532523 |
| GAGGGGLTAGAN | TCSAENSGNTLY | 0.6810492 |
| FVDALVNM | TCSAENSGNTLY | 0.4725882 |
| FLTVAPEL | TCSAENSGNTLY | 0.40363103 |
| FETLTAL | TCSAENSGNTLY | 0.83532286 |
| DVINPIEL | TCSAENSGNTLY | 0.44609702 |
| APVSGAPV | TCSAENSGNTLY | 0.43963253 |
| ALVGGLAW | TCSAENSGNTLY | 0.64858925 |
| ALAAAAATAI | TCSAENSGNTLY | 0.8054533 |
| AFGLGLLL | TCSAENSGNTLY | 0.80657077 |
| AFFAAASL | TCSAENSGNTLY | 0.5272943 |
| AAAALPAFSPP | TCSAENSGNTLY | 0.87673986 |
| VVLAPAL | TCSAENSGNTLY | 0.84998345 |
| TIGHVDHGKTTL | TCSAENSGNTLY | 0.867834 |
| LISNASDALDK | TCSAENSGNTLY | 0.7354331 |
| LISNASDAL | TCSAENSGNTLY | 0.4437031 |
| ISNASDALDK | TCSAENSGNTLY | 0.8440319 |
| NQAETLVYQTE | TCSAENSGNTLY | 0.67460203 |
| TGGTGGTAGAG | TCSAENSGNTLY | 0.73348004 |
| TGSTAGGAGGA | TCSAENSGNTLY | 0.81453216 |
| DPLAVPVAAGPV | TCSAENSGNTLY | 0.68421566 |
| GGTGGTPGNGGN | TCSAENSGNTLY | 0.5254748 |
| PAGLAGVDPATL | TCSAENSGNTLY | 0.74505126 |
| GNNLVVEAPHAGASV | TCSAENSGNTLY | 0.88119316 |
| KVTRIMSASTLEQPA | TCSAENSGNTLY | 0.54372597 |
| LGFALPIALPRRTVV | TCSAENSGNTLY | 0.4481967 |
| TRLGNLTARKLVVLV | TCSAENSGNTLY | 0.68940675 |
| WSQQRRLIYFPSAGP | TCSAENSGNTLY | 0.66330254 |
| GPNGNPDPMAAAVDIR | TCSAENSGNTLY | 0.83330286 |
| TQRCVSVADLRAAASR | TCSAENSGNTLY | 0.81458133 |
| QFVLDVAEHPVEVLTETV | TCSAENSGNTLY | 0.7150758 |
| GGAGGTGGAGGNGGTQAGDG | TCSAENSGNTLY | 0.79562914 |
| PLLATLRAIAVDVDAKPAQV | TCSAENSGNTLY | 0.81092435 |
| APKFRTGGYMIDFWGVGYQVAK | TCSAENSGNTLY | 0.76474726 |
| GAGGAGGAGGAAGGPGGTGGTG | TCSAENSGNTLY | 0.78255045 |
| IGALGLLATDVAPAGSVAFGVA | TCSAENSGNTLY | 0.6160584 |
| AGGAGGIGGNGGAGGAGGVLMG | TCSAENSGNTLY | 0.87696254 |
| ADELIGLL | TCSAENSGNTLY | 0.8225404 |
| ADRALKA | TCSAENSGNTLY | 0.84880686 |

| | | |
|---|---|---|
| AGAVGQVGGAGGA | TCSAENSGNTLY | 0.8704338 |
| AGGGGGLGGAP | TCSAENSGNTLY | 0.72775805 |
| AGGLLYGNGGAG | TCSAENSGNTLY | 0.80218697 |
| AGLESIGL | TCSAENSGNTLY | 0.36309543 |
| AGLKTGII | TCSAENSGNTLY | 0.5571709 |
| ASLAALLL | TCSAENSGNTLY | 0.74183035 |
| ASVINISSI | TCSAENSGNTLY | 0.8218037 |
| ATPPGPA | TCSAENSGNTLY | 0.8454093 |
| ATVASALV | TCSAENSGNTLY | 0.83423734 |
| ATVASLIL | TCSAENSGNTLY | 0.61224896 |
| AVNGPVLIRH | TCSAENSGNTLY | 0.8616241 |
| DGAAGAVGI | TCSAENSGNTLY | 0.86096203 |
| DPLLHGI | TCSAENSGNTLY | 0.46477413 |
| ELGAALGAL | TCSAENSGNTLY | 0.68295294 |
| ELGRTQAV | TCSAENSGNTLY | 0.8766649 |
| FVAGGYLS | TCSAENSGNTLY | 0.8106382 |
| GASGGAGGAGGW | TCSAENSGNTLY | 0.6542163 |
| GGAGAGGDGGAGGI | TCSAENSGNTLY | 0.69816685 |
| GTGHNGSQPGMGGNG | TCSAENSGNTLY | 0.8661245 |
| GYAAASASAS | TCSAENSGNTLY | 0.75911856 |
| HQVDVAAVQQG | TCSAENSGNTLY | 0.8734343 |
| IGAAIAHT | TCSAENSGNTLY | 0.24258068 |
| IGMIEVAQL | TCSAENSGNTLY | 0.772805 |
| IRLAPEHTSAMD | TCSAENSGNTLY | 0.7195431 |
| IYNVSVL | TCSAENSGNTLY | 0.21814434 |
| KQGFLGM | TCSAENSGNTLY | 0.829903 |
| LESVITQL | TCSAENSGNTLY | 0.66257036 |
| LGLGALVVV | TCSAENSGNTLY | 0.78577113 |
| LIGNGGNGGSGGI | TCSAENSGNTLY | 0.66481143 |
| LNAINAPTL | TCSAENSGNTLY | 0.84918296 |
| LQLLNTGV | TCSAENSGNTLY | 0.79190636 |
| LQTSIEF | TCSAENSGNTLY | 0.6516458 |
| MSFAVSEL | TCSAENSGNTLY | 0.49285597 |
| NTGALGL | TCSAENSGNTLY | 0.7720294 |
| PPGAGGAGGAGTT | TCSAENSGNTLY | 0.87185425 |
| QLVGLGI | TCSAENSGNTLY | 0.8699544 |
| RFLEQVHQL | TCSAENSGNTLY | 0.27119058 |
| RGVGYVLR | TCSAENSGNTLY | 0.8255457 |
| SATGVFAGVI | TCSAENSGNTLY | 0.69053507 |
| SLPAITLP | TCSAENSGNTLY | 0.2594155 |
| STALAFTV | TCSAENSGNTLY | 0.2796719 |
| SVLTVGLGK | TCSAENSGNTLY | 0.78960294 |
| TALAGAGSGI | TCSAENSGNTLY | 0.76282847 |
| TGLPFTGI | TCSAENSGNTLY | 0.7966534 |
| VAEHVTV | TCSAENSGNTLY | 0.7913173 |
| VDIVLNSL | TCSAENSGNTLY | 0.5908923 |
| VLTGGGAVL | TCSAENSGNTLY | 0.80507535 |
| VNVGLAYGI | TCSAENSGNTLY | 0.7288655 |
| VSSAAVGAV | TCSAENSGNTLY | 0.096139476 |
| VTSVYEAGV | TCSAENSGNTLY | 0.5980302 |
| WVALDRGA | TCSAENSGNTLY | 0.3581726 |

| | | |
|---|---|---|
| YLVGSVVI | TCSAENSGNTLY | 0.87798876 |
| GPPGTGKTLL | TCSAENSGNTLY | 0.5450245 |
| GVGMTKF | TCSAENSGNTLY | 0.65294504 |
| IGIGGSDL | TCSAENSGNTLY | 0.67100006 |
| IGTLLAEL | TCSAENSGNTLY | 0.8160478 |
| IYTSGTT | TCSAENSGNTLY | 0.7363498 |
| LSFPAGV | TCSAENSGNTLY | 0.6121088 |
| LYTSGTT | TCSAENSGNTLY | 0.27096 |
| VGAFPFGF | TCSAENSGNTLY | 0.75967693 |
| VGAGTVEF | TCSAENSGNTLY | 0.527549 |
| VGKSTLLL | TCSAENSGNTLY | 0.47379467 |
| AALAHAGGGIV | TCSAENSGNTLY | 0.52546334 |
| AALVAVSLL | TCSAENSGNTLY | 0.61528885 |
| AAVSRAVAAL | TCSAENSGNTLY | 0.33045006 |
| AGGSSGIGGF | TCSAENSGNTLY | 0.8561974 |
| AGLTVANANG | TCSAENSGNTLY | 0.55981183 |
| AGLVGAVL | TCSAENSGNTLY | 0.65299976 |
| ALSQAFTV | TCSAENSGNTLY | 0.8785273 |
| ALTFSVSL | TCSAENSGNTLY | 0.76791674 |
| ALTGAALVAL | TCSAENSGNTLY | 0.69207186 |
| ASFSIGAGVL | TCSAENSGNTLY | 0.52526563 |
| DGSVYKI | TCSAENSGNTLY | 0.5538608 |
| EQLAEIV | TCSAENSGNTLY | 0.53256917 |
| FAVSELTVM | TCSAENSGNTLY | 0.5571059 |
| GAAPLAAGL | TCSAENSGNTLY | 0.8485519 |
| GADGGDGGSGGAGG | TCSAENSGNTLY | 0.804994 |
| GFGLDTGGAGG | TCSAENSGNTLY | 0.46037167 |
| GLASGGLL | TCSAENSGNTLY | 0.8573526 |
| GRIAGAAVLL | TCSAENSGNTLY | 0.8756106 |
| GSAGSLNTSF | TCSAENSGNTLY | 0.8821841 |
| GTGQIGI | TCSAENSGNTLY | 0.52547956 |
| GTLIALIL | TCSAENSGNTLY | 0.6306999 |
| IAATMGFVVG | TCSAENSGNTLY | 0.66885036 |
| IFAQALL | TCSAENSGNTLY | 0.7122124 |
| IGAAVALGF | TCSAENSGNTLY | 0.47856668 |
| IGLTPLAL | TCSAENSGNTLY | 0.46369186 |
| ILGSNTSTL | TCSAENSGNTLY | 0.8624843 |
| IVFSLTVL | TCSAENSGNTLY | 0.8796654 |
| LASKATGF | TCSAENSGNTLY | 0.7039702 |
| LASVALIL | TCSAENSGNTLY | 0.2746408 |
| LFAVGLTL | TCSAENSGNTLY | 0.6626463 |
| LFNGGIGTYI | TCSAENSGNTLY | 0.56168723 |
| LTLGAGII | TCSAENSGNTLY | 0.5369945 |
| LVAAALGDI | TCSAENSGNTLY | 0.58500683 |
| NTAFVQLGI | TCSAENSGNTLY | 0.36619833 |
| QLFSGAEL | TCSAENSGNTLY | 0.7224983 |
| SAATFITL | TCSAENSGNTLY | 0.87479377 |
| SNYGLANLGV | TCSAENSGNTLY | 0.70198035 |
| SYGGAYAVM | TCSAENSGNTLY | 0.8128536 |
| TTGVGGLTLG | TCSAENSGNTLY | 0.8668034 |
| TTQTIYI | TCSAENSGNTLY | 0.8333705 |

| | | |
|---|---|---|
| VAAGAGLGAVI | TCSAENSGNTLY | 0.8620109 |
| VAAVVTLL | TCSAENSGNTLY | 0.8524984 |
| VGFLGGLI | TCSAENSGNTLY | 0.7318943 |
| VGVIPILL | TCSAENSGNTLY | 0.79168975 |
| VIFASSLI | TCSAENSGNTLY | 0.85451084 |
| VKGGDGGAAAG | TCSAENSGNTLY | 0.6355296 |
| VTKTFTNI | TCSAENSGNTLY | 0.7300357 |
| VVAAAILL | TCSAENSGNTLY | 0.757517 |
| YAALDVEL | TCSAENSGNTLY | 0.8279643 |
| LASVALLL | TCSAENSGNTLY | 0.72138095 |
| LDAVHRV | TCSAENSGNTLY | 0.87328184 |
| QGLSSSIF | TCSAENSGNTLY | 0.25080028 |
| RGTFANIRL | TCSAENSGNTLY | 0.42579445 |
| RQLWWGHRI | TCSAENSGNTLY | 0.8645705 |
| YGPPGTGKTLL | TCSAENSGNTLY | 0.13423653 |
| AALVGAPANVA | TCSAENSGNTLY | 0.3425012 |
| AATFRRTE | TCSAENSGNTLY | 0.81538045 |
| AGALGGGI | TCSAENSGNTLY | 0.863605 |
| AGGAGGAGGIGAG | TCSAENSGNTLY | 0.15840356 |
| AGSGALGL | TCSAENSGNTLY | 0.561854 |
| AQIADQI | TCSAENSGNTLY | 0.3562063 |
| ASAVAVTYL | TCSAENSGNTLY | 0.8507149 |
| ASTVLGGTGG | TCSAENSGNTLY | 0.6790169 |
| AVGGEPAK | TCSAENSGNTLY | 0.32240826 |
| CEQGTFGGTC | TCSAENSGNTLY | 0.8284754 |
| DEGKAPAG | TCSAENSGNTLY | 0.37522924 |
| DTVVFPAL | TCSAENSGNTLY | 0.7199297 |
| FEGITMAG | TCSAENSGNTLY | 0.8445393 |
| FPLVVQD | TCSAENSGNTLY | 0.8759791 |
| GAFTTPQ | TCSAENSGNTLY | 0.41276038 |
| GFGGFGTAGGVGGA | TCSAENSGNTLY | 0.7862917 |
| GFLAQFL | TCSAENSGNTLY | 0.7256044 |
| GGAGGAGGNAGVA | TCSAENSGNTLY | 0.8835965 |
| GGVGTAGAGGAGGAGG | TCSAENSGNTLY | 0.6347004 |
| GNGGGGDAGNAGSGGN | TCSAENSGNTLY | 0.39331463 |
| GTGGAGGAGNDAGS | TCSAENSGNTLY | 0.85436916 |
| GTTNTGWFNS | TCSAENSGNTLY | 0.8579598 |
| IFMGPLFTL | TCSAENSGNTLY | 0.6595641 |
| ILAHAIEAI | TCSAENSGNTLY | 0.8540029 |
| IRSAIREAL | TCSAENSGNTLY | 0.7618313 |
| ITFAGGG | TCSAENSGNTLY | 0.6938394 |
| ITGMSGHQEV | TCSAENSGNTLY | 0.82452285 |
| IVAGLACTL | TCSAENSGNTLY | 0.8260796 |
| IVGGGLAGA | TCSAENSGNTLY | 0.53873885 |
| IVTGVLNI | TCSAENSGNTLY | 0.8655635 |
| LAQQGCI | TCSAENSGNTLY | 0.69962597 |
| LCLHVEN | TCSAENSGNTLY | 0.43109393 |
| LDAISDGS | TCSAENSGNTLY | 0.886847 |
| LGGFAAML | TCSAENSGNTLY | 0.7592282 |
| LGGKNPMI | TCSAENSGNTLY | 0.8404225 |
| LGPKGVAAIL | TCSAENSGNTLY | 0.8246572 |

| | | |
|---|---|---|
| LMVGVSI | TCSAENSGNTLY | 0.86257553 |
| LVGAAFVL | TCSAENSGNTLY | 0.44937703 |
| LVLAVAPL | TCSAENSGNTLY | 0.7257806 |
| MAAIAVIDAA | TCSAENSGNTLY | 0.6994642 |
| NNGINAELQ | TCSAENSGNTLY | 0.8396706 |
| RGEPLVLVHG | TCSAENSGNTLY | 0.76789016 |
| RGGAGGAAGLI | TCSAENSGNTLY | 0.70342106 |
| RSYRMHPAV | TCSAENSGNTLY | 0.8454493 |
| SAAHLLAGQLA | TCSAENSGNTLY | 0.8597956 |
| SAAVGSY | TCSAENSGNTLY | 0.795516 |
| SPHAVLHRI | TCSAENSGNTLY | 0.75607836 |
| SQIAKVL | TCSAENSGNTLY | 0.8684706 |
| TALGHLVP | TCSAENSGNTLY | 0.5414643 |
| TAPTTSGAA | TCSAENSGNTLY | 0.69307363 |
| TDGDDGNGGSPGT | TCSAENSGNTLY | 0.8588241 |
| TDGLGEN | TCSAENSGNTLY | 0.853943 |
| TGGPPLPGQA | TCSAENSGNTLY | 0.7723576 |
| TIPAITI | TCSAENSGNTLY | 0.8661843 |
| TIPGFTI | TCSAENSGNTLY | 0.589406 |
| TSIASGALA | TCSAENSGNTLY | 0.77496314 |
| TVTLPGAAI | TCSAENSGNTLY | 0.7369676 |
| VASGLSAALP | TCSAENSGNTLY | 0.5874376 |
| VGAGVIGI | TCSAENSGNTLY | 0.5485401 |
| VGPKRFGV | TCSAENSGNTLY | 0.72897434 |
| VNVAFPD | TCSAENSGNTLY | 0.84151804 |
| VSMTNSGVS | TCSAENSGNTLY | 0.7707689 |
| VTDVLGDLSV | TCSAENSGNTLY | 0.24518539 |
| VVVVGAGFAGL | TCSAENSGNTLY | 0.7902994 |
| VYTVGTVV | TCSAENSGNTLY | 0.7718416 |
| WTTFSGP | TCSAENSGNTLY | 0.88328695 |
| YDELVEAF | TCSAENSGNTLY | 0.7364701 |
| YGTGGAGGNGG | TCSAENSGNTLY | 0.82726324 |
| YTNVMAV | TCSAENSGNTLY | 0.55028427 |
| FLEQVHQL | TCSAENSGNTLY | 0.77934426 |
| GLSLLAVAV | TCSAENSGNTLY | 0.3783928 |
| LEQVHQL | TCSAENSGNTLY | 0.53475046 |
| LNAAGAI | TCSAENSGNTLY | 0.6495025 |
| PAVHASL | TCSAENSGNTLY | 0.5682024 |
| TELEDDA | TCSAENSGNTLY | 0.8359729 |
| TPPGPPN | TCSAENSGNTLY | 0.84024197 |
| TPPPPPPP | TCSAENSGNTLY | 0.832381 |
| TQSPALI | ASRGPVDTNYAEQF | 0.8591895 |
| EFAAAFPQ | ASRGPVDTNYAEQF | 0.87614316 |
| LLSGNLDVL | ASRGPVDTNYAEQF | 0.8588548 |
| GPANLGNYNI | ASRGPVDTNYAEQF | 0.60419136 |
| TLGPTLP | ASRGPVDTNYAEQF | 0.66248727 |
| VYAYSSCCK | ASRGPVDTNYAEQF | 0.36492276 |
| APAPAPAPNI | ASRGPVDTNYAEQF | 0.6838215 |
| PGGPVPIP | ASRGPVDTNYAEQF | 0.66604877 |
| FLTVALEP | ASRGPVDTNYAEQF | 0.7128651 |
| SQYFGANL | ASRGPVDTNYAEQF | 0.14931405 |

| | | |
|---|---|---|
| GGVKPAA | ASRGPVDTNYAEQF | 0.8043561 |
| GGTGGAGGAGGW | ASRGPVDTNYAEQF | 0.85792047 |
| GVGGVAGLGGAGG | ASRGPVDTNYAEQF | 0.6606384 |
| VLLVLATTI | ASRGPVDTNYAEQF | 0.79258 |
| IGIFAAML | ASRGPVDTNYAEQF | 0.8329028 |
| AVGDLID | ASRGPVDTNYAEQF | 0.7205863 |
| NISPLPL | ASRGPVDTNYAEQF | 0.81511915 |
| VTAVFVAI | ASRGPVDTNYAEQF | 0.64123464 |
| SGDDNDAAL | ASRGPVDTNYAEQF | 0.8462336 |
| AVAALAHS | ASRGPVDTNYAEQF | 0.8660919 |
| GGVTVMATP | ASRGPVDTNYAEQF | 0.8079846 |
| AGLAANVRD | ASRGPVDTNYAEQF | 0.7643776 |
| TNRPDLIDPALLRPG | ASRGPVDTNYAEQF | 0.7680732 |
| LANLGVTAL | ASRGPVDTNYAEQF | 0.39299423 |
| TGERIEL | ASRGPVDTNYAEQF | 0.7853817 |
| NGEELAA | ASRGPVDTNYAEQF | 0.18900493 |
| VLSLVWG | ASRGPVDTNYAEQF | 0.7745174 |
| IGYALWL | ASRGPVDTNYAEQF | 0.5695416 |
| LSLLLAPL | ASRGPVDTNYAEQF | 0.76149607 |
| TLDEALK | ASRGPVDTNYAEQF | 0.72099245 |
| TDGAGTAEGG | ASRGPVDTNYAEQF | 0.8209877 |
| NAPGGAGGNGG | ASRGPVDTNYAEQF | 0.49564636 |
| VSVFMGMGGNF | ASRGPVDTNYAEQF | 0.822742 |
| LASYLAAL | ASRGPVDTNYAEQF | 0.7702033 |
| LVGGTPAL | ASRGPVDTNYAEQF | 0.3823793 |
| VRNADVL | ASRGPVDTNYAEQF | 0.8615949 |
| TCSSPFLVSAM | ASRGPVDTNYAEQF | 0.7933811 |
| VHITDAE | ASRGPVDTNYAEQF | 0.75132227 |
| EYAAPSF | ASRGPVDTNYAEQF | 0.84271944 |
| VVDAFHGGL | ASRGPVDTNYAEQF | 0.73978996 |
| ASVVDPQ | ASRGPVDTNYAEQF | 0.42135146 |
| SYNEFGGLS | ASRGPVDTNYAEQF | 0.6640339 |
| DPDGSVY | ASRGPVDTNYAEQF | 0.7927782 |
| MVSTIHATE | ASRGPVDTNYAEQF | 0.6697439 |
| LAAGVADPV | ASRGPVDTNYAEQF | 0.15462631 |
| TKEQIAA | ASRGPVDTNYAEQF | 0.8388871 |
| SLASPAAL | ASRGPVDTNYAEQF | 0.43087327 |
| IGLVFYGV | ASRGPVDTNYAEQF | 0.6353955 |
| FLLDASNL | ASRGPVDTNYAEQF | 0.678337 |
| YQVGNFP | ASRGPVDTNYAEQF | 0.8239199 |
| ESLLSQST | ASRGPVDTNYAEQF | 0.67684245 |
| LFTVGFAAVG | ASRGPVDTNYAEQF | 0.632713 |
| ATPPTPT | ASRGPVDTNYAEQF | 0.81705326 |
| TDLATLG | ASRGPVDTNYAEQF | 0.7181983 |
| SAGGTGR | ASRGPVDTNYAEQF | 0.8205111 |
| GFSNSGS | ASRGPVDTNYAEQF | 0.7131419 |
| GGNGGSGGTSV | ASRGPVDTNYAEQF | 0.79395247 |
| GTSVATGGAGN | ASRGPVDTNYAEQF | 0.7858956 |
| NGGNGGSGGTS | ASRGPVDTNYAEQF | 0.5494107 |
| AVVKFPH | ASRGPVDTNYAEQF | 0.59464717 |
| TPPPPPP | ASRGPVDTNYAEQF | 0.5359621 |

| | | |
|---|---|---|
| AAPAAAAAPA | ASRGPVDTNYAEQF | 0.4302264 |
| IALNEDL | ASRGPVDTNYAEQF | 0.4978295 |
| IVNVSSM | ASRGPVDTNYAEQF | 0.78135735 |
| TGIPPAPR | ASRGPVDTNYAEQF | 0.8755924 |
| FDLGGGTF | ASRGPVDTNYAEQF | 0.78368795 |
| FDLGGGTFDVS | ASRGPVDTNYAEQF | 0.38708174 |
| NEPTAAA | ASRGPVDTNYAEQF | 0.74352145 |
| NEPTAAALA | ASRGPVDTNYAEQF | 0.69629335 |
| GSIINISSI | ASRGPVDTNYAEQF | 0.8177748 |
| VAPGNAGTA | ASRGPVDTNYAEQF | 0.8512974 |
| TGGGASLE | ASRGPVDTNYAEQF | 0.87228835 |
| VAVYNHYKRI | ASRGPVDTNYAEQF | 0.8150219 |
| TDPAPATT | ASRGPVDTNYAEQF | 0.61332935 |
| TNRPDILDPALLRPG | ASRGPVDTNYAEQF | 0.83194035 |
| LTTTTTS | ASRGPVDTNYAEQF | 0.7141465 |
| PTGSGKTL | ASRGPVDTNYAEQF | 0.7919565 |
| TGSGKTL | ASRGPVDTNYAEQF | 0.8019208 |
| TIGHVDHGK | ASRGPVDTNYAEQF | 0.84051204 |
| TIGHVDHGKTT | ASRGPVDTNYAEQF | 0.62962997 |
| GHVDHGKTT | ASRGPVDTNYAEQF | 0.69676876 |
| AQFEHTLL | ASRGPVDTNYAEQF | 0.32942772 |
| IVNQDEL | ASRGPVDTNYAEQF | 0.8532125 |
| TNGSQFF | ASRGPVDTNYAEQF | 0.43694586 |
| TGKGIVE | ASRGPVDTNYAEQF | 0.57849574 |
| LISNASDA | ASRGPVDTNYAEQF | 0.66647243 |
| AELRTLGELTKIAKAHGAQ | ASRGPVDTNYAEQF | 0.8701617 |
| APNAVAAAEPSW | ASRGPVDTNYAEQF | 0.85219747 |
| APQVAGAATAVL | ASRGPVDTNYAEQF | 0.64115965 |
| ARRDPAAAAAILPTDA | ASRGPVDTNYAEQF | 0.5613042 |
| DAAARDVLAEAGLAEYF | ASRGPVDTNYAEQF | 0.8153862 |
| DDQIVLSAKANLAKVEQ | ASRGPVDTNYAEQF | 0.53850996 |
| DELLPDTGEDSDEDGH | ASRGPVDTNYAEQF | 0.8513793 |
| ELPEAPANRATVPQD | ASRGPVDTNYAEQF | 0.7320466 |
| FGQSKIERVTGIG | ASRGPVDTNYAEQF | 0.7256217 |
| GAGGTGSTAGGAGGAGGAGGLY | ASRGPVDTNYAEQF | 0.6730486 |
| GHGGTGAAGGNGGTGGAGGSGI | ASRGPVDTNYAEQF | 0.8406586 |
| GNGGNGGAGGEGGPGGAGGAGA | ASRGPVDTNYAEQF | 0.6553401 |
| GSTAGNGGNGGNATNGTIAGQP | ASRGPVDTNYAEQF | 0.81827486 |
| GTGGDGGNGGNGGTGG | ASRGPVDTNYAEQF | 0.18620989 |
| GVRRPAGIPADAPV | ASRGPVDTNYAEQF | 0.8451737 |
| GYRQVTDAHLLTLARR | ASRGPVDTNYAEQF | 0.4716259 |
| IAKMGQDGHDRG | ASRGPVDTNYAEQF | 0.55663335 |
| KAEPSIEIQYT | ASRGPVDTNYAEQF | 0.24023291 |
| KPVRIRMAADEDTVAVT | ASRGPVDTNYAEQF | 0.67876947 |
| MPPRIAGMRLLVIKPEP | ASRGPVDTNYAEQF | 0.8091912 |
| MKFYAVALFRSRGVPPTR | ASRGPVDTNYAEQF | 0.8678688 |
| NAGGFGILGAVAHSP | ASRGPVDTNYAEQF | 0.77478004 |
| NEGREITLAMAEM | ASRGPVDTNYAEQF | 0.8025242 |
| NELVEWSHPLGHRWRWE | ASRGPVDTNYAEQF | 0.84872365 |
| NLGAGGTGGNGGNPG | ASRGPVDTNYAEQF | 0.6416407 |
| NTEWPKWETVMAADPVD | ASRGPVDTNYAEQF | 0.2580818 |

| | | |
|---|---|---|
| PASAAPAETQP | ASRGPVDTNYAEQF | 0.7360071 |
| QQSSDGKNGVRAAEITGE | ASRGPVDTNYAEQF | 0.7804965 |
| RAPDAVTTSWWKE | ASRGPVDTNYAEQF | 0.8673819 |
| SGFGNVGAHTSGWFNQ | ASRGPVDTNYAEQF | 0.7511401 |
| SGVAGAVGQAASVGGLKV | ASRGPVDTNYAEQF | 0.40576595 |
| SNGGAGTGSAGGNGGT | ASRGPVDTNYAEQF | 0.8260201 |
| TGGAGGEGGAAG | ASRGPVDTNYAEQF | 0.7860782 |
| TGGDGGAAGAGGAPGHG | ASRGPVDTNYAEQF | 0.4478637 |
| TMPPLLAAIANAGRHGV | ASRGPVDTNYAEQF | 0.57807064 |
| TSSTAGNSGGA | ASRGPVDTNYAEQF | 0.5677774 |
| VPATVPVGILC | ASRGPVDTNYAEQF | 0.3673364 |
| VQRPDSGGVVLTGRIS | ASRGPVDTNYAEQF | 0.80238116 |
| WTPNPVPATKGVPAGLVQAT | ASRGPVDTNYAEQF | 0.69247234 |
| YALKRALAKFSDRFDVVII | ASRGPVDTNYAEQF | 0.5685365 |
| YLIGSVIV | ASRGPVDTNYAEQF | 0.73799515 |
| YGQLVAVID | ASRGPVDTNYAEQF | 0.58154625 |
| VTVGGMTLPA | ASRGPVDTNYAEQF | 0.50322497 |
| VLALPAV | ASRGPVDTNYAEQF | 0.6001801 |
| VIPPTLNYET | ASRGPVDTNYAEQF | 0.7824813 |
| VGMGGMHPSSG | ASRGPVDTNYAEQF | 0.8521703 |
| VGLAGGAATA | ASRGPVDTNYAEQF | 0.70965827 |
| VGGAGGAGGSGP | ASRGPVDTNYAEQF | 0.76141787 |
| VFAAFMV | ASRGPVDTNYAEQF | 0.8534619 |
| TLYDKLR | ASRGPVDTNYAEQF | 0.34438246 |
| TLAVFTVL | ASRGPVDTNYAEQF | 0.55836916 |
| TGSVQPARV | ASRGPVDTNYAEQF | 0.82267463 |
| TGGGAGSGGT | ASRGPVDTNYAEQF | 0.65886915 |
| SHLETSL | ASRGPVDTNYAEQF | 0.8556802 |
| SFTTGMT | ASRGPVDTNYAEQF | 0.7521337 |
| QVEVAKRMGVRQPTVS | ASRGPVDTNYAEQF | 0.7613828 |
| QGSGGPALAM | ASRGPVDTNYAEQF | 0.76308775 |
| MLAPSNP | ASRGPVDTNYAEQF | 0.8851813 |
| LVNNAAI | ASRGPVDTNYAEQF | 0.79317737 |
| LTGEINTLA | ASRGPVDTNYAEQF | 0.7476131 |
| LSIFGPV | ASRGPVDTNYAEQF | 0.8518045 |
| LLALLANL | ASRGPVDTNYAEQF | 0.40816402 |
| LISNASDALDKLR | ASRGPVDTNYAEQF | 0.8400154 |
| LINEAVP | ASRGPVDTNYAEQF | 0.8427671 |
| LGYAFSGL | ASRGPVDTNYAEQF | 0.49622083 |
| LDFVAVAL | ASRGPVDTNYAEQF | 0.70891875 |
| LAQPGAE | ASRGPVDTNYAEQF | 0.8807541 |
| ISNLGAVTGF | ASRGPVDTNYAEQF | 0.85137486 |
| HQEDPGRI | ASRGPVDTNYAEQF | 0.5555018 |
| GTPGEPTP | ASRGPVDTNYAEQF | 0.5865912 |
| GFFNSGS | ASRGPVDTNYAEQF | 0.4690825 |
| GAGGGGLTAGAN | ASRGPVDTNYAEQF | 0.67938066 |
| FVDALVNM | ASRGPVDTNYAEQF | 0.33238974 |
| FLTVAPEL | ASRGPVDTNYAEQF | 0.8192482 |
| FETLTAL | ASRGPVDTNYAEQF | 0.61111885 |
| DVINPIEL | ASRGPVDTNYAEQF | 0.82913625 |
| APVSGAPV | ASRGPVDTNYAEQF | 0.77420664 |

| | | |
|---|---|---|
| ALVGGLAW | ASRGPVDTNYAEQF | 0.8370948 |
| ALAAAAATAI | ASRGPVDTNYAEQF | 0.68876624 |
| AFGLGLLL | ASRGPVDTNYAEQF | 0.63586044 |
| AFFAAASL | ASRGPVDTNYAEQF | 0.39903086 |
| AAAALPAFSPP | ASRGPVDTNYAEQF | 0.3417428 |
| VVLAPAL | ASRGPVDTNYAEQF | 0.6934806 |
| TIGHVDHGKTTL | ASRGPVDTNYAEQF | 0.66092277 |
| LISNASDALDK | ASRGPVDTNYAEQF | 0.24353367 |
| LISNASDAL | ASRGPVDTNYAEQF | 0.8744588 |
| ISNASDALDK | ASRGPVDTNYAEQF | 0.66839063 |
| NQAETLVYQTE | ASRGPVDTNYAEQF | 0.7779043 |
| TGGTGGTAGAG | ASRGPVDTNYAEQF | 0.5884354 |
| TGSTAGGAGGA | ASRGPVDTNYAEQF | 0.4125201 |
| DPLAVPVAAGPV | ASRGPVDTNYAEQF | 0.623032 |
| GGTGGTPGNGGN | ASRGPVDTNYAEQF | 0.8098068 |
| PAGLAGVDPATL | ASRGPVDTNYAEQF | 0.50746137 |
| GNNLVVEAPHAGASV | ASRGPVDTNYAEQF | 0.25732574 |
| KVTRIMSASTLEQPA | ASRGPVDTNYAEQF | 0.47802848 |
| LGFALPIALPRRTVV | ASRGPVDTNYAEQF | 0.5181595 |
| TRLGNLTARKLVVLV | ASRGPVDTNYAEQF | 0.8869507 |
| WSQQRRLIYFPSAGP | ASRGPVDTNYAEQF | 0.8600504 |
| GPNGNPDPMAAAVDIR | ASRGPVDTNYAEQF | 0.7824912 |
| TQRCVSVADLRAAASR | ASRGPVDTNYAEQF | 0.88521385 |
| QFVLDVAEHPVEVLTETV | ASRGPVDTNYAEQF | 0.17280118 |
| GGAGGTGGAGGNGGTQAGDG | ASRGPVDTNYAEQF | 0.63503647 |
| PLLATLRAIAVDVDAKPAQV | ASRGPVDTNYAEQF | 0.8464831 |
| APKFRTGGYMIDFWGVGYQVAK | ASRGPVDTNYAEQF | 0.8074142 |
| GAGGAGGAGGAAGGPGGTGGTG | ASRGPVDTNYAEQF | 0.6555755 |
| IGALGLLATDVAPAGSVAFGVA | ASRGPVDTNYAEQF | 0.840559 |
| AGGAGGIGGNGGAGGAGGVLMG | ASRGPVDTNYAEQF | 0.52784085 |
| ADELIGLL | ASRGPVDTNYAEQF | 0.6430355 |
| ADRALKA | ASRGPVDTNYAEQF | 0.70597816 |
| AGAVGQVGGAGGA | ASRGPVDTNYAEQF | 0.8527263 |
| AGGGGGLGGAP | ASRGPVDTNYAEQF | 0.81977546 |
| AGGLLYGNGGAG | ASRGPVDTNYAEQF | 0.36390394 |
| AGLESIGL | ASRGPVDTNYAEQF | 0.7995148 |
| AGLKTGII | ASRGPVDTNYAEQF | 0.8879978 |
| ASLAALLL | ASRGPVDTNYAEQF | 0.6070791 |
| ASVINISSI | ASRGPVDTNYAEQF | 0.7967774 |
| ATPPGPA | ASRGPVDTNYAEQF | 0.73372185 |
| ATVASALV | ASRGPVDTNYAEQF | 0.48732293 |
| ATVASLIL | ASRGPVDTNYAEQF | 0.5642665 |
| AVNGPVLIRH | ASRGPVDTNYAEQF | 0.4999503 |
| DGAAGAVGI | ASRGPVDTNYAEQF | 0.818995 |
| DPLLHGI | ASRGPVDTNYAEQF | 0.4771087 |
| ELGAALGAL | ASRGPVDTNYAEQF | 0.7696061 |
| ELGRTQAV | ASRGPVDTNYAEQF | 0.7305959 |
| FVAGGYLS | ASRGPVDTNYAEQF | 0.8914633 |
| GASGGAGGAGGW | ASRGPVDTNYAEQF | 0.8135445 |
| GGAGAGGDGGAGGI | ASRGPVDTNYAEQF | 0.78583455 |
| GTGHNGSQPGMGGNG | ASRGPVDTNYAEQF | 0.5302465 |

| | | |
|---|---|---|
| GYAAASASAS | ASRGPVDTNYAEQF | 0.84285724 |
| HQVDVAAVQQG | ASRGPVDTNYAEQF | 0.8287218 |
| IGAAIAHT | ASRGPVDTNYAEQF | 0.35678208 |
| IGMIEVAQL | ASRGPVDTNYAEQF | 0.84754926 |
| IRLAPEHTSAMD | ASRGPVDTNYAEQF | 0.41697282 |
| IYNVSVL | ASRGPVDTNYAEQF | 0.8016868 |
| KQGFLGM | ASRGPVDTNYAEQF | 0.75431967 |
| LESVITQL | ASRGPVDTNYAEQF | 0.8473824 |
| LGLGALVVV | ASRGPVDTNYAEQF | 0.74651676 |
| LIGNGGNGGSGGI | ASRGPVDTNYAEQF | 0.80316186 |
| LNAINAPTL | ASRGPVDTNYAEQF | 0.48608112 |
| LQLLNTGV | ASRGPVDTNYAEQF | 0.8500694 |
| LQTSIEF | ASRGPVDTNYAEQF | 0.8388008 |
| MSFAVSEL | ASRGPVDTNYAEQF | 0.7433954 |
| NTGALGL | ASRGPVDTNYAEQF | 0.8815551 |
| PPGAGGAGGAGTT | ASRGPVDTNYAEQF | 0.8146 |
| QLVGLGI | ASRGPVDTNYAEQF | 0.2810865 |
| RFLEQVHQL | ASRGPVDTNYAEQF | 0.7014314 |
| RGVGYVLR | ASRGPVDTNYAEQF | 0.8566762 |
| SATGVFAGVI | ASRGPVDTNYAEQF | 0.87524366 |
| SLPAITLP | ASRGPVDTNYAEQF | 0.811414 |
| STALAFTV | ASRGPVDTNYAEQF | 0.82894623 |
| SVLTVGLGK | ASRGPVDTNYAEQF | 0.720713 |
| TALAGAGSGI | ASRGPVDTNYAEQF | 0.86989254 |
| TGLPFTGI | ASRGPVDTNYAEQF | 0.89450145 |
| VAEHVTV | ASRGPVDTNYAEQF | 0.79458064 |
| VDIVLNSL | ASRGPVDTNYAEQF | 0.5154805 |
| VLTGGGAVL | ASRGPVDTNYAEQF | 0.54256666 |
| VNVGLAYGI | ASRGPVDTNYAEQF | 0.840643 |
| VSSAAVGAV | ASRGPVDTNYAEQF | 0.85916376 |
| VTSVYEAGV | ASRGPVDTNYAEQF | 0.52184665 |
| WVALDRGA | ASRGPVDTNYAEQF | 0.69041276 |
| YLVGSVVI | ASRGPVDTNYAEQF | 0.7383535 |
| GPPGTGKTLL | ASRGPVDTNYAEQF | 0.23051879 |
| GVGMTKF | ASRGPVDTNYAEQF | 0.80182457 |
| IGIGGSDL | ASRGPVDTNYAEQF | 0.8744807 |
| IGTLLAEL | ASRGPVDTNYAEQF | 0.83074045 |
| IYTSGTT | ASRGPVDTNYAEQF | 0.4915371 |
| LSFPAGV | ASRGPVDTNYAEQF | 0.8557707 |
| LYTSGTT | ASRGPVDTNYAEQF | 0.86693025 |
| VGAFPFGF | ASRGPVDTNYAEQF | 0.88352 |
| VGAGTVEF | ASRGPVDTNYAEQF | 0.29464844 |
| VGKSTLLL | ASRGPVDTNYAEQF | 0.18323708 |
| AALAHAGGGIV | ASRGPVDTNYAEQF | 0.7298329 |
| AALVAVSLL | ASRGPVDTNYAEQF | 0.82219243 |
| AAVSRAVAAL | ASRGPVDTNYAEQF | 0.6464217 |
| AGGSSGIGGF | ASRGPVDTNYAEQF | 0.5750923 |
| AGLTVANANG | ASRGPVDTNYAEQF | 0.77805686 |
| AGLVGAVL | ASRGPVDTNYAEQF | 0.8491584 |
| ALSQAFTV | ASRGPVDTNYAEQF | 0.5101348 |
| ALTFSVSL | ASRGPVDTNYAEQF | 0.43552268 |

| | | |
|---|---|---|
| ALTGAALVAL | ASRGPVDTNYAEQF | 0.84134305 |
| ASFSIGAGVL | ASRGPVDTNYAEQF | 0.8473511 |
| DGSVYKI | ASRGPVDTNYAEQF | 0.5377791 |
| EQLAEIV | ASRGPVDTNYAEQF | 0.70340234 |
| FAVSELTVM | ASRGPVDTNYAEQF | 0.73755 |
| GAAPLAAGL | ASRGPVDTNYAEQF | 0.8054626 |
| GADGGDGGSGGAGG | ASRGPVDTNYAEQF | 0.7849498 |
| GFGLDTGGAGG | ASRGPVDTNYAEQF | 0.40673178 |
| GLASGGLL | ASRGPVDTNYAEQF | 0.84320307 |
| GRIAGAAVLL | ASRGPVDTNYAEQF | 0.4046319 |
| GSAGSLNTSF | ASRGPVDTNYAEQF | 0.8537994 |
| GTGQIGI | ASRGPVDTNYAEQF | 0.7301885 |
| GTLIALIL | ASRGPVDTNYAEQF | 0.81725377 |
| IAATMGFVVG | ASRGPVDTNYAEQF | 0.67141056 |
| IFAQALL | ASRGPVDTNYAEQF | 0.39410067 |
| IGAAVALGF | ASRGPVDTNYAEQF | 0.53513503 |
| IGLTPLAL | ASRGPVDTNYAEQF | 0.4444586 |
| ILGSNTSTL | ASRGPVDTNYAEQF | 0.8539576 |
| IVFSLTVL | ASRGPVDTNYAEQF | 0.27432704 |
| LASKATGF | ASRGPVDTNYAEQF | 0.8407718 |
| LASVALIL | ASRGPVDTNYAEQF | 0.6961161 |
| LFAVGLTL | ASRGPVDTNYAEQF | 0.4682795 |
| LFNGGIGTYI | ASRGPVDTNYAEQF | 0.83900094 |
| LTLGAGII | ASRGPVDTNYAEQF | 0.41894788 |
| LVAAALGDI | ASRGPVDTNYAEQF | 0.7975401 |
| NTAFVQLGI | ASRGPVDTNYAEQF | 0.69352514 |
| QLFSGAEL | ASRGPVDTNYAEQF | 0.8202785 |
| SAATFITL | ASRGPVDTNYAEQF | 0.5504607 |
| SNYGLANLGV | ASRGPVDTNYAEQF | 0.8409219 |
| SYGGAYAVM | ASRGPVDTNYAEQF | 0.81073403 |
| TTGVGGLTLG | ASRGPVDTNYAEQF | 0.45691168 |
| TTQTIYI | ASRGPVDTNYAEQF | 0.8300425 |
| VAAGAGLGAVI | ASRGPVDTNYAEQF | 0.8310997 |
| VAAVVTLL | ASRGPVDTNYAEQF | 0.80122447 |
| VGFLGGLI | ASRGPVDTNYAEQF | 0.33615136 |
| VGVIPILL | ASRGPVDTNYAEQF | 0.77200353 |
| VIFASSLI | ASRGPVDTNYAEQF | 0.5686357 |
| VKGGDGGAAAG | ASRGPVDTNYAEQF | 0.7943833 |
| VTKTFTNI | ASRGPVDTNYAEQF | 0.75626487 |
| VVAAAILL | ASRGPVDTNYAEQF | 0.58534616 |
| YAALDVEL | ASRGPVDTNYAEQF | 0.8313122 |
| LASVALLL | ASRGPVDTNYAEQF | 0.31554547 |
| LDAVHRV | ASRGPVDTNYAEQF | 0.6388142 |
| QGLSSSIF | ASRGPVDTNYAEQF | 0.66549784 |
| RGTFANIRL | ASRGPVDTNYAEQF | 0.7331295 |
| RQLWWGHRI | ASRGPVDTNYAEQF | 0.8007733 |
| YGPPGTGKTLL | ASRGPVDTNYAEQF | 0.46020874 |
| AALVGAPANVA | ASRGPVDTNYAEQF | 0.7695972 |
| AATFRRTE | ASRGPVDTNYAEQF | 0.70241344 |
| AGALGGGI | ASRGPVDTNYAEQF | 0.47030967 |
| AGGAGGAGGIGAG | ASRGPVDTNYAEQF | 0.8169737 |

| | | |
|---|---|---|
| AGSGALGL | ASRGPVDTNYAEQF | 0.65470517 |
| AQIADQI | ASRGPVDTNYAEQF | 0.38094956 |
| ASAVAVTYL | ASRGPVDTNYAEQF | 0.8437891 |
| ASTVLGGTGG | ASRGPVDTNYAEQF | 0.52780026 |
| AVGGEPAK | ASRGPVDTNYAEQF | 0.77425015 |
| CEQGTFGGTC | ASRGPVDTNYAEQF | 0.88657546 |
| DEGKAPAG | ASRGPVDTNYAEQF | 0.35286248 |
| DTVVFPAL | ASRGPVDTNYAEQF | 0.8531211 |
| FEGITMAG | ASRGPVDTNYAEQF | 0.7724916 |
| FPLVVQD | ASRGPVDTNYAEQF | 0.7708273 |
| GAFTTPQ | ASRGPVDTNYAEQF | 0.6659718 |
| GFGGFGTAGGVGGA | ASRGPVDTNYAEQF | 0.5765922 |
| GFLAQFL | ASRGPVDTNYAEQF | 0.8641928 |
| GGAGGAGGNAGVA | ASRGPVDTNYAEQF | 0.764683 |
| GGVGTAGAGGAGGAGG | ASRGPVDTNYAEQF | 0.7475587 |
| GNGGGGDAGNAGSGGN | ASRGPVDTNYAEQF | 0.83102614 |
| GTGGAGGAGNDAGS | ASRGPVDTNYAEQF | 0.8402226 |
| GTTNTGWFNS | ASRGPVDTNYAEQF | 0.829531 |
| IFMGPLFTL | ASRGPVDTNYAEQF | 0.5287117 |
| ILAHAIEAI | ASRGPVDTNYAEQF | 0.35233283 |
| IRSAIREAL | ASRGPVDTNYAEQF | 0.6713574 |
| ITFAGGG | ASRGPVDTNYAEQF | 0.70301056 |
| ITGMSGHQEV | ASRGPVDTNYAEQF | 0.5639642 |
| IVAGLACTL | ASRGPVDTNYAEQF | 0.3458771 |
| IVGGGLAGA | ASRGPVDTNYAEQF | 0.8203237 |
| IVTGVLNI | ASRGPVDTNYAEQF | 0.8866415 |
| LAQQGCI | ASRGPVDTNYAEQF | 0.5440346 |
| LCLHVEN | ASRGPVDTNYAEQF | 0.8943131 |
| LDAISDGS | ASRGPVDTNYAEQF | 0.4683838 |
| LGGFAAML | ASRGPVDTNYAEQF | 0.80454284 |
| LGGKNPMI | ASRGPVDTNYAEQF | 0.8272437 |
| LGPKGVAAIL | ASRGPVDTNYAEQF | 0.42159384 |
| LMVGVSI | ASRGPVDTNYAEQF | 0.7450061 |
| LVGAAFVL | ASRGPVDTNYAEQF | 0.4904864 |
| LVLAVAPL | ASRGPVDTNYAEQF | 0.5656185 |
| MAAIAVIDAA | ASRGPVDTNYAEQF | 0.85016656 |
| NNGINAELQ | ASRGPVDTNYAEQF | 0.5734241 |
| RGEPLVLVHG | ASRGPVDTNYAEQF | 0.82494295 |
| RGGAGGAAGLI | ASRGPVDTNYAEQF | 0.6179461 |
| RSYRMHPAV | ASRGPVDTNYAEQF | 0.5147651 |
| SAAHLLAGQLA | ASRGPVDTNYAEQF | 0.8050934 |
| SAAVGSY | ASRGPVDTNYAEQF | 0.8147496 |
| SPHAVLHRI | ASRGPVDTNYAEQF | 0.7801479 |
| SQIAKVL | ASRGPVDTNYAEQF | 0.8928983 |
| TALGHLVP | ASRGPVDTNYAEQF | 0.52667946 |
| TAPTTSGAA | ASRGPVDTNYAEQF | 0.87245023 |
| TDGDDGNGGSPGT | ASRGPVDTNYAEQF | 0.8715455 |
| TDGLGEN | ASRGPVDTNYAEQF | 0.76594794 |
| TGGPPLPGQA | ASRGPVDTNYAEQF | 0.6784816 |
| TIPAITI | ASRGPVDTNYAEQF | 0.8159094 |
| TIPGFTI | ASRGPVDTNYAEQF | 0.6099124 |

| | | |
|---|---|---|
| TSIASGALA | ASRGPVDTNYAEQF | 0.7675185 |
| TVTLPGAAI | ASRGPVDTNYAEQF | 0.88009405 |
| VASGLSAALP | ASRGPVDTNYAEQF | 0.6923059 |
| VGAGVIGI | ASRGPVDTNYAEQF | 0.78896224 |
| VGPKRFGV | ASRGPVDTNYAEQF | 0.7682579 |
| VNVAFPD | ASRGPVDTNYAEQF | 0.77071756 |
| VSMTNSGVS | ASRGPVDTNYAEQF | 0.6923512 |
| VTDVLGDLSV | ASRGPVDTNYAEQF | 0.52682155 |
| VVVVGAGFAGL | ASRGPVDTNYAEQF | 0.7714283 |
| VYTVGTVV | ASRGPVDTNYAEQF | 0.599982 |
| WTTFSGP | ASRGPVDTNYAEQF | 0.81130993 |
| YDELVEAF | ASRGPVDTNYAEQF | 0.79962134 |
| YGTGGAGGNGG | ASRGPVDTNYAEQF | 0.6806766 |
| YTNVMAV | ASRGPVDTNYAEQF | 0.80390775 |
| FLEQVHQL | ASRGPVDTNYAEQF | 0.78983104 |
| GLSLLAVAV | ASRGPVDTNYAEQF | 0.64440393 |
| LEQVHQL | ASRGPVDTNYAEQF | 0.76188624 |
| LNAAGAI | ASRGPVDTNYAEQF | 0.7331939 |
| PAVHASL | ASRGPVDTNYAEQF | 0.74739087 |
| TELEDDA | ASRGPVDTNYAEQF | 0.6431043 |
| TPPGPPN | ASRGPVDTNYAEQF | 0.7755736 |
| TPPPPPPP | ASRGPVDTNYAEQF | 0.7526256 |
| TQSPALI | ASSQDTGAEEQY | 0.45178652 |
| EFAAAFPQ | ASSQDTGAEEQY | 0.7945243 |
| LLSGNLDVL | ASSQDTGAEEQY | 0.73785716 |
| GPANLGNYNI | ASSQDTGAEEQY | 0.73207784 |
| TLGPTLP | ASSQDTGAEEQY | 0.7898769 |
| VYAYSSCCK | ASSQDTGAEEQY | 0.65734446 |
| APAPAPAPNI | ASSQDTGAEEQY | 0.6847464 |
| PGGPVPIP | ASSQDTGAEEQY | 0.7845141 |
| FLTVALEP | ASSQDTGAEEQY | 0.80352443 |
| SQYFGANL | ASSQDTGAEEQY | 0.5729195 |
| GGVKPAA | ASSQDTGAEEQY | 0.7631059 |
| GGTGGAGGAGGW | ASSQDTGAEEQY | 0.80537033 |
| GVGGVAGLGGAGG | ASSQDTGAEEQY | 0.7671695 |
| VLLVLATTI | ASSQDTGAEEQY | 0.8065671 |
| IGIFAAML | ASSQDTGAEEQY | 0.8090012 |
| AVGDLID | ASSQDTGAEEQY | 0.7233189 |
| NISPLPL | ASSQDTGAEEQY | 0.7999059 |
| VTAVFVAI | ASSQDTGAEEQY | 0.81340194 |
| SGDDNDAAL | ASSQDTGAEEQY | 0.7201109 |
| AVAALAHS | ASSQDTGAEEQY | 0.7496436 |
| GGVTVMATP | ASSQDTGAEEQY | 0.6997485 |
| AGLAANVRD | ASSQDTGAEEQY | 0.7766714 |
| TNRPDLIDPALLRPG | ASSQDTGAEEQY | 0.3941055 |
| LANLGVTAL | ASSQDTGAEEQY | 0.76622516 |
| TGERIEL | ASSQDTGAEEQY | 0.8012094 |
| NGEELAA | ASSQDTGAEEQY | 0.7431314 |
| VLSLVWG | ASSQDTGAEEQY | 0.635592 |
| IGYALWL | ASSQDTGAEEQY | 0.6527635 |
| LSLLLAPL | ASSQDTGAEEQY | 0.7196066 |

| | | |
|---|---|---|
| TLDEALK | ASSQDTGAEEQY | 0.6529852 |
| TDGAGTAEGG | ASSQDTGAEEQY | 0.473292 |
| NAPGGAGGNGG | ASSQDTGAEEQY | 0.7198718 |
| VSVFMGMGGNF | ASSQDTGAEEQY | 0.7950748 |
| LASYLAAL | ASSQDTGAEEQY | 0.7966883 |
| LVGGTPAL | ASSQDTGAEEQY | 0.6018748 |
| VRNADVL | ASSQDTGAEEQY | 0.8009576 |
| TCSSPFLVSAM | ASSQDTGAEEQY | 0.8023577 |
| VHITDAE | ASSQDTGAEEQY | 0.79481304 |
| EYAAPSF | ASSQDTGAEEQY | 0.75374997 |
| VVDAFHGGL | ASSQDTGAEEQY | 0.65325034 |
| ASVVDPQ | ASSQDTGAEEQY | 0.80652165 |
| SYNEFGGLS | ASSQDTGAEEQY | 0.7251397 |
| DPDGSVY | ASSQDTGAEEQY | 0.77153826 |
| MVSTIHATE | ASSQDTGAEEQY | 0.7995608 |
| LAAGVADPV | ASSQDTGAEEQY | 0.732661 |
| TKEQIAA | ASSQDTGAEEQY | 0.66924286 |
| SLASPAAL | ASSQDTGAEEQY | 0.76731586 |
| IGLVFYGV | ASSQDTGAEEQY | 0.80710816 |
| FLLDASNL | ASSQDTGAEEQY | 0.67354 |
| YQVGNFP | ASSQDTGAEEQY | 0.541662 |
| ESLLSQST | ASSQDTGAEEQY | 0.7002911 |
| LFTVGFAAVG | ASSQDTGAEEQY | 0.7429948 |
| ATPPTPT | ASSQDTGAEEQY | 0.801668 |
| TDLATLG | ASSQDTGAEEQY | 0.78176093 |
| SAGGTGR | ASSQDTGAEEQY | 0.7449155 |
| GFSNSGS | ASSQDTGAEEQY | 0.78820384 |
| GGNGGSGGTSV | ASSQDTGAEEQY | 0.77204245 |
| GTSVATGGAGN | ASSQDTGAEEQY | 0.7528166 |
| NGGNGGSGGTS | ASSQDTGAEEQY | 0.80459034 |
| AVVKFPH | ASSQDTGAEEQY | 0.6719289 |
| TPPPPPP | ASSQDTGAEEQY | 0.8014883 |
| AAPAAAAAPA | ASSQDTGAEEQY | 0.7864151 |
| IALNEDL | ASSQDTGAEEQY | 0.79113865 |
| IVNVSSM | ASSQDTGAEEQY | 0.8046788 |
| TGIPPAPR | ASSQDTGAEEQY | 0.7538706 |
| FDLGGGTF | ASSQDTGAEEQY | 0.7716398 |
| FDLGGGTFDVS | ASSQDTGAEEQY | 0.56076574 |
| NEPTAAA | ASSQDTGAEEQY | 0.75487226 |
| NEPTAAALA | ASSQDTGAEEQY | 0.7882015 |
| GSIINISSI | ASSQDTGAEEQY | 0.76690114 |
| VAPGNAGTA | ASSQDTGAEEQY | 0.8004315 |
| TGGGASLE | ASSQDTGAEEQY | 0.77900773 |
| VAVYNHYKRI | ASSQDTGAEEQY | 0.7265035 |
| TDPAPATT | ASSQDTGAEEQY | 0.80656165 |
| TNRPDILDPALLRPG | ASSQDTGAEEQY | 0.80385745 |
| LTTTTTS | ASSQDTGAEEQY | 0.5876926 |
| PTGSGKTL | ASSQDTGAEEQY | 0.73715496 |
| TGSGKTL | ASSQDTGAEEQY | 0.8054418 |
| TIGHVDHGK | ASSQDTGAEEQY | 0.7930549 |
| TIGHVDHGKTT | ASSQDTGAEEQY | 0.7468428 |

| | | |
|---|---|---|
| GHVDHGKTT | ASSQDTGAEEQY | 0.75693834 |
| AQFEHTLL | ASSQDTGAEEQY | 0.8162921 |
| IVNQDEL | ASSQDTGAEEQY | 0.7521995 |
| TNGSQFF | ASSQDTGAEEQY | 0.8112669 |
| TGKGIVE | ASSQDTGAEEQY | 0.38429946 |
| LISNASDA | ASSQDTGAEEQY | 0.72548604 |
| AELRTLGELTKIAKAHGAQ | ASSQDTGAEEQY | 0.64826894 |
| APNAVAAAEPSW | ASSQDTGAEEQY | 0.3422102 |
| APQVAGAATAVL | ASSQDTGAEEQY | 0.7902492 |
| ARRDPAAAAAILPTDA | ASSQDTGAEEQY | 0.7321342 |
| DAAARDVLAEAGLAEYF | ASSQDTGAEEQY | 0.79144394 |
| DDQIVLSAKANLAKVEQ | ASSQDTGAEEQY | 0.7793983 |
| DELLPDTGEDSDEDGH | ASSQDTGAEEQY | 0.7922932 |
| ELPEAPANRATVPQD | ASSQDTGAEEQY | 0.7586553 |
| FGQSKIERVTGIG | ASSQDTGAEEQY | 0.781667 |
| GAGGTGSTAGGAGGAGGAGGLY | ASSQDTGAEEQY | 0.7189283 |
| GHGGTGAAGGNGGTGGAGGSGI | ASSQDTGAEEQY | 0.78126264 |
| GNGGNGGAGGEGGPGGAGGAGA | ASSQDTGAEEQY | 0.80955255 |
| GSTAGNGGNGGNATNGTIAGQP | ASSQDTGAEEQY | 0.7986408 |
| GTGGDGGNGGNGGTGG | ASSQDTGAEEQY | 0.45407718 |
| GVRRPAGIPADAPV | ASSQDTGAEEQY | 0.77409613 |
| GYRQVTDAHLLTLARR | ASSQDTGAEEQY | 0.7405155 |
| IAKMGQDGHDRG | ASSQDTGAEEQY | 0.44267172 |
| KAEPSIEIQYT | ASSQDTGAEEQY | 0.51220655 |
| KPVRIRMAADEDTVAVT | ASSQDTGAEEQY | 0.69906485 |
| MPPRIAGMRLLVIKPEP | ASSQDTGAEEQY | 0.7077058 |
| MKFYAVALFRSRGVPPTR | ASSQDTGAEEQY | 0.79553235 |
| NAGGFGILGAVAHSP | ASSQDTGAEEQY | 0.79488736 |
| NEGREITLAMAEM | ASSQDTGAEEQY | 0.7205545 |
| NELVEWSHPLGHRWRWE | ASSQDTGAEEQY | 0.79232603 |
| NLGAGGTGGNGGNPG | ASSQDTGAEEQY | 0.7591058 |
| NTEWPKWETVMAADPVD | ASSQDTGAEEQY | 0.16700943 |
| PASAAPAETQP | ASSQDTGAEEQY | 0.72532225 |
| QQSSDGKNGVRAAEITGE | ASSQDTGAEEQY | 0.6556561 |
| RAPDAVTTSWWKE | ASSQDTGAEEQY | 0.8022938 |
| SGFGNVGAHTSGWFNQ | ASSQDTGAEEQY | 0.7278091 |
| SGVAGAVGQAASVGGLKV | ASSQDTGAEEQY | 0.736568 |
| SNGGAGTGSAGGNGGT | ASSQDTGAEEQY | 0.74586874 |
| TGGAGGEGGAAG | ASSQDTGAEEQY | 0.78372777 |
| TGGDGGAAGAGGAPGHG | ASSQDTGAEEQY | 0.7581192 |
| TMPPLLAAIANAGRHGV | ASSQDTGAEEQY | 0.6980042 |
| TSSTAGNSGGA | ASSQDTGAEEQY | 0.39805835 |
| VPATVPVGILC | ASSQDTGAEEQY | 0.7870506 |
| VQRPDSGGVVLTGRIS | ASSQDTGAEEQY | 0.66804135 |
| WTPNPVPATKGVPAGLVQAT | ASSQDTGAEEQY | 0.6514747 |
| YALKRALAKFSDRFDVVII | ASSQDTGAEEQY | 0.7066525 |
| YLIGSVIV | ASSQDTGAEEQY | 0.7664083 |
| YGQLVAVID | ASSQDTGAEEQY | 0.4881029 |
| VTVGGMTLPA | ASSQDTGAEEQY | 0.80355024 |
| VLALPAV | ASSQDTGAEEQY | 0.69601834 |
| VIPPTLNYET | ASSQDTGAEEQY | 0.73540634 |

| | | |
|---|---|---|
| VGMGGMHPSSG | ASSQDTGAEEQY | 0.80249643 |
| VGLAGGAATA | ASSQDTGAEEQY | 0.781005 |
| VGGAGGAGGSGP | ASSQDTGAEEQY | 0.73914844 |
| VFAAFMV | ASSQDTGAEEQY | 0.71328497 |
| TLYDKLR | ASSQDTGAEEQY | 0.6903627 |
| TLAVFTVL | ASSQDTGAEEQY | 0.706275 |
| TGSVQPARV | ASSQDTGAEEQY | 0.68584967 |
| TGGGAGSGGT | ASSQDTGAEEQY | 0.80789846 |
| SHLETSL | ASSQDTGAEEQY | 0.72728956 |
| SFTTGMT | ASSQDTGAEEQY | 0.6441776 |
| QVEVAKRMGVRQPTVS | ASSQDTGAEEQY | 0.7275492 |
| QGSGGPALAM | ASSQDTGAEEQY | 0.80560505 |
| MLAPSNP | ASSQDTGAEEQY | 0.8099699 |
| LVNNAAI | ASSQDTGAEEQY | 0.74332666 |
| LTGEINTLA | ASSQDTGAEEQY | 0.71688384 |
| LSIFGPV | ASSQDTGAEEQY | 0.73588383 |
| LLALLANL | ASSQDTGAEEQY | 0.7426588 |
| LISNASDALDKLR | ASSQDTGAEEQY | 0.73148286 |
| LINEAVP | ASSQDTGAEEQY | 0.5734948 |
| LGYAFSGL | ASSQDTGAEEQY | 0.8007579 |
| LDFVAVAL | ASSQDTGAEEQY | 0.8189331 |
| LAQPGAE | ASSQDTGAEEQY | 0.73683023 |
| ISNLGAVTGF | ASSQDTGAEEQY | 0.70283663 |
| HQEDPGRI | ASSQDTGAEEQY | 0.7572974 |
| GTPGEPTP | ASSQDTGAEEQY | 0.67863965 |
| GFFNSGS | ASSQDTGAEEQY | 0.69345284 |
| GAGGGGLTAGAN | ASSQDTGAEEQY | 0.7403995 |
| FVDALVNM | ASSQDTGAEEQY | 0.5236668 |
| FLTVAPEL | ASSQDTGAEEQY | 0.7328321 |
| FETLTAL | ASSQDTGAEEQY | 0.8041743 |
| DVINPIEL | ASSQDTGAEEQY | 0.75387776 |
| APVSGAPV | ASSQDTGAEEQY | 0.7748296 |
| ALVGGLAW | ASSQDTGAEEQY | 0.8012328 |
| ALAAAAATAI | ASSQDTGAEEQY | 0.78037834 |
| AFGLGLLL | ASSQDTGAEEQY | 0.7963629 |
| AFFAAASL | ASSQDTGAEEQY | 0.80862266 |
| AAAALPAFSPP | ASSQDTGAEEQY | 0.77474046 |
| VVLAPAL | ASSQDTGAEEQY | 0.76261383 |
| TIGHVDHGKTTL | ASSQDTGAEEQY | 0.7959397 |
| LISNASDALDK | ASSQDTGAEEQY | 0.76068306 |
| LISNASDAL | ASSQDTGAEEQY | 0.7655027 |
| ISNASDALDK | ASSQDTGAEEQY | 0.74451065 |
| NQAETLVYQTE | ASSQDTGAEEQY | 0.7926172 |
| TGGTGGTAGAG | ASSQDTGAEEQY | 0.75145465 |
| TGSTAGGAGGA | ASSQDTGAEEQY | 0.80813664 |
| DPLAVPVAAGPV | ASSQDTGAEEQY | 0.40969038 |
| GGTGGTPGNGGN | ASSQDTGAEEQY | 0.6062902 |
| PAGLAGVDPATL | ASSQDTGAEEQY | 0.7825341 |
| GNNLVVEAPHAGASV | ASSQDTGAEEQY | 0.27700216 |
| KVTRIMSASTLEQPA | ASSQDTGAEEQY | 0.5474781 |
| LGFALPIALPRRTVV | ASSQDTGAEEQY | 0.7826265 |

| | | |
|---|---|---|
| TRLGNLTARKLVVLV | ASSQDTGAEEQY | 0.7918278 |
| WSQQRRLIYFPSAGP | ASSQDTGAEEQY | 0.24261963 |
| GPNGNPDPMAAAVDIR | ASSQDTGAEEQY | 0.6815344 |
| TQRCVSVADLRAAASR | ASSQDTGAEEQY | 0.5994949 |
| QFVLDVAEHPVEVLTETV | ASSQDTGAEEQY | 0.74875486 |
| GGAGGTGGAGGNGGTQAGDG | ASSQDTGAEEQY | 0.75175405 |
| PLLATLRAIAVDVDAKPAQV | ASSQDTGAEEQY | 0.49521106 |
| APKFRTGGYMIDFWGVGYQVAK | ASSQDTGAEEQY | 0.7841928 |
| GAGGAGGAGGAAGGPGGTGGTG | ASSQDTGAEEQY | 0.6484301 |
| IGALGLLATDVAPAGSVAFGVA | ASSQDTGAEEQY | 0.7311883 |
| AGGAGGIGGNGGAGGAGGVLMG | ASSQDTGAEEQY | 0.79943645 |
| ADELIGLL | ASSQDTGAEEQY | 0.81445193 |
| ADRALKA | ASSQDTGAEEQY | 0.70638674 |
| AGAVGQVGGAGGA | ASSQDTGAEEQY | 0.789446 |
| AGGGGGLGGAP | ASSQDTGAEEQY | 0.72205466 |
| AGGLLYGNGGAG | ASSQDTGAEEQY | 0.8146092 |
| AGLESIGL | ASSQDTGAEEQY | 0.7243735 |
| AGLKTGII | ASSQDTGAEEQY | 0.52674913 |
| ASLAALLL | ASSQDTGAEEQY | 0.783333 |
| ASVINISSI | ASSQDTGAEEQY | 0.77024114 |
| ATPPGPA | ASSQDTGAEEQY | 0.7666834 |
| ATVASALV | ASSQDTGAEEQY | 0.79771745 |
| ATVASLIL | ASSQDTGAEEQY | 0.5930673 |
| AVNGPVLIRH | ASSQDTGAEEQY | 0.73372823 |
| DGAAGAVGI | ASSQDTGAEEQY | 0.79093146 |
| DPLLHGI | ASSQDTGAEEQY | 0.79365027 |
| ELGAALGAL | ASSQDTGAEEQY | 0.534816 |
| ELGRTQAV | ASSQDTGAEEQY | 0.80450654 |
| FVAGGYLS | ASSQDTGAEEQY | 0.7577908 |
| GASGGAGGAGGW | ASSQDTGAEEQY | 0.71277153 |
| GGAGAGGDGGAGGI | ASSQDTGAEEQY | 0.8025427 |
| GTGHNGSQPGMGGNG | ASSQDTGAEEQY | 0.77797204 |
| GYAAASASAS | ASSQDTGAEEQY | 0.7916562 |
| HQVDVAAVQQG | ASSQDTGAEEQY | 0.7784023 |
| IGAAIAHT | ASSQDTGAEEQY | 0.81639725 |
| IGMIEVAQL | ASSQDTGAEEQY | 0.64793456 |
| IRLAPEHTSAMD | ASSQDTGAEEQY | 0.7941306 |
| IYNVSVL | ASSQDTGAEEQY | 0.77442074 |
| KQGFLGM | ASSQDTGAEEQY | 0.7977582 |
| LESVITQL | ASSQDTGAEEQY | 0.74704194 |
| LGLGALVVV | ASSQDTGAEEQY | 0.75030684 |
| LIGNGGNGGSGGI | ASSQDTGAEEQY | 0.7860439 |
| LNAINAPTL | ASSQDTGAEEQY | 0.7996454 |
| LQLLNTGV | ASSQDTGAEEQY | 0.782464 |
| LQTSIEF | ASSQDTGAEEQY | 0.7803281 |
| MSFAVSEL | ASSQDTGAEEQY | 0.80295956 |
| NTGALGL | ASSQDTGAEEQY | 0.7378186 |
| PPGAGGAGGAGTT | ASSQDTGAEEQY | 0.74581444 |
| QLVGLGI | ASSQDTGAEEQY | 0.79187006 |
| RFLEQVHQL | ASSQDTGAEEQY | 0.80635756 |
| RGVGYVLR | ASSQDTGAEEQY | 0.7339176 |

| | | |
|---|---|---|
| SATGVFAGVI | ASSQDTGAEEQY | 0.8100822 |
| SLPAITLP | ASSQDTGAEEQY | 0.7291893 |
| STALAFTV | ASSQDTGAEEQY | 0.794768 |
| SVLTVGLGK | ASSQDTGAEEQY | 0.69178003 |
| TALAGAGSGI | ASSQDTGAEEQY | 0.76826805 |
| TGLPFTGI | ASSQDTGAEEQY | 0.649771 |
| VAEHVTV | ASSQDTGAEEQY | 0.78899837 |
| VDIVLNSL | ASSQDTGAEEQY | 0.7917596 |
| VLTGGGAVL | ASSQDTGAEEQY | 0.795607 |
| VNVGLAYGI | ASSQDTGAEEQY | 0.52582556 |
| VSSAAVGAV | ASSQDTGAEEQY | 0.77565503 |
| VTSVYEAGV | ASSQDTGAEEQY | 0.7961446 |
| WVALDRGA | ASSQDTGAEEQY | 0.81478393 |
| YLVGSVVI | ASSQDTGAEEQY | 0.75926995 |
| GPPGTGKTLL | ASSQDTGAEEQY | 0.7864049 |
| GVGMTKF | ASSQDTGAEEQY | 0.74317706 |
| IGIGGSDL | ASSQDTGAEEQY | 0.751914 |
| IGTLLAEL | ASSQDTGAEEQY | 0.6607993 |
| IYTSGTT | ASSQDTGAEEQY | 0.7468548 |
| LSFPAGV | ASSQDTGAEEQY | 0.73465276 |
| LYTSGTT | ASSQDTGAEEQY | 0.6936462 |
| VGAFPFGF | ASSQDTGAEEQY | 0.7995049 |
| VGAGTVEF | ASSQDTGAEEQY | 0.80878216 |
| VGKSTLLL | ASSQDTGAEEQY | 0.7879787 |
| AALAHAGGGIV | ASSQDTGAEEQY | 0.79959595 |
| AALVAVSLL | ASSQDTGAEEQY | 0.8028635 |
| AAVSRAVAAL | ASSQDTGAEEQY | 0.7988581 |
| AGGSSGIGGF | ASSQDTGAEEQY | 0.70777035 |
| AGLTVANANG | ASSQDTGAEEQY | 0.7778884 |
| AGLVGAVL | ASSQDTGAEEQY | 0.59879255 |
| ALSQAFTV | ASSQDTGAEEQY | 0.73383045 |
| ALTFSVSL | ASSQDTGAEEQY | 0.6853344 |
| ALTGAALVAL | ASSQDTGAEEQY | 0.7625081 |
| ASFSIGAGVL | ASSQDTGAEEQY | 0.19421154 |
| DGSVYKI | ASSQDTGAEEQY | 0.7804321 |
| EQLAEIV | ASSQDTGAEEQY | 0.80810285 |
| FAVSELTVM | ASSQDTGAEEQY | 0.7284274 |
| GAAPLAAGL | ASSQDTGAEEQY | 0.78458536 |
| GADGGDGGSGGAGG | ASSQDTGAEEQY | 0.77911764 |
| GFGLDTGGAGG | ASSQDTGAEEQY | 0.7155827 |
| GLASGGLL | ASSQDTGAEEQY | 0.79050624 |
| GRIAGAAVLL | ASSQDTGAEEQY | 0.6886568 |
| GSAGSLNTSF | ASSQDTGAEEQY | 0.789542 |
| GTGQIGI | ASSQDTGAEEQY | 0.8018775 |
| GTLIALIL | ASSQDTGAEEQY | 0.7770517 |
| IAATMGFVVG | ASSQDTGAEEQY | 0.7657515 |
| IFAQALL | ASSQDTGAEEQY | 0.76173085 |
| IGAAVALGF | ASSQDTGAEEQY | 0.59639657 |
| IGLTPLAL | ASSQDTGAEEQY | 0.7800325 |
| ILGSNTSTL | ASSQDTGAEEQY | 0.6951659 |
| IVFSLTVL | ASSQDTGAEEQY | 0.7580699 |

| | | |
|---|---|---|
| LASKATGF | ASSQDTGAEEQY | 0.6705064 |
| LASVALIL | ASSQDTGAEEQY | 0.78569394 |
| LFAVGLTL | ASSQDTGAEEQY | 0.7487556 |
| LFNGGIGTYI | ASSQDTGAEEQY | 0.7934052 |
| LTLGAGII | ASSQDTGAEEQY | 0.7139932 |
| LVAAALGDI | ASSQDTGAEEQY | 0.7618254 |
| NTAFVQLGI | ASSQDTGAEEQY | 0.75746214 |
| QLFSGAEL | ASSQDTGAEEQY | 0.57726115 |
| SAATFITL | ASSQDTGAEEQY | 0.8004774 |
| SNYGLANLGV | ASSQDTGAEEQY | 0.75774413 |
| SYGGAYAVM | ASSQDTGAEEQY | 0.71433234 |
| TTGVGGLTLG | ASSQDTGAEEQY | 0.79408914 |
| TTQTIYI | ASSQDTGAEEQY | 0.7661513 |
| VAAGAGLGAVI | ASSQDTGAEEQY | 0.6116793 |
| VAAVVTLL | ASSQDTGAEEQY | 0.73789203 |
| VGFLGGLI | ASSQDTGAEEQY | 0.7666655 |
| VGVIPILL | ASSQDTGAEEQY | 0.73524225 |
| VIFASSLI | ASSQDTGAEEQY | 0.30007124 |
| VKGGDGGAAAG | ASSQDTGAEEQY | 0.7899581 |
| VTKTFTNI | ASSQDTGAEEQY | 0.7063135 |
| VVAAAILL | ASSQDTGAEEQY | 0.67895615 |
| YAALDVEL | ASSQDTGAEEQY | 0.7161031 |
| LASVALLL | ASSQDTGAEEQY | 0.7922089 |
| LDAVHRV | ASSQDTGAEEQY | 0.752604 |
| QGLSSSIF | ASSQDTGAEEQY | 0.711694 |
| RGTFANIRL | ASSQDTGAEEQY | 0.8038789 |
| RQLWWGHRI | ASSQDTGAEEQY | 0.7799015 |
| YGPPGTGKTLL | ASSQDTGAEEQY | 0.79860216 |
| AALVGAPANVA | ASSQDTGAEEQY | 0.7814641 |
| AATFRRTE | ASSQDTGAEEQY | 0.7445427 |
| AGALGGGI | ASSQDTGAEEQY | 0.7880174 |
| AGGAGGAGGIGAG | ASSQDTGAEEQY | 0.7568912 |
| AGSGALGL | ASSQDTGAEEQY | 0.70291936 |
| AQIADQI | ASSQDTGAEEQY | 0.6487764 |
| ASAVAVTYL | ASSQDTGAEEQY | 0.74531233 |
| ASTVLGGTGG | ASSQDTGAEEQY | 0.70346177 |
| AVGGEPAK | ASSQDTGAEEQY | 0.770405 |
| CEQGTFGGTC | ASSQDTGAEEQY | 0.8038254 |
| DEGKAPAG | ASSQDTGAEEQY | 0.71045 |
| DTVVFPAL | ASSQDTGAEEQY | 0.6326556 |
| FEGITMAG | ASSQDTGAEEQY | 0.7461723 |
| FPLVVQD | ASSQDTGAEEQY | 0.7294251 |
| GAFTTPQ | ASSQDTGAEEQY | 0.78980577 |
| GFGGFGTAGGVGGA | ASSQDTGAEEQY | 0.76265836 |
| GFLAQFL | ASSQDTGAEEQY | 0.81911755 |
| GGAGGAGGNAGVA | ASSQDTGAEEQY | 0.7951616 |
| GGVGTAGAGGAGGAGG | ASSQDTGAEEQY | 0.7078563 |
| GNGGGGDAGNAGSGGN | ASSQDTGAEEQY | 0.79108346 |
| GTGGAGGAGNDAGS | ASSQDTGAEEQY | 0.7084667 |
| GTTNTGWFNS | ASSQDTGAEEQY | 0.77666605 |
| IFMGPLFTL | ASSQDTGAEEQY | 0.7874923 |

| | | |
|---|---|---|
| ILAHAIEAI | ASSQDTGAEEQY | 0.7605884 |
| IRSAIREAL | ASSQDTGAEEQY | 0.7571083 |
| ITFAGGG | ASSQDTGAEEQY | 0.7824673 |
| ITGMSGHQEV | ASSQDTGAEEQY | 0.5901766 |
| IVAGLACTL | ASSQDTGAEEQY | 0.8033526 |
| IVGGGLAGA | ASSQDTGAEEQY | 0.66536355 |
| IVTGVLNI | ASSQDTGAEEQY | 0.5896316 |
| LAQQGCI | ASSQDTGAEEQY | 0.7363639 |
| LCLHVEN | ASSQDTGAEEQY | 0.8089486 |
| LDAISDGS | ASSQDTGAEEQY | 0.8050554 |
| LGGFAAML | ASSQDTGAEEQY | 0.7060832 |
| LGGKNPMI | ASSQDTGAEEQY | 0.67338574 |
| LGPKGVAAIL | ASSQDTGAEEQY | 0.7895774 |
| LMVGVSI | ASSQDTGAEEQY | 0.62501454 |
| LVGAAFVL | ASSQDTGAEEQY | 0.77456045 |
| LVLAVAPL | ASSQDTGAEEQY | 0.7340524 |
| MAAIAVIDAA | ASSQDTGAEEQY | 0.71454 |
| NNGINAELQ | ASSQDTGAEEQY | 0.6973026 |
| RGEPLVLVHG | ASSQDTGAEEQY | 0.7795699 |
| RGGAGGAAGLI | ASSQDTGAEEQY | 0.72058797 |
| RSYRMHPAV | ASSQDTGAEEQY | 0.7979369 |
| SAAHLLAGQLA | ASSQDTGAEEQY | 0.35705668 |
| SAAVGSY | ASSQDTGAEEQY | 0.7804841 |
| SPHAVLHRI | ASSQDTGAEEQY | 0.68692994 |
| SQIAKVL | ASSQDTGAEEQY | 0.6892834 |
| TALGHLVP | ASSQDTGAEEQY | 0.5139687 |
| TAPTTSGAA | ASSQDTGAEEQY | 0.76912236 |
| TDGDDGNGGSPGT | ASSQDTGAEEQY | 0.79204535 |
| TDGLGEN | ASSQDTGAEEQY | 0.79575276 |
| TGGPPLPGQA | ASSQDTGAEEQY | 0.76672924 |
| TIPAITI | ASSQDTGAEEQY | 0.7967383 |
| TIPGFTI | ASSQDTGAEEQY | 0.80386806 |
| TSIASGALA | ASSQDTGAEEQY | 0.7276611 |
| TVTLPGAAI | ASSQDTGAEEQY | 0.57796025 |
| VASGLSAALP | ASSQDTGAEEQY | 0.75263 |
| VGAGVIGI | ASSQDTGAEEQY | 0.78295416 |
| VGPKRFGV | ASSQDTGAEEQY | 0.7810595 |
| VNVAFPD | ASSQDTGAEEQY | 0.79811275 |
| VSMTNSGVS | ASSQDTGAEEQY | 0.6499766 |
| VTDVLGDLSV | ASSQDTGAEEQY | 0.77564687 |
| VVVVGAGFAGL | ASSQDTGAEEQY | 0.78406584 |
| VYTVGTVV | ASSQDTGAEEQY | 0.64150935 |
| WTTFSGP | ASSQDTGAEEQY | 0.7534832 |
| YDELVEAF | ASSQDTGAEEQY | 0.7147167 |
| YGTGGAGGNGG | ASSQDTGAEEQY | 0.5369744 |
| YTNVMAV | ASSQDTGAEEQY | 0.779866 |
| FLEQVHQL | ASSQDTGAEEQY | 0.72272456 |
| GLSLLAVAV | ASSQDTGAEEQY | 0.70837176 |
| LEQVHQL | ASSQDTGAEEQY | 0.7331036 |
| LNAAGAI | ASSQDTGAEEQY | 0.70535606 |
| PAVHASL | ASSQDTGAEEQY | 0.68560493 |

| | | |
|---|---|---|
| TELEDDA | ASSQDTGAEEQY | 0.6819726 |
| TPPGPPN | ASSQDTGAEEQY | 0.76591474 |
| TPPPPPPP | ASSQDTGAEEQY | 0.78579545 |
| TQSPALI | ASSTDRLSGNTLY | 0.7344477 |
| EFAAAFPQ | ASSTDRLSGNTLY | 0.7690877 |
| LLSGNLDVL | ASSTDRLSGNTLY | 0.80318713 |
| GPANLGNYNI | ASSTDRLSGNTLY | 0.5110254 |
| TLGPTLP | ASSTDRLSGNTLY | 0.64921916 |
| VYAYSSCCK | ASSTDRLSGNTLY | 0.7738849 |
| APAPAPAPNI | ASSTDRLSGNTLY | 0.638818 |
| PGGPVPIP | ASSTDRLSGNTLY | 0.79381263 |
| FLTVALEP | ASSTDRLSGNTLY | 0.7643248 |
| SQYFGANL | ASSTDRLSGNTLY | 0.7724309 |
| GGVKPAA | ASSTDRLSGNTLY | 0.785789 |
| GGTGGAGGAGGW | ASSTDRLSGNTLY | 0.8105581 |
| GVGGVAGLGGAGG | ASSTDRLSGNTLY | 0.77149534 |
| VLLVLATTI | ASSTDRLSGNTLY | 0.73827255 |
| IGIFAAML | ASSTDRLSGNTLY | 0.7982818 |
| AVGDLID | ASSTDRLSGNTLY | 0.6474858 |
| NISPLPL | ASSTDRLSGNTLY | 0.7971246 |
| VTAVFVAI | ASSTDRLSGNTLY | 0.7924826 |
| SGDDNDAAL | ASSTDRLSGNTLY | 0.6299528 |
| AVAALAHS | ASSTDRLSGNTLY | 0.73767024 |
| GGVTVMATP | ASSTDRLSGNTLY | 0.8091354 |
| AGLAANVRD | ASSTDRLSGNTLY | 0.8034701 |
| TNRPDLIDPALLRPG | ASSTDRLSGNTLY | 0.65495443 |
| LANLGVTAL | ASSTDRLSGNTLY | 0.71587515 |
| TGERIEL | ASSTDRLSGNTLY | 0.6413054 |
| NGEELAA | ASSTDRLSGNTLY | 0.72145563 |
| VLSLVWG | ASSTDRLSGNTLY | 0.52211297 |
| IGYALWL | ASSTDRLSGNTLY | 0.7891737 |
| LSLLLAPL | ASSTDRLSGNTLY | 0.7498965 |
| TLDEALK | ASSTDRLSGNTLY | 0.8058531 |
| TDGAGTAEGG | ASSTDRLSGNTLY | 0.7682369 |
| NAPGGAGGNGG | ASSTDRLSGNTLY | 0.8004046 |
| VSVFMGMGGNF | ASSTDRLSGNTLY | 0.73645425 |
| LASYLAAL | ASSTDRLSGNTLY | 0.734498 |
| LVGGTPAL | ASSTDRLSGNTLY | 0.6006409 |
| VRNADVL | ASSTDRLSGNTLY | 0.6238003 |
| TCSSPFLVSAM | ASSTDRLSGNTLY | 0.7339745 |
| VHITDAE | ASSTDRLSGNTLY | 0.7074902 |
| EYAAPSF | ASSTDRLSGNTLY | 0.46744072 |
| VVDAFHGGL | ASSTDRLSGNTLY | 0.79185224 |
| ASVVDPQ | ASSTDRLSGNTLY | 0.74242115 |
| SYNEFGGLS | ASSTDRLSGNTLY | 0.7901039 |
| DPDGSVY | ASSTDRLSGNTLY | 0.7248562 |
| MVSTIHATE | ASSTDRLSGNTLY | 0.7398494 |
| LAAGVADPV | ASSTDRLSGNTLY | 0.6737566 |
| TKEQIAA | ASSTDRLSGNTLY | 0.7923948 |
| SLASPAAL | ASSTDRLSGNTLY | 0.6968102 |
| IGLVFYGV | ASSTDRLSGNTLY | 0.56628835 |

| | | |
|---|---|---|
| FLLDASNL | ASSTDRLSGNTLY | 0.75391304 |
| YQVGNFP | ASSTDRLSGNTLY | 0.7344991 |
| ESLLSQST | ASSTDRLSGNTLY | 0.80365914 |
| LFTVGFAAVG | ASSTDRLSGNTLY | 0.79594564 |
| ATPPTPT | ASSTDRLSGNTLY | 0.763962 |
| TDLATLG | ASSTDRLSGNTLY | 0.80786836 |
| SAGGTGR | ASSTDRLSGNTLY | 0.25088465 |
| GFSNSGS | ASSTDRLSGNTLY | 0.72747517 |
| GGNGGSGGTSV | ASSTDRLSGNTLY | 0.80534226 |
| GTSVATGGAGN | ASSTDRLSGNTLY | 0.7732661 |
| NGGNGGSGGTS | ASSTDRLSGNTLY | 0.76360583 |
| AVVKFPH | ASSTDRLSGNTLY | 0.79955405 |
| TPPPPPP | ASSTDRLSGNTLY | 0.6801497 |
| AAPAAAAAPA | ASSTDRLSGNTLY | 0.7528372 |
| IALNEDL | ASSTDRLSGNTLY | 0.7543531 |
| IVNVSSM | ASSTDRLSGNTLY | 0.8024551 |
| TGIPPAPR | ASSTDRLSGNTLY | 0.77027655 |
| FDLGGGTF | ASSTDRLSGNTLY | 0.7202114 |
| FDLGGGTFDVS | ASSTDRLSGNTLY | 0.77762645 |
| NEPTAAA | ASSTDRLSGNTLY | 0.81534874 |
| NEPTAAALA | ASSTDRLSGNTLY | 0.7393608 |
| GSIINISSI | ASSTDRLSGNTLY | 0.79127693 |
| VAPGNAGTA | ASSTDRLSGNTLY | 0.7710198 |
| TGGGASLE | ASSTDRLSGNTLY | 0.7741592 |
| VAVYNHYKRI | ASSTDRLSGNTLY | 0.69774497 |
| TDPAPATT | ASSTDRLSGNTLY | 0.71426517 |
| TNRPDILDPALLRPG | ASSTDRLSGNTLY | 0.77121854 |
| LTTTTTS | ASSTDRLSGNTLY | 0.6473422 |
| PTGSGKTL | ASSTDRLSGNTLY | 0.7682098 |
| TGSGKTL | ASSTDRLSGNTLY | 0.7176901 |
| TIGHVDHGK | ASSTDRLSGNTLY | 0.81685734 |
| TIGHVDHGKTT | ASSTDRLSGNTLY | 0.7886646 |
| GHVDHGKTT | ASSTDRLSGNTLY | 0.7691548 |
| AQFEHTLL | ASSTDRLSGNTLY | 0.6847223 |
| IVNQDEL | ASSTDRLSGNTLY | 0.78164506 |
| TNGSQFF | ASSTDRLSGNTLY | 0.761476 |
| TGKGIVE | ASSTDRLSGNTLY | 0.55616474 |
| LISNASDA | ASSTDRLSGNTLY | 0.8017942 |
| AELRTLGELTKIAKAHGAQ | ASSTDRLSGNTLY | 0.64365405 |
| APNAVAAAEPSW | ASSTDRLSGNTLY | 0.6996974 |
| APQVAGAATAVL | ASSTDRLSGNTLY | 0.77511287 |
| ARRDPAAAAAILPTDA | ASSTDRLSGNTLY | 0.8134621 |
| DAAARDVLAEAGLAEYF | ASSTDRLSGNTLY | 0.7339356 |
| DDQIVLSAKANLAKVEQ | ASSTDRLSGNTLY | 0.7737857 |
| DELLPDTGEDSDEDGH | ASSTDRLSGNTLY | 0.65268636 |
| ELPEAPANRATVPQD | ASSTDRLSGNTLY | 0.78841877 |
| FGQSKIERVTGIG | ASSTDRLSGNTLY | 0.79770637 |
| GAGGTGSTAGGAGGAGGAGGLY | ASSTDRLSGNTLY | 0.78931975 |
| GHGGTGAAGGNGGTGGAGGSGI | ASSTDRLSGNTLY | 0.81715274 |
| GNGGNGGAGGEGGPGGAGGAGA | ASSTDRLSGNTLY | 0.7417859 |
| GSTAGNGGNGGNATNGTIAGQP | ASSTDRLSGNTLY | 0.4916829 |

| | | |
|---|---|---|
| GTGGDGGNGGNGGTGG | ASSTDRLSGNTLY | 0.7698388 |
| GVRRPAGIPADAPV | ASSTDRLSGNTLY | 0.79770243 |
| GYRQVTDAHLLTLARR | ASSTDRLSGNTLY | 0.800141 |
| IAKMGQDGHDRG | ASSTDRLSGNTLY | 0.805181 |
| KAEPSIEIQYT | ASSTDRLSGNTLY | 0.79849654 |
| KPVRIRMAADEDTVAVT | ASSTDRLSGNTLY | 0.74005973 |
| MPPRIAGMRLLVIKPEP | ASSTDRLSGNTLY | 0.7996732 |
| MKFYAVALFRSRGVPPTR | ASSTDRLSGNTLY | 0.80812526 |
| NAGGFGILGAVAHSP | ASSTDRLSGNTLY | 0.78256977 |
| NEGREITLAMAEM | ASSTDRLSGNTLY | 0.72638905 |
| NELVEWSHPLGHRWRWE | ASSTDRLSGNTLY | 0.671537 |
| NLGAGGTGGNGGNPG | ASSTDRLSGNTLY | 0.77713263 |
| NTEWPKWETVMAADPVD | ASSTDRLSGNTLY | 0.7669141 |
| PASAAPAETQP | ASSTDRLSGNTLY | 0.6522591 |
| QQSSDGKNGVRAAEITGE | ASSTDRLSGNTLY | 0.7511894 |
| RAPDAVTTSWWKE | ASSTDRLSGNTLY | 0.76511955 |
| SGFGNVGAHTSGWFNQ | ASSTDRLSGNTLY | 0.55620146 |
| SGVAGAVGQAASVGGLKV | ASSTDRLSGNTLY | 0.7719399 |
| SNGGAGTGSAGGNGGT | ASSTDRLSGNTLY | 0.8081893 |
| TGGAGGEGGAAG | ASSTDRLSGNTLY | 0.7515957 |
| TGGDGGAAGAGGAPGHG | ASSTDRLSGNTLY | 0.65535456 |
| TMPPLLAAIANAGRHGV | ASSTDRLSGNTLY | 0.80050004 |
| TSSTAGNSGGA | ASSTDRLSGNTLY | 0.8140292 |
| VPATVPVGILC | ASSTDRLSGNTLY | 0.7953675 |
| VQRPDSGGVVLTGRIS | ASSTDRLSGNTLY | 0.5232742 |
| WTPNPVPATKGVPAGLVQAT | ASSTDRLSGNTLY | 0.30239487 |
| YALKRALAKFSDRFDVVII | ASSTDRLSGNTLY | 0.75386465 |
| YLIGSVIV | ASSTDRLSGNTLY | 0.8005488 |
| YGQLVAVID | ASSTDRLSGNTLY | 0.7166153 |
| VTVGGMTLPA | ASSTDRLSGNTLY | 0.67925525 |
| VLALPAV | ASSTDRLSGNTLY | 0.75772226 |
| VIPPTLNYET | ASSTDRLSGNTLY | 0.7849671 |
| VGMGGMHPSSG | ASSTDRLSGNTLY | 0.7196874 |
| VGLAGGAATA | ASSTDRLSGNTLY | 0.6527976 |
| VGGAGGAGGSGP | ASSTDRLSGNTLY | 0.7885906 |
| VFAAFMV | ASSTDRLSGNTLY | 0.79612887 |
| TLYDKLR | ASSTDRLSGNTLY | 0.70473206 |
| TLAVFTVL | ASSTDRLSGNTLY | 0.73747253 |
| TGSVQPARV | ASSTDRLSGNTLY | 0.75185233 |
| TGGGAGSGGT | ASSTDRLSGNTLY | 0.6991867 |
| SHLETSL | ASSTDRLSGNTLY | 0.75267994 |
| SFTTGMT | ASSTDRLSGNTLY | 0.5857518 |
| QVEVAKRMGVRQPTVS | ASSTDRLSGNTLY | 0.7964796 |
| QGSGGPALAM | ASSTDRLSGNTLY | 0.5936372 |
| MLAPSNP | ASSTDRLSGNTLY | 0.8014015 |
| LVNNAAI | ASSTDRLSGNTLY | 0.7830548 |
| LTGEINTLA | ASSTDRLSGNTLY | 0.7899234 |
| LSIFGPV | ASSTDRLSGNTLY | 0.7264993 |
| LLALLANL | ASSTDRLSGNTLY | 0.71253747 |
| LISNASDALDKLR | ASSTDRLSGNTLY | 0.6285505 |
| LINEAVP | ASSTDRLSGNTLY | 0.6387846 |

| | | |
|---|---|---|
| LGYAFSGL | ASSTDRLSGNTLY | 0.80480325 |
| LDFVAVAL | ASSTDRLSGNTLY | 0.73022217 |
| LAQPGAE | ASSTDRLSGNTLY | 0.8000674 |
| ISNLGAVTGF | ASSTDRLSGNTLY | 0.77530575 |
| HQEDPGRI | ASSTDRLSGNTLY | 0.71016836 |
| GTPGEPTP | ASSTDRLSGNTLY | 0.78601277 |
| GFFNSGS | ASSTDRLSGNTLY | 0.5356535 |
| GAGGGGLTAGAN | ASSTDRLSGNTLY | 0.77669036 |
| FVDALVNM | ASSTDRLSGNTLY | 0.7557599 |
| FLTVAPEL | ASSTDRLSGNTLY | 0.7702402 |
| FETLTAL | ASSTDRLSGNTLY | 0.73888737 |
| DVINPIEL | ASSTDRLSGNTLY | 0.77836734 |
| APVSGAPV | ASSTDRLSGNTLY | 0.77935827 |
| ALVGGLAW | ASSTDRLSGNTLY | 0.72595584 |
| ALAAAAATAI | ASSTDRLSGNTLY | 0.74762964 |
| AFGLGLLL | ASSTDRLSGNTLY | 0.7926028 |
| AFFAAASL | ASSTDRLSGNTLY | 0.781994 |
| AAAALPAFSPP | ASSTDRLSGNTLY | 0.68140197 |
| VVLAPAL | ASSTDRLSGNTLY | 0.7692603 |
| TIGHVDHGKTTL | ASSTDRLSGNTLY | 0.719746 |
| LISNASDALDK | ASSTDRLSGNTLY | 0.7690761 |
| LISNASDAL | ASSTDRLSGNTLY | 0.77682406 |
| ISNASDALDK | ASSTDRLSGNTLY | 0.74752104 |
| NQAETLVYQTE | ASSTDRLSGNTLY | 0.77888423 |
| TGGTGGTAGAG | ASSTDRLSGNTLY | 0.56796575 |
| TGSTAGGAGGA | ASSTDRLSGNTLY | 0.7150088 |
| DPLAVPVAAGPV | ASSTDRLSGNTLY | 0.7398342 |
| GGTGGTPGNGGN | ASSTDRLSGNTLY | 0.75671566 |
| PAGLAGVDPATL | ASSTDRLSGNTLY | 0.7769953 |
| GNNLVVEAPHAGASV | ASSTDRLSGNTLY | 0.6112641 |
| KVTRIMSASTLEQPA | ASSTDRLSGNTLY | 0.77697384 |
| LGFALPIALPRRTVV | ASSTDRLSGNTLY | 0.7323526 |
| TRLGNLTARKLVVLV | ASSTDRLSGNTLY | 0.66880834 |
| WSQQRRLIYFPSAGP | ASSTDRLSGNTLY | 0.7248001 |
| GPNGNPDPMAAAVDIR | ASSTDRLSGNTLY | 0.75317717 |
| TQRCVSVADLRAAASR | ASSTDRLSGNTLY | 0.6222849 |
| QFVLDVAEHPVEVLTETV | ASSTDRLSGNTLY | 0.8005221 |
| GGAGGTGGAGGNGGTQAGDG | ASSTDRLSGNTLY | 0.67855036 |
| PLLATLRAIAVDVDAKPAQV | ASSTDRLSGNTLY | 0.7724216 |
| APKFRTGGYMIDFWGVGYQVAK | ASSTDRLSGNTLY | 0.7624663 |
| GAGGAGGAGGAAGGPGGTGGTG | ASSTDRLSGNTLY | 0.73806536 |
| IGALGLLATDVAPAGSVAFGVA | ASSTDRLSGNTLY | 0.7960168 |
| AGGAGGIGGNGGAGGAGGVLMG | ASSTDRLSGNTLY | 0.73570836 |
| ADELIGLL | ASSTDRLSGNTLY | 0.79107 |
| ADRALKA | ASSTDRLSGNTLY | 0.7197149 |
| AGAVGQVGGAGGA | ASSTDRLSGNTLY | 0.70797706 |
| AGGGGGLGGAP | ASSTDRLSGNTLY | 0.7870781 |
| AGGLLYGNGGAG | ASSTDRLSGNTLY | 0.77978575 |
| AGLESIGL | ASSTDRLSGNTLY | 0.797328 |
| AGLKTGII | ASSTDRLSGNTLY | 0.75904226 |
| ASLAALLL | ASSTDRLSGNTLY | 0.805713 |

| | | |
|---|---|---|
| ASVINISSI | ASSTDRLSGNTLY | 0.787132 |
| ATPPGPA | ASSTDRLSGNTLY | 0.7053532 |
| ATVASALV | ASSTDRLSGNTLY | 0.48320422 |
| ATVASLIL | ASSTDRLSGNTLY | 0.7291646 |
| AVNGPVLIRH | ASSTDRLSGNTLY | 0.6998507 |
| DGAAGAVGI | ASSTDRLSGNTLY | 0.7413652 |
| DPLLHGI | ASSTDRLSGNTLY | 0.486939 |
| ELGAALGAL | ASSTDRLSGNTLY | 0.7904149 |
| ELGRTQAV | ASSTDRLSGNTLY | 0.81602865 |
| FVAGGYLS | ASSTDRLSGNTLY | 0.7035444 |
| GASGGAGGAGGW | ASSTDRLSGNTLY | 0.8071197 |
| GGAGAGGDGGAGGI | ASSTDRLSGNTLY | 0.55745816 |
| GTGHNGSQPGMGGNG | ASSTDRLSGNTLY | 0.78377604 |
| GYAAASASAS | ASSTDRLSGNTLY | 0.8004368 |
| HQVDVAAVQQG | ASSTDRLSGNTLY | 0.6419255 |
| IGAAIAHT | ASSTDRLSGNTLY | 0.7706146 |
| IGMIEVAQL | ASSTDRLSGNTLY | 0.7367258 |
| IRLAPEHTSAMD | ASSTDRLSGNTLY | 0.7174927 |
| IYNVSVL | ASSTDRLSGNTLY | 0.80312455 |
| KQGFLGM | ASSTDRLSGNTLY | 0.68917906 |
| LESVITQL | ASSTDRLSGNTLY | 0.79350686 |
| LGLGALVVV | ASSTDRLSGNTLY | 0.739092 |
| LIGNGGNGGSGGI | ASSTDRLSGNTLY | 0.6504247 |
| LNAINAPTL | ASSTDRLSGNTLY | 0.79494715 |
| LQLLNTGV | ASSTDRLSGNTLY | 0.80446565 |
| LQTSIEF | ASSTDRLSGNTLY | 0.7538214 |
| MSFAVSEL | ASSTDRLSGNTLY | 0.807829 |
| NTGALGL | ASSTDRLSGNTLY | 0.7404666 |
| PPGAGGAGGAGTT | ASSTDRLSGNTLY | 0.8092898 |
| QLVGLGI | ASSTDRLSGNTLY | 0.8027117 |
| RFLEQVHQL | ASSTDRLSGNTLY | 0.75186116 |
| RGVGYVLR | ASSTDRLSGNTLY | 0.74773735 |
| SATGVFAGVI | ASSTDRLSGNTLY | 0.7989279 |
| SLPAITLP | ASSTDRLSGNTLY | 0.76748496 |
| STALAFTV | ASSTDRLSGNTLY | 0.79619217 |
| SVLTVGLGK | ASSTDRLSGNTLY | 0.80204767 |
| TALAGAGSGI | ASSTDRLSGNTLY | 0.65674686 |
| TGLPFTGI | ASSTDRLSGNTLY | 0.78839266 |
| VAEHVTV | ASSTDRLSGNTLY | 0.7460702 |
| VDIVLNSL | ASSTDRLSGNTLY | 0.76681185 |
| VLTGGGAVL | ASSTDRLSGNTLY | 0.6871699 |
| VNVGLAYGI | ASSTDRLSGNTLY | 0.5751146 |
| VSSAAVGAV | ASSTDRLSGNTLY | 0.7538326 |
| VTSVYEAGV | ASSTDRLSGNTLY | 0.63599765 |
| WVALDRGA | ASSTDRLSGNTLY | 0.78975964 |
| YLVGSVVI | ASSTDRLSGNTLY | 0.7868645 |
| GPPGTGKTLL | ASSTDRLSGNTLY | 0.68773484 |
| GVGMTKF | ASSTDRLSGNTLY | 0.7874388 |
| IGIGGSDL | ASSTDRLSGNTLY | 0.7956679 |
| IGTLLAEL | ASSTDRLSGNTLY | 0.5738985 |
| IYTSGTT | ASSTDRLSGNTLY | 0.71212995 |

| | | |
|---|---|---|
| LSFPAGV | ASSTDRLSGNTLY | 0.72947705 |
| LYTSGTT | ASSTDRLSGNTLY | 0.7340969 |
| VGAFPFGF | ASSTDRLSGNTLY | 0.5084801 |
| VGAGTVEF | ASSTDRLSGNTLY | 0.78303885 |
| VGKSTLLL | ASSTDRLSGNTLY | 0.67657435 |
| AALAHAGGGIV | ASSTDRLSGNTLY | 0.32035777 |
| AALVAVSLL | ASSTDRLSGNTLY | 0.76705337 |
| AAVSRAVAAL | ASSTDRLSGNTLY | 0.7354455 |
| AGGSSGIGGF | ASSTDRLSGNTLY | 0.71986645 |
| AGLTVANANG | ASSTDRLSGNTLY | 0.7819431 |
| AGLVGAVL | ASSTDRLSGNTLY | 0.6552057 |
| ALSQAFTV | ASSTDRLSGNTLY | 0.6581738 |
| ALTFSVSL | ASSTDRLSGNTLY | 0.76233685 |
| ALTGAALVAL | ASSTDRLSGNTLY | 0.79386914 |
| ASFSIGAGVL | ASSTDRLSGNTLY | 0.6081097 |
| DGSVYKI | ASSTDRLSGNTLY | 0.747113 |
| EQLAEIV | ASSTDRLSGNTLY | 0.79294467 |
| FAVSELTVM | ASSTDRLSGNTLY | 0.76255715 |
| GAAPLAAGL | ASSTDRLSGNTLY | 0.79644716 |
| GADGGDGGSGGAGG | ASSTDRLSGNTLY | 0.80338573 |
| GFGLDTGGAGG | ASSTDRLSGNTLY | 0.71184325 |
| GLASGGLL | ASSTDRLSGNTLY | 0.7899295 |
| GRIAGAAVLL | ASSTDRLSGNTLY | 0.7977563 |
| GSAGSLNTSF | ASSTDRLSGNTLY | 0.7143613 |
| GTGQIGI | ASSTDRLSGNTLY | 0.736043 |
| GTLIALIL | ASSTDRLSGNTLY | 0.6517973 |
| IAATMGFVVG | ASSTDRLSGNTLY | 0.7398126 |
| IFAQALL | ASSTDRLSGNTLY | 0.492908 |
| IGAAVALGF | ASSTDRLSGNTLY | 0.74022365 |
| IGLTPLAL | ASSTDRLSGNTLY | 0.79479027 |
| ILGSNTSTL | ASSTDRLSGNTLY | 0.7137619 |
| IVFSLTVL | ASSTDRLSGNTLY | 0.6033585 |
| LASKATGF | ASSTDRLSGNTLY | 0.58516467 |
| LASVALIL | ASSTDRLSGNTLY | 0.71666694 |
| LFAVGLTL | ASSTDRLSGNTLY | 0.685249 |
| LFNGGIGTYI | ASSTDRLSGNTLY | 0.5311363 |
| LTLGAGII | ASSTDRLSGNTLY | 0.70513135 |
| LVAAALGDI | ASSTDRLSGNTLY | 0.7898915 |
| NTAFVQLGI | ASSTDRLSGNTLY | 0.79236555 |
| QLFSGAEL | ASSTDRLSGNTLY | 0.55755556 |
| SAATFITL | ASSTDRLSGNTLY | 0.801085 |
| SNYGLANLGV | ASSTDRLSGNTLY | 0.78119045 |
| SYGGAYAVM | ASSTDRLSGNTLY | 0.7978505 |
| TTGVGGLTLG | ASSTDRLSGNTLY | 0.7597407 |
| TTQTIYI | ASSTDRLSGNTLY | 0.6328 |
| VAAGAGLGAVI | ASSTDRLSGNTLY | 0.79001 |
| VAAVVTLL | ASSTDRLSGNTLY | 0.677311 |
| VGFLGGLI | ASSTDRLSGNTLY | 0.702326 |
| VGVIPILL | ASSTDRLSGNTLY | 0.78771317 |
| VIFASSLI | ASSTDRLSGNTLY | 0.73367 |
| VKGGDGGAAAG | ASSTDRLSGNTLY | 0.64522624 |

| | | |
|---|---|---|
| VTKTFTNI | ASSTDRLSGNTLY | 0.7049509 |
| VVAAAILL | ASSTDRLSGNTLY | 0.79748553 |
| YAALDVEL | ASSTDRLSGNTLY | 0.60228795 |
| LASVALLL | ASSTDRLSGNTLY | 0.61238956 |
| LDAVHRV | ASSTDRLSGNTLY | 0.6819702 |
| QGLSSSIF | ASSTDRLSGNTLY | 0.72928 |
| RGTFANIRL | ASSTDRLSGNTLY | 0.7886137 |
| RQLWWGHRI | ASSTDRLSGNTLY | 0.78105104 |
| YGPPGTGKTLL | ASSTDRLSGNTLY | 0.6716677 |
| AALVGAPANVA | ASSTDRLSGNTLY | 0.7565471 |
| AATFRRTE | ASSTDRLSGNTLY | 0.7731221 |
| AGALGGGI | ASSTDRLSGNTLY | 0.75841904 |
| AGGAGGAGGIGAG | ASSTDRLSGNTLY | 0.77026266 |
| AGSGALGL | ASSTDRLSGNTLY | 0.63207567 |
| AQIADQI | ASSTDRLSGNTLY | 0.7905648 |
| ASAVAVTYL | ASSTDRLSGNTLY | 0.7807127 |
| ASTVLGGTGG | ASSTDRLSGNTLY | 0.7900398 |
| AVGGEPAK | ASSTDRLSGNTLY | 0.79363143 |
| CEQGTFGGTC | ASSTDRLSGNTLY | 0.7328562 |
| DEGKAPAG | ASSTDRLSGNTLY | 0.7576567 |
| DTVVFPAL | ASSTDRLSGNTLY | 0.5058935 |
| FEGITMAG | ASSTDRLSGNTLY | 0.72452486 |
| FPLVVQD | ASSTDRLSGNTLY | 0.7480812 |
| GAFTTPQ | ASSTDRLSGNTLY | 0.7464097 |
| GFGGFGTAGGVGGA | ASSTDRLSGNTLY | 0.7834797 |
| GFLAQFL | ASSTDRLSGNTLY | 0.7850771 |
| GGAGGAGGNAGVA | ASSTDRLSGNTLY | 0.6946666 |
| GGVGTAGAGGAGGAGG | ASSTDRLSGNTLY | 0.80227643 |
| GNGGGGDAGNAGSGGN | ASSTDRLSGNTLY | 0.79149705 |
| GTGGAGGAGNDAGS | ASSTDRLSGNTLY | 0.5667172 |
| GTTNTGWFNS | ASSTDRLSGNTLY | 0.7120701 |
| IFMGPLFTL | ASSTDRLSGNTLY | 0.8039871 |
| ILAHAIEAI | ASSTDRLSGNTLY | 0.76696396 |
| IRSAIREAL | ASSTDRLSGNTLY | 0.7040893 |
| ITFAGGG | ASSTDRLSGNTLY | 0.7273142 |
| ITGMSGHQEV | ASSTDRLSGNTLY | 0.81007487 |
| IVAGLACTL | ASSTDRLSGNTLY | 0.6983393 |
| IVGGGLAGA | ASSTDRLSGNTLY | 0.8015124 |
| IVTGVLNI | ASSTDRLSGNTLY | 0.4322904 |
| LAQQGCI | ASSTDRLSGNTLY | 0.7309705 |
| LCLHVEN | ASSTDRLSGNTLY | 0.67404765 |
| LDAISDGS | ASSTDRLSGNTLY | 0.39858052 |
| LGGFAAML | ASSTDRLSGNTLY | 0.7873719 |
| LGGKNPMI | ASSTDRLSGNTLY | 0.7530613 |
| LGPKGVAAIL | ASSTDRLSGNTLY | 0.7740509 |
| LMVGVSI | ASSTDRLSGNTLY | 0.74687254 |
| LVGAAFVL | ASSTDRLSGNTLY | 0.7917297 |
| LVLAVAPL | ASSTDRLSGNTLY | 0.72600377 |
| MAAIAVIDAA | ASSTDRLSGNTLY | 0.75961477 |
| NNGINAELQ | ASSTDRLSGNTLY | 0.66499317 |
| RGEPLVLVHG | ASSTDRLSGNTLY | 0.7793927 |

| | | |
|---|---|---|
| RGGAGGAAGLI | ASSTDRLSGNTLY | 0.8069557 |
| RSYRMHPAV | ASSTDRLSGNTLY | 0.7884923 |
| SAAHLLAGQLA | ASSTDRLSGNTLY | 0.5041996 |
| SAAVGSY | ASSTDRLSGNTLY | 0.71751904 |
| SPHAVLHRI | ASSTDRLSGNTLY | 0.74794716 |
| SQIAKVL | ASSTDRLSGNTLY | 0.44574577 |
| TALGHLVP | ASSTDRLSGNTLY | 0.37153298 |
| TAPTTSGAA | ASSTDRLSGNTLY | 0.6898464 |
| TDGDDGNGGSPGT | ASSTDRLSGNTLY | 0.6669894 |
| TDGLGEN | ASSTDRLSGNTLY | 0.7810474 |
| TGGPPLPGQA | ASSTDRLSGNTLY | 0.7636676 |
| TIPAITI | ASSTDRLSGNTLY | 0.7177155 |
| TIPGFTI | ASSTDRLSGNTLY | 0.75944173 |
| TSIASGALA | ASSTDRLSGNTLY | 0.7327424 |
| TVTLPGAAI | ASSTDRLSGNTLY | 0.21179555 |
| VASGLSAALP | ASSTDRLSGNTLY | 0.6820835 |
| VGAGVIGI | ASSTDRLSGNTLY | 0.58430195 |
| VGPKRFGV | ASSTDRLSGNTLY | 0.8057846 |
| VNVAFPD | ASSTDRLSGNTLY | 0.6807898 |
| VSMTNSGVS | ASSTDRLSGNTLY | 0.6587113 |
| VTDVLGDLSV | ASSTDRLSGNTLY | 0.72919905 |
| VVVVGAGFAGL | ASSTDRLSGNTLY | 0.7621071 |
| VYTVGTVV | ASSTDRLSGNTLY | 0.7335733 |
| WTTFSGP | ASSTDRLSGNTLY | 0.6286763 |
| YDELVEAF | ASSTDRLSGNTLY | 0.3832203 |
| YGTGGAGGNGG | ASSTDRLSGNTLY | 0.7831807 |
| YTNVMAV | ASSTDRLSGNTLY | 0.6182261 |
| FLEQVHQL | ASSTDRLSGNTLY | 0.6351557 |
| GLSLLAVAV | ASSTDRLSGNTLY | 0.55014753 |
| LEQVHQL | ASSTDRLSGNTLY | 0.6343752 |
| LNAAGAI | ASSTDRLSGNTLY | 0.5588894 |
| PAVHASL | ASSTDRLSGNTLY | 0.79167795 |
| TELEDDA | ASSTDRLSGNTLY | 0.6911052 |
| TPPGPPN | ASSTDRLSGNTLY | 0.70371807 |
| TPPPPPPP | ASSTDRLSGNTLY | 0.80569494 |
| TQSPALI | ASSPGTGGAETLY | 0.7406058 |
| EFAAAFPQ | ASSPGTGGAETLY | 0.70292443 |
| LLSGNLDVL | ASSPGTGGAETLY | 0.67329687 |
| GPANLGNYNI | ASSPGTGGAETLY | 0.6339205 |
| TLGPTLP | ASSPGTGGAETLY | 0.66583014 |
| VYAYSSCCK | ASSPGTGGAETLY | 0.6911764 |
| APAPAPAPNI | ASSPGTGGAETLY | 0.80408883 |
| PGGPVPIP | ASSPGTGGAETLY | 0.74217296 |
| FLTVALEP | ASSPGTGGAETLY | 0.6786668 |
| SQYFGANL | ASSPGTGGAETLY | 0.7639924 |
| GGVKPAA | ASSPGTGGAETLY | 0.8011502 |
| GGTGGAGGAGGW | ASSPGTGGAETLY | 0.80580795 |
| GVGGVAGLGGAGG | ASSPGTGGAETLY | 0.64408493 |
| VLLVLATTI | ASSPGTGGAETLY | 0.7256873 |
| IGIFAAML | ASSPGTGGAETLY | 0.7207465 |
| AVGDLID | ASSPGTGGAETLY | 0.7121436 |

| | | |
|---|---|---|
| NISPLPL | ASSPGTGGAETLY | 0.6925818 |
| VTAVFVAI | ASSPGTGGAETLY | 0.6467402 |
| SGDDNDAAL | ASSPGTGGAETLY | 0.78194666 |
| AVAALAHS | ASSPGTGGAETLY | 0.8147856 |
| GGVTVMATP | ASSPGTGGAETLY | 0.72613645 |
| AGLAANVRD | ASSPGTGGAETLY | 0.6252921 |
| TNRPDLIDPALLRPG | ASSPGTGGAETLY | 0.7179277 |
| LANLGVTAL | ASSPGTGGAETLY | 0.62095416 |
| TGERIEL | ASSPGTGGAETLY | 0.66623616 |
| NGEELAA | ASSPGTGGAETLY | 0.7309829 |
| VLSLVWG | ASSPGTGGAETLY | 0.5434077 |
| IGYALWL | ASSPGTGGAETLY | 0.7379074 |
| LSLLLAPL | ASSPGTGGAETLY | 0.8043082 |
| TLDEALK | ASSPGTGGAETLY | 0.7061331 |
| TDGAGTAEGG | ASSPGTGGAETLY | 0.7712761 |
| NAPGGAGGNGG | ASSPGTGGAETLY | 0.7930373 |
| VSVFMGMGGNF | ASSPGTGGAETLY | 0.76846766 |
| LASYLAAL | ASSPGTGGAETLY | 0.794599 |
| LVGGTPAL | ASSPGTGGAETLY | 0.7887517 |
| VRNADVL | ASSPGTGGAETLY | 0.76730585 |
| TCSSPFLVSAM | ASSPGTGGAETLY | 0.73855543 |
| VHITDAE | ASSPGTGGAETLY | 0.77420866 |
| EYAAPSF | ASSPGTGGAETLY | 0.72921014 |
| VVDAFHGGL | ASSPGTGGAETLY | 0.7283596 |
| ASVVDPQ | ASSPGTGGAETLY | 0.69123995 |
| SYNEFGGLS | ASSPGTGGAETLY | 0.7881173 |
| DPDGSVY | ASSPGTGGAETLY | 0.71695113 |
| MVSTIHATE | ASSPGTGGAETLY | 0.8033875 |
| LAAGVADPV | ASSPGTGGAETLY | 0.5084409 |
| TKEQIAA | ASSPGTGGAETLY | 0.64109564 |
| SLASPAAL | ASSPGTGGAETLY | 0.78040445 |
| IGLVFYGV | ASSPGTGGAETLY | 0.2275282 |
| FLLDASNL | ASSPGTGGAETLY | 0.40337893 |
| YQVGNFP | ASSPGTGGAETLY | 0.7384867 |
| ESLLSQST | ASSPGTGGAETLY | 0.7891786 |
| LFTVGFAAVG | ASSPGTGGAETLY | 0.24477051 |
| ATPPTPT | ASSPGTGGAETLY | 0.6530447 |
| TDLATLG | ASSPGTGGAETLY | 0.5568592 |
| SAGGTGR | ASSPGTGGAETLY | 0.77846134 |
| GFSNSGS | ASSPGTGGAETLY | 0.7249766 |
| GGNGGSGGTSV | ASSPGTGGAETLY | 0.50895196 |
| GTSVATGGAGN | ASSPGTGGAETLY | 0.78204274 |
| NGGNGGSGGTS | ASSPGTGGAETLY | 0.5429578 |
| AVVKFPH | ASSPGTGGAETLY | 0.6586826 |
| TPPPPPP | ASSPGTGGAETLY | 0.7871957 |
| AAPAAAAAPA | ASSPGTGGAETLY | 0.8011226 |
| IALNEDL | ASSPGTGGAETLY | 0.6391859 |
| IVNVSSM | ASSPGTGGAETLY | 0.76869357 |
| TGIPPAPR | ASSPGTGGAETLY | 0.6911392 |
| FDLGGGTF | ASSPGTGGAETLY | 0.8090118 |
| FDLGGGTFDVS | ASSPGTGGAETLY | 0.71647537 |

| | | |
|---|---|---|
| NEPTAAA | ASSPGTGGAETLY | 0.58637 |
| NEPTAAALA | ASSPGTGGAETLY | 0.79033905 |
| GSIINISSI | ASSPGTGGAETLY | 0.7745359 |
| VAPGNAGTA | ASSPGTGGAETLY | 0.7453812 |
| TGGGASLE | ASSPGTGGAETLY | 0.7827699 |
| VAVYNHYKRI | ASSPGTGGAETLY | 0.70052934 |
| TDPAPATT | ASSPGTGGAETLY | 0.7042552 |
| TNRPDILDPALLRPG | ASSPGTGGAETLY | 0.7853515 |
| LTTTTTS | ASSPGTGGAETLY | 0.7827145 |
| PTGSGKTL | ASSPGTGGAETLY | 0.6466525 |
| TGSGKTL | ASSPGTGGAETLY | 0.79536545 |
| TIGHVDHGK | ASSPGTGGAETLY | 0.72344166 |
| TIGHVDHGKTT | ASSPGTGGAETLY | 0.66363525 |
| GHVDHGKTT | ASSPGTGGAETLY | 0.80774933 |
| AQFEHTLL | ASSPGTGGAETLY | 0.7490481 |
| IVNQDEL | ASSPGTGGAETLY | 0.7781577 |
| TNGSQFF | ASSPGTGGAETLY | 0.79061675 |
| TGKGIVE | ASSPGTGGAETLY | 0.8059792 |
| LISNASDA | ASSPGTGGAETLY | 0.6090171 |
| AELRTLGELTKIAKAHGAQ | ASSPGTGGAETLY | 0.7713109 |
| APNAVAAAEPSW | ASSPGTGGAETLY | 0.7261336 |
| APQVAGAATAVL | ASSPGTGGAETLY | 0.78336155 |
| ARRDPAAAAAILPTDA | ASSPGTGGAETLY | 0.75125736 |
| DAAARDVLAEAGLAEYF | ASSPGTGGAETLY | 0.70244825 |
| DDQIVLSAKANLAKVEQ | ASSPGTGGAETLY | 0.78096414 |
| DELLPDTGEDSDEDGH | ASSPGTGGAETLY | 0.7981014 |
| ELPEAPANRATVPQD | ASSPGTGGAETLY | 0.74498427 |
| FGQSKIERVTGIG | ASSPGTGGAETLY | 0.766624 |
| GAGGTGSTAGGAGGAGGAGGLY | ASSPGTGGAETLY | 0.7992449 |
| GHGGTGAAGGNGGTGGAGGSGI | ASSPGTGGAETLY | 0.6857209 |
| GNGGNGGAGGEGGPGGAGGAGA | ASSPGTGGAETLY | 0.7321062 |
| GSTAGNGGNGGNATNGTIAGQP | ASSPGTGGAETLY | 0.77990043 |
| GTGGDGGNGGNGGTGG | ASSPGTGGAETLY | 0.7990651 |
| GVRRPAGIPADAPV | ASSPGTGGAETLY | 0.736842 |
| GYRQVTDAHLLTLARR | ASSPGTGGAETLY | 0.79541624 |
| IAKMGQDGHDRG | ASSPGTGGAETLY | 0.6632761 |
| KAEPSIEIQYT | ASSPGTGGAETLY | 0.7641223 |
| KPVRIRMAADEDTVAVT | ASSPGTGGAETLY | 0.6565536 |
| MPPRIAGMRLLVIKPEP | ASSPGTGGAETLY | 0.73966634 |
| MKFYAVALFRSRGVPPTR | ASSPGTGGAETLY | 0.6091024 |
| NAGGFGILGAVAHSP | ASSPGTGGAETLY | 0.76086235 |
| NEGREITLAMAEM | ASSPGTGGAETLY | 0.7728677 |
| NELVEWSHPLGHRWRWE | ASSPGTGGAETLY | 0.770054 |
| NLGAGGTGGNGGNPG | ASSPGTGGAETLY | 0.44063216 |
| NTEWPKWETVMAADPVD | ASSPGTGGAETLY | 0.7662567 |
| PASAAPAETQP | ASSPGTGGAETLY | 0.77232647 |
| QQSSDGKNGVRAAEITGE | ASSPGTGGAETLY | 0.8101288 |
| RAPDAVTTSWWKE | ASSPGTGGAETLY | 0.7204821 |
| SGFGNVGAHTSGWFNQ | ASSPGTGGAETLY | 0.77694726 |
| SGVAGAVGQAASVGGLKV | ASSPGTGGAETLY | 0.67598593 |
| SNGGAGTGSAGGNGGT | ASSPGTGGAETLY | 0.73730785 |

| | | |
|---|---|---|
| TGGAGGEGGAAG | ASSPGTGGAETLY | 0.56063986 |
| TGGDGGAAGAGGAPGHG | ASSPGTGGAETLY | 0.6878915 |
| TMPPLLAAIANAGRHGV | ASSPGTGGAETLY | 0.6677674 |
| TSSTAGNSGGA | ASSPGTGGAETLY | 0.6349532 |
| VPATVPVGILC | ASSPGTGGAETLY | 0.7817273 |
| VQRPDSGGVVLTGRIS | ASSPGTGGAETLY | 0.79059654 |
| WTPNPVPATKGVPAGLVQAT | ASSPGTGGAETLY | 0.78517866 |
| YALKRALAKFSDRFDVVII | ASSPGTGGAETLY | 0.78648144 |
| YLIGSVIV | ASSPGTGGAETLY | 0.7967957 |
| YGQLVAVID | ASSPGTGGAETLY | 0.7958493 |
| VTVGGMTLPA | ASSPGTGGAETLY | 0.67390597 |
| VLALPAV | ASSPGTGGAETLY | 0.73267496 |
| VIPPTLNYET | ASSPGTGGAETLY | 0.5831132 |
| VGMGGMHPSSG | ASSPGTGGAETLY | 0.7263825 |
| VGLAGGAATA | ASSPGTGGAETLY | 0.6886705 |
| VGGAGGAGGSGP | ASSPGTGGAETLY | 0.75455594 |
| VFAAFMV | ASSPGTGGAETLY | 0.23809174 |
| TLYDKLR | ASSPGTGGAETLY | 0.7409438 |
| TLAVFTVL | ASSPGTGGAETLY | 0.80335075 |
| TGSVQPARV | ASSPGTGGAETLY | 0.6270511 |
| TGGGAGSGGT | ASSPGTGGAETLY | 0.76683336 |
| SHLETSL | ASSPGTGGAETLY | 0.7544771 |
| SFTTGMT | ASSPGTGGAETLY | 0.66043925 |
| QVEVAKRMGVRQPTVS | ASSPGTGGAETLY | 0.76799077 |
| QGSGGPALAM | ASSPGTGGAETLY | 0.6663871 |
| MLAPSNP | ASSPGTGGAETLY | 0.7802013 |
| LVNNAAI | ASSPGTGGAETLY | 0.80423766 |
| LTGEINTLA | ASSPGTGGAETLY | 0.75792575 |
| LSIFGPV | ASSPGTGGAETLY | 0.7485827 |
| LLALLANL | ASSPGTGGAETLY | 0.7513554 |
| LISNASDALDKLR | ASSPGTGGAETLY | 0.56101555 |
| LINEAVP | ASSPGTGGAETLY | 0.7631756 |
| LGYAFSGL | ASSPGTGGAETLY | 0.33624497 |
| LDFVAVAL | ASSPGTGGAETLY | 0.73448145 |
| LAQPGAE | ASSPGTGGAETLY | 0.6477121 |
| ISNLGAVTGF | ASSPGTGGAETLY | 0.75116116 |
| HQEDPGRI | ASSPGTGGAETLY | 0.7287091 |
| GTPGEPTP | ASSPGTGGAETLY | 0.75279546 |
| GFFNSGS | ASSPGTGGAETLY | 0.6342351 |
| GAGGGGLTAGAN | ASSPGTGGAETLY | 0.7350307 |
| FVDALVNM | ASSPGTGGAETLY | 0.71328354 |
| FLTVAPEL | ASSPGTGGAETLY | 0.6046157 |
| FETLTAL | ASSPGTGGAETLY | 0.7839122 |
| DVINPIEL | ASSPGTGGAETLY | 0.7401335 |
| APVSGAPV | ASSPGTGGAETLY | 0.6944086 |
| ALVGGLAW | ASSPGTGGAETLY | 0.7921584 |
| ALAAAAATAI | ASSPGTGGAETLY | 0.74836373 |
| AFGLGLLL | ASSPGTGGAETLY | 0.5412766 |
| AFFAAASL | ASSPGTGGAETLY | 0.6925907 |
| AAAALPAFSPP | ASSPGTGGAETLY | 0.7382853 |
| VVLAPAL | ASSPGTGGAETLY | 0.7092427 |

| | | |
|---|---|---|
| TIGHVDHGKTTL | ASSPGTGGAETLY | 0.28527513 |
| LISNASDALDK | ASSPGTGGAETLY | 0.76803064 |
| LISNASDAL | ASSPGTGGAETLY | 0.66518253 |
| ISNASDALDK | ASSPGTGGAETLY | 0.6081009 |
| NQAETLVYQTE | ASSPGTGGAETLY | 0.6998904 |
| TGGTGGTAGAG | ASSPGTGGAETLY | 0.7783009 |
| TGSTAGGAGGA | ASSPGTGGAETLY | 0.6451429 |
| DPLAVPVAAGPV | ASSPGTGGAETLY | 0.73873276 |
| GGTGGTPGNGGN | ASSPGTGGAETLY | 0.7711445 |
| PAGLAGVDPATL | ASSPGTGGAETLY | 0.76626563 |
| GNNLVVEAPHAGASV | ASSPGTGGAETLY | 0.7922628 |
| KVTRIMSASTLEQPA | ASSPGTGGAETLY | 0.76323044 |
| LGFALPIALPRRTVV | ASSPGTGGAETLY | 0.7463955 |
| TRLGNLTARKLVVLV | ASSPGTGGAETLY | 0.78203154 |
| WSQQRRLIYFPSAGP | ASSPGTGGAETLY | 0.735616 |
| GPNGNPDPMAAAVDIR | ASSPGTGGAETLY | 0.61046875 |
| TQRCVSVADLRAAASR | ASSPGTGGAETLY | 0.5396469 |
| QFVLDVAEHPVEVLTETV | ASSPGTGGAETLY | 0.7101839 |
| GGAGGTGGAGGNGGTQAGDG | ASSPGTGGAETLY | 0.5374609 |
| PLLATLRAIAVDVDAKPAQV | ASSPGTGGAETLY | 0.7522676 |
| APKFRTGGYMIDFWGVGYQVAK | ASSPGTGGAETLY | 0.8031608 |
| GAGGAGGAGGAAGGPGGTGGTG | ASSPGTGGAETLY | 0.73797995 |
| IGALGLLATDVAPAGSVAFGVA | ASSPGTGGAETLY | 0.5743333 |
| AGGAGGIGGNGGAGGAGGVLMG | ASSPGTGGAETLY | 0.7046298 |
| ADELIGLL | ASSPGTGGAETLY | 0.6892884 |
| ADRALKA | ASSPGTGGAETLY | 0.77262545 |
| AGAVGQVGGAGGA | ASSPGTGGAETLY | 0.7897916 |
| AGGGGGLGGAP | ASSPGTGGAETLY | 0.81189555 |
| AGGLLYGNGGAG | ASSPGTGGAETLY | 0.7800819 |
| AGLESIGL | ASSPGTGGAETLY | 0.6646638 |
| AGLKTGII | ASSPGTGGAETLY | 0.7758234 |
| ASLAALLL | ASSPGTGGAETLY | 0.7277813 |
| ASVINISSI | ASSPGTGGAETLY | 0.7480805 |
| ATPPGPA | ASSPGTGGAETLY | 0.7566469 |
| ATVASALV | ASSPGTGGAETLY | 0.7484677 |
| ATVASLIL | ASSPGTGGAETLY | 0.7258147 |
| AVNGPVLIRH | ASSPGTGGAETLY | 0.7244839 |
| DGAAGAVGI | ASSPGTGGAETLY | 0.5006862 |
| DPLLHGI | ASSPGTGGAETLY | 0.7810211 |
| ELGAALGAL | ASSPGTGGAETLY | 0.72950065 |
| ELGRTQAV | ASSPGTGGAETLY | 0.56003994 |
| FVAGGYLS | ASSPGTGGAETLY | 0.7150725 |
| GASGGAGGAGGW | ASSPGTGGAETLY | 0.80385625 |
| GGAGAGGDGGAGGI | ASSPGTGGAETLY | 0.79475605 |
| GTGHNGSQPGMGGNG | ASSPGTGGAETLY | 0.6523895 |
| GYAAASASAS | ASSPGTGGAETLY | 0.6556032 |
| HQVDVAAVQQG | ASSPGTGGAETLY | 0.74413675 |
| IGAAIAHT | ASSPGTGGAETLY | 0.5592613 |
| IGMIEVAQL | ASSPGTGGAETLY | 0.7736524 |
| IRLAPEHTSAMD | ASSPGTGGAETLY | 0.7023003 |
| IYNVSVL | ASSPGTGGAETLY | 0.68863297 |

| | | |
|---|---|---|
| KQGFLGM | ASSPGTGGAETLY | 0.69874334 |
| LESVITQL | ASSPGTGGAETLY | 0.74960524 |
| LGLGALVVV | ASSPGTGGAETLY | 0.66003597 |
| LIGNGGNGGSGGI | ASSPGTGGAETLY | 0.7836923 |
| LNAINAPTL | ASSPGTGGAETLY | 0.29804015 |
| LQLLNTGV | ASSPGTGGAETLY | 0.78684044 |
| LQTSIEF | ASSPGTGGAETLY | 0.6221496 |
| MSFAVSEL | ASSPGTGGAETLY | 0.663851 |
| NTGALGL | ASSPGTGGAETLY | 0.4256062 |
| PPGAGGAGGAGTT | ASSPGTGGAETLY | 0.75819933 |
| QLVGLGI | ASSPGTGGAETLY | 0.7711148 |
| RFLEQVHQL | ASSPGTGGAETLY | 0.7823596 |
| RGVGYVLR | ASSPGTGGAETLY | 0.67485774 |
| SATGVFAGVI | ASSPGTGGAETLY | 0.78682864 |
| SLPAITLP | ASSPGTGGAETLY | 0.78539115 |
| STALAFTV | ASSPGTGGAETLY | 0.69238746 |
| SVLTVGLGK | ASSPGTGGAETLY | 0.50832194 |
| TALAGAGSGI | ASSPGTGGAETLY | 0.763956 |
| TGLPFTGI | ASSPGTGGAETLY | 0.7376824 |
| VAEHVTV | ASSPGTGGAETLY | 0.7816448 |
| VDIVLNSL | ASSPGTGGAETLY | 0.7603124 |
| VLTGGGAVL | ASSPGTGGAETLY | 0.55100954 |
| VNVGLAYGI | ASSPGTGGAETLY | 0.79120916 |
| VSSAAVGAV | ASSPGTGGAETLY | 0.78970873 |
| VTSVYEAGV | ASSPGTGGAETLY | 0.60810375 |
| WVALDRGA | ASSPGTGGAETLY | 0.6948049 |
| YLVGSVVI | ASSPGTGGAETLY | 0.71618104 |
| GPPGTGKTLL | ASSPGTGGAETLY | 0.53805 |
| GVGMTKF | ASSPGTGGAETLY | 0.7773646 |
| IGIGGSDL | ASSPGTGGAETLY | 0.6661577 |
| IGTLLAEL | ASSPGTGGAETLY | 0.6924355 |
| IYTSGTT | ASSPGTGGAETLY | 0.73498553 |
| LSFPAGV | ASSPGTGGAETLY | 0.70891356 |
| LYTSGTT | ASSPGTGGAETLY | 0.7083554 |
| VGAFPFGF | ASSPGTGGAETLY | 0.712186 |
| VGAGTVEF | ASSPGTGGAETLY | 0.7602371 |
| VGKSTLLL | ASSPGTGGAETLY | 0.7609649 |
| AALAHAGGGIV | ASSPGTGGAETLY | 0.73030895 |
| AALVAVSLL | ASSPGTGGAETLY | 0.7635473 |
| AAVSRAVAAL | ASSPGTGGAETLY | 0.79229784 |
| AGGSSGIGGF | ASSPGTGGAETLY | 0.54624236 |
| AGLTVANANG | ASSPGTGGAETLY | 0.6416515 |
| AGLVGAVL | ASSPGTGGAETLY | 0.7648896 |
| ALSQAFTV | ASSPGTGGAETLY | 0.62907636 |
| ALTFSVSL | ASSPGTGGAETLY | 0.7823101 |
| ALTGAALVAL | ASSPGTGGAETLY | 0.73640496 |
| ASFSIGAGVL | ASSPGTGGAETLY | 0.7610726 |
| DGSVYKI | ASSPGTGGAETLY | 0.74564016 |
| EQLAEIV | ASSPGTGGAETLY | 0.80604374 |
| FAVSELTVM | ASSPGTGGAETLY | 0.77991724 |
| GAAPLAAGL | ASSPGTGGAETLY | 0.76104015 |

| | | |
|---|---|---|
| GADGGDGGSGGAGG | ASSPGTGGAETLY | 0.7838812 |
| GFGLDTGGAGG | ASSPGTGGAETLY | 0.6285123 |
| GLASGGLL | ASSPGTGGAETLY | 0.79041606 |
| GRIAGAAVLL | ASSPGTGGAETLY | 0.7778936 |
| GSAGSLNTSF | ASSPGTGGAETLY | 0.5932568 |
| GTGQIGI | ASSPGTGGAETLY | 0.71034884 |
| GTLIALIL | ASSPGTGGAETLY | 0.7959572 |
| IAATMGFVVG | ASSPGTGGAETLY | 0.7915398 |
| IFAQALL | ASSPGTGGAETLY | 0.5852153 |
| IGAAVALGF | ASSPGTGGAETLY | 0.6864956 |
| IGLTPLAL | ASSPGTGGAETLY | 0.5219336 |
| ILGSNTSTL | ASSPGTGGAETLY | 0.6432353 |
| IVFSLTVL | ASSPGTGGAETLY | 0.5626377 |
| LASKATGF | ASSPGTGGAETLY | 0.76888585 |
| LASVALIL | ASSPGTGGAETLY | 0.69014114 |
| LFAVGLTL | ASSPGTGGAETLY | 0.7816063 |
| LFNGGIGTYI | ASSPGTGGAETLY | 0.75535715 |
| LTLGAGII | ASSPGTGGAETLY | 0.7996167 |
| LVAAALGDI | ASSPGTGGAETLY | 0.67632276 |
| NTAFVQLGI | ASSPGTGGAETLY | 0.7166884 |
| QLFSGAEL | ASSPGTGGAETLY | 0.5016775 |
| SAATFITL | ASSPGTGGAETLY | 0.59570897 |
| SNYGLANLGV | ASSPGTGGAETLY | 0.6828997 |
| SYGGAYAVM | ASSPGTGGAETLY | 0.7149128 |
| TTGVGGLTLG | ASSPGTGGAETLY | 0.3480764 |
| TTQTIYI | ASSPGTGGAETLY | 0.79685104 |
| VAAGAGLGAVI | ASSPGTGGAETLY | 0.73268247 |
| VAAVVTLL | ASSPGTGGAETLY | 0.7833553 |
| VGFLGGLI | ASSPGTGGAETLY | 0.6766424 |
| VGVIPILL | ASSPGTGGAETLY | 0.7224662 |
| VIFASSLI | ASSPGTGGAETLY | 0.6676575 |
| VKGGDGGAAAG | ASSPGTGGAETLY | 0.7674704 |
| VTKTFTNI | ASSPGTGGAETLY | 0.64960665 |
| VVAAAILL | ASSPGTGGAETLY | 0.4822146 |
| YAALDVEL | ASSPGTGGAETLY | 0.68768764 |
| LASVALLL | ASSPGTGGAETLY | 0.6250545 |
| LDAVHRV | ASSPGTGGAETLY | 0.8028858 |
| QGLSSSIF | ASSPGTGGAETLY | 0.7894922 |
| RGTFANIRL | ASSPGTGGAETLY | 0.74938536 |
| RQLWWGHRI | ASSPGTGGAETLY | 0.7975703 |
| YGPPGTGKTLL | ASSPGTGGAETLY | 0.30053282 |
| AALVGAPANVA | ASSPGTGGAETLY | 0.6871026 |
| AATFRRTE | ASSPGTGGAETLY | 0.79849696 |
| AGALGGGI | ASSPGTGGAETLY | 0.7670765 |
| AGGAGGAGGIGAG | ASSPGTGGAETLY | 0.71776915 |
| AGSGALGL | ASSPGTGGAETLY | 0.7896379 |
| AQIADQI | ASSPGTGGAETLY | 0.6393278 |
| ASAVAVTYL | ASSPGTGGAETLY | 0.7047062 |
| ASTVLGGTGG | ASSPGTGGAETLY | 0.7399785 |
| AVGGEPAK | ASSPGTGGAETLY | 0.7909514 |
| CEQGTFGGTC | ASSPGTGGAETLY | 0.7656645 |

| | | |
|---|---|---|
| DEGKAPAG | ASSPGTGGAETLY | 0.64376134 |
| DTVVFPAL | ASSPGTGGAETLY | 0.77419275 |
| FEGITMAG | ASSPGTGGAETLY | 0.80845344 |
| FPLVVQD | ASSPGTGGAETLY | 0.6789718 |
| GAFTTPQ | ASSPGTGGAETLY | 0.77298176 |
| GFGGFGTAGGVGGA | ASSPGTGGAETLY | 0.7554804 |
| GFLAQFL | ASSPGTGGAETLY | 0.74259233 |
| GGAGGAGGNAGVA | ASSPGTGGAETLY | 0.64324003 |
| GGVGTAGAGGAGGAGG | ASSPGTGGAETLY | 0.6170101 |
| GNGGGGDAGNAGSGGN | ASSPGTGGAETLY | 0.74440074 |
| GTGGAGGAGNDAGS | ASSPGTGGAETLY | 0.59518707 |
| GTTNTGWFNS | ASSPGTGGAETLY | 0.7358273 |
| IFMGPLFTL | ASSPGTGGAETLY | 0.736395 |
| ILAHAIEAI | ASSPGTGGAETLY | 0.8138846 |
| IRSAIREAL | ASSPGTGGAETLY | 0.7646947 |
| ITFAGGG | ASSPGTGGAETLY | 0.754879 |
| ITGMSGHQEV | ASSPGTGGAETLY | 0.6245912 |
| IVAGLACTL | ASSPGTGGAETLY | 0.755108 |
| IVGGGLAGA | ASSPGTGGAETLY | 0.75956625 |
| IVTGVLNI | ASSPGTGGAETLY | 0.5491674 |
| LAQQGCI | ASSPGTGGAETLY | 0.79118204 |
| LCLHVEN | ASSPGTGGAETLY | 0.63623655 |
| LDAISDGS | ASSPGTGGAETLY | 0.70437574 |
| LGGFAAML | ASSPGTGGAETLY | 0.7732835 |
| LGGKNPMI | ASSPGTGGAETLY | 0.80259514 |
| LGPKGVAAIL | ASSPGTGGAETLY | 0.7315529 |
| LMVGVSI | ASSPGTGGAETLY | 0.736529 |
| LVGAAFVL | ASSPGTGGAETLY | 0.62680674 |
| LVLAVAPL | ASSPGTGGAETLY | 0.78630495 |
| MAAIAVIDAA | ASSPGTGGAETLY | 0.7867024 |
| NNGINAELQ | ASSPGTGGAETLY | 0.7613971 |
| RGEPLVLVHG | ASSPGTGGAETLY | 0.8069828 |
| RGGAGGAAGLI | ASSPGTGGAETLY | 0.78236157 |
| RSYRMHPAV | ASSPGTGGAETLY | 0.43752924 |
| SAAHLLAGQLA | ASSPGTGGAETLY | 0.7454086 |
| SAAVGSY | ASSPGTGGAETLY | 0.79376936 |
| SPHAVLHRI | ASSPGTGGAETLY | 0.7828135 |
| SQIAKVL | ASSPGTGGAETLY | 0.8046905 |
| TALGHLVP | ASSPGTGGAETLY | 0.78265697 |
| TAPTTSGAA | ASSPGTGGAETLY | 0.69023156 |
| TDGDDGNGGSPGT | ASSPGTGGAETLY | 0.7977467 |
| TDGLGEN | ASSPGTGGAETLY | 0.81376845 |
| TGGPPLPGQA | ASSPGTGGAETLY | 0.7803509 |
| TIPAITI | ASSPGTGGAETLY | 0.66922945 |
| TIPGFTI | ASSPGTGGAETLY | 0.6325178 |
| TSIASGALA | ASSPGTGGAETLY | 0.7652217 |
| TVTLPGAAI | ASSPGTGGAETLY | 0.77804565 |
| VASGLSAALP | ASSPGTGGAETLY | 0.61746764 |
| VGAGVIGI | ASSPGTGGAETLY | 0.73487866 |
| VGPKRFGV | ASSPGTGGAETLY | 0.7489429 |
| VNVAFPD | ASSPGTGGAETLY | 0.4644274 |

| | | |
|---|---|---|
| VSMTNSGVS | ASSPGTGGAETLY | 0.7761133 |
| VTDVLGDLSV | ASSPGTGGAETLY | 0.7909298 |
| VVVVGAGFAGL | ASSPGTGGAETLY | 0.7652443 |
| VYTVGTVV | ASSPGTGGAETLY | 0.6757246 |
| WTTFSGP | ASSPGTGGAETLY | 0.78659946 |
| YDELVEAF | ASSPGTGGAETLY | 0.80891454 |
| YGTGGAGGNGG | ASSPGTGGAETLY | 0.8031435 |
| YTNVMAV | ASSPGTGGAETLY | 0.49014938 |
| FLEQVHQL | ASSPGTGGAETLY | 0.3591417 |
| GLSLLAVAV | ASSPGTGGAETLY | 0.7222054 |
| LEQVHQL | ASSPGTGGAETLY | 0.7809016 |
| LNAAGAI | ASSPGTGGAETLY | 0.71538454 |
| PAVHASL | ASSPGTGGAETLY | 0.67596394 |
| TELEDDA | ASSPGTGGAETLY | 0.75111187 |
| TPPGPPN | ASSPGTGGAETLY | 0.79025304 |
| TPPPPPPP | ASSPGTGGAETLY | 0.66858184 |
| TQSPALI | ASSLGSSGNTLY | 0.5331391 |
| EFAAAFPQ | ASSLGSSGNTLY | 0.78111386 |
| LLSGNLDVL | ASSLGSSGNTLY | 0.7803377 |
| GPANLGNYNI | ASSLGSSGNTLY | 0.62215906 |
| TLGPTLP | ASSLGSSGNTLY | 0.69673455 |
| VYAYSSCCK | ASSLGSSGNTLY | 0.7251816 |
| APAPAPAPNI | ASSLGSSGNTLY | 0.7088468 |
| PGGPVPIP | ASSLGSSGNTLY | 0.73915154 |
| FLTVALEP | ASSLGSSGNTLY | 0.5872271 |
| SQYFGANL | ASSLGSSGNTLY | 0.7780137 |
| GGVKPAA | ASSLGSSGNTLY | 0.5986727 |
| GGTGGAGGAGGW | ASSLGSSGNTLY | 0.7694135 |
| GVGGVAGLGGAGG | ASSLGSSGNTLY | 0.7532759 |
| VLLVLATTI | ASSLGSSGNTLY | 0.7784926 |
| IGIFAAML | ASSLGSSGNTLY | 0.6983092 |
| AVGDLID | ASSLGSSGNTLY | 0.6591436 |
| NISPLPL | ASSLGSSGNTLY | 0.6325573 |
| VTAVFVAI | ASSLGSSGNTLY | 0.48844612 |
| SGDDNDAAL | ASSLGSSGNTLY | 0.7937192 |
| AVAALAHS | ASSLGSSGNTLY | 0.59674984 |
| GGVTVMATP | ASSLGSSGNTLY | 0.7937268 |
| AGLAANVRD | ASSLGSSGNTLY | 0.7703279 |
| TNRPDLIDPALLRPG | ASSLGSSGNTLY | 0.64763486 |
| LANLGVTAL | ASSLGSSGNTLY | 0.7655093 |
| TGERIEL | ASSLGSSGNTLY | 0.5799657 |
| NGEELAA | ASSLGSSGNTLY | 0.732074 |
| VLSLVWG | ASSLGSSGNTLY | 0.7499325 |
| IGYALWL | ASSLGSSGNTLY | 0.7824994 |
| LSLLLAPL | ASSLGSSGNTLY | 0.65407455 |
| TLDEALK | ASSLGSSGNTLY | 0.7703105 |
| TDGAGTAEGG | ASSLGSSGNTLY | 0.7627565 |
| NAPGGAGGNGG | ASSLGSSGNTLY | 0.67845654 |
| VSVFMGMGGNF | ASSLGSSGNTLY | 0.7655756 |
| LASYLAAL | ASSLGSSGNTLY | 0.7851167 |
| LVGGTPAL | ASSLGSSGNTLY | 0.773875 |

| | | |
|---|---|---|
| VRNADVL | ASSLGSSGNTLY | 0.63676286 |
| TCSSPFLVSAM | ASSLGSSGNTLY | 0.76500607 |
| VHITDAE | ASSLGSSGNTLY | 0.63412845 |
| EYAAPSF | ASSLGSSGNTLY | 0.7478416 |
| VVDAFHGGL | ASSLGSSGNTLY | 0.7421384 |
| ASVVDPQ | ASSLGSSGNTLY | 0.6948974 |
| SYNEFGGLS | ASSLGSSGNTLY | 0.7844974 |
| DPDGSVY | ASSLGSSGNTLY | 0.6121081 |
| MVSTIHATE | ASSLGSSGNTLY | 0.66830814 |
| LAAGVADPV | ASSLGSSGNTLY | 0.7108164 |
| TKEQIAA | ASSLGSSGNTLY | 0.7486026 |
| SLASPAAL | ASSLGSSGNTLY | 0.7772625 |
| IGLVFYGV | ASSLGSSGNTLY | 0.63414323 |
| FLLDASNL | ASSLGSSGNTLY | 0.7416484 |
| YQVGNFP | ASSLGSSGNTLY | 0.7045981 |
| ESLLSQST | ASSLGSSGNTLY | 0.62657106 |
| LFTVGFAAVG | ASSLGSSGNTLY | 0.7671554 |
| ATPPTPT | ASSLGSSGNTLY | 0.74048793 |
| TDLATLG | ASSLGSSGNTLY | 0.5907155 |
| SAGGTGR | ASSLGSSGNTLY | 0.7824913 |
| GFSNSGS | ASSLGSSGNTLY | 0.69195604 |
| GGNGGSGGTSV | ASSLGSSGNTLY | 0.76403743 |
| GTSVATGGAGN | ASSLGSSGNTLY | 0.8081814 |
| NGGNGGSGGTS | ASSLGSSGNTLY | 0.66592 |
| AVVKFPH | ASSLGSSGNTLY | 0.7706626 |
| TPPPPPP | ASSLGSSGNTLY | 0.7460835 |
| AAPAAAAAPA | ASSLGSSGNTLY | 0.76062274 |
| IALNEDL | ASSLGSSGNTLY | 0.72392744 |
| IVNVSSM | ASSLGSSGNTLY | 0.6816677 |
| TGIPPAPR | ASSLGSSGNTLY | 0.7994962 |
| FDLGGGTF | ASSLGSSGNTLY | 0.7681366 |
| FDLGGGTFDVS | ASSLGSSGNTLY | 0.7834766 |
| NEPTAAA | ASSLGSSGNTLY | 0.76919234 |
| NEPTAAALA | ASSLGSSGNTLY | 0.79245484 |
| GSIINISSI | ASSLGSSGNTLY | 0.7795547 |
| VAPGNAGTA | ASSLGSSGNTLY | 0.65395063 |
| TGGGASLE | ASSLGSSGNTLY | 0.5155476 |
| VAVYNHYKRI | ASSLGSSGNTLY | 0.70538867 |
| TDPAPATT | ASSLGSSGNTLY | 0.7228278 |
| TNRPDILDPALLRPG | ASSLGSSGNTLY | 0.73455274 |
| LTTTTTS | ASSLGSSGNTLY | 0.45201167 |
| PTGSGKTL | ASSLGSSGNTLY | 0.7804612 |
| TGSGKTL | ASSLGSSGNTLY | 0.80961525 |
| TIGHVDHGK | ASSLGSSGNTLY | 0.6500077 |
| TIGHVDHGKTT | ASSLGSSGNTLY | 0.809371 |
| GHVDHGKTT | ASSLGSSGNTLY | 0.63719696 |
| AQFEHTLL | ASSLGSSGNTLY | 0.77024746 |
| IVNQDEL | ASSLGSSGNTLY | 0.7878767 |
| TNGSQFF | ASSLGSSGNTLY | 0.5701892 |
| TGKGIVE | ASSLGSSGNTLY | 0.7796718 |
| LISNASDA | ASSLGSSGNTLY | 0.70394397 |

| | | |
|---|---|---|
| AELRTLGELTKIAKAHGAQ | ASSLGSSGNTLY | 0.6928357 |
| APNAVAAAEPSW | ASSLGSSGNTLY | 0.7939184 |
| APQVAGAATAVL | ASSLGSSGNTLY | 0.70882237 |
| ARRDPAAAAAILPTDA | ASSLGSSGNTLY | 0.77623165 |
| DAAARDVLAEAGLAEYF | ASSLGSSGNTLY | 0.73803854 |
| DDQIVLSAKANLAKVEQ | ASSLGSSGNTLY | 0.6475029 |
| DELLPDTGEDSDEDGH | ASSLGSSGNTLY | 0.7723797 |
| ELPEAPANRATVPQD | ASSLGSSGNTLY | 0.7919947 |
| FGQSKIERVTGIG | ASSLGSSGNTLY | 0.7023212 |
| GAGGTGSTAGGAGGAGGAGGLY | ASSLGSSGNTLY | 0.8119396 |
| GHGGTGAAGGNGGTGGAGGSGI | ASSLGSSGNTLY | 0.64676535 |
| GNGGNGGAGGEGGPGGAGGAGA | ASSLGSSGNTLY | 0.8083076 |
| GSTAGNGGNGGNATNGTIAGQP | ASSLGSSGNTLY | 0.77836585 |
| GTGGDGGNGGNGGTGG | ASSLGSSGNTLY | 0.7248468 |
| GVRRPAGIPADAPV | ASSLGSSGNTLY | 0.7099633 |
| GYRQVTDAHLLTLARR | ASSLGSSGNTLY | 0.7800886 |
| IAKMGQDGHDRG | ASSLGSSGNTLY | 0.7097274 |
| KAEPSIEIQYT | ASSLGSSGNTLY | 0.7744781 |
| KPVRIRMAADEDTVAVT | ASSLGSSGNTLY | 0.80284536 |
| MPPRIAGMRLLVIKPEP | ASSLGSSGNTLY | 0.44995552 |
| MKFYAVALFRSRGVPPTR | ASSLGSSGNTLY | 0.671329 |
| NAGGFGILGAVAHSP | ASSLGSSGNTLY | 0.5934052 |
| NEGREITLAMAEM | ASSLGSSGNTLY | 0.5930865 |
| NELVEWSHPLGHRWRWE | ASSLGSSGNTLY | 0.50438267 |
| NLGAGGTGGNGGNPG | ASSLGSSGNTLY | 0.4354081 |
| NTEWPKWETVMAADPVD | ASSLGSSGNTLY | 0.5695155 |
| PASAAPAETQP | ASSLGSSGNTLY | 0.4597643 |
| QQSSDGKNGVRAAEITGE | ASSLGSSGNTLY | 0.678773 |
| RAPDAVTTSWWKE | ASSLGSSGNTLY | 0.6422733 |
| SGFGNVGAHTSGWFNQ | ASSLGSSGNTLY | 0.4930883 |
| SGVAGAVGQAASVGGLKV | ASSLGSSGNTLY | 0.68021226 |
| SNGGAGTGSAGGNGGT | ASSLGSSGNTLY | 0.6895415 |
| TGGAGGEGGAAG | ASSLGSSGNTLY | 0.37858397 |
| TGGDGGAAGAGGAPGHG | ASSLGSSGNTLY | 0.5267385 |
| TMPPLLAAIANAGRHGV | ASSLGSSGNTLY | 0.47299507 |
| TSSTAGNSGGA | ASSLGSSGNTLY | 0.53891546 |
| VPATVPVGILC | ASSLGSSGNTLY | 0.33261746 |
| VQRPDSGGVVLTGRIS | ASSLGSSGNTLY | 0.6337061 |
| WTPNPVPATKGVPAGLVQAT | ASSLGSSGNTLY | 0.4532227 |
| YALKRALAKFSDRFDVVII | ASSLGSSGNTLY | 0.29527858 |
| YLIGSVIV | ASSLGSSGNTLY | 0.5882163 |
| YGQLVAVID | ASSLGSSGNTLY | 0.5583195 |
| VTVGGMTLPA | ASSLGSSGNTLY | 0.5581796 |
| VLALPAV | ASSLGSSGNTLY | 0.6434142 |
| VIPPTLNYET | ASSLGSSGNTLY | 0.49750975 |
| VGMGGMHPSSG | ASSLGSSGNTLY | 0.5184217 |
| VGLAGGAATA | ASSLGSSGNTLY | 0.621569 |
| VGGAGGAGGSGP | ASSLGSSGNTLY | 0.6719981 |
| VFAAFMV | ASSLGSSGNTLY | 0.48899144 |
| TLYDKLR | ASSLGSSGNTLY | 0.5793201 |
| TLAVFTVL | ASSLGSSGNTLY | 0.68542117 |

| | | |
|---|---|---|
| TGSVQPARV | ASSLGSSGNTLY | 0.5844835 |
| TGGGAGSGGT | ASSLGSSGNTLY | 0.6813583 |
| SHLETSL | ASSLGSSGNTLY | 0.68989015 |
| SFTTGMT | ASSLGSSGNTLY | 0.5018482 |
| QVEVAKRMGVRQPTVS | ASSLGSSGNTLY | 0.6685341 |
| QGSGGPALAM | ASSLGSSGNTLY | 0.6915056 |
| MLAPSNP | ASSLGSSGNTLY | 0.55637515 |
| LVNNAAI | ASSLGSSGNTLY | 0.56645423 |
| LTGEINTLA | ASSLGSSGNTLY | 0.4366528 |
| LSIFGPV | ASSLGSSGNTLY | 0.5657468 |
| LLALLANL | ASSLGSSGNTLY | 0.39323977 |
| LISNASDALDKLR | ASSLGSSGNTLY | 0.58743167 |
| LINEAVP | ASSLGSSGNTLY | 0.68391806 |
| LGYAFSGL | ASSLGSSGNTLY | 0.5245801 |
| LDFVAVAL | ASSLGSSGNTLY | 0.39082533 |
| LAQPGAE | ASSLGSSGNTLY | 0.41896778 |
| ISNLGAVTGF | ASSLGSSGNTLY | 0.5598115 |
| HQEDPGRI | ASSLGSSGNTLY | 0.47275445 |
| GTPGEPTP | ASSLGSSGNTLY | 0.39003378 |
| GFFNSGS | ASSLGSSGNTLY | 0.45260286 |
| GAGGGGLTAGAN | ASSLGSSGNTLY | 0.6650383 |
| FVDALVNM | ASSLGSSGNTLY | 0.66725534 |
| FLTVAPEL | ASSLGSSGNTLY | 0.42245737 |
| FETLTAL | ASSLGSSGNTLY | 0.6841014 |
| DVINPIEL | ASSLGSSGNTLY | 0.6189835 |
| APVSGAPV | ASSLGSSGNTLY | 0.69799197 |
| ALVGGLAW | ASSLGSSGNTLY | 0.5932251 |
| ALAAAAATAI | ASSLGSSGNTLY | 0.4551784 |
| AFGLGLLL | ASSLGSSGNTLY | 0.644489 |
| AFFAAASL | ASSLGSSGNTLY | 0.5114443 |
| AAAALPAFSPP | ASSLGSSGNTLY | 0.580291 |
| VVLAPAL | ASSLGSSGNTLY | 0.6438905 |
| TIGHVDHGKTTL | ASSLGSSGNTLY | 0.5875564 |
| LISNASDALDK | ASSLGSSGNTLY | 0.46332714 |
| LISNASDAL | ASSLGSSGNTLY | 0.5282811 |
| ISNASDALDK | ASSLGSSGNTLY | 0.6549927 |
| NQAETLVYQTE | ASSLGSSGNTLY | 0.40779704 |
| TGGTGGTAGAG | ASSLGSSGNTLY | 0.48241693 |
| TGSTAGGAGGA | ASSLGSSGNTLY | 0.54026186 |
| DPLAVPVAAGPV | ASSLGSSGNTLY | 0.585155 |
| GGTGGTPGNGGN | ASSLGSSGNTLY | 0.6684289 |
| PAGLAGVDPATL | ASSLGSSGNTLY | 0.63261366 |
| GNNLVVEAPHAGASV | ASSLGSSGNTLY | 0.504331 |
| KVTRIMSASTLEQPA | ASSLGSSGNTLY | 0.6253766 |
| LGFALPIALPRRTVV | ASSLGSSGNTLY | 0.6355244 |
| TRLGNLTARKLVVLV | ASSLGSSGNTLY | 0.5936308 |
| WSQQRRLIYFPSAGP | ASSLGSSGNTLY | 0.6334863 |
| GPNGNPDPMAAAVDIR | ASSLGSSGNTLY | 0.45462942 |
| TQRCVSVADLRAAASR | ASSLGSSGNTLY | 0.6408444 |
| QFVLDVAEHPVEVLTETV | ASSLGSSGNTLY | 0.62223333 |
| GGAGGTGGAGGNGGTQAGDG | ASSLGSSGNTLY | 0.65025336 |

| | | |
|---|---|---|
| PLLATLRAIAVDVDAKPAQV | ASSLGSSGNTLY | 0.66674405 |
| APKFRTGGYMIDFWGVGYQVAK | ASSLGSSGNTLY | 0.6165434 |
| GAGGAGGAGGAAGGPGGTGGTG | ASSLGSSGNTLY | 0.5796989 |
| IGALGLLATDVAPAGSVAFGVA | ASSLGSSGNTLY | 0.36695135 |
| AGGAGGIGGNGGAGGAGGVLMG | ASSLGSSGNTLY | 0.49031973 |
| ADELIGLL | ASSLGSSGNTLY | 0.5965351 |
| ADRALKA | ASSLGSSGNTLY | 0.5715355 |
| AGAVGQVGGAGGA | ASSLGSSGNTLY | 0.6519777 |
| AGGGGGLGGAP | ASSLGSSGNTLY | 0.65607363 |
| AGGLLYGNGGAG | ASSLGSSGNTLY | 0.47956973 |
| AGLESIGL | ASSLGSSGNTLY | 0.6917291 |
| AGLKTGII | ASSLGSSGNTLY | 0.67663056 |
| ASLAALLL | ASSLGSSGNTLY | 0.35778025 |
| ASVINISSI | ASSLGSSGNTLY | 0.4806453 |
| ATPPGPA | ASSLGSSGNTLY | 0.6962737 |
| ATVASALV | ASSLGSSGNTLY | 0.59148777 |
| ATVASLIL | ASSLGSSGNTLY | 0.54523724 |
| AVNGPVLIRH | ASSLGSSGNTLY | 0.5555982 |
| DGAAGAVGI | ASSLGSSGNTLY | 0.69922185 |
| DPLLHGI | ASSLGSSGNTLY | 0.5418864 |
| ELGAALGAL | ASSLGSSGNTLY | 0.6919895 |
| ELGRTQAV | ASSLGSSGNTLY | 0.33373266 |
| FVAGGYLS | ASSLGSSGNTLY | 0.5182667 |
| GASGGAGGAGGW | ASSLGSSGNTLY | 0.46686932 |
| GGAGAGGDGGAGGI | ASSLGSSGNTLY | 0.31476924 |
| GTGHNGSQPGMGGNG | ASSLGSSGNTLY | 0.64248943 |
| GYAAASASAS | ASSLGSSGNTLY | 0.56228983 |
| HQVDVAAVQQG | ASSLGSSGNTLY | 0.65439683 |
| IGAAIAHT | ASSLGSSGNTLY | 0.5656074 |
| IGMIEVAQL | ASSLGSSGNTLY | 0.6576388 |
| IRLAPEHTSAMD | ASSLGSSGNTLY | 0.57264495 |
| IYNVSVL | ASSLGSSGNTLY | 0.60172564 |
| KQGFLGM | ASSLGSSGNTLY | 0.49945936 |
| LESVITQL | ASSLGSSGNTLY | 0.6331643 |
| LGLGALVVV | ASSLGSSGNTLY | 0.6922215 |
| LIGNGGNGGSGGI | ASSLGSSGNTLY | 0.63717556 |
| LNAINAPTL | ASSLGSSGNTLY | 0.4317848 |
| LQLLNTGV | ASSLGSSGNTLY | 0.57334054 |
| LQTSIEF | ASSLGSSGNTLY | 0.5820266 |
| MSFAVSEL | ASSLGSSGNTLY | 0.42309612 |
| NTGALGL | ASSLGSSGNTLY | 0.37081367 |
| PPGAGGAGGAGTT | ASSLGSSGNTLY | 0.48668963 |
| QLVGLGI | ASSLGSSGNTLY | 0.47453967 |
| RFLEQVHQL | ASSLGSSGNTLY | 0.6498378 |
| RGVGYVLR | ASSLGSSGNTLY | 0.6266348 |
| SATGVFAGVI | ASSLGSSGNTLY | 0.5156133 |
| SLPAITLP | ASSLGSSGNTLY | 0.62142384 |
| STALAFTV | ASSLGSSGNTLY | 0.6101593 |
| SVLTVGLGK | ASSLGSSGNTLY | 0.2507228 |
| TALAGAGSGI | ASSLGSSGNTLY | 0.5112654 |
| TGLPFTGI | ASSLGSSGNTLY | 0.39055312 |

| | | |
|---|---|---|
| VAEHVTV | ASSLGSSGNTLY | 0.69758886 |
| VDIVLNSL | ASSLGSSGNTLY | 0.42624792 |
| VLTGGGAVL | ASSLGSSGNTLY | 0.42821595 |
| VNVGLAYGI | ASSLGSSGNTLY | 0.57071376 |
| VSSAAVGAV | ASSLGSSGNTLY | 0.5776599 |
| VTSVYEAGV | ASSLGSSGNTLY | 0.5667318 |
| WVALDRGA | ASSLGSSGNTLY | 0.44316468 |
| YLVGSVVI | ASSLGSSGNTLY | 0.34266853 |
| GPPGTGKTLL | ASSLGSSGNTLY | 0.6474231 |
| GVGMTKF | ASSLGSSGNTLY | 0.50931215 |
| IGIGGSDL | ASSLGSSGNTLY | 0.49277976 |
| IGTLLAEL | ASSLGSSGNTLY | 0.40015945 |
| IYTSGTT | ASSLGSSGNTLY | 0.5412247 |
| LSFPAGV | ASSLGSSGNTLY | 0.4005749 |
| LYTSGTT | ASSLGSSGNTLY | 0.66281456 |
| VGAFPFGF | ASSLGSSGNTLY | 0.48642716 |
| VGAGTVEF | ASSLGSSGNTLY | 0.51770097 |
| VGKSTLLL | ASSLGSSGNTLY | 0.69359064 |
| AALAHAGGGIV | ASSLGSSGNTLY | 0.59255236 |
| AALVAVSLL | ASSLGSSGNTLY | 0.50842524 |
| AAVSRAVAAL | ASSLGSSGNTLY | 0.5617154 |
| AGGSSGIGGF | ASSLGSSGNTLY | 0.4221329 |
| AGLTVANANG | ASSLGSSGNTLY | 0.49792194 |
| AGLVGAVL | ASSLGSSGNTLY | 0.53575486 |
| ALSQAFTV | ASSLGSSGNTLY | 0.69671327 |
| ALTFSVSL | ASSLGSSGNTLY | 0.5537027 |
| ALTGAALVAL | ASSLGSSGNTLY | 0.4316867 |
| ASFSIGAGVL | ASSLGSSGNTLY | 0.5895224 |
| DGSVYKI | ASSLGSSGNTLY | 0.69353724 |
| EQLAEIV | ASSLGSSGNTLY | 0.7061975 |
| FAVSELTVM | ASSLGSSGNTLY | 0.44613597 |
| GAAPLAAGL | ASSLGSSGNTLY | 0.609794 |
| GADGGDGGSGGAGG | ASSLGSSGNTLY | 0.54073507 |
| GFGLDTGGAGG | ASSLGSSGNTLY | 0.53757066 |
| GLASGGLL | ASSLGSSGNTLY | 0.509325 |
| GRIAGAAVLL | ASSLGSSGNTLY | 0.4719379 |
| GSAGSLNTSF | ASSLGSSGNTLY | 0.6218148 |
| GTGQIGI | ASSLGSSGNTLY | 0.71112967 |
| GTLIALIL | ASSLGSSGNTLY | 0.5297461 |
| IAATMGFVVG | ASSLGSSGNTLY | 0.36801577 |
| IFAQALL | ASSLGSSGNTLY | 0.554688 |
| IGAAVALGF | ASSLGSSGNTLY | 0.43249267 |
| IGLTPLAL | ASSLGSSGNTLY | 0.48307273 |
| ILGSNTSTL | ASSLGSSGNTLY | 0.5653602 |
| IVFSLTVL | ASSLGSSGNTLY | 0.32976875 |
| LASKATGF | ASSLGSSGNTLY | 0.57624817 |
| LASVALIL | ASSLGSSGNTLY | 0.70031077 |
| LFAVGLTL | ASSLGSSGNTLY | 0.52282774 |
| LFNGGIGTYI | ASSLGSSGNTLY | 0.6144971 |
| LTLGAGII | ASSLGSSGNTLY | 0.6680432 |
| LVAAALGDI | ASSLGSSGNTLY | 0.61549425 |

| | | |
|---|---|---|
| NTAFVQLGI | ASSLGSSGNTLY | 0.6562181 |
| QLFSGAEL | ASSLGSSGNTLY | 0.651561 |
| SAATFITL | ASSLGSSGNTLY | 0.60336006 |
| SNYGLANLGV | ASSLGSSGNTLY | 0.5762862 |
| SYGGAYAVM | ASSLGSSGNTLY | 0.6524428 |
| TTGVGGLTLG | ASSLGSSGNTLY | 0.57918906 |
| TTQTIYI | ASSLGSSGNTLY | 0.5484551 |
| VAAGAGLGAVI | ASSLGSSGNTLY | 0.48257416 |
| VAAVVTLL | ASSLGSSGNTLY | 0.66288936 |
| VGFLGGLI | ASSLGSSGNTLY | 0.52825713 |
| VGVIPILL | ASSLGSSGNTLY | 0.6755384 |
| VIFASSLI | ASSLGSSGNTLY | 0.3798844 |
| VKGGDGGAAAG | ASSLGSSGNTLY | 0.4725142 |
| VTKTFTNI | ASSLGSSGNTLY | 0.6443522 |
| VVAAAILL | ASSLGSSGNTLY | 0.29366815 |
| YAALDVEL | ASSLGSSGNTLY | 0.37715587 |
| LASVALLL | ASSLGSSGNTLY | 0.5313729 |
| LDAVHRV | ASSLGSSGNTLY | 0.67819834 |
| QGLSSSIF | ASSLGSSGNTLY | 0.28614697 |
| RGTFANIRL | ASSLGSSGNTLY | 0.49387345 |
| RQLWWGHRI | ASSLGSSGNTLY | 0.4075133 |
| YGPPGTGKTLL | ASSLGSSGNTLY | 0.657894 |
| AALVGAPANVA | ASSLGSSGNTLY | 0.5778731 |
| AATFRRTE | ASSLGSSGNTLY | 0.35741416 |
| AGALGGGI | ASSLGSSGNTLY | 0.6381439 |
| AGGAGGAGGIGAG | ASSLGSSGNTLY | 0.33843914 |
| AGSGALGL | ASSLGSSGNTLY | 0.5597148 |
| AQIADQI | ASSLGSSGNTLY | 0.64458835 |
| ASAVAVTYL | ASSLGSSGNTLY | 0.68283606 |
| ASTVLGGTGG | ASSLGSSGNTLY | 0.40178126 |
| AVGGEPAK | ASSLGSSGNTLY | 0.6413542 |
| CEQGTFGGTC | ASSLGSSGNTLY | 0.47489536 |
| DEGKAPAG | ASSLGSSGNTLY | 0.7146907 |
| DTVVFPAL | ASSLGSSGNTLY | 0.54316115 |
| FEGITMAG | ASSLGSSGNTLY | 0.40293872 |
| FPLVVQD | ASSLGSSGNTLY | 0.65498906 |
| GAFTTPQ | ASSLGSSGNTLY | 0.61577684 |
| GFGGFGTAGGVGGA | ASSLGSSGNTLY | 0.6024318 |
| GFLAQFL | ASSLGSSGNTLY | 0.6136111 |
| GGAGGAGGNAGVA | ASSLGSSGNTLY | 0.514614 |
| GGVGTAGAGGAGGAGG | ASSLGSSGNTLY | 0.54197663 |
| GNGGGGDAGNAGSGGN | ASSLGSSGNTLY | 0.6547825 |
| GTGGAGGAGNDAGS | ASSLGSSGNTLY | 0.6294167 |
| GTTNTGWFNS | ASSLGSSGNTLY | 0.4985077 |
| IFMGPLFTL | ASSLGSSGNTLY | 0.6626704 |
| ILAHAIEAI | ASSLGSSGNTLY | 0.56624067 |
| IRSAIREAL | ASSLGSSGNTLY | 0.5034204 |
| ITFAGGG | ASSLGSSGNTLY | 0.68506646 |
| ITGMSGHQEV | ASSLGSSGNTLY | 0.5587865 |
| IVAGLACTL | ASSLGSSGNTLY | 0.63967955 |
| IVGGGLAGA | ASSLGSSGNTLY | 0.6480824 |

| | | |
|---|---|---|
| IVTGVLNI | ASSLGSSGNTLY | 0.70957196 |
| LAQQGCI | ASSLGSSGNTLY | 0.41861397 |
| LCLHVEN | ASSLGSSGNTLY | 0.6394815 |
| LDAISDGS | ASSLGSSGNTLY | 0.5800303 |
| LGGFAAML | ASSLGSSGNTLY | 0.66053027 |
| LGGKNPMI | ASSLGSSGNTLY | 0.5811007 |
| LGPKGVAAIL | ASSLGSSGNTLY | 0.58183515 |
| LMVGVSI | ASSLGSSGNTLY | 0.6229523 |
| LVGAAFVL | ASSLGSSGNTLY | 0.67799807 |
| LVLAVAPL | ASSLGSSGNTLY | 0.54730666 |
| MAAIAVIDAA | ASSLGSSGNTLY | 0.6118968 |
| NNGINAELQ | ASSLGSSGNTLY | 0.6745191 |
| RGEPLVLVHG | ASSLGSSGNTLY | 0.49072474 |
| RGGAGGAAGLI | ASSLGSSGNTLY | 0.5770766 |
| RSYRMHPAV | ASSLGSSGNTLY | 0.626921 |
| SAAHLLAGQLA | ASSLGSSGNTLY | 0.6779522 |
| SAAVGSY | ASSLGSSGNTLY | 0.54380375 |
| SPHAVLHRI | ASSLGSSGNTLY | 0.67313504 |
| SQIAKVL | ASSLGSSGNTLY | 0.44222942 |
| TALGHLVP | ASSLGSSGNTLY | 0.6372081 |
| TAPTTSGAA | ASSLGSSGNTLY | 0.49547276 |
| TDGDDGNGGSPGT | ASSLGSSGNTLY | 0.6018414 |
| TDGLGEN | ASSLGSSGNTLY | 0.47434467 |
| TGGPPLPGQA | ASSLGSSGNTLY | 0.6370342 |
| TIPAITI | ASSLGSSGNTLY | 0.6452183 |
| TIPGFTI | ASSLGSSGNTLY | 0.6599785 |
| TSIASGALA | ASSLGSSGNTLY | 0.35907146 |
| TVTLPGAAI | ASSLGSSGNTLY | 0.61807686 |
| VASGLSAALP | ASSLGSSGNTLY | 0.6704289 |
| VGAGVIGI | ASSLGSSGNTLY | 0.68383086 |
| VGPKRFGV | ASSLGSSGNTLY | 0.55012715 |
| VNVAFPD | ASSLGSSGNTLY | 0.632139 |
| VSMTNSGVS | ASSLGSSGNTLY | 0.5473918 |
| VTDVLGDLSV | ASSLGSSGNTLY | 0.5524708 |
| VVVVGAGFAGL | ASSLGSSGNTLY | 0.40188304 |
| VYTVGTVV | ASSLGSSGNTLY | 0.5487485 |
| WTTFSGP | ASSLGSSGNTLY | 0.49779847 |
| YDELVEAF | ASSLGSSGNTLY | 0.44223392 |
| YGTGGAGGNGG | ASSLGSSGNTLY | 0.6329343 |
| YTNVMAV | ASSLGSSGNTLY | 0.6621098 |
| FLEQVHQL | ASSLGSSGNTLY | 0.66117126 |
| GLSLLAVAV | ASSLGSSGNTLY | 0.67638206 |
| LEQVHQL | ASSLGSSGNTLY | 0.6745491 |
| LNAAGAI | ASSLGSSGNTLY | 0.66098475 |
| PAVHASL | ASSLGSSGNTLY | 0.5074608 |
| TELEDDA | ASSLGSSGNTLY | 0.5611434 |
| TPPGPPN | ASSLGSSGNTLY | 0.38771987 |
| TPPPPPPP | ASSLGSSGNTLY | 0.51748693 |
| TQSPALI | ASSLSASRNTLY | 0.50081986 |
| EFAAAFPQ | ASSLSASRNTLY | 0.5813621 |
| LLSGNLDVL | ASSLSASRNTLY | 0.286213 |

| | | |
|---|---|---|
| GPANLGNYNI | ASSLSASRNTLY | 0.55404615 |
| TLGPTLP | ASSLSASRNTLY | 0.69504243 |
| VYAYSSCCK | ASSLSASRNTLY | 0.53822637 |
| APAPAPAPNI | ASSLSASRNTLY | 0.63981825 |
| PGGPVPIP | ASSLSASRNTLY | 0.60830945 |
| FLTVALEP | ASSLSASRNTLY | 0.4449565 |
| SQYFGANL | ASSLSASRNTLY | 0.65126103 |
| GGVKPAA | ASSLSASRNTLY | 0.47476685 |
| GGTGGAGGAGGW | ASSLSASRNTLY | 0.63860166 |
| GVGGVAGLGGAGG | ASSLSASRNTLY | 0.68710595 |
| VLLVLATTI | ASSLSASRNTLY | 0.6085762 |
| IGIFAAML | ASSLSASRNTLY | 0.5641062 |
| AVGDLID | ASSLSASRNTLY | 0.5880718 |
| NISPLPL | ASSLSASRNTLY | 0.40471613 |
| VTAVFVAI | ASSLSASRNTLY | 0.6000485 |
| SGDDNDAAL | ASSLSASRNTLY | 0.4378419 |
| AVAALAHS | ASSLSASRNTLY | 0.5380079 |
| GGVTVMATP | ASSLSASRNTLY | 0.46538877 |
| AGLAANVRD | ASSLSASRNTLY | 0.62240684 |
| TNRPDLIDPALLRPG | ASSLSASRNTLY | 0.59874666 |
| LANLGVTAL | ASSLSASRNTLY | 0.5742127 |
| TGERIEL | ASSLSASRNTLY | 0.47440982 |
| NGEELAA | ASSLSASRNTLY | 0.59748673 |
| VLSLVWG | ASSLSASRNTLY | 0.5785488 |
| IGYALWL | ASSLSASRNTLY | 0.45806637 |
| LSLLLAPL | ASSLSASRNTLY | 0.65261763 |
| TLDEALK | ASSLSASRNTLY | 0.536486 |
| TDGAGTAEGG | ASSLSASRNTLY | 0.4933989 |
| NAPGGAGGNGG | ASSLSASRNTLY | 0.6702401 |
| VSVFMGMGGNF | ASSLSASRNTLY | 0.59225947 |
| LASYLAAL | ASSLSASRNTLY | 0.35445935 |
| LVGGTPAL | ASSLSASRNTLY | 0.54256684 |
| VRNADVL | ASSLSASRNTLY | 0.58608663 |
| TCSSPFLVSAM | ASSLSASRNTLY | 0.5481754 |
| VHITDAE | ASSLSASRNTLY | 0.30955997 |
| EYAAPSF | ASSLSASRNTLY | 0.60925716 |
| VVDAFHGGL | ASSLSASRNTLY | 0.5114739 |
| ASVVDPQ | ASSLSASRNTLY | 0.4065205 |
| SYNEFGGLS | ASSLSASRNTLY | 0.4661686 |
| DPDGSVY | ASSLSASRNTLY | 0.6398392 |
| MVSTIHATE | ASSLSASRNTLY | 0.5724048 |
| LAAGVADPV | ASSLSASRNTLY | 0.6174648 |
| TKEQIAA | ASSLSASRNTLY | 0.5996961 |
| SLASPAAL | ASSLSASRNTLY | 0.65675753 |
| IGLVFYGV | ASSLSASRNTLY | 0.67877203 |
| FLLDASNL | ASSLSASRNTLY | 0.64532644 |
| YQVGNFP | ASSLSASRNTLY | 0.5532203 |
| ESLLSQST | ASSLSASRNTLY | 0.642426 |
| LFTVGFAAVG | ASSLSASRNTLY | 0.56152517 |
| ATPPTPT | ASSLSASRNTLY | 0.41493446 |
| TDLATLG | ASSLSASRNTLY | 0.37659746 |

| | | |
|---|---|---|
| SAGGTGR | ASSLSASRNTLY | 0.583728 |
| GFSNSGS | ASSLSASRNTLY | 0.4428051 |
| GGNGGSGGTSV | ASSLSASRNTLY | 0.61032325 |
| GTSVATGGAGN | ASSLSASRNTLY | 0.68205285 |
| NGGNGGSGGTS | ASSLSASRNTLY | 0.5422022 |
| AVVKFPH | ASSLSASRNTLY | 0.41667706 |
| TPPPPPP | ASSLSASRNTLY | 0.48470134 |
| AAPAAAAAPA | ASSLSASRNTLY | 0.4575374 |
| IALNEDL | ASSLSASRNTLY | 0.6329247 |
| IVNVSSM | ASSLSASRNTLY | 0.66010314 |
| TGIPPAPR | ASSLSASRNTLY | 0.700871 |
| FDLGGGTF | ASSLSASRNTLY | 0.6547348 |
| FDLGGGTFDVS | ASSLSASRNTLY | 0.5213648 |
| NEPTAAA | ASSLSASRNTLY | 0.61922127 |
| NEPTAAALA | ASSLSASRNTLY | 0.47391072 |
| GSIINISSI | ASSLSASRNTLY | 0.607636 |
| VAPGNAGTA | ASSLSASRNTLY | 0.5483463 |
| TGGGASLE | ASSLSASRNTLY | 0.5705845 |
| VAVYNHYKRI | ASSLSASRNTLY | 0.56910646 |
| TDPAPATT | ASSLSASRNTLY | 0.5397829 |
| TNRPDILDPALLRPG | ASSLSASRNTLY | 0.39717546 |
| LTTTTTS | ASSLSASRNTLY | 0.62089634 |
| PTGSGKTL | ASSLSASRNTLY | 0.55454445 |
| TGSGKTL | ASSLSASRNTLY | 0.40133196 |
| TIGHVDHGK | ASSLSASRNTLY | 0.5243227 |
| TIGHVDHGKTT | ASSLSASRNTLY | 0.6824274 |
| GHVDHGKTT | ASSLSASRNTLY | 0.6876333 |
| AQFEHTLL | ASSLSASRNTLY | 0.46701318 |
| IVNQDEL | ASSLSASRNTLY | 0.4886113 |
| TNGSQFF | ASSLSASRNTLY | 0.5712518 |
| TGKGIVE | ASSLSASRNTLY | 0.39458165 |
| LISNASDA | ASSLSASRNTLY | 0.60277593 |
| AELRTLGELTKIAKAHGAQ | ASSLSASRNTLY | 0.52477765 |
| APNAVAAAEPSW | ASSLSASRNTLY | 0.54077256 |
| APQVAGAATAVL | ASSLSASRNTLY | 0.5104559 |
| ARRDPAAAAAILPTDA | ASSLSASRNTLY | 0.59810966 |
| DAAARDVLAEAGLAEYF | ASSLSASRNTLY | 0.53619003 |
| DDQIVLSAKANLAKVEQ | ASSLSASRNTLY | 0.63521844 |
| DELLPDTGEDSDEDGH | ASSLSASRNTLY | 0.287343 |
| ELPEAPANRATVPQD | ASSLSASRNTLY | 0.622686 |
| FGQSKIERVTGIG | ASSLSASRNTLY | 0.43684316 |
| GAGGTGSTAGGAGGAGGAGGLY | ASSLSASRNTLY | 0.4834771 |
| GHGGTGAAGGNGGTGGAGGSGI | ASSLSASRNTLY | 0.33234167 |
| GNGGNGGAGGEGGPGGAGGAGA | ASSLSASRNTLY | 0.5911074 |
| GSTAGNGGNGGNATNGTIAGQP | ASSLSASRNTLY | 0.6594353 |
| GTGGDGGNGGNGGTGG | ASSLSASRNTLY | 0.6424209 |
| GVRRPAGIPADAPV | ASSLSASRNTLY | 0.4922291 |
| GYRQVTDAHLLTLARR | ASSLSASRNTLY | 0.65037465 |
| IAKMGQDGHDRG | ASSLSASRNTLY | 0.6849308 |
| KAEPSIEIQYT | ASSLSASRNTLY | 0.47610843 |
| KPVRIRMAADEDTVAVT | ASSLSASRNTLY | 0.37243629 |

| | | |
|---|---|---|
| MPPRIAGMRLLVIKPEP | ASSLSASRNTLY | 0.6164674 |
| MKFYAVALFRSRGVPPTR | ASSLSASRNTLY | 0.5842037 |
| NAGGFGILGAVAHSP | ASSLSASRNTLY | 0.6581786 |
| NEGREITLAMAEM | ASSLSASRNTLY | 0.6663591 |
| NELVEWSHPLGHRWRWE | ASSLSASRNTLY | 0.3720464 |
| NLGAGGTGGNGGNPG | ASSLSASRNTLY | 0.67116493 |
| NTEWPKWETVMAADPVD | ASSLSASRNTLY | 0.6485944 |
| PASAAPAETQP | ASSLSASRNTLY | 0.41324762 |
| QQSSDGKNGVRAAEITGE | ASSLSASRNTLY | 0.4576294 |
| RAPDAVTTSWWKE | ASSLSASRNTLY | 0.54849577 |
| SGFGNVGAHTSGWFNQ | ASSLSASRNTLY | 0.37284604 |
| SGVAGAVGQAASVGGLKV | ASSLSASRNTLY | 0.65730906 |
| SNGGAGTGSAGGNGGT | ASSLSASRNTLY | 0.47708595 |
| TGGAGGEGGAAG | ASSLSASRNTLY | 0.5610931 |
| TGGDGGAAGAGGAPGHG | ASSLSASRNTLY | 0.5669727 |
| TMPPLLAAIANAGRHGV | ASSLSASRNTLY | 0.559187 |
| TSSTAGNSGGA | ASSLSASRNTLY | 0.5439643 |
| VPATVPVGILC | ASSLSASRNTLY | 0.55495983 |
| VQRPDSGGVVLTGRIS | ASSLSASRNTLY | 0.6296492 |
| WTPNPVPATKGVPAGLVQAT | ASSLSASRNTLY | 0.60291576 |
| YALKRALAKFSDRFDVVII | ASSLSASRNTLY | 0.55651355 |
| YLIGSVIV | ASSLSASRNTLY | 0.60793585 |
| YGQLVAVID | ASSLSASRNTLY | 0.67497617 |
| VTVGGMTLPA | ASSLSASRNTLY | 0.44565374 |
| VLALPAV | ASSLSASRNTLY | 0.5196284 |
| VIPPTLNYET | ASSLSASRNTLY | 0.6158684 |
| VGMGGMHPSSG | ASSLSASRNTLY | 0.4601041 |
| VGLAGGAATA | ASSLSASRNTLY | 0.64735615 |
| VGGAGGAGGSGP | ASSLSASRNTLY | 0.6080412 |
| VFAAFMV | ASSLSASRNTLY | 0.5890782 |
| TLYDKLR | ASSLSASRNTLY | 0.5596002 |
| TLAVFTVL | ASSLSASRNTLY | 0.6891043 |
| TGSVQPARV | ASSLSASRNTLY | 0.62745374 |
| TGGGAGSGGT | ASSLSASRNTLY | 0.5671698 |
| SHLETSL | ASSLSASRNTLY | 0.64314425 |
| SFTTGMT | ASSLSASRNTLY | 0.5175854 |
| QVEVAKRMGVRQPTVS | ASSLSASRNTLY | 0.6565906 |
| QGSGGPALAM | ASSLSASRNTLY | 0.63972586 |
| MLAPSNP | ASSLSASRNTLY | 0.40047744 |
| LVNNAAI | ASSLSASRNTLY | 0.54620004 |
| LTGEINTLA | ASSLSASRNTLY | 0.6668943 |
| LSIFGPV | ASSLSASRNTLY | 0.65329343 |
| LLALLANL | ASSLSASRNTLY | 0.45587444 |
| LISNASDALDKLR | ASSLSASRNTLY | 0.4857148 |
| LINEAVP | ASSLSASRNTLY | 0.42253205 |
| LGYAFSGL | ASSLSASRNTLY | 0.470844 |
| LDFVAVAL | ASSLSASRNTLY | 0.48486006 |
| LAQPGAE | ASSLSASRNTLY | 0.6112858 |
| ISNLGAVTGF | ASSLSASRNTLY | 0.54045624 |
| HQEDPGRI | ASSLSASRNTLY | 0.6519962 |
| GTPGEPTP | ASSLSASRNTLY | 0.58981925 |

| | | |
|---|---|---|
| GFFNSGS | ASSLSASRNTLY | 0.66668755 |
| GAGGGGLTAGAN | ASSLSASRNTLY | 0.4961936 |
| FVDALVNM | ASSLSASRNTLY | 0.59895825 |
| FLTVAPEL | ASSLSASRNTLY | 0.37948 |
| FETLTAL | ASSLSASRNTLY | 0.59856725 |
| DVINPIEL | ASSLSASRNTLY | 0.46905813 |
| APVSGAPV | ASSLSASRNTLY | 0.582099 |
| ALVGGLAW | ASSLSASRNTLY | 0.365738 |
| ALAAAAATAI | ASSLSASRNTLY | 0.68928564 |
| AFGLGLLL | ASSLSASRNTLY | 0.5371647 |
| AFFAAASL | ASSLSASRNTLY | 0.6311772 |
| AAAALPAFSPP | ASSLSASRNTLY | 0.5195262 |
| VVLAPAL | ASSLSASRNTLY | 0.5156317 |
| TIGHVDHGKTTL | ASSLSASRNTLY | 0.48232263 |
| LISNASDALDK | ASSLSASRNTLY | 0.6400906 |
| LISNASDAL | ASSLSASRNTLY | 0.39818463 |
| ISNASDALDK | ASSLSASRNTLY | 0.34061798 |
| NQAETLVYQTE | ASSLSASRNTLY | 0.52927506 |
| TGGTGGTAGAG | ASSLSASRNTLY | 0.3765457 |
| TGSTAGGAGGA | ASSLSASRNTLY | 0.67805165 |
| DPLAVPVAAGPV | ASSLSASRNTLY | 0.6729176 |
| GGTGGTPGNGGN | ASSLSASRNTLY | 0.5815371 |
| PAGLAGVDPATL | ASSLSASRNTLY | 0.66851443 |
| GNNLVVEAPHAGASV | ASSLSASRNTLY | 0.3004049 |
| KVTRIMSASTLEQPA | ASSLSASRNTLY | 0.50372744 |
| LGFALPIALPRRTVV | ASSLSASRNTLY | 0.68874 |
| TRLGNLTARKLVVLV | ASSLSASRNTLY | 0.6083394 |
| WSQQRRLIYFPSAGP | ASSLSASRNTLY | 0.5731679 |
| GPNGNPDPMAAAVDIR | ASSLSASRNTLY | 0.6633113 |
| TQRCVSVADLRAAASR | ASSLSASRNTLY | 0.5108424 |
| QFVLDVAEHPVEVLTETV | ASSLSASRNTLY | 0.56928617 |
| GGAGGTGGAGGNGGTQAGDG | ASSLSASRNTLY | 0.5833493 |
| PLLATLRAIAVDVDAKPAQV | ASSLSASRNTLY | 0.6204775 |
| APKFRTGGYMIDFWGVGYQVAK | ASSLSASRNTLY | 0.6009405 |
| GAGGAGGAGGAAGGPGGTGGTG | ASSLSASRNTLY | 0.42602822 |
| IGALGLLATDVAPAGSVAFGVA | ASSLSASRNTLY | 0.61498785 |
| AGGAGGIGGNGGAGGAGGVLMG | ASSLSASRNTLY | 0.70471 |
| ADELIGLL | ASSLSASRNTLY | 0.5168302 |
| ADRALKA | ASSLSASRNTLY | 0.6554098 |
| AGAVGQVGGAGGA | ASSLSASRNTLY | 0.6027496 |
| AGGGGGLGGAP | ASSLSASRNTLY | 0.6227451 |
| AGGLLYGNGGAG | ASSLSASRNTLY | 0.4595117 |
| AGLESIGL | ASSLSASRNTLY | 0.4094963 |
| AGLKTGII | ASSLSASRNTLY | 0.61103624 |
| ASLAALLL | ASSLSASRNTLY | 0.40193143 |
| ASVINISSI | ASSLSASRNTLY | 0.57627 |
| ATPPGPA | ASSLSASRNTLY | 0.508904 |
| ATVASALV | ASSLSASRNTLY | 0.7183496 |
| ATVASLIL | ASSLSASRNTLY | 0.6340684 |
| AVNGPVLIRH | ASSLSASRNTLY | 0.601558 |
| DGAAGAVGI | ASSLSASRNTLY | 0.46239626 |

| | | |
|---|---|---|
| DPLLHGI | ASSLSASRNTLY | 0.567167 |
| ELGAALGAL | ASSLSASRNTLY | 0.6044202 |
| ELGRTQAV | ASSLSASRNTLY | 0.40679863 |
| FVAGGYLS | ASSLSASRNTLY | 0.6724591 |
| GASGGAGGAGGW | ASSLSASRNTLY | 0.46279076 |
| GGAGAGGDGGAGGI | ASSLSASRNTLY | 0.5256067 |
| GTGHNGSQPGMGGNG | ASSLSASRNTLY | 0.6490853 |
| GYAAASASAS | ASSLSASRNTLY | 0.6892599 |
| HQVDVAAVQQG | ASSLSASRNTLY | 0.53009677 |
| IGAAIAHT | ASSLSASRNTLY | 0.59008354 |
| IGMIEVAQL | ASSLSASRNTLY | 0.47701985 |
| IRLAPEHTSAMD | ASSLSASRNTLY | 0.63053703 |
| IYNVSVL | ASSLSASRNTLY | 0.6598262 |
| KQGFLGM | ASSLSASRNTLY | 0.60696536 |
| LESVITQL | ASSLSASRNTLY | 0.6839322 |
| LGLGALVVV | ASSLSASRNTLY | 0.63993657 |
| LIGNGGNGGSGGI | ASSLSASRNTLY | 0.4204399 |
| LNAINAPTL | ASSLSASRNTLY | 0.57498693 |
| LQLLNTGV | ASSLSASRNTLY | 0.6441076 |
| LQTSIEF | ASSLSASRNTLY | 0.6561612 |
| MSFAVSEL | ASSLSASRNTLY | 0.705489 |
| NTGALGL | ASSLSASRNTLY | 0.6403587 |
| PPGAGGAGGAGTT | ASSLSASRNTLY | 0.49912292 |
| QLVGLGI | ASSLSASRNTLY | 0.6785873 |
| RFLEQVHQL | ASSLSASRNTLY | 0.71368396 |
| RGVGYVLR | ASSLSASRNTLY | 0.62333864 |
| SATGVFAGVI | ASSLSASRNTLY | 0.47007072 |
| SLPAITLP | ASSLSASRNTLY | 0.47382325 |
| STALAFTV | ASSLSASRNTLY | 0.6274785 |
| SVLTVGLGK | ASSLSASRNTLY | 0.62107885 |
| TALAGAGSGI | ASSLSASRNTLY | 0.44206056 |
| TGLPFTGI | ASSLSASRNTLY | 0.58506376 |
| VAEHVTV | ASSLSASRNTLY | 0.6094275 |
| VDIVLNSL | ASSLSASRNTLY | 0.39663562 |
| VLTGGGAVL | ASSLSASRNTLY | 0.63159937 |
| VNVGLAYGI | ASSLSASRNTLY | 0.6574006 |
| VSSAAVGAV | ASSLSASRNTLY | 0.57528883 |
| VTSVYEAGV | ASSLSASRNTLY | 0.46468076 |
| WVALDRGA | ASSLSASRNTLY | 0.6648241 |
| YLVGSVVI | ASSLSASRNTLY | 0.6947242 |
| GPPGTGKTLL | ASSLSASRNTLY | 0.6851836 |
| GVGMTKF | ASSLSASRNTLY | 0.4087811 |
| IGIGGSDL | ASSLSASRNTLY | 0.29748031 |
| IGTLLAEL | ASSLSASRNTLY | 0.5309496 |
| IYTSGTT | ASSLSASRNTLY | 0.657617 |
| LSFPAGV | ASSLSASRNTLY | 0.5861151 |
| LYTSGTT | ASSLSASRNTLY | 0.5034782 |
| VGAFPFGF | ASSLSASRNTLY | 0.587558 |
| VGAGTVEF | ASSLSASRNTLY | 0.6619152 |
| VGKSTLLL | ASSLSASRNTLY | 0.49192974 |
| AALAHAGGGIV | ASSLSASRNTLY | 0.42109594 |

| | | |
|---|---|---|
| AALVAVSLL | ASSLSASRNTLY | 0.64912254 |
| AAVSRAVAAL | ASSLSASRNTLY | 0.6364713 |
| AGGSSGIGGF | ASSLSASRNTLY | 0.44107687 |
| AGLTVANANG | ASSLSASRNTLY | 0.50932646 |
| AGLVGAVL | ASSLSASRNTLY | 0.5767639 |
| ALSQAFTV | ASSLSASRNTLY | 0.5661186 |
| ALTFSVSL | ASSLSASRNTLY | 0.5808235 |
| ALTGAALVAL | ASSLSASRNTLY | 0.42841402 |
| ASFSIGAGVL | ASSLSASRNTLY | 0.66075945 |
| DGSVYKI | ASSLSASRNTLY | 0.38498607 |
| EQLAEIV | ASSLSASRNTLY | 0.5929816 |
| FAVSELTVM | ASSLSASRNTLY | 0.6137866 |
| GAAPLAAGL | ASSLSASRNTLY | 0.60202813 |
| GADGGDGGSGGAGG | ASSLSASRNTLY | 0.51725394 |
| GFGLDTGGAGG | ASSLSASRNTLY | 0.46922985 |
| GLASGGLL | ASSLSASRNTLY | 0.39631858 |
| GRIAGAAVLL | ASSLSASRNTLY | 0.37483343 |
| GSAGSLNTSF | ASSLSASRNTLY | 0.66694933 |
| GTGQIGI | ASSLSASRNTLY | 0.5454772 |
| GTLIALIL | ASSLSASRNTLY | 0.6479331 |
| IAATMGFVVG | ASSLSASRNTLY | 0.61766213 |
| IFAQALL | ASSLSASRNTLY | 0.47743312 |
| IGAAVALGF | ASSLSASRNTLY | 0.59902716 |
| IGLTPLAL | ASSLSASRNTLY | 0.4502269 |
| ILGSNTSTL | ASSLSASRNTLY | 0.5568161 |
| IVFSLTVL | ASSLSASRNTLY | 0.6151513 |
| LASKATGF | ASSLSASRNTLY | 0.6299983 |
| LASVALIL | ASSLSASRNTLY | 0.46116957 |
| LFAVGLTL | ASSLSASRNTLY | 0.6048885 |
| LFNGGIGTYI | ASSLSASRNTLY | 0.60788673 |
| LTLGAGII | ASSLSASRNTLY | 0.50722414 |
| LVAAALGDI | ASSLSASRNTLY | 0.6023911 |
| NTAFVQLGI | ASSLSASRNTLY | 0.6673386 |
| QLFSGAEL | ASSLSASRNTLY | 0.5985243 |
| SAATFITL | ASSLSASRNTLY | 0.40564522 |
| SNYGLANLGV | ASSLSASRNTLY | 0.6258786 |
| SYGGAYAVM | ASSLSASRNTLY | 0.4397523 |
| TTGVGGLTLG | ASSLSASRNTLY | 0.59543765 |
| TTQTIYI | ASSLSASRNTLY | 0.5538855 |
| VAAGAGLGAVI | ASSLSASRNTLY | 0.5901749 |
| VAAVVTLL | ASSLSASRNTLY | 0.6370543 |
| VGFLGGLI | ASSLSASRNTLY | 0.5109916 |
| VGVIPILL | ASSLSASRNTLY | 0.457648 |
| VIFASSLI | ASSLSASRNTLY | 0.5056776 |
| VKGGDGGAAAG | ASSLSASRNTLY | 0.61616063 |
| VTKTFTNI | ASSLSASRNTLY | 0.63374776 |
| VVAAAILL | ASSLSASRNTLY | 0.42209518 |
| YAALDVEL | ASSLSASRNTLY | 0.5878241 |
| LASVALLL | ASSLSASRNTLY | 0.56871426 |
| LDAVHRV | ASSLSASRNTLY | 0.49141333 |
| QGLSSSIF | ASSLSASRNTLY | 0.56358397 |

| | | |
|---|---|---|
| RGTFANIRL | ASSLSASRNTLY | 0.5567404 |
| RQLWWGHRI | ASSLSASRNTLY | 0.39931992 |
| YGPPGTGKTLL | ASSLSASRNTLY | 0.63361704 |
| AALVGAPANVA | ASSLSASRNTLY | 0.5173301 |
| AATFRRTE | ASSLSASRNTLY | 0.6175511 |
| AGALGGGI | ASSLSASRNTLY | 0.6296838 |
| AGGAGGAGGIGAG | ASSLSASRNTLY | 0.4300483 |
| AGSGALGL | ASSLSASRNTLY | 0.58425397 |
| AQIADQI | ASSLSASRNTLY | 0.58041847 |
| ASAVAVTYL | ASSLSASRNTLY | 0.6049343 |
| ASTVLGGTGG | ASSLSASRNTLY | 0.5677637 |
| AVGGEPAK | ASSLSASRNTLY | 0.4803595 |
| CEQGTFGGTC | ASSLSASRNTLY | 0.6500473 |
| DEGKAPAG | ASSLSASRNTLY | 0.63867337 |
| DTVVFPAL | ASSLSASRNTLY | 0.64970714 |
| FEGITMAG | ASSLSASRNTLY | 0.6328635 |
| FPLVVQD | ASSLSASRNTLY | 0.64348865 |
| GAFTTPQ | ASSLSASRNTLY | 0.6355434 |
| GFGGFGTAGGVGGA | ASSLSASRNTLY | 0.49646395 |
| GFLAQFL | ASSLSASRNTLY | 0.36052638 |
| GGAGGAGGNAGVA | ASSLSASRNTLY | 0.5702936 |
| GGVGTAGAGGAGGAGG | ASSLSASRNTLY | 0.5532076 |
| GNGGGGDAGNAGSGGN | ASSLSASRNTLY | 0.51707405 |
| GTGGAGGAGNDAGS | ASSLSASRNTLY | 0.40742937 |
| GTTNTGWFNS | ASSLSASRNTLY | 0.66550875 |
| IFMGPLFTL | ASSLSASRNTLY | 0.70607424 |
| ILAHAIEAI | ASSLSASRNTLY | 0.51148564 |
| IRSAIREAL | ASSLSASRNTLY | 0.70237255 |
| ITFAGGG | ASSLSASRNTLY | 0.3824929 |
| ITGMSGHQEV | ASSLSASRNTLY | 0.6437261 |
| IVAGLACTL | ASSLSASRNTLY | 0.6764164 |
| IVGGGLAGA | ASSLSASRNTLY | 0.39798808 |
| IVTGVLNI | ASSLSASRNTLY | 0.6595819 |
| LAQQGCI | ASSLSASRNTLY | 0.44377938 |
| LCLHVEN | ASSLSASRNTLY | 0.48484114 |
| LDAISDGS | ASSLSASRNTLY | 0.67286015 |
| LGGFAAML | ASSLSASRNTLY | 0.5808152 |
| LGGKNPMI | ASSLSASRNTLY | 0.6439788 |
| LGPKGVAAIL | ASSLSASRNTLY | 0.56581676 |
| LMVGVSI | ASSLSASRNTLY | 0.47690144 |
| LVGAAFVL | ASSLSASRNTLY | 0.584679 |
| LVLAVAPL | ASSLSASRNTLY | 0.6873565 |
| MAAIAVIDAA | ASSLSASRNTLY | 0.509916 |
| NNGINAELQ | ASSLSASRNTLY | 0.71161324 |
| RGEPLVLVHG | ASSLSASRNTLY | 0.46632704 |
| RGGAGGAAGLI | ASSLSASRNTLY | 0.6912555 |
| RSYRMHPAV | ASSLSASRNTLY | 0.6187008 |
| SAAHLLAGQLA | ASSLSASRNTLY | 0.54885364 |
| SAAVGSY | ASSLSASRNTLY | 0.52817476 |
| SPHAVLHRI | ASSLSASRNTLY | 0.6579925 |
| SQIAKVL | ASSLSASRNTLY | 0.5552824 |

| | | |
|---|---|---|
| TALGHLVP | ASSLSASRNTLY | 0.6195026 |
| TAPTTSGAA | ASSLSASRNTLY | 0.6758883 |
| TDGDDGNGGSPGT | ASSLSASRNTLY | 0.5123028 |
| TDGLGEN | ASSLSASRNTLY | 0.8419912 |
| TGGPPLPGQA | ASSLSASRNTLY | 0.74498516 |
| TIPAITI | ASSLSASRNTLY | 0.78006434 |
| TIPGFTI | ASSLSASRNTLY | 0.58331954 |
| TSIASGALA | ASSLSASRNTLY | 0.42350286 |
| TVTLPGAAI | ASSLSASRNTLY | 0.6822121 |
| VASGLSAALP | ASSLSASRNTLY | 0.47286522 |
| VGAGVIGI | ASSLSASRNTLY | 0.7603204 |
| VGPKRFGV | ASSLSASRNTLY | 0.8042164 |
| VNVAFPD | ASSLSASRNTLY | 0.5493642 |
| VSMTNSGVS | ASSLSASRNTLY | 0.83900887 |
| VTDVLGDLSV | ASSLSASRNTLY | 0.84474957 |
| VVVVGAGFAGL | ASSLSASRNTLY | 0.37522662 |
| VYTVGTVV | ASSLSASRNTLY | 0.6351094 |
| WTTFSGP | ASSLSASRNTLY | 0.5774305 |
| YDELVEAF | ASSLSASRNTLY | 0.6879982 |
| YGTGGAGGNGG | ASSLSASRNTLY | 0.2509748 |
| YTNVMAV | ASSLSASRNTLY | 0.8036224 |
| FLEQVHQL | ASSLSASRNTLY | 0.53778154 |
| GLSLLAVAV | ASSLSASRNTLY | 0.16523226 |
| LEQVHQL | ASSLSASRNTLY | 0.7616257 |
| LNAAGAI | ASSLSASRNTLY | 0.6716813 |
| PAVHASL | ASSLSASRNTLY | 0.6463228 |
| TELEDDA | ASSLSASRNTLY | 0.82378674 |
| TPPGPPN | ASSLSASRNTLY | 0.51180977 |
| TPPPPPPP | ASSLSASRNTLY | 0.49584138 |
| TQSPALI | ASSLWGAAETLY | 0.7798646 |
| EFAAAFPQ | ASSLWGAAETLY | 0.825019 |
| LLSGNLDVL | ASSLWGAAETLY | 0.47022316 |
| GPANLGNYNI | ASSLWGAAETLY | 0.7310139 |
| TLGPTLP | ASSLWGAAETLY | 0.8335996 |
| VYAYSSCCK | ASSLWGAAETLY | 0.7431621 |
| APAPAPAPNI | ASSLWGAAETLY | 0.83074343 |
| PGGPVPIP | ASSLWGAAETLY | 0.8471438 |
| FLTVALEP | ASSLWGAAETLY | 0.6533514 |
| SQYFGANL | ASSLWGAAETLY | 0.81873775 |
| GGVKPAA | ASSLWGAAETLY | 0.8579222 |
| GGTGGAGGAGGW | ASSLWGAAETLY | 0.6500418 |
| GVGGVAGLGGAGG | ASSLWGAAETLY | 0.7257912 |
| VLLVLATTI | ASSLWGAAETLY | 0.4761761 |
| IGIFAAML | ASSLWGAAETLY | 0.7016336 |
| AVGDLID | ASSLWGAAETLY | 0.36732727 |
| NISPLPL | ASSLWGAAETLY | 0.7327398 |
| VTAVFVAI | ASSLWGAAETLY | 0.84319806 |
| SGDDNDAAL | ASSLWGAAETLY | 0.65941805 |
| AVAALAHS | ASSLWGAAETLY | 0.44428027 |
| GGVTVMATP | ASSLWGAAETLY | 0.375771 |
| AGLAANVRD | ASSLWGAAETLY | 0.7202927 |

| | | |
|---|---|---|
| TNRPDLIDPALLRPG | ASSLWGAAETLY | 0.6018111 |
| LANLGVTAL | ASSLWGAAETLY | 0.33019972 |
| TGERIEL | ASSLWGAAETLY | 0.7211498 |
| NGEELAA | ASSLWGAAETLY | 0.8298237 |
| VLSLVWG | ASSLWGAAETLY | 0.8281725 |
| IGYALWL | ASSLWGAAETLY | 0.44228798 |
| LSLLLAPL | ASSLWGAAETLY | 0.8605397 |
| TLDEALK | ASSLWGAAETLY | 0.75257826 |
| TDGAGTAEGG | ASSLWGAAETLY | 0.8342423 |
| NAPGGAGGNGG | ASSLWGAAETLY | 0.67148226 |
| VSVFMGMGGNF | ASSLWGAAETLY | 0.48790485 |
| LASYLAAL | ASSLWGAAETLY | 0.8276149 |
| LVGGTPAL | ASSLWGAAETLY | 0.66442037 |
| VRNADVL | ASSLWGAAETLY | 0.65281975 |
| TCSSPFLVSAM | ASSLWGAAETLY | 0.82664335 |
| VHITDAE | ASSLWGAAETLY | 0.6891396 |
| EYAAPSF | ASSLWGAAETLY | 0.5565932 |
| VVDAFHGGL | ASSLWGAAETLY | 0.61965746 |
| ASVVDPQ | ASSLWGAAETLY | 0.8215399 |
| SYNEFGGLS | ASSLWGAAETLY | 0.49228394 |
| DPDGSVY | ASSLWGAAETLY | 0.5024413 |
| MVSTIHATE | ASSLWGAAETLY | 0.6206589 |
| LAAGVADPV | ASSLWGAAETLY | 0.694811 |
| TKEQIAA | ASSLWGAAETLY | 0.82273 |
| SLASPAAL | ASSLWGAAETLY | 0.7681924 |
| IGLVFYGV | ASSLWGAAETLY | 0.6507652 |
| FLLDASNL | ASSLWGAAETLY | 0.75712323 |
| YQVGNFP | ASSLWGAAETLY | 0.78984 |
| ESLLSQST | ASSLWGAAETLY | 0.74161065 |
| LFTVGFAAVG | ASSLWGAAETLY | 0.78282547 |
| ATPPTPT | ASSLWGAAETLY | 0.54721004 |
| TDLATLG | ASSLWGAAETLY | 0.8231456 |
| SAGGTGR | ASSLWGAAETLY | 0.8143745 |
| GFSNSGS | ASSLWGAAETLY | 0.8136027 |
| GGNGGSGGTSV | ASSLWGAAETLY | 0.82262075 |
| GTSVATGGAGN | ASSLWGAAETLY | 0.7635237 |
| NGGNGGSGGTS | ASSLWGAAETLY | 0.7374369 |
| AVVKFPH | ASSLWGAAETLY | 0.28005716 |
| TPPPPPP | ASSLWGAAETLY | 0.660251 |
| AAPAAAAAPA | ASSLWGAAETLY | 0.74493474 |
| IALNEDL | ASSLWGAAETLY | 0.74886465 |
| IVNVSSM | ASSLWGAAETLY | 0.7973882 |
| TGIPPAPR | ASSLWGAAETLY | 0.8063414 |
| FDLGGGTF | ASSLWGAAETLY | 0.59088904 |
| FDLGGGTFDVS | ASSLWGAAETLY | 0.83033615 |
| NEPTAAA | ASSLWGAAETLY | 0.8325388 |
| NEPTAAALA | ASSLWGAAETLY | 0.36829698 |
| GSIINISSI | ASSLWGAAETLY | 0.6531036 |
| VAPGNAGTA | ASSLWGAAETLY | 0.87053585 |
| TGGGASLE | ASSLWGAAETLY | 0.7492963 |
| VAVYNHYKRI | ASSLWGAAETLY | 0.65505195 |

| | | |
|---|---|---|
| TDPAPATT | ASSLWGAAETLY | 0.69643456 |
| TNRPDILDPALLRPG | ASSLWGAAETLY | 0.84453607 |
| LTTTTTS | ASSLWGAAETLY | 0.6694214 |
| PTGSGKTL | ASSLWGAAETLY | 0.84997696 |
| TGSGKTL | ASSLWGAAETLY | 0.2510964 |
| TIGHVDHGK | ASSLWGAAETLY | 0.63248825 |
| TIGHVDHGKTT | ASSLWGAAETLY | 0.5221628 |
| GHVDHGKTT | ASSLWGAAETLY | 0.24694696 |
| AQFEHTLL | ASSLWGAAETLY | 0.8130032 |
| IVNQDEL | ASSLWGAAETLY | 0.6755521 |
| TNGSQFF | ASSLWGAAETLY | 0.81914914 |
| TGKGIVE | ASSLWGAAETLY | 0.6847057 |
| LISNASDA | ASSLWGAAETLY | 0.826589 |
| AELRTLGELTKIAKAHGAQ | ASSLWGAAETLY | 0.7255667 |
| APNAVAAAEPSW | ASSLWGAAETLY | 0.7558474 |
| APQVAGAATAVL | ASSLWGAAETLY | 0.57596356 |
| ARRDPAAAAAILPTDA | ASSLWGAAETLY | 0.7979884 |
| DAAARDVLAEAGLAEYF | ASSLWGAAETLY | 0.84816444 |
| DDQIVLSAKANLAKVEQ | ASSLWGAAETLY | 0.8081446 |
| DELLPDTGEDSDEDGH | ASSLWGAAETLY | 0.3618871 |
| ELPEAPANRATVPQD | ASSLWGAAETLY | 0.6711428 |
| FGQSKIERVTGIG | ASSLWGAAETLY | 0.68051577 |
| GAGGTGSTAGGAGGAGGAGGLY | ASSLWGAAETLY | 0.27956137 |
| GHGGTGAAGGNGGTGGAGGSGI | ASSLWGAAETLY | 0.24677119 |
| GNGGNGGAGGEGGPGGAGGAGA | ASSLWGAAETLY | 0.63174015 |
| GSTAGNGGNGGNATNGTIAGQP | ASSLWGAAETLY | 0.6402435 |
| GTGGDGGNGGNGGTGG | ASSLWGAAETLY | 0.80756176 |
| GVRRPAGIPADAPV | ASSLWGAAETLY | 0.77506554 |
| GYRQVTDAHLLTLARR | ASSLWGAAETLY | 0.65785855 |
| IAKMGQDGHDRG | ASSLWGAAETLY | 0.7749989 |
| KAEPSIEIQYT | ASSLWGAAETLY | 0.75636303 |
| KPVRIRMAADEDTVAVT | ASSLWGAAETLY | 0.11039317 |
| MPPRIAGMRLLVIKPEP | ASSLWGAAETLY | 0.6379448 |
| MKFYAVALFRSRGVPPTR | ASSLWGAAETLY | 0.39945468 |
| NAGGFGILGAVAHSP | ASSLWGAAETLY | 0.85594726 |
| NEGREITLAMAEM | ASSLWGAAETLY | 0.53525484 |
| NELVEWSHPLGHRWRWE | ASSLWGAAETLY | 0.57109636 |
| NLGAGGTGGNGGNPG | ASSLWGAAETLY | 0.70787346 |
| NTEWPKWETVMAADPVD | ASSLWGAAETLY | 0.7388923 |
| PASAAPAETQP | ASSLWGAAETLY | 0.7196565 |
| QQSSDGKNGVRAAEITGE | ASSLWGAAETLY | 0.4597568 |
| RAPDAVTTSWWKE | ASSLWGAAETLY | 0.22455719 |
| SGFGNVGAHTSGWFNQ | ASSLWGAAETLY | 0.81570935 |
| SGVAGAVGQAASVGGLKV | ASSLWGAAETLY | 0.45846438 |
| SNGGAGTGSAGGNGGT | ASSLWGAAETLY | 0.5031757 |
| TGGAGGEGGAAG | ASSLWGAAETLY | 0.38341862 |
| TGGDGGAAGAGGAPGHG | ASSLWGAAETLY | 0.28696376 |
| TMPPLLAAIANAGRHGV | ASSLWGAAETLY | 0.3632767 |
| TSSTAGNSGGA | ASSLWGAAETLY | 0.8169347 |
| VPATVPVGILC | ASSLWGAAETLY | 0.6211546 |
| VQRPDSGGVVLTGRIS | ASSLWGAAETLY | 0.66752297 |

| | | |
|---|---|---|
| WTPNPVPATKGVPAGLVQAT | ASSLWGAAETLY | 0.86678296 |
| YALKRALAKFSDRFDVVII | ASSLWGAAETLY | 0.7050924 |
| YLIGSVIV | ASSLWGAAETLY | 0.6933732 |
| YGQLVAVID | ASSLWGAAETLY | 0.6057495 |
| VTVGGMTLPA | ASSLWGAAETLY | 0.47063607 |
| VLALPAV | ASSLWGAAETLY | 0.5176969 |
| VIPPTLNYET | ASSLWGAAETLY | 0.59332824 |
| VGMGGMHPSSG | ASSLWGAAETLY | 0.859216 |
| VGLAGGAATA | ASSLWGAAETLY | 0.6640608 |
| VGGAGGAGGSGP | ASSLWGAAETLY | 0.46521258 |
| VFAAFMV | ASSLWGAAETLY | 0.739733 |
| TLYDKLR | ASSLWGAAETLY | 0.8486192 |
| TLAVFTVL | ASSLWGAAETLY | 0.8639371 |
| TGSVQPARV | ASSLWGAAETLY | 0.50968987 |
| TGGGAGSGGT | ASSLWGAAETLY | 0.6623902 |
| SHLETSL | ASSLWGAAETLY | 0.6599499 |
| SFTTGMT | ASSLWGAAETLY | 0.6681498 |
| QVEVAKRMGVRQPTVS | ASSLWGAAETLY | 0.6234914 |
| QGSGGPALAM | ASSLWGAAETLY | 0.4935711 |
| MLAPSNP | ASSLWGAAETLY | 0.79669654 |
| LVNNAAI | ASSLWGAAETLY | 0.86183876 |
| LTGEINTLA | ASSLWGAAETLY | 0.68265086 |
| LSIFGPV | ASSLWGAAETLY | 0.2785055 |
| LLALLANL | ASSLWGAAETLY | 0.64425933 |
| LISNASDALDKLR | ASSLWGAAETLY | 0.47324944 |
| LINEAVP | ASSLWGAAETLY | 0.5428872 |
| LGYAFSGL | ASSLWGAAETLY | 0.66152644 |
| LDFVAVAL | ASSLWGAAETLY | 0.3580124 |
| LAQPGAE | ASSLWGAAETLY | 0.7089896 |
| ISNLGAVTGF | ASSLWGAAETLY | 0.8593766 |
| HQEDPGRI | ASSLWGAAETLY | 0.62964547 |
| GTPGEPTP | ASSLWGAAETLY | 0.7983463 |
| GFFNSGS | ASSLWGAAETLY | 0.832585 |
| GAGGGGLTAGAN | ASSLWGAAETLY | 0.7952368 |
| FVDALVNM | ASSLWGAAETLY | 0.836877 |
| FLTVAPEL | ASSLWGAAETLY | 0.77302414 |
| FETLTAL | ASSLWGAAETLY | 0.76123136 |
| DVINPIEL | ASSLWGAAETLY | 0.7276716 |
| APVSGAPV | ASSLWGAAETLY | 0.7773318 |
| ALVGGLAW | ASSLWGAAETLY | 0.6827714 |
| ALAAAAATAI | ASSLWGAAETLY | 0.6846163 |
| AFGLGLLL | ASSLWGAAETLY | 0.6330806 |
| AFFAAASL | ASSLWGAAETLY | 0.8154808 |
| AAAALPAFSPP | ASSLWGAAETLY | 0.6270498 |
| VVLAPAL | ASSLWGAAETLY | 0.8255871 |
| TIGHVDHGKTTL | ASSLWGAAETLY | 0.36963478 |
| LISNASDALDK | ASSLWGAAETLY | 0.49507546 |
| LISNASDAL | ASSLWGAAETLY | 0.82089394 |
| ISNASDALDK | ASSLWGAAETLY | 0.1415345 |
| NQAETLVYQTE | ASSLWGAAETLY | 0.25867984 |
| TGGTGGTAGAG | ASSLWGAAETLY | 0.6809404 |

| | | |
|---|---|---|
| TGSTAGGAGGA | ASSLWGAAETLY | 0.8385015 |
| DPLAVPVAAGPV | ASSLWGAAETLY | 0.15298067 |
| GGTGGTPGNGGN | ASSLWGAAETLY | 0.52180684 |
| PAGLAGVDPATL | ASSLWGAAETLY | 0.40996212 |
| GNNLVVEAPHAGASV | ASSLWGAAETLY | 0.81403553 |
| KVTRIMSASTLEQPA | ASSLWGAAETLY | 0.7062176 |
| LGFALPIALPRRTVV | ASSLWGAAETLY | 0.3816751 |
| TRLGNLTARKLVVLV | ASSLWGAAETLY | 0.80666584 |
| WSQQRRLIYFPSAGP | ASSLWGAAETLY | 0.3072468 |
| GPNGNPDPMAAAVDIR | ASSLWGAAETLY | 0.5475521 |
| TQRCVSVADLRAAASR | ASSLWGAAETLY | 0.80336094 |
| QFVLDVAEHPVEVLTETV | ASSLWGAAETLY | 0.83089626 |
| GGAGGTGGAGGNGGTQAGDG | ASSLWGAAETLY | 0.44947347 |
| PLLATLRAIAVDVDAKPAQV | ASSLWGAAETLY | 0.783085 |
| APKFRTGGYMIDFWGVGYQVAK | ASSLWGAAETLY | 0.55823964 |
| GAGGAGGAGGAAGGPGGTGGTG | ASSLWGAAETLY | 0.87027985 |
| IGALGLLATDVAPAGSVAFGVA | ASSLWGAAETLY | 0.6506696 |
| AGGAGGIGGNGGAGGAGGVLMG | ASSLWGAAETLY | 0.37424415 |
| ADELIGLL | ASSLWGAAETLY | 0.8155358 |
| ADRALKA | ASSLWGAAETLY | 0.81242573 |
| AGAVGQVGGAGGA | ASSLWGAAETLY | 0.71064246 |
| AGGGGGLGGAP | ASSLWGAAETLY | 0.776211 |
| AGGLLYGNGGAG | ASSLWGAAETLY | 0.60842943 |
| AGLESIGL | ASSLWGAAETLY | 0.6483104 |
| AGLKTGII | ASSLWGAAETLY | 0.8283013 |
| ASLAALLL | ASSLWGAAETLY | 0.80647445 |
| ASVINISSI | ASSLWGAAETLY | 0.52577025 |
| ATPPGPA | ASSLWGAAETLY | 0.83415806 |
| ATVASALV | ASSLWGAAETLY | 0.7166648 |
| ATVASLIL | ASSLWGAAETLY | 0.51462877 |
| AVNGPVLIRH | ASSLWGAAETLY | 0.85923326 |
| DGAAGAVGI | ASSLWGAAETLY | 0.71129644 |
| DPLLHGI | ASSLWGAAETLY | 0.8045391 |
| ELGAALGAL | ASSLWGAAETLY | 0.8004165 |
| ELGRTQAV | ASSLWGAAETLY | 0.79818606 |
| FVAGGYLS | ASSLWGAAETLY | 0.50473934 |
| GASGGAGGAGGW | ASSLWGAAETLY | 0.7897331 |
| GGAGAGGDGGAGGI | ASSLWGAAETLY | 0.6940019 |
| GTGHNGSQPGMGGNG | ASSLWGAAETLY | 0.803298 |
| GYAAASASAS | ASSLWGAAETLY | 0.7099961 |
| HQVDVAAVQQG | ASSLWGAAETLY | 0.5821068 |
| IGAAIAHT | ASSLWGAAETLY | 0.79771405 |
| IGMIEVAQL | ASSLWGAAETLY | 0.8484402 |
| IRLAPEHTSAMD | ASSLWGAAETLY | 0.72362185 |
| IYNVSVL | ASSLWGAAETLY | 0.75132143 |
| KQGFLGM | ASSLWGAAETLY | 0.8520242 |
| LESVITQL | ASSLWGAAETLY | 0.51403123 |
| LGLGALVVV | ASSLWGAAETLY | 0.71546644 |
| LIGNGGNGGSGGI | ASSLWGAAETLY | 0.7958989 |
| LNAINAPTL | ASSLWGAAETLY | 0.83608854 |
| LQLLNTGV | ASSLWGAAETLY | 0.68382764 |

| | | |
|---|---|---|
| LQTSIEF | ASSLWGAAETLY | 0.83375925 |
| MSFAVSEL | ASSLWGAAETLY | 0.5697043 |
| NTGALGL | ASSLWGAAETLY | 0.794041 |
| PPGAGGAGGAGTT | ASSLWGAAETLY | 0.5741709 |
| QLVGLGI | ASSLWGAAETLY | 0.74293786 |
| RFLEQVHQL | ASSLWGAAETLY | 0.515689 |
| RGVGYVLR | ASSLWGAAETLY | 0.7739868 |
| SATGVFAGVI | ASSLWGAAETLY | 0.79542184 |
| SLPAITLP | ASSLWGAAETLY | 0.81217587 |
| STALAFTV | ASSLWGAAETLY | 0.24978161 |
| SVLTVGLGK | ASSLWGAAETLY | 0.78671265 |
| TALAGAGSGI | ASSLWGAAETLY | 0.8072345 |
| TGLPFTGI | ASSLWGAAETLY | 0.86812 |
| VAEHVTV | ASSLWGAAETLY | 0.6755794 |
| VDIVLNSL | ASSLWGAAETLY | 0.7974843 |
| VLTGGGAVL | ASSLWGAAETLY | 0.577658 |
| VNVGLAYGI | ASSLWGAAETLY | 0.6921777 |
| VSSAAVGAV | ASSLWGAAETLY | 0.42081648 |
| VTSVYEAGV | ASSLWGAAETLY | 0.6077751 |
| WVALDRGA | ASSLWGAAETLY | 0.60972726 |
| YLVGSVVI | ASSLWGAAETLY | 0.50803614 |
| GPPGTGKTLL | ASSLWGAAETLY | 0.7989383 |
| GVGMTKF | ASSLWGAAETLY | 0.8150897 |
| IGIGGSDL | ASSLWGAAETLY | 0.82613707 |
| IGTLLAEL | ASSLWGAAETLY | 0.8188976 |
| IYTSGTT | ASSLWGAAETLY | 0.84105986 |
| LSFPAGV | ASSLWGAAETLY | 0.8207129 |
| LYTSGTT | ASSLWGAAETLY | 0.58162576 |
| VGAFPFGF | ASSLWGAAETLY | 0.68898964 |
| VGAGTVEF | ASSLWGAAETLY | 0.3748642 |
| VGKSTLLL | ASSLWGAAETLY | 0.66359895 |
| AALAHAGGGIV | ASSLWGAAETLY | 0.61957103 |
| AALVAVSLL | ASSLWGAAETLY | 0.7447906 |
| AAVSRAVAAL | ASSLWGAAETLY | 0.13668352 |
| AGGSSGIGGF | ASSLWGAAETLY | 0.7065312 |
| AGLTVANANG | ASSLWGAAETLY | 0.8620279 |
| AGLVGAVL | ASSLWGAAETLY | 0.63860714 |
| ALSQAFTV | ASSLWGAAETLY | 0.7794586 |
| ALTFSVSL | ASSLWGAAETLY | 0.77879715 |
| ALTGAALVAL | ASSLWGAAETLY | 0.5911916 |
| ASFSIGAGVL | ASSLWGAAETLY | 0.78011024 |
| DGSVYKI | ASSLWGAAETLY | 0.56623864 |
| EQLAEIV | ASSLWGAAETLY | 0.8018565 |
| FAVSELTVM | ASSLWGAAETLY | 0.8596363 |
| GAAPLAAGL | ASSLWGAAETLY | 0.755638 |
| GADGGDGGSGGAGG | ASSLWGAAETLY | 0.69824004 |
| GFGLDTGGAGG | ASSLWGAAETLY | 0.7386078 |
| GLASGGLL | ASSLWGAAETLY | 0.43113795 |
| GRIAGAAVLL | ASSLWGAAETLY | 0.7321687 |
| GSAGSLNTSF | ASSLWGAAETLY | 0.15543383 |
| GTGQIGI | ASSLWGAAETLY | 0.69087386 |

| | | |
|---|---|---|
| GTLIALIL | ASSLWGAAETLY | 0.53862685 |
| IAATMGFVVG | ASSLWGAAETLY | 0.7421501 |
| IFAQALL | ASSLWGAAETLY | 0.7383179 |
| IGAAVALGF | ASSLWGAAETLY | 0.74483657 |
| IGLTPLAL | ASSLWGAAETLY | 0.48737386 |
| ILGSNTSTL | ASSLWGAAETLY | 0.7404554 |
| IVFSLTVL | ASSLWGAAETLY | 0.6644571 |
| LASKATGF | ASSLWGAAETLY | 0.4245001 |
| LASVALIL | ASSLWGAAETLY | 0.8096687 |
| LFAVGLTL | ASSLWGAAETLY | 0.66844094 |
| LFNGGIGTYI | ASSLWGAAETLY | 0.68817204 |
| LTLGAGII | ASSLWGAAETLY | 0.8422053 |
| LVAAALGDI | ASSLWGAAETLY | 0.7460408 |
| NTAFVQLGI | ASSLWGAAETLY | 0.33856302 |
| QLFSGAEL | ASSLWGAAETLY | 0.6600946 |
| SAATFITL | ASSLWGAAETLY | 0.7039878 |
| SNYGLANLGV | ASSLWGAAETLY | 0.66664577 |
| SYGGAYAVM | ASSLWGAAETLY | 0.14567728 |
| TTGVGGLTLG | ASSLWGAAETLY | 0.8118484 |
| TTQTIYI | ASSLWGAAETLY | 0.5346409 |
| VAAGAGLGAVI | ASSLWGAAETLY | 0.4818921 |
| VAAVVTLL | ASSLWGAAETLY | 0.59673613 |
| VGFLGGLI | ASSLWGAAETLY | 0.8127544 |
| VGVIPILL | ASSLWGAAETLY | 0.6365153 |
| VIFASSLI | ASSLWGAAETLY | 0.69222015 |
| VKGGDGGAAAG | ASSLWGAAETLY | 0.7742865 |
| VTKTFTNI | ASSLWGAAETLY | 0.77832323 |
| VVAAAILL | ASSLWGAAETLY | 0.8460022 |
| YAALDVEL | ASSLWGAAETLY | 0.76257896 |
| LASVALLL | ASSLWGAAETLY | 0.6900133 |
| LDAVHRV | ASSLWGAAETLY | 0.80582446 |
| QGLSSSIF | ASSLWGAAETLY | 0.6770389 |
| RGTFANIRL | ASSLWGAAETLY | 0.42491502 |
| RQLWWGHRI | ASSLWGAAETLY | 0.33821127 |
| YGPPGTGKTLL | ASSLWGAAETLY | 0.6553385 |
| AALVGAPANVA | ASSLWGAAETLY | 0.37222123 |
| AATFRRTE | ASSLWGAAETLY | 0.7443888 |
| AGALGGGI | ASSLWGAAETLY | 0.8418983 |
| AGGAGGAGGIGAG | ASSLWGAAETLY | 0.6877063 |
| AGSGALGL | ASSLWGAAETLY | 0.40521684 |
| AQIADQI | ASSLWGAAETLY | 0.6003054 |
| ASAVAVTYL | ASSLWGAAETLY | 0.6757637 |
| ASTVLGGTGG | ASSLWGAAETLY | 0.7951587 |
| AVGGEPAK | ASSLWGAAETLY | 0.7990151 |
| CEQGTFGGTC | ASSLWGAAETLY | 0.8467736 |
| DEGKAPAG | ASSLWGAAETLY | 0.7964551 |
| DTVVFPAL | ASSLWGAAETLY | 0.62162995 |
| FEGITMAG | ASSLWGAAETLY | 0.7879493 |
| FPLVVQD | ASSLWGAAETLY | 0.67892504 |
| GAFTTPQ | ASSLWGAAETLY | 0.7375765 |
| GFGGFGTAGGVGGA | ASSLWGAAETLY | 0.7367938 |

| | | |
|---|---|---|
| GFLAQFL | ASSLWGAAETLY | 0.7354299 |
| GGAGGAGGNAGVA | ASSLWGAAETLY | 0.6747463 |
| GGVGTAGAGGAGGAGG | ASSLWGAAETLY | 0.6812974 |
| GNGGGGDAGNAGSGGN | ASSLWGAAETLY | 0.34315193 |
| GTGGAGGAGNDAGS | ASSLWGAAETLY | 0.7463949 |
| GTTNTGWFNS | ASSLWGAAETLY | 0.7344583 |
| IFMGPLFTL | ASSLWGAAETLY | 0.3924178 |
| ILAHAIEAI | ASSLWGAAETLY | 0.6705012 |
| IRSAIREAL | ASSLWGAAETLY | 0.85077065 |
| ITFAGGG | ASSLWGAAETLY | 0.83061266 |
| ITGMSGHQEV | ASSLWGAAETLY | 0.55077416 |
| IVAGLACTL | ASSLWGAAETLY | 0.56432104 |
| IVGGGLAGA | ASSLWGAAETLY | 0.7244098 |
| IVTGVLNI | ASSLWGAAETLY | 0.37578967 |
| LAQQGCI | ASSLWGAAETLY | 0.77729905 |
| LCLHVEN | ASSLWGAAETLY | 0.68459845 |
| LDAISDGS | ASSLWGAAETLY | 0.70731246 |
| LGGFAAML | ASSLWGAAETLY | 0.62873244 |
| LGGKNPMI | ASSLWGAAETLY | 0.7484163 |
| LGPKGVAAIL | ASSLWGAAETLY | 0.6481765 |
| LMVGVSI | ASSLWGAAETLY | 0.79061466 |
| LVGAAFVL | ASSLWGAAETLY | 0.15134953 |
| LVLAVAPL | ASSLWGAAETLY | 0.7972107 |
| MAAIAVIDAA | ASSLWGAAETLY | 0.51188636 |
| NNGINAELQ | ASSLWGAAETLY | 0.5097805 |
| RGEPLVLVHG | ASSLWGAAETLY | 0.26802486 |
| RGGAGGAAGLI | ASSLWGAAETLY | 0.7128545 |
| RSYRMHPAV | ASSLWGAAETLY | 0.78380936 |
| SAAHLLAGQLA | ASSLWGAAETLY | 0.8155477 |
| SAAVGSY | ASSLWGAAETLY | 0.61985373 |
| SPHAVLHRI | ASSLWGAAETLY | 0.80767584 |
| SQIAKVL | ASSLWGAAETLY | 0.8318097 |
| TALGHLVP | ASSLWGAAETLY | 0.6505261 |
| TAPTTSGAA | ASSLWGAAETLY | 0.34533355 |
| TDGDDGNGGSPGT | ASSLWGAAETLY | 0.7723355 |
| TDGLGEN | ASSLWGAAETLY | 0.6459013 |
| TGGPPLPGQA | ASSLWGAAETLY | 0.8085495 |
| TIPAITI | ASSLWGAAETLY | 0.80255765 |
| TIPGFTI | ASSLWGAAETLY | 0.39735273 |
| TSIASGALA | ASSLWGAAETLY | 0.81520134 |
| TVTLPGAAI | ASSLWGAAETLY | 0.83188885 |
| VASGLSAALP | ASSLWGAAETLY | 0.51487076 |
| VGAGVIGI | ASSLWGAAETLY | 0.52481294 |
| VGPKRFGV | ASSLWGAAETLY | 0.65772593 |
| VNVAFPD | ASSLWGAAETLY | 0.31227493 |
| VSMTNSGVS | ASSLWGAAETLY | 0.7943531 |
| VTDVLGDLSV | ASSLWGAAETLY | 0.626235 |
| VVVVGAGFAGL | ASSLWGAAETLY | 0.5913797 |
| VYTVGTVV | ASSLWGAAETLY | 0.7385533 |
| WTTFSGP | ASSLWGAAETLY | 0.608942 |
| YDELVEAF | ASSLWGAAETLY | 0.6275157 |

| | | |
|---|---|---|
| YGTGGAGGNGG | ASSLWGAAETLY | 0.6603382 |
| YTNVMAV | ASSLWGAAETLY | 0.72810966 |
| FLEQVHQL | ASSLWGAAETLY | 0.7618895 |
| GLSLLAVAV | ASSLWGAAETLY | 0.6892228 |
| LEQVHQL | ASSLWGAAETLY | 0.75194156 |
| LNAAGAI | ASSLWGAAETLY | 0.8490174 |
| PAVHASL | ASSLWGAAETLY | 0.41420552 |
| TELEDDA | ASSLWGAAETLY | 0.5841876 |
| TPPGPPN | ASSLWGAAETLY | 0.7774259 |
| TPPPPPPP | ASSLWGAAETLY | 0.50798047 |
| TQSPALI | ASSQGAPL | 0.8061571 |
| EFAAAFPQ | ASSQGAPL | 0.7515433 |
| LLSGNLDVL | ASSQGAPL | 0.7701902 |
| GPANLGNYNI | ASSQGAPL | 0.65720534 |
| TLGPTLP | ASSQGAPL | 0.86718845 |
| VYAYSSCCK | ASSQGAPL | 0.8038776 |
| APAPAPAPNI | ASSQGAPL | 0.7384232 |
| PGGPVPIP | ASSQGAPL | 0.7684897 |
| FLTVALEP | ASSQGAPL | 0.52902997 |
| SQYFGANL | ASSQGAPL | 0.8152786 |
| GGVKPAA | ASSQGAPL | 0.77602243 |
| GGTGGAGGAGGW | ASSQGAPL | 0.44674116 |
| GVGGVAGLGGAGG | ASSQGAPL | 0.6486909 |
| VLLVLATTI | ASSQGAPL | 0.8320589 |
| IGIFAAML | ASSQGAPL | 0.83944416 |
| AVGDLID | ASSQGAPL | 0.42660937 |
| NISPLPL | ASSQGAPL | 0.6189144 |
| VTAVFVAI | ASSQGAPL | 0.39802593 |
| SGDDNDAAL | ASSQGAPL | 0.56483495 |
| AVAALAHS | ASSQGAPL | 0.4718496 |
| GGVTVMATP | ASSQGAPL | 0.76629835 |
| AGLAANVRD | ASSQGAPL | 0.6304666 |
| TNRPDLIDPALLRPG | ASSQGAPL | 0.7728123 |
| LANLGVTAL | ASSQGAPL | 0.73251706 |
| TGERIEL | ASSQGAPL | 0.8346739 |
| NGEELAA | ASSQGAPL | 0.54150677 |
| VLSLVWG | ASSQGAPL | 0.688873 |
| IGYALWL | ASSQGAPL | 0.36452198 |
| LSLLLAPL | ASSQGAPL | 0.503695 |
| TLDEALK | ASSQGAPL | 0.5150958 |
| TDGAGTAEGG | ASSQGAPL | 0.67100096 |
| NAPGGAGGNGG | ASSQGAPL | 0.25121275 |
| VSVFMGMGGNF | ASSQGAPL | 0.8574468 |
| LASYLAAL | ASSQGAPL | 0.69644076 |
| LVGGTPAL | ASSQGAPL | 0.80608684 |
| VRNADVL | ASSQGAPL | 0.6004371 |
| TCSSPFLVSAM | ASSQGAPL | 0.596673 |
| VHITDAE | ASSQGAPL | 0.5572302 |
| EYAAPSF | ASSQGAPL | 0.7625903 |
| VVDAFHGGL | ASSQGAPL | 0.47240773 |
| ASVVDPQ | ASSQGAPL | 0.2699825 |

| | | |
|---|---|---|
| SYNEFGGLS | ASSQGAPL | 0.5653056 |
| DPDGSVY | ASSQGAPL | 0.36672083 |
| MVSTIHATE | ASSQGAPL | 0.8518013 |
| LAAGVADPV | ASSQGAPL | 0.82166886 |
| TKEQIAA | ASSQGAPL | 0.7410517 |
| SLASPAAL | ASSQGAPL | 0.8400508 |
| IGLVFYGV | ASSQGAPL | 0.194354 |
| FLLDASNL | ASSQGAPL | 0.6230794 |
| YQVGNFP | ASSQGAPL | 0.8528496 |
| ESLLSQST | ASSQGAPL | 0.75842804 |
| LFTVGFAAVG | ASSQGAPL | 0.6953331 |
| ATPPTPT | ASSQGAPL | 0.8285153 |
| TDLATLG | ASSQGAPL | 0.5594344 |
| SAGGTGR | ASSQGAPL | 0.6788381 |
| GFSNSGS | ASSQGAPL | 0.6897477 |
| GGNGGSGGTSV | ASSQGAPL | 0.81582576 |
| GTSVATGGAGN | ASSQGAPL | 0.73483807 |
| NGGNGGSGGTS | ASSQGAPL | 0.45739982 |
| AVVKFPH | ASSQGAPL | 0.7860755 |
| TPPPPPP | ASSQGAPL | 0.87038386 |
| AAPAAAAAPA | ASSQGAPL | 0.5992302 |
| IALNEDL | ASSQGAPL | 0.80547357 |
| IVNVSSM | ASSQGAPL | 0.7649541 |
| TGIPPAPR | ASSQGAPL | 0.7479147 |
| FDLGGGTF | ASSQGAPL | 0.5041731 |
| FDLGGGTFDVS | ASSQGAPL | 0.516338 |
| NEPTAAA | ASSQGAPL | 0.7472839 |
| NEPTAAALA | ASSQGAPL | 0.47503334 |
| GSIINISSI | ASSQGAPL | 0.6965133 |
| VAPGNAGTA | ASSQGAPL | 0.68197715 |
| TGGGASLE | ASSQGAPL | 0.87818754 |
| VAVYNHYKRI | ASSQGAPL | 0.7849311 |
| TDPAPATT | ASSQGAPL | 0.7802348 |
| TNRPDILDPALLRPG | ASSQGAPL | 0.45453042 |
| LTTTTTS | ASSQGAPL | 0.7394401 |
| PTGSGKTL | ASSQGAPL | 0.77154195 |
| TGSGKTL | ASSQGAPL | 0.394893 |
| TIGHVDHGK | ASSQGAPL | 0.8398038 |
| TIGHVDHGKTT | ASSQGAPL | 0.5381998 |
| GHVDHGKTT | ASSQGAPL | 0.68363494 |
| AQFEHTLL | ASSQGAPL | 0.8107612 |
| IVNQDEL | ASSQGAPL | 0.81664014 |
| TNGSQFF | ASSQGAPL | 0.6790549 |
| TGKGIVE | ASSQGAPL | 0.74047893 |
| LISNASDA | ASSQGAPL | 0.52861476 |
| AELRTLGELTKIAKAHGAQ | ASSQGAPL | 0.8016742 |
| APNAVAAAEPSW | ASSQGAPL | 0.8245095 |
| APQVAGAATAVL | ASSQGAPL | 0.64549613 |
| ARRDPAAAAAILPTDA | ASSQGAPL | 0.8547219 |
| DAAARDVLAEAGLAEYF | ASSQGAPL | 0.77347255 |
| DDQIVLSAKANLAKVEQ | ASSQGAPL | 0.37215886 |

| | | |
|---|---|---|
| DELLPDTGEDSDEDGH | ASSQGAPL | 0.7018485 |
| ELPEAPANRATVPQD | ASSQGAPL | 0.8315102 |
| FGQSKIERVTGIG | ASSQGAPL | 0.8211316 |
| GAGGTGSTAGGAGGAGGAGGLY | ASSQGAPL | 0.84165895 |
| GHGGTGAAGGNGGTGGAGGSGI | ASSQGAPL | 0.78228015 |
| GNGGNGGAGGEGGPGGAGGAGA | ASSQGAPL | 0.58537793 |
| GSTAGNGGNGGNATNGTIAGQP | ASSQGAPL | 0.8453369 |
| GTGGDGGNGGNGGTGG | ASSQGAPL | 0.8676944 |
| GVRRPAGIPADAPV | ASSQGAPL | 0.8013848 |
| GYRQVTDAHLLTLARR | ASSQGAPL | 0.5421832 |
| IAKMGQDGHDRG | ASSQGAPL | 0.54846936 |
| KAEPSIEIQYT | ASSQGAPL | 0.75478244 |
| KPVRIRMAADEDTVAVT | ASSQGAPL | 0.8105592 |
| MPPRIAGMRLLVIKPEP | ASSQGAPL | 0.44754744 |
| MKFYAVALFRSRGVPPTR | ASSQGAPL | 0.7544061 |
| NAGGFGILGAVAHSP | ASSQGAPL | 0.75282073 |
| NEGREITLAMAEM | ASSQGAPL | 0.3087841 |
| NELVEWSHPLGHRWRWE | ASSQGAPL | 0.7918674 |
| NLGAGGTGGNGGNPG | ASSQGAPL | 0.8418222 |
| NTEWPKWETVMAADPVD | ASSQGAPL | 0.7590984 |
| PASAAPAETQP | ASSQGAPL | 0.5577062 |
| QQSSDGKNGVRAAEITGE | ASSQGAPL | 0.8249054 |
| RAPDAVTTSWWKE | ASSQGAPL | 0.8617266 |
| SGFGNVGAHTSGWFNQ | ASSQGAPL | 0.8483693 |
| SGVAGAVGQAASVGGLKV | ASSQGAPL | 0.35054976 |
| SNGGAGTGSAGGNGGT | ASSQGAPL | 0.199391 |
| TGGAGGEGGAAG | ASSQGAPL | 0.6614668 |
| TGGDGGAAGAGGAPGHG | ASSQGAPL | 0.79166865 |
| TMPPLLAAIANAGRHGV | ASSQGAPL | 0.67186236 |
| TSSTAGNSGGA | ASSQGAPL | 0.5563617 |
| VPATVPVGILC | ASSQGAPL | 0.7639352 |
| VQRPDSGGVVLTGRIS | ASSQGAPL | 0.832652 |
| WTPNPVPATKGVPAGLVQAT | ASSQGAPL | 0.5267253 |
| YALKRALAKFSDRFDVVII | ASSQGAPL | 0.38494244 |
| YLIGSVIV | ASSQGAPL | 0.7950052 |
| YGQLVAVID | ASSQGAPL | 0.7859897 |
| VTVGGMTLPA | ASSQGAPL | 0.54606795 |
| VLALPAV | ASSQGAPL | 0.62044704 |
| VIPPTLNYET | ASSQGAPL | 0.6896526 |
| VGMGGMHPSSG | ASSQGAPL | 0.64184785 |
| VGLAGGAATA | ASSQGAPL | 0.7496761 |
| VGGAGGAGGSGP | ASSQGAPL | 0.37093884 |
| VFAAFMV | ASSQGAPL | 0.8338466 |
| TLYDKLR | ASSQGAPL | 0.39116216 |
| TLAVFTVL | ASSQGAPL | 0.7851721 |
| TGSVQPARV | ASSQGAPL | 0.75491625 |
| TGGGAGSGGT | ASSQGAPL | 0.779485 |
| SHLETSL | ASSQGAPL | 0.6327516 |
| SFTTGMT | ASSQGAPL | 0.5169853 |
| QVEVAKRMGVRQPTVS | ASSQGAPL | 0.49184176 |
| QGSGGPALAM | ASSQGAPL | 0.31049082 |

| | | |
|---|---|---|
| MLAPSNP | ASSQGAPL | 0.8298017 |
| LVNNAAI | ASSQGAPL | 0.28631565 |
| LTGEINTLA | ASSQGAPL | 0.8337779 |
| LSIFGPV | ASSQGAPL | 0.76628387 |
| LLALLANL | ASSQGAPL | 0.55821365 |
| LISNASDALDKLR | ASSQGAPL | 0.7612944 |
| LINEAVP | ASSQGAPL | 0.42003995 |
| LGYAFSGL | ASSQGAPL | 0.66960454 |
| LDFVAVAL | ASSQGAPL | 0.7852744 |
| LAQPGAE | ASSQGAPL | 0.815755 |
| ISNLGAVTGF | ASSQGAPL | 0.5023315 |
| HQEDPGRI | ASSQGAPL | 0.77494085 |
| GTPGEPTP | ASSQGAPL | 0.76466095 |
| GFFNSGS | ASSQGAPL | 0.5424566 |
| GAGGGGLTAGAN | ASSQGAPL | 0.7654561 |
| FVDALVNM | ASSQGAPL | 0.81708694 |
| FLTVAPEL | ASSQGAPL | 0.7714324 |
| FETLTAL | ASSQGAPL | 0.40249676 |
| DVINPIEL | ASSQGAPL | 0.77709615 |
| APVSGAPV | ASSQGAPL | 0.5342051 |
| ALVGGLAW | ASSQGAPL | 0.75953597 |
| ALAAAAATAI | ASSQGAPL | 0.7469795 |
| AFGLGLLL | ASSQGAPL | 0.6526835 |
| AFFAAASL | ASSQGAPL | 0.81656337 |
| AAAALPAFSPP | ASSQGAPL | 0.5078758 |
| VVLAPAL | ASSQGAPL | 0.5178211 |
| TIGHVDHGKTTL | ASSQGAPL | 0.66852564 |
| LISNASDALDK | ASSQGAPL | 0.71065617 |
| LISNASDAL | ASSQGAPL | 0.8081515 |
| ISNASDALDK | ASSQGAPL | 0.50145406 |
| NQAETLVYQTE | ASSQGAPL | 0.7217874 |
| TGGTGGTAGAG | ASSQGAPL | 0.68892324 |
| TGSTAGGAGGA | ASSQGAPL | 0.53550863 |
| DPLAVPVAAGPV | ASSQGAPL | 0.72159743 |
| GGTGGTPGNGGN | ASSQGAPL | 0.6770169 |
| PAGLAGVDPATL | ASSQGAPL | 0.43060207 |
| GNNLVVEAPHAGASV | ASSQGAPL | 0.8140274 |
| KVTRIMSASTLEQPA | ASSQGAPL | 0.5703292 |
| LGFALPIALPRRTVV | ASSQGAPL | 0.7609443 |
| TRLGNLTARKLVVLV | ASSQGAPL | 0.8558674 |
| WSQQRRLIYFPSAGP | ASSQGAPL | 0.4888298 |
| GPNGNPDPMAAAVDIR | ASSQGAPL | 0.7719425 |
| TQRCVSVADLRAAASR | ASSQGAPL | 0.7382781 |
| QFVLDVAEHPVEVLTETV | ASSQGAPL | 0.76787007 |
| GGAGGTGGAGGNGGTQAGDG | ASSQGAPL | 0.6861297 |
| PLLATLRAIAVDVDAKPAQV | ASSQGAPL | 0.59940785 |
| APKFRTGGYMIDFWGVGYQVAK | ASSQGAPL | 0.84449977 |
| GAGGAGGAGGAAGGPGGTGGTG | ASSQGAPL | 0.7909347 |
| IGALGLLATDVAPAGSVAFGVA | ASSQGAPL | 0.80418706 |
| AGGAGGIGGNGGAGGAGGVLMG | ASSQGAPL | 0.8160076 |
| ADELIGLL | ASSQGAPL | 0.78215635 |

| | | |
|---|---|---|
| ADRALKA | ASSQGAPL | 0.80745983 |
| AGAVGQVGGAGGA | ASSQGAPL | 0.5738667 |
| AGGGGGLGGAP | ASSQGAPL | 0.35030794 |
| AGGLLYGNGGAG | ASSQGAPL | 0.6523623 |
| AGLESIGL | ASSQGAPL | 0.7137706 |
| AGLKTGII | ASSQGAPL | 0.60672915 |
| ASLAALLL | ASSQGAPL | 0.3488584 |
| ASVINISSI | ASSQGAPL | 0.8267868 |
| ATPPGPA | ASSQGAPL | 0.8655279 |
| ATVASALV | ASSQGAPL | 0.52823025 |
| ATVASLIL | ASSQGAPL | 0.86380476 |
| AVNGPVLIRH | ASSQGAPL | 0.42673627 |
| DGAAGAVGI | ASSQGAPL | 0.79096043 |
| DPLLHGI | ASSQGAPL | 0.83703685 |
| ELGAALGAL | ASSQGAPL | 0.45759588 |
| ELGRTQAV | ASSQGAPL | 0.7957572 |
| FVAGGYLS | ASSQGAPL | 0.55173844 |
| GASGGAGGAGGW | ASSQGAPL | 0.635265 |
| GGAGAGGDGGAGGI | ASSQGAPL | 0.8368478 |
| GTGHNGSQPGMGGNG | ASSQGAPL | 0.64822024 |
| GYAAASASAS | ASSQGAPL | 0.8004106 |
| HQVDVAAVQQG | ASSQGAPL | 0.65957075 |
| IGAAIAHT | ASSQGAPL | 0.5295767 |
| IGMIEVAQL | ASSQGAPL | 0.7755244 |
| IRLAPEHTSAMD | ASSQGAPL | 0.83332837 |
| IYNVSVL | ASSQGAPL | 0.6386589 |
| KQGFLGM | ASSQGAPL | 0.85978556 |
| LESVITQL | ASSQGAPL | 0.5339649 |
| LGLGALVVV | ASSQGAPL | 0.8580153 |
| LIGNGGNGGSGGI | ASSQGAPL | 0.8182038 |
| LNAINAPTL | ASSQGAPL | 0.70205104 |
| LQLLNTGV | ASSQGAPL | 0.656398 |
| LQTSIEF | ASSQGAPL | 0.80553377 |
| MSFAVSEL | ASSQGAPL | 0.61979675 |
| NTGALGL | ASSQGAPL | 0.76961726 |
| PPGAGGAGGAGTT | ASSQGAPL | 0.844016 |
| QLVGLGI | ASSQGAPL | 0.35517183 |
| RFLEQVHQL | ASSQGAPL | 0.8411316 |
| RGVGYVLR | ASSQGAPL | 0.69603956 |
| SATGVFAGVI | ASSQGAPL | 0.7266492 |
| SLPAITLP | ASSQGAPL | 0.4504857 |
| STALAFTV | ASSQGAPL | 0.30267 |
| SVLTVGLGK | ASSQGAPL | 0.673096 |
| TALAGAGSGI | ASSQGAPL | 0.31889245 |
| TGLPFTGI | ASSQGAPL | 0.8040197 |
| VAEHVTV | ASSQGAPL | 0.7999257 |
| VDIVLNSL | ASSQGAPL | 0.46851307 |
| VLTGGGAVL | ASSQGAPL | 0.76031053 |
| VNVGLAYGI | ASSQGAPL | 0.7993206 |
| VSSAAVGAV | ASSQGAPL | 0.32507226 |
| VTSVYEAGV | ASSQGAPL | 0.59630716 |

| | | |
|---|---|---|
| WVALDRGA | ASSQGAPL | 0.5121777 |
| YLVGSVVI | ASSQGAPL | 0.6054221 |
| GPPGTGKTLL | ASSQGAPL | 0.2396272 |
| GVGMTKF | ASSQGAPL | 0.7439332 |
| IGIGGSDL | ASSQGAPL | 0.48494732 |
| IGTLLAEL | ASSQGAPL | 0.14132795 |
| IYTSGTT | ASSQGAPL | 0.7579852 |
| LSFPAGV | ASSQGAPL | 0.565355 |
| LYTSGTT | ASSQGAPL | 0.48507485 |
| VGAFPFGF | ASSQGAPL | 0.7873405 |
| VGAGTVEF | ASSQGAPL | 0.6342598 |
| VGKSTLLL | ASSQGAPL | 0.39847285 |
| AALAHAGGGIV | ASSQGAPL | 0.730883 |
| AALVAVSLL | ASSQGAPL | 0.8250622 |
| AAVSRAVAAL | ASSQGAPL | 0.3217434 |
| AGGSSGIGGF | ASSQGAPL | 0.6943894 |
| AGLTVANANG | ASSQGAPL | 0.85467595 |
| AGLVGAVL | ASSQGAPL | 0.6383589 |
| ALSQAFTV | ASSQGAPL | 0.8611592 |
| ALTFSVSL | ASSQGAPL | 0.86827904 |
| ALTGAALVAL | ASSQGAPL | 0.58059275 |
| ASFSIGAGVL | ASSQGAPL | 0.8145927 |
| DGSVYKI | ASSQGAPL | 0.8159968 |
| EQLAEIV | ASSQGAPL | 0.6233215 |
| FAVSELTVM | ASSQGAPL | 0.6321132 |
| GAAPLAAGL | ASSQGAPL | 0.3789484 |
| GADGGDGGSGGAGG | ASSQGAPL | 0.69873977 |
| GFGLDTGGAGG | ASSQGAPL | 0.27671963 |
| GLASGGLL | ASSQGAPL | 0.71109337 |
| GRIAGAAVLL | ASSQGAPL | 0.83937275 |
| GSAGSLNTSF | ASSQGAPL | 0.5907614 |
| GTGQIGI | ASSQGAPL | 0.30194098 |
| GTLIALIL | ASSQGAPL | 0.30740404 |
| IAATMGFVVG | ASSQGAPL | 0.7433777 |
| IFAQALL | ASSQGAPL | 0.3792327 |
| IGAAVALGF | ASSQGAPL | 0.34035504 |
| IGLTPLAL | ASSQGAPL | 0.57327783 |
| ILGSNTSTL | ASSQGAPL | 0.7913359 |
| IVFSLTVL | ASSQGAPL | 0.8183975 |
| LASKATGF | ASSQGAPL | 0.31666583 |
| LASVALIL | ASSQGAPL | 0.8404813 |
| LFAVGLTL | ASSQGAPL | 0.8159164 |
| LFNGGIGTYI | ASSQGAPL | 0.8564712 |
| LTLGAGII | ASSQGAPL | 0.6824171 |
| LVAAALGDI | ASSQGAPL | 0.39100507 |
| NTAFVQLGI | ASSQGAPL | 0.8137957 |
| QLFSGAEL | ASSQGAPL | 0.50050974 |
| SAATFITL | ASSQGAPL | 0.70784956 |
| SNYGLANLGV | ASSQGAPL | 0.80696845 |
| SYGGAYAVM | ASSQGAPL | 0.44834977 |
| TTGVGGLTLG | ASSQGAPL | 0.37537307 |

| | | |
|---|---|---|
| TTQTIYI | ASSQGAPL | 0.6882478 |
| VAAGAGLGAVI | ASSQGAPL | 0.8630618 |
| VAAVVTLL | ASSQGAPL | 0.35957333 |
| VGFLGGLI | ASSQGAPL | 0.383096 |
| VGVIPILL | ASSQGAPL | 0.6082306 |
| VIFASSLI | ASSQGAPL | 0.6151649 |
| VKGGDGGAAAG | ASSQGAPL | 0.8315171 |
| VTKTFTNI | ASSQGAPL | 0.76176906 |
| VVAAAILL | ASSQGAPL | 0.6705538 |
| YAALDVEL | ASSQGAPL | 0.75419474 |
| LASVALLL | ASSQGAPL | 0.7880345 |
| LDAVHRV | ASSQGAPL | 0.68965214 |
| QGLSSSIF | ASSQGAPL | 0.7281747 |
| RGTFANIRL | ASSQGAPL | 0.38665342 |
| RQLWWGHRI | ASSQGAPL | 0.8592502 |
| YGPPGTGKTLL | ASSQGAPL | 0.7675895 |
| AALVGAPANVA | ASSQGAPL | 0.81501377 |
| AATFRRTE | ASSQGAPL | 0.85401314 |
| AGALGGGI | ASSQGAPL | 0.7400251 |
| AGGAGGAGGIGAG | ASSQGAPL | 0.7358594 |
| AGSGALGL | ASSQGAPL | 0.28604722 |
| AQIADQI | ASSQGAPL | 0.52769697 |
| ASAVAVTYL | ASSQGAPL | 0.655177 |
| ASTVLGGTGG | ASSQGAPL | 0.6979219 |
| AVGGEPAK | ASSQGAPL | 0.7778946 |
| CEQGTFGGTC | ASSQGAPL | 0.8112157 |
| DEGKAPAG | ASSQGAPL | 0.5651058 |
| DTVVFPAL | ASSQGAPL | 0.8346156 |
| FEGITMAG | ASSQGAPL | 0.83508307 |
| FPLVVQD | ASSQGAPL | 0.31430122 |
| GAFTTPQ | ASSQGAPL | 0.630152 |
| GFGGFGTAGGVGGA | ASSQGAPL | 0.864511 |
| GFLAQFL | ASSQGAPL | 0.76399493 |
| GGAGGAGGNAGVA | ASSQGAPL | 0.53198063 |
| GGVGTAGAGGAGGAGG | ASSQGAPL | 0.6341933 |
| GNGGGGDAGNAGSGGN | ASSQGAPL | 0.8495897 |
| GTGGAGGAGNDAGS | ASSQGAPL | 0.64904976 |
| GTTNTGWFNS | ASSQGAPL | 0.8448572 |
| IFMGPLFTL | ASSQGAPL | 0.19821873 |
| ILAHAIEAI | ASSQGAPL | 0.63950706 |
| IRSAIREAL | ASSQGAPL | 0.5498512 |
| ITFAGGG | ASSQGAPL | 0.16678475 |
| ITGMSGHQEV | ASSQGAPL | 0.79823947 |
| IVAGLACTL | ASSQGAPL | 0.5521371 |
| IVGGGLAGA | ASSQGAPL | 0.7978889 |
| IVTGVLNI | ASSQGAPL | 0.49682727 |
| LAQQGCI | ASSQGAPL | 0.8196034 |
| LCLHVEN | ASSQGAPL | 0.71644044 |
| LDAISDGS | ASSQGAPL | 0.68915313 |
| LGGFAAML | ASSQGAPL | 0.53837675 |
| LGGKNPMI | ASSQGAPL | 0.6660569 |

| | | |
|---|---|---|
| LGPKGVAAIL | ASSQGAPL | 0.84275496 |
| LMVGVSI | ASSQGAPL | 0.8604288 |
| LVGAAFVL | ASSQGAPL | 0.24566802 |
| LVLAVAPL | ASSQGAPL | 0.74052763 |
| MAAIAVIDAA | ASSQGAPL | 0.5928886 |
| NNGINAELQ | ASSQGAPL | 0.21243449 |
| RGEPLVLVHG | ASSQGAPL | 0.21701415 |
| RGGAGGAAGLI | ASSQGAPL | 0.510671 |
| RSYRMHPAV | ASSQGAPL | 0.6440948 |
| SAAHLLAGQLA | ASSQGAPL | 0.7355758 |
| SAAVGSY | ASSQGAPL | 0.7689965 |
| SPHAVLHRI | ASSQGAPL | 0.4458353 |
| SQIAKVL | ASSQGAPL | 0.76723385 |
| TALGHLVP | ASSQGAPL | 0.6214365 |
| TAPTTSGAA | ASSQGAPL | 0.078722596 |
| TDGDDGNGGSPGT | ASSQGAPL | 0.56386375 |
| TDGLGEN | ASSQGAPL | 0.31777057 |
| TGGPPLPGQA | ASSQGAPL | 0.8595135 |
| TIPAITI | ASSQGAPL | 0.39997232 |
| TIPGFTI | ASSQGAPL | 0.5296768 |
| TSIASGALA | ASSQGAPL | 0.5789252 |
| TVTLPGAAI | ASSQGAPL | 0.74571383 |
| VASGLSAALP | ASSQGAPL | 0.6975641 |
| VGAGVIGI | ASSQGAPL | 0.4759946 |
| VGPKRFGV | ASSQGAPL | 0.16951677 |
| VNVAFPD | ASSQGAPL | 0.7309489 |
| VSMTNSGVS | ASSQGAPL | 0.52315116 |
| VTDVLGDLSV | ASSQGAPL | 0.42881456 |
| VVVVGAGFAGL | ASSQGAPL | 0.44196242 |
| VYTVGTVV | ASSQGAPL | 0.39135712 |
| WTTFSGP | ASSQGAPL | 0.26041842 |
| YDELVEAF | ASSQGAPL | 0.8415242 |
| YGTGGAGGNGG | ASSQGAPL | 0.48797256 |
| YTNVMAV | ASSQGAPL | 0.5039375 |
| FLEQVHQL | ASSQGAPL | 0.8574859 |
| GLSLLAVAV | ASSQGAPL | 0.6956255 |
| LEQVHQL | ASSQGAPL | 0.5770445 |
| LNAAGAI | ASSQGAPL | 0.47137517 |
| PAVHASL | ASSQGAPL | 0.43997955 |
| TELEDDA | ASSQGAPL | 0.39329344 |
| TPPGPPN | ASSQGAPL | 0.4216696 |
| TPPPPPPP | ASSQGAPL | 0.8251221 |
| TQSPALI | ASSGWVGAETLY | 0.69341207 |
| EFAAAFPQ | ASSGWVGAETLY | 0.37970966 |
| LLSGNLDVL | ASSGWVGAETLY | 0.83488035 |
| GPANLGNYNI | ASSGWVGAETLY | 0.82540053 |
| TLGPTLP | ASSGWVGAETLY | 0.8585738 |
| VYAYSSCCK | ASSGWVGAETLY | 0.46186602 |
| APAPAPAPNI | ASSGWVGAETLY | 0.59412855 |
| PGGPVPIP | ASSGWVGAETLY | 0.6574606 |
| FLTVALEP | ASSGWVGAETLY | 0.61128694 |

| | | |
|---|---|---|
| SQYFGANL | ASSGWVGAETLY | 0.53296936 |
| GGVKPAA | ASSGWVGAETLY | 0.36013067 |
| GGTGGAGGAGGW | ASSGWVGAETLY | 0.84304166 |
| GVGGVAGLGGAGG | ASSGWVGAETLY | 0.88031507 |
| VLLVLATTI | ASSGWVGAETLY | 0.6374443 |
| IGIFAAML | ASSGWVGAETLY | 0.22435147 |
| AVGDLID | ASSGWVGAETLY | 0.54805285 |
| NISPLPL | ASSGWVGAETLY | 0.38127962 |
| VTAVFVAI | ASSGWVGAETLY | 0.4645087 |
| SGDDNDAAL | ASSGWVGAETLY | 0.6436838 |
| AVAALAHS | ASSGWVGAETLY | 0.2822627 |
| GGVTVMATP | ASSGWVGAETLY | 0.606447 |
| AGLAANVRD | ASSGWVGAETLY | 0.84984994 |
| TNRPDLIDPALLRPG | ASSGWVGAETLY | 0.5791641 |
| LANLGVTAL | ASSGWVGAETLY | 0.74216473 |
| TGERIEL | ASSGWVGAETLY | 0.80969524 |
| NGEELAA | ASSGWVGAETLY | 0.7605283 |
| VLSLVWG | ASSGWVGAETLY | 0.84537005 |
| IGYALWL | ASSGWVGAETLY | 0.83155656 |
| LSLLLAPL | ASSGWVGAETLY | 0.7227745 |
| TLDEALK | ASSGWVGAETLY | 0.72299755 |
| TDGAGTAEGG | ASSGWVGAETLY | 0.78159213 |
| NAPGGAGGNGG | ASSGWVGAETLY | 0.612242 |
| VSVFMGMGGNF | ASSGWVGAETLY | 0.6229961 |
| LASYLAAL | ASSGWVGAETLY | 0.6630688 |
| LVGGTPAL | ASSGWVGAETLY | 0.7424278 |
| VRNADVL | ASSGWVGAETLY | 0.67735064 |
| TCSSPFLVSAM | ASSGWVGAETLY | 0.8442771 |
| VHITDAE | ASSGWVGAETLY | 0.20067996 |
| EYAAPSF | ASSGWVGAETLY | 0.35764143 |
| VVDAFHGGL | ASSGWVGAETLY | 0.82430947 |
| ASVVDPQ | ASSGWVGAETLY | 0.14621024 |
| SYNEFGGLS | ASSGWVGAETLY | 0.31367674 |
| DPDGSVY | ASSGWVGAETLY | 0.7305199 |
| MVSTIHATE | ASSGWVGAETLY | 0.8151543 |
| LAAGVADPV | ASSGWVGAETLY | 0.15920961 |
| TKEQIAA | ASSGWVGAETLY | 0.5315888 |
| SLASPAAL | ASSGWVGAETLY | 0.34862262 |
| IGLVFYGV | ASSGWVGAETLY | 0.85178924 |
| FLLDASNL | ASSGWVGAETLY | 0.5621532 |
| YQVGNFP | ASSGWVGAETLY | 0.27013952 |
| ESLLSQST | ASSGWVGAETLY | 0.79197526 |
| LFTVGFAAVG | ASSGWVGAETLY | 0.29785722 |
| ATPPTPT | ASSGWVGAETLY | 0.6324277 |
| TDLATLG | ASSGWVGAETLY | 0.8181029 |
| SAGGTGR | ASSGWVGAETLY | 0.8755721 |
| GFSNSGS | ASSGWVGAETLY | 0.34788057 |
| GGNGGSGGTSV | ASSGWVGAETLY | 0.79100585 |
| GTSVATGGAGN | ASSGWVGAETLY | 0.5515536 |
| NGGNGGSGGTS | ASSGWVGAETLY | 0.86716694 |
| AVVKFPH | ASSGWVGAETLY | 0.5167641 |

| | | |
|---|---|---|
| TPPPPPP | ASSGWVGAETLY | 0.28573063 |
| AAPAAAAAPA | ASSGWVGAETLY | 0.8002597 |
| IALNEDL | ASSGWVGAETLY | 0.806343 |
| IVNVSSM | ASSGWVGAETLY | 0.6496026 |
| TGIPPAPR | ASSGWVGAETLY | 0.80469894 |
| FDLGGGTF | ASSGWVGAETLY | 0.68641984 |
| FDLGGGTFDVS | ASSGWVGAETLY | 0.5700153 |
| NEPTAAA | ASSGWVGAETLY | 0.7786379 |
| NEPTAAALA | ASSGWVGAETLY | 0.76440656 |
| GSIINISSI | ASSGWVGAETLY | 0.39105812 |
| VAPGNAGTA | ASSGWVGAETLY | 0.834125 |
| TGGGASLE | ASSGWVGAETLY | 0.57445323 |
| VAVYNHYKRI | ASSGWVGAETLY | 0.4334509 |
| TDPAPATT | ASSGWVGAETLY | 0.8648732 |
| TNRPDILDPALLRPG | ASSGWVGAETLY | 0.7230931 |
| LTTTTTS | ASSGWVGAETLY | 0.78289926 |
| PTGSGKTL | ASSGWVGAETLY | 0.74040335 |
| TGSGKTL | ASSGWVGAETLY | 0.8590772 |
| TIGHVDHGK | ASSGWVGAETLY | 0.34197557 |
| TIGHVDHGKTT | ASSGWVGAETLY | 0.70532787 |
| GHVDHGKTT | ASSGWVGAETLY | 0.6745662 |
| AQFEHTLL | ASSGWVGAETLY | 0.8218591 |
| IVNQDEL | ASSGWVGAETLY | 0.7034412 |
| TNGSQFF | ASSGWVGAETLY | 0.5126593 |
| TGKGIVE | ASSGWVGAETLY | 0.771963 |
| LISNASDA | ASSGWVGAETLY | 0.82012796 |
| AELRTLGELTKIAKAHGAQ | ASSGWVGAETLY | 0.7038485 |
| APNAVAAAEPSW | ASSGWVGAETLY | 0.6774075 |
| APQVAGAATAVL | ASSGWVGAETLY | 0.8673757 |
| ARRDPAAAAAILPTDA | ASSGWVGAETLY | 0.39429742 |
| DAAARDVLAEAGLAEYF | ASSGWVGAETLY | 0.593944 |
| DDQIVLSAKANLAKVEQ | ASSGWVGAETLY | 0.8313135 |
| DELLPDTGEDSDEDGH | ASSGWVGAETLY | 0.8296977 |
| ELPEAPANRATVPQD | ASSGWVGAETLY | 0.66257423 |
| FGQSKIERVTGIG | ASSGWVGAETLY | 0.83777916 |
| GAGGTGSTAGGAGGAGGAGGLY | ASSGWVGAETLY | 0.61669457 |
| GHGGTGAAGGNGGTGGAGGSGI | ASSGWVGAETLY | 0.7493281 |
| GNGGNGGAGGEGGPGGAGGAGA | ASSGWVGAETLY | 0.6155498 |
| GSTAGNGGNGGNATNGTIAGQP | ASSGWVGAETLY | 0.5983827 |
| GTGGDGGNGGNGGTGG | ASSGWVGAETLY | 0.33335644 |
| GVRRPAGIPADAPV | ASSGWVGAETLY | 0.6729797 |
| GYRQVTDAHLLTLARR | ASSGWVGAETLY | 0.7633637 |
| IAKMGQDGHDRG | ASSGWVGAETLY | 0.71912456 |
| KAEPSIEIQYT | ASSGWVGAETLY | 0.21349886 |
| KPVRIRMAADEDTVAVT | ASSGWVGAETLY | 0.72551084 |
| MPPRIAGMRLLVIKPEP | ASSGWVGAETLY | 0.7764616 |
| MKFYAVALFRSRGVPPTR | ASSGWVGAETLY | 0.87581766 |
| NAGGFGILGAVAHSP | ASSGWVGAETLY | 0.69028914 |
| NEGREITLAMAEM | ASSGWVGAETLY | 0.81454504 |
| NELVEWSHPLGHRWRWE | ASSGWVGAETLY | 0.5566663 |
| NLGAGGTGGNGGNPG | ASSGWVGAETLY | 0.73665106 |

| | | |
|---|---|---|
| NTEWPKWETVMAADPVD | ASSGWVGAETLY | 0.31152928 |
| PASAAPAETQP | ASSGWVGAETLY | 0.50015146 |
| QQSSDGKNGVRAAEITGE | ASSGWVGAETLY | 0.5758443 |
| RAPDAVTTSWWKE | ASSGWVGAETLY | 0.42204696 |
| SGFGNVGAHTSGWFNQ | ASSGWVGAETLY | 0.7710576 |
| SGVAGAVGQAASVGGLKV | ASSGWVGAETLY | 0.81311184 |
| SNGGAGTGSAGGNGGT | ASSGWVGAETLY | 0.7735108 |
| TGGAGGEGGAAG | ASSGWVGAETLY | 0.81155324 |
| TGGDGGAAGAGGAPGHG | ASSGWVGAETLY | 0.839387 |
| TMPPLLAAIANAGRHGV | ASSGWVGAETLY | 0.8396262 |
| TSSTAGNSGGA | ASSGWVGAETLY | 0.43567288 |
| VPATVPVGILC | ASSGWVGAETLY | 0.5699901 |
| VQRPDSGGVVLTGRIS | ASSGWVGAETLY | 0.26159653 |
| WTPNPVPATKGVPAGLVQAT | ASSGWVGAETLY | 0.5357812 |
| YALKRALAKFSDRFDVVII | ASSGWVGAETLY | 0.45671397 |
| YLIGSVIV | ASSGWVGAETLY | 0.65365714 |
| YGQLVAVID | ASSGWVGAETLY | 0.13057956 |
| VTVGGMTLPA | ASSGWVGAETLY | 0.7123195 |
| VLALPAV | ASSGWVGAETLY | 0.8172815 |
| VIPPTLNYET | ASSGWVGAETLY | 0.4587134 |
| VGMGGMHPSSG | ASSGWVGAETLY | 0.769431 |
| VGLAGGAATA | ASSGWVGAETLY | 0.7417593 |
| VGGAGGAGGSGP | ASSGWVGAETLY | 0.48949647 |
| VFAAFMV | ASSGWVGAETLY | 0.75551605 |
| TLYDKLR | ASSGWVGAETLY | 0.49059618 |
| TLAVFTVL | ASSGWVGAETLY | 0.8142507 |
| TGSVQPARV | ASSGWVGAETLY | 0.8331143 |
| TGGGAGSGGT | ASSGWVGAETLY | 0.72598386 |
| SHLETSL | ASSGWVGAETLY | 0.60800064 |
| SFTTGMT | ASSGWVGAETLY | 0.7248744 |
| QVEVAKRMGVRQPTVS | ASSGWVGAETLY | 0.3414295 |
| QGSGGPALAM | ASSGWVGAETLY | 0.6543162 |
| MLAPSNP | ASSGWVGAETLY | 0.17381383 |
| LVNNAAI | ASSGWVGAETLY | 0.6845317 |
| LTGEINTLA | ASSGWVGAETLY | 0.4456669 |
| LSIFGPV | ASSGWVGAETLY | 0.74000883 |
| LLALLANL | ASSGWVGAETLY | 0.5682211 |
| LISNASDALDKLR | ASSGWVGAETLY | 0.7778389 |
| LINEAVP | ASSGWVGAETLY | 0.39228213 |
| LGYAFSGL | ASSGWVGAETLY | 0.76303947 |
| LDFVAVAL | ASSGWVGAETLY | 0.72068083 |
| LAQPGAE | ASSGWVGAETLY | 0.28700587 |
| ISNLGAVTGF | ASSGWVGAETLY | 0.799628 |
| HQEDPGRI | ASSGWVGAETLY | 0.6980748 |
| GTPGEPTP | ASSGWVGAETLY | 0.61865735 |
| GFFNSGS | ASSGWVGAETLY | 0.8138714 |
| GAGGGGLTAGAN | ASSGWVGAETLY | 0.6906924 |
| FVDALVNM | ASSGWVGAETLY | 0.38704368 |
| FLTVAPEL | ASSGWVGAETLY | 0.51334333 |
| FETLTAL | ASSGWVGAETLY | 0.6741626 |
| DVINPIEL | ASSGWVGAETLY | 0.5380055 |

| | | |
|---|---|---|
| APVSGAPV | ASSGWVGAETLY | 0.14218426 |
| ALVGGLAW | ASSGWVGAETLY | 0.80977654 |
| ALAAAAATAI | ASSGWVGAETLY | 0.440702 |
| AFGLGLLL | ASSGWVGAETLY | 0.46442366 |
| AFFAAASL | ASSGWVGAETLY | 0.5334779 |
| AAAALPAFSPP | ASSGWVGAETLY | 0.80909884 |
| VVLAPAL | ASSGWVGAETLY | 0.6554263 |
| TIGHVDHGKTTL | ASSGWVGAETLY | 0.64156866 |
| LISNASDALDK | ASSGWVGAETLY | 0.7534733 |
| LISNASDAL | ASSGWVGAETLY | 0.6893418 |
| ISNASDALDK | ASSGWVGAETLY | 0.79667044 |
| NQAETLVYQTE | ASSGWVGAETLY | 0.6906738 |
| TGGTGGTAGAG | ASSGWVGAETLY | 0.7094976 |
| TGSTAGGAGGA | ASSGWVGAETLY | 0.79581225 |
| DPLAVPVAAGPV | ASSGWVGAETLY | 0.6179991 |
| GGTGGTPGNGGN | ASSGWVGAETLY | 0.4779287 |
| PAGLAGVDPATL | ASSGWVGAETLY | 0.48180565 |
| GNNLVVEAPHAGASV | ASSGWVGAETLY | 0.47681037 |
| KVTRIMSASTLEQPA | ASSGWVGAETLY | 0.4165444 |
| LGFALPIALPRRTVV | ASSGWVGAETLY | 0.7058543 |
| TRLGNLTARKLVVLV | ASSGWVGAETLY | 0.8728472 |
| WSQQRRLIYFPSAGP | ASSGWVGAETLY | 0.7489213 |
| GPNGNPDPMAAAVDIR | ASSGWVGAETLY | 0.31928828 |
| TQRCVSVADLRAAASR | ASSGWVGAETLY | 0.55751705 |
| QFVLDVAEHPVEVLTETV | ASSGWVGAETLY | 0.5368621 |
| GGAGGTGGAGGNGGTQAGDG | ASSGWVGAETLY | 0.769325 |
| PLLATLRAIAVDVDAKPAQV | ASSGWVGAETLY | 0.804343 |
| APKFRTGGYMIDFWGVGYQVAK | ASSGWVGAETLY | 0.8626406 |
| GAGGAGGAGGAAGGPGGTGGTG | ASSGWVGAETLY | 0.77734834 |
| IGALGLLATDVAPAGSVAFGVA | ASSGWVGAETLY | 0.44637638 |
| AGGAGGIGGNGGAGGAGGVLMG | ASSGWVGAETLY | 0.79624903 |
| ADELIGLL | ASSGWVGAETLY | 0.5996589 |
| ADRALKA | ASSGWVGAETLY | 0.69897103 |
| AGAVGQVGGAGGA | ASSGWVGAETLY | 0.7207705 |
| AGGGGGLGGAP | ASSGWVGAETLY | 0.79977226 |
| AGGLLYGNGGAG | ASSGWVGAETLY | 0.566605 |
| AGLESIGL | ASSGWVGAETLY | 0.6529256 |
| AGLKTGII | ASSGWVGAETLY | 0.3200242 |
| ASLAALLL | ASSGWVGAETLY | 0.82304454 |
| ASVINISSI | ASSGWVGAETLY | 0.31737632 |
| ATPPGPA | ASSGWVGAETLY | 0.50698435 |
| ATVASALV | ASSGWVGAETLY | 0.6368158 |
| ATVASLIL | ASSGWVGAETLY | 0.84620434 |
| AVNGPVLIRH | ASSGWVGAETLY | 0.84441566 |
| DGAAGAVGI | ASSGWVGAETLY | 0.48559278 |
| DPLLHGI | ASSGWVGAETLY | 0.43238118 |
| ELGAALGAL | ASSGWVGAETLY | 0.75199866 |
| ELGRTQAV | ASSGWVGAETLY | 0.3762101 |
| FVAGGYLS | ASSGWVGAETLY | 0.79701453 |
| GASGGAGGAGGW | ASSGWVGAETLY | 0.61493504 |
| GGAGAGGDGGAGGI | ASSGWVGAETLY | 0.5806178 |

| | | |
|---|---|---|
| GTGHNGSQPGMGGNG | ASSGWVGAETLY | 0.5524477 |
| GYAAASASAS | ASSGWVGAETLY | 0.7619914 |
| HQVDVAAVQQG | ASSGWVGAETLY | 0.55468184 |
| IGAAIAHT | ASSGWVGAETLY | 0.80556464 |
| IGMIEVAQL | ASSGWVGAETLY | 0.1558847 |
| IRLAPEHTSAMD | ASSGWVGAETLY | 0.8135711 |
| IYNVSVL | ASSGWVGAETLY | 0.3579768 |
| KQGFLGM | ASSGWVGAETLY | 0.52281064 |
| LESVITQL | ASSGWVGAETLY | 0.20791051 |
| LGLGALVVV | ASSGWVGAETLY | 0.6839221 |
| LIGNGGNGGSGGI | ASSGWVGAETLY | 0.73905873 |
| LNAINAPTL | ASSGWVGAETLY | 0.8325584 |
| LQLLNTGV | ASSGWVGAETLY | 0.714223 |
| LQTSIEF | ASSGWVGAETLY | 0.7300023 |
| MSFAVSEL | ASSGWVGAETLY | 0.8263401 |
| NTGALGL | ASSGWVGAETLY | 0.48750055 |
| PPGAGGAGGAGTT | ASSGWVGAETLY | 0.2424498 |
| QLVGLGI | ASSGWVGAETLY | 0.6904422 |
| RFLEQVHQL | ASSGWVGAETLY | 0.6926955 |
| RGVGYVLR | ASSGWVGAETLY | 0.84728086 |
| SATGVFAGVI | ASSGWVGAETLY | 0.77602553 |
| SLPAITLP | ASSGWVGAETLY | 0.29246154 |
| STALAFTV | ASSGWVGAETLY | 0.74909395 |
| SVLTVGLGK | ASSGWVGAETLY | 0.81549674 |
| TALAGAGSGI | ASSGWVGAETLY | 0.40211883 |
| TGLPFTGI | ASSGWVGAETLY | 0.5082351 |
| VAEHVTV | ASSGWVGAETLY | 0.5835062 |
| VDIVLNSL | ASSGWVGAETLY | 0.1917904 |
| VLTGGGAVL | ASSGWVGAETLY | 0.74098706 |
| VNVGLAYGI | ASSGWVGAETLY | 0.49084038 |
| VSSAAVGAV | ASSGWVGAETLY | 0.52147967 |
| VTSVYEAGV | ASSGWVGAETLY | 0.60682786 |
| WVALDRGA | ASSGWVGAETLY | 0.4685201 |
| YLVGSVVI | ASSGWVGAETLY | 0.47897452 |
| GPPGTGKTLL | ASSGWVGAETLY | 0.4595529 |
| GVGMTKF | ASSGWVGAETLY | 0.715446 |
| IGIGGSDL | ASSGWVGAETLY | 0.77648234 |
| IGTLLAEL | ASSGWVGAETLY | 0.57754385 |
| IYTSGTT | ASSGWVGAETLY | 0.7562429 |
| LSFPAGV | ASSGWVGAETLY | 0.7938489 |
| LYTSGTT | ASSGWVGAETLY | 0.28893095 |
| VGAFPFGF | ASSGWVGAETLY | 0.43588513 |
| VGAGTVEF | ASSGWVGAETLY | 0.78011775 |
| VGKSTLLL | ASSGWVGAETLY | 0.4482359 |
| AALAHAGGGIV | ASSGWVGAETLY | 0.83434796 |
| AALVAVSLL | ASSGWVGAETLY | 0.6398867 |
| AAVSRAVAAL | ASSGWVGAETLY | 0.6795654 |
| AGGSSGIGGF | ASSGWVGAETLY | 0.7094615 |
| AGLTVANANG | ASSGWVGAETLY | 0.87080556 |
| AGLVGAVL | ASSGWVGAETLY | 0.76081145 |
| ALSQAFTV | ASSGWVGAETLY | 0.6318462 |

| | | |
|---|---|---|
| ALTFSVSL | ASSGWVGAETLY | 0.7367077 |
| ALTGAALVAL | ASSGWVGAETLY | 0.42752397 |
| ASFSIGAGVL | ASSGWVGAETLY | 0.7669381 |
| DGSVYKI | ASSGWVGAETLY | 0.795997 |
| EQLAEIV | ASSGWVGAETLY | 0.305022 |
| FAVSELTVM | ASSGWVGAETLY | 0.570966 |
| GAAPLAAGL | ASSGWVGAETLY | 0.8628344 |
| GADGGDGGSGGAGG | ASSGWVGAETLY | 0.82859355 |
| GFGLDTGGAGG | ASSGWVGAETLY | 0.46804866 |
| GLASGGLL | ASSGWVGAETLY | 0.52628416 |
| GRIAGAAVLL | ASSGWVGAETLY | 0.462599 |
| GSAGSLNTSF | ASSGWVGAETLY | 0.61296767 |
| GTGQIGI | ASSGWVGAETLY | 0.35103443 |
| GTLIALIL | ASSGWVGAETLY | 0.7761736 |
| IAATMGFVVG | ASSGWVGAETLY | 0.5439262 |
| IFAQALL | ASSGWVGAETLY | 0.80901337 |
| IGAAVALGF | ASSGWVGAETLY | 0.6750705 |
| IGLTPLAL | ASSGWVGAETLY | 0.8083417 |
| ILGSNTSTL | ASSGWVGAETLY | 0.5677638 |
| IVFSLTVL | ASSGWVGAETLY | 0.5008426 |
| LASKATGF | ASSGWVGAETLY | 0.30695003 |
| LASVALIL | ASSGWVGAETLY | 0.28789383 |
| LFAVGLTL | ASSGWVGAETLY | 0.48192698 |
| LFNGGIGTYI | ASSGWVGAETLY | 0.50418437 |
| LTLGAGII | ASSGWVGAETLY | 0.19219422 |
| LVAAALGDI | ASSGWVGAETLY | 0.85691214 |
| NTAFVQLGI | ASSGWVGAETLY | 0.6254101 |
| QLFSGAEL | ASSGWVGAETLY | 0.71270144 |
| SAATFITL | ASSGWVGAETLY | 0.5955112 |
| SNYGLANLGV | ASSGWVGAETLY | 0.34664363 |
| SYGGAYAVM | ASSGWVGAETLY | 0.40446222 |
| TTGVGGLTLG | ASSGWVGAETLY | 0.7832905 |
| TTQTIYI | ASSGWVGAETLY | 0.35445344 |
| VAAGAGLGAVI | ASSGWVGAETLY | 0.27598363 |
| VAAVVTLL | ASSGWVGAETLY | 0.5212661 |
| VGFLGGLI | ASSGWVGAETLY | 0.59386706 |
| VGVIPILL | ASSGWVGAETLY | 0.8658285 |
| VIFASSLI | ASSGWVGAETLY | 0.8014338 |
| VKGGDGGAAAG | ASSGWVGAETLY | 0.7157234 |
| VTKTFTNI | ASSGWVGAETLY | 0.8686733 |
| VVAAAILL | ASSGWVGAETLY | 0.1587226 |
| YAALDVEL | ASSGWVGAETLY | 0.58282346 |
| LASVALLL | ASSGWVGAETLY | 0.8456989 |
| LDAVHRV | ASSGWVGAETLY | 0.72858006 |
| QGLSSSIF | ASSGWVGAETLY | 0.56472874 |
| RGTFANIRL | ASSGWVGAETLY | 0.8185075 |
| RQLWWGHRI | ASSGWVGAETLY | 0.3975104 |
| YGPPGTGKTLL | ASSGWVGAETLY | 0.57935834 |
| AALVGAPANVA | ASSGWVGAETLY | 0.62993544 |
| AATFRRTE | ASSGWVGAETLY | 0.8169068 |
| AGALGGGI | ASSGWVGAETLY | 0.7799421 |

| | | |
|---|---|---|
| AGGAGGAGGIGAG | ASSGWVGAETLY | 0.32703435 |
| AGSGALGL | ASSGWVGAETLY | 0.7808467 |
| AQIADQI | ASSGWVGAETLY | 0.8631313 |
| ASAVAVTYL | ASSGWVGAETLY | 0.53593326 |
| ASTVLGGTGG | ASSGWVGAETLY | 0.77421224 |
| AVGGEPAK | ASSGWVGAETLY | 0.7108028 |
| CEQGTFGGTC | ASSGWVGAETLY | 0.5260886 |
| DEGKAPAG | ASSGWVGAETLY | 0.43200567 |
| DTVVFPAL | ASSGWVGAETLY | 0.378442 |
| FEGITMAG | ASSGWVGAETLY | 0.7755034 |
| FPLVVQD | ASSGWVGAETLY | 0.36161804 |
| GAFTTPQ | ASSGWVGAETLY | 0.6891413 |
| GFGGFGTAGGVGGA | ASSGWVGAETLY | 0.631799 |
| GFLAQFL | ASSGWVGAETLY | 0.8830544 |
| GGAGGAGGNAGVA | ASSGWVGAETLY | 0.75335705 |
| GGVGTAGAGGAGGAGG | ASSGWVGAETLY | 0.75714403 |
| GNGGGGDAGNAGSGGN | ASSGWVGAETLY | 0.40458882 |
| GTGGAGGAGNDAGS | ASSGWVGAETLY | 0.76726454 |
| GTTNTGWFNS | ASSGWVGAETLY | 0.7292853 |
| IFMGPLFTL | ASSGWVGAETLY | 0.34788987 |
| ILAHAIEAI | ASSGWVGAETLY | 0.819984 |
| IRSAIREAL | ASSGWVGAETLY | 0.329253 |
| ITFAGGG | ASSGWVGAETLY | 0.7209387 |
| ITGMSGHQEV | ASSGWVGAETLY | 0.7653886 |
| IVAGLACTL | ASSGWVGAETLY | 0.82182795 |
| IVGGGLAGA | ASSGWVGAETLY | 0.6082847 |
| IVTGVLNI | ASSGWVGAETLY | 0.756117 |
| LAQQGCI | ASSGWVGAETLY | 0.3690794 |
| LCLHVEN | ASSGWVGAETLY | 0.81529236 |
| LDAISDGS | ASSGWVGAETLY | 0.82800823 |
| LGGFAAML | ASSGWVGAETLY | 0.7301626 |
| LGGKNPMI | ASSGWVGAETLY | 0.8762856 |
| LGPKGVAAIL | ASSGWVGAETLY | 0.80764765 |
| LMVGVSI | ASSGWVGAETLY | 0.31026185 |
| LVGAAFVL | ASSGWVGAETLY | 0.66955495 |
| LVLAVAPL | ASSGWVGAETLY | 0.8291743 |
| MAAIAVIDAA | ASSGWVGAETLY | 0.82512933 |
| NNGINAELQ | ASSGWVGAETLY | 0.78500533 |
| RGEPLVLVHG | ASSGWVGAETLY | 0.78443897 |
| RGGAGGAAGLI | ASSGWVGAETLY | 0.5947329 |
| RSYRMHPAV | ASSGWVGAETLY | 0.82993996 |
| SAAHLLAGQLA | ASSGWVGAETLY | 0.8988982 |
| SAAVGSY | ASSGWVGAETLY | 0.78009534 |
| SPHAVLHRI | ASSGWVGAETLY | 0.5222216 |
| SQIAKVL | ASSGWVGAETLY | 0.5384823 |
| TALGHLVP | ASSGWVGAETLY | 0.7591747 |
| TAPTTSGAA | ASSGWVGAETLY | 0.80324316 |
| TDGDDGNGGSPGT | ASSGWVGAETLY | 0.39123422 |
| TDGLGEN | ASSGWVGAETLY | 0.60688263 |
| TGGPPLPGQA | ASSGWVGAETLY | 0.6373634 |
| TIPAITI | ASSGWVGAETLY | 0.21462806 |

| | | |
|---|---|---|
| TIPGFTI | ASSGWVGAETLY | 0.79530114 |
| TSIASGALA | ASSGWVGAETLY | 0.83835125 |
| TVTLPGAAI | ASSGWVGAETLY | 0.7209902 |
| VASGLSAALP | ASSGWVGAETLY | 0.36809802 |
| VGAGVIGI | ASSGWVGAETLY | 0.8130065 |
| VGPKRFGV | ASSGWVGAETLY | 0.8263892 |
| VNVAFPD | ASSGWVGAETLY | 0.85122395 |
| VSMTNSGVS | ASSGWVGAETLY | 0.2863126 |
| VTDVLGDLSV | ASSGWVGAETLY | 0.15352191 |
| VVVVGAGFAGL | ASSGWVGAETLY | 0.6325575 |
| VYTVGTVV | ASSGWVGAETLY | 0.78902125 |
| WTTFSGP | ASSGWVGAETLY | 0.5066139 |
| YDELVEAF | ASSGWVGAETLY | 0.41820908 |
| YGTGGAGGNGG | ASSGWVGAETLY | 0.6116482 |
| YTNVMAV | ASSGWVGAETLY | 0.7876743 |
| FLEQVHQL | ASSGWVGAETLY | 0.38018486 |
| GLSLLAVAV | ASSGWVGAETLY | 0.38916624 |
| LEQVHQL | ASSGWVGAETLY | 0.71027374 |
| LNAAGAI | ASSGWVGAETLY | 0.82684064 |
| PAVHASL | ASSGWVGAETLY | 0.41867816 |
| TELEDDA | ASSGWVGAETLY | 0.5461893 |
| TPPGPPN | ASSGWVGAETLY | 0.6278792 |
| TPPPPPPP | ASSGWVGAETLY | 0.7680646 |
| TQSPALI | ASSQEGDRGYEQY | 0.65905255 |
| EFAAAFPQ | ASSQEGDRGYEQY | 0.35237604 |
| LLSGNLDVL | ASSQEGDRGYEQY | 0.82043505 |
| GPANLGNYNI | ASSQEGDRGYEQY | 0.32329294 |
| TLGPTLP | ASSQEGDRGYEQY | 0.822041 |
| VYAYSSCCK | ASSQEGDRGYEQY | 0.67616534 |
| APAPAPAPNI | ASSQEGDRGYEQY | 0.77118397 |
| PGGPVPIP | ASSQEGDRGYEQY | 0.50017464 |
| FLTVALEP | ASSQEGDRGYEQY | 0.41691157 |
| SQYFGANL | ASSQEGDRGYEQY | 0.35422313 |
| GGVKPAA | ASSQEGDRGYEQY | 0.3766334 |
| GGTGGAGGAGGW | ASSQEGDRGYEQY | 0.818472 |
| GVGGVAGLGGAGG | ASSQEGDRGYEQY | 0.20041943 |
| VLLVLATTI | ASSQEGDRGYEQY | 0.8017305 |
| IGIFAAML | ASSQEGDRGYEQY | 0.6846787 |
| AVGDLID | ASSQEGDRGYEQY | 0.47235984 |
| NISPLPL | ASSQEGDRGYEQY | 0.78804004 |
| VTAVFVAI | ASSQEGDRGYEQY | 0.29634532 |
| SGDDNDAAL | ASSQEGDRGYEQY | 0.77371955 |
| AVAALAHS | ASSQEGDRGYEQY | 0.7161245 |
| GGVTVMATP | ASSQEGDRGYEQY | 0.77773285 |
| AGLAANVRD | ASSQEGDRGYEQY | 0.54987454 |
| TNRPDLIDPALLRPG | ASSQEGDRGYEQY | 0.7677934 |
| LANLGVTAL | ASSQEGDRGYEQY | 0.7419256 |
| TGERIEL | ASSQEGDRGYEQY | 0.4426855 |
| NGEELAA | ASSQEGDRGYEQY | 0.7069712 |
| VLSLVWG | ASSQEGDRGYEQY | 0.7934729 |
| IGYALWL | ASSQEGDRGYEQY | 0.73650056 |

| | | |
|---|---|---|
| LSLLLAPL | ASSQEGDRGYEQY | 0.2771878 |
| TLDEALK | ASSQEGDRGYEQY | 0.69622976 |
| TDGAGTAEGG | ASSQEGDRGYEQY | 0.49313414 |
| NAPGGAGGNGG | ASSQEGDRGYEQY | 0.71252906 |
| VSVFMGMGGNF | ASSQEGDRGYEQY | 0.7199249 |
| LASYLAAL | ASSQEGDRGYEQY | 0.63983655 |
| LVGGTPAL | ASSQEGDRGYEQY | 0.7778022 |
| VRNADVL | ASSQEGDRGYEQY | 0.40365288 |
| TCSSPFLVSAM | ASSQEGDRGYEQY | 0.539639 |
| VHITDAE | ASSQEGDRGYEQY | 0.6013731 |
| EYAAPSF | ASSQEGDRGYEQY | 0.6437754 |
| VVDAFHGGL | ASSQEGDRGYEQY | 0.78151107 |
| ASVVDPQ | ASSQEGDRGYEQY | 0.365165 |
| SYNEFGGLS | ASSQEGDRGYEQY | 0.7297713 |
| DPDGSVY | ASSQEGDRGYEQY | 0.5864835 |
| MVSTIHATE | ASSQEGDRGYEQY | 0.39899555 |
| LAAGVADPV | ASSQEGDRGYEQY | 0.70498705 |
| TKEQIAA | ASSQEGDRGYEQY | 0.5645114 |
| SLASPAAL | ASSQEGDRGYEQY | 0.3542844 |
| IGLVFYGV | ASSQEGDRGYEQY | 0.8090167 |
| FLLDASNL | ASSQEGDRGYEQY | 0.40798002 |
| YQVGNFP | ASSQEGDRGYEQY | 0.7398845 |
| ESLLSQST | ASSQEGDRGYEQY | 0.8555794 |
| LFTVGFAAVG | ASSQEGDRGYEQY | 0.31434852 |
| ATPPTPT | ASSQEGDRGYEQY | 0.8173732 |
| TDLATLG | ASSQEGDRGYEQY | 0.69686055 |
| SAGGTGR | ASSQEGDRGYEQY | 0.69012547 |
| GFSNSGS | ASSQEGDRGYEQY | 0.4768229 |
| GGNGGSGGTSV | ASSQEGDRGYEQY | 0.46128032 |
| GTSVATGGAGN | ASSQEGDRGYEQY | 0.8339942 |
| NGGNGGSGGTS | ASSQEGDRGYEQY | 0.7135297 |
| AVVKFPH | ASSQEGDRGYEQY | 0.7412653 |
| TPPPPPP | ASSQEGDRGYEQY | 0.7902487 |
| AAPAAAAAPA | ASSQEGDRGYEQY | 0.7835742 |
| IALNEDL | ASSQEGDRGYEQY | 0.81609535 |
| IVNVSSM | ASSQEGDRGYEQY | 0.49244544 |
| TGIPPAPR | ASSQEGDRGYEQY | 0.26873118 |
| FDLGGGTF | ASSQEGDRGYEQY | 0.5128643 |
| FDLGGGTFDVS | ASSQEGDRGYEQY | 0.48765612 |
| NEPTAAA | ASSQEGDRGYEQY | 0.5154475 |
| NEPTAAALA | ASSQEGDRGYEQY | 0.3504489 |
| GSIINISSI | ASSQEGDRGYEQY | 0.78262043 |
| VAPGNAGTA | ASSQEGDRGYEQY | 0.8841335 |
| TGGGASLE | ASSQEGDRGYEQY | 0.4811488 |
| VAVYNHYKRI | ASSQEGDRGYEQY | 0.8802201 |
| TDPAPATT | ASSQEGDRGYEQY | 0.30942675 |
| TNRPDILDPALLRPG | ASSQEGDRGYEQY | 0.7487234 |
| LTTTTTS | ASSQEGDRGYEQY | 0.8149788 |
| PTGSGKTL | ASSQEGDRGYEQY | 0.3194465 |
| TGSGKTL | ASSQEGDRGYEQY | 0.6946709 |
| TIGHVDHGK | ASSQEGDRGYEQY | 0.36530483 |

| | | |
|---|---|---|
| TIGHVDHGKTT | ASSQEGDRGYEQY | 0.49561715 |
| GHVDHGKTT | ASSQEGDRGYEQY | 0.8013995 |
| AQFEHTLL | ASSQEGDRGYEQY | 0.49055964 |
| IVNQDEL | ASSQEGDRGYEQY | 0.7825891 |
| TNGSQFF | ASSQEGDRGYEQY | 0.5278329 |
| TGKGIVE | ASSQEGDRGYEQY | 0.41192803 |
| LISNASDA | ASSQEGDRGYEQY | 0.7581777 |
| AELRTLGELTKIAKAHGAQ | ASSQEGDRGYEQY | 0.79781663 |
| APNAVAAAEPSW | ASSQEGDRGYEQY | 0.7384006 |
| APQVAGAATAVL | ASSQEGDRGYEQY | 0.8949367 |
| ARRDPAAAAAILPTDA | ASSQEGDRGYEQY | 0.5177087 |
| DAAARDVLAEAGLAEYF | ASSQEGDRGYEQY | 0.8370793 |
| DDQIVLSAKANLAKVEQ | ASSQEGDRGYEQY | 0.85305583 |
| DELLPDTGEDSDEDGH | ASSQEGDRGYEQY | 0.67230964 |
| ELPEAPANRATVPQD | ASSQEGDRGYEQY | 0.637995 |
| FGQSKIERVTGIG | ASSQEGDRGYEQY | 0.79784966 |
| GAGGTGSTAGGAGGAGGAGGLY | ASSQEGDRGYEQY | 0.62813914 |
| GHGGTGAAGGNGGTGGAGGSGI | ASSQEGDRGYEQY | 0.7182753 |
| GNGGNGGAGGEGGPGGAGGAGA | ASSQEGDRGYEQY | 0.8519733 |
| GSTAGNGGNGGNATNGTIAGQP | ASSQEGDRGYEQY | 0.43861413 |
| GTGGDGGNGGNGGTGG | ASSQEGDRGYEQY | 0.87453246 |
| GVRRPAGIPADAPV | ASSQEGDRGYEQY | 0.7925485 |
| GYRQVTDAHLLTLARR | ASSQEGDRGYEQY | 0.709107 |
| IAKMGQDGHDRG | ASSQEGDRGYEQY | 0.5822245 |
| KAEPSIEIQYT | ASSQEGDRGYEQY | 0.2314633 |
| KPVRIRMAADEDTVAVT | ASSQEGDRGYEQY | 0.670629 |
| MPPRIAGMRLLVIKPEP | ASSQEGDRGYEQY | 0.48541957 |
| MKFYAVALFRSRGVPPTR | ASSQEGDRGYEQY | 0.8347236 |
| NAGGFGILGAVAHSP | ASSQEGDRGYEQY | 0.7551303 |
| NEGREITLAMAEM | ASSQEGDRGYEQY | 0.5410968 |
| NELVEWSHPLGHRWRWE | ASSQEGDRGYEQY | 0.8551041 |
| NLGAGGTGGNGGNPG | ASSQEGDRGYEQY | 0.8566579 |
| NTEWPKWETVMAADPVD | ASSQEGDRGYEQY | 0.36337793 |
| PASAAPAETQP | ASSQEGDRGYEQY | 0.6550311 |
| QQSSDGKNGVRAAEITGE | ASSQEGDRGYEQY | 0.5262035 |
| RAPDAVTTSWWKE | ASSQEGDRGYEQY | 0.71103567 |
| SGFGNVGAHTSGWFNQ | ASSQEGDRGYEQY | 0.1954469 |
| SGVAGAVGQAASVGGLKV | ASSQEGDRGYEQY | 0.8146509 |
| SNGGAGTGSAGGNGGT | ASSQEGDRGYEQY | 0.55848813 |
| TGGAGGEGGAAG | ASSQEGDRGYEQY | 0.121754184 |
| TGGDGGAAGAGGAPGHG | ASSQEGDRGYEQY | 0.73890364 |
| TMPPLLAAIANAGRHGV | ASSQEGDRGYEQY | 0.6779504 |
| TSSTAGNSGGA | ASSQEGDRGYEQY | 0.67432636 |
| VPATVPVGILC | ASSQEGDRGYEQY | 0.86320555 |
| VQRPDSGGVVLTGRIS | ASSQEGDRGYEQY | 0.39405218 |
| WTPNPVPATKGVPAGLVQAT | ASSQEGDRGYEQY | 0.40415773 |
| YALKRALAKFSDRFDVVII | ASSQEGDRGYEQY | 0.84669393 |
| YLIGSVIV | ASSQEGDRGYEQY | 0.86573267 |
| YGQLVAVID | ASSQEGDRGYEQY | 0.31213742 |
| VTVGGMTLPA | ASSQEGDRGYEQY | 0.8253464 |
| VLALPAV | ASSQEGDRGYEQY | 0.8623565 |

| | | |
|---|---|---|
| VIPPTLNYET | ASSQEGDRGYEQY | 0.73617446 |
| VGMGGMHPSSG | ASSQEGDRGYEQY | 0.839053 |
| VGLAGGAATA | ASSQEGDRGYEQY | 0.8448044 |
| VGGAGGAGGSGP | ASSQEGDRGYEQY | 0.33130336 |
| VFAAFMV | ASSQEGDRGYEQY | 0.86264145 |
| TLYDKLR | ASSQEGDRGYEQY | 0.87135786 |
| TLAVFTVL | ASSQEGDRGYEQY | 0.6030382 |
| TGSVQPARV | ASSQEGDRGYEQY | 0.75288236 |
| TGGGAGSGGT | ASSQEGDRGYEQY | 0.4202299 |
| SHLETSL | ASSQEGDRGYEQY | 0.63131577 |
| SFTTGMT | ASSQEGDRGYEQY | 0.21226296 |
| QVEVAKRMGVRQPTVS | ASSQEGDRGYEQY | 0.83577275 |
| QGSGGPALAM | ASSQEGDRGYEQY | 0.8368155 |
| MLAPSNP | ASSQEGDRGYEQY | 0.7317554 |
| LVNNAAI | ASSQEGDRGYEQY | 0.4594169 |
| LTGEINTLA | ASSQEGDRGYEQY | 0.44895983 |
| LSIFGPV | ASSQEGDRGYEQY | 0.64625126 |
| LLALLANL | ASSQEGDRGYEQY | 0.5285889 |
| LISNASDALDKLR | ASSQEGDRGYEQY | 0.18927397 |
| LINEAVP | ASSQEGDRGYEQY | 0.67235506 |
| LGYAFSGL | ASSQEGDRGYEQY | 0.84359396 |
| LDFVAVAL | ASSQEGDRGYEQY | 0.8415617 |
| LAQPGAE | ASSQEGDRGYEQY | 0.37340948 |
| ISNLGAVTGF | ASSQEGDRGYEQY | 0.88223314 |
| HQEDPGRI | ASSQEGDRGYEQY | 0.8074155 |
| GTPGEPTP | ASSQEGDRGYEQY | 0.85777575 |
| GFFNSGS | ASSQEGDRGYEQY | 0.68039423 |
| GAGGGGLTAGAN | ASSQEGDRGYEQY | 0.4997856 |
| FVDALVNM | ASSQEGDRGYEQY | 0.857931 |
| FLTVAPEL | ASSQEGDRGYEQY | 0.7271226 |
| FETLTAL | ASSQEGDRGYEQY | 0.456279 |
| DVINPIEL | ASSQEGDRGYEQY | 0.84990525 |
| APVSGAPV | ASSQEGDRGYEQY | 0.58340216 |
| ALVGGLAW | ASSQEGDRGYEQY | 0.55879307 |
| ALAAAAATAI | ASSQEGDRGYEQY | 0.759901 |
| AFGLGLLL | ASSQEGDRGYEQY | 0.80519116 |
| AFFAAASL | ASSQEGDRGYEQY | 0.5277295 |
| AAAALPAFSPP | ASSQEGDRGYEQY | 0.34388146 |
| VVLAPAL | ASSQEGDRGYEQY | 0.44592994 |
| TIGHVDHGKTTL | ASSQEGDRGYEQY | 0.71906763 |
| LISNASDALDK | ASSQEGDRGYEQY | 0.86084473 |
| LISNASDAL | ASSQEGDRGYEQY | 0.7230997 |
| ISNASDALDK | ASSQEGDRGYEQY | 0.6264472 |
| NQAETLVYQTE | ASSQEGDRGYEQY | 0.83620644 |
| TGGTGGTAGAG | ASSQEGDRGYEQY | 0.79079294 |
| TGSTAGGAGGA | ASSQEGDRGYEQY | 0.7355372 |
| DPLAVPVAAGPV | ASSQEGDRGYEQY | 0.720937 |
| GGTGGTPGNGGN | ASSQEGDRGYEQY | 0.6635066 |
| PAGLAGVDPATL | ASSQEGDRGYEQY | 0.83286357 |
| GNNLVVEAPHAGASV | ASSQEGDRGYEQY | 0.8053982 |
| KVTRIMSASTLEQPA | ASSQEGDRGYEQY | 0.7968629 |

| | | |
|---|---|---|
| LGFALPIALPRRTVV | ASSQEGDRGYEQY | 0.81130856 |
| TRLGNLTARKLVVLV | ASSQEGDRGYEQY | 0.6765271 |
| WSQQRRLIYFPSAGP | ASSQEGDRGYEQY | 0.74444216 |
| GPNGNPDPMAAAVDIR | ASSQEGDRGYEQY | 0.23560753 |
| TQRCVSVADLRAAASR | ASSQEGDRGYEQY | 0.45866022 |
| QFVLDVAEHPVEVLTETV | ASSQEGDRGYEQY | 0.77048004 |
| GGAGGTGGAGGNGGTQAGDG | ASSQEGDRGYEQY | 0.8064835 |
| PLLATLRAIAVDVDAKPAQV | ASSQEGDRGYEQY | 0.8318759 |
| APKFRTGGYMIDFWGVGYQVAK | ASSQEGDRGYEQY | 0.73509324 |
| GAGGAGGAGGAAGGPGGTGGTG | ASSQEGDRGYEQY | 0.5234691 |
| IGALGLLATDVAPAGSVAFGVA | ASSQEGDRGYEQY | 0.74373525 |
| AGGAGGIGGNGGAGGAGGVLMG | ASSQEGDRGYEQY | 0.86470795 |
| ADELIGLL | ASSQEGDRGYEQY | 0.4001985 |
| ADRALKA | ASSQEGDRGYEQY | 0.5898072 |
| AGAVGQVGGAGGA | ASSQEGDRGYEQY | 0.88948214 |
| AGGGGGLGGAP | ASSQEGDRGYEQY | 0.810486 |
| AGGLLYGNGGAG | ASSQEGDRGYEQY | 0.6634278 |
| AGLESIGL | ASSQEGDRGYEQY | 0.5244802 |
| AGLKTGII | ASSQEGDRGYEQY | 0.8674116 |
| ASLAALLL | ASSQEGDRGYEQY | 0.43933564 |
| ASVINISSI | ASSQEGDRGYEQY | 0.8596796 |
| ATPPGPA | ASSQEGDRGYEQY | 0.17904335 |
| ATVASALV | ASSQEGDRGYEQY | 0.5562475 |
| ATVASLIL | ASSQEGDRGYEQY | 0.34165883 |
| AVNGPVLIRH | ASSQEGDRGYEQY | 0.19390105 |
| DGAAGAVGI | ASSQEGDRGYEQY | 0.82060367 |
| DPLLHGI | ASSQEGDRGYEQY | 0.5137433 |
| ELGAALGAL | ASSQEGDRGYEQY | 0.8429767 |
| ELGRTQAV | ASSQEGDRGYEQY | 0.66297114 |
| FVAGGYLS | ASSQEGDRGYEQY | 0.8335607 |
| GASGGAGGAGGW | ASSQEGDRGYEQY | 0.8363521 |
| GGAGAGGDGGAGGI | ASSQEGDRGYEQY | 0.69101226 |
| GTGHNGSQPGMGGNG | ASSQEGDRGYEQY | 0.4575026 |
| GYAAASASAS | ASSQEGDRGYEQY | 0.76472276 |
| HQVDVAAVQQG | ASSQEGDRGYEQY | 0.7840934 |
| IGAAIAHT | ASSQEGDRGYEQY | 0.78385437 |
| IGMIEVAQL | ASSQEGDRGYEQY | 0.22166196 |
| IRLAPEHTSAMD | ASSQEGDRGYEQY | 0.6250727 |
| IYNVSVL | ASSQEGDRGYEQY | 0.7060152 |
| KQGFLGM | ASSQEGDRGYEQY | 0.16599558 |
| LESVITQL | ASSQEGDRGYEQY | 0.13507377 |
| LGLGALVVV | ASSQEGDRGYEQY | 0.46360856 |
| LIGNGGNGGSGGI | ASSQEGDRGYEQY | 0.67711294 |
| LNAINAPTL | ASSQEGDRGYEQY | 0.85137695 |
| LQLLNTGV | ASSQEGDRGYEQY | 0.85950327 |
| LQTSIEF | ASSQEGDRGYEQY | 0.6761693 |
| MSFAVSEL | ASSQEGDRGYEQY | 0.8170565 |
| NTGALGL | ASSQEGDRGYEQY | 0.7261815 |
| PPGAGGAGGAGTT | ASSQEGDRGYEQY | 0.09499664 |
| QLVGLGI | ASSQEGDRGYEQY | 0.58117616 |
| RFLEQVHQL | ASSQEGDRGYEQY | 0.38487127 |

| | | |
|---|---|---|
| RGVGYVLR | ASSQEGDRGYEQY | 0.8226038 |
| SATGVFAGVI | ASSQEGDRGYEQY | 0.477309 |
| SLPAITLP | ASSQEGDRGYEQY | 0.60580134 |
| STALAFTV | ASSQEGDRGYEQY | 0.7806568 |
| SVLTVGLGK | ASSQEGDRGYEQY | 0.8108435 |
| TALAGAGSGI | ASSQEGDRGYEQY | 0.75890267 |
| TGLPFTGI | ASSQEGDRGYEQY | 0.4994384 |
| VAEHVTV | ASSQEGDRGYEQY | 0.13672753 |
| VDIVLNSL | ASSQEGDRGYEQY | 0.7962774 |
| VLTGGGAVL | ASSQEGDRGYEQY | 0.30235463 |
| VNVGLAYGI | ASSQEGDRGYEQY | 0.38276997 |
| VSSAAVGAV | ASSQEGDRGYEQY | 0.18342653 |
| VTSVYEAGV | ASSQEGDRGYEQY | 0.18918034 |
| WVALDRGA | ASSQEGDRGYEQY | 0.255148 |
| YLVGSVVI | ASSQEGDRGYEQY | 0.80260456 |
| GPPGTGKTLL | ASSQEGDRGYEQY | 0.75141215 |
| GVGMTKF | ASSQEGDRGYEQY | 0.6903697 |
| IGIGGSDL | ASSQEGDRGYEQY | 0.8840659 |
| IGTLLAEL | ASSQEGDRGYEQY | 0.81849265 |
| IYTSGTT | ASSQEGDRGYEQY | 0.73055327 |
| LSFPAGV | ASSQEGDRGYEQY | 0.44789782 |
| LYTSGTT | ASSQEGDRGYEQY | 0.5014757 |
| VGAFPFGF | ASSQEGDRGYEQY | 0.44067097 |
| VGAGTVEF | ASSQEGDRGYEQY | 0.43693227 |
| VGKSTLLL | ASSQEGDRGYEQY | 0.8728808 |
| AALAHAGGGIV | ASSQEGDRGYEQY | 0.5427689 |
| AALVAVSLL | ASSQEGDRGYEQY | 0.3836801 |
| AAVSRAVAAL | ASSQEGDRGYEQY | 0.68376637 |
| AGGSSGIGGF | ASSQEGDRGYEQY | 0.8715668 |
| AGLTVANANG | ASSQEGDRGYEQY | 0.8896514 |
| AGLVGAVL | ASSQEGDRGYEQY | 0.48195356 |
| ALSQAFTV | ASSQEGDRGYEQY | 0.58419204 |
| ALTFSVSL | ASSQEGDRGYEQY | 0.5988162 |
| ALTGAALVAL | ASSQEGDRGYEQY | 0.7800389 |
| ASFSIGAGVL | ASSQEGDRGYEQY | 0.43241256 |
| DGSVYKI | ASSQEGDRGYEQY | 0.3501908 |
| EQLAEIV | ASSQEGDRGYEQY | 0.81916213 |
| FAVSELTVM | ASSQEGDRGYEQY | 0.87800467 |
| GAAPLAAGL | ASSQEGDRGYEQY | 0.58691555 |
| GADGGDGGSGGAGG | ASSQEGDRGYEQY | 0.32665604 |
| GFGLDTGGAGG | ASSQEGDRGYEQY | 0.7082337 |
| GLASGGLL | ASSQEGDRGYEQY | 0.4890671 |
| GRIAGAAVLL | ASSQEGDRGYEQY | 0.6786234 |
| GSAGSLNTSF | ASSQEGDRGYEQY | 0.63084996 |
| GTGQIGI | ASSQEGDRGYEQY | 0.42614546 |
| GTLIALIL | ASSQEGDRGYEQY | 0.7065686 |
| IAATMGFVVG | ASSQEGDRGYEQY | 0.8840427 |
| IFAQALL | ASSQEGDRGYEQY | 0.76008546 |
| IGAAVALGF | ASSQEGDRGYEQY | 0.82564795 |
| IGLTPLAL | ASSQEGDRGYEQY | 0.87206554 |
| ILGSNTSTL | ASSQEGDRGYEQY | 0.8457599 |

| | | |
|---|---|---|
| IVFSLTVL | ASSQEGDRGYEQY | 0.88163453 |
| LASKATGF | ASSQEGDRGYEQY | 0.8250366 |
| LASVALIL | ASSQEGDRGYEQY | 0.6494183 |
| LFAVGLTL | ASSQEGDRGYEQY | 0.67280304 |
| LFNGGIGTYI | ASSQEGDRGYEQY | 0.8478466 |
| LTLGAGII | ASSQEGDRGYEQY | 0.7454034 |
| LVAAALGDI | ASSQEGDRGYEQY | 0.74681425 |
| NTAFVQLGI | ASSQEGDRGYEQY | 0.54056466 |
| QLFSGAEL | ASSQEGDRGYEQY | 0.8637152 |
| SAATFITL | ASSQEGDRGYEQY | 0.66902924 |
| SNYGLANLGV | ASSQEGDRGYEQY | 0.84135133 |
| SYGGAYAVM | ASSQEGDRGYEQY | 0.22506891 |
| TTGVGGLTLG | ASSQEGDRGYEQY | 0.35515732 |
| TTQTIYI | ASSQEGDRGYEQY | 0.8638642 |
| VAAGAGLGAVI | ASSQEGDRGYEQY | 0.11558493 |
| VAAVVTLL | ASSQEGDRGYEQY | 0.15319909 |
| VGFLGGLI | ASSQEGDRGYEQY | 0.7431466 |
| VGVIPILL | ASSQEGDRGYEQY | 0.8656187 |
| VIFASSLI | ASSQEGDRGYEQY | 0.11308694 |
| VKGGDGGAAAG | ASSQEGDRGYEQY | 0.36479586 |
| VTKTFTNI | ASSQEGDRGYEQY | 0.33707255 |
| VVAAAILL | ASSQEGDRGYEQY | 0.69533086 |
| YAALDVEL | ASSQEGDRGYEQY | 0.70384336 |
| LASVALLL | ASSQEGDRGYEQY | 0.23476812 |
| LDAVHRV | ASSQEGDRGYEQY | 0.8307346 |
| QGLSSSIF | ASSQEGDRGYEQY | 0.31382254 |
| RGTFANIRL | ASSQEGDRGYEQY | 0.45221883 |
| RQLWWGHRI | ASSQEGDRGYEQY | 0.8149721 |
| YGPPGTGKTLL | ASSQEGDRGYEQY | 0.86032116 |
| AALVGAPANVA | ASSQEGDRGYEQY | 0.35668486 |
| AATFRRTE | ASSQEGDRGYEQY | 0.8149452 |
| AGALGGGI | ASSQEGDRGYEQY | 0.57190716 |
| AGGAGGAGGIGAG | ASSQEGDRGYEQY | 0.89270467 |
| AGSGALGL | ASSQEGDRGYEQY | 0.5974221 |
| AQIADQI | ASSQEGDRGYEQY | 0.28041014 |
| ASAVAVTYL | ASSQEGDRGYEQY | 0.84709257 |
| ASTVLGGTGG | ASSQEGDRGYEQY | 0.8514478 |
| AVGGEPAK | ASSQEGDRGYEQY | 0.5667037 |
| CEQGTFGGTC | ASSQEGDRGYEQY | 0.84134996 |
| DEGKAPAG | ASSQEGDRGYEQY | 0.48580486 |
| DTVVFPAL | ASSQEGDRGYEQY | 0.78351825 |
| FEGITMAG | ASSQEGDRGYEQY | 0.8676585 |
| FPLVVQD | ASSQEGDRGYEQY | 0.8315749 |
| GAFTTPQ | ASSQEGDRGYEQY | 0.33503637 |
| GFGGFGTAGGVGGA | ASSQEGDRGYEQY | 0.76607215 |
| GFLAQFL | ASSQEGDRGYEQY | 0.71302515 |
| GGAGGAGGNAGVA | ASSQEGDRGYEQY | 0.47835687 |
| GGVGTAGAGGAGGAGG | ASSQEGDRGYEQY | 0.8701533 |
| GNGGGGDAGNAGSGGN | ASSQEGDRGYEQY | 0.65820193 |
| GTGGAGGAGNDAGS | ASSQEGDRGYEQY | 0.84627795 |
| GTTNTGWFNS | ASSQEGDRGYEQY | 0.7802701 |

| | | |
|---|---|---|
| IFMGPLFTL | ASSQEGDRGYEQY | 0.8668467 |
| ILAHAIEAI | ASSQEGDRGYEQY | 0.28258345 |
| IRSAIREAL | ASSQEGDRGYEQY | 0.7367666 |
| ITFAGGG | ASSQEGDRGYEQY | 0.7669027 |
| ITGMSGHQEV | ASSQEGDRGYEQY | 0.83954734 |
| IVAGLACTL | ASSQEGDRGYEQY | 0.6183386 |
| IVGGGLAGA | ASSQEGDRGYEQY | 0.4518028 |
| IVTGVLNI | ASSQEGDRGYEQY | 0.78796995 |
| LAQQGCI | ASSQEGDRGYEQY | 0.8668635 |
| LCLHVEN | ASSQEGDRGYEQY | 0.6573067 |
| LDAISDGS | ASSQEGDRGYEQY | 0.70115674 |
| LGGFAAML | ASSQEGDRGYEQY | 0.8891123 |
| LGGKNPMI | ASSQEGDRGYEQY | 0.44300038 |
| LGPKGVAAIL | ASSQEGDRGYEQY | 0.69468945 |
| LMVGVSI | ASSQEGDRGYEQY | 0.8082987 |
| LVGAAFVL | ASSQEGDRGYEQY | 0.83418167 |
| LVLAVAPL | ASSQEGDRGYEQY | 0.6692762 |
| MAAIAVIDAA | ASSQEGDRGYEQY | 0.8369664 |
| NNGINAELQ | ASSQEGDRGYEQY | 0.41575673 |
| RGEPLVLVHG | ASSQEGDRGYEQY | 0.8282492 |
| RGGAGGAAGLI | ASSQEGDRGYEQY | 0.5468304 |
| RSYRMHPAV | ASSQEGDRGYEQY | 0.76961374 |
| SAAHLLAGQLA | ASSQEGDRGYEQY | 0.41039455 |
| SAAVGSY | ASSQEGDRGYEQY | 0.81759584 |
| SPHAVLHRI | ASSQEGDRGYEQY | 0.8580381 |
| SQIAKVL | ASSQEGDRGYEQY | 0.8439487 |
| TALGHLVP | ASSQEGDRGYEQY | 0.14386645 |
| TAPTTSGAA | ASSQEGDRGYEQY | 0.85633063 |
| TDGDDGNGGSPGT | ASSQEGDRGYEQY | 0.8425869 |
| TDGLGEN | ASSQEGDRGYEQY | 0.88593155 |
| TGGPPLPGQA | ASSQEGDRGYEQY | 0.65877306 |
| TIPAITI | ASSQEGDRGYEQY | 0.79806244 |
| TIPGFTI | ASSQEGDRGYEQY | 0.5038701 |
| TSIASGALA | ASSQEGDRGYEQY | 0.68449306 |
| TVTLPGAAI | ASSQEGDRGYEQY | 0.5016355 |
| VASGLSAALP | ASSQEGDRGYEQY | 0.56478876 |
| VGAGVIGI | ASSQEGDRGYEQY | 0.7310685 |
| VGPKRFGV | ASSQEGDRGYEQY | 0.65877 |
| VNVAFPD | ASSQEGDRGYEQY | 0.84073997 |
| VSMTNSGVS | ASSQEGDRGYEQY | 0.84470755 |
| VTDVLGDLSV | ASSQEGDRGYEQY | 0.8371193 |
| VVVVGAGFAGL | ASSQEGDRGYEQY | 0.8705342 |
| VYTVGTVV | ASSQEGDRGYEQY | 0.881814 |
| WTTFSGP | ASSQEGDRGYEQY | 0.797899 |
| YDELVEAF | ASSQEGDRGYEQY | 0.5548207 |
| YGTGGAGGNGG | ASSQEGDRGYEQY | 0.58291525 |
| YTNVMAV | ASSQEGDRGYEQY | 0.42748672 |
| FLEQVHQL | ASSQEGDRGYEQY | 0.67708886 |
| GLSLLAVAV | ASSQEGDRGYEQY | 0.62140477 |
| LEQVHQL | ASSQEGDRGYEQY | 0.80949265 |
| LNAAGAI | ASSQEGDRGYEQY | 0.10315886 |

| | | |
|---|---|---|
| PAVHASL | ASSQEGDRGYEQY | 0.80508924 |
| TELEDDA | ASSQEGDRGYEQY | 0.8740988 |
| TPPGPPN | ASSQEGDRGYEQY | 0.6011339 |
| TPPPPPPP | ASSQEGDRGYEQY | 0.7936337 |
| TQSPALI | ASPGGNQDTQY | 0.83358896 |
| EFAAAFPQ | ASPGGNQDTQY | 0.57866466 |
| LLSGNLDVL | ASPGGNQDTQY | 0.83954954 |
| GPANLGNYNI | ASPGGNQDTQY | 0.46791 |
| TLGPTLP | ASPGGNQDTQY | 0.81531173 |
| VYAYSSCCK | ASPGGNQDTQY | 0.86431915 |
| APAPAPAPNI | ASPGGNQDTQY | 0.8006835 |
| PGGPVPIP | ASPGGNQDTQY | 0.58394307 |
| FLTVALEP | ASPGGNQDTQY | 0.6112046 |
| SQYFGANL | ASPGGNQDTQY | 0.23768404 |
| GGVKPAA | ASPGGNQDTQY | 0.6921716 |
| GGTGGAGGAGGW | ASPGGNQDTQY | 0.11850353 |
| GVGGVAGLGGAGG | ASPGGNQDTQY | 0.80054116 |
| VLLVLATTI | ASPGGNQDTQY | 0.40744117 |
| IGIFAAML | ASPGGNQDTQY | 0.80323124 |
| AVGDLID | ASPGGNQDTQY | 0.74634117 |
| NISPLPL | ASPGGNQDTQY | 0.7494307 |
| VTAVFVAI | ASPGGNQDTQY | 0.5559344 |
| SGDDNDAAL | ASPGGNQDTQY | 0.72297114 |
| AVAALAHS | ASPGGNQDTQY | 0.55898887 |
| GGVTVMATP | ASPGGNQDTQY | 0.3043435 |
| AGLAANVRD | ASPGGNQDTQY | 0.84872377 |
| TNRPDLIDPALLRPG | ASPGGNQDTQY | 0.5937995 |
| LANLGVTAL | ASPGGNQDTQY | 0.77533644 |
| TGERIEL | ASPGGNQDTQY | 0.8424355 |
| NGEELAA | ASPGGNQDTQY | 0.7272565 |
| VLSLVWG | ASPGGNQDTQY | 0.24799438 |
| IGYALWL | ASPGGNQDTQY | 0.7637148 |
| LSLLLAPL | ASPGGNQDTQY | 0.7469532 |
| TLDEALK | ASPGGNQDTQY | 0.67150015 |
| TDGAGTAEGG | ASPGGNQDTQY | 0.1098829 |
| NAPGGAGGNGG | ASPGGNQDTQY | 0.7920103 |
| VSVFMGMGGNF | ASPGGNQDTQY | 0.47572318 |
| LASYLAAL | ASPGGNQDTQY | 0.62718016 |
| LVGGTPAL | ASPGGNQDTQY | 0.61396646 |
| VRNADVL | ASPGGNQDTQY | 0.8720787 |
| TCSSPFLVSAM | ASPGGNQDTQY | 0.48315066 |
| VHITDAE | ASPGGNQDTQY | 0.4746276 |
| EYAAPSF | ASPGGNQDTQY | 0.8345405 |
| VVDAFHGGL | ASPGGNQDTQY | 0.7123635 |
| ASVVDPQ | ASPGGNQDTQY | 0.8676417 |
| SYNEFGGLS | ASPGGNQDTQY | 0.66752636 |
| DPDGSVY | ASPGGNQDTQY | 0.5230336 |
| MVSTIHATE | ASPGGNQDTQY | 0.7250725 |
| LAAGVADPV | ASPGGNQDTQY | 0.50413924 |
| TKEQIAA | ASPGGNQDTQY | 0.24001075 |
| SLASPAAL | ASPGGNQDTQY | 0.20899782 |

| | | |
|---|---|---|
| IGLVFYGV | ASPGGNQDTQY | 0.7640288 |
| FLLDASNL | ASPGGNQDTQY | 0.20559806 |
| YQVGNFP | ASPGGNQDTQY | 0.7322477 |
| ESLLSQST | ASPGGNQDTQY | 0.8143903 |
| LFTVGFAAVG | ASPGGNQDTQY | 0.5672385 |
| ATPPTPT | ASPGGNQDTQY | 0.26872665 |
| TDLATLG | ASPGGNQDTQY | 0.55904037 |
| SAGGTGR | ASPGGNQDTQY | 0.6091618 |
| GFSNSGS | ASPGGNQDTQY | 0.8453176 |
| GGNGGSGGTSV | ASPGGNQDTQY | 0.70903283 |
| GTSVATGGAGN | ASPGGNQDTQY | 0.8674749 |
| NGGNGGSGGTS | ASPGGNQDTQY | 0.8416463 |
| AVVKFPH | ASPGGNQDTQY | 0.6091022 |
| TPPPPPP | ASPGGNQDTQY | 0.84964365 |
| AAPAAAAAPA | ASPGGNQDTQY | 0.67032164 |
| IALNEDL | ASPGGNQDTQY | 0.6679983 |
| IVNVSSM | ASPGGNQDTQY | 0.66955596 |
| TGIPPAPR | ASPGGNQDTQY | 0.7436988 |
| FDLGGGTF | ASPGGNQDTQY | 0.5880131 |
| FDLGGGTFDVS | ASPGGNQDTQY | 0.59590983 |
| NEPTAAA | ASPGGNQDTQY | 0.28858298 |
| NEPTAAALA | ASPGGNQDTQY | 0.6795119 |
| GSIINISSI | ASPGGNQDTQY | 0.6033112 |
| VAPGNAGTA | ASPGGNQDTQY | 0.21195146 |
| TGGGASLE | ASPGGNQDTQY | 0.498052 |
| VAVYNHYKRI | ASPGGNQDTQY | 0.8903438 |
| TDPAPATT | ASPGGNQDTQY | 0.8674785 |
| TNRPDILDPALLRPG | ASPGGNQDTQY | 0.63521135 |
| LTTTTTS | ASPGGNQDTQY | 0.50158644 |
| PTGSGKTL | ASPGGNQDTQY | 0.8116041 |
| TGSGKTL | ASPGGNQDTQY | 0.31671005 |
| TIGHVDHGK | ASPGGNQDTQY | 0.7373722 |
| TIGHVDHGKTT | ASPGGNQDTQY | 0.7917162 |
| GHVDHGKTT | ASPGGNQDTQY | 0.7954663 |
| AQFEHTLL | ASPGGNQDTQY | 0.43940043 |
| IVNQDEL | ASPGGNQDTQY | 0.7943252 |
| TNGSQFF | ASPGGNQDTQY | 0.69276476 |
| TGKGIVE | ASPGGNQDTQY | 0.7235447 |
| LISNASDA | ASPGGNQDTQY | 0.11353783 |
| AELRTLGELTKIAKAHGAQ | ASPGGNQDTQY | 0.73164666 |
| APNAVAAAEPSW | ASPGGNQDTQY | 0.5061469 |
| APQVAGAATAVL | ASPGGNQDTQY | 0.4366763 |
| ARRDPAAAAAILPTDA | ASPGGNQDTQY | 0.18453799 |
| DAAARDVLAEAGLAEYF | ASPGGNQDTQY | 0.68620896 |
| DDQIVLSAKANLAKVEQ | ASPGGNQDTQY | 0.86430514 |
| DELLPDTGEDSDEDGH | ASPGGNQDTQY | 0.8663293 |
| ELPEAPANRATVPQD | ASPGGNQDTQY | 0.45448616 |
| FGQSKIERVTGIG | ASPGGNQDTQY | 0.843921 |
| GAGGTGSTAGGAGGAGGAGGLY | ASPGGNQDTQY | 0.87179047 |
| GHGGTGAAGGNGGTGGAGGSGI | ASPGGNQDTQY | 0.7053602 |
| GNGGNGGAGGEGGPGGAGGAGA | ASPGGNQDTQY | 0.19059782 |

| | | |
|---|---|---|
| GSTAGNGGNGGNATNGTIAGQP | ASPGGNQDTQY | 0.7858965 |
| GTGGDGGNGGNGGTGG | ASPGGNQDTQY | 0.6608057 |
| GVRRPAGIPADAPV | ASPGGNQDTQY | 0.8357327 |
| GYRQVTDAHLLTLARR | ASPGGNQDTQY | 0.83173287 |
| IAKMGQDGHDRG | ASPGGNQDTQY | 0.5239239 |
| KAEPSIEIQYT | ASPGGNQDTQY | 0.7659348 |
| KPVRIRMAADEDTVAVT | ASPGGNQDTQY | 0.8605928 |
| MPPRIAGMRLLVIKPEP | ASPGGNQDTQY | 0.46110827 |
| MKFYAVALFRSRGVPPTR | ASPGGNQDTQY | 0.39839077 |
| NAGGFGILGAVAHSP | ASPGGNQDTQY | 0.50799775 |
| NEGREITLAMAEM | ASPGGNQDTQY | 0.22913426 |
| NELVEWSHPLGHRWRWE | ASPGGNQDTQY | 0.71005976 |
| NLGAGGTGGNGGNPG | ASPGGNQDTQY | 0.67840934 |
| NTEWPKWETVMAADPVD | ASPGGNQDTQY | 0.4778389 |
| PASAAPAETQP | ASPGGNQDTQY | 0.78835714 |
| QQSSDGKNGVRAAEITGE | ASPGGNQDTQY | 0.41602212 |
| RAPDAVTTSWWKE | ASPGGNQDTQY | 0.53647244 |
| SGFGNVGAHTSGWFNQ | ASPGGNQDTQY | 0.5650547 |
| SGVAGAVGQAASVGGLKV | ASPGGNQDTQY | 0.6458216 |
| SNGGAGTGSAGGNGGT | ASPGGNQDTQY | 0.818663 |
| TGGAGGEGGAAG | ASPGGNQDTQY | 0.7478112 |
| TGGDGGAAGAGGAPGHG | ASPGGNQDTQY | 0.78758395 |
| TMPPLLAAIANAGRHGV | ASPGGNQDTQY | 0.8795407 |
| TSSTAGNSGGA | ASPGGNQDTQY | 0.26561606 |
| VPATVPVGILC | ASPGGNQDTQY | 0.4527057 |
| VQRPDSGGVVLTGRIS | ASPGGNQDTQY | 0.8277498 |
| WTPNPVPATKGVPAGLVQAT | ASPGGNQDTQY | 0.36411923 |
| YALKRALAKFSDRFDVVII | ASPGGNQDTQY | 0.84297323 |
| YLIGSVIV | ASPGGNQDTQY | 0.67675126 |
| YGQLVAVID | ASPGGNQDTQY | 0.81836784 |
| VTVGGMTLPA | ASPGGNQDTQY | 0.6228312 |
| VLALPAV | ASPGGNQDTQY | 0.88548887 |
| VIPPTLNYET | ASPGGNQDTQY | 0.85378075 |
| VGMGGMHPSSG | ASPGGNQDTQY | 0.6603937 |
| VGLAGGAATA | ASPGGNQDTQY | 0.79534006 |
| VGGAGGAGGSGP | ASPGGNQDTQY | 0.33897448 |
| VFAAFMV | ASPGGNQDTQY | 0.8249015 |
| TLYDKLR | ASPGGNQDTQY | 0.8024379 |
| TLAVFTVL | ASPGGNQDTQY | 0.5810547 |
| TGSVQPARV | ASPGGNQDTQY | 0.7805331 |
| TGGGAGSGGT | ASPGGNQDTQY | 0.8468356 |
| SHLETSL | ASPGGNQDTQY | 0.8756257 |
| SFTTGMT | ASPGGNQDTQY | 0.2510692 |
| QVEVAKRMGVRQPTVS | ASPGGNQDTQY | 0.7095101 |
| QGSGGPALAM | ASPGGNQDTQY | 0.23939714 |
| MLAPSNP | ASPGGNQDTQY | 0.41143465 |
| LVNNAAI | ASPGGNQDTQY | 0.29986826 |
| LTGEINTLA | ASPGGNQDTQY | 0.8252248 |
| LSIFGPV | ASPGGNQDTQY | 0.65231866 |
| LLALLANL | ASPGGNQDTQY | 0.8224959 |
| LISNASDALDKLR | ASPGGNQDTQY | 0.786796 |

| | | |
|---|---|---|
| LINEAVP | ASPGGNQDTQY | 0.8557714 |
| LGYAFSGL | ASPGGNQDTQY | 0.52663195 |
| LDFVAVAL | ASPGGNQDTQY | 0.57814336 |
| LAQPGAE | ASPGGNQDTQY | 0.25655186 |
| ISNLGAVTGF | ASPGGNQDTQY | 0.23180631 |
| HQEDPGRI | ASPGGNQDTQY | 0.34812814 |
| GTPGEPTP | ASPGGNQDTQY | 0.6331333 |
| GFFNSGS | ASPGGNQDTQY | 0.14920463 |
| GAGGGGLTAGAN | ASPGGNQDTQY | 0.86737263 |
| FVDALVNM | ASPGGNQDTQY | 0.7748774 |
| FLTVAPEL | ASPGGNQDTQY | 0.80582774 |
| FETLTAL | ASPGGNQDTQY | 0.40320623 |
| DVINPIEL | ASPGGNQDTQY | 0.48647198 |
| APVSGAPV | ASPGGNQDTQY | 0.6081344 |
| ALVGGLAW | ASPGGNQDTQY | 0.82817465 |
| ALAAAAATAI | ASPGGNQDTQY | 0.4678272 |
| AFGLGLLL | ASPGGNQDTQY | 0.23368591 |
| AFFAAASL | ASPGGNQDTQY | 0.44470167 |
| AAAALPAFSPP | ASPGGNQDTQY | 0.41660166 |
| VVLAPAL | ASPGGNQDTQY | 0.8787585 |
| TIGHVDHGKTTL | ASPGGNQDTQY | 0.8571522 |
| LISNASDALDK | ASPGGNQDTQY | 0.8288436 |
| LISNASDAL | ASPGGNQDTQY | 0.84279597 |
| ISNASDALDK | ASPGGNQDTQY | 0.13633683 |
| NQAETLVYQTE | ASPGGNQDTQY | 0.46718404 |
| TGGTGGTAGAG | ASPGGNQDTQY | 0.8808681 |
| TGSTAGGAGGA | ASPGGNQDTQY | 0.804705 |
| DPLAVPVAAGPV | ASPGGNQDTQY | 0.7019584 |
| GGTGGTPGNGGN | ASPGGNQDTQY | 0.85877645 |
| PAGLAGVDPATL | ASPGGNQDTQY | 0.4868305 |
| GNNLVVEAPHAGASV | ASPGGNQDTQY | 0.6447276 |
| KVTRIMSASTLEQPA | ASPGGNQDTQY | 0.74398434 |
| LGFALPIALPRRTVV | ASPGGNQDTQY | 0.7517299 |
| TRLGNLTARKLVVLV | ASPGGNQDTQY | 0.68744713 |
| WSQQRRLIYFPSAGP | ASPGGNQDTQY | 0.3436616 |
| GPNGNPDPMAAAVDIR | ASPGGNQDTQY | 0.7616297 |
| TQRCVSVADLRAAASR | ASPGGNQDTQY | 0.88591456 |
| QFVLDVAEHPVEVLTETV | ASPGGNQDTQY | 0.7600299 |
| GGAGGTGGAGGNGGTQAGDG | ASPGGNQDTQY | 0.86170775 |
| PLLATLRAIAVDVDAKPAQV | ASPGGNQDTQY | 0.77494395 |
| APKFRTGGYMIDFWGVGYQVAK | ASPGGNQDTQY | 0.8144619 |
| GAGGAGGAGGAAGGPGGTGGTG | ASPGGNQDTQY | 0.59654725 |
| IGALGLLATDVAPAGSVAFGVA | ASPGGNQDTQY | 0.6009216 |
| AGGAGGIGGNGGAGGAGGVLMG | ASPGGNQDTQY | 0.6840447 |
| ADELIGLL | ASPGGNQDTQY | 0.5020726 |
| ADRALKA | ASPGGNQDTQY | 0.69070965 |
| AGAVGQVGGAGGA | ASPGGNQDTQY | 0.64969397 |
| AGGGGGLGGAP | ASPGGNQDTQY | 0.8913107 |
| AGGLLYGNGGAG | ASPGGNQDTQY | 0.8127914 |
| AGLESIGL | ASPGGNQDTQY | 0.80818117 |
| AGLKTGII | ASPGGNQDTQY | 0.4702063 |

| | | |
|---|---|---|
| ASLAALLL | ASPGGNQDTQY | 0.6944264 |
| ASVINISSI | ASPGGNQDTQY | 0.81412745 |
| ATPPGPA | ASPGGNQDTQY | 0.3415569 |
| ATVASALV | ASPGGNQDTQY | 0.8652481 |
| ATVASLIL | ASPGGNQDTQY | 0.44375432 |
| AVNGPVLIRH | ASPGGNQDTQY | 0.7113164 |
| DGAAGAVGI | ASPGGNQDTQY | 0.80794626 |
| DPLLHGI | ASPGGNQDTQY | 0.8532069 |
| ELGAALGAL | ASPGGNQDTQY | 0.6602594 |
| ELGRTQAV | ASPGGNQDTQY | 0.6796069 |
| FVAGGYLS | ASPGGNQDTQY | 0.56471366 |
| GASGGAGGAGGW | ASPGGNQDTQY | 0.8418696 |
| GGAGAGGDGGAGGI | ASPGGNQDTQY | 0.8774904 |
| GTGHNGSQPGMGGNG | ASPGGNQDTQY | 0.73940575 |
| GYAAASASAS | ASPGGNQDTQY | 0.8772323 |
| HQVDVAAVQQG | ASPGGNQDTQY | 0.69848365 |
| IGAAIAHT | ASPGGNQDTQY | 0.23744367 |
| IGMIEVAQL | ASPGGNQDTQY | 0.6733457 |
| IRLAPEHTSAMD | ASPGGNQDTQY | 0.85627854 |
| IYNVSVL | ASPGGNQDTQY | 0.864194 |
| KQGFLGM | ASPGGNQDTQY | 0.8055161 |
| LESVITQL | ASPGGNQDTQY | 0.6800907 |
| LGLGALVVV | ASPGGNQDTQY | 0.6455506 |
| LIGNGGNGGSGGI | ASPGGNQDTQY | 0.8755839 |
| LNAINAPTL | ASPGGNQDTQY | 0.8773807 |
| LQLLNTGV | ASPGGNQDTQY | 0.83731955 |
| LQTSIEF | ASPGGNQDTQY | 0.56367874 |
| MSFAVSEL | ASPGGNQDTQY | 0.42848226 |
| NTGALGL | ASPGGNQDTQY | 0.7614976 |
| PPGAGGAGGAGTT | ASPGGNQDTQY | 0.8559257 |
| QLVGLGI | ASPGGNQDTQY | 0.3679727 |
| RFLEQVHQL | ASPGGNQDTQY | 0.7914505 |
| RGVGYVLR | ASPGGNQDTQY | 0.7426436 |
| SATGVFAGVI | ASPGGNQDTQY | 0.23858368 |
| SLPAITLP | ASPGGNQDTQY | 0.78726894 |
| STALAFTV | ASPGGNQDTQY | 0.8694874 |
| SVLTVGLGK | ASPGGNQDTQY | 0.6840998 |
| TALAGAGSGI | ASPGGNQDTQY | 0.47048363 |
| TGLPFTGI | ASPGGNQDTQY | 0.8617822 |
| VAEHVTV | ASPGGNQDTQY | 0.8892152 |
| VDIVLNSL | ASPGGNQDTQY | 0.862688 |
| VLTGGGAVL | ASPGGNQDTQY | 0.2126722 |
| VNVGLAYGI | ASPGGNQDTQY | 0.16674934 |
| VSSAAVGAV | ASPGGNQDTQY | 0.65307194 |
| VTSVYEAGV | ASPGGNQDTQY | 0.829363 |
| WVALDRGA | ASPGGNQDTQY | 0.5339097 |
| YLVGSVVI | ASPGGNQDTQY | 0.48938364 |
| GPPGTGKTLL | ASPGGNQDTQY | 0.8290004 |
| GVGMTKF | ASPGGNQDTQY | 0.86333144 |
| IGIGGSDL | ASPGGNQDTQY | 0.3895164 |
| IGTLLAEL | ASPGGNQDTQY | 0.27446833 |

| | | |
|---|---|---|
| IYTSGTT | ASPGGNQDTQY | 0.8154541 |
| LSFPAGV | ASPGGNQDTQY | 0.844463 |
| LYTSGTT | ASPGGNQDTQY | 0.5649185 |
| VGAFPFGF | ASPGGNQDTQY | 0.76096404 |
| VGAGTVEF | ASPGGNQDTQY | 0.6355991 |
| VGKSTLLL | ASPGGNQDTQY | 0.55040324 |
| AALAHAGGGIV | ASPGGNQDTQY | 0.80013967 |
| AALVAVSLL | ASPGGNQDTQY | 0.41795218 |
| AAVSRAVAAL | ASPGGNQDTQY | 0.8703653 |
| AGGSSGIGGF | ASPGGNQDTQY | 0.4391137 |
| AGLTVANANG | ASPGGNQDTQY | 0.7976176 |
| AGLVGAVL | ASPGGNQDTQY | 0.7782502 |
| ALSQAFTV | ASPGGNQDTQY | 0.77444273 |
| ALTFSVSL | ASPGGNQDTQY | 0.67396414 |
| ALTGAALVAL | ASPGGNQDTQY | 0.5871204 |
| ASFSIGAGVL | ASPGGNQDTQY | 0.39372844 |
| DGSVYKI | ASPGGNQDTQY | 0.22171852 |
| EQLAEIV | ASPGGNQDTQY | 0.8581244 |
| FAVSELTVM | ASPGGNQDTQY | 0.21617481 |
| GAAPLAAGL | ASPGGNQDTQY | 0.8586816 |
| GADGGDGGSGGAGG | ASPGGNQDTQY | 0.7749933 |
| GFGLDTGGAGG | ASPGGNQDTQY | 0.51471794 |
| GLASGGLL | ASPGGNQDTQY | 0.8011612 |
| GRIAGAAVLL | ASPGGNQDTQY | 0.32698298 |
| GSAGSLNTSF | ASPGGNQDTQY | 0.48854205 |
| GTGQIGI | ASPGGNQDTQY | 0.8221805 |
| GTLIALIL | ASPGGNQDTQY | 0.82750666 |
| IAATMGFVVG | ASPGGNQDTQY | 0.47093686 |
| IFAQALL | ASPGGNQDTQY | 0.7553292 |
| IGAAVALGF | ASPGGNQDTQY | 0.81764776 |
| IGLTPLAL | ASPGGNQDTQY | 0.4058649 |
| ILGSNTSTL | ASPGGNQDTQY | 0.61764514 |
| IVFSLTVL | ASPGGNQDTQY | 0.8548558 |
| LASKATGF | ASPGGNQDTQY | 0.67558146 |
| LASVALIL | ASPGGNQDTQY | 0.44443497 |
| LFAVGLTL | ASPGGNQDTQY | 0.7967981 |
| LFNGGIGTYI | ASPGGNQDTQY | 0.62152135 |
| LTLGAGII | ASPGGNQDTQY | 0.8376813 |
| LVAAALGDI | ASPGGNQDTQY | 0.76315755 |
| NTAFVQLGI | ASPGGNQDTQY | 0.58179426 |
| QLFSGAEL | ASPGGNQDTQY | 0.8278837 |
| SAATFITL | ASPGGNQDTQY | 0.29512262 |
| SNYGLANLGV | ASPGGNQDTQY | 0.55512273 |
| SYGGAYAVM | ASPGGNQDTQY | 0.58418703 |
| TTGVGGLTLG | ASPGGNQDTQY | 0.6634389 |
| TTQTIYI | ASPGGNQDTQY | 0.8472052 |
| VAAGAGLGAVI | ASPGGNQDTQY | 0.5144663 |
| VAAVVTLL | ASPGGNQDTQY | 0.79262435 |
| VGFLGGLI | ASPGGNQDTQY | 0.5941 |
| VGVIPILL | ASPGGNQDTQY | 0.4148826 |
| VIFASSLI | ASPGGNQDTQY | 0.64519095 |

| | | |
|---|---|---|
| VKGGDGGAAAG | ASPGGNQDTQY | 0.502367 |
| VTKTFTNI | ASPGGNQDTQY | 0.40203494 |
| VVAAAILL | ASPGGNQDTQY | 0.80626065 |
| YAALDVEL | ASPGGNQDTQY | 0.40538794 |
| LASVALLL | ASPGGNQDTQY | 0.63191384 |
| LDAVHRV | ASPGGNQDTQY | 0.85509276 |
| QGLSSSIF | ASPGGNQDTQY | 0.42684296 |
| RGTFANIRL | ASPGGNQDTQY | 0.80592924 |
| RQLWWGHRI | ASPGGNQDTQY | 0.812161 |
| YGPPGTGKTLL | ASPGGNQDTQY | 0.813671 |
| AALVGAPANVA | ASPGGNQDTQY | 0.5906178 |
| AATFRRTE | ASPGGNQDTQY | 0.4837377 |
| AGALGGGI | ASPGGNQDTQY | 0.8759044 |
| AGGAGGAGGIGAG | ASPGGNQDTQY | 0.8103261 |
| AGSGALGL | ASPGGNQDTQY | 0.7951447 |
| AQIADQI | ASPGGNQDTQY | 0.828756 |
| ASAVAVTYL | ASPGGNQDTQY | 0.8293891 |
| ASTVLGGTGG | ASPGGNQDTQY | 0.8491279 |
| AVGGEPAK | ASPGGNQDTQY | 0.6612627 |
| CEQGTFGGTC | ASPGGNQDTQY | 0.28363037 |
| DEGKAPAG | ASPGGNQDTQY | 0.5693855 |
| DTVVFPAL | ASPGGNQDTQY | 0.61562973 |
| FEGITMAG | ASPGGNQDTQY | 0.5626409 |
| FPLVVQD | ASPGGNQDTQY | 0.22963184 |
| GAFTTPQ | ASPGGNQDTQY | 0.8667567 |
| GFGGFGTAGGVGGA | ASPGGNQDTQY | 0.8945061 |
| GFLAQFL | ASPGGNQDTQY | 0.39250147 |
| GGAGGAGGNAGVA | ASPGGNQDTQY | 0.8853493 |
| GGVGTAGAGGAGGAGG | ASPGGNQDTQY | 0.3304302 |
| GNGGGGDAGNAGSGGN | ASPGGNQDTQY | 0.76808524 |
| GTGGAGGAGNDAGS | ASPGGNQDTQY | 0.85747755 |
| GTTNTGWFNS | ASPGGNQDTQY | 0.47006318 |
| IFMGPLFTL | ASPGGNQDTQY | 0.75171304 |
| ILAHAIEAI | ASPGGNQDTQY | 0.4429084 |
| IRSAIREAL | ASPGGNQDTQY | 0.5761364 |
| ITFAGGG | ASPGGNQDTQY | 0.8608111 |
| ITGMSGHQEV | ASPGGNQDTQY | 0.44555342 |
| IVAGLACTL | ASPGGNQDTQY | 0.71590936 |
| IVGGGLAGA | ASPGGNQDTQY | 0.6509757 |
| IVTGVLNI | ASPGGNQDTQY | 0.45879063 |
| LAQQGCI | ASPGGNQDTQY | 0.7698419 |
| LCLHVEN | ASPGGNQDTQY | 0.8223996 |
| LDAISDGS | ASPGGNQDTQY | 0.47019762 |
| LGGFAAML | ASPGGNQDTQY | 0.87051445 |
| LGGKNPMI | ASPGGNQDTQY | 0.4026463 |
| LGPKGVAAIL | ASPGGNQDTQY | 0.891137 |
| LMVGVSI | ASPGGNQDTQY | 0.8528124 |
| LVGAAFVL | ASPGGNQDTQY | 0.80726683 |
| LVLAVAPL | ASPGGNQDTQY | 0.61384255 |
| MAAIAVIDAA | ASPGGNQDTQY | 0.864694 |
| NNGINAELQ | ASPGGNQDTQY | 0.6451372 |

| | | |
|---|---|---|
| RGEPLVLVHG | ASPGGNQDTQY | 0.7264545 |
| RGGAGGAAGLI | ASPGGNQDTQY | 0.8114363 |
| RSYRMHPAV | ASPGGNQDTQY | 0.71156055 |
| SAAHLLAGQLA | ASPGGNQDTQY | 0.86022663 |
| SAAVGSY | ASPGGNQDTQY | 0.81395036 |
| SPHAVLHRI | ASPGGNQDTQY | 0.8217272 |
| SQIAKVL | ASPGGNQDTQY | 0.74384016 |
| TALGHLVP | ASPGGNQDTQY | 0.6415404 |
| TAPTTSGAA | ASPGGNQDTQY | 0.80087674 |
| TDGDDGNGGSPGT | ASPGGNQDTQY | 0.6883173 |
| TDGLGEN | ASPGGNQDTQY | 0.8417542 |
| TGGPPLPGQA | ASPGGNQDTQY | 0.8703904 |
| TIPAITI | ASPGGNQDTQY | 0.72079206 |
| TIPGFTI | ASPGGNQDTQY | 0.8471704 |
| TSIASGALA | ASPGGNQDTQY | 0.84255964 |
| TVTLPGAAI | ASPGGNQDTQY | 0.59870285 |
| VASGLSAALP | ASPGGNQDTQY | 0.7838974 |
| VGAGVIGI | ASPGGNQDTQY | 0.76343596 |
| VGPKRFGV | ASPGGNQDTQY | 0.80612725 |
| VNVAFPD | ASPGGNQDTQY | 0.49345154 |
| VSMTNSGVS | ASPGGNQDTQY | 0.8525348 |
| VTDVLGDLSV | ASPGGNQDTQY | 0.6738831 |
| VVVVGAGFAGL | ASPGGNQDTQY | 0.37378967 |
| VYTVGTVV | ASPGGNQDTQY | 0.843526 |
| WTTFSGP | ASPGGNQDTQY | 0.77047765 |
| YDELVEAF | ASPGGNQDTQY | 0.78187084 |
| YGTGGAGGNGG | ASPGGNQDTQY | 0.8596935 |
| YTNVMAV | ASPGGNQDTQY | 0.691519 |
| FLEQVHQL | ASPGGNQDTQY | 0.68850994 |
| GLSLLAVAV | ASPGGNQDTQY | 0.8189741 |
| LEQVHQL | ASPGGNQDTQY | 0.86726594 |
| LNAAGAI | ASPGGNQDTQY | 0.65797037 |
| PAVHASL | ASPGGNQDTQY | 0.80877626 |
| TELEDDA | ASPGGNQDTQY | 0.8619038 |
| TPPGPPN | ASPGGNQDTQY | 0.8280588 |
| TPPPPPPP | ASPGGNQDTQY | 0.8820282 |
| TQSPALI | ASSQEPSGTGNENTLY | 0.8899016 |
| EFAAAFPQ | ASSQEPSGTGNENTLY | 0.72620326 |
| LLSGNLDVL | ASSQEPSGTGNENTLY | 0.86454153 |
| GPANLGNYNI | ASSQEPSGTGNENTLY | 0.86336094 |
| TLGPTLP | ASSQEPSGTGNENTLY | 0.77848774 |
| VYAYSSCCK | ASSQEPSGTGNENTLY | 0.7961955 |
| APAPAPAPNI | ASSQEPSGTGNENTLY | 0.68576145 |
| PGGPVPIP | ASSQEPSGTGNENTLY | 0.7857896 |
| FLTVALEP | ASSQEPSGTGNENTLY | 0.6178175 |
| SQYFGANL | ASSQEPSGTGNENTLY | 0.8020549 |
| GGVKPAA | ASSQEPSGTGNENTLY | 0.87820315 |
| GGTGGAGGAGGW | ASSQEPSGTGNENTLY | 0.7566834 |
| GVGGVAGLGGAGG | ASSQEPSGTGNENTLY | 0.632949 |
| VLLVLATTI | ASSQEPSGTGNENTLY | 0.5921818 |
| IGIFAAML | ASSQEPSGTGNENTLY | 0.8145973 |

| | | |
|---|---|---|
| AVGDLID | ASSQEPSGTGNENTLY | 0.7266043 |
| NISPLPL | ASSQEPSGTGNENTLY | 0.6148952 |
| VTAVFVAI | ASSQEPSGTGNENTLY | 0.7355467 |
| SGDDNDAAL | ASSQEPSGTGNENTLY | 0.84421617 |
| AVAALAHS | ASSQEPSGTGNENTLY | 0.84755266 |
| GGVTVMATP | ASSQEPSGTGNENTLY | 0.6529647 |
| AGLAANVRD | ASSQEPSGTGNENTLY | 0.86997545 |
| TNRPDLIDPALLRPG | ASSQEPSGTGNENTLY | 0.8297382 |
| LANLGVTAL | ASSQEPSGTGNENTLY | 0.86046445 |
| TGERIEL | ASSQEPSGTGNENTLY | 0.8113735 |
| NGEELAA | ASSQEPSGTGNENTLY | 0.69851756 |
| VLSLVWG | ASSQEPSGTGNENTLY | 0.835134 |
| IGYALWL | ASSQEPSGTGNENTLY | 0.72072816 |
| LSLLLAPL | ASSQEPSGTGNENTLY | 0.8161185 |
| TLDEALK | ASSQEPSGTGNENTLY | 0.85336614 |
| TDGAGTAEGG | ASSQEPSGTGNENTLY | 0.7692293 |
| NAPGGAGGNGG | ASSQEPSGTGNENTLY | 0.7170973 |
| VSVFMGMGGNF | ASSQEPSGTGNENTLY | 0.68855715 |
| LASYLAAL | ASSQEPSGTGNENTLY | 0.88785857 |
| LVGGTPAL | ASSQEPSGTGNENTLY | 0.6416068 |
| VRNADVL | ASSQEPSGTGNENTLY | 0.63387895 |
| TCSSPFLVSAM | ASSQEPSGTGNENTLY | 0.74322957 |
| VHITDAE | ASSQEPSGTGNENTLY | 0.77060854 |
| EYAAPSF | ASSQEPSGTGNENTLY | 0.86575764 |
| VVDAFHGGL | ASSQEPSGTGNENTLY | 0.85167044 |
| ASVVDPQ | ASSQEPSGTGNENTLY | 0.7571378 |
| SYNEFGGLS | ASSQEPSGTGNENTLY | 0.82706696 |
| DPDGSVY | ASSQEPSGTGNENTLY | 0.852244 |
| MVSTIHATE | ASSQEPSGTGNENTLY | 0.81340325 |
| LAAGVADPV | ASSQEPSGTGNENTLY | 0.8460985 |
| TKEQIAA | ASSQEPSGTGNENTLY | 0.67521536 |
| SLASPAAL | ASSQEPSGTGNENTLY | 0.88128513 |
| IGLVFYGV | ASSQEPSGTGNENTLY | 0.8498162 |
| FLLDASNL | ASSQEPSGTGNENTLY | 0.86640584 |
| YQVGNFP | ASSQEPSGTGNENTLY | 0.87561095 |
| ESLLSQST | ASSQEPSGTGNENTLY | 0.80433375 |
| LFTVGFAAVG | ASSQEPSGTGNENTLY | 0.82370937 |
| ATPPTPT | ASSQEPSGTGNENTLY | 0.5988969 |
| TDLATLG | ASSQEPSGTGNENTLY | 0.713874 |
| SAGGTGR | ASSQEPSGTGNENTLY | 0.7984113 |
| GFSNSGS | ASSQEPSGTGNENTLY | 0.8201066 |
| GGNGGSGGTSV | ASSQEPSGTGNENTLY | 0.84621936 |
| GTSVATGGAGN | ASSQEPSGTGNENTLY | 0.8506292 |
| NGGNGGSGGTS | ASSQEPSGTGNENTLY | 0.692765 |
| AVVKFPH | ASSQEPSGTGNENTLY | 0.86736095 |
| TPPPPPP | ASSQEPSGTGNENTLY | 0.8558382 |
| AAPAAAAAPA | ASSQEPSGTGNENTLY | 0.6210278 |
| IALNEDL | ASSQEPSGTGNENTLY | 0.784371 |
| IVNVSSM | ASSQEPSGTGNENTLY | 0.88000375 |
| TGIPPAPR | ASSQEPSGTGNENTLY | 0.795694 |
| FDLGGGTF | ASSQEPSGTGNENTLY | 0.75789666 |

| | | |
|---|---|---|
| FDLGGGTFDVS | ASSQEPSGTGNENTLY | 0.7663383 |
| NEPTAAA | ASSQEPSGTGNENTLY | 0.8546449 |
| NEPTAAALA | ASSQEPSGTGNENTLY | 0.80069715 |
| GSIINISSI | ASSQEPSGTGNENTLY | 0.87542087 |
| VAPGNAGTA | ASSQEPSGTGNENTLY | 0.48063418 |
| TGGGASLE | ASSQEPSGTGNENTLY | 0.8020826 |
| VAVYNHYKRI | ASSQEPSGTGNENTLY | 0.7724832 |
| TDPAPATT | ASSQEPSGTGNENTLY | 0.41005406 |
| TNRPDILDPALLRPG | ASSQEPSGTGNENTLY | 0.86146355 |
| LTTTTTS | ASSQEPSGTGNENTLY | 0.7702806 |
| PTGSGKTL | ASSQEPSGTGNENTLY | 0.8266251 |
| TGSGKTL | ASSQEPSGTGNENTLY | 0.76004016 |
| TIGHVDHGK | ASSQEPSGTGNENTLY | 0.872754 |
| TIGHVDHGKTT | ASSQEPSGTGNENTLY | 0.77413964 |
| GHVDHGKTT | ASSQEPSGTGNENTLY | 0.77146554 |
| AQFEHTLL | ASSQEPSGTGNENTLY | 0.7675535 |
| IVNQDEL | ASSQEPSGTGNENTLY | 0.8456161 |
| TNGSQFF | ASSQEPSGTGNENTLY | 0.87496054 |
| TGKGIVE | ASSQEPSGTGNENTLY | 0.880637 |
| LISNASDA | ASSQEPSGTGNENTLY | 0.52236366 |
| AELRTLGELTKIAKAHGAQ | ASSQEPSGTGNENTLY | 0.81920445 |
| APNAVAAAEPSW | ASSQEPSGTGNENTLY | 0.79388416 |
| APQVAGAATAVL | ASSQEPSGTGNENTLY | 0.46185067 |
| ARRDPAAAAAILPTDA | ASSQEPSGTGNENTLY | 0.5351535 |
| DAAARDVLAEAGLAEYF | ASSQEPSGTGNENTLY | 0.75344884 |
| DDQIVLSAKANLAKVEQ | ASSQEPSGTGNENTLY | 0.7208526 |
| DELLPDTGEDSDEDGH | ASSQEPSGTGNENTLY | 0.8251716 |
| ELPEAPANRATVPQD | ASSQEPSGTGNENTLY | 0.8093078 |
| FGQSKIERVTGIG | ASSQEPSGTGNENTLY | 0.7556397 |
| GAGGTGSTAGGAGGAGGAGGLY | ASSQEPSGTGNENTLY | 0.8169658 |
| GHGGTGAAGGNGGTGGAGGSGI | ASSQEPSGTGNENTLY | 0.80061173 |
| GNGGNGGAGGEGGPGGAGGAGA | ASSQEPSGTGNENTLY | 0.101090245 |
| GSTAGNGGNGGNATNGTIAGQP | ASSQEPSGTGNENTLY | 0.71603453 |
| GTGGDGGNGGNGGTGG | ASSQEPSGTGNENTLY | 0.591027 |
| GVRRPAGIPADAPV | ASSQEPSGTGNENTLY | 0.88198745 |
| GYRQVTDAHLLTLARR | ASSQEPSGTGNENTLY | 0.74296325 |
| IAKMGQDGHDRG | ASSQEPSGTGNENTLY | 0.70575583 |
| KAEPSIEIQYT | ASSQEPSGTGNENTLY | 0.77279425 |
| KPVRIRMAADEDTVAVT | ASSQEPSGTGNENTLY | 0.7672912 |
| MPPRIAGMRLLVIKPEP | ASSQEPSGTGNENTLY | 0.7534801 |
| MKFYAVALFRSRGVPPTR | ASSQEPSGTGNENTLY | 0.70244956 |
| NAGGFGILGAVAHSP | ASSQEPSGTGNENTLY | 0.4915187 |
| NEGREITLAMAEM | ASSQEPSGTGNENTLY | 0.8495116 |
| NELVEWSHPLGHRWRWE | ASSQEPSGTGNENTLY | 0.7144512 |
| NLGAGGTGGNGGNPG | ASSQEPSGTGNENTLY | 0.68804777 |
| NTEWPKWETVMAADPVD | ASSQEPSGTGNENTLY | 0.71015966 |
| PASAAPAETQP | ASSQEPSGTGNENTLY | 0.7670773 |
| QQSSDGKNGVRAAEITGE | ASSQEPSGTGNENTLY | 0.56681454 |
| RAPDAVTTSWWKE | ASSQEPSGTGNENTLY | 0.8747096 |
| SGFGNVGAHTSGWFNQ | ASSQEPSGTGNENTLY | 0.6846207 |
| SGVAGAVGQAASVGGLKV | ASSQEPSGTGNENTLY | 0.7452966 |

| | | |
|---|---|---|
| SNGGAGTGSAGGNGGT | ASSQEPSGTGNENTLY | 0.87450296 |
| TGGAGGEGGAAG | ASSQEPSGTGNENTLY | 0.738546 |
| TGGDGGAAGAGGAPGHG | ASSQEPSGTGNENTLY | 0.749807 |
| TMPPLLAAIANAGRHGV | ASSQEPSGTGNENTLY | 0.7680779 |
| TSSTAGNSGGA | ASSQEPSGTGNENTLY | 0.6862012 |
| VPATVPVGILC | ASSQEPSGTGNENTLY | 0.7292634 |
| VQRPDSGGVVLTGRIS | ASSQEPSGTGNENTLY | 0.736904 |
| WTPNPVPATKGVPAGLVQAT | ASSQEPSGTGNENTLY | 0.88115513 |
| YALKRALAKFSDRFDVVII | ASSQEPSGTGNENTLY | 0.8249053 |
| YLIGSVIV | ASSQEPSGTGNENTLY | 0.72297555 |
| YGQLVAVID | ASSQEPSGTGNENTLY | 0.84479284 |
| VTVGGMTLPA | ASSQEPSGTGNENTLY | 0.8821387 |
| VLALPAV | ASSQEPSGTGNENTLY | 0.8800552 |
| VIPPTLNYET | ASSQEPSGTGNENTLY | 0.6729052 |
| VGMGGMHPSSG | ASSQEPSGTGNENTLY | 0.7648351 |
| VGLAGGAATA | ASSQEPSGTGNENTLY | 0.7604184 |
| VGGAGGAGGSGP | ASSQEPSGTGNENTLY | 0.7293174 |
| VFAAFMV | ASSQEPSGTGNENTLY | 0.7408748 |
| TLYDKLR | ASSQEPSGTGNENTLY | 0.69893837 |
| TLAVFTVL | ASSQEPSGTGNENTLY | 0.87015164 |
| TGSVQPARV | ASSQEPSGTGNENTLY | 0.87217695 |
| TGGGAGSGGT | ASSQEPSGTGNENTLY | 0.7965611 |
| SHLETSL | ASSQEPSGTGNENTLY | 0.57546806 |
| SFTTGMT | ASSQEPSGTGNENTLY | 0.7526401 |
| QVEVAKRMGVRQPTVS | ASSQEPSGTGNENTLY | 0.6664249 |
| QGSGGPALAM | ASSQEPSGTGNENTLY | 0.6928698 |
| MLAPSNP | ASSQEPSGTGNENTLY | 0.7357988 |
| LVNNAAI | ASSQEPSGTGNENTLY | 0.58024186 |
| LTGEINTLA | ASSQEPSGTGNENTLY | 0.7849349 |
| LSIFGPV | ASSQEPSGTGNENTLY | 0.875892 |
| LLALLANL | ASSQEPSGTGNENTLY | 0.69136965 |
| LISNASDALDKLR | ASSQEPSGTGNENTLY | 0.8211365 |
| LINEAVP | ASSQEPSGTGNENTLY | 0.86619973 |
| LGYAFSGL | ASSQEPSGTGNENTLY | 0.84015834 |
| LDFVAVAL | ASSQEPSGTGNENTLY | 0.8683509 |
| LAQPGAE | ASSQEPSGTGNENTLY | 0.8472575 |
| ISNLGAVTGF | ASSQEPSGTGNENTLY | 0.8485584 |
| HQEDPGRI | ASSQEPSGTGNENTLY | 0.8099015 |
| GTPGEPTP | ASSQEPSGTGNENTLY | 0.83864623 |
| GFFNSGS | ASSQEPSGTGNENTLY | 0.76102 |
| GAGGGGLTAGAN | ASSQEPSGTGNENTLY | 0.7797253 |
| FVDALVNM | ASSQEPSGTGNENTLY | 0.7574418 |
| FLTVAPEL | ASSQEPSGTGNENTLY | 0.83724296 |
| FETLTAL | ASSQEPSGTGNENTLY | 0.7919911 |
| DVINPIEL | ASSQEPSGTGNENTLY | 0.86541766 |
| APVSGAPV | ASSQEPSGTGNENTLY | 0.4841941 |
| ALVGGLAW | ASSQEPSGTGNENTLY | 0.6863127 |
| ALAAAAATAI | ASSQEPSGTGNENTLY | 0.85690916 |
| AFGLGLLL | ASSQEPSGTGNENTLY | 0.20807901 |
| AFFAAASL | ASSQEPSGTGNENTLY | 0.58779943 |
| AAAALPAFSPP | ASSQEPSGTGNENTLY | 0.8133851 |

| | | |
|---|---|---|
| VVLAPAL | ASSQEPSGTGNENTLY | 0.8492451 |
| TIGHVDHGKTTL | ASSQEPSGTGNENTLY | 0.22139624 |
| LISNASDALDK | ASSQEPSGTGNENTLY | 0.7281529 |
| LISNASDAL | ASSQEPSGTGNENTLY | 0.57706356 |
| ISNASDALDK | ASSQEPSGTGNENTLY | 0.8700324 |
| NQAETLVYQTE | ASSQEPSGTGNENTLY | 0.77573395 |
| TGGTGGTAGAG | ASSQEPSGTGNENTLY | 0.5842953 |
| TGSTAGGAGGA | ASSQEPSGTGNENTLY | 0.86404514 |
| DPLAVPVAAGPV | ASSQEPSGTGNENTLY | 0.584234 |
| GGTGGTPGNGGN | ASSQEPSGTGNENTLY | 0.7625644 |
| PAGLAGVDPATL | ASSQEPSGTGNENTLY | 0.87144685 |
| GNNLVVEAPHAGASV | ASSQEPSGTGNENTLY | 0.8717376 |
| KVTRIMSASTLEQPA | ASSQEPSGTGNENTLY | 0.6779347 |
| LGFALPIALPRRTVV | ASSQEPSGTGNENTLY | 0.8273681 |
| TRLGNLTARKLVVLV | ASSQEPSGTGNENTLY | 0.7262212 |
| WSQQRRLIYFPSAGP | ASSQEPSGTGNENTLY | 0.88678473 |
| GPNGNPDPMAAAVDIR | ASSQEPSGTGNENTLY | 0.76423186 |
| TQRCVSVADLRAAASR | ASSQEPSGTGNENTLY | 0.64818704 |
| QFVLDVAEHPVEVLTETV | ASSQEPSGTGNENTLY | 0.8542152 |
| GGAGGTGGAGGNGGTQAGDG | ASSQEPSGTGNENTLY | 0.8381922 |
| PLLATLRAIAVDVDAKPAQV | ASSQEPSGTGNENTLY | 0.81096643 |
| APKFRTGGYMIDFWGVGYQVAK | ASSQEPSGTGNENTLY | 0.8329599 |
| GAGGAGGAGGAAGGPGGTGGTG | ASSQEPSGTGNENTLY | 0.78518355 |
| IGALGLLATDVAPAGSVAFGVA | ASSQEPSGTGNENTLY | 0.73874885 |
| AGGAGGIGGNGGAGGAGGVLMG | ASSQEPSGTGNENTLY | 0.83941865 |
| ADELIGLL | ASSQEPSGTGNENTLY | 0.85655 |
| ADRALKA | ASSQEPSGTGNENTLY | 0.71643656 |
| AGAVGQVGGAGGA | ASSQEPSGTGNENTLY | 0.8773662 |
| AGGGGGLGGAP | ASSQEPSGTGNENTLY | 0.7785087 |
| AGGLLYGNGGAG | ASSQEPSGTGNENTLY | 0.7194646 |
| AGLESIGL | ASSQEPSGTGNENTLY | 0.8873092 |
| AGLKTGII | ASSQEPSGTGNENTLY | 0.81048644 |
| ASLAALLL | ASSQEPSGTGNENTLY | 0.8501682 |
| ASVINISSI | ASSQEPSGTGNENTLY | 0.85984766 |
| ATPPGPA | ASSQEPSGTGNENTLY | 0.83665156 |
| ATVASALV | ASSQEPSGTGNENTLY | 0.70681965 |
| ATVASLIL | ASSQEPSGTGNENTLY | 0.8294668 |
| AVNGPVLIRH | ASSQEPSGTGNENTLY | 0.77140576 |
| DGAAGAVGI | ASSQEPSGTGNENTLY | 0.85674417 |
| DPLLHGI | ASSQEPSGTGNENTLY | 0.81240296 |
| ELGAALGAL | ASSQEPSGTGNENTLY | 0.73176813 |
| ELGRTQAV | ASSQEPSGTGNENTLY | 0.85532236 |
| FVAGGYLS | ASSQEPSGTGNENTLY | 0.87632024 |
| GASGGAGGAGGW | ASSQEPSGTGNENTLY | 0.84269786 |
| GGAGAGGDGGAGGI | ASSQEPSGTGNENTLY | 0.8299644 |
| GTGHNGSQPGMGGNG | ASSQEPSGTGNENTLY | 0.8746128 |
| GYAAASASAS | ASSQEPSGTGNENTLY | 0.7391701 |
| HQVDVAAVQQG | ASSQEPSGTGNENTLY | 0.7979579 |
| IGAAIAHT | ASSQEPSGTGNENTLY | 0.8637371 |
| IGMIEVAQL | ASSQEPSGTGNENTLY | 0.87082255 |
| IRLAPEHTSAMD | ASSQEPSGTGNENTLY | 0.8105165 |

| | | |
|---|---|---|
| IYNVSVL | ASSQEPSGTGNENTLY | 0.86553144 |
| KQGFLGM | ASSQEPSGTGNENTLY | 0.6760864 |
| LESVITQL | ASSQEPSGTGNENTLY | 0.784597 |
| LGLGALVVV | ASSQEPSGTGNENTLY | 0.61613667 |
| LIGNGGNGGSGGI | ASSQEPSGTGNENTLY | 0.7622748 |
| LNAINAPTL | ASSQEPSGTGNENTLY | 0.6478153 |
| LQLLNTGV | ASSQEPSGTGNENTLY | 0.785184 |
| LQTSIEF | ASSQEPSGTGNENTLY | 0.8380463 |
| MSFAVSEL | ASSQEPSGTGNENTLY | 0.80169976 |
| NTGALGL | ASSQEPSGTGNENTLY | 0.53359187 |
| PPGAGGAGGAGTT | ASSQEPSGTGNENTLY | 0.82697225 |
| QLVGLGI | ASSQEPSGTGNENTLY | 0.7788406 |
| RFLEQVHQL | ASSQEPSGTGNENTLY | 0.87976277 |
| RGVGYVLR | ASSQEPSGTGNENTLY | 0.7829509 |
| SATGVFAGVI | ASSQEPSGTGNENTLY | 0.8307432 |
| SLPAITLP | ASSQEPSGTGNENTLY | 0.7075716 |
| STALAFTV | ASSQEPSGTGNENTLY | 0.82727623 |
| SVLTVGLGK | ASSQEPSGTGNENTLY | 0.5861731 |
| TALAGAGSGI | ASSQEPSGTGNENTLY | 0.7151212 |
| TGLPFTGI | ASSQEPSGTGNENTLY | 0.6977602 |
| VAEHVTV | ASSQEPSGTGNENTLY | 0.6603919 |
| VDIVLNSL | ASSQEPSGTGNENTLY | 0.83279175 |
| VLTGGGAVL | ASSQEPSGTGNENTLY | 0.8690289 |
| VNVGLAYGI | ASSQEPSGTGNENTLY | 0.8406359 |
| VSSAAVGAV | ASSQEPSGTGNENTLY | 0.85767424 |
| VTSVYEAGV | ASSQEPSGTGNENTLY | 0.8666667 |
| WVALDRGA | ASSQEPSGTGNENTLY | 0.87942994 |
| YLVGSVVI | ASSQEPSGTGNENTLY | 0.7270781 |
| GPPGTGKTLL | ASSQEPSGTGNENTLY | 0.76178735 |
| GVGMTKF | ASSQEPSGTGNENTLY | 0.62367755 |
| IGIGGSDL | ASSQEPSGTGNENTLY | 0.76987374 |
| IGTLLAEL | ASSQEPSGTGNENTLY | 0.75640416 |
| IYTSGTT | ASSQEPSGTGNENTLY | 0.8043176 |
| LSFPAGV | ASSQEPSGTGNENTLY | 0.17675135 |
| LYTSGTT | ASSQEPSGTGNENTLY | 0.76955163 |
| VGAFPFGF | ASSQEPSGTGNENTLY | 0.88323474 |
| VGAGTVEF | ASSQEPSGTGNENTLY | 0.6927308 |
| VGKSTLLL | ASSQEPSGTGNENTLY | 0.81111884 |
| AALAHAGGGIV | ASSQEPSGTGNENTLY | 0.8176168 |
| AALVAVSLL | ASSQEPSGTGNENTLY | 0.74601865 |
| AAVSRAVAAL | ASSQEPSGTGNENTLY | 0.8269025 |
| AGGSSGIGGF | ASSQEPSGTGNENTLY | 0.74540377 |
| AGLTVANANG | ASSQEPSGTGNENTLY | 0.85916483 |
| AGLVGAVL | ASSQEPSGTGNENTLY | 0.8748273 |
| ALSQAFTV | ASSQEPSGTGNENTLY | 0.8090672 |
| ALTFSVSL | ASSQEPSGTGNENTLY | 0.7772868 |
| ALTGAALVAL | ASSQEPSGTGNENTLY | 0.8069994 |
| ASFSIGAGVL | ASSQEPSGTGNENTLY | 0.66796225 |
| DGSVYKI | ASSQEPSGTGNENTLY | 0.8103983 |
| EQLAEIV | ASSQEPSGTGNENTLY | 0.594393 |
| FAVSELTVM | ASSQEPSGTGNENTLY | 0.7538088 |

| | | |
|---|---|---|
| GAAPLAAGL | ASSQEPSGTGNENTLY | 0.7501241 |
| GADGGDGGSGGAGG | ASSQEPSGTGNENTLY | 0.81552815 |
| GFGLDTGGAGG | ASSQEPSGTGNENTLY | 0.7998978 |
| GLASGGLL | ASSQEPSGTGNENTLY | 0.8371696 |
| GRIAGAAVLL | ASSQEPSGTGNENTLY | 0.63557833 |
| GSAGSLNTSF | ASSQEPSGTGNENTLY | 0.84518266 |
| GTGQIGI | ASSQEPSGTGNENTLY | 0.7952783 |
| GTLIALIL | ASSQEPSGTGNENTLY | 0.62966835 |
| IAATMGFVVG | ASSQEPSGTGNENTLY | 0.86147696 |
| IFAQALL | ASSQEPSGTGNENTLY | 0.83427215 |
| IGAAVALGF | ASSQEPSGTGNENTLY | 0.7266041 |
| IGLTPLAL | ASSQEPSGTGNENTLY | 0.8508662 |
| ILGSNTSTL | ASSQEPSGTGNENTLY | 0.8191054 |
| IVFSLTVL | ASSQEPSGTGNENTLY | 0.62999004 |
| LASKATGF | ASSQEPSGTGNENTLY | 0.6678616 |
| LASVALIL | ASSQEPSGTGNENTLY | 0.78920275 |
| LFAVGLTL | ASSQEPSGTGNENTLY | 0.7782898 |
| LFNGGIGTYI | ASSQEPSGTGNENTLY | 0.309811 |
| LTLGAGII | ASSQEPSGTGNENTLY | 0.8457161 |
| LVAAALGDI | ASSQEPSGTGNENTLY | 0.7261754 |
| NTAFVQLGI | ASSQEPSGTGNENTLY | 0.6443194 |
| QLFSGAEL | ASSQEPSGTGNENTLY | 0.7827375 |
| SAATFITL | ASSQEPSGTGNENTLY | 0.84911877 |
| SNYGLANLGV | ASSQEPSGTGNENTLY | 0.7228116 |
| SYGGAYAVM | ASSQEPSGTGNENTLY | 0.73396075 |
| TTGVGGLTLG | ASSQEPSGTGNENTLY | 0.8248285 |
| TTQTIYI | ASSQEPSGTGNENTLY | 0.83016217 |
| VAAGAGLGAVI | ASSQEPSGTGNENTLY | 0.86022544 |
| VAAVVTLL | ASSQEPSGTGNENTLY | 0.8282319 |
| VGFLGGLI | ASSQEPSGTGNENTLY | 0.81064194 |
| VGVIPILL | ASSQEPSGTGNENTLY | 0.82491815 |
| VIFASSLI | ASSQEPSGTGNENTLY | 0.7921616 |
| VKGGDGGAAAG | ASSQEPSGTGNENTLY | 0.7126522 |
| VTKTFTNI | ASSQEPSGTGNENTLY | 0.6513761 |
| VVAAAILL | ASSQEPSGTGNENTLY | 0.7032716 |
| YAALDVEL | ASSQEPSGTGNENTLY | 0.68662775 |
| LASVALLL | ASSQEPSGTGNENTLY | 0.802775 |
| LDAVHRV | ASSQEPSGTGNENTLY | 0.88342214 |
| QGLSSSIF | ASSQEPSGTGNENTLY | 0.7778261 |
| RGTFANIRL | ASSQEPSGTGNENTLY | 0.59213334 |
| RQLWWGHRI | ASSQEPSGTGNENTLY | 0.7762051 |
| YGPPGTGKTLL | ASSQEPSGTGNENTLY | 0.771702 |
| AALVGAPANVA | ASSQEPSGTGNENTLY | 0.83833563 |
| AATFRRTE | ASSQEPSGTGNENTLY | 0.85517484 |
| AGALGGGI | ASSQEPSGTGNENTLY | 0.87104726 |
| AGGAGGAGGIGAG | ASSQEPSGTGNENTLY | 0.84289205 |
| AGSGALGL | ASSQEPSGTGNENTLY | 0.7220414 |
| AQIADQI | ASSQEPSGTGNENTLY | 0.84612554 |
| ASAVAVTYL | ASSQEPSGTGNENTLY | 0.76909494 |
| ASTVLGGTGG | ASSQEPSGTGNENTLY | 0.8095213 |
| AVGGEPAK | ASSQEPSGTGNENTLY | 0.82868195 |

| | | |
|---|---|---|
| CEQGTFGGTC | ASSQEPSGTGNENTLY | 0.81835663 |
| DEGKAPAG | ASSQEPSGTGNENTLY | 0.79629004 |
| DTVVFPAL | ASSQEPSGTGNENTLY | 0.7902149 |
| FEGITMAG | ASSQEPSGTGNENTLY | 0.60200816 |
| FPLVVQD | ASSQEPSGTGNENTLY | 0.8454376 |
| GAFTTPQ | ASSQEPSGTGNENTLY | 0.70231307 |
| GFGGFGTAGGVGGA | ASSQEPSGTGNENTLY | 0.70357865 |
| GFLAQFL | ASSQEPSGTGNENTLY | 0.7451412 |
| GGAGGAGGNAGVA | ASSQEPSGTGNENTLY | 0.8777565 |
| GGVGTAGAGGAGGAGG | ASSQEPSGTGNENTLY | 0.85566777 |
| GNGGGGDAGNAGSGGN | ASSQEPSGTGNENTLY | 0.6844108 |
| GTGGAGGAGNDAGS | ASSQEPSGTGNENTLY | 0.73812073 |
| GTTNTGWFNS | ASSQEPSGTGNENTLY | 0.78063506 |
| IFMGPLFTL | ASSQEPSGTGNENTLY | 0.6573796 |
| ILAHAIEAI | ASSQEPSGTGNENTLY | 0.86071503 |
| IRSAIREAL | ASSQEPSGTGNENTLY | 0.7511066 |
| ITFAGGG | ASSQEPSGTGNENTLY | 0.7571937 |
| ITGMSGHQEV | ASSQEPSGTGNENTLY | 0.73690164 |
| IVAGLACTL | ASSQEPSGTGNENTLY | 0.83294964 |
| IVGGGLAGA | ASSQEPSGTGNENTLY | 0.7278947 |
| IVTGVLNI | ASSQEPSGTGNENTLY | 0.8625057 |
| LAQQGCI | ASSQEPSGTGNENTLY | 0.2578851 |
| LCLHVEN | ASSQEPSGTGNENTLY | 0.87896127 |
| LDAISDGS | ASSQEPSGTGNENTLY | 0.6380752 |
| LGGFAAML | ASSQEPSGTGNENTLY | 0.72068465 |
| LGGKNPMI | ASSQEPSGTGNENTLY | 0.42037565 |
| LGPKGVAAIL | ASSQEPSGTGNENTLY | 0.8282378 |
| LMVGVSI | ASSQEPSGTGNENTLY | 0.80763435 |
| LVGAAFVL | ASSQEPSGTGNENTLY | 0.8648565 |
| LVLAVAPL | ASSQEPSGTGNENTLY | 0.7824708 |
| MAAIAVIDAA | ASSQEPSGTGNENTLY | 0.84542847 |
| NNGINAELQ | ASSQEPSGTGNENTLY | 0.83835334 |
| RGEPLVLVHG | ASSQEPSGTGNENTLY | 0.7319582 |
| RGGAGGAAGLI | ASSQEPSGTGNENTLY | 0.54570425 |
| RSYRMHPAV | ASSQEPSGTGNENTLY | 0.8231251 |
| SAAHLLAGQLA | ASSQEPSGTGNENTLY | 0.74874604 |
| SAAVGSY | ASSQEPSGTGNENTLY | 0.8642911 |
| SPHAVLHRI | ASSQEPSGTGNENTLY | 0.7955495 |
| SQIAKVL | ASSQEPSGTGNENTLY | 0.57883084 |
| TALGHLVP | ASSQEPSGTGNENTLY | 0.83111525 |
| TAPTTSGAA | ASSQEPSGTGNENTLY | 0.82383263 |
| TDGDDGNGGSPGT | ASSQEPSGTGNENTLY | 0.61851245 |
| TDGLGEN | ASSQEPSGTGNENTLY | 0.758917 |
| TGGPPLPGQA | ASSQEPSGTGNENTLY | 0.8013144 |
| TIPAITI | ASSQEPSGTGNENTLY | 0.58801186 |
| TIPGFTI | ASSQEPSGTGNENTLY | 0.8341671 |
| TSIASGALA | ASSQEPSGTGNENTLY | 0.72164875 |
| TVTLPGAAI | ASSQEPSGTGNENTLY | 0.71739745 |
| VASGLSAALP | ASSQEPSGTGNENTLY | 0.80010605 |
| VGAGVIGI | ASSQEPSGTGNENTLY | 0.7524017 |
| VGPKRFGV | ASSQEPSGTGNENTLY | 0.76459795 |

| | | |
|---|---|---|
| VNVAFPD | ASSQEPSGTGNENTLY | 0.7644025 |
| VSMTNSGVS | ASSQEPSGTGNENTLY | 0.8474579 |
| VTDVLGDLSV | ASSQEPSGTGNENTLY | 0.8453822 |
| VVVVGAGFAGL | ASSQEPSGTGNENTLY | 0.75704753 |
| VYTVGTVV | ASSQEPSGTGNENTLY | 0.7930621 |
| WTTFSGP | ASSQEPSGTGNENTLY | 0.8643829 |
| YDELVEAF | ASSQEPSGTGNENTLY | 0.58886874 |
| YGTGGAGGNGG | ASSQEPSGTGNENTLY | 0.7206328 |
| YTNVMAV | ASSQEPSGTGNENTLY | 0.8312546 |
| FLEQVHQL | ASSQEPSGTGNENTLY | 0.7353374 |
| GLSLLAVAV | ASSQEPSGTGNENTLY | 0.8567657 |
| LEQVHQL | ASSQEPSGTGNENTLY | 0.8064512 |
| LNAAGAI | ASSQEPSGTGNENTLY | 0.81177187 |
| PAVHASL | ASSQEPSGTGNENTLY | 0.7419121 |
| TELEDDA | ASSQEPSGTGNENTLY | 0.87510014 |
| TPPGPPN | ASSQEPSGTGNENTLY | 0.84405816 |
| TPPPPPPP | ASSQEPSGTGNENTLY | 0.8243075 |
| TQSPALI | ASSLSEGPNSDYT | 0.8301898 |
| EFAAAFPQ | ASSLSEGPNSDYT | 0.7032793 |
| LLSGNLDVL | ASSLSEGPNSDYT | 0.85623264 |
| GPANLGNYNI | ASSLSEGPNSDYT | 0.8444544 |
| TLGPTLP | ASSLSEGPNSDYT | 0.63375366 |
| VYAYSSCCK | ASSLSEGPNSDYT | 0.7460053 |
| APAPAPAPNI | ASSLSEGPNSDYT | 0.8879024 |
| PGGPVPIP | ASSLSEGPNSDYT | 0.85659754 |
| FLTVALEP | ASSLSEGPNSDYT | 0.74982846 |
| SQYFGANL | ASSLSEGPNSDYT | 0.71660006 |
| GGVKPAA | ASSLSEGPNSDYT | 0.68897176 |
| GGTGGAGGAGGW | ASSLSEGPNSDYT | 0.7590293 |
| GVGGVAGLGGAGG | ASSLSEGPNSDYT | 0.64533496 |
| VLLVLATTI | ASSLSEGPNSDYT | 0.83120793 |
| IGIFAAML | ASSLSEGPNSDYT | 0.7354368 |
| AVGDLID | ASSLSEGPNSDYT | 0.8085543 |
| NISPLPL | ASSLSEGPNSDYT | 0.80926085 |
| VTAVFVAI | ASSLSEGPNSDYT | 0.86964047 |
| SGDDNDAAL | ASSLSEGPNSDYT | 0.7356899 |
| AVAALAHS | ASSLSEGPNSDYT | 0.7808319 |
| GGVTVMATP | ASSLSEGPNSDYT | 0.570013 |
| AGLAANVRD | ASSLSEGPNSDYT | 0.67287433 |
| TNRPDLIDPALLRPG | ASSLSEGPNSDYT | 0.7635105 |
| LANLGVTAL | ASSLSEGPNSDYT | 0.74965036 |
| TGERIEL | ASSLSEGPNSDYT | 0.38741922 |
| NGEELAA | ASSLSEGPNSDYT | 0.8722416 |
| VLSLVWG | ASSLSEGPNSDYT | 0.78054297 |
| IGYALWL | ASSLSEGPNSDYT | 0.84564334 |
| LSLLLAPL | ASSLSEGPNSDYT | 0.73991054 |
| TLDEALK | ASSLSEGPNSDYT | 0.7690568 |
| TDGAGTAEGG | ASSLSEGPNSDYT | 0.7319604 |
| NAPGGAGGNGG | ASSLSEGPNSDYT | 0.8351291 |
| VSVFMGMGGNF | ASSLSEGPNSDYT | 0.68236095 |
| LASYLAAL | ASSLSEGPNSDYT | 0.48021168 |

| | | |
|---|---|---|
| LVGGTPAL | ASSLSEGPNSDYT | 0.7519311 |
| VRNADVL | ASSLSEGPNSDYT | 0.6718015 |
| TCSSPFLVSAM | ASSLSEGPNSDYT | 0.8762082 |
| VHITDAE | ASSLSEGPNSDYT | 0.8627962 |
| EYAAPSF | ASSLSEGPNSDYT | 0.8007382 |
| VVDAFHGGL | ASSLSEGPNSDYT | 0.8807663 |
| ASVVDPQ | ASSLSEGPNSDYT | 0.24093772 |
| SYNEFGGLS | ASSLSEGPNSDYT | 0.72923493 |
| DPDGSVY | ASSLSEGPNSDYT | 0.8549832 |
| MVSTIHATE | ASSLSEGPNSDYT | 0.8396746 |
| LAAGVADPV | ASSLSEGPNSDYT | 0.7730199 |
| TKEQIAA | ASSLSEGPNSDYT | 0.8714464 |
| SLASPAAL | ASSLSEGPNSDYT | 0.6681043 |
| IGLVFYGV | ASSLSEGPNSDYT | 0.77084136 |
| FLLDASNL | ASSLSEGPNSDYT | 0.8082815 |
| YQVGNFP | ASSLSEGPNSDYT | 0.8703643 |
| ESLLSQST | ASSLSEGPNSDYT | 0.82687294 |
| LFTVGFAAVG | ASSLSEGPNSDYT | 0.68926376 |
| ATPPTPT | ASSLSEGPNSDYT | 0.8259666 |
| TDLATLG | ASSLSEGPNSDYT | 0.8844433 |
| SAGGTGR | ASSLSEGPNSDYT | 0.70633435 |
| GFSNSGS | ASSLSEGPNSDYT | 0.8354676 |
| GGNGGSGGTSV | ASSLSEGPNSDYT | 0.8110797 |
| GTSVATGGAGN | ASSLSEGPNSDYT | 0.67901844 |
| NGGNGGSGGTS | ASSLSEGPNSDYT | 0.66576195 |
| AVVKFPH | ASSLSEGPNSDYT | 0.6596636 |
| TPPPPPP | ASSLSEGPNSDYT | 0.8465852 |
| AAPAAAAAPA | ASSLSEGPNSDYT | 0.61719954 |
| IALNEDL | ASSLSEGPNSDYT | 0.804366 |
| IVNVSSM | ASSLSEGPNSDYT | 0.78836834 |
| TGIPPAPR | ASSLSEGPNSDYT | 0.89099896 |
| FDLGGGTF | ASSLSEGPNSDYT | 0.82614756 |
| FDLGGGTFDVS | ASSLSEGPNSDYT | 0.84606767 |
| NEPTAAA | ASSLSEGPNSDYT | 0.67449474 |
| NEPTAAALA | ASSLSEGPNSDYT | 0.84681225 |
| GSIINISSI | ASSLSEGPNSDYT | 0.829538 |
| VAPGNAGTA | ASSLSEGPNSDYT | 0.65286106 |
| TGGGASLE | ASSLSEGPNSDYT | 0.8648622 |
| VAVYNHYKRI | ASSLSEGPNSDYT | 0.7012186 |
| TDPAPATT | ASSLSEGPNSDYT | 0.74865055 |
| TNRPDILDPALLRPG | ASSLSEGPNSDYT | 0.84063315 |
| LTTTTTS | ASSLSEGPNSDYT | 0.83538246 |
| PTGSGKTL | ASSLSEGPNSDYT | 0.80030394 |
| TGSGKTL | ASSLSEGPNSDYT | 0.84489685 |
| TIGHVDHGK | ASSLSEGPNSDYT | 0.69868594 |
| TIGHVDHGKTT | ASSLSEGPNSDYT | 0.85095966 |
| GHVDHGKTT | ASSLSEGPNSDYT | 0.8510444 |
| AQFEHTLL | ASSLSEGPNSDYT | 0.7476111 |
| IVNQDEL | ASSLSEGPNSDYT | 0.8848677 |
| TNGSQFF | ASSLSEGPNSDYT | 0.8610147 |
| TGKGIVE | ASSLSEGPNSDYT | 0.5322292 |

| | | |
|---|---|---|
| LISNASDA | ASSLSEGPNSDYT | 0.80466974 |
| AELRTLGELTKIAKAHGAQ | ASSLSEGPNSDYT | 0.87354434 |
| APNAVAAAEPSW | ASSLSEGPNSDYT | 0.8625771 |
| APQVAGAATAVL | ASSLSEGPNSDYT | 0.8723664 |
| ARRDPAAAAAILPTDA | ASSLSEGPNSDYT | 0.8325754 |
| DAAARDVLAEAGLAEYF | ASSLSEGPNSDYT | 0.7094167 |
| DDQIVLSAKANLAKVEQ | ASSLSEGPNSDYT | 0.873325 |
| DELLPDTGEDSDEDGH | ASSLSEGPNSDYT | 0.8929967 |
| ELPEAPANRATVPQD | ASSLSEGPNSDYT | 0.8402114 |
| FGQSKIERVTGIG | ASSLSEGPNSDYT | 0.6985823 |
| GAGGTGSTAGGAGGAGGAGGLY | ASSLSEGPNSDYT | 0.71452284 |
| GHGGTGAAGGNGGTGGAGGSGI | ASSLSEGPNSDYT | 0.84392035 |
| GNGGNGGAGGEGGPGGAGGAGA | ASSLSEGPNSDYT | 0.85766494 |
| GSTAGNGGNGGNATNGTIAGQP | ASSLSEGPNSDYT | 0.69824165 |
| GTGGDGGNGGNGGTGG | ASSLSEGPNSDYT | 0.79260397 |
| GVRRPAGIPADAPV | ASSLSEGPNSDYT | 0.8018071 |
| GYRQVTDAHLLTLARR | ASSLSEGPNSDYT | 0.48903114 |
| IAKMGQDGHDRG | ASSLSEGPNSDYT | 0.8116182 |
| KAEPSIEIQYT | ASSLSEGPNSDYT | 0.8585637 |
| KPVRIRMAADEDTVAVT | ASSLSEGPNSDYT | 0.8521042 |
| MPPRIAGMRLLVIKPEP | ASSLSEGPNSDYT | 0.6733155 |
| MKFYAVALFRSRGVPPTR | ASSLSEGPNSDYT | 0.86214435 |
| NAGGFGILGAVAHSP | ASSLSEGPNSDYT | 0.85881364 |
| NEGREITLAMAEM | ASSLSEGPNSDYT | 0.8897701 |
| NELVEWSHPLGHRWRWE | ASSLSEGPNSDYT | 0.5674767 |
| NLGAGGTGGNGGNPG | ASSLSEGPNSDYT | 0.3402198 |
| NTEWPKWETVMAADPVD | ASSLSEGPNSDYT | 0.8099563 |
| PASAAPAETQP | ASSLSEGPNSDYT | 0.7930339 |
| QQSSDGKNGVRAAEITGE | ASSLSEGPNSDYT | 0.76517177 |
| RAPDAVTTSWWKE | ASSLSEGPNSDYT | 0.7228982 |
| SGFGNVGAHTSGWFNQ | ASSLSEGPNSDYT | 0.7996542 |
| SGVAGAVGQAASVGGLKV | ASSLSEGPNSDYT | 0.8604579 |
| SNGGAGTGSAGGNGGT | ASSLSEGPNSDYT | 0.7099662 |
| TGGAGGEGGAAG | ASSLSEGPNSDYT | 0.64441335 |
| TGGDGGAAGAGGAPGHG | ASSLSEGPNSDYT | 0.85329986 |
| TMPPLLAAIANAGRHGV | ASSLSEGPNSDYT | 0.8492023 |
| TSSTAGNSGGA | ASSLSEGPNSDYT | 0.66302335 |
| VPATVPVGILC | ASSLSEGPNSDYT | 0.7446301 |
| VQRPDSGGVVLTGRIS | ASSLSEGPNSDYT | 0.7987235 |
| WTPNPVPATKGVPAGLVQAT | ASSLSEGPNSDYT | 0.78288746 |
| YALKRALAKFSDRFDVVII | ASSLSEGPNSDYT | 0.8039092 |
| YLIGSVIV | ASSLSEGPNSDYT | 0.6139571 |
| YGQLVAVID | ASSLSEGPNSDYT | 0.85094666 |
| VTVGGMTLPA | ASSLSEGPNSDYT | 0.6289997 |
| VLALPAV | ASSLSEGPNSDYT | 0.85373795 |
| VIPPTLNYET | ASSLSEGPNSDYT | 0.8151691 |
| VGMGGMHPSSG | ASSLSEGPNSDYT | 0.8399107 |
| VGLAGGAATA | ASSLSEGPNSDYT | 0.74712706 |
| VGGAGGAGGSGP | ASSLSEGPNSDYT | 0.6236408 |
| VFAAFMV | ASSLSEGPNSDYT | 0.7076134 |
| TLYDKLR | ASSLSEGPNSDYT | 0.5867997 |

| | | |
|---|---|---|
| TLAVFTVL | ASSLSEGPNSDYT | 0.86915237 |
| TGSVQPARV | ASSLSEGPNSDYT | 0.7184837 |
| TGGGAGSGGT | ASSLSEGPNSDYT | 0.8636397 |
| SHLETSL | ASSLSEGPNSDYT | 0.8353753 |
| SFTTGMT | ASSLSEGPNSDYT | 0.7424799 |
| QVEVAKRMGVRQPTVS | ASSLSEGPNSDYT | 0.85021526 |
| QGSGGPALAM | ASSLSEGPNSDYT | 0.58972883 |
| MLAPSNP | ASSLSEGPNSDYT | 0.8221922 |
| LVNNAAI | ASSLSEGPNSDYT | 0.7598689 |
| LTGEINTLA | ASSLSEGPNSDYT | 0.8589976 |
| LSIFGPV | ASSLSEGPNSDYT | 0.7017167 |
| LLALLANL | ASSLSEGPNSDYT | 0.85698664 |
| LISNASDALDKLR | ASSLSEGPNSDYT | 0.8158747 |
| LINEAVP | ASSLSEGPNSDYT | 0.66400206 |
| LGYAFSGL | ASSLSEGPNSDYT | 0.8335949 |
| LDFVAVAL | ASSLSEGPNSDYT | 0.8404163 |
| LAQPGAE | ASSLSEGPNSDYT | 0.8561732 |
| ISNLGAVTGF | ASSLSEGPNSDYT | 0.5893276 |
| HQEDPGRI | ASSLSEGPNSDYT | 0.81230295 |
| GTPGEPTP | ASSLSEGPNSDYT | 0.65670085 |
| GFFNSGS | ASSLSEGPNSDYT | 0.8023547 |
| GAGGGGLTAGAN | ASSLSEGPNSDYT | 0.8250377 |
| FVDALVNM | ASSLSEGPNSDYT | 0.7466283 |
| FLTVAPEL | ASSLSEGPNSDYT | 0.8507762 |
| FETLTAL | ASSLSEGPNSDYT | 0.6216632 |
| DVINPIEL | ASSLSEGPNSDYT | 0.6581091 |
| APVSGAPV | ASSLSEGPNSDYT | 0.75104606 |
| ALVGGLAW | ASSLSEGPNSDYT | 0.8169204 |
| ALAAAAATAI | ASSLSEGPNSDYT | 0.8430908 |
| AFGLGLLL | ASSLSEGPNSDYT | 0.6353704 |
| AFFAAASL | ASSLSEGPNSDYT | 0.77628946 |
| AAAALPAFSPP | ASSLSEGPNSDYT | 0.7872571 |
| VVLAPAL | ASSLSEGPNSDYT | 0.67472196 |
| TIGHVDHGKTTL | ASSLSEGPNSDYT | 0.8376258 |
| LISNASDALDK | ASSLSEGPNSDYT | 0.8014934 |
| LISNASDAL | ASSLSEGPNSDYT | 0.5429107 |
| ISNASDALDK | ASSLSEGPNSDYT | 0.8626272 |
| NQAETLVYQTE | ASSLSEGPNSDYT | 0.73296916 |
| TGGTGGTAGAG | ASSLSEGPNSDYT | 0.83645606 |
| TGSTAGGAGGA | ASSLSEGPNSDYT | 0.87974405 |
| DPLAVPVAAGPV | ASSLSEGPNSDYT | 0.6007992 |
| GGTGGTPGNGGN | ASSLSEGPNSDYT | 0.85787046 |
| PAGLAGVDPATL | ASSLSEGPNSDYT | 0.80797356 |
| GNNLVVEAPHAGASV | ASSLSEGPNSDYT | 0.80545807 |
| KVTRIMSASTLEQPA | ASSLSEGPNSDYT | 0.78180075 |
| LGFALPIALPRRTVV | ASSLSEGPNSDYT | 0.7314292 |
| TRLGNLTARKLVVLV | ASSLSEGPNSDYT | 0.87861884 |
| WSQQRRLIYFPSAGP | ASSLSEGPNSDYT | 0.83805203 |
| GPNGNPDPMAAAVDIR | ASSLSEGPNSDYT | 0.82271373 |
| TQRCVSVADLRAAASR | ASSLSEGPNSDYT | 0.83818626 |
| QFVLDVAEHPVEVLTETV | ASSLSEGPNSDYT | 0.8408641 |

| | | |
|---|---|---|
| GGAGGTGGAGGNGGTQAGDG | ASSLSEGPNSDYT | 0.8533144 |
| PLLATLRAIAVDVDAKPAQV | ASSLSEGPNSDYT | 0.66368026 |
| APKFRTGGYMIDFWGVGYQVAK | ASSLSEGPNSDYT | 0.56497395 |
| GAGGAGGAGGAAGGPGGTGGTG | ASSLSEGPNSDYT | 0.7667394 |
| IGALGLLATDVAPAGSVAFGVA | ASSLSEGPNSDYT | 0.73544705 |
| AGGAGGIGGNGGAGGAGGVLMG | ASSLSEGPNSDYT | 0.7887521 |
| ADELIGLL | ASSLSEGPNSDYT | 0.59373426 |
| ADRALKA | ASSLSEGPNSDYT | 0.83354723 |
| AGAVGQVGGAGGA | ASSLSEGPNSDYT | 0.88935024 |
| AGGGGGLGGAP | ASSLSEGPNSDYT | 0.6917185 |
| AGGLLYGNGGAG | ASSLSEGPNSDYT | 0.8890307 |
| AGLESIGL | ASSLSEGPNSDYT | 0.66918623 |
| AGLKTGII | ASSLSEGPNSDYT | 0.84441125 |
| ASLAALLL | ASSLSEGPNSDYT | 0.8537377 |
| ASVINISSI | ASSLSEGPNSDYT | 0.63711506 |
| ATPPGPA | ASSLSEGPNSDYT | 0.8296641 |
| ATVASALV | ASSLSEGPNSDYT | 0.7158994 |
| ATVASLIL | ASSLSEGPNSDYT | 0.77910465 |
| AVNGPVLIRH | ASSLSEGPNSDYT | 0.86679137 |
| DGAAGAVGI | ASSLSEGPNSDYT | 0.73409814 |
| DPLLHGI | ASSLSEGPNSDYT | 0.8419887 |
| ELGAALGAL | ASSLSEGPNSDYT | 0.7965238 |
| ELGRTQAV | ASSLSEGPNSDYT | 0.7373349 |
| FVAGGYLS | ASSLSEGPNSDYT | 0.8578029 |
| GASGGAGGAGGW | ASSLSEGPNSDYT | 0.8556391 |
| GGAGAGGDGGAGGI | ASSLSEGPNSDYT | 0.7890156 |
| GTGHNGSQPGMGGNG | ASSLSEGPNSDYT | 0.8879125 |
| GYAAASASAS | ASSLSEGPNSDYT | 0.6979412 |
| HQVDVAAVQQG | ASSLSEGPNSDYT | 0.87982005 |
| IGAAIAHT | ASSLSEGPNSDYT | 0.8477037 |
| IGMIEVAQL | ASSLSEGPNSDYT | 0.77211183 |
| IRLAPEHTSAMD | ASSLSEGPNSDYT | 0.7976589 |
| IYNVSVL | ASSLSEGPNSDYT | 0.81926996 |
| KQGFLGM | ASSLSEGPNSDYT | 0.736254 |
| LESVITQL | ASSLSEGPNSDYT | 0.8446252 |
| LGLGALVVV | ASSLSEGPNSDYT | 0.88702416 |
| LIGNGGNGGSGGI | ASSLSEGPNSDYT | 0.58047587 |
| LNAINAPTL | ASSLSEGPNSDYT | 0.89368033 |
| LQLLNTGV | ASSLSEGPNSDYT | 0.841377 |
| LQTSIEF | ASSLSEGPNSDYT | 0.7178505 |
| MSFAVSEL | ASSLSEGPNSDYT | 0.63983583 |
| NTGALGL | ASSLSEGPNSDYT | 0.24856508 |
| PPGAGGAGGAGTT | ASSLSEGPNSDYT | 0.66649747 |
| QLVGLGI | ASSLSEGPNSDYT | 0.527433 |
| RFLEQVHQL | ASSLSEGPNSDYT | 0.8441244 |
| RGVGYVLR | ASSLSEGPNSDYT | 0.8219068 |
| SATGVFAGVI | ASSLSEGPNSDYT | 0.5765315 |
| SLPAITLP | ASSLSEGPNSDYT | 0.89416695 |
| STALAFTV | ASSLSEGPNSDYT | 0.8598313 |
| SVLTVGLGK | ASSLSEGPNSDYT | 0.34787124 |
| TALAGAGSGI | ASSLSEGPNSDYT | 0.72998 |

| | | |
|---|---|---|
| TGLPFTGI | ASSLSEGPNSDYT | 0.53204215 |
| VAEHVTV | ASSLSEGPNSDYT | 0.718941 |
| VDIVLNSL | ASSLSEGPNSDYT | 0.3280955 |
| VLTGGGAVL | ASSLSEGPNSDYT | 0.8471983 |
| VNVGLAYGI | ASSLSEGPNSDYT | 0.6360204 |
| VSSAAVGAV | ASSLSEGPNSDYT | 0.11133923 |
| VTSVYEAGV | ASSLSEGPNSDYT | 0.7720239 |
| WVALDRGA | ASSLSEGPNSDYT | 0.74920464 |
| YLVGSVVI | ASSLSEGPNSDYT | 0.6554613 |
| GPPGTGKTLL | ASSLSEGPNSDYT | 0.87969923 |
| GVGMTKF | ASSLSEGPNSDYT | 0.38043064 |
| IGIGGSDL | ASSLSEGPNSDYT | 0.51303446 |
| IGTLLAEL | ASSLSEGPNSDYT | 0.8753177 |
| IYTSGTT | ASSLSEGPNSDYT | 0.8864505 |
| LSFPAGV | ASSLSEGPNSDYT | 0.3902516 |
| LYTSGTT | ASSLSEGPNSDYT | 0.8430674 |
| VGAFPFGF | ASSLSEGPNSDYT | 0.87458956 |
| VGAGTVEF | ASSLSEGPNSDYT | 0.8315515 |
| VGKSTLLL | ASSLSEGPNSDYT | 0.83774114 |
| AALAHAGGGIV | ASSLSEGPNSDYT | 0.8697846 |
| AALVAVSLL | ASSLSEGPNSDYT | 0.59664255 |
| AAVSRAVAAL | ASSLSEGPNSDYT | 0.88065827 |
| AGGSSGIGGF | ASSLSEGPNSDYT | 0.89619076 |
| AGLTVANANG | ASSLSEGPNSDYT | 0.66735244 |
| AGLVGAVL | ASSLSEGPNSDYT | 0.7671977 |
| ALSQAFTV | ASSLSEGPNSDYT | 0.476142 |
| ALTFSVSL | ASSLSEGPNSDYT | 0.71457267 |
| ALTGAALVAL | ASSLSEGPNSDYT | 0.2596814 |
| ASFSIGAGVL | ASSLSEGPNSDYT | 0.8490908 |
| DGSVYKI | ASSLSEGPNSDYT | 0.8598609 |
| EQLAEIV | ASSLSEGPNSDYT | 0.723517 |
| FAVSELTVM | ASSLSEGPNSDYT | 0.5763693 |
| GAAPLAAGL | ASSLSEGPNSDYT | 0.54272974 |
| GADGGDGGSGGAGG | ASSLSEGPNSDYT | 0.77804196 |
| GFGLDTGGAGG | ASSLSEGPNSDYT | 0.6694777 |
| GLASGGLL | ASSLSEGPNSDYT | 0.23468766 |
| GRIAGAAVLL | ASSLSEGPNSDYT | 0.69030386 |
| GSAGSLNTSF | ASSLSEGPNSDYT | 0.8736656 |
| GTGQIGI | ASSLSEGPNSDYT | 0.86239195 |
| GTLIALIL | ASSLSEGPNSDYT | 0.35598695 |
| IAATMGFVVG | ASSLSEGPNSDYT | 0.89472246 |
| IFAQALL | ASSLSEGPNSDYT | 0.8528123 |
| IGAAVALGF | ASSLSEGPNSDYT | 0.8893534 |
| IGLTPLAL | ASSLSEGPNSDYT | 0.6485232 |
| ILGSNTSTL | ASSLSEGPNSDYT | 0.5824787 |
| IVFSLTVL | ASSLSEGPNSDYT | 0.8683541 |
| LASKATGF | ASSLSEGPNSDYT | 0.77918947 |
| LASVALIL | ASSLSEGPNSDYT | 0.47408485 |
| LFAVGLTL | ASSLSEGPNSDYT | 0.8552244 |
| LFNGGIGTYI | ASSLSEGPNSDYT | 0.70473313 |
| LTLGAGII | ASSLSEGPNSDYT | 0.632259 |

| | | |
|---|---|---|
| LVAAALGDI | ASSLSEGPNSDYT | 0.73283154 |
| NTAFVQLGI | ASSLSEGPNSDYT | 0.85904586 |
| QLFSGAEL | ASSLSEGPNSDYT | 0.55140543 |
| SAATFITL | ASSLSEGPNSDYT | 0.43731117 |
| SNYGLANLGV | ASSLSEGPNSDYT | 0.49644855 |
| SYGGAYAVM | ASSLSEGPNSDYT | 0.79177076 |
| TTGVGGLTLG | ASSLSEGPNSDYT | 0.8668039 |
| TTQTIYI | ASSLSEGPNSDYT | 0.78588235 |
| VAAGAGLGAVI | ASSLSEGPNSDYT | 0.7245666 |
| VAAVVTLL | ASSLSEGPNSDYT | 0.8442843 |
| VGFLGGLI | ASSLSEGPNSDYT | 0.7902334 |
| VGVIPILL | ASSLSEGPNSDYT | 0.7792073 |
| VIFASSLI | ASSLSEGPNSDYT | 0.855773 |
| VKGGDGGAAAG | ASSLSEGPNSDYT | 0.6853637 |
| VTKTFTNI | ASSLSEGPNSDYT | 0.8655678 |
| VVAAAILL | ASSLSEGPNSDYT | 0.8055686 |
| YAALDVEL | ASSLSEGPNSDYT | 0.79698503 |
| LASVALLL | ASSLSEGPNSDYT | 0.85316324 |
| LDAVHRV | ASSLSEGPNSDYT | 0.83329046 |
| QGLSSSIF | ASSLSEGPNSDYT | 0.79222995 |
| RGTFANIRL | ASSLSEGPNSDYT | 0.22836271 |
| RQLWWGHRI | ASSLSEGPNSDYT | 0.54056925 |
| YGPPGTGKTLL | ASSLSEGPNSDYT | 0.8515484 |
| AALVGAPANVA | ASSLSEGPNSDYT | 0.82908905 |
| AATFRRTE | ASSLSEGPNSDYT | 0.8635258 |
| AGALGGGI | ASSLSEGPNSDYT | 0.69118226 |
| AGGAGGAGGIGAG | ASSLSEGPNSDYT | 0.65914726 |
| AGSGALGL | ASSLSEGPNSDYT | 0.8550521 |
| AQIADQI | ASSLSEGPNSDYT | 0.8767165 |
| ASAVAVTYL | ASSLSEGPNSDYT | 0.3077376 |
| ASTVLGGTGG | ASSLSEGPNSDYT | 0.68323815 |
| AVGGEPAK | ASSLSEGPNSDYT | 0.9043566 |
| CEQGTFGGTC | ASSLSEGPNSDYT | 0.827435 |
| DEGKAPAG | ASSLSEGPNSDYT | 0.706983 |
| DTVVFPAL | ASSLSEGPNSDYT | 0.5607985 |
| FEGITMAG | ASSLSEGPNSDYT | 0.8872598 |
| FPLVVQD | ASSLSEGPNSDYT | 0.587725 |
| GAFTTPQ | ASSLSEGPNSDYT | 0.89921767 |
| GFGGFGTAGGVGGA | ASSLSEGPNSDYT | 0.14401051 |
| GFLAQFL | ASSLSEGPNSDYT | 0.56843513 |
| GGAGGAGGNAGVA | ASSLSEGPNSDYT | 0.47267205 |
| GGVGTAGAGGAGGAGG | ASSLSEGPNSDYT | 0.26421535 |
| GNGGGGDAGNAGSGGN | ASSLSEGPNSDYT | 0.86025244 |
| GTGGAGGAGNDAGS | ASSLSEGPNSDYT | 0.5710549 |
| GTTNTGWFNS | ASSLSEGPNSDYT | 0.8558639 |
| IFMGPLFTL | ASSLSEGPNSDYT | 0.76237816 |
| ILAHAIEAI | ASSLSEGPNSDYT | 0.86000335 |
| IRSAIREAL | ASSLSEGPNSDYT | 0.832283 |
| ITFAGGG | ASSLSEGPNSDYT | 0.80362165 |
| ITGMSGHQEV | ASSLSEGPNSDYT | 0.47187585 |
| IVAGLACTL | ASSLSEGPNSDYT | 0.85539305 |

| | | |
|---|---|---|
| IVGGGLAGA | ASSLSEGPNSDYT | 0.87627363 |
| IVTGVLNI | ASSLSEGPNSDYT | 0.7865772 |
| LAQQGCI | ASSLSEGPNSDYT | 0.29610747 |
| LCLHVEN | ASSLSEGPNSDYT | 0.76714945 |
| LDAISDGS | ASSLSEGPNSDYT | 0.7635329 |
| LGGFAAML | ASSLSEGPNSDYT | 0.19029292 |
| LGGKNPMI | ASSLSEGPNSDYT | 0.16443928 |
| LGPKGVAAIL | ASSLSEGPNSDYT | 0.7525703 |
| LMVGVSI | ASSLSEGPNSDYT | 0.66289 |
| LVGAAFVL | ASSLSEGPNSDYT | 0.8823125 |
| LVLAVAPL | ASSLSEGPNSDYT | 0.8684955 |
| MAAIAVIDAA | ASSLSEGPNSDYT | 0.7437216 |
| NNGINAELQ | ASSLSEGPNSDYT | 0.8649978 |
| RGEPLVLVHG | ASSLSEGPNSDYT | 0.8267536 |
| RGGAGGAAGLI | ASSLSEGPNSDYT | 0.08863447 |
| RSYRMHPAV | ASSLSEGPNSDYT | 0.5933926 |
| SAAHLLAGQLA | ASSLSEGPNSDYT | 0.33226013 |
| SAAVGSY | ASSLSEGPNSDYT | 0.8511318 |
| SPHAVLHRI | ASSLSEGPNSDYT | 0.5586206 |
| SQIAKVL | ASSLSEGPNSDYT | 0.6189786 |
| TALGHLVP | ASSLSEGPNSDYT | 0.7753295 |
| TAPTTSGAA | ASSLSEGPNSDYT | 0.8149102 |
| TDGDDGNGGSPGT | ASSLSEGPNSDYT | 0.7540681 |
| TDGLGEN | ASSLSEGPNSDYT | 0.52111804 |
| TGGPPLPGQA | ASSLSEGPNSDYT | 0.14134946 |
| TIPAITI | ASSLSEGPNSDYT | 0.85470766 |
| TIPGFTI | ASSLSEGPNSDYT | 0.29354322 |
| TSIASGALA | ASSLSEGPNSDYT | 0.44027358 |
| TVTLPGAAI | ASSLSEGPNSDYT | 0.18969336 |
| VASGLSAALP | ASSLSEGPNSDYT | 0.16825764 |
| VGAGVIGI | ASSLSEGPNSDYT | 0.23598084 |
| VGPKRFGV | ASSLSEGPNSDYT | 0.81162226 |
| VNVAFPD | ASSLSEGPNSDYT | 0.74952143 |
| VSMTNSGVS | ASSLSEGPNSDYT | 0.73758507 |
| VTDVLGDLSV | ASSLSEGPNSDYT | 0.89959717 |
| VVVVGAGFAGL | ASSLSEGPNSDYT | 0.8107408 |
| VYTVGTVV | ASSLSEGPNSDYT | 0.7594812 |
| WTTFSGP | ASSLSEGPNSDYT | 0.59339064 |
| YDELVEAF | ASSLSEGPNSDYT | 0.4954812 |
| YGTGGAGGNGG | ASSLSEGPNSDYT | 0.5118247 |
| YTNVMAV | ASSLSEGPNSDYT | 0.47136268 |
| FLEQVHQL | ASSLSEGPNSDYT | 0.8931805 |
| GLSLLAVAV | ASSLSEGPNSDYT | 0.57691383 |
| LEQVHQL | ASSLSEGPNSDYT | 0.31357837 |
| LNAAGAI | ASSLSEGPNSDYT | 0.64681864 |
| PAVHASL | ASSLSEGPNSDYT | 0.8894689 |
| TELEDDA | ASSLSEGPNSDYT | 0.90467477 |
| TPPGPPN | ASSLSEGPNSDYT | 0.69157356 |
| TPPPPPPP | ASSLSEGPNSDYT | 0.591249 |
| TQSPALI | ASSDGTGGGTEVF | 0.64266145 |
| EFAAAFPQ | ASSDGTGGGTEVF | 0.80763555 |

| | | |
|---|---|---|
| LLSGNLDVL | ASSDGTGGGTEVF | 0.6018481 |
| GPANLGNYNI | ASSDGTGGGTEVF | 0.3375109 |
| TLGPTLP | ASSDGTGGGTEVF | 0.8410618 |
| VYAYSSCCK | ASSDGTGGGTEVF | 0.89255893 |
| APAPAPAPNI | ASSDGTGGGTEVF | 0.6569163 |
| PGGPVPIP | ASSDGTGGGTEVF | 0.26042306 |
| FLTVALEP | ASSDGTGGGTEVF | 0.7674401 |
| SQYFGANL | ASSDGTGGGTEVF | 0.57914925 |
| GGVKPAA | ASSDGTGGGTEVF | 0.7037812 |
| GGTGGAGGAGGW | ASSDGTGGGTEVF | 0.77287173 |
| GVGGVAGLGGAGG | ASSDGTGGGTEVF | 0.31920236 |
| VLLVLATTI | ASSDGTGGGTEVF | 0.7069942 |
| IGIFAAML | ASSDGTGGGTEVF | 0.90273786 |
| AVGDLID | ASSDGTGGGTEVF | 0.7550998 |
| NISPLPL | ASSDGTGGGTEVF | 0.84555244 |
| VTAVFVAI | ASSDGTGGGTEVF | 0.8886312 |
| SGDDNDAAL | ASSDGTGGGTEVF | 0.8654804 |
| AVAALAHS | ASSDGTGGGTEVF | 0.8908849 |
| GGVTVMATP | ASSDGTGGGTEVF | 0.8617142 |
| AGLAANVRD | ASSDGTGGGTEVF | 0.6250856 |
| TNRPDLIDPALLRPG | ASSDGTGGGTEVF | 0.7963518 |
| LANLGVTAL | ASSDGTGGGTEVF | 0.8366839 |
| TGERIEL | ASSDGTGGGTEVF | 0.78868955 |
| NGEELAA | ASSDGTGGGTEVF | 0.7458848 |
| VLSLVWG | ASSDGTGGGTEVF | 0.5301206 |
| IGYALWL | ASSDGTGGGTEVF | 0.87823874 |
| LSLLLAPL | ASSDGTGGGTEVF | 0.73341775 |
| TLDEALK | ASSDGTGGGTEVF | 0.85985553 |
| TDGAGTAEGG | ASSDGTGGGTEVF | 0.33498526 |
| NAPGGAGGNGG | ASSDGTGGGTEVF | 0.40424797 |
| VSVFMGMGGNF | ASSDGTGGGTEVF | 0.8759637 |
| LASYLAAL | ASSDGTGGGTEVF | 0.112207055 |
| LVGGTPAL | ASSDGTGGGTEVF | 0.15648234 |
| VRNADVL | ASSDGTGGGTEVF | 0.78161764 |
| TCSSPFLVSAM | ASSDGTGGGTEVF | 0.8919753 |
| VHITDAE | ASSDGTGGGTEVF | 0.122443594 |
| EYAAPSF | ASSDGTGGGTEVF | 0.40609783 |
| VVDAFHGGL | ASSDGTGGGTEVF | 0.40231127 |
| ASVVDPQ | ASSDGTGGGTEVF | 0.76089203 |
| SYNEFGGLS | ASSDGTGGGTEVF | 0.77657115 |
| DPDGSVY | ASSDGTGGGTEVF | 0.2546551 |
| MVSTIHATE | ASSDGTGGGTEVF | 0.823925 |
| LAAGVADPV | ASSDGTGGGTEVF | 0.2649968 |
| TKEQIAA | ASSDGTGGGTEVF | 0.42339867 |
| SLASPAAL | ASSDGTGGGTEVF | 0.849945 |
| IGLVFYGV | ASSDGTGGGTEVF | 0.8797002 |
| FLLDASNL | ASSDGTGGGTEVF | 0.3993479 |
| YQVGNFP | ASSDGTGGGTEVF | 0.8566761 |
| ESLLSQST | ASSDGTGGGTEVF | 0.6067717 |
| LFTVGFAAVG | ASSDGTGGGTEVF | 0.90546274 |
| ATPPTPT | ASSDGTGGGTEVF | 0.69501215 |

| | | |
|---|---|---|
| TDLATLG | ASSDGTGGGTEVF | 0.28468627 |
| SAGGTGR | ASSDGTGGGTEVF | 0.8638681 |
| GFSNSGS | ASSDGTGGGTEVF | 0.85383856 |
| GGNGGSGGTSV | ASSDGTGGGTEVF | 0.7454766 |
| GTSVATGGAGN | ASSDGTGGGTEVF | 0.8605999 |
| NGGNGGSGGTS | ASSDGTGGGTEVF | 0.5135648 |
| AVVKFPH | ASSDGTGGGTEVF | 0.756124 |
| TPPPPPP | ASSDGTGGGTEVF | 0.8848927 |
| AAPAAAAAPA | ASSDGTGGGTEVF | 0.8804498 |
| IALNEDL | ASSDGTGGGTEVF | 0.40547985 |
| IVNVSSM | ASSDGTGGGTEVF | 0.84963375 |
| TGIPPAPR | ASSDGTGGGTEVF | 0.7736368 |
| FDLGGGTF | ASSDGTGGGTEVF | 0.56597203 |
| FDLGGGTFDVS | ASSDGTGGGTEVF | 0.8942852 |
| NEPTAAA | ASSDGTGGGTEVF | 0.6322645 |
| NEPTAAALA | ASSDGTGGGTEVF | 0.8734888 |
| GSIINISSI | ASSDGTGGGTEVF | 0.8338134 |
| VAPGNAGTA | ASSDGTGGGTEVF | 0.84184587 |
| TGGGASLE | ASSDGTGGGTEVF | 0.31267524 |
| VAVYNHYKRI | ASSDGTGGGTEVF | 0.82989943 |
| TDPAPATT | ASSDGTGGGTEVF | 0.8070762 |
| TNRPDILDPALLRPG | ASSDGTGGGTEVF | 0.8314225 |
| LTTTTTS | ASSDGTGGGTEVF | 0.72471064 |
| PTGSGKTL | ASSDGTGGGTEVF | 0.4975691 |
| TGSGKTL | ASSDGTGGGTEVF | 0.8123567 |
| TIGHVDHGK | ASSDGTGGGTEVF | 0.884089 |
| TIGHVDHGKTT | ASSDGTGGGTEVF | 0.744208 |
| GHVDHGKTT | ASSDGTGGGTEVF | 0.7749636 |
| AQFEHTLL | ASSDGTGGGTEVF | 0.8942503 |
| IVNQDEL | ASSDGTGGGTEVF | 0.5492969 |
| TNGSQFF | ASSDGTGGGTEVF | 0.70076 |
| TGKGIVE | ASSDGTGGGTEVF | 0.82439595 |
| LISNASDA | ASSDGTGGGTEVF | 0.8759027 |
| AELRTLGELTKIAKAHGAQ | ASSDGTGGGTEVF | 0.6509464 |
| APNAVAAAEPSW | ASSDGTGGGTEVF | 0.8928846 |
| APQVAGAATAVL | ASSDGTGGGTEVF | 0.5842979 |
| ARRDPAAAAAILPTDA | ASSDGTGGGTEVF | 0.87736696 |
| DAAARDVLAEAGLAEYF | ASSDGTGGGTEVF | 0.7277557 |
| DDQIVLSAKANLAKVEQ | ASSDGTGGGTEVF | 0.8264392 |
| DELLPDTGEDSDEDGH | ASSDGTGGGTEVF | 0.64392537 |
| ELPEAPANRATVPQD | ASSDGTGGGTEVF | 0.8754022 |
| FGQSKIERVTGIG | ASSDGTGGGTEVF | 0.8758588 |
| GAGGTGSTAGGAGGAGGAGGLY | ASSDGTGGGTEVF | 0.8821796 |
| GHGGTGAAGGNGGTGGAGGSGI | ASSDGTGGGTEVF | 0.15003842 |
| GNGGNGGAGGEGGPGGAGGAGA | ASSDGTGGGTEVF | 0.86715794 |
| GSTAGNGGNGGNATNGTIAGQP | ASSDGTGGGTEVF | 0.86936724 |
| GTGGDGGNGGNGGTGG | ASSDGTGGGTEVF | 0.9029666 |
| GVRRPAGIPADAPV | ASSDGTGGGTEVF | 0.7839 |
| GYRQVTDAHLLTLARR | ASSDGTGGGTEVF | 0.8030159 |
| IAKMGQDGHDRG | ASSDGTGGGTEVF | 0.7616904 |
| KAEPSIEIQYT | ASSDGTGGGTEVF | 0.64400065 |

| | | |
|---|---|---|
| KPVRIRMAADEDTVAVT | ASSDGTGGGTEVF | 0.51829636 |
| MPPRIAGMRLLVIKPEP | ASSDGTGGGTEVF | 0.7609612 |
| MKFYAVALFRSRGVPPTR | ASSDGTGGGTEVF | 0.72464526 |
| NAGGFGILGAVAHSP | ASSDGTGGGTEVF | 0.6601043 |
| NEGREITLAMAEM | ASSDGTGGGTEVF | 0.8649489 |
| NELVEWSHPLGHRWRWE | ASSDGTGGGTEVF | 0.87956697 |
| NLGAGGTGGNGGNPG | ASSDGTGGGTEVF | 0.8826335 |
| NTEWPKWETVMAADPVD | ASSDGTGGGTEVF | 0.88102716 |
| PASAAPAETQP | ASSDGTGGGTEVF | 0.88923013 |
| QQSSDGKNGVRAAEITGE | ASSDGTGGGTEVF | 0.8038946 |
| RAPDAVTTSWWKE | ASSDGTGGGTEVF | 0.63735414 |
| SGFGNVGAHTSGWFNQ | ASSDGTGGGTEVF | 0.6492598 |
| SGVAGAVGQAASVGGLKV | ASSDGTGGGTEVF | 0.43516213 |
| SNGGAGTGSAGGNGGT | ASSDGTGGGTEVF | 0.72735655 |
| TGGAGGEGGAAG | ASSDGTGGGTEVF | 0.6419015 |
| TGGDGGAAGAGGAPGHG | ASSDGTGGGTEVF | 0.8133078 |
| TMPPLLAAIANAGRHGV | ASSDGTGGGTEVF | 0.10179099 |
| TSSTAGNSGGA | ASSDGTGGGTEVF | 0.8247315 |
| VPATVPVGILC | ASSDGTGGGTEVF | 0.89616555 |
| VQRPDSGGVVLTGRIS | ASSDGTGGGTEVF | 0.7138246 |
| WTPNPVPATKGVPAGLVQAT | ASSDGTGGGTEVF | 0.80002904 |
| YALKRALAKFSDRFDVVII | ASSDGTGGGTEVF | 0.8196952 |
| YLIGSVIV | ASSDGTGGGTEVF | 0.65994537 |
| YGQLVAVID | ASSDGTGGGTEVF | 0.85624266 |
| VTVGGMTLPA | ASSDGTGGGTEVF | 0.48340294 |
| VLALPAV | ASSDGTGGGTEVF | 0.81234825 |
| VIPPTLNYET | ASSDGTGGGTEVF | 0.8905016 |
| VGMGGMHPSSG | ASSDGTGGGTEVF | 0.8033715 |
| VGLAGGAATA | ASSDGTGGGTEVF | 0.7978103 |
| VGGAGGAGGSGP | ASSDGTGGGTEVF | 0.67592084 |
| VFAAFMV | ASSDGTGGGTEVF | 0.2556119 |
| TLYDKLR | ASSDGTGGGTEVF | 0.8321193 |
| TLAVFTVL | ASSDGTGGGTEVF | 0.09425379 |
| TGSVQPARV | ASSDGTGGGTEVF | 0.78934216 |
| TGGGAGSGGT | ASSDGTGGGTEVF | 0.46127266 |
| SHLETSL | ASSDGTGGGTEVF | 0.8079065 |
| SFTTGMT | ASSDGTGGGTEVF | 0.8128014 |
| QVEVAKRMGVRQPTVS | ASSDGTGGGTEVF | 0.81808734 |
| QGSGGPALAM | ASSDGTGGGTEVF | 0.5165281 |
| MLAPSNP | ASSDGTGGGTEVF | 0.7532027 |
| LVNNAAI | ASSDGTGGGTEVF | 0.67193764 |
| LTGEINTLA | ASSDGTGGGTEVF | 0.37767175 |
| LSIFGPV | ASSDGTGGGTEVF | 0.8747701 |
| LLALLANL | ASSDGTGGGTEVF | 0.5972281 |
| LISNASDALDKLR | ASSDGTGGGTEVF | 0.78047824 |
| LINEAVP | ASSDGTGGGTEVF | 0.88067406 |
| LGYAFSGL | ASSDGTGGGTEVF | 0.7834618 |
| LDFVAVAL | ASSDGTGGGTEVF | 0.250586 |
| LAQPGAE | ASSDGTGGGTEVF | 0.81016856 |
| ISNLGAVTGF | ASSDGTGGGTEVF | 0.7717756 |
| HQEDPGRI | ASSDGTGGGTEVF | 0.7417233 |

| | | |
|---|---|---|
| GTPGEPTP | ASSDGTGGGTEVF | 0.09471062 |
| GFFNSGS | ASSDGTGGGTEVF | 0.84708834 |
| GAGGGGLTAGAN | ASSDGTGGGTEVF | 0.55899227 |
| FVDALVNM | ASSDGTGGGTEVF | 0.62900144 |
| FLTVAPEL | ASSDGTGGGTEVF | 0.5794519 |
| FETLTAL | ASSDGTGGGTEVF | 0.8666615 |
| DVINPIEL | ASSDGTGGGTEVF | 0.6634487 |
| APVSGAPV | ASSDGTGGGTEVF | 0.6878804 |
| ALVGGLAW | ASSDGTGGGTEVF | 0.86553156 |
| ALAAAAATAI | ASSDGTGGGTEVF | 0.82616174 |
| AFGLGLLL | ASSDGTGGGTEVF | 0.89375305 |
| AFFAAASL | ASSDGTGGGTEVF | 0.7821022 |
| AAAALPAFSPP | ASSDGTGGGTEVF | 0.66622233 |
| VVLAPAL | ASSDGTGGGTEVF | 0.7975599 |
| TIGHVDHGKTTL | ASSDGTGGGTEVF | 0.6331405 |
| LISNASDALDK | ASSDGTGGGTEVF | 0.26522213 |
| LISNASDAL | ASSDGTGGGTEVF | 0.19531427 |
| ISNASDALDK | ASSDGTGGGTEVF | 0.82485354 |
| NQAETLVYQTE | ASSDGTGGGTEVF | 0.21802747 |
| TGGTGGTAGAG | ASSDGTGGGTEVF | 0.79022586 |
| TGSTAGGAGGA | ASSDGTGGGTEVF | 0.82827055 |
| DPLAVPVAAGPV | ASSDGTGGGTEVF | 0.70428824 |
| GGTGGTPGNGGN | ASSDGTGGGTEVF | 0.38250828 |
| PAGLAGVDPATL | ASSDGTGGGTEVF | 0.70035386 |
| GNNLVVEAPHAGASV | ASSDGTGGGTEVF | 0.58701116 |
| KVTRIMSASTLEQPA | ASSDGTGGGTEVF | 0.8634599 |
| LGFALPIALPRRTVV | ASSDGTGGGTEVF | 0.8405696 |
| TRLGNLTARKLVVLV | ASSDGTGGGTEVF | 0.8949819 |
| WSQQRRLIYFPSAGP | ASSDGTGGGTEVF | 0.87702906 |
| GPNGNPDPMAAAVDIR | ASSDGTGGGTEVF | 0.7147084 |
| TQRCVSVADLRAAASR | ASSDGTGGGTEVF | 0.86176 |
| QFVLDVAEHPVEVLTETV | ASSDGTGGGTEVF | 0.69818103 |
| GGAGGTGGAGGNGGTQAGDG | ASSDGTGGGTEVF | 0.76699024 |
| PLLATLRAIAVDVDAKPAQV | ASSDGTGGGTEVF | 0.7179658 |
| APKFRTGGYMIDFWGVGYQVAK | ASSDGTGGGTEVF | 0.7990644 |
| GAGGAGGAGGAAGGPGGTGGTG | ASSDGTGGGTEVF | 0.71420586 |
| IGALGLLATDVAPAGSVAFGVA | ASSDGTGGGTEVF | 0.66446996 |
| AGGAGGIGGNGGAGGAGGVLMG | ASSDGTGGGTEVF | 0.2579578 |
| ADELIGLL | ASSDGTGGGTEVF | 0.8155877 |
| ADRALKA | ASSDGTGGGTEVF | 0.8728359 |
| AGAVGQVGGAGGA | ASSDGTGGGTEVF | 0.2764783 |
| AGGGGGLGGAP | ASSDGTGGGTEVF | 0.6358214 |
| AGGLLYGNGGAG | ASSDGTGGGTEVF | 0.90006495 |
| AGLESIGL | ASSDGTGGGTEVF | 0.8811854 |
| AGLKTGII | ASSDGTGGGTEVF | 0.6536711 |
| ASLAALLL | ASSDGTGGGTEVF | 0.542889 |
| ASVINISSI | ASSDGTGGGTEVF | 0.81796324 |
| ATPPGPA | ASSDGTGGGTEVF | 0.2679737 |
| ATVASALV | ASSDGTGGGTEVF | 0.78068125 |
| ATVASLIL | ASSDGTGGGTEVF | 0.80614495 |
| AVNGPVLIRH | ASSDGTGGGTEVF | 0.8146001 |

| | | |
|---|---|---|
| DGAAGAVGI | ASSDGTGGGTEVF | 0.5148854 |
| DPLLHGI | ASSDGTGGGTEVF | 0.81031585 |
| ELGAALGAL | ASSDGTGGGTEVF | 0.7561405 |
| ELGRTQAV | ASSDGTGGGTEVF | 0.7608749 |
| FVAGGYLS | ASSDGTGGGTEVF | 0.11917708 |
| GASGGAGGAGGW | ASSDGTGGGTEVF | 0.7593012 |
| GGAGAGGDGGAGGI | ASSDGTGGGTEVF | 0.5764232 |
| GTGHNGSQPGMGGNG | ASSDGTGGGTEVF | 0.55231154 |
| GYAAASASAS | ASSDGTGGGTEVF | 0.26836988 |
| HQVDVAAVQQG | ASSDGTGGGTEVF | 0.6971351 |
| IGAAIAHT | ASSDGTGGGTEVF | 0.8789488 |
| IGMIEVAQL | ASSDGTGGGTEVF | 0.8762731 |
| IRLAPEHTSAMD | ASSDGTGGGTEVF | 0.52584094 |
| IYNVSVL | ASSDGTGGGTEVF | 0.8723576 |
| KQGFLGM | ASSDGTGGGTEVF | 0.9021946 |
| LESVITQL | ASSDGTGGGTEVF | 0.7305548 |
| LGLGALVVV | ASSDGTGGGTEVF | 0.24539898 |
| LIGNGGNGGSGGI | ASSDGTGGGTEVF | 0.7995461 |
| LNAINAPTL | ASSDGTGGGTEVF | 0.7441542 |
| LQLLNTGV | ASSDGTGGGTEVF | 0.82859826 |
| LQTSIEF | ASSDGTGGGTEVF | 0.87202394 |
| MSFAVSEL | ASSDGTGGGTEVF | 0.50091934 |
| NTGALGL | ASSDGTGGGTEVF | 0.81384534 |
| PPGAGGAGGAGTT | ASSDGTGGGTEVF | 0.8812044 |
| QLVGLGI | ASSDGTGGGTEVF | 0.6250232 |
| RFLEQVHQL | ASSDGTGGGTEVF | 0.5170839 |
| RGVGYVLR | ASSDGTGGGTEVF | 0.49879026 |
| SATGVFAGVI | ASSDGTGGGTEVF | 0.25937673 |
| SLPAITLP | ASSDGTGGGTEVF | 0.84370327 |
| STALAFTV | ASSDGTGGGTEVF | 0.70374453 |
| SVLTVGLGK | ASSDGTGGGTEVF | 0.49210793 |
| TALAGAGSGI | ASSDGTGGGTEVF | 0.7908774 |
| TGLPFTGI | ASSDGTGGGTEVF | 0.5030062 |
| VAEHVTV | ASSDGTGGGTEVF | 0.52325106 |
| VDIVLNSL | ASSDGTGGGTEVF | 0.650933 |
| VLTGGGAVL | ASSDGTGGGTEVF | 0.733795 |
| VNVGLAYGI | ASSDGTGGGTEVF | 0.83979934 |
| VSSAAVGAV | ASSDGTGGGTEVF | 0.7657683 |
| VTSVYEAGV | ASSDGTGGGTEVF | 0.72112 |
| WVALDRGA | ASSDGTGGGTEVF | 0.89249 |
| YLVGSVVI | ASSDGTGGGTEVF | 0.3925761 |
| GPPGTGKTLL | ASSDGTGGGTEVF | 0.52280307 |
| GVGMTKF | ASSDGTGGGTEVF | 0.8378021 |
| IGIGGSDL | ASSDGTGGGTEVF | 0.36081025 |
| IGTLLAEL | ASSDGTGGGTEVF | 0.855307 |
| IYTSGTT | ASSDGTGGGTEVF | 0.81639075 |
| LSFPAGV | ASSDGTGGGTEVF | 0.8329213 |
| LYTSGTT | ASSDGTGGGTEVF | 0.7650318 |
| VGAFPFGF | ASSDGTGGGTEVF | 0.9038807 |
| VGAGTVEF | ASSDGTGGGTEVF | 0.8598942 |
| VGKSTLLL | ASSDGTGGGTEVF | 0.6516515 |

| | | |
|---|---|---|
| AALAHAGGGIV | ASSDGTGGGTEVF | 0.8332852 |
| AALVAVSLL | ASSDGTGGGTEVF | 0.3959608 |
| AAVSRAVAAL | ASSDGTGGGTEVF | 0.86867285 |
| AGGSSGIGGF | ASSDGTGGGTEVF | 0.82508814 |
| AGLTVANANG | ASSDGTGGGTEVF | 0.5515117 |
| AGLVGAVL | ASSDGTGGGTEVF | 0.77046645 |
| ALSQAFTV | ASSDGTGGGTEVF | 0.88174164 |
| ALTFSVSL | ASSDGTGGGTEVF | 0.88626635 |
| ALTGAALVAL | ASSDGTGGGTEVF | 0.2437916 |
| ASFSIGAGVL | ASSDGTGGGTEVF | 0.7090073 |
| DGSVYKI | ASSDGTGGGTEVF | 0.27336743 |
| EQLAEIV | ASSDGTGGGTEVF | 0.65991116 |
| FAVSELTVM | ASSDGTGGGTEVF | 0.39127135 |
| GAAPLAAGL | ASSDGTGGGTEVF | 0.85121596 |
| GADGGDGGSGGAGG | ASSDGTGGGTEVF | 0.6691907 |
| GFGLDTGGAGG | ASSDGTGGGTEVF | 0.8360019 |
| GLASGGLL | ASSDGTGGGTEVF | 0.7927064 |
| GRIAGAAVLL | ASSDGTGGGTEVF | 0.87838155 |
| GSAGSLNTSF | ASSDGTGGGTEVF | 0.62460625 |
| GTGQIGI | ASSDGTGGGTEVF | 0.7411515 |
| GTLIALIL | ASSDGTGGGTEVF | 0.3097771 |
| IAATMGFVVG | ASSDGTGGGTEVF | 0.39456788 |
| IFAQALL | ASSDGTGGGTEVF | 0.540075 |
| IGAAVALGF | ASSDGTGGGTEVF | 0.628546 |
| IGLTPLAL | ASSDGTGGGTEVF | 0.17775513 |
| ILGSNTSTL | ASSDGTGGGTEVF | 0.8840035 |
| IVFSLTVL | ASSDGTGGGTEVF | 0.80499625 |
| LASKATGF | ASSDGTGGGTEVF | 0.8473842 |
| LASVALIL | ASSDGTGGGTEVF | 0.6059542 |
| LFAVGLTL | ASSDGTGGGTEVF | 0.6217555 |
| LFNGGIGTYI | ASSDGTGGGTEVF | 0.6547361 |
| LTLGAGII | ASSDGTGGGTEVF | 0.8519313 |
| LVAAALGDI | ASSDGTGGGTEVF | 0.5779433 |
| NTAFVQLGI | ASSDGTGGGTEVF | 0.22073252 |
| QLFSGAEL | ASSDGTGGGTEVF | 0.52002835 |
| SAATFITL | ASSDGTGGGTEVF | 0.37536916 |
| SNYGLANLGV | ASSDGTGGGTEVF | 0.8937525 |
| SYGGAYAVM | ASSDGTGGGTEVF | 0.87426054 |
| TTGVGGLTLG | ASSDGTGGGTEVF | 0.83079803 |
| TTQTIYI | ASSDGTGGGTEVF | 0.8687902 |
| VAAGAGLGAVI | ASSDGTGGGTEVF | 0.17389642 |
| VAAVVTLL | ASSDGTGGGTEVF | 0.65587366 |
| VGFLGGLI | ASSDGTGGGTEVF | 0.88622236 |
| VGVIPILL | ASSDGTGGGTEVF | 0.81967753 |
| VIFASSLI | ASSDGTGGGTEVF | 0.72072744 |
| VKGGDGGAAAG | ASSDGTGGGTEVF | 0.8823733 |
| VTKTFTNI | ASSDGTGGGTEVF | 0.6525538 |
| VVAAAILL | ASSDGTGGGTEVF | 0.7095996 |
| YAALDVEL | ASSDGTGGGTEVF | 0.7304276 |
| LASVALLL | ASSDGTGGGTEVF | 0.81921506 |
| LDAVHRV | ASSDGTGGGTEVF | 0.7957244 |

| | | |
|---|---|---|
| QGLSSSIF | ASSDGTGGGTEVF | 0.49104857 |
| RGTFANIRL | ASSDGTGGGTEVF | 0.76467204 |
| RQLWWGHRI | ASSDGTGGGTEVF | 0.9064745 |
| YGPPGTGKTLL | ASSDGTGGGTEVF | 0.73836225 |
| AALVGAPANVA | ASSDGTGGGTEVF | 0.8781799 |
| AATFRRTE | ASSDGTGGGTEVF | 0.82790315 |
| AGALGGGI | ASSDGTGGGTEVF | 0.8544791 |
| AGGAGGAGGIGAG | ASSDGTGGGTEVF | 0.5831175 |
| AGSGALGL | ASSDGTGGGTEVF | 0.6996369 |
| AQIADQI | ASSDGTGGGTEVF | 0.75818145 |
| ASAVAVTYL | ASSDGTGGGTEVF | 0.61448085 |
| ASTVLGGTGG | ASSDGTGGGTEVF | 0.6759961 |
| AVGGEPAK | ASSDGTGGGTEVF | 0.605615 |
| CEQGTFGGTC | ASSDGTGGGTEVF | 0.9093405 |
| DEGKAPAG | ASSDGTGGGTEVF | 0.85441244 |
| DTVVFPAL | ASSDGTGGGTEVF | 0.8267797 |
| FEGITMAG | ASSDGTGGGTEVF | 0.57764614 |
| FPLVVQD | ASSDGTGGGTEVF | 0.79023993 |
| GAFTTPQ | ASSDGTGGGTEVF | 0.8419764 |
| GFGGFGTAGGVGGA | ASSDGTGGGTEVF | 0.29811123 |
| GFLAQFL | ASSDGTGGGTEVF | 0.89071584 |
| GGAGGAGGNAGVA | ASSDGTGGGTEVF | 0.547752 |
| GGVGTAGAGGAGGAGG | ASSDGTGGGTEVF | 0.753 |
| GNGGGGDAGNAGSGGN | ASSDGTGGGTEVF | 0.8203616 |
| GTGGAGGAGNDAGS | ASSDGTGGGTEVF | 0.8788154 |
| GTTNTGWFNS | ASSDGTGGGTEVF | 0.7006786 |
| IFMGPLFTL | ASSDGTGGGTEVF | 0.7046294 |
| ILAHAIEAI | ASSDGTGGGTEVF | 0.5509508 |
| IRSAIREAL | ASSDGTGGGTEVF | 0.85793906 |
| ITFAGGG | ASSDGTGGGTEVF | 0.8784541 |
| ITGMSGHQEV | ASSDGTGGGTEVF | 0.7445973 |
| IVAGLACTL | ASSDGTGGGTEVF | 0.89545643 |
| IVGGGLAGA | ASSDGTGGGTEVF | 0.61566746 |
| IVTGVLNI | ASSDGTGGGTEVF | 0.31470448 |
| LAQQGCI | ASSDGTGGGTEVF | 0.71771884 |
| LCLHVEN | ASSDGTGGGTEVF | 0.8618982 |
| LDAISDGS | ASSDGTGGGTEVF | 0.8846885 |
| LGGFAAML | ASSDGTGGGTEVF | 0.88484484 |
| LGGKNPMI | ASSDGTGGGTEVF | 0.8461485 |
| LGPKGVAAIL | ASSDGTGGGTEVF | 0.7393952 |
| LMVGVSI | ASSDGTGGGTEVF | 0.88842213 |
| LVGAAFVL | ASSDGTGGGTEVF | 0.89865345 |
| LVLAVAPL | ASSDGTGGGTEVF | 0.83388734 |
| MAAIAVIDAA | ASSDGTGGGTEVF | 0.6121986 |
| NNGINAELQ | ASSDGTGGGTEVF | 0.49405313 |
| RGEPLVLVHG | ASSDGTGGGTEVF | 0.784809 |
| RGGAGGAAGLI | ASSDGTGGGTEVF | 0.8777647 |
| RSYRMHPAV | ASSDGTGGGTEVF | 0.40758893 |
| SAAHLLAGQLA | ASSDGTGGGTEVF | 0.82179415 |
| SAAVGSY | ASSDGTGGGTEVF | 0.81455433 |
| SPHAVLHRI | ASSDGTGGGTEVF | 0.30997688 |

| | | |
|---|---|---|
| SQIAKVL | ASSDGTGGGTEVF | 0.85033154 |
| TALGHLVP | ASSDGTGGGTEVF | 0.8926524 |
| TAPTTSGAA | ASSDGTGGGTEVF | 0.6924542 |
| TDGDDGNGGSPGT | ASSDGTGGGTEVF | 0.61208606 |
| TDGLGEN | ASSDGTGGGTEVF | 0.887904 |
| TGGPPLPGQA | ASSDGTGGGTEVF | 0.9085536 |
| TIPAITI | ASSDGTGGGTEVF | 0.87794995 |
| TIPGFTI | ASSDGTGGGTEVF | 0.25207967 |
| TSIASGALA | ASSDGTGGGTEVF | 0.16892567 |
| TVTLPGAAI | ASSDGTGGGTEVF | 0.7116292 |
| VASGLSAALP | ASSDGTGGGTEVF | 0.853549 |
| VGAGVIGI | ASSDGTGGGTEVF | 0.60505325 |
| VGPKRFGV | ASSDGTGGGTEVF | 0.56180143 |
| VNVAFPD | ASSDGTGGGTEVF | 0.8541027 |
| VSMTNSGVS | ASSDGTGGGTEVF | 0.87562 |
| VTDVLGDLSV | ASSDGTGGGTEVF | 0.53676844 |
| VVVVGAGFAGL | ASSDGTGGGTEVF | 0.31971544 |
| VYTVGTVV | ASSDGTGGGTEVF | 0.8366684 |
| WTTFSGP | ASSDGTGGGTEVF | 0.83052194 |
| YDELVEAF | ASSDGTGGGTEVF | 0.64958924 |
| YGTGGAGGNGG | ASSDGTGGGTEVF | 0.78274274 |
| YTNVMAV | ASSDGTGGGTEVF | 0.6989472 |
| FLEQVHQL | ASSDGTGGGTEVF | 0.59962624 |
| GLSLLAVAV | ASSDGTGGGTEVF | 0.83700585 |
| LEQVHQL | ASSDGTGGGTEVF | 0.47513855 |
| LNAAGAI | ASSDGTGGGTEVF | 0.89790237 |
| PAVHASL | ASSDGTGGGTEVF | 0.46352574 |
| TELEDDA | ASSDGTGGGTEVF | 0.8638178 |
| TPPGPPN | ASSDGTGGGTEVF | 0.8290246 |
| TPPPPPPP | ASSDGTGGGTEVF | 0.81364024 |
| TQSPALI | ASSTTASQNTLY | 0.7252734 |
| EFAAAFPQ | ASSTTASQNTLY | 0.6155939 |
| LLSGNLDVL | ASSTTASQNTLY | 0.42227766 |
| GPANLGNYNI | ASSTTASQNTLY | 0.23089983 |
| TLGPTLP | ASSTTASQNTLY | 0.8859309 |
| VYAYSSCCK | ASSTTASQNTLY | 0.15880753 |
| APAPAPAPNI | ASSTTASQNTLY | 0.8838471 |
| PGGPVPIP | ASSTTASQNTLY | 0.83598804 |
| FLTVALEP | ASSTTASQNTLY | 0.5354699 |
| SQYFGANL | ASSTTASQNTLY | 0.80929065 |
| GGVKPAA | ASSTTASQNTLY | 0.49754208 |
| GGTGGAGGAGGW | ASSTTASQNTLY | 0.5601208 |
| GVGGVAGLGGAGG | ASSTTASQNTLY | 0.8776835 |
| VLLVLATTI | ASSTTASQNTLY | 0.80729556 |
| IGIFAAML | ASSTTASQNTLY | 0.539394 |
| AVGDLID | ASSTTASQNTLY | 0.7678763 |
| NISPLPL | ASSTTASQNTLY | 0.82265323 |
| VTAVFVAI | ASSTTASQNTLY | 0.5400401 |
| SGDDNDAAL | ASSTTASQNTLY | 0.66494036 |
| AVAALAHS | ASSTTASQNTLY | 0.8828568 |
| GGVTVMATP | ASSTTASQNTLY | 0.7454541 |

| | | |
|---|---|---|
| AGLAANVRD | ASSTTASQNTLY | 0.5106313 |
| TNRPDLIDPALLRPG | ASSTTASQNTLY | 0.85767245 |
| LANLGVTAL | ASSTTASQNTLY | 0.6038736 |
| TGERIEL | ASSTTASQNTLY | 0.8429738 |
| NGEELAA | ASSTTASQNTLY | 0.77840495 |
| VLSLVWG | ASSTTASQNTLY | 0.6680567 |
| IGYALWL | ASSTTASQNTLY | 0.8640387 |
| LSLLLAPL | ASSTTASQNTLY | 0.33115166 |
| TLDEALK | ASSTTASQNTLY | 0.6252786 |
| TDGAGTAEGG | ASSTTASQNTLY | 0.73735154 |
| NAPGGAGGNGG | ASSTTASQNTLY | 0.77459294 |
| VSVFMGMGGNF | ASSTTASQNTLY | 0.8607335 |
| LASYLAAL | ASSTTASQNTLY | 0.57616 |
| LVGGTPAL | ASSTTASQNTLY | 0.79923445 |
| VRNADVL | ASSTTASQNTLY | 0.6168563 |
| TCSSPFLVSAM | ASSTTASQNTLY | 0.46843594 |
| VHITDAE | ASSTTASQNTLY | 0.6068869 |
| EYAAPSF | ASSTTASQNTLY | 0.64194965 |
| VVDAFHGGL | ASSTTASQNTLY | 0.5289976 |
| ASVVDPQ | ASSTTASQNTLY | 0.8314756 |
| SYNEFGGLS | ASSTTASQNTLY | 0.48369962 |
| DPDGSVY | ASSTTASQNTLY | 0.81536317 |
| MVSTIHATE | ASSTTASQNTLY | 0.89766276 |
| LAAGVADPV | ASSTTASQNTLY | 0.60024583 |
| TKEQIAA | ASSTTASQNTLY | 0.8364006 |
| SLASPAAL | ASSTTASQNTLY | 0.8296037 |
| IGLVFYGV | ASSTTASQNTLY | 0.8604332 |
| FLLDASNL | ASSTTASQNTLY | 0.657521 |
| YQVGNFP | ASSTTASQNTLY | 0.5745541 |
| ESLLSQST | ASSTTASQNTLY | 0.87962407 |
| LFTVGFAAVG | ASSTTASQNTLY | 0.8357518 |
| ATPPTPT | ASSTTASQNTLY | 0.8192151 |
| TDLATLG | ASSTTASQNTLY | 0.85514385 |
| SAGGTGR | ASSTTASQNTLY | 0.8390177 |
| GFSNSGS | ASSTTASQNTLY | 0.8445216 |
| GGNGGSGGTSV | ASSTTASQNTLY | 0.6848272 |
| GTSVATGGAGN | ASSTTASQNTLY | 0.34805807 |
| NGGNGGSGGTS | ASSTTASQNTLY | 0.72638875 |
| AVVKFPH | ASSTTASQNTLY | 0.8124379 |
| TPPPPPP | ASSTTASQNTLY | 0.5171149 |
| AAPAAAAAPA | ASSTTASQNTLY | 0.2339682 |
| IALNEDL | ASSTTASQNTLY | 0.8804894 |
| IVNVSSM | ASSTTASQNTLY | 0.90593845 |
| TGIPPAPR | ASSTTASQNTLY | 0.4894147 |
| FDLGGGTF | ASSTTASQNTLY | 0.89965534 |
| FDLGGGTFDVS | ASSTTASQNTLY | 0.3701957 |
| NEPTAAA | ASSTTASQNTLY | 0.8499477 |
| NEPTAAALA | ASSTTASQNTLY | 0.88374114 |
| GSIINISSI | ASSTTASQNTLY | 0.51638925 |
| VAPGNAGTA | ASSTTASQNTLY | 0.8276005 |
| TGGGASLE | ASSTTASQNTLY | 0.54766953 |

| | | |
|---|---|---|
| VAVYNHYKRI | ASSTTASQNTLY | 0.61453116 |
| TDPAPATT | ASSTTASQNTLY | 0.8921318 |
| TNRPDILDPALLRPG | ASSTTASQNTLY | 0.5869376 |
| LTTTTTS | ASSTTASQNTLY | 0.79404664 |
| PTGSGKTL | ASSTTASQNTLY | 0.6036125 |
| TGSGKTL | ASSTTASQNTLY | 0.49473462 |
| TIGHVDHGK | ASSTTASQNTLY | 0.7776174 |
| TIGHVDHGKTT | ASSTTASQNTLY | 0.86431915 |
| GHVDHGKTT | ASSTTASQNTLY | 0.5958592 |
| AQFEHTLL | ASSTTASQNTLY | 0.8944287 |
| IVNQDEL | ASSTTASQNTLY | 0.6024602 |
| TNGSQFF | ASSTTASQNTLY | 0.9030806 |
| TGKGIVE | ASSTTASQNTLY | 0.8625244 |
| LISNASDA | ASSTTASQNTLY | 0.8128429 |
| AELRTLGELTKIAKAHGAQ | ASSTTASQNTLY | 0.6438631 |
| APNAVAAAEPSW | ASSTTASQNTLY | 0.87912464 |
| APQVAGAATAVL | ASSTTASQNTLY | 0.67966354 |
| ARRDPAAAAAILPTDA | ASSTTASQNTLY | 0.84117067 |
| DAAARDVLAEAGLAEYF | ASSTTASQNTLY | 0.8598469 |
| DDQIVLSAKANLAKVEQ | ASSTTASQNTLY | 0.7242062 |
| DELLPDTGEDSDEDGH | ASSTTASQNTLY | 0.77966905 |
| ELPEAPANRATVPQD | ASSTTASQNTLY | 0.747511 |
| FGQSKIERVTGIG | ASSTTASQNTLY | 0.7487281 |
| GAGGTGSTAGGAGGAGGAGGLY | ASSTTASQNTLY | 0.72864723 |
| GHGGTGAAGGNGGTGGAGGSGI | ASSTTASQNTLY | 0.5760173 |
| GNGGNGGAGGEGGPGGAGGAGA | ASSTTASQNTLY | 0.7500621 |
| GSTAGNGGNGGNATNGTIAGQP | ASSTTASQNTLY | 0.6968412 |
| GTGGDGGNGGNGGTGG | ASSTTASQNTLY | 0.7627871 |
| GVRRPAGIPADAPV | ASSTTASQNTLY | 0.77491426 |
| GYRQVTDAHLLTLARR | ASSTTASQNTLY | 0.7154752 |
| IAKMGQDGHDRG | ASSTTASQNTLY | 0.7867751 |
| KAEPSIEIQYT | ASSTTASQNTLY | 0.7820703 |
| KPVRIRMAADEDTVAVT | ASSTTASQNTLY | 0.5994078 |
| MPPRIAGMRLLVIKPEP | ASSTTASQNTLY | 0.7317493 |
| MKFYAVALFRSRGVPPTR | ASSTTASQNTLY | 0.69113857 |
| NAGGFGILGAVAHSP | ASSTTASQNTLY | 0.7442316 |
| NEGREITLAMAEM | ASSTTASQNTLY | 0.42653012 |
| NELVEWSHPLGHRWRWE | ASSTTASQNTLY | 0.7784107 |
| NLGAGGTGGNGGNPG | ASSTTASQNTLY | 0.68955284 |
| NTEWPKWETVMAADPVD | ASSTTASQNTLY | 0.25861135 |
| PASAAPAETQP | ASSTTASQNTLY | 0.75571656 |
| QQSSDGKNGVRAAEITGE | ASSTTASQNTLY | 0.7536383 |
| RAPDAVTTSWWKE | ASSTTASQNTLY | 0.74458086 |
| SGFGNVGAHTSGWFNQ | ASSTTASQNTLY | 0.7709403 |
| SGVAGAVGQAASVGGLKV | ASSTTASQNTLY | 0.45778462 |
| SNGGAGTGSAGGNGGT | ASSTTASQNTLY | 0.7031404 |
| TGGAGGEGGAAG | ASSTTASQNTLY | 0.7769779 |
| TGGDGGAAGAGGAPGHG | ASSTTASQNTLY | 0.7846128 |
| TMPPLLAAIANAGRHGV | ASSTTASQNTLY | 0.6563778 |
| TSSTAGNSGGA | ASSTTASQNTLY | 0.7565707 |
| VPATVPVGILC | ASSTTASQNTLY | 0.77947307 |

| | | |
|---|---|---|
| VQRPDSGGVVLTGRIS | ASSTTASQNTLY | 0.7620667 |
| WTPNPVPATKGVPAGLVQAT | ASSTTASQNTLY | 0.7794825 |
| YALKRALAKFSDRFDVVII | ASSTTASQNTLY | 0.78652936 |
| YLIGSVIV | ASSTTASQNTLY | 0.6442923 |
| YGQLVAVID | ASSTTASQNTLY | 0.7833036 |
| VTVGGMTLPA | ASSTTASQNTLY | 0.78282636 |
| VLALPAV | ASSTTASQNTLY | 0.7433117 |
| VIPPTLNYET | ASSTTASQNTLY | 0.7550231 |
| VGMGGMHPSSG | ASSTTASQNTLY | 0.67833257 |
| VGLAGGAATA | ASSTTASQNTLY | 0.7246362 |
| VGGAGGAGGSGP | ASSTTASQNTLY | 0.48436248 |
| VFAAFMV | ASSTTASQNTLY | 0.7563424 |
| TLYDKLR | ASSTTASQNTLY | 0.77792543 |
| TLAVFTVL | ASSTTASQNTLY | 0.7239083 |
| TGSVQPARV | ASSTTASQNTLY | 0.6435411 |
| TGGGAGSGGT | ASSTTASQNTLY | 0.7009954 |
| SHLETSL | ASSTTASQNTLY | 0.7038646 |
| SFTTGMT | ASSTTASQNTLY | 0.7303513 |
| QVEVAKRMGVRQPTVS | ASSTTASQNTLY | 0.48547706 |
| QGSGGPALAM | ASSTTASQNTLY | 0.6782678 |
| MLAPSNP | ASSTTASQNTLY | 0.7826711 |
| LVNNAAI | ASSTTASQNTLY | 0.78199804 |
| LTGEINTLA | ASSTTASQNTLY | 0.5455834 |
| LSIFGPV | ASSTTASQNTLY | 0.7809803 |
| LLALLANL | ASSTTASQNTLY | 0.77856207 |
| LISNASDALDKLR | ASSTTASQNTLY | 0.7829332 |
| LINEAVP | ASSTTASQNTLY | 0.74943566 |
| LGYAFSGL | ASSTTASQNTLY | 0.67589366 |
| LDFVAVAL | ASSTTASQNTLY | 0.7689289 |
| LAQPGAE | ASSTTASQNTLY | 0.70108914 |
| ISNLGAVTGF | ASSTTASQNTLY | 0.6367568 |
| HQEDPGRI | ASSTTASQNTLY | 0.7779282 |
| GTPGEPTP | ASSTTASQNTLY | 0.7332283 |
| GFFNSGS | ASSTTASQNTLY | 0.6823974 |
| GAGGGGLTAGAN | ASSTTASQNTLY | 0.6740842 |
| FVDALVNM | ASSTTASQNTLY | 0.77823246 |
| FLTVAPEL | ASSTTASQNTLY | 0.6328201 |
| FETLTAL | ASSTTASQNTLY | 0.65148693 |
| DVINPIEL | ASSTTASQNTLY | 0.66651094 |
| APVSGAPV | ASSTTASQNTLY | 0.7473203 |
| ALVGGLAW | ASSTTASQNTLY | 0.77891695 |
| ALAAAAATAI | ASSTTASQNTLY | 0.7726294 |
| AFGLGLLL | ASSTTASQNTLY | 0.66234285 |
| AFFAAASL | ASSTTASQNTLY | 0.76121235 |
| AAAALPAFSPP | ASSTTASQNTLY | 0.7658762 |
| VVLAPAL | ASSTTASQNTLY | 0.75545263 |
| TIGHVDHGKTTL | ASSTTASQNTLY | 0.75778866 |
| LISNASDALDK | ASSTTASQNTLY | 0.66945964 |
| LISNASDAL | ASSTTASQNTLY | 0.7805481 |
| ISNASDALDK | ASSTTASQNTLY | 0.762233 |
| NQAETLVYQTE | ASSTTASQNTLY | 0.78052104 |

| | | |
|---|---|---|
| TGGTGGTAGAG | ASSTTASQNTLY | 0.7814432 |
| TGSTAGGAGGA | ASSTTASQNTLY | 0.7436756 |
| DPLAVPVAAGPV | ASSTTASQNTLY | 0.74949324 |
| GGTGGTPGNGGN | ASSTTASQNTLY | 0.517824 |
| PAGLAGVDPATL | ASSTTASQNTLY | 0.656481 |
| GNNLVVEAPHAGASV | ASSTTASQNTLY | 0.7574594 |
| KVTRIMSASTLEQPA | ASSTTASQNTLY | 0.74313116 |
| LGFALPIALPRRTVV | ASSTTASQNTLY | 0.7753302 |
| TRLGNLTARKLVVLV | ASSTTASQNTLY | 0.77177083 |
| WSQQRRLIYFPSAGP | ASSTTASQNTLY | 0.6913177 |
| GPNGNPDPMAAAVDIR | ASSTTASQNTLY | 0.77880824 |
| TQRCVSVADLRAAASR | ASSTTASQNTLY | 0.7767459 |
| QFVLDVAEHPVEVLTETV | ASSTTASQNTLY | 0.5905895 |
| GGAGGTGGAGGNGGTQAGDG | ASSTTASQNTLY | 0.718531 |
| PLLATLRAIAVDVDAKPAQV | ASSTTASQNTLY | 0.7888913 |
| APKFRTGGYMIDFWGVGYQVAK | ASSTTASQNTLY | 0.75068873 |
| GAGGAGGAGGAAGGPGGTGGTG | ASSTTASQNTLY | 0.72885084 |
| IGALGLLATDVAPAGSVAFGVA | ASSTTASQNTLY | 0.7169848 |
| AGGAGGIGGNGGAGGAGGVLMG | ASSTTASQNTLY | 0.7884526 |
| ADELIGLL | ASSTTASQNTLY | 0.70376825 |
| ADRALKA | ASSTTASQNTLY | 0.7874896 |
| AGAVGQVGGAGGA | ASSTTASQNTLY | 0.37089697 |
| AGGGGGLGGAP | ASSTTASQNTLY | 0.72150993 |
| AGGLLYGNGGAG | ASSTTASQNTLY | 0.69010377 |
| AGLESIGL | ASSTTASQNTLY | 0.5135856 |
| AGLKTGII | ASSTTASQNTLY | 0.7809465 |
| ASLAALLL | ASSTTASQNTLY | 0.7404052 |
| ASVINISSI | ASSTTASQNTLY | 0.7717667 |
| ATPPGPA | ASSTTASQNTLY | 0.73910606 |
| ATVASALV | ASSTTASQNTLY | 0.7843824 |
| ATVASLIL | ASSTTASQNTLY | 0.7373047 |
| AVNGPVLIRH | ASSTTASQNTLY | 0.74881774 |
| DGAAGAVGI | ASSTTASQNTLY | 0.6570074 |
| DPLLHGI | ASSTTASQNTLY | 0.77807224 |
| ELGAALGAL | ASSTTASQNTLY | 0.7841363 |
| ELGRTQAV | ASSTTASQNTLY | 0.77346116 |
| FVAGGYLS | ASSTTASQNTLY | 0.62327325 |
| GASGGAGGAGGW | ASSTTASQNTLY | 0.7000171 |
| GGAGAGGDGGAGGI | ASSTTASQNTLY | 0.76891524 |
| GTGHNGSQPGMGGNG | ASSTTASQNTLY | 0.33686355 |
| GYAAASASAS | ASSTTASQNTLY | 0.39807037 |
| HQVDVAAVQQG | ASSTTASQNTLY | 0.7117859 |
| IGAAIAHT | ASSTTASQNTLY | 0.6424109 |
| IGMIEVAQL | ASSTTASQNTLY | 0.78300095 |
| IRLAPEHTSAMD | ASSTTASQNTLY | 0.77153146 |
| IYNVSVL | ASSTTASQNTLY | 0.7377112 |
| KQGFLGM | ASSTTASQNTLY | 0.75989187 |
| LESVITQL | ASSTTASQNTLY | 0.73765385 |
| LGLGALVVV | ASSTTASQNTLY | 0.21253306 |
| LIGNGGNGGSGGI | ASSTTASQNTLY | 0.68362033 |
| LNAINAPTL | ASSTTASQNTLY | 0.62385726 |

| | | |
|---|---|---|
| LQLLNTGV | ASSTTASQNTLY | 0.7896934 |
| LQTSIEF | ASSTTASQNTLY | 0.6900141 |
| MSFAVSEL | ASSTTASQNTLY | 0.6415845 |
| NTGALGL | ASSTTASQNTLY | 0.74248123 |
| PPGAGGAGGAGTT | ASSTTASQNTLY | 0.74724793 |
| QLVGLGI | ASSTTASQNTLY | 0.7342463 |
| RFLEQVHQL | ASSTTASQNTLY | 0.65922797 |
| RGVGYVLR | ASSTTASQNTLY | 0.28428984 |
| SATGVFAGVI | ASSTTASQNTLY | 0.7790439 |
| SLPAITLP | ASSTTASQNTLY | 0.5160811 |
| STALAFTV | ASSTTASQNTLY | 0.6983415 |
| SVLTVGLGK | ASSTTASQNTLY | 0.36683 |
| TALAGAGSGI | ASSTTASQNTLY | 0.37345374 |
| TGLPFTGI | ASSTTASQNTLY | 0.5687438 |
| VAEHVTV | ASSTTASQNTLY | 0.77497935 |
| VDIVLNSL | ASSTTASQNTLY | 0.7186174 |
| VLTGGGAVL | ASSTTASQNTLY | 0.7071336 |
| VNVGLAYGI | ASSTTASQNTLY | 0.7885895 |
| VSSAAVGAV | ASSTTASQNTLY | 0.74611235 |
| VTSVYEAGV | ASSTTASQNTLY | 0.6997092 |
| WVALDRGA | ASSTTASQNTLY | 0.71370006 |
| YLVGSVVI | ASSTTASQNTLY | 0.67975736 |
| GPPGTGKTLL | ASSTTASQNTLY | 0.6710619 |
| GVGMTKF | ASSTTASQNTLY | 0.70196164 |
| IGIGGSDL | ASSTTASQNTLY | 0.7881031 |
| IGTLLAEL | ASSTTASQNTLY | 0.72843987 |
| IYTSGTT | ASSTTASQNTLY | 0.60906327 |
| LSFPAGV | ASSTTASQNTLY | 0.7432877 |
| LYTSGTT | ASSTTASQNTLY | 0.7864225 |
| VGAFPFGF | ASSTTASQNTLY | 0.7878815 |
| VGAGTVEF | ASSTTASQNTLY | 0.66446364 |
| VGKSTLLL | ASSTTASQNTLY | 0.73016983 |
| AALAHAGGGIV | ASSTTASQNTLY | 0.72325754 |
| AALVAVSLL | ASSTTASQNTLY | 0.7412887 |
| AAVSRAVAAL | ASSTTASQNTLY | 0.6935075 |
| AGGSSGIGGF | ASSTTASQNTLY | 0.6526729 |
| AGLTVANANG | ASSTTASQNTLY | 0.767645 |
| AGLVGAVL | ASSTTASQNTLY | 0.79347754 |
| ALSQAFTV | ASSTTASQNTLY | 0.7066007 |
| ALTFSVSL | ASSTTASQNTLY | 0.548189 |
| ALTGAALVAL | ASSTTASQNTLY | 0.7375134 |
| ASFSIGAGVL | ASSTTASQNTLY | 0.6708368 |
| DGSVYKI | ASSTTASQNTLY | 0.70759225 |
| EQLAEIV | ASSTTASQNTLY | 0.7358774 |
| FAVSELTVM | ASSTTASQNTLY | 0.5774162 |
| GAAPLAAGL | ASSTTASQNTLY | 0.73174095 |
| GADGGDGGSGGAGG | ASSTTASQNTLY | 0.78896314 |
| GFGLDTGGAGG | ASSTTASQNTLY | 0.7240367 |
| GLASGGLL | ASSTTASQNTLY | 0.75376546 |
| GRIAGAAVLL | ASSTTASQNTLY | 0.78182137 |
| GSAGSLNTSF | ASSTTASQNTLY | 0.76186347 |

| | | |
|---|---|---|
| GTGQIGI | ASSTTASQNTLY | 0.780396 |
| GTLIALIL | ASSTTASQNTLY | 0.78025866 |
| IAATMGFVVG | ASSTTASQNTLY | 0.7558341 |
| IFAQALL | ASSTTASQNTLY | 0.75132626 |
| IGAAVALGF | ASSTTASQNTLY | 0.7496736 |
| IGLTPLAL | ASSTTASQNTLY | 0.7477893 |
| ILGSNTSTL | ASSTTASQNTLY | 0.7200546 |
| IVFSLTVL | ASSTTASQNTLY | 0.68720555 |
| LASKATGF | ASSTTASQNTLY | 0.7847537 |
| LASVALIL | ASSTTASQNTLY | 0.7345865 |
| LFAVGLTL | ASSTTASQNTLY | 0.78184235 |
| LFNGGIGTYI | ASSTTASQNTLY | 0.640221 |
| LTLGAGII | ASSTTASQNTLY | 0.6362426 |
| LVAAALGDI | ASSTTASQNTLY | 0.7676878 |
| NTAFVQLGI | ASSTTASQNTLY | 0.23481736 |
| QLFSGAEL | ASSTTASQNTLY | 0.3769123 |
| SAATFITL | ASSTTASQNTLY | 0.74982697 |
| SNYGLANLGV | ASSTTASQNTLY | 0.7789 |
| SYGGAYAVM | ASSTTASQNTLY | 0.24565297 |
| TTGVGGLTLG | ASSTTASQNTLY | 0.6697355 |
| TTQTIYI | ASSTTASQNTLY | 0.6163949 |
| VAAGAGLGAVI | ASSTTASQNTLY | 0.76318145 |
| VAAVVTLL | ASSTTASQNTLY | 0.7570642 |
| VGFLGGLI | ASSTTASQNTLY | 0.5929662 |
| VGVIPILL | ASSTTASQNTLY | 0.7751806 |
| VIFASSLI | ASSTTASQNTLY | 0.47839767 |
| VKGGDGGAAAG | ASSTTASQNTLY | 0.5598711 |
| VTKTFTNI | ASSTTASQNTLY | 0.78052294 |
| VVAAAILL | ASSTTASQNTLY | 0.7840371 |
| YAALDVEL | ASSTTASQNTLY | 0.644871 |
| LASVALLL | ASSTTASQNTLY | 0.7764547 |
| LDAVHRV | ASSTTASQNTLY | 0.6823492 |
| QGLSSSIF | ASSTTASQNTLY | 0.78888404 |
| RGTFANIRL | ASSTTASQNTLY | 0.73063415 |
| RQLWWGHRI | ASSTTASQNTLY | 0.5921047 |
| YGPPGTGKTLL | ASSTTASQNTLY | 0.7765193 |
| AALVGAPANVA | ASSTTASQNTLY | 0.75960684 |
| AATFRRTE | ASSTTASQNTLY | 0.7608969 |
| AGALGGGI | ASSTTASQNTLY | 0.7765852 |
| AGGAGGAGGIGAG | ASSTTASQNTLY | 0.65808105 |
| AGSGALGL | ASSTTASQNTLY | 0.7336684 |
| AQIADQI | ASSTTASQNTLY | 0.7861873 |
| ASAVAVTYL | ASSTTASQNTLY | 0.7769372 |
| ASTVLGGTGG | ASSTTASQNTLY | 0.6761142 |
| AVGGEPAK | ASSTTASQNTLY | 0.77802634 |
| CEQGTFGGTC | ASSTTASQNTLY | 0.7182132 |
| DEGKAPAG | ASSTTASQNTLY | 0.7084497 |
| DTVVFPAL | ASSTTASQNTLY | 0.7865137 |
| FEGITMAG | ASSTTASQNTLY | 0.73962134 |
| FPLVVQD | ASSTTASQNTLY | 0.7705711 |
| GAFTTPQ | ASSTTASQNTLY | 0.782237 |

| | | |
|---|---|---|
| GFGGFGTAGGVGGA | ASSTTASQNTLY | 0.77963465 |
| GFLAQFL | ASSTTASQNTLY | 0.5935493 |
| GGAGGAGGNAGVA | ASSTTASQNTLY | 0.73861253 |
| GGVGTAGAGGAGGAGG | ASSTTASQNTLY | 0.73502475 |
| GNGGGGDAGNAGSGGN | ASSTTASQNTLY | 0.7781478 |
| GTGGAGGAGNDAGS | ASSTTASQNTLY | 0.7479466 |
| GTTNTGWFNS | ASSTTASQNTLY | 0.5989073 |
| IFMGPLFTL | ASSTTASQNTLY | 0.76465845 |
| ILAHAIEAI | ASSTTASQNTLY | 0.78014976 |
| IRSAIREAL | ASSTTASQNTLY | 0.75135565 |
| ITFAGGG | ASSTTASQNTLY | 0.74899566 |
| ITGMSGHQEV | ASSTTASQNTLY | 0.7842226 |
| IVAGLACTL | ASSTTASQNTLY | 0.7237276 |
| IVGGGLAGA | ASSTTASQNTLY | 0.7290379 |
| IVTGVLNI | ASSTTASQNTLY | 0.7790146 |
| LAQQGCI | ASSTTASQNTLY | 0.77809346 |
| LCLHVEN | ASSTTASQNTLY | 0.7416698 |
| LDAISDGS | ASSTTASQNTLY | 0.7787198 |
| LGGFAAML | ASSTTASQNTLY | 0.64527434 |
| LGGKNPMI | ASSTTASQNTLY | 0.7699118 |
| LGPKGVAAIL | ASSTTASQNTLY | 0.64023757 |
| LMVGVSI | ASSTTASQNTLY | 0.7671418 |
| LVGAAFVL | ASSTTASQNTLY | 0.6621728 |
| LVLAVAPL | ASSTTASQNTLY | 0.7761156 |
| MAAIAVIDAA | ASSTTASQNTLY | 0.7675474 |
| NNGINAELQ | ASSTTASQNTLY | 0.7777202 |
| RGEPLVLVHG | ASSTTASQNTLY | 0.33827317 |
| RGGAGGAAGLI | ASSTTASQNTLY | 0.77472794 |
| RSYRMHPAV | ASSTTASQNTLY | 0.7810379 |
| SAAHLLAGQLA | ASSTTASQNTLY | 0.79157543 |
| SAAVGSY | ASSTTASQNTLY | 0.743348 |
| SPHAVLHRI | ASSTTASQNTLY | 0.7632061 |
| SQIAKVL | ASSTTASQNTLY | 0.6986949 |
| TALGHLVP | ASSTTASQNTLY | 0.72318757 |
| TAPTTSGAA | ASSTTASQNTLY | 0.6232681 |
| TDGDDGNGGSPGT | ASSTTASQNTLY | 0.7119448 |
| TDGLGEN | ASSTTASQNTLY | 0.68095016 |
| TGGPPLPGQA | ASSTTASQNTLY | 0.6839415 |
| TIPAITI | ASSTTASQNTLY | 0.77059555 |
| TIPGFTI | ASSTTASQNTLY | 0.7749877 |
| TSIASGALA | ASSTTASQNTLY | 0.7765261 |
| TVTLPGAAI | ASSTTASQNTLY | 0.7835124 |
| VASGLSAALP | ASSTTASQNTLY | 0.7765774 |
| VGAGVIGI | ASSTTASQNTLY | 0.77999115 |
| VGPKRFGV | ASSTTASQNTLY | 0.7071819 |
| VNVAFPD | ASSTTASQNTLY | 0.7190213 |
| VSMTNSGVS | ASSTTASQNTLY | 0.628274 |
| VTDVLGDLSV | ASSTTASQNTLY | 0.7381945 |
| VVVVGAGFAGL | ASSTTASQNTLY | 0.7120401 |
| VYTVGTVV | ASSTTASQNTLY | 0.7613202 |
| WTTFSGP | ASSTTASQNTLY | 0.23101665 |

| | | |
|---|---|---|
| YDELVEAF | ASSTTASQNTLY | 0.73579574 |
| YGTGGAGGNGG | ASSTTASQNTLY | 0.78584 |
| YTNVMAV | ASSTTASQNTLY | 0.6132936 |
| FLEQVHQL | ASSTTASQNTLY | 0.762277 |
| GLSLLAVAV | ASSTTASQNTLY | 0.735648 |
| LEQVHQL | ASSTTASQNTLY | 0.6874992 |
| LNAAGAI | ASSTTASQNTLY | 0.77253187 |
| PAVHASL | ASSTTASQNTLY | 0.6835399 |
| TELEDDA | ASSTTASQNTLY | 0.7638711 |
| TPPGPPN | ASSTTASQNTLY | 0.7855544 |
| TPPPPPPP | ASSTTASQNTLY | 0.7543937 |
| TQSPALI | ASGEGGQDTQY | 0.74506545 |
| EFAAAFPQ | ASGEGGQDTQY | 0.745528 |
| LLSGNLDVL | ASGEGGQDTQY | 0.5428599 |
| GPANLGNYNI | ASGEGGQDTQY | 0.7628598 |
| TLGPTLP | ASGEGGQDTQY | 0.21719605 |
| VYAYSSCCK | ASGEGGQDTQY | 0.739362 |
| APAPAPAPNI | ASGEGGQDTQY | 0.6735819 |
| PGGPVPIP | ASGEGGQDTQY | 0.76072085 |
| FLTVALEP | ASGEGGQDTQY | 0.7371844 |
| SQYFGANL | ASGEGGQDTQY | 0.7633101 |
| GGVKPAA | ASGEGGQDTQY | 0.6786793 |
| GGTGGAGGAGGW | ASGEGGQDTQY | 0.7176162 |
| GVGGVAGLGGAGG | ASGEGGQDTQY | 0.66267866 |
| VLLVLATTI | ASGEGGQDTQY | 0.663223 |
| IGIFAAML | ASGEGGQDTQY | 0.7627599 |
| AVGDLID | ASGEGGQDTQY | 0.74105096 |
| NISPLPL | ASGEGGQDTQY | 0.70578027 |
| VTAVFVAI | ASGEGGQDTQY | 0.78535473 |
| SGDDNDAAL | ASGEGGQDTQY | 0.7580733 |
| AVAALAHS | ASGEGGQDTQY | 0.47917157 |
| GGVTVMATP | ASGEGGQDTQY | 0.7205191 |
| AGLAANVRD | ASGEGGQDTQY | 0.7408206 |
| TNRPDLIDPALLRPG | ASGEGGQDTQY | 0.7364582 |
| LANLGVTAL | ASGEGGQDTQY | 0.21665901 |
| TGERIEL | ASGEGGQDTQY | 0.748039 |
| NGEELAA | ASGEGGQDTQY | 0.7058415 |
| VLSLVWG | ASGEGGQDTQY | 0.6448456 |
| IGYALWL | ASGEGGQDTQY | 0.70082104 |
| LSLLLAPL | ASGEGGQDTQY | 0.76979685 |
| TLDEALK | ASGEGGQDTQY | 0.5156589 |
| TDGAGTAEGG | ASGEGGQDTQY | 0.7530718 |
| NAPGGAGGNGG | ASGEGGQDTQY | 0.766271 |
| VSVFMGMGGNF | ASGEGGQDTQY | 0.77587533 |
| LASYLAAL | ASGEGGQDTQY | 0.78565705 |
| LVGGTPAL | ASGEGGQDTQY | 0.7546072 |
| VRNADVL | ASGEGGQDTQY | 0.7343375 |
| TCSSPFLVSAM | ASGEGGQDTQY | 0.7699665 |
| VHITDAE | ASGEGGQDTQY | 0.72403014 |
| EYAAPSF | ASGEGGQDTQY | 0.5698526 |
| VVDAFHGGL | ASGEGGQDTQY | 0.41462743 |

| | | |
|---|---|---|
| ASVVDPQ | ASGEGGQDTQY | 0.7543144 |
| SYNEFGGLS | ASGEGGQDTQY | 0.46867362 |
| DPDGSVY | ASGEGGQDTQY | 0.75656855 |
| MVSTIHATE | ASGEGGQDTQY | 0.7861023 |
| LAAGVADPV | ASGEGGQDTQY | 0.71271527 |
| TKEQIAA | ASGEGGQDTQY | 0.60830206 |
| SLASPAAL | ASGEGGQDTQY | 0.7190556 |
| IGLVFYGV | ASGEGGQDTQY | 0.72304034 |
| FLLDASNL | ASGEGGQDTQY | 0.76746666 |
| YQVGNFP | ASGEGGQDTQY | 0.77380365 |
| ESLLSQST | ASGEGGQDTQY | 0.7914475 |
| LFTVGFAAVG | ASGEGGQDTQY | 0.7799019 |
| ATPPTPT | ASGEGGQDTQY | 0.7027134 |
| TDLATLG | ASGEGGQDTQY | 0.7592307 |
| SAGGTGR | ASGEGGQDTQY | 0.72950053 |
| GFSNSGS | ASGEGGQDTQY | 0.71921796 |
| GGNGGSGGTSV | ASGEGGQDTQY | 0.7452101 |
| GTSVATGGAGN | ASGEGGQDTQY | 0.7500375 |
| NGGNGGSGGTS | ASGEGGQDTQY | 0.7382973 |
| AVVKFPH | ASGEGGQDTQY | 0.7285056 |
| TPPPPPP | ASGEGGQDTQY | 0.55932474 |
| AAPAAAAAPA | ASGEGGQDTQY | 0.7764238 |
| IALNEDL | ASGEGGQDTQY | 0.7559432 |
| IVNVSSM | ASGEGGQDTQY | 0.5183601 |
| TGIPPAPR | ASGEGGQDTQY | 0.71847236 |
| FDLGGGTF | ASGEGGQDTQY | 0.7880577 |
| FDLGGGTFDVS | ASGEGGQDTQY | 0.78418624 |
| NEPTAAA | ASGEGGQDTQY | 0.67926025 |
| NEPTAAALA | ASGEGGQDTQY | 0.70392275 |
| GSIINISSI | ASGEGGQDTQY | 0.73502845 |
| VAPGNAGTA | ASGEGGQDTQY | 0.5099162 |
| TGGGASLE | ASGEGGQDTQY | 0.7569591 |
| VAVYNHYKRI | ASGEGGQDTQY | 0.73434556 |
| TDPAPATT | ASGEGGQDTQY | 0.71276534 |
| TNRPDILDPALLRPG | ASGEGGQDTQY | 0.7022277 |
| LTTTTTS | ASGEGGQDTQY | 0.740515 |
| PTGSGKTL | ASGEGGQDTQY | 0.6806463 |
| TGSGKTL | ASGEGGQDTQY | 0.7667385 |
| TIGHVDHGK | ASGEGGQDTQY | 0.23792058 |
| TIGHVDHGKTT | ASGEGGQDTQY | 0.7669205 |
| GHVDHGKTT | ASGEGGQDTQY | 0.64417267 |
| AQFEHTLL | ASGEGGQDTQY | 0.66941524 |
| IVNQDEL | ASGEGGQDTQY | 0.4576859 |
| TNGSQFF | ASGEGGQDTQY | 0.7383944 |
| TGKGIVE | ASGEGGQDTQY | 0.77298963 |
| LISNASDA | ASGEGGQDTQY | 0.77077466 |
| AELRTLGELTKIAKAHGAQ | ASGEGGQDTQY | 0.61177117 |
| APNAVAAAEPSW | ASGEGGQDTQY | 0.77878237 |
| APQVAGAATAVL | ASGEGGQDTQY | 0.78295183 |
| ARRDPAAAAAILPTDA | ASGEGGQDTQY | 0.7030841 |
| DAAARDVLAEAGLAEYF | ASGEGGQDTQY | 0.47222587 |

| | | |
|---|---|---|
| DDQIVLSAKANLAKVEQ | ASGEGGQDTQY | 0.76781154 |
| DELLPDTGEDSDEDGH | ASGEGGQDTQY | 0.7278181 |
| ELPEAPANRATVPQD | ASGEGGQDTQY | 0.7692218 |
| FGQSKIERVTGIG | ASGEGGQDTQY | 0.7502197 |
| GAGGTGSTAGGAGGAGGAGGLY | ASGEGGQDTQY | 0.6286665 |
| GHGGTGAAGGNGGTGGAGGSGI | ASGEGGQDTQY | 0.7782325 |
| GNGGNGGAGGEGGPGGAGGAGA | ASGEGGQDTQY | 0.7791616 |
| GSTAGNGGNGGNATNGTIAGQP | ASGEGGQDTQY | 0.66728956 |
| GTGGDGGNGGNGGTGG | ASGEGGQDTQY | 0.6512788 |
| GVRRPAGIPADAPV | ASGEGGQDTQY | 0.67602 |
| GYRQVTDAHLLTLARR | ASGEGGQDTQY | 0.41651723 |
| IAKMGQDGHDRG | ASGEGGQDTQY | 0.7782223 |
| KAEPSIEIQYT | ASGEGGQDTQY | 0.67481136 |
| KPVRIRMAADEDTVAVT | ASGEGGQDTQY | 0.7107036 |
| MPPRIAGMRLLVIKPEP | ASGEGGQDTQY | 0.75519776 |
| MKFYAVALFRSRGVPPTR | ASGEGGQDTQY | 0.72206855 |
| NAGGFGILGAVAHSP | ASGEGGQDTQY | 0.7248398 |
| NEGREITLAMAEM | ASGEGGQDTQY | 0.72977173 |
| NELVEWSHPLGHRWRWE | ASGEGGQDTQY | 0.74939257 |
| NLGAGGTGGNGGNPG | ASGEGGQDTQY | 0.7429914 |
| NTEWPKWETVMAADPVD | ASGEGGQDTQY | 0.7350141 |
| PASAAPAETQP | ASGEGGQDTQY | 0.74472684 |
| QQSSDGKNGVRAAEITGE | ASGEGGQDTQY | 0.78381443 |
| RAPDAVTTSWWKE | ASGEGGQDTQY | 0.6302084 |
| SGFGNVGAHTSGWFNQ | ASGEGGQDTQY | 0.6961833 |
| SGVAGAVGQAASVGGLKV | ASGEGGQDTQY | 0.74902004 |
| SNGGAGTGSAGGNGGT | ASGEGGQDTQY | 0.58113337 |
| TGGAGGEGGAAG | ASGEGGQDTQY | 0.7633279 |
| TGGDGGAAGAGGAPGHG | ASGEGGQDTQY | 0.7356471 |
| TMPPLLAAIANAGRHGV | ASGEGGQDTQY | 0.7542591 |
| TSSTAGNSGGA | ASGEGGQDTQY | 0.73750424 |
| VPATVPVGILC | ASGEGGQDTQY | 0.7893511 |
| VQRPDSGGVVLTGRIS | ASGEGGQDTQY | 0.77676046 |
| WTPNPVPATKGVPAGLVQAT | ASGEGGQDTQY | 0.7516229 |
| YALKRALAKFSDRFDVVII | ASGEGGQDTQY | 0.7523444 |
| YLIGSVIV | ASGEGGQDTQY | 0.61885554 |
| YGQLVAVID | ASGEGGQDTQY | 0.7794064 |
| VTVGGMTLPA | ASGEGGQDTQY | 0.7634629 |
| VLALPAV | ASGEGGQDTQY | 0.6467271 |
| VIPPTLNYET | ASGEGGQDTQY | 0.74021804 |
| VGMGGMHPSSG | ASGEGGQDTQY | 0.78438973 |
| VGLAGGAATA | ASGEGGQDTQY | 0.77948755 |
| VGGAGGAGGSGP | ASGEGGQDTQY | 0.54227364 |
| VFAAFMV | ASGEGGQDTQY | 0.71741843 |
| TLYDKLR | ASGEGGQDTQY | 0.47932094 |
| TLAVFTVL | ASGEGGQDTQY | 0.6803973 |
| TGSVQPARV | ASGEGGQDTQY | 0.649264 |
| TGGGAGSGGT | ASGEGGQDTQY | 0.76009893 |
| SHLETSL | ASGEGGQDTQY | 0.712337 |
| SFTTGMT | ASGEGGQDTQY | 0.7599926 |
| QVEVAKRMGVRQPTVS | ASGEGGQDTQY | 0.7552018 |

| | | |
|---|---|---|
| QGSGGPALAM | ASGEGGQDTQY | 0.78666043 |
| MLAPSNP | ASGEGGQDTQY | 0.7055011 |
| LVNNAAI | ASGEGGQDTQY | 0.7401222 |
| LTGEINTLA | ASGEGGQDTQY | 0.53836465 |
| LSIFGPV | ASGEGGQDTQY | 0.57806665 |
| LLALLANL | ASGEGGQDTQY | 0.6866097 |
| LISNASDALDKLR | ASGEGGQDTQY | 0.7281865 |
| LINEAVP | ASGEGGQDTQY | 0.3039811 |
| LGYAFSGL | ASGEGGQDTQY | 0.7791458 |
| LDFVAVAL | ASGEGGQDTQY | 0.7572653 |
| LAQPGAE | ASGEGGQDTQY | 0.7738666 |
| ISNLGAVTGF | ASGEGGQDTQY | 0.71271825 |
| HQEDPGRI | ASGEGGQDTQY | 0.73086494 |
| GTPGEPTP | ASGEGGQDTQY | 0.6945403 |
| GFFNSGS | ASGEGGQDTQY | 0.7677679 |
| GAGGGGLTAGAN | ASGEGGQDTQY | 0.679332 |
| FVDALVNM | ASGEGGQDTQY | 0.5430285 |
| FLTVAPEL | ASGEGGQDTQY | 0.6928496 |
| FETLTAL | ASGEGGQDTQY | 0.5580958 |
| DVINPIEL | ASGEGGQDTQY | 0.78008074 |
| APVSGAPV | ASGEGGQDTQY | 0.77774096 |
| ALVGGLAW | ASGEGGQDTQY | 0.76299775 |
| ALAAAAATAI | ASGEGGQDTQY | 0.78069186 |
| AFGLGLLL | ASGEGGQDTQY | 0.33961886 |
| AFFAAASL | ASGEGGQDTQY | 0.70686173 |
| AAAALPAFSPP | ASGEGGQDTQY | 0.78747845 |
| VVLAPAL | ASGEGGQDTQY | 0.7601516 |
| TIGHVDHGKTTL | ASGEGGQDTQY | 0.7252779 |
| LISNASDALDK | ASGEGGQDTQY | 0.7803128 |
| LISNASDAL | ASGEGGQDTQY | 0.68894035 |
| ISNASDALDK | ASGEGGQDTQY | 0.7408075 |
| NQAETLVYQTE | ASGEGGQDTQY | 0.7322131 |
| TGGTGGTAGAG | ASGEGGQDTQY | 0.7721672 |
| TGSTAGGAGGA | ASGEGGQDTQY | 0.7487564 |
| DPLAVPVAAGPV | ASGEGGQDTQY | 0.6838076 |
| GGTGGTPGNGGN | ASGEGGQDTQY | 0.76563203 |
| PAGLAGVDPATL | ASGEGGQDTQY | 0.79009485 |
| GNNLVVEAPHAGASV | ASGEGGQDTQY | 0.7234856 |
| KVTRIMSASTLEQPA | ASGEGGQDTQY | 0.767421 |
| LGFALPIALPRRTVV | ASGEGGQDTQY | 0.7649617 |
| TRLGNLTARKLVVLV | ASGEGGQDTQY | 0.7638978 |
| WSQQRRLIYFPSAGP | ASGEGGQDTQY | 0.69092715 |
| GPNGNPDPMAAAVDIR | ASGEGGQDTQY | 0.6967307 |
| TQRCVSVADLRAAASR | ASGEGGQDTQY | 0.7314836 |
| QFVLDVAEHPVEVLTETV | ASGEGGQDTQY | 0.62324023 |
| GGAGGTGGAGGNGGTQAGDG | ASGEGGQDTQY | 0.73671603 |
| PLLATLRAIAVDVDAKPAQV | ASGEGGQDTQY | 0.7261733 |
| APKFRTGGYMIDFWGVGYQVAK | ASGEGGQDTQY | 0.7940347 |
| GAGGAGGAGGAAGGPGGTGGTG | ASGEGGQDTQY | 0.7621804 |
| IGALGLLATDVAPAGSVAFGVA | ASGEGGQDTQY | 0.75556993 |
| AGGAGGIGGNGGAGGAGGVLMG | ASGEGGQDTQY | 0.66651577 |

| | | |
|---|---|---|
| ADELIGLL | ASGEGGQDTQY | 0.7651216 |
| ADRALKA | ASGEGGQDTQY | 0.7621199 |
| AGAVGQVGGAGGA | ASGEGGQDTQY | 0.5990537 |
| AGGGGGLGGAP | ASGEGGQDTQY | 0.7831614 |
| AGGLLYGNGGAG | ASGEGGQDTQY | 0.6901405 |
| AGLESIGL | ASGEGGQDTQY | 0.7070038 |
| AGLKTGII | ASGEGGQDTQY | 0.774263 |
| ASLAALLL | ASGEGGQDTQY | 0.7892858 |
| ASVINISSI | ASGEGGQDTQY | 0.7240659 |
| ATPPGPA | ASGEGGQDTQY | 0.69981915 |
| ATVASALV | ASGEGGQDTQY | 0.68473446 |
| ATVASLIL | ASGEGGQDTQY | 0.7739393 |
| AVNGPVLIRH | ASGEGGQDTQY | 0.780817 |
| DGAAGAVGI | ASGEGGQDTQY | 0.7329142 |
| DPLLHGI | ASGEGGQDTQY | 0.7867927 |
| ELGAALGAL | ASGEGGQDTQY | 0.76381963 |
| ELGRTQAV | ASGEGGQDTQY | 0.55436766 |
| FVAGGYLS | ASGEGGQDTQY | 0.73868036 |
| GASGGAGGAGGW | ASGEGGQDTQY | 0.77670115 |
| GGAGAGGDGGAGGI | ASGEGGQDTQY | 0.7643268 |
| GTGHNGSQPGMGGNG | ASGEGGQDTQY | 0.7898041 |
| GYAAASASAS | ASGEGGQDTQY | 0.7672026 |
| HQVDVAAVQQG | ASGEGGQDTQY | 0.71224415 |
| IGAAIAHT | ASGEGGQDTQY | 0.78510267 |
| IGMIEVAQL | ASGEGGQDTQY | 0.7898185 |
| IRLAPEHTSAMD | ASGEGGQDTQY | 0.77643275 |
| IYNVSVL | ASGEGGQDTQY | 0.71099925 |
| KQGFLGM | ASGEGGQDTQY | 0.69422853 |
| LESVITQL | ASGEGGQDTQY | 0.7546879 |
| LGLGALVVV | ASGEGGQDTQY | 0.7676929 |
| LIGNGGNGGSGGI | ASGEGGQDTQY | 0.6539581 |
| LNAINAPTL | ASGEGGQDTQY | 0.74762756 |
| LQLLNTGV | ASGEGGQDTQY | 0.7556017 |
| LQTSIEF | ASGEGGQDTQY | 0.57880205 |
| MSFAVSEL | ASGEGGQDTQY | 0.76649046 |
| NTGALGL | ASGEGGQDTQY | 0.77992237 |
| PPGAGGAGGAGTT | ASGEGGQDTQY | 0.744611 |
| QLVGLGI | ASGEGGQDTQY | 0.7213799 |
| RFLEQVHQL | ASGEGGQDTQY | 0.7781487 |
| RGVGYVLR | ASGEGGQDTQY | 0.79389966 |
| SATGVFAGVI | ASGEGGQDTQY | 0.78386843 |
| SLPAITLP | ASGEGGQDTQY | 0.57072705 |
| STALAFTV | ASGEGGQDTQY | 0.37653193 |
| SVLTVGLGK | ASGEGGQDTQY | 0.69719917 |
| TALAGAGSGI | ASGEGGQDTQY | 0.77002764 |
| TGLPFTGI | ASGEGGQDTQY | 0.7349657 |
| VAEHVTV | ASGEGGQDTQY | 0.72692263 |
| VDIVLNSL | ASGEGGQDTQY | 0.7480889 |
| VLTGGGAVL | ASGEGGQDTQY | 0.7828546 |
| VNVGLAYGI | ASGEGGQDTQY | 0.7245338 |
| VSSAAVGAV | ASGEGGQDTQY | 0.54884076 |

| | | |
|---|---|---|
| VTSVYEAGV | ASGEGGQDTQY | 0.7523413 |
| WVALDRGA | ASGEGGQDTQY | 0.7697904 |
| YLVGSVVI | ASGEGGQDTQY | 0.6888964 |
| GPPGTGKTLL | ASGEGGQDTQY | 0.7257608 |
| GVGMTKF | ASGEGGQDTQY | 0.6855229 |
| IGIGGSDL | ASGEGGQDTQY | 0.58502734 |
| IGTLLAEL | ASGEGGQDTQY | 0.7482705 |
| IYTSGTT | ASGEGGQDTQY | 0.6838223 |
| LSFPAGV | ASGEGGQDTQY | 0.77828264 |
| LYTSGTT | ASGEGGQDTQY | 0.6842567 |
| VGAFPFGF | ASGEGGQDTQY | 0.75884104 |
| VGAGTVEF | ASGEGGQDTQY | 0.75967145 |
| VGKSTLLL | ASGEGGQDTQY | 0.77083397 |
| AALAHAGGGIV | ASGEGGQDTQY | 0.71519494 |
| AALVAVSLL | ASGEGGQDTQY | 0.71512115 |
| AAVSRAVAAL | ASGEGGQDTQY | 0.6455174 |
| AGGSSGIGGF | ASGEGGQDTQY | 0.4188568 |
| AGLTVANANG | ASGEGGQDTQY | 0.77937466 |
| AGLVGAVL | ASGEGGQDTQY | 0.26368195 |
| ALSQAFTV | ASGEGGQDTQY | 0.7826565 |
| ALTFSVSL | ASGEGGQDTQY | 0.7759616 |
| ALTGAALVAL | ASGEGGQDTQY | 0.6751589 |
| ASFSIGAGVL | ASGEGGQDTQY | 0.7623255 |
| DGSVYKI | ASGEGGQDTQY | 0.65483624 |
| EQLAEIV | ASGEGGQDTQY | 0.6796405 |
| FAVSELTVM | ASGEGGQDTQY | 0.7690868 |
| GAAPLAAGL | ASGEGGQDTQY | 0.77152467 |
| GADGGDGGSGGAGG | ASGEGGQDTQY | 0.6699206 |
| GFGLDTGGAGG | ASGEGGQDTQY | 0.7506507 |
| GLASGGLL | ASGEGGQDTQY | 0.75066 |
| GRIAGAAVLL | ASGEGGQDTQY | 0.6641106 |
| GSAGSLNTSF | ASGEGGQDTQY | 0.7537791 |
| GTGQIGI | ASGEGGQDTQY | 0.7816987 |
| GTLIALIL | ASGEGGQDTQY | 0.75887734 |
| IAATMGFVVG | ASGEGGQDTQY | 0.6990545 |
| IFAQALL | ASGEGGQDTQY | 0.7682827 |
| IGAAVALGF | ASGEGGQDTQY | 0.64021057 |
| IGLTPLAL | ASGEGGQDTQY | 0.76025 |
| ILGSNTSTL | ASGEGGQDTQY | 0.72785354 |
| IVFSLTVL | ASGEGGQDTQY | 0.72628784 |
| LASKATGF | ASGEGGQDTQY | 0.7701905 |
| LASVALIL | ASGEGGQDTQY | 0.48316145 |
| LFAVGLTL | ASGEGGQDTQY | 0.66192454 |
| LFNGGIGTYI | ASGEGGQDTQY | 0.7288101 |
| LTLGAGII | ASGEGGQDTQY | 0.7605802 |
| LVAAALGDI | ASGEGGQDTQY | 0.7761207 |
| NTAFVQLGI | ASGEGGQDTQY | 0.68825644 |
| QLFSGAEL | ASGEGGQDTQY | 0.7464717 |
| SAATFITL | ASGEGGQDTQY | 0.7245244 |
| SNYGLANLGV | ASGEGGQDTQY | 0.6883457 |
| SYGGAYAVM | ASGEGGQDTQY | 0.7508043 |

| | | |
|---|---|---|
| TTGVGGLTLG | ASGEGGQDTQY | 0.7347471 |
| TTQTIYI | ASGEGGQDTQY | 0.654531 |
| VAAGAGLGAVI | ASGEGGQDTQY | 0.76716423 |
| VAAVVTLL | ASGEGGQDTQY | 0.7023787 |
| VGFLGGLI | ASGEGGQDTQY | 0.7686484 |
| VGVIPILL | ASGEGGQDTQY | 0.7841715 |
| VIFASSLI | ASGEGGQDTQY | 0.70456755 |
| VKGGDGGAAAG | ASGEGGQDTQY | 0.76436037 |
| VTKTFTNI | ASGEGGQDTQY | 0.75863874 |
| VVAAAILL | ASGEGGQDTQY | 0.760453 |
| YAALDVEL | ASGEGGQDTQY | 0.7312686 |
| LASVALLL | ASGEGGQDTQY | 0.7012675 |
| LDAVHRV | ASGEGGQDTQY | 0.7833951 |
| QGLSSSIF | ASGEGGQDTQY | 0.77149034 |
| RGTFANIRL | ASGEGGQDTQY | 0.7667991 |
| RQLWWGHRI | ASGEGGQDTQY | 0.76202327 |
| YGPPGTGKTLL | ASGEGGQDTQY | 0.7718826 |
| AALVGAPANVA | ASGEGGQDTQY | 0.7673113 |
| AATFRRTE | ASGEGGQDTQY | 0.68258405 |
| AGALGGGI | ASGEGGQDTQY | 0.57707936 |
| AGGAGGAGGIGAG | ASGEGGQDTQY | 0.73644066 |
| AGSGALGL | ASGEGGQDTQY | 0.7532151 |
| AQIADQI | ASGEGGQDTQY | 0.7276825 |
| ASAVAVTYL | ASGEGGQDTQY | 0.48356247 |
| ASTVLGGTGG | ASGEGGQDTQY | 0.7816534 |
| AVGGEPAK | ASGEGGQDTQY | 0.7916118 |
| CEQGTFGGTC | ASGEGGQDTQY | 0.6859592 |
| DEGKAPAG | ASGEGGQDTQY | 0.7858847 |
| DTVVFPAL | ASGEGGQDTQY | 0.6285112 |
| FEGITMAG | ASGEGGQDTQY | 0.77055657 |
| FPLVVQD | ASGEGGQDTQY | 0.7808229 |
| GAFTTPQ | ASGEGGQDTQY | 0.64973307 |
| GFGGFGTAGGVGGA | ASGEGGQDTQY | 0.7799207 |
| GFLAQFL | ASGEGGQDTQY | 0.6895411 |
| GGAGGAGGNAGVA | ASGEGGQDTQY | 0.73094136 |
| GGVGTAGAGGAGGAGG | ASGEGGQDTQY | 0.7847511 |
| GNGGGGDAGNAGSGGN | ASGEGGQDTQY | 0.72604 |
| GTGGAGGAGNDAGS | ASGEGGQDTQY | 0.759312 |
| GTTNTGWFNS | ASGEGGQDTQY | 0.75280255 |
| IFMGPLFTL | ASGEGGQDTQY | 0.6849274 |
| ILAHAIEAI | ASGEGGQDTQY | 0.7648405 |
| IRSAIREAL | ASGEGGQDTQY | 0.77984107 |
| ITFAGGG | ASGEGGQDTQY | 0.6832414 |
| ITGMSGHQEV | ASGEGGQDTQY | 0.78901803 |
| IVAGLACTL | ASGEGGQDTQY | 0.6804458 |
| IVGGGLAGA | ASGEGGQDTQY | 0.7926035 |
| IVTGVLNI | ASGEGGQDTQY | 0.76285076 |
| LAQQGCI | ASGEGGQDTQY | 0.73896194 |
| LCLHVEN | ASGEGGQDTQY | 0.7091737 |
| LDAISDGS | ASGEGGQDTQY | 0.7824437 |
| LGGFAAML | ASGEGGQDTQY | 0.7319256 |

| | | |
|---|---|---|
| LGGKNPMI | ASGEGGQDTQY | 0.7602859 |
| LGPKGVAAIL | ASGEGGQDTQY | 0.78761697 |
| LMVGVSI | ASGEGGQDTQY | 0.44370952 |
| LVGAAFVL | ASGEGGQDTQY | 0.77421755 |
| LVLAVAPL | ASGEGGQDTQY | 0.6237097 |
| MAAIAVIDAA | ASGEGGQDTQY | 0.47633544 |
| NNGINAELQ | ASGEGGQDTQY | 0.41473776 |
| RGEPLVLVHG | ASGEGGQDTQY | 0.27196506 |
| RGGAGGAAGLI | ASGEGGQDTQY | 0.4772639 |
| RSYRMHPAV | ASGEGGQDTQY | 0.3726839 |
| SAAHLLAGQLA | ASGEGGQDTQY | 0.61438835 |
| SAAVGSY | ASGEGGQDTQY | 0.5698811 |
| SPHAVLHRI | ASGEGGQDTQY | 0.4156798 |
| SQIAKVL | ASGEGGQDTQY | 0.8496262 |
| TALGHLVP | ASGEGGQDTQY | 0.7328143 |
| TAPTTSGAA | ASGEGGQDTQY | 0.2963452 |
| TDGDDGNGGSPGT | ASGEGGQDTQY | 0.48416728 |
| TDGLGEN | ASGEGGQDTQY | 0.42657784 |
| TGGPPLPGQA | ASGEGGQDTQY | 0.48107988 |
| TIPAITI | ASGEGGQDTQY | 0.24467422 |
| TIPGFTI | ASGEGGQDTQY | 0.6143676 |
| TSIASGALA | ASGEGGQDTQY | 0.36190134 |
| TVTLPGAAI | ASGEGGQDTQY | 0.10943056 |
| VASGLSAALP | ASGEGGQDTQY | 0.55282855 |
| VGAGVIGI | ASGEGGQDTQY | 0.47278526 |
| VGPKRFGV | ASGEGGQDTQY | 0.4892286 |
| VNVAFPD | ASGEGGQDTQY | 0.78205884 |
| VSMTNSGVS | ASGEGGQDTQY | 0.1646464 |
| VTDVLGDLSV | ASGEGGQDTQY | 0.39116424 |
| VVVVGAGFAGL | ASGEGGQDTQY | 0.80834424 |
| VYTVGTVV | ASGEGGQDTQY | 0.81802773 |
| WTTFSGP | ASGEGGQDTQY | 0.33898902 |
| YDELVEAF | ASGEGGQDTQY | 0.60820866 |
| YGTGGAGGNGG | ASGEGGQDTQY | 0.75939524 |
| YTNVMAV | ASGEGGQDTQY | 0.6072746 |
| FLEQVHQL | ASGEGGQDTQY | 0.59284496 |
| GLSLLAVAV | ASGEGGQDTQY | 0.64264375 |
| LEQVHQL | ASGEGGQDTQY | 0.37697956 |
| LNAAGAI | ASGEGGQDTQY | 0.7743859 |
| PAVHASL | ASGEGGQDTQY | 0.85906625 |
| TELEDDA | ASGEGGQDTQY | 0.45323527 |
| TPPGPPN | ASGEGGQDTQY | 0.5835692 |
| TPPPPPPP | ASGEGGQDTQY | 0.44592202 |
| TQSPALI | ASRDNSNERLF | 0.40612465 |
| EFAAAFPQ | ASRDNSNERLF | 0.14406595 |
| LLSGNLDVL | ASRDNSNERLF | 0.63619024 |
| GPANLGNYNI | ASRDNSNERLF | 0.7024025 |
| TLGPTLP | ASRDNSNERLF | 0.47710142 |
| VYAYSSCCK | ASRDNSNERLF | 0.34944215 |
| APAPAPAPNI | ASRDNSNERLF | 0.36567035 |
| PGGPVPIP | ASRDNSNERLF | 0.3595983 |

| | | |
|---|---|---|
| FLTVALEP | ASRDNSNERLF | 0.5045498 |
| SQYFGANL | ASRDNSNERLF | 0.15022823 |
| GGVKPAA | ASRDNSNERLF | 0.40384007 |
| GGTGGAGGAGGW | ASRDNSNERLF | 0.80363774 |
| GVGGVAGLGGAGG | ASRDNSNERLF | 0.68574727 |
| VLLVLATTI | ASRDNSNERLF | 0.18224744 |
| IGIFAAML | ASRDNSNERLF | 0.8268224 |
| AVGDLID | ASRDNSNERLF | 0.6043215 |
| NISPLPL | ASRDNSNERLF | 0.69480747 |
| VTAVFVAI | ASRDNSNERLF | 0.4310492 |
| SGDDNDAAL | ASRDNSNERLF | 0.42085734 |
| AVAALAHS | ASRDNSNERLF | 0.6919173 |
| GGVTVMATP | ASRDNSNERLF | 0.5826919 |
| AGLAANVRD | ASRDNSNERLF | 0.23291299 |
| TNRPDLIDPALLRPG | ASRDNSNERLF | 0.7279663 |
| LANLGVTAL | ASRDNSNERLF | 0.48112363 |
| TGERIEL | ASRDNSNERLF | 0.4277973 |
| NGEELAA | ASRDNSNERLF | 0.35601097 |
| VLSLVWG | ASRDNSNERLF | 0.5825519 |
| IGYALWL | ASRDNSNERLF | 0.3207875 |
| LSLLLAPL | ASRDNSNERLF | 0.40439376 |
| TLDEALK | ASRDNSNERLF | 0.20454778 |
| TDGAGTAEGG | ASRDNSNERLF | 0.5593434 |
| NAPGGAGGNGG | ASRDNSNERLF | 0.7955688 |
| VSVFMGMGGNF | ASRDNSNERLF | 0.51112986 |
| LASYLAAL | ASRDNSNERLF | 0.38070652 |
| LVGGTPAL | ASRDNSNERLF | 0.729009 |
| VRNADVL | ASRDNSNERLF | 0.4901844 |
| TCSSPFLVSAM | ASRDNSNERLF | 0.49008048 |
| VHITDAE | ASRDNSNERLF | 0.69739246 |
| EYAAPSF | ASRDNSNERLF | 0.3972969 |
| VVDAFHGGL | ASRDNSNERLF | 0.601517 |
| ASVVDPQ | ASRDNSNERLF | 0.5230933 |
| SYNEFGGLS | ASRDNSNERLF | 0.5701272 |
| DPDGSVY | ASRDNSNERLF | 0.58021444 |
| MVSTIHATE | ASRDNSNERLF | 0.49301705 |
| LAAGVADPV | ASRDNSNERLF | 0.43634292 |
| TKEQIAA | ASRDNSNERLF | 0.16183177 |
| SLASPAAL | ASRDNSNERLF | 0.40041563 |
| IGLVFYGV | ASRDNSNERLF | 0.71155787 |
| FLLDASNL | ASRDNSNERLF | 0.60585076 |
| YQVGNFP | ASRDNSNERLF | 0.66498727 |
| ESLLSQST | ASRDNSNERLF | 0.51495194 |
| LFTVGFAAVG | ASRDNSNERLF | 0.3506 |
| ATPPTPT | ASRDNSNERLF | 0.59957594 |
| TDLATLG | ASRDNSNERLF | 0.6989558 |
| SAGGTGR | ASRDNSNERLF | 0.2723993 |
| GFSNSGS | ASRDNSNERLF | 0.46477526 |
| GGNGGSGGTSV | ASRDNSNERLF | 0.8393241 |
| GTSVATGGAGN | ASRDNSNERLF | 0.5587196 |
| NGGNGGSGGTS | ASRDNSNERLF | 0.5222206 |

| | | |
|---|---|---|
| AVVKFPH | ASRDNSNERLF | 0.39669263 |
| TPPPPPP | ASRDNSNERLF | 0.79333436 |
| AAPAAAAAPA | ASRDNSNERLF | 0.25972426 |
| IALNEDL | ASRDNSNERLF | 0.854992 |
| IVNVSSM | ASRDNSNERLF | 0.13030116 |
| TGIPPAPR | ASRDNSNERLF | 0.3473911 |
| FDLGGGTF | ASRDNSNERLF | 0.3387029 |
| FDLGGGTFDVS | ASRDNSNERLF | 0.31021982 |
| NEPTAAA | ASRDNSNERLF | 0.7179393 |
| NEPTAAALA | ASRDNSNERLF | 0.40664917 |
| GSIINISSI | ASRDNSNERLF | 0.747142 |
| VAPGNAGTA | ASRDNSNERLF | 0.60654455 |
| TGGGASLE | ASRDNSNERLF | 0.68646353 |
| VAVYNHYKRI | ASRDNSNERLF | 0.5930538 |
| TDPAPATT | ASRDNSNERLF | 0.567564 |
| TNRPDILDPALLRPG | ASRDNSNERLF | 0.39401644 |
| LTTTTTS | ASRDNSNERLF | 0.7158833 |
| PTGSGKTL | ASRDNSNERLF | 0.6874601 |
| TGSGKTL | ASRDNSNERLF | 0.49110138 |
| TIGHVDHGK | ASRDNSNERLF | 0.33981112 |
| TIGHVDHGKTT | ASRDNSNERLF | 0.27730283 |
| GHVDHGKTT | ASRDNSNERLF | 0.60917985 |
| AQFEHTLL | ASRDNSNERLF | 0.15504453 |
| IVNQDEL | ASRDNSNERLF | 0.14455524 |
| TNGSQFF | ASRDNSNERLF | 0.36771592 |
| TGKGIVE | ASRDNSNERLF | 0.2885107 |
| LISNASDA | ASRDNSNERLF | 0.8376988 |
| AELRTLGELTKIAKAHGAQ | ASRDNSNERLF | 0.82083946 |
| APNAVAAAEPSW | ASRDNSNERLF | 0.5347279 |
| APQVAGAATAVL | ASRDNSNERLF | 0.608539 |
| ARRDPAAAAAILPTDA | ASRDNSNERLF | 0.69795823 |
| DAAARDVLAEAGLAEYF | ASRDNSNERLF | 0.09960657 |
| DDQIVLSAKANLAKVEQ | ASRDNSNERLF | 0.38661486 |
| DELLPDTGEDSDEDGH | ASRDNSNERLF | 0.34084716 |
| ELPEAPANRATVPQD | ASRDNSNERLF | 0.65348846 |
| FGQSKIERVTGIG | ASRDNSNERLF | 0.43229887 |
| GAGGTGSTAGGAGGAGGAGGLY | ASRDNSNERLF | 0.37528545 |
| GHGGTGAAGGNGGTGGAGGSGI | ASRDNSNERLF | 0.6470955 |
| GNGGNGGAGGEGGPGGAGGAGA | ASRDNSNERLF | 0.5787312 |
| GSTAGNGGNGGNATNGTIAGQP | ASRDNSNERLF | 0.42375726 |
| GTGGDGGNGGNGGTGG | ASRDNSNERLF | 0.26768142 |
| GVRRPAGIPADAPV | ASRDNSNERLF | 0.10688642 |
| GYRQVTDAHLLTLARR | ASRDNSNERLF | 0.75508434 |
| IAKMGQDGHDRG | ASRDNSNERLF | 0.1777325 |
| KAEPSIEIQYT | ASRDNSNERLF | 0.39397338 |
| KPVRIRMAADEDTVAVT | ASRDNSNERLF | 0.15693247 |
| MPPRIAGMRLLVIKPEP | ASRDNSNERLF | 0.17827308 |
| MKFYAVALFRSRGVPPTR | ASRDNSNERLF | 0.2741701 |
| NAGGFGILGAVAHSP | ASRDNSNERLF | 0.5901252 |
| NEGREITLAMAEM | ASRDNSNERLF | 0.49389258 |
| NELVEWSHPLGHRWRWE | ASRDNSNERLF | 0.49549168 |

| | | |
|---|---|---|
| NLGAGGTGGNGGNPG | ASRDNSNERLF | 0.8388267 |
| NTEWPKWETVMAADPVD | ASRDNSNERLF | 0.6805742 |
| PASAAPAETQP | ASRDNSNERLF | 0.49949774 |
| QQSSDGKNGVRAAEITGE | ASRDNSNERLF | 0.44007218 |
| RAPDAVTTSWWKE | ASRDNSNERLF | 0.3655651 |
| SGFGNVGAHTSGWFNQ | ASRDNSNERLF | 0.39459506 |
| SGVAGAVGQAASVGGLKV | ASRDNSNERLF | 0.385633 |
| SNGGAGTGSAGGNGGT | ASRDNSNERLF | 0.8302473 |
| TGGAGGEGGAAG | ASRDNSNERLF | 0.3896491 |
| TGGDGGAAGAGGAPGHG | ASRDNSNERLF | 0.34415284 |
| TMPPLLAAIANAGRHGV | ASRDNSNERLF | 0.38994303 |
| TSSTAGNSGGA | ASRDNSNERLF | 0.74759024 |
| VPATVPVGILC | ASRDNSNERLF | 0.8520525 |
| VQRPDSGGVVLTGRIS | ASRDNSNERLF | 0.47196984 |
| WTPNPVPATKGVPAGLVQAT | ASRDNSNERLF | 0.41920403 |
| YALKRALAKFSDRFDVVII | ASRDNSNERLF | 0.41109705 |
| YLIGSVIV | ASRDNSNERLF | 0.6680226 |
| YGQLVAVID | ASRDNSNERLF | 0.42681777 |
| VTVGGMTLPA | ASRDNSNERLF | 0.34728163 |
| VLALPAV | ASRDNSNERLF | 0.5229912 |
| VIPPTLNYET | ASRDNSNERLF | 0.78987753 |
| VGMGGMHPSSG | ASRDNSNERLF | 0.40197814 |
| VGLAGGAATA | ASRDNSNERLF | 0.306303 |
| VGGAGGAGGSGP | ASRDNSNERLF | 0.61822796 |
| VFAAFMV | ASRDNSNERLF | 0.3355597 |
| TLYDKLR | ASRDNSNERLF | 0.5106641 |
| TLAVFTVL | ASRDNSNERLF | 0.51012313 |
| TGSVQPARV | ASRDNSNERLF | 0.24048637 |
| TGGGAGSGGT | ASRDNSNERLF | 0.36491767 |
| SHLETSL | ASRDNSNERLF | 0.7984978 |
| SFTTGMT | ASRDNSNERLF | 0.48049584 |
| QVEVAKRMGVRQPTVS | ASRDNSNERLF | 0.5613381 |
| QGSGGPALAM | ASRDNSNERLF | 0.80718726 |
| MLAPSNP | ASRDNSNERLF | 0.70917755 |
| LVNNAAI | ASRDNSNERLF | 0.79927576 |
| LTGEINTLA | ASRDNSNERLF | 0.61927843 |
| LSIFGPV | ASRDNSNERLF | 0.4912474 |
| LLALLANL | ASRDNSNERLF | 0.4772615 |
| LISNASDALDKLR | ASRDNSNERLF | 0.5782086 |
| LINEAVP | ASRDNSNERLF | 0.7096902 |
| LGYAFSGL | ASRDNSNERLF | 0.5052882 |
| LDFVAVAL | ASRDNSNERLF | 0.34836394 |
| LAQPGAE | ASRDNSNERLF | 0.8150089 |
| ISNLGAVTGF | ASRDNSNERLF | 0.41170463 |
| HQEDPGRI | ASRDNSNERLF | 0.55496895 |
| GTPGEPTP | ASRDNSNERLF | 0.3875345 |
| GFFNSGS | ASRDNSNERLF | 0.35804805 |
| GAGGGGLTAGAN | ASRDNSNERLF | 0.68430525 |
| FVDALVNM | ASRDNSNERLF | 0.10833415 |
| FLTVAPEL | ASRDNSNERLF | 0.12342888 |
| FETLTAL | ASRDNSNERLF | 0.5009812 |

| | | |
|---|---|---|
| DVINPIEL | ASRDNSNERLF | 0.74669385 |
| APVSGAPV | ASRDNSNERLF | 0.11533049 |
| ALVGGLAW | ASRDNSNERLF | 0.3405525 |
| ALAAAAATAI | ASRDNSNERLF | 0.24442157 |
| AFGLGLLL | ASRDNSNERLF | 0.43885905 |
| AFFAAASL | ASRDNSNERLF | 0.581423 |
| AAAALPAFSPP | ASRDNSNERLF | 0.23009872 |
| VVLAPAL | ASRDNSNERLF | 0.547877 |
| TIGHVDHGKTTL | ASRDNSNERLF | 0.24146688 |
| LISNASDALDK | ASRDNSNERLF | 0.1882897 |
| LISNASDAL | ASRDNSNERLF | 0.63964534 |
| ISNASDALDK | ASRDNSNERLF | 0.7203282 |
| NQAETLVYQTE | ASRDNSNERLF | 0.37253743 |
| TGGTGGTAGAG | ASRDNSNERLF | 0.715209 |
| TGSTAGGAGGA | ASRDNSNERLF | 0.28924596 |
| DPLAVPVAAGPV | ASRDNSNERLF | 0.84167063 |
| GGTGGTPGNGGN | ASRDNSNERLF | 0.45266747 |
| PAGLAGVDPATL | ASRDNSNERLF | 0.2849025 |
| GNNLVVEAPHAGASV | ASRDNSNERLF | 0.6457261 |
| KVTRIMSASTLEQPA | ASRDNSNERLF | 0.55326533 |
| LGFALPIALPRRTVV | ASRDNSNERLF | 0.54749066 |
| TRLGNLTARKLVVLV | ASRDNSNERLF | 0.65845937 |
| WSQQRRLIYFPSAGP | ASRDNSNERLF | 0.33272833 |
| GPNGNPDPMAAAVDIR | ASRDNSNERLF | 0.55081666 |
| TQRCVSVADLRAAASR | ASRDNSNERLF | 0.7995826 |
| QFVLDVAEHPVEVLTETV | ASRDNSNERLF | 0.7638341 |
| GGAGGTGGAGGNGGTQAGDG | ASRDNSNERLF | 0.3654322 |
| PLLATLRAIAVDVDAKPAQV | ASRDNSNERLF | 0.6272016 |
| APKFRTGGYMIDFWGVGYQVAK | ASRDNSNERLF | 0.6249869 |
| GAGGAGGAGGAAGGPGGTGGTG | ASRDNSNERLF | 0.46362305 |
| IGALGLLATDVAPAGSVAFGVA | ASRDNSNERLF | 0.7187789 |
| AGGAGGIGGNGGAGGAGGVLMG | ASRDNSNERLF | 0.48191673 |
| ADELIGLL | ASRDNSNERLF | 0.6620481 |
| ADRALKA | ASRDNSNERLF | 0.58606994 |
| AGAVGQVGGAGGA | ASRDNSNERLF | 0.7303506 |
| AGGGGGLGGAP | ASRDNSNERLF | 0.32905674 |
| AGGLLYGNGGAG | ASRDNSNERLF | 0.6684653 |
| AGLESIGL | ASRDNSNERLF | 0.52257496 |
| AGLKTGII | ASRDNSNERLF | 0.61984974 |
| ASLAALLL | ASRDNSNERLF | 0.4488166 |
| ASVINISSI | ASRDNSNERLF | 0.21992084 |
| ATPPGPA | ASRDNSNERLF | 0.52110296 |
| ATVASALV | ASRDNSNERLF | 0.7277614 |
| ATVASLIL | ASRDNSNERLF | 0.53478616 |
| AVNGPVLIRH | ASRDNSNERLF | 0.49546593 |
| DGAAGAVGI | ASRDNSNERLF | 0.82007337 |
| DPLLHGI | ASRDNSNERLF | 0.44409508 |
| ELGAALGAL | ASRDNSNERLF | 0.52246964 |
| ELGRTQAV | ASRDNSNERLF | 0.59189045 |
| FVAGGYLS | ASRDNSNERLF | 0.69702446 |
| GASGGAGGAGGW | ASRDNSNERLF | 0.4197113 |

| | | |
|---|---|---|
| GGAGAGGDGGAGGI | ASRDNSNERLF | 0.77650464 |
| GTGHNGSQPGMGGNG | ASRDNSNERLF | 0.28573534 |
| GYAAASASAS | ASRDNSNERLF | 0.80351293 |
| HQVDVAAVQQG | ASRDNSNERLF | 0.34359846 |
| IGAAIAHT | ASRDNSNERLF | 0.69226557 |
| IGMIEVAQL | ASRDNSNERLF | 0.4164043 |
| IRLAPEHTSAMD | ASRDNSNERLF | 0.8369707 |
| IYNVSVL | ASRDNSNERLF | 0.76905024 |
| KQGFLGM | ASRDNSNERLF | 0.8253895 |
| LESVITQL | ASRDNSNERLF | 0.1285592 |
| LGLGALVVV | ASRDNSNERLF | 0.7333277 |
| LIGNGGNGGSGGI | ASRDNSNERLF | 0.77284616 |
| LNAINAPTL | ASRDNSNERLF | 0.84935665 |
| LQLLNTGV | ASRDNSNERLF | 0.44317904 |
| LQTSIEF | ASRDNSNERLF | 0.53604686 |
| MSFAVSEL | ASRDNSNERLF | 0.582981 |
| NTGALGL | ASRDNSNERLF | 0.40667382 |
| PPGAGGAGGAGTT | ASRDNSNERLF | 0.34221342 |
| QLVGLGI | ASRDNSNERLF | 0.5873852 |
| RFLEQVHQL | ASRDNSNERLF | 0.514301 |
| RGVGYVLR | ASRDNSNERLF | 0.4376855 |
| SATGVFAGVI | ASRDNSNERLF | 0.74838734 |
| SLPAITLP | ASRDNSNERLF | 0.75804687 |
| STALAFTV | ASRDNSNERLF | 0.72101456 |
| SVLTVGLGK | ASRDNSNERLF | 0.7511923 |
| TALAGAGSGI | ASRDNSNERLF | 0.7386139 |
| TGLPFTGI | ASRDNSNERLF | 0.54829264 |
| VAEHVTV | ASRDNSNERLF | 0.42118457 |
| VDIVLNSL | ASRDNSNERLF | 0.49279037 |
| VLTGGGAVL | ASRDNSNERLF | 0.36481038 |
| VNVGLAYGI | ASRDNSNERLF | 0.4053915 |
| VSSAAVGAV | ASRDNSNERLF | 0.43215075 |
| VTSVYEAGV | ASRDNSNERLF | 0.5602294 |
| WVALDRGA | ASRDNSNERLF | 0.0973677 |
| YLVGSVVI | ASRDNSNERLF | 0.5210401 |
| GPPGTGKTLL | ASRDNSNERLF | 0.83631104 |
| GVGMTKF | ASRDNSNERLF | 0.3767505 |
| IGIGGSDL | ASRDNSNERLF | 0.50376326 |
| IGTLLAEL | ASRDNSNERLF | 0.46835625 |
| IYTSGTT | ASRDNSNERLF | 0.3816396 |
| LSFPAGV | ASRDNSNERLF | 0.72411895 |
| LYTSGTT | ASRDNSNERLF | 0.3304604 |
| VGAFPFGF | ASRDNSNERLF | 0.5134232 |
| VGAGTVEF | ASRDNSNERLF | 0.7734555 |
| VGKSTLLL | ASRDNSNERLF | 0.52666944 |
| AALAHAGGGIV | ASRDNSNERLF | 0.50895 |
| AALVAVSLL | ASRDNSNERLF | 0.4392268 |
| AAVSRAVAAL | ASRDNSNERLF | 0.29775628 |
| AGGSSGIGGF | ASRDNSNERLF | 0.5909447 |
| AGLTVANANG | ASRDNSNERLF | 0.14506443 |
| AGLVGAVL | ASRDNSNERLF | 0.4559353 |

| | | |
|---|---|---|
| ALSQAFTV | ASRDNSNERLF | 0.354905 |
| ALTFSVSL | ASRDNSNERLF | 0.7297015 |
| ALTGAALVAL | ASRDNSNERLF | 0.7037189 |
| ASFSIGAGVL | ASRDNSNERLF | 0.5876915 |
| DGSVYKI | ASRDNSNERLF | 0.40911657 |
| EQLAEIV | ASRDNSNERLF | 0.3339303 |
| FAVSELTVM | ASRDNSNERLF | 0.24975449 |
| GAAPLAAGL | ASRDNSNERLF | 0.32955348 |
| GADGGDGGSGGAGG | ASRDNSNERLF | 0.7257651 |
| GFGLDTGGAGG | ASRDNSNERLF | 0.46145618 |
| GLASGGLL | ASRDNSNERLF | 0.38797453 |
| GRIAGAAVLL | ASRDNSNERLF | 0.78364784 |
| GSAGSLNTSF | ASRDNSNERLF | 0.54564995 |
| GTGQIGI | ASRDNSNERLF | 0.32414475 |
| GTLIALIL | ASRDNSNERLF | 0.5984504 |
| IAATMGFVVG | ASRDNSNERLF | 0.51839536 |
| IFAQALL | ASRDNSNERLF | 0.46630883 |
| IGAAVALGF | ASRDNSNERLF | 0.104953006 |
| IGLTPLAL | ASRDNSNERLF | 0.51001376 |
| ILGSNTSTL | ASRDNSNERLF | 0.48391438 |
| IVFSLTVL | ASRDNSNERLF | 0.3478656 |
| LASKATGF | ASRDNSNERLF | 0.4193756 |
| LASVALIL | ASRDNSNERLF | 0.76794696 |
| LFAVGLTL | ASRDNSNERLF | 0.2592228 |
| LFNGGIGTYI | ASRDNSNERLF | 0.4784168 |
| LTLGAGII | ASRDNSNERLF | 0.6882146 |
| LVAAALGDI | ASRDNSNERLF | 0.74339885 |
| NTAFVQLGI | ASRDNSNERLF | 0.82987857 |
| QLFSGAEL | ASRDNSNERLF | 0.5987372 |
| SAATFITL | ASRDNSNERLF | 0.40244237 |
| SNYGLANLGV | ASRDNSNERLF | 0.55171967 |
| SYGGAYAVM | ASRDNSNERLF | 0.4911806 |
| TTGVGGLTLG | ASRDNSNERLF | 0.16329776 |
| TTQTIYI | ASRDNSNERLF | 0.12641738 |
| VAAGAGLGAVI | ASRDNSNERLF | 0.69436806 |
| VAAVVTLL | ASRDNSNERLF | 0.14527129 |
| VGFLGGLI | ASRDNSNERLF | 0.5321295 |
| VGVIPILL | ASRDNSNERLF | 0.5586927 |
| VIFASSLI | ASRDNSNERLF | 0.36868703 |
| VKGGDGGAAAG | ASRDNSNERLF | 0.37272492 |
| VTKTFTNI | ASRDNSNERLF | 0.43719673 |
| VVAAAILL | ASRDNSNERLF | 0.4295432 |
| YAALDVEL | ASRDNSNERLF | 0.70242023 |
| LASVALLL | ASRDNSNERLF | 0.51745766 |
| LDAVHRV | ASRDNSNERLF | 0.81795895 |
| QGLSSSIF | ASRDNSNERLF | 0.8105617 |
| RGTFANIRL | ASRDNSNERLF | 0.556002 |
| RQLWWGHRI | ASRDNSNERLF | 0.68544996 |
| YGPPGTGKTLL | ASRDNSNERLF | 0.3840444 |
| AALVGAPANVA | ASRDNSNERLF | 0.49639472 |
| AATFRRTE | ASRDNSNERLF | 0.48030618 |

| | | |
|---|---|---|
| AGALGGGI | ASRDNSNERLF | 0.48819283 |
| AGGAGGAGGIGAG | ASRDNSNERLF | 0.45433983 |
| AGSGALGL | ASRDNSNERLF | 0.4669999 |
| AQIADQI | ASRDNSNERLF | 0.2512535 |
| ASAVAVTYL | ASRDNSNERLF | 0.5036253 |
| ASTVLGGTGG | ASRDNSNERLF | 0.6000836 |
| AVGGEPAK | ASRDNSNERLF | 0.15461549 |
| CEQGTFGGTC | ASRDNSNERLF | 0.37306648 |
| DEGKAPAG | ASRDNSNERLF | 0.86449826 |
| DTVVFPAL | ASRDNSNERLF | 0.7498543 |
| FEGITMAG | ASRDNSNERLF | 0.32467404 |
| FPLVVQD | ASRDNSNERLF | 0.38950512 |
| GAFTTPQ | ASRDNSNERLF | 0.5315564 |
| GFGGFGTAGGVGGA | ASRDNSNERLF | 0.15747042 |
| GFLAQFL | ASRDNSNERLF | 0.48787072 |
| GGAGGAGGNAGVA | ASRDNSNERLF | 0.563995 |
| GGVGTAGAGGAGGAGG | ASRDNSNERLF | 0.62344384 |
| GNGGGGDAGNAGSGGN | ASRDNSNERLF | 0.36821377 |
| GTGGAGGAGNDAGS | ASRDNSNERLF | 0.55930966 |
| GTTNTGWFNS | ASRDNSNERLF | 0.50775313 |
| IFMGPLFTL | ASRDNSNERLF | 0.52677965 |
| ILAHAIEAI | ASRDNSNERLF | 0.11567382 |
| IRSAIREAL | ASRDNSNERLF | 0.5349517 |
| ITFAGGG | ASRDNSNERLF | 0.40522626 |
| ITGMSGHQEV | ASRDNSNERLF | 0.3553644 |
| IVAGLACTL | ASRDNSNERLF | 0.30437165 |
| IVGGGLAGA | ASRDNSNERLF | 0.5263169 |
| IVTGVLNI | ASRDNSNERLF | 0.7896317 |
| LAQQGCI | ASRDNSNERLF | 0.7298777 |
| LCLHVEN | ASRDNSNERLF | 0.19488172 |
| LDAISDGS | ASRDNSNERLF | 0.7921101 |
| LGGFAAML | ASRDNSNERLF | 0.84857917 |
| LGGKNPMI | ASRDNSNERLF | 0.38962984 |
| LGPKGVAAIL | ASRDNSNERLF | 0.32687134 |
| LMVGVSI | ASRDNSNERLF | 0.59525245 |
| LVGAAFVL | ASRDNSNERLF | 0.45364362 |
| LVLAVAPL | ASRDNSNERLF | 0.5216763 |
| MAAIAVIDAA | ASRDNSNERLF | 0.72247124 |
| NNGINAELQ | ASRDNSNERLF | 0.33054864 |
| RGEPLVLVHG | ASRDNSNERLF | 0.605084 |
| RGGAGGAAGLI | ASRDNSNERLF | 0.6357205 |
| RSYRMHPAV | ASRDNSNERLF | 0.36372983 |
| SAAHLLAGQLA | ASRDNSNERLF | 0.45397416 |
| SAAVGSY | ASRDNSNERLF | 0.43248355 |
| SPHAVLHRI | ASRDNSNERLF | 0.15277417 |
| SQIAKVL | ASRDNSNERLF | 0.6741631 |
| TALGHLVP | ASRDNSNERLF | 0.40755475 |
| TAPTTSGAA | ASRDNSNERLF | 0.39811355 |
| TDGDDGNGGSPGT | ASRDNSNERLF | 0.63482296 |
| TDGLGEN | ASRDNSNERLF | 0.4231473 |
| TGGPPLPGQA | ASRDNSNERLF | 0.45427236 |

| | | |
|---|---|---|
| TIPAITI | ASRDNSNERLF | 0.44935572 |
| TIPGFTI | ASRDNSNERLF | 0.5678292 |
| TSIASGALA | ASRDNSNERLF | 0.5636486 |
| TVTLPGAAI | ASRDNSNERLF | 0.49311164 |
| VASGLSAALP | ASRDNSNERLF | 0.4439256 |
| VGAGVIGI | ASRDNSNERLF | 0.8516593 |
| VGPKRFGV | ASRDNSNERLF | 0.30537352 |
| VNVAFPD | ASRDNSNERLF | 0.41948068 |
| VSMTNSGVS | ASRDNSNERLF | 0.5107893 |
| VTDVLGDLSV | ASRDNSNERLF | 0.1975614 |
| VVVVGAGFAGL | ASRDNSNERLF | 0.58897316 |
| VYTVGTVV | ASRDNSNERLF | 0.6785829 |
| WTTFSGP | ASRDNSNERLF | 0.57032555 |
| YDELVEAF | ASRDNSNERLF | 0.36522815 |
| YGTGGAGGNGG | ASRDNSNERLF | 0.8530317 |
| YTNVMAV | ASRDNSNERLF | 0.6716577 |
| FLEQVHQL | ASRDNSNERLF | 0.49698973 |
| GLSLLAVAV | ASRDNSNERLF | 0.6155463 |
| LEQVHQL | ASRDNSNERLF | 0.39767647 |
| LNAAGAI | ASRDNSNERLF | 0.7096111 |
| PAVHASL | ASRDNSNERLF | 0.5086131 |
| TELEDDA | ASRDNSNERLF | 0.3314432 |
| TPPGPPN | ASRDNSNERLF | 0.56553674 |
| TPPPPPPP | ASRDNSNERLF | 0.6748866 |
| TQSPALI | ASSLSRGQGNQAPL | 0.804007 |
| EFAAAFPQ | ASSLSRGQGNQAPL | 0.15788063 |
| LLSGNLDVL | ASSLSRGQGNQAPL | 0.52025586 |
| GPANLGNYNI | ASSLSRGQGNQAPL | 0.15869272 |
| TLGPTLP | ASSLSRGQGNQAPL | 0.38154903 |
| VYAYSSCCK | ASSLSRGQGNQAPL | 0.46971393 |
| APAPAPAPNI | ASSLSRGQGNQAPL | 0.5628748 |
| PGGPVPIP | ASSLSRGQGNQAPL | 0.5052277 |
| FLTVALEP | ASSLSRGQGNQAPL | 0.6101804 |
| SQYFGANL | ASSLSRGQGNQAPL | 0.5569328 |
| GGVKPAA | ASSLSRGQGNQAPL | 0.767008 |
| GGTGGAGGAGGW | ASSLSRGQGNQAPL | 0.35220343 |
| GVGGVAGLGGAGG | ASSLSRGQGNQAPL | 0.5979903 |
| VLLVLATTI | ASSLSRGQGNQAPL | 0.2873675 |
| IGIFAAML | ASSLSRGQGNQAPL | 0.21893844 |
| AVGDLID | ASSLSRGQGNQAPL | 0.40743086 |
| NISPLPL | ASSLSRGQGNQAPL | 0.44878042 |
| VTAVFVAI | ASSLSRGQGNQAPL | 0.14796472 |
| SGDDNDAAL | ASSLSRGQGNQAPL | 0.60560215 |
| AVAALAHS | ASSLSRGQGNQAPL | 0.5794153 |
| GGVTVMATP | ASSLSRGQGNQAPL | 0.6998082 |
| AGLAANVRD | ASSLSRGQGNQAPL | 0.41944304 |
| TNRPDLIDPALLRPG | ASSLSRGQGNQAPL | 0.531286 |
| LANLGVTAL | ASSLSRGQGNQAPL | 0.41066173 |
| TGERIEL | ASSLSRGQGNQAPL | 0.71431696 |
| NGEELAA | ASSLSRGQGNQAPL | 0.40788108 |
| VLSLVWG | ASSLSRGQGNQAPL | 0.2975091 |

| | | |
|---|---|---|
| IGYALWL | ASSLSRGQGNQAPL | 0.44407922 |
| LSLLLAPL | ASSLSRGQGNQAPL | 0.32836708 |
| TLDEALK | ASSLSRGQGNQAPL | 0.72886354 |
| TDGAGTAEGG | ASSLSRGQGNQAPL | 0.67028046 |
| NAPGGAGGNGG | ASSLSRGQGNQAPL | 0.6171852 |
| VSVFMGMGGNF | ASSLSRGQGNQAPL | 0.55674046 |
| LASYLAAL | ASSLSRGQGNQAPL | 0.21537593 |
| LVGGTPAL | ASSLSRGQGNQAPL | 0.4576527 |
| VRNADVL | ASSLSRGQGNQAPL | 0.84821606 |
| TCSSPFLVSAM | ASSLSRGQGNQAPL | 0.5394571 |
| VHITDAE | ASSLSRGQGNQAPL | 0.52145225 |
| EYAAPSF | ASSLSRGQGNQAPL | 0.7561764 |
| VVDAFHGGL | ASSLSRGQGNQAPL | 0.5080446 |
| ASVVDPQ | ASSLSRGQGNQAPL | 0.47884923 |
| SYNEFGGLS | ASSLSRGQGNQAPL | 0.51929766 |
| DPDGSVY | ASSLSRGQGNQAPL | 0.5354736 |
| MVSTIHATE | ASSLSRGQGNQAPL | 0.5154824 |
| LAAGVADPV | ASSLSRGQGNQAPL | 0.42829436 |
| TKEQIAA | ASSLSRGQGNQAPL | 0.47454256 |
| SLASPAAL | ASSLSRGQGNQAPL | 0.8718759 |
| IGLVFYGV | ASSLSRGQGNQAPL | 0.63104975 |
| FLLDASNL | ASSLSRGQGNQAPL | 0.7618983 |
| YQVGNFP | ASSLSRGQGNQAPL | 0.61752725 |
| ESLLSQST | ASSLSRGQGNQAPL | 0.6575929 |
| LFTVGFAAVG | ASSLSRGQGNQAPL | 0.37241095 |
| ATPPTPT | ASSLSRGQGNQAPL | 0.4730125 |
| TDLATLG | ASSLSRGQGNQAPL | 0.3398218 |
| SAGGTGR | ASSLSRGQGNQAPL | 0.36985922 |
| GFSNSGS | ASSLSRGQGNQAPL | 0.44891417 |
| GGNGGSGGTSV | ASSLSRGQGNQAPL | 0.39872673 |
| GTSVATGGAGN | ASSLSRGQGNQAPL | 0.87968564 |
| NGGNGGSGGTS | ASSLSRGQGNQAPL | 0.642438 |
| AVVKFPH | ASSLSRGQGNQAPL | 0.58075386 |
| TPPPPPP | ASSLSRGQGNQAPL | 0.37061954 |
| AAPAAAAAPA | ASSLSRGQGNQAPL | 0.51489025 |
| IALNEDL | ASSLSRGQGNQAPL | 0.5647366 |
| IVNVSSM | ASSLSRGQGNQAPL | 0.31914118 |
| TGIPPAPR | ASSLSRGQGNQAPL | 0.7773144 |
| FDLGGGTF | ASSLSRGQGNQAPL | 0.41536787 |
| FDLGGGTFDVS | ASSLSRGQGNQAPL | 0.300246 |
| NEPTAAA | ASSLSRGQGNQAPL | 0.599912 |
| NEPTAAALA | ASSLSRGQGNQAPL | 0.7904254 |
| GSIINISSI | ASSLSRGQGNQAPL | 0.4042404 |
| VAPGNAGTA | ASSLSRGQGNQAPL | 0.2986479 |
| TGGGASLE | ASSLSRGQGNQAPL | 0.47746256 |
| VAVYNHYKRI | ASSLSRGQGNQAPL | 0.5341374 |
| TDPAPATT | ASSLSRGQGNQAPL | 0.80820435 |
| TNRPDILDPALLRPG | ASSLSRGQGNQAPL | 0.3346917 |
| LTTTTTS | ASSLSRGQGNQAPL | 0.8045342 |
| PTGSGKTL | ASSLSRGQGNQAPL | 0.4426079 |
| TGSGKTL | ASSLSRGQGNQAPL | 0.35042173 |

| | | |
|---|---|---|
| TIGHVDHGK | ASSLSRGQGNQAPL | 0.47276717 |
| TIGHVDHGKTT | ASSLSRGQGNQAPL | 0.6792789 |
| GHVDHGKTT | ASSLSRGQGNQAPL | 0.7775297 |
| AQFEHTLL | ASSLSRGQGNQAPL | 0.84127307 |
| IVNQDEL | ASSLSRGQGNQAPL | 0.5647756 |
| TNGSQFF | ASSLSRGQGNQAPL | 0.40158954 |
| TGKGIVE | ASSLSRGQGNQAPL | 0.73295 |
| LISNASDA | ASSLSRGQGNQAPL | 0.62837666 |
| AELRTLGELTKIAKAHGAQ | ASSLSRGQGNQAPL | 0.621777 |
| APNAVAAAEPSW | ASSLSRGQGNQAPL | 0.38218004 |
| APQVAGAATAVL | ASSLSRGQGNQAPL | 0.31624812 |
| ARRDPAAAAAILPTDA | ASSLSRGQGNQAPL | 0.4670006 |
| DAAARDVLAEAGLAEYF | ASSLSRGQGNQAPL | 0.7364593 |
| DDQIVLSAKANLAKVEQ | ASSLSRGQGNQAPL | 0.34086788 |
| DELLPDTGEDSDEDGH | ASSLSRGQGNQAPL | 0.53284955 |
| ELPEAPANRATVPQD | ASSLSRGQGNQAPL | 0.6391828 |
| FGQSKIERVTGIG | ASSLSRGQGNQAPL | 0.3641557 |
| GAGGTGSTAGGAGGAGGAGGLY | ASSLSRGQGNQAPL | 0.51500726 |
| GHGGTGAAGGNGGTGGAGGSGI | ASSLSRGQGNQAPL | 0.8176398 |
| GNGGNGGAGGEGGPGGAGGAGA | ASSLSRGQGNQAPL | 0.42878824 |
| GSTAGNGGNGGNATNGTIAGQP | ASSLSRGQGNQAPL | 0.40546265 |
| GTGGDGGNGGNGGTGG | ASSLSRGQGNQAPL | 0.7478012 |
| GVRRPAGIPADAPV | ASSLSRGQGNQAPL | 0.88420075 |
| GYRQVTDAHLLTLARR | ASSLSRGQGNQAPL | 0.5845078 |
| IAKMGQDGHDRG | ASSLSRGQGNQAPL | 0.3321274 |
| KAEPSIEIQYT | ASSLSRGQGNQAPL | 0.1921386 |
| KPVRIRMAADEDTVAVT | ASSLSRGQGNQAPL | 0.3110042 |
| MPPRIAGMRLLVIKPEP | ASSLSRGQGNQAPL | 0.6243915 |
| MKFYAVALFRSRGVPPTR | ASSLSRGQGNQAPL | 0.44197395 |
| NAGGFGILGAVAHSP | ASSLSRGQGNQAPL | 0.41452104 |
| NEGREITLAMAEM | ASSLSRGQGNQAPL | 0.7002487 |
| NELVEWSHPLGHRWRWE | ASSLSRGQGNQAPL | 0.7073047 |
| NLGAGGTGGNGGNPG | ASSLSRGQGNQAPL | 0.4347734 |
| NTEWPKWETVMAADPVD | ASSLSRGQGNQAPL | 0.3188187 |
| PASAAPAETQP | ASSLSRGQGNQAPL | 0.6048428 |
| QQSSDGKNGVRAAEITGE | ASSLSRGQGNQAPL | 0.52632356 |
| RAPDAVTTSWWKE | ASSLSRGQGNQAPL | 0.47034162 |
| SGFGNVGAHTSGWFNQ | ASSLSRGQGNQAPL | 0.5623609 |
| SGVAGAVGQAASVGGLKV | ASSLSRGQGNQAPL | 0.3133583 |
| SNGGAGTGSAGGNGGT | ASSLSRGQGNQAPL | 0.17962861 |
| TGGAGGEGGAAG | ASSLSRGQGNQAPL | 0.62953436 |
| TGGDGGAAGAGGAPGHG | ASSLSRGQGNQAPL | 0.31410405 |
| TMPPLLAAIANAGRHGV | ASSLSRGQGNQAPL | 0.8401154 |
| TSSTAGNSGGA | ASSLSRGQGNQAPL | 0.38687992 |
| VPATVPVGILC | ASSLSRGQGNQAPL | 0.6045558 |
| VQRPDSGGVVLTGRIS | ASSLSRGQGNQAPL | 0.74729276 |
| WTPNPVPATKGVPAGLVQAT | ASSLSRGQGNQAPL | 0.5781821 |
| YALKRALAKFSDRFDVVII | ASSLSRGQGNQAPL | 0.4647878 |
| YLIGSVIV | ASSLSRGQGNQAPL | 0.38098443 |
| YGQLVAVID | ASSLSRGQGNQAPL | 0.31720394 |
| VTVGGMTLPA | ASSLSRGQGNQAPL | 0.13063236 |

| | | |
|---|---|---|
| VLALPAV | ASSLSRGQGNQAPL | 0.7718568 |
| VIPPTLNYET | ASSLSRGQGNQAPL | 0.17840837 |
| VGMGGMHPSSG | ASSLSRGQGNQAPL | 0.73624086 |
| VGLAGGAATA | ASSLSRGQGNQAPL | 0.6754387 |
| VGGAGGAGGSGP | ASSLSRGQGNQAPL | 0.41527447 |
| VFAAFMV | ASSLSRGQGNQAPL | 0.56179464 |
| TLYDKLR | ASSLSRGQGNQAPL | 0.36446998 |
| TLAVFTVL | ASSLSRGQGNQAPL | 0.2381505 |
| TGSVQPARV | ASSLSRGQGNQAPL | 0.71201843 |
| TGGGAGSGGT | ASSLSRGQGNQAPL | 0.56042254 |
| SHLETSL | ASSLSRGQGNQAPL | 0.39775786 |
| SFTTGMT | ASSLSRGQGNQAPL | 0.5078833 |
| QVEVAKRMGVRQPTVS | ASSLSRGQGNQAPL | 0.5448495 |
| QGSGGPALAM | ASSLSRGQGNQAPL | 0.34792972 |
| MLAPSNP | ASSLSRGQGNQAPL | 0.4543092 |
| LVNNAAI | ASSLSRGQGNQAPL | 0.7931757 |
| LTGEINTLA | ASSLSRGQGNQAPL | 0.48013076 |
| LSIFGPV | ASSLSRGQGNQAPL | 0.43688425 |
| LLALLANL | ASSLSRGQGNQAPL | 0.6015538 |
| LISNASDALDKLR | ASSLSRGQGNQAPL | 0.30774093 |
| LINEAVP | ASSLSRGQGNQAPL | 0.7152067 |
| LGYAFSGL | ASSLSRGQGNQAPL | 0.52359957 |
| LDFVAVAL | ASSLSRGQGNQAPL | 0.48020777 |
| LAQPGAE | ASSLSRGQGNQAPL | 0.60194635 |
| ISNLGAVTGF | ASSLSRGQGNQAPL | 0.35761526 |
| HQEDPGRI | ASSLSRGQGNQAPL | 0.31860787 |
| GTPGEPTP | ASSLSRGQGNQAPL | 0.41850144 |
| GFFNSGS | ASSLSRGQGNQAPL | 0.6341497 |
| GAGGGGLTAGAN | ASSLSRGQGNQAPL | 0.6610088 |
| FVDALVNM | ASSLSRGQGNQAPL | 0.39539158 |
| FLTVAPEL | ASSLSRGQGNQAPL | 0.48896685 |
| FETLTAL | ASSLSRGQGNQAPL | 0.4489546 |
| DVINPIEL | ASSLSRGQGNQAPL | 0.36510783 |
| APVSGAPV | ASSLSRGQGNQAPL | 0.43886542 |
| ALVGGLAW | ASSLSRGQGNQAPL | 0.46814004 |
| ALAAAAATAI | ASSLSRGQGNQAPL | 0.38073304 |
| AFGLGLLL | ASSLSRGQGNQAPL | 0.58054894 |
| AFFAAASL | ASSLSRGQGNQAPL | 0.4020799 |
| AAAALPAFSPP | ASSLSRGQGNQAPL | 0.52082837 |
| VVLAPAL | ASSLSRGQGNQAPL | 0.6117091 |
| TIGHVDHGKTTL | ASSLSRGQGNQAPL | 0.4960937 |
| LISNASDALDK | ASSLSRGQGNQAPL | 0.56162924 |
| LISNASDAL | ASSLSRGQGNQAPL | 0.6011025 |
| ISNASDALDK | ASSLSRGQGNQAPL | 0.70574594 |
| NQAETLVYQTE | ASSLSRGQGNQAPL | 0.45800143 |
| TGGTGGTAGAG | ASSLSRGQGNQAPL | 0.4539904 |
| TGSTAGGAGGA | ASSLSRGQGNQAPL | 0.72814065 |
| DPLAVPVAAGPV | ASSLSRGQGNQAPL | 0.6767875 |
| GGTGGTPGNGGN | ASSLSRGQGNQAPL | 0.65048105 |
| PAGLAGVDPATL | ASSLSRGQGNQAPL | 0.57963777 |
| GNNLVVEAPHAGASV | ASSLSRGQGNQAPL | 0.6492833 |

| | | |
|---|---|---|
| KVTRIMSASTLEQPA | ASSLSRGQGNQAPL | 0.571804 |
| LGFALPIALPRRTVV | ASSLSRGQGNQAPL | 0.5390632 |
| TRLGNLTARKLVVLV | ASSLSRGQGNQAPL | 0.26234645 |
| WSQQRRLIYFPSAGP | ASSLSRGQGNQAPL | 0.48223677 |
| GPNGNPDPMAAAVDIR | ASSLSRGQGNQAPL | 0.5407152 |
| TQRCVSVADLRAAASR | ASSLSRGQGNQAPL | 0.47331375 |
| QFVLDVAEHPVEVLTETV | ASSLSRGQGNQAPL | 0.21630909 |
| GGAGGTGGAGGNGGTQAGDG | ASSLSRGQGNQAPL | 0.8122176 |
| PLLATLRAIAVDVDAKPAQV | ASSLSRGQGNQAPL | 0.8723064 |
| APKFRTGGYMIDFWGVGYQVAK | ASSLSRGQGNQAPL | 0.3785407 |
| GAGGAGGAGGAAGGPGGTGGTG | ASSLSRGQGNQAPL | 0.74151444 |
| IGALGLLATDVAPAGSVAFGVA | ASSLSRGQGNQAPL | 0.28132126 |
| AGGAGGIGGNGGAGGAGGVLMG | ASSLSRGQGNQAPL | 0.6283196 |
| ADELIGLL | ASSLSRGQGNQAPL | 0.7823021 |
| ADRALKA | ASSLSRGQGNQAPL | 0.33921966 |
| AGAVGQVGGAGGA | ASSLSRGQGNQAPL | 0.67148876 |
| AGGGGGLGGAP | ASSLSRGQGNQAPL | 0.40688962 |
| AGGLLYGNGGAG | ASSLSRGQGNQAPL | 0.4785665 |
| AGLESIGL | ASSLSRGQGNQAPL | 0.80046785 |
| AGLKTGII | ASSLSRGQGNQAPL | 0.4749474 |
| ASLAALLL | ASSLSRGQGNQAPL | 0.55405146 |
| ASVINISSI | ASSLSRGQGNQAPL | 0.48699173 |
| ATPPGPA | ASSLSRGQGNQAPL | 0.41633078 |
| ATVASALV | ASSLSRGQGNQAPL | 0.5246253 |
| ATVASLIL | ASSLSRGQGNQAPL | 0.7651171 |
| AVNGPVLIRH | ASSLSRGQGNQAPL | 0.21498184 |
| DGAAGAVGI | ASSLSRGQGNQAPL | 0.60726357 |
| DPLLHGI | ASSLSRGQGNQAPL | 0.4438158 |
| ELGAALGAL | ASSLSRGQGNQAPL | 0.87554395 |
| ELGRTQAV | ASSLSRGQGNQAPL | 0.6934517 |
| FVAGGYLS | ASSLSRGQGNQAPL | 0.6260209 |
| GASGGAGGAGGW | ASSLSRGQGNQAPL | 0.42187932 |
| GGAGAGGDGGAGGI | ASSLSRGQGNQAPL | 0.8341455 |
| GTGHNGSQPGMGGNG | ASSLSRGQGNQAPL | 0.5629672 |
| GYAAASASAS | ASSLSRGQGNQAPL | 0.6449788 |
| HQVDVAAVQQG | ASSLSRGQGNQAPL | 0.59156775 |
| IGAAIAHT | ASSLSRGQGNQAPL | 0.34909594 |
| IGMIEVAQL | ASSLSRGQGNQAPL | 0.71126854 |
| IRLAPEHTSAMD | ASSLSRGQGNQAPL | 0.5098517 |
| IYNVSVL | ASSLSRGQGNQAPL | 0.6207094 |
| KQGFLGM | ASSLSRGQGNQAPL | 0.43626523 |
| LESVITQL | ASSLSRGQGNQAPL | 0.32079837 |
| LGLGALVVV | ASSLSRGQGNQAPL | 0.47924894 |
| LIGNGGNGGSGGI | ASSLSRGQGNQAPL | 0.349799 |
| LNAINAPTL | ASSLSRGQGNQAPL | 0.7493585 |
| LQLLNTGV | ASSLSRGQGNQAPL | 0.643727 |
| LQTSIEF | ASSLSRGQGNQAPL | 0.49396828 |
| MSFAVSEL | ASSLSRGQGNQAPL | 0.7042825 |
| NTGALGL | ASSLSRGQGNQAPL | 0.6962638 |
| PPGAGGAGGAGTT | ASSLSRGQGNQAPL | 0.31031176 |
| QLVGLGI | ASSLSRGQGNQAPL | 0.42378253 |

| | | |
|---|---|---|
| RFLEQVHQL | ASSLSRGQGNQAPL | 0.49710795 |
| RGVGYVLR | ASSLSRGQGNQAPL | 0.49842647 |
| SATGVFAGVI | ASSLSRGQGNQAPL | 0.22029547 |
| SLPAITLP | ASSLSRGQGNQAPL | 0.6228371 |
| STALAFTV | ASSLSRGQGNQAPL | 0.32692224 |
| SVLTVGLGK | ASSLSRGQGNQAPL | 0.18187958 |
| TALAGAGSGI | ASSLSRGQGNQAPL | 0.5613644 |
| TGLPFTGI | ASSLSRGQGNQAPL | 0.46339697 |
| VAEHVTV | ASSLSRGQGNQAPL | 0.44517022 |
| VDIVLNSL | ASSLSRGQGNQAPL | 0.6860614 |
| VLTGGGAVL | ASSLSRGQGNQAPL | 0.35860935 |
| VNVGLAYGI | ASSLSRGQGNQAPL | 0.37289417 |
| VSSAAVGAV | ASSLSRGQGNQAPL | 0.6178032 |
| VTSVYEAGV | ASSLSRGQGNQAPL | 0.6927093 |
| WVALDRGA | ASSLSRGQGNQAPL | 0.34825936 |
| YLVGSVVI | ASSLSRGQGNQAPL | 0.5394076 |
| GPPGTGKTLL | ASSLSRGQGNQAPL | 0.7201223 |
| GVGMTKF | ASSLSRGQGNQAPL | 0.55235434 |
| IGIGGSDL | ASSLSRGQGNQAPL | 0.6918083 |
| IGTLLAEL | ASSLSRGQGNQAPL | 0.7049565 |
| IYTSGTT | ASSLSRGQGNQAPL | 0.42568156 |
| LSFPAGV | ASSLSRGQGNQAPL | 0.7182599 |
| LYTSGTT | ASSLSRGQGNQAPL | 0.7638427 |
| VGAFPFGF | ASSLSRGQGNQAPL | 0.49608976 |
| VGAGTVEF | ASSLSRGQGNQAPL | 0.5333932 |
| VGKSTLLL | ASSLSRGQGNQAPL | 0.3911041 |
| AALAHAGGGIV | ASSLSRGQGNQAPL | 0.5271673 |
| AALVAVSLL | ASSLSRGQGNQAPL | 0.33559832 |
| AAVSRAVAAL | ASSLSRGQGNQAPL | 0.55645066 |
| AGGSSGIGGF | ASSLSRGQGNQAPL | 0.7077652 |
| AGLTVANANG | ASSLSRGQGNQAPL | 0.47961858 |
| AGLVGAVL | ASSLSRGQGNQAPL | 0.2845788 |
| ALSQAFTV | ASSLSRGQGNQAPL | 0.28004947 |
| ALTFSVSL | ASSLSRGQGNQAPL | 0.505926 |
| ALTGAALVAL | ASSLSRGQGNQAPL | 0.40614378 |
| ASFSIGAGVL | ASSLSRGQGNQAPL | 0.29860914 |
| DGSVYKI | ASSLSRGQGNQAPL | 0.40275168 |
| EQLAEIV | ASSLSRGQGNQAPL | 0.681537 |
| FAVSELTVM | ASSLSRGQGNQAPL | 0.6745082 |
| GAAPLAAGL | ASSLSRGQGNQAPL | 0.37310034 |
| GADGGDGGSGGAGG | ASSLSRGQGNQAPL | 0.74311984 |
| GFGLDTGGAGG | ASSLSRGQGNQAPL | 0.6133339 |
| GLASGGLL | ASSLSRGQGNQAPL | 0.72625756 |
| GRIAGAAVLL | ASSLSRGQGNQAPL | 0.5409063 |
| GSAGSLNTSF | ASSLSRGQGNQAPL | 0.34322494 |
| GTGQIGI | ASSLSRGQGNQAPL | 0.74977815 |
| GTLIALIL | ASSLSRGQGNQAPL | 0.44255257 |
| IAATMGFVVG | ASSLSRGQGNQAPL | 0.59524995 |
| IFAQALL | ASSLSRGQGNQAPL | 0.66071385 |
| IGAAVALGF | ASSLSRGQGNQAPL | 0.5119441 |
| IGLTPLAL | ASSLSRGQGNQAPL | 0.35667694 |

| | | |
|---|---|---|
| ILGSNTSTL | ASSLSRGQGNQAPL | 0.5026593 |
| IVFSLTVL | ASSLSRGQGNQAPL | 0.64301646 |
| LASKATGF | ASSLSRGQGNQAPL | 0.3225506 |
| LASVALIL | ASSLSRGQGNQAPL | 0.39622918 |
| LFAVGLTL | ASSLSRGQGNQAPL | 0.4535574 |
| LFNGGIGTYI | ASSLSRGQGNQAPL | 0.4763316 |
| LTLGAGII | ASSLSRGQGNQAPL | 0.7005789 |
| LVAAALGDI | ASSLSRGQGNQAPL | 0.5495335 |
| NTAFVQLGI | ASSLSRGQGNQAPL | 0.40431947 |
| QLFSGAEL | ASSLSRGQGNQAPL | 0.59955925 |
| SAATFITL | ASSLSRGQGNQAPL | 0.60874337 |
| SNYGLANLGV | ASSLSRGQGNQAPL | 0.53391254 |
| SYGGAYAVM | ASSLSRGQGNQAPL | 0.4632284 |
| TTGVGGLTLG | ASSLSRGQGNQAPL | 0.38627458 |
| TTQTIYI | ASSLSRGQGNQAPL | 0.6274941 |
| VAAGAGLGAVI | ASSLSRGQGNQAPL | 0.687528 |
| VAAVVTLL | ASSLSRGQGNQAPL | 0.63517267 |
| VGFLGGLI | ASSLSRGQGNQAPL | 0.6664417 |
| VGVIPILL | ASSLSRGQGNQAPL | 0.5439137 |
| VIFASSLI | ASSLSRGQGNQAPL | 0.5398465 |
| VKGGDGGAAAG | ASSLSRGQGNQAPL | 0.25822917 |
| VTKTFTNI | ASSLSRGQGNQAPL | 0.49700147 |
| VVAAAILL | ASSLSRGQGNQAPL | 0.48953858 |
| YAALDVEL | ASSLSRGQGNQAPL | 0.5451178 |
| LASVALLL | ASSLSRGQGNQAPL | 0.657457 |
| LDAVHRV | ASSLSRGQGNQAPL | 0.60154915 |
| QGLSSSIF | ASSLSRGQGNQAPL | 0.38905048 |
| RGTFANIRL | ASSLSRGQGNQAPL | 0.5764109 |
| RQLWWGHRI | ASSLSRGQGNQAPL | 0.71788484 |
| YGPPGTGKTLL | ASSLSRGQGNQAPL | 0.31062734 |
| AALVGAPANVA | ASSLSRGQGNQAPL | 0.43183023 |
| AATFRRTE | ASSLSRGQGNQAPL | 0.76205415 |
| AGALGGGI | ASSLSRGQGNQAPL | 0.61060137 |
| AGGAGGAGGIGAG | ASSLSRGQGNQAPL | 0.52154505 |
| AGSGALGL | ASSLSRGQGNQAPL | 0.53012455 |
| AQIADQI | ASSLSRGQGNQAPL | 0.74249876 |
| ASAVAVTYL | ASSLSRGQGNQAPL | 0.4918156 |
| ASTVLGGTGG | ASSLSRGQGNQAPL | 0.718406 |
| AVGGEPAK | ASSLSRGQGNQAPL | 0.23696408 |
| CEQGTFGGTC | ASSLSRGQGNQAPL | 0.47589958 |
| DEGKAPAG | ASSLSRGQGNQAPL | 0.34564897 |
| DTVVFPAL | ASSLSRGQGNQAPL | 0.17481388 |
| FEGITMAG | ASSLSRGQGNQAPL | 0.68494195 |
| FPLVVQD | ASSLSRGQGNQAPL | 0.47606662 |
| GAFTTPQ | ASSLSRGQGNQAPL | 0.6041953 |
| GFGGFGTAGGVGGA | ASSLSRGQGNQAPL | 0.49652407 |
| GFLAQFL | ASSLSRGQGNQAPL | 0.6367042 |
| GGAGGAGGNAGVA | ASSLSRGQGNQAPL | 0.5577501 |
| GGVGTAGAGGAGGAGG | ASSLSRGQGNQAPL | 0.4684604 |
| GNGGGGDAGNAGSGGN | ASSLSRGQGNQAPL | 0.4037362 |
| GTGGAGGAGNDAGS | ASSLSRGQGNQAPL | 0.6097822 |

| | | |
|---|---|---|
| GTTNTGWFNS | ASSLSRGQGNQAPL | 0.6157596 |
| IFMGPLFTL | ASSLSRGQGNQAPL | 0.6178011 |
| ILAHAIEAI | ASSLSRGQGNQAPL | 0.2763243 |
| IRSAIREAL | ASSLSRGQGNQAPL | 0.5093363 |
| ITFAGGG | ASSLSRGQGNQAPL | 0.47808722 |
| ITGMSGHQEV | ASSLSRGQGNQAPL | 0.31840304 |
| IVAGLACTL | ASSLSRGQGNQAPL | 0.22928895 |
| IVGGGLAGA | ASSLSRGQGNQAPL | 0.35216635 |
| IVTGVLNI | ASSLSRGQGNQAPL | 0.44919062 |
| LAQQGCI | ASSLSRGQGNQAPL | 0.6229315 |
| LCLHVEN | ASSLSRGQGNQAPL | 0.6487972 |
| LDAISDGS | ASSLSRGQGNQAPL | 0.47235063 |
| LGGFAAML | ASSLSRGQGNQAPL | 0.6272855 |
| LGGKNPMI | ASSLSRGQGNQAPL | 0.52799845 |
| LGPKGVAAIL | ASSLSRGQGNQAPL | 0.112921156 |
| LMVGVSI | ASSLSRGQGNQAPL | 0.44422063 |
| LVGAAFVL | ASSLSRGQGNQAPL | 0.34084705 |
| LVLAVAPL | ASSLSRGQGNQAPL | 0.7208541 |
| MAAIAVIDAA | ASSLSRGQGNQAPL | 0.29993182 |
| NNGINAELQ | ASSLSRGQGNQAPL | 0.47622108 |
| RGEPLVLVHG | ASSLSRGQGNQAPL | 0.5336608 |
| RGGAGGAAGLI | ASSLSRGQGNQAPL | 0.6305406 |
| RSYRMHPAV | ASSLSRGQGNQAPL | 0.5217469 |
| SAAHLLAGQLA | ASSLSRGQGNQAPL | 0.28729695 |
| SAAVGSY | ASSLSRGQGNQAPL | 0.21692637 |
| SPHAVLHRI | ASSLSRGQGNQAPL | 0.60983527 |
| SQIAKVL | ASSLSRGQGNQAPL | 0.4355625 |
| TALGHLVP | ASSLSRGQGNQAPL | 0.36651447 |
| TAPTTSGAA | ASSLSRGQGNQAPL | 0.27655426 |
| TDGDDGNGGSPGT | ASSLSRGQGNQAPL | 0.5388876 |
| TDGLGEN | ASSLSRGQGNQAPL | 0.29165298 |
| TGGPPLPGQA | ASSLSRGQGNQAPL | 0.6575524 |
| TIPAITI | ASSLSRGQGNQAPL | 0.40798396 |
| TIPGFTI | ASSLSRGQGNQAPL | 0.4617054 |
| TSIASGALA | ASSLSRGQGNQAPL | 0.769634 |
| TVTLPGAAI | ASSLSRGQGNQAPL | 0.6003483 |
| VASGLSAALP | ASSLSRGQGNQAPL | 0.50897133 |
| VGAGVIGI | ASSLSRGQGNQAPL | 0.42518178 |
| VGPKRFGV | ASSLSRGQGNQAPL | 0.41215643 |
| VNVAFPD | ASSLSRGQGNQAPL | 0.36701316 |
| VSMTNSGVS | ASSLSRGQGNQAPL | 0.41692153 |
| VTDVLGDLSV | ASSLSRGQGNQAPL | 0.7130993 |
| VVVVGAGFAGL | ASSLSRGQGNQAPL | 0.55490553 |
| VYTVGTVV | ASSLSRGQGNQAPL | 0.34750447 |
| WTTFSGP | ASSLSRGQGNQAPL | 0.6398559 |
| YDELVEAF | ASSLSRGQGNQAPL | 0.7556172 |
| YGTGGAGGNGG | ASSLSRGQGNQAPL | 0.779954 |
| YTNVMAV | ASSLSRGQGNQAPL | 0.3383916 |
| FLEQVHQL | ASSLSRGQGNQAPL | 0.4573075 |
| GLSLLAVAV | ASSLSRGQGNQAPL | 0.47546053 |
| LEQVHQL | ASSLSRGQGNQAPL | 0.47045046 |

| | | |
|---|---|---|
| LNAAGAI | ASSLSRGQGNQAPL | 0.41589886 |
| PAVHASL | ASSLSRGQGNQAPL | 0.3753034 |
| TELEDDA | ASSLSRGQGNQAPL | 0.6018619 |
| TPPGPPN | ASSLSRGQGNQAPL | 0.81781507 |
| TPPPPPPP | ASSLSRGQGNQAPL | 0.4822445 |
| TQSPALI | ASSRDHGAPNTEVF | 0.27621138 |
| EFAAAFPQ | ASSRDHGAPNTEVF | 0.49266484 |
| LLSGNLDVL | ASSRDHGAPNTEVF | 0.32612982 |
| GPANLGNYNI | ASSRDHGAPNTEVF | 0.37791568 |
| TLGPTLP | ASSRDHGAPNTEVF | 0.49902284 |
| VYAYSSCCK | ASSRDHGAPNTEVF | 0.260314 |
| APAPAPAPNI | ASSRDHGAPNTEVF | 0.51496685 |
| PGGPVPIP | ASSRDHGAPNTEVF | 0.7357828 |
| FLTVALEP | ASSRDHGAPNTEVF | 0.48164585 |
| SQYFGANL | ASSRDHGAPNTEVF | 0.6522822 |
| GGVKPAA | ASSRDHGAPNTEVF | 0.74395394 |
| GGTGGAGGAGGW | ASSRDHGAPNTEVF | 0.64763725 |
| GVGGVAGLGGAGG | ASSRDHGAPNTEVF | 0.74165815 |
| VLLVLATTI | ASSRDHGAPNTEVF | 0.6543516 |
| IGIFAAML | ASSRDHGAPNTEVF | 0.5086501 |
| AVGDLID | ASSRDHGAPNTEVF | 0.44207227 |
| NISPLPL | ASSRDHGAPNTEVF | 0.73529667 |
| VTAVFVAI | ASSRDHGAPNTEVF | 0.51107794 |
| SGDDNDAAL | ASSRDHGAPNTEVF | 0.5900936 |
| AVAALAHS | ASSRDHGAPNTEVF | 0.5072192 |
| GGVTVMATP | ASSRDHGAPNTEVF | 0.6839088 |
| AGLAANVRD | ASSRDHGAPNTEVF | 0.42383963 |
| TNRPDLIDPALLRPG | ASSRDHGAPNTEVF | 0.77249247 |
| LANLGVTAL | ASSRDHGAPNTEVF | 0.2552454 |
| TGERIEL | ASSRDHGAPNTEVF | 0.3674495 |
| NGEELAA | ASSRDHGAPNTEVF | 0.71122044 |
| VLSLVWG | ASSRDHGAPNTEVF | 0.1483715 |
| IGYALWL | ASSRDHGAPNTEVF | 0.25985402 |
| LSLLLAPL | ASSRDHGAPNTEVF | 0.48108763 |
| TLDEALK | ASSRDHGAPNTEVF | 0.701952 |
| TDGAGTAEGG | ASSRDHGAPNTEVF | 0.15450297 |
| NAPGGAGGNGG | ASSRDHGAPNTEVF | 0.44337898 |
| VSVFMGMGGNF | ASSRDHGAPNTEVF | 0.35171813 |
| LASYLAAL | ASSRDHGAPNTEVF | 0.5822832 |
| LVGGTPAL | ASSRDHGAPNTEVF | 0.5219508 |
| VRNADVL | ASSRDHGAPNTEVF | 0.29636565 |
| TCSSPFLVSAM | ASSRDHGAPNTEVF | 0.63903534 |
| VHITDAE | ASSRDHGAPNTEVF | 0.32474756 |
| EYAAPSF | ASSRDHGAPNTEVF | 0.531137 |
| VVDAFHGGL | ASSRDHGAPNTEVF | 0.6439515 |
| ASVVDPQ | ASSRDHGAPNTEVF | 0.7670809 |
| SYNEFGGLS | ASSRDHGAPNTEVF | 0.34422153 |
| DPDGSVY | ASSRDHGAPNTEVF | 0.6071506 |
| MVSTIHATE | ASSRDHGAPNTEVF | 0.36993033 |
| LAAGVADPV | ASSRDHGAPNTEVF | 0.8198206 |
| TKEQIAA | ASSRDHGAPNTEVF | 0.48650795 |

| | | |
|---|---|---|
| SLASPAAL | ASSRDHGAPNTEVF | 0.2990642 |
| IGLVFYGV | ASSRDHGAPNTEVF | 0.6561764 |
| FLLDASNL | ASSRDHGAPNTEVF | 0.6486001 |
| YQVGNFP | ASSRDHGAPNTEVF | 0.525759 |
| ESLLSQST | ASSRDHGAPNTEVF | 0.6465522 |
| LFTVGFAAVG | ASSRDHGAPNTEVF | 0.45377317 |
| ATPPTPT | ASSRDHGAPNTEVF | 0.5054564 |
| TDLATLG | ASSRDHGAPNTEVF | 0.68915576 |
| SAGGTGR | ASSRDHGAPNTEVF | 0.5596246 |
| GFSNSGS | ASSRDHGAPNTEVF | 0.41027486 |
| GGNGGSGGTSV | ASSRDHGAPNTEVF | 0.6679688 |
| GTSVATGGAGN | ASSRDHGAPNTEVF | 0.5585291 |
| NGGNGGSGGTS | ASSRDHGAPNTEVF | 0.3791215 |
| AVVKFPH | ASSRDHGAPNTEVF | 0.76777136 |
| TPPPPPP | ASSRDHGAPNTEVF | 0.51771295 |
| AAPAAAAAPA | ASSRDHGAPNTEVF | 0.6831643 |
| IALNEDL | ASSRDHGAPNTEVF | 0.61681354 |
| IVNVSSM | ASSRDHGAPNTEVF | 0.7565176 |
| TGIPPAPR | ASSRDHGAPNTEVF | 0.3487223 |
| FDLGGGTF | ASSRDHGAPNTEVF | 0.5107015 |
| FDLGGGTFDVS | ASSRDHGAPNTEVF | 0.5276994 |
| NEPTAAA | ASSRDHGAPNTEVF | 0.6125967 |
| NEPTAAALA | ASSRDHGAPNTEVF | 0.5060618 |
| GSIINISSI | ASSRDHGAPNTEVF | 0.5513914 |
| VAPGNAGTA | ASSRDHGAPNTEVF | 0.64402324 |
| TGGGASLE | ASSRDHGAPNTEVF | 0.7414618 |
| VAVYNHYKRI | ASSRDHGAPNTEVF | 0.45359996 |
| TDPAPATT | ASSRDHGAPNTEVF | 0.454148 |
| TNRPDILDPALLRPG | ASSRDHGAPNTEVF | 0.78071177 |
| LTTTTTS | ASSRDHGAPNTEVF | 0.4003746 |
| PTGSGKTL | ASSRDHGAPNTEVF | 0.5621647 |
| TGSGKTL | ASSRDHGAPNTEVF | 0.61408895 |
| TIGHVDHGK | ASSRDHGAPNTEVF | 0.64641595 |
| TIGHVDHGKTT | ASSRDHGAPNTEVF | 0.49883148 |
| GHVDHGKTT | ASSRDHGAPNTEVF | 0.6712943 |
| AQFEHTLL | ASSRDHGAPNTEVF | 0.40962976 |
| IVNQDEL | ASSRDHGAPNTEVF | 0.6519002 |
| TNGSQFF | ASSRDHGAPNTEVF | 0.45978147 |
| TGKGIVE | ASSRDHGAPNTEVF | 0.49340528 |
| LISNASDA | ASSRDHGAPNTEVF | 0.31234792 |
| AELRTLGELTKIAKAHGAQ | ASSRDHGAPNTEVF | 0.6148343 |
| APNAVAAAEPSW | ASSRDHGAPNTEVF | 0.706685 |
| APQVAGAATAVL | ASSRDHGAPNTEVF | 0.6463448 |
| ARRDPAAAAAILPTDA | ASSRDHGAPNTEVF | 0.22931871 |
| DAAARDVLAEAGLAEYF | ASSRDHGAPNTEVF | 0.64787626 |
| DDQIVLSAKANLAKVEQ | ASSRDHGAPNTEVF | 0.6347443 |
| DELLPDTGEDSDEDGH | ASSRDHGAPNTEVF | 0.80731505 |
| ELPEAPANRATVPQD | ASSRDHGAPNTEVF | 0.42089996 |
| FGQSKIERVTGIG | ASSRDHGAPNTEVF | 0.6453734 |
| GAGGTGSTAGGAGGAGGAGGLY | ASSRDHGAPNTEVF | 0.47234043 |
| GHGGTGAAGGNGGTGGAGGSGI | ASSRDHGAPNTEVF | 0.5193472 |

| | | |
|---|---|---|
| GNGGNGGAGGEGGPGGAGGAGA | ASSRDHGAPNTEVF | 0.25322872 |
| GSTAGNGGNGGNATNGTIAGQP | ASSRDHGAPNTEVF | 0.4061947 |
| GTGGDGGNGGNGGTGG | ASSRDHGAPNTEVF | 0.4964105 |
| GVRRPAGIPADAPV | ASSRDHGAPNTEVF | 0.3584269 |
| GYRQVTDAHLLTLARR | ASSRDHGAPNTEVF | 0.69625306 |
| IAKMGQDGHDRG | ASSRDHGAPNTEVF | 0.6228504 |
| KAEPSIEIQYT | ASSRDHGAPNTEVF | 0.6915499 |
| KPVRIRMAADEDTVAVT | ASSRDHGAPNTEVF | 0.7191598 |
| MPPRIAGMRLLVIKPEP | ASSRDHGAPNTEVF | 0.7701115 |
| MKFYAVALFRSRGVPPTR | ASSRDHGAPNTEVF | 0.6376499 |
| NAGGFGILGAVAHSP | ASSRDHGAPNTEVF | 0.39817125 |
| NEGREITLAMAEM | ASSRDHGAPNTEVF | 0.553185 |
| NELVEWSHPLGHRWRWE | ASSRDHGAPNTEVF | 0.26675543 |
| NLGAGGTGGNGGNPG | ASSRDHGAPNTEVF | 0.49201816 |
| NTEWPKWETVMAADPVD | ASSRDHGAPNTEVF | 0.44103312 |
| PASAAPAETQP | ASSRDHGAPNTEVF | 0.57369655 |
| QQSSDGKNGVRAAEITGE | ASSRDHGAPNTEVF | 0.1423442 |
| RAPDAVTTSWWKE | ASSRDHGAPNTEVF | 0.60788083 |
| SGFGNVGAHTSGWFNQ | ASSRDHGAPNTEVF | 0.7338873 |
| SGVAGAVGQAASVGGLKV | ASSRDHGAPNTEVF | 0.42136806 |
| SNGGAGTGSAGGNGGT | ASSRDHGAPNTEVF | 0.6322906 |
| TGGAGGEGGAAG | ASSRDHGAPNTEVF | 0.67702365 |
| TGGDGGAAGAGGAPGHG | ASSRDHGAPNTEVF | 0.319302 |
| TMPPLLAAIANAGRHGV | ASSRDHGAPNTEVF | 0.6374011 |
| TSSTAGNSGGA | ASSRDHGAPNTEVF | 0.4005575 |
| VPATVPVGILC | ASSRDHGAPNTEVF | 0.6648703 |
| VQRPDSGGVVLTGRIS | ASSRDHGAPNTEVF | 0.7301601 |
| WTPNPVPATKGVPAGLVQAT | ASSRDHGAPNTEVF | 0.6193604 |
| YALKRALAKFSDRFDVVII | ASSRDHGAPNTEVF | 0.41432723 |
| YLIGSVIV | ASSRDHGAPNTEVF | 0.5604004 |
| YGQLVAVID | ASSRDHGAPNTEVF | 0.34717828 |
| VTVGGMTLPA | ASSRDHGAPNTEVF | 0.4194604 |
| VLALPAV | ASSRDHGAPNTEVF | 0.32702833 |
| VIPPTLNYET | ASSRDHGAPNTEVF | 0.5603458 |
| VGMGGMHPSSG | ASSRDHGAPNTEVF | 0.40015322 |
| VGLAGGAATA | ASSRDHGAPNTEVF | 0.5802512 |
| VGGAGGAGGSGP | ASSRDHGAPNTEVF | 0.55350834 |
| VFAAFMV | ASSRDHGAPNTEVF | 0.48425797 |
| TLYDKLR | ASSRDHGAPNTEVF | 0.39554137 |
| TLAVFTVL | ASSRDHGAPNTEVF | 0.60761905 |
| TGSVQPARV | ASSRDHGAPNTEVF | 0.5586318 |
| TGGGAGSGGT | ASSRDHGAPNTEVF | 0.30283296 |
| SHLETSL | ASSRDHGAPNTEVF | 0.611784 |
| SFTTGMT | ASSRDHGAPNTEVF | 0.47929692 |
| QVEVAKRMGVRQPTVS | ASSRDHGAPNTEVF | 0.48739278 |
| QGSGGPALAM | ASSRDHGAPNTEVF | 0.69567186 |
| MLAPSNP | ASSRDHGAPNTEVF | 0.54786146 |
| LVNNAAI | ASSRDHGAPNTEVF | 0.3301619 |
| LTGEINTLA | ASSRDHGAPNTEVF | 0.45036453 |
| LSIFGPV | ASSRDHGAPNTEVF | 0.6020292 |
| LLALLANL | ASSRDHGAPNTEVF | 0.44766235 |

| | | |
|---|---|---|
| LISNASDALDKLR | ASSRDHGAPNTEVF | 0.18105693 |
| LINEAVP | ASSRDHGAPNTEVF | 0.60945 |
| LGYAFSGL | ASSRDHGAPNTEVF | 0.38856345 |
| LDFVAVAL | ASSRDHGAPNTEVF | 0.4034479 |
| LAQPGAE | ASSRDHGAPNTEVF | 0.43909204 |
| ISNLGAVTGF | ASSRDHGAPNTEVF | 0.6957518 |
| HQEDPGRI | ASSRDHGAPNTEVF | 0.4180413 |
| GTPGEPTP | ASSRDHGAPNTEVF | 0.50791407 |
| GFFNSGS | ASSRDHGAPNTEVF | 0.60392666 |
| GAGGGGLTAGAN | ASSRDHGAPNTEVF | 0.60665816 |
| FVDALVNM | ASSRDHGAPNTEVF | 0.69462454 |
| FLTVAPEL | ASSRDHGAPNTEVF | 0.57505405 |
| FETLTAL | ASSRDHGAPNTEVF | 0.43560752 |
| DVINPIEL | ASSRDHGAPNTEVF | 0.5852951 |
| APVSGAPV | ASSRDHGAPNTEVF | 0.48757958 |
| ALVGGLAW | ASSRDHGAPNTEVF | 0.37499735 |
| ALAAAAATAI | ASSRDHGAPNTEVF | 0.3022106 |
| AFGLGLLL | ASSRDHGAPNTEVF | 0.4440402 |
| AFFAAASL | ASSRDHGAPNTEVF | 0.3440992 |
| AAAALPAFSPP | ASSRDHGAPNTEVF | 0.6042508 |
| VVLAPAL | ASSRDHGAPNTEVF | 0.713406 |
| TIGHVDHGKTTL | ASSRDHGAPNTEVF | 0.4215659 |
| LISNASDALDK | ASSRDHGAPNTEVF | 0.31914523 |
| LISNASDAL | ASSRDHGAPNTEVF | 0.3985669 |
| ISNASDALDK | ASSRDHGAPNTEVF | 0.48414835 |
| NQAETLVYQTE | ASSRDHGAPNTEVF | 0.6608422 |
| TGGTGGTAGAG | ASSRDHGAPNTEVF | 0.5848278 |
| TGSTAGGAGGA | ASSRDHGAPNTEVF | 0.7779802 |
| DPLAVPVAAGPV | ASSRDHGAPNTEVF | 0.6305285 |
| GGTGGTPGNGGN | ASSRDHGAPNTEVF | 0.42603293 |
| PAGLAGVDPATL | ASSRDHGAPNTEVF | 0.6645593 |
| GNNLVVEAPHAGASV | ASSRDHGAPNTEVF | 0.46403846 |
| KVTRIMSASTLEQPA | ASSRDHGAPNTEVF | 0.52148426 |
| LGFALPIALPRRTVV | ASSRDHGAPNTEVF | 0.5209224 |
| TRLGNLTARKLVVLV | ASSRDHGAPNTEVF | 0.5094674 |
| WSQQRRLIYFPSAGP | ASSRDHGAPNTEVF | 0.38903838 |
| GPNGNPDPMAAAVDIR | ASSRDHGAPNTEVF | 0.45665547 |
| TQRCVSVADLRAAASR | ASSRDHGAPNTEVF | 0.3136449 |
| QFVLDVAEHPVEVLTETV | ASSRDHGAPNTEVF | 0.56403166 |
| GGAGGTGGAGGNGGTQAGDG | ASSRDHGAPNTEVF | 0.43285826 |
| PLLATLRAIAVDVDAKPAQV | ASSRDHGAPNTEVF | 0.3262249 |
| APKFRTGGYMIDFWGVGYQVAK | ASSRDHGAPNTEVF | 0.4439375 |
| GAGGAGGAGGAAGGPGGTGGTG | ASSRDHGAPNTEVF | 0.76106435 |
| IGALGLLATDVAPAGSVAFGVA | ASSRDHGAPNTEVF | 0.740731 |
| AGGAGGIGGNGGAGGAGGVLMG | ASSRDHGAPNTEVF | 0.38117716 |
| ADELIGLL | ASSRDHGAPNTEVF | 0.38292208 |
| ADRALKA | ASSRDHGAPNTEVF | 0.6173606 |
| AGAVGQVGGAGGA | ASSRDHGAPNTEVF | 0.35024834 |
| AGGGGGLGGAP | ASSRDHGAPNTEVF | 0.594595 |
| AGGLLYGNGGAG | ASSRDHGAPNTEVF | 0.50028193 |
| AGLESIGL | ASSRDHGAPNTEVF | 0.51695895 |

| | | |
|---|---|---|
| AGLKTGII | ASSRDHGAPNTEVF | 0.4111052 |
| ASLAALLL | ASSRDHGAPNTEVF | 0.58208704 |
| ASVINISSI | ASSRDHGAPNTEVF | 0.45517862 |
| ATPPGPA | ASSRDHGAPNTEVF | 0.56823075 |
| ATVASALV | ASSRDHGAPNTEVF | 0.19121835 |
| ATVASLIL | ASSRDHGAPNTEVF | 0.6179422 |
| AVNGPVLIRH | ASSRDHGAPNTEVF | 0.36474636 |
| DGAAGAVGI | ASSRDHGAPNTEVF | 0.33672816 |
| DPLLHGI | ASSRDHGAPNTEVF | 0.21648331 |
| ELGAALGAL | ASSRDHGAPNTEVF | 0.47978285 |
| ELGRTQAV | ASSRDHGAPNTEVF | 0.681002 |
| FVAGGYLS | ASSRDHGAPNTEVF | 0.70855814 |
| GASGGAGGAGGW | ASSRDHGAPNTEVF | 0.47803396 |
| GGAGAGGDGGAGGI | ASSRDHGAPNTEVF | 0.61470306 |
| GTGHNGSQPGMGGNG | ASSRDHGAPNTEVF | 0.71581197 |
| GYAAASASAS | ASSRDHGAPNTEVF | 0.46178472 |
| HQVDVAAVQQG | ASSRDHGAPNTEVF | 0.2417799 |
| IGAAIAHT | ASSRDHGAPNTEVF | 0.60187864 |
| IGMIEVAQL | ASSRDHGAPNTEVF | 0.43877757 |
| IRLAPEHTSAMD | ASSRDHGAPNTEVF | 0.6525946 |
| IYNVSVL | ASSRDHGAPNTEVF | 0.67225385 |
| KQGFLGM | ASSRDHGAPNTEVF | 0.28203565 |
| LESVITQL | ASSRDHGAPNTEVF | 0.6552326 |
| LGLGALVVV | ASSRDHGAPNTEVF | 0.6770691 |
| LIGNGGNGGSGGI | ASSRDHGAPNTEVF | 0.2903521 |
| LNAINAPTL | ASSRDHGAPNTEVF | 0.39207417 |
| LQLLNTGV | ASSRDHGAPNTEVF | 0.414198 |
| LQTSIEF | ASSRDHGAPNTEVF | 0.2156096 |
| MSFAVSEL | ASSRDHGAPNTEVF | 0.6167999 |
| NTGALGL | ASSRDHGAPNTEVF | 0.45381206 |
| PPGAGGAGGAGTT | ASSRDHGAPNTEVF | 0.4568498 |
| QLVGLGI | ASSRDHGAPNTEVF | 0.5107198 |
| RFLEQVHQL | ASSRDHGAPNTEVF | 0.43284863 |
| RGVGYVLR | ASSRDHGAPNTEVF | 0.47653612 |
| SATGVFAGVI | ASSRDHGAPNTEVF | 0.49971306 |
| SLPAITLP | ASSRDHGAPNTEVF | 0.6090161 |
| STALAFTV | ASSRDHGAPNTEVF | 0.60119164 |
| SVLTVGLGK | ASSRDHGAPNTEVF | 0.4778052 |
| TALAGAGSGI | ASSRDHGAPNTEVF | 0.55288035 |
| TGLPFTGI | ASSRDHGAPNTEVF | 0.75799763 |
| VAEHVTV | ASSRDHGAPNTEVF | 0.30623248 |
| VDIVLNSL | ASSRDHGAPNTEVF | 0.38631484 |
| VLTGGGAVL | ASSRDHGAPNTEVF | 0.5819495 |
| VNVGLAYGI | ASSRDHGAPNTEVF | 0.4187131 |
| VSSAAVGAV | ASSRDHGAPNTEVF | 0.6984799 |
| VTSVYEAGV | ASSRDHGAPNTEVF | 0.57721114 |
| WVALDRGA | ASSRDHGAPNTEVF | 0.62260973 |
| YLVGSVVI | ASSRDHGAPNTEVF | 0.45992008 |
| GPPGTGKTLL | ASSRDHGAPNTEVF | 0.7877066 |
| GVGMTKF | ASSRDHGAPNTEVF | 0.6263432 |
| IGIGGSDL | ASSRDHGAPNTEVF | 0.48477402 |

| | | |
|---|---|---|
| IGTLLAEL | ASSRDHGAPNTEVF | 0.6558775 |
| IYTSGTT | ASSRDHGAPNTEVF | 0.3329512 |
| LSFPAGV | ASSRDHGAPNTEVF | 0.6179229 |
| LYTSGTT | ASSRDHGAPNTEVF | 0.63814086 |
| VGAFPFGF | ASSRDHGAPNTEVF | 0.33829236 |
| VGAGTVEF | ASSRDHGAPNTEVF | 0.5120864 |
| VGKSTLLL | ASSRDHGAPNTEVF | 0.6723156 |
| AALAHAGGGIV | ASSRDHGAPNTEVF | 0.69597745 |
| AALVAVSLL | ASSRDHGAPNTEVF | 0.36567646 |
| AAVSRAVAAL | ASSRDHGAPNTEVF | 0.39691907 |
| AGGSSGIGGF | ASSRDHGAPNTEVF | 0.31145477 |
| AGLTVANANG | ASSRDHGAPNTEVF | 0.32785687 |
| AGLVGAVL | ASSRDHGAPNTEVF | 0.36499578 |
| ALSQAFTV | ASSRDHGAPNTEVF | 0.60522634 |
| ALTFSVSL | ASSRDHGAPNTEVF | 0.52175534 |
| ALTGAALVAL | ASSRDHGAPNTEVF | 0.71263015 |
| ASFSIGAGVL | ASSRDHGAPNTEVF | 0.60042095 |
| DGSVYKI | ASSRDHGAPNTEVF | 0.6984143 |
| EQLAEIV | ASSRDHGAPNTEVF | 0.4330119 |
| FAVSELTVM | ASSRDHGAPNTEVF | 0.5059044 |
| GAAPLAAGL | ASSRDHGAPNTEVF | 0.29380482 |
| GADGGDGGSGGAGG | ASSRDHGAPNTEVF | 0.36824083 |
| GFGLDTGGAGG | ASSRDHGAPNTEVF | 0.36888134 |
| GLASGGLL | ASSRDHGAPNTEVF | 0.48854285 |
| GRIAGAAVLL | ASSRDHGAPNTEVF | 0.25414506 |
| GSAGSLNTSF | ASSRDHGAPNTEVF | 0.7558638 |
| GTGQIGI | ASSRDHGAPNTEVF | 0.5093143 |
| GTLIALIL | ASSRDHGAPNTEVF | 0.6365039 |
| IAATMGFVVG | ASSRDHGAPNTEVF | 0.5035696 |
| IFAQALL | ASSRDHGAPNTEVF | 0.39056677 |
| IGAAVALGF | ASSRDHGAPNTEVF | 0.42298913 |
| IGLTPLAL | ASSRDHGAPNTEVF | 0.6112971 |
| ILGSNTSTL | ASSRDHGAPNTEVF | 0.27859163 |
| IVFSLTVL | ASSRDHGAPNTEVF | 0.28473166 |
| LASKATGF | ASSRDHGAPNTEVF | 0.40293863 |
| LASVALIL | ASSRDHGAPNTEVF | 0.38839018 |
| LFAVGLTL | ASSRDHGAPNTEVF | 0.7966213 |
| LFNGGIGTYI | ASSRDHGAPNTEVF | 0.7302291 |
| LTLGAGII | ASSRDHGAPNTEVF | 0.5819379 |
| LVAAALGDI | ASSRDHGAPNTEVF | 0.8012683 |
| NTAFVQLGI | ASSRDHGAPNTEVF | 0.1807329 |
| QLFSGAEL | ASSRDHGAPNTEVF | 0.5077234 |
| SAATFITL | ASSRDHGAPNTEVF | 0.74240786 |
| SNYGLANLGV | ASSRDHGAPNTEVF | 0.55311024 |
| SYGGAYAVM | ASSRDHGAPNTEVF | 0.56338024 |
| TTGVGGLTLG | ASSRDHGAPNTEVF | 0.6899939 |
| TTQTIYI | ASSRDHGAPNTEVF | 0.40155435 |
| VAAGAGLGAVI | ASSRDHGAPNTEVF | 0.45713666 |
| VAAVVTLL | ASSRDHGAPNTEVF | 0.4707789 |
| VGFLGGLI | ASSRDHGAPNTEVF | 0.58703125 |
| VGVIPILL | ASSRDHGAPNTEVF | 0.4942986 |

| | | |
|---|---|---|
| VIFASSLI | ASSRDHGAPNTEVF | 0.2941462 |
| VKGGDGGAAAG | ASSRDHGAPNTEVF | 0.6347825 |
| VTKTFTNI | ASSRDHGAPNTEVF | 0.81280005 |
| VVAAAILL | ASSRDHGAPNTEVF | 0.41891903 |
| YAALDVEL | ASSRDHGAPNTEVF | 0.6784484 |
| LASVALLL | ASSRDHGAPNTEVF | 0.563447 |
| LDAVHRV | ASSRDHGAPNTEVF | 0.47140864 |
| QGLSSSIF | ASSRDHGAPNTEVF | 0.42012298 |
| RGTFANIRL | ASSRDHGAPNTEVF | 0.30187213 |
| RQLWWGHRI | ASSRDHGAPNTEVF | 0.610042 |
| YGPPGTGKTLL | ASSRDHGAPNTEVF | 0.32456005 |
| AALVGAPANVA | ASSRDHGAPNTEVF | 0.51243186 |
| AATFRRTE | ASSRDHGAPNTEVF | 0.5345438 |
| AGALGGGI | ASSRDHGAPNTEVF | 0.81393313 |
| AGGAGGAGGIGAG | ASSRDHGAPNTEVF | 0.62895215 |
| AGSGALGL | ASSRDHGAPNTEVF | 0.59505475 |
| AQIADQI | ASSRDHGAPNTEVF | 0.31182632 |
| ASAVAVTYL | ASSRDHGAPNTEVF | 0.496523 |
| ASTVLGGTGG | ASSRDHGAPNTEVF | 0.5720105 |
| AVGGEPAK | ASSRDHGAPNTEVF | 0.34223577 |
| CEQGTFGGTC | ASSRDHGAPNTEVF | 0.7122929 |
| DEGKAPAG | ASSRDHGAPNTEVF | 0.3690802 |
| DTVVFPAL | ASSRDHGAPNTEVF | 0.50869215 |
| FEGITMAG | ASSRDHGAPNTEVF | 0.5713633 |
| FPLVVQD | ASSRDHGAPNTEVF | 0.7463824 |
| GAFTTPQ | ASSRDHGAPNTEVF | 0.5189194 |
| GFGGFGTAGGVGGA | ASSRDHGAPNTEVF | 0.6393928 |
| GFLAQFL | ASSRDHGAPNTEVF | 0.35732102 |
| GGAGGAGGNAGVA | ASSRDHGAPNTEVF | 0.65283495 |
| GGVGTAGAGGAGGAGG | ASSRDHGAPNTEVF | 0.7398821 |
| GNGGGGDAGNAGSGGN | ASSRDHGAPNTEVF | 0.5005355 |
| GTGGAGGAGNDAGS | ASSRDHGAPNTEVF | 0.78947246 |
| GTTNTGWFNS | ASSRDHGAPNTEVF | 0.56918216 |
| IFMGPLFTL | ASSRDHGAPNTEVF | 0.2842407 |
| ILAHAIEAI | ASSRDHGAPNTEVF | 0.5300288 |
| IRSAIREAL | ASSRDHGAPNTEVF | 0.65708715 |
| ITFAGGG | ASSRDHGAPNTEVF | 0.71999574 |
| ITGMSGHQEV | ASSRDHGAPNTEVF | 0.66156536 |
| IVAGLACTL | ASSRDHGAPNTEVF | 0.582379 |
| IVGGGLAGA | ASSRDHGAPNTEVF | 0.42221677 |
| IVTGVLNI | ASSRDHGAPNTEVF | 0.74856114 |
| LAQQGCI | ASSRDHGAPNTEVF | 0.8021172 |
| LCLHVEN | ASSRDHGAPNTEVF | 0.65887165 |
| LDAISDGS | ASSRDHGAPNTEVF | 0.3748531 |
| LGGFAAML | ASSRDHGAPNTEVF | 0.35633987 |
| LGGKNPMI | ASSRDHGAPNTEVF | 0.6333873 |
| LGPKGVAAIL | ASSRDHGAPNTEVF | 0.6810861 |
| LMVGVSI | ASSRDHGAPNTEVF | 0.35768625 |
| LVGAAFVL | ASSRDHGAPNTEVF | 0.5569132 |
| LVLAVAPL | ASSRDHGAPNTEVF | 0.56073207 |
| MAAIAVIDAA | ASSRDHGAPNTEVF | 0.2529832 |

| | | |
|---|---|---|
| NNGINAELQ | ASSRDHGAPNTEVF | 0.6836714 |
| RGEPLVLVHG | ASSRDHGAPNTEVF | 0.7479786 |
| RGGAGGAAGLI | ASSRDHGAPNTEVF | 0.6011275 |
| RSYRMHPAV | ASSRDHGAPNTEVF | 0.33632192 |
| SAAHLLAGQLA | ASSRDHGAPNTEVF | 0.72366774 |
| SAAVGSY | ASSRDHGAPNTEVF | 0.7705441 |
| SPHAVLHRI | ASSRDHGAPNTEVF | 0.7598032 |
| SQIAKVL | ASSRDHGAPNTEVF | 0.29825777 |
| TALGHLVP | ASSRDHGAPNTEVF | 0.18918347 |
| TAPTTSGAA | ASSRDHGAPNTEVF | 0.4797927 |
| TDGDDGNGGSPGT | ASSRDHGAPNTEVF | 0.6756034 |
| TDGLGEN | ASSRDHGAPNTEVF | 0.4662478 |
| TGGPPLPGQA | ASSRDHGAPNTEVF | 0.36685985 |
| TIPAITI | ASSRDHGAPNTEVF | 0.5336859 |
| TIPGFTI | ASSRDHGAPNTEVF | 0.70742863 |
| TSIASGALA | ASSRDHGAPNTEVF | 0.39197183 |
| TVTLPGAAI | ASSRDHGAPNTEVF | 0.26867265 |
| VASGLSAALP | ASSRDHGAPNTEVF | 0.6155795 |
| VGAGVIGI | ASSRDHGAPNTEVF | 0.6844467 |
| VGPKRFGV | ASSRDHGAPNTEVF | 0.30580688 |
| VNVAFPD | ASSRDHGAPNTEVF | 0.40626746 |
| VSMTNSGVS | ASSRDHGAPNTEVF | 0.5913552 |
| VTDVLGDLSV | ASSRDHGAPNTEVF | 0.50549513 |
| VVVVGAGFAGL | ASSRDHGAPNTEVF | 0.5090811 |
| VYTVGTVV | ASSRDHGAPNTEVF | 0.3025098 |
| WTTFSGP | ASSRDHGAPNTEVF | 0.66911036 |
| YDELVEAF | ASSRDHGAPNTEVF | 0.34608346 |
| YGTGGAGGNGG | ASSRDHGAPNTEVF | 0.56980175 |
| YTNVMAV | ASSRDHGAPNTEVF | 0.5135665 |
| FLEQVHQL | ASSRDHGAPNTEVF | 0.5773656 |
| GLSLLAVAV | ASSRDHGAPNTEVF | 0.49951327 |
| LEQVHQL | ASSRDHGAPNTEVF | 0.3774017 |
| LNAAGAI | ASSRDHGAPNTEVF | 0.34628698 |
| PAVHASL | ASSRDHGAPNTEVF | 0.23028724 |
| TELEDDA | ASSRDHGAPNTEVF | 0.6473647 |
| TPPGPPN | ASSRDHGAPNTEVF | 0.48828816 |
| TPPPPPP | ASSRDHGAPNTEVF | 0.67612416 |
| TQSPALI | ASSPSGDSDYT | 0.573219 |
| EFAAAFPQ | ASSPSGDSDYT | 0.38318086 |
| LLSGNLDVL | ASSPSGDSDYT | 0.5533013 |
| GPANLGNYNI | ASSPSGDSDYT | 0.29143068 |
| TLGPTLP | ASSPSGDSDYT | 0.55575883 |
| VYAYSSCCK | ASSPSGDSDYT | 0.5023596 |
| APAPAPAPNI | ASSPSGDSDYT | 0.5988873 |
| PGGPVPIP | ASSPSGDSDYT | 0.46792755 |
| FLTVALEP | ASSPSGDSDYT | 0.6263417 |
| SQYFGANL | ASSPSGDSDYT | 0.60603356 |
| GGVKPAA | ASSPSGDSDYT | 0.35538384 |
| GGTGGAGGAGGW | ASSPSGDSDYT | 0.6026606 |
| GVGGVAGLGGAGG | ASSPSGDSDYT | 0.68018305 |
| VLLVLATTI | ASSPSGDSDYT | 0.6039281 |

| | | |
|---|---|---|
| IGIFAAML | ASSPSGDSDYT | 0.29450503 |
| AVGDLID | ASSPSGDSDYT | 0.6042002 |
| NISPLPL | ASSPSGDSDYT | 0.39798892 |
| VTAVFVAI | ASSPSGDSDYT | 0.5937348 |
| SGDDNDAAL | ASSPSGDSDYT | 0.51883894 |
| AVAALAHS | ASSPSGDSDYT | 0.4715898 |
| GGVTVMATP | ASSPSGDSDYT | 0.60340995 |
| AGLAANVRD | ASSPSGDSDYT | 0.36864287 |
| TNRPDLIDPALLRPG | ASSPSGDSDYT | 0.39290878 |
| LANLGVTAL | ASSPSGDSDYT | 0.4798177 |
| TGERIEL | ASSPSGDSDYT | 0.59791774 |
| NGEELAA | ASSPSGDSDYT | 0.64241374 |
| VLSLVWG | ASSPSGDSDYT | 0.3264073 |
| IGYALWL | ASSPSGDSDYT | 0.5547365 |
| LSLLLAPL | ASSPSGDSDYT | 0.5051419 |
| TLDEALK | ASSPSGDSDYT | 0.37794936 |
| TDGAGTAEGG | ASSPSGDSDYT | 0.56685966 |
| NAPGGAGGNGG | ASSPSGDSDYT | 0.47132027 |
| VSVFMGMGGNF | ASSPSGDSDYT | 0.3291112 |
| LASYLAAL | ASSPSGDSDYT | 0.60903376 |
| LVGGTPAL | ASSPSGDSDYT | 0.4116223 |
| VRNADVL | ASSPSGDSDYT | 0.55515075 |
| TCSSPFLVSAM | ASSPSGDSDYT | 0.49088433 |
| VHITDAE | ASSPSGDSDYT | 0.29855734 |
| EYAAPSF | ASSPSGDSDYT | 0.57179326 |
| VVDAFHGGL | ASSPSGDSDYT | 0.54522884 |
| ASVVDPQ | ASSPSGDSDYT | 0.53194463 |
| SYNEFGGLS | ASSPSGDSDYT | 0.52490133 |
| DPDGSVY | ASSPSGDSDYT | 0.41845912 |
| MVSTIHATE | ASSPSGDSDYT | 0.7071197 |
| LAAGVADPV | ASSPSGDSDYT | 0.6475111 |
| TKEQIAA | ASSPSGDSDYT | 0.6044071 |
| SLASPAAL | ASSPSGDSDYT | 0.5773854 |
| IGLVFYGV | ASSPSGDSDYT | 0.6307893 |
| FLLDASNL | ASSPSGDSDYT | 0.6478798 |
| YQVGNFP | ASSPSGDSDYT | 0.39261243 |
| ESLLSQST | ASSPSGDSDYT | 0.2647689 |
| LFTVGFAAVG | ASSPSGDSDYT | 0.49591875 |
| ATPPTPT | ASSPSGDSDYT | 0.48370752 |
| TDLATLG | ASSPSGDSDYT | 0.4539341 |
| SAGGTGR | ASSPSGDSDYT | 0.25988513 |
| GFSNSGS | ASSPSGDSDYT | 0.6624404 |
| GGNGGSGGTSV | ASSPSGDSDYT | 0.80983835 |
| GTSVATGGAGN | ASSPSGDSDYT | 0.37054053 |
| NGGNGGSGGTS | ASSPSGDSDYT | 0.80912894 |
| AVVKFPH | ASSPSGDSDYT | 0.3347459 |
| TPPPPPP | ASSPSGDSDYT | 0.6053362 |
| AAPAAAAAPA | ASSPSGDSDYT | 0.6860196 |
| IALNEDL | ASSPSGDSDYT | 0.29832172 |
| IVNVSSM | ASSPSGDSDYT | 0.58573663 |
| TGIPPAPR | ASSPSGDSDYT | 0.45165747 |

| | | |
|---|---|---|
| FDLGGGTF | ASSPSGDSDYT | 0.3857814 |
| FDLGGGTFDVS | ASSPSGDSDYT | 0.70818293 |
| NEPTAAA | ASSPSGDSDYT | 0.4908718 |
| NEPTAAALA | ASSPSGDSDYT | 0.58376324 |
| GSIINISSI | ASSPSGDSDYT | 0.4620901 |
| VAPGNAGTA | ASSPSGDSDYT | 0.38214436 |
| TGGGASLE | ASSPSGDSDYT | 0.55797786 |
| VAVYNHYKRI | ASSPSGDSDYT | 0.6416389 |
| TDPAPATT | ASSPSGDSDYT | 0.48260358 |
| TNRPDILDPALLRPG | ASSPSGDSDYT | 0.79964614 |
| LTTTTTS | ASSPSGDSDYT | 0.41499132 |
| PTGSGKTL | ASSPSGDSDYT | 0.8060048 |
| TGSGKTL | ASSPSGDSDYT | 0.7713782 |
| TIGHVDHGK | ASSPSGDSDYT | 0.5384103 |
| TIGHVDHGKTT | ASSPSGDSDYT | 0.43852854 |
| GHVDHGKTT | ASSPSGDSDYT | 0.7085744 |
| AQFEHTLL | ASSPSGDSDYT | 0.44621554 |
| IVNQDEL | ASSPSGDSDYT | 0.5123372 |
| TNGSQFF | ASSPSGDSDYT | 0.68107635 |
| TGKGIVE | ASSPSGDSDYT | 0.20292908 |
| LISNASDA | ASSPSGDSDYT | 0.75036424 |
| AELRTLGELTKIAKAHGAQ | ASSPSGDSDYT | 0.48877236 |
| APNAVAAAEPSW | ASSPSGDSDYT | 0.42258012 |
| APQVAGAATAVL | ASSPSGDSDYT | 0.2231987 |
| ARRDPAAAAAILPTDA | ASSPSGDSDYT | 0.15311122 |
| DAAARDVLAEAGLAEYF | ASSPSGDSDYT | 0.3258298 |
| DDQIVLSAKANLAKVEQ | ASSPSGDSDYT | 0.20744097 |
| DELLPDTGEDSDEDGH | ASSPSGDSDYT | 0.71606994 |
| ELPEAPANRATVPQD | ASSPSGDSDYT | 0.48475713 |
| FGQSKIERVTGIG | ASSPSGDSDYT | 0.22316714 |
| GAGGTGSTAGGAGGAGGAGGLY | ASSPSGDSDYT | 0.7048801 |
| GHGGTGAAGGNGGTGGAGGSGI | ASSPSGDSDYT | 0.59470075 |
| GNGGNGGAGGEGGPGGAGGAGA | ASSPSGDSDYT | 0.16921742 |
| GSTAGNGGNGGNATNGTIAGQP | ASSPSGDSDYT | 0.31389907 |
| GTGGDGGNGGNGGTGG | ASSPSGDSDYT | 0.21807921 |
| GVRRPAGIPADAPV | ASSPSGDSDYT | 0.37573296 |
| GYRQVTDAHLLTLARR | ASSPSGDSDYT | 0.15511839 |
| IAKMGQDGHDRG | ASSPSGDSDYT | 0.47976935 |
| KAEPSIEIQYT | ASSPSGDSDYT | 0.26259762 |
| KPVRIRMAADEDTVAVT | ASSPSGDSDYT | 0.13600077 |
| MPPRIAGMRLLVIKPEP | ASSPSGDSDYT | 0.3552605 |
| MKFYAVALFRSRGVPPTR | ASSPSGDSDYT | 0.32146114 |
| NAGGFGILGAVAHSP | ASSPSGDSDYT | 0.3194958 |
| NEGREITLAMAEM | ASSPSGDSDYT | 0.7120161 |
| NELVEWSHPLGHRWRWE | ASSPSGDSDYT | 0.21728587 |
| NLGAGGTGGNGGNPG | ASSPSGDSDYT | 0.20747264 |
| NTEWPKWETVMAADPVD | ASSPSGDSDYT | 0.6187458 |
| PASAAPAETQP | ASSPSGDSDYT | 0.6267869 |
| QQSSDGKNGVRAAEITGE | ASSPSGDSDYT | 0.1954635 |
| RAPDAVTTSWWKE | ASSPSGDSDYT | 0.5498985 |
| SGFGNVGAHTSGWFNQ | ASSPSGDSDYT | 0.6523632 |

| | | |
|---|---|---|
| SGVAGAVGQAASVGGLKV | ASSPSGDSDYT | 0.34412348 |
| SNGGAGTGSAGGNGGT | ASSPSGDSDYT | 0.6530031 |
| TGGAGGEGGAAG | ASSPSGDSDYT | 0.64162445 |
| TGGDGGAAGAGGAPGHG | ASSPSGDSDYT | 0.1776298 |
| TMPPLLAAIANAGRHGV | ASSPSGDSDYT | 0.6695135 |
| TSSTAGNSGGA | ASSPSGDSDYT | 0.779353 |
| VPATVPVGILC | ASSPSGDSDYT | 0.3197303 |
| VQRPDSGGVVLTGRIS | ASSPSGDSDYT | 0.38899845 |
| WTPNPVPATKGVPAGLVQAT | ASSPSGDSDYT | 0.18601526 |
| YALKRALAKFSDRFDVVII | ASSPSGDSDYT | 0.2121603 |
| YLIGSVIV | ASSPSGDSDYT | 0.16065176 |
| YGQLVAVID | ASSPSGDSDYT | 0.59366107 |
| VTVGGMTLPA | ASSPSGDSDYT | 0.566566 |
| VLALPAV | ASSPSGDSDYT | 0.35745478 |
| VIPPTLNYET | ASSPSGDSDYT | 0.2115378 |
| VGMGGMHPSSG | ASSPSGDSDYT | 0.17804407 |
| VGLAGGAATA | ASSPSGDSDYT | 0.34851047 |
| VGGAGGAGGSGP | ASSPSGDSDYT | 0.24988651 |
| VFAAFMV | ASSPSGDSDYT | 0.16609667 |
| TLYDKLR | ASSPSGDSDYT | 0.18766259 |
| TLAVFTVL | ASSPSGDSDYT | 0.46298862 |
| TGSVQPARV | ASSPSGDSDYT | 0.5285578 |
| TGGGAGSGGT | ASSPSGDSDYT | 0.21029237 |
| SHLETSL | ASSPSGDSDYT | 0.77748644 |
| SFTTGMT | ASSPSGDSDYT | 0.58848536 |
| QVEVAKRMGVRQPTVS | ASSPSGDSDYT | 0.62640315 |
| QGSGGPALAM | ASSPSGDSDYT | 0.35574245 |
| MLAPSNP | ASSPSGDSDYT | 0.20619984 |
| LVNNAAI | ASSPSGDSDYT | 0.784803 |
| LTGEINTLA | ASSPSGDSDYT | 0.30905342 |
| LSIFGPV | ASSPSGDSDYT | 0.3466422 |
| LLALLANL | ASSPSGDSDYT | 0.32795182 |
| LISNASDALDKLR | ASSPSGDSDYT | 0.29310626 |
| LINEAVP | ASSPSGDSDYT | 0.26966083 |
| LGYAFSGL | ASSPSGDSDYT | 0.6176275 |
| LDFVAVAL | ASSPSGDSDYT | 0.5086957 |
| LAQPGAE | ASSPSGDSDYT | 0.23361431 |
| ISNLGAVTGF | ASSPSGDSDYT | 0.19358155 |
| HQEDPGRI | ASSPSGDSDYT | 0.31084043 |
| GTPGEPTP | ASSPSGDSDYT | 0.4377787 |
| GFFNSGS | ASSPSGDSDYT | 0.6079575 |
| GAGGGGLTAGAN | ASSPSGDSDYT | 0.38791412 |
| FVDALVNM | ASSPSGDSDYT | 0.38049102 |
| FLTVAPEL | ASSPSGDSDYT | 0.6418884 |
| FETLTAL | ASSPSGDSDYT | 0.52068114 |
| DVINPIEL | ASSPSGDSDYT | 0.34725535 |
| APVSGAPV | ASSPSGDSDYT | 0.3107812 |
| ALVGGLAW | ASSPSGDSDYT | 0.3150918 |
| ALAAAAATAI | ASSPSGDSDYT | 0.5474393 |
| AFGLGLLL | ASSPSGDSDYT | 0.56989 |
| AFFAAASL | ASSPSGDSDYT | 0.50198317 |

| | | |
|---|---|---|
| AAAALPAFSPP | ASSPSGDSDYT | 0.5557011 |
| VVLAPAL | ASSPSGDSDYT | 0.35928926 |
| TIGHVDHGKTTL | ASSPSGDSDYT | 0.29935223 |
| LISNASDALDK | ASSPSGDSDYT | 0.16617149 |
| LISNASDAL | ASSPSGDSDYT | 0.19762036 |
| ISNASDALDK | ASSPSGDSDYT | 0.35237777 |
| NQAETLVYQTE | ASSPSGDSDYT | 0.50172055 |
| TGGTGGTAGAG | ASSPSGDSDYT | 0.49676716 |
| TGSTAGGAGGA | ASSPSGDSDYT | 0.35055375 |
| DPLAVPVAAGPV | ASSPSGDSDYT | 0.26056296 |
| GGTGGTPGNGGN | ASSPSGDSDYT | 0.32489127 |
| PAGLAGVDPATL | ASSPSGDSDYT | 0.6961169 |
| GNNLVVEAPHAGASV | ASSPSGDSDYT | 0.17451207 |
| KVTRIMSASTLEQPA | ASSPSGDSDYT | 0.26485506 |
| LGFALPIALPRRTVV | ASSPSGDSDYT | 0.7885388 |
| TRLGNLTARKLVVLV | ASSPSGDSDYT | 0.6383147 |
| WSQQRRLIYFPSAGP | ASSPSGDSDYT | 0.28432986 |
| GPNGNPDPMAAAVDIR | ASSPSGDSDYT | 0.36952007 |
| TQRCVSVADLRAAASR | ASSPSGDSDYT | 0.787758 |
| QFVLDVAEHPVEVLTETV | ASSPSGDSDYT | 0.31417763 |
| GGAGGTGGAGGNGGTQAGDG | ASSPSGDSDYT | 0.6842632 |
| PLLATLRAIAVDVDAKPAQV | ASSPSGDSDYT | 0.15549141 |
| APKFRTGGYMIDFWGVGYQVAK | ASSPSGDSDYT | 0.3218912 |
| GAGGAGGAGGAAGGPGGTGGTG | ASSPSGDSDYT | 0.20654501 |
| IGALGLLATDVAPAGSVAFGVA | ASSPSGDSDYT | 0.14714618 |
| AGGAGGIGGNGGAGGAGGVLMG | ASSPSGDSDYT | 0.55289817 |
| ADELIGLL | ASSPSGDSDYT | 0.252387 |
| ADRALKA | ASSPSGDSDYT | 0.66375494 |
| AGAVGQVGGAGGA | ASSPSGDSDYT | 0.28274977 |
| AGGGGGLGGAP | ASSPSGDSDYT | 0.5818784 |
| AGGLLYGNGGAG | ASSPSGDSDYT | 0.5474475 |
| AGLESIGL | ASSPSGDSDYT | 0.24734318 |
| AGLKTGII | ASSPSGDSDYT | 0.21766646 |
| ASLAALLL | ASSPSGDSDYT | 0.41792178 |
| ASVINISSI | ASSPSGDSDYT | 0.39984414 |
| ATPPGPA | ASSPSGDSDYT | 0.3915839 |
| ATVASALV | ASSPSGDSDYT | 0.15503243 |
| ATVASLIL | ASSPSGDSDYT | 0.41676843 |
| AVNGPVLIRH | ASSPSGDSDYT | 0.29543668 |
| DGAAGAVGI | ASSPSGDSDYT | 0.17130044 |
| DPLLHGI | ASSPSGDSDYT | 0.13729674 |
| ELGAALGAL | ASSPSGDSDYT | 0.20302328 |
| ELGRTQAV | ASSPSGDSDYT | 0.31975484 |
| FVAGGYLS | ASSPSGDSDYT | 0.6468017 |
| GASGGAGGAGGW | ASSPSGDSDYT | 0.6661864 |
| GGAGAGGDGGAGGI | ASSPSGDSDYT | 0.32024333 |
| GTGHNGSQPGMGGNG | ASSPSGDSDYT | 0.5952215 |
| GYAAASASAS | ASSPSGDSDYT | 0.3093711 |
| HQVDVAAVQQG | ASSPSGDSDYT | 0.124147095 |
| IGAAIAHT | ASSPSGDSDYT | 0.2593388 |
| IGMIEVAQL | ASSPSGDSDYT | 0.17440423 |

| | | |
|---|---|---|
| IRLAPEHTSAMD | ASSPSGDSDYT | 0.55018926 |
| IYNVSVL | ASSPSGDSDYT | 0.19667634 |
| KQGFLGM | ASSPSGDSDYT | 0.38016787 |
| LESVITQL | ASSPSGDSDYT | 0.34488022 |
| LGLGALVVV | ASSPSGDSDYT | 0.726372 |
| LIGNGGNGGSGGI | ASSPSGDSDYT | 0.37823942 |
| LNAINAPTL | ASSPSGDSDYT | 0.21821217 |
| LQLLNTGV | ASSPSGDSDYT | 0.13396353 |
| LQTSIEF | ASSPSGDSDYT | 0.39820153 |
| MSFAVSEL | ASSPSGDSDYT | 0.22494958 |
| NTGALGL | ASSPSGDSDYT | 0.20448123 |
| PPGAGGAGGAGTT | ASSPSGDSDYT | 0.18485732 |
| QLVGLGI | ASSPSGDSDYT | 0.27653566 |
| RFLEQVHQL | ASSPSGDSDYT | 0.17317493 |
| RGVGYVLR | ASSPSGDSDYT | 0.529953 |
| SATGVFAGVI | ASSPSGDSDYT | 0.26998675 |
| SLPAITLP | ASSPSGDSDYT | 0.41171318 |
| STALAFTV | ASSPSGDSDYT | 0.71110296 |
| SVLTVGLGK | ASSPSGDSDYT | 0.70349556 |
| TALAGAGSGI | ASSPSGDSDYT | 0.32168418 |
| TGLPFTGI | ASSPSGDSDYT | 0.2377295 |
| VAEHVTV | ASSPSGDSDYT | 0.20835139 |
| VDIVLNSL | ASSPSGDSDYT | 0.2123376 |
| VLTGGGAVL | ASSPSGDSDYT | 0.2325348 |
| VNVGLAYGI | ASSPSGDSDYT | 0.65538776 |
| VSSAAVGAV | ASSPSGDSDYT | 0.35711488 |
| VTSVYEAGV | ASSPSGDSDYT | 0.22062704 |
| WVALDRGA | ASSPSGDSDYT | 0.48012567 |
| YLVGSVVI | ASSPSGDSDYT | 0.71679044 |
| GPPGTGKTLL | ASSPSGDSDYT | 0.7439201 |
| GVGMTKF | ASSPSGDSDYT | 0.19068967 |
| IGIGGSDL | ASSPSGDSDYT | 0.26202068 |
| IGTLLAEL | ASSPSGDSDYT | 0.3292888 |
| IYTSGTT | ASSPSGDSDYT | 0.42592293 |
| LSFPAGV | ASSPSGDSDYT | 0.27650917 |
| LYTSGTT | ASSPSGDSDYT | 0.21496767 |
| VGAFPFGF | ASSPSGDSDYT | 0.48226997 |
| VGAGTVEF | ASSPSGDSDYT | 0.8153213 |
| VGKSTLLL | ASSPSGDSDYT | 0.28000617 |
| AALAHAGGGIV | ASSPSGDSDYT | 0.15907463 |
| AALVAVSLL | ASSPSGDSDYT | 0.2741898 |
| AAVSRAVAAL | ASSPSGDSDYT | 0.22888413 |
| AGGSSGIGGF | ASSPSGDSDYT | 0.23839323 |
| AGLTVANANG | ASSPSGDSDYT | 0.3369756 |
| AGLVGAVL | ASSPSGDSDYT | 0.1982708 |
| ALSQAFTV | ASSPSGDSDYT | 0.36644953 |
| ALTFSVSL | ASSPSGDSDYT | 0.7147364 |
| ALTGAALVAL | ASSPSGDSDYT | 0.25541514 |
| ASFSIGAGVL | ASSPSGDSDYT | 0.57410926 |
| DGSVYKI | ASSPSGDSDYT | 0.7571812 |
| EQLAEIV | ASSPSGDSDYT | 0.58204657 |

| | | |
|---|---|---|
| FAVSELTVM | ASSPSGDSDYT | 0.77523774 |
| GAAPLAAGL | ASSPSGDSDYT | 0.67407596 |
| GADGGDGGSGGAGG | ASSPSGDSDYT | 0.24174961 |
| GFGLDTGGAGG | ASSPSGDSDYT | 0.3082185 |
| GLASGGLL | ASSPSGDSDYT | 0.566653 |
| GRIAGAAVLL | ASSPSGDSDYT | 0.27185082 |
| GSAGSLNTSF | ASSPSGDSDYT | 0.33991426 |
| GTGQIGI | ASSPSGDSDYT | 0.27511075 |
| GTLIALIL | ASSPSGDSDYT | 0.7073927 |
| IAATMGFVVG | ASSPSGDSDYT | 0.32523108 |
| IFAQALL | ASSPSGDSDYT | 0.71751595 |
| IGAAVALGF | ASSPSGDSDYT | 0.1588046 |
| IGLTPLAL | ASSPSGDSDYT | 0.20434107 |
| ILGSNTSTL | ASSPSGDSDYT | 0.7451943 |
| IVFSLTVL | ASSPSGDSDYT | 0.130375 |
| LASKATGF | ASSPSGDSDYT | 0.15018749 |
| LASVALIL | ASSPSGDSDYT | 0.3148281 |
| LFAVGLTL | ASSPSGDSDYT | 0.74624026 |
| LFNGGIGTYI | ASSPSGDSDYT | 0.13264306 |
| LTLGAGII | ASSPSGDSDYT | 0.21673934 |
| LVAAALGDI | ASSPSGDSDYT | 0.19226196 |
| NTAFVQLGI | ASSPSGDSDYT | 0.39680666 |
| QLFSGAEL | ASSPSGDSDYT | 0.34730968 |
| SAATFITL | ASSPSGDSDYT | 0.14431514 |
| SNYGLANLGV | ASSPSGDSDYT | 0.5925263 |
| SYGGAYAVM | ASSPSGDSDYT | 0.1932547 |
| TTGVGGLTLG | ASSPSGDSDYT | 0.31130448 |
| TTQTIYI | ASSPSGDSDYT | 0.56120783 |
| VAAGAGLGAVI | ASSPSGDSDYT | 0.7750051 |
| VAAVVTLL | ASSPSGDSDYT | 0.18522194 |
| VGFLGGLI | ASSPSGDSDYT | 0.51051426 |
| VGVIPILL | ASSPSGDSDYT | 0.27483284 |
| VIFASSLI | ASSPSGDSDYT | 0.80230975 |
| VKGGDGGAAAG | ASSPSGDSDYT | 0.23875974 |
| VTKTFTNI | ASSPSGDSDYT | 0.17956688 |
| VVAAAILL | ASSPSGDSDYT | 0.6395769 |
| YAALDVEL | ASSPSGDSDYT | 0.6030327 |
| LASVALLL | ASSPSGDSDYT | 0.28393686 |
| LDAVHRV | ASSPSGDSDYT | 0.7132902 |
| QGLSSSIF | ASSPSGDSDYT | 0.3060626 |
| RGTFANIRL | ASSPSGDSDYT | 0.33210942 |
| RQLWWGHRI | ASSPSGDSDYT | 0.7128424 |
| YGPPGTGKTLL | ASSPSGDSDYT | 0.47598836 |
| AALVGAPANVA | ASSPSGDSDYT | 0.21470937 |
| AATFRRTE | ASSPSGDSDYT | 0.5010184 |
| AGALGGGI | ASSPSGDSDYT | 0.35557684 |
| AGGAGGAGGIGAG | ASSPSGDSDYT | 0.18317318 |
| AGSGALGL | ASSPSGDSDYT | 0.727384 |
| AQIADQI | ASSPSGDSDYT | 0.32882452 |
| ASAVAVTYL | ASSPSGDSDYT | 0.6586676 |
| ASTVLGGTGG | ASSPSGDSDYT | 0.41090626 |

| | | |
|---|---|---|
| AVGGEPAK | ASSPSGDSDYT | 0.7443003 |
| CEQGTFGGTC | ASSPSGDSDYT | 0.1569848 |
| DEGKAPAG | ASSPSGDSDYT | 0.38631612 |
| DTVVFPAL | ASSPSGDSDYT | 0.3192451 |
| FEGITMAG | ASSPSGDSDYT | 0.5067836 |
| FPLVVQD | ASSPSGDSDYT | 0.32292116 |
| GAFTTPQ | ASSPSGDSDYT | 0.28474724 |
| GFGGFGTAGGVGGA | ASSPSGDSDYT | 0.34508628 |
| GFLAQFL | ASSPSGDSDYT | 0.7047286 |
| GGAGGAGGNAGVA | ASSPSGDSDYT | 0.29077938 |
| GGVGTAGAGGAGGAGG | ASSPSGDSDYT | 0.28944814 |
| GNGGGGDAGNAGSGGN | ASSPSGDSDYT | 0.79260063 |
| GTGGAGGAGNDAGS | ASSPSGDSDYT | 0.18630171 |
| GTTNTGWFNS | ASSPSGDSDYT | 0.32178655 |
| IFMGPLFTL | ASSPSGDSDYT | 0.43361282 |
| ILAHAIEAI | ASSPSGDSDYT | 0.47662508 |
| IRSAIREAL | ASSPSGDSDYT | 0.41614068 |
| ITFAGGG | ASSPSGDSDYT | 0.6170765 |
| ITGMSGHQEV | ASSPSGDSDYT | 0.169641 |
| IVAGLACTL | ASSPSGDSDYT | 0.6527733 |
| IVGGGLAGA | ASSPSGDSDYT | 0.26012582 |
| IVTGVLNI | ASSPSGDSDYT | 0.4860283 |
| LAQQGCI | ASSPSGDSDYT | 0.18643992 |
| LCLHVEN | ASSPSGDSDYT | 0.47986308 |
| LDAISDGS | ASSPSGDSDYT | 0.691854 |
| LGGFAAML | ASSPSGDSDYT | 0.6607133 |
| LGGKNPMI | ASSPSGDSDYT | 0.15623747 |
| LGPKGVAAIL | ASSPSGDSDYT | 0.61174166 |
| LMVGVSI | ASSPSGDSDYT | 0.64339924 |
| LVGAAFVL | ASSPSGDSDYT | 0.8238162 |
| LVLAVAPL | ASSPSGDSDYT | 0.28063017 |
| MAAIAVIDAA | ASSPSGDSDYT | 0.51456493 |
| NNGINAELQ | ASSPSGDSDYT | 0.27612242 |
| RGEPLVLVHG | ASSPSGDSDYT | 0.34688506 |
| RGGAGGAAGLI | ASSPSGDSDYT | 0.19873707 |
| RSYRMHPAV | ASSPSGDSDYT | 0.2020736 |
| SAAHLLAGQLA | ASSPSGDSDYT | 0.37302 |
| SAAVGSY | ASSPSGDSDYT | 0.2592434 |
| SPHAVLHRI | ASSPSGDSDYT | 0.7557572 |
| SQIAKVL | ASSPSGDSDYT | 0.5635215 |
| TALGHLVP | ASSPSGDSDYT | 0.5621542 |
| TAPTTSGAA | ASSPSGDSDYT | 0.7110486 |
| TDGDDGNGGSPGT | ASSPSGDSDYT | 0.7489612 |
| TDGLGEN | ASSPSGDSDYT | 0.55551195 |
| TGGPPLPGQA | ASSPSGDSDYT | 0.22435102 |
| TIPAITI | ASSPSGDSDYT | 0.26376364 |
| TIPGFTI | ASSPSGDSDYT | 0.20458026 |
| TSIASGALA | ASSPSGDSDYT | 0.33741266 |
| TVTLPGAAI | ASSPSGDSDYT | 0.26617643 |
| VASGLSAALP | ASSPSGDSDYT | 0.467485 |
| VGAGVIGI | ASSPSGDSDYT | 0.13016738 |

| | | |
|---|---|---|
| VGPKRFGV | ASSPSGDSDYT | 0.7170775 |
| VNVAFPD | ASSPSGDSDYT | 0.7161125 |
| VSMTNSGVS | ASSPSGDSDYT | 0.2141828 |
| VTDVLGDLSV | ASSPSGDSDYT | 0.48154914 |
| VVVVGAGFAGL | ASSPSGDSDYT | 0.66065294 |
| VYTVGTVV | ASSPSGDSDYT | 0.26983845 |
| WTTFSGP | ASSPSGDSDYT | 0.5743185 |
| YDELVEAF | ASSPSGDSDYT | 0.25931555 |
| YGTGGAGGNGG | ASSPSGDSDYT | 0.6443605 |
| YTNVMAV | ASSPSGDSDYT | 0.65638024 |
| FLEQVHQL | ASSPSGDSDYT | 0.39228502 |
| GLSLLAVAV | ASSPSGDSDYT | 0.19130099 |
| LEQVHQL | ASSPSGDSDYT | 0.3841267 |
| LNAAGAI | ASSPSGDSDYT | 0.15185486 |
| PAVHASL | ASSPSGDSDYT | 0.21132456 |
| TELEDDA | ASSPSGDSDYT | 0.13698635 |
| TPPGPPN | ASSPSGDSDYT | 0.40130877 |
| TPPPPPPP | ASSPSGDSDYT | 0.20219594 |
| TQSPALI | ASSQGGDGEQY | 0.4286706 |
| EFAAAFPQ | ASSQGGDGEQY | 0.3568216 |
| LLSGNLDVL | ASSQGGDGEQY | 0.3315783 |
| GPANLGNYNI | ASSQGGDGEQY | 0.18986861 |
| TLGPTLP | ASSQGGDGEQY | 0.5647119 |
| VYAYSSCCK | ASSQGGDGEQY | 0.46851158 |
| APAPAPAPNI | ASSQGGDGEQY | 0.17730911 |
| PGGPVPIP | ASSQGGDGEQY | 0.67459375 |
| FLTVALEP | ASSQGGDGEQY | 0.2800581 |
| SQYFGANL | ASSQGGDGEQY | 0.314206 |
| GGVKPAA | ASSQGGDGEQY | 0.5129125 |
| GGTGGAGGAGGW | ASSQGGDGEQY | 0.3620873 |
| GVGGVAGLGGAGG | ASSQGGDGEQY | 0.16145132 |
| VLLVLATTI | ASSQGGDGEQY | 0.36011636 |
| IGIFAAML | ASSQGGDGEQY | 0.3370365 |
| AVGDLID | ASSQGGDGEQY | 0.22554019 |
| NISPLPL | ASSQGGDGEQY | 0.13353597 |
| VTAVFVAI | ASSQGGDGEQY | 0.41313303 |
| SGDDNDAAL | ASSQGGDGEQY | 0.18248396 |
| AVAALAHS | ASSQGGDGEQY | 0.20464373 |
| GGVTVMATP | ASSQGGDGEQY | 0.31184 |
| AGLAANVRD | ASSQGGDGEQY | 0.716185 |
| TNRPDLIDPALLRPG | ASSQGGDGEQY | 0.23223615 |
| LANLGVTAL | ASSQGGDGEQY | 0.3404593 |
| TGERIEL | ASSQGGDGEQY | 0.4998884 |
| NGEELAA | ASSQGGDGEQY | 0.3257007 |
| VLSLVWG | ASSQGGDGEQY | 0.7254199 |
| IGYALWL | ASSQGGDGEQY | 0.35741892 |
| LSLLLAPL | ASSQGGDGEQY | 0.26312515 |
| TLDEALK | ASSQGGDGEQY | 0.35413292 |
| TDGAGTAEGG | ASSQGGDGEQY | 0.32515013 |
| NAPGGAGGNGG | ASSQGGDGEQY | 0.1868597 |
| VSVFMGMGGNF | ASSQGGDGEQY | 0.17306602 |

| | | |
|---|---|---|
| LASYLAAL | ASSQGGDGEQY | 0.29670927 |
| LVGGTPAL | ASSQGGDGEQY | 0.1644479 |
| VRNADVL | ASSQGGDGEQY | 0.30367434 |
| TCSSPFLVSAM | ASSQGGDGEQY | 0.5635333 |
| VHITDAE | ASSQGGDGEQY | 0.23934063 |
| EYAAPSF | ASSQGGDGEQY | 0.16279438 |
| VVDAFHGGL | ASSQGGDGEQY | 0.2567212 |
| ASVVDPQ | ASSQGGDGEQY | 0.17808603 |
| SYNEFGGLS | ASSQGGDGEQY | 0.55019474 |
| DPDGSVY | ASSQGGDGEQY | 0.29075912 |
| MVSTIHATE | ASSQGGDGEQY | 0.8016752 |
| LAAGVADPV | ASSQGGDGEQY | 0.6141838 |
| TKEQIAA | ASSQGGDGEQY | 0.26485044 |
| SLASPAAL | ASSQGGDGEQY | 0.66974187 |
| IGLVFYGV | ASSQGGDGEQY | 0.280319 |
| FLLDASNL | ASSQGGDGEQY | 0.37392086 |
| YQVGNFP | ASSQGGDGEQY | 0.2633662 |
| ESLLSQST | ASSQGGDGEQY | 0.34515667 |
| LFTVGFAAVG | ASSQGGDGEQY | 0.24096504 |
| ATPPTPT | ASSQGGDGEQY | 0.22845206 |
| TDLATLG | ASSQGGDGEQY | 0.16172776 |
| SAGGTGR | ASSQGGDGEQY | 0.35183507 |
| GFSNSGS | ASSQGGDGEQY | 0.23393081 |
| GGNGGSGGTSV | ASSQGGDGEQY | 0.18318544 |
| GTSVATGGAGN | ASSQGGDGEQY | 0.23590991 |
| NGGNGGSGGTS | ASSQGGDGEQY | 0.77618116 |
| AVVKFPH | ASSQGGDGEQY | 0.7370248 |
| TPPPPPP | ASSQGGDGEQY | 0.25755507 |
| AAPAAAAAPA | ASSQGGDGEQY | 0.2214101 |
| IALNEDL | ASSQGGDGEQY | 0.73022634 |
| IVNVSSM | ASSQGGDGEQY | 0.19151324 |
| TGIPPAPR | ASSQGGDGEQY | 0.5357584 |
| FDLGGGTF | ASSQGGDGEQY | 0.4822342 |
| FDLGGGTFDVS | ASSQGGDGEQY | 0.35653162 |
| NEPTAAA | ASSQGGDGEQY | 0.22395377 |
| NEPTAAALA | ASSQGGDGEQY | 0.48834234 |
| GSIINISSI | ASSQGGDGEQY | 0.27137363 |
| VAPGNAGTA | ASSQGGDGEQY | 0.4120987 |
| TGGGASLE | ASSQGGDGEQY | 0.1342059 |
| VAVYNHYKRI | ASSQGGDGEQY | 0.45178586 |
| TDPAPATT | ASSQGGDGEQY | 0.27692664 |
| TNRPDILDPALLRPG | ASSQGGDGEQY | 0.20456038 |
| LTTTTTS | ASSQGGDGEQY | 0.14825599 |
| PTGSGKTL | ASSQGGDGEQY | 0.30702496 |
| TGSGKTL | ASSQGGDGEQY | 0.62910557 |
| TIGHVDHGK | ASSQGGDGEQY | 0.7005732 |
| TIGHVDHGKTT | ASSQGGDGEQY | 0.19620484 |
| GHVDHGKTT | ASSQGGDGEQY | 0.5098847 |
| AQFEHTLL | ASSQGGDGEQY | 0.65263265 |
| IVNQDEL | ASSQGGDGEQY | 0.430229 |
| TNGSQFF | ASSQGGDGEQY | 0.14537358 |

| | | |
|---|---|---|
| TGKGIVE | ASSQGGDGEQY | 0.46905267 |
| LISNASDA | ASSQGGDGEQY | 0.39217216 |
| AELRTLGELTKIAKAHGAQ | ASSQGGDGEQY | 0.5706442 |
| APNAVAAAEPSW | ASSQGGDGEQY | 0.427041 |
| APQVAGAATAVL | ASSQGGDGEQY | 0.17573215 |
| ARRDPAAAAAILPTDA | ASSQGGDGEQY | 0.42268997 |
| DAAARDVLAEAGLAEYF | ASSQGGDGEQY | 0.6753906 |
| DDQIVLSAKANLAKVEQ | ASSQGGDGEQY | 0.18352287 |
| DELLPDTGEDSDEDGH | ASSQGGDGEQY | 0.23052487 |
| ELPEAPANRATVPQD | ASSQGGDGEQY | 0.23844604 |
| FGQSKIERVTGIG | ASSQGGDGEQY | 0.17157993 |
| GAGGTGSTAGGAGGAGGAGGLY | ASSQGGDGEQY | 0.33554864 |
| GHGGTGAAGGNGGTGGAGGSGI | ASSQGGDGEQY | 0.37534785 |
| GNGGNGGAGGEGGPGGAGGAGA | ASSQGGDGEQY | 0.2537985 |
| GSTAGNGGNGGNATNGTIAGQP | ASSQGGDGEQY | 0.35083333 |
| GTGGDGGNGGNGGTGG | ASSQGGDGEQY | 0.22039106 |
| GVRRPAGIPADAPV | ASSQGGDGEQY | 0.25882745 |
| GYRQVTDAHLLTLARR | ASSQGGDGEQY | 0.28024462 |
| IAKMGQDGHDRG | ASSQGGDGEQY | 0.3228323 |
| KAEPSIEIQYT | ASSQGGDGEQY | 0.5477497 |
| KPVRIRMAADEDTVAVT | ASSQGGDGEQY | 0.39767092 |
| MPPRIAGMRLLVIKPEP | ASSQGGDGEQY | 0.43138713 |
| MKFYAVALFRSRGVPPTR | ASSQGGDGEQY | 0.78703976 |
| NAGGFGILGAVAHSP | ASSQGGDGEQY | 0.17891179 |
| NEGREITLAMAEM | ASSQGGDGEQY | 0.20607138 |
| NELVEWSHPLGHRWRWE | ASSQGGDGEQY | 0.55597407 |
| NLGAGGTGGNGGNPG | ASSQGGDGEQY | 0.2423746 |
| NTEWPKWETVMAADPVD | ASSQGGDGEQY | 0.69252086 |
| PASAAPAETQP | ASSQGGDGEQY | 0.33820683 |
| QQSSDGKNGVRAAEITGE | ASSQGGDGEQY | 0.5400412 |
| RAPDAVTTSWWKE | ASSQGGDGEQY | 0.34190857 |
| SGFGNVGAHTSGWFNQ | ASSQGGDGEQY | 0.77111965 |
| SGVAGAVGQAASVGGLKV | ASSQGGDGEQY | 0.6357191 |
| SNGGAGTGSAGGNGGT | ASSQGGDGEQY | 0.34441143 |
| TGGAGGEGGAAG | ASSQGGDGEQY | 0.6720039 |
| TGGDGGAAGAGGAPGHG | ASSQGGDGEQY | 0.18504846 |
| TMPPLLAAIANAGRHGV | ASSQGGDGEQY | 0.47956175 |
| TSSTAGNSGGA | ASSQGGDGEQY | 0.5557096 |
| VPATVPVGILC | ASSQGGDGEQY | 0.17920849 |
| VQRPDSGGVVLTGRIS | ASSQGGDGEQY | 0.32829326 |
| WTPNPVPATKGVPAGLVQAT | ASSQGGDGEQY | 0.592492 |
| YALKRALAKFSDRFDVVII | ASSQGGDGEQY | 0.75310975 |
| YLIGSVIV | ASSQGGDGEQY | 0.17629634 |
| YGQLVAVID | ASSQGGDGEQY | 0.30220577 |
| VTVGGMTLPA | ASSQGGDGEQY | 0.18865679 |
| VLALPAV | ASSQGGDGEQY | 0.19805758 |
| VIPPTLNYET | ASSQGGDGEQY | 0.17015845 |
| VGMGGMHPSSG | ASSQGGDGEQY | 0.48089886 |
| VGLAGGAATA | ASSQGGDGEQY | 0.29000968 |
| VGGAGGAGGSGP | ASSQGGDGEQY | 0.7493519 |
| VFAAFMV | ASSQGGDGEQY | 0.3048349 |

| | | |
|---|---|---|
| TLYDKLR | ASSQGGDGEQY | 0.5028451 |
| TLAVFTVL | ASSQGGDGEQY | 0.23320818 |
| TGSVQPARV | ASSQGGDGEQY | 0.293618 |
| TGGGAGSGGT | ASSQGGDGEQY | 0.17006797 |
| SHLETSL | ASSQGGDGEQY | 0.17236838 |
| SFTTGMT | ASSQGGDGEQY | 0.1913466 |
| QVEVAKRMGVRQPTVS | ASSQGGDGEQY | 0.31094822 |
| QGSGGPALAM | ASSQGGDGEQY | 0.15164247 |
| MLAPSNP | ASSQGGDGEQY | 0.7665939 |
| LVNNAAI | ASSQGGDGEQY | 0.44363883 |
| LTGEINTLA | ASSQGGDGEQY | 0.43695834 |
| LSIFGPV | ASSQGGDGEQY | 0.2427158 |
| LLALLANL | ASSQGGDGEQY | 0.23501001 |
| LISNASDALDKLR | ASSQGGDGEQY | 0.25545013 |
| LINEAVP | ASSQGGDGEQY | 0.61563134 |
| LGYAFSGL | ASSQGGDGEQY | 0.16638193 |
| LDFVAVAL | ASSQGGDGEQY | 0.16537513 |
| LAQPGAE | ASSQGGDGEQY | 0.24812554 |
| ISNLGAVTGF | ASSQGGDGEQY | 0.21612278 |
| HQEDPGRI | ASSQGGDGEQY | 0.66947305 |
| GTPGEPTP | ASSQGGDGEQY | 0.67220026 |
| GFFNSGS | ASSQGGDGEQY | 0.4702887 |
| GAGGGGLTAGAN | ASSQGGDGEQY | 0.71652687 |
| FVDALVNM | ASSQGGDGEQY | 0.13890064 |
| FLTVAPEL | ASSQGGDGEQY | 0.2574968 |
| FETLTAL | ASSQGGDGEQY | 0.7233943 |
| DVINPIEL | ASSQGGDGEQY | 0.34173045 |
| APVSGAPV | ASSQGGDGEQY | 0.30848294 |
| ALVGGLAW | ASSQGGDGEQY | 0.6953175 |
| ALAAAAATAI | ASSQGGDGEQY | 0.21978952 |
| AFGLGLLL | ASSQGGDGEQY | 0.2224337 |
| AFFAAASL | ASSQGGDGEQY | 0.42294854 |
| AAAALPAFSPP | ASSQGGDGEQY | 0.3407786 |
| VVLAPAL | ASSQGGDGEQY | 0.30969816 |
| TIGHVDHGKTTL | ASSQGGDGEQY | 0.16265179 |
| LISNASDALDK | ASSQGGDGEQY | 0.47999245 |
| LISNASDAL | ASSQGGDGEQY | 0.81915724 |
| ISNASDALDK | ASSQGGDGEQY | 0.34869647 |
| NQAETLVYQTE | ASSQGGDGEQY | 0.5140463 |
| TGGTGGTAGAG | ASSQGGDGEQY | 0.46669906 |
| TGSTAGGAGGA | ASSQGGDGEQY | 0.53342086 |
| DPLAVPVAAGPV | ASSQGGDGEQY | 0.2267224 |
| GGTGGTPGNGGN | ASSQGGDGEQY | 0.20409706 |
| PAGLAGVDPATL | ASSQGGDGEQY | 0.49809846 |
| GNNLVVEAPHAGASV | ASSQGGDGEQY | 0.23507628 |
| KVTRIMSASTLEQPA | ASSQGGDGEQY | 0.4109919 |
| LGFALPIALPRRTVV | ASSQGGDGEQY | 0.35737517 |
| TRLGNLTARKLVVLV | ASSQGGDGEQY | 0.82634896 |
| WSQQRRLIYFPSAGP | ASSQGGDGEQY | 0.568082 |
| GPNGNPDPMAAAVDIR | ASSQGGDGEQY | 0.45559967 |
| TQRCVSVADLRAAASR | ASSQGGDGEQY | 0.20146097 |

| | | |
|---|---|---|
| QFVLDVAEHPVEVLTETV | ASSQGGDGEQY | 0.33181533 |
| GGAGGTGGAGGNGGTQAGDG | ASSQGGDGEQY | 0.57869816 |
| PLLATLRAIAVDVDAKPAQV | ASSQGGDGEQY | 0.17287806 |
| APKFRTGGYMIDFWGVGYQVAK | ASSQGGDGEQY | 0.6736314 |
| GAGGAGGAGGAAGGPGGTGGTG | ASSQGGDGEQY | 0.23730654 |
| IGALGLLATDVAPAGSVAFGVA | ASSQGGDGEQY | 0.43886757 |
| AGGAGGIGGNGGAGGAGGVLMG | ASSQGGDGEQY | 0.5441564 |
| ADELIGLL | ASSQGGDGEQY | 0.7887907 |
| ADRALKA | ASSQGGDGEQY | 0.3715424 |
| AGAVGQVGGAGGA | ASSQGGDGEQY | 0.5374067 |
| AGGGGGLGGAP | ASSQGGDGEQY | 0.1998449 |
| AGGLLYGNGGAG | ASSQGGDGEQY | 0.5070466 |
| AGLESIGL | ASSQGGDGEQY | 0.71081483 |
| AGLKTGII | ASSQGGDGEQY | 0.28556955 |
| ASLAALLL | ASSQGGDGEQY | 0.75232065 |
| ASVINISSI | ASSQGGDGEQY | 0.40497762 |
| ATPPGPA | ASSQGGDGEQY | 0.15931654 |
| ATVASALV | ASSQGGDGEQY | 0.25696245 |
| ATVASLIL | ASSQGGDGEQY | 0.6290945 |
| AVNGPVLIRH | ASSQGGDGEQY | 0.7391447 |
| DGAAGAVGI | ASSQGGDGEQY | 0.5312863 |
| DPLLHGI | ASSQGGDGEQY | 0.32718435 |
| ELGAALGAL | ASSQGGDGEQY | 0.20307961 |
| ELGRTQAV | ASSQGGDGEQY | 0.7282813 |
| FVAGGYLS | ASSQGGDGEQY | 0.7727772 |
| GASGGAGGAGGW | ASSQGGDGEQY | 0.53840387 |
| GGAGAGGDGGAGGI | ASSQGGDGEQY | 0.24130999 |
| GTGHNGSQPGMGGNG | ASSQGGDGEQY | 0.21159421 |
| GYAAASASAS | ASSQGGDGEQY | 0.41234714 |
| HQVDVAAVQQG | ASSQGGDGEQY | 0.72537225 |
| IGAAIAHT | ASSQGGDGEQY | 0.20790437 |
| IGMIEVAQL | ASSQGGDGEQY | 0.4947247 |
| IRLAPEHTSAMD | ASSQGGDGEQY | 0.44207346 |
| IYNVSVL | ASSQGGDGEQY | 0.14740227 |
| KQGFLGM | ASSQGGDGEQY | 0.46212757 |
| LESVITQL | ASSQGGDGEQY | 0.7235757 |
| LGLGALVVV | ASSQGGDGEQY | 0.4316002 |
| LIGNGGNGGSGGI | ASSQGGDGEQY | 0.20765789 |
| LNAINAPTL | ASSQGGDGEQY | 0.7222798 |
| LQLLNTGV | ASSQGGDGEQY | 0.8047928 |
| LQTSIEF | ASSQGGDGEQY | 0.7249371 |
| MSFAVSEL | ASSQGGDGEQY | 0.1688842 |
| NTGALGL | ASSQGGDGEQY | 0.14516804 |
| PPGAGGAGGAGTT | ASSQGGDGEQY | 0.37375885 |
| QLVGLGI | ASSQGGDGEQY | 0.72879136 |
| RFLEQVHQL | ASSQGGDGEQY | 0.26607502 |
| RGVGYVLR | ASSQGGDGEQY | 0.23842312 |
| SATGVFAGVI | ASSQGGDGEQY | 0.6338184 |
| SLPAITLP | ASSQGGDGEQY | 0.6975099 |
| STALAFTV | ASSQGGDGEQY | 0.17778137 |
| SVLTVGLGK | ASSQGGDGEQY | 0.17617308 |

| | | |
|---|---|---|
| TALAGAGSGI | ASSQGGDGEQY | 0.6349752 |
| TGLPFTGI | ASSQGGDGEQY | 0.6378404 |
| VAEHVTV | ASSQGGDGEQY | 0.22351325 |
| VDIVLNSL | ASSQGGDGEQY | 0.31794658 |
| VLTGGGAVL | ASSQGGDGEQY | 0.3604585 |
| VNVGLAYGI | ASSQGGDGEQY | 0.33522648 |
| VSSAAVGAV | ASSQGGDGEQY | 0.45055693 |
| VTSVYEAGV | ASSQGGDGEQY | 0.17740378 |
| WVALDRGA | ASSQGGDGEQY | 0.67042017 |
| YLVGSVVI | ASSQGGDGEQY | 0.20007294 |
| GPPGTGKTLL | ASSQGGDGEQY | 0.5049931 |
| GVGMTKF | ASSQGGDGEQY | 0.35798252 |
| IGIGGSDL | ASSQGGDGEQY | 0.29297864 |
| IGTLLAEL | ASSQGGDGEQY | 0.33712086 |
| IYTSGTT | ASSQGGDGEQY | 0.2583085 |
| LSFPAGV | ASSQGGDGEQY | 0.17144707 |
| LYTSGTT | ASSQGGDGEQY | 0.17388624 |
| VGAFPFGF | ASSQGGDGEQY | 0.5535111 |
| VGAGTVEF | ASSQGGDGEQY | 0.1833287 |
| VGKSTLLL | ASSQGGDGEQY | 0.5632582 |
| AALAHAGGGIV | ASSQGGDGEQY | 0.3997736 |
| AALVAVSLL | ASSQGGDGEQY | 0.1964103 |
| AAVSRAVAAL | ASSQGGDGEQY | 0.46628797 |
| AGGSSGIGGF | ASSQGGDGEQY | 0.18104368 |
| AGLTVANANG | ASSQGGDGEQY | 0.20813353 |
| AGLVGAVL | ASSQGGDGEQY | 0.33433387 |
| ALSQAFTV | ASSQGGDGEQY | 0.5875313 |
| ALTFSVSL | ASSQGGDGEQY | 0.21679728 |
| ALTGAALVAL | ASSQGGDGEQY | 0.52086127 |
| ASFSIGAGVL | ASSQGGDGEQY | 0.37280232 |
| DGSVYKI | ASSQGGDGEQY | 0.19546232 |
| EQLAEIV | ASSQGGDGEQY | 0.39631334 |
| FAVSELTVM | ASSQGGDGEQY | 0.70584476 |
| GAAPLAAGL | ASSQGGDGEQY | 0.42988056 |
| GADGGDGGSGGAGG | ASSQGGDGEQY | 0.17122488 |
| GFGLDTGGAGG | ASSQGGDGEQY | 0.4855777 |
| GLASGGLL | ASSQGGDGEQY | 0.22826976 |
| GRIAGAAVLL | ASSQGGDGEQY | 0.47207028 |
| GSAGSLNTSF | ASSQGGDGEQY | 0.31643 |
| GTGQIGI | ASSQGGDGEQY | 0.2640749 |
| GTLIALIL | ASSQGGDGEQY | 0.5184469 |
| IAATMGFVVG | ASSQGGDGEQY | 0.212991 |
| IFAQALL | ASSQGGDGEQY | 0.20450255 |
| IGAAVALGF | ASSQGGDGEQY | 0.28191015 |
| IGLTPLAL | ASSQGGDGEQY | 0.35826814 |
| ILGSNTSTL | ASSQGGDGEQY | 0.596264 |
| IVFSLTVL | ASSQGGDGEQY | 0.21274388 |
| LASKATGF | ASSQGGDGEQY | 0.31648368 |
| LASVALIL | ASSQGGDGEQY | 0.2269354 |
| LFAVGLTL | ASSQGGDGEQY | 0.19268198 |
| LFNGGIGTYI | ASSQGGDGEQY | 0.43512684 |

| | | |
|---|---|---|
| LTLGAGII | ASSQGGDGEQY | 0.32953626 |
| LVAAALGDI | ASSQGGDGEQY | 0.21668157 |
| NTAFVQLGI | ASSQGGDGEQY | 0.3237841 |
| QLFSGAEL | ASSQGGDGEQY | 0.2373507 |
| SAATFITL | ASSQGGDGEQY | 0.32600403 |
| SNYGLANLGV | ASSQGGDGEQY | 0.75463474 |
| SYGGAYAVM | ASSQGGDGEQY | 0.17382064 |
| TTGVGGLTLG | ASSQGGDGEQY | 0.45786595 |
| TTQTIYI | ASSQGGDGEQY | 0.39631763 |
| VAAGAGLGAVI | ASSQGGDGEQY | 0.45385477 |
| VAAVVTLL | ASSQGGDGEQY | 0.2939512 |
| VGFLGGLI | ASSQGGDGEQY | 0.19509804 |
| VGVIPILL | ASSQGGDGEQY | 0.6939349 |
| VIFASSLI | ASSQGGDGEQY | 0.4527911 |
| VKGGDGGAAAG | ASSQGGDGEQY | 0.41070965 |
| VTKTFTNI | ASSQGGDGEQY | 0.52025956 |
| VVAAAILL | ASSQGGDGEQY | 0.71326005 |
| YAALDVEL | ASSQGGDGEQY | 0.6290543 |
| LASVALLL | ASSQGGDGEQY | 0.27088195 |
| LDAVHRV | ASSQGGDGEQY | 0.17594057 |
| QGLSSSIF | ASSQGGDGEQY | 0.31027287 |
| RGTFANIRL | ASSQGGDGEQY | 0.38832492 |
| RQLWWGHRI | ASSQGGDGEQY | 0.28818902 |
| YGPPGTGKTLL | ASSQGGDGEQY | 0.17048559 |
| AALVGAPANVA | ASSQGGDGEQY | 0.7152577 |
| AATFRRTE | ASSQGGDGEQY | 0.81096315 |
| AGALGGGI | ASSQGGDGEQY | 0.21315639 |
| AGGAGGAGGIGAG | ASSQGGDGEQY | 0.7591814 |
| AGSGALGL | ASSQGGDGEQY | 0.21960187 |
| AQIADQI | ASSQGGDGEQY | 0.41559303 |
| ASAVAVTYL | ASSQGGDGEQY | 0.7498911 |
| ASTVLGGTGG | ASSQGGDGEQY | 0.21675798 |
| AVGGEPAK | ASSQGGDGEQY | 0.4027754 |
| CEQGTFGGTC | ASSQGGDGEQY | 0.17809017 |
| DEGKAPAG | ASSQGGDGEQY | 0.21818906 |
| DTVVFPAL | ASSQGGDGEQY | 0.70716655 |
| FEGITMAG | ASSQGGDGEQY | 0.24538596 |
| FPLVVQD | ASSQGGDGEQY | 0.28780645 |
| GAFTTPQ | ASSQGGDGEQY | 0.33699274 |
| GFGGFGTAGGVGGA | ASSQGGDGEQY | 0.20885266 |
| GFLAQFL | ASSQGGDGEQY | 0.31246385 |
| GGAGGAGGNAGVA | ASSQGGDGEQY | 0.58176666 |
| GGVGTAGAGGAGGAGG | ASSQGGDGEQY | 0.26328325 |
| GNGGGGDAGNAGSGGN | ASSQGGDGEQY | 0.7527064 |
| GTGGAGGAGNDAGS | ASSQGGDGEQY | 0.19415836 |
| GTTNTGWFNS | ASSQGGDGEQY | 0.79156905 |
| IFMGPLFTL | ASSQGGDGEQY | 0.7861843 |
| ILAHAIEAI | ASSQGGDGEQY | 0.47655076 |
| IRSAIREAL | ASSQGGDGEQY | 0.27660823 |
| ITFAGGG | ASSQGGDGEQY | 0.7413603 |
| ITGMSGHQEV | ASSQGGDGEQY | 0.33082846 |

| | | |
|---|---|---|
| IVAGLACTL | ASSQGGDGEQY | 0.3229918 |
| IVGGGLAGA | ASSQGGDGEQY | 0.5770943 |
| IVTGVLNI | ASSQGGDGEQY | 0.7034999 |
| LAQQGCI | ASSQGGDGEQY | 0.88511527 |
| LCLHVEN | ASSQGGDGEQY | 0.8566814 |
| LDAISDGS | ASSQGGDGEQY | 0.85474914 |
| LGGFAAML | ASSQGGDGEQY | 0.79230595 |
| LGGKNPMI | ASSQGGDGEQY | 0.36145788 |
| LGPKGVAAIL | ASSQGGDGEQY | 0.8004773 |
| LMVGVSI | ASSQGGDGEQY | 0.7441621 |
| LVGAAFVL | ASSQGGDGEQY | 0.83658063 |
| LVLAVAPL | ASSQGGDGEQY | 0.8517531 |
| MAAIAVIDAA | ASSQGGDGEQY | 0.77107185 |
| NNGINAELQ | ASSQGGDGEQY | 0.881557 |
| RGEPLVLVHG | ASSQGGDGEQY | 0.8836454 |
| RGGAGGAAGLI | ASSQGGDGEQY | 0.57377255 |
| RSYRMHPAV | ASSQGGDGEQY | 0.8154861 |
| SAAHLLAGQLA | ASSQGGDGEQY | 0.74148095 |
| SAAVGSY | ASSQGGDGEQY | 0.8347541 |
| SPHAVLHRI | ASSQGGDGEQY | 0.33565253 |
| SQIAKVL | ASSQGGDGEQY | 0.8711524 |
| TALGHLVP | ASSQGGDGEQY | 0.7514695 |
| TAPTTSGAA | ASSQGGDGEQY | 0.15108591 |
| TDGDDGNGGSPGT | ASSQGGDGEQY | 0.81215507 |
| TDGLGEN | ASSQGGDGEQY | 0.8152903 |
| TGGPPLPGQA | ASSQGGDGEQY | 0.82095957 |
| TIPAITI | ASSQGGDGEQY | 0.8728596 |
| TIPGFTI | ASSQGGDGEQY | 0.5329125 |
| TSIASGALA | ASSQGGDGEQY | 0.70832086 |
| TVTLPGAAI | ASSQGGDGEQY | 0.8689234 |
| VASGLSAALP | ASSQGGDGEQY | 0.88490313 |
| VGAGVIGI | ASSQGGDGEQY | 0.6338284 |
| VGPKRFGV | ASSQGGDGEQY | 0.85144 |
| VNVAFPD | ASSQGGDGEQY | 0.8871519 |
| VSMTNSGVS | ASSQGGDGEQY | 0.853058 |
| VTDVLGDLSV | ASSQGGDGEQY | 0.873322 |
| VVVVGAGFAGL | ASSQGGDGEQY | 0.88327205 |
| VYTVGTVV | ASSQGGDGEQY | 0.6606412 |
| WTTFSGP | ASSQGGDGEQY | 0.87801516 |
| YDELVEAF | ASSQGGDGEQY | 0.87604034 |
| YGTGGAGGNGG | ASSQGGDGEQY | 0.82394314 |
| YTNVMAV | ASSQGGDGEQY | 0.8416765 |
| FLEQVHQL | ASSQGGDGEQY | 0.6630311 |
| GLSLLAVAV | ASSQGGDGEQY | 0.77885395 |
| LEQVHQL | ASSQGGDGEQY | 0.43912208 |
| LNAAGAI | ASSQGGDGEQY | 0.8481166 |
| PAVHASL | ASSQGGDGEQY | 0.8770629 |
| TELEDDA | ASSQGGDGEQY | 0.81323576 |
| TPPGPPN | ASSQGGDGEQY | 0.6918428 |
| TPPPPPPP | ASSQGGDGEQY | 0.7058307 |
| TQSPALI | ASSQDGDWGINQDTQY | 0.80955327 |

| | | |
|---|---|---|
| EFAAAFPQ | ASSQDGDWGINQDTQY | 0.81235015 |
| LLSGNLDVL | ASSQDGDWGINQDTQY | 0.3791032 |
| GPANLGNYNI | ASSQDGDWGINQDTQY | 0.75814164 |
| TLGPTLP | ASSQDGDWGINQDTQY | 0.880914 |
| VYAYSSCCK | ASSQDGDWGINQDTQY | 0.8855218 |
| APAPAPAPNI | ASSQDGDWGINQDTQY | 0.513981 |
| PGGPVPIP | ASSQDGDWGINQDTQY | 0.8789468 |
| FLTVALEP | ASSQDGDWGINQDTQY | 0.8504684 |
| SQYFGANL | ASSQDGDWGINQDTQY | 0.8838347 |
| GGVKPAA | ASSQDGDWGINQDTQY | 0.82813907 |
| GGTGGAGGAGGW | ASSQDGDWGINQDTQY | 0.74363315 |
| GVGGVAGLGGAGG | ASSQDGDWGINQDTQY | 0.86787385 |
| VLLVLATTI | ASSQDGDWGINQDTQY | 0.8110971 |
| IGIFAAML | ASSQDGDWGINQDTQY | 0.6520802 |
| AVGDLID | ASSQDGDWGINQDTQY | 0.87425476 |
| NISPLPL | ASSQDGDWGINQDTQY | 0.8137305 |
| VTAVFVAI | ASSQDGDWGINQDTQY | 0.7561898 |
| SGDDNDAAL | ASSQDGDWGINQDTQY | 0.77917254 |
| AVAALAHS | ASSQDGDWGINQDTQY | 0.87314296 |
| GGVTVMATP | ASSQDGDWGINQDTQY | 0.69004625 |
| AGLAANVRD | ASSQDGDWGINQDTQY | 0.6665907 |
| TNRPDLIDPALLRPG | ASSQDGDWGINQDTQY | 0.72521013 |
| LANLGVTAL | ASSQDGDWGINQDTQY | 0.8402087 |
| TGERIEL | ASSQDGDWGINQDTQY | 0.88014174 |
| NGEELAA | ASSQDGDWGINQDTQY | 0.84987736 |
| VLSLVWG | ASSQDGDWGINQDTQY | 0.7907749 |
| IGYALWL | ASSQDGDWGINQDTQY | 0.86644524 |
| LSLLLAPL | ASSQDGDWGINQDTQY | 0.8669834 |
| TLDEALK | ASSQDGDWGINQDTQY | 0.8481388 |
| TDGAGTAEGG | ASSQDGDWGINQDTQY | 0.8647835 |
| NAPGGAGGNGG | ASSQDGDWGINQDTQY | 0.7720518 |
| VSVFMGMGGNF | ASSQDGDWGINQDTQY | 0.87245774 |
| LASYLAAL | ASSQDGDWGINQDTQY | 0.8715134 |
| LVGGTPAL | ASSQDGDWGINQDTQY | 0.8713224 |
| VRNADVL | ASSQDGDWGINQDTQY | 0.8686429 |
| TCSSPFLVSAM | ASSQDGDWGINQDTQY | 0.8552207 |
| VHITDAE | ASSQDGDWGINQDTQY | 0.85042536 |
| EYAAPSF | ASSQDGDWGINQDTQY | 0.38255006 |
| VVDAFHGGL | ASSQDGDWGINQDTQY | 0.75322723 |
| ASVVDPQ | ASSQDGDWGINQDTQY | 0.8500904 |
| SYNEFGGLS | ASSQDGDWGINQDTQY | 0.84767354 |
| DPDGSVY | ASSQDGDWGINQDTQY | 0.85607666 |
| MVSTIHATE | ASSQDGDWGINQDTQY | 0.8538933 |
| LAAGVADPV | ASSQDGDWGINQDTQY | 0.77617 |
| TKEQIAA | ASSQDGDWGINQDTQY | 0.8715369 |
| SLASPAAL | ASSQDGDWGINQDTQY | 0.8763831 |
| IGLVFYGV | ASSQDGDWGINQDTQY | 0.55664307 |
| FLLDASNL | ASSQDGDWGINQDTQY | 0.80241716 |
| YQVGNFP | ASSQDGDWGINQDTQY | 0.887258 |
| ESLLSQST | ASSQDGDWGINQDTQY | 0.83640444 |
| LFTVGFAAVG | ASSQDGDWGINQDTQY | 0.802738 |

| | | |
|---|---|---|
| ATPPTPT | ASSQDGDWGINQDTQY | 0.79443204 |
| TDLATLG | ASSQDGDWGINQDTQY | 0.87455684 |
| SAGGTGR | ASSQDGDWGINQDTQY | 0.75122106 |
| GFSNSGS | ASSQDGDWGINQDTQY | 0.8842254 |
| GGNGGSGGTSV | ASSQDGDWGINQDTQY | 0.32909507 |
| GTSVATGGAGN | ASSQDGDWGINQDTQY | 0.8077306 |
| NGGNGGSGGTS | ASSQDGDWGINQDTQY | 0.7071303 |
| AVVKFPH | ASSQDGDWGINQDTQY | 0.41349733 |
| TPPPPPP | ASSQDGDWGINQDTQY | 0.8758316 |
| AAPAAAAAPA | ASSQDGDWGINQDTQY | 0.76728576 |
| IALNEDL | ASSQDGDWGINQDTQY | 0.87717265 |
| IVNVSSM | ASSQDGDWGINQDTQY | 0.8085091 |
| TGIPPAPR | ASSQDGDWGINQDTQY | 0.8700569 |
| FDLGGGTF | ASSQDGDWGINQDTQY | 0.8357926 |
| FDLGGGTFDVS | ASSQDGDWGINQDTQY | 0.83521724 |
| NEPTAAA | ASSQDGDWGINQDTQY | 0.68947875 |
| NEPTAAALA | ASSQDGDWGINQDTQY | 0.86873 |
| GSIINISSI | ASSQDGDWGINQDTQY | 0.8715571 |
| VAPGNAGTA | ASSQDGDWGINQDTQY | 0.8617371 |
| TGGGASLE | ASSQDGDWGINQDTQY | 0.47878087 |
| VAVYNHYKRI | ASSQDGDWGINQDTQY | 0.7791127 |
| TDPAPATT | ASSQDGDWGINQDTQY | 0.85052896 |
| TNRPDILDPALLRPG | ASSQDGDWGINQDTQY | 0.22066705 |
| LTTTTTS | ASSQDGDWGINQDTQY | 0.20140933 |
| PTGSGKTL | ASSQDGDWGINQDTQY | 0.76683354 |
| TGSGKTL | ASSQDGDWGINQDTQY | 0.72385937 |
| TIGHVDHGK | ASSQDGDWGINQDTQY | 0.8792046 |
| TIGHVDHGKTT | ASSQDGDWGINQDTQY | 0.87013626 |
| GHVDHGKTT | ASSQDGDWGINQDTQY | 0.8298764 |
| AQFEHTLL | ASSQDGDWGINQDTQY | 0.8649863 |
| IVNQDEL | ASSQDGDWGINQDTQY | 0.84473246 |
| TNGSQFF | ASSQDGDWGINQDTQY | 0.10544912 |
| TGKGIVE | ASSQDGDWGINQDTQY | 0.79500437 |
| LISNASDA | ASSQDGDWGINQDTQY | 0.61106205 |
| AELRTLGELTKIAKAHGAQ | ASSQDGDWGINQDTQY | 0.8801218 |
| APNAVAAAEPSW | ASSQDGDWGINQDTQY | 0.6919153 |
| APQVAGAATAVL | ASSQDGDWGINQDTQY | 0.7231693 |
| ARRDPAAAAAILPTDA | ASSQDGDWGINQDTQY | 0.8418216 |
| DAAARDVLAEAGLAEYF | ASSQDGDWGINQDTQY | 0.828761 |
| DDQIVLSAKANLAKVEQ | ASSQDGDWGINQDTQY | 0.8313271 |
| DELLPDTGEDSDEDGH | ASSQDGDWGINQDTQY | 0.70202124 |
| ELPEAPANRATVPQD | ASSQDGDWGINQDTQY | 0.17517218 |
| FGQSKIERVTGIG | ASSQDGDWGINQDTQY | 0.8753923 |
| GAGGTGSTAGGAGGAGGAGGLY | ASSQDGDWGINQDTQY | 0.48312765 |
| GHGGTGAAGGNGGTGGAGGSGI | ASSQDGDWGINQDTQY | 0.7073495 |
| GNGGNGGAGGEGGPGGAGGAGA | ASSQDGDWGINQDTQY | 0.2676167 |
| GSTAGNGGNGGNATNGTIAGQP | ASSQDGDWGINQDTQY | 0.15383956 |
| GTGGDGGNGGNGGTGG | ASSQDGDWGINQDTQY | 0.4856998 |
| GVRRPAGIPADAPV | ASSQDGDWGINQDTQY | 0.86720943 |
| GYRQVTDAHLLTLARR | ASSQDGDWGINQDTQY | 0.8038236 |
| IAKMGQDGHDRG | ASSQDGDWGINQDTQY | 0.78446484 |

| | | |
|---|---|---|
| KAEPSIEIQYT | ASSQDGDWGINQDTQY | 0.8842275 |
| KPVRIRMAADEDTVAVT | ASSQDGDWGINQDTQY | 0.8359413 |
| MPPRIAGMRLLVIKPEP | ASSQDGDWGINQDTQY | 0.81034374 |
| MKFYAVALFRSRGVPPTR | ASSQDGDWGINQDTQY | 0.77267355 |
| NAGGFGILGAVAHSP | ASSQDGDWGINQDTQY | 0.71487105 |
| NEGREITLAMAEM | ASSQDGDWGINQDTQY | 0.7124734 |
| NELVEWSHPLGHRWRWE | ASSQDGDWGINQDTQY | 0.72258276 |
| NLGAGGTGGNGGNPG | ASSQDGDWGINQDTQY | 0.8872735 |
| NTEWPKWETVMAADPVD | ASSQDGDWGINQDTQY | 0.7990689 |
| PASAAPAETQP | ASSQDGDWGINQDTQY | 0.5197127 |
| QQSSDGKNGVRAAEITGE | ASSQDGDWGINQDTQY | 0.84905946 |
| RAPDAVTTSWWKE | ASSQDGDWGINQDTQY | 0.8855145 |
| SGFGNVGAHTSGWFNQ | ASSQDGDWGINQDTQY | 0.8902352 |
| SGVAGAVGQAASVGGLKV | ASSQDGDWGINQDTQY | 0.72528595 |
| SNGGAGTGSAGGNGGT | ASSQDGDWGINQDTQY | 0.82422704 |
| TGGAGGEGGAAG | ASSQDGDWGINQDTQY | 0.80409217 |
| TGGDGGAAGAGGAPGHG | ASSQDGDWGINQDTQY | 0.8299633 |
| TMPPLLAAIANAGRHGV | ASSQDGDWGINQDTQY | 0.7706851 |
| TSSTAGNSGGA | ASSQDGDWGINQDTQY | 0.56983685 |
| VPATVPVGILC | ASSQDGDWGINQDTQY | 0.8696612 |
| VQRPDSGGVVLTGRIS | ASSQDGDWGINQDTQY | 0.8891972 |
| WTPNPVPATKGVPAGLVQAT | ASSQDGDWGINQDTQY | 0.7908137 |
| YALKRALAKFSDRFDVVII | ASSQDGDWGINQDTQY | 0.46114805 |
| YLIGSVIV | ASSQDGDWGINQDTQY | 0.8247779 |
| YGQLVAVID | ASSQDGDWGINQDTQY | 0.72953856 |
| VTVGGMTLPA | ASSQDGDWGINQDTQY | 0.8052686 |
| VLALPAV | ASSQDGDWGINQDTQY | 0.8358357 |
| VIPPTLNYET | ASSQDGDWGINQDTQY | 0.5637917 |
| VGMGGMHPSSG | ASSQDGDWGINQDTQY | 0.78941 |
| VGLAGGAATA | ASSQDGDWGINQDTQY | 0.89200807 |
| VGGAGGAGGSGP | ASSQDGDWGINQDTQY | 0.8194574 |
| VFAAFMV | ASSQDGDWGINQDTQY | 0.8518581 |
| TLYDKLR | ASSQDGDWGINQDTQY | 0.8782192 |
| TLAVFTVL | ASSQDGDWGINQDTQY | 0.8586318 |
| TGSVQPARV | ASSQDGDWGINQDTQY | 0.88318676 |
| TGGGAGSGGT | ASSQDGDWGINQDTQY | 0.86174965 |
| SHLETSL | ASSQDGDWGINQDTQY | 0.77810633 |
| SFTTGMT | ASSQDGDWGINQDTQY | 0.8192872 |
| QVEVAKRMGVRQPTVS | ASSQDGDWGINQDTQY | 0.82678485 |
| QGSGGPALAM | ASSQDGDWGINQDTQY | 0.84186137 |
| MLAPSNP | ASSQDGDWGINQDTQY | 0.8316542 |
| LVNNAAI | ASSQDGDWGINQDTQY | 0.72709537 |
| LTGEINTLA | ASSQDGDWGINQDTQY | 0.88319933 |
| LSIFGPV | ASSQDGDWGINQDTQY | 0.83662605 |
| LLALLANL | ASSQDGDWGINQDTQY | 0.8694091 |
| LISNASDALDKLR | ASSQDGDWGINQDTQY | 0.60005784 |
| LINEAVP | ASSQDGDWGINQDTQY | 0.68896496 |
| LGYAFSGL | ASSQDGDWGINQDTQY | 0.8719504 |
| LDFVAVAL | ASSQDGDWGINQDTQY | 0.16479328 |
| LAQPGAE | ASSQDGDWGINQDTQY | 0.2941206 |
| ISNLGAVTGF | ASSQDGDWGINQDTQY | 0.83492434 |

| | | |
|---|---|---|
| HQEDPGRI | ASSQDGDWGINQDTQY | 0.8821544 |
| GTPGEPTP | ASSQDGDWGINQDTQY | 0.15418355 |
| GFFNSGS | ASSQDGDWGINQDTQY | 0.70283765 |
| GAGGGGLTAGAN | ASSQDGDWGINQDTQY | 0.63977563 |
| FVDALVNM | ASSQDGDWGINQDTQY | 0.85639775 |
| FLTVAPEL | ASSQDGDWGINQDTQY | 0.8380161 |
| FETLTAL | ASSQDGDWGINQDTQY | 0.4736995 |
| DVINPIEL | ASSQDGDWGINQDTQY | 0.87278485 |
| APVSGAPV | ASSQDGDWGINQDTQY | 0.47281775 |
| ALVGGLAW | ASSQDGDWGINQDTQY | 0.6121783 |
| ALAAAAATAI | ASSQDGDWGINQDTQY | 0.8616922 |
| AFGLGLLL | ASSQDGDWGINQDTQY | 0.8758817 |
| AFFAAASL | ASSQDGDWGINQDTQY | 0.61741054 |
| AAAALPAFSPP | ASSQDGDWGINQDTQY | 0.869663 |
| VVLAPAL | ASSQDGDWGINQDTQY | 0.7804129 |
| TIGHVDHGKTTL | ASSQDGDWGINQDTQY | 0.8904233 |
| LISNASDALDK | ASSQDGDWGINQDTQY | 0.79632896 |
| LISNASDAL | ASSQDGDWGINQDTQY | 0.4670093 |
| ISNASDALDK | ASSQDGDWGINQDTQY | 0.8722273 |
| NQAETLVYQTE | ASSQDGDWGINQDTQY | 0.8566469 |
| TGGTGGTAGAG | ASSQDGDWGINQDTQY | 0.8297418 |
| TGSTAGGAGGA | ASSQDGDWGINQDTQY | 0.86108243 |
| DPLAVPVAAGPV | ASSQDGDWGINQDTQY | 0.7818942 |
| GGTGGTPGNGGN | ASSQDGDWGINQDTQY | 0.82298267 |
| PAGLAGVDPATL | ASSQDGDWGINQDTQY | 0.8812443 |
| GNNLVVEAPHAGASV | ASSQDGDWGINQDTQY | 0.8678033 |
| KVTRIMSASTLEQPA | ASSQDGDWGINQDTQY | 0.64184296 |
| LGFALPIALPRRTVV | ASSQDGDWGINQDTQY | 0.85519516 |
| TRLGNLTARKLVVLV | ASSQDGDWGINQDTQY | 0.8326838 |
| WSQQRRLIYFPSAGP | ASSQDGDWGINQDTQY | 0.70785224 |
| GPNGNPDPMAAAVDIR | ASSQDGDWGINQDTQY | 0.88803434 |
| TQRCVSVADLRAAASR | ASSQDGDWGINQDTQY | 0.82763106 |
| QFVLDVAEHPVEVLTETV | ASSQDGDWGINQDTQY | 0.8742428 |
| GGAGGTGGAGGNGGTQAGDG | ASSQDGDWGINQDTQY | 0.8706535 |
| PLLATLRAIAVDVDAKPAQV | ASSQDGDWGINQDTQY | 0.8667581 |
| APKFRTGGYMIDFWGVGYQVAK | ASSQDGDWGINQDTQY | 0.4789614 |
| GAGGAGGAGGAAGGPGGTGGTG | ASSQDGDWGINQDTQY | 0.8448533 |
| IGALGLLATDVAPAGSVAFGVA | ASSQDGDWGINQDTQY | 0.8420504 |
| AGGAGGIGGNGGAGGAGGVLMG | ASSQDGDWGINQDTQY | 0.86884975 |
| ADELIGLL | ASSQDGDWGINQDTQY | 0.83070225 |
| ADRALKA | ASSQDGDWGINQDTQY | 0.64879626 |
| AGAVGQVGGAGGA | ASSQDGDWGINQDTQY | 0.8545213 |
| AGGGGGLGGAP | ASSQDGDWGINQDTQY | 0.8817767 |
| AGGLLYGNGGAG | ASSQDGDWGINQDTQY | 0.81718093 |
| AGLESIGL | ASSQDGDWGINQDTQY | 0.84578 |
| AGLKTGII | ASSQDGDWGINQDTQY | 0.8905196 |
| ASLAALLL | ASSQDGDWGINQDTQY | 0.70866704 |
| ASVINISSI | ASSQDGDWGINQDTQY | 0.82148623 |
| ATPPGPA | ASSQDGDWGINQDTQY | 0.86549497 |
| ATVASALV | ASSQDGDWGINQDTQY | 0.8703801 |
| ATVASLIL | ASSQDGDWGINQDTQY | 0.83027995 |

| | | |
|---|---|---|
| AVNGPVLIRH | ASSQDGDWGINQDTQY | 0.880992 |
| DGAAGAVGI | ASSQDGDWGINQDTQY | 0.69439185 |
| DPLLHGI | ASSQDGDWGINQDTQY | 0.8713512 |
| ELGAALGAL | ASSQDGDWGINQDTQY | 0.71688163 |
| ELGRTQAV | ASSQDGDWGINQDTQY | 0.8559174 |
| FVAGGYLS | ASSQDGDWGINQDTQY | 0.71020156 |
| GASGGAGGAGGW | ASSQDGDWGINQDTQY | 0.86586815 |
| GGAGAGGDGGAGGI | ASSQDGDWGINQDTQY | 0.8699907 |
| GTGHNGSQPGMGGNG | ASSQDGDWGINQDTQY | 0.8698438 |
| GYAAASASAS | ASSQDGDWGINQDTQY | 0.20593342 |
| HQVDVAAVQQG | ASSQDGDWGINQDTQY | 0.87323785 |
| IGAAIAHT | ASSQDGDWGINQDTQY | 0.87710625 |
| IGMIEVAQL | ASSQDGDWGINQDTQY | 0.890183 |
| IRLAPEHTSAMD | ASSQDGDWGINQDTQY | 0.82904845 |
| IYNVSVL | ASSQDGDWGINQDTQY | 0.86629796 |
| KQGFLGM | ASSQDGDWGINQDTQY | 0.80195385 |
| LESVITQL | ASSQDGDWGINQDTQY | 0.8185953 |
| LGLGALVVV | ASSQDGDWGINQDTQY | 0.7319001 |
| LIGNGGNGGSGGI | ASSQDGDWGINQDTQY | 0.8023809 |
| LNAINAPTL | ASSQDGDWGINQDTQY | 0.7898117 |
| LQLLNTGV | ASSQDGDWGINQDTQY | 0.7742224 |
| LQTSIEF | ASSQDGDWGINQDTQY | 0.8674979 |
| MSFAVSEL | ASSQDGDWGINQDTQY | 0.8688104 |
| NTGALGL | ASSQDGDWGINQDTQY | 0.874229 |
| PPGAGGAGGAGTT | ASSQDGDWGINQDTQY | 0.8841255 |
| QLVGLGI | ASSQDGDWGINQDTQY | 0.8783173 |
| RFLEQVHQL | ASSQDGDWGINQDTQY | 0.8758719 |
| RGVGYVLR | ASSQDGDWGINQDTQY | 0.795989 |
| SATGVFAGVI | ASSQDGDWGINQDTQY | 0.7980984 |
| SLPAITLP | ASSQDGDWGINQDTQY | 0.6565228 |
| STALAFTV | ASSQDGDWGINQDTQY | 0.7963381 |
| SVLTVGLGK | ASSQDGDWGINQDTQY | 0.7961955 |
| TALAGAGSGI | ASSQDGDWGINQDTQY | 0.85602105 |
| TGLPFTGI | ASSQDGDWGINQDTQY | 0.113447145 |
| VAEHVTV | ASSQDGDWGINQDTQY | 0.8104093 |
| VDIVLNSL | ASSQDGDWGINQDTQY | 0.88345754 |
| VLTGGGAVL | ASSQDGDWGINQDTQY | 0.74328303 |
| VNVGLAYGI | ASSQDGDWGINQDTQY | 0.86775327 |
| VSSAAVGAV | ASSQDGDWGINQDTQY | 0.81371117 |
| VTSVYEAGV | ASSQDGDWGINQDTQY | 0.7554089 |
| WVALDRGA | ASSQDGDWGINQDTQY | 0.8752923 |
| YLVGSVVI | ASSQDGDWGINQDTQY | 0.7134707 |
| GPPGTGKTLL | ASSQDGDWGINQDTQY | 0.870448 |
| GVGMTKF | ASSQDGDWGINQDTQY | 0.88557863 |
| IGIGGSDL | ASSQDGDWGINQDTQY | 0.8525569 |
| IGTLLAEL | ASSQDGDWGINQDTQY | 0.827274 |
| IYTSGTT | ASSQDGDWGINQDTQY | 0.82807803 |
| LSFPAGV | ASSQDGDWGINQDTQY | 0.39587826 |
| LYTSGTT | ASSQDGDWGINQDTQY | 0.8401949 |
| VGAFPFGF | ASSQDGDWGINQDTQY | 0.104414865 |
| VGAGTVEF | ASSQDGDWGINQDTQY | 0.8374938 |

| | | |
|---|---|---|
| VGKSTLLL | ASSQDGDWGINQDTQY | 0.65814054 |
| AALAHAGGGIV | ASSQDGDWGINQDTQY | 0.8578726 |
| AALVAVSLL | ASSQDGDWGINQDTQY | 0.8327124 |
| AAVSRAVAAL | ASSQDGDWGINQDTQY | 0.8482506 |
| AGGSSGIGGF | ASSQDGDWGINQDTQY | 0.7389767 |
| AGLTVANANG | ASSQDGDWGINQDTQY | 0.81067383 |
| AGLVGAVL | ASSQDGDWGINQDTQY | 0.76161695 |
| ALSQAFTV | ASSQDGDWGINQDTQY | 0.6200756 |
| ALTFSVSL | ASSQDGDWGINQDTQY | 0.84992194 |
| ALTGAALVAL | ASSQDGDWGINQDTQY | 0.79322684 |
| ASFSIGAGVL | ASSQDGDWGINQDTQY | 0.7907438 |
| DGSVYKI | ASSQDGDWGINQDTQY | 0.87964416 |
| EQLAEIV | ASSQDGDWGINQDTQY | 0.83625585 |
| FAVSELTVM | ASSQDGDWGINQDTQY | 0.3700841 |
| GAAPLAAGL | ASSQDGDWGINQDTQY | 0.8224108 |
| GADGGDGGSGGAGG | ASSQDGDWGINQDTQY | 0.8424574 |
| GFGLDTGGAGG | ASSQDGDWGINQDTQY | 0.8238462 |
| GLASGGLL | ASSQDGDWGINQDTQY | 0.10977137 |
| GRIAGAAVLL | ASSQDGDWGINQDTQY | 0.84652364 |
| GSAGSLNTSF | ASSQDGDWGINQDTQY | 0.7205517 |
| GTGQIGI | ASSQDGDWGINQDTQY | 0.73379076 |
| GTLIALIL | ASSQDGDWGINQDTQY | 0.7526029 |
| IAATMGFVVG | ASSQDGDWGINQDTQY | 0.87680566 |
| IFAQALL | ASSQDGDWGINQDTQY | 0.53662694 |
| IGAAVALGF | ASSQDGDWGINQDTQY | 0.8495088 |
| IGLTPLAL | ASSQDGDWGINQDTQY | 0.8644369 |
| ILGSNTSTL | ASSQDGDWGINQDTQY | 0.8594345 |
| IVFSLTVL | ASSQDGDWGINQDTQY | 0.88550836 |
| LASKATGF | ASSQDGDWGINQDTQY | 0.86253536 |
| LASVALIL | ASSQDGDWGINQDTQY | 0.7876923 |
| LFAVGLTL | ASSQDGDWGINQDTQY | 0.84968495 |
| LFNGGIGTYI | ASSQDGDWGINQDTQY | 0.81877875 |
| LTLGAGII | ASSQDGDWGINQDTQY | 0.42284465 |
| LVAAALGDI | ASSQDGDWGINQDTQY | 0.40035427 |
| NTAFVQLGI | ASSQDGDWGINQDTQY | 0.83778685 |
| QLFSGAEL | ASSQDGDWGINQDTQY | 0.33267695 |
| SAATFITL | ASSQDGDWGINQDTQY | 0.85165435 |
| SNYGLANLGV | ASSQDGDWGINQDTQY | 0.87869537 |
| SYGGAYAVM | ASSQDGDWGINQDTQY | 0.8176659 |
| TTGVGGLTLG | ASSQDGDWGINQDTQY | 0.48229998 |
| TTQTIYI | ASSQDGDWGINQDTQY | 0.7976022 |
| VAAGAGLGAVI | ASSQDGDWGINQDTQY | 0.75162905 |
| VAAVVTLL | ASSQDGDWGINQDTQY | 0.8701918 |
| VGFLGGLI | ASSQDGDWGINQDTQY | 0.8652985 |
| VGVIPILL | ASSQDGDWGINQDTQY | 0.8806946 |
| VIFASSLI | ASSQDGDWGINQDTQY | 0.8774674 |
| VKGGDGGAAAG | ASSQDGDWGINQDTQY | 0.80476534 |
| VTKTFTNI | ASSQDGDWGINQDTQY | 0.8703271 |
| VVAAAILL | ASSQDGDWGINQDTQY | 0.80785155 |
| YAALDVEL | ASSQDGDWGINQDTQY | 0.77068615 |
| LASVALLL | ASSQDGDWGINQDTQY | 0.7985304 |

| | | |
|---|---|---|
| LDAVHRV | ASSQDGDWGINQDTQY | 0.83043087 |
| QGLSSSIF | ASSQDGDWGINQDTQY | 0.8209303 |
| RGTFANIRL | ASSQDGDWGINQDTQY | 0.76295745 |
| RQLWWGHRI | ASSQDGDWGINQDTQY | 0.5042663 |
| YGPPGTGKTLL | ASSQDGDWGINQDTQY | 0.829771 |
| AALVGAPANVA | ASSQDGDWGINQDTQY | 0.85419166 |
| AATFRRTE | ASSQDGDWGINQDTQY | 0.52030337 |
| AGALGGGI | ASSQDGDWGINQDTQY | 0.777754 |
| AGGAGGAGGIGAG | ASSQDGDWGINQDTQY | 0.8894915 |
| AGSGALGL | ASSQDGDWGINQDTQY | 0.88815206 |
| AQIADQI | ASSQDGDWGINQDTQY | 0.75596094 |
| ASAVAVTYL | ASSQDGDWGINQDTQY | 0.74520063 |
| ASTVLGGTGG | ASSQDGDWGINQDTQY | 0.8184329 |
| AVGGEPAK | ASSQDGDWGINQDTQY | 0.5050269 |
| CEQGTFGGTC | ASSQDGDWGINQDTQY | 0.85737133 |
| DEGKAPAG | ASSQDGDWGINQDTQY | 0.82649577 |
| DTVVFPAL | ASSQDGDWGINQDTQY | 0.82071453 |
| FEGITMAG | ASSQDGDWGINQDTQY | 0.7307838 |
| FPLVVQD | ASSQDGDWGINQDTQY | 0.8367878 |
| GAFTTPQ | ASSQDGDWGINQDTQY | 0.79357195 |
| GFGGFGTAGGVGGA | ASSQDGDWGINQDTQY | 0.8624885 |
| GFLAQFL | ASSQDGDWGINQDTQY | 0.14281635 |
| GGAGGAGGNAGVA | ASSQDGDWGINQDTQY | 0.8407535 |
| GGVGTAGAGGAGGAGG | ASSQDGDWGINQDTQY | 0.71896076 |
| GNGGGGDAGNAGSGGN | ASSQDGDWGINQDTQY | 0.7436929 |
| GTGGAGGAGNDAGS | ASSQDGDWGINQDTQY | 0.40675184 |
| GTTNTGWFNS | ASSQDGDWGINQDTQY | 0.805676 |
| IFMGPLFTL | ASSQDGDWGINQDTQY | 0.8723587 |
| ILAHAIEAI | ASSQDGDWGINQDTQY | 0.8770267 |
| IRSAIREAL | ASSQDGDWGINQDTQY | 0.6643329 |
| ITFAGGG | ASSQDGDWGINQDTQY | 0.8711045 |
| ITGMSGHQEV | ASSQDGDWGINQDTQY | 0.8827745 |
| IVAGLACTL | ASSQDGDWGINQDTQY | 0.7858635 |
| IVGGGLAGA | ASSQDGDWGINQDTQY | 0.29024208 |
| IVTGVLNI | ASSQDGDWGINQDTQY | 0.8614126 |
| LAQQGCI | ASSQDGDWGINQDTQY | 0.757152 |
| LCLHVEN | ASSQDGDWGINQDTQY | 0.8644652 |
| LDAISDGS | ASSQDGDWGINQDTQY | 0.8701289 |
| LGGFAAML | ASSQDGDWGINQDTQY | 0.6769307 |
| LGGKNPMI | ASSQDGDWGINQDTQY | 0.8760723 |
| LGPKGVAAIL | ASSQDGDWGINQDTQY | 0.88235366 |
| LMVGVSI | ASSQDGDWGINQDTQY | 0.7500284 |
| LVGAAFVL | ASSQDGDWGINQDTQY | 0.6514615 |
| LVLAVAPL | ASSQDGDWGINQDTQY | 0.6378827 |
| MAAIAVIDAA | ASSQDGDWGINQDTQY | 0.3663226 |
| NNGINAELQ | ASSQDGDWGINQDTQY | 0.8519629 |
| RGEPLVLVHG | ASSQDGDWGINQDTQY | 0.7645235 |
| RGGAGGAAGLI | ASSQDGDWGINQDTQY | 0.7771898 |
| RSYRMHPAV | ASSQDGDWGINQDTQY | 0.85206753 |
| SAAHLLAGQLA | ASSQDGDWGINQDTQY | 0.7168951 |
| SAAVGSY | ASSQDGDWGINQDTQY | 0.80475986 |

| | | |
|---|---|---|
| SPHAVLHRI | ASSQDGDWGINQDTQY | 0.8047081 |
| SQIAKVL | ASSQDGDWGINQDTQY | 0.81560034 |
| TALGHLVP | ASSQDGDWGINQDTQY | 0.84693277 |
| TAPTTSGAA | ASSQDGDWGINQDTQY | 0.8224265 |
| TDGDDGNGGSPGT | ASSQDGDWGINQDTQY | 0.8307355 |
| TDGLGEN | ASSQDGDWGINQDTQY | 0.8842846 |
| TGGPPLPGQA | ASSQDGDWGINQDTQY | 0.60280466 |
| TIPAITI | ASSQDGDWGINQDTQY | 0.7123983 |
| TIPGFTI | ASSQDGDWGINQDTQY | 0.86603063 |
| TSIASGALA | ASSQDGDWGINQDTQY | 0.6330579 |
| TVTLPGAAI | ASSQDGDWGINQDTQY | 0.86584604 |
| VASGLSAALP | ASSQDGDWGINQDTQY | 0.83607745 |
| VGAGVIGI | ASSQDGDWGINQDTQY | 0.8593208 |
| VGPKRFGV | ASSQDGDWGINQDTQY | 0.8005353 |
| VNVAFPD | ASSQDGDWGINQDTQY | 0.8833363 |
| VSMTNSGVS | ASSQDGDWGINQDTQY | 0.8776574 |
| VTDVLGDLSV | ASSQDGDWGINQDTQY | 0.7856179 |
| VVVVGAGFAGL | ASSQDGDWGINQDTQY | 0.7908024 |
| VYTVGTVV | ASSQDGDWGINQDTQY | 0.56919503 |
| WTTFSGP | ASSQDGDWGINQDTQY | 0.8683729 |
| YDELVEAF | ASSQDGDWGINQDTQY | 0.8681632 |
| YGTGGAGGNGG | ASSQDGDWGINQDTQY | 0.70229244 |
| YTNVMAV | ASSQDGDWGINQDTQY | 0.82489735 |
| FLEQVHQL | ASSQDGDWGINQDTQY | 0.87757826 |
| GLSLLAVAV | ASSQDGDWGINQDTQY | 0.87827206 |
| LEQVHQL | ASSQDGDWGINQDTQY | 0.4063691 |
| LNAAGAI | ASSQDGDWGINQDTQY | 0.8060038 |
| PAVHASL | ASSQDGDWGINQDTQY | 0.41758993 |
| TELEDDA | ASSQDGDWGINQDTQY | 0.7857685 |
| TPPGPPN | ASSQDGDWGINQDTQY | 0.51078385 |
| TPPPPPPP | ASSQDGDWGINQDTQY | 0.85432434 |
| TQSPALI | ASSLSHSGNTLY | 0.7920785 |
| EFAAAFPQ | ASSLSHSGNTLY | 0.8585672 |
| LLSGNLDVL | ASSLSHSGNTLY | 0.85656476 |
| GPANLGNYNI | ASSLSHSGNTLY | 0.8863614 |
| TLGPTLP | ASSLSHSGNTLY | 0.777419 |
| VYAYSSCCK | ASSLSHSGNTLY | 0.8257685 |
| APAPAPAPNI | ASSLSHSGNTLY | 0.5042966 |
| PGGPVPIP | ASSLSHSGNTLY | 0.4760929 |
| FLTVALEP | ASSLSHSGNTLY | 0.68181276 |
| SQYFGANL | ASSLSHSGNTLY | 0.82847345 |
| GGVKPAA | ASSLSHSGNTLY | 0.22732177 |
| GGTGGAGGAGGW | ASSLSHSGNTLY | 0.8828535 |
| GVGGVAGLGGAGG | ASSLSHSGNTLY | 0.8522066 |
| VLLVLATTI | ASSLSHSGNTLY | 0.87613857 |
| IGIFAAML | ASSLSHSGNTLY | 0.71775043 |
| AVGDLID | ASSLSHSGNTLY | 0.7625497 |
| NISPLPL | ASSLSHSGNTLY | 0.7863068 |
| VTAVFVAI | ASSLSHSGNTLY | 0.87464523 |
| SGDDNDAAL | ASSLSHSGNTLY | 0.6998335 |
| AVAALAHS | ASSLSHSGNTLY | 0.39037243 |

| | | |
|---|---|---|
| GGVTVMATP | ASSLSHSGNTLY | 0.7466023 |
| AGLAANVRD | ASSLSHSGNTLY | 0.5730907 |
| TNRPDLIDPALLRPG | ASSLSHSGNTLY | 0.87895083 |
| LANLGVTAL | ASSLSHSGNTLY | 0.87672234 |
| TGERIEL | ASSLSHSGNTLY | 0.8616021 |
| NGEELAA | ASSLSHSGNTLY | 0.87783384 |
| VLSLVWG | ASSLSHSGNTLY | 0.21043204 |
| IGYALWL | ASSLSHSGNTLY | 0.7387223 |
| LSLLLAPL | ASSLSHSGNTLY | 0.88952106 |
| TLDEALK | ASSLSHSGNTLY | 0.8605219 |
| TDGAGTAEGG | ASSLSHSGNTLY | 0.8125 |
| NAPGGAGGNGG | ASSLSHSGNTLY | 0.8818422 |
| VSVFMGMGGNF | ASSLSHSGNTLY | 0.76040673 |
| LASYLAAL | ASSLSHSGNTLY | 0.82439524 |
| LVGGTPAL | ASSLSHSGNTLY | 0.82950395 |
| VRNADVL | ASSLSHSGNTLY | 0.8428258 |
| TCSSPFLVSAM | ASSLSHSGNTLY | 0.81989247 |
| VHITDAE | ASSLSHSGNTLY | 0.6391095 |
| EYAAPSF | ASSLSHSGNTLY | 0.8614153 |
| VVDAFHGGL | ASSLSHSGNTLY | 0.8838558 |
| ASVVDPQ | ASSLSHSGNTLY | 0.82237375 |
| SYNEFGGLS | ASSLSHSGNTLY | 0.8724873 |
| DPDGSVY | ASSLSHSGNTLY | 0.8522773 |
| MVSTIHATE | ASSLSHSGNTLY | 0.8613641 |
| LAAGVADPV | ASSLSHSGNTLY | 0.7579025 |
| TKEQIAA | ASSLSHSGNTLY | 0.7156297 |
| SLASPAAL | ASSLSHSGNTLY | 0.80722976 |
| IGLVFYGV | ASSLSHSGNTLY | 0.7221831 |
| FLLDASNL | ASSLSHSGNTLY | 0.83193356 |
| YQVGNFP | ASSLSHSGNTLY | 0.8091785 |
| ESLLSQST | ASSLSHSGNTLY | 0.894085 |
| LFTVGFAAVG | ASSLSHSGNTLY | 0.8683637 |
| ATPPTPT | ASSLSHSGNTLY | 0.85122 |
| TDLATLG | ASSLSHSGNTLY | 0.7246206 |
| SAGGTGR | ASSLSHSGNTLY | 0.83120686 |
| GFSNSGS | ASSLSHSGNTLY | 0.8518816 |
| GGNGGSGGTSV | ASSLSHSGNTLY | 0.49609655 |
| GTSVATGGAGN | ASSLSHSGNTLY | 0.88531595 |
| NGGNGGSGGTS | ASSLSHSGNTLY | 0.7213307 |
| AVVKFPH | ASSLSHSGNTLY | 0.7962688 |
| TPPPPPP | ASSLSHSGNTLY | 0.87240714 |
| AAPAAAAAPA | ASSLSHSGNTLY | 0.8769388 |
| IALNEDL | ASSLSHSGNTLY | 0.7959732 |
| IVNVSSM | ASSLSHSGNTLY | 0.78688985 |
| TGIPPAPR | ASSLSHSGNTLY | 0.75137985 |
| FDLGGGTF | ASSLSHSGNTLY | 0.87262106 |
| FDLGGGTFDVS | ASSLSHSGNTLY | 0.8832048 |
| NEPTAAA | ASSLSHSGNTLY | 0.75257856 |
| NEPTAAALA | ASSLSHSGNTLY | 0.8804501 |
| GSIINISSI | ASSLSHSGNTLY | 0.84396493 |
| VAPGNAGTA | ASSLSHSGNTLY | 0.42854738 |

| | | |
|---|---|---|
| TGGGASLE | ASSLSHSGNTLY | 0.83813465 |
| VAVYNHYKRI | ASSLSHSGNTLY | 0.8676117 |
| TDPAPATT | ASSLSHSGNTLY | 0.8657769 |
| TNRPDILDPALLRPG | ASSLSHSGNTLY | 0.8827028 |
| LTTTTTS | ASSLSHSGNTLY | 0.8370413 |
| PTGSGKTL | ASSLSHSGNTLY | 0.78856784 |
| TGSGKTL | ASSLSHSGNTLY | 0.8852814 |
| TIGHVDHGK | ASSLSHSGNTLY | 0.8901788 |
| TIGHVDHGKTT | ASSLSHSGNTLY | 0.8756437 |
| GHVDHGKTT | ASSLSHSGNTLY | 0.81274694 |
| AQFEHTLL | ASSLSHSGNTLY | 0.76843953 |
| IVNQDEL | ASSLSHSGNTLY | 0.8473393 |
| TNGSQFF | ASSLSHSGNTLY | 0.87032527 |
| TGKGIVE | ASSLSHSGNTLY | 0.6837668 |
| LISNASDA | ASSLSHSGNTLY | 0.8461326 |
| AELRTLGELTKIAKAHGAQ | ASSLSHSGNTLY | 0.85457575 |
| APNAVAAAEPSW | ASSLSHSGNTLY | 0.41887826 |
| APQVAGAATAVL | ASSLSHSGNTLY | 0.8595978 |
| ARRDPAAAAAILPTDA | ASSLSHSGNTLY | 0.8670715 |
| DAAARDVLAEAGLAEYF | ASSLSHSGNTLY | 0.8190707 |
| DDQIVLSAKANLAKVEQ | ASSLSHSGNTLY | 0.78948915 |
| DELLPDTGEDSDEDGH | ASSLSHSGNTLY | 0.8785207 |
| ELPEAPANRATVPQD | ASSLSHSGNTLY | 0.892663 |
| FGQSKIERVTGIG | ASSLSHSGNTLY | 0.885543 |
| GAGGTGSTAGGAGGAGGAGGLY | ASSLSHSGNTLY | 0.40985698 |
| GHGGTGAAGGNGGTGGAGGSGI | ASSLSHSGNTLY | 0.31395873 |
| GNGGNGGAGGEGGPGGAGGAGA | ASSLSHSGNTLY | 0.7708024 |
| GSTAGNGGNGGNATNGTIAGQP | ASSLSHSGNTLY | 0.86379683 |
| GTGGDGGNGGNGGTGG | ASSLSHSGNTLY | 0.78770304 |
| GVRRPAGIPADAPV | ASSLSHSGNTLY | 0.7767849 |
| GYRQVTDAHLLTLARR | ASSLSHSGNTLY | 0.83438325 |
| IAKMGQDGHDRG | ASSLSHSGNTLY | 0.88187563 |
| KAEPSIEIQYT | ASSLSHSGNTLY | 0.6733388 |
| KPVRIRMAADEDTVAVT | ASSLSHSGNTLY | 0.52765685 |
| MPPRIAGMRLLVIKPEP | ASSLSHSGNTLY | 0.8255725 |
| MKFYAVALFRSRGVPPTR | ASSLSHSGNTLY | 0.8776162 |
| NAGGFGILGAVAHSP | ASSLSHSGNTLY | 0.73939997 |
| NEGREITLAMAEM | ASSLSHSGNTLY | 0.8234235 |
| NELVEWSHPLGHRWRWE | ASSLSHSGNTLY | 0.7497219 |
| NLGAGGTGGNGGNPG | ASSLSHSGNTLY | 0.68847 |
| NTEWPKWETVMAADPVD | ASSLSHSGNTLY | 0.8431742 |
| PASAAPAETQP | ASSLSHSGNTLY | 0.71054727 |
| QQSSDGKNGVRAAEITGE | ASSLSHSGNTLY | 0.8745312 |
| RAPDAVTTSWWKE | ASSLSHSGNTLY | 0.7043222 |
| SGFGNVGAHTSGWFNQ | ASSLSHSGNTLY | 0.86565053 |
| SGVAGAVGQAASVGGLKV | ASSLSHSGNTLY | 0.8524134 |
| SNGGAGTGSAGGNGGT | ASSLSHSGNTLY | 0.85785335 |
| TGGAGGEGGAAG | ASSLSHSGNTLY | 0.8063432 |
| TGGDGGAAGAGGAPGHG | ASSLSHSGNTLY | 0.765838 |
| TMPPLLAAIANAGRHGV | ASSLSHSGNTLY | 0.6311451 |
| TSSTAGNSGGA | ASSLSHSGNTLY | 0.4471373 |

| | | |
|---|---|---|
| VPATVPVGILC | ASSLSHSGNTLY | 0.87054455 |
| VQRPDSGGVVLTGRIS | ASSLSHSGNTLY | 0.15081383 |
| WTPNPVPATKGVPAGLVQAT | ASSLSHSGNTLY | 0.87882715 |
| YALKRALAKFSDRFDVVII | ASSLSHSGNTLY | 0.8678782 |
| YLIGSVIV | ASSLSHSGNTLY | 0.6720843 |
| YGQLVAVID | ASSLSHSGNTLY | 0.8551984 |
| VTVGGMTLPA | ASSLSHSGNTLY | 0.6944138 |
| VLALPAV | ASSLSHSGNTLY | 0.71785873 |
| VIPPTLNYET | ASSLSHSGNTLY | 0.86852264 |
| VGMGGMHPSSG | ASSLSHSGNTLY | 0.868835 |
| VGLAGGAATA | ASSLSHSGNTLY | 0.697737 |
| VGGAGGAGGSGP | ASSLSHSGNTLY | 0.8514763 |
| VFAAFMV | ASSLSHSGNTLY | 0.8533977 |
| TLYDKLR | ASSLSHSGNTLY | 0.7621927 |
| TLAVFTVL | ASSLSHSGNTLY | 0.8153459 |
| TGSVQPARV | ASSLSHSGNTLY | 0.88027656 |
| TGGGAGSGGT | ASSLSHSGNTLY | 0.8352703 |
| SHLETSL | ASSLSHSGNTLY | 0.7485439 |
| SFTTGMT | ASSLSHSGNTLY | 0.8631299 |
| QVEVAKRMGVRQPTVS | ASSLSHSGNTLY | 0.728819 |
| QGSGGPALAM | ASSLSHSGNTLY | 0.86041915 |
| MLAPSNP | ASSLSHSGNTLY | 0.8186381 |
| LVNNAAI | ASSLSHSGNTLY | 0.83116806 |
| LTGEINTLA | ASSLSHSGNTLY | 0.8689338 |
| LSIFGPV | ASSLSHSGNTLY | 0.5853032 |
| LLALLANL | ASSLSHSGNTLY | 0.74749327 |
| LISNASDALDKLR | ASSLSHSGNTLY | 0.8011764 |
| LINEAVP | ASSLSHSGNTLY | 0.8493693 |
| LGYAFSGL | ASSLSHSGNTLY | 0.8744386 |
| LDFVAVAL | ASSLSHSGNTLY | 0.7470107 |
| LAQPGAE | ASSLSHSGNTLY | 0.8537517 |
| ISNLGAVTGF | ASSLSHSGNTLY | 0.8094233 |
| HQEDPGRI | ASSLSHSGNTLY | 0.7276696 |
| GTPGEPTP | ASSLSHSGNTLY | 0.83445454 |
| GFFNSGS | ASSLSHSGNTLY | 0.7802036 |
| GAGGGGLTAGAN | ASSLSHSGNTLY | 0.709998 |
| FVDALVNM | ASSLSHSGNTLY | 0.8524927 |
| FLTVAPEL | ASSLSHSGNTLY | 0.72184515 |
| FETLTAL | ASSLSHSGNTLY | 0.83682364 |
| DVINPIEL | ASSLSHSGNTLY | 0.8693937 |
| APVSGAPV | ASSLSHSGNTLY | 0.7951627 |
| ALVGGLAW | ASSLSHSGNTLY | 0.8608612 |
| ALAAAAATAI | ASSLSHSGNTLY | 0.8512045 |
| AFGLGLLL | ASSLSHSGNTLY | 0.8491914 |
| AFFAAASL | ASSLSHSGNTLY | 0.8100635 |
| AAAALPAFSPP | ASSLSHSGNTLY | 0.7704842 |
| VVLAPAL | ASSLSHSGNTLY | 0.8797202 |
| TIGHVDHGKTTL | ASSLSHSGNTLY | 0.86835647 |
| LISNASDALDK | ASSLSHSGNTLY | 0.86833644 |
| LISNASDAL | ASSLSHSGNTLY | 0.85412765 |
| ISNASDALDK | ASSLSHSGNTLY | 0.8460112 |

| | | |
|---|---|---|
| NQAETLVYQTE | ASSLSHSGNTLY | 0.87422466 |
| TGGTGGTAGAG | ASSLSHSGNTLY | 0.792078 |
| TGSTAGGAGGA | ASSLSHSGNTLY | 0.5975381 |
| DPLAVPVAAGPV | ASSLSHSGNTLY | 0.8112507 |
| GGTGGTPGNGGN | ASSLSHSGNTLY | 0.8242965 |
| PAGLAGVDPATL | ASSLSHSGNTLY | 0.7297634 |
| GNNLVVEAPHAGASV | ASSLSHSGNTLY | 0.4140424 |
| KVTRIMSASTLEQPA | ASSLSHSGNTLY | 0.87899256 |
| LGFALPIALPRRTVV | ASSLSHSGNTLY | 0.89224505 |
| TRLGNLTARKLVVLV | ASSLSHSGNTLY | 0.6322588 |
| WSQQRRLIYFPSAGP | ASSLSHSGNTLY | 0.88271874 |
| GPNGNPDPMAAAVDIR | ASSLSHSGNTLY | 0.62365365 |
| TQRCVSVADLRAAASR | ASSLSHSGNTLY | 0.8615813 |
| QFVLDVAEHPVEVLTETV | ASSLSHSGNTLY | 0.88049614 |
| GGAGGTGGAGGNGGTQAGDG | ASSLSHSGNTLY | 0.68183434 |
| PLLATLRAIAVDVDAKPAQV | ASSLSHSGNTLY | 0.8646386 |
| APKFRTGGYMIDFWGVGYQVAK | ASSLSHSGNTLY | 0.73198545 |
| GAGGAGGAGGAAGGPGGTGGTG | ASSLSHSGNTLY | 0.73852324 |
| IGALGLLATDVAPAGSVAFGVA | ASSLSHSGNTLY | 0.8828858 |
| AGGAGGIGGNGGAGGAGGVLMG | ASSLSHSGNTLY | 0.79222846 |
| ADELIGLL | ASSLSHSGNTLY | 0.8180849 |
| ADRALKA | ASSLSHSGNTLY | 0.8248529 |
| AGAVGQVGGAGGA | ASSLSHSGNTLY | 0.714472 |
| AGGGGGLGGAP | ASSLSHSGNTLY | 0.83466077 |
| AGGLLYGNGGAG | ASSLSHSGNTLY | 0.87970316 |
| AGLESIGL | ASSLSHSGNTLY | 0.73975337 |
| AGLKTGII | ASSLSHSGNTLY | 0.8880465 |
| ASLAALLL | ASSLSHSGNTLY | 0.66170615 |
| ASVINISSI | ASSLSHSGNTLY | 0.8897487 |
| ATPPGPA | ASSLSHSGNTLY | 0.8653518 |
| ATVASALV | ASSLSHSGNTLY | 0.8404534 |
| ATVASLIL | ASSLSHSGNTLY | 0.79087377 |
| AVNGPVLIRH | ASSLSHSGNTLY | 0.8828277 |
| DGAAGAVGI | ASSLSHSGNTLY | 0.8346594 |
| DPLLHGI | ASSLSHSGNTLY | 0.8521489 |
| ELGAALGAL | ASSLSHSGNTLY | 0.8803835 |
| ELGRTQAV | ASSLSHSGNTLY | 0.543429 |
| FVAGGYLS | ASSLSHSGNTLY | 0.71839833 |
| GASGGAGGAGGW | ASSLSHSGNTLY | 0.66478086 |
| GGAGAGGDGGAGGI | ASSLSHSGNTLY | 0.6958963 |
| GTGHNGSQPGMGGNG | ASSLSHSGNTLY | 0.52137774 |
| GYAAASASAS | ASSLSHSGNTLY | 0.44992614 |
| HQVDVAAVQQG | ASSLSHSGNTLY | 0.6726886 |
| IGAAIAHT | ASSLSHSGNTLY | 0.49150056 |
| IGMIEVAQL | ASSLSHSGNTLY | 0.7255057 |
| IRLAPEHTSAMD | ASSLSHSGNTLY | 0.7356912 |
| IYNVSVL | ASSLSHSGNTLY | 0.5138373 |
| KQGFLGM | ASSLSHSGNTLY | 0.6932198 |
| LESVITQL | ASSLSHSGNTLY | 0.72236687 |
| LGLGALVVV | ASSLSHSGNTLY | 0.4663991 |
| LIGNGGNGGSGGI | ASSLSHSGNTLY | 0.6313866 |

| | | |
|---|---|---|
| LNAINAPTL | ASSLSHSGNTLY | 0.5531002 |
| LQLLNTGV | ASSLSHSGNTLY | 0.6255012 |
| LQTSIEF | ASSLSHSGNTLY | 0.34790558 |
| MSFAVSEL | ASSLSHSGNTLY | 0.709417 |
| NTGALGL | ASSLSHSGNTLY | 0.523966 |
| PPGAGGAGGAGTT | ASSLSHSGNTLY | 0.29264644 |
| QLVGLGI | ASSLSHSGNTLY | 0.6826102 |
| RFLEQVHQL | ASSLSHSGNTLY | 0.5443483 |
| RGVGYVLR | ASSLSHSGNTLY | 0.6883437 |
| SATGVFAGVI | ASSLSHSGNTLY | 0.74473155 |
| SLPAITLP | ASSLSHSGNTLY | 0.5735879 |
| STALAFTV | ASSLSHSGNTLY | 0.5327307 |
| SVLTVGLGK | ASSLSHSGNTLY | 0.67164546 |
| TALAGAGSGI | ASSLSHSGNTLY | 0.7498012 |
| TGLPFTGI | ASSLSHSGNTLY | 0.4819507 |
| VAEHVTV | ASSLSHSGNTLY | 0.66767144 |
| VDIVLNSL | ASSLSHSGNTLY | 0.73896927 |
| VLTGGGAVL | ASSLSHSGNTLY | 0.63099736 |
| VNVGLAYGI | ASSLSHSGNTLY | 0.7586156 |
| VSSAAVGAV | ASSLSHSGNTLY | 0.76522666 |
| VTSVYEAGV | ASSLSHSGNTLY | 0.51680636 |
| WVALDRGA | ASSLSHSGNTLY | 0.7339607 |
| YLVGSVVI | ASSLSHSGNTLY | 0.7184833 |
| GPPGTGKTLL | ASSLSHSGNTLY | 0.6096344 |
| GVGMTKF | ASSLSHSGNTLY | 0.5937778 |
| IGIGGSDL | ASSLSHSGNTLY | 0.55066484 |
| IGTLLAEL | ASSLSHSGNTLY | 0.6661429 |
| IYTSGTT | ASSLSHSGNTLY | 0.44436634 |
| LSFPAGV | ASSLSHSGNTLY | 0.6144906 |
| LYTSGTT | ASSLSHSGNTLY | 0.73829615 |
| VGAFPFGF | ASSLSHSGNTLY | 0.5621954 |
| VGAGTVEF | ASSLSHSGNTLY | 0.45819137 |
| VGKSTLLL | ASSLSHSGNTLY | 0.39875337 |
| AALAHAGGGIV | ASSLSHSGNTLY | 0.64285636 |
| AALVAVSLL | ASSLSHSGNTLY | 0.58547896 |
| AAVSRAVAAL | ASSLSHSGNTLY | 0.4287397 |
| AGGSSGIGGF | ASSLSHSGNTLY | 0.56375104 |
| AGLTVANANG | ASSLSHSGNTLY | 0.7151224 |
| AGLVGAVL | ASSLSHSGNTLY | 0.7243773 |
| ALSQAFTV | ASSLSHSGNTLY | 0.4183265 |
| ALTFSVSL | ASSLSHSGNTLY | 0.7543647 |
| ALTGAALVAL | ASSLSHSGNTLY | 0.72271645 |
| ASFSIGAGVL | ASSLSHSGNTLY | 0.72449684 |
| DGSVYKI | ASSLSHSGNTLY | 0.6482188 |
| EQLAEIV | ASSLSHSGNTLY | 0.5270205 |
| FAVSELTVM | ASSLSHSGNTLY | 0.70357955 |
| GAAPLAAGL | ASSLSHSGNTLY | 0.50260186 |
| GADGGDGGSGGAGG | ASSLSHSGNTLY | 0.6539736 |
| GFGLDTGGAGG | ASSLSHSGNTLY | 0.71741635 |
| GLASGGLL | ASSLSHSGNTLY | 0.56554466 |
| GRIAGAAVLL | ASSLSHSGNTLY | 0.47375432 |

| | | |
|---|---|---|
| GSAGSLNTSF | ASSLSHSGNTLY | 0.59344715 |
| GTGQIGI | ASSLSHSGNTLY | 0.7492632 |
| GTLIALIL | ASSLSHSGNTLY | 0.44212297 |
| IAATMGFVVG | ASSLSHSGNTLY | 0.5426216 |
| IFAQALL | ASSLSHSGNTLY | 0.6108532 |
| IGAAVALGF | ASSLSHSGNTLY | 0.5951841 |
| IGLTPLAL | ASSLSHSGNTLY | 0.74919516 |
| ILGSNTSTL | ASSLSHSGNTLY | 0.71851325 |
| IVFSLTVL | ASSLSHSGNTLY | 0.57925695 |
| LASKATGF | ASSLSHSGNTLY | 0.72303516 |
| LASVALIL | ASSLSHSGNTLY | 0.67612886 |
| LFAVGLTL | ASSLSHSGNTLY | 0.62474936 |
| LFNGGIGTYI | ASSLSHSGNTLY | 0.6940256 |
| LTLGAGII | ASSLSHSGNTLY | 0.46544266 |
| LVAAALGDI | ASSLSHSGNTLY | 0.741495 |
| NTAFVQLGI | ASSLSHSGNTLY | 0.66712195 |
| QLFSGAEL | ASSLSHSGNTLY | 0.7502954 |
| SAATFITL | ASSLSHSGNTLY | 0.75004077 |
| SNYGLANLGV | ASSLSHSGNTLY | 0.67460936 |
| SYGGAYAVM | ASSLSHSGNTLY | 0.655823 |
| TTGVGGLTLG | ASSLSHSGNTLY | 0.37475276 |
| TTQTIYI | ASSLSHSGNTLY | 0.575439 |
| VAAGAGLGAVI | ASSLSHSGNTLY | 0.6644257 |
| VAAVVTLL | ASSLSHSGNTLY | 0.6624817 |
| VGFLGGLI | ASSLSHSGNTLY | 0.71724033 |
| VGVIPILL | ASSLSHSGNTLY | 0.74710155 |
| VIFASSLI | ASSLSHSGNTLY | 0.5350416 |
| VKGGDGGAAAG | ASSLSHSGNTLY | 0.74592817 |
| VTKTFTNI | ASSLSHSGNTLY | 0.7316859 |
| VVAAAILL | ASSLSHSGNTLY | 0.4073891 |
| YAALDVEL | ASSLSHSGNTLY | 0.60584724 |
| LASVALLL | ASSLSHSGNTLY | 0.74544394 |
| LDAVHRV | ASSLSHSGNTLY | 0.6895231 |
| QGLSSSIF | ASSLSHSGNTLY | 0.5250876 |
| RGTFANIRL | ASSLSHSGNTLY | 0.6406302 |
| RQLWWGHRI | ASSLSHSGNTLY | 0.73732287 |
| YGPPGTGKTLL | ASSLSHSGNTLY | 0.6436798 |
| AALVGAPANVA | ASSLSHSGNTLY | 0.74109596 |
| AATFRRTE | ASSLSHSGNTLY | 0.31140882 |
| AGALGGGI | ASSLSHSGNTLY | 0.6533066 |
| AGGAGGAGGIGAG | ASSLSHSGNTLY | 0.5429356 |
| AGSGALGL | ASSLSHSGNTLY | 0.35679075 |
| AQIADQI | ASSLSHSGNTLY | 0.7064385 |
| ASAVAVTYL | ASSLSHSGNTLY | 0.63976777 |
| ASTVLGGTGG | ASSLSHSGNTLY | 0.7318728 |
| AVGGEPAK | ASSLSHSGNTLY | 0.552226 |
| CEQGTFGGTC | ASSLSHSGNTLY | 0.7313746 |
| DEGKAPAG | ASSLSHSGNTLY | 0.6012234 |
| DTVVFPAL | ASSLSHSGNTLY | 0.6580122 |
| FEGITMAG | ASSLSHSGNTLY | 0.57774854 |
| FPLVVQD | ASSLSHSGNTLY | 0.6389166 |

| | | |
|---|---|---|
| GAFTTPQ | ASSLSHSGNTLY | 0.75227314 |
| GFGGFGTAGGVGGA | ASSLSHSGNTLY | 0.7560688 |
| GFLAQFL | ASSLSHSGNTLY | 0.41081718 |
| GGAGGAGGNAGVA | ASSLSHSGNTLY | 0.66592884 |
| GGVGTAGAGGAGGAGG | ASSLSHSGNTLY | 0.60381395 |
| GNGGGGDAGNAGSGGN | ASSLSHSGNTLY | 0.40165913 |
| GTGGAGGAGNDAGS | ASSLSHSGNTLY | 0.3784398 |
| GTTNTGWFNS | ASSLSHSGNTLY | 0.6128882 |
| IFMGPLFTL | ASSLSHSGNTLY | 0.5559347 |
| ILAHAIEAI | ASSLSHSGNTLY | 0.66188157 |
| IRSAIREAL | ASSLSHSGNTLY | 0.692116 |
| ITFAGGG | ASSLSHSGNTLY | 0.4895029 |
| ITGMSGHQEV | ASSLSHSGNTLY | 0.6764697 |
| IVAGLACTL | ASSLSHSGNTLY | 0.6102111 |
| IVGGGLAGA | ASSLSHSGNTLY | 0.21293819 |
| IVTGVLNI | ASSLSHSGNTLY | 0.54543895 |
| LAQQGCI | ASSLSHSGNTLY | 0.48281962 |
| LCLHVEN | ASSLSHSGNTLY | 0.76949924 |
| LDAISDGS | ASSLSHSGNTLY | 0.54489404 |
| LGGFAAML | ASSLSHSGNTLY | 0.5086794 |
| LGGKNPMI | ASSLSHSGNTLY | 0.5829206 |
| LGPKGVAAIL | ASSLSHSGNTLY | 0.64770633 |
| LMVGVSI | ASSLSHSGNTLY | 0.58263606 |
| LVGAAFVL | ASSLSHSGNTLY | 0.48698846 |
| LVLAVAPL | ASSLSHSGNTLY | 0.33786336 |
| MAAIAVIDAA | ASSLSHSGNTLY | 0.68239397 |
| NNGINAELQ | ASSLSHSGNTLY | 0.5634301 |
| RGEPLVLVHG | ASSLSHSGNTLY | 0.62633467 |
| RGGAGGAAGLI | ASSLSHSGNTLY | 0.49240547 |
| RSYRMHPAV | ASSLSHSGNTLY | 0.596137 |
| SAAHLLAGQLA | ASSLSHSGNTLY | 0.39875296 |
| SAAVGSY | ASSLSHSGNTLY | 0.7527008 |
| SPHAVLHRI | ASSLSHSGNTLY | 0.6133084 |
| SQIAKVL | ASSLSHSGNTLY | 0.5512556 |
| TALGHLVP | ASSLSHSGNTLY | 0.74923664 |
| TAPTTSGAA | ASSLSHSGNTLY | 0.6287818 |
| TDGDDGNGGSPGT | ASSLSHSGNTLY | 0.54279244 |
| TDGLGEN | ASSLSHSGNTLY | 0.5364547 |
| TGGPPLPGQA | ASSLSHSGNTLY | 0.5016313 |
| TIPAITI | ASSLSHSGNTLY | 0.51137435 |
| TIPGFTI | ASSLSHSGNTLY | 0.5449611 |
| TSIASGALA | ASSLSHSGNTLY | 0.74317414 |
| TVTLPGAAI | ASSLSHSGNTLY | 0.68527246 |
| VASGLSAALP | ASSLSHSGNTLY | 0.5315923 |
| VGAGVIGI | ASSLSHSGNTLY | 0.73926705 |
| VGPKRFGV | ASSLSHSGNTLY | 0.7312069 |
| VNVAFPD | ASSLSHSGNTLY | 0.7454959 |
| VSMTNSGVS | ASSLSHSGNTLY | 0.5499143 |
| VTDVLGDLSV | ASSLSHSGNTLY | 0.6190979 |
| VVVVGAGFAGL | ASSLSHSGNTLY | 0.64503384 |
| VYTVGTVV | ASSLSHSGNTLY | 0.5523103 |

| | | |
|---|---|---|
| WTTFSGP | ASSLSHSGNTLY | 0.60870415 |
| YDELVEAF | ASSLSHSGNTLY | 0.51967096 |
| YGTGGAGGNGG | ASSLSHSGNTLY | 0.7329503 |
| YTNVMAV | ASSLSHSGNTLY | 0.7795975 |
| FLEQVHQL | ASSLSHSGNTLY | 0.5823414 |
| GLSLLAVAV | ASSLSHSGNTLY | 0.41791588 |
| LEQVHQL | ASSLSHSGNTLY | 0.50740284 |
| LNAAGAI | ASSLSHSGNTLY | 0.44421756 |
| PAVHASL | ASSLSHSGNTLY | 0.48586997 |
| TELEDDA | ASSLSHSGNTLY | 0.67046374 |
| TPPGPPN | ASSLSHSGNTLY | 0.43325883 |
| TPPPPPPP | ASSLSHSGNTLY | 0.60599625 |
| TQSPALI | ASSAGTTSQNTLY | 0.7451123 |
| EFAAAFPQ | ASSAGTTSQNTLY | 0.5601028 |
| LLSGNLDVL | ASSAGTTSQNTLY | 0.706493 |
| GPANLGNYNI | ASSAGTTSQNTLY | 0.721076 |
| TLGPTLP | ASSAGTTSQNTLY | 0.699802 |
| VYAYSSCCK | ASSAGTTSQNTLY | 0.7592619 |
| APAPAPAPNI | ASSAGTTSQNTLY | 0.7400247 |
| PGGPVPIP | ASSAGTTSQNTLY | 0.66577715 |
| FLTVALEP | ASSAGTTSQNTLY | 0.619031 |
| SQYFGANL | ASSAGTTSQNTLY | 0.71470785 |
| GGVKPAA | ASSAGTTSQNTLY | 0.573477 |
| GGTGGAGGAGGW | ASSAGTTSQNTLY | 0.55660415 |
| GVGGVAGLGGAGG | ASSAGTTSQNTLY | 0.55160487 |
| VLLVLATTI | ASSAGTTSQNTLY | 0.67677987 |
| IGIFAAML | ASSAGTTSQNTLY | 0.5640323 |
| AVGDLID | ASSAGTTSQNTLY | 0.7596394 |
| NISPLPL | ASSAGTTSQNTLY | 0.3998899 |
| VTAVFVAI | ASSAGTTSQNTLY | 0.49329084 |
| SGDDNDAAL | ASSAGTTSQNTLY | 0.7278104 |
| AVAALAHS | ASSAGTTSQNTLY | 0.25022262 |
| GGVTVMATP | ASSAGTTSQNTLY | 0.44956425 |
| AGLAANVRD | ASSAGTTSQNTLY | 0.6558403 |
| TNRPDLIDPALLRPG | ASSAGTTSQNTLY | 0.6924445 |
| LANLGVTAL | ASSAGTTSQNTLY | 0.28627154 |
| TGERIEL | ASSAGTTSQNTLY | 0.563619 |
| NGEELAA | ASSAGTTSQNTLY | 0.5295245 |
| VLSLVWG | ASSAGTTSQNTLY | 0.7535135 |
| IGYALWL | ASSAGTTSQNTLY | 0.6142019 |
| LSLLLAPL | ASSAGTTSQNTLY | 0.40579695 |
| TLDEALK | ASSAGTTSQNTLY | 0.74685323 |
| TDGAGTAEGG | ASSAGTTSQNTLY | 0.40082604 |
| NAPGGAGGNGG | ASSAGTTSQNTLY | 0.60263765 |
| VSVFMGMGGNF | ASSAGTTSQNTLY | 0.7315674 |
| LASYLAAL | ASSAGTTSQNTLY | 0.7782589 |
| LVGGTPAL | ASSAGTTSQNTLY | 0.51554316 |
| VRNADVL | ASSAGTTSQNTLY | 0.719365 |
| TCSSPFLVSAM | ASSAGTTSQNTLY | 0.5316516 |
| VHITDAE | ASSAGTTSQNTLY | 0.76286924 |
| EYAAPSF | ASSAGTTSQNTLY | 0.56003034 |

| | | |
|---|---|---|
| VVDAFHGGL | ASSAGTTSQNTLY | 0.45175412 |
| ASVVDPQ | ASSAGTTSQNTLY | 0.74335426 |
| SYNEFGGLS | ASSAGTTSQNTLY | 0.737172 |
| DPDGSVY | ASSAGTTSQNTLY | 0.72904825 |
| MVSTIHATE | ASSAGTTSQNTLY | 0.72652644 |
| LAAGVADPV | ASSAGTTSQNTLY | 0.6886369 |
| TKEQIAA | ASSAGTTSQNTLY | 0.5575556 |
| SLASPAAL | ASSAGTTSQNTLY | 0.68235874 |
| IGLVFYGV | ASSAGTTSQNTLY | 0.6986697 |
| FLLDASNL | ASSAGTTSQNTLY | 0.55504024 |
| YQVGNFP | ASSAGTTSQNTLY | 0.7476162 |
| ESLLSQST | ASSAGTTSQNTLY | 0.561728 |
| LFTVGFAAVG | ASSAGTTSQNTLY | 0.5799943 |
| ATPPTPT | ASSAGTTSQNTLY | 0.74699455 |
| TDLATLG | ASSAGTTSQNTLY | 0.69632226 |
| SAGGTGR | ASSAGTTSQNTLY | 0.68644464 |
| GFSNSGS | ASSAGTTSQNTLY | 0.6911267 |
| GGNGGSGGTSV | ASSAGTTSQNTLY | 0.76110303 |
| GTSVATGGAGN | ASSAGTTSQNTLY | 0.50288904 |
| NGGNGGSGGTS | ASSAGTTSQNTLY | 0.680774 |
| AVVKFPH | ASSAGTTSQNTLY | 0.6044773 |
| TPPPPPP | ASSAGTTSQNTLY | 0.74573135 |
| AAPAAAAAPA | ASSAGTTSQNTLY | 0.6743252 |
| IALNEDL | ASSAGTTSQNTLY | 0.5953237 |
| IVNVSSM | ASSAGTTSQNTLY | 0.71689934 |
| TGIPPAPR | ASSAGTTSQNTLY | 0.73216945 |
| FDLGGGTF | ASSAGTTSQNTLY | 0.6631662 |
| FDLGGGTFDVS | ASSAGTTSQNTLY | 0.6836549 |
| NEPTAAA | ASSAGTTSQNTLY | 0.76858705 |
| NEPTAAALA | ASSAGTTSQNTLY | 0.55897564 |
| GSIINISSI | ASSAGTTSQNTLY | 0.613813 |
| VAPGNAGTA | ASSAGTTSQNTLY | 0.7367154 |
| TGGGASLE | ASSAGTTSQNTLY | 0.75265646 |
| VAVYNHYKRI | ASSAGTTSQNTLY | 0.68595535 |
| TDPAPATT | ASSAGTTSQNTLY | 0.7302549 |
| TNRPDILDPALLRPG | ASSAGTTSQNTLY | 0.54690105 |
| LTTTTTS | ASSAGTTSQNTLY | 0.6629862 |
| PTGSGKTL | ASSAGTTSQNTLY | 0.52074546 |
| TGSGKTL | ASSAGTTSQNTLY | 0.5877124 |
| TIGHVDHGK | ASSAGTTSQNTLY | 0.48051453 |
| TIGHVDHGKTT | ASSAGTTSQNTLY | 0.65272796 |
| GHVDHGKTT | ASSAGTTSQNTLY | 0.6906414 |
| AQFEHTLL | ASSAGTTSQNTLY | 0.6479188 |
| IVNQDEL | ASSAGTTSQNTLY | 0.3863054 |
| TNGSQFF | ASSAGTTSQNTLY | 0.67728925 |
| TGKGIVE | ASSAGTTSQNTLY | 0.68178535 |
| LISNASDA | ASSAGTTSQNTLY | 0.7818604 |
| AELRTLGELTKIAKAHGAQ | ASSAGTTSQNTLY | 0.58060575 |
| APNAVAAAEPSW | ASSAGTTSQNTLY | 0.72613895 |
| APQVAGAATAVL | ASSAGTTSQNTLY | 0.5972093 |
| ARRDPAAAAAILPTDA | ASSAGTTSQNTLY | 0.67483383 |

| | | |
|---|---|---|
| DAAARDVLAEAGLAEYF | ASSAGTTSQNTLY | 0.37161714 |
| DDQIVLSAKANLAKVEQ | ASSAGTTSQNTLY | 0.6040913 |
| DELLPDTGEDSDEDGH | ASSAGTTSQNTLY | 0.5431701 |
| ELPEAPANRATVPQD | ASSAGTTSQNTLY | 0.48564044 |
| FGQSKIERVTGIG | ASSAGTTSQNTLY | 0.6816726 |
| GAGGTGSTAGGAGGAGGAGGLY | ASSAGTTSQNTLY | 0.7376809 |
| GHGGTGAAGGNGGTGGAGGSGI | ASSAGTTSQNTLY | 0.7182482 |
| GNGGNGGAGGEGGPGGAGGAGA | ASSAGTTSQNTLY | 0.7165718 |
| GSTAGNGGNGGNATNGTIAGQP | ASSAGTTSQNTLY | 0.73285127 |
| GTGGDGGNGGNGGTGG | ASSAGTTSQNTLY | 0.76527727 |
| GVRRPAGIPADAPV | ASSAGTTSQNTLY | 0.53449684 |
| GYRQVTDAHLLTLARR | ASSAGTTSQNTLY | 0.5948886 |
| IAKMGQDGHDRG | ASSAGTTSQNTLY | 0.4026802 |
| KAEPSIEIQYT | ASSAGTTSQNTLY | 0.5466732 |
| KPVRIRMAADEDTVAVT | ASSAGTTSQNTLY | 0.5387971 |
| MPPRIAGMRLLVIKPEP | ASSAGTTSQNTLY | 0.62344176 |
| MKFYAVALFRSRGVPPTR | ASSAGTTSQNTLY | 0.28623888 |
| NAGGFGILGAVAHSP | ASSAGTTSQNTLY | 0.6733569 |
| NEGREITLAMAEM | ASSAGTTSQNTLY | 0.7463546 |
| NELVEWSHPLGHRWRWE | ASSAGTTSQNTLY | 0.45886406 |
| NLGAGGTGGNGGNPG | ASSAGTTSQNTLY | 0.6763888 |
| NTEWPKWETVMAADPVD | ASSAGTTSQNTLY | 0.62600094 |
| PASAAPAETQP | ASSAGTTSQNTLY | 0.52584666 |
| QQSSDGKNGVRAAEITGE | ASSAGTTSQNTLY | 0.66552675 |
| RAPDAVTTSWWKE | ASSAGTTSQNTLY | 0.5425949 |
| SGFGNVGAHTSGWFNQ | ASSAGTTSQNTLY | 0.7393676 |
| SGVAGAVGQAASVGGLKV | ASSAGTTSQNTLY | 0.73773307 |
| SNGGAGTGSAGGNGGT | ASSAGTTSQNTLY | 0.66429055 |
| TGGAGGEGGAAG | ASSAGTTSQNTLY | 0.62151575 |
| TGGDGGAAGAGGAPGHG | ASSAGTTSQNTLY | 0.6850575 |
| TMPPLLAAIANAGRHGV | ASSAGTTSQNTLY | 0.46400428 |
| TSSTAGNSGGA | ASSAGTTSQNTLY | 0.65601575 |
| VPATVPVGILC | ASSAGTTSQNTLY | 0.4366276 |
| VQRPDSGGVVLTGRIS | ASSAGTTSQNTLY | 0.6091359 |
| WTPNPVPATKGVPAGLVQAT | ASSAGTTSQNTLY | 0.50843406 |
| YALKRALAKFSDRFDVVII | ASSAGTTSQNTLY | 0.6922219 |
| YLIGSVIV | ASSAGTTSQNTLY | 0.6151307 |
| YGQLVAVID | ASSAGTTSQNTLY | 0.6958958 |
| VTVGGMTLPA | ASSAGTTSQNTLY | 0.43047035 |
| VLALPAV | ASSAGTTSQNTLY | 0.7020044 |
| VIPPTLNYET | ASSAGTTSQNTLY | 0.64967334 |
| VGMGGMHPSSG | ASSAGTTSQNTLY | 0.46212596 |
| VGLAGGAATA | ASSAGTTSQNTLY | 0.7132431 |
| VGGAGGAGGSGP | ASSAGTTSQNTLY | 0.6558167 |
| VFAAFMV | ASSAGTTSQNTLY | 0.5812571 |
| TLYDKLR | ASSAGTTSQNTLY | 0.7211908 |
| TLAVFTVL | ASSAGTTSQNTLY | 0.72542894 |
| TGSVQPARV | ASSAGTTSQNTLY | 0.46305132 |
| TGGGAGSGGT | ASSAGTTSQNTLY | 0.5374669 |
| SHLETSL | ASSAGTTSQNTLY | 0.63274103 |
| SFTTGMT | ASSAGTTSQNTLY | 0.5516119 |

| | | |
|---|---|---|
| QVEVAKRMGVRQPTVS | ASSAGTTSQNTLY | 0.3156363 |
| QGSGGPALAM | ASSAGTTSQNTLY | 0.70165956 |
| MLAPSNP | ASSAGTTSQNTLY | 0.582905 |
| LVNNAAI | ASSAGTTSQNTLY | 0.46474844 |
| LTGEINTLA | ASSAGTTSQNTLY | 0.5967573 |
| LSIFGPV | ASSAGTTSQNTLY | 0.7511443 |
| LLALLANL | ASSAGTTSQNTLY | 0.5697065 |
| LISNASDALDKLR | ASSAGTTSQNTLY | 0.7365837 |
| LINEAVP | ASSAGTTSQNTLY | 0.7036713 |
| LGYAFSGL | ASSAGTTSQNTLY | 0.6853756 |
| LDFVAVAL | ASSAGTTSQNTLY | 0.6971968 |
| LAQPGAE | ASSAGTTSQNTLY | 0.68705857 |
| ISNLGAVTGF | ASSAGTTSQNTLY | 0.645195 |
| HQEDPGRI | ASSAGTTSQNTLY | 0.7101103 |
| GTPGEPTP | ASSAGTTSQNTLY | 0.63147426 |
| GFFNSGS | ASSAGTTSQNTLY | 0.56006867 |
| GAGGGGLTAGAN | ASSAGTTSQNTLY | 0.4913193 |
| FVDALVNM | ASSAGTTSQNTLY | 0.49577537 |
| FLTVAPEL | ASSAGTTSQNTLY | 0.4464534 |
| FETLTAL | ASSAGTTSQNTLY | 0.63589436 |
| DVINPIEL | ASSAGTTSQNTLY | 0.7713348 |
| APVSGAPV | ASSAGTTSQNTLY | 0.62756455 |
| ALVGGLAW | ASSAGTTSQNTLY | 0.46205908 |
| ALAAAAATAI | ASSAGTTSQNTLY | 0.609809 |
| AFGLGLLL | ASSAGTTSQNTLY | 0.5921811 |
| AFFAAASL | ASSAGTTSQNTLY | 0.7296384 |
| AAAALPAFSPP | ASSAGTTSQNTLY | 0.7092665 |
| VVLAPAL | ASSAGTTSQNTLY | 0.7686795 |
| TIGHVDHGKTTL | ASSAGTTSQNTLY | 0.70127505 |
| LISNASDALDK | ASSAGTTSQNTLY | 0.50942117 |
| LISNASDAL | ASSAGTTSQNTLY | 0.73232394 |
| ISNASDALDK | ASSAGTTSQNTLY | 0.5817921 |
| NQAETLVYQTE | ASSAGTTSQNTLY | 0.64014757 |
| TGGTGGTAGAG | ASSAGTTSQNTLY | 0.6673236 |
| TGSTAGGAGGA | ASSAGTTSQNTLY | 0.6891333 |
| DPLAVPVAAGPV | ASSAGTTSQNTLY | 0.6221341 |
| GGTGGTPGNGGN | ASSAGTTSQNTLY | 0.67250127 |
| PAGLAGVDPATL | ASSAGTTSQNTLY | 0.40458456 |
| GNNLVVEAPHAGASV | ASSAGTTSQNTLY | 0.7392074 |
| KVTRIMSASTLEQPA | ASSAGTTSQNTLY | 0.41769043 |
| LGFALPIALPRRTVV | ASSAGTTSQNTLY | 0.48853904 |
| TRLGNLTARKLVVLV | ASSAGTTSQNTLY | 0.57459027 |
| WSQQRRLIYFPSAGP | ASSAGTTSQNTLY | 0.7585728 |
| GPNGNPDPMAAAVDIR | ASSAGTTSQNTLY | 0.7415074 |
| TQRCVSVADLRAAASR | ASSAGTTSQNTLY | 0.49551076 |
| QFVLDVAEHPVEVLTETV | ASSAGTTSQNTLY | 0.501139 |
| GGAGGTGGAGGNGGTQAGDG | ASSAGTTSQNTLY | 0.6711482 |
| PLLATLRAIAVDVDAKPAQV | ASSAGTTSQNTLY | 0.416986 |
| APKFRTGGYMIDFWGVGYQVAK | ASSAGTTSQNTLY | 0.72787774 |
| GAGGAGGAGGAAGGPGGTGGTG | ASSAGTTSQNTLY | 0.5797661 |
| IGALGLLATDVAPAGSVAFGVA | ASSAGTTSQNTLY | 0.58507097 |

| | | |
|---|---|---|
| AGGAGGIGGNGGAGGAGGVLMG | ASSAGTTSQNTLY | 0.5572894 |
| ADELIGLL | ASSAGTTSQNTLY | 0.6972555 |
| ADRALKA | ASSAGTTSQNTLY | 0.5714194 |
| AGAVGQVGGAGGA | ASSAGTTSQNTLY | 0.7406474 |
| AGGGGGLGGAP | ASSAGTTSQNTLY | 0.2867015 |
| AGGLLYGNGGAG | ASSAGTTSQNTLY | 0.7349838 |
| AGLESIGL | ASSAGTTSQNTLY | 0.4404901 |
| AGLKTGII | ASSAGTTSQNTLY | 0.49861234 |
| ASLAALLL | ASSAGTTSQNTLY | 0.36058658 |
| ASVINISSI | ASSAGTTSQNTLY | 0.6777795 |
| ATPPGPA | ASSAGTTSQNTLY | 0.68564135 |
| ATVASALV | ASSAGTTSQNTLY | 0.7413601 |
| ATVASLIL | ASSAGTTSQNTLY | 0.64219916 |
| AVNGPVLIRH | ASSAGTTSQNTLY | 0.7081373 |
| DGAAGAVGI | ASSAGTTSQNTLY | 0.70898366 |
| DPLLHGI | ASSAGTTSQNTLY | 0.518247 |
| ELGAALGAL | ASSAGTTSQNTLY | 0.4352934 |
| ELGRTQAV | ASSAGTTSQNTLY | 0.7072173 |
| FVAGGYLS | ASSAGTTSQNTLY | 0.62942827 |
| GASGGAGGAGGW | ASSAGTTSQNTLY | 0.7528207 |
| GGAGAGGDGGAGGI | ASSAGTTSQNTLY | 0.6781651 |
| GTGHNGSQPGMGGNG | ASSAGTTSQNTLY | 0.3828654 |
| GYAAASASAS | ASSAGTTSQNTLY | 0.739787 |
| HQVDVAAVQQG | ASSAGTTSQNTLY | 0.73204714 |
| IGAAIAHT | ASSAGTTSQNTLY | 0.38539338 |
| IGMIEVAQL | ASSAGTTSQNTLY | 0.5737991 |
| IRLAPEHTSAMD | ASSAGTTSQNTLY | 0.5846572 |
| IYNVSVL | ASSAGTTSQNTLY | 0.42234778 |
| KQGFLGM | ASSAGTTSQNTLY | 0.6931051 |
| LESVITQL | ASSAGTTSQNTLY | 0.5133937 |
| LGLGALVVV | ASSAGTTSQNTLY | 0.60024315 |
| LIGNGGNGGSGGI | ASSAGTTSQNTLY | 0.6214207 |
| LNAINAPTL | ASSAGTTSQNTLY | 0.5666813 |
| LQLLNTGV | ASSAGTTSQNTLY | 0.5631403 |
| LQTSIEF | ASSAGTTSQNTLY | 0.58785546 |
| MSFAVSEL | ASSAGTTSQNTLY | 0.6775742 |
| NTGALGL | ASSAGTTSQNTLY | 0.7005085 |
| PPGAGGAGGAGTT | ASSAGTTSQNTLY | 0.56720924 |
| QLVGLGI | ASSAGTTSQNTLY | 0.67994225 |
| RFLEQVHQL | ASSAGTTSQNTLY | 0.7179283 |
| RGVGYVLR | ASSAGTTSQNTLY | 0.45245916 |
| SATGVFAGVI | ASSAGTTSQNTLY | 0.47410443 |
| SLPAITLP | ASSAGTTSQNTLY | 0.7285398 |
| STALAFTV | ASSAGTTSQNTLY | 0.5189246 |
| SVLTVGLGK | ASSAGTTSQNTLY | 0.73714375 |
| TALAGAGSGI | ASSAGTTSQNTLY | 0.6026793 |
| TGLPFTGI | ASSAGTTSQNTLY | 0.6140509 |
| VAEHVTV | ASSAGTTSQNTLY | 0.6114598 |
| VDIVLNSL | ASSAGTTSQNTLY | 0.7573269 |
| VLTGGGAVL | ASSAGTTSQNTLY | 0.74658597 |
| VNVGLAYGI | ASSAGTTSQNTLY | 0.62357295 |

| | | |
|---|---|---|
| VSSAAVGAV | ASSAGTTSQNTLY | 0.73115635 |
| VTSVYEAGV | ASSAGTTSQNTLY | 0.46984503 |
| WVALDRGA | ASSAGTTSQNTLY | 0.70700943 |
| YLVGSVVI | ASSAGTTSQNTLY | 0.7301616 |
| GPPGTGKTLL | ASSAGTTSQNTLY | 0.43029755 |
| GVGMTKF | ASSAGTTSQNTLY | 0.55453235 |
| IGIGGSDL | ASSAGTTSQNTLY | 0.7450609 |
| IGTLLAEL | ASSAGTTSQNTLY | 0.7460263 |
| IYTSGTT | ASSAGTTSQNTLY | 0.5301193 |
| LSFPAGV | ASSAGTTSQNTLY | 0.55555075 |
| LYTSGTT | ASSAGTTSQNTLY | 0.45982295 |
| VGAFPFGF | ASSAGTTSQNTLY | 0.535857 |
| VGAGTVEF | ASSAGTTSQNTLY | 0.47735104 |
| VGKSTLLL | ASSAGTTSQNTLY | 0.67861784 |
| AALAHAGGGIV | ASSAGTTSQNTLY | 0.5146286 |
| AALVAVSLL | ASSAGTTSQNTLY | 0.72443753 |
| AAVSRAVAAL | ASSAGTTSQNTLY | 0.64204615 |
| AGGSSGIGGF | ASSAGTTSQNTLY | 0.7212924 |
| AGLTVANANG | ASSAGTTSQNTLY | 0.52013665 |
| AGLVGAVL | ASSAGTTSQNTLY | 0.60663843 |
| ALSQAFTV | ASSAGTTSQNTLY | 0.45230368 |
| ALTFSVSL | ASSAGTTSQNTLY | 0.49707416 |
| ALTGAALVAL | ASSAGTTSQNTLY | 0.515182 |
| ASFSIGAGVL | ASSAGTTSQNTLY | 0.5891173 |
| DGSVYKI | ASSAGTTSQNTLY | 0.34376803 |
| EQLAEIV | ASSAGTTSQNTLY | 0.72951 |
| FAVSELTVM | ASSAGTTSQNTLY | 0.67822653 |
| GAAPLAAGL | ASSAGTTSQNTLY | 0.65400034 |
| GADGGDGGSGGAGG | ASSAGTTSQNTLY | 0.62066597 |
| GFGLDTGGAGG | ASSAGTTSQNTLY | 0.5808253 |
| GLASGGLL | ASSAGTTSQNTLY | 0.53939366 |
| GRIAGAAVLL | ASSAGTTSQNTLY | 0.7255801 |
| GSAGSLNTSF | ASSAGTTSQNTLY | 0.49464682 |
| GTGQIGI | ASSAGTTSQNTLY | 0.38326827 |
| GTLIALIL | ASSAGTTSQNTLY | 0.5447834 |
| IAATMGFVVG | ASSAGTTSQNTLY | 0.5205599 |
| IFAQALL | ASSAGTTSQNTLY | 0.7484596 |
| IGAAVALGF | ASSAGTTSQNTLY | 0.7046795 |
| IGLTPLAL | ASSAGTTSQNTLY | 0.68480134 |
| ILGSNTSTL | ASSAGTTSQNTLY | 0.7578721 |
| IVFSLTVL | ASSAGTTSQNTLY | 0.29704753 |
| LASKATGF | ASSAGTTSQNTLY | 0.63035387 |
| LASVALIL | ASSAGTTSQNTLY | 0.7304215 |
| LFAVGLTL | ASSAGTTSQNTLY | 0.66297317 |
| LFNGGIGTYI | ASSAGTTSQNTLY | 0.53886217 |
| LTLGAGII | ASSAGTTSQNTLY | 0.74352646 |
| LVAAALGDI | ASSAGTTSQNTLY | 0.51801467 |
| NTAFVQLGI | ASSAGTTSQNTLY | 0.5523918 |
| QLFSGAEL | ASSAGTTSQNTLY | 0.72210836 |
| SAATFITL | ASSAGTTSQNTLY | 0.725319 |
| SNYGLANLGV | ASSAGTTSQNTLY | 0.6742715 |

| | | |
|---|---|---|
| SYGGAYAVM | ASSAGTTSQNTLY | 0.4411891 |
| TTGVGGLTLG | ASSAGTTSQNTLY | 0.7257802 |
| TTQTIYI | ASSAGTTSQNTLY | 0.74528813 |
| VAAGAGLGAVI | ASSAGTTSQNTLY | 0.611568 |
| VAAVVTLL | ASSAGTTSQNTLY | 0.6931632 |
| VGFLGGLI | ASSAGTTSQNTLY | 0.6882572 |
| VGVIPILL | ASSAGTTSQNTLY | 0.5398799 |
| VIFASSLI | ASSAGTTSQNTLY | 0.4644696 |
| VKGGDGGAAAG | ASSAGTTSQNTLY | 0.43548635 |
| VTKTFTNI | ASSAGTTSQNTLY | 0.69853634 |
| VVAAAILL | ASSAGTTSQNTLY | 0.42142683 |
| YAALDVEL | ASSAGTTSQNTLY | 0.6720756 |
| LASVALLL | ASSAGTTSQNTLY | 0.6030199 |
| LDAVHRV | ASSAGTTSQNTLY | 0.7517246 |
| QGLSSSIF | ASSAGTTSQNTLY | 0.66346 |
| RGTFANIRL | ASSAGTTSQNTLY | 0.70722437 |
| RQLWWGHRI | ASSAGTTSQNTLY | 0.4576789 |
| YGPPGTGKTLL | ASSAGTTSQNTLY | 0.66861135 |
| AALVGAPANVA | ASSAGTTSQNTLY | 0.64739287 |
| AATFRRTE | ASSAGTTSQNTLY | 0.45224446 |
| AGALGGGI | ASSAGTTSQNTLY | 0.7128487 |
| AGGAGGAGGIGAG | ASSAGTTSQNTLY | 0.43839052 |
| AGSGALGL | ASSAGTTSQNTLY | 0.60201615 |
| AQIADQI | ASSAGTTSQNTLY | 0.65694505 |
| ASAVAVTYL | ASSAGTTSQNTLY | 0.7547831 |
| ASTVLGGTGG | ASSAGTTSQNTLY | 0.55923736 |
| AVGGEPAK | ASSAGTTSQNTLY | 0.65708524 |
| CEQGTFGGTC | ASSAGTTSQNTLY | 0.48735163 |
| DEGKAPAG | ASSAGTTSQNTLY | 0.70745575 |
| DTVVFPAL | ASSAGTTSQNTLY | 0.74199224 |
| FEGITMAG | ASSAGTTSQNTLY | 0.64878047 |
| FPLVVQD | ASSAGTTSQNTLY | 0.77782965 |
| GAFTTPQ | ASSAGTTSQNTLY | 0.7401954 |
| GFGGFGTAGGVGGA | ASSAGTTSQNTLY | 0.43422347 |
| GFLAQFL | ASSAGTTSQNTLY | 0.70702356 |
| GGAGGAGGNAGVA | ASSAGTTSQNTLY | 0.7343071 |
| GGVGTAGAGGAGGAGG | ASSAGTTSQNTLY | 0.7283101 |
| GNGGGGDAGNAGSGGN | ASSAGTTSQNTLY | 0.71701825 |
| GTGGAGGAGNDAGS | ASSAGTTSQNTLY | 0.7063724 |
| GTTNTGWFNS | ASSAGTTSQNTLY | 0.5581338 |
| IFMGPLFTL | ASSAGTTSQNTLY | 0.7277216 |
| ILAHAIEAI | ASSAGTTSQNTLY | 0.7667514 |
| IRSAIREAL | ASSAGTTSQNTLY | 0.7206855 |
| ITFAGGG | ASSAGTTSQNTLY | 0.51738375 |
| ITGMSGHQEV | ASSAGTTSQNTLY | 0.4900791 |
| IVAGLACTL | ASSAGTTSQNTLY | 0.6896916 |
| IVGGGLAGA | ASSAGTTSQNTLY | 0.7290743 |
| IVTGVLNI | ASSAGTTSQNTLY | 0.5212886 |
| LAQQGCI | ASSAGTTSQNTLY | 0.5673629 |
| LCLHVEN | ASSAGTTSQNTLY | 0.59196967 |
| LDAISDGS | ASSAGTTSQNTLY | 0.41757527 |

| | | |
|---|---|---|
| LGGFAAML | ASSAGTTSQNTLY | 0.6877344 |
| LGGKNPMI | ASSAGTTSQNTLY | 0.73393106 |
| LGPKGVAAIL | ASSAGTTSQNTLY | 0.6894658 |
| LMVGVSI | ASSAGTTSQNTLY | 0.5048294 |
| LVGAAFVL | ASSAGTTSQNTLY | 0.7188858 |
| LVLAVAPL | ASSAGTTSQNTLY | 0.73674935 |
| MAAIAVIDAA | ASSAGTTSQNTLY | 0.7671399 |
| NNGINAELQ | ASSAGTTSQNTLY | 0.42918798 |
| RGEPLVLVHG | ASSAGTTSQNTLY | 0.26554847 |
| RGGAGGAAGLI | ASSAGTTSQNTLY | 0.64960194 |
| RSYRMHPAV | ASSAGTTSQNTLY | 0.68729633 |
| SAAHLLAGQLA | ASSAGTTSQNTLY | 0.5888624 |
| SAAVGSY | ASSAGTTSQNTLY | 0.61898947 |
| SPHAVLHRI | ASSAGTTSQNTLY | 0.67886686 |
| SQIAKVL | ASSAGTTSQNTLY | 0.7012724 |
| TALGHLVP | ASSAGTTSQNTLY | 0.54577667 |
| TAPTTSGAA | ASSAGTTSQNTLY | 0.45924392 |
| TDGDDGNGGSPGT | ASSAGTTSQNTLY | 0.68683237 |
| TDGLGEN | ASSAGTTSQNTLY | 0.7484466 |
| TGGPPLPGQA | ASSAGTTSQNTLY | 0.46413085 |
| TIPAITI | ASSAGTTSQNTLY | 0.5920955 |
| TIPGFTI | ASSAGTTSQNTLY | 0.6238191 |
| TSIASGALA | ASSAGTTSQNTLY | 0.61596 |
| TVTLPGAAI | ASSAGTTSQNTLY | 0.5791346 |
| VASGLSAALP | ASSAGTTSQNTLY | 0.4022621 |
| VGAGVIGI | ASSAGTTSQNTLY | 0.68608195 |
| VGPKRFGV | ASSAGTTSQNTLY | 0.41651362 |
| VNVAFPD | ASSAGTTSQNTLY | 0.70494175 |
| VSMTNSGVS | ASSAGTTSQNTLY | 0.6946011 |
| VTDVLGDLSV | ASSAGTTSQNTLY | 0.7083069 |
| VVVVGAGFAGL | ASSAGTTSQNTLY | 0.51782954 |
| VYTVGTVV | ASSAGTTSQNTLY | 0.45238638 |
| WTTFSGP | ASSAGTTSQNTLY | 0.43138966 |
| YDELVEAF | ASSAGTTSQNTLY | 0.40319565 |
| YGTGGAGGNGG | ASSAGTTSQNTLY | 0.74292725 |
| YTNVMAV | ASSAGTTSQNTLY | 0.59955275 |
| FLEQVHQL | ASSAGTTSQNTLY | 0.7445382 |
| GLSLLAVAV | ASSAGTTSQNTLY | 0.6602427 |
| LEQVHQL | ASSAGTTSQNTLY | 0.5303719 |
| LNAAGAI | ASSAGTTSQNTLY | 0.69917905 |
| PAVHASL | ASSAGTTSQNTLY | 0.55405694 |
| TELEDDA | ASSAGTTSQNTLY | 0.6561085 |
| TPPGPPN | ASSAGTTSQNTLY | 0.6139674 |
| TPPPPPPP | ASSAGTTSQNTLY | 0.7385578 |
| TQSPALI | ASSHTGASQNTLY | 0.48846686 |
| EFAAAFPQ | ASSHTGASQNTLY | 0.7227936 |
| LLSGNLDVL | ASSHTGASQNTLY | 0.67276347 |
| GPANLGNYNI | ASSHTGASQNTLY | 0.49755532 |
| TLGPTLP | ASSHTGASQNTLY | 0.6845859 |
| VYAYSSCCK | ASSHTGASQNTLY | 0.70741904 |
| APAPAPAPNI | ASSHTGASQNTLY | 0.6906973 |

| | | |
|---|---|---|
| PGGPVPIP | ASSHTGASQNTLY | 0.44245934 |
| FLTVALEP | ASSHTGASQNTLY | 0.6865115 |
| SQYFGANL | ASSHTGASQNTLY | 0.4955066 |
| GGVKPAA | ASSHTGASQNTLY | 0.6272632 |
| GGTGGAGGAGGW | ASSHTGASQNTLY | 0.6390319 |
| GVGGVAGLGGAGG | ASSHTGASQNTLY | 0.5657694 |
| VLLVLATTI | ASSHTGASQNTLY | 0.75886226 |
| IGIFAAML | ASSHTGASQNTLY | 0.5249198 |
| AVGDLID | ASSHTGASQNTLY | 0.516413 |
| NISPLPL | ASSHTGASQNTLY | 0.5471807 |
| VTAVFVAI | ASSHTGASQNTLY | 0.6488064 |
| SGDDNDAAL | ASSHTGASQNTLY | 0.7213622 |
| AVAALAHS | ASSHTGASQNTLY | 0.4491939 |
| GGVTVMATP | ASSHTGASQNTLY | 0.6575811 |
| AGLAANVRD | ASSHTGASQNTLY | 0.5984058 |
| TNRPDLIDPALLRPG | ASSHTGASQNTLY | 0.46430868 |
| LANLGVTAL | ASSHTGASQNTLY | 0.7095235 |
| TGERIEL | ASSHTGASQNTLY | 0.588713 |
| NGEELAA | ASSHTGASQNTLY | 0.55389696 |
| VLSLVWG | ASSHTGASQNTLY | 0.7206603 |
| IGYALWL | ASSHTGASQNTLY | 0.662528 |
| LSLLLAPL | ASSHTGASQNTLY | 0.69785297 |
| TLDEALK | ASSHTGASQNTLY | 0.7324432 |
| TDGAGTAEGG | ASSHTGASQNTLY | 0.49532348 |
| NAPGGAGGNGG | ASSHTGASQNTLY | 0.7033325 |
| VSVFMGMGGNF | ASSHTGASQNTLY | 0.6706882 |
| LASYLAAL | ASSHTGASQNTLY | 0.67943853 |
| LVGGTPAL | ASSHTGASQNTLY | 0.559792 |
| VRNADVL | ASSHTGASQNTLY | 0.47881877 |
| TCSSPFLVSAM | ASSHTGASQNTLY | 0.7487217 |
| VHITDAE | ASSHTGASQNTLY | 0.7018481 |
| EYAAPSF | ASSHTGASQNTLY | 0.6986559 |
| VVDAFHGGL | ASSHTGASQNTLY | 0.7122936 |
| ASVVDPQ | ASSHTGASQNTLY | 0.72181636 |
| SYNEFGGLS | ASSHTGASQNTLY | 0.7496418 |
| DPDGSVY | ASSHTGASQNTLY | 0.51133883 |
| MVSTIHATE | ASSHTGASQNTLY | 0.4643243 |
| LAAGVADPV | ASSHTGASQNTLY | 0.60379523 |
| TKEQIAA | ASSHTGASQNTLY | 0.52602404 |
| SLASPAAL | ASSHTGASQNTLY | 0.62399286 |
| IGLVFYGV | ASSHTGASQNTLY | 0.42622238 |
| FLLDASNL | ASSHTGASQNTLY | 0.6814969 |
| YQVGNFP | ASSHTGASQNTLY | 0.77136904 |
| ESLLSQST | ASSHTGASQNTLY | 0.54090506 |
| LFTVGFAAVG | ASSHTGASQNTLY | 0.7692551 |
| ATPPTPT | ASSHTGASQNTLY | 0.44490513 |
| TDLATLG | ASSHTGASQNTLY | 0.68136585 |
| SAGGTGR | ASSHTGASQNTLY | 0.6948183 |
| GFSNSGS | ASSHTGASQNTLY | 0.41490138 |
| GGNGGSGGTSV | ASSHTGASQNTLY | 0.68434525 |
| GTSVATGGAGN | ASSHTGASQNTLY | 0.48559746 |

| | | |
|---|---|---|
| NGGNGGSGGTS | ASSHTGASQNTLY | 0.5559037 |
| AVVKFPH | ASSHTGASQNTLY | 0.73048824 |
| TPPPPPP | ASSHTGASQNTLY | 0.5725085 |
| AAPAAAAAPA | ASSHTGASQNTLY | 0.6985528 |
| IALNEDL | ASSHTGASQNTLY | 0.69844306 |
| IVNVSSM | ASSHTGASQNTLY | 0.5311835 |
| TGIPPAPR | ASSHTGASQNTLY | 0.699242 |
| FDLGGGTF | ASSHTGASQNTLY | 0.7237803 |
| FDLGGGTFDVS | ASSHTGASQNTLY | 0.6180581 |
| NEPTAAA | ASSHTGASQNTLY | 0.7681881 |
| NEPTAAALA | ASSHTGASQNTLY | 0.50901276 |
| GSIINISSI | ASSHTGASQNTLY | 0.7466774 |
| VAPGNAGTA | ASSHTGASQNTLY | 0.7501348 |
| TGGGASLE | ASSHTGASQNTLY | 0.6083281 |
| VAVYNHYKRI | ASSHTGASQNTLY | 0.5738588 |
| TDPAPATT | ASSHTGASQNTLY | 0.7074362 |
| TNRPDILDPALLRPG | ASSHTGASQNTLY | 0.67489046 |
| LTTTTTS | ASSHTGASQNTLY | 0.7016853 |
| PTGSGKTL | ASSHTGASQNTLY | 0.7615095 |
| TGSGKTL | ASSHTGASQNTLY | 0.64029515 |
| TIGHVDHGK | ASSHTGASQNTLY | 0.7171964 |
| TIGHVDHGKTT | ASSHTGASQNTLY | 0.70203495 |
| GHVDHGKTT | ASSHTGASQNTLY | 0.68621933 |
| AQFEHTLL | ASSHTGASQNTLY | 0.67815995 |
| IVNQDEL | ASSHTGASQNTLY | 0.47548747 |
| TNGSQFF | ASSHTGASQNTLY | 0.681813 |
| TGKGIVE | ASSHTGASQNTLY | 0.6671113 |
| LISNASDA | ASSHTGASQNTLY | 0.70413613 |
| AELRTLGELTKIAKAHGAQ | ASSHTGASQNTLY | 0.70232254 |
| APNAVAAAEPSW | ASSHTGASQNTLY | 0.66823816 |
| APQVAGAATAVL | ASSHTGASQNTLY | 0.7195488 |
| ARRDPAAAAAILPTDA | ASSHTGASQNTLY | 0.72103214 |
| DAAARDVLAEAGLAEYF | ASSHTGASQNTLY | 0.5878452 |
| DDQIVLSAKANLAKVEQ | ASSHTGASQNTLY | 0.6835171 |
| DELLPDTGEDSDEDGH | ASSHTGASQNTLY | 0.63774586 |
| ELPEAPANRATVPQD | ASSHTGASQNTLY | 0.6811409 |
| FGQSKIERVTGIG | ASSHTGASQNTLY | 0.45773375 |
| GAGGTGSTAGGAGGAGGAGGLY | ASSHTGASQNTLY | 0.70904493 |
| GHGGTGAAGGNGGTGGAGGSGI | ASSHTGASQNTLY | 0.6559797 |
| GNGGNGGAGGEGGPGGAGGAGA | ASSHTGASQNTLY | 0.33384928 |
| GSTAGNGGNGGNATNGTIAGQP | ASSHTGASQNTLY | 0.6935039 |
| GTGGDGGNGGNGGTGG | ASSHTGASQNTLY | 0.69601333 |
| GVRRPAGIPADAPV | ASSHTGASQNTLY | 0.69448525 |
| GYRQVTDAHLLTLARR | ASSHTGASQNTLY | 0.7130648 |
| IAKMGQDGHDRG | ASSHTGASQNTLY | 0.4765222 |
| KAEPSIEIQYT | ASSHTGASQNTLY | 0.65026486 |
| KPVRIRMAADEDTVAVT | ASSHTGASQNTLY | 0.70850885 |
| MPPRIAGMRLLVIKPEP | ASSHTGASQNTLY | 0.7178807 |
| MKFYAVALFRSRGVPPTR | ASSHTGASQNTLY | 0.620947 |
| NAGGFGILGAVAHSP | ASSHTGASQNTLY | 0.69561744 |
| NEGREITLAMAEM | ASSHTGASQNTLY | 0.71371853 |

| | | |
|---|---|---|
| NELVEWSHPLGHRWRWE | ASSHTGASQNTLY | 0.69078684 |
| NLGAGGTGGNGGNPG | ASSHTGASQNTLY | 0.70812595 |
| NTEWPKWETVMAADPVD | ASSHTGASQNTLY | 0.7159965 |
| PASAAPAETQP | ASSHTGASQNTLY | 0.61173964 |
| QQSSDGKNGVRAAEITGE | ASSHTGASQNTLY | 0.7161927 |
| RAPDAVTTSWWKE | ASSHTGASQNTLY | 0.7192539 |
| SGFGNVGAHTSGWFNQ | ASSHTGASQNTLY | 0.6886665 |
| SGVAGAVGQAASVGGLKV | ASSHTGASQNTLY | 0.69349086 |
| SNGGAGTGSAGGNGGT | ASSHTGASQNTLY | 0.5903861 |
| TGGAGGEGGAAG | ASSHTGASQNTLY | 0.6706106 |
| TGGDGGAAGAGGAPGHG | ASSHTGASQNTLY | 0.44688928 |
| TMPPLLAAIANAGRHGV | ASSHTGASQNTLY | 0.7027496 |
| TSSTAGNSGGA | ASSHTGASQNTLY | 0.71360993 |
| VPATVPVGILC | ASSHTGASQNTLY | 0.6756265 |
| VQRPDSGGVVLTGRIS | ASSHTGASQNTLY | 0.61440194 |
| WTPNPVPATKGVPAGLVQAT | ASSHTGASQNTLY | 0.6239966 |
| YALKRALAKFSDRFDVVII | ASSHTGASQNTLY | 0.65784466 |
| YLIGSVIV | ASSHTGASQNTLY | 0.6838974 |
| YGQLVAVID | ASSHTGASQNTLY | 0.39152548 |
| VTVGGMTLPA | ASSHTGASQNTLY | 0.6464379 |
| VLALPAV | ASSHTGASQNTLY | 0.71612537 |
| VIPPTLNYET | ASSHTGASQNTLY | 0.71635866 |
| VGMGGMHPSSG | ASSHTGASQNTLY | 0.5698642 |
| VGLAGGAATA | ASSHTGASQNTLY | 0.71901095 |
| VGGAGGAGGSGP | ASSHTGASQNTLY | 0.70111954 |
| VFAAFMV | ASSHTGASQNTLY | 0.7134203 |
| TLYDKLR | ASSHTGASQNTLY | 0.6778041 |
| TLAVFTVL | ASSHTGASQNTLY | 0.6175129 |
| TGSVQPARV | ASSHTGASQNTLY | 0.70766085 |
| TGGGAGSGGT | ASSHTGASQNTLY | 0.6761976 |
| SHLETSL | ASSHTGASQNTLY | 0.6452116 |
| SFTTGMT | ASSHTGASQNTLY | 0.7129127 |
| QVEVAKRMGVRQPTVS | ASSHTGASQNTLY | 0.6892083 |
| QGSGGPALAM | ASSHTGASQNTLY | 0.6699065 |
| MLAPSNP | ASSHTGASQNTLY | 0.6476224 |
| LVNNAAI | ASSHTGASQNTLY | 0.6993725 |
| LTGEINTLA | ASSHTGASQNTLY | 0.61455804 |
| LSIFGPV | ASSHTGASQNTLY | 0.59417814 |
| LLALLANL | ASSHTGASQNTLY | 0.6336663 |
| LISNASDALDKLR | ASSHTGASQNTLY | 0.68895817 |
| LINEAVP | ASSHTGASQNTLY | 0.7147949 |
| LGYAFSGL | ASSHTGASQNTLY | 0.6995213 |
| LDFVAVAL | ASSHTGASQNTLY | 0.6460984 |
| LAQPGAE | ASSHTGASQNTLY | 0.70610833 |
| ISNLGAVTGF | ASSHTGASQNTLY | 0.7017282 |
| HQEDPGRI | ASSHTGASQNTLY | 0.69126374 |
| GTPGEPTP | ASSHTGASQNTLY | 0.70698845 |
| GFFNSGS | ASSHTGASQNTLY | 0.66466695 |
| GAGGGGLTAGAN | ASSHTGASQNTLY | 0.70680106 |
| FVDALVNM | ASSHTGASQNTLY | 0.7015977 |
| FLTVAPEL | ASSHTGASQNTLY | 0.70890754 |

| | | |
|---|---|---|
| FETLTAL | ASSHTGASQNTLY | 0.70523846 |
| DVINPIEL | ASSHTGASQNTLY | 0.6996888 |
| APVSGAPV | ASSHTGASQNTLY | 0.67901087 |
| ALVGGLAW | ASSHTGASQNTLY | 0.38035035 |
| ALAAAAATAI | ASSHTGASQNTLY | 0.6346638 |
| AFGLGLLL | ASSHTGASQNTLY | 0.7047285 |
| AFFAAASL | ASSHTGASQNTLY | 0.6990225 |
| AAAALPAFSPP | ASSHTGASQNTLY | 0.71235615 |
| VVLAPAL | ASSHTGASQNTLY | 0.69981664 |
| TIGHVDHGKTTL | ASSHTGASQNTLY | 0.6578148 |
| LISNASDALDK | ASSHTGASQNTLY | 0.70700383 |
| LISNASDAL | ASSHTGASQNTLY | 0.7093867 |
| ISNASDALDK | ASSHTGASQNTLY | 0.50961167 |
| NQAETLVYQTE | ASSHTGASQNTLY | 0.6808344 |
| TGGTGGTAGAG | ASSHTGASQNTLY | 0.72380924 |
| TGSTAGGAGGA | ASSHTGASQNTLY | 0.6838876 |
| DPLAVPVAAGPV | ASSHTGASQNTLY | 0.68615687 |
| GGTGGTPGNGGN | ASSHTGASQNTLY | 0.6348188 |
| PAGLAGVDPATL | ASSHTGASQNTLY | 0.7203362 |
| GNNLVVEAPHAGASV | ASSHTGASQNTLY | 0.6523705 |
| KVTRIMSASTLEQPA | ASSHTGASQNTLY | 0.7169101 |
| LGFALPIALPRRTVV | ASSHTGASQNTLY | 0.38095394 |
| TRLGNLTARKLVVLV | ASSHTGASQNTLY | 0.6124807 |
| WSQQRRLIYFPSAGP | ASSHTGASQNTLY | 0.52860576 |
| GPNGNPDPMAAAVDIR | ASSHTGASQNTLY | 0.4759186 |
| TQRCVSVADLRAAASR | ASSHTGASQNTLY | 0.71064526 |
| QFVLDVAEHPVEVLTETV | ASSHTGASQNTLY | 0.6695349 |
| GGAGGTGGAGGNGGTQAGDG | ASSHTGASQNTLY | 0.7110508 |
| PLLATLRAIAVDVDAKPAQV | ASSHTGASQNTLY | 0.69397163 |
| APKFRTGGYMIDFWGVGYQVAK | ASSHTGASQNTLY | 0.7141203 |
| GAGGAGGAGGAAGGPGGTGGTG | ASSHTGASQNTLY | 0.6849984 |
| IGALGLLATDVAPAGSVAFGVA | ASSHTGASQNTLY | 0.7025646 |
| AGGAGGIGGNGGAGGAGGVLMG | ASSHTGASQNTLY | 0.583071 |
| ADELIGLL | ASSHTGASQNTLY | 0.7140646 |
| ADRALKA | ASSHTGASQNTLY | 0.7135862 |
| AGAVGQVGGAGGA | ASSHTGASQNTLY | 0.6651702 |
| AGGGGGLGGAP | ASSHTGASQNTLY | 0.61474717 |
| AGGLLYGNGGAG | ASSHTGASQNTLY | 0.6576954 |
| AGLESIGL | ASSHTGASQNTLY | 0.6987587 |
| AGLKTGII | ASSHTGASQNTLY | 0.32840598 |
| ASLAALLL | ASSHTGASQNTLY | 0.3702509 |
| ASVINISSI | ASSHTGASQNTLY | 0.6535932 |
| ATPPGPA | ASSHTGASQNTLY | 0.6145575 |
| ATVASALV | ASSHTGASQNTLY | 0.7183641 |
| ATVASLIL | ASSHTGASQNTLY | 0.7091046 |
| AVNGPVLIRH | ASSHTGASQNTLY | 0.68689036 |
| DGAAGAVGI | ASSHTGASQNTLY | 0.7011653 |
| DPLLHGI | ASSHTGASQNTLY | 0.7013933 |
| ELGAALGAL | ASSHTGASQNTLY | 0.25980917 |
| ELGRTQAV | ASSHTGASQNTLY | 0.6399542 |
| FVAGGYLS | ASSHTGASQNTLY | 0.6162226 |

| | | |
|---|---|---|
| GASGGAGGAGGW | ASSHTGASQNTLY | 0.71960735 |
| GGAGAGGDGGAGGI | ASSHTGASQNTLY | 0.65793335 |
| GTGHNGSQPGMGGNG | ASSHTGASQNTLY | 0.64603937 |
| GYAAASASAS | ASSHTGASQNTLY | 0.69340575 |
| HQVDVAAVQQG | ASSHTGASQNTLY | 0.68500435 |
| IGAAIAHT | ASSHTGASQNTLY | 0.6782781 |
| IGMIEVAQL | ASSHTGASQNTLY | 0.6104164 |
| IRLAPEHTSAMD | ASSHTGASQNTLY | 0.2972704 |
| IYNVSVL | ASSHTGASQNTLY | 0.71212566 |
| KQGFLGM | ASSHTGASQNTLY | 0.47397453 |
| LESVITQL | ASSHTGASQNTLY | 0.6569934 |
| LGLGALVVV | ASSHTGASQNTLY | 0.37646955 |
| LIGNGGNGGSGGI | ASSHTGASQNTLY | 0.51790005 |
| LNAINAPTL | ASSHTGASQNTLY | 0.5206864 |
| LQLLNTGV | ASSHTGASQNTLY | 0.707741 |
| LQTSIEF | ASSHTGASQNTLY | 0.67817914 |
| MSFAVSEL | ASSHTGASQNTLY | 0.6650413 |
| NTGALGL | ASSHTGASQNTLY | 0.71948516 |
| PPGAGGAGGAGTT | ASSHTGASQNTLY | 0.69453 |
| QLVGLGI | ASSHTGASQNTLY | 0.67250204 |
| RFLEQVHQL | ASSHTGASQNTLY | 0.6853869 |
| RGVGYVLR | ASSHTGASQNTLY | 0.6416813 |
| SATGVFAGVI | ASSHTGASQNTLY | 0.634943 |
| SLPAITLP | ASSHTGASQNTLY | 0.6592345 |
| STALAFTV | ASSHTGASQNTLY | 0.7224107 |
| SVLTVGLGK | ASSHTGASQNTLY | 0.6155159 |
| TALAGAGSGI | ASSHTGASQNTLY | 0.5580371 |
| TGLPFTGI | ASSHTGASQNTLY | 0.655288 |
| VAEHVTV | ASSHTGASQNTLY | 0.72064006 |
| VDIVLNSL | ASSHTGASQNTLY | 0.72203195 |
| VLTGGGAVL | ASSHTGASQNTLY | 0.64556384 |
| VNVGLAYGI | ASSHTGASQNTLY | 0.65439594 |
| VSSAAVGAV | ASSHTGASQNTLY | 0.6433614 |
| VTSVYEAGV | ASSHTGASQNTLY | 0.6926836 |
| WVALDRGA | ASSHTGASQNTLY | 0.6124472 |
| YLVGSVVI | ASSHTGASQNTLY | 0.5606177 |
| GPPGTGKTLL | ASSHTGASQNTLY | 0.6934279 |
| GVGMTKF | ASSHTGASQNTLY | 0.72133815 |
| IGIGGSDL | ASSHTGASQNTLY | 0.6667167 |
| IGTLLAEL | ASSHTGASQNTLY | 0.52165294 |
| IYTSGTT | ASSHTGASQNTLY | 0.69296265 |
| LSFPAGV | ASSHTGASQNTLY | 0.6493641 |
| LYTSGTT | ASSHTGASQNTLY | 0.6749568 |
| VGAFPFGF | ASSHTGASQNTLY | 0.69238997 |
| VGAGTVEF | ASSHTGASQNTLY | 0.4702736 |
| VGKSTLLL | ASSHTGASQNTLY | 0.66320044 |
| AALAHAGGGIV | ASSHTGASQNTLY | 0.72133803 |
| AALVAVSLL | ASSHTGASQNTLY | 0.6730845 |
| AAVSRAVAAL | ASSHTGASQNTLY | 0.6983892 |
| AGGSSGIGGF | ASSHTGASQNTLY | 0.7155619 |
| AGLTVANANG | ASSHTGASQNTLY | 0.706866 |

| | | |
|---|---|---|
| AGLVGAVL | ASSHTGASQNTLY | 0.7167487 |
| ALSQAFTV | ASSHTGASQNTLY | 0.7049296 |
| ALTFSVSL | ASSHTGASQNTLY | 0.6793356 |
| ALTGAALVAL | ASSHTGASQNTLY | 0.67484146 |
| ASFSIGAGVL | ASSHTGASQNTLY | 0.7027613 |
| DGSVYKI | ASSHTGASQNTLY | 0.7036555 |
| EQLAEIV | ASSHTGASQNTLY | 0.6871069 |
| FAVSELTVM | ASSHTGASQNTLY | 0.5970974 |
| GAAPLAAGL | ASSHTGASQNTLY | 0.71656746 |
| GADGGDGGSGGAGG | ASSHTGASQNTLY | 0.67344975 |
| GFGLDTGGAGG | ASSHTGASQNTLY | 0.7084862 |
| GLASGGLL | ASSHTGASQNTLY | 0.6337249 |
| GRIAGAAVLL | ASSHTGASQNTLY | 0.6581322 |
| GSAGSLNTSF | ASSHTGASQNTLY | 0.71142054 |
| GTGQIGI | ASSHTGASQNTLY | 0.2640087 |
| GTLIALIL | ASSHTGASQNTLY | 0.37638032 |
| IAATMGFVVG | ASSHTGASQNTLY | 0.67800605 |
| IFAQALL | ASSHTGASQNTLY | 0.7145904 |
| IGAAVALGF | ASSHTGASQNTLY | 0.27048478 |
| IGLTPLAL | ASSHTGASQNTLY | 0.6434492 |
| ILGSNTSTL | ASSHTGASQNTLY | 0.51473945 |
| IVFSLTVL | ASSHTGASQNTLY | 0.69349885 |
| LASKATGF | ASSHTGASQNTLY | 0.6993577 |
| LASVALIL | ASSHTGASQNTLY | 0.5501885 |
| LFAVGLTL | ASSHTGASQNTLY | 0.7078248 |
| LFNGGIGTYI | ASSHTGASQNTLY | 0.56510985 |
| LTLGAGII | ASSHTGASQNTLY | 0.46039402 |
| LVAAALGDI | ASSHTGASQNTLY | 0.7099805 |
| NTAFVQLGI | ASSHTGASQNTLY | 0.71255857 |
| QLFSGAEL | ASSHTGASQNTLY | 0.5932498 |
| SAATFITL | ASSHTGASQNTLY | 0.7079356 |
| SNYGLANLGV | ASSHTGASQNTLY | 0.5997075 |
| SYGGAYAVM | ASSHTGASQNTLY | 0.72232085 |
| TTGVGGLTLG | ASSHTGASQNTLY | 0.67856646 |
| TTQTIYI | ASSHTGASQNTLY | 0.43111882 |
| VAAGAGLGAVI | ASSHTGASQNTLY | 0.71048915 |
| VAAVVTLL | ASSHTGASQNTLY | 0.696041 |
| VGFLGGLI | ASSHTGASQNTLY | 0.69864076 |
| VGVIPILL | ASSHTGASQNTLY | 0.7052057 |
| VIFASSLI | ASSHTGASQNTLY | 0.5616669 |
| VKGGDGGAAAG | ASSHTGASQNTLY | 0.6853596 |
| VTKTFTNI | ASSHTGASQNTLY | 0.7193381 |
| VVAAAILL | ASSHTGASQNTLY | 0.7138179 |
| YAALDVEL | ASSHTGASQNTLY | 0.54669 |
| LASVALLL | ASSHTGASQNTLY | 0.70276165 |
| LDAVHRV | ASSHTGASQNTLY | 0.6947331 |
| QGLSSSIF | ASSHTGASQNTLY | 0.64628124 |
| RGTFANIRL | ASSHTGASQNTLY | 0.71498704 |
| RQLWWGHRI | ASSHTGASQNTLY | 0.66242635 |
| YGPPGTGKTLL | ASSHTGASQNTLY | 0.71176 |
| AALVGAPANVA | ASSHTGASQNTLY | 0.7042035 |

| | | |
|---|---|---|
| AATFRRTE | ASSHTGASQNTLY | 0.7134288 |
| AGALGGGI | ASSHTGASQNTLY | 0.55485713 |
| AGGAGGAGGIGAG | ASSHTGASQNTLY | 0.7007524 |
| AGSGALGL | ASSHTGASQNTLY | 0.683777 |
| AQIADQI | ASSHTGASQNTLY | 0.7094277 |
| ASAVAVTYL | ASSHTGASQNTLY | 0.6799905 |
| ASTVLGGTGG | ASSHTGASQNTLY | 0.607371 |
| AVGGEPAK | ASSHTGASQNTLY | 0.69201934 |
| CEQGTFGGTC | ASSHTGASQNTLY | 0.7165576 |
| DEGKAPAG | ASSHTGASQNTLY | 0.6730075 |
| DTVVFPAL | ASSHTGASQNTLY | 0.6935562 |
| FEGITMAG | ASSHTGASQNTLY | 0.72012985 |
| FPLVVQD | ASSHTGASQNTLY | 0.6363381 |
| GAFTTPQ | ASSHTGASQNTLY | 0.6876236 |
| GFGGFGTAGGVGGA | ASSHTGASQNTLY | 0.70426834 |
| GFLAQFL | ASSHTGASQNTLY | 0.7129421 |
| GGAGGAGGNAGVA | ASSHTGASQNTLY | 0.6570151 |
| GGVGTAGAGGAGGAGG | ASSHTGASQNTLY | 0.71465373 |
| GNGGGGDAGNAGSGGN | ASSHTGASQNTLY | 0.62453884 |
| GTGGAGGAGNDAGS | ASSHTGASQNTLY | 0.7123233 |
| GTTNTGWFNS | ASSHTGASQNTLY | 0.629813 |
| IFMGPLFTL | ASSHTGASQNTLY | 0.70241594 |
| ILAHAIEAI | ASSHTGASQNTLY | 0.6640392 |
| IRSAIREAL | ASSHTGASQNTLY | 0.7085222 |
| ITFAGGG | ASSHTGASQNTLY | 0.71151465 |
| ITGMSGHQEV | ASSHTGASQNTLY | 0.713964 |
| IVAGLACTL | ASSHTGASQNTLY | 0.37667188 |
| IVGGGLAGA | ASSHTGASQNTLY | 0.7147211 |
| IVTGVLNI | ASSHTGASQNTLY | 0.71337926 |
| LAQQGCI | ASSHTGASQNTLY | 0.721404 |
| LCLHVEN | ASSHTGASQNTLY | 0.68021715 |
| LDAISDGS | ASSHTGASQNTLY | 0.69647074 |
| LGGFAAML | ASSHTGASQNTLY | 0.6852671 |
| LGGKNPMI | ASSHTGASQNTLY | 0.65371644 |
| LGPKGVAAIL | ASSHTGASQNTLY | 0.64166963 |
| LMVGVSI | ASSHTGASQNTLY | 0.6712892 |
| LVGAAFVL | ASSHTGASQNTLY | 0.6671747 |
| LVLAVAPL | ASSHTGASQNTLY | 0.6643861 |
| MAAIAVIDAA | ASSHTGASQNTLY | 0.71041 |
| NNGINAELQ | ASSHTGASQNTLY | 0.71137214 |
| RGEPLVLVHG | ASSHTGASQNTLY | 0.71235937 |
| RGGAGGAAGLI | ASSHTGASQNTLY | 0.7164643 |
| RSYRMHPAV | ASSHTGASQNTLY | 0.7163911 |
| SAAHLLAGQLA | ASSHTGASQNTLY | 0.70491827 |
| SAAVGSY | ASSHTGASQNTLY | 0.66721654 |
| SPHAVLHRI | ASSHTGASQNTLY | 0.6854562 |
| SQIAKVL | ASSHTGASQNTLY | 0.6488094 |
| TALGHLVP | ASSHTGASQNTLY | 0.6656741 |
| TAPTTSGAA | ASSHTGASQNTLY | 0.6768998 |
| TDGDDGNGGSPGT | ASSHTGASQNTLY | 0.7011523 |
| TDGLGEN | ASSHTGASQNTLY | 0.2686532 |

| | | |
|---|---|---|
| TGGPPLPGQA | ASSHTGASQNTLY | 0.676509 |
| TIPAITI | ASSHTGASQNTLY | 0.72106767 |
| TIPGFTI | ASSHTGASQNTLY | 0.63465863 |
| TSIASGALA | ASSHTGASQNTLY | 0.7018722 |
| TVTLPGAAI | ASSHTGASQNTLY | 0.68705124 |
| VASGLSAALP | ASSHTGASQNTLY | 0.6475854 |
| VGAGVIGI | ASSHTGASQNTLY | 0.71312654 |
| VGPKRFGV | ASSHTGASQNTLY | 0.6354941 |
| VNVAFPD | ASSHTGASQNTLY | 0.69618654 |
| VSMTNSGVS | ASSHTGASQNTLY | 0.71641946 |
| VTDVLGDLSV | ASSHTGASQNTLY | 0.69167525 |
| VVVVGAGFAGL | ASSHTGASQNTLY | 0.6793245 |
| VYTVGTVV | ASSHTGASQNTLY | 0.65023607 |
| WTTFSGP | ASSHTGASQNTLY | 0.53183204 |
| YDELVEAF | ASSHTGASQNTLY | 0.6949271 |
| YGTGGAGGNGG | ASSHTGASQNTLY | 0.4079417 |
| YTNVMAV | ASSHTGASQNTLY | 0.68426657 |
| FLEQVHQL | ASSHTGASQNTLY | 0.61413074 |
| GLSLLAVAV | ASSHTGASQNTLY | 0.7068751 |
| LEQVHQL | ASSHTGASQNTLY | 0.69737256 |
| LNAAGAI | ASSHTGASQNTLY | 0.6884769 |
| PAVHASL | ASSHTGASQNTLY | 0.65861356 |
| TELEDDA | ASSHTGASQNTLY | 0.6780428 |
| TPPGPPN | ASSHTGASQNTLY | 0.6552174 |
| TPPPPPPP | ASSHTGASQNTLY | 0.59645355 |
| TQSPALI | ASSHLGGWAETLY | 0.704049 |
| EFAAAFPQ | ASSHLGGWAETLY | 0.64843583 |
| LLSGNLDVL | ASSHLGGWAETLY | 0.6610593 |
| GPANLGNYNI | ASSHLGGWAETLY | 0.71690845 |
| TLGPTLP | ASSHLGGWAETLY | 0.69685304 |
| VYAYSSCCK | ASSHLGGWAETLY | 0.5312593 |
| APAPAPAPNI | ASSHLGGWAETLY | 0.67983687 |
| PGGPVPIP | ASSHLGGWAETLY | 0.6899151 |
| FLTVALEP | ASSHLGGWAETLY | 0.68432677 |
| SQYFGANL | ASSHLGGWAETLY | 0.27014053 |
| GGVKPAA | ASSHLGGWAETLY | 0.7033484 |
| GGTGGAGGAGGW | ASSHLGGWAETLY | 0.65641403 |
| GVGGVAGLGGAGG | ASSHLGGWAETLY | 0.6297796 |
| VLLVLATTI | ASSHLGGWAETLY | 0.6422421 |
| IGIFAAML | ASSHLGGWAETLY | 0.71046746 |
| AVGDLID | ASSHLGGWAETLY | 0.5128953 |
| NISPLPL | ASSHLGGWAETLY | 0.6827059 |
| VTAVFVAI | ASSHLGGWAETLY | 0.7070932 |
| SGDDNDAAL | ASSHLGGWAETLY | 0.71779 |
| AVAALAHS | ASSHLGGWAETLY | 0.718537 |
| GGVTVMATP | ASSHLGGWAETLY | 0.7117471 |
| AGLAANVRD | ASSHLGGWAETLY | 0.6038465 |
| TNRPDLIDPALLRPG | ASSHLGGWAETLY | 0.70743346 |
| LANLGVTAL | ASSHLGGWAETLY | 0.67618924 |
| TGERIEL | ASSHLGGWAETLY | 0.5489248 |
| NGEELAA | ASSHLGGWAETLY | 0.45893133 |

| | | |
|---|---|---|
| VLSLVWG | ASSHLGGWAETLY | 0.69773257 |
| IGYALWL | ASSHLGGWAETLY | 0.53967386 |
| LSLLLAPL | ASSHLGGWAETLY | 0.6948465 |
| TLDEALK | ASSHLGGWAETLY | 0.7105504 |
| TDGAGTAEGG | ASSHLGGWAETLY | 0.6685847 |
| NAPGGAGGNGG | ASSHLGGWAETLY | 0.51533854 |
| VSVFMGMGGNF | ASSHLGGWAETLY | 0.6675379 |
| LASYLAAL | ASSHLGGWAETLY | 0.67248154 |
| LVGGTPAL | ASSHLGGWAETLY | 0.7116132 |
| VRNADVL | ASSHLGGWAETLY | 0.69581103 |
| TCSSPFLVSAM | ASSHLGGWAETLY | 0.7178273 |
| VHITDAE | ASSHLGGWAETLY | 0.7165016 |
| EYAAPSF | ASSHLGGWAETLY | 0.6781732 |
| VVDAFHGGL | ASSHLGGWAETLY | 0.70400184 |
| ASVVDPQ | ASSHLGGWAETLY | 0.6285246 |
| SYNEFGGLS | ASSHLGGWAETLY | 0.6692884 |
| DPDGSVY | ASSHLGGWAETLY | 0.6543164 |
| MVSTIHATE | ASSHLGGWAETLY | 0.6829845 |
| LAAGVADPV | ASSHLGGWAETLY | 0.69736946 |
| TKEQIAA | ASSHLGGWAETLY | 0.66547346 |
| SLASPAAL | ASSHLGGWAETLY | 0.49358884 |
| IGLVFYGV | ASSHLGGWAETLY | 0.70457315 |
| FLLDASNL | ASSHLGGWAETLY | 0.6998632 |
| YQVGNFP | ASSHLGGWAETLY | 0.40077275 |
| ESLLSQST | ASSHLGGWAETLY | 0.66564196 |
| LFTVGFAAVG | ASSHLGGWAETLY | 0.72073746 |
| ATPPTPT | ASSHLGGWAETLY | 0.71647704 |
| TDLATLG | ASSHLGGWAETLY | 0.64601296 |
| SAGGTGR | ASSHLGGWAETLY | 0.66537297 |
| GFSNSGS | ASSHLGGWAETLY | 0.6790296 |
| GGNGGSGGTSV | ASSHLGGWAETLY | 0.5270794 |
| GTSVATGGAGN | ASSHLGGWAETLY | 0.6749866 |
| NGGNGGSGGTS | ASSHLGGWAETLY | 0.6910456 |
| AVVKFPH | ASSHLGGWAETLY | 0.6857005 |
| TPPPPPP | ASSHLGGWAETLY | 0.5994091 |
| AAPAAAAAPA | ASSHLGGWAETLY | 0.6857591 |
| IALNEDL | ASSHLGGWAETLY | 0.6731583 |
| IVNVSSM | ASSHLGGWAETLY | 0.69513994 |
| TGIPPAPR | ASSHLGGWAETLY | 0.26967895 |
| FDLGGGTF | ASSHLGGWAETLY | 0.70302314 |
| FDLGGGTFDVS | ASSHLGGWAETLY | 0.64474416 |
| NEPTAAA | ASSHLGGWAETLY | 0.6033039 |
| NEPTAAALA | ASSHLGGWAETLY | 0.38375646 |
| GSIINISSI | ASSHLGGWAETLY | 0.6713773 |
| VAPGNAGTA | ASSHLGGWAETLY | 0.70871204 |
| TGGGASLE | ASSHLGGWAETLY | 0.71175593 |
| VAVYNHYKRI | ASSHLGGWAETLY | 0.5911891 |
| TDPAPATT | ASSHLGGWAETLY | 0.71339047 |
| TNRPDILDPALLRPG | ASSHLGGWAETLY | 0.71601677 |
| LTTTTTS | ASSHLGGWAETLY | 0.6661068 |
| PTGSGKTL | ASSHLGGWAETLY | 0.48731935 |

| | | |
|---|---|---|
| TGSGKTL | ASSHLGGWAETLY | 0.7119563 |
| TIGHVDHGK | ASSHLGGWAETLY | 0.65936625 |
| TIGHVDHGKTT | ASSHLGGWAETLY | 0.705186 |
| GHVDHGKTT | ASSHLGGWAETLY | 0.70497584 |
| AQFEHTLL | ASSHLGGWAETLY | 0.63858366 |
| IVNQDEL | ASSHLGGWAETLY | 0.7166917 |
| TNGSQFF | ASSHLGGWAETLY | 0.71570754 |
| TGKGIVE | ASSHLGGWAETLY | 0.6475674 |
| LISNASDA | ASSHLGGWAETLY | 0.5651915 |
| AELRTLGELTKIAKAHGAQ | ASSHLGGWAETLY | 0.6512735 |
| APNAVAAAEPSW | ASSHLGGWAETLY | 0.35649145 |
| APQVAGAATAVL | ASSHLGGWAETLY | 0.71521294 |
| ARRDPAAAAAILPTDA | ASSHLGGWAETLY | 0.6569935 |
| DAAARDVLAEAGLAEYF | ASSHLGGWAETLY | 0.6663492 |
| DDQIVLSAKANLAKVEQ | ASSHLGGWAETLY | 0.70037234 |
| DELLPDTGEDSDEDGH | ASSHLGGWAETLY | 0.66926956 |
| ELPEAPANRATVPQD | ASSHLGGWAETLY | 0.6743295 |
| FGQSKIERVTGIG | ASSHLGGWAETLY | 0.67822444 |
| GAGGTGSTAGGAGGAGGAGGLY | ASSHLGGWAETLY | 0.698363 |
| GHGGTGAAGGNGGTGGAGGSGI | ASSHLGGWAETLY | 0.6932311 |
| GNGGNGGAGGEGGPGGAGGAGA | ASSHLGGWAETLY | 0.684552 |
| GSTAGNGGNGGNATNGTIAGQP | ASSHLGGWAETLY | 0.6787481 |
| GTGGDGGNGGNGGTGG | ASSHLGGWAETLY | 0.7179742 |
| GVRRPAGIPADAPV | ASSHLGGWAETLY | 0.5983249 |
| GYRQVTDAHLLTLARR | ASSHLGGWAETLY | 0.65994954 |
| IAKMGQDGHDRG | ASSHLGGWAETLY | 0.69762594 |
| KAEPSIEIQYT | ASSHLGGWAETLY | 0.582824 |
| KPVRIRMAADEDTVAVT | ASSHLGGWAETLY | 0.70158494 |
| MPPRIAGMRLLVIKPEP | ASSHLGGWAETLY | 0.6964874 |
| MKFYAVALFRSRGVPPTR | ASSHLGGWAETLY | 0.702627 |
| NAGGFGILGAVAHSP | ASSHLGGWAETLY | 0.67136383 |
| NEGREITLAMAEM | ASSHLGGWAETLY | 0.72055244 |
| NELVEWSHPLGHRWRWE | ASSHLGGWAETLY | 0.71243685 |
| NLGAGGTGGNGGNPG | ASSHLGGWAETLY | 0.66762424 |
| NTEWPKWETVMAADPVD | ASSHLGGWAETLY | 0.7082181 |
| PASAAPAETQP | ASSHLGGWAETLY | 0.60507786 |
| QQSSDGKNGVRAAEITGE | ASSHLGGWAETLY | 0.7107247 |
| RAPDAVTTSWWKE | ASSHLGGWAETLY | 0.7115903 |
| SGFGNVGAHTSGWFNQ | ASSHLGGWAETLY | 0.6192877 |
| SGVAGAVGQAASVGGLKV | ASSHLGGWAETLY | 0.68638366 |
| SNGGAGTGSAGGNGGT | ASSHLGGWAETLY | 0.7112694 |
| TGGAGGEGGAAG | ASSHLGGWAETLY | 0.7137073 |
| TGGDGGAAGAGGAPGHG | ASSHLGGWAETLY | 0.5258973 |
| TMPPLLAAIANAGRHGV | ASSHLGGWAETLY | 0.6783972 |
| TSSTAGNSGGA | ASSHLGGWAETLY | 0.424323 |
| VPATVPVGILC | ASSHLGGWAETLY | 0.5651937 |
| VQRPDSGGVVLTGRIS | ASSHLGGWAETLY | 0.6305539 |
| WTPNPVPATKGVPAGLVQAT | ASSHLGGWAETLY | 0.7013585 |
| YALKRALAKFSDRFDVVII | ASSHLGGWAETLY | 0.6783832 |
| YLIGSVIV | ASSHLGGWAETLY | 0.700579 |
| YGQLVAVID | ASSHLGGWAETLY | 0.6992064 |

| | | |
|---|---|---|
| VTVGGMTLPA | ASSHLGGWAETLY | 0.7199644 |
| VLALPAV | ASSHLGGWAETLY | 0.656428 |
| VIPPTLNYET | ASSHLGGWAETLY | 0.70101017 |
| VGMGGMHPSSG | ASSHLGGWAETLY | 0.461536 |
| VGLAGGAATA | ASSHLGGWAETLY | 0.48318815 |
| VGGAGGAGGSGP | ASSHLGGWAETLY | 0.5570335 |
| VFAAFMV | ASSHLGGWAETLY | 0.6897909 |
| TLYDKLR | ASSHLGGWAETLY | 0.31428787 |
| TLAVFTVL | ASSHLGGWAETLY | 0.71659714 |
| TGSVQPARV | ASSHLGGWAETLY | 0.6923095 |
| TGGGAGSGGT | ASSHLGGWAETLY | 0.71281004 |
| SHLETSL | ASSHLGGWAETLY | 0.67690754 |
| SFTTGMT | ASSHLGGWAETLY | 0.6745608 |
| QVEVAKRMGVRQPTVS | ASSHLGGWAETLY | 0.67570454 |
| QGSGGPALAM | ASSHLGGWAETLY | 0.7059238 |
| MLAPSNP | ASSHLGGWAETLY | 0.648792 |
| LVNNAAI | ASSHLGGWAETLY | 0.44423532 |
| LTGEINTLA | ASSHLGGWAETLY | 0.64359725 |
| LSIFGPV | ASSHLGGWAETLY | 0.60406923 |
| LLALLANL | ASSHLGGWAETLY | 0.71700037 |
| LISNASDALDKLR | ASSHLGGWAETLY | 0.7138027 |
| LINEAVP | ASSHLGGWAETLY | 0.7051857 |
| LGYAFSGL | ASSHLGGWAETLY | 0.70701385 |
| LDFVAVAL | ASSHLGGWAETLY | 0.30435792 |
| LAQPGAE | ASSHLGGWAETLY | 0.6622087 |
| ISNLGAVTGF | ASSHLGGWAETLY | 0.7208997 |
| HQEDPGRI | ASSHLGGWAETLY | 0.6909703 |
| GTPGEPTP | ASSHLGGWAETLY | 0.68453574 |
| GFFNSGS | ASSHLGGWAETLY | 0.7178216 |
| GAGGGGLTAGAN | ASSHLGGWAETLY | 0.6714938 |
| FVDALVNM | ASSHLGGWAETLY | 0.6857076 |
| FLTVAPEL | ASSHLGGWAETLY | 0.68619835 |
| FETLTAL | ASSHLGGWAETLY | 0.6978879 |
| DVINPIEL | ASSHLGGWAETLY | 0.6735783 |
| APVSGAPV | ASSHLGGWAETLY | 0.6438981 |
| ALVGGLAW | ASSHLGGWAETLY | 0.68746674 |
| ALAAAAATAI | ASSHLGGWAETLY | 0.7218598 |
| AFGLGLLL | ASSHLGGWAETLY | 0.68908507 |
| AFFAAASL | ASSHLGGWAETLY | 0.71337223 |
| AAAALPAFSPP | ASSHLGGWAETLY | 0.7017281 |
| VVLAPAL | ASSHLGGWAETLY | 0.7062439 |
| TIGHVDHGKTTL | ASSHLGGWAETLY | 0.66125774 |
| LISNASDALDK | ASSHLGGWAETLY | 0.6761014 |
| LISNASDAL | ASSHLGGWAETLY | 0.6815699 |
| ISNASDALDK | ASSHLGGWAETLY | 0.63650316 |
| NQAETLVYQTE | ASSHLGGWAETLY | 0.67750025 |
| TGGTGGTAGAG | ASSHLGGWAETLY | 0.64345324 |
| TGSTAGGAGGA | ASSHLGGWAETLY | 0.7240118 |
| DPLAVPVAAGPV | ASSHLGGWAETLY | 0.70620847 |
| GGTGGTPGNGGN | ASSHLGGWAETLY | 0.69833636 |
| PAGLAGVDPATL | ASSHLGGWAETLY | 0.66230756 |

| | | |
|---|---|---|
| GNNLVVEAPHAGASV | ASSHLGGWAETLY | 0.67402756 |
| KVTRIMSASTLEQPA | ASSHLGGWAETLY | 0.7060666 |
| LGFALPIALPRRTVV | ASSHLGGWAETLY | 0.51685935 |
| TRLGNLTARKLVVLV | ASSHLGGWAETLY | 0.7174156 |
| WSQQRRLIYFPSAGP | ASSHLGGWAETLY | 0.66329974 |
| GPNGNPDPMAAAVDIR | ASSHLGGWAETLY | 0.65152526 |
| TQRCVSVADLRAAASR | ASSHLGGWAETLY | 0.71348125 |
| QFVLDVAEHPVEVLTETV | ASSHLGGWAETLY | 0.7170223 |
| GGAGGTGGAGGNGGTQAGDG | ASSHLGGWAETLY | 0.6486071 |
| PLLATLRAIAVDVDAKPAQV | ASSHLGGWAETLY | 0.662755 |
| APKFRTGGYMIDFWGVGYQVAK | ASSHLGGWAETLY | 0.6710697 |
| GAGGAGGAGGAAGGPGGTGGTG | ASSHLGGWAETLY | 0.70549047 |
| IGALGLLATDVAPAGSVAFGVA | ASSHLGGWAETLY | 0.7184087 |
| AGGAGGIGGNGGAGGAGGVLMG | ASSHLGGWAETLY | 0.67890984 |
| ADELIGLL | ASSHLGGWAETLY | 0.7210389 |
| ADRALKA | ASSHLGGWAETLY | 0.69001967 |
| AGAVGQVGGAGGA | ASSHLGGWAETLY | 0.50693643 |
| AGGGGGLGGAP | ASSHLGGWAETLY | 0.684538 |
| AGGLLYGNGGAG | ASSHLGGWAETLY | 0.7074317 |
| AGLESIGL | ASSHLGGWAETLY | 0.71173555 |
| AGLKTGII | ASSHLGGWAETLY | 0.7228404 |
| ASLAALLL | ASSHLGGWAETLY | 0.6974195 |
| ASVINISSI | ASSHLGGWAETLY | 0.64053917 |
| ATPPGPA | ASSHLGGWAETLY | 0.71640307 |
| ATVASALV | ASSHLGGWAETLY | 0.71846914 |
| ATVASLIL | ASSHLGGWAETLY | 0.70979774 |
| AVNGPVLIRH | ASSHLGGWAETLY | 0.64786744 |
| DGAAGAVGI | ASSHLGGWAETLY | 0.64014024 |
| DPLLHGI | ASSHLGGWAETLY | 0.6819318 |
| ELGAALGAL | ASSHLGGWAETLY | 0.7113391 |
| ELGRTQAV | ASSHLGGWAETLY | 0.5830551 |
| FVAGGYLS | ASSHLGGWAETLY | 0.69544536 |
| GASGGAGGAGGW | ASSHLGGWAETLY | 0.7011357 |
| GGAGAGGDGGAGGI | ASSHLGGWAETLY | 0.58292776 |
| GTGHNGSQPGMGGNG | ASSHLGGWAETLY | 0.68987274 |
| GYAAASASAS | ASSHLGGWAETLY | 0.7145934 |
| HQVDVAAVQQG | ASSHLGGWAETLY | 0.61904013 |
| IGAAIAHT | ASSHLGGWAETLY | 0.657035 |
| IGMIEVAQL | ASSHLGGWAETLY | 0.71193033 |
| IRLAPEHTSAMD | ASSHLGGWAETLY | 0.7243857 |
| IYNVSVL | ASSHLGGWAETLY | 0.71162885 |
| KQGFLGM | ASSHLGGWAETLY | 0.5432521 |
| LESVITQL | ASSHLGGWAETLY | 0.4277807 |
| LGLGALVVV | ASSHLGGWAETLY | 0.63959295 |
| LIGNGGNGGSGGI | ASSHLGGWAETLY | 0.7101084 |
| LNAINAPTL | ASSHLGGWAETLY | 0.68445766 |
| LQLLNTGV | ASSHLGGWAETLY | 0.65462977 |
| LQTSIEF | ASSHLGGWAETLY | 0.7029508 |
| MSFAVSEL | ASSHLGGWAETLY | 0.71479154 |
| NTGALGL | ASSHLGGWAETLY | 0.640592 |
| PPGAGGAGGAGTT | ASSHLGGWAETLY | 0.47460023 |

| | | |
|---|---|---|
| QLVGLGI | ASSHLGGWAETLY | 0.69446236 |
| RFLEQVHQL | ASSHLGGWAETLY | 0.70796275 |
| RGVGYVLR | ASSHLGGWAETLY | 0.6512102 |
| SATGVFAGVI | ASSHLGGWAETLY | 0.6820692 |
| SLPAITLP | ASSHLGGWAETLY | 0.63068426 |
| STALAFTV | ASSHLGGWAETLY | 0.5161651 |
| SVLTVGLGK | ASSHLGGWAETLY | 0.7007196 |
| TALAGAGSGI | ASSHLGGWAETLY | 0.65058404 |
| TGLPFTGI | ASSHLGGWAETLY | 0.7161406 |
| VAEHVTV | ASSHLGGWAETLY | 0.6307838 |
| VDIVLNSL | ASSHLGGWAETLY | 0.69572854 |
| VLTGGGAVL | ASSHLGGWAETLY | 0.70789003 |
| VNVGLAYGI | ASSHLGGWAETLY | 0.70652705 |
| VSSAAVGAV | ASSHLGGWAETLY | 0.67463624 |
| VTSVYEAGV | ASSHLGGWAETLY | 0.64950025 |
| WVALDRGA | ASSHLGGWAETLY | 0.62198687 |
| YLVGSVVI | ASSHLGGWAETLY | 0.48742968 |
| GPPGTGKTLL | ASSHLGGWAETLY | 0.7131268 |
| GVGMTKF | ASSHLGGWAETLY | 0.50085664 |
| IGIGGSDL | ASSHLGGWAETLY | 0.71891975 |
| IGTLLAEL | ASSHLGGWAETLY | 0.70756304 |
| IYTSGTT | ASSHLGGWAETLY | 0.6547419 |
| LSFPAGV | ASSHLGGWAETLY | 0.70035064 |
| LYTSGTT | ASSHLGGWAETLY | 0.59756756 |
| VGAFPFGF | ASSHLGGWAETLY | 0.6420627 |
| VGAGTVEF | ASSHLGGWAETLY | 0.7044916 |
| VGKSTLLL | ASSHLGGWAETLY | 0.7026386 |
| AALAHAGGGIV | ASSHLGGWAETLY | 0.6525225 |
| AALVAVSLL | ASSHLGGWAETLY | 0.68138313 |
| AAVSRAVAAL | ASSHLGGWAETLY | 0.6916519 |
| AGGSSGIGGF | ASSHLGGWAETLY | 0.63012344 |
| AGLTVANANG | ASSHLGGWAETLY | 0.6463082 |
| AGLVGAVL | ASSHLGGWAETLY | 0.7133892 |
| ALSQAFTV | ASSHLGGWAETLY | 0.66825753 |
| ALTFSVSL | ASSHLGGWAETLY | 0.664961 |
| ALTGAALVAL | ASSHLGGWAETLY | 0.7067651 |
| ASFSIGAGVL | ASSHLGGWAETLY | 0.62671643 |
| DGSVYKI | ASSHLGGWAETLY | 0.707813 |
| EQLAEIV | ASSHLGGWAETLY | 0.6877924 |
| FAVSELTVM | ASSHLGGWAETLY | 0.6786485 |
| GAAPLAAGL | ASSHLGGWAETLY | 0.70418286 |
| GADGGDGGSGGAGG | ASSHLGGWAETLY | 0.5557822 |
| GFGLDTGGAGG | ASSHLGGWAETLY | 0.6133182 |
| GLASGGLL | ASSHLGGWAETLY | 0.6714953 |
| GRIAGAAVLL | ASSHLGGWAETLY | 0.7056403 |
| GSAGSLNTSF | ASSHLGGWAETLY | 0.7135081 |
| GTGQIGI | ASSHLGGWAETLY | 0.6651274 |
| GTLIALIL | ASSHLGGWAETLY | 0.6902922 |
| IAATMGFVVG | ASSHLGGWAETLY | 0.6665467 |
| IFAQALL | ASSHLGGWAETLY | 0.65129006 |
| IGAAVALGF | ASSHLGGWAETLY | 0.63883865 |

| | | |
|---|---|---|
| IGLTPLAL | ASSHLGGWAETLY | 0.65268534 |
| ILGSNTSTL | ASSHLGGWAETLY | 0.6093947 |
| IVFSLTVL | ASSHLGGWAETLY | 0.7044834 |
| LASKATGF | ASSHLGGWAETLY | 0.60209525 |
| LASVALIL | ASSHLGGWAETLY | 0.69459105 |
| LFAVGLTL | ASSHLGGWAETLY | 0.7018127 |
| LFNGGIGTYI | ASSHLGGWAETLY | 0.6712528 |
| LTLGAGII | ASSHLGGWAETLY | 0.6936418 |
| LVAAALGDI | ASSHLGGWAETLY | 0.7004024 |
| NTAFVQLGI | ASSHLGGWAETLY | 0.70541143 |
| QLFSGAEL | ASSHLGGWAETLY | 0.691077 |
| SAATFITL | ASSHLGGWAETLY | 0.67291415 |
| SNYGLANLGV | ASSHLGGWAETLY | 0.7176036 |
| SYGGAYAVM | ASSHLGGWAETLY | 0.71003115 |
| TTGVGGLTLG | ASSHLGGWAETLY | 0.7099736 |
| TTQTIYI | ASSHLGGWAETLY | 0.71082675 |
| VAAGAGLGAVI | ASSHLGGWAETLY | 0.70066273 |
| VAAVVTLL | ASSHLGGWAETLY | 0.70093143 |
| VGFLGGLI | ASSHLGGWAETLY | 0.68416667 |
| VGVIPILL | ASSHLGGWAETLY | 0.50269496 |
| VIFASSLI | ASSHLGGWAETLY | 0.688347 |
| VKGGDGGAAAG | ASSHLGGWAETLY | 0.68768275 |
| VTKTFTNI | ASSHLGGWAETLY | 0.6431879 |
| VVAAAILL | ASSHLGGWAETLY | 0.4609984 |
| YAALDVEL | ASSHLGGWAETLY | 0.7168458 |
| LASVALLL | ASSHLGGWAETLY | 0.723329 |
| LDAVHRV | ASSHLGGWAETLY | 0.6402898 |
| QGLSSSIF | ASSHLGGWAETLY | 0.72112346 |
| RGTFANIRL | ASSHLGGWAETLY | 0.5179532 |
| RQLWWGHRI | ASSHLGGWAETLY | 0.70769674 |
| YGPPGTGKTLL | ASSHLGGWAETLY | 0.7146041 |
| AALVGAPANVA | ASSHLGGWAETLY | 0.64964086 |
| AATFRRTE | ASSHLGGWAETLY | 0.7128433 |
| AGALGGGI | ASSHLGGWAETLY | 0.65280807 |
| AGGAGGAGGIGAG | ASSHLGGWAETLY | 0.6712798 |
| AGSGALGL | ASSHLGGWAETLY | 0.72072005 |
| AQIADQI | ASSHLGGWAETLY | 0.69145674 |
| ASAVAVTYL | ASSHLGGWAETLY | 0.69411623 |
| ASTVLGGTGG | ASSHLGGWAETLY | 0.6945541 |
| AVGGEPAK | ASSHLGGWAETLY | 0.66930306 |
| CEQGTFGGTC | ASSHLGGWAETLY | 0.689721 |
| DEGKAPAG | ASSHLGGWAETLY | 0.71845454 |
| DTVVFPAL | ASSHLGGWAETLY | 0.6290853 |
| FEGITMAG | ASSHLGGWAETLY | 0.7176074 |
| FPLVVQD | ASSHLGGWAETLY | 0.6604724 |
| GAFTTPQ | ASSHLGGWAETLY | 0.72408617 |
| GFGGFGTAGGVGGA | ASSHLGGWAETLY | 0.6994333 |
| GFLAQFL | ASSHLGGWAETLY | 0.6941129 |
| GGAGGAGGNAGVA | ASSHLGGWAETLY | 0.65407324 |
| GGVGTAGAGGAGGAGG | ASSHLGGWAETLY | 0.7160041 |
| GNGGGGDAGNAGSGGN | ASSHLGGWAETLY | 0.69372046 |

| | | |
|---|---|---|
| GTGGAGGAGNDAGS | ASSHLGGWAETLY | 0.69971174 |
| GTTNTGWFNS | ASSHLGGWAETLY | 0.71424186 |
| IFMGPLFTL | ASSHLGGWAETLY | 0.61589736 |
| ILAHAIEAI | ASSHLGGWAETLY | 0.69547904 |
| IRSAIREAL | ASSHLGGWAETLY | 0.6626179 |
| ITFAGGG | ASSHLGGWAETLY | 0.6761864 |
| ITGMSGHQEV | ASSHLGGWAETLY | 0.6201258 |
| IVAGLACTL | ASSHLGGWAETLY | 0.5720184 |
| IVGGGLAGA | ASSHLGGWAETLY | 0.6729763 |
| IVTGVLNI | ASSHLGGWAETLY | 0.6122625 |
| LAQQGCI | ASSHLGGWAETLY | 0.68201846 |
| LCLHVEN | ASSHLGGWAETLY | 0.69882834 |
| LDAISDGS | ASSHLGGWAETLY | 0.6189326 |
| LGGFAAML | ASSHLGGWAETLY | 0.69905937 |
| LGGKNPMI | ASSHLGGWAETLY | 0.70331156 |
| LGPKGVAAIL | ASSHLGGWAETLY | 0.52282846 |
| LMVGVSI | ASSHLGGWAETLY | 0.63128924 |
| LVGAAFVL | ASSHLGGWAETLY | 0.6338924 |
| LVLAVAPL | ASSHLGGWAETLY | 0.64889675 |
| MAAIAVIDAA | ASSHLGGWAETLY | 0.46494466 |
| NNGINAELQ | ASSHLGGWAETLY | 0.6952368 |
| RGEPLVLVHG | ASSHLGGWAETLY | 0.6098281 |
| RGGAGGAAGLI | ASSHLGGWAETLY | 0.36707026 |
| RSYRMHPAV | ASSHLGGWAETLY | 0.6702709 |
| SAAHLLAGQLA | ASSHLGGWAETLY | 0.6476939 |
| SAAVGSY | ASSHLGGWAETLY | 0.66542685 |
| SPHAVLHRI | ASSHLGGWAETLY | 0.69008213 |
| SQIAKVL | ASSHLGGWAETLY | 0.56470525 |
| TALGHLVP | ASSHLGGWAETLY | 0.63085884 |
| TAPTTSGAA | ASSHLGGWAETLY | 0.6771952 |
| TDGDDGNGGSPGT | ASSHLGGWAETLY | 0.7036454 |
| TDGLGEN | ASSHLGGWAETLY | 0.6186634 |
| TGGPPLPGQA | ASSHLGGWAETLY | 0.66454935 |
| TIPAITI | ASSHLGGWAETLY | 0.70581436 |
| TIPGFTI | ASSHLGGWAETLY | 0.6801745 |
| TSIASGALA | ASSHLGGWAETLY | 0.70638394 |
| TVTLPGAAI | ASSHLGGWAETLY | 0.71143585 |
| VASGLSAALP | ASSHLGGWAETLY | 0.6131834 |
| VGAGVIGI | ASSHLGGWAETLY | 0.6943723 |
| VGPKRFGV | ASSHLGGWAETLY | 0.70160353 |
| VNVAFPD | ASSHLGGWAETLY | 0.6446906 |
| VSMTNSGVS | ASSHLGGWAETLY | 0.6636019 |
| VTDVLGDLSV | ASSHLGGWAETLY | 0.6318733 |
| VVVVGAGFAGL | ASSHLGGWAETLY | 0.64882904 |
| VYTVGTVV | ASSHLGGWAETLY | 0.4401034 |
| WTTFSGP | ASSHLGGWAETLY | 0.66159505 |
| YDELVEAF | ASSHLGGWAETLY | 0.70181715 |
| YGTGGAGGNGG | ASSHLGGWAETLY | 0.62937367 |
| YTNVMAV | ASSHLGGWAETLY | 0.5774324 |
| FLEQVHQL | ASSHLGGWAETLY | 0.57119614 |
| GLSLLAVAV | ASSHLGGWAETLY | 0.6393386 |

| | | |
|---|---|---|
| LEQVHQL | ASSHLGGWAETLY | 0.6366371 |
| LNAAGAI | ASSHLGGWAETLY | 0.56124634 |
| PAVHASL | ASSHLGGWAETLY | 0.5842067 |
| TELEDDA | ASSHLGGWAETLY | 0.6990859 |
| TPPGPPN | ASSHLGGWAETLY | 0.702881 |
| TPPPPPPP | ASSHLGGWAETLY | 0.49337417 |
| TQSPALI | ASSLNGNERLF | 0.6926912 |
| EFAAAFPQ | ASSLNGNERLF | 0.68961835 |
| LLSGNLDVL | ASSLNGNERLF | 0.7079163 |
| GPANLGNYNI | ASSLNGNERLF | 0.67913866 |
| TLGPTLP | ASSLNGNERLF | 0.5909479 |
| VYAYSSCCK | ASSLNGNERLF | 0.6810914 |
| APAPAPAPNI | ASSLNGNERLF | 0.6082948 |
| PGGPVPIP | ASSLNGNERLF | 0.66441566 |
| FLTVALEP | ASSLNGNERLF | 0.69567275 |
| SQYFGANL | ASSLNGNERLF | 0.65869963 |
| GGVKPAA | ASSLNGNERLF | 0.600478 |
| GGTGGAGGAGGW | ASSLNGNERLF | 0.61453307 |
| GVGGVAGLGGAGG | ASSLNGNERLF | 0.70898736 |
| VLLVLATTI | ASSLNGNERLF | 0.53979164 |
| IGIFAAML | ASSLNGNERLF | 0.6177045 |
| AVGDLID | ASSLNGNERLF | 0.6284438 |
| NISPLPL | ASSLNGNERLF | 0.6616214 |
| VTAVFVAI | ASSLNGNERLF | 0.6975223 |
| SGDDNDAAL | ASSLNGNERLF | 0.6908215 |
| AVAALAHS | ASSLNGNERLF | 0.61025643 |
| GGVTVMATP | ASSLNGNERLF | 0.6806333 |
| AGLAANVRD | ASSLNGNERLF | 0.6892294 |
| TNRPDLIDPALLRPG | ASSLNGNERLF | 0.67488456 |
| LANLGVTAL | ASSLNGNERLF | 0.6964247 |
| TGERIEL | ASSLNGNERLF | 0.57478935 |
| NGEELAA | ASSLNGNERLF | 0.7036005 |
| VLSLVWG | ASSLNGNERLF | 0.68186665 |
| IGYALWL | ASSLNGNERLF | 0.7017139 |
| LSLLLAPL | ASSLNGNERLF | 0.6961462 |
| TLDEALK | ASSLNGNERLF | 0.67930424 |
| TDGAGTAEGG | ASSLNGNERLF | 0.6722601 |
| NAPGGAGGNGG | ASSLNGNERLF | 0.4684843 |
| VSVFMGMGGNF | ASSLNGNERLF | 0.64134157 |
| LASYLAAL | ASSLNGNERLF | 0.6697659 |
| LVGGTPAL | ASSLNGNERLF | 0.6622633 |
| VRNADVL | ASSLNGNERLF | 0.6882789 |
| TCSSPFLVSAM | ASSLNGNERLF | 0.701985 |
| VHITDAE | ASSLNGNERLF | 0.6128537 |
| EYAAPSF | ASSLNGNERLF | 0.7093593 |
| VVDAFHGGL | ASSLNGNERLF | 0.6900649 |
| ASVVDPQ | ASSLNGNERLF | 0.5129686 |
| SYNEFGGLS | ASSLNGNERLF | 0.64088595 |
| DPDGSVY | ASSLNGNERLF | 0.6985673 |
| MVSTIHATE | ASSLNGNERLF | 0.6670726 |
| LAAGVADPV | ASSLNGNERLF | 0.6347148 |

| | | |
|---|---|---|
| TKEQIAA | ASSLNGNERLF | 0.65959305 |
| SLASPAAL | ASSLNGNERLF | 0.7077662 |
| IGLVFYGV | ASSLNGNERLF | 0.67426115 |
| FLLDASNL | ASSLNGNERLF | 0.70835924 |
| YQVGNFP | ASSLNGNERLF | 0.3628435 |
| ESLLSQST | ASSLNGNERLF | 0.6467049 |
| LFTVGFAAVG | ASSLNGNERLF | 0.62516916 |
| ATPPTPT | ASSLNGNERLF | 0.5018285 |
| TDLATLG | ASSLNGNERLF | 0.69818115 |
| SAGGTGR | ASSLNGNERLF | 0.6578795 |
| GFSNSGS | ASSLNGNERLF | 0.68824863 |
| GGNGGSGGTSV | ASSLNGNERLF | 0.6574894 |
| GTSVATGGAGN | ASSLNGNERLF | 0.698954 |
| NGGNGGSGGTS | ASSLNGNERLF | 0.6418338 |
| AVVKFPH | ASSLNGNERLF | 0.6630664 |
| TPPPPPP | ASSLNGNERLF | 0.6236328 |
| AAPAAAAAPA | ASSLNGNERLF | 0.69158936 |
| IALNEDL | ASSLNGNERLF | 0.7116322 |
| IVNVSSM | ASSLNGNERLF | 0.6989588 |
| TGIPPAPR | ASSLNGNERLF | 0.5857736 |
| FDLGGGTF | ASSLNGNERLF | 0.6525042 |
| FDLGGGTFDVS | ASSLNGNERLF | 0.6743754 |
| NEPTAAA | ASSLNGNERLF | 0.41198477 |
| NEPTAAALA | ASSLNGNERLF | 0.46911913 |
| GSIINISSI | ASSLNGNERLF | 0.6278404 |
| VAPGNAGTA | ASSLNGNERLF | 0.5706236 |
| TGGGASLE | ASSLNGNERLF | 0.69424987 |
| VAVYNHYKRI | ASSLNGNERLF | 0.6796783 |
| TDPAPATT | ASSLNGNERLF | 0.63484013 |
| TNRPDILDPALLRPG | ASSLNGNERLF | 0.68269515 |
| LTTTTTS | ASSLNGNERLF | 0.6652185 |
| PTGSGKTL | ASSLNGNERLF | 0.27054054 |
| TGSGKTL | ASSLNGNERLF | 0.63266516 |
| TIGHVDHGK | ASSLNGNERLF | 0.5516367 |
| TIGHVDHGKTT | ASSLNGNERLF | 0.71033907 |
| GHVDHGKTT | ASSLNGNERLF | 0.623847 |
| AQFEHTLL | ASSLNGNERLF | 0.57368636 |
| IVNQDEL | ASSLNGNERLF | 0.6470515 |
| TNGSQFF | ASSLNGNERLF | 0.64699507 |
| TGKGIVE | ASSLNGNERLF | 0.6482804 |
| LISNASDA | ASSLNGNERLF | 0.59927696 |
| AELRTLGELTKIAKAHGAQ | ASSLNGNERLF | 0.33284506 |
| APNAVAAAEPSW | ASSLNGNERLF | 0.6952348 |
| APQVAGAATAVL | ASSLNGNERLF | 0.53824 |
| ARRDPAAAAAILPTDA | ASSLNGNERLF | 0.63150084 |
| DAAARDVLAEAGLAEYF | ASSLNGNERLF | 0.4518421 |
| DDQIVLSAKANLAKVEQ | ASSLNGNERLF | 0.59887516 |
| DELLPDTGEDSDEDGH | ASSLNGNERLF | 0.5689198 |
| ELPEAPANRATVPQD | ASSLNGNERLF | 0.70346165 |
| FGQSKIERVTGIG | ASSLNGNERLF | 0.59470844 |
| GAGGTGSTAGGAGGAGGAGGLY | ASSLNGNERLF | 0.612985 |

| | | |
|---|---|---|
| GHGGTGAAGGNGGTGGAGGSGI | ASSLNGNERLF | 0.6976598 |
| GNGGNGGAGGEGGPGGAGGAGA | ASSLNGNERLF | 0.64942163 |
| GSTAGNGGNGGNATNGTIAGQP | ASSLNGNERLF | 0.61239886 |
| GTGGDGGNGGNGGTGG | ASSLNGNERLF | 0.633669 |
| GVRRPAGIPADAPV | ASSLNGNERLF | 0.57977796 |
| GYRQVTDAHLLTLARR | ASSLNGNERLF | 0.63668966 |
| IAKMGQDGHDRG | ASSLNGNERLF | 0.64057106 |
| KAEPSIEIQYT | ASSLNGNERLF | 0.71079516 |
| KPVRIRMAADEDTVAVT | ASSLNGNERLF | 0.6473662 |
| MPPRIAGMRLLVIKPEP | ASSLNGNERLF | 0.59016204 |
| MKFYAVALFRSRGVPPTR | ASSLNGNERLF | 0.67015064 |
| NAGGFGILGAVAHSP | ASSLNGNERLF | 0.7083163 |
| NEGREITLAMAEM | ASSLNGNERLF | 0.71237624 |
| NELVEWSHPLGHRWRWE | ASSLNGNERLF | 0.59275895 |
| NLGAGGTGGNGGNPG | ASSLNGNERLF | 0.6715147 |
| NTEWPKWETVMAADPVD | ASSLNGNERLF | 0.6500877 |
| PASAAPAETQP | ASSLNGNERLF | 0.6405098 |
| QQSSDGKNGVRAAEITGE | ASSLNGNERLF | 0.65007645 |
| RAPDAVTTSWWKE | ASSLNGNERLF | 0.59043926 |
| SGFGNVGAHTSGWFNQ | ASSLNGNERLF | 0.6961243 |
| SGVAGAVGQAASVGGLKV | ASSLNGNERLF | 0.7188077 |
| SNGGAGTGSAGGNGGT | ASSLNGNERLF | 0.65275204 |
| TGGAGGEGGAAG | ASSLNGNERLF | 0.58489347 |
| TGGDGGAAGAGGAPGHG | ASSLNGNERLF | 0.63823473 |
| TMPPLLAAIANAGRHGV | ASSLNGNERLF | 0.57604 |
| TSSTAGNSGGA | ASSLNGNERLF | 0.61204904 |
| VPATVPVGILC | ASSLNGNERLF | 0.6778067 |
| VQRPDSGGVVLTGRIS | ASSLNGNERLF | 0.49901837 |
| WTPNPVPATKGVPAGLVQAT | ASSLNGNERLF | 0.6399305 |
| YALKRALAKFSDRFDVVII | ASSLNGNERLF | 0.712939 |
| YLIGSVIV | ASSLNGNERLF | 0.59876525 |
| YGQLVAVID | ASSLNGNERLF | 0.6485065 |
| VTVGGMTLPA | ASSLNGNERLF | 0.6943003 |
| VLALPAV | ASSLNGNERLF | 0.67205846 |
| VIPPTLNYET | ASSLNGNERLF | 0.70077574 |
| VGMGGMHPSSG | ASSLNGNERLF | 0.6950934 |
| VGLAGGAATA | ASSLNGNERLF | 0.67442966 |
| VGGAGGAGGSGP | ASSLNGNERLF | 0.6673056 |
| VFAAFMV | ASSLNGNERLF | 0.64784616 |
| TLYDKLR | ASSLNGNERLF | 0.6484592 |
| TLAVFTVL | ASSLNGNERLF | 0.6360145 |
| TGSVQPARV | ASSLNGNERLF | 0.6034982 |
| TGGGAGSGGT | ASSLNGNERLF | 0.69899 |
| SHLETSL | ASSLNGNERLF | 0.66758037 |
| SFTTGMT | ASSLNGNERLF | 0.6982652 |
| QVEVAKRMGVRQPTVS | ASSLNGNERLF | 0.58502245 |
| QGSGGPALAM | ASSLNGNERLF | 0.61238134 |
| MLAPSNP | ASSLNGNERLF | 0.6869774 |
| LVNNAAI | ASSLNGNERLF | 0.33073848 |
| LTGEINTLA | ASSLNGNERLF | 0.44095767 |
| LSIFGPV | ASSLNGNERLF | 0.6645243 |

| | | |
|---|---|---|
| LLALLANL | ASSLNGNERLF | 0.70159614 |
| LISNASDALDKLR | ASSLNGNERLF | 0.31898117 |
| LINEAVP | ASSLNGNERLF | 0.62392545 |
| LGYAFSGL | ASSLNGNERLF | 0.53495675 |
| LDFVAVAL | ASSLNGNERLF | 0.69057906 |
| LAQPGAE | ASSLNGNERLF | 0.66773456 |
| ISNLGAVTGF | ASSLNGNERLF | 0.5191774 |
| HQEDPGRI | ASSLNGNERLF | 0.6899302 |
| GTPGEPTP | ASSLNGNERLF | 0.50611323 |
| GFFNSGS | ASSLNGNERLF | 0.5304855 |
| GAGGGGLTAGAN | ASSLNGNERLF | 0.69725686 |
| FVDALVNM | ASSLNGNERLF | 0.70281094 |
| FLTVAPEL | ASSLNGNERLF | 0.5971298 |
| FETLTAL | ASSLNGNERLF | 0.69400036 |
| DVINPIEL | ASSLNGNERLF | 0.62599397 |
| APVSGAPV | ASSLNGNERLF | 0.7131476 |
| ALVGGLAW | ASSLNGNERLF | 0.6288939 |
| ALAAAAATAI | ASSLNGNERLF | 0.5582638 |
| AFGLGLLL | ASSLNGNERLF | 0.6905344 |
| AFFAAASL | ASSLNGNERLF | 0.65559137 |
| AAAALPAFSPP | ASSLNGNERLF | 0.6979846 |
| VVLAPAL | ASSLNGNERLF | 0.69005406 |
| TIGHVDHGKTTL | ASSLNGNERLF | 0.62713647 |
| LISNASDALDK | ASSLNGNERLF | 0.6236847 |
| LISNASDAL | ASSLNGNERLF | 0.6896708 |
| ISNASDALDK | ASSLNGNERLF | 0.6917149 |
| NQAETLVYQTE | ASSLNGNERLF | 0.5731561 |
| TGGTGGTAGAG | ASSLNGNERLF | 0.7033533 |
| TGSTAGGAGGA | ASSLNGNERLF | 0.64146996 |
| DPLAVPVAAGPV | ASSLNGNERLF | 0.6268339 |
| GGTGGTPGNGGN | ASSLNGNERLF | 0.7090342 |
| PAGLAGVDPATL | ASSLNGNERLF | 0.68204594 |
| GNNLVVEAPHAGASV | ASSLNGNERLF | 0.68635 |
| KVTRIMSASTLEQPA | ASSLNGNERLF | 0.700634 |
| LGFALPIALPRRTVV | ASSLNGNERLF | 0.7100257 |
| TRLGNLTARKLVVLV | ASSLNGNERLF | 0.57625026 |
| WSQQRRLIYFPSAGP | ASSLNGNERLF | 0.6740292 |
| GPNGNPDPMAAAVDIR | ASSLNGNERLF | 0.6605718 |
| TQRCVSVADLRAAASR | ASSLNGNERLF | 0.6926879 |
| QFVLDVAEHPVEVLTETV | ASSLNGNERLF | 0.66014665 |
| GGAGGTGGAGGNGGTQAGDG | ASSLNGNERLF | 0.57577837 |
| PLLATLRAIAVDVDAKPAQV | ASSLNGNERLF | 0.68424565 |
| APKFRTGGYMIDFWGVGYQVAK | ASSLNGNERLF | 0.70104474 |
| GAGGAGGAGGAAGGPGGTGGTG | ASSLNGNERLF | 0.67509085 |
| IGALGLLATDVAPAGSVAFGVA | ASSLNGNERLF | 0.68418986 |
| AGGAGGIGGNGGAGGAGGVLMG | ASSLNGNERLF | 0.70521784 |
| ADELIGLL | ASSLNGNERLF | 0.6313288 |
| ADRALKA | ASSLNGNERLF | 0.65182567 |
| AGAVGQVGGAGGA | ASSLNGNERLF | 0.6938665 |
| AGGGGGLGGAP | ASSLNGNERLF | 0.70436436 |
| AGGLLYGNGGAG | ASSLNGNERLF | 0.65444815 |

| | | |
|---|---|---|
| AGLESIGL | ASSLNGNERLF | 0.70270985 |
| AGLKTGII | ASSLNGNERLF | 0.61402535 |
| ASLAALLL | ASSLNGNERLF | 0.6777595 |
| ASVINISSI | ASSLNGNERLF | 0.59213686 |
| ATPPGPA | ASSLNGNERLF | 0.6688911 |
| ATVASALV | ASSLNGNERLF | 0.5804063 |
| ATVASLIL | ASSLNGNERLF | 0.67114526 |
| AVNGPVLIRH | ASSLNGNERLF | 0.68815213 |
| DGAAGAVGI | ASSLNGNERLF | 0.6827333 |
| DPLLHGI | ASSLNGNERLF | 0.48367527 |
| ELGAALGAL | ASSLNGNERLF | 0.68549514 |
| ELGRTQAV | ASSLNGNERLF | 0.68357086 |
| FVAGGYLS | ASSLNGNERLF | 0.7129209 |
| GASGGAGGAGGW | ASSLNGNERLF | 0.6631629 |
| GGAGAGGDGGAGGI | ASSLNGNERLF | 0.6895991 |
| GTGHNGSQPGMGGNG | ASSLNGNERLF | 0.6381117 |
| GYAAASASAS | ASSLNGNERLF | 0.6606544 |
| HQVDVAAVQQG | ASSLNGNERLF | 0.5590119 |
| IGAAIAHT | ASSLNGNERLF | 0.6394496 |
| IGMIEVAQL | ASSLNGNERLF | 0.61332035 |
| IRLAPEHTSAMD | ASSLNGNERLF | 0.5761963 |
| IYNVSVL | ASSLNGNERLF | 0.68321645 |
| KQGFLGM | ASSLNGNERLF | 0.70433044 |
| LESVITQL | ASSLNGNERLF | 0.69161844 |
| LGLGALVVV | ASSLNGNERLF | 0.697277 |
| LIGNGGNGGSGGI | ASSLNGNERLF | 0.69312966 |
| LNAINAPTL | ASSLNGNERLF | 0.70590806 |
| LQLLNTGV | ASSLNGNERLF | 0.6391007 |
| LQTSIEF | ASSLNGNERLF | 0.6422254 |
| MSFAVSEL | ASSLNGNERLF | 0.56844014 |
| NTGALGL | ASSLNGNERLF | 0.6312711 |
| PPGAGGAGGAGTT | ASSLNGNERLF | 0.6375415 |
| QLVGLGI | ASSLNGNERLF | 0.6654023 |
| RFLEQVHQL | ASSLNGNERLF | 0.29336897 |
| RGVGYVLR | ASSLNGNERLF | 0.65211755 |
| SATGVFAGVI | ASSLNGNERLF | 0.7076993 |
| SLPAITLP | ASSLNGNERLF | 0.56766236 |
| STALAFTV | ASSLNGNERLF | 0.6877685 |
| SVLTVGLGK | ASSLNGNERLF | 0.63596106 |
| TALAGAGSGI | ASSLNGNERLF | 0.60292345 |
| TGLPFTGI | ASSLNGNERLF | 0.68809736 |
| VAEHVTV | ASSLNGNERLF | 0.6062194 |
| VDIVLNSL | ASSLNGNERLF | 0.68855774 |
| VLTGGGAVL | ASSLNGNERLF | 0.7077816 |
| VNVGLAYGI | ASSLNGNERLF | 0.6673368 |
| VSSAAVGAV | ASSLNGNERLF | 0.6719805 |
| VTSVYEAGV | ASSLNGNERLF | 0.6778503 |
| WVALDRGA | ASSLNGNERLF | 0.54266584 |
| YLVGSVVI | ASSLNGNERLF | 0.6785449 |
| GPPGTGKTLL | ASSLNGNERLF | 0.40237612 |
| GVGMTKF | ASSLNGNERLF | 0.6332663 |

| | | |
|---|---|---|
| IGIGGSDL | ASSLNGNERLF | 0.6083423 |
| IGTLLAEL | ASSLNGNERLF | 0.67343503 |
| IYTSGTT | ASSLNGNERLF | 0.6545687 |
| LSFPAGV | ASSLNGNERLF | 0.6823342 |
| LYTSGTT | ASSLNGNERLF | 0.5671183 |
| VGAFPFGF | ASSLNGNERLF | 0.6811253 |
| VGAGTVEF | ASSLNGNERLF | 0.6549178 |
| VGKSTLLL | ASSLNGNERLF | 0.59353924 |
| AALAHAGGGIV | ASSLNGNERLF | 0.68874896 |
| AALVAVSLL | ASSLNGNERLF | 0.67107904 |
| AAVSRAVAAL | ASSLNGNERLF | 0.6063893 |
| AGGSSGIGGF | ASSLNGNERLF | 0.7007928 |
| AGLTVANANG | ASSLNGNERLF | 0.6797056 |
| AGLVGAVL | ASSLNGNERLF | 0.5556847 |
| ALSQAFTV | ASSLNGNERLF | 0.62421626 |
| ALTFSVSL | ASSLNGNERLF | 0.6518876 |
| ALTGAALVAL | ASSLNGNERLF | 0.64425415 |
| ASFSIGAGVL | ASSLNGNERLF | 0.33723044 |
| DGSVYKI | ASSLNGNERLF | 0.66488194 |
| EQLAEIV | ASSLNGNERLF | 0.62783587 |
| FAVSELTVM | ASSLNGNERLF | 0.54851305 |
| GAAPLAAGL | ASSLNGNERLF | 0.6337445 |
| GADGGDGGSGGAGG | ASSLNGNERLF | 0.68686956 |
| GFGLDTGGAGG | ASSLNGNERLF | 0.6152277 |
| GLASGGLL | ASSLNGNERLF | 0.6896245 |
| GRIAGAAVLL | ASSLNGNERLF | 0.68969405 |
| GSAGSLNTSF | ASSLNGNERLF | 0.6985081 |
| GTGQIGI | ASSLNGNERLF | 0.70533985 |
| GTLIALIL | ASSLNGNERLF | 0.6870688 |
| IAATMGFVVG | ASSLNGNERLF | 0.6590376 |
| IFAQALL | ASSLNGNERLF | 0.6840971 |
| IGAAVALGF | ASSLNGNERLF | 0.6821624 |
| IGLTPLAL | ASSLNGNERLF | 0.5394584 |
| ILGSNTSTL | ASSLNGNERLF | 0.54419875 |
| IVFSLTVL | ASSLNGNERLF | 0.651869 |
| LASKATGF | ASSLNGNERLF | 0.48759386 |
| LASVALIL | ASSLNGNERLF | 0.66291314 |
| LFAVGLTL | ASSLNGNERLF | 0.7094502 |
| LFNGGIGTYI | ASSLNGNERLF | 0.6539656 |
| LTLGAGII | ASSLNGNERLF | 0.5891188 |
| LVAAALGDI | ASSLNGNERLF | 0.6373995 |
| NTAFVQLGI | ASSLNGNERLF | 0.6225265 |
| QLFSGAEL | ASSLNGNERLF | 0.6880362 |
| SAATFITL | ASSLNGNERLF | 0.69699144 |
| SNYGLANLGV | ASSLNGNERLF | 0.7085887 |
| SYGGAYAVM | ASSLNGNERLF | 0.69161904 |
| TTGVGGLTLG | ASSLNGNERLF | 0.61618996 |
| TTQTIYI | ASSLNGNERLF | 0.6783442 |
| VAAGAGLGAVI | ASSLNGNERLF | 0.6294385 |
| VAAVVTLL | ASSLNGNERLF | 0.6404544 |
| VGFLGGLI | ASSLNGNERLF | 0.6625668 |

| | | |
|---|---|---|
| VGVIPILL | ASSLNGNERLF | 0.655567 |
| VIFASSLI | ASSLNGNERLF | 0.66690004 |
| VKGGDGGAAAG | ASSLNGNERLF | 0.6342452 |
| VTKTFTNI | ASSLNGNERLF | 0.51882774 |
| VVAAAILL | ASSLNGNERLF | 0.68601674 |
| YAALDVEL | ASSLNGNERLF | 0.64310014 |
| LASVALLL | ASSLNGNERLF | 0.5944747 |
| LDAVHRV | ASSLNGNERLF | 0.6227009 |
| QGLSSSIF | ASSLNGNERLF | 0.7060356 |
| RGTFANIRL | ASSLNGNERLF | 0.70728207 |
| RQLWWGHRI | ASSLNGNERLF | 0.58891314 |
| YGPPGTGKTLL | ASSLNGNERLF | 0.6270518 |
| AALVGAPANVA | ASSLNGNERLF | 0.63817763 |
| AATFRRTE | ASSLNGNERLF | 0.47420886 |
| AGALGGGI | ASSLNGNERLF | 0.6814027 |
| AGGAGGAGGIGAG | ASSLNGNERLF | 0.6287093 |
| AGSGALGL | ASSLNGNERLF | 0.6242756 |
| AQIADQI | ASSLNGNERLF | 0.6394558 |
| ASAVAVTYL | ASSLNGNERLF | 0.6571851 |
| ASTVLGGTGG | ASSLNGNERLF | 0.60858786 |
| AVGGEPAK | ASSLNGNERLF | 0.7009575 |
| CEQGTFGGTC | ASSLNGNERLF | 0.31017312 |
| DEGKAPAG | ASSLNGNERLF | 0.6901708 |
| DTVVFPAL | ASSLNGNERLF | 0.56218135 |
| FEGITMAG | ASSLNGNERLF | 0.63736135 |
| FPLVVQD | ASSLNGNERLF | 0.5091595 |
| GAFTTPQ | ASSLNGNERLF | 0.66584766 |
| GFGGFGTAGGVGGA | ASSLNGNERLF | 0.6883877 |
| GFLAQFL | ASSLNGNERLF | 0.6906833 |
| GGAGGAGGNAGVA | ASSLNGNERLF | 0.59544545 |
| GGVGTAGAGGAGGAGG | ASSLNGNERLF | 0.6921092 |
| GNGGGGDAGNAGSGGN | ASSLNGNERLF | 0.7102065 |
| GTGGAGGAGNDAGS | ASSLNGNERLF | 0.6091498 |
| GTTNTGWFNS | ASSLNGNERLF | 0.51120573 |
| IFMGPLFTL | ASSLNGNERLF | 0.6887448 |
| ILAHAIEAI | ASSLNGNERLF | 0.6388159 |
| IRSAIREAL | ASSLNGNERLF | 0.6857585 |
| ITFAGGG | ASSLNGNERLF | 0.67549765 |
| ITGMSGHQEV | ASSLNGNERLF | 0.52325505 |
| IVAGLACTL | ASSLNGNERLF | 0.70943373 |
| IVGGGLAGA | ASSLNGNERLF | 0.6860023 |
| IVTGVLNI | ASSLNGNERLF | 0.58371794 |
| LAQQGCI | ASSLNGNERLF | 0.64779556 |
| LCLHVEN | ASSLNGNERLF | 0.62907654 |
| LDAISDGS | ASSLNGNERLF | 0.44809216 |
| LGGFAAML | ASSLNGNERLF | 0.70182467 |
| LGGKNPMI | ASSLNGNERLF | 0.5864156 |
| LGPKGVAAIL | ASSLNGNERLF | 0.6584929 |
| LMVGVSI | ASSLNGNERLF | 0.660519 |
| LVGAAFVL | ASSLNGNERLF | 0.6663866 |
| LVLAVAPL | ASSLNGNERLF | 0.65136975 |

| | | |
|---|---|---|
| MAAIAVIDAA | ASSLNGNERLF | 0.65370476 |
| NNGINAELQ | ASSLNGNERLF | 0.67727244 |
| RGEPLVLVHG | ASSLNGNERLF | 0.66649175 |
| RGGAGGAAGLI | ASSLNGNERLF | 0.6435058 |
| RSYRMHPAV | ASSLNGNERLF | 0.6663989 |
| SAAHLLAGQLA | ASSLNGNERLF | 0.70023054 |
| SAAVGSY | ASSLNGNERLF | 0.56301004 |
| SPHAVLHRI | ASSLNGNERLF | 0.6322827 |
| SQIAKVL | ASSLNGNERLF | 0.67607117 |
| TALGHLVP | ASSLNGNERLF | 0.59443676 |
| TAPTTSGAA | ASSLNGNERLF | 0.6846784 |
| TDGDDGNGGSPGT | ASSLNGNERLF | 0.6574694 |
| TDGLGEN | ASSLNGNERLF | 0.66226554 |
| TGGPPLPGQA | ASSLNGNERLF | 0.657892 |
| TIPAITI | ASSLNGNERLF | 0.70667595 |
| TIPGFTI | ASSLNGNERLF | 0.69152254 |
| TSIASGALA | ASSLNGNERLF | 0.6700176 |
| TVTLPGAAI | ASSLNGNERLF | 0.672039 |
| VASGLSAALP | ASSLNGNERLF | 0.61369824 |
| VGAGVIGI | ASSLNGNERLF | 0.69350255 |
| VGPKRFGV | ASSLNGNERLF | 0.68477464 |
| VNVAFPD | ASSLNGNERLF | 0.5511209 |
| VSMTNSGVS | ASSLNGNERLF | 0.6293 |
| VTDVLGDLSV | ASSLNGNERLF | 0.7093173 |
| VVVVGAGFAGL | ASSLNGNERLF | 0.6900356 |
| VYTVGTVV | ASSLNGNERLF | 0.57836014 |
| WTTFSGP | ASSLNGNERLF | 0.61951506 |
| YDELVEAF | ASSLNGNERLF | 0.5147957 |
| YGTGGAGGNGG | ASSLNGNERLF | 0.6449916 |
| YTNVMAV | ASSLNGNERLF | 0.58611214 |
| FLEQVHQL | ASSLNGNERLF | 0.6812749 |
| GLSLLAVAV | ASSLNGNERLF | 0.6225618 |
| LEQVHQL | ASSLNGNERLF | 0.6859828 |
| LNAAGAI | ASSLNGNERLF | 0.6607639 |
| PAVHASL | ASSLNGNERLF | 0.70606107 |
| TELEDDA | ASSLNGNERLF | 0.6106487 |
| TPPGPPN | ASSLNGNERLF | 0.6492006 |
| TPPPPPPP | ASSLNGNERLF | 0.5160173 |
| TQSPALI | ASSPRDRGHEQY | 0.58738697 |
| EFAAAFPQ | ASSPRDRGHEQY | 0.632604 |
| LLSGNLDVL | ASSPRDRGHEQY | 0.66795564 |
| GPANLGNYNI | ASSPRDRGHEQY | 0.38339072 |
| TLGPTLP | ASSPRDRGHEQY | 0.6947185 |
| VYAYSSCCK | ASSPRDRGHEQY | 0.65788144 |
| APAPAPAPNI | ASSPRDRGHEQY | 0.6892371 |
| PGGPVPIP | ASSPRDRGHEQY | 0.6543607 |
| FLTVALEP | ASSPRDRGHEQY | 0.66054773 |
| SQYFGANL | ASSPRDRGHEQY | 0.6299306 |
| GGVKPAA | ASSPRDRGHEQY | 0.6865995 |
| GGTGGAGGAGGW | ASSPRDRGHEQY | 0.60405016 |
| GVGGVAGLGGAGG | ASSPRDRGHEQY | 0.4970311 |

| | | |
|---|---|---|
| VLLVLATTI | ASSPRDRGHEQY | 0.6534763 |
| IGIFAAML | ASSPRDRGHEQY | 0.57005525 |
| AVGDLID | ASSPRDRGHEQY | 0.7037609 |
| NISPLPL | ASSPRDRGHEQY | 0.69550836 |
| VTAVFVAI | ASSPRDRGHEQY | 0.662551 |
| SGDDNDAAL | ASSPRDRGHEQY | 0.6922498 |
| AVAALAHS | ASSPRDRGHEQY | 0.3899462 |
| GGVTVMATP | ASSPRDRGHEQY | 0.64983916 |
| AGLAANVRD | ASSPRDRGHEQY | 0.7074142 |
| TNRPDLIDPALLRPG | ASSPRDRGHEQY | 0.677118 |
| LANLGVTAL | ASSPRDRGHEQY | 0.6346897 |
| TGERIEL | ASSPRDRGHEQY | 0.7007147 |
| NGEELAA | ASSPRDRGHEQY | 0.61740226 |
| VLSLVWG | ASSPRDRGHEQY | 0.64653003 |
| IGYALWL | ASSPRDRGHEQY | 0.67862594 |
| LSLLLAPL | ASSPRDRGHEQY | 0.6892799 |
| TLDEALK | ASSPRDRGHEQY | 0.68330276 |
| TDGAGTAEGG | ASSPRDRGHEQY | 0.6017816 |
| NAPGGAGGNGG | ASSPRDRGHEQY | 0.688849 |
| VSVFMGMGGNF | ASSPRDRGHEQY | 0.7119359 |
| LASYLAAL | ASSPRDRGHEQY | 0.62311095 |
| LVGGTPAL | ASSPRDRGHEQY | 0.6879851 |
| VRNADVL | ASSPRDRGHEQY | 0.6783446 |
| TCSSPFLVSAM | ASSPRDRGHEQY | 0.6557761 |
| VHITDAE | ASSPRDRGHEQY | 0.5817459 |
| EYAAPSF | ASSPRDRGHEQY | 0.5995819 |
| VVDAFHGGL | ASSPRDRGHEQY | 0.68574333 |
| ASVVDPQ | ASSPRDRGHEQY | 0.56605 |
| SYNEFGGLS | ASSPRDRGHEQY | 0.65967846 |
| DPDGSVY | ASSPRDRGHEQY | 0.6388979 |
| MVSTIHATE | ASSPRDRGHEQY | 0.7170232 |
| LAAGVADPV | ASSPRDRGHEQY | 0.67672235 |
| TKEQIAA | ASSPRDRGHEQY | 0.68027663 |
| SLASPAAL | ASSPRDRGHEQY | 0.60510504 |
| IGLVFYGV | ASSPRDRGHEQY | 0.6782036 |
| FLLDASNL | ASSPRDRGHEQY | 0.6723219 |
| YQVGNFP | ASSPRDRGHEQY | 0.54274404 |
| ESLLSQST | ASSPRDRGHEQY | 0.70257425 |
| LFTVGFAAVG | ASSPRDRGHEQY | 0.6005143 |
| ATPPTPT | ASSPRDRGHEQY | 0.60813147 |
| TDLATLG | ASSPRDRGHEQY | 0.68770474 |
| SAGGTGR | ASSPRDRGHEQY | 0.70635957 |
| GFSNSGS | ASSPRDRGHEQY | 0.6371139 |
| GGNGGSGGTSV | ASSPRDRGHEQY | 0.66910076 |
| GTSVATGGAGN | ASSPRDRGHEQY | 0.61106306 |
| NGGNGGSGGTS | ASSPRDRGHEQY | 0.6950451 |
| AVVKFPH | ASSPRDRGHEQY | 0.6948498 |
| TPPPPPP | ASSPRDRGHEQY | 0.66186684 |
| AAPAAAAAPA | ASSPRDRGHEQY | 0.70290077 |
| IALNEDL | ASSPRDRGHEQY | 0.69560707 |
| IVNVSSM | ASSPRDRGHEQY | 0.52711725 |

| | | |
|---|---|---|
| TGIPPAPR | ASSPRDRGHEQY | 0.6655332 |
| FDLGGGTF | ASSPRDRGHEQY | 0.6923611 |
| FDLGGGTFDVS | ASSPRDRGHEQY | 0.6901109 |
| NEPTAAA | ASSPRDRGHEQY | 0.71018463 |
| NEPTAAALA | ASSPRDRGHEQY | 0.6803236 |
| GSIINISSI | ASSPRDRGHEQY | 0.610937 |
| VAPGNAGTA | ASSPRDRGHEQY | 0.70180845 |
| TGGGASLE | ASSPRDRGHEQY | 0.7181368 |
| VAVYNHYKRI | ASSPRDRGHEQY | 0.6914527 |
| TDPAPATT | ASSPRDRGHEQY | 0.6351124 |
| TNRPDILDPALLRPG | ASSPRDRGHEQY | 0.63140655 |
| LTTTTTS | ASSPRDRGHEQY | 0.68325806 |
| PTGSGKTL | ASSPRDRGHEQY | 0.683582 |
| TGSGKTL | ASSPRDRGHEQY | 0.5939278 |
| TIGHVDHGK | ASSPRDRGHEQY | 0.65428877 |
| TIGHVDHGKTT | ASSPRDRGHEQY | 0.6693486 |
| GHVDHGKTT | ASSPRDRGHEQY | 0.5392996 |
| AQFEHTLL | ASSPRDRGHEQY | 0.67483425 |
| IVNQDEL | ASSPRDRGHEQY | 0.6934666 |
| TNGSQFF | ASSPRDRGHEQY | 0.66635007 |
| TGKGIVE | ASSPRDRGHEQY | 0.62338006 |
| LISNASDA | ASSPRDRGHEQY | 0.69443065 |
| AELRTLGELTKIAKAHGAQ | ASSPRDRGHEQY | 0.71128505 |
| APNAVAAAEPSW | ASSPRDRGHEQY | 0.7065481 |
| APQVAGAATAVL | ASSPRDRGHEQY | 0.5633534 |
| ARRDPAAAAAILPTDA | ASSPRDRGHEQY | 0.43839252 |
| DAAARDVLAEAGLAEYF | ASSPRDRGHEQY | 0.6215008 |
| DDQIVLSAKANLAKVEQ | ASSPRDRGHEQY | 0.6889816 |
| DELLPDTGEDSDEDGH | ASSPRDRGHEQY | 0.66750467 |
| ELPEAPANRATVPQD | ASSPRDRGHEQY | 0.64048713 |
| FGQSKIERVTGIG | ASSPRDRGHEQY | 0.6570804 |
| GAGGTGSTAGGAGGAGGAGGLY | ASSPRDRGHEQY | 0.69534993 |
| GHGGTGAAGGNGGTGGAGGSGI | ASSPRDRGHEQY | 0.6235874 |
| GNGGNGGAGGEGGPGGAGGAGA | ASSPRDRGHEQY | 0.578392 |
| GSTAGNGGNGGNATNGTIAGQP | ASSPRDRGHEQY | 0.67844266 |
| GTGGDGGNGGNGGTGG | ASSPRDRGHEQY | 0.6876483 |
| GVRRPAGIPADAPV | ASSPRDRGHEQY | 0.5686163 |
| GYRQVTDAHLLTLARR | ASSPRDRGHEQY | 0.6284882 |
| IAKMGQDGHDRG | ASSPRDRGHEQY | 0.6241896 |
| KAEPSIEIQYT | ASSPRDRGHEQY | 0.61342984 |
| KPVRIRMAADEDTVAVT | ASSPRDRGHEQY | 0.65339065 |
| MPPRIAGMRLLVIKPEP | ASSPRDRGHEQY | 0.58112454 |
| MKFYAVALFRSRGVPPTR | ASSPRDRGHEQY | 0.680197 |
| NAGGFGILGAVAHSP | ASSPRDRGHEQY | 0.5769496 |
| NEGREITLAMAEM | ASSPRDRGHEQY | 0.69459987 |
| NELVEWSHPLGHRWRWE | ASSPRDRGHEQY | 0.67639273 |
| NLGAGGTGGNGGNPG | ASSPRDRGHEQY | 0.68429995 |
| NTEWPKWETVMAADPVD | ASSPRDRGHEQY | 0.6217254 |
| PASAAPAETQP | ASSPRDRGHEQY | 0.5953253 |
| QQSSDGKNGVRAAEITGE | ASSPRDRGHEQY | 0.5624175 |
| RAPDAVTTSWWKE | ASSPRDRGHEQY | 0.5251757 |

| | | |
|---|---|---|
| SGFGNVGAHTSGWFNQ | ASSPRDRGHEQY | 0.70561063 |
| SGVAGAVGQAASVGGLKV | ASSPRDRGHEQY | 0.5838637 |
| SNGGAGTGSAGGNGGT | ASSPRDRGHEQY | 0.6977509 |
| TGGAGGEGGAAG | ASSPRDRGHEQY | 0.69123304 |
| TGGDGGAAGAGGAPGHG | ASSPRDRGHEQY | 0.5745459 |
| TMPPLLAAIANAGRHGV | ASSPRDRGHEQY | 0.6799071 |
| TSSTAGNSGGA | ASSPRDRGHEQY | 0.58966386 |
| VPATVPVGILC | ASSPRDRGHEQY | 0.61316144 |
| VQRPDSGGVVLTGRIS | ASSPRDRGHEQY | 0.67970026 |
| WTPNPVPATKGVPAGLVQAT | ASSPRDRGHEQY | 0.6884148 |
| YALKRALAKFSDRFDVVII | ASSPRDRGHEQY | 0.5964859 |
| YLIGSVIV | ASSPRDRGHEQY | 0.67630243 |
| YGQLVAVID | ASSPRDRGHEQY | 0.667519 |
| VTVGGMTLPA | ASSPRDRGHEQY | 0.6282497 |
| VLALPAV | ASSPRDRGHEQY | 0.6781747 |
| VIPPTLNYET | ASSPRDRGHEQY | 0.690972 |
| VGMGGMHPSSG | ASSPRDRGHEQY | 0.6865229 |
| VGLAGGAATA | ASSPRDRGHEQY | 0.5968138 |
| VGGAGGAGGSGP | ASSPRDRGHEQY | 0.68686014 |
| VFAAFMV | ASSPRDRGHEQY | 0.5640852 |
| TLYDKLR | ASSPRDRGHEQY | 0.6617541 |
| TLAVFTVL | ASSPRDRGHEQY | 0.6493207 |
| TGSVQPARV | ASSPRDRGHEQY | 0.656726 |
| TGGGAGSGGT | ASSPRDRGHEQY | 0.6837126 |
| SHLETSL | ASSPRDRGHEQY | 0.61663276 |
| SFTTGMT | ASSPRDRGHEQY | 0.6045606 |
| QVEVAKRMGVRQPTVS | ASSPRDRGHEQY | 0.63099587 |
| QGSGGPALAM | ASSPRDRGHEQY | 0.6728613 |
| MLAPSNP | ASSPRDRGHEQY | 0.6875328 |
| LVNNAAI | ASSPRDRGHEQY | 0.575785 |
| LTGEINTLA | ASSPRDRGHEQY | 0.66359925 |
| LSIFGPV | ASSPRDRGHEQY | 0.63765657 |
| LLALLANL | ASSPRDRGHEQY | 0.59239304 |
| LISNASDALDKLR | ASSPRDRGHEQY | 0.6746106 |
| LINEAVP | ASSPRDRGHEQY | 0.6641587 |
| LGYAFSGL | ASSPRDRGHEQY | 0.6106316 |
| LDFVAVAL | ASSPRDRGHEQY | 0.67795265 |
| LAQPGAE | ASSPRDRGHEQY | 0.6323984 |
| ISNLGAVTGF | ASSPRDRGHEQY | 0.6926403 |
| HQEDPGRI | ASSPRDRGHEQY | 0.70685333 |
| GTPGEPTP | ASSPRDRGHEQY | 0.6425594 |
| GFFNSGS | ASSPRDRGHEQY | 0.69276416 |
| GAGGGGLTAGAN | ASSPRDRGHEQY | 0.6628766 |
| FVDALVNM | ASSPRDRGHEQY | 0.67244506 |
| FLTVAPEL | ASSPRDRGHEQY | 0.66008544 |
| FETLTAL | ASSPRDRGHEQY | 0.61376405 |
| DVINPIEL | ASSPRDRGHEQY | 0.6937478 |
| APVSGAPV | ASSPRDRGHEQY | 0.6943897 |
| ALVGGLAW | ASSPRDRGHEQY | 0.69822556 |
| ALAAAAATAI | ASSPRDRGHEQY | 0.661811 |
| AFGLGLLL | ASSPRDRGHEQY | 0.6842773 |

| | | |
|---|---|---|
| AFFAAASL | ASSPRDRGHEQY | 0.68631876 |
| AAAALPAFSPP | ASSPRDRGHEQY | 0.5939584 |
| VVLAPAL | ASSPRDRGHEQY | 0.52266127 |
| TIGHVDHGKTTL | ASSPRDRGHEQY | 0.6515411 |
| LISNASDALDK | ASSPRDRGHEQY | 0.58065456 |
| LISNASDAL | ASSPRDRGHEQY | 0.66393405 |
| ISNASDALDK | ASSPRDRGHEQY | 0.5348517 |
| NQAETLVYQTE | ASSPRDRGHEQY | 0.69024104 |
| TGGTGGTAGAG | ASSPRDRGHEQY | 0.7142235 |
| TGSTAGGAGGA | ASSPRDRGHEQY | 0.64755815 |
| DPLAVPVAAGPV | ASSPRDRGHEQY | 0.70900893 |
| GGTGGTPGNGGN | ASSPRDRGHEQY | 0.57768583 |
| PAGLAGVDPATL | ASSPRDRGHEQY | 0.69778085 |
| GNNLVVEAPHAGASV | ASSPRDRGHEQY | 0.69950306 |
| KVTRIMSASTLEQPA | ASSPRDRGHEQY | 0.56913453 |
| LGFALPIALPRRTVV | ASSPRDRGHEQY | 0.69279873 |
| TRLGNLTARKLVVLV | ASSPRDRGHEQY | 0.6190939 |
| WSQQRRLIYFPSAGP | ASSPRDRGHEQY | 0.65592086 |
| GPNGNPDPMAAAVDIR | ASSPRDRGHEQY | 0.7020945 |
| TQRCVSVADLRAAASR | ASSPRDRGHEQY | 0.6607909 |
| QFVLDVAEHPVEVLTETV | ASSPRDRGHEQY | 0.6800049 |
| GGAGGTGGAGGNGGTQAGDG | ASSPRDRGHEQY | 0.6765371 |
| PLLATLRAIAVDVDAKPAQV | ASSPRDRGHEQY | 0.626848 |
| APKFRTGGYMIDFWGVGYQVAK | ASSPRDRGHEQY | 0.6704409 |
| GAGGAGGAGGAAGGPGGTGGTG | ASSPRDRGHEQY | 0.70213556 |
| IGALGLLATDVAPAGSVAFGVA | ASSPRDRGHEQY | 0.6460861 |
| AGGAGGIGGNGGAGGAGGVLMG | ASSPRDRGHEQY | 0.7171774 |
| ADELIGLL | ASSPRDRGHEQY | 0.5956795 |
| ADRALKA | ASSPRDRGHEQY | 0.70375323 |
| AGAVGQVGGAGGA | ASSPRDRGHEQY | 0.67139304 |
| AGGGGGLGGAP | ASSPRDRGHEQY | 0.6441538 |
| AGGLLYGNGGAG | ASSPRDRGHEQY | 0.6407071 |
| AGLESIGL | ASSPRDRGHEQY | 0.6967446 |
| AGLKTGII | ASSPRDRGHEQY | 0.64202666 |
| ASLAALLL | ASSPRDRGHEQY | 0.6968201 |
| ASVINISSI | ASSPRDRGHEQY | 0.7081336 |
| ATPPGPA | ASSPRDRGHEQY | 0.49257684 |
| ATVASALV | ASSPRDRGHEQY | 0.6873642 |
| ATVASLIL | ASSPRDRGHEQY | 0.66215944 |
| AVNGPVLIRH | ASSPRDRGHEQY | 0.65944546 |
| DGAAGAVGI | ASSPRDRGHEQY | 0.58689654 |
| DPLLHGI | ASSPRDRGHEQY | 0.35617617 |
| ELGAALGAL | ASSPRDRGHEQY | 0.5223905 |
| ELGRTQAV | ASSPRDRGHEQY | 0.554649 |
| FVAGGYLS | ASSPRDRGHEQY | 0.69852483 |
| GASGGAGGAGGW | ASSPRDRGHEQY | 0.67355144 |
| GGAGAGGDGGAGGI | ASSPRDRGHEQY | 0.51700604 |
| GTGHNGSQPGMGGNG | ASSPRDRGHEQY | 0.6858798 |
| GYAAASASAS | ASSPRDRGHEQY | 0.67167187 |
| HQVDVAAVQQG | ASSPRDRGHEQY | 0.40962055 |
| IGAAIAHT | ASSPRDRGHEQY | 0.6098976 |

| | | |
|---|---|---|
| IGMIEVAQL | ASSPRDRGHEQY | 0.44360477 |
| IRLAPEHTSAMD | ASSPRDRGHEQY | 0.64801466 |
| IYNVSVL | ASSPRDRGHEQY | 0.30565834 |
| KQGFLGM | ASSPRDRGHEQY | 0.6775529 |
| LESVITQL | ASSPRDRGHEQY | 0.5138342 |
| LGLGALVVV | ASSPRDRGHEQY | 0.2782233 |
| LIGNGGNGGSGGI | ASSPRDRGHEQY | 0.6391039 |
| LNAINAPTL | ASSPRDRGHEQY | 0.61004746 |
| LQLLNTGV | ASSPRDRGHEQY | 0.6622555 |
| LQTSIEF | ASSPRDRGHEQY | 0.68462944 |
| MSFAVSEL | ASSPRDRGHEQY | 0.49098408 |
| NTGALGL | ASSPRDRGHEQY | 0.4688552 |
| PPGAGGAGGAGTT | ASSPRDRGHEQY | 0.66600716 |
| QLVGLGI | ASSPRDRGHEQY | 0.6963401 |
| RFLEQVHQL | ASSPRDRGHEQY | 0.48193577 |
| RGVGYVLR | ASSPRDRGHEQY | 0.67089057 |
| SATGVFAGVI | ASSPRDRGHEQY | 0.676276 |
| SLPAITLP | ASSPRDRGHEQY | 0.62943333 |
| STALAFTV | ASSPRDRGHEQY | 0.6910995 |
| SVLTVGLGK | ASSPRDRGHEQY | 0.69962573 |
| TALAGAGSGI | ASSPRDRGHEQY | 0.50509095 |
| TGLPFTGI | ASSPRDRGHEQY | 0.69345975 |
| VAEHVTV | ASSPRDRGHEQY | 0.68986154 |
| VDIVLNSL | ASSPRDRGHEQY | 0.58441985 |
| VLTGGGAVL | ASSPRDRGHEQY | 0.61403525 |
| VNVGLAYGI | ASSPRDRGHEQY | 0.39938322 |
| VSSAAVGAV | ASSPRDRGHEQY | 0.615357 |
| VTSVYEAGV | ASSPRDRGHEQY | 0.44418186 |
| WVALDRGA | ASSPRDRGHEQY | 0.6485111 |
| YLVGSVVI | ASSPRDRGHEQY | 0.68716455 |
| GPPGTGKTLL | ASSPRDRGHEQY | 0.61669624 |
| GVGMTKF | ASSPRDRGHEQY | 0.4252325 |
| IGIGGSDL | ASSPRDRGHEQY | 0.44056368 |
| IGTLLAEL | ASSPRDRGHEQY | 0.5921782 |
| IYTSGTT | ASSPRDRGHEQY | 0.6001749 |
| LSFPAGV | ASSPRDRGHEQY | 0.35401845 |
| LYTSGTT | ASSPRDRGHEQY | 0.57524395 |
| VGAFPFGF | ASSPRDRGHEQY | 0.68152523 |
| VGAGTVEF | ASSPRDRGHEQY | 0.6835459 |
| VGKSTLLL | ASSPRDRGHEQY | 0.4487108 |
| AALAHAGGGIV | ASSPRDRGHEQY | 0.69767165 |
| AALVAVSLL | ASSPRDRGHEQY | 0.68340814 |
| AAVSRAVAAL | ASSPRDRGHEQY | 0.6921104 |
| AGGSSGIGGF | ASSPRDRGHEQY | 0.53496593 |
| AGLTVANANG | ASSPRDRGHEQY | 0.4493827 |
| AGLVGAVL | ASSPRDRGHEQY | 0.68567467 |
| ALSQAFTV | ASSPRDRGHEQY | 0.61040664 |
| ALTFSVSL | ASSPRDRGHEQY | 0.6168872 |
| ALTGAALVAL | ASSPRDRGHEQY | 0.6810665 |
| ASFSIGAGVL | ASSPRDRGHEQY | 0.5712837 |
| DGSVYKI | ASSPRDRGHEQY | 0.5082232 |

| | | |
|---|---|---|
| EQLAEIV | ASSPRDRGHEQY | 0.64656377 |
| FAVSELTVM | ASSPRDRGHEQY | 0.6827123 |
| GAAPLAAGL | ASSPRDRGHEQY | 0.46931827 |
| GADGGDGGSGGAGG | ASSPRDRGHEQY | 0.51571614 |
| GFGLDTGGAGG | ASSPRDRGHEQY | 0.5754357 |
| GLASGGLL | ASSPRDRGHEQY | 0.6263553 |
| GRIAGAAVLL | ASSPRDRGHEQY | 0.69339067 |
| GSAGSLNTSF | ASSPRDRGHEQY | 0.6538516 |
| GTGQIGI | ASSPRDRGHEQY | 0.5721669 |
| GTLIALIL | ASSPRDRGHEQY | 0.6817105 |
| IAATMGFVVG | ASSPRDRGHEQY | 0.6614611 |
| IFAQALL | ASSPRDRGHEQY | 0.64633083 |
| IGAAVALGF | ASSPRDRGHEQY | 0.64460623 |
| IGLTPLAL | ASSPRDRGHEQY | 0.5494667 |
| ILGSNTSTL | ASSPRDRGHEQY | 0.6754572 |
| IVFSLTVL | ASSPRDRGHEQY | 0.66628855 |
| LASKATGF | ASSPRDRGHEQY | 0.6854174 |
| LASVALIL | ASSPRDRGHEQY | 0.68160915 |
| LFAVGLTL | ASSPRDRGHEQY | 0.6357861 |
| LFNGGIGTYI | ASSPRDRGHEQY | 0.63294995 |
| LTLGAGII | ASSPRDRGHEQY | 0.37875766 |
| LVAAALGDI | ASSPRDRGHEQY | 0.4464206 |
| NTAFVQLGI | ASSPRDRGHEQY | 0.657598 |
| QLFSGAEL | ASSPRDRGHEQY | 0.64376163 |
| SAATFITL | ASSPRDRGHEQY | 0.6894069 |
| SNYGLANLGV | ASSPRDRGHEQY | 0.64887166 |
| SYGGAYAVM | ASSPRDRGHEQY | 0.55785054 |
| TTGVGGLTLG | ASSPRDRGHEQY | 0.66600335 |
| TTQTIYI | ASSPRDRGHEQY | 0.68466556 |
| VAAGAGLGAVI | ASSPRDRGHEQY | 0.39131075 |
| VAAVVTLL | ASSPRDRGHEQY | 0.60497034 |
| VGFLGGLI | ASSPRDRGHEQY | 0.7039684 |
| VGVIPILL | ASSPRDRGHEQY | 0.67045873 |
| VIFASSLI | ASSPRDRGHEQY | 0.5539947 |
| VKGGDGGAAAG | ASSPRDRGHEQY | 0.6505476 |
| VTKTFTNI | ASSPRDRGHEQY | 0.706401 |
| VVAAAILL | ASSPRDRGHEQY | 0.6498966 |
| YAALDVEL | ASSPRDRGHEQY | 0.6901655 |
| LASVALLL | ASSPRDRGHEQY | 0.3404866 |
| LDAVHRV | ASSPRDRGHEQY | 0.61177325 |
| QGLSSSIF | ASSPRDRGHEQY | 0.46844685 |
| RGTFANIRL | ASSPRDRGHEQY | 0.3653773 |
| RQLWWGHRI | ASSPRDRGHEQY | 0.6759633 |
| YGPPGTGKTLL | ASSPRDRGHEQY | 0.5693461 |
| AALVGAPANVA | ASSPRDRGHEQY | 0.68550307 |
| AATFRRTE | ASSPRDRGHEQY | 0.55710244 |
| AGALGGGI | ASSPRDRGHEQY | 0.6936126 |
| AGGAGGAGGIGAG | ASSPRDRGHEQY | 0.6324799 |
| AGSGALGL | ASSPRDRGHEQY | 0.65600455 |
| AQIADQI | ASSPRDRGHEQY | 0.5202451 |
| ASAVAVTYL | ASSPRDRGHEQY | 0.66537124 |

| | | |
|---|---|---|
| ASTVLGGTGG | ASSPRDRGHEQY | 0.6855669 |
| AVGGEPAK | ASSPRDRGHEQY | 0.6680862 |
| CEQGTFGGTC | ASSPRDRGHEQY | 0.37208188 |
| DEGKAPAG | ASSPRDRGHEQY | 0.6494781 |
| DTVVFPAL | ASSPRDRGHEQY | 0.6405617 |
| FEGITMAG | ASSPRDRGHEQY | 0.31294882 |
| FPLVVQD | ASSPRDRGHEQY | 0.3259059 |
| GAFTTPQ | ASSPRDRGHEQY | 0.5301253 |
| GFGGFGTAGGVGGA | ASSPRDRGHEQY | 0.5418853 |
| GFLAQFL | ASSPRDRGHEQY | 0.6768329 |
| GGAGGAGGNAGVA | ASSPRDRGHEQY | 0.6607964 |
| GGVGTAGAGGAGGAGG | ASSPRDRGHEQY | 0.60986125 |
| GNGGGGDAGNAGSGGN | ASSPRDRGHEQY | 0.6439458 |
| GTGGAGGAGNDAGS | ASSPRDRGHEQY | 0.6438403 |
| GTTNTGWFNS | ASSPRDRGHEQY | 0.26566702 |
| IFMGPLFTL | ASSPRDRGHEQY | 0.44174653 |
| ILAHAIEAI | ASSPRDRGHEQY | 0.41125333 |
| IRSAIREAL | ASSPRDRGHEQY | 0.70032954 |
| ITFAGGG | ASSPRDRGHEQY | 0.5559361 |
| ITGMSGHQEV | ASSPRDRGHEQY | 0.48777074 |
| IVAGLACTL | ASSPRDRGHEQY | 0.6342155 |
| IVGGGLAGA | ASSPRDRGHEQY | 0.67195237 |
| IVTGVLNI | ASSPRDRGHEQY | 0.6230533 |
| LAQQGCI | ASSPRDRGHEQY | 0.49484837 |
| LCLHVEN | ASSPRDRGHEQY | 0.33309692 |
| LDAISDGS | ASSPRDRGHEQY | 0.6628412 |
| LGGFAAML | ASSPRDRGHEQY | 0.40803242 |
| LGGKNPMI | ASSPRDRGHEQY | 0.55702376 |
| LGPKGVAAIL | ASSPRDRGHEQY | 0.35390195 |
| LMVGVSI | ASSPRDRGHEQY | 0.5925034 |
| LVGAAFVL | ASSPRDRGHEQY | 0.3658554 |
| LVLAVAPL | ASSPRDRGHEQY | 0.67980677 |
| MAAIAVIDAA | ASSPRDRGHEQY | 0.59402514 |
| NNGINAELQ | ASSPRDRGHEQY | 0.57931316 |
| RGEPLVLVHG | ASSPRDRGHEQY | 0.69736654 |
| RGGAGGAAGLI | ASSPRDRGHEQY | 0.67497456 |
| RSYRMHPAV | ASSPRDRGHEQY | 0.5991646 |
| SAAHLLAGQLA | ASSPRDRGHEQY | 0.5703255 |
| SAAVGSY | ASSPRDRGHEQY | 0.47599158 |
| SPHAVLHRI | ASSPRDRGHEQY | 0.51287436 |
| SQIAKVL | ASSPRDRGHEQY | 0.5598749 |
| TALGHLVP | ASSPRDRGHEQY | 0.69061935 |
| TAPTTSGAA | ASSPRDRGHEQY | 0.6611495 |
| TDGDDGNGGSPGT | ASSPRDRGHEQY | 0.3857902 |
| TDGLGEN | ASSPRDRGHEQY | 0.65425074 |
| TGGPPLPGQA | ASSPRDRGHEQY | 0.69721293 |
| TIPAITI | ASSPRDRGHEQY | 0.6982769 |
| TIPGFTI | ASSPRDRGHEQY | 0.5651804 |
| TSIASGALA | ASSPRDRGHEQY | 0.4785015 |
| TVTLPGAAI | ASSPRDRGHEQY | 0.57118016 |
| VASGLSAALP | ASSPRDRGHEQY | 0.62229186 |

| | | |
|---|---|---|
| VGAGVIGI | ASSPRDRGHEQY | 0.49191636 |
| VGPKRFGV | ASSPRDRGHEQY | 0.4824649 |
| VNVAFPD | ASSPRDRGHEQY | 0.6683229 |
| VSMTNSGVS | ASSPRDRGHEQY | 0.71404004 |
| VTDVLGDLSV | ASSPRDRGHEQY | 0.59643817 |
| VVVVGAGFAGL | ASSPRDRGHEQY | 0.30224115 |
| VYTVGTVV | ASSPRDRGHEQY | 0.6159083 |
| WTTFSGP | ASSPRDRGHEQY | 0.4992047 |
| YDELVEAF | ASSPRDRGHEQY | 0.5690813 |
| YGTGGAGGNGG | ASSPRDRGHEQY | 0.59525263 |
| YTNVMAV | ASSPRDRGHEQY | 0.37879354 |
| FLEQVHQL | ASSPRDRGHEQY | 0.61857224 |
| GLSLLAVAV | ASSPRDRGHEQY | 0.7007048 |
| LEQVHQL | ASSPRDRGHEQY | 0.6287456 |
| LNAAGAI | ASSPRDRGHEQY | 0.67042816 |
| PAVHASL | ASSPRDRGHEQY | 0.69214785 |
| TELEDDA | ASSPRDRGHEQY | 0.6728127 |
| TPPGPPN | ASSPRDRGHEQY | 0.6966866 |
| TPPPPPPP | ASSPRDRGHEQY | 0.6906903 |
| TQSPALI | ASSLSIGANTEVF | 0.64598197 |
| EFAAAFPQ | ASSLSIGANTEVF | 0.6052995 |
| LLSGNLDVL | ASSLSIGANTEVF | 0.6657084 |
| GPANLGNYNI | ASSLSIGANTEVF | 0.627319 |
| TLGPTLP | ASSLSIGANTEVF | 0.6022241 |
| VYAYSSCCK | ASSLSIGANTEVF | 0.5675953 |
| APAPAPAPNI | ASSLSIGANTEVF | 0.68735826 |
| PGGPVPIP | ASSLSIGANTEVF | 0.507912 |
| FLTVALEP | ASSLSIGANTEVF | 0.70432234 |
| SQYFGANL | ASSLSIGANTEVF | 0.3949699 |
| GGVKPAA | ASSLSIGANTEVF | 0.48954964 |
| GGTGGAGGAGGW | ASSLSIGANTEVF | 0.6885254 |
| GVGGVAGLGGAGG | ASSLSIGANTEVF | 0.2676971 |
| VLLVLATTI | ASSLSIGANTEVF | 0.35828567 |
| IGIFAAML | ASSLSIGANTEVF | 0.5842416 |
| AVGDLID | ASSLSIGANTEVF | 0.6909274 |
| NISPLPL | ASSLSIGANTEVF | 0.27559665 |
| VTAVFVAI | ASSLSIGANTEVF | 0.43294156 |
| SGDDNDAAL | ASSLSIGANTEVF | 0.38557398 |
| AVAALAHS | ASSLSIGANTEVF | 0.6607497 |
| GGVTVMATP | ASSLSIGANTEVF | 0.63095546 |
| AGLAANVRD | ASSLSIGANTEVF | 0.33298445 |
| TNRPDLIDPALLRPG | ASSLSIGANTEVF | 0.69541866 |
| LANLGVTAL | ASSLSIGANTEVF | 0.31854302 |
| TGERIEL | ASSLSIGANTEVF | 0.59551305 |
| NGEELAA | ASSLSIGANTEVF | 0.67887604 |
| VLSLVWG | ASSLSIGANTEVF | 0.70319355 |
| IGYALWL | ASSLSIGANTEVF | 0.41539955 |
| LSLLLAPL | ASSLSIGANTEVF | 0.6680223 |
| TLDEALK | ASSLSIGANTEVF | 0.56587434 |
| TDGAGTAEGG | ASSLSIGANTEVF | 0.7029152 |
| NAPGGAGGNGG | ASSLSIGANTEVF | 0.5962626 |

| | | |
|---|---|---|
| VSVFMGMGGNF | ASSLSIGANTEVF | 0.3778619 |
| LASYLAAL | ASSLSIGANTEVF | 0.6926334 |
| LVGGTPAL | ASSLSIGANTEVF | 0.68272793 |
| VRNADVL | ASSLSIGANTEVF | 0.60967934 |
| TCSSPFLVSAM | ASSLSIGANTEVF | 0.68640435 |
| VHITDAE | ASSLSIGANTEVF | 0.60258865 |
| EYAAPSF | ASSLSIGANTEVF | 0.6365043 |
| VVDAFHGGL | ASSLSIGANTEVF | 0.6932205 |
| ASVVDPQ | ASSLSIGANTEVF | 0.68092686 |
| SYNEFGGLS | ASSLSIGANTEVF | 0.4726514 |
| DPDGSVY | ASSLSIGANTEVF | 0.6825044 |
| MVSTIHATE | ASSLSIGANTEVF | 0.61815166 |
| LAAGVADPV | ASSLSIGANTEVF | 0.47681832 |
| TKEQIAA | ASSLSIGANTEVF | 0.69749856 |
| SLASPAAL | ASSLSIGANTEVF | 0.596012 |
| IGLVFYGV | ASSLSIGANTEVF | 0.6742941 |
| FLLDASNL | ASSLSIGANTEVF | 0.6815591 |
| YQVGNFP | ASSLSIGANTEVF | 0.7053093 |
| ESLLSQST | ASSLSIGANTEVF | 0.35724834 |
| LFTVGFAAVG | ASSLSIGANTEVF | 0.59775347 |
| ATPPTPT | ASSLSIGANTEVF | 0.60546017 |
| TDLATLG | ASSLSIGANTEVF | 0.680831 |
| SAGGTGR | ASSLSIGANTEVF | 0.6205513 |
| GFSNSGS | ASSLSIGANTEVF | 0.5422536 |
| GGNGGSGGTSV | ASSLSIGANTEVF | 0.68033695 |
| GTSVATGGAGN | ASSLSIGANTEVF | 0.6807188 |
| NGGNGGSGGTS | ASSLSIGANTEVF | 0.5989152 |
| AVVKFPH | ASSLSIGANTEVF | 0.61686325 |
| TPPPPPP | ASSLSIGANTEVF | 0.7026961 |
| AAPAAAAAPA | ASSLSIGANTEVF | 0.4961909 |
| IALNEDL | ASSLSIGANTEVF | 0.6181915 |
| IVNVSSM | ASSLSIGANTEVF | 0.6676003 |
| TGIPPAPR | ASSLSIGANTEVF | 0.6888012 |
| FDLGGGTF | ASSLSIGANTEVF | 0.6405724 |
| FDLGGGTFDVS | ASSLSIGANTEVF | 0.68926823 |
| NEPTAAA | ASSLSIGANTEVF | 0.47434083 |
| NEPTAAALA | ASSLSIGANTEVF | 0.6792611 |
| GSIINISSI | ASSLSIGANTEVF | 0.57887644 |
| VAPGNAGTA | ASSLSIGANTEVF | 0.6382886 |
| TGGGASLE | ASSLSIGANTEVF | 0.5015178 |
| VAVYNHYKRI | ASSLSIGANTEVF | 0.6582729 |
| TDPAPATT | ASSLSIGANTEVF | 0.666522 |
| TNRPDILDPALLRPG | ASSLSIGANTEVF | 0.67827415 |
| LTTTTTS | ASSLSIGANTEVF | 0.32496172 |
| PTGSGKTL | ASSLSIGANTEVF | 0.6714232 |
| TGSGKTL | ASSLSIGANTEVF | 0.67864573 |
| TIGHVDHGK | ASSLSIGANTEVF | 0.7094057 |
| TIGHVDHGKTT | ASSLSIGANTEVF | 0.59465504 |
| GHVDHGKTT | ASSLSIGANTEVF | 0.66201335 |
| AQFEHTLL | ASSLSIGANTEVF | 0.56742007 |
| IVNQDEL | ASSLSIGANTEVF | 0.6229635 |

| | | |
|---|---|---|
| TNGSQFF | ASSLSIGANTEVF | 0.44946754 |
| TGKGIVE | ASSLSIGANTEVF | 0.533147 |
| LISNASDA | ASSLSIGANTEVF | 0.5714012 |
| AELRTLGELTKIAKAHGAQ | ASSLSIGANTEVF | 0.5719231 |
| APNAVAAAEPSW | ASSLSIGANTEVF | 0.69160306 |
| APQVAGAATAVL | ASSLSIGANTEVF | 0.6843925 |
| ARRDPAAAAAILPTDA | ASSLSIGANTEVF | 0.6718364 |
| DAAARDVLAEAGLAEYF | ASSLSIGANTEVF | 0.6797564 |
| DDQIVLSAKANLAKVEQ | ASSLSIGANTEVF | 0.6875322 |
| DELLPDTGEDSDEDGH | ASSLSIGANTEVF | 0.6946559 |
| ELPEAPANRATVPQD | ASSLSIGANTEVF | 0.60585684 |
| FGQSKIERVTGIG | ASSLSIGANTEVF | 0.5387337 |
| GAGGTGSTAGGAGGAGGAGGLY | ASSLSIGANTEVF | 0.41655576 |
| GHGGTGAAGGNGGTGGAGGSGI | ASSLSIGANTEVF | 0.5339031 |
| GNGGNGGAGGEGGPGGAGGAGA | ASSLSIGANTEVF | 0.5905129 |
| GSTAGNGGNGGNATNGTIAGQP | ASSLSIGANTEVF | 0.63418317 |
| GTGGDGGNGGNGGTGG | ASSLSIGANTEVF | 0.2805714 |
| GVRRPAGIPADAPV | ASSLSIGANTEVF | 0.65829384 |
| GYRQVTDAHLLTLARR | ASSLSIGANTEVF | 0.6963274 |
| IAKMGQDGHDRG | ASSLSIGANTEVF | 0.60099876 |
| KAEPSIEIQYT | ASSLSIGANTEVF | 0.6690737 |
| KPVRIRMAADEDTVAVT | ASSLSIGANTEVF | 0.6284988 |
| MPPRIAGMRLLVIKPEP | ASSLSIGANTEVF | 0.55501616 |
| MKFYAVALFRSRGVPPTR | ASSLSIGANTEVF | 0.64734656 |
| NAGGFGILGAVAHSP | ASSLSIGANTEVF | 0.5367433 |
| NEGREITLAMAEM | ASSLSIGANTEVF | 0.69138604 |
| NELVEWSHPLGHRWRWE | ASSLSIGANTEVF | 0.69775546 |
| NLGAGGTGGNGGNPG | ASSLSIGANTEVF | 0.66265976 |
| NTEWPKWETVMAADPVD | ASSLSIGANTEVF | 0.56528366 |
| PASAAPAETQP | ASSLSIGANTEVF | 0.6595142 |
| QQSSDGKNGVRAAEITGE | ASSLSIGANTEVF | 0.33707908 |
| RAPDAVTTSWWKE | ASSLSIGANTEVF | 0.6009756 |
| SGFGNVGAHTSGWFNQ | ASSLSIGANTEVF | 0.4023814 |
| SGVAGAVGQAASVGGLKV | ASSLSIGANTEVF | 0.6296452 |
| SNGGAGTGSAGGNGGT | ASSLSIGANTEVF | 0.5446712 |
| TGGAGGEGGAAG | ASSLSIGANTEVF | 0.6719065 |
| TGGDGGAAGAGGAPGHG | ASSLSIGANTEVF | 0.6157849 |
| TMPPLLAAIANAGRHGV | ASSLSIGANTEVF | 0.6296294 |
| TSSTAGNSGGA | ASSLSIGANTEVF | 0.46749467 |
| VPATVPVGILC | ASSLSIGANTEVF | 0.67928445 |
| VQRPDSGGVVLTGRIS | ASSLSIGANTEVF | 0.6565748 |
| WTPNPVPATKGVPAGLVQAT | ASSLSIGANTEVF | 0.44269258 |
| YALKRALAKFSDRFDVVII | ASSLSIGANTEVF | 0.67182887 |
| YLIGSVIV | ASSLSIGANTEVF | 0.57570565 |
| YGQLVAVID | ASSLSIGANTEVF | 0.5970346 |
| VTVGGMTLPA | ASSLSIGANTEVF | 0.68706024 |
| VLALPAV | ASSLSIGANTEVF | 0.65653527 |
| VIPPTLNYET | ASSLSIGANTEVF | 0.31629872 |
| VGMGGMHPSSG | ASSLSIGANTEVF | 0.608613 |
| VGLAGGAATA | ASSLSIGANTEVF | 0.6080548 |
| VGGAGGAGGSGP | ASSLSIGANTEVF | 0.62958926 |

| | | |
|---|---|---|
| VFAAFMV | ASSLSIGANTEVF | 0.28573444 |
| TLYDKLR | ASSLSIGANTEVF | 0.6764045 |
| TLAVFTVL | ASSLSIGANTEVF | 0.4793669 |
| TGSVQPARV | ASSLSIGANTEVF | 0.50142115 |
| TGGGAGSGGT | ASSLSIGANTEVF | 0.5851097 |
| SHLETSL | ASSLSIGANTEVF | 0.69245136 |
| SFTTGMT | ASSLSIGANTEVF | 0.60270214 |
| QVEVAKRMGVRQPTVS | ASSLSIGANTEVF | 0.665021 |
| QGSGGPALAM | ASSLSIGANTEVF | 0.6662142 |
| MLAPSNP | ASSLSIGANTEVF | 0.61795175 |
| LVNNAAI | ASSLSIGANTEVF | 0.6916538 |
| LTGEINTLA | ASSLSIGANTEVF | 0.6515163 |
| LSIFGPV | ASSLSIGANTEVF | 0.55343324 |
| LLALLANL | ASSLSIGANTEVF | 0.658365 |
| LISNASDALDKLR | ASSLSIGANTEVF | 0.42581993 |
| LINEAVP | ASSLSIGANTEVF | 0.46702796 |
| LGYAFSGL | ASSLSIGANTEVF | 0.50781494 |
| LDFVAVAL | ASSLSIGANTEVF | 0.56338966 |
| LAQPGAE | ASSLSIGANTEVF | 0.39514408 |
| ISNLGAVTGF | ASSLSIGANTEVF | 0.6556847 |
| HQEDPGRI | ASSLSIGANTEVF | 0.6918466 |
| GTPGEPTP | ASSLSIGANTEVF | 0.5888621 |
| GFFNSGS | ASSLSIGANTEVF | 0.36214072 |
| GAGGGGLTAGAN | ASSLSIGANTEVF | 0.5402086 |
| FVDALVNM | ASSLSIGANTEVF | 0.40302244 |
| FLTVAPEL | ASSLSIGANTEVF | 0.68391275 |
| FETLTAL | ASSLSIGANTEVF | 0.6742822 |
| DVINPIEL | ASSLSIGANTEVF | 0.7072332 |
| APVSGAPV | ASSLSIGANTEVF | 0.68045807 |
| ALVGGLAW | ASSLSIGANTEVF | 0.5930934 |
| ALAAAAATAI | ASSLSIGANTEVF | 0.6844475 |
| AFGLGLLL | ASSLSIGANTEVF | 0.58937466 |
| AFFAAASL | ASSLSIGANTEVF | 0.575173 |
| AAAALPAFSPP | ASSLSIGANTEVF | 0.6309227 |
| VVLAPAL | ASSLSIGANTEVF | 0.6436533 |
| TIGHVDHGKTTL | ASSLSIGANTEVF | 0.6101587 |
| LISNASDALDK | ASSLSIGANTEVF | 0.64451075 |
| LISNASDAL | ASSLSIGANTEVF | 0.35193467 |
| ISNASDALDK | ASSLSIGANTEVF | 0.66401124 |
| NQAETLVYQTE | ASSLSIGANTEVF | 0.56662714 |
| TGGTGGTAGAG | ASSLSIGANTEVF | 0.41644388 |
| TGSTAGGAGGA | ASSLSIGANTEVF | 0.6109023 |
| DPLAVPVAAGPV | ASSLSIGANTEVF | 0.6990136 |
| GGTGGTPGNGGN | ASSLSIGANTEVF | 0.6773033 |
| PAGLAGVDPATL | ASSLSIGANTEVF | 0.5784538 |
| GNNLVVEAPHAGASV | ASSLSIGANTEVF | 0.5539895 |
| KVTRIMSASTLEQPA | ASSLSIGANTEVF | 0.6453643 |
| LGFALPIALPRRTVV | ASSLSIGANTEVF | 0.4755604 |
| TRLGNLTARKLVVLV | ASSLSIGANTEVF | 0.6785915 |
| WSQQRRLIYFPSAGP | ASSLSIGANTEVF | 0.6322824 |
| GPNGNPDPMAAAVDIR | ASSLSIGANTEVF | 0.61614656 |

| | | |
|---|---|---|
| TQRCVSVADLRAAASR | ASSLSIGANTEVF | 0.52885973 |
| QFVLDVAEHPVEVLTETV | ASSLSIGANTEVF | 0.65496653 |
| GGAGGTGGAGGNGGTQAGDG | ASSLSIGANTEVF | 0.5989536 |
| PLLATLRAIAVDVDAKPAQV | ASSLSIGANTEVF | 0.65786207 |
| APKFRTGGYMIDFWGVGYQVAK | ASSLSIGANTEVF | 0.2715342 |
| GAGGAGGAGGAAGGPGGTGGTG | ASSLSIGANTEVF | 0.670792 |
| IGALGLLATDVAPAGSVAFGVA | ASSLSIGANTEVF | 0.49280667 |
| AGGAGGIGGNGGAGGAGGVLMG | ASSLSIGANTEVF | 0.45555276 |
| ADELIGLL | ASSLSIGANTEVF | 0.3054809 |
| ADRALKA | ASSLSIGANTEVF | 0.5471929 |
| AGAVGQVGGAGGA | ASSLSIGANTEVF | 0.6584106 |
| AGGGGGLGGAP | ASSLSIGANTEVF | 0.6828622 |
| AGGLLYGNGGAG | ASSLSIGANTEVF | 0.60199904 |
| AGLESIGL | ASSLSIGANTEVF | 0.67737424 |
| AGLKTGII | ASSLSIGANTEVF | 0.68254185 |
| ASLAALLL | ASSLSIGANTEVF | 0.5401822 |
| ASVINISSI | ASSLSIGANTEVF | 0.32637548 |
| ATPPGPA | ASSLSIGANTEVF | 0.66539884 |
| ATVASALV | ASSLSIGANTEVF | 0.59437764 |
| ATVASLIL | ASSLSIGANTEVF | 0.68708646 |
| AVNGPVLIRH | ASSLSIGANTEVF | 0.67003775 |
| DGAAGAVGI | ASSLSIGANTEVF | 0.4203583 |
| DPLLHGI | ASSLSIGANTEVF | 0.6783786 |
| ELGAALGAL | ASSLSIGANTEVF | 0.68962175 |
| ELGRTQAV | ASSLSIGANTEVF | 0.46895432 |
| FVAGGYLS | ASSLSIGANTEVF | 0.390609 |
| GASGGAGGAGGW | ASSLSIGANTEVF | 0.44034386 |
| GGAGAGGDGGAGGI | ASSLSIGANTEVF | 0.38603315 |
| GTGHNGSQPGMGGNG | ASSLSIGANTEVF | 0.63605297 |
| GYAAASASAS | ASSLSIGANTEVF | 0.56347924 |
| HQVDVAAVQQG | ASSLSIGANTEVF | 0.54966915 |
| IGAAIAHT | ASSLSIGANTEVF | 0.6265862 |
| IGMIEVAQL | ASSLSIGANTEVF | 0.5594915 |
| IRLAPEHTSAMD | ASSLSIGANTEVF | 0.5831839 |
| IYNVSVL | ASSLSIGANTEVF | 0.5911883 |
| KQGFLGM | ASSLSIGANTEVF | 0.627944 |
| LESVITQL | ASSLSIGANTEVF | 0.65827334 |
| LGLGALVVV | ASSLSIGANTEVF | 0.5819086 |
| LIGNGGNGGSGGI | ASSLSIGANTEVF | 0.5984454 |
| LNAINAPTL | ASSLSIGANTEVF | 0.6915312 |
| LQLLNTGV | ASSLSIGANTEVF | 0.42171323 |
| LQTSIEF | ASSLSIGANTEVF | 0.5071713 |
| MSFAVSEL | ASSLSIGANTEVF | 0.6773474 |
| NTGALGL | ASSLSIGANTEVF | 0.5900729 |
| PPGAGGAGGAGTT | ASSLSIGANTEVF | 0.68738705 |
| QLVGLGI | ASSLSIGANTEVF | 0.6260332 |
| RFLEQVHQL | ASSLSIGANTEVF | 0.64036375 |
| RGVGYVLR | ASSLSIGANTEVF | 0.52161264 |
| SATGVFAGVI | ASSLSIGANTEVF | 0.69718474 |
| SLPAITLP | ASSLSIGANTEVF | 0.6892902 |
| STALAFTV | ASSLSIGANTEVF | 0.5890883 |

| | | |
|---|---|---|
| SVLTVGLGK | ASSLSIGANTEVF | 0.67558146 |
| TALAGAGSGI | ASSLSIGANTEVF | 0.36555877 |
| TGLPFTGI | ASSLSIGANTEVF | 0.6806959 |
| VAEHVTV | ASSLSIGANTEVF | 0.68069804 |
| VDIVLNSL | ASSLSIGANTEVF | 0.49391228 |
| VLTGGGAVL | ASSLSIGANTEVF | 0.63715345 |
| VNVGLAYGI | ASSLSIGANTEVF | 0.68810654 |
| VSSAAVGAV | ASSLSIGANTEVF | 0.69743115 |
| VTSVYEAGV | ASSLSIGANTEVF | 0.54962337 |
| WVALDRGA | ASSLSIGANTEVF | 0.58038837 |
| YLVGSVVI | ASSLSIGANTEVF | 0.34833133 |
| GPPGTGKTLL | ASSLSIGANTEVF | 0.4260741 |
| GVGMTKF | ASSLSIGANTEVF | 0.37684408 |
| IGIGGSDL | ASSLSIGANTEVF | 0.6590658 |
| IGTLLAEL | ASSLSIGANTEVF | 0.54425204 |
| IYTSGTT | ASSLSIGANTEVF | 0.6976919 |
| LSFPAGV | ASSLSIGANTEVF | 0.65618277 |
| LYTSGTT | ASSLSIGANTEVF | 0.6934451 |
| VGAFPFGF | ASSLSIGANTEVF | 0.51249635 |
| VGAGTVEF | ASSLSIGANTEVF | 0.6164621 |
| VGKSTLLL | ASSLSIGANTEVF | 0.44880396 |
| AALAHAGGGIV | ASSLSIGANTEVF | 0.5011305 |
| AALVAVSLL | ASSLSIGANTEVF | 0.4500246 |
| AAVSRAVAAL | ASSLSIGANTEVF | 0.6236263 |
| AGGSSGIGGF | ASSLSIGANTEVF | 0.29470098 |
| AGLTVANANG | ASSLSIGANTEVF | 0.6965409 |
| AGLVGAVL | ASSLSIGANTEVF | 0.6573458 |
| ALSQAFTV | ASSLSIGANTEVF | 0.6641396 |
| ALTFSVSL | ASSLSIGANTEVF | 0.47557634 |
| ALTGAALVAL | ASSLSIGANTEVF | 0.5095097 |
| ASFSIGAGVL | ASSLSIGANTEVF | 0.5879047 |
| DGSVYKI | ASSLSIGANTEVF | 0.6839496 |
| EQLAEIV | ASSLSIGANTEVF | 0.504145 |
| FAVSELTVM | ASSLSIGANTEVF | 0.31047577 |
| GAAPLAAGL | ASSLSIGANTEVF | 0.4051127 |
| GADGGDGGSGGAGG | ASSLSIGANTEVF | 0.32470667 |
| GFGLDTGGAGG | ASSLSIGANTEVF | 0.69817525 |
| GLASGGLL | ASSLSIGANTEVF | 0.6860801 |
| GRIAGAAVLL | ASSLSIGANTEVF | 0.66638756 |
| GSAGSLNTSF | ASSLSIGANTEVF | 0.70138216 |
| GTGQIGI | ASSLSIGANTEVF | 0.2777147 |
| GTLIALIL | ASSLSIGANTEVF | 0.47442415 |
| IAATMGFVVG | ASSLSIGANTEVF | 0.69435483 |
| IFAQALL | ASSLSIGANTEVF | 0.65231353 |
| IGAAVALGF | ASSLSIGANTEVF | 0.5858122 |
| IGLTPLAL | ASSLSIGANTEVF | 0.6937336 |
| ILGSNTSTL | ASSLSIGANTEVF | 0.5792464 |
| IVFSLTVL | ASSLSIGANTEVF | 0.6368985 |
| LASKATGF | ASSLSIGANTEVF | 0.65325826 |
| LASVALIL | ASSLSIGANTEVF | 0.6781907 |
| LFAVGLTL | ASSLSIGANTEVF | 0.6030036 |

| | | |
|---|---|---|
| LFNGGIGTYI | ASSLSIGANTEVF | 0.45930094 |
| LTLGAGII | ASSLSIGANTEVF | 0.6688529 |
| LVAAALGDI | ASSLSIGANTEVF | 0.69298893 |
| NTAFVQLGI | ASSLSIGANTEVF | 0.6356254 |
| QLFSGAEL | ASSLSIGANTEVF | 0.6527803 |
| SAATFITL | ASSLSIGANTEVF | 0.65652394 |
| SNYGLANLGV | ASSLSIGANTEVF | 0.66507185 |
| SYGGAYAVM | ASSLSIGANTEVF | 0.530059 |
| TTGVGGLTLG | ASSLSIGANTEVF | 0.48671412 |
| TTQTIYI | ASSLSIGANTEVF | 0.6809257 |
| VAAGAGLGAVI | ASSLSIGANTEVF | 0.48341465 |
| VAAVVTLL | ASSLSIGANTEVF | 0.62196505 |
| VGFLGGLI | ASSLSIGANTEVF | 0.57050586 |
| VGVIPILL | ASSLSIGANTEVF | 0.7090682 |
| VIFASSLI | ASSLSIGANTEVF | 0.6644527 |
| VKGGDGGAAAG | ASSLSIGANTEVF | 0.6447215 |
| VTKTFTNI | ASSLSIGANTEVF | 0.53582954 |
| VVAAAILL | ASSLSIGANTEVF | 0.63518137 |
| YAALDVEL | ASSLSIGANTEVF | 0.66191316 |
| LASVALLL | ASSLSIGANTEVF | 0.35358918 |
| LDAVHRV | ASSLSIGANTEVF | 0.67203265 |
| QGLSSSIF | ASSLSIGANTEVF | 0.47685486 |
| RGTFANIRL | ASSLSIGANTEVF | 0.6322107 |
| RQLWWGHRI | ASSLSIGANTEVF | 0.6683476 |
| YGPPGTGKTLL | ASSLSIGANTEVF | 0.7073539 |
| AALVGAPANVA | ASSLSIGANTEVF | 0.595312 |
| AATFRRTE | ASSLSIGANTEVF | 0.6511756 |
| AGALGGGI | ASSLSIGANTEVF | 0.48331383 |
| AGGAGGAGGIGAG | ASSLSIGANTEVF | 0.67227376 |
| AGSGALGL | ASSLSIGANTEVF | 0.69690317 |
| AQIADQI | ASSLSIGANTEVF | 0.6655811 |
| ASAVAVTYL | ASSLSIGANTEVF | 0.7078066 |
| ASTVLGGTGG | ASSLSIGANTEVF | 0.64966726 |
| AVGGEPAK | ASSLSIGANTEVF | 0.33419597 |
| CEQGTFGGTC | ASSLSIGANTEVF | 0.6446711 |
| DEGKAPAG | ASSLSIGANTEVF | 0.68407583 |
| DTVVFPAL | ASSLSIGANTEVF | 0.6775367 |
| FEGITMAG | ASSLSIGANTEVF | 0.69492924 |
| FPLVVQD | ASSLSIGANTEVF | 0.66654915 |
| GAFTTPQ | ASSLSIGANTEVF | 0.5840514 |
| GFGGFGTAGGVGGA | ASSLSIGANTEVF | 0.69252884 |
| GFLAQFL | ASSLSIGANTEVF | 0.7015887 |
| GGAGGAGGNAGVA | ASSLSIGANTEVF | 0.67496806 |
| GGVGTAGAGGAGGAGG | ASSLSIGANTEVF | 0.545032 |
| GNGGGGDAGNAGSGGN | ASSLSIGANTEVF | 0.47450763 |
| GTGGAGGAGNDAGS | ASSLSIGANTEVF | 0.6687666 |
| GTTNTGWFNS | ASSLSIGANTEVF | 0.69263935 |
| IFMGPLFTL | ASSLSIGANTEVF | 0.46502757 |
| ILAHAIEAI | ASSLSIGANTEVF | 0.6450157 |
| IRSAIREAL | ASSLSIGANTEVF | 0.63745904 |
| ITFAGGG | ASSLSIGANTEVF | 0.3301033 |

| | | |
|---|---|---|
| ITGMSGHQEV | ASSLSIGANTEVF | 0.6468797 |
| IVAGLACTL | ASSLSIGANTEVF | 0.6779423 |
| IVGGGLAGA | ASSLSIGANTEVF | 0.6601675 |
| IVTGVLNI | ASSLSIGANTEVF | 0.53485894 |
| LAQQGCI | ASSLSIGANTEVF | 0.67326087 |
| LCLHVEN | ASSLSIGANTEVF | 0.70201194 |
| LDAISDGS | ASSLSIGANTEVF | 0.7044009 |
| LGGFAAML | ASSLSIGANTEVF | 0.3616077 |
| LGGKNPMI | ASSLSIGANTEVF | 0.28208813 |
| LGPKGVAAIL | ASSLSIGANTEVF | 0.639343 |
| LMVGVSI | ASSLSIGANTEVF | 0.6878412 |
| LVGAAFVL | ASSLSIGANTEVF | 0.5908607 |
| LVLAVAPL | ASSLSIGANTEVF | 0.62590665 |
| MAAIAVIDAA | ASSLSIGANTEVF | 0.6556906 |
| NNGINAELQ | ASSLSIGANTEVF | 0.6860737 |
| RGEPLVLVHG | ASSLSIGANTEVF | 0.4905883 |
| RGGAGGAAGLI | ASSLSIGANTEVF | 0.38596573 |
| RSYRMHPAV | ASSLSIGANTEVF | 0.6500933 |
| SAAHLLAGQLA | ASSLSIGANTEVF | 0.6873818 |
| SAAVGSY | ASSLSIGANTEVF | 0.5560341 |
| SPHAVLHRI | ASSLSIGANTEVF | 0.6276344 |
| SQIAKVL | ASSLSIGANTEVF | 0.59388936 |
| TALGHLVP | ASSLSIGANTEVF | 0.5936812 |
| TAPTTSGAA | ASSLSIGANTEVF | 0.62826145 |
| TDGDDGNGGSPGT | ASSLSIGANTEVF | 0.46321827 |
| TDGLGEN | ASSLSIGANTEVF | 0.67887306 |
| TGGPPLPGQA | ASSLSIGANTEVF | 0.4161117 |
| TIPAITI | ASSLSIGANTEVF | 0.66035914 |
| TIPGFTI | ASSLSIGANTEVF | 0.66933733 |
| TSIASGALA | ASSLSIGANTEVF | 0.667426 |
| TVTLPGAAI | ASSLSIGANTEVF | 0.5735444 |
| VASGLSAALP | ASSLSIGANTEVF | 0.49317142 |
| VGAGVIGI | ASSLSIGANTEVF | 0.40655765 |
| VGPKRFGV | ASSLSIGANTEVF | 0.43165106 |
| VNVAFPD | ASSLSIGANTEVF | 0.68810153 |
| VSMTNSGVS | ASSLSIGANTEVF | 0.57874936 |
| VTDVLGDLSV | ASSLSIGANTEVF | 0.69497764 |
| VVVVGAGFAGL | ASSLSIGANTEVF | 0.66315866 |
| VYTVGTVV | ASSLSIGANTEVF | 0.5029151 |
| WTTFSGP | ASSLSIGANTEVF | 0.6600349 |
| YDELVEAF | ASSLSIGANTEVF | 0.51725996 |
| YGTGGAGGNGG | ASSLSIGANTEVF | 0.5963214 |
| YTNVMAV | ASSLSIGANTEVF | 0.65961146 |
| FLEQVHQL | ASSLSIGANTEVF | 0.69131374 |
| GLSLLAVAV | ASSLSIGANTEVF | 0.4090356 |
| LEQVHQL | ASSLSIGANTEVF | 0.6649784 |
| LNAAGAI | ASSLSIGANTEVF | 0.6527968 |
| PAVHASL | ASSLSIGANTEVF | 0.3962769 |
| TELEDDA | ASSLSIGANTEVF | 0.67691267 |
| TPPGPPN | ASSLSIGANTEVF | 0.68786657 |
| TPPPPPPP | ASSLSIGANTEVF | 0.6707103 |

| | | |
|---|---|---|
| TQSPALI | ASGEGGNQDTQY | 0.45343184 |
| EFAAAFPQ | ASGEGGNQDTQY | 0.64370036 |
| LLSGNLDVL | ASGEGGNQDTQY | 0.5459065 |
| GPANLGNYNI | ASGEGGNQDTQY | 0.656443 |
| TLGPTLP | ASGEGGNQDTQY | 0.624468 |
| VYAYSSCCK | ASGEGGNQDTQY | 0.52907133 |
| APAPAPAPNI | ASGEGGNQDTQY | 0.67718554 |
| PGGPVPIP | ASGEGGNQDTQY | 0.34682298 |
| FLTVALEP | ASGEGGNQDTQY | 0.49439 |
| SQYFGANL | ASGEGGNQDTQY | 0.6207352 |
| GGVKPAA | ASGEGGNQDTQY | 0.6371887 |
| GGTGGAGGAGGW | ASGEGGNQDTQY | 0.6754173 |
| GVGGVAGLGGAGG | ASGEGGNQDTQY | 0.5341094 |
| VLLVLATTI | ASGEGGNQDTQY | 0.62841904 |
| IGIFAAML | ASGEGGNQDTQY | 0.60265756 |
| AVGDLID | ASGEGGNQDTQY | 0.55329686 |
| NISPLPL | ASGEGGNQDTQY | 0.66436636 |
| VTAVFVAI | ASGEGGNQDTQY | 0.50646585 |
| SGDDNDAAL | ASGEGGNQDTQY | 0.45289117 |
| AVAALAHS | ASGEGGNQDTQY | 0.67479384 |
| GGVTVMATP | ASGEGGNQDTQY | 0.54723096 |
| AGLAANVRD | ASGEGGNQDTQY | 0.6109074 |
| TNRPDLIDPALLRPG | ASGEGGNQDTQY | 0.68937624 |
| LANLGVTAL | ASGEGGNQDTQY | 0.45086998 |
| TGERIEL | ASGEGGNQDTQY | 0.64347005 |
| NGEELAA | ASGEGGNQDTQY | 0.6568836 |
| VLSLVWG | ASGEGGNQDTQY | 0.6688508 |
| IGYALWL | ASGEGGNQDTQY | 0.60667574 |
| LSLLLAPL | ASGEGGNQDTQY | 0.48236078 |
| TLDEALK | ASGEGGNQDTQY | 0.6902709 |
| TDGAGTAEGG | ASGEGGNQDTQY | 0.6560725 |
| NAPGGAGGNGG | ASGEGGNQDTQY | 0.63714004 |
| VSVFMGMGGNF | ASGEGGNQDTQY | 0.6806561 |
| LASYLAAL | ASGEGGNQDTQY | 0.67216575 |
| LVGGTPAL | ASGEGGNQDTQY | 0.6841847 |
| VRNADVL | ASGEGGNQDTQY | 0.5798216 |
| TCSSPFLVSAM | ASGEGGNQDTQY | 0.44970453 |
| VHITDAE | ASGEGGNQDTQY | 0.6241226 |
| EYAAPSF | ASGEGGNQDTQY | 0.580245 |
| VVDAFHGGL | ASGEGGNQDTQY | 0.48896274 |
| ASVVDPQ | ASGEGGNQDTQY | 0.33999923 |
| SYNEFGGLS | ASGEGGNQDTQY | 0.68813586 |
| DPDGSVY | ASGEGGNQDTQY | 0.70726776 |
| MVSTIHATE | ASGEGGNQDTQY | 0.45082235 |
| LAAGVADPV | ASGEGGNQDTQY | 0.7052857 |
| TKEQIAA | ASGEGGNQDTQY | 0.42816988 |
| SLASPAAL | ASGEGGNQDTQY | 0.66795576 |
| IGLVFYGV | ASGEGGNQDTQY | 0.68796766 |
| FLLDASNL | ASGEGGNQDTQY | 0.4495775 |
| YQVGNFP | ASGEGGNQDTQY | 0.658803 |
| ESLLSQST | ASGEGGNQDTQY | 0.44651663 |

| | | |
|---|---|---|
| LFTVGFAAVG | ASGEGGNQDTQY | 0.47489738 |
| ATPPTPT | ASGEGGNQDTQY | 0.69549406 |
| TDLATLG | ASGEGGNQDTQY | 0.54499257 |
| SAGGTGR | ASGEGGNQDTQY | 0.6597962 |
| GFSNSGS | ASGEGGNQDTQY | 0.67596287 |
| GGNGGSGGTSV | ASGEGGNQDTQY | 0.5475431 |
| GTSVATGGAGN | ASGEGGNQDTQY | 0.6733015 |
| NGGNGGSGGTS | ASGEGGNQDTQY | 0.67445624 |
| AVVKFPH | ASGEGGNQDTQY | 0.5877656 |
| TPPPPPP | ASGEGGNQDTQY | 0.69950664 |
| AAPAAAAAPA | ASGEGGNQDTQY | 0.43938458 |
| IALNEDL | ASGEGGNQDTQY | 0.69488657 |
| IVNVSSM | ASGEGGNQDTQY | 0.67781985 |
| TGIPPAPR | ASGEGGNQDTQY | 0.6446635 |
| FDLGGGTF | ASGEGGNQDTQY | 0.57707214 |
| FDLGGGTFDVS | ASGEGGNQDTQY | 0.68367726 |
| NEPTAAA | ASGEGGNQDTQY | 0.6528833 |
| NEPTAAALA | ASGEGGNQDTQY | 0.62338054 |
| GSIINISSI | ASGEGGNQDTQY | 0.6985487 |
| VAPGNAGTA | ASGEGGNQDTQY | 0.57769513 |
| TGGGASLE | ASGEGGNQDTQY | 0.685482 |
| VAVYNHYKRI | ASGEGGNQDTQY | 0.65895754 |
| TDPAPATT | ASGEGGNQDTQY | 0.63564515 |
| TNRPDILDPALLRPG | ASGEGGNQDTQY | 0.6305139 |
| LTTTTTS | ASGEGGNQDTQY | 0.47641185 |
| PTGSGKTL | ASGEGGNQDTQY | 0.59659445 |
| TGSGKTL | ASGEGGNQDTQY | 0.6096556 |
| TIGHVDHGK | ASGEGGNQDTQY | 0.67606026 |
| TIGHVDHGKTT | ASGEGGNQDTQY | 0.674103 |
| GHVDHGKTT | ASGEGGNQDTQY | 0.61865795 |
| AQFEHTLL | ASGEGGNQDTQY | 0.6790504 |
| IVNQDEL | ASGEGGNQDTQY | 0.66480416 |
| TNGSQFF | ASGEGGNQDTQY | 0.5282456 |
| TGKGIVE | ASGEGGNQDTQY | 0.636758 |
| LISNASDA | ASGEGGNQDTQY | 0.575045 |
| AELRTLGELTKIAKAHGAQ | ASGEGGNQDTQY | 0.6457263 |
| APNAVAAAEPSW | ASGEGGNQDTQY | 0.43676132 |
| APQVAGAATAVL | ASGEGGNQDTQY | 0.67478764 |
| ARRDPAAAAAILPTDA | ASGEGGNQDTQY | 0.5952071 |
| DAAARDVLAEAGLAEYF | ASGEGGNQDTQY | 0.35376456 |
| DDQIVLSAKANLAKVEQ | ASGEGGNQDTQY | 0.6561767 |
| DELLPDTGEDSDEDGH | ASGEGGNQDTQY | 0.65226233 |
| ELPEAPANRATVPQD | ASGEGGNQDTQY | 0.653472 |
| FGQSKIERVTGIG | ASGEGGNQDTQY | 0.67034596 |
| GAGGTGSTAGGAGGAGGAGGLY | ASGEGGNQDTQY | 0.5830898 |
| GHGGTGAAGGNGGTGGAGGSGI | ASGEGGNQDTQY | 0.5882892 |
| GNGGNGGAGGEGGPGGAGGAGA | ASGEGGNQDTQY | 0.6729629 |
| GSTAGNGGNGGNATNGTIAGQP | ASGEGGNQDTQY | 0.68193084 |
| GTGGDGGNGGNGGTGG | ASGEGGNQDTQY | 0.56151736 |
| GVRRPAGIPADAPV | ASGEGGNQDTQY | 0.66001487 |
| GYRQVTDAHLLTLARR | ASGEGGNQDTQY | 0.6767438 |

| | | |
|---|---|---|
| IAKMGQDGHDRG | ASGEGGNQDTQY | 0.6606455 |
| KAEPSIEIQYT | ASGEGGNQDTQY | 0.6760118 |
| KPVRIRMAADEDTVAVT | ASGEGGNQDTQY | 0.68296564 |
| MPPRIAGMRLLVIKPEP | ASGEGGNQDTQY | 0.57439756 |
| MKFYAVALFRSRGVPPTR | ASGEGGNQDTQY | 0.6819331 |
| NAGGFGILGAVAHSP | ASGEGGNQDTQY | 0.68038213 |
| NEGREITLAMAEM | ASGEGGNQDTQY | 0.6436999 |
| NELVEWSHPLGHRWRWE | ASGEGGNQDTQY | 0.6471275 |
| NLGAGGTGGNGGNPG | ASGEGGNQDTQY | 0.5323869 |
| NTEWPKWETVMAADPVD | ASGEGGNQDTQY | 0.6319525 |
| PASAAPAETQP | ASGEGGNQDTQY | 0.536926 |
| QQSSDGKNGVRAAEITGE | ASGEGGNQDTQY | 0.6582561 |
| RAPDAVTTSWWKE | ASGEGGNQDTQY | 0.6844316 |
| SGFGNVGAHTSGWFNQ | ASGEGGNQDTQY | 0.6331843 |
| SGVAGAVGQAASVGGLKV | ASGEGGNQDTQY | 0.57587016 |
| SNGGAGTGSAGGNGGT | ASGEGGNQDTQY | 0.5854442 |
| TGGAGGEGGAAG | ASGEGGNQDTQY | 0.6526767 |
| TGGDGGAAGAGGAPGHG | ASGEGGNQDTQY | 0.5947609 |
| TMPPLLAAIANAGRHGV | ASGEGGNQDTQY | 0.44872475 |
| TSSTAGNSGGA | ASGEGGNQDTQY | 0.6260545 |
| VPATVPVGILC | ASGEGGNQDTQY | 0.67756647 |
| VQRPDSGGVVLTGRIS | ASGEGGNQDTQY | 0.6771462 |
| WTPNPVPATKGVPAGLVQAT | ASGEGGNQDTQY | 0.56647444 |
| YALKRALAKFSDRFDVVII | ASGEGGNQDTQY | 0.68391514 |
| YLIGSVIV | ASGEGGNQDTQY | 0.6659344 |
| YGQLVAVID | ASGEGGNQDTQY | 0.6832558 |
| VTVGGMTLPA | ASGEGGNQDTQY | 0.62540674 |
| VLALPAV | ASGEGGNQDTQY | 0.5838943 |
| VIPPTLNYET | ASGEGGNQDTQY | 0.66979706 |
| VGMGGMHPSSG | ASGEGGNQDTQY | 0.62808746 |
| VGLAGGAATA | ASGEGGNQDTQY | 0.63419694 |
| VGGAGGAGGSGP | ASGEGGNQDTQY | 0.6720211 |
| VFAAFMV | ASGEGGNQDTQY | 0.6455919 |
| TLYDKLR | ASGEGGNQDTQY | 0.613679 |
| TLAVFTVL | ASGEGGNQDTQY | 0.6194096 |
| TGSVQPARV | ASGEGGNQDTQY | 0.68252575 |
| TGGGAGSGGT | ASGEGGNQDTQY | 0.5759521 |
| SHLETSL | ASGEGGNQDTQY | 0.62426305 |
| SFTTGMT | ASGEGGNQDTQY | 0.62111914 |
| QVEVAKRMGVRQPTVS | ASGEGGNQDTQY | 0.64967954 |
| QGSGGPALAM | ASGEGGNQDTQY | 0.67996013 |
| MLAPSNP | ASGEGGNQDTQY | 0.6659241 |
| LVNNAAI | ASGEGGNQDTQY | 0.62496024 |
| LTGEINTLA | ASGEGGNQDTQY | 0.66322714 |
| LSIFGPV | ASGEGGNQDTQY | 0.67426574 |
| LLALLANL | ASGEGGNQDTQY | 0.6601579 |
| LISNASDALDKLR | ASGEGGNQDTQY | 0.656382 |
| LINEAVP | ASGEGGNQDTQY | 0.61729413 |
| LGYAFSGL | ASGEGGNQDTQY | 0.68060696 |
| LDFVAVAL | ASGEGGNQDTQY | 0.6652496 |
| LAQPGAE | ASGEGGNQDTQY | 0.6712657 |

| | | |
|---|---|---|
| ISNLGAVTGF | ASGEGGNQDTQY | 0.6760647 |
| HQEDPGRI | ASGEGGNQDTQY | 0.67130053 |
| GTPGEPTP | ASGEGGNQDTQY | 0.65343106 |
| GFFNSGS | ASGEGGNQDTQY | 0.48947585 |
| GAGGGGLTAGAN | ASGEGGNQDTQY | 0.5183769 |
| FVDALVNM | ASGEGGNQDTQY | 0.65479755 |
| FLTVAPEL | ASGEGGNQDTQY | 0.6552648 |
| FETLTAL | ASGEGGNQDTQY | 0.6730207 |
| DVINPIEL | ASGEGGNQDTQY | 0.6496631 |
| APVSGAPV | ASGEGGNQDTQY | 0.61594 |
| ALVGGLAW | ASGEGGNQDTQY | 0.6697521 |
| ALAAAAATAI | ASGEGGNQDTQY | 0.6777879 |
| AFGLGLLL | ASGEGGNQDTQY | 0.5261561 |
| AFFAAASL | ASGEGGNQDTQY | 0.61773217 |
| AAAALPAFSPP | ASGEGGNQDTQY | 0.6909219 |
| VVLAPAL | ASGEGGNQDTQY | 0.64536864 |
| TIGHVDHGKTTL | ASGEGGNQDTQY | 0.63170254 |
| LISNASDALDK | ASGEGGNQDTQY | 0.6103684 |
| LISNASDAL | ASGEGGNQDTQY | 0.6841884 |
| ISNASDALDK | ASGEGGNQDTQY | 0.63017434 |
| NQAETLVYQTE | ASGEGGNQDTQY | 0.6841669 |
| TGGTGGTAGAG | ASGEGGNQDTQY | 0.39420006 |
| TGSTAGGAGGA | ASGEGGNQDTQY | 0.6247079 |
| DPLAVPVAAGPV | ASGEGGNQDTQY | 0.60414517 |
| GGTGGTPGNGGN | ASGEGGNQDTQY | 0.46068007 |
| PAGLAGVDPATL | ASGEGGNQDTQY | 0.6749321 |
| GNNLVVEAPHAGASV | ASGEGGNQDTQY | 0.57842606 |
| KVTRIMSASTLEQPA | ASGEGGNQDTQY | 0.6642966 |
| LGFALPIALPRRTVV | ASGEGGNQDTQY | 0.6299118 |
| TRLGNLTARKLVVLV | ASGEGGNQDTQY | 0.6788667 |
| WSQQRRLIYFPSAGP | ASGEGGNQDTQY | 0.64713085 |
| GPNGNPDPMAAAVDIR | ASGEGGNQDTQY | 0.6515098 |
| TQRCVSVADLRAAASR | ASGEGGNQDTQY | 0.6128092 |
| QFVLDVAEHPVEVLTETV | ASGEGGNQDTQY | 0.66715115 |
| GGAGGTGGAGGNGGTQAGDG | ASGEGGNQDTQY | 0.67640096 |
| PLLATLRAIAVDVDAKPAQV | ASGEGGNQDTQY | 0.6734162 |
| APKFRTGGYMIDFWGVGYQVAK | ASGEGGNQDTQY | 0.47976106 |
| GAGGAGGAGGAAGGPGGTGGTG | ASGEGGNQDTQY | 0.6495293 |
| IGALGLLATDVAPAGSVAFGVA | ASGEGGNQDTQY | 0.6530224 |
| AGGAGGIGGNGGAGGAGGVLMG | ASGEGGNQDTQY | 0.48994774 |
| ADELIGLL | ASGEGGNQDTQY | 0.4139384 |
| ADRALKA | ASGEGGNQDTQY | 0.6294311 |
| AGAVGQVGGAGGA | ASGEGGNQDTQY | 0.62289155 |
| AGGGGGLGGAP | ASGEGGNQDTQY | 0.67287016 |
| AGGLLYGNGGAG | ASGEGGNQDTQY | 0.669876 |
| AGLESIGL | ASGEGGNQDTQY | 0.63353384 |
| AGLKTGII | ASGEGGNQDTQY | 0.6645179 |
| ASLAALLL | ASGEGGNQDTQY | 0.6542214 |
| ASVINISSI | ASGEGGNQDTQY | 0.2974749 |
| ATPPGPA | ASGEGGNQDTQY | 0.59269094 |
| ATVASALV | ASGEGGNQDTQY | 0.53701985 |

| | | |
|---|---|---|
| ATVASLIL | ASGEGGNQDTQY | 0.6815588 |
| AVNGPVLIRH | ASGEGGNQDTQY | 0.6093099 |
| DGAAGAVGI | ASGEGGNQDTQY | 0.58045805 |
| DPLLHGI | ASGEGGNQDTQY | 0.6529389 |
| ELGAALGAL | ASGEGGNQDTQY | 0.64880043 |
| ELGRTQAV | ASGEGGNQDTQY | 0.6368753 |
| FVAGGYLS | ASGEGGNQDTQY | 0.60421884 |
| GASGGAGGAGGW | ASGEGGNQDTQY | 0.43688813 |
| GGAGAGGDGGAGGI | ASGEGGNQDTQY | 0.6714973 |
| GTGHNGSQPGMGGNG | ASGEGGNQDTQY | 0.55904466 |
| GYAAASASAS | ASGEGGNQDTQY | 0.59486115 |
| HQVDVAAVQQG | ASGEGGNQDTQY | 0.3980037 |
| IGAAIAHT | ASGEGGNQDTQY | 0.6124259 |
| IGMIEVAQL | ASGEGGNQDTQY | 0.44257534 |
| IRLAPEHTSAMD | ASGEGGNQDTQY | 0.6678548 |
| IYNVSVL | ASGEGGNQDTQY | 0.62560123 |
| KQGFLGM | ASGEGGNQDTQY | 0.6289262 |
| LESVITQL | ASGEGGNQDTQY | 0.6889279 |
| LGLGALVVV | ASGEGGNQDTQY | 0.64817595 |
| LIGNGGNGGSGGI | ASGEGGNQDTQY | 0.6256801 |
| LNAINAPTL | ASGEGGNQDTQY | 0.62341654 |
| LQLLNTGV | ASGEGGNQDTQY | 0.57792383 |
| LQTSIEF | ASGEGGNQDTQY | 0.6143695 |
| MSFAVSEL | ASGEGGNQDTQY | 0.6197833 |
| NTGALGL | ASGEGGNQDTQY | 0.6810696 |
| PPGAGGAGGAGTT | ASGEGGNQDTQY | 0.6404091 |
| QLVGLGI | ASGEGGNQDTQY | 0.4662423 |
| RFLEQVHQL | ASGEGGNQDTQY | 0.6458645 |
| RGVGYVLR | ASGEGGNQDTQY | 0.6856249 |
| SATGVFAGVI | ASGEGGNQDTQY | 0.6885782 |
| SLPAITLP | ASGEGGNQDTQY | 0.60890573 |
| STALAFTV | ASGEGGNQDTQY | 0.58245516 |
| SVLTVGLGK | ASGEGGNQDTQY | 0.58857065 |
| TALAGAGSGI | ASGEGGNQDTQY | 0.63843477 |
| TGLPFTGI | ASGEGGNQDTQY | 0.5899307 |
| VAEHVTV | ASGEGGNQDTQY | 0.57656604 |
| VDIVLNSL | ASGEGGNQDTQY | 0.67409796 |
| VLTGGGAVL | ASGEGGNQDTQY | 0.68854773 |
| VNVGLAYGI | ASGEGGNQDTQY | 0.64187944 |
| VSSAAVGAV | ASGEGGNQDTQY | 0.41379845 |
| VTSVYEAGV | ASGEGGNQDTQY | 0.64444226 |
| WVALDRGA | ASGEGGNQDTQY | 0.5893606 |
| YLVGSVVI | ASGEGGNQDTQY | 0.6281514 |
| GPPGTGKTLL | ASGEGGNQDTQY | 0.62110317 |
| GVGMTKF | ASGEGGNQDTQY | 0.47704694 |
| IGIGGSDL | ASGEGGNQDTQY | 0.65105724 |
| IGTLLAEL | ASGEGGNQDTQY | 0.6894263 |
| IYTSGTT | ASGEGGNQDTQY | 0.62774277 |
| LSFPAGV | ASGEGGNQDTQY | 0.6552755 |
| LYTSGTT | ASGEGGNQDTQY | 0.6839462 |
| VGAFPFGF | ASGEGGNQDTQY | 0.67070085 |

| | | |
|---|---|---|
| VGAGTVEF | ASGEGGNQDTQY | 0.67938495 |
| VGKSTLLL | ASGEGGNQDTQY | 0.67074126 |
| AALAHAGGGIV | ASGEGGNQDTQY | 0.6580609 |
| AALVAVSLL | ASGEGGNQDTQY | 0.6503712 |
| AAVSRAVAAL | ASGEGGNQDTQY | 0.6747076 |
| AGGSSGIGGF | ASGEGGNQDTQY | 0.64843833 |
| AGLTVANANG | ASGEGGNQDTQY | 0.6412636 |
| AGLVGAVL | ASGEGGNQDTQY | 0.5904731 |
| ALSQAFTV | ASGEGGNQDTQY | 0.67789924 |
| ALTFSVSL | ASGEGGNQDTQY | 0.6305435 |
| ALTGAALVAL | ASGEGGNQDTQY | 0.67984235 |
| ASFSIGAGVL | ASGEGGNQDTQY | 0.42960227 |
| DGSVYKI | ASGEGGNQDTQY | 0.5794331 |
| EQLAEIV | ASGEGGNQDTQY | 0.67350924 |
| FAVSELTVM | ASGEGGNQDTQY | 0.31502423 |
| GAAPLAAGL | ASGEGGNQDTQY | 0.41120327 |
| GADGGDGGSGGAGG | ASGEGGNQDTQY | 0.649052 |
| GFGLDTGGAGG | ASGEGGNQDTQY | 0.6800248 |
| GLASGGLL | ASGEGGNQDTQY | 0.33340713 |
| GRIAGAAVLL | ASGEGGNQDTQY | 0.559292 |
| GSAGSLNTSF | ASGEGGNQDTQY | 0.49254173 |
| GTGQIGI | ASGEGGNQDTQY | 0.67052865 |
| GTLIALIL | ASGEGGNQDTQY | 0.6523608 |
| IAATMGFVVG | ASGEGGNQDTQY | 0.47943366 |
| IFAQALL | ASGEGGNQDTQY | 0.67411834 |
| IGAAVALGF | ASGEGGNQDTQY | 0.43821985 |
| IGLTPLAL | ASGEGGNQDTQY | 0.64234436 |
| ILGSNTSTL | ASGEGGNQDTQY | 0.67101216 |
| IVFSLTVL | ASGEGGNQDTQY | 0.6800809 |
| LASKATGF | ASGEGGNQDTQY | 0.5227477 |
| LASVALIL | ASGEGGNQDTQY | 0.66622233 |
| LFAVGLTL | ASGEGGNQDTQY | 0.61377424 |
| LFNGGIGTYI | ASGEGGNQDTQY | 0.690882 |
| LTLGAGII | ASGEGGNQDTQY | 0.6336683 |
| LVAAALGDI | ASGEGGNQDTQY | 0.5107867 |
| NTAFVQLGI | ASGEGGNQDTQY | 0.6759181 |
| QLFSGAEL | ASGEGGNQDTQY | 0.66957885 |
| SAATFITL | ASGEGGNQDTQY | 0.6459147 |
| SNYGLANLGV | ASGEGGNQDTQY | 0.66941154 |
| SYGGAYAVM | ASGEGGNQDTQY | 0.62375003 |
| TTGVGGLTLG | ASGEGGNQDTQY | 0.64878917 |
| TTQTIYI | ASGEGGNQDTQY | 0.67614377 |
| VAAGAGLGAVI | ASGEGGNQDTQY | 0.6721468 |
| VAAVVTLL | ASGEGGNQDTQY | 0.59473217 |
| VGFLGGLI | ASGEGGNQDTQY | 0.67450947 |
| VGVIPILL | ASGEGGNQDTQY | 0.6413244 |
| VIFASSLI | ASGEGGNQDTQY | 0.5737277 |
| VKGGDGGAAAG | ASGEGGNQDTQY | 0.6860625 |
| VTKTFTNI | ASGEGGNQDTQY | 0.62642527 |
| VVAAAILL | ASGEGGNQDTQY | 0.6719079 |
| YAALDVEL | ASGEGGNQDTQY | 0.67644906 |

| | | |
|---|---|---|
| LASVALLL | ASGEGGNQDTQY | 0.6796509 |
| LDAVHRV | ASGEGGNQDTQY | 0.5027551 |
| QGLSSSIF | ASGEGGNQDTQY | 0.63156986 |
| RGTFANIRL | ASGEGGNQDTQY | 0.64222616 |
| RQLWWGHRI | ASGEGGNQDTQY | 0.6744541 |
| YGPPGTGKTLL | ASGEGGNQDTQY | 0.6401246 |
| AALVGAPANVA | ASGEGGNQDTQY | 0.6228206 |
| AATFRRTE | ASGEGGNQDTQY | 0.6702333 |
| AGALGGGI | ASGEGGNQDTQY | 0.6803043 |
| AGGAGGAGGIGAG | ASGEGGNQDTQY | 0.65249574 |
| AGSGALGL | ASGEGGNQDTQY | 0.6504669 |
| AQIADQI | ASGEGGNQDTQY | 0.68411034 |
| ASAVAVTYL | ASGEGGNQDTQY | 0.60109735 |
| ASTVLGGTGG | ASGEGGNQDTQY | 0.6422542 |
| AVGGEPAK | ASGEGGNQDTQY | 0.6724855 |
| CEQGTFGGTC | ASGEGGNQDTQY | 0.6757694 |
| DEGKAPAG | ASGEGGNQDTQY | 0.6415847 |
| DTVVFPAL | ASGEGGNQDTQY | 0.6806917 |
| FEGITMAG | ASGEGGNQDTQY | 0.58198 |
| FPLVVQD | ASGEGGNQDTQY | 0.6684342 |
| GAFTTPQ | ASGEGGNQDTQY | 0.611121 |
| GFGGFGTAGGVGGA | ASGEGGNQDTQY | 0.6425344 |
| GFLAQFL | ASGEGGNQDTQY | 0.6145137 |
| GGAGGAGGNAGVA | ASGEGGNQDTQY | 0.6664983 |
| GGVGTAGAGGAGGAGG | ASGEGGNQDTQY | 0.6700048 |
| GNGGGGDAGNAGSGGN | ASGEGGNQDTQY | 0.67212725 |
| GTGGAGGAGNDAGS | ASGEGGNQDTQY | 0.3560167 |
| GTTNTGWFNS | ASGEGGNQDTQY | 0.6770056 |
| IFMGPLFTL | ASGEGGNQDTQY | 0.67357373 |
| ILAHAIEAI | ASGEGGNQDTQY | 0.68404365 |
| IRSAIREAL | ASGEGGNQDTQY | 0.64152074 |
| ITFAGGG | ASGEGGNQDTQY | 0.65089715 |
| ITGMSGHQEV | ASGEGGNQDTQY | 0.62546074 |
| IVAGLACTL | ASGEGGNQDTQY | 0.6521703 |
| IVGGGLAGA | ASGEGGNQDTQY | 0.5650233 |
| IVTGVLNI | ASGEGGNQDTQY | 0.62665117 |
| LAQQGCI | ASGEGGNQDTQY | 0.6169545 |
| LCLHVEN | ASGEGGNQDTQY | 0.6144788 |
| LDAISDGS | ASGEGGNQDTQY | 0.6698901 |
| LGGFAAML | ASGEGGNQDTQY | 0.67618597 |
| LGGKNPMI | ASGEGGNQDTQY | 0.6749829 |
| LGPKGVAAIL | ASGEGGNQDTQY | 0.6793678 |
| LMVGVSI | ASGEGGNQDTQY | 0.6821624 |
| LVGAAFVL | ASGEGGNQDTQY | 0.6775546 |
| LVLAVAPL | ASGEGGNQDTQY | 0.627941 |
| MAAIAVIDAA | ASGEGGNQDTQY | 0.6113121 |
| NNGINAELQ | ASGEGGNQDTQY | 0.5537983 |
| RGEPLVLVHG | ASGEGGNQDTQY | 0.63752675 |
| RGGAGGAAGLI | ASGEGGNQDTQY | 0.6361613 |
| RSYRMHPAV | ASGEGGNQDTQY | 0.6609308 |
| SAAHLLAGQLA | ASGEGGNQDTQY | 0.31822774 |

| | | |
|---|---|---|
| SAAVGSY | ASGEGGNQDTQY | 0.6457866 |
| SPHAVLHRI | ASGEGGNQDTQY | 0.6813326 |
| SQIAKVL | ASGEGGNQDTQY | 0.6246624 |
| TALGHLVP | ASGEGGNQDTQY | 0.65767676 |
| TAPTTSGAA | ASGEGGNQDTQY | 0.6569698 |
| TDGDDGNGGSPGT | ASGEGGNQDTQY | 0.6123925 |
| TDGLGEN | ASGEGGNQDTQY | 0.66613424 |
| TGGPPLPGQA | ASGEGGNQDTQY | 0.6040387 |
| TIPAITI | ASGEGGNQDTQY | 0.66642725 |
| TIPGFTI | ASGEGGNQDTQY | 0.68368614 |
| TSIASGALA | ASGEGGNQDTQY | 0.66077644 |
| TVTLPGAAI | ASGEGGNQDTQY | 0.6194645 |
| VASGLSAALP | ASGEGGNQDTQY | 0.6294846 |
| VGAGVIGI | ASGEGGNQDTQY | 0.44064978 |
| VGPKRFGV | ASGEGGNQDTQY | 0.6541903 |
| VNVAFPD | ASGEGGNQDTQY | 0.5596443 |
| VSMTNSGVS | ASGEGGNQDTQY | 0.6452933 |
| VTDVLGDLSV | ASGEGGNQDTQY | 0.61070514 |
| VVVVGAGFAGL | ASGEGGNQDTQY | 0.6657787 |
| VYTVGTVV | ASGEGGNQDTQY | 0.6554293 |
| WTTFSGP | ASGEGGNQDTQY | 0.65650094 |
| YDELVEAF | ASGEGGNQDTQY | 0.5843082 |
| YGTGGAGGNGG | ASGEGGNQDTQY | 0.66368306 |
| YTNVMAV | ASGEGGNQDTQY | 0.6447214 |
| FLEQVHQL | ASGEGGNQDTQY | 0.56275254 |
| GLSLLAVAV | ASGEGGNQDTQY | 0.66873133 |
| LEQVHQL | ASGEGGNQDTQY | 0.6403928 |
| LNAAGAI | ASGEGGNQDTQY | 0.63056594 |
| PAVHASL | ASGEGGNQDTQY | 0.6798451 |
| TELEDDA | ASGEGGNQDTQY | 0.6586964 |
| TPPGPPN | ASGEGGNQDTQY | 0.42347637 |
| TPPPPPPP | ASGEGGNQDTQY | 0.63434315 |
| TQSPALI | ASSLADQDTQY | 0.6493728 |
| EFAAAFPQ | ASSLADQDTQY | 0.64824545 |
| LLSGNLDVL | ASSLADQDTQY | 0.32126456 |
| GPANLGNYNI | ASSLADQDTQY | 0.6712957 |
| TLGPTLP | ASSLADQDTQY | 0.56803775 |
| VYAYSSCCK | ASSLADQDTQY | 0.5892694 |
| APAPAPAPNI | ASSLADQDTQY | 0.615945 |
| PGGPVPIP | ASSLADQDTQY | 0.6717834 |
| FLTVALEP | ASSLADQDTQY | 0.64048594 |
| SQYFGANL | ASSLADQDTQY | 0.65063536 |
| GGVKPAA | ASSLADQDTQY | 0.6599901 |
| GGTGGAGGAGGW | ASSLADQDTQY | 0.65971047 |
| GVGGVAGLGGAGG | ASSLADQDTQY | 0.6786002 |
| VLLVLATTI | ASSLADQDTQY | 0.6725905 |
| IGIFAAML | ASSLADQDTQY | 0.64680684 |
| AVGDLID | ASSLADQDTQY | 0.65352845 |
| NISPLPL | ASSLADQDTQY | 0.55786043 |
| VTAVFVAI | ASSLADQDTQY | 0.58210015 |
| SGDDNDAAL | ASSLADQDTQY | 0.5209411 |

| | | |
|---|---|---|
| AVAALAHS | ASSLADQDTQY | 0.64459085 |
| GGVTVMATP | ASSLADQDTQY | 0.5357818 |
| AGLAANVRD | ASSLADQDTQY | 0.6560644 |
| TNRPDLIDPALLRPG | ASSLADQDTQY | 0.67916006 |
| LANLGVTAL | ASSLADQDTQY | 0.6378526 |
| TGERIEL | ASSLADQDTQY | 0.48532975 |
| NGEELAA | ASSLADQDTQY | 0.61748296 |
| VLSLVWG | ASSLADQDTQY | 0.59434575 |
| IGYALWL | ASSLADQDTQY | 0.66792077 |
| LSLLLAPL | ASSLADQDTQY | 0.67838895 |
| TLDEALK | ASSLADQDTQY | 0.688928 |
| TDGAGTAEGG | ASSLADQDTQY | 0.6801804 |
| NAPGGAGGNGG | ASSLADQDTQY | 0.6250236 |
| VSVFMGMGGNF | ASSLADQDTQY | 0.66944695 |
| LASYLAAL | ASSLADQDTQY | 0.62135184 |
| LVGGTPAL | ASSLADQDTQY | 0.62141645 |
| VRNADVL | ASSLADQDTQY | 0.64330375 |
| TCSSPFLVSAM | ASSLADQDTQY | 0.6619835 |
| VHITDAE | ASSLADQDTQY | 0.6447404 |
| EYAAPSF | ASSLADQDTQY | 0.63387865 |
| VVDAFHGGL | ASSLADQDTQY | 0.45932248 |
| ASVVDPQ | ASSLADQDTQY | 0.6679194 |
| SYNEFGGLS | ASSLADQDTQY | 0.6326283 |
| DPDGSVY | ASSLADQDTQY | 0.48884305 |
| MVSTIHATE | ASSLADQDTQY | 0.6155616 |
| LAAGVADPV | ASSLADQDTQY | 0.686394 |
| TKEQIAA | ASSLADQDTQY | 0.68030405 |
| SLASPAAL | ASSLADQDTQY | 0.6146456 |
| IGLVFYGV | ASSLADQDTQY | 0.6174388 |
| FLLDASNL | ASSLADQDTQY | 0.64536595 |
| YQVGNFP | ASSLADQDTQY | 0.54675865 |
| ESLLSQST | ASSLADQDTQY | 0.6660158 |
| LFTVGFAAVG | ASSLADQDTQY | 0.6416773 |
| ATPPTPT | ASSLADQDTQY | 0.64964926 |
| TDLATLG | ASSLADQDTQY | 0.54210716 |
| SAGGTGR | ASSLADQDTQY | 0.6592336 |
| GFSNSGS | ASSLADQDTQY | 0.6431829 |
| GGNGGSGGTSV | ASSLADQDTQY | 0.66084564 |
| GTSVATGGAGN | ASSLADQDTQY | 0.34561235 |
| NGGNGGSGGTS | ASSLADQDTQY | 0.66911554 |
| AVVKFPH | ASSLADQDTQY | 0.58515614 |
| TPPPPPP | ASSLADQDTQY | 0.6006814 |
| AAPAAAAAPA | ASSLADQDTQY | 0.42324245 |
| IALNEDL | ASSLADQDTQY | 0.6048806 |
| IVNVSSM | ASSLADQDTQY | 0.67049354 |
| TGIPPAPR | ASSLADQDTQY | 0.6765735 |
| FDLGGGTF | ASSLADQDTQY | 0.62662673 |
| FDLGGGTFDVS | ASSLADQDTQY | 0.6711401 |
| NEPTAAA | ASSLADQDTQY | 0.6835934 |
| NEPTAAALA | ASSLADQDTQY | 0.6132292 |
| GSIINISSI | ASSLADQDTQY | 0.4152286 |

| | | |
|---|---|---|
| VAPGNAGTA | ASSLADQDTQY | 0.65518296 |
| TGGGASLE | ASSLADQDTQY | 0.6191764 |
| VAVYNHYKRI | ASSLADQDTQY | 0.6748401 |
| TDPAPATT | ASSLADQDTQY | 0.665509 |
| TNRPDILDPALLRPG | ASSLADQDTQY | 0.5643163 |
| LTTTTTS | ASSLADQDTQY | 0.6624327 |
| PTGSGKTL | ASSLADQDTQY | 0.67833054 |
| TGSGKTL | ASSLADQDTQY | 0.5823174 |
| TIGHVDHGK | ASSLADQDTQY | 0.49541348 |
| TIGHVDHGKTT | ASSLADQDTQY | 0.5516457 |
| GHVDHGKTT | ASSLADQDTQY | 0.45309073 |
| AQFEHTLL | ASSLADQDTQY | 0.66579354 |
| IVNQDEL | ASSLADQDTQY | 0.61147225 |
| TNGSQFF | ASSLADQDTQY | 0.61135364 |
| TGKGIVE | ASSLADQDTQY | 0.6544929 |
| LISNASDA | ASSLADQDTQY | 0.61752105 |
| AELRTLGELTKIAKAHGAQ | ASSLADQDTQY | 0.60412043 |
| APNAVAAAEPSW | ASSLADQDTQY | 0.634433 |
| APQVAGAATAVL | ASSLADQDTQY | 0.66618466 |
| ARRDPAAAAAILPTDA | ASSLADQDTQY | 0.6660734 |
| DAAARDVLAEAGLAEYF | ASSLADQDTQY | 0.63827825 |
| DDQIVLSAKANLAKVEQ | ASSLADQDTQY | 0.6164349 |
| DELLPDTGEDSDEDGH | ASSLADQDTQY | 0.6822435 |
| ELPEAPANRATVPQD | ASSLADQDTQY | 0.5659188 |
| FGQSKIERVTGIG | ASSLADQDTQY | 0.6137368 |
| GAGGTGSTAGGAGGAGGAGGLY | ASSLADQDTQY | 0.6682446 |
| GHGGTGAAGGNGGTGGAGGSGI | ASSLADQDTQY | 0.5938487 |
| GNGGNGGAGGEGGPGGAGGAGA | ASSLADQDTQY | 0.6713878 |
| GSTAGNGGNGGNATNGTIAGQP | ASSLADQDTQY | 0.6512624 |
| GTGGDGGNGGNGGTGG | ASSLADQDTQY | 0.6613971 |
| GVRRPAGIPADAPV | ASSLADQDTQY | 0.63708925 |
| GYRQVTDAHLLTLARR | ASSLADQDTQY | 0.68447584 |
| IAKMGQDGHDRG | ASSLADQDTQY | 0.66891354 |
| KAEPSIEIQYT | ASSLADQDTQY | 0.6225008 |
| KPVRIRMAADEDTVAVT | ASSLADQDTQY | 0.6705431 |
| MPPRIAGMRLLVIKPEP | ASSLADQDTQY | 0.48359904 |
| MKFYAVALFRSRGVPPTR | ASSLADQDTQY | 0.6720985 |
| NAGGFGILGAVAHSP | ASSLADQDTQY | 0.6615925 |
| NEGREITLAMAEM | ASSLADQDTQY | 0.5927243 |
| NELVEWSHPLGHRWRWE | ASSLADQDTQY | 0.649862 |
| NLGAGGTGGNGGNPG | ASSLADQDTQY | 0.68201846 |
| NTEWPKWETVMAADPVD | ASSLADQDTQY | 0.6791538 |
| PASAAPAETQP | ASSLADQDTQY | 0.5851384 |
| QQSSDGKNGVRAAEITGE | ASSLADQDTQY | 0.6221935 |
| RAPDAVTTSWWKE | ASSLADQDTQY | 0.5150941 |
| SGFGNVGAHTSGWFNQ | ASSLADQDTQY | 0.5934555 |
| SGVAGAVGQAASVGGLKV | ASSLADQDTQY | 0.49130988 |
| SNGGAGTGSAGGNGGT | ASSLADQDTQY | 0.6643808 |
| TGGAGGEGGAAG | ASSLADQDTQY | 0.6227447 |
| TGGDGGAAGAGGAPGHG | ASSLADQDTQY | 0.66423213 |
| TMPPLLAAIANAGRHGV | ASSLADQDTQY | 0.6597775 |

| | | |
|---|---|---|
| TSSTAGNSGGA | ASSLADQDTQY | 0.6824131 |
| VPATVPVGILC | ASSLADQDTQY | 0.620406 |
| VQRPDSGGVVLTGRIS | ASSLADQDTQY | 0.6459298 |
| WTPNPVPATKGVPAGLVQAT | ASSLADQDTQY | 0.49114865 |
| YALKRALAKFSDRFDVVII | ASSLADQDTQY | 0.6195861 |
| YLIGSVIV | ASSLADQDTQY | 0.59774226 |
| YGQLVAVID | ASSLADQDTQY | 0.63847554 |
| VTVGGMTLPA | ASSLADQDTQY | 0.4235763 |
| VLALPAV | ASSLADQDTQY | 0.6872523 |
| VIPPTLNYET | ASSLADQDTQY | 0.6440809 |
| VGMGGMHPSSG | ASSLADQDTQY | 0.66902965 |
| VGLAGGAATA | ASSLADQDTQY | 0.5952893 |
| VGGAGGAGGSGP | ASSLADQDTQY | 0.5922482 |
| VFAAFMV | ASSLADQDTQY | 0.6327293 |
| TLYDKLR | ASSLADQDTQY | 0.66820514 |
| TLAVFTVL | ASSLADQDTQY | 0.6045164 |
| TGSVQPARV | ASSLADQDTQY | 0.3931314 |
| TGGGAGSGGT | ASSLADQDTQY | 0.5326549 |
| SHLETSL | ASSLADQDTQY | 0.47567302 |
| SFTTGMT | ASSLADQDTQY | 0.68356943 |
| QVEVAKRMGVRQPTVS | ASSLADQDTQY | 0.6795232 |
| QGSGGPALAM | ASSLADQDTQY | 0.6614536 |
| MLAPSNP | ASSLADQDTQY | 0.68130434 |
| LVNNAAI | ASSLADQDTQY | 0.33207014 |
| LTGEINTLA | ASSLADQDTQY | 0.6038559 |
| LSIFGPV | ASSLADQDTQY | 0.67996037 |
| LLALLANL | ASSLADQDTQY | 0.6691355 |
| LISNASDALDKLR | ASSLADQDTQY | 0.63839746 |
| LINEAVP | ASSLADQDTQY | 0.677799 |
| LGYAFSGL | ASSLADQDTQY | 0.6025558 |
| LDFVAVAL | ASSLADQDTQY | 0.6473572 |
| LAQPGAE | ASSLADQDTQY | 0.65186936 |
| ISNLGAVTGF | ASSLADQDTQY | 0.67086875 |
| HQEDPGRI | ASSLADQDTQY | 0.65701944 |
| GTPGEPTP | ASSLADQDTQY | 0.55657417 |
| GFFNSGS | ASSLADQDTQY | 0.64540774 |
| GAGGGGLTAGAN | ASSLADQDTQY | 0.690109 |
| FVDALVNM | ASSLADQDTQY | 0.63360286 |
| FLTVAPEL | ASSLADQDTQY | 0.6750388 |
| FETLTAL | ASSLADQDTQY | 0.6536968 |
| DVINPIEL | ASSLADQDTQY | 0.6553513 |
| APVSGAPV | ASSLADQDTQY | 0.605807 |
| ALVGGLAW | ASSLADQDTQY | 0.6016648 |
| ALAAAAATAI | ASSLADQDTQY | 0.6645633 |
| AFGLGLLL | ASSLADQDTQY | 0.5759946 |
| AFFAAASL | ASSLADQDTQY | 0.6309231 |
| AAAALPAFSPP | ASSLADQDTQY | 0.62184954 |
| VVLAPAL | ASSLADQDTQY | 0.69365275 |
| TIGHVDHGKTTL | ASSLADQDTQY | 0.6609564 |
| LISNASDALDK | ASSLADQDTQY | 0.6696381 |
| LISNASDAL | ASSLADQDTQY | 0.60670054 |

| | | |
|---|---|---|
| ISNASDALDK | ASSLADQDTQY | 0.66050553 |
| NQAETLVYQTE | ASSLADQDTQY | 0.66347605 |
| TGGTGGTAGAG | ASSLADQDTQY | 0.48827347 |
| TGSTAGGAGGA | ASSLADQDTQY | 0.67709875 |
| DPLAVPVAAGPV | ASSLADQDTQY | 0.57406604 |
| GGTGGTPGNGGN | ASSLADQDTQY | 0.6442548 |
| PAGLAGVDPATL | ASSLADQDTQY | 0.66944087 |
| GNNLVVEAPHAGASV | ASSLADQDTQY | 0.6824952 |
| KVTRIMSASTLEQPA | ASSLADQDTQY | 0.640147 |
| LGFALPIALPRRTVV | ASSLADQDTQY | 0.65139455 |
| TRLGNLTARKLVVLV | ASSLADQDTQY | 0.61012864 |
| WSQQRRLIYFPSAGP | ASSLADQDTQY | 0.674024 |
| GPNGNPDPMAAAVDIR | ASSLADQDTQY | 0.68062687 |
| TQRCVSVADLRAAASR | ASSLADQDTQY | 0.6464294 |
| QFVLDVAEHPVEVLTETV | ASSLADQDTQY | 0.68856066 |
| GGAGGTGGAGGNGGTQAGDG | ASSLADQDTQY | 0.65800625 |
| PLLATLRAIAVDVDAKPAQV | ASSLADQDTQY | 0.43344703 |
| APKFRTGGYMIDFWGVGYQVAK | ASSLADQDTQY | 0.6491654 |
| GAGGAGGAGGAAGGPGGTGGTG | ASSLADQDTQY | 0.67327046 |
| IGALGLLATDVAPAGSVAFGVA | ASSLADQDTQY | 0.67545325 |
| AGGAGGIGGNGGAGGAGGVLMG | ASSLADQDTQY | 0.6859439 |
| ADELIGLL | ASSLADQDTQY | 0.6643613 |
| ADRALKA | ASSLADQDTQY | 0.6310562 |
| AGAVGQVGGAGGA | ASSLADQDTQY | 0.6841032 |
| AGGGGGLGGAP | ASSLADQDTQY | 0.6894641 |
| AGGLLYGNGGAG | ASSLADQDTQY | 0.6631875 |
| AGLESIGL | ASSLADQDTQY | 0.5982881 |
| AGLKTGII | ASSLADQDTQY | 0.58304715 |
| ASLAALLL | ASSLADQDTQY | 0.66192585 |
| ASVINISSI | ASSLADQDTQY | 0.67737913 |
| ATPPGPA | ASSLADQDTQY | 0.53302395 |
| ATVASALV | ASSLADQDTQY | 0.65471506 |
| ATVASLIL | ASSLADQDTQY | 0.65613264 |
| AVNGPVLIRH | ASSLADQDTQY | 0.44180712 |
| DGAAGAVGI | ASSLADQDTQY | 0.6726949 |
| DPLLHGI | ASSLADQDTQY | 0.6762463 |
| ELGAALGAL | ASSLADQDTQY | 0.6529658 |
| ELGRTQAV | ASSLADQDTQY | 0.5832535 |
| FVAGGYLS | ASSLADQDTQY | 0.67772394 |
| GASGGAGGAGGW | ASSLADQDTQY | 0.6896162 |
| GGAGAGGDGGAGGI | ASSLADQDTQY | 0.6852185 |
| GTGHNGSQPGMGGNG | ASSLADQDTQY | 0.4434644 |
| GYAAASASAS | ASSLADQDTQY | 0.36540776 |
| HQVDVAAVQQG | ASSLADQDTQY | 0.63691986 |
| IGAAIAHT | ASSLADQDTQY | 0.66505206 |
| IGMIEVAQL | ASSLADQDTQY | 0.6403022 |
| IRLAPEHTSAMD | ASSLADQDTQY | 0.6279007 |
| IYNVSVL | ASSLADQDTQY | 0.65503824 |
| KQGFLGM | ASSLADQDTQY | 0.6805539 |
| LESVITQL | ASSLADQDTQY | 0.59865004 |
| LGLGALVVV | ASSLADQDTQY | 0.5583992 |

| | | |
|---|---|---|
| LIGNGGNGGSGGI | ASSLADQDTQY | 0.65388894 |
| LNAINAPTL | ASSLADQDTQY | 0.6692307 |
| LQLLNTGV | ASSLADQDTQY | 0.59138364 |
| LQTSIEF | ASSLADQDTQY | 0.64457715 |
| MSFAVSEL | ASSLADQDTQY | 0.6454164 |
| NTGALGL | ASSLADQDTQY | 0.6256912 |
| PPGAGGAGGAGTT | ASSLADQDTQY | 0.6513096 |
| QLVGLGI | ASSLADQDTQY | 0.59400344 |
| RFLEQVHQL | ASSLADQDTQY | 0.6828357 |
| RGVGYVLR | ASSLADQDTQY | 0.5595281 |
| SATGVFAGVI | ASSLADQDTQY | 0.6646422 |
| SLPAITLP | ASSLADQDTQY | 0.65752244 |
| STALAFTV | ASSLADQDTQY | 0.6635887 |
| SVLTVGLGK | ASSLADQDTQY | 0.63154036 |
| TALAGAGSGI | ASSLADQDTQY | 0.60042584 |
| TGLPFTGI | ASSLADQDTQY | 0.5571787 |
| VAEHVTV | ASSLADQDTQY | 0.5144366 |
| VDIVLNSL | ASSLADQDTQY | 0.67643446 |
| VLTGGGAVL | ASSLADQDTQY | 0.6215993 |
| VNVGLAYGI | ASSLADQDTQY | 0.67876434 |
| VSSAAVGAV | ASSLADQDTQY | 0.6649314 |
| VTSVYEAGV | ASSLADQDTQY | 0.6145168 |
| WVALDRGA | ASSLADQDTQY | 0.6670682 |
| YLVGSVVI | ASSLADQDTQY | 0.5975177 |
| GPPGTGKTLL | ASSLADQDTQY | 0.6413424 |
| GVGMTKF | ASSLADQDTQY | 0.661695 |
| IGIGGSDL | ASSLADQDTQY | 0.6656716 |
| IGTLLAEL | ASSLADQDTQY | 0.57506144 |
| IYTSGTT | ASSLADQDTQY | 0.6585828 |
| LSFPAGV | ASSLADQDTQY | 0.6628747 |
| LYTSGTT | ASSLADQDTQY | 0.5325211 |
| VGAFPFGF | ASSLADQDTQY | 0.65266067 |
| VGAGTVEF | ASSLADQDTQY | 0.67760885 |
| VGKSTLLL | ASSLADQDTQY | 0.65408146 |
| AALAHAGGGIV | ASSLADQDTQY | 0.5794767 |
| AALVAVSLL | ASSLADQDTQY | 0.6638362 |
| AAVSRAVAAL | ASSLADQDTQY | 0.59931946 |
| AGGSSGIGGF | ASSLADQDTQY | 0.6595421 |
| AGLTVANANG | ASSLADQDTQY | 0.6558603 |
| AGLVGAVL | ASSLADQDTQY | 0.6013728 |
| ALSQAFTV | ASSLADQDTQY | 0.6748505 |
| ALTFSVSL | ASSLADQDTQY | 0.47726145 |
| ALTGAALVAL | ASSLADQDTQY | 0.59903026 |
| ASFSIGAGVL | ASSLADQDTQY | 0.62409043 |
| DGSVYKI | ASSLADQDTQY | 0.6560655 |
| EQLAEIV | ASSLADQDTQY | 0.6710429 |
| FAVSELTVM | ASSLADQDTQY | 0.5982294 |
| GAAPLAAGL | ASSLADQDTQY | 0.6517708 |
| GADGGDGGSGGAGG | ASSLADQDTQY | 0.6362126 |
| GFGLDTGGAGG | ASSLADQDTQY | 0.5934533 |
| GLASGGLL | ASSLADQDTQY | 0.64673585 |

| | | |
|---|---|---|
| GRIAGAAVLL | ASSLADQDTQY | 0.61283344 |
| GSAGSLNTSF | ASSLADQDTQY | 0.53404343 |
| GTGQIGI | ASSLADQDTQY | 0.66566265 |
| GTLIALIL | ASSLADQDTQY | 0.6142785 |
| IAATMGFVVG | ASSLADQDTQY | 0.6629076 |
| IFAQALL | ASSLADQDTQY | 0.6670847 |
| IGAAVALGF | ASSLADQDTQY | 0.5386845 |
| IGLTPLAL | ASSLADQDTQY | 0.66707695 |
| ILGSNTSTL | ASSLADQDTQY | 0.66294783 |
| IVFSLTVL | ASSLADQDTQY | 0.66155976 |
| LASKATGF | ASSLADQDTQY | 0.6390171 |
| LASVALIL | ASSLADQDTQY | 0.59877837 |
| LFAVGLTL | ASSLADQDTQY | 0.68033093 |
| LFNGGIGTYI | ASSLADQDTQY | 0.67124903 |
| LTLGAGII | ASSLADQDTQY | 0.6325583 |
| LVAAALGDI | ASSLADQDTQY | 0.6711248 |
| NTAFVQLGI | ASSLADQDTQY | 0.6540244 |
| QLFSGAEL | ASSLADQDTQY | 0.6700065 |
| SAATFITL | ASSLADQDTQY | 0.6108199 |
| SNYGLANLGV | ASSLADQDTQY | 0.49778104 |
| SYGGAYAVM | ASSLADQDTQY | 0.6467853 |
| TTGVGGLTLG | ASSLADQDTQY | 0.63393867 |
| TTQTIYI | ASSLADQDTQY | 0.5622908 |
| VAAGAGLGAVI | ASSLADQDTQY | 0.48576114 |
| VAAVVTLL | ASSLADQDTQY | 0.6749109 |
| VGFLGGLI | ASSLADQDTQY | 0.6914325 |
| VGVIPILL | ASSLADQDTQY | 0.58408535 |
| VIFASSLI | ASSLADQDTQY | 0.6899132 |
| VKGGDGGAAAG | ASSLADQDTQY | 0.5323975 |
| VTKTFTNI | ASSLADQDTQY | 0.67278486 |
| VVAAAILL | ASSLADQDTQY | 0.67452043 |
| YAALDVEL | ASSLADQDTQY | 0.57969445 |
| LASVALLL | ASSLADQDTQY | 0.6692449 |
| LDAVHRV | ASSLADQDTQY | 0.56184864 |
| QGLSSSIF | ASSLADQDTQY | 0.60300654 |
| RGTFANIRL | ASSLADQDTQY | 0.67707115 |
| RQLWWGHRI | ASSLADQDTQY | 0.6064137 |
| YGPPGTGKTLL | ASSLADQDTQY | 0.66066444 |
| AALVGAPANVA | ASSLADQDTQY | 0.64726746 |
| AATFRRTE | ASSLADQDTQY | 0.6163987 |
| AGALGGGI | ASSLADQDTQY | 0.6652138 |
| AGGAGGAGGIGAG | ASSLADQDTQY | 0.6654351 |
| AGSGALGL | ASSLADQDTQY | 0.63154745 |
| AQIADQI | ASSLADQDTQY | 0.68723285 |
| ASAVAVTYL | ASSLADQDTQY | 0.5798814 |
| ASTVLGGTGG | ASSLADQDTQY | 0.6881274 |
| AVGGEPAK | ASSLADQDTQY | 0.6648146 |
| CEQGTFGGTC | ASSLADQDTQY | 0.6523955 |
| DEGKAPAG | ASSLADQDTQY | 0.62539065 |
| DTVVFPAL | ASSLADQDTQY | 0.67386466 |
| FEGITMAG | ASSLADQDTQY | 0.637743 |

| | | |
|---|---|---|
| FPLVVQD | ASSLADQDTQY | 0.6557611 |
| GAFTTPQ | ASSLADQDTQY | 0.6825183 |
| GFGGFGTAGGVGGA | ASSLADQDTQY | 0.5336357 |
| GFLAQFL | ASSLADQDTQY | 0.67451674 |
| GGAGGAGGNAGVA | ASSLADQDTQY | 0.64370215 |
| GGVGTAGAGGAGGAGG | ASSLADQDTQY | 0.62841916 |
| GNGGGGDAGNAGSGGN | ASSLADQDTQY | 0.56532156 |
| GTGGAGGAGNDAGS | ASSLADQDTQY | 0.47429144 |
| GTTNTGWFNS | ASSLADQDTQY | 0.62597585 |
| IFMGPLFTL | ASSLADQDTQY | 0.5201647 |
| ILAHAIEAI | ASSLADQDTQY | 0.66135657 |
| IRSAIREAL | ASSLADQDTQY | 0.6630789 |
| ITFAGGG | ASSLADQDTQY | 0.54899335 |
| ITGMSGHQEV | ASSLADQDTQY | 0.66409963 |
| IVAGLACTL | ASSLADQDTQY | 0.6651163 |
| IVGGGLAGA | ASSLADQDTQY | 0.47501236 |
| IVTGVLNI | ASSLADQDTQY | 0.5972075 |
| LAQQGCI | ASSLADQDTQY | 0.61007184 |
| LCLHVEN | ASSLADQDTQY | 0.5999254 |
| LDAISDGS | ASSLADQDTQY | 0.45464978 |
| LGGFAAML | ASSLADQDTQY | 0.6501118 |
| LGGKNPMI | ASSLADQDTQY | 0.5744192 |
| LGPKGVAAIL | ASSLADQDTQY | 0.3158624 |
| LMVGVSI | ASSLADQDTQY | 0.6231683 |
| LVGAAFVL | ASSLADQDTQY | 0.59409094 |
| LVLAVAPL | ASSLADQDTQY | 0.59379995 |
| MAAIAVIDAA | ASSLADQDTQY | 0.6588043 |
| NNGINAELQ | ASSLADQDTQY | 0.5492795 |
| RGEPLVLVHG | ASSLADQDTQY | 0.5728216 |
| RGGAGGAAGLI | ASSLADQDTQY | 0.64916736 |
| RSYRMHPAV | ASSLADQDTQY | 0.6650253 |
| SAAHLLAGQLA | ASSLADQDTQY | 0.505998 |
| SAAVGSY | ASSLADQDTQY | 0.62655246 |
| SPHAVLHRI | ASSLADQDTQY | 0.66892344 |
| SQIAKVL | ASSLADQDTQY | 0.6400069 |
| TALGHLVP | ASSLADQDTQY | 0.6770451 |
| TAPTTSGAA | ASSLADQDTQY | 0.6809751 |
| TDGDDGNGGSPGT | ASSLADQDTQY | 0.5805808 |
| TDGLGEN | ASSLADQDTQY | 0.66442204 |
| TGGPPLPGQA | ASSLADQDTQY | 0.6856754 |
| TIPAITI | ASSLADQDTQY | 0.6091322 |
| TIPGFTI | ASSLADQDTQY | 0.6092816 |
| TSIASGALA | ASSLADQDTQY | 0.53496236 |
| TVTLPGAAI | ASSLADQDTQY | 0.6099075 |
| VASGLSAALP | ASSLADQDTQY | 0.4437387 |
| VGAGVIGI | ASSLADQDTQY | 0.62383443 |
| VGPKRFGV | ASSLADQDTQY | 0.67356455 |
| VNVAFPD | ASSLADQDTQY | 0.5790783 |
| VSMTNSGVS | ASSLADQDTQY | 0.4979112 |
| VTDVLGDLSV | ASSLADQDTQY | 0.5213364 |
| VVVVGAGFAGL | ASSLADQDTQY | 0.6249449 |

| | | |
|---|---|---|
| VYTVGTVV | ASSLADQDTQY | 0.56537485 |
| WTTFSGP | ASSLADQDTQY | 0.4316102 |
| YDELVEAF | ASSLADQDTQY | 0.58371717 |
| YGTGGAGGNGG | ASSLADQDTQY | 0.6642107 |
| YTNVMAV | ASSLADQDTQY | 0.6644845 |
| FLEQVHQL | ASSLADQDTQY | 0.43814144 |
| GLSLLAVAV | ASSLADQDTQY | 0.67615616 |
| LEQVHQL | ASSLADQDTQY | 0.66143304 |
| LNAAGAI | ASSLADQDTQY | 0.683989 |
| PAVHASL | ASSLADQDTQY | 0.6405173 |
| TELEDDA | ASSLADQDTQY | 0.5300698 |
| TPPGPPN | ASSLADQDTQY | 0.656492 |
| TPPPPPPP | ASSLADQDTQY | 0.5595356 |
| TQSPALI | APPRGRDTEVF | 0.5636261 |
| EFAAAFPQ | APPRGRDTEVF | 0.6547473 |
| LLSGNLDVL | APPRGRDTEVF | 0.6133951 |
| GPANLGNYNI | APPRGRDTEVF | 0.5412486 |
| TLGPTLP | APPRGRDTEVF | 0.57088566 |
| VYAYSSCCK | APPRGRDTEVF | 0.67996883 |
| APAPAPAPNI | APPRGRDTEVF | 0.51949346 |
| PGGPVPIP | APPRGRDTEVF | 0.53423584 |
| FLTVALEP | APPRGRDTEVF | 0.5312735 |
| SQYFGANL | APPRGRDTEVF | 0.6023612 |
| GGVKPAA | APPRGRDTEVF | 0.6621039 |
| GGTGGAGGAGGW | APPRGRDTEVF | 0.65879244 |
| GVGGVAGLGGAGG | APPRGRDTEVF | 0.5914998 |
| VLLVLATTI | APPRGRDTEVF | 0.6329998 |
| IGIFAAML | APPRGRDTEVF | 0.6522335 |
| AVGDLID | APPRGRDTEVF | 0.6234815 |
| NISPLPL | APPRGRDTEVF | 0.6462892 |
| VTAVFVAI | APPRGRDTEVF | 0.5261354 |
| SGDDNDAAL | APPRGRDTEVF | 0.6762689 |
| AVAALAHS | APPRGRDTEVF | 0.6452802 |
| GGVTVMATP | APPRGRDTEVF | 0.66974956 |
| AGLAANVRD | APPRGRDTEVF | 0.67256093 |
| TNRPDLIDPALLRPG | APPRGRDTEVF | 0.637619 |
| LANLGVTAL | APPRGRDTEVF | 0.6349127 |
| TGERIEL | APPRGRDTEVF | 0.43273777 |
| NGEELAA | APPRGRDTEVF | 0.60615045 |
| VLSLVWG | APPRGRDTEVF | 0.62423134 |
| IGYALWL | APPRGRDTEVF | 0.62946653 |
| LSLLLAPL | APPRGRDTEVF | 0.65326023 |
| TLDEALK | APPRGRDTEVF | 0.6651593 |
| TDGAGTAEGG | APPRGRDTEVF | 0.56614393 |
| NAPGGAGGNGG | APPRGRDTEVF | 0.67002547 |
| VSVFMGMGGNF | APPRGRDTEVF | 0.6640726 |
| LASYLAAL | APPRGRDTEVF | 0.4691569 |
| LVGGTPAL | APPRGRDTEVF | 0.5813972 |
| VRNADVL | APPRGRDTEVF | 0.6805179 |
| TCSSPFLVSAM | APPRGRDTEVF | 0.6272237 |
| VHITDAE | APPRGRDTEVF | 0.58604765 |

| | | |
|---|---|---|
| EYAAPSF | APPRGRDTEVF | 0.60173166 |
| VVDAFHGGL | APPRGRDTEVF | 0.6832776 |
| ASVVDPQ | APPRGRDTEVF | 0.5666101 |
| SYNEFGGLS | APPRGRDTEVF | 0.67733264 |
| DPDGSVY | APPRGRDTEVF | 0.359605 |
| MVSTIHATE | APPRGRDTEVF | 0.61096585 |
| LAAGVADPV | APPRGRDTEVF | 0.58141655 |
| TKEQIAA | APPRGRDTEVF | 0.36279577 |
| SLASPAAL | APPRGRDTEVF | 0.6629114 |
| IGLVFYGV | APPRGRDTEVF | 0.60981566 |
| FLLDASNL | APPRGRDTEVF | 0.6509438 |
| YQVGNFP | APPRGRDTEVF | 0.62654054 |
| ESLLSQST | APPRGRDTEVF | 0.6667833 |
| LFTVGFAAVG | APPRGRDTEVF | 0.6282986 |
| ATPPTPT | APPRGRDTEVF | 0.62470007 |
| TDLATLG | APPRGRDTEVF | 0.55154276 |
| SAGGTGR | APPRGRDTEVF | 0.655683 |
| GFSNSGS | APPRGRDTEVF | 0.6757958 |
| GGNGGSGGTSV | APPRGRDTEVF | 0.67275643 |
| GTSVATGGAGN | APPRGRDTEVF | 0.46636838 |
| NGGNGGSGGTS | APPRGRDTEVF | 0.5913375 |
| AVVKFPH | APPRGRDTEVF | 0.6266942 |
| TPPPPPP | APPRGRDTEVF | 0.39840698 |
| AAPAAAAAPA | APPRGRDTEVF | 0.33615252 |
| IALNEDL | APPRGRDTEVF | 0.5941904 |
| IVNVSSM | APPRGRDTEVF | 0.57305926 |
| TGIPPAPR | APPRGRDTEVF | 0.65225554 |
| FDLGGGTF | APPRGRDTEVF | 0.6516336 |
| FDLGGGTFDVS | APPRGRDTEVF | 0.5769325 |
| NEPTAAA | APPRGRDTEVF | 0.6474693 |
| NEPTAAALA | APPRGRDTEVF | 0.6063046 |
| GSIINISSI | APPRGRDTEVF | 0.28069556 |
| VAPGNAGTA | APPRGRDTEVF | 0.6028342 |
| TGGGASLE | APPRGRDTEVF | 0.47285682 |
| VAVYNHYKRI | APPRGRDTEVF | 0.6845705 |
| TDPAPATT | APPRGRDTEVF | 0.55261594 |
| TNRPDILDPALLRPG | APPRGRDTEVF | 0.5464108 |
| LTTTTTS | APPRGRDTEVF | 0.5957254 |
| PTGSGKTL | APPRGRDTEVF | 0.62567335 |
| TGSGKTL | APPRGRDTEVF | 0.6035201 |
| TIGHVDHGK | APPRGRDTEVF | 0.55592424 |
| TIGHVDHGKTT | APPRGRDTEVF | 0.35210463 |
| GHVDHGKTT | APPRGRDTEVF | 0.66000366 |
| AQFEHTLL | APPRGRDTEVF | 0.5518609 |
| IVNQDEL | APPRGRDTEVF | 0.5407894 |
| TNGSQFF | APPRGRDTEVF | 0.42021573 |
| TGKGIVE | APPRGRDTEVF | 0.48606792 |
| LISNASDA | APPRGRDTEVF | 0.47568446 |
| AELRTLGELTKIAKAHGAQ | APPRGRDTEVF | 0.67102677 |
| APNAVAAAEPSW | APPRGRDTEVF | 0.5902613 |
| APQVAGAATAVL | APPRGRDTEVF | 0.5788152 |

| | | |
|---|---|---|
| ARRDPAAAAAILPTDA | APPRGRDTEVF | 0.68233395 |
| DAAARDVLAEAGLAEYF | APPRGRDTEVF | 0.6066178 |
| DDQIVLSAKANLAKVEQ | APPRGRDTEVF | 0.5678151 |
| DELLPDTGEDSDEDGH | APPRGRDTEVF | 0.5702359 |
| ELPEAPANRATVPQD | APPRGRDTEVF | 0.5107929 |
| FGQSKIERVTGIG | APPRGRDTEVF | 0.5546016 |
| GAGGTGSTAGGAGGAGGAGGLY | APPRGRDTEVF | 0.5616399 |
| GHGGTGAAGGNGGTGGAGGSGI | APPRGRDTEVF | 0.67600745 |
| GNGGNGGAGGEGGPGGAGGAGA | APPRGRDTEVF | 0.6132537 |
| GSTAGNGGNGGNATNGTIAGQP | APPRGRDTEVF | 0.55830616 |
| GTGGDGGNGGNGGTGG | APPRGRDTEVF | 0.6550665 |
| GVRRPAGIPADAPV | APPRGRDTEVF | 0.6750676 |
| GYRQVTDAHLLTLARR | APPRGRDTEVF | 0.68878514 |
| IAKMGQDGHDRG | APPRGRDTEVF | 0.5329492 |
| KAEPSIEIQYT | APPRGRDTEVF | 0.62774956 |
| KPVRIRMAADEDTVAVT | APPRGRDTEVF | 0.6005438 |
| MPPRIAGMRLLVIKPEP | APPRGRDTEVF | 0.5916173 |
| MKFYAVALFRSRGVPPTR | APPRGRDTEVF | 0.59680325 |
| NAGGFGILGAVAHSP | APPRGRDTEVF | 0.49253944 |
| NEGREITLAMAEM | APPRGRDTEVF | 0.6675172 |
| NELVEWSHPLGHRWRWE | APPRGRDTEVF | 0.688583 |
| NLGAGGTGGNGGNPG | APPRGRDTEVF | 0.59545076 |
| NTEWPKWETVMAADPVD | APPRGRDTEVF | 0.55623555 |
| PASAAPAETQP | APPRGRDTEVF | 0.5809342 |
| QQSSDGKNGVRAAEITGE | APPRGRDTEVF | 0.5385352 |
| RAPDAVTTSWWKE | APPRGRDTEVF | 0.5377809 |
| SGFGNVGAHTSGWFNQ | APPRGRDTEVF | 0.6277642 |
| SGVAGAVGQAASVGGLKV | APPRGRDTEVF | 0.46551892 |
| SNGGAGTGSAGGNGGT | APPRGRDTEVF | 0.606058 |
| TGGAGGEGGAAG | APPRGRDTEVF | 0.6852184 |
| TGGDGGAAGAGGAPGHG | APPRGRDTEVF | 0.58730745 |
| TMPPLLAAIANAGRHGV | APPRGRDTEVF | 0.6458774 |
| TSSTAGNSGGA | APPRGRDTEVF | 0.6649289 |
| VPATVPVGILC | APPRGRDTEVF | 0.64175177 |
| VQRPDSGGVVLTGRIS | APPRGRDTEVF | 0.66955304 |
| WTPNPVPATKGVPAGLVQAT | APPRGRDTEVF | 0.66504526 |
| YALKRALAKFSDRFDVVII | APPRGRDTEVF | 0.62192464 |
| YLIGSVIV | APPRGRDTEVF | 0.62475497 |
| YGQLVAVID | APPRGRDTEVF | 0.6390215 |
| VTVGGMTLPA | APPRGRDTEVF | 0.5935643 |
| VLALPAV | APPRGRDTEVF | 0.5903093 |
| VIPPTLNYET | APPRGRDTEVF | 0.5733987 |
| VGMGGMHPSSG | APPRGRDTEVF | 0.6541997 |
| VGLAGGAATA | APPRGRDTEVF | 0.6201445 |
| VGGAGGAGGSGP | APPRGRDTEVF | 0.67227954 |
| VFAAFMV | APPRGRDTEVF | 0.42607027 |
| TLYDKLR | APPRGRDTEVF | 0.49474487 |
| TLAVFTVL | APPRGRDTEVF | 0.65924823 |
| TGSVQPARV | APPRGRDTEVF | 0.30797076 |
| TGGGAGSGGT | APPRGRDTEVF | 0.37560755 |
| SHLETSL | APPRGRDTEVF | 0.6391636 |

| | | |
|---|---|---|
| SFTTGMT | APPRGRDTEVF | 0.6710906 |
| QVEVAKRMGVRQPTVS | APPRGRDTEVF | 0.3275755 |
| QGSGGPALAM | APPRGRDTEVF | 0.5453439 |
| MLAPSNP | APPRGRDTEVF | 0.5054941 |
| LVNNAAI | APPRGRDTEVF | 0.6682001 |
| LTGEINTLA | APPRGRDTEVF | 0.612527 |
| LSIFGPV | APPRGRDTEVF | 0.4238315 |
| LLALLANL | APPRGRDTEVF | 0.65565026 |
| LISNASDALDKLR | APPRGRDTEVF | 0.45798498 |
| LINEAVP | APPRGRDTEVF | 0.5310445 |
| LGYAFSGL | APPRGRDTEVF | 0.665686 |
| LDFVAVAL | APPRGRDTEVF | 0.6780389 |
| LAQPGAE | APPRGRDTEVF | 0.5173663 |
| ISNLGAVTGF | APPRGRDTEVF | 0.6530286 |
| HQEDPGRI | APPRGRDTEVF | 0.575489 |
| GTPGEPTP | APPRGRDTEVF | 0.6925334 |
| GFFNSGS | APPRGRDTEVF | 0.5852778 |
| GAGGGGLTAGAN | APPRGRDTEVF | 0.46648878 |
| FVDALVNM | APPRGRDTEVF | 0.6612803 |
| FLTVAPEL | APPRGRDTEVF | 0.65188646 |
| FETLTAL | APPRGRDTEVF | 0.62980556 |
| DVINPIEL | APPRGRDTEVF | 0.65957385 |
| APVSGAPV | APPRGRDTEVF | 0.5949278 |
| ALVGGLAW | APPRGRDTEVF | 0.58602625 |
| ALAAAAATAI | APPRGRDTEVF | 0.6596215 |
| AFGLGLLL | APPRGRDTEVF | 0.65380406 |
| AFFAAASL | APPRGRDTEVF | 0.5023349 |
| AAAALPAFSPP | APPRGRDTEVF | 0.6699508 |
| VVLAPAL | APPRGRDTEVF | 0.58831775 |
| TIGHVDHGKTTL | APPRGRDTEVF | 0.5412413 |
| LISNASDALDK | APPRGRDTEVF | 0.687953 |
| LISNASDAL | APPRGRDTEVF | 0.6411308 |
| ISNASDALDK | APPRGRDTEVF | 0.6514968 |
| NQAETLVYQTE | APPRGRDTEVF | 0.6622013 |
| TGGTGGTAGAG | APPRGRDTEVF | 0.68756354 |
| TGSTAGGAGGA | APPRGRDTEVF | 0.4982638 |
| DPLAVPVAAGPV | APPRGRDTEVF | 0.6528808 |
| GGTGGTPGNGGN | APPRGRDTEVF | 0.6102492 |
| PAGLAGVDPATL | APPRGRDTEVF | 0.664815 |
| GNNLVVEAPHAGASV | APPRGRDTEVF | 0.65196663 |
| KVTRIMSASTLEQPA | APPRGRDTEVF | 0.6016039 |
| LGFALPIALPRRTVV | APPRGRDTEVF | 0.6485102 |
| TRLGNLTARKLVVLV | APPRGRDTEVF | 0.67156476 |
| WSQQRRLIYFPSAGP | APPRGRDTEVF | 0.63867724 |
| GPNGNPDPMAAAVDIR | APPRGRDTEVF | 0.63978595 |
| TQRCVSVADLRAAASR | APPRGRDTEVF | 0.6735404 |
| QFVLDVAEHPVEVLTETV | APPRGRDTEVF | 0.55913854 |
| GGAGGTGGAGGNGGTQAGDG | APPRGRDTEVF | 0.5928755 |
| PLLATLRAIAVDVDAKPAQV | APPRGRDTEVF | 0.6695303 |
| APKFRTGGYMIDFWGVGYQVAK | APPRGRDTEVF | 0.66531646 |
| GAGGAGGAGGAAGGPGGTGGTG | APPRGRDTEVF | 0.616905 |

| | | |
|---|---|---|
| IGALGLLATDVAPAGSVAFGVA | APPRGRDTEVF | 0.67218447 |
| AGGAGGIGGNGGAGGAGGVLMG | APPRGRDTEVF | 0.56122625 |
| ADELIGLL | APPRGRDTEVF | 0.64384186 |
| ADRALKA | APPRGRDTEVF | 0.5778614 |
| AGAVGQVGGAGGA | APPRGRDTEVF | 0.63160354 |
| AGGGGGLGGAP | APPRGRDTEVF | 0.50072914 |
| AGGLLYGNGGAG | APPRGRDTEVF | 0.63310575 |
| AGLESIGL | APPRGRDTEVF | 0.65722007 |
| AGLKTGII | APPRGRDTEVF | 0.6474131 |
| ASLAALLL | APPRGRDTEVF | 0.36196914 |
| ASVINISSI | APPRGRDTEVF | 0.6508884 |
| ATPPGPA | APPRGRDTEVF | 0.6482193 |
| ATVASALV | APPRGRDTEVF | 0.68421495 |
| ATVASLIL | APPRGRDTEVF | 0.61007667 |
| AVNGPVLIRH | APPRGRDTEVF | 0.64647245 |
| DGAAGAVGI | APPRGRDTEVF | 0.59746796 |
| DPLLHGI | APPRGRDTEVF | 0.6189058 |
| ELGAALGAL | APPRGRDTEVF | 0.4761271 |
| ELGRTQAV | APPRGRDTEVF | 0.5882413 |
| FVAGGYLS | APPRGRDTEVF | 0.56488055 |
| GASGGAGGAGGW | APPRGRDTEVF | 0.5188276 |
| GGAGAGGDGGAGGI | APPRGRDTEVF | 0.65033656 |
| GTGHNGSQPGMGGNG | APPRGRDTEVF | 0.66235226 |
| GYAAASASAS | APPRGRDTEVF | 0.66332227 |
| HQVDVAAVQQG | APPRGRDTEVF | 0.6671586 |
| IGAAIAHT | APPRGRDTEVF | 0.6737584 |
| IGMIEVAQL | APPRGRDTEVF | 0.675992 |
| IRLAPEHTSAMD | APPRGRDTEVF | 0.5543879 |
| IYNVSVL | APPRGRDTEVF | 0.61596435 |
| KQGFLGM | APPRGRDTEVF | 0.46322966 |
| LESVITQL | APPRGRDTEVF | 0.59815264 |
| LGLGALVVV | APPRGRDTEVF | 0.57111543 |
| LIGNGGNGGSGGI | APPRGRDTEVF | 0.61758137 |
| LNAINAPTL | APPRGRDTEVF | 0.30660462 |
| LQLLNTGV | APPRGRDTEVF | 0.6184267 |
| LQTSIEF | APPRGRDTEVF | 0.6763473 |
| MSFAVSEL | APPRGRDTEVF | 0.5567069 |
| NTGALGL | APPRGRDTEVF | 0.6393318 |
| PPGAGGAGGAGTT | APPRGRDTEVF | 0.6316175 |
| QLVGLGI | APPRGRDTEVF | 0.5657085 |
| RFLEQVHQL | APPRGRDTEVF | 0.64902234 |
| RGVGYVLR | APPRGRDTEVF | 0.542601 |
| SATGVFAGVI | APPRGRDTEVF | 0.6530324 |
| SLPAITLP | APPRGRDTEVF | 0.68455994 |
| STALAFTV | APPRGRDTEVF | 0.63298744 |
| SVLTVGLGK | APPRGRDTEVF | 0.6207659 |
| TALAGAGSGI | APPRGRDTEVF | 0.62609947 |
| TGLPFTGI | APPRGRDTEVF | 0.44787842 |
| VAEHVTV | APPRGRDTEVF | 0.6282929 |
| VDIVLNSL | APPRGRDTEVF | 0.33001807 |
| VLTGGGAVL | APPRGRDTEVF | 0.6052531 |

| | | |
|---|---|---|
| VNVGLAYGI | APPRGRDTEVF | 0.5526799 |
| VSSAAVGAV | APPRGRDTEVF | 0.6222336 |
| VTSVYEAGV | APPRGRDTEVF | 0.6058676 |
| WVALDRGA | APPRGRDTEVF | 0.6480471 |
| YLVGSVVI | APPRGRDTEVF | 0.5395235 |
| GPPGTGKTLL | APPRGRDTEVF | 0.6216086 |
| GVGMTKF | APPRGRDTEVF | 0.58653086 |
| IGIGGSDL | APPRGRDTEVF | 0.51503277 |
| IGTLLAEL | APPRGRDTEVF | 0.66414905 |
| IYTSGTT | APPRGRDTEVF | 0.6407515 |
| LSFPAGV | APPRGRDTEVF | 0.59894973 |
| LYTSGTT | APPRGRDTEVF | 0.67044747 |
| VGAFPFGF | APPRGRDTEVF | 0.63386816 |
| VGAGTVEF | APPRGRDTEVF | 0.4812364 |
| VGKSTLLL | APPRGRDTEVF | 0.59960407 |
| AALAHAGGGIV | APPRGRDTEVF | 0.6123315 |
| AALVAVSLL | APPRGRDTEVF | 0.5943711 |
| AAVSRAVAAL | APPRGRDTEVF | 0.31627846 |
| AGGSSGIGGF | APPRGRDTEVF | 0.6533181 |
| AGLTVANANG | APPRGRDTEVF | 0.5434664 |
| AGLVGAVL | APPRGRDTEVF | 0.5426229 |
| ALSQAFTV | APPRGRDTEVF | 0.568422 |
| ALTFSVSL | APPRGRDTEVF | 0.6527259 |
| ALTGAALVAL | APPRGRDTEVF | 0.5941862 |
| ASFSIGAGVL | APPRGRDTEVF | 0.63121057 |
| DGSVYKI | APPRGRDTEVF | 0.6447401 |
| EQLAEIV | APPRGRDTEVF | 0.6491467 |
| FAVSELTVM | APPRGRDTEVF | 0.6650041 |
| GAAPLAAGL | APPRGRDTEVF | 0.64222765 |
| GADGGDGGSGGAGG | APPRGRDTEVF | 0.62924564 |
| GFGLDTGGAGG | APPRGRDTEVF | 0.6412909 |
| GLASGGLL | APPRGRDTEVF | 0.6312171 |
| GRIAGAAVLL | APPRGRDTEVF | 0.43544525 |
| GSAGSLNTSF | APPRGRDTEVF | 0.41846853 |
| GTGQIGI | APPRGRDTEVF | 0.62635535 |
| GTLIALIL | APPRGRDTEVF | 0.42833263 |
| IAATMGFVVG | APPRGRDTEVF | 0.6245475 |
| IFAQALL | APPRGRDTEVF | 0.68281144 |
| IGAAVALGF | APPRGRDTEVF | 0.63049334 |
| IGLTPLAL | APPRGRDTEVF | 0.49509168 |
| ILGSNTSTL | APPRGRDTEVF | 0.59612364 |
| IVFSLTVL | APPRGRDTEVF | 0.5994216 |
| LASKATGF | APPRGRDTEVF | 0.64909685 |
| LASVALIL | APPRGRDTEVF | 0.6760871 |
| LFAVGLTL | APPRGRDTEVF | 0.6796585 |
| LFNGGIGTYI | APPRGRDTEVF | 0.652297 |
| LTLGAGII | APPRGRDTEVF | 0.5474671 |
| LVAAALGDI | APPRGRDTEVF | 0.64255834 |
| NTAFVQLGI | APPRGRDTEVF | 0.5928974 |
| QLFSGAEL | APPRGRDTEVF | 0.63124907 |
| SAATFITL | APPRGRDTEVF | 0.6282364 |

| | | |
|---|---|---|
| SNYGLANLGV | APPRGRDTEVF | 0.64726377 |
| SYGGAYAVM | APPRGRDTEVF | 0.6059178 |
| TTGVGGLTLG | APPRGRDTEVF | 0.5942718 |
| TTQTIYI | APPRGRDTEVF | 0.48426253 |
| VAAGAGLGAVI | APPRGRDTEVF | 0.66691107 |
| VAAVVTLL | APPRGRDTEVF | 0.649331 |
| VGFLGGLI | APPRGRDTEVF | 0.4749818 |
| VGVIPILL | APPRGRDTEVF | 0.5770091 |
| VIFASSLI | APPRGRDTEVF | 0.6816104 |
| VKGGDGGAAAG | APPRGRDTEVF | 0.6714531 |
| VTKTFTNI | APPRGRDTEVF | 0.53230894 |
| VVAAAILL | APPRGRDTEVF | 0.5554163 |
| YAALDVEL | APPRGRDTEVF | 0.6177743 |
| LASVALLL | APPRGRDTEVF | 0.43347308 |
| LDAVHRV | APPRGRDTEVF | 0.6480768 |
| QGLSSSIF | APPRGRDTEVF | 0.59929425 |
| RGTFANIRL | APPRGRDTEVF | 0.59818894 |
| RQLWWGHRI | APPRGRDTEVF | 0.5867573 |
| YGPPGTGKTLL | APPRGRDTEVF | 0.62974614 |
| AALVGAPANVA | APPRGRDTEVF | 0.5778346 |
| AATFRRTE | APPRGRDTEVF | 0.66382784 |
| AGALGGGI | APPRGRDTEVF | 0.3306069 |
| AGGAGGAGGIGAG | APPRGRDTEVF | 0.65928614 |
| AGSGALGL | APPRGRDTEVF | 0.5133987 |
| AQIADQI | APPRGRDTEVF | 0.5690347 |
| ASAVAVTYL | APPRGRDTEVF | 0.45264414 |
| ASTVLGGTGG | APPRGRDTEVF | 0.61942816 |
| AVGGEPAK | APPRGRDTEVF | 0.66976887 |
| CEQGTFGGTC | APPRGRDTEVF | 0.6605545 |
| DEGKAPAG | APPRGRDTEVF | 0.53854066 |
| DTVVFPAL | APPRGRDTEVF | 0.65672183 |
| FEGITMAG | APPRGRDTEVF | 0.6843716 |
| FPLVVQD | APPRGRDTEVF | 0.57010496 |
| GAFTTPQ | APPRGRDTEVF | 0.4040873 |
| GFGGFGTAGGVGGA | APPRGRDTEVF | 0.6416031 |
| GFLAQFL | APPRGRDTEVF | 0.6176367 |
| GGAGGAGGNAGVA | APPRGRDTEVF | 0.6693114 |
| GGVGTAGAGGAGGAGG | APPRGRDTEVF | 0.65543497 |
| GNGGGGDAGNAGSGGN | APPRGRDTEVF | 0.4536395 |
| GTGGAGGAGNDAGS | APPRGRDTEVF | 0.66083765 |
| GTTNTGWFNS | APPRGRDTEVF | 0.67217016 |
| IFMGPLFTL | APPRGRDTEVF | 0.5103063 |
| ILAHAIEAI | APPRGRDTEVF | 0.5817315 |
| IRSAIREAL | APPRGRDTEVF | 0.58278733 |
| ITFAGGG | APPRGRDTEVF | 0.48976105 |
| ITGMSGHQEV | APPRGRDTEVF | 0.6600288 |
| IVAGLACTL | APPRGRDTEVF | 0.55969507 |
| IVGGGLAGA | APPRGRDTEVF | 0.5806169 |
| IVTGVLNI | APPRGRDTEVF | 0.60934186 |
| LAQQGCI | APPRGRDTEVF | 0.5755639 |
| LCLHVEN | APPRGRDTEVF | 0.5803371 |

| | | |
|---|---|---|
| LDAISDGS | APPRGRDTEVF | 0.5959822 |
| LGGFAAML | APPRGRDTEVF | 0.6474514 |
| LGGKNPMI | APPRGRDTEVF | 0.6422602 |
| LGPKGVAAIL | APPRGRDTEVF | 0.6227945 |
| LMVGVSI | APPRGRDTEVF | 0.63823164 |
| LVGAAFVL | APPRGRDTEVF | 0.6683805 |
| LVLAVAPL | APPRGRDTEVF | 0.49120802 |
| MAAIAVIDAA | APPRGRDTEVF | 0.5311287 |
| NNGINAELQ | APPRGRDTEVF | 0.6472016 |
| RGEPLVLVHG | APPRGRDTEVF | 0.49154618 |
| RGGAGGAAGLI | APPRGRDTEVF | 0.65654325 |
| RSYRMHPAV | APPRGRDTEVF | 0.6060111 |
| SAAHLLAGQLA | APPRGRDTEVF | 0.6240634 |
| SAAVGSY | APPRGRDTEVF | 0.62579477 |
| SPHAVLHRI | APPRGRDTEVF | 0.68467504 |
| SQIAKVL | APPRGRDTEVF | 0.65775746 |
| TALGHLVP | APPRGRDTEVF | 0.6282452 |
| TAPTTSGAA | APPRGRDTEVF | 0.65771705 |
| TDGDDGNGGSPGT | APPRGRDTEVF | 0.53653616 |
| TDGLGEN | APPRGRDTEVF | 0.6623905 |
| TGGPPLPGQA | APPRGRDTEVF | 0.6518371 |
| TIPAITI | APPRGRDTEVF | 0.50598586 |
| TIPGFTI | APPRGRDTEVF | 0.5894454 |
| TSIASGALA | APPRGRDTEVF | 0.6773435 |
| TVTLPGAAI | APPRGRDTEVF | 0.6665795 |
| VASGLSAALP | APPRGRDTEVF | 0.4658562 |
| VGAGVIGI | APPRGRDTEVF | 0.5668871 |
| VGPKRFGV | APPRGRDTEVF | 0.43665662 |
| VNVAFPD | APPRGRDTEVF | 0.5811208 |
| VSMTNSGVS | APPRGRDTEVF | 0.48081166 |
| VTDVLGDLSV | APPRGRDTEVF | 0.64012986 |
| VVVVGAGFAGL | APPRGRDTEVF | 0.57492983 |
| VYTVGTVV | APPRGRDTEVF | 0.6451987 |
| WTTFSGP | APPRGRDTEVF | 0.6207359 |
| YDELVEAF | APPRGRDTEVF | 0.66891617 |
| YGTGGAGGNGG | APPRGRDTEVF | 0.56529236 |
| YTNVMAV | APPRGRDTEVF | 0.5977971 |
| FLEQVHQL | APPRGRDTEVF | 0.45793554 |
| GLSLLAVAV | APPRGRDTEVF | 0.58196485 |
| LEQVHQL | APPRGRDTEVF | 0.59449404 |
| LNAAGAI | APPRGRDTEVF | 0.6032827 |
| PAVHASL | APPRGRDTEVF | 0.3928277 |
| TELEDDA | APPRGRDTEVF | 0.6752955 |
| TPPGPPN | APPRGRDTEVF | 0.6160475 |
| TPPPPPPP | APPRGRDTEVF | 0.6539912 |
| TQSPALI | ASSSDRGRTEVF | 0.595401 |
| EFAAAFPQ | ASSSDRGRTEVF | 0.5969368 |
| LLSGNLDVL | ASSSDRGRTEVF | 0.54705364 |
| GPANLGNYNI | ASSSDRGRTEVF | 0.63877964 |
| TLGPTLP | ASSSDRGRTEVF | 0.52001214 |
| VYAYSSCCK | ASSSDRGRTEVF | 0.4845194 |

| | | |
|---|---|---|
| APAPAPAPNI | ASSSDRGRTEVF | 0.5991455 |
| PGGPVPIP | ASSSDRGRTEVF | 0.5425353 |
| FLTVALEP | ASSSDRGRTEVF | 0.6805554 |
| SQYFGANL | ASSSDRGRTEVF | 0.6666494 |
| GGVKPAA | ASSSDRGRTEVF | 0.6227591 |
| GGTGGAGGAGGW | ASSSDRGRTEVF | 0.67554224 |
| GVGGVAGLGGAGG | ASSSDRGRTEVF | 0.34458193 |
| VLLVLATTI | ASSSDRGRTEVF | 0.60658926 |
| IGIFAAML | ASSSDRGRTEVF | 0.6703297 |
| AVGDLID | ASSSDRGRTEVF | 0.64528 |
| NISPLPL | ASSSDRGRTEVF | 0.5958493 |
| VTAVFVAI | ASSSDRGRTEVF | 0.66759235 |
| SGDDNDAAL | ASSSDRGRTEVF | 0.5294 |
| AVAALAHS | ASSSDRGRTEVF | 0.59510916 |
| GGVTVMATP | ASSSDRGRTEVF | 0.6217427 |
| AGLAANVRD | ASSSDRGRTEVF | 0.65827703 |
| TNRPDLIDPALLRPG | ASSSDRGRTEVF | 0.6626153 |
| LANLGVTAL | ASSSDRGRTEVF | 0.49660107 |
| TGERIEL | ASSSDRGRTEVF | 0.64679235 |
| NGEELAA | ASSSDRGRTEVF | 0.68574524 |
| VLSLVWG | ASSSDRGRTEVF | 0.5755038 |
| IGYALWL | ASSSDRGRTEVF | 0.6662905 |
| LSLLLAPL | ASSSDRGRTEVF | 0.636754 |
| TLDEALK | ASSSDRGRTEVF | 0.6562139 |
| TDGAGTAEGG | ASSSDRGRTEVF | 0.52989894 |
| NAPGGAGGNGG | ASSSDRGRTEVF | 0.5357684 |
| VSVFMGMGGNF | ASSSDRGRTEVF | 0.61860764 |
| LASYLAAL | ASSSDRGRTEVF | 0.51563746 |
| LVGGTPAL | ASSSDRGRTEVF | 0.62699604 |
| VRNADVL | ASSSDRGRTEVF | 0.6028875 |
| TCSSPFLVSAM | ASSSDRGRTEVF | 0.6905349 |
| VHITDAE | ASSSDRGRTEVF | 0.6360979 |
| EYAAPSF | ASSSDRGRTEVF | 0.6438721 |
| VVDAFHGGL | ASSSDRGRTEVF | 0.5040272 |
| ASVVDPQ | ASSSDRGRTEVF | 0.64677125 |
| SYNEFGGLS | ASSSDRGRTEVF | 0.6335454 |
| DPDGSVY | ASSSDRGRTEVF | 0.5040287 |
| MVSTIHATE | ASSSDRGRTEVF | 0.66688985 |
| LAAGVADPV | ASSSDRGRTEVF | 0.52655053 |
| TKEQIAA | ASSSDRGRTEVF | 0.6010283 |
| SLASPAAL | ASSSDRGRTEVF | 0.65347004 |
| IGLVFYGV | ASSSDRGRTEVF | 0.6673727 |
| FLLDASNL | ASSSDRGRTEVF | 0.58301616 |
| YQVGNFP | ASSSDRGRTEVF | 0.60925674 |
| ESLLSQST | ASSSDRGRTEVF | 0.49833232 |
| LFTVGFAAVG | ASSSDRGRTEVF | 0.6602673 |
| ATPPTPT | ASSSDRGRTEVF | 0.6563401 |
| TDLATLG | ASSSDRGRTEVF | 0.64335907 |
| SAGGTGR | ASSSDRGRTEVF | 0.6842092 |
| GFSNSGS | ASSSDRGRTEVF | 0.66625565 |
| GGNGGSGGTSV | ASSSDRGRTEVF | 0.42227235 |

| | | |
|---|---|---|
| GTSVATGGAGN | ASSSDRGRTEVF | 0.6170075 |
| NGGNGGSGGTS | ASSSDRGRTEVF | 0.66728795 |
| AVVKFPH | ASSSDRGRTEVF | 0.6684685 |
| TPPPPPP | ASSSDRGRTEVF | 0.688033 |
| AAPAAAAAPA | ASSSDRGRTEVF | 0.66537213 |
| IALNEDL | ASSSDRGRTEVF | 0.5697451 |
| IVNVSSM | ASSSDRGRTEVF | 0.6708337 |
| TGIPPAPR | ASSSDRGRTEVF | 0.6947546 |
| FDLGGGTF | ASSSDRGRTEVF | 0.65173614 |
| FDLGGGTFDVS | ASSSDRGRTEVF | 0.569972 |
| NEPTAAA | ASSSDRGRTEVF | 0.55265594 |
| NEPTAAALA | ASSSDRGRTEVF | 0.6409317 |
| GSIINISSI | ASSSDRGRTEVF | 0.65899694 |
| VAPGNAGTA | ASSSDRGRTEVF | 0.5346692 |
| TGGGASLE | ASSSDRGRTEVF | 0.6100501 |
| VAVYNHYKRI | ASSSDRGRTEVF | 0.61942524 |
| TDPAPATT | ASSSDRGRTEVF | 0.43509874 |
| TNRPDILDPALLRPG | ASSSDRGRTEVF | 0.6685638 |
| LTTTTTS | ASSSDRGRTEVF | 0.6700747 |
| PTGSGKTL | ASSSDRGRTEVF | 0.6321023 |
| TGSGKTL | ASSSDRGRTEVF | 0.5692718 |
| TIGHVDHGK | ASSSDRGRTEVF | 0.6597847 |
| TIGHVDHGKTT | ASSSDRGRTEVF | 0.6852434 |
| GHVDHGKTT | ASSSDRGRTEVF | 0.67889553 |
| AQFEHTLL | ASSSDRGRTEVF | 0.4559608 |
| IVNQDEL | ASSSDRGRTEVF | 0.3446606 |
| TNGSQFF | ASSSDRGRTEVF | 0.5643909 |
| TGKGIVE | ASSSDRGRTEVF | 0.6570833 |
| LISNASDA | ASSSDRGRTEVF | 0.60258055 |
| AELRTLGELTKIAKAHGAQ | ASSSDRGRTEVF | 0.5779666 |
| APNAVAAAEPSW | ASSSDRGRTEVF | 0.63897496 |
| APQVAGAATAVL | ASSSDRGRTEVF | 0.66665995 |
| ARRDPAAAAAILPTDA | ASSSDRGRTEVF | 0.5360765 |
| DAAARDVLAEAGLAEYF | ASSSDRGRTEVF | 0.5070139 |
| DDQIVLSAKANLAKVEQ | ASSSDRGRTEVF | 0.6653942 |
| DELLPDTGEDSDEDGH | ASSSDRGRTEVF | 0.6545046 |
| ELPEAPANRATVPQD | ASSSDRGRTEVF | 0.5239007 |
| FGQSKIERVTGIG | ASSSDRGRTEVF | 0.5759993 |
| GAGGTGSTAGGAGGAGGAGGLY | ASSSDRGRTEVF | 0.5903147 |
| GHGGTGAAGGNGGTGGAGGSGI | ASSSDRGRTEVF | 0.60624284 |
| GNGGNGGAGGEGGPGGAGGAGA | ASSSDRGRTEVF | 0.61747336 |
| GSTAGNGGNGGNATNGTIAGQP | ASSSDRGRTEVF | 0.4866036 |
| GTGGDGGNGGNGGTGG | ASSSDRGRTEVF | 0.6649476 |
| GVRRPAGIPADAPV | ASSSDRGRTEVF | 0.49039242 |
| GYRQVTDAHLLTLARR | ASSSDRGRTEVF | 0.6602503 |
| IAKMGQDGHDRG | ASSSDRGRTEVF | 0.62273157 |
| KAEPSIEIQYT | ASSSDRGRTEVF | 0.6494024 |
| KPVRIRMAADEDTVAVT | ASSSDRGRTEVF | 0.57061464 |
| MPPRIAGMRLLVIKPEP | ASSSDRGRTEVF | 0.56746995 |
| MKFYAVALFRSRGVPPTR | ASSSDRGRTEVF | 0.5041673 |
| NAGGFGILGAVAHSP | ASSSDRGRTEVF | 0.397854 |

| | | |
|---|---|---|
| NEGREITLAMAEM | ASSSDRGRTEVF | 0.6656679 |
| NELVEWSHPLGHRWRWE | ASSSDRGRTEVF | 0.425573 |
| NLGAGGTGGNGGNPG | ASSSDRGRTEVF | 0.6649389 |
| NTEWPKWETVMAADPVD | ASSSDRGRTEVF | 0.65023404 |
| PASAAPAETQP | ASSSDRGRTEVF | 0.51702183 |
| QQSSDGKNGVRAAEITGE | ASSSDRGRTEVF | 0.64893866 |
| RAPDAVTTSWWKE | ASSSDRGRTEVF | 0.48874357 |
| SGFGNVGAHTSGWFNQ | ASSSDRGRTEVF | 0.5850959 |
| SGVAGAVGQAASVGGLKV | ASSSDRGRTEVF | 0.6530241 |
| SNGGAGTGSAGGNGGT | ASSSDRGRTEVF | 0.6561502 |
| TGGAGGEGGAAG | ASSSDRGRTEVF | 0.5446351 |
| TGGDGGAAGAGGAPGHG | ASSSDRGRTEVF | 0.64921105 |
| TMPPLLAAIANAGRHGV | ASSSDRGRTEVF | 0.6305711 |
| TSSTAGNSGGA | ASSSDRGRTEVF | 0.5837803 |
| VPATVPVGILC | ASSSDRGRTEVF | 0.634347 |
| VQRPDSGGVVLTGRIS | ASSSDRGRTEVF | 0.65859985 |
| WTPNPVPATKGVPAGLVQAT | ASSSDRGRTEVF | 0.6408534 |
| YALKRALAKFSDRFDVVII | ASSSDRGRTEVF | 0.52096796 |
| YLIGSVIV | ASSSDRGRTEVF | 0.65338373 |
| YGQLVAVID | ASSSDRGRTEVF | 0.52049303 |
| VTVGGMTLPA | ASSSDRGRTEVF | 0.6252934 |
| VLALPAV | ASSSDRGRTEVF | 0.6067928 |
| VIPPTLNYET | ASSSDRGRTEVF | 0.58990586 |
| VGMGGMHPSSG | ASSSDRGRTEVF | 0.66954935 |
| VGLAGGAATA | ASSSDRGRTEVF | 0.5829712 |
| VGGAGGAGGSGP | ASSSDRGRTEVF | 0.5509314 |
| VFAAFMV | ASSSDRGRTEVF | 0.5688364 |
| TLYDKLR | ASSSDRGRTEVF | 0.62359244 |
| TLAVFTVL | ASSSDRGRTEVF | 0.6551208 |
| TGSVQPARV | ASSSDRGRTEVF | 0.50949854 |
| TGGGAGSGGT | ASSSDRGRTEVF | 0.6237248 |
| SHLETSL | ASSSDRGRTEVF | 0.60057414 |
| SFTTGMT | ASSSDRGRTEVF | 0.52117455 |
| QVEVAKRMGVRQPTVS | ASSSDRGRTEVF | 0.62947 |
| QGSGGPALAM | ASSSDRGRTEVF | 0.6252632 |
| MLAPSNP | ASSSDRGRTEVF | 0.5520104 |
| LVNNAAI | ASSSDRGRTEVF | 0.65197265 |
| LTGEINTLA | ASSSDRGRTEVF | 0.56930476 |
| LSIFGPV | ASSSDRGRTEVF | 0.64890206 |
| LLALLANL | ASSSDRGRTEVF | 0.6896665 |
| LISNASDALDKLR | ASSSDRGRTEVF | 0.5308052 |
| LINEAVP | ASSSDRGRTEVF | 0.6608311 |
| LGYAFSGL | ASSSDRGRTEVF | 0.64028907 |
| LDFVAVAL | ASSSDRGRTEVF | 0.6377975 |
| LAQPGAE | ASSSDRGRTEVF | 0.60135335 |
| ISNLGAVTGF | ASSSDRGRTEVF | 0.5744298 |
| HQEDPGRI | ASSSDRGRTEVF | 0.67656916 |
| GTPGEPTP | ASSSDRGRTEVF | 0.6511215 |
| GFFNSGS | ASSSDRGRTEVF | 0.65295243 |
| GAGGGGLTAGAN | ASSSDRGRTEVF | 0.6497544 |
| FVDALVNM | ASSSDRGRTEVF | 0.6534992 |

| | | |
|---|---|---|
| FLTVAPEL | ASSSDRGRTEVF | 0.6487977 |
| FETLTAL | ASSSDRGRTEVF | 0.53577614 |
| DVINPIEL | ASSSDRGRTEVF | 0.4599893 |
| APVSGAPV | ASSSDRGRTEVF | 0.58109313 |
| ALVGGLAW | ASSSDRGRTEVF | 0.5934863 |
| ALAAAAATAI | ASSSDRGRTEVF | 0.61219114 |
| AFGLGLLL | ASSSDRGRTEVF | 0.42684662 |
| AFFAAASL | ASSSDRGRTEVF | 0.6550212 |
| AAAALPAFSPP | ASSSDRGRTEVF | 0.6871753 |
| VVLAPAL | ASSSDRGRTEVF | 0.5719373 |
| TIGHVDHGKTTL | ASSSDRGRTEVF | 0.6889914 |
| LISNASDALDK | ASSSDRGRTEVF | 0.5083143 |
| LISNASDAL | ASSSDRGRTEVF | 0.6471813 |
| ISNASDALDK | ASSSDRGRTEVF | 0.6648817 |
| NQAETLVYQTE | ASSSDRGRTEVF | 0.50681037 |
| TGGTGGTAGAG | ASSSDRGRTEVF | 0.65189916 |
| TGSTAGGAGGA | ASSSDRGRTEVF | 0.58602107 |
| DPLAVPVAAGPV | ASSSDRGRTEVF | 0.5981908 |
| GGTGGTPGNGGN | ASSSDRGRTEVF | 0.663476 |
| PAGLAGVDPATL | ASSSDRGRTEVF | 0.5912219 |
| GNNLVVEAPHAGASV | ASSSDRGRTEVF | 0.64615184 |
| KVTRIMSASTLEQPA | ASSSDRGRTEVF | 0.6051771 |
| LGFALPIALPRRTVV | ASSSDRGRTEVF | 0.52086186 |
| TRLGNLTARKLVVLV | ASSSDRGRTEVF | 0.6352909 |
| WSQQRRLIYFPSAGP | ASSSDRGRTEVF | 0.6629338 |
| GPNGNPDPMAAAVDIR | ASSSDRGRTEVF | 0.5768715 |
| TQRCVSVADLRAAASR | ASSSDRGRTEVF | 0.6934738 |
| QFVLDVAEHPVEVLTETV | ASSSDRGRTEVF | 0.5434571 |
| GGAGGTGGAGGNGGTQAGDG | ASSSDRGRTEVF | 0.6863905 |
| PLLATLRAIAVDVDAKPAQV | ASSSDRGRTEVF | 0.65394133 |
| APKFRTGGYMIDFWGVGYQVAK | ASSSDRGRTEVF | 0.5990012 |
| GAGGAGGAGGAAGGPGGTGGTG | ASSSDRGRTEVF | 0.5808897 |
| IGALGLLATDVAPAGSVAFGVA | ASSSDRGRTEVF | 0.6569591 |
| AGGAGGIGGNGGAGGAGGVLMG | ASSSDRGRTEVF | 0.58924514 |
| ADELIGLL | ASSSDRGRTEVF | 0.64980835 |
| ADRALKA | ASSSDRGRTEVF | 0.68209577 |
| AGAVGQVGGAGGA | ASSSDRGRTEVF | 0.71088487 |
| AGGGGGLGGAP | ASSSDRGRTEVF | 0.85116553 |
| AGGLLYGNGGAG | ASSSDRGRTEVF | 0.7640895 |
| AGLESIGL | ASSSDRGRTEVF | 0.79602915 |
| AGLKTGII | ASSSDRGRTEVF | 0.7209232 |
| ASLAALLL | ASSSDRGRTEVF | 0.53073263 |
| ASVINISSI | ASSSDRGRTEVF | 0.80220366 |
| ATPPGPA | ASSSDRGRTEVF | 0.65569735 |
| ATVASALV | ASSSDRGRTEVF | 0.80858564 |
| ATVASLIL | ASSSDRGRTEVF | 0.83658373 |
| AVNGPVLIRH | ASSSDRGRTEVF | 0.7266634 |
| DGAAGAVGI | ASSSDRGRTEVF | 0.8484999 |
| DPLLHGI | ASSSDRGRTEVF | 0.8567077 |
| ELGAALGAL | ASSSDRGRTEVF | 0.56367326 |
| ELGRTQAV | ASSSDRGRTEVF | 0.7157151 |

| | | |
|---|---|---|
| FVAGGYLS | ASSSDRGRTEVF | 0.74835443 |
| GASGGAGGAGGW | ASSSDRGRTEVF | 0.7657397 |
| GGAGAGGDGGAGGI | ASSSDRGRTEVF | 0.5068422 |
| GTGHNGSQPGMGGNG | ASSSDRGRTEVF | 0.828771 |
| GYAAASASAS | ASSSDRGRTEVF | 0.66351855 |
| HQVDVAAVQQG | ASSSDRGRTEVF | 0.22449356 |
| IGAAIAHT | ASSSDRGRTEVF | 0.79004616 |
| IGMIEVAQL | ASSSDRGRTEVF | 0.7751025 |
| IRLAPEHTSAMD | ASSSDRGRTEVF | 0.7487912 |
| IYNVSVL | ASSSDRGRTEVF | 0.83811224 |
| KQGFLGM | ASSSDRGRTEVF | 0.58411205 |
| LESVITQL | ASSSDRGRTEVF | 0.703037 |
| LGLGALVVV | ASSSDRGRTEVF | 0.8053212 |
| LIGNGGNGGSGGI | ASSSDRGRTEVF | 0.8546006 |
| LNAINAPTL | ASSSDRGRTEVF | 0.6395896 |
| LQLLNTGV | ASSSDRGRTEVF | 0.7747284 |
| LQTSIEF | ASSSDRGRTEVF | 0.8517686 |
| MSFAVSEL | ASSSDRGRTEVF | 0.8143011 |
| NTGALGL | ASSSDRGRTEVF | 0.8529693 |
| PPGAGGAGGAGTT | ASSSDRGRTEVF | 0.8626301 |
| QLVGLGI | ASSSDRGRTEVF | 0.65922743 |
| RFLEQVHQL | ASSSDRGRTEVF | 0.8371469 |
| RGVGYVLR | ASSSDRGRTEVF | 0.8620896 |
| SATGVFAGVI | ASSSDRGRTEVF | 0.76311535 |
| SLPAITLP | ASSSDRGRTEVF | 0.786503 |
| STALAFTV | ASSSDRGRTEVF | 0.7022412 |
| SVLTVGLGK | ASSSDRGRTEVF | 0.727966 |
| TALAGAGSGI | ASSSDRGRTEVF | 0.36267334 |
| TGLPFTGI | ASSSDRGRTEVF | 0.7849234 |
| VAEHVTV | ASSSDRGRTEVF | 0.8464055 |
| VDIVLNSL | ASSSDRGRTEVF | 0.7380766 |
| VLTGGGAVL | ASSSDRGRTEVF | 0.5895316 |
| VNVGLAYGI | ASSSDRGRTEVF | 0.5962655 |
| VSSAAVGAV | ASSSDRGRTEVF | 0.7303529 |
| VTSVYEAGV | ASSSDRGRTEVF | 0.73052204 |
| WVALDRGA | ASSSDRGRTEVF | 0.43862036 |
| YLVGSVVI | ASSSDRGRTEVF | 0.6882921 |
| GPPGTGKTLL | ASSSDRGRTEVF | 0.8523206 |
| GVGMTKF | ASSSDRGRTEVF | 0.85224074 |
| IGIGGSDL | ASSSDRGRTEVF | 0.47476435 |
| IGTLLAEL | ASSSDRGRTEVF | 0.85392755 |
| IYTSGTT | ASSSDRGRTEVF | 0.79555744 |
| LSFPAGV | ASSSDRGRTEVF | 0.8521006 |
| LYTSGTT | ASSSDRGRTEVF | 0.7994042 |
| VGAFPFGF | ASSSDRGRTEVF | 0.6929435 |
| VGAGTVEF | ASSSDRGRTEVF | 0.82167315 |
| VGKSTLLL | ASSSDRGRTEVF | 0.70138645 |
| AALAHAGGGIV | ASSSDRGRTEVF | 0.7151115 |
| AALVAVSLL | ASSSDRGRTEVF | 0.84413505 |
| AAVSRAVAAL | ASSSDRGRTEVF | 0.7496877 |
| AGGSSGIGGF | ASSSDRGRTEVF | 0.69583315 |

| | | |
|---|---|---|
| AGLTVANANG | ASSSDRGRTEVF | 0.64552504 |
| AGLVGAVL | ASSSDRGRTEVF | 0.8471292 |
| ALSQAFTV | ASSSDRGRTEVF | 0.56852263 |
| ALTFSVSL | ASSSDRGRTEVF | 0.6628038 |
| ALTGAALVAL | ASSSDRGRTEVF | 0.5961678 |
| ASFSIGAGVL | ASSSDRGRTEVF | 0.7652132 |
| DGSVYKI | ASSSDRGRTEVF | 0.847731 |
| EQLAEIV | ASSSDRGRTEVF | 0.8276774 |
| FAVSELTVM | ASSSDRGRTEVF | 0.71780443 |
| GAAPLAAGL | ASSSDRGRTEVF | 0.80094737 |
| GADGGDGGSGGAGG | ASSSDRGRTEVF | 0.8287095 |
| GFGLDTGGAGG | ASSSDRGRTEVF | 0.80316085 |
| GLASGGLL | ASSSDRGRTEVF | 0.82242703 |
| GRIAGAAVLL | ASSSDRGRTEVF | 0.6578468 |
| GSAGSLNTSF | ASSSDRGRTEVF | 0.84534943 |
| GTGQIGI | ASSSDRGRTEVF | 0.8232868 |
| GTLIALIL | ASSSDRGRTEVF | 0.84827423 |
| IAATMGFVVG | ASSSDRGRTEVF | 0.8341005 |
| IFAQALL | ASSSDRGRTEVF | 0.7894625 |
| IGAAVALGF | ASSSDRGRTEVF | 0.7709409 |
| IGLTPLAL | ASSSDRGRTEVF | 0.39788067 |
| ILGSNTSTL | ASSSDRGRTEVF | 0.71753526 |
| IVFSLTVL | ASSSDRGRTEVF | 0.7718048 |
| LASKATGF | ASSSDRGRTEVF | 0.77135324 |
| LASVALIL | ASSSDRGRTEVF | 0.7978299 |
| LFAVGLTL | ASSSDRGRTEVF | 0.83879507 |
| LFNGGIGTYI | ASSSDRGRTEVF | 0.6653308 |
| LTLGAGII | ASSSDRGRTEVF | 0.8310554 |
| LVAAALGDI | ASSSDRGRTEVF | 0.82539606 |
| NTAFVQLGI | ASSSDRGRTEVF | 0.5412115 |
| QLFSGAEL | ASSSDRGRTEVF | 0.7290299 |
| SAATFITL | ASSSDRGRTEVF | 0.86630005 |
| SNYGLANLGV | ASSSDRGRTEVF | 0.7800848 |
| SYGGAYAVM | ASSSDRGRTEVF | 0.74544287 |
| TTGVGGLTLG | ASSSDRGRTEVF | 0.74159586 |
| TTQTIYI | ASSSDRGRTEVF | 0.8569512 |
| VAAGAGLGAVI | ASSSDRGRTEVF | 0.6958872 |
| VAAVVTLL | ASSSDRGRTEVF | 0.8451444 |
| VGFLGGLI | ASSSDRGRTEVF | 0.29178643 |
| VGVIPILL | ASSSDRGRTEVF | 0.7145158 |
| VIFASSLI | ASSSDRGRTEVF | 0.6817266 |
| VKGGDGGAAAG | ASSSDRGRTEVF | 0.42873576 |
| VTKTFTNI | ASSSDRGRTEVF | 0.8466252 |
| VVAAAILL | ASSSDRGRTEVF | 0.75434166 |
| YAALDVEL | ASSSDRGRTEVF | 0.8175698 |
| LASVALLL | ASSSDRGRTEVF | 0.7750343 |
| LDAVHRV | ASSSDRGRTEVF | 0.8470454 |
| QGLSSSIF | ASSSDRGRTEVF | 0.76615965 |
| RGTFANIRL | ASSSDRGRTEVF | 0.7923128 |
| RQLWWGHRI | ASSSDRGRTEVF | 0.72197545 |
| YGPPGTGKTLL | ASSSDRGRTEVF | 0.8308051 |

| | | |
|---|---|---|
| AALVGAPANVA | ASSSDRGRTEVF | 0.838055 |
| AATFRRTE | ASSSDRGRTEVF | 0.8303672 |
| AGALGGGI | ASSSDRGRTEVF | 0.55431145 |
| AGGAGGAGGIGAG | ASSSDRGRTEVF | 0.6747928 |
| AGSGALGL | ASSSDRGRTEVF | 0.7998568 |
| AQIADQI | ASSSDRGRTEVF | 0.43861356 |
| ASAVAVTYL | ASSSDRGRTEVF | 0.32943162 |
| ASTVLGGTGG | ASSSDRGRTEVF | 0.62916386 |
| AVGGEPAK | ASSSDRGRTEVF | 0.65333396 |
| CEQGTFGGTC | ASSSDRGRTEVF | 0.83253884 |
| DEGKAPAG | ASSSDRGRTEVF | 0.8335203 |
| DTVVFPAL | ASSSDRGRTEVF | 0.7507311 |
| FEGITMAG | ASSSDRGRTEVF | 0.8151689 |
| FPLVVQD | ASSSDRGRTEVF | 0.7790747 |
| GAFTTPQ | ASSSDRGRTEVF | 0.12452464 |
| GFGGFGTAGGVGGA | ASSSDRGRTEVF | 0.74933887 |
| GFLAQFL | ASSSDRGRTEVF | 0.5700263 |
| GGAGGAGGNAGVA | ASSSDRGRTEVF | 0.8630257 |
| GGVGTAGAGGAGGAGG | ASSSDRGRTEVF | 0.66601956 |
| GNGGGGDAGNAGSGGN | ASSSDRGRTEVF | 0.66496986 |
| GTGGAGGAGNDAGS | ASSSDRGRTEVF | 0.77651614 |
| GTTNTGWFNS | ASSSDRGRTEVF | 0.75265557 |
| IFMGPLFTL | ASSSDRGRTEVF | 0.7605866 |
| ILAHAIEAI | ASSSDRGRTEVF | 0.62771773 |
| IRSAIREAL | ASSSDRGRTEVF | 0.237461 |
| ITFAGGG | ASSSDRGRTEVF | 0.84675926 |
| ITGMSGHQEV | ASSSDRGRTEVF | 0.5619882 |
| IVAGLACTL | ASSSDRGRTEVF | 0.6507313 |
| IVGGGLAGA | ASSSDRGRTEVF | 0.42129657 |
| IVTGVLNI | ASSSDRGRTEVF | 0.58573633 |
| LAQQGCI | ASSSDRGRTEVF | 0.5548129 |
| LCLHVEN | ASSSDRGRTEVF | 0.8533032 |
| LDAISDGS | ASSSDRGRTEVF | 0.686197 |
| LGGFAAML | ASSSDRGRTEVF | 0.71782815 |
| LGGKNPMI | ASSSDRGRTEVF | 0.8498811 |
| LGPKGVAAIL | ASSSDRGRTEVF | 0.7402866 |
| LMVGVSI | ASSSDRGRTEVF | 0.72921324 |
| LVGAAFVL | ASSSDRGRTEVF | 0.7036503 |
| LVLAVAPL | ASSSDRGRTEVF | 0.66848195 |
| MAAIAVIDAA | ASSSDRGRTEVF | 0.694617 |
| NNGINAELQ | ASSSDRGRTEVF | 0.671253 |
| RGEPLVLVHG | ASSSDRGRTEVF | 0.8636147 |
| RGGAGGAAGLI | ASSSDRGRTEVF | 0.749735 |
| RSYRMHPAV | ASSSDRGRTEVF | 0.66463894 |
| SAAHLLAGQLA | ASSSDRGRTEVF | 0.80781955 |
| SAAVGSY | ASSSDRGRTEVF | 0.85166264 |
| SPHAVLHRI | ASSSDRGRTEVF | 0.8696399 |
| SQIAKVL | ASSSDRGRTEVF | 0.66472065 |
| TALGHLVP | ASSSDRGRTEVF | 0.7589185 |
| TAPTTSGAA | ASSSDRGRTEVF | 0.74360436 |
| TDGDDGNGGSPGT | ASSSDRGRTEVF | 0.75192505 |

| | | |
|---|---|---|
| TDGLGEN | ASSSDRGRTEVF | 0.7350573 |
| TGGPPLPGQA | ASSSDRGRTEVF | 0.619279 |
| TIPAITI | ASSSDRGRTEVF | 0.826379 |
| TIPGFTI | ASSSDRGRTEVF | 0.86434805 |
| TSIASGALA | ASSSDRGRTEVF | 0.7555499 |
| TVTLPGAAI | ASSSDRGRTEVF | 0.65871155 |
| VASGLSAALP | ASSSDRGRTEVF | 0.73944724 |
| VGAGVIGI | ASSSDRGRTEVF | 0.6251314 |
| VGPKRFGV | ASSSDRGRTEVF | 0.7085227 |
| VNVAFPD | ASSSDRGRTEVF | 0.79193366 |
| VSMTNSGVS | ASSSDRGRTEVF | 0.5374869 |
| VTDVLGDLSV | ASSSDRGRTEVF | 0.7331987 |
| VVVVGAGFAGL | ASSSDRGRTEVF | 0.8660872 |
| VYTVGTVV | ASSSDRGRTEVF | 0.6907403 |
| WTTFSGP | ASSSDRGRTEVF | 0.77274597 |
| YDELVEAF | ASSSDRGRTEVF | 0.83562297 |
| YGTGGAGGNGG | ASSSDRGRTEVF | 0.8024239 |
| YTNVMAV | ASSSDRGRTEVF | 0.85215634 |
| FLEQVHQL | ASSSDRGRTEVF | 0.808303 |
| GLSLLAVAV | ASSSDRGRTEVF | 0.8050438 |
| LEQVHQL | ASSSDRGRTEVF | 0.76902735 |
| LNAAGAI | ASSSDRGRTEVF | 0.7536321 |
| PAVHASL | ASSSDRGRTEVF | 0.76664805 |
| TELEDDA | ASSSDRGRTEVF | 0.755615 |
| TPPGPPN | ASSSDRGRTEVF | 0.6934909 |
| TPPPPPPP | ASSSDRGRTEVF | 0.8291441 |
| TQSPALI | ASSRDRGNYAEQF | 0.7907738 |
| EFAAAFPQ | ASSRDRGNYAEQF | 0.82905996 |
| LLSGNLDVL | ASSRDRGNYAEQF | 0.5609673 |
| GPANLGNYNI | ASSRDRGNYAEQF | 0.64996904 |
| TLGPTLP | ASSRDRGNYAEQF | 0.82577634 |
| VYAYSSCCK | ASSRDRGNYAEQF | 0.2222194 |
| APAPAPAPNI | ASSRDRGNYAEQF | 0.36269385 |
| PGGPVPIP | ASSRDRGNYAEQF | 0.76721823 |
| FLTVALEP | ASSRDRGNYAEQF | 0.8477539 |
| SQYFGANL | ASSRDRGNYAEQF | 0.23679908 |
| GGVKPAA | ASSRDRGNYAEQF | 0.68391156 |
| GGTGGAGGAGGW | ASSRDRGNYAEQF | 0.5542146 |
| GVGGVAGLGGAGG | ASSRDRGNYAEQF | 0.8348211 |
| VLLVLATTI | ASSRDRGNYAEQF | 0.7883733 |
| IGIFAAML | ASSRDRGNYAEQF | 0.51647717 |
| AVGDLID | ASSRDRGNYAEQF | 0.82813454 |
| NISPLPL | ASSRDRGNYAEQF | 0.54637265 |
| VTAVFVAI | ASSRDRGNYAEQF | 0.59073794 |
| SGDDNDAAL | ASSRDRGNYAEQF | 0.8440349 |
| AVAALAHS | ASSRDRGNYAEQF | 0.8422306 |
| GGVTVMATP | ASSRDRGNYAEQF | 0.6418484 |
| AGLAANVRD | ASSRDRGNYAEQF | 0.8390674 |
| TNRPDLIDPALLRPG | ASSRDRGNYAEQF | 0.6706364 |
| LANLGVTAL | ASSRDRGNYAEQF | 0.8678562 |
| TGERIEL | ASSRDRGNYAEQF | 0.7382505 |

| | | |
|---|---|---|
| NGEELAA | ASSRDRGNYAEQF | 0.5121282 |
| VLSLVWG | ASSRDRGNYAEQF | 0.82240784 |
| IGYALWL | ASSRDRGNYAEQF | 0.79504067 |
| LSLLLAPL | ASSRDRGNYAEQF | 0.81506646 |
| TLDEALK | ASSRDRGNYAEQF | 0.80567116 |
| TDGAGTAEGG | ASSRDRGNYAEQF | 0.7238516 |
| NAPGGAGGNGG | ASSRDRGNYAEQF | 0.725584 |
| VSVFMGMGGNF | ASSRDRGNYAEQF | 0.84560525 |
| LASYLAAL | ASSRDRGNYAEQF | 0.8279994 |
| LVGGTPAL | ASSRDRGNYAEQF | 0.6168767 |
| VRNADVL | ASSRDRGNYAEQF | 0.8461334 |
| TCSSPFLVSAM | ASSRDRGNYAEQF | 0.757641 |
| VHITDAE | ASSRDRGNYAEQF | 0.70769036 |
| EYAAPSF | ASSRDRGNYAEQF | 0.86182135 |
| VVDAFHGGL | ASSRDRGNYAEQF | 0.80048066 |
| ASVVDPQ | ASSRDRGNYAEQF | 0.8222093 |
| SYNEFGGLS | ASSRDRGNYAEQF | 0.83335495 |
| DPDGSVY | ASSRDRGNYAEQF | 0.84366894 |
| MVSTIHATE | ASSRDRGNYAEQF | 0.57073826 |
| LAAGVADPV | ASSRDRGNYAEQF | 0.8012088 |
| TKEQIAA | ASSRDRGNYAEQF | 0.75597423 |
| SLASPAAL | ASSRDRGNYAEQF | 0.8361496 |
| IGLVFYGV | ASSRDRGNYAEQF | 0.7744694 |
| FLLDASNL | ASSRDRGNYAEQF | 0.6183867 |
| YQVGNFP | ASSRDRGNYAEQF | 0.7986487 |
| ESLLSQST | ASSRDRGNYAEQF | 0.8482002 |
| LFTVGFAAVG | ASSRDRGNYAEQF | 0.79412454 |
| ATPPTPT | ASSRDRGNYAEQF | 0.79778343 |
| TDLATLG | ASSRDRGNYAEQF | 0.8592596 |
| SAGGTGR | ASSRDRGNYAEQF | 0.69026273 |
| GFSNSGS | ASSRDRGNYAEQF | 0.771492 |
| GGNGGSGGTSV | ASSRDRGNYAEQF | 0.8279836 |
| GTSVATGGAGN | ASSRDRGNYAEQF | 0.8355272 |
| NGGNGGSGGTS | ASSRDRGNYAEQF | 0.7780391 |
| AVVKFPH | ASSRDRGNYAEQF | 0.8446546 |
| TPPPPPP | ASSRDRGNYAEQF | 0.67310566 |
| AAPAAAAAPA | ASSRDRGNYAEQF | 0.805435 |
| IALNEDL | ASSRDRGNYAEQF | 0.58806163 |
| IVNVSSM | ASSRDRGNYAEQF | 0.8033092 |
| TGIPPAPR | ASSRDRGNYAEQF | 0.59065795 |
| FDLGGGTF | ASSRDRGNYAEQF | 0.80123764 |
| FDLGGGTFDVS | ASSRDRGNYAEQF | 0.8309546 |
| NEPTAAA | ASSRDRGNYAEQF | 0.82391596 |
| NEPTAAALA | ASSRDRGNYAEQF | 0.35769442 |
| GSIINISSI | ASSRDRGNYAEQF | 0.83876646 |
| VAPGNAGTA | ASSRDRGNYAEQF | 0.8199959 |
| TGGGASLE | ASSRDRGNYAEQF | 0.8669575 |
| VAVYNHYKRI | ASSRDRGNYAEQF | 0.76673084 |
| TDPAPATT | ASSRDRGNYAEQF | 0.8184911 |
| TNRPDILDPALLRPG | ASSRDRGNYAEQF | 0.72954786 |
| LTTTTS | ASSRDRGNYAEQF | 0.7785637 |

| | | |
|---|---|---|
| PTGSGKTL | ASSRDRGNYAEQF | 0.5763019 |
| TGSGKTL | ASSRDRGNYAEQF | 0.69994783 |
| TIGHVDHGK | ASSRDRGNYAEQF | 0.69663054 |
| TIGHVDHGKTT | ASSRDRGNYAEQF | 0.6523516 |
| GHVDHGKTT | ASSRDRGNYAEQF | 0.828843 |
| AQFEHTLL | ASSRDRGNYAEQF | 0.8356924 |
| IVNQDEL | ASSRDRGNYAEQF | 0.8325317 |
| TNGSQFF | ASSRDRGNYAEQF | 0.83060837 |
| TGKGIVE | ASSRDRGNYAEQF | 0.836564 |
| LISNASDA | ASSRDRGNYAEQF | 0.85221756 |
| AELRTLGELTKIAKAHGAQ | ASSRDRGNYAEQF | 0.70436 |
| APNAVAAAEPSW | ASSRDRGNYAEQF | 0.7523698 |
| APQVAGAATAVL | ASSRDRGNYAEQF | 0.6196172 |
| ARRDPAAAAAILPTDA | ASSRDRGNYAEQF | 0.7388903 |
| DAAARDVLAEAGLAEYF | ASSRDRGNYAEQF | 0.70927227 |
| DDQIVLSAKANLAKVEQ | ASSRDRGNYAEQF | 0.8044089 |
| DELLPDTGEDSDEDGH | ASSRDRGNYAEQF | 0.20538577 |
| ELPEAPANRATVPQD | ASSRDRGNYAEQF | 0.7310185 |
| FGQSKIERVTGIG | ASSRDRGNYAEQF | 0.86056006 |
| GAGGTGSTAGGAGGAGGAGGLY | ASSRDRGNYAEQF | 0.60550857 |
| GHGGTGAAGGNGGTGGAGGSGI | ASSRDRGNYAEQF | 0.8094488 |
| GNGGNGGAGGEGGPGGAGGAGA | ASSRDRGNYAEQF | 0.7470459 |
| GSTAGNGGNGGNATNGTIAGQP | ASSRDRGNYAEQF | 0.6707472 |
| GTGGDGGNGGNGGTGG | ASSRDRGNYAEQF | 0.8161541 |
| GVRRPAGIPADAPV | ASSRDRGNYAEQF | 0.6495573 |
| GYRQVTDAHLLTLARR | ASSRDRGNYAEQF | 0.82134944 |
| IAKMGQDGHDRG | ASSRDRGNYAEQF | 0.8593843 |
| KAEPSIEIQYT | ASSRDRGNYAEQF | 0.77576077 |
| KPVRIRMAADEDTVAVT | ASSRDRGNYAEQF | 0.7380712 |
| MPPRIAGMRLLVIKPEP | ASSRDRGNYAEQF | 0.7792212 |
| MKFYAVALFRSRGVPPTR | ASSRDRGNYAEQF | 0.55404806 |
| NAGGFGILGAVAHSP | ASSRDRGNYAEQF | 0.7576937 |
| NEGREITLAMAEM | ASSRDRGNYAEQF | 0.26821238 |
| NELVEWSHPLGHRWRWE | ASSRDRGNYAEQF | 0.74040824 |
| NLGAGGTGGNGGNPG | ASSRDRGNYAEQF | 0.6431092 |
| NTEWPKWETVMAADPVD | ASSRDRGNYAEQF | 0.78514284 |
| PASAAPAETQP | ASSRDRGNYAEQF | 0.7839803 |
| QQSSDGKNGVRAAEITGE | ASSRDRGNYAEQF | 0.8092609 |
| RAPDAVTTSWWKE | ASSRDRGNYAEQF | 0.6202173 |
| SGFGNVGAHTSGWFNQ | ASSRDRGNYAEQF | 0.75489944 |
| SGVAGAVGQAASVGGLKV | ASSRDRGNYAEQF | 0.6872593 |
| SNGGAGTGSAGGNGGT | ASSRDRGNYAEQF | 0.58393556 |
| TGGAGGEGGAAG | ASSRDRGNYAEQF | 0.8239347 |
| TGGDGGAAGAGGAPGHG | ASSRDRGNYAEQF | 0.7690762 |
| TMPPLLAAIANAGRHGV | ASSRDRGNYAEQF | 0.72399276 |
| TSSTAGNSGGA | ASSRDRGNYAEQF | 0.8468275 |
| VPATVPVGILC | ASSRDRGNYAEQF | 0.806072 |
| VQRPDSGGVVLTGRIS | ASSRDRGNYAEQF | 0.6093214 |
| WTPNPVPATKGVPAGLVQAT | ASSRDRGNYAEQF | 0.707065 |
| YALKRALAKFSDRFDVVII | ASSRDRGNYAEQF | 0.7391885 |
| YLIGSVIV | ASSRDRGNYAEQF | 0.73957366 |

| | | |
|---|---|---|
| YGQLVAVID | ASSRDRGNYAEQF | 0.24481612 |
| VTVGGMTLPA | ASSRDRGNYAEQF | 0.8042772 |
| VLALPAV | ASSRDRGNYAEQF | 0.7101543 |
| VIPPTLNYET | ASSRDRGNYAEQF | 0.6108991 |
| VGMGGMHPSSG | ASSRDRGNYAEQF | 0.72253245 |
| VGLAGGAATA | ASSRDRGNYAEQF | 0.82772267 |
| VGGAGGAGGSGP | ASSRDRGNYAEQF | 0.69417846 |
| VFAAFMV | ASSRDRGNYAEQF | 0.77272284 |
| TLYDKLR | ASSRDRGNYAEQF | 0.8136352 |
| TLAVFTVL | ASSRDRGNYAEQF | 0.8406264 |
| TGSVQPARV | ASSRDRGNYAEQF | 0.8527211 |
| TGGGAGSGGT | ASSRDRGNYAEQF | 0.81525666 |
| SHLETSL | ASSRDRGNYAEQF | 0.7569956 |
| SFTTGMT | ASSRDRGNYAEQF | 0.8234954 |
| QVEVAKRMGVRQPTVS | ASSRDRGNYAEQF | 0.81594115 |
| QGSGGPALAM | ASSRDRGNYAEQF | 0.47857457 |
| MLAPSNP | ASSRDRGNYAEQF | 0.43137282 |
| LVNNAAI | ASSRDRGNYAEQF | 0.7822948 |
| LTGEINTLA | ASSRDRGNYAEQF | 0.44933277 |
| LSIFGPV | ASSRDRGNYAEQF | 0.7938715 |
| LLALLANL | ASSRDRGNYAEQF | 0.8606379 |
| LISNASDALDKLR | ASSRDRGNYAEQF | 0.7502184 |
| LINEAVP | ASSRDRGNYAEQF | 0.5996464 |
| LGYAFSGL | ASSRDRGNYAEQF | 0.70321983 |
| LDFVAVAL | ASSRDRGNYAEQF | 0.7429676 |
| LAQPGAE | ASSRDRGNYAEQF | 0.82811534 |
| ISNLGAVTGF | ASSRDRGNYAEQF | 0.81820536 |
| HQEDPGRI | ASSRDRGNYAEQF | 0.8504437 |
| GTPGEPTP | ASSRDRGNYAEQF | 0.8409282 |
| GFFNSGS | ASSRDRGNYAEQF | 0.69728893 |
| GAGGGGLTAGAN | ASSRDRGNYAEQF | 0.8183168 |
| FVDALVNM | ASSRDRGNYAEQF | 0.72982776 |
| FLTVAPEL | ASSRDRGNYAEQF | 0.74114513 |
| FETLTAL | ASSRDRGNYAEQF | 0.768268 |
| DVINPIEL | ASSRDRGNYAEQF | 0.78726184 |
| APVSGAPV | ASSRDRGNYAEQF | 0.7476326 |
| ALVGGLAW | ASSRDRGNYAEQF | 0.7165716 |
| ALAAAAATAI | ASSRDRGNYAEQF | 0.5889931 |
| AFGLGLLL | ASSRDRGNYAEQF | 0.80959904 |
| AFFAAASL | ASSRDRGNYAEQF | 0.7598274 |
| AAAALPAFSPP | ASSRDRGNYAEQF | 0.492447 |
| VVLAPAL | ASSRDRGNYAEQF | 0.6957462 |
| TIGHVDHGKTTL | ASSRDRGNYAEQF | 0.8625554 |
| LISNASDALDK | ASSRDRGNYAEQF | 0.85289097 |
| LISNASDAL | ASSRDRGNYAEQF | 0.6126059 |
| ISNASDALDK | ASSRDRGNYAEQF | 0.6770592 |
| NQAETLVYQTE | ASSRDRGNYAEQF | 0.7398147 |
| TGGTGGTAGAG | ASSRDRGNYAEQF | 0.4503444 |
| TGSTAGGAGGA | ASSRDRGNYAEQF | 0.81452703 |
| DPLAVPVAAGPV | ASSRDRGNYAEQF | 0.7502143 |
| GGTGGTPGNGGN | ASSRDRGNYAEQF | 0.74730575 |

| | | |
|---|---|---|
| PAGLAGVDPATL | ASSRDRGNYAEQF | 0.67853767 |
| GNNLVVEAPHAGASV | ASSRDRGNYAEQF | 0.7574184 |
| KVTRIMSASTLEQPA | ASSRDRGNYAEQF | 0.7045672 |
| LGFALPIALPRRTVV | ASSRDRGNYAEQF | 0.8389257 |
| TRLGNLTARKLVVLV | ASSRDRGNYAEQF | 0.22338673 |
| WSQQRRLIYFPSAGP | ASSRDRGNYAEQF | 0.82867783 |
| GPNGNPDPMAAAVDIR | ASSRDRGNYAEQF | 0.63466734 |
| TQRCVSVADLRAAASR | ASSRDRGNYAEQF | 0.73006475 |
| QFVLDVAEHPVEVLTETV | ASSRDRGNYAEQF | 0.4603855 |
| GGAGGTGGAGGNGGTQAGDG | ASSRDRGNYAEQF | 0.7992382 |
| PLLATLRAIAVDVDAKPAQV | ASSRDRGNYAEQF | 0.8252268 |
| APKFRTGGYMIDFWGVGYQVAK | ASSRDRGNYAEQF | 0.8318004 |
| GAGGAGGAGGAAGGPGGTGGTG | ASSRDRGNYAEQF | 0.59399414 |
| IGALGLLATDVAPAGSVAFGVA | ASSRDRGNYAEQF | 0.82266265 |
| AGGAGGIGGNGGAGGAGGVLMG | ASSRDRGNYAEQF | 0.8574572 |
| ADELIGLL | ASSRDRGNYAEQF | 0.7189296 |
| ADRALKA | ASSRDRGNYAEQF | 0.48868585 |
| AGAVGQVGGAGGA | ASSRDRGNYAEQF | 0.8222425 |
| AGGGGGLGGAP | ASSRDRGNYAEQF | 0.7506201 |
| AGGLLYGNGGAG | ASSRDRGNYAEQF | 0.83369577 |
| AGLESIGL | ASSRDRGNYAEQF | 0.7995036 |
| AGLKTGII | ASSRDRGNYAEQF | 0.5666989 |
| ASLAALLL | ASSRDRGNYAEQF | 0.8460493 |
| ASVINISSI | ASSRDRGNYAEQF | 0.8281474 |
| ATPPGPA | ASSRDRGNYAEQF | 0.6094038 |
| ATVASALV | ASSRDRGNYAEQF | 0.7423677 |
| ATVASLIL | ASSRDRGNYAEQF | 0.7699264 |
| AVNGPVLIRH | ASSRDRGNYAEQF | 0.38492954 |
| DGAAGAVGI | ASSRDRGNYAEQF | 0.8300764 |
| DPLLHGI | ASSRDRGNYAEQF | 0.6625442 |
| ELGAALGAL | ASSRDRGNYAEQF | 0.7236343 |
| ELGRTQAV | ASSRDRGNYAEQF | 0.78360474 |
| FVAGGYLS | ASSRDRGNYAEQF | 0.7132706 |
| GASGGAGGAGGW | ASSRDRGNYAEQF | 0.7518686 |
| GGAGAGGDGGAGGI | ASSRDRGNYAEQF | 0.7476326 |
| GTGHNGSQPGMGGNG | ASSRDRGNYAEQF | 0.8039892 |
| GYAAASASAS | ASSRDRGNYAEQF | 0.78002065 |
| HQVDVAAVQQG | ASSRDRGNYAEQF | 0.7785786 |
| IGAAIAHT | ASSRDRGNYAEQF | 0.80425024 |
| IGMIEVAQL | ASSRDRGNYAEQF | 0.85210097 |
| IRLAPEHTSAMD | ASSRDRGNYAEQF | 0.57784873 |
| IYNVSVL | ASSRDRGNYAEQF | 0.68992424 |
| KQGFLGM | ASSRDRGNYAEQF | 0.8107817 |
| LESVITQL | ASSRDRGNYAEQF | 0.5271273 |
| LGLGALVVV | ASSRDRGNYAEQF | 0.8191671 |
| LIGNGGNGGSGGI | ASSRDRGNYAEQF | 0.77215385 |
| LNAINAPTL | ASSRDRGNYAEQF | 0.8010974 |
| LQLLNTGV | ASSRDRGNYAEQF | 0.75663275 |
| LQTSIEF | ASSRDRGNYAEQF | 0.86183274 |
| MSFAVSEL | ASSRDRGNYAEQF | 0.832425 |
| NTGALGL | ASSRDRGNYAEQF | 0.7881802 |

| | | |
|---|---|---|
| PPGAGGAGGAGTT | ASSRDRGNYAEQF | 0.8173872 |
| QLVGLGI | ASSRDRGNYAEQF | 0.70191544 |
| RFLEQVHQL | ASSRDRGNYAEQF | 0.8205503 |
| RGVGYVLR | ASSRDRGNYAEQF | 0.8157228 |
| SATGVFAGVI | ASSRDRGNYAEQF | 0.60364413 |
| SLPAITLP | ASSRDRGNYAEQF | 0.7319227 |
| STALAFTV | ASSRDRGNYAEQF | 0.8526381 |
| SVLTVGLGK | ASSRDRGNYAEQF | 0.8411827 |
| TALAGAGSGI | ASSRDRGNYAEQF | 0.49323863 |
| TGLPFTGI | ASSRDRGNYAEQF | 0.739155 |
| VAEHVTV | ASSRDRGNYAEQF | 0.46596855 |
| VDIVLNSL | ASSRDRGNYAEQF | 0.7116167 |
| VLTGGGAVL | ASSRDRGNYAEQF | 0.6507298 |
| VNVGLAYGI | ASSRDRGNYAEQF | 0.796719 |
| VSSAAVGAV | ASSRDRGNYAEQF | 0.72887844 |
| VTSVYEAGV | ASSRDRGNYAEQF | 0.7818275 |
| WVALDRGA | ASSRDRGNYAEQF | 0.79962975 |
| YLVGSVVI | ASSRDRGNYAEQF | 0.8581057 |
| GPPGTGKTLL | ASSRDRGNYAEQF | 0.64785767 |
| GVGMTKF | ASSRDRGNYAEQF | 0.7517001 |
| IGIGGSDL | ASSRDRGNYAEQF | 0.5102458 |
| IGTLLAEL | ASSRDRGNYAEQF | 0.53860587 |
| IYTSGTT | ASSRDRGNYAEQF | 0.6722266 |
| LSFPAGV | ASSRDRGNYAEQF | 0.7505643 |
| LYTSGTT | ASSRDRGNYAEQF | 0.38403752 |
| VGAFPFGF | ASSRDRGNYAEQF | 0.85459954 |
| VGAGTVEF | ASSRDRGNYAEQF | 0.77769977 |
| VGKSTLLL | ASSRDRGNYAEQF | 0.83799905 |
| AALAHAGGGIV | ASSRDRGNYAEQF | 0.73921084 |
| AALVAVSLL | ASSRDRGNYAEQF | 0.7512189 |
| AAVSRAVAAL | ASSRDRGNYAEQF | 0.6993968 |
| AGGSSGIGGF | ASSRDRGNYAEQF | 0.7981714 |
| AGLTVANANG | ASSRDRGNYAEQF | 0.6209402 |
| AGLVGAVL | ASSRDRGNYAEQF | 0.5572678 |
| ALSQAFTV | ASSRDRGNYAEQF | 0.76904887 |
| ALTFSVSL | ASSRDRGNYAEQF | 0.6542149 |
| ALTGAALVAL | ASSRDRGNYAEQF | 0.8452565 |
| ASFSIGAGVL | ASSRDRGNYAEQF | 0.83024347 |
| DGSVYKI | ASSRDRGNYAEQF | 0.7773009 |
| EQLAEIV | ASSRDRGNYAEQF | 0.8312085 |
| FAVSELTVM | ASSRDRGNYAEQF | 0.33587196 |
| GAAPLAAGL | ASSRDRGNYAEQF | 0.7536356 |
| GADGGDGGSGGAGG | ASSRDRGNYAEQF | 0.8600455 |
| GFGLDTGGAGG | ASSRDRGNYAEQF | 0.7990031 |
| GLASGGLL | ASSRDRGNYAEQF | 0.75417036 |
| GRIAGAAVLL | ASSRDRGNYAEQF | 0.8568311 |
| GSAGSLNTSF | ASSRDRGNYAEQF | 0.65174127 |
| GTGQIGI | ASSRDRGNYAEQF | 0.7438848 |
| GTLIALIL | ASSRDRGNYAEQF | 0.7863718 |
| IAATMGFVVG | ASSRDRGNYAEQF | 0.81591225 |
| IFAQALL | ASSRDRGNYAEQF | 0.79710025 |

| | | |
|---|---|---|
| IGAAVALGF | ASSRDRGNYAEQF | 0.66606486 |
| IGLTPLAL | ASSRDRGNYAEQF | 0.8172042 |
| ILGSNTSTL | ASSRDRGNYAEQF | 0.8653179 |
| IVFSLTVL | ASSRDRGNYAEQF | 0.7102407 |
| LASKATGF | ASSRDRGNYAEQF | 0.8384349 |
| LASVALIL | ASSRDRGNYAEQF | 0.81637275 |
| LFAVGLTL | ASSRDRGNYAEQF | 0.7485693 |
| LFNGGIGTYI | ASSRDRGNYAEQF | 0.66124403 |
| LTLGAGII | ASSRDRGNYAEQF | 0.68705523 |
| LVAAALGDI | ASSRDRGNYAEQF | 0.75758815 |
| NTAFVQLGI | ASSRDRGNYAEQF | 0.61980355 |
| QLFSGAEL | ASSRDRGNYAEQF | 0.77269876 |
| SAATFITL | ASSRDRGNYAEQF | 0.76236945 |
| SNYGLANLGV | ASSRDRGNYAEQF | 0.8705369 |
| SYGGAYAVM | ASSRDRGNYAEQF | 0.7983195 |
| TTGVGGLTLG | ASSRDRGNYAEQF | 0.8281114 |
| TTQTIYI | ASSRDRGNYAEQF | 0.6373045 |
| VAAGAGLGAVI | ASSRDRGNYAEQF | 0.7869536 |
| VAAVVTLL | ASSRDRGNYAEQF | 0.7922019 |
| VGFLGGLI | ASSRDRGNYAEQF | 0.6454208 |
| VGVIPILL | ASSRDRGNYAEQF | 0.8515477 |
| VIFASSLI | ASSRDRGNYAEQF | 0.6997625 |
| VKGGDGGAAAG | ASSRDRGNYAEQF | 0.7104385 |
| VTKTFTNI | ASSRDRGNYAEQF | 0.8363053 |
| VVAAAILL | ASSRDRGNYAEQF | 0.8274751 |
| YAALDVEL | ASSRDRGNYAEQF | 0.75059676 |
| LASVALLL | ASSRDRGNYAEQF | 0.7583705 |
| LDAVHRV | ASSRDRGNYAEQF | 0.66701466 |
| QGLSSSIF | ASSRDRGNYAEQF | 0.8251983 |
| RGTFANIRL | ASSRDRGNYAEQF | 0.8436173 |
| RQLWWGHRI | ASSRDRGNYAEQF | 0.75667226 |
| YGPPGTGKTLL | ASSRDRGNYAEQF | 0.85248804 |
| AALVGAPANVA | ASSRDRGNYAEQF | 0.8359235 |
| AATFRRTE | ASSRDRGNYAEQF | 0.5535953 |
| AGALGGGI | ASSRDRGNYAEQF | 0.7714986 |
| AGGAGGAGGIGAG | ASSRDRGNYAEQF | 0.83807516 |
| AGSGALGL | ASSRDRGNYAEQF | 0.82695925 |
| AQIADQI | ASSRDRGNYAEQF | 0.86231446 |
| ASAVAVTYL | ASSRDRGNYAEQF | 0.79507995 |
| ASTVLGGTGG | ASSRDRGNYAEQF | 0.66833997 |
| AVGGEPAK | ASSRDRGNYAEQF | 0.8464458 |
| CEQGTFGGTC | ASSRDRGNYAEQF | 0.8685863 |
| DEGKAPAG | ASSRDRGNYAEQF | 0.8365425 |
| DTVVFPAL | ASSRDRGNYAEQF | 0.7190013 |
| FEGITMAG | ASSRDRGNYAEQF | 0.6982457 |
| FPLVVQD | ASSRDRGNYAEQF | 0.78247523 |
| GAFTTPQ | ASSRDRGNYAEQF | 0.83134687 |
| GFGGFGTAGGVGGA | ASSRDRGNYAEQF | 0.6459257 |
| GFLAQFL | ASSRDRGNYAEQF | 0.7902935 |
| GGAGGAGGNAGVA | ASSRDRGNYAEQF | 0.78755903 |
| GGVGTAGAGGAGGAGG | ASSRDRGNYAEQF | 0.52081764 |

| | | |
|---|---|---|
| GNGGGGDAGNAGSGGN | ASSRDRGNYAEQF | 0.8080207 |
| GTGGAGGAGNDAGS | ASSRDRGNYAEQF | 0.837353 |
| GTTNTGWFNS | ASSRDRGNYAEQF | 0.7647122 |
| IFMGPLFTL | ASSRDRGNYAEQF | 0.6687906 |
| ILAHAIEAI | ASSRDRGNYAEQF | 0.8347366 |
| IRSAIREAL | ASSRDRGNYAEQF | 0.8638389 |
| ITFAGGG | ASSRDRGNYAEQF | 0.84549004 |
| ITGMSGHQEV | ASSRDRGNYAEQF | 0.56903386 |
| IVAGLACTL | ASSRDRGNYAEQF | 0.34556568 |
| IVGGGLAGA | ASSRDRGNYAEQF | 0.6532116 |
| IVTGVLNI | ASSRDRGNYAEQF | 0.80148345 |
| LAQQGCI | ASSRDRGNYAEQF | 0.7390848 |
| LCLHVEN | ASSRDRGNYAEQF | 0.6687795 |
| LDAISDGS | ASSRDRGNYAEQF | 0.7791682 |
| LGGFAAML | ASSRDRGNYAEQF | 0.84801155 |
| LGGKNPMI | ASSRDRGNYAEQF | 0.67658174 |
| LGPKGVAAIL | ASSRDRGNYAEQF | 0.62698734 |
| LMVGVSI | ASSRDRGNYAEQF | 0.8239355 |
| LVGAAFVL | ASSRDRGNYAEQF | 0.8308344 |
| LVLAVAPL | ASSRDRGNYAEQF | 0.61985075 |
| MAAIAVIDAA | ASSRDRGNYAEQF | 0.7001509 |
| NNGINAELQ | ASSRDRGNYAEQF | 0.68616533 |
| RGEPLVLVHG | ASSRDRGNYAEQF | 0.6516366 |
| RGGAGGAAGLI | ASSRDRGNYAEQF | 0.7800705 |
| RSYRMHPAV | ASSRDRGNYAEQF | 0.62501645 |
| SAAHLLAGQLA | ASSRDRGNYAEQF | 0.84799933 |
| SAAVGSY | ASSRDRGNYAEQF | 0.6499685 |
| SPHAVLHRI | ASSRDRGNYAEQF | 0.82798564 |
| SQIAKVL | ASSRDRGNYAEQF | 0.77606153 |
| TALGHLVP | ASSRDRGNYAEQF | 0.80808276 |
| TAPTTSGAA | ASSRDRGNYAEQF | 0.72216785 |
| TDGDDGNGGSPGT | ASSRDRGNYAEQF | 0.6960161 |
| TDGLGEN | ASSRDRGNYAEQF | 0.5892616 |
| TGGPPLPGQA | ASSRDRGNYAEQF | 0.37265772 |
| TIPAITI | ASSRDRGNYAEQF | 0.84046096 |
| TIPGFTI | ASSRDRGNYAEQF | 0.51205873 |
| TSIASGALA | ASSRDRGNYAEQF | 0.832801 |
| TVTLPGAAI | ASSRDRGNYAEQF | 0.832538 |
| VASGLSAALP | ASSRDRGNYAEQF | 0.6881983 |
| VGAGVIGI | ASSRDRGNYAEQF | 0.80973387 |
| VGPKRFGV | ASSRDRGNYAEQF | 0.5909769 |
| VNVAFPD | ASSRDRGNYAEQF | 0.6770845 |
| VSMTNSGVS | ASSRDRGNYAEQF | 0.8044639 |
| VTDVLGDLSV | ASSRDRGNYAEQF | 0.82465315 |
| VVVVGAGFAGL | ASSRDRGNYAEQF | 0.6722468 |
| VYTVGTVV | ASSRDRGNYAEQF | 0.81418335 |
| WTTFSGP | ASSRDRGNYAEQF | 0.78284883 |
| YDELVEAF | ASSRDRGNYAEQF | 0.68327546 |
| YGTGGAGGNGG | ASSRDRGNYAEQF | 0.7669824 |
| YTNVMAV | ASSRDRGNYAEQF | 0.8386238 |
| FLEQVHQL | ASSRDRGNYAEQF | 0.8085675 |

| | | |
|---|---|---|
| GLSLLAVAV | ASSRDRGNYAEQF | 0.6356262 |
| LEQVHQL | ASSRDRGNYAEQF | 0.8233361 |
| LNAAGAI | ASSRDRGNYAEQF | 0.6241126 |
| PAVHASL | ASSRDRGNYAEQF | 0.7945425 |
| TELEDDA | ASSRDRGNYAEQF | 0.79116285 |
| TPPGPPN | ASSRDRGNYAEQF | 0.7449546 |
| TPPPPPPP | ASSRDRGNYAEQF | 0.8291192 |
| TQSPALI | ASGDRSPYEQY | 0.6990827 |
| EFAAAFPQ | ASGDRSPYEQY | 0.63999045 |
| LLSGNLDVL | ASGDRSPYEQY | 0.7234677 |
| GPANLGNYNI | ASGDRSPYEQY | 0.8003002 |
| TLGPTLP | ASGDRSPYEQY | 0.83910334 |
| VYAYSSCCK | ASGDRSPYEQY | 0.63554984 |
| APAPAPAPNI | ASGDRSPYEQY | 0.7773858 |
| PGGPVPIP | ASGDRSPYEQY | 0.72109205 |
| FLTVALEP | ASGDRSPYEQY | 0.6403023 |
| SQYFGANL | ASGDRSPYEQY | 0.7896577 |
| GGVKPAA | ASGDRSPYEQY | 0.7589612 |
| GGTGGAGGAGGW | ASGDRSPYEQY | 0.6891351 |
| GVGGVAGLGGAGG | ASGDRSPYEQY | 0.80713296 |
| VLLVLATTI | ASGDRSPYEQY | 0.6672058 |
| IGIFAAML | ASGDRSPYEQY | 0.7999201 |
| AVGDLID | ASGDRSPYEQY | 0.8663133 |
| NISPLPL | ASGDRSPYEQY | 0.69181526 |
| VTAVFVAI | ASGDRSPYEQY | 0.82687366 |
| SGDDNDAAL | ASGDRSPYEQY | 0.81177294 |
| AVAALAHS | ASGDRSPYEQY | 0.7865908 |
| GGVTVMATP | ASGDRSPYEQY | 0.7516799 |
| AGLAANVRD | ASGDRSPYEQY | 0.69090146 |
| TNRPDLIDPALLRPG | ASGDRSPYEQY | 0.85419524 |
| LANLGVTAL | ASGDRSPYEQY | 0.8358713 |
| TGERIEL | ASGDRSPYEQY | 0.83625126 |
| NGEELAA | ASGDRSPYEQY | 0.8118722 |
| VLSLVWG | ASGDRSPYEQY | 0.8220242 |
| IGYALWL | ASGDRSPYEQY | 0.82133687 |
| LSLLLAPL | ASGDRSPYEQY | 0.6769607 |
| TLDEALK | ASGDRSPYEQY | 0.5031491 |
| TDGAGTAEGG | ASGDRSPYEQY | 0.7380854 |
| NAPGGAGGNGG | ASGDRSPYEQY | 0.70268726 |
| VSVFMGMGGNF | ASGDRSPYEQY | 0.7798416 |
| LASYLAAL | ASGDRSPYEQY | 0.5356926 |
| LVGGTPAL | ASGDRSPYEQY | 0.83976376 |
| VRNADVL | ASGDRSPYEQY | 0.87189156 |
| TCSSPFLVSAM | ASGDRSPYEQY | 0.71145844 |
| VHITDAE | ASGDRSPYEQY | 0.85523605 |
| EYAAPSF | ASGDRSPYEQY | 0.55846345 |
| VVDAFHGGL | ASGDRSPYEQY | 0.8203255 |
| ASVVDPQ | ASGDRSPYEQY | 0.84762263 |
| SYNEFGGLS | ASGDRSPYEQY | 0.6317658 |
| DPDGSVY | ASGDRSPYEQY | 0.83299553 |
| MVSTIHATE | ASGDRSPYEQY | 0.6748769 |

| | | |
|---|---|---|
| LAAGVADPV | ASGDRSPYEQY | 0.76611465 |
| TKEQIAA | ASGDRSPYEQY | 0.847406 |
| SLASPAAL | ASGDRSPYEQY | 0.73560584 |
| IGLVFYGV | ASGDRSPYEQY | 0.8062248 |
| FLLDASNL | ASGDRSPYEQY | 0.7622449 |
| YQVGNFP | ASGDRSPYEQY | 0.6790974 |
| ESLLSQST | ASGDRSPYEQY | 0.8053735 |
| LFTVGFAAVG | ASGDRSPYEQY | 0.84081376 |
| ATPPTPT | ASGDRSPYEQY | 0.65042377 |
| TDLATLG | ASGDRSPYEQY | 0.86664635 |
| SAGGTGR | ASGDRSPYEQY | 0.6685571 |
| GFSNSGS | ASGDRSPYEQY | 0.8624137 |
| GGNGGSGGTSV | ASGDRSPYEQY | 0.8149684 |
| GTSVATGGAGN | ASGDRSPYEQY | 0.75561535 |
| NGGNGGSGGTS | ASGDRSPYEQY | 0.7404364 |
| AVVKFPH | ASGDRSPYEQY | 0.8389753 |
| TPPPPPP | ASGDRSPYEQY | 0.71221495 |
| AAPAAAAAPA | ASGDRSPYEQY | 0.8170904 |
| IALNEDL | ASGDRSPYEQY | 0.8557601 |
| IVNVSSM | ASGDRSPYEQY | 0.20783697 |
| TGIPPAPR | ASGDRSPYEQY | 0.45011318 |
| FDLGGGTF | ASGDRSPYEQY | 0.36252832 |
| FDLGGGTFDVS | ASGDRSPYEQY | 0.36387062 |
| NEPTAAA | ASGDRSPYEQY | 0.24360982 |
| NEPTAAALA | ASGDRSPYEQY | 0.21979907 |
| GSIINISSI | ASGDRSPYEQY | 0.267961 |
| VAPGNAGTA | ASGDRSPYEQY | 0.23607527 |
| TGGGASLE | ASGDRSPYEQY | 0.60876364 |
| VAVYNHYKRI | ASGDRSPYEQY | 0.34805614 |
| TDPAPATT | ASGDRSPYEQY | 0.25282744 |
| TNRPDILDPALLRPG | ASGDRSPYEQY | 0.46731812 |
| LTTTTTS | ASGDRSPYEQY | 0.38168985 |
| PTGSGKTL | ASGDRSPYEQY | 0.21676028 |
| TGSGKTL | ASGDRSPYEQY | 0.3136289 |
| TIGHVDHGK | ASGDRSPYEQY | 0.2887141 |
| TIGHVDHGKTT | ASGDRSPYEQY | 0.25005722 |
| GHVDHGKTT | ASGDRSPYEQY | 0.19921987 |
| AQFEHTLL | ASGDRSPYEQY | 0.30192357 |
| IVNQDEL | ASGDRSPYEQY | 0.37970155 |
| TNGSQFF | ASGDRSPYEQY | 0.19953582 |
| TGKGIVE | ASGDRSPYEQY | 0.39581722 |
| LISNASDA | ASGDRSPYEQY | 0.23448952 |
| AELRTLGELTKIAKAHGAQ | ASGDRSPYEQY | 0.25678214 |
| APNAVAAAEPSW | ASGDRSPYEQY | 0.40706244 |
| APQVAGAATAVL | ASGDRSPYEQY | 0.24512167 |
| ARRDPAAAAAILPTDA | ASGDRSPYEQY | 0.29035726 |
| DAAARDVLAEAGLAEYF | ASGDRSPYEQY | 0.36884892 |
| DDQIVLSAKANLAKVEQ | ASGDRSPYEQY | 0.50987387 |
| DELLPDTGEDSDEDGH | ASGDRSPYEQY | 0.2508626 |
| ELPEAPANRATVPQD | ASGDRSPYEQY | 0.27336347 |
| FGQSKIERVTGIG | ASGDRSPYEQY | 0.6100034 |

| | | |
|---|---|---|
| GAGGTGSTAGGAGGAGGAGGLY | ASGDRSPYEQY | 0.31301025 |
| GHGGTGAAGGNGGTGGAGGSGI | ASGDRSPYEQY | 0.5316085 |
| GNGGNGGAGGEGGPGGAGGAGA | ASGDRSPYEQY | 0.5534805 |
| GSTAGNGGNGGNATNGTIAGQP | ASGDRSPYEQY | 0.22969408 |
| GTGGDGGNGGNGGTGG | ASGDRSPYEQY | 0.43184182 |
| GVRRPAGIPADAPV | ASGDRSPYEQY | 0.36482975 |
| GYRQVTDAHLLTLARR | ASGDRSPYEQY | 0.32268313 |
| IAKMGQDGHDRG | ASGDRSPYEQY | 0.29739565 |
| KAEPSIEIQYT | ASGDRSPYEQY | 0.21472527 |
| KPVRIRMAADEDTVAVT | ASGDRSPYEQY | 0.2866124 |
| MPPRIAGMRLLVIKPEP | ASGDRSPYEQY | 0.21288495 |
| MKFYAVALFRSRGVPPTR | ASGDRSPYEQY | 0.35196924 |
| NAGGFGILGAVAHSP | ASGDRSPYEQY | 0.5131788 |
| NEGREITLAMAEM | ASGDRSPYEQY | 0.33886874 |
| NELVEWSHPLGHRWRWE | ASGDRSPYEQY | 0.20335783 |
| NLGAGGTGGNGGNPG | ASGDRSPYEQY | 0.21369624 |
| NTEWPKWETVMAADPVD | ASGDRSPYEQY | 0.2582171 |
| PASAAPAETQP | ASGDRSPYEQY | 0.21572286 |
| QQSSDGKNGVRAAEITGE | ASGDRSPYEQY | 0.22067833 |
| RAPDAVTTSWWKE | ASGDRSPYEQY | 0.23331437 |
| SGFGNVGAHTSGWFNQ | ASGDRSPYEQY | 0.5283749 |
| SGVAGAVGQAASVGGLKV | ASGDRSPYEQY | 0.5007546 |
| SNGGAGTGSAGGNGGT | ASGDRSPYEQY | 0.27272713 |
| TGGAGGEGGAAG | ASGDRSPYEQY | 0.41420114 |
| TGGDGGAAGAGGAPGHG | ASGDRSPYEQY | 0.6337901 |
| TMPPLLAAIANAGRHGV | ASGDRSPYEQY | 0.4024398 |
| TSSTAGNSGGA | ASGDRSPYEQY | 0.31470504 |
| VPATVPVGILC | ASGDRSPYEQY | 0.20826934 |
| VQRPDSGGVVLTGRIS | ASGDRSPYEQY | 0.6493317 |
| WTPNPVPATKGVPAGLVQAT | ASGDRSPYEQY | 0.30535084 |
| YALKRALAKFSDRFDVVII | ASGDRSPYEQY | 0.4288717 |
| YLIGSVIV | ASGDRSPYEQY | 0.48017162 |
| YGQLVAVID | ASGDRSPYEQY | 0.34934938 |
| VTVGGMTLPA | ASGDRSPYEQY | 0.23325668 |
| VLALPAV | ASGDRSPYEQY | 0.6039183 |
| VIPPTLNYET | ASGDRSPYEQY | 0.45183724 |
| VGMGGMHPSSG | ASGDRSPYEQY | 0.21970312 |
| VGLAGGAATA | ASGDRSPYEQY | 0.2139639 |
| VGGAGGAGGSGP | ASGDRSPYEQY | 0.3181474 |
| VFAAFMV | ASGDRSPYEQY | 0.33715126 |
| TLYDKLR | ASGDRSPYEQY | 0.5343697 |
| TLAVFTVL | ASGDRSPYEQY | 0.28035676 |
| TGSVQPARV | ASGDRSPYEQY | 0.26607907 |
| TGGGAGSGGT | ASGDRSPYEQY | 0.4502554 |
| SHLETSL | ASGDRSPYEQY | 0.38986278 |
| SFTTGMT | ASGDRSPYEQY | 0.28129512 |
| QVEVAKRMGVRQPTVS | ASGDRSPYEQY | 0.36528918 |
| QGSGGPALAM | ASGDRSPYEQY | 0.2638171 |
| MLAPSNP | ASGDRSPYEQY | 0.43119597 |
| LVNNAAI | ASGDRSPYEQY | 0.45524538 |
| LTGEINTLA | ASGDRSPYEQY | 0.36151716 |

| | | |
|---|---|---|
| LSIFGPV | ASGDRSPYEQY | 0.48736066 |
| LLALLANL | ASGDRSPYEQY | 0.3297806 |
| LISNASDALDKLR | ASGDRSPYEQY | 0.31294227 |
| LINEAVP | ASGDRSPYEQY | 0.24112138 |
| LGYAFSGL | ASGDRSPYEQY | 0.29995343 |
| LDFVAVAL | ASGDRSPYEQY | 0.31975198 |
| LAQPGAE | ASGDRSPYEQY | 0.37599254 |
| ISNLGAVTGF | ASGDRSPYEQY | 0.41504624 |
| HQEDPGRI | ASGDRSPYEQY | 0.394533 |
| GTPGEPTP | ASGDRSPYEQY | 0.25300604 |
| GFFNSGS | ASGDRSPYEQY | 0.4912137 |
| GAGGGGLTAGAN | ASGDRSPYEQY | 0.5357522 |
| FVDALVNM | ASGDRSPYEQY | 0.22739266 |
| FLTVAPEL | ASGDRSPYEQY | 0.28019863 |
| FETLTAL | ASGDRSPYEQY | 0.4111275 |
| DVINPIEL | ASGDRSPYEQY | 0.5756833 |
| APVSGAPV | ASGDRSPYEQY | 0.30162695 |
| ALVGGLAW | ASGDRSPYEQY | 0.32089674 |
| ALAAAAATAI | ASGDRSPYEQY | 0.64060616 |
| AFGLGLLL | ASGDRSPYEQY | 0.32921165 |
| AFFAAASL | ASGDRSPYEQY | 0.5347873 |
| AAAALPAFSPP | ASGDRSPYEQY | 0.21755561 |
| VVLAPAL | ASGDRSPYEQY | 0.279397 |
| TIGHVDHGKTTL | ASGDRSPYEQY | 0.24330008 |
| LISNASDALDK | ASGDRSPYEQY | 0.19650236 |
| LISNASDAL | ASGDRSPYEQY | 0.45118433 |
| ISNASDALDK | ASGDRSPYEQY | 0.24178615 |
| NQAETLVYQTE | ASGDRSPYEQY | 0.37590754 |
| TGGTGGTAGAG | ASGDRSPYEQY | 0.29692727 |
| TGSTAGGAGGA | ASGDRSPYEQY | 0.39211267 |
| DPLAVPVAAGPV | ASGDRSPYEQY | 0.3394633 |
| GGTGGTPGNGGN | ASGDRSPYEQY | 0.2649535 |
| PAGLAGVDPATL | ASGDRSPYEQY | 0.22310276 |
| GNNLVVEAPHAGASV | ASGDRSPYEQY | 0.39230803 |
| KVTRIMSASTLEQPA | ASGDRSPYEQY | 0.4574487 |
| LGFALPIALPRRTVV | ASGDRSPYEQY | 0.43921068 |
| TRLGNLTARKLVVLV | ASGDRSPYEQY | 0.22068956 |
| WSQQRRLIYFPSAGP | ASGDRSPYEQY | 0.42957866 |
| GPNGNPDPMAAAVDIR | ASGDRSPYEQY | 0.30720198 |
| TQRCVSVADLRAAASR | ASGDRSPYEQY | 0.20926246 |
| QFVLDVAEHPVEVLTETV | ASGDRSPYEQY | 0.20777652 |
| GGAGGTGGAGGNGGTQAGDG | ASGDRSPYEQY | 0.22906235 |
| PLLATLRAIAVDVDAKPAQV | ASGDRSPYEQY | 0.2488629 |
| APKFRTGGYMIDFWGVGYQVAK | ASGDRSPYEQY | 0.40610015 |
| GAGGAGGAGGAAGGPGGTGGTG | ASGDRSPYEQY | 0.5605862 |
| IGALGLLATDVAPAGSVAFGVA | ASGDRSPYEQY | 0.24930954 |
| AGGAGGIGGNGGAGGAGGVLMG | ASGDRSPYEQY | 0.44431323 |
| ADELIGLL | ASGDRSPYEQY | 0.3331608 |
| ADRALKA | ASGDRSPYEQY | 0.19431952 |
| AGAVGQVGGAGGA | ASGDRSPYEQY | 0.26190904 |
| AGGGGGLGGAP | ASGDRSPYEQY | 0.22016284 |

| | | |
|---|---|---|
| AGGLLYGNGGAG | ASGDRSPYEQY | 0.43623856 |
| AGLESIGL | ASGDRSPYEQY | 0.21308321 |
| AGLKTGII | ASGDRSPYEQY | 0.3154468 |
| ASLAALLL | ASGDRSPYEQY | 0.30453846 |
| ASVINISSI | ASGDRSPYEQY | 0.5838371 |
| ATPPGPA | ASGDRSPYEQY | 0.40386748 |
| ATVASALV | ASGDRSPYEQY | 0.22616395 |
| ATVASLIL | ASGDRSPYEQY | 0.21387583 |
| AVNGPVLIRH | ASGDRSPYEQY | 0.41029274 |
| DGAAGAVGI | ASGDRSPYEQY | 0.30053005 |
| DPLLHGI | ASGDRSPYEQY | 0.24631266 |
| ELGAALGAL | ASGDRSPYEQY | 0.24492331 |
| ELGRTQAV | ASGDRSPYEQY | 0.4106531 |
| FVAGGYLS | ASGDRSPYEQY | 0.21172942 |
| GASGGAGGAGGW | ASGDRSPYEQY | 0.4878139 |
| GGAGAGGDGGAGGI | ASGDRSPYEQY | 0.23558995 |
| GTGHNGSQPGMGGNG | ASGDRSPYEQY | 0.30665523 |
| GYAAASASAS | ASGDRSPYEQY | 0.45526132 |
| HQVDVAAVQQG | ASGDRSPYEQY | 0.60371375 |
| IGAAIAHT | ASGDRSPYEQY | 0.31232062 |
| IGMIEVAQL | ASGDRSPYEQY | 0.30594826 |
| IRLAPEHTSAMD | ASGDRSPYEQY | 0.22373927 |
| IYNVSVL | ASGDRSPYEQY | 0.26483867 |
| KQGFLGM | ASGDRSPYEQY | 0.26419926 |
| LESVITQL | ASGDRSPYEQY | 0.49157917 |
| LGLGALVVV | ASGDRSPYEQY | 0.28199753 |
| LIGNGGNGGSGGI | ASGDRSPYEQY | 0.21081765 |
| LNAINAPTL | ASGDRSPYEQY | 0.3148883 |
| LQLLNTGV | ASGDRSPYEQY | 0.49697006 |
| LQTSIEF | ASGDRSPYEQY | 0.50679106 |
| MSFAVSEL | ASGDRSPYEQY | 0.22536655 |
| NTGALGL | ASGDRSPYEQY | 0.2539767 |
| PPGAGGAGGAGTT | ASGDRSPYEQY | 0.28420407 |
| QLVGLGI | ASGDRSPYEQY | 0.27112392 |
| RFLEQVHQL | ASGDRSPYEQY | 0.23097149 |
| RGVGYVLR | ASGDRSPYEQY | 0.24022602 |
| SATGVFAGVI | ASGDRSPYEQY | 0.45766836 |
| SLPAITLP | ASGDRSPYEQY | 0.7081654 |
| STALAFTV | ASGDRSPYEQY | 0.2548736 |
| SVLTVGLGK | ASGDRSPYEQY | 0.20895576 |
| TALAGAGSGI | ASGDRSPYEQY | 0.3073244 |
| TGLPFTGI | ASGDRSPYEQY | 0.22803193 |
| VAEHVTV | ASGDRSPYEQY | 0.24651664 |
| VDIVLNSL | ASGDRSPYEQY | 0.31363395 |
| VLTGGGAVL | ASGDRSPYEQY | 0.20629008 |
| VNVGLAYGI | ASGDRSPYEQY | 0.2758312 |
| VSSAAVGAV | ASGDRSPYEQY | 0.5619623 |
| VTSVYEAGV | ASGDRSPYEQY | 0.3895977 |
| WVALDRGA | ASGDRSPYEQY | 0.35977292 |
| YLVGSVVI | ASGDRSPYEQY | 0.4784246 |
| GPPGTGKTLL | ASGDRSPYEQY | 0.33619067 |

| | | |
|---|---|---|
| GVGMTKF | ASGDRSPYEQY | 0.48818102 |
| IGIGGSDL | ASGDRSPYEQY | 0.641696 |
| IGTLLAEL | ASGDRSPYEQY | 0.29104066 |
| IYTSGTT | ASGDRSPYEQY | 0.3042949 |
| LSFPAGV | ASGDRSPYEQY | 0.4391849 |
| LYTSGTT | ASGDRSPYEQY | 0.31067464 |
| VGAFPFGF | ASGDRSPYEQY | 0.35287938 |
| VGAGTVEF | ASGDRSPYEQY | 0.307536 |
| VGKSTLLL | ASGDRSPYEQY | 0.43838358 |
| AALAHAGGGIV | ASGDRSPYEQY | 0.3394868 |
| AALVAVSLL | ASGDRSPYEQY | 0.61982167 |
| AAVSRAVAAL | ASGDRSPYEQY | 0.20294759 |
| AGGSSGIGGF | ASGDRSPYEQY | 0.2511498 |
| AGLTVANANG | ASGDRSPYEQY | 0.4694268 |
| AGLVGAVL | ASGDRSPYEQY | 0.20461814 |
| ALSQAFTV | ASGDRSPYEQY | 0.22929785 |
| ALTFSVSL | ASGDRSPYEQY | 0.3605783 |
| ALTGAALVAL | ASGDRSPYEQY | 0.49492645 |
| ASFSIGAGVL | ASGDRSPYEQY | 0.19747537 |
| DGSVYKI | ASGDRSPYEQY | 0.35655877 |
| EQLAEIV | ASGDRSPYEQY | 0.21630204 |
| FAVSELTVM | ASGDRSPYEQY | 0.336015 |
| GAAPLAAGL | ASGDRSPYEQY | 0.2864862 |
| GADGGDGGSGGAGG | ASGDRSPYEQY | 0.21432522 |
| GFGLDTGGAGG | ASGDRSPYEQY | 0.34310472 |
| GLASGGLL | ASGDRSPYEQY | 0.21665451 |
| GRIAGAAVLL | ASGDRSPYEQY | 0.33299094 |
| GSAGSLNTSF | ASGDRSPYEQY | 0.46243572 |
| GTGQIGI | ASGDRSPYEQY | 0.6807939 |
| GTLIALIL | ASGDRSPYEQY | 0.2163455 |
| IAATMGFVVG | ASGDRSPYEQY | 0.42500734 |
| IFAQALL | ASGDRSPYEQY | 0.22683248 |
| IGAAVALGF | ASGDRSPYEQY | 0.6311768 |
| IGLTPLAL | ASGDRSPYEQY | 0.28885803 |
| ILGSNTSTL | ASGDRSPYEQY | 0.20627768 |
| IVFSLTVL | ASGDRSPYEQY | 0.400384 |
| LASKATGF | ASGDRSPYEQY | 0.28662688 |
| LASVALIL | ASGDRSPYEQY | 0.27987817 |
| LFAVGLTL | ASGDRSPYEQY | 0.62294835 |
| LFNGGIGTYI | ASGDRSPYEQY | 0.23937508 |
| LTLGAGII | ASGDRSPYEQY | 0.25836322 |
| LVAAALGDI | ASGDRSPYEQY | 0.37651503 |
| NTAFVQLGI | ASGDRSPYEQY | 0.32918644 |
| QLFSGAEL | ASGDRSPYEQY | 0.25491837 |
| SAATFITL | ASGDRSPYEQY | 0.40769836 |
| SNYGLANLGV | ASGDRSPYEQY | 0.36876348 |
| SYGGAYAVM | ASGDRSPYEQY | 0.23262423 |
| TTGVGGLTLG | ASGDRSPYEQY | 0.5294501 |
| TTQTIYI | ASGDRSPYEQY | 0.2741141 |
| VAAGAGLGAVI | ASGDRSPYEQY | 0.40257078 |
| VAAVVTLL | ASGDRSPYEQY | 0.32686913 |

| | | |
|---|---|---|
| VGFLGGLI | ASGDRSPYEQY | 0.6957314 |
| VGVIPILL | ASGDRSPYEQY | 0.26021525 |
| VIFASSLI | ASGDRSPYEQY | 0.25795764 |
| VKGGDGGAAAG | ASGDRSPYEQY | 0.38840067 |
| VTKTFTNI | ASGDRSPYEQY | 0.40247673 |
| VVAAAILL | ASGDRSPYEQY | 0.22581616 |
| YAALDVEL | ASGDRSPYEQY | 0.43764028 |
| LASVALLL | ASGDRSPYEQY | 0.33296224 |
| LDAVHRV | ASGDRSPYEQY | 0.5210768 |
| QGLSSSIF | ASGDRSPYEQY | 0.2704618 |
| RGTFANIRL | ASGDRSPYEQY | 0.31460053 |
| RQLWWGHRI | ASGDRSPYEQY | 0.58599913 |
| YGPPGTGKTLL | ASGDRSPYEQY | 0.22394557 |
| AALVGAPANVA | ASGDRSPYEQY | 0.35725158 |
| AATFRRTE | ASGDRSPYEQY | 0.46499723 |
| AGALGGGI | ASGDRSPYEQY | 0.5022525 |
| AGGAGGAGGIGAG | ASGDRSPYEQY | 0.25349674 |
| AGSGALGL | ASGDRSPYEQY | 0.4564147 |
| AQIADQI | ASGDRSPYEQY | 0.28510207 |
| ASAVAVTYL | ASGDRSPYEQY | 0.3845542 |
| ASTVLGGTGG | ASGDRSPYEQY | 0.26377007 |
| AVGGEPAK | ASGDRSPYEQY | 0.29548225 |
| CEQGTFGGTC | ASGDRSPYEQY | 0.23004493 |
| DEGKAPAG | ASGDRSPYEQY | 0.32578915 |
| DTVVFPAL | ASGDRSPYEQY | 0.42828697 |
| FEGITMAG | ASGDRSPYEQY | 0.40376383 |
| FPLVVQD | ASGDRSPYEQY | 0.21813685 |
| GAFTTPQ | ASGDRSPYEQY | 0.43229362 |
| GFGGFGTAGGVGGA | ASGDRSPYEQY | 0.4814176 |
| GFLAQFL | ASGDRSPYEQY | 0.68872404 |
| GGAGGAGGNAGVA | ASGDRSPYEQY | 0.32994962 |
| GGVGTAGAGGAGGAGG | ASGDRSPYEQY | 0.43898138 |
| GNGGGGDAGNAGSGGN | ASGDRSPYEQY | 0.39015985 |
| GTGGAGGAGNDAGS | ASGDRSPYEQY | 0.28932613 |
| GTTNTGWFNS | ASGDRSPYEQY | 0.21663223 |
| IFMGPLFTL | ASGDRSPYEQY | 0.23933044 |
| ILAHAIEAI | ASGDRSPYEQY | 0.33922884 |
| IRSAIREAL | ASGDRSPYEQY | 0.2418404 |
| ITFAGGG | ASGDRSPYEQY | 0.59903806 |
| ITGMSGHQEV | ASGDRSPYEQY | 0.51175135 |
| IVAGLACTL | ASGDRSPYEQY | 0.44414887 |
| IVGGGLAGA | ASGDRSPYEQY | 0.57161945 |
| IVTGVLNI | ASGDRSPYEQY | 0.47447795 |
| LAQQGCI | ASGDRSPYEQY | 0.38574994 |
| LCLHVEN | ASGDRSPYEQY | 0.25674972 |
| LDAISDGS | ASGDRSPYEQY | 0.34088665 |
| LGGFAAML | ASGDRSPYEQY | 0.20481013 |
| LGGKNPMI | ASGDRSPYEQY | 0.23659131 |
| LGPKGVAAIL | ASGDRSPYEQY | 0.24770966 |
| LMVGVSI | ASGDRSPYEQY | 0.3112136 |
| LVGAAFVL | ASGDRSPYEQY | 0.19659464 |

| | | |
|---|---|---|
| LVLAVAPL | ASGDRSPYEQY | 0.6025743 |
| MAAIAVIDAA | ASGDRSPYEQY | 0.50693476 |
| NNGINAELQ | ASGDRSPYEQY | 0.3513835 |
| RGEPLVLVHG | ASGDRSPYEQY | 0.4505062 |
| RGGAGGAAGLI | ASGDRSPYEQY | 0.3429469 |
| RSYRMHPAV | ASGDRSPYEQY | 0.2235666 |
| SAAHLLAGQLA | ASGDRSPYEQY | 0.4830377 |
| SAAVGSY | ASGDRSPYEQY | 0.2630872 |
| SPHAVLHRI | ASGDRSPYEQY | 0.45940626 |
| SQIAKVL | ASGDRSPYEQY | 0.41950387 |
| TALGHLVP | ASGDRSPYEQY | 0.33832368 |
| TAPTTSGAA | ASGDRSPYEQY | 0.27324286 |
| TDGDDGNGGSPGT | ASGDRSPYEQY | 0.3618572 |
| TDGLGEN | ASGDRSPYEQY | 0.25819635 |
| TGGPPLPGQA | ASGDRSPYEQY | 0.28113142 |
| TIPAITI | ASGDRSPYEQY | 0.27658355 |
| TIPGFTI | ASGDRSPYEQY | 0.3582369 |
| TSIASGALA | ASGDRSPYEQY | 0.21963547 |
| TVTLPGAAI | ASGDRSPYEQY | 0.46585953 |
| VASGLSAALP | ASGDRSPYEQY | 0.36772856 |
| VGAGVIGI | ASGDRSPYEQY | 0.36800522 |
| VGPKRFGV | ASGDRSPYEQY | 0.24782339 |
| VNVAFPD | ASGDRSPYEQY | 0.3897515 |
| VSMTNSGVS | ASGDRSPYEQY | 0.41077358 |
| VTDVLGDLSV | ASGDRSPYEQY | 0.22728883 |
| VVVVGAGFAGL | ASGDRSPYEQY | 0.30516997 |
| VYTVGTVV | ASGDRSPYEQY | 0.28607213 |
| WTTFSGP | ASGDRSPYEQY | 0.2846893 |
| YDELVEAF | ASGDRSPYEQY | 0.46171755 |
| YGTGGAGGNGG | ASGDRSPYEQY | 0.31344965 |
| YTNVMAV | ASGDRSPYEQY | 0.23380178 |
| FLEQVHQL | ASGDRSPYEQY | 0.2787737 |
| GLSLLAVAV | ASGDRSPYEQY | 0.36282885 |
| LEQVHQL | ASGDRSPYEQY | 0.28954142 |
| LNAAGAI | ASGDRSPYEQY | 0.1981451 |
| PAVHASL | ASGDRSPYEQY | 0.27869907 |
| TELEDDA | ASGDRSPYEQY | 0.23332001 |
| TPPGPPN | ASGDRSPYEQY | 0.2532818 |
| TPPPPPPP | ASGDRSPYEQY | 0.23327155 |
| TQSPALI | ASSFGGRNTEVF | 0.49938455 |
| EFAAAFPQ | ASSFGGRNTEVF | 0.4711547 |
| LLSGNLDVL | ASSFGGRNTEVF | 0.32709366 |
| GPANLGNYNI | ASSFGGRNTEVF | 0.4109899 |
| TLGPTLP | ASSFGGRNTEVF | 0.39457607 |
| VYAYSSCCK | ASSFGGRNTEVF | 0.46018496 |
| APAPAPAPNI | ASSFGGRNTEVF | 0.41868138 |
| PGGPVPIP | ASSFGGRNTEVF | 0.30682203 |
| FLTVALEP | ASSFGGRNTEVF | 0.32906947 |
| SQYFGANL | ASSFGGRNTEVF | 0.27530327 |
| GGVKPAA | ASSFGGRNTEVF | 0.22563386 |
| GGTGGAGGAGGW | ASSFGGRNTEVF | 0.2499784 |

| | | |
|---|---|---|
| GVGGVAGLGGAGG | ASSFGGRNTEVF | 0.24521047 |
| VLLVLATTI | ASSFGGRNTEVF | 0.2593625 |
| IGIFAAML | ASSFGGRNTEVF | 0.38342023 |
| AVGDLID | ASSFGGRNTEVF | 0.48259106 |
| NISPLPL | ASSFGGRNTEVF | 0.27328092 |
| VTAVFVAI | ASSFGGRNTEVF | 0.20551117 |
| SGDDNDAAL | ASSFGGRNTEVF | 0.2695672 |
| AVAALAHS | ASSFGGRNTEVF | 0.27714363 |
| GGVTVMATP | ASSFGGRNTEVF | 0.42526138 |
| AGLAANVRD | ASSFGGRNTEVF | 0.29994217 |
| TNRPDLIDPALLRPG | ASSFGGRNTEVF | 0.68847734 |
| LANLGVTAL | ASSFGGRNTEVF | 0.530134 |
| TGERIEL | ASSFGGRNTEVF | 0.27994668 |
| NGEELAA | ASSFGGRNTEVF | 0.4116376 |
| VLSLVWG | ASSFGGRNTEVF | 0.24589674 |
| IGYALWL | ASSFGGRNTEVF | 0.2691214 |
| LSLLLAPL | ASSFGGRNTEVF | 0.28467163 |
| TLDEALK | ASSFGGRNTEVF | 0.2988676 |
| TDGAGTAEGG | ASSFGGRNTEVF | 0.27335817 |
| NAPGGAGGNGG | ASSFGGRNTEVF | 0.2600657 |
| VSVFMGMGGNF | ASSFGGRNTEVF | 0.2168164 |
| LASYLAAL | ASSFGGRNTEVF | 0.46295685 |
| LVGGTPAL | ASSFGGRNTEVF | 0.25399938 |
| VRNADVL | ASSFGGRNTEVF | 0.2350463 |
| TCSSPFLVSAM | ASSFGGRNTEVF | 0.29416284 |
| VHITDAE | ASSFGGRNTEVF | 0.4524817 |
| EYAAPSF | ASSFGGRNTEVF | 0.64706284 |
| VVDAFHGGL | ASSFGGRNTEVF | 0.25813708 |
| ASVVDPQ | ASSFGGRNTEVF | 0.24471548 |
| SYNEFGGLS | ASSFGGRNTEVF | 0.62876314 |
| DPDGSVY | ASSFGGRNTEVF | 0.2757213 |
| MVSTIHATE | ASSFGGRNTEVF | 0.3020599 |
| LAAGVADPV | ASSFGGRNTEVF | 0.34225702 |
| TKEQIAA | ASSFGGRNTEVF | 0.2870835 |
| SLASPAAL | ASSFGGRNTEVF | 0.23864391 |
| IGLVFYGV | ASSFGGRNTEVF | 0.43006438 |
| FLLDASNL | ASSFGGRNTEVF | 0.28486767 |
| YQVGNFP | ASSFGGRNTEVF | 0.34872937 |
| ESLLSQST | ASSFGGRNTEVF | 0.20226406 |
| LFTVGFAAVG | ASSFGGRNTEVF | 0.30185318 |
| ATPPTPT | ASSFGGRNTEVF | 0.24048723 |
| TDLATLG | ASSFGGRNTEVF | 0.23817927 |
| SAGGTGR | ASSFGGRNTEVF | 0.20046562 |
| GFSNSGS | ASSFGGRNTEVF | 0.2611642 |
| GGNGGSGGTSV | ASSFGGRNTEVF | 0.4220284 |
| GTSVATGGAGN | ASSFGGRNTEVF | 0.58524996 |
| NGGNGGSGGTS | ASSFGGRNTEVF | 0.33428204 |
| AVVKFPH | ASSFGGRNTEVF | 0.3560973 |
| TPPPPPP | ASSFGGRNTEVF | 0.5316691 |
| AAPAAAAAPA | ASSFGGRNTEVF | 0.2628019 |
| IALNEDL | ASSFGGRNTEVF | 0.21830167 |

| | | |
|---|---|---|
| IVNVSSM | ASSFGGRNTEVF | 0.32266963 |
| TGIPPAPR | ASSFGGRNTEVF | 0.28042644 |
| FDLGGGTF | ASSFGGRNTEVF | 0.37901604 |
| FDLGGGTFDVS | ASSFGGRNTEVF | 0.4858119 |
| NEPTAAA | ASSFGGRNTEVF | 0.21050698 |
| NEPTAAALA | ASSFGGRNTEVF | 0.36092627 |
| GSIINISSI | ASSFGGRNTEVF | 0.44017667 |
| VAPGNAGTA | ASSFGGRNTEVF | 0.22005276 |
| TGGGASLE | ASSFGGRNTEVF | 0.23127687 |
| VAVYNHYKRI | ASSFGGRNTEVF | 0.22446641 |
| TDPAPATT | ASSFGGRNTEVF | 0.21261771 |
| TNRPDILDPALLRPG | ASSFGGRNTEVF | 0.31585163 |
| LTTTTTS | ASSFGGRNTEVF | 0.26044288 |
| PTGSGKTL | ASSFGGRNTEVF | 0.37761933 |
| TGSGKTL | ASSFGGRNTEVF | 0.23897257 |
| TIGHVDHGK | ASSFGGRNTEVF | 0.26846653 |
| TIGHVDHGKTT | ASSFGGRNTEVF | 0.31020576 |
| GHVDHGKTT | ASSFGGRNTEVF | 0.32890025 |
| AQFEHTLL | ASSFGGRNTEVF | 0.30909866 |
| IVNQDEL | ASSFGGRNTEVF | 0.44835085 |
| TNGSQFF | ASSFGGRNTEVF | 0.2269051 |
| TGKGIVE | ASSFGGRNTEVF | 0.2720447 |
| LISNASDA | ASSFGGRNTEVF | 0.5929817 |
| AELRTLGELTKIAKAHGAQ | ASSFGGRNTEVF | 0.22576016 |
| APNAVAAAEPSW | ASSFGGRNTEVF | 0.2387218 |
| APQVAGAATAVL | ASSFGGRNTEVF | 0.3575582 |
| ARRDPAAAAAILPTDA | ASSFGGRNTEVF | 0.24711171 |
| DAAARDVLAEAGLAEYF | ASSFGGRNTEVF | 0.4735004 |
| DDQIVLSAKANLAKVEQ | ASSFGGRNTEVF | 0.37035626 |
| DELLPDTGEDSDEDGH | ASSFGGRNTEVF | 0.3436906 |
| ELPEAPANRATVPQD | ASSFGGRNTEVF | 0.5345945 |
| FGQSKIERVTGIG | ASSFGGRNTEVF | 0.50200266 |
| GAGGTGSTAGGAGGAGGAGGLY | ASSFGGRNTEVF | 0.3780208 |
| GHGGTGAAGGNGGTGGAGGSGI | ASSFGGRNTEVF | 0.23085608 |
| GNGGNGGAGGEGGPGGAGGAGA | ASSFGGRNTEVF | 0.31304234 |
| GSTAGNGGNGGNATNGTIAGQP | ASSFGGRNTEVF | 0.21988708 |
| GTGGDGGNGGNGGTGG | ASSFGGRNTEVF | 0.3569466 |
| GVRRPAGIPADAPV | ASSFGGRNTEVF | 0.45157564 |
| GYRQVTDAHLLTLARR | ASSFGGRNTEVF | 0.24229649 |
| IAKMGQDGHDRG | ASSFGGRNTEVF | 0.26314205 |
| KAEPSIEIQYT | ASSFGGRNTEVF | 0.55320346 |
| KPVRIRMAADEDTVAVT | ASSFGGRNTEVF | 0.4379483 |
| MPPRIAGMRLLVIKPEP | ASSFGGRNTEVF | 0.2428132 |
| MKFYAVALFRSRGVPPTR | ASSFGGRNTEVF | 0.22610141 |
| NAGGFGILGAVAHSP | ASSFGGRNTEVF | 0.33414176 |
| NEGREITLAMAEM | ASSFGGRNTEVF | 0.27775773 |
| NELVEWSHPLGHRWRWE | ASSFGGRNTEVF | 0.21230488 |
| NLGAGGTGGNGGNPG | ASSFGGRNTEVF | 0.4096631 |
| NTEWPKWETVMAADPVD | ASSFGGRNTEVF | 0.31754988 |
| PASAAPAETQP | ASSFGGRNTEVF | 0.6615256 |
| QQSSDGKNGVRAAEITGE | ASSFGGRNTEVF | 0.38547012 |

| | | |
|---|---|---|
| RAPDAVTTSWWKE | ASSFGGRNTEVF | 0.49919987 |
| SGFGNVGAHTSGWFNQ | ASSFGGRNTEVF | 0.27157927 |
| SGVAGAVGQAASVGGLKV | ASSFGGRNTEVF | 0.32091957 |
| SNGGAGTGSAGGNGGT | ASSFGGRNTEVF | 0.2088332 |
| TGGAGGEGGAAG | ASSFGGRNTEVF | 0.24460281 |
| TGGDGGAAGAGGAPGHG | ASSFGGRNTEVF | 0.24183457 |
| TMPPLLAAIANAGRHGV | ASSFGGRNTEVF | 0.33528858 |
| TSSTAGNSGGA | ASSFGGRNTEVF | 0.20635606 |
| VPATVPVGILC | ASSFGGRNTEVF | 0.44842517 |
| VQRPDSGGVVLTGRIS | ASSFGGRNTEVF | 0.2518283 |
| WTPNPVPATKGVPAGLVQAT | ASSFGGRNTEVF | 0.43208584 |
| YALKRALAKFSDRFDVVII | ASSFGGRNTEVF | 0.29266825 |
| YLIGSVIV | ASSFGGRNTEVF | 0.22449343 |
| YGQLVAVID | ASSFGGRNTEVF | 0.2505057 |
| VTVGGMTLPA | ASSFGGRNTEVF | 0.53590906 |
| VLALPAV | ASSFGGRNTEVF | 0.21482188 |
| VIPPTLNYET | ASSFGGRNTEVF | 0.20589593 |
| VGMGGMHPSSG | ASSFGGRNTEVF | 0.2441281 |
| VGLAGGAATA | ASSFGGRNTEVF | 0.31914207 |
| VGGAGGAGGSGP | ASSFGGRNTEVF | 0.49479336 |
| VFAAFMV | ASSFGGRNTEVF | 0.4606536 |
| TLYDKLR | ASSFGGRNTEVF | 0.27735984 |
| TLAVFTVL | ASSFGGRNTEVF | 0.6513126 |
| TGSVQPARV | ASSFGGRNTEVF | 0.19741452 |
| TGGGAGSGGT | ASSFGGRNTEVF | 0.2749607 |
| SHLETSL | ASSFGGRNTEVF | 0.6016225 |
| SFTTGMT | ASSFGGRNTEVF | 0.3005012 |
| QVEVAKRMGVRQPTVS | ASSFGGRNTEVF | 0.34517413 |
| QGSGGPALAM | ASSFGGRNTEVF | 0.49397483 |
| MLAPSNP | ASSFGGRNTEVF | 0.26545653 |
| LVNNAAI | ASSFGGRNTEVF | 0.30044144 |
| LTGEINTLA | ASSFGGRNTEVF | 0.33818433 |
| LSIFGPV | ASSFGGRNTEVF | 0.31738818 |
| LLALLANL | ASSFGGRNTEVF | 0.2714179 |
| LISNASDALDKLR | ASSFGGRNTEVF | 0.21133642 |
| LINEAVP | ASSFGGRNTEVF | 0.41679657 |
| LGYAFSGL | ASSFGGRNTEVF | 0.612427 |
| LDFVAVAL | ASSFGGRNTEVF | 0.30488262 |
| LAQPGAE | ASSFGGRNTEVF | 0.41815054 |
| ISNLGAVTGF | ASSFGGRNTEVF | 0.30874583 |
| HQEDPGRI | ASSFGGRNTEVF | 0.5112389 |
| GTPGEPTP | ASSFGGRNTEVF | 0.24489093 |
| GFFNSGS | ASSFGGRNTEVF | 0.21333681 |
| GAGGGGLTAGAN | ASSFGGRNTEVF | 0.38461936 |
| FVDALVNM | ASSFGGRNTEVF | 0.22992826 |
| FLTVAPEL | ASSFGGRNTEVF | 0.35652357 |
| FETLTAL | ASSFGGRNTEVF | 0.27740026 |
| DVINPIEL | ASSFGGRNTEVF | 0.6726155 |
| APVSGAPV | ASSFGGRNTEVF | 0.3913013 |
| ALVGGLAW | ASSFGGRNTEVF | 0.33473098 |
| ALAAAAATAI | ASSFGGRNTEVF | 0.24883765 |

| | | |
|---|---|---|
| AFGLGLLL | ASSFGGRNTEVF | 0.31351393 |
| AFFAAASL | ASSFGGRNTEVF | 0.3977099 |
| AAAALPAFSPP | ASSFGGRNTEVF | 0.21497945 |
| VVLAPAL | ASSFGGRNTEVF | 0.51794046 |
| TIGHVDHGKTTL | ASSFGGRNTEVF | 0.24102549 |
| LISNASDALDK | ASSFGGRNTEVF | 0.2632004 |
| LISNASDAL | ASSFGGRNTEVF | 0.36191306 |
| ISNASDALDK | ASSFGGRNTEVF | 0.674602 |
| NQAETLVYQTE | ASSFGGRNTEVF | 0.28090256 |
| TGGTGGTAGAG | ASSFGGRNTEVF | 0.44640613 |
| TGSTAGGAGGA | ASSFGGRNTEVF | 0.23283443 |
| DPLAVPVAAGPV | ASSFGGRNTEVF | 0.4309255 |
| GGTGGTPGNGGN | ASSFGGRNTEVF | 0.59611166 |
| PAGLAGVDPATL | ASSFGGRNTEVF | 0.3907585 |
| GNNLVVEAPHAGASV | ASSFGGRNTEVF | 0.6789872 |
| KVTRIMSASTLEQPA | ASSFGGRNTEVF | 0.27132824 |
| LGFALPIALPRRTVV | ASSFGGRNTEVF | 0.21279131 |
| TRLGNLTARKLVVLV | ASSFGGRNTEVF | 0.28546986 |
| WSQQRRLIYFPSAGP | ASSFGGRNTEVF | 0.34739387 |
| GPNGNPDPMAAAVDIR | ASSFGGRNTEVF | 0.5650899 |
| TQRCVSVADLRAAASR | ASSFGGRNTEVF | 0.35134196 |
| QFVLDVAEHPVEVLTETV | ASSFGGRNTEVF | 0.3196834 |
| GGAGGTGGAGGNGGTQAGDG | ASSFGGRNTEVF | 0.25900608 |
| PLLATLRAIAVDVDAKPAQV | ASSFGGRNTEVF | 0.5131643 |
| APKFRTGGYMIDFWGVGYQVAK | ASSFGGRNTEVF | 0.56902623 |
| GAGGAGGAGGAAGGPGGTGGTG | ASSFGGRNTEVF | 0.4135576 |
| IGALGLLATDVAPAGSVAFGVA | ASSFGGRNTEVF | 0.24674708 |
| AGGAGGIGGNGGAGGAGGVLMG | ASSFGGRNTEVF | 0.25390142 |
| ADELIGLL | ASSFGGRNTEVF | 0.36489737 |
| ADRALKA | ASSFGGRNTEVF | 0.33458927 |
| AGAVGQVGGAGGA | ASSFGGRNTEVF | 0.23939073 |
| AGGGGGLGGAP | ASSFGGRNTEVF | 0.27411216 |
| AGGLLYGNGGAG | ASSFGGRNTEVF | 0.37538597 |
| AGLESIGL | ASSFGGRNTEVF | 0.20609711 |
| AGLKTGII | ASSFGGRNTEVF | 0.30076027 |
| ASLAALLL | ASSFGGRNTEVF | 0.51695585 |
| ASVINISSI | ASSFGGRNTEVF | 0.296072 |
| ATPPGPA | ASSFGGRNTEVF | 0.21758918 |
| ATVASALV | ASSFGGRNTEVF | 0.5484752 |
| ATVASLIL | ASSFGGRNTEVF | 0.62227976 |
| AVNGPVLIRH | ASSFGGRNTEVF | 0.44182348 |
| DGAAGAVGI | ASSFGGRNTEVF | 0.2308721 |
| DPLLHGI | ASSFGGRNTEVF | 0.1974594 |
| ELGAALGAL | ASSFGGRNTEVF | 0.4659931 |
| ELGRTQAV | ASSFGGRNTEVF | 0.56585044 |
| FVAGGYLS | ASSFGGRNTEVF | 0.30628395 |
| GASGGAGGAGGW | ASSFGGRNTEVF | 0.2268106 |
| GGAGAGGDGGAGGI | ASSFGGRNTEVF | 0.23733671 |
| GTGHNGSQPGMGGNG | ASSFGGRNTEVF | 0.39725325 |
| GYAAASASAS | ASSFGGRNTEVF | 0.22269517 |
| HQVDVAAVQQG | ASSFGGRNTEVF | 0.2220291 |

| | | |
|---|---|---|
| IGAAIAHT | ASSFGGRNTEVF | 0.25329155 |
| IGMIEVAQL | ASSFGGRNTEVF | 0.48976925 |
| IRLAPEHTSAMD | ASSFGGRNTEVF | 0.21967292 |
| IYNVSVL | ASSFGGRNTEVF | 0.24633852 |
| KQGFLGM | ASSFGGRNTEVF | 0.49789196 |
| LESVITQL | ASSFGGRNTEVF | 0.38082892 |
| LGLGALVVV | ASSFGGRNTEVF | 0.31825858 |
| LIGNGGNGGSGGI | ASSFGGRNTEVF | 0.22806032 |
| LNAINAPTL | ASSFGGRNTEVF | 0.3641966 |
| LQLLNTGV | ASSFGGRNTEVF | 0.22604528 |
| LQTSIEF | ASSFGGRNTEVF | 0.46120948 |
| MSFAVSEL | ASSFGGRNTEVF | 0.34410718 |
| NTGALGL | ASSFGGRNTEVF | 0.30812073 |
| PPGAGGAGGAGTT | ASSFGGRNTEVF | 0.28672284 |
| QLVGLGI | ASSFGGRNTEVF | 0.22848427 |
| RFLEQVHQL | ASSFGGRNTEVF | 0.22996674 |
| RGVGYVLR | ASSFGGRNTEVF | 0.25560012 |
| SATGVFAGVI | ASSFGGRNTEVF | 0.5027166 |
| SLPAITLP | ASSFGGRNTEVF | 0.34142378 |
| STALAFTV | ASSFGGRNTEVF | 0.37779623 |
| SVLTVGLGK | ASSFGGRNTEVF | 0.37463534 |
| TALAGAGSGI | ASSFGGRNTEVF | 0.23709668 |
| TGLPFTGI | ASSFGGRNTEVF | 0.33555022 |
| VAEHVTV | ASSFGGRNTEVF | 0.2088634 |
| VDIVLNSL | ASSFGGRNTEVF | 0.33202136 |
| VLTGGGAVL | ASSFGGRNTEVF | 0.25156733 |
| VNVGLAYGI | ASSFGGRNTEVF | 0.3118825 |
| VSSAAVGAV | ASSFGGRNTEVF | 0.29288062 |
| VTSVYEAGV | ASSFGGRNTEVF | 0.3350641 |
| WVALDRGA | ASSFGGRNTEVF | 0.36348414 |
| YLVGSVVI | ASSFGGRNTEVF | 0.27749884 |
| GPPGTGKTLL | ASSFGGRNTEVF | 0.28202477 |
| GVGMTKF | ASSFGGRNTEVF | 0.44665605 |
| IGIGGSDL | ASSFGGRNTEVF | 0.35429466 |
| IGTLLAEL | ASSFGGRNTEVF | 0.21375239 |
| IYTSGTT | ASSFGGRNTEVF | 0.31749898 |
| LSFPAGV | ASSFGGRNTEVF | 0.22783993 |
| LYTSGTT | ASSFGGRNTEVF | 0.33457214 |
| VGAFPFGF | ASSFGGRNTEVF | 0.31688863 |
| VGAGTVEF | ASSFGGRNTEVF | 0.34165388 |
| VGKSTLLL | ASSFGGRNTEVF | 0.3277556 |
| AALAHAGGGIV | ASSFGGRNTEVF | 0.2085507 |
| AALVAVSLL | ASSFGGRNTEVF | 0.25238007 |
| AAVSRAVAAL | ASSFGGRNTEVF | 0.26033095 |
| AGGSSGIGGF | ASSFGGRNTEVF | 0.42932618 |
| AGLTVANANG | ASSFGGRNTEVF | 0.33874995 |
| AGLVGAVL | ASSFGGRNTEVF | 0.2150681 |
| ALSQAFTV | ASSFGGRNTEVF | 0.3800525 |
| ALTFSVSL | ASSFGGRNTEVF | 0.28817716 |
| ALTGAALVAL | ASSFGGRNTEVF | 0.24072747 |
| ASFSIGAGVL | ASSFGGRNTEVF | 0.27145 |

| | | |
|---|---|---|
| DGSVYKI | ASSFGGRNTEVF | 0.2430365 |
| EQLAEIV | ASSFGGRNTEVF | 0.2172768 |
| FAVSELTVM | ASSFGGRNTEVF | 0.35337067 |
| GAAPLAAGL | ASSFGGRNTEVF | 0.25291753 |
| GADGGDGGSGGAGG | ASSFGGRNTEVF | 0.34474012 |
| GFGLDTGGAGG | ASSFGGRNTEVF | 0.24574524 |
| GLASGGLL | ASSFGGRNTEVF | 0.22369744 |
| GRIAGAAVLL | ASSFGGRNTEVF | 0.43105146 |
| GSAGSLNTSF | ASSFGGRNTEVF | 0.24529955 |
| GTGQIGI | ASSFGGRNTEVF | 0.33197564 |
| GTLIALIL | ASSFGGRNTEVF | 0.34161007 |
| IAATMGFVVG | ASSFGGRNTEVF | 0.23878872 |
| IFAQALL | ASSFGGRNTEVF | 0.28362104 |
| IGAAVALGF | ASSFGGRNTEVF | 0.45028472 |
| IGLTPLAL | ASSFGGRNTEVF | 0.37223515 |
| ILGSNTSTL | ASSFGGRNTEVF | 0.26687747 |
| IVFSLTVL | ASSFGGRNTEVF | 0.5545937 |
| LASKATGF | ASSFGGRNTEVF | 0.4201646 |
| LASVALIL | ASSFGGRNTEVF | 0.27630103 |
| LFAVGLTL | ASSFGGRNTEVF | 0.20538257 |
| LFNGGIGTYI | ASSFGGRNTEVF | 0.31676447 |
| LTLGAGII | ASSFGGRNTEVF | 0.30986375 |
| LVAAALGDI | ASSFGGRNTEVF | 0.22924271 |
| NTAFVQLGI | ASSFGGRNTEVF | 0.2189664 |
| QLFSGAEL | ASSFGGRNTEVF | 0.40898806 |
| SAATFITL | ASSFGGRNTEVF | 0.57361805 |
| SNYGLANLGV | ASSFGGRNTEVF | 0.34759712 |
| SYGGAYAVM | ASSFGGRNTEVF | 0.53638184 |
| TTGVGGLTLG | ASSFGGRNTEVF | 0.20681162 |
| TTQTIYI | ASSFGGRNTEVF | 0.4248551 |
| VAAGAGLGAVI | ASSFGGRNTEVF | 0.50770366 |
| VAAVVTLL | ASSFGGRNTEVF | 0.2328147 |
| VGFLGGLI | ASSFGGRNTEVF | 0.31429085 |
| VGVIPILL | ASSFGGRNTEVF | 0.28212082 |
| VIFASSLI | ASSFGGRNTEVF | 0.23306286 |
| VKGGDGGAAAG | ASSFGGRNTEVF | 0.44566566 |
| VTKTFTNI | ASSFGGRNTEVF | 0.3560115 |
| VVAAAILL | ASSFGGRNTEVF | 0.34377858 |
| YAALDVEL | ASSFGGRNTEVF | 0.26743248 |
| LASVALLL | ASSFGGRNTEVF | 0.23194006 |
| LDAVHRV | ASSFGGRNTEVF | 0.3023442 |
| QGLSSSIF | ASSFGGRNTEVF | 0.44360906 |
| RGTFANIRL | ASSFGGRNTEVF | 0.39144373 |
| RQLWWGHRI | ASSFGGRNTEVF | 0.5332283 |
| YGPPGTGKTLL | ASSFGGRNTEVF | 0.2644424 |
| AALVGAPANVA | ASSFGGRNTEVF | 0.5237366 |
| AATFRRTE | ASSFGGRNTEVF | 0.6792773 |
| AGALGGGI | ASSFGGRNTEVF | 0.33433196 |
| AGGAGGAGGIGAG | ASSFGGRNTEVF | 0.25453362 |
| AGSGALGL | ASSFGGRNTEVF | 0.5962812 |
| AQIADQI | ASSFGGRNTEVF | 0.36719108 |

| | | |
|---|---|---|
| ASAVAVTYL | ASSFGGRNTEVF | 0.39545327 |
| ASTVLGGTGG | ASSFGGRNTEVF | 0.41896302 |
| AVGGEPAK | ASSFGGRNTEVF | 0.38424793 |
| CEQGTFGGTC | ASSFGGRNTEVF | 0.65769774 |
| DEGKAPAG | ASSFGGRNTEVF | 0.5667312 |
| DTVVFPAL | ASSFGGRNTEVF | 0.5710853 |
| FEGITMAG | ASSFGGRNTEVF | 0.4567319 |
| FPLVVQD | ASSFGGRNTEVF | 0.3809052 |
| GAFTTPQ | ASSFGGRNTEVF | 0.5026579 |
| GFGGFGTAGGVGGA | ASSFGGRNTEVF | 0.42009026 |
| GFLAQFL | ASSFGGRNTEVF | 0.6439403 |
| GGAGGAGGNAGVA | ASSFGGRNTEVF | 0.6044951 |
| GGVGTAGAGGAGGAGG | ASSFGGRNTEVF | 0.46898112 |
| GNGGGGDAGNAGSGGN | ASSFGGRNTEVF | 0.62183154 |
| GTGGAGGAGNDAGS | ASSFGGRNTEVF | 0.6448462 |
| GTTNTGWFNS | ASSFGGRNTEVF | 0.34734774 |
| IFMGPLFTL | ASSFGGRNTEVF | 0.51805997 |
| ILAHAIEAI | ASSFGGRNTEVF | 0.42782038 |
| IRSAIREAL | ASSFGGRNTEVF | 0.47800666 |
| ITFAGGG | ASSFGGRNTEVF | 0.2838103 |
| ITGMSGHQEV | ASSFGGRNTEVF | 0.56767 |
| IVAGLACTL | ASSFGGRNTEVF | 0.44175386 |
| IVGGGLAGA | ASSFGGRNTEVF | 0.26277095 |
| IVTGVLNI | ASSFGGRNTEVF | 0.54736775 |
| LAQQGCI | ASSFGGRNTEVF | 0.48208094 |
| LCLHVEN | ASSFGGRNTEVF | 0.49902296 |
| LDAISDGS | ASSFGGRNTEVF | 0.6173756 |
| LGGFAAML | ASSFGGRNTEVF | 0.44340327 |
| LGGKNPMI | ASSFGGRNTEVF | 0.44026154 |
| LGPKGVAAIL | ASSFGGRNTEVF | 0.5631341 |
| LMVGVSI | ASSFGGRNTEVF | 0.64706063 |
| LVGAAFVL | ASSFGGRNTEVF | 0.4208004 |
| LVLAVAPL | ASSFGGRNTEVF | 0.53007996 |
| MAAIAVIDAA | ASSFGGRNTEVF | 0.6690768 |
| NNGINAELQ | ASSFGGRNTEVF | 0.5217578 |
| RGEPLVLVHG | ASSFGGRNTEVF | 0.63964146 |
| RGGAGGAAGLI | ASSFGGRNTEVF | 0.66117245 |
| RSYRMHPAV | ASSFGGRNTEVF | 0.4262866 |
| SAAHLLAGQLA | ASSFGGRNTEVF | 0.63525265 |
| SAAVGSY | ASSFGGRNTEVF | 0.66093755 |
| SPHAVLHRI | ASSFGGRNTEVF | 0.524066 |
| SQIAKVL | ASSFGGRNTEVF | 0.48787233 |
| TALGHLVP | ASSFGGRNTEVF | 0.40272766 |
| TAPTTSGAA | ASSFGGRNTEVF | 0.50197005 |
| TDGDDGNGGSPGT | ASSFGGRNTEVF | 0.3548056 |
| TDGLGEN | ASSFGGRNTEVF | 0.5280087 |
| TGGPPLPGQA | ASSFGGRNTEVF | 0.6518728 |
| TIPAITI | ASSFGGRNTEVF | 0.47426385 |
| TIPGFTI | ASSFGGRNTEVF | 0.3293074 |
| TSIASGALA | ASSFGGRNTEVF | 0.34181795 |
| TVTLPGAAI | ASSFGGRNTEVF | 0.5596791 |

| | | |
|---|---|---|
| VASGLSAALP | ASSFGGRNTEVF | 0.41636974 |
| VGAGVIGI | ASSFGGRNTEVF | 0.347066 |
| VGPKRFGV | ASSFGGRNTEVF | 0.4458951 |
| VNVAFPD | ASSFGGRNTEVF | 0.62377167 |
| VSMTNSGVS | ASSFGGRNTEVF | 0.62892747 |
| VTDVLGDLSV | ASSFGGRNTEVF | 0.4019411 |
| VVVVGAGFAGL | ASSFGGRNTEVF | 0.66879296 |
| VYTVGTVV | ASSFGGRNTEVF | 0.610613 |
| WTTFSGP | ASSFGGRNTEVF | 0.6763825 |
| YDELVEAF | ASSFGGRNTEVF | 0.538483 |
| YGTGGAGGNGG | ASSFGGRNTEVF | 0.39436555 |
| YTNVMAV | ASSFGGRNTEVF | 0.644364 |
| FLEQVHQL | ASSFGGRNTEVF | 0.4680571 |
| GLSLLAVAV | ASSFGGRNTEVF | 0.50304085 |
| LEQVHQL | ASSFGGRNTEVF | 0.60570365 |
| LNAAGAI | ASSFGGRNTEVF | 0.5544141 |
| PAVHASL | ASSFGGRNTEVF | 0.39845505 |
| TELEDDA | ASSFGGRNTEVF | 0.52014285 |
| TPPGPPN | ASSFGGRNTEVF | 0.6422195 |
| TPPPPPPP | ASSFGGRNTEVF | 0.35228908 |
| TQSPALI | ASSPTGYQDTQY | 0.44369522 |
| EFAAAFPQ | ASSPTGYQDTQY | 0.4604202 |
| LLSGNLDVL | ASSPTGYQDTQY | 0.51186097 |
| GPANLGNYNI | ASSPTGYQDTQY | 0.64644897 |
| TLGPTLP | ASSPTGYQDTQY | 0.5694798 |
| VYAYSSCCK | ASSPTGYQDTQY | 0.47405016 |
| APAPAPAPNI | ASSPTGYQDTQY | 0.5910686 |
| PGGPVPIP | ASSPTGYQDTQY | 0.6118478 |
| FLTVALEP | ASSPTGYQDTQY | 0.526078 |
| SQYFGANL | ASSPTGYQDTQY | 0.5687306 |
| GGVKPAA | ASSPTGYQDTQY | 0.406563 |
| GGTGGAGGAGGW | ASSPTGYQDTQY | 0.6290191 |
| GVGGVAGLGGAGG | ASSPTGYQDTQY | 0.60034007 |
| VLLVLATTI | ASSPTGYQDTQY | 0.6029791 |
| IGIFAAML | ASSPTGYQDTQY | 0.6287518 |
| AVGDLID | ASSPTGYQDTQY | 0.6021863 |
| NISPLPL | ASSPTGYQDTQY | 0.5493159 |
| VTAVFVAI | ASSPTGYQDTQY | 0.34313577 |
| SGDDNDAAL | ASSPTGYQDTQY | 0.4659515 |
| AVAALAHS | ASSPTGYQDTQY | 0.55173117 |
| GGVTVMATP | ASSPTGYQDTQY | 0.55030465 |
| AGLAANVRD | ASSPTGYQDTQY | 0.6102553 |
| TNRPDLIDPALLRPG | ASSPTGYQDTQY | 0.60068846 |
| LANLGVTAL | ASSPTGYQDTQY | 0.41841733 |
| TGERIEL | ASSPTGYQDTQY | 0.6474897 |
| NGEELAA | ASSPTGYQDTQY | 0.66294837 |
| VLSLVWG | ASSPTGYQDTQY | 0.33541083 |
| IGYALWL | ASSPTGYQDTQY | 0.47205096 |
| LSLLLAPL | ASSPTGYQDTQY | 0.6719538 |
| TLDEALK | ASSPTGYQDTQY | 0.58295095 |
| TDGAGTAEGG | ASSPTGYQDTQY | 0.48857576 |

| | | |
|---|---|---|
| NAPGGAGGNGG | ASSPTGYQDTQY | 0.50025433 |
| VSVFMGMGGNF | ASSPTGYQDTQY | 0.68308914 |
| LASYLAAL | ASSPTGYQDTQY | 0.46477318 |
| LVGGTPAL | ASSPTGYQDTQY | 0.6661811 |
| VRNADVL | ASSPTGYQDTQY | 0.30132276 |
| TCSSPFLVSAM | ASSPTGYQDTQY | 0.5128507 |
| VHITDAE | ASSPTGYQDTQY | 0.42034414 |
| EYAAPSF | ASSPTGYQDTQY | 0.28327975 |
| VVDAFHGGL | ASSPTGYQDTQY | 0.5924431 |
| ASVVDPQ | ASSPTGYQDTQY | 0.49871483 |
| SYNEFGGLS | ASSPTGYQDTQY | 0.6032826 |
| DPDGSVY | ASSPTGYQDTQY | 0.47184667 |
| MVSTIHATE | ASSPTGYQDTQY | 0.609764 |
| LAAGVADPV | ASSPTGYQDTQY | 0.53077877 |
| TKEQIAA | ASSPTGYQDTQY | 0.5180467 |
| SLASPAAL | ASSPTGYQDTQY | 0.42961004 |
| IGLVFYGV | ASSPTGYQDTQY | 0.575705 |
| FLLDASNL | ASSPTGYQDTQY | 0.6547886 |
| YQVGNFP | ASSPTGYQDTQY | 0.62755 |
| ESLLSQST | ASSPTGYQDTQY | 0.35446453 |
| LFTVGFAAVG | ASSPTGYQDTQY | 0.5104753 |
| ATPPTPT | ASSPTGYQDTQY | 0.5029742 |
| TDLATLG | ASSPTGYQDTQY | 0.34082896 |
| SAGGTGR | ASSPTGYQDTQY | 0.29386187 |
| GFSNSGS | ASSPTGYQDTQY | 0.46781054 |
| GGNGGSGGTSV | ASSPTGYQDTQY | 0.46601838 |
| GTSVATGGAGN | ASSPTGYQDTQY | 0.59741557 |
| NGGNGGSGGTS | ASSPTGYQDTQY | 0.5890149 |
| AVVKFPH | ASSPTGYQDTQY | 0.46203592 |
| TPPPPPP | ASSPTGYQDTQY | 0.60367215 |
| AAPAAAAAPA | ASSPTGYQDTQY | 0.5573211 |
| IALNEDL | ASSPTGYQDTQY | 0.23855232 |
| IVNVSSM | ASSPTGYQDTQY | 0.44462067 |
| TGIPPAPR | ASSPTGYQDTQY | 0.34499058 |
| FDLGGGTF | ASSPTGYQDTQY | 0.64680856 |
| FDLGGGTFDVS | ASSPTGYQDTQY | 0.38544387 |
| NEPTAAA | ASSPTGYQDTQY | 0.39815968 |
| NEPTAAALA | ASSPTGYQDTQY | 0.51323336 |
| GSIINISSI | ASSPTGYQDTQY | 0.58814204 |
| VAPGNAGTA | ASSPTGYQDTQY | 0.51087356 |
| TGGGASLE | ASSPTGYQDTQY | 0.39978918 |
| VAVYNHYKRI | ASSPTGYQDTQY | 0.30191368 |
| TDPAPATT | ASSPTGYQDTQY | 0.59017605 |
| TNRPDILDPALLRPG | ASSPTGYQDTQY | 0.4506997 |
| LTTTTTS | ASSPTGYQDTQY | 0.43102655 |
| PTGSGKTL | ASSPTGYQDTQY | 0.34002858 |
| TGSGKTL | ASSPTGYQDTQY | 0.46953923 |
| TIGHVDHGK | ASSPTGYQDTQY | 0.34751266 |
| TIGHVDHGKTT | ASSPTGYQDTQY | 0.61877537 |
| GHVDHGKTT | ASSPTGYQDTQY | 0.44107953 |
| AQFEHTLL | ASSPTGYQDTQY | 0.45751652 |

| | | |
|---|---|---|
| IVNQDEL | ASSPTGYQDTQY | 0.6664931 |
| TNGSQFF | ASSPTGYQDTQY | 0.57045615 |
| TGKGIVE | ASSPTGYQDTQY | 0.46581468 |
| LISNASDA | ASSPTGYQDTQY | 0.4957787 |
| AELRTLGELTKIAKAHGAQ | ASSPTGYQDTQY | 0.38697374 |
| APNAVAAAEPSW | ASSPTGYQDTQY | 0.44433206 |
| APQVAGAATAVL | ASSPTGYQDTQY | 0.4615426 |
| ARRDPAAAAAILPTDA | ASSPTGYQDTQY | 0.66193616 |
| DAAARDVLAEAGLAEYF | ASSPTGYQDTQY | 0.5034697 |
| DDQIVLSAKANLAKVEQ | ASSPTGYQDTQY | 0.38647664 |
| DELLPDTGEDSDEDGH | ASSPTGYQDTQY | 0.5925551 |
| ELPEAPANRATVPQD | ASSPTGYQDTQY | 0.6594948 |
| FGQSKIERVTGIG | ASSPTGYQDTQY | 0.69016 |
| GAGGTGSTAGGAGGAGGAGGLY | ASSPTGYQDTQY | 0.3847665 |
| GHGGTGAAGGNGGTGGAGGSGI | ASSPTGYQDTQY | 0.4836815 |
| GNGGNGGAGGEGGPGGAGGAGA | ASSPTGYQDTQY | 0.50196505 |
| GSTAGNGGNGGNATNGTIAGQP | ASSPTGYQDTQY | 0.46080106 |
| GTGGDGGNGGNGGTGG | ASSPTGYQDTQY | 0.45351502 |
| GVRRPAGIPADAPV | ASSPTGYQDTQY | 0.4285071 |
| GYRQVTDAHLLTLARR | ASSPTGYQDTQY | 0.6182002 |
| IAKMGQDGHDRG | ASSPTGYQDTQY | 0.7063999 |
| KAEPSIEIQYT | ASSPTGYQDTQY | 0.4827973 |
| KPVRIRMAADEDTVAVT | ASSPTGYQDTQY | 0.3133122 |
| MPPRIAGMRLLVIKPEP | ASSPTGYQDTQY | 0.5045201 |
| MKFYAVALFRSRGVPPTR | ASSPTGYQDTQY | 0.37303004 |
| NAGGFGILGAVAHSP | ASSPTGYQDTQY | 0.4141338 |
| NEGREITLAMAEM | ASSPTGYQDTQY | 0.5148126 |
| NELVEWSHPLGHRWRWE | ASSPTGYQDTQY | 0.31879508 |
| NLGAGGTGGNGGNPG | ASSPTGYQDTQY | 0.49981305 |
| NTEWPKWETVMAADPVD | ASSPTGYQDTQY | 0.6728915 |
| PASAAPAETQP | ASSPTGYQDTQY | 0.49372578 |
| QQSSDGKNGVRAAEITGE | ASSPTGYQDTQY | 0.6186734 |
| RAPDAVTTSWWKE | ASSPTGYQDTQY | 0.6392035 |
| SGFGNVGAHTSGWFNQ | ASSPTGYQDTQY | 0.57229674 |
| SGVAGAVGQAASVGGLKV | ASSPTGYQDTQY | 0.638273 |
| SNGGAGTGSAGGNGGT | ASSPTGYQDTQY | 0.6296479 |
| TGGAGGEGGAAG | ASSPTGYQDTQY | 0.53138113 |
| TGGDGGAAGAGGAPGHG | ASSPTGYQDTQY | 0.5009309 |
| TMPPLLAAIANAGRHGV | ASSPTGYQDTQY | 0.62496614 |
| TSSTAGNSGGA | ASSPTGYQDTQY | 0.5050456 |
| VPATVPVGILC | ASSPTGYQDTQY | 0.54159415 |
| VQRPDSGGVVLTGRIS | ASSPTGYQDTQY | 0.4455006 |
| WTPNPVPATKGVPAGLVQAT | ASSPTGYQDTQY | 0.61592877 |
| YALKRALAKFSDRFDVVII | ASSPTGYQDTQY | 0.50497717 |
| YLIGSVIV | ASSPTGYQDTQY | 0.6771092 |
| YGQLVAVID | ASSPTGYQDTQY | 0.34104446 |
| VTVGGMTLPA | ASSPTGYQDTQY | 0.42394775 |
| VLALPAV | ASSPTGYQDTQY | 0.62428606 |
| VIPPTLNYET | ASSPTGYQDTQY | 0.27771002 |
| VGMGGMHPSSG | ASSPTGYQDTQY | 0.3194422 |
| VGLAGGAATA | ASSPTGYQDTQY | 0.54042983 |

| | | |
|---|---|---|
| VGGAGGAGGSGP | ASSPTGYQDTQY | 0.6500902 |
| VFAAFMV | ASSPTGYQDTQY | 0.27360755 |
| TLYDKLR | ASSPTGYQDTQY | 0.4693045 |
| TLAVFTVL | ASSPTGYQDTQY | 0.3891641 |
| TGSVQPARV | ASSPTGYQDTQY | 0.6157124 |
| TGGGAGSGGT | ASSPTGYQDTQY | 0.50761306 |
| SHLETSL | ASSPTGYQDTQY | 0.35562754 |
| SFTTGMT | ASSPTGYQDTQY | 0.60569483 |
| QVEVAKRMGVRQPTVS | ASSPTGYQDTQY | 0.31349733 |
| QGSGGPALAM | ASSPTGYQDTQY | 0.51214826 |
| MLAPSNP | ASSPTGYQDTQY | 0.59978133 |
| LVNNAAI | ASSPTGYQDTQY | 0.67255354 |
| LTGEINTLA | ASSPTGYQDTQY | 0.37440658 |
| LSIFGPV | ASSPTGYQDTQY | 0.59730405 |
| LLALLANL | ASSPTGYQDTQY | 0.40587088 |
| LISNASDALDKLR | ASSPTGYQDTQY | 0.7056488 |
| LINEAVP | ASSPTGYQDTQY | 0.49106598 |
| LGYAFSGL | ASSPTGYQDTQY | 0.35267612 |
| LDFVAVAL | ASSPTGYQDTQY | 0.6202836 |
| LAQPGAE | ASSPTGYQDTQY | 0.6123593 |
| ISNLGAVTGF | ASSPTGYQDTQY | 0.5406367 |
| HQEDPGRI | ASSPTGYQDTQY | 0.6111583 |
| GTPGEPTP | ASSPTGYQDTQY | 0.50208443 |
| GFFNSGS | ASSPTGYQDTQY | 0.48753554 |
| GAGGGGLTAGAN | ASSPTGYQDTQY | 0.60903764 |
| FVDALVNM | ASSPTGYQDTQY | 0.5872223 |
| FLTVAPEL | ASSPTGYQDTQY | 0.45086586 |
| FETLTAL | ASSPTGYQDTQY | 0.6339384 |
| DVINPIEL | ASSPTGYQDTQY | 0.536596 |
| APVSGAPV | ASSPTGYQDTQY | 0.43739185 |
| ALVGGLAW | ASSPTGYQDTQY | 0.668939 |
| ALAAAAATAI | ASSPTGYQDTQY | 0.52975637 |
| AFGLGLLL | ASSPTGYQDTQY | 0.61501795 |
| AFFAAASL | ASSPTGYQDTQY | 0.59129494 |
| AAAALPAFSPP | ASSPTGYQDTQY | 0.69105077 |
| VVLAPAL | ASSPTGYQDTQY | 0.39829534 |
| TIGHVDHGKTTL | ASSPTGYQDTQY | 0.588241 |
| LISNASDALDK | ASSPTGYQDTQY | 0.5424385 |
| LISNASDAL | ASSPTGYQDTQY | 0.61377305 |
| ISNASDALDK | ASSPTGYQDTQY | 0.49937916 |
| NQAETLVYQTE | ASSPTGYQDTQY | 0.5046356 |
| TGGTGGTAGAG | ASSPTGYQDTQY | 0.5769714 |
| TGSTAGGAGGA | ASSPTGYQDTQY | 0.65270114 |
| DPLAVPVAAGPV | ASSPTGYQDTQY | 0.5155996 |
| GGTGGTPGNGGN | ASSPTGYQDTQY | 0.55228734 |
| PAGLAGVDPATL | ASSPTGYQDTQY | 0.6640351 |
| GNNLVVEAPHAGASV | ASSPTGYQDTQY | 0.4483884 |
| KVTRIMSASTLEQPA | ASSPTGYQDTQY | 0.5395577 |
| LGFALPIALPRRTVV | ASSPTGYQDTQY | 0.61360025 |
| TRLGNLTARKLVVLV | ASSPTGYQDTQY | 0.65052783 |
| WSQQRRLIYFPSAGP | ASSPTGYQDTQY | 0.4952472 |

| | | |
|---|---|---|
| GPNGNPDPMAAAVDIR | ASSPTGYQDTQY | 0.64659166 |
| TQRCVSVADLRAAASR | ASSPTGYQDTQY | 0.4274439 |
| QFVLDVAEHPVEVLTETV | ASSPTGYQDTQY | 0.61166555 |
| GGAGGTGGAGGNGGTQAGDG | ASSPTGYQDTQY | 0.4455232 |
| PLLATLRAIAVDVDAKPAQV | ASSPTGYQDTQY | 0.49581325 |
| APKFRTGGYMIDFWGVGYQVAK | ASSPTGYQDTQY | 0.41708595 |
| GAGGAGGAGGAAGGPGGTGGTG | ASSPTGYQDTQY | 0.5533697 |
| IGALGLLATDVAPAGSVAFGVA | ASSPTGYQDTQY | 0.6110608 |
| AGGAGGIGGNGGAGGAGGVLMG | ASSPTGYQDTQY | 0.5979352 |
| ADELIGLL | ASSPTGYQDTQY | 0.30419612 |
| ADRALKA | ASSPTGYQDTQY | 0.57567143 |
| AGAVGQVGGAGGA | ASSPTGYQDTQY | 0.61210424 |
| AGGGGGLGGAP | ASSPTGYQDTQY | 0.6830121 |
| AGGLLYGNGGAG | ASSPTGYQDTQY | 0.48648354 |
| AGLESIGL | ASSPTGYQDTQY | 0.6194941 |
| AGLKTGII | ASSPTGYQDTQY | 0.48652384 |
| ASLAALLL | ASSPTGYQDTQY | 0.5191394 |
| ASVINISSI | ASSPTGYQDTQY | 0.33333766 |
| ATPPGPA | ASSPTGYQDTQY | 0.49802285 |
| ATVASALV | ASSPTGYQDTQY | 0.43324715 |
| ATVASLIL | ASSPTGYQDTQY | 0.39102897 |
| AVNGPVLIRH | ASSPTGYQDTQY | 0.63036925 |
| DGAAGAVGI | ASSPTGYQDTQY | 0.6237043 |
| DPLLHGI | ASSPTGYQDTQY | 0.63029975 |
| ELGAALGAL | ASSPTGYQDTQY | 0.6608698 |
| ELGRTQAV | ASSPTGYQDTQY | 0.67084897 |
| FVAGGYLS | ASSPTGYQDTQY | 0.63385594 |
| GASGGAGGAGGW | ASSPTGYQDTQY | 0.4470832 |
| GGAGAGGDGGAGGI | ASSPTGYQDTQY | 0.50255924 |
| GTGHNGSQPGMGGNG | ASSPTGYQDTQY | 0.33493355 |
| GYAAASASAS | ASSPTGYQDTQY | 0.44880912 |
| HQVDVAAVQQG | ASSPTGYQDTQY | 0.4662426 |
| IGAAIAHT | ASSPTGYQDTQY | 0.51820976 |
| IGMIEVAQL | ASSPTGYQDTQY | 0.27603903 |
| IRLAPEHTSAMD | ASSPTGYQDTQY | 0.54797554 |
| IYNVSVL | ASSPTGYQDTQY | 0.66960764 |
| KQGFLGM | ASSPTGYQDTQY | 0.50898874 |
| LESVITQL | ASSPTGYQDTQY | 0.5944861 |
| LGLGALVVV | ASSPTGYQDTQY | 0.586199 |
| LIGNGGNGGSGGI | ASSPTGYQDTQY | 0.4008626 |
| LNAINAPTL | ASSPTGYQDTQY | 0.6227023 |
| LQLLNTGV | ASSPTGYQDTQY | 0.41190192 |
| LQTSIEF | ASSPTGYQDTQY | 0.60307664 |
| MSFAVSEL | ASSPTGYQDTQY | 0.6615664 |
| NTGALGL | ASSPTGYQDTQY | 0.5683123 |
| PPGAGGAGGAGTT | ASSPTGYQDTQY | 0.5094801 |
| QLVGLGI | ASSPTGYQDTQY | 0.5359784 |
| RFLEQVHQL | ASSPTGYQDTQY | 0.36537695 |
| RGVGYVLR | ASSPTGYQDTQY | 0.5395498 |
| SATGVFAGVI | ASSPTGYQDTQY | 0.40645522 |
| SLPAITLP | ASSPTGYQDTQY | 0.5264216 |

| | | |
|---|---|---|
| STALAFTV | ASSPTGYQDTQY | 0.39836362 |
| SVLTVGLGK | ASSPTGYQDTQY | 0.5788642 |
| TALAGAGSGI | ASSPTGYQDTQY | 0.5411446 |
| TGLPFTGI | ASSPTGYQDTQY | 0.567535 |
| VAEHVTV | ASSPTGYQDTQY | 0.39336774 |
| VDIVLNSL | ASSPTGYQDTQY | 0.543595 |
| VLTGGGAVL | ASSPTGYQDTQY | 0.50798804 |
| VNVGLAYGI | ASSPTGYQDTQY | 0.37629136 |
| VSSAAVGAV | ASSPTGYQDTQY | 0.6203539 |
| VTSVYEAGV | ASSPTGYQDTQY | 0.511426 |
| WVALDRGA | ASSPTGYQDTQY | 0.47529107 |
| YLVGSVVI | ASSPTGYQDTQY | 0.6296159 |
| GPPGTGKTLL | ASSPTGYQDTQY | 0.5466366 |
| GVGMTKF | ASSPTGYQDTQY | 0.33587617 |
| IGIGGSDL | ASSPTGYQDTQY | 0.49808556 |
| IGTLLAEL | ASSPTGYQDTQY | 0.5429276 |
| IYTSGTT | ASSPTGYQDTQY | 0.49481353 |
| LSFPAGV | ASSPTGYQDTQY | 0.27846187 |
| LYTSGTT | ASSPTGYQDTQY | 0.57731473 |
| VGAFPFGF | ASSPTGYQDTQY | 0.443154 |
| VGAGTVEF | ASSPTGYQDTQY | 0.38990754 |
| VGKSTLLL | ASSPTGYQDTQY | 0.45861742 |
| AALAHAGGGIV | ASSPTGYQDTQY | 0.6340413 |
| AALVAVSLL | ASSPTGYQDTQY | 0.55772686 |
| AAVSRAVAAL | ASSPTGYQDTQY | 0.54097545 |
| AGGSSGIGGF | ASSPTGYQDTQY | 0.580483 |
| AGLTVANANG | ASSPTGYQDTQY | 0.57340324 |
| AGLVGAVL | ASSPTGYQDTQY | 0.63989913 |
| ALSQAFTV | ASSPTGYQDTQY | 0.6096365 |
| ALTFSVSL | ASSPTGYQDTQY | 0.506644 |
| ALTGAALVAL | ASSPTGYQDTQY | 0.57232606 |
| ASFSIGAGVL | ASSPTGYQDTQY | 0.49015674 |
| DGSVYKI | ASSPTGYQDTQY | 0.33808607 |
| EQLAEIV | ASSPTGYQDTQY | 0.3433286 |
| FAVSELTVM | ASSPTGYQDTQY | 0.44805297 |
| GAAPLAAGL | ASSPTGYQDTQY | 0.34109366 |
| GADGGDGGSGGAGG | ASSPTGYQDTQY | 0.572191 |
| GFGLDTGGAGG | ASSPTGYQDTQY | 0.6564665 |
| GLASGGLL | ASSPTGYQDTQY | 0.54366785 |
| GRIAGAAVLL | ASSPTGYQDTQY | 0.3601968 |
| GSAGSLNTSF | ASSPTGYQDTQY | 0.45492953 |
| GTGQIGI | ASSPTGYQDTQY | 0.43442848 |
| GTLIALIL | ASSPTGYQDTQY | 0.59964585 |
| IAATMGFVVG | ASSPTGYQDTQY | 0.6107218 |
| IFAQALL | ASSPTGYQDTQY | 0.68358076 |
| IGAAVALGF | ASSPTGYQDTQY | 0.61197525 |
| IGLTPLAL | ASSPTGYQDTQY | 0.45996025 |
| ILGSNTSTL | ASSPTGYQDTQY | 0.60967606 |
| IVFSLTVL | ASSPTGYQDTQY | 0.44212317 |
| LASKATGF | ASSPTGYQDTQY | 0.52169055 |
| LASVALIL | ASSPTGYQDTQY | 0.5036106 |

| | | |
|---|---|---|
| LFAVGLTL | ASSPTGYQDTQY | 0.564606 |
| LFNGGIGTYI | ASSPTGYQDTQY | 0.5363066 |
| LTLGAGII | ASSPTGYQDTQY | 0.47352275 |
| LVAAALGDI | ASSPTGYQDTQY | 0.34469348 |
| NTAFVQLGI | ASSPTGYQDTQY | 0.61824536 |
| QLFSGAEL | ASSPTGYQDTQY | 0.5125419 |
| SAATFITL | ASSPTGYQDTQY | 0.36049652 |
| SNYGLANLGV | ASSPTGYQDTQY | 0.45017493 |
| SYGGAYAVM | ASSPTGYQDTQY | 0.6818109 |
| TTGVGGLTLG | ASSPTGYQDTQY | 0.6797405 |
| TTQTIYI | ASSPTGYQDTQY | 0.4141562 |
| VAAGAGLGAVI | ASSPTGYQDTQY | 0.4207837 |
| VAAVVTLL | ASSPTGYQDTQY | 0.5792033 |
| VGFLGGLI | ASSPTGYQDTQY | 0.4017957 |
| VGVIPILL | ASSPTGYQDTQY | 0.5727691 |
| VIFASSLI | ASSPTGYQDTQY | 0.47951728 |
| VKGGDGGAAAG | ASSPTGYQDTQY | 0.5108413 |
| VTKTFTNI | ASSPTGYQDTQY | 0.46599966 |
| VVAAAILL | ASSPTGYQDTQY | 0.57614917 |
| YAALDVEL | ASSPTGYQDTQY | 0.48990977 |
| LASVALLL | ASSPTGYQDTQY | 0.5951034 |
| LDAVHRV | ASSPTGYQDTQY | 0.28160977 |
| QGLSSSIF | ASSPTGYQDTQY | 0.58940756 |
| RGTFANIRL | ASSPTGYQDTQY | 0.37880751 |
| RQLWWGHRI | ASSPTGYQDTQY | 0.420605 |
| YGPPGTGKTLL | ASSPTGYQDTQY | 0.3180247 |
| AALVGAPANVA | ASSPTGYQDTQY | 0.4967354 |
| AATFRRTE | ASSPTGYQDTQY | 0.6281042 |
| AGALGGGI | ASSPTGYQDTQY | 0.64228696 |
| AGGAGGAGGIGAG | ASSPTGYQDTQY | 0.43539795 |
| AGSGALGL | ASSPTGYQDTQY | 0.6050167 |
| AQIADQI | ASSPTGYQDTQY | 0.67317927 |
| ASAVAVTYL | ASSPTGYQDTQY | 0.41038814 |
| ASTVLGGTGG | ASSPTGYQDTQY | 0.3325905 |
| AVGGEPAK | ASSPTGYQDTQY | 0.5699825 |
| CEQGTFGGTC | ASSPTGYQDTQY | 0.5223033 |
| DEGKAPAG | ASSPTGYQDTQY | 0.6224438 |
| DTVVFPAL | ASSPTGYQDTQY | 0.65159416 |
| FEGITMAG | ASSPTGYQDTQY | 0.32762092 |
| FPLVVQD | ASSPTGYQDTQY | 0.6144041 |
| GAFTTPQ | ASSPTGYQDTQY | 0.6404662 |
| GFGGFGTAGGVGGA | ASSPTGYQDTQY | 0.3802015 |
| GFLAQFL | ASSPTGYQDTQY | 0.39654192 |
| GGAGGAGGNAGVA | ASSPTGYQDTQY | 0.4377485 |
| GGVGTAGAGGAGGAGG | ASSPTGYQDTQY | 0.32814643 |
| GNGGGGDAGNAGSGGN | ASSPTGYQDTQY | 0.59450036 |
| GTGGAGGAGNDAGS | ASSPTGYQDTQY | 0.4347065 |
| GTTNTGWFNS | ASSPTGYQDTQY | 0.50333834 |
| IFMGPLFTL | ASSPTGYQDTQY | 0.48101306 |
| ILAHAIEAI | ASSPTGYQDTQY | 0.49197996 |
| IRSAIREAL | ASSPTGYQDTQY | 0.49749464 |

| | | |
|---|---|---|
| ITFAGGG | ASSPTGYQDTQY | 0.5171625 |
| ITGMSGHQEV | ASSPTGYQDTQY | 0.5497402 |
| IVAGLACTL | ASSPTGYQDTQY | 0.5898713 |
| IVGGGLAGA | ASSPTGYQDTQY | 0.4637465 |
| IVTGVLNI | ASSPTGYQDTQY | 0.56428397 |
| LAQQGCI | ASSPTGYQDTQY | 0.65526843 |
| LCLHVEN | ASSPTGYQDTQY | 0.39980847 |
| LDAISDGS | ASSPTGYQDTQY | 0.43348843 |
| LGGFAAML | ASSPTGYQDTQY | 0.61403793 |
| LGGKNPMI | ASSPTGYQDTQY | 0.4159842 |
| LGPKGVAAIL | ASSPTGYQDTQY | 0.6228749 |
| LMVGVSI | ASSPTGYQDTQY | 0.56810063 |
| LVGAAFVL | ASSPTGYQDTQY | 0.5562506 |
| LVLAVAPL | ASSPTGYQDTQY | 0.5417699 |
| MAAIAVIDAA | ASSPTGYQDTQY | 0.68661815 |
| NNGINAELQ | ASSPTGYQDTQY | 0.57949936 |
| RGEPLVLVHG | ASSPTGYQDTQY | 0.48835608 |
| RGGAGGAAGLI | ASSPTGYQDTQY | 0.62381184 |
| RSYRMHPAV | ASSPTGYQDTQY | 0.41832057 |
| SAAHLLAGQLA | ASSPTGYQDTQY | 0.613716 |
| SAAVGSY | ASSPTGYQDTQY | 0.62672013 |
| SPHAVLHRI | ASSPTGYQDTQY | 0.36989367 |
| SQIAKVL | ASSPTGYQDTQY | 0.49109325 |
| TALGHLVP | ASSPTGYQDTQY | 0.654226 |
| TAPTTSGAA | ASSPTGYQDTQY | 0.6357671 |
| TDGDDGNGGSPGT | ASSPTGYQDTQY | 0.37788245 |
| TDGLGEN | ASSPTGYQDTQY | 0.4102815 |
| TGGPPLPGQA | ASSPTGYQDTQY | 0.40415892 |
| TIPAITI | ASSPTGYQDTQY | 0.46978027 |
| TIPGFTI | ASSPTGYQDTQY | 0.40570518 |
| TSIASGALA | ASSPTGYQDTQY | 0.5874035 |
| TVTLPGAAI | ASSPTGYQDTQY | 0.48261034 |
| VASGLSAALP | ASSPTGYQDTQY | 0.6351843 |
| VGAGVIGI | ASSPTGYQDTQY | 0.5663091 |
| VGPKRFGV | ASSPTGYQDTQY | 0.63380593 |
| VNVAFPD | ASSPTGYQDTQY | 0.45627883 |
| VSMTNSGVS | ASSPTGYQDTQY | 0.5174569 |
| VTDVLGDLSV | ASSPTGYQDTQY | 0.35091177 |
| VVVVGAGFAGL | ASSPTGYQDTQY | 0.45974907 |
| VYTVGTVV | ASSPTGYQDTQY | 0.4155825 |
| WTTFSGP | ASSPTGYQDTQY | 0.52981776 |
| YDELVEAF | ASSPTGYQDTQY | 0.31189167 |
| YGTGGAGGNGG | ASSPTGYQDTQY | 0.66934466 |
| YTNVMAV | ASSPTGYQDTQY | 0.47318467 |
| FLEQVHQL | ASSPTGYQDTQY | 0.59360063 |
| GLSLLAVAV | ASSPTGYQDTQY | 0.46563786 |
| LEQVHQL | ASSPTGYQDTQY | 0.4464567 |
| LNAAGAI | ASSPTGYQDTQY | 0.45999202 |
| PAVHASL | ASSPTGYQDTQY | 0.61977 |
| TELEDDA | ASSPTGYQDTQY | 0.3681106 |
| TPPGPPN | ASSPTGYQDTQY | 0.3100819 |

| | | |
|---|---|---|
| TPPPPPPP | ASSPTGYQDTQY | 0.44644845 |
| TQSPALI | ASSLTAYRNSDYT | 0.3245213 |
| EFAAAFPQ | ASSLTAYRNSDYT | 0.6826009 |
| LLSGNLDVL | ASSLTAYRNSDYT | 0.63631344 |
| GPANLGNYNI | ASSLTAYRNSDYT | 0.52744335 |
| TLGPTLP | ASSLTAYRNSDYT | 0.69439584 |
| VYAYSSCCK | ASSLTAYRNSDYT | 0.27590135 |
| APAPAPAPNI | ASSLTAYRNSDYT | 0.47625545 |
| PGGPVPIP | ASSLTAYRNSDYT | 0.6612801 |
| FLTVALEP | ASSLTAYRNSDYT | 0.55230194 |
| SQYFGANL | ASSLTAYRNSDYT | 0.5304404 |
| GGVKPAA | ASSLTAYRNSDYT | 0.61644053 |
| GGTGGAGGAGGW | ASSLTAYRNSDYT | 0.4425007 |
| GVGGVAGLGGAGG | ASSLTAYRNSDYT | 0.50662726 |
| VLLVLATTI | ASSLTAYRNSDYT | 0.519778 |
| IGIFAAML | ASSLTAYRNSDYT | 0.5776557 |
| AVGDLID | ASSLTAYRNSDYT | 0.5567313 |
| NISPLPL | ASSLTAYRNSDYT | 0.36245337 |
| VTAVFVAI | ASSLTAYRNSDYT | 0.59447414 |
| SGDDNDAAL | ASSLTAYRNSDYT | 0.69116133 |
| AVAALAHS | ASSLTAYRNSDYT | 0.46475372 |
| GGVTVMATP | ASSLTAYRNSDYT | 0.6015993 |
| AGLAANVRD | ASSLTAYRNSDYT | 0.5543285 |
| TNRPDLIDPALLRPG | ASSLTAYRNSDYT | 0.5611769 |
| LANLGVTAL | ASSLTAYRNSDYT | 0.41875255 |
| TGERIEL | ASSLTAYRNSDYT | 0.35398778 |
| NGEELAA | ASSLTAYRNSDYT | 0.5322162 |
| VLSLVWG | ASSLTAYRNSDYT | 0.34314966 |
| IGYALWL | ASSLTAYRNSDYT | 0.52977306 |
| LSLLLAPL | ASSLTAYRNSDYT | 0.4723016 |
| TLDEALK | ASSLTAYRNSDYT | 0.71173453 |
| TDGAGTAEGG | ASSLTAYRNSDYT | 0.5881274 |
| NAPGGAGGNGG | ASSLTAYRNSDYT | 0.5750784 |
| VSVFMGMGGNF | ASSLTAYRNSDYT | 0.4149369 |
| LASYLAAL | ASSLTAYRNSDYT | 0.536018 |
| LVGGTPAL | ASSLTAYRNSDYT | 0.5404053 |
| VRNADVL | ASSLTAYRNSDYT | 0.3526843 |
| TCSSPFLVSAM | ASSLTAYRNSDYT | 0.6448546 |
| VHITDAE | ASSLTAYRNSDYT | 0.4040655 |
| EYAAPSF | ASSLTAYRNSDYT | 0.51127446 |
| VVDAFHGGL | ASSLTAYRNSDYT | 0.5954526 |
| ASVVDPQ | ASSLTAYRNSDYT | 0.64536965 |
| SYNEFGGLS | ASSLTAYRNSDYT | 0.48415926 |
| DPDGSVY | ASSLTAYRNSDYT | 0.5389261 |
| MVSTIHATE | ASSLTAYRNSDYT | 0.42235953 |
| LAAGVADPV | ASSLTAYRNSDYT | 0.6212381 |
| TKEQIAA | ASSLTAYRNSDYT | 0.6405884 |
| SLASPAAL | ASSLTAYRNSDYT | 0.59590125 |
| IGLVFYGV | ASSLTAYRNSDYT | 0.68846387 |
| FLLDASNL | ASSLTAYRNSDYT | 0.56910473 |
| YQVGNFP | ASSLTAYRNSDYT | 0.3792218 |

| | | |
|---|---|---|
| ESLLSQST | ASSLTAYRNSDYT | 0.54283017 |
| LFTVGFAAVG | ASSLTAYRNSDYT | 0.62351733 |
| ATPPTPT | ASSLTAYRNSDYT | 0.64907897 |
| TDLATLG | ASSLTAYRNSDYT | 0.6435922 |
| SAGGTGR | ASSLTAYRNSDYT | 0.6170039 |
| GFSNSGS | ASSLTAYRNSDYT | 0.43743125 |
| GGNGGSGGTSV | ASSLTAYRNSDYT | 0.65813875 |
| GTSVATGGAGN | ASSLTAYRNSDYT | 0.7052785 |
| NGGNGGSGGTS | ASSLTAYRNSDYT | 0.610579 |
| AVVKFPH | ASSLTAYRNSDYT | 0.4369033 |
| TPPPPPP | ASSLTAYRNSDYT | 0.45827907 |
| AAPAAAAAPA | ASSLTAYRNSDYT | 0.5692001 |
| IALNEDL | ASSLTAYRNSDYT | 0.6146356 |
| IVNVSSM | ASSLTAYRNSDYT | 0.4132546 |
| TGIPPAPR | ASSLTAYRNSDYT | 0.5295491 |
| FDLGGGTF | ASSLTAYRNSDYT | 0.5571048 |
| FDLGGGTFDVS | ASSLTAYRNSDYT | 0.337068 |
| NEPTAAA | ASSLTAYRNSDYT | 0.6102464 |
| NEPTAAALA | ASSLTAYRNSDYT | 0.65272087 |
| GSIINISSI | ASSLTAYRNSDYT | 0.5379226 |
| VAPGNAGTA | ASSLTAYRNSDYT | 0.40655166 |
| TGGGASLE | ASSLTAYRNSDYT | 0.6479259 |
| VAVYNHYKRI | ASSLTAYRNSDYT | 0.6851982 |
| TDPAPATT | ASSLTAYRNSDYT | 0.64481777 |
| TNRPDILDPALLRPG | ASSLTAYRNSDYT | 0.34541497 |
| LTTTTTS | ASSLTAYRNSDYT | 0.2782644 |
| PTGSGKTL | ASSLTAYRNSDYT | 0.5175676 |
| TGSGKTL | ASSLTAYRNSDYT | 0.6353986 |
| TIGHVDHGK | ASSLTAYRNSDYT | 0.5089423 |
| TIGHVDHGKTT | ASSLTAYRNSDYT | 0.44090846 |
| GHVDHGKTT | ASSLTAYRNSDYT | 0.54469407 |
| AQFEHTLL | ASSLTAYRNSDYT | 0.6292452 |
| IVNQDEL | ASSLTAYRNSDYT | 0.44211024 |
| TNGSQFF | ASSLTAYRNSDYT | 0.39535552 |
| TGKGIVE | ASSLTAYRNSDYT | 0.57612425 |
| LISNASDA | ASSLTAYRNSDYT | 0.6415333 |
| AELRTLGELTKIAKAHGAQ | ASSLTAYRNSDYT | 0.36780378 |
| APNAVAAAEPSW | ASSLTAYRNSDYT | 0.4448419 |
| APQVAGAATAVL | ASSLTAYRNSDYT | 0.56054705 |
| ARRDPAAAAAILPTDA | ASSLTAYRNSDYT | 0.53311485 |
| DAAARDVLAEAGLAEYF | ASSLTAYRNSDYT | 0.5143821 |
| DDQIVLSAKANLAKVEQ | ASSLTAYRNSDYT | 0.39274755 |
| DELLPDTGEDSDEDGH | ASSLTAYRNSDYT | 0.62015975 |
| ELPEAPANRATVPQD | ASSLTAYRNSDYT | 0.34430316 |
| FGQSKIERVTGIG | ASSLTAYRNSDYT | 0.5983834 |
| GAGGTGSTAGGAGGAGGAGGLY | ASSLTAYRNSDYT | 0.56625825 |
| GHGGTGAAGGNGGTGGAGGSGI | ASSLTAYRNSDYT | 0.5628166 |
| GNGGNGGAGGEGGPGGAGGAGA | ASSLTAYRNSDYT | 0.48093304 |
| GSTAGNGGNGGNATNGTIAGQP | ASSLTAYRNSDYT | 0.39018914 |
| GTGGDGGNGGNGGTGG | ASSLTAYRNSDYT | 0.38820896 |
| GVRRPAGIPADAPV | ASSLTAYRNSDYT | 0.35793933 |

| | | |
|---|---|---|
| GYRQVTDAHLLTLARR | ASSLTAYRNSDYT | 0.6330527 |
| IAKMGQDGHDRG | ASSLTAYRNSDYT | 0.49956062 |
| KAEPSIEIQYT | ASSLTAYRNSDYT | 0.63127095 |
| KPVRIRMAADEDTVAVT | ASSLTAYRNSDYT | 0.53550076 |
| MPPRIAGMRLLVIKPEP | ASSLTAYRNSDYT | 0.4112112 |
| MKFYAVALFRSRGVPPTR | ASSLTAYRNSDYT | 0.5709691 |
| NAGGFGILGAVAHSP | ASSLTAYRNSDYT | 0.38828224 |
| NEGREITLAMAEM | ASSLTAYRNSDYT | 0.4667979 |
| NELVEWSHPLGHRWRWE | ASSLTAYRNSDYT | 0.6019638 |
| NLGAGGTGGNGGNPG | ASSLTAYRNSDYT | 0.58811015 |
| NTEWPKWETVMAADPVD | ASSLTAYRNSDYT | 0.42258418 |
| PASAAPAETQP | ASSLTAYRNSDYT | 0.5852007 |
| QQSSDGKNGVRAAEITGE | ASSLTAYRNSDYT | 0.56399834 |
| RAPDAVTTSWWKE | ASSLTAYRNSDYT | 0.43101737 |
| SGFGNVGAHTSGWFNQ | ASSLTAYRNSDYT | 0.5462149 |
| SGVAGAVGQAASVGGLKV | ASSLTAYRNSDYT | 0.62396204 |
| SNGGAGTGSAGGNGGT | ASSLTAYRNSDYT | 0.5539814 |
| TGGAGGEGGAAG | ASSLTAYRNSDYT | 0.34843603 |
| TGGDGGAAGAGGAPGHG | ASSLTAYRNSDYT | 0.55757505 |
| TMPPLLAAIANAGRHGV | ASSLTAYRNSDYT | 0.39409718 |
| TSSTAGNSGGA | ASSLTAYRNSDYT | 0.55329823 |
| VPATVPVGILC | ASSLTAYRNSDYT | 0.5158341 |
| VQRPDSGGVVLTGRIS | ASSLTAYRNSDYT | 0.51840633 |
| WTPNPVPATKGVPAGLVQAT | ASSLTAYRNSDYT | 0.60971826 |
| YALKRALAKFSDRFDVVII | ASSLTAYRNSDYT | 0.39250562 |
| YLIGSVIV | ASSLTAYRNSDYT | 0.4137546 |
| YGQLVAVID | ASSLTAYRNSDYT | 0.4550725 |
| VTVGGMTLPA | ASSLTAYRNSDYT | 0.57514375 |
| VLALPAV | ASSLTAYRNSDYT | 0.5861616 |
| VIPPTLNYET | ASSLTAYRNSDYT | 0.35514054 |
| VGMGGMHPSSG | ASSLTAYRNSDYT | 0.5535521 |
| VGLAGGAATA | ASSLTAYRNSDYT | 0.49073607 |
| VGGAGGAGGSGP | ASSLTAYRNSDYT | 0.41290203 |
| VFAAFMV | ASSLTAYRNSDYT | 0.5319677 |
| TLYDKLR | ASSLTAYRNSDYT | 0.4657216 |
| TLAVFTVL | ASSLTAYRNSDYT | 0.36778963 |
| TGSVQPARV | ASSLTAYRNSDYT | 0.5897496 |
| TGGGAGSGGT | ASSLTAYRNSDYT | 0.4727018 |
| SHLETSL | ASSLTAYRNSDYT | 0.5647167 |
| SFTTGMT | ASSLTAYRNSDYT | 0.5355262 |
| QVEVAKRMGVRQPTVS | ASSLTAYRNSDYT | 0.40026146 |
| QGSGGPALAM | ASSLTAYRNSDYT | 0.58682525 |
| MLAPSNP | ASSLTAYRNSDYT | 0.54188174 |
| LVNNAAI | ASSLTAYRNSDYT | 0.56861556 |
| LTGEINTLA | ASSLTAYRNSDYT | 0.5329598 |
| LSIFGPV | ASSLTAYRNSDYT | 0.40772364 |
| LLALLANL | ASSLTAYRNSDYT | 0.6265802 |
| LISNASDALDKLR | ASSLTAYRNSDYT | 0.59626114 |
| LINEAVP | ASSLTAYRNSDYT | 0.61458737 |
| LGYAFSGL | ASSLTAYRNSDYT | 0.5858207 |
| LDFVAVAL | ASSLTAYRNSDYT | 0.59918004 |

| | | |
|---|---|---|
| LAQPGAE | ASSLTAYRNSDYT | 0.6059646 |
| ISNLGAVTGF | ASSLTAYRNSDYT | 0.42384374 |
| HQEDPGRI | ASSLTAYRNSDYT | 0.33574447 |
| GTPGEPTP | ASSLTAYRNSDYT | 0.50865865 |
| GFFNSGS | ASSLTAYRNSDYT | 0.5171129 |
| GAGGGGLTAGAN | ASSLTAYRNSDYT | 0.4593774 |
| FVDALVNM | ASSLTAYRNSDYT | 0.3768498 |
| FLTVAPEL | ASSLTAYRNSDYT | 0.6127752 |
| FETLTAL | ASSLTAYRNSDYT | 0.7040545 |
| DVINPIEL | ASSLTAYRNSDYT | 0.47691607 |
| APVSGAPV | ASSLTAYRNSDYT | 0.6974591 |
| ALVGGLAW | ASSLTAYRNSDYT | 0.34348655 |
| ALAAAAATAI | ASSLTAYRNSDYT | 0.5894805 |
| AFGLGLLL | ASSLTAYRNSDYT | 0.6360298 |
| AFFAAASL | ASSLTAYRNSDYT | 0.37200534 |
| AAAALPAFSPP | ASSLTAYRNSDYT | 0.59042066 |
| VVLAPAL | ASSLTAYRNSDYT | 0.3964146 |
| TIGHVDHGKTTL | ASSLTAYRNSDYT | 0.43750608 |
| LISNASDALDK | ASSLTAYRNSDYT | 0.64436984 |
| LISNASDAL | ASSLTAYRNSDYT | 0.523626 |
| ISNASDALDK | ASSLTAYRNSDYT | 0.5779414 |
| NQAETLVYQTE | ASSLTAYRNSDYT | 0.51191384 |
| TGGTGGTAGAG | ASSLTAYRNSDYT | 0.43152234 |
| TGSTAGGAGGA | ASSLTAYRNSDYT | 0.5478624 |
| DPLAVPVAAGPV | ASSLTAYRNSDYT | 0.6446162 |
| GGTGGTPGNGGN | ASSLTAYRNSDYT | 0.48439074 |
| PAGLAGVDPATL | ASSLTAYRNSDYT | 0.7043259 |
| GNNLVVEAPHAGASV | ASSLTAYRNSDYT | 0.39672217 |
| KVTRIMSASTLEQPA | ASSLTAYRNSDYT | 0.67223084 |
| LGFALPIALPRRTVV | ASSLTAYRNSDYT | 0.6645857 |
| TRLGNLTARKLVVLV | ASSLTAYRNSDYT | 0.52052176 |
| WSQQRRLIYFPSAGP | ASSLTAYRNSDYT | 0.45872742 |
| GPNGNPDPMAAAVDIR | ASSLTAYRNSDYT | 0.62258416 |
| TQRCVSVADLRAAASR | ASSLTAYRNSDYT | 0.51350754 |
| QFVLDVAEHPVEVLTETV | ASSLTAYRNSDYT | 0.56798923 |
| GGAGGTGGAGGNGGTQAGDG | ASSLTAYRNSDYT | 0.63991284 |
| PLLATLRAIAVDVDAKPAQV | ASSLTAYRNSDYT | 0.2581856 |
| APKFRTGGYMIDFWGVGYQVAK | ASSLTAYRNSDYT | 0.5811034 |
| GAGGAGGAGGAAGGPGGTGGTG | ASSLTAYRNSDYT | 0.42973122 |
| IGALGLLATDVAPAGSVAFGVA | ASSLTAYRNSDYT | 0.4397934 |
| AGGAGGIGGNGGAGGAGGVLMG | ASSLTAYRNSDYT | 0.32618302 |
| ADELIGLL | ASSLTAYRNSDYT | 0.24478929 |
| ADRALKA | ASSLTAYRNSDYT | 0.32466042 |
| AGAVGQVGGAGGA | ASSLTAYRNSDYT | 0.2844437 |
| AGGGGGLGGAP | ASSLTAYRNSDYT | 0.6365278 |
| AGGLLYGNGGAG | ASSLTAYRNSDYT | 0.45830563 |
| AGLESIGL | ASSLTAYRNSDYT | 0.3302003 |
| AGLKTGII | ASSLTAYRNSDYT | 0.60507226 |
| ASLAALLL | ASSLTAYRNSDYT | 0.5544752 |
| ASVINISSI | ASSLTAYRNSDYT | 0.22057584 |
| ATPPGPA | ASSLTAYRNSDYT | 0.32233167 |

| | | |
|---|---|---|
| ATVASALV | ASSLTAYRNSDYT | 0.2914932 |
| ATVASLIL | ASSLTAYRNSDYT | 0.3487811 |
| AVNGPVLIRH | ASSLTAYRNSDYT | 0.17324238 |
| DGAAGAVGI | ASSLTAYRNSDYT | 0.4638237 |
| DPLLHGI | ASSLTAYRNSDYT | 0.27557823 |
| ELGAALGAL | ASSLTAYRNSDYT | 0.17528746 |
| ELGRTQAV | ASSLTAYRNSDYT | 0.3496557 |
| FVAGGYLS | ASSLTAYRNSDYT | 0.30834723 |
| GASGGAGGAGGW | ASSLTAYRNSDYT | 0.3500129 |
| GGAGAGGDGGAGGI | ASSLTAYRNSDYT | 0.5884743 |
| GTGHNGSQPGMGGNG | ASSLTAYRNSDYT | 0.262602 |
| GYAAASASAS | ASSLTAYRNSDYT | 0.28244254 |
| HQVDVAAVQQG | ASSLTAYRNSDYT | 0.51166284 |
| IGAAIAHT | ASSLTAYRNSDYT | 0.6497234 |
| IGMIEVAQL | ASSLTAYRNSDYT | 0.28779382 |
| IRLAPEHTSAMD | ASSLTAYRNSDYT | 0.39911476 |
| IYNVSVL | ASSLTAYRNSDYT | 0.68956375 |
| KQGFLGM | ASSLTAYRNSDYT | 0.4371131 |
| LESVITQL | ASSLTAYRNSDYT | 0.5180643 |
| LGLGALVVV | ASSLTAYRNSDYT | 0.56617343 |
| LIGNGGNGGSGGI | ASSLTAYRNSDYT | 0.26246107 |
| LNAINAPTL | ASSLTAYRNSDYT | 0.5762571 |
| LQLLNTGV | ASSLTAYRNSDYT | 0.6595162 |
| LQTSIEF | ASSLTAYRNSDYT | 0.35655758 |
| MSFAVSEL | ASSLTAYRNSDYT | 0.39326853 |
| NTGALGL | ASSLTAYRNSDYT | 0.27925104 |
| PPGAGGAGGAGTT | ASSLTAYRNSDYT | 0.30802998 |
| QLVGLGI | ASSLTAYRNSDYT | 0.22004887 |
| RFLEQVHQL | ASSLTAYRNSDYT | 0.44000232 |
| RGVGYVLR | ASSLTAYRNSDYT | 0.6028297 |
| SATGVFAGVI | ASSLTAYRNSDYT | 0.3566065 |
| SLPAITLP | ASSLTAYRNSDYT | 0.21524642 |
| STALAFTV | ASSLTAYRNSDYT | 0.22381431 |
| SVLTVGLGK | ASSLTAYRNSDYT | 0.29195407 |
| TALAGAGSGI | ASSLTAYRNSDYT | 0.30904093 |
| TGLPFTGI | ASSLTAYRNSDYT | 0.23091166 |
| VAEHVTV | ASSLTAYRNSDYT | 0.24075273 |
| VDIVLNSL | ASSLTAYRNSDYT | 0.6034014 |
| VLTGGGAVL | ASSLTAYRNSDYT | 0.5902759 |
| VNVGLAYGI | ASSLTAYRNSDYT | 0.26405066 |
| VSSAAVGAV | ASSLTAYRNSDYT | 0.6239199 |
| VTSVYEAGV | ASSLTAYRNSDYT | 0.54333276 |
| WVALDRGA | ASSLTAYRNSDYT | 0.5814715 |
| YLVGSVVI | ASSLTAYRNSDYT | 0.37640303 |
| GPPGTGKTLL | ASSLTAYRNSDYT | 0.24585554 |
| GVGMTKF | ASSLTAYRNSDYT | 0.65589523 |
| IGIGGSDL | ASSLTAYRNSDYT | 0.3841332 |
| IGTLLAEL | ASSLTAYRNSDYT | 0.4617175 |
| IYTSGTT | ASSLTAYRNSDYT | 0.4956908 |
| LSFPAGV | ASSLTAYRNSDYT | 0.40186808 |
| LYTSGTT | ASSLTAYRNSDYT | 0.30949077 |

| | | |
|---|---|---|
| VGAFPFGF | ASSLTAYRNSDYT | 0.4751349 |
| VGAGTVEF | ASSLTAYRNSDYT | 0.46215993 |
| VGKSTLLL | ASSLTAYRNSDYT | 0.22639501 |
| AALAHAGGGIV | ASSLTAYRNSDYT | 0.29656696 |
| AALVAVSLL | ASSLTAYRNSDYT | 0.32020432 |
| AAVSRAVAAL | ASSLTAYRNSDYT | 0.39464098 |
| AGGSSGIGGF | ASSLTAYRNSDYT | 0.61887515 |
| AGLTVANANG | ASSLTAYRNSDYT | 0.40470958 |
| AGLVGAVL | ASSLTAYRNSDYT | 0.321728 |
| ALSQAFTV | ASSLTAYRNSDYT | 0.5136958 |
| ALTFSVSL | ASSLTAYRNSDYT | 0.47586304 |
| ALTGAALVAL | ASSLTAYRNSDYT | 0.3820587 |
| ASFSIGAGVL | ASSLTAYRNSDYT | 0.3477995 |
| DGSVYKI | ASSLTAYRNSDYT | 0.2770963 |
| EQLAEIV | ASSLTAYRNSDYT | 0.46859735 |
| FAVSELTVM | ASSLTAYRNSDYT | 0.49382484 |
| GAAPLAAGL | ASSLTAYRNSDYT | 0.47560072 |
| GADGGDGGSGGAGG | ASSLTAYRNSDYT | 0.46474737 |
| GFGLDTGGAGG | ASSLTAYRNSDYT | 0.45865324 |
| GLASGGLL | ASSLTAYRNSDYT | 0.37293366 |
| GRIAGAAVLL | ASSLTAYRNSDYT | 0.22965464 |
| GSAGSLNTSF | ASSLTAYRNSDYT | 0.2659344 |
| GTGQIGI | ASSLTAYRNSDYT | 0.35425764 |
| GTLIALIL | ASSLTAYRNSDYT | 0.42047578 |
| IAATMGFVVG | ASSLTAYRNSDYT | 0.4163768 |
| IFAQALL | ASSLTAYRNSDYT | 0.48624197 |
| IGAAVALGF | ASSLTAYRNSDYT | 0.27511793 |
| IGLTPLAL | ASSLTAYRNSDYT | 0.43146178 |
| ILGSNTSTL | ASSLTAYRNSDYT | 0.61972535 |
| IVFSLTVL | ASSLTAYRNSDYT | 0.22852151 |
| LASKATGF | ASSLTAYRNSDYT | 0.3058046 |
| LASVALIL | ASSLTAYRNSDYT | 0.6049113 |
| LFAVGLTL | ASSLTAYRNSDYT | 0.5440893 |
| LFNGGIGTYI | ASSLTAYRNSDYT | 0.36646536 |
| LTLGAGII | ASSLTAYRNSDYT | 0.3858699 |
| LVAAALGDI | ASSLTAYRNSDYT | 0.71306974 |
| NTAFVQLGI | ASSLTAYRNSDYT | 0.3399523 |
| QLFSGAEL | ASSLTAYRNSDYT | 0.57153875 |
| SAATFITL | ASSLTAYRNSDYT | 0.20499258 |
| SNYGLANLGV | ASSLTAYRNSDYT | 0.34436953 |
| SYGGAYAVM | ASSLTAYRNSDYT | 0.2659409 |
| TTGVGGLTLG | ASSLTAYRNSDYT | 0.1831502 |
| TTQTIYI | ASSLTAYRNSDYT | 0.54023045 |
| VAAGAGLGAVI | ASSLTAYRNSDYT | 0.29606903 |
| VAAVVTLL | ASSLTAYRNSDYT | 0.5254068 |
| VGFLGGLI | ASSLTAYRNSDYT | 0.3118674 |
| VGVIPILL | ASSLTAYRNSDYT | 0.49003592 |
| VIFASSLI | ASSLTAYRNSDYT | 0.4159892 |
| VKGGDGGAAAG | ASSLTAYRNSDYT | 0.3438656 |
| VTKTFTNI | ASSLTAYRNSDYT | 0.27677783 |
| VVAAAILL | ASSLTAYRNSDYT | 0.51211965 |

| | | |
|---|---|---|
| YAALDVEL | ASSLTAYRNSDYT | 0.42418426 |
| LASVALLL | ASSLTAYRNSDYT | 0.47553498 |
| LDAVHRV | ASSLTAYRNSDYT | 0.24229023 |
| QGLSSSIF | ASSLTAYRNSDYT | 0.30123353 |
| RGTFANIRL | ASSLTAYRNSDYT | 0.4224117 |
| RQLWWGHRI | ASSLTAYRNSDYT | 0.23597372 |
| YGPPGTGKTLL | ASSLTAYRNSDYT | 0.19386113 |
| AALVGAPANVA | ASSLTAYRNSDYT | 0.25171924 |
| AATFRRTE | ASSLTAYRNSDYT | 0.24862434 |
| AGALGGGI | ASSLTAYRNSDYT | 0.5082498 |
| AGGAGGAGGIGAG | ASSLTAYRNSDYT | 0.5544782 |
| AGSGALGL | ASSLTAYRNSDYT | 0.33870628 |
| AQIADQI | ASSLTAYRNSDYT | 0.5397813 |
| ASAVAVTYL | ASSLTAYRNSDYT | 0.46304578 |
| ASTVLGGTGG | ASSLTAYRNSDYT | 0.15505007 |
| AVGGEPAK | ASSLTAYRNSDYT | 0.2991776 |
| CEQGTFGGTC | ASSLTAYRNSDYT | 0.23016344 |
| DEGKAPAG | ASSLTAYRNSDYT | 0.53189105 |
| DTVVFPAL | ASSLTAYRNSDYT | 0.22844665 |
| FEGITMAG | ASSLTAYRNSDYT | 0.27123943 |
| FPLVVQD | ASSLTAYRNSDYT | 0.39778012 |
| GAFTTPQ | ASSLTAYRNSDYT | 0.58400685 |
| GFGGFGTAGGVGGA | ASSLTAYRNSDYT | 0.42385808 |
| GFLAQFL | ASSLTAYRNSDYT | 0.25637212 |
| GGAGGAGGNAGVA | ASSLTAYRNSDYT | 0.1812939 |
| GGVGTAGAGGAGGAGG | ASSLTAYRNSDYT | 0.54687095 |
| GNGGGGDAGNAGSGGN | ASSLTAYRNSDYT | 0.28663972 |
| GTGGAGGAGNDAGS | ASSLTAYRNSDYT | 0.3004816 |
| GTTNTGWFNS | ASSLTAYRNSDYT | 0.19749644 |
| IFMGPLFTL | ASSLTAYRNSDYT | 0.29883373 |
| ILAHAIEAI | ASSLTAYRNSDYT | 0.23040716 |
| IRSAIREAL | ASSLTAYRNSDYT | 0.52878565 |
| ITFAGGG | ASSLTAYRNSDYT | 0.2979624 |
| ITGMSGHQEV | ASSLTAYRNSDYT | 0.33342704 |
| IVAGLACTL | ASSLTAYRNSDYT | 0.6547675 |
| IVGGGLAGA | ASSLTAYRNSDYT | 0.60207266 |
| IVTGVLNI | ASSLTAYRNSDYT | 0.34213462 |
| LAQQGCI | ASSLTAYRNSDYT | 0.34348306 |
| LCLHVEN | ASSLTAYRNSDYT | 0.26076016 |
| LDAISDGS | ASSLTAYRNSDYT | 0.28665674 |
| LGGFAAML | ASSLTAYRNSDYT | 0.31984714 |
| LGGKNPMI | ASSLTAYRNSDYT | 0.5686443 |
| LGPKGVAAIL | ASSLTAYRNSDYT | 0.362084 |
| LMVGVSI | ASSLTAYRNSDYT | 0.22361465 |
| LVGAAFVL | ASSLTAYRNSDYT | 0.43977943 |
| LVLAVAPL | ASSLTAYRNSDYT | 0.6406656 |
| MAAIAVIDAA | ASSLTAYRNSDYT | 0.70437765 |
| NNGINAELQ | ASSLTAYRNSDYT | 0.24950479 |
| RGEPLVLVHG | ASSLTAYRNSDYT | 0.35203928 |
| RGGAGGAAGLI | ASSLTAYRNSDYT | 0.34830153 |
| RSYRMHPAV | ASSLTAYRNSDYT | 0.37316483 |

| | | |
|---|---|---|
| SAAHLLAGQLA | ASSLTAYRNSDYT | 0.30790666 |
| SAAVGSY | ASSLTAYRNSDYT | 0.30077922 |
| SPHAVLHRI | ASSLTAYRNSDYT | 0.45214337 |
| SQIAKVL | ASSLTAYRNSDYT | 0.75492084 |
| TALGHLVP | ASSLTAYRNSDYT | 0.30219007 |
| TAPTTSGAA | ASSLTAYRNSDYT | 0.19889164 |
| TDGDDGNGGSPGT | ASSLTAYRNSDYT | 0.35083726 |
| TDGLGEN | ASSLTAYRNSDYT | 0.2453172 |
| TGGPPLPGQA | ASSLTAYRNSDYT | 0.2893272 |
| TIPAITI | ASSLTAYRNSDYT | 0.4205385 |
| TIPGFTI | ASSLTAYRNSDYT | 0.20910564 |
| TSIASGALA | ASSLTAYRNSDYT | 0.33241913 |
| TVTLPGAAI | ASSLTAYRNSDYT | 0.69628954 |
| VASGLSAALP | ASSLTAYRNSDYT | 0.3828158 |
| VGAGVIGI | ASSLTAYRNSDYT | 0.43010217 |
| VGPKRFGV | ASSLTAYRNSDYT | 0.60145813 |
| VNVAFPD | ASSLTAYRNSDYT | 0.45793763 |
| VSMTNSGVS | ASSLTAYRNSDYT | 0.63940454 |
| VTDVLGDLSV | ASSLTAYRNSDYT | 0.56566066 |
| VVVVGAGFAGL | ASSLTAYRNSDYT | 0.35705787 |
| VYTVGTVV | ASSLTAYRNSDYT | 0.35455135 |
| WTTFSGP | ASSLTAYRNSDYT | 0.4647974 |
| YDELVEAF | ASSLTAYRNSDYT | 0.3846758 |
| YGTGGAGGNGG | ASSLTAYRNSDYT | 0.40359503 |
| YTNVMAV | ASSLTAYRNSDYT | 0.3111642 |
| FLEQVHQL | ASSLTAYRNSDYT | 0.5633358 |
| GLSLLAVAV | ASSLTAYRNSDYT | 0.34365118 |
| LEQVHQL | ASSLTAYRNSDYT | 0.6259491 |
| LNAAGAI | ASSLTAYRNSDYT | 0.22575526 |
| PAVHASL | ASSLTAYRNSDYT | 0.2913359 |
| TELEDDA | ASSLTAYRNSDYT | 0.57166 |
| TPPGPPN | ASSLTAYRNSDYT | 0.17342423 |
| TPPPPPPP | ASSLTAYRNSDYT | 0.21823633 |
| TQSPALI | ASRQDGDYEQY | 0.34989637 |
| EFAAAFPQ | ASRQDGDYEQY | 0.6395855 |
| LLSGNLDVL | ASRQDGDYEQY | 0.18226565 |
| GPANLGNYNI | ASRQDGDYEQY | 0.29178032 |
| TLGPTLP | ASRQDGDYEQY | 0.25337762 |
| VYAYSSCCK | ASRQDGDYEQY | 0.43023837 |
| APAPAPAPNI | ASRQDGDYEQY | 0.4490062 |
| PGGPVPIP | ASRQDGDYEQY | 0.20428926 |
| FLTVALEP | ASRQDGDYEQY | 0.46078074 |
| SQYFGANL | ASRQDGDYEQY | 0.19820333 |
| GGVKPAA | ASRQDGDYEQY | 0.32555357 |
| GGTGGAGGAGGW | ASRQDGDYEQY | 0.46738598 |
| GVGGVAGLGGAGG | ASRQDGDYEQY | 0.65850043 |
| VLLVLATTI | ASRQDGDYEQY | 0.23356868 |
| IGIFAAML | ASRQDGDYEQY | 0.55912256 |
| AVGDLID | ASRQDGDYEQY | 0.2556048 |
| NISPLPL | ASRQDGDYEQY | 0.735533 |
| VTAVFVAI | ASRQDGDYEQY | 0.3414429 |

| | | |
|---|---|---|
| SGDDNDAAL | ASRQDGDYEQY | 0.22446373 |
| AVAALAHS | ASRQDGDYEQY | 0.48417115 |
| GGVTVMATP | ASRQDGDYEQY | 0.46112412 |
| AGLAANVRD | ASRQDGDYEQY | 0.43495715 |
| TNRPDLIDPALLRPG | ASRQDGDYEQY | 0.55540895 |
| LANLGVTAL | ASRQDGDYEQY | 0.3393606 |
| TGERIEL | ASRQDGDYEQY | 0.34040833 |
| NGEELAA | ASRQDGDYEQY | 0.54371524 |
| VLSLVWG | ASRQDGDYEQY | 0.38872135 |
| IGYALWL | ASRQDGDYEQY | 0.29131863 |
| LSLLLAPL | ASRQDGDYEQY | 0.47883686 |
| TLDEALK | ASRQDGDYEQY | 0.4345315 |
| TDGAGTAEGG | ASRQDGDYEQY | 0.28805482 |
| NAPGGAGGNGG | ASRQDGDYEQY | 0.64928436 |
| VSVFMGMGGNF | ASRQDGDYEQY | 0.3936894 |
| LASYLAAL | ASRQDGDYEQY | 0.5742898 |
| LVGGTPAL | ASRQDGDYEQY | 0.48749882 |
| VRNADVL | ASRQDGDYEQY | 0.7270958 |
| TCSSPFLVSAM | ASRQDGDYEQY | 0.24003837 |
| VHITDAE | ASRQDGDYEQY | 0.35537207 |
| EYAAPSF | ASRQDGDYEQY | 0.39053243 |
| VVDAFHGGL | ASRQDGDYEQY | 0.48657444 |
| ASVVDPQ | ASRQDGDYEQY | 0.3510206 |
| SYNEFGGLS | ASRQDGDYEQY | 0.3370168 |
| DPDGSVY | ASRQDGDYEQY | 0.4269862 |
| MVSTIHATE | ASRQDGDYEQY | 0.645622 |
| LAAGVADPV | ASRQDGDYEQY | 0.34436786 |
| TKEQIAA | ASRQDGDYEQY | 0.31605896 |
| SLASPAAL | ASRQDGDYEQY | 0.68620384 |
| IGLVFYGV | ASRQDGDYEQY | 0.27978006 |
| FLLDASNL | ASRQDGDYEQY | 0.42967954 |
| YQVGNFP | ASRQDGDYEQY | 0.46340078 |
| ESLLSQST | ASRQDGDYEQY | 0.45686132 |
| LFTVGFAAVG | ASRQDGDYEQY | 0.3706668 |
| ATPPTPT | ASRQDGDYEQY | 0.47314698 |
| TDLATLG | ASRQDGDYEQY | 0.20652328 |
| SAGGTGR | ASRQDGDYEQY | 0.51052624 |
| GFSNSGS | ASRQDGDYEQY | 0.24318597 |
| GGNGGSGGTSV | ASRQDGDYEQY | 0.4151738 |
| GTSVATGGAGN | ASRQDGDYEQY | 0.23873104 |
| NGGNGGSGGTS | ASRQDGDYEQY | 0.4227699 |
| AVVKFPH | ASRQDGDYEQY | 0.57852787 |
| TPPPPPP | ASRQDGDYEQY | 0.55845183 |
| AAPAAAAAPA | ASRQDGDYEQY | 0.19156235 |
| IALNEDL | ASRQDGDYEQY | 0.5226947 |
| IVNVSSM | ASRQDGDYEQY | 0.53516304 |
| TGIPPAPR | ASRQDGDYEQY | 0.73948175 |
| FDLGGGTF | ASRQDGDYEQY | 0.35848525 |
| FDLGGGTFDVS | ASRQDGDYEQY | 0.4894409 |
| NEPTAAA | ASRQDGDYEQY | 0.34605387 |
| NEPTAAALA | ASRQDGDYEQY | 0.33957976 |

| | | |
|---|---|---|
| GSIINISSI | ASRQDGDYEQY | 0.2283217 |
| VAPGNAGTA | ASRQDGDYEQY | 0.31085774 |
| TGGGASLE | ASRQDGDYEQY | 0.36520892 |
| VAVYNHYKRI | ASRQDGDYEQY | 0.26073727 |
| TDPAPATT | ASRQDGDYEQY | 0.6501799 |
| TNRPDILDPALLRPG | ASRQDGDYEQY | 0.47434664 |
| LTTTTTS | ASRQDGDYEQY | 0.6081307 |
| PTGSGKTL | ASRQDGDYEQY | 0.61205137 |
| TGSGKTL | ASRQDGDYEQY | 0.6450227 |
| TIGHVDHGK | ASRQDGDYEQY | 0.47086963 |
| TIGHVDHGKTT | ASRQDGDYEQY | 0.2948947 |
| GHVDHGKTT | ASRQDGDYEQY | 0.3213671 |
| AQFEHTLL | ASRQDGDYEQY | 0.22611055 |
| IVNQDEL | ASRQDGDYEQY | 0.30610436 |
| TNGSQFF | ASRQDGDYEQY | 0.32338884 |
| TGKGIVE | ASRQDGDYEQY | 0.4061162 |
| LISNASDA | ASRQDGDYEQY | 0.18405949 |
| AELRTLGELTKIAKAHGAQ | ASRQDGDYEQY | 0.5165743 |
| APNAVAAAEPSW | ASRQDGDYEQY | 0.5929483 |
| APQVAGAATAVL | ASRQDGDYEQY | 0.41490325 |
| ARRDPAAAAAILPTDA | ASRQDGDYEQY | 0.49374944 |
| DAAARDVLAEAGLAEYF | ASRQDGDYEQY | 0.38853282 |
| DDQIVLSAKANLAKVEQ | ASRQDGDYEQY | 0.25152788 |
| DELLPDTGEDSDEDGH | ASRQDGDYEQY | 0.54807836 |
| ELPEAPANRATVPQD | ASRQDGDYEQY | 0.28246787 |
| FGQSKIERVTGIG | ASRQDGDYEQY | 0.52039725 |
| GAGGTGSTAGGAGGAGGAGGLY | ASRQDGDYEQY | 0.65521014 |
| GHGGTGAAGGNGGTGGAGGSGI | ASRQDGDYEQY | 0.41946912 |
| GNGGNGGAGGEGGPGGAGGAGA | ASRQDGDYEQY | 0.27538237 |
| GSTAGNGGNGGNATNGTIAGQP | ASRQDGDYEQY | 0.40726474 |
| GTGGDGGNGGNGGTGG | ASRQDGDYEQY | 0.23631126 |
| GVRRPAGIPADAPV | ASRQDGDYEQY | 0.29157874 |
| GYRQVTDAHLLTLARR | ASRQDGDYEQY | 0.21935154 |
| IAKMGQDGHDRG | ASRQDGDYEQY | 0.38724613 |
| KAEPSIEIQYT | ASRQDGDYEQY | 0.2546185 |
| KPVRIRMAADEDTVAVT | ASRQDGDYEQY | 0.47927064 |
| MPPRIAGMRLLVIKPEP | ASRQDGDYEQY | 0.43813366 |
| MKFYAVALFRSRGVPPTR | ASRQDGDYEQY | 0.38755342 |
| NAGGFGILGAVAHSP | ASRQDGDYEQY | 0.26821625 |
| NEGREITLAMAEM | ASRQDGDYEQY | 0.43774498 |
| NELVEWSHPLGHRWRWE | ASRQDGDYEQY | 0.41666603 |
| NLGAGGTGGNGGNPG | ASRQDGDYEQY | 0.2483668 |
| NTEWPKWETVMAADPVD | ASRQDGDYEQY | 0.4115109 |
| PASAAPAETQP | ASRQDGDYEQY | 0.3563821 |
| QQSSDGKNGVRAAEITGE | ASRQDGDYEQY | 0.31554866 |
| RAPDAVTTSWWKE | ASRQDGDYEQY | 0.59832275 |
| SGFGNVGAHTSGWFNQ | ASRQDGDYEQY | 0.45024344 |
| SGVAGAVGQAASVGGLKV | ASRQDGDYEQY | 0.2115471 |
| SNGGAGTGSAGGNGGT | ASRQDGDYEQY | 0.3836035 |
| TGGAGGEGGAAG | ASRQDGDYEQY | 0.39929208 |
| TGGDGGAAGAGGAPGHG | ASRQDGDYEQY | 0.35525292 |

| | | |
|---|---|---|
| TMPPLLAAIANAGRHGV | ASRQDGDYEQY | 0.16934034 |
| TSSTAGNSGGA | ASRQDGDYEQY | 0.33514932 |
| VPATVPVGILC | ASRQDGDYEQY | 0.2959627 |
| VQRPDSGGVVLTGRIS | ASRQDGDYEQY | 0.23219715 |
| WTPNPVPATKGVPAGLVQAT | ASRQDGDYEQY | 0.29013973 |
| YALKRALAKFSDRFDVVII | ASRQDGDYEQY | 0.57885396 |
| YLIGSVIV | ASRQDGDYEQY | 0.47388023 |
| YGQLVAVID | ASRQDGDYEQY | 0.3991195 |
| VTVGGMTLPA | ASRQDGDYEQY | 0.42203024 |
| VLALPAV | ASRQDGDYEQY | 0.5387245 |
| VIPPTLNYET | ASRQDGDYEQY | 0.6469983 |
| VGMGGMHPSSG | ASRQDGDYEQY | 0.43371806 |
| VGLAGGAATA | ASRQDGDYEQY | 0.31769305 |
| VGGAGGAGGSGP | ASRQDGDYEQY | 0.3935585 |
| VFAAFMV | ASRQDGDYEQY | 0.35033792 |
| TLYDKLR | ASRQDGDYEQY | 0.23801506 |
| TLAVFTVL | ASRQDGDYEQY | 0.23559096 |
| TGSVQPARV | ASRQDGDYEQY | 0.37445882 |
| TGGGAGSGGT | ASRQDGDYEQY | 0.24905561 |
| SHLETSL | ASRQDGDYEQY | 0.44131222 |
| SFTTGMT | ASRQDGDYEQY | 0.5315149 |
| QVEVAKRMGVRQPTVS | ASRQDGDYEQY | 0.29701802 |
| QGSGGPALAM | ASRQDGDYEQY | 0.23278145 |
| MLAPSNP | ASRQDGDYEQY | 0.28556985 |
| LVNNAAI | ASRQDGDYEQY | 0.29034537 |
| LTGEINTLA | ASRQDGDYEQY | 0.5581541 |
| LSIFGPV | ASRQDGDYEQY | 0.40132475 |
| LLALLANL | ASRQDGDYEQY | 0.71157694 |
| LISNASDALDKLR | ASRQDGDYEQY | 0.54814947 |
| LINEAVP | ASRQDGDYEQY | 0.33606577 |
| LGYAFSGL | ASRQDGDYEQY | 0.5016981 |
| LDFVAVAL | ASRQDGDYEQY | 0.3225576 |
| LAQPGAE | ASRQDGDYEQY | 0.28851476 |
| ISNLGAVTGF | ASRQDGDYEQY | 0.33809453 |
| HQEDPGRI | ASRQDGDYEQY | 0.3533327 |
| GTPGEPTP | ASRQDGDYEQY | 0.2936772 |
| GFFNSGS | ASRQDGDYEQY | 0.3140135 |
| GAGGGGLTAGAN | ASRQDGDYEQY | 0.22941403 |
| FVDALVNM | ASRQDGDYEQY | 0.4142329 |
| FLTVAPEL | ASRQDGDYEQY | 0.44317618 |
| FETLTAL | ASRQDGDYEQY | 0.2721776 |
| DVINPIEL | ASRQDGDYEQY | 0.30036902 |
| APVSGAPV | ASRQDGDYEQY | 0.6946974 |
| ALVGGLAW | ASRQDGDYEQY | 0.6741273 |
| ALAAAAATAI | ASRQDGDYEQY | 0.27432847 |
| AFGLGLLL | ASRQDGDYEQY | 0.2967342 |
| AFFAAASL | ASRQDGDYEQY | 0.5149782 |
| AAAALPAFSPP | ASRQDGDYEQY | 0.25685677 |
| VVLAPAL | ASRQDGDYEQY | 0.41820028 |
| TIGHVDHGKTTL | ASRQDGDYEQY | 0.44171917 |
| LISNASDALDK | ASRQDGDYEQY | 0.39054048 |

| | | |
|---|---|---|
| LISNASDAL | ASRQDGDYEQY | 0.29937017 |
| ISNASDALDK | ASRQDGDYEQY | 0.4154513 |
| NQAETLVYQTE | ASRQDGDYEQY | 0.3223223 |
| TGGTGGTAGAG | ASRQDGDYEQY | 0.4668043 |
| TGSTAGGAGGA | ASRQDGDYEQY | 0.18774712 |
| DPLAVPVAAGPV | ASRQDGDYEQY | 0.38197917 |
| GGTGGTPGNGGN | ASRQDGDYEQY | 0.27200714 |
| PAGLAGVDPATL | ASRQDGDYEQY | 0.280342 |
| GNNLVVEAPHAGASV | ASRQDGDYEQY | 0.20812555 |
| KVTRIMSASTLEQPA | ASRQDGDYEQY | 0.31022668 |
| LGFALPIALPRRTVV | ASRQDGDYEQY | 0.5626729 |
| TRLGNLTARKLVVLV | ASRQDGDYEQY | 0.58543736 |
| WSQQRRLIYFPSAGP | ASRQDGDYEQY | 0.27932853 |
| GPNGNPDPMAAAVDIR | ASRQDGDYEQY | 0.4250976 |
| TQRCVSVADLRAAASR | ASRQDGDYEQY | 0.7084104 |
| QFVLDVAEHPVEVLTETV | ASRQDGDYEQY | 0.30777025 |
| GGAGGTGGAGGNGGTQAGDG | ASRQDGDYEQY | 0.20504606 |
| PLLATLRAIAVDVDAKPAQV | ASRQDGDYEQY | 0.5175413 |
| APKFRTGGYMIDFWGVGYQVAK | ASRQDGDYEQY | 0.2730482 |
| GAGGAGGAGGAAGGPGGTGGTG | ASRQDGDYEQY | 0.42979148 |
| IGALGLLATDVAPAGSVAFGVA | ASRQDGDYEQY | 0.6103616 |
| AGGAGGIGGNGGAGGAGGVLMG | ASRQDGDYEQY | 0.21029651 |
| ADELIGLL | ASRQDGDYEQY | 0.54090774 |
| ADRALKA | ASRQDGDYEQY | 0.5336868 |
| AGAVGQVGGAGGA | ASRQDGDYEQY | 0.25227812 |
| AGGGGGLGGAP | ASRQDGDYEQY | 0.2385639 |
| AGGLLYGNGGAG | ASRQDGDYEQY | 0.26098895 |
| AGLESIGL | ASRQDGDYEQY | 0.19292456 |
| AGLKTGII | ASRQDGDYEQY | 0.5064891 |
| ASLAALLL | ASRQDGDYEQY | 0.2982173 |
| ASVINISSI | ASRQDGDYEQY | 0.3603211 |
| ATPPGPA | ASRQDGDYEQY | 0.3697434 |
| ATVASALV | ASRQDGDYEQY | 0.3251154 |
| ATVASLIL | ASRQDGDYEQY | 0.3915496 |
| AVNGPVLIRH | ASRQDGDYEQY | 0.36753306 |
| DGAAGAVGI | ASRQDGDYEQY | 0.36424136 |
| DPLLHGI | ASRQDGDYEQY | 0.4172393 |
| ELGAALGAL | ASRQDGDYEQY | 0.31548375 |
| ELGRTQAV | ASRQDGDYEQY | 0.34743467 |
| FVAGGYLS | ASRQDGDYEQY | 0.66856855 |
| GASGGAGGAGGW | ASRQDGDYEQY | 0.2501002 |
| GGAGAGGDGGAGGI | ASRQDGDYEQY | 0.30879706 |
| GTGHNGSQPGMGGNG | ASRQDGDYEQY | 0.44484618 |
| GYAAASASAS | ASRQDGDYEQY | 0.2903166 |
| HQVDVAAVQQG | ASRQDGDYEQY | 0.55308795 |
| IGAAIAHT | ASRQDGDYEQY | 0.42948806 |
| IGMIEVAQL | ASRQDGDYEQY | 0.45076808 |
| IRLAPEHTSAMD | ASRQDGDYEQY | 0.3729566 |
| IYNVSVL | ASRQDGDYEQY | 0.6447029 |
| KQGFLGM | ASRQDGDYEQY | 0.510105 |
| LESVITQL | ASRQDGDYEQY | 0.31252763 |

| | | |
|---|---|---|
| LGLGALVVV | ASRQDGDYEQY | 0.58203495 |
| LIGNGGNGGSGGI | ASRQDGDYEQY | 0.24478655 |
| LNAINAPTL | ASRQDGDYEQY | 0.39549133 |
| LQLLNTGV | ASRQDGDYEQY | 0.4638836 |
| LQTSIEF | ASRQDGDYEQY | 0.24510957 |
| MSFAVSEL | ASRQDGDYEQY | 0.3394574 |
| NTGALGL | ASRQDGDYEQY | 0.5227072 |
| PPGAGGAGGAGTT | ASRQDGDYEQY | 0.5216934 |
| QLVGLGI | ASRQDGDYEQY | 0.26160857 |
| RFLEQVHQL | ASRQDGDYEQY | 0.29427564 |
| RGVGYVLR | ASRQDGDYEQY | 0.2550519 |
| SATGVFAGVI | ASRQDGDYEQY | 0.31349766 |
| SLPAITLP | ASRQDGDYEQY | 0.2765738 |
| STALAFTV | ASRQDGDYEQY | 0.4734376 |
| SVLTVGLGK | ASRQDGDYEQY | 0.36744916 |
| TALAGAGSGI | ASRQDGDYEQY | 0.6556699 |
| TGLPFTGI | ASRQDGDYEQY | 0.44406226 |
| VAEHVTV | ASRQDGDYEQY | 0.5738424 |
| VDIVLNSL | ASRQDGDYEQY | 0.29157615 |
| VLTGGGAVL | ASRQDGDYEQY | 0.3863462 |
| VNVGLAYGI | ASRQDGDYEQY | 0.24055852 |
| VSSAAVGAV | ASRQDGDYEQY | 0.3389408 |
| VTSVYEAGV | ASRQDGDYEQY | 0.2668281 |
| WVALDRGA | ASRQDGDYEQY | 0.3805466 |
| YLVGSVVI | ASRQDGDYEQY | 0.21264306 |
| GPPGTGKTLL | ASRQDGDYEQY | 0.60645497 |
| GVGMTKF | ASRQDGDYEQY | 0.40252665 |
| IGIGGSDL | ASRQDGDYEQY | 0.5511496 |
| IGTLLAEL | ASRQDGDYEQY | 0.3319589 |
| IYTSGTT | ASRQDGDYEQY | 0.28321126 |
| LSFPAGV | ASRQDGDYEQY | 0.31580317 |
| LYTSGTT | ASRQDGDYEQY | 0.52703923 |
| VGAFPFGF | ASRQDGDYEQY | 0.22168104 |
| VGAGTVEF | ASRQDGDYEQY | 0.19711478 |
| VGKSTLLL | ASRQDGDYEQY | 0.2967596 |
| AALAHAGGGIV | ASRQDGDYEQY | 0.20298366 |
| AALVAVSLL | ASRQDGDYEQY | 0.6463069 |
| AAVSRAVAAL | ASRQDGDYEQY | 0.6126691 |
| AGGSSGIGGF | ASRQDGDYEQY | 0.39199692 |
| AGLTVANANG | ASRQDGDYEQY | 0.6018139 |
| AGLVGAVL | ASRQDGDYEQY | 0.18104033 |
| ALSQAFTV | ASRQDGDYEQY | 0.33607116 |
| ALTFSVSL | ASRQDGDYEQY | 0.684498 |
| ALTGAALVAL | ASRQDGDYEQY | 0.4126216 |
| ASFSIGAGVL | ASRQDGDYEQY | 0.39861965 |
| DGSVYKI | ASRQDGDYEQY | 0.61163604 |
| EQLAEIV | ASRQDGDYEQY | 0.3124554 |
| FAVSELTVM | ASRQDGDYEQY | 0.35892344 |
| GAAPLAAGL | ASRQDGDYEQY | 0.37748432 |
| GADGGDGGSGGAGG | ASRQDGDYEQY | 0.38623583 |
| GFGLDTGGAGG | ASRQDGDYEQY | 0.36613643 |

| | | |
|---|---|---|
| GLASGGLL | ASRQDGDYEQY | 0.23231994 |
| GRIAGAAVLL | ASRQDGDYEQY | 0.48552892 |
| GSAGSLNTSF | ASRQDGDYEQY | 0.71535975 |
| GTGQIGI | ASRQDGDYEQY | 0.32701614 |
| GTLIALIL | ASRQDGDYEQY | 0.60001504 |
| IAATMGFVVG | ASRQDGDYEQY | 0.4702996 |
| IFAQALL | ASRQDGDYEQY | 0.6167416 |
| IGAAVALGF | ASRQDGDYEQY | 0.29107848 |
| IGLTPLAL | ASRQDGDYEQY | 0.25616446 |
| ILGSNTSTL | ASRQDGDYEQY | 0.44026068 |
| IVFSLTVL | ASRQDGDYEQY | 0.24663906 |
| LASKATGF | ASRQDGDYEQY | 0.40506378 |
| LASVALIL | ASRQDGDYEQY | 0.33431074 |
| LFAVGLTL | ASRQDGDYEQY | 0.73998606 |
| LFNGGIGTYI | ASRQDGDYEQY | 0.5363995 |
| LTLGAGII | ASRQDGDYEQY | 0.47234738 |
| LVAAALGDI | ASRQDGDYEQY | 0.26298875 |
| NTAFVQLGI | ASRQDGDYEQY | 0.34867704 |
| QLFSGAEL | ASRQDGDYEQY | 0.47147653 |
| SAATFITL | ASRQDGDYEQY | 0.24093074 |
| SNYGLANLGV | ASRQDGDYEQY | 0.6508044 |
| SYGGAYAVM | ASRQDGDYEQY | 0.3095382 |
| TTGVGGLTLG | ASRQDGDYEQY | 0.29575446 |
| TTQTIYI | ASRQDGDYEQY | 0.45917892 |
| VAAGAGLGAVI | ASRQDGDYEQY | 0.697863 |
| VAAVVTLL | ASRQDGDYEQY | 0.3381183 |
| VGFLGGLI | ASRQDGDYEQY | 0.50232995 |
| VGVIPILL | ASRQDGDYEQY | 0.2885157 |
| VIFASSLI | ASRQDGDYEQY | 0.5351515 |
| VKGGDGGAAAG | ASRQDGDYEQY | 0.6234326 |
| VTKTFTNI | ASRQDGDYEQY | 0.48909694 |
| VVAAAILL | ASRQDGDYEQY | 0.6955992 |
| YAALDVEL | ASRQDGDYEQY | 0.40574333 |
| LASVALLL | ASRQDGDYEQY | 0.24712744 |
| LDAVHRV | ASRQDGDYEQY | 0.39276114 |
| QGLSSSIF | ASRQDGDYEQY | 0.46331516 |
| RGTFANIRL | ASRQDGDYEQY | 0.6030445 |
| RQLWWGHRI | ASRQDGDYEQY | 0.5282361 |
| YGPPGTGKTLL | ASRQDGDYEQY | 0.3414943 |
| AALVGAPANVA | ASRQDGDYEQY | 0.26107413 |
| AATFRRTE | ASRQDGDYEQY | 0.63389254 |
| AGALGGGI | ASRQDGDYEQY | 0.633839 |
| AGGAGGAGGIGAG | ASRQDGDYEQY | 0.538114 |
| AGSGALGL | ASRQDGDYEQY | 0.30133682 |
| AQIADQI | ASRQDGDYEQY | 0.29074967 |
| ASAVAVTYL | ASRQDGDYEQY | 0.39554954 |
| ASTVLGGTGG | ASRQDGDYEQY | 0.55318767 |
| AVGGEPAK | ASRQDGDYEQY | 0.28482482 |
| CEQGTFGGTC | ASRQDGDYEQY | 0.43642473 |
| DEGKAPAG | ASRQDGDYEQY | 0.45260128 |
| DTVVFPAL | ASRQDGDYEQY | 0.2126625 |

| | | |
|---|---|---|
| FEGITMAG | ASRQDGDYEQY | 0.38600078 |
| FPLVVQD | ASRQDGDYEQY | 0.526847 |
| GAFTTPQ | ASRQDGDYEQY | 0.37196288 |
| GFGGFGTAGGVGGA | ASRQDGDYEQY | 0.2825985 |
| GFLAQFL | ASRQDGDYEQY | 0.6321134 |
| GGAGGAGGNAGVA | ASRQDGDYEQY | 0.71154964 |
| GGVGTAGAGGAGGAGG | ASRQDGDYEQY | 0.5321718 |
| GNGGGGDAGNAGSGGN | ASRQDGDYEQY | 0.23905571 |
| GTGGAGGAGNDAGS | ASRQDGDYEQY | 0.18248637 |
| GTTNTGWFNS | ASRQDGDYEQY | 0.3092129 |
| IFMGPLFTL | ASRQDGDYEQY | 0.6834183 |
| ILAHAIEAI | ASRQDGDYEQY | 0.3641125 |
| IRSAIREAL | ASRQDGDYEQY | 0.2929856 |
| ITFAGGG | ASRQDGDYEQY | 0.4426156 |
| ITGMSGHQEV | ASRQDGDYEQY | 0.5976645 |
| IVAGLACTL | ASRQDGDYEQY | 0.27875972 |
| IVGGGLAGA | ASRQDGDYEQY | 0.2335497 |
| IVTGVLNI | ASRQDGDYEQY | 0.37446678 |
| LAQQGCI | ASRQDGDYEQY | 0.5272388 |
| LCLHVEN | ASRQDGDYEQY | 0.24547453 |
| LDAISDGS | ASRQDGDYEQY | 0.30893496 |
| LGGFAAML | ASRQDGDYEQY | 0.44067475 |
| LGGKNPMI | ASRQDGDYEQY | 0.32242504 |
| LGPKGVAAIL | ASRQDGDYEQY | 0.44470006 |
| LMVGVSI | ASRQDGDYEQY | 0.26176715 |
| LVGAAFVL | ASRQDGDYEQY | 0.5225173 |
| LVLAVAPL | ASRQDGDYEQY | 0.2307295 |
| MAAIAVIDAA | ASRQDGDYEQY | 0.4581712 |
| NNGINAELQ | ASRQDGDYEQY | 0.35731298 |
| RGEPLVLVHG | ASRQDGDYEQY | 0.39756685 |
| RGGAGGAAGLI | ASRQDGDYEQY | 0.3419165 |
| RSYRMHPAV | ASRQDGDYEQY | 0.2861193 |
| SAAHLLAGQLA | ASRQDGDYEQY | 0.23455545 |
| SAAVGSY | ASRQDGDYEQY | 0.2134414 |
| SPHAVLHRI | ASRQDGDYEQY | 0.46576127 |
| SQIAKVL | ASRQDGDYEQY | 0.31343958 |
| TALGHLVP | ASRQDGDYEQY | 0.45918012 |
| TAPTTSGAA | ASRQDGDYEQY | 0.48561087 |
| TDGDDGNGGSPGT | ASRQDGDYEQY | 0.2606113 |
| TDGLGEN | ASRQDGDYEQY | 0.4076687 |
| TGGPPLPGQA | ASRQDGDYEQY | 0.23825742 |
| TIPAITI | ASRQDGDYEQY | 0.35438982 |
| TIPGFTI | ASRQDGDYEQY | 0.5406071 |
| TSIASGALA | ASRQDGDYEQY | 0.4630977 |
| TVTLPGAAI | ASRQDGDYEQY | 0.28113785 |
| VASGLSAALP | ASRQDGDYEQY | 0.4494023 |
| VGAGVIGI | ASRQDGDYEQY | 0.41594547 |
| VGPKRFGV | ASRQDGDYEQY | 0.27482128 |
| VNVAFPD | ASRQDGDYEQY | 0.39233708 |
| VSMTNSGVS | ASRQDGDYEQY | 0.5558529 |
| VTDVLGDLSV | ASRQDGDYEQY | 0.4072362 |

| | | |
|---|---|---|
| VVVVGAGFAGL | ASRQDGDYEQY | 0.24884342 |
| VYTVGTVV | ASRQDGDYEQY | 0.5229179 |
| WTTFSGP | ASRQDGDYEQY | 0.26632085 |
| YDELVEAF | ASRQDGDYEQY | 0.46884337 |
| YGTGGAGGNGG | ASRQDGDYEQY | 0.33668226 |
| YTNVMAV | ASRQDGDYEQY | 0.37126204 |
| FLEQVHQL | ASRQDGDYEQY | 0.5136199 |
| GLSLLAVAV | ASRQDGDYEQY | 0.24963252 |
| LEQVHQL | ASRQDGDYEQY | 0.24267437 |
| LNAAGAI | ASRQDGDYEQY | 0.3525672 |
| PAVHASL | ASRQDGDYEQY | 0.46000478 |
| TELEDDA | ASRQDGDYEQY | 0.55056304 |
| TPPGPPN | ASRQDGDYEQY | 0.26506227 |
| TPPPPPPP | ASRQDGDYEQY | 0.41868654 |
| TQSPALI | ASSSGGQDTQY | 0.35940006 |
| EFAAAFPQ | ASSSGGQDTQY | 0.2934464 |
| LLSGNLDVL | ASSSGGQDTQY | 0.39213645 |
| GPANLGNYNI | ASSSGGQDTQY | 0.32311493 |
| TLGPTLP | ASSSGGQDTQY | 0.28761724 |
| VYAYSSCCK | ASSSGGQDTQY | 0.41388956 |
| APAPAPAPNI | ASSSGGQDTQY | 0.31928074 |
| PGGPVPIP | ASSSGGQDTQY | 0.45783302 |
| FLTVALEP | ASSSGGQDTQY | 0.2911257 |
| SQYFGANL | ASSSGGQDTQY | 0.24909656 |
| GGVKPAA | ASSSGGQDTQY | 0.43495923 |
| GGTGGAGGAGGW | ASSSGGQDTQY | 0.39880836 |
| GVGGVAGLGGAGG | ASSSGGQDTQY | 0.38571036 |
| VLLVLATTI | ASSSGGQDTQY | 0.38686845 |
| IGIFAAML | ASSSGGQDTQY | 0.2701793 |
| AVGDLID | ASSSGGQDTQY | 0.51264846 |
| NISPLPL | ASSSGGQDTQY | 0.58501697 |
| VTAVFVAI | ASSSGGQDTQY | 0.43038636 |
| SGDDNDAAL | ASSSGGQDTQY | 0.3967865 |
| AVAALAHS | ASSSGGQDTQY | 0.4620004 |
| GGVTVMATP | ASSSGGQDTQY | 0.49421406 |
| AGLAANVRD | ASSSGGQDTQY | 0.30329207 |
| TNRPDLIDPALLRPG | ASSSGGQDTQY | 0.2219012 |
| LANLGVTAL | ASSSGGQDTQY | 0.3738913 |
| TGERIEL | ASSSGGQDTQY | 0.35319608 |
| NGEELAA | ASSSGGQDTQY | 0.30526525 |
| VLSLVWG | ASSSGGQDTQY | 0.24303442 |
| IGYALWL | ASSSGGQDTQY | 0.59568405 |
| LSLLLAPL | ASSSGGQDTQY | 0.7311436 |
| TLDEALK | ASSSGGQDTQY | 0.31438956 |
| TDGAGTAEGG | ASSSGGQDTQY | 0.6291589 |
| NAPGGAGGNGG | ASSSGGQDTQY | 0.23009813 |
| VSVFMGMGGNF | ASSSGGQDTQY | 0.5715445 |
| LASYLAAL | ASSSGGQDTQY | 0.59786415 |
| LVGGTPAL | ASSSGGQDTQY | 0.27090693 |
| VRNADVL | ASSSGGQDTQY | 0.4491076 |
| TCSSPFLVSAM | ASSSGGQDTQY | 0.26619422 |

| | | |
|---|---|---|
| VHITDAE | ASSSGGQDTQY | 0.30219215 |
| EYAAPSF | ASSSGGQDTQY | 0.555279 |
| VVDAFHGGL | ASSSGGQDTQY | 0.38003245 |
| ASVVDPQ | ASSSGGQDTQY | 0.41702318 |
| SYNEFGGLS | ASSSGGQDTQY | 0.37840375 |
| DPDGSVY | ASSSGGQDTQY | 0.29372615 |
| MVSTIHATE | ASSSGGQDTQY | 0.36849648 |
| LAAGVADPV | ASSSGGQDTQY | 0.4631114 |
| TKEQIAA | ASSSGGQDTQY | 0.34001347 |
| SLASPAAL | ASSSGGQDTQY | 0.6427803 |
| IGLVFYGV | ASSSGGQDTQY | 0.24985033 |
| FLLDASNL | ASSSGGQDTQY | 0.6511958 |
| YQVGNFP | ASSSGGQDTQY | 0.6263708 |
| ESLLSQST | ASSSGGQDTQY | 0.4017254 |
| LFTVGFAAVG | ASSSGGQDTQY | 0.31793582 |
| ATPPTPT | ASSSGGQDTQY | 0.6287564 |
| TDLATLG | ASSSGGQDTQY | 0.34864306 |
| SAGGTGR | ASSSGGQDTQY | 0.35156053 |
| GFSNSGS | ASSSGGQDTQY | 0.4788801 |
| GGNGGSGGTSV | ASSSGGQDTQY | 0.7230618 |
| GTSVATGGAGN | ASSSGGQDTQY | 0.9014115 |
| NGGNGGSGGTS | ASSSGGQDTQY | 0.8670989 |
| AVVKFPH | ASSSGGQDTQY | 0.853575 |
| TPPPPPP | ASSSGGQDTQY | 0.8132607 |
| AAPAAAAAPA | ASSSGGQDTQY | 0.4144155 |
| IALNEDL | ASSSGGQDTQY | 0.7737174 |
| IVNVSSM | ASSSGGQDTQY | 0.7565657 |
| TGIPPAPR | ASSSGGQDTQY | 0.839314 |
| FDLGGGTF | ASSSGGQDTQY | 0.8874242 |
| FDLGGGTFDVS | ASSSGGQDTQY | 0.7808055 |
| NEPTAAA | ASSSGGQDTQY | 0.89248824 |
| NEPTAAALA | ASSSGGQDTQY | 0.8754345 |
| GSIINISSI | ASSSGGQDTQY | 0.5930509 |
| VAPGNAGTA | ASSSGGQDTQY | 0.8282004 |
| TGGGASLE | ASSSGGQDTQY | 0.70030284 |
| VAVYNHYKRI | ASSSGGQDTQY | 0.84118354 |
| TDPAPATT | ASSSGGQDTQY | 0.33923316 |
| TNRPDILDPALLRPG | ASSSGGQDTQY | 0.8847002 |
| LTTTTTS | ASSSGGQDTQY | 0.7347076 |
| PTGSGKTL | ASSSGGQDTQY | 0.149423 |
| TGSGKTL | ASSSGGQDTQY | 0.86289763 |
| TIGHVDHGK | ASSSGGQDTQY | 0.81619227 |
| TIGHVDHGKTT | ASSSGGQDTQY | 0.81741595 |
| GHVDHGKTT | ASSSGGQDTQY | 0.8905435 |
| AQFEHTLL | ASSSGGQDTQY | 0.55365264 |
| IVNQDEL | ASSSGGQDTQY | 0.6871619 |
| TNGSQFF | ASSSGGQDTQY | 0.8745358 |
| TGKGIVE | ASSSGGQDTQY | 0.89976126 |
| LISNASDA | ASSSGGQDTQY | 0.6525146 |
| AELRTLGELTKIAKAHGAQ | ASSSGGQDTQY | 0.86649966 |
| APNAVAAAEPSW | ASSSGGQDTQY | 0.89510024 |

| | | |
|---|---|---|
| APQVAGAATAVL | ASSSGGQDTQY | 0.87503284 |
| ARRDPAAAAAILPTDA | ASSSGGQDTQY | 0.8922006 |
| DAAARDVLAEAGLAEYF | ASSSGGQDTQY | 0.9031277 |
| DDQIVLSAKANLAKVEQ | ASSSGGQDTQY | 0.7022054 |
| DELLPDTGEDSDEDGH | ASSSGGQDTQY | 0.89494693 |
| ELPEAPANRATVPQD | ASSSGGQDTQY | 0.90348756 |
| FGQSKIERVTGIG | ASSSGGQDTQY | 0.7865606 |
| GAGGTGSTAGGAGGAGGAGGLY | ASSSGGQDTQY | 0.85246027 |
| GHGGTGAAGGNGGTGGAGGSGI | ASSSGGQDTQY | 0.7223064 |
| GNGGNGGAGGEGGPGGAGGAGA | ASSSGGQDTQY | 0.8090966 |
| GSTAGNGGNGGNATNGTIAGQP | ASSSGGQDTQY | 0.41002846 |
| GTGGDGGNGGNGGTGG | ASSSGGQDTQY | 0.8572368 |
| GVRRPAGIPADAPV | ASSSGGQDTQY | 0.903309 |
| GYRQVTDAHLLTLARR | ASSSGGQDTQY | 0.8311121 |
| IAKMGQDGHDRG | ASSSGGQDTQY | 0.6926217 |
| KAEPSIEIQYT | ASSSGGQDTQY | 0.6838236 |
| KPVRIRMAADEDTVAVT | ASSSGGQDTQY | 0.8560177 |
| MPPRIAGMRLLVIKPEP | ASSSGGQDTQY | 0.804327 |
| MKFYAVALFRSRGVPPTR | ASSSGGQDTQY | 0.38908988 |
| NAGGFGILGAVAHSP | ASSSGGQDTQY | 0.745324 |
| NEGREITLAMAEM | ASSSGGQDTQY | 0.89152145 |
| NELVEWSHPLGHRWRWE | ASSSGGQDTQY | 0.8913303 |
| NLGAGGTGGNGGNPG | ASSSGGQDTQY | 0.64918685 |
| NTEWPKWETVMAADPVD | ASSSGGQDTQY | 0.9053415 |
| PASAAPAETQP | ASSSGGQDTQY | 0.8672638 |
| QQSSDGKNGVRAAEITGE | ASSSGGQDTQY | 0.9026847 |
| RAPDAVTTSWWKE | ASSSGGQDTQY | 0.8005772 |
| SGFGNVGAHTSGWFNQ | ASSSGGQDTQY | 0.7512243 |
| SGVAGAVGQAASVGGLKV | ASSSGGQDTQY | 0.87107897 |
| SNGGAGTGSAGGNGGT | ASSSGGQDTQY | 0.81496775 |
| TGGAGGEGGAAG | ASSSGGQDTQY | 0.7681691 |
| TGGDGGAAGAGGAPGHG | ASSSGGQDTQY | 0.8892821 |
| TMPPLLAAIANAGRHGV | ASSSGGQDTQY | 0.8272618 |
| TSSTAGNSGGA | ASSSGGQDTQY | 0.7613755 |
| VPATVPVGILC | ASSSGGQDTQY | 0.76888716 |
| VQRPDSGGVVLTGRIS | ASSSGGQDTQY | 0.9091488 |
| WTPNPVPATKGVPAGLVQAT | ASSSGGQDTQY | 0.69784504 |
| YALKRALAKFSDRFDVVII | ASSSGGQDTQY | 0.7368115 |
| YLIGSVIV | ASSSGGQDTQY | 0.6806721 |
| YGQLVAVID | ASSSGGQDTQY | 0.82352805 |
| VTVGGMTLPA | ASSSGGQDTQY | 0.8969091 |
| VLALPAV | ASSSGGQDTQY | 0.864426 |
| VIPPTLNYET | ASSSGGQDTQY | 0.8080152 |
| VGMGGMHPSSG | ASSSGGQDTQY | 0.8643733 |
| VGLAGGAATA | ASSSGGQDTQY | 0.880962 |
| VGGAGGAGGSGP | ASSSGGQDTQY | 0.8540473 |
| VFAAFMV | ASSSGGQDTQY | 0.8791394 |
| TLYDKLR | ASSSGGQDTQY | 0.75555336 |
| TLAVFTVL | ASSSGGQDTQY | 0.9045721 |
| TGSVQPARV | ASSSGGQDTQY | 0.8863126 |
| TGGGAGSGGT | ASSSGGQDTQY | 0.8874533 |

| | | |
|---|---|---|
| SHLETSL | ASSSGGQDTQY | 0.8868571 |
| SFTTGMT | ASSSGGQDTQY | 0.8545928 |
| QVEVAKRMGVRQPTVS | ASSSGGQDTQY | 0.85951626 |
| QGSGGPALAM | ASSSGGQDTQY | 0.5629835 |
| MLAPSNP | ASSSGGQDTQY | 0.58084786 |
| LVNNAAI | ASSSGGQDTQY | 0.85693645 |
| LTGEINTLA | ASSSGGQDTQY | 0.8593118 |
| LSIFGPV | ASSSGGQDTQY | 0.87728333 |
| LLALLANL | ASSSGGQDTQY | 0.8594039 |
| LISNASDALDKLR | ASSSGGQDTQY | 0.7800461 |
| LINEAVP | ASSSGGQDTQY | 0.88782144 |
| LGYAFSGL | ASSSGGQDTQY | 0.8874452 |
| LDFVAVAL | ASSSGGQDTQY | 0.6375964 |
| LAQPGAE | ASSSGGQDTQY | 0.81130767 |
| ISNLGAVTGF | ASSSGGQDTQY | 0.9071348 |
| HQEDPGRI | ASSSGGQDTQY | 0.85398257 |
| GTPGEPTP | ASSSGGQDTQY | 0.81783557 |
| GFFNSGS | ASSSGGQDTQY | 0.80949986 |
| GAGGGGLTAGAN | ASSSGGQDTQY | 0.8942834 |
| FVDALVNM | ASSSGGQDTQY | 0.74758554 |
| FLTVAPEL | ASSSGGQDTQY | 0.90661234 |
| FETLTAL | ASSSGGQDTQY | 0.33175588 |
| DVINPIEL | ASSSGGQDTQY | 0.8178476 |
| APVSGAPV | ASSSGGQDTQY | 0.8033288 |
| ALVGGLAW | ASSSGGQDTQY | 0.46209887 |
| ALAAAAATAI | ASSSGGQDTQY | 0.8951795 |
| AFGLGLLL | ASSSGGQDTQY | 0.72599185 |
| AFFAAASL | ASSSGGQDTQY | 0.8721338 |
| AAAALPAFSPP | ASSSGGQDTQY | 0.7928458 |
| VVLAPAL | ASSSGGQDTQY | 0.8973671 |
| TIGHVDHGKTTL | ASSSGGQDTQY | 0.8570681 |
| LISNASDALDK | ASSSGGQDTQY | 0.8211343 |
| LISNASDAL | ASSSGGQDTQY | 0.7061159 |
| ISNASDALDK | ASSSGGQDTQY | 0.88768154 |
| NQAETLVYQTE | ASSSGGQDTQY | 0.8954679 |
| TGGTGGTAGAG | ASSSGGQDTQY | 0.8986548 |
| TGSTAGGAGGA | ASSSGGQDTQY | 0.4542628 |
| DPLAVPVAAGPV | ASSSGGQDTQY | 0.8190678 |
| GGTGGTPGNGGN | ASSSGGQDTQY | 0.8529035 |
| PAGLAGVDPATL | ASSSGGQDTQY | 0.28286135 |
| GNNLVVEAPHAGASV | ASSSGGQDTQY | 0.26642108 |
| KVTRIMSASTLEQPA | ASSSGGQDTQY | 0.80857277 |
| LGFALPIALPRRTVV | ASSSGGQDTQY | 0.79055023 |
| TRLGNLTARKLVVLV | ASSSGGQDTQY | 0.8859043 |
| WSQQRRLIYFPSAGP | ASSSGGQDTQY | 0.8759658 |
| GPNGNPDPMAAAVDIR | ASSSGGQDTQY | 0.83701897 |
| TQRCVSVADLRAAASR | ASSSGGQDTQY | 0.87020457 |
| QFVLDVAEHPVEVLTETV | ASSSGGQDTQY | 0.8520056 |
| GGAGGTGGAGGNGGTQAGDG | ASSSGGQDTQY | 0.082466066 |
| PLLATLRAIAVDVDAKPAQV | ASSSGGQDTQY | 0.71829504 |
| APKFRTGGYMIDFWGVGYQVAK | ASSSGGQDTQY | 0.60828847 |

| | | |
|---|---|---|
| GAGGAGGAGGAAGGPGGTGGTG | ASSSGGQDTQY | 0.8928516 |
| IGALGLLATDVAPAGSVAFGVA | ASSSGGQDTQY | 0.7304379 |
| AGGAGGIGGNGGAGGAGGVLMG | ASSSGGQDTQY | 0.7425868 |
| ADELIGLL | ASSSGGQDTQY | 0.85226417 |
| ADRALKA | ASSSGGQDTQY | 0.83295125 |
| AGAVGQVGGAGGA | ASSSGGQDTQY | 0.82165384 |
| AGGGGGLGGAP | ASSSGGQDTQY | 0.7566359 |
| AGGLLYGNGGAG | ASSSGGQDTQY | 0.2579186 |
| AGLESIGL | ASSSGGQDTQY | 0.8927137 |
| AGLKTGII | ASSSGGQDTQY | 0.47221267 |
| ASLAALLL | ASSSGGQDTQY | 0.6646258 |
| ASVINISSI | ASSSGGQDTQY | 0.37442672 |
| ATPPGPA | ASSSGGQDTQY | 0.36403054 |
| ATVASALV | ASSSGGQDTQY | 0.36065775 |
| ATVASLIL | ASSSGGQDTQY | 0.8897424 |
| AVNGPVLIRH | ASSSGGQDTQY | 0.774946 |
| DGAAGAVGI | ASSSGGQDTQY | 0.8078159 |
| DPLLHGI | ASSSGGQDTQY | 0.9083574 |
| ELGAALGAL | ASSSGGQDTQY | 0.79892963 |
| ELGRTQAV | ASSSGGQDTQY | 0.8280151 |
| FVAGGYLS | ASSSGGQDTQY | 0.7576914 |
| GASGGAGGAGGW | ASSSGGQDTQY | 0.75171125 |
| GGAGAGGDGGAGGI | ASSSGGQDTQY | 0.7884096 |
| GTGHNGSQPGMGGNG | ASSSGGQDTQY | 0.745484 |
| GYAAASASAS | ASSSGGQDTQY | 0.90187466 |
| HQVDVAAVQQG | ASSSGGQDTQY | 0.84151113 |
| IGAAIAHT | ASSSGGQDTQY | 0.50787985 |
| IGMIEVAQL | ASSSGGQDTQY | 0.8569933 |
| IRLAPEHTSAMD | ASSSGGQDTQY | 0.9033629 |
| IYNVSVL | ASSSGGQDTQY | 0.9130238 |
| KQGFLGM | ASSSGGQDTQY | 0.7516761 |
| LESVITQL | ASSSGGQDTQY | 0.7584036 |
| LGLGALVVV | ASSSGGQDTQY | 0.7844525 |
| LIGNGGNGGSGGI | ASSSGGQDTQY | 0.8249737 |
| LNAINAPTL | ASSSGGQDTQY | 0.72022796 |
| LQLLNTGV | ASSSGGQDTQY | 0.6397284 |
| LQTSIEF | ASSSGGQDTQY | 0.8953117 |
| MSFAVSEL | ASSSGGQDTQY | 0.89871323 |
| NTGALGL | ASSSGGQDTQY | 0.8243081 |
| PPGAGGAGGAGTT | ASSSGGQDTQY | 0.39978176 |
| QLVGLGI | ASSSGGQDTQY | 0.8224089 |
| RFLEQVHQL | ASSSGGQDTQY | 0.74620515 |
| RGVGYVLR | ASSSGGQDTQY | 0.8007561 |
| SATGVFAGVI | ASSSGGQDTQY | 0.815307 |
| SLPAITLP | ASSSGGQDTQY | 0.5980938 |
| STALAFTV | ASSSGGQDTQY | 0.8275688 |
| SVLTVGLGK | ASSSGGQDTQY | 0.9141737 |
| TALAGAGSGI | ASSSGGQDTQY | 0.76989377 |
| TGLPFTGI | ASSSGGQDTQY | 0.8607032 |
| VAEHVTV | ASSSGGQDTQY | 0.90182185 |
| VDIVLNSL | ASSSGGQDTQY | 0.8853551 |

| | | |
|---|---|---|
| VLTGGGAVL | ASSSGGQDTQY | 0.8997547 |
| VNVGLAYGI | ASSSGGQDTQY | 0.86006117 |
| VSSAAVGAV | ASSSGGQDTQY | 0.8462846 |
| VTSVYEAGV | ASSSGGQDTQY | 0.85002816 |
| WVALDRGA | ASSSGGQDTQY | 0.847664 |
| YLVGSVVI | ASSSGGQDTQY | 0.8298187 |
| GPPGTGKTLL | ASSSGGQDTQY | 0.8431893 |
| GVGMTKF | ASSSGGQDTQY | 0.79889536 |
| IGIGGSDL | ASSSGGQDTQY | 0.8853406 |
| IGTLLAEL | ASSSGGQDTQY | 0.8362054 |
| IYTSGTT | ASSSGGQDTQY | 0.8902632 |
| LSFPAGV | ASSSGGQDTQY | 0.45129776 |
| LYTSGTT | ASSSGGQDTQY | 0.7046394 |
| VGAFPFGF | ASSSGGQDTQY | 0.89072627 |
| VGAGTVEF | ASSSGGQDTQY | 0.16659643 |
| VGKSTLLL | ASSSGGQDTQY | 0.2784319 |
| AALAHAGGGIV | ASSSGGQDTQY | 0.8606186 |
| AALVAVSLL | ASSSGGQDTQY | 0.90225726 |
| AAVSRAVAAL | ASSSGGQDTQY | 0.15341027 |
| AGGSSGIGGF | ASSSGGQDTQY | 0.6124717 |
| AGLTVANANG | ASSSGGQDTQY | 0.6383563 |
| AGLVGAVL | ASSSGGQDTQY | 0.8679384 |
| ALSQAFTV | ASSSGGQDTQY | 0.83614343 |
| ALTFSVSL | ASSSGGQDTQY | 0.40987808 |
| ALTGAALVAL | ASSSGGQDTQY | 0.8903166 |
| ASFSIGAGVL | ASSSGGQDTQY | 0.5000203 |
| DGSVYKI | ASSSGGQDTQY | 0.7052487 |
| EQLAEIV | ASSSGGQDTQY | 0.8928151 |
| FAVSELTVM | ASSSGGQDTQY | 0.89058745 |
| GAAPLAAGL | ASSSGGQDTQY | 0.65258193 |
| GADGGDGGSGGAGG | ASSSGGQDTQY | 0.88305676 |
| GFGLDTGGAGG | ASSSGGQDTQY | 0.76979387 |
| GLASGGLL | ASSSGGQDTQY | 0.91375387 |
| GRIAGAAVLL | ASSSGGQDTQY | 0.8151425 |
| GSAGSLNTSF | ASSSGGQDTQY | 0.5061949 |
| GTGQIGI | ASSSGGQDTQY | 0.8900864 |
| GTLIALIL | ASSSGGQDTQY | 0.8812711 |
| IAATMGFVVG | ASSSGGQDTQY | 0.842546 |
| IFAQALL | ASSSGGQDTQY | 0.87532425 |
| IGAAVALGF | ASSSGGQDTQY | 0.80438864 |
| IGLTPLAL | ASSSGGQDTQY | 0.8165573 |
| ILGSNTSTL | ASSSGGQDTQY | 0.8865757 |
| IVFSLTVL | ASSSGGQDTQY | 0.88873696 |
| LASKATGF | ASSSGGQDTQY | 0.6399225 |
| LASVALIL | ASSSGGQDTQY | 0.9039531 |
| LFAVGLTL | ASSSGGQDTQY | 0.83687186 |
| LFNGGIGTYI | ASSSGGQDTQY | 0.7335179 |
| LTLGAGII | ASSSGGQDTQY | 0.912303 |
| LVAAALGDI | ASSSGGQDTQY | 0.8390418 |
| NTAFVQLGI | ASSSGGQDTQY | 0.89409375 |
| QLFSGAEL | ASSSGGQDTQY | 0.8957473 |

| | | |
|---|---|---|
| SAATFITL | ASSSGGQDTQY | 0.8823128 |
| SNYGLANLGV | ASSSGGQDTQY | 0.48010933 |
| SYGGAYAVM | ASSSGGQDTQY | 0.80873203 |
| TTGVGGLTLG | ASSSGGQDTQY | 0.8298124 |
| TTQTIYI | ASSSGGQDTQY | 0.8753933 |
| VAAGAGLGAVI | ASSSGGQDTQY | 0.8063735 |
| VAAVVTLL | ASSSGGQDTQY | 0.687056 |
| VGFLGGLI | ASSSGGQDTQY | 0.8821645 |
| VGVIPILL | ASSSGGQDTQY | 0.9044957 |
| VIFASSLI | ASSSGGQDTQY | 0.8703017 |
| VKGGDGGAAAG | ASSSGGQDTQY | 0.8575449 |
| VTKTFTNI | ASSSGGQDTQY | 0.90855527 |
| VVAAAILL | ASSSGGQDTQY | 0.7679764 |
| YAALDVEL | ASSSGGQDTQY | 0.8430191 |
| LASVALLL | ASSSGGQDTQY | 0.8938694 |
| LDAVHRV | ASSSGGQDTQY | 0.89069414 |
| QGLSSSIF | ASSSGGQDTQY | 0.83807325 |
| RGTFANIRL | ASSSGGQDTQY | 0.89786136 |
| RQLWWGHRI | ASSSGGQDTQY | 0.70804226 |
| YGPPGTGKTLL | ASSSGGQDTQY | 0.86032724 |
| AALVGAPANVA | ASSSGGQDTQY | 0.7179954 |
| AATFRRTE | ASSSGGQDTQY | 0.8374716 |
| AGALGGGI | ASSSGGQDTQY | 0.71866226 |
| AGGAGGAGGIGAG | ASSSGGQDTQY | 0.870276 |
| AGSGALGL | ASSSGGQDTQY | 0.88692135 |
| AQIADQI | ASSSGGQDTQY | 0.87443084 |
| ASAVAVTYL | ASSSGGQDTQY | 0.2058245 |
| ASTVLGGTGG | ASSSGGQDTQY | 0.8819926 |
| AVGGEPAK | ASSSGGQDTQY | 0.8524679 |
| CEQGTFGGTC | ASSSGGQDTQY | 0.9096173 |
| DEGKAPAG | ASSSGGQDTQY | 0.8406156 |
| DTVVFPAL | ASSSGGQDTQY | 0.8668962 |
| FEGITMAG | ASSSGGQDTQY | 0.7907101 |
| FPLVVQD | ASSSGGQDTQY | 0.8552307 |
| GAFTTPQ | ASSSGGQDTQY | 0.6656202 |
| GFGGFGTAGGVGGA | ASSSGGQDTQY | 0.77645576 |
| GFLAQFL | ASSSGGQDTQY | 0.7721176 |
| GGAGGAGGNAGVA | ASSSGGQDTQY | 0.75899744 |
| GGVGTAGAGGAGGAGG | ASSSGGQDTQY | 0.87501454 |
| GNGGGGDAGNAGSGGN | ASSSGGQDTQY | 0.8941339 |
| GTGGAGGAGNDAGS | ASSSGGQDTQY | 0.88924783 |
| GTTNTGWFNS | ASSSGGQDTQY | 0.8889692 |
| IFMGPLFTL | ASSSGGQDTQY | 0.90273213 |
| ILAHAIEAI | ASSSGGQDTQY | 0.89436316 |
| IRSAIREAL | ASSSGGQDTQY | 0.8139035 |
| ITFAGGG | ASSSGGQDTQY | 0.71161675 |
| ITGMSGHQEV | ASSSGGQDTQY | 0.67463833 |
| IVAGLACTL | ASSSGGQDTQY | 0.7864437 |
| IVGGGLAGA | ASSSGGQDTQY | 0.8115543 |
| IVTGVLNI | ASSSGGQDTQY | 0.854898 |
| LAQQGCI | ASSSGGQDTQY | 0.12076319 |

| | | |
|---|---|---|
| LCLHVEN | ASSSGGQDTQY | 0.82113767 |
| LDAISDGS | ASSSGGQDTQY | 0.903782 |
| LGGFAAML | ASSSGGQDTQY | 0.7658925 |
| LGGKNPMI | ASSSGGQDTQY | 0.8543382 |
| LGPKGVAAIL | ASSSGGQDTQY | 0.8406992 |
| LMVGVSI | ASSSGGQDTQY | 0.79868263 |
| LVGAAFVL | ASSSGGQDTQY | 0.87023425 |
| LVLAVAPL | ASSSGGQDTQY | 0.7832371 |
| MAAIAVIDAA | ASSSGGQDTQY | 0.8906187 |
| NNGINAELQ | ASSSGGQDTQY | 0.91054523 |
| RGEPLVLVHG | ASSSGGQDTQY | 0.8570874 |
| RGGAGGAAGLI | ASSSGGQDTQY | 0.8302293 |
| RSYRMHPAV | ASSSGGQDTQY | 0.84752774 |
| SAAHLLAGQLA | ASSSGGQDTQY | 0.37988165 |
| SAAVGSY | ASSSGGQDTQY | 0.8440221 |
| SPHAVLHRI | ASSSGGQDTQY | 0.12069656 |
| SQIAKVL | ASSSGGQDTQY | 0.82440865 |
| TALGHLVP | ASSSGGQDTQY | 0.8037244 |
| TAPTTSGAA | ASSSGGQDTQY | 0.85817844 |
| TDGDDGNGGSPGT | ASSSGGQDTQY | 0.8458466 |
| TDGLGEN | ASSSGGQDTQY | 0.8816633 |
| TGGPPLPGQA | ASSSGGQDTQY | 0.71204287 |
| TIPAITI | ASSSGGQDTQY | 0.83448327 |
| TIPGFTI | ASSSGGQDTQY | 0.7817582 |
| TSIASGALA | ASSSGGQDTQY | 0.5747074 |
| TVTLPGAAI | ASSSGGQDTQY | 0.88146365 |
| VASGLSAALP | ASSSGGQDTQY | 0.8620089 |
| VGAGVIGI | ASSSGGQDTQY | 0.82551 |
| VGPKRFGV | ASSSGGQDTQY | 0.8944514 |
| VNVAFPD | ASSSGGQDTQY | 0.86926746 |
| VSMTNSGVS | ASSSGGQDTQY | 0.4319362 |
| VTDVLGDLSV | ASSSGGQDTQY | 0.7968646 |
| VVVVGAGFAGL | ASSSGGQDTQY | 0.8391932 |
| VYTVGTVV | ASSSGGQDTQY | 0.8495914 |
| WTTFSGP | ASSSGGQDTQY | 0.10836543 |
| YDELVEAF | ASSSGGQDTQY | 0.873916 |
| YGTGGAGGNGG | ASSSGGQDTQY | 0.7344936 |
| YTNVMAV | ASSSGGQDTQY | 0.7179564 |
| FLEQVHQL | ASSSGGQDTQY | 0.81545496 |
| GLSLLAVAV | ASSSGGQDTQY | 0.8890719 |
| LEQVHQL | ASSSGGQDTQY | 0.79324085 |
| LNAAGAI | ASSSGGQDTQY | 0.76482296 |
| PAVHASL | ASSSGGQDTQY | 0.88398755 |
| TELEDDA | ASSSGGQDTQY | 0.86706424 |
| TPPGPPN | ASSSGGQDTQY | 0.8965411 |
| TPPPPPPP | ASSSGGQDTQY | 0.8600913 |
| TQSPALI | ASSRREVTEVF | 0.8519697 |
| EFAAAFPQ | ASSRREVTEVF | 0.8528804 |
| LLSGNLDVL | ASSRREVTEVF | 0.72513634 |
| GPANLGNYNI | ASSRREVTEVF | 0.5550482 |
| TLGPTLP | ASSRREVTEVF | 0.5412313 |

| | | |
|---|---|---|
| VYAYSSCCK | ASSRREVTEVF | 0.8224714 |
| APAPAPAPNI | ASSRREVTEVF | 0.48335433 |
| PGGPVPIP | ASSRREVTEVF | 0.8601984 |
| FLTVALEP | ASSRREVTEVF | 0.8953195 |
| SQYFGANL | ASSRREVTEVF | 0.78953457 |
| GGVKPAA | ASSRREVTEVF | 0.6194172 |
| GGTGGAGGAGGW | ASSRREVTEVF | 0.821285 |
| GVGGVAGLGGAGG | ASSRREVTEVF | 0.71632814 |
| VLLVLATTI | ASSRREVTEVF | 0.8835944 |
| IGIFAAML | ASSRREVTEVF | 0.88025475 |
| AVGDLID | ASSRREVTEVF | 0.89405954 |
| NISPLPL | ASSRREVTEVF | 0.88416195 |
| VTAVFVAI | ASSRREVTEVF | 0.7997775 |
| SGDDNDAAL | ASSRREVTEVF | 0.8909258 |
| AVAALAHS | ASSRREVTEVF | 0.8183836 |
| GGVTVMATP | ASSRREVTEVF | 0.7717215 |
| AGLAANVRD | ASSRREVTEVF | 0.8547852 |
| TNRPDLIDPALLRPG | ASSRREVTEVF | 0.8748964 |
| LANLGVTAL | ASSRREVTEVF | 0.8288461 |
| TGERIEL | ASSRREVTEVF | 0.8122977 |
| NGEELAA | ASSRREVTEVF | 0.48125106 |
| VLSLVWG | ASSRREVTEVF | 0.8699608 |
| IGYALWL | ASSRREVTEVF | 0.8371093 |
| LSLLLAPL | ASSRREVTEVF | 0.51454973 |
| TLDEALK | ASSRREVTEVF | 0.7929747 |
| TDGAGTAEGG | ASSRREVTEVF | 0.90624154 |
| NAPGGAGGNGG | ASSRREVTEVF | 0.89167213 |
| VSVFMGMGGNF | ASSRREVTEVF | 0.7597656 |
| LASYLAAL | ASSRREVTEVF | 0.78506786 |
| LVGGTPAL | ASSRREVTEVF | 0.8241737 |
| VRNADVL | ASSRREVTEVF | 0.66916347 |
| TCSSPFLVSAM | ASSRREVTEVF | 0.88315403 |
| VHITDAE | ASSRREVTEVF | 0.82208574 |
| EYAAPSF | ASSRREVTEVF | 0.82973754 |
| VVDAFHGGL | ASSRREVTEVF | 0.7476156 |
| ASVVDPQ | ASSRREVTEVF | 0.8617451 |
| SYNEFGGLS | ASSRREVTEVF | 0.80588233 |
| DPDGSVY | ASSRREVTEVF | 0.88196087 |
| MVSTIHATE | ASSRREVTEVF | 0.1594165 |
| LAAGVADPV | ASSRREVTEVF | 0.86574566 |
| TKEQIAA | ASSRREVTEVF | 0.72200227 |
| SLASPAAL | ASSRREVTEVF | 0.7861929 |
| IGLVFYGV | ASSRREVTEVF | 0.42148942 |
| FLLDASNL | ASSRREVTEVF | 0.79089504 |
| YQVGNFP | ASSRREVTEVF | 0.87419164 |
| ESLLSQST | ASSRREVTEVF | 0.8945192 |
| LFTVGFAAVG | ASSRREVTEVF | 0.76065266 |
| ATPPTPT | ASSRREVTEVF | 0.8767824 |
| TDLATLG | ASSRREVTEVF | 0.9058697 |
| SAGGTGR | ASSRREVTEVF | 0.80056167 |
| GFSNSGS | ASSRREVTEVF | 0.3347345 |

| | | |
|---|---|---|
| GGNGGSGGTSV | ASSRREVTEVF | 0.8597076 |
| GTSVATGGAGN | ASSRREVTEVF | 0.766572 |
| NGGNGGSGGTS | ASSRREVTEVF | 0.88403815 |
| AVVKFPH | ASSRREVTEVF | 0.8751476 |
| TPPPPPP | ASSRREVTEVF | 0.66571915 |
| AAPAAAAAPA | ASSRREVTEVF | 0.8658825 |
| IALNEDL | ASSRREVTEVF | 0.8821423 |
| IVNVSSM | ASSRREVTEVF | 0.7296902 |
| TGIPPAPR | ASSRREVTEVF | 0.72610533 |
| FDLGGGTF | ASSRREVTEVF | 0.64667404 |
| FDLGGGTFDVS | ASSRREVTEVF | 0.4045945 |
| NEPTAAA | ASSRREVTEVF | 0.8722022 |
| NEPTAAALA | ASSRREVTEVF | 0.7841393 |
| GSIINISSI | ASSRREVTEVF | 0.7026972 |
| VAPGNAGTA | ASSRREVTEVF | 0.8558102 |
| TGGGASLE | ASSRREVTEVF | 0.769521 |
| VAVYNHYKRI | ASSRREVTEVF | 0.77268684 |
| TDPAPATT | ASSRREVTEVF | 0.7976512 |
| TNRPDILDPALLRPG | ASSRREVTEVF | 0.8043009 |
| LTTTTTS | ASSRREVTEVF | 0.8762852 |
| PTGSGKTL | ASSRREVTEVF | 0.79298806 |
| TGSGKTL | ASSRREVTEVF | 0.80697167 |
| TIGHVDHGK | ASSRREVTEVF | 0.9026551 |
| TIGHVDHGKTT | ASSRREVTEVF | 0.5238506 |
| GHVDHGKTT | ASSRREVTEVF | 0.7064477 |
| AQFEHTLL | ASSRREVTEVF | 0.8783395 |
| IVNQDEL | ASSRREVTEVF | 0.70916796 |
| TNGSQFF | ASSRREVTEVF | 0.8857982 |
| TGKGIVE | ASSRREVTEVF | 0.8488859 |
| LISNASDA | ASSRREVTEVF | 0.8688735 |
| AELRTLGELTKIAKAHGAQ | ASSRREVTEVF | 0.7604877 |
| APNAVAAAEPSW | ASSRREVTEVF | 0.9090092 |
| APQVAGAATAVL | ASSRREVTEVF | 0.8900182 |
| ARRDPAAAAAILPTDA | ASSRREVTEVF | 0.79737955 |
| DAAARDVLAEAGLAEYF | ASSRREVTEVF | 0.8247163 |
| DDQIVLSAKANLAKVEQ | ASSRREVTEVF | 0.53303355 |
| DELLPDTGEDSDEDGH | ASSRREVTEVF | 0.879475 |
| ELPEAPANRATVPQD | ASSRREVTEVF | 0.86857814 |
| FGQSKIERVTGIG | ASSRREVTEVF | 0.7345784 |
| GAGGTGSTAGGAGGAGGAGGLY | ASSRREVTEVF | 0.81675184 |
| GHGGTGAAGGNGGTGGAGGSGI | ASSRREVTEVF | 0.9093536 |
| GNGGNGGAGGEGGPGGAGGAGA | ASSRREVTEVF | 0.8973961 |
| GSTAGNGGNGGNATNGTIAGQP | ASSRREVTEVF | 0.64778185 |
| GTGGDGGNGGNGGTGG | ASSRREVTEVF | 0.7984648 |
| GVRRPAGIPADAPV | ASSRREVTEVF | 0.57395416 |
| GYRQVTDAHLLTLARR | ASSRREVTEVF | 0.81529915 |
| IAKMGQDGHDRG | ASSRREVTEVF | 0.5349875 |
| KAEPSIEIQYT | ASSRREVTEVF | 0.87551725 |
| KPVRIRMAADEDTVAVT | ASSRREVTEVF | 0.8002099 |
| MPPRIAGMRLLVIKPEP | ASSRREVTEVF | 0.85117376 |
| MKFYAVALFRSRGVPPTR | ASSRREVTEVF | 0.866043 |

| | | |
|---|---|---|
| NAGGFGILGAVAHSP | ASSRREVTEVF | 0.90716624 |
| NEGREITLAMAEM | ASSRREVTEVF | 0.7895521 |
| NELVEWSHPLGHRWRWE | ASSRREVTEVF | 0.81455994 |
| NLGAGGTGGNGGNPG | ASSRREVTEVF | 0.5995171 |
| NTEWPKWETVMAADPVD | ASSRREVTEVF | 0.5734745 |
| PASAAPAETQP | ASSRREVTEVF | 0.7741573 |
| QQSSDGKNGVRAAEITGE | ASSRREVTEVF | 0.7744436 |
| RAPDAVTTSWWKE | ASSRREVTEVF | 0.21412338 |
| SGFGNVGAHTSGWFNQ | ASSRREVTEVF | 0.90303755 |
| SGVAGAVGQAASVGGLKV | ASSRREVTEVF | 0.84935135 |
| SNGGAGTGSAGGNGGT | ASSRREVTEVF | 0.89364135 |
| TGGAGGEGGAAG | ASSRREVTEVF | 0.7439577 |
| TGGDGGAAGAGGAPGHG | ASSRREVTEVF | 0.7609267 |
| TMPPLLAAIANAGRHGV | ASSRREVTEVF | 0.80472785 |
| TSSTAGNSGGA | ASSRREVTEVF | 0.8706857 |
| VPATVPVGILC | ASSRREVTEVF | 0.70895094 |
| VQRPDSGGVVLTGRIS | ASSRREVTEVF | 0.40289515 |
| WTPNPVPATKGVPAGLVQAT | ASSRREVTEVF | 0.67800474 |
| YALKRALAKFSDRFDVVII | ASSRREVTEVF | 0.5495479 |
| YLIGSVIV | ASSRREVTEVF | 0.909636 |
| YGQLVAVID | ASSRREVTEVF | 0.89779556 |
| VTVGGMTLPA | ASSRREVTEVF | 0.85932684 |
| VLALPAV | ASSRREVTEVF | 0.905007 |
| VIPPTLNYET | ASSRREVTEVF | 0.20519558 |
| VGMGGMHPSSG | ASSRREVTEVF | 0.7322105 |
| VGLAGGAATA | ASSRREVTEVF | 0.8958759 |
| VGGAGGAGGSGP | ASSRREVTEVF | 0.88020426 |
| VFAAFMV | ASSRREVTEVF | 0.82496405 |
| TLYDKLR | ASSRREVTEVF | 0.90350044 |
| TLAVFTVL | ASSRREVTEVF | 0.74957764 |
| TGSVQPARV | ASSRREVTEVF | 0.8446015 |
| TGGGAGSGGT | ASSRREVTEVF | 0.83172965 |
| SHLETSL | ASSRREVTEVF | 0.88838696 |
| SFTTGMT | ASSRREVTEVF | 0.8578983 |
| QVEVAKRMGVRQPTVS | ASSRREVTEVF | 0.6743523 |
| QGSGGPALAM | ASSRREVTEVF | 0.86098105 |
| MLAPSNP | ASSRREVTEVF | 0.9136818 |
| LVNNAAI | ASSRREVTEVF | 0.80622214 |
| LTGEINTLA | ASSRREVTEVF | 0.890617 |
| LSIFGPV | ASSRREVTEVF | 0.8683171 |
| LLALLANL | ASSRREVTEVF | 0.82287526 |
| LISNASDALDKLR | ASSRREVTEVF | 0.75221604 |
| LINEAVP | ASSRREVTEVF | 0.7663361 |
| LGYAFSGL | ASSRREVTEVF | 0.8313621 |
| LDFVAVAL | ASSRREVTEVF | 0.66586053 |
| LAQPGAE | ASSRREVTEVF | 0.8273252 |
| ISNLGAVTGF | ASSRREVTEVF | 0.79758006 |
| HQEDPGRI | ASSRREVTEVF | 0.92065215 |
| GTPGEPTP | ASSRREVTEVF | 0.8771877 |
| GFFNSGS | ASSRREVTEVF | 0.87901855 |
| GAGGGGLTAGAN | ASSRREVTEVF | 0.7333336 |

| | | |
|---|---|---|
| FVDALVNM | ASSRREVTEVF | 0.8766979 |
| FLTVAPEL | ASSRREVTEVF | 0.87051374 |
| FETLTAL | ASSRREVTEVF | 0.5372257 |
| DVINPIEL | ASSRREVTEVF | 0.89823866 |
| APVSGAPV | ASSRREVTEVF | 0.75733256 |
| ALVGGLAW | ASSRREVTEVF | 0.8317218 |
| ALAAAAATAI | ASSRREVTEVF | 0.8607763 |
| AFGLGLLL | ASSRREVTEVF | 0.87862134 |
| AFFAAASL | ASSRREVTEVF | 0.83663166 |
| AAAALPAFSPP | ASSRREVTEVF | 0.8335957 |
| VVLAPAL | ASSRREVTEVF | 0.74430704 |
| TIGHVDHGKTTL | ASSRREVTEVF | 0.88657963 |
| LISNASDALDK | ASSRREVTEVF | 0.89749825 |
| LISNASDAL | ASSRREVTEVF | 0.77409124 |
| ISNASDALDK | ASSRREVTEVF | 0.9060123 |
| NQAETLVYQTE | ASSRREVTEVF | 0.7846047 |
| TGGTGGTAGAG | ASSRREVTEVF | 0.43376753 |
| TGSTAGGAGGA | ASSRREVTEVF | 0.83627903 |
| DPLAVPVAAGPV | ASSRREVTEVF | 0.89234436 |
| GGTGGTPGNGGN | ASSRREVTEVF | 0.8956076 |
| PAGLAGVDPATL | ASSRREVTEVF | 0.9018856 |
| GNNLVVEAPHAGASV | ASSRREVTEVF | 0.8546858 |
| KVTRIMSASTLEQPA | ASSRREVTEVF | 0.7928154 |
| LGFALPIALPRRTVV | ASSRREVTEVF | 0.9014747 |
| TRLGNLTARKLVVLV | ASSRREVTEVF | 0.91435003 |
| WSQQRRLIYFPSAGP | ASSRREVTEVF | 0.88369244 |
| GPNGNPDPMAAAVDIR | ASSRREVTEVF | 0.77572656 |
| TQRCVSVADLRAAASR | ASSRREVTEVF | 0.6914729 |
| QFVLDVAEHPVEVLTETV | ASSRREVTEVF | 0.8716907 |
| GGAGGTGGAGGNGGTQAGDG | ASSRREVTEVF | 0.89552236 |
| PLLATLRAIAVDVDAKPAQV | ASSRREVTEVF | 0.7190705 |
| APKFRTGGYMIDFWGVGYQVAK | ASSRREVTEVF | 0.8489796 |
| GAGGAGGAGGAAGGPGGTGGTG | ASSRREVTEVF | 0.8544395 |
| IGALGLLATDVAPAGSVAFGVA | ASSRREVTEVF | 0.44383585 |
| AGGAGGIGGNGGAGGAGGVLMG | ASSRREVTEVF | 0.84286255 |
| ADELIGLL | ASSRREVTEVF | 0.8932884 |
| ADRALKA | ASSRREVTEVF | 0.86948997 |
| AGAVGQVGGAGGA | ASSRREVTEVF | 0.70067525 |
| AGGGGGLGGAP | ASSRREVTEVF | 0.89341664 |
| AGGLLYGNGGAG | ASSRREVTEVF | 0.91456974 |
| AGLESIGL | ASSRREVTEVF | 0.906174 |
| AGLKTGII | ASSRREVTEVF | 0.433881 |
| ASLAALLL | ASSRREVTEVF | 0.3013193 |
| ASVINISSI | ASSRREVTEVF | 0.81319124 |
| ATPPGPA | ASSRREVTEVF | 0.85536796 |
| ATVASALV | ASSRREVTEVF | 0.78086805 |
| ATVASLIL | ASSRREVTEVF | 0.7640977 |
| AVNGPVLIRH | ASSRREVTEVF | 0.84283847 |
| DGAAGAVGI | ASSRREVTEVF | 0.89624274 |
| DPLLHGI | ASSRREVTEVF | 0.7138787 |
| ELGAALGAL | ASSRREVTEVF | 0.67027056 |

| | | |
|---|---|---|
| ELGRTQAV | ASSRREVTEVF | 0.8468503 |
| FVAGGYLS | ASSRREVTEVF | 0.8842401 |
| GASGGAGGAGGW | ASSRREVTEVF | 0.7562119 |
| GGAGAGGDGGAGGI | ASSRREVTEVF | 0.83189106 |
| GTGHNGSQPGMGGNG | ASSRREVTEVF | 0.82573235 |
| GYAAASASAS | ASSRREVTEVF | 0.8032862 |
| HQVDVAAVQQG | ASSRREVTEVF | 0.8559612 |
| IGAAIAHT | ASSRREVTEVF | 0.6468106 |
| IGMIEVAQL | ASSRREVTEVF | 0.90070844 |
| IRLAPEHTSAMD | ASSRREVTEVF | 0.70477307 |
| IYNVSVL | ASSRREVTEVF | 0.8855227 |
| KQGFLGM | ASSRREVTEVF | 0.8598571 |
| LESVITQL | ASSRREVTEVF | 0.8741518 |
| LGLGALVVV | ASSRREVTEVF | 0.80578107 |
| LIGNGGNGGSGGI | ASSRREVTEVF | 0.712407 |
| LNAINAPTL | ASSRREVTEVF | 0.66530484 |
| LQLLNTGV | ASSRREVTEVF | 0.54709697 |
| LQTSIEF | ASSRREVTEVF | 0.8900941 |
| MSFAVSEL | ASSRREVTEVF | 0.2365334 |
| NTGALGL | ASSRREVTEVF | 0.8947555 |
| PPGAGGAGGAGTT | ASSRREVTEVF | 0.8873986 |
| QLVGLGI | ASSRREVTEVF | 0.7548247 |
| RFLEQVHQL | ASSRREVTEVF | 0.8842825 |
| RGVGYVLR | ASSRREVTEVF | 0.6334254 |
| SATGVFAGVI | ASSRREVTEVF | 0.7928355 |
| SLPAITLP | ASSRREVTEVF | 0.87106776 |
| STALAFTV | ASSRREVTEVF | 0.8828688 |
| SVLTVGLGK | ASSRREVTEVF | 0.735574 |
| TALAGAGSGI | ASSRREVTEVF | 0.8799156 |
| TGLPFTGI | ASSRREVTEVF | 0.8606016 |
| VAEHVTV | ASSRREVTEVF | 0.62489545 |
| VDIVLNSL | ASSRREVTEVF | 0.85667515 |
| VLTGGGAVL | ASSRREVTEVF | 0.8860549 |
| VNVGLAYGI | ASSRREVTEVF | 0.8817712 |
| VSSAAVGAV | ASSRREVTEVF | 0.6806127 |
| VTSVYEAGV | ASSRREVTEVF | 0.8754131 |
| WVALDRGA | ASSRREVTEVF | 0.7310746 |
| YLVGSVVI | ASSRREVTEVF | 0.85935223 |
| GPPGTGKTLL | ASSRREVTEVF | 0.857249 |
| GVGMTKF | ASSRREVTEVF | 0.790216 |
| IGIGGSDL | ASSRREVTEVF | 0.8935294 |
| IGTLLAEL | ASSRREVTEVF | 0.37053454 |
| IYTSGTT | ASSRREVTEVF | 0.7558156 |
| LSFPAGV | ASSRREVTEVF | 0.81637764 |
| LYTSGTT | ASSRREVTEVF | 0.85214734 |
| VGAFPFGF | ASSRREVTEVF | 0.8840351 |
| VGAGTVEF | ASSRREVTEVF | 0.7144464 |
| VGKSTLLL | ASSRREVTEVF | 0.83740926 |
| AALAHAGGGIV | ASSRREVTEVF | 0.8258 |
| AALVAVSLL | ASSRREVTEVF | 0.7425704 |
| AAVSRAVAAL | ASSRREVTEVF | 0.8624061 |

| | | |
|---|---|---|
| AGGSSGIGGF | ASSRREVTEVF | 0.7678919 |
| AGLTVANANG | ASSRREVTEVF | 0.62716895 |
| AGLVGAVL | ASSRREVTEVF | 0.8802617 |
| ALSQAFTV | ASSRREVTEVF | 0.7402158 |
| ALTFSVSL | ASSRREVTEVF | 0.86682427 |
| ALTGAALVAL | ASSRREVTEVF | 0.88897306 |
| ASFSIGAGVL | ASSRREVTEVF | 0.7401967 |
| DGSVYKI | ASSRREVTEVF | 0.89059705 |
| EQLAEIV | ASSRREVTEVF | 0.86460274 |
| FAVSELTVM | ASSRREVTEVF | 0.8645497 |
| GAAPLAAGL | ASSRREVTEVF | 0.8106267 |
| GADGGDGGSGGAGG | ASSRREVTEVF | 0.7117643 |
| GFGLDTGGAGG | ASSRREVTEVF | 0.8973619 |
| GLASGGLL | ASSRREVTEVF | 0.88197374 |
| GRIAGAAVLL | ASSRREVTEVF | 0.85331786 |
| GSAGSLNTSF | ASSRREVTEVF | 0.8669945 |
| GTGQIGI | ASSRREVTEVF | 0.8417815 |
| GTLIALIL | ASSRREVTEVF | 0.88674355 |
| IAATMGFVVG | ASSRREVTEVF | 0.77662265 |
| IFAQALL | ASSRREVTEVF | 0.6693486 |
| IGAAVALGF | ASSRREVTEVF | 0.81298804 |
| IGLTPLAL | ASSRREVTEVF | 0.79634756 |
| ILGSNTSTL | ASSRREVTEVF | 0.75173116 |
| IVFSLTVL | ASSRREVTEVF | 0.41771838 |
| LASKATGF | ASSRREVTEVF | 0.8810797 |
| LASVALIL | ASSRREVTEVF | 0.9158325 |
| LFAVGLTL | ASSRREVTEVF | 0.66545534 |
| LFNGGIGTYI | ASSRREVTEVF | 0.90890694 |
| LTLGAGII | ASSRREVTEVF | 0.6359611 |
| LVAAALGDI | ASSRREVTEVF | 0.8843818 |
| NTAFVQLGI | ASSRREVTEVF | 0.88693386 |
| QLFSGAEL | ASSRREVTEVF | 0.71506387 |
| SAATFITL | ASSRREVTEVF | 0.8497302 |
| SNYGLANLGV | ASSRREVTEVF | 0.69281876 |
| SYGGAYAVM | ASSRREVTEVF | 0.7884308 |
| TTGVGGLTLG | ASSRREVTEVF | 0.8985282 |
| TTQTIYI | ASSRREVTEVF | 0.7361224 |
| VAAGAGLGAVI | ASSRREVTEVF | 0.86409664 |
| VAAVVTLL | ASSRREVTEVF | 0.820277 |
| VGFLGGLI | ASSRREVTEVF | 0.7825167 |
| VGVIPILL | ASSRREVTEVF | 0.8787444 |
| VIFASSLI | ASSRREVTEVF | 0.8888336 |
| VKGGDGGAAAG | ASSRREVTEVF | 0.7767528 |
| VTKTFTNI | ASSRREVTEVF | 0.91294646 |
| VVAAAILL | ASSRREVTEVF | 0.7794509 |
| YAALDVEL | ASSRREVTEVF | 0.9033419 |
| LASVALLL | ASSRREVTEVF | 0.8845353 |
| LDAVHRV | ASSRREVTEVF | 0.84039664 |
| QGLSSSIF | ASSRREVTEVF | 0.83447635 |
| RGTFANIRL | ASSRREVTEVF | 0.88475156 |
| RQLWWGHRI | ASSRREVTEVF | 0.7845613 |

| | | |
|---|---|---|
| YGPPGTGKTLL | ASSRREVTEVF | 0.87044954 |
| AALVGAPANVA | ASSRREVTEVF | 0.9019404 |
| AATFRRTE | ASSRREVTEVF | 0.40934175 |
| AGALGGGI | ASSRREVTEVF | 0.8665687 |
| AGGAGGAGGIGAG | ASSRREVTEVF | 0.8216681 |
| AGSGALGL | ASSRREVTEVF | 0.81316984 |
| AQIADQI | ASSRREVTEVF | 0.6955932 |
| ASAVAVTYL | ASSRREVTEVF | 0.30578923 |
| ASTVLGGTGG | ASSRREVTEVF | 0.66319686 |
| AVGGEPAK | ASSRREVTEVF | 0.62524843 |
| CEQGTFGGTC | ASSRREVTEVF | 0.84038883 |
| DEGKAPAG | ASSRREVTEVF | 0.8508525 |
| DTVVFPAL | ASSRREVTEVF | 0.624226 |
| FEGITMAG | ASSRREVTEVF | 0.81461424 |
| FPLVVQD | ASSRREVTEVF | 0.7727934 |
| GAFTTPQ | ASSRREVTEVF | 0.45390525 |
| GFGGFGTAGGVGGA | ASSRREVTEVF | 0.7776945 |
| GFLAQFL | ASSRREVTEVF | 0.6822476 |
| GGAGGAGGNAGVA | ASSRREVTEVF | 0.7496716 |
| GGVGTAGAGGAGGAGG | ASSRREVTEVF | 0.34584796 |
| GNGGGGDAGNAGSGGN | ASSRREVTEVF | 0.8247907 |
| GTGGAGGAGNDAGS | ASSRREVTEVF | 0.67343116 |
| GTTNTGWFNS | ASSRREVTEVF | 0.15873441 |
| IFMGPLFTL | ASSRREVTEVF | 0.8197255 |
| ILAHAIEAI | ASSRREVTEVF | 0.609702 |
| IRSAIREAL | ASSRREVTEVF | 0.65600693 |
| ITFAGGG | ASSRREVTEVF | 0.83242154 |
| ITGMSGHQEV | ASSRREVTEVF | 0.6270848 |
| IVAGLACTL | ASSRREVTEVF | 0.56455016 |
| IVGGGLAGA | ASSRREVTEVF | 0.81209385 |
| IVTGVLNI | ASSRREVTEVF | 0.8440988 |
| LAQQGCI | ASSRREVTEVF | 0.45930082 |
| LCLHVEN | ASSRREVTEVF | 0.7813035 |
| LDAISDGS | ASSRREVTEVF | 0.8718317 |
| LGGFAAML | ASSRREVTEVF | 0.79445904 |
| LGGKNPMI | ASSRREVTEVF | 0.87752044 |
| LGPKGVAAIL | ASSRREVTEVF | 0.88359255 |
| LMVGVSI | ASSRREVTEVF | 0.6541141 |
| LVGAAFVL | ASSRREVTEVF | 0.8542074 |
| LVLAVAPL | ASSRREVTEVF | 0.86376053 |
| MAAIAVIDAA | ASSRREVTEVF | 0.6869782 |
| NNGINAELQ | ASSRREVTEVF | 0.7422701 |
| RGEPLVLVHG | ASSRREVTEVF | 0.4450752 |
| RGGAGGAAGLI | ASSRREVTEVF | 0.71404946 |
| RSYRMHPAV | ASSRREVTEVF | 0.493379 |
| SAAHLLAGQLA | ASSRREVTEVF | 0.7954271 |
| SAAVGSY | ASSRREVTEVF | 0.8732474 |
| SPHAVLHRI | ASSRREVTEVF | 0.7460332 |
| SQIAKVL | ASSRREVTEVF | 0.4661645 |
| TALGHLVP | ASSRREVTEVF | 0.4746558 |
| TAPTTSGAA | ASSRREVTEVF | 0.85116017 |

| | | |
|---|---|---|
| TDGDDGNGGSPGT | ASSRREVTEVF | 0.44078103 |
| TDGLGEN | ASSRREVTEVF | 0.46203408 |
| TGGPPLPGQA | ASSRREVTEVF | 0.6093774 |
| TIPAITI | ASSRREVTEVF | 0.8316953 |
| TIPGFTI | ASSRREVTEVF | 0.84022474 |
| TSIASGALA | ASSRREVTEVF | 0.61855245 |
| TVTLPGAAI | ASSRREVTEVF | 0.87286526 |
| VASGLSAALP | ASSRREVTEVF | 0.8450308 |
| VGAGVIGI | ASSRREVTEVF | 0.88343537 |
| VGPKRFGV | ASSRREVTEVF | 0.7573601 |
| VNVAFPD | ASSRREVTEVF | 0.47763914 |
| VSMTNSGVS | ASSRREVTEVF | 0.8320622 |
| VTDVLGDLSV | ASSRREVTEVF | 0.69890225 |
| VVVVGAGFAGL | ASSRREVTEVF | 0.8138116 |
| VYTVGTVV | ASSRREVTEVF | 0.8316394 |
| WTTFSGP | ASSRREVTEVF | 0.6910188 |
| YDELVEAF | ASSRREVTEVF | 0.58704364 |
| YGTGGAGGNGG | ASSRREVTEVF | 0.7424141 |
| YTNVMAV | ASSRREVTEVF | 0.9000304 |
| FLEQVHQL | ASSRREVTEVF | 0.58506453 |
| GLSLLAVAV | ASSRREVTEVF | 0.46862108 |
| LEQVHQL | ASSRREVTEVF | 0.668581 |
| LNAAGAI | ASSRREVTEVF | 0.68677026 |
| PAVHASL | ASSRREVTEVF | 0.8588631 |
| TELEDDA | ASSRREVTEVF | 0.8159272 |
| TPPGPPN | ASSRREVTEVF | 0.7620299 |
| TPPPPPPP | ASSRREVTEVF | 0.7867216 |
| TQSPALI | ASGDKGANTEFF | 0.84053326 |
| EFAAAFPQ | ASGDKGANTEFF | 0.76220965 |
| LLSGNLDVL | ASGDKGANTEFF | 0.79311484 |
| GPANLGNYNI | ASGDKGANTEFF | 0.65631115 |
| TLGPTLP | ASGDKGANTEFF | 0.8911359 |
| VYAYSSCCK | ASGDKGANTEFF | 0.8482753 |
| APAPAPAPNI | ASGDKGANTEFF | 0.8457917 |
| PGGPVPIP | ASGDKGANTEFF | 0.86646724 |
| FLTVALEP | ASGDKGANTEFF | 0.7540745 |
| SQYFGANL | ASGDKGANTEFF | 0.8084544 |
| GGVKPAA | ASGDKGANTEFF | 0.54184806 |
| GGTGGAGGAGGW | ASGDKGANTEFF | 0.5299307 |
| GVGGVAGLGGAGG | ASGDKGANTEFF | 0.73286605 |
| VLLVLATTI | ASGDKGANTEFF | 0.81776834 |
| IGIFAAML | ASGDKGANTEFF | 0.827706 |
| AVGDLID | ASGDKGANTEFF | 0.80456877 |
| NISPLPL | ASGDKGANTEFF | 0.65411204 |
| VTAVFVAI | ASGDKGANTEFF | 0.85923654 |
| SGDDNDAAL | ASGDKGANTEFF | 0.86050135 |
| AVAALAHS | ASGDKGANTEFF | 0.5813137 |
| GGVTVMATP | ASGDKGANTEFF | 0.72993004 |
| AGLAANVRD | ASGDKGANTEFF | 0.87119234 |
| TNRPDLIDPALLRPG | ASGDKGANTEFF | 0.7998768 |
| LANLGVTAL | ASGDKGANTEFF | 0.7159257 |

| | | |
|---|---|---|
| TGERIEL | ASGDKGANTEFF | 0.73184395 |
| NGEELAA | ASGDKGANTEFF | 0.8582976 |
| VLSLVWG | ASGDKGANTEFF | 0.7147559 |
| IGYALWL | ASGDKGANTEFF | 0.8813996 |
| LSLLLAPL | ASGDKGANTEFF | 0.31181818 |
| TLDEALK | ASGDKGANTEFF | 0.79031706 |
| TDGAGTAEGG | ASGDKGANTEFF | 0.7679986 |
| NAPGGAGGNGG | ASGDKGANTEFF | 0.1910887 |
| VSVFMGMGGNF | ASGDKGANTEFF | 0.85482377 |
| LASYLAAL | ASGDKGANTEFF | 0.5260368 |
| LVGGTPAL | ASGDKGANTEFF | 0.7817182 |
| VRNADVL | ASGDKGANTEFF | 0.62345725 |
| TCSSPFLVSAM | ASGDKGANTEFF | 0.8621503 |
| VHITDAE | ASGDKGANTEFF | 0.80602556 |
| EYAAPSF | ASGDKGANTEFF | 0.61839986 |
| VVDAFHGGL | ASGDKGANTEFF | 0.5634283 |
| ASVVDPQ | ASGDKGANTEFF | 0.812675 |
| SYNEFGGLS | ASGDKGANTEFF | 0.8657108 |
| DPDGSVY | ASGDKGANTEFF | 0.8971851 |
| MVSTIHATE | ASGDKGANTEFF | 0.31000698 |
| LAAGVADPV | ASGDKGANTEFF | 0.83290046 |
| TKEQIAA | ASGDKGANTEFF | 0.7400762 |
| SLASPAAL | ASGDKGANTEFF | 0.33010238 |
| IGLVFYGV | ASGDKGANTEFF | 0.24173847 |
| FLLDASNL | ASGDKGANTEFF | 0.74838495 |
| YQVGNFP | ASGDKGANTEFF | 0.76962733 |
| ESLLSQST | ASGDKGANTEFF | 0.7966307 |
| LFTVGFAAVG | ASGDKGANTEFF | 0.8388909 |
| ATPPTPT | ASGDKGANTEFF | 0.69322133 |
| TDLATLG | ASGDKGANTEFF | 0.8155536 |
| SAGGTGR | ASGDKGANTEFF | 0.73053193 |
| GFSNSGS | ASGDKGANTEFF | 0.07878874 |
| GGNGGSGGTSV | ASGDKGANTEFF | 0.61076105 |
| GTSVATGGAGN | ASGDKGANTEFF | 0.408339 |
| NGGNGGSGGTS | ASGDKGANTEFF | 0.86751604 |
| AVVKFPH | ASGDKGANTEFF | 0.56471246 |
| TPPPPPP | ASGDKGANTEFF | 0.7233081 |
| AAPAAAAAPA | ASGDKGANTEFF | 0.7513313 |
| IALNEDL | ASGDKGANTEFF | 0.8011265 |
| IVNVSSM | ASGDKGANTEFF | 0.7513627 |
| TGIPPAPR | ASGDKGANTEFF | 0.7095618 |
| FDLGGGTF | ASGDKGANTEFF | 0.3878333 |
| FDLGGGTFDVS | ASGDKGANTEFF | 0.82140696 |
| NEPTAAA | ASGDKGANTEFF | 0.6820656 |
| NEPTAAALA | ASGDKGANTEFF | 0.48764646 |
| GSIINISSI | ASGDKGANTEFF | 0.58966345 |
| VAPGNAGTA | ASGDKGANTEFF | 0.6279649 |
| TGGGASLE | ASGDKGANTEFF | 0.2705702 |
| VAVYNHYKRI | ASGDKGANTEFF | 0.8657306 |
| TDPAPATT | ASGDKGANTEFF | 0.47038376 |
| TNRPDILDPALLRPG | ASGDKGANTEFF | 0.70579004 |

| | | |
|---|---|---|
| LTTTTTS | ASGDKGANTEFF | 0.87686723 |
| PTGSGKTL | ASGDKGANTEFF | 0.73628116 |
| TGSGKTL | ASGDKGANTEFF | 0.7484994 |
| TIGHVDHGK | ASGDKGANTEFF | 0.6323235 |
| TIGHVDHGKTT | ASGDKGANTEFF | 0.62327313 |
| GHVDHGKTT | ASGDKGANTEFF | 0.6643472 |
| AQFEHTLL | ASGDKGANTEFF | 0.5300405 |
| IVNQDEL | ASGDKGANTEFF | 0.84532225 |
| TNGSQFF | ASGDKGANTEFF | 0.8346405 |
| TGKGIVE | ASGDKGANTEFF | 0.45227826 |
| LISNASDA | ASGDKGANTEFF | 0.87316835 |
| AELRTLGELTKIAKAHGAQ | ASGDKGANTEFF | 0.8742874 |
| APNAVAAAEPSW | ASGDKGANTEFF | 0.8882998 |
| APQVAGAATAVL | ASGDKGANTEFF | 0.5820223 |
| ARRDPAAAAAILPTDA | ASGDKGANTEFF | 0.6591669 |
| DAAARDVLAEAGLAEYF | ASGDKGANTEFF | 0.69002223 |
| DDQIVLSAKANLAKVEQ | ASGDKGANTEFF | 0.7137703 |
| DELLPDTGEDSDEDGH | ASGDKGANTEFF | 0.5062814 |
| ELPEAPANRATVPQD | ASGDKGANTEFF | 0.51248443 |
| FGQSKIERVTGIG | ASGDKGANTEFF | 0.88376105 |
| GAGGTGSTAGGAGGAGGAGGLY | ASGDKGANTEFF | 0.8772329 |
| GHGGTGAAGGNGGTGGAGGSGI | ASGDKGANTEFF | 0.7260585 |
| GNGGNGGAGGEGGPGGAGGAGA | ASGDKGANTEFF | 0.34967986 |
| GSTAGNGGNGGNATNGTIAGQP | ASGDKGANTEFF | 0.7133167 |
| GTGGDGGNGGNGGTGG | ASGDKGANTEFF | 0.61136156 |
| GVRRPAGIPADAPV | ASGDKGANTEFF | 0.63847756 |
| GYRQVTDAHLLTLARR | ASGDKGANTEFF | 0.5837331 |
| IAKMGQDGHDRG | ASGDKGANTEFF | 0.43519443 |
| KAEPSIEIQYT | ASGDKGANTEFF | 0.714423 |
| KPVRIRMAADEDTVAVT | ASGDKGANTEFF | 0.8856809 |
| MPPRIAGMRLLVIKPEP | ASGDKGANTEFF | 0.6979591 |
| MKFYAVALFRSRGVPPTR | ASGDKGANTEFF | 0.771154 |
| NAGGFGILGAVAHSP | ASGDKGANTEFF | 0.87085325 |
| NEGREITLAMAEM | ASGDKGANTEFF | 0.8334384 |
| NELVEWSHPLGHRWRWE | ASGDKGANTEFF | 0.85677433 |
| NLGAGGTGGNGGNPG | ASGDKGANTEFF | 0.8336922 |
| NTEWPKWETVMAADPVD | ASGDKGANTEFF | 0.7600988 |
| PASAAPAETQP | ASGDKGANTEFF | 0.7868934 |
| QQSSDGKNGVRAAEITGE | ASGDKGANTEFF | 0.8401121 |
| RAPDAVTTSWWKE | ASGDKGANTEFF | 0.6728029 |
| SGFGNVGAHTSGWFNQ | ASGDKGANTEFF | 0.7882893 |
| SGVAGAVGQAASVGGLKV | ASGDKGANTEFF | 0.78324664 |
| SNGGAGTGSAGGNGGT | ASGDKGANTEFF | 0.7998471 |
| TGGAGGEGGAAG | ASGDKGANTEFF | 0.7686707 |
| TGGDGGAAGAGGAPGHG | ASGDKGANTEFF | 0.8816359 |
| TMPPLLAAIANAGRHGV | ASGDKGANTEFF | 0.18323235 |
| TSSTAGNSGGA | ASGDKGANTEFF | 0.52124596 |
| VPATVPVGILC | ASGDKGANTEFF | 0.8629599 |
| VQRPDSGGVVLTGRIS | ASGDKGANTEFF | 0.160976 |
| WTPNPVPATKGVPAGLVQAT | ASGDKGANTEFF | 0.36331904 |
| YALKRALAKFSDRFDVVII | ASGDKGANTEFF | 0.8232473 |

| | | |
|---|---|---|
| YLIGSVIV | ASGDKGANTEFF | 0.8642901 |
| YGQLVAVID | ASGDKGANTEFF | 0.1594038 |
| VTVGGMTLPA | ASGDKGANTEFF | 0.6571654 |
| VLALPAV | ASGDKGANTEFF | 0.39250922 |
| VIPPTLNYET | ASGDKGANTEFF | 0.86793387 |
| VGMGGMHPSSG | ASGDKGANTEFF | 0.6939507 |
| VGLAGGAATA | ASGDKGANTEFF | 0.3470898 |
| VGGAGGAGGSGP | ASGDKGANTEFF | 0.8456993 |
| VFAAFMV | ASGDKGANTEFF | 0.60127306 |
| TLYDKLR | ASGDKGANTEFF | 0.80520034 |
| TLAVFTVL | ASGDKGANTEFF | 0.86293757 |
| TGSVQPARV | ASGDKGANTEFF | 0.87979364 |
| TGGGAGSGGT | ASGDKGANTEFF | 0.49133348 |
| SHLETSL | ASGDKGANTEFF | 0.80665004 |
| SFTTGMT | ASGDKGANTEFF | 0.70616907 |
| QVEVAKRMGVRQPTVS | ASGDKGANTEFF | 0.9010774 |
| QGSGGPALAM | ASGDKGANTEFF | 0.6846553 |
| MLAPSNP | ASGDKGANTEFF | 0.39818144 |
| LVNNAAI | ASGDKGANTEFF | 0.84571576 |
| LTGEINTLA | ASGDKGANTEFF | 0.8448286 |
| LSIFGPV | ASGDKGANTEFF | 0.6825316 |
| LLALLANL | ASGDKGANTEFF | 0.8423743 |
| LISNASDALDKLR | ASGDKGANTEFF | 0.811066 |
| LINEAVP | ASGDKGANTEFF | 0.7007773 |
| LGYAFSGL | ASGDKGANTEFF | 0.8111017 |
| LDFVAVAL | ASGDKGANTEFF | 0.818507 |
| LAQPGAE | ASGDKGANTEFF | 0.6084472 |
| ISNLGAVTGF | ASGDKGANTEFF | 0.8795253 |
| HQEDPGRI | ASGDKGANTEFF | 0.75225055 |
| GTPGEPTP | ASGDKGANTEFF | 0.4794879 |
| GFFNSGS | ASGDKGANTEFF | 0.89503264 |
| GAGGGGLTAGAN | ASGDKGANTEFF | 0.7784276 |
| FVDALVNM | ASGDKGANTEFF | 0.8492327 |
| FLTVAPEL | ASGDKGANTEFF | 0.8445107 |
| FETLTAL | ASGDKGANTEFF | 0.8775878 |
| DVINPIEL | ASGDKGANTEFF | 0.50747806 |
| APVSGAPV | ASGDKGANTEFF | 0.7350801 |
| ALVGGLAW | ASGDKGANTEFF | 0.7427778 |
| ALAAAAATAI | ASGDKGANTEFF | 0.83168185 |
| AFGLGLLL | ASGDKGANTEFF | 0.7309227 |
| AFFAAASL | ASGDKGANTEFF | 0.7846304 |
| AAAALPAFSPP | ASGDKGANTEFF | 0.8432939 |
| VVLAPAL | ASGDKGANTEFF | 0.8735838 |
| TIGHVDHGKTTL | ASGDKGANTEFF | 0.82288426 |
| LISNASDALDK | ASGDKGANTEFF | 0.7658503 |
| LISNASDAL | ASGDKGANTEFF | 0.87653345 |
| ISNASDALDK | ASGDKGANTEFF | 0.5379312 |
| NQAETLVYQTE | ASGDKGANTEFF | 0.7676774 |
| TGGTGGTAGAG | ASGDKGANTEFF | 0.86495584 |
| TGSTAGGAGGA | ASGDKGANTEFF | 0.84525216 |
| DPLAVPVAAGPV | ASGDKGANTEFF | 0.7949449 |

| | | |
|---|---|---|
| GGTGGTPGNGGN | ASGDKGANTEFF | 0.860927 |
| PAGLAGVDPATL | ASGDKGANTEFF | 0.6250564 |
| GNNLVVEAPHAGASV | ASGDKGANTEFF | 0.7754913 |
| KVTRIMSASTLEQPA | ASGDKGANTEFF | 0.6957887 |
| LGFALPIALPRRTVV | ASGDKGANTEFF | 0.65361583 |
| TRLGNLTARKLVVLV | ASGDKGANTEFF | 0.5948767 |
| WSQQRRLIYFPSAGP | ASGDKGANTEFF | 0.76486975 |
| GPNGNPDPMAAAVDIR | ASGDKGANTEFF | 0.84099376 |
| TQRCVSVADLRAAASR | ASGDKGANTEFF | 0.7684562 |
| QFVLDVAEHPVEVLTETV | ASGDKGANTEFF | 0.2410102 |
| GGAGGTGGAGGNGGTQAGDG | ASGDKGANTEFF | 0.7955477 |
| PLLATLRAIAVDVDAKPAQV | ASGDKGANTEFF | 0.765309 |
| APKFRTGGYMIDFWGVGYQVAK | ASGDKGANTEFF | 0.87970877 |
| GAGGAGGAGGAAGGPGGTGGTG | ASGDKGANTEFF | 0.7707746 |
| IGALGLLATDVAPAGSVAFGVA | ASGDKGANTEFF | 0.82681066 |
| AGGAGGIGGNGGAGGAGGVLMG | ASGDKGANTEFF | 0.7278632 |
| ADELIGLL | ASGDKGANTEFF | 0.828837 |
| ADRALKA | ASGDKGANTEFF | 0.4389488 |
| AGAVGQVGGAGGA | ASGDKGANTEFF | 0.504066 |
| AGGGGGLGGAP | ASGDKGANTEFF | 0.7004743 |
| AGGLLYGNGGAG | ASGDKGANTEFF | 0.6242273 |
| AGLESIGL | ASGDKGANTEFF | 0.81999874 |
| AGLKTGII | ASGDKGANTEFF | 0.8527554 |
| ASLAALLL | ASGDKGANTEFF | 0.8270626 |
| ASVINISSI | ASGDKGANTEFF | 0.85633814 |
| ATPPGPA | ASGDKGANTEFF | 0.880772 |
| ATVASALV | ASGDKGANTEFF | 0.8751168 |
| ATVASLIL | ASGDKGANTEFF | 0.706833 |
| AVNGPVLIRH | ASGDKGANTEFF | 0.6163664 |
| DGAAGAVGI | ASGDKGANTEFF | 0.44055155 |
| DPLLHGI | ASGDKGANTEFF | 0.61330825 |
| ELGAALGAL | ASGDKGANTEFF | 0.71513397 |
| ELGRTQAV | ASGDKGANTEFF | 0.7249606 |
| FVAGGYLS | ASGDKGANTEFF | 0.14104375 |
| GASGGAGGAGGW | ASGDKGANTEFF | 0.78351736 |
| GGAGAGGDGGAGGI | ASGDKGANTEFF | 0.8526952 |
| GTGHNGSQPGMGGNG | ASGDKGANTEFF | 0.70841014 |
| GYAAASASAS | ASGDKGANTEFF | 0.7934104 |
| HQVDVAAVQQG | ASGDKGANTEFF | 0.8069122 |
| IGAAIAHT | ASGDKGANTEFF | 0.7331457 |
| IGMIEVAQL | ASGDKGANTEFF | 0.81029433 |
| IRLAPEHTSAMD | ASGDKGANTEFF | 0.71553683 |
| IYNVSVL | ASGDKGANTEFF | 0.86347604 |
| KQGFLGM | ASGDKGANTEFF | 0.8789225 |
| LESVITQL | ASGDKGANTEFF | 0.804351 |
| LGLGALVVV | ASGDKGANTEFF | 0.6938734 |
| LIGNGGNGGSGGI | ASGDKGANTEFF | 0.7742948 |
| LNAINAPTL | ASGDKGANTEFF | 0.5622139 |
| LQLLNTGV | ASGDKGANTEFF | 0.7295723 |
| LQTSIEF | ASGDKGANTEFF | 0.21370766 |
| MSFAVSEL | ASGDKGANTEFF | 0.76398855 |

| | | |
|---|---|---|
| NTGALGL | ASGDKGANTEFF | 0.6901907 |
| PPGAGGAGGAGTT | ASGDKGANTEFF | 0.79224414 |
| QLVGLGI | ASGDKGANTEFF | 0.7470989 |
| RFLEQVHQL | ASGDKGANTEFF | 0.8459437 |
| RGVGYVLR | ASGDKGANTEFF | 0.5888438 |
| SATGVFAGVI | ASGDKGANTEFF | 0.83104104 |
| SLPAITLP | ASGDKGANTEFF | 0.77802294 |
| STALAFTV | ASGDKGANTEFF | 0.35944825 |
| SVLTVGLGK | ASGDKGANTEFF | 0.8533175 |
| TALAGAGSGI | ASGDKGANTEFF | 0.8134749 |
| TGLPFTGI | ASGDKGANTEFF | 0.75569725 |
| VAEHVTV | ASGDKGANTEFF | 0.8376354 |
| VDIVLNSL | ASGDKGANTEFF | 0.76391786 |
| VLTGGGAVL | ASGDKGANTEFF | 0.60497445 |
| VNVGLAYGI | ASGDKGANTEFF | 0.6447377 |
| VSSAAVGAV | ASGDKGANTEFF | 0.7852582 |
| VTSVYEAGV | ASGDKGANTEFF | 0.75567377 |
| WVALDRGA | ASGDKGANTEFF | 0.18261328 |
| YLVGSVVI | ASGDKGANTEFF | 0.83449113 |
| GPPGTGKTLL | ASGDKGANTEFF | 0.48127216 |
| GVGMTKF | ASGDKGANTEFF | 0.66618925 |
| IGIGGSDL | ASGDKGANTEFF | 0.75136817 |
| IGTLLAEL | ASGDKGANTEFF | 0.8424537 |
| IYTSGTT | ASGDKGANTEFF | 0.77966964 |
| LSFPAGV | ASGDKGANTEFF | 0.53694916 |
| LYTSGTT | ASGDKGANTEFF | 0.8130923 |
| VGAFPFGF | ASGDKGANTEFF | 0.76337874 |
| VGAGTVEF | ASGDKGANTEFF | 0.8143528 |
| VGKSTLLL | ASGDKGANTEFF | 0.75876445 |
| AALAHAGGGIV | ASGDKGANTEFF | 0.8165741 |
| AALVAVSLL | ASGDKGANTEFF | 0.7631093 |
| AAVSRAVAAL | ASGDKGANTEFF | 0.6057187 |
| AGGSSGIGGF | ASGDKGANTEFF | 0.600472 |
| AGLTVANANG | ASGDKGANTEFF | 0.6530589 |
| AGLVGAVL | ASGDKGANTEFF | 0.6326074 |
| ALSQAFTV | ASGDKGANTEFF | 0.53943706 |
| ALTFSVSL | ASGDKGANTEFF | 0.79004955 |
| ALTGAALVAL | ASGDKGANTEFF | 0.8837526 |
| ASFSIGAGVL | ASGDKGANTEFF | 0.7798104 |
| DGSVYKI | ASGDKGANTEFF | 0.3569008 |
| EQLAEIV | ASGDKGANTEFF | 0.74405944 |
| FAVSELTVM | ASGDKGANTEFF | 0.7327348 |
| GAAPLAAGL | ASGDKGANTEFF | 0.81836903 |
| GADGGDGGSGGAGG | ASGDKGANTEFF | 0.86158955 |
| GFGLDTGGAGG | ASGDKGANTEFF | 0.8533524 |
| GLASGGLL | ASGDKGANTEFF | 0.81864893 |
| GRIAGAAVLL | ASGDKGANTEFF | 0.6513177 |
| GSAGSLNTSF | ASGDKGANTEFF | 0.8552705 |
| GTGQIGI | ASGDKGANTEFF | 0.73991704 |
| GTLIALIL | ASGDKGANTEFF | 0.7385305 |
| IAATMGFVVG | ASGDKGANTEFF | 0.80348784 |

| | | |
|---|---|---|
| IFAQALL | ASGDKGANTEFF | 0.8504408 |
| IGAAVALGF | ASGDKGANTEFF | 0.6375153 |
| IGLTPLAL | ASGDKGANTEFF | 0.7169766 |
| ILGSNTSTL | ASGDKGANTEFF | 0.48399138 |
| IVFSLTVL | ASGDKGANTEFF | 0.8544054 |
| LASKATGF | ASGDKGANTEFF | 0.44336683 |
| LASVALIL | ASGDKGANTEFF | 0.62672 |
| LFAVGLTL | ASGDKGANTEFF | 0.7049897 |
| LFNGGIGTYI | ASGDKGANTEFF | 0.8712497 |
| LTLGAGII | ASGDKGANTEFF | 0.85791576 |
| LVAAALGDI | ASGDKGANTEFF | 0.6799488 |
| NTAFVQLGI | ASGDKGANTEFF | 0.6803826 |
| QLFSGAEL | ASGDKGANTEFF | 0.80721474 |
| SAATFITL | ASGDKGANTEFF | 0.6833175 |
| SNYGLANLGV | ASGDKGANTEFF | 0.87079036 |
| SYGGAYAVM | ASGDKGANTEFF | 0.729387 |
| TTGVGGLTLG | ASGDKGANTEFF | 0.72874 |
| TTQTIYI | ASGDKGANTEFF | 0.62924844 |
| VAAGAGLGAVI | ASGDKGANTEFF | 0.8472885 |
| VAAVVTLL | ASGDKGANTEFF | 0.70509434 |
| VGFLGGLI | ASGDKGANTEFF | 0.84522665 |
| VGVIPILL | ASGDKGANTEFF | 0.33946228 |
| VIFASSLI | ASGDKGANTEFF | 0.8563227 |
| VKGGDGGAAAG | ASGDKGANTEFF | 0.5681723 |
| VTKTFTNI | ASGDKGANTEFF | 0.65982056 |
| VVAAAILL | ASGDKGANTEFF | 0.26789904 |
| YAALDVEL | ASGDKGANTEFF | 0.71052366 |
| LASVALLL | ASGDKGANTEFF | 0.81939733 |
| LDAVHRV | ASGDKGANTEFF | 0.87722737 |
| QGLSSSIF | ASGDKGANTEFF | 0.80636144 |
| RGTFANIRL | ASGDKGANTEFF | 0.7670145 |
| RQLWWGHRI | ASGDKGANTEFF | 0.86070454 |
| YGPPGTGKTLL | ASGDKGANTEFF | 0.68818164 |
| AALVGAPANVA | ASGDKGANTEFF | 0.25593787 |
| AATFRRTE | ASGDKGANTEFF | 0.7410022 |
| AGALGGGI | ASGDKGANTEFF | 0.7432203 |
| AGGAGGAGGIGAG | ASGDKGANTEFF | 0.8745091 |
| AGSGALGL | ASGDKGANTEFF | 0.8085134 |
| AQIADQI | ASGDKGANTEFF | 0.45816815 |
| ASAVAVTYL | ASGDKGANTEFF | 0.7612718 |
| ASTVLGGTGG | ASGDKGANTEFF | 0.7781951 |
| AVGGEPAK | ASGDKGANTEFF | 0.42881542 |
| CEQGTFGGTC | ASGDKGANTEFF | 0.65895295 |
| DEGKAPAG | ASGDKGANTEFF | 0.52486163 |
| DTVVFPAL | ASGDKGANTEFF | 0.4504222 |
| FEGITMAG | ASGDKGANTEFF | 0.7826296 |
| FPLVVQD | ASGDKGANTEFF | 0.7096151 |
| GAFTTPQ | ASGDKGANTEFF | 0.6143709 |
| GFGGFGTAGGVGGA | ASGDKGANTEFF | 0.72076637 |
| GFLAQFL | ASGDKGANTEFF | 0.5545275 |
| GGAGGAGGNAGVA | ASGDKGANTEFF | 0.6400964 |

| | | |
|---|---|---|
| GGVGTAGAGGAGGAGG | ASGDKGANTEFF | 0.6486497 |
| GNGGGGDAGNAGSGGN | ASGDKGANTEFF | 0.76259196 |
| GTGGAGGAGNDAGS | ASGDKGANTEFF | 0.8642571 |
| GTTNTGWFNS | ASGDKGANTEFF | 0.709939 |
| IFMGPLFTL | ASGDKGANTEFF | 0.7318529 |
| ILAHAIEAI | ASGDKGANTEFF | 0.85498583 |
| IRSAIREAL | ASGDKGANTEFF | 0.366717 |
| ITFAGGG | ASGDKGANTEFF | 0.5568189 |
| ITGMSGHQEV | ASGDKGANTEFF | 0.83007 |
| IVAGLACTL | ASGDKGANTEFF | 0.72477186 |
| IVGGGLAGA | ASGDKGANTEFF | 0.8567382 |
| IVTGVLNI | ASGDKGANTEFF | 0.77463746 |
| LAQQGCI | ASGDKGANTEFF | 0.78226 |
| LCLHVEN | ASGDKGANTEFF | 0.73286134 |
| LDAISDGS | ASGDKGANTEFF | 0.87719476 |
| LGGFAAML | ASGDKGANTEFF | 0.8026954 |
| LGGKNPMI | ASGDKGANTEFF | 0.6963167 |
| LGPKGVAAIL | ASGDKGANTEFF | 0.82184434 |
| LMVGVSI | ASGDKGANTEFF | 0.4042686 |
| LVGAAFVL | ASGDKGANTEFF | 0.8120245 |
| LVLAVAPL | ASGDKGANTEFF | 0.8412012 |
| MAAIAVIDAA | ASGDKGANTEFF | 0.6023241 |
| NNGINAELQ | ASGDKGANTEFF | 0.6981504 |
| RGEPLVLVHG | ASGDKGANTEFF | 0.89484304 |
| RGGAGGAAGLI | ASGDKGANTEFF | 0.8476438 |
| RSYRMHPAV | ASGDKGANTEFF | 0.7043618 |
| SAAHLLAGQLA | ASGDKGANTEFF | 0.62566817 |
| SAAVGSY | ASGDKGANTEFF | 0.6096901 |
| SPHAVLHRI | ASGDKGANTEFF | 0.75280476 |
| SQIAKVL | ASGDKGANTEFF | 0.3203051 |
| TALGHLVP | ASGDKGANTEFF | 0.828426 |
| TAPTTSGAA | ASGDKGANTEFF | 0.70391846 |
| TDGDDGNGGSPGT | ASGDKGANTEFF | 0.8325006 |
| TDGLGEN | ASGDKGANTEFF | 0.8087831 |
| TGGPPLPGQA | ASGDKGANTEFF | 0.8663753 |
| TIPAITI | ASGDKGANTEFF | 0.71446264 |
| TIPGFTI | ASGDKGANTEFF | 0.7095963 |
| TSIASGALA | ASGDKGANTEFF | 0.40453744 |
| TVTLPGAAI | ASGDKGANTEFF | 0.42842358 |
| VASGLSAALP | ASGDKGANTEFF | 0.7000476 |
| VGAGVIGI | ASGDKGANTEFF | 0.65210545 |
| VGPKRFGV | ASGDKGANTEFF | 0.30146325 |
| VNVAFPD | ASGDKGANTEFF | 0.8841889 |
| VSMTNSGVS | ASGDKGANTEFF | 0.63500285 |
| VTDVLGDLSV | ASGDKGANTEFF | 0.8410834 |
| VVVVGAGFAGL | ASGDKGANTEFF | 0.6657337 |
| VYTVGTVV | ASGDKGANTEFF | 0.45007586 |
| WTTFSGP | ASGDKGANTEFF | 0.70955724 |
| YDELVEAF | ASGDKGANTEFF | 0.8055117 |
| YGTGGAGGNGG | ASGDKGANTEFF | 0.4928759 |
| YTNVMAV | ASGDKGANTEFF | 0.36843032 |

| | | |
|---|---|---|
| FLEQVHQL | ASGDKGANTEFF | 0.57005024 |
| GLSLLAVAV | ASGDKGANTEFF | 0.7686053 |
| LEQVHQL | ASGDKGANTEFF | 0.8957361 |
| LNAAGAI | ASGDKGANTEFF | 0.8648 |
| PAVHASL | ASGDKGANTEFF | 0.7746983 |
| TELEDDA | ASGDKGANTEFF | 0.9013709 |
| TPPGPPN | ASGDKGANTEFF | 0.13488011 |
| TPPPPPPP | ASGDKGANTEFF | 0.6706234 |
| TQSPALI | AWSRTGSNERLF | 0.85159487 |
| EFAAAFPQ | AWSRTGSNERLF | 0.83477485 |
| LLSGNLDVL | AWSRTGSNERLF | 0.7271249 |
| GPANLGNYNI | AWSRTGSNERLF | 0.86371577 |
| TLGPTLP | AWSRTGSNERLF | 0.57547617 |
| VYAYSSCCK | AWSRTGSNERLF | 0.7338416 |
| APAPAPAPNI | AWSRTGSNERLF | 0.73292685 |
| PGGPVPIP | AWSRTGSNERLF | 0.8584744 |
| FLTVALEP | AWSRTGSNERLF | 0.8374041 |
| SQYFGANL | AWSRTGSNERLF | 0.36443117 |
| GGVKPAA | AWSRTGSNERLF | 0.8111167 |
| GGTGGAGGAGGW | AWSRTGSNERLF | 0.8915 |
| GVGGVAGLGGAGG | AWSRTGSNERLF | 0.6630645 |
| VLLVLATTI | AWSRTGSNERLF | 0.8558557 |
| IGIFAAML | AWSRTGSNERLF | 0.7388162 |
| AVGDLID | AWSRTGSNERLF | 0.71645606 |
| NISPLPL | AWSRTGSNERLF | 0.63253057 |
| VTAVFVAI | AWSRTGSNERLF | 0.5478854 |
| SGDDNDAAL | AWSRTGSNERLF | 0.8202059 |
| AVAALAHS | AWSRTGSNERLF | 0.49748525 |
| GGVTVMATP | AWSRTGSNERLF | 0.772774 |
| AGLAANVRD | AWSRTGSNERLF | 0.7409229 |
| TNRPDLIDPALLRPG | AWSRTGSNERLF | 0.89932454 |
| LANLGVTAL | AWSRTGSNERLF | 0.8101574 |
| TGERIEL | AWSRTGSNERLF | 0.80958056 |
| NGEELAA | AWSRTGSNERLF | 0.62707096 |
| VLSLVWG | AWSRTGSNERLF | 0.8494083 |
| IGYALWL | AWSRTGSNERLF | 0.8276719 |
| LSLLLAPL | AWSRTGSNERLF | 0.57257235 |
| TLDEALK | AWSRTGSNERLF | 0.8545333 |
| TDGAGTAEGG | AWSRTGSNERLF | 0.5175915 |
| NAPGGAGGNGG | AWSRTGSNERLF | 0.80739594 |
| VSVFMGMGGNF | AWSRTGSNERLF | 0.7493644 |
| LASYLAAL | AWSRTGSNERLF | 0.83548164 |
| LVGGTPAL | AWSRTGSNERLF | 0.77538246 |
| VRNADVL | AWSRTGSNERLF | 0.8397658 |
| TCSSPFLVSAM | AWSRTGSNERLF | 0.5770155 |
| VHITDAE | AWSRTGSNERLF | 0.8539033 |
| EYAAPSF | AWSRTGSNERLF | 0.85177326 |
| VVDAFHGGL | AWSRTGSNERLF | 0.76195556 |
| ASVVDPQ | AWSRTGSNERLF | 0.8886975 |
| SYNEFGGLS | AWSRTGSNERLF | 0.77765775 |
| DPDGSVY | AWSRTGSNERLF | 0.29115304 |

| | | |
|---|---|---|
| MVSTIHATE | AWSRTGSNERLF | 0.73360884 |
| LAAGVADPV | AWSRTGSNERLF | 0.86552554 |
| TKEQIAA | AWSRTGSNERLF | 0.88247216 |
| SLASPAAL | AWSRTGSNERLF | 0.8246677 |
| IGLVFYGV | AWSRTGSNERLF | 0.80165833 |
| FLLDASNL | AWSRTGSNERLF | 0.7043371 |
| YQVGNFP | AWSRTGSNERLF | 0.8746903 |
| ESLLSQST | AWSRTGSNERLF | 0.91152465 |
| LFTVGFAAVG | AWSRTGSNERLF | 0.8033551 |
| ATPPTPT | AWSRTGSNERLF | 0.60961676 |
| TDLATLG | AWSRTGSNERLF | 0.53443134 |
| SAGGTGR | AWSRTGSNERLF | 0.8420191 |
| GFSNSGS | AWSRTGSNERLF | 0.8555789 |
| GGNGGSGGTSV | AWSRTGSNERLF | 0.56280184 |
| GTSVATGGAGN | AWSRTGSNERLF | 0.7056035 |
| NGGNGGSGGTS | AWSRTGSNERLF | 0.75378764 |
| AVVKFPH | AWSRTGSNERLF | 0.2242862 |
| TPPPPPP | AWSRTGSNERLF | 0.8107997 |
| AAPAAAAAPA | AWSRTGSNERLF | 0.8559389 |
| IALNEDL | AWSRTGSNERLF | 0.8514557 |
| IVNVSSM | AWSRTGSNERLF | 0.3686674 |
| TGIPPAPR | AWSRTGSNERLF | 0.8527783 |
| FDLGGGTF | AWSRTGSNERLF | 0.8398802 |
| FDLGGGTFDVS | AWSRTGSNERLF | 0.89110434 |
| NEPTAAA | AWSRTGSNERLF | 0.35614562 |
| NEPTAAALA | AWSRTGSNERLF | 0.18553257 |
| GSIINISSI | AWSRTGSNERLF | 0.77367806 |
| VAPGNAGTA | AWSRTGSNERLF | 0.80059004 |
| TGGGASLE | AWSRTGSNERLF | 0.57248545 |
| VAVYNHYKRI | AWSRTGSNERLF | 0.49748674 |
| TDPAPATT | AWSRTGSNERLF | 0.7196406 |
| TNRPDILDPALLRPG | AWSRTGSNERLF | 0.85606885 |
| LTTTTTS | AWSRTGSNERLF | 0.42413938 |
| PTGSGKTL | AWSRTGSNERLF | 0.6124133 |
| TGSGKTL | AWSRTGSNERLF | 0.80877316 |
| TIGHVDHGK | AWSRTGSNERLF | 0.862602 |
| TIGHVDHGKTT | AWSRTGSNERLF | 0.59616673 |
| GHVDHGKTT | AWSRTGSNERLF | 0.7186142 |
| AQFEHTLL | AWSRTGSNERLF | 0.81069475 |
| IVNQDEL | AWSRTGSNERLF | 0.8360063 |
| TNGSQFF | AWSRTGSNERLF | 0.7887344 |
| TGKGIVE | AWSRTGSNERLF | 0.42492527 |
| LISNASDA | AWSRTGSNERLF | 0.8520988 |
| AELRTLGELTKIAKAHGAQ | AWSRTGSNERLF | 0.53715473 |
| APNAVAAAEPSW | AWSRTGSNERLF | 0.8526938 |
| APQVAGAATAVL | AWSRTGSNERLF | 0.73480886 |
| ARRDPAAAAAILPTDA | AWSRTGSNERLF | 0.8140453 |
| DAAARDVLAEAGLAEYF | AWSRTGSNERLF | 0.7083261 |
| DDQIVLSAKANLAKVEQ | AWSRTGSNERLF | 0.4219381 |
| DELLPDTGEDSDEDGH | AWSRTGSNERLF | 0.55962634 |
| ELPEAPANRATVPQD | AWSRTGSNERLF | 0.62142694 |

| | | |
|---|---|---|
| FGQSKIERVTGIG | AWSRTGSNERLF | 0.8433591 |
| GAGGTGSTAGGAGGAGGAGGLY | AWSRTGSNERLF | 0.41144824 |
| GHGGTGAAGGNGGTGGAGGSGI | AWSRTGSNERLF | 0.84325325 |
| GNGGNGGAGGEGGPGGAGGAGA | AWSRTGSNERLF | 0.7871941 |
| GSTAGNGGNGGNATNGTIAGQP | AWSRTGSNERLF | 0.59851515 |
| GTGGDGGNGGNGGTGG | AWSRTGSNERLF | 0.8583951 |
| GVRRPAGIPADAPV | AWSRTGSNERLF | 0.37843847 |
| GYRQVTDAHLLTLARR | AWSRTGSNERLF | 0.8202419 |
| IAKMGQDGHDRG | AWSRTGSNERLF | 0.73518115 |
| KAEPSIEIQYT | AWSRTGSNERLF | 0.81834936 |
| KPVRIRMAADEDTVAVT | AWSRTGSNERLF | 0.6740701 |
| MPPRIAGMRLLVIKPEP | AWSRTGSNERLF | 0.85996187 |
| MKFYAVALFRSRGVPPTR | AWSRTGSNERLF | 0.8066133 |
| NAGGFGILGAVAHSP | AWSRTGSNERLF | 0.45105773 |
| NEGREITLAMAEM | AWSRTGSNERLF | 0.8127979 |
| NELVEWSHPLGHRWRWE | AWSRTGSNERLF | 0.83020335 |
| NLGAGGTGGNGGNPG | AWSRTGSNERLF | 0.8321768 |
| NTEWPKWETVMAADPVD | AWSRTGSNERLF | 0.33624214 |
| PASAAPAETQP | AWSRTGSNERLF | 0.76268923 |
| QQSSDGKNGVRAAEITGE | AWSRTGSNERLF | 0.6209319 |
| RAPDAVTTSWWKE | AWSRTGSNERLF | 0.7872431 |
| SGFGNVGAHTSGWFNQ | AWSRTGSNERLF | 0.78832364 |
| SGVAGAVGQAASVGGLKV | AWSRTGSNERLF | 0.63477415 |
| SNGGAGTGSAGGNGGT | AWSRTGSNERLF | 0.82985044 |
| TGGAGGEGGAAG | AWSRTGSNERLF | 0.42361945 |
| TGGDGGAAGAGGAPGHG | AWSRTGSNERLF | 0.6473694 |
| TMPPLLAAIANAGRHGV | AWSRTGSNERLF | 0.69663846 |
| TSSTAGNSGGA | AWSRTGSNERLF | 0.7927742 |
| VPATVPVGILC | AWSRTGSNERLF | 0.80459833 |
| VQRPDSGGVVLTGRIS | AWSRTGSNERLF | 0.47623435 |
| WTPNPVPATKGVPAGLVQAT | AWSRTGSNERLF | 0.7662015 |
| YALKRALAKFSDRFDVVII | AWSRTGSNERLF | 0.7417196 |
| YLIGSVIV | AWSRTGSNERLF | 0.5650089 |
| YGQLVAVID | AWSRTGSNERLF | 0.83036315 |
| VTVGGMTLPA | AWSRTGSNERLF | 0.67419773 |
| VLALPAV | AWSRTGSNERLF | 0.3039204 |
| VIPPTLNYET | AWSRTGSNERLF | 0.8426703 |
| VGMGGMHPSSG | AWSRTGSNERLF | 0.51330316 |
| VGLAGGAATA | AWSRTGSNERLF | 0.78630745 |
| VGGAGGAGGSGP | AWSRTGSNERLF | 0.8563521 |
| VFAAFMV | AWSRTGSNERLF | 0.361063 |
| TLYDKLR | AWSRTGSNERLF | 0.86505467 |
| TLAVFTVL | AWSRTGSNERLF | 0.73111737 |
| TGSVQPARV | AWSRTGSNERLF | 0.7488328 |
| TGGGAGSGGT | AWSRTGSNERLF | 0.711864 |
| SHLETSL | AWSRTGSNERLF | 0.57283247 |
| SFTTGMT | AWSRTGSNERLF | 0.8662126 |
| QVEVAKRMGVRQPTVS | AWSRTGSNERLF | 0.8100781 |
| QGSGGPALAM | AWSRTGSNERLF | 0.6953765 |
| MLAPSNP | AWSRTGSNERLF | 0.7877794 |
| LVNNAAI | AWSRTGSNERLF | 0.8064349 |

| | | |
|---|---|---|
| LTGEINTLA | AWSRTGSNERLF | 0.8492862 |
| LSIFGPV | AWSRTGSNERLF | 0.599292 |
| LLALLANL | AWSRTGSNERLF | 0.41118264 |
| LISNASDALDKLR | AWSRTGSNERLF | 0.72190094 |
| LINEAVP | AWSRTGSNERLF | 0.6787855 |
| LGYAFSGL | AWSRTGSNERLF | 0.60842055 |
| LDFVAVAL | AWSRTGSNERLF | 0.4219662 |
| LAQPGAE | AWSRTGSNERLF | 0.80613244 |
| ISNLGAVTGF | AWSRTGSNERLF | 0.896406 |
| HQEDPGRI | AWSRTGSNERLF | 0.6060908 |
| GTPGEPTP | AWSRTGSNERLF | 0.9010644 |
| GFFNSGS | AWSRTGSNERLF | 0.5114838 |
| GAGGGGLTAGAN | AWSRTGSNERLF | 0.82381094 |
| FVDALVNM | AWSRTGSNERLF | 0.82911074 |
| FLTVAPEL | AWSRTGSNERLF | 0.538937 |
| FETLTAL | AWSRTGSNERLF | 0.72438943 |
| DVINPIEL | AWSRTGSNERLF | 0.5587455 |
| APVSGAPV | AWSRTGSNERLF | 0.6180108 |
| ALVGGLAW | AWSRTGSNERLF | 0.8338553 |
| ALAAAAATAI | AWSRTGSNERLF | 0.57692516 |
| AFGLGLLL | AWSRTGSNERLF | 0.81090534 |
| AFFAAASL | AWSRTGSNERLF | 0.64833903 |
| AAAALPAFSPP | AWSRTGSNERLF | 0.64482826 |
| VVLAPAL | AWSRTGSNERLF | 0.83880186 |
| TIGHVDHGKTTL | AWSRTGSNERLF | 0.8075118 |
| LISNASDALDK | AWSRTGSNERLF | 0.80657244 |
| LISNASDAL | AWSRTGSNERLF | 0.9097681 |
| ISNASDALDK | AWSRTGSNERLF | 0.67053735 |
| NQAETLVYQTE | AWSRTGSNERLF | 0.8803855 |
| TGGTGGTAGAG | AWSRTGSNERLF | 0.8746984 |
| TGSTAGGAGGA | AWSRTGSNERLF | 0.7732908 |
| DPLAVPVAAGPV | AWSRTGSNERLF | 0.7546663 |
| GGTGGTPGNGGN | AWSRTGSNERLF | 0.8135663 |
| PAGLAGVDPATL | AWSRTGSNERLF | 0.6374829 |
| GNNLVVEAPHAGASV | AWSRTGSNERLF | 0.7915065 |
| KVTRIMSASTLEQPA | AWSRTGSNERLF | 0.8838526 |
| LGFALPIALPRRTVV | AWSRTGSNERLF | 0.5257417 |
| TRLGNLTARKLVVLV | AWSRTGSNERLF | 0.70808935 |
| WSQQRRLIYFPSAGP | AWSRTGSNERLF | 0.67408156 |
| GPNGNPDPMAAAVDIR | AWSRTGSNERLF | 0.6442169 |
| TQRCVSVADLRAAASR | AWSRTGSNERLF | 0.57104826 |
| QFVLDVAEHPVEVLTETV | AWSRTGSNERLF | 0.41082847 |
| GGAGGTGGAGGNGGTQAGDG | AWSRTGSNERLF | 0.59322 |
| PLLATLRAIAVDVDAKPAQV | AWSRTGSNERLF | 0.5575906 |
| APKFRTGGYMIDFWGVGYQVAK | AWSRTGSNERLF | 0.6856872 |
| GAGGAGGAGGAAGGPGGTGGTG | AWSRTGSNERLF | 0.6843532 |
| IGALGLLATDVAPAGSVAFGVA | AWSRTGSNERLF | 0.5752314 |
| AGGAGGIGGNGGAGGAGGVLMG | AWSRTGSNERLF | 0.70494246 |
| ADELIGLL | AWSRTGSNERLF | 0.7052691 |
| ADRALKA | AWSRTGSNERLF | 0.40678397 |
| AGAVGQVGGAGGA | AWSRTGSNERLF | 0.6226827 |

| | | |
|---|---|---|
| AGGGGGLGGAP | AWSRTGSNERLF | 0.5431607 |
| AGGLLYGNGGAG | AWSRTGSNERLF | 0.6166847 |
| AGLESIGL | AWSRTGSNERLF | 0.37504426 |
| AGLKTGII | AWSRTGSNERLF | 0.6869985 |
| ASLAALLL | AWSRTGSNERLF | 0.57964945 |
| ASVINISSI | AWSRTGSNERLF | 0.28481793 |
| ATPPGPA | AWSRTGSNERLF | 0.6299238 |
| ATVASALV | AWSRTGSNERLF | 0.65141946 |
| ATVASLIL | AWSRTGSNERLF | 0.6332422 |
| AVNGPVLIRH | AWSRTGSNERLF | 0.6961028 |
| DGAAGAVGI | AWSRTGSNERLF | 0.45307225 |
| DPLLHGI | AWSRTGSNERLF | 0.55155957 |
| ELGAALGAL | AWSRTGSNERLF | 0.6884115 |
| ELGRTQAV | AWSRTGSNERLF | 0.7051463 |
| FVAGGYLS | AWSRTGSNERLF | 0.5060887 |
| GASGGAGGAGGW | AWSRTGSNERLF | 0.67369914 |
| GGAGAGGDGGAGGI | AWSRTGSNERLF | 0.7064054 |
| GTGHNGSQPGMGGNG | AWSRTGSNERLF | 0.66408217 |
| GYAAASASAS | AWSRTGSNERLF | 0.69563377 |
| HQVDVAAVQQG | AWSRTGSNERLF | 0.7052885 |
| IGAAIAHT | AWSRTGSNERLF | 0.49123508 |
| IGMIEVAQL | AWSRTGSNERLF | 0.69956505 |
| IRLAPEHTSAMD | AWSRTGSNERLF | 0.7042155 |
| IYNVSVL | AWSRTGSNERLF | 0.58952624 |
| KQGFLGM | AWSRTGSNERLF | 0.6613556 |
| LESVITQL | AWSRTGSNERLF | 0.48419514 |
| LGLGALVVV | AWSRTGSNERLF | 0.6068504 |
| LIGNGGNGGSGGI | AWSRTGSNERLF | 0.33600616 |
| LNAINAPTL | AWSRTGSNERLF | 0.6846568 |
| LQLLNTGV | AWSRTGSNERLF | 0.6989013 |
| LQTSIEF | AWSRTGSNERLF | 0.6299179 |
| MSFAVSEL | AWSRTGSNERLF | 0.5112945 |
| NTGALGL | AWSRTGSNERLF | 0.5297389 |
| PPGAGGAGGAGTT | AWSRTGSNERLF | 0.6370608 |
| QLVGLGI | AWSRTGSNERLF | 0.598703 |
| RFLEQVHQL | AWSRTGSNERLF | 0.350905 |
| RGVGYVLR | AWSRTGSNERLF | 0.61614496 |
| SATGVFAGVI | AWSRTGSNERLF | 0.69988096 |
| SLPAITLP | AWSRTGSNERLF | 0.69941825 |
| STALAFTV | AWSRTGSNERLF | 0.41979176 |
| SVLTVGLGK | AWSRTGSNERLF | 0.70636487 |
| TALAGAGSGI | AWSRTGSNERLF | 0.6838874 |
| TGLPFTGI | AWSRTGSNERLF | 0.6998956 |
| VAEHVTV | AWSRTGSNERLF | 0.62918174 |
| VDIVLNSL | AWSRTGSNERLF | 0.5602878 |
| VLTGGGAVL | AWSRTGSNERLF | 0.6950962 |
| VNVGLAYGI | AWSRTGSNERLF | 0.6374573 |
| VSSAAVGAV | AWSRTGSNERLF | 0.5245288 |
| VTSVYEAGV | AWSRTGSNERLF | 0.69545615 |
| WVALDRGA | AWSRTGSNERLF | 0.6347047 |
| YLVGSVVI | AWSRTGSNERLF | 0.60851574 |

| | | |
|---|---|---|
| GPPGTGKTLL | AWSRTGSNERLF | 0.6254035 |
| GVGMTKF | AWSRTGSNERLF | 0.6938343 |
| IGIGGSDL | AWSRTGSNERLF | 0.5138057 |
| IGTLLAEL | AWSRTGSNERLF | 0.538185 |
| IYTSGTT | AWSRTGSNERLF | 0.5006857 |
| LSFPAGV | AWSRTGSNERLF | 0.66141504 |
| LYTSGTT | AWSRTGSNERLF | 0.7037542 |
| VGAFPFGF | AWSRTGSNERLF | 0.6695712 |
| VGAGTVEF | AWSRTGSNERLF | 0.62104356 |
| VGKSTLLL | AWSRTGSNERLF | 0.67872715 |
| AALAHAGGGIV | AWSRTGSNERLF | 0.68363833 |
| AALVAVSLL | AWSRTGSNERLF | 0.65888035 |
| AAVSRAVAAL | AWSRTGSNERLF | 0.69701385 |
| AGGSSGIGGF | AWSRTGSNERLF | 0.6232073 |
| AGLTVANANG | AWSRTGSNERLF | 0.6926707 |
| AGLVGAVL | AWSRTGSNERLF | 0.68251294 |
| ALSQAFTV | AWSRTGSNERLF | 0.68862915 |
| ALTFSVSL | AWSRTGSNERLF | 0.69152886 |
| ALTGAALVAL | AWSRTGSNERLF | 0.6856046 |
| ASFSIGAGVL | AWSRTGSNERLF | 0.6285588 |
| DGSVYKI | AWSRTGSNERLF | 0.34169194 |
| EQLAEIV | AWSRTGSNERLF | 0.5520641 |
| FAVSELTVM | AWSRTGSNERLF | 0.6757267 |
| GAAPLAAGL | AWSRTGSNERLF | 0.6702205 |
| GADGGDGGSGGAGG | AWSRTGSNERLF | 0.687488 |
| GFGLDTGGAGG | AWSRTGSNERLF | 0.66765726 |
| GLASGGLL | AWSRTGSNERLF | 0.5454402 |
| GRIAGAAVLL | AWSRTGSNERLF | 0.7009896 |
| GSAGSLNTSF | AWSRTGSNERLF | 0.6948662 |
| GTGQIGI | AWSRTGSNERLF | 0.4126937 |
| GTLIALIL | AWSRTGSNERLF | 0.6277962 |
| IAATMGFVVG | AWSRTGSNERLF | 0.71325636 |
| IFAQALL | AWSRTGSNERLF | 0.6708399 |
| IGAAVALGF | AWSRTGSNERLF | 0.6005026 |
| IGLTPLAL | AWSRTGSNERLF | 0.5703972 |
| ILGSNTSTL | AWSRTGSNERLF | 0.7059213 |
| IVFSLTVL | AWSRTGSNERLF | 0.51116705 |
| LASKATGF | AWSRTGSNERLF | 0.70690215 |
| LASVALIL | AWSRTGSNERLF | 0.29911244 |
| LFAVGLTL | AWSRTGSNERLF | 0.51340014 |
| LFNGGIGTYI | AWSRTGSNERLF | 0.49502677 |
| LTLGAGII | AWSRTGSNERLF | 0.399455 |
| LVAAALGDI | AWSRTGSNERLF | 0.6909604 |
| NTAFVQLGI | AWSRTGSNERLF | 0.50100434 |
| QLFSGAEL | AWSRTGSNERLF | 0.68517315 |
| SAATFITL | AWSRTGSNERLF | 0.62882924 |
| SNYGLANLGV | AWSRTGSNERLF | 0.69977725 |
| SYGGAYAVM | AWSRTGSNERLF | 0.66677225 |
| TTGVGGLTLG | AWSRTGSNERLF | 0.6559397 |
| TTQTIYI | AWSRTGSNERLF | 0.5296464 |
| VAAGAGLGAVI | AWSRTGSNERLF | 0.6841766 |

| | | |
|---|---|---|
| VAAVVTLL | AWSRTGSNERLF | 0.7025664 |
| VGFLGGLI | AWSRTGSNERLF | 0.6626458 |
| VGVIPILL | AWSRTGSNERLF | 0.45209005 |
| VIFASSLI | AWSRTGSNERLF | 0.6081285 |
| VKGGDGGAAAG | AWSRTGSNERLF | 0.6560512 |
| VTKTFTNI | AWSRTGSNERLF | 0.3681149 |
| VVAAAILL | AWSRTGSNERLF | 0.32157734 |
| YAALDVEL | AWSRTGSNERLF | 0.5587745 |
| LASVALLL | AWSRTGSNERLF | 0.61646855 |
| LDAVHRV | AWSRTGSNERLF | 0.6987761 |
| QGLSSSIF | AWSRTGSNERLF | 0.6952723 |
| RGTFANIRL | AWSRTGSNERLF | 0.6426259 |
| RQLWWGHRI | AWSRTGSNERLF | 0.6874135 |
| YGPPGTGKTLL | AWSRTGSNERLF | 0.6747868 |
| AALVGAPANVA | AWSRTGSNERLF | 0.26662835 |
| AATFRRTE | AWSRTGSNERLF | 0.5702423 |
| AGALGGGI | AWSRTGSNERLF | 0.4238906 |
| AGGAGGAGGIGAG | AWSRTGSNERLF | 0.6999131 |
| AGSGALGL | AWSRTGSNERLF | 0.5472262 |
| AQIADQI | AWSRTGSNERLF | 0.5601704 |
| ASAVAVTYL | AWSRTGSNERLF | 0.67016995 |
| ASTVLGGTGG | AWSRTGSNERLF | 0.6560559 |
| AVGGEPAK | AWSRTGSNERLF | 0.6494691 |
| CEQGTFGGTC | AWSRTGSNERLF | 0.56253886 |
| DEGKAPAG | AWSRTGSNERLF | 0.30361524 |
| DTVVFPAL | AWSRTGSNERLF | 0.6908153 |
| FEGITMAG | AWSRTGSNERLF | 0.44888946 |
| FPLVVQD | AWSRTGSNERLF | 0.51481175 |
| GAFTTPQ | AWSRTGSNERLF | 0.33860925 |
| GFGGFGTAGGVGGA | AWSRTGSNERLF | 0.57769275 |
| GFLAQFL | AWSRTGSNERLF | 0.41328514 |
| GGAGGAGGNAGVA | AWSRTGSNERLF | 0.68943685 |
| GGVGTAGAGGAGGAGG | AWSRTGSNERLF | 0.6066786 |
| GNGGGGDAGNAGSGGN | AWSRTGSNERLF | 0.62138104 |
| GTGGAGGAGNDAGS | AWSRTGSNERLF | 0.7057353 |
| GTTNTGWFNS | AWSRTGSNERLF | 0.66597486 |
| IFMGPLFTL | AWSRTGSNERLF | 0.6430018 |
| ILAHAIEAI | AWSRTGSNERLF | 0.5637385 |
| IRSAIREAL | AWSRTGSNERLF | 0.4987594 |
| ITFAGGG | AWSRTGSNERLF | 0.5483843 |
| ITGMSGHQEV | AWSRTGSNERLF | 0.55194867 |
| IVAGLACTL | AWSRTGSNERLF | 0.71167797 |
| IVGGGLAGA | AWSRTGSNERLF | 0.54755545 |
| IVTGVLNI | AWSRTGSNERLF | 0.42408192 |
| LAQQGCI | AWSRTGSNERLF | 0.5943714 |
| LCLHVEN | AWSRTGSNERLF | 0.7073339 |
| LDAISDGS | AWSRTGSNERLF | 0.7121422 |
| LGGFAAML | AWSRTGSNERLF | 0.54415023 |
| LGGKNPMI | AWSRTGSNERLF | 0.6125158 |
| LGPKGVAAIL | AWSRTGSNERLF | 0.57255363 |
| LMVGVSI | AWSRTGSNERLF | 0.6607901 |

| | | |
|---|---|---|
| LVGAAFVL | AWSRTGSNERLF | 0.5754202 |
| LVLAVAPL | AWSRTGSNERLF | 0.46748674 |
| MAAIAVIDAA | AWSRTGSNERLF | 0.68332565 |
| NNGINAELQ | AWSRTGSNERLF | 0.711833 |
| RGEPLVLVHG | AWSRTGSNERLF | 0.59161365 |
| RGGAGGAAGLI | AWSRTGSNERLF | 0.41455647 |
| RSYRMHPAV | AWSRTGSNERLF | 0.65908754 |
| SAAHLLAGQLA | AWSRTGSNERLF | 0.56909275 |
| SAAVGSY | AWSRTGSNERLF | 0.6369362 |
| SPHAVLHRI | AWSRTGSNERLF | 0.64583313 |
| SQIAKVL | AWSRTGSNERLF | 0.40655723 |
| TALGHLVP | AWSRTGSNERLF | 0.6118779 |
| TAPTTSGAA | AWSRTGSNERLF | 0.71238154 |
| TDGDDGNGGSPGT | AWSRTGSNERLF | 0.5987852 |
| TDGLGEN | AWSRTGSNERLF | 0.64526385 |
| TGGPPLPGQA | AWSRTGSNERLF | 0.70163035 |
| TIPAITI | AWSRTGSNERLF | 0.6834229 |
| TIPGFTI | AWSRTGSNERLF | 0.70290864 |
| TSIASGALA | AWSRTGSNERLF | 0.6917305 |
| TVTLPGAAI | AWSRTGSNERLF | 0.6271026 |
| VASGLSAALP | AWSRTGSNERLF | 0.6514957 |
| VGAGVIGI | AWSRTGSNERLF | 0.67241216 |
| VGPKRFGV | AWSRTGSNERLF | 0.6709537 |
| VNVAFPD | AWSRTGSNERLF | 0.6354202 |
| VSMTNSGVS | AWSRTGSNERLF | 0.5438353 |
| VTDVLGDLSV | AWSRTGSNERLF | 0.6984631 |
| VVVVGAGFAGL | AWSRTGSNERLF | 0.6520112 |
| VYTVGTVV | AWSRTGSNERLF | 0.68924075 |
| WTTFSGP | AWSRTGSNERLF | 0.43395668 |
| YDELVEAF | AWSRTGSNERLF | 0.5171448 |
| YGTGGAGGNGG | AWSRTGSNERLF | 0.69667214 |
| YTNVMAV | AWSRTGSNERLF | 0.29257128 |
| FLEQVHQL | AWSRTGSNERLF | 0.32787505 |
| GLSLLAVAV | AWSRTGSNERLF | 0.62061965 |
| LEQVHQL | AWSRTGSNERLF | 0.70445824 |
| LNAAGAI | AWSRTGSNERLF | 0.2963468 |
| PAVHASL | AWSRTGSNERLF | 0.5430707 |
| TELEDDA | AWSRTGSNERLF | 0.42350549 |
| TPPGPPN | AWSRTGSNERLF | 0.6844703 |
| TPPPPPPP | AWSRTGSNERLF | 0.6652442 |
| TQSPALI | ASSSGQGAGEQY | 0.42701864 |
| EFAAAFPQ | ASSSGQGAGEQY | 0.6852399 |
| LLSGNLDVL | ASSSGQGAGEQY | 0.4128171 |
| GPANLGNYNI | ASSSGQGAGEQY | 0.55948824 |
| TLGPTLP | ASSSGQGAGEQY | 0.6934167 |
| VYAYSSCCK | ASSSGQGAGEQY | 0.7054342 |
| APAPAPAPNI | ASSSGQGAGEQY | 0.46037757 |
| PGGPVPIP | ASSSGQGAGEQY | 0.69526684 |
| FLTVALEP | ASSSGQGAGEQY | 0.5915371 |
| SQYFGANL | ASSSGQGAGEQY | 0.71203244 |
| GGVKPAA | ASSSGQGAGEQY | 0.59076595 |

| | | |
|---|---|---|
| GGTGGAGGAGGW | ASSSGQGAGEQY | 0.39827958 |
| GVGGVAGLGGAGG | ASSSGQGAGEQY | 0.6908784 |
| VLLVLATTI | ASSSGQGAGEQY | 0.6652427 |
| IGIFAAML | ASSSGQGAGEQY | 0.6296821 |
| AVGDLID | ASSSGQGAGEQY | 0.6895259 |
| NISPLPL | ASSSGQGAGEQY | 0.5751014 |
| VTAVFVAI | ASSSGQGAGEQY | 0.6432226 |
| SGDDNDAAL | ASSSGQGAGEQY | 0.7020836 |
| AVAALAHS | ASSSGQGAGEQY | 0.6977357 |
| GGVTVMATP | ASSSGQGAGEQY | 0.48645395 |
| AGLAANVRD | ASSSGQGAGEQY | 0.69240177 |
| TNRPDLIDPALLRPG | ASSSGQGAGEQY | 0.660465 |
| LANLGVTAL | ASSSGQGAGEQY | 0.52609026 |
| TGERIEL | ASSSGQGAGEQY | 0.70646584 |
| NGEELAA | ASSSGQGAGEQY | 0.6348845 |
| VLSLVWG | ASSSGQGAGEQY | 0.6924466 |
| IGYALWL | ASSSGQGAGEQY | 0.67762065 |
| LSLLLAPL | ASSSGQGAGEQY | 0.7036932 |
| TLDEALK | ASSSGQGAGEQY | 0.4401613 |
| TDGAGTAEGG | ASSSGQGAGEQY | 0.6511748 |
| NAPGGAGGNGG | ASSSGQGAGEQY | 0.645318 |
| VSVFMGMGGNF | ASSSGQGAGEQY | 0.69401556 |
| LASYLAAL | ASSSGQGAGEQY | 0.61292446 |
| LVGGTPAL | ASSSGQGAGEQY | 0.5622231 |
| VRNADVL | ASSSGQGAGEQY | 0.65648806 |
| TCSSPFLVSAM | ASSSGQGAGEQY | 0.70303947 |
| VHITDAE | ASSSGQGAGEQY | 0.6434396 |
| EYAAPSF | ASSSGQGAGEQY | 0.6813277 |
| VVDAFHGGL | ASSSGQGAGEQY | 0.70958495 |
| ASVVDPQ | ASSSGQGAGEQY | 0.48141468 |
| SYNEFGGLS | ASSSGQGAGEQY | 0.6384567 |
| DPDGSVY | ASSSGQGAGEQY | 0.68026483 |
| MVSTIHATE | ASSSGQGAGEQY | 0.6994418 |
| LAAGVADPV | ASSSGQGAGEQY | 0.61660284 |
| TKEQIAA | ASSSGQGAGEQY | 0.7041702 |
| SLASPAAL | ASSSGQGAGEQY | 0.54874146 |
| IGLVFYGV | ASSSGQGAGEQY | 0.6863488 |
| FLLDASNL | ASSSGQGAGEQY | 0.5440467 |
| YQVGNFP | ASSSGQGAGEQY | 0.6699457 |
| ESLLSQST | ASSSGQGAGEQY | 0.5912655 |
| LFTVGFAAVG | ASSSGQGAGEQY | 0.6825336 |
| ATPPTPT | ASSSGQGAGEQY | 0.6953714 |
| TDLATLG | ASSSGQGAGEQY | 0.6961043 |
| SAGGTGR | ASSSGQGAGEQY | 0.30574086 |
| GFSNSGS | ASSSGQGAGEQY | 0.70000696 |
| GGNGGSGGTSV | ASSSGQGAGEQY | 0.6952572 |
| GTSVATGGAGN | ASSSGQGAGEQY | 0.711871 |
| NGGNGGSGGTS | ASSSGQGAGEQY | 0.65071565 |
| AVVKFPH | ASSSGQGAGEQY | 0.66622263 |
| TPPPPPP | ASSSGQGAGEQY | 0.6104106 |
| AAPAAAAPA | ASSSGQGAGEQY | 0.62973845 |

| | | |
|---|---|---|
| IALNEDL | ASSSGQGAGEQY | 0.5487917 |
| IVNVSSM | ASSSGQGAGEQY | 0.56846935 |
| TGIPPAPR | ASSSGQGAGEQY | 0.64036167 |
| FDLGGGTF | ASSSGQGAGEQY | 0.58858943 |
| FDLGGGTFDVS | ASSSGQGAGEQY | 0.6907171 |
| NEPTAAA | ASSSGQGAGEQY | 0.7007737 |
| NEPTAAALA | ASSSGQGAGEQY | 0.69784224 |
| GSIINISSI | ASSSGQGAGEQY | 0.7031691 |
| VAPGNAGTA | ASSSGQGAGEQY | 0.7046684 |
| TGGGASLE | ASSSGQGAGEQY | 0.68954074 |
| VAVYNHYKRI | ASSSGQGAGEQY | 0.61152935 |
| TDPAPATT | ASSSGQGAGEQY | 0.61736387 |
| TNRPDILDPALLRPG | ASSSGQGAGEQY | 0.50659186 |
| LTTTTTS | ASSSGQGAGEQY | 0.59769297 |
| PTGSGKTL | ASSSGQGAGEQY | 0.6140542 |
| TGSGKTL | ASSSGQGAGEQY | 0.6806154 |
| TIGHVDHGK | ASSSGQGAGEQY | 0.28688732 |
| TIGHVDHGKTT | ASSSGQGAGEQY | 0.663296 |
| GHVDHGKTT | ASSSGQGAGEQY | 0.7091565 |
| AQFEHTLL | ASSSGQGAGEQY | 0.60002345 |
| IVNQDEL | ASSSGQGAGEQY | 0.66920394 |
| TNGSQFF | ASSSGQGAGEQY | 0.65739834 |
| TGKGIVE | ASSSGQGAGEQY | 0.6104556 |
| LISNASDA | ASSSGQGAGEQY | 0.6934267 |
| AELRTLGELTKIAKAHGAQ | ASSSGQGAGEQY | 0.54027426 |
| APNAVAAAEPSW | ASSSGQGAGEQY | 0.677909 |
| APQVAGAATAVL | ASSSGQGAGEQY | 0.7055133 |
| ARRDPAAAAAILPTDA | ASSSGQGAGEQY | 0.6463964 |
| DAAARDVLAEAGLAEYF | ASSSGQGAGEQY | 0.662189 |
| DDQIVLSAKANLAKVEQ | ASSSGQGAGEQY | 0.6272057 |
| DELLPDTGEDSDEDGH | ASSSGQGAGEQY | 0.40443653 |
| ELPEAPANRATVPQD | ASSSGQGAGEQY | 0.67495096 |
| FGQSKIERVTGIG | ASSSGQGAGEQY | 0.37528074 |
| GAGGTGSTAGGAGGAGGAGGLY | ASSSGQGAGEQY | 0.635382 |
| GHGGTGAAGGNGGTGGAGGSGI | ASSSGQGAGEQY | 0.48110712 |
| GNGGNGGAGGEGGPGGAGGAGA | ASSSGQGAGEQY | 0.6691989 |
| GSTAGNGGNGGNATNGTIAGQP | ASSSGQGAGEQY | 0.66840667 |
| GTGGDGGNGGNGGTGG | ASSSGQGAGEQY | 0.67642 |
| GVRRPAGIPADAPV | ASSSGQGAGEQY | 0.48823023 |
| GYRQVTDAHLLTLARR | ASSSGQGAGEQY | 0.627131 |
| IAKMGQDGHDRG | ASSSGQGAGEQY | 0.5251024 |
| KAEPSIEIQYT | ASSSGQGAGEQY | 0.47133934 |
| KPVRIRMAADEDTVAVT | ASSSGQGAGEQY | 0.6839088 |
| MPPRIAGMRLLVIKPEP | ASSSGQGAGEQY | 0.57373834 |
| MKFYAVALFRSRGVPPTR | ASSSGQGAGEQY | 0.65364444 |
| NAGGFGILGAVAHSP | ASSSGQGAGEQY | 0.69612885 |
| NEGREITLAMAEM | ASSSGQGAGEQY | 0.6594198 |
| NELVEWSHPLGHRWRWE | ASSSGQGAGEQY | 0.4128151 |
| NLGAGGTGGNGGNPG | ASSSGQGAGEQY | 0.63777316 |
| NTEWPKWETVMAADPVD | ASSSGQGAGEQY | 0.61467075 |
| PASAAPAETQP | ASSSGQGAGEQY | 0.6182988 |

| | | |
|---|---|---|
| QQSSDGKNGVRAAEITGE | ASSSGQGAGEQY | 0.2923678 |
| RAPDAVTTSWWKE | ASSSGQGAGEQY | 0.6768968 |
| SGFGNVGAHTSGWFNQ | ASSSGQGAGEQY | 0.52702975 |
| SGVAGAVGQAASVGGLKV | ASSSGQGAGEQY | 0.5727269 |
| SNGGAGTGSAGGNGGT | ASSSGQGAGEQY | 0.5963931 |
| TGGAGGEGGAAG | ASSSGQGAGEQY | 0.6946082 |
| TGGDGGAAGAGGAPGHG | ASSSGQGAGEQY | 0.6274749 |
| TMPPLLAAIANAGRHGV | ASSSGQGAGEQY | 0.63568044 |
| TSSTAGNSGGA | ASSSGQGAGEQY | 0.68956023 |
| VPATVPVGILC | ASSSGQGAGEQY | 0.6921427 |
| VQRPDSGGVVLTGRIS | ASSSGQGAGEQY | 0.7057055 |
| WTPNPVPATKGVPAGLVQAT | ASSSGQGAGEQY | 0.6800128 |
| YALKRALAKFSDRFDVVII | ASSSGQGAGEQY | 0.6102033 |
| YLIGSVIV | ASSSGQGAGEQY | 0.66546255 |
| YGQLVAVID | ASSSGQGAGEQY | 0.6409396 |
| VTVGGMTLPA | ASSSGQGAGEQY | 0.36720088 |
| VLALPAV | ASSSGQGAGEQY | 0.33233714 |
| VIPPTLNYET | ASSSGQGAGEQY | 0.66187894 |
| VGMGGMHPSSG | ASSSGQGAGEQY | 0.35025218 |
| VGLAGGAATA | ASSSGQGAGEQY | 0.6444221 |
| VGGAGGAGGSGP | ASSSGQGAGEQY | 0.6927999 |
| VFAAFMV | ASSSGQGAGEQY | 0.63657486 |
| TLYDKLR | ASSSGQGAGEQY | 0.40836918 |
| TLAVFTVL | ASSSGQGAGEQY | 0.5994635 |
| TGSVQPARV | ASSSGQGAGEQY | 0.59457636 |
| TGGGAGSGGT | ASSSGQGAGEQY | 0.6908152 |
| SHLETSL | ASSSGQGAGEQY | 0.67443776 |
| SFTTGMT | ASSSGQGAGEQY | 0.707553 |
| QVEVAKRMGVRQPTVS | ASSSGQGAGEQY | 0.70211905 |
| QGSGGPALAM | ASSSGQGAGEQY | 0.6209587 |
| MLAPSNP | ASSSGQGAGEQY | 0.6903627 |
| LVNNAAI | ASSSGQGAGEQY | 0.5857041 |
| LTGEINTLA | ASSSGQGAGEQY | 0.5913096 |
| LSIFGPV | ASSSGQGAGEQY | 0.59681016 |
| LLALLANL | ASSSGQGAGEQY | 0.6660455 |
| LISNASDALDKLR | ASSSGQGAGEQY | 0.6520892 |
| LINEAVP | ASSSGQGAGEQY | 0.5634978 |
| LGYAFSGL | ASSSGQGAGEQY | 0.39799234 |
| LDFVAVAL | ASSSGQGAGEQY | 0.66500974 |
| LAQPGAE | ASSSGQGAGEQY | 0.63656676 |
| ISNLGAVTGF | ASSSGQGAGEQY | 0.37133622 |
| HQEDPGRI | ASSSGQGAGEQY | 0.5807142 |
| GTPGEPTP | ASSSGQGAGEQY | 0.7084865 |
| GFFNSGS | ASSSGQGAGEQY | 0.70756316 |
| GAGGGGLTAGAN | ASSSGQGAGEQY | 0.588749 |
| FVDALVNM | ASSSGQGAGEQY | 0.5472507 |
| FLTVAPEL | ASSSGQGAGEQY | 0.6685978 |
| FETLTAL | ASSSGQGAGEQY | 0.38210773 |
| DVINPIEL | ASSSGQGAGEQY | 0.65278727 |
| APVSGAPV | ASSSGQGAGEQY | 0.6662837 |
| ALVGGLAW | ASSSGQGAGEQY | 0.6724498 |

| | | |
|---|---|---|
| ALAAAAATAI | ASSSGQGAGEQY | 0.51794523 |
| AFGLGLLL | ASSSGQGAGEQY | 0.66586244 |
| AFFAAASL | ASSSGQGAGEQY | 0.6497334 |
| AAAALPAFSPP | ASSSGQGAGEQY | 0.6813174 |
| VVLAPAL | ASSSGQGAGEQY | 0.3071365 |
| TIGHVDHGKTTL | ASSSGQGAGEQY | 0.6902589 |
| LISNASDALDK | ASSSGQGAGEQY | 0.55243397 |
| LISNASDAL | ASSSGQGAGEQY | 0.5655584 |
| ISNASDALDK | ASSSGQGAGEQY | 0.364407 |
| NQAETLVYQTE | ASSSGQGAGEQY | 0.62589306 |
| TGGTGGTAGAG | ASSSGQGAGEQY | 0.6901279 |
| TGSTAGGAGGA | ASSSGQGAGEQY | 0.6987779 |
| DPLAVPVAAGPV | ASSSGQGAGEQY | 0.4384682 |
| GGTGGTPGNGGN | ASSSGQGAGEQY | 0.6982976 |
| PAGLAGVDPATL | ASSSGQGAGEQY | 0.7107042 |
| GNNLVVEAPHAGASV | ASSSGQGAGEQY | 0.6185266 |
| KVTRIMSASTLEQPA | ASSSGQGAGEQY | 0.3890289 |
| LGFALPIALPRRTVV | ASSSGQGAGEQY | 0.6788306 |
| TRLGNLTARKLVVLV | ASSSGQGAGEQY | 0.6229615 |
| WSQQRRLIYFPSAGP | ASSSGQGAGEQY | 0.6971368 |
| GPNGNPDPMAAAVDIR | ASSSGQGAGEQY | 0.6821361 |
| TQRCVSVADLRAAASR | ASSSGQGAGEQY | 0.5197957 |
| QFVLDVAEHPVEVLTETV | ASSSGQGAGEQY | 0.68113935 |
| GGAGGTGGAGGNGGTQAGDG | ASSSGQGAGEQY | 0.6917782 |
| PLLATLRAIAVDVDAKPAQV | ASSSGQGAGEQY | 0.5509262 |
| APKFRTGGYMIDFWGVGYQVAK | ASSSGQGAGEQY | 0.5116308 |
| GAGGAGGAGGAAGGPGGTGGTG | ASSSGQGAGEQY | 0.52757955 |
| IGALGLLATDVAPAGSVAFGVA | ASSSGQGAGEQY | 0.3538472 |
| AGGAGGIGGNGGAGGAGGVLMG | ASSSGQGAGEQY | 0.68844163 |
| ADELIGLL | ASSSGQGAGEQY | 0.59490746 |
| ADRALKA | ASSSGQGAGEQY | 0.5913195 |
| AGAVGQVGGAGGA | ASSSGQGAGEQY | 0.66112745 |
| AGGGGGLGGAP | ASSSGQGAGEQY | 0.5681165 |
| AGGLLYGNGGAG | ASSSGQGAGEQY | 0.5934516 |
| AGLESIGL | ASSSGQGAGEQY | 0.5965687 |
| AGLKTGII | ASSSGQGAGEQY | 0.6723487 |
| ASLAALLL | ASSSGQGAGEQY | 0.677289 |
| ASVINISSI | ASSSGQGAGEQY | 0.6420509 |
| ATPPGPA | ASSSGQGAGEQY | 0.6380969 |
| ATVASALV | ASSSGQGAGEQY | 0.70641506 |
| ATVASLIL | ASSSGQGAGEQY | 0.50251186 |
| AVNGPVLIRH | ASSSGQGAGEQY | 0.5968151 |
| DGAAGAVGI | ASSSGQGAGEQY | 0.6859081 |
| DPLLHGI | ASSSGQGAGEQY | 0.3981546 |
| ELGAALGAL | ASSSGQGAGEQY | 0.68514395 |
| ELGRTQAV | ASSSGQGAGEQY | 0.6669687 |
| FVAGGYLS | ASSSGQGAGEQY | 0.68415964 |
| GASGGAGGAGGW | ASSSGQGAGEQY | 0.6421052 |
| GGAGAGGDGGAGGI | ASSSGQGAGEQY | 0.7120911 |
| GTGHNGSQPGMGGNG | ASSSGQGAGEQY | 0.6894159 |
| GYAAASASAS | ASSSGQGAGEQY | 0.5839126 |

| | | |
|---|---|---|
| HQVDVAAVQQG | ASSSGQGAGEQY | 0.6876197 |
| IGAAIAHT | ASSSGQGAGEQY | 0.48149094 |
| IGMIEVAQL | ASSSGQGAGEQY | 0.69803727 |
| IRLAPEHTSAMD | ASSSGQGAGEQY | 0.6687963 |
| IYNVSVL | ASSSGQGAGEQY | 0.54944515 |
| KQGFLGM | ASSSGQGAGEQY | 0.6485459 |
| LESVITQL | ASSSGQGAGEQY | 0.7022995 |
| LGLGALVVV | ASSSGQGAGEQY | 0.69728976 |
| LIGNGGNGGSGGI | ASSSGQGAGEQY | 0.37259448 |
| LNAINAPTL | ASSSGQGAGEQY | 0.6264775 |
| LQLLNTGV | ASSSGQGAGEQY | 0.38396358 |
| LQTSIEF | ASSSGQGAGEQY | 0.57077813 |
| MSFAVSEL | ASSSGQGAGEQY | 0.45158336 |
| NTGALGL | ASSSGQGAGEQY | 0.6815178 |
| PPGAGGAGGAGTT | ASSSGQGAGEQY | 0.57941353 |
| QLVGLGI | ASSSGQGAGEQY | 0.6838581 |
| RFLEQVHQL | ASSSGQGAGEQY | 0.6644239 |
| RGVGYVLR | ASSSGQGAGEQY | 0.70477515 |
| SATGVFAGVI | ASSSGQGAGEQY | 0.5190195 |
| SLPAITLP | ASSSGQGAGEQY | 0.63775194 |
| STALAFTV | ASSSGQGAGEQY | 0.3920614 |
| SVLTVGLGK | ASSSGQGAGEQY | 0.48644683 |
| TALAGAGSGI | ASSSGQGAGEQY | 0.5182252 |
| TGLPFTGI | ASSSGQGAGEQY | 0.6570572 |
| VAEHVTV | ASSSGQGAGEQY | 0.3443558 |
| VDIVLNSL | ASSSGQGAGEQY | 0.7080045 |
| VLTGGGAVL | ASSSGQGAGEQY | 0.6563437 |
| VNVGLAYGI | ASSSGQGAGEQY | 0.68739897 |
| VSSAAVGAV | ASSSGQGAGEQY | 0.5717703 |
| VTSVYEAGV | ASSSGQGAGEQY | 0.6051929 |
| WVALDRGA | ASSSGQGAGEQY | 0.61811596 |
| YLVGSVVI | ASSSGQGAGEQY | 0.6840377 |
| GPPGTGKTLL | ASSSGQGAGEQY | 0.5337583 |
| GVGMTKF | ASSSGQGAGEQY | 0.3519595 |
| IGIGGSDL | ASSSGQGAGEQY | 0.5903199 |
| IGTLLAEL | ASSSGQGAGEQY | 0.48711255 |
| IYTSGTT | ASSSGQGAGEQY | 0.7035873 |
| LSFPAGV | ASSSGQGAGEQY | 0.69647944 |
| LYTSGTT | ASSSGQGAGEQY | 0.67025506 |
| VGAFPFGF | ASSSGQGAGEQY | 0.6886102 |
| VGAGTVEF | ASSSGQGAGEQY | 0.31359562 |
| VGKSTLLL | ASSSGQGAGEQY | 0.6046028 |
| AALAHAGGGIV | ASSSGQGAGEQY | 0.7100929 |
| AALVAVSLL | ASSSGQGAGEQY | 0.6735302 |
| AAVSRAVAAL | ASSSGQGAGEQY | 0.60445267 |
| AGGSSGIGGF | ASSSGQGAGEQY | 0.70349956 |
| AGLTVANANG | ASSSGQGAGEQY | 0.58515847 |
| AGLVGAVL | ASSSGQGAGEQY | 0.61483836 |
| ALSQAFTV | ASSSGQGAGEQY | 0.656523 |
| ALTFSVSL | ASSSGQGAGEQY | 0.6577039 |
| ALTGAALVAL | ASSSGQGAGEQY | 0.62217605 |

| | | |
|---|---|---|
| ASFSIGAGVL | ASSSGQGAGEQY | 0.51041317 |
| DGSVYKI | ASSSGQGAGEQY | 0.64112794 |
| EQLAEIV | ASSSGQGAGEQY | 0.7156031 |
| FAVSELTVM | ASSSGQGAGEQY | 0.6257881 |
| GAAPLAAGL | ASSSGQGAGEQY | 0.7006466 |
| GADGGDGGSGGAGG | ASSSGQGAGEQY | 0.67580914 |
| GFGLDTGGAGG | ASSSGQGAGEQY | 0.6681864 |
| GLASGGLL | ASSSGQGAGEQY | 0.5283424 |
| GRIAGAAVLL | ASSSGQGAGEQY | 0.6055754 |
| GSAGSLNTSF | ASSSGQGAGEQY | 0.6253072 |
| GTGQIGI | ASSSGQGAGEQY | 0.57988006 |
| GTLIALIL | ASSSGQGAGEQY | 0.5861454 |
| IAATMGFVVG | ASSSGQGAGEQY | 0.57415503 |
| IFAQALL | ASSSGQGAGEQY | 0.71652305 |
| IGAAVALGF | ASSSGQGAGEQY | 0.681095 |
| IGLTPLAL | ASSSGQGAGEQY | 0.68810713 |
| ILGSNTSTL | ASSSGQGAGEQY | 0.5844496 |
| IVFSLTVL | ASSSGQGAGEQY | 0.6480886 |
| LASKATGF | ASSSGQGAGEQY | 0.68645704 |
| LASVALIL | ASSSGQGAGEQY | 0.41909778 |
| LFAVGLTL | ASSSGQGAGEQY | 0.70664465 |
| LFNGGIGTYI | ASSSGQGAGEQY | 0.5755244 |
| LTLGAGII | ASSSGQGAGEQY | 0.6413378 |
| LVAAALGDI | ASSSGQGAGEQY | 0.6911048 |
| NTAFVQLGI | ASSSGQGAGEQY | 0.7012087 |
| QLFSGAEL | ASSSGQGAGEQY | 0.5907254 |
| SAATFITL | ASSSGQGAGEQY | 0.6076537 |
| SNYGLANLGV | ASSSGQGAGEQY | 0.57915556 |
| SYGGAYAVM | ASSSGQGAGEQY | 0.69031596 |
| TTGVGGLTLG | ASSSGQGAGEQY | 0.7006883 |
| TTQTIYI | ASSSGQGAGEQY | 0.6567049 |
| VAAGAGLGAVI | ASSSGQGAGEQY | 0.7091421 |
| VAAVVTLL | ASSSGQGAGEQY | 0.67649865 |
| VGFLGGLI | ASSSGQGAGEQY | 0.40006015 |
| VGVIPILL | ASSSGQGAGEQY | 0.61676323 |
| VIFASSLI | ASSSGQGAGEQY | 0.69613975 |
| VKGGDGGAAAG | ASSSGQGAGEQY | 0.6991999 |
| VTKTFTNI | ASSSGQGAGEQY | 0.7071799 |
| VVAAAILL | ASSSGQGAGEQY | 0.6596415 |
| YAALDVEL | ASSSGQGAGEQY | 0.5982615 |
| LASVALLL | ASSSGQGAGEQY | 0.7061235 |
| LDAVHRV | ASSSGQGAGEQY | 0.71117616 |
| QGLSSSIF | ASSSGQGAGEQY | 0.6931766 |
| RGTFANIRL | ASSSGQGAGEQY | 0.6174474 |
| RQLWWGHRI | ASSSGQGAGEQY | 0.55639035 |
| YGPPGTGKTLL | ASSSGQGAGEQY | 0.6510925 |
| AALVGAPANVA | ASSSGQGAGEQY | 0.7004943 |
| AATFRRTE | ASSSGQGAGEQY | 0.4484725 |
| AGALGGGI | ASSSGQGAGEQY | 0.6657181 |
| AGGAGGAGGIGAG | ASSSGQGAGEQY | 0.6720027 |
| AGSGALGL | ASSSGQGAGEQY | 0.38361868 |

| | | |
|---|---|---|
| AQIADQI | ASSSGQGAGEQY | 0.65046537 |
| ASAVAVTYL | ASSSGQGAGEQY | 0.70361495 |
| ASTVLGGTGG | ASSSGQGAGEQY | 0.60089 |
| AVGGEPAK | ASSSGQGAGEQY | 0.5692927 |
| CEQGTFGGTC | ASSSGQGAGEQY | 0.69985837 |
| DEGKAPAG | ASSSGQGAGEQY | 0.7144778 |
| DTVVFPAL | ASSSGQGAGEQY | 0.6976079 |
| FEGITMAG | ASSSGQGAGEQY | 0.41300932 |
| FPLVVQD | ASSSGQGAGEQY | 0.32385638 |
| GAFTTPQ | ASSSGQGAGEQY | 0.59862715 |
| GFGGFGTAGGVGGA | ASSSGQGAGEQY | 0.6879171 |
| GFLAQFL | ASSSGQGAGEQY | 0.6204946 |
| GGAGGAGGNAGVA | ASSSGQGAGEQY | 0.58374894 |
| GGVGTAGAGGAGGAGG | ASSSGQGAGEQY | 0.67240644 |
| GNGGGGDAGNAGSGGN | ASSSGQGAGEQY | 0.70124894 |
| GTGGAGGAGNDAGS | ASSSGQGAGEQY | 0.46232593 |
| GTTNTGWFNS | ASSSGQGAGEQY | 0.4815231 |
| IFMGPLFTL | ASSSGQGAGEQY | 0.6506599 |
| ILAHAIEAI | ASSSGQGAGEQY | 0.682642 |
| IRSAIREAL | ASSSGQGAGEQY | 0.5717033 |
| ITFAGGG | ASSSGQGAGEQY | 0.6300668 |
| ITGMSGHQEV | ASSSGQGAGEQY | 0.55683017 |
| IVAGLACTL | ASSSGQGAGEQY | 0.52989453 |
| IVGGGLAGA | ASSSGQGAGEQY | 0.68065023 |
| IVTGVLNI | ASSSGQGAGEQY | 0.54908687 |
| LAQQGCI | ASSSGQGAGEQY | 0.7054981 |
| LCLHVEN | ASSSGQGAGEQY | 0.5199417 |
| LDAISDGS | ASSSGQGAGEQY | 0.686115 |
| LGGFAAML | ASSSGQGAGEQY | 0.6728059 |
| LGGKNPMI | ASSSGQGAGEQY | 0.6546677 |
| LGPKGVAAIL | ASSSGQGAGEQY | 0.60107195 |
| LMVGVSI | ASSSGQGAGEQY | 0.571169 |
| LVGAAFVL | ASSSGQGAGEQY | 0.45228237 |
| LVLAVAPL | ASSSGQGAGEQY | 0.35746416 |
| MAAIAVIDAA | ASSSGQGAGEQY | 0.70067513 |
| NNGINAELQ | ASSSGQGAGEQY | 0.5694663 |
| RGEPLVLVHG | ASSSGQGAGEQY | 0.70040476 |
| RGGAGGAAGLI | ASSSGQGAGEQY | 0.6782657 |
| RSYRMHPAV | ASSSGQGAGEQY | 0.54608315 |
| SAAHLLAGQLA | ASSSGQGAGEQY | 0.67977977 |
| SAAVGSY | ASSSGQGAGEQY | 0.546942 |
| SPHAVLHRI | ASSSGQGAGEQY | 0.46132004 |
| SQIAKVL | ASSSGQGAGEQY | 0.6814813 |
| TALGHLVP | ASSSGQGAGEQY | 0.6808162 |
| TAPTTSGAA | ASSSGQGAGEQY | 0.50262684 |
| TDGDDGNGGSPGT | ASSSGQGAGEQY | 0.65287554 |
| TDGLGEN | ASSSGQGAGEQY | 0.66040194 |
| TGGPPLPGQA | ASSSGQGAGEQY | 0.5191829 |
| TIPAITI | ASSSGQGAGEQY | 0.59837025 |
| TIPGFTI | ASSSGQGAGEQY | 0.70194507 |
| TSIASGALA | ASSSGQGAGEQY | 0.6371602 |

| | | |
|---|---|---|
| TVTLPGAAI | ASSSGQGAGEQY | 0.5228957 |
| VASGLSAALP | ASSSGQGAGEQY | 0.6877848 |
| VGAGVIGI | ASSSGQGAGEQY | 0.53818357 |
| VGPKRFGV | ASSSGQGAGEQY | 0.6820142 |
| VNVAFPD | ASSSGQGAGEQY | 0.6484165 |
| VSMTNSGVS | ASSSGQGAGEQY | 0.60724586 |
| VTDVLGDLSV | ASSSGQGAGEQY | 0.68980545 |
| VVVVGAGFAGL | ASSSGQGAGEQY | 0.514397 |
| VYTVGTVV | ASSSGQGAGEQY | 0.5441299 |
| WTTFSGP | ASSSGQGAGEQY | 0.6077328 |
| YDELVEAF | ASSSGQGAGEQY | 0.65292287 |
| YGTGGAGGNGG | ASSSGQGAGEQY | 0.693426 |
| YTNVMAV | ASSSGQGAGEQY | 0.5588076 |
| FLEQVHQL | ASSSGQGAGEQY | 0.6543527 |
| GLSLLAVAV | ASSSGQGAGEQY | 0.55976075 |
| LEQVHQL | ASSSGQGAGEQY | 0.49581712 |
| LNAAGAI | ASSSGQGAGEQY | 0.58300674 |
| PAVHASL | ASSSGQGAGEQY | 0.5946301 |
| TELEDDA | ASSSGQGAGEQY | 0.51017463 |
| TPPGPPN | ASSSGQGAGEQY | 0.6692433 |
| TPPPPPPP | ASSSGQGAGEQY | 0.5331198 |
| TQSPALI | ASSMISGGPSQNTLY | 0.66252613 |
| EFAAAFPQ | ASSMISGGPSQNTLY | 0.70234156 |
| LLSGNLDVL | ASSMISGGPSQNTLY | 0.62290686 |
| GPANLGNYNI | ASSMISGGPSQNTLY | 0.679548 |
| TLGPTLP | ASSMISGGPSQNTLY | 0.6855759 |
| VYAYSSCCK | ASSMISGGPSQNTLY | 0.679441 |
| APAPAPAPNI | ASSMISGGPSQNTLY | 0.61211497 |
| PGGPVPIP | ASSMISGGPSQNTLY | 0.56997985 |
| FLTVALEP | ASSMISGGPSQNTLY | 0.70127845 |
| SQYFGANL | ASSMISGGPSQNTLY | 0.6947733 |
| GGVKPAA | ASSMISGGPSQNTLY | 0.6931255 |
| GGTGGAGGAGGW | ASSMISGGPSQNTLY | 0.68987155 |
| GVGGVAGLGGAGG | ASSMISGGPSQNTLY | 0.67254806 |
| VLLVLATTI | ASSMISGGPSQNTLY | 0.6893674 |
| IGIFAAML | ASSMISGGPSQNTLY | 0.64131975 |
| AVGDLID | ASSMISGGPSQNTLY | 0.44422784 |
| NISPLPL | ASSMISGGPSQNTLY | 0.63320684 |
| VTAVFVAI | ASSMISGGPSQNTLY | 0.58771956 |
| SGDDNDAAL | ASSMISGGPSQNTLY | 0.54461175 |
| AVAALAHS | ASSMISGGPSQNTLY | 0.4319454 |
| GGVTVMATP | ASSMISGGPSQNTLY | 0.7005698 |
| AGLAANVRD | ASSMISGGPSQNTLY | 0.7149833 |
| TNRPDLIDPALLRPG | ASSMISGGPSQNTLY | 0.56397617 |
| LANLGVTAL | ASSMISGGPSQNTLY | 0.70613146 |
| TGERIEL | ASSMISGGPSQNTLY | 0.44119212 |
| NGEELAA | ASSMISGGPSQNTLY | 0.69066155 |
| VLSLVWG | ASSMISGGPSQNTLY | 0.69934416 |
| IGYALWL | ASSMISGGPSQNTLY | 0.5722775 |
| LSLLLAPL | ASSMISGGPSQNTLY | 0.6910638 |
| TLDEALK | ASSMISGGPSQNTLY | 0.5213361 |

| | | |
|---|---|---|
| TDGAGTAEGG | ASSMISGGPSQNTLY | 0.6113919 |
| NAPGGAGGNGG | ASSMISGGPSQNTLY | 0.7040551 |
| VSVFMGMGGNF | ASSMISGGPSQNTLY | 0.6016064 |
| LASYLAAL | ASSMISGGPSQNTLY | 0.6527201 |
| LVGGTPAL | ASSMISGGPSQNTLY | 0.6216022 |
| VRNADVL | ASSMISGGPSQNTLY | 0.5527803 |
| TCSSPFLVSAM | ASSMISGGPSQNTLY | 0.6531592 |
| VHITDAE | ASSMISGGPSQNTLY | 0.7018455 |
| EYAAPSF | ASSMISGGPSQNTLY | 0.5104675 |
| VVDAFHGGL | ASSMISGGPSQNTLY | 0.70953214 |
| ASVVDPQ | ASSMISGGPSQNTLY | 0.5359206 |
| SYNEFGGLS | ASSMISGGPSQNTLY | 0.7085757 |
| DPDGSVY | ASSMISGGPSQNTLY | 0.69366276 |
| MVSTIHATE | ASSMISGGPSQNTLY | 0.66657853 |
| LAAGVADPV | ASSMISGGPSQNTLY | 0.60415965 |
| TKEQIAA | ASSMISGGPSQNTLY | 0.700281 |
| SLASPAAL | ASSMISGGPSQNTLY | 0.5984686 |
| IGLVFYGV | ASSMISGGPSQNTLY | 0.6882003 |
| FLLDASNL | ASSMISGGPSQNTLY | 0.7002278 |
| YQVGNFP | ASSMISGGPSQNTLY | 0.5751351 |
| ESLLSQST | ASSMISGGPSQNTLY | 0.8240973 |
| LFTVGFAAVG | ASSMISGGPSQNTLY | 0.7724103 |
| ATPPTPT | ASSMISGGPSQNTLY | 0.7625116 |
| TDLATLG | ASSMISGGPSQNTLY | 0.59108603 |
| SAGGTGR | ASSMISGGPSQNTLY | 0.32425624 |
| GFSNSGS | ASSMISGGPSQNTLY | 0.7174212 |
| GGNGGSGGTSV | ASSMISGGPSQNTLY | 0.48856598 |
| GTSVATGGAGN | ASSMISGGPSQNTLY | 0.7907447 |
| NGGNGGSGGTS | ASSMISGGPSQNTLY | 0.7959738 |
| AVVKFPH | ASSMISGGPSQNTLY | 0.62206435 |
| TPPPPPP | ASSMISGGPSQNTLY | 0.81739914 |
| AAPAAAAAPA | ASSMISGGPSQNTLY | 0.8206851 |
| IALNEDL | ASSMISGGPSQNTLY | 0.36790377 |
| IVNVSSM | ASSMISGGPSQNTLY | 0.74170053 |
| TGIPPAPR | ASSMISGGPSQNTLY | 0.6119982 |
| FDLGGGTF | ASSMISGGPSQNTLY | 0.71839035 |
| FDLGGGTFDVS | ASSMISGGPSQNTLY | 0.34984672 |
| NEPTAAA | ASSMISGGPSQNTLY | 0.78710747 |
| NEPTAAALA | ASSMISGGPSQNTLY | 0.68827593 |
| GSIINISSI | ASSMISGGPSQNTLY | 0.17956161 |
| VAPGNAGTA | ASSMISGGPSQNTLY | 0.75233173 |
| TGGGASLE | ASSMISGGPSQNTLY | 0.74083304 |
| VAVYNHYKRI | ASSMISGGPSQNTLY | 0.6526992 |
| TDPAPATT | ASSMISGGPSQNTLY | 0.80948013 |
| TNRPDILDPALLRPG | ASSMISGGPSQNTLY | 0.5416293 |
| LTTTTTS | ASSMISGGPSQNTLY | 0.53684723 |
| PTGSGKTL | ASSMISGGPSQNTLY | 0.80291045 |
| TGSGKTL | ASSMISGGPSQNTLY | 0.8178246 |
| TIGHVDHGK | ASSMISGGPSQNTLY | 0.41372252 |
| TIGHVDHGKTT | ASSMISGGPSQNTLY | 0.7762946 |
| GHVDHGKTT | ASSMISGGPSQNTLY | 0.8218243 |

| | | |
|---|---|---|
| AQFEHTLL | ASSMISGGPSQNTLY | 0.73660797 |
| IVNQDEL | ASSMISGGPSQNTLY | 0.8103827 |
| TNGSQFF | ASSMISGGPSQNTLY | 0.8197204 |
| TGKGIVE | ASSMISGGPSQNTLY | 0.619336 |
| LISNASDA | ASSMISGGPSQNTLY | 0.8119587 |
| AELRTLGELTKIAKAHGAQ | ASSMISGGPSQNTLY | 0.8201273 |
| APNAVAAAEPSW | ASSMISGGPSQNTLY | 0.6658623 |
| APQVAGAATAVL | ASSMISGGPSQNTLY | 0.75046325 |
| ARRDPAAAAAILPTDA | ASSMISGGPSQNTLY | 0.53767484 |
| DAAARDVLAEAGLAEYF | ASSMISGGPSQNTLY | 0.73725474 |
| DDQIVLSAKANLAKVEQ | ASSMISGGPSQNTLY | 0.23863384 |
| DELLPDTGEDSDEDGH | ASSMISGGPSQNTLY | 0.79095757 |
| ELPEAPANRATVPQD | ASSMISGGPSQNTLY | 0.80493104 |
| FGQSKIERVTGIG | ASSMISGGPSQNTLY | 0.708838 |
| GAGGTGSTAGGAGGAGGAGGLY | ASSMISGGPSQNTLY | 0.5307154 |
| GHGGTGAAGGNGGTGGAGGSGI | ASSMISGGPSQNTLY | 0.5248134 |
| GNGGNGGAGGEGGPGGAGGAGA | ASSMISGGPSQNTLY | 0.72858113 |
| GSTAGNGGNGGNATNGTIAGQP | ASSMISGGPSQNTLY | 0.6630451 |
| GTGGDGGNGGNGGTGG | ASSMISGGPSQNTLY | 0.25795156 |
| GVRRPAGIPADAPV | ASSMISGGPSQNTLY | 0.7454007 |
| GYRQVTDAHLLTLARR | ASSMISGGPSQNTLY | 0.7994461 |
| IAKMGQDGHDRG | ASSMISGGPSQNTLY | 0.8015903 |
| KAEPSIEIQYT | ASSMISGGPSQNTLY | 0.4232909 |
| KPVRIRMAADEDTVAVT | ASSMISGGPSQNTLY | 0.8243841 |
| MPPRIAGMRLLVIKPEP | ASSMISGGPSQNTLY | 0.8052033 |
| MKFYAVALFRSRGVPPTR | ASSMISGGPSQNTLY | 0.815585 |
| NAGGFGILGAVAHSP | ASSMISGGPSQNTLY | 0.742253 |
| NEGREITLAMAEM | ASSMISGGPSQNTLY | 0.5914899 |
| NELVEWSHPLGHRWRWE | ASSMISGGPSQNTLY | 0.8151407 |
| NLGAGGTGGNGGNPG | ASSMISGGPSQNTLY | 0.72219175 |
| NTEWPKWETVMAADPVD | ASSMISGGPSQNTLY | 0.61544204 |
| PASAAPAETQP | ASSMISGGPSQNTLY | 0.81298345 |
| QQSSDGKNGVRAAEITGE | ASSMISGGPSQNTLY | 0.6429074 |
| RAPDAVTTSWWKE | ASSMISGGPSQNTLY | 0.63738 |
| SGFGNVGAHTSGWFNQ | ASSMISGGPSQNTLY | 0.7406857 |
| SGVAGAVGQAASVGGLKV | ASSMISGGPSQNTLY | 0.8087534 |
| SNGGAGTGSAGGNGGT | ASSMISGGPSQNTLY | 0.5864225 |
| TGGAGGEGGAAG | ASSMISGGPSQNTLY | 0.45428652 |
| TGGDGGAAGAGGAPGHG | ASSMISGGPSQNTLY | 0.56764823 |
| TMPPLLAAIANAGRHGV | ASSMISGGPSQNTLY | 0.7397511 |
| TSSTAGNSGGA | ASSMISGGPSQNTLY | 0.81696624 |
| VPATVPVGILC | ASSMISGGPSQNTLY | 0.76636875 |
| VQRPDSGGVVLTGRIS | ASSMISGGPSQNTLY | 0.7250322 |
| WTPNPVPATKGVPAGLVQAT | ASSMISGGPSQNTLY | 0.7918413 |
| YALKRALAKFSDRFDVVII | ASSMISGGPSQNTLY | 0.7723603 |
| YLIGSVIV | ASSMISGGPSQNTLY | 0.76044774 |
| YGQLVAVID | ASSMISGGPSQNTLY | 0.81848085 |
| VTVGGMTLPA | ASSMISGGPSQNTLY | 0.67985237 |
| VLALPAV | ASSMISGGPSQNTLY | 0.8096435 |
| VIPPTLNYET | ASSMISGGPSQNTLY | 0.79928136 |
| VGMGGMHPSSG | ASSMISGGPSQNTLY | 0.80005807 |

| | | |
|---|---|---|
| VGLAGGAATA | ASSMISGGPSQNTLY | 0.813606 |
| VGGAGGAGGSGP | ASSMISGGPSQNTLY | 0.7726996 |
| VFAAFMV | ASSMISGGPSQNTLY | 0.7543193 |
| TLYDKLR | ASSMISGGPSQNTLY | 0.28929216 |
| TLAVFTVL | ASSMISGGPSQNTLY | 0.70006114 |
| TGSVQPARV | ASSMISGGPSQNTLY | 0.7940535 |
| TGGGAGSGGT | ASSMISGGPSQNTLY | 0.7848454 |
| SHLETSL | ASSMISGGPSQNTLY | 0.80154085 |
| SFTTGMT | ASSMISGGPSQNTLY | 0.7687795 |
| QVEVAKRMGVRQPTVS | ASSMISGGPSQNTLY | 0.6172633 |
| QGSGGPALAM | ASSMISGGPSQNTLY | 0.8205334 |
| MLAPSNP | ASSMISGGPSQNTLY | 0.8144518 |
| LVNNAAI | ASSMISGGPSQNTLY | 0.39092213 |
| LTGEINTLA | ASSMISGGPSQNTLY | 0.72268975 |
| LSIFGPV | ASSMISGGPSQNTLY | 0.82880783 |
| LLALLANL | ASSMISGGPSQNTLY | 0.79062647 |
| LISNASDALDKLR | ASSMISGGPSQNTLY | 0.70067424 |
| LINEAVP | ASSMISGGPSQNTLY | 0.63446236 |
| LGYAFSGL | ASSMISGGPSQNTLY | 0.81954753 |
| LDFVAVAL | ASSMISGGPSQNTLY | 0.59373516 |
| LAQPGAE | ASSMISGGPSQNTLY | 0.82376206 |
| ISNLGAVTGF | ASSMISGGPSQNTLY | 0.22436999 |
| HQEDPGRI | ASSMISGGPSQNTLY | 0.6096426 |
| GTPGEPTP | ASSMISGGPSQNTLY | 0.48481333 |
| GFFNSGS | ASSMISGGPSQNTLY | 0.25654122 |
| GAGGGGLTAGAN | ASSMISGGPSQNTLY | 0.78729934 |
| FVDALVNM | ASSMISGGPSQNTLY | 0.6117291 |
| FLTVAPEL | ASSMISGGPSQNTLY | 0.7982362 |
| FETLTAL | ASSMISGGPSQNTLY | 0.7346791 |
| DVINPIEL | ASSMISGGPSQNTLY | 0.8116518 |
| APVSGAPV | ASSMISGGPSQNTLY | 0.78925455 |
| ALVGGLAW | ASSMISGGPSQNTLY | 0.7731087 |
| ALAAAAATAI | ASSMISGGPSQNTLY | 0.52559614 |
| AFGLGLLL | ASSMISGGPSQNTLY | 0.77526677 |
| AFFAAASL | ASSMISGGPSQNTLY | 0.8229619 |
| AAAALPAFSPP | ASSMISGGPSQNTLY | 0.76618075 |
| VVLAPAL | ASSMISGGPSQNTLY | 0.31548735 |
| TIGHVDHGKTTL | ASSMISGGPSQNTLY | 0.7370764 |
| LISNASDALDK | ASSMISGGPSQNTLY | 0.67613524 |
| LISNASDAL | ASSMISGGPSQNTLY | 0.23930086 |
| ISNASDALDK | ASSMISGGPSQNTLY | 0.20820898 |
| NQAETLVYQTE | ASSMISGGPSQNTLY | 0.6199442 |
| TGGTGGTAGAG | ASSMISGGPSQNTLY | 0.6864397 |
| TGSTAGGAGGA | ASSMISGGPSQNTLY | 0.81180555 |
| DPLAVPVAAGPV | ASSMISGGPSQNTLY | 0.8112751 |
| GGTGGTPGNGGN | ASSMISGGPSQNTLY | 0.71506196 |
| PAGLAGVDPATL | ASSMISGGPSQNTLY | 0.8085755 |
| GNNLVVEAPHAGASV | ASSMISGGPSQNTLY | 0.7594466 |
| KVTRIMSASTLEQPA | ASSMISGGPSQNTLY | 0.15198345 |
| LGFALPIALPRRTVV | ASSMISGGPSQNTLY | 0.6823518 |
| TRLGNLTARKLVVLV | ASSMISGGPSQNTLY | 0.38523647 |

| | | |
|---|---|---|
| WSQQRRLIYFPSAGP | ASSMISGGPSQNTLY | 0.8173736 |
| GPNGNPDPMAAAVDIR | ASSMISGGPSQNTLY | 0.6372075 |
| TQRCVSVADLRAAASR | ASSMISGGPSQNTLY | 0.6558453 |
| QFVLDVAEHPVEVLTETV | ASSMISGGPSQNTLY | 0.7532774 |
| GGAGGTGGAGGNGGTQAGDG | ASSMISGGPSQNTLY | 0.7786639 |
| PLLATLRAIAVDVDAKPAQV | ASSMISGGPSQNTLY | 0.7590212 |
| APKFRTGGYMIDFWGVGYQVAK | ASSMISGGPSQNTLY | 0.57877004 |
| GAGGAGGAGGAAGGPGGTGGTG | ASSMISGGPSQNTLY | 0.19114296 |
| IGALGLLATDVAPAGSVAFGVA | ASSMISGGPSQNTLY | 0.78075516 |
| AGGAGGIGGNGGAGGAGGVLMG | ASSMISGGPSQNTLY | 0.43578324 |
| ADELIGLL | ASSMISGGPSQNTLY | 0.46442616 |
| ADRALKA | ASSMISGGPSQNTLY | 0.3358798 |
| AGAVGQVGGAGGA | ASSMISGGPSQNTLY | 0.23891445 |
| AGGGGGLGGAP | ASSMISGGPSQNTLY | 0.33152226 |
| AGGLLYGNGGAG | ASSMISGGPSQNTLY | 0.8035179 |
| AGLESIGL | ASSMISGGPSQNTLY | 0.72978574 |
| AGLKTGII | ASSMISGGPSQNTLY | 0.7165238 |
| ASLAALLL | ASSMISGGPSQNTLY | 0.8236582 |
| ASVINISSI | ASSMISGGPSQNTLY | 0.7766547 |
| ATPPGPA | ASSMISGGPSQNTLY | 0.7469264 |
| ATVASALV | ASSMISGGPSQNTLY | 0.5386638 |
| ATVASLIL | ASSMISGGPSQNTLY | 0.57265997 |
| AVNGPVLIRH | ASSMISGGPSQNTLY | 0.5169631 |
| DGAAGAVGI | ASSMISGGPSQNTLY | 0.5115669 |
| DPLLHGI | ASSMISGGPSQNTLY | 0.82870567 |
| ELGAALGAL | ASSMISGGPSQNTLY | 0.61952037 |
| ELGRTQAV | ASSMISGGPSQNTLY | 0.43965608 |
| FVAGGYLS | ASSMISGGPSQNTLY | 0.71902716 |
| GASGGAGGAGGW | ASSMISGGPSQNTLY | 0.82271945 |
| GGAGAGGDGGAGGI | ASSMISGGPSQNTLY | 0.83080983 |
| GTGHNGSQPGMGGNG | ASSMISGGPSQNTLY | 0.60405236 |
| GYAAASASAS | ASSMISGGPSQNTLY | 0.7182379 |
| HQVDVAAVQQG | ASSMISGGPSQNTLY | 0.7038723 |
| IGAAIAHT | ASSMISGGPSQNTLY | 0.76303595 |
| IGMIEVAQL | ASSMISGGPSQNTLY | 0.6697675 |
| IRLAPEHTSAMD | ASSMISGGPSQNTLY | 0.40076184 |
| IYNVSVL | ASSMISGGPSQNTLY | 0.7974359 |
| KQGFLGM | ASSMISGGPSQNTLY | 0.826645 |
| LESVITQL | ASSMISGGPSQNTLY | 0.68873584 |
| LGLGALVVV | ASSMISGGPSQNTLY | 0.36866686 |
| LIGNGGNGGSGGI | ASSMISGGPSQNTLY | 0.7430375 |
| LNAINAPTL | ASSMISGGPSQNTLY | 0.613698 |
| LQLLNTGV | ASSMISGGPSQNTLY | 0.67218685 |
| LQTSIEF | ASSMISGGPSQNTLY | 0.72190845 |
| MSFAVSEL | ASSMISGGPSQNTLY | 0.42454097 |
| NTGALGL | ASSMISGGPSQNTLY | 0.7047448 |
| PPGAGGAGGAGTT | ASSMISGGPSQNTLY | 0.82727355 |
| QLVGLGI | ASSMISGGPSQNTLY | 0.7163386 |
| RFLEQVHQL | ASSMISGGPSQNTLY | 0.7882722 |
| RGVGYVLR | ASSMISGGPSQNTLY | 0.81383926 |
| SATGVFAGVI | ASSMISGGPSQNTLY | 0.7924133 |

| | | |
|---|---|---|
| SLPAITLP | ASSMISGGPSQNTLY | 0.81900156 |
| STALAFTV | ASSMISGGPSQNTLY | 0.8112483 |
| SVLTVGLGK | ASSMISGGPSQNTLY | 0.7251141 |
| TALAGAGSGI | ASSMISGGPSQNTLY | 0.7587718 |
| TGLPFTGI | ASSMISGGPSQNTLY | 0.77972 |
| VAEHVTV | ASSMISGGPSQNTLY | 0.7577013 |
| VDIVLNSL | ASSMISGGPSQNTLY | 0.7463935 |
| VLTGGGAVL | ASSMISGGPSQNTLY | 0.67989546 |
| VNVGLAYGI | ASSMISGGPSQNTLY | 0.8101555 |
| VSSAAVGAV | ASSMISGGPSQNTLY | 0.7583531 |
| VTSVYEAGV | ASSMISGGPSQNTLY | 0.80196506 |
| WVALDRGA | ASSMISGGPSQNTLY | 0.36273998 |
| YLVGSVVI | ASSMISGGPSQNTLY | 0.41966474 |
| GPPGTGKTLL | ASSMISGGPSQNTLY | 0.8087319 |
| GVGMTKF | ASSMISGGPSQNTLY | 0.19406123 |
| IGIGGSDL | ASSMISGGPSQNTLY | 0.23022619 |
| IGTLLAEL | ASSMISGGPSQNTLY | 0.7592231 |
| IYTSGTT | ASSMISGGPSQNTLY | 0.82134765 |
| LSFPAGV | ASSMISGGPSQNTLY | 0.1856365 |
| LYTSGTT | ASSMISGGPSQNTLY | 0.51207614 |
| VGAFPFGF | ASSMISGGPSQNTLY | 0.46321452 |
| VGAGTVEF | ASSMISGGPSQNTLY | 0.77533776 |
| VGKSTLLL | ASSMISGGPSQNTLY | 0.73836213 |
| AALAHAGGGIV | ASSMISGGPSQNTLY | 0.3672994 |
| AALVAVSLL | ASSMISGGPSQNTLY | 0.79602706 |
| AAVSRAVAAL | ASSMISGGPSQNTLY | 0.4012881 |
| AGGSSGIGGF | ASSMISGGPSQNTLY | 0.5517492 |
| AGLTVANANG | ASSMISGGPSQNTLY | 0.8075044 |
| AGLVGAVL | ASSMISGGPSQNTLY | 0.82209903 |
| ALSQAFTV | ASSMISGGPSQNTLY | 0.5324484 |
| ALTFSVSL | ASSMISGGPSQNTLY | 0.80479604 |
| ALTGAALVAL | ASSMISGGPSQNTLY | 0.6823394 |
| ASFSIGAGVL | ASSMISGGPSQNTLY | 0.8298539 |
| DGSVYKI | ASSMISGGPSQNTLY | 0.6407149 |
| EQLAEIV | ASSMISGGPSQNTLY | 0.31570408 |
| FAVSELTVM | ASSMISGGPSQNTLY | 0.80314636 |
| GAAPLAAGL | ASSMISGGPSQNTLY | 0.7933769 |
| GADGGDGGSGGAGG | ASSMISGGPSQNTLY | 0.7254708 |
| GFGLDTGGAGG | ASSMISGGPSQNTLY | 0.8037372 |
| GLASGGLL | ASSMISGGPSQNTLY | 0.59845716 |
| GRIAGAAVLL | ASSMISGGPSQNTLY | 0.71797085 |
| GSAGSLNTSF | ASSMISGGPSQNTLY | 0.81640095 |
| GTGQIGI | ASSMISGGPSQNTLY | 0.8188044 |
| GTLIALIL | ASSMISGGPSQNTLY | 0.41208297 |
| IAATMGFVVG | ASSMISGGPSQNTLY | 0.80010265 |
| IFAQALL | ASSMISGGPSQNTLY | 0.7326808 |
| IGAAVALGF | ASSMISGGPSQNTLY | 0.6280997 |
| IGLTPLAL | ASSMISGGPSQNTLY | 0.8215342 |
| ILGSNTSTL | ASSMISGGPSQNTLY | 0.72611094 |
| IVFSLTVL | ASSMISGGPSQNTLY | 0.8048255 |
| LASKATGF | ASSMISGGPSQNTLY | 0.76497436 |

| | | |
|---|---|---|
| LASVALIL | ASSMISGGPSQNTLY | 0.8158649 |
| LFAVGLTL | ASSMISGGPSQNTLY | 0.40261292 |
| LFNGGIGTYI | ASSMISGGPSQNTLY | 0.7814288 |
| LTLGAGII | ASSMISGGPSQNTLY | 0.75430596 |
| LVAAALGDI | ASSMISGGPSQNTLY | 0.8134233 |
| NTAFVQLGI | ASSMISGGPSQNTLY | 0.70719016 |
| QLFSGAEL | ASSMISGGPSQNTLY | 0.49864447 |
| SAATFITL | ASSMISGGPSQNTLY | 0.76757264 |
| SNYGLANLGV | ASSMISGGPSQNTLY | 0.8177202 |
| SYGGAYAVM | ASSMISGGPSQNTLY | 0.76391983 |
| TTGVGGLTLG | ASSMISGGPSQNTLY | 0.7993456 |
| TTQTIYI | ASSMISGGPSQNTLY | 0.8253652 |
| VAAGAGLGAVI | ASSMISGGPSQNTLY | 0.51351255 |
| VAAVVTLL | ASSMISGGPSQNTLY | 0.69572294 |
| VGFLGGLI | ASSMISGGPSQNTLY | 0.80228347 |
| VGVIPILL | ASSMISGGPSQNTLY | 0.81850004 |
| VIFASSLI | ASSMISGGPSQNTLY | 0.7058364 |
| VKGGDGGAAAG | ASSMISGGPSQNTLY | 0.8245443 |
| VTKTFTNI | ASSMISGGPSQNTLY | 0.6831287 |
| VVAAAILL | ASSMISGGPSQNTLY | 0.8003345 |
| YAALDVEL | ASSMISGGPSQNTLY | 0.66132593 |
| LASVALLL | ASSMISGGPSQNTLY | 0.7774943 |
| LDAVHRV | ASSMISGGPSQNTLY | 0.56231594 |
| QGLSSSIF | ASSMISGGPSQNTLY | 0.79708856 |
| RGTFANIRL | ASSMISGGPSQNTLY | 0.81197786 |
| RQLWWGHRI | ASSMISGGPSQNTLY | 0.80628306 |
| YGPPGTGKTLL | ASSMISGGPSQNTLY | 0.21928683 |
| AALVGAPANVA | ASSMISGGPSQNTLY | 0.8058121 |
| AATFRRTE | ASSMISGGPSQNTLY | 0.79831225 |
| AGALGGGI | ASSMISGGPSQNTLY | 0.8284881 |
| AGGAGGAGGIGAG | ASSMISGGPSQNTLY | 0.7487179 |
| AGSGALGL | ASSMISGGPSQNTLY | 0.7922758 |
| AQIADQI | ASSMISGGPSQNTLY | 0.62274003 |
| ASAVAVTYL | ASSMISGGPSQNTLY | 0.67913795 |
| ASTVLGGTGG | ASSMISGGPSQNTLY | 0.5673262 |
| AVGGEPAK | ASSMISGGPSQNTLY | 0.6550682 |
| CEQGTFGGTC | ASSMISGGPSQNTLY | 0.72908473 |
| DEGKAPAG | ASSMISGGPSQNTLY | 0.6223761 |
| DTVVFPAL | ASSMISGGPSQNTLY | 0.8031036 |
| FEGITMAG | ASSMISGGPSQNTLY | 0.8182956 |
| FPLVVQD | ASSMISGGPSQNTLY | 0.81330967 |
| GAFTTPQ | ASSMISGGPSQNTLY | 0.81894433 |
| GFGGFGTAGGVGGA | ASSMISGGPSQNTLY | 0.82150376 |
| GFLAQFL | ASSMISGGPSQNTLY | 0.795043 |
| GGAGGAGGNAGVA | ASSMISGGPSQNTLY | 0.6025562 |
| GGVGTAGAGGAGGAGG | ASSMISGGPSQNTLY | 0.7338332 |
| GNGGGGDAGNAGSGGN | ASSMISGGPSQNTLY | 0.46665448 |
| GTGGAGGAGNDAGS | ASSMISGGPSQNTLY | 0.70874584 |
| GTTNTGWFNS | ASSMISGGPSQNTLY | 0.6202803 |
| IFMGPLFTL | ASSMISGGPSQNTLY | 0.7789681 |
| ILAHAIEAI | ASSMISGGPSQNTLY | 0.16683179 |

| | | |
|---|---|---|
| IRSAIREAL | ASSMISGGPSQNTLY | 0.7756779 |
| ITFAGGG | ASSMISGGPSQNTLY | 0.826509 |
| ITGMSGHQEV | ASSMISGGPSQNTLY | 0.6873357 |
| IVAGLACTL | ASSMISGGPSQNTLY | 0.77680707 |
| IVGGGLAGA | ASSMISGGPSQNTLY | 0.7884613 |
| IVTGVLNI | ASSMISGGPSQNTLY | 0.69088244 |
| LAQQGCI | ASSMISGGPSQNTLY | 0.80603564 |
| LCLHVEN | ASSMISGGPSQNTLY | 0.5773612 |
| LDAISDGS | ASSMISGGPSQNTLY | 0.7779746 |
| LGGFAAML | ASSMISGGPSQNTLY | 0.8172806 |
| LGGKNPMI | ASSMISGGPSQNTLY | 0.7601734 |
| LGPKGVAAIL | ASSMISGGPSQNTLY | 0.7881316 |
| LMVGVSI | ASSMISGGPSQNTLY | 0.7133635 |
| LVGAAFVL | ASSMISGGPSQNTLY | 0.31905615 |
| LVLAVAPL | ASSMISGGPSQNTLY | 0.79678595 |
| MAAIAVIDAA | ASSMISGGPSQNTLY | 0.16763286 |
| NNGINAELQ | ASSMISGGPSQNTLY | 0.76306915 |
| RGEPLVLVHG | ASSMISGGPSQNTLY | 0.45376647 |
| RGGAGGAAGLI | ASSMISGGPSQNTLY | 0.7547537 |
| RSYRMHPAV | ASSMISGGPSQNTLY | 0.7423396 |
| SAAHLLAGQLA | ASSMISGGPSQNTLY | 0.80209476 |
| SAAVGSY | ASSMISGGPSQNTLY | 0.5727361 |
| SPHAVLHRI | ASSMISGGPSQNTLY | 0.7228619 |
| SQIAKVL | ASSMISGGPSQNTLY | 0.6363678 |
| TALGHLVP | ASSMISGGPSQNTLY | 0.38076636 |
| TAPTTSGAA | ASSMISGGPSQNTLY | 0.80636525 |
| TDGDDGNGGSPGT | ASSMISGGPSQNTLY | 0.64668494 |
| TDGLGEN | ASSMISGGPSQNTLY | 0.7569592 |
| TGGPPLPGQA | ASSMISGGPSQNTLY | 0.80026454 |
| TIPAITI | ASSMISGGPSQNTLY | 0.754778 |
| TIPGFTI | ASSMISGGPSQNTLY | 0.3243575 |
| TSIASGALA | ASSMISGGPSQNTLY | 0.7571868 |
| TVTLPGAAI | ASSMISGGPSQNTLY | 0.742835 |
| VASGLSAALP | ASSMISGGPSQNTLY | 0.6515018 |
| VGAGVIGI | ASSMISGGPSQNTLY | 0.16970733 |
| VGPKRFGV | ASSMISGGPSQNTLY | 0.79310393 |
| VNVAFPD | ASSMISGGPSQNTLY | 0.6245496 |
| VSMTNSGVS | ASSMISGGPSQNTLY | 0.6726955 |
| VTDVLGDLSV | ASSMISGGPSQNTLY | 0.69150555 |
| VVVVGAGFAGL | ASSMISGGPSQNTLY | 0.80862755 |
| VYTVGTVV | ASSMISGGPSQNTLY | 0.6656236 |
| WTTFSGP | ASSMISGGPSQNTLY | 0.5319933 |
| YDELVEAF | ASSMISGGPSQNTLY | 0.8086675 |
| YGTGGAGGNGG | ASSMISGGPSQNTLY | 0.7724293 |
| YTNVMAV | ASSMISGGPSQNTLY | 0.81929195 |
| FLEQVHQL | ASSMISGGPSQNTLY | 0.7632244 |
| GLSLLAVAV | ASSMISGGPSQNTLY | 0.6900194 |
| LEQVHQL | ASSMISGGPSQNTLY | 0.79008335 |
| LNAAGAI | ASSMISGGPSQNTLY | 0.7250716 |
| PAVHASL | ASSMISGGPSQNTLY | 0.32107833 |
| TELEDDA | ASSMISGGPSQNTLY | 0.27010566 |

| | | |
|---|---|---|
| TPPGPPN | ASSMISGGPSQNTLY | 0.7648885 |
| TPPPPPPP | ASSMISGGPSQNTLY | 0.31087765 |
| TQSPALI | ASGERVGASQNTLY | 0.7371716 |
| EFAAAFPQ | ASGERVGASQNTLY | 0.809077 |
| LLSGNLDVL | ASGERVGASQNTLY | 0.75062215 |
| GPANLGNYNI | ASGERVGASQNTLY | 0.41789705 |
| TLGPTLP | ASGERVGASQNTLY | 0.7161617 |
| VYAYSSCCK | ASGERVGASQNTLY | 0.66213816 |
| APAPAPAPNI | ASGERVGASQNTLY | 0.78958607 |
| PGGPVPIP | ASGERVGASQNTLY | 0.7439297 |
| FLTVALEP | ASGERVGASQNTLY | 0.8217498 |
| SQYFGANL | ASGERVGASQNTLY | 0.8116569 |
| GGVKPAA | ASGERVGASQNTLY | 0.64997745 |
| GGTGGAGGAGGW | ASGERVGASQNTLY | 0.8083824 |
| GVGGVAGLGGAGG | ASGERVGASQNTLY | 0.6654167 |
| VLLVLATTI | ASGERVGASQNTLY | 0.7400665 |
| IGIFAAML | ASGERVGASQNTLY | 0.73681843 |
| AVGDLID | ASGERVGASQNTLY | 0.78232443 |
| NISPLPL | ASGERVGASQNTLY | 0.7411653 |
| VTAVFVAI | ASGERVGASQNTLY | 0.7035202 |
| SGDDNDAAL | ASGERVGASQNTLY | 0.36108616 |
| AVAALAHS | ASGERVGASQNTLY | 0.78880763 |
| GGVTVMATP | ASGERVGASQNTLY | 0.73849404 |
| AGLAANVRD | ASGERVGASQNTLY | 0.30839312 |
| TNRPDLIDPALLRPG | ASGERVGASQNTLY | 0.68073905 |
| LANLGVTAL | ASGERVGASQNTLY | 0.82745504 |
| TGERIEL | ASGERVGASQNTLY | 0.82410514 |
| NGEELAA | ASGERVGASQNTLY | 0.6135024 |
| VLSLVWG | ASGERVGASQNTLY | 0.5118722 |
| IGYALWL | ASGERVGASQNTLY | 0.7946441 |
| LSLLLAPL | ASGERVGASQNTLY | 0.38005942 |
| TLDEALK | ASGERVGASQNTLY | 0.75257564 |
| TDGAGTAEGG | ASGERVGASQNTLY | 0.7688339 |
| NAPGGAGGNGG | ASGERVGASQNTLY | 0.7710901 |
| VSVFMGMGGNF | ASGERVGASQNTLY | 0.6331811 |
| LASYLAAL | ASGERVGASQNTLY | 0.7812931 |
| LVGGTPAL | ASGERVGASQNTLY | 0.717643 |
| VRNADVL | ASGERVGASQNTLY | 0.7907568 |
| TCSSPFLVSAM | ASGERVGASQNTLY | 0.1849277 |
| VHITDAE | ASGERVGASQNTLY | 0.78065807 |
| EYAAPSF | ASGERVGASQNTLY | 0.5948682 |
| VVDAFHGGL | ASGERVGASQNTLY | 0.61684716 |
| ASVVDPQ | ASGERVGASQNTLY | 0.3044143 |
| SYNEFGGLS | ASGERVGASQNTLY | 0.74879825 |
| DPDGSVY | ASGERVGASQNTLY | 0.80987775 |
| MVSTIHATE | ASGERVGASQNTLY | 0.8173194 |
| LAAGVADPV | ASGERVGASQNTLY | 0.57564384 |
| TKEQIAA | ASGERVGASQNTLY | 0.81500506 |
| SLASPAAL | ASGERVGASQNTLY | 0.8264202 |
| IGLVFYGV | ASGERVGASQNTLY | 0.72026944 |
| FLLDASNL | ASGERVGASQNTLY | 0.28779697 |

| | | |
|---|---|---|
| YQVGNFP | ASGERVGASQNTLY | 0.76157975 |
| ESLLSQST | ASGERVGASQNTLY | 0.782989 |
| LFTVGFAAVG | ASGERVGASQNTLY | 0.8065924 |
| ATPPTPT | ASGERVGASQNTLY | 0.8041838 |
| TDLATLG | ASGERVGASQNTLY | 0.5450161 |
| SAGGTGR | ASGERVGASQNTLY | 0.78466046 |
| GFSNSGS | ASGERVGASQNTLY | 0.80945307 |
| GGNGGSGGTSV | ASGERVGASQNTLY | 0.61243296 |
| GTSVATGGAGN | ASGERVGASQNTLY | 0.6468817 |
| NGGNGGSGGTS | ASGERVGASQNTLY | 0.6239213 |
| AVVKFPH | ASGERVGASQNTLY | 0.27179334 |
| TPPPPPP | ASGERVGASQNTLY | 0.78929305 |
| AAPAAAAAPA | ASGERVGASQNTLY | 0.69331706 |
| IALNEDL | ASGERVGASQNTLY | 0.55478096 |
| IVNVSSM | ASGERVGASQNTLY | 0.7427516 |
| TGIPPAPR | ASGERVGASQNTLY | 0.54674804 |
| FDLGGGTF | ASGERVGASQNTLY | 0.5923579 |
| FDLGGGTFDVS | ASGERVGASQNTLY | 0.6070212 |
| NEPTAAA | ASGERVGASQNTLY | 0.7139658 |
| NEPTAAALA | ASGERVGASQNTLY | 0.79493904 |
| GSIINISSI | ASGERVGASQNTLY | 0.7516218 |
| VAPGNAGTA | ASGERVGASQNTLY | 0.7839346 |
| TGGGASLE | ASGERVGASQNTLY | 0.8213663 |
| VAVYNHYKRI | ASGERVGASQNTLY | 0.3985437 |
| TDPAPATT | ASGERVGASQNTLY | 0.59177405 |
| TNRPDILDPALLRPG | ASGERVGASQNTLY | 0.79338527 |
| LTTTTTS | ASGERVGASQNTLY | 0.3946817 |
| PTGSGKTL | ASGERVGASQNTLY | 0.7998596 |
| TGSGKTL | ASGERVGASQNTLY | 0.75592816 |
| TIGHVDHGK | ASGERVGASQNTLY | 0.79135257 |
| TIGHVDHGKTT | ASGERVGASQNTLY | 0.7543138 |
| GHVDHGKTT | ASGERVGASQNTLY | 0.8308677 |
| AQFEHTLL | ASGERVGASQNTLY | 0.7896732 |
| IVNQDEL | ASGERVGASQNTLY | 0.678198 |
| TNGSQFF | ASGERVGASQNTLY | 0.7646424 |
| TGKGIVE | ASGERVGASQNTLY | 0.51040196 |
| LISNASDA | ASGERVGASQNTLY | 0.815428 |
| AELRTLGELTKIAKAHGAQ | ASGERVGASQNTLY | 0.80007994 |
| APNAVAAAEPSW | ASGERVGASQNTLY | 0.5633175 |
| APQVAGAATAVL | ASGERVGASQNTLY | 0.7284113 |
| ARRDPAAAAAILPTDA | ASGERVGASQNTLY | 0.82113993 |
| DAAARDVLAEAGLAEYF | ASGERVGASQNTLY | 0.81584775 |
| DDQIVLSAKANLAKVEQ | ASGERVGASQNTLY | 0.31580228 |
| DELLPDTGEDSDEDGH | ASGERVGASQNTLY | 0.68421245 |
| ELPEAPANRATVPQD | ASGERVGASQNTLY | 0.39858973 |
| FGQSKIERVTGIG | ASGERVGASQNTLY | 0.6670974 |
| GAGGTGSTAGGAGGAGGAGGLY | ASGERVGASQNTLY | 0.41929704 |
| GHGGTGAAGGNGGTGGAGGSGI | ASGERVGASQNTLY | 0.7977626 |
| GNGGNGGAGGEGGPGGAGGAGA | ASGERVGASQNTLY | 0.67678016 |
| GSTAGNGGNGGNATNGTIAGQP | ASGERVGASQNTLY | 0.79474044 |
| GTGGDGGNGGNGGTGG | ASGERVGASQNTLY | 0.7540802 |

| | | |
|---|---|---|
| GVRRPAGIPADAPV | ASGERVGASQNTLY | 0.80979526 |
| GYRQVTDAHLLTLARR | ASGERVGASQNTLY | 0.6308832 |
| IAKMGQDGHDRG | ASGERVGASQNTLY | 0.6383014 |
| KAEPSIEIQYT | ASGERVGASQNTLY | 0.3725945 |
| KPVRIRMAADEDTVAVT | ASGERVGASQNTLY | 0.2957608 |
| MPPRIAGMRLLVIKPEP | ASGERVGASQNTLY | 0.582247 |
| MKFYAVALFRSRGVPPTR | ASGERVGASQNTLY | 0.68899286 |
| NAGGFGILGAVAHSP | ASGERVGASQNTLY | 0.21991839 |
| NEGREITLAMAEM | ASGERVGASQNTLY | 0.8177472 |
| NELVEWSHPLGHRWRWE | ASGERVGASQNTLY | 0.7532156 |
| NLGAGGTGGNGGNPG | ASGERVGASQNTLY | 0.78151673 |
| NTEWPKWETVMAADPVD | ASGERVGASQNTLY | 0.56671846 |
| PASAAPAETQP | ASGERVGASQNTLY | 0.67484105 |
| QQSSDGKNGVRAAEITGE | ASGERVGASQNTLY | 0.6208569 |
| RAPDAVTTSWWKE | ASGERVGASQNTLY | 0.7995373 |
| SGFGNVGAHTSGWFNQ | ASGERVGASQNTLY | 0.57952166 |
| SGVAGAVGQAASVGGLKV | ASGERVGASQNTLY | 0.31933302 |
| SNGGAGTGSAGGNGGT | ASGERVGASQNTLY | 0.64590734 |
| TGGAGGEGGAAG | ASGERVGASQNTLY | 0.5359814 |
| TGGDGGAAGAGGAPGHG | ASGERVGASQNTLY | 0.8236238 |
| TMPPLLAAIANAGRHGV | ASGERVGASQNTLY | 0.80612695 |
| TSSTAGNSGGA | ASGERVGASQNTLY | 0.7637117 |
| VPATVPVGILC | ASGERVGASQNTLY | 0.80941665 |
| VQRPDSGGVVLTGRIS | ASGERVGASQNTLY | 0.22707811 |
| WTPNPVPATKGVPAGLVQAT | ASGERVGASQNTLY | 0.64046 |
| YALKRALAKFSDRFDVVII | ASGERVGASQNTLY | 0.8262818 |
| YLIGSVIV | ASGERVGASQNTLY | 0.7658386 |
| YGQLVAVID | ASGERVGASQNTLY | 0.7179494 |
| VTVGGMTLPA | ASGERVGASQNTLY | 0.81039447 |
| VLALPAV | ASGERVGASQNTLY | 0.5397028 |
| VIPPTLNYET | ASGERVGASQNTLY | 0.64136666 |
| VGMGGMHPSSG | ASGERVGASQNTLY | 0.7083558 |
| VGLAGGAATA | ASGERVGASQNTLY | 0.78978443 |
| VGGAGGAGGSGP | ASGERVGASQNTLY | 0.71810293 |
| VFAAFMV | ASGERVGASQNTLY | 0.5185324 |
| TLYDKLR | ASGERVGASQNTLY | 0.76875573 |
| TLAVFTVL | ASGERVGASQNTLY | 0.8306105 |
| TGSVQPARV | ASGERVGASQNTLY | 0.68894047 |
| TGGGAGSGGT | ASGERVGASQNTLY | 0.8133444 |
| SHLETSL | ASGERVGASQNTLY | 0.7679909 |
| SFTTGMT | ASGERVGASQNTLY | 0.7450979 |
| QVEVAKRMGVRQPTVS | ASGERVGASQNTLY | 0.5921111 |
| QGSGGPALAM | ASGERVGASQNTLY | 0.66759646 |
| MLAPSNP | ASGERVGASQNTLY | 0.73812616 |
| LVNNAAI | ASGERVGASQNTLY | 0.6183457 |
| LTGEINTLA | ASGERVGASQNTLY | 0.73971355 |
| LSIFGPV | ASGERVGASQNTLY | 0.6798537 |
| LLALLANL | ASGERVGASQNTLY | 0.8325946 |
| LISNASDALDKLR | ASGERVGASQNTLY | 0.79291046 |
| LINEAVP | ASGERVGASQNTLY | 0.78108096 |
| LGYAFSGL | ASGERVGASQNTLY | 0.54662836 |

| | | |
|---|---|---|
| LDFVAVAL | ASGERVGASQNTLY | 0.75655234 |
| LAQPGAE | ASGERVGASQNTLY | 0.78934145 |
| ISNLGAVTGF | ASGERVGASQNTLY | 0.38988695 |
| HQEDPGRI | ASGERVGASQNTLY | 0.82475775 |
| GTPGEPTP | ASGERVGASQNTLY | 0.5633411 |
| GFFNSGS | ASGERVGASQNTLY | 0.7373451 |
| GAGGGGLTAGAN | ASGERVGASQNTLY | 0.7957753 |
| FVDALVNM | ASGERVGASQNTLY | 0.81366336 |
| FLTVAPEL | ASGERVGASQNTLY | 0.68942505 |
| FETLTAL | ASGERVGASQNTLY | 0.7200837 |
| DVINPIEL | ASGERVGASQNTLY | 0.5226551 |
| APVSGAPV | ASGERVGASQNTLY | 0.8050834 |
| ALVGGLAW | ASGERVGASQNTLY | 0.8117698 |
| ALAAAAATAI | ASGERVGASQNTLY | 0.7354097 |
| AFGLGLLL | ASGERVGASQNTLY | 0.8269061 |
| AFFAAASL | ASGERVGASQNTLY | 0.7449231 |
| AAAALPAFSPP | ASGERVGASQNTLY | 0.34563166 |
| VVLAPAL | ASGERVGASQNTLY | 0.7286749 |
| TIGHVDHGKTTL | ASGERVGASQNTLY | 0.81708455 |
| LISNASDALDK | ASGERVGASQNTLY | 0.81637347 |
| LISNASDAL | ASGERVGASQNTLY | 0.8012655 |
| ISNASDALDK | ASGERVGASQNTLY | 0.77981603 |
| NQAETLVYQTE | ASGERVGASQNTLY | 0.7128136 |
| TGGTGGTAGAG | ASGERVGASQNTLY | 0.81895214 |
| TGSTAGGAGGA | ASGERVGASQNTLY | 0.82906425 |
| DPLAVPVAAGPV | ASGERVGASQNTLY | 0.8020373 |
| GGTGGTPGNGGN | ASGERVGASQNTLY | 0.70125765 |
| PAGLAGVDPATL | ASGERVGASQNTLY | 0.60011137 |
| GNNLVVEAPHAGASV | ASGERVGASQNTLY | 0.7493609 |
| KVTRIMSASTLEQPA | ASGERVGASQNTLY | 0.8050282 |
| LGFALPIALPRRTVV | ASGERVGASQNTLY | 0.5228611 |
| TRLGNLTARKLVVLV | ASGERVGASQNTLY | 0.7690456 |
| WSQQRRLIYFPSAGP | ASGERVGASQNTLY | 0.7647052 |
| GPNGNPDPMAAAVDIR | ASGERVGASQNTLY | 0.35941383 |
| TQRCVSVADLRAAASR | ASGERVGASQNTLY | 0.78114367 |
| QFVLDVAEHPVEVLTETV | ASGERVGASQNTLY | 0.82421577 |
| GGAGGTGGAGGNGGTQAGDG | ASGERVGASQNTLY | 0.68719053 |
| PLLATLRAIAVDVDAKPAQV | ASGERVGASQNTLY | 0.6013389 |
| APKFRTGGYMIDFWGVGYQVAK | ASGERVGASQNTLY | 0.8190175 |
| GAGGAGGAGGAAGGPGGTGGTG | ASGERVGASQNTLY | 0.8309933 |
| IGALGLLATDVAPAGSVAFGVA | ASGERVGASQNTLY | 0.8044437 |
| AGGAGGIGGNGGAGGAGGVLMG | ASGERVGASQNTLY | 0.30348435 |
| ADELIGLL | ASGERVGASQNTLY | 0.23822635 |
| ADRALKA | ASGERVGASQNTLY | 0.7030728 |
| AGAVGQVGGAGGA | ASGERVGASQNTLY | 0.79767275 |
| AGGGGGLGGAP | ASGERVGASQNTLY | 0.64239043 |
| AGGLLYGNGGAG | ASGERVGASQNTLY | 0.5598562 |
| AGLESIGL | ASGERVGASQNTLY | 0.7746631 |
| AGLKTGII | ASGERVGASQNTLY | 0.8129171 |
| ASLAALLL | ASGERVGASQNTLY | 0.50594366 |
| ASVINISSI | ASGERVGASQNTLY | 0.43966115 |

| | | |
|---|---|---|
| ATPPGPA | ASGERVGASQNTLY | 0.7762103 |
| ATVASALV | ASGERVGASQNTLY | 0.8069669 |
| ATVASLIL | ASGERVGASQNTLY | 0.611737 |
| AVNGPVLIRH | ASGERVGASQNTLY | 0.7079167 |
| DGAAGAVGI | ASGERVGASQNTLY | 0.6696385 |
| DPLLHGI | ASGERVGASQNTLY | 0.63257605 |
| ELGAALGAL | ASGERVGASQNTLY | 0.7851294 |
| ELGRTQAV | ASGERVGASQNTLY | 0.50028145 |
| FVAGGYLS | ASGERVGASQNTLY | 0.8237851 |
| GASGGAGGAGGW | ASGERVGASQNTLY | 0.49454555 |
| GGAGAGGDGGAGGI | ASGERVGASQNTLY | 0.8126636 |
| GTGHNGSQPGMGGNG | ASGERVGASQNTLY | 0.7790276 |
| GYAAASASAS | ASGERVGASQNTLY | 0.77183294 |
| HQVDVAAVQQG | ASGERVGASQNTLY | 0.69277954 |
| IGAAIAHT | ASGERVGASQNTLY | 0.6546666 |
| IGMIEVAQL | ASGERVGASQNTLY | 0.49463052 |
| IRLAPEHTSAMD | ASGERVGASQNTLY | 0.33147025 |
| IYNVSVL | ASGERVGASQNTLY | 0.8195922 |
| KQGFLGM | ASGERVGASQNTLY | 0.1915161 |
| LESVITQL | ASGERVGASQNTLY | 0.8176296 |
| LGLGALVVV | ASGERVGASQNTLY | 0.74293244 |
| LIGNGGNGGSGGI | ASGERVGASQNTLY | 0.5429158 |
| LNAINAPTL | ASGERVGASQNTLY | 0.7904625 |
| LQLLNTGV | ASGERVGASQNTLY | 0.4500808 |
| LQTSIEF | ASGERVGASQNTLY | 0.59972453 |
| MSFAVSEL | ASGERVGASQNTLY | 0.7993866 |
| NTGALGL | ASGERVGASQNTLY | 0.7895577 |
| PPGAGGAGGAGTT | ASGERVGASQNTLY | 0.5901568 |
| QLVGLGI | ASGERVGASQNTLY | 0.7639627 |
| RFLEQVHQL | ASGERVGASQNTLY | 0.77720046 |
| RGVGYVLR | ASGERVGASQNTLY | 0.6160282 |
| SATGVFAGVI | ASGERVGASQNTLY | 0.6845699 |
| SLPAITLP | ASGERVGASQNTLY | 0.8148606 |
| STALAFTV | ASGERVGASQNTLY | 0.75753504 |
| SVLTVGLGK | ASGERVGASQNTLY | 0.49583447 |
| TALAGAGSGI | ASGERVGASQNTLY | 0.7878891 |
| TGLPFTGI | ASGERVGASQNTLY | 0.64860946 |
| VAEHVTV | ASGERVGASQNTLY | 0.78859544 |
| VDIVLNSL | ASGERVGASQNTLY | 0.7667382 |
| VLTGGGAVL | ASGERVGASQNTLY | 0.6901723 |
| VNVGLAYGI | ASGERVGASQNTLY | 0.79722977 |
| VSSAAVGAV | ASGERVGASQNTLY | 0.44952583 |
| VTSVYEAGV | ASGERVGASQNTLY | 0.6159345 |
| WVALDRGA | ASGERVGASQNTLY | 0.6866833 |
| YLVGSVVI | ASGERVGASQNTLY | 0.6637696 |
| GPPGTGKTLL | ASGERVGASQNTLY | 0.7955704 |
| GVGMTKF | ASGERVGASQNTLY | 0.52786195 |
| IGIGGSDL | ASGERVGASQNTLY | 0.7682375 |
| IGTLLAEL | ASGERVGASQNTLY | 0.6197723 |
| IYTSGTT | ASGERVGASQNTLY | 0.45475602 |
| LSFPAGV | ASGERVGASQNTLY | 0.6782161 |

| | | |
|---|---|---|
| LYTSGTT | ASGERVGASQNTLY | 0.6525752 |
| VGAFPFGF | ASGERVGASQNTLY | 0.5828446 |
| VGAGTVEF | ASGERVGASQNTLY | 0.7961223 |
| VGKSTLLL | ASGERVGASQNTLY | 0.43886343 |
| AALAHAGGGIV | ASGERVGASQNTLY | 0.7487124 |
| AALVAVSLL | ASGERVGASQNTLY | 0.8258241 |
| AAVSRAVAAL | ASGERVGASQNTLY | 0.6966552 |
| AGGSSGIGGF | ASGERVGASQNTLY | 0.8019742 |
| AGLTVANANG | ASGERVGASQNTLY | 0.7962676 |
| AGLVGAVL | ASGERVGASQNTLY | 0.7997133 |
| ALSQAFTV | ASGERVGASQNTLY | 0.63515663 |
| ALTFSVSL | ASGERVGASQNTLY | 0.60303265 |
| ALTGAALVAL | ASGERVGASQNTLY | 0.819744 |
| ASFSIGAGVL | ASGERVGASQNTLY | 0.78969663 |
| DGSVYKI | ASGERVGASQNTLY | 0.8105056 |
| EQLAEIV | ASGERVGASQNTLY | 0.79681504 |
| FAVSELTVM | ASGERVGASQNTLY | 0.8035791 |
| GAAPLAAGL | ASGERVGASQNTLY | 0.8022949 |
| GADGGDGGSGGAGG | ASGERVGASQNTLY | 0.7016145 |
| GFGLDTGGAGG | ASGERVGASQNTLY | 0.42391855 |
| GLASGGLL | ASGERVGASQNTLY | 0.6356151 |
| GRIAGAAVLL | ASGERVGASQNTLY | 0.707417 |
| GSAGSLNTSF | ASGERVGASQNTLY | 0.6192944 |
| GTGQIGI | ASGERVGASQNTLY | 0.34885174 |
| GTLIALIL | ASGERVGASQNTLY | 0.8141426 |
| IAATMGFVVG | ASGERVGASQNTLY | 0.83205664 |
| IFAQALL | ASGERVGASQNTLY | 0.5702323 |
| IGAAVALGF | ASGERVGASQNTLY | 0.8222515 |
| IGLTPLAL | ASGERVGASQNTLY | 0.41735485 |
| ILGSNTSTL | ASGERVGASQNTLY | 0.7790104 |
| IVFSLTVL | ASGERVGASQNTLY | 0.81141084 |
| LASKATGF | ASGERVGASQNTLY | 0.56765616 |
| LASVALIL | ASGERVGASQNTLY | 0.77886295 |
| LFAVGLTL | ASGERVGASQNTLY | 0.55856526 |
| LFNGGIGTYI | ASGERVGASQNTLY | 0.673723 |
| LTLGAGII | ASGERVGASQNTLY | 0.81881833 |
| LVAAALGDI | ASGERVGASQNTLY | 0.59940016 |
| NTAFVQLGI | ASGERVGASQNTLY | 0.77493703 |
| QLFSGAEL | ASGERVGASQNTLY | 0.6467114 |
| SAATFITL | ASGERVGASQNTLY | 0.50832224 |
| SNYGLANLGV | ASGERVGASQNTLY | 0.77523285 |
| SYGGAYAVM | ASGERVGASQNTLY | 0.8187337 |
| TTGVGGLTLG | ASGERVGASQNTLY | 0.6373316 |
| TTQTIYI | ASGERVGASQNTLY | 0.82592136 |
| VAAGAGLGAVI | ASGERVGASQNTLY | 0.61382747 |
| VAAVVTLL | ASGERVGASQNTLY | 0.8294016 |
| VGFLGGLI | ASGERVGASQNTLY | 0.81824625 |
| VGVIPILL | ASGERVGASQNTLY | 0.76735103 |
| VIFASSLI | ASGERVGASQNTLY | 0.69573104 |
| VKGGDGGAAAG | ASGERVGASQNTLY | 0.81086844 |
| VTKTFTNI | ASGERVGASQNTLY | 0.670965 |

| | | |
|---|---|---|
| VVAAAILL | ASGERVGASQNTLY | 0.8097755 |
| YAALDVEL | ASGERVGASQNTLY | 0.8143827 |
| LASVALLL | ASGERVGASQNTLY | 0.424548 |
| LDAVHRV | ASGERVGASQNTLY | 0.72670305 |
| QGLSSSIF | ASGERVGASQNTLY | 0.6308018 |
| RGTFANIRL | ASGERVGASQNTLY | 0.50787485 |
| RQLWWGHRI | ASGERVGASQNTLY | 0.46248192 |
| YGPPGTGKTLL | ASGERVGASQNTLY | 0.37515002 |
| AALVGAPANVA | ASGERVGASQNTLY | 0.5432662 |
| AATFRRTE | ASGERVGASQNTLY | 0.41075855 |
| AGALGGGI | ASGERVGASQNTLY | 0.67125636 |
| AGGAGGAGGIGAG | ASGERVGASQNTLY | 0.6611348 |
| AGSGALGL | ASGERVGASQNTLY | 0.419977 |
| AQIADQI | ASGERVGASQNTLY | 0.71103245 |
| ASAVAVTYL | ASGERVGASQNTLY | 0.6333532 |
| ASTVLGGTGG | ASGERVGASQNTLY | 0.35250312 |
| AVGGEPAK | ASGERVGASQNTLY | 0.48288682 |
| CEQGTFGGTC | ASGERVGASQNTLY | 0.5843686 |
| DEGKAPAG | ASGERVGASQNTLY | 0.5113647 |
| DTVVFPAL | ASGERVGASQNTLY | 0.31948486 |
| FEGITMAG | ASGERVGASQNTLY | 0.6395961 |
| FPLVVQD | ASGERVGASQNTLY | 0.47965914 |
| GAFTTPQ | ASGERVGASQNTLY | 0.22281897 |
| GFGGFGTAGGVGGA | ASGERVGASQNTLY | 0.595448 |
| GFLAQFL | ASGERVGASQNTLY | 0.5291112 |
| GGAGGAGGNAGVA | ASGERVGASQNTLY | 0.5448964 |
| GGVGTAGAGGAGGAGG | ASGERVGASQNTLY | 0.6967895 |
| GNGGGGDAGNAGSGGN | ASGERVGASQNTLY | 0.4223378 |
| GTGGAGGAGNDAGS | ASGERVGASQNTLY | 0.42695904 |
| GTTNTGWFNS | ASGERVGASQNTLY | 0.6769748 |
| IFMGPLFTL | ASGERVGASQNTLY | 0.7059983 |
| ILAHAIEAI | ASGERVGASQNTLY | 0.3709194 |
| IRSAIREAL | ASGERVGASQNTLY | 0.6063654 |
| ITFAGGG | ASGERVGASQNTLY | 0.6611101 |
| ITGMSGHQEV | ASGERVGASQNTLY | 0.62523866 |
| IVAGLACTL | ASGERVGASQNTLY | 0.7174815 |
| IVGGGLAGA | ASGERVGASQNTLY | 0.7199417 |
| IVTGVLNI | ASGERVGASQNTLY | 0.39953044 |
| LAQQGCI | ASGERVGASQNTLY | 0.70722824 |
| LCLHVEN | ASGERVGASQNTLY | 0.7385982 |
| LDAISDGS | ASGERVGASQNTLY | 0.51037824 |
| LGGFAAML | ASGERVGASQNTLY | 0.53603554 |
| LGGKNPMI | ASGERVGASQNTLY | 0.430523 |
| LGPKGVAAIL | ASGERVGASQNTLY | 0.50372124 |
| LMVGVSI | ASGERVGASQNTLY | 0.3348496 |
| LVGAAFVL | ASGERVGASQNTLY | 0.63875157 |
| LVLAVAPL | ASGERVGASQNTLY | 0.7153718 |
| MAAIAVIDAA | ASGERVGASQNTLY | 0.5190495 |
| NNGINAELQ | ASGERVGASQNTLY | 0.40105984 |
| RGEPLVLVHG | ASGERVGASQNTLY | 0.34809983 |
| RGGAGGAAGLI | ASGERVGASQNTLY | 0.5522229 |

| | | |
|---|---|---|
| RSYRMHPAV | ASGERVGASQNTLY | 0.3995748 |
| SAAHLLAGQLA | ASGERVGASQNTLY | 0.3076227 |
| SAAVGSY | ASGERVGASQNTLY | 0.5582508 |
| SPHAVLHRI | ASGERVGASQNTLY | 0.67913735 |
| SQIAKVL | ASGERVGASQNTLY | 0.6817424 |
| TALGHLVP | ASGERVGASQNTLY | 0.29304993 |
| TAPTTSGAA | ASGERVGASQNTLY | 0.73127645 |
| TDGDDGNGGSPGT | ASGERVGASQNTLY | 0.6238043 |
| TDGLGEN | ASGERVGASQNTLY | 0.69663405 |
| TGGPPLPGQA | ASGERVGASQNTLY | 0.5678133 |
| TIPAITI | ASGERVGASQNTLY | 0.45005387 |
| TIPGFTI | ASGERVGASQNTLY | 0.6706876 |
| TSIASGALA | ASGERVGASQNTLY | 0.51013863 |
| TVTLPGAAI | ASGERVGASQNTLY | 0.4881287 |
| VASGLSAALP | ASGERVGASQNTLY | 0.65493786 |
| VGAGVIGI | ASGERVGASQNTLY | 0.58309054 |
| VGPKRFGV | ASGERVGASQNTLY | 0.44368738 |
| VNVAFPD | ASGERVGASQNTLY | 0.4976929 |
| VSMTNSGVS | ASGERVGASQNTLY | 0.68000203 |
| VTDVLGDLSV | ASGERVGASQNTLY | 0.38027665 |
| VVVVGAGFAGL | ASGERVGASQNTLY | 0.43172795 |
| VYTVGTVV | ASGERVGASQNTLY | 0.39176145 |
| WTTFSGP | ASGERVGASQNTLY | 0.5368676 |
| YDELVEAF | ASGERVGASQNTLY | 0.69885665 |
| YGTGGAGGNGG | ASGERVGASQNTLY | 0.63258636 |
| YTNVMAV | ASGERVGASQNTLY | 0.4847083 |
| FLEQVHQL | ASGERVGASQNTLY | 0.6356878 |
| GLSLLAVAV | ASGERVGASQNTLY | 0.63174593 |
| LEQVHQL | ASGERVGASQNTLY | 0.5582524 |
| LNAAGAI | ASGERVGASQNTLY | 0.5852912 |
| PAVHASL | ASGERVGASQNTLY | 0.46340537 |
| TELEDDA | ASGERVGASQNTLY | 0.6940733 |
| TPPGPPN | ASGERVGASQNTLY | 0.6223662 |
| TPPPPPPP | ASGERVGASQNTLY | 0.6626747 |
| TQSPALI | ASSQGGAPDYT | 0.6975052 |
| EFAAAFPQ | ASSQGGAPDYT | 0.6206864 |
| LLSGNLDVL | ASSQGGAPDYT | 0.5339798 |
| GPANLGNYNI | ASSQGGAPDYT | 0.27389345 |
| TLGPTLP | ASSQGGAPDYT | 0.50264215 |
| VYAYSSCCK | ASSQGGAPDYT | 0.5942943 |
| APAPAPAPNI | ASSQGGAPDYT | 0.6181408 |
| PGGPVPIP | ASSQGGAPDYT | 0.6557653 |
| FLTVALEP | ASSQGGAPDYT | 0.6208628 |
| SQYFGANL | ASSQGGAPDYT | 0.45783457 |
| GGVKPAA | ASSQGGAPDYT | 0.6199252 |
| GGTGGAGGAGGW | ASSQGGAPDYT | 0.6725702 |
| GVGGVAGLGGAGG | ASSQGGAPDYT | 0.32548746 |
| VLLVLATTI | ASSQGGAPDYT | 0.5037521 |
| IGIFAAML | ASSQGGAPDYT | 0.75996876 |
| AVGDLID | ASSQGGAPDYT | 0.5998808 |
| NISPLPL | ASSQGGAPDYT | 0.49740177 |

| | | |
|---|---|---|
| VTAVFVAI | ASSQGGAPDYT | 0.42771992 |
| SGDDNDAAL | ASSQGGAPDYT | 0.70913243 |
| AVAALAHS | ASSQGGAPDYT | 0.44838315 |
| GGVTVMATP | ASSQGGAPDYT | 0.70257646 |
| AGLAANVRD | ASSQGGAPDYT | 0.23365244 |
| TNRPDLIDPALLRPG | ASSQGGAPDYT | 0.46123618 |
| LANLGVTAL | ASSQGGAPDYT | 0.40357348 |
| TGERIEL | ASSQGGAPDYT | 0.2815504 |
| NGEELAA | ASSQGGAPDYT | 0.6864491 |
| VLSLVWG | ASSQGGAPDYT | 0.4457778 |
| IGYALWL | ASSQGGAPDYT | 0.60786766 |
| LSLLLAPL | ASSQGGAPDYT | 0.6172899 |
| TLDEALK | ASSQGGAPDYT | 0.6939284 |
| TDGAGTAEGG | ASSQGGAPDYT | 0.5828519 |
| NAPGGAGGNGG | ASSQGGAPDYT | 0.5221262 |
| VSVFMGMGGNF | ASSQGGAPDYT | 0.4904856 |
| LASYLAAL | ASSQGGAPDYT | 0.64496005 |
| LVGGTPAL | ASSQGGAPDYT | 0.65723896 |
| VRNADVL | ASSQGGAPDYT | 0.64601445 |
| TCSSPFLVSAM | ASSQGGAPDYT | 0.31016654 |
| VHITDAE | ASSQGGAPDYT | 0.5223248 |
| EYAAPSF | ASSQGGAPDYT | 0.5372014 |
| VVDAFHGGL | ASSQGGAPDYT | 0.32750696 |
| ASVVDPQ | ASSQGGAPDYT | 0.24763763 |
| SYNEFGGLS | ASSQGGAPDYT | 0.47659865 |
| DPDGSVY | ASSQGGAPDYT | 0.5104191 |
| MVSTIHATE | ASSQGGAPDYT | 0.6718925 |
| LAAGVADPV | ASSQGGAPDYT | 0.68586665 |
| TKEQIAA | ASSQGGAPDYT | 0.4903694 |
| SLASPAAL | ASSQGGAPDYT | 0.6458768 |
| IGLVFYGV | ASSQGGAPDYT | 0.5241296 |
| FLLDASNL | ASSQGGAPDYT | 0.19145828 |
| YQVGNFP | ASSQGGAPDYT | 0.4672205 |
| ESLLSQST | ASSQGGAPDYT | 0.34874165 |
| LFTVGFAAVG | ASSQGGAPDYT | 0.70708776 |
| ATPPTPT | ASSQGGAPDYT | 0.46384263 |
| TDLATLG | ASSQGGAPDYT | 0.47812322 |
| SAGGTGR | ASSQGGAPDYT | 0.526683 |
| GFSNSGS | ASSQGGAPDYT | 0.5944015 |
| GGNGGSGGTSV | ASSQGGAPDYT | 0.53981906 |
| GTSVATGGAGN | ASSQGGAPDYT | 0.39976394 |
| NGGNGGSGGTS | ASSQGGAPDYT | 0.2592919 |
| AVVKFPH | ASSQGGAPDYT | 0.6463791 |
| TPPPPPP | ASSQGGAPDYT | 0.4759105 |
| AAPAAAAAPA | ASSQGGAPDYT | 0.3822701 |
| IALNEDL | ASSQGGAPDYT | 0.28307864 |
| IVNVSSM | ASSQGGAPDYT | 0.5915279 |
| TGIPPAPR | ASSQGGAPDYT | 0.32219845 |
| FDLGGGTF | ASSQGGAPDYT | 0.69004977 |
| FDLGGGTFDVS | ASSQGGAPDYT | 0.48392475 |
| NEPTAAA | ASSQGGAPDYT | 0.5294915 |

| | | |
|---|---|---|
| NEPTAAALA | ASSQGGAPDYT | 0.75505495 |
| GSIINISSI | ASSQGGAPDYT | 0.571489 |
| VAPGNAGTA | ASSQGGAPDYT | 0.49792403 |
| TGGGASLE | ASSQGGAPDYT | 0.44230852 |
| VAVYNHYKRI | ASSQGGAPDYT | 0.41778615 |
| TDPAPATT | ASSQGGAPDYT | 0.45972386 |
| TNRPDILDPALLRPG | ASSQGGAPDYT | 0.44404116 |
| LTTTTTS | ASSQGGAPDYT | 0.72294825 |
| PTGSGKTL | ASSQGGAPDYT | 0.48382202 |
| TGSGKTL | ASSQGGAPDYT | 0.43183273 |
| TIGHVDHGK | ASSQGGAPDYT | 0.5386169 |
| TIGHVDHGKTT | ASSQGGAPDYT | 0.71918654 |
| GHVDHGKTT | ASSQGGAPDYT | 0.7466152 |
| AQFEHTLL | ASSQGGAPDYT | 0.43167862 |
| IVNQDEL | ASSQGGAPDYT | 0.47876102 |
| TNGSQFF | ASSQGGAPDYT | 0.4241944 |
| TGKGIVE | ASSQGGAPDYT | 0.5365619 |
| LISNASDA | ASSQGGAPDYT | 0.45746216 |
| AELRTLGELTKIAKAHGAQ | ASSQGGAPDYT | 0.4048689 |
| APNAVAAAEPSW | ASSQGGAPDYT | 0.64972186 |
| APQVAGAATAVL | ASSQGGAPDYT | 0.7224066 |
| ARRDPAAAAAILPTDA | ASSQGGAPDYT | 0.49758315 |
| DAAARDVLAEAGLAEYF | ASSQGGAPDYT | 0.46139145 |
| DDQIVLSAKANLAKVEQ | ASSQGGAPDYT | 0.4960932 |
| DELLPDTGEDSDEDGH | ASSQGGAPDYT | 0.4112379 |
| ELPEAPANRATVPQD | ASSQGGAPDYT | 0.44477034 |
| FGQSKIERVTGIG | ASSQGGAPDYT | 0.54981667 |
| GAGGTGSTAGGAGGAGGAGGLY | ASSQGGAPDYT | 0.32178947 |
| GHGGTGAAGGNGGTGGAGGSGI | ASSQGGAPDYT | 0.5425625 |
| GNGGNGGAGGEGGPGGAGGAGA | ASSQGGAPDYT | 0.73656595 |
| GSTAGNGGNGGNATNGTIAGQP | ASSQGGAPDYT | 0.52274346 |
| GTGGDGGNGGNGGTGG | ASSQGGAPDYT | 0.6121836 |
| GVRRPAGIPADAPV | ASSQGGAPDYT | 0.726449 |
| GYRQVTDAHLLTLARR | ASSQGGAPDYT | 0.65368605 |
| IAKMGQDGHDRG | ASSQGGAPDYT | 0.7192952 |
| KAEPSIEIQYT | ASSQGGAPDYT | 0.66958976 |
| KPVRIRMAADEDTVAVT | ASSQGGAPDYT | 0.53988 |
| MPPRIAGMRLLVIKPEP | ASSQGGAPDYT | 0.52057135 |
| MKFYAVALFRSRGVPPTR | ASSQGGAPDYT | 0.69565165 |
| NAGGFGILGAVAHSP | ASSQGGAPDYT | 0.50666857 |
| NEGREITLAMAEM | ASSQGGAPDYT | 0.52878344 |
| NELVEWSHPLGHRWRWE | ASSQGGAPDYT | 0.42769665 |
| NLGAGGTGGNGGNPG | ASSQGGAPDYT | 0.6740937 |
| NTEWPKWETVMAADPVD | ASSQGGAPDYT | 0.538717 |
| PASAAPAETQP | ASSQGGAPDYT | 0.6755564 |
| QQSSDGKNGVRAAEITGE | ASSQGGAPDYT | 0.32033694 |
| RAPDAVTTSWWKE | ASSQGGAPDYT | 0.4033268 |
| SGFGNVGAHTSGWFNQ | ASSQGGAPDYT | 0.6908145 |
| SGVAGAVGQAASVGGLKV | ASSQGGAPDYT | 0.2033283 |
| SNGGAGTGSAGGNGGT | ASSQGGAPDYT | 0.2505761 |
| TGGAGGEGGAAG | ASSQGGAPDYT | 0.560056 |

| | | |
|---|---|---|
| TGGDGGAAGAGGAPGHG | ASSQGGAPDYT | 0.6742447 |
| TMPPLLAAIANAGRHGV | ASSQGGAPDYT | 0.22701856 |
| TSSTAGNSGGA | ASSQGGAPDYT | 0.40683025 |
| VPATVPVGILC | ASSQGGAPDYT | 0.34003028 |
| VQRPDSGGVVLTGRIS | ASSQGGAPDYT | 0.6251495 |
| WTPNPVPATKGVPAGLVQAT | ASSQGGAPDYT | 0.57719773 |
| YALKRALAKFSDRFDVVII | ASSQGGAPDYT | 0.32540336 |
| YLIGSVIV | ASSQGGAPDYT | 0.6564684 |
| YGQLVAVID | ASSQGGAPDYT | 0.34599483 |
| VTVGGMTLPA | ASSQGGAPDYT | 0.4761579 |
| VLALPAV | ASSQGGAPDYT | 0.69195485 |
| VIPPTLNYET | ASSQGGAPDYT | 0.71459 |
| VGMGGMHPSSG | ASSQGGAPDYT | 0.4114138 |
| VGLAGGAATA | ASSQGGAPDYT | 0.6495122 |
| VGGAGGAGGSGP | ASSQGGAPDYT | 0.43567654 |
| VFAAFMV | ASSQGGAPDYT | 0.76205033 |
| TLYDKLR | ASSQGGAPDYT | 0.485164 |
| TLAVFTVL | ASSQGGAPDYT | 0.3561515 |
| TGSVQPARV | ASSQGGAPDYT | 0.6879083 |
| TGGGAGSGGT | ASSQGGAPDYT | 0.6194804 |
| SHLETSL | ASSQGGAPDYT | 0.51187354 |
| SFTTGMT | ASSQGGAPDYT | 0.6526685 |
| QVEVAKRMGVRQPTVS | ASSQGGAPDYT | 0.46340454 |
| QGSGGPALAM | ASSQGGAPDYT | 0.50808704 |
| MLAPSNP | ASSQGGAPDYT | 0.70991486 |
| LVNNAAI | ASSQGGAPDYT | 0.6660373 |
| LTGEINTLA | ASSQGGAPDYT | 0.44959491 |
| LSIFGPV | ASSQGGAPDYT | 0.6856429 |
| LLALLANL | ASSQGGAPDYT | 0.511444 |
| LISNASDALDKLR | ASSQGGAPDYT | 0.40486467 |
| LINEAVP | ASSQGGAPDYT | 0.73908883 |
| LGYAFSGL | ASSQGGAPDYT | 0.5769825 |
| LDFVAVAL | ASSQGGAPDYT | 0.6663596 |
| LAQPGAE | ASSQGGAPDYT | 0.63232315 |
| ISNLGAVTGF | ASSQGGAPDYT | 0.65055007 |
| HQEDPGRI | ASSQGGAPDYT | 0.44374943 |
| GTPGEPTP | ASSQGGAPDYT | 0.58554643 |
| GFFNSGS | ASSQGGAPDYT | 0.52744365 |
| GAGGGGLTAGAN | ASSQGGAPDYT | 0.68047345 |
| FVDALVNM | ASSQGGAPDYT | 0.5603767 |
| FLTVAPEL | ASSQGGAPDYT | 0.6103851 |
| FETLTAL | ASSQGGAPDYT | 0.6217797 |
| DVINPIEL | ASSQGGAPDYT | 0.72413045 |
| APVSGAPV | ASSQGGAPDYT | 0.5420265 |
| ALVGGLAW | ASSQGGAPDYT | 0.552458 |
| ALAAAAATAI | ASSQGGAPDYT | 0.73187935 |
| AFGLGLLL | ASSQGGAPDYT | 0.45470178 |
| AFFAAASL | ASSQGGAPDYT | 0.490968 |
| AAAALPAFSPP | ASSQGGAPDYT | 0.6386233 |
| VVLAPAL | ASSQGGAPDYT | 0.6709415 |
| TIGHVDHGKTTL | ASSQGGAPDYT | 0.53747046 |

| | | |
|---|---|---|
| LISNASDALDK | ASSQGGAPDYT | 0.7098193 |
| LISNASDAL | ASSQGGAPDYT | 0.4690455 |
| ISNASDALDK | ASSQGGAPDYT | 0.67122436 |
| NQAETLVYQTE | ASSQGGAPDYT | 0.5031216 |
| TGGTGGTAGAG | ASSQGGAPDYT | 0.57659036 |
| TGSTAGGAGGA | ASSQGGAPDYT | 0.4522545 |
| DPLAVPVAAGPV | ASSQGGAPDYT | 0.642195 |
| GGTGGTPGNGGN | ASSQGGAPDYT | 0.69947714 |
| PAGLAGVDPATL | ASSQGGAPDYT | 0.6629296 |
| GNNLVVEAPHAGASV | ASSQGGAPDYT | 0.25629216 |
| KVTRIMSASTLEQPA | ASSQGGAPDYT | 0.69392204 |
| LGFALPIALPRRTVV | ASSQGGAPDYT | 0.66544783 |
| TRLGNLTARKLVVLV | ASSQGGAPDYT | 0.73510283 |
| WSQQRRLIYFPSAGP | ASSQGGAPDYT | 0.5059284 |
| GPNGNPDPMAAAVDIR | ASSQGGAPDYT | 0.550496 |
| TQRCVSVADLRAAASR | ASSQGGAPDYT | 0.48941657 |
| QFVLDVAEHPVEVLTETV | ASSQGGAPDYT | 0.56580025 |
| GGAGGTGGAGGNGGTQAGDG | ASSQGGAPDYT | 0.31273386 |
| PLLATLRAIAVDVDAKPAQV | ASSQGGAPDYT | 0.48584947 |
| APKFRTGGYMIDFWGVGYQVAK | ASSQGGAPDYT | 0.5051874 |
| GAGGAGGAGGAAGGPGGTGGTG | ASSQGGAPDYT | 0.44266915 |
| IGALGLLATDVAPAGSVAFGVA | ASSQGGAPDYT | 0.6547297 |
| AGGAGGIGGNGGAGGAGGVLMG | ASSQGGAPDYT | 0.69741 |
| ADELIGLL | ASSQGGAPDYT | 0.69328976 |
| ADRALKA | ASSQGGAPDYT | 0.70425266 |
| AGAVGQVGGAGGA | ASSQGGAPDYT | 0.7193295 |
| AGGGGGLGGAP | ASSQGGAPDYT | 0.688843 |
| AGGLLYGNGGAG | ASSQGGAPDYT | 0.4485857 |
| AGLESIGL | ASSQGGAPDYT | 0.5824681 |
| AGLKTGII | ASSQGGAPDYT | 0.38524866 |
| ASLAALLL | ASSQGGAPDYT | 0.53892595 |
| ASVINISSI | ASSQGGAPDYT | 0.4908328 |
| ATPPGPA | ASSQGGAPDYT | 0.61294013 |
| ATVASALV | ASSQGGAPDYT | 0.20865242 |
| ATVASLIL | ASSQGGAPDYT | 0.57947904 |
| AVNGPVLIRH | ASSQGGAPDYT | 0.72841233 |
| DGAAGAVGI | ASSQGGAPDYT | 0.47451735 |
| DPLLHGI | ASSQGGAPDYT | 0.5880877 |
| ELGAALGAL | ASSQGGAPDYT | 0.6299422 |
| ELGRTQAV | ASSQGGAPDYT | 0.4459235 |
| FVAGGYLS | ASSQGGAPDYT | 0.64270854 |
| GASGGAGGAGGW | ASSQGGAPDYT | 0.4209259 |
| GGAGAGGDGGAGGI | ASSQGGAPDYT | 0.6441782 |
| GTGHNGSQPGMGGNG | ASSQGGAPDYT | 0.7195562 |
| GYAAASASAS | ASSQGGAPDYT | 0.5880345 |
| HQVDVAAVQQG | ASSQGGAPDYT | 0.4667129 |
| IGAAIAHT | ASSQGGAPDYT | 0.4900712 |
| IGMIEVAQL | ASSQGGAPDYT | 0.38691807 |
| IRLAPEHTSAMD | ASSQGGAPDYT | 0.5284465 |
| IYNVSVL | ASSQGGAPDYT | 0.33790165 |
| KQGFLGM | ASSQGGAPDYT | 0.5770544 |

| | | |
|---|---|---|
| LESVITQL | ASSQGGAPDYT | 0.42631623 |
| LGLGALVVV | ASSQGGAPDYT | 0.58900803 |
| LIGNGGNGGSGGI | ASSQGGAPDYT | 0.5664965 |
| LNAINAPTL | ASSQGGAPDYT | 0.5525977 |
| LQLLNTGV | ASSQGGAPDYT | 0.37631044 |
| LQTSIEF | ASSQGGAPDYT | 0.58670604 |
| MSFAVSEL | ASSQGGAPDYT | 0.50190383 |
| NTGALGL | ASSQGGAPDYT | 0.3556545 |
| PPGAGGAGGAGTT | ASSQGGAPDYT | 0.69221985 |
| QLVGLGI | ASSQGGAPDYT | 0.53255796 |
| RFLEQVHQL | ASSQGGAPDYT | 0.5519751 |
| RGVGYVLR | ASSQGGAPDYT | 0.6863446 |
| SATGVFAGVI | ASSQGGAPDYT | 0.5930329 |
| SLPAITLP | ASSQGGAPDYT | 0.38831183 |
| STALAFTV | ASSQGGAPDYT | 0.5298697 |
| SVLTVGLGK | ASSQGGAPDYT | 0.52739614 |
| TALAGAGSGI | ASSQGGAPDYT | 0.5218623 |
| TGLPFTGI | ASSQGGAPDYT | 0.23455748 |
| VAEHVTV | ASSQGGAPDYT | 0.6577823 |
| VDIVLNSL | ASSQGGAPDYT | 0.4068049 |
| VLTGGGAVL | ASSQGGAPDYT | 0.41110355 |
| VNVGLAYGI | ASSQGGAPDYT | 0.4769376 |
| VSSAAVGAV | ASSQGGAPDYT | 0.68579954 |
| VTSVYEAGV | ASSQGGAPDYT | 0.48031145 |
| WVALDRGA | ASSQGGAPDYT | 0.48598236 |
| YLVGSVVI | ASSQGGAPDYT | 0.6658716 |
| GPPGTGKTLL | ASSQGGAPDYT | 0.61201864 |
| GVGMTKF | ASSQGGAPDYT | 0.6977552 |
| IGIGGSDL | ASSQGGAPDYT | 0.61031264 |
| IGTLLAEL | ASSQGGAPDYT | 0.52025586 |
| IYTSGTT | ASSQGGAPDYT | 0.5302361 |
| LSFPAGV | ASSQGGAPDYT | 0.5247846 |
| LYTSGTT | ASSQGGAPDYT | 0.34114805 |
| VGAFPFGF | ASSQGGAPDYT | 0.2834988 |
| VGAGTVEF | ASSQGGAPDYT | 0.5466811 |
| VGKSTLLL | ASSQGGAPDYT | 0.2782504 |
| AALAHAGGGIV | ASSQGGAPDYT | 0.55499554 |
| AALVAVSLL | ASSQGGAPDYT | 0.70431805 |
| AAVSRAVAAL | ASSQGGAPDYT | 0.52855206 |
| AGGSSGIGGF | ASSQGGAPDYT | 0.36055797 |
| AGLTVANANG | ASSQGGAPDYT | 0.47968876 |
| AGLVGAVL | ASSQGGAPDYT | 0.58614767 |
| ALSQAFTV | ASSQGGAPDYT | 0.6746506 |
| ALTFSVSL | ASSQGGAPDYT | 0.6152607 |
| ALTGAALVAL | ASSQGGAPDYT | 0.733878 |
| ASFSIGAGVL | ASSQGGAPDYT | 0.68588114 |
| DGSVYKI | ASSQGGAPDYT | 0.43154877 |
| EQLAEIV | ASSQGGAPDYT | 0.6542803 |
| FAVSELTVM | ASSQGGAPDYT | 0.49872756 |
| GAAPLAAGL | ASSQGGAPDYT | 0.56180793 |
| GADGGDGGSGGAGG | ASSQGGAPDYT | 0.5587833 |

| | | |
|---|---|---|
| GFGLDTGGAGG | ASSQGGAPDYT | 0.5787269 |
| GLASGGLL | ASSQGGAPDYT | 0.5214037 |
| GRIAGAAVLL | ASSQGGAPDYT | 0.48123443 |
| GSAGSLNTSF | ASSQGGAPDYT | 0.34248853 |
| GTGQIGI | ASSQGGAPDYT | 0.5851455 |
| GTLIALIL | ASSQGGAPDYT | 0.45799604 |
| IAATMGFVVG | ASSQGGAPDYT | 0.30722427 |
| IFAQALL | ASSQGGAPDYT | 0.41154966 |
| IGAAVALGF | ASSQGGAPDYT | 0.7419063 |
| IGLTPLAL | ASSQGGAPDYT | 0.71504265 |
| ILGSNTSTL | ASSQGGAPDYT | 0.44287795 |
| IVFSLTVL | ASSQGGAPDYT | 0.4301956 |
| LASKATGF | ASSQGGAPDYT | 0.6013472 |
| LASVALIL | ASSQGGAPDYT | 0.2954768 |
| LFAVGLTL | ASSQGGAPDYT | 0.618396 |
| LFNGGIGTYI | ASSQGGAPDYT | 0.57813066 |
| LTLGAGII | ASSQGGAPDYT | 0.5502411 |
| LVAAALGDI | ASSQGGAPDYT | 0.36290336 |
| NTAFVQLGI | ASSQGGAPDYT | 0.6074623 |
| QLFSGAEL | ASSQGGAPDYT | 0.49764985 |
| SAATFITL | ASSQGGAPDYT | 0.6380646 |
| SNYGLANLGV | ASSQGGAPDYT | 0.27130675 |
| SYGGAYAVM | ASSQGGAPDYT | 0.6737341 |
| TTGVGGLTLG | ASSQGGAPDYT | 0.3903091 |
| TTQTIYI | ASSQGGAPDYT | 0.45295784 |
| VAAGAGLGAVI | ASSQGGAPDYT | 0.3016435 |
| VAAVVTLL | ASSQGGAPDYT | 0.5257381 |
| VGFLGGLI | ASSQGGAPDYT | 0.69075197 |
| VGVIPILL | ASSQGGAPDYT | 0.7052924 |
| VIFASSLI | ASSQGGAPDYT | 0.436625 |
| VKGGDGGAAAG | ASSQGGAPDYT | 0.6703226 |
| VTKTFTNI | ASSQGGAPDYT | 0.722162 |
| VVAAAILL | ASSQGGAPDYT | 0.5399518 |
| YAALDVEL | ASSQGGAPDYT | 0.36477938 |
| LASVALLL | ASSQGGAPDYT | 0.62734926 |
| LDAVHRV | ASSQGGAPDYT | 0.5216123 |
| QGLSSSIF | ASSQGGAPDYT | 0.6960939 |
| RGTFANIRL | ASSQGGAPDYT | 0.6142394 |
| RQLWWGHRI | ASSQGGAPDYT | 0.35266396 |
| YGPPGTGKTLL | ASSQGGAPDYT | 0.54603225 |
| AALVGAPANVA | ASSQGGAPDYT | 0.6834758 |
| AATFRRTE | ASSQGGAPDYT | 0.35282767 |
| AGALGGGI | ASSQGGAPDYT | 0.43034366 |
| AGGAGGAGGIGAG | ASSQGGAPDYT | 0.47724673 |
| AGSGALGL | ASSQGGAPDYT | 0.3532055 |
| AQIADQI | ASSQGGAPDYT | 0.60082775 |
| ASAVAVTYL | ASSQGGAPDYT | 0.48281562 |
| ASTVLGGTGG | ASSQGGAPDYT | 0.47324383 |
| AVGGEPAK | ASSQGGAPDYT | 0.5342966 |
| CEQGTFGGTC | ASSQGGAPDYT | 0.44906056 |
| DEGKAPAG | ASSQGGAPDYT | 0.46739584 |

| | | |
|---|---|---|
| DTVVFPAL | ASSQGGAPDYT | 0.52084315 |
| FEGITMAG | ASSQGGAPDYT | 0.64441043 |
| FPLVVQD | ASSQGGAPDYT | 0.6534211 |
| GAFTTPQ | ASSQGGAPDYT | 0.57705224 |
| GFGGFGTAGGVGGA | ASSQGGAPDYT | 0.5364714 |
| GFLAQFL | ASSQGGAPDYT | 0.7325778 |
| GGAGGAGGNAGVA | ASSQGGAPDYT | 0.3932882 |
| GGVGTAGAGGAGGAGG | ASSQGGAPDYT | 0.3701861 |
| GNGGGGDAGNAGSGGN | ASSQGGAPDYT | 0.6447089 |
| GTGGAGGAGNDAGS | ASSQGGAPDYT | 0.3784025 |
| GTTNTGWFNS | ASSQGGAPDYT | 0.68079084 |
| IFMGPLFTL | ASSQGGAPDYT | 0.54331875 |
| ILAHAIEAI | ASSQGGAPDYT | 0.6226787 |
| IRSAIREAL | ASSQGGAPDYT | 0.6013893 |
| ITFAGGG | ASSQGGAPDYT | 0.74113834 |
| ITGMSGHQEV | ASSQGGAPDYT | 0.688082 |
| IVAGLACTL | ASSQGGAPDYT | 0.5100404 |
| IVGGGLAGA | ASSQGGAPDYT | 0.6606874 |
| IVTGVLNI | ASSQGGAPDYT | 0.36818168 |
| LAQQGCI | ASSQGGAPDYT | 0.6648896 |
| LCLHVEN | ASSQGGAPDYT | 0.61644995 |
| LDAISDGS | ASSQGGAPDYT | 0.38937318 |
| LGGFAAML | ASSQGGAPDYT | 0.5093828 |
| LGGKNPMI | ASSQGGAPDYT | 0.6999286 |
| LGPKGVAAIL | ASSQGGAPDYT | 0.7120945 |
| LMVGVSI | ASSQGGAPDYT | 0.33287266 |
| LVGAAFVL | ASSQGGAPDYT | 0.4808546 |
| LVLAVAPL | ASSQGGAPDYT | 0.3100369 |
| MAAIAVIDAA | ASSQGGAPDYT | 0.3897052 |
| NNGINAELQ | ASSQGGAPDYT | 0.37495777 |
| RGEPLVLVHG | ASSQGGAPDYT | 0.6479938 |
| RGGAGGAAGLI | ASSQGGAPDYT | 0.47562808 |
| RSYRMHPAV | ASSQGGAPDYT | 0.60478914 |
| SAAHLLAGQLA | ASSQGGAPDYT | 0.5945041 |
| SAAVGSY | ASSQGGAPDYT | 0.7034125 |
| SPHAVLHRI | ASSQGGAPDYT | 0.40437663 |
| SQIAKVL | ASSQGGAPDYT | 0.51158226 |
| TALGHLVP | ASSQGGAPDYT | 0.31111327 |
| TAPTTSGAA | ASSQGGAPDYT | 0.41899437 |
| TDGDDGNGGSPGT | ASSQGGAPDYT | 0.46155387 |
| TDGLGEN | ASSQGGAPDYT | 0.50345504 |
| TGGPPLPGQA | ASSQGGAPDYT | 0.32755378 |
| TIPAITI | ASSQGGAPDYT | 0.7186194 |
| TIPGFTI | ASSQGGAPDYT | 0.5637319 |
| TSIASGALA | ASSQGGAPDYT | 0.6571531 |
| TVTLPGAAI | ASSQGGAPDYT | 0.46004838 |
| VASGLSAALP | ASSQGGAPDYT | 0.50149626 |
| VGAGVIGI | ASSQGGAPDYT | 0.48107284 |
| VGPKRFGV | ASSQGGAPDYT | 0.6126598 |
| VNVAFPD | ASSQGGAPDYT | 0.4412616 |
| VSMTNSGVS | ASSQGGAPDYT | 0.30143878 |

| | | |
|---|---|---|
| VTDVLGDLSV | ASSQGGAPDYT | 0.468902 |
| VVVVGAGFAGL | ASSQGGAPDYT | 0.45941928 |
| VYTVGTVV | ASSQGGAPDYT | 0.7165071 |
| WTTFSGP | ASSQGGAPDYT | 0.70383215 |
| YDELVEAF | ASSQGGAPDYT | 0.6055031 |
| YGTGGAGGNGG | ASSQGGAPDYT | 0.6827073 |
| YTNVMAV | ASSQGGAPDYT | 0.2514881 |
| FLEQVHQL | ASSQGGAPDYT | 0.5026834 |
| GLSLLAVAV | ASSQGGAPDYT | 0.7035723 |
| LEQVHQL | ASSQGGAPDYT | 0.64382637 |
| LNAAGAI | ASSQGGAPDYT | 0.5100932 |
| PAVHASL | ASSQGGAPDYT | 0.7200812 |
| TELEDDA | ASSQGGAPDYT | 0.36330247 |
| TPPGPPN | ASSQGGAPDYT | 0.47963336 |
| TPPPPPPP | ASSQGGAPDYT | 0.6192311 |
| TQSPALI | ATGQGPNSDYT | 0.62107235 |
| EFAAAFPQ | ATGQGPNSDYT | 0.6081184 |
| LLSGNLDVL | ATGQGPNSDYT | 0.4138178 |
| GPANLGNYNI | ATGQGPNSDYT | 0.5612509 |
| TLGPTLP | ATGQGPNSDYT | 0.7679857 |
| VYAYSSCCK | ATGQGPNSDYT | 0.5138705 |
| APAPAPAPNI | ATGQGPNSDYT | 0.70787036 |
| PGGPVPIP | ATGQGPNSDYT | 0.6058655 |
| FLTVALEP | ATGQGPNSDYT | 0.6006406 |
| SQYFGANL | ATGQGPNSDYT | 0.39932087 |
| GGVKPAA | ATGQGPNSDYT | 0.43466866 |
| GGTGGAGGAGGW | ATGQGPNSDYT | 0.56014127 |
| GVGGVAGLGGAGG | ATGQGPNSDYT | 0.3924415 |
| VLLVLATTI | ATGQGPNSDYT | 0.51632446 |
| IGIFAAML | ATGQGPNSDYT | 0.505913 |
| AVGDLID | ATGQGPNSDYT | 0.76019144 |
| NISPLPL | ATGQGPNSDYT | 0.59856594 |
| VTAVFVAI | ATGQGPNSDYT | 0.66879296 |
| SGDDNDAAL | ATGQGPNSDYT | 0.39665943 |
| AVAALAHS | ATGQGPNSDYT | 0.5640784 |
| GGVTVMATP | ATGQGPNSDYT | 0.6510977 |
| AGLAANVRD | ATGQGPNSDYT | 0.40114996 |
| TNRPDLIDPALLRPG | ATGQGPNSDYT | 0.69820017 |
| LANLGVTAL | ATGQGPNSDYT | 0.4157894 |
| TGERIEL | ATGQGPNSDYT | 0.5396158 |
| NGEELAA | ATGQGPNSDYT | 0.6221969 |
| VLSLVWG | ATGQGPNSDYT | 0.6788645 |
| IGYALWL | ATGQGPNSDYT | 0.5000027 |
| LSLLLAPL | ATGQGPNSDYT | 0.53432345 |
| TLDEALK | ATGQGPNSDYT | 0.4134629 |
| TDGAGTAEGG | ATGQGPNSDYT | 0.6592541 |
| NAPGGAGGNGG | ATGQGPNSDYT | 0.70991576 |
| VSVFMGMGGNF | ATGQGPNSDYT | 0.5382053 |
| LASYLAAL | ATGQGPNSDYT | 0.7414508 |
| LVGGTPAL | ATGQGPNSDYT | 0.6346623 |
| VRNADVL | ATGQGPNSDYT | 0.3122043 |

| | | |
|---|---|---|
| TCSSPFLVSAM | ATGQGPNSDYT | 0.5228739 |
| VHITDAE | ATGQGPNSDYT | 0.7014976 |
| EYAAPSF | ATGQGPNSDYT | 0.7147064 |
| VVDAFHGGL | ATGQGPNSDYT | 0.7243323 |
| ASVVDPQ | ATGQGPNSDYT | 0.6290252 |
| SYNEFGGLS | ATGQGPNSDYT | 0.450866 |
| DPDGSVY | ATGQGPNSDYT | 0.7200279 |
| MVSTIHATE | ATGQGPNSDYT | 0.7459565 |
| LAAGVADPV | ATGQGPNSDYT | 0.6607841 |
| TKEQIAA | ATGQGPNSDYT | 0.42010796 |
| SLASPAAL | ATGQGPNSDYT | 0.41329834 |
| IGLVFYGV | ATGQGPNSDYT | 0.59051394 |
| FLLDASNL | ATGQGPNSDYT | 0.6826732 |
| YQVGNFP | ATGQGPNSDYT | 0.36700228 |
| ESLLSQST | ATGQGPNSDYT | 0.58641005 |
| LFTVGFAAVG | ATGQGPNSDYT | 0.5762945 |
| ATPPTPT | ATGQGPNSDYT | 0.3032076 |
| TDLATLG | ATGQGPNSDYT | 0.6645665 |
| SAGGTGR | ATGQGPNSDYT | 0.71633756 |
| GFSNSGS | ATGQGPNSDYT | 0.55789655 |
| GGNGGSGGTSV | ATGQGPNSDYT | 0.41702658 |
| GTSVATGGAGN | ATGQGPNSDYT | 0.7025696 |
| NGGNGGSGGTS | ATGQGPNSDYT | 0.7557387 |
| AVVKFPH | ATGQGPNSDYT | 0.71989506 |
| TPPPPPP | ATGQGPNSDYT | 0.30932406 |
| AAPAAAAAPA | ATGQGPNSDYT | 0.23113447 |
| IALNEDL | ATGQGPNSDYT | 0.5004079 |
| IVNVSSM | ATGQGPNSDYT | 0.64617693 |
| TGIPPAPR | ATGQGPNSDYT | 0.4793896 |
| FDLGGGTF | ATGQGPNSDYT | 0.44619483 |
| FDLGGGTFDVS | ATGQGPNSDYT | 0.6452157 |
| NEPTAAA | ATGQGPNSDYT | 0.6945231 |
| NEPTAAALA | ATGQGPNSDYT | 0.41321117 |
| GSIINISSI | ATGQGPNSDYT | 0.37233776 |
| VAPGNAGTA | ATGQGPNSDYT | 0.67250544 |
| TGGGASLE | ATGQGPNSDYT | 0.65389824 |
| VAVYNHYKRI | ATGQGPNSDYT | 0.38784993 |
| TDPAPATT | ATGQGPNSDYT | 0.512691 |
| TNRPDILDPALLRPG | ATGQGPNSDYT | 0.5180003 |
| LTTTTTS | ATGQGPNSDYT | 0.5111682 |
| PTGSGKTL | ATGQGPNSDYT | 0.6140576 |
| TGSGKTL | ATGQGPNSDYT | 0.32923084 |
| TIGHVDHGK | ATGQGPNSDYT | 0.72466516 |
| TIGHVDHGKTT | ATGQGPNSDYT | 0.37391856 |
| GHVDHGKTT | ATGQGPNSDYT | 0.6530845 |
| AQFEHTLL | ATGQGPNSDYT | 0.56962305 |
| IVNQDEL | ATGQGPNSDYT | 0.6104141 |
| TNGSQFF | ATGQGPNSDYT | 0.48880938 |
| TGKGIVE | ATGQGPNSDYT | 0.44585487 |
| LISNASDA | ATGQGPNSDYT | 0.39656007 |
| AELRTLGELTKIAKAHGAQ | ATGQGPNSDYT | 0.26284572 |

| | | |
|---|---|---|
| APNAVAAAEPSW | ATGQGPNSDYT | 0.69705766 |
| APQVAGAATAVL | ATGQGPNSDYT | 0.56929 |
| ARRDPAAAAAILPTDA | ATGQGPNSDYT | 0.6975762 |
| DAAARDVLAEAGLAEYF | ATGQGPNSDYT | 0.64502805 |
| DDQIVLSAKANLAKVEQ | ATGQGPNSDYT | 0.45159453 |
| DELLPDTGEDSDEDGH | ATGQGPNSDYT | 0.62736315 |
| ELPEAPANRATVPQD | ATGQGPNSDYT | 0.40997717 |
| FGQSKIERVTGIG | ATGQGPNSDYT | 0.46045446 |
| GAGGTGSTAGGAGGAGGAGGLY | ATGQGPNSDYT | 0.5544927 |
| GHGGTGAAGGNGGTGGAGGSGI | ATGQGPNSDYT | 0.62557614 |
| GNGGNGGAGGEGGPGGAGGAGA | ATGQGPNSDYT | 0.4110549 |
| GSTAGNGGNGGNATNGTIAGQP | ATGQGPNSDYT | 0.6158868 |
| GTGGDGGNGGNGGTGG | ATGQGPNSDYT | 0.60892916 |
| GVRRPAGIPADAPV | ATGQGPNSDYT | 0.40561914 |
| GYRQVTDAHLLTLARR | ATGQGPNSDYT | 0.50976986 |
| IAKMGQDGHDRG | ATGQGPNSDYT | 0.6894547 |
| KAEPSIEIQYT | ATGQGPNSDYT | 0.5691484 |
| KPVRIRMAADEDTVAVT | ATGQGPNSDYT | 0.44016188 |
| MPPRIAGMRLLVIKPEP | ATGQGPNSDYT | 0.65817344 |
| MKFYAVALFRSRGVPPTR | ATGQGPNSDYT | 0.38899624 |
| NAGGFGILGAVAHSP | ATGQGPNSDYT | 0.62113357 |
| NEGREITLAMAEM | ATGQGPNSDYT | 0.6098596 |
| NELVEWSHPLGHRWRWE | ATGQGPNSDYT | 0.42961693 |
| NLGAGGTGGNGGNPG | ATGQGPNSDYT | 0.6760793 |
| NTEWPKWETVMAADPVD | ATGQGPNSDYT | 0.42740974 |
| PASAAPAETQP | ATGQGPNSDYT | 0.4250178 |
| QQSSDGKNGVRAAEITGE | ATGQGPNSDYT | 0.45140234 |
| RAPDAVTTSWWKE | ATGQGPNSDYT | 0.57645243 |
| SGFGNVGAHTSGWFNQ | ATGQGPNSDYT | 0.6910376 |
| SGVAGAVGQAASVGGLKV | ATGQGPNSDYT | 0.3931871 |
| SNGGAGTGSAGGNGGT | ATGQGPNSDYT | 0.56765556 |
| TGGAGGEGGAAG | ATGQGPNSDYT | 0.45891038 |
| TGGDGGAAGAGGAPGHG | ATGQGPNSDYT | 0.36899385 |
| TMPPLLAAIANAGRHGV | ATGQGPNSDYT | 0.52415067 |
| TSSTAGNSGGA | ATGQGPNSDYT | 0.52791923 |
| VPATVPVGILC | ATGQGPNSDYT | 0.4135166 |
| VQRPDSGGVVLTGRIS | ATGQGPNSDYT | 0.61863077 |
| WTPNPVPATKGVPAGLVQAT | ATGQGPNSDYT | 0.47285667 |
| YALKRALAKFSDRFDVVII | ATGQGPNSDYT | 0.5744725 |
| YLIGSVIV | ATGQGPNSDYT | 0.7208613 |
| YGQLVAVID | ATGQGPNSDYT | 0.40574044 |
| VTVGGMTLPA | ATGQGPNSDYT | 0.6534336 |
| VLALPAV | ATGQGPNSDYT | 0.6284879 |
| VIPPTLNYET | ATGQGPNSDYT | 0.6372267 |
| VGMGGMHPSSG | ATGQGPNSDYT | 0.52174646 |
| VGLAGGAATA | ATGQGPNSDYT | 0.45012328 |
| VGGAGGAGGSGP | ATGQGPNSDYT | 0.7367055 |
| VFAAFMV | ATGQGPNSDYT | 0.6605229 |
| TLYDKLR | ATGQGPNSDYT | 0.55827177 |
| TLAVFTVL | ATGQGPNSDYT | 0.647509 |
| TGSVQPARV | ATGQGPNSDYT | 0.64296067 |

| | | |
|---|---|---|
| TGGGAGSGGT | ATGQGPNSDYT | 0.6565316 |
| SHLETSL | ATGQGPNSDYT | 0.44448453 |
| SFTTGMT | ATGQGPNSDYT | 0.3000003 |
| QVEVAKRMGVRQPTVS | ATGQGPNSDYT | 0.5212627 |
| QGSGGPALAM | ATGQGPNSDYT | 0.49440643 |
| MLAPSNP | ATGQGPNSDYT | 0.52881026 |
| LVNNAAI | ATGQGPNSDYT | 0.32007706 |
| LTGEINTLA | ATGQGPNSDYT | 0.68129915 |
| LSIFGPV | ATGQGPNSDYT | 0.75616705 |
| LLALLANL | ATGQGPNSDYT | 0.44197497 |
| LISNASDALDKLR | ATGQGPNSDYT | 0.7514551 |
| LINEAVP | ATGQGPNSDYT | 0.36252373 |
| LGYAFSGL | ATGQGPNSDYT | 0.64661354 |
| LDFVAVAL | ATGQGPNSDYT | 0.67879224 |
| LAQPGAE | ATGQGPNSDYT | 0.38343522 |
| ISNLGAVTGF | ATGQGPNSDYT | 0.62929356 |
| HQEDPGRI | ATGQGPNSDYT | 0.5221422 |
| GTPGEPTP | ATGQGPNSDYT | 0.552006 |
| GFFNSGS | ATGQGPNSDYT | 0.7055781 |
| GAGGGGLTAGAN | ATGQGPNSDYT | 0.44369 |
| FVDALVNM | ATGQGPNSDYT | 0.5669986 |
| FLTVAPEL | ATGQGPNSDYT | 0.5324432 |
| FETLTAL | ATGQGPNSDYT | 0.4250573 |
| DVINPIEL | ATGQGPNSDYT | 0.615229 |
| APVSGAPV | ATGQGPNSDYT | 0.6195674 |
| ALVGGLAW | ATGQGPNSDYT | 0.4313757 |
| ALAAAAATAI | ATGQGPNSDYT | 0.73381114 |
| AFGLGLLL | ATGQGPNSDYT | 0.42130306 |
| AFFAAASL | ATGQGPNSDYT | 0.7576472 |
| AAAALPAFSPP | ATGQGPNSDYT | 0.66982853 |
| VVLAPAL | ATGQGPNSDYT | 0.58521867 |
| TIGHVDHGKTTL | ATGQGPNSDYT | 0.50267327 |
| LISNASDALDK | ATGQGPNSDYT | 0.7017669 |
| LISNASDAL | ATGQGPNSDYT | 0.427041 |
| ISNASDALDK | ATGQGPNSDYT | 0.5993876 |
| NQAETLVYQTE | ATGQGPNSDYT | 0.71254337 |
| TGGTGGTAGAG | ATGQGPNSDYT | 0.68524766 |
| TGSTAGGAGGA | ATGQGPNSDYT | 0.8731587 |
| DPLAVPVAAGPV | ATGQGPNSDYT | 0.84074247 |
| GGTGGTPGNGGN | ATGQGPNSDYT | 0.82227886 |
| PAGLAGVDPATL | ATGQGPNSDYT | 0.71896166 |
| GNNLVVEAPHAGASV | ATGQGPNSDYT | 0.40215838 |
| KVTRIMSASTLEQPA | ATGQGPNSDYT | 0.81460667 |
| LGFALPIALPRRTVV | ATGQGPNSDYT | 0.6445915 |
| TRLGNLTARKLVVLV | ATGQGPNSDYT | 0.84917843 |
| WSQQRRLIYFPSAGP | ATGQGPNSDYT | 0.85413945 |
| GPNGNPDPMAAAVDIR | ATGQGPNSDYT | 0.7224937 |
| TQRCVSVADLRAAASR | ATGQGPNSDYT | 0.86767685 |
| QFVLDVAEHPVEVLTETV | ATGQGPNSDYT | 0.87085795 |
| GGAGGTGGAGGNGGTQAGDG | ATGQGPNSDYT | 0.5090758 |
| PLLATLRAIAVDVDAKPAQV | ATGQGPNSDYT | 0.77100325 |

| | | |
|---|---|---|
| APKFRTGGYMIDFWGVGYQVAK | ATGQGPNSDYT | 0.7397524 |
| GAGGAGGAGGAAGGPGGTGGTG | ATGQGPNSDYT | 0.80999446 |
| IGALGLLATDVAPAGSVAFGVA | ATGQGPNSDYT | 0.46051133 |
| AGGAGGIGGNGGAGGAGGVLMG | ATGQGPNSDYT | 0.8601717 |
| ADELIGLL | ATGQGPNSDYT | 0.7690383 |
| ADRALKA | ATGQGPNSDYT | 0.18890747 |
| AGAVGQVGGAGGA | ATGQGPNSDYT | 0.8205918 |
| AGGGGGLGGAP | ATGQGPNSDYT | 0.7684863 |
| AGGLLYGNGGAG | ATGQGPNSDYT | 0.75722873 |
| AGLESIGL | ATGQGPNSDYT | 0.8665019 |
| AGLKTGII | ATGQGPNSDYT | 0.58135796 |
| ASLAALLL | ATGQGPNSDYT | 0.6625143 |
| ASVINISSI | ATGQGPNSDYT | 0.8511021 |
| ATPPGPA | ATGQGPNSDYT | 0.86603546 |
| ATVASALV | ATGQGPNSDYT | 0.5249624 |
| ATVASLIL | ATGQGPNSDYT | 0.84544814 |
| AVNGPVLIRH | ATGQGPNSDYT | 0.87579286 |
| DGAAGAVGI | ATGQGPNSDYT | 0.8434956 |
| DPLLHGI | ATGQGPNSDYT | 0.8666287 |
| ELGAALGAL | ATGQGPNSDYT | 0.8749887 |
| ELGRTQAV | ATGQGPNSDYT | 0.7769767 |
| FVAGGYLS | ATGQGPNSDYT | 0.86496216 |
| GASGGAGGAGGW | ATGQGPNSDYT | 0.8772037 |
| GGAGAGGDGGAGGI | ATGQGPNSDYT | 0.74010324 |
| GTGHNGSQPGMGGNG | ATGQGPNSDYT | 0.79932356 |
| GYAAASASAS | ATGQGPNSDYT | 0.73229015 |
| HQVDVAAVQQG | ATGQGPNSDYT | 0.81149375 |
| IGAAIAHT | ATGQGPNSDYT | 0.41641068 |
| IGMIEVAQL | ATGQGPNSDYT | 0.8320507 |
| IRLAPEHTSAMD | ATGQGPNSDYT | 0.87004966 |
| IYNVSVL | ATGQGPNSDYT | 0.7662517 |
| KQGFLGM | ATGQGPNSDYT | 0.69972193 |
| LESVITQL | ATGQGPNSDYT | 0.61134636 |
| LGLGALVVV | ATGQGPNSDYT | 0.83652663 |
| LIGNGGNGGSGGI | ATGQGPNSDYT | 0.75443316 |
| LNAINAPTL | ATGQGPNSDYT | 0.46279225 |
| LQLLNTGV | ATGQGPNSDYT | 0.73719287 |
| LQTSIEF | ATGQGPNSDYT | 0.85892737 |
| MSFAVSEL | ATGQGPNSDYT | 0.8640771 |
| NTGALGL | ATGQGPNSDYT | 0.46792966 |
| PPGAGGAGGAGTT | ATGQGPNSDYT | 0.87352324 |
| QLVGLGI | ATGQGPNSDYT | 0.8569533 |
| RFLEQVHQL | ATGQGPNSDYT | 0.8815901 |
| RGVGYVLR | ATGQGPNSDYT | 0.83450484 |
| SATGVFAGVI | ATGQGPNSDYT | 0.7051475 |
| SLPAITLP | ATGQGPNSDYT | 0.86167204 |
| STALAFTV | ATGQGPNSDYT | 0.77565944 |
| SVLTVGLGK | ATGQGPNSDYT | 0.6421869 |
| TALAGAGSGI | ATGQGPNSDYT | 0.85747445 |
| TGLPFTGI | ATGQGPNSDYT | 0.8064431 |
| VAEHVTV | ATGQGPNSDYT | 0.6782497 |

| | | |
|---|---|---|
| VDIVLNSL | ATGQGPNSDYT | 0.7893335 |
| VLTGGGAVL | ATGQGPNSDYT | 0.8791884 |
| VNVGLAYGI | ATGQGPNSDYT | 0.67092633 |
| VSSAAVGAV | ATGQGPNSDYT | 0.6548354 |
| VTSVYEAGV | ATGQGPNSDYT | 0.7273123 |
| WVALDRGA | ATGQGPNSDYT | 0.81386447 |
| YLVGSVVI | ATGQGPNSDYT | 0.8528136 |
| GPPGTGKTLL | ATGQGPNSDYT | 0.8395917 |
| GVGMTKF | ATGQGPNSDYT | 0.78092957 |
| IGIGGSDL | ATGQGPNSDYT | 0.8410801 |
| IGTLLAEL | ATGQGPNSDYT | 0.8479826 |
| IYTSGTT | ATGQGPNSDYT | 0.8340969 |
| LSFPAGV | ATGQGPNSDYT | 0.85318136 |
| LYTSGTT | ATGQGPNSDYT | 0.7124535 |
| VGAFPFGF | ATGQGPNSDYT | 0.87497175 |
| VGAGTVEF | ATGQGPNSDYT | 0.845315 |
| VGKSTLLL | ATGQGPNSDYT | 0.8546858 |
| AALAHAGGGIV | ATGQGPNSDYT | 0.8638879 |
| AALVAVSLL | ATGQGPNSDYT | 0.81956375 |
| AAVSRAVAAL | ATGQGPNSDYT | 0.85066885 |
| AGGSSGIGGF | ATGQGPNSDYT | 0.41107726 |
| AGLTVANANG | ATGQGPNSDYT | 0.76179355 |
| AGLVGAVL | ATGQGPNSDYT | 0.82963026 |
| ALSQAFTV | ATGQGPNSDYT | 0.8372766 |
| ALTFSVSL | ATGQGPNSDYT | 0.85132176 |
| ALTGAALVAL | ATGQGPNSDYT | 0.83109915 |
| ASFSIGAGVL | ATGQGPNSDYT | 0.77327317 |
| DGSVYKI | ATGQGPNSDYT | 0.87005734 |
| EQLAEIV | ATGQGPNSDYT | 0.86835825 |
| FAVSELTVM | ATGQGPNSDYT | 0.49687773 |
| GAAPLAAGL | ATGQGPNSDYT | 0.77128935 |
| GADGGDGGSGGAGG | ATGQGPNSDYT | 0.88153064 |
| GFGLDTGGAGG | ATGQGPNSDYT | 0.8378396 |
| GLASGGLL | ATGQGPNSDYT | 0.78483164 |
| GRIAGAAVLL | ATGQGPNSDYT | 0.79538995 |
| GSAGSLNTSF | ATGQGPNSDYT | 0.8762362 |
| GTGQIGI | ATGQGPNSDYT | 0.7503185 |
| GTLIALIL | ATGQGPNSDYT | 0.88020295 |
| IAATMGFVVG | ATGQGPNSDYT | 0.28333828 |
| IFAQALL | ATGQGPNSDYT | 0.79682857 |
| IGAAVALGF | ATGQGPNSDYT | 0.79119307 |
| IGLTPLAL | ATGQGPNSDYT | 0.40086365 |
| ILGSNTSTL | ATGQGPNSDYT | 0.8630382 |
| IVFSLTVL | ATGQGPNSDYT | 0.79730004 |
| LASKATGF | ATGQGPNSDYT | 0.8655317 |
| LASVALIL | ATGQGPNSDYT | 0.8223182 |
| LFAVGLTL | ATGQGPNSDYT | 0.8556087 |
| LFNGGIGTYI | ATGQGPNSDYT | 0.818455 |
| LTLGAGII | ATGQGPNSDYT | 0.83341944 |
| LVAAALGDI | ATGQGPNSDYT | 0.6582097 |
| NTAFVQLGI | ATGQGPNSDYT | 0.8556502 |

| | | |
|---|---|---|
| QLFSGAEL | ATGQGPNSDYT | 0.8707912 |
| SAATFITL | ATGQGPNSDYT | 0.8604992 |
| SNYGLANLGV | ATGQGPNSDYT | 0.40155047 |
| SYGGAYAVM | ATGQGPNSDYT | 0.8022494 |
| TTGVGGLTLG | ATGQGPNSDYT | 0.80216587 |
| TTQTIYI | ATGQGPNSDYT | 0.2929249 |
| VAAGAGLGAVI | ATGQGPNSDYT | 0.27361444 |
| VAAVVTLL | ATGQGPNSDYT | 0.7942337 |
| VGFLGGLI | ATGQGPNSDYT | 0.7602824 |
| VGVIPILL | ATGQGPNSDYT | 0.8616023 |
| VIFASSLI | ATGQGPNSDYT | 0.8459699 |
| VKGGDGGAAAG | ATGQGPNSDYT | 0.77701664 |
| VTKTFTNI | ATGQGPNSDYT | 0.8567807 |
| VVAAAILL | ATGQGPNSDYT | 0.81505764 |
| YAALDVEL | ATGQGPNSDYT | 0.11004899 |
| LASVALLL | ATGQGPNSDYT | 0.7827781 |
| LDAVHRV | ATGQGPNSDYT | 0.5356367 |
| QGLSSSIF | ATGQGPNSDYT | 0.874052 |
| RGTFANIRL | ATGQGPNSDYT | 0.7203802 |
| RQLWWGHRI | ATGQGPNSDYT | 0.7272122 |
| YGPPGTGKTLL | ATGQGPNSDYT | 0.79193443 |
| AALVGAPANVA | ATGQGPNSDYT | 0.8260407 |
| AATFRRTE | ATGQGPNSDYT | 0.8163557 |
| AGALGGGI | ATGQGPNSDYT | 0.6807369 |
| AGGAGGAGGIGAG | ATGQGPNSDYT | 0.21441886 |
| AGSGALGL | ATGQGPNSDYT | 0.85158867 |
| AQIADQI | ATGQGPNSDYT | 0.47720647 |
| ASAVAVTYL | ATGQGPNSDYT | 0.5929365 |
| ASTVLGGTGG | ATGQGPNSDYT | 0.5104481 |
| AVGGEPAK | ATGQGPNSDYT | 0.288566 |
| CEQGTFGGTC | ATGQGPNSDYT | 0.46446455 |
| DEGKAPAG | ATGQGPNSDYT | 0.8609798 |
| DTVVFPAL | ATGQGPNSDYT | 0.7856693 |
| FEGITMAG | ATGQGPNSDYT | 0.75739944 |
| FPLVVQD | ATGQGPNSDYT | 0.8756354 |
| GAFTTPQ | ATGQGPNSDYT | 0.81181854 |
| GFGGFGTAGGVGGA | ATGQGPNSDYT | 0.7690946 |
| GFLAQFL | ATGQGPNSDYT | 0.68558174 |
| GGAGGAGGNAGVA | ATGQGPNSDYT | 0.6856469 |
| GGVGTAGAGGAGGAGG | ATGQGPNSDYT | 0.7465856 |
| GNGGGGDAGNAGSGGN | ATGQGPNSDYT | 0.69443345 |
| GTGGAGGAGNDAGS | ATGQGPNSDYT | 0.8782972 |
| GTTNTGWFNS | ATGQGPNSDYT | 0.8119135 |
| IFMGPLFTL | ATGQGPNSDYT | 0.64879435 |
| ILAHAIEAI | ATGQGPNSDYT | 0.85695446 |
| IRSAIREAL | ATGQGPNSDYT | 0.8777176 |
| ITFAGGG | ATGQGPNSDYT | 0.8869934 |
| ITGMSGHQEV | ATGQGPNSDYT | 0.715472 |
| IVAGLACTL | ATGQGPNSDYT | 0.8162197 |
| IVGGGLAGA | ATGQGPNSDYT | 0.799039 |
| IVTGVLNI | ATGQGPNSDYT | 0.80176246 |

| | | |
|---|---|---|
| LAQQGCI | ATGQGPNSDYT | 0.7730129 |
| LCLHVEN | ATGQGPNSDYT | 0.53257114 |
| LDAISDGS | ATGQGPNSDYT | 0.86958086 |
| LGGFAAML | ATGQGPNSDYT | 0.887411 |
| LGGKNPMI | ATGQGPNSDYT | 0.7986598 |
| LGPKGVAAIL | ATGQGPNSDYT | 0.6795594 |
| LMVGVSI | ATGQGPNSDYT | 0.7801969 |
| LVGAAFVL | ATGQGPNSDYT | 0.66993594 |
| LVLAVAPL | ATGQGPNSDYT | 0.7437295 |
| MAAIAVIDAA | ATGQGPNSDYT | 0.8170788 |
| NNGINAELQ | ATGQGPNSDYT | 0.47642052 |
| RGEPLVLVHG | ATGQGPNSDYT | 0.7884502 |
| RGGAGGAAGLI | ATGQGPNSDYT | 0.8845556 |
| RSYRMHPAV | ATGQGPNSDYT | 0.785174 |
| SAAHLLAGQLA | ATGQGPNSDYT | 0.8477312 |
| SAAVGSY | ATGQGPNSDYT | 0.8693445 |
| SPHAVLHRI | ATGQGPNSDYT | 0.85416913 |
| SQIAKVL | ATGQGPNSDYT | 0.87674737 |
| TALGHLVP | ATGQGPNSDYT | 0.8599703 |
| TAPTTSGAA | ATGQGPNSDYT | 0.7655511 |
| TDGDDGNGGSPGT | ATGQGPNSDYT | 0.82392555 |
| TDGLGEN | ATGQGPNSDYT | 0.83221805 |
| TGGPPLPGQA | ATGQGPNSDYT | 0.7994618 |
| TIPAITI | ATGQGPNSDYT | 0.79024655 |
| TIPGFTI | ATGQGPNSDYT | 0.7657754 |
| TSIASGALA | ATGQGPNSDYT | 0.86510456 |
| TVTLPGAAI | ATGQGPNSDYT | 0.8273634 |
| VASGLSAALP | ATGQGPNSDYT | 0.87210524 |
| VGAGVIGI | ATGQGPNSDYT | 0.4192056 |
| VGPKRFGV | ATGQGPNSDYT | 0.62096816 |
| VNVAFPD | ATGQGPNSDYT | 0.86708474 |
| VSMTNSGVS | ATGQGPNSDYT | 0.16459318 |
| VTDVLGDLSV | ATGQGPNSDYT | 0.31790572 |
| VVVVGAGFAGL | ATGQGPNSDYT | 0.8407886 |
| VYTVGTVV | ATGQGPNSDYT | 0.876455 |
| WTTFSGP | ATGQGPNSDYT | 0.17405173 |
| YDELVEAF | ATGQGPNSDYT | 0.55723417 |
| YGTGGAGGNGG | ATGQGPNSDYT | 0.5245505 |
| YTNVMAV | ATGQGPNSDYT | 0.8564749 |
| FLEQVHQL | ATGQGPNSDYT | 0.7780552 |
| GLSLLAVAV | ATGQGPNSDYT | 0.3203091 |
| LEQVHQL | ATGQGPNSDYT | 0.8487866 |
| LNAAGAI | ATGQGPNSDYT | 0.54312825 |
| PAVHASL | ATGQGPNSDYT | 0.622829 |
| TELEDDA | ATGQGPNSDYT | 0.8600995 |
| TPPGPPN | ATGQGPNSDYT | 0.872087 |
| TPPPPPPP | ATGQGPNSDYT | 0.6103944 |
| TQSPALI | ASSYQGGNERLF | 0.8458823 |
| EFAAAFPQ | ASSYQGGNERLF | 0.7856137 |
| LLSGNLDVL | ASSYQGGNERLF | 0.8860942 |
| GPANLGNYNI | ASSYQGGNERLF | 0.7536453 |

| | | |
|---|---|---|
| TLGPTLP | ASSYQGGNERLF | 0.5025971 |
| VYAYSSCCK | ASSYQGGNERLF | 0.86427844 |
| APAPAPAPNI | ASSYQGGNERLF | 0.85612583 |
| PGGPVPIP | ASSYQGGNERLF | 0.80012864 |
| FLTVALEP | ASSYQGGNERLF | 0.8615385 |
| SQYFGANL | ASSYQGGNERLF | 0.80367124 |
| GGVKPAA | ASSYQGGNERLF | 0.7831595 |
| GGTGGAGGAGGW | ASSYQGGNERLF | 0.8623206 |
| GVGGVAGLGGAGG | ASSYQGGNERLF | 0.8542327 |
| VLLVLATTI | ASSYQGGNERLF | 0.56143814 |
| IGIFAAML | ASSYQGGNERLF | 0.8608779 |
| AVGDLID | ASSYQGGNERLF | 0.79318464 |
| NISPLPL | ASSYQGGNERLF | 0.67247736 |
| VTAVFVAI | ASSYQGGNERLF | 0.88567454 |
| SGDDNDAAL | ASSYQGGNERLF | 0.81603885 |
| AVAALAHS | ASSYQGGNERLF | 0.86464095 |
| GGVTVMATP | ASSYQGGNERLF | 0.8640586 |
| AGLAANVRD | ASSYQGGNERLF | 0.8717563 |
| TNRPDLIDPALLRPG | ASSYQGGNERLF | 0.45148838 |
| LANLGVTAL | ASSYQGGNERLF | 0.84511197 |
| TGERIEL | ASSYQGGNERLF | 0.82237226 |
| NGEELAA | ASSYQGGNERLF | 0.8482846 |
| VLSLVWG | ASSYQGGNERLF | 0.8300291 |
| IGYALWL | ASSYQGGNERLF | 0.7129576 |
| LSLLLAPL | ASSYQGGNERLF | 0.8559885 |
| TLDEALK | ASSYQGGNERLF | 0.8731149 |
| TDGAGTAEGG | ASSYQGGNERLF | 0.8303354 |
| NAPGGAGGNGG | ASSYQGGNERLF | 0.8304731 |
| VSVFMGMGGNF | ASSYQGGNERLF | 0.8789847 |
| LASYLAAL | ASSYQGGNERLF | 0.70702696 |
| LVGGTPAL | ASSYQGGNERLF | 0.7666446 |
| VRNADVL | ASSYQGGNERLF | 0.86514616 |
| TCSSPFLVSAM | ASSYQGGNERLF | 0.8632474 |
| VHITDAE | ASSYQGGNERLF | 0.806216 |
| EYAAPSF | ASSYQGGNERLF | 0.87848204 |
| VVDAFHGGL | ASSYQGGNERLF | 0.6551025 |
| ASVVDPQ | ASSYQGGNERLF | 0.8543706 |
| SYNEFGGLS | ASSYQGGNERLF | 0.755564 |
| DPDGSVY | ASSYQGGNERLF | 0.8382089 |
| MVSTIHATE | ASSYQGGNERLF | 0.67876 |
| LAAGVADPV | ASSYQGGNERLF | 0.8483622 |
| TKEQIAA | ASSYQGGNERLF | 0.86376745 |
| SLASPAAL | ASSYQGGNERLF | 0.85369587 |
| IGLVFYGV | ASSYQGGNERLF | 0.2856932 |
| FLLDASNL | ASSYQGGNERLF | 0.8494347 |
| YQVGNFP | ASSYQGGNERLF | 0.8547226 |
| ESLLSQST | ASSYQGGNERLF | 0.8852259 |
| LFTVGFAAVG | ASSYQGGNERLF | 0.8110676 |
| ATPPTPT | ASSYQGGNERLF | 0.85498536 |
| TDLATLG | ASSYQGGNERLF | 0.78031945 |
| SAGGTGR | ASSYQGGNERLF | 0.81592 |

| | | |
|---|---|---|
| GFSNSGS | ASSYQGGNERLF | 0.66035724 |
| GGNGGSGGTSV | ASSYQGGNERLF | 0.8074547 |
| GTSVATGGAGN | ASSYQGGNERLF | 0.72996324 |
| NGGNGGSGGTS | ASSYQGGNERLF | 0.69833994 |
| AVVKFPH | ASSYQGGNERLF | 0.8566815 |
| TPPPPPP | ASSYQGGNERLF | 0.86510265 |
| AAPAAAAAPA | ASSYQGGNERLF | 0.8607564 |
| IALNEDL | ASSYQGGNERLF | 0.8696978 |
| IVNVSSM | ASSYQGGNERLF | 0.8744664 |
| TGIPPAPR | ASSYQGGNERLF | 0.8591316 |
| FDLGGGTF | ASSYQGGNERLF | 0.7523371 |
| FDLGGGTFDVS | ASSYQGGNERLF | 0.79546404 |
| NEPTAAA | ASSYQGGNERLF | 0.6467409 |
| NEPTAAALA | ASSYQGGNERLF | 0.78854954 |
| GSIINISSI | ASSYQGGNERLF | 0.7617239 |
| VAPGNAGTA | ASSYQGGNERLF | 0.8129693 |
| TGGGASLE | ASSYQGGNERLF | 0.13199098 |
| VAVYNHYKRI | ASSYQGGNERLF | 0.8204478 |
| TDPAPATT | ASSYQGGNERLF | 0.8767699 |
| TNRPDILDPALLRPG | ASSYQGGNERLF | 0.7420083 |
| LTTTTTS | ASSYQGGNERLF | 0.8528981 |
| PTGSGKTL | ASSYQGGNERLF | 0.8235881 |
| TGSGKTL | ASSYQGGNERLF | 0.76406443 |
| TIGHVDHGK | ASSYQGGNERLF | 0.853869 |
| TIGHVDHGKTT | ASSYQGGNERLF | 0.6678766 |
| GHVDHGKTT | ASSYQGGNERLF | 0.8584163 |
| AQFEHTLL | ASSYQGGNERLF | 0.88341767 |
| IVNQDEL | ASSYQGGNERLF | 0.83777624 |
| TNGSQFF | ASSYQGGNERLF | 0.82021964 |
| TGKGIVE | ASSYQGGNERLF | 0.81003606 |
| LISNASDA | ASSYQGGNERLF | 0.36167344 |
| AELRTLGELTKIAKAHGAQ | ASSYQGGNERLF | 0.82512116 |
| APNAVAAAEPSW | ASSYQGGNERLF | 0.124403074 |
| APQVAGAATAVL | ASSYQGGNERLF | 0.81625485 |
| ARRDPAAAAAILPTDA | ASSYQGGNERLF | 0.7231577 |
| DAAARDVLAEAGLAEYF | ASSYQGGNERLF | 0.8151729 |
| DDQIVLSAKANLAKVEQ | ASSYQGGNERLF | 0.8011484 |
| DELLPDTGEDSDEDGH | ASSYQGGNERLF | 0.8493626 |
| ELPEAPANRATVPQD | ASSYQGGNERLF | 0.70559025 |
| FGQSKIERVTGIG | ASSYQGGNERLF | 0.79729235 |
| GAGGTGSTAGGAGGAGGAGGLY | ASSYQGGNERLF | 0.78816867 |
| GHGGTGAAGGNGGTGGAGGSGI | ASSYQGGNERLF | 0.5768271 |
| GNGGNGGAGGEGGPGGAGGAGA | ASSYQGGNERLF | 0.86672884 |
| GSTAGNGGNGGNATNGTIAGQP | ASSYQGGNERLF | 0.82449365 |
| GTGGDGGNGGNGGTGG | ASSYQGGNERLF | 0.7906301 |
| GVRRPAGIPADAPV | ASSYQGGNERLF | 0.869307 |
| GYRQVTDAHLLTLARR | ASSYQGGNERLF | 0.799825 |
| IAKMGQDGHDRG | ASSYQGGNERLF | 0.48221147 |
| KAEPSIEIQYT | ASSYQGGNERLF | 0.7973424 |
| KPVRIRMAADEDTVAVT | ASSYQGGNERLF | 0.8261305 |
| MPPRIAGMRLLVIKPEP | ASSYQGGNERLF | 0.78251576 |

| | | |
|---|---|---|
| MKFYAVALFRSRGVPPTR | ASSYQGGNERLF | 0.14130764 |
| NAGGFGILGAVAHSP | ASSYQGGNERLF | 0.84887004 |
| NEGREITLAMAEM | ASSYQGGNERLF | 0.6726643 |
| NELVEWSHPLGHRWRWE | ASSYQGGNERLF | 0.7024514 |
| NLGAGGTGGNGGNPG | ASSYQGGNERLF | 0.75879526 |
| NTEWPKWETVMAADPVD | ASSYQGGNERLF | 0.85621023 |
| PASAAPAETQP | ASSYQGGNERLF | 0.59742725 |
| QQSSDGKNGVRAAEITGE | ASSYQGGNERLF | 0.8229957 |
| RAPDAVTTSWWKE | ASSYQGGNERLF | 0.8575492 |
| SGFGNVGAHTSGWFNQ | ASSYQGGNERLF | 0.81394196 |
| SGVAGAVGQAASVGGLKV | ASSYQGGNERLF | 0.8750293 |
| SNGGAGTGSAGGNGGT | ASSYQGGNERLF | 0.8511339 |
| TGGAGGEGGAAG | ASSYQGGNERLF | 0.82949656 |
| TGGDGGAAGAGGAPGHG | ASSYQGGNERLF | 0.84204894 |
| TMPPLLAAIANAGRHGV | ASSYQGGNERLF | 0.82163644 |
| TSSTAGNSGGA | ASSYQGGNERLF | 0.460177 |
| VPATVPVGILC | ASSYQGGNERLF | 0.5057856 |
| VQRPDSGGVVLTGRIS | ASSYQGGNERLF | 0.82372355 |
| WTPNPVPATKGVPAGLVQAT | ASSYQGGNERLF | 0.43173784 |
| YALKRALAKFSDRFDVVII | ASSYQGGNERLF | 0.8245129 |
| YLIGSVIV | ASSYQGGNERLF | 0.8696784 |
| YGQLVAVID | ASSYQGGNERLF | 0.81945324 |
| VTVGGMTLPA | ASSYQGGNERLF | 0.64776 |
| VLALPAV | ASSYQGGNERLF | 0.7964233 |
| VIPPTLNYET | ASSYQGGNERLF | 0.6457257 |
| VGMGGMHPSSG | ASSYQGGNERLF | 0.8600888 |
| VGLAGGAATA | ASSYQGGNERLF | 0.8666992 |
| VGGAGGAGGSGP | ASSYQGGNERLF | 0.87790716 |
| VFAAFMV | ASSYQGGNERLF | 0.8604313 |
| TLYDKLR | ASSYQGGNERLF | 0.7470738 |
| TLAVFTVL | ASSYQGGNERLF | 0.8588786 |
| TGSVQPARV | ASSYQGGNERLF | 0.78875065 |
| TGGGAGSGGT | ASSYQGGNERLF | 0.82818437 |
| SHLETSL | ASSYQGGNERLF | 0.8230059 |
| SFTTGMT | ASSYQGGNERLF | 0.8420538 |
| QVEVAKRMGVRQPTVS | ASSYQGGNERLF | 0.75736463 |
| QGSGGPALAM | ASSYQGGNERLF | 0.76303303 |
| MLAPSNP | ASSYQGGNERLF | 0.54805017 |
| LVNNAAI | ASSYQGGNERLF | 0.85368127 |
| LTGEINTLA | ASSYQGGNERLF | 0.8193151 |
| LSIFGPV | ASSYQGGNERLF | 0.52648246 |
| LLALLANL | ASSYQGGNERLF | 0.74302727 |
| LISNASDALDKLR | ASSYQGGNERLF | 0.8775115 |
| LINEAVP | ASSYQGGNERLF | 0.8733226 |
| LGYAFSGL | ASSYQGGNERLF | 0.6861973 |
| LDFVAVAL | ASSYQGGNERLF | 0.6894465 |
| LAQPGAE | ASSYQGGNERLF | 0.81795424 |
| ISNLGAVTGF | ASSYQGGNERLF | 0.44807553 |
| HQEDPGRI | ASSYQGGNERLF | 0.85656965 |
| GTPGEPTP | ASSYQGGNERLF | 0.78747594 |
| GFFNSGS | ASSYQGGNERLF | 0.7834838 |

| | | |
|---|---|---|
| GAGGGGLTAGAN | ASSYQGGNERLF | 0.75009716 |
| FVDALVNM | ASSYQGGNERLF | 0.8333969 |
| FLTVAPEL | ASSYQGGNERLF | 0.76995206 |
| FETLTAL | ASSYQGGNERLF | 0.85478127 |
| DVINPIEL | ASSYQGGNERLF | 0.179197 |
| APVSGAPV | ASSYQGGNERLF | 0.83884263 |
| ALVGGLAW | ASSYQGGNERLF | 0.69800735 |
| ALAAAAATAI | ASSYQGGNERLF | 0.7431302 |
| AFGLGLLL | ASSYQGGNERLF | 0.40916365 |
| AFFAAASL | ASSYQGGNERLF | 0.8221391 |
| AAAALPAFSPP | ASSYQGGNERLF | 0.8660414 |
| VVLAPAL | ASSYQGGNERLF | 0.8685782 |
| TIGHVDHGKTTL | ASSYQGGNERLF | 0.7125446 |
| LISNASDALDK | ASSYQGGNERLF | 0.8621185 |
| LISNASDAL | ASSYQGGNERLF | 0.88010526 |
| ISNASDALDK | ASSYQGGNERLF | 0.7664299 |
| NQAETLVYQTE | ASSYQGGNERLF | 0.33952308 |
| TGGTGGTAGAG | ASSYQGGNERLF | 0.8297876 |
| TGSTAGGAGGA | ASSYQGGNERLF | 0.8068645 |
| DPLAVPVAAGPV | ASSYQGGNERLF | 0.856109 |
| GGTGGTPGNGGN | ASSYQGGNERLF | 0.8600483 |
| PAGLAGVDPATL | ASSYQGGNERLF | 0.581988 |
| GNNLVVEAPHAGASV | ASSYQGGNERLF | 0.85816777 |
| KVTRIMSASTLEQPA | ASSYQGGNERLF | 0.8658832 |
| LGFALPIALPRRTVV | ASSYQGGNERLF | 0.6806886 |
| TRLGNLTARKLVVLV | ASSYQGGNERLF | 0.79364145 |
| WSQQRRLIYFPSAGP | ASSYQGGNERLF | 0.7510574 |
| GPNGNPDPMAAAVDIR | ASSYQGGNERLF | 0.4257425 |
| TQRCVSVADLRAAASR | ASSYQGGNERLF | 0.8321948 |
| QFVLDVAEHPVEVLTETV | ASSYQGGNERLF | 0.7490182 |
| GGAGGTGGAGGNGGTQAGDG | ASSYQGGNERLF | 0.67294264 |
| PLLATLRAIAVDVDAKPAQV | ASSYQGGNERLF | 0.81791484 |
| APKFRTGGYMIDFWGVGYQVAK | ASSYQGGNERLF | 0.67691636 |
| GAGGAGGAGGAAGGPGGTGGTG | ASSYQGGNERLF | 0.6599849 |
| IGALGLLATDVAPAGSVAFGVA | ASSYQGGNERLF | 0.721645 |
| AGGAGGIGGNGGAGGAGGVLMG | ASSYQGGNERLF | 0.82922983 |
| ADELIGLL | ASSYQGGNERLF | 0.84931123 |
| ADRALKA | ASSYQGGNERLF | 0.8048587 |
| AGAVGQVGGAGGA | ASSYQGGNERLF | 0.83626235 |
| AGGGGGLGGAP | ASSYQGGNERLF | 0.8715682 |
| AGGLLYGNGGAG | ASSYQGGNERLF | 0.48659706 |
| AGLESIGL | ASSYQGGNERLF | 0.5826492 |
| AGLKTGII | ASSYQGGNERLF | 0.85059524 |
| ASLAALLL | ASSYQGGNERLF | 0.6157119 |
| ASVINISSI | ASSYQGGNERLF | 0.86177105 |
| ATPPGPA | ASSYQGGNERLF | 0.80959076 |
| ATVASALV | ASSYQGGNERLF | 0.829923 |
| ATVASLIL | ASSYQGGNERLF | 0.8154616 |
| AVNGPVLIRH | ASSYQGGNERLF | 0.8859675 |
| DGAAGAVGI | ASSYQGGNERLF | 0.85710263 |
| DPLLHGI | ASSYQGGNERLF | 0.82537496 |

| | | |
|---|---|---|
| ELGAALGAL | ASSYQGGNERLF | 0.84081495 |
| ELGRTQAV | ASSYQGGNERLF | 0.66645235 |
| FVAGGYLS | ASSYQGGNERLF | 0.8655371 |
| GASGGAGGAGGW | ASSYQGGNERLF | 0.861784 |
| GGAGAGGDGGAGGI | ASSYQGGNERLF | 0.61036897 |
| GTGHNGSQPGMGGNG | ASSYQGGNERLF | 0.7896521 |
| GYAAASASAS | ASSYQGGNERLF | 0.8784781 |
| HQVDVAAVQQG | ASSYQGGNERLF | 0.87341404 |
| IGAAIAHT | ASSYQGGNERLF | 0.47350618 |
| IGMIEVAQL | ASSYQGGNERLF | 0.74901974 |
| IRLAPEHTSAMD | ASSYQGGNERLF | 0.40899107 |
| IYNVSVL | ASSYQGGNERLF | 0.79826725 |
| KQGFLGM | ASSYQGGNERLF | 0.5226504 |
| LESVITQL | ASSYQGGNERLF | 0.8502711 |
| LGLGALVVV | ASSYQGGNERLF | 0.7637522 |
| LIGNGGNGGSGGI | ASSYQGGNERLF | 0.8535584 |
| LNAINAPTL | ASSYQGGNERLF | 0.81311846 |
| LQLLNTGV | ASSYQGGNERLF | 0.86931765 |
| LQTSIEF | ASSYQGGNERLF | 0.7811354 |
| MSFAVSEL | ASSYQGGNERLF | 0.77649057 |
| NTGALGL | ASSYQGGNERLF | 0.53710914 |
| PPGAGGAGGAGTT | ASSYQGGNERLF | 0.52338535 |
| QLVGLGI | ASSYQGGNERLF | 0.768455 |
| RFLEQVHQL | ASSYQGGNERLF | 0.76634896 |
| RGVGYVLR | ASSYQGGNERLF | 0.2726943 |
| SATGVFAGVI | ASSYQGGNERLF | 0.87657845 |
| SLPAITLP | ASSYQGGNERLF | 0.8300077 |
| STALAFTV | ASSYQGGNERLF | 0.8535497 |
| SVLTVGLGK | ASSYQGGNERLF | 0.6678296 |
| TALAGAGSGI | ASSYQGGNERLF | 0.7390596 |
| TGLPFTGI | ASSYQGGNERLF | 0.74027914 |
| VAEHVTV | ASSYQGGNERLF | 0.85204554 |
| VDIVLNSL | ASSYQGGNERLF | 0.67122215 |
| VLTGGGAVL | ASSYQGGNERLF | 0.5274669 |
| VNVGLAYGI | ASSYQGGNERLF | 0.763458 |
| VSSAAVGAV | ASSYQGGNERLF | 0.65487117 |
| VTSVYEAGV | ASSYQGGNERLF | 0.88192534 |
| WVALDRGA | ASSYQGGNERLF | 0.8692303 |
| YLVGSVVI | ASSYQGGNERLF | 0.83209884 |
| GPPGTGKTLL | ASSYQGGNERLF | 0.876917 |
| GVGMTKF | ASSYQGGNERLF | 0.26793277 |
| IGIGGSDL | ASSYQGGNERLF | 0.6937757 |
| IGTLLAEL | ASSYQGGNERLF | 0.87649024 |
| IYTSGTT | ASSYQGGNERLF | 0.85420346 |
| LSFPAGV | ASSYQGGNERLF | 0.7844051 |
| LYTSGTT | ASSYQGGNERLF | 0.8707528 |
| VGAFPFGF | ASSYQGGNERLF | 0.67104095 |
| VGAGTVEF | ASSYQGGNERLF | 0.7739469 |
| VGKSTLLL | ASSYQGGNERLF | 0.8185744 |
| AALAHAGGGIV | ASSYQGGNERLF | 0.8529861 |
| AALVAVSLL | ASSYQGGNERLF | 0.84912187 |

| | | |
|---|---|---|
| AAVSRAVAAL | ASSYQGGNERLF | 0.6420207 |
| AGGSSGIGGF | ASSYQGGNERLF | 0.83518285 |
| AGLTVANANG | ASSYQGGNERLF | 0.88482976 |
| AGLVGAVL | ASSYQGGNERLF | 0.7592077 |
| ALSQAFTV | ASSYQGGNERLF | 0.85556066 |
| ALTFSVSL | ASSYQGGNERLF | 0.8373981 |
| ALTGAALVAL | ASSYQGGNERLF | 0.837013 |
| ASFSIGAGVL | ASSYQGGNERLF | 0.6894871 |
| DGSVYKI | ASSYQGGNERLF | 0.7020774 |
| EQLAEIV | ASSYQGGNERLF | 0.81215286 |
| FAVSELTVM | ASSYQGGNERLF | 0.6274606 |
| GAAPLAAGL | ASSYQGGNERLF | 0.8241467 |
| GADGGDGGSGGAGG | ASSYQGGNERLF | 0.7701563 |
| GFGLDTGGAGG | ASSYQGGNERLF | 0.8905273 |
| GLASGGLL | ASSYQGGNERLF | 0.85686433 |
| GRIAGAAVLL | ASSYQGGNERLF | 0.8307606 |
| GSAGSLNTSF | ASSYQGGNERLF | 0.66995144 |
| GTGQIGI | ASSYQGGNERLF | 0.8427223 |
| GTLIALIL | ASSYQGGNERLF | 0.8367889 |
| IAATMGFVVG | ASSYQGGNERLF | 0.5495079 |
| IFAQALL | ASSYQGGNERLF | 0.8747412 |
| IGAAVALGF | ASSYQGGNERLF | 0.5963726 |
| IGLTPLAL | ASSYQGGNERLF | 0.8017123 |
| ILGSNTSTL | ASSYQGGNERLF | 0.8542948 |
| IVFSLTVL | ASSYQGGNERLF | 0.8707806 |
| LASKATGF | ASSYQGGNERLF | 0.7949871 |
| LASVALIL | ASSYQGGNERLF | 0.7868961 |
| LFAVGLTL | ASSYQGGNERLF | 0.6313422 |
| LFNGGIGTYI | ASSYQGGNERLF | 0.86882365 |
| LTLGAGII | ASSYQGGNERLF | 0.86565024 |
| LVAAALGDI | ASSYQGGNERLF | 0.8208117 |
| NTAFVQLGI | ASSYQGGNERLF | 0.8795781 |
| QLFSGAEL | ASSYQGGNERLF | 0.84581566 |
| SAATFITL | ASSYQGGNERLF | 0.41213265 |
| SNYGLANLGV | ASSYQGGNERLF | 0.8167424 |
| SYGGAYAVM | ASSYQGGNERLF | 0.85908055 |
| TTGVGGLTLG | ASSYQGGNERLF | 0.87147653 |
| TTQTIYI | ASSYQGGNERLF | 0.87746257 |
| VAAGAGLGAVI | ASSYQGGNERLF | 0.8510835 |
| VAAVVTLL | ASSYQGGNERLF | 0.78338087 |
| VGFLGGLI | ASSYQGGNERLF | 0.87499434 |
| VGVIPILL | ASSYQGGNERLF | 0.8904947 |
| VIFASSLI | ASSYQGGNERLF | 0.8537437 |
| VKGGDGGAAAG | ASSYQGGNERLF | 0.7341521 |
| VTKTFTNI | ASSYQGGNERLF | 0.6773287 |
| VVAAAILL | ASSYQGGNERLF | 0.8485007 |
| YAALDVEL | ASSYQGGNERLF | 0.8618397 |
| LASVALLL | ASSYQGGNERLF | 0.6965092 |
| LDAVHRV | ASSYQGGNERLF | 0.8179759 |
| QGLSSSIF | ASSYQGGNERLF | 0.83109856 |
| RGTFANIRL | ASSYQGGNERLF | 0.47192803 |

| | | |
|---|---|---|
| RQLWWGHRI | ASSYQGGNERLF | 0.8524606 |
| YGPPGTGKTLL | ASSYQGGNERLF | 0.8747047 |
| AALVGAPANVA | ASSYQGGNERLF | 0.8204708 |
| AATFRRTE | ASSYQGGNERLF | 0.7337016 |
| AGALGGGI | ASSYQGGNERLF | 0.8693443 |
| AGGAGGAGGIGAG | ASSYQGGNERLF | 0.8868773 |
| AGSGALGL | ASSYQGGNERLF | 0.8764365 |
| AQIADQI | ASSYQGGNERLF | 0.3804829 |
| ASAVAVTYL | ASSYQGGNERLF | 0.34789732 |
| ASTVLGGTGG | ASSYQGGNERLF | 0.7425388 |
| AVGGEPAK | ASSYQGGNERLF | 0.854694 |
| CEQGTFGGTC | ASSYQGGNERLF | 0.76644826 |
| DEGKAPAG | ASSYQGGNERLF | 0.73468924 |
| DTVVFPAL | ASSYQGGNERLF | 0.8362206 |
| FEGITMAG | ASSYQGGNERLF | 0.8681904 |
| FPLVVQD | ASSYQGGNERLF | 0.68632007 |
| GAFTTPQ | ASSYQGGNERLF | 0.5986395 |
| GFGGFGTAGGVGGA | ASSYQGGNERLF | 0.86057335 |
| GFLAQFL | ASSYQGGNERLF | 0.86053395 |
| GGAGGAGGNAGVA | ASSYQGGNERLF | 0.66219884 |
| GGVGTAGAGGAGGAGG | ASSYQGGNERLF | 0.8031752 |
| GNGGGGDAGNAGSGGN | ASSYQGGNERLF | 0.73939264 |
| GTGGAGGAGNDAGS | ASSYQGGNERLF | 0.7069874 |
| GTTNTGWFNS | ASSYQGGNERLF | 0.82095855 |
| IFMGPLFTL | ASSYQGGNERLF | 0.5480032 |
| ILAHAIEAI | ASSYQGGNERLF | 0.86442107 |
| IRSAIREAL | ASSYQGGNERLF | 0.60959685 |
| ITFAGGG | ASSYQGGNERLF | 0.8663614 |
| ITGMSGHQEV | ASSYQGGNERLF | 0.82044643 |
| IVAGLACTL | ASSYQGGNERLF | 0.8418294 |
| IVGGGLAGA | ASSYQGGNERLF | 0.77666605 |
| IVTGVLNI | ASSYQGGNERLF | 0.72113216 |
| LAQQGCI | ASSYQGGNERLF | 0.6406566 |
| LCLHVEN | ASSYQGGNERLF | 0.4781843 |
| LDAISDGS | ASSYQGGNERLF | 0.86636245 |
| LGGFAAML | ASSYQGGNERLF | 0.26241708 |
| LGGKNPMI | ASSYQGGNERLF | 0.8611998 |
| LGPKGVAAIL | ASSYQGGNERLF | 0.8486301 |
| LMVGVSI | ASSYQGGNERLF | 0.61453843 |
| LVGAAFVL | ASSYQGGNERLF | 0.8492527 |
| LVLAVAPL | ASSYQGGNERLF | 0.63034356 |
| MAAIAVIDAA | ASSYQGGNERLF | 0.7830759 |
| NNGINAELQ | ASSYQGGNERLF | 0.86012423 |
| RGEPLVLVHG | ASSYQGGNERLF | 0.8481866 |
| RGGAGGAAGLI | ASSYQGGNERLF | 0.7076305 |
| RSYRMHPAV | ASSYQGGNERLF | 0.8510261 |
| SAAHLLAGQLA | ASSYQGGNERLF | 0.84673274 |
| SAAVGSY | ASSYQGGNERLF | 0.77844566 |
| SPHAVLHRI | ASSYQGGNERLF | 0.8191922 |
| SQIAKVL | ASSYQGGNERLF | 0.85992974 |
| TALGHLVP | ASSYQGGNERLF | 0.84372836 |

| | | |
|---|---|---|
| TAPTTSGAA | ASSYQGGNERLF | 0.68261075 |
| TDGDDGNGGSPGT | ASSYQGGNERLF | 0.84883094 |
| TDGLGEN | ASSYQGGNERLF | 0.7033806 |
| TGGPPLPGQA | ASSYQGGNERLF | 0.83033276 |
| TIPAITI | ASSYQGGNERLF | 0.81087494 |
| TIPGFTI | ASSYQGGNERLF | 0.79851496 |
| TSIASGALA | ASSYQGGNERLF | 0.85200346 |
| TVTLPGAAI | ASSYQGGNERLF | 0.67853785 |
| VASGLSAALP | ASSYQGGNERLF | 0.74627906 |
| VGAGVIGI | ASSYQGGNERLF | 0.7786368 |
| VGPKRFGV | ASSYQGGNERLF | 0.7882571 |
| VNVAFPD | ASSYQGGNERLF | 0.8531124 |
| VSMTNSGVS | ASSYQGGNERLF | 0.6580676 |
| VTDVLGDLSV | ASSYQGGNERLF | 0.83109957 |
| VVVVGAGFAGL | ASSYQGGNERLF | 0.7802553 |
| VYTVGTVV | ASSYQGGNERLF | 0.6318188 |
| WTTFSGP | ASSYQGGNERLF | 0.8272051 |
| YDELVEAF | ASSYQGGNERLF | 0.80490154 |
| YGTGGAGGNGG | ASSYQGGNERLF | 0.67925376 |
| YTNVMAV | ASSYQGGNERLF | 0.8488461 |
| FLEQVHQL | ASSYQGGNERLF | 0.7071204 |
| GLSLLAVAV | ASSYQGGNERLF | 0.8260628 |
| LEQVHQL | ASSYQGGNERLF | 0.8790163 |
| LNAAGAI | ASSYQGGNERLF | 0.7140303 |
| PAVHASL | ASSYQGGNERLF | 0.8642471 |
| TELEDDA | ASSYQGGNERLF | 0.83297217 |
| TPPGPPN | ASSYQGGNERLF | 0.8348112 |
| TPPPPPPP | ASSYQGGNERLF | 0.7539799 |
| TQSPALI | ASSDAGLGASAETLY | 0.7388871 |
| EFAAAFPQ | ASSDAGLGASAETLY | 0.8667829 |
| LLSGNLDVL | ASSDAGLGASAETLY | 0.81739044 |
| GPANLGNYNI | ASSDAGLGASAETLY | 0.8624687 |
| TLGPTLP | ASSDAGLGASAETLY | 0.84622985 |
| VYAYSSCCK | ASSDAGLGASAETLY | 0.8594614 |
| APAPAPAPNI | ASSDAGLGASAETLY | 0.8492995 |
| PGGPVPIP | ASSDAGLGASAETLY | 0.69830215 |
| FLTVALEP | ASSDAGLGASAETLY | 0.5268749 |
| SQYFGANL | ASSDAGLGASAETLY | 0.76740074 |
| GGVKPAA | ASSDAGLGASAETLY | 0.7825578 |
| GGTGGAGGAGGW | ASSDAGLGASAETLY | 0.7813835 |
| GVGGVAGLGGAGG | ASSDAGLGASAETLY | 0.36999786 |
| VLLVLATTI | ASSDAGLGASAETLY | 0.862661 |
| IGIFAAML | ASSDAGLGASAETLY | 0.88478476 |
| AVGDLID | ASSDAGLGASAETLY | 0.6208761 |
| NISPLPL | ASSDAGLGASAETLY | 0.8830788 |
| VTAVFVAI | ASSDAGLGASAETLY | 0.6732575 |
| SGDDNDAAL | ASSDAGLGASAETLY | 0.83746946 |
| AVAALAHS | ASSDAGLGASAETLY | 0.8712558 |
| GGVTVMATP | ASSDAGLGASAETLY | 0.5389269 |
| AGLAANVRD | ASSDAGLGASAETLY | 0.8412343 |
| TNRPDLIDPALLRPG | ASSDAGLGASAETLY | 0.6988675 |

| | | |
|---|---|---|
| LANLGVTAL | ASSDAGLGASAETLY | 0.7417486 |
| TGERIEL | ASSDAGLGASAETLY | 0.87117124 |
| NGEELAA | ASSDAGLGASAETLY | 0.7269167 |
| VLSLVWG | ASSDAGLGASAETLY | 0.83663505 |
| IGYALWL | ASSDAGLGASAETLY | 0.75442123 |
| LSLLLAPL | ASSDAGLGASAETLY | 0.679492 |
| TLDEALK | ASSDAGLGASAETLY | 0.8204528 |
| TDGAGTAEGG | ASSDAGLGASAETLY | 0.8693254 |
| NAPGGAGGNGG | ASSDAGLGASAETLY | 0.74741757 |
| VSVFMGMGGNF | ASSDAGLGASAETLY | 0.8889086 |
| LASYLAAL | ASSDAGLGASAETLY | 0.70154786 |
| LVGGTPAL | ASSDAGLGASAETLY | 0.8791101 |
| VRNADVL | ASSDAGLGASAETLY | 0.86321497 |
| TCSSPFLVSAM | ASSDAGLGASAETLY | 0.8112115 |
| VHITDAE | ASSDAGLGASAETLY | 0.78760135 |
| EYAAPSF | ASSDAGLGASAETLY | 0.86319065 |
| VVDAFHGGL | ASSDAGLGASAETLY | 0.7851771 |
| ASVVDPQ | ASSDAGLGASAETLY | 0.8493331 |
| SYNEFGGLS | ASSDAGLGASAETLY | 0.86910903 |
| DPDGSVY | ASSDAGLGASAETLY | 0.32290837 |
| MVSTIHATE | ASSDAGLGASAETLY | 0.8690665 |
| LAAGVADPV | ASSDAGLGASAETLY | 0.72504926 |
| TKEQIAA | ASSDAGLGASAETLY | 0.6709707 |
| SLASPAAL | ASSDAGLGASAETLY | 0.42810863 |
| IGLVFYGV | ASSDAGLGASAETLY | 0.28889096 |
| FLLDASNL | ASSDAGLGASAETLY | 0.5712015 |
| YQVGNFP | ASSDAGLGASAETLY | 0.31834835 |
| ESLLSQST | ASSDAGLGASAETLY | 0.8315783 |
| LFTVGFAAVG | ASSDAGLGASAETLY | 0.73711735 |
| ATPPTPT | ASSDAGLGASAETLY | 0.46466285 |
| TDLATLG | ASSDAGLGASAETLY | 0.7174903 |
| SAGGTGR | ASSDAGLGASAETLY | 0.8015382 |
| GFSNSGS | ASSDAGLGASAETLY | 0.3159984 |
| GGNGGSGGTSV | ASSDAGLGASAETLY | 0.563069 |
| GTSVATGGAGN | ASSDAGLGASAETLY | 0.5262868 |
| NGGNGGSGGTS | ASSDAGLGASAETLY | 0.5764599 |
| AVVKFPH | ASSDAGLGASAETLY | 0.24809 |
| TPPPPPP | ASSDAGLGASAETLY | 0.7106827 |
| AAPAAAAAPA | ASSDAGLGASAETLY | 0.45441204 |
| IALNEDL | ASSDAGLGASAETLY | 0.14604633 |
| IVNVSSM | ASSDAGLGASAETLY | 0.69825643 |
| TGIPPAPR | ASSDAGLGASAETLY | 0.5023457 |
| FDLGGGTF | ASSDAGLGASAETLY | 0.43518186 |
| FDLGGGTFDVS | ASSDAGLGASAETLY | 0.7432399 |
| NEPTAAA | ASSDAGLGASAETLY | 0.66807806 |
| NEPTAAALA | ASSDAGLGASAETLY | 0.3341719 |
| GSIINISSI | ASSDAGLGASAETLY | 0.7595842 |
| VAPGNAGTA | ASSDAGLGASAETLY | 0.792762 |
| TGGGASLE | ASSDAGLGASAETLY | 0.26768294 |
| VAVYNHYKRI | ASSDAGLGASAETLY | 0.6572337 |
| TDPAPATT | ASSDAGLGASAETLY | 0.84948474 |

| | | |
|---|---|---|
| TNRPDILDPALLRPG | ASSDAGLGASAETLY | 0.5684806 |
| LTTTTTS | ASSDAGLGASAETLY | 0.82895964 |
| PTGSGKTL | ASSDAGLGASAETLY | 0.8391443 |
| TGSGKTL | ASSDAGLGASAETLY | 0.57543474 |
| TIGHVDHGK | ASSDAGLGASAETLY | 0.7630694 |
| TIGHVDHGKTT | ASSDAGLGASAETLY | 0.8260922 |
| GHVDHGKTT | ASSDAGLGASAETLY | 0.6938045 |
| AQFEHTLL | ASSDAGLGASAETLY | 0.5801369 |
| IVNQDEL | ASSDAGLGASAETLY | 0.33510882 |
| TNGSQFF | ASSDAGLGASAETLY | 0.65964043 |
| TGKGIVE | ASSDAGLGASAETLY | 0.38680375 |
| LISNASDA | ASSDAGLGASAETLY | 0.67960477 |
| AELRTLGELTKIAKAHGAQ | ASSDAGLGASAETLY | 0.83117104 |
| APNAVAAAEPSW | ASSDAGLGASAETLY | 0.51740265 |
| APQVAGAATAVL | ASSDAGLGASAETLY | 0.25362802 |
| ARRDPAAAAAILPTDA | ASSDAGLGASAETLY | 0.34809452 |
| DAAARDVLAEAGLAEYF | ASSDAGLGASAETLY | 0.7870928 |
| DDQIVLSAKANLAKVEQ | ASSDAGLGASAETLY | 0.30787233 |
| DELLPDTGEDSDEDGH | ASSDAGLGASAETLY | 0.3527817 |
| ELPEAPANRATVPQD | ASSDAGLGASAETLY | 0.6196579 |
| FGQSKIERVTGIG | ASSDAGLGASAETLY | 0.75185144 |
| GAGGTGSTAGGAGGAGGAGGLY | ASSDAGLGASAETLY | 0.8083422 |
| GHGGTGAAGGNGGTGGAGGSGI | ASSDAGLGASAETLY | 0.3112637 |
| GNGGNGGAGGEGGPGGAGGAGA | ASSDAGLGASAETLY | 0.8294269 |
| GSTAGNGGNGGNATNGTIAGQP | ASSDAGLGASAETLY | 0.79767346 |
| GTGGDGGNGGNGGTGG | ASSDAGLGASAETLY | 0.8780303 |
| GVRRPAGIPADAPV | ASSDAGLGASAETLY | 0.7223567 |
| GYRQVTDAHLLTLARR | ASSDAGLGASAETLY | 0.3485149 |
| IAKMGQDGHDRG | ASSDAGLGASAETLY | 0.7961174 |
| KAEPSIEIQYT | ASSDAGLGASAETLY | 0.44234228 |
| KPVRIRMAADEDTVAVT | ASSDAGLGASAETLY | 0.61735 |
| MPPRIAGMRLLVIKPEP | ASSDAGLGASAETLY | 0.7718613 |
| MKFYAVALFRSRGVPPTR | ASSDAGLGASAETLY | 0.40509626 |
| NAGGFGILGAVAHSP | ASSDAGLGASAETLY | 0.30988982 |
| NEGREITLAMAEM | ASSDAGLGASAETLY | 0.6804944 |
| NELVEWSHPLGHRWRWE | ASSDAGLGASAETLY | 0.8418565 |
| NLGAGGTGGNGGNPG | ASSDAGLGASAETLY | 0.36179745 |
| NTEWPKWETVMAADPVD | ASSDAGLGASAETLY | 0.40571484 |
| PASAAPAETQP | ASSDAGLGASAETLY | 0.611676 |
| QQSSDGKNGVRAAEITGE | ASSDAGLGASAETLY | 0.6509874 |
| RAPDAVTTSWWKE | ASSDAGLGASAETLY | 0.77830315 |
| SGFGNVGAHTSGWFNQ | ASSDAGLGASAETLY | 0.75962186 |
| SGVAGAVGQAASVGGLKV | ASSDAGLGASAETLY | 0.66519916 |
| SNGGAGTGSAGGNGGT | ASSDAGLGASAETLY | 0.71970797 |
| TGGAGGEGGAAG | ASSDAGLGASAETLY | 0.8129034 |
| TGGDGGAAGAGGAPGHG | ASSDAGLGASAETLY | 0.69974566 |
| TMPPLLAAIANAGRHGV | ASSDAGLGASAETLY | 0.7751602 |
| TSSTAGNSGGA | ASSDAGLGASAETLY | 0.37200865 |
| VPATVPVGILC | ASSDAGLGASAETLY | 0.8250019 |
| VQRPDSGGVVLTGRIS | ASSDAGLGASAETLY | 0.70675504 |
| WTPNPVPATKGVPAGLVQAT | ASSDAGLGASAETLY | 0.7378429 |

| | | |
|---|---|---|
| YALKRALAKFSDRFDVVII | ASSDAGLGASAETLY | 0.8181496 |
| YLIGSVIV | ASSDAGLGASAETLY | 0.79138684 |
| YGQLVAVID | ASSDAGLGASAETLY | 0.74567986 |
| VTVGGMTLPA | ASSDAGLGASAETLY | 0.26543075 |
| VLALPAV | ASSDAGLGASAETLY | 0.59212863 |
| VIPPTLNYET | ASSDAGLGASAETLY | 0.6458703 |
| VGMGGMHPSSG | ASSDAGLGASAETLY | 0.64269865 |
| VGLAGGAATA | ASSDAGLGASAETLY | 0.75816727 |
| VGGAGGAGGSGP | ASSDAGLGASAETLY | 0.7539295 |
| VFAAFMV | ASSDAGLGASAETLY | 0.533658 |
| TLYDKLR | ASSDAGLGASAETLY | 0.8200262 |
| TLAVFTVL | ASSDAGLGASAETLY | 0.8411335 |
| TGSVQPARV | ASSDAGLGASAETLY | 0.2802625 |
| TGGGAGSGGT | ASSDAGLGASAETLY | 0.56602985 |
| SHLETSL | ASSDAGLGASAETLY | 0.8756324 |
| SFTTGMT | ASSDAGLGASAETLY | 0.727774 |
| QVEVAKRMGVRQPTVS | ASSDAGLGASAETLY | 0.49514863 |
| QGSGGPALAM | ASSDAGLGASAETLY | 0.56614983 |
| MLAPSNP | ASSDAGLGASAETLY | 0.85304147 |
| LVNNAAI | ASSDAGLGASAETLY | 0.56494164 |
| LTGEINTLA | ASSDAGLGASAETLY | 0.84461105 |
| LSIFGPV | ASSDAGLGASAETLY | 0.18351783 |
| LLALLANL | ASSDAGLGASAETLY | 0.6346022 |
| LISNASDALDKLR | ASSDAGLGASAETLY | 0.62108666 |
| LINEAVP | ASSDAGLGASAETLY | 0.16406268 |
| LGYAFSGL | ASSDAGLGASAETLY | 0.7528311 |
| LDFVAVAL | ASSDAGLGASAETLY | 0.5275788 |
| LAQPGAE | ASSDAGLGASAETLY | 0.7624143 |
| ISNLGAVTGF | ASSDAGLGASAETLY | 0.58644277 |
| HQEDPGRI | ASSDAGLGASAETLY | 0.76747954 |
| GTPGEPTP | ASSDAGLGASAETLY | 0.7227967 |
| GFFNSGS | ASSDAGLGASAETLY | 0.70534265 |
| GAGGGGLTAGAN | ASSDAGLGASAETLY | 0.4841206 |
| FVDALVNM | ASSDAGLGASAETLY | 0.5683895 |
| FLTVAPEL | ASSDAGLGASAETLY | 0.8115289 |
| FETLTAL | ASSDAGLGASAETLY | 0.82319224 |
| DVINPIEL | ASSDAGLGASAETLY | 0.19166559 |
| APVSGAPV | ASSDAGLGASAETLY | 0.6923717 |
| ALVGGLAW | ASSDAGLGASAETLY | 0.49281386 |
| ALAAAAATAI | ASSDAGLGASAETLY | 0.25131097 |
| AFGLGLLL | ASSDAGLGASAETLY | 0.16301195 |
| AFFAAASL | ASSDAGLGASAETLY | 0.43320802 |
| AAAALPAFSPP | ASSDAGLGASAETLY | 0.69565403 |
| VVLAPAL | ASSDAGLGASAETLY | 0.68578064 |
| TIGHVDHGKTTL | ASSDAGLGASAETLY | 0.7554079 |
| LISNASDALDK | ASSDAGLGASAETLY | 0.38432586 |
| LISNASDAL | ASSDAGLGASAETLY | 0.79842615 |
| ISNASDALDK | ASSDAGLGASAETLY | 0.56232905 |
| NQAETLVYQTE | ASSDAGLGASAETLY | 0.07620175 |
| TGGTGGTAGAG | ASSDAGLGASAETLY | 0.65796995 |
| TGSTAGGAGGA | ASSDAGLGASAETLY | 0.27555418 |

| | | |
|---|---|---|
| DPLAVPVAAGPV | ASSDAGLGASAETLY | 0.81259024 |
| GGTGGTPGNGGN | ASSDAGLGASAETLY | 0.4269591 |
| PAGLAGVDPATL | ASSDAGLGASAETLY | 0.46162295 |
| GNNLVVEAPHAGASV | ASSDAGLGASAETLY | 0.53892505 |
| KVTRIMSASTLEQPA | ASSDAGLGASAETLY | 0.74227804 |
| LGFALPIALPRRTVV | ASSDAGLGASAETLY | 0.6839453 |
| TRLGNLTARKLVVLV | ASSDAGLGASAETLY | 0.53211963 |
| WSQQRRLIYFPSAGP | ASSDAGLGASAETLY | 0.19361983 |
| GPNGNPDPMAAAVDIR | ASSDAGLGASAETLY | 0.63591146 |
| TQRCVSVADLRAAASR | ASSDAGLGASAETLY | 0.63607156 |
| QFVLDVAEHPVEVLTETV | ASSDAGLGASAETLY | 0.3381363 |
| GGAGGTGGAGGNGGTQAGDG | ASSDAGLGASAETLY | 0.34794933 |
| PLLATLRAIAVDVDAKPAQV | ASSDAGLGASAETLY | 0.38898852 |
| APKFRTGGYMIDFWGVGYQVAK | ASSDAGLGASAETLY | 0.31014508 |
| GAGGAGGAGGAAGGPGGTGGTG | ASSDAGLGASAETLY | 0.80181897 |
| IGALGLLATDVAPAGSVAFGVA | ASSDAGLGASAETLY | 0.39742398 |
| AGGAGGIGGNGGAGGAGGVLMG | ASSDAGLGASAETLY | 0.4123013 |
| ADELIGLL | ASSDAGLGASAETLY | 0.8555075 |
| ADRALKA | ASSDAGLGASAETLY | 0.6883698 |
| AGAVGQVGGAGGA | ASSDAGLGASAETLY | 0.49523252 |
| AGGGGGLGGAP | ASSDAGLGASAETLY | 0.52978504 |
| AGGLLYGNGGAG | ASSDAGLGASAETLY | 0.41250956 |
| AGLESIGL | ASSDAGLGASAETLY | 0.39659745 |
| AGLKTGII | ASSDAGLGASAETLY | 0.41295925 |
| ASLAALLL | ASSDAGLGASAETLY | 0.80222666 |
| ASVINISSI | ASSDAGLGASAETLY | 0.6302274 |
| ATPPGPA | ASSDAGLGASAETLY | 0.37696844 |
| ATVASALV | ASSDAGLGASAETLY | 0.8245419 |
| ATVASLIL | ASSDAGLGASAETLY | 0.8402299 |
| AVNGPVLIRH | ASSDAGLGASAETLY | 0.8839042 |
| DGAAGAVGI | ASSDAGLGASAETLY | 0.44794482 |
| DPLLHGI | ASSDAGLGASAETLY | 0.6838598 |
| ELGAALGAL | ASSDAGLGASAETLY | 0.567395 |
| ELGRTQAV | ASSDAGLGASAETLY | 0.54755557 |
| FVAGGYLS | ASSDAGLGASAETLY | 0.49901584 |
| GASGGAGGAGGW | ASSDAGLGASAETLY | 0.30132172 |
| GGAGAGGDGGAGGI | ASSDAGLGASAETLY | 0.8105484 |
| GTGHNGSQPGMGGNG | ASSDAGLGASAETLY | 0.8770256 |
| GYAAASASAS | ASSDAGLGASAETLY | 0.68643546 |
| HQVDVAAVQQG | ASSDAGLGASAETLY | 0.23947543 |
| IGAAIAHT | ASSDAGLGASAETLY | 0.4607976 |
| IGMIEVAQL | ASSDAGLGASAETLY | 0.3955745 |
| IRLAPEHTSAMD | ASSDAGLGASAETLY | 0.37912935 |
| IYNVSVL | ASSDAGLGASAETLY | 0.6385542 |
| KQGFLGM | ASSDAGLGASAETLY | 0.24200906 |
| LESVITQL | ASSDAGLGASAETLY | 0.6268002 |
| LGLGALVVV | ASSDAGLGASAETLY | 0.87039816 |
| LIGNGGNGGSGGI | ASSDAGLGASAETLY | 0.59147334 |
| LNAINAPTL | ASSDAGLGASAETLY | 0.70334285 |
| LQLLNTGV | ASSDAGLGASAETLY | 0.81611943 |
| LQTSIEF | ASSDAGLGASAETLY | 0.69773626 |

| | | |
|---|---|---|
| MSFAVSEL | ASSDAGLGASAETLY | 0.8338405 |
| NTGALGL | ASSDAGLGASAETLY | 0.81637 |
| PPGAGGAGGAGTT | ASSDAGLGASAETLY | 0.67635965 |
| QLVGLGI | ASSDAGLGASAETLY | 0.7387012 |
| RFLEQVHQL | ASSDAGLGASAETLY | 0.8149851 |
| RGVGYVLR | ASSDAGLGASAETLY | 0.5571217 |
| SATGVFAGVI | ASSDAGLGASAETLY | 0.5701162 |
| SLPAITLP | ASSDAGLGASAETLY | 0.6112776 |
| STALAFTV | ASSDAGLGASAETLY | 0.7621355 |
| SVLTVGLGK | ASSDAGLGASAETLY | 0.72514725 |
| TALAGAGSGI | ASSDAGLGASAETLY | 0.8530698 |
| TGLPFTGI | ASSDAGLGASAETLY | 0.19488743 |
| VAEHVTV | ASSDAGLGASAETLY | 0.2785457 |
| VDIVLNSL | ASSDAGLGASAETLY | 0.7979504 |
| VLTGGGAVL | ASSDAGLGASAETLY | 0.16582999 |
| VNVGLAYGI | ASSDAGLGASAETLY | 0.24028978 |
| VSSAAVGAV | ASSDAGLGASAETLY | 0.6880111 |
| VTSVYEAGV | ASSDAGLGASAETLY | 0.8486053 |
| WVALDRGA | ASSDAGLGASAETLY | 0.16029699 |
| YLVGSVVI | ASSDAGLGASAETLY | 0.42237514 |
| GPPGTGKTLL | ASSDAGLGASAETLY | 0.27534768 |
| GVGMTKF | ASSDAGLGASAETLY | 0.8366864 |
| IGIGGSDL | ASSDAGLGASAETLY | 0.5356206 |
| IGTLLAEL | ASSDAGLGASAETLY | 0.2642894 |
| IYTSGTT | ASSDAGLGASAETLY | 0.7498255 |
| LSFPAGV | ASSDAGLGASAETLY | 0.29448664 |
| LYTSGTT | ASSDAGLGASAETLY | 0.6741656 |
| VGAFPFGF | ASSDAGLGASAETLY | 0.76728547 |
| VGAGTVEF | ASSDAGLGASAETLY | 0.86081296 |
| VGKSTLLL | ASSDAGLGASAETLY | 0.31184545 |
| AALAHAGGGIV | ASSDAGLGASAETLY | 0.7038199 |
| AALVAVSLL | ASSDAGLGASAETLY | 0.53440726 |
| AAVSRAVAAL | ASSDAGLGASAETLY | 0.8903908 |
| AGGSSGIGGF | ASSDAGLGASAETLY | 0.45087558 |
| AGLTVANANG | ASSDAGLGASAETLY | 0.2776843 |
| AGLVGAVL | ASSDAGLGASAETLY | 0.7732729 |
| ALSQAFTV | ASSDAGLGASAETLY | 0.80572695 |
| ALTFSVSL | ASSDAGLGASAETLY | 0.63518333 |
| ALTGAALVAL | ASSDAGLGASAETLY | 0.7925662 |
| ASFSIGAGVL | ASSDAGLGASAETLY | 0.6814028 |
| DGSVYKI | ASSDAGLGASAETLY | 0.5936611 |
| EQLAEIV | ASSDAGLGASAETLY | 0.775501 |
| FAVSELTVM | ASSDAGLGASAETLY | 0.74263847 |
| GAAPLAAGL | ASSDAGLGASAETLY | 0.37523928 |
| GADGGDGGSGGAGG | ASSDAGLGASAETLY | 0.7981416 |
| GFGLDTGGAGG | ASSDAGLGASAETLY | 0.5160454 |
| GLASGGLL | ASSDAGLGASAETLY | 0.36727828 |
| GRIAGAAVLL | ASSDAGLGASAETLY | 0.8585135 |
| GSAGSLNTSF | ASSDAGLGASAETLY | 0.6453788 |
| GTGQIGI | ASSDAGLGASAETLY | 0.7895688 |
| GTLIALIL | ASSDAGLGASAETLY | 0.7473808 |

| | | |
|---|---|---|
| IAATMGFVVG | ASSDAGLGASAETLY | 0.8816277 |
| IFAQALL | ASSDAGLGASAETLY | 0.40226245 |
| IGAAVALGF | ASSDAGLGASAETLY | 0.7683068 |
| IGLTPLAL | ASSDAGLGASAETLY | 0.7255046 |
| ILGSNTSTL | ASSDAGLGASAETLY | 0.76901054 |
| IVFSLTVL | ASSDAGLGASAETLY | 0.6834898 |
| LASKATGF | ASSDAGLGASAETLY | 0.58385634 |
| LASVALIL | ASSDAGLGASAETLY | 0.7441567 |
| LFAVGLTL | ASSDAGLGASAETLY | 0.8320521 |
| LFNGGIGTYI | ASSDAGLGASAETLY | 0.639519 |
| LTLGAGII | ASSDAGLGASAETLY | 0.70437074 |
| LVAAALGDI | ASSDAGLGASAETLY | 0.85352874 |
| NTAFVQLGI | ASSDAGLGASAETLY | 0.35275197 |
| QLFSGAEL | ASSDAGLGASAETLY | 0.55394757 |
| SAATFITL | ASSDAGLGASAETLY | 0.8031724 |
| SNYGLANLGV | ASSDAGLGASAETLY | 0.7977555 |
| SYGGAYAVM | ASSDAGLGASAETLY | 0.6039195 |
| TTGVGGLTLG | ASSDAGLGASAETLY | 0.8319617 |
| TTQTIYI | ASSDAGLGASAETLY | 0.6850538 |
| VAAGAGLGAVI | ASSDAGLGASAETLY | 0.7155261 |
| VAAVVTLL | ASSDAGLGASAETLY | 0.67289275 |
| VGFLGGLI | ASSDAGLGASAETLY | 0.569126 |
| VGVIPILL | ASSDAGLGASAETLY | 0.31339088 |
| VIFASSLI | ASSDAGLGASAETLY | 0.6591277 |
| VKGGDGGAAAG | ASSDAGLGASAETLY | 0.78491557 |
| VTKTFTNI | ASSDAGLGASAETLY | 0.66596955 |
| VVAAAILL | ASSDAGLGASAETLY | 0.21407722 |
| YAALDVEL | ASSDAGLGASAETLY | 0.7276174 |
| LASVALLL | ASSDAGLGASAETLY | 0.7316699 |
| LDAVHRV | ASSDAGLGASAETLY | 0.8658452 |
| QGLSSSIF | ASSDAGLGASAETLY | 0.7197596 |
| RGTFANIRL | ASSDAGLGASAETLY | 0.75869036 |
| RQLWWGHRI | ASSDAGLGASAETLY | 0.5212271 |
| YGPPGTGKTLL | ASSDAGLGASAETLY | 0.689268 |
| AALVGAPANVA | ASSDAGLGASAETLY | 0.32346395 |
| AATFRRTE | ASSDAGLGASAETLY | 0.4795692 |
| AGALGGGI | ASSDAGLGASAETLY | 0.5508831 |
| AGGAGGAGGIGAG | ASSDAGLGASAETLY | 0.36272764 |
| AGSGALGL | ASSDAGLGASAETLY | 0.77805364 |
| AQIADQI | ASSDAGLGASAETLY | 0.7955455 |
| ASAVAVTYL | ASSDAGLGASAETLY | 0.761876 |
| ASTVLGGTGG | ASSDAGLGASAETLY | 0.8260491 |
| AVGGEPAK | ASSDAGLGASAETLY | 0.8513483 |
| CEQGTFGGTC | ASSDAGLGASAETLY | 0.78926516 |
| DEGKAPAG | ASSDAGLGASAETLY | 0.41293472 |
| DTVVFPAL | ASSDAGLGASAETLY | 0.6129656 |
| FEGITMAG | ASSDAGLGASAETLY | 0.22702743 |
| FPLVVQD | ASSDAGLGASAETLY | 0.57048225 |
| GAFTTPQ | ASSDAGLGASAETLY | 0.49328148 |
| GFGGFGTAGGVGGA | ASSDAGLGASAETLY | 0.60502416 |
| GFLAQFL | ASSDAGLGASAETLY | 0.12626755 |

| | | |
|---|---|---|
| GGAGGAGGNAGVA | ASSDAGLGASAETLY | 0.7428372 |
| GGVGTAGAGGAGGAGG | ASSDAGLGASAETLY | 0.79887795 |
| GNGGGGDAGNAGSGGN | ASSDAGLGASAETLY | 0.5304457 |
| GTGGAGGAGNDAGS | ASSDAGLGASAETLY | 0.7848485 |
| GTTNTGWFNS | ASSDAGLGASAETLY | 0.75612706 |
| IFMGPLFTL | ASSDAGLGASAETLY | 0.4475875 |
| ILAHAIEAI | ASSDAGLGASAETLY | 0.75747716 |
| IRSAIREAL | ASSDAGLGASAETLY | 0.51107466 |
| ITFAGGG | ASSDAGLGASAETLY | 0.77130246 |
| ITGMSGHQEV | ASSDAGLGASAETLY | 0.81549346 |
| IVAGLACTL | ASSDAGLGASAETLY | 0.7455152 |
| IVGGGLAGA | ASSDAGLGASAETLY | 0.6155198 |
| IVTGVLNI | ASSDAGLGASAETLY | 0.64196 |
| LAQQGCI | ASSDAGLGASAETLY | 0.31412116 |
| LCLHVEN | ASSDAGLGASAETLY | 0.624792 |
| LDAISDGS | ASSDAGLGASAETLY | 0.14804357 |
| LGGFAAML | ASSDAGLGASAETLY | 0.7032566 |
| LGGKNPMI | ASSDAGLGASAETLY | 0.46528304 |
| LGPKGVAAIL | ASSDAGLGASAETLY | 0.6632339 |
| LMVGVSI | ASSDAGLGASAETLY | 0.5228381 |
| LVGAAFVL | ASSDAGLGASAETLY | 0.7188153 |
| LVLAVAPL | ASSDAGLGASAETLY | 0.36463964 |
| MAAIAVIDAA | ASSDAGLGASAETLY | 0.68998724 |
| NNGINAELQ | ASSDAGLGASAETLY | 0.7275874 |
| RGEPLVLVHG | ASSDAGLGASAETLY | 0.28384262 |
| RGGAGGAAGLI | ASSDAGLGASAETLY | 0.81784356 |
| RSYRMHPAV | ASSDAGLGASAETLY | 0.6485353 |
| SAAHLLAGQLA | ASSDAGLGASAETLY | 0.64039457 |
| SAAVGSY | ASSDAGLGASAETLY | 0.79801154 |
| SPHAVLHRI | ASSDAGLGASAETLY | 0.62532306 |
| SQIAKVL | ASSDAGLGASAETLY | 0.3517756 |
| TALGHLVP | ASSDAGLGASAETLY | 0.617118 |
| TAPTTSGAA | ASSDAGLGASAETLY | 0.70934093 |
| TDGDDGNGGSPGT | ASSDAGLGASAETLY | 0.56970257 |
| TDGLGEN | ASSDAGLGASAETLY | 0.14768626 |
| TGGPPLPGQA | ASSDAGLGASAETLY | 0.8158995 |
| TIPAITI | ASSDAGLGASAETLY | 0.3645629 |
| TIPGFTI | ASSDAGLGASAETLY | 0.5008862 |
| TSIASGALA | ASSDAGLGASAETLY | 0.47587532 |
| TVTLPGAAI | ASSDAGLGASAETLY | 0.7803557 |
| VASGLSAALP | ASSDAGLGASAETLY | 0.64231133 |
| VGAGVIGI | ASSDAGLGASAETLY | 0.5907956 |
| VGPKRFGV | ASSDAGLGASAETLY | 0.747556 |
| VNVAFPD | ASSDAGLGASAETLY | 0.6168529 |
| VSMTNSGVS | ASSDAGLGASAETLY | 0.7808377 |
| VTDVLGDLSV | ASSDAGLGASAETLY | 0.7452241 |
| VVVVGAGFAGL | ASSDAGLGASAETLY | 0.74287796 |
| VYTVGTVV | ASSDAGLGASAETLY | 0.7722367 |
| WTTFSGP | ASSDAGLGASAETLY | 0.64225876 |
| YDELVEAF | ASSDAGLGASAETLY | 0.3180452 |
| YGTGGAGGNGG | ASSDAGLGASAETLY | 0.38695705 |

| | | |
|---|---|---|
| YTNVMAV | ASSDAGLGASAETLY | 0.5986397 |
| FLEQVHQL | ASSDAGLGASAETLY | 0.31412584 |
| GLSLLAVAV | ASSDAGLGASAETLY | 0.6721153 |
| LEQVHQL | ASSDAGLGASAETLY | 0.8398864 |
| LNAAGAI | ASSDAGLGASAETLY | 0.7898064 |
| PAVHASL | ASSDAGLGASAETLY | 0.27270615 |
| TELEDDA | ASSDAGLGASAETLY | 0.47860426 |
| TPPGPPN | ASSDAGLGASAETLY | 0.5804408 |
| TPPPPPPP | ASSDAGLGASAETLY | 0.7298389 |
| TQSPALI | ASSQGQTEVF | 0.8415909 |
| EFAAAFPQ | ASSQGQTEVF | 0.8600693 |
| LLSGNLDVL | ASSQGQTEVF | 0.77319074 |
| GPANLGNYNI | ASSQGQTEVF | 0.41096538 |
| TLGPTLP | ASSQGQTEVF | 0.7579347 |
| VYAYSSCCK | ASSQGQTEVF | 0.5780248 |
| APAPAPAPNI | ASSQGQTEVF | 0.6617756 |
| PGGPVPIP | ASSQGQTEVF | 0.6451787 |
| FLTVALEP | ASSQGQTEVF | 0.7757635 |
| SQYFGANL | ASSQGQTEVF | 0.5955892 |
| GGVKPAA | ASSQGQTEVF | 0.54999304 |
| GGTGGAGGAGGW | ASSQGQTEVF | 0.32783434 |
| GVGGVAGLGGAGG | ASSQGQTEVF | 0.7519827 |
| VLLVLATTI | ASSQGQTEVF | 0.3510085 |
| IGIFAAML | ASSQGQTEVF | 0.40168047 |
| AVGDLID | ASSQGQTEVF | 0.53566825 |
| NISPLPL | ASSQGQTEVF | 0.85581243 |
| VTAVFVAI | ASSQGQTEVF | 0.8505478 |
| SGDDNDAAL | ASSQGQTEVF | 0.49141958 |
| AVAALAHS | ASSQGQTEVF | 0.45813993 |
| GGVTVMATP | ASSQGQTEVF | 0.7532265 |
| AGLAANVRD | ASSQGQTEVF | 0.34871972 |
| TNRPDLIDPALLRPG | ASSQGQTEVF | 0.7523184 |
| LANLGVTAL | ASSQGQTEVF | 0.57428896 |
| TGERIEL | ASSQGQTEVF | 0.6016129 |
| NGEELAA | ASSQGQTEVF | 0.50894237 |
| VLSLVWG | ASSQGQTEVF | 0.7176606 |
| IGYALWL | ASSQGQTEVF | 0.5415021 |
| LSLLLAPL | ASSQGQTEVF | 0.7374331 |
| TLDEALK | ASSQGQTEVF | 0.19100349 |
| TDGAGTAEGG | ASSQGQTEVF | 0.7910918 |
| NAPGGAGGNGG | ASSQGQTEVF | 0.27754048 |
| VSVFMGMGGNF | ASSQGQTEVF | 0.56318617 |
| LASYLAAL | ASSQGQTEVF | 0.21206723 |
| LVGGTPAL | ASSQGQTEVF | 0.6216818 |
| VRNADVL | ASSQGQTEVF | 0.8043233 |
| TCSSPFLVSAM | ASSQGQTEVF | 0.8088371 |
| VHITDAE | ASSQGQTEVF | 0.60360134 |
| EYAAPSF | ASSQGQTEVF | 0.71113384 |
| VVDAFHGGL | ASSQGQTEVF | 0.8233162 |
| ASVVDPQ | ASSQGQTEVF | 0.41293377 |
| SYNEFGGLS | ASSQGQTEVF | 0.22063352 |

| | | |
|---|---|---|
| DPDGSVY | ASSQGQTEVF | 0.6740764 |
| MVSTIHATE | ASSQGQTEVF | 0.7104554 |
| LAAGVADPV | ASSQGQTEVF | 0.8254893 |
| TKEQIAA | ASSQGQTEVF | 0.8082094 |
| SLASPAAL | ASSQGQTEVF | 0.25988355 |
| IGLVFYGV | ASSQGQTEVF | 0.7349303 |
| FLLDASNL | ASSQGQTEVF | 0.7790096 |
| YQVGNFP | ASSQGQTEVF | 0.3478608 |
| ESLLSQST | ASSQGQTEVF | 0.48700994 |
| LFTVGFAAVG | ASSQGQTEVF | 0.5421786 |
| ATPPTPT | ASSQGQTEVF | 0.2573117 |
| TDLATLG | ASSQGQTEVF | 0.7217233 |
| SAGGTGR | ASSQGQTEVF | 0.4935295 |
| GFSNSGS | ASSQGQTEVF | 0.43300557 |
| GGNGGSGGTSV | ASSQGQTEVF | 0.5409349 |
| GTSVATGGAGN | ASSQGQTEVF | 0.42874634 |
| NGGNGGSGGTS | ASSQGQTEVF | 0.3681922 |
| AVVKFPH | ASSQGQTEVF | 0.39943102 |
| TPPPPPP | ASSQGQTEVF | 0.7532749 |
| AAPAAAAAPA | ASSQGQTEVF | 0.74365306 |
| IALNEDL | ASSQGQTEVF | 0.7067085 |
| IVNVSSM | ASSQGQTEVF | 0.7528758 |
| TGIPPAPR | ASSQGQTEVF | 0.77498335 |
| FDLGGGTF | ASSQGQTEVF | 0.27570593 |
| FDLGGGTFDVS | ASSQGQTEVF | 0.37088788 |
| NEPTAAA | ASSQGQTEVF | 0.7740724 |
| NEPTAAALA | ASSQGQTEVF | 0.5105089 |
| GSIINISSI | ASSQGQTEVF | 0.80711156 |
| VAPGNAGTA | ASSQGQTEVF | 0.5533662 |
| TGGGASLE | ASSQGQTEVF | 0.62162983 |
| VAVYNHYKRI | ASSQGQTEVF | 0.7338705 |
| TDPAPATT | ASSQGQTEVF | 0.8757465 |
| TNRPDILDPALLRPG | ASSQGQTEVF | 0.715617 |
| LTTTTTS | ASSQGQTEVF | 0.583511 |
| PTGSGKTL | ASSQGQTEVF | 0.74600834 |
| TGSGKTL | ASSQGQTEVF | 0.4467082 |
| TIGHVDHGK | ASSQGQTEVF | 0.76060325 |
| TIGHVDHGKTT | ASSQGQTEVF | 0.7741572 |
| GHVDHGKTT | ASSQGQTEVF | 0.30789053 |
| AQFEHTLL | ASSQGQTEVF | 0.5990147 |
| IVNQDEL | ASSQGQTEVF | 0.8484334 |
| TNGSQFF | ASSQGQTEVF | 0.8034589 |
| TGKGIVE | ASSQGQTEVF | 0.42672783 |
| LISNASDA | ASSQGQTEVF | 0.4748268 |
| AELRTLGELTKIAKAHGAQ | ASSQGQTEVF | 0.49063262 |
| APNAVAAAEPSW | ASSQGQTEVF | 0.66853565 |
| APQVAGAATAVL | ASSQGQTEVF | 0.3026456 |
| ARRDPAAAAAILPTDA | ASSQGQTEVF | 0.7670928 |
| DAAARDVLAEAGLAEYF | ASSQGQTEVF | 0.50684494 |
| DDQIVLSAKANLAKVEQ | ASSQGQTEVF | 0.80905306 |
| DELLPDTGEDSDEDGH | ASSQGQTEVF | 0.6466911 |

| | | |
|---|---|---|
| ELPEAPANRATVPQD | ASSQGQTEVF | 0.7832485 |
| FGQSKIERVTGIG | ASSQGQTEVF | 0.61385816 |
| GAGGTGSTAGGAGGAGGAGGLY | ASSQGQTEVF | 0.4769115 |
| GHGGTGAAGGNGGTGGAGGSGI | ASSQGQTEVF | 0.28350705 |
| GNGGNGGAGGEGGPGGAGGAGA | ASSQGQTEVF | 0.26249442 |
| GSTAGNGGNGGNATNGTIAGQP | ASSQGQTEVF | 0.43154165 |
| GTGGDGGNGGNGGTGG | ASSQGQTEVF | 0.4487818 |
| GVRRPAGIPADAPV | ASSQGQTEVF | 0.27810088 |
| GYRQVTDAHLLTLARR | ASSQGQTEVF | 0.8695297 |
| IAKMGQDGHDRG | ASSQGQTEVF | 0.53019124 |
| KAEPSIEIQYT | ASSQGQTEVF | 0.62587875 |
| KPVRIRMAADEDTVAVT | ASSQGQTEVF | 0.5437628 |
| MPPRIAGMRLLVIKPEP | ASSQGQTEVF | 0.34524193 |
| MKFYAVALFRSRGVPPTR | ASSQGQTEVF | 0.40581584 |
| NAGGFGILGAVAHSP | ASSQGQTEVF | 0.76126283 |
| NEGREITLAMAEM | ASSQGQTEVF | 0.39102846 |
| NELVEWSHPLGHRWRWE | ASSQGQTEVF | 0.23337682 |
| NLGAGGTGGNGGNPG | ASSQGQTEVF | 0.5330798 |
| NTEWPKWETVMAADPVD | ASSQGQTEVF | 0.56327665 |
| PASAAPAETQP | ASSQGQTEVF | 0.87953603 |
| QQSSDGKNGVRAAEITGE | ASSQGQTEVF | 0.81439996 |
| RAPDAVTTSWWKE | ASSQGQTEVF | 0.63343287 |
| SGFGNVGAHTSGWFNQ | ASSQGQTEVF | 0.8727007 |
| SGVAGAVGQAASVGGLKV | ASSQGQTEVF | 0.14369588 |
| SNGGAGTGSAGGNGGT | ASSQGQTEVF | 0.54650736 |
| TGGAGGEGGAAG | ASSQGQTEVF | 0.83142686 |
| TGGDGGAAGAGGAPGHG | ASSQGQTEVF | 0.75184506 |
| TMPPLLAAIANAGRHGV | ASSQGQTEVF | 0.51411146 |
| TSSTAGNSGGA | ASSQGQTEVF | 0.79782563 |
| VPATVPVGILC | ASSQGQTEVF | 0.3498898 |
| VQRPDSGGVVLTGRIS | ASSQGQTEVF | 0.5940403 |
| WTPNPVPATKGVPAGLVQAT | ASSQGQTEVF | 0.56262404 |
| YALKRALAKFSDRFDVVII | ASSQGQTEVF | 0.74867857 |
| YLIGSVIV | ASSQGQTEVF | 0.75503117 |
| YGQLVAVID | ASSQGQTEVF | 0.3159783 |
| VTVGGMTLPA | ASSQGQTEVF | 0.70795834 |
| VLALPAV | ASSQGQTEVF | 0.8556421 |
| VIPPTLNYET | ASSQGQTEVF | 0.47355336 |
| VGMGGMHPSSG | ASSQGQTEVF | 0.7999977 |
| VGLAGGAATA | ASSQGQTEVF | 0.65734917 |
| VGGAGGAGGSGP | ASSQGQTEVF | 0.59410536 |
| VFAAFMV | ASSQGQTEVF | 0.4150396 |
| TLYDKLR | ASSQGQTEVF | 0.30714658 |
| TLAVFTVL | ASSQGQTEVF | 0.6854842 |
| TGSVQPARV | ASSQGQTEVF | 0.31519395 |
| TGGGAGSGGT | ASSQGQTEVF | 0.6601645 |
| SHLETSL | ASSQGQTEVF | 0.60476685 |
| SFTTGMT | ASSQGQTEVF | 0.8880972 |
| QVEVAKRMGVRQPTVS | ASSQGQTEVF | 0.7547792 |
| QGSGGPALAM | ASSQGQTEVF | 0.68637586 |
| MLAPSNP | ASSQGQTEVF | 0.3255992 |

| | | |
|---|---|---|
| LVNNAAI | ASSQGQTEVF | 0.7196628 |
| LTGEINTLA | ASSQGQTEVF | 0.74072737 |
| LSIFGPV | ASSQGQTEVF | 0.33947378 |
| LLALLANL | ASSQGQTEVF | 0.82765174 |
| LISNASDALDKLR | ASSQGQTEVF | 0.25229794 |
| LINEAVP | ASSQGQTEVF | 0.7828872 |
| LGYAFSGL | ASSQGQTEVF | 0.7935237 |
| LDFVAVAL | ASSQGQTEVF | 0.822493 |
| LAQPGAE | ASSQGQTEVF | 0.5734195 |
| ISNLGAVTGF | ASSQGQTEVF | 0.7013074 |
| HQEDPGRI | ASSQGQTEVF | 0.32072842 |
| GTPGEPTP | ASSQGQTEVF | 0.8125266 |
| GFFNSGS | ASSQGQTEVF | 0.8065479 |
| GAGGGGLTAGAN | ASSQGQTEVF | 0.7185293 |
| FVDALVNM | ASSQGQTEVF | 0.86361957 |
| FLTVAPEL | ASSQGQTEVF | 0.79214054 |
| FETLTAL | ASSQGQTEVF | 0.28324854 |
| DVINPIEL | ASSQGQTEVF | 0.57957494 |
| APVSGAPV | ASSQGQTEVF | 0.8122982 |
| ALVGGLAW | ASSQGQTEVF | 0.80837643 |
| ALAAAAATAI | ASSQGQTEVF | 0.80595946 |
| AFGLGLLL | ASSQGQTEVF | 0.7777822 |
| AFFAAASL | ASSQGQTEVF | 0.6126596 |
| AAAALPAFSPP | ASSQGQTEVF | 0.8323823 |
| VVLAPAL | ASSQGQTEVF | 0.9093032 |
| TIGHVDHGKTTL | ASSQGQTEVF | 0.7396693 |
| LISNASDALDK | ASSQGQTEVF | 0.51786184 |
| LISNASDAL | ASSQGQTEVF | 0.5855425 |
| ISNASDALDK | ASSQGQTEVF | 0.7841001 |
| NQAETLVYQTE | ASSQGQTEVF | 0.8102385 |
| TGGTGGTAGAG | ASSQGQTEVF | 0.3568135 |
| TGSTAGGAGGA | ASSQGQTEVF | 0.55593944 |
| DPLAVPVAAGPV | ASSQGQTEVF | 0.6294842 |
| GGTGGTPGNGGN | ASSQGQTEVF | 0.18386628 |
| PAGLAGVDPATL | ASSQGQTEVF | 0.8366293 |
| GNNLVVEAPHAGASV | ASSQGQTEVF | 0.84557647 |
| KVTRIMSASTLEQPA | ASSQGQTEVF | 0.68362373 |
| LGFALPIALPRRTVV | ASSQGQTEVF | 0.2983505 |
| TRLGNLTARKLVVLV | ASSQGQTEVF | 0.8296819 |
| WSQQRRLIYFPSAGP | ASSQGQTEVF | 0.82097584 |
| GPNGNPDPMAAAVDIR | ASSQGQTEVF | 0.83558524 |
| TQRCVSVADLRAAASR | ASSQGQTEVF | 0.22801019 |
| QFVLDVAEHPVEVLTETV | ASSQGQTEVF | 0.14063741 |
| GGAGGTGGAGGNGGTQAGDG | ASSQGQTEVF | 0.599298 |
| PLLATLRAIAVDVDAKPAQV | ASSQGQTEVF | 0.79890335 |
| APKFRTGGYMIDFWGVGYQVAK | ASSQGQTEVF | 0.50593144 |
| GAGGAGGAGGAAGGPGGTGGTG | ASSQGQTEVF | 0.33414388 |
| IGALGLLATDVAPAGSVAFGVA | ASSQGQTEVF | 0.6566637 |
| AGGAGGIGGNGGAGGAGGVLMG | ASSQGQTEVF | 0.77874565 |
| ADELIGLL | ASSQGQTEVF | 0.3475837 |
| ADRALKA | ASSQGQTEVF | 0.42937392 |

| | | |
|---|---|---|
| AGAVGQVGGAGGA | ASSQGQTEVF | 0.71671283 |
| AGGGGGLGGAP | ASSQGQTEVF | 0.80831146 |
| AGGLLYGNGGAG | ASSQGQTEVF | 0.32998163 |
| AGLESIGL | ASSQGQTEVF | 0.4631346 |
| AGLKTGII | ASSQGQTEVF | 0.60026217 |
| ASLAALLL | ASSQGQTEVF | 0.7948241 |
| ASVINISSI | ASSQGQTEVF | 0.6410443 |
| ATPPGPA | ASSQGQTEVF | 0.37373418 |
| ATVASALV | ASSQGQTEVF | 0.82106483 |
| ATVASLIL | ASSQGQTEVF | 0.30396593 |
| AVNGPVLIRH | ASSQGQTEVF | 0.8071919 |
| DGAAGAVGI | ASSQGQTEVF | 0.58949006 |
| DPLLHGI | ASSQGQTEVF | 0.7067708 |
| ELGAALGAL | ASSQGQTEVF | 0.51488614 |
| ELGRTQAV | ASSQGQTEVF | 0.369231 |
| FVAGGYLS | ASSQGQTEVF | 0.3871686 |
| GASGGAGGAGGW | ASSQGQTEVF | 0.35372078 |
| GGAGAGGDGGAGGI | ASSQGQTEVF | 0.7966456 |
| GTGHNGSQPGMGGNG | ASSQGQTEVF | 0.40807557 |
| GYAAASASAS | ASSQGQTEVF | 0.771647 |
| HQVDVAAVQQG | ASSQGQTEVF | 0.6100391 |
| IGAAIAHT | ASSQGQTEVF | 0.39897713 |
| IGMIEVAQL | ASSQGQTEVF | 0.7546378 |
| IRLAPEHTSAMD | ASSQGQTEVF | 0.27708748 |
| IYNVSVL | ASSQGQTEVF | 0.6797488 |
| KQGFLGM | ASSQGQTEVF | 0.75698704 |
| LESVITQL | ASSQGQTEVF | 0.7365729 |
| LGLGALVVV | ASSQGQTEVF | 0.5295789 |
| LIGNGGNGGSGGI | ASSQGQTEVF | 0.70735925 |
| LNAINAPTL | ASSQGQTEVF | 0.7432256 |
| LQLLNTGV | ASSQGQTEVF | 0.48955306 |
| LQTSIEF | ASSQGQTEVF | 0.65687704 |
| MSFAVSEL | ASSQGQTEVF | 0.8012837 |
| NTGALGL | ASSQGQTEVF | 0.6925864 |
| PPGAGGAGGAGTT | ASSQGQTEVF | 0.2689408 |
| QLVGLGI | ASSQGQTEVF | 0.7135643 |
| RFLEQVHQL | ASSQGQTEVF | 0.46234006 |
| RGVGYVLR | ASSQGQTEVF | 0.6940727 |
| SATGVFAGVI | ASSQGQTEVF | 0.6775874 |
| SLPAITLP | ASSQGQTEVF | 0.67382795 |
| STALAFTV | ASSQGQTEVF | 0.75340855 |
| SVLTVGLGK | ASSQGQTEVF | 0.54908204 |
| TALAGAGSGI | ASSQGQTEVF | 0.57351756 |
| TGLPFTGI | ASSQGQTEVF | 0.54188484 |
| VAEHVTV | ASSQGQTEVF | 0.5893228 |
| VDIVLNSL | ASSQGQTEVF | 0.7446226 |
| VLTGGGAVL | ASSQGQTEVF | 0.3123902 |
| VNVGLAYGI | ASSQGQTEVF | 0.7131921 |
| VSSAAVGAV | ASSQGQTEVF | 0.6241807 |
| VTSVYEAGV | ASSQGQTEVF | 0.33764556 |
| WVALDRGA | ASSQGQTEVF | 0.6329512 |

| | | |
|---|---|---|
| YLVGSVVI | ASSQGQTEVF | 0.62064314 |
| GPPGTGKTLL | ASSQGQTEVF | 0.324599 |
| GVGMTKF | ASSQGQTEVF | 0.74056035 |
| IGIGGSDL | ASSQGQTEVF | 0.36691487 |
| IGTLLAEL | ASSQGQTEVF | 0.76068145 |
| IYTSGTT | ASSQGQTEVF | 0.843123 |
| LSFPAGV | ASSQGQTEVF | 0.33082336 |
| LYTSGTT | ASSQGQTEVF | 0.7922414 |
| VGAFPFGF | ASSQGQTEVF | 0.6685069 |
| VGAGTVEF | ASSQGQTEVF | 0.6840164 |
| VGKSTLLL | ASSQGQTEVF | 0.39647198 |
| AALAHAGGGIV | ASSQGQTEVF | 0.5211339 |
| AALVAVSLL | ASSQGQTEVF | 0.83439124 |
| AAVSRAVAAL | ASSQGQTEVF | 0.6757963 |
| AGGSSGIGGF | ASSQGQTEVF | 0.74109584 |
| AGLTVANANG | ASSQGQTEVF | 0.7931758 |
| AGLVGAVL | ASSQGQTEVF | 0.78415334 |
| ALSQAFTV | ASSQGQTEVF | 0.771737 |
| ALTFSVSL | ASSQGQTEVF | 0.35629964 |
| ALTGAALVAL | ASSQGQTEVF | 0.23090672 |
| ASFSIGAGVL | ASSQGQTEVF | 0.4885156 |
| DGSVYKI | ASSQGQTEVF | 0.4982413 |
| EQLAEIV | ASSQGQTEVF | 0.45274305 |
| FAVSELTVM | ASSQGQTEVF | 0.35087982 |
| GAAPLAAGL | ASSQGQTEVF | 0.7767977 |
| GADGGDGGSGGAGG | ASSQGQTEVF | 0.87845945 |
| GFGLDTGGAGG | ASSQGQTEVF | 0.37093836 |
| GLASGGLL | ASSQGQTEVF | 0.88228273 |
| GRIAGAAVLL | ASSQGQTEVF | 0.36270255 |
| GSAGSLNTSF | ASSQGQTEVF | 0.7663851 |
| GTGQIGI | ASSQGQTEVF | 0.8036393 |
| GTLIALIL | ASSQGQTEVF | 0.25234327 |
| IAATMGFVVG | ASSQGQTEVF | 0.6841196 |
| IFAQALL | ASSQGQTEVF | 0.4807763 |
| IGAAVALGF | ASSQGQTEVF | 0.4255625 |
| IGLTPLAL | ASSQGQTEVF | 0.7916581 |
| ILGSNTSTL | ASSQGQTEVF | 0.39385578 |
| IVFSLTVL | ASSQGQTEVF | 0.7316019 |
| LASKATGF | ASSQGQTEVF | 0.47410864 |
| LASVALIL | ASSQGQTEVF | 0.36607152 |
| LFAVGLTL | ASSQGQTEVF | 0.7104299 |
| LFNGGIGTYI | ASSQGQTEVF | 0.768346 |
| LTLGAGII | ASSQGQTEVF | 0.678996 |
| LVAAALGDI | ASSQGQTEVF | 0.90438354 |
| NTAFVQLGI | ASSQGQTEVF | 0.417136 |
| QLFSGAEL | ASSQGQTEVF | 0.8729056 |
| SAATFITL | ASSQGQTEVF | 0.8389837 |
| SNYGLANLGV | ASSQGQTEVF | 0.6193101 |
| SYGGAYAVM | ASSQGQTEVF | 0.583617 |
| TTGVGGLTLG | ASSQGQTEVF | 0.78853667 |
| TTQTIYI | ASSQGQTEVF | 0.57077014 |

| | | |
|---|---|---|
| VAAGAGLGAVI | ASSQGQTEVF | 0.74387217 |
| VAAVVTLL | ASSQGQTEVF | 0.81579715 |
| VGFLGGLI | ASSQGQTEVF | 0.73963225 |
| VGVIPILL | ASSQGQTEVF | 0.81699747 |
| VIFASSLI | ASSQGQTEVF | 0.7952418 |
| VKGGDGGAAAG | ASSQGQTEVF | 0.79958534 |
| VTKTFTNI | ASSQGQTEVF | 0.769649 |
| VVAAAILL | ASSQGQTEVF | 0.5557446 |
| YAALDVEL | ASSQGQTEVF | 0.7646493 |
| LASVALLL | ASSQGQTEVF | 0.7406362 |
| LDAVHRV | ASSQGQTEVF | 0.8061333 |
| QGLSSSIF | ASSQGQTEVF | 0.8158894 |
| RGTFANIRL | ASSQGQTEVF | 0.75970054 |
| RQLWWGHRI | ASSQGQTEVF | 0.8144951 |
| YGPPGTGKTLL | ASSQGQTEVF | 0.81110036 |
| AALVGAPANVA | ASSQGQTEVF | 0.63398224 |
| AATFRRTE | ASSQGQTEVF | 0.77555656 |
| AGALGGGI | ASSQGQTEVF | 0.7401601 |
| AGGAGGAGGIGAG | ASSQGQTEVF | 0.7932995 |
| AGSGALGL | ASSQGQTEVF | 0.41904226 |
| AQIADQI | ASSQGQTEVF | 0.8178687 |
| ASAVAVTYL | ASSQGQTEVF | 0.741575 |
| ASTVLGGTGG | ASSQGQTEVF | 0.2651284 |
| AVGGEPAK | ASSQGQTEVF | 0.79639965 |
| CEQGTFGGTC | ASSQGQTEVF | 0.78953373 |
| DEGKAPAG | ASSQGQTEVF | 0.7839746 |
| DTVVFPAL | ASSQGQTEVF | 0.81451416 |
| FEGITMAG | ASSQGQTEVF | 0.63761383 |
| FPLVVQD | ASSQGQTEVF | 0.7081678 |
| GAFTTPQ | ASSQGQTEVF | 0.8045515 |
| GFGGFGTAGGVGGA | ASSQGQTEVF | 0.82136667 |
| GFLAQFL | ASSQGQTEVF | 0.67183596 |
| GGAGGAGGNAGVA | ASSQGQTEVF | 0.7997137 |
| GGVGTAGAGGAGGAGG | ASSQGQTEVF | 0.81856203 |
| GNGGGGDAGNAGSGGN | ASSQGQTEVF | 0.80222166 |
| GTGGAGGAGNDAGS | ASSQGQTEVF | 0.8196691 |
| GTTNTGWFNS | ASSQGQTEVF | 0.8257593 |
| IFMGPLFTL | ASSQGQTEVF | 0.7135453 |
| ILAHAIEAI | ASSQGQTEVF | 0.8198684 |
| IRSAIREAL | ASSQGQTEVF | 0.8122736 |
| ITFAGGG | ASSQGQTEVF | 0.7683336 |
| ITGMSGHQEV | ASSQGQTEVF | 0.789039 |
| IVAGLACTL | ASSQGQTEVF | 0.70672977 |
| IVGGGLAGA | ASSQGQTEVF | 0.77204704 |
| IVTGVLNI | ASSQGQTEVF | 0.6012225 |
| LAQQGCI | ASSQGQTEVF | 0.79638916 |
| LCLHVEN | ASSQGQTEVF | 0.82259345 |
| LDAISDGS | ASSQGQTEVF | 0.77535313 |
| LGGFAAML | ASSQGQTEVF | 0.68997896 |
| LGGKNPMI | ASSQGQTEVF | 0.7227772 |
| LGPKGVAAIL | ASSQGQTEVF | 0.77289623 |

| | | |
|---|---|---|
| LMVGVSI | ASSQGQTEVF | 0.772333 |
| LVGAAFVL | ASSQGQTEVF | 0.5585426 |
| LVLAVAPL | ASSQGQTEVF | 0.73692757 |
| MAAIAVIDAA | ASSQGQTEVF | 0.81492245 |
| NNGINAELQ | ASSQGQTEVF | 0.8177768 |
| RGEPLVLVHG | ASSQGQTEVF | 0.6692393 |
| RGGAGGAAGLI | ASSQGQTEVF | 0.81817573 |
| RSYRMHPAV | ASSQGQTEVF | 0.8062713 |
| SAAHLLAGQLA | ASSQGQTEVF | 0.82255995 |
| SAAVGSY | ASSQGQTEVF | 0.78268963 |
| SPHAVLHRI | ASSQGQTEVF | 0.7150699 |
| SQIAKVL | ASSQGQTEVF | 0.80766577 |
| TALGHLVP | ASSQGQTEVF | 0.76211315 |
| TAPTTSGAA | ASSQGQTEVF | 0.7424398 |
| TDGDDGNGGSPGT | ASSQGQTEVF | 0.8143826 |
| TDGLGEN | ASSQGQTEVF | 0.77102935 |
| TGGPPLPGQA | ASSQGQTEVF | 0.7533933 |
| TIPAITI | ASSQGQTEVF | 0.75430906 |
| TIPGFTI | ASSQGQTEVF | 0.82445794 |
| TSIASGALA | ASSQGQTEVF | 0.6958523 |
| TVTLPGAAI | ASSQGQTEVF | 0.7531123 |
| VASGLSAALP | ASSQGQTEVF | 0.74198383 |
| VGAGVIGI | ASSQGQTEVF | 0.78506064 |
| VGPKRFGV | ASSQGQTEVF | 0.8180671 |
| VNVAFPD | ASSQGQTEVF | 0.8126919 |
| VSMTNSGVS | ASSQGQTEVF | 0.7601129 |
| VTDVLGDLSV | ASSQGQTEVF | 0.8000375 |
| VVVVGAGFAGL | ASSQGQTEVF | 0.81393254 |
| VYTVGTVV | ASSQGQTEVF | 0.8017489 |
| WTTFSGP | ASSQGQTEVF | 0.7995665 |
| YDELVEAF | ASSQGQTEVF | 0.7323835 |
| YGTGGAGGNGG | ASSQGQTEVF | 0.82177705 |
| YTNVMAV | ASSQGQTEVF | 0.8120127 |
| FLEQVHQL | ASSQGQTEVF | 0.81949866 |
| GLSLLAVAV | ASSQGQTEVF | 0.8170059 |
| LEQVHQL | ASSQGQTEVF | 0.7893591 |
| LNAAGAI | ASSQGQTEVF | 0.7992834 |
| PAVHASL | ASSQGQTEVF | 0.6153368 |
| TELEDDA | ASSQGQTEVF | 0.67507017 |
| TPPGPPN | ASSQGQTEVF | 0.7954253 |
| TPPPPPPP | ASSQGQTEVF | 0.7965673 |
| TQSPALI | ASGRDLSYEQY | 0.8111464 |
| EFAAAFPQ | ASGRDLSYEQY | 0.8028616 |
| LLSGNLDVL | ASGRDLSYEQY | 0.75023687 |
| GPANLGNYNI | ASGRDLSYEQY | 0.8172883 |
| TLGPTLP | ASGRDLSYEQY | 0.8177691 |
| VYAYSSCCK | ASGRDLSYEQY | 0.6510952 |
| APAPAPAPNI | ASGRDLSYEQY | 0.7544043 |
| PGGPVPIP | ASGRDLSYEQY | 0.8252746 |
| FLTVALEP | ASGRDLSYEQY | 0.7923721 |
| SQYFGANL | ASGRDLSYEQY | 0.7682859 |

| | | |
|---|---|---|
| GGVKPAA | ASGRDLSYEQY | 0.7757779 |
| GGTGGAGGAGGW | ASGRDLSYEQY | 0.82409716 |
| GVGGVAGLGGAGG | ASGRDLSYEQY | 0.76303697 |
| VLLVLATTI | ASGRDLSYEQY | 0.8247778 |
| IGIFAAML | ASGRDLSYEQY | 0.43254995 |
| AVGDLID | ASGRDLSYEQY | 0.78305054 |
| NISPLPL | ASGRDLSYEQY | 0.7230016 |
| VTAVFVAI | ASGRDLSYEQY | 0.5772165 |
| SGDDNDAAL | ASGRDLSYEQY | 0.8166311 |
| AVAALAHS | ASGRDLSYEQY | 0.7757838 |
| GGVTVMATP | ASGRDLSYEQY | 0.8009968 |
| AGLAANVRD | ASGRDLSYEQY | 0.7774529 |
| TNRPDLIDPALLRPG | ASGRDLSYEQY | 0.8218087 |
| LANLGVTAL | ASGRDLSYEQY | 0.7847807 |
| TGERIEL | ASGRDLSYEQY | 0.78694546 |
| NGEELAA | ASGRDLSYEQY | 0.7570839 |
| VLSLVWG | ASGRDLSYEQY | 0.8109126 |
| IGYALWL | ASGRDLSYEQY | 0.82166356 |
| LSLLLAPL | ASGRDLSYEQY | 0.81923914 |
| TLDEALK | ASGRDLSYEQY | 0.5400015 |
| TDGAGTAEGG | ASGRDLSYEQY | 0.77466357 |
| NAPGGAGGNGG | ASGRDLSYEQY | 0.79783875 |
| VSVFMGMGGNF | ASGRDLSYEQY | 0.37592772 |
| LASYLAAL | ASGRDLSYEQY | 0.38865516 |
| LVGGTPAL | ASGRDLSYEQY | 0.75140953 |
| VRNADVL | ASGRDLSYEQY | 0.7394955 |
| TCSSPFLVSAM | ASGRDLSYEQY | 0.8126097 |
| VHITDAE | ASGRDLSYEQY | 0.8011578 |
| EYAAPSF | ASGRDLSYEQY | 0.78503025 |
| VVDAFHGGL | ASGRDLSYEQY | 0.8062327 |
| ASVVDPQ | ASGRDLSYEQY | 0.7851385 |
| SYNEFGGLS | ASGRDLSYEQY | 0.17858747 |
| DPDGSVY | ASGRDLSYEQY | 0.72573674 |
| MVSTIHATE | ASGRDLSYEQY | 0.6460637 |
| LAAGVADPV | ASGRDLSYEQY | 0.8243782 |
| TKEQIAA | ASGRDLSYEQY | 0.72507286 |
| SLASPAAL | ASGRDLSYEQY | 0.71646154 |
| IGLVFYGV | ASGRDLSYEQY | 0.79353595 |
| FLLDASNL | ASGRDLSYEQY | 0.78993845 |
| YQVGNFP | ASGRDLSYEQY | 0.77727526 |
| ESLLSQST | ASGRDLSYEQY | 0.72990465 |
| LFTVGFAAVG | ASGRDLSYEQY | 0.35028177 |
| ATPPTPT | ASGRDLSYEQY | 0.81198424 |
| TDLATLG | ASGRDLSYEQY | 0.60347295 |
| SAGGTGR | ASGRDLSYEQY | 0.67953783 |
| GFSNSGS | ASGRDLSYEQY | 0.47380796 |
| GGNGGSGGTSV | ASGRDLSYEQY | 0.7134737 |
| GTSVATGGAGN | ASGRDLSYEQY | 0.54830074 |
| NGGNGGSGGTS | ASGRDLSYEQY | 0.8178227 |
| AVVKFPH | ASGRDLSYEQY | 0.7503357 |
| TPPPPPP | ASGRDLSYEQY | 0.75742203 |

| | | |
|---|---|---|
| AAPAAAAAPA | ASGRDLSYEQY | 0.82093483 |
| IALNEDL | ASGRDLSYEQY | 0.78409827 |
| IVNVSSM | ASGRDLSYEQY | 0.77020955 |
| TGIPPAPR | ASGRDLSYEQY | 0.74040407 |
| FDLGGGTF | ASGRDLSYEQY | 0.7402141 |
| FDLGGGTFDVS | ASGRDLSYEQY | 0.73700035 |
| NEPTAAA | ASGRDLSYEQY | 0.7427825 |
| NEPTAAALA | ASGRDLSYEQY | 0.82196873 |
| GSIINISSI | ASGRDLSYEQY | 0.7889612 |
| VAPGNAGTA | ASGRDLSYEQY | 0.6408225 |
| TGGGASLE | ASGRDLSYEQY | 0.794724 |
| VAVYNHYKRI | ASGRDLSYEQY | 0.8241278 |
| TDPAPATT | ASGRDLSYEQY | 0.8282146 |
| TNRPDILDPALLRPG | ASGRDLSYEQY | 0.7330351 |
| LTTTTTS | ASGRDLSYEQY | 0.7460969 |
| PTGSGKTL | ASGRDLSYEQY | 0.76560414 |
| TGSGKTL | ASGRDLSYEQY | 0.7780514 |
| TIGHVDHGK | ASGRDLSYEQY | 0.73947114 |
| TIGHVDHGKTT | ASGRDLSYEQY | 0.70648694 |
| GHVDHGKTT | ASGRDLSYEQY | 0.81630397 |
| AQFEHTLL | ASGRDLSYEQY | 0.8298383 |
| IVNQDEL | ASGRDLSYEQY | 0.7771967 |
| TNGSQFF | ASGRDLSYEQY | 0.55553675 |
| TGKGIVE | ASGRDLSYEQY | 0.7812105 |
| LISNASDA | ASGRDLSYEQY | 0.73535085 |
| AELRTLGELTKIAKAHGAQ | ASGRDLSYEQY | 0.76460487 |
| APNAVAAAEPSW | ASGRDLSYEQY | 0.78251046 |
| APQVAGAATAVL | ASGRDLSYEQY | 0.61844206 |
| ARRDPAAAAAILPTDA | ASGRDLSYEQY | 0.77848 |
| DAAARDVLAEAGLAEYF | ASGRDLSYEQY | 0.82599723 |
| DDQIVLSAKANLAKVEQ | ASGRDLSYEQY | 0.7547145 |
| DELLPDTGEDSDEDGH | ASGRDLSYEQY | 0.791196 |
| ELPEAPANRATVPQD | ASGRDLSYEQY | 0.8207365 |
| FGQSKIERVTGIG | ASGRDLSYEQY | 0.80885994 |
| GAGGTGSTAGGAGGAGGAGGLY | ASGRDLSYEQY | 0.8210016 |
| GHGGTGAAGGNGGTGGAGGSGI | ASGRDLSYEQY | 0.8121358 |
| GNGGNGGAGGEGGPGGAGGAGA | ASGRDLSYEQY | 0.80055046 |
| GSTAGNGGNGGNATNGTIAGQP | ASGRDLSYEQY | 0.79107445 |
| GTGGDGGNGGNGGTGG | ASGRDLSYEQY | 0.793813 |
| GVRRPAGIPADAPV | ASGRDLSYEQY | 0.7867529 |
| GYRQVTDAHLLTLARR | ASGRDLSYEQY | 0.78146136 |
| IAKMGQDGHDRG | ASGRDLSYEQY | 0.75621283 |
| KAEPSIEIQYT | ASGRDLSYEQY | 0.8150099 |
| KPVRIRMAADEDTVAVT | ASGRDLSYEQY | 0.7785988 |
| MPPRIAGMRLLVIKPEP | ASGRDLSYEQY | 0.8240753 |
| MKFYAVALFRSRGVPPTR | ASGRDLSYEQY | 0.6241434 |
| NAGGFGILGAVAHSP | ASGRDLSYEQY | 0.7158959 |
| NEGREITLAMAEM | ASGRDLSYEQY | 0.81596094 |
| NELVEWSHPLGHRWRWE | ASGRDLSYEQY | 0.20925105 |
| NLGAGGTGGNGGNPG | ASGRDLSYEQY | 0.39418775 |
| NTEWPKWETVMAADPVD | ASGRDLSYEQY | 0.7921742 |

| | | |
|---|---|---|
| PASAAPAETQP | ASGRDLSYEQY | 0.82119095 |
| QQSSDGKNGVRAAEITGE | ASGRDLSYEQY | 0.22963524 |
| RAPDAVTTSWWKE | ASGRDLSYEQY | 0.6845801 |
| SGFGNVGAHTSGWFNQ | ASGRDLSYEQY | 0.67474085 |
| SGVAGAVGQAASVGGLKV | ASGRDLSYEQY | 0.80447465 |
| SNGGAGTGSAGGNGGT | ASGRDLSYEQY | 0.7853542 |
| TGGAGGEGGAAG | ASGRDLSYEQY | 0.5498947 |
| TGGDGGAAGAGGAPGHG | ASGRDLSYEQY | 0.81823695 |
| TMPPLLAAIANAGRHGV | ASGRDLSYEQY | 0.56527984 |
| TSSTAGNSGGA | ASGRDLSYEQY | 0.7250383 |
| VPATVPVGILC | ASGRDLSYEQY | 0.81578815 |
| VQRPDSGGVVLTGRIS | ASGRDLSYEQY | 0.82165706 |
| WTPNPVPATKGVPAGLVQAT | ASGRDLSYEQY | 0.66602707 |
| YALKRALAKFSDRFDVVII | ASGRDLSYEQY | 0.8116103 |
| YLIGSVIV | ASGRDLSYEQY | 0.7442856 |
| YGQLVAVID | ASGRDLSYEQY | 0.82859635 |
| VTVGGMTLPA | ASGRDLSYEQY | 0.7670763 |
| VLALPAV | ASGRDLSYEQY | 0.63632524 |
| VIPPTLNYET | ASGRDLSYEQY | 0.81521094 |
| VGMGGMHPSSG | ASGRDLSYEQY | 0.80545175 |
| VGLAGGAATA | ASGRDLSYEQY | 0.78592706 |
| VGGAGGAGGSGP | ASGRDLSYEQY | 0.8100009 |
| VFAAFMV | ASGRDLSYEQY | 0.7581601 |
| TLYDKLR | ASGRDLSYEQY | 0.7799959 |
| TLAVFTVL | ASGRDLSYEQY | 0.8190988 |
| TGSVQPARV | ASGRDLSYEQY | 0.8125074 |
| TGGGAGSGGT | ASGRDLSYEQY | 0.6835847 |
| SHLETSL | ASGRDLSYEQY | 0.8223036 |
| SFTTGMT | ASGRDLSYEQY | 0.77965915 |
| QVEVAKRMGVRQPTVS | ASGRDLSYEQY | 0.7134493 |
| QGSGGPALAM | ASGRDLSYEQY | 0.826066 |
| MLAPSNP | ASGRDLSYEQY | 0.79305255 |
| LVNNAAI | ASGRDLSYEQY | 0.81465626 |
| LTGEINTLA | ASGRDLSYEQY | 0.82038456 |
| LSIFGPV | ASGRDLSYEQY | 0.8199671 |
| LLALLANL | ASGRDLSYEQY | 0.61630946 |
| LISNASDALDKLR | ASGRDLSYEQY | 0.76694024 |
| LINEAVP | ASGRDLSYEQY | 0.777223 |
| LGYAFSGL | ASGRDLSYEQY | 0.8145925 |
| LDFVAVAL | ASGRDLSYEQY | 0.77353346 |
| LAQPGAE | ASGRDLSYEQY | 0.6832803 |
| ISNLGAVTGF | ASGRDLSYEQY | 0.8136621 |
| HQEDPGRI | ASGRDLSYEQY | 0.81895417 |
| GTPGEPTP | ASGRDLSYEQY | 0.78917766 |
| GFFNSGS | ASGRDLSYEQY | 0.78891677 |
| GAGGGGLTAGAN | ASGRDLSYEQY | 0.82452345 |
| FVDALVNM | ASGRDLSYEQY | 0.75447434 |
| FLTVAPEL | ASGRDLSYEQY | 0.7845644 |
| FETLTAL | ASGRDLSYEQY | 0.8111171 |
| DVINPIEL | ASGRDLSYEQY | 0.81519043 |
| APVSGAPV | ASGRDLSYEQY | 0.7922854 |

| | | |
|---|---|---|
| ALVGGLAW | ASGRDLSYEQY | 0.8240047 |
| ALAAAAATAI | ASGRDLSYEQY | 0.6980553 |
| AFGLGLLL | ASGRDLSYEQY | 0.81071436 |
| AFFAAASL | ASGRDLSYEQY | 0.7166456 |
| AAAALPAFSPP | ASGRDLSYEQY | 0.7866298 |
| VVLAPAL | ASGRDLSYEQY | 0.7069073 |
| TIGHVDHGKTTL | ASGRDLSYEQY | 0.8003133 |
| LISNASDALDK | ASGRDLSYEQY | 0.81161726 |
| LISNASDAL | ASGRDLSYEQY | 0.8059497 |
| ISNASDALDK | ASGRDLSYEQY | 0.34757203 |
| NQAETLVYQTE | ASGRDLSYEQY | 0.81192017 |
| TGGTGGTAGAG | ASGRDLSYEQY | 0.80625045 |
| TGSTAGGAGGA | ASGRDLSYEQY | 0.8276726 |
| DPLAVPVAAGPV | ASGRDLSYEQY | 0.78617704 |
| GGTGGTPGNGGN | ASGRDLSYEQY | 0.8072959 |
| PAGLAGVDPATL | ASGRDLSYEQY | 0.74492985 |
| GNNLVVEAPHAGASV | ASGRDLSYEQY | 0.79455024 |
| KVTRIMSASTLEQPA | ASGRDLSYEQY | 0.69863844 |
| LGFALPIALPRRTVV | ASGRDLSYEQY | 0.73634624 |
| TRLGNLTARKLVVLV | ASGRDLSYEQY | 0.74481463 |
| WSQQRRLIYFPSAGP | ASGRDLSYEQY | 0.7382136 |
| GPNGNPDPMAAAVDIR | ASGRDLSYEQY | 0.80996 |
| TQRCVSVADLRAAASR | ASGRDLSYEQY | 0.81535494 |
| QFVLDVAEHPVEVLTETV | ASGRDLSYEQY | 0.8074415 |
| GGAGGTGGAGGNGGTQAGDG | ASGRDLSYEQY | 0.817045 |
| PLLATLRAIAVDVDAKPAQV | ASGRDLSYEQY | 0.8180678 |
| APKFRTGGYMIDFWGVGYQVAK | ASGRDLSYEQY | 0.82368374 |
| GAGGAGGAGGAAGGPGGTGGTG | ASGRDLSYEQY | 0.7649589 |
| IGALGLLATDVAPAGSVAFGVA | ASGRDLSYEQY | 0.75185525 |
| AGGAGGIGGNGGAGGAGGVLMG | ASGRDLSYEQY | 0.6941217 |
| ADELIGLL | ASGRDLSYEQY | 0.76853824 |
| ADRALKA | ASGRDLSYEQY | 0.76629716 |
| AGAVGQVGGAGGA | ASGRDLSYEQY | 0.7986051 |
| AGGGGGLGGAP | ASGRDLSYEQY | 0.20481518 |
| AGGLLYGNGGAG | ASGRDLSYEQY | 0.78075546 |
| AGLESIGL | ASGRDLSYEQY | 0.8211731 |
| AGLKTGII | ASGRDLSYEQY | 0.74209446 |
| ASLAALLL | ASGRDLSYEQY | 0.79779136 |
| ASVINISSI | ASGRDLSYEQY | 0.7832873 |
| ATPPGPA | ASGRDLSYEQY | 0.7576467 |
| ATVASALV | ASGRDLSYEQY | 0.8048147 |
| ATVASLIL | ASGRDLSYEQY | 0.7591777 |
| AVNGPVLIRH | ASGRDLSYEQY | 0.81528366 |
| DGAAGAVGI | ASGRDLSYEQY | 0.8223096 |
| DPLLHGI | ASGRDLSYEQY | 0.7929347 |
| ELGAALGAL | ASGRDLSYEQY | 0.78414047 |
| ELGRTQAV | ASGRDLSYEQY | 0.7980837 |
| FVAGGYLS | ASGRDLSYEQY | 0.5438382 |
| GASGGAGGAGGW | ASGRDLSYEQY | 0.7925091 |
| GGAGAGGDGGAGGI | ASGRDLSYEQY | 0.4161805 |
| GTGHNGSQPGMGGNG | ASGRDLSYEQY | 0.7786139 |

| | | |
|---|---|---|
| GYAAASASAS | ASGRDLSYEQY | 0.7614351 |
| HQVDVAAVQQG | ASGRDLSYEQY | 0.79503006 |
| IGAAIAHT | ASGRDLSYEQY | 0.7823914 |
| IGMIEVAQL | ASGRDLSYEQY | 0.8001734 |
| IRLAPEHTSAMD | ASGRDLSYEQY | 0.7047514 |
| IYNVSVL | ASGRDLSYEQY | 0.7921896 |
| KQGFLGM | ASGRDLSYEQY | 0.7634635 |
| LESVITQL | ASGRDLSYEQY | 0.65515864 |
| LGLGALVVV | ASGRDLSYEQY | 0.8045528 |
| LIGNGGNGGSGGI | ASGRDLSYEQY | 0.7830066 |
| LNAINAPTL | ASGRDLSYEQY | 0.76128304 |
| LQLLNTGV | ASGRDLSYEQY | 0.81757724 |
| LQTSIEF | ASGRDLSYEQY | 0.79846686 |
| MSFAVSEL | ASGRDLSYEQY | 0.5378903 |
| NTGALGL | ASGRDLSYEQY | 0.76088905 |
| PPGAGGAGGAGTT | ASGRDLSYEQY | 0.78802395 |
| QLVGLGI | ASGRDLSYEQY | 0.783158 |
| RFLEQVHQL | ASGRDLSYEQY | 0.23370525 |
| RGVGYVLR | ASGRDLSYEQY | 0.7993885 |
| SATGVFAGVI | ASGRDLSYEQY | 0.7116382 |
| SLPAITLP | ASGRDLSYEQY | 0.68801975 |
| STALAFTV | ASGRDLSYEQY | 0.75636554 |
| SVLTVGLGK | ASGRDLSYEQY | 0.81176215 |
| TALAGAGSGI | ASGRDLSYEQY | 0.752772 |
| TGLPFTGI | ASGRDLSYEQY | 0.77005094 |
| VAEHVTV | ASGRDLSYEQY | 0.80543363 |
| VDIVLNSL | ASGRDLSYEQY | 0.79957956 |
| VLTGGGAVL | ASGRDLSYEQY | 0.820462 |
| VNVGLAYGI | ASGRDLSYEQY | 0.7962103 |
| VSSAAVGAV | ASGRDLSYEQY | 0.7851614 |
| VTSVYEAGV | ASGRDLSYEQY | 0.8013928 |
| WVALDRGA | ASGRDLSYEQY | 0.7498579 |
| YLVGSVVI | ASGRDLSYEQY | 0.6298249 |
| GPPGTGKTLL | ASGRDLSYEQY | 0.57465017 |
| GVGMTKF | ASGRDLSYEQY | 0.7853018 |
| IGIGGSDL | ASGRDLSYEQY | 0.561398 |
| IGTLLAEL | ASGRDLSYEQY | 0.79418623 |
| IYTSGTT | ASGRDLSYEQY | 0.8237002 |
| LSFPAGV | ASGRDLSYEQY | 0.7603294 |
| LYTSGTT | ASGRDLSYEQY | 0.63236624 |
| VGAFPFGF | ASGRDLSYEQY | 0.7648635 |
| VGAGTVEF | ASGRDLSYEQY | 0.74398595 |
| VGKSTLLL | ASGRDLSYEQY | 0.81428206 |
| AALAHAGGGIV | ASGRDLSYEQY | 0.81657904 |
| AALVAVSLL | ASGRDLSYEQY | 0.82714486 |
| AAVSRAVAAL | ASGRDLSYEQY | 0.8124634 |
| AGGSSGIGGF | ASGRDLSYEQY | 0.7542081 |
| AGLTVANANG | ASGRDLSYEQY | 0.81186336 |
| AGLVGAVL | ASGRDLSYEQY | 0.77733886 |
| ALSQAFTV | ASGRDLSYEQY | 0.73616177 |
| ALTFSVSL | ASGRDLSYEQY | 0.7953073 |

| | | |
|---|---|---|
| ALTGAALVAL | ASGRDLSYEQY | 0.79551387 |
| ASFSIGAGVL | ASGRDLSYEQY | 0.78382313 |
| DGSVYKI | ASGRDLSYEQY | 0.7664739 |
| EQLAEIV | ASGRDLSYEQY | 0.60458493 |
| FAVSELTVM | ASGRDLSYEQY | 0.81427765 |
| GAAPLAAGL | ASGRDLSYEQY | 0.79008305 |
| GADGGDGGSGGAGG | ASGRDLSYEQY | 0.6471782 |
| GFGLDTGGAGG | ASGRDLSYEQY | 0.7671354 |
| GLASGGLL | ASGRDLSYEQY | 0.8230014 |
| GRIAGAAVLL | ASGRDLSYEQY | 0.82047594 |
| GSAGSLNTSF | ASGRDLSYEQY | 0.7463505 |
| GTGQIGI | ASGRDLSYEQY | 0.76013553 |
| GTLIALIL | ASGRDLSYEQY | 0.77546644 |
| IAATMGFVVG | ASGRDLSYEQY | 0.66652584 |
| IFAQALL | ASGRDLSYEQY | 0.8124698 |
| IGAAVALGF | ASGRDLSYEQY | 0.7779921 |
| IGLTPLAL | ASGRDLSYEQY | 0.76507473 |
| ILGSNTSTL | ASGRDLSYEQY | 0.7494012 |
| IVFSLTVL | ASGRDLSYEQY | 0.7940045 |
| LASKATGF | ASGRDLSYEQY | 0.74389267 |
| LASVALIL | ASGRDLSYEQY | 0.8165258 |
| LFAVGLTL | ASGRDLSYEQY | 0.2507057 |
| LFNGGIGTYI | ASGRDLSYEQY | 0.80174285 |
| LTLGAGII | ASGRDLSYEQY | 0.71252906 |
| LVAAALGDI | ASGRDLSYEQY | 0.75912607 |
| NTAFVQLGI | ASGRDLSYEQY | 0.5312013 |
| QLFSGAEL | ASGRDLSYEQY | 0.7661967 |
| SAATFITL | ASGRDLSYEQY | 0.80639577 |
| SNYGLANLGV | ASGRDLSYEQY | 0.8155618 |
| SYGGAYAVM | ASGRDLSYEQY | 0.72986543 |
| TTGVGGLTLG | ASGRDLSYEQY | 0.80826676 |
| TTQTIYI | ASGRDLSYEQY | 0.8238324 |
| VAAGAGLGAVI | ASGRDLSYEQY | 0.75272256 |
| VAAVVTLL | ASGRDLSYEQY | 0.4490296 |
| VGFLGGLI | ASGRDLSYEQY | 0.80694234 |
| VGVIPILL | ASGRDLSYEQY | 0.74358463 |
| VIFASSLI | ASGRDLSYEQY | 0.8113617 |
| VKGGDGGAAAG | ASGRDLSYEQY | 0.8024819 |
| VTKTFTNI | ASGRDLSYEQY | 0.6852883 |
| VVAAAILL | ASGRDLSYEQY | 0.81728876 |
| YAALDVEL | ASGRDLSYEQY | 0.8107268 |
| LASVALLL | ASGRDLSYEQY | 0.7249613 |
| LDAVHRV | ASGRDLSYEQY | 0.67718923 |
| QGLSSSIF | ASGRDLSYEQY | 0.6929399 |
| RGTFANIRL | ASGRDLSYEQY | 0.49869722 |
| RQLWWGHRI | ASGRDLSYEQY | 0.8083204 |
| YGPPGTGKTLL | ASGRDLSYEQY | 0.74667513 |
| AALVGAPANVA | ASGRDLSYEQY | 0.7378988 |
| AATFRRTE | ASGRDLSYEQY | 0.79574263 |
| AGALGGGI | ASGRDLSYEQY | 0.74746835 |
| AGGAGGAGGIGAG | ASGRDLSYEQY | 0.76154053 |

| | | |
|---|---|---|
| AGSGALGL | ASGRDLSYEQY | 0.7619648 |
| AQIADQI | ASGRDLSYEQY | 0.77270746 |
| ASAVAVTYL | ASGRDLSYEQY | 0.80390626 |
| ASTVLGGTGG | ASGRDLSYEQY | 0.76258147 |
| AVGGEPAK | ASGRDLSYEQY | 0.7780715 |
| CEQGTFGGTC | ASGRDLSYEQY | 0.81907254 |
| DEGKAPAG | ASGRDLSYEQY | 0.6222964 |
| DTVVFPAL | ASGRDLSYEQY | 0.7316648 |
| FEGITMAG | ASGRDLSYEQY | 0.8051096 |
| FPLVVQD | ASGRDLSYEQY | 0.7051895 |
| GAFTTPQ | ASGRDLSYEQY | 0.81018245 |
| GFGGFGTAGGVGGA | ASGRDLSYEQY | 0.7821342 |
| GFLAQFL | ASGRDLSYEQY | 0.80165374 |
| GGAGGAGGNAGVA | ASGRDLSYEQY | 0.7720587 |
| GGVGTAGAGGAGGAGG | ASGRDLSYEQY | 0.82303524 |
| GNGGGGDAGNAGSGGN | ASGRDLSYEQY | 0.8122071 |
| GTGGAGGAGNDAGS | ASGRDLSYEQY | 0.7750412 |
| GTTNTGWFNS | ASGRDLSYEQY | 0.80112594 |
| IFMGPLFTL | ASGRDLSYEQY | 0.6232238 |
| ILAHAIEAI | ASGRDLSYEQY | 0.814657 |
| IRSAIREAL | ASGRDLSYEQY | 0.8102733 |
| ITFAGGG | ASGRDLSYEQY | 0.7078383 |
| ITGMSGHQEV | ASGRDLSYEQY | 0.78370637 |
| IVAGLACTL | ASGRDLSYEQY | 0.8237539 |
| IVGGGLAGA | ASGRDLSYEQY | 0.81964076 |
| IVTGVLNI | ASGRDLSYEQY | 0.6855749 |
| LAQQGCI | ASGRDLSYEQY | 0.7629489 |
| LCLHVEN | ASGRDLSYEQY | 0.580215 |
| LDAISDGS | ASGRDLSYEQY | 0.72951376 |
| LGGFAAML | ASGRDLSYEQY | 0.5918795 |
| LGGKNPMI | ASGRDLSYEQY | 0.8055097 |
| LGPKGVAAIL | ASGRDLSYEQY | 0.7556536 |
| LMVGVSI | ASGRDLSYEQY | 0.80494475 |
| LVGAAFVL | ASGRDLSYEQY | 0.79783714 |
| LVLAVAPL | ASGRDLSYEQY | 0.8239598 |
| MAAIAVIDAA | ASGRDLSYEQY | 0.75782514 |
| NNGINAELQ | ASGRDLSYEQY | 0.7637807 |
| RGEPLVLVHG | ASGRDLSYEQY | 0.64093953 |
| RGGAGGAAGLI | ASGRDLSYEQY | 0.6362736 |
| RSYRMHPAV | ASGRDLSYEQY | 0.742659 |
| SAAHLLAGQLA | ASGRDLSYEQY | 0.7597167 |
| SAAVGSY | ASGRDLSYEQY | 0.3961847 |
| SPHAVLHRI | ASGRDLSYEQY | 0.8212844 |
| SQIAKVL | ASGRDLSYEQY | 0.79249346 |
| TALGHLVP | ASGRDLSYEQY | 0.8134328 |
| TAPTTSGAA | ASGRDLSYEQY | 0.73333406 |
| TDGDDGNGGSPGT | ASGRDLSYEQY | 0.7569548 |
| TDGLGEN | ASGRDLSYEQY | 0.762568 |
| TGGPPLPGQA | ASGRDLSYEQY | 0.80677617 |
| TIPAITI | ASGRDLSYEQY | 0.716418 |
| TIPGFTI | ASGRDLSYEQY | 0.51254773 |

| | | |
|---|---|---|
| TSIASGALA | ASGRDLSYEQY | 0.7093759 |
| TVTLPGAAI | ASGRDLSYEQY | 0.61792946 |
| VASGLSAALP | ASGRDLSYEQY | 0.824771 |
| VGAGVIGI | ASGRDLSYEQY | 0.81958973 |
| VGPKRFGV | ASGRDLSYEQY | 0.7984894 |
| VNVAFPD | ASGRDLSYEQY | 0.8251958 |
| VSMTNSGVS | ASGRDLSYEQY | 0.30149496 |
| VTDVLGDLSV | ASGRDLSYEQY | 0.7265387 |
| VVVVGAGFAGL | ASGRDLSYEQY | 0.81934726 |
| VYTVGTVV | ASGRDLSYEQY | 0.80846703 |
| WTTFSGP | ASGRDLSYEQY | 0.76984906 |
| YDELVEAF | ASGRDLSYEQY | 0.8227347 |
| YGTGGAGGNGG | ASGRDLSYEQY | 0.72250915 |
| YTNVMAV | ASGRDLSYEQY | 0.7812213 |
| FLEQVHQL | ASGRDLSYEQY | 0.78556824 |
| GLSLLAVAV | ASGRDLSYEQY | 0.8163382 |
| LEQVHQL | ASGRDLSYEQY | 0.7981681 |
| LNAAGAI | ASGRDLSYEQY | 0.71396947 |
| PAVHASL | ASGRDLSYEQY | 0.8071625 |
| TELEDDA | ASGRDLSYEQY | 0.8245572 |
| TPPGPPN | ASGRDLSYEQY | 0.7573211 |
| TPPPPPPP | ASGRDLSYEQY | 0.8142259 |
| TQSPALI | ASSGHLNQDTQY | 0.80377316 |
| EFAAAFPQ | ASSGHLNQDTQY | 0.79054236 |
| LLSGNLDVL | ASSGHLNQDTQY | 0.7379149 |
| GPANLGNYNI | ASSGHLNQDTQY | 0.7417503 |
| TLGPTLP | ASSGHLNQDTQY | 0.79345477 |
| VYAYSSCCK | ASSGHLNQDTQY | 0.70329887 |
| APAPAPAPNI | ASSGHLNQDTQY | 0.7791288 |
| PGGPVPIP | ASSGHLNQDTQY | 0.77645415 |
| FLTVALEP | ASSGHLNQDTQY | 0.8319733 |
| SQYFGANL | ASSGHLNQDTQY | 0.80560887 |
| GGVKPAA | ASSGHLNQDTQY | 0.8067164 |
| GGTGGAGGAGGW | ASSGHLNQDTQY | 0.7351615 |
| GVGGVAGLGGAGG | ASSGHLNQDTQY | 0.8017237 |
| VLLVLATTI | ASSGHLNQDTQY | 0.8048718 |
| IGIFAAML | ASSGHLNQDTQY | 0.64258564 |
| AVGDLID | ASSGHLNQDTQY | 0.82017374 |
| NISPLPL | ASSGHLNQDTQY | 0.7404154 |
| VTAVFVAI | ASSGHLNQDTQY | 0.76647687 |
| SGDDNDAAL | ASSGHLNQDTQY | 0.8084562 |
| AVAALAHS | ASSGHLNQDTQY | 0.81855315 |
| GGVTVMATP | ASSGHLNQDTQY | 0.7849771 |
| AGLAANVRD | ASSGHLNQDTQY | 0.77994794 |
| TNRPDLIDPALLRPG | ASSGHLNQDTQY | 0.7257308 |
| LANLGVTAL | ASSGHLNQDTQY | 0.8171254 |
| TGERIEL | ASSGHLNQDTQY | 0.8185189 |
| NGEELAA | ASSGHLNQDTQY | 0.78238034 |
| VLSLVWG | ASSGHLNQDTQY | 0.8276382 |
| IGYALWL | ASSGHLNQDTQY | 0.7869465 |
| LSLLLAPL | ASSGHLNQDTQY | 0.49552113 |

| | | |
|---|---|---|
| TLDEALK | ASSGHLNQDTQY | 0.79433715 |
| TDGAGTAEGG | ASSGHLNQDTQY | 0.8144952 |
| NAPGGAGGNGG | ASSGHLNQDTQY | 0.81526756 |
| VSVFMGMGGNF | ASSGHLNQDTQY | 0.82339686 |
| LASYLAAL | ASSGHLNQDTQY | 0.79424095 |
| LVGGTPAL | ASSGHLNQDTQY | 0.7630308 |
| VRNADVL | ASSGHLNQDTQY | 0.8236399 |
| TCSSPFLVSAM | ASSGHLNQDTQY | 0.83011186 |
| VHITDAE | ASSGHLNQDTQY | 0.81577295 |
| EYAAPSF | ASSGHLNQDTQY | 0.7656181 |
| VVDAFHGGL | ASSGHLNQDTQY | 0.74029124 |
| ASVVDPQ | ASSGHLNQDTQY | 0.8003595 |
| SYNEFGGLS | ASSGHLNQDTQY | 0.81332755 |
| DPDGSVY | ASSGHLNQDTQY | 0.7196101 |
| MVSTIHATE | ASSGHLNQDTQY | 0.7947606 |
| LAAGVADPV | ASSGHLNQDTQY | 0.7955493 |
| TKEQIAA | ASSGHLNQDTQY | 0.5023786 |
| SLASPAAL | ASSGHLNQDTQY | 0.8003689 |
| IGLVFYGV | ASSGHLNQDTQY | 0.8129827 |
| FLLDASNL | ASSGHLNQDTQY | 0.8008314 |
| YQVGNFP | ASSGHLNQDTQY | 0.73547745 |
| ESLLSQST | ASSGHLNQDTQY | 0.81590116 |
| LFTVGFAAVG | ASSGHLNQDTQY | 0.8284037 |
| ATPPTPT | ASSGHLNQDTQY | 0.82734704 |
| TDLATLG | ASSGHLNQDTQY | 0.5167794 |
| SAGGTGR | ASSGHLNQDTQY | 0.44091314 |
| GFSNSGS | ASSGHLNQDTQY | 0.77193254 |
| GGNGGSGGTSV | ASSGHLNQDTQY | 0.80232954 |
| GTSVATGGAGN | ASSGHLNQDTQY | 0.7673458 |
| NGGNGGSGGTS | ASSGHLNQDTQY | 0.75885165 |
| AVVKFPH | ASSGHLNQDTQY | 0.78511727 |
| TPPPPPP | ASSGHLNQDTQY | 0.8205425 |
| AAPAAAAAPA | ASSGHLNQDTQY | 0.73574436 |
| IALNEDL | ASSGHLNQDTQY | 0.6731919 |
| IVNVSSM | ASSGHLNQDTQY | 0.7822778 |
| TGIPPAPR | ASSGHLNQDTQY | 0.81427294 |
| FDLGGGTF | ASSGHLNQDTQY | 0.7245165 |
| FDLGGGTFDVS | ASSGHLNQDTQY | 0.7766731 |
| NEPTAAA | ASSGHLNQDTQY | 0.7577088 |
| NEPTAAALA | ASSGHLNQDTQY | 0.7583716 |
| GSIINISSI | ASSGHLNQDTQY | 0.79468125 |
| VAPGNAGTA | ASSGHLNQDTQY | 0.71842504 |
| TGGGASLE | ASSGHLNQDTQY | 0.81171143 |
| VAVYNHYKRI | ASSGHLNQDTQY | 0.71237314 |
| TDPAPATT | ASSGHLNQDTQY | 0.80873466 |
| TNRPDILDPALLRPG | ASSGHLNQDTQY | 0.7939594 |
| LTTTTTS | ASSGHLNQDTQY | 0.8077098 |
| PTGSGKTL | ASSGHLNQDTQY | 0.76460004 |
| TGSGKTL | ASSGHLNQDTQY | 0.73907125 |
| TIGHVDHGK | ASSGHLNQDTQY | 0.6966052 |
| TIGHVDHGKTT | ASSGHLNQDTQY | 0.5927714 |

| | | |
|---|---|---|
| GHVDHGKTT | ASSGHLNQDTQY | 0.8163154 |
| AQFEHTLL | ASSGHLNQDTQY | 0.7001681 |
| IVNQDEL | ASSGHLNQDTQY | 0.82132864 |
| TNGSQFF | ASSGHLNQDTQY | 0.80999625 |
| TGKGIVE | ASSGHLNQDTQY | 0.7382301 |
| LISNASDA | ASSGHLNQDTQY | 0.80866015 |
| AELRTLGELTKIAKAHGAQ | ASSGHLNQDTQY | 0.6884931 |
| APNAVAAAEPSW | ASSGHLNQDTQY | 0.7701247 |
| APQVAGAATAVL | ASSGHLNQDTQY | 0.8016599 |
| ARRDPAAAAAILPTDA | ASSGHLNQDTQY | 0.8142425 |
| DAAARDVLAEAGLAEYF | ASSGHLNQDTQY | 0.71925956 |
| DDQIVLSAKANLAKVEQ | ASSGHLNQDTQY | 0.8079774 |
| DELLPDTGEDSDEDGH | ASSGHLNQDTQY | 0.7984333 |
| ELPEAPANRATVPQD | ASSGHLNQDTQY | 0.6987374 |
| FGQSKIERVTGIG | ASSGHLNQDTQY | 0.79979813 |
| GAGGTGSTAGGAGGAGGAGGLY | ASSGHLNQDTQY | 0.81942767 |
| GHGGTGAAGGNGGTGGAGGSGI | ASSGHLNQDTQY | 0.8089855 |
| GNGGNGGAGGEGGPGGAGGAGA | ASSGHLNQDTQY | 0.71572006 |
| GSTAGNGGNGGNATNGTIAGQP | ASSGHLNQDTQY | 0.79828334 |
| GTGGDGGNGGNGGTGG | ASSGHLNQDTQY | 0.7203183 |
| GVRRPAGIPADAPV | ASSGHLNQDTQY | 0.79779017 |
| GYRQVTDAHLLTLARR | ASSGHLNQDTQY | 0.79249024 |
| IAKMGQDGHDRG | ASSGHLNQDTQY | 0.74788517 |
| KAEPSIEIQYT | ASSGHLNQDTQY | 0.8067912 |
| KPVRIRMAADEDTVAVT | ASSGHLNQDTQY | 0.45524973 |
| MPPRIAGMRLLVIKPEP | ASSGHLNQDTQY | 0.7283572 |
| MKFYAVALFRSRGVPPTR | ASSGHLNQDTQY | 0.7804415 |
| NAGGFGILGAVAHSP | ASSGHLNQDTQY | 0.7862476 |
| NEGREITLAMAEM | ASSGHLNQDTQY | 0.81554806 |
| NELVEWSHPLGHRWRWE | ASSGHLNQDTQY | 0.72572625 |
| NLGAGGTGGNGGNPG | ASSGHLNQDTQY | 0.7915586 |
| NTEWPKWETVMAADPVD | ASSGHLNQDTQY | 0.76811755 |
| PASAAPAETQP | ASSGHLNQDTQY | 0.7235075 |
| QQSSDGKNGVRAAEITGE | ASSGHLNQDTQY | 0.8044212 |
| RAPDAVTTSWWKE | ASSGHLNQDTQY | 0.7722317 |
| SGFGNVGAHTSGWFNQ | ASSGHLNQDTQY | 0.6994401 |
| SGVAGAVGQAASVGGLKV | ASSGHLNQDTQY | 0.81012475 |
| SNGGAGTGSAGGNGGT | ASSGHLNQDTQY | 0.696233 |
| TGGAGGEGGAAG | ASSGHLNQDTQY | 0.8035023 |
| TGGDGGAAGAGGAPGHG | ASSGHLNQDTQY | 0.82419354 |
| TMPPLLAAIANAGRHGV | ASSGHLNQDTQY | 0.731634 |
| TSSTAGNSGGA | ASSGHLNQDTQY | 0.8093282 |
| VPATVPVGILC | ASSGHLNQDTQY | 0.79715836 |
| VQRPDSGGVVLTGRIS | ASSGHLNQDTQY | 0.80177665 |
| WTPNPVPATKGVPAGLVQAT | ASSGHLNQDTQY | 0.76744384 |
| YALKRALAKFSDRFDVVII | ASSGHLNQDTQY | 0.7350023 |
| YLIGSVIV | ASSGHLNQDTQY | 0.8171455 |
| YGQLVAVID | ASSGHLNQDTQY | 0.8060297 |
| VTVGGMTLPA | ASSGHLNQDTQY | 0.7955401 |
| VLALPAV | ASSGHLNQDTQY | 0.79766095 |
| VIPPTLNYET | ASSGHLNQDTQY | 0.79482764 |

| | | |
|---|---|---|
| VGMGGMHPSSG | ASSGHLNQDTQY | 0.8117583 |
| VGLAGGAATA | ASSGHLNQDTQY | 0.7512709 |
| VGGAGGAGGSGP | ASSGHLNQDTQY | 0.66971976 |
| VFAAFMV | ASSGHLNQDTQY | 0.7744061 |
| TLYDKLR | ASSGHLNQDTQY | 0.7541356 |
| TLAVFTVL | ASSGHLNQDTQY | 0.7626798 |
| TGSVQPARV | ASSGHLNQDTQY | 0.5766153 |
| TGGGAGSGGT | ASSGHLNQDTQY | 0.81307554 |
| SHLETSL | ASSGHLNQDTQY | 0.82893157 |
| SFTTGMT | ASSGHLNQDTQY | 0.69674313 |
| QVEVAKRMGVRQPTVS | ASSGHLNQDTQY | 0.8283864 |
| QGSGGPALAM | ASSGHLNQDTQY | 0.6853609 |
| MLAPSNP | ASSGHLNQDTQY | 0.81043386 |
| LVNNAAI | ASSGHLNQDTQY | 0.81519103 |
| LTGEINTLA | ASSGHLNQDTQY | 0.71674216 |
| LSIFGPV | ASSGHLNQDTQY | 0.8025389 |
| LLALLANL | ASSGHLNQDTQY | 0.7377846 |
| LISNASDALDKLR | ASSGHLNQDTQY | 0.7581339 |
| LINEAVP | ASSGHLNQDTQY | 0.81864 |
| LGYAFSGL | ASSGHLNQDTQY | 0.7414533 |
| LDFVAVAL | ASSGHLNQDTQY | 0.80053186 |
| LAQPGAE | ASSGHLNQDTQY | 0.78995466 |
| ISNLGAVTGF | ASSGHLNQDTQY | 0.7607836 |
| HQEDPGRI | ASSGHLNQDTQY | 0.80991244 |
| GTPGEPTP | ASSGHLNQDTQY | 0.81220925 |
| GFFNSGS | ASSGHLNQDTQY | 0.76092947 |
| GAGGGGLTAGAN | ASSGHLNQDTQY | 0.8291577 |
| FVDALVNM | ASSGHLNQDTQY | 0.7107177 |
| FLTVAPEL | ASSGHLNQDTQY | 0.822878 |
| FETLTAL | ASSGHLNQDTQY | 0.8076517 |
| DVINPIEL | ASSGHLNQDTQY | 0.78964144 |
| APVSGAPV | ASSGHLNQDTQY | 0.78058606 |
| ALVGGLAW | ASSGHLNQDTQY | 0.8127321 |
| ALAAAAATAI | ASSGHLNQDTQY | 0.76798594 |
| AFGLGLLL | ASSGHLNQDTQY | 0.79551625 |
| AFFAAASL | ASSGHLNQDTQY | 0.8262782 |
| AAAALPAFSPP | ASSGHLNQDTQY | 0.43886235 |
| VVLAPAL | ASSGHLNQDTQY | 0.6969659 |
| TIGHVDHGKTTL | ASSGHLNQDTQY | 0.60456604 |
| LISNASDALDK | ASSGHLNQDTQY | 0.65629816 |
| LISNASDAL | ASSGHLNQDTQY | 0.5381484 |
| ISNASDALDK | ASSGHLNQDTQY | 0.50589 |
| NQAETLVYQTE | ASSGHLNQDTQY | 0.5787068 |
| TGGTGGTAGAG | ASSGHLNQDTQY | 0.50817144 |
| TGSTAGGAGGA | ASSGHLNQDTQY | 0.66921854 |
| DPLAVPVAAGPV | ASSGHLNQDTQY | 0.6771029 |
| GGTGGTPGNGGN | ASSGHLNQDTQY | 0.52443033 |
| PAGLAGVDPATL | ASSGHLNQDTQY | 0.6726792 |
| GNNLVVEAPHAGASV | ASSGHLNQDTQY | 0.6921777 |
| KVTRIMSASTLEQPA | ASSGHLNQDTQY | 0.4010461 |
| LGFALPIALPRRTVV | ASSGHLNQDTQY | 0.5893782 |

| | | |
|---|---|---|
| TRLGNLTARKLVVLV | ASSGHLNQDTQY | 0.59934676 |
| WSQQRRLIYFPSAGP | ASSGHLNQDTQY | 0.5634607 |
| GPNGNPDPMAAAVDIR | ASSGHLNQDTQY | 0.33185214 |
| TQRCVSVADLRAAASR | ASSGHLNQDTQY | 0.65434104 |
| QFVLDVAEHPVEVLTETV | ASSGHLNQDTQY | 0.4359063 |
| GGAGGTGGAGGNGGTQAGDG | ASSGHLNQDTQY | 0.3049271 |
| PLLATLRAIAVDVDAKPAQV | ASSGHLNQDTQY | 0.63998747 |
| APKFRTGGYMIDFWGVGYQVAK | ASSGHLNQDTQY | 0.54341143 |
| GAGGAGGAGGAAGGPGGTGGTG | ASSGHLNQDTQY | 0.60246605 |
| IGALGLLATDVAPAGSVAFGVA | ASSGHLNQDTQY | 0.6681502 |
| AGGAGGIGGNGGAGGAGGVLMG | ASSGHLNQDTQY | 0.5796247 |
| ADELIGLL | ASSGHLNQDTQY | 0.57128006 |
| ADRALKA | ASSGHLNQDTQY | 0.61999655 |
| AGAVGQVGGAGGA | ASSGHLNQDTQY | 0.68689555 |
| AGGGGGLGGAP | ASSGHLNQDTQY | 0.54082644 |
| AGGLLYGNGGAG | ASSGHLNQDTQY | 0.5484359 |
| AGLESIGL | ASSGHLNQDTQY | 0.7089893 |
| AGLKTGII | ASSGHLNQDTQY | 0.6151952 |
| ASLAALLL | ASSGHLNQDTQY | 0.70996577 |
| ASVINISSI | ASSGHLNQDTQY | 0.7180197 |
| ATPPGPA | ASSGHLNQDTQY | 0.49792123 |
| ATVASALV | ASSGHLNQDTQY | 0.67906237 |
| ATVASLIL | ASSGHLNQDTQY | 0.6847946 |
| AVNGPVLIRH | ASSGHLNQDTQY | 0.60329276 |
| DGAAGAVGI | ASSGHLNQDTQY | 0.5656763 |
| DPLLHGI | ASSGHLNQDTQY | 0.40956315 |
| ELGAALGAL | ASSGHLNQDTQY | 0.57562363 |
| ELGRTQAV | ASSGHLNQDTQY | 0.4996942 |
| FVAGGYLS | ASSGHLNQDTQY | 0.5969297 |
| GASGGAGGAGGW | ASSGHLNQDTQY | 0.71373624 |
| GGAGAGGDGGAGGI | ASSGHLNQDTQY | 0.5561191 |
| GTGHNGSQPGMGGNG | ASSGHLNQDTQY | 0.3826102 |
| GYAAASASAS | ASSGHLNQDTQY | 0.36670578 |
| HQVDVAAVQQG | ASSGHLNQDTQY | 0.5952507 |
| IGAAIAHT | ASSGHLNQDTQY | 0.41059634 |
| IGMIEVAQL | ASSGHLNQDTQY | 0.4546932 |
| IRLAPEHTSAMD | ASSGHLNQDTQY | 0.49644518 |
| IYNVSVL | ASSGHLNQDTQY | 0.69889516 |
| KQGFLGM | ASSGHLNQDTQY | 0.7094681 |
| LESVITQL | ASSGHLNQDTQY | 0.49376345 |
| LGLGALVVV | ASSGHLNQDTQY | 0.68800586 |
| LIGNGGNGGSGGI | ASSGHLNQDTQY | 0.6273252 |
| LNAINAPTL | ASSGHLNQDTQY | 0.6881228 |
| LQLLNTGV | ASSGHLNQDTQY | 0.6178245 |
| LQTSIEF | ASSGHLNQDTQY | 0.46391228 |
| MSFAVSEL | ASSGHLNQDTQY | 0.6680807 |
| NTGALGL | ASSGHLNQDTQY | 0.5074185 |
| PPGAGGAGGAGTT | ASSGHLNQDTQY | 0.69905764 |
| QLVGLGI | ASSGHLNQDTQY | 0.6809671 |
| RFLEQVHQL | ASSGHLNQDTQY | 0.63591963 |
| RGVGYVLR | ASSGHLNQDTQY | 0.5020691 |

| | | |
|---|---|---|
| SATGVFAGVI | ASSGHLNQDTQY | 0.5456268 |
| SLPAITLP | ASSGHLNQDTQY | 0.7109846 |
| STALAFTV | ASSGHLNQDTQY | 0.4167195 |
| SVLTVGLGK | ASSGHLNQDTQY | 0.5071656 |
| TALAGAGSGI | ASSGHLNQDTQY | 0.6078168 |
| TGLPFTGI | ASSGHLNQDTQY | 0.5861914 |
| VAEHVTV | ASSGHLNQDTQY | 0.70902824 |
| VDIVLNSL | ASSGHLNQDTQY | 0.6622723 |
| VLTGGGAVL | ASSGHLNQDTQY | 0.54112184 |
| VNVGLAYGI | ASSGHLNQDTQY | 0.6251681 |
| VSSAAVGAV | ASSGHLNQDTQY | 0.67315894 |
| VTSVYEAGV | ASSGHLNQDTQY | 0.620466 |
| WVALDRGA | ASSGHLNQDTQY | 0.61960936 |
| YLVGSVVI | ASSGHLNQDTQY | 0.48452872 |
| GPPGTGKTLL | ASSGHLNQDTQY | 0.69883204 |
| GVGMTKF | ASSGHLNQDTQY | 0.67918575 |
| IGIGGSDL | ASSGHLNQDTQY | 0.6754466 |
| IGTLLAEL | ASSGHLNQDTQY | 0.68760526 |
| IYTSGTT | ASSGHLNQDTQY | 0.65034634 |
| LSFPAGV | ASSGHLNQDTQY | 0.6065497 |
| LYTSGTT | ASSGHLNQDTQY | 0.4571838 |
| VGAFPFGF | ASSGHLNQDTQY | 0.51437 |
| VGAGTVEF | ASSGHLNQDTQY | 0.5455244 |
| VGKSTLLL | ASSGHLNQDTQY | 0.61215794 |
| AALAHAGGGIV | ASSGHLNQDTQY | 0.6346675 |
| AALVAVSLL | ASSGHLNQDTQY | 0.69010824 |
| AAVSRAVAAL | ASSGHLNQDTQY | 0.48764586 |
| AGGSSGIGGF | ASSGHLNQDTQY | 0.703512 |
| AGLTVANANG | ASSGHLNQDTQY | 0.6786858 |
| AGLVGAVL | ASSGHLNQDTQY | 0.41134295 |
| ALSQAFTV | ASSGHLNQDTQY | 0.5623733 |
| ALTFSVSL | ASSGHLNQDTQY | 0.71258825 |
| ALTGAALVAL | ASSGHLNQDTQY | 0.6154769 |
| ASFSIGAGVL | ASSGHLNQDTQY | 0.5625447 |
| DGSVYKI | ASSGHLNQDTQY | 0.5906435 |
| EQLAEIV | ASSGHLNQDTQY | 0.6969374 |
| FAVSELTVM | ASSGHLNQDTQY | 0.6338953 |
| GAAPLAAGL | ASSGHLNQDTQY | 0.69700086 |
| GADGGDGGSGGAGG | ASSGHLNQDTQY | 0.36994317 |
| GFGLDTGGAGG | ASSGHLNQDTQY | 0.5532819 |
| GLASGGLL | ASSGHLNQDTQY | 0.515343 |
| GRIAGAAVLL | ASSGHLNQDTQY | 0.29378027 |
| GSAGSLNTSF | ASSGHLNQDTQY | 0.6960449 |
| GTGQIGI | ASSGHLNQDTQY | 0.5388048 |
| GTLIALIL | ASSGHLNQDTQY | 0.6317576 |
| IAATMGFVVG | ASSGHLNQDTQY | 0.5257121 |
| IFAQALL | ASSGHLNQDTQY | 0.6889433 |
| IGAAVALGF | ASSGHLNQDTQY | 0.5645916 |
| IGLTPLAL | ASSGHLNQDTQY | 0.5846209 |
| ILGSNTSTL | ASSGHLNQDTQY | 0.5538703 |
| IVFSLTVL | ASSGHLNQDTQY | 0.6609828 |

| | | |
|---|---|---|
| LASKATGF | ASSGHLNQDTQY | 0.7052094 |
| LASVALIL | ASSGHLNQDTQY | 0.7039553 |
| LFAVGLTL | ASSGHLNQDTQY | 0.43851176 |
| LFNGGIGTYI | ASSGHLNQDTQY | 0.66562843 |
| LTLGAGII | ASSGHLNQDTQY | 0.603895 |
| LVAAALGDI | ASSGHLNQDTQY | 0.47505993 |
| NTAFVQLGI | ASSGHLNQDTQY | 0.3962181 |
| QLFSGAEL | ASSGHLNQDTQY | 0.5824525 |
| SAATFITL | ASSGHLNQDTQY | 0.5220718 |
| SNYGLANLGV | ASSGHLNQDTQY | 0.64508384 |
| SYGGAYAVM | ASSGHLNQDTQY | 0.6678957 |
| TTGVGGLTLG | ASSGHLNQDTQY | 0.5091217 |
| TTQTIYI | ASSGHLNQDTQY | 0.62769717 |
| VAAGAGLGAVI | ASSGHLNQDTQY | 0.5911762 |
| VAAVVTLL | ASSGHLNQDTQY | 0.2026528 |
| VGFLGGLI | ASSGHLNQDTQY | 0.5186131 |
| VGVIPILL | ASSGHLNQDTQY | 0.39329112 |
| VIFASSLI | ASSGHLNQDTQY | 0.71888745 |
| VKGGDGGAAAG | ASSGHLNQDTQY | 0.47054407 |
| VTKTFTNI | ASSGHLNQDTQY | 0.52894837 |
| VVAAAILL | ASSGHLNQDTQY | 0.5737899 |
| YAALDVEL | ASSGHLNQDTQY | 0.59741986 |
| LASVALLL | ASSGHLNQDTQY | 0.58142084 |
| LDAVHRV | ASSGHLNQDTQY | 0.4948842 |
| QGLSSSIF | ASSGHLNQDTQY | 0.4295569 |
| RGTFANIRL | ASSGHLNQDTQY | 0.6684466 |
| RQLWWGHRI | ASSGHLNQDTQY | 0.6236159 |
| YGPPGTGKTLL | ASSGHLNQDTQY | 0.55520725 |
| AALVGAPANVA | ASSGHLNQDTQY | 0.5000138 |
| AATFRRTE | ASSGHLNQDTQY | 0.6670622 |
| AGALGGGI | ASSGHLNQDTQY | 0.41814122 |
| AGGAGGAGGIGAG | ASSGHLNQDTQY | 0.6853722 |
| AGSGALGL | ASSGHLNQDTQY | 0.44445854 |
| AQIADQI | ASSGHLNQDTQY | 0.5298494 |
| ASAVAVTYL | ASSGHLNQDTQY | 0.71089697 |
| ASTVLGGTGG | ASSGHLNQDTQY | 0.5981442 |
| AVGGEPAK | ASSGHLNQDTQY | 0.5536386 |
| CEQGTFGGTC | ASSGHLNQDTQY | 0.6003915 |
| DEGKAPAG | ASSGHLNQDTQY | 0.46950498 |
| DTVVFPAL | ASSGHLNQDTQY | 0.55371106 |
| FEGITMAG | ASSGHLNQDTQY | 0.5832261 |
| FPLVVQD | ASSGHLNQDTQY | 0.71194625 |
| GAFTTPQ | ASSGHLNQDTQY | 0.59891057 |
| GFGGFGTAGGVGGA | ASSGHLNQDTQY | 0.44973084 |
| GFLAQFL | ASSGHLNQDTQY | 0.66577375 |
| GGAGGAGGNAGVA | ASSGHLNQDTQY | 0.7069286 |
| GGVGTAGAGGAGGAGG | ASSGHLNQDTQY | 0.70563847 |
| GNGGGGDAGNAGSGGN | ASSGHLNQDTQY | 0.4554408 |
| GTGGAGGAGNDAGS | ASSGHLNQDTQY | 0.57025445 |
| GTTNTGWFNS | ASSGHLNQDTQY | 0.5713562 |
| IFMGPLFTL | ASSGHLNQDTQY | 0.5333612 |

| | | |
|---|---|---|
| ILAHAIEAI | ASSGHLNQDTQY | 0.49283418 |
| IRSAIREAL | ASSGHLNQDTQY | 0.55207103 |
| ITFAGGG | ASSGHLNQDTQY | 0.68246794 |
| ITGMSGHQEV | ASSGHLNQDTQY | 0.73520243 |
| IVAGLACTL | ASSGHLNQDTQY | 0.57828784 |
| IVGGGLAGA | ASSGHLNQDTQY | 0.36472493 |
| IVTGVLNI | ASSGHLNQDTQY | 0.557266 |
| LAQQGCI | ASSGHLNQDTQY | 0.45200688 |
| LCLHVEN | ASSGHLNQDTQY | 0.46942016 |
| LDAISDGS | ASSGHLNQDTQY | 0.5575048 |
| LGGFAAML | ASSGHLNQDTQY | 0.34640104 |
| LGGKNPMI | ASSGHLNQDTQY | 0.581663 |
| LGPKGVAAIL | ASSGHLNQDTQY | 0.7111397 |
| LMVGVSI | ASSGHLNQDTQY | 0.5145152 |
| LVGAAFVL | ASSGHLNQDTQY | 0.60735965 |
| LVLAVAPL | ASSGHLNQDTQY | 0.68658745 |
| MAAIAVIDAA | ASSGHLNQDTQY | 0.6224218 |
| NNGINAELQ | ASSGHLNQDTQY | 0.67201185 |
| RGEPLVLVHG | ASSGHLNQDTQY | 0.64732194 |
| RGGAGGAAGLI | ASSGHLNQDTQY | 0.6586756 |
| RSYRMHPAV | ASSGHLNQDTQY | 0.58503157 |
| SAAHLLAGQLA | ASSGHLNQDTQY | 0.6500318 |
| SAAVGSY | ASSGHLNQDTQY | 0.5424428 |
| SPHAVLHRI | ASSGHLNQDTQY | 0.60866004 |
| SQIAKVL | ASSGHLNQDTQY | 0.5359514 |
| TALGHLVP | ASSGHLNQDTQY | 0.65407026 |
| TAPTTSGAA | ASSGHLNQDTQY | 0.57167834 |
| TDGDDGNGGSPGT | ASSGHLNQDTQY | 0.69526565 |
| TDGLGEN | ASSGHLNQDTQY | 0.32302374 |
| TGGPPLPGQA | ASSGHLNQDTQY | 0.548178 |
| TIPAITI | ASSGHLNQDTQY | 0.6681775 |
| TIPGFTI | ASSGHLNQDTQY | 0.28162733 |
| TSIASGALA | ASSGHLNQDTQY | 0.4314066 |
| TVTLPGAAI | ASSGHLNQDTQY | 0.5712003 |
| VASGLSAALP | ASSGHLNQDTQY | 0.67943245 |
| VGAGVIGI | ASSGHLNQDTQY | 0.28540343 |
| VGPKRFGV | ASSGHLNQDTQY | 0.6183833 |
| VNVAFPD | ASSGHLNQDTQY | 0.41264337 |
| VSMTNSGVS | ASSGHLNQDTQY | 0.7097682 |
| VTDVLGDLSV | ASSGHLNQDTQY | 0.60510695 |
| VVVVGAGFAGL | ASSGHLNQDTQY | 0.41859487 |
| VYTVGTVV | ASSGHLNQDTQY | 0.66701126 |
| WTTFSGP | ASSGHLNQDTQY | 0.42551088 |
| YDELVEAF | ASSGHLNQDTQY | 0.65181303 |
| YGTGGAGGNGG | ASSGHLNQDTQY | 0.6715301 |
| YTNVMAV | ASSGHLNQDTQY | 0.7179843 |
| FLEQVHQL | ASSGHLNQDTQY | 0.4099753 |
| GLSLLAVAV | ASSGHLNQDTQY | 0.65237314 |
| LEQVHQL | ASSGHLNQDTQY | 0.5009917 |
| LNAAGAI | ASSGHLNQDTQY | 0.72332835 |
| PAVHASL | ASSGHLNQDTQY | 0.56551015 |

| | | |
|---|---|---|
| TELEDDA | ASSGHLNQDTQY | 0.3784731 |
| TPPGPPN | ASSGHLNQDTQY | 0.6689159 |
| TPPPPPPP | ASSGHLNQDTQY | 0.63199055 |
| TQSPALI | ASGAMTGGAYEQY | 0.6450983 |
| EFAAAFPQ | ASGAMTGGAYEQY | 0.624876 |
| LLSGNLDVL | ASGAMTGGAYEQY | 0.57140064 |
| GPANLGNYNI | ASGAMTGGAYEQY | 0.5119738 |
| TLGPTLP | ASGAMTGGAYEQY | 0.6513796 |
| VYAYSSCCK | ASGAMTGGAYEQY | 0.6184393 |
| APAPAPAPNI | ASGAMTGGAYEQY | 0.5886396 |
| PGGPVPIP | ASGAMTGGAYEQY | 0.70779604 |
| FLTVALEP | ASGAMTGGAYEQY | 0.611619 |
| SQYFGANL | ASGAMTGGAYEQY | 0.48275712 |
| GGVKPAA | ASGAMTGGAYEQY | 0.7176604 |
| GGTGGAGGAGGW | ASGAMTGGAYEQY | 0.5791551 |
| GVGGVAGLGGAGG | ASGAMTGGAYEQY | 0.64785486 |
| VLLVLATTI | ASGAMTGGAYEQY | 0.66781354 |
| IGIFAAML | ASGAMTGGAYEQY | 0.6929211 |
| AVGDLID | ASGAMTGGAYEQY | 0.55689377 |
| NISPLPL | ASGAMTGGAYEQY | 0.5821596 |
| VTAVFVAI | ASGAMTGGAYEQY | 0.5797094 |
| SGDDNDAAL | ASGAMTGGAYEQY | 0.67875665 |
| AVAALAHS | ASGAMTGGAYEQY | 0.5128262 |
| GGVTVMATP | ASGAMTGGAYEQY | 0.6433619 |
| AGLAANVRD | ASGAMTGGAYEQY | 0.6513322 |
| TNRPDLIDPALLRPG | ASGAMTGGAYEQY | 0.7113708 |
| LANLGVTAL | ASGAMTGGAYEQY | 0.5851811 |
| TGERIEL | ASGAMTGGAYEQY | 0.6013147 |
| NGEELAA | ASGAMTGGAYEQY | 0.7093911 |
| VLSLVWG | ASGAMTGGAYEQY | 0.47272882 |
| IGYALWL | ASGAMTGGAYEQY | 0.6129138 |
| LSLLLAPL | ASGAMTGGAYEQY | 0.6613388 |
| TLDEALK | ASGAMTGGAYEQY | 0.6918372 |
| TDGAGTAEGG | ASGAMTGGAYEQY | 0.5828335 |
| NAPGGAGGNGG | ASGAMTGGAYEQY | 0.6763333 |
| VSVFMGMGGNF | ASGAMTGGAYEQY | 0.5306238 |
| LASYLAAL | ASGAMTGGAYEQY | 0.6318168 |
| LVGGTPAL | ASGAMTGGAYEQY | 0.452754 |
| VRNADVL | ASGAMTGGAYEQY | 0.54264313 |
| TCSSPFLVSAM | ASGAMTGGAYEQY | 0.5015136 |
| VHITDAE | ASGAMTGGAYEQY | 0.5844242 |
| EYAAPSF | ASGAMTGGAYEQY | 0.65468144 |
| VVDAFHGGL | ASGAMTGGAYEQY | 0.6330877 |
| ASVVDPQ | ASGAMTGGAYEQY | 0.4083603 |
| SYNEFGGLS | ASGAMTGGAYEQY | 0.6425948 |
| DPDGSVY | ASGAMTGGAYEQY | 0.6624957 |
| MVSTIHATE | ASGAMTGGAYEQY | 0.70849276 |
| LAAGVADPV | ASGAMTGGAYEQY | 0.55747914 |
| TKEQIAA | ASGAMTGGAYEQY | 0.67932373 |
| SLASPAAL | ASGAMTGGAYEQY | 0.5842645 |
| IGLVFYGV | ASGAMTGGAYEQY | 0.6086535 |

| | | |
|---|---|---|
| FLLDASNL | ASGAMTGGAYEQY | 0.3669128 |
| YQVGNFP | ASGAMTGGAYEQY | 0.44548303 |
| ESLLSQST | ASGAMTGGAYEQY | 0.53432983 |
| LFTVGFAAVG | ASGAMTGGAYEQY | 0.47399414 |
| ATPPTPT | ASGAMTGGAYEQY | 0.64494807 |
| TDLATLG | ASGAMTGGAYEQY | 0.665685 |
| SAGGTGR | ASGAMTGGAYEQY | 0.68036073 |
| GFSNSGS | ASGAMTGGAYEQY | 0.69136447 |
| GGNGGSGGTSV | ASGAMTGGAYEQY | 0.6783475 |
| GTSVATGGAGN | ASGAMTGGAYEQY | 0.69506013 |
| NGGNGGSGGTS | ASGAMTGGAYEQY | 0.55790156 |
| AVVKFPH | ASGAMTGGAYEQY | 0.564962 |
| TPPPPPP | ASGAMTGGAYEQY | 0.38300568 |
| AAPAAAAAPA | ASGAMTGGAYEQY | 0.5060479 |
| IALNEDL | ASGAMTGGAYEQY | 0.5642891 |
| IVNVSSM | ASGAMTGGAYEQY | 0.6077709 |
| TGIPPAPR | ASGAMTGGAYEQY | 0.27595633 |
| FDLGGGTF | ASGAMTGGAYEQY | 0.5625191 |
| FDLGGGTFDVS | ASGAMTGGAYEQY | 0.70862323 |
| NEPTAAA | ASGAMTGGAYEQY | 0.556543 |
| NEPTAAALA | ASGAMTGGAYEQY | 0.6599785 |
| GSIINISSI | ASGAMTGGAYEQY | 0.61864674 |
| VAPGNAGTA | ASGAMTGGAYEQY | 0.4754691 |
| TGGGASLE | ASGAMTGGAYEQY | 0.6587919 |
| VAVYNHYKRI | ASGAMTGGAYEQY | 0.52395666 |
| TDPAPATT | ASGAMTGGAYEQY | 0.66379535 |
| TNRPDILDPALLRPG | ASGAMTGGAYEQY | 0.6992951 |
| LTTTTTS | ASGAMTGGAYEQY | 0.6209785 |
| PTGSGKTL | ASGAMTGGAYEQY | 0.51813185 |
| TGSGKTL | ASGAMTGGAYEQY | 0.629557 |
| TIGHVDHGK | ASGAMTGGAYEQY | 0.5609125 |
| TIGHVDHGKTT | ASGAMTGGAYEQY | 0.559556 |
| GHVDHGKTT | ASGAMTGGAYEQY | 0.56108207 |
| AQFEHTLL | ASGAMTGGAYEQY | 0.56188166 |
| IVNQDEL | ASGAMTGGAYEQY | 0.49720186 |
| TNGSQFF | ASGAMTGGAYEQY | 0.6536758 |
| TGKGIVE | ASGAMTGGAYEQY | 0.61568475 |
| LISNASDA | ASGAMTGGAYEQY | 0.6187143 |
| AELRTLGELTKIAKAHGAQ | ASGAMTGGAYEQY | 0.44386667 |
| APNAVAAAEPSW | ASGAMTGGAYEQY | 0.6890415 |
| APQVAGAATAVL | ASGAMTGGAYEQY | 0.6427741 |
| ARRDPAAAAAILPTDA | ASGAMTGGAYEQY | 0.4793755 |
| DAAARDVLAEAGLAEYF | ASGAMTGGAYEQY | 0.62380266 |
| DDQIVLSAKANLAKVEQ | ASGAMTGGAYEQY | 0.57209694 |
| DELLPDTGEDSDEDGH | ASGAMTGGAYEQY | 0.5226164 |
| ELPEAPANRATVPQD | ASGAMTGGAYEQY | 0.6918488 |
| FGQSKIERVTGIG | ASGAMTGGAYEQY | 0.61600065 |
| GAGGTGSTAGGAGGAGGAGGLY | ASGAMTGGAYEQY | 0.4799648 |
| GHGGTGAAGGNGGTGGAGGSGI | ASGAMTGGAYEQY | 0.48901862 |
| GNGGNGGAGGEGGPGGAGGAGA | ASGAMTGGAYEQY | 0.5919231 |
| GSTAGNGGNGGNATNGTIAGQP | ASGAMTGGAYEQY | 0.56944895 |

| | | |
|---|---|---|
| GTGGDGGNGGNGGTGG | ASGAMTGGAYEQY | 0.32444292 |
| GVRRPAGIPADAPV | ASGAMTGGAYEQY | 0.6424299 |
| GYRQVTDAHLLTLARR | ASGAMTGGAYEQY | 0.48545963 |
| IAKMGGQDGHDRG | ASGAMTGGAYEQY | 0.45389456 |
| KAEPSIEIQYT | ASGAMTGGAYEQY | 0.5507554 |
| KPVRIRMAADEDTVAVT | ASGAMTGGAYEQY | 0.66881466 |
| MPPRIAGMRLLVIKPEP | ASGAMTGGAYEQY | 0.6312264 |
| MKFYAVALFRSRGVPPTR | ASGAMTGGAYEQY | 0.62373656 |
| NAGGFGILGAVAHSP | ASGAMTGGAYEQY | 0.5957226 |
| NEGREITLAMAEM | ASGAMTGGAYEQY | 0.660594 |
| NELVEWSHPLGHRWRWE | ASGAMTGGAYEQY | 0.6677841 |
| NLGAGGTGGNGGNPG | ASGAMTGGAYEQY | 0.6868827 |
| NTEWPKWETVMAADPVD | ASGAMTGGAYEQY | 0.62219304 |
| PASAAPAETQP | ASGAMTGGAYEQY | 0.66382605 |
| QQSSDGKNGVRAAEITGE | ASGAMTGGAYEQY | 0.5599064 |
| RAPDAVTTSWWKE | ASGAMTGGAYEQY | 0.50761116 |
| SGFGNVGAHTSGWFNQ | ASGAMTGGAYEQY | 0.5083381 |
| SGVAGAVGQAASVGGLKV | ASGAMTGGAYEQY | 0.47326022 |
| SNGGAGTGSAGGNGGT | ASGAMTGGAYEQY | 0.5194861 |
| TGGAGGEGGAAG | ASGAMTGGAYEQY | 0.6446446 |
| TGGDGGAAGAGGAPGHG | ASGAMTGGAYEQY | 0.721509 |
| TMPPLLAAIANAGRHGV | ASGAMTGGAYEQY | 0.537917 |
| TSSTAGNSGGA | ASGAMTGGAYEQY | 0.37383342 |
| VPATVPVGILC | ASGAMTGGAYEQY | 0.5342101 |
| VQRPDSGGVVLTGRIS | ASGAMTGGAYEQY | 0.5869006 |
| WTPNPVPATKGVPAGLVQAT | ASGAMTGGAYEQY | 0.6448488 |
| YALKRALAKFSDRFDVVII | ASGAMTGGAYEQY | 0.6756906 |
| YLIGSVIV | ASGAMTGGAYEQY | 0.71340334 |
| YGQLVAVID | ASGAMTGGAYEQY | 0.67614 |
| VTVGGMTLPA | ASGAMTGGAYEQY | 0.5324389 |
| VLALPAV | ASGAMTGGAYEQY | 0.66496366 |
| VIPPTLNYET | ASGAMTGGAYEQY | 0.49373543 |
| VGMGGMHPSSG | ASGAMTGGAYEQY | 0.586865 |
| VGLAGGAATA | ASGAMTGGAYEQY | 0.5898973 |
| VGGAGGAGGSGP | ASGAMTGGAYEQY | 0.5885059 |
| VFAAFMV | ASGAMTGGAYEQY | 0.5766996 |
| TLYDKLR | ASGAMTGGAYEQY | 0.5714139 |
| TLAVFTVL | ASGAMTGGAYEQY | 0.43786612 |
| TGSVQPARV | ASGAMTGGAYEQY | 0.6595142 |
| TGGGAGSGGT | ASGAMTGGAYEQY | 0.48650652 |
| SHLETSL | ASGAMTGGAYEQY | 0.49682835 |
| SFTTGMT | ASGAMTGGAYEQY | 0.51213706 |
| QVEVAKRMGVRQPTVS | ASGAMTGGAYEQY | 0.6903963 |
| QGSGGPALAM | ASGAMTGGAYEQY | 0.71685576 |
| MLAPSNP | ASGAMTGGAYEQY | 0.5044424 |
| LVNNAAI | ASGAMTGGAYEQY | 0.5463171 |
| LTGEINTLA | ASGAMTGGAYEQY | 0.619291 |
| LSIFGPV | ASGAMTGGAYEQY | 0.51149637 |
| LLALLANL | ASGAMTGGAYEQY | 0.65503347 |
| LISNASDALDKLR | ASGAMTGGAYEQY | 0.533094 |
| LINEAVP | ASGAMTGGAYEQY | 0.5362855 |

| | | |
|---|---|---|
| LGYAFSGL | ASGAMTGGAYEQY | 0.50999194 |
| LDFVAVAL | ASGAMTGGAYEQY | 0.6462722 |
| LAQPGAE | ASGAMTGGAYEQY | 0.5686637 |
| ISNLGAVTGF | ASGAMTGGAYEQY | 0.6733967 |
| HQEDPGRI | ASGAMTGGAYEQY | 0.34647557 |
| GTPGEPTP | ASGAMTGGAYEQY | 0.68113494 |
| GFFNSGS | ASGAMTGGAYEQY | 0.45271963 |
| GAGGGGLTAGAN | ASGAMTGGAYEQY | 0.49082428 |
| FVDALVNM | ASGAMTGGAYEQY | 0.30150568 |
| FLTVAPEL | ASGAMTGGAYEQY | 0.61196667 |
| FETLTAL | ASGAMTGGAYEQY | 0.60189044 |
| DVINPIEL | ASGAMTGGAYEQY | 0.6998042 |
| APVSGAPV | ASGAMTGGAYEQY | 0.6255522 |
| ALVGGLAW | ASGAMTGGAYEQY | 0.6182954 |
| ALAAAAATAI | ASGAMTGGAYEQY | 0.6722361 |
| AFGLGLLL | ASGAMTGGAYEQY | 0.5105776 |
| AFFAAASL | ASGAMTGGAYEQY | 0.41961634 |
| AAAALPAFSPP | ASGAMTGGAYEQY | 0.64160997 |
| VVLAPAL | ASGAMTGGAYEQY | 0.49109566 |
| TIGHVDHGKTTL | ASGAMTGGAYEQY | 0.69643503 |
| LISNASDALDK | ASGAMTGGAYEQY | 0.62489474 |
| LISNASDAL | ASGAMTGGAYEQY | 0.35321313 |
| ISNASDALDK | ASGAMTGGAYEQY | 0.6829722 |
| NQAETLVYQTE | ASGAMTGGAYEQY | 0.64197266 |
| TGGTGGTAGAG | ASGAMTGGAYEQY | 0.3941998 |
| TGSTAGGAGGA | ASGAMTGGAYEQY | 0.46926 |
| DPLAVPVAAGPV | ASGAMTGGAYEQY | 0.5460844 |
| GGTGGTPGNGGN | ASGAMTGGAYEQY | 0.4102946 |
| PAGLAGVDPATL | ASGAMTGGAYEQY | 0.6534255 |
| GNNLVVEAPHAGASV | ASGAMTGGAYEQY | 0.51565474 |
| KVTRIMSASTLEQPA | ASGAMTGGAYEQY | 0.62997013 |
| LGFALPIALPRRTVV | ASGAMTGGAYEQY | 0.55975366 |
| TRLGNLTARKLVVLV | ASGAMTGGAYEQY | 0.5973944 |
| WSQQRRLIYFPSAGP | ASGAMTGGAYEQY | 0.6009625 |
| GPNGNPDPMAAAVDIR | ASGAMTGGAYEQY | 0.6083631 |
| TQRCVSVADLRAAASR | ASGAMTGGAYEQY | 0.6803676 |
| QFVLDVAEHPVEVLTETV | ASGAMTGGAYEQY | 0.6587078 |
| GGAGGTGGAGGNGGTQAGDG | ASGAMTGGAYEQY | 0.5249938 |
| PLLATLRAIAVDVDAKPAQV | ASGAMTGGAYEQY | 0.5868056 |
| APKFRTGGYMIDFWGVGYQVAK | ASGAMTGGAYEQY | 0.6903615 |
| GAGGAGGAGGAAGGPGGTGGTG | ASGAMTGGAYEQY | 0.49235475 |
| IGALGLLATDVAPAGSVAFGVA | ASGAMTGGAYEQY | 0.50970304 |
| AGGAGGIGGNGGAGGAGGVLMG | ASGAMTGGAYEQY | 0.65321565 |
| ADELIGLL | ASGAMTGGAYEQY | 0.5725978 |
| ADRALKA | ASGAMTGGAYEQY | 0.65816075 |
| AGAVGQVGGAGGA | ASGAMTGGAYEQY | 0.6402443 |
| AGGGGGLGGAP | ASGAMTGGAYEQY | 0.6266375 |
| AGGLLYGNGGAG | ASGAMTGGAYEQY | 0.5530641 |
| AGLESIGL | ASGAMTGGAYEQY | 0.6795427 |
| AGLKTGII | ASGAMTGGAYEQY | 0.6455644 |
| ASLAALLL | ASGAMTGGAYEQY | 0.55713063 |

| | | |
|---|---|---|
| ASVINISSI | ASGAMTGGAYEQY | 0.60790956 |
| ATPPGPA | ASGAMTGGAYEQY | 0.46357578 |
| ATVASALV | ASGAMTGGAYEQY | 0.6344935 |
| ATVASLIL | ASGAMTGGAYEQY | 0.66802263 |
| AVNGPVLIRH | ASGAMTGGAYEQY | 0.41389945 |
| DGAAGAVGI | ASGAMTGGAYEQY | 0.5166868 |
| DPLLHGI | ASGAMTGGAYEQY | 0.70245016 |
| ELGAALGAL | ASGAMTGGAYEQY | 0.66310906 |
| ELGRTQAV | ASGAMTGGAYEQY | 0.5889256 |
| FVAGGYLS | ASGAMTGGAYEQY | 0.48690227 |
| GASGGAGGAGGW | ASGAMTGGAYEQY | 0.53712803 |
| GGAGAGGDGGAGGI | ASGAMTGGAYEQY | 0.51798874 |
| GTGHNGSQPGMGGNG | ASGAMTGGAYEQY | 0.43759102 |
| GYAAASASAS | ASGAMTGGAYEQY | 0.6443588 |
| HQVDVAAVQQG | ASGAMTGGAYEQY | 0.54940176 |
| IGAAIAHT | ASGAMTGGAYEQY | 0.6430777 |
| IGMIEVAQL | ASGAMTGGAYEQY | 0.6443288 |
| IRLAPEHTSAMD | ASGAMTGGAYEQY | 0.7112017 |
| IYNVSVL | ASGAMTGGAYEQY | 0.51970047 |
| KQGFLGM | ASGAMTGGAYEQY | 0.60928166 |
| LESVITQL | ASGAMTGGAYEQY | 0.3621109 |
| LGLGALVVV | ASGAMTGGAYEQY | 0.58125836 |
| LIGNGGNGGSGGI | ASGAMTGGAYEQY | 0.48732352 |
| LNAINAPTL | ASGAMTGGAYEQY | 0.6271286 |
| LQLLNTGV | ASGAMTGGAYEQY | 0.45020673 |
| LQTSIEF | ASGAMTGGAYEQY | 0.71213466 |
| MSFAVSEL | ASGAMTGGAYEQY | 0.50179154 |
| NTGALGL | ASGAMTGGAYEQY | 0.675354 |
| PPGAGGAGGAGTT | ASGAMTGGAYEQY | 0.5683992 |
| QLVGLGI | ASGAMTGGAYEQY | 0.45275444 |
| RFLEQVHQL | ASGAMTGGAYEQY | 0.5465079 |
| RGVGYVLR | ASGAMTGGAYEQY | 0.6486627 |
| SATGVFAGVI | ASGAMTGGAYEQY | 0.4372144 |
| SLPAITLP | ASGAMTGGAYEQY | 0.35166708 |
| STALAFTV | ASGAMTGGAYEQY | 0.5022577 |
| SVLTVGLGK | ASGAMTGGAYEQY | 0.5630362 |
| TALAGAGSGI | ASGAMTGGAYEQY | 0.69495237 |
| TGLPFTGI | ASGAMTGGAYEQY | 0.6862811 |
| VAEHVTV | ASGAMTGGAYEQY | 0.5734441 |
| VDIVLNSL | ASGAMTGGAYEQY | 0.70404375 |
| VLTGGGAVL | ASGAMTGGAYEQY | 0.26161876 |
| VNVGLAYGI | ASGAMTGGAYEQY | 0.59648633 |
| VSSAAVGAV | ASGAMTGGAYEQY | 0.71064997 |
| VTSVYEAGV | ASGAMTGGAYEQY | 0.62342596 |
| WVALDRGA | ASGAMTGGAYEQY | 0.5818311 |
| YLVGSVVI | ASGAMTGGAYEQY | 0.6814626 |
| GPPGTGKTLL | ASGAMTGGAYEQY | 0.5128281 |
| GVGMTKF | ASGAMTGGAYEQY | 0.57427686 |
| IGIGGSDL | ASGAMTGGAYEQY | 0.64075166 |
| IGTLLAEL | ASGAMTGGAYEQY | 0.677414 |
| IYTSGTT | ASGAMTGGAYEQY | 0.59597695 |

| | | |
|---|---|---|
| LSFPAGV | ASGAMTGGAYEQY | 0.39841416 |
| LYTSGTT | ASGAMTGGAYEQY | 0.65846145 |
| VGAFPFGF | ASGAMTGGAYEQY | 0.71752596 |
| VGAGTVEF | ASGAMTGGAYEQY | 0.50996405 |
| VGKSTLLL | ASGAMTGGAYEQY | 0.6797892 |
| AALAHAGGGIV | ASGAMTGGAYEQY | 0.60052675 |
| AALVAVSLL | ASGAMTGGAYEQY | 0.5757016 |
| AAVSRAVAAL | ASGAMTGGAYEQY | 0.48735017 |
| AGGSSGIGGF | ASGAMTGGAYEQY | 0.4456062 |
| AGLTVANANG | ASGAMTGGAYEQY | 0.6870517 |
| AGLVGAVL | ASGAMTGGAYEQY | 0.42202598 |
| ALSQAFTV | ASGAMTGGAYEQY | 0.5879962 |
| ALTFSVSL | ASGAMTGGAYEQY | 0.581646 |
| ALTGAALVAL | ASGAMTGGAYEQY | 0.7327109 |
| ASFSIGAGVL | ASGAMTGGAYEQY | 0.60615486 |
| DGSVYKI | ASGAMTGGAYEQY | 0.6492313 |
| EQLAEIV | ASGAMTGGAYEQY | 0.51964885 |
| FAVSELTVM | ASGAMTGGAYEQY | 0.5958919 |
| GAAPLAAGL | ASGAMTGGAYEQY | 0.6169687 |
| GADGGDGGSGGAGG | ASGAMTGGAYEQY | 0.4583413 |
| GFGLDTGGAGG | ASGAMTGGAYEQY | 0.6877849 |
| GLASGGLL | ASGAMTGGAYEQY | 0.46717924 |
| GRIAGAAVLL | ASGAMTGGAYEQY | 0.5632916 |
| GSAGSLNTSF | ASGAMTGGAYEQY | 0.6591243 |
| GTGQIGI | ASGAMTGGAYEQY | 0.68175995 |
| GTLIALIL | ASGAMTGGAYEQY | 0.5566503 |
| IAATMGFVVG | ASGAMTGGAYEQY | 0.6969842 |
| IFAQALL | ASGAMTGGAYEQY | 0.519713 |
| IGAAVALGF | ASGAMTGGAYEQY | 0.6620555 |
| IGLTPLAL | ASGAMTGGAYEQY | 0.69239366 |
| ILGSNTSTL | ASGAMTGGAYEQY | 0.53480244 |
| IVFSLTVL | ASGAMTGGAYEQY | 0.7207304 |
| LASKATGF | ASGAMTGGAYEQY | 0.67288256 |
| LASVALIL | ASGAMTGGAYEQY | 0.39975697 |
| LFAVGLTL | ASGAMTGGAYEQY | 0.60962725 |
| LFNGGIGTYI | ASGAMTGGAYEQY | 0.6750373 |
| LTLGAGII | ASGAMTGGAYEQY | 0.6822097 |
| LVAAALGDI | ASGAMTGGAYEQY | 0.7073282 |
| NTAFVQLGI | ASGAMTGGAYEQY | 0.61368525 |
| QLFSGAEL | ASGAMTGGAYEQY | 0.49454814 |
| SAATFITL | ASGAMTGGAYEQY | 0.699638 |
| SNYGLANLGV | ASGAMTGGAYEQY | 0.750368 |
| SYGGAYAVM | ASGAMTGGAYEQY | 0.6498079 |
| TTGVGGLTLG | ASGAMTGGAYEQY | 0.47803855 |
| TTQTIYI | ASGAMTGGAYEQY | 0.5293549 |
| VAAGAGLGAVI | ASGAMTGGAYEQY | 0.6426778 |
| VAAVVTLL | ASGAMTGGAYEQY | 0.6632925 |
| VGFLGGLI | ASGAMTGGAYEQY | 0.4273839 |
| VGVIPILL | ASGAMTGGAYEQY | 0.56398636 |
| VIFASSLI | ASGAMTGGAYEQY | 0.63345087 |
| VKGGDGGAAAG | ASGAMTGGAYEQY | 0.3345054 |

| | | |
|---|---|---|
| VTKTFTNI | ASGAMTGGAYEQY | 0.5866065 |
| VVAAAILL | ASGAMTGGAYEQY | 0.6555541 |
| YAALDVEL | ASGAMTGGAYEQY | 0.6294531 |
| LASVALLL | ASGAMTGGAYEQY | 0.4063886 |
| LDAVHRV | ASGAMTGGAYEQY | 0.68198735 |
| QGLSSSIF | ASGAMTGGAYEQY | 0.6853228 |
| RGTFANIRL | ASGAMTGGAYEQY | 0.7031339 |
| RQLWWGHRI | ASGAMTGGAYEQY | 0.4524919 |
| YGPPGTGKTLL | ASGAMTGGAYEQY | 0.26180094 |
| AALVGAPANVA | ASGAMTGGAYEQY | 0.6246306 |
| AATFRRTE | ASGAMTGGAYEQY | 0.63165694 |
| AGALGGGI | ASGAMTGGAYEQY | 0.60942286 |
| AGGAGGAGGIGAG | ASGAMTGGAYEQY | 0.5016439 |
| AGSGALGL | ASGAMTGGAYEQY | 0.53442544 |
| AQIADQI | ASGAMTGGAYEQY | 0.68305165 |
| ASAVAVTYL | ASGAMTGGAYEQY | 0.46487248 |
| ASTVLGGTGG | ASGAMTGGAYEQY | 0.49417457 |
| AVGGEPAK | ASGAMTGGAYEQY | 0.6259413 |
| CEQGTFGGTC | ASGAMTGGAYEQY | 0.6811604 |
| DEGKAPAG | ASGAMTGGAYEQY | 0.4088416 |
| DTVVFPAL | ASGAMTGGAYEQY | 0.4975702 |
| FEGITMAG | ASGAMTGGAYEQY | 0.65360343 |
| FPLVVQD | ASGAMTGGAYEQY | 0.66017807 |
| GAFTTPQ | ASGAMTGGAYEQY | 0.5957394 |
| GFGGFGTAGGVGGA | ASGAMTGGAYEQY | 0.41477543 |
| GFLAQFL | ASGAMTGGAYEQY | 0.6498693 |
| GGAGGAGGNAGVA | ASGAMTGGAYEQY | 0.40812206 |
| GGVGTAGAGGAGGAGG | ASGAMTGGAYEQY | 0.6401637 |
| GNGGGGDAGNAGSGGN | ASGAMTGGAYEQY | 0.61337477 |
| GTGGAGGAGNDAGS | ASGAMTGGAYEQY | 0.6346518 |
| GTTNTGWFNS | ASGAMTGGAYEQY | 0.53224385 |
| IFMGPLFTL | ASGAMTGGAYEQY | 0.41409078 |
| ILAHAIEAI | ASGAMTGGAYEQY | 0.47128248 |
| IRSAIREAL | ASGAMTGGAYEQY | 0.46217668 |
| ITFAGGG | ASGAMTGGAYEQY | 0.6706254 |
| ITGMSGHQEV | ASGAMTGGAYEQY | 0.6409509 |
| IVAGLACTL | ASGAMTGGAYEQY | 0.6531657 |
| IVGGGLAGA | ASGAMTGGAYEQY | 0.63384515 |
| IVTGVLNI | ASGAMTGGAYEQY | 0.52507824 |
| LAQQGCI | ASGAMTGGAYEQY | 0.6406646 |
| LCLHVEN | ASGAMTGGAYEQY | 0.46334815 |
| LDAISDGS | ASGAMTGGAYEQY | 0.6321116 |
| LGGFAAML | ASGAMTGGAYEQY | 0.47600442 |
| LGGKNPMI | ASGAMTGGAYEQY | 0.64363265 |
| LGPKGVAAIL | ASGAMTGGAYEQY | 0.5273951 |
| LMVGVSI | ASGAMTGGAYEQY | 0.6544515 |
| LVGAAFVL | ASGAMTGGAYEQY | 0.63006103 |
| LVLAVAPL | ASGAMTGGAYEQY | 0.4907275 |
| MAAIAVIDAA | ASGAMTGGAYEQY | 0.6285417 |
| NNGINAELQ | ASGAMTGGAYEQY | 0.6809975 |
| RGEPLVLVHG | ASGAMTGGAYEQY | 0.6518562 |

| | | |
|---|---|---|
| RGGAGGAAGLI | ASGAMTGGAYEQY | 0.35653687 |
| RSYRMHPAV | ASGAMTGGAYEQY | 0.63797814 |
| SAAHLLAGQLA | ASGAMTGGAYEQY | 0.44419777 |
| SAAVGSY | ASGAMTGGAYEQY | 0.602456 |
| SPHAVLHRI | ASGAMTGGAYEQY | 0.60990644 |
| SQIAKVL | ASGAMTGGAYEQY | 0.59425706 |
| TALGHLVP | ASGAMTGGAYEQY | 0.65507674 |
| TAPTTSGAA | ASGAMTGGAYEQY | 0.4540337 |
| TDGDDGNGGSPGT | ASGAMTGGAYEQY | 0.4784611 |
| TDGLGEN | ASGAMTGGAYEQY | 0.5095819 |
| TGGPPLPGQA | ASGAMTGGAYEQY | 0.6481105 |
| TIPAITI | ASGAMTGGAYEQY | 0.6565247 |
| TIPGFTI | ASGAMTGGAYEQY | 0.4167591 |
| TSIASGALA | ASGAMTGGAYEQY | 0.61552846 |
| TVTLPGAAI | ASGAMTGGAYEQY | 0.5764495 |
| VASGLSAALP | ASGAMTGGAYEQY | 0.50164926 |
| VGAGVIGI | ASGAMTGGAYEQY | 0.61230665 |
| VGPKRFGV | ASGAMTGGAYEQY | 0.5272592 |
| VNVAFPD | ASGAMTGGAYEQY | 0.40538275 |
| VSMTNSGVS | ASGAMTGGAYEQY | 0.65662426 |
| VTDVLGDLSV | ASGAMTGGAYEQY | 0.55973303 |
| VVVVGAGFAGL | ASGAMTGGAYEQY | 0.6477958 |
| VYTVGTVV | ASGAMTGGAYEQY | 0.6389305 |
| WTTFSGP | ASGAMTGGAYEQY | 0.37553298 |
| YDELVEAF | ASGAMTGGAYEQY | 0.6321862 |
| YGTGGAGGNGG | ASGAMTGGAYEQY | 0.5684947 |
| YTNVMAV | ASGAMTGGAYEQY | 0.56915903 |
| FLEQVHQL | ASGAMTGGAYEQY | 0.63942707 |
| GLSLLAVAV | ASGAMTGGAYEQY | 0.4739974 |
| LEQVHQL | ASGAMTGGAYEQY | 0.67853725 |
| LNAAGAI | ASGAMTGGAYEQY | 0.6771833 |
| PAVHASL | ASGAMTGGAYEQY | 0.633238 |
| TELEDDA | ASGAMTGGAYEQY | 0.62838775 |
| TPPGPPN | ASGAMTGGAYEQY | 0.6259995 |
| TPPPPPPP | ASGAMTGGAYEQY | 0.6751914 |
| TQSPALI | ASSADWGERTGQLY | 0.4753338 |
| EFAAAFPQ | ASSADWGERTGQLY | 0.3549465 |
| LLSGNLDVL | ASSADWGERTGQLY | 0.6265171 |
| GPANLGNYNI | ASSADWGERTGQLY | 0.5682523 |
| TLGPTLP | ASSADWGERTGQLY | 0.5021451 |
| VYAYSSCCK | ASSADWGERTGQLY | 0.43451363 |
| APAPAPAPNI | ASSADWGERTGQLY | 0.6661105 |
| PGGPVPIP | ASSADWGERTGQLY | 0.7272898 |
| FLTVALEP | ASSADWGERTGQLY | 0.5975922 |
| SQYFGANL | ASSADWGERTGQLY | 0.7279573 |
| GGVKPAA | ASSADWGERTGQLY | 0.38894346 |
| GGTGGAGGAGGW | ASSADWGERTGQLY | 0.67338324 |
| GVGGVAGLGGAGG | ASSADWGERTGQLY | 0.6773327 |
| VLLVLATTI | ASSADWGERTGQLY | 0.44781983 |
| IGIFAAML | ASSADWGERTGQLY | 0.6679759 |
| AVGDLID | ASSADWGERTGQLY | 0.5027509 |

| | | |
|---|---|---|
| NISPLPL | ASSADWGERTGQLY | 0.49877048 |
| VTAVFVAI | ASSADWGERTGQLY | 0.6705691 |
| SGDDNDAAL | ASSADWGERTGQLY | 0.6217679 |
| AVAALAHS | ASSADWGERTGQLY | 0.656498 |
| GGVTVMATP | ASSADWGERTGQLY | 0.5976439 |
| AGLAANVRD | ASSADWGERTGQLY | 0.5460342 |
| TNRPDLIDPALLRPG | ASSADWGERTGQLY | 0.6455647 |
| LANLGVTAL | ASSADWGERTGQLY | 0.67142415 |
| TGERIEL | ASSADWGERTGQLY | 0.6240524 |
| NGEELAA | ASSADWGERTGQLY | 0.74718326 |
| VLSLVWG | ASSADWGERTGQLY | 0.46661314 |
| IGYALWL | ASSADWGERTGQLY | 0.7151955 |
| LSLLLAPL | ASSADWGERTGQLY | 0.67898524 |
| TLDEALK | ASSADWGERTGQLY | 0.5929638 |
| TDGAGTAEGG | ASSADWGERTGQLY | 0.5743208 |
| NAPGGAGGNGG | ASSADWGERTGQLY | 0.6731912 |
| VSVFMGMGGNF | ASSADWGERTGQLY | 0.56914794 |
| LASYLAAL | ASSADWGERTGQLY | 0.60617447 |
| LVGGTPAL | ASSADWGERTGQLY | 0.71176726 |
| VRNADVL | ASSADWGERTGQLY | 0.2279518 |
| TCSSPFLVSAM | ASSADWGERTGQLY | 0.70894015 |
| VHITDAE | ASSADWGERTGQLY | 0.56672317 |
| EYAAPSF | ASSADWGERTGQLY | 0.4992072 |
| VVDAFHGGL | ASSADWGERTGQLY | 0.411587 |
| ASVVDPQ | ASSADWGERTGQLY | 0.24905458 |
| SYNEFGGLS | ASSADWGERTGQLY | 0.3434843 |
| DPDGSVY | ASSADWGERTGQLY | 0.41563642 |
| MVSTIHATE | ASSADWGERTGQLY | 0.8440767 |
| LAAGVADPV | ASSADWGERTGQLY | 0.6046331 |
| TKEQIAA | ASSADWGERTGQLY | 0.38934594 |
| SLASPAAL | ASSADWGERTGQLY | 0.77698076 |
| IGLVFYGV | ASSADWGERTGQLY | 0.73716664 |
| FLLDASNL | ASSADWGERTGQLY | 0.19698185 |
| YQVGNFP | ASSADWGERTGQLY | 0.5373886 |
| ESLLSQST | ASSADWGERTGQLY | 0.2225927 |
| LFTVGFAAVG | ASSADWGERTGQLY | 0.3754372 |
| ATPPTPT | ASSADWGERTGQLY | 0.10964188 |
| TDLATLG | ASSADWGERTGQLY | 0.5624342 |
| SAGGTGR | ASSADWGERTGQLY | 0.64506984 |
| GFSNSGS | ASSADWGERTGQLY | 0.14591312 |
| GGNGGSGGTSV | ASSADWGERTGQLY | 0.6096891 |
| GTSVATGGAGN | ASSADWGERTGQLY | 0.5455088 |
| NGGNGGSGGTS | ASSADWGERTGQLY | 0.59299725 |
| AVVKFPH | ASSADWGERTGQLY | 0.63622105 |
| TPPPPPP | ASSADWGERTGQLY | 0.3197454 |
| AAPAAAAAPA | ASSADWGERTGQLY | 0.43979993 |
| IALNEDL | ASSADWGERTGQLY | 0.6759076 |
| IVNVSSM | ASSADWGERTGQLY | 0.76107085 |
| TGIPPAPR | ASSADWGERTGQLY | 0.3439172 |
| FDLGGGTF | ASSADWGERTGQLY | 0.4459353 |
| FDLGGGTFDVS | ASSADWGERTGQLY | 0.74776065 |

| | | |
|---|---|---|
| NEPTAAA | ASSADWGERTGQLY | 0.50269467 |
| NEPTAAALA | ASSADWGERTGQLY | 0.68679607 |
| GSIINISSI | ASSADWGERTGQLY | 0.7425182 |
| VAPGNAGTA | ASSADWGERTGQLY | 0.22982079 |
| TGGGASLE | ASSADWGERTGQLY | 0.77202314 |
| VAVYNHYKRI | ASSADWGERTGQLY | 0.686456 |
| TDPAPATT | ASSADWGERTGQLY | 0.4710928 |
| TNRPDILDPALLRPG | ASSADWGERTGQLY | 0.46917367 |
| LTTTTTS | ASSADWGERTGQLY | 0.20453338 |
| PTGSGKTL | ASSADWGERTGQLY | 0.4487626 |
| TGSGKTL | ASSADWGERTGQLY | 0.2464143 |
| TIGHVDHGK | ASSADWGERTGQLY | 0.68480027 |
| TIGHVDHGKTT | ASSADWGERTGQLY | 0.7387444 |
| GHVDHGKTT | ASSADWGERTGQLY | 0.5676155 |
| AQFEHTLL | ASSADWGERTGQLY | 0.1642592 |
| IVNQDEL | ASSADWGERTGQLY | 0.2113697 |
| TNGSQFF | ASSADWGERTGQLY | 0.26864198 |
| TGKGIVE | ASSADWGERTGQLY | 0.20316193 |
| LISNASDA | ASSADWGERTGQLY | 0.20310274 |
| AELRTLGELTKIAKAHGAQ | ASSADWGERTGQLY | 0.23471394 |
| APNAVAAAEPSW | ASSADWGERTGQLY | 0.7755045 |
| APQVAGAATAVL | ASSADWGERTGQLY | 0.7629037 |
| ARRDPAAAAAILPTDA | ASSADWGERTGQLY | 0.55987436 |
| DAAARDVLAEAGLAEYF | ASSADWGERTGQLY | 0.68328285 |
| DDQIVLSAKANLAKVEQ | ASSADWGERTGQLY | 0.8146407 |
| DELLPDTGEDSDEDGH | ASSADWGERTGQLY | 0.4137895 |
| ELPEAPANRATVPQD | ASSADWGERTGQLY | 0.42402133 |
| FGQSKIERVTGIG | ASSADWGERTGQLY | 0.20606449 |
| GAGGTGSTAGGAGGAGGAGGLY | ASSADWGERTGQLY | 0.85369956 |
| GHGGTGAAGGNGGTGGAGGSGI | ASSADWGERTGQLY | 0.5439491 |
| GNGGNGGAGGEGGPGGAGGAGA | ASSADWGERTGQLY | 0.734144 |
| GSTAGNGGNGGNATNGTIAGQP | ASSADWGERTGQLY | 0.69182575 |
| GTGGDGGNGGNGGTGG | ASSADWGERTGQLY | 0.6048444 |
| GVRRPAGIPADAPV | ASSADWGERTGQLY | 0.33849904 |
| GYRQVTDAHLLTLARR | ASSADWGERTGQLY | 0.72721845 |
| IAKMGQDGHDRG | ASSADWGERTGQLY | 0.5320629 |
| KAEPSIEIQYT | ASSADWGERTGQLY | 0.32557565 |
| KPVRIRMAADEDTVAVT | ASSADWGERTGQLY | 0.18956251 |
| MPPRIAGMRLLVIKPEP | ASSADWGERTGQLY | 0.6878797 |
| MKFYAVALFRSRGVPPTR | ASSADWGERTGQLY | 0.59232277 |
| NAGGFGILGAVAHSP | ASSADWGERTGQLY | 0.7891365 |
| NEGREITLAMAEM | ASSADWGERTGQLY | 0.4402992 |
| NELVEWSHPLGHRWRWE | ASSADWGERTGQLY | 0.24275655 |
| NLGAGGTGGNGGNPG | ASSADWGERTGQLY | 0.6974786 |
| NTEWPKWETVMAADPVD | ASSADWGERTGQLY | 0.5622712 |
| PASAAPAETQP | ASSADWGERTGQLY | 0.47585368 |
| QQSSDGKNGVRAAEITGE | ASSADWGERTGQLY | 0.66726 |
| RAPDAVTTSWWKE | ASSADWGERTGQLY | 0.53548074 |
| SGFGNVGAHTSGWFNQ | ASSADWGERTGQLY | 0.5485651 |
| SGVAGAVGQAASVGGLKV | ASSADWGERTGQLY | 0.66836333 |
| SNGGAGTGSAGGNGGT | ASSADWGERTGQLY | 0.50121135 |

| | | |
|---|---|---|
| TGGAGGEGGAAG | ASSADWGERTGQLY | 0.68333805 |
| TGGDGGAAGAGGAPGHG | ASSADWGERTGQLY | 0.42905238 |
| TMPPLLAAIANAGRHGV | ASSADWGERTGQLY | 0.394565 |
| TSSTAGNSGGA | ASSADWGERTGQLY | 0.3056693 |
| VPATVPVGILC | ASSADWGERTGQLY | 0.34953266 |
| VQRPDSGGVVLTGRIS | ASSADWGERTGQLY | 0.6006936 |
| WTPNPVPATKGVPAGLVQAT | ASSADWGERTGQLY | 0.57352036 |
| YALKRALAKFSDRFDVVII | ASSADWGERTGQLY | 0.7354629 |
| YLIGSVIV | ASSADWGERTGQLY | 0.45981452 |
| YGQLVAVID | ASSADWGERTGQLY | 0.31330025 |
| VTVGGMTLPA | ASSADWGERTGQLY | 0.74798983 |
| VLALPAV | ASSADWGERTGQLY | 0.69392145 |
| VIPPTLNYET | ASSADWGERTGQLY | 0.21395782 |
| VGMGGMHPSSG | ASSADWGERTGQLY | 0.5100252 |
| VGLAGGAATA | ASSADWGERTGQLY | 0.769534 |
| VGGAGGAGGSGP | ASSADWGERTGQLY | 0.7678695 |
| VFAAFMV | ASSADWGERTGQLY | 0.53966784 |
| TLYDKLR | ASSADWGERTGQLY | 0.39054203 |
| TLAVFTVL | ASSADWGERTGQLY | 0.86458737 |
| TGSVQPARV | ASSADWGERTGQLY | 0.62824893 |
| TGGGAGSGGT | ASSADWGERTGQLY | 0.79209244 |
| SHLETSL | ASSADWGERTGQLY | 0.19489071 |
| SFTTGMT | ASSADWGERTGQLY | 0.2985389 |
| QVEVAKRMGVRQPTVS | ASSADWGERTGQLY | 0.23156995 |
| QGSGGPALAM | ASSADWGERTGQLY | 0.104426354 |
| MLAPSNP | ASSADWGERTGQLY | 0.759869 |
| LVNNAAI | ASSADWGERTGQLY | 0.2795713 |
| LTGEINTLA | ASSADWGERTGQLY | 0.6015292 |
| LSIFGPV | ASSADWGERTGQLY | 0.48404127 |
| LLALLANL | ASSADWGERTGQLY | 0.7268096 |
| LISNASDALDKLR | ASSADWGERTGQLY | 0.3629759 |
| LINEAVP | ASSADWGERTGQLY | 0.39753073 |
| LGYAFSGL | ASSADWGERTGQLY | 0.24110328 |
| LDFVAVAL | ASSADWGERTGQLY | 0.7116968 |
| LAQPGAE | ASSADWGERTGQLY | 0.7514318 |
| ISNLGAVTGF | ASSADWGERTGQLY | 0.4334072 |
| HQEDPGRI | ASSADWGERTGQLY | 0.19747251 |
| GTPGEPTP | ASSADWGERTGQLY | 0.7245103 |
| GFFNSGS | ASSADWGERTGQLY | 0.5618793 |
| GAGGGGLTAGAN | ASSADWGERTGQLY | 0.17177276 |
| FVDALVNM | ASSADWGERTGQLY | 0.18672945 |
| FLTVAPEL | ASSADWGERTGQLY | 0.22523001 |
| FETLTAL | ASSADWGERTGQLY | 0.25446963 |
| DVINPIEL | ASSADWGERTGQLY | 0.7634583 |
| APVSGAPV | ASSADWGERTGQLY | 0.75310564 |
| ALVGGLAW | ASSADWGERTGQLY | 0.4517366 |
| ALAAAAATAI | ASSADWGERTGQLY | 0.5844813 |
| AFGLGLLL | ASSADWGERTGQLY | 0.63643503 |
| AFFAAASL | ASSADWGERTGQLY | 0.081539266 |
| AAAALPAFSPP | ASSADWGERTGQLY | 0.31093866 |
| VVLAPAL | ASSADWGERTGQLY | 0.21826756 |

| | | |
|---|---|---|
| TIGHVDHGKTTL | ASSADWGERTGQLY | 0.70861936 |
| LISNASDALDK | ASSADWGERTGQLY | 0.3150363 |
| LISNASDAL | ASSADWGERTGQLY | 0.5109538 |
| ISNASDALDK | ASSADWGERTGQLY | 0.59148407 |
| NQAETLVYQTE | ASSADWGERTGQLY | 0.7928483 |
| TGGTGGTAGAG | ASSADWGERTGQLY | 0.62184274 |
| TGSTAGGAGGA | ASSADWGERTGQLY | 0.20829438 |
| DPLAVPVAAGPV | ASSADWGERTGQLY | 0.16374269 |
| GGTGGTPGNGGN | ASSADWGERTGQLY | 0.6671939 |
| PAGLAGVDPATL | ASSADWGERTGQLY | 0.37464327 |
| GNNLVVEAPHAGASV | ASSADWGERTGQLY | 0.3714615 |
| KVTRIMSASTLEQPA | ASSADWGERTGQLY | 0.29181647 |
| LGFALPIALPRRTVV | ASSADWGERTGQLY | 0.69421685 |
| TRLGNLTARKLVVLV | ASSADWGERTGQLY | 0.1810277 |
| WSQQRRLIYFPSAGP | ASSADWGERTGQLY | 0.54025674 |
| GPNGNPDPMAAAVDIR | ASSADWGERTGQLY | 0.2363675 |
| TQRCVSVADLRAAASR | ASSADWGERTGQLY | 0.62314504 |
| QFVLDVAEHPVEVLTETV | ASSADWGERTGQLY | 0.7683636 |
| GGAGGTGGAGGNGGTQAGDG | ASSADWGERTGQLY | 0.8091478 |
| PLLATLRAIAVDVDAKPAQV | ASSADWGERTGQLY | 0.43799978 |
| APKFRTGGYMIDFWGVGYQVAK | ASSADWGERTGQLY | 0.46891862 |
| GAGGAGGAGGAAGGPGGTGGTG | ASSADWGERTGQLY | 0.27711946 |
| IGALGLLATDVAPAGSVAFGVA | ASSADWGERTGQLY | 0.3552812 |
| AGGAGGIGGNGGAGGAGGVLMG | ASSADWGERTGQLY | 0.4455508 |
| ADELIGLL | ASSADWGERTGQLY | 0.7863858 |
| ADRALKA | ASSADWGERTGQLY | 0.3237203 |
| AGAVGQVGGAGGA | ASSADWGERTGQLY | 0.20324129 |
| AGGGGGLGGAP | ASSADWGERTGQLY | 0.34837973 |
| AGGLLYGNGGAG | ASSADWGERTGQLY | 0.8051601 |
| AGLESIGL | ASSADWGERTGQLY | 0.69583786 |
| AGLKTGII | ASSADWGERTGQLY | 0.25750762 |
| ASLAALLL | ASSADWGERTGQLY | 0.28795683 |
| ASVINISSI | ASSADWGERTGQLY | 0.32229835 |
| ATPPGPA | ASSADWGERTGQLY | 0.5920615 |
| ATVASALV | ASSADWGERTGQLY | 0.22632697 |
| ATVASLIL | ASSADWGERTGQLY | 0.359644 |
| AVNGPVLIRH | ASSADWGERTGQLY | 0.46091524 |
| DGAAGAVGI | ASSADWGERTGQLY | 0.88261425 |
| DPLLHGI | ASSADWGERTGQLY | 0.3718094 |
| ELGAALGAL | ASSADWGERTGQLY | 0.11569489 |
| ELGRTQAV | ASSADWGERTGQLY | 0.663726 |
| FVAGGYLS | ASSADWGERTGQLY | 0.2859047 |
| GASGGAGGAGGW | ASSADWGERTGQLY | 0.35344404 |
| GGAGAGGDGGAGGI | ASSADWGERTGQLY | 0.3367623 |
| GTGHNGSQPGMGGNG | ASSADWGERTGQLY | 0.16263518 |
| GYAAASASAS | ASSADWGERTGQLY | 0.5457287 |
| HQVDVAAVQQG | ASSADWGERTGQLY | 0.7098416 |
| IGAAIAHT | ASSADWGERTGQLY | 0.7309313 |
| IGMIEVAQL | ASSADWGERTGQLY | 0.3892559 |
| IRLAPEHTSAMD | ASSADWGERTGQLY | 0.7856143 |
| IYNVSVL | ASSADWGERTGQLY | 0.60344726 |

| | | |
|---|---|---|
| KQGFLGM | ASSADWGERTGQLY | 0.74400616 |
| LESVITQL | ASSADWGERTGQLY | 0.8348582 |
| LGLGALVVV | ASSADWGERTGQLY | 0.41512567 |
| LIGNGGNGGSGGI | ASSADWGERTGQLY | 0.38448435 |
| LNAINAPTL | ASSADWGERTGQLY | 0.7229733 |
| LQLLNTGV | ASSADWGERTGQLY | 0.63517606 |
| LQTSIEF | ASSADWGERTGQLY | 0.6403692 |
| MSFAVSEL | ASSADWGERTGQLY | 0.3476827 |
| NTGALGL | ASSADWGERTGQLY | 0.8053194 |
| PPGAGGAGGAGTT | ASSADWGERTGQLY | 0.40095425 |
| QLVGLGI | ASSADWGERTGQLY | 0.84638745 |
| RFLEQVHQL | ASSADWGERTGQLY | 0.14632586 |
| RGVGYVLR | ASSADWGERTGQLY | 0.32294047 |
| SATGVFAGVI | ASSADWGERTGQLY | 0.6977212 |
| SLPAITLP | ASSADWGERTGQLY | 0.09979453 |
| STALAFTV | ASSADWGERTGQLY | 0.19957341 |
| SVLTVGLGK | ASSADWGERTGQLY | 0.3766531 |
| TALAGAGSGI | ASSADWGERTGQLY | 0.5162978 |
| TGLPFTGI | ASSADWGERTGQLY | 0.09682832 |
| VAEHVTV | ASSADWGERTGQLY | 0.5461632 |
| VDIVLNSL | ASSADWGERTGQLY | 0.18089223 |
| VLTGGGAVL | ASSADWGERTGQLY | 0.32079273 |
| VNVGLAYGI | ASSADWGERTGQLY | 0.54179114 |
| VSSAAVGAV | ASSADWGERTGQLY | 0.20435801 |
| VTSVYEAGV | ASSADWGERTGQLY | 0.6259788 |
| WVALDRGA | ASSADWGERTGQLY | 0.1290345 |
| YLVGSVVI | ASSADWGERTGQLY | 0.5627049 |
| GPPGTGKTLL | ASSADWGERTGQLY | 0.6674943 |
| GVGMTKF | ASSADWGERTGQLY | 0.86907405 |
| IGIGGSDL | ASSADWGERTGQLY | 0.23799123 |
| IGTLLAEL | ASSADWGERTGQLY | 0.6391343 |
| IYTSGTT | ASSADWGERTGQLY | 0.24438092 |
| LSFPAGV | ASSADWGERTGQLY | 0.83757114 |
| LYTSGTT | ASSADWGERTGQLY | 0.54642344 |
| VGAFPFGF | ASSADWGERTGQLY | 0.1585118 |
| VGAGTVEF | ASSADWGERTGQLY | 0.71553886 |
| VGKSTLLL | ASSADWGERTGQLY | 0.3171521 |
| AALAHAGGGIV | ASSADWGERTGQLY | 0.38829574 |
| AALVAVSLL | ASSADWGERTGQLY | 0.8125885 |
| AAVSRAVAAL | ASSADWGERTGQLY | 0.18567318 |
| AGGSSGIGGF | ASSADWGERTGQLY | 0.5746922 |
| AGLTVANANG | ASSADWGERTGQLY | 0.7502286 |
| AGLVGAVL | ASSADWGERTGQLY | 0.69626087 |
| ALSQAFTV | ASSADWGERTGQLY | 0.31578028 |
| ALTFSVSL | ASSADWGERTGQLY | 0.5706104 |
| ALTGAALVAL | ASSADWGERTGQLY | 0.67134786 |
| ASFSIGAGVL | ASSADWGERTGQLY | 0.26680964 |
| DGSVYKI | ASSADWGERTGQLY | 0.7815974 |
| EQLAEIV | ASSADWGERTGQLY | 0.27038527 |
| FAVSELTVM | ASSADWGERTGQLY | 0.6449562 |
| GAAPLAAGL | ASSADWGERTGQLY | 0.6354525 |

| | | |
|---|---|---|
| GADGGDGGSGGAGG | ASSADWGERTGQLY | 0.86359894 |
| GFGLDTGGAGG | ASSADWGERTGQLY | 0.34654158 |
| GLASGGLL | ASSADWGERTGQLY | 0.36830068 |
| GRIAGAAVLL | ASSADWGERTGQLY | 0.5748782 |
| GSAGSLNTSF | ASSADWGERTGQLY | 0.66004026 |
| GTGQIGI | ASSADWGERTGQLY | 0.24856488 |
| GTLIALIL | ASSADWGERTGQLY | 0.5568883 |
| IAATMGFVVG | ASSADWGERTGQLY | 0.49175563 |
| IFAQALL | ASSADWGERTGQLY | 0.7755666 |
| IGAAVALGF | ASSADWGERTGQLY | 0.39773497 |
| IGLTPLAL | ASSADWGERTGQLY | 0.6095387 |
| ILGSNTSTL | ASSADWGERTGQLY | 0.8137452 |
| IVFSLTVL | ASSADWGERTGQLY | 0.2651717 |
| LASKATGF | ASSADWGERTGQLY | 0.55117524 |
| LASVALIL | ASSADWGERTGQLY | 0.6358979 |
| LFAVGLTL | ASSADWGERTGQLY | 0.7598824 |
| LFNGGIGTYI | ASSADWGERTGQLY | 0.32871455 |
| LTLGAGII | ASSADWGERTGQLY | 0.68945515 |
| LVAAALGDI | ASSADWGERTGQLY | 0.40285772 |
| NTAFVQLGI | ASSADWGERTGQLY | 0.6248933 |
| QLFSGAEL | ASSADWGERTGQLY | 0.19610822 |
| SAATFITL | ASSADWGERTGQLY | 0.40286043 |
| SNYGLANLGV | ASSADWGERTGQLY | 0.34530216 |
| SYGGAYAVM | ASSADWGERTGQLY | 0.6069291 |
| TTGVGGLTLG | ASSADWGERTGQLY | 0.6688874 |
| TTQTIYI | ASSADWGERTGQLY | 0.7058636 |
| VAAGAGLGAVI | ASSADWGERTGQLY | 0.19727817 |
| VAAVVTLL | ASSADWGERTGQLY | 0.72943026 |
| VGFLGGLI | ASSADWGERTGQLY | 0.7013639 |
| VGVIPILL | ASSADWGERTGQLY | 0.8745898 |
| VIFASSLI | ASSADWGERTGQLY | 0.37287003 |
| VKGGDGGAAAG | ASSADWGERTGQLY | 0.5687423 |
| VTKTFTNI | ASSADWGERTGQLY | 0.4739298 |
| VVAAAILL | ASSADWGERTGQLY | 0.3506511 |
| YAALDVEL | ASSADWGERTGQLY | 0.1842207 |
| LASVALLL | ASSADWGERTGQLY | 0.21477719 |
| LDAVHRV | ASSADWGERTGQLY | 0.51384735 |
| QGLSSSIF | ASSADWGERTGQLY | 0.43580627 |
| RGTFANIRL | ASSADWGERTGQLY | 0.84170306 |
| RQLWWGHRI | ASSADWGERTGQLY | 0.7749541 |
| YGPPGTGKTLL | ASSADWGERTGQLY | 0.68781894 |
| AALVGAPANVA | ASSADWGERTGQLY | 0.81476635 |
| AATFRRTE | ASSADWGERTGQLY | 0.603229 |
| AGALGGGI | ASSADWGERTGQLY | 0.53912526 |
| AGGAGGAGGIGAG | ASSADWGERTGQLY | 0.46132982 |
| AGSGALGL | ASSADWGERTGQLY | 0.49915904 |
| AQIADQI | ASSADWGERTGQLY | 0.19741623 |
| ASAVAVTYL | ASSADWGERTGQLY | 0.4552465 |
| ASTVLGGTGG | ASSADWGERTGQLY | 0.43469003 |
| AVGGEPAK | ASSADWGERTGQLY | 0.65804327 |
| CEQGTFGGTC | ASSADWGERTGQLY | 0.09684164 |

| | | |
|---|---|---|
| DEGKAPAG | ASSADWGERTGQLY | 0.7828916 |
| DTVVFPAL | ASSADWGERTGQLY | 0.7938311 |
| FEGITMAG | ASSADWGERTGQLY | 0.57390755 |
| FPLVVQD | ASSADWGERTGQLY | 0.6532708 |
| GAFTTPQ | ASSADWGERTGQLY | 0.5909988 |
| GFGGFGTAGGVGGA | ASSADWGERTGQLY | 0.26157022 |
| GFLAQFL | ASSADWGERTGQLY | 0.70696497 |
| GGAGGAGGNAGVA | ASSADWGERTGQLY | 0.33808905 |
| GGVGTAGAGGAGGAGG | ASSADWGERTGQLY | 0.6424485 |
| GNGGGGDAGNAGSGGN | ASSADWGERTGQLY | 0.71008825 |
| GTGGAGGAGNDAGS | ASSADWGERTGQLY | 0.5803777 |
| GTTNTGWFNS | ASSADWGERTGQLY | 0.49765664 |
| IFMGPLFTL | ASSADWGERTGQLY | 0.4136331 |
| ILAHAIEAI | ASSADWGERTGQLY | 0.3258227 |
| IRSAIREAL | ASSADWGERTGQLY | 0.5254421 |
| ITFAGGG | ASSADWGERTGQLY | 0.2940771 |
| ITGMSGHQEV | ASSADWGERTGQLY | 0.58307683 |
| IVAGLACTL | ASSADWGERTGQLY | 0.22656475 |
| IVGGGLAGA | ASSADWGERTGQLY | 0.7366332 |
| IVTGVLNI | ASSADWGERTGQLY | 0.68857753 |
| LAQQGCI | ASSADWGERTGQLY | 0.47395805 |
| LCLHVEN | ASSADWGERTGQLY | 0.36533985 |
| LDAISDGS | ASSADWGERTGQLY | 0.74902 |
| LGGFAAML | ASSADWGERTGQLY | 0.720871 |
| LGGKNPMI | ASSADWGERTGQLY | 0.26672512 |
| LGPKGVAAIL | ASSADWGERTGQLY | 0.361292 |
| LMVGVSI | ASSADWGERTGQLY | 0.31530845 |
| LVGAAFVL | ASSADWGERTGQLY | 0.30094647 |
| LVLAVAPL | ASSADWGERTGQLY | 0.73890704 |
| MAAIAVIDAA | ASSADWGERTGQLY | 0.5294342 |
| NNGINAELQ | ASSADWGERTGQLY | 0.13953969 |
| RGEPLVLVHG | ASSADWGERTGQLY | 0.40596443 |
| RGGAGGAAGLI | ASSADWGERTGQLY | 0.6405374 |
| RSYRMHPAV | ASSADWGERTGQLY | 0.44823194 |
| SAAHLLAGQLA | ASSADWGERTGQLY | 0.10650702 |
| SAAVGSY | ASSADWGERTGQLY | 0.3942253 |
| SPHAVLHRI | ASSADWGERTGQLY | 0.2611641 |
| SQIAKVL | ASSADWGERTGQLY | 0.26020783 |
| TALGHLVP | ASSADWGERTGQLY | 0.39244035 |
| TAPTTSGAA | ASSADWGERTGQLY | 0.76213187 |
| TDGDDGNGGSPGT | ASSADWGERTGQLY | 0.4664846 |
| TDGLGEN | ASSADWGERTGQLY | 0.3563659 |
| TGGPPLPGQA | ASSADWGERTGQLY | 0.65543526 |
| TIPAITI | ASSADWGERTGQLY | 0.6478793 |
| TIPGFTI | ASSADWGERTGQLY | 0.75241023 |
| TSIASGALA | ASSADWGERTGQLY | 0.6656281 |
| TVTLPGAAI | ASSADWGERTGQLY | 0.34222198 |
| VASGLSAALP | ASSADWGERTGQLY | 0.53040266 |
| VGAGVIGI | ASSADWGERTGQLY | 0.29963827 |
| VGPKRFGV | ASSADWGERTGQLY | 0.52479124 |
| VNVAFPD | ASSADWGERTGQLY | 0.31588227 |

| | | |
|---|---|---|
| VSMTNSGVS | ASSADWGERTGQLY | 0.23222062 |
| VTDVLGDLSV | ASSADWGERTGQLY | 0.48202428 |
| VVVVGAGFAGL | ASSADWGERTGQLY | 0.6772147 |
| VYTVGTVV | ASSADWGERTGQLY | 0.58271873 |
| WTTFSGP | ASSADWGERTGQLY | 0.25701916 |
| YDELVEAF | ASSADWGERTGQLY | 0.1746829 |
| YGTGGAGGNGG | ASSADWGERTGQLY | 0.41835743 |
| YTNVMAV | ASSADWGERTGQLY | 0.36738437 |
| FLEQVHQL | ASSADWGERTGQLY | 0.6844765 |
| GLSLLAVAV | ASSADWGERTGQLY | 0.4292075 |
| LEQVHQL | ASSADWGERTGQLY | 0.87371504 |
| LNAAGAI | ASSADWGERTGQLY | 0.7888178 |
| PAVHASL | ASSADWGERTGQLY | 0.51544785 |
| TELEDDA | ASSADWGERTGQLY | 0.66136116 |
| TPPGPPN | ASSADWGERTGQLY | 0.28831744 |
| TPPPPPPP | ASSADWGERTGQLY | 0.28138408 |
| TQSPALI | ASSKLGADTQY | 0.36224923 |
| EFAAAFPQ | ASSKLGADTQY | 0.29131663 |
| LLSGNLDVL | ASSKLGADTQY | 0.47264987 |
| GPANLGNYNI | ASSKLGADTQY | 0.32941887 |
| TLGPTLP | ASSKLGADTQY | 0.14898783 |
| VYAYSSCCK | ASSKLGADTQY | 0.7558484 |
| APAPAPAPNI | ASSKLGADTQY | 0.22262442 |
| PGGPVPIP | ASSKLGADTQY | 0.19363296 |
| FLTVALEP | ASSKLGADTQY | 0.36763155 |
| SQYFGANL | ASSKLGADTQY | 0.8069499 |
| GGVKPAA | ASSKLGADTQY | 0.80673206 |
| GGTGGAGGAGGW | ASSKLGADTQY | 0.43378207 |
| GVGGVAGLGGAGG | ASSKLGADTQY | 0.3287735 |
| VLLVLATTI | ASSKLGADTQY | 0.80192035 |
| IGIFAAML | ASSKLGADTQY | 0.4706868 |
| AVGDLID | ASSKLGADTQY | 0.36675286 |
| NISPLPL | ASSKLGADTQY | 0.611163 |
| VTAVFVAI | ASSKLGADTQY | 0.4851884 |
| SGDDNDAAL | ASSKLGADTQY | 0.25092596 |
| AVAALAHS | ASSKLGADTQY | 0.673456 |
| GGVTVMATP | ASSKLGADTQY | 0.42938137 |
| AGLAANVRD | ASSKLGADTQY | 0.36467358 |
| TNRPDLIDPALLRPG | ASSKLGADTQY | 0.09973691 |
| LANLGVTAL | ASSKLGADTQY | 0.6094117 |
| TGERIEL | ASSKLGADTQY | 0.38126746 |
| NGEELAA | ASSKLGADTQY | 0.20808017 |
| VLSLVWG | ASSKLGADTQY | 0.11646108 |
| IGYALWL | ASSKLGADTQY | 0.35901132 |
| LSLLLAPL | ASSKLGADTQY | 0.5692551 |
| TLDEALK | ASSKLGADTQY | 0.79438496 |
| TDGAGTAEGG | ASSKLGADTQY | 0.5363218 |
| NAPGGAGGNGG | ASSKLGADTQY | 0.7167325 |
| VSVFMGMGGNF | ASSKLGADTQY | 0.43710762 |
| LASYLAAL | ASSKLGADTQY | 0.49568474 |
| LVGGTPAL | ASSKLGADTQY | 0.19977568 |

| | | |
|---|---|---|
| VRNADVL | ASSKLGADTQY | 0.54307646 |
| TCSSPFLVSAM | ASSKLGADTQY | 0.68986005 |
| VHITDAE | ASSKLGADTQY | 0.5576869 |
| EYAAPSF | ASSKLGADTQY | 0.58986676 |
| VVDAFHGGL | ASSKLGADTQY | 0.27291393 |
| ASVVDPQ | ASSKLGADTQY | 0.4907967 |
| SYNEFGGLS | ASSKLGADTQY | 0.50055784 |
| DPDGSVY | ASSKLGADTQY | 0.17957363 |
| MVSTIHATE | ASSKLGADTQY | 0.24569702 |
| LAAGVADPV | ASSKLGADTQY | 0.29911625 |
| TKEQIAA | ASSKLGADTQY | 0.18526582 |
| SLASPAAL | ASSKLGADTQY | 0.60697436 |
| IGLVFYGV | ASSKLGADTQY | 0.50521374 |
| FLLDASNL | ASSKLGADTQY | 0.5925219 |
| YQVGNFP | ASSKLGADTQY | 0.4070416 |
| ESLLSQST | ASSKLGADTQY | 0.41682962 |
| LFTVGFAAVG | ASSKLGADTQY | 0.49192932 |
| ATPPTPT | ASSKLGADTQY | 0.62951493 |
| TDLATLG | ASSKLGADTQY | 0.53850317 |
| SAGGTGR | ASSKLGADTQY | 0.63527906 |
| GFSNSGS | ASSKLGADTQY | 0.3502088 |
| GGNGGSGGTSV | ASSKLGADTQY | 0.27244365 |
| GTSVATGGAGN | ASSKLGADTQY | 0.8279918 |
| NGGNGGSGGTS | ASSKLGADTQY | 0.24639647 |
| AVVKFPH | ASSKLGADTQY | 0.33526927 |
| TPPPPPP | ASSKLGADTQY | 0.46588492 |
| AAPAAAAAPA | ASSKLGADTQY | 0.53545713 |
| IALNEDL | ASSKLGADTQY | 0.5805268 |
| IVNVSSM | ASSKLGADTQY | 0.6938148 |
| TGIPPAPR | ASSKLGADTQY | 0.6752739 |
| FDLGGGTF | ASSKLGADTQY | 0.7640718 |
| FDLGGGTFDVS | ASSKLGADTQY | 0.7957531 |
| NEPTAAA | ASSKLGADTQY | 0.67515683 |
| NEPTAAALA | ASSKLGADTQY | 0.3017737 |
| GSIINISSI | ASSKLGADTQY | 0.80590165 |
| VAPGNAGTA | ASSKLGADTQY | 0.22154573 |
| TGGGASLE | ASSKLGADTQY | 0.6639374 |
| VAVYNHYKRI | ASSKLGADTQY | 0.61937237 |
| TDPAPATT | ASSKLGADTQY | 0.34263945 |
| TNRPDILDPALLRPG | ASSKLGADTQY | 0.5859041 |
| LTTTTTS | ASSKLGADTQY | 0.73233086 |
| PTGSGKTL | ASSKLGADTQY | 0.7494792 |
| TGSGKTL | ASSKLGADTQY | 0.4069078 |
| TIGHVDHGK | ASSKLGADTQY | 0.32711995 |
| TIGHVDHGKTT | ASSKLGADTQY | 0.3901872 |
| GHVDHGKTT | ASSKLGADTQY | 0.22819845 |
| AQFEHTLL | ASSKLGADTQY | 0.20686117 |
| IVNQDEL | ASSKLGADTQY | 0.54424214 |
| TNGSQFF | ASSKLGADTQY | 0.5441207 |
| TGKGIVE | ASSKLGADTQY | 0.8272131 |
| LISNASDA | ASSKLGADTQY | 0.690016 |

| | | |
|---|---|---|
| AELRTLGELTKIAKAHGAQ | ASSKLGADTQY | 0.797325 |
| APNAVAAAEPSW | ASSKLGADTQY | 0.36749062 |
| APQVAGAATAVL | ASSKLGADTQY | 0.6335665 |
| ARRDPAAAAAILPTDA | ASSKLGADTQY | 0.13434896 |
| DAAARDVLAEAGLAEYF | ASSKLGADTQY | 0.3482511 |
| DDQIVLSAKANLAKVEQ | ASSKLGADTQY | 0.23408797 |
| DELLPDTGEDSDEDGH | ASSKLGADTQY | 0.64944166 |
| ELPEAPANRATVPQD | ASSKLGADTQY | 0.15132406 |
| FGQSKIERVTGIG | ASSKLGADTQY | 0.64118123 |
| GAGGTGSTAGGAGGAGGAGGLY | ASSKLGADTQY | 0.37439448 |
| GHGGTGAAGGNGGTGGAGGSGI | ASSKLGADTQY | 0.710054 |
| GNGGNGGAGGEGGPGGAGGAGA | ASSKLGADTQY | 0.30722147 |
| GSTAGNGGNGGNATNGTIAGQP | ASSKLGADTQY | 0.26940253 |
| GTGGDGGNGGNGGTGG | ASSKLGADTQY | 0.45912898 |
| GVRRPAGIPADAPV | ASSKLGADTQY | 0.74699044 |
| GYRQVTDAHLLTLARR | ASSKLGADTQY | 0.28315014 |
| IAKMGQDGHDRG | ASSKLGADTQY | 0.113759466 |
| KAEPSIEIQYT | ASSKLGADTQY | 0.23206475 |
| KPVRIRMAADEDTVAVT | ASSKLGADTQY | 0.18715356 |
| MPPRIAGMRLLVIKPEP | ASSKLGADTQY | 0.7283881 |
| MKFYAVALFRSRGVPPTR | ASSKLGADTQY | 0.6973206 |
| NAGGFGILGAVAHSP | ASSKLGADTQY | 0.60436285 |
| NEGREITLAMAEM | ASSKLGADTQY | 0.85981774 |
| NELVEWSHPLGHRWRWE | ASSKLGADTQY | 0.09772829 |
| NLGAGGTGGNGGNPG | ASSKLGADTQY | 0.34684697 |
| NTEWPKWETVMAADPVD | ASSKLGADTQY | 0.8047935 |
| PASAAPAETQP | ASSKLGADTQY | 0.49868527 |
| QQSSDGKNGVRAAEITGE | ASSKLGADTQY | 0.59280074 |
| RAPDAVTTSWWKE | ASSKLGADTQY | 0.7113812 |
| SGFGNVGAHTSGWFNQ | ASSKLGADTQY | 0.41714573 |
| SGVAGAVGQAASVGGLKV | ASSKLGADTQY | 0.49550506 |
| SNGGAGTGSAGGNGGT | ASSKLGADTQY | 0.5735439 |
| TGGAGGEGGAAG | ASSKLGADTQY | 0.4656104 |
| TGGDGGAAGAGGAPGHG | ASSKLGADTQY | 0.22098102 |
| TMPPLLAAIANAGRHGV | ASSKLGADTQY | 0.21491098 |
| TSSTAGNSGGA | ASSKLGADTQY | 0.4815694 |
| VPATVPVGILC | ASSKLGADTQY | 0.86468154 |
| VQRPDSGGVVLTGRIS | ASSKLGADTQY | 0.6129587 |
| WTPNPVPATKGVPAGLVQAT | ASSKLGADTQY | 0.39005095 |
| YALKRALAKFSDRFDVVII | ASSKLGADTQY | 0.47648138 |
| YLIGSVIV | ASSKLGADTQY | 0.60075504 |
| YGQLVAVID | ASSKLGADTQY | 0.43868697 |
| VTVGGMTLPA | ASSKLGADTQY | 0.24958888 |
| VLALPAV | ASSKLGADTQY | 0.7761454 |
| VIPPTLNYET | ASSKLGADTQY | 0.19758733 |
| VGMGGMHPSSG | ASSKLGADTQY | 0.41615522 |
| VGLAGGAATA | ASSKLGADTQY | 0.33621785 |
| VGGAGGAGGSGP | ASSKLGADTQY | 0.8476734 |
| VFAAFMV | ASSKLGADTQY | 0.62410986 |
| TLYDKLR | ASSKLGADTQY | 0.5507787 |
| TLAVFTVL | ASSKLGADTQY | 0.4785872 |

| | | |
|---|---|---|
| TGSVQPARV | ASSKLGADTQY | 0.4329282 |
| TGGGAGSGGT | ASSKLGADTQY | 0.66802657 |
| SHLETSL | ASSKLGADTQY | 0.15398975 |
| SFTTGMT | ASSKLGADTQY | 0.7624855 |
| QVEVAKRMGVRQPTVS | ASSKLGADTQY | 0.28710794 |
| QGSGGPALAM | ASSKLGADTQY | 0.37986177 |
| MLAPSNP | ASSKLGADTQY | 0.5821331 |
| LVNNAAI | ASSKLGADTQY | 0.8572187 |
| LTGEINTLA | ASSKLGADTQY | 0.40033454 |
| LSIFGPV | ASSKLGADTQY | 0.7381591 |
| LLALLANL | ASSKLGADTQY | 0.4623366 |
| LISNASDALDKLR | ASSKLGADTQY | 0.5267832 |
| LINEAVP | ASSKLGADTQY | 0.82892585 |
| LGYAFSGL | ASSKLGADTQY | 0.65548825 |
| LDFVAVAL | ASSKLGADTQY | 0.8683127 |
| LAQPGAE | ASSKLGADTQY | 0.39718938 |
| ISNLGAVTGF | ASSKLGADTQY | 0.17023847 |
| HQEDPGRI | ASSKLGADTQY | 0.32051468 |
| GTPGEPTP | ASSKLGADTQY | 0.63980067 |
| GFFNSGS | ASSKLGADTQY | 0.6685939 |
| GAGGGGLTAGAN | ASSKLGADTQY | 0.6203041 |
| FVDALVNM | ASSKLGADTQY | 0.5014354 |
| FLTVAPEL | ASSKLGADTQY | 0.3258518 |
| FETLTAL | ASSKLGADTQY | 0.7445239 |
| DVINPIEL | ASSKLGADTQY | 0.71219075 |
| APVSGAPV | ASSKLGADTQY | 0.6117091 |
| ALVGGLAW | ASSKLGADTQY | 0.23199114 |
| ALAAAAATAI | ASSKLGADTQY | 0.273253 |
| AFGLGLLL | ASSKLGADTQY | 0.5839556 |
| AFFAAASL | ASSKLGADTQY | 0.54913986 |
| AAAALPAFSPP | ASSKLGADTQY | 0.19784707 |
| VVLAPAL | ASSKLGADTQY | 0.6073402 |
| TIGHVDHGKTTL | ASSKLGADTQY | 0.6772246 |
| LISNASDALDK | ASSKLGADTQY | 0.16233905 |
| LISNASDAL | ASSKLGADTQY | 0.31757155 |
| ISNASDALDK | ASSKLGADTQY | 0.7435399 |
| NQAETLVYQTE | ASSKLGADTQY | 0.39059216 |
| TGGTGGTAGAG | ASSKLGADTQY | 0.23352511 |
| TGSTAGGAGGA | ASSKLGADTQY | 0.77934206 |
| DPLAVPVAAGPV | ASSKLGADTQY | 0.8374479 |
| GGTGGTPGNGGN | ASSKLGADTQY | 0.741683 |
| PAGLAGVDPATL | ASSKLGADTQY | 0.20494294 |
| GNNLVVEAPHAGASV | ASSKLGADTQY | 0.097451694 |
| KVTRIMSASTLEQPA | ASSKLGADTQY | 0.717343 |
| LGFALPIALPRRTVV | ASSKLGADTQY | 0.81917995 |
| TRLGNLTARKLVVLV | ASSKLGADTQY | 0.5543395 |
| WSQQRRLIYFPSAGP | ASSKLGADTQY | 0.35832724 |
| GPNGNPDPMAAAVDIR | ASSKLGADTQY | 0.3519354 |
| TQRCVSVADLRAAASR | ASSKLGADTQY | 0.71599615 |
| QFVLDVAEHPVEVLTETV | ASSKLGADTQY | 0.25254852 |
| GGAGGTGGAGGNGGTQAGDG | ASSKLGADTQY | 0.18999684 |

| | | |
|---|---|---|
| PLLATLRAIAVDVDAKPAQV | ASSKLGADTQY | 0.3871776 |
| APKFRTGGYMIDFWGVGYQVAK | ASSKLGADTQY | 0.677114 |
| GAGGAGGAGGAAGGPGGTGGTG | ASSKLGADTQY | 0.24214846 |
| IGALGLLATDVAPAGSVAFGVA | ASSKLGADTQY | 0.36771578 |
| AGGAGGIGGNGGAGGAGGVLMG | ASSKLGADTQY | 0.76948094 |
| ADELIGLL | ASSKLGADTQY | 0.48648483 |
| ADRALKA | ASSKLGADTQY | 0.5741074 |
| AGAVGQVGGAGGA | ASSKLGADTQY | 0.30513608 |
| AGGGGGLGGAP | ASSKLGADTQY | 0.6414257 |
| AGGLLYGNGGAG | ASSKLGADTQY | 0.24556592 |
| AGLESIGL | ASSKLGADTQY | 0.45480698 |
| AGLKTGII | ASSKLGADTQY | 0.6074544 |
| ASLAALLL | ASSKLGADTQY | 0.46887314 |
| ASVINISSI | ASSKLGADTQY | 0.5388335 |
| ATPPGPA | ASSKLGADTQY | 0.27838117 |
| ATVASALV | ASSKLGADTQY | 0.2646155 |
| ATVASLIL | ASSKLGADTQY | 0.30272752 |
| AVNGPVLIRH | ASSKLGADTQY | 0.7724993 |
| DGAAGAVGI | ASSKLGADTQY | 0.5741042 |
| DPLLHGI | ASSKLGADTQY | 0.7153014 |
| ELGAALGAL | ASSKLGADTQY | 0.62987256 |
| ELGRTQAV | ASSKLGADTQY | 0.34207767 |
| FVAGGYLS | ASSKLGADTQY | 0.5421431 |
| GASGGAGGAGGW | ASSKLGADTQY | 0.1955226 |
| GGAGAGGDGGAGGI | ASSKLGADTQY | 0.62145746 |
| GTGHNGSQPGMGGNG | ASSKLGADTQY | 0.24579003 |
| GYAAASASAS | ASSKLGADTQY | 0.520481 |
| HQVDVAAVQQG | ASSKLGADTQY | 0.37282228 |
| IGAAIAHT | ASSKLGADTQY | 0.391745 |
| IGMIEVAQL | ASSKLGADTQY | 0.54793596 |
| IRLAPEHTSAMD | ASSKLGADTQY | 0.28977948 |
| IYNVSVL | ASSKLGADTQY | 0.4078608 |
| KQGFLGM | ASSKLGADTQY | 0.76025474 |
| LESVITQL | ASSKLGADTQY | 0.4871929 |
| LGLGALVVV | ASSKLGADTQY | 0.2562654 |
| LIGNGGNGGSGGI | ASSKLGADTQY | 0.5950537 |
| LNAINAPTL | ASSKLGADTQY | 0.29320908 |
| LQLLNTGV | ASSKLGADTQY | 0.66100985 |
| LQTSIEF | ASSKLGADTQY | 0.47160274 |
| MSFAVSEL | ASSKLGADTQY | 0.42650557 |
| NTGALGL | ASSKLGADTQY | 0.4462183 |
| PPGAGGAGGAGTT | ASSKLGADTQY | 0.15673879 |
| QLVGLGI | ASSKLGADTQY | 0.2712186 |
| RFLEQVHQL | ASSKLGADTQY | 0.33034834 |
| RGVGYVLR | ASSKLGADTQY | 0.71515286 |
| SATGVFAGVI | ASSKLGADTQY | 0.6465156 |
| SLPAITLP | ASSKLGADTQY | 0.30218112 |
| STALAFTV | ASSKLGADTQY | 0.49476212 |
| SVLTVGLGK | ASSKLGADTQY | 0.44750172 |
| TALAGAGSGI | ASSKLGADTQY | 0.33473462 |
| TGLPFTGI | ASSKLGADTQY | 0.37344044 |

| | | |
|---|---|---|
| VAEHVTV | ASSKLGADTQY | 0.28486443 |
| VDIVLNSL | ASSKLGADTQY | 0.15650624 |
| VLTGGGAVL | ASSKLGADTQY | 0.51865447 |
| VNVGLAYGI | ASSKLGADTQY | 0.32843858 |
| VSSAAVGAV | ASSKLGADTQY | 0.4793812 |
| VTSVYEAGV | ASSKLGADTQY | 0.60172385 |
| WVALDRGA | ASSKLGADTQY | 0.22343032 |
| YLVGSVVI | ASSKLGADTQY | 0.47302324 |
| GPPGTGKTLL | ASSKLGADTQY | 0.39992702 |
| GVGMTKF | ASSKLGADTQY | 0.60924697 |
| IGIGGSDL | ASSKLGADTQY | 0.6977534 |
| IGTLLAEL | ASSKLGADTQY | 0.32571787 |
| IYTSGTT | ASSKLGADTQY | 0.7416612 |
| LSFPAGV | ASSKLGADTQY | 0.6966421 |
| LYTSGTT | ASSKLGADTQY | 0.62512803 |
| VGAFPFGF | ASSKLGADTQY | 0.43588305 |
| VGAGTVEF | ASSKLGADTQY | 0.8131094 |
| VGKSTLLL | ASSKLGADTQY | 0.61057836 |
| AALAHAGGGIV | ASSKLGADTQY | 0.4176265 |
| AALVAVSLL | ASSKLGADTQY | 0.12598139 |
| AAVSRAVAAL | ASSKLGADTQY | 0.6594868 |
| AGGSSGIGGF | ASSKLGADTQY | 0.62666696 |
| AGLTVANANG | ASSKLGADTQY | 0.25231487 |
| AGLVGAVL | ASSKLGADTQY | 0.15511955 |
| ALSQAFTV | ASSKLGADTQY | 0.72553337 |
| ALTFSVSL | ASSKLGADTQY | 0.8026238 |
| ALTGAALVAL | ASSKLGADTQY | 0.5577042 |
| ASFSIGAGVL | ASSKLGADTQY | 0.5953814 |
| DGSVYKI | ASSKLGADTQY | 0.16431355 |
| EQLAEIV | ASSKLGADTQY | 0.6819627 |
| FAVSELTVM | ASSKLGADTQY | 0.7354587 |
| GAAPLAAGL | ASSKLGADTQY | 0.31787592 |
| GADGGDGGSGGAGG | ASSKLGADTQY | 0.6528602 |
| GFGLDTGGAGG | ASSKLGADTQY | 0.3132292 |
| GLASGGLL | ASSKLGADTQY | 0.32829148 |
| GRIAGAAVLL | ASSKLGADTQY | 0.77929425 |
| GSAGSLNTSF | ASSKLGADTQY | 0.55780774 |
| GTGQIGI | ASSKLGADTQY | 0.50213724 |
| GTLIALIL | ASSKLGADTQY | 0.5368874 |
| IAATMGFVVG | ASSKLGADTQY | 0.34859872 |
| IFAQALL | ASSKLGADTQY | 0.4764146 |
| IGAAVALGF | ASSKLGADTQY | 0.7281383 |
| IGLTPLAL | ASSKLGADTQY | 0.6148858 |
| ILGSNTSTL | ASSKLGADTQY | 0.5778062 |
| IVFSLTVL | ASSKLGADTQY | 0.32799512 |
| LASKATGF | ASSKLGADTQY | 0.8196261 |
| LASVALIL | ASSKLGADTQY | 0.84724736 |
| LFAVGLTL | ASSKLGADTQY | 0.62297535 |
| LFNGGIGTYI | ASSKLGADTQY | 0.2654829 |
| LTLGAGII | ASSKLGADTQY | 0.81672895 |
| LVAAALGDI | ASSKLGADTQY | 0.46268603 |

| | | |
|---|---|---|
| NTAFVQLGI | ASSKLGADTQY | 0.63546765 |
| QLFSGAEL | ASSKLGADTQY | 0.644093 |
| SAATFITL | ASSKLGADTQY | 0.33439544 |
| SNYGLANLGV | ASSKLGADTQY | 0.60693145 |
| SYGGAYAVM | ASSKLGADTQY | 0.38935032 |
| TTGVGGLTLG | ASSKLGADTQY | 0.4058628 |
| TTQTIYI | ASSKLGADTQY | 0.33356625 |
| VAAGAGLGAVI | ASSKLGADTQY | 0.27961963 |
| VAAVVTLL | ASSKLGADTQY | 0.41257495 |
| VGFLGGLI | ASSKLGADTQY | 0.32130936 |
| VGVIPILL | ASSKLGADTQY | 0.53748167 |
| VIFASSLI | ASSKLGADTQY | 0.43617535 |
| VKGGDGGAAAG | ASSKLGADTQY | 0.32730433 |
| VTKTFTNI | ASSKLGADTQY | 0.6338384 |
| VVAAAILL | ASSKLGADTQY | 0.64635754 |
| YAALDVEL | ASSKLGADTQY | 0.28689015 |
| LASVALLL | ASSKLGADTQY | 0.35112008 |
| LDAVHRV | ASSKLGADTQY | 0.35460874 |
| QGLSSSIF | ASSKLGADTQY | 0.354271 |
| RGTFANIRL | ASSKLGADTQY | 0.26809567 |
| RQLWWGHRI | ASSKLGADTQY | 0.45891646 |
| YGPPGTGKTLL | ASSKLGADTQY | 0.28809804 |
| AALVGAPANVA | ASSKLGADTQY | 0.24081878 |
| AATFRRTE | ASSKLGADTQY | 0.413952 |
| AGALGGGI | ASSKLGADTQY | 0.39622265 |
| AGGAGGAGGIGAG | ASSKLGADTQY | 0.35164183 |
| AGSGALGL | ASSKLGADTQY | 0.5844696 |
| AQIADQI | ASSKLGADTQY | 0.26867846 |
| ASAVAVTYL | ASSKLGADTQY | 0.3217358 |
| ASTVLGGTGG | ASSKLGADTQY | 0.5421609 |
| AVGGEPAK | ASSKLGADTQY | 0.6125557 |
| CEQGTFGGTC | ASSKLGADTQY | 0.30370367 |
| DEGKAPAG | ASSKLGADTQY | 0.47245404 |
| DTVVFPAL | ASSKLGADTQY | 0.6287421 |
| FEGITMAG | ASSKLGADTQY | 0.46447247 |
| FPLVVQD | ASSKLGADTQY | 0.5061416 |
| GAFTTPQ | ASSKLGADTQY | 0.54483134 |
| GFGGFGTAGGVGGA | ASSKLGADTQY | 0.3173986 |
| GFLAQFL | ASSKLGADTQY | 0.5465074 |
| GGAGGAGGNAGVA | ASSKLGADTQY | 0.65153706 |
| GGVGTAGAGGAGGAGG | ASSKLGADTQY | 0.3636927 |
| GNGGGGDAGNAGSGGN | ASSKLGADTQY | 0.41246966 |
| GTGGAGGAGNDAGS | ASSKLGADTQY | 0.3331805 |
| GTTNTGWFNS | ASSKLGADTQY | 0.33753958 |
| IFMGPLFTL | ASSKLGADTQY | 0.25476867 |
| ILAHAIEAI | ASSKLGADTQY | 0.522317 |
| IRSAIREAL | ASSKLGADTQY | 0.48739243 |
| ITFAGGG | ASSKLGADTQY | 0.3592971 |
| ITGMSGHQEV | ASSKLGADTQY | 0.2928529 |
| IVAGLACTL | ASSKLGADTQY | 0.28416646 |
| IVGGGLAGA | ASSKLGADTQY | 0.3537789 |

| | | |
|---|---|---|
| IVTGVLNI | ASSKLGADTQY | 0.33933717 |
| LAQQGCI | ASSKLGADTQY | 0.26024318 |
| LCLHVEN | ASSKLGADTQY | 0.32450598 |
| LDAISDGS | ASSKLGADTQY | 0.569565 |
| LGGFAAML | ASSKLGADTQY | 0.56107694 |
| LGGKNPMI | ASSKLGADTQY | 0.26757273 |
| LGPKGVAAIL | ASSKLGADTQY | 0.6360745 |
| LMVGVSI | ASSKLGADTQY | 0.47954035 |
| LVGAAFVL | ASSKLGADTQY | 0.5702356 |
| LVLAVAPL | ASSKLGADTQY | 0.37827083 |
| MAAIAVIDAA | ASSKLGADTQY | 0.3403324 |
| NNGINAELQ | ASSKLGADTQY | 0.5829757 |
| RGEPLVLVHG | ASSKLGADTQY | 0.38029543 |
| RGGAGGAAGLI | ASSKLGADTQY | 0.29666403 |
| RSYRMHPAV | ASSKLGADTQY | 0.5089619 |
| SAAHLLAGQLA | ASSKLGADTQY | 0.3589992 |
| SAAVGSY | ASSKLGADTQY | 0.34368023 |
| SPHAVLHRI | ASSKLGADTQY | 0.41594398 |
| SQIAKVL | ASSKLGADTQY | 0.4563594 |
| TALGHLVP | ASSKLGADTQY | 0.2879361 |
| TAPTTSGAA | ASSKLGADTQY | 0.3124877 |
| TDGDDGNGGSPGT | ASSKLGADTQY | 0.2805466 |
| TDGLGEN | ASSKLGADTQY | 0.42744645 |
| TGGPPLPGQA | ASSKLGADTQY | 0.58231354 |
| TIPAITI | ASSKLGADTQY | 0.40570775 |
| TIPGFTI | ASSKLGADTQY | 0.3250755 |
| TSIASGALA | ASSKLGADTQY | 0.5134654 |
| TVTLPGAAI | ASSKLGADTQY | 0.41115683 |
| VASGLSAALP | ASSKLGADTQY | 0.41546115 |
| VGAGVIGI | ASSKLGADTQY | 0.5760093 |
| VGPKRFGV | ASSKLGADTQY | 0.35496065 |
| VNVAFPD | ASSKLGADTQY | 0.46181828 |
| VSMTNSGVS | ASSKLGADTQY | 0.44965386 |
| VTDVLGDLSV | ASSKLGADTQY | 0.44351354 |
| VVVVGAGFAGL | ASSKLGADTQY | 0.4910757 |
| VYTVGTVV | ASSKLGADTQY | 0.40842915 |
| WTTFSGP | ASSKLGADTQY | 0.37291482 |
| YDELVEAF | ASSKLGADTQY | 0.2584576 |
| YGTGGAGGNGG | ASSKLGADTQY | 0.3618651 |
| YTNVMAV | ASSKLGADTQY | 0.47802472 |
| FLEQVHQL | ASSKLGADTQY | 0.4398259 |
| GLSLLAVAV | ASSKLGADTQY | 0.4898345 |
| LEQVHQL | ASSKLGADTQY | 0.43788114 |
| LNAAGAI | ASSKLGADTQY | 0.31964052 |
| PAVHASL | ASSKLGADTQY | 0.5652111 |
| TELEDDA | ASSKLGADTQY | 0.5714339 |
| TPPGPPN | ASSKLGADTQY | 0.28302103 |
| TPPPPPPP | ASSKLGADTQY | 0.36940885 |
| TQSPALI | ASSRGATEVF | 0.6621733 |
| EFAAAFPQ | ASSRGATEVF | 0.47743195 |
| LLSGNLDVL | ASSRGATEVF | 0.34677455 |

| | | |
|---|---|---|
| GPANLGNYNI | ASSRGATEVF | 0.334052 |
| TLGPTLP | ASSRGATEVF | 0.64855283 |
| VYAYSSCCK | ASSRGATEVF | 0.2985866 |
| APAPAPAPNI | ASSRGATEVF | 0.62270284 |
| PGGPVPIP | ASSRGATEVF | 0.25018024 |
| FLTVALEP | ASSRGATEVF | 0.32170114 |
| SQYFGANL | ASSRGATEVF | 0.32577118 |
| GGVKPAA | ASSRGATEVF | 0.26469833 |
| GGTGGAGGAGGW | ASSRGATEVF | 0.52218306 |
| GVGGVAGLGGAGG | ASSRGATEVF | 0.36028415 |
| VLLVLATTI | ASSRGATEVF | 0.55052555 |
| IGIFAAML | ASSRGATEVF | 0.3887205 |
| AVGDLID | ASSRGATEVF | 0.53065485 |
| NISPLPL | ASSRGATEVF | 0.4616977 |
| VTAVFVAI | ASSRGATEVF | 0.4251924 |
| SGDDNDAAL | ASSRGATEVF | 0.33105165 |
| AVAALAHS | ASSRGATEVF | 0.49977446 |
| GGVTVMATP | ASSRGATEVF | 0.57164866 |
| AGLAANVRD | ASSRGATEVF | 0.41709542 |
| TNRPDLIDPALLRPG | ASSRGATEVF | 0.2854988 |
| LANLGVTAL | ASSRGATEVF | 0.33272037 |
| TGERIEL | ASSRGATEVF | 0.42295468 |
| NGEELAA | ASSRGATEVF | 0.2575469 |
| VLSLVWG | ASSRGATEVF | 0.2472946 |
| IGYALWL | ASSRGATEVF | 0.33763513 |
| LSLLLAPL | ASSRGATEVF | 0.30813432 |
| TLDEALK | ASSRGATEVF | 0.58626026 |
| TDGAGTAEGG | ASSRGATEVF | 0.5683514 |
| NAPGGAGGNGG | ASSRGATEVF | 0.36548918 |
| VSVFMGMGGNF | ASSRGATEVF | 0.5238443 |
| LASYLAAL | ASSRGATEVF | 0.48680913 |
| LVGGTPAL | ASSRGATEVF | 0.23688045 |
| VRNADVL | ASSRGATEVF | 0.34771052 |
| TCSSPFLVSAM | ASSRGATEVF | 0.2886494 |
| VHITDAE | ASSRGATEVF | 0.5358984 |
| EYAAPSF | ASSRGATEVF | 0.31928727 |
| VVDAFHGGL | ASSRGATEVF | 0.34456104 |
| ASVVDPQ | ASSRGATEVF | 0.40209523 |
| SYNEFGGLS | ASSRGATEVF | 0.5231744 |
| DPDGSVY | ASSRGATEVF | 0.40612572 |
| MVSTIHATE | ASSRGATEVF | 0.29903173 |
| LAAGVADPV | ASSRGATEVF | 0.24210268 |
| TKEQIAA | ASSRGATEVF | 0.5140569 |
| SLASPAAL | ASSRGATEVF | 0.27849355 |
| IGLVFYGV | ASSRGATEVF | 0.31455696 |
| FLLDASNL | ASSRGATEVF | 0.25883302 |
| YQVGNFP | ASSRGATEVF | 0.24407476 |
| ESLLSQST | ASSRGATEVF | 0.2847279 |
| LFTVGFAAVG | ASSRGATEVF | 0.46459532 |
| ATPPTPT | ASSRGATEVF | 0.3395806 |
| TDLATLG | ASSRGATEVF | 0.38627347 |

| | | |
|---|---|---|
| SAGGTGR | ASSRGATEVF | 0.6434881 |
| GFSNSGS | ASSRGATEVF | 0.5525124 |
| GGNGGSGGTSV | ASSRGATEVF | 0.36094248 |
| GTSVATGGAGN | ASSRGATEVF | 0.3397143 |
| NGGNGGSGGTS | ASSRGATEVF | 0.29890975 |
| AVVKFPH | ASSRGATEVF | 0.314651 |
| TPPPPPP | ASSRGATEVF | 0.30608395 |
| AAPAAAAAPA | ASSRGATEVF | 0.6566059 |
| IALNEDL | ASSRGATEVF | 0.3290442 |
| IVNVSSM | ASSRGATEVF | 0.31808355 |
| TGIPPAPR | ASSRGATEVF | 0.37619233 |
| FDLGGGTF | ASSRGATEVF | 0.59558094 |
| FDLGGGTFDVS | ASSRGATEVF | 0.6580503 |
| NEPTAAA | ASSRGATEVF | 0.31587377 |
| NEPTAAALA | ASSRGATEVF | 0.36345497 |
| GSIINISSI | ASSRGATEVF | 0.36463535 |
| VAPGNAGTA | ASSRGATEVF | 0.4717389 |
| TGGGASLE | ASSRGATEVF | 0.3292094 |
| VAVYNHYKRI | ASSRGATEVF | 0.31467476 |
| TDPAPATT | ASSRGATEVF | 0.42564076 |
| TNRPDILDPALLRPG | ASSRGATEVF | 0.65885 |
| LTTTTTS | ASSRGATEVF | 0.3442615 |
| PTGSGKTL | ASSRGATEVF | 0.2899412 |
| TGSGKTL | ASSRGATEVF | 0.38543463 |
| TIGHVDHGK | ASSRGATEVF | 0.29986873 |
| TIGHVDHGKTT | ASSRGATEVF | 0.3414795 |
| GHVDHGKTT | ASSRGATEVF | 0.4179654 |
| AQFEHTLL | ASSRGATEVF | 0.2798813 |
| IVNQDEL | ASSRGATEVF | 0.3486835 |
| TNGSQFF | ASSRGATEVF | 0.64642555 |
| TGKGIVE | ASSRGATEVF | 0.34932622 |
| LISNASDA | ASSRGATEVF | 0.44724345 |
| AELRTLGELTKIAKAHGAQ | ASSRGATEVF | 0.6017972 |
| APNAVAAAEPSW | ASSRGATEVF | 0.47807333 |
| APQVAGAATAVL | ASSRGATEVF | 0.62326574 |
| ARRDPAAAAAILPTDA | ASSRGATEVF | 0.5433223 |
| DAAARDVLAEAGLAEYF | ASSRGATEVF | 0.38102704 |
| DDQIVLSAKANLAKVEQ | ASSRGATEVF | 0.38144895 |
| DELLPDTGEDSDEDGH | ASSRGATEVF | 0.43157604 |
| ELPEAPANRATVPQD | ASSRGATEVF | 0.41384587 |
| FGQSKIERVTGIG | ASSRGATEVF | 0.3655059 |
| GAGGTGSTAGGAGGAGGAGGLY | ASSRGATEVF | 0.30862567 |
| GHGGTGAAGGNGGTGGAGGSGI | ASSRGATEVF | 0.59849083 |
| GNGGNGGAGGEGGPGGAGGAGA | ASSRGATEVF | 0.38826424 |
| GSTAGNGGNGGNATNGTIAGQP | ASSRGATEVF | 0.46187827 |
| GTGGDGGNGGNGGTGG | ASSRGATEVF | 0.28328303 |
| GVRRPAGIPADAPV | ASSRGATEVF | 0.2968136 |
| GYRQVTDAHLLTLARR | ASSRGATEVF | 0.5266337 |
| IAKMGQDGHDRG | ASSRGATEVF | 0.2424248 |
| KAEPSIEIQYT | ASSRGATEVF | 0.24964882 |
| KPVRIRMAADEDTVAVT | ASSRGATEVF | 0.36384666 |

| | | |
|---|---|---|
| MPPRIAGMRLLVIKPEP | ASSRGATEVF | 0.61627996 |
| MKFYAVALFRSRGVPPTR | ASSRGATEVF | 0.24353406 |
| NAGGFGILGAVAHSP | ASSRGATEVF | 0.32537273 |
| NEGREITLAMAEM | ASSRGATEVF | 0.26831496 |
| NELVEWSHPLGHRWRWE | ASSRGATEVF | 0.4482695 |
| NLGAGGTGGNGGNPG | ASSRGATEVF | 0.43683636 |
| NTEWPKWETVMAADPVD | ASSRGATEVF | 0.27119932 |
| PASAAPAETQP | ASSRGATEVF | 0.4540136 |
| QQSSDGKNGVRAAEITGE | ASSRGATEVF | 0.26934412 |
| RAPDAVTTSWWKE | ASSRGATEVF | 0.2743265 |
| SGFGNVGAHTSGWFNQ | ASSRGATEVF | 0.4914897 |
| SGVAGAVGQAASVGGLKV | ASSRGATEVF | 0.5986991 |
| SNGGAGTGSAGGNGGT | ASSRGATEVF | 0.3005443 |
| TGGAGGEGGAAG | ASSRGATEVF | 0.51856756 |
| TGGDGGAAGAGGAPGHG | ASSRGATEVF | 0.3266299 |
| TMPPLLAAIANAGRHGV | ASSRGATEVF | 0.6624573 |
| TSSTAGNSGGA | ASSRGATEVF | 0.353874 |
| VPATVPVGILC | ASSRGATEVF | 0.27484953 |
| VQRPDSGGVVLTGRIS | ASSRGATEVF | 0.5084558 |
| WTPNPVPATKGVPAGLVQAT | ASSRGATEVF | 0.42560872 |
| YALKRALAKFSDRFDVVII | ASSRGATEVF | 0.41388243 |
| YLIGSVIV | ASSRGATEVF | 0.5059842 |
| YGQLVAVID | ASSRGATEVF | 0.3285487 |
| VTVGGMTLPA | ASSRGATEVF | 0.35965744 |
| VLALPAV | ASSRGATEVF | 0.6160538 |
| VIPPTLNYET | ASSRGATEVF | 0.57699424 |
| VGMGGMHPSSG | ASSRGATEVF | 0.32648078 |
| VGLAGGAATA | ASSRGATEVF | 0.47643903 |
| VGGAGGAGGSGP | ASSRGATEVF | 0.4310293 |
| VFAAFMV | ASSRGATEVF | 0.33509776 |
| TLYDKLR | ASSRGATEVF | 0.60822725 |
| TLAVFTVL | ASSRGATEVF | 0.38435256 |
| TGSVQPARV | ASSRGATEVF | 0.5057818 |
| TGGGAGSGGT | ASSRGATEVF | 0.43724716 |
| SHLETSL | ASSRGATEVF | 0.57047844 |
| SFTTGMT | ASSRGATEVF | 0.2980197 |
| QVEVAKRMGVRQPTVS | ASSRGATEVF | 0.5159616 |
| QGSGGPALAM | ASSRGATEVF | 0.41983306 |
| MLAPSNP | ASSRGATEVF | 0.52170956 |
| LVNNAAI | ASSRGATEVF | 0.36330062 |
| LTGEINTLA | ASSRGATEVF | 0.28757125 |
| LSIFGPV | ASSRGATEVF | 0.38389868 |
| LLALLANL | ASSRGATEVF | 0.5765653 |
| LISNASDALDKLR | ASSRGATEVF | 0.37051767 |
| LINEAVP | ASSRGATEVF | 0.50463504 |
| LGYAFSGL | ASSRGATEVF | 0.6426358 |
| LDFVAVAL | ASSRGATEVF | 0.3197442 |
| LAQPGAE | ASSRGATEVF | 0.38490355 |
| ISNLGAVTGF | ASSRGATEVF | 0.42690313 |
| HQEDPGRI | ASSRGATEVF | 0.6040743 |
| GTPGEPTP | ASSRGATEVF | 0.35271022 |

| | | |
|---|---|---|
| GFFNSGS | ASSRGATEVF | 0.5964549 |
| GAGGGGLTAGAN | ASSRGATEVF | 0.3169058 |
| FVDALVNM | ASSRGATEVF | 0.55370545 |
| FLTVAPEL | ASSRGATEVF | 0.30729 |
| FETLTAL | ASSRGATEVF | 0.50080925 |
| DVINPIEL | ASSRGATEVF | 0.30732128 |
| APVSGAPV | ASSRGATEVF | 0.5579886 |
| ALVGGLAW | ASSRGATEVF | 0.5716438 |
| ALAAAAATAI | ASSRGATEVF | 0.6042371 |
| AFGLGLLL | ASSRGATEVF | 0.24998464 |
| AFFAAASL | ASSRGATEVF | 0.5530932 |
| AAAALPAFSPP | ASSRGATEVF | 0.59008914 |
| VVLAPAL | ASSRGATEVF | 0.6689475 |
| TIGHVDHGKTTL | ASSRGATEVF | 0.37639725 |
| LISNASDALDK | ASSRGATEVF | 0.49569905 |
| LISNASDAL | ASSRGATEVF | 0.36868823 |
| ISNASDALDK | ASSRGATEVF | 0.33764157 |
| NQAETLVYQTE | ASSRGATEVF | 0.29424205 |
| TGGTGGTAGAG | ASSRGATEVF | 0.3742499 |
| TGSTAGGAGGA | ASSRGATEVF | 0.39776158 |
| DPLAVPVAAGPV | ASSRGATEVF | 0.31347156 |
| GGTGGTPGNGGN | ASSRGATEVF | 0.56156945 |
| PAGLAGVDPATL | ASSRGATEVF | 0.611415 |
| GNNLVVEAPHAGASV | ASSRGATEVF | 0.577543 |
| KVTRIMSASTLEQPA | ASSRGATEVF | 0.5865925 |
| LGFALPIALPRRTVV | ASSRGATEVF | 0.59593916 |
| TRLGNLTARKLVVLV | ASSRGATEVF | 0.44085097 |
| WSQQRRLIYFPSAGP | ASSRGATEVF | 0.327207 |
| GPNGNPDPMAAAVDIR | ASSRGATEVF | 0.37161848 |
| TQRCVSVADLRAAASR | ASSRGATEVF | 0.2968923 |
| QFVLDVAEHPVEVLTETV | ASSRGATEVF | 0.36852586 |
| GGAGGTGGAGGNGGTQAGDG | ASSRGATEVF | 0.32648063 |
| PLLATLRAIAVDVDAKPAQV | ASSRGATEVF | 0.4595452 |
| APKFRTGGYMIDFWGVGYQVAK | ASSRGATEVF | 0.24066618 |
| GAGGAGGAGGAAGGPGGTGGTG | ASSRGATEVF | 0.490452 |
| IGALGLLATDVAPAGSVAFGVA | ASSRGATEVF | 0.6509494 |
| AGGAGGIGGNGGAGGAGGVLMG | ASSRGATEVF | 0.30230907 |
| ADELIGLL | ASSRGATEVF | 0.44161123 |
| ADRALKA | ASSRGATEVF | 0.41811413 |
| AGAVGQVGGAGGA | ASSRGATEVF | 0.30811897 |
| AGGGGGLGGAP | ASSRGATEVF | 0.525792 |
| AGGLLYGNGGAG | ASSRGATEVF | 0.29377136 |
| AGLESIGL | ASSRGATEVF | 0.43409735 |
| AGLKTGII | ASSRGATEVF | 0.61067027 |
| ASLAALLL | ASSRGATEVF | 0.3941406 |
| ASVINISSI | ASSRGATEVF | 0.43291438 |
| ATPPGPA | ASSRGATEVF | 0.3697707 |
| ATVASALV | ASSRGATEVF | 0.28155896 |
| ATVASLIL | ASSRGATEVF | 0.49575716 |
| AVNGPVLIRH | ASSRGATEVF | 0.23732851 |
| DGAAGAVGI | ASSRGATEVF | 0.39519563 |

| | | |
|---|---|---|
| DPLLHGI | ASSRGATEVF | 0.30042353 |
| ELGAALGAL | ASSRGATEVF | 0.4322469 |
| ELGRTQAV | ASSRGATEVF | 0.45432633 |
| FVAGGYLS | ASSRGATEVF | 0.41839945 |
| GASGGAGGAGGW | ASSRGATEVF | 0.2988501 |
| GGAGAGGDGGAGGI | ASSRGATEVF | 0.34267133 |
| GTGHNGSQPGMGGNG | ASSRGATEVF | 0.29472008 |
| GYAAASASAS | ASSRGATEVF | 0.29176667 |
| HQVDVAAVQQG | ASSRGATEVF | 0.55639774 |
| IGAAIAHT | ASSRGATEVF | 0.35713467 |
| IGMIEVAQL | ASSRGATEVF | 0.370079 |
| IRLAPEHTSAMD | ASSRGATEVF | 0.55543816 |
| IYNVSVL | ASSRGATEVF | 0.3984529 |
| KQGFLGM | ASSRGATEVF | 0.28360173 |
| LESVITQL | ASSRGATEVF | 0.42629558 |
| LGLGALVVV | ASSRGATEVF | 0.36854845 |
| LIGNGGNGGSGGI | ASSRGATEVF | 0.33736098 |
| LNAINAPTL | ASSRGATEVF | 0.2387301 |
| LQLLNTGV | ASSRGATEVF | 0.43000346 |
| LQTSIEF | ASSRGATEVF | 0.33669052 |
| MSFAVSEL | ASSRGATEVF | 0.32087836 |
| NTGALGL | ASSRGATEVF | 0.35193932 |
| PPGAGGAGGAGTT | ASSRGATEVF | 0.56180036 |
| QLVGLGI | ASSRGATEVF | 0.27362606 |
| RFLEQVHQL | ASSRGATEVF | 0.38798115 |
| RGVGYVLR | ASSRGATEVF | 0.5605355 |
| SATGVFAGVI | ASSRGATEVF | 0.51291114 |
| SLPAITLP | ASSRGATEVF | 0.6418544 |
| STALAFTV | ASSRGATEVF | 0.45962307 |
| SVLTVGLGK | ASSRGATEVF | 0.3580018 |
| TALAGAGSGI | ASSRGATEVF | 0.462236 |
| TGLPFTGI | ASSRGATEVF | 0.40346575 |
| VAEHVTV | ASSRGATEVF | 0.25371888 |
| VDIVLNSL | ASSRGATEVF | 0.24975207 |
| VLTGGGAVL | ASSRGATEVF | 0.38404346 |
| VNVGLAYGI | ASSRGATEVF | 0.25364864 |
| VSSAAVGAV | ASSRGATEVF | 0.4063558 |
| VTSVYEAGV | ASSRGATEVF | 0.498716 |
| WVALDRGA | ASSRGATEVF | 0.3625737 |
| YLVGSVVI | ASSRGATEVF | 0.29909384 |
| GPPGTGKTLL | ASSRGATEVF | 0.33723664 |
| GVGMTKF | ASSRGATEVF | 0.34867385 |
| IGIGGSDL | ASSRGATEVF | 0.50223553 |
| IGTLLAEL | ASSRGATEVF | 0.39676905 |
| IYTSGTT | ASSRGATEVF | 0.6435782 |
| LSFPAGV | ASSRGATEVF | 0.6042011 |
| LYTSGTT | ASSRGATEVF | 0.35340053 |
| VGAFPFGF | ASSRGATEVF | 0.53411186 |
| VGAGTVEF | ASSRGATEVF | 0.33679977 |
| VGKSTLLL | ASSRGATEVF | 0.39981768 |
| AALAHAGGGIV | ASSRGATEVF | 0.3578769 |

| | | |
|---|---|---|
| AALVAVSLL | ASSRGATEVF | 0.3849705 |
| AAVSRAVAAL | ASSRGATEVF | 0.36963138 |
| AGGSSGIGGF | ASSRGATEVF | 0.3219246 |
| AGLTVANANG | ASSRGATEVF | 0.2732618 |
| AGLVGAVL | ASSRGATEVF | 0.3910411 |
| ALSQAFTV | ASSRGATEVF | 0.44029793 |
| ALTFSVSL | ASSRGATEVF | 0.25948074 |
| ALTGAALVAL | ASSRGATEVF | 0.32128477 |
| ASFSIGAGVL | ASSRGATEVF | 0.6702608 |
| DGSVYKI | ASSRGATEVF | 0.6154606 |
| EQLAEIV | ASSRGATEVF | 0.3102155 |
| FAVSELTVM | ASSRGATEVF | 0.3131491 |
| GAAPLAAGL | ASSRGATEVF | 0.52050614 |
| GADGGDGGSGGAGG | ASSRGATEVF | 0.26280284 |
| GFGLDTGGAGG | ASSRGATEVF | 0.39490017 |
| GLASGGLL | ASSRGATEVF | 0.46780837 |
| GRIAGAAVLL | ASSRGATEVF | 0.41861552 |
| GSAGSLNTSF | ASSRGATEVF | 0.32281998 |
| GTGQIGI | ASSRGATEVF | 0.39064872 |
| GTLIALIL | ASSRGATEVF | 0.36827767 |
| IAATMGFVVG | ASSRGATEVF | 0.4429409 |
| IFAQALL | ASSRGATEVF | 0.24211003 |
| IGAAVALGF | ASSRGATEVF | 0.43522364 |
| IGLTPLAL | ASSRGATEVF | 0.3116873 |
| ILGSNTSTL | ASSRGATEVF | 0.32991233 |
| IVFSLTVL | ASSRGATEVF | 0.26886883 |
| LASKATGF | ASSRGATEVF | 0.442741 |
| LASVALIL | ASSRGATEVF | 0.567728 |
| LFAVGLTL | ASSRGATEVF | 0.55212456 |
| LFNGGIGTYI | ASSRGATEVF | 0.26625696 |
| LTLGAGII | ASSRGATEVF | 0.59957266 |
| LVAAALGDI | ASSRGATEVF | 0.6377934 |
| NTAFVQLGI | ASSRGATEVF | 0.36369854 |
| QLFSGAEL | ASSRGATEVF | 0.28321055 |
| SAATFITL | ASSRGATEVF | 0.47473896 |
| SNYGLANLGV | ASSRGATEVF | 0.4249028 |
| SYGGAYAVM | ASSRGATEVF | 0.46904817 |
| TTGVGGLTLG | ASSRGATEVF | 0.5492938 |
| TTQTIYI | ASSRGATEVF | 0.28120747 |
| VAAGAGLGAVI | ASSRGATEVF | 0.52340907 |
| VAAVVTLL | ASSRGATEVF | 0.52304494 |
| VGFLGGLI | ASSRGATEVF | 0.2927175 |
| VGVIPILL | ASSRGATEVF | 0.34850407 |
| VIFASSLI | ASSRGATEVF | 0.3647924 |
| VKGGDGGAAAG | ASSRGATEVF | 0.25408754 |
| VTKTFTNI | ASSRGATEVF | 0.4983543 |
| VVAAAILL | ASSRGATEVF | 0.3300781 |
| YAALDVEL | ASSRGATEVF | 0.3383732 |
| LASVALLL | ASSRGATEVF | 0.40016362 |
| LDAVHRV | ASSRGATEVF | 0.32741386 |
| QGLSSSIF | ASSRGATEVF | 0.35144305 |

| | | |
|---|---|---|
| RGTFANIRL | ASSRGATEVF | 0.35837975 |
| RQLWWGHRI | ASSRGATEVF | 0.3974323 |
| YGPPGTGKTLL | ASSRGATEVF | 0.43601894 |
| AALVGAPANVA | ASSRGATEVF | 0.41994688 |
| AATFRRTE | ASSRGATEVF | 0.4547428 |
| AGALGGGI | ASSRGATEVF | 0.63522506 |
| AGGAGGAGGIGAG | ASSRGATEVF | 0.2896241 |
| AGSGALGL | ASSRGATEVF | 0.3290566 |
| AQIADQI | ASSRGATEVF | 0.47759658 |
| ASAVAVTYL | ASSRGATEVF | 0.26890555 |
| ASTVLGGTGG | ASSRGATEVF | 0.49476922 |
| AVGGEPAK | ASSRGATEVF | 0.47261027 |
| CEQGTFGGTC | ASSRGATEVF | 0.47837287 |
| DEGKAPAG | ASSRGATEVF | 0.36338207 |
| DTVVFPAL | ASSRGATEVF | 0.6749846 |
| FEGITMAG | ASSRGATEVF | 0.50318867 |
| FPLVVQD | ASSRGATEVF | 0.38289696 |
| GAFTTPQ | ASSRGATEVF | 0.4661857 |
| GFGGFGTAGGVGGA | ASSRGATEVF | 0.3320502 |
| GFLAQFL | ASSRGATEVF | 0.55619276 |
| GGAGGAGGNAGVA | ASSRGATEVF | 0.45821327 |
| GGVGTAGAGGAGGAGG | ASSRGATEVF | 0.28839394 |
| GNGGGGDAGNAGSGGN | ASSRGATEVF | 0.36851537 |
| GTGGAGGAGNDAGS | ASSRGATEVF | 0.49712184 |
| GTTNTGWFNS | ASSRGATEVF | 0.6031506 |
| IFMGPLFTL | ASSRGATEVF | 0.25916988 |
| ILAHAIEAI | ASSRGATEVF | 0.3687088 |
| IRSAIREAL | ASSRGATEVF | 0.26627782 |
| ITFAGGG | ASSRGATEVF | 0.3434952 |
| ITGMSGHQEV | ASSRGATEVF | 0.32428625 |
| IVAGLACTL | ASSRGATEVF | 0.4813143 |
| IVGGGLAGA | ASSRGATEVF | 0.3360409 |
| IVTGVLNI | ASSRGATEVF | 0.52236843 |
| LAQQGCI | ASSRGATEVF | 0.40279067 |
| LCLHVEN | ASSRGATEVF | 0.56558764 |
| LDAISDGS | ASSRGATEVF | 0.30421308 |
| LGGFAAML | ASSRGATEVF | 0.39030325 |
| LGGKNPMI | ASSRGATEVF | 0.27364275 |
| LGPKGVAAIL | ASSRGATEVF | 0.27254903 |
| LMVGVSI | ASSRGATEVF | 0.29991978 |
| LVGAAFVL | ASSRGATEVF | 0.35755745 |
| LVLAVAPL | ASSRGATEVF | 0.25109443 |
| MAAIAVIDAA | ASSRGATEVF | 0.5630692 |
| NNGINAELQ | ASSRGATEVF | 0.41980654 |
| RGEPLVLVHG | ASSRGATEVF | 0.5095335 |
| RGGAGGAAGLI | ASSRGATEVF | 0.3523442 |
| RSYRMHPAV | ASSRGATEVF | 0.3566266 |
| SAAHLLAGQLA | ASSRGATEVF | 0.3394321 |
| SAAVGSY | ASSRGATEVF | 0.52718574 |
| SPHAVLHRI | ASSRGATEVF | 0.31121024 |
| SQIAKVL | ASSRGATEVF | 0.26744053 |

| | | |
|---|---|---|
| TALGHLVP | ASSRGATEVF | 0.34606203 |
| TAPTTSGAA | ASSRGATEVF | 0.30911747 |
| TDGDDGNGGSPGT | ASSRGATEVF | 0.57641363 |
| TDGLGEN | ASSRGATEVF | 0.48461694 |
| TGGPPLPGQA | ASSRGATEVF | 0.41585347 |
| TIPAITI | ASSRGATEVF | 0.4567791 |
| TIPGFTI | ASSRGATEVF | 0.2609332 |
| TSIASGALA | ASSRGATEVF | 0.37281072 |
| TVTLPGAAI | ASSRGATEVF | 0.6645962 |
| VASGLSAALP | ASSRGATEVF | 0.39979967 |
| VGAGVIGI | ASSRGATEVF | 0.35685238 |
| VGPKRFGV | ASSRGATEVF | 0.5532086 |
| VNVAFPD | ASSRGATEVF | 0.33069193 |
| VSMTNSGVS | ASSRGATEVF | 0.33992893 |
| VTDVLGDLSV | ASSRGATEVF | 0.41102076 |
| VVVVGAGFAGL | ASSRGATEVF | 0.40983266 |
| VYTVGTVV | ASSRGATEVF | 0.36734998 |
| WTTFSGP | ASSRGATEVF | 0.30240357 |
| YDELVEAF | ASSRGATEVF | 0.43183506 |
| YGTGGAGGNGG | ASSRGATEVF | 0.6837782 |
| YTNVMAV | ASSRGATEVF | 0.35279977 |
| FLEQVHQL | ASSRGATEVF | 0.54286426 |
| GLSLLAVAV | ASSRGATEVF | 0.46274868 |
| LEQVHQL | ASSRGATEVF | 0.53485346 |
| LNAAGAI | ASSRGATEVF | 0.30855322 |
| PAVHASL | ASSRGATEVF | 0.3277348 |
| TELEDDA | ASSRGATEVF | 0.34425315 |
| TPPGPPN | ASSRGATEVF | 0.31038758 |
| TPPPPPPP | ASSRGATEVF | 0.36599484 |
| TQSPALI | ASSQHWGTSQNTLY | 0.3661333 |
| EFAAAFPQ | ASSQHWGTSQNTLY | 0.6933986 |
| LLSGNLDVL | ASSQHWGTSQNTLY | 0.47748226 |
| GPANLGNYNI | ASSQHWGTSQNTLY | 0.46060327 |
| TLGPTLP | ASSQHWGTSQNTLY | 0.3181432 |
| VYAYSSCCK | ASSQHWGTSQNTLY | 0.3670946 |
| APAPAPAPNI | ASSQHWGTSQNTLY | 0.42844242 |
| PGGPVPIP | ASSQHWGTSQNTLY | 0.2882096 |
| FLTVALEP | ASSQHWGTSQNTLY | 0.61007893 |
| SQYFGANL | ASSQHWGTSQNTLY | 0.3168918 |
| GGVKPAA | ASSQHWGTSQNTLY | 0.3383007 |
| GGTGGAGGAGGW | ASSQHWGTSQNTLY | 0.49738723 |
| GVGGVAGLGGAGG | ASSQHWGTSQNTLY | 0.6230353 |
| VLLVLATTI | ASSQHWGTSQNTLY | 0.34318373 |
| IGIFAAML | ASSQHWGTSQNTLY | 0.33468288 |
| AVGDLID | ASSQHWGTSQNTLY | 0.32194436 |
| NISPLPL | ASSQHWGTSQNTLY | 0.48062715 |
| VTAVFVAI | ASSQHWGTSQNTLY | 0.5784828 |
| SGDDNDAAL | ASSQHWGTSQNTLY | 0.33669215 |
| AVAALAHS | ASSQHWGTSQNTLY | 0.60685974 |
| GGVTVMATP | ASSQHWGTSQNTLY | 0.4169671 |
| AGLAANVRD | ASSQHWGTSQNTLY | 0.29447782 |

| | | |
|---|---|---|
| TNRPDLIDPALLRPG | ASSQHWGTSQNTLY | 0.35051212 |
| LANLGVTAL | ASSQHWGTSQNTLY | 0.54112566 |
| TGERIEL | ASSQHWGTSQNTLY | 0.59170336 |
| NGEELAA | ASSQHWGTSQNTLY | 0.57873386 |
| VLSLVWG | ASSQHWGTSQNTLY | 0.45485085 |
| IGYALWL | ASSQHWGTSQNTLY | 0.3268889 |
| LSLLLAPL | ASSQHWGTSQNTLY | 0.5857581 |
| TLDEALK | ASSQHWGTSQNTLY | 0.64045537 |
| TDGAGTAEGG | ASSQHWGTSQNTLY | 0.54041344 |
| NAPGGAGGNGG | ASSQHWGTSQNTLY | 0.33204162 |
| VSVFMGMGGNF | ASSQHWGTSQNTLY | 0.32045677 |
| LASYLAAL | ASSQHWGTSQNTLY | 0.36154744 |
| LVGGTPAL | ASSQHWGTSQNTLY | 0.54292583 |
| VRNADVL | ASSQHWGTSQNTLY | 0.29369646 |
| TCSSPFLVSAM | ASSQHWGTSQNTLY | 0.42975375 |
| VHITDAE | ASSQHWGTSQNTLY | 0.44284177 |
| EYAAPSF | ASSQHWGTSQNTLY | 0.2798611 |
| VVDAFHGGL | ASSQHWGTSQNTLY | 0.44672966 |
| ASVVDPQ | ASSQHWGTSQNTLY | 0.64091617 |
| SYNEFGGLS | ASSQHWGTSQNTLY | 0.35268676 |
| DPDGSVY | ASSQHWGTSQNTLY | 0.31638765 |
| MVSTIHATE | ASSQHWGTSQNTLY | 0.60387444 |
| LAAGVADPV | ASSQHWGTSQNTLY | 0.68489784 |
| TKEQIAA | ASSQHWGTSQNTLY | 0.50476074 |
| SLASPAAL | ASSQHWGTSQNTLY | 0.28474948 |
| IGLVFYGV | ASSQHWGTSQNTLY | 0.25210467 |
| FLLDASNL | ASSQHWGTSQNTLY | 0.3163805 |
| YQVGNFP | ASSQHWGTSQNTLY | 0.5334816 |
| ESLLSQST | ASSQHWGTSQNTLY | 0.34263387 |
| LFTVGFAAVG | ASSQHWGTSQNTLY | 0.3152661 |
| ATPPTPT | ASSQHWGTSQNTLY | 0.4586866 |
| TDLATLG | ASSQHWGTSQNTLY | 0.5653672 |
| SAGGTGR | ASSQHWGTSQNTLY | 0.32114398 |
| GFSNSGS | ASSQHWGTSQNTLY | 0.2959641 |
| GGNGGSGGTSV | ASSQHWGTSQNTLY | 0.50181156 |
| GTSVATGGAGN | ASSQHWGTSQNTLY | 0.4761505 |
| NGGNGGSGGTS | ASSQHWGTSQNTLY | 0.30537707 |
| AVVKFPH | ASSQHWGTSQNTLY | 0.36984602 |
| TPPPPPP | ASSQHWGTSQNTLY | 0.33800992 |
| AAPAAAAAPA | ASSQHWGTSQNTLY | 0.2713834 |
| IALNEDL | ASSQHWGTSQNTLY | 0.4720959 |
| IVNVSSM | ASSQHWGTSQNTLY | 0.29128462 |
| TGIPPAPR | ASSQHWGTSQNTLY | 0.6298892 |
| FDLGGGTF | ASSQHWGTSQNTLY | 0.29516897 |
| FDLGGGTFDVS | ASSQHWGTSQNTLY | 0.4573918 |
| NEPTAAA | ASSQHWGTSQNTLY | 0.46727443 |
| NEPTAAALA | ASSQHWGTSQNTLY | 0.41584426 |
| GSIINISSI | ASSQHWGTSQNTLY | 0.3469477 |
| VAPGNAGTA | ASSQHWGTSQNTLY | 0.33039907 |
| TGGGASLE | ASSQHWGTSQNTLY | 0.29874766 |
| VAVYNHYKRI | ASSQHWGTSQNTLY | 0.24902697 |

| | | |
|---|---|---|
| TDPAPATT | ASSQHWGTSQNTLY | 0.61778873 |
| TNRPDILDPALLRPG | ASSQHWGTSQNTLY | 0.2425423 |
| LTTTTTS | ASSQHWGTSQNTLY | 0.5602803 |
| PTGSGKTL | ASSQHWGTSQNTLY | 0.45832783 |
| TGSGKTL | ASSQHWGTSQNTLY | 0.31593406 |
| TIGHVDHGK | ASSQHWGTSQNTLY | 0.3909964 |
| TIGHVDHGKTT | ASSQHWGTSQNTLY | 0.30919576 |
| GHVDHGKTT | ASSQHWGTSQNTLY | 0.27637607 |
| AQFEHTLL | ASSQHWGTSQNTLY | 0.50157654 |
| IVNQDEL | ASSQHWGTSQNTLY | 0.47309816 |
| TNGSQFF | ASSQHWGTSQNTLY | 0.31116673 |
| TGKGIVE | ASSQHWGTSQNTLY | 0.38353348 |
| LISNASDA | ASSQHWGTSQNTLY | 0.42863634 |
| AELRTLGELTKIAKAHGAQ | ASSQHWGTSQNTLY | 0.3121496 |
| APNAVAAAEPSW | ASSQHWGTSQNTLY | 0.36201808 |
| APQVAGAATAVL | ASSQHWGTSQNTLY | 0.6071065 |
| ARRDPAAAAAILPTDA | ASSQHWGTSQNTLY | 0.38541752 |
| DAAARDVLAEAGLAEYF | ASSQHWGTSQNTLY | 0.2979707 |
| DDQIVLSAKANLAKVEQ | ASSQHWGTSQNTLY | 0.49129942 |
| DELLPDTGEDSDEDGH | ASSQHWGTSQNTLY | 0.30755544 |
| ELPEAPANRATVPQD | ASSQHWGTSQNTLY | 0.47463566 |
| FGQSKIERVTGIG | ASSQHWGTSQNTLY | 0.3871693 |
| GAGGTGSTAGGAGGAGGAGGLY | ASSQHWGTSQNTLY | 0.3566753 |
| GHGGTGAAGGNGGTGGAGGSGI | ASSQHWGTSQNTLY | 0.4573848 |
| GNGGNGGAGGEGGPGGAGGAGA | ASSQHWGTSQNTLY | 0.31828785 |
| GSTAGNGGNGGNATNGTIAGQP | ASSQHWGTSQNTLY | 0.30456576 |
| GTGGDGGNGGNGGTGG | ASSQHWGTSQNTLY | 0.34157807 |
| GVRRPAGIPADAPV | ASSQHWGTSQNTLY | 0.42472196 |
| GYRQVTDAHLLTLARR | ASSQHWGTSQNTLY | 0.55757374 |
| IAKMGQDGHDRG | ASSQHWGTSQNTLY | 0.3081905 |
| KAEPSIEIQYT | ASSQHWGTSQNTLY | 0.39545503 |
| KPVRIRMAADEDTVAVT | ASSQHWGTSQNTLY | 0.32568476 |
| MPPRIAGMRLLVIKPEP | ASSQHWGTSQNTLY | 0.29843202 |
| MKFYAVALFRSRGVPPTR | ASSQHWGTSQNTLY | 0.3574491 |
| NAGGFGILGAVAHSP | ASSQHWGTSQNTLY | 0.34015706 |
| NEGREITLAMAEM | ASSQHWGTSQNTLY | 0.31584665 |
| NELVEWSHPLGHRWRWE | ASSQHWGTSQNTLY | 0.428988 |
| NLGAGGTGGNGGNPG | ASSQHWGTSQNTLY | 0.31712043 |
| NTEWPKWETVMAADPVD | ASSQHWGTSQNTLY | 0.41401148 |
| PASAAPAETQP | ASSQHWGTSQNTLY | 0.65758693 |
| QQSSDGKNGVRAAEITGE | ASSQHWGTSQNTLY | 0.3448262 |
| RAPDAVTTSWWKE | ASSQHWGTSQNTLY | 0.4206113 |
| SGFGNVGAHTSGWFNQ | ASSQHWGTSQNTLY | 0.45735055 |
| SGVAGAVGQAASVGGLKV | ASSQHWGTSQNTLY | 0.5018071 |
| SNGGAGTGSAGGNGGT | ASSQHWGTSQNTLY | 0.3556577 |
| TGGAGGEGGAAG | ASSQHWGTSQNTLY | 0.31331885 |
| TGGDGGAAGAGGAPGHG | ASSQHWGTSQNTLY | 0.59507674 |
| TMPPLLAAIANAGRHGV | ASSQHWGTSQNTLY | 0.5065646 |
| TSSTAGNSGGA | ASSQHWGTSQNTLY | 0.5941556 |
| VPATVPVGILC | ASSQHWGTSQNTLY | 0.48583543 |
| VQRPDSGGVVLTGRIS | ASSQHWGTSQNTLY | 0.5522803 |

| | | |
|---|---|---|
| WTPNPVPATKGVPAGLVQAT | ASSQHWGTSQNTLY | 0.5162809 |
| YALKRALAKFSDRFDVVII | ASSQHWGTSQNTLY | 0.34643573 |
| YLIGSVIV | ASSQHWGTSQNTLY | 0.27242452 |
| YGQLVAVID | ASSQHWGTSQNTLY | 0.36218697 |
| VTVGGMTLPA | ASSQHWGTSQNTLY | 0.45243347 |
| VLALPAV | ASSQHWGTSQNTLY | 0.34885433 |
| VIPPTLNYET | ASSQHWGTSQNTLY | 0.27361017 |
| VGMGGMHPSSG | ASSQHWGTSQNTLY | 0.6326048 |
| VGLAGGAATA | ASSQHWGTSQNTLY | 0.68511236 |
| VGGAGGAGGSGP | ASSQHWGTSQNTLY | 0.32622868 |
| VFAAFMV | ASSQHWGTSQNTLY | 0.5851463 |
| TLYDKLR | ASSQHWGTSQNTLY | 0.27930084 |
| TLAVFTVL | ASSQHWGTSQNTLY | 0.48118162 |
| TGSVQPARV | ASSQHWGTSQNTLY | 0.6198086 |
| TGGGAGSGGT | ASSQHWGTSQNTLY | 0.29959077 |
| SHLETSL | ASSQHWGTSQNTLY | 0.53816783 |
| SFTTGMT | ASSQHWGTSQNTLY | 0.30839258 |
| QVEVAKRMGVRQPTVS | ASSQHWGTSQNTLY | 0.36699805 |
| QGSGGPALAM | ASSQHWGTSQNTLY | 0.62453896 |
| MLAPSNP | ASSQHWGTSQNTLY | 0.3623502 |
| LVNNAAI | ASSQHWGTSQNTLY | 0.41181886 |
| LTGEINTLA | ASSQHWGTSQNTLY | 0.3566905 |
| LSIFGPV | ASSQHWGTSQNTLY | 0.31152615 |
| LLALLANL | ASSQHWGTSQNTLY | 0.38963026 |
| LISNASDALDKLR | ASSQHWGTSQNTLY | 0.60012144 |
| LINEAVP | ASSQHWGTSQNTLY | 0.28206405 |
| LGYAFSGL | ASSQHWGTSQNTLY | 0.61806226 |
| LDFVAVAL | ASSQHWGTSQNTLY | 0.30434746 |
| LAQPGAE | ASSQHWGTSQNTLY | 0.67540014 |
| ISNLGAVTGF | ASSQHWGTSQNTLY | 0.5773032 |
| HQEDPGRI | ASSQHWGTSQNTLY | 0.44429758 |
| GTPGEPTP | ASSQHWGTSQNTLY | 0.33334005 |
| GFFNSGS | ASSQHWGTSQNTLY | 0.59900016 |
| GAGGGGLTAGAN | ASSQHWGTSQNTLY | 0.33698258 |
| FVDALVNM | ASSQHWGTSQNTLY | 0.55054015 |
| FLTVAPEL | ASSQHWGTSQNTLY | 0.48864245 |
| FETLTAL | ASSQHWGTSQNTLY | 0.3568923 |
| DVINPIEL | ASSQHWGTSQNTLY | 0.87308794 |
| APVSGAPV | ASSQHWGTSQNTLY | 0.7697787 |
| ALVGGLAW | ASSQHWGTSQNTLY | 0.7948743 |
| ALAAAAATAI | ASSQHWGTSQNTLY | 0.503088 |
| AFGLGLLL | ASSQHWGTSQNTLY | 0.27546087 |
| AFFAAASL | ASSQHWGTSQNTLY | 0.5753577 |
| AAAALPAFSPP | ASSQHWGTSQNTLY | 0.39031368 |
| VVLAPAL | ASSQHWGTSQNTLY | 0.8355037 |
| TIGHVDHGKTTL | ASSQHWGTSQNTLY | 0.83426523 |
| LISNASDALDK | ASSQHWGTSQNTLY | 0.5035809 |
| LISNASDAL | ASSQHWGTSQNTLY | 0.8231816 |
| ISNASDALDK | ASSQHWGTSQNTLY | 0.87604153 |
| NQAETLVYQTE | ASSQHWGTSQNTLY | 0.29048246 |
| TGGTGGTAGAG | ASSQHWGTSQNTLY | 0.71325886 |

| | | |
|---|---|---|
| TGSTAGGAGGA | ASSQHWGTSQNTLY | 0.37004435 |
| DPLAVPVAAGPV | ASSQHWGTSQNTLY | 0.6024622 |
| GGTGGTPGNGGN | ASSQHWGTSQNTLY | 0.20654213 |
| PAGLAGVDPATL | ASSQHWGTSQNTLY | 0.8232471 |
| GNNLVVEAPHAGASV | ASSQHWGTSQNTLY | 0.61315346 |
| KVTRIMSASTLEQPA | ASSQHWGTSQNTLY | 0.13333005 |
| LGFALPIALPRRTVV | ASSQHWGTSQNTLY | 0.74664354 |
| TRLGNLTARKLVVLV | ASSQHWGTSQNTLY | 0.6060997 |
| WSQQRRLIYFPSAGP | ASSQHWGTSQNTLY | 0.50641763 |
| GPNGNPDPMAAAVDIR | ASSQHWGTSQNTLY | 0.8251258 |
| TQRCVSVADLRAAASR | ASSQHWGTSQNTLY | 0.48487175 |
| QFVLDVAEHPVEVLTETV | ASSQHWGTSQNTLY | 0.3938188 |
| GGAGGTGGAGGNGGTQAGDG | ASSQHWGTSQNTLY | 0.8197787 |
| PLLATLRAIAVDVDAKPAQV | ASSQHWGTSQNTLY | 0.84542185 |
| APKFRTGGYMIDFWGVGYQVAK | ASSQHWGTSQNTLY | 0.3093824 |
| GAGGAGGAGGAAGGPGGTGGTG | ASSQHWGTSQNTLY | 0.7806539 |
| IGALGLLATDVAPAGSVAFGVA | ASSQHWGTSQNTLY | 0.87907696 |
| AGGAGGIGGNGGAGGAGGVLMG | ASSQHWGTSQNTLY | 0.66559494 |
| ADELIGLL | ASSQHWGTSQNTLY | 0.853919 |
| ADRALKA | ASSQHWGTSQNTLY | 0.8715843 |
| AGAVGQVGGAGGA | ASSQHWGTSQNTLY | 0.7290087 |
| AGGGGGLGGAP | ASSQHWGTSQNTLY | 0.8480062 |
| AGGLLYGNGGAG | ASSQHWGTSQNTLY | 0.83662844 |
| AGLESIGL | ASSQHWGTSQNTLY | 0.5740738 |
| AGLKTGII | ASSQHWGTSQNTLY | 0.6913388 |
| ASLAALLL | ASSQHWGTSQNTLY | 0.3479576 |
| ASVINISSI | ASSQHWGTSQNTLY | 0.74273205 |
| ATPPGPA | ASSQHWGTSQNTLY | 0.26730245 |
| ATVASALV | ASSQHWGTSQNTLY | 0.7925648 |
| ATVASLIL | ASSQHWGTSQNTLY | 0.8524063 |
| AVNGPVLIRH | ASSQHWGTSQNTLY | 0.6400599 |
| DGAAGAVGI | ASSQHWGTSQNTLY | 0.36522394 |
| DPLLHGI | ASSQHWGTSQNTLY | 0.30901876 |
| ELGAALGAL | ASSQHWGTSQNTLY | 0.8006711 |
| ELGRTQAV | ASSQHWGTSQNTLY | 0.38770682 |
| FVAGGYLS | ASSQHWGTSQNTLY | 0.27409247 |
| GASGGAGGAGGW | ASSQHWGTSQNTLY | 0.5680715 |
| GGAGAGGDGGAGGI | ASSQHWGTSQNTLY | 0.8199054 |
| GTGHNGSQPGMGGNG | ASSQHWGTSQNTLY | 0.8079354 |
| GYAAASASAS | ASSQHWGTSQNTLY | 0.44572142 |
| HQVDVAAVQQG | ASSQHWGTSQNTLY | 0.8791724 |
| IGAAIAHT | ASSQHWGTSQNTLY | 0.84863794 |
| IGMIEVAQL | ASSQHWGTSQNTLY | 0.8611547 |
| IRLAPEHTSAMD | ASSQHWGTSQNTLY | 0.66083086 |
| IYNVSVL | ASSQHWGTSQNTLY | 0.35153714 |
| KQGFLGM | ASSQHWGTSQNTLY | 0.86311287 |
| LESVITQL | ASSQHWGTSQNTLY | 0.6267704 |
| LGLGALVVV | ASSQHWGTSQNTLY | 0.717158 |
| LIGNGGNGGSGGI | ASSQHWGTSQNTLY | 0.8260943 |
| LNAINAPTL | ASSQHWGTSQNTLY | 0.525365 |
| LQLLNTGV | ASSQHWGTSQNTLY | 0.5115868 |

| | | |
|---|---|---|
| LQTSIEF | ASSQHWGTSQNTLY | 0.73247516 |
| MSFAVSEL | ASSQHWGTSQNTLY | 0.878574 |
| NTGALGL | ASSQHWGTSQNTLY | 0.48078865 |
| PPGAGGAGGAGTT | ASSQHWGTSQNTLY | 0.32634974 |
| QLVGLGI | ASSQHWGTSQNTLY | 0.61662614 |
| RFLEQVHQL | ASSQHWGTSQNTLY | 0.63184756 |
| RGVGYVLR | ASSQHWGTSQNTLY | 0.84258306 |
| SATGVFAGVI | ASSQHWGTSQNTLY | 0.7514297 |
| SLPAITLP | ASSQHWGTSQNTLY | 0.68827343 |
| STALAFTV | ASSQHWGTSQNTLY | 0.7728428 |
| SVLTVGLGK | ASSQHWGTSQNTLY | 0.78364336 |
| TALAGAGSGI | ASSQHWGTSQNTLY | 0.7468037 |
| TGLPFTGI | ASSQHWGTSQNTLY | 0.84109116 |
| VAEHVTV | ASSQHWGTSQNTLY | 0.5595598 |
| VDIVLNSL | ASSQHWGTSQNTLY | 0.87384206 |
| VLTGGGAVL | ASSQHWGTSQNTLY | 0.82736444 |
| VNVGLAYGI | ASSQHWGTSQNTLY | 0.80652505 |
| VSSAAVGAV | ASSQHWGTSQNTLY | 0.86561906 |
| VTSVYEAGV | ASSQHWGTSQNTLY | 0.7475456 |
| WVALDRGA | ASSQHWGTSQNTLY | 0.76611006 |
| YLVGSVVI | ASSQHWGTSQNTLY | 0.24748549 |
| GPPGTGKTLL | ASSQHWGTSQNTLY | 0.69373274 |
| GVGMTKF | ASSQHWGTSQNTLY | 0.766754 |
| IGIGGSDL | ASSQHWGTSQNTLY | 0.78947306 |
| IGTLLAEL | ASSQHWGTSQNTLY | 0.83560395 |
| IYTSGTT | ASSQHWGTSQNTLY | 0.8163215 |
| LSFPAGV | ASSQHWGTSQNTLY | 0.5728377 |
| LYTSGTT | ASSQHWGTSQNTLY | 0.88098276 |
| VGAFPFGF | ASSQHWGTSQNTLY | 0.86460435 |
| VGAGTVEF | ASSQHWGTSQNTLY | 0.35967243 |
| VGKSTLLL | ASSQHWGTSQNTLY | 0.6662797 |
| AALAHAGGGIV | ASSQHWGTSQNTLY | 0.87808084 |
| AALVAVSLL | ASSQHWGTSQNTLY | 0.8035804 |
| AAVSRAVAAL | ASSQHWGTSQNTLY | 0.58421636 |
| AGGSSGIGGF | ASSQHWGTSQNTLY | 0.6521187 |
| AGLTVANANG | ASSQHWGTSQNTLY | 0.86910224 |
| AGLVGAVL | ASSQHWGTSQNTLY | 0.68325543 |
| ALSQAFTV | ASSQHWGTSQNTLY | 0.88135743 |
| ALTFSVSL | ASSQHWGTSQNTLY | 0.16247822 |
| ALTGAALVAL | ASSQHWGTSQNTLY | 0.6318028 |
| ASFSIGAGVL | ASSQHWGTSQNTLY | 0.5846974 |
| DGSVYKI | ASSQHWGTSQNTLY | 0.15368272 |
| EQLAEIV | ASSQHWGTSQNTLY | 0.81631476 |
| FAVSELTVM | ASSQHWGTSQNTLY | 0.5916116 |
| GAAPLAAGL | ASSQHWGTSQNTLY | 0.79476845 |
| GADGGDGGSGGAGG | ASSQHWGTSQNTLY | 0.6397394 |
| GFGLDTGGAGG | ASSQHWGTSQNTLY | 0.84294176 |
| GLASGGLL | ASSQHWGTSQNTLY | 0.7820145 |
| GRIAGAAVLL | ASSQHWGTSQNTLY | 0.651966 |
| GSAGSLNTSF | ASSQHWGTSQNTLY | 0.37837464 |
| GTGQIGI | ASSQHWGTSQNTLY | 0.7278446 |

| | | |
|---|---|---|
| GTLIALIL | ASSQHWGTSQNTLY | 0.88711166 |
| IAATMGFVVG | ASSQHWGTSQNTLY | 0.84737504 |
| IFAQALL | ASSQHWGTSQNTLY | 0.20980817 |
| IGAAVALGF | ASSQHWGTSQNTLY | 0.775367 |
| IGLTPLAL | ASSQHWGTSQNTLY | 0.60458934 |
| ILGSNTSTL | ASSQHWGTSQNTLY | 0.20497963 |
| IVFSLTVL | ASSQHWGTSQNTLY | 0.1658209 |
| LASKATGF | ASSQHWGTSQNTLY | 0.64793026 |
| LASVALIL | ASSQHWGTSQNTLY | 0.7063328 |
| LFAVGLTL | ASSQHWGTSQNTLY | 0.7925829 |
| LFNGGIGTYI | ASSQHWGTSQNTLY | 0.81838006 |
| LTLGAGII | ASSQHWGTSQNTLY | 0.5464283 |
| LVAAALGDI | ASSQHWGTSQNTLY | 0.82709515 |
| NTAFVQLGI | ASSQHWGTSQNTLY | 0.7593572 |
| QLFSGAEL | ASSQHWGTSQNTLY | 0.072601944 |
| SAATFITL | ASSQHWGTSQNTLY | 0.6093698 |
| SNYGLANLGV | ASSQHWGTSQNTLY | 0.2691717 |
| SYGGAYAVM | ASSQHWGTSQNTLY | 0.87569773 |
| TTGVGGLTLG | ASSQHWGTSQNTLY | 0.47208518 |
| TTQTIYI | ASSQHWGTSQNTLY | 0.5880882 |
| VAAGAGLGAVI | ASSQHWGTSQNTLY | 0.64321196 |
| VAAVVTLL | ASSQHWGTSQNTLY | 0.8017902 |
| VGFLGGLI | ASSQHWGTSQNTLY | 0.6904583 |
| VGVIPILL | ASSQHWGTSQNTLY | 0.562623 |
| VIFASSLI | ASSQHWGTSQNTLY | 0.15884161 |
| VKGGDGGAAAG | ASSQHWGTSQNTLY | 0.7602732 |
| VTKTFTNI | ASSQHWGTSQNTLY | 0.412215 |
| VVAAAILL | ASSQHWGTSQNTLY | 0.36019486 |
| YAALDVEL | ASSQHWGTSQNTLY | 0.36947757 |
| LASVALLL | ASSQHWGTSQNTLY | 0.28110492 |
| LDAVHRV | ASSQHWGTSQNTLY | 0.23138353 |
| QGLSSSIF | ASSQHWGTSQNTLY | 0.83630216 |
| RGTFANIRL | ASSQHWGTSQNTLY | 0.46678358 |
| RQLWWGHRI | ASSQHWGTSQNTLY | 0.62660474 |
| YGPPGTGKTLL | ASSQHWGTSQNTLY | 0.88635725 |
| AALVGAPANVA | ASSQHWGTSQNTLY | 0.77731514 |
| AATFRRTE | ASSQHWGTSQNTLY | 0.67104197 |
| AGALGGGI | ASSQHWGTSQNTLY | 0.3928631 |
| AGGAGGAGGIGAG | ASSQHWGTSQNTLY | 0.4112719 |
| AGSGALGL | ASSQHWGTSQNTLY | 0.39089713 |
| AQIADQI | ASSQHWGTSQNTLY | 0.45723423 |
| ASAVAVTYL | ASSQHWGTSQNTLY | 0.87490433 |
| ASTVLGGTGG | ASSQHWGTSQNTLY | 0.6665546 |
| AVGGEPAK | ASSQHWGTSQNTLY | 0.37103227 |
| CEQGTFGGTC | ASSQHWGTSQNTLY | 0.78977025 |
| DEGKAPAG | ASSQHWGTSQNTLY | 0.8792692 |
| DTVVFPAL | ASSQHWGTSQNTLY | 0.887233 |
| FEGITMAG | ASSQHWGTSQNTLY | 0.41021258 |
| FPLVVQD | ASSQHWGTSQNTLY | 0.57455826 |
| GAFTTPQ | ASSQHWGTSQNTLY | 0.6719827 |
| GFGGFGTAGGVGGA | ASSQHWGTSQNTLY | 0.7218617 |

| | | |
|---|---|---|
| GFLAQFL | ASSQHWGTSQNTLY | 0.4754222 |
| GGAGGAGGNAGVA | ASSQHWGTSQNTLY | 0.36132863 |
| GGVGTAGAGGAGGAGG | ASSQHWGTSQNTLY | 0.8482115 |
| GNGGGGDAGNAGSGGN | ASSQHWGTSQNTLY | 0.8918003 |
| GTGGAGGAGNDAGS | ASSQHWGTSQNTLY | 0.6680707 |
| GTTNTGWFNS | ASSQHWGTSQNTLY | 0.19299653 |
| IFMGPLFTL | ASSQHWGTSQNTLY | 0.6158393 |
| ILAHAIEAI | ASSQHWGTSQNTLY | 0.4866591 |
| IRSAIREAL | ASSQHWGTSQNTLY | 0.49758297 |
| ITFAGGG | ASSQHWGTSQNTLY | 0.65815353 |
| ITGMSGHQEV | ASSQHWGTSQNTLY | 0.3060069 |
| IVAGLACTL | ASSQHWGTSQNTLY | 0.68117356 |
| IVGGGLAGA | ASSQHWGTSQNTLY | 0.8816118 |
| IVTGVLNI | ASSQHWGTSQNTLY | 0.57779634 |
| LAQQGCI | ASSQHWGTSQNTLY | 0.78530335 |
| LCLHVEN | ASSQHWGTSQNTLY | 0.8646438 |
| LDAISDGS | ASSQHWGTSQNTLY | 0.8110839 |
| LGGFAAML | ASSQHWGTSQNTLY | 0.8657248 |
| LGGKNPMI | ASSQHWGTSQNTLY | 0.8500514 |
| LGPKGVAAIL | ASSQHWGTSQNTLY | 0.6536409 |
| LMVGVSI | ASSQHWGTSQNTLY | 0.77873194 |
| LVGAAFVL | ASSQHWGTSQNTLY | 0.83198774 |
| LVLAVAPL | ASSQHWGTSQNTLY | 0.5913865 |
| MAAIAVIDAA | ASSQHWGTSQNTLY | 0.69731367 |
| NNGINAELQ | ASSQHWGTSQNTLY | 0.6813055 |
| RGEPLVLVHG | ASSQHWGTSQNTLY | 0.8255764 |
| RGGAGGAAGLI | ASSQHWGTSQNTLY | 0.6955637 |
| RSYRMHPAV | ASSQHWGTSQNTLY | 0.87138414 |
| SAAHLLAGQLA | ASSQHWGTSQNTLY | 0.20244245 |
| SAAVGSY | ASSQHWGTSQNTLY | 0.3084643 |
| SPHAVLHRI | ASSQHWGTSQNTLY | 0.8591026 |
| SQIAKVL | ASSQHWGTSQNTLY | 0.11869389 |
| TALGHLVP | ASSQHWGTSQNTLY | 0.18675011 |
| TAPTTSGAA | ASSQHWGTSQNTLY | 0.76268315 |
| TDGDDGNGGSPGT | ASSQHWGTSQNTLY | 0.8666584 |
| TDGLGEN | ASSQHWGTSQNTLY | 0.122307636 |
| TGGPPLPGQA | ASSQHWGTSQNTLY | 0.52151954 |
| TIPAITI | ASSQHWGTSQNTLY | 0.27585262 |
| TIPGFTI | ASSQHWGTSQNTLY | 0.8214438 |
| TSIASGALA | ASSQHWGTSQNTLY | 0.6410185 |
| TVTLPGAAI | ASSQHWGTSQNTLY | 0.27429342 |
| VASGLSAALP | ASSQHWGTSQNTLY | 0.80735195 |
| VGAGVIGI | ASSQHWGTSQNTLY | 0.25368023 |
| VGPKRFGV | ASSQHWGTSQNTLY | 0.650939 |
| VNVAFPD | ASSQHWGTSQNTLY | 0.8223853 |
| VSMTNSGVS | ASSQHWGTSQNTLY | 0.8736795 |
| VTDVLGDLSV | ASSQHWGTSQNTLY | 0.39822987 |
| VVVVGAGFAGL | ASSQHWGTSQNTLY | 0.783319 |
| VYTVGTVV | ASSQHWGTSQNTLY | 0.6729552 |
| WTTFSGP | ASSQHWGTSQNTLY | 0.893574 |
| YDELVEAF | ASSQHWGTSQNTLY | 0.5168294 |

| | | |
|---|---|---|
| YGTGGAGGNGG | ASSQHWGTSQNTLY | 0.27941477 |
| YTNVMAV | ASSQHWGTSQNTLY | 0.844033 |
| FLEQVHQL | ASSQHWGTSQNTLY | 0.8305304 |
| GLSLLAVAV | ASSQHWGTSQNTLY | 0.66728014 |
| LEQVHQL | ASSQHWGTSQNTLY | 0.8435439 |
| LNAAGAI | ASSQHWGTSQNTLY | 0.6241118 |
| PAVHASL | ASSQHWGTSQNTLY | 0.6348979 |
| TELEDDA | ASSQHWGTSQNTLY | 0.8227121 |
| TPPGPPN | ASSQHWGTSQNTLY | 0.84538305 |
| TPPPPPPP | ASSQHWGTSQNTLY | 0.3582175 |
| TQSPALI | ASSQDNAEQF | 0.8607142 |
| EFAAAFPQ | ASSQDNAEQF | 0.6912495 |
| LLSGNLDVL | ASSQDNAEQF | 0.41037878 |
| GPANLGNYNI | ASSQDNAEQF | 0.89211977 |
| TLGPTLP | ASSQDNAEQF | 0.69390255 |
| VYAYSSCCK | ASSQDNAEQF | 0.8257008 |
| APAPAPAPNI | ASSQDNAEQF | 0.79283845 |
| PGGPVPIP | ASSQDNAEQF | 0.865173 |
| FLTVALEP | ASSQDNAEQF | 0.33238226 |
| SQYFGANL | ASSQDNAEQF | 0.7456409 |
| GGVKPAA | ASSQDNAEQF | 0.70475286 |
| GGTGGAGGAGGW | ASSQDNAEQF | 0.82193226 |
| GVGGVAGLGGAGG | ASSQDNAEQF | 0.6480131 |
| VLLVLATTI | ASSQDNAEQF | 0.4655136 |
| IGIFAAML | ASSQDNAEQF | 0.81531227 |
| AVGDLID | ASSQDNAEQF | 0.85890234 |
| NISPLPL | ASSQDNAEQF | 0.75415266 |
| VTAVFVAI | ASSQDNAEQF | 0.82705337 |
| SGDDNDAAL | ASSQDNAEQF | 0.8799696 |
| AVAALAHS | ASSQDNAEQF | 0.377905 |
| GGVTVMATP | ASSQDNAEQF | 0.66600657 |
| AGLAANVRD | ASSQDNAEQF | 0.8426628 |
| TNRPDLIDPALLRPG | ASSQDNAEQF | 0.8766185 |
| LANLGVTAL | ASSQDNAEQF | 0.6755158 |
| TGERIEL | ASSQDNAEQF | 0.8769289 |
| NGEELAA | ASSQDNAEQF | 0.6753 |
| VLSLVWG | ASSQDNAEQF | 0.78513306 |
| IGYALWL | ASSQDNAEQF | 0.71536547 |
| LSLLLAPL | ASSQDNAEQF | 0.6058025 |
| TLDEALK | ASSQDNAEQF | 0.34398913 |
| TDGAGTAEGG | ASSQDNAEQF | 0.73114884 |
| NAPGGAGGNGG | ASSQDNAEQF | 0.83121365 |
| VSVFMGMGGNF | ASSQDNAEQF | 0.77424765 |
| LASYLAAL | ASSQDNAEQF | 0.19110495 |
| LVGGTPAL | ASSQDNAEQF | 0.780414 |
| VRNADVL | ASSQDNAEQF | 0.7849794 |
| TCSSPFLVSAM | ASSQDNAEQF | 0.89163077 |
| VHITDAE | ASSQDNAEQF | 0.6972275 |
| EYAAPSF | ASSQDNAEQF | 0.83913994 |
| VVDAFHGGL | ASSQDNAEQF | 0.4896739 |
| ASVVDPQ | ASSQDNAEQF | 0.69656146 |

| | | |
|---|---|---|
| SYNEFGGLS | ASSQDNAEQF | 0.3637596 |
| DPDGSVY | ASSQDNAEQF | 0.50806457 |
| MVSTIHATE | ASSQDNAEQF | 0.6704267 |
| LAAGVADPV | ASSQDNAEQF | 0.46569747 |
| TKEQIAA | ASSQDNAEQF | 0.83627546 |
| SLASPAAL | ASSQDNAEQF | 0.8572567 |
| IGLVFYGV | ASSQDNAEQF | 0.8357624 |
| FLLDASNL | ASSQDNAEQF | 0.8615018 |
| YQVGNFP | ASSQDNAEQF | 0.8755587 |
| ESLLSQST | ASSQDNAEQF | 0.820179 |
| LFTVGFAAVG | ASSQDNAEQF | 0.46790338 |
| ATPPTPT | ASSQDNAEQF | 0.69743955 |
| TDLATLG | ASSQDNAEQF | 0.27938968 |
| SAGGTGR | ASSQDNAEQF | 0.6692727 |
| GFSNSGS | ASSQDNAEQF | 0.56220454 |
| GGNGGSGGTSV | ASSQDNAEQF | 0.7060457 |
| GTSVATGGAGN | ASSQDNAEQF | 0.10507614 |
| NGGNGGSGGTS | ASSQDNAEQF | 0.8042456 |
| AVVKFPH | ASSQDNAEQF | 0.8744687 |
| TPPPPPP | ASSQDNAEQF | 0.584725 |
| AAPAAAAAPA | ASSQDNAEQF | 0.76233625 |
| IALNEDL | ASSQDNAEQF | 0.82356083 |
| IVNVSSM | ASSQDNAEQF | 0.6299515 |
| TGIPPAPR | ASSQDNAEQF | 0.8124053 |
| FDLGGGTF | ASSQDNAEQF | 0.5330456 |
| FDLGGGTFDVS | ASSQDNAEQF | 0.8270261 |
| NEPTAAA | ASSQDNAEQF | 0.8522934 |
| NEPTAAALA | ASSQDNAEQF | 0.7843946 |
| GSIINISSI | ASSQDNAEQF | 0.8086629 |
| VAPGNAGTA | ASSQDNAEQF | 0.737957 |
| TGGGASLE | ASSQDNAEQF | 0.2747049 |
| VAVYNHYKRI | ASSQDNAEQF | 0.8136455 |
| TDPAPATT | ASSQDNAEQF | 0.09729501 |
| TNRPDILDPALLRPG | ASSQDNAEQF | 0.7403592 |
| LTTTTTS | ASSQDNAEQF | 0.4322452 |
| PTGSGKTL | ASSQDNAEQF | 0.73115146 |
| TGSGKTL | ASSQDNAEQF | 0.6996287 |
| TIGHVDHGK | ASSQDNAEQF | 0.81831014 |
| TIGHVDHGKTT | ASSQDNAEQF | 0.38523987 |
| GHVDHGKTT | ASSQDNAEQF | 0.7744343 |
| AQFEHTLL | ASSQDNAEQF | 0.7409836 |
| IVNQDEL | ASSQDNAEQF | 0.3011881 |
| TNGSQFF | ASSQDNAEQF | 0.8368113 |
| TGKGIVE | ASSQDNAEQF | 0.69916356 |
| LISNASDA | ASSQDNAEQF | 0.70102555 |
| AELRTLGELTKIAKAHGAQ | ASSQDNAEQF | 0.84655464 |
| APNAVAAAEPSW | ASSQDNAEQF | 0.68750584 |
| APQVAGAATAVL | ASSQDNAEQF | 0.23716307 |
| ARRDPAAAAAILPTDA | ASSQDNAEQF | 0.6200057 |
| DAAARDVLAEAGLAEYF | ASSQDNAEQF | 0.74800545 |
| DDQIVLSAKANLAKVEQ | ASSQDNAEQF | 0.5293118 |

| | | |
|---|---|---|
| DELLPDTGEDSDEDGH | ASSQDNAEQF | 0.09408498 |
| ELPEAPANRATVPQD | ASSQDNAEQF | 0.8217495 |
| FGQSKIERVTGIG | ASSQDNAEQF | 0.41283032 |
| GAGGTGSTAGGAGGAGGAGGLY | ASSQDNAEQF | 0.60219264 |
| GHGGTGAAGGNGGTGGAGGSGI | ASSQDNAEQF | 0.5956466 |
| GNGGNGGAGGEGGPGGAGGAGA | ASSQDNAEQF | 0.8276949 |
| GSTAGNGGNGGNATNGTIAGQP | ASSQDNAEQF | 0.59216326 |
| GTGGDGGNGGNGGTGG | ASSQDNAEQF | 0.50237787 |
| GVRRPAGIPADAPV | ASSQDNAEQF | 0.8207567 |
| GYRQVTDAHLLTLARR | ASSQDNAEQF | 0.7457511 |
| IAKMGQDGHDRG | ASSQDNAEQF | 0.8481202 |
| KAEPSIEIQYT | ASSQDNAEQF | 0.64138836 |
| KPVRIRMAADEDTVAVT | ASSQDNAEQF | 0.74896646 |
| MPPRIAGMRLLVIKPEP | ASSQDNAEQF | 0.80695873 |
| MKFYAVALFRSRGVPPTR | ASSQDNAEQF | 0.56133693 |
| NAGGFGILGAVAHSP | ASSQDNAEQF | 0.39471805 |
| NEGREITLAMAEM | ASSQDNAEQF | 0.31153005 |
| NELVEWSHPLGHRWRWE | ASSQDNAEQF | 0.37812304 |
| NLGAGGTGGNGGNPG | ASSQDNAEQF | 0.32606283 |
| NTEWPKWETVMAADPVD | ASSQDNAEQF | 0.71563166 |
| PASAAPAETQP | ASSQDNAEQF | 0.86866224 |
| QQSSDGKNGVRAAEITGE | ASSQDNAEQF | 0.77544236 |
| RAPDAVTTSWWKE | ASSQDNAEQF | 0.31151587 |
| SGFGNVGAHTSGWFNQ | ASSQDNAEQF | 0.696527 |
| SGVAGAVGQAASVGGLKV | ASSQDNAEQF | 0.6139803 |
| SNGGAGTGSAGGNGGT | ASSQDNAEQF | 0.78758776 |
| TGGAGGEGGAAG | ASSQDNAEQF | 0.81784403 |
| TGGDGGAAGAGGAPGHG | ASSQDNAEQF | 0.88707304 |
| TMPPLLAAIANAGRHGV | ASSQDNAEQF | 0.8263188 |
| TSSTAGNSGGA | ASSQDNAEQF | 0.53100777 |
| VPATVPVGILC | ASSQDNAEQF | 0.83069146 |
| VQRPDSGGVVLTGRIS | ASSQDNAEQF | 0.6289927 |
| WTPNPVPATKGVPAGLVQAT | ASSQDNAEQF | 0.73481977 |
| YALKRALAKFSDRFDVVII | ASSQDNAEQF | 0.739867 |
| YLIGSVIV | ASSQDNAEQF | 0.81318754 |
| YGQLVAVID | ASSQDNAEQF | 0.65871406 |
| VTVGGMTLPA | ASSQDNAEQF | 0.55886924 |
| VLALPAV | ASSQDNAEQF | 0.22935754 |
| VIPPTLNYET | ASSQDNAEQF | 0.83120215 |
| VGMGGMHPSSG | ASSQDNAEQF | 0.2966037 |
| VGLAGGAATA | ASSQDNAEQF | 0.38352543 |
| VGGAGGAGGSGP | ASSQDNAEQF | 0.58519864 |
| VFAAFMV | ASSQDNAEQF | 0.87657046 |
| TLYDKLR | ASSQDNAEQF | 0.8658396 |
| TLAVFTVL | ASSQDNAEQF | 0.6053219 |
| TGSVQPARV | ASSQDNAEQF | 0.446765 |
| TGGGAGSGGT | ASSQDNAEQF | 0.8264899 |
| SHLETSL | ASSQDNAEQF | 0.39142868 |
| SFTTGMT | ASSQDNAEQF | 0.8097503 |
| QVEVAKRMGVRQPTVS | ASSQDNAEQF | 0.72724533 |
| QGSGGPALAM | ASSQDNAEQF | 0.74895716 |

| | | |
|---|---|---|
| MLAPSNP | ASSQDNAEQF | 0.6233797 |
| LVNNAAI | ASSQDNAEQF | 0.81386244 |
| LTGEINTLA | ASSQDNAEQF | 0.5896162 |
| LSIFGPV | ASSQDNAEQF | 0.79116285 |
| LLALLANL | ASSQDNAEQF | 0.14230374 |
| LISNASDALDKLR | ASSQDNAEQF | 0.8337456 |
| LINEAVP | ASSQDNAEQF | 0.43834794 |
| LGYAFSGL | ASSQDNAEQF | 0.5068164 |
| LDFVAVAL | ASSQDNAEQF | 0.20043841 |
| LAQPGAE | ASSQDNAEQF | 0.68991995 |
| ISNLGAVTGF | ASSQDNAEQF | 0.78667897 |
| HQEDPGRI | ASSQDNAEQF | 0.8650876 |
| GTPGEPTP | ASSQDNAEQF | 0.67088014 |
| GFFNSGS | ASSQDNAEQF | 0.8231839 |
| GAGGGGLTAGAN | ASSQDNAEQF | 0.84298944 |
| FVDALVNM | ASSQDNAEQF | 0.5794562 |
| FLTVAPEL | ASSQDNAEQF | 0.19721784 |
| FETLTAL | ASSQDNAEQF | 0.7526734 |
| DVINPIEL | ASSQDNAEQF | 0.7332111 |
| APVSGAPV | ASSQDNAEQF | 0.8447869 |
| ALVGGLAW | ASSQDNAEQF | 0.7948103 |
| ALAAAAATAI | ASSQDNAEQF | 0.32723707 |
| AFGLGLLL | ASSQDNAEQF | 0.85891086 |
| AFFAAASL | ASSQDNAEQF | 0.76411235 |
| AAAALPAFSPP | ASSQDNAEQF | 0.43732452 |
| VVLAPAL | ASSQDNAEQF | 0.5319309 |
| TIGHVDHGKTTL | ASSQDNAEQF | 0.5086168 |
| LISNASDALDK | ASSQDNAEQF | 0.1893431 |
| LISNASDAL | ASSQDNAEQF | 0.8300278 |
| ISNASDALDK | ASSQDNAEQF | 0.6298445 |
| NQAETLVYQTE | ASSQDNAEQF | 0.4883081 |
| TGGTGGTAGAG | ASSQDNAEQF | 0.7089455 |
| TGSTAGGAGGA | ASSQDNAEQF | 0.49674445 |
| DPLAVPVAAGPV | ASSQDNAEQF | 0.4509552 |
| GGTGGTPGNGGN | ASSQDNAEQF | 0.50259626 |
| PAGLAGVDPATL | ASSQDNAEQF | 0.63687265 |
| GNNLVVEAPHAGASV | ASSQDNAEQF | 0.8365247 |
| KVTRIMSASTLEQPA | ASSQDNAEQF | 0.69792455 |
| LGFALPIALPRRTVV | ASSQDNAEQF | 0.7716376 |
| TRLGNLTARKLVVLV | ASSQDNAEQF | 0.8571953 |
| WSQQRRLIYFPSAGP | ASSQDNAEQF | 0.25387722 |
| GPNGNPDPMAAAVDIR | ASSQDNAEQF | 0.49758178 |
| TQRCVSVADLRAAASR | ASSQDNAEQF | 0.8075906 |
| QFVLDVAEHPVEVLTETV | ASSQDNAEQF | 0.49683526 |
| GGAGGTGGAGGNGGTQAGDG | ASSQDNAEQF | 0.85428154 |
| PLLATLRAIAVDVDAKPAQV | ASSQDNAEQF | 0.7457669 |
| APKFRTGGYMIDFWGVGYQVAK | ASSQDNAEQF | 0.7563028 |
| GAGGAGGAGGAAGGPGGTGGTG | ASSQDNAEQF | 0.7621236 |
| IGALGLLATDVAPAGSVAFGVA | ASSQDNAEQF | 0.89487326 |
| AGGAGGIGGNGGAGGAGGVLMG | ASSQDNAEQF | 0.7614095 |
| ADELIGLL | ASSQDNAEQF | 0.67466295 |

| | | |
|---|---|---|
| ADRALKA | ASSQDNAEQF | 0.78159356 |
| AGAVGQVGGAGGA | ASSQDNAEQF | 0.37766343 |
| AGGGGGLGGAP | ASSQDNAEQF | 0.847659 |
| AGGLLYGNGGAG | ASSQDNAEQF | 0.8024658 |
| AGLESIGL | ASSQDNAEQF | 0.40548754 |
| AGLKTGII | ASSQDNAEQF | 0.64955425 |
| ASLAALLL | ASSQDNAEQF | 0.8814873 |
| ASVINISSI | ASSQDNAEQF | 0.8409719 |
| ATPPGPA | ASSQDNAEQF | 0.38131124 |
| ATVASALV | ASSQDNAEQF | 0.56032526 |
| ATVASLIL | ASSQDNAEQF | 0.42545202 |
| AVNGPVLIRH | ASSQDNAEQF | 0.66593003 |
| DGAAGAVGI | ASSQDNAEQF | 0.28963357 |
| DPLLHGI | ASSQDNAEQF | 0.80926704 |
| ELGAALGAL | ASSQDNAEQF | 0.55713105 |
| ELGRTQAV | ASSQDNAEQF | 0.82823205 |
| FVAGGYLS | ASSQDNAEQF | 0.73518455 |
| GASGGAGGAGGW | ASSQDNAEQF | 0.8373493 |
| GGAGAGGDGGAGGI | ASSQDNAEQF | 0.60858196 |
| GTGHNGSQPGMGGNG | ASSQDNAEQF | 0.5572436 |
| GYAAASASAS | ASSQDNAEQF | 0.24772969 |
| HQVDVAAVQQG | ASSQDNAEQF | 0.24025533 |
| IGAAIAHT | ASSQDNAEQF | 0.5896577 |
| IGMIEVAQL | ASSQDNAEQF | 0.6219404 |
| IRLAPEHTSAMD | ASSQDNAEQF | 0.15584454 |
| IYNVSVL | ASSQDNAEQF | 0.8751216 |
| KQGFLGM | ASSQDNAEQF | 0.59647644 |
| LESVITQL | ASSQDNAEQF | 0.76961505 |
| LGLGALVVV | ASSQDNAEQF | 0.60116684 |
| LIGNGGNGGSGGI | ASSQDNAEQF | 0.4154408 |
| LNAINAPTL | ASSQDNAEQF | 0.51188743 |
| LQLLNTGV | ASSQDNAEQF | 0.80509794 |
| LQTSIEF | ASSQDNAEQF | 0.4428725 |
| MSFAVSEL | ASSQDNAEQF | 0.17396174 |
| NTGALGL | ASSQDNAEQF | 0.4223625 |
| PPGAGGAGGAGTT | ASSQDNAEQF | 0.39158034 |
| QLVGLGI | ASSQDNAEQF | 0.8915814 |
| RFLEQVHQL | ASSQDNAEQF | 0.85168535 |
| RGVGYVLR | ASSQDNAEQF | 0.7441797 |
| SATGVFAGVI | ASSQDNAEQF | 0.88799495 |
| SLPAITLP | ASSQDNAEQF | 0.14652774 |
| STALAFTV | ASSQDNAEQF | 0.6355332 |
| SVLTVGLGK | ASSQDNAEQF | 0.87769604 |
| TALAGAGSGI | ASSQDNAEQF | 0.7624543 |
| TGLPFTGI | ASSQDNAEQF | 0.60169953 |
| VAEHVTV | ASSQDNAEQF | 0.820625 |
| VDIVLNSL | ASSQDNAEQF | 0.45761526 |
| VLTGGGAVL | ASSQDNAEQF | 0.4972209 |
| VNVGLAYGI | ASSQDNAEQF | 0.5994583 |
| VSSAAVGAV | ASSQDNAEQF | 0.82524085 |
| VTSVYEAGV | ASSQDNAEQF | 0.75587153 |

| | | |
|---|---|---|
| WVALDRGA | ASSQDNAEQF | 0.29871848 |
| YLVGSVVI | ASSQDNAEQF | 0.7876896 |
| GPPGTGKTLL | ASSQDNAEQF | 0.89478713 |
| GVGMTKF | ASSQDNAEQF | 0.50551254 |
| IGIGGSDL | ASSQDNAEQF | 0.7749392 |
| IGTLLAEL | ASSQDNAEQF | 0.7228385 |
| IYTSGTT | ASSQDNAEQF | 0.4032067 |
| LSFPAGV | ASSQDNAEQF | 0.4630862 |
| LYTSGTT | ASSQDNAEQF | 0.44993815 |
| VGAFPFGF | ASSQDNAEQF | 0.788504 |
| VGAGTVEF | ASSQDNAEQF | 0.42627627 |
| VGKSTLLL | ASSQDNAEQF | 0.6798907 |
| AALAHAGGGIV | ASSQDNAEQF | 0.6856904 |
| AALVAVSLL | ASSQDNAEQF | 0.90076053 |
| AAVSRAVAAL | ASSQDNAEQF | 0.79509914 |
| AGGSSGIGGF | ASSQDNAEQF | 0.75800735 |
| AGLTVANANG | ASSQDNAEQF | 0.47395325 |
| AGLVGAVL | ASSQDNAEQF | 0.8078988 |
| ALSQAFTV | ASSQDNAEQF | 0.81499004 |
| ALTFSVSL | ASSQDNAEQF | 0.2491621 |
| ALTGAALVAL | ASSQDNAEQF | 0.8725771 |
| ASFSIGAGVL | ASSQDNAEQF | 0.3575328 |
| DGSVYKI | ASSQDNAEQF | 0.782124 |
| EQLAEIV | ASSQDNAEQF | 0.7613074 |
| FAVSELTVM | ASSQDNAEQF | 0.85825443 |
| GAAPLAAGL | ASSQDNAEQF | 0.6650003 |
| GADGGDGGSGGAGG | ASSQDNAEQF | 0.78176194 |
| GFGLDTGGAGG | ASSQDNAEQF | 0.37699467 |
| GLASGGLL | ASSQDNAEQF | 0.84564954 |
| GRIAGAAVLL | ASSQDNAEQF | 0.8347181 |
| GSAGSLNTSF | ASSQDNAEQF | 0.6501546 |
| GTGQIGI | ASSQDNAEQF | 0.8891647 |
| GTLIALIL | ASSQDNAEQF | 0.7622324 |
| IAATMGFVVG | ASSQDNAEQF | 0.22799861 |
| IFAQALL | ASSQDNAEQF | 0.60836434 |
| IGAAVALGF | ASSQDNAEQF | 0.8637004 |
| IGLTPLAL | ASSQDNAEQF | 0.866841 |
| ILGSNTSTL | ASSQDNAEQF | 0.7370316 |
| IVFSLTVL | ASSQDNAEQF | 0.8199712 |
| LASKATGF | ASSQDNAEQF | 0.6851867 |
| LASVALIL | ASSQDNAEQF | 0.87243915 |
| LFAVGLTL | ASSQDNAEQF | 0.9057169 |
| LFNGGIGTYI | ASSQDNAEQF | 0.8122957 |
| LTLGAGII | ASSQDNAEQF | 0.51688623 |
| LVAAALGDI | ASSQDNAEQF | 0.4883791 |
| NTAFVQLGI | ASSQDNAEQF | 0.80989003 |
| QLFSGAEL | ASSQDNAEQF | 0.8496735 |
| SAATFITL | ASSQDNAEQF | 0.34260827 |
| SNYGLANLGV | ASSQDNAEQF | 0.6708284 |
| SYGGAYAVM | ASSQDNAEQF | 0.6937589 |
| TTGVGGLTLG | ASSQDNAEQF | 0.19232258 |

| | | |
|---|---|---|
| TTQTIYI | ASSQDNAEQF | 0.81644285 |
| VAAGAGLGAVI | ASSQDNAEQF | 0.8820454 |
| VAAVVTLL | ASSQDNAEQF | 0.7729306 |
| VGFLGGLI | ASSQDNAEQF | 0.44825286 |
| VGVIPILL | ASSQDNAEQF | 0.86873186 |
| VIFASSLI | ASSQDNAEQF | 0.8689412 |
| VKGGDGGAAAG | ASSQDNAEQF | 0.8750037 |
| VTKTFTNI | ASSQDNAEQF | 0.22464173 |
| VVAAAILL | ASSQDNAEQF | 0.1432117 |
| YAALDVEL | ASSQDNAEQF | 0.7161648 |
| LASVALLL | ASSQDNAEQF | 0.8218583 |
| LDAVHRV | ASSQDNAEQF | 0.558957 |
| QGLSSSIF | ASSQDNAEQF | 0.44867915 |
| RGTFANIRL | ASSQDNAEQF | 0.70552576 |
| RQLWWGHRI | ASSQDNAEQF | 0.8524505 |
| YGPPGTGKTLL | ASSQDNAEQF | 0.37442258 |
| AALVGAPANVA | ASSQDNAEQF | 0.31881052 |
| AATFRRTE | ASSQDNAEQF | 0.7929578 |
| AGALGGGI | ASSQDNAEQF | 0.82972205 |
| AGGAGGAGGIGAG | ASSQDNAEQF | 0.48498014 |
| AGSGALGL | ASSQDNAEQF | 0.6257118 |
| AQIADQI | ASSQDNAEQF | 0.71960276 |
| ASAVAVTYL | ASSQDNAEQF | 0.73075664 |
| ASTVLGGTGG | ASSQDNAEQF | 0.7809015 |
| AVGGEPAK | ASSQDNAEQF | 0.31380472 |
| CEQGTFGGTC | ASSQDNAEQF | 0.8480323 |
| DEGKAPAG | ASSQDNAEQF | 0.29956928 |
| DTVVFPAL | ASSQDNAEQF | 0.85517585 |
| FEGITMAG | ASSQDNAEQF | 0.7521305 |
| FPLVVQD | ASSQDNAEQF | 0.7923216 |
| GAFTTPQ | ASSQDNAEQF | 0.5878081 |
| GFGGFGTAGGVGGA | ASSQDNAEQF | 0.34180307 |
| GFLAQFL | ASSQDNAEQF | 0.42146537 |
| GGAGGAGGNAGVA | ASSQDNAEQF | 0.32863665 |
| GGVGTAGAGGAGGAGG | ASSQDNAEQF | 0.86878407 |
| GNGGGGDAGNAGSGGN | ASSQDNAEQF | 0.13207604 |
| GTGGAGGAGNDAGS | ASSQDNAEQF | 0.84547526 |
| GTTNTGWFNS | ASSQDNAEQF | 0.6214988 |
| IFMGPLFTL | ASSQDNAEQF | 0.35376272 |
| ILAHAIEAI | ASSQDNAEQF | 0.8049104 |
| IRSAIREAL | ASSQDNAEQF | 0.27094516 |
| ITFAGGG | ASSQDNAEQF | 0.71397233 |
| ITGMSGHQEV | ASSQDNAEQF | 0.7568761 |
| IVAGLACTL | ASSQDNAEQF | 0.8007116 |
| IVGGGLAGA | ASSQDNAEQF | 0.47362357 |
| IVTGVLNI | ASSQDNAEQF | 0.7762575 |
| LAQQGCI | ASSQDNAEQF | 0.7945814 |
| LCLHVEN | ASSQDNAEQF | 0.48950702 |
| LDAISDGS | ASSQDNAEQF | 0.7871392 |
| LGGFAAML | ASSQDNAEQF | 0.837402 |
| LGGKNPMI | ASSQDNAEQF | 0.7609767 |

| | | |
|---|---|---|
| LGPKGVAAIL | ASSQDNAEQF | 0.26879454 |
| LMVGVSI | ASSQDNAEQF | 0.74796295 |
| LVGAAFVL | ASSQDNAEQF | 0.5001666 |
| LVLAVAPL | ASSQDNAEQF | 0.7888174 |
| MAAIAVIDAA | ASSQDNAEQF | 0.68527544 |
| NNGINAELQ | ASSQDNAEQF | 0.5821876 |
| RGEPLVLVHG | ASSQDNAEQF | 0.7985089 |
| RGGAGGAAGLI | ASSQDNAEQF | 0.3136199 |
| RSYRMHPAV | ASSQDNAEQF | 0.60504925 |
| SAAHLLAGQLA | ASSQDNAEQF | 0.6063084 |
| SAAVGSY | ASSQDNAEQF | 0.6534679 |
| SPHAVLHRI | ASSQDNAEQF | 0.8092083 |
| SQIAKVL | ASSQDNAEQF | 0.40214336 |
| TALGHLVP | ASSQDNAEQF | 0.75747395 |
| TAPTTSGAA | ASSQDNAEQF | 0.5687804 |
| TDGDDGNGGSPGT | ASSQDNAEQF | 0.3262981 |
| TDGLGEN | ASSQDNAEQF | 0.72271705 |
| TGGPPLPGQA | ASSQDNAEQF | 0.60527354 |
| TIPAITI | ASSQDNAEQF | 0.36065573 |
| TIPGFTI | ASSQDNAEQF | 0.82290196 |
| TSIASGALA | ASSQDNAEQF | 0.36890462 |
| TVTLPGAAI | ASSQDNAEQF | 0.78709215 |
| VASGLSAALP | ASSQDNAEQF | 0.88320506 |
| VGAGVIGI | ASSQDNAEQF | 0.4791541 |
| VGPKRFGV | ASSQDNAEQF | 0.83528787 |
| VNVAFPD | ASSQDNAEQF | 0.74132264 |
| VSMTNSGVS | ASSQDNAEQF | 0.7802861 |
| VTDVLGDLSV | ASSQDNAEQF | 0.56357014 |
| VVVVGAGFAGL | ASSQDNAEQF | 0.52723616 |
| VYTVGTVV | ASSQDNAEQF | 0.87398124 |
| WTTFSGP | ASSQDNAEQF | 0.7693615 |
| YDELVEAF | ASSQDNAEQF | 0.8342747 |
| YGTGGAGGNGG | ASSQDNAEQF | 0.82306385 |
| YTNVMAV | ASSQDNAEQF | 0.8145206 |
| FLEQVHQL | ASSQDNAEQF | 0.8287691 |
| GLSLLAVAV | ASSQDNAEQF | 0.51730525 |
| LEQVHQL | ASSQDNAEQF | 0.24551146 |
| LNAAGAI | ASSQDNAEQF | 0.5481131 |
| PAVHASL | ASSQDNAEQF | 0.5868925 |
| TELEDDA | ASSQDNAEQF | 0.4430551 |
| TPPGPPN | ASSQDNAEQF | 0.28655532 |
| TPPPPPPP | ASSQDNAEQF | 0.83719045 |
| TQSPALI | ASSRTGGRNTLY | 0.8983587 |
| EFAAAFPQ | ASSRTGGRNTLY | 0.50327647 |
| LLSGNLDVL | ASSRTGGRNTLY | 0.89698935 |
| GPANLGNYNI | ASSRTGGRNTLY | 0.32956326 |
| TLGPTLP | ASSRTGGRNTLY | 0.78748214 |
| VYAYSSCCK | ASSRTGGRNTLY | 0.8397815 |
| APAPAPAPNI | ASSRTGGRNTLY | 0.3741197 |
| PGGPVPIP | ASSRTGGRNTLY | 0.7633704 |
| FLTVALEP | ASSRTGGRNTLY | 0.3952613 |

| | | |
|---|---|---|
| SQYFGANL | ASSRTGGRNTLY | 0.5158514 |
| GGVKPAA | ASSRTGGRNTLY | 0.85301614 |
| GGTGGAGGAGGW | ASSRTGGRNTLY | 0.52705455 |
| GVGGVAGLGGAGG | ASSRTGGRNTLY | 0.81727874 |
| VLLVLATTI | ASSRTGGRNTLY | 0.51024723 |
| IGIFAAML | ASSRTGGRNTLY | 0.38381758 |
| AVGDLID | ASSRTGGRNTLY | 0.7508701 |
| NISPLPL | ASSRTGGRNTLY | 0.8402089 |
| VTAVFVAI | ASSRTGGRNTLY | 0.7270265 |
| SGDDNDAAL | ASSRTGGRNTLY | 0.9006654 |
| AVAALAHS | ASSRTGGRNTLY | 0.5150506 |
| GGVTVMATP | ASSRTGGRNTLY | 0.8774637 |
| AGLAANVRD | ASSRTGGRNTLY | 0.8756056 |
| TNRPDLIDPALLRPG | ASSRTGGRNTLY | 0.7561458 |
| LANLGVTAL | ASSRTGGRNTLY | 0.5878285 |
| TGERIEL | ASSRTGGRNTLY | 0.82366556 |
| NGEELAA | ASSRTGGRNTLY | 0.52702516 |
| VLSLVWG | ASSRTGGRNTLY | 0.84222627 |
| IGYALWL | ASSRTGGRNTLY | 0.8669874 |
| LSLLLAPL | ASSRTGGRNTLY | 0.30171072 |
| TLDEALK | ASSRTGGRNTLY | 0.8182963 |
| TDGAGTAEGG | ASSRTGGRNTLY | 0.7548698 |
| NAPGGAGGNGG | ASSRTGGRNTLY | 0.7511016 |
| VSVFMGMGGNF | ASSRTGGRNTLY | 0.41817817 |
| LASYLAAL | ASSRTGGRNTLY | 0.27458858 |
| LVGGTPAL | ASSRTGGRNTLY | 0.64200664 |
| VRNADVL | ASSRTGGRNTLY | 0.3119006 |
| TCSSPFLVSAM | ASSRTGGRNTLY | 0.86667335 |
| VHITDAE | ASSRTGGRNTLY | 0.78385425 |
| EYAAPSF | ASSRTGGRNTLY | 0.46488938 |
| VVDAFHGGL | ASSRTGGRNTLY | 0.7971051 |
| ASVVDPQ | ASSRTGGRNTLY | 0.8490685 |
| SYNEFGGLS | ASSRTGGRNTLY | 0.2287761 |
| DPDGSVY | ASSRTGGRNTLY | 0.6343782 |
| MVSTIHATE | ASSRTGGRNTLY | 0.63452506 |
| LAAGVADPV | ASSRTGGRNTLY | 0.5290935 |
| TKEQIAA | ASSRTGGRNTLY | 0.23107177 |
| SLASPAAL | ASSRTGGRNTLY | 0.76114297 |
| IGLVFYGV | ASSRTGGRNTLY | 0.54806197 |
| FLLDASNL | ASSRTGGRNTLY | 0.17261285 |
| YQVGNFP | ASSRTGGRNTLY | 0.7527441 |
| ESLLSQST | ASSRTGGRNTLY | 0.5057008 |
| LFTVGFAAVG | ASSRTGGRNTLY | 0.4640545 |
| ATPPTPT | ASSRTGGRNTLY | 0.8039954 |
| TDLATLG | ASSRTGGRNTLY | 0.72227764 |
| SAGGTGR | ASSRTGGRNTLY | 0.48401254 |
| GFSNSGS | ASSRTGGRNTLY | 0.7687485 |
| GGNGGSGGTSV | ASSRTGGRNTLY | 0.8299494 |
| GTSVATGGAGN | ASSRTGGRNTLY | 0.36238444 |
| NGGNGGSGGTS | ASSRTGGRNTLY | 0.716455 |
| AVVKFPH | ASSRTGGRNTLY | 0.8680258 |

| | | |
|---|---|---|
| TPPPPPP | ASSRTGGRNTLY | 0.63954055 |
| AAPAAAAAPA | ASSRTGGRNTLY | 0.864166 |
| IALNEDL | ASSRTGGRNTLY | 0.87274456 |
| IVNVSSM | ASSRTGGRNTLY | 0.62794477 |
| TGIPPAPR | ASSRTGGRNTLY | 0.81527376 |
| FDLGGGTF | ASSRTGGRNTLY | 0.84676546 |
| FDLGGGTFDVS | ASSRTGGRNTLY | 0.46088326 |
| NEPTAAA | ASSRTGGRNTLY | 0.6536779 |
| NEPTAAALA | ASSRTGGRNTLY | 0.304811 |
| GSIINISSI | ASSRTGGRNTLY | 0.74391973 |
| VAPGNAGTA | ASSRTGGRNTLY | 0.3199814 |
| TGGGASLE | ASSRTGGRNTLY | 0.7646992 |
| VAVYNHYKRI | ASSRTGGRNTLY | 0.8473184 |
| TDPAPATT | ASSRTGGRNTLY | 0.619044 |
| TNRPDILDPALLRPG | ASSRTGGRNTLY | 0.26690054 |
| LTTTTTS | ASSRTGGRNTLY | 0.3418563 |
| PTGSGKTL | ASSRTGGRNTLY | 0.67320234 |
| TGSGKTL | ASSRTGGRNTLY | 0.3511079 |
| TIGHVDHGK | ASSRTGGRNTLY | 0.3244483 |
| TIGHVDHGKTT | ASSRTGGRNTLY | 0.4541487 |
| GHVDHGKTT | ASSRTGGRNTLY | 0.81277156 |
| AQFEHTLL | ASSRTGGRNTLY | 0.8139441 |
| IVNQDEL | ASSRTGGRNTLY | 0.4155987 |
| TNGSQFF | ASSRTGGRNTLY | 0.8468915 |
| TGKGIVE | ASSRTGGRNTLY | 0.87536824 |
| LISNASDA | ASSRTGGRNTLY | 0.87247705 |
| AELRTLGELTKIAKAHGAQ | ASSRTGGRNTLY | 0.69610727 |
| APNAVAAAEPSW | ASSRTGGRNTLY | 0.26407185 |
| APQVAGAATAVL | ASSRTGGRNTLY | 0.85534656 |
| ARRDPAAAAAILPTDA | ASSRTGGRNTLY | 0.5455051 |
| DAAARDVLAEAGLAEYF | ASSRTGGRNTLY | 0.8166399 |
| DDQIVLSAKANLAKVEQ | ASSRTGGRNTLY | 0.81666464 |
| DELLPDTGEDSDEDGH | ASSRTGGRNTLY | 0.5722297 |
| ELPEAPANRATVPQD | ASSRTGGRNTLY | 0.4183671 |
| FGQSKIERVTGIG | ASSRTGGRNTLY | 0.79209065 |
| GAGGTGSTAGGAGGAGGAGGLY | ASSRTGGRNTLY | 0.8668723 |
| GHGGTGAAGGNGGTGGAGGSGI | ASSRTGGRNTLY | 0.4258728 |
| GNGGNGGAGGEGGPGGAGGAGA | ASSRTGGRNTLY | 0.2580983 |
| GSTAGNGGNGGNATNGTIAGQP | ASSRTGGRNTLY | 0.72016054 |
| GTGGDGGNGGNGGTGG | ASSRTGGRNTLY | 0.66756976 |
| GVRRPAGIPADAPV | ASSRTGGRNTLY | 0.82885844 |
| GYRQVTDAHLLTLARR | ASSRTGGRNTLY | 0.73079157 |
| IAKMGQDGHDRG | ASSRTGGRNTLY | 0.6929816 |
| KAEPSIEIQYT | ASSRTGGRNTLY | 0.778859 |
| KPVRIRMAADEDTVAVT | ASSRTGGRNTLY | 0.7436732 |
| MPPRIAGMRLLVIKPEP | ASSRTGGRNTLY | 0.6961249 |
| MKFYAVALFRSRGVPPTR | ASSRTGGRNTLY | 0.8459479 |
| NAGGFGILGAVAHSP | ASSRTGGRNTLY | 0.3788572 |
| NEGREITLAMAEM | ASSRTGGRNTLY | 0.8593254 |
| NELVEWSHPLGHRWRWE | ASSRTGGRNTLY | 0.81611913 |
| NLGAGGTGGNGGNPG | ASSRTGGRNTLY | 0.8141524 |

| | | |
|---|---|---|
| NTEWPKWETVMAADPVD | ASSRTGGRNTLY | 0.86168945 |
| PASAAPAETQP | ASSRTGGRNTLY | 0.6362526 |
| QQSSDGKNGVRAAEITGE | ASSRTGGRNTLY | 0.747092 |
| RAPDAVTTSWWKE | ASSRTGGRNTLY | 0.30582446 |
| SGFGNVGAHTSGWFNQ | ASSRTGGRNTLY | 0.67020905 |
| SGVAGAVGQAASVGGLKV | ASSRTGGRNTLY | 0.7359478 |
| SNGGAGTGSAGGNGGT | ASSRTGGRNTLY | 0.7490468 |
| TGGAGGEGGAAG | ASSRTGGRNTLY | 0.82204837 |
| TGGDGGAAGAGGAPGHG | ASSRTGGRNTLY | 0.81854415 |
| TMPPLLAAIANAGRHGV | ASSRTGGRNTLY | 0.5324105 |
| TSSTAGNSGGA | ASSRTGGRNTLY | 0.87796557 |
| VPATVPVGILC | ASSRTGGRNTLY | 0.85233104 |
| VQRPDSGGVVLTGRIS | ASSRTGGRNTLY | 0.33123434 |
| WTPNPVPATKGVPAGLVQAT | ASSRTGGRNTLY | 0.54725283 |
| YALKRALAKFSDRFDVVII | ASSRTGGRNTLY | 0.8173934 |
| YLIGSVIV | ASSRTGGRNTLY | 0.8455225 |
| YGQLVAVID | ASSRTGGRNTLY | 0.5914608 |
| VTVGGMTLPA | ASSRTGGRNTLY | 0.6782875 |
| VLALPAV | ASSRTGGRNTLY | 0.8783729 |
| VIPPTLNYET | ASSRTGGRNTLY | 0.7522025 |
| VGMGGMHPSSG | ASSRTGGRNTLY | 0.8589659 |
| VGLAGGAATA | ASSRTGGRNTLY | 0.22109497 |
| VGGAGGAGGSGP | ASSRTGGRNTLY | 0.6072699 |
| VFAAFMV | ASSRTGGRNTLY | 0.58151066 |
| TLYDKLR | ASSRTGGRNTLY | 0.13419129 |
| TLAVFTVL | ASSRTGGRNTLY | 0.7989179 |
| TGSVQPARV | ASSRTGGRNTLY | 0.5192267 |
| TGGGAGSGGT | ASSRTGGRNTLY | 0.80186063 |
| SHLETSL | ASSRTGGRNTLY | 0.5213348 |
| SFTTGMT | ASSRTGGRNTLY | 0.81482583 |
| QVEVAKRMGVRQPTVS | ASSRTGGRNTLY | 0.76454353 |
| QGSGGPALAM | ASSRTGGRNTLY | 0.72469807 |
| MLAPSNP | ASSRTGGRNTLY | 0.34187838 |
| LVNNAAI | ASSRTGGRNTLY | 0.70069236 |
| LTGEINTLA | ASSRTGGRNTLY | 0.87683666 |
| LSIFGPV | ASSRTGGRNTLY | 0.8541049 |
| LLALLANL | ASSRTGGRNTLY | 0.2497122 |
| LISNASDALDKLR | ASSRTGGRNTLY | 0.8163926 |
| LINEAVP | ASSRTGGRNTLY | 0.56548375 |
| LGYAFSGL | ASSRTGGRNTLY | 0.21478324 |
| LDFVAVAL | ASSRTGGRNTLY | 0.23375264 |
| LAQPGAE | ASSRTGGRNTLY | 0.52441776 |
| ISNLGAVTGF | ASSRTGGRNTLY | 0.56263006 |
| HQEDPGRI | ASSRTGGRNTLY | 0.7558293 |
| GTPGEPTP | ASSRTGGRNTLY | 0.80569893 |
| GFFNSGS | ASSRTGGRNTLY | 0.48049623 |
| GAGGGGLTAGAN | ASSRTGGRNTLY | 0.8074316 |
| FVDALVNM | ASSRTGGRNTLY | 0.6509205 |
| FLTVAPEL | ASSRTGGRNTLY | 0.08752178 |
| FETLTAL | ASSRTGGRNTLY | 0.49133745 |
| DVINPIEL | ASSRTGGRNTLY | 0.23401415 |

| | | |
|---|---|---|
| APVSGAPV | ASSRTGGRNTLY | 0.87251186 |
| ALVGGLAW | ASSRTGGRNTLY | 0.3223894 |
| ALAAAAATAI | ASSRTGGRNTLY | 0.59211826 |
| AFGLGLLL | ASSRTGGRNTLY | 0.5952136 |
| AFFAAASL | ASSRTGGRNTLY | 0.8226893 |
| AAAALPAFSPP | ASSRTGGRNTLY | 0.7374519 |
| VVLAPAL | ASSRTGGRNTLY | 0.47120264 |
| TIGHVDHGKTTL | ASSRTGGRNTLY | 0.32059535 |
| LISNASDALDK | ASSRTGGRNTLY | 0.7344067 |
| LISNASDAL | ASSRTGGRNTLY | 0.64068377 |
| ISNASDALDK | ASSRTGGRNTLY | 0.4028167 |
| NQAETLVYQTE | ASSRTGGRNTLY | 0.5313336 |
| TGGTGGTAGAG | ASSRTGGRNTLY | 0.81723195 |
| TGSTAGGAGGA | ASSRTGGRNTLY | 0.22255065 |
| DPLAVPVAAGPV | ASSRTGGRNTLY | 0.8500053 |
| GGTGGTPGNGGN | ASSRTGGRNTLY | 0.37329373 |
| PAGLAGVDPATL | ASSRTGGRNTLY | 0.5768964 |
| GNNLVVEAPHAGASV | ASSRTGGRNTLY | 0.86245155 |
| KVTRIMSASTLEQPA | ASSRTGGRNTLY | 0.7930273 |
| LGFALPIALPRRTVV | ASSRTGGRNTLY | 0.6314316 |
| TRLGNLTARKLVVLV | ASSRTGGRNTLY | 0.36471802 |
| WSQQRRLIYFPSAGP | ASSRTGGRNTLY | 0.32113105 |
| GPNGNPDPMAAAVDIR | ASSRTGGRNTLY | 0.38267726 |
| TQRCVSVADLRAAASR | ASSRTGGRNTLY | 0.41036713 |
| QFVLDVAEHPVEVLTETV | ASSRTGGRNTLY | 0.8614255 |
| GGAGGTGGAGGNGGTQAGDG | ASSRTGGRNTLY | 0.68507487 |
| PLLATLRAIAVDVDAKPAQV | ASSRTGGRNTLY | 0.31253797 |
| APKFRTGGYMIDFWGVGYQVAK | ASSRTGGRNTLY | 0.77790636 |
| GAGGAGGAGGAAGGPGGTGGTG | ASSRTGGRNTLY | 0.8635744 |
| IGALGLLATDVAPAGSVAFGVA | ASSRTGGRNTLY | 0.8688208 |
| AGGAGGIGGNGGAGGAGGVLMG | ASSRTGGRNTLY | 0.36846155 |
| ADELIGLL | ASSRTGGRNTLY | 0.71551716 |
| ADRALKA | ASSRTGGRNTLY | 0.6540084 |
| AGAVGQVGGAGGA | ASSRTGGRNTLY | 0.646211 |
| AGGGGGLGGAP | ASSRTGGRNTLY | 0.52938306 |
| AGGLLYGNGGAG | ASSRTGGRNTLY | 0.33453226 |
| AGLESIGL | ASSRTGGRNTLY | 0.8381394 |
| AGLKTGII | ASSRTGGRNTLY | 0.89757735 |
| ASLAALLL | ASSRTGGRNTLY | 0.5716434 |
| ASVINISSI | ASSRTGGRNTLY | 0.23311843 |
| ATPPGPA | ASSRTGGRNTLY | 0.6090932 |
| ATVASALV | ASSRTGGRNTLY | 0.4221378 |
| ATVASLIL | ASSRTGGRNTLY | 0.41182274 |
| AVNGPVLIRH | ASSRTGGRNTLY | 0.5768199 |
| DGAAGAVGI | ASSRTGGRNTLY | 0.22105478 |
| DPLLHGI | ASSRTGGRNTLY | 0.62568486 |
| ELGAALGAL | ASSRTGGRNTLY | 0.86373925 |
| ELGRTQAV | ASSRTGGRNTLY | 0.6471348 |
| FVAGGYLS | ASSRTGGRNTLY | 0.72690487 |
| GASGGAGGAGGW | ASSRTGGRNTLY | 0.83439887 |
| GGAGAGGDGGAGGI | ASSRTGGRNTLY | 0.77277213 |

| | | |
|---|---|---|
| GTGHNGSQPGMGGNG | ASSRTGGRNTLY | 0.8411044 |
| GYAAASASAS | ASSRTGGRNTLY | 0.8673591 |
| HQVDVAAVQQG | ASSRTGGRNTLY | 0.6189873 |
| IGAAIAHT | ASSRTGGRNTLY | 0.7256069 |
| IGMIEVAQL | ASSRTGGRNTLY | 0.80616355 |
| IRLAPEHTSAMD | ASSRTGGRNTLY | 0.5961583 |
| IYNVSVL | ASSRTGGRNTLY | 0.69137913 |
| KQGFLGM | ASSRTGGRNTLY | 0.6436854 |
| LESVITQL | ASSRTGGRNTLY | 0.7883994 |
| LGLGALVVV | ASSRTGGRNTLY | 0.6390398 |
| LIGNGGNGGSGGI | ASSRTGGRNTLY | 0.8771032 |
| LNAINAPTL | ASSRTGGRNTLY | 0.17821878 |
| LQLLNTGV | ASSRTGGRNTLY | 0.34873712 |
| LQTSIEF | ASSRTGGRNTLY | 0.83507025 |
| MSFAVSEL | ASSRTGGRNTLY | 0.14998943 |
| NTGALGL | ASSRTGGRNTLY | 0.27237242 |
| PPGAGGAGGAGTT | ASSRTGGRNTLY | 0.73326147 |
| QLVGLGI | ASSRTGGRNTLY | 0.85276973 |
| RFLEQVHQL | ASSRTGGRNTLY | 0.13915278 |
| RGVGYVLR | ASSRTGGRNTLY | 0.61375636 |
| SATGVFAGVI | ASSRTGGRNTLY | 0.2684041 |
| SLPAITLP | ASSRTGGRNTLY | 0.81461203 |
| STALAFTV | ASSRTGGRNTLY | 0.55436873 |
| SVLTVGLGK | ASSRTGGRNTLY | 0.24512702 |
| TALAGAGSGI | ASSRTGGRNTLY | 0.79189956 |
| TGLPFTGI | ASSRTGGRNTLY | 0.35876352 |
| VAEHVTV | ASSRTGGRNTLY | 0.6811391 |
| VDIVLNSL | ASSRTGGRNTLY | 0.8080133 |
| VLTGGGAVL | ASSRTGGRNTLY | 0.8817084 |
| VNVGLAYGI | ASSRTGGRNTLY | 0.29446495 |
| VSSAAVGAV | ASSRTGGRNTLY | 0.76690054 |
| VTSVYEAGV | ASSRTGGRNTLY | 0.6200218 |
| WVALDRGA | ASSRTGGRNTLY | 0.8851568 |
| YLVGSVVI | ASSRTGGRNTLY | 0.5307134 |
| GPPGTGKTLL | ASSRTGGRNTLY | 0.24047382 |
| GVGMTKF | ASSRTGGRNTLY | 0.8219867 |
| IGIGGSDL | ASSRTGGRNTLY | 0.7708944 |
| IGTLLAEL | ASSRTGGRNTLY | 0.6141094 |
| IYTSGTT | ASSRTGGRNTLY | 0.84821427 |
| LSFPAGV | ASSRTGGRNTLY | 0.58146894 |
| LYTSGTT | ASSRTGGRNTLY | 0.568044 |
| VGAFPFGF | ASSRTGGRNTLY | 0.7816262 |
| VGAGTVEF | ASSRTGGRNTLY | 0.80829334 |
| VGKSTLLL | ASSRTGGRNTLY | 0.36846307 |
| AALAHAGGGIV | ASSRTGGRNTLY | 0.8171851 |
| AALVAVSLL | ASSRTGGRNTLY | 0.64444816 |
| AAVSRAVAAL | ASSRTGGRNTLY | 0.42374706 |
| AGGSSGIGGF | ASSRTGGRNTLY | 0.8676249 |
| AGLTVANANG | ASSRTGGRNTLY | 0.67927 |
| AGLVGAVL | ASSRTGGRNTLY | 0.7951584 |
| ALSQAFTV | ASSRTGGRNTLY | 0.74624133 |

| | | |
|---|---|---|
| ALTFSVSL | ASSRTGGRNTLY | 0.8881691 |
| ALTGAALVAL | ASSRTGGRNTLY | 0.42143583 |
| ASFSIGAGVL | ASSRTGGRNTLY | 0.805813 |
| DGSVYKI | ASSRTGGRNTLY | 0.7316923 |
| EQLAEIV | ASSRTGGRNTLY | 0.83148634 |
| FAVSELTVM | ASSRTGGRNTLY | 0.7393407 |
| GAAPLAAGL | ASSRTGGRNTLY | 0.7376118 |
| GADGGDGGSGGAGG | ASSRTGGRNTLY | 0.7794548 |
| GFGLDTGGAGG | ASSRTGGRNTLY | 0.8397016 |
| GLASGGLL | ASSRTGGRNTLY | 0.7243442 |
| GRIAGAAVLL | ASSRTGGRNTLY | 0.79627776 |
| GSAGSLNTSF | ASSRTGGRNTLY | 0.8594619 |
| GTGQIGI | ASSRTGGRNTLY | 0.34803182 |
| GTLIALIL | ASSRTGGRNTLY | 0.6367036 |
| IAATMGFVVG | ASSRTGGRNTLY | 0.83669376 |
| IFAQALL | ASSRTGGRNTLY | 0.8600956 |
| IGAAVALGF | ASSRTGGRNTLY | 0.6099821 |
| IGLTPLAL | ASSRTGGRNTLY | 0.86309993 |
| ILGSNTSTL | ASSRTGGRNTLY | 0.6017388 |
| IVFSLTVL | ASSRTGGRNTLY | 0.7599968 |
| LASKATGF | ASSRTGGRNTLY | 0.7005267 |
| LASVALIL | ASSRTGGRNTLY | 0.5876866 |
| LFAVGLTL | ASSRTGGRNTLY | 0.2846391 |
| LFNGGIGTYI | ASSRTGGRNTLY | 0.6787558 |
| LTLGAGII | ASSRTGGRNTLY | 0.81019884 |
| LVAAALGDI | ASSRTGGRNTLY | 0.74975723 |
| NTAFVQLGI | ASSRTGGRNTLY | 0.2711353 |
| QLFSGAEL | ASSRTGGRNTLY | 0.70684415 |
| SAATFITL | ASSRTGGRNTLY | 0.77879643 |
| SNYGLANLGV | ASSRTGGRNTLY | 0.8859298 |
| SYGGAYAVM | ASSRTGGRNTLY | 0.69030106 |
| TTGVGGLTLG | ASSRTGGRNTLY | 0.82573783 |
| TTQTIYI | ASSRTGGRNTLY | 0.59748256 |
| VAAGAGLGAVI | ASSRTGGRNTLY | 0.6650742 |
| VAAVVTLL | ASSRTGGRNTLY | 0.31808725 |
| VGFLGGLI | ASSRTGGRNTLY | 0.4754738 |
| VGVIPILL | ASSRTGGRNTLY | 0.681846 |
| VIFASSLI | ASSRTGGRNTLY | 0.37200367 |
| VKGGDGGAAAG | ASSRTGGRNTLY | 0.8188875 |
| VTKTFTNI | ASSRTGGRNTLY | 0.8639835 |
| VVAAAILL | ASSRTGGRNTLY | 0.8027267 |
| YAALDVEL | ASSRTGGRNTLY | 0.843951 |
| LASVALLL | ASSRTGGRNTLY | 0.84761465 |
| LDAVHRV | ASSRTGGRNTLY | 0.83584684 |
| QGLSSSIF | ASSRTGGRNTLY | 0.41577023 |
| RGTFANIRL | ASSRTGGRNTLY | 0.6401276 |
| RQLWWGHRI | ASSRTGGRNTLY | 0.19867155 |
| YGPPGTGKTLL | ASSRTGGRNTLY | 0.51464427 |
| AALVGAPANVA | ASSRTGGRNTLY | 0.43121305 |
| AATFRRTE | ASSRTGGRNTLY | 0.6651475 |
| AGALGGGI | ASSRTGGRNTLY | 0.11015919 |

| | | |
|---|---|---|
| AGGAGGAGGIGAG | ASSRTGGRNTLY | 0.8000984 |
| AGSGALGL | ASSRTGGRNTLY | 0.8568343 |
| AQIADQI | ASSRTGGRNTLY | 0.6407229 |
| ASAVAVTYL | ASSRTGGRNTLY | 0.7989186 |
| ASTVLGGTGG | ASSRTGGRNTLY | 0.7710612 |
| AVGGEPAK | ASSRTGGRNTLY | 0.5173437 |
| CEQGTFGGTC | ASSRTGGRNTLY | 0.80067575 |
| DEGKAPAG | ASSRTGGRNTLY | 0.5233078 |
| DTVVFPAL | ASSRTGGRNTLY | 0.8281962 |
| FEGITMAG | ASSRTGGRNTLY | 0.82376313 |
| FPLVVQD | ASSRTGGRNTLY | 0.7578288 |
| GAFTTPQ | ASSRTGGRNTLY | 0.777194 |
| GFGGFGTAGGVGGA | ASSRTGGRNTLY | 0.7560931 |
| GFLAQFL | ASSRTGGRNTLY | 0.45406595 |
| GGAGGAGGNAGVA | ASSRTGGRNTLY | 0.7934027 |
| GGVGTAGAGGAGGAGG | ASSRTGGRNTLY | 0.5918672 |
| GNGGGGDAGNAGSGGN | ASSRTGGRNTLY | 0.7110516 |
| GTGGAGGAGNDAGS | ASSRTGGRNTLY | 0.37819612 |
| GTTNTGWFNS | ASSRTGGRNTLY | 0.74614215 |
| IFMGPLFTL | ASSRTGGRNTLY | 0.6228108 |
| ILAHAIEAI | ASSRTGGRNTLY | 0.8035329 |
| IRSAIREAL | ASSRTGGRNTLY | 0.47247672 |
| ITFAGGG | ASSRTGGRNTLY | 0.8104906 |
| ITGMSGHQEV | ASSRTGGRNTLY | 0.81078184 |
| IVAGLACTL | ASSRTGGRNTLY | 0.28953767 |
| IVGGGLAGA | ASSRTGGRNTLY | 0.83772093 |
| IVTGVLNI | ASSRTGGRNTLY | 0.6878878 |
| LAQQGCI | ASSRTGGRNTLY | 0.57834876 |
| LCLHVEN | ASSRTGGRNTLY | 0.82168204 |
| LDAISDGS | ASSRTGGRNTLY | 0.61754507 |
| LGGFAAML | ASSRTGGRNTLY | 0.3031453 |
| LGGKNPMI | ASSRTGGRNTLY | 0.59911335 |
| LGPKGVAAIL | ASSRTGGRNTLY | 0.7273081 |
| LMVGVSI | ASSRTGGRNTLY | 0.54537857 |
| LVGAAFVL | ASSRTGGRNTLY | 0.19040608 |
| LVLAVAPL | ASSRTGGRNTLY | 0.74291515 |
| MAAIAVIDAA | ASSRTGGRNTLY | 0.42621124 |
| NNGINAELQ | ASSRTGGRNTLY | 0.43911892 |
| RGEPLVLVHG | ASSRTGGRNTLY | 0.45260698 |
| RGGAGGAAGLI | ASSRTGGRNTLY | 0.8120126 |
| RSYRMHPAV | ASSRTGGRNTLY | 0.7888165 |
| SAAHLLAGQLA | ASSRTGGRNTLY | 0.56042504 |
| SAAVGSY | ASSRTGGRNTLY | 0.79930794 |
| SPHAVLHRI | ASSRTGGRNTLY | 0.726163 |
| SQIAKVL | ASSRTGGRNTLY | 0.8216663 |
| TALGHLVP | ASSRTGGRNTLY | 0.64404535 |
| TAPTTSGAA | ASSRTGGRNTLY | 0.735216 |
| TDGDDGNGGSPGT | ASSRTGGRNTLY | 0.81325835 |
| TDGLGEN | ASSRTGGRNTLY | 0.5959463 |
| TGGPPLPGQA | ASSRTGGRNTLY | 0.5067887 |
| TIPAITI | ASSRTGGRNTLY | 0.5307025 |

| | | |
|---|---|---|
| TIPGFTI | ASSRTGGRNTLY | 0.56018084 |
| TSIASGALA | ASSRTGGRNTLY | 0.49161422 |
| TVTLPGAAI | ASSRTGGRNTLY | 0.7244035 |
| VASGLSAALP | ASSRTGGRNTLY | 0.86590743 |
| VGAGVIGI | ASSRTGGRNTLY | 0.72159517 |
| VGPKRFGV | ASSRTGGRNTLY | 0.26164883 |
| VNVAFPD | ASSRTGGRNTLY | 0.6053066 |
| VSMTNSGVS | ASSRTGGRNTLY | 0.60206497 |
| VTDVLGDLSV | ASSRTGGRNTLY | 0.7831651 |
| VVVVGAGFAGL | ASSRTGGRNTLY | 0.84168047 |
| VYTVGTVV | ASSRTGGRNTLY | 0.8855212 |
| WTTFSGP | ASSRTGGRNTLY | 0.7857032 |
| YDELVEAF | ASSRTGGRNTLY | 0.4541359 |
| YGTGGAGGNGG | ASSRTGGRNTLY | 0.7913526 |
| YTNVMAV | ASSRTGGRNTLY | 0.589493 |
| FLEQVHQL | ASSRTGGRNTLY | 0.76574254 |
| GLSLLAVAV | ASSRTGGRNTLY | 0.71159005 |
| LEQVHQL | ASSRTGGRNTLY | 0.79451686 |
| LNAAGAI | ASSRTGGRNTLY | 0.5404943 |
| PAVHASL | ASSRTGGRNTLY | 0.6108998 |
| TELEDDA | ASSRTGGRNTLY | 0.2673092 |
| TPPGPPN | ASSRTGGRNTLY | 0.84254146 |
| TPPPPPPP | ASSRTGGRNTLY | 0.47078836 |
| TQSPALI | ASSQDYGREQY | 0.52810353 |
| EFAAAFPQ | ASSQDYGREQY | 0.6228421 |
| LLSGNLDVL | ASSQDYGREQY | 0.8580601 |
| GPANLGNYNI | ASSQDYGREQY | 0.8664322 |
| TLGPTLP | ASSQDYGREQY | 0.5544262 |
| VYAYSSCCK | ASSQDYGREQY | 0.39103362 |
| APAPAPAPNI | ASSQDYGREQY | 0.8154828 |
| PGGPVPIP | ASSQDYGREQY | 0.43607906 |
| FLTVALEP | ASSQDYGREQY | 0.7760502 |
| SQYFGANL | ASSQDYGREQY | 0.67645746 |
| GGVKPAA | ASSQDYGREQY | 0.57597303 |
| GGTGGAGGAGGW | ASSQDYGREQY | 0.6081817 |
| GVGGVAGLGGAGG | ASSQDYGREQY | 0.7914099 |
| VLLVLATTI | ASSQDYGREQY | 0.4523591 |
| IGIFAAML | ASSQDYGREQY | 0.81809664 |
| AVGDLID | ASSQDYGREQY | 0.17976263 |
| NISPLPL | ASSQDYGREQY | 0.7870046 |
| VTAVFVAI | ASSQDYGREQY | 0.3882385 |
| SGDDNDAAL | ASSQDYGREQY | 0.41654438 |
| AVAALAHS | ASSQDYGREQY | 0.20457965 |
| GGVTVMATP | ASSQDYGREQY | 0.6814742 |
| AGLAANVRD | ASSQDYGREQY | 0.79643726 |
| TNRPDLIDPALLRPG | ASSQDYGREQY | 0.84343684 |
| LANLGVTAL | ASSQDYGREQY | 0.7704316 |
| TGERIEL | ASSQDYGREQY | 0.80245245 |
| NGEELAA | ASSQDYGREQY | 0.85617524 |
| VLSLVWG | ASSQDYGREQY | 0.51157105 |
| IGYALWL | ASSQDYGREQY | 0.20581824 |

| | | |
|---|---|---|
| LSLLLAPL | ASSQDYGREQY | 0.64694947 |
| TLDEALK | ASSQDYGREQY | 0.77836025 |
| TDGAGTAEGG | ASSQDYGREQY | 0.8372169 |
| NAPGGAGGNGG | ASSQDYGREQY | 0.8301199 |
| VSVFMGMGGNF | ASSQDYGREQY | 0.22719857 |
| LASYLAAL | ASSQDYGREQY | 0.78415036 |
| LVGGTPAL | ASSQDYGREQY | 0.75575364 |
| VRNADVL | ASSQDYGREQY | 0.3878159 |
| TCSSPFLVSAM | ASSQDYGREQY | 0.5500901 |
| VHITDAE | ASSQDYGREQY | 0.43367356 |
| EYAAPSF | ASSQDYGREQY | 0.42013466 |
| VVDAFHGGL | ASSQDYGREQY | 0.7756323 |
| ASVVDPQ | ASSQDYGREQY | 0.5426257 |
| SYNEFGGLS | ASSQDYGREQY | 0.63285017 |
| DPDGSVY | ASSQDYGREQY | 0.55588025 |
| MVSTIHATE | ASSQDYGREQY | 0.46219492 |
| LAAGVADPV | ASSQDYGREQY | 0.5460401 |
| TKEQIAA | ASSQDYGREQY | 0.54768705 |
| SLASPAAL | ASSQDYGREQY | 0.7118615 |
| IGLVFYGV | ASSQDYGREQY | 0.80662113 |
| FLLDASNL | ASSQDYGREQY | 0.6229451 |
| YQVGNFP | ASSQDYGREQY | 0.7731577 |
| ESLLSQST | ASSQDYGREQY | 0.8255446 |
| LFTVGFAAVG | ASSQDYGREQY | 0.25843117 |
| ATPPTPT | ASSQDYGREQY | 0.43065912 |
| TDLATLG | ASSQDYGREQY | 0.7622927 |
| SAGGTGR | ASSQDYGREQY | 0.55065596 |
| GFSNSGS | ASSQDYGREQY | 0.8487927 |
| GGNGGSGGTSV | ASSQDYGREQY | 0.6660674 |
| GTSVATGGAGN | ASSQDYGREQY | 0.7026076 |
| NGGNGGSGGTS | ASSQDYGREQY | 0.7501818 |
| AVVKFPH | ASSQDYGREQY | 0.8690431 |
| TPPPPPP | ASSQDYGREQY | 0.7351092 |
| AAPAAAAAPA | ASSQDYGREQY | 0.60947424 |
| IALNEDL | ASSQDYGREQY | 0.85036874 |
| IVNVSSM | ASSQDYGREQY | 0.34080413 |
| TGIPPAPR | ASSQDYGREQY | 0.8304018 |
| FDLGGGTF | ASSQDYGREQY | 0.8070787 |
| FDLGGGTFDVS | ASSQDYGREQY | 0.32147652 |
| NEPTAAA | ASSQDYGREQY | 0.58183604 |
| NEPTAAALA | ASSQDYGREQY | 0.8762429 |
| GSIINISSI | ASSQDYGREQY | 0.8167969 |
| VAPGNAGTA | ASSQDYGREQY | 0.58922744 |
| TGGGASLE | ASSQDYGREQY | 0.44465834 |
| VAVYNHYKRI | ASSQDYGREQY | 0.5018085 |
| TDPAPATT | ASSQDYGREQY | 0.5986641 |
| TNRPDILDPALLRPG | ASSQDYGREQY | 0.25356722 |
| LTTTTTS | ASSQDYGREQY | 0.7989594 |
| PTGSGKTL | ASSQDYGREQY | 0.5689719 |
| TGSGKTL | ASSQDYGREQY | 0.86353683 |
| TIGHVDHGK | ASSQDYGREQY | 0.70622945 |

| | | |
|---|---|---|
| TIGHVDHGKTT | ASSQDYGREQY | 0.81173503 |
| GHVDHGKTT | ASSQDYGREQY | 0.57604307 |
| AQFEHTLL | ASSQDYGREQY | 0.53586376 |
| IVNQDEL | ASSQDYGREQY | 0.23291919 |
| TNGSQFF | ASSQDYGREQY | 0.4766231 |
| TGKGIVE | ASSQDYGREQY | 0.5601295 |
| LISNASDA | ASSQDYGREQY | 0.6413741 |
| AELRTLGELTKIAKAHGAQ | ASSQDYGREQY | 0.19469792 |
| APNAVAAAEPSW | ASSQDYGREQY | 0.8398744 |
| APQVAGAATAVL | ASSQDYGREQY | 0.5839449 |
| ARRDPAAAAAILPTDA | ASSQDYGREQY | 0.73377776 |
| DAAARDVLAEAGLAEYF | ASSQDYGREQY | 0.5120102 |
| DDQIVLSAKANLAKVEQ | ASSQDYGREQY | 0.44113487 |
| DELLPDTGEDSDEDGH | ASSQDYGREQY | 0.38817275 |
| ELPEAPANRATVPQD | ASSQDYGREQY | 0.8101262 |
| FGQSKIERVTGIG | ASSQDYGREQY | 0.2671966 |
| GAGGTGSTAGGAGGAGGAGGLY | ASSQDYGREQY | 0.23617318 |
| GHGGTGAAGGNGGTGGAGGSGI | ASSQDYGREQY | 0.4867027 |
| GNGGNGGAGGEGGPGGAGGAGA | ASSQDYGREQY | 0.67321897 |
| GSTAGNGGNGGNATNGTIAGQP | ASSQDYGREQY | 0.8697142 |
| GTGGDGGNGGNGGTGG | ASSQDYGREQY | 0.8354552 |
| GVRRPAGIPADAPV | ASSQDYGREQY | 0.6844242 |
| GYRQVTDAHLLTLARR | ASSQDYGREQY | 0.87240314 |
| IAKMGQDGHDRG | ASSQDYGREQY | 0.128325 |
| KAEPSIEIQYT | ASSQDYGREQY | 0.546993 |
| KPVRIRMAADEDTVAVT | ASSQDYGREQY | 0.86361974 |
| MPPRIAGMRLLVIKPEP | ASSQDYGREQY | 0.7658715 |
| MKFYAVALFRSRGVPPTR | ASSQDYGREQY | 0.61057323 |
| NAGGFGILGAVAHSP | ASSQDYGREQY | 0.8212187 |
| NEGREITLAMAEM | ASSQDYGREQY | 0.39514858 |
| NELVEWSHPLGHRWRWE | ASSQDYGREQY | 0.5807156 |
| NLGAGGTGGNGGNPG | ASSQDYGREQY | 0.63727826 |
| NTEWPKWETVMAADPVD | ASSQDYGREQY | 0.81330407 |
| PASAAPAETQP | ASSQDYGREQY | 0.81874 |
| QQSSDGKNGVRAAEITGE | ASSQDYGREQY | 0.27750102 |
| RAPDAVTTSWWKE | ASSQDYGREQY | 0.788832 |
| SGFGNVGAHTSGWFNQ | ASSQDYGREQY | 0.8707047 |
| SGVAGAVGQAASVGGLKV | ASSQDYGREQY | 0.4951409 |
| SNGGAGTGSAGGNGGT | ASSQDYGREQY | 0.8361418 |
| TGGAGGEGGAAG | ASSQDYGREQY | 0.6674041 |
| TGGDGGAAGAGGAPGHG | ASSQDYGREQY | 0.7610798 |
| TMPPLLAAIANAGRHGV | ASSQDYGREQY | 0.45607156 |
| TSSTAGNSGGA | ASSQDYGREQY | 0.31715465 |
| VPATVPVGILC | ASSQDYGREQY | 0.83188164 |
| VQRPDSGGVVLTGRIS | ASSQDYGREQY | 0.42594135 |
| WTPNPVPATKGVPAGLVQAT | ASSQDYGREQY | 0.68231463 |
| YALKRALAKFSDRFDVVII | ASSQDYGREQY | 0.6393422 |
| YLIGSVIV | ASSQDYGREQY | 0.8819727 |
| YGQLVAVID | ASSQDYGREQY | 0.78161085 |
| VTVGGMTLPA | ASSQDYGREQY | 0.6784901 |
| VLALPAV | ASSQDYGREQY | 0.42013174 |

| | | |
|---|---|---|
| VIPPTLNYET | ASSQDYGREQY | 0.78931564 |
| VGMGGMHPSSG | ASSQDYGREQY | 0.7881515 |
| VGLAGGAATA | ASSQDYGREQY | 0.28140974 |
| VGGAGGAGGSGP | ASSQDYGREQY | 0.8447962 |
| VFAAFMV | ASSQDYGREQY | 0.3292898 |
| TLYDKLR | ASSQDYGREQY | 0.66437626 |
| TLAVFTVL | ASSQDYGREQY | 0.69370663 |
| TGSVQPARV | ASSQDYGREQY | 0.8774458 |
| TGGGAGSGGT | ASSQDYGREQY | 0.6546185 |
| SHLETSL | ASSQDYGREQY | 0.8471813 |
| SFTTGMT | ASSQDYGREQY | 0.30638662 |
| QVEVAKRMGVRQPTVS | ASSQDYGREQY | 0.82736766 |
| QGSGGPALAM | ASSQDYGREQY | 0.81937456 |
| MLAPSNP | ASSQDYGREQY | 0.86138123 |
| LVNNAAI | ASSQDYGREQY | 0.8803943 |
| LTGEINTLA | ASSQDYGREQY | 0.8032539 |
| LSIFGPV | ASSQDYGREQY | 0.20759913 |
| LLALLANL | ASSQDYGREQY | 0.65313566 |
| LISNASDALDKLR | ASSQDYGREQY | 0.83076906 |
| LINEAVP | ASSQDYGREQY | 0.8493114 |
| LGYAFSGL | ASSQDYGREQY | 0.83888614 |
| LDFVAVAL | ASSQDYGREQY | 0.83149093 |
| LAQPGAE | ASSQDYGREQY | 0.64472806 |
| ISNLGAVTGF | ASSQDYGREQY | 0.85137534 |
| HQEDPGRI | ASSQDYGREQY | 0.8955527 |
| GTPGEPTP | ASSQDYGREQY | 0.7540957 |
| GFFNSGS | ASSQDYGREQY | 0.49136445 |
| GAGGGGLTAGAN | ASSQDYGREQY | 0.42370123 |
| FVDALVNM | ASSQDYGREQY | 0.80192685 |
| FLTVAPEL | ASSQDYGREQY | 0.77848625 |
| FETLTAL | ASSQDYGREQY | 0.3450656 |
| DVINPIEL | ASSQDYGREQY | 0.6223613 |
| APVSGAPV | ASSQDYGREQY | 0.6997268 |
| ALVGGLAW | ASSQDYGREQY | 0.18987742 |
| ALAAAAATAI | ASSQDYGREQY | 0.7903861 |
| AFGLGLLL | ASSQDYGREQY | 0.85407025 |
| AFFAAASL | ASSQDYGREQY | 0.74108136 |
| AAAALPAFSPP | ASSQDYGREQY | 0.37732685 |
| VVLAPAL | ASSQDYGREQY | 0.84343773 |
| TIGHVDHGKTTL | ASSQDYGREQY | 0.84017336 |
| LISNASDALDK | ASSQDYGREQY | 0.85913026 |
| LISNASDAL | ASSQDYGREQY | 0.27921504 |
| ISNASDALDK | ASSQDYGREQY | 0.13892399 |
| NQAETLVYQTE | ASSQDYGREQY | 0.7600862 |
| TGGTGGTAGAG | ASSQDYGREQY | 0.83182776 |
| TGSTAGGAGGA | ASSQDYGREQY | 0.5432461 |
| DPLAVPVAAGPV | ASSQDYGREQY | 0.39761347 |
| GGTGGTPGNGGN | ASSQDYGREQY | 0.72082675 |
| PAGLAGVDPATL | ASSQDYGREQY | 0.80858946 |
| GNNLVVEAPHAGASV | ASSQDYGREQY | 0.35462934 |
| KVTRIMSASTLEQPA | ASSQDYGREQY | 0.30794537 |

| | | |
|---|---|---|
| LGFALPIALPRRTVV | ASSQDYGREQY | 0.7823355 |
| TRLGNLTARKLVVLV | ASSQDYGREQY | 0.8214973 |
| WSQQRRLIYFPSAGP | ASSQDYGREQY | 0.39122558 |
| GPNGNPDPMAAAVDIR | ASSQDYGREQY | 0.5226907 |
| TQRCVSVADLRAAASR | ASSQDYGREQY | 0.7703223 |
| QFVLDVAEHPVEVLTETV | ASSQDYGREQY | 0.8041623 |
| GGAGGTGGAGGNGGTQAGDG | ASSQDYGREQY | 0.7185986 |
| PLLATLRAIAVDVDAKPAQV | ASSQDYGREQY | 0.2748214 |
| APKFRTGGYMIDFWGVGYQVAK | ASSQDYGREQY | 0.75794315 |
| GAGGAGGAGGAAGGPGGTGGTG | ASSQDYGREQY | 0.29001647 |
| IGALGLLATDVAPAGSVAFGVA | ASSQDYGREQY | 0.84997284 |
| AGGAGGIGGNGGAGGAGGVLMG | ASSQDYGREQY | 0.69156206 |
| ADELIGLL | ASSQDYGREQY | 0.7678229 |
| ADRALKA | ASSQDYGREQY | 0.572526 |
| AGAVGQVGGAGGA | ASSQDYGREQY | 0.40205467 |
| AGGGGGLGGAP | ASSQDYGREQY | 0.3116408 |
| AGGLLYGNGGAG | ASSQDYGREQY | 0.41870463 |
| AGLESIGL | ASSQDYGREQY | 0.85937357 |
| AGLKTGII | ASSQDYGREQY | 0.7862047 |
| ASLAALLL | ASSQDYGREQY | 0.8160519 |
| ASVINISSI | ASSQDYGREQY | 0.6421777 |
| ATPPGPA | ASSQDYGREQY | 0.307011 |
| ATVASALV | ASSQDYGREQY | 0.7934169 |
| ATVASLIL | ASSQDYGREQY | 0.23268911 |
| AVNGPVLIRH | ASSQDYGREQY | 0.77775764 |
| DGAAGAVGI | ASSQDYGREQY | 0.7159156 |
| DPLLHGI | ASSQDYGREQY | 0.77128613 |
| ELGAALGAL | ASSQDYGREQY | 0.5056751 |
| ELGRTQAV | ASSQDYGREQY | 0.7713437 |
| FVAGGYLS | ASSQDYGREQY | 0.7778213 |
| GASGGAGGAGGW | ASSQDYGREQY | 0.4916641 |
| GGAGAGGDGGAGGI | ASSQDYGREQY | 0.75600827 |
| GTGHNGSQPGMGGNG | ASSQDYGREQY | 0.8045636 |
| GYAAASASAS | ASSQDYGREQY | 0.77771175 |
| HQVDVAAVQQG | ASSQDYGREQY | 0.20356707 |
| IGAAIAHT | ASSQDYGREQY | 0.67897594 |
| IGMIEVAQL | ASSQDYGREQY | 0.44697654 |
| IRLAPEHTSAMD | ASSQDYGREQY | 0.7197511 |
| IYNVSVL | ASSQDYGREQY | 0.6031805 |
| KQGFLGM | ASSQDYGREQY | 0.6535344 |
| LESVITQL | ASSQDYGREQY | 0.7331032 |
| LGLGALVVV | ASSQDYGREQY | 0.3348186 |
| LIGNGGNGGSGGI | ASSQDYGREQY | 0.54175127 |
| LNAINAPTL | ASSQDYGREQY | 0.59837395 |
| LQLLNTGV | ASSQDYGREQY | 0.67897487 |
| LQTSIEF | ASSQDYGREQY | 0.74553037 |
| MSFAVSEL | ASSQDYGREQY | 0.35649785 |
| NTGALGL | ASSQDYGREQY | 0.7526831 |
| PPGAGGAGGAGTT | ASSQDYGREQY | 0.5968147 |
| QLVGLGI | ASSQDYGREQY | 0.38641927 |
| RFLEQVHQL | ASSQDYGREQY | 0.70386887 |

| | | |
|---|---|---|
| RGVGYVLR | ASSQDYGREQY | 0.57928157 |
| SATGVFAGVI | ASSQDYGREQY | 0.282058 |
| SLPAITLP | ASSQDYGREQY | 0.82337916 |
| STALAFTV | ASSQDYGREQY | 0.3447305 |
| SVLTVGLGK | ASSQDYGREQY | 0.73948836 |
| TALAGAGSGI | ASSQDYGREQY | 0.87954754 |
| TGLPFTGI | ASSQDYGREQY | 0.3985579 |
| VAEHVTV | ASSQDYGREQY | 0.8283671 |
| VDIVLNSL | ASSQDYGREQY | 0.6075623 |
| VLTGGGAVL | ASSQDYGREQY | 0.7537811 |
| VNVGLAYGI | ASSQDYGREQY | 0.595377 |
| VSSAAVGAV | ASSQDYGREQY | 0.43416816 |
| VTSVYEAGV | ASSQDYGREQY | 0.8255505 |
| WVALDRGA | ASSQDYGREQY | 0.74683213 |
| YLVGSVVI | ASSQDYGREQY | 0.8172989 |
| GPPGTGKTLL | ASSQDYGREQY | 0.7055614 |
| GVGMTKF | ASSQDYGREQY | 0.85106933 |
| IGIGGSDL | ASSQDYGREQY | 0.8216522 |
| IGTLLAEL | ASSQDYGREQY | 0.54340124 |
| IYTSGTT | ASSQDYGREQY | 0.21463908 |
| LSFPAGV | ASSQDYGREQY | 0.52806354 |
| LYTSGTT | ASSQDYGREQY | 0.554304 |
| VGAFPFGF | ASSQDYGREQY | 0.47553128 |
| VGAGTVEF | ASSQDYGREQY | 0.2513287 |
| VGKSTLLL | ASSQDYGREQY | 0.8019576 |
| AALAHAGGGIV | ASSQDYGREQY | 0.87742126 |
| AALVAVSLL | ASSQDYGREQY | 0.5685112 |
| AAVSRAVAAL | ASSQDYGREQY | 0.8782205 |
| AGGSSGIGGF | ASSQDYGREQY | 0.3047242 |
| AGLTVANANG | ASSQDYGREQY | 0.7685373 |
| AGLVGAVL | ASSQDYGREQY | 0.834339 |
| ALSQAFTV | ASSQDYGREQY | 0.2998919 |
| ALTFSVSL | ASSQDYGREQY | 0.73894244 |
| ALTGAALVAL | ASSQDYGREQY | 0.46381858 |
| ASFSIGAGVL | ASSQDYGREQY | 0.40830165 |
| DGSVYKI | ASSQDYGREQY | 0.83254206 |
| EQLAEIV | ASSQDYGREQY | 0.5783511 |
| FAVSELTVM | ASSQDYGREQY | 0.81927896 |
| GAAPLAAGL | ASSQDYGREQY | 0.49950004 |
| GADGGDGGSGGAGG | ASSQDYGREQY | 0.30032617 |
| GFGLDTGGAGG | ASSQDYGREQY | 0.71355844 |
| GLASGGLL | ASSQDYGREQY | 0.8367916 |
| GRIAGAAVLL | ASSQDYGREQY | 0.7805535 |
| GSAGSLNTSF | ASSQDYGREQY | 0.8932182 |
| GTGQIGI | ASSQDYGREQY | 0.4651938 |
| GTLIALIL | ASSQDYGREQY | 0.8678797 |
| IAATMGFVVG | ASSQDYGREQY | 0.8580443 |
| IFAQALL | ASSQDYGREQY | 0.70189714 |
| IGAAVALGF | ASSQDYGREQY | 0.6113012 |
| IGLTPLAL | ASSQDYGREQY | 0.8240194 |
| ILGSNTSTL | ASSQDYGREQY | 0.5952714 |

| | | |
|---|---|---|
| IVFSLTVL | ASSQDYGREQY | 0.8325388 |
| LASKATGF | ASSQDYGREQY | 0.85839415 |
| LASVALIL | ASSQDYGREQY | 0.5151258 |
| LFAVGLTL | ASSQDYGREQY | 0.7748357 |
| LFNGGIGTYI | ASSQDYGREQY | 0.7181351 |
| LTLGAGII | ASSQDYGREQY | 0.6808518 |
| LVAAALGDI | ASSQDYGREQY | 0.48845637 |
| NTAFVQLGI | ASSQDYGREQY | 0.28902313 |
| QLFSGAEL | ASSQDYGREQY | 0.6946882 |
| SAATFITL | ASSQDYGREQY | 0.52209646 |
| SNYGLANLGV | ASSQDYGREQY | 0.7693206 |
| SYGGAYAVM | ASSQDYGREQY | 0.74532217 |
| TTGVGGLTLG | ASSQDYGREQY | 0.5559969 |
| TTQTIYI | ASSQDYGREQY | 0.7814209 |
| VAAGAGLGAVI | ASSQDYGREQY | 0.78631645 |
| VAAVVTLL | ASSQDYGREQY | 0.37863886 |
| VGFLGGLI | ASSQDYGREQY | 0.6921172 |
| VGVIPILL | ASSQDYGREQY | 0.43505973 |
| VIFASSLI | ASSQDYGREQY | 0.60389555 |
| VKGGDGGAAAG | ASSQDYGREQY | 0.31661832 |
| VTKTFTNI | ASSQDYGREQY | 0.73729104 |
| VVAAAILL | ASSQDYGREQY | 0.6603618 |
| YAALDVEL | ASSQDYGREQY | 0.24362919 |
| LASVALLL | ASSQDYGREQY | 0.71167326 |
| LDAVHRV | ASSQDYGREQY | 0.6947628 |
| QGLSSSIF | ASSQDYGREQY | 0.69547176 |
| RGTFANIRL | ASSQDYGREQY | 0.7684337 |
| RQLWWGHRI | ASSQDYGREQY | 0.5270618 |
| YGPPGTGKTLL | ASSQDYGREQY | 0.63554406 |
| AALVGAPANVA | ASSQDYGREQY | 0.75907505 |
| AATFRRTE | ASSQDYGREQY | 0.7749167 |
| AGALGGGI | ASSQDYGREQY | 0.54693305 |
| AGGAGGAGGIGAG | ASSQDYGREQY | 0.72541595 |
| AGSGALGL | ASSQDYGREQY | 0.777586 |
| AQIADQI | ASSQDYGREQY | 0.6379157 |
| ASAVAVTYL | ASSQDYGREQY | 0.75222826 |
| ASTVLGGTGG | ASSQDYGREQY | 0.765795 |
| AVGGEPAK | ASSQDYGREQY | 0.46087992 |
| CEQGTFGGTC | ASSQDYGREQY | 0.7714567 |
| DEGKAPAG | ASSQDYGREQY | 0.7892748 |
| DTVVFPAL | ASSQDYGREQY | 0.6040378 |
| FEGITMAG | ASSQDYGREQY | 0.66476667 |
| FPLVVQD | ASSQDYGREQY | 0.47381443 |
| GAFTTPQ | ASSQDYGREQY | 0.64514315 |
| GFGGFGTAGGVGGA | ASSQDYGREQY | 0.300174 |
| GFLAQFL | ASSQDYGREQY | 0.7381307 |
| GGAGGAGGNAGVA | ASSQDYGREQY | 0.75401956 |
| GGVGTAGAGGAGGAGG | ASSQDYGREQY | 0.69028515 |
| GNGGGGDAGNAGSGGN | ASSQDYGREQY | 0.4815937 |
| GTGGAGGAGNDAGS | ASSQDYGREQY | 0.45480037 |
| GTTNTGWFNS | ASSQDYGREQY | 0.67947024 |

| | | |
|---|---|---|
| IFMGPLFTL | ASSQDYGREQY | 0.5579883 |
| ILAHAIEAI | ASSQDYGREQY | 0.35899118 |
| IRSAIREAL | ASSQDYGREQY | 0.35363078 |
| ITFAGGG | ASSQDYGREQY | 0.7555152 |
| ITGMSGHQEV | ASSQDYGREQY | 0.7321514 |
| IVAGLACTL | ASSQDYGREQY | 0.59719 |
| IVGGGLAGA | ASSQDYGREQY | 0.78180957 |
| IVTGVLNI | ASSQDYGREQY | 0.7678068 |
| LAQQGCI | ASSQDYGREQY | 0.7541065 |
| LCLHVEN | ASSQDYGREQY | 0.6731024 |
| LDAISDGS | ASSQDYGREQY | 0.49698323 |
| LGGFAAML | ASSQDYGREQY | 0.776939 |
| LGGKNPMI | ASSQDYGREQY | 0.6419064 |
| LGPKGVAAIL | ASSQDYGREQY | 0.647272 |
| LMVGVSI | ASSQDYGREQY | 0.7040407 |
| LVGAAFVL | ASSQDYGREQY | 0.7171918 |
| LVLAVAPL | ASSQDYGREQY | 0.5931014 |
| MAAIAVIDAA | ASSQDYGREQY | 0.7248186 |
| NNGINAELQ | ASSQDYGREQY | 0.7017336 |
| RGEPLVLVHG | ASSQDYGREQY | 0.5368875 |
| RGGAGGAAGLI | ASSQDYGREQY | 0.3533205 |
| RSYRMHPAV | ASSQDYGREQY | 0.65310645 |
| SAAHLLAGQLA | ASSQDYGREQY | 0.7298214 |
| SAAVGSY | ASSQDYGREQY | 0.7711485 |
| SPHAVLHRI | ASSQDYGREQY | 0.68204415 |
| SQIAKVL | ASSQDYGREQY | 0.5502331 |
| TALGHLVP | ASSQDYGREQY | 0.7603514 |
| TAPTTSGAA | ASSQDYGREQY | 0.6824655 |
| TDGDDGNGGSPGT | ASSQDYGREQY | 0.64366615 |
| TDGLGEN | ASSQDYGREQY | 0.76963514 |
| TGGPPLPGQA | ASSQDYGREQY | 0.65255165 |
| TIPAITI | ASSQDYGREQY | 0.7041552 |
| TIPGFTI | ASSQDYGREQY | 0.7048558 |
| TSIASGALA | ASSQDYGREQY | 0.712181 |
| TVTLPGAAI | ASSQDYGREQY | 0.7586563 |
| VASGLSAALP | ASSQDYGREQY | 0.6714156 |
| VGAGVIGI | ASSQDYGREQY | 0.5907562 |
| VGPKRFGV | ASSQDYGREQY | 0.33291954 |
| VNVAFPD | ASSQDYGREQY | 0.5523119 |
| VSMTNSGVS | ASSQDYGREQY | 0.75419235 |
| VTDVLGDLSV | ASSQDYGREQY | 0.7299031 |
| VVVVGAGFAGL | ASSQDYGREQY | 0.7736453 |
| VYTVGTVV | ASSQDYGREQY | 0.7153243 |
| WTTFSGP | ASSQDYGREQY | 0.6174418 |
| YDELVEAF | ASSQDYGREQY | 0.7838998 |
| YGTGGAGGNGG | ASSQDYGREQY | 0.76898813 |
| YTNVMAV | ASSQDYGREQY | 0.34301534 |
| FLEQVHQL | ASSQDYGREQY | 0.5930955 |
| GLSLLAVAV | ASSQDYGREQY | 0.7866931 |
| LEQVHQL | ASSQDYGREQY | 0.76034075 |
| LNAAGAI | ASSQDYGREQY | 0.60341287 |

| | | |
|---|---|---|
| PAVHASL | ASSQDYGREQY | 0.5768614 |
| TELEDDA | ASSQDYGREQY | 0.79076076 |
| TPPGPPN | ASSQDYGREQY | 0.648281 |
| TPPPPPPP | ASSQDYGREQY | 0.78778 |
| TQSPALI | ASSRTGDPEVF | 0.28125253 |
| EFAAAFPQ | ASSRTGDPEVF | 0.52535063 |
| LLSGNLDVL | ASSRTGDPEVF | 0.3910846 |
| GPANLGNYNI | ASSRTGDPEVF | 0.2784027 |
| TLGPTLP | ASSRTGDPEVF | 0.73279655 |
| VYAYSSCCK | ASSRTGDPEVF | 0.6756921 |
| APAPAPAPNI | ASSRTGDPEVF | 0.764773 |
| PGGPVPIP | ASSRTGDPEVF | 0.62403744 |
| FLTVALEP | ASSRTGDPEVF | 0.75854254 |
| SQYFGANL | ASSRTGDPEVF | 0.717015 |
| GGVKPAA | ASSRTGDPEVF | 0.6358254 |
| GGTGGAGGAGGW | ASSRTGDPEVF | 0.3856784 |
| GVGGVAGLGGAGG | ASSRTGDPEVF | 0.7288525 |
| VLLVLATTI | ASSRTGDPEVF | 0.78915805 |
| IGIFAAML | ASSRTGDPEVF | 0.6173428 |
| AVGDLID | ASSRTGDPEVF | 0.41524172 |
| NISPLPL | ASSRTGDPEVF | 0.70346993 |
| VTAVFVAI | ASSRTGDPEVF | 0.6311079 |
| SGDDNDAAL | ASSRTGDPEVF | 0.27503178 |
| AVAALAHS | ASSRTGDPEVF | 0.255755 |
| GGVTVMATP | ASSRTGDPEVF | 0.6248149 |
| AGLAANVRD | ASSRTGDPEVF | 0.5999758 |
| TNRPDLIDPALLRPG | ASSRTGDPEVF | 0.77283037 |
| LANLGVTAL | ASSRTGDPEVF | 0.7654365 |
| TGERIEL | ASSRTGDPEVF | 0.66451705 |
| NGEELAA | ASSRTGDPEVF | 0.74633443 |
| VLSLVWG | ASSRTGDPEVF | 0.7405123 |
| IGYALWL | ASSRTGDPEVF | 0.19264877 |
| LSLLLAPL | ASSRTGDPEVF | 0.435692 |
| TLDEALK | ASSRTGDPEVF | 0.39151958 |
| TDGAGTAEGG | ASSRTGDPEVF | 0.7610823 |
| NAPGGAGGNGG | ASSRTGDPEVF | 0.51317674 |
| VSVFMGMGGNF | ASSRTGDPEVF | 0.6423112 |
| LASYLAAL | ASSRTGDPEVF | 0.7029259 |
| LVGGTPAL | ASSRTGDPEVF | 0.7605399 |
| VRNADVL | ASSRTGDPEVF | 0.6538507 |
| TCSSPFLVSAM | ASSRTGDPEVF | 0.40187243 |
| VHITDAE | ASSRTGDPEVF | 0.24692853 |
| EYAAPSF | ASSRTGDPEVF | 0.7134236 |
| VVDAFHGGL | ASSRTGDPEVF | 0.43800712 |
| ASVVDPQ | ASSRTGDPEVF | 0.5459134 |
| SYNEFGGLS | ASSRTGDPEVF | 0.3941236 |
| DPDGSVY | ASSRTGDPEVF | 0.6581677 |
| MVSTIHATE | ASSRTGDPEVF | 0.46144295 |
| LAAGVADPV | ASSRTGDPEVF | 0.74275786 |
| TKEQIAA | ASSRTGDPEVF | 0.4915066 |
| SLASPAAL | ASSRTGDPEVF | 0.7006042 |

| | | |
|---|---|---|
| IGLVFYGV | ASSRTGDPEVF | 0.78163093 |
| FLLDASNL | ASSRTGDPEVF | 0.75661016 |
| YQVGNFP | ASSRTGDPEVF | 0.5998178 |
| ESLLSQST | ASSRTGDPEVF | 0.59500414 |
| LFTVGFAAVG | ASSRTGDPEVF | 0.42572117 |
| ATPPTPT | ASSRTGDPEVF | 0.58761406 |
| TDLATLG | ASSRTGDPEVF | 0.62250555 |
| SAGGTGR | ASSRTGDPEVF | 0.78666794 |
| GFSNSGS | ASSRTGDPEVF | 0.5230396 |
| GGNGGSGGTSV | ASSRTGDPEVF | 0.52075493 |
| GTSVATGGAGN | ASSRTGDPEVF | 0.5726571 |
| NGGNGGSGGTS | ASSRTGDPEVF | 0.7765018 |
| AVVKFPH | ASSRTGDPEVF | 0.77977407 |
| TPPPPPP | ASSRTGDPEVF | 0.53502655 |
| AAPAAAAAPA | ASSRTGDPEVF | 0.56593853 |
| IALNEDL | ASSRTGDPEVF | 0.6306759 |
| IVNVSSM | ASSRTGDPEVF | 0.71973884 |
| TGIPPAPR | ASSRTGDPEVF | 0.4916855 |
| FDLGGGTF | ASSRTGDPEVF | 0.47461897 |
| FDLGGGTFDVS | ASSRTGDPEVF | 0.6725954 |
| NEPTAAA | ASSRTGDPEVF | 0.7914455 |
| NEPTAAALA | ASSRTGDPEVF | 0.6088321 |
| GSIINISSI | ASSRTGDPEVF | 0.3397905 |
| VAPGNAGTA | ASSRTGDPEVF | 0.7088021 |
| TGGGASLE | ASSRTGDPEVF | 0.5525333 |
| VAVYNHYKRI | ASSRTGDPEVF | 0.58592415 |
| TDPAPATT | ASSRTGDPEVF | 0.7008029 |
| TNRPDILDPALLRPG | ASSRTGDPEVF | 0.40616092 |
| LTTTTTS | ASSRTGDPEVF | 0.68231755 |
| PTGSGKTL | ASSRTGDPEVF | 0.7825235 |
| TGSGKTL | ASSRTGDPEVF | 0.6863084 |
| TIGHVDHGK | ASSRTGDPEVF | 0.7444655 |
| TIGHVDHGKTT | ASSRTGDPEVF | 0.7793937 |
| GHVDHGKTT | ASSRTGDPEVF | 0.746632 |
| AQFEHTLL | ASSRTGDPEVF | 0.77546763 |
| IVNQDEL | ASSRTGDPEVF | 0.7754778 |
| TNGSQFF | ASSRTGDPEVF | 0.60737973 |
| TGKGIVE | ASSRTGDPEVF | 0.62290096 |
| LISNASDA | ASSRTGDPEVF | 0.7578925 |
| AELRTLGELTKIAKAHGAQ | ASSRTGDPEVF | 0.70683146 |
| APNAVAAAEPSW | ASSRTGDPEVF | 0.6728212 |
| APQVAGAATAVL | ASSRTGDPEVF | 0.4761651 |
| ARRDPAAAAAILPTDA | ASSRTGDPEVF | 0.78063273 |
| DAAARDVLAEAGLAEYF | ASSRTGDPEVF | 0.55845475 |
| DDQIVLSAKANLAKVEQ | ASSRTGDPEVF | 0.77741563 |
| DELLPDTGEDSDEDGH | ASSRTGDPEVF | 0.3064402 |
| ELPEAPANRATVPQD | ASSRTGDPEVF | 0.51863927 |
| FGQSKIERVTGIG | ASSRTGDPEVF | 0.7606033 |
| GAGGTGSTAGGAGGAGGAGGLY | ASSRTGDPEVF | 0.20606606 |
| GHGGTGAAGGNGGTGGAGGSGI | ASSRTGDPEVF | 0.25253123 |
| GNGGNGGAGGEGGPGGAGGAGA | ASSRTGDPEVF | 0.6601602 |

| | | |
|---|---|---|
| GSTAGNGGNGGNATNGTIAGQP | ASSRTGDPEVF | 0.768214 |
| GTGGDGGNGGNGGTGG | ASSRTGDPEVF | 0.20557027 |
| GVRRPAGIPADAPV | ASSRTGDPEVF | 0.59577787 |
| GYRQVTDAHLLTLARR | ASSRTGDPEVF | 0.33813536 |
| IAKMGQDGHDRG | ASSRTGDPEVF | 0.6925628 |
| KAEPSIEIQYT | ASSRTGDPEVF | 0.70130825 |
| KPVRIRMAADEDTVAVT | ASSRTGDPEVF | 0.27678835 |
| MPPRIAGMRLLVIKPEP | ASSRTGDPEVF | 0.7333807 |
| MKFYAVALFRSRGVPPTR | ASSRTGDPEVF | 0.3518324 |
| NAGGFGILGAVAHSP | ASSRTGDPEVF | 0.52966034 |
| NEGREITLAMAEM | ASSRTGDPEVF | 0.75175214 |
| NELVEWSHPLGHRWRWE | ASSRTGDPEVF | 0.7852947 |
| NLGAGGTGGNGGNPG | ASSRTGDPEVF | 0.41117284 |
| NTEWPKWETVMAADPVD | ASSRTGDPEVF | 0.74721247 |
| PASAAPAETQP | ASSRTGDPEVF | 0.5831224 |
| QQSSDGKNGVRAAEITGE | ASSRTGDPEVF | 0.7859812 |
| RAPDAVTTSWWKE | ASSRTGDPEVF | 0.5325187 |
| SGFGNVGAHTSGWFNQ | ASSRTGDPEVF | 0.3233397 |
| SGVAGAVGQAASVGGLKV | ASSRTGDPEVF | 0.7708952 |
| SNGGAGTGSAGGNGGT | ASSRTGDPEVF | 0.6937251 |
| TGGAGGEGGAAG | ASSRTGDPEVF | 0.68507135 |
| TGGDGGAAGAGGAPGHG | ASSRTGDPEVF | 0.7712703 |
| TMPPLLAAIANAGRHGV | ASSRTGDPEVF | 0.45429236 |
| TSSTAGNSGGA | ASSRTGDPEVF | 0.6809002 |
| VPATVPVGILC | ASSRTGDPEVF | 0.7723408 |
| VQRPDSGGVVLTGRIS | ASSRTGDPEVF | 0.7673524 |
| WTPNPVPATKGVPAGLVQAT | ASSRTGDPEVF | 0.4453289 |
| YALKRALAKFSDRFDVVII | ASSRTGDPEVF | 0.75658494 |
| YLIGSVIV | ASSRTGDPEVF | 0.7353777 |
| YGQLVAVID | ASSRTGDPEVF | 0.51439637 |
| VTVGGMTLPA | ASSRTGDPEVF | 0.7636385 |
| VLALPAV | ASSRTGDPEVF | 0.5938333 |
| VIPPTLNYET | ASSRTGDPEVF | 0.75223166 |
| VGMGGMHPSSG | ASSRTGDPEVF | 0.7104753 |
| VGLAGGAATA | ASSRTGDPEVF | 0.78363013 |
| VGGAGGAGGSGP | ASSRTGDPEVF | 0.29634553 |
| VFAAFMV | ASSRTGDPEVF | 0.73926145 |
| TLYDKLR | ASSRTGDPEVF | 0.67982167 |
| TLAVFTVL | ASSRTGDPEVF | 0.75683033 |
| TGSVQPARV | ASSRTGDPEVF | 0.5980543 |
| TGGGAGSGGT | ASSRTGDPEVF | 0.6107551 |
| SHLETSL | ASSRTGDPEVF | 0.6538079 |
| SFTTGMT | ASSRTGDPEVF | 0.77983385 |
| QVEVAKRMGVRQPTVS | ASSRTGDPEVF | 0.5449961 |
| QGSGGPALAM | ASSRTGDPEVF | 0.76304233 |
| MLAPSNP | ASSRTGDPEVF | 0.7835424 |
| LVNNAAI | ASSRTGDPEVF | 0.5136284 |
| LTGEINTLA | ASSRTGDPEVF | 0.7139881 |
| LSIFGPV | ASSRTGDPEVF | 0.7338491 |
| LLALLANL | ASSRTGDPEVF | 0.786399 |
| LISNASDALDKLR | ASSRTGDPEVF | 0.57350147 |

| | | |
|---|---|---|
| LINEAVP | ASSRTGDPEVF | 0.77769023 |
| LGYAFSGL | ASSRTGDPEVF | 0.52987266 |
| LDFVAVAL | ASSRTGDPEVF | 0.76269853 |
| LAQPGAE | ASSRTGDPEVF | 0.51217335 |
| ISNLGAVTGF | ASSRTGDPEVF | 0.7182659 |
| HQEDPGRI | ASSRTGDPEVF | 0.5975569 |
| GTPGEPTP | ASSRTGDPEVF | 0.76171654 |
| GFFNSGS | ASSRTGDPEVF | 0.7674227 |
| GAGGGGLTAGAN | ASSRTGDPEVF | 0.7720514 |
| FVDALVNM | ASSRTGDPEVF | 0.34372345 |
| FLTVAPEL | ASSRTGDPEVF | 0.7609415 |
| FETLTAL | ASSRTGDPEVF | 0.76513255 |
| DVINPIEL | ASSRTGDPEVF | 0.7921046 |
| APVSGAPV | ASSRTGDPEVF | 0.5763147 |
| ALVGGLAW | ASSRTGDPEVF | 0.7500547 |
| ALAAAAATAI | ASSRTGDPEVF | 0.6758031 |
| AFGLGLLL | ASSRTGDPEVF | 0.5782614 |
| AFFAAASL | ASSRTGDPEVF | 0.47841305 |
| AAAALPAFSPP | ASSRTGDPEVF | 0.6416236 |
| VVLAPAL | ASSRTGDPEVF | 0.65867436 |
| TIGHVDHGKTTL | ASSRTGDPEVF | 0.5869446 |
| LISNASDALDK | ASSRTGDPEVF | 0.77586967 |
| LISNASDAL | ASSRTGDPEVF | 0.77987033 |
| ISNASDALDK | ASSRTGDPEVF | 0.76573354 |
| NQAETLVYQTE | ASSRTGDPEVF | 0.7783076 |
| TGGTGGTAGAG | ASSRTGDPEVF | 0.77973175 |
| TGSTAGGAGGA | ASSRTGDPEVF | 0.7210548 |
| DPLAVPVAAGPV | ASSRTGDPEVF | 0.6254153 |
| GGTGGTPGNGGN | ASSRTGDPEVF | 0.59017634 |
| PAGLAGVDPATL | ASSRTGDPEVF | 0.5106394 |
| GNNLVVEAPHAGASV | ASSRTGDPEVF | 0.64775586 |
| KVTRIMSASTLEQPA | ASSRTGDPEVF | 0.61048996 |
| LGFALPIALPRRTVV | ASSRTGDPEVF | 0.717435 |
| TRLGNLTARKLVVLV | ASSRTGDPEVF | 0.20247458 |
| WSQQRRLIYFPSAGP | ASSRTGDPEVF | 0.74970126 |
| GPNGNPDPMAAAVDIR | ASSRTGDPEVF | 0.78829646 |
| TQRCVSVADLRAAASR | ASSRTGDPEVF | 0.6416651 |
| QFVLDVAEHPVEVLTETV | ASSRTGDPEVF | 0.7185806 |
| GGAGGTGGAGGNGGTQAGDG | ASSRTGDPEVF | 0.73968816 |
| PLLATLRAIAVDVDAKPAQV | ASSRTGDPEVF | 0.5354821 |
| APKFRTGGYMIDFWGVGYQVAK | ASSRTGDPEVF | 0.76072013 |
| GAGGAGGAGGAAGGPGGTGGTG | ASSRTGDPEVF | 0.53465164 |
| IGALGLLATDVAPAGSVAFGVA | ASSRTGDPEVF | 0.71591514 |
| AGGAGGIGGNGGAGGAGGVLMG | ASSRTGDPEVF | 0.7645334 |
| ADELIGLL | ASSRTGDPEVF | 0.6926004 |
| ADRALKA | ASSRTGDPEVF | 0.7559291 |
| AGAVGQVGGAGGA | ASSRTGDPEVF | 0.62210596 |
| AGGGGGLGGAP | ASSRTGDPEVF | 0.26369125 |
| AGGLLYGNGGAG | ASSRTGDPEVF | 0.7662945 |
| AGLESIGL | ASSRTGDPEVF | 0.43360516 |
| AGLKTGII | ASSRTGDPEVF | 0.66632974 |

| | | |
|---|---|---|
| ASLAALLL | ASSRTGDPEVF | 0.37503624 |
| ASVINISSI | ASSRTGDPEVF | 0.73831224 |
| ATPPGPA | ASSRTGDPEVF | 0.7436987 |
| ATVASALV | ASSRTGDPEVF | 0.68319225 |
| ATVASLIL | ASSRTGDPEVF | 0.48931265 |
| AVNGPVLIRH | ASSRTGDPEVF | 0.70513165 |
| DGAAGAVGI | ASSRTGDPEVF | 0.68886817 |
| DPLLHGI | ASSRTGDPEVF | 0.49488008 |
| ELGAALGAL | ASSRTGDPEVF | 0.76314604 |
| ELGRTQAV | ASSRTGDPEVF | 0.51564026 |
| FVAGGYLS | ASSRTGDPEVF | 0.60800564 |
| GASGGAGGAGGW | ASSRTGDPEVF | 0.76207703 |
| GGAGAGGDGGAGGI | ASSRTGDPEVF | 0.6982307 |
| GTGHNGSQPGMGGNG | ASSRTGDPEVF | 0.31026247 |
| GYAAASASAS | ASSRTGDPEVF | 0.67603517 |
| HQVDVAAVQQG | ASSRTGDPEVF | 0.7077467 |
| IGAAIAHT | ASSRTGDPEVF | 0.51525986 |
| IGMIEVAQL | ASSRTGDPEVF | 0.2441912 |
| IRLAPEHTSAMD | ASSRTGDPEVF | 0.67460537 |
| IYNVSVL | ASSRTGDPEVF | 0.50847566 |
| KQGFLGM | ASSRTGDPEVF | 0.46444526 |
| LESVITQL | ASSRTGDPEVF | 0.65713465 |
| LGLGALVVV | ASSRTGDPEVF | 0.7727214 |
| LIGNGGNGGSGGI | ASSRTGDPEVF | 0.60175925 |
| LNAINAPTL | ASSRTGDPEVF | 0.6717212 |
| LQLLNTGV | ASSRTGDPEVF | 0.7648296 |
| LQTSIEF | ASSRTGDPEVF | 0.74052 |
| MSFAVSEL | ASSRTGDPEVF | 0.7801168 |
| NTGALGL | ASSRTGDPEVF | 0.7567256 |
| PPGAGGAGGAGTT | ASSRTGDPEVF | 0.5261405 |
| QLVGLGI | ASSRTGDPEVF | 0.7154783 |
| RFLEQVHQL | ASSRTGDPEVF | 0.57220143 |
| RGVGYVLR | ASSRTGDPEVF | 0.4162215 |
| SATGVFAGVI | ASSRTGDPEVF | 0.36405382 |
| SLPAITLP | ASSRTGDPEVF | 0.6039847 |
| STALAFTV | ASSRTGDPEVF | 0.36037177 |
| SVLTVGLGK | ASSRTGDPEVF | 0.65850645 |
| TALAGAGSGI | ASSRTGDPEVF | 0.75529057 |
| TGLPFTGI | ASSRTGDPEVF | 0.51169705 |
| VAEHVTV | ASSRTGDPEVF | 0.37969083 |
| VDIVLNSL | ASSRTGDPEVF | 0.6563043 |
| VLTGGGAVL | ASSRTGDPEVF | 0.3312804 |
| VNVGLAYGI | ASSRTGDPEVF | 0.735367 |
| VSSAAVGAV | ASSRTGDPEVF | 0.6071598 |
| VTSVYEAGV | ASSRTGDPEVF | 0.7918699 |
| WVALDRGA | ASSRTGDPEVF | 0.7745154 |
| YLVGSVVI | ASSRTGDPEVF | 0.6928554 |
| GPPGTGKTLL | ASSRTGDPEVF | 0.75905013 |
| GVGMTKF | ASSRTGDPEVF | 0.49745077 |
| IGIGGSDL | ASSRTGDPEVF | 0.71667755 |
| IGTLLAEL | ASSRTGDPEVF | 0.6100725 |

| | | |
|---|---|---|
| IYTSGTT | ASSRTGDPEVF | 0.68303883 |
| LSFPAGV | ASSRTGDPEVF | 0.71811104 |
| LYTSGTT | ASSRTGDPEVF | 0.5061969 |
| VGAFPFGF | ASSRTGDPEVF | 0.26908767 |
| VGAGTVEF | ASSRTGDPEVF | 0.7605598 |
| VGKSTLLL | ASSRTGDPEVF | 0.6489345 |
| AALAHAGGGIV | ASSRTGDPEVF | 0.31205893 |
| AALVAVSLL | ASSRTGDPEVF | 0.5186894 |
| AAVSRAVAAL | ASSRTGDPEVF | 0.7869363 |
| AGGSSGIGGF | ASSRTGDPEVF | 0.7839962 |
| AGLTVANANG | ASSRTGDPEVF | 0.58497626 |
| AGLVGAVL | ASSRTGDPEVF | 0.5226231 |
| ALSQAFTV | ASSRTGDPEVF | 0.756974 |
| ALTFSVSL | ASSRTGDPEVF | 0.4576748 |
| ALTGAALVAL | ASSRTGDPEVF | 0.63464576 |
| ASFSIGAGVL | ASSRTGDPEVF | 0.6971693 |
| DGSVYKI | ASSRTGDPEVF | 0.69014215 |
| EQLAEIV | ASSRTGDPEVF | 0.5091452 |
| FAVSELTVM | ASSRTGDPEVF | 0.7285117 |
| GAAPLAAGL | ASSRTGDPEVF | 0.71240586 |
| GADGGDGGSGGAGG | ASSRTGDPEVF | 0.69938326 |
| GFGLDTGGAGG | ASSRTGDPEVF | 0.24128275 |
| GLASGGLL | ASSRTGDPEVF | 0.70414674 |
| GRIAGAAVLL | ASSRTGDPEVF | 0.59069324 |
| GSAGSLNTSF | ASSRTGDPEVF | 0.42674905 |
| GTGQIGI | ASSRTGDPEVF | 0.2534432 |
| GTLIALIL | ASSRTGDPEVF | 0.7088691 |
| IAATMGFVVG | ASSRTGDPEVF | 0.76685345 |
| IFAQALL | ASSRTGDPEVF | 0.7710913 |
| IGAAVALGF | ASSRTGDPEVF | 0.4847014 |
| IGLTPLAL | ASSRTGDPEVF | 0.77708554 |
| ILGSNTSTL | ASSRTGDPEVF | 0.7535619 |
| IVFSLTVL | ASSRTGDPEVF | 0.60006166 |
| LASKATGF | ASSRTGDPEVF | 0.26023754 |
| LASVALIL | ASSRTGDPEVF | 0.74711496 |
| LFAVGLTL | ASSRTGDPEVF | 0.75725895 |
| LFNGGIGTYI | ASSRTGDPEVF | 0.73399645 |
| LTLGAGII | ASSRTGDPEVF | 0.7441083 |
| LVAAALGDI | ASSRTGDPEVF | 0.47268975 |
| NTAFVQLGI | ASSRTGDPEVF | 0.7748039 |
| QLFSGAEL | ASSRTGDPEVF | 0.74260926 |
| SAATFITL | ASSRTGDPEVF | 0.43565494 |
| SNYGLANLGV | ASSRTGDPEVF | 0.57597804 |
| SYGGAYAVM | ASSRTGDPEVF | 0.6219722 |
| TTGVGGLTLG | ASSRTGDPEVF | 0.37081325 |
| TTQTIYI | ASSRTGDPEVF | 0.76174706 |
| VAAGAGLGAVI | ASSRTGDPEVF | 0.64207566 |
| VAAVVTLL | ASSRTGDPEVF | 0.6506345 |
| VGFLGGLI | ASSRTGDPEVF | 0.68482685 |
| VGVIPILL | ASSRTGDPEVF | 0.66838396 |
| VIFASSLI | ASSRTGDPEVF | 0.61028737 |

| | | |
|---|---|---|
| VKGGDGGAAAG | ASSRTGDPEVF | 0.67822605 |
| VTKTFTNI | ASSRTGDPEVF | 0.6819707 |
| VVAAAILL | ASSRTGDPEVF | 0.740031 |
| YAALDVEL | ASSRTGDPEVF | 0.61071116 |
| LASVALLL | ASSRTGDPEVF | 0.7364027 |
| LDAVHRV | ASSRTGDPEVF | 0.7841631 |
| QGLSSSIF | ASSRTGDPEVF | 0.47365505 |
| RGTFANIRL | ASSRTGDPEVF | 0.5886766 |
| RQLWWGHRI | ASSRTGDPEVF | 0.74553865 |
| YGPPGTGKTLL | ASSRTGDPEVF | 0.4632913 |
| AALVGAPANVA | ASSRTGDPEVF | 0.7530468 |
| AATFRRTE | ASSRTGDPEVF | 0.7495889 |
| AGALGGGI | ASSRTGDPEVF | 0.7382604 |
| AGGAGGAGGIGAG | ASSRTGDPEVF | 0.6932374 |
| AGSGALGL | ASSRTGDPEVF | 0.7895365 |
| AQIADQI | ASSRTGDPEVF | 0.7583416 |
| ASAVAVTYL | ASSRTGDPEVF | 0.6738552 |
| ASTVLGGTGG | ASSRTGDPEVF | 0.78177106 |
| AVGGEPAK | ASSRTGDPEVF | 0.5613266 |
| CEQGTFGGTC | ASSRTGDPEVF | 0.77337766 |
| DEGKAPAG | ASSRTGDPEVF | 0.7534501 |
| DTVVFPAL | ASSRTGDPEVF | 0.4035824 |
| FEGITMAG | ASSRTGDPEVF | 0.69281006 |
| FPLVVQD | ASSRTGDPEVF | 0.74799573 |
| GAFTTPQ | ASSRTGDPEVF | 0.77851653 |
| GFGGFGTAGGVGGA | ASSRTGDPEVF | 0.32258424 |
| GFLAQFL | ASSRTGDPEVF | 0.6213676 |
| GGAGGAGGNAGVA | ASSRTGDPEVF | 0.39728644 |
| GGVGTAGAGGAGGAGG | ASSRTGDPEVF | 0.403997 |
| GNGGGGDAGNAGSGGN | ASSRTGDPEVF | 0.40693662 |
| GTGGAGGAGNDAGS | ASSRTGDPEVF | 0.71564645 |
| GTTNTGWFNS | ASSRTGDPEVF | 0.636551 |
| IFMGPLFTL | ASSRTGDPEVF | 0.76893425 |
| ILAHAIEAI | ASSRTGDPEVF | 0.65004 |
| IRSAIREAL | ASSRTGDPEVF | 0.7558242 |
| ITFAGGG | ASSRTGDPEVF | 0.5692605 |
| ITGMSGHQEV | ASSRTGDPEVF | 0.70590496 |
| IVAGLACTL | ASSRTGDPEVF | 0.3066694 |
| IVGGGLAGA | ASSRTGDPEVF | 0.35227653 |
| IVTGVLNI | ASSRTGDPEVF | 0.3591729 |
| LAQQGCI | ASSRTGDPEVF | 0.6881855 |
| LCLHVEN | ASSRTGDPEVF | 0.24936774 |
| LDAISDGS | ASSRTGDPEVF | 0.75763214 |
| LGGFAAML | ASSRTGDPEVF | 0.6854217 |
| LGGKNPMI | ASSRTGDPEVF | 0.716966 |
| LGPKGVAAIL | ASSRTGDPEVF | 0.607234 |
| LMVGVSI | ASSRTGDPEVF | 0.66717017 |
| LVGAAFVL | ASSRTGDPEVF | 0.6115619 |
| LVLAVAPL | ASSRTGDPEVF | 0.7667929 |
| MAAIAVIDAA | ASSRTGDPEVF | 0.4109503 |
| NNGINAELQ | ASSRTGDPEVF | 0.2942861 |

| | | |
|---|---|---|
| RGEPLVLVHG | ASSRTGDPEVF | 0.5345771 |
| RGGAGGAAGLI | ASSRTGDPEVF | 0.48086035 |
| RSYRMHPAV | ASSRTGDPEVF | 0.7728423 |
| SAAHLLAGQLA | ASSRTGDPEVF | 0.75709677 |
| SAAVGSY | ASSRTGDPEVF | 0.71274066 |
| SPHAVLHRI | ASSRTGDPEVF | 0.7788158 |
| SQIAKVL | ASSRTGDPEVF | 0.2281925 |
| TALGHLVP | ASSRTGDPEVF | 0.6273098 |
| TAPTTSGAA | ASSRTGDPEVF | 0.7876822 |
| TDGDDGNGGSPGT | ASSRTGDPEVF | 0.68125045 |
| TDGLGEN | ASSRTGDPEVF | 0.66353357 |
| TGGPPLPGQA | ASSRTGDPEVF | 0.7637787 |
| TIPAITI | ASSRTGDPEVF | 0.6650815 |
| TIPGFTI | ASSRTGDPEVF | 0.51421154 |
| TSIASGALA | ASSRTGDPEVF | 0.7284002 |
| TVTLPGAAI | ASSRTGDPEVF | 0.69527096 |
| VASGLSAALP | ASSRTGDPEVF | 0.6283654 |
| VGAGVIGI | ASSRTGDPEVF | 0.36535826 |
| VGPKRFGV | ASSRTGDPEVF | 0.69598985 |
| VNVAFPD | ASSRTGDPEVF | 0.7931944 |
| VSMTNSGVS | ASSRTGDPEVF | 0.6961191 |
| VTDVLGDLSV | ASSRTGDPEVF | 0.7158323 |
| VVVVGAGFAGL | ASSRTGDPEVF | 0.7314682 |
| VYTVGTVV | ASSRTGDPEVF | 0.7314631 |
| WTTFSGP | ASSRTGDPEVF | 0.54405874 |
| YDELVEAF | ASSRTGDPEVF | 0.5297263 |
| YGTGGAGGNGG | ASSRTGDPEVF | 0.7049062 |
| YTNVMAV | ASSRTGDPEVF | 0.49580142 |
| FLEQVHQL | ASSRTGDPEVF | 0.6567478 |
| GLSLLAVAV | ASSRTGDPEVF | 0.5473299 |
| LEQVHQL | ASSRTGDPEVF | 0.79401386 |
| LNAAGAI | ASSRTGDPEVF | 0.7433269 |
| PAVHASL | ASSRTGDPEVF | 0.73166776 |
| TELEDDA | ASSRTGDPEVF | 0.6367638 |
| TPPGPPN | ASSRTGDPEVF | 0.5923791 |
| TPPPPPPP | ASSRTGDPEVF | 0.72879374 |
| TQSPALI | ASSPTEDQDTQY | 0.28543344 |
| EFAAAFPQ | ASSPTEDQDTQY | 0.76368326 |
| LLSGNLDVL | ASSPTEDQDTQY | 0.552545 |
| GPANLGNYNI | ASSPTEDQDTQY | 0.5663917 |
| TLGPTLP | ASSPTEDQDTQY | 0.74427235 |
| VYAYSSCCK | ASSPTEDQDTQY | 0.7858602 |
| APAPAPAPNI | ASSPTEDQDTQY | 0.5988089 |
| PGGPVPIP | ASSPTEDQDTQY | 0.69545984 |
| FLTVALEP | ASSPTEDQDTQY | 0.600709 |
| SQYFGANL | ASSPTEDQDTQY | 0.70761955 |
| GGVKPAA | ASSPTEDQDTQY | 0.7769331 |
| GGTGGAGGAGGW | ASSPTEDQDTQY | 0.7364523 |
| GVGGVAGLGGAGG | ASSPTEDQDTQY | 0.78540516 |
| VLLVLATTI | ASSPTEDQDTQY | 0.6687876 |
| IGIFAAML | ASSPTEDQDTQY | 0.31968042 |

| | | |
|---|---|---|
| AVGDLID | ASSPTEDQDTQY | 0.5312055 |
| NISPLPL | ASSPTEDQDTQY | 0.747306 |
| VTAVFVAI | ASSPTEDQDTQY | 0.7700498 |
| SGDDNDAAL | ASSPTEDQDTQY | 0.77434444 |
| AVAALAHS | ASSPTEDQDTQY | 0.7651617 |
| GGVTVMATP | ASSPTEDQDTQY | 0.5309955 |
| AGLAANVRD | ASSPTEDQDTQY | 0.76558167 |
| TNRPDLIDPALLRPG | ASSPTEDQDTQY | 0.7823856 |
| LANLGVTAL | ASSPTEDQDTQY | 0.755718 |
| TGERIEL | ASSPTEDQDTQY | 0.44488993 |
| NGEELAA | ASSPTEDQDTQY | 0.40980622 |
| VLSLVWG | ASSPTEDQDTQY | 0.6883754 |
| IGYALWL | ASSPTEDQDTQY | 0.76239955 |
| LSLLLAPL | ASSPTEDQDTQY | 0.42576438 |
| TLDEALK | ASSPTEDQDTQY | 0.69366443 |
| TDGAGTAEGG | ASSPTEDQDTQY | 0.74008256 |
| NAPGGAGGNGG | ASSPTEDQDTQY | 0.33844054 |
| VSVFMGMGGNF | ASSPTEDQDTQY | 0.603978 |
| LASYLAAL | ASSPTEDQDTQY | 0.7816927 |
| LVGGTPAL | ASSPTEDQDTQY | 0.5957256 |
| VRNADVL | ASSPTEDQDTQY | 0.48882404 |
| TCSSPFLVSAM | ASSPTEDQDTQY | 0.7771529 |
| VHITDAE | ASSPTEDQDTQY | 0.79091555 |
| EYAAPSF | ASSPTEDQDTQY | 0.75791657 |
| VVDAFHGGL | ASSPTEDQDTQY | 0.3886984 |
| ASVVDPQ | ASSPTEDQDTQY | 0.24563533 |
| SYNEFGGLS | ASSPTEDQDTQY | 0.7050301 |
| DPDGSVY | ASSPTEDQDTQY | 0.77657515 |
| MVSTIHATE | ASSPTEDQDTQY | 0.7055055 |
| LAAGVADPV | ASSPTEDQDTQY | 0.63343126 |
| TKEQIAA | ASSPTEDQDTQY | 0.72560614 |
| SLASPAAL | ASSPTEDQDTQY | 0.75364006 |
| IGLVFYGV | ASSPTEDQDTQY | 0.47210932 |
| FLLDASNL | ASSPTEDQDTQY | 0.33025524 |
| YQVGNFP | ASSPTEDQDTQY | 0.7469006 |
| ESLLSQST | ASSPTEDQDTQY | 0.75262594 |
| LFTVGFAAVG | ASSPTEDQDTQY | 0.5555299 |
| ATPPTPT | ASSPTEDQDTQY | 0.67219335 |
| TDLATLG | ASSPTEDQDTQY | 0.68327415 |
| SAGGTGR | ASSPTEDQDTQY | 0.53181136 |
| GFSNSGS | ASSPTEDQDTQY | 0.71112734 |
| GGNGGSGGTSV | ASSPTEDQDTQY | 0.48859495 |
| GTSVATGGAGN | ASSPTEDQDTQY | 0.7748287 |
| NGGNGGSGGTS | ASSPTEDQDTQY | 0.44446814 |
| AVVKFPH | ASSPTEDQDTQY | 0.70615256 |
| TPPPPPP | ASSPTEDQDTQY | 0.7599971 |
| AAPAAAAAPA | ASSPTEDQDTQY | 0.6955367 |
| IALNEDL | ASSPTEDQDTQY | 0.62705356 |
| IVNVSSM | ASSPTEDQDTQY | 0.54897064 |
| TGIPPAPR | ASSPTEDQDTQY | 0.39632887 |
| FDLGGGTF | ASSPTEDQDTQY | 0.42535168 |

| | | |
|---|---|---|
| FDLGGGTFDVS | ASSPTEDQDTQY | 0.7831503 |
| NEPTAAA | ASSPTEDQDTQY | 0.60378826 |
| NEPTAAALA | ASSPTEDQDTQY | 0.7776382 |
| GSIINISSI | ASSPTEDQDTQY | 0.6835871 |
| VAPGNAGTA | ASSPTEDQDTQY | 0.4846632 |
| TGGGASLE | ASSPTEDQDTQY | 0.6822109 |
| VAVYNHYKRI | ASSPTEDQDTQY | 0.52737665 |
| TDPAPATT | ASSPTEDQDTQY | 0.4318179 |
| TNRPDILDPALLRPG | ASSPTEDQDTQY | 0.68091184 |
| LTTTTTS | ASSPTEDQDTQY | 0.7223444 |
| PTGSGKTL | ASSPTEDQDTQY | 0.503641 |
| TGSGKTL | ASSPTEDQDTQY | 0.6706946 |
| TIGHVDHGK | ASSPTEDQDTQY | 0.6838223 |
| TIGHVDHGKTT | ASSPTEDQDTQY | 0.39245373 |
| GHVDHGKTT | ASSPTEDQDTQY | 0.6320146 |
| AQFEHTLL | ASSPTEDQDTQY | 0.7719265 |
| IVNQDEL | ASSPTEDQDTQY | 0.67758936 |
| TNGSQFF | ASSPTEDQDTQY | 0.34123874 |
| TGKGIVE | ASSPTEDQDTQY | 0.743031 |
| LISNASDA | ASSPTEDQDTQY | 0.46780664 |
| AELRTLGELTKIAKAHGAQ | ASSPTEDQDTQY | 0.72891665 |
| APNAVAAAEPSW | ASSPTEDQDTQY | 0.6652143 |
| APQVAGAATAVL | ASSPTEDQDTQY | 0.59763443 |
| ARRDPAAAAAILPTDA | ASSPTEDQDTQY | 0.73685074 |
| DAAARDVLAEAGLAEYF | ASSPTEDQDTQY | 0.33704185 |
| DDQIVLSAKANLAKVEQ | ASSPTEDQDTQY | 0.55280447 |
| DELLPDTGEDSDEDGH | ASSPTEDQDTQY | 0.56332874 |
| ELPEAPANRATVPQD | ASSPTEDQDTQY | 0.71642566 |
| FGQSKIERVTGIG | ASSPTEDQDTQY | 0.7531416 |
| GAGGTGSTAGGAGGAGGAGGLY | ASSPTEDQDTQY | 0.48649147 |
| GHGGTGAAGGNGGTGGAGGSGI | ASSPTEDQDTQY | 0.6465732 |
| GNGGNGGAGGEGGPGGAGGAGA | ASSPTEDQDTQY | 0.50099576 |
| GSTAGNGGNGGNATNGTIAGQP | ASSPTEDQDTQY | 0.38588053 |
| GTGGDGGNGGNGGTGG | ASSPTEDQDTQY | 0.59397995 |
| GVRRPAGIPADAPV | ASSPTEDQDTQY | 0.5484173 |
| GYRQVTDAHLLTLARR | ASSPTEDQDTQY | 0.42607796 |
| IAKMGQDGHDRG | ASSPTEDQDTQY | 0.70938766 |
| KAEPSIEIQYT | ASSPTEDQDTQY | 0.57267356 |
| KPVRIRMAADEDTVAVT | ASSPTEDQDTQY | 0.7028355 |
| MPPRIAGMRLLVIKPEP | ASSPTEDQDTQY | 0.7639563 |
| MKFYAVALFRSRGVPPTR | ASSPTEDQDTQY | 0.57866037 |
| NAGGFGILGAVAHSP | ASSPTEDQDTQY | 0.6566222 |
| NEGREITLAMAEM | ASSPTEDQDTQY | 0.71296585 |
| NELVEWSHPLGHRWRWE | ASSPTEDQDTQY | 0.7555474 |
| NLGAGGTGGNGGNPG | ASSPTEDQDTQY | 0.6882595 |
| NTEWPKWETVMAADPVD | ASSPTEDQDTQY | 0.44270742 |
| PASAAPAETQP | ASSPTEDQDTQY | 0.757769 |
| QQSSDGKNGVRAAEITGE | ASSPTEDQDTQY | 0.7433988 |
| RAPDAVTTSWWKE | ASSPTEDQDTQY | 0.77939737 |
| SGFGNVGAHTSGWFNQ | ASSPTEDQDTQY | 0.7489647 |
| SGVAGAVGQAASVGGLKV | ASSPTEDQDTQY | 0.77616173 |

| | | |
|---|---|---|
| SNGGAGTGSAGGNGGT | ASSPTEDQDTQY | 0.747362 |
| TGGAGGEGGAAG | ASSPTEDQDTQY | 0.6545099 |
| TGGDGGAAGAGGAPGHG | ASSPTEDQDTQY | 0.3237867 |
| TMPPLLAAIANAGRHGV | ASSPTEDQDTQY | 0.6934824 |
| TSSTAGNSGGA | ASSPTEDQDTQY | 0.69984484 |
| VPATVPVGILC | ASSPTEDQDTQY | 0.58339304 |
| VQRPDSGGVVLTGRIS | ASSPTEDQDTQY | 0.29230735 |
| WTPNPVPATKGVPAGLVQAT | ASSPTEDQDTQY | 0.77399004 |
| YALKRALAKFSDRFDVVII | ASSPTEDQDTQY | 0.79309744 |
| YLIGSVIV | ASSPTEDQDTQY | 0.6342906 |
| YGQLVAVID | ASSPTEDQDTQY | 0.77724093 |
| VTVGGMTLPA | ASSPTEDQDTQY | 0.3813173 |
| VLALPAV | ASSPTEDQDTQY | 0.7439624 |
| VIPPTLNYET | ASSPTEDQDTQY | 0.7763326 |
| VGMGGMHPSSG | ASSPTEDQDTQY | 0.51074517 |
| VGLAGGAATA | ASSPTEDQDTQY | 0.75696194 |
| VGGAGGAGGSGP | ASSPTEDQDTQY | 0.316978 |
| VFAAFMV | ASSPTEDQDTQY | 0.4552645 |
| TLYDKLR | ASSPTEDQDTQY | 0.7842263 |
| TLAVFTVL | ASSPTEDQDTQY | 0.69898486 |
| TGSVQPARV | ASSPTEDQDTQY | 0.67263496 |
| TGGGAGSGGT | ASSPTEDQDTQY | 0.69689584 |
| SHLETSL | ASSPTEDQDTQY | 0.51283133 |
| SFTTGMT | ASSPTEDQDTQY | 0.6638763 |
| QVEVAKRMGVRQPTVS | ASSPTEDQDTQY | 0.78494376 |
| QGSGGPALAM | ASSPTEDQDTQY | 0.52918917 |
| MLAPSNP | ASSPTEDQDTQY | 0.78100705 |
| LVNNAAI | ASSPTEDQDTQY | 0.5140339 |
| LTGEINTLA | ASSPTEDQDTQY | 0.78913534 |
| LSIFGPV | ASSPTEDQDTQY | 0.7821253 |
| LLALLANL | ASSPTEDQDTQY | 0.7265834 |
| LISNASDALDKLR | ASSPTEDQDTQY | 0.49837968 |
| LINEAVP | ASSPTEDQDTQY | 0.7742319 |
| LGYAFSGL | ASSPTEDQDTQY | 0.6657554 |
| LDFVAVAL | ASSPTEDQDTQY | 0.7704232 |
| LAQPGAE | ASSPTEDQDTQY | 0.746186 |
| ISNLGAVTGF | ASSPTEDQDTQY | 0.14496091 |
| HQEDPGRI | ASSPTEDQDTQY | 0.7300259 |
| GTPGEPTP | ASSPTEDQDTQY | 0.49448374 |
| GFFNSGS | ASSPTEDQDTQY | 0.38381934 |
| GAGGGGLTAGAN | ASSPTEDQDTQY | 0.22317421 |
| FVDALVNM | ASSPTEDQDTQY | 0.1725081 |
| FLTVAPEL | ASSPTEDQDTQY | 0.24205288 |
| FETLTAL | ASSPTEDQDTQY | 0.18243788 |
| DVINPIEL | ASSPTEDQDTQY | 0.71791834 |
| APVSGAPV | ASSPTEDQDTQY | 0.47276837 |
| ALVGGLAW | ASSPTEDQDTQY | 0.22672041 |
| ALAAAAATAI | ASSPTEDQDTQY | 0.3855145 |
| AFGLGLLL | ASSPTEDQDTQY | 0.4573307 |
| AFFAAASL | ASSPTEDQDTQY | 0.13059059 |
| AAAALPAFSPP | ASSPTEDQDTQY | 0.32360125 |

| | | |
|---|---|---|
| VVLAPAL | ASSPTEDQDTQY | 0.2321445 |
| TIGHVDHGKTTL | ASSPTEDQDTQY | 0.22393982 |
| LISNASDALDK | ASSPTEDQDTQY | 0.10733079 |
| LISNASDAL | ASSPTEDQDTQY | 0.54431033 |
| ISNASDALDK | ASSPTEDQDTQY | 0.24542889 |
| NQAETLVYQTE | ASSPTEDQDTQY | 0.09890453 |
| TGGTGGTAGAG | ASSPTEDQDTQY | 0.31987214 |
| TGSTAGGAGGA | ASSPTEDQDTQY | 0.21171692 |
| DPLAVPVAAGPV | ASSPTEDQDTQY | 0.20944901 |
| GGTGGTPGNGGN | ASSPTEDQDTQY | 0.38200778 |
| PAGLAGVDPATL | ASSPTEDQDTQY | 0.41856772 |
| GNNLVVEAPHAGASV | ASSPTEDQDTQY | 0.20040332 |
| KVTRIMSASTLEQPA | ASSPTEDQDTQY | 0.46937925 |
| LGFALPIALPRRTVV | ASSPTEDQDTQY | 0.5378251 |
| TRLGNLTARKLVVLV | ASSPTEDQDTQY | 0.19682315 |
| WSQQRRLIYFPSAGP | ASSPTEDQDTQY | 0.3331809 |
| GPNGNPDPMAAAVDIR | ASSPTEDQDTQY | 0.7522568 |
| TQRCVSVADLRAAASR | ASSPTEDQDTQY | 0.2833126 |
| QFVLDVAEHPVEVLTETV | ASSPTEDQDTQY | 0.64738536 |
| GGAGGTGGAGGNGGTQAGDG | ASSPTEDQDTQY | 0.69965374 |
| PLLATLRAIAVDVDAKPAQV | ASSPTEDQDTQY | 0.27111492 |
| APKFRTGGYMIDFWGVGYQVAK | ASSPTEDQDTQY | 0.5389362 |
| GAGGAGGAGGAAGGPGGTGGTG | ASSPTEDQDTQY | 0.6427454 |
| IGALGLLATDVAPAGSVAFGVA | ASSPTEDQDTQY | 0.3294807 |
| AGGAGGIGGNGGAGGAGGVLMG | ASSPTEDQDTQY | 0.2539689 |
| ADELIGLL | ASSPTEDQDTQY | 0.1600656 |
| ADRALKA | ASSPTEDQDTQY | 0.30622488 |
| AGAVGQVGGAGGA | ASSPTEDQDTQY | 0.1910646 |
| AGGGGGLGGAP | ASSPTEDQDTQY | 0.46049672 |
| AGGLLYGNGGAG | ASSPTEDQDTQY | 0.70215786 |
| AGLESIGL | ASSPTEDQDTQY | 0.23966679 |
| AGLKTGII | ASSPTEDQDTQY | 0.12289525 |
| ASLAALLL | ASSPTEDQDTQY | 0.13197051 |
| ASVINISSI | ASSPTEDQDTQY | 0.5585879 |
| ATPPGPA | ASSPTEDQDTQY | 0.15352136 |
| ATVASALV | ASSPTEDQDTQY | 0.16917595 |
| ATVASLIL | ASSPTEDQDTQY | 0.24131875 |
| AVNGPVLIRH | ASSPTEDQDTQY | 0.47316676 |
| DGAAGAVGI | ASSPTEDQDTQY | 0.52680975 |
| DPLLHGI | ASSPTEDQDTQY | 0.15874356 |
| ELGAALGAL | ASSPTEDQDTQY | 0.6888101 |
| ELGRTQAV | ASSPTEDQDTQY | 0.7130961 |
| FVAGGYLS | ASSPTEDQDTQY | 0.8004676 |
| GASGGAGGAGGW | ASSPTEDQDTQY | 0.37811098 |
| GGAGAGGDGGAGGI | ASSPTEDQDTQY | 0.15491158 |
| GTGHNGSQPGMGGNG | ASSPTEDQDTQY | 0.7300592 |
| GYAAASASAS | ASSPTEDQDTQY | 0.22155753 |
| HQVDVAAVQQG | ASSPTEDQDTQY | 0.383465 |
| IGAAIAHT | ASSPTEDQDTQY | 0.39549023 |
| IGMIEVAQL | ASSPTEDQDTQY | 0.301624 |
| IRLAPEHTSAMD | ASSPTEDQDTQY | 0.18552157 |

| | | |
|---|---|---|
| IYNVSVL | ASSPTEDQDTQY | 0.6313945 |
| KQGFLGM | ASSPTEDQDTQY | 0.6718079 |
| LESVITQL | ASSPTEDQDTQY | 0.15257056 |
| LGLGALVVV | ASSPTEDQDTQY | 0.21099938 |
| LIGNGGNGGSGGI | ASSPTEDQDTQY | 0.30506986 |
| LNAINAPTL | ASSPTEDQDTQY | 0.2667127 |
| LQLLNTGV | ASSPTEDQDTQY | 0.50794536 |
| LQTSIEF | ASSPTEDQDTQY | 0.40825862 |
| MSFAVSEL | ASSPTEDQDTQY | 0.41691947 |
| NTGALGL | ASSPTEDQDTQY | 0.38959587 |
| PPGAGGAGGAGTT | ASSPTEDQDTQY | 0.56169415 |
| QLVGLGI | ASSPTEDQDTQY | 0.33574465 |
| RFLEQVHQL | ASSPTEDQDTQY | 0.4106655 |
| RGVGYVLR | ASSPTEDQDTQY | 0.17475647 |
| SATGVFAGVI | ASSPTEDQDTQY | 0.62762463 |
| SLPAITLP | ASSPTEDQDTQY | 0.5270413 |
| STALAFTV | ASSPTEDQDTQY | 0.5102369 |
| SVLTVGLGK | ASSPTEDQDTQY | 0.6752206 |
| TALAGAGSGI | ASSPTEDQDTQY | 0.54008734 |
| TGLPFTGI | ASSPTEDQDTQY | 0.45510972 |
| VAEHVTV | ASSPTEDQDTQY | 0.13041045 |
| VDIVLNSL | ASSPTEDQDTQY | 0.2697707 |
| VLTGGGAVL | ASSPTEDQDTQY | 0.22534765 |
| VNVGLAYGI | ASSPTEDQDTQY | 0.46076688 |
| VSSAAVGAV | ASSPTEDQDTQY | 0.41981736 |
| VTSVYEAGV | ASSPTEDQDTQY | 0.6240562 |
| WVALDRGA | ASSPTEDQDTQY | 0.23091304 |
| YLVGSVVI | ASSPTEDQDTQY | 0.60965276 |
| GPPGTGKTLL | ASSPTEDQDTQY | 0.6951777 |
| GVGMTKF | ASSPTEDQDTQY | 0.15203576 |
| IGIGGSDL | ASSPTEDQDTQY | 0.22524907 |
| IGTLLAEL | ASSPTEDQDTQY | 0.6981523 |
| IYTSGTT | ASSPTEDQDTQY | 0.43765792 |
| LSFPAGV | ASSPTEDQDTQY | 0.20939551 |
| LYTSGTT | ASSPTEDQDTQY | 0.2702679 |
| VGAFPFGF | ASSPTEDQDTQY | 0.79570806 |
| VGAGTVEF | ASSPTEDQDTQY | 0.30757385 |
| VGKSTLLL | ASSPTEDQDTQY | 0.7113134 |
| AALAHAGGGIV | ASSPTEDQDTQY | 0.118161954 |
| AALVAVSLL | ASSPTEDQDTQY | 0.3659034 |
| AAVSRAVAAL | ASSPTEDQDTQY | 0.31600326 |
| AGGSSGIGGF | ASSPTEDQDTQY | 0.100405335 |
| AGLTVANANG | ASSPTEDQDTQY | 0.47094047 |
| AGLVGAVL | ASSPTEDQDTQY | 0.19931026 |
| ALSQAFTV | ASSPTEDQDTQY | 0.33572242 |
| ALTFSVSL | ASSPTEDQDTQY | 0.22794712 |
| ALTGAALVAL | ASSPTEDQDTQY | 0.52526283 |
| ASFSIGAGVL | ASSPTEDQDTQY | 0.4205053 |
| DGSVYKI | ASSPTEDQDTQY | 0.22073056 |
| EQLAEIV | ASSPTEDQDTQY | 0.20235145 |
| FAVSELTVM | ASSPTEDQDTQY | 0.33032334 |

| | | |
|---|---|---|
| GAAPLAAGL | ASSPTEDQDTQY | 0.56956625 |
| GADGGDGGSGGAGG | ASSPTEDQDTQY | 0.6463768 |
| GFGLDTGGAGG | ASSPTEDQDTQY | 0.14031492 |
| GLASGGLL | ASSPTEDQDTQY | 0.34354392 |
| GRIAGAAVLL | ASSPTEDQDTQY | 0.27181607 |
| GSAGSLNTSF | ASSPTEDQDTQY | 0.13984266 |
| GTGQIGI | ASSPTEDQDTQY | 0.09901167 |
| GTLIALIL | ASSPTEDQDTQY | 0.34316105 |
| IAATMGFVVG | ASSPTEDQDTQY | 0.3366976 |
| IFAQALL | ASSPTEDQDTQY | 0.3159392 |
| IGAAVALGF | ASSPTEDQDTQY | 0.4271698 |
| IGLTPLAL | ASSPTEDQDTQY | 0.1855048 |
| ILGSNTSTL | ASSPTEDQDTQY | 0.5563483 |
| IVFSLTVL | ASSPTEDQDTQY | 0.3618593 |
| LASKATGF | ASSPTEDQDTQY | 0.08038978 |
| LASVALIL | ASSPTEDQDTQY | 0.23975015 |
| LFAVGLTL | ASSPTEDQDTQY | 0.12186147 |
| LFNGGIGTYI | ASSPTEDQDTQY | 0.7093415 |
| LTLGAGII | ASSPTEDQDTQY | 0.14457086 |
| LVAAALGDI | ASSPTEDQDTQY | 0.20139644 |
| NTAFVQLGI | ASSPTEDQDTQY | 0.23013845 |
| QLFSGAEL | ASSPTEDQDTQY | 0.72138816 |
| SAATFITL | ASSPTEDQDTQY | 0.39593467 |
| SNYGLANLGV | ASSPTEDQDTQY | 0.2374017 |
| SYGGAYAVM | ASSPTEDQDTQY | 0.12206016 |
| TTGVGGLTLG | ASSPTEDQDTQY | 0.3445236 |
| TTQTIYI | ASSPTEDQDTQY | 0.43281868 |
| VAAGAGLGAVI | ASSPTEDQDTQY | 0.20644334 |
| VAAVVTLL | ASSPTEDQDTQY | 0.14221142 |
| VGFLGGLI | ASSPTEDQDTQY | 0.18270858 |
| VGVIPILL | ASSPTEDQDTQY | 0.19213504 |
| VIFASSLI | ASSPTEDQDTQY | 0.6010467 |
| VKGGDGGAAAG | ASSPTEDQDTQY | 0.1591831 |
| VTKTFTNI | ASSPTEDQDTQY | 0.19028717 |
| VVAAAILL | ASSPTEDQDTQY | 0.7498833 |
| YAALDVEL | ASSPTEDQDTQY | 0.63406503 |
| LASVALLL | ASSPTEDQDTQY | 0.22407213 |
| LDAVHRV | ASSPTEDQDTQY | 0.23733842 |
| QGLSSSIF | ASSPTEDQDTQY | 0.1624113 |
| RGTFANIRL | ASSPTEDQDTQY | 0.19212186 |
| RQLWWGHRI | ASSPTEDQDTQY | 0.23674801 |
| YGPPGTGKTLL | ASSPTEDQDTQY | 0.41254792 |
| AALVGAPANVA | ASSPTEDQDTQY | 0.36847386 |
| AATFRRTE | ASSPTEDQDTQY | 0.16249971 |
| AGALGGGI | ASSPTEDQDTQY | 0.76944935 |
| AGGAGGAGGIGAG | ASSPTEDQDTQY | 0.71001667 |
| AGSGALGL | ASSPTEDQDTQY | 0.7752524 |
| AQIADQI | ASSPTEDQDTQY | 0.14518946 |
| ASAVAVTYL | ASSPTEDQDTQY | 0.34767702 |
| ASTVLGGTGG | ASSPTEDQDTQY | 0.2583605 |
| AVGGEPAK | ASSPTEDQDTQY | 0.23088303 |

| | | |
|---|---|---|
| CEQGTFGGTC | ASSPTEDQDTQY | 0.19568504 |
| DEGKAPAG | ASSPTEDQDTQY | 0.20277125 |
| DTVVFPAL | ASSPTEDQDTQY | 0.6416657 |
| FEGITMAG | ASSPTEDQDTQY | 0.8343515 |
| FPLVVQD | ASSPTEDQDTQY | 0.2867282 |
| GAFTTPQ | ASSPTEDQDTQY | 0.12425023 |
| GFGGFGTAGGVGGA | ASSPTEDQDTQY | 0.20871359 |
| GFLAQFL | ASSPTEDQDTQY | 0.20031938 |
| GGAGGAGGNAGVA | ASSPTEDQDTQY | 0.15357788 |
| GGVGTAGAGGAGGAGG | ASSPTEDQDTQY | 0.27542034 |
| GNGGGGDAGNAGSGGN | ASSPTEDQDTQY | 0.123759635 |
| GTGGAGGAGNDAGS | ASSPTEDQDTQY | 0.28401256 |
| GTTNTGWFNS | ASSPTEDQDTQY | 0.78817505 |
| IFMGPLFTL | ASSPTEDQDTQY | 0.31294376 |
| ILAHAIEAI | ASSPTEDQDTQY | 0.36435258 |
| IRSAIREAL | ASSPTEDQDTQY | 0.6543186 |
| ITFAGGG | ASSPTEDQDTQY | 0.44222808 |
| ITGMSGHQEV | ASSPTEDQDTQY | 0.5725473 |
| IVAGLACTL | ASSPTEDQDTQY | 0.7257571 |
| IVGGGLAGA | ASSPTEDQDTQY | 0.30344117 |
| IVTGVLNI | ASSPTEDQDTQY | 0.4744842 |
| LAQQGCI | ASSPTEDQDTQY | 0.50056446 |
| LCLHVEN | ASSPTEDQDTQY | 0.20381786 |
| LDAISDGS | ASSPTEDQDTQY | 0.28627685 |
| LGGFAAML | ASSPTEDQDTQY | 0.3540893 |
| LGGKNPMI | ASSPTEDQDTQY | 0.43893903 |
| LGPKGVAAIL | ASSPTEDQDTQY | 0.42606792 |
| LMVGVSI | ASSPTEDQDTQY | 0.7573819 |
| LVGAAFVL | ASSPTEDQDTQY | 0.11360771 |
| LVLAVAPL | ASSPTEDQDTQY | 0.17779925 |
| MAAIAVIDAA | ASSPTEDQDTQY | 0.65045524 |
| NNGINAELQ | ASSPTEDQDTQY | 0.114295594 |
| RGEPLVLVHG | ASSPTEDQDTQY | 0.14520967 |
| RGGAGGAAGLI | ASSPTEDQDTQY | 0.3937202 |
| RSYRMHPAV | ASSPTEDQDTQY | 0.7651967 |
| SAAHLLAGQLA | ASSPTEDQDTQY | 0.10927872 |
| SAAVGSY | ASSPTEDQDTQY | 0.28072572 |
| SPHAVLHRI | ASSPTEDQDTQY | 0.15166879 |
| SQIAKVL | ASSPTEDQDTQY | 0.7443022 |
| TALGHLVP | ASSPTEDQDTQY | 0.2936849 |
| TAPTTSGAA | ASSPTEDQDTQY | 0.14858077 |
| TDGDDGNGGSPGT | ASSPTEDQDTQY | 0.46262997 |
| TDGLGEN | ASSPTEDQDTQY | 0.12829998 |
| TGGPPLPGQA | ASSPTEDQDTQY | 0.30141646 |
| TIPAITI | ASSPTEDQDTQY | 0.5024331 |
| TIPGFTI | ASSPTEDQDTQY | 0.7711626 |
| TSIASGALA | ASSPTEDQDTQY | 0.13493541 |
| TVTLPGAAI | ASSPTEDQDTQY | 0.4402929 |
| VASGLSAALP | ASSPTEDQDTQY | 0.28901574 |
| VGAGVIGI | ASSPTEDQDTQY | 0.81924653 |
| VGPKRFGV | ASSPTEDQDTQY | 0.22785713 |

| | | |
|---|---|---|
| VNVAFPD | ASSPTEDQDTQY | 0.14778465 |
| VSMTNSGVS | ASSPTEDQDTQY | 0.5422979 |
| VTDVLGDLSV | ASSPTEDQDTQY | 0.47055525 |
| VVVVGAGFAGL | ASSPTEDQDTQY | 0.32842702 |
| VYTVGTVV | ASSPTEDQDTQY | 0.6818532 |
| WTTFSGP | ASSPTEDQDTQY | 0.5242468 |
| YDELVEAF | ASSPTEDQDTQY | 0.24439535 |
| YGTGGAGGNGG | ASSPTEDQDTQY | 0.406542 |
| YTNVMAV | ASSPTEDQDTQY | 0.38331968 |
| FLEQVHQL | ASSPTEDQDTQY | 0.22574191 |
| GLSLLAVAV | ASSPTEDQDTQY | 0.5941007 |
| LEQVHQL | ASSPTEDQDTQY | 0.2977897 |
| LNAAGAI | ASSPTEDQDTQY | 0.15745892 |
| PAVHASL | ASSPTEDQDTQY | 0.7872463 |
| TELEDDA | ASSPTEDQDTQY | 0.35239545 |
| TPPGPPN | ASSPTEDQDTQY | 0.51596934 |
| TPPPPPPP | ASSPTEDQDTQY | 0.4682486 |
| TQSPALI | ASSLGTVANTEVF | 0.79236746 |
| EFAAAFPQ | ASSLGTVANTEVF | 0.18669541 |
| LLSGNLDVL | ASSLGTVANTEVF | 0.39315695 |
| GPANLGNYNI | ASSLGTVANTEVF | 0.31696782 |
| TLGPTLP | ASSLGTVANTEVF | 0.5640277 |
| VYAYSSCCK | ASSLGTVANTEVF | 0.39400733 |
| APAPAPAPNI | ASSLGTVANTEVF | 0.4787475 |
| PGGPVPIP | ASSLGTVANTEVF | 0.4912817 |
| FLTVALEP | ASSLGTVANTEVF | 0.6062771 |
| SQYFGANL | ASSLGTVANTEVF | 0.33382374 |
| GGVKPAA | ASSLGTVANTEVF | 0.35753867 |
| GGTGGAGGAGGW | ASSLGTVANTEVF | 0.6794741 |
| GVGGVAGLGGAGG | ASSLGTVANTEVF | 0.16048926 |
| VLLVLATTI | ASSLGTVANTEVF | 0.31282097 |
| IGIFAAML | ASSLGTVANTEVF | 0.5867941 |
| AVGDLID | ASSLGTVANTEVF | 0.4806286 |
| NISPLPL | ASSLGTVANTEVF | 0.33483768 |
| VTAVFVAI | ASSLGTVANTEVF | 0.5600107 |
| SGDDNDAAL | ASSLGTVANTEVF | 0.26517957 |
| AVAALAHS | ASSLGTVANTEVF | 0.29836988 |
| GGVTVMATP | ASSLGTVANTEVF | 0.37121558 |
| AGLAANVRD | ASSLGTVANTEVF | 0.20209284 |
| TNRPDLIDPALLRPG | ASSLGTVANTEVF | 0.15880352 |
| LANLGVTAL | ASSLGTVANTEVF | 0.23253298 |
| TGERIEL | ASSLGTVANTEVF | 0.548661 |
| NGEELAA | ASSLGTVANTEVF | 0.32162464 |
| VLSLVWG | ASSLGTVANTEVF | 0.10852753 |
| IGYALWL | ASSLGTVANTEVF | 0.3920061 |
| LSLLLAPL | ASSLGTVANTEVF | 0.38638785 |
| TLDEALK | ASSLGTVANTEVF | 0.78509104 |
| TDGAGTAEGG | ASSLGTVANTEVF | 0.4294346 |
| NAPGGAGGNGG | ASSLGTVANTEVF | 0.52591896 |
| VSVFMGMGGNF | ASSLGTVANTEVF | 0.21471772 |
| LASYLAAL | ASSLGTVANTEVF | 0.4271773 |

| | | |
|---|---|---|
| LVGGTPAL | ASSLGTVANTEVF | 0.1323256 |
| VRNADVL | ASSLGTVANTEVF | 0.21494727 |
| TCSSPFLVSAM | ASSLGTVANTEVF | 0.25018817 |
| VHITDAE | ASSLGTVANTEVF | 0.1462399 |
| EYAAPSF | ASSLGTVANTEVF | 0.7241425 |
| VVDAFHGGL | ASSLGTVANTEVF | 0.5243349 |
| ASVVDPQ | ASSLGTVANTEVF | 0.61693513 |
| SYNEFGGLS | ASSLGTVANTEVF | 0.69997793 |
| DPDGSVY | ASSLGTVANTEVF | 0.6999244 |
| MVSTIHATE | ASSLGTVANTEVF | 0.4904703 |
| LAAGVADPV | ASSLGTVANTEVF | 0.21994707 |
| TKEQIAA | ASSLGTVANTEVF | 0.25894648 |
| SLASPAAL | ASSLGTVANTEVF | 0.11828743 |
| IGLVFYGV | ASSLGTVANTEVF | 0.24918386 |
| FLLDASNL | ASSLGTVANTEVF | 0.2805822 |
| YQVGNFP | ASSLGTVANTEVF | 0.267281 |
| ESLLSQST | ASSLGTVANTEVF | 0.09974344 |
| LFTVGFAAVG | ASSLGTVANTEVF | 0.6577962 |
| ATPPTPT | ASSLGTVANTEVF | 0.40957177 |
| TDLATLG | ASSLGTVANTEVF | 0.35254675 |
| SAGGTGR | ASSLGTVANTEVF | 0.48636767 |
| GFSNSGS | ASSLGTVANTEVF | 0.53044635 |
| GGNGGSGGTSV | ASSLGTVANTEVF | 0.21584144 |
| GTSVATGGAGN | ASSLGTVANTEVF | 0.5434152 |
| NGGNGGSGGTS | ASSLGTVANTEVF | 0.26406014 |
| AVVKFPH | ASSLGTVANTEVF | 0.55166066 |
| TPPPPPP | ASSLGTVANTEVF | 0.6553509 |
| AAPAAAAAPA | ASSLGTVANTEVF | 0.44038403 |
| IALNEDL | ASSLGTVANTEVF | 0.28367066 |
| IVNVSSM | ASSLGTVANTEVF | 0.3711242 |
| TGIPPAPR | ASSLGTVANTEVF | 0.17645866 |
| FDLGGGTF | ASSLGTVANTEVF | 0.2930795 |
| FDLGGGTFDVS | ASSLGTVANTEVF | 0.09513373 |
| NEPTAAA | ASSLGTVANTEVF | 0.44601625 |
| NEPTAAALA | ASSLGTVANTEVF | 0.20985924 |
| GSIINISSI | ASSLGTVANTEVF | 0.31529903 |
| VAPGNAGTA | ASSLGTVANTEVF | 0.2666231 |
| TGGGASLE | ASSLGTVANTEVF | 0.46775782 |
| VAVYNHYKRI | ASSLGTVANTEVF | 0.14479 |
| TDPAPATT | ASSLGTVANTEVF | 0.4516536 |
| TNRPDILDPALLRPG | ASSLGTVANTEVF | 0.4730219 |
| LTTTTTS | ASSLGTVANTEVF | 0.1678807 |
| PTGSGKTL | ASSLGTVANTEVF | 0.4960484 |
| TGSGKTL | ASSLGTVANTEVF | 0.35750824 |
| TIGHVDHGK | ASSLGTVANTEVF | 0.3639791 |
| TIGHVDHGKTT | ASSLGTVANTEVF | 0.6161177 |
| GHVDHGKTT | ASSLGTVANTEVF | 0.32504788 |
| AQFEHTLL | ASSLGTVANTEVF | 0.17186205 |
| IVNQDEL | ASSLGTVANTEVF | 0.24586672 |
| TNGSQFF | ASSLGTVANTEVF | 0.38124698 |
| TGKGIVE | ASSLGTVANTEVF | 0.25845656 |

| | | |
|---|---|---|
| LISNASDA | ASSLGTVANTEVF | 0.09919438 |
| AELRTLGELTKIAKAHGAQ | ASSLGTVANTEVF | 0.48558903 |
| APNAVAAAEPSW | ASSLGTVANTEVF | 0.1606152 |
| APQVAGAATAVL | ASSLGTVANTEVF | 0.22021976 |
| ARRDPAAAAAILPTDA | ASSLGTVANTEVF | 0.21210836 |
| DAAARDVLAEAGLAEYF | ASSLGTVANTEVF | 0.4697142 |
| DDQIVLSAKANLAKVEQ | ASSLGTVANTEVF | 0.6484504 |
| DELLPDTGEDSDEDGH | ASSLGTVANTEVF | 0.3991379 |
| ELPEAPANRATVPQD | ASSLGTVANTEVF | 0.3689388 |
| FGQSKIERVTGIG | ASSLGTVANTEVF | 0.38706863 |
| GAGGTGSTAGGAGGAGGAGGLY | ASSLGTVANTEVF | 0.49076098 |
| GHGGTGAAGGNGGTGGAGGSGI | ASSLGTVANTEVF | 0.42271376 |
| GNGGNGGAGGEGGPGGAGGAGA | ASSLGTVANTEVF | 0.45333457 |
| GSTAGNGGNGGNATNGTIAGQP | ASSLGTVANTEVF | 0.38569066 |
| GTGGDGGNGGNGGTGG | ASSLGTVANTEVF | 0.28822637 |
| GVRRPAGIPADAPV | ASSLGTVANTEVF | 0.13177985 |
| GYRQVTDAHLLTLARR | ASSLGTVANTEVF | 0.20626208 |
| IAKMGQDGHDRG | ASSLGTVANTEVF | 0.20270982 |
| KAEPSIEIQYT | ASSLGTVANTEVF | 0.1445412 |
| KPVRIRMAADEDTVAVT | ASSLGTVANTEVF | 0.4096777 |
| MPPRIAGMRLLVIKPEP | ASSLGTVANTEVF | 0.7103758 |
| MKFYAVALFRSRGVPPTR | ASSLGTVANTEVF | 0.6147927 |
| NAGGFGILGAVAHSP | ASSLGTVANTEVF | 0.13752922 |
| NEGREITLAMAEM | ASSLGTVANTEVF | 0.26825184 |
| NELVEWSHPLGHRWRWE | ASSLGTVANTEVF | 0.2798177 |
| NLGAGGTGGNGGNPG | ASSLGTVANTEVF | 0.33884507 |
| NTEWPKWETVMAADPVD | ASSLGTVANTEVF | 0.6943379 |
| PASAAPAETQP | ASSLGTVANTEVF | 0.800226 |
| QQSSDGKNGVRAAEITGE | ASSLGTVANTEVF | 0.5413873 |
| RAPDAVTTSWWKE | ASSLGTVANTEVF | 0.1996896 |
| SGFGNVGAHTSGWFNQ | ASSLGTVANTEVF | 0.41033113 |
| SGVAGAVGQAASVGGLKV | ASSLGTVANTEVF | 0.2952103 |
| SNGGAGTGSAGGNGGT | ASSLGTVANTEVF | 0.2544805 |
| TGGAGGEGGAAG | ASSLGTVANTEVF | 0.33091855 |
| TGGDGGAAGAGGAPGHG | ASSLGTVANTEVF | 0.5913093 |
| TMPPLLAAIANAGRHGV | ASSLGTVANTEVF | 0.26323926 |
| TSSTAGNSGGA | ASSLGTVANTEVF | 0.20205885 |
| VPATVPVGILC | ASSLGTVANTEVF | 0.16907597 |
| VQRPDSGGVVLTGRIS | ASSLGTVANTEVF | 0.52382815 |
| WTPNPVPATKGVPAGLVQAT | ASSLGTVANTEVF | 0.2720445 |
| YALKRALAKFSDRFDVVII | ASSLGTVANTEVF | 0.21940911 |
| YLIGSVIV | ASSLGTVANTEVF | 0.2542229 |
| YGQLVAVID | ASSLGTVANTEVF | 0.7356876 |
| VTVGGMTLPA | ASSLGTVANTEVF | 0.7418624 |
| VLALPAV | ASSLGTVANTEVF | 0.2371436 |
| VIPPTLNYET | ASSLGTVANTEVF | 0.22694169 |
| VGMGGMHPSSG | ASSLGTVANTEVF | 0.6724433 |
| VGLAGGAATA | ASSLGTVANTEVF | 0.18962443 |
| VGGAGGAGGSGP | ASSLGTVANTEVF | 0.5318156 |
| VFAAFMV | ASSLGTVANTEVF | 0.39508682 |
| TLYDKLR | ASSLGTVANTEVF | 0.26410553 |

| | | |
|---|---|---|
| TLAVFTVL | ASSLGTVANTEVF | 0.23131391 |
| TGSVQPARV | ASSLGTVANTEVF | 0.5528962 |
| TGGGAGSGGT | ASSLGTVANTEVF | 0.23722677 |
| SHLETSL | ASSLGTVANTEVF | 0.5236346 |
| SFTTGMT | ASSLGTVANTEVF | 0.1259793 |
| QVEVAKRMGVRQPTVS | ASSLGTVANTEVF | 0.6268289 |
| QGSGGPALAM | ASSLGTVANTEVF | 0.13733783 |
| MLAPSNP | ASSLGTVANTEVF | 0.26171765 |
| LVNNAAI | ASSLGTVANTEVF | 0.114070244 |
| LTGEINTLA | ASSLGTVANTEVF | 0.2248003 |
| LSIFGPV | ASSLGTVANTEVF | 0.5435849 |
| LLALLANL | ASSLGTVANTEVF | 0.70727885 |
| LISNASDALDKLR | ASSLGTVANTEVF | 0.2656551 |
| LINEAVP | ASSLGTVANTEVF | 0.28420314 |
| LGYAFSGL | ASSLGTVANTEVF | 0.7728129 |
| LDFVAVAL | ASSLGTVANTEVF | 0.17045815 |
| LAQPGAE | ASSLGTVANTEVF | 0.12239572 |
| ISNLGAVTGF | ASSLGTVANTEVF | 0.3403434 |
| HQEDPGRI | ASSLGTVANTEVF | 0.5252893 |
| GTPGEPTP | ASSLGTVANTEVF | 0.7088591 |
| GFFNSGS | ASSLGTVANTEVF | 0.69068617 |
| GAGGGGLTAGAN | ASSLGTVANTEVF | 0.11214448 |
| FVDALVNM | ASSLGTVANTEVF | 0.5732974 |
| FLTVAPEL | ASSLGTVANTEVF | 0.54299897 |
| FETLTAL | ASSLGTVANTEVF | 0.16317867 |
| DVINPIEL | ASSLGTVANTEVF | 0.17851901 |
| APVSGAPV | ASSLGTVANTEVF | 0.2076509 |
| ALVGGLAW | ASSLGTVANTEVF | 0.12209229 |
| ALAAAAATAI | ASSLGTVANTEVF | 0.4810471 |
| AFGLGLLL | ASSLGTVANTEVF | 0.22916904 |
| AFFAAASL | ASSLGTVANTEVF | 0.34317616 |
| AAAALPAFSPP | ASSLGTVANTEVF | 0.25660658 |
| VVLAPAL | ASSLGTVANTEVF | 0.27408865 |
| TIGHVDHGKTTL | ASSLGTVANTEVF | 0.25502878 |
| LISNASDALDK | ASSLGTVANTEVF | 0.24960962 |
| LISNASDAL | ASSLGTVANTEVF | 0.5199975 |
| ISNASDALDK | ASSLGTVANTEVF | 0.61115515 |
| NQAETLVYQTE | ASSLGTVANTEVF | 0.3985629 |
| TGGTGGTAGAG | ASSLGTVANTEVF | 0.3523715 |
| TGSTAGGAGGA | ASSLGTVANTEVF | 0.5632467 |
| DPLAVPVAAGPV | ASSLGTVANTEVF | 0.17986625 |
| GGTGGTPGNGGN | ASSLGTVANTEVF | 0.20914781 |
| PAGLAGVDPATL | ASSLGTVANTEVF | 0.5471773 |
| GNNLVVEAPHAGASV | ASSLGTVANTEVF | 0.26013532 |
| KVTRIMSASTLEQPA | ASSLGTVANTEVF | 0.63826686 |
| LGFALPIALPRRTVV | ASSLGTVANTEVF | 0.33511993 |
| TRLGNLTARKLVVLV | ASSLGTVANTEVF | 0.33217406 |
| WSQQRRLIYFPSAGP | ASSLGTVANTEVF | 0.58450335 |
| GPNGNPDPMAAAVDIR | ASSLGTVANTEVF | 0.6808795 |
| TQRCVSVADLRAAASR | ASSLGTVANTEVF | 0.34601024 |
| QFVLDVAEHPVEVLTETV | ASSLGTVANTEVF | 0.23789299 |

| | | |
|---|---|---|
| GGAGGTGGAGGNGGTQAGDG | ASSLGTVANTEVF | 0.6240449 |
| PLLATLRAIAVDVDAKPAQV | ASSLGTVANTEVF | 0.19118226 |
| APKFRTGGYMIDFWGVGYQVAK | ASSLGTVANTEVF | 0.38430384 |
| GAGGAGGAGGAAGGPGGTGGTG | ASSLGTVANTEVF | 0.55747354 |
| IGALGLLATDVAPAGSVAFGVA | ASSLGTVANTEVF | 0.143814 |
| AGGAGGIGGNGGAGGAGGVLMG | ASSLGTVANTEVF | 0.25150806 |
| ADELIGLL | ASSLGTVANTEVF | 0.69821405 |
| ADRALKA | ASSLGTVANTEVF | 0.3829958 |
| AGAVGQVGGAGGA | ASSLGTVANTEVF | 0.20530061 |
| AGGGGGLGGAP | ASSLGTVANTEVF | 0.17043264 |
| AGGLLYGNGGAG | ASSLGTVANTEVF | 0.22389095 |
| AGLESIGL | ASSLGTVANTEVF | 0.44633925 |
| AGLKTGII | ASSLGTVANTEVF | 0.15527555 |
| ASLAALLL | ASSLGTVANTEVF | 0.51960546 |
| ASVINISSI | ASSLGTVANTEVF | 0.20839077 |
| ATPPGPA | ASSLGTVANTEVF | 0.7455481 |
| ATVASALV | ASSLGTVANTEVF | 0.37104636 |
| ATVASLIL | ASSLGTVANTEVF | 0.51272744 |
| AVNGPVLIRH | ASSLGTVANTEVF | 0.2614017 |
| DGAAGAVGI | ASSLGTVANTEVF | 0.2393837 |
| DPLLHGI | ASSLGTVANTEVF | 0.14820799 |
| ELGAALGAL | ASSLGTVANTEVF | 0.19429207 |
| ELGRTQAV | ASSLGTVANTEVF | 0.19643404 |
| FVAGGYLS | ASSLGTVANTEVF | 0.3322276 |
| GASGGAGGAGGW | ASSLGTVANTEVF | 0.15398107 |
| GGAGAGGDGGAGGI | ASSLGTVANTEVF | 0.7778692 |
| GTGHNGSQPGMGGNG | ASSLGTVANTEVF | 0.20515674 |
| GYAAASASAS | ASSLGTVANTEVF | 0.37553203 |
| HQVDVAAVQQG | ASSLGTVANTEVF | 0.31355256 |
| IGAAIAHT | ASSLGTVANTEVF | 0.1517398 |
| IGMIEVAQL | ASSLGTVANTEVF | 0.21297272 |
| IRLAPEHTSAMD | ASSLGTVANTEVF | 0.52127874 |
| IYNVSVL | ASSLGTVANTEVF | 0.13700865 |
| KQGFLGM | ASSLGTVANTEVF | 0.117568836 |
| LESVITQL | ASSLGTVANTEVF | 0.22381845 |
| LGLGALVVV | ASSLGTVANTEVF | 0.17602614 |
| LIGNGGNGGSGGI | ASSLGTVANTEVF | 0.764916 |
| LNAINAPTL | ASSLGTVANTEVF | 0.6474576 |
| LQLLNTGV | ASSLGTVANTEVF | 0.34309945 |
| LQTSIEF | ASSLGTVANTEVF | 0.7702434 |
| MSFAVSEL | ASSLGTVANTEVF | 0.10181018 |
| NTGALGL | ASSLGTVANTEVF | 0.2872536 |
| PPGAGGAGGAGTT | ASSLGTVANTEVF | 0.65916324 |
| QLVGLGI | ASSLGTVANTEVF | 0.47965986 |
| RFLEQVHQL | ASSLGTVANTEVF | 0.22932193 |
| RGVGYVLR | ASSLGTVANTEVF | 0.49299043 |
| SATGVFAGVI | ASSLGTVANTEVF | 0.19324806 |
| SLPAITLP | ASSLGTVANTEVF | 0.25933784 |
| STALAFTV | ASSLGTVANTEVF | 0.25065222 |
| SVLTVGLGK | ASSLGTVANTEVF | 0.43123427 |
| TALAGAGSGI | ASSLGTVANTEVF | 0.4197243 |

| | | |
|---|---|---|
| TGLPFTGI | ASSLGTVANTEVF | 0.12763792 |
| VAEHVTV | ASSLGTVANTEVF | 0.4669812 |
| VDIVLNSL | ASSLGTVANTEVF | 0.7891094 |
| VLTGGGAVL | ASSLGTVANTEVF | 0.17433205 |
| VNVGLAYGI | ASSLGTVANTEVF | 0.5888968 |
| VSSAAVGAV | ASSLGTVANTEVF | 0.3068205 |
| VTSVYEAGV | ASSLGTVANTEVF | 0.59170616 |
| WVALDRGA | ASSLGTVANTEVF | 0.1837923 |
| YLVGSVVI | ASSLGTVANTEVF | 0.13202113 |
| GPPGTGKTLL | ASSLGTVANTEVF | 0.4745636 |
| GVGMTKF | ASSLGTVANTEVF | 0.14531371 |
| IGIGGSDL | ASSLGTVANTEVF | 0.38788798 |
| IGTLLAEL | ASSLGTVANTEVF | 0.32916692 |
| IYTSGTT | ASSLGTVANTEVF | 0.82825255 |
| LSFPAGV | ASSLGTVANTEVF | 0.5118863 |
| LYTSGTT | ASSLGTVANTEVF | 0.34127092 |
| VGAFPFGF | ASSLGTVANTEVF | 0.16134185 |
| VGAGTVEF | ASSLGTVANTEVF | 0.40537983 |
| VGKSTLLL | ASSLGTVANTEVF | 0.59025174 |
| AALAHAGGGIV | ASSLGTVANTEVF | 0.17710881 |
| AALVAVSLL | ASSLGTVANTEVF | 0.6335648 |
| AAVSRAVAAL | ASSLGTVANTEVF | 0.16960411 |
| AGGSSGIGGF | ASSLGTVANTEVF | 0.54049706 |
| AGLTVANANG | ASSLGTVANTEVF | 0.44564947 |
| AGLVGAVL | ASSLGTVANTEVF | 0.77342236 |
| ALSQAFTV | ASSLGTVANTEVF | 0.26464763 |
| ALTFSVSL | ASSLGTVANTEVF | 0.46988612 |
| ALTGAALVAL | ASSLGTVANTEVF | 0.15167478 |
| ASFSIGAGVL | ASSLGTVANTEVF | 0.6460737 |
| DGSVYKI | ASSLGTVANTEVF | 0.5588341 |
| EQLAEIV | ASSLGTVANTEVF | 0.62155426 |
| FAVSELTVM | ASSLGTVANTEVF | 0.7742504 |
| GAAPLAAGL | ASSLGTVANTEVF | 0.7075987 |
| GADGGDGGSGGAGG | ASSLGTVANTEVF | 0.15961532 |
| GFGLDTGGAGG | ASSLGTVANTEVF | 0.26067635 |
| GLASGGLL | ASSLGTVANTEVF | 0.6040343 |
| GRIAGAAVLL | ASSLGTVANTEVF | 0.7229213 |
| GSAGSLNTSF | ASSLGTVANTEVF | 0.55380315 |
| GTGQIGI | ASSLGTVANTEVF | 0.3699589 |
| GTLIALIL | ASSLGTVANTEVF | 0.28373468 |
| IAATMGFVVG | ASSLGTVANTEVF | 0.712849 |
| IFAQALL | ASSLGTVANTEVF | 0.8649268 |
| IGAAVALGF | ASSLGTVANTEVF | 0.44147104 |
| IGLTPLAL | ASSLGTVANTEVF | 0.2230863 |
| ILGSNTSTL | ASSLGTVANTEVF | 0.30493474 |
| IVFSLTVL | ASSLGTVANTEVF | 0.46899262 |
| LASKATGF | ASSLGTVANTEVF | 0.6176133 |
| LASVALIL | ASSLGTVANTEVF | 0.18490155 |
| LFAVGLTL | ASSLGTVANTEVF | 0.29091442 |
| LFNGGIGTYI | ASSLGTVANTEVF | 0.30725315 |
| LTLGAGII | ASSLGTVANTEVF | 0.11124989 |

| | | |
|---|---|---|
| LVAAALGDI | ASSLGTVANTEVF | 0.56547916 |
| NTAFVQLGI | ASSLGTVANTEVF | 0.6334282 |
| QLFSGAEL | ASSLGTVANTEVF | 0.36812332 |
| SAATFITL | ASSLGTVANTEVF | 0.15493879 |
| SNYGLANLGV | ASSLGTVANTEVF | 0.6456324 |
| SYGGAYAVM | ASSLGTVANTEVF | 0.65477467 |
| TTGVGGLTLG | ASSLGTVANTEVF | 0.6649676 |
| TTQTIYI | ASSLGTVANTEVF | 0.1562359 |
| VAAGAGLGAVI | ASSLGTVANTEVF | 0.100613765 |
| VAAVVTLL | ASSLGTVANTEVF | 0.22091068 |
| VGFLGGLI | ASSLGTVANTEVF | 0.753574 |
| VGVIPILL | ASSLGTVANTEVF | 0.33184132 |
| VIFASSLI | ASSLGTVANTEVF | 0.20494877 |
| VKGGDGGAAAG | ASSLGTVANTEVF | 0.28744704 |
| VTKTFTNI | ASSLGTVANTEVF | 0.56534404 |
| VVAAAILL | ASSLGTVANTEVF | 0.16323723 |
| YAALDVEL | ASSLGTVANTEVF | 0.19170584 |
| LASVALLL | ASSLGTVANTEVF | 0.32300717 |
| LDAVHRV | ASSLGTVANTEVF | 0.63003993 |
| QGLSSSIF | ASSLGTVANTEVF | 0.13771747 |
| RGTFANIRL | ASSLGTVANTEVF | 0.20087598 |
| RQLWWGHRI | ASSLGTVANTEVF | 0.3192499 |
| YGPPGTGKTLL | ASSLGTVANTEVF | 0.6039324 |
| AALVGAPANVA | ASSLGTVANTEVF | 0.45154893 |
| AATFRRTE | ASSLGTVANTEVF | 0.17205487 |
| AGALGGGI | ASSLGTVANTEVF | 0.5795085 |
| AGGAGGAGGIGAG | ASSLGTVANTEVF | 0.13335079 |
| AGSGALGL | ASSLGTVANTEVF | 0.6101759 |
| AQIADQI | ASSLGTVANTEVF | 0.23463853 |
| ASAVAVTYL | ASSLGTVANTEVF | 0.39289135 |
| ASTVLGGTGG | ASSLGTVANTEVF | 0.20985734 |
| AVGGEPAK | ASSLGTVANTEVF | 0.15844615 |
| CEQGTFGGTC | ASSLGTVANTEVF | 0.18201108 |
| DEGKAPAG | ASSLGTVANTEVF | 0.19127363 |
| DTVVFPAL | ASSLGTVANTEVF | 0.50768995 |
| FEGITMAG | ASSLGTVANTEVF | 0.19871879 |
| FPLVVQD | ASSLGTVANTEVF | 0.38355488 |
| GAFTTPQ | ASSLGTVANTEVF | 0.27862656 |
| GFGGFGTAGGVGGA | ASSLGTVANTEVF | 0.1626096 |
| GFLAQFL | ASSLGTVANTEVF | 0.4599956 |
| GGAGGAGGNAGVA | ASSLGTVANTEVF | 0.14823505 |
| GGVGTAGAGGAGGAGG | ASSLGTVANTEVF | 0.33000147 |
| GNGGGGDAGNAGSGGN | ASSLGTVANTEVF | 0.5721705 |
| GTGGAGGAGNDAGS | ASSLGTVANTEVF | 0.40538806 |
| GTTNTGWFNS | ASSLGTVANTEVF | 0.19593859 |
| IFMGPLFTL | ASSLGTVANTEVF | 0.4340049 |
| ILAHAIEAI | ASSLGTVANTEVF | 0.48587322 |
| IRSAIREAL | ASSLGTVANTEVF | 0.19694161 |
| ITFAGGG | ASSLGTVANTEVF | 0.37514913 |
| ITGMSGHQEV | ASSLGTVANTEVF | 0.5635853 |
| IVAGLACTL | ASSLGTVANTEVF | 0.35859588 |

| | | |
|---|---|---|
| IVGGGLAGA | ASSLGTVANTEVF | 0.11632374 |
| IVTGVLNI | ASSLGTVANTEVF | 0.4393989 |
| LAQQGCI | ASSLGTVANTEVF | 0.16282934 |
| LCLHVEN | ASSLGTVANTEVF | 0.38753325 |
| LDAISDGS | ASSLGTVANTEVF | 0.2657419 |
| LGGFAAML | ASSLGTVANTEVF | 0.2979481 |
| LGGKNPMI | ASSLGTVANTEVF | 0.5389586 |
| LGPKGVAAIL | ASSLGTVANTEVF | 0.24603643 |
| LMVGVSI | ASSLGTVANTEVF | 0.24266872 |
| LVGAAFVL | ASSLGTVANTEVF | 0.25967824 |
| LVLAVAPL | ASSLGTVANTEVF | 0.33335593 |
| MAAIAVIDAA | ASSLGTVANTEVF | 0.41119838 |
| NNGINAELQ | ASSLGTVANTEVF | 0.14465603 |
| RGEPLVLVHG | ASSLGTVANTEVF | 0.46191034 |
| RGGAGGAAGLI | ASSLGTVANTEVF | 0.30059028 |
| RSYRMHPAV | ASSLGTVANTEVF | 0.17722066 |
| SAAHLLAGQLA | ASSLGTVANTEVF | 0.37312156 |
| SAAVGSY | ASSLGTVANTEVF | 0.25033075 |
| SPHAVLHRI | ASSLGTVANTEVF | 0.15052341 |
| SQIAKVL | ASSLGTVANTEVF | 0.4990015 |
| TALGHLVP | ASSLGTVANTEVF | 0.22529748 |
| TAPTTSGAA | ASSLGTVANTEVF | 0.5173601 |
| TDGDDGNGGSPGT | ASSLGTVANTEVF | 0.7891933 |
| TDGLGEN | ASSLGTVANTEVF | 0.14691769 |
| TGGPPLPGQA | ASSLGTVANTEVF | 0.5316137 |
| TIPAITI | ASSLGTVANTEVF | 0.37112555 |
| TIPGFTI | ASSLGTVANTEVF | 0.25186396 |
| TSIASGALA | ASSLGTVANTEVF | 0.2846619 |
| TVTLPGAAI | ASSLGTVANTEVF | 0.20430492 |
| VASGLSAALP | ASSLGTVANTEVF | 0.63889146 |
| VGAGVIGI | ASSLGTVANTEVF | 0.42035514 |
| VGPKRFGV | ASSLGTVANTEVF | 0.31605676 |
| VNVAFPD | ASSLGTVANTEVF | 0.4719818 |
| VSMTNSGVS | ASSLGTVANTEVF | 0.45662937 |
| VTDVLGDLSV | ASSLGTVANTEVF | 0.45188412 |
| VVVVGAGFAGL | ASSLGTVANTEVF | 0.16205871 |
| VYTVGTVV | ASSLGTVANTEVF | 0.14192729 |
| WTTFSGP | ASSLGTVANTEVF | 0.25712037 |
| YDELVEAF | ASSLGTVANTEVF | 0.26350403 |
| YGTGGAGGNGG | ASSLGTVANTEVF | 0.1876197 |
| YTNVMAV | ASSLGTVANTEVF | 0.20613681 |
| FLEQVHQL | ASSLGTVANTEVF | 0.45496014 |
| GLSLLAVAV | ASSLGTVANTEVF | 0.7978703 |
| LEQVHQL | ASSLGTVANTEVF | 0.27895978 |
| LNAAGAI | ASSLGTVANTEVF | 0.79526174 |
| PAVHASL | ASSLGTVANTEVF | 0.16102779 |
| TELEDDA | ASSLGTVANTEVF | 0.54435277 |
| TPPGPPN | ASSLGTVANTEVF | 0.5284606 |
| TPPPPPPP | ASSLGTVANTEVF | 0.16372782 |
| TQSPALI | ASSQEWGEDEQY | 0.3437223 |
| EFAAAFPQ | ASSQEWGEDEQY | 0.16718046 |

| | | |
|---|---|---|
| LLSGNLDVL | ASSQEWGEDEQY | 0.18893036 |
| GPANLGNYNI | ASSQEWGEDEQY | 0.375408 |
| TLGPTLP | ASSQEWGEDEQY | 0.31959012 |
| VYAYSSCCK | ASSQEWGEDEQY | 0.45306623 |
| APAPAPAPNI | ASSQEWGEDEQY | 0.24051756 |
| PGGPVPIP | ASSQEWGEDEQY | 0.18189692 |
| FLTVALEP | ASSQEWGEDEQY | 0.3178859 |
| SQYFGANL | ASSQEWGEDEQY | 0.35884506 |
| GGVKPAA | ASSQEWGEDEQY | 0.40348876 |
| GGTGGAGGAGGW | ASSQEWGEDEQY | 0.8557582 |
| GVGGVAGLGGAGG | ASSQEWGEDEQY | 0.15327325 |
| VLLVLATTI | ASSQEWGEDEQY | 0.7452065 |
| IGIFAAML | ASSQEWGEDEQY | 0.7602186 |
| AVGDLID | ASSQEWGEDEQY | 0.3587395 |
| NISPLPL | ASSQEWGEDEQY | 0.27462786 |
| VTAVFVAI | ASSQEWGEDEQY | 0.5811511 |
| SGDDNDAAL | ASSQEWGEDEQY | 0.24193905 |
| AVAALAHS | ASSQEWGEDEQY | 0.38371992 |
| GGVTVMATP | ASSQEWGEDEQY | 0.6593894 |
| AGLAANVRD | ASSQEWGEDEQY | 0.45274872 |
| TNRPDLIDPALLRPG | ASSQEWGEDEQY | 0.8552712 |
| LANLGVTAL | ASSQEWGEDEQY | 0.77075726 |
| TGERIEL | ASSQEWGEDEQY | 0.8059077 |
| NGEELAA | ASSQEWGEDEQY | 0.619647 |
| VLSLVWG | ASSQEWGEDEQY | 0.36816818 |
| IGYALWL | ASSQEWGEDEQY | 0.72910684 |
| LSLLLAPL | ASSQEWGEDEQY | 0.5094443 |
| TLDEALK | ASSQEWGEDEQY | 0.8419535 |
| TDGAGTAEGG | ASSQEWGEDEQY | 0.8444146 |
| NAPGGAGGNGG | ASSQEWGEDEQY | 0.6190649 |
| VSVFMGMGGNF | ASSQEWGEDEQY | 0.83412635 |
| LASYLAAL | ASSQEWGEDEQY | 0.86740744 |
| LVGGTPAL | ASSQEWGEDEQY | 0.3685032 |
| VRNADVL | ASSQEWGEDEQY | 0.7034441 |
| TCSSPFLVSAM | ASSQEWGEDEQY | 0.52212113 |
| VHITDAE | ASSQEWGEDEQY | 0.6640258 |
| EYAAPSF | ASSQEWGEDEQY | 0.22062528 |
| VVDAFHGGL | ASSQEWGEDEQY | 0.8321158 |
| ASVVDPQ | ASSQEWGEDEQY | 0.618454 |
| SYNEFGGLS | ASSQEWGEDEQY | 0.19267657 |
| DPDGSVY | ASSQEWGEDEQY | 0.789827 |
| MVSTIHATE | ASSQEWGEDEQY | 0.68746656 |
| LAAGVADPV | ASSQEWGEDEQY | 0.6111691 |
| TKEQIAA | ASSQEWGEDEQY | 0.82573193 |
| SLASPAAL | ASSQEWGEDEQY | 0.6264859 |
| IGLVFYGV | ASSQEWGEDEQY | 0.54946554 |
| FLLDASNL | ASSQEWGEDEQY | 0.80689585 |
| YQVGNFP | ASSQEWGEDEQY | 0.8544819 |
| ESLLSQST | ASSQEWGEDEQY | 0.44088924 |
| LFTVGFAAVG | ASSQEWGEDEQY | 0.7760315 |
| ATPPTPT | ASSQEWGEDEQY | 0.8698312 |

| | | |
|---|---|---|
| TDLATLG | ASSQEWGEDEQY | 0.75891376 |
| SAGGTGR | ASSQEWGEDEQY | 0.8562463 |
| GFSNSGS | ASSQEWGEDEQY | 0.8701892 |
| GGNGGSGGTSV | ASSQEWGEDEQY | 0.73919 |
| GTSVATGGAGN | ASSQEWGEDEQY | 0.84410036 |
| NGGNGGSGGTS | ASSQEWGEDEQY | 0.8496939 |
| AVVKFPH | ASSQEWGEDEQY | 0.668792 |
| TPPPPPP | ASSQEWGEDEQY | 0.7458128 |
| AAPAAAAAPA | ASSQEWGEDEQY | 0.461307 |
| IALNEDL | ASSQEWGEDEQY | 0.7623266 |
| IVNVSSM | ASSQEWGEDEQY | 0.46038812 |
| TGIPPAPR | ASSQEWGEDEQY | 0.7739943 |
| FDLGGGTF | ASSQEWGEDEQY | 0.8556862 |
| FDLGGGTFDVS | ASSQEWGEDEQY | 0.7025163 |
| NEPTAAA | ASSQEWGEDEQY | 0.44353214 |
| NEPTAAALA | ASSQEWGEDEQY | 0.44595534 |
| GSIINISSI | ASSQEWGEDEQY | 0.7721429 |
| VAPGNAGTA | ASSQEWGEDEQY | 0.5436256 |
| TGGGASLE | ASSQEWGEDEQY | 0.41858935 |
| VAVYNHYKRI | ASSQEWGEDEQY | 0.5975876 |
| TDPAPATT | ASSQEWGEDEQY | 0.8468744 |
| TNRPDILDPALLRPG | ASSQEWGEDEQY | 0.8339008 |
| LTTTTTS | ASSQEWGEDEQY | 0.5133316 |
| PTGSGKTL | ASSQEWGEDEQY | 0.868703 |
| TGSGKTL | ASSQEWGEDEQY | 0.83985263 |
| TIGHVDHGK | ASSQEWGEDEQY | 0.8596331 |
| TIGHVDHGKTT | ASSQEWGEDEQY | 0.75239134 |
| GHVDHGKTT | ASSQEWGEDEQY | 0.40680397 |
| AQFEHTLL | ASSQEWGEDEQY | 0.85810727 |
| IVNQDEL | ASSQEWGEDEQY | 0.68453896 |
| TNGSQFF | ASSQEWGEDEQY | 0.73067194 |
| TGKGIVE | ASSQEWGEDEQY | 0.8318236 |
| LISNASDA | ASSQEWGEDEQY | 0.6290443 |
| AELRTLGELTKIAKAHGAQ | ASSQEWGEDEQY | 0.59244883 |
| APNAVAAAEPSW | ASSQEWGEDEQY | 0.7302294 |
| APQVAGAATAVL | ASSQEWGEDEQY | 0.8579856 |
| ARRDPAAAAAILPTDA | ASSQEWGEDEQY | 0.5688286 |
| DAAARDVLAEAGLAEYF | ASSQEWGEDEQY | 0.44254613 |
| DDQIVLSAKANLAKVEQ | ASSQEWGEDEQY | 0.7044169 |
| DELLPDTGEDSDEDGH | ASSQEWGEDEQY | 0.7358358 |
| ELPEAPANRATVPQD | ASSQEWGEDEQY | 0.85102713 |
| FGQSKIERVTGIG | ASSQEWGEDEQY | 0.7934584 |
| GAGGTGSTAGGAGGAGGAGGLY | ASSQEWGEDEQY | 0.67983484 |
| GHGGTGAAGGNGGTGGAGGSGI | ASSQEWGEDEQY | 0.7902012 |
| GNGGNGGAGGEGGPGGAGGAGA | ASSQEWGEDEQY | 0.8077346 |
| GSTAGNGGNGGNATNGTIAGQP | ASSQEWGEDEQY | 0.7728559 |
| GTGGDGGNGGNGGTGG | ASSQEWGEDEQY | 0.8303561 |
| GVRRPAGIPADAPV | ASSQEWGEDEQY | 0.5743095 |
| GYRQVTDAHLLTLARR | ASSQEWGEDEQY | 0.8508949 |
| IAKMGQDGHDRG | ASSQEWGEDEQY | 0.82732195 |
| KAEPSIEIQYT | ASSQEWGEDEQY | 0.83881634 |

| | | |
|---|---|---|
| KPVRIRMAADEDTVAVT | ASSQEWGEDEQY | 0.8610751 |
| MPPRIAGMRLLVIKPEP | ASSQEWGEDEQY | 0.7800288 |
| MKFYAVALFRSRGVPPTR | ASSQEWGEDEQY | 0.78408647 |
| NAGGFGILGAVAHSP | ASSQEWGEDEQY | 0.35431778 |
| NEGREITLAMAEM | ASSQEWGEDEQY | 0.75005835 |
| NELVEWSHPLGHRWRWE | ASSQEWGEDEQY | 0.77136433 |
| NLGAGGTGGNGGNPG | ASSQEWGEDEQY | 0.77246356 |
| NTEWPKWETVMAADPVD | ASSQEWGEDEQY | 0.82970166 |
| PASAAPAETQP | ASSQEWGEDEQY | 0.8490819 |
| QQSSDGKNGVRAAEITGE | ASSQEWGEDEQY | 0.63651663 |
| RAPDAVTTSWWKE | ASSQEWGEDEQY | 0.8706113 |
| SGFGNVGAHTSGWFNQ | ASSQEWGEDEQY | 0.8561561 |
| SGVAGAVGQAASVGGLKV | ASSQEWGEDEQY | 0.46710545 |
| SNGGAGTGSAGGNGGT | ASSQEWGEDEQY | 0.710315 |
| TGGAGGEGGAAG | ASSQEWGEDEQY | 0.8654946 |
| TGGDGGAAGAGGAPGHG | ASSQEWGEDEQY | 0.8088274 |
| TMPPLLAAIANAGRHGV | ASSQEWGEDEQY | 0.67337537 |
| TSSTAGNSGGA | ASSQEWGEDEQY | 0.70898783 |
| VPATVPVGILC | ASSQEWGEDEQY | 0.87863374 |
| VQRPDSGGVVLTGRIS | ASSQEWGEDEQY | 0.7392638 |
| WTPNPVPATKGVPAGLVQAT | ASSQEWGEDEQY | 0.8668792 |
| YALKRALAKFSDRFDVVII | ASSQEWGEDEQY | 0.24296364 |
| YLIGSVIV | ASSQEWGEDEQY | 0.70714766 |
| YGQLVAVID | ASSQEWGEDEQY | 0.6334541 |
| VTVGGMTLPA | ASSQEWGEDEQY | 0.18902248 |
| VLALPAV | ASSQEWGEDEQY | 0.82871914 |
| VIPPTLNYET | ASSQEWGEDEQY | 0.7205523 |
| VGMGGMHPSSG | ASSQEWGEDEQY | 0.8088137 |
| VGLAGGAATA | ASSQEWGEDEQY | 0.7046071 |
| VGGAGGAGGSGP | ASSQEWGEDEQY | 0.84479946 |
| VFAAFMV | ASSQEWGEDEQY | 0.76421535 |
| TLYDKLR | ASSQEWGEDEQY | 0.70925426 |
| TLAVFTVL | ASSQEWGEDEQY | 0.4543516 |
| TGSVQPARV | ASSQEWGEDEQY | 0.78380394 |
| TGGGAGSGGT | ASSQEWGEDEQY | 0.87808335 |
| SHLETSL | ASSQEWGEDEQY | 0.83837986 |
| SFTTGMT | ASSQEWGEDEQY | 0.32339954 |
| QVEVAKRMGVRQPTVS | ASSQEWGEDEQY | 0.7533436 |
| QGSGGPALAM | ASSQEWGEDEQY | 0.71994567 |
| MLAPSNP | ASSQEWGEDEQY | 0.2706375 |
| LVNNAAI | ASSQEWGEDEQY | 0.22943452 |
| LTGEINTLA | ASSQEWGEDEQY | 0.64560676 |
| LSIFGPV | ASSQEWGEDEQY | 0.6786417 |
| LLALLANL | ASSQEWGEDEQY | 0.81302124 |
| LISNASDALDKLR | ASSQEWGEDEQY | 0.81011665 |
| LINEAVP | ASSQEWGEDEQY | 0.6314797 |
| LGYAFSGL | ASSQEWGEDEQY | 0.8136873 |
| LDFVAVAL | ASSQEWGEDEQY | 0.7797997 |
| LAQPGAE | ASSQEWGEDEQY | 0.09116367 |
| ISNLGAVTGF | ASSQEWGEDEQY | 0.6469109 |
| HQEDPGRI | ASSQEWGEDEQY | 0.38019148 |

| | | |
|---|---|---|
| GTPGEPTP | ASSQEWGEDEQY | 0.87613547 |
| GFFNSGS | ASSQEWGEDEQY | 0.553815 |
| GAGGGGLTAGAN | ASSQEWGEDEQY | 0.63740754 |
| FVDALVNM | ASSQEWGEDEQY | 0.71335924 |
| FLTVAPEL | ASSQEWGEDEQY | 0.80554056 |
| FETLTAL | ASSQEWGEDEQY | 0.73294413 |
| DVINPIEL | ASSQEWGEDEQY | 0.54060596 |
| APVSGAPV | ASSQEWGEDEQY | 0.23023105 |
| ALVGGLAW | ASSQEWGEDEQY | 0.81177825 |
| ALAAAAATAI | ASSQEWGEDEQY | 0.5542983 |
| AFGLGLLL | ASSQEWGEDEQY | 0.5201551 |
| AFFAAASL | ASSQEWGEDEQY | 0.4531114 |
| AAAALPAFSPP | ASSQEWGEDEQY | 0.31524965 |
| VVLAPAL | ASSQEWGEDEQY | 0.338146 |
| TIGHVDHGKTTL | ASSQEWGEDEQY | 0.8443101 |
| LISNASDALDK | ASSQEWGEDEQY | 0.5265341 |
| LISNASDAL | ASSQEWGEDEQY | 0.6923609 |
| ISNASDALDK | ASSQEWGEDEQY | 0.8773619 |
| NQAETLVYQTE | ASSQEWGEDEQY | 0.7716824 |
| TGGTGGTAGAG | ASSQEWGEDEQY | 0.69439065 |
| TGSTAGGAGGA | ASSQEWGEDEQY | 0.5389761 |
| DPLAVPVAAGPV | ASSQEWGEDEQY | 0.47945476 |
| GGTGGTPGNGGN | ASSQEWGEDEQY | 0.50597835 |
| PAGLAGVDPATL | ASSQEWGEDEQY | 0.57865906 |
| GNNLVVEAPHAGASV | ASSQEWGEDEQY | 0.8681855 |
| KVTRIMSASTLEQPA | ASSQEWGEDEQY | 0.72746265 |
| LGFALPIALPRRTVV | ASSQEWGEDEQY | 0.47752202 |
| TRLGNLTARKLVVLV | ASSQEWGEDEQY | 0.80996615 |
| WSQQRRLIYFPSAGP | ASSQEWGEDEQY | 0.87103844 |
| GPNGNPDPMAAAVDIR | ASSQEWGEDEQY | 0.87448645 |
| TQRCVSVADLRAAASR | ASSQEWGEDEQY | 0.47789693 |
| QFVLDVAEHPVEVLTETV | ASSQEWGEDEQY | 0.697074 |
| GGAGGTGGAGGNGGTQAGDG | ASSQEWGEDEQY | 0.736544 |
| PLLATLRAIAVDVDAKPAQV | ASSQEWGEDEQY | 0.729439 |
| APKFRTGGYMIDFWGVGYQVAK | ASSQEWGEDEQY | 0.58543724 |
| GAGGAGGAGGAAGGPGGTGGTG | ASSQEWGEDEQY | 0.4866693 |
| IGALGLLATDVAPAGSVAFGVA | ASSQEWGEDEQY | 0.82588506 |
| AGGAGGIGGNGGAGGAGGVLMG | ASSQEWGEDEQY | 0.8886555 |
| ADELIGLL | ASSQEWGEDEQY | 0.72658336 |
| ADRALKA | ASSQEWGEDEQY | 0.2783107 |
| AGAVGQVGGAGGA | ASSQEWGEDEQY | 0.68598163 |
| AGGGGGLGGAP | ASSQEWGEDEQY | 0.58357406 |
| AGGLLYGNGGAG | ASSQEWGEDEQY | 0.5880994 |
| AGLESIGL | ASSQEWGEDEQY | 0.7308783 |
| AGLKTGII | ASSQEWGEDEQY | 0.38093477 |
| ASLAALLL | ASSQEWGEDEQY | 0.7195226 |
| ASVINISSI | ASSQEWGEDEQY | 0.87622786 |
| ATPPGPA | ASSQEWGEDEQY | 0.69000363 |
| ATVASALV | ASSQEWGEDEQY | 0.78458124 |
| ATVASLIL | ASSQEWGEDEQY | 0.85164213 |
| AVNGPVLIRH | ASSQEWGEDEQY | 0.7962462 |

| | | |
|---|---|---|
| DGAAGAVGI | ASSQEWGEDEQY | 0.8554357 |
| DPLLHGI | ASSQEWGEDEQY | 0.85231256 |
| ELGAALGAL | ASSQEWGEDEQY | 0.7448801 |
| ELGRTQAV | ASSQEWGEDEQY | 0.772828 |
| FVAGGYLS | ASSQEWGEDEQY | 0.8272245 |
| GASGGAGGAGGW | ASSQEWGEDEQY | 0.7166767 |
| GGAGAGGDGGAGGI | ASSQEWGEDEQY | 0.7258581 |
| GTGHNGSQPGMGGNG | ASSQEWGEDEQY | 0.69604826 |
| GYAAASASAS | ASSQEWGEDEQY | 0.8357023 |
| HQVDVAAVQQG | ASSQEWGEDEQY | 0.6794684 |
| IGAAIAHT | ASSQEWGEDEQY | 0.87349075 |
| IGMIEVAQL | ASSQEWGEDEQY | 0.26254028 |
| IRLAPEHTSAMD | ASSQEWGEDEQY | 0.40946347 |
| IYNVSVL | ASSQEWGEDEQY | 0.84101117 |
| KQGFLGM | ASSQEWGEDEQY | 0.1343995 |
| LESVITQL | ASSQEWGEDEQY | 0.24658966 |
| LGLGALVVV | ASSQEWGEDEQY | 0.75081694 |
| LIGNGGNGGSGGI | ASSQEWGEDEQY | 0.8541089 |
| LNAINAPTL | ASSQEWGEDEQY | 0.13949586 |
| LQLLNTGV | ASSQEWGEDEQY | 0.6574382 |
| LQTSIEF | ASSQEWGEDEQY | 0.34939533 |
| MSFAVSEL | ASSQEWGEDEQY | 0.8465115 |
| NTGALGL | ASSQEWGEDEQY | 0.7123828 |
| PPGAGGAGGAGTT | ASSQEWGEDEQY | 0.39004296 |
| QLVGLGI | ASSQEWGEDEQY | 0.8254625 |
| RFLEQVHQL | ASSQEWGEDEQY | 0.30957392 |
| RGVGYVLR | ASSQEWGEDEQY | 0.6907213 |
| SATGVFAGVI | ASSQEWGEDEQY | 0.8360721 |
| SLPAITLP | ASSQEWGEDEQY | 0.8706969 |
| STALAFTV | ASSQEWGEDEQY | 0.49968302 |
| SVLTVGLGK | ASSQEWGEDEQY | 0.8078182 |
| TALAGAGSGI | ASSQEWGEDEQY | 0.6849972 |
| TGLPFTGI | ASSQEWGEDEQY | 0.8839526 |
| VAEHVTV | ASSQEWGEDEQY | 0.64748156 |
| VDIVLNSL | ASSQEWGEDEQY | 0.3968143 |
| VLTGGGAVL | ASSQEWGEDEQY | 0.84031254 |
| VNVGLAYGI | ASSQEWGEDEQY | 0.81927073 |
| VSSAAVGAV | ASSQEWGEDEQY | 0.79200095 |
| VTSVYEAGV | ASSQEWGEDEQY | 0.83825123 |
| WVALDRGA | ASSQEWGEDEQY | 0.67083967 |
| YLVGSVVI | ASSQEWGEDEQY | 0.6917198 |
| GPPGTGKTLL | ASSQEWGEDEQY | 0.82333124 |
| GVGMTKF | ASSQEWGEDEQY | 0.82797045 |
| IGIGGSDL | ASSQEWGEDEQY | 0.46902788 |
| IGTLLAEL | ASSQEWGEDEQY | 0.8640125 |
| IYTSGTT | ASSQEWGEDEQY | 0.73243636 |
| LSFPAGV | ASSQEWGEDEQY | 0.551327 |
| LYTSGTT | ASSQEWGEDEQY | 0.8801702 |
| VGAFPFGF | ASSQEWGEDEQY | 0.7314285 |
| VGAGTVEF | ASSQEWGEDEQY | 0.82213646 |
| VGKSTLLL | ASSQEWGEDEQY | 0.82378197 |

| | | |
|---|---|---|
| AALAHAGGGIV | ASSQEWGEDEQY | 0.87214017 |
| AALVAVSLL | ASSQEWGEDEQY | 0.48586112 |
| AAVSRAVAAL | ASSQEWGEDEQY | 0.811794 |
| AGGSSGIGGF | ASSQEWGEDEQY | 0.763601 |
| AGLTVANANG | ASSQEWGEDEQY | 0.8328134 |
| AGLVGAVL | ASSQEWGEDEQY | 0.7505106 |
| ALSQAFTV | ASSQEWGEDEQY | 0.62985766 |
| ALTFSVSL | ASSQEWGEDEQY | 0.8217492 |
| ALTGAALVAL | ASSQEWGEDEQY | 0.8563136 |
| ASFSIGAGVL | ASSQEWGEDEQY | 0.7500975 |
| DGSVYKI | ASSQEWGEDEQY | 0.82333857 |
| EQLAEIV | ASSQEWGEDEQY | 0.8670387 |
| FAVSELTVM | ASSQEWGEDEQY | 0.49343902 |
| GAAPLAAGL | ASSQEWGEDEQY | 0.744164 |
| GADGGDGGSGGAGG | ASSQEWGEDEQY | 0.839809 |
| GFGLDTGGAGG | ASSQEWGEDEQY | 0.86614096 |
| GLASGGLL | ASSQEWGEDEQY | 0.7229575 |
| GRIAGAAVLL | ASSQEWGEDEQY | 0.8533317 |
| GSAGSLNTSF | ASSQEWGEDEQY | 0.7096296 |
| GTGQIGI | ASSQEWGEDEQY | 0.79401773 |
| GTLIALIL | ASSQEWGEDEQY | 0.7001083 |
| IAATMGFVVG | ASSQEWGEDEQY | 0.70342433 |
| IFAQALL | ASSQEWGEDEQY | 0.41378668 |
| IGAAVALGF | ASSQEWGEDEQY | 0.76962274 |
| IGLTPLAL | ASSQEWGEDEQY | 0.82249737 |
| ILGSNTSTL | ASSQEWGEDEQY | 0.7973527 |
| IVFSLTVL | ASSQEWGEDEQY | 0.2523052 |
| LASKATGF | ASSQEWGEDEQY | 0.8058684 |
| LASVALIL | ASSQEWGEDEQY | 0.7958776 |
| LFAVGLTL | ASSQEWGEDEQY | 0.88474596 |
| LFNGGIGTYI | ASSQEWGEDEQY | 0.7167957 |
| LTLGAGII | ASSQEWGEDEQY | 0.84325683 |
| LVAAALGDI | ASSQEWGEDEQY | 0.6015795 |
| NTAFVQLGI | ASSQEWGEDEQY | 0.73491144 |
| QLFSGAEL | ASSQEWGEDEQY | 0.44198418 |
| SAATFITL | ASSQEWGEDEQY | 0.6402559 |
| SNYGLANLGV | ASSQEWGEDEQY | 0.69702554 |
| SYGGAYAVM | ASSQEWGEDEQY | 0.5482658 |
| TTGVGGLTLG | ASSQEWGEDEQY | 0.8382169 |
| TTQTIYI | ASSQEWGEDEQY | 0.84440863 |
| VAAGAGLGAVI | ASSQEWGEDEQY | 0.8459376 |
| VAAVVTLL | ASSQEWGEDEQY | 0.84741044 |
| VGFLGGLI | ASSQEWGEDEQY | 0.8645638 |
| VGVIPILL | ASSQEWGEDEQY | 0.84525037 |
| VIFASSLI | ASSQEWGEDEQY | 0.60514957 |
| VKGGDGGAAAG | ASSQEWGEDEQY | 0.7348889 |
| VTKTFTNI | ASSQEWGEDEQY | 0.36620283 |
| VVAAAILL | ASSQEWGEDEQY | 0.7160534 |
| YAALDVEL | ASSQEWGEDEQY | 0.6344894 |
| LASVALLL | ASSQEWGEDEQY | 0.76065814 |
| LDAVHRV | ASSQEWGEDEQY | 0.13331282 |

| | | |
|---|---|---|
| QGLSSSIF | ASSQEWGEDEQY | 0.80122983 |
| RGTFANIRL | ASSQEWGEDEQY | 0.8686589 |
| RQLWWGHRI | ASSQEWGEDEQY | 0.63987076 |
| YGPPGTGKTLL | ASSQEWGEDEQY | 0.7939813 |
| AALVGAPANVA | ASSQEWGEDEQY | 0.8113945 |
| AATFRRTE | ASSQEWGEDEQY | 0.61287 |
| AGALGGGI | ASSQEWGEDEQY | 0.81450653 |
| AGGAGGAGGIGAG | ASSQEWGEDEQY | 0.62533617 |
| AGSGALGL | ASSQEWGEDEQY | 0.82624626 |
| AQIADQI | ASSQEWGEDEQY | 0.8578547 |
| ASAVAVTYL | ASSQEWGEDEQY | 0.7937724 |
| ASTVLGGTGG | ASSQEWGEDEQY | 0.80305296 |
| AVGGEPAK | ASSQEWGEDEQY | 0.78203 |
| CEQGTFGGTC | ASSQEWGEDEQY | 0.35836077 |
| DEGKAPAG | ASSQEWGEDEQY | 0.8098545 |
| DTVVFPAL | ASSQEWGEDEQY | 0.12668976 |
| FEGITMAG | ASSQEWGEDEQY | 0.748469 |
| FPLVVQD | ASSQEWGEDEQY | 0.57966393 |
| GAFTTPQ | ASSQEWGEDEQY | 0.770201 |
| GFGGFGTAGGVGGA | ASSQEWGEDEQY | 0.7528212 |
| GFLAQFL | ASSQEWGEDEQY | 0.8016607 |
| GGAGGAGGNAGVA | ASSQEWGEDEQY | 0.5591273 |
| GGVGTAGAGGAGGAGG | ASSQEWGEDEQY | 0.7882817 |
| GNGGGGDAGNAGSGGN | ASSQEWGEDEQY | 0.74548656 |
| GTGGAGGAGNDAGS | ASSQEWGEDEQY | 0.44931737 |
| GTTNTGWFNS | ASSQEWGEDEQY | 0.8344484 |
| IFMGPLFTL | ASSQEWGEDEQY | 0.73048526 |
| ILAHAIEAI | ASSQEWGEDEQY | 0.70793486 |
| IRSAIREAL | ASSQEWGEDEQY | 0.8508897 |
| ITFAGGG | ASSQEWGEDEQY | 0.75368255 |
| ITGMSGHQEV | ASSQEWGEDEQY | 0.29781762 |
| IVAGLACTL | ASSQEWGEDEQY | 0.6804644 |
| IVGGGLAGA | ASSQEWGEDEQY | 0.7761636 |
| IVTGVLNI | ASSQEWGEDEQY | 0.6785665 |
| LAQQGCI | ASSQEWGEDEQY | 0.13838683 |
| LCLHVEN | ASSQEWGEDEQY | 0.807962 |
| LDAISDGS | ASSQEWGEDEQY | 0.5575844 |
| LGGFAAML | ASSQEWGEDEQY | 0.59034497 |
| LGGKNPMI | ASSQEWGEDEQY | 0.64597875 |
| LGPKGVAAIL | ASSQEWGEDEQY | 0.8276186 |
| LMVGVSI | ASSQEWGEDEQY | 0.706573 |
| LVGAAFVL | ASSQEWGEDEQY | 0.6674804 |
| LVLAVAPL | ASSQEWGEDEQY | 0.80363244 |
| MAAIAVIDAA | ASSQEWGEDEQY | 0.8110663 |
| NNGINAELQ | ASSQEWGEDEQY | 0.8509252 |
| RGEPLVLVHG | ASSQEWGEDEQY | 0.7694278 |
| RGGAGGAAGLI | ASSQEWGEDEQY | 0.76573265 |
| RSYRMHPAV | ASSQEWGEDEQY | 0.8351786 |
| SAAHLLAGQLA | ASSQEWGEDEQY | 0.66565764 |
| SAAVGSY | ASSQEWGEDEQY | 0.47376236 |
| SPHAVLHRI | ASSQEWGEDEQY | 0.3849996 |

| | | |
|---|---|---|
| SQIAKVL | ASSQEWGEDEQY | 0.51404506 |
| TALGHLVP | ASSQEWGEDEQY | 0.39035696 |
| TAPTTSGAA | ASSQEWGEDEQY | 0.78166825 |
| TDGDDGNGGSPGT | ASSQEWGEDEQY | 0.8714359 |
| TDGLGEN | ASSQEWGEDEQY | 0.7498711 |
| TGGPPLPGQA | ASSQEWGEDEQY | 0.40393937 |
| TIPAITI | ASSQEWGEDEQY | 0.71355283 |
| TIPGFTI | ASSQEWGEDEQY | 0.67860234 |
| TSIASGALA | ASSQEWGEDEQY | 0.7951343 |
| TVTLPGAAI | ASSQEWGEDEQY | 0.8409518 |
| VASGLSAALP | ASSQEWGEDEQY | 0.8825252 |
| VGAGVIGI | ASSQEWGEDEQY | 0.8218846 |
| VGPKRFGV | ASSQEWGEDEQY | 0.63631415 |
| VNVAFPD | ASSQEWGEDEQY | 0.8168677 |
| VSMTNSGVS | ASSQEWGEDEQY | 0.6936301 |
| VTDVLGDLSV | ASSQEWGEDEQY | 0.771543 |
| VVVVGAGFAGL | ASSQEWGEDEQY | 0.76167846 |
| VYTVGTVV | ASSQEWGEDEQY | 0.78257823 |
| WTTFSGP | ASSQEWGEDEQY | 0.72607595 |
| YDELVEAF | ASSQEWGEDEQY | 0.6269609 |
| YGTGGAGGNGG | ASSQEWGEDEQY | 0.29329544 |
| YTNVMAV | ASSQEWGEDEQY | 0.82922626 |
| FLEQVHQL | ASSQEWGEDEQY | 0.46255106 |
| GLSLLAVAV | ASSQEWGEDEQY | 0.45074013 |
| LEQVHQL | ASSQEWGEDEQY | 0.70749056 |
| LNAAGAI | ASSQEWGEDEQY | 0.869648 |
| PAVHASL | ASSQEWGEDEQY | 0.8635101 |
| TELEDDA | ASSQEWGEDEQY | 0.67045057 |
| TPPGPPN | ASSQEWGEDEQY | 0.5724769 |
| TPPPPPPP | ASSQEWGEDEQY | 0.81087255 |
| TQSPALI | ASSQETGNQDTQY | 0.49873608 |
| EFAAAFPQ | ASSQETGNQDTQY | 0.81487155 |
| LLSGNLDVL | ASSQETGNQDTQY | 0.736084 |
| GPANLGNYNI | ASSQETGNQDTQY | 0.7278557 |
| TLGPTLP | ASSQETGNQDTQY | 0.69436 |
| VYAYSSCCK | ASSQETGNQDTQY | 0.79625136 |
| APAPAPAPNI | ASSQETGNQDTQY | 0.6706072 |
| PGGPVPIP | ASSQETGNQDTQY | 0.82499254 |
| FLTVALEP | ASSQETGNQDTQY | 0.17090763 |
| SQYFGANL | ASSQETGNQDTQY | 0.8453017 |
| GGVKPAA | ASSQETGNQDTQY | 0.5489926 |
| GGTGGAGGAGGW | ASSQETGNQDTQY | 0.5054101 |
| GVGGVAGLGGAGG | ASSQETGNQDTQY | 0.23686768 |
| VLLVLATTI | ASSQETGNQDTQY | 0.76657414 |
| IGIFAAML | ASSQETGNQDTQY | 0.80535555 |
| AVGDLID | ASSQETGNQDTQY | 0.84340703 |
| NISPLPL | ASSQETGNQDTQY | 0.67830205 |
| VTAVFVAI | ASSQETGNQDTQY | 0.83084935 |
| SGDDNDAAL | ASSQETGNQDTQY | 0.83514214 |
| AVAALAHS | ASSQETGNQDTQY | 0.64096963 |
| GGVTVMATP | ASSQETGNQDTQY | 0.24645503 |

| | | |
|---|---|---|
| AGLAANVRD | ASSQETGNQDTQY | 0.77881014 |
| TNRPDLIDPALLRPG | ASSQETGNQDTQY | 0.7823206 |
| LANLGVTAL | ASSQETGNQDTQY | 0.8415177 |
| TGERIEL | ASSQETGNQDTQY | 0.8115003 |
| NGEELAA | ASSQETGNQDTQY | 0.39940834 |
| VLSLVWG | ASSQETGNQDTQY | 0.8608614 |
| IGYALWL | ASSQETGNQDTQY | 0.81784904 |
| LSLLLAPL | ASSQETGNQDTQY | 0.53524184 |
| TLDEALK | ASSQETGNQDTQY | 0.62386715 |
| TDGAGTAEGG | ASSQETGNQDTQY | 0.65390354 |
| NAPGGAGGNGG | ASSQETGNQDTQY | 0.3676451 |
| VSVFMGMGGNF | ASSQETGNQDTQY | 0.85023165 |
| LASYLAAL | ASSQETGNQDTQY | 0.6647662 |
| LVGGTPAL | ASSQETGNQDTQY | 0.64645314 |
| VRNADVL | ASSQETGNQDTQY | 0.7362587 |
| TCSSPFLVSAM | ASSQETGNQDTQY | 0.6439233 |
| VHITDAE | ASSQETGNQDTQY | 0.5775429 |
| EYAAPSF | ASSQETGNQDTQY | 0.64048797 |
| VVDAFHGGL | ASSQETGNQDTQY | 0.7714615 |
| ASVVDPQ | ASSQETGNQDTQY | 0.81150424 |
| SYNEFGGLS | ASSQETGNQDTQY | 0.730445 |
| DPDGSVY | ASSQETGNQDTQY | 0.7924451 |
| MVSTIHATE | ASSQETGNQDTQY | 0.85120744 |
| LAAGVADPV | ASSQETGNQDTQY | 0.3708369 |
| TKEQIAA | ASSQETGNQDTQY | 0.6343639 |
| SLASPAAL | ASSQETGNQDTQY | 0.7991885 |
| IGLVFYGV | ASSQETGNQDTQY | 0.56192195 |
| FLLDASNL | ASSQETGNQDTQY | 0.8455787 |
| YQVGNFP | ASSQETGNQDTQY | 0.7792975 |
| ESLLSQST | ASSQETGNQDTQY | 0.77871084 |
| LFTVGFAAVG | ASSQETGNQDTQY | 0.7552521 |
| ATPPTPT | ASSQETGNQDTQY | 0.87881213 |
| TDLATLG | ASSQETGNQDTQY | 0.80069864 |
| SAGGTGR | ASSQETGNQDTQY | 0.7658157 |
| GFSNSGS | ASSQETGNQDTQY | 0.8172033 |
| GGNGGSGGTSV | ASSQETGNQDTQY | 0.5637664 |
| GTSVATGGAGN | ASSQETGNQDTQY | 0.83990145 |
| NGGNGGSGGTS | ASSQETGNQDTQY | 0.81181943 |
| AVVKFPH | ASSQETGNQDTQY | 0.48431367 |
| TPPPPPP | ASSQETGNQDTQY | 0.7047836 |
| AAPAAAAAPA | ASSQETGNQDTQY | 0.8616176 |
| IALNEDL | ASSQETGNQDTQY | 0.8339349 |
| IVNVSSM | ASSQETGNQDTQY | 0.5061063 |
| TGIPPAPR | ASSQETGNQDTQY | 0.5884399 |
| FDLGGGTF | ASSQETGNQDTQY | 0.45666564 |
| FDLGGGTFDVS | ASSQETGNQDTQY | 0.60846895 |
| NEPTAAA | ASSQETGNQDTQY | 0.40534925 |
| NEPTAAALA | ASSQETGNQDTQY | 0.797521 |
| GSIINISSI | ASSQETGNQDTQY | 0.650753 |
| VAPGNAGTA | ASSQETGNQDTQY | 0.8362248 |
| TGGGASLE | ASSQETGNQDTQY | 0.7766199 |

| | | |
|---|---|---|
| VAVYNHYKRI | ASSQETGNQDTQY | 0.84513295 |
| TDPAPATT | ASSQETGNQDTQY | 0.6538327 |
| TNRPDILDPALLRPG | ASSQETGNQDTQY | 0.6623278 |
| LTTTTTS | ASSQETGNQDTQY | 0.31757492 |
| PTGSGKTL | ASSQETGNQDTQY | 0.3769021 |
| TGSGKTL | ASSQETGNQDTQY | 0.5806594 |
| TIGHVDHGK | ASSQETGNQDTQY | 0.68829334 |
| TIGHVDHGKTT | ASSQETGNQDTQY | 0.2130824 |
| GHVDHGKTT | ASSQETGNQDTQY | 0.86435986 |
| AQFEHTLL | ASSQETGNQDTQY | 0.7107111 |
| IVNQDEL | ASSQETGNQDTQY | 0.794029 |
| TNGSQFF | ASSQETGNQDTQY | 0.7014164 |
| TGKGIVE | ASSQETGNQDTQY | 0.5947728 |
| LISNASDA | ASSQETGNQDTQY | 0.6047473 |
| AELRTLGELTKIAKAHGAQ | ASSQETGNQDTQY | 0.8164409 |
| APNAVAAAEPSW | ASSQETGNQDTQY | 0.48388216 |
| APQVAGAATAVL | ASSQETGNQDTQY | 0.23503129 |
| ARRDPAAAAAILPTDA | ASSQETGNQDTQY | 0.5661639 |
| DAAARDVLAEAGLAEYF | ASSQETGNQDTQY | 0.45417643 |
| DDQIVLSAKANLAKVEQ | ASSQETGNQDTQY | 0.87948155 |
| DELLPDTGEDSDEDGH | ASSQETGNQDTQY | 0.84348595 |
| ELPEAPANRATVPQD | ASSQETGNQDTQY | 0.759575 |
| FGQSKIERVTGIG | ASSQETGNQDTQY | 0.8783978 |
| GAGGTGSTAGGAGGAGGAGGLY | ASSQETGNQDTQY | 0.16821732 |
| GHGGTGAAGGNGGTGGAGGSGI | ASSQETGNQDTQY | 0.70327854 |
| GNGGNGGAGGEGGPGGAGGAGA | ASSQETGNQDTQY | 0.87027615 |
| GSTAGNGGNGGNATNGTIAGQP | ASSQETGNQDTQY | 0.8000605 |
| GTGGDGGNGGNGGTGG | ASSQETGNQDTQY | 0.6972959 |
| GVRRPAGIPADAPV | ASSQETGNQDTQY | 0.83441573 |
| GYRQVTDAHLLTLARR | ASSQETGNQDTQY | 0.60069907 |
| IAKMGQDGHDRG | ASSQETGNQDTQY | 0.6783829 |
| KAEPSIEIQYT | ASSQETGNQDTQY | 0.70222354 |
| KPVRIRMAADEDTVAVT | ASSQETGNQDTQY | 0.8291375 |
| MPPRIAGMRLLVIKPEP | ASSQETGNQDTQY | 0.7583437 |
| MKFYAVALFRSRGVPPTR | ASSQETGNQDTQY | 0.38731682 |
| NAGGFGILGAVAHSP | ASSQETGNQDTQY | 0.82080376 |
| NEGREITLAMAEM | ASSQETGNQDTQY | 0.8777236 |
| NELVEWSHPLGHRWRWE | ASSQETGNQDTQY | 0.6374383 |
| NLGAGGTGGNGGNPG | ASSQETGNQDTQY | 0.79779005 |
| NTEWPKWETVMAADPVD | ASSQETGNQDTQY | 0.77302647 |
| PASAAPAETQP | ASSQETGNQDTQY | 0.6407237 |
| QQSSDGKNGVRAAEITGE | ASSQETGNQDTQY | 0.5660915 |
| RAPDAVTTSWWKE | ASSQETGNQDTQY | 0.48601288 |
| SGFGNVGAHTSGWFNQ | ASSQETGNQDTQY | 0.7929734 |
| SGVAGAVGQAASVGGLKV | ASSQETGNQDTQY | 0.54485047 |
| SNGGAGTGSAGGNGGT | ASSQETGNQDTQY | 0.7348466 |
| TGGAGGEGGAAG | ASSQETGNQDTQY | 0.7368898 |
| TGGDGGAAGAGGAPGHG | ASSQETGNQDTQY | 0.8866513 |
| TMPPLLAAIANAGRHGV | ASSQETGNQDTQY | 0.8025497 |
| TSSTAGNSGGA | ASSQETGNQDTQY | 0.78323495 |
| VPATVPVGILC | ASSQETGNQDTQY | 0.54988897 |

| | | |
|---|---|---|
| VQRPDSGGVVLTGRIS | ASSQETGNQDTQY | 0.79275656 |
| WTPNPVPATKGVPAGLVQAT | ASSQETGNQDTQY | 0.8062823 |
| YALKRALAKFSDRFDVVII | ASSQETGNQDTQY | 0.32364452 |
| YLIGSVIV | ASSQETGNQDTQY | 0.86279756 |
| YGQLVAVID | ASSQETGNQDTQY | 0.48633218 |
| VTVGGMTLPA | ASSQETGNQDTQY | 0.7589164 |
| VLALPAV | ASSQETGNQDTQY | 0.8006046 |
| VIPPTLNYET | ASSQETGNQDTQY | 0.8623198 |
| VGMGGMHPSSG | ASSQETGNQDTQY | 0.72461224 |
| VGLAGGAATA | ASSQETGNQDTQY | 0.79292893 |
| VGGAGGAGGSGP | ASSQETGNQDTQY | 0.44729266 |
| VFAAFMV | ASSQETGNQDTQY | 0.82866603 |
| TLYDKLR | ASSQETGNQDTQY | 0.83388984 |
| TLAVFTVL | ASSQETGNQDTQY | 0.75279546 |
| TGSVQPARV | ASSQETGNQDTQY | 0.8791863 |
| TGGGAGSGGT | ASSQETGNQDTQY | 0.83245 |
| SHLETSL | ASSQETGNQDTQY | 0.30211812 |
| SFTTGMT | ASSQETGNQDTQY | 0.7064316 |
| QVEVAKRMGVRQPTVS | ASSQETGNQDTQY | 0.8602538 |
| QGSGGPALAM | ASSQETGNQDTQY | 0.8449186 |
| MLAPSNP | ASSQETGNQDTQY | 0.8472931 |
| LVNNAAI | ASSQETGNQDTQY | 0.8276857 |
| LTGEINTLA | ASSQETGNQDTQY | 0.691679 |
| LSIFGPV | ASSQETGNQDTQY | 0.86132 |
| LLALLANL | ASSQETGNQDTQY | 0.89120555 |
| LISNASDALDKLR | ASSQETGNQDTQY | 0.8290863 |
| LINEAVP | ASSQETGNQDTQY | 0.5425683 |
| LGYAFSGL | ASSQETGNQDTQY | 0.52265143 |
| LDFVAVAL | ASSQETGNQDTQY | 0.81114423 |
| LAQPGAE | ASSQETGNQDTQY | 0.83449197 |
| ISNLGAVTGF | ASSQETGNQDTQY | 0.44398513 |
| HQEDPGRI | ASSQETGNQDTQY | 0.74927235 |
| GTPGEPTP | ASSQETGNQDTQY | 0.75639075 |
| GFFNSGS | ASSQETGNQDTQY | 0.27392474 |
| GAGGGGLTAGAN | ASSQETGNQDTQY | 0.8155594 |
| FVDALVNM | ASSQETGNQDTQY | 0.85983074 |
| FLTVAPEL | ASSQETGNQDTQY | 0.79767644 |
| FETLTAL | ASSQETGNQDTQY | 0.5812534 |
| DVINPIEL | ASSQETGNQDTQY | 0.85581225 |
| APVSGAPV | ASSQETGNQDTQY | 0.8600717 |
| ALVGGLAW | ASSQETGNQDTQY | 0.872101 |
| ALAAAAATAI | ASSQETGNQDTQY | 0.30336693 |
| AFGLGLLL | ASSQETGNQDTQY | 0.17917873 |
| AFFAAASL | ASSQETGNQDTQY | 0.72977984 |
| AAAALPAFSPP | ASSQETGNQDTQY | 0.84228724 |
| VVLAPAL | ASSQETGNQDTQY | 0.7033411 |
| TIGHVDHGKTTL | ASSQETGNQDTQY | 0.6015679 |
| LISNASDALDK | ASSQETGNQDTQY | 0.7612997 |
| LISNASDAL | ASSQETGNQDTQY | 0.84459805 |
| ISNASDALDK | ASSQETGNQDTQY | 0.50004625 |
| NQAETLVYQTE | ASSQETGNQDTQY | 0.39433402 |

| | | |
|---|---|---|
| TGGTGGTAGAG | ASSQETGNQDTQY | 0.8201954 |
| TGSTAGGAGGA | ASSQETGNQDTQY | 0.8314514 |
| DPLAVPVAAGPV | ASSQETGNQDTQY | 0.5293895 |
| GGTGGTPGNGGN | ASSQETGNQDTQY | 0.6655559 |
| PAGLAGVDPATL | ASSQETGNQDTQY | 0.7535621 |
| GNNLVVEAPHAGASV | ASSQETGNQDTQY | 0.7453393 |
| KVTRIMSASTLEQPA | ASSQETGNQDTQY | 0.7808336 |
| LGFALPIALPRRTVV | ASSQETGNQDTQY | 0.4316732 |
| TRLGNLTARKLVVLV | ASSQETGNQDTQY | 0.82972085 |
| WSQQRRLIYFPSAGP | ASSQETGNQDTQY | 0.4170909 |
| GPNGNPDPMAAAVDIR | ASSQETGNQDTQY | 0.82792497 |
| TQRCVSVADLRAAASR | ASSQETGNQDTQY | 0.7791983 |
| QFVLDVAEHPVEVLTETV | ASSQETGNQDTQY | 0.8088333 |
| GGAGGTGGAGGNGGTQAGDG | ASSQETGNQDTQY | 0.6774348 |
| PLLATLRAIAVDVDAKPAQV | ASSQETGNQDTQY | 0.5087882 |
| APKFRTGGYMIDFWGVGYQVAK | ASSQETGNQDTQY | 0.5293432 |
| GAGGAGGAGGAAGGPGGTGGTG | ASSQETGNQDTQY | 0.36203858 |
| IGALGLLATDVAPAGSVAFGVA | ASSQETGNQDTQY | 0.8556886 |
| AGGAGGIGGNGGAGGAGGVLMG | ASSQETGNQDTQY | 0.21078233 |
| ADELIGLL | ASSQETGNQDTQY | 0.84111315 |
| ADRALKA | ASSQETGNQDTQY | 0.729396 |
| AGAVGQVGGAGGA | ASSQETGNQDTQY | 0.4543038 |
| AGGGGGLGGAP | ASSQETGNQDTQY | 0.7853489 |
| AGGLLYGNGGAG | ASSQETGNQDTQY | 0.38032603 |
| AGLESIGL | ASSQETGNQDTQY | 0.7412728 |
| AGLKTGII | ASSQETGNQDTQY | 0.78227586 |
| ASLAALLL | ASSQETGNQDTQY | 0.8187145 |
| ASVINISSI | ASSQETGNQDTQY | 0.57777345 |
| ATPPGPA | ASSQETGNQDTQY | 0.79434335 |
| ATVASALV | ASSQETGNQDTQY | 0.804863 |
| ATVASLIL | ASSQETGNQDTQY | 0.58162385 |
| AVNGPVLIRH | ASSQETGNQDTQY | 0.811429 |
| DGAAGAVGI | ASSQETGNQDTQY | 0.82444406 |
| DPLLHGI | ASSQETGNQDTQY | 0.7973347 |
| ELGAALGAL | ASSQETGNQDTQY | 0.3513138 |
| ELGRTQAV | ASSQETGNQDTQY | 0.7964395 |
| FVAGGYLS | ASSQETGNQDTQY | 0.5727307 |
| GASGGAGGAGGW | ASSQETGNQDTQY | 0.7868283 |
| GGAGAGGDGGAGGI | ASSQETGNQDTQY | 0.70334935 |
| GTGHNGSQPGMGGNG | ASSQETGNQDTQY | 0.6617012 |
| GYAAASASAS | ASSQETGNQDTQY | 0.82209 |
| HQVDVAAVQQG | ASSQETGNQDTQY | 0.44074816 |
| IGAAIAHT | ASSQETGNQDTQY | 0.6457422 |
| IGMIEVAQL | ASSQETGNQDTQY | 0.7114064 |
| IRLAPEHTSAMD | ASSQETGNQDTQY | 0.74003017 |
| IYNVSVL | ASSQETGNQDTQY | 0.8183106 |
| KQGFLGM | ASSQETGNQDTQY | 0.5059172 |
| LESVITQL | ASSQETGNQDTQY | 0.76990604 |
| LGLGALVVV | ASSQETGNQDTQY | 0.6933082 |
| LIGNGGNGGSGGI | ASSQETGNQDTQY | 0.5007206 |
| LNAINAPTL | ASSQETGNQDTQY | 0.7775518 |

| | | |
|---|---|---|
| LQLLNTGV | ASSQETGNQDTQY | 0.6942076 |
| LQTSIEF | ASSQETGNQDTQY | 0.4338204 |
| MSFAVSEL | ASSQETGNQDTQY | 0.8327631 |
| NTGALGL | ASSQETGNQDTQY | 0.48217505 |
| PPGAGGAGGAGTT | ASSQETGNQDTQY | 0.81979764 |
| QLVGLGI | ASSQETGNQDTQY | 0.8769257 |
| RFLEQVHQL | ASSQETGNQDTQY | 0.5501882 |
| RGVGYVLR | ASSQETGNQDTQY | 0.81657743 |
| SATGVFAGVI | ASSQETGNQDTQY | 0.77918315 |
| SLPAITLP | ASSQETGNQDTQY | 0.7824341 |
| STALAFTV | ASSQETGNQDTQY | 0.66401196 |
| SVLTVGLGK | ASSQETGNQDTQY | 0.62118435 |
| TALAGAGSGI | ASSQETGNQDTQY | 0.86430126 |
| TGLPFTGI | ASSQETGNQDTQY | 0.81533384 |
| VAEHVTV | ASSQETGNQDTQY | 0.84173036 |
| VDIVLNSL | ASSQETGNQDTQY | 0.8238285 |
| VLTGGGAVL | ASSQETGNQDTQY | 0.83255994 |
| VNVGLAYGI | ASSQETGNQDTQY | 0.83361495 |
| VSSAAVGAV | ASSQETGNQDTQY | 0.6073046 |
| VTSVYEAGV | ASSQETGNQDTQY | 0.32581562 |
| WVALDRGA | ASSQETGNQDTQY | 0.6300466 |
| YLVGSVVI | ASSQETGNQDTQY | 0.6675727 |
| GPPGTGKTLL | ASSQETGNQDTQY | 0.6104634 |
| GVGMTKF | ASSQETGNQDTQY | 0.32386023 |
| IGIGGSDL | ASSQETGNQDTQY | 0.8274925 |
| IGTLLAEL | ASSQETGNQDTQY | 0.8833792 |
| IYTSGTT | ASSQETGNQDTQY | 0.6103978 |
| LSFPAGV | ASSQETGNQDTQY | 0.88391525 |
| LYTSGTT | ASSQETGNQDTQY | 0.49065205 |
| VGAFPFGF | ASSQETGNQDTQY | 0.8226994 |
| VGAGTVEF | ASSQETGNQDTQY | 0.8356411 |
| VGKSTLLL | ASSQETGNQDTQY | 0.47233987 |
| AALAHAGGGIV | ASSQETGNQDTQY | 0.8158396 |
| AALVAVSLL | ASSQETGNQDTQY | 0.5257939 |
| AAVSRAVAAL | ASSQETGNQDTQY | 0.63109076 |
| AGGSSGIGGF | ASSQETGNQDTQY | 0.8478708 |
| AGLTVANANG | ASSQETGNQDTQY | 0.67852217 |
| AGLVGAVL | ASSQETGNQDTQY | 0.8265478 |
| ALSQAFTV | ASSQETGNQDTQY | 0.6456063 |
| ALTFSVSL | ASSQETGNQDTQY | 0.49070823 |
| ALTGAALVAL | ASSQETGNQDTQY | 0.76415956 |
| ASFSIGAGVL | ASSQETGNQDTQY | 0.8519782 |
| DGSVYKI | ASSQETGNQDTQY | 0.72359234 |
| EQLAEIV | ASSQETGNQDTQY | 0.8886042 |
| FAVSELTVM | ASSQETGNQDTQY | 0.6098436 |
| GAAPLAAGL | ASSQETGNQDTQY | 0.8677888 |
| GADGGDGGSGGAGG | ASSQETGNQDTQY | 0.8623935 |
| GFGLDTGGAGG | ASSQETGNQDTQY | 0.7515906 |
| GLASGGLL | ASSQETGNQDTQY | 0.6439054 |
| GRIAGAAVLL | ASSQETGNQDTQY | 0.8250214 |
| GSAGSLNTSF | ASSQETGNQDTQY | 0.64149237 |

| | | |
|---|---|---|
| GTGQIGI | ASSQETGNQDTQY | 0.83830595 |
| GTLIALIL | ASSQETGNQDTQY | 0.8663199 |
| IAATMGFVVG | ASSQETGNQDTQY | 0.6087474 |
| IFAQALL | ASSQETGNQDTQY | 0.7692861 |
| IGAAVALGF | ASSQETGNQDTQY | 0.73090994 |
| IGLTPLAL | ASSQETGNQDTQY | 0.72221136 |
| ILGSNTSTL | ASSQETGNQDTQY | 0.48972365 |
| IVFSLTVL | ASSQETGNQDTQY | 0.3164572 |
| LASKATGF | ASSQETGNQDTQY | 0.7103278 |
| LASVALIL | ASSQETGNQDTQY | 0.48846388 |
| LFAVGLTL | ASSQETGNQDTQY | 0.7624532 |
| LFNGGIGTYI | ASSQETGNQDTQY | 0.75792265 |
| LTLGAGII | ASSQETGNQDTQY | 0.54600203 |
| LVAAALGDI | ASSQETGNQDTQY | 0.7747301 |
| NTAFVQLGI | ASSQETGNQDTQY | 0.7551446 |
| QLFSGAEL | ASSQETGNQDTQY | 0.41569096 |
| SAATFITL | ASSQETGNQDTQY | 0.70073485 |
| SNYGLANLGV | ASSQETGNQDTQY | 0.49823174 |
| SYGGAYAVM | ASSQETGNQDTQY | 0.696918 |
| TTGVGGLTLG | ASSQETGNQDTQY | 0.48245418 |
| TTQTIYI | ASSQETGNQDTQY | 0.7520982 |
| VAAGAGLGAVI | ASSQETGNQDTQY | 0.64911556 |
| VAAVVTLL | ASSQETGNQDTQY | 0.2917945 |
| VGFLGGLI | ASSQETGNQDTQY | 0.73781717 |
| VGVIPILL | ASSQETGNQDTQY | 0.7193235 |
| VIFASSLI | ASSQETGNQDTQY | 0.66266847 |
| VKGGDGGAAAG | ASSQETGNQDTQY | 0.75775325 |
| VTKTFTNI | ASSQETGNQDTQY | 0.50528014 |
| VVAAAILL | ASSQETGNQDTQY | 0.60811377 |
| YAALDVEL | ASSQETGNQDTQY | 0.75778604 |
| LASVALLL | ASSQETGNQDTQY | 0.7635048 |
| LDAVHRV | ASSQETGNQDTQY | 0.51430005 |
| QGLSSSIF | ASSQETGNQDTQY | 0.72463757 |
| RGTFANIRL | ASSQETGNQDTQY | 0.76860183 |
| RQLWWGHRI | ASSQETGNQDTQY | 0.66545373 |
| YGPPGTGKTLL | ASSQETGNQDTQY | 0.7652261 |
| AALVGAPANVA | ASSQETGNQDTQY | 0.76833236 |
| AATFRRTE | ASSQETGNQDTQY | 0.5397558 |
| AGALGGGI | ASSQETGNQDTQY | 0.76375294 |
| AGGAGGAGGIGAG | ASSQETGNQDTQY | 0.7778343 |
| AGSGALGL | ASSQETGNQDTQY | 0.6627481 |
| AQIADQI | ASSQETGNQDTQY | 0.7147108 |
| ASAVAVTYL | ASSQETGNQDTQY | 0.47039855 |
| ASTVLGGTGG | ASSQETGNQDTQY | 0.69894946 |
| AVGGEPAK | ASSQETGNQDTQY | 0.27903023 |
| CEQGTFGGTC | ASSQETGNQDTQY | 0.7403196 |
| DEGKAPAG | ASSQETGNQDTQY | 0.7579335 |
| DTVVFPAL | ASSQETGNQDTQY | 0.68695295 |
| FEGITMAG | ASSQETGNQDTQY | 0.5939555 |
| FPLVVQD | ASSQETGNQDTQY | 0.55401564 |
| GAFTTPQ | ASSQETGNQDTQY | 0.6729417 |

| | | |
|---|---|---|
| GFGGFGTAGGVGGA | ASSQETGNQDTQY | 0.54789245 |
| GFLAQFL | ASSQETGNQDTQY | 0.45850918 |
| GGAGGAGGNAGVA | ASSQETGNQDTQY | 0.5645271 |
| GGVGTAGAGGAGGAGG | ASSQETGNQDTQY | 0.748114 |
| GNGGGGDAGNAGSGGN | ASSQETGNQDTQY | 0.7580627 |
| GTGGAGGAGNDAGS | ASSQETGNQDTQY | 0.5254985 |
| GTTNTGWFNS | ASSQETGNQDTQY | 0.7722028 |
| IFMGPLFTL | ASSQETGNQDTQY | 0.75670695 |
| ILAHAIEAI | ASSQETGNQDTQY | 0.76206964 |
| IRSAIREAL | ASSQETGNQDTQY | 0.71274155 |
| ITFAGGG | ASSQETGNQDTQY | 0.5247237 |
| ITGMSGHQEV | ASSQETGNQDTQY | 0.7676646 |
| IVAGLACTL | ASSQETGNQDTQY | 0.63664925 |
| IVGGGLAGA | ASSQETGNQDTQY | 0.7032722 |
| IVTGVLNI | ASSQETGNQDTQY | 0.7447239 |
| LAQQGCI | ASSQETGNQDTQY | 0.69059587 |
| LCLHVEN | ASSQETGNQDTQY | 0.5717247 |
| LDAISDGS | ASSQETGNQDTQY | 0.7125164 |
| LGGFAAML | ASSQETGNQDTQY | 0.7535405 |
| LGGKNPMI | ASSQETGNQDTQY | 0.5698153 |
| LGPKGVAAIL | ASSQETGNQDTQY | 0.3544021 |
| LMVGVSI | ASSQETGNQDTQY | 0.67509806 |
| LVGAAFVL | ASSQETGNQDTQY | 0.71652114 |
| LVLAVAPL | ASSQETGNQDTQY | 0.760846 |
| MAAIAVIDAA | ASSQETGNQDTQY | 0.75038135 |
| NNGINAELQ | ASSQETGNQDTQY | 0.6603576 |
| RGEPLVLVHG | ASSQETGNQDTQY | 0.750852 |
| RGGAGGAAGLI | ASSQETGNQDTQY | 0.73377055 |
| RSYRMHPAV | ASSQETGNQDTQY | 0.6946816 |
| SAAHLLAGQLA | ASSQETGNQDTQY | 0.7480042 |
| SAAVGSY | ASSQETGNQDTQY | 0.65228987 |
| SPHAVLHRI | ASSQETGNQDTQY | 0.7487407 |
| SQIAKVL | ASSQETGNQDTQY | 0.7437659 |
| TALGHLVP | ASSQETGNQDTQY | 0.7353971 |
| TAPTTSGAA | ASSQETGNQDTQY | 0.764216 |
| TDGDDGNGGSPGT | ASSQETGNQDTQY | 0.6845126 |
| TDGLGEN | ASSQETGNQDTQY | 0.6989165 |
| TGGPPLPGQA | ASSQETGNQDTQY | 0.2891188 |
| TIPAITI | ASSQETGNQDTQY | 0.5002052 |
| TIPGFTI | ASSQETGNQDTQY | 0.737931 |
| TSIASGALA | ASSQETGNQDTQY | 0.7229223 |
| TVTLPGAAI | ASSQETGNQDTQY | 0.76153946 |
| VASGLSAALP | ASSQETGNQDTQY | 0.7340702 |
| VGAGVIGI | ASSQETGNQDTQY | 0.62488043 |
| VGPKRFGV | ASSQETGNQDTQY | 0.76627743 |
| VNVAFPD | ASSQETGNQDTQY | 0.7693418 |
| VSMTNSGVS | ASSQETGNQDTQY | 0.38048476 |
| VTDVLGDLSV | ASSQETGNQDTQY | 0.62388325 |
| VVVVGAGFAGL | ASSQETGNQDTQY | 0.7799708 |
| VYTVGTVV | ASSQETGNQDTQY | 0.74507207 |
| WTTFSGP | ASSQETGNQDTQY | 0.64766765 |

| | | |
|---|---|---|
| YDELVEAF | ASSQETGNQDTQY | 0.6989687 |
| YGTGGAGGNGG | ASSQETGNQDTQY | 0.7784094 |
| YTNVMAV | ASSQETGNQDTQY | 0.6870781 |
| FLEQVHQL | ASSQETGNQDTQY | 0.779425 |
| GLSLLAVAV | ASSQETGNQDTQY | 0.28220013 |
| LEQVHQL | ASSQETGNQDTQY | 0.6851329 |
| LNAAGAI | ASSQETGNQDTQY | 0.515429 |
| PAVHASL | ASSQETGNQDTQY | 0.28453904 |
| TELEDDA | ASSQETGNQDTQY | 0.7386634 |
| TPPGPPN | ASSQETGNQDTQY | 0.68368125 |
| TPPPPPPP | ASSQETGNQDTQY | 0.7556144 |
| TQSPALI | ASSQEGGANTEVF | 0.682788 |
| EFAAAFPQ | ASSQEGGANTEVF | 0.7596947 |
| LLSGNLDVL | ASSQEGGANTEVF | 0.7282873 |
| GPANLGNYNI | ASSQEGGANTEVF | 0.7063123 |
| TLGPTLP | ASSQEGGANTEVF | 0.5480887 |
| VYAYSSCCK | ASSQEGGANTEVF | 0.72064686 |
| APAPAPAPNI | ASSQEGGANTEVF | 0.7712238 |
| PGGPVPIP | ASSQEGGANTEVF | 0.7450576 |
| FLTVALEP | ASSQEGGANTEVF | 0.35703695 |
| SQYFGANL | ASSQEGGANTEVF | 0.73607165 |
| GGVKPAA | ASSQEGGANTEVF | 0.63315123 |
| GGTGGAGGAGGW | ASSQEGGANTEVF | 0.2906174 |
| GVGGVAGLGGAGG | ASSQEGGANTEVF | 0.32322 |
| VLLVLATTI | ASSQEGGANTEVF | 0.57250845 |
| IGIFAAML | ASSQEGGANTEVF | 0.65459955 |
| AVGDLID | ASSQEGGANTEVF | 0.758829 |
| NISPLPL | ASSQEGGANTEVF | 0.7593187 |
| VTAVFVAI | ASSQEGGANTEVF | 0.66886985 |
| SGDDNDAAL | ASSQEGGANTEVF | 0.74381983 |
| AVAALAHS | ASSQEGGANTEVF | 0.71729696 |
| GGVTVMATP | ASSQEGGANTEVF | 0.20081489 |
| AGLAANVRD | ASSQEGGANTEVF | 0.5317711 |
| TNRPDLIDPALLRPG | ASSQEGGANTEVF | 0.44154894 |
| LANLGVTAL | ASSQEGGANTEVF | 0.7634031 |
| TGERIEL | ASSQEGGANTEVF | 0.5843933 |
| NGEELAA | ASSQEGGANTEVF | 0.6631978 |
| VLSLVWG | ASSQEGGANTEVF | 0.69811404 |
| IGYALWL | ASSQEGGANTEVF | 0.74526656 |
| LSLLLAPL | ASSQEGGANTEVF | 0.7066776 |
| TLDEALK | ASSQEGGANTEVF | 0.54993755 |
| TDGAGTAEGG | ASSQEGGANTEVF | 0.2866064 |
| NAPGGAGGNGG | ASSQEGGANTEVF | 0.72331303 |
| VSVFMGMGGNF | ASSQEGGANTEVF | 0.47304994 |
| LASYLAAL | ASSQEGGANTEVF | 0.560377 |
| LVGGTPAL | ASSQEGGANTEVF | 0.44945288 |
| VRNADVL | ASSQEGGANTEVF | 0.6963438 |
| TCSSPFLVSAM | ASSQEGGANTEVF | 0.40629324 |
| VHITDAE | ASSQEGGANTEVF | 0.76075494 |
| EYAAPSF | ASSQEGGANTEVF | 0.61063516 |
| VVDAFHGGL | ASSQEGGANTEVF | 0.70570487 |

| | | |
|---|---|---|
| ASVVDPQ | ASSQEGGANTEVF | 0.7782415 |
| SYNEFGGLS | ASSQEGGANTEVF | 0.73886764 |
| DPDGSVY | ASSQEGGANTEVF | 0.63691485 |
| MVSTIHATE | ASSQEGGANTEVF | 0.619772 |
| LAAGVADPV | ASSQEGGANTEVF | 0.52812874 |
| TKEQIAA | ASSQEGGANTEVF | 0.5641911 |
| SLASPAAL | ASSQEGGANTEVF | 0.6368172 |
| IGLVFYGV | ASSQEGGANTEVF | 0.7730945 |
| FLLDASNL | ASSQEGGANTEVF | 0.67751086 |
| YQVGNFP | ASSQEGGANTEVF | 0.487895 |
| ESLLSQST | ASSQEGGANTEVF | 0.690829 |
| LFTVGFAAVG | ASSQEGGANTEVF | 0.77282345 |
| ATPPTPT | ASSQEGGANTEVF | 0.7725908 |
| TDLATLG | ASSQEGGANTEVF | 0.5853866 |
| SAGGTGR | ASSQEGGANTEVF | 0.66587067 |
| GFSNSGS | ASSQEGGANTEVF | 0.67587954 |
| GGNGGSGGTSV | ASSQEGGANTEVF | 0.71177006 |
| GTSVATGGAGN | ASSQEGGANTEVF | 0.6507568 |
| NGGNGGSGGTS | ASSQEGGANTEVF | 0.49686092 |
| AVVKFPH | ASSQEGGANTEVF | 0.73858786 |
| TPPPPPP | ASSQEGGANTEVF | 0.7795732 |
| AAPAAAAAPA | ASSQEGGANTEVF | 0.653449 |
| IALNEDL | ASSQEGGANTEVF | 0.38445622 |
| IVNVSSM | ASSQEGGANTEVF | 0.7033482 |
| TGIPPAPR | ASSQEGGANTEVF | 0.5737327 |
| FDLGGGTF | ASSQEGGANTEVF | 0.59492105 |
| FDLGGGTFDVS | ASSQEGGANTEVF | 0.6226573 |
| NEPTAAA | ASSQEGGANTEVF | 0.45481724 |
| NEPTAAALA | ASSQEGGANTEVF | 0.7102964 |
| GSIINISSI | ASSQEGGANTEVF | 0.7739484 |
| VAPGNAGTA | ASSQEGGANTEVF | 0.63980436 |
| TGGGASLE | ASSQEGGANTEVF | 0.74642456 |
| VAVYNHYKRI | ASSQEGGANTEVF | 0.773187 |
| TDPAPATT | ASSQEGGANTEVF | 0.7451632 |
| TNRPDILDPALLRPG | ASSQEGGANTEVF | 0.76613355 |
| LTTTTTS | ASSQEGGANTEVF | 0.76525795 |
| PTGSGKTL | ASSQEGGANTEVF | 0.6855086 |
| TGSGKTL | ASSQEGGANTEVF | 0.7170707 |
| TIGHVDHGK | ASSQEGGANTEVF | 0.7568949 |
| TIGHVDHGKTT | ASSQEGGANTEVF | 0.70002294 |
| GHVDHGKTT | ASSQEGGANTEVF | 0.68690693 |
| AQFEHTLL | ASSQEGGANTEVF | 0.6518097 |
| IVNQDEL | ASSQEGGANTEVF | 0.76762575 |
| TNGSQFF | ASSQEGGANTEVF | 0.6468294 |
| TGKGIVE | ASSQEGGANTEVF | 0.766743 |
| LISNASDA | ASSQEGGANTEVF | 0.3394167 |
| AELRTLGELTKIAKAHGAQ | ASSQEGGANTEVF | 0.48246294 |
| APNAVAAAEPSW | ASSQEGGANTEVF | 0.7540593 |
| APQVAGAATAVL | ASSQEGGANTEVF | 0.2130235 |
| ARRDPAAAAAILPTDA | ASSQEGGANTEVF | 0.35799342 |
| DAAARDVLAEAGLAEYF | ASSQEGGANTEVF | 0.705423 |

| | | |
|---|---|---|
| DDQIVLSAKANLAKVEQ | ASSQEGGANTEVF | 0.76471305 |
| DELLPDTGEDSDEDGH | ASSQEGGANTEVF | 0.22254969 |
| ELPEAPANRATVPQD | ASSQEGGANTEVF | 0.5774768 |
| FGQSKIERVTGIG | ASSQEGGANTEVF | 0.4480511 |
| GAGGTGSTAGGAGGAGGAGGLY | ASSQEGGANTEVF | 0.7247938 |
| GHGGTGAAGGNGGTGGAGGSGI | ASSQEGGANTEVF | 0.7007358 |
| GNGGNGGAGGEGGPGGAGGAGA | ASSQEGGANTEVF | 0.27503675 |
| GSTAGNGGNGGNATNGTIAGQP | ASSQEGGANTEVF | 0.74526465 |
| GTGGDGGNGGNGGTGG | ASSQEGGANTEVF | 0.39165616 |
| GVRRPAGIPADAPV | ASSQEGGANTEVF | 0.5673226 |
| GYRQVTDAHLLTLARR | ASSQEGGANTEVF | 0.76035845 |
| IAKMGQDGHDRG | ASSQEGGANTEVF | 0.77203 |
| KAEPSIEIQYT | ASSQEGGANTEVF | 0.5240878 |
| KPVRIRMAADEDTVAVT | ASSQEGGANTEVF | 0.74279237 |
| MPPRIAGMRLLVIKPEP | ASSQEGGANTEVF | 0.6580505 |
| MKFYAVALFRSRGVPPTR | ASSQEGGANTEVF | 0.77834094 |
| NAGGFGILGAVAHSP | ASSQEGGANTEVF | 0.576675 |
| NEGREITLAMAEM | ASSQEGGANTEVF | 0.502931 |
| NELVEWSHPLGHRWRWE | ASSQEGGANTEVF | 0.76253736 |
| NLGAGGTGGNGGNPG | ASSQEGGANTEVF | 0.7370968 |
| NTEWPKWETVMAADPVD | ASSQEGGANTEVF | 0.6786872 |
| PASAAPAETQP | ASSQEGGANTEVF | 0.75834405 |
| QQSSDGKNGVRAAEITGE | ASSQEGGANTEVF | 0.55108213 |
| RAPDAVTTSWWKE | ASSQEGGANTEVF | 0.6823205 |
| SGFGNVGAHTSGWFNQ | ASSQEGGANTEVF | 0.7694609 |
| SGVAGAVGQAASVGGLKV | ASSQEGGANTEVF | 0.75819206 |
| SNGGAGTGSAGGNGGT | ASSQEGGANTEVF | 0.4656334 |
| TGGAGGEGGAAG | ASSQEGGANTEVF | 0.76157993 |
| TGGDGGAAGAGGAPGHG | ASSQEGGANTEVF | 0.7012357 |
| TMPPLLAAIANAGRHGV | ASSQEGGANTEVF | 0.5680029 |
| TSSTAGNSGGA | ASSQEGGANTEVF | 0.76480424 |
| VPATVPVGILC | ASSQEGGANTEVF | 0.64681184 |
| VQRPDSGGVVLTGRIS | ASSQEGGANTEVF | 0.7578217 |
| WTPNPVPATKGVPAGLVQAT | ASSQEGGANTEVF | 0.7313132 |
| YALKRALAKFSDRFDVVII | ASSQEGGANTEVF | 0.7694824 |
| YLIGSVIV | ASSQEGGANTEVF | 0.3025975 |
| YGQLVAVID | ASSQEGGANTEVF | 0.7364609 |
| VTVGGMTLPA | ASSQEGGANTEVF | 0.6833701 |
| VLALPAV | ASSQEGGANTEVF | 0.75284195 |
| VIPPTLNYET | ASSQEGGANTEVF | 0.69870055 |
| VGMGGMHPSSG | ASSQEGGANTEVF | 0.5860392 |
| VGLAGGAATA | ASSQEGGANTEVF | 0.735579 |
| VGGAGGAGGSGP | ASSQEGGANTEVF | 0.76933104 |
| VFAAFMV | ASSQEGGANTEVF | 0.6828144 |
| TLYDKLR | ASSQEGGANTEVF | 0.73561275 |
| TLAVFTVL | ASSQEGGANTEVF | 0.77036077 |
| TGSVQPARV | ASSQEGGANTEVF | 0.5708399 |
| TGGGAGSGGT | ASSQEGGANTEVF | 0.68316114 |
| SHLETSL | ASSQEGGANTEVF | 0.7418238 |
| SFTTGMT | ASSQEGGANTEVF | 0.76680183 |
| QVEVAKRMGVRQPTVS | ASSQEGGANTEVF | 0.6823753 |

| | | |
|---|---|---|
| QGSGGPALAM | ASSQEGGANTEVF | 0.77112794 |
| MLAPSNP | ASSQEGGANTEVF | 0.5178418 |
| LVNNAAI | ASSQEGGANTEVF | 0.75825036 |
| LTGEINTLA | ASSQEGGANTEVF | 0.6775303 |
| LSIFGPV | ASSQEGGANTEVF | 0.719435 |
| LLALLANL | ASSQEGGANTEVF | 0.5340168 |
| LISNASDALDKLR | ASSQEGGANTEVF | 0.75364375 |
| LINEAVP | ASSQEGGANTEVF | 0.75486803 |
| LGYAFSGL | ASSQEGGANTEVF | 0.76419467 |
| LDFVAVAL | ASSQEGGANTEVF | 0.34865305 |
| LAQPGAE | ASSQEGGANTEVF | 0.75182635 |
| ISNLGAVTGF | ASSQEGGANTEVF | 0.7646781 |
| HQEDPGRI | ASSQEGGANTEVF | 0.7789519 |
| GTPGEPTP | ASSQEGGANTEVF | 0.6710596 |
| GFFNSGS | ASSQEGGANTEVF | 0.7427673 |
| GAGGGGLTAGAN | ASSQEGGANTEVF | 0.65298283 |
| FVDALVNM | ASSQEGGANTEVF | 0.6626324 |
| FLTVAPEL | ASSQEGGANTEVF | 0.4967933 |
| FETLTAL | ASSQEGGANTEVF | 0.62052387 |
| DVINPIEL | ASSQEGGANTEVF | 0.6835859 |
| APVSGAPV | ASSQEGGANTEVF | 0.62836254 |
| ALVGGLAW | ASSQEGGANTEVF | 0.76472175 |
| ALAAAAATAI | ASSQEGGANTEVF | 0.76842076 |
| AFGLGLLL | ASSQEGGANTEVF | 0.75738215 |
| AFFAAASL | ASSQEGGANTEVF | 0.7657347 |
| AAAALPAFSPP | ASSQEGGANTEVF | 0.76982415 |
| VVLAPAL | ASSQEGGANTEVF | 0.7462312 |
| TIGHVDHGKTTL | ASSQEGGANTEVF | 0.5981047 |
| LISNASDALDK | ASSQEGGANTEVF | 0.62354505 |
| LISNASDAL | ASSQEGGANTEVF | 0.49339676 |
| ISNASDALDK | ASSQEGGANTEVF | 0.7121162 |
| NQAETLVYQTE | ASSQEGGANTEVF | 0.58431315 |
| TGGTGGTAGAG | ASSQEGGANTEVF | 0.7385355 |
| TGSTAGGAGGA | ASSQEGGANTEVF | 0.21186274 |
| DPLAVPVAAGPV | ASSQEGGANTEVF | 0.7421167 |
| GGTGGTPGNGGN | ASSQEGGANTEVF | 0.7749913 |
| PAGLAGVDPATL | ASSQEGGANTEVF | 0.5710656 |
| GNNLVVEAPHAGASV | ASSQEGGANTEVF | 0.7382577 |
| KVTRIMSASTLEQPA | ASSQEGGANTEVF | 0.74892306 |
| LGFALPIALPRRTVV | ASSQEGGANTEVF | 0.6331279 |
| TRLGNLTARKLVVLV | ASSQEGGANTEVF | 0.75056255 |
| WSQQRRLIYFPSAGP | ASSQEGGANTEVF | 0.6026561 |
| GPNGNPDPMAAAVDIR | ASSQEGGANTEVF | 0.7571702 |
| TQRCVSVADLRAAASR | ASSQEGGANTEVF | 0.7604542 |
| QFVLDVAEHPVEVLTETV | ASSQEGGANTEVF | 0.736338 |
| GGAGGTGGAGGNGGTQAGDG | ASSQEGGANTEVF | 0.70414376 |
| PLLATLRAIAVDVDAKPAQV | ASSQEGGANTEVF | 0.71594065 |
| APKFRTGGYMIDFWGVGYQVAK | ASSQEGGANTEVF | 0.27653882 |
| GAGGAGGAGGAAGGPGGTGGTG | ASSQEGGANTEVF | 0.7282275 |
| IGALGLLATDVAPAGSVAFGVA | ASSQEGGANTEVF | 0.31386393 |
| AGGAGGIGGNGGAGGAGGVLMG | ASSQEGGANTEVF | 0.70151603 |

| | | |
|---|---|---|
| ADELIGLL | ASSQEGGANTEVF | 0.43600294 |
| ADRALKA | ASSQEGGANTEVF | 0.7347777 |
| AGAVGQVGGAGGA | ASSQEGGANTEVF | 0.7140691 |
| AGGGGGLGGAP | ASSQEGGANTEVF | 0.7207054 |
| AGGLLYGNGGAG | ASSQEGGANTEVF | 0.5293485 |
| AGLESIGL | ASSQEGGANTEVF | 0.7263689 |
| AGLKTGII | ASSQEGGANTEVF | 0.72701794 |
| ASLAALLL | ASSQEGGANTEVF | 0.49051234 |
| ASVINISSI | ASSQEGGANTEVF | 0.7626729 |
| ATPPGPA | ASSQEGGANTEVF | 0.6366048 |
| ATVASALV | ASSQEGGANTEVF | 0.6543081 |
| ATVASLIL | ASSQEGGANTEVF | 0.7569684 |
| AVNGPVLIRH | ASSQEGGANTEVF | 0.6930584 |
| DGAAGAVGI | ASSQEGGANTEVF | 0.2945726 |
| DPLLHGI | ASSQEGGANTEVF | 0.6731049 |
| ELGAALGAL | ASSQEGGANTEVF | 0.71895945 |
| ELGRTQAV | ASSQEGGANTEVF | 0.5254628 |
| FVAGGYLS | ASSQEGGANTEVF | 0.24434562 |
| GASGGAGGAGGW | ASSQEGGANTEVF | 0.74052346 |
| GGAGAGGDGGAGGI | ASSQEGGANTEVF | 0.5736233 |
| GTGHNGSQPGMGGNG | ASSQEGGANTEVF | 0.54828197 |
| GYAAASASAS | ASSQEGGANTEVF | 0.6504714 |
| HQVDVAAVQQG | ASSQEGGANTEVF | 0.75610316 |
| IGAAIAHT | ASSQEGGANTEVF | 0.43614602 |
| IGMIEVAQL | ASSQEGGANTEVF | 0.66823214 |
| IRLAPEHTSAMD | ASSQEGGANTEVF | 0.74632466 |
| IYNVSVL | ASSQEGGANTEVF | 0.7420425 |
| KQGFLGM | ASSQEGGANTEVF | 0.76871085 |
| LESVITQL | ASSQEGGANTEVF | 0.70932066 |
| LGLGALVVV | ASSQEGGANTEVF | 0.6445347 |
| LIGNGGNGGSGGI | ASSQEGGANTEVF | 0.7464278 |
| LNAINAPTL | ASSQEGGANTEVF | 0.6831105 |
| LQLLNTGV | ASSQEGGANTEVF | 0.64070857 |
| LQTSIEF | ASSQEGGANTEVF | 0.4766521 |
| MSFAVSEL | ASSQEGGANTEVF | 0.67658013 |
| NTGALGL | ASSQEGGANTEVF | 0.52633935 |
| PPGAGGAGGAGTT | ASSQEGGANTEVF | 0.67121774 |
| QLVGLGI | ASSQEGGANTEVF | 0.76351666 |
| RFLEQVHQL | ASSQEGGANTEVF | 0.5896854 |
| RGVGYVLR | ASSQEGGANTEVF | 0.43976772 |
| SATGVFAGVI | ASSQEGGANTEVF | 0.68141973 |
| SLPAITLP | ASSQEGGANTEVF | 0.37637284 |
| STALAFTV | ASSQEGGANTEVF | 0.7415749 |
| SVLTVGLGK | ASSQEGGANTEVF | 0.69831276 |
| TALAGAGSGI | ASSQEGGANTEVF | 0.7812103 |
| TGLPFTGI | ASSQEGGANTEVF | 0.76707417 |
| VAEHVTV | ASSQEGGANTEVF | 0.6551702 |
| VDIVLNSL | ASSQEGGANTEVF | 0.75285804 |
| VLTGGGAVL | ASSQEGGANTEVF | 0.6302123 |
| VNVGLAYGI | ASSQEGGANTEVF | 0.741624 |
| VSSAAVGAV | ASSQEGGANTEVF | 0.71221936 |

| | | |
|---|---|---|
| VTSVYEAGV | ASSQEGGANTEVF | 0.7080907 |
| WVALDRGA | ASSQEGGANTEVF | 0.6608922 |
| YLVGSVVI | ASSQEGGANTEVF | 0.69497347 |
| GPPGTGKTLL | ASSQEGGANTEVF | 0.29369077 |
| GVGMTKF | ASSQEGGANTEVF | 0.756582 |
| IGIGGSDL | ASSQEGGANTEVF | 0.5327848 |
| IGTLLAEL | ASSQEGGANTEVF | 0.41286463 |
| IYTSGTT | ASSQEGGANTEVF | 0.62861586 |
| LSFPAGV | ASSQEGGANTEVF | 0.7738832 |
| LYTSGTT | ASSQEGGANTEVF | 0.76925814 |
| VGAFPFGF | ASSQEGGANTEVF | 0.6363392 |
| VGAGTVEF | ASSQEGGANTEVF | 0.5257951 |
| VGKSTLLL | ASSQEGGANTEVF | 0.74728084 |
| AALAHAGGGIV | ASSQEGGANTEVF | 0.40926546 |
| AALVAVSLL | ASSQEGGANTEVF | 0.71885526 |
| AAVSRAVAAL | ASSQEGGANTEVF | 0.698117 |
| AGGSSGIGGF | ASSQEGGANTEVF | 0.67678595 |
| AGLTVANANG | ASSQEGGANTEVF | 0.6413189 |
| AGLVGAVL | ASSQEGGANTEVF | 0.7380581 |
| ALSQAFTV | ASSQEGGANTEVF | 0.6276585 |
| ALTFSVSL | ASSQEGGANTEVF | 0.72078 |
| ALTGAALVAL | ASSQEGGANTEVF | 0.22373702 |
| ASFSIGAGVL | ASSQEGGANTEVF | 0.7353332 |
| DGSVYKI | ASSQEGGANTEVF | 0.597049 |
| EQLAEIV | ASSQEGGANTEVF | 0.5167151 |
| FAVSELTVM | ASSQEGGANTEVF | 0.31390882 |
| GAAPLAAGL | ASSQEGGANTEVF | 0.70984495 |
| GADGGDGGSGGAGG | ASSQEGGANTEVF | 0.7516563 |
| GFGLDTGGAGG | ASSQEGGANTEVF | 0.75713146 |
| GLASGGLL | ASSQEGGANTEVF | 0.63572454 |
| GRIAGAAVLL | ASSQEGGANTEVF | 0.7633163 |
| GSAGSLNTSF | ASSQEGGANTEVF | 0.76062256 |
| GTGQIGI | ASSQEGGANTEVF | 0.6854328 |
| GTLIALIL | ASSQEGGANTEVF | 0.2817614 |
| IAATMGFVVG | ASSQEGGANTEVF | 0.69696045 |
| IFAQALL | ASSQEGGANTEVF | 0.74184203 |
| IGAAVALGF | ASSQEGGANTEVF | 0.74645174 |
| IGLTPLAL | ASSQEGGANTEVF | 0.7273141 |
| ILGSNTSTL | ASSQEGGANTEVF | 0.48879236 |
| IVFSLTVL | ASSQEGGANTEVF | 0.7390489 |
| LASKATGF | ASSQEGGANTEVF | 0.76165986 |
| LASVALIL | ASSQEGGANTEVF | 0.51684225 |
| LFAVGLTL | ASSQEGGANTEVF | 0.636727 |
| LFNGGIGTYI | ASSQEGGANTEVF | 0.6461151 |
| LTLGAGII | ASSQEGGANTEVF | 0.42704964 |
| LVAAALGDI | ASSQEGGANTEVF | 0.7456923 |
| NTAFVQLGI | ASSQEGGANTEVF | 0.6641489 |
| QLFSGAEL | ASSQEGGANTEVF | 0.6572375 |
| SAATFITL | ASSQEGGANTEVF | 0.6903358 |
| SNYGLANLGV | ASSQEGGANTEVF | 0.6705419 |
| SYGGAYAVM | ASSQEGGANTEVF | 0.60096526 |

| | | |
|---|---|---|
| TTGVGGLTLG | ASSQEGGANTEVF | 0.6631192 |
| TTQTIYI | ASSQEGGANTEVF | 0.68365276 |
| VAAGAGLGAVI | ASSQEGGANTEVF | 0.7387731 |
| VAAVVTLL | ASSQEGGANTEVF | 0.71404207 |
| VGFLGGLI | ASSQEGGANTEVF | 0.73297846 |
| VGVIPILL | ASSQEGGANTEVF | 0.7740685 |
| VIFASSLI | ASSQEGGANTEVF | 0.4568981 |
| VKGGDGGAAAG | ASSQEGGANTEVF | 0.57225335 |
| VTKTFTNI | ASSQEGGANTEVF | 0.7332295 |
| VVAAAILL | ASSQEGGANTEVF | 0.5103513 |
| YAALDVEL | ASSQEGGANTEVF | 0.7584458 |
| LASVALLL | ASSQEGGANTEVF | 0.73088855 |
| LDAVHRV | ASSQEGGANTEVF | 0.7417323 |
| QGLSSSIF | ASSQEGGANTEVF | 0.72046083 |
| RGTFANIRL | ASSQEGGANTEVF | 0.7775805 |
| RQLWWGHRI | ASSQEGGANTEVF | 0.7496072 |
| YGPPGTGKTLL | ASSQEGGANTEVF | 0.7158104 |
| AALVGAPANVA | ASSQEGGANTEVF | 0.76835114 |
| AATFRRTE | ASSQEGGANTEVF | 0.49680966 |
| AGALGGGI | ASSQEGGANTEVF | 0.76461965 |
| AGGAGGAGGIGAG | ASSQEGGANTEVF | 0.7503022 |
| AGSGALGL | ASSQEGGANTEVF | 0.47022116 |
| AQIADQI | ASSQEGGANTEVF | 0.6924492 |
| ASAVAVTYL | ASSQEGGANTEVF | 0.7670957 |
| ASTVLGGTGG | ASSQEGGANTEVF | 0.7697545 |
| AVGGEPAK | ASSQEGGANTEVF | 0.42584997 |
| CEQGTFGGTC | ASSQEGGANTEVF | 0.651423 |
| DEGKAPAG | ASSQEGGANTEVF | 0.45720872 |
| DTVVFPAL | ASSQEGGANTEVF | 0.54590905 |
| FEGITMAG | ASSQEGGANTEVF | 0.41364637 |
| FPLVVQD | ASSQEGGANTEVF | 0.7217361 |
| GAFTTPQ | ASSQEGGANTEVF | 0.666366 |
| GFGGFGTAGGVGGA | ASSQEGGANTEVF | 0.76047313 |
| GFLAQFL | ASSQEGGANTEVF | 0.65588653 |
| GGAGGAGGNAGVA | ASSQEGGANTEVF | 0.7542125 |
| GGVGTAGAGGAGGAGG | ASSQEGGANTEVF | 0.6377743 |
| GNGGGGDAGNAGSGGN | ASSQEGGANTEVF | 0.6902642 |
| GTGGAGGAGNDAGS | ASSQEGGANTEVF | 0.37634844 |
| GTTNTGWFNS | ASSQEGGANTEVF | 0.3563491 |
| IFMGPLFTL | ASSQEGGANTEVF | 0.5678358 |
| ILAHAIEAI | ASSQEGGANTEVF | 0.657498 |
| IRSAIREAL | ASSQEGGANTEVF | 0.25604248 |
| ITFAGGG | ASSQEGGANTEVF | 0.7708291 |
| ITGMSGHQEV | ASSQEGGANTEVF | 0.69532806 |
| IVAGLACTL | ASSQEGGANTEVF | 0.7258184 |
| IVGGGLAGA | ASSQEGGANTEVF | 0.6239984 |
| IVTGVLNI | ASSQEGGANTEVF | 0.665275 |
| LAQQGCI | ASSQEGGANTEVF | 0.53891605 |
| LCLHVEN | ASSQEGGANTEVF | 0.7539356 |
| LDAISDGS | ASSQEGGANTEVF | 0.47651398 |
| LGGFAAML | ASSQEGGANTEVF | 0.34136882 |

| | | |
|---|---|---|
| LGGKNPMI | ASSQEGGANTEVF | 0.6791078 |
| LGPKGVAAIL | ASSQEGGANTEVF | 0.5403668 |
| LMVGVSI | ASSQEGGANTEVF | 0.76881593 |
| LVGAAFVL | ASSQEGGANTEVF | 0.76363844 |
| LVLAVAPL | ASSQEGGANTEVF | 0.7326313 |
| MAAIAVIDAA | ASSQEGGANTEVF | 0.76514614 |
| NNGINAELQ | ASSQEGGANTEVF | 0.23066403 |
| RGEPLVLVHG | ASSQEGGANTEVF | 0.6720753 |
| RGGAGGAAGLI | ASSQEGGANTEVF | 0.7733483 |
| RSYRMHPAV | ASSQEGGANTEVF | 0.7087158 |
| SAAHLLAGQLA | ASSQEGGANTEVF | 0.68088526 |
| SAAVGSY | ASSQEGGANTEVF | 0.765361 |
| SPHAVLHRI | ASSQEGGANTEVF | 0.64785564 |
| SQIAKVL | ASSQEGGANTEVF | 0.55051476 |
| TALGHLVP | ASSQEGGANTEVF | 0.70170105 |
| TAPTTSGAA | ASSQEGGANTEVF | 0.75157 |
| TDGDDGNGGSPGT | ASSQEGGANTEVF | 0.7399156 |
| TDGLGEN | ASSQEGGANTEVF | 0.47477895 |
| TGGPPLPGQA | ASSQEGGANTEVF | 0.7349339 |
| TIPAITI | ASSQEGGANTEVF | 0.78253657 |
| TIPGFTI | ASSQEGGANTEVF | 0.6828206 |
| TSIASGALA | ASSQEGGANTEVF | 0.72572476 |
| TVTLPGAAI | ASSQEGGANTEVF | 0.7156286 |
| VASGLSAALP | ASSQEGGANTEVF | 0.69491255 |
| VGAGVIGI | ASSQEGGANTEVF | 0.5898253 |
| VGPKRFGV | ASSQEGGANTEVF | 0.5782227 |
| VNVAFPD | ASSQEGGANTEVF | 0.73501736 |
| VSMTNSGVS | ASSQEGGANTEVF | 0.5540447 |
| VTDVLGDLSV | ASSQEGGANTEVF | 0.71075034 |
| VVVVGAGFAGL | ASSQEGGANTEVF | 0.6434802 |
| VYTVGTVV | ASSQEGGANTEVF | 0.7811676 |
| WTTFSGP | ASSQEGGANTEVF | 0.7484163 |
| YDELVEAF | ASSQEGGANTEVF | 0.71643996 |
| YGTGGAGGNGG | ASSQEGGANTEVF | 0.54532146 |
| YTNVMAV | ASSQEGGANTEVF | 0.7171339 |
| FLEQVHQL | ASSQEGGANTEVF | 0.7460599 |
| GLSLLAVAV | ASSQEGGANTEVF | 0.30573577 |
| LEQVHQL | ASSQEGGANTEVF | 0.76638436 |
| LNAAGAI | ASSQEGGANTEVF | 0.5451896 |
| PAVHASL | ASSQEGGANTEVF | 0.696388 |
| TELEDDA | ASSQEGGANTEVF | 0.744868 |
| TPPGPPN | ASSQEGGANTEVF | 0.77685094 |
| TPPPPPPP | ASSQEGGANTEVF | 0.69843525 |
| TQSPALI | ASSFIANTEVF | 0.7312403 |
| EFAAAFPQ | ASSFIANTEVF | 0.5446354 |
| LLSGNLDVL | ASSFIANTEVF | 0.7217017 |
| GPANLGNYNI | ASSFIANTEVF | 0.7625793 |
| TLGPTLP | ASSFIANTEVF | 0.69653213 |
| VYAYSSCCK | ASSFIANTEVF | 0.77441186 |
| APAPAPAPNI | ASSFIANTEVF | 0.68851256 |
| PGGPVPIP | ASSFIANTEVF | 0.32671875 |

| | | |
|---|---|---|
| FLTVALEP | ASSFIANTEVF | 0.62421596 |
| SQYFGANL | ASSFIANTEVF | 0.76017475 |
| GGVKPAA | ASSFIANTEVF | 0.7596936 |
| GGTGGAGGAGGW | ASSFIANTEVF | 0.76910216 |
| GVGGVAGLGGAGG | ASSFIANTEVF | 0.7572844 |
| VLLVLATTI | ASSFIANTEVF | 0.64031786 |
| IGIFAAML | ASSFIANTEVF | 0.769567 |
| AVGDLID | ASSFIANTEVF | 0.77611685 |
| NISPLPL | ASSFIANTEVF | 0.7213995 |
| VTAVFVAI | ASSFIANTEVF | 0.54442406 |
| SGDDNDAAL | ASSFIANTEVF | 0.47334546 |
| AVAALAHS | ASSFIANTEVF | 0.7431618 |
| GGVTVMATP | ASSFIANTEVF | 0.7484162 |
| AGLAANVRD | ASSFIANTEVF | 0.5130077 |
| TNRPDLIDPALLRPG | ASSFIANTEVF | 0.72533584 |
| LANLGVTAL | ASSFIANTEVF | 0.7300452 |
| TGERIEL | ASSFIANTEVF | 0.37301466 |
| NGEELAA | ASSFIANTEVF | 0.7336517 |
| VLSLVWG | ASSFIANTEVF | 0.77151006 |
| IGYALWL | ASSFIANTEVF | 0.73250496 |
| LSLLLAPL | ASSFIANTEVF | 0.5639236 |
| TLDEALK | ASSFIANTEVF | 0.7710657 |
| TDGAGTAEGG | ASSFIANTEVF | 0.7774807 |
| NAPGGAGGNGG | ASSFIANTEVF | 0.7696185 |
| VSVFMGMGGNF | ASSFIANTEVF | 0.41877216 |
| LASYLAAL | ASSFIANTEVF | 0.27692997 |
| LVGGTPAL | ASSFIANTEVF | 0.70370436 |
| VRNADVL | ASSFIANTEVF | 0.7645798 |
| TCSSPFLVSAM | ASSFIANTEVF | 0.67716026 |
| VHITDAE | ASSFIANTEVF | 0.58262837 |
| EYAAPSF | ASSFIANTEVF | 0.7412679 |
| VVDAFHGGL | ASSFIANTEVF | 0.7616395 |
| ASVVDPQ | ASSFIANTEVF | 0.56809807 |
| SYNEFGGLS | ASSFIANTEVF | 0.35845256 |
| DPDGSVY | ASSFIANTEVF | 0.7624844 |
| MVSTIHATE | ASSFIANTEVF | 0.7488582 |
| LAAGVADPV | ASSFIANTEVF | 0.63705397 |
| TKEQIAA | ASSFIANTEVF | 0.66610014 |
| SLASPAAL | ASSFIANTEVF | 0.6995146 |
| IGLVFYGV | ASSFIANTEVF | 0.59832454 |
| FLLDASNL | ASSFIANTEVF | 0.72052586 |
| YQVGNFP | ASSFIANTEVF | 0.4811812 |
| ESLLSQST | ASSFIANTEVF | 0.7715335 |
| LFTVGFAAVG | ASSFIANTEVF | 0.48586294 |
| ATPPTPT | ASSFIANTEVF | 0.7427216 |
| TDLATLG | ASSFIANTEVF | 0.7309137 |
| SAGGTGR | ASSFIANTEVF | 0.7280352 |
| GFSNSGS | ASSFIANTEVF | 0.67448574 |
| GGNGGSGGTSV | ASSFIANTEVF | 0.59187627 |
| GTSVATGGAGN | ASSFIANTEVF | 0.37873125 |
| NGGNGGSGGTS | ASSFIANTEVF | 0.42828724 |

| | | |
|---|---|---|
| AVVKFPH | ASSFIANTEVF | 0.7672067 |
| TPPPPPP | ASSFIANTEVF | 0.5378074 |
| AAPAAAAAPA | ASSFIANTEVF | 0.76495314 |
| IALNEDL | ASSFIANTEVF | 0.694762 |
| IVNVSSM | ASSFIANTEVF | 0.53168154 |
| TGIPPAPR | ASSFIANTEVF | 0.7328931 |
| FDLGGGTF | ASSFIANTEVF | 0.3933298 |
| FDLGGGTFDVS | ASSFIANTEVF | 0.68755734 |
| NEPTAAA | ASSFIANTEVF | 0.6985866 |
| NEPTAAALA | ASSFIANTEVF | 0.7421073 |
| GSIINISSI | ASSFIANTEVF | 0.5771683 |
| VAPGNAGTA | ASSFIANTEVF | 0.7255579 |
| TGGGASLE | ASSFIANTEVF | 0.7258574 |
| VAVYNHYKRI | ASSFIANTEVF | 0.52544045 |
| TDPAPATT | ASSFIANTEVF | 0.7350973 |
| TNRPDILDPALLRPG | ASSFIANTEVF | 0.7696016 |
| LTTTTTS | ASSFIANTEVF | 0.7376496 |
| PTGSGKTL | ASSFIANTEVF | 0.43575037 |
| TGSGKTL | ASSFIANTEVF | 0.7422607 |
| TIGHVDHGK | ASSFIANTEVF | 0.5712844 |
| TIGHVDHGKTT | ASSFIANTEVF | 0.727329 |
| GHVDHGKTT | ASSFIANTEVF | 0.68909 |
| AQFEHTLL | ASSFIANTEVF | 0.6479971 |
| IVNQDEL | ASSFIANTEVF | 0.74167144 |
| TNGSQFF | ASSFIANTEVF | 0.45800433 |
| TGKGIVE | ASSFIANTEVF | 0.61116165 |
| LISNASDA | ASSFIANTEVF | 0.6456248 |
| AELRTLGELTKIAKAHGAQ | ASSFIANTEVF | 0.7273011 |
| APNAVAAAEPSW | ASSFIANTEVF | 0.7413876 |
| APQVAGAATAVL | ASSFIANTEVF | 0.5506209 |
| ARRDPAAAAAILPTDA | ASSFIANTEVF | 0.70071054 |
| DAAARDVLAEAGLAEYF | ASSFIANTEVF | 0.5449859 |
| DDQIVLSAKANLAKVEQ | ASSFIANTEVF | 0.44248563 |
| DELLPDTGEDSDEDGH | ASSFIANTEVF | 0.71887213 |
| ELPEAPANRATVPQD | ASSFIANTEVF | 0.72719574 |
| FGQSKIERVTGIG | ASSFIANTEVF | 0.47289625 |
| GAGGTGSTAGGAGGAGGAGGLY | ASSFIANTEVF | 0.74306667 |
| GHGGTGAAGGNGGTGGAGGSGI | ASSFIANTEVF | 0.57020044 |
| GNGGNGGAGGEGGPGGAGGAGA | ASSFIANTEVF | 0.7339736 |
| GSTAGNGGNGGNATNGTIAGQP | ASSFIANTEVF | 0.7701805 |
| GTGGDGGNGGNGGTGG | ASSFIANTEVF | 0.47043735 |
| GVRRPAGIPADAPV | ASSFIANTEVF | 0.7238547 |
| GYRQVTDAHLLTLARR | ASSFIANTEVF | 0.70277923 |
| IAKMGQDGHDRG | ASSFIANTEVF | 0.745142 |
| KAEPSIEIQYT | ASSFIANTEVF | 0.6696541 |
| KPVRIRMAADEDTVAVT | ASSFIANTEVF | 0.4902459 |
| MPPRIAGMRLLVIKPEP | ASSFIANTEVF | 0.76494217 |
| MKFYAVALFRSRGVPPTR | ASSFIANTEVF | 0.7427004 |
| NAGGFGILGAVAHSP | ASSFIANTEVF | 0.7365766 |
| NEGREITLAMAEM | ASSFIANTEVF | 0.7370924 |
| NELVEWSHPLGHRWRWE | ASSFIANTEVF | 0.7691661 |

| | | |
|---|---|---|
| NLGAGGTGGNGGNPG | ASSFIANTEVF | 0.7516471 |
| NTEWPKWETVMAADPVD | ASSFIANTEVF | 0.6288074 |
| PASAAPAETQP | ASSFIANTEVF | 0.36247933 |
| QQSSDGKNGVRAAEITGE | ASSFIANTEVF | 0.7007728 |
| RAPDAVTTSWWKE | ASSFIANTEVF | 0.6721813 |
| SGFGNVGAHTSGWFNQ | ASSFIANTEVF | 0.6453092 |
| SGVAGAVGQAASVGGLKV | ASSFIANTEVF | 0.32446852 |
| SNGGAGTGSAGGNGGT | ASSFIANTEVF | 0.7648424 |
| TGGAGGEGGAAG | ASSFIANTEVF | 0.7798643 |
| TGGDGGAAGAGGAPGHG | ASSFIANTEVF | 0.62160504 |
| TMPPLLAAIANAGRHGV | ASSFIANTEVF | 0.77386856 |
| TSSTAGNSGGA | ASSFIANTEVF | 0.51120365 |
| VPATVPVGILC | ASSFIANTEVF | 0.74492824 |
| VQRPDSGGVVLTGRIS | ASSFIANTEVF | 0.77046967 |
| WTPNPVPATKGVPAGLVQAT | ASSFIANTEVF | 0.50094587 |
| YALKRALAKFSDRFDVVII | ASSFIANTEVF | 0.74132615 |
| YLIGSVIV | ASSFIANTEVF | 0.44376385 |
| YGQLVAVID | ASSFIANTEVF | 0.524153 |
| VTVGGMTLPA | ASSFIANTEVF | 0.77347803 |
| VLALPAV | ASSFIANTEVF | 0.63831365 |
| VIPPTLNYET | ASSFIANTEVF | 0.7492902 |
| VGMGGMHPSSG | ASSFIANTEVF | 0.6604532 |
| VGLAGGAATA | ASSFIANTEVF | 0.4844304 |
| VGGAGGAGGSGP | ASSFIANTEVF | 0.7296858 |
| VFAAFMV | ASSFIANTEVF | 0.7618129 |
| TLYDKLR | ASSFIANTEVF | 0.60911345 |
| TLAVFTVL | ASSFIANTEVF | 0.77620316 |
| TGSVQPARV | ASSFIANTEVF | 0.59392595 |
| TGGGAGSGGT | ASSFIANTEVF | 0.77827185 |
| SHLETSL | ASSFIANTEVF | 0.7617332 |
| SFTTGMT | ASSFIANTEVF | 0.7024231 |
| QVEVAKRMGVRQPTVS | ASSFIANTEVF | 0.6033438 |
| QGSGGPALAM | ASSFIANTEVF | 0.7627083 |
| MLAPSNP | ASSFIANTEVF | 0.66648906 |
| LVNNAAI | ASSFIANTEVF | 0.7461485 |
| LTGEINTLA | ASSFIANTEVF | 0.7658508 |
| LSIFGPV | ASSFIANTEVF | 0.5208422 |
| LLALLANL | ASSFIANTEVF | 0.7583166 |
| LISNASDALDKLR | ASSFIANTEVF | 0.6747081 |
| LINEAVP | ASSFIANTEVF | 0.67151535 |
| LGYAFSGL | ASSFIANTEVF | 0.5566759 |
| LDFVAVAL | ASSFIANTEVF | 0.39125168 |
| LAQPGAE | ASSFIANTEVF | 0.6523082 |
| ISNLGAVTGF | ASSFIANTEVF | 0.52352387 |
| HQEDPGRI | ASSFIANTEVF | 0.74404633 |
| GTPGEPTP | ASSFIANTEVF | 0.7302387 |
| GFFNSGS | ASSFIANTEVF | 0.55431503 |
| GAGGGGLTAGAN | ASSFIANTEVF | 0.76428103 |
| FVDALVNM | ASSFIANTEVF | 0.76630855 |
| FLTVAPEL | ASSFIANTEVF | 0.40013465 |
| FETLTAL | ASSFIANTEVF | 0.63435197 |

| | | |
|---|---|---|
| DVINPIEL | ASSFIANTEVF | 0.5513976 |
| APVSGAPV | ASSFIANTEVF | 0.6128616 |
| ALVGGLAW | ASSFIANTEVF | 0.31300128 |
| ALAAAAATAI | ASSFIANTEVF | 0.72561663 |
| AFGLGLLL | ASSFIANTEVF | 0.5665886 |
| AFFAAASL | ASSFIANTEVF | 0.24633358 |
| AAAALPAFSPP | ASSFIANTEVF | 0.6841183 |
| VVLAPAL | ASSFIANTEVF | 0.6733953 |
| TIGHVDHGKTTL | ASSFIANTEVF | 0.63173544 |
| LISNASDALDK | ASSFIANTEVF | 0.74841243 |
| LISNASDAL | ASSFIANTEVF | 0.55798084 |
| ISNASDALDK | ASSFIANTEVF | 0.55016077 |
| NQAETLVYQTE | ASSFIANTEVF | 0.7381867 |
| TGGTGGTAGAG | ASSFIANTEVF | 0.76255727 |
| TGSTAGGAGGA | ASSFIANTEVF | 0.4598217 |
| DPLAVPVAAGPV | ASSFIANTEVF | 0.6976458 |
| GGTGGTPGNGGN | ASSFIANTEVF | 0.76014745 |
| PAGLAGVDPATL | ASSFIANTEVF | 0.6777475 |
| GNNLVVEAPHAGASV | ASSFIANTEVF | 0.7465558 |
| KVTRIMSASTLEQPA | ASSFIANTEVF | 0.75930893 |
| LGFALPIALPRRTVV | ASSFIANTEVF | 0.56183696 |
| TRLGNLTARKLVVLV | ASSFIANTEVF | 0.7580035 |
| WSQQRRLIYFPSAGP | ASSFIANTEVF | 0.76579815 |
| GPNGNPDPMAAAVDIR | ASSFIANTEVF | 0.62846714 |
| TQRCVSVADLRAAASR | ASSFIANTEVF | 0.6601495 |
| QFVLDVAEHPVEVLTETV | ASSFIANTEVF | 0.5058691 |
| GGAGGTGGAGGNGGTQAGDG | ASSFIANTEVF | 0.67210364 |
| PLLATLRAIAVDVDAKPAQV | ASSFIANTEVF | 0.3506381 |
| APKFRTGGYMIDFWGVGYQVAK | ASSFIANTEVF | 0.71539664 |
| GAGGAGGAGGAAGGPGGTGGTG | ASSFIANTEVF | 0.75068665 |
| IGALGLLATDVAPAGSVAFGVA | ASSFIANTEVF | 0.6385212 |
| AGGAGGIGGNGGAGGAGGVLMG | ASSFIANTEVF | 0.46135813 |
| ADELIGLL | ASSFIANTEVF | 0.4750523 |
| ADRALKA | ASSFIANTEVF | 0.6553781 |
| AGAVGQVGGAGGA | ASSFIANTEVF | 0.5377236 |
| AGGGGGLGGAP | ASSFIANTEVF | 0.3507014 |
| AGGLLYGNGGAG | ASSFIANTEVF | 0.5934101 |
| AGLESIGL | ASSFIANTEVF | 0.75432646 |
| AGLKTGII | ASSFIANTEVF | 0.7519121 |
| ASLAALLL | ASSFIANTEVF | 0.4263885 |
| ASVINISSI | ASSFIANTEVF | 0.7716428 |
| ATPPGPA | ASSFIANTEVF | 0.73152673 |
| ATVASALV | ASSFIANTEVF | 0.73159903 |
| ATVASLIL | ASSFIANTEVF | 0.6692445 |
| AVNGPVLIRH | ASSFIANTEVF | 0.5036283 |
| DGAAGAVGI | ASSFIANTEVF | 0.7556765 |
| DPLLHGI | ASSFIANTEVF | 0.64233863 |
| ELGAALGAL | ASSFIANTEVF | 0.5627202 |
| ELGRTQAV | ASSFIANTEVF | 0.7381725 |
| FVAGGYLS | ASSFIANTEVF | 0.6565328 |
| GASGGAGGAGGW | ASSFIANTEVF | 0.57694095 |

| | | |
|---|---|---|
| GGAGAGGDGGAGGI | ASSFIANTEVF | 0.63777554 |
| GTGHNGSQPGMGGNG | ASSFIANTEVF | 0.7369691 |
| GYAAASASAS | ASSFIANTEVF | 0.5243403 |
| HQVDVAAVQQG | ASSFIANTEVF | 0.4357539 |
| IGAAIAHT | ASSFIANTEVF | 0.5979787 |
| IGMIEVAQL | ASSFIANTEVF | 0.6840335 |
| IRLAPEHTSAMD | ASSFIANTEVF | 0.7599351 |
| IYNVSVL | ASSFIANTEVF | 0.6971214 |
| KQGFLGM | ASSFIANTEVF | 0.5883359 |
| LESVITQL | ASSFIANTEVF | 0.71601856 |
| LGLGALVVV | ASSFIANTEVF | 0.69439316 |
| LIGNGGNGGSGGI | ASSFIANTEVF | 0.68512964 |
| LNAINAPTL | ASSFIANTEVF | 0.72945285 |
| LQLLNTGV | ASSFIANTEVF | 0.5852881 |
| LQTSIEF | ASSFIANTEVF | 0.73831904 |
| MSFAVSEL | ASSFIANTEVF | 0.7163265 |
| NTGALGL | ASSFIANTEVF | 0.72708356 |
| PPGAGGAGGAGTT | ASSFIANTEVF | 0.7480229 |
| QLVGLGI | ASSFIANTEVF | 0.6927477 |
| RFLEQVHQL | ASSFIANTEVF | 0.6813649 |
| RGVGYVLR | ASSFIANTEVF | 0.33741635 |
| SATGVFAGVI | ASSFIANTEVF | 0.58473265 |
| SLPAITLP | ASSFIANTEVF | 0.69135517 |
| STALAFTV | ASSFIANTEVF | 0.7060766 |
| SVLTVGLGK | ASSFIANTEVF | 0.7388699 |
| TALAGAGSGI | ASSFIANTEVF | 0.714541 |
| TGLPFTGI | ASSFIANTEVF | 0.5849905 |
| VAEHVTV | ASSFIANTEVF | 0.75467974 |
| VDIVLNSL | ASSFIANTEVF | 0.7562003 |
| VLTGGGAVL | ASSFIANTEVF | 0.39091846 |
| VNVGLAYGI | ASSFIANTEVF | 0.6361013 |
| VSSAAVGAV | ASSFIANTEVF | 0.7743052 |
| VTSVYEAGV | ASSFIANTEVF | 0.7117306 |
| WVALDRGA | ASSFIANTEVF | 0.63471806 |
| YLVGSVVI | ASSFIANTEVF | 0.56872606 |
| GPPGTGKTLL | ASSFIANTEVF | 0.76858747 |
| GVGMTKF | ASSFIANTEVF | 0.60230184 |
| IGIGGSDL | ASSFIANTEVF | 0.764574 |
| IGTLLAEL | ASSFIANTEVF | 0.28558877 |
| IYTSGTT | ASSFIANTEVF | 0.5680248 |
| LSFPAGV | ASSFIANTEVF | 0.49695697 |
| LYTSGTT | ASSFIANTEVF | 0.28703752 |
| VGAFPFGF | ASSFIANTEVF | 0.73591167 |
| VGAGTVEF | ASSFIANTEVF | 0.5955405 |
| VGKSTLLL | ASSFIANTEVF | 0.73210216 |
| AALAHAGGGIV | ASSFIANTEVF | 0.64668834 |
| AALVAVSLL | ASSFIANTEVF | 0.74369895 |
| AAVSRAVAAL | ASSFIANTEVF | 0.67298853 |
| AGGSSGIGGF | ASSFIANTEVF | 0.6914915 |
| AGLTVANANG | ASSFIANTEVF | 0.48279378 |
| AGLVGAVL | ASSFIANTEVF | 0.7322172 |

| | | |
|---|---|---|
| ALSQAFTV | ASSFIANTEVF | 0.760372 |
| ALTFSVSL | ASSFIANTEVF | 0.6854705 |
| ALTGAALVAL | ASSFIANTEVF | 0.38496113 |
| ASFSIGAGVL | ASSFIANTEVF | 0.64394045 |
| DGSVYKI | ASSFIANTEVF | 0.6741241 |
| EQLAEIV | ASSFIANTEVF | 0.3152582 |
| FAVSELTVM | ASSFIANTEVF | 0.31071517 |
| GAAPLAAGL | ASSFIANTEVF | 0.56639457 |
| GADGGDGGSGGAGG | ASSFIANTEVF | 0.5986646 |
| GFGLDTGGAGG | ASSFIANTEVF | 0.75373936 |
| GLASGGLL | ASSFIANTEVF | 0.74299294 |
| GRIAGAAVLL | ASSFIANTEVF | 0.616458 |
| GSAGSLNTSF | ASSFIANTEVF | 0.72670656 |
| GTGQIGI | ASSFIANTEVF | 0.6925886 |
| GTLIALIL | ASSFIANTEVF | 0.19761337 |
| IAATMGFVVG | ASSFIANTEVF | 0.6019207 |
| IFAQALL | ASSFIANTEVF | 0.4278012 |
| IGAAVALGF | ASSFIANTEVF | 0.7652326 |
| IGLTPLAL | ASSFIANTEVF | 0.54869777 |
| ILGSNTSTL | ASSFIANTEVF | 0.5926068 |
| IVFSLTVL | ASSFIANTEVF | 0.6709306 |
| LASKATGF | ASSFIANTEVF | 0.70519996 |
| LASVALIL | ASSFIANTEVF | 0.6822573 |
| LFAVGLTL | ASSFIANTEVF | 0.5107247 |
| LFNGGIGTYI | ASSFIANTEVF | 0.28229332 |
| LTLGAGII | ASSFIANTEVF | 0.73167324 |
| LVAAALGDI | ASSFIANTEVF | 0.4974039 |
| NTAFVQLGI | ASSFIANTEVF | 0.50724053 |
| QLFSGAEL | ASSFIANTEVF | 0.37037188 |
| SAATFITL | ASSFIANTEVF | 0.6594619 |
| SNYGLANLGV | ASSFIANTEVF | 0.39187497 |
| SYGGAYAVM | ASSFIANTEVF | 0.7235302 |
| TTGVGGLTLG | ASSFIANTEVF | 0.62842524 |
| TTQTIYI | ASSFIANTEVF | 0.6612599 |
| VAAGAGLGAVI | ASSFIANTEVF | 0.7760932 |
| VAAVVTLL | ASSFIANTEVF | 0.6975665 |
| VGFLGGLI | ASSFIANTEVF | 0.6336633 |
| VGVIPILL | ASSFIANTEVF | 0.54500186 |
| VIFASSLI | ASSFIANTEVF | 0.47048593 |
| VKGGDGGAAAG | ASSFIANTEVF | 0.5241099 |
| VTKTFTNI | ASSFIANTEVF | 0.5480126 |
| VVAAAILL | ASSFIANTEVF | 0.77405906 |
| YAALDVEL | ASSFIANTEVF | 0.58896714 |
| LASVALLL | ASSFIANTEVF | 0.4583586 |
| LDAVHRV | ASSFIANTEVF | 0.6314732 |
| QGLSSSIF | ASSFIANTEVF | 0.7679986 |
| RGTFANIRL | ASSFIANTEVF | 0.76974046 |
| RQLWWGHRI | ASSFIANTEVF | 0.5106932 |
| YGPPGTGKTLL | ASSFIANTEVF | 0.63283175 |
| AALVGAPANVA | ASSFIANTEVF | 0.6239505 |
| AATFRRTE | ASSFIANTEVF | 0.67811704 |

| | | |
|---|---|---|
| AGALGGGI | ASSFIANTEVF | 0.5188457 |
| AGGAGGAGGIGAG | ASSFIANTEVF | 0.46888313 |
| AGSGALGL | ASSFIANTEVF | 0.7201359 |
| AQIADQI | ASSFIANTEVF | 0.7778879 |
| ASAVAVTYL | ASSFIANTEVF | 0.6135799 |
| ASTVLGGTGG | ASSFIANTEVF | 0.37871286 |
| AVGGEPAK | ASSFIANTEVF | 0.6720909 |
| CEQGTFGGTC | ASSFIANTEVF | 0.5397823 |
| DEGKAPAG | ASSFIANTEVF | 0.5831716 |
| DTVVFPAL | ASSFIANTEVF | 0.6553271 |
| FEGITMAG | ASSFIANTEVF | 0.37801236 |
| FPLVVQD | ASSFIANTEVF | 0.6512388 |
| GAFTTPQ | ASSFIANTEVF | 0.7675451 |
| GFGGFGTAGGVGGA | ASSFIANTEVF | 0.6576129 |
| GFLAQFL | ASSFIANTEVF | 0.7062688 |
| GGAGGAGGNAGVA | ASSFIANTEVF | 0.7567043 |
| GGVGTAGAGGAGGAGG | ASSFIANTEVF | 0.72935575 |
| GNGGGGDAGNAGSGGN | ASSFIANTEVF | 0.7564031 |
| GTGGAGGAGNDAGS | ASSFIANTEVF | 0.74566543 |
| GTTNTGWFNS | ASSFIANTEVF | 0.6334942 |
| IFMGPLFTL | ASSFIANTEVF | 0.67577237 |
| ILAHAIEAI | ASSFIANTEVF | 0.7496091 |
| IRSAIREAL | ASSFIANTEVF | 0.6901152 |
| ITFAGGG | ASSFIANTEVF | 0.64723307 |
| ITGMSGHQEV | ASSFIANTEVF | 0.5427798 |
| IVAGLACTL | ASSFIANTEVF | 0.7591239 |
| IVGGGLAGA | ASSFIANTEVF | 0.636634 |
| IVTGVLNI | ASSFIANTEVF | 0.7520505 |
| LAQQGCI | ASSFIANTEVF | 0.36969233 |
| LCLHVEN | ASSFIANTEVF | 0.46630841 |
| LDAISDGS | ASSFIANTEVF | 0.7460108 |
| LGGFAAML | ASSFIANTEVF | 0.22353187 |
| LGGKNPMI | ASSFIANTEVF | 0.30161616 |
| LGPKGVAAIL | ASSFIANTEVF | 0.6447874 |
| LMVGVSI | ASSFIANTEVF | 0.7514795 |
| LVGAAFVL | ASSFIANTEVF | 0.22314563 |
| LVLAVAPL | ASSFIANTEVF | 0.5560998 |
| MAAIAVIDAA | ASSFIANTEVF | 0.3904418 |
| NNGINAELQ | ASSFIANTEVF | 0.69228256 |
| RGEPLVLVHG | ASSFIANTEVF | 0.65932333 |
| RGGAGGAAGLI | ASSFIANTEVF | 0.36574316 |
| RSYRMHPAV | ASSFIANTEVF | 0.7259973 |
| SAAHLLAGQLA | ASSFIANTEVF | 0.36192912 |
| SAAVGSY | ASSFIANTEVF | 0.58951265 |
| SPHAVLHRI | ASSFIANTEVF | 0.7372589 |
| SQIAKVL | ASSFIANTEVF | 0.7624885 |
| TALGHLVP | ASSFIANTEVF | 0.46617177 |
| TAPTTSGAA | ASSFIANTEVF | 0.72900784 |
| TDGDDGNGGSPGT | ASSFIANTEVF | 0.607682 |
| TDGLGEN | ASSFIANTEVF | 0.7713307 |
| TGGPPLPGQA | ASSFIANTEVF | 0.6355136 |

| | | |
|---|---|---|
| TIPAITI | ASSFIANTEVF | 0.38547027 |
| TIPGFTI | ASSFIANTEVF | 0.73890173 |
| TSIASGALA | ASSFIANTEVF | 0.7080803 |
| TVTLPGAAI | ASSFIANTEVF | 0.6653464 |
| VASGLSAALP | ASSFIANTEVF | 0.7306894 |
| VGAGVIGI | ASSFIANTEVF | 0.5453162 |
| VGPKRFGV | ASSFIANTEVF | 0.654721 |
| VNVAFPD | ASSFIANTEVF | 0.7582958 |
| VSMTNSGVS | ASSFIANTEVF | 0.73447454 |
| VTDVLGDLSV | ASSFIANTEVF | 0.46718258 |
| VVVVGAGFAGL | ASSFIANTEVF | 0.7374829 |
| VYTVGTVV | ASSFIANTEVF | 0.6684725 |
| WTTFSGP | ASSFIANTEVF | 0.55562645 |
| YDELVEAF | ASSFIANTEVF | 0.7658691 |
| YGTGGAGGNGG | ASSFIANTEVF | 0.6323333 |
| YTNVMAV | ASSFIANTEVF | 0.7336161 |
| FLEQVHQL | ASSFIANTEVF | 0.7153769 |
| GLSLLAVAV | ASSFIANTEVF | 0.76016074 |
| LEQVHQL | ASSFIANTEVF | 0.4426893 |
| LNAAGAI | ASSFIANTEVF | 0.71966004 |
| PAVHASL | ASSFIANTEVF | 0.6805971 |
| TELEDDA | ASSFIANTEVF | 0.7448374 |
| TPPGPPN | ASSFIANTEVF | 0.61976266 |
| TPPPPPPP | ASSFIANTEVF | 0.642223 |
| TQSPALI | TCSADWGYNYAEQF | 0.70657885 |
| EFAAAFPQ | TCSADWGYNYAEQF | 0.7599221 |
| LLSGNLDVL | TCSADWGYNYAEQF | 0.64569646 |
| GPANLGNYNI | TCSADWGYNYAEQF | 0.7109313 |
| TLGPTLP | TCSADWGYNYAEQF | 0.7696456 |
| VYAYSSCCK | TCSADWGYNYAEQF | 0.5263013 |
| APAPAPAPNI | TCSADWGYNYAEQF | 0.636788 |
| PGGPVPIP | TCSADWGYNYAEQF | 0.72053814 |
| FLTVALEP | TCSADWGYNYAEQF | 0.75807965 |
| SQYFGANL | TCSADWGYNYAEQF | 0.6237649 |
| GGVKPAA | TCSADWGYNYAEQF | 0.75610816 |
| GGTGGAGGAGGW | TCSADWGYNYAEQF | 0.5657111 |
| GVGGVAGLGGAGG | TCSADWGYNYAEQF | 0.7310618 |
| VLLVLATTI | TCSADWGYNYAEQF | 0.56513095 |
| IGIFAAML | TCSADWGYNYAEQF | 0.70502794 |
| AVGDLID | TCSADWGYNYAEQF | 0.5221378 |
| NISPLPL | TCSADWGYNYAEQF | 0.7284179 |
| VTAVFVAI | TCSADWGYNYAEQF | 0.7425078 |
| SGDDNDAAL | TCSADWGYNYAEQF | 0.7446207 |
| AVAALAHS | TCSADWGYNYAEQF | 0.2878236 |
| GGVTVMATP | TCSADWGYNYAEQF | 0.7373667 |
| AGLAANVRD | TCSADWGYNYAEQF | 0.7481985 |
| TNRPDLIDPALLRPG | TCSADWGYNYAEQF | 0.77740264 |
| LANLGVTAL | TCSADWGYNYAEQF | 0.64274466 |
| TGERIEL | TCSADWGYNYAEQF | 0.7305121 |
| NGEELAA | TCSADWGYNYAEQF | 0.6141225 |
| VLSLVWG | TCSADWGYNYAEQF | 0.61066115 |

| | | |
|---|---|---|
| IGYALWL | TCSADWGYNYAEQF | 0.4724686 |
| LSLLLAPL | TCSADWGYNYAEQF | 0.6589873 |
| TLDEALK | TCSADWGYNYAEQF | 0.6239131 |
| TDGAGTAEGG | TCSADWGYNYAEQF | 0.5607867 |
| NAPGGAGGNGG | TCSADWGYNYAEQF | 0.74229777 |
| VSVFMGMGGNF | TCSADWGYNYAEQF | 0.7484281 |
| LASYLAAL | TCSADWGYNYAEQF | 0.75029576 |
| LVGGTPAL | TCSADWGYNYAEQF | 0.76105595 |
| VRNADVL | TCSADWGYNYAEQF | 0.7638762 |
| TCSSPFLVSAM | TCSADWGYNYAEQF | 0.7224704 |
| VHITDAE | TCSADWGYNYAEQF | 0.5589068 |
| EYAAPSF | TCSADWGYNYAEQF | 0.650566 |
| VVDAFHGGL | TCSADWGYNYAEQF | 0.46440518 |
| ASVVDPQ | TCSADWGYNYAEQF | 0.6584063 |
| SYNEFGGLS | TCSADWGYNYAEQF | 0.5921115 |
| DPDGSVY | TCSADWGYNYAEQF | 0.7088848 |
| MVSTIHATE | TCSADWGYNYAEQF | 0.2320217 |
| LAAGVADPV | TCSADWGYNYAEQF | 0.69187236 |
| TKEQIAA | TCSADWGYNYAEQF | 0.7740582 |
| SLASPAAL | TCSADWGYNYAEQF | 0.57309914 |
| IGLVFYGV | TCSADWGYNYAEQF | 0.71237254 |
| FLLDASNL | TCSADWGYNYAEQF | 0.71950495 |
| YQVGNFP | TCSADWGYNYAEQF | 0.5780389 |
| ESLLSQST | TCSADWGYNYAEQF | 0.7414468 |
| LFTVGFAAVG | TCSADWGYNYAEQF | 0.52231973 |
| ATPPTPT | TCSADWGYNYAEQF | 0.70624244 |
| TDLATLG | TCSADWGYNYAEQF | 0.7646222 |
| SAGGTGR | TCSADWGYNYAEQF | 0.6912693 |
| GFSNSGS | TCSADWGYNYAEQF | 0.68054783 |
| GGNGGSGGTSV | TCSADWGYNYAEQF | 0.622772 |
| GTSVATGGAGN | TCSADWGYNYAEQF | 0.3885851 |
| NGGNGGSGGTS | TCSADWGYNYAEQF | 0.7100656 |
| AVVKFPH | TCSADWGYNYAEQF | 0.43766975 |
| TPPPPPP | TCSADWGYNYAEQF | 0.67706716 |
| AAPAAAAAPA | TCSADWGYNYAEQF | 0.48059812 |
| IALNEDL | TCSADWGYNYAEQF | 0.7090882 |
| IVNVSSM | TCSADWGYNYAEQF | 0.69071287 |
| TGIPPAPR | TCSADWGYNYAEQF | 0.6886054 |
| FDLGGGTF | TCSADWGYNYAEQF | 0.52503294 |
| FDLGGGTFDVS | TCSADWGYNYAEQF | 0.6672219 |
| NEPTAAA | TCSADWGYNYAEQF | 0.6027024 |
| NEPTAAALA | TCSADWGYNYAEQF | 0.46495134 |
| GSIINISSI | TCSADWGYNYAEQF | 0.73389715 |
| VAPGNAGTA | TCSADWGYNYAEQF | 0.5884303 |
| TGGGASLE | TCSADWGYNYAEQF | 0.6378622 |
| VAVYNHYKRI | TCSADWGYNYAEQF | 0.7564384 |
| TDPAPATT | TCSADWGYNYAEQF | 0.6610807 |
| TNRPDILDPALLRPG | TCSADWGYNYAEQF | 0.35399398 |
| LTTTTTS | TCSADWGYNYAEQF | 0.6456201 |
| PTGSGKTL | TCSADWGYNYAEQF | 0.67495 |
| TGSGKTL | TCSADWGYNYAEQF | 0.5992277 |

| | | |
|---|---|---|
| TIGHVDHGK | TCSADWGYNYAEQF | 0.2312442 |
| TIGHVDHGKTT | TCSADWGYNYAEQF | 0.7136103 |
| GHVDHGKTT | TCSADWGYNYAEQF | 0.55021423 |
| AQFEHTLL | TCSADWGYNYAEQF | 0.521661 |
| IVNQDEL | TCSADWGYNYAEQF | 0.56181353 |
| TNGSQFF | TCSADWGYNYAEQF | 0.7433957 |
| TGKGIVE | TCSADWGYNYAEQF | 0.60637474 |
| LISNASDA | TCSADWGYNYAEQF | 0.6126326 |
| AELRTLGELTKIAKAHGAQ | TCSADWGYNYAEQF | 0.72870755 |
| APNAVAAAEPSW | TCSADWGYNYAEQF | 0.7219243 |
| APQVAGAATAVL | TCSADWGYNYAEQF | 0.76404 |
| ARRDPAAAAAILPTDA | TCSADWGYNYAEQF | 0.7066978 |
| DAAARDVLAEAGLAEYF | TCSADWGYNYAEQF | 0.64612323 |
| DDQIVLSAKANLAKVEQ | TCSADWGYNYAEQF | 0.730384 |
| DELLPDTGEDSDEDGH | TCSADWGYNYAEQF | 0.61831033 |
| ELPEAPANRATVPQD | TCSADWGYNYAEQF | 0.44386506 |
| FGQSKIERVTGIG | TCSADWGYNYAEQF | 0.3449206 |
| GAGGTGSTAGGAGGAGGAGGLY | TCSADWGYNYAEQF | 0.6174666 |
| GHGGTGAAGGNGGTGGAGGSGI | TCSADWGYNYAEQF | 0.35830364 |
| GNGGNGGAGGEGGPGGAGGAGA | TCSADWGYNYAEQF | 0.6980435 |
| GSTAGNGGNGGNATNGTIAGQP | TCSADWGYNYAEQF | 0.7459121 |
| GTGGDGGNGGNGGTGG | TCSADWGYNYAEQF | 0.6106524 |
| GVRRPAGIPADAPV | TCSADWGYNYAEQF | 0.44986367 |
| GYRQVTDAHLLTLARR | TCSADWGYNYAEQF | 0.6226836 |
| IAKMGQDGHDRG | TCSADWGYNYAEQF | 0.5896026 |
| KAEPSIEIQYT | TCSADWGYNYAEQF | 0.7299339 |
| KPVRIRMAADEDTVAVT | TCSADWGYNYAEQF | 0.6951058 |
| MPPRIAGMRLLVIKPEP | TCSADWGYNYAEQF | 0.77483225 |
| MKFYAVALFRSRGVPPTR | TCSADWGYNYAEQF | 0.7509235 |
| NAGGFGILGAVAHSP | TCSADWGYNYAEQF | 0.58718437 |
| NEGREITLAMAEM | TCSADWGYNYAEQF | 0.7278217 |
| NELVEWSHPLGHRWRWE | TCSADWGYNYAEQF | 0.5939052 |
| NLGAGGTGGNGGNPG | TCSADWGYNYAEQF | 0.68810874 |
| NTEWPKWETVMAADPVD | TCSADWGYNYAEQF | 0.64953667 |
| PASAAPAETQP | TCSADWGYNYAEQF | 0.6568599 |
| QQSSDGKNGVRAAEITGE | TCSADWGYNYAEQF | 0.6276555 |
| RAPDAVTTSWWKE | TCSADWGYNYAEQF | 0.59062207 |
| SGFGNVGAHTSGWFNQ | TCSADWGYNYAEQF | 0.3848769 |
| SGVAGAVGQAASVGGLKV | TCSADWGYNYAEQF | 0.7200588 |
| SNGGAGTGSAGGNGGT | TCSADWGYNYAEQF | 0.5773939 |
| TGGAGGEGGAAG | TCSADWGYNYAEQF | 0.36605492 |
| TGGDGGAAGAGGAPGHG | TCSADWGYNYAEQF | 0.59213936 |
| TMPPLLAAIANAGRHGV | TCSADWGYNYAEQF | 0.77476555 |
| TSSTAGNSGGA | TCSADWGYNYAEQF | 0.7660111 |
| VPATVPVGILC | TCSADWGYNYAEQF | 0.59752595 |
| VQRPDSGGVVLTGRIS | TCSADWGYNYAEQF | 0.5432025 |
| WTPNPVPATKGVPAGLVQAT | TCSADWGYNYAEQF | 0.7141533 |
| YALKRALAKFSDRFDVVII | TCSADWGYNYAEQF | 0.3799387 |
| YLIGSVIV | TCSADWGYNYAEQF | 0.6825852 |
| YGQLVAVID | TCSADWGYNYAEQF | 0.68618715 |
| VTVGGMTLPA | TCSADWGYNYAEQF | 0.6534499 |

| | | |
|---|---|---|
| VLALPAV | TCSADWGYNYAEQF | 0.5699765 |
| VIPPTLNYET | TCSADWGYNYAEQF | 0.6979085 |
| VGMGGMHPSSG | TCSADWGYNYAEQF | 0.6249517 |
| VGLAGGAATA | TCSADWGYNYAEQF | 0.7057513 |
| VGGAGGAGGSGP | TCSADWGYNYAEQF | 0.27252835 |
| VFAAFMV | TCSADWGYNYAEQF | 0.7204309 |
| TLYDKLR | TCSADWGYNYAEQF | 0.54407406 |
| TLAVFTVL | TCSADWGYNYAEQF | 0.50786805 |
| TGSVQPARV | TCSADWGYNYAEQF | 0.28848836 |
| TGGGAGSGGT | TCSADWGYNYAEQF | 0.6676482 |
| SHLETSL | TCSADWGYNYAEQF | 0.726614 |
| SFTTGMT | TCSADWGYNYAEQF | 0.7501595 |
| QVEVAKRMGVRQPTVS | TCSADWGYNYAEQF | 0.5414628 |
| QGSGGPALAM | TCSADWGYNYAEQF | 0.74858975 |
| MLAPSNP | TCSADWGYNYAEQF | 0.7520112 |
| LVNNAAI | TCSADWGYNYAEQF | 0.63970387 |
| LTGEINTLA | TCSADWGYNYAEQF | 0.35564655 |
| LSIFGPV | TCSADWGYNYAEQF | 0.7045499 |
| LLALLANL | TCSADWGYNYAEQF | 0.67672604 |
| LISNASDALDKLR | TCSADWGYNYAEQF | 0.74342644 |
| LINEAVP | TCSADWGYNYAEQF | 0.7133042 |
| LGYAFSGL | TCSADWGYNYAEQF | 0.46195602 |
| LDFVAVAL | TCSADWGYNYAEQF | 0.74493945 |
| LAQPGAE | TCSADWGYNYAEQF | 0.7364428 |
| ISNLGAVTGF | TCSADWGYNYAEQF | 0.47200757 |
| HQEDPGRI | TCSADWGYNYAEQF | 0.51383823 |
| GTPGEPTP | TCSADWGYNYAEQF | 0.58330417 |
| GFFNSGS | TCSADWGYNYAEQF | 0.33729744 |
| GAGGGGLTAGAN | TCSADWGYNYAEQF | 0.72124815 |
| FVDALVNM | TCSADWGYNYAEQF | 0.6114595 |
| FLTVAPEL | TCSADWGYNYAEQF | 0.563715 |
| FETLTAL | TCSADWGYNYAEQF | 0.6633877 |
| DVINPIEL | TCSADWGYNYAEQF | 0.560149 |
| APVSGAPV | TCSADWGYNYAEQF | 0.56071496 |
| ALVGGLAW | TCSADWGYNYAEQF | 0.6127786 |
| ALAAAAATAI | TCSADWGYNYAEQF | 0.67911804 |
| AFGLGLLL | TCSADWGYNYAEQF | 0.71723133 |
| AFFAAASL | TCSADWGYNYAEQF | 0.6482247 |
| AAAALPAFSPP | TCSADWGYNYAEQF | 0.6880816 |
| VVLAPAL | TCSADWGYNYAEQF | 0.76593924 |
| TIGHVDHGKTTL | TCSADWGYNYAEQF | 0.42792606 |
| LISNASDALDK | TCSADWGYNYAEQF | 0.5282912 |
| LISNASDAL | TCSADWGYNYAEQF | 0.7044734 |
| ISNASDALDK | TCSADWGYNYAEQF | 0.44832656 |
| NQAETLVYQTE | TCSADWGYNYAEQF | 0.7318754 |
| TGGTGGTAGAG | TCSADWGYNYAEQF | 0.68964505 |
| TGSTAGGAGGA | TCSADWGYNYAEQF | 0.7147496 |
| DPLAVPVAAGPV | TCSADWGYNYAEQF | 0.66957515 |
| GGTGGTPGNGGN | TCSADWGYNYAEQF | 0.7745745 |
| PAGLAGVDPATL | TCSADWGYNYAEQF | 0.73196685 |
| GNNLVVEAPHAGASV | TCSADWGYNYAEQF | 0.6411567 |

| | | |
|---|---|---|
| KVTRIMSASTLEQPA | TCSADWGYNYAEQF | 0.7367516 |
| LGFALPIALPRRTVV | TCSADWGYNYAEQF | 0.5243437 |
| TRLGNLTARKLVVLV | TCSADWGYNYAEQF | 0.74669343 |
| WSQQRRLIYFPSAGP | TCSADWGYNYAEQF | 0.7253332 |
| GPNGNPDPMAAAVDIR | TCSADWGYNYAEQF | 0.4822318 |
| TQRCVSVADLRAAASR | TCSADWGYNYAEQF | 0.66716456 |
| QFVLDVAEHPVEVLTETV | TCSADWGYNYAEQF | 0.7506647 |
| GGAGGTGGAGGNGGTQAGDG | TCSADWGYNYAEQF | 0.749912 |
| PLLATLRAIAVDVDAKPAQV | TCSADWGYNYAEQF | 0.37576523 |
| APKFRTGGYMIDFWGVGYQVAK | TCSADWGYNYAEQF | 0.5875929 |
| GAGGAGGAGGAAGGPGGTGGTG | TCSADWGYNYAEQF | 0.39815456 |
| IGALGLLATDVAPAGSVAFGVA | TCSADWGYNYAEQF | 0.5486331 |
| AGGAGGIGGNGGAGGAGGVLMG | TCSADWGYNYAEQF | 0.4486096 |
| ADELIGLL | TCSADWGYNYAEQF | 0.72178 |
| ADRALKA | TCSADWGYNYAEQF | 0.6012237 |
| AGAVGQVGGAGGA | TCSADWGYNYAEQF | 0.724802 |
| AGGGGGLGGAP | TCSADWGYNYAEQF | 0.6921848 |
| AGGLLYGNGGAG | TCSADWGYNYAEQF | 0.7551419 |
| AGLESIGL | TCSADWGYNYAEQF | 0.5617023 |
| AGLKTGII | TCSADWGYNYAEQF | 0.65590644 |
| ASLAALLL | TCSADWGYNYAEQF | 0.34982502 |
| ASVINISSI | TCSADWGYNYAEQF | 0.5028105 |
| ATPPGPA | TCSADWGYNYAEQF | 0.46519265 |
| ATVASALV | TCSADWGYNYAEQF | 0.6418583 |
| ATVASLIL | TCSADWGYNYAEQF | 0.3105766 |
| AVNGPVLIRH | TCSADWGYNYAEQF | 0.75678504 |
| DGAAGAVGI | TCSADWGYNYAEQF | 0.67295784 |
| DPLLHGI | TCSADWGYNYAEQF | 0.72154737 |
| ELGAALGAL | TCSADWGYNYAEQF | 0.5616725 |
| ELGRTQAV | TCSADWGYNYAEQF | 0.54807997 |
| FVAGGYLS | TCSADWGYNYAEQF | 0.5789899 |
| GASGGAGGAGGW | TCSADWGYNYAEQF | 0.7310504 |
| GGAGAGGDGGAGGI | TCSADWGYNYAEQF | 0.49320516 |
| GTGHNGSQPGMGGNG | TCSADWGYNYAEQF | 0.32888627 |
| GYAAASASAS | TCSADWGYNYAEQF | 0.51934296 |
| HQVDVAAVQQG | TCSADWGYNYAEQF | 0.4818997 |
| IGAAIAHT | TCSADWGYNYAEQF | 0.7615851 |
| IGMIEVAQL | TCSADWGYNYAEQF | 0.74415827 |
| IRLAPEHTSAMD | TCSADWGYNYAEQF | 0.700288 |
| IYNVSVL | TCSADWGYNYAEQF | 0.7522439 |
| KQGFLGM | TCSADWGYNYAEQF | 0.2731347 |
| LESVITQL | TCSADWGYNYAEQF | 0.59524417 |
| LGLGALVVV | TCSADWGYNYAEQF | 0.7726344 |
| LIGNGGNGGSGGI | TCSADWGYNYAEQF | 0.69669795 |
| LNAINAPTL | TCSADWGYNYAEQF | 0.6461171 |
| LQLLNTGV | TCSADWGYNYAEQF | 0.749688 |
| LQTSIEF | TCSADWGYNYAEQF | 0.52902216 |
| MSFAVSEL | TCSADWGYNYAEQF | 0.6179361 |
| NTGALGL | TCSADWGYNYAEQF | 0.6698987 |
| PPGAGGAGGAGTT | TCSADWGYNYAEQF | 0.7169938 |
| QLVGLGI | TCSADWGYNYAEQF | 0.6374955 |

| | | |
|---|---|---|
| RFLEQVHQL | TCSADWGYNYAEQF | 0.43176275 |
| RGVGYVLR | TCSADWGYNYAEQF | 0.6987964 |
| SATGVFAGVI | TCSADWGYNYAEQF | 0.7786772 |
| SLPAITLP | TCSADWGYNYAEQF | 0.63146806 |
| STALAFTV | TCSADWGYNYAEQF | 0.7316662 |
| SVLTVGLGK | TCSADWGYNYAEQF | 0.6993325 |
| TALAGAGSGI | TCSADWGYNYAEQF | 0.7031237 |
| TGLPFTGI | TCSADWGYNYAEQF | 0.5380072 |
| VAEHVTV | TCSADWGYNYAEQF | 0.573526 |
| VDIVLNSL | TCSADWGYNYAEQF | 0.6771596 |
| VLTGGGAVL | TCSADWGYNYAEQF | 0.48473117 |
| VNVGLAYGI | TCSADWGYNYAEQF | 0.63947415 |
| VSSAAVGAV | TCSADWGYNYAEQF | 0.6027221 |
| VTSVYEAGV | TCSADWGYNYAEQF | 0.781364 |
| WVALDRGA | TCSADWGYNYAEQF | 0.71724874 |
| YLVGSVVI | TCSADWGYNYAEQF | 0.69064724 |
| GPPGTGKTLL | TCSADWGYNYAEQF | 0.5405371 |
| GVGMTKF | TCSADWGYNYAEQF | 0.6609475 |
| IGIGGSDL | TCSADWGYNYAEQF | 0.719918 |
| IGTLLAEL | TCSADWGYNYAEQF | 0.406512 |
| IYTSGTT | TCSADWGYNYAEQF | 0.7637274 |
| LSFPAGV | TCSADWGYNYAEQF | 0.52427185 |
| LYTSGTT | TCSADWGYNYAEQF | 0.63525 |
| VGAFPFGF | TCSADWGYNYAEQF | 0.72714585 |
| VGAGTVEF | TCSADWGYNYAEQF | 0.75888306 |
| VGKSTLLL | TCSADWGYNYAEQF | 0.5928575 |
| AALAHAGGGIV | TCSADWGYNYAEQF | 0.6466609 |
| AALVAVSLL | TCSADWGYNYAEQF | 0.52786237 |
| AAVSRAVAAL | TCSADWGYNYAEQF | 0.72880346 |
| AGGSSGIGGF | TCSADWGYNYAEQF | 0.75691926 |
| AGLTVANANG | TCSADWGYNYAEQF | 0.6758375 |
| AGLVGAVL | TCSADWGYNYAEQF | 0.7721304 |
| ALSQAFTV | TCSADWGYNYAEQF | 0.6800041 |
| ALTFSVSL | TCSADWGYNYAEQF | 0.38175362 |
| ALTGAALVAL | TCSADWGYNYAEQF | 0.64533603 |
| ASFSIGAGVL | TCSADWGYNYAEQF | 0.7461165 |
| DGSVYKI | TCSADWGYNYAEQF | 0.7521823 |
| EQLAEIV | TCSADWGYNYAEQF | 0.75952834 |
| FAVSELTVM | TCSADWGYNYAEQF | 0.70978457 |
| GAAPLAAGL | TCSADWGYNYAEQF | 0.61778915 |
| GADGGDGGSGGAGG | TCSADWGYNYAEQF | 0.7615516 |
| GFGLDTGGAGG | TCSADWGYNYAEQF | 0.7729212 |
| GLASGGLL | TCSADWGYNYAEQF | 0.73616934 |
| GRIAGAAVLL | TCSADWGYNYAEQF | 0.54416525 |
| GSAGSLNTSF | TCSADWGYNYAEQF | 0.530109 |
| GTGQIGI | TCSADWGYNYAEQF | 0.6807807 |
| GTLIALIL | TCSADWGYNYAEQF | 0.7314429 |
| IAATMGFVVG | TCSADWGYNYAEQF | 0.4404978 |
| IFAQALL | TCSADWGYNYAEQF | 0.6938288 |
| IGAAVALGF | TCSADWGYNYAEQF | 0.70982635 |
| IGLTPLAL | TCSADWGYNYAEQF | 0.3848098 |

| | | |
|---|---|---|
| ILGSNTSTL | TCSADWGYNYAEQF | 0.68596816 |
| IVFSLTVL | TCSADWGYNYAEQF | 0.7615674 |
| LASKATGF | TCSADWGYNYAEQF | 0.6458205 |
| LASVALIL | TCSADWGYNYAEQF | 0.5464017 |
| LFAVGLTL | TCSADWGYNYAEQF | 0.75769866 |
| LFNGGIGTYI | TCSADWGYNYAEQF | 0.7804321 |
| LTLGAGII | TCSADWGYNYAEQF | 0.7570043 |
| LVAAALGDI | TCSADWGYNYAEQF | 0.3524465 |
| NTAFVQLGI | TCSADWGYNYAEQF | 0.26466805 |
| QLFSGAEL | TCSADWGYNYAEQF | 0.6118018 |
| SAATFITL | TCSADWGYNYAEQF | 0.73871493 |
| SNYGLANLGV | TCSADWGYNYAEQF | 0.6284398 |
| SYGGAYAVM | TCSADWGYNYAEQF | 0.57586646 |
| TTGVGGLTLG | TCSADWGYNYAEQF | 0.7086541 |
| TTQTIYI | TCSADWGYNYAEQF | 0.7494634 |
| VAAGAGLGAVI | TCSADWGYNYAEQF | 0.5357453 |
| VAAVVTLL | TCSADWGYNYAEQF | 0.41473466 |
| VGFLGGLI | TCSADWGYNYAEQF | 0.7260673 |
| VGVIPILL | TCSADWGYNYAEQF | 0.7301084 |
| VIFASSLI | TCSADWGYNYAEQF | 0.5207093 |
| VKGGDGGAAAG | TCSADWGYNYAEQF | 0.6298251 |
| VTKTFTNI | TCSADWGYNYAEQF | 0.6471062 |
| VVAAAILL | TCSADWGYNYAEQF | 0.61077756 |
| YAALDVEL | TCSADWGYNYAEQF | 0.71423006 |
| LASVALLL | TCSADWGYNYAEQF | 0.4838609 |
| LDAVHRV | TCSADWGYNYAEQF | 0.7563579 |
| QGLSSSIF | TCSADWGYNYAEQF | 0.44619772 |
| RGTFANIRL | TCSADWGYNYAEQF | 0.7083441 |
| RQLWWGHRI | TCSADWGYNYAEQF | 0.69787765 |
| YGPPGTGKTLL | TCSADWGYNYAEQF | 0.7038629 |
| AALVGAPANVA | TCSADWGYNYAEQF | 0.6100173 |
| AATFRRTE | TCSADWGYNYAEQF | 0.57730746 |
| AGALGGGI | TCSADWGYNYAEQF | 0.46329033 |
| AGGAGGAGGIGAG | TCSADWGYNYAEQF | 0.35353696 |
| AGSGALGL | TCSADWGYNYAEQF | 0.7546545 |
| AQIADQI | TCSADWGYNYAEQF | 0.6006493 |
| ASAVAVTYL | TCSADWGYNYAEQF | 0.7481489 |
| ASTVLGGTGG | TCSADWGYNYAEQF | 0.7036197 |
| AVGGEPAK | TCSADWGYNYAEQF | 0.50772506 |
| CEQGTFGGTC | TCSADWGYNYAEQF | 0.6940077 |
| DEGKAPAG | TCSADWGYNYAEQF | 0.46114045 |
| DTVVFPAL | TCSADWGYNYAEQF | 0.5723878 |
| FEGITMAG | TCSADWGYNYAEQF | 0.7035099 |
| FPLVVQD | TCSADWGYNYAEQF | 0.7231083 |
| GAFTTPQ | TCSADWGYNYAEQF | 0.52381533 |
| GFGGFGTAGGVGGA | TCSADWGYNYAEQF | 0.69025433 |
| GFLAQFL | TCSADWGYNYAEQF | 0.71630394 |
| GGAGGAGGNAGVA | TCSADWGYNYAEQF | 0.5579251 |
| GGVGTAGAGGAGGAGG | TCSADWGYNYAEQF | 0.656417 |
| GNGGGGDAGNAGSGGN | TCSADWGYNYAEQF | 0.7547301 |
| GTGGAGGAGNDAGS | TCSADWGYNYAEQF | 0.6607988 |

| | | |
|---|---|---|
| GTTNTGWFNS | TCSADWGYNYAEQF | 0.45093817 |
| IFMGPLFTL | TCSADWGYNYAEQF | 0.7137238 |
| ILAHAIEAI | TCSADWGYNYAEQF | 0.537361 |
| IRSAIREAL | TCSADWGYNYAEQF | 0.7185396 |
| ITFAGGG | TCSADWGYNYAEQF | 0.6578404 |
| ITGMSGHQEV | TCSADWGYNYAEQF | 0.6277071 |
| IVAGLACTL | TCSADWGYNYAEQF | 0.7150374 |
| IVGGGLAGA | TCSADWGYNYAEQF | 0.42184156 |
| IVTGVLNI | TCSADWGYNYAEQF | 0.53220147 |
| LAQQGCI | TCSADWGYNYAEQF | 0.6000617 |
| LCLHVEN | TCSADWGYNYAEQF | 0.67953527 |
| LDAISDGS | TCSADWGYNYAEQF | 0.7347064 |
| LGGFAAML | TCSADWGYNYAEQF | 0.51044416 |
| LGGKNPMI | TCSADWGYNYAEQF | 0.6977246 |
| LGPKGVAAIL | TCSADWGYNYAEQF | 0.5885245 |
| LMVGVSI | TCSADWGYNYAEQF | 0.49874753 |
| LVGAAFVL | TCSADWGYNYAEQF | 0.623399 |
| LVLAVAPL | TCSADWGYNYAEQF | 0.5842092 |
| MAAIAVIDAA | TCSADWGYNYAEQF | 0.479764 |
| NNGINAELQ | TCSADWGYNYAEQF | 0.7217387 |
| RGEPLVLVHG | TCSADWGYNYAEQF | 0.52724636 |
| RGGAGGAAGLI | TCSADWGYNYAEQF | 0.69304544 |
| RSYRMHPAV | TCSADWGYNYAEQF | 0.7593292 |
| SAAHLLAGQLA | TCSADWGYNYAEQF | 0.5203427 |
| SAAVGSY | TCSADWGYNYAEQF | 0.6953416 |
| SPHAVLHRI | TCSADWGYNYAEQF | 0.7067244 |
| SQIAKVL | TCSADWGYNYAEQF | 0.70969105 |
| TALGHLVP | TCSADWGYNYAEQF | 0.6309207 |
| TAPTTSGAA | TCSADWGYNYAEQF | 0.53902423 |
| TDGDDGNGGSPGT | TCSADWGYNYAEQF | 0.76049113 |
| TDGLGEN | TCSADWGYNYAEQF | 0.7204295 |
| TGGPPLPGQA | TCSADWGYNYAEQF | 0.7380662 |
| TIPAITI | TCSADWGYNYAEQF | 0.72840184 |
| TIPGFTI | TCSADWGYNYAEQF | 0.7439252 |
| TSIASGALA | TCSADWGYNYAEQF | 0.72909355 |
| TVTLPGAAI | TCSADWGYNYAEQF | 0.61627716 |
| VASGLSAALP | TCSADWGYNYAEQF | 0.36295038 |
| VGAGVIGI | TCSADWGYNYAEQF | 0.62691927 |
| VGPKRFGV | TCSADWGYNYAEQF | 0.6580073 |
| VNVAFPD | TCSADWGYNYAEQF | 0.547016 |
| VSMTNSGVS | TCSADWGYNYAEQF | 0.35517704 |
| VTDVLGDLSV | TCSADWGYNYAEQF | 0.75621694 |
| VVVVGAGFAGL | TCSADWGYNYAEQF | 0.7812823 |
| VYTVGTVV | TCSADWGYNYAEQF | 0.55769897 |
| WTTFSGP | TCSADWGYNYAEQF | 0.7709074 |
| YDELVEAF | TCSADWGYNYAEQF | 0.4083423 |
| YGTGGAGGNGG | TCSADWGYNYAEQF | 0.71828055 |
| YTNVMAV | TCSADWGYNYAEQF | 0.7542399 |
| FLEQVHQL | TCSADWGYNYAEQF | 0.4930138 |
| GLSLLAVAV | TCSADWGYNYAEQF | 0.73817885 |
| LEQVHQL | TCSADWGYNYAEQF | 0.48365963 |

| | | |
|---|---|---|
| LNAAGAI | TCSADWGYNYAEQF | 0.5870295 |
| PAVHASL | TCSADWGYNYAEQF | 0.75946677 |
| TELEDDA | TCSADWGYNYAEQF | 0.6216887 |
| TPPGPPN | TCSADWGYNYAEQF | 0.7077588 |
| TPPPPPPP | TCSADWGYNYAEQF | 0.62178123 |
| TQSPALI | ASAGQNNQAPL | 0.51151574 |
| EFAAAFPQ | ASAGQNNQAPL | 0.67888117 |
| LLSGNLDVL | ASAGQNNQAPL | 0.75159156 |
| GPANLGNYNI | ASAGQNNQAPL | 0.57771266 |
| TLGPTLP | ASAGQNNQAPL | 0.76760936 |
| VYAYSSCCK | ASAGQNNQAPL | 0.52973485 |
| APAPAPAPNI | ASAGQNNQAPL | 0.7758219 |
| PGGPVPIP | ASAGQNNQAPL | 0.74694455 |
| FLTVALEP | ASAGQNNQAPL | 0.68805695 |
| SQYFGANL | ASAGQNNQAPL | 0.57610327 |
| GGVKPAA | ASAGQNNQAPL | 0.7552073 |
| GGTGGAGGAGGW | ASAGQNNQAPL | 0.61067736 |
| GVGGVAGLGGAGG | ASAGQNNQAPL | 0.71954226 |
| VLLVLATTI | ASAGQNNQAPL | 0.7522341 |
| IGIFAAML | ASAGQNNQAPL | 0.47403425 |
| AVGDLID | ASAGQNNQAPL | 0.71935934 |
| NISPLPL | ASAGQNNQAPL | 0.67177844 |
| VTAVFVAI | ASAGQNNQAPL | 0.6824781 |
| SGDDNDAAL | ASAGQNNQAPL | 0.55391216 |
| AVAALAHS | ASAGQNNQAPL | 0.4190547 |
| GGVTVMATP | ASAGQNNQAPL | 0.6013123 |
| AGLAANVRD | ASAGQNNQAPL | 0.5225065 |
| TNRPDLIDPALLRPG | ASAGQNNQAPL | 0.72607636 |
| LANLGVTAL | ASAGQNNQAPL | 0.7144196 |
| TGERIEL | ASAGQNNQAPL | 0.61264694 |
| NGEELAA | ASAGQNNQAPL | 0.721504 |
| VLSLVWG | ASAGQNNQAPL | 0.6934892 |
| IGYALWL | ASAGQNNQAPL | 0.44848895 |
| LSLLLAPL | ASAGQNNQAPL | 0.6550577 |
| TLDEALK | ASAGQNNQAPL | 0.5438176 |
| TDGAGTAEGG | ASAGQNNQAPL | 0.6299316 |
| NAPGGAGGNGG | ASAGQNNQAPL | 0.39227623 |
| VSVFMGMGGNF | ASAGQNNQAPL | 0.6905876 |
| LASYLAAL | ASAGQNNQAPL | 0.61030525 |
| LVGGTPAL | ASAGQNNQAPL | 0.3951645 |
| VRNADVL | ASAGQNNQAPL | 0.6881865 |
| TCSSPFLVSAM | ASAGQNNQAPL | 0.65699613 |
| VHITDAE | ASAGQNNQAPL | 0.61741304 |
| EYAAPSF | ASAGQNNQAPL | 0.7078835 |
| VVDAFHGGL | ASAGQNNQAPL | 0.48684803 |
| ASVVDPQ | ASAGQNNQAPL | 0.5921261 |
| SYNEFGGLS | ASAGQNNQAPL | 0.7070474 |
| DPDGSVY | ASAGQNNQAPL | 0.7269922 |
| MVSTIHATE | ASAGQNNQAPL | 0.54085565 |
| LAAGVADPV | ASAGQNNQAPL | 0.67085695 |
| TKEQIAA | ASAGQNNQAPL | 0.7206126 |

| | | |
|---|---|---|
| SLASPAAL | ASAGQNNQAPL | 0.65434307 |
| IGLVFYGV | ASAGQNNQAPL | 0.72229606 |
| FLLDASNL | ASAGQNNQAPL | 0.727297 |
| YQVGNFP | ASAGQNNQAPL | 0.5518862 |
| ESLLSQST | ASAGQNNQAPL | 0.72664106 |
| LFTVGFAAVG | ASAGQNNQAPL | 0.7305323 |
| ATPPTPT | ASAGQNNQAPL | 0.6388194 |
| TDLATLG | ASAGQNNQAPL | 0.66706526 |
| SAGGTGR | ASAGQNNQAPL | 0.4566995 |
| GFSNSGS | ASAGQNNQAPL | 0.61465776 |
| GGNGGSGGTSV | ASAGQNNQAPL | 0.4047075 |
| GTSVATGGAGN | ASAGQNNQAPL | 0.686974 |
| NGGNGGSGGTS | ASAGQNNQAPL | 0.7196878 |
| AVVKFPH | ASAGQNNQAPL | 0.6433706 |
| TPPPPPP | ASAGQNNQAPL | 0.49578053 |
| AAPAAAAAPA | ASAGQNNQAPL | 0.5113369 |
| IALNEDL | ASAGQNNQAPL | 0.62483823 |
| IVNVSSM | ASAGQNNQAPL | 0.48943907 |
| TGIPPAPR | ASAGQNNQAPL | 0.40587324 |
| FDLGGGTF | ASAGQNNQAPL | 0.49068123 |
| FDLGGGTFDVS | ASAGQNNQAPL | 0.70655906 |
| NEPTAAA | ASAGQNNQAPL | 0.7097761 |
| NEPTAAALA | ASAGQNNQAPL | 0.5891943 |
| GSIINISSI | ASAGQNNQAPL | 0.7242569 |
| VAPGNAGTA | ASAGQNNQAPL | 0.70437825 |
| TGGGASLE | ASAGQNNQAPL | 0.6899065 |
| VAVYNHYKRI | ASAGQNNQAPL | 0.663023 |
| TDPAPATT | ASAGQNNQAPL | 0.44870433 |
| TNRPDILDPALLRPG | ASAGQNNQAPL | 0.72622454 |
| LTTTTTS | ASAGQNNQAPL | 0.63466084 |
| PTGSGKTL | ASAGQNNQAPL | 0.6921911 |
| TGSGKTL | ASAGQNNQAPL | 0.7063279 |
| TIGHVDHGK | ASAGQNNQAPL | 0.6421618 |
| TIGHVDHGKTT | ASAGQNNQAPL | 0.60160357 |
| GHVDHGKTT | ASAGQNNQAPL | 0.64943016 |
| AQFEHTLL | ASAGQNNQAPL | 0.70726097 |
| IVNQDEL | ASAGQNNQAPL | 0.5786284 |
| TNGSQFF | ASAGQNNQAPL | 0.37603775 |
| TGKGIVE | ASAGQNNQAPL | 0.6589656 |
| LISNASDA | ASAGQNNQAPL | 0.6676359 |
| AELRTLGELTKIAKAHGAQ | ASAGQNNQAPL | 0.72679365 |
| APNAVAAAEPSW | ASAGQNNQAPL | 0.64547247 |
| APQVAGAATAVL | ASAGQNNQAPL | 0.6142777 |
| ARRDPAAAAAILPTDA | ASAGQNNQAPL | 0.70273155 |
| DAAARDVLAEAGLAEYF | ASAGQNNQAPL | 0.6851202 |
| DDQIVLSAKANLAKVEQ | ASAGQNNQAPL | 0.66060555 |
| DELLPDTGEDSDEDGH | ASAGQNNQAPL | 0.6114734 |
| ELPEAPANRATVPQD | ASAGQNNQAPL | 0.61133003 |
| FGQSKIERVTGIG | ASAGQNNQAPL | 0.7021775 |
| GAGGTGSTAGGAGGAGGAGGLY | ASAGQNNQAPL | 0.70655143 |
| GHGGTGAAGGNGGTGGAGGSGI | ASAGQNNQAPL | 0.67787385 |

| | | |
|---|---|---|
| GNGGNGGAGGEGGPGGAGGAGA | ASAGQNNQAPL | 0.7222036 |
| GSTAGNGGNGGNATNGTIAGQP | ASAGQNNQAPL | 0.6518553 |
| GTGGDGGNGGNGGTGG | ASAGQNNQAPL | 0.66414165 |
| GVRRPAGIPADAPV | ASAGQNNQAPL | 0.4065899 |
| GYRQVTDAHLLTLARR | ASAGQNNQAPL | 0.5490865 |
| IAKMGQDGHDRG | ASAGQNNQAPL | 0.6688812 |
| KAEPSIEIQYT | ASAGQNNQAPL | 0.6825706 |
| KPVRIRMAADEDTVAVT | ASAGQNNQAPL | 0.71003854 |
| MPPRIAGMRLLVIKPEP | ASAGQNNQAPL | 0.6806767 |
| MKFYAVALFRSRGVPPTR | ASAGQNNQAPL | 0.6082548 |
| NAGGFGILGAVAHSP | ASAGQNNQAPL | 0.7033895 |
| NEGREITLAMAEM | ASAGQNNQAPL | 0.7226577 |
| NELVEWSHPLGHRWRWE | ASAGQNNQAPL | 0.48593885 |
| NLGAGGTGGNGGNPG | ASAGQNNQAPL | 0.6359381 |
| NTEWPKWETVMAADPVD | ASAGQNNQAPL | 0.7323187 |
| PASAAPAETQP | ASAGQNNQAPL | 0.68566704 |
| QQSSDGKNGVRAAEITGE | ASAGQNNQAPL | 0.65692437 |
| RAPDAVTTSWWKE | ASAGQNNQAPL | 0.6484024 |
| SGFGNVGAHTSGWFNQ | ASAGQNNQAPL | 0.731537 |
| SGVAGAVGQAASVGGLKV | ASAGQNNQAPL | 0.64885545 |
| SNGGAGTGSAGGNGGT | ASAGQNNQAPL | 0.732533 |
| TGGAGGEGGAAG | ASAGQNNQAPL | 0.39062193 |
| TGGDGGAAGAGGAPGHG | ASAGQNNQAPL | 0.5967272 |
| TMPPLLAAIANAGRHGV | ASAGQNNQAPL | 0.39707968 |
| TSSTAGNSGGA | ASAGQNNQAPL | 0.2896039 |
| VPATVPVGILC | ASAGQNNQAPL | 0.705864 |
| VQRPDSGGVVLTGRIS | ASAGQNNQAPL | 0.5708419 |
| WTPNPVPATKGVPAGLVQAT | ASAGQNNQAPL | 0.68350464 |
| YALKRALAKFSDRFDVVII | ASAGQNNQAPL | 0.6510054 |
| YLIGSVIV | ASAGQNNQAPL | 0.71592224 |
| YGQLVAVID | ASAGQNNQAPL | 0.6617907 |
| VTVGGMTLPA | ASAGQNNQAPL | 0.6147534 |
| VLALPAV | ASAGQNNQAPL | 0.5131989 |
| VIPPTLNYET | ASAGQNNQAPL | 0.6824587 |
| VGMGGMHPSSG | ASAGQNNQAPL | 0.71656895 |
| VGLAGGAATA | ASAGQNNQAPL | 0.6913716 |
| VGGAGGAGGSGP | ASAGQNNQAPL | 0.43293726 |
| VFAAFMV | ASAGQNNQAPL | 0.6805383 |
| TLYDKLR | ASAGQNNQAPL | 0.6240673 |
| TLAVFTVL | ASAGQNNQAPL | 0.4151597 |
| TGSVQPARV | ASAGQNNQAPL | 0.38176462 |
| TGGGAGSGGT | ASAGQNNQAPL | 0.54332525 |
| SHLETSL | ASAGQNNQAPL | 0.5915713 |
| SFTTGMT | ASAGQNNQAPL | 0.7114774 |
| QVEVAKRMGVRQPTVS | ASAGQNNQAPL | 0.7100296 |
| QGSGGPALAM | ASAGQNNQAPL | 0.6287523 |
| MLAPSNP | ASAGQNNQAPL | 0.69749963 |
| LVNNAAI | ASAGQNNQAPL | 0.6774374 |
| LTGEINTLA | ASAGQNNQAPL | 0.23772669 |
| LSIFGPV | ASAGQNNQAPL | 0.58871293 |
| LLALLANL | ASAGQNNQAPL | 0.46985573 |

| | | |
|---|---|---|
| LISNASDALDKLR | ASAGQNNQAPL | 0.7263464 |
| LINEAVP | ASAGQNNQAPL | 0.56187963 |
| LGYAFSGL | ASAGQNNQAPL | 0.6503642 |
| LDFVAVAL | ASAGQNNQAPL | 0.6629612 |
| LAQPGAE | ASAGQNNQAPL | 0.692453 |
| ISNLGAVTGF | ASAGQNNQAPL | 0.6709122 |
| HQEDPGRI | ASAGQNNQAPL | 0.54591393 |
| GTPGEPTP | ASAGQNNQAPL | 0.33121136 |
| GFFNSGS | ASAGQNNQAPL | 0.6935557 |
| GAGGGGLTAGAN | ASAGQNNQAPL | 0.55104035 |
| FVDALVNM | ASAGQNNQAPL | 0.5592171 |
| FLTVAPEL | ASAGQNNQAPL | 0.48783726 |
| FETLTAL | ASAGQNNQAPL | 0.6700523 |
| DVINPIEL | ASAGQNNQAPL | 0.49143326 |
| APVSGAPV | ASAGQNNQAPL | 0.70529723 |
| ALVGGLAW | ASAGQNNQAPL | 0.5077859 |
| ALAAAAATAI | ASAGQNNQAPL | 0.6587765 |
| AFGLGLLL | ASAGQNNQAPL | 0.72877777 |
| AFFAAASL | ASAGQNNQAPL | 0.68887794 |
| AAAALPAFSPP | ASAGQNNQAPL | 0.6292851 |
| VVLAPAL | ASAGQNNQAPL | 0.6244215 |
| TIGHVDHGKTTL | ASAGQNNQAPL | 0.52444905 |
| LISNASDALDK | ASAGQNNQAPL | 0.5561546 |
| LISNASDAL | ASAGQNNQAPL | 0.608649 |
| ISNASDALDK | ASAGQNNQAPL | 0.7274508 |
| NQAETLVYQTE | ASAGQNNQAPL | 0.6168207 |
| TGGTGGTAGAG | ASAGQNNQAPL | 0.4512432 |
| TGSTAGGAGGA | ASAGQNNQAPL | 0.62780017 |
| DPLAVPVAAGPV | ASAGQNNQAPL | 0.7342691 |
| GGTGGTPGNGGN | ASAGQNNQAPL | 0.72603756 |
| PAGLAGVDPATL | ASAGQNNQAPL | 0.51934755 |
| GNNLVVEAPHAGASV | ASAGQNNQAPL | 0.55864817 |
| KVTRIMSASTLEQPA | ASAGQNNQAPL | 0.61944515 |
| LGFALPIALPRRTVV | ASAGQNNQAPL | 0.65002054 |
| TRLGNLTARKLVVLV | ASAGQNNQAPL | 0.49881983 |
| WSQQRRLIYFPSAGP | ASAGQNNQAPL | 0.5390235 |
| GPNGNPDPMAAAVDIR | ASAGQNNQAPL | 0.69379145 |
| TQRCVSVADLRAAASR | ASAGQNNQAPL | 0.7383392 |
| QFVLDVAEHPVEVLTETV | ASAGQNNQAPL | 0.6250874 |
| GGAGGTGGAGGNGGTQAGDG | ASAGQNNQAPL | 0.42117405 |
| PLLATLRAIAVDVDAKPAQV | ASAGQNNQAPL | 0.6721953 |
| APKFRTGGYMIDFWGVGYQVAK | ASAGQNNQAPL | 0.5710847 |
| GAGGAGGAGGAAGGPGGTGGTG | ASAGQNNQAPL | 0.5923375 |
| IGALGLLATDVAPAGSVAFGVA | ASAGQNNQAPL | 0.5705068 |
| AGGAGGIGGNGGAGGAGGVLMG | ASAGQNNQAPL | 0.45192528 |
| ADELIGLL | ASAGQNNQAPL | 0.66777873 |
| ADRALKA | ASAGQNNQAPL | 0.7158185 |
| AGAVGQVGGAGGA | ASAGQNNQAPL | 0.6346041 |
| AGGGGGLGGAP | ASAGQNNQAPL | 0.6652672 |
| AGGLLYGNGGAG | ASAGQNNQAPL | 0.7268156 |
| AGLESIGL | ASAGQNNQAPL | 0.69261664 |

| | | |
|---|---|---|
| AGLKTGII | ASAGQNNQAPL | 0.7216536 |
| ASLAALLL | ASAGQNNQAPL | 0.7195278 |
| ASVINISSI | ASAGQNNQAPL | 0.6392479 |
| ATPPGPA | ASAGQNNQAPL | 0.61940765 |
| ATVASALV | ASAGQNNQAPL | 0.7223378 |
| ATVASLIL | ASAGQNNQAPL | 0.67691654 |
| AVNGPVLIRH | ASAGQNNQAPL | 0.68105185 |
| DGAAGAVGI | ASAGQNNQAPL | 0.6195687 |
| DPLLHGI | ASAGQNNQAPL | 0.72290933 |
| ELGAALGAL | ASAGQNNQAPL | 0.5782882 |
| ELGRTQAV | ASAGQNNQAPL | 0.7251177 |
| FVAGGYLS | ASAGQNNQAPL | 0.3300367 |
| GASGGAGGAGGW | ASAGQNNQAPL | 0.5204512 |
| GGAGAGGDGGAGGI | ASAGQNNQAPL | 0.71032935 |
| GTGHNGSQPGMGGNG | ASAGQNNQAPL | 0.24840672 |
| GYAAASASAS | ASAGQNNQAPL | 0.43516842 |
| HQVDVAAVQQG | ASAGQNNQAPL | 0.6466347 |
| IGAAIAHT | ASAGQNNQAPL | 0.70761377 |
| IGMIEVAQL | ASAGQNNQAPL | 0.2643019 |
| IRLAPEHTSAMD | ASAGQNNQAPL | 0.6249044 |
| IYNVSVL | ASAGQNNQAPL | 0.4081644 |
| KQGFLGM | ASAGQNNQAPL | 0.64128923 |
| LESVITQL | ASAGQNNQAPL | 0.6605023 |
| LGLGALVVV | ASAGQNNQAPL | 0.43688965 |
| LIGNGGNGGSGGI | ASAGQNNQAPL | 0.69629526 |
| LNAINAPTL | ASAGQNNQAPL | 0.43030638 |
| LQLLNTGV | ASAGQNNQAPL | 0.6526568 |
| LQTSIEF | ASAGQNNQAPL | 0.7103535 |
| MSFAVSEL | ASAGQNNQAPL | 0.7319157 |
| NTGALGL | ASAGQNNQAPL | 0.53551704 |
| PPGAGGAGGAGTT | ASAGQNNQAPL | 0.70137554 |
| QLVGLGI | ASAGQNNQAPL | 0.6036545 |
| RFLEQVHQL | ASAGQNNQAPL | 0.73756903 |
| RGVGYVLR | ASAGQNNQAPL | 0.5945559 |
| SATGVFAGVI | ASAGQNNQAPL | 0.4229884 |
| SLPAITLP | ASAGQNNQAPL | 0.70563674 |
| STALAFTV | ASAGQNNQAPL | 0.6413069 |
| SVLTVGLGK | ASAGQNNQAPL | 0.6215556 |
| TALAGAGSGI | ASAGQNNQAPL | 0.7041395 |
| TGLPFTGI | ASAGQNNQAPL | 0.47263414 |
| VAEHVTV | ASAGQNNQAPL | 0.66418195 |
| VDIVLNSL | ASAGQNNQAPL | 0.7084657 |
| VLTGGGAVL | ASAGQNNQAPL | 0.7064446 |
| VNVGLAYGI | ASAGQNNQAPL | 0.50831604 |
| VSSAAVGAV | ASAGQNNQAPL | 0.708434 |
| VTSVYEAGV | ASAGQNNQAPL | 0.6678796 |
| WVALDRGA | ASAGQNNQAPL | 0.544322 |
| YLVGSVVI | ASAGQNNQAPL | 0.7232434 |
| GPPGTGKTLL | ASAGQNNQAPL | 0.5768341 |
| GVGMTKF | ASAGQNNQAPL | 0.7099679 |
| IGIGGSDL | ASAGQNNQAPL | 0.67867684 |

| | | |
|---|---|---|
| IGTLLAEL | ASAGQNNQAPL | 0.7188153 |
| IYTSGTT | ASAGQNNQAPL | 0.48435396 |
| LSFPAGV | ASAGQNNQAPL | 0.64470303 |
| LYTSGTT | ASAGQNNQAPL | 0.6694977 |
| VGAFPFGF | ASAGQNNQAPL | 0.71353054 |
| VGAGTVEF | ASAGQNNQAPL | 0.6006031 |
| VGKSTLLL | ASAGQNNQAPL | 0.62410176 |
| AALAHAGGGIV | ASAGQNNQAPL | 0.68941486 |
| AALVAVSLL | ASAGQNNQAPL | 0.72374 |
| AAVSRAVAAL | ASAGQNNQAPL | 0.63193697 |
| AGGSSGIGGF | ASAGQNNQAPL | 0.65859425 |
| AGLTVANANG | ASAGQNNQAPL | 0.7280711 |
| AGLVGAVL | ASAGQNNQAPL | 0.5103947 |
| ALSQAFTV | ASAGQNNQAPL | 0.65828156 |
| ALTFSVSL | ASAGQNNQAPL | 0.6910385 |
| ALTGAALVAL | ASAGQNNQAPL | 0.7133139 |
| ASFSIGAGVL | ASAGQNNQAPL | 0.6182544 |
| DGSVYKI | ASAGQNNQAPL | 0.7167165 |
| EQLAEIV | ASAGQNNQAPL | 0.5761125 |
| FAVSELTVM | ASAGQNNQAPL | 0.7064929 |
| GAAPLAAGL | ASAGQNNQAPL | 0.59605205 |
| GADGGDGGSGGAGG | ASAGQNNQAPL | 0.6375725 |
| GFGLDTGGAGG | ASAGQNNQAPL | 0.5318929 |
| GLASGGLL | ASAGQNNQAPL | 0.6976242 |
| GRIAGAAVLL | ASAGQNNQAPL | 0.7095795 |
| GSAGSLNTSF | ASAGQNNQAPL | 0.7095547 |
| GTGQIGI | ASAGQNNQAPL | 0.3868872 |
| GTLIALIL | ASAGQNNQAPL | 0.7023704 |
| IAATMGFVVG | ASAGQNNQAPL | 0.71474653 |
| IFAQALL | ASAGQNNQAPL | 0.73597795 |
| IGAAVALGF | ASAGQNNQAPL | 0.57258344 |
| IGLTPLAL | ASAGQNNQAPL | 0.695992 |
| ILGSNTSTL | ASAGQNNQAPL | 0.6142175 |
| IVFSLTVL | ASAGQNNQAPL | 0.63483834 |
| LASKATGF | ASAGQNNQAPL | 0.49711692 |
| LASVALIL | ASAGQNNQAPL | 0.5091374 |
| LFAVGLTL | ASAGQNNQAPL | 0.6606592 |
| LFNGGIGTYI | ASAGQNNQAPL | 0.5771979 |
| LTLGAGII | ASAGQNNQAPL | 0.71650696 |
| LVAAALGDI | ASAGQNNQAPL | 0.71564496 |
| NTAFVQLGI | ASAGQNNQAPL | 0.703961 |
| QLFSGAEL | ASAGQNNQAPL | 0.72674125 |
| SAATFITL | ASAGQNNQAPL | 0.72307265 |
| SNYGLANLGV | ASAGQNNQAPL | 0.69774616 |
| SYGGAYAVM | ASAGQNNQAPL | 0.5586021 |
| TTGVGGLTLG | ASAGQNNQAPL | 0.66768193 |
| TTQTIYI | ASAGQNNQAPL | 0.47414353 |
| VAAGAGLGAVI | ASAGQNNQAPL | 0.6675602 |
| VAAVVTLL | ASAGQNNQAPL | 0.5894426 |
| VGFLGGLI | ASAGQNNQAPL | 0.694038 |
| VGVIPILL | ASAGQNNQAPL | 0.2527979 |

| | | |
|---|---|---|
| VIFASSLI | ASAGQNNQAPL | 0.68942046 |
| VKGGDGGAAAG | ASAGQNNQAPL | 0.72945845 |
| VTKTFTNI | ASAGQNNQAPL | 0.6031003 |
| VVAAAILL | ASAGQNNQAPL | 0.697773 |
| YAALDVEL | ASAGQNNQAPL | 0.705762 |
| LASVALLL | ASAGQNNQAPL | 0.5799166 |
| LDAVHRV | ASAGQNNQAPL | 0.70875967 |
| QGLSSSIF | ASAGQNNQAPL | 0.60375696 |
| RGTFANIRL | ASAGQNNQAPL | 0.70561326 |
| RQLWWGHRI | ASAGQNNQAPL | 0.70946836 |
| YGPPGTGKTLL | ASAGQNNQAPL | 0.6945232 |
| AALVGAPANVA | ASAGQNNQAPL | 0.5342632 |
| AATFRRTE | ASAGQNNQAPL | 0.6664884 |
| AGALGGGI | ASAGQNNQAPL | 0.40639177 |
| AGGAGGAGGIGAG | ASAGQNNQAPL | 0.63670623 |
| AGSGALGL | ASAGQNNQAPL | 0.4209727 |
| AQIADQI | ASAGQNNQAPL | 0.67141616 |
| ASAVAVTYL | ASAGQNNQAPL | 0.47315168 |
| ASTVLGGTGG | ASAGQNNQAPL | 0.70232546 |
| AVGGEPAK | ASAGQNNQAPL | 0.6790023 |
| CEQGTFGGTC | ASAGQNNQAPL | 0.67017066 |
| DEGKAPAG | ASAGQNNQAPL | 0.58689564 |
| DTVVFPAL | ASAGQNNQAPL | 0.69322515 |
| FEGITMAG | ASAGQNNQAPL | 0.684326 |
| FPLVVQD | ASAGQNNQAPL | 0.49656904 |
| GAFTTPQ | ASAGQNNQAPL | 0.678218 |
| GFGGFGTAGGVGGA | ASAGQNNQAPL | 0.59812075 |
| GFLAQFL | ASAGQNNQAPL | 0.5903511 |
| GGAGGAGGNAGVA | ASAGQNNQAPL | 0.7105857 |
| GGVGTAGAGGAGGAGG | ASAGQNNQAPL | 0.6618553 |
| GNGGGGDAGNAGSGGN | ASAGQNNQAPL | 0.3787577 |
| GTGGAGGAGNDAGS | ASAGQNNQAPL | 0.61614823 |
| GTTNTGWFNS | ASAGQNNQAPL | 0.6944088 |
| IFMGPLFTL | ASAGQNNQAPL | 0.57309645 |
| ILAHAIEAI | ASAGQNNQAPL | 0.27650452 |
| IRSAIREAL | ASAGQNNQAPL | 0.66507214 |
| ITFAGGG | ASAGQNNQAPL | 0.54228884 |
| ITGMSGHQEV | ASAGQNNQAPL | 0.56649536 |
| IVAGLACTL | ASAGQNNQAPL | 0.6237962 |
| IVGGGLAGA | ASAGQNNQAPL | 0.71309376 |
| IVTGVLNI | ASAGQNNQAPL | 0.5657394 |
| LAQQGCI | ASAGQNNQAPL | 0.55893695 |
| LCLHVEN | ASAGQNNQAPL | 0.68685496 |
| LDAISDGS | ASAGQNNQAPL | 0.696678 |
| LGGFAAML | ASAGQNNQAPL | 0.7144104 |
| LGGKNPMI | ASAGQNNQAPL | 0.6677382 |
| LGPKGVAAIL | ASAGQNNQAPL | 0.52067924 |
| LMVGVSI | ASAGQNNQAPL | 0.6543691 |
| LVGAAFVL | ASAGQNNQAPL | 0.55734015 |
| LVLAVAPL | ASAGQNNQAPL | 0.606344 |
| MAAIAVIDAA | ASAGQNNQAPL | 0.519485 |

| | | |
|---|---|---|
| NNGINAELQ | ASAGQNNQAPL | 0.598002 |
| RGEPLVLVHG | ASAGQNNQAPL | 0.6210269 |
| RGGAGGAAGLI | ASAGQNNQAPL | 0.67413497 |
| RSYRMHPAV | ASAGQNNQAPL | 0.7196872 |
| SAAHLLAGQLA | ASAGQNNQAPL | 0.5537781 |
| SAAVGSY | ASAGQNNQAPL | 0.40305555 |
| SPHAVLHRI | ASAGQNNQAPL | 0.61599743 |
| SQIAKVL | ASAGQNNQAPL | 0.5832077 |
| TALGHLVP | ASAGQNNQAPL | 0.7027446 |
| TAPTTSGAA | ASAGQNNQAPL | 0.6102928 |
| TDGDDGNGGSPGT | ASAGQNNQAPL | 0.7391572 |
| TDGLGEN | ASAGQNNQAPL | 0.7129601 |
| TGGPPLPGQA | ASAGQNNQAPL | 0.6243073 |
| TIPAITI | ASAGQNNQAPL | 0.70690763 |
| TIPGFTI | ASAGQNNQAPL | 0.584895 |
| TSIASGALA | ASAGQNNQAPL | 0.6258657 |
| TVTLPGAAI | ASAGQNNQAPL | 0.64191335 |
| VASGLSAALP | ASAGQNNQAPL | 0.5947014 |
| VGAGVIGI | ASAGQNNQAPL | 0.591678 |
| VGPKRFGV | ASAGQNNQAPL | 0.62397695 |
| VNVAFPD | ASAGQNNQAPL | 0.3965681 |
| VSMTNSGVS | ASAGQNNQAPL | 0.6977763 |
| VTDVLGDLSV | ASAGQNNQAPL | 0.4758823 |
| VVVVGAGFAGL | ASAGQNNQAPL | 0.45630053 |
| VYTVGTVV | ASAGQNNQAPL | 0.6096032 |
| WTTFSGP | ASAGQNNQAPL | 0.73580027 |
| YDELVEAF | ASAGQNNQAPL | 0.7242309 |
| YGTGGAGGNGG | ASAGQNNQAPL | 0.61674684 |
| YTNVMAV | ASAGQNNQAPL | 0.555478 |
| FLEQVHQL | ASAGQNNQAPL | 0.70031464 |
| GLSLLAVAV | ASAGQNNQAPL | 0.54555035 |
| LEQVHQL | ASAGQNNQAPL | 0.6687051 |
| LNAAGAI | ASAGQNNQAPL | 0.65450525 |
| PAVHASL | ASAGQNNQAPL | 0.63281274 |
| TELEDDA | ASAGQNNQAPL | 0.56263024 |
| TPPGPPN | ASAGQNNQAPL | 0.6984726 |
| TPPPPPPP | ASAGQNNQAPL | 0.6195798 |
| TQSPALI | ASSPIRDWGNYAEQF | 0.6368109 |
| EFAAAFPQ | ASSPIRDWGNYAEQF | 0.32557452 |
| LLSGNLDVL | ASSPIRDWGNYAEQF | 0.69837284 |
| GPANLGNYNI | ASSPIRDWGNYAEQF | 0.5909711 |
| TLGPTLP | ASSPIRDWGNYAEQF | 0.471339 |
| VYAYSSCCK | ASSPIRDWGNYAEQF | 0.30485994 |
| APAPAPAPNI | ASSPIRDWGNYAEQF | 0.6307329 |
| PGGPVPIP | ASSPIRDWGNYAEQF | 0.69069743 |
| FLTVALEP | ASSPIRDWGNYAEQF | 0.71810853 |
| SQYFGANL | ASSPIRDWGNYAEQF | 0.63864726 |
| GGVKPAA | ASSPIRDWGNYAEQF | 0.7082736 |
| GGTGGAGGAGGW | ASSPIRDWGNYAEQF | 0.71358204 |
| GVGGVAGLGGAGG | ASSPIRDWGNYAEQF | 0.6489288 |
| VLLVLATTI | ASSPIRDWGNYAEQF | 0.3390615 |

| | | |
|---|---|---|
| IGIFAAML | ASSPIRDWGNYAEQF | 0.6654633 |
| AVGDLID | ASSPIRDWGNYAEQF | 0.6575243 |
| NISPLPL | ASSPIRDWGNYAEQF | 0.6929271 |
| VTAVFVAI | ASSPIRDWGNYAEQF | 0.69328487 |
| SGDDNDAAL | ASSPIRDWGNYAEQF | 0.51090395 |
| AVAALAHS | ASSPIRDWGNYAEQF | 0.68996763 |
| GGVTVMATP | ASSPIRDWGNYAEQF | 0.6963952 |
| AGLAANVRD | ASSPIRDWGNYAEQF | 0.4880712 |
| TNRPDLIDPALLRPG | ASSPIRDWGNYAEQF | 0.51787853 |
| LANLGVTAL | ASSPIRDWGNYAEQF | 0.5571644 |
| TGERIEL | ASSPIRDWGNYAEQF | 0.35891154 |
| NGEELAA | ASSPIRDWGNYAEQF | 0.69950914 |
| VLSLVWG | ASSPIRDWGNYAEQF | 0.6401633 |
| IGYALWL | ASSPIRDWGNYAEQF | 0.644112 |
| LSLLLAPL | ASSPIRDWGNYAEQF | 0.6341578 |
| TLDEALK | ASSPIRDWGNYAEQF | 0.6194029 |
| TDGAGTAEGG | ASSPIRDWGNYAEQF | 0.6159958 |
| NAPGGAGGNGG | ASSPIRDWGNYAEQF | 0.63708323 |
| VSVFMGMGGNF | ASSPIRDWGNYAEQF | 0.6891923 |
| LASYLAAL | ASSPIRDWGNYAEQF | 0.70136714 |
| LVGGTPAL | ASSPIRDWGNYAEQF | 0.64872086 |
| VRNADVL | ASSPIRDWGNYAEQF | 0.6184305 |
| TCSSPFLVSAM | ASSPIRDWGNYAEQF | 0.7317649 |
| VHITDAE | ASSPIRDWGNYAEQF | 0.47520244 |
| EYAAPSF | ASSPIRDWGNYAEQF | 0.62254727 |
| VVDAFHGGL | ASSPIRDWGNYAEQF | 0.67915595 |
| ASVVDPQ | ASSPIRDWGNYAEQF | 0.5776 |
| SYNEFGGLS | ASSPIRDWGNYAEQF | 0.71062815 |
| DPDGSVY | ASSPIRDWGNYAEQF | 0.68741953 |
| MVSTIHATE | ASSPIRDWGNYAEQF | 0.6974786 |
| LAAGVADPV | ASSPIRDWGNYAEQF | 0.6741433 |
| TKEQIAA | ASSPIRDWGNYAEQF | 0.734859 |
| SLASPAAL | ASSPIRDWGNYAEQF | 0.69295466 |
| IGLVFYGV | ASSPIRDWGNYAEQF | 0.61213696 |
| FLLDASNL | ASSPIRDWGNYAEQF | 0.71612036 |
| YQVGNFP | ASSPIRDWGNYAEQF | 0.44152564 |
| ESLLSQST | ASSPIRDWGNYAEQF | 0.71280193 |
| LFTVGFAAVG | ASSPIRDWGNYAEQF | 0.7011603 |
| ATPPTPT | ASSPIRDWGNYAEQF | 0.520017 |
| TDLATLG | ASSPIRDWGNYAEQF | 0.66773975 |
| SAGGTGR | ASSPIRDWGNYAEQF | 0.719315 |
| GFSNSGS | ASSPIRDWGNYAEQF | 0.71780497 |
| GGNGGSGGTSV | ASSPIRDWGNYAEQF | 0.54513526 |
| GTSVATGGAGN | ASSPIRDWGNYAEQF | 0.5855478 |
| NGGNGGSGGTS | ASSPIRDWGNYAEQF | 0.4874028 |
| AVVKFPH | ASSPIRDWGNYAEQF | 0.4072533 |
| TPPPPPP | ASSPIRDWGNYAEQF | 0.44193354 |
| AAPAAAAAPA | ASSPIRDWGNYAEQF | 0.6907786 |
| IALNEDL | ASSPIRDWGNYAEQF | 0.66180485 |
| IVNVSSM | ASSPIRDWGNYAEQF | 0.70963216 |
| TGIPPAPR | ASSPIRDWGNYAEQF | 0.6733901 |

| | | |
|---|---|---|
| FDLGGGTF | ASSPIRDWGNYAEQF | 0.7103492 |
| FDLGGGTFDVS | ASSPIRDWGNYAEQF | 0.62099695 |
| NEPTAAA | ASSPIRDWGNYAEQF | 0.665066 |
| NEPTAAALA | ASSPIRDWGNYAEQF | 0.36500004 |
| GSIINISSI | ASSPIRDWGNYAEQF | 0.4323417 |
| VAPGNAGTA | ASSPIRDWGNYAEQF | 0.5203945 |
| TGGGASLE | ASSPIRDWGNYAEQF | 0.65043294 |
| VAVYNHYKRI | ASSPIRDWGNYAEQF | 0.33277875 |
| TDPAPATT | ASSPIRDWGNYAEQF | 0.7191157 |
| TNRPDILDPALLRPG | ASSPIRDWGNYAEQF | 0.6257306 |
| LTTTTTS | ASSPIRDWGNYAEQF | 0.6922299 |
| PTGSGKTL | ASSPIRDWGNYAEQF | 0.61864805 |
| TGSGKTL | ASSPIRDWGNYAEQF | 0.4947094 |
| TIGHVDHGK | ASSPIRDWGNYAEQF | 0.57582664 |
| TIGHVDHGKTT | ASSPIRDWGNYAEQF | 0.70893073 |
| GHVDHGKTT | ASSPIRDWGNYAEQF | 0.50540614 |
| AQFEHTLL | ASSPIRDWGNYAEQF | 0.32764715 |
| IVNQDEL | ASSPIRDWGNYAEQF | 0.5601281 |
| TNGSQFF | ASSPIRDWGNYAEQF | 0.56503254 |
| TGKGIVE | ASSPIRDWGNYAEQF | 0.7310618 |
| LISNASDA | ASSPIRDWGNYAEQF | 0.7201007 |
| AELRTLGELTKIAKAHGAQ | ASSPIRDWGNYAEQF | 0.6785726 |
| APNAVAAAEPSW | ASSPIRDWGNYAEQF | 0.72906053 |
| APQVAGAATAVL | ASSPIRDWGNYAEQF | 0.27002725 |
| ARRDPAAAAAILPTDA | ASSPIRDWGNYAEQF | 0.6370913 |
| DAAARDVLAEAGLAEYF | ASSPIRDWGNYAEQF | 0.72138083 |
| DDQIVLSAKANLAKVEQ | ASSPIRDWGNYAEQF | 0.66092694 |
| DELLPDTGEDSDEDGH | ASSPIRDWGNYAEQF | 0.6761453 |
| ELPEAPANRATVPQD | ASSPIRDWGNYAEQF | 0.7157644 |
| FGQSKIERVTGIG | ASSPIRDWGNYAEQF | 0.58870435 |
| GAGGTGSTAGGAGGAGGAGGLY | ASSPIRDWGNYAEQF | 0.6030442 |
| GHGGTGAAGGNGGTGGAGGSGI | ASSPIRDWGNYAEQF | 0.64821786 |
| GNGGNGGAGGEGGPGGAGGAGA | ASSPIRDWGNYAEQF | 0.68675524 |
| GSTAGNGGNGGNATNGTIAGQP | ASSPIRDWGNYAEQF | 0.55646265 |
| GTGGDGGNGGNGGTGG | ASSPIRDWGNYAEQF | 0.4478532 |
| GVRRPAGIPADAPV | ASSPIRDWGNYAEQF | 0.687979 |
| GYRQVTDAHLLTLARR | ASSPIRDWGNYAEQF | 0.7445702 |
| IAKMGQDGHDRG | ASSPIRDWGNYAEQF | 0.63694394 |
| KAEPSIEIQYT | ASSPIRDWGNYAEQF | 0.68361837 |
| KPVRIRMAADEDTVAVT | ASSPIRDWGNYAEQF | 0.66672003 |
| MPPRIAGMRLLVIKPEP | ASSPIRDWGNYAEQF | 0.61345637 |
| MKFYAVALFRSRGVPPTR | ASSPIRDWGNYAEQF | 0.60929704 |
| NAGGFGILGAVAHSP | ASSPIRDWGNYAEQF | 0.5321338 |
| NEGREITLAMAEM | ASSPIRDWGNYAEQF | 0.69891214 |
| NELVEWSHPLGHRWRWE | ASSPIRDWGNYAEQF | 0.5536042 |
| NLGAGGTGGNGGNPG | ASSPIRDWGNYAEQF | 0.6641384 |
| NTEWPKWETVMAADPVD | ASSPIRDWGNYAEQF | 0.63146883 |
| PASAAPAETQP | ASSPIRDWGNYAEQF | 0.74156076 |
| QQSSDGKNGVRAAEITGE | ASSPIRDWGNYAEQF | 0.69783473 |
| RAPDAVTTSWWKE | ASSPIRDWGNYAEQF | 0.68607295 |
| SGFGNVGAHTSGWFNQ | ASSPIRDWGNYAEQF | 0.58377516 |

| | | |
|---|---|---|
| SGVAGAVGQAASVGGLKV | ASSPIRDWGNYAEQF | 0.65759194 |
| SNGGAGTGSAGGNGGT | ASSPIRDWGNYAEQF | 0.69341946 |
| TGGAGGEGGAAG | ASSPIRDWGNYAEQF | 0.4012748 |
| TGGDGGAAGAGGAPGHG | ASSPIRDWGNYAEQF | 0.7248485 |
| TMPPLLAAIANAGRHGV | ASSPIRDWGNYAEQF | 0.5502818 |
| TSSTAGNSGGA | ASSPIRDWGNYAEQF | 0.6210705 |
| VPATVPVGILC | ASSPIRDWGNYAEQF | 0.68477297 |
| VQRPDSGGVVLTGRIS | ASSPIRDWGNYAEQF | 0.72880685 |
| WTPNPVPATKGVPAGLVQAT | ASSPIRDWGNYAEQF | 0.6576293 |
| YALKRALAKFSDRFDVVII | ASSPIRDWGNYAEQF | 0.6997818 |
| YLIGSVIV | ASSPIRDWGNYAEQF | 0.5766334 |
| YGQLVAVID | ASSPIRDWGNYAEQF | 0.6935622 |
| VTVGGMTLPA | ASSPIRDWGNYAEQF | 0.7194356 |
| VLALPAV | ASSPIRDWGNYAEQF | 0.6453544 |
| VIPPTLNYET | ASSPIRDWGNYAEQF | 0.7364528 |
| VGMGGMHPSSG | ASSPIRDWGNYAEQF | 0.6274103 |
| VGLAGGAATA | ASSPIRDWGNYAEQF | 0.36059618 |
| VGGAGGAGGSGP | ASSPIRDWGNYAEQF | 0.6184356 |
| VFAAFMV | ASSPIRDWGNYAEQF | 0.7191386 |
| TLYDKLR | ASSPIRDWGNYAEQF | 0.719965 |
| TLAVFTVL | ASSPIRDWGNYAEQF | 0.71496856 |
| TGSVQPARV | ASSPIRDWGNYAEQF | 0.6869575 |
| TGGGAGSGGT | ASSPIRDWGNYAEQF | 0.5852751 |
| SHLETSL | ASSPIRDWGNYAEQF | 0.7279718 |
| SFTTGMT | ASSPIRDWGNYAEQF | 0.7290654 |
| QVEVAKRMGVRQPTVS | ASSPIRDWGNYAEQF | 0.69089633 |
| QGSGGPALAM | ASSPIRDWGNYAEQF | 0.5329635 |
| MLAPSNP | ASSPIRDWGNYAEQF | 0.47179884 |
| LVNNAAI | ASSPIRDWGNYAEQF | 0.69972456 |
| LTGEINTLA | ASSPIRDWGNYAEQF | 0.6902834 |
| LSIFGPV | ASSPIRDWGNYAEQF | 0.47259855 |
| LLALLANL | ASSPIRDWGNYAEQF | 0.6832801 |
| LISNASDALDKLR | ASSPIRDWGNYAEQF | 0.684955 |
| LINEAVP | ASSPIRDWGNYAEQF | 0.38124242 |
| LGYAFSGL | ASSPIRDWGNYAEQF | 0.6764803 |
| LDFVAVAL | ASSPIRDWGNYAEQF | 0.7240368 |
| LAQPGAE | ASSPIRDWGNYAEQF | 0.67490935 |
| ISNLGAVTGF | ASSPIRDWGNYAEQF | 0.5519662 |
| HQEDPGRI | ASSPIRDWGNYAEQF | 0.72542703 |
| GTPGEPTP | ASSPIRDWGNYAEQF | 0.7320173 |
| GFFNSGS | ASSPIRDWGNYAEQF | 0.7267648 |
| GAGGGGLTAGAN | ASSPIRDWGNYAEQF | 0.43066692 |
| FVDALVNM | ASSPIRDWGNYAEQF | 0.29760134 |
| FLTVAPEL | ASSPIRDWGNYAEQF | 0.6633016 |
| FETLTAL | ASSPIRDWGNYAEQF | 0.70954883 |
| DVINPIEL | ASSPIRDWGNYAEQF | 0.64283955 |
| APVSGAPV | ASSPIRDWGNYAEQF | 0.54178846 |
| ALVGGLAW | ASSPIRDWGNYAEQF | 0.6621938 |
| ALAAAAATAI | ASSPIRDWGNYAEQF | 0.7147685 |
| AFGLGLLL | ASSPIRDWGNYAEQF | 0.50294864 |
| AFFAAASL | ASSPIRDWGNYAEQF | 0.3696471 |

| | | |
|---|---|---|
| AAAALPAFSPP | ASSPIRDWGNYAEQF | 0.68052804 |
| VVLAPAL | ASSPIRDWGNYAEQF | 0.70916903 |
| TIGHVDHGKTTL | ASSPIRDWGNYAEQF | 0.56325614 |
| LISNASDALDK | ASSPIRDWGNYAEQF | 0.6192472 |
| LISNASDAL | ASSPIRDWGNYAEQF | 0.67959094 |
| ISNASDALDK | ASSPIRDWGNYAEQF | 0.60632944 |
| NQAETLVYQTE | ASSPIRDWGNYAEQF | 0.6759981 |
| TGGTGGTAGAG | ASSPIRDWGNYAEQF | 0.5222385 |
| TGSTAGGAGGA | ASSPIRDWGNYAEQF | 0.7116045 |
| DPLAVPVAAGPV | ASSPIRDWGNYAEQF | 0.49778146 |
| GGTGGTPGNGGN | ASSPIRDWGNYAEQF | 0.6852461 |
| PAGLAGVDPATL | ASSPIRDWGNYAEQF | 0.6754248 |
| GNNLVVEAPHAGASV | ASSPIRDWGNYAEQF | 0.6720623 |
| KVTRIMSASTLEQPA | ASSPIRDWGNYAEQF | 0.65707946 |
| LGFALPIALPRRTVV | ASSPIRDWGNYAEQF | 0.582284 |
| TRLGNLTARKLVVLV | ASSPIRDWGNYAEQF | 0.52193993 |
| WSQQRRLIYFPSAGP | ASSPIRDWGNYAEQF | 0.49498367 |
| GPNGNPDPMAAAVDIR | ASSPIRDWGNYAEQF | 0.7178838 |
| TQRCVSVADLRAAASR | ASSPIRDWGNYAEQF | 0.60893714 |
| QFVLDVAEHPVEVLTETV | ASSPIRDWGNYAEQF | 0.7119002 |
| GGAGGTGGAGGNGGTQAGDG | ASSPIRDWGNYAEQF | 0.66368586 |
| PLLATLRAIAVDVDAKPAQV | ASSPIRDWGNYAEQF | 0.57222205 |
| APKFRTGGYMIDFWGVGYQVAK | ASSPIRDWGNYAEQF | 0.67684186 |
| GAGGAGGAGGAAGGPGGTGGTG | ASSPIRDWGNYAEQF | 0.4078378 |
| IGALGLLATDVAPAGSVAFGVA | ASSPIRDWGNYAEQF | 0.68388104 |
| AGGAGGIGGNGGAGGAGGVLMG | ASSPIRDWGNYAEQF | 0.58070457 |
| ADELIGLL | ASSPIRDWGNYAEQF | 0.68775594 |
| ADRALKA | ASSPIRDWGNYAEQF | 0.6293118 |
| AGAVGQVGGAGGA | ASSPIRDWGNYAEQF | 0.6723769 |
| AGGGGGLGGAP | ASSPIRDWGNYAEQF | 0.69230425 |
| AGGLLYGNGGAG | ASSPIRDWGNYAEQF | 0.5282936 |
| AGLESIGL | ASSPIRDWGNYAEQF | 0.6793442 |
| AGLKTGII | ASSPIRDWGNYAEQF | 0.7114315 |
| ASLAALLL | ASSPIRDWGNYAEQF | 0.68190193 |
| ASVINISSI | ASSPIRDWGNYAEQF | 0.47881198 |
| ATPPGPA | ASSPIRDWGNYAEQF | 0.68180096 |
| ATVASALV | ASSPIRDWGNYAEQF | 0.57988435 |
| ATVASLIL | ASSPIRDWGNYAEQF | 0.68646777 |
| AVNGPVLIRH | ASSPIRDWGNYAEQF | 0.6494628 |
| DGAAGAVGI | ASSPIRDWGNYAEQF | 0.61842626 |
| DPLLHGI | ASSPIRDWGNYAEQF | 0.66371465 |
| ELGAALGAL | ASSPIRDWGNYAEQF | 0.3885042 |
| ELGRTQAV | ASSPIRDWGNYAEQF | 0.52496564 |
| FVAGGYLS | ASSPIRDWGNYAEQF | 0.5944823 |
| GASGGAGGAGGW | ASSPIRDWGNYAEQF | 0.68549275 |
| GGAGAGGDGGAGGI | ASSPIRDWGNYAEQF | 0.6962377 |
| GTGHNGSQPGMGGNG | ASSPIRDWGNYAEQF | 0.5049271 |
| GYAAASASAS | ASSPIRDWGNYAEQF | 0.68248105 |
| HQVDVAAVQQG | ASSPIRDWGNYAEQF | 0.61551017 |
| IGAAIAHT | ASSPIRDWGNYAEQF | 0.5099803 |
| IGMIEVAQL | ASSPIRDWGNYAEQF | 0.615399 |

| | | |
|---|---|---|
| IRLAPEHTSAMD | ASSPIRDWGNYAEQF | 0.5974417 |
| IYNVSVL | ASSPIRDWGNYAEQF | 0.40711227 |
| KQGFLGM | ASSPIRDWGNYAEQF | 0.6963022 |
| LESVITQL | ASSPIRDWGNYAEQF | 0.49387455 |
| LGLGALVVV | ASSPIRDWGNYAEQF | 0.67742753 |
| LIGNGGNGGSGGI | ASSPIRDWGNYAEQF | 0.6475839 |
| LNAINAPTL | ASSPIRDWGNYAEQF | 0.40894455 |
| LQLLNTGV | ASSPIRDWGNYAEQF | 0.66991353 |
| LQTSIEF | ASSPIRDWGNYAEQF | 0.63585985 |
| MSFAVSEL | ASSPIRDWGNYAEQF | 0.6828728 |
| NTGALGL | ASSPIRDWGNYAEQF | 0.6636262 |
| PPGAGGAGGAGTT | ASSPIRDWGNYAEQF | 0.60993135 |
| QLVGLGI | ASSPIRDWGNYAEQF | 0.72330755 |
| RFLEQVHQL | ASSPIRDWGNYAEQF | 0.6991718 |
| RGVGYVLR | ASSPIRDWGNYAEQF | 0.60929674 |
| SATGVFAGVI | ASSPIRDWGNYAEQF | 0.6507616 |
| SLPAITLP | ASSPIRDWGNYAEQF | 0.7104087 |
| STALAFTV | ASSPIRDWGNYAEQF | 0.70502764 |
| SVLTVGLGK | ASSPIRDWGNYAEQF | 0.6050644 |
| TALAGAGSGI | ASSPIRDWGNYAEQF | 0.34983486 |
| TGLPFTGI | ASSPIRDWGNYAEQF | 0.6507838 |
| VAEHVTV | ASSPIRDWGNYAEQF | 0.6453215 |
| VDIVLNSL | ASSPIRDWGNYAEQF | 0.57029355 |
| VLTGGGAVL | ASSPIRDWGNYAEQF | 0.38722146 |
| VNVGLAYGI | ASSPIRDWGNYAEQF | 0.70905364 |
| VSSAAVGAV | ASSPIRDWGNYAEQF | 0.7421514 |
| VTSVYEAGV | ASSPIRDWGNYAEQF | 0.6283375 |
| WVALDRGA | ASSPIRDWGNYAEQF | 0.7356722 |
| YLVGSVVI | ASSPIRDWGNYAEQF | 0.45318955 |
| GPPGTGKTLL | ASSPIRDWGNYAEQF | 0.6955172 |
| GVGMTKF | ASSPIRDWGNYAEQF | 0.71864957 |
| IGIGGSDL | ASSPIRDWGNYAEQF | 0.5484003 |
| IGTLLAEL | ASSPIRDWGNYAEQF | 0.691182 |
| IYTSGTT | ASSPIRDWGNYAEQF | 0.55226797 |
| LSFPAGV | ASSPIRDWGNYAEQF | 0.57462835 |
| LYTSGTT | ASSPIRDWGNYAEQF | 0.7124473 |
| VGAFPFGF | ASSPIRDWGNYAEQF | 0.6309167 |
| VGAGTVEF | ASSPIRDWGNYAEQF | 0.690427 |
| VGKSTLLL | ASSPIRDWGNYAEQF | 0.6136583 |
| AALAHAGGGIV | ASSPIRDWGNYAEQF | 0.502385 |
| AALVAVSLL | ASSPIRDWGNYAEQF | 0.66545045 |
| AAVSRAVAAL | ASSPIRDWGNYAEQF | 0.70018137 |
| AGGSSGIGGF | ASSPIRDWGNYAEQF | 0.6506496 |
| AGLTVANANG | ASSPIRDWGNYAEQF | 0.7271471 |
| AGLVGAVL | ASSPIRDWGNYAEQF | 0.5809521 |
| ALSQAFTV | ASSPIRDWGNYAEQF | 0.7376156 |
| ALTFSVSL | ASSPIRDWGNYAEQF | 0.7239311 |
| ALTGAALVAL | ASSPIRDWGNYAEQF | 0.66578645 |
| ASFSIGAGVL | ASSPIRDWGNYAEQF | 0.58609086 |
| DGSVYKI | ASSPIRDWGNYAEQF | 0.7183814 |
| EQLAEIV | ASSPIRDWGNYAEQF | 0.64221287 |

| | | |
|---|---|---|
| FAVSELTVM | ASSPIRDWGNYAEQF | 0.65844226 |
| GAAPLAAGL | ASSPIRDWGNYAEQF | 0.7173033 |
| GADGGDGGSGGAGG | ASSPIRDWGNYAEQF | 0.82864934 |
| GFGLDTGGAGG | ASSPIRDWGNYAEQF | 0.90806216 |
| GLASGGLL | ASSPIRDWGNYAEQF | 0.87761843 |
| GRIAGAAVLL | ASSPIRDWGNYAEQF | 0.87662053 |
| GSAGSLNTSF | ASSPIRDWGNYAEQF | 0.84045 |
| GTGQIGI | ASSPIRDWGNYAEQF | 0.6174975 |
| GTLIALIL | ASSPIRDWGNYAEQF | 0.85569316 |
| IAATMGFVVG | ASSPIRDWGNYAEQF | 0.8043577 |
| IFAQALL | ASSPIRDWGNYAEQF | 0.8449428 |
| IGAAVALGF | ASSPIRDWGNYAEQF | 0.8980378 |
| IGLTPLAL | ASSPIRDWGNYAEQF | 0.837736 |
| ILGSNTSTL | ASSPIRDWGNYAEQF | 0.90787923 |
| IVFSLTVL | ASSPIRDWGNYAEQF | 0.9060886 |
| LASKATGF | ASSPIRDWGNYAEQF | 0.7369033 |
| LASVALIL | ASSPIRDWGNYAEQF | 0.8474691 |
| LFAVGLTL | ASSPIRDWGNYAEQF | 0.820395 |
| LFNGGIGTYI | ASSPIRDWGNYAEQF | 0.8678559 |
| LTLGAGII | ASSPIRDWGNYAEQF | 0.56446767 |
| LVAAALGDI | ASSPIRDWGNYAEQF | 0.9009018 |
| NTAFVQLGI | ASSPIRDWGNYAEQF | 0.7965425 |
| QLFSGAEL | ASSPIRDWGNYAEQF | 0.20688894 |
| SAATFITL | ASSPIRDWGNYAEQF | 0.87588704 |
| SNYGLANLGV | ASSPIRDWGNYAEQF | 0.8548063 |
| SYGGAYAVM | ASSPIRDWGNYAEQF | 0.8600638 |
| TTGVGGLTLG | ASSPIRDWGNYAEQF | 0.89761996 |
| TTQTIYI | ASSPIRDWGNYAEQF | 0.45910287 |
| VAAGAGLGAVI | ASSPIRDWGNYAEQF | 0.824617 |
| VAAVVTLL | ASSPIRDWGNYAEQF | 0.89594746 |
| VGFLGGLI | ASSPIRDWGNYAEQF | 0.909286 |
| VGVIPILL | ASSPIRDWGNYAEQF | 0.7776952 |
| VIFASSLI | ASSPIRDWGNYAEQF | 0.8828939 |
| VKGGDGGAAAG | ASSPIRDWGNYAEQF | 0.90629953 |
| VTKTFTNI | ASSPIRDWGNYAEQF | 0.8885437 |
| VVAAAILL | ASSPIRDWGNYAEQF | 0.90291226 |
| YAALDVEL | ASSPIRDWGNYAEQF | 0.9107528 |
| LASVALLL | ASSPIRDWGNYAEQF | 0.80165386 |
| LDAVHRV | ASSPIRDWGNYAEQF | 0.9048549 |
| QGLSSSIF | ASSPIRDWGNYAEQF | 0.90613425 |
| RGTFANIRL | ASSPIRDWGNYAEQF | 0.8659164 |
| RQLWWGHRI | ASSPIRDWGNYAEQF | 0.8750882 |
| YGPPGTGKTLL | ASSPIRDWGNYAEQF | 0.82132226 |
| AALVGAPANVA | ASSPIRDWGNYAEQF | 0.83226186 |
| AATFRRTE | ASSPIRDWGNYAEQF | 0.4320649 |
| AGALGGGI | ASSPIRDWGNYAEQF | 0.8758794 |
| AGGAGGAGGIGAG | ASSPIRDWGNYAEQF | 0.9064914 |
| AGSGALGL | ASSPIRDWGNYAEQF | 0.8389853 |
| AQIADQI | ASSPIRDWGNYAEQF | 0.7526206 |
| ASAVAVTYL | ASSPIRDWGNYAEQF | 0.7896532 |
| ASTVLGGTGG | ASSPIRDWGNYAEQF | 0.8521069 |

| | | |
|---|---|---|
| AVGGEPAK | ASSPIRDWGNYAEQF | 0.8609544 |
| CEQGTFGGTC | ASSPIRDWGNYAEQF | 0.5338717 |
| DEGKAPAG | ASSPIRDWGNYAEQF | 0.77060413 |
| DTVVFPAL | ASSPIRDWGNYAEQF | 0.9065949 |
| FEGITMAG | ASSPIRDWGNYAEQF | 0.9073449 |
| FPLVVQD | ASSPIRDWGNYAEQF | 0.5923252 |
| GAFTTPQ | ASSPIRDWGNYAEQF | 0.90546083 |
| GFGGFGTAGGVGGA | ASSPIRDWGNYAEQF | 0.88674694 |
| GFLAQFL | ASSPIRDWGNYAEQF | 0.9057836 |
| GGAGGAGGNAGVA | ASSPIRDWGNYAEQF | 0.8731663 |
| GGVGTAGAGGAGGAGG | ASSPIRDWGNYAEQF | 0.8145189 |
| GNGGGGDAGNAGSGGN | ASSPIRDWGNYAEQF | 0.87629557 |
| GTGGAGGAGNDAGS | ASSPIRDWGNYAEQF | 0.823684 |
| GTTNTGWFNS | ASSPIRDWGNYAEQF | 0.7498098 |
| IFMGPLFTL | ASSPIRDWGNYAEQF | 0.90243363 |
| ILAHAIEAI | ASSPIRDWGNYAEQF | 0.85978806 |
| IRSAIREAL | ASSPIRDWGNYAEQF | 0.8047878 |
| ITFAGGG | ASSPIRDWGNYAEQF | 0.75150084 |
| ITGMSGHQEV | ASSPIRDWGNYAEQF | 0.9108211 |
| IVAGLACTL | ASSPIRDWGNYAEQF | 0.7182291 |
| IVGGGLAGA | ASSPIRDWGNYAEQF | 0.80725026 |
| IVTGVLNI | ASSPIRDWGNYAEQF | 0.741673 |
| LAQQGCI | ASSPIRDWGNYAEQF | 0.86016774 |
| LCLHVEN | ASSPIRDWGNYAEQF | 0.9040518 |
| LDAISDGS | ASSPIRDWGNYAEQF | 0.8945041 |
| LGGFAAML | ASSPIRDWGNYAEQF | 0.82393944 |
| LGGKNPMI | ASSPIRDWGNYAEQF | 0.88340324 |
| LGPKGVAAIL | ASSPIRDWGNYAEQF | 0.8961723 |
| LMVGVSI | ASSPIRDWGNYAEQF | 0.8803987 |
| LVGAAFVL | ASSPIRDWGNYAEQF | 0.9002812 |
| LVLAVAPL | ASSPIRDWGNYAEQF | 0.77556294 |
| MAAIAVIDAA | ASSPIRDWGNYAEQF | 0.9092945 |
| NNGINAELQ | ASSPIRDWGNYAEQF | 0.89294946 |
| RGEPLVLVHG | ASSPIRDWGNYAEQF | 0.904958 |
| RGGAGGAAGLI | ASSPIRDWGNYAEQF | 0.9007921 |
| RSYRMHPAV | ASSPIRDWGNYAEQF | 0.88514835 |
| SAAHLLAGQLA | ASSPIRDWGNYAEQF | 0.8787339 |
| SAAVGSY | ASSPIRDWGNYAEQF | 0.5493351 |
| SPHAVLHRI | ASSPIRDWGNYAEQF | 0.8009053 |
| SQIAKVL | ASSPIRDWGNYAEQF | 0.8802866 |
| TALGHLVP | ASSPIRDWGNYAEQF | 0.8714057 |
| TAPTTSGAA | ASSPIRDWGNYAEQF | 0.89260364 |
| TDGDDGNGGSPGT | ASSPIRDWGNYAEQF | 0.89251566 |
| TDGLGEN | ASSPIRDWGNYAEQF | 0.81625843 |
| TGGPPLPGQA | ASSPIRDWGNYAEQF | 0.90308 |
| TIPAITI | ASSPIRDWGNYAEQF | 0.8965808 |
| TIPGFTI | ASSPIRDWGNYAEQF | 0.691648 |
| TSIASGALA | ASSPIRDWGNYAEQF | 0.8443662 |
| TVTLPGAAI | ASSPIRDWGNYAEQF | 0.9129356 |
| VASGLSAALP | ASSPIRDWGNYAEQF | 0.8577105 |
| VGAGVIGI | ASSPIRDWGNYAEQF | 0.8506494 |

| | | |
|---|---|---|
| VGPKRFGV | ASSPIRDWGNYAEQF | 0.83735347 |
| VNVAFPD | ASSPIRDWGNYAEQF | 0.9055103 |
| VSMTNSGVS | ASSPIRDWGNYAEQF | 0.7977499 |
| VTDVLGDLSV | ASSPIRDWGNYAEQF | 0.914685 |
| VVVVGAGFAGL | ASSPIRDWGNYAEQF | 0.32408988 |
| VYTVGTVV | ASSPIRDWGNYAEQF | 0.8470573 |
| WTTFSGP | ASSPIRDWGNYAEQF | 0.8281859 |
| YDELVEAF | ASSPIRDWGNYAEQF | 0.70005786 |
| YGTGGAGGNGG | ASSPIRDWGNYAEQF | 0.90625936 |
| YTNVMAV | ASSPIRDWGNYAEQF | 0.84936416 |
| FLEQVHQL | ASSPIRDWGNYAEQF | 0.89641637 |
| GLSLLAVAV | ASSPIRDWGNYAEQF | 0.85230273 |
| LEQVHQL | ASSPIRDWGNYAEQF | 0.9034511 |
| LNAAGAI | ASSPIRDWGNYAEQF | 0.8616709 |
| PAVHASL | ASSPIRDWGNYAEQF | 0.8750976 |
| TELEDDA | ASSPIRDWGNYAEQF | 0.8281872 |
| TPPGPPN | ASSPIRDWGNYAEQF | 0.9018248 |
| TPPPPPPP | ASSPIRDWGNYAEQF | 0.9084258 |
| TQSPALI | ASRQY | 0.9012656 |
| EFAAAFPQ | ASRQY | 0.7259224 |
| LLSGNLDVL | ASRQY | 0.8104701 |
| GPANLGNYNI | ASRQY | 0.88741183 |
| TLGPTLP | ASRQY | 0.29648873 |
| VYAYSSCCK | ASRQY | 0.36014247 |
| APAPAPAPNI | ASRQY | 0.8290069 |
| PGGPVPIP | ASRQY | 0.7500887 |
| FLTVALEP | ASRQY | 0.9068357 |
| SQYFGANL | ASRQY | 0.8969644 |
| GGVKPAA | ASRQY | 0.85885465 |
| GGTGGAGGAGGW | ASRQY | 0.89108485 |
| GVGGVAGLGGAGG | ASRQY | 0.87045234 |
| VLLVLATTI | ASRQY | 0.10003533 |
| IGIFAAML | ASRQY | 0.8411362 |
| AVGDLID | ASRQY | 0.7394886 |
| NISPLPL | ASRQY | 0.9096227 |
| VTAVFVAI | ASRQY | 0.8102931 |
| SGDDNDAAL | ASRQY | 0.7554716 |
| AVAALAHS | ASRQY | 0.86444014 |
| GGVTVMATP | ASRQY | 0.84829843 |
| AGLAANVRD | ASRQY | 0.8428863 |
| TNRPDLIDPALLRPG | ASRQY | 0.7934183 |
| LANLGVTAL | ASRQY | 0.1801785 |
| TGERIEL | ASRQY | 0.9036231 |
| NGEELAA | ASRQY | 0.5167788 |
| VLSLVWG | ASRQY | 0.815281 |
| IGYALWL | ASRQY | 0.45529234 |
| LSLLLAPL | ASRQY | 0.24357742 |
| TLDEALK | ASRQY | 0.6570536 |
| TDGAGTAEGG | ASRQY | 0.90203226 |
| NAPGGAGGNGG | ASRQY | 0.8131693 |
| VSVFMGMGGNF | ASRQY | 0.80182225 |

| | | |
|---|---|---|
| LASYLAAL | ASRQY | 0.90857613 |
| LVGGTPAL | ASRQY | 0.8372911 |
| VRNADVL | ASRQY | 0.8274332 |
| TCSSPFLVSAM | ASRQY | 0.843276 |
| VHITDAE | ASRQY | 0.8006694 |
| EYAAPSF | ASRQY | 0.8285126 |
| VVDAFHGGL | ASRQY | 0.819711 |
| ASVVDPQ | ASRQY | 0.9135065 |
| SYNEFGGLS | ASRQY | 0.84707385 |
| DPDGSVY | ASRQY | 0.75558096 |
| MVSTIHATE | ASRQY | 0.8666929 |
| LAAGVADPV | ASRQY | 0.90790695 |
| TKEQIAA | ASRQY | 0.9156002 |
| SLASPAAL | ASRQY | 0.8085415 |
| IGLVFYGV | ASRQY | 0.83972156 |
| FLLDASNL | ASRQY | 0.8457285 |
| YQVGNFP | ASRQY | 0.8531154 |
| ESLLSQST | ASRQY | 0.8317362 |
| LFTVGFAAVG | ASRQY | 0.74024963 |
| ATPPTPT | ASRQY | 0.9023974 |
| TDLATLG | ASRQY | 0.9125871 |
| SAGGTGR | ASRQY | 0.86095625 |
| GFSNSGS | ASRQY | 0.6961078 |
| GGNGGSGGTSV | ASRQY | 0.8553047 |
| GTSVATGGAGN | ASRQY | 0.78475076 |
| NGGNGGSGGTS | ASRQY | 0.8394046 |
| AVVKFPH | ASRQY | 0.8818003 |
| TPPPPPP | ASRQY | 0.65930104 |
| AAPAAAAAPA | ASRQY | 0.8233521 |
| IALNEDL | ASRQY | 0.9154912 |
| IVNVSSM | ASRQY | 0.79867077 |
| TGIPPAPR | ASRQY | 0.86626995 |
| FDLGGGTF | ASRQY | 0.902948 |
| FDLGGGTFDVS | ASRQY | 0.88737595 |
| NEPTAAA | ASRQY | 0.90798503 |
| NEPTAAALA | ASRQY | 0.8905827 |
| GSIINISSI | ASRQY | 0.8746463 |
| VAPGNAGTA | ASRQY | 0.8722446 |
| TGGGASLE | ASRQY | 0.8400972 |
| VAVYNHYKRI | ASRQY | 0.867384 |
| TDPAPATT | ASRQY | 0.8529051 |
| TNRPDILDPALLRPG | ASRQY | 0.81506693 |
| LTTTTTS | ASRQY | 0.9025079 |
| PTGSGKTL | ASRQY | 0.87561095 |
| TGSGKTL | ASRQY | 0.89321774 |
| TIGHVDHGK | ASRQY | 0.7629537 |
| TIGHVDHGKTT | ASRQY | 0.7969225 |
| GHVDHGKTT | ASRQY | 0.8964107 |
| AQFEHTLL | ASRQY | 0.21110463 |
| IVNQDEL | ASRQY | 0.36744633 |
| TNGSQFF | ASRQY | 0.88090485 |

| | | |
|---|---|---|
| TGKGIVE | ASRQY | 0.90568554 |
| LISNASDA | ASRQY | 0.19782975 |
| AELRTLGELTKIAKAHGAQ | ASRQY | 0.81095576 |
| APNAVAAAEPSW | ASRQY | 0.69102776 |
| APQVAGAATAVL | ASRQY | 0.8926445 |
| ARRDPAAAAAILPTDA | ASRQY | 0.8696245 |
| DAAARDVLAEAGLAEYF | ASRQY | 0.6547874 |
| DDQIVLSAKANLAKVEQ | ASRQY | 0.89574105 |
| DELLPDTGEDSDEDGH | ASRQY | 0.6163955 |
| ELPEAPANRATVPQD | ASRQY | 0.6037355 |
| FGQSKIERVTGIG | ASRQY | 0.90313005 |
| GAGGTGSTAGGAGGAGGAGGLY | ASRQY | 0.9025217 |
| GHGGTGAAGGNGGTGGAGGSGI | ASRQY | 0.7856766 |
| GNGGNGGAGGEGGPGGAGGAGA | ASRQY | 0.90159523 |
| GSTAGNGGNGGNATNGTIAGQP | ASRQY | 0.80158967 |
| GTGGDGGNGGNGGTGG | ASRQY | 0.9121565 |
| GVRRPAGIPADAPV | ASRQY | 0.83861434 |
| GYRQVTDAHLLTLARR | ASRQY | 0.6886341 |
| IAKMGQDGHDRG | ASRQY | 0.8967744 |
| KAEPSIEIQYT | ASRQY | 0.87904704 |
| KPVRIRMAADEDTVAVT | ASRQY | 0.88581204 |
| MPPRIAGMRLLVIKPEP | ASRQY | 0.88703287 |
| MKFYAVALFRSRGVPPTR | ASRQY | 0.8345646 |
| NAGGFGILGAVAHSP | ASRQY | 0.83723 |
| NEGREITLAMAEM | ASRQY | 0.904795 |
| NELVEWSHPLGHRWRWE | ASRQY | 0.90059936 |
| NLGAGGTGGNGGNPG | ASRQY | 0.7753757 |
| NTEWPKWETVMAADPVD | ASRQY | 0.9016656 |
| PASAAPAETQP | ASRQY | 0.8558616 |
| QQSSDGKNGVRAAEITGE | ASRQY | 0.8140248 |
| RAPDAVTTSWWKE | ASRQY | 0.913693 |
| SGFGNVGAHTSGWFNQ | ASRQY | 0.88275063 |
| SGVAGAVGQAASVGGLKV | ASRQY | 0.89775574 |
| SNGGAGTGSAGGNGGT | ASRQY | 0.9051136 |
| TGGAGGEGGAAG | ASRQY | 0.8974873 |
| TGGDGGAAGAGGAPGHG | ASRQY | 0.69253856 |
| TMPPLLAAIANAGRHGV | ASRQY | 0.8668593 |
| TSSTAGNSGGA | ASRQY | 0.862553 |
| VPATVPVGILC | ASRQY | 0.8936459 |
| VQRPDSGGVVLTGRIS | ASRQY | 0.8672825 |
| WTPNPVPATKGVPAGLVQAT | ASRQY | 0.6304045 |
| YALKRALAKFSDRFDVVII | ASRQY | 0.8845526 |
| YLIGSVIV | ASRQY | 0.9061915 |
| YGQLVAVID | ASRQY | 0.88564295 |
| VTVGGMTLPA | ASRQY | 0.88357234 |
| VLALPAV | ASRQY | 0.9129816 |
| VIPPTLNYET | ASRQY | 0.83655083 |
| VGMGGMHPSSG | ASRQY | 0.85906994 |
| VGLAGGAATA | ASRQY | 0.903721 |
| VGGAGGAGGSGP | ASRQY | 0.89926386 |
| VFAAFMV | ASRQY | 0.86056274 |

| | | |
|---|---|---|
| TLYDKLR | ASRQY | 0.9112328 |
| TLAVFTVL | ASRQY | 0.7709156 |
| TGSVQPARV | ASRQY | 0.8952882 |
| TGGGAGSGGT | ASRQY | 0.73715776 |
| SHLETSL | ASRQY | 0.88380206 |
| SFTTGMT | ASRQY | 0.77069974 |
| QVEVAKRMGVRQPTVS | ASRQY | 0.89401233 |
| QGSGGPALAM | ASRQY | 0.8968598 |
| MLAPSNP | ASRQY | 0.8953464 |
| LVNNAAI | ASRQY | 0.3200622 |
| LTGEINTLA | ASRQY | 0.9011227 |
| LSIFGPV | ASRQY | 0.89635146 |
| LLALLANL | ASRQY | 0.91501385 |
| LISNASDALDKLR | ASRQY | 0.8694612 |
| LINEAVP | ASRQY | 0.889197 |
| LGYAFSGL | ASRQY | 0.8480495 |
| LDFVAVAL | ASRQY | 0.8678692 |
| LAQPGAE | ASRQY | 0.74952054 |
| ISNLGAVTGF | ASRQY | 0.8306948 |
| HQEDPGRI | ASRQY | 0.8149682 |
| GTPGEPTP | ASRQY | 0.79014164 |
| GFFNSGS | ASRQY | 0.8901048 |
| GAGGGGLTAGAN | ASRQY | 0.90138996 |
| FVDALVNM | ASRQY | 0.8992678 |
| FLTVAPEL | ASRQY | 0.9045013 |
| FETLTAL | ASRQY | 0.901639 |
| DVINPIEL | ASRQY | 0.90735257 |
| APVSGAPV | ASRQY | 0.8316016 |
| ALVGGLAW | ASRQY | 0.8382826 |
| ALAAAAATAI | ASRQY | 0.7610166 |
| AFGLGLLL | ASRQY | 0.8375947 |
| AFFAAASL | ASRQY | 0.8357005 |
| AAAALPAFSPP | ASRQY | 0.87704134 |
| VVLAPAL | ASRQY | 0.13976382 |
| TIGHVDHGKTTL | ASRQY | 0.82462466 |
| LISNASDALDK | ASRQY | 0.9101766 |
| LISNASDAL | ASRQY | 0.72422516 |
| ISNASDALDK | ASRQY | 0.88164616 |
| NQAETLVYQTE | ASRQY | 0.82231426 |
| TGGTGGTAGAG | ASRQY | 0.8222829 |
| TGSTAGGAGGA | ASRQY | 0.8957714 |
| DPLAVPVAAGPV | ASRQY | 0.80248797 |
| GGTGGTPGNGGN | ASRQY | 0.89095473 |
| PAGLAGVDPATL | ASRQY | 0.91230583 |
| GNNLVVEAPHAGASV | ASRQY | 0.8703782 |
| KVTRIMSASTLEQPA | ASRQY | 0.8728225 |
| LGFALPIALPRRTVV | ASRQY | 0.8662791 |
| TRLGNLTARKLVVLV | ASRQY | 0.6242797 |
| WSQQRRLIYFPSAGP | ASRQY | 0.8811908 |
| GPNGNPDPMAAAVDIR | ASRQY | 0.10924333 |
| TQRCVSVADLRAAASR | ASRQY | 0.8490861 |

| | | |
|---|---|---|
| QFVLDVAEHPVEVLTETV | ASRQY | 0.8224472 |
| GGAGGTGGAGGNGGTQAGDG | ASRQY | 0.8814011 |
| PLLATLRAIAVDVDAKPAQV | ASRQY | 0.86411333 |
| APKFRTGGYMIDFWGVGYQVAK | ASRQY | 0.8948717 |
| GAGGAGGAGGAAGGPGGTGGTG | ASRQY | 0.76676685 |
| IGALGLLATDVAPAGSVAFGVA | ASRQY | 0.8172102 |
| AGGAGGIGGNGGAGGAGGVLMG | ASRQY | 0.7571087 |
| ADELIGLL | ASRQY | 0.7699008 |
| ADRALKA | ASRQY | 0.88937294 |
| AGAVGQVGGAGGA | ASRQY | 0.8828426 |
| AGGGGGLGGAP | ASRQY | 0.80768573 |
| AGGLLYGNGGAG | ASRQY | 0.90775096 |
| AGLESIGL | ASRQY | 0.87976265 |
| AGLKTGII | ASRQY | 0.71030194 |
| ASLAALLL | ASRQY | 0.8244777 |
| ASVINISSI | ASRQY | 0.86514413 |
| ATPPGPA | ASRQY | 0.85526264 |
| ATVASALV | ASRQY | 0.121481314 |
| ATVASLIL | ASRQY | 0.87989366 |
| AVNGPVLIRH | ASRQY | 0.81741273 |
| DGAAGAVGI | ASRQY | 0.7456883 |
| DPLLHGI | ASRQY | 0.8219583 |
| ELGAALGAL | ASRQY | 0.8938004 |
| ELGRTQAV | ASRQY | 0.65744317 |
| FVAGGYLS | ASRQY | 0.8773111 |
| GASGGAGGAGGW | ASRQY | 0.8968392 |
| GGAGAGGDGGAGGI | ASRQY | 0.90110457 |
| GTGHNGSQPGMGGNG | ASRQY | 0.9096692 |
| GYAAASASAS | ASRQY | 0.89014167 |
| HQVDVAAVQQG | ASRQY | 0.86972994 |
| IGAAIAHT | ASRQY | 0.88647115 |
| IGMIEVAQL | ASRQY | 0.8713331 |
| IRLAPEHTSAMD | ASRQY | 0.5545317 |
| IYNVSVL | ASRQY | 0.5197371 |
| KQGFLGM | ASRQY | 0.8673313 |
| LESVITQL | ASRQY | 0.433565 |
| LGLGALVVV | ASRQY | 0.8719945 |
| LIGNGGNGGSGGI | ASRQY | 0.90810823 |
| LNAINAPTL | ASRQY | 0.8557016 |
| LQLLNTGV | ASRQY | 0.7606464 |
| LQTSIEF | ASRQY | 0.84577084 |
| MSFAVSEL | ASRQY | 0.8295393 |
| NTGALGL | ASRQY | 0.8947073 |
| PPGAGGAGGAGTT | ASRQY | 0.8969944 |
| QLVGLGI | ASRQY | 0.90772635 |
| RFLEQVHQL | ASRQY | 0.90096366 |
| RGVGYVLR | ASRQY | 0.83104265 |
| SATGVFAGVI | ASRQY | 0.89252067 |
| SLPAITLP | ASRQY | 0.835904 |
| STALAFTV | ASRQY | 0.82873404 |
| SVLTVGLGK | ASRQY | 0.87219566 |

| | | |
|---|---|---|
| TALAGAGSGI | ASRQY | 0.8840769 |
| TGLPFTGI | ASRQY | 0.87154496 |
| VAEHVTV | ASRQY | 0.8409655 |
| VDIVLNSL | ASRQY | 0.679149 |
| VLTGGGAVL | ASRQY | 0.89121664 |
| VNVGLAYGI | ASRQY | 0.85302055 |
| VSSAAVGAV | ASRQY | 0.611743 |
| VTSVYEAGV | ASRQY | 0.82275724 |
| WVALDRGA | ASRQY | 0.9142498 |
| YLVGSVVI | ASRQY | 0.90972674 |
| GPPGTGKTLL | ASRQY | 0.7801329 |
| GVGMTKF | ASRQY | 0.8283936 |
| IGIGGSDL | ASRQY | 0.83443606 |
| IGTLLAEL | ASRQY | 0.57672465 |
| IYTSGTT | ASRQY | 0.89119595 |
| LSFPAGV | ASRQY | 0.8432422 |
| LYTSGTT | ASRQY | 0.83576477 |
| VGAFPFGF | ASRQY | 0.8182974 |
| VGAGTVEF | ASRQY | 0.8620919 |
| VGKSTLLL | ASRQY | 0.8162137 |
| AALAHAGGGIV | ASRQY | 0.9010097 |
| AALVAVSLL | ASRQY | 0.18539345 |
| AAVSRAVAAL | ASRQY | 0.8902271 |
| AGGSSGIGGF | ASRQY | 0.7456383 |
| AGLTVANANG | ASRQY | 0.8430437 |
| AGLVGAVL | ASRQY | 0.6234858 |
| ALSQAFTV | ASRQY | 0.8538206 |
| ALTFSVSL | ASRQY | 0.8938787 |
| ALTGAALVAL | ASRQY | 0.9007422 |
| ASFSIGAGVL | ASRQY | 0.6663288 |
| DGSVYKI | ASRQY | 0.8999187 |
| EQLAEIV | ASRQY | 0.9155282 |
| FAVSELTVM | ASRQY | 0.80226266 |
| GAAPLAAGL | ASRQY | 0.5837782 |
| GADGGDGGSGGAGG | ASRQY | 0.8907771 |
| GFGLDTGGAGG | ASRQY | 0.8130354 |
| GLASGGLL | ASRQY | 0.89227104 |
| GRIAGAAVLL | ASRQY | 0.87389845 |
| GSAGSLNTSF | ASRQY | 0.70988166 |
| GTGQIGI | ASRQY | 0.9015329 |
| GTLIALIL | ASRQY | 0.9016249 |
| IAATMGFVVG | ASRQY | 0.80347514 |
| IFAQALL | ASRQY | 0.8275633 |
| IGAAVALGF | ASRQY | 0.8119941 |
| IGLTPLAL | ASRQY | 0.42781854 |
| ILGSNTSTL | ASRQY | 0.900595 |
| IVFSLTVL | ASRQY | 0.77052796 |
| LASKATGF | ASRQY | 0.83464813 |
| LASVALIL | ASRQY | 0.873683 |
| LFAVGLTL | ASRQY | 0.8415355 |
| LFNGGIGTYI | ASRQY | 0.845387 |

| | | |
|---|---|---|
| LTLGAGII | ASRQY | 0.8494724 |
| LVAAALGDI | ASRQY | 0.8675231 |
| NTAFVQLGI | ASRQY | 0.87761045 |
| QLFSGAEL | ASRQY | 0.8560799 |
| SAATFITL | ASRQY | 0.8519203 |
| SNYGLANLGV | ASRQY | 0.9069727 |
| SYGGAYAVM | ASRQY | 0.74460727 |
| TTGVGGLTLG | ASRQY | 0.80067766 |
| TTQTIYI | ASRQY | 0.88734305 |
| VAAGAGLGAVI | ASRQY | 0.67160904 |
| VAAVVTLL | ASRQY | 0.88961947 |
| VGFLGGLI | ASRQY | 0.86778015 |
| VGVIPILL | ASRQY | 0.8775054 |
| VIFASSLI | ASRQY | 0.8391977 |
| VKGGDGGAAAG | ASRQY | 0.91598254 |
| VTKTFTNI | ASRQY | 0.9008088 |
| VVAAAILL | ASRQY | 0.86433387 |
| YAALDVEL | ASRQY | 0.8351437 |
| LASVALLL | ASRQY | 0.7955621 |
| LDAVHRV | ASRQY | 0.90241635 |
| QGLSSSIF | ASRQY | 0.8897752 |
| RGTFANIRL | ASRQY | 0.742226 |
| RQLWWGHRI | ASRQY | 0.8426423 |
| YGPPGTGKTLL | ASRQY | 0.9109057 |
| AALVGAPANVA | ASRQY | 0.90350854 |
| AATFRRTE | ASRQY | 0.60841423 |
| AGALGGGI | ASRQY | 0.839027 |
| AGGAGGAGGIGAG | ASRQY | 0.5931169 |
| AGSGALGL | ASRQY | 0.85688347 |
| AQIADQI | ASRQY | 0.7726408 |
| ASAVAVTYL | ASRQY | 0.88677263 |
| ASTVLGGTGG | ASRQY | 0.83374876 |
| AVGGEPAK | ASRQY | 0.8689531 |
| CEQGTFGGTC | ASRQY | 0.8773913 |
| DEGKAPAG | ASRQY | 0.9135305 |
| DTVVFPAL | ASRQY | 0.82853615 |
| FEGITMAG | ASRQY | 0.85719776 |
| FPLVVQD | ASRQY | 0.70579803 |
| GAFTTPQ | ASRQY | 0.5764981 |
| GFGGFGTAGGVGGA | ASRQY | 0.8287921 |
| GFLAQFL | ASRQY | 0.85005784 |
| GGAGGAGGNAGVA | ASRQY | 0.30810916 |
| GGVGTAGAGGAGGAGG | ASRQY | 0.9055239 |
| GNGGGGDAGNAGSGGN | ASRQY | 0.88176787 |
| GTGGAGGAGNDAGS | ASRQY | 0.90237534 |
| GTTNTGWFNS | ASRQY | 0.8476821 |
| IFMGPLFTL | ASRQY | 0.849916 |
| ILAHAIEAI | ASRQY | 0.82424784 |
| IRSAIREAL | ASRQY | 0.889385 |
| ITFAGGG | ASRQY | 0.7976931 |
| ITGMSGHQEV | ASRQY | 0.68292695 |

| | | |
|---|---|---|
| IVAGLACTL | ASRQY | 0.83120096 |
| IVGGGLAGA | ASRQY | 0.7297845 |
| IVTGVLNI | ASRQY | 0.9041359 |
| LAQQGCI | ASRQY | 0.90274745 |
| LCLHVEN | ASRQY | 0.88276744 |
| LDAISDGS | ASRQY | 0.9037035 |
| LGGFAAML | ASRQY | 0.43729594 |
| LGGKNPMI | ASRQY | 0.8364291 |
| LGPKGVAAIL | ASRQY | 0.91145766 |
| LMVGVSI | ASRQY | 0.8905537 |
| LVGAAFVL | ASRQY | 0.8419543 |
| LVLAVAPL | ASRQY | 0.90806115 |
| MAAIAVIDAA | ASRQY | 0.80599976 |
| NNGINAELQ | ASRQY | 0.855697 |
| RGEPLVLVHG | ASRQY | 0.8632241 |
| RGGAGGAAGLI | ASRQY | 0.89403003 |
| RSYRMHPAV | ASRQY | 0.87938654 |
| SAAHLLAGQLA | ASRQY | 0.8011742 |
| SAAVGSY | ASRQY | 0.88396555 |
| SPHAVLHRI | ASRQY | 0.9145448 |
| SQIAKVL | ASRQY | 0.84439284 |
| TALGHLVP | ASRQY | 0.8979762 |
| TAPTTSGAA | ASRQY | 0.8873049 |
| TDGDDGNGGSPGT | ASRQY | 0.8606274 |
| TDGLGEN | ASRQY | 0.7936393 |
| TGGPPLPGQA | ASRQY | 0.8255285 |
| TIPAITI | ASRQY | 0.8268455 |
| TIPGFTI | ASRQY | 0.7557428 |
| TSIASGALA | ASRQY | 0.85541856 |
| TVTLPGAAI | ASRQY | 0.84906805 |
| VASGLSAALP | ASRQY | 0.9197029 |
| VGAGVIGI | ASRQY | 0.8856211 |
| VGPKRFGV | ASRQY | 0.8889955 |
| VNVAFPD | ASRQY | 0.7872849 |
| VSMTNSGVS | ASRQY | 0.8888098 |
| VTDVLGDLSV | ASRQY | 0.88475287 |
| VVVVGAGFAGL | ASRQY | 0.70088303 |
| VYTVGTVV | ASRQY | 0.91012824 |
| WTTFSGP | ASRQY | 0.8082037 |
| YDELVEAF | ASRQY | 0.81101155 |
| YGTGGAGGNGG | ASRQY | 0.8985969 |
| YTNVMAV | ASRQY | 0.90104103 |
| FLEQVHQL | ASRQY | 0.83940274 |
| GLSLLAVAV | ASRQY | 0.8023859 |
| LEQVHQL | ASRQY | 0.7972826 |
| LNAAGAI | ASRQY | 0.90289855 |
| PAVHASL | ASRQY | 0.9065362 |
| TELEDDA | ASRQY | 0.8076886 |
| TPPGPPN | ASRQY | 0.9086628 |
| TPPPPPPP | ASRQY | 0.8856716 |
| TQSPALI | AWSPQDWGEGAETLY | 0.68878233 |

| | | |
|---|---|---|
| EFAAAFPQ | AWSPQDWGEGAETLY | 0.8703251 |
| LLSGNLDVL | AWSPQDWGEGAETLY | 0.8971537 |
| GPANLGNYNI | AWSPQDWGEGAETLY | 0.8950883 |
| TLGPTLP | AWSPQDWGEGAETLY | 0.91440535 |
| VYAYSSCCK | AWSPQDWGEGAETLY | 0.8840933 |
| APAPAPAPNI | AWSPQDWGEGAETLY | 0.8262528 |
| PGGPVPIP | AWSPQDWGEGAETLY | 0.9084769 |
| FLTVALEP | AWSPQDWGEGAETLY | 0.9158884 |
| SQYFGANL | AWSPQDWGEGAETLY | 0.90298605 |
| GGVKPAA | AWSPQDWGEGAETLY | 0.85325325 |
| GGTGGAGGAGGW | AWSPQDWGEGAETLY | 0.834844 |
| GVGGVAGLGGAGG | AWSPQDWGEGAETLY | 0.8712943 |
| VLLVLATTI | AWSPQDWGEGAETLY | 0.8979134 |
| IGIFAAML | AWSPQDWGEGAETLY | 0.7867806 |
| AVGDLID | AWSPQDWGEGAETLY | 0.8531449 |
| NISPLPL | AWSPQDWGEGAETLY | 0.87746733 |
| VTAVFVAI | AWSPQDWGEGAETLY | 0.68986773 |
| SGDDNDAAL | AWSPQDWGEGAETLY | 0.88658714 |
| AVAALAHS | AWSPQDWGEGAETLY | 0.9064474 |
| GGVTVMATP | AWSPQDWGEGAETLY | 0.8609106 |
| AGLAANVRD | AWSPQDWGEGAETLY | 0.8262806 |
| TNRPDLIDPALLRPG | AWSPQDWGEGAETLY | 0.90192306 |
| LANLGVTAL | AWSPQDWGEGAETLY | 0.91980875 |
| TGERIEL | AWSPQDWGEGAETLY | 0.9088137 |
| NGEELAA | AWSPQDWGEGAETLY | 0.68959033 |
| VLSLVWG | AWSPQDWGEGAETLY | 0.51966584 |
| IGYALWL | AWSPQDWGEGAETLY | 0.78730434 |
| LSLLLAPL | AWSPQDWGEGAETLY | 0.8744065 |
| TLDEALK | AWSPQDWGEGAETLY | 0.84671926 |
| TDGAGTAEGG | AWSPQDWGEGAETLY | 0.8428525 |
| NAPGGAGGNGG | AWSPQDWGEGAETLY | 0.8560694 |
| VSVFMGMGGNF | AWSPQDWGEGAETLY | 0.90540755 |
| LASYLAAL | AWSPQDWGEGAETLY | 0.8192737 |
| LVGGTPAL | AWSPQDWGEGAETLY | 0.773909 |
| VRNADVL | AWSPQDWGEGAETLY | 0.8636466 |
| TCSSPFLVSAM | AWSPQDWGEGAETLY | 0.8949096 |
| VHITDAE | AWSPQDWGEGAETLY | 0.78177595 |
| EYAAPSF | AWSPQDWGEGAETLY | 0.85208786 |
| VVDAFHGGL | AWSPQDWGEGAETLY | 0.7748341 |
| ASVVDPQ | AWSPQDWGEGAETLY | 0.6352964 |
| SYNEFGGLS | AWSPQDWGEGAETLY | 0.8708439 |
| DPDGSVY | AWSPQDWGEGAETLY | 0.7945127 |
| MVSTIHATE | AWSPQDWGEGAETLY | 0.9077115 |
| LAAGVADPV | AWSPQDWGEGAETLY | 0.796579 |
| TKEQIAA | AWSPQDWGEGAETLY | 0.89936674 |
| SLASPAAL | AWSPQDWGEGAETLY | 0.88451064 |
| IGLVFYGV | AWSPQDWGEGAETLY | 0.89225805 |
| FLLDASNL | AWSPQDWGEGAETLY | 0.8290578 |
| YQVGNFP | AWSPQDWGEGAETLY | 0.79474545 |
| ESLLSQST | AWSPQDWGEGAETLY | 0.7604834 |
| LFTVGFAAVG | AWSPQDWGEGAETLY | 0.5738129 |

| | | |
|---|---|---|
| ATPPTPT | AWSPQDWGEGAETLY | 0.90101457 |
| TDLATLG | AWSPQDWGEGAETLY | 0.17538363 |
| SAGGTGR | AWSPQDWGEGAETLY | 0.90688705 |
| GFSNSGS | AWSPQDWGEGAETLY | 0.90048826 |
| GGNGGSGGTSV | AWSPQDWGEGAETLY | 0.8042039 |
| GTSVATGGAGN | AWSPQDWGEGAETLY | 0.89199984 |
| NGGNGGSGGTS | AWSPQDWGEGAETLY | 0.77356386 |
| AVVKFPH | AWSPQDWGEGAETLY | 0.77182364 |
| TPPPPPP | AWSPQDWGEGAETLY | 0.89558256 |
| AAPAAAAAPA | AWSPQDWGEGAETLY | 0.8916854 |
| IALNEDL | AWSPQDWGEGAETLY | 0.8037547 |
| IVNVSSM | AWSPQDWGEGAETLY | 0.88419867 |
| TGIPPAPR | AWSPQDWGEGAETLY | 0.87309384 |
| FDLGGGTF | AWSPQDWGEGAETLY | 0.8134365 |
| FDLGGGTFDVS | AWSPQDWGEGAETLY | 0.8659813 |
| NEPTAAA | AWSPQDWGEGAETLY | 0.90226305 |
| NEPTAAALA | AWSPQDWGEGAETLY | 0.88453865 |
| GSIINISSI | AWSPQDWGEGAETLY | 0.81832266 |
| VAPGNAGTA | AWSPQDWGEGAETLY | 0.89031065 |
| TGGGASLE | AWSPQDWGEGAETLY | 0.73203725 |
| VAVYNHYKRI | AWSPQDWGEGAETLY | 0.8816848 |
| TDPAPATT | AWSPQDWGEGAETLY | 0.86950403 |
| TNRPDILDPALLRPG | AWSPQDWGEGAETLY | 0.85437644 |
| LTTTTTS | AWSPQDWGEGAETLY | 0.89031327 |
| PTGSGKTL | AWSPQDWGEGAETLY | 0.7038311 |
| TGSGKTL | AWSPQDWGEGAETLY | 0.80019057 |
| TIGHVDHGK | AWSPQDWGEGAETLY | 0.8445382 |
| TIGHVDHGKTT | AWSPQDWGEGAETLY | 0.88547444 |
| GHVDHGKTT | AWSPQDWGEGAETLY | 0.89971757 |
| AQFEHTLL | AWSPQDWGEGAETLY | 0.7848764 |
| IVNQDEL | AWSPQDWGEGAETLY | 0.874174 |
| TNGSQFF | AWSPQDWGEGAETLY | 0.8404342 |
| TGKGIVE | AWSPQDWGEGAETLY | 0.78781176 |
| LISNASDA | AWSPQDWGEGAETLY | 0.86910427 |
| AELRTLGELTKIAKAHGAQ | AWSPQDWGEGAETLY | 0.85755575 |
| APNAVAAAEPSW | AWSPQDWGEGAETLY | 0.8206558 |
| APQVAGAATAVL | AWSPQDWGEGAETLY | 0.8880887 |
| ARRDPAAAAAILPTDA | AWSPQDWGEGAETLY | 0.8135173 |
| DAAARDVLAEAGLAEYF | AWSPQDWGEGAETLY | 0.8951661 |
| DDQIVLSAKANLAKVEQ | AWSPQDWGEGAETLY | 0.9082502 |
| DELLPDTGEDSDEDGH | AWSPQDWGEGAETLY | 0.8487468 |
| ELPEAPANRATVPQD | AWSPQDWGEGAETLY | 0.8993541 |
| FGQSKIERVTGIG | AWSPQDWGEGAETLY | 0.88473135 |
| GAGGTGSTAGGAGGAGGAGGLY | AWSPQDWGEGAETLY | 0.879363 |
| GHGGTGAAGGNGGTGGAGGSGI | AWSPQDWGEGAETLY | 0.85534644 |
| GNGGNGGAGGEGGPGGAGGAGA | AWSPQDWGEGAETLY | 0.81330454 |
| GSTAGNGGNGGNATNGTIAGQP | AWSPQDWGEGAETLY | 0.9048227 |
| GTGGDGGNGGNGGTGG | AWSPQDWGEGAETLY | 0.8956953 |
| GVRRPAGIPADAPV | AWSPQDWGEGAETLY | 0.8960538 |
| GYRQVTDAHLLTLARR | AWSPQDWGEGAETLY | 0.8830064 |
| IAKMGQDGHDRG | AWSPQDWGEGAETLY | 0.87200844 |

| | | |
|---|---|---|
| KAEPSIEIQYT | AWSPQDWGEGAETLY | 0.89261895 |
| KPVRIRMAADEDTVAVT | AWSPQDWGEGAETLY | 0.80837977 |
| MPPRIAGMRLLVIKPEP | AWSPQDWGEGAETLY | 0.7066752 |
| MKFYAVALFRSRGVPPTR | AWSPQDWGEGAETLY | 0.8479501 |
| NAGGFGILGAVAHSP | AWSPQDWGEGAETLY | 0.8450912 |
| NEGREITLAMAEM | AWSPQDWGEGAETLY | 0.8558297 |
| NELVEWSHPLGHRWRWE | AWSPQDWGEGAETLY | 0.6736471 |
| NLGAGGTGGNGGNPG | AWSPQDWGEGAETLY | 0.90111 |
| NTEWPKWETVMAADPVD | AWSPQDWGEGAETLY | 0.9189113 |
| PASAAPAETQP | AWSPQDWGEGAETLY | 0.83981323 |
| QQSSDGKNGVRAAEITGE | AWSPQDWGEGAETLY | 0.91332483 |
| RAPDAVTTSWWKE | AWSPQDWGEGAETLY | 0.72512627 |
| SGFGNVGAHTSGWFNQ | AWSPQDWGEGAETLY | 0.8957181 |
| SGVAGAVGQAASVGGLKV | AWSPQDWGEGAETLY | 0.9039731 |
| SNGGAGTGSAGGNGGT | AWSPQDWGEGAETLY | 0.77565 |
| TGGAGGEGGAAG | AWSPQDWGEGAETLY | 0.89383996 |
| TGGDGGAAGAGGAPGHG | AWSPQDWGEGAETLY | 0.78987455 |
| TMPPLLAAIANAGRHGV | AWSPQDWGEGAETLY | 0.8538611 |
| TSSTAGNSGGA | AWSPQDWGEGAETLY | 0.9072373 |
| VPATVPVGILC | AWSPQDWGEGAETLY | 0.84797126 |
| VQRPDSGGVVLTGRIS | AWSPQDWGEGAETLY | 0.8746706 |
| WTPNPVPATKGVPAGLVQAT | AWSPQDWGEGAETLY | 0.86828446 |
| YALKRALAKFSDRFDVVII | AWSPQDWGEGAETLY | 0.8193495 |
| YLIGSVIV | AWSPQDWGEGAETLY | 0.8847573 |
| YGQLVAVID | AWSPQDWGEGAETLY | 0.9073497 |
| VTVGGMTLPA | AWSPQDWGEGAETLY | 0.76585084 |
| VLALPAV | AWSPQDWGEGAETLY | 0.9139291 |
| VIPPTLNYET | AWSPQDWGEGAETLY | 0.8129206 |
| VGMGGMHPSSG | AWSPQDWGEGAETLY | 0.9143115 |
| VGLAGGAATA | AWSPQDWGEGAETLY | 0.89073443 |
| VGGAGGAGGSGP | AWSPQDWGEGAETLY | 0.8627092 |
| VFAAFMV | AWSPQDWGEGAETLY | 0.84699553 |
| TLYDKLR | AWSPQDWGEGAETLY | 0.9046676 |
| TLAVFTVL | AWSPQDWGEGAETLY | 0.852893 |
| TGSVQPARV | AWSPQDWGEGAETLY | 0.8942045 |
| TGGGAGSGGT | AWSPQDWGEGAETLY | 0.9117574 |
| SHLETSL | AWSPQDWGEGAETLY | 0.636014 |
| SFTTGMT | AWSPQDWGEGAETLY | 0.6813754 |
| QVEVAKRMGVRQPTVS | AWSPQDWGEGAETLY | 0.6639365 |
| QGSGGPALAM | AWSPQDWGEGAETLY | 0.66320944 |
| MLAPSNP | AWSPQDWGEGAETLY | 0.64197993 |
| LVNNAAI | AWSPQDWGEGAETLY | 0.57822156 |
| LTGEINTLA | AWSPQDWGEGAETLY | 0.6637321 |
| LSIFGPV | AWSPQDWGEGAETLY | 0.6186668 |
| LLALLANL | AWSPQDWGEGAETLY | 0.6813611 |
| LISNASDALDKLR | AWSPQDWGEGAETLY | 0.6831691 |
| LINEAVP | AWSPQDWGEGAETLY | 0.63493305 |
| LGYAFSGL | AWSPQDWGEGAETLY | 0.6837417 |
| LDFVAVAL | AWSPQDWGEGAETLY | 0.68037754 |
| LAQPGAE | AWSPQDWGEGAETLY | 0.59677434 |
| ISNLGAVTGF | AWSPQDWGEGAETLY | 0.6518369 |

| | | |
|---|---|---|
| HQEDPGRI | AWSPQDWGEGAETLY | 0.648169 |
| GTPGEPTP | AWSPQDWGEGAETLY | 0.66091263 |
| GFFNSGS | AWSPQDWGEGAETLY | 0.5365858 |
| GAGGGGLTAGAN | AWSPQDWGEGAETLY | 0.6799127 |
| FVDALVNM | AWSPQDWGEGAETLY | 0.6367884 |
| FLTVAPEL | AWSPQDWGEGAETLY | 0.4444211 |
| FETLTAL | AWSPQDWGEGAETLY | 0.67026794 |
| DVINPIEL | AWSPQDWGEGAETLY | 0.66635025 |
| APVSGAPV | AWSPQDWGEGAETLY | 0.6605488 |
| ALVGGLAW | AWSPQDWGEGAETLY | 0.67939436 |
| ALAAAAATAI | AWSPQDWGEGAETLY | 0.62048525 |
| AFGLGLLL | AWSPQDWGEGAETLY | 0.6214185 |
| AFFAAASL | AWSPQDWGEGAETLY | 0.67473173 |
| AAAALPAFSPP | AWSPQDWGEGAETLY | 0.68256533 |
| VVLAPAL | AWSPQDWGEGAETLY | 0.61369586 |
| TIGHVDHGKTTL | AWSPQDWGEGAETLY | 0.66772676 |
| LISNASDALDK | AWSPQDWGEGAETLY | 0.6812279 |
| LISNASDAL | AWSPQDWGEGAETLY | 0.67455035 |
| ISNASDALDK | AWSPQDWGEGAETLY | 0.6848306 |
| NQAETLVYQTE | AWSPQDWGEGAETLY | 0.68718445 |
| TGGTGGTAGAG | AWSPQDWGEGAETLY | 0.63808376 |
| TGSTAGGAGGA | AWSPQDWGEGAETLY | 0.6823288 |
| DPLAVPVAAGPV | AWSPQDWGEGAETLY | 0.68450326 |
| GGTGGTPGNGGN | AWSPQDWGEGAETLY | 0.6550097 |
| PAGLAGVDPATL | AWSPQDWGEGAETLY | 0.6611788 |
| GNNLVVEAPHAGASV | AWSPQDWGEGAETLY | 0.61516905 |
| KVTRIMSASTLEQPA | AWSPQDWGEGAETLY | 0.6604315 |
| LGFALPIALPRRTVV | AWSPQDWGEGAETLY | 0.5814391 |
| TRLGNLTARKLVVLV | AWSPQDWGEGAETLY | 0.66944396 |
| WSQQRRLIYFPSAGP | AWSPQDWGEGAETLY | 0.685424 |
| GPNGNPDPMAAAVDIR | AWSPQDWGEGAETLY | 0.65511 |
| TQRCVSVADLRAAASR | AWSPQDWGEGAETLY | 0.6156229 |
| QFVLDVAEHPVEVLTETV | AWSPQDWGEGAETLY | 0.61610365 |
| GGAGGTGGAGGNGGTQAGDG | AWSPQDWGEGAETLY | 0.6549443 |
| PLLATLRAIAVDVDAKPAQV | AWSPQDWGEGAETLY | 0.64579576 |
| APKFRTGGYMIDFWGVGYQVAK | AWSPQDWGEGAETLY | 0.58117586 |
| GAGGAGGAGGAAGGPGGTGGTG | AWSPQDWGEGAETLY | 0.6461127 |
| IGALGLLATDVAPAGSVAFGVA | AWSPQDWGEGAETLY | 0.68118346 |
| AGGAGGIGGNGGAGGAGGVLMG | AWSPQDWGEGAETLY | 0.68238825 |
| ADELIGLL | AWSPQDWGEGAETLY | 0.5869685 |
| ADRALKA | AWSPQDWGEGAETLY | 0.68365955 |
| AGAVGQVGGAGGA | AWSPQDWGEGAETLY | 0.67959535 |
| AGGGGGLGGAP | AWSPQDWGEGAETLY | 0.6833502 |
| AGGLLYGNGGAG | AWSPQDWGEGAETLY | 0.666803 |
| AGLESIGL | AWSPQDWGEGAETLY | 0.61276656 |
| AGLKTGII | AWSPQDWGEGAETLY | 0.67998123 |
| ASLAALLL | AWSPQDWGEGAETLY | 0.63915807 |
| ASVINISSI | AWSPQDWGEGAETLY | 0.65641326 |
| ATPPGPA | AWSPQDWGEGAETLY | 0.67997265 |
| ATVASALV | AWSPQDWGEGAETLY | 0.6539745 |
| ATVASLIL | AWSPQDWGEGAETLY | 0.6339067 |

| | | |
|---|---|---|
| AVNGPVLIRH | AWSPQDWGEGAETLY | 0.6496028 |
| DGAAGAVGI | AWSPQDWGEGAETLY | 0.6862565 |
| DPLLHGI | AWSPQDWGEGAETLY | 0.6086377 |
| ELGAALGAL | AWSPQDWGEGAETLY | 0.62129605 |
| ELGRTQAV | AWSPQDWGEGAETLY | 0.64414394 |
| FVAGGYLS | AWSPQDWGEGAETLY | 0.6586108 |
| GASGGAGGAGGW | AWSPQDWGEGAETLY | 0.68114734 |
| GGAGAGGDGGAGGI | AWSPQDWGEGAETLY | 0.6810011 |
| GTGHNGSQPGMGGNG | AWSPQDWGEGAETLY | 0.64210284 |
| GYAAASASAS | AWSPQDWGEGAETLY | 0.6690664 |
| HQVDVAAVQQG | AWSPQDWGEGAETLY | 0.6764115 |
| IGAAIAHT | AWSPQDWGEGAETLY | 0.668584 |
| IGMIEVAQL | AWSPQDWGEGAETLY | 0.67330456 |
| IRLAPEHTSAMD | AWSPQDWGEGAETLY | 0.6259011 |
| IYNVSVL | AWSPQDWGEGAETLY | 0.6838949 |
| KQGFLGM | AWSPQDWGEGAETLY | 0.67398804 |
| LESVITQL | AWSPQDWGEGAETLY | 0.6831444 |
| LGLGALVVV | AWSPQDWGEGAETLY | 0.6836446 |
| LIGNGGNGGSGGI | AWSPQDWGEGAETLY | 0.662909 |
| LNAINAPTL | AWSPQDWGEGAETLY | 0.66799927 |
| LQLLNTGV | AWSPQDWGEGAETLY | 0.529453 |
| LQTSIEF | AWSPQDWGEGAETLY | 0.64936197 |
| MSFAVSEL | AWSPQDWGEGAETLY | 0.6666872 |
| NTGALGL | AWSPQDWGEGAETLY | 0.66670513 |
| PPGAGGAGGAGTT | AWSPQDWGEGAETLY | 0.6801194 |
| QLVGLGI | AWSPQDWGEGAETLY | 0.6782912 |
| RFLEQVHQL | AWSPQDWGEGAETLY | 0.64367247 |
| RGVGYVLR | AWSPQDWGEGAETLY | 0.6835233 |
| SATGVFAGVI | AWSPQDWGEGAETLY | 0.681463 |
| SLPAITLP | AWSPQDWGEGAETLY | 0.5811479 |
| STALAFTV | AWSPQDWGEGAETLY | 0.643308 |
| SVLTVGLGK | AWSPQDWGEGAETLY | 0.6868756 |
| TALAGAGSGI | AWSPQDWGEGAETLY | 0.66687226 |
| TGLPFTGI | AWSPQDWGEGAETLY | 0.6497146 |
| VAEHVTV | AWSPQDWGEGAETLY | 0.6533804 |
| VDIVLNSL | AWSPQDWGEGAETLY | 0.68939126 |
| VLTGGGAVL | AWSPQDWGEGAETLY | 0.66757107 |
| VNVGLAYGI | AWSPQDWGEGAETLY | 0.685372 |
| VSSAAVGAV | AWSPQDWGEGAETLY | 0.50019175 |
| VTSVYEAGV | AWSPQDWGEGAETLY | 0.6573232 |
| WVALDRGA | AWSPQDWGEGAETLY | 0.60924375 |
| YLVGSVVI | AWSPQDWGEGAETLY | 0.49157086 |
| GPPGTGKTLL | AWSPQDWGEGAETLY | 0.6807644 |
| GVGMTKF | AWSPQDWGEGAETLY | 0.65845484 |
| IGIGGSDL | AWSPQDWGEGAETLY | 0.67517114 |
| IGTLLAEL | AWSPQDWGEGAETLY | 0.6638526 |
| IYTSGTT | AWSPQDWGEGAETLY | 0.68330705 |
| LSFPAGV | AWSPQDWGEGAETLY | 0.653248 |
| LYTSGTT | AWSPQDWGEGAETLY | 0.6696912 |
| VGAFPFGF | AWSPQDWGEGAETLY | 0.6484649 |
| VGAGTVEF | AWSPQDWGEGAETLY | 0.67729795 |

| | | |
|---|---|---|
| VGKSTLLL | AWSPQDWGEGAETLY | 0.6860794 |
| AALAHAGGGIV | AWSPQDWGEGAETLY | 0.68256783 |
| AALVAVSLL | AWSPQDWGEGAETLY | 0.560903 |
| AAVSRAVAAL | AWSPQDWGEGAETLY | 0.6662211 |
| AGGSSGIGGF | AWSPQDWGEGAETLY | 0.6647124 |
| AGLTVANANG | AWSPQDWGEGAETLY | 0.4455514 |
| AGLVGAVL | AWSPQDWGEGAETLY | 0.54003465 |
| ALSQAFTV | AWSPQDWGEGAETLY | 0.64461285 |
| ALTFSVSL | AWSPQDWGEGAETLY | 0.6345307 |
| ALTGAALVAL | AWSPQDWGEGAETLY | 0.67815876 |
| ASFSIGAGVL | AWSPQDWGEGAETLY | 0.67065334 |
| DGSVYKI | AWSPQDWGEGAETLY | 0.6546775 |
| EQLAEIV | AWSPQDWGEGAETLY | 0.6718267 |
| FAVSELTVM | AWSPQDWGEGAETLY | 0.66006416 |
| GAAPLAAGL | AWSPQDWGEGAETLY | 0.30266836 |
| GADGGDGGSGGAGG | AWSPQDWGEGAETLY | 0.63927513 |
| GFGLDTGGAGG | AWSPQDWGEGAETLY | 0.60239875 |
| GLASGGLL | AWSPQDWGEGAETLY | 0.6876625 |
| GRIAGAAVLL | AWSPQDWGEGAETLY | 0.63738143 |
| GSAGSLNTSF | AWSPQDWGEGAETLY | 0.6254577 |
| GTGQIGI | AWSPQDWGEGAETLY | 0.6595366 |
| GTLIALIL | AWSPQDWGEGAETLY | 0.669474 |
| IAATMGFVVG | AWSPQDWGEGAETLY | 0.658759 |
| IFAQALL | AWSPQDWGEGAETLY | 0.62811273 |
| IGAAVALGF | AWSPQDWGEGAETLY | 0.45370096 |
| IGLTPLAL | AWSPQDWGEGAETLY | 0.67947423 |
| ILGSNTSTL | AWSPQDWGEGAETLY | 0.6057018 |
| IVFSLTVL | AWSPQDWGEGAETLY | 0.61420983 |
| LASKATGF | AWSPQDWGEGAETLY | 0.5492819 |
| LASVALIL | AWSPQDWGEGAETLY | 0.64472866 |
| LFAVGLTL | AWSPQDWGEGAETLY | 0.5446055 |
| LFNGGIGTYI | AWSPQDWGEGAETLY | 0.6824578 |
| LTLGAGII | AWSPQDWGEGAETLY | 0.6398164 |
| LVAAALGDI | AWSPQDWGEGAETLY | 0.6469532 |
| NTAFVQLGI | AWSPQDWGEGAETLY | 0.68693376 |
| QLFSGAEL | AWSPQDWGEGAETLY | 0.66533947 |
| SAATFITL | AWSPQDWGEGAETLY | 0.6484474 |
| SNYGLANLGV | AWSPQDWGEGAETLY | 0.63721734 |
| SYGGAYAVM | AWSPQDWGEGAETLY | 0.6259806 |
| TTGVGGLTLG | AWSPQDWGEGAETLY | 0.62822294 |
| TTQTIYI | AWSPQDWGEGAETLY | 0.64040977 |
| VAAGAGLGAVI | AWSPQDWGEGAETLY | 0.68613136 |
| VAAVVTLL | AWSPQDWGEGAETLY | 0.6622666 |
| VGFLGGLI | AWSPQDWGEGAETLY | 0.6122165 |
| VGVIPILL | AWSPQDWGEGAETLY | 0.6668213 |
| VIFASSLI | AWSPQDWGEGAETLY | 0.68600553 |
| VKGGDGGAAAG | AWSPQDWGEGAETLY | 0.68598974 |
| VTKTFTNI | AWSPQDWGEGAETLY | 0.63440835 |
| VVAAAILL | AWSPQDWGEGAETLY | 0.6477685 |
| YAALDVEL | AWSPQDWGEGAETLY | 0.6381253 |
| LASVALLL | AWSPQDWGEGAETLY | 0.6515816 |

| | | |
|---|---|---|
| LDAVHRV | AWSPQDWGEGAETLY | 0.6461051 |
| QGLSSSIF | AWSPQDWGEGAETLY | 0.60163486 |
| RGTFANIRL | AWSPQDWGEGAETLY | 0.6808795 |
| RQLWWGHRI | AWSPQDWGEGAETLY | 0.6915087 |
| YGPPGTGKTLL | AWSPQDWGEGAETLY | 0.6544388 |
| AALVGAPANVA | AWSPQDWGEGAETLY | 0.5661714 |
| AATFRRTE | AWSPQDWGEGAETLY | 0.65399015 |
| AGALGGGI | AWSPQDWGEGAETLY | 0.6243306 |
| AGGAGGAGGIGAG | AWSPQDWGEGAETLY | 0.6423424 |
| AGSGALGL | AWSPQDWGEGAETLY | 0.6516978 |
| AQIADQI | AWSPQDWGEGAETLY | 0.5785277 |
| ASAVAVTYL | AWSPQDWGEGAETLY | 0.66260284 |
| ASTVLGGTGG | AWSPQDWGEGAETLY | 0.68595695 |
| AVGGEPAK | AWSPQDWGEGAETLY | 0.63962936 |
| CEQGTFGGTC | AWSPQDWGEGAETLY | 0.6688938 |
| DEGKAPAG | AWSPQDWGEGAETLY | 0.6845591 |
| DTVVFPAL | AWSPQDWGEGAETLY | 0.67588496 |
| FEGITMAG | AWSPQDWGEGAETLY | 0.68286765 |
| FPLVVQD | AWSPQDWGEGAETLY | 0.6834171 |
| GAFTTPQ | AWSPQDWGEGAETLY | 0.67287123 |
| GFGGFGTAGGVGGA | AWSPQDWGEGAETLY | 0.6665422 |
| GFLAQFL | AWSPQDWGEGAETLY | 0.674039 |
| GGAGGAGGNAGVA | AWSPQDWGEGAETLY | 0.65841293 |
| GGVGTAGAGGAGGAGG | AWSPQDWGEGAETLY | 0.6554916 |
| GNGGGGDAGNAGSGGN | AWSPQDWGEGAETLY | 0.6350899 |
| GTGGAGGAGNDAGS | AWSPQDWGEGAETLY | 0.68043625 |
| GTTNTGWFNS | AWSPQDWGEGAETLY | 0.6509475 |
| IFMGPLFTL | AWSPQDWGEGAETLY | 0.685831 |
| ILAHAIEAI | AWSPQDWGEGAETLY | 0.56667036 |
| IRSAIREAL | AWSPQDWGEGAETLY | 0.6257919 |
| ITFAGGG | AWSPQDWGEGAETLY | 0.67915606 |
| ITGMSGHQEV | AWSPQDWGEGAETLY | 0.31329438 |
| IVAGLACTL | AWSPQDWGEGAETLY | 0.53543943 |
| IVGGGLAGA | AWSPQDWGEGAETLY | 0.66352516 |
| IVTGVLNI | AWSPQDWGEGAETLY | 0.6797419 |
| LAQQGCI | AWSPQDWGEGAETLY | 0.3311865 |
| LCLHVEN | AWSPQDWGEGAETLY | 0.6184696 |
| LDAISDGS | AWSPQDWGEGAETLY | 0.60645336 |
| LGGFAAML | AWSPQDWGEGAETLY | 0.6762271 |
| LGGKNPMI | AWSPQDWGEGAETLY | 0.6646072 |
| LGPKGVAAIL | AWSPQDWGEGAETLY | 0.5717613 |
| LMVGVSI | AWSPQDWGEGAETLY | 0.67899996 |
| LVGAAFVL | AWSPQDWGEGAETLY | 0.5545777 |
| LVLAVAPL | AWSPQDWGEGAETLY | 0.640297 |
| MAAIAVIDAA | AWSPQDWGEGAETLY | 0.6797252 |
| NNGINAELQ | AWSPQDWGEGAETLY | 0.68629265 |
| RGEPLVLVHG | AWSPQDWGEGAETLY | 0.6153646 |
| RGGAGGAAGLI | AWSPQDWGEGAETLY | 0.6761731 |
| RSYRMHPAV | AWSPQDWGEGAETLY | 0.6430192 |
| SAAHLLAGQLA | AWSPQDWGEGAETLY | 0.687726 |
| SAAVGSY | AWSPQDWGEGAETLY | 0.65009284 |

| | | |
|---|---|---|
| SPHAVLHRI | AWSPQDWGEGAETLY | 0.61035746 |
| SQIAKVL | AWSPQDWGEGAETLY | 0.6799673 |
| TALGHLVP | AWSPQDWGEGAETLY | 0.6708952 |
| TAPTTSGAA | AWSPQDWGEGAETLY | 0.6644637 |
| TDGDDGNGGSPGT | AWSPQDWGEGAETLY | 0.68095016 |
| TDGLGEN | AWSPQDWGEGAETLY | 0.6473025 |
| TGGPPLPGQA | AWSPQDWGEGAETLY | 0.6550141 |
| TIPAITI | AWSPQDWGEGAETLY | 0.68348205 |
| TIPGFTI | AWSPQDWGEGAETLY | 0.6784832 |
| TSIASGALA | AWSPQDWGEGAETLY | 0.6121014 |
| TVTLPGAAI | AWSPQDWGEGAETLY | 0.6857473 |
| VASGLSAALP | AWSPQDWGEGAETLY | 0.6538553 |
| VGAGVIGI | AWSPQDWGEGAETLY | 0.62387955 |
| VGPKRFGV | AWSPQDWGEGAETLY | 0.6864798 |
| VNVAFPD | AWSPQDWGEGAETLY | 0.67240274 |
| VSMTNSGVS | AWSPQDWGEGAETLY | 0.677255 |
| VTDVLGDLSV | AWSPQDWGEGAETLY | 0.682236 |
| VVVVGAGFAGL | AWSPQDWGEGAETLY | 0.6869216 |
| VYTVGTVV | AWSPQDWGEGAETLY | 0.60911083 |
| WTTFSGP | AWSPQDWGEGAETLY | 0.66327924 |
| YDELVEAF | AWSPQDWGEGAETLY | 0.6516167 |
| YGTGGAGGNGG | AWSPQDWGEGAETLY | 0.6811789 |
| YTNVMAV | AWSPQDWGEGAETLY | 0.66375136 |
| FLEQVHQL | AWSPQDWGEGAETLY | 0.63199824 |
| GLSLLAVAV | AWSPQDWGEGAETLY | 0.6780882 |
| LEQVHQL | AWSPQDWGEGAETLY | 0.683254 |
| LNAAGAI | AWSPQDWGEGAETLY | 0.6676621 |
| PAVHASL | AWSPQDWGEGAETLY | 0.6718824 |
| TELEDDA | AWSPQDWGEGAETLY | 0.68502486 |
| TPPGPPN | AWSPQDWGEGAETLY | 0.6438217 |
| TPPPPPPP | AWSPQDWGEGAETLY | 0.65824807 |
| TQSPALI | ASSLDWGGSSYEQY | 0.6751265 |
| EFAAAFPQ | ASSLDWGGSSYEQY | 0.6825042 |
| LLSGNLDVL | ASSLDWGGSSYEQY | 0.66371095 |
| GPANLGNYNI | ASSLDWGGSSYEQY | 0.6852988 |
| TLGPTLP | ASSLDWGGSSYEQY | 0.62537986 |
| VYAYSSCCK | ASSLDWGGSSYEQY | 0.67710865 |
| APAPAPAPNI | ASSLDWGGSSYEQY | 0.61823237 |
| PGGPVPIP | ASSLDWGGSSYEQY | 0.6657409 |
| FLTVALEP | ASSLDWGGSSYEQY | 0.61606973 |
| SQYFGANL | ASSLDWGGSSYEQY | 0.67125475 |
| GGVKPAA | ASSLDWGGSSYEQY | 0.6765021 |
| GGTGGAGGAGGW | ASSLDWGGSSYEQY | 0.6776442 |
| GVGGVAGLGGAGG | ASSLDWGGSSYEQY | 0.49065757 |
| VLLVLATTI | ASSLDWGGSSYEQY | 0.67860186 |
| IGIFAAML | ASSLDWGGSSYEQY | 0.6794636 |
| AVGDLID | ASSLDWGGSSYEQY | 0.68858486 |
| NISPLPL | ASSLDWGGSSYEQY | 0.65806067 |
| VTAVFVAI | ASSLDWGGSSYEQY | 0.6702999 |
| SGDDNDAAL | ASSLDWGGSSYEQY | 0.632519 |
| AVAALAHS | ASSLDWGGSSYEQY | 0.6669865 |

| | | |
|---|---|---|
| GGVTVMATP | ASSLDWGGSSYEQY | 0.5943005 |
| AGLAANVRD | ASSLDWGGSSYEQY | 0.62811303 |
| TNRPDLIDPALLRPG | ASSLDWGGSSYEQY | 0.63617516 |
| LANLGVTAL | ASSLDWGGSSYEQY | 0.63253033 |
| TGERIEL | ASSLDWGGSSYEQY | 0.67671514 |
| NGEELAA | ASSLDWGGSSYEQY | 0.682531 |
| VLSLVWG | ASSLDWGGSSYEQY | 0.6765738 |
| IGYALWL | ASSLDWGGSSYEQY | 0.68137443 |
| LSLLLAPL | ASSLDWGGSSYEQY | 0.68174833 |
| TLDEALK | ASSLDWGGSSYEQY | 0.6860877 |
| TDGAGTAEGG | ASSLDWGGSSYEQY | 0.6409228 |
| NAPGGAGGNGG | ASSLDWGGSSYEQY | 0.66127896 |
| VSVFMGMGGNF | ASSLDWGGSSYEQY | 0.602019 |
| LASYLAAL | ASSLDWGGSSYEQY | 0.6585057 |
| LVGGTPAL | ASSLDWGGSSYEQY | 0.64347655 |
| VRNADVL | ASSLDWGGSSYEQY | 0.6686489 |
| TCSSPFLVSAM | ASSLDWGGSSYEQY | 0.3277446 |
| VHITDAE | ASSLDWGGSSYEQY | 0.66730523 |
| EYAAPSF | ASSLDWGGSSYEQY | 0.6861342 |
| VVDAFHGGL | ASSLDWGGSSYEQY | 0.6172401 |
| ASVVDPQ | ASSLDWGGSSYEQY | 0.67075324 |
| SYNEFGGLS | ASSLDWGGSSYEQY | 0.666168 |
| DPDGSVY | ASSLDWGGSSYEQY | 0.63758594 |
| MVSTIHATE | ASSLDWGGSSYEQY | 0.6734383 |
| LAAGVADPV | ASSLDWGGSSYEQY | 0.6408764 |
| TKEQIAA | ASSLDWGGSSYEQY | 0.6767587 |
| SLASPAAL | ASSLDWGGSSYEQY | 0.6849562 |
| IGLVFYGV | ASSLDWGGSSYEQY | 0.66777885 |
| FLLDASNL | ASSLDWGGSSYEQY | 0.65917397 |
| YQVGNFP | ASSLDWGGSSYEQY | 0.6638954 |
| ESLLSQST | ASSLDWGGSSYEQY | 0.57783973 |
| LFTVGFAAVG | ASSLDWGGSSYEQY | 0.66897166 |
| ATPPTPT | ASSLDWGGSSYEQY | 0.5323183 |
| TDLATLG | ASSLDWGGSSYEQY | 0.6586697 |
| SAGGTGR | ASSLDWGGSSYEQY | 0.6396271 |
| GFSNSGS | ASSLDWGGSSYEQY | 0.6680685 |
| GGNGGSGGTSV | ASSLDWGGSSYEQY | 0.658172 |
| GTSVATGGAGN | ASSLDWGGSSYEQY | 0.6668544 |
| NGGNGGSGGTS | ASSLDWGGSSYEQY | 0.6114744 |
| AVVKFPH | ASSLDWGGSSYEQY | 0.6760043 |
| TPPPPPP | ASSLDWGGSSYEQY | 0.6585663 |
| AAPAAAAAPA | ASSLDWGGSSYEQY | 0.58722776 |
| IALNEDL | ASSLDWGGSSYEQY | 0.67708117 |
| IVNVSSM | ASSLDWGGSSYEQY | 0.6601819 |
| TGIPPAPR | ASSLDWGGSSYEQY | 0.6457553 |
| FDLGGGTF | ASSLDWGGSSYEQY | 0.6836618 |
| FDLGGGTFDVS | ASSLDWGGSSYEQY | 0.66900826 |
| NEPTAAA | ASSLDWGGSSYEQY | 0.5437465 |
| NEPTAAALA | ASSLDWGGSSYEQY | 0.64094555 |
| GSIINISSI | ASSLDWGGSSYEQY | 0.65822345 |
| VAPGNAGTA | ASSLDWGGSSYEQY | 0.6564275 |

| | | |
|---|---|---|
| TGGGASLE | ASSLDWGGSSYEQY | 0.37077773 |
| VAVYNHYKRI | ASSLDWGGSSYEQY | 0.67347586 |
| TDPAPATT | ASSLDWGGSSYEQY | 0.62696517 |
| TNRPDILDPALLRPG | ASSLDWGGSSYEQY | 0.61041635 |
| LTTTTTS | ASSLDWGGSSYEQY | 0.64384115 |
| PTGSGKTL | ASSLDWGGSSYEQY | 0.67749727 |
| TGSGKTL | ASSLDWGGSSYEQY | 0.63704926 |
| TIGHVDHGK | ASSLDWGGSSYEQY | 0.6604757 |
| TIGHVDHGKTT | ASSLDWGGSSYEQY | 0.6740383 |
| GHVDHGKTT | ASSLDWGGSSYEQY | 0.67653555 |
| AQFEHTLL | ASSLDWGGSSYEQY | 0.6837559 |
| IVNQDEL | ASSLDWGGSSYEQY | 0.6666674 |
| TNGSQFF | ASSLDWGGSSYEQY | 0.65069103 |
| TGKGIVE | ASSLDWGGSSYEQY | 0.6724788 |
| LISNASDA | ASSLDWGGSSYEQY | 0.65222025 |
| AELRTLGELTKIAKAHGAQ | ASSLDWGGSSYEQY | 0.6368204 |
| APNAVAAAEPSW | ASSLDWGGSSYEQY | 0.6105131 |
| APQVAGAATAVL | ASSLDWGGSSYEQY | 0.6478021 |
| ARRDPAAAAAILPTDA | ASSLDWGGSSYEQY | 0.6016642 |
| DAAARDVLAEAGLAEYF | ASSLDWGGSSYEQY | 0.66377175 |
| DDQIVLSAKANLAKVEQ | ASSLDWGGSSYEQY | 0.6866696 |
| DELLPDTGEDSDEDGH | ASSLDWGGSSYEQY | 0.6556239 |
| ELPEAPANRATVPQD | ASSLDWGGSSYEQY | 0.5739465 |
| FGQSKIERVTGIG | ASSLDWGGSSYEQY | 0.644449 |
| GAGGTGSTAGGAGGAGGAGGLY | ASSLDWGGSSYEQY | 0.66137284 |
| GHGGTGAAGGNGGTGGAGGSGI | ASSLDWGGSSYEQY | 0.6765452 |
| GNGGNGGAGGEGGPGGAGGAGA | ASSLDWGGSSYEQY | 0.6793386 |
| GSTAGNGGNGGNATNGTIAGQP | ASSLDWGGSSYEQY | 0.6905862 |
| GTGGDGGNGGNGGTGG | ASSLDWGGSSYEQY | 0.6796082 |
| GVRRPAGIPADAPV | ASSLDWGGSSYEQY | 0.6353327 |
| GYRQVTDAHLLTLARR | ASSLDWGGSSYEQY | 0.6757501 |
| IAKMGQDGHDRG | ASSLDWGGSSYEQY | 0.65756726 |
| KAEPSIEIQYT | ASSLDWGGSSYEQY | 0.6554547 |
| KPVRIRMAADEDTVAVT | ASSLDWGGSSYEQY | 0.6701796 |
| MPPRIAGMRLLVIKPEP | ASSLDWGGSSYEQY | 0.6652591 |
| MKFYAVALFRSRGVPPTR | ASSLDWGGSSYEQY | 0.65735054 |
| NAGGFGILGAVAHSP | ASSLDWGGSSYEQY | 0.65407026 |
| NEGREITLAMAEM | ASSLDWGGSSYEQY | 0.5450977 |
| NELVEWSHPLGHRWRWE | ASSLDWGGSSYEQY | 0.6799977 |
| NLGAGGTGGNGGNPG | ASSLDWGGSSYEQY | 0.6552139 |
| NTEWPKWETVMAADPVD | ASSLDWGGSSYEQY | 0.63152826 |
| PASAAPAETQP | ASSLDWGGSSYEQY | 0.6502405 |
| QQSSDGKNGVRAAEITGE | ASSLDWGGSSYEQY | 0.68451226 |
| RAPDAVTTSWWKE | ASSLDWGGSSYEQY | 0.6833129 |
| SGFGNVGAHTSGWFNQ | ASSLDWGGSSYEQY | 0.6308857 |
| SGVAGAVGQAASVGGLKV | ASSLDWGGSSYEQY | 0.636075 |
| SNGGAGTGSAGGNGGT | ASSLDWGGSSYEQY | 0.66042626 |
| TGGAGGEGGAAG | ASSLDWGGSSYEQY | 0.5773108 |
| TGGDGGAAGAGGAPGHG | ASSLDWGGSSYEQY | 0.67703354 |
| TMPPLLAAIANAGRHGV | ASSLDWGGSSYEQY | 0.6578474 |
| TSSTAGNSGGA | ASSLDWGGSSYEQY | 0.6460751 |

| | | |
|---|---|---|
| VPATVPVGILC | ASSLDWGGSSYEQY | 0.63847136 |
| VQRPDSGGVVLTGRIS | ASSLDWGGSSYEQY | 0.6659133 |
| WTPNPVPATKGVPAGLVQAT | ASSLDWGGSSYEQY | 0.6256785 |
| YALKRALAKFSDRFDVVII | ASSLDWGGSSYEQY | 0.6818984 |
| YLIGSVIV | ASSLDWGGSSYEQY | 0.35732365 |
| YGQLVAVID | ASSLDWGGSSYEQY | 0.6783304 |
| VTVGGMTLPA | ASSLDWGGSSYEQY | 0.61559105 |
| VLALPAV | ASSLDWGGSSYEQY | 0.6272866 |
| VIPPTLNYET | ASSLDWGGSSYEQY | 0.49442196 |
| VGMGGMHPSSG | ASSLDWGGSSYEQY | 0.66128945 |
| VGLAGGAATA | ASSLDWGGSSYEQY | 0.67134 |
| VGGAGGAGGSGP | ASSLDWGGSSYEQY | 0.67732215 |
| VFAAFMV | ASSLDWGGSSYEQY | 0.65403146 |
| TLYDKLR | ASSLDWGGSSYEQY | 0.6801355 |
| TLAVFTVL | ASSLDWGGSSYEQY | 0.68103707 |
| TGSVQPARV | ASSLDWGGSSYEQY | 0.6450019 |
| TGGGAGSGGT | ASSLDWGGSSYEQY | 0.5307343 |
| SHLETSL | ASSLDWGGSSYEQY | 0.6740583 |
| SFTTGMT | ASSLDWGGSSYEQY | 0.652014 |
| QVEVAKRMGVRQPTVS | ASSLDWGGSSYEQY | 0.67909145 |
| QGSGGPALAM | ASSLDWGGSSYEQY | 0.66685194 |
| MLAPSNP | ASSLDWGGSSYEQY | 0.5997449 |
| LVNNAAI | ASSLDWGGSSYEQY | 0.6786555 |
| LTGEINTLA | ASSLDWGGSSYEQY | 0.68075013 |
| LSIFGPV | ASSLDWGGSSYEQY | 0.61518294 |
| LLALLANL | ASSLDWGGSSYEQY | 0.6195708 |
| LISNASDALDKLR | ASSLDWGGSSYEQY | 0.6362619 |
| LINEAVP | ASSLDWGGSSYEQY | 0.55431753 |
| LGYAFSGL | ASSLDWGGSSYEQY | 0.67977023 |
| LDFVAVAL | ASSLDWGGSSYEQY | 0.6370512 |
| LAQPGAE | ASSLDWGGSSYEQY | 0.64407897 |
| ISNLGAVTGF | ASSLDWGGSSYEQY | 0.6646373 |
| HQEDPGRI | ASSLDWGGSSYEQY | 0.65008825 |
| GTPGEPTP | ASSLDWGGSSYEQY | 0.6514863 |
| GFFNSGS | ASSLDWGGSSYEQY | 0.64990544 |
| GAGGGGLTAGAN | ASSLDWGGSSYEQY | 0.6633433 |
| FVDALVNM | ASSLDWGGSSYEQY | 0.6725451 |
| FLTVAPEL | ASSLDWGGSSYEQY | 0.65124106 |
| FETLTAL | ASSLDWGGSSYEQY | 0.6677567 |
| DVINPIEL | ASSLDWGGSSYEQY | 0.68328094 |
| APVSGAPV | ASSLDWGGSSYEQY | 0.57716 |
| ALVGGLAW | ASSLDWGGSSYEQY | 0.63107175 |
| ALAAAAATAI | ASSLDWGGSSYEQY | 0.671515 |
| AFGLGLLL | ASSLDWGGSSYEQY | 0.6321279 |
| AFFAAASL | ASSLDWGGSSYEQY | 0.678333 |
| AAAALPAFSPP | ASSLDWGGSSYEQY | 0.659636 |
| VVLAPAL | ASSLDWGGSSYEQY | 0.67029274 |
| TIGHVDHGKTTL | ASSLDWGGSSYEQY | 0.6584355 |
| LISNASDALDK | ASSLDWGGSSYEQY | 0.6871798 |
| LISNASDAL | ASSLDWGGSSYEQY | 0.6769855 |
| ISNASDALDK | ASSLDWGGSSYEQY | 0.6665257 |

| | | |
|---|---|---|
| NQAETLVYQTE | ASSLDWGGSSYEQY | 0.67757845 |
| TGGTGGTAGAG | ASSLDWGGSSYEQY | 0.5730728 |
| TGSTAGGAGGA | ASSLDWGGSSYEQY | 0.6828865 |
| DPLAVPVAAGPV | ASSLDWGGSSYEQY | 0.6746818 |
| GGTGGTPGNGGN | ASSLDWGGSSYEQY | 0.61350435 |
| PAGLAGVDPATL | ASSLDWGGSSYEQY | 0.6563883 |
| GNNLVVEAPHAGASV | ASSLDWGGSSYEQY | 0.68472993 |
| KVTRIMSASTLEQPA | ASSLDWGGSSYEQY | 0.6829784 |
| LGFALPIALPRRTVV | ASSLDWGGSSYEQY | 0.6286081 |
| TRLGNLTARKLVVLV | ASSLDWGGSSYEQY | 0.65065837 |
| WSQQRRLIYFPSAGP | ASSLDWGGSSYEQY | 0.5593457 |
| GPNGNPDPMAAAVDIR | ASSLDWGGSSYEQY | 0.600376 |
| TQRCVSVADLRAAASR | ASSLDWGGSSYEQY | 0.5626151 |
| QFVLDVAEHPVEVLTETV | ASSLDWGGSSYEQY | 0.6746975 |
| GGAGGTGGAGGNGGTQAGDG | ASSLDWGGSSYEQY | 0.6415431 |
| PLLATLRAIAVDVDAKPAQV | ASSLDWGGSSYEQY | 0.67739296 |
| APKFRTGGYMIDFWGVGYQVAK | ASSLDWGGSSYEQY | 0.6641271 |
| GAGGAGGAGGAAGGPGGTGGTG | ASSLDWGGSSYEQY | 0.6852994 |
| IGALGLLATDVAPAGSVAFGVA | ASSLDWGGSSYEQY | 0.63780636 |
| AGGAGGIGGNGGAGGAGGVLMG | ASSLDWGGSSYEQY | 0.65785515 |
| ADELIGLL | ASSLDWGGSSYEQY | 0.57640356 |
| ADRALKA | ASSLDWGGSSYEQY | 0.6287958 |
| AGAVGQVGGAGGA | ASSLDWGGSSYEQY | 0.64075965 |
| AGGGGGLGGAP | ASSLDWGGSSYEQY | 0.65162754 |
| AGGLLYGNGGAG | ASSLDWGGSSYEQY | 0.4598475 |
| AGLESIGL | ASSLDWGGSSYEQY | 0.6834451 |
| AGLKTGII | ASSLDWGGSSYEQY | 0.66032755 |
| ASLAALLL | ASSLDWGGSSYEQY | 0.6789338 |
| ASVINISSI | ASSLDWGGSSYEQY | 0.6482584 |
| ATPPGPA | ASSLDWGGSSYEQY | 0.6559317 |
| ATVASALV | ASSLDWGGSSYEQY | 0.63788664 |
| ATVASLIL | ASSLDWGGSSYEQY | 0.671824 |
| AVNGPVLIRH | ASSLDWGGSSYEQY | 0.6244687 |
| DGAAGAVGI | ASSLDWGGSSYEQY | 0.49473953 |
| DPLLHGI | ASSLDWGGSSYEQY | 0.63677394 |
| ELGAALGAL | ASSLDWGGSSYEQY | 0.59242773 |
| ELGRTQAV | ASSLDWGGSSYEQY | 0.6841319 |
| FVAGGYLS | ASSLDWGGSSYEQY | 0.68261385 |
| GASGGAGGAGGW | ASSLDWGGSSYEQY | 0.6679425 |
| GGAGAGGDGGAGGI | ASSLDWGGSSYEQY | 0.68368137 |
| GTGHNGSQPGMGGNG | ASSLDWGGSSYEQY | 0.3630409 |
| GYAAASASAS | ASSLDWGGSSYEQY | 0.65157527 |
| HQVDVAAVQQG | ASSLDWGGSSYEQY | 0.684746 |
| IGAAIAHT | ASSLDWGGSSYEQY | 0.674407 |
| IGMIEVAQL | ASSLDWGGSSYEQY | 0.6542637 |
| IRLAPEHTSAMD | ASSLDWGGSSYEQY | 0.68285894 |
| IYNVSVL | ASSLDWGGSSYEQY | 0.6225308 |
| KQGFLGM | ASSLDWGGSSYEQY | 0.655114 |
| LESVITQL | ASSLDWGGSSYEQY | 0.66296947 |
| LGLGALVVV | ASSLDWGGSSYEQY | 0.68120563 |
| LIGNGGNGGSGGI | ASSLDWGGSSYEQY | 0.66931486 |

| | | |
|---|---|---|
| LNAINAPTL | ASSLDWGGSSYEQY | 0.6104683 |
| LQLLNTGV | ASSLDWGGSSYEQY | 0.67274976 |
| LQTSIEF | ASSLDWGGSSYEQY | 0.6882072 |
| MSFAVSEL | ASSLDWGGSSYEQY | 0.6419206 |
| NTGALGL | ASSLDWGGSSYEQY | 0.6752617 |
| PPGAGGAGGAGTT | ASSLDWGGSSYEQY | 0.6708368 |
| QLVGLGI | ASSLDWGGSSYEQY | 0.66712755 |
| RFLEQVHQL | ASSLDWGGSSYEQY | 0.62560904 |
| RGVGYVLR | ASSLDWGGSSYEQY | 0.6330588 |
| SATGVFAGVI | ASSLDWGGSSYEQY | 0.6745738 |
| SLPAITLP | ASSLDWGGSSYEQY | 0.61245656 |
| STALAFTV | ASSLDWGGSSYEQY | 0.6570493 |
| SVLTVGLGK | ASSLDWGGSSYEQY | 0.6541238 |
| TALAGAGSGI | ASSLDWGGSSYEQY | 0.6900401 |
| TGLPFTGI | ASSLDWGGSSYEQY | 0.67131174 |
| VAEHVTV | ASSLDWGGSSYEQY | 0.67623985 |
| VDIVLNSL | ASSLDWGGSSYEQY | 0.6185609 |
| VLTGGGAVL | ASSLDWGGSSYEQY | 0.6712917 |
| VNVGLAYGI | ASSLDWGGSSYEQY | 0.6733165 |
| VSSAAVGAV | ASSLDWGGSSYEQY | 0.57423806 |
| VTSVYEAGV | ASSLDWGGSSYEQY | 0.6823712 |
| WVALDRGA | ASSLDWGGSSYEQY | 0.6337564 |
| YLVGSVVI | ASSLDWGGSSYEQY | 0.6446332 |
| GPPGTGKTLL | ASSLDWGGSSYEQY | 0.67651486 |
| GVGMTKF | ASSLDWGGSSYEQY | 0.6878714 |
| IGIGGSDL | ASSLDWGGSSYEQY | 0.65711653 |
| IGTLLAEL | ASSLDWGGSSYEQY | 0.669345 |
| IYTSGTT | ASSLDWGGSSYEQY | 0.61981744 |
| LSFPAGV | ASSLDWGGSSYEQY | 0.6802408 |
| LYTSGTT | ASSLDWGGSSYEQY | 0.6803472 |
| VGAFPFGF | ASSLDWGGSSYEQY | 0.665299 |
| VGAGTVEF | ASSLDWGGSSYEQY | 0.68849444 |
| VGKSTLLL | ASSLDWGGSSYEQY | 0.67686766 |
| AALAHAGGGIV | ASSLDWGGSSYEQY | 0.488752 |
| AALVAVSLL | ASSLDWGGSSYEQY | 0.6633435 |
| AAVSRAVAAL | ASSLDWGGSSYEQY | 0.6813246 |
| AGGSSGIGGF | ASSLDWGGSSYEQY | 0.6785969 |
| AGLTVANANG | ASSLDWGGSSYEQY | 0.6859736 |
| AGLVGAVL | ASSLDWGGSSYEQY | 0.67488384 |
| ALSQAFTV | ASSLDWGGSSYEQY | 0.6354791 |
| ALTFSVSL | ASSLDWGGSSYEQY | 0.68525785 |
| ALTGAALVAL | ASSLDWGGSSYEQY | 0.6894616 |
| ASFSIGAGVL | ASSLDWGGSSYEQY | 0.6767023 |
| DGSVYKI | ASSLDWGGSSYEQY | 0.6371198 |
| EQLAEIV | ASSLDWGGSSYEQY | 0.62047297 |
| FAVSELTVM | ASSLDWGGSSYEQY | 0.669888 |
| GAAPLAAGL | ASSLDWGGSSYEQY | 0.6752868 |
| GADGGDGGSGGAGG | ASSLDWGGSSYEQY | 0.61287355 |
| GFGLDTGGAGG | ASSLDWGGSSYEQY | 0.66570604 |
| GLASGGLL | ASSLDWGGSSYEQY | 0.6656945 |
| GRIAGAAVLL | ASSLDWGGSSYEQY | 0.5211919 |

| | | |
|---|---|---|
| GSAGSLNTSF | ASSLDWGGSSYEQY | 0.6761745 |
| GTGQIGI | ASSLDWGGSSYEQY | 0.6817492 |
| GTLIALIL | ASSLDWGGSSYEQY | 0.67054373 |
| IAATMGFVVG | ASSLDWGGSSYEQY | 0.63504547 |
| IFAQALL | ASSLDWGGSSYEQY | 0.67938405 |
| IGAAVALGF | ASSLDWGGSSYEQY | 0.6873807 |
| IGLTPLAL | ASSLDWGGSSYEQY | 0.6863059 |
| ILGSNTSTL | ASSLDWGGSSYEQY | 0.56025887 |
| IVFSLTVL | ASSLDWGGSSYEQY | 0.47608358 |
| LASKATGF | ASSLDWGGSSYEQY | 0.6556802 |
| LASVALIL | ASSLDWGGSSYEQY | 0.67563057 |
| LFAVGLTL | ASSLDWGGSSYEQY | 0.6542298 |
| LFNGGIGTYI | ASSLDWGGSSYEQY | 0.64148027 |
| LTLGAGII | ASSLDWGGSSYEQY | 0.6647581 |
| LVAAALGDI | ASSLDWGGSSYEQY | 0.6828228 |
| NTAFVQLGI | ASSLDWGGSSYEQY | 0.6327057 |
| QLFSGAEL | ASSLDWGGSSYEQY | 0.56903446 |
| SAATFITL | ASSLDWGGSSYEQY | 0.6724446 |
| SNYGLANLGV | ASSLDWGGSSYEQY | 0.6755105 |
| SYGGAYAVM | ASSLDWGGSSYEQY | 0.629331 |
| TTGVGGLTLG | ASSLDWGGSSYEQY | 0.6525934 |
| TTQTIYI | ASSLDWGGSSYEQY | 0.6550222 |
| VAAGAGLGAVI | ASSLDWGGSSYEQY | 0.6524422 |
| VAAVVTLL | ASSLDWGGSSYEQY | 0.66344357 |
| VGFLGGLI | ASSLDWGGSSYEQY | 0.61047447 |
| VGVIPILL | ASSLDWGGSSYEQY | 0.6760107 |
| VIFASSLI | ASSLDWGGSSYEQY | 0.6126349 |
| VKGGDGGAAAG | ASSLDWGGSSYEQY | 0.67543244 |
| VTKTFTNI | ASSLDWGGSSYEQY | 0.66773087 |
| VVAAAILL | ASSLDWGGSSYEQY | 0.67505974 |
| YAALDVEL | ASSLDWGGSSYEQY | 0.64521104 |
| LASVALLL | ASSLDWGGSSYEQY | 0.6369269 |
| LDAVHRV | ASSLDWGGSSYEQY | 0.61125594 |
| QGLSSSIF | ASSLDWGGSSYEQY | 0.58667 |
| RGTFANIRL | ASSLDWGGSSYEQY | 0.6826327 |
| RQLWWGHRI | ASSLDWGGSSYEQY | 0.63324547 |
| YGPPGTGKTLL | ASSLDWGGSSYEQY | 0.6831119 |
| AALVGAPANVA | ASSLDWGGSSYEQY | 0.6753138 |
| AATFRRTE | ASSLDWGGSSYEQY | 0.6354086 |
| AGALGGGI | ASSLDWGGSSYEQY | 0.6738356 |
| AGGAGGAGGIGAG | ASSLDWGGSSYEQY | 0.6028145 |
| AGSGALGL | ASSLDWGGSSYEQY | 0.6621009 |
| AQIADQI | ASSLDWGGSSYEQY | 0.6635306 |
| ASAVAVTYL | ASSLDWGGSSYEQY | 0.6781095 |
| ASTVLGGTGG | ASSLDWGGSSYEQY | 0.6227337 |
| AVGGEPAK | ASSLDWGGSSYEQY | 0.674428 |
| CEQGTFGGTC | ASSLDWGGSSYEQY | 0.6702905 |
| DEGKAPAG | ASSLDWGGSSYEQY | 0.62172496 |
| DTVVFPAL | ASSLDWGGSSYEQY | 0.6700327 |
| FEGITMAG | ASSLDWGGSSYEQY | 0.6818955 |
| FPLVVQD | ASSLDWGGSSYEQY | 0.67320275 |

| | | |
|---|---|---|
| GAFTTPQ | ASSLDWGGSSYEQY | 0.6152717 |
| GFGGFGTAGGVGGA | ASSLDWGGSSYEQY | 0.6713693 |
| GFLAQFL | ASSLDWGGSSYEQY | 0.6280158 |
| GGAGGAGGNAGVA | ASSLDWGGSSYEQY | 0.6677548 |
| GGVGTAGAGGAGGAGG | ASSLDWGGSSYEQY | 0.6669337 |
| GNGGGGDAGNAGSGGN | ASSLDWGGSSYEQY | 0.6426673 |
| GTGGAGGAGNDAGS | ASSLDWGGSSYEQY | 0.6785977 |
| GTTNTGWFNS | ASSLDWGGSSYEQY | 0.55343115 |
| IFMGPLFTL | ASSLDWGGSSYEQY | 0.62003815 |
| ILAHAIEAI | ASSLDWGGSSYEQY | 0.65112305 |
| IRSAIREAL | ASSLDWGGSSYEQY | 0.6648082 |
| ITFAGGG | ASSLDWGGSSYEQY | 0.6795498 |
| ITGMSGHQEV | ASSLDWGGSSYEQY | 0.62477136 |
| IVAGLACTL | ASSLDWGGSSYEQY | 0.6656431 |
| IVGGGLAGA | ASSLDWGGSSYEQY | 0.6525594 |
| IVTGVLNI | ASSLDWGGSSYEQY | 0.62580746 |
| LAQQGCI | ASSLDWGGSSYEQY | 0.6721244 |
| LCLHVEN | ASSLDWGGSSYEQY | 0.6642389 |
| LDAISDGS | ASSLDWGGSSYEQY | 0.60351026 |
| LGGFAAML | ASSLDWGGSSYEQY | 0.6773968 |
| LGGKNPMI | ASSLDWGGSSYEQY | 0.6298314 |
| LGPKGVAAIL | ASSLDWGGSSYEQY | 0.67307085 |
| LMVGVSI | ASSLDWGGSSYEQY | 0.68674695 |
| LVGAAFVL | ASSLDWGGSSYEQY | 0.62171507 |
| LVLAVAPL | ASSLDWGGSSYEQY | 0.6758406 |
| MAAIAVIDAA | ASSLDWGGSSYEQY | 0.6671634 |
| NNGINAELQ | ASSLDWGGSSYEQY | 0.6728405 |
| RGEPLVLVHG | ASSLDWGGSSYEQY | 0.6530063 |
| RGGAGGAAGLI | ASSLDWGGSSYEQY | 0.6396833 |
| RSYRMHPAV | ASSLDWGGSSYEQY | 0.68249583 |
| SAAHLLAGQLA | ASSLDWGGSSYEQY | 0.6763235 |
| SAAVGSY | ASSLDWGGSSYEQY | 0.6736331 |
| SPHAVLHRI | ASSLDWGGSSYEQY | 0.67239785 |
| SQIAKVL | ASSLDWGGSSYEQY | 0.6777245 |
| TALGHLVP | ASSLDWGGSSYEQY | 0.67627966 |
| TAPTTSGAA | ASSLDWGGSSYEQY | 0.63833153 |
| TDGDDGNGGSPGT | ASSLDWGGSSYEQY | 0.5797172 |
| TDGLGEN | ASSLDWGGSSYEQY | 0.6555835 |
| TGGPPLPGQA | ASSLDWGGSSYEQY | 0.65447545 |
| TIPAITI | ASSLDWGGSSYEQY | 0.66039217 |
| TIPGFTI | ASSLDWGGSSYEQY | 0.5523332 |
| TSIASGALA | ASSLDWGGSSYEQY | 0.67942536 |
| TVTLPGAAI | ASSLDWGGSSYEQY | 0.68793064 |
| VASGLSAALP | ASSLDWGGSSYEQY | 0.63287085 |
| VGAGVIGI | ASSLDWGGSSYEQY | 0.6891568 |
| VGPKRFGV | ASSLDWGGSSYEQY | 0.61179584 |
| VNVAFPD | ASSLDWGGSSYEQY | 0.6764276 |
| VSMTNSGVS | ASSLDWGGSSYEQY | 0.68149316 |
| VTDVLGDLSV | ASSLDWGGSSYEQY | 0.6112645 |
| VVVVGAGFAGL | ASSLDWGGSSYEQY | 0.680583 |
| VYTVGTVV | ASSLDWGGSSYEQY | 0.6423812 |

| | | |
|---|---|---|
| WTTFSGP | ASSLDWGGSSYEQY | 0.6515263 |
| YDELVEAF | ASSLDWGGSSYEQY | 0.68275243 |
| YGTGGAGGNGG | ASSLDWGGSSYEQY | 0.6463492 |
| YTNVMAV | ASSLDWGGSSYEQY | 0.67465377 |
| FLEQVHQL | ASSLDWGGSSYEQY | 0.6641063 |
| GLSLLAVAV | ASSLDWGGSSYEQY | 0.6289339 |
| LEQVHQL | ASSLDWGGSSYEQY | 0.67610455 |
| LNAAGAI | ASSLDWGGSSYEQY | 0.6793395 |
| PAVHASL | ASSLDWGGSSYEQY | 0.6615553 |
| TELEDDA | ASSLDWGGSSYEQY | 0.68856955 |
| TPPGPPN | ASSLDWGGSSYEQY | 0.61943156 |
| TPPPPPPP | ASSLDWGGSSYEQY | 0.6865469 |
| TQSPALI | ASSQETGGLNYAEQF | 0.67517555 |
| EFAAAFPQ | ASSQETGGLNYAEQF | 0.6621081 |
| LLSGNLDVL | ASSQETGGLNYAEQF | 0.6515062 |
| GPANLGNYNI | ASSQETGGLNYAEQF | 0.6785148 |
| TLGPTLP | ASSQETGGLNYAEQF | 0.6503954 |
| VYAYSSCCK | ASSQETGGLNYAEQF | 0.67350376 |
| APAPAPAPNI | ASSQETGGLNYAEQF | 0.6881611 |
| PGGPVPIP | ASSQETGGLNYAEQF | 0.6761092 |
| FLTVALEP | ASSQETGGLNYAEQF | 0.74998695 |
| SQYFGANL | ASSQETGGLNYAEQF | 0.7068567 |
| GGVKPAA | ASSQETGGLNYAEQF | 0.73017967 |
| GGTGGAGGAGGW | ASSQETGGLNYAEQF | 0.6734225 |
| GVGGVAGLGGAGG | ASSQETGGLNYAEQF | 0.5336494 |
| VLLVLATTI | ASSQETGGLNYAEQF | 0.71476305 |
| IGIFAAML | ASSQETGGLNYAEQF | 0.64721614 |
| AVGDLID | ASSQETGGLNYAEQF | 0.6927227 |
| NISPLPL | ASSQETGGLNYAEQF | 0.7447398 |
| VTAVFVAI | ASSQETGGLNYAEQF | 0.6835303 |
| SGDDNDAAL | ASSQETGGLNYAEQF | 0.7530477 |
| AVAALAHS | ASSQETGGLNYAEQF | 0.751317 |
| GGVTVMATP | ASSQETGGLNYAEQF | 0.55294275 |
| AGLAANVRD | ASSQETGGLNYAEQF | 0.696434 |
| TNRPDLIDPALLRPG | ASSQETGGLNYAEQF | 0.6729393 |
| LANLGVTAL | ASSQETGGLNYAEQF | 0.7156395 |
| TGERIEL | ASSQETGGLNYAEQF | 0.5660362 |
| NGEELAA | ASSQETGGLNYAEQF | 0.7430931 |
| VLSLVWG | ASSQETGGLNYAEQF | 0.6368468 |
| IGYALWL | ASSQETGGLNYAEQF | 0.357346 |
| LSLLLAPL | ASSQETGGLNYAEQF | 0.7351309 |
| TLDEALK | ASSQETGGLNYAEQF | 0.7138127 |
| TDGAGTAEGG | ASSQETGGLNYAEQF | 0.7016769 |
| NAPGGAGGNGG | ASSQETGGLNYAEQF | 0.7442927 |
| VSVFMGMGGNF | ASSQETGGLNYAEQF | 0.54562855 |
| LASYLAAL | ASSQETGGLNYAEQF | 0.65773565 |
| LVGGTPAL | ASSQETGGLNYAEQF | 0.73135555 |
| VRNADVL | ASSQETGGLNYAEQF | 0.7509494 |
| TCSSPFLVSAM | ASSQETGGLNYAEQF | 0.6609496 |
| VHITDAE | ASSQETGGLNYAEQF | 0.7126428 |
| EYAAPSF | ASSQETGGLNYAEQF | 0.7488303 |

| | | |
|---|---|---|
| VVDAFHGGL | ASSQETGGLNYAEQF | 0.7267616 |
| ASVVDPQ | ASSQETGGLNYAEQF | 0.7511076 |
| SYNEFGGLS | ASSQETGGLNYAEQF | 0.754233 |
| DPDGSVY | ASSQETGGLNYAEQF | 0.66172874 |
| MVSTIHATE | ASSQETGGLNYAEQF | 0.7427392 |
| LAAGVADPV | ASSQETGGLNYAEQF | 0.7527185 |
| TKEQIAA | ASSQETGGLNYAEQF | 0.7137284 |
| SLASPAAL | ASSQETGGLNYAEQF | 0.7309579 |
| IGLVFYGV | ASSQETGGLNYAEQF | 0.64773387 |
| FLLDASNL | ASSQETGGLNYAEQF | 0.685603 |
| YQVGNFP | ASSQETGGLNYAEQF | 0.39352542 |
| ESLLSQST | ASSQETGGLNYAEQF | 0.7211221 |
| LFTVGFAAVG | ASSQETGGLNYAEQF | 0.7498273 |
| ATPPTPT | ASSQETGGLNYAEQF | 0.6961832 |
| TDLATLG | ASSQETGGLNYAEQF | 0.6525463 |
| SAGGTGR | ASSQETGGLNYAEQF | 0.57623637 |
| GFSNSGS | ASSQETGGLNYAEQF | 0.71455735 |
| GGNGGSGGTSV | ASSQETGGLNYAEQF | 0.69923735 |
| GTSVATGGAGN | ASSQETGGLNYAEQF | 0.5185859 |
| NGGNGGSGGTS | ASSQETGGLNYAEQF | 0.6526692 |
| AVVKFPH | ASSQETGGLNYAEQF | 0.74500805 |
| TPPPPPP | ASSQETGGLNYAEQF | 0.7441126 |
| AAPAAAAAPA | ASSQETGGLNYAEQF | 0.5993645 |
| IALNEDL | ASSQETGGLNYAEQF | 0.75115925 |
| IVNVSSM | ASSQETGGLNYAEQF | 0.7116106 |
| TGIPPAPR | ASSQETGGLNYAEQF | 0.72630423 |
| FDLGGGTF | ASSQETGGLNYAEQF | 0.7252865 |
| FDLGGGTFDVS | ASSQETGGLNYAEQF | 0.66969764 |
| NEPTAAA | ASSQETGGLNYAEQF | 0.72563326 |
| NEPTAAALA | ASSQETGGLNYAEQF | 0.6806356 |
| GSIINISSI | ASSQETGGLNYAEQF | 0.67934847 |
| VAPGNAGTA | ASSQETGGLNYAEQF | 0.7444213 |
| TGGGASLE | ASSQETGGLNYAEQF | 0.6864028 |
| VAVYNHYKRI | ASSQETGGLNYAEQF | 0.67318106 |
| TDPAPATT | ASSQETGGLNYAEQF | 0.6285319 |
| TNRPDILDPALLRPG | ASSQETGGLNYAEQF | 0.748118 |
| LTTTTTS | ASSQETGGLNYAEQF | 0.59403336 |
| PTGSGKTL | ASSQETGGLNYAEQF | 0.603839 |
| TGSGKTL | ASSQETGGLNYAEQF | 0.64501595 |
| TIGHVDHGK | ASSQETGGLNYAEQF | 0.71441364 |
| TIGHVDHGKTT | ASSQETGGLNYAEQF | 0.7475671 |
| GHVDHGKTT | ASSQETGGLNYAEQF | 0.732515 |
| AQFEHTLL | ASSQETGGLNYAEQF | 0.70796883 |
| IVNQDEL | ASSQETGGLNYAEQF | 0.7358563 |
| TNGSQFF | ASSQETGGLNYAEQF | 0.74179745 |
| TGKGIVE | ASSQETGGLNYAEQF | 0.72570205 |
| LISNASDA | ASSQETGGLNYAEQF | 0.7294594 |
| AELRTLGELTKIAKAHGAQ | ASSQETGGLNYAEQF | 0.6628956 |
| APNAVAAAEPSW | ASSQETGGLNYAEQF | 0.7472243 |
| APQVAGAATAVL | ASSQETGGLNYAEQF | 0.7371652 |
| ARRDPAAAAAILPTDA | ASSQETGGLNYAEQF | 0.74473 |

| | | |
|---|---|---|
| DAAARDVLAEAGLAEYF | ASSQETGGLNYAEQF | 0.74507856 |
| DDQIVLSAKANLAKVEQ | ASSQETGGLNYAEQF | 0.7251702 |
| DELLPDTGEDSDEDGH | ASSQETGGLNYAEQF | 0.71238947 |
| ELPEAPANRATVPQD | ASSQETGGLNYAEQF | 0.42833948 |
| FGQSKIERVTGIG | ASSQETGGLNYAEQF | 0.6778956 |
| GAGGTGSTAGGAGGAGGAGGLY | ASSQETGGLNYAEQF | 0.7208462 |
| GHGGTGAAGGNGGTGGAGGSGI | ASSQETGGLNYAEQF | 0.71809256 |
| GNGGNGGAGGEGGPGGAGGAGA | ASSQETGGLNYAEQF | 0.7325283 |
| GSTAGNGGNGGNATNGTIAGQP | ASSQETGGLNYAEQF | 0.7449395 |
| GTGGDGGNGGNGGTGG | ASSQETGGLNYAEQF | 0.6573949 |
| GVRRPAGIPADAPV | ASSQETGGLNYAEQF | 0.7460825 |
| GYRQVTDAHLLTLARR | ASSQETGGLNYAEQF | 0.74755234 |
| IAKMGQDGHDRG | ASSQETGGLNYAEQF | 0.5812914 |
| KAEPSIEIQYT | ASSQETGGLNYAEQF | 0.6720795 |
| KPVRIRMAADEDTVAVT | ASSQETGGLNYAEQF | 0.75601596 |
| MPPRIAGMRLLVIKPEP | ASSQETGGLNYAEQF | 0.70624113 |
| MKFYAVALFRSRGVPPTR | ASSQETGGLNYAEQF | 0.7000519 |
| NAGGFGILGAVAHSP | ASSQETGGLNYAEQF | 0.7137252 |
| NEGREITLAMAEM | ASSQETGGLNYAEQF | 0.74010885 |
| NELVEWSHPLGHRWRWE | ASSQETGGLNYAEQF | 0.66632074 |
| NLGAGGTGGNGGNPG | ASSQETGGLNYAEQF | 0.75694907 |
| NTEWPKWETVMAADPVD | ASSQETGGLNYAEQF | 0.40766987 |
| PASAAPAETQP | ASSQETGGLNYAEQF | 0.7052358 |
| QQSSDGKNGVRAAEITGE | ASSQETGGLNYAEQF | 0.5835686 |
| RAPDAVTTSWWKE | ASSQETGGLNYAEQF | 0.51652473 |
| SGFGNVGAHTSGWFNQ | ASSQETGGLNYAEQF | 0.7394148 |
| SGVAGAVGQAASVGGLKV | ASSQETGGLNYAEQF | 0.7155501 |
| SNGGAGTGSAGGNGGT | ASSQETGGLNYAEQF | 0.74040633 |
| TGGAGGEGGAAG | ASSQETGGLNYAEQF | 0.7070251 |
| TGGDGGAAGAGGAPGHG | ASSQETGGLNYAEQF | 0.7457542 |
| TMPPLLAAIANAGRHGV | ASSQETGGLNYAEQF | 0.71066916 |
| TSSTAGNSGGA | ASSQETGGLNYAEQF | 0.7222621 |
| VPATVPVGILC | ASSQETGGLNYAEQF | 0.68592983 |
| VQRPDSGGVVLTGRIS | ASSQETGGLNYAEQF | 0.7329365 |
| WTPNPVPATKGVPAGLVQAT | ASSQETGGLNYAEQF | 0.7510903 |
| YALKRALAKFSDRFDVVII | ASSQETGGLNYAEQF | 0.7458549 |
| YLIGSVIV | ASSQETGGLNYAEQF | 0.58929044 |
| YGQLVAVID | ASSQETGGLNYAEQF | 0.6754027 |
| VTVGGMTLPA | ASSQETGGLNYAEQF | 0.7089286 |
| VLALPAV | ASSQETGGLNYAEQF | 0.3886708 |
| VIPPTLNYET | ASSQETGGLNYAEQF | 0.4551255 |
| VGMGGMHPSSG | ASSQETGGLNYAEQF | 0.63996804 |
| VGLAGGAATA | ASSQETGGLNYAEQF | 0.6765188 |
| VGGAGGAGGSGP | ASSQETGGLNYAEQF | 0.74287385 |
| VFAAFMV | ASSQETGGLNYAEQF | 0.7367011 |
| TLYDKLR | ASSQETGGLNYAEQF | 0.70465076 |
| TLAVFTVL | ASSQETGGLNYAEQF | 0.7338539 |
| TGSVQPARV | ASSQETGGLNYAEQF | 0.72257006 |
| TGGGAGSGGT | ASSQETGGLNYAEQF | 0.23283829 |
| SHLETSL | ASSQETGGLNYAEQF | 0.6886287 |
| SFTTGMT | ASSQETGGLNYAEQF | 0.61048836 |

| | | |
|---|---|---|
| QVEVAKRMGVRQPTVS | ASSQETGGLNYAEQF | 0.75265425 |
| QGSGGPALAM | ASSQETGGLNYAEQF | 0.65732396 |
| MLAPSNP | ASSQETGGLNYAEQF | 0.6686113 |
| LVNNAAI | ASSQETGGLNYAEQF | 0.7172995 |
| LTGEINTLA | ASSQETGGLNYAEQF | 0.6852299 |
| LSIFGPV | ASSQETGGLNYAEQF | 0.6956421 |
| LLALLANL | ASSQETGGLNYAEQF | 0.64830405 |
| LISNASDALDKLR | ASSQETGGLNYAEQF | 0.3042065 |
| LINEAVP | ASSQETGGLNYAEQF | 0.74057156 |
| LGYAFSGL | ASSQETGGLNYAEQF | 0.5463499 |
| LDFVAVAL | ASSQETGGLNYAEQF | 0.66379446 |
| LAQPGAE | ASSQETGGLNYAEQF | 0.54503775 |
| ISNLGAVTGF | ASSQETGGLNYAEQF | 0.60769624 |
| HQEDPGRI | ASSQETGGLNYAEQF | 0.54916453 |
| GTPGEPTP | ASSQETGGLNYAEQF | 0.75378823 |
| GFFNSGS | ASSQETGGLNYAEQF | 0.64654166 |
| GAGGGGLTAGAN | ASSQETGGLNYAEQF | 0.6845369 |
| FVDALVNM | ASSQETGGLNYAEQF | 0.75374866 |
| FLTVAPEL | ASSQETGGLNYAEQF | 0.68819404 |
| FETLTAL | ASSQETGGLNYAEQF | 0.69719696 |
| DVINPIEL | ASSQETGGLNYAEQF | 0.7067685 |
| APVSGAPV | ASSQETGGLNYAEQF | 0.6477708 |
| ALVGGLAW | ASSQETGGLNYAEQF | 0.66349405 |
| ALAAAAATAI | ASSQETGGLNYAEQF | 0.6714244 |
| AFGLGLLL | ASSQETGGLNYAEQF | 0.75581205 |
| AFFAAASL | ASSQETGGLNYAEQF | 0.70871043 |
| AAAALPAFSPP | ASSQETGGLNYAEQF | 0.6170765 |
| VVLAPAL | ASSQETGGLNYAEQF | 0.71886003 |
| TIGHVDHGKTTL | ASSQETGGLNYAEQF | 0.75336623 |
| LISNASDALDK | ASSQETGGLNYAEQF | 0.74903536 |
| LISNASDAL | ASSQETGGLNYAEQF | 0.65005517 |
| ISNASDALDK | ASSQETGGLNYAEQF | 0.6992115 |
| NQAETLVYQTE | ASSQETGGLNYAEQF | 0.70802927 |
| TGGTGGTAGAG | ASSQETGGLNYAEQF | 0.7004583 |
| TGSTAGGAGGA | ASSQETGGLNYAEQF | 0.71215576 |
| DPLAVPVAAGPV | ASSQETGGLNYAEQF | 0.6601871 |
| GGTGGTPGNGGN | ASSQETGGLNYAEQF | 0.7398053 |
| PAGLAGVDPATL | ASSQETGGLNYAEQF | 0.745828 |
| GNNLVVEAPHAGASV | ASSQETGGLNYAEQF | 0.71205145 |
| KVTRIMSASTLEQPA | ASSQETGGLNYAEQF | 0.55414814 |
| LGFALPIALPRRTVV | ASSQETGGLNYAEQF | 0.69524723 |
| TRLGNLTARKLVVLV | ASSQETGGLNYAEQF | 0.60696054 |
| WSQQRRLIYFPSAGP | ASSQETGGLNYAEQF | 0.6808615 |
| GPNGNPDPMAAAVDIR | ASSQETGGLNYAEQF | 0.712838 |
| TQRCVSVADLRAAASR | ASSQETGGLNYAEQF | 0.60527056 |
| QFVLDVAEHPVEVLTETV | ASSQETGGLNYAEQF | 0.7023605 |
| GGAGGTGGAGGNGGTQAGDG | ASSQETGGLNYAEQF | 0.7479711 |
| PLLATLRAIAVDVDAKPAQV | ASSQETGGLNYAEQF | 0.61983275 |
| APKFRTGGYMIDFWGVGYQVAK | ASSQETGGLNYAEQF | 0.72690976 |
| GAGGAGGAGGAAGGPGGTGGTG | ASSQETGGLNYAEQF | 0.7465329 |
| IGALGLLATDVAPAGSVAFGVA | ASSQETGGLNYAEQF | 0.7330018 |

| | | |
|---|---|---|
| AGGAGGIGGNGGAGGAGGVLMG | ASSQETGGLNYAEQF | 0.7491098 |
| ADELIGLL | ASSQETGGLNYAEQF | 0.71410394 |
| ADRALKA | ASSQETGGLNYAEQF | 0.7354159 |
| AGAVGQVGGAGGA | ASSQETGGLNYAEQF | 0.7148718 |
| AGGGGGLGGAP | ASSQETGGLNYAEQF | 0.70640796 |
| AGGLLYGNGGAG | ASSQETGGLNYAEQF | 0.7116193 |
| AGLESIGL | ASSQETGGLNYAEQF | 0.7018633 |
| AGLKTGII | ASSQETGGLNYAEQF | 0.686079 |
| ASLAALLL | ASSQETGGLNYAEQF | 0.7440856 |
| ASVINISSI | ASSQETGGLNYAEQF | 0.71170706 |
| ATPPGPA | ASSQETGGLNYAEQF | 0.7293234 |
| ATVASALV | ASSQETGGLNYAEQF | 0.6371757 |
| ATVASLIL | ASSQETGGLNYAEQF | 0.6596644 |
| AVNGPVLIRH | ASSQETGGLNYAEQF | 0.73813015 |
| DGAAGAVGI | ASSQETGGLNYAEQF | 0.27173066 |
| DPLLHGI | ASSQETGGLNYAEQF | 0.4464228 |
| ELGAALGAL | ASSQETGGLNYAEQF | 0.7215308 |
| ELGRTQAV | ASSQETGGLNYAEQF | 0.7435967 |
| FVAGGYLS | ASSQETGGLNYAEQF | 0.28944254 |
| GASGGAGGAGGW | ASSQETGGLNYAEQF | 0.66097033 |
| GGAGAGGDGGAGGI | ASSQETGGLNYAEQF | 0.59028006 |
| GTGHNGSQPGMGGNG | ASSQETGGLNYAEQF | 0.7461508 |
| GYAAASASAS | ASSQETGGLNYAEQF | 0.71816844 |
| HQVDVAAVQQG | ASSQETGGLNYAEQF | 0.51237726 |
| IGAAIAHT | ASSQETGGLNYAEQF | 0.739391 |
| IGMIEVAQL | ASSQETGGLNYAEQF | 0.5686518 |
| IRLAPEHTSAMD | ASSQETGGLNYAEQF | 0.58967376 |
| IYNVSVL | ASSQETGGLNYAEQF | 0.743665 |
| KQGFLGM | ASSQETGGLNYAEQF | 0.73685306 |
| LESVITQL | ASSQETGGLNYAEQF | 0.6590561 |
| LGLGALVVV | ASSQETGGLNYAEQF | 0.7435287 |
| LIGNGGNGGSGGI | ASSQETGGLNYAEQF | 0.66916656 |
| LNAINAPTL | ASSQETGGLNYAEQF | 0.75013775 |
| LQLLNTGV | ASSQETGGLNYAEQF | 0.68558776 |
| LQTSIEF | ASSQETGGLNYAEQF | 0.61583245 |
| MSFAVSEL | ASSQETGGLNYAEQF | 0.7359444 |
| NTGALGL | ASSQETGGLNYAEQF | 0.72025275 |
| PPGAGGAGGAGTT | ASSQETGGLNYAEQF | 0.7244602 |
| QLVGLGI | ASSQETGGLNYAEQF | 0.7181187 |
| RFLEQVHQL | ASSQETGGLNYAEQF | 0.6884436 |
| RGVGYVLR | ASSQETGGLNYAEQF | 0.69648206 |
| SATGVFAGVI | ASSQETGGLNYAEQF | 0.74490064 |
| SLPAITLP | ASSQETGGLNYAEQF | 0.74369717 |
| STALAFTV | ASSQETGGLNYAEQF | 0.6379446 |
| SVLTVGLGK | ASSQETGGLNYAEQF | 0.7466718 |
| TALAGAGSGI | ASSQETGGLNYAEQF | 0.7079787 |
| TGLPFTGI | ASSQETGGLNYAEQF | 0.66937953 |
| VAEHVTV | ASSQETGGLNYAEQF | 0.74708396 |
| VDIVLNSL | ASSQETGGLNYAEQF | 0.72034895 |
| VLTGGGAVL | ASSQETGGLNYAEQF | 0.7399123 |
| VNVGLAYGI | ASSQETGGLNYAEQF | 0.7342565 |

| | | |
|---|---|---|
| VSSAAVGAV | ASSQETGGLNYAEQF | 0.72814626 |
| VTSVYEAGV | ASSQETGGLNYAEQF | 0.57900035 |
| WVALDRGA | ASSQETGGLNYAEQF | 0.7145764 |
| YLVGSVVI | ASSQETGGLNYAEQF | 0.7094461 |
| GPPGTGKTLL | ASSQETGGLNYAEQF | 0.74070436 |
| GVGMTKF | ASSQETGGLNYAEQF | 0.71084076 |
| IGIGGSDL | ASSQETGGLNYAEQF | 0.5614929 |
| IGTLLAEL | ASSQETGGLNYAEQF | 0.72694033 |
| IYTSGTT | ASSQETGGLNYAEQF | 0.75131977 |
| LSFPAGV | ASSQETGGLNYAEQF | 0.72969615 |
| LYTSGTT | ASSQETGGLNYAEQF | 0.7293777 |
| VGAFPFGF | ASSQETGGLNYAEQF | 0.7505039 |
| VGAGTVEF | ASSQETGGLNYAEQF | 0.66849804 |
| VGKSTLLL | ASSQETGGLNYAEQF | 0.72129697 |
| AALAHAGGGIV | ASSQETGGLNYAEQF | 0.7448124 |
| AALVAVSLL | ASSQETGGLNYAEQF | 0.7460016 |
| AAVSRAVAAL | ASSQETGGLNYAEQF | 0.7199869 |
| AGGSSGIGGF | ASSQETGGLNYAEQF | 0.7462249 |
| AGLTVANANG | ASSQETGGLNYAEQF | 0.5993556 |
| AGLVGAVL | ASSQETGGLNYAEQF | 0.73894095 |
| ALSQAFTV | ASSQETGGLNYAEQF | 0.61811346 |
| ALTFSVSL | ASSQETGGLNYAEQF | 0.73176765 |
| ALTGAALVAL | ASSQETGGLNYAEQF | 0.6091393 |
| ASFSIGAGVL | ASSQETGGLNYAEQF | 0.7364761 |
| DGSVYKI | ASSQETGGLNYAEQF | 0.7357366 |
| EQLAEIV | ASSQETGGLNYAEQF | 0.74597275 |
| FAVSELTVM | ASSQETGGLNYAEQF | 0.4642405 |
| GAAPLAAGL | ASSQETGGLNYAEQF | 0.73868084 |
| GADGGDGGSGGAGG | ASSQETGGLNYAEQF | 0.73996305 |
| GFGLDTGGAGG | ASSQETGGLNYAEQF | 0.7524405 |
| GLASGGLL | ASSQETGGLNYAEQF | 0.6975538 |
| GRIAGAAVLL | ASSQETGGLNYAEQF | 0.7360278 |
| GSAGSLNTSF | ASSQETGGLNYAEQF | 0.6848612 |
| GTGQIGI | ASSQETGGLNYAEQF | 0.7088922 |
| GTLIALIL | ASSQETGGLNYAEQF | 0.6389818 |
| IAATMGFVVG | ASSQETGGLNYAEQF | 0.65318066 |
| IFAQALL | ASSQETGGLNYAEQF | 0.6910727 |
| IGAAVALGF | ASSQETGGLNYAEQF | 0.6513202 |
| IGLTPLAL | ASSQETGGLNYAEQF | 0.7297956 |
| ILGSNTSTL | ASSQETGGLNYAEQF | 0.7431272 |
| IVFSLTVL | ASSQETGGLNYAEQF | 0.7371831 |
| LASKATGF | ASSQETGGLNYAEQF | 0.7457345 |
| LASVALIL | ASSQETGGLNYAEQF | 0.7463819 |
| LFAVGLTL | ASSQETGGLNYAEQF | 0.7484358 |
| LFNGGIGTYI | ASSQETGGLNYAEQF | 0.679228 |
| LTLGAGII | ASSQETGGLNYAEQF | 0.697013 |
| LVAAALGDI | ASSQETGGLNYAEQF | 0.6298287 |
| NTAFVQLGI | ASSQETGGLNYAEQF | 0.7024597 |
| QLFSGAEL | ASSQETGGLNYAEQF | 0.6897473 |
| SAATFITL | ASSQETGGLNYAEQF | 0.72697103 |
| SNYGLANLGV | ASSQETGGLNYAEQF | 0.25410703 |

| | | |
|---|---|---|
| SYGGAYAVM | ASSQETGGLNYAEQF | 0.6875986 |
| TTGVGGLTLG | ASSQETGGLNYAEQF | 0.7554664 |
| TTQTIYI | ASSQETGGLNYAEQF | 0.57876396 |
| VAAGAGLGAVI | ASSQETGGLNYAEQF | 0.72859824 |
| VAAVVTLL | ASSQETGGLNYAEQF | 0.701124 |
| VGFLGGLI | ASSQETGGLNYAEQF | 0.6778462 |
| VGVIPILL | ASSQETGGLNYAEQF | 0.739562 |
| VIFASSLI | ASSQETGGLNYAEQF | 0.67480266 |
| VKGGDGGAAAG | ASSQETGGLNYAEQF | 0.7412546 |
| VTKTFTNI | ASSQETGGLNYAEQF | 0.74635243 |
| VVAAAILL | ASSQETGGLNYAEQF | 0.7220322 |
| YAALDVEL | ASSQETGGLNYAEQF | 0.6989114 |
| LASVALLL | ASSQETGGLNYAEQF | 0.7282895 |
| LDAVHRV | ASSQETGGLNYAEQF | 0.5067501 |
| QGLSSSIF | ASSQETGGLNYAEQF | 0.7212837 |
| RGTFANIRL | ASSQETGGLNYAEQF | 0.34139073 |
| RQLWWGHRI | ASSQETGGLNYAEQF | 0.68205845 |
| YGPPGTGKTLL | ASSQETGGLNYAEQF | 0.6395273 |
| AALVGAPANVA | ASSQETGGLNYAEQF | 0.72208077 |
| AATFRRTE | ASSQETGGLNYAEQF | 0.7218047 |
| AGALGGGI | ASSQETGGLNYAEQF | 0.7312518 |
| AGGAGGAGGIGAG | ASSQETGGLNYAEQF | 0.60781205 |
| AGSGALGL | ASSQETGGLNYAEQF | 0.6925111 |
| AQIADQI | ASSQETGGLNYAEQF | 0.65782374 |
| ASAVAVTYL | ASSQETGGLNYAEQF | 0.60316813 |
| ASTVLGGTGG | ASSQETGGLNYAEQF | 0.72586685 |
| AVGGEPAK | ASSQETGGLNYAEQF | 0.72554815 |
| CEQGTFGGTC | ASSQETGGLNYAEQF | 0.6645888 |
| DEGKAPAG | ASSQETGGLNYAEQF | 0.746372 |
| DTVVFPAL | ASSQETGGLNYAEQF | 0.7254965 |
| FEGITMAG | ASSQETGGLNYAEQF | 0.5629872 |
| FPLVVQD | ASSQETGGLNYAEQF | 0.65583766 |
| GAFTTPQ | ASSQETGGLNYAEQF | 0.7127316 |
| GFGGFGTAGGVGGA | ASSQETGGLNYAEQF | 0.6895503 |
| GFLAQFL | ASSQETGGLNYAEQF | 0.27127996 |
| GGAGGAGGNAGVA | ASSQETGGLNYAEQF | 0.72043383 |
| GGVGTAGAGGAGGAGG | ASSQETGGLNYAEQF | 0.6711609 |
| GNGGGGDAGNAGSGGN | ASSQETGGLNYAEQF | 0.60147524 |
| GTGGAGGAGNDAGS | ASSQETGGLNYAEQF | 0.6896907 |
| GTTNTGWFNS | ASSQETGGLNYAEQF | 0.7378684 |
| IFMGPLFTL | ASSQETGGLNYAEQF | 0.58825624 |
| ILAHAIEAI | ASSQETGGLNYAEQF | 0.68971884 |
| IRSAIREAL | ASSQETGGLNYAEQF | 0.7202077 |
| ITFAGGG | ASSQETGGLNYAEQF | 0.7421845 |
| ITGMSGHQEV | ASSQETGGLNYAEQF | 0.753538 |
| IVAGLACTL | ASSQETGGLNYAEQF | 0.72721577 |
| IVGGGLAGA | ASSQETGGLNYAEQF | 0.6844245 |
| IVTGVLNI | ASSQETGGLNYAEQF | 0.7368034 |
| LAQQGCI | ASSQETGGLNYAEQF | 0.73168314 |
| LCLHVEN | ASSQETGGLNYAEQF | 0.57775885 |
| LDAISDGS | ASSQETGGLNYAEQF | 0.50052524 |

| | | |
|---|---|---|
| LGGFAAML | ASSQETGGLNYAEQF | 0.678921 |
| LGGKNPMI | ASSQETGGLNYAEQF | 0.5630928 |
| LGPKGVAAIL | ASSQETGGLNYAEQF | 0.7158122 |
| LMVGVSI | ASSQETGGLNYAEQF | 0.7530172 |
| LVGAAFVL | ASSQETGGLNYAEQF | 0.6824474 |
| LVLAVAPL | ASSQETGGLNYAEQF | 0.60690236 |
| MAAIAVIDAA | ASSQETGGLNYAEQF | 0.6868898 |
| NNGINAELQ | ASSQETGGLNYAEQF | 0.65338945 |
| RGEPLVLVHG | ASSQETGGLNYAEQF | 0.73574513 |
| RGGAGGAAGLI | ASSQETGGLNYAEQF | 0.69033575 |
| RSYRMHPAV | ASSQETGGLNYAEQF | 0.74739575 |
| SAAHLLAGQLA | ASSQETGGLNYAEQF | 0.7395208 |
| SAAVGSY | ASSQETGGLNYAEQF | 0.6844157 |
| SPHAVLHRI | ASSQETGGLNYAEQF | 0.73784244 |
| SQIAKVL | ASSQETGGLNYAEQF | 0.678185 |
| TALGHLVP | ASSQETGGLNYAEQF | 0.69867176 |
| TAPTTSGAA | ASSQETGGLNYAEQF | 0.71407825 |
| TDGDDGNGGSPGT | ASSQETGGLNYAEQF | 0.72029054 |
| TDGLGEN | ASSQETGGLNYAEQF | 0.71460253 |
| TGGPPLPGQA | ASSQETGGLNYAEQF | 0.7100625 |
| TIPAITI | ASSQETGGLNYAEQF | 0.4904846 |
| TIPGFTI | ASSQETGGLNYAEQF | 0.7284108 |
| TSIASGALA | ASSQETGGLNYAEQF | 0.62582284 |
| TVTLPGAAI | ASSQETGGLNYAEQF | 0.5801916 |
| VASGLSAALP | ASSQETGGLNYAEQF | 0.66385776 |
| VGAGVIGI | ASSQETGGLNYAEQF | 0.7525829 |
| VGPKRFGV | ASSQETGGLNYAEQF | 0.7483234 |
| VNVAFPD | ASSQETGGLNYAEQF | 0.6402757 |
| VSMTNSGVS | ASSQETGGLNYAEQF | 0.64719605 |
| VTDVLGDLSV | ASSQETGGLNYAEQF | 0.6926997 |
| VVVVGAGFAGL | ASSQETGGLNYAEQF | 0.56798446 |
| VYTVGTVV | ASSQETGGLNYAEQF | 0.7344898 |
| WTTFSGP | ASSQETGGLNYAEQF | 0.69470644 |
| YDELVEAF | ASSQETGGLNYAEQF | 0.7056921 |
| YGTGGAGGNGG | ASSQETGGLNYAEQF | 0.6942725 |
| YTNVMAV | ASSQETGGLNYAEQF | 0.7195918 |
| FLEQVHQL | ASSQETGGLNYAEQF | 0.6765643 |
| GLSLLAVAV | ASSQETGGLNYAEQF | 0.74116176 |
| LEQVHQL | ASSQETGGLNYAEQF | 0.28742304 |
| LNAAGAI | ASSQETGGLNYAEQF | 0.729738 |
| PAVHASL | ASSQETGGLNYAEQF | 0.6610794 |
| TELEDDA | ASSQETGGLNYAEQF | 0.66327596 |
| TPPGPPN | ASSQETGGLNYAEQF | 0.54643065 |
| TPPPPPPP | ASSQETGGLNYAEQF | 0.72015136 |
| TQSPALI | ASSGQYEQY | 0.7173917 |
| EFAAAFPQ | ASSGQYEQY | 0.7397105 |
| LLSGNLDVL | ASSGQYEQY | 0.604246 |
| GPANLGNYNI | ASSGQYEQY | 0.7420649 |
| TLGPTLP | ASSGQYEQY | 0.7462565 |
| VYAYSSCCK | ASSGQYEQY | 0.6905329 |
| APAPAPAPNI | ASSGQYEQY | 0.47049916 |

| | | |
|---|---|---|
| PGGPVPIP | ASSGQYEQY | 0.7324941 |
| FLTVALEP | ASSGQYEQY | 0.67203444 |
| SQYFGANL | ASSGQYEQY | 0.7343691 |
| GGVKPAA | ASSGQYEQY | 0.70372784 |
| GGTGGAGGAGGW | ASSGQYEQY | 0.6096387 |
| GVGGVAGLGGAGG | ASSGQYEQY | 0.7443266 |
| VLLVLATTI | ASSGQYEQY | 0.7424235 |
| IGIFAAML | ASSGQYEQY | 0.64667153 |
| AVGDLID | ASSGQYEQY | 0.68529516 |
| NISPLPL | ASSGQYEQY | 0.70271385 |
| VTAVFVAI | ASSGQYEQY | 0.46612138 |
| SGDDNDAAL | ASSGQYEQY | 0.73948854 |
| AVAALAHS | ASSGQYEQY | 0.6581455 |
| GGVTVMATP | ASSGQYEQY | 0.69029975 |
| AGLAANVRD | ASSGQYEQY | 0.7273302 |
| TNRPDLIDPALLRPG | ASSGQYEQY | 0.7019628 |
| LANLGVTAL | ASSGQYEQY | 0.7046901 |
| TGERIEL | ASSGQYEQY | 0.6979101 |
| NGEELAA | ASSGQYEQY | 0.70538026 |
| VLSLVWG | ASSGQYEQY | 0.72153413 |
| IGYALWL | ASSGQYEQY | 0.7125524 |
| LSLLLAPL | ASSGQYEQY | 0.71980625 |
| TLDEALK | ASSGQYEQY | 0.7534383 |
| TDGAGTAEGG | ASSGQYEQY | 0.57590437 |
| NAPGGAGGNGG | ASSGQYEQY | 0.6964258 |
| VSVFMGMGGNF | ASSGQYEQY | 0.729145 |
| LASYLAAL | ASSGQYEQY | 0.57857233 |
| LVGGTPAL | ASSGQYEQY | 0.7411177 |
| VRNADVL | ASSGQYEQY | 0.7186359 |
| TCSSPFLVSAM | ASSGQYEQY | 0.7249443 |
| VHITDAE | ASSGQYEQY | 0.6691614 |
| EYAAPSF | ASSGQYEQY | 0.7552844 |
| VVDAFHGGL | ASSGQYEQY | 0.74051756 |
| ASVVDPQ | ASSGQYEQY | 0.72327954 |
| SYNEFGGLS | ASSGQYEQY | 0.73124593 |
| DPDGSVY | ASSGQYEQY | 0.6797793 |
| MVSTIHATE | ASSGQYEQY | 0.7442497 |
| LAAGVADPV | ASSGQYEQY | 0.7337566 |
| TKEQIAA | ASSGQYEQY | 0.59417474 |
| SLASPAAL | ASSGQYEQY | 0.6962484 |
| IGLVFYGV | ASSGQYEQY | 0.7512742 |
| FLLDASNL | ASSGQYEQY | 0.7459255 |
| YQVGNFP | ASSGQYEQY | 0.5004672 |
| ESLLSQST | ASSGQYEQY | 0.694136 |
| LFTVGFAAVG | ASSGQYEQY | 0.5737707 |
| ATPPTPT | ASSGQYEQY | 0.6552415 |
| TDLATLG | ASSGQYEQY | 0.65383375 |
| SAGGTGR | ASSGQYEQY | 0.72417736 |
| GFSNSGS | ASSGQYEQY | 0.69832087 |
| GGNGGSGGTSV | ASSGQYEQY | 0.7088158 |
| GTSVATGGAGN | ASSGQYEQY | 0.7175938 |

| | | |
|---|---|---|
| NGGNGGSGGTS | ASSGQYEQY | 0.7447907 |
| AVVKFPH | ASSGQYEQY | 0.669611 |
| TPPPPPP | ASSGQYEQY | 0.70893687 |
| AAPAAAAAPA | ASSGQYEQY | 0.56608677 |
| IALNEDL | ASSGQYEQY | 0.42421153 |
| IVNVSSM | ASSGQYEQY | 0.6836659 |
| TGIPPAPR | ASSGQYEQY | 0.6860787 |
| FDLGGGTF | ASSGQYEQY | 0.35813525 |
| FDLGGGTFDVS | ASSGQYEQY | 0.75101984 |
| NEPTAAA | ASSGQYEQY | 0.7211107 |
| NEPTAAALA | ASSGQYEQY | 0.73052764 |
| GSIINISSI | ASSGQYEQY | 0.7045554 |
| VAPGNAGTA | ASSGQYEQY | 0.7193708 |
| TGGGASLE | ASSGQYEQY | 0.6778205 |
| VAVYNHYKRI | ASSGQYEQY | 0.7358405 |
| TDPAPATT | ASSGQYEQY | 0.6275375 |
| TNRPDILDPALLRPG | ASSGQYEQY | 0.57039946 |
| LTTTTTS | ASSGQYEQY | 0.6998545 |
| PTGSGKTL | ASSGQYEQY | 0.62792397 |
| TGSGKTL | ASSGQYEQY | 0.7403813 |
| TIGHVDHGK | ASSGQYEQY | 0.7414331 |
| TIGHVDHGKTT | ASSGQYEQY | 0.72238845 |
| GHVDHGKTT | ASSGQYEQY | 0.7391158 |
| AQFEHTLL | ASSGQYEQY | 0.35760498 |
| IVNQDEL | ASSGQYEQY | 0.7063649 |
| TNGSQFF | ASSGQYEQY | 0.75174767 |
| TGKGIVE | ASSGQYEQY | 0.723245 |
| LISNASDA | ASSGQYEQY | 0.7082572 |
| AELRTLGELTKIAKAHGAQ | ASSGQYEQY | 0.75140256 |
| APNAVAAAEPSW | ASSGQYEQY | 0.67041016 |
| APQVAGAATAVL | ASSGQYEQY | 0.68915683 |
| ARRDPAAAAAILPTDA | ASSGQYEQY | 0.7203419 |
| DAAARDVLAEAGLAEYF | ASSGQYEQY | 0.73744535 |
| DDQIVLSAKANLAKVEQ | ASSGQYEQY | 0.7253621 |
| DELLPDTGEDSDEDGH | ASSGQYEQY | 0.6352304 |
| ELPEAPANRATVPQD | ASSGQYEQY | 0.7359401 |
| FGQSKIERVTGIG | ASSGQYEQY | 0.7585405 |
| GAGGTGSTAGGAGGAGGAGGLY | ASSGQYEQY | 0.6760787 |
| GHGGTGAAGGNGGTGGAGGSGI | ASSGQYEQY | 0.73177624 |
| GNGGNGGAGGEGGPGGAGGAGA | ASSGQYEQY | 0.7357651 |
| GSTAGNGGNGGNATNGTIAGQP | ASSGQYEQY | 0.60597277 |
| GTGGDGGNGGNGGTGG | ASSGQYEQY | 0.6527643 |
| GVRRPAGIPADAPV | ASSGQYEQY | 0.67403424 |
| GYRQVTDAHLLTLARR | ASSGQYEQY | 0.6901909 |
| IAKMGQDGHDRG | ASSGQYEQY | 0.6599655 |
| KAEPSIEIQYT | ASSGQYEQY | 0.7223045 |
| KPVRIRMAADEDTVAVT | ASSGQYEQY | 0.70945096 |
| MPPRIAGMRLLVIKPEP | ASSGQYEQY | 0.756497 |
| MKFYAVALFRSRGVPPTR | ASSGQYEQY | 0.73063874 |
| NAGGFGILGAVAHSP | ASSGQYEQY | 0.73455876 |
| NEGREITLAMAEM | ASSGQYEQY | 0.64078367 |

| | | |
|---|---|---|
| NELVEWSHPLGHRWRWE | ASSGQYEQY | 0.725887 |
| NLGAGGTGGNGGNPG | ASSGQYEQY | 0.723452 |
| NTEWPKWETVMAADPVD | ASSGQYEQY | 0.5368937 |
| PASAAPAETQP | ASSGQYEQY | 0.7519841 |
| QQSSDGKNGVRAAEITGE | ASSGQYEQY | 0.6823714 |
| RAPDAVTTSWWKE | ASSGQYEQY | 0.6731856 |
| SGFGNVGAHTSGWFNQ | ASSGQYEQY | 0.74494696 |
| SGVAGAVGQAASVGGLKV | ASSGQYEQY | 0.73398304 |
| SNGGAGTGSAGGNGGT | ASSGQYEQY | 0.72088516 |
| TGGAGGEGGAAG | ASSGQYEQY | 0.6959526 |
| TGGDGGAAGAGGAPGHG | ASSGQYEQY | 0.6775805 |
| TMPPLLAAIANAGRHGV | ASSGQYEQY | 0.7370875 |
| TSSTAGNSGGA | ASSGQYEQY | 0.742024 |
| VPATVPVGILC | ASSGQYEQY | 0.63270646 |
| VQRPDSGGVVLTGRIS | ASSGQYEQY | 0.7398255 |
| WTPNPVPATKGVPAGLVQAT | ASSGQYEQY | 0.73082405 |
| YALKRALAKFSDRFDVVII | ASSGQYEQY | 0.57027483 |
| YLIGSVIV | ASSGQYEQY | 0.70697975 |
| YGQLVAVID | ASSGQYEQY | 0.73944646 |
| VTVGGMTLPA | ASSGQYEQY | 0.7346859 |
| VLALPAV | ASSGQYEQY | 0.7493323 |
| VIPPTLNYET | ASSGQYEQY | 0.7333014 |
| VGMGGMHPSSG | ASSGQYEQY | 0.61617744 |
| VGLAGGAATA | ASSGQYEQY | 0.7511893 |
| VGGAGGAGGSGP | ASSGQYEQY | 0.7501128 |
| VFAAFMV | ASSGQYEQY | 0.7398559 |
| TLYDKLR | ASSGQYEQY | 0.6743458 |
| TLAVFTVL | ASSGQYEQY | 0.66344625 |
| TGSVQPARV | ASSGQYEQY | 0.73054844 |
| TGGGAGSGGT | ASSGQYEQY | 0.7335802 |
| SHLETSL | ASSGQYEQY | 0.6705815 |
| SFTTGMT | ASSGQYEQY | 0.72137034 |
| QVEVAKRMGVRQPTVS | ASSGQYEQY | 0.71910036 |
| QGSGGPALAM | ASSGQYEQY | 0.5653986 |
| MLAPSNP | ASSGQYEQY | 0.7259551 |
| LVNNAAI | ASSGQYEQY | 0.7463722 |
| LTGEINTLA | ASSGQYEQY | 0.7252458 |
| LSIFGPV | ASSGQYEQY | 0.6821184 |
| LLALLANL | ASSGQYEQY | 0.7474915 |
| LISNASDALDKLR | ASSGQYEQY | 0.75184417 |
| LINEAVP | ASSGQYEQY | 0.7498186 |
| LGYAFSGL | ASSGQYEQY | 0.61814183 |
| LDFVAVAL | ASSGQYEQY | 0.48402566 |
| LAQPGAE | ASSGQYEQY | 0.68764716 |
| ISNLGAVTGF | ASSGQYEQY | 0.71882254 |
| HQEDPGRI | ASSGQYEQY | 0.6992583 |
| GTPGEPTP | ASSGQYEQY | 0.66577524 |
| GFFNSGS | ASSGQYEQY | 0.7183864 |
| GAGGGGLTAGAN | ASSGQYEQY | 0.7492594 |
| FVDALVNM | ASSGQYEQY | 0.66000867 |
| FLTVAPEL | ASSGQYEQY | 0.52247924 |

| | | |
|---|---|---|
| FETLTAL | ASSGQYEQY | 0.72938704 |
| DVINPIEL | ASSGQYEQY | 0.73711383 |
| APVSGAPV | ASSGQYEQY | 0.6727773 |
| ALVGGLAW | ASSGQYEQY | 0.6760998 |
| ALAAAAATAI | ASSGQYEQY | 0.691535 |
| AFGLGLLL | ASSGQYEQY | 0.63174 |
| AFFAAASL | ASSGQYEQY | 0.7156361 |
| AAAALPAFSPP | ASSGQYEQY | 0.635934 |
| VVLAPAL | ASSGQYEQY | 0.74923825 |
| TIGHVDHGKTTL | ASSGQYEQY | 0.6600773 |
| LISNASDALDK | ASSGQYEQY | 0.7407882 |
| LISNASDAL | ASSGQYEQY | 0.7281023 |
| ISNASDALDK | ASSGQYEQY | 0.7260669 |
| NQAETLVYQTE | ASSGQYEQY | 0.6961243 |
| TGGTGGTAGAG | ASSGQYEQY | 0.65887034 |
| TGSTAGGAGGA | ASSGQYEQY | 0.5973393 |
| DPLAVPVAAGPV | ASSGQYEQY | 0.5328966 |
| GGTGGTPGNGGN | ASSGQYEQY | 0.7490371 |
| PAGLAGVDPATL | ASSGQYEQY | 0.45453125 |
| GNNLVVEAPHAGASV | ASSGQYEQY | 0.74731046 |
| KVTRIMSASTLEQPA | ASSGQYEQY | 0.7354573 |
| LGFALPIALPRRTVV | ASSGQYEQY | 0.67909145 |
| TRLGNLTARKLVVLV | ASSGQYEQY | 0.7334119 |
| WSQQRRLIYFPSAGP | ASSGQYEQY | 0.59693795 |
| GPNGNPDPMAAAVDIR | ASSGQYEQY | 0.63208205 |
| TQRCVSVADLRAAASR | ASSGQYEQY | 0.7147257 |
| QFVLDVAEHPVEVLTETV | ASSGQYEQY | 0.73852324 |
| GGAGGTGGAGGNGGTQAGDG | ASSGQYEQY | 0.6749803 |
| PLLATLRAIAVDVDAKPAQV | ASSGQYEQY | 0.7365193 |
| APKFRTGGYMIDFWGVGYQVAK | ASSGQYEQY | 0.7186411 |
| GAGGAGGAGGAAGGPGGTGGTG | ASSGQYEQY | 0.66282207 |
| IGALGLLATDVAPAGSVAFGVA | ASSGQYEQY | 0.72860014 |
| AGGAGGIGGNGGAGGAGGVLMG | ASSGQYEQY | 0.7444283 |
| ADELIGLL | ASSGQYEQY | 0.73904157 |
| ADRALKA | ASSGQYEQY | 0.63607216 |
| AGAVGQVGGAGGA | ASSGQYEQY | 0.7347143 |
| AGGGGGLGGAP | ASSGQYEQY | 0.6474784 |
| AGGLLYGNGGAG | ASSGQYEQY | 0.7227793 |
| AGLESIGL | ASSGQYEQY | 0.72269183 |
| AGLKTGII | ASSGQYEQY | 0.7127665 |
| ASLAALLL | ASSGQYEQY | 0.7357044 |
| ASVINISSI | ASSGQYEQY | 0.6213191 |
| ATPPGPA | ASSGQYEQY | 0.6156468 |
| ATVASALV | ASSGQYEQY | 0.69869417 |
| ATVASLIL | ASSGQYEQY | 0.7256625 |
| AVNGPVLIRH | ASSGQYEQY | 0.7419232 |
| DGAAGAVGI | ASSGQYEQY | 0.6361275 |
| DPLLHGI | ASSGQYEQY | 0.7132659 |
| ELGAALGAL | ASSGQYEQY | 0.69347966 |
| ELGRTQAV | ASSGQYEQY | 0.6290724 |
| FVAGGYLS | ASSGQYEQY | 0.73178947 |

| | | |
|---|---|---|
| GASGGAGGAGGW | ASSGQYEQY | 0.7088245 |
| GGAGAGGDGGAGGI | ASSGQYEQY | 0.6608156 |
| GTGHNGSQPGMGGNG | ASSGQYEQY | 0.7378619 |
| GYAAASASAS | ASSGQYEQY | 0.65695864 |
| HQVDVAAVQQG | ASSGQYEQY | 0.7337164 |
| IGAAIAHT | ASSGQYEQY | 0.7499435 |
| IGMIEVAQL | ASSGQYEQY | 0.6576999 |
| IRLAPEHTSAMD | ASSGQYEQY | 0.73964417 |
| IYNVSVL | ASSGQYEQY | 0.71418244 |
| KQGFLGM | ASSGQYEQY | 0.7277082 |
| LESVITQL | ASSGQYEQY | 0.6952918 |
| LGLGALVVV | ASSGQYEQY | 0.6535078 |
| LIGNGGNGGSGGI | ASSGQYEQY | 0.74700683 |
| LNAINAPTL | ASSGQYEQY | 0.74162066 |
| LQLLNTGV | ASSGQYEQY | 0.7449236 |
| LQTSIEF | ASSGQYEQY | 0.7280211 |
| MSFAVSEL | ASSGQYEQY | 0.7291872 |
| NTGALGL | ASSGQYEQY | 0.7403383 |
| PPGAGGAGGAGTT | ASSGQYEQY | 0.6683639 |
| QLVGLGI | ASSGQYEQY | 0.57611716 |
| RFLEQVHQL | ASSGQYEQY | 0.69319534 |
| RGVGYVLR | ASSGQYEQY | 0.69890237 |
| SATGVFAGVI | ASSGQYEQY | 0.71742344 |
| SLPAITLP | ASSGQYEQY | 0.59762836 |
| STALAFTV | ASSGQYEQY | 0.7462635 |
| SVLTVGLGK | ASSGQYEQY | 0.75679314 |
| TALAGAGSGI | ASSGQYEQY | 0.68286717 |
| TGLPFTGI | ASSGQYEQY | 0.7510807 |
| VAEHVTV | ASSGQYEQY | 0.6152942 |
| VDIVLNSL | ASSGQYEQY | 0.7385465 |
| VLTGGGAVL | ASSGQYEQY | 0.7527218 |
| VNVGLAYGI | ASSGQYEQY | 0.6412076 |
| VSSAAVGAV | ASSGQYEQY | 0.7379447 |
| VTSVYEAGV | ASSGQYEQY | 0.6677745 |
| WVALDRGA | ASSGQYEQY | 0.7017961 |
| YLVGSVVI | ASSGQYEQY | 0.7490266 |
| GPPGTGKTLL | ASSGQYEQY | 0.69406354 |
| GVGMTKF | ASSGQYEQY | 0.7367338 |
| IGIGGSDL | ASSGQYEQY | 0.70490634 |
| IGTLLAEL | ASSGQYEQY | 0.66158 |
| IYTSGTT | ASSGQYEQY | 0.7318077 |
| LSFPAGV | ASSGQYEQY | 0.7509726 |
| LYTSGTT | ASSGQYEQY | 0.6393971 |
| VGAFPFGF | ASSGQYEQY | 0.75103533 |
| VGAGTVEF | ASSGQYEQY | 0.68840086 |
| VGKSTLLL | ASSGQYEQY | 0.75628835 |
| AALAHAGGGIV | ASSGQYEQY | 0.7401528 |
| AALVAVSLL | ASSGQYEQY | 0.7010809 |
| AAVSRAVAAL | ASSGQYEQY | 0.70463216 |
| AGGSSGIGGF | ASSGQYEQY | 0.7448374 |
| AGLTVANANG | ASSGQYEQY | 0.6760684 |

| | | |
|---|---|---|
| AGLVGAVL | ASSGQYEQY | 0.73722994 |
| ALSQAFTV | ASSGQYEQY | 0.7491313 |
| ALTFSVSL | ASSGQYEQY | 0.37004974 |
| ALTGAALVAL | ASSGQYEQY | 0.63600254 |
| ASFSIGAGVL | ASSGQYEQY | 0.5871351 |
| DGSVYKI | ASSGQYEQY | 0.622013 |
| EQLAEIV | ASSGQYEQY | 0.46793535 |
| FAVSELTVM | ASSGQYEQY | 0.38394007 |
| GAAPLAAGL | ASSGQYEQY | 0.535956 |
| GADGGDGGSGGAGG | ASSGQYEQY | 0.42526624 |
| GFGLDTGGAGG | ASSGQYEQY | 0.65608424 |
| GLASGGLL | ASSGQYEQY | 0.6297803 |
| GRIAGAAVLL | ASSGQYEQY | 0.44237188 |
| GSAGSLNTSF | ASSGQYEQY | 0.5823313 |
| GTGQIGI | ASSGQYEQY | 0.63539577 |
| GTLIALIL | ASSGQYEQY | 0.3653607 |
| IAATMGFVVG | ASSGQYEQY | 0.53466415 |
| IFAQALL | ASSGQYEQY | 0.51677275 |
| IGAAVALGF | ASSGQYEQY | 0.49676913 |
| IGLTPLAL | ASSGQYEQY | 0.32786575 |
| ILGSNTSTL | ASSGQYEQY | 0.60886765 |
| IVFSLTVL | ASSGQYEQY | 0.4778037 |
| LASKATGF | ASSGQYEQY | 0.31613156 |
| LASVALIL | ASSGQYEQY | 0.57755125 |
| LFAVGLTL | ASSGQYEQY | 0.46239918 |
| LFNGGIGTYI | ASSGQYEQY | 0.48790666 |
| LTLGAGII | ASSGQYEQY | 0.60783213 |
| LVAAALGDI | ASSGQYEQY | 0.60284734 |
| NTAFVQLGI | ASSGQYEQY | 0.47526133 |
| QLFSGAEL | ASSGQYEQY | 0.57745576 |
| SAATFITL | ASSGQYEQY | 0.6444566 |
| SNYGLANLGV | ASSGQYEQY | 0.42761123 |
| SYGGAYAVM | ASSGQYEQY | 0.54379433 |
| TTGVGGLTLG | ASSGQYEQY | 0.66954076 |
| TTQTIYI | ASSGQYEQY | 0.55599296 |
| VAAGAGLGAVI | ASSGQYEQY | 0.66609484 |
| VAAVVTLL | ASSGQYEQY | 0.6754437 |
| VGFLGGLI | ASSGQYEQY | 0.51382565 |
| VGVIPILL | ASSGQYEQY | 0.6349294 |
| VIFASSLI | ASSGQYEQY | 0.631606 |
| VKGGDGGAAAG | ASSGQYEQY | 0.54092777 |
| VTKTFTNI | ASSGQYEQY | 0.53076315 |
| VVAAAILL | ASSGQYEQY | 0.36785215 |
| YAALDVEL | ASSGQYEQY | 0.53276634 |
| LASVALLL | ASSGQYEQY | 0.42843288 |
| LDAVHRV | ASSGQYEQY | 0.5646701 |
| QGLSSSIF | ASSGQYEQY | 0.6545562 |
| RGTFANIRL | ASSGQYEQY | 0.5323298 |
| RQLWWGHRI | ASSGQYEQY | 0.33721036 |
| YGPPGTGKTLL | ASSGQYEQY | 0.36967683 |
| AALVGAPANVA | ASSGQYEQY | 0.57553583 |

| | | |
|---|---|---|
| AATFRRTE | ASSGQYEQY | 0.40032512 |
| AGALGGGI | ASSGQYEQY | 0.4295853 |
| AGGAGGAGGIGAG | ASSGQYEQY | 0.4269954 |
| AGSGALGL | ASSGQYEQY | 0.63301146 |
| AQIADQI | ASSGQYEQY | 0.64172536 |
| ASAVAVTYL | ASSGQYEQY | 0.46622747 |
| ASTVLGGTGG | ASSGQYEQY | 0.6410098 |
| AVGGEPAK | ASSGQYEQY | 0.65930676 |
| CEQGTFGGTC | ASSGQYEQY | 0.67018056 |
| DEGKAPAG | ASSGQYEQY | 0.57984376 |
| DTVVFPAL | ASSGQYEQY | 0.36593196 |
| FEGITMAG | ASSGQYEQY | 0.64536935 |
| FPLVVQD | ASSGQYEQY | 0.46835867 |
| GAFTTPQ | ASSGQYEQY | 0.6259059 |
| GFGGFGTAGGVGGA | ASSGQYEQY | 0.6128111 |
| GFLAQFL | ASSGQYEQY | 0.50260794 |
| GGAGGAGGNAGVA | ASSGQYEQY | 0.44358784 |
| GGVGTAGAGGAGGAGG | ASSGQYEQY | 0.60422724 |
| GNGGGGDAGNAGSGGN | ASSGQYEQY | 0.66951597 |
| GTGGAGGAGNDAGS | ASSGQYEQY | 0.39577335 |
| GTTNTGWFNS | ASSGQYEQY | 0.40884984 |
| IFMGPLFTL | ASSGQYEQY | 0.56516016 |
| ILAHAIEAI | ASSGQYEQY | 0.53152853 |
| IRSAIREAL | ASSGQYEQY | 0.6516157 |
| ITFAGGG | ASSGQYEQY | 0.61202437 |
| ITGMSGHQEV | ASSGQYEQY | 0.5570827 |
| IVAGLACTL | ASSGQYEQY | 0.58395106 |
| IVGGGLAGA | ASSGQYEQY | 0.62684137 |
| IVTGVLNI | ASSGQYEQY | 0.56088877 |
| LAQQGCI | ASSGQYEQY | 0.57344234 |
| LCLHVEN | ASSGQYEQY | 0.43246445 |
| LDAISDGS | ASSGQYEQY | 0.6564746 |
| LGGFAAML | ASSGQYEQY | 0.6343665 |
| LGGKNPMI | ASSGQYEQY | 0.6443336 |
| LGPKGVAAIL | ASSGQYEQY | 0.660813 |
| LMVGVSI | ASSGQYEQY | 0.5935013 |
| LVGAAFVL | ASSGQYEQY | 0.5868694 |
| LVLAVAPL | ASSGQYEQY | 0.42685184 |
| MAAIAVIDAA | ASSGQYEQY | 0.51743084 |
| NNGINAELQ | ASSGQYEQY | 0.51574206 |
| RGEPLVLVHG | ASSGQYEQY | 0.5741001 |
| RGGAGGAAGLI | ASSGQYEQY | 0.61511624 |
| RSYRMHPAV | ASSGQYEQY | 0.6510692 |
| SAAHLLAGQLA | ASSGQYEQY | 0.4609499 |
| SAAVGSY | ASSGQYEQY | 0.6513317 |
| SPHAVLHRI | ASSGQYEQY | 0.6535318 |
| SQIAKVL | ASSGQYEQY | 0.42071778 |
| TALGHLVP | ASSGQYEQY | 0.49714977 |
| TAPTTSGAA | ASSGQYEQY | 0.63339347 |
| TDGDDGNGGSPGT | ASSGQYEQY | 0.62128437 |
| TDGLGEN | ASSGQYEQY | 0.47522035 |

| | | |
|---|---|---|
| TGGPPLPGQA | ASSGQYEQY | 0.559561 |
| TIPAITI | ASSGQYEQY | 0.6641532 |
| TIPGFTI | ASSGQYEQY | 0.60175365 |
| TSIASGALA | ASSGQYEQY | 0.64605045 |
| TVTLPGAAI | ASSGQYEQY | 0.36910105 |
| VASGLSAALP | ASSGQYEQY | 0.5583914 |
| VGAGVIGI | ASSGQYEQY | 0.503284 |
| VGPKRFGV | ASSGQYEQY | 0.29856607 |
| VNVAFPD | ASSGQYEQY | 0.62391996 |
| VSMTNSGVS | ASSGQYEQY | 0.49475604 |
| VTDVLGDLSV | ASSGQYEQY | 0.59720397 |
| VVVVGAGFAGL | ASSGQYEQY | 0.45808005 |
| VYTVGTVV | ASSGQYEQY | 0.644883 |
| WTTFSGP | ASSGQYEQY | 0.576893 |
| YDELVEAF | ASSGQYEQY | 0.5346829 |
| YGTGGAGGNGG | ASSGQYEQY | 0.4518979 |
| YTNVMAV | ASSGQYEQY | 0.58072907 |
| FLEQVHQL | ASSGQYEQY | 0.6509025 |
| GLSLLAVAV | ASSGQYEQY | 0.6761504 |
| LEQVHQL | ASSGQYEQY | 0.37402058 |
| LNAAGAI | ASSGQYEQY | 0.6245123 |
| PAVHASL | ASSGQYEQY | 0.5552495 |
| TELEDDA | ASSGQYEQY | 0.40082112 |
| TPPGPPN | ASSGQYEQY | 0.33712295 |
| TPPPPPPP | ASSGQYEQY | 0.4569437 |
| TQSPALI | ASSETAESSYEQY | 0.48312712 |
| EFAAAFPQ | ASSETAESSYEQY | 0.55639976 |
| LLSGNLDVL | ASSETAESSYEQY | 0.6224442 |
| GPANLGNYNI | ASSETAESSYEQY | 0.47404554 |
| TLGPTLP | ASSETAESSYEQY | 0.60077846 |
| VYAYSSCCK | ASSETAESSYEQY | 0.5355936 |
| APAPAPAPNI | ASSETAESSYEQY | 0.27110788 |
| PGGPVPIP | ASSETAESSYEQY | 0.4742666 |
| FLTVALEP | ASSETAESSYEQY | 0.35337916 |
| SQYFGANL | ASSETAESSYEQY | 0.67568225 |
| GGVKPAA | ASSETAESSYEQY | 0.39175558 |
| GGTGGAGGAGGW | ASSETAESSYEQY | 0.48397756 |
| GVGGVAGLGGAGG | ASSETAESSYEQY | 0.5292738 |
| VLLVLATTI | ASSETAESSYEQY | 0.62779856 |
| IGIFAAML | ASSETAESSYEQY | 0.5525594 |
| AVGDLID | ASSETAESSYEQY | 0.47504485 |
| NISPLPL | ASSETAESSYEQY | 0.36038935 |
| VTAVFVAI | ASSETAESSYEQY | 0.6020262 |
| SGDDNDAAL | ASSETAESSYEQY | 0.5889312 |
| AVAALAHS | ASSETAESSYEQY | 0.4506544 |
| GGVTVMATP | ASSETAESSYEQY | 0.4736873 |
| AGLAANVRD | ASSETAESSYEQY | 0.5265139 |
| TNRPDLIDPALLRPG | ASSETAESSYEQY | 0.3533319 |
| LANLGVTAL | ASSETAESSYEQY | 0.6603294 |
| TGERIEL | ASSETAESSYEQY | 0.39891517 |
| NGEELAA | ASSETAESSYEQY | 0.47021657 |

| | | |
|---|---|---|
| VLSLVWG | ASSETAESSYEQY | 0.65583235 |
| IGYALWL | ASSETAESSYEQY | 0.60341024 |
| LSLLLAPL | ASSETAESSYEQY | 0.5016119 |
| TLDEALK | ASSETAESSYEQY | 0.48844266 |
| TDGAGTAEGG | ASSETAESSYEQY | 0.40552932 |
| NAPGGAGGNGG | ASSETAESSYEQY | 0.44112226 |
| VSVFMGMGGNF | ASSETAESSYEQY | 0.45732993 |
| LASYLAAL | ASSETAESSYEQY | 0.63765883 |
| LVGGTPAL | ASSETAESSYEQY | 0.5928438 |
| VRNADVL | ASSETAESSYEQY | 0.3966715 |
| TCSSPFLVSAM | ASSETAESSYEQY | 0.6480997 |
| VHITDAE | ASSETAESSYEQY | 0.6462572 |
| EYAAPSF | ASSETAESSYEQY | 0.6637435 |
| VVDAFHGGL | ASSETAESSYEQY | 0.4124467 |
| ASVVDPQ | ASSETAESSYEQY | 0.5444959 |
| SYNEFGGLS | ASSETAESSYEQY | 0.5496551 |
| DPDGSVY | ASSETAESSYEQY | 0.5051822 |
| MVSTIHATE | ASSETAESSYEQY | 0.47904357 |
| LAAGVADPV | ASSETAESSYEQY | 0.4487463 |
| TKEQIAA | ASSETAESSYEQY | 0.6540787 |
| SLASPAAL | ASSETAESSYEQY | 0.6952132 |
| IGLVFYGV | ASSETAESSYEQY | 0.52949834 |
| FLLDASNL | ASSETAESSYEQY | 0.32921126 |
| YQVGNFP | ASSETAESSYEQY | 0.50928104 |
| ESLLSQST | ASSETAESSYEQY | 0.43442756 |
| LFTVGFAAVG | ASSETAESSYEQY | 0.43098235 |
| ATPPTPT | ASSETAESSYEQY | 0.53737974 |
| TDLATLG | ASSETAESSYEQY | 0.3486038 |
| SAGGTGR | ASSETAESSYEQY | 0.52027136 |
| GFSNSGS | ASSETAESSYEQY | 0.66414285 |
| GGNGGSGGTSV | ASSETAESSYEQY | 0.520965 |
| GTSVATGGAGN | ASSETAESSYEQY | 0.5679922 |
| NGGNGGSGGTS | ASSETAESSYEQY | 0.62777877 |
| AVVKFPH | ASSETAESSYEQY | 0.57943505 |
| TPPPPPP | ASSETAESSYEQY | 0.6388339 |
| AAPAAAAAPA | ASSETAESSYEQY | 0.66245985 |
| IALNEDL | ASSETAESSYEQY | 0.56188637 |
| IVNVSSM | ASSETAESSYEQY | 0.56734574 |
| TGIPPAPR | ASSETAESSYEQY | 0.60574424 |
| FDLGGGTF | ASSETAESSYEQY | 0.49931383 |
| FDLGGGTFDVS | ASSETAESSYEQY | 0.54505324 |
| NEPTAAA | ASSETAESSYEQY | 0.55039424 |
| NEPTAAALA | ASSETAESSYEQY | 0.5702087 |
| GSIINISSI | ASSETAESSYEQY | 0.5455164 |
| VAPGNAGTA | ASSETAESSYEQY | 0.68292344 |
| TGGGASLE | ASSETAESSYEQY | 0.31564003 |
| VAVYNHYKRI | ASSETAESSYEQY | 0.4345193 |
| TDPAPATT | ASSETAESSYEQY | 0.63901174 |
| TNRPDILDPALLRPG | ASSETAESSYEQY | 0.31255502 |
| LTTTTTS | ASSETAESSYEQY | 0.4006204 |
| PTGSGKTL | ASSETAESSYEQY | 0.5696726 |

| | | |
|---|---|---|
| TGSGKTL | ASSETAESSYEQY | 0.6436559 |
| TIGHVDHGK | ASSETAESSYEQY | 0.3124298 |
| TIGHVDHGKTT | ASSETAESSYEQY | 0.55096006 |
| GHVDHGKTT | ASSETAESSYEQY | 0.38216674 |
| AQFEHTLL | ASSETAESSYEQY | 0.66423666 |
| IVNQDEL | ASSETAESSYEQY | 0.53594375 |
| TNGSQFF | ASSETAESSYEQY | 0.38682044 |
| TGKGIVE | ASSETAESSYEQY | 0.62931275 |
| LISNASDA | ASSETAESSYEQY | 0.38672698 |
| AELRTLGELTKIAKAHGAQ | ASSETAESSYEQY | 0.60984516 |
| APNAVAAAEPSW | ASSETAESSYEQY | 0.6319326 |
| APQVAGAATAVL | ASSETAESSYEQY | 0.68647975 |
| ARRDPAAAAAILPTDA | ASSETAESSYEQY | 0.37360042 |
| DAAARDVLAEAGLAEYF | ASSETAESSYEQY | 0.6232802 |
| DDQIVLSAKANLAKVEQ | ASSETAESSYEQY | 0.5028575 |
| DELLPDTGEDSDEDGH | ASSETAESSYEQY | 0.6773283 |
| ELPEAPANRATVPQD | ASSETAESSYEQY | 0.4858554 |
| FGQSKIERVTGIG | ASSETAESSYEQY | 0.36890987 |
| GAGGTGSTAGGAGGAGGAGGLY | ASSETAESSYEQY | 0.6269032 |
| GHGGTGAAGGNGGTGGAGGSGI | ASSETAESSYEQY | 0.5884602 |
| GNGGNGGAGGEGGPGGAGGAGA | ASSETAESSYEQY | 0.54840297 |
| GSTAGNGGNGGNATNGTIAGQP | ASSETAESSYEQY | 0.64385647 |
| GTGGDGGNGGNGGTGG | ASSETAESSYEQY | 0.5779792 |
| GVRRPAGIPADAPV | ASSETAESSYEQY | 0.5061147 |
| GYRQVTDAHLLTLARR | ASSETAESSYEQY | 0.59947115 |
| IAKMGQDGHDRG | ASSETAESSYEQY | 0.5889735 |
| KAEPSIEIQYT | ASSETAESSYEQY | 0.45887202 |
| KPVRIRMAADEDTVAVT | ASSETAESSYEQY | 0.6571686 |
| MPPRIAGMRLLVIKPEP | ASSETAESSYEQY | 0.54689467 |
| MKFYAVALFRSRGVPPTR | ASSETAESSYEQY | 0.4030603 |
| NAGGFGILGAVAHSP | ASSETAESSYEQY | 0.66240025 |
| NEGREITLAMAEM | ASSETAESSYEQY | 0.58647984 |
| NELVEWSHPLGHRWRWE | ASSETAESSYEQY | 0.60414755 |
| NLGAGGTGGNGGNPG | ASSETAESSYEQY | 0.6113953 |
| NTEWPKWETVMAADPVD | ASSETAESSYEQY | 0.6950518 |
| PASAAPAETQP | ASSETAESSYEQY | 0.45021218 |
| QQSSDGKNGVRAAEITGE | ASSETAESSYEQY | 0.570322 |
| RAPDAVTTSWWKE | ASSETAESSYEQY | 0.54945445 |
| SGFGNVGAHTSGWFNQ | ASSETAESSYEQY | 0.6455108 |
| SGVAGAVGQAASVGGLKV | ASSETAESSYEQY | 0.5809241 |
| SNGGAGTGSAGGNGGT | ASSETAESSYEQY | 0.5922096 |
| TGGAGGEGGAAG | ASSETAESSYEQY | 0.6046247 |
| TGGDGGAAGAGGAPGHG | ASSETAESSYEQY | 0.6398508 |
| TMPPLLAAIANAGRHGV | ASSETAESSYEQY | 0.5505634 |
| TSSTAGNSGGA | ASSETAESSYEQY | 0.55879885 |
| VPATVPVGILC | ASSETAESSYEQY | 0.6590924 |
| VQRPDSGGVVLTGRIS | ASSETAESSYEQY | 0.40047047 |
| WTPNPVPATKGVPAGLVQAT | ASSETAESSYEQY | 0.5660994 |
| YALKRALAKFSDRFDVVII | ASSETAESSYEQY | 0.64648396 |
| YLIGSVIV | ASSETAESSYEQY | 0.63842356 |
| YGQLVAVID | ASSETAESSYEQY | 0.55580986 |

| | | |
|---|---|---|
| VTVGGMTLPA | ASSETAESSYEQY | 0.64624906 |
| VLALPAV | ASSETAESSYEQY | 0.49377346 |
| VIPPTLNYET | ASSETAESSYEQY | 0.56983143 |
| VGMGGMHPSSG | ASSETAESSYEQY | 0.52568066 |
| VGLAGGAATA | ASSETAESSYEQY | 0.47760472 |
| VGGAGGAGGSGP | ASSETAESSYEQY | 0.3788823 |
| VFAAFMV | ASSETAESSYEQY | 0.4908666 |
| TLYDKLR | ASSETAESSYEQY | 0.60352546 |
| TLAVFTVL | ASSETAESSYEQY | 0.5472445 |
| TGSVQPARV | ASSETAESSYEQY | 0.3544798 |
| TGGGAGSGGT | ASSETAESSYEQY | 0.56552994 |
| SHLETSL | ASSETAESSYEQY | 0.58494246 |
| SFTTGMT | ASSETAESSYEQY | 0.6762104 |
| QVEVAKRMGVRQPTVS | ASSETAESSYEQY | 0.55401534 |
| QGSGGPALAM | ASSETAESSYEQY | 0.6453339 |
| MLAPSNP | ASSETAESSYEQY | 0.5314742 |
| LVNNAAI | ASSETAESSYEQY | 0.5892361 |
| LTGEINTLA | ASSETAESSYEQY | 0.35666296 |
| LSIFGPV | ASSETAESSYEQY | 0.40894786 |
| LLALLANL | ASSETAESSYEQY | 0.530437 |
| LISNASDALDKLR | ASSETAESSYEQY | 0.4032674 |
| LINEAVP | ASSETAESSYEQY | 0.6261488 |
| LGYAFSGL | ASSETAESSYEQY | 0.6190875 |
| LDFVAVAL | ASSETAESSYEQY | 0.6273452 |
| LAQPGAE | ASSETAESSYEQY | 0.64110595 |
| ISNLGAVTGF | ASSETAESSYEQY | 0.6411217 |
| HQEDPGRI | ASSETAESSYEQY | 0.66221565 |
| GTPGEPTP | ASSETAESSYEQY | 0.45623487 |
| GFFNSGS | ASSETAESSYEQY | 0.51758444 |
| GAGGGGLTAGAN | ASSETAESSYEQY | 0.3372742 |
| FVDALVNM | ASSETAESSYEQY | 0.467101 |
| FLTVAPEL | ASSETAESSYEQY | 0.46509364 |
| FETLTAL | ASSETAESSYEQY | 0.5334065 |
| DVINPIEL | ASSETAESSYEQY | 0.30169147 |
| APVSGAPV | ASSETAESSYEQY | 0.61097145 |
| ALVGGLAW | ASSETAESSYEQY | 0.63384783 |
| ALAAAAATAI | ASSETAESSYEQY | 0.5472341 |
| AFGLGLLL | ASSETAESSYEQY | 0.60429597 |
| AFFAAASL | ASSETAESSYEQY | 0.5680718 |
| AAAALPAFSPP | ASSETAESSYEQY | 0.4450116 |
| VVLAPAL | ASSETAESSYEQY | 0.60179687 |
| TIGHVDHGKTTL | ASSETAESSYEQY | 0.50941986 |
| LISNASDALDK | ASSETAESSYEQY | 0.6437393 |
| LISNASDAL | ASSETAESSYEQY | 0.64304954 |
| ISNASDALDK | ASSETAESSYEQY | 0.5865345 |
| NQAETLVYQTE | ASSETAESSYEQY | 0.51589006 |
| TGGTGGTAGAG | ASSETAESSYEQY | 0.6055904 |
| TGSTAGGAGGA | ASSETAESSYEQY | 0.46152797 |
| DPLAVPVAAGPV | ASSETAESSYEQY | 0.54004425 |
| GGTGGTPGNGGN | ASSETAESSYEQY | 0.3736784 |
| PAGLAGVDPATL | ASSETAESSYEQY | 0.5552163 |

| | | |
|---|---|---|
| GNNLVVEAPHAGASV | ASSETAESSYEQY | 0.4296606 |
| KVTRIMSASTLEQPA | ASSETAESSYEQY | 0.57944775 |
| LGFALPIALPRRTVV | ASSETAESSYEQY | 0.5297358 |
| TRLGNLTARKLVVLV | ASSETAESSYEQY | 0.6066792 |
| WSQQRRLIYFPSAGP | ASSETAESSYEQY | 0.40470886 |
| GPNGNPDPMAAAVDIR | ASSETAESSYEQY | 0.64149654 |
| TQRCVSVADLRAAASR | ASSETAESSYEQY | 0.6134074 |
| QFVLDVAEHPVEVLTETV | ASSETAESSYEQY | 0.3913051 |
| GGAGGTGGAGGNGGTQAGDG | ASSETAESSYEQY | 0.60985726 |
| PLLATLRAIAVDVDAKPAQV | ASSETAESSYEQY | 0.56135523 |
| APKFRTGGYMIDFWGVGYQVAK | ASSETAESSYEQY | 0.48872858 |
| GAGGAGGAGGAAGGPGGTGGTG | ASSETAESSYEQY | 0.63532865 |
| IGALGLLATDVAPAGSVAFGVA | ASSETAESSYEQY | 0.5565535 |
| AGGAGGIGGNGGAGGAGGVLMG | ASSETAESSYEQY | 0.4170598 |
| ADELIGLL | ASSETAESSYEQY | 0.47180235 |
| ADRALKA | ASSETAESSYEQY | 0.56737787 |
| AGAVGQVGGAGGA | ASSETAESSYEQY | 0.5084859 |
| AGGGGGLGGAP | ASSETAESSYEQY | 0.33871016 |
| AGGLLYGNGGAG | ASSETAESSYEQY | 0.5836165 |
| AGLESIGL | ASSETAESSYEQY | 0.4064628 |
| AGLKTGII | ASSETAESSYEQY | 0.42661715 |
| ASLAALLL | ASSETAESSYEQY | 0.4650424 |
| ASVINISSI | ASSETAESSYEQY | 0.62243026 |
| ATPPGPA | ASSETAESSYEQY | 0.65631986 |
| ATVASALV | ASSETAESSYEQY | 0.53224516 |
| ATVASLIL | ASSETAESSYEQY | 0.5644705 |
| AVNGPVLIRH | ASSETAESSYEQY | 0.5923477 |
| DGAAGAVGI | ASSETAESSYEQY | 0.6213439 |
| DPLLHGI | ASSETAESSYEQY | 0.5651362 |
| ELGAALGAL | ASSETAESSYEQY | 0.57480365 |
| ELGRTQAV | ASSETAESSYEQY | 0.6114565 |
| FVAGGYLS | ASSETAESSYEQY | 0.5459012 |
| GASGGAGGAGGW | ASSETAESSYEQY | 0.4462903 |
| GGAGAGGDGGAGGI | ASSETAESSYEQY | 0.4833123 |
| GTGHNGSQPGMGGNG | ASSETAESSYEQY | 0.4468491 |
| GYAAASASAS | ASSETAESSYEQY | 0.44983408 |
| HQVDVAAVQQG | ASSETAESSYEQY | 0.58259374 |
| IGAAIAHT | ASSETAESSYEQY | 0.6825867 |
| IGMIEVAQL | ASSETAESSYEQY | 0.58951247 |
| IRLAPEHTSAMD | ASSETAESSYEQY | 0.35171232 |
| IYNVSVL | ASSETAESSYEQY | 0.49138862 |
| KQGFLGM | ASSETAESSYEQY | 0.5417031 |
| LESVITQL | ASSETAESSYEQY | 0.5893806 |
| LGLGALVVV | ASSETAESSYEQY | 0.6536011 |
| LIGNGGNGGSGGI | ASSETAESSYEQY | 0.6809015 |
| LNAINAPTL | ASSETAESSYEQY | 0.60331154 |
| LQLLNTGV | ASSETAESSYEQY | 0.46285075 |
| LQTSIEF | ASSETAESSYEQY | 0.61475784 |
| MSFAVSEL | ASSETAESSYEQY | 0.50938606 |
| NTGALGL | ASSETAESSYEQY | 0.5079128 |
| PPGAGGAGGAGTT | ASSETAESSYEQY | 0.5538959 |

| | | |
|---|---|---|
| QLVGLGI | ASSETAESSYEQY | 0.61277264 |
| RFLEQVHQL | ASSETAESSYEQY | 0.49859226 |
| RGVGYVLR | ASSETAESSYEQY | 0.50821847 |
| SATGVFAGVI | ASSETAESSYEQY | 0.39841253 |
| SLPAITLP | ASSETAESSYEQY | 0.6590248 |
| STALAFTV | ASSETAESSYEQY | 0.43458968 |
| SVLTVGLGK | ASSETAESSYEQY | 0.53349507 |
| TALAGAGSGI | ASSETAESSYEQY | 0.51234585 |
| TGLPFTGI | ASSETAESSYEQY | 0.65315473 |
| VAEHVTV | ASSETAESSYEQY | 0.6589287 |
| VDIVLNSL | ASSETAESSYEQY | 0.49713606 |
| VLTGGGAVL | ASSETAESSYEQY | 0.4503755 |
| VNVGLAYGI | ASSETAESSYEQY | 0.61754704 |
| VSSAAVGAV | ASSETAESSYEQY | 0.4928356 |
| VTSVYEAGV | ASSETAESSYEQY | 0.63502634 |
| WVALDRGA | ASSETAESSYEQY | 0.54950494 |
| YLVGSVVI | ASSETAESSYEQY | 0.50173736 |
| GPPGTGKTLL | ASSETAESSYEQY | 0.50813925 |
| GVGMTKF | ASSETAESSYEQY | 0.6150615 |
| IGIGGSDL | ASSETAESSYEQY | 0.48576662 |
| IGTLLAEL | ASSETAESSYEQY | 0.6438162 |
| IYTSGTT | ASSETAESSYEQY | 0.38949317 |
| LSFPAGV | ASSETAESSYEQY | 0.63542414 |
| LYTSGTT | ASSETAESSYEQY | 0.3927496 |
| VGAFPFGF | ASSETAESSYEQY | 0.4899022 |
| VGAGTVEF | ASSETAESSYEQY | 0.32865945 |
| VGKSTLLL | ASSETAESSYEQY | 0.53733385 |
| AALAHAGGGIV | ASSETAESSYEQY | 0.5930957 |
| AALVAVSLL | ASSETAESSYEQY | 0.64818406 |
| AAVSRAVAAL | ASSETAESSYEQY | 0.5886506 |
| AGGSSGIGGF | ASSETAESSYEQY | 0.5761654 |
| AGLTVANANG | ASSETAESSYEQY | 0.6582353 |
| AGLVGAVL | ASSETAESSYEQY | 0.42661083 |
| ALSQAFTV | ASSETAESSYEQY | 0.3319086 |
| ALTFSVSL | ASSETAESSYEQY | 0.58441097 |
| ALTGAALVAL | ASSETAESSYEQY | 0.59536743 |
| ASFSIGAGVL | ASSETAESSYEQY | 0.6531005 |
| DGSVYKI | ASSETAESSYEQY | 0.6065048 |
| EQLAEIV | ASSETAESSYEQY | 0.33123964 |
| FAVSELTVM | ASSETAESSYEQY | 0.63008773 |
| GAAPLAAGL | ASSETAESSYEQY | 0.6050124 |
| GADGGDGGSGGAGG | ASSETAESSYEQY | 0.38644958 |
| GFGLDTGGAGG | ASSETAESSYEQY | 0.4218515 |
| GLASGGLL | ASSETAESSYEQY | 0.45843163 |
| GRIAGAAVLL | ASSETAESSYEQY | 0.39354968 |
| GSAGSLNTSF | ASSETAESSYEQY | 0.60565233 |
| GTGQIGI | ASSETAESSYEQY | 0.46132165 |
| GTLIALIL | ASSETAESSYEQY | 0.55482024 |
| IAATMGFVVG | ASSETAESSYEQY | 0.4985693 |
| IFAQALL | ASSETAESSYEQY | 0.47478205 |
| IGAAVALGF | ASSETAESSYEQY | 0.5339566 |

| | | |
|---|---|---|
| IGLTPLAL | ASSETAESSYEQY | 0.50603735 |
| ILGSNTSTL | ASSETAESSYEQY | 0.59749645 |
| IVFSLTVL | ASSETAESSYEQY | 0.6228203 |
| LASKATGF | ASSETAESSYEQY | 0.51604235 |
| LASVALIL | ASSETAESSYEQY | 0.58875525 |
| LFAVGLTL | ASSETAESSYEQY | 0.615497 |
| LFNGGIGTYI | ASSETAESSYEQY | 0.39667988 |
| LTLGAGII | ASSETAESSYEQY | 0.45623374 |
| LVAAALGDI | ASSETAESSYEQY | 0.6126561 |
| NTAFVQLGI | ASSETAESSYEQY | 0.5093297 |
| QLFSGAEL | ASSETAESSYEQY | 0.6464381 |
| SAATFITL | ASSETAESSYEQY | 0.5651873 |
| SNYGLANLGV | ASSETAESSYEQY | 0.5630601 |
| SYGGAYAVM | ASSETAESSYEQY | 0.59235495 |
| TTGVGGLTLG | ASSETAESSYEQY | 0.648712 |
| TTQTIYI | ASSETAESSYEQY | 0.5829159 |
| VAAGAGLGAVI | ASSETAESSYEQY | 0.5062953 |
| VAAVVTLL | ASSETAESSYEQY | 0.60657984 |
| VGFLGGLI | ASSETAESSYEQY | 0.40926054 |
| VGVIPILL | ASSETAESSYEQY | 0.60494816 |
| VIFASSLI | ASSETAESSYEQY | 0.62904316 |
| VKGGDGGAAAG | ASSETAESSYEQY | 0.38786554 |
| VTKTFTNI | ASSETAESSYEQY | 0.50639015 |
| VVAAAILL | ASSETAESSYEQY | 0.66372496 |
| YAALDVEL | ASSETAESSYEQY | 0.6139005 |
| LASVALLL | ASSETAESSYEQY | 0.50121176 |
| LDAVHRV | ASSETAESSYEQY | 0.44773695 |
| QGLSSSIF | ASSETAESSYEQY | 0.4919905 |
| RGTFANIRL | ASSETAESSYEQY | 0.5040132 |
| RQLWWGHRI | ASSETAESSYEQY | 0.36388218 |
| YGPPGTGKTLL | ASSETAESSYEQY | 0.6024298 |
| AALVGAPANVA | ASSETAESSYEQY | 0.48453838 |
| AATFRRTE | ASSETAESSYEQY | 0.66229516 |
| AGALGGGI | ASSETAESSYEQY | 0.5761258 |
| AGGAGGAGGIGAG | ASSETAESSYEQY | 0.63823575 |
| AGSGALGL | ASSETAESSYEQY | 0.47914127 |
| AQIADQI | ASSETAESSYEQY | 0.51869154 |
| ASAVAVTYL | ASSETAESSYEQY | 0.37116703 |
| ASTVLGGTGG | ASSETAESSYEQY | 0.44956 |
| AVGGEPAK | ASSETAESSYEQY | 0.46390525 |
| CEQGTFGGTC | ASSETAESSYEQY | 0.53504175 |
| DEGKAPAG | ASSETAESSYEQY | 0.3596491 |
| DTVVFPAL | ASSETAESSYEQY | 0.65577006 |
| FEGITMAG | ASSETAESSYEQY | 0.48484802 |
| FPLVVQD | ASSETAESSYEQY | 0.6103122 |
| GAFTTPQ | ASSETAESSYEQY | 0.52905077 |
| GFGGFGTAGGVGGA | ASSETAESSYEQY | 0.4044444 |
| GFLAQFL | ASSETAESSYEQY | 0.4629287 |
| GGAGGAGGNAGVA | ASSETAESSYEQY | 0.61278284 |
| GGVGTAGAGGAGGAGG | ASSETAESSYEQY | 0.36780357 |
| GNGGGGDAGNAGSGGN | ASSETAESSYEQY | 0.34573826 |

| | | |
|---|---|---|
| GTGGAGGAGNDAGS | ASSETAESSYEQY | 0.48744503 |
| GTTNTGWFNS | ASSETAESSYEQY | 0.5320463 |
| IFMGPLFTL | ASSETAESSYEQY | 0.66654384 |
| ILAHAIEAI | ASSETAESSYEQY | 0.63509476 |
| IRSAIREAL | ASSETAESSYEQY | 0.54839057 |
| ITFAGGG | ASSETAESSYEQY | 0.6855618 |
| ITGMSGHQEV | ASSETAESSYEQY | 0.29708514 |
| IVAGLACTL | ASSETAESSYEQY | 0.54557294 |
| IVGGGLAGA | ASSETAESSYEQY | 0.65679735 |
| IVTGVLNI | ASSETAESSYEQY | 0.5825192 |
| LAQQGCI | ASSETAESSYEQY | 0.49527895 |
| LCLHVEN | ASSETAESSYEQY | 0.63484013 |
| LDAISDGS | ASSETAESSYEQY | 0.44784907 |
| LGGFAAML | ASSETAESSYEQY | 0.51527 |
| LGGKNPMI | ASSETAESSYEQY | 0.5383208 |
| LGPKGVAAIL | ASSETAESSYEQY | 0.6157135 |
| LMVGVSI | ASSETAESSYEQY | 0.61216676 |
| LVGAAFVL | ASSETAESSYEQY | 0.3564943 |
| LVLAVAPL | ASSETAESSYEQY | 0.64034456 |
| MAAIAVIDAA | ASSETAESSYEQY | 0.64761925 |
| NNGINAELQ | ASSETAESSYEQY | 0.45516816 |
| RGEPLVLVHG | ASSETAESSYEQY | 0.62976587 |
| RGGAGGAAGLI | ASSETAESSYEQY | 0.56559664 |
| RSYRMHPAV | ASSETAESSYEQY | 0.57160544 |
| SAAHLLAGQLA | ASSETAESSYEQY | 0.43596894 |
| SAAVGSY | ASSETAESSYEQY | 0.36814052 |
| SPHAVLHRI | ASSETAESSYEQY | 0.64400065 |
| SQIAKVL | ASSETAESSYEQY | 0.4241017 |
| TALGHLVP | ASSETAESSYEQY | 0.57501066 |
| TAPTTSGAA | ASSETAESSYEQY | 0.5639492 |
| TDGDDGNGGSPGT | ASSETAESSYEQY | 0.66923326 |
| TDGLGEN | ASSETAESSYEQY | 0.5822661 |
| TGGPPLPGQA | ASSETAESSYEQY | 0.5782952 |
| TIPAITI | ASSETAESSYEQY | 0.4542238 |
| TIPGFTI | ASSETAESSYEQY | 0.5940622 |
| TSIASGALA | ASSETAESSYEQY | 0.6067342 |
| TVTLPGAAI | ASSETAESSYEQY | 0.41721675 |
| VASGLSAALP | ASSETAESSYEQY | 0.63306344 |
| VGAGVIGI | ASSETAESSYEQY | 0.4195421 |
| VGPKRFGV | ASSETAESSYEQY | 0.5715804 |
| VNVAFPD | ASSETAESSYEQY | 0.60014606 |
| VSMTNSGVS | ASSETAESSYEQY | 0.66916376 |
| VTDVLGDLSV | ASSETAESSYEQY | 0.529005 |
| VVVVGAGFAGL | ASSETAESSYEQY | 0.64340574 |
| VYTVGTVV | ASSETAESSYEQY | 0.4102499 |
| WTTFSGP | ASSETAESSYEQY | 0.6375081 |
| YDELVEAF | ASSETAESSYEQY | 0.6296099 |
| YGTGGAGGNGG | ASSETAESSYEQY | 0.6248018 |
| YTNVMAV | ASSETAESSYEQY | 0.68150574 |
| FLEQVHQL | ASSETAESSYEQY | 0.64065504 |
| GLSLLAVAV | ASSETAESSYEQY | 0.35631394 |

| | | |
|---|---|---|
| LEQVHQL | ASSETAESSYEQY | 0.55278623 |
| LNAAGAI | ASSETAESSYEQY | 0.6446878 |
| PAVHASL | ASSETAESSYEQY | 0.6442086 |
| TELEDDA | ASSETAESSYEQY | 0.63786715 |
| TPPGPPN | ASSETAESSYEQY | 0.6093659 |
| TPPPPPPP | ASSETAESSYEQY | 0.5040141 |
| TQSPALI | ASSRDRVNTEVF | 0.6516973 |
| EFAAAFPQ | ASSRDRVNTEVF | 0.69680154 |
| LLSGNLDVL | ASSRDRVNTEVF | 0.61566013 |
| GPANLGNYNI | ASSRDRVNTEVF | 0.48518738 |
| TLGPTLP | ASSRDRVNTEVF | 0.47455055 |
| VYAYSSCCK | ASSRDRVNTEVF | 0.6036634 |
| APAPAPAPNI | ASSRDRVNTEVF | 0.6072857 |
| PGGPVPIP | ASSRDRVNTEVF | 0.41419852 |
| FLTVALEP | ASSRDRVNTEVF | 0.51952195 |
| SQYFGANL | ASSRDRVNTEVF | 0.55532074 |
| GGVKPAA | ASSRDRVNTEVF | 0.30915713 |
| GGTGGAGGAGGW | ASSRDRVNTEVF | 0.6121239 |
| GVGGVAGLGGAGG | ASSRDRVNTEVF | 0.6229212 |
| VLLVLATTI | ASSRDRVNTEVF | 0.59966373 |
| IGIFAAML | ASSRDRVNTEVF | 0.39537823 |
| AVGDLID | ASSRDRVNTEVF | 0.6305345 |
| NISPLPL | ASSRDRVNTEVF | 0.60794455 |
| VTAVFVAI | ASSRDRVNTEVF | 0.66762125 |
| SGDDNDAAL | ASSRDRVNTEVF | 0.39214087 |
| AVAALAHS | ASSRDRVNTEVF | 0.29914168 |
| GGVTVMATP | ASSRDRVNTEVF | 0.5873654 |
| AGLAANVRD | ASSRDRVNTEVF | 0.62746847 |
| TNRPDLIDPALLRPG | ASSRDRVNTEVF | 0.50303686 |
| LANLGVTAL | ASSRDRVNTEVF | 0.43489802 |
| TGERIEL | ASSRDRVNTEVF | 0.5079503 |
| NGEELAA | ASSRDRVNTEVF | 0.6302639 |
| VLSLVWG | ASSRDRVNTEVF | 0.39110032 |
| IGYALWL | ASSRDRVNTEVF | 0.4174757 |
| LSLLLAPL | ASSRDRVNTEVF | 0.5614033 |
| TLDEALK | ASSRDRVNTEVF | 0.64704955 |
| TDGAGTAEGG | ASSRDRVNTEVF | 0.4138277 |
| NAPGGAGGNGG | ASSRDRVNTEVF | 0.45519426 |
| VSVFMGMGGNF | ASSRDRVNTEVF | 0.6015567 |
| LASYLAAL | ASSRDRVNTEVF | 0.6545473 |
| LVGGTPAL | ASSRDRVNTEVF | 0.5655426 |
| VRNADVL | ASSRDRVNTEVF | 0.4110859 |
| TCSSPFLVSAM | ASSRDRVNTEVF | 0.57586557 |
| VHITDAE | ASSRDRVNTEVF | 0.38859808 |
| EYAAPSF | ASSRDRVNTEVF | 0.6339315 |
| VVDAFHGGL | ASSRDRVNTEVF | 0.54311466 |
| ASVVDPQ | ASSRDRVNTEVF | 0.58196914 |
| SYNEFGGLS | ASSRDRVNTEVF | 0.4717559 |
| DPDGSVY | ASSRDRVNTEVF | 0.40016583 |
| MVSTIHATE | ASSRDRVNTEVF | 0.40467566 |
| LAAGVADPV | ASSRDRVNTEVF | 0.4411661 |

| | | |
|---|---|---|
| TKEQIAA | ASSRDRVNTEVF | 0.62295777 |
| SLASPAAL | ASSRDRVNTEVF | 0.49654233 |
| IGLVFYGV | ASSRDRVNTEVF | 0.6036217 |
| FLLDASNL | ASSRDRVNTEVF | 0.55558497 |
| YQVGNFP | ASSRDRVNTEVF | 0.41169995 |
| ESLLSQST | ASSRDRVNTEVF | 0.6093185 |
| LFTVGFAAVG | ASSRDRVNTEVF | 0.35811937 |
| ATPPTPT | ASSRDRVNTEVF | 0.6221781 |
| TDLATLG | ASSRDRVNTEVF | 0.56367624 |
| SAGGTGR | ASSRDRVNTEVF | 0.62388134 |
| GFSNSGS | ASSRDRVNTEVF | 0.45453456 |
| GGNGGSGGTSV | ASSRDRVNTEVF | 0.62066096 |
| GTSVATGGAGN | ASSRDRVNTEVF | 0.5987009 |
| NGGNGGSGGTS | ASSRDRVNTEVF | 0.4407274 |
| AVVKFPH | ASSRDRVNTEVF | 0.59142405 |
| TPPPPPP | ASSRDRVNTEVF | 0.6223334 |
| AAPAAAAAPA | ASSRDRVNTEVF | 0.6023568 |
| IALNEDL | ASSRDRVNTEVF | 0.33396128 |
| IVNVSSM | ASSRDRVNTEVF | 0.57362545 |
| TGIPPAPR | ASSRDRVNTEVF | 0.43760043 |
| FDLGGGTF | ASSRDRVNTEVF | 0.5704761 |
| FDLGGGTFDVS | ASSRDRVNTEVF | 0.52721953 |
| NEPTAAA | ASSRDRVNTEVF | 0.49664116 |
| NEPTAAALA | ASSRDRVNTEVF | 0.6065521 |
| GSIINISSI | ASSRDRVNTEVF | 0.4523994 |
| VAPGNAGTA | ASSRDRVNTEVF | 0.45870522 |
| TGGGASLE | ASSRDRVNTEVF | 0.52131844 |
| VAVYNHYKRI | ASSRDRVNTEVF | 0.5752531 |
| TDPAPATT | ASSRDRVNTEVF | 0.6080517 |
| TNRPDILDPALLRPG | ASSRDRVNTEVF | 0.39418843 |
| LTTTTTS | ASSRDRVNTEVF | 0.59929836 |
| PTGSGKTL | ASSRDRVNTEVF | 0.51530755 |
| TGSGKTL | ASSRDRVNTEVF | 0.42969984 |
| TIGHVDHGK | ASSRDRVNTEVF | 0.59054744 |
| TIGHVDHGKTT | ASSRDRVNTEVF | 0.5256629 |
| GHVDHGKTT | ASSRDRVNTEVF | 0.37981498 |
| AQFEHTLL | ASSRDRVNTEVF | 0.61913514 |
| IVNQDEL | ASSRDRVNTEVF | 0.41399717 |
| TNGSQFF | ASSRDRVNTEVF | 0.6113821 |
| TGKGIVE | ASSRDRVNTEVF | 0.66725576 |
| LISNASDA | ASSRDRVNTEVF | 0.36944175 |
| AELRTLGELTKIAKAHGAQ | ASSRDRVNTEVF | 0.6268015 |
| APNAVAAAEPSW | ASSRDRVNTEVF | 0.53077245 |
| APQVAGAATAVL | ASSRDRVNTEVF | 0.536875 |
| ARRDPAAAAAILPTDA | ASSRDRVNTEVF | 0.5314121 |
| DAAARDVLAEAGLAEYF | ASSRDRVNTEVF | 0.45289758 |
| DDQIVLSAKANLAKVEQ | ASSRDRVNTEVF | 0.6387427 |
| DELLPDTGEDSDEDGH | ASSRDRVNTEVF | 0.61392784 |
| ELPEAPANRATVPQD | ASSRDRVNTEVF | 0.5934234 |
| FGQSKIERVTGIG | ASSRDRVNTEVF | 0.56689465 |
| GAGGTGSTAGGAGGAGGAGGLY | ASSRDRVNTEVF | 0.63776255 |

| | | |
|---|---|---|
| GHGGTGAAGGNGGTGGAGGSGI | ASSRDRVNTEVF | 0.63588166 |
| GNGGNGGAGGEGGPGGAGGAGA | ASSRDRVNTEVF | 0.46163177 |
| GSTAGNGGNGGNATNGTIAGQP | ASSRDRVNTEVF | 0.36122736 |
| GTGGDGGNGGNGGTGG | ASSRDRVNTEVF | 0.5221176 |
| GVRRPAGIPADAPV | ASSRDRVNTEVF | 0.51513946 |
| GYRQVTDAHLLTLARR | ASSRDRVNTEVF | 0.46430066 |
| IAKMGQDGHDRG | ASSRDRVNTEVF | 0.41249856 |
| KAEPSIEIQYT | ASSRDRVNTEVF | 0.60830694 |
| KPVRIRMAADEDTVAVT | ASSRDRVNTEVF | 0.6724376 |
| MPPRIAGMRLLVIKPEP | ASSRDRVNTEVF | 0.5300981 |
| MKFYAVALFRSRGVPPTR | ASSRDRVNTEVF | 0.6850481 |
| NAGGFGILGAVAHSP | ASSRDRVNTEVF | 0.39285937 |
| NEGREITLAMAEM | ASSRDRVNTEVF | 0.6231964 |
| NELVEWSHPLGHRWRWE | ASSRDRVNTEVF | 0.62281406 |
| NLGAGGTGGNGGNPG | ASSRDRVNTEVF | 0.41674435 |
| NTEWPKWETVMAADPVD | ASSRDRVNTEVF | 0.5943235 |
| PASAAPAETQP | ASSRDRVNTEVF | 0.4232703 |
| QQSSDGKNGVRAAEITGE | ASSRDRVNTEVF | 0.45004213 |
| RAPDAVTTSWWKE | ASSRDRVNTEVF | 0.62082803 |
| SGFGNVGAHTSGWFNQ | ASSRDRVNTEVF | 0.5446733 |
| SGVAGAVGQAASVGGLKV | ASSRDRVNTEVF | 0.60095334 |
| SNGGAGTGSAGGNGGT | ASSRDRVNTEVF | 0.51046705 |
| TGGAGGEGGAAG | ASSRDRVNTEVF | 0.43220535 |
| TGGDGGAAGAGGAPGHG | ASSRDRVNTEVF | 0.58764774 |
| TMPPLLAAIANAGRHGV | ASSRDRVNTEVF | 0.605427 |
| TSSTAGNSGGA | ASSRDRVNTEVF | 0.61814785 |
| VPATVPVGILC | ASSRDRVNTEVF | 0.69440305 |
| VQRPDSGGVVLTGRIS | ASSRDRVNTEVF | 0.4065247 |
| WTPNPVPATKGVPAGLVQAT | ASSRDRVNTEVF | 0.6443469 |
| YALKRALAKFSDRFDVVII | ASSRDRVNTEVF | 0.6651401 |
| YLIGSVIV | ASSRDRVNTEVF | 0.5604954 |
| YGQLVAVID | ASSRDRVNTEVF | 0.5196 |
| VTVGGMTLPA | ASSRDRVNTEVF | 0.61552227 |
| VLALPAV | ASSRDRVNTEVF | 0.5188563 |
| VIPPTLNYET | ASSRDRVNTEVF | 0.57107097 |
| VGMGGMHPSSG | ASSRDRVNTEVF | 0.67272013 |
| VGLAGGAATA | ASSRDRVNTEVF | 0.3153105 |
| VGGAGGAGGSGP | ASSRDRVNTEVF | 0.6180749 |
| VFAAFMV | ASSRDRVNTEVF | 0.5505234 |
| TLYDKLR | ASSRDRVNTEVF | 0.56054133 |
| TLAVFTVL | ASSRDRVNTEVF | 0.40821466 |
| TGSVQPARV | ASSRDRVNTEVF | 0.33359164 |
| TGGGAGSGGT | ASSRDRVNTEVF | 0.467067 |
| SHLETSL | ASSRDRVNTEVF | 0.37478885 |
| SFTTGMT | ASSRDRVNTEVF | 0.65032434 |
| QVEVAKRMGVRQPTVS | ASSRDRVNTEVF | 0.5831554 |
| QGSGGPALAM | ASSRDRVNTEVF | 0.39237273 |
| MLAPSNP | ASSRDRVNTEVF | 0.555645 |
| LVNNAAI | ASSRDRVNTEVF | 0.5737496 |
| LTGEINTLA | ASSRDRVNTEVF | 0.3288635 |
| LSIFGPV | ASSRDRVNTEVF | 0.46496928 |

| | | |
|---|---|---|
| LLALLANL | ASSRDRVNTEVF | 0.45063806 |
| LISNASDALDKLR | ASSRDRVNTEVF | 0.43890092 |
| LINEAVP | ASSRDRVNTEVF | 0.30017793 |
| LGYAFSGL | ASSRDRVNTEVF | 0.5697628 |
| LDFVAVAL | ASSRDRVNTEVF | 0.39415976 |
| LAQPGAE | ASSRDRVNTEVF | 0.29524624 |
| ISNLGAVTGF | ASSRDRVNTEVF | 0.5286641 |
| HQEDPGRI | ASSRDRVNTEVF | 0.40581965 |
| GTPGEPTP | ASSRDRVNTEVF | 0.42311403 |
| GFFNSGS | ASSRDRVNTEVF | 0.5678421 |
| GAGGGGLTAGAN | ASSRDRVNTEVF | 0.5844438 |
| FVDALVNM | ASSRDRVNTEVF | 0.43034106 |
| FLTVAPEL | ASSRDRVNTEVF | 0.55768603 |
| FETLTAL | ASSRDRVNTEVF | 0.6232565 |
| DVINPIEL | ASSRDRVNTEVF | 0.3757765 |
| APVSGAPV | ASSRDRVNTEVF | 0.4959823 |
| ALVGGLAW | ASSRDRVNTEVF | 0.6507803 |
| ALAAAAATAI | ASSRDRVNTEVF | 0.514366 |
| AFGLGLLL | ASSRDRVNTEVF | 0.64142513 |
| AFFAAASL | ASSRDRVNTEVF | 0.65143484 |
| AAAALPAFSPP | ASSRDRVNTEVF | 0.43587068 |
| VVLAPAL | ASSRDRVNTEVF | 0.6047104 |
| TIGHVDHGKTTL | ASSRDRVNTEVF | 0.60827315 |
| LISNASDALDK | ASSRDRVNTEVF | 0.47116917 |
| LISNASDAL | ASSRDRVNTEVF | 0.4756778 |
| ISNASDALDK | ASSRDRVNTEVF | 0.31390268 |
| NQAETLVYQTE | ASSRDRVNTEVF | 0.45902726 |
| TGGTGGTAGAG | ASSRDRVNTEVF | 0.3862543 |
| TGSTAGGAGGA | ASSRDRVNTEVF | 0.55424386 |
| DPLAVPVAAGPV | ASSRDRVNTEVF | 0.6343858 |
| GGTGGTPGNGGN | ASSRDRVNTEVF | 0.5048037 |
| PAGLAGVDPATL | ASSRDRVNTEVF | 0.30229825 |
| GNNLVVEAPHAGASV | ASSRDRVNTEVF | 0.33767965 |
| KVTRIMSASTLEQPA | ASSRDRVNTEVF | 0.5066626 |
| LGFALPIALPRRTVV | ASSRDRVNTEVF | 0.337308 |
| TRLGNLTARKLVVLV | ASSRDRVNTEVF | 0.35570943 |
| WSQQRRLIYFPSAGP | ASSRDRVNTEVF | 0.37622613 |
| GPNGNPDPMAAAVDIR | ASSRDRVNTEVF | 0.6231704 |
| TQRCVSVADLRAAASR | ASSRDRVNTEVF | 0.6279603 |
| QFVLDVAEHPVEVLTETV | ASSRDRVNTEVF | 0.42403376 |
| GGAGGTGGAGGNGGTQAGDG | ASSRDRVNTEVF | 0.6231246 |
| PLLATLRAIAVDVDAKPAQV | ASSRDRVNTEVF | 0.6441415 |
| APKFRTGGYMIDFWGVGYQVAK | ASSRDRVNTEVF | 0.65587276 |
| GAGGAGGAGGAAGGPGGTGGTG | ASSRDRVNTEVF | 0.5140524 |
| IGALGLLATDVAPAGSVAFGVA | ASSRDRVNTEVF | 0.31761277 |
| AGGAGGIGGNGGAGGAGGVLMG | ASSRDRVNTEVF | 0.6321236 |
| ADELIGLL | ASSRDRVNTEVF | 0.41423574 |
| ADRALKA | ASSRDRVNTEVF | 0.62542766 |
| AGAVGQVGGAGGA | ASSRDRVNTEVF | 0.5778413 |
| AGGGGGLGGAP | ASSRDRVNTEVF | 0.45402527 |
| AGGLLYGNGGAG | ASSRDRVNTEVF | 0.39067352 |

| | | |
|---|---|---|
| AGLESIGL | ASSRDRVNTEVF | 0.6141199 |
| AGLKTGII | ASSRDRVNTEVF | 0.6268071 |
| ASLAALLL | ASSRDRVNTEVF | 0.3601401 |
| ASVINISSI | ASSRDRVNTEVF | 0.3664955 |
| ATPPGPA | ASSRDRVNTEVF | 0.50932413 |
| ATVASALV | ASSRDRVNTEVF | 0.49594796 |
| ATVASLIL | ASSRDRVNTEVF | 0.6268821 |
| AVNGPVLIRH | ASSRDRVNTEVF | 0.5466578 |
| DGAAGAVGI | ASSRDRVNTEVF | 0.49769396 |
| DPLLHGI | ASSRDRVNTEVF | 0.5518844 |
| ELGAALGAL | ASSRDRVNTEVF | 0.5851507 |
| ELGRTQAV | ASSRDRVNTEVF | 0.5153551 |
| FVAGGYLS | ASSRDRVNTEVF | 0.49200615 |
| GASGGAGGAGGW | ASSRDRVNTEVF | 0.38347602 |
| GGAGAGGDGGAGGI | ASSRDRVNTEVF | 0.60020626 |
| GTGHNGSQPGMGGNG | ASSRDRVNTEVF | 0.6127002 |
| GYAAASASAS | ASSRDRVNTEVF | 0.5994431 |
| HQVDVAAVQQG | ASSRDRVNTEVF | 0.6382228 |
| IGAAIAHT | ASSRDRVNTEVF | 0.57622504 |
| IGMIEVAQL | ASSRDRVNTEVF | 0.5247429 |
| IRLAPEHTSAMD | ASSRDRVNTEVF | 0.39029187 |
| IYNVSVL | ASSRDRVNTEVF | 0.45235234 |
| KQGFLGM | ASSRDRVNTEVF | 0.44746634 |
| LESVITQL | ASSRDRVNTEVF | 0.530565 |
| LGLGALVVV | ASSRDRVNTEVF | 0.5952014 |
| LIGNGGNGGSGGI | ASSRDRVNTEVF | 0.5838829 |
| LNAINAPTL | ASSRDRVNTEVF | 0.41383916 |
| LQLLNTGV | ASSRDRVNTEVF | 0.61765426 |
| LQTSIEF | ASSRDRVNTEVF | 0.6270651 |
| MSFAVSEL | ASSRDRVNTEVF | 0.38567975 |
| NTGALGL | ASSRDRVNTEVF | 0.42000228 |
| PPGAGGAGGAGTT | ASSRDRVNTEVF | 0.617393 |
| QLVGLGI | ASSRDRVNTEVF | 0.5986289 |
| RFLEQVHQL | ASSRDRVNTEVF | 0.43659645 |
| RGVGYVLR | ASSRDRVNTEVF | 0.5005393 |
| SATGVFAGVI | ASSRDRVNTEVF | 0.6595862 |
| SLPAITLP | ASSRDRVNTEVF | 0.5783456 |
| STALAFTV | ASSRDRVNTEVF | 0.61817193 |
| SVLTVGLGK | ASSRDRVNTEVF | 0.34262782 |
| TALAGAGSGI | ASSRDRVNTEVF | 0.46389467 |
| TGLPFTGI | ASSRDRVNTEVF | 0.40476435 |
| VAEHVTV | ASSRDRVNTEVF | 0.28195074 |
| VDIVLNSL | ASSRDRVNTEVF | 0.6111356 |
| VLTGGGAVL | ASSRDRVNTEVF | 0.40821844 |
| VNVGLAYGI | ASSRDRVNTEVF | 0.5416606 |
| VSSAAVGAV | ASSRDRVNTEVF | 0.38815796 |
| VTSVYEAGV | ASSRDRVNTEVF | 0.62498075 |
| WVALDRGA | ASSRDRVNTEVF | 0.5696715 |
| YLVGSVVI | ASSRDRVNTEVF | 0.46370262 |
| GPPGTGKTLL | ASSRDRVNTEVF | 0.38287792 |
| GVGMTKF | ASSRDRVNTEVF | 0.54831624 |

| | | |
|---|---|---|
| IGIGGSDL | ASSRDRVNTEVF | 0.6129916 |
| IGTLLAEL | ASSRDRVNTEVF | 0.63044524 |
| IYTSGTT | ASSRDRVNTEVF | 0.34451386 |
| LSFPAGV | ASSRDRVNTEVF | 0.58100754 |
| LYTSGTT | ASSRDRVNTEVF | 0.5126731 |
| VGAFPFGF | ASSRDRVNTEVF | 0.3358449 |
| VGAGTVEF | ASSRDRVNTEVF | 0.2992627 |
| VGKSTLLL | ASSRDRVNTEVF | 0.37802473 |
| AALAHAGGGIV | ASSRDRVNTEVF | 0.4280337 |
| AALVAVSLL | ASSRDRVNTEVF | 0.52603436 |
| AAVSRAVAAL | ASSRDRVNTEVF | 0.61066043 |
| AGGSSGIGGF | ASSRDRVNTEVF | 0.42188895 |
| AGLTVANANG | ASSRDRVNTEVF | 0.57526684 |
| AGLVGAVL | ASSRDRVNTEVF | 0.4832193 |
| ALSQAFTV | ASSRDRVNTEVF | 0.27235794 |
| ALTFSVSL | ASSRDRVNTEVF | 0.41382557 |
| ALTGAALVAL | ASSRDRVNTEVF | 0.32053512 |
| ASFSIGAGVL | ASSRDRVNTEVF | 0.6483488 |
| DGSVYKI | ASSRDRVNTEVF | 0.3321648 |
| EQLAEIV | ASSRDRVNTEVF | 0.4474624 |
| FAVSELTVM | ASSRDRVNTEVF | 0.49297178 |
| GAAPLAAGL | ASSRDRVNTEVF | 0.6306902 |
| GADGGDGGSGGAGG | ASSRDRVNTEVF | 0.5320935 |
| GFGLDTGGAGG | ASSRDRVNTEVF | 0.41104192 |
| GLASGGLL | ASSRDRVNTEVF | 0.33498648 |
| GRIAGAAVLL | ASSRDRVNTEVF | 0.58042336 |
| GSAGSLNTSF | ASSRDRVNTEVF | 0.602076 |
| GTGQIGI | ASSRDRVNTEVF | 0.38550025 |
| GTLIALIL | ASSRDRVNTEVF | 0.38691378 |
| IAATMGFVVG | ASSRDRVNTEVF | 0.48191184 |
| IFAQALL | ASSRDRVNTEVF | 0.30991945 |
| IGAAVALGF | ASSRDRVNTEVF | 0.62439126 |
| IGLTPLAL | ASSRDRVNTEVF | 0.35341212 |
| ILGSNTSTL | ASSRDRVNTEVF | 0.4322155 |
| IVFSLTVL | ASSRDRVNTEVF | 0.64027286 |
| LASKATGF | ASSRDRVNTEVF | 0.6163215 |
| LASVALIL | ASSRDRVNTEVF | 0.4524127 |
| LFAVGLTL | ASSRDRVNTEVF | 0.4138232 |
| LFNGGIGTYI | ASSRDRVNTEVF | 0.35741156 |
| LTLGAGII | ASSRDRVNTEVF | 0.3872105 |
| LVAAALGDI | ASSRDRVNTEVF | 0.3973139 |
| NTAFVQLGI | ASSRDRVNTEVF | 0.59485817 |
| QLFSGAEL | ASSRDRVNTEVF | 0.5156697 |
| SAATFITL | ASSRDRVNTEVF | 0.34684882 |
| SNYGLANLGV | ASSRDRVNTEVF | 0.60831356 |
| SYGGAYAVM | ASSRDRVNTEVF | 0.6187187 |
| TTGVGGLTLG | ASSRDRVNTEVF | 0.6448858 |
| TTQTIYI | ASSRDRVNTEVF | 0.3670983 |
| VAAGAGLGAVI | ASSRDRVNTEVF | 0.48393053 |
| VAAVVTLL | ASSRDRVNTEVF | 0.47034758 |
| VGFLGGLI | ASSRDRVNTEVF | 0.46227163 |

| | | |
|---|---|---|
| VGVIPILL | ASSRDRVNTEVF | 0.39527002 |
| VIFASSLI | ASSRDRVNTEVF | 0.3895049 |
| VKGGDGGAAAG | ASSRDRVNTEVF | 0.603763 |
| VTKTFTNI | ASSRDRVNTEVF | 0.68234277 |
| VVAAAILL | ASSRDRVNTEVF | 0.477692 |
| YAALDVEL | ASSRDRVNTEVF | 0.29995894 |
| LASVALLL | ASSRDRVNTEVF | 0.47136217 |
| LDAVHRV | ASSRDRVNTEVF | 0.39404345 |
| QGLSSSIF | ASSRDRVNTEVF | 0.39068094 |
| RGTFANIRL | ASSRDRVNTEVF | 0.48366666 |
| RQLWWGHRI | ASSRDRVNTEVF | 0.31961942 |
| YGPPGTGKTLL | ASSRDRVNTEVF | 0.47357476 |
| AALVGAPANVA | ASSRDRVNTEVF | 0.65437067 |
| AATFRRTE | ASSRDRVNTEVF | 0.50353426 |
| AGALGGGI | ASSRDRVNTEVF | 0.4857614 |
| AGGAGGAGGIGAG | ASSRDRVNTEVF | 0.60938823 |
| AGSGALGL | ASSRDRVNTEVF | 0.5408809 |
| AQIADQI | ASSRDRVNTEVF | 0.6144692 |
| ASAVAVTYL | ASSRDRVNTEVF | 0.64839756 |
| ASTVLGGTGG | ASSRDRVNTEVF | 0.48840588 |
| AVGGEPAK | ASSRDRVNTEVF | 0.5066198 |
| CEQGTFGGTC | ASSRDRVNTEVF | 0.58351505 |
| DEGKAPAG | ASSRDRVNTEVF | 0.45980942 |
| DTVVFPAL | ASSRDRVNTEVF | 0.52020574 |
| FEGITMAG | ASSRDRVNTEVF | 0.46851158 |
| FPLVVQD | ASSRDRVNTEVF | 0.55600685 |
| GAFTTPQ | ASSRDRVNTEVF | 0.47242513 |
| GFGGFGTAGGVGGA | ASSRDRVNTEVF | 0.6596395 |
| GFLAQFL | ASSRDRVNTEVF | 0.2888503 |
| GGAGGAGGNAGVA | ASSRDRVNTEVF | 0.37941054 |
| GGVGTAGAGGAGGAGG | ASSRDRVNTEVF | 0.61632603 |
| GNGGGGDAGNAGSGGN | ASSRDRVNTEVF | 0.29455757 |
| GTGGAGGAGNDAGS | ASSRDRVNTEVF | 0.33680734 |
| GTTNTGWFNS | ASSRDRVNTEVF | 0.49092478 |
| IFMGPLFTL | ASSRDRVNTEVF | 0.6401778 |
| ILAHAIEAI | ASSRDRVNTEVF | 0.29684278 |
| IRSAIREAL | ASSRDRVNTEVF | 0.5100732 |
| ITFAGGG | ASSRDRVNTEVF | 0.32847697 |
| ITGMSGHQEV | ASSRDRVNTEVF | 0.629552 |
| IVAGLACTL | ASSRDRVNTEVF | 0.5010043 |
| IVGGGLAGA | ASSRDRVNTEVF | 0.3437941 |
| IVTGVLNI | ASSRDRVNTEVF | 0.5900632 |
| LAQQGCI | ASSRDRVNTEVF | 0.34220102 |
| LCLHVEN | ASSRDRVNTEVF | 0.58011353 |
| LDAISDGS | ASSRDRVNTEVF | 0.6099973 |
| LGGFAAML | ASSRDRVNTEVF | 0.6685936 |
| LGGKNPMI | ASSRDRVNTEVF | 0.31917933 |
| LGPKGVAAIL | ASSRDRVNTEVF | 0.5998781 |
| LMVGVSI | ASSRDRVNTEVF | 0.4244405 |
| LVGAAFVL | ASSRDRVNTEVF | 0.6623866 |
| LVLAVAPL | ASSRDRVNTEVF | 0.4339819 |

| | | |
|---|---|---|
| MAAIAVIDAA | ASSRDRVNTEVF | 0.3143554 |
| NNGINAELQ | ASSRDRVNTEVF | 0.59188306 |
| RGEPLVLVHG | ASSRDRVNTEVF | 0.5430622 |
| RGGAGGAAGLI | ASSRDRVNTEVF | 0.47124636 |
| RSYRMHPAV | ASSRDRVNTEVF | 0.62181693 |
| SAAHLLAGQLA | ASSRDRVNTEVF | 0.5091577 |
| SAAVGSY | ASSRDRVNTEVF | 0.46554312 |
| SPHAVLHRI | ASSRDRVNTEVF | 0.58011216 |
| SQIAKVL | ASSRDRVNTEVF | 0.5532602 |
| TALGHLVP | ASSRDRVNTEVF | 0.4138968 |
| TAPTTSGAA | ASSRDRVNTEVF | 0.6237135 |
| TDGDDGNGGSPGT | ASSRDRVNTEVF | 0.5077364 |
| TDGLGEN | ASSRDRVNTEVF | 0.34993368 |
| TGGPPLPGQA | ASSRDRVNTEVF | 0.65096563 |
| TIPAITI | ASSRDRVNTEVF | 0.5149188 |
| TIPGFTI | ASSRDRVNTEVF | 0.5753609 |
| TSIASGALA | ASSRDRVNTEVF | 0.5708141 |
| TVTLPGAAI | ASSRDRVNTEVF | 0.68806976 |
| VASGLSAALP | ASSRDRVNTEVF | 0.40295392 |
| VGAGVIGI | ASSRDRVNTEVF | 0.5075979 |
| VGPKRFGV | ASSRDRVNTEVF | 0.5097161 |
| VNVAFPD | ASSRDRVNTEVF | 0.61893874 |
| VSMTNSGVS | ASSRDRVNTEVF | 0.5385304 |
| VTDVLGDLSV | ASSRDRVNTEVF | 0.57436156 |
| VVVVGAGFAGL | ASSRDRVNTEVF | 0.5491903 |
| VYTVGTVV | ASSRDRVNTEVF | 0.61119115 |
| WTTFSGP | ASSRDRVNTEVF | 0.46524096 |
| YDELVEAF | ASSRDRVNTEVF | 0.48152152 |
| YGTGGAGGNGG | ASSRDRVNTEVF | 0.6422764 |
| YTNVMAV | ASSRDRVNTEVF | 0.33397108 |
| FLEQVHQL | ASSRDRVNTEVF | 0.5222312 |
| GLSLLAVAV | ASSRDRVNTEVF | 0.60344946 |
| LEQVHQL | ASSRDRVNTEVF | 0.59954906 |
| LNAAGAI | ASSRDRVNTEVF | 0.4576828 |
| PAVHASL | ASSRDRVNTEVF | 0.610643 |
| TELEDDA | ASSRDRVNTEVF | 0.4349067 |
| TPPGPPN | ASSRDRVNTEVF | 0.5356194 |
| TPPPPPPP | ASSRDRVNTEVF | 0.5107992 |
| TQSPALI | ASSLVGNYAEQF | 0.42207742 |
| EFAAAFPQ | ASSLVGNYAEQF | 0.34200427 |
| LLSGNLDVL | ASSLVGNYAEQF | 0.44117495 |
| GPANLGNYNI | ASSLVGNYAEQF | 0.58433264 |
| TLGPTLP | ASSLVGNYAEQF | 0.5246034 |
| VYAYSSCCK | ASSLVGNYAEQF | 0.3271849 |
| APAPAPAPNI | ASSLVGNYAEQF | 0.5382819 |
| PGGPVPIP | ASSLVGNYAEQF | 0.5634426 |
| FLTVALEP | ASSLVGNYAEQF | 0.654683 |
| SQYFGANL | ASSLVGNYAEQF | 0.49716687 |
| GGVKPAA | ASSLVGNYAEQF | 0.6013636 |
| GGTGGAGGAGGW | ASSLVGNYAEQF | 0.48497957 |
| GVGGVAGLGGAGG | ASSLVGNYAEQF | 0.528865 |

| | | |
|---|---|---|
| VLLVLATTI | ASSLVGNYAEQF | 0.32225835 |
| IGIFAAML | ASSLVGNYAEQF | 0.3459408 |
| AVGDLID | ASSLVGNYAEQF | 0.5061885 |
| NISPLPL | ASSLVGNYAEQF | 0.35915142 |
| VTAVFVAI | ASSLVGNYAEQF | 0.6178924 |
| SGDDNDAAL | ASSLVGNYAEQF | 0.5927209 |
| AVAALAHS | ASSLVGNYAEQF | 0.6088729 |
| GGVTVMATP | ASSLVGNYAEQF | 0.62288404 |
| AGLAANVRD | ASSLVGNYAEQF | 0.6251551 |
| TNRPDLIDPALLRPG | ASSLVGNYAEQF | 0.6191183 |
| LANLGVTAL | ASSLVGNYAEQF | 0.39886138 |
| TGERIEL | ASSLVGNYAEQF | 0.4482223 |
| NGEELAA | ASSLVGNYAEQF | 0.3037382 |
| VLSLVWG | ASSLVGNYAEQF | 0.4020256 |
| IGYALWL | ASSLVGNYAEQF | 0.39930218 |
| LSLLLAPL | ASSLVGNYAEQF | 0.48397976 |
| TLDEALK | ASSLVGNYAEQF | 0.2923368 |
| TDGAGTAEGG | ASSLVGNYAEQF | 0.6010127 |
| NAPGGAGGNGG | ASSLVGNYAEQF | 0.590944 |
| VSVFMGMGGNF | ASSLVGNYAEQF | 0.5228286 |
| LASYLAAL | ASSLVGNYAEQF | 0.5889176 |
| LVGGTPAL | ASSLVGNYAEQF | 0.5363648 |
| VRNADVL | ASSLVGNYAEQF | 0.3755713 |
| TCSSPFLVSAM | ASSLVGNYAEQF | 0.5775517 |
| VHITDAE | ASSLVGNYAEQF | 0.45764753 |
| EYAAPSF | ASSLVGNYAEQF | 0.6158184 |
| VVDAFHGGL | ASSLVGNYAEQF | 0.61932856 |
| ASVVDPQ | ASSLVGNYAEQF | 0.55745584 |
| SYNEFGGLS | ASSLVGNYAEQF | 0.43522906 |
| DPDGSVY | ASSLVGNYAEQF | 0.563589 |
| MVSTIHATE | ASSLVGNYAEQF | 0.4250849 |
| LAAGVADPV | ASSLVGNYAEQF | 0.47128487 |
| TKEQIAA | ASSLVGNYAEQF | 0.33234933 |
| SLASPAAL | ASSLVGNYAEQF | 0.5406092 |
| IGLVFYGV | ASSLVGNYAEQF | 0.37405485 |
| FLLDASNL | ASSLVGNYAEQF | 0.5488438 |
| YQVGNFP | ASSLVGNYAEQF | 0.47521937 |
| ESLLSQST | ASSLVGNYAEQF | 0.5533238 |
| LFTVGFAAVG | ASSLVGNYAEQF | 0.38641724 |
| ATPPTPT | ASSLVGNYAEQF | 0.61781305 |
| TDLATLG | ASSLVGNYAEQF | 0.613907 |
| SAGGTGR | ASSLVGNYAEQF | 0.3418619 |
| GFSNSGS | ASSLVGNYAEQF | 0.57779247 |
| GGNGGSGGTSV | ASSLVGNYAEQF | 0.46189958 |
| GTSVATGGAGN | ASSLVGNYAEQF | 0.45998785 |
| NGGNGGSGGTS | ASSLVGNYAEQF | 0.61720085 |
| AVVKFPH | ASSLVGNYAEQF | 0.5089182 |
| TPPPPPP | ASSLVGNYAEQF | 0.34921733 |
| AAPAAAAAPA | ASSLVGNYAEQF | 0.44646707 |
| IALNEDL | ASSLVGNYAEQF | 0.5271421 |
| IVNVSSM | ASSLVGNYAEQF | 0.45923674 |

| | | |
|---|---|---|
| TGIPPAPR | ASSLVGNYAEQF | 0.32534724 |
| FDLGGGTF | ASSLVGNYAEQF | 0.5503506 |
| FDLGGGTFDVS | ASSLVGNYAEQF | 0.35099602 |
| NEPTAAA | ASSLVGNYAEQF | 0.41486484 |
| NEPTAAALA | ASSLVGNYAEQF | 0.39343727 |
| GSIINISSI | ASSLVGNYAEQF | 0.60263836 |
| VAPGNAGTA | ASSLVGNYAEQF | 0.6563617 |
| TGGGASLE | ASSLVGNYAEQF | 0.4638224 |
| VAVYNHYKRI | ASSLVGNYAEQF | 0.5076737 |
| TDPAPATT | ASSLVGNYAEQF | 0.5536907 |
| TNRPDILDPALLRPG | ASSLVGNYAEQF | 0.58856905 |
| LTTTTTS | ASSLVGNYAEQF | 0.50430876 |
| PTGSGKTL | ASSLVGNYAEQF | 0.49526578 |
| TGSGKTL | ASSLVGNYAEQF | 0.5460298 |
| TIGHVDHGK | ASSLVGNYAEQF | 0.47166154 |
| TIGHVDHGKTT | ASSLVGNYAEQF | 0.36720303 |
| GHVDHGKTT | ASSLVGNYAEQF | 0.46508858 |
| AQFEHTLL | ASSLVGNYAEQF | 0.4087002 |
| IVNQDEL | ASSLVGNYAEQF | 0.400154 |
| TNGSQFF | ASSLVGNYAEQF | 0.55381095 |
| TGKGIVE | ASSLVGNYAEQF | 0.6550401 |
| LISNASDA | ASSLVGNYAEQF | 0.55267656 |
| AELRTLGELTKIAKAHGAQ | ASSLVGNYAEQF | 0.30357477 |
| APNAVAAAEPSW | ASSLVGNYAEQF | 0.403035 |
| APQVAGAATAVL | ASSLVGNYAEQF | 0.5078757 |
| ARRDPAAAAAILPTDA | ASSLVGNYAEQF | 0.55627406 |
| DAAARDVLAEAGLAEYF | ASSLVGNYAEQF | 0.6241558 |
| DDQIVLSAKANLAKVEQ | ASSLVGNYAEQF | 0.664019 |
| DELLPDTGEDSDEDGH | ASSLVGNYAEQF | 0.5869207 |
| ELPEAPANRATVPQD | ASSLVGNYAEQF | 0.4137771 |
| FGQSKIERVTGIG | ASSLVGNYAEQF | 0.58815306 |
| GAGGTGSTAGGAGGAGGAGGLY | ASSLVGNYAEQF | 0.4489723 |
| GHGGTGAAGGNGGTGGAGGSGI | ASSLVGNYAEQF | 0.43154433 |
| GNGGNGGAGGEGGPGGAGGAGA | ASSLVGNYAEQF | 0.4912681 |
| GSTAGNGGNGGNATNGTIAGQP | ASSLVGNYAEQF | 0.57857823 |
| GTGGDGGNGGNGGTGG | ASSLVGNYAEQF | 0.4151308 |
| GVRRPAGIPADAPV | ASSLVGNYAEQF | 0.42665863 |
| GYRQVTDAHLLTLARR | ASSLVGNYAEQF | 0.34123623 |
| IAKMGQDGHDRG | ASSLVGNYAEQF | 0.59743434 |
| KAEPSIEIQYT | ASSLVGNYAEQF | 0.4578485 |
| KPVRIRMAADEDTVAVT | ASSLVGNYAEQF | 0.428264 |
| MPPRIAGMRLLVIKPEP | ASSLVGNYAEQF | 0.4737702 |
| MKFYAVALFRSRGVPPTR | ASSLVGNYAEQF | 0.6379554 |
| NAGGFGILGAVAHSP | ASSLVGNYAEQF | 0.64687526 |
| NEGREITLAMAEM | ASSLVGNYAEQF | 0.4421435 |
| NELVEWSHPLGHRWRWE | ASSLVGNYAEQF | 0.38531768 |
| NLGAGGTGGNGGNPG | ASSLVGNYAEQF | 0.59994197 |
| NTEWPKWETVMAADPVD | ASSLVGNYAEQF | 0.45042223 |
| PASAAPAETQP | ASSLVGNYAEQF | 0.57082283 |
| QQSSDGKNGVRAAEITGE | ASSLVGNYAEQF | 0.54511255 |
| RAPDAVTTSWWKE | ASSLVGNYAEQF | 0.47372124 |

| | | |
|---|---|---|
| SGFGNVGAHTSGWFNQ | ASSLVGNYAEQF | 0.42487764 |
| SGVAGAVGQAASVGGLKV | ASSLVGNYAEQF | 0.57431823 |
| SNGGAGTGSAGGNGGT | ASSLVGNYAEQF | 0.44409907 |
| TGGAGGEGGAAG | ASSLVGNYAEQF | 0.590708 |
| TGGDGGAAGAGGAPGHG | ASSLVGNYAEQF | 0.3569421 |
| TMPPLLAAIANAGRHGV | ASSLVGNYAEQF | 0.61668754 |
| TSSTAGNSGGA | ASSLVGNYAEQF | 0.35096 |
| VPATVPVGILC | ASSLVGNYAEQF | 0.42221135 |
| VQRPDSGGVVLTGRIS | ASSLVGNYAEQF | 0.29338598 |
| WTPNPVPATKGVPAGLVQAT | ASSLVGNYAEQF | 0.46418902 |
| YALKRALAKFSDRFDVVII | ASSLVGNYAEQF | 0.570597 |
| YLIGSVIV | ASSLVGNYAEQF | 0.62301356 |
| YGQLVAVID | ASSLVGNYAEQF | 0.53886354 |
| VTVGGMTLPA | ASSLVGNYAEQF | 0.53452736 |
| VLALPAV | ASSLVGNYAEQF | 0.64449716 |
| VIPPTLNYET | ASSLVGNYAEQF | 0.3733622 |
| VGMGGMHPSSG | ASSLVGNYAEQF | 0.30551016 |
| VGLAGGAATA | ASSLVGNYAEQF | 0.54327774 |
| VGGAGGAGGSGP | ASSLVGNYAEQF | 0.5679233 |
| VFAAFMV | ASSLVGNYAEQF | 0.6336312 |
| TLYDKLR | ASSLVGNYAEQF | 0.5974879 |
| TLAVFTVL | ASSLVGNYAEQF | 0.3074514 |
| TGSVQPARV | ASSLVGNYAEQF | 0.5407888 |
| TGGGAGSGGT | ASSLVGNYAEQF | 0.56829005 |
| SHLETSL | ASSLVGNYAEQF | 0.33933213 |
| SFTTGMT | ASSLVGNYAEQF | 0.35417366 |
| QVEVAKRMGVRQPTVS | ASSLVGNYAEQF | 0.37322155 |
| QGSGGPALAM | ASSLVGNYAEQF | 0.35273376 |
| MLAPSNP | ASSLVGNYAEQF | 0.5693527 |
| LVNNAAI | ASSLVGNYAEQF | 0.41194767 |
| LTGEINTLA | ASSLVGNYAEQF | 0.5326231 |
| LSIFGPV | ASSLVGNYAEQF | 0.4328004 |
| LLALLANL | ASSLVGNYAEQF | 0.40746915 |
| LISNASDALDKLR | ASSLVGNYAEQF | 0.49894914 |
| LINEAVP | ASSLVGNYAEQF | 0.45835626 |
| LGYAFSGL | ASSLVGNYAEQF | 0.5675098 |
| LDFVAVAL | ASSLVGNYAEQF | 0.5872617 |
| LAQPGAE | ASSLVGNYAEQF | 0.50588435 |
| ISNLGAVTGF | ASSLVGNYAEQF | 0.5571145 |
| HQEDPGRI | ASSLVGNYAEQF | 0.59720004 |
| GTPGEPTP | ASSLVGNYAEQF | 0.3582248 |
| GFFNSGS | ASSLVGNYAEQF | 0.39922166 |
| GAGGGGLTAGAN | ASSLVGNYAEQF | 0.5764151 |
| FVDALVNM | ASSLVGNYAEQF | 0.48637003 |
| FLTVAPEL | ASSLVGNYAEQF | 0.62619996 |
| FETLTAL | ASSLVGNYAEQF | 0.51490766 |
| DVINPIEL | ASSLVGNYAEQF | 0.52637 |
| APVSGAPV | ASSLVGNYAEQF | 0.5365076 |
| ALVGGLAW | ASSLVGNYAEQF | 0.6243571 |
| ALAAAAATAI | ASSLVGNYAEQF | 0.5380869 |
| AFGLGLLL | ASSLVGNYAEQF | 0.42298692 |

| | | |
|---|---|---|
| AFFAAASL | ASSLVGNYAEQF | 0.60854834 |
| AAAALPAFSPP | ASSLVGNYAEQF | 0.3365531 |
| VVLAPAL | ASSLVGNYAEQF | 0.57158935 |
| TIGHVDHGKTTL | ASSLVGNYAEQF | 0.5868362 |
| LISNASDALDK | ASSLVGNYAEQF | 0.3613661 |
| LISNASDAL | ASSLVGNYAEQF | 0.46650508 |
| ISNASDALDK | ASSLVGNYAEQF | 0.62964916 |
| NQAETLVYQTE | ASSLVGNYAEQF | 0.58090436 |
| TGGTGGTAGAG | ASSLVGNYAEQF | 0.45944524 |
| TGSTAGGAGGA | ASSLVGNYAEQF | 0.4057418 |
| DPLAVPVAAGPV | ASSLVGNYAEQF | 0.41910285 |
| GGTGGTPGNGGN | ASSLVGNYAEQF | 0.39244592 |
| PAGLAGVDPATL | ASSLVGNYAEQF | 0.3152887 |
| GNNLVVEAPHAGASV | ASSLVGNYAEQF | 0.5728637 |
| KVTRIMSASTLEQPA | ASSLVGNYAEQF | 0.44298825 |
| LGFALPIALPRRTVV | ASSLVGNYAEQF | 0.6490888 |
| TRLGNLTARKLVVLV | ASSLVGNYAEQF | 0.5448561 |
| WSQQRRLIYFPSAGP | ASSLVGNYAEQF | 0.62174654 |
| GPNGNPDPMAAAVDIR | ASSLVGNYAEQF | 0.43050084 |
| TQRCVSVADLRAAASR | ASSLVGNYAEQF | 0.46361208 |
| QFVLDVAEHPVEVLTETV | ASSLVGNYAEQF | 0.3197699 |
| GGAGGTGGAGGNGGTQAGDG | ASSLVGNYAEQF | 0.45107585 |
| PLLATLRAIAVDVDAKPAQV | ASSLVGNYAEQF | 0.36968118 |
| APKFRTGGYMIDFWGVGYQVAK | ASSLVGNYAEQF | 0.5167232 |
| GAGGAGGAGGAAGGPGGTGGTG | ASSLVGNYAEQF | 0.3338132 |
| IGALGLLATDVAPAGSVAFGVA | ASSLVGNYAEQF | 0.6453091 |
| AGGAGGIGGNGGAGGAGGVLMG | ASSLVGNYAEQF | 0.40950868 |
| ADELIGLL | ASSLVGNYAEQF | 0.5905775 |
| ADRALKA | ASSLVGNYAEQF | 0.48048395 |
| AGAVGQVGGAGGA | ASSLVGNYAEQF | 0.33361977 |
| AGGGGGLGGAP | ASSLVGNYAEQF | 0.4105508 |
| AGGLLYGNGGAG | ASSLVGNYAEQF | 0.58892167 |
| AGLESIGL | ASSLVGNYAEQF | 0.32251576 |
| AGLKTGII | ASSLVGNYAEQF | 0.30886427 |
| ASLAALLL | ASSLVGNYAEQF | 0.4142769 |
| ASVINISSI | ASSLVGNYAEQF | 0.49234995 |
| ATPPGPA | ASSLVGNYAEQF | 0.6495577 |
| ATVASALV | ASSLVGNYAEQF | 0.61398387 |
| ATVASLIL | ASSLVGNYAEQF | 0.47699067 |
| AVNGPVLIRH | ASSLVGNYAEQF | 0.6586446 |
| DGAAGAVGI | ASSLVGNYAEQF | 0.28051704 |
| DPLLHGI | ASSLVGNYAEQF | 0.5139607 |
| ELGAALGAL | ASSLVGNYAEQF | 0.6429553 |
| ELGRTQAV | ASSLVGNYAEQF | 0.55142075 |
| FVAGGYLS | ASSLVGNYAEQF | 0.45465997 |
| GASGGAGGAGGW | ASSLVGNYAEQF | 0.61005944 |
| GGAGAGGDGGAGGI | ASSLVGNYAEQF | 0.40993348 |
| GTGHNGSQPGMGGNG | ASSLVGNYAEQF | 0.46905163 |
| GYAAASASAS | ASSLVGNYAEQF | 0.4917606 |
| HQVDVAAVQQG | ASSLVGNYAEQF | 0.5510904 |
| IGAAIAHT | ASSLVGNYAEQF | 0.55527914 |

| | | |
|---|---|---|
| IGMIEVAQL | ASSLVGNYAEQF | 0.31760848 |
| IRLAPEHTSAMD | ASSLVGNYAEQF | 0.6027553 |
| IYNVSVL | ASSLVGNYAEQF | 0.63577557 |
| KQGFLGM | ASSLVGNYAEQF | 0.41472661 |
| LESVITQL | ASSLVGNYAEQF | 0.6245096 |
| LGLGALVVV | ASSLVGNYAEQF | 0.51667637 |
| LIGNGGNGGSGGI | ASSLVGNYAEQF | 0.59251994 |
| LNAINAPTL | ASSLVGNYAEQF | 0.40053284 |
| LQLLNTGV | ASSLVGNYAEQF | 0.32762045 |
| LQTSIEF | ASSLVGNYAEQF | 0.6200937 |
| MSFAVSEL | ASSLVGNYAEQF | 0.3858921 |
| NTGALGL | ASSLVGNYAEQF | 0.49914372 |
| PPGAGGAGGAGTT | ASSLVGNYAEQF | 0.519669 |
| QLVGLGI | ASSLVGNYAEQF | 0.6558689 |
| RFLEQVHQL | ASSLVGNYAEQF | 0.55082333 |
| RGVGYVLR | ASSLVGNYAEQF | 0.5340695 |
| SATGVFAGVI | ASSLVGNYAEQF | 0.4193912 |
| SLPAITLP | ASSLVGNYAEQF | 0.51595664 |
| STALAFTV | ASSLVGNYAEQF | 0.588976 |
| SVLTVGLGK | ASSLVGNYAEQF | 0.35119486 |
| TALAGAGSGI | ASSLVGNYAEQF | 0.6111022 |
| TGLPFTGI | ASSLVGNYAEQF | 0.36386803 |
| VAEHVTV | ASSLVGNYAEQF | 0.54572195 |
| VDIVLNSL | ASSLVGNYAEQF | 0.54416156 |
| VLTGGGAVL | ASSLVGNYAEQF | 0.6611547 |
| VNVGLAYGI | ASSLVGNYAEQF | 0.47743434 |
| VSSAAVGAV | ASSLVGNYAEQF | 0.6343437 |
| VTSVYEAGV | ASSLVGNYAEQF | 0.3609471 |
| WVALDRGA | ASSLVGNYAEQF | 0.60243237 |
| YLVGSVVI | ASSLVGNYAEQF | 0.6130234 |
| GPPGTGKTLL | ASSLVGNYAEQF | 0.6359397 |
| GVGMTKF | ASSLVGNYAEQF | 0.6581218 |
| IGIGGSDL | ASSLVGNYAEQF | 0.6088141 |
| IGTLLAEL | ASSLVGNYAEQF | 0.30654097 |
| IYTSGTT | ASSLVGNYAEQF | 0.47684464 |
| LSFPAGV | ASSLVGNYAEQF | 0.62987614 |
| LYTSGTT | ASSLVGNYAEQF | 0.6291255 |
| VGAFPFGF | ASSLVGNYAEQF | 0.6094018 |
| VGAGTVEF | ASSLVGNYAEQF | 0.5506445 |
| VGKSTLLL | ASSLVGNYAEQF | 0.45895988 |
| AALAHAGGGIV | ASSLVGNYAEQF | 0.62823 |
| AALVAVSLL | ASSLVGNYAEQF | 0.68308055 |
| AAVSRAVAAL | ASSLVGNYAEQF | 0.5833921 |
| AGGSSGIGGF | ASSLVGNYAEQF | 0.4276691 |
| AGLTVANANG | ASSLVGNYAEQF | 0.3820566 |
| AGLVGAVL | ASSLVGNYAEQF | 0.56263137 |
| ALSQAFTV | ASSLVGNYAEQF | 0.5770675 |
| ALTFSVSL | ASSLVGNYAEQF | 0.3418783 |
| ALTGAALVAL | ASSLVGNYAEQF | 0.46963286 |
| ASFSIGAGVL | ASSLVGNYAEQF | 0.5116999 |
| DGSVYKI | ASSLVGNYAEQF | 0.28851107 |

| | | |
|---|---|---|
| EQLAEIV | ASSLVGNYAEQF | 0.59377897 |
| FAVSELTVM | ASSLVGNYAEQF | 0.59486216 |
| GAAPLAAGL | ASSLVGNYAEQF | 0.5336054 |
| GADGGDGGSGGAGG | ASSLVGNYAEQF | 0.33014557 |
| GFGLDTGGAGG | ASSLVGNYAEQF | 0.61294484 |
| GLASGGLL | ASSLVGNYAEQF | 0.59105206 |
| GRIAGAAVLL | ASSLVGNYAEQF | 0.63594526 |
| GSAGSLNTSF | ASSLVGNYAEQF | 0.35518178 |
| GTGQIGI | ASSLVGNYAEQF | 0.28358945 |
| GTLIALIL | ASSLVGNYAEQF | 0.5561801 |
| IAATMGFVVG | ASSLVGNYAEQF | 0.638041 |
| IFAQALL | ASSLVGNYAEQF | 0.45318723 |
| IGAAVALGF | ASSLVGNYAEQF | 0.3695965 |
| IGLTPLAL | ASSLVGNYAEQF | 0.45181108 |
| ILGSNTSTL | ASSLVGNYAEQF | 0.6070555 |
| IVFSLTVL | ASSLVGNYAEQF | 0.32361382 |
| LASKATGF | ASSLVGNYAEQF | 0.35542232 |
| LASVALIL | ASSLVGNYAEQF | 0.4916835 |
| LFAVGLTL | ASSLVGNYAEQF | 0.61374307 |
| LFNGGIGTYI | ASSLVGNYAEQF | 0.37457997 |
| LTLGAGII | ASSLVGNYAEQF | 0.40231013 |
| LVAAALGDI | ASSLVGNYAEQF | 0.59377444 |
| NTAFVQLGI | ASSLVGNYAEQF | 0.64593166 |
| QLFSGAEL | ASSLVGNYAEQF | 0.5270009 |
| SAATFITL | ASSLVGNYAEQF | 0.35446346 |
| SNYGLANLGV | ASSLVGNYAEQF | 0.5468377 |
| SYGGAYAVM | ASSLVGNYAEQF | 0.3508066 |
| TTGVGGLTLG | ASSLVGNYAEQF | 0.5939572 |
| TTQTIYI | ASSLVGNYAEQF | 0.48099756 |
| VAAGAGLGAVI | ASSLVGNYAEQF | 0.5275537 |
| VAAVVTLL | ASSLVGNYAEQF | 0.43162557 |
| VGFLGGLI | ASSLVGNYAEQF | 0.36189854 |
| VGVIPILL | ASSLVGNYAEQF | 0.3442501 |
| VIFASSLI | ASSLVGNYAEQF | 0.4099009 |
| VKGGDGGAAAG | ASSLVGNYAEQF | 0.5981684 |
| VTKTFTNI | ASSLVGNYAEQF | 0.45558405 |
| VVAAAILL | ASSLVGNYAEQF | 0.54472107 |
| YAALDVEL | ASSLVGNYAEQF | 0.5188483 |
| LASVALLL | ASSLVGNYAEQF | 0.3568043 |
| LDAVHRV | ASSLVGNYAEQF | 0.5551529 |
| QGLSSSIF | ASSLVGNYAEQF | 0.326908 |
| RGTFANIRL | ASSLVGNYAEQF | 0.5596073 |
| RQLWWGHRI | ASSLVGNYAEQF | 0.50243 |
| YGPPGTGKTLL | ASSLVGNYAEQF | 0.5731152 |
| AALVGAPANVA | ASSLVGNYAEQF | 0.40917903 |
| AATFRRTE | ASSLVGNYAEQF | 0.54678965 |
| AGALGGGI | ASSLVGNYAEQF | 0.56214035 |
| AGGAGGAGGIGAG | ASSLVGNYAEQF | 0.36996812 |
| AGSGALGL | ASSLVGNYAEQF | 0.5181551 |
| AQIADQI | ASSLVGNYAEQF | 0.60980356 |
| ASAVAVTYL | ASSLVGNYAEQF | 0.54516774 |

| | | |
|---|---|---|
| ASTVLGGTGG | ASSLVGNYAEQF | 0.30113629 |
| AVGGEPAK | ASSLVGNYAEQF | 0.5490491 |
| CEQGTFGGTC | ASSLVGNYAEQF | 0.3875172 |
| DEGKAPAG | ASSLVGNYAEQF | 0.53344935 |
| DTVVFPAL | ASSLVGNYAEQF | 0.4706574 |
| FEGITMAG | ASSLVGNYAEQF | 0.42388016 |
| FPLVVQD | ASSLVGNYAEQF | 0.5695138 |
| GAFTTPQ | ASSLVGNYAEQF | 0.38126853 |
| GFGGFGTAGGVGGA | ASSLVGNYAEQF | 0.40536046 |
| GFLAQFL | ASSLVGNYAEQF | 0.46857154 |
| GGAGGAGGNAGVA | ASSLVGNYAEQF | 0.5324988 |
| GGVGTAGAGGAGGAGG | ASSLVGNYAEQF | 0.5675202 |
| GNGGGGDAGNAGSGGN | ASSLVGNYAEQF | 0.35185018 |
| GTGGAGGAGNDAGS | ASSLVGNYAEQF | 0.56491715 |
| GTTNTGWFNS | ASSLVGNYAEQF | 0.46543252 |
| IFMGPLFTL | ASSLVGNYAEQF | 0.38315868 |
| ILAHAIEAI | ASSLVGNYAEQF | 0.49448088 |
| IRSAIREAL | ASSLVGNYAEQF | 0.4462402 |
| ITFAGGG | ASSLVGNYAEQF | 0.3230938 |
| ITGMSGHQEV | ASSLVGNYAEQF | 0.57601416 |
| IVAGLACTL | ASSLVGNYAEQF | 0.3495125 |
| IVGGGLAGA | ASSLVGNYAEQF | 0.5658352 |
| IVTGVLNI | ASSLVGNYAEQF | 0.6532148 |
| LAQQGCI | ASSLVGNYAEQF | 0.31590816 |
| LCLHVEN | ASSLVGNYAEQF | 0.5763734 |
| LDAISDGS | ASSLVGNYAEQF | 0.48011678 |
| LGGFAAML | ASSLVGNYAEQF | 0.48243022 |
| LGGKNPMI | ASSLVGNYAEQF | 0.48360485 |
| LGPKGVAAIL | ASSLVGNYAEQF | 0.3914645 |
| LMVGVSI | ASSLVGNYAEQF | 0.616022 |
| LVGAAFVL | ASSLVGNYAEQF | 0.5845809 |
| LVLAVAPL | ASSLVGNYAEQF | 0.50761247 |
| MAAIAVIDAA | ASSLVGNYAEQF | 0.49857458 |
| NNGINAELQ | ASSLVGNYAEQF | 0.6009624 |
| RGEPLVLVHG | ASSLVGNYAEQF | 0.6071428 |
| RGGAGGAAGLI | ASSLVGNYAEQF | 0.43185097 |
| RSYRMHPAV | ASSLVGNYAEQF | 0.32285795 |
| SAAHLLAGQLA | ASSLVGNYAEQF | 0.48047352 |
| SAAVGSY | ASSLVGNYAEQF | 0.47906277 |
| SPHAVLHRI | ASSLVGNYAEQF | 0.3896941 |
| SQIAKVL | ASSLVGNYAEQF | 0.3361535 |
| TALGHLVP | ASSLVGNYAEQF | 0.5859508 |
| TAPTTSGAA | ASSLVGNYAEQF | 0.65815246 |
| TDGDDGNGGSPGT | ASSLVGNYAEQF | 0.49803877 |
| TDGLGEN | ASSLVGNYAEQF | 0.66875184 |
| TGGPPLPGQA | ASSLVGNYAEQF | 0.33054897 |
| TIPAITI | ASSLVGNYAEQF | 0.6055701 |
| TIPGFTI | ASSLVGNYAEQF | 0.60062474 |
| TSIASGALA | ASSLVGNYAEQF | 0.3683188 |
| TVTLPGAAI | ASSLVGNYAEQF | 0.5638575 |
| VASGLSAALP | ASSLVGNYAEQF | 0.36752093 |

| | | |
|---|---|---|
| VGAGVIGI | ASSLVGNYAEQF | 0.38014555 |
| VGPKRFGV | ASSLVGNYAEQF | 0.58112967 |
| VNVAFPD | ASSLVGNYAEQF | 0.5062137 |
| VSMTNSGVS | ASSLVGNYAEQF | 0.53671277 |
| VTDVLGDLSV | ASSLVGNYAEQF | 0.45092526 |
| VVVVGAGFAGL | ASSLVGNYAEQF | 0.37300172 |
| VYTVGTVV | ASSLVGNYAEQF | 0.53561026 |
| WTTFSGP | ASSLVGNYAEQF | 0.5371716 |
| YDELVEAF | ASSLVGNYAEQF | 0.5657502 |
| YGTGGAGGNGG | ASSLVGNYAEQF | 0.6786778 |
| YTNVMAV | ASSLVGNYAEQF | 0.35783502 |
| FLEQVHQL | ASSLVGNYAEQF | 0.6356995 |
| GLSLLAVAV | ASSLVGNYAEQF | 0.63837874 |
| LEQVHQL | ASSLVGNYAEQF | 0.53963435 |
| LNAAGAI | ASSLVGNYAEQF | 0.46690506 |
| PAVHASL | ASSLVGNYAEQF | 0.6186719 |
| TELEDDA | ASSLVGNYAEQF | 0.47755164 |
| TPPGPPN | ASSLVGNYAEQF | 0.48748013 |
| TPPPPPPP | ASSLVGNYAEQF | 0.64272153 |

PanPep_cluster15

| Peptide | CDR3 | Score |
| --- | --- | --- |
| TQSPALI | ASGELGGSSYEQY | 0.6759249 |
| EFAAAFPQ | ASGELGGSSYEQY | 0.7171448 |
| LLSGNLDVL | ASGELGGSSYEQY | 0.5961815 |
| GPANLGNYNI | ASGELGGSSYEQY | 0.5356473 |
| TLGPTLP | ASGELGGSSYEQY | 0.65381336 |
| VYAYSSCCK | ASGELGGSSYEQY | 0.68268 |
| APAPAPAPNI | ASGELGGSSYEQY | 0.7155036 |
| PGGPVPIP | ASGELGGSSYEQY | 0.6864308 |
| FLTVALEP | ASGELGGSSYEQY | 0.5076953 |
| SQYFGANL | ASGELGGSSYEQY | 0.6648581 |
| GGVKPAA | ASGELGGSSYEQY | 0.6038815 |
| GGTGGAGGAGGW | ASGELGGSSYEQY | 0.6369971 |
| GVGGVAGLGGAGG | ASGELGGSSYEQY | 0.65658027 |
| VLLVLATTI | ASGELGGSSYEQY | 0.631425 |
| IGIFAAML | ASGELGGSSYEQY | 0.62358475 |
| AVGDLID | ASGELGGSSYEQY | 0.57269853 |
| NISPLPL | ASGELGGSSYEQY | 0.6624925 |
| VTAVFVAI | ASGELGGSSYEQY | 0.68347955 |
| SGDDNDAAL | ASGELGGSSYEQY | 0.682304 |
| AVAALAHS | ASGELGGSSYEQY | 0.70183986 |
| GGVTVMATP | ASGELGGSSYEQY | 0.6973531 |
| AGLAANVRD | ASGELGGSSYEQY | 0.6539254 |
| TNRPDLIDPALLRPG | ASGELGGSSYEQY | 0.2780354 |
| LANLGVTAL | ASGELGGSSYEQY | 0.34008244 |
| TGERIEL | ASGELGGSSYEQY | 0.56993234 |
| NGEELAA | ASGELGGSSYEQY | 0.5783057 |
| VLSLVWG | ASGELGGSSYEQY | 0.33822635 |
| IGYALWL | ASGELGGSSYEQY | 0.64419794 |
| LSLLLAPL | ASGELGGSSYEQY | 0.61624056 |
| TLDEALK | ASGELGGSSYEQY | 0.5723225 |
| TDGAGTAEGG | ASGELGGSSYEQY | 0.70076734 |
| NAPGGAGGNGG | ASGELGGSSYEQY | 0.6168668 |
| VSVFMGMGGNF | ASGELGGSSYEQY | 0.49777126 |
| LASYLAAL | ASGELGGSSYEQY | 0.523142 |
| LVGGTPAL | ASGELGGSSYEQY | 0.664073 |
| VRNADVL | ASGELGGSSYEQY | 0.4388168 |
| TCSSPFLVSAM | ASGELGGSSYEQY | 0.67916906 |
| VHITDAE | ASGELGGSSYEQY | 0.46849486 |
| EYAAPSF | ASGELGGSSYEQY | 0.68018174 |
| VVDAFHGGL | ASGELGGSSYEQY | 0.4374906 |
| ASVVDPQ | ASGELGGSSYEQY | 0.6787029 |
| SYNEFGGLS | ASGELGGSSYEQY | 0.63421226 |
| DPDGSVY | ASGELGGSSYEQY | 0.728996 |
| MVSTIHATE | ASGELGGSSYEQY | 0.5880126 |
| LAAGVADPV | ASGELGGSSYEQY | 0.6452134 |
| TKEQIAA | ASGELGGSSYEQY | 0.50820434 |
| SLASPAAL | ASGELGGSSYEQY | 0.7113445 |
| IGLVFYGV | ASGELGGSSYEQY | 0.4676926 |
| FLLDASNL | ASGELGGSSYEQY | 0.6193912 |

| | | |
|---|---|---|
| YQVGNFP | ASGELGGSSYEQY | 0.36777276 |
| ESLLSQST | ASGELGGSSYEQY | 0.65627 |
| LFTVGFAAVG | ASGELGGSSYEQY | 0.6777487 |
| ATPPTPT | ASGELGGSSYEQY | 0.42211998 |
| TDLATLG | ASGELGGSSYEQY | 0.7069544 |
| SAGGTGR | ASGELGGSSYEQY | 0.6196387 |
| GFSNSGS | ASGELGGSSYEQY | 0.543461 |
| GGNGGSGGTSV | ASGELGGSSYEQY | 0.70509124 |
| GTSVATGGAGN | ASGELGGSSYEQY | 0.60881305 |
| NGGNGGSGGTS | ASGELGGSSYEQY | 0.7143199 |
| AVVKFPH | ASGELGGSSYEQY | 0.5976421 |
| TPPPPPP | ASGELGGSSYEQY | 0.60166985 |
| AAPAAAAAPA | ASGELGGSSYEQY | 0.6507252 |
| IALNEDL | ASGELGGSSYEQY | 0.59889513 |
| IVNVSSM | ASGELGGSSYEQY | 0.52656275 |
| TGIPPAPR | ASGELGGSSYEQY | 0.70209295 |
| FDLGGGTF | ASGELGGSSYEQY | 0.8573632 |
| FDLGGGTFDVS | ASGELGGSSYEQY | 0.85335463 |
| NEPTAAA | ASGELGGSSYEQY | 0.69441015 |
| NEPTAAALA | ASGELGGSSYEQY | 0.6887909 |
| GSIINISSI | ASGELGGSSYEQY | 0.7770362 |
| VAPGNAGTA | ASGELGGSSYEQY | 0.8396752 |
| TGGGASLE | ASGELGGSSYEQY | 0.88836294 |
| VAVYNHYKRI | ASGELGGSSYEQY | 0.8355307 |
| TDPAPATT | ASGELGGSSYEQY | 0.67971355 |
| TNRPDILDPALLRPG | ASGELGGSSYEQY | 0.8583324 |
| LTTTTTS | ASGELGGSSYEQY | 0.81862915 |
| PTGSGKTL | ASGELGGSSYEQY | 0.7960743 |
| TGSGKTL | ASGELGGSSYEQY | 0.852952 |
| TIGHVDHGK | ASGELGGSSYEQY | 0.82306206 |
| TIGHVDHGKTT | ASGELGGSSYEQY | 0.7517328 |
| GHVDHGKTT | ASGELGGSSYEQY | 0.7224124 |
| AQFEHTLL | ASGELGGSSYEQY | 0.8513397 |
| IVNQDEL | ASGELGGSSYEQY | 0.87120855 |
| TNGSQFF | ASGELGGSSYEQY | 0.8636012 |
| TGKGIVE | ASGELGGSSYEQY | 0.8796552 |
| LISNASDA | ASGELGGSSYEQY | 0.8673164 |
| AELRTLGELTKIAKAHGAQ | ASGELGGSSYEQY | 0.8462416 |
| APNAVAAAEPSW | ASGELGGSSYEQY | 0.34653744 |
| APQVAGAATAVL | ASGELGGSSYEQY | 0.5630412 |
| ARRDPAAAAAILPTDA | ASGELGGSSYEQY | 0.7167666 |
| DAAARDVLAEAGLAEYF | ASGELGGSSYEQY | 0.7619904 |
| DDQIVLSAKANLAKVEQ | ASGELGGSSYEQY | 0.34099746 |
| DELLPDTGEDSDEDGH | ASGELGGSSYEQY | 0.8182892 |
| ELPEAPANRATVPQD | ASGELGGSSYEQY | 0.80178595 |
| FGQSKIERVTGIG | ASGELGGSSYEQY | 0.7137245 |
| GAGGTGSTAGGAGGAGGAGGI | ASGELGGSSYEQY | 0.8698364 |
| GHGGTGAAGGNGGTGGAGGS( | ASGELGGSSYEQY | 0.76415807 |
| GNGGNGGAGGEGGPGGAGGA( | ASGELGGSSYEQY | 0.6371534 |
| GSTAGNGGNGGNATNGTIAG( | ASGELGGSSYEQY | 0.7616954 |
| GTGGDGGNGGNGGTGG | ASGELGGSSYEQY | 0.8768631 |

| | | |
|---|---|---|
| GVRRPAGIPADAPV | ASGELGGSSYEQY | 0.7066145 |
| GYRQVTDAHLLTLARR | ASGELGGSSYEQY | 0.8795733 |
| IAKMGQDGHDRG | ASGELGGSSYEQY | 0.7336253 |
| KAEPSIEIQYT | ASGELGGSSYEQY | 0.88143754 |
| KPVRIRMAADEDTVAVT | ASGELGGSSYEQY | 0.60490364 |
| MPPRIAGMRLLVIKPEP | ASGELGGSSYEQY | 0.8640988 |
| MKFYAVALFRSRGVPPTR | ASGELGGSSYEQY | 0.8071181 |
| NAGGFGILGAVAHSP | ASGELGGSSYEQY | 0.8992108 |
| NEGREITLAMAEM | ASGELGGSSYEQY | 0.80376434 |
| NELVEWSHPLGHRWRWE | ASGELGGSSYEQY | 0.8694681 |
| NLGAGGTGGNGGNPG | ASGELGGSSYEQY | 0.8382913 |
| NTEWPKWETVMAADPVD | ASGELGGSSYEQY | 0.8768378 |
| PASAAPAETQP | ASGELGGSSYEQY | 0.64356434 |
| QQSSDGKNGVRAAEITGE | ASGELGGSSYEQY | 0.80207616 |
| RAPDAVTTSWWKE | ASGELGGSSYEQY | 0.6438755 |
| SGFGNVGAHTSGWFNQ | ASGELGGSSYEQY | 0.8092553 |
| SGVAGAVGQAASVGGLKV | ASGELGGSSYEQY | 0.84903497 |
| SNGGAGTGSAGGNGGT | ASGELGGSSYEQY | 0.24347113 |
| TGGAGGEGGAAG | ASGELGGSSYEQY | 0.88427365 |
| TGGDGGAAGAGGAPGHG | ASGELGGSSYEQY | 0.55939794 |
| TMPPLLAAIANAGRHGV | ASGELGGSSYEQY | 0.6764305 |
| TSSTAGNSGGA | ASGELGGSSYEQY | 0.8665968 |
| VPATVPVGILC | ASGELGGSSYEQY | 0.8061519 |
| VQRPDSGGVVLTGRIS | ASGELGGSSYEQY | 0.86638737 |
| WTPNPVPATKGVPAGLVQAT | ASGELGGSSYEQY | 0.7577207 |
| YALKRALAKFSDRFDVVII | ASGELGGSSYEQY | 0.8282973 |
| YLIGSVIV | ASGELGGSSYEQY | 0.82881963 |
| YGQLVAVID | ASGELGGSSYEQY | 0.5144826 |
| VTVGGMTLPA | ASGELGGSSYEQY | 0.62889177 |
| VLALPAV | ASGELGGSSYEQY | 0.8806102 |
| VIPPTLNYET | ASGELGGSSYEQY | 0.64844 |
| VGMGGMHPSSG | ASGELGGSSYEQY | 0.6910795 |
| VGLAGGAATA | ASGELGGSSYEQY | 0.59306425 |
| VGGAGGAGGSGP | ASGELGGSSYEQY | 0.5262803 |
| VFAAFMV | ASGELGGSSYEQY | 0.5963479 |
| TLYDKLR | ASGELGGSSYEQY | 0.67848015 |
| TLAVFTVL | ASGELGGSSYEQY | 0.69381255 |
| TGSVQPARV | ASGELGGSSYEQY | 0.66803384 |
| TGGGAGSGGT | ASGELGGSSYEQY | 0.48813576 |
| SHLETSL | ASGELGGSSYEQY | 0.67105436 |
| SFTTGMT | ASGELGGSSYEQY | 0.6161414 |
| QVEVAKRMGVRQPTVS | ASGELGGSSYEQY | 0.6121515 |
| QGSGGPALAM | ASGELGGSSYEQY | 0.6364397 |
| MLAPSNP | ASGELGGSSYEQY | 0.6408221 |
| LVNNAAI | ASGELGGSSYEQY | 0.5758089 |
| LTGEINTLA | ASGELGGSSYEQY | 0.54536414 |
| LSIFGPV | ASGELGGSSYEQY | 0.6562295 |
| LLALLANL | ASGELGGSSYEQY | 0.67185354 |
| LISNASDALDKLR | ASGELGGSSYEQY | 0.67547274 |
| LINEAVP | ASGELGGSSYEQY | 0.7023078 |
| LGYAFSGL | ASGELGGSSYEQY | 0.68538296 |

| | | |
|---|---|---|
| LDFVAVAL | ASGELGGSSYEQY | 0.67660487 |
| LAQPGAE | ASGELGGSSYEQY | 0.34518862 |
| ISNLGAVTGF | ASGELGGSSYEQY | 0.42152628 |
| HQEDPGRI | ASGELGGSSYEQY | 0.5612578 |
| GTPGEPTP | ASGELGGSSYEQY | 0.5987636 |
| GFFNSGS | ASGELGGSSYEQY | 0.36343184 |
| GAGGGGLTAGAN | ASGELGGSSYEQY | 0.6470556 |
| FVDALVNM | ASGELGGSSYEQY | 0.5832022 |
| FLTVAPEL | ASGELGGSSYEQY | 0.55627215 |
| FETLTAL | ASGELGGSSYEQY | 0.682171 |
| DVINPIEL | ASGELGGSSYEQY | 0.5888402 |
| APVSGAPV | ASGELGGSSYEQY | 0.5254615 |
| ALVGGLAW | ASGELGGSSYEQY | 0.5330229 |
| ALAAAAATAI | ASGELGGSSYEQY | 0.66712505 |
| AFGLGLLL | ASGELGGSSYEQY | 0.4660921 |
| AFFAAASL | ASGELGGSSYEQY | 0.6714042 |
| AAAALPAFSPP | ASGELGGSSYEQY | 0.4890643 |
| VVLAPAL | ASGELGGSSYEQY | 0.68955326 |
| TIGHVDHGKTTL | ASGELGGSSYEQY | 0.45459086 |
| LISNASDALDK | ASGELGGSSYEQY | 0.67709094 |
| LISNASDAL | ASGELGGSSYEQY | 0.6247588 |
| ISNASDALDK | ASGELGGSSYEQY | 0.7159968 |
| NQAETLVYQTE | ASGELGGSSYEQY | 0.6002543 |
| TGGTGGTAGAG | ASGELGGSSYEQY | 0.6451512 |
| TGSTAGGAGGA | ASGELGGSSYEQY | 0.5585809 |
| DPLAVPVAAGPV | ASGELGGSSYEQY | 0.6859846 |
| GGTGGTPGNGGN | ASGELGGSSYEQY | 0.42348403 |
| PAGLAGVDPATL | ASGELGGSSYEQY | 0.6062934 |
| GNNLVVEAPHAGASV | ASGELGGSSYEQY | 0.44670463 |
| KVTRIMSASTLEQPA | ASGELGGSSYEQY | 0.64628786 |
| LGFALPIALPRRTVV | ASGELGGSSYEQY | 0.66370046 |
| TRLGNLTARKLVVLV | ASGELGGSSYEQY | 0.3899907 |
| WSQQRRLIYFPSAGP | ASGELGGSSYEQY | 0.6767258 |
| GPNGNPDPMAAAVDIR | ASGELGGSSYEQY | 0.57752836 |
| TQRCVSVADLRAAASR | ASGELGGSSYEQY | 0.54105425 |
| QFVLDVAEHPVEVLTETV | ASGELGGSSYEQY | 0.692176 |
| GGAGGTGGAGGNGGTQAGDG | ASGELGGSSYEQY | 0.649912 |
| PLLATLRAIAVDVDAKPAQV | ASGELGGSSYEQY | 0.68428695 |
| APKFRTGGYMIDFWGVGYQV | ASGELGGSSYEQY | 0.5796566 |
| GAGGAGGAGGAAGGPGGTGG | ASGELGGSSYEQY | 0.58665407 |
| IGALGLLATDVAPAGSVAFG | ASGELGGSSYEQY | 0.64761215 |
| AGGAGGIGGNGGAGGAGGVL | ASGELGGSSYEQY | 0.59597445 |
| ADELIGLL | ASGELGGSSYEQY | 0.54887456 |
| ADRALKA | ASGELGGSSYEQY | 0.6899879 |
| AGAVGQVGGAGGA | ASGELGGSSYEQY | 0.78730106 |
| AGGGGGLGGAP | ASGELGGSSYEQY | 0.79730946 |
| AGGLLYGNGGAG | ASGELGGSSYEQY | 0.72569454 |
| AGLESIGL | ASGELGGSSYEQY | 0.7396352 |
| AGLKTGII | ASGELGGSSYEQY | 0.7601129 |
| ASLAALLL | ASGELGGSSYEQY | 0.80076325 |
| ASVINISSI | ASGELGGSSYEQY | 0.7959901 |

| | | |
|---|---|---|
| ATPPGPA | ASGELGGSSYEQY | 0.7933221 |
| ATVASALV | ASGELGGSSYEQY | 0.74678916 |
| ATVASLIL | ASGELGGSSYEQY | 0.78384745 |
| AVNGPVLIRH | ASGELGGSSYEQY | 0.7636118 |
| DGAAGAVGI | ASGELGGSSYEQY | 0.77023304 |
| DPLLHGI | ASGELGGSSYEQY | 0.7888522 |
| ELGAALGAL | ASGELGGSSYEQY | 0.78237504 |
| ELGRTQAV | ASGELGGSSYEQY | 0.7606105 |
| FVAGGYLS | ASGELGGSSYEQY | 0.7613349 |
| GASGGAGGAGGW | ASGELGGSSYEQY | 0.7823928 |
| GGAGAGGDGGAGGI | ASGELGGSSYEQY | 0.79277 |
| GTGHNGSQPGMGGNG | ASGELGGSSYEQY | 0.7862972 |
| GYAAASASAS | ASGELGGSSYEQY | 0.8040296 |
| HQVDVAAVQQG | ASGELGGSSYEQY | 0.7916132 |
| IGAAIAHT | ASGELGGSSYEQY | 0.7844068 |
| IGMIEVAQL | ASGELGGSSYEQY | 0.4618693 |
| IRLAPEHTSAMD | ASGELGGSSYEQY | 0.6938093 |
| IYNVSVL | ASGELGGSSYEQY | 0.7346697 |
| KQGFLGM | ASGELGGSSYEQY | 0.7698945 |
| LESVITQL | ASGELGGSSYEQY | 0.4302985 |
| LGLGALVVV | ASGELGGSSYEQY | 0.79353964 |
| LIGNGGNGGSGGI | ASGELGGSSYEQY | 0.77154315 |
| LNAINAPTL | ASGELGGSSYEQY | 0.73371613 |
| LQLLNTGV | ASGELGGSSYEQY | 0.79066706 |
| LQTSIEF | ASGELGGSSYEQY | 0.75669503 |
| MSFAVSEL | ASGELGGSSYEQY | 0.7423243 |
| NTGALGL | ASGELGGSSYEQY | 0.7701225 |
| PPGAGGAGGAGTT | ASGELGGSSYEQY | 0.7964246 |
| QLVGLGI | ASGELGGSSYEQY | 0.6986567 |
| RFLEQVHQL | ASGELGGSSYEQY | 0.7953477 |
| RGVGYVLR | ASGELGGSSYEQY | 0.77940536 |
| SATGVFAGVI | ASGELGGSSYEQY | 0.7998049 |
| SLPAITLP | ASGELGGSSYEQY | 0.71414983 |
| STALAFTV | ASGELGGSSYEQY | 0.7925686 |
| SVLTVGLGK | ASGELGGSSYEQY | 0.7750815 |
| TALAGAGSGI | ASGELGGSSYEQY | 0.8053682 |
| TGLPFTGI | ASGELGGSSYEQY | 0.77038455 |
| VAEHVTV | ASGELGGSSYEQY | 0.7938493 |
| VDIVLNSL | ASGELGGSSYEQY | 0.78760505 |
| VLTGGGAVL | ASGELGGSSYEQY | 0.79073274 |
| VNVGLAYGI | ASGELGGSSYEQY | 0.67696846 |
| VSSAAVGAV | ASGELGGSSYEQY | 0.7670547 |
| VTSVYEAGV | ASGELGGSSYEQY | 0.7239281 |
| WVALDRGA | ASGELGGSSYEQY | 0.7808776 |
| YLVGSVVI | ASGELGGSSYEQY | 0.7931581 |
| GPPGTGKTLL | ASGELGGSSYEQY | 0.51593816 |
| GVGMTKF | ASGELGGSSYEQY | 0.7891644 |
| IGIGGSDL | ASGELGGSSYEQY | 0.74704146 |
| IGTLLAEL | ASGELGGSSYEQY | 0.76066375 |
| IYTSGTT | ASGELGGSSYEQY | 0.79363066 |
| LSFPAGV | ASGELGGSSYEQY | 0.76344335 |

| | | |
|---|---|---|
| LYTSGTT | ASGELGGSSYEQY | 0.78939575 |
| VGAFPFGF | ASGELGGSSYEQY | 0.75712514 |
| VGAGTVEF | ASGELGGSSYEQY | 0.77096534 |
| VGKSTLLL | ASGELGGSSYEQY | 0.78649104 |
| AALAHAGGGIV | ASGELGGSSYEQY | 0.57206494 |
| AALVAVSLL | ASGELGGSSYEQY | 0.73241067 |
| AAVSRAVAAL | ASGELGGSSYEQY | 0.8009317 |
| AGGSSGIGGF | ASGELGGSSYEQY | 0.69741756 |
| AGLTVANANG | ASGELGGSSYEQY | 0.7306535 |
| AGLVGAVL | ASGELGGSSYEQY | 0.57444966 |
| ALSQAFTV | ASGELGGSSYEQY | 0.5330366 |
| ALTFSVSL | ASGELGGSSYEQY | 0.6028844 |
| ALTGAALVAL | ASGELGGSSYEQY | 0.72855234 |
| ASFSIGAGVL | ASGELGGSSYEQY | 0.7467778 |
| DGSVYKI | ASGELGGSSYEQY | 0.70136905 |
| EQLAEIV | ASGELGGSSYEQY | 0.58439577 |
| FAVSELTVM | ASGELGGSSYEQY | 0.66704655 |
| GAAPLAAGL | ASGELGGSSYEQY | 0.6163825 |
| GADGGDGGSGGAGG | ASGELGGSSYEQY | 0.63319427 |
| GFGLDTGGAGG | ASGELGGSSYEQY | 0.69708014 |
| GLASGGLL | ASGELGGSSYEQY | 0.66806555 |
| GRIAGAAVLL | ASGELGGSSYEQY | 0.59192467 |
| GSAGSLNTSF | ASGELGGSSYEQY | 0.6040712 |
| GTGQIGI | ASGELGGSSYEQY | 0.6534003 |
| GTLIALIL | ASGELGGSSYEQY | 0.7099682 |
| IAATMGFVVG | ASGELGGSSYEQY | 0.6947212 |
| IFAQALL | ASGELGGSSYEQY | 0.7534167 |
| IGAAVALGF | ASGELGGSSYEQY | 0.71818215 |
| IGLTPLAL | ASGELGGSSYEQY | 0.72866964 |
| ILGSNTSTL | ASGELGGSSYEQY | 0.27640945 |
| IVFSLTVL | ASGELGGSSYEQY | 0.40580684 |
| LASKATGF | ASGELGGSSYEQY | 0.64430594 |
| LASVALIL | ASGELGGSSYEQY | 0.63170916 |
| LFAVGLTL | ASGELGGSSYEQY | 0.34606305 |
| LFNGGIGTYI | ASGELGGSSYEQY | 0.7096013 |
| LTLGAGII | ASGELGGSSYEQY | 0.619964 |
| LVAAALGDI | ASGELGGSSYEQY | 0.54484904 |
| NTAFVQLGI | ASGELGGSSYEQY | 0.6983731 |
| QLFSGAEL | ASGELGGSSYEQY | 0.60425735 |
| SAATFITL | ASGELGGSSYEQY | 0.5625294 |
| SNYGLANLGV | ASGELGGSSYEQY | 0.60073394 |
| SYGGAYAVM | ASGELGGSSYEQY | 0.7255254 |
| TTGVGGLTLG | ASGELGGSSYEQY | 0.44828218 |
| TTQTIYI | ASGELGGSSYEQY | 0.70610714 |
| VAAGAGLGAVI | ASGELGGSSYEQY | 0.5569589 |
| VAAVVTLL | ASGELGGSSYEQY | 0.72847307 |
| VGFLGGLI | ASGELGGSSYEQY | 0.49418652 |
| VGVIPILL | ASGELGGSSYEQY | 0.7038097 |
| VIFASSLI | ASGELGGSSYEQY | 0.6314566 |
| VKGGDGGAAAG | ASGELGGSSYEQY | 0.7569674 |
| VTKTFTNI | ASGELGGSSYEQY | 0.61378837 |

| | | |
|---|---|---|
| VVAAAILL | ASGELGGSSYEQY | 0.6901263 |
| YAALDVEL | ASGELGGSSYEQY | 0.64854383 |
| LASVALLL | ASGELGGSSYEQY | 0.7286267 |
| LDAVHRV | ASGELGGSSYEQY | 0.42363173 |
| QGLSSSIF | ASGELGGSSYEQY | 0.62972695 |
| RGTFANIRL | ASGELGGSSYEQY | 0.42494535 |
| RQLWWGHRI | ASGELGGSSYEQY | 0.68282735 |
| YGPPGTGKTLL | ASGELGGSSYEQY | 0.69483995 |
| AALVGAPANVA | ASGELGGSSYEQY | 0.3913956 |
| AATFRRTE | ASGELGGSSYEQY | 0.70840204 |
| AGALGGGI | ASGELGGSSYEQY | 0.6268487 |
| AGGAGGAGGIGAG | ASGELGGSSYEQY | 0.6006105 |
| AGSGALGL | ASGELGGSSYEQY | 0.7283913 |
| AQIADQI | ASGELGGSSYEQY | 0.6304085 |
| ASAVAVTYL | ASGELGGSSYEQY | 0.7282052 |
| ASTVLGGTGG | ASGELGGSSYEQY | 0.61889875 |
| AVGGEPAK | ASGELGGSSYEQY | 0.5942885 |
| CEQGTFGGTC | ASGELGGSSYEQY | 0.7055676 |
| DEGKAPAG | ASGELGGSSYEQY | 0.39419436 |
| DTVVFPAL | ASGELGGSSYEQY | 0.5320619 |
| FEGITMAG | ASGELGGSSYEQY | 0.73062843 |
| FPLVVQD | ASGELGGSSYEQY | 0.6566684 |
| GAFTTPQ | ASGELGGSSYEQY | 0.79712415 |
| GFGGFGTAGGVGGA | ASGELGGSSYEQY | 0.40581167 |
| GFLAQFL | ASGELGGSSYEQY | 0.45877108 |
| GGAGGAGGNAGVA | ASGELGGSSYEQY | 0.63055515 |
| GGVGTAGAGGAGGAGG | ASGELGGSSYEQY | 0.72194266 |
| GNGGGGDAGNAGSGGN | ASGELGGSSYEQY | 0.793216 |
| GTGGAGGAGNDAGS | ASGELGGSSYEQY | 0.69402176 |
| GTTNTGWFNS | ASGELGGSSYEQY | 0.4850543 |
| IFMGPLFTL | ASGELGGSSYEQY | 0.69476867 |
| ILAHAIEAI | ASGELGGSSYEQY | 0.51423174 |
| IRSAIREAL | ASGELGGSSYEQY | 0.61418074 |
| ITFAGGG | ASGELGGSSYEQY | 0.7306266 |
| ITGMSGHQEV | ASGELGGSSYEQY | 0.61156213 |
| IVAGLACTL | ASGELGGSSYEQY | 0.46957412 |
| IVGGGLAGA | ASGELGGSSYEQY | 0.5261656 |
| IVTGVLNI | ASGELGGSSYEQY | 0.74153364 |
| LAQQGCI | ASGELGGSSYEQY | 0.7575454 |
| LCLHVEN | ASGELGGSSYEQY | 0.741547 |
| LDAISDGS | ASGELGGSSYEQY | 0.8014767 |
| LGGFAAML | ASGELGGSSYEQY | 0.7331995 |
| LGGKNPMI | ASGELGGSSYEQY | 0.68892217 |
| LGPKGVAAIL | ASGELGGSSYEQY | 0.2490146 |
| LMVGVSI | ASGELGGSSYEQY | 0.3890002 |
| LVGAAFVL | ASGELGGSSYEQY | 0.49505574 |
| LVLAVAPL | ASGELGGSSYEQY | 0.6486236 |
| MAAIAVIDAA | ASGELGGSSYEQY | 0.24088608 |
| NNGINAELQ | ASGELGGSSYEQY | 0.7242155 |
| RGEPLVLVHG | ASGELGGSSYEQY | 0.54881144 |
| RGGAGGAAGLI | ASGELGGSSYEQY | 0.45536083 |

| | | |
|---|---|---|
| RSYRMHPAV | ASGELGGSSYEQY | 0.7772026 |
| SAAHLLAGQLA | ASGELGGSSYEQY | 0.49028075 |
| SAAVGSY | ASGELGGSSYEQY | 0.46751362 |
| SPHAVLHRI | ASGELGGSSYEQY | 0.52466273 |
| SQIAKVL | ASGELGGSSYEQY | 0.79532874 |
| TALGHLVP | ASGELGGSSYEQY | 0.37952465 |
| TAPTTSGAA | ASGELGGSSYEQY | 0.73794127 |
| TDGDDGNGGSPGT | ASGELGGSSYEQY | 0.6715369 |
| TDGLGEN | ASGELGGSSYEQY | 0.7562025 |
| TGGPPLPGQA | ASGELGGSSYEQY | 0.36092317 |
| TIPAITI | ASGELGGSSYEQY | 0.7226426 |
| TIPGFTI | ASGELGGSSYEQY | 0.5701927 |
| TSIASGALA | ASGELGGSSYEQY | 0.8187829 |
| TVTLPGAAI | ASGELGGSSYEQY | 0.5888723 |
| VASGLSAALP | ASGELGGSSYEQY | 0.73568845 |
| VGAGVIGI | ASGELGGSSYEQY | 0.67671394 |
| VGPKRFGV | ASGELGGSSYEQY | 0.7792523 |
| VNVAFPD | ASGELGGSSYEQY | 0.35396966 |
| VSMTNSGVS | ASGELGGSSYEQY | 0.5769682 |
| VTDVLGDLSV | ASGELGGSSYEQY | 0.49045765 |
| VVVVGAGFAGL | ASGELGGSSYEQY | 0.6300752 |
| VYTVGTVV | ASGELGGSSYEQY | 0.688336 |
| WTTFSGP | ASGELGGSSYEQY | 0.36050335 |
| YDELVEAF | ASGELGGSSYEQY | 0.76563525 |
| YGTGGAGGNGG | ASGELGGSSYEQY | 0.65873206 |
| YTNVMAV | ASGELGGSSYEQY | 0.446091 |
| FLEQVHQL | ASGELGGSSYEQY | 0.7384335 |
| GLSLLAVAV | ASGELGGSSYEQY | 0.5900092 |
| LEQVHQL | ASGELGGSSYEQY | 0.70194566 |
| LNAAGAI | ASGELGGSSYEQY | 0.48642173 |
| PAVHASL | ASGELGGSSYEQY | 0.6578998 |
| TELEDDA | ASGELGGSSYEQY | 0.6841085 |
| TPPGPPN | ASGELGGSSYEQY | 0.42970714 |
| TPPPPPPP | ASGELGGSSYEQY | 0.44581735 |
| TQSPALI | AWSLGGAETLY | 0.7543733 |
| EFAAAFPQ | AWSLGGAETLY | 0.7170559 |
| LLSGNLDVL | AWSLGGAETLY | 0.7348478 |
| GPANLGNYNI | AWSLGGAETLY | 0.60151136 |
| TLGPTLP | AWSLGGAETLY | 0.59081185 |
| VYAYSSCCK | AWSLGGAETLY | 0.64553237 |
| APAPAPAPNI | AWSLGGAETLY | 0.7145263 |
| PGGPVPIP | AWSLGGAETLY | 0.73492265 |
| FLTVALEP | AWSLGGAETLY | 0.7056658 |
| SQYFGANL | AWSLGGAETLY | 0.5328977 |
| GGVKPAA | AWSLGGAETLY | 0.6860498 |
| GGTGGAGGAGGW | AWSLGGAETLY | 0.66669786 |
| GVGGVAGLGGAGG | AWSLGGAETLY | 0.69608605 |
| VLLVLATTI | AWSLGGAETLY | 0.64418626 |
| IGIFAAML | AWSLGGAETLY | 0.6704186 |
| AVGDLID | AWSLGGAETLY | 0.5427755 |
| NISPLPL | AWSLGGAETLY | 0.5038372 |

| | | |
|---|---|---|
| VTAVFVAI | AWSLGGAETLY | 0.6982434 |
| SGDDNDAAL | AWSLGGAETLY | 0.72823566 |
| AVAALAHS | AWSLGGAETLY | 0.7217651 |
| GGVTVMATP | AWSLGGAETLY | 0.7214037 |
| AGLAANVRD | AWSLGGAETLY | 0.7315917 |
| TNRPDLIDPALLRPG | AWSLGGAETLY | 0.6543316 |
| LANLGVTAL | AWSLGGAETLY | 0.32853293 |
| TGERIEL | AWSLGGAETLY | 0.418128 |
| NGEELAA | AWSLGGAETLY | 0.47361428 |
| VLSLVWG | AWSLGGAETLY | 0.65322447 |
| IGYALWL | AWSLGGAETLY | 0.25505278 |
| LSLLLAPL | AWSLGGAETLY | 0.6650482 |
| TLDEALK | AWSLGGAETLY | 0.5655662 |
| TDGAGTAEGG | AWSLGGAETLY | 0.5869409 |
| NAPGGAGGNGG | AWSLGGAETLY | 0.7227955 |
| VSVFMGMGGNF | AWSLGGAETLY | 0.65618885 |
| LASYLAAL | AWSLGGAETLY | 0.4775341 |
| LVGGTPAL | AWSLGGAETLY | 0.46357882 |
| VRNADVL | AWSLGGAETLY | 0.6803169 |
| TCSSPFLVSAM | AWSLGGAETLY | 0.42355335 |
| VHITDAE | AWSLGGAETLY | 0.7334179 |
| EYAAPSF | AWSLGGAETLY | 0.39062643 |
| VVDAFHGGL | AWSLGGAETLY | 0.7354697 |
| ASVVDPQ | AWSLGGAETLY | 0.4953081 |
| SYNEFGGLS | AWSLGGAETLY | 0.72371995 |
| DPDGSVY | AWSLGGAETLY | 0.709913 |
| MVSTIHATE | AWSLGGAETLY | 0.7483419 |
| LAAGVADPV | AWSLGGAETLY | 0.6745479 |
| TKEQIAA | AWSLGGAETLY | 0.72056174 |
| SLASPAAL | AWSLGGAETLY | 0.6719129 |
| IGLVFYGV | AWSLGGAETLY | 0.74138725 |
| FLLDASNL | AWSLGGAETLY | 0.52310187 |
| YQVGNFP | AWSLGGAETLY | 0.6549805 |
| ESLLSQST | AWSLGGAETLY | 0.347277 |
| LFTVGFAAVG | AWSLGGAETLY | 0.65861905 |
| ATPPTPT | AWSLGGAETLY | 0.71817905 |
| TDLATLG | AWSLGGAETLY | 0.28167486 |
| SAGGTGR | AWSLGGAETLY | 0.64138275 |
| GFSNSGS | AWSLGGAETLY | 0.57890165 |
| GGNGGSGGTSV | AWSLGGAETLY | 0.5970551 |
| GTSVATGGAGN | AWSLGGAETLY | 0.7044814 |
| NGGNGGSGGTS | AWSLGGAETLY | 0.6736917 |
| AVVKFPH | AWSLGGAETLY | 0.676674 |
| TPPPPPP | AWSLGGAETLY | 0.44986808 |
| AAPAAAAAPA | AWSLGGAETLY | 0.53818583 |
| IALNEDL | AWSLGGAETLY | 0.63656455 |
| IVNVSSM | AWSLGGAETLY | 0.4687038 |
| TGIPPAPR | AWSLGGAETLY | 0.3908596 |
| FDLGGGTF | AWSLGGAETLY | 0.7395636 |
| FDLGGGTFDVS | AWSLGGAETLY | 0.6507982 |
| NEPTAAA | AWSLGGAETLY | 0.78856295 |

| | | |
|---|---|---|
| NEPTAAALA | AWSLGGAETLY | 0.44363618 |
| GSIINISSI | AWSLGGAETLY | 0.42393923 |
| VAPGNAGTA | AWSLGGAETLY | 0.5563357 |
| TGGGASLE | AWSLGGAETLY | 0.7728876 |
| VAVYNHYKRI | AWSLGGAETLY | 0.77189314 |
| TDPAPATT | AWSLGGAETLY | 0.679011 |
| TNRPDILDPALLRPG | AWSLGGAETLY | 0.4532454 |
| LTTTTTS | AWSLGGAETLY | 0.5176933 |
| PTGSGKTL | AWSLGGAETLY | 0.49801487 |
| TGSGKTL | AWSLGGAETLY | 0.5545837 |
| TIGHVDHGK | AWSLGGAETLY | 0.65226007 |
| TIGHVDHGKTT | AWSLGGAETLY | 0.6255274 |
| GHVDHGKTT | AWSLGGAETLY | 0.48294616 |
| AQFEHTLL | AWSLGGAETLY | 0.52312976 |
| IVNQDEL | AWSLGGAETLY | 0.6182555 |
| TNGSQFF | AWSLGGAETLY | 0.65794885 |
| TGKGIVE | AWSLGGAETLY | 0.5543967 |
| LISNASDA | AWSLGGAETLY | 0.800318 |
| AELRTLGELTKIAKAHGAQ | AWSLGGAETLY | 0.6982763 |
| APNAVAAAEPSW | AWSLGGAETLY | 0.68824375 |
| APQVAGAATAVL | AWSLGGAETLY | 0.17878775 |
| ARRDPAAAAAILPTDA | AWSLGGAETLY | 0.24496175 |
| DAAARDVLAEAGLAEYF | AWSLGGAETLY | 0.54511344 |
| DDQIVLSAKANLAKVEQ | AWSLGGAETLY | 0.55169797 |
| DELLPDTGEDSDEDGH | AWSLGGAETLY | 0.26123196 |
| ELPEAPANRATVPQD | AWSLGGAETLY | 0.7168899 |
| FGQSKIERVTGIG | AWSLGGAETLY | 0.51436406 |
| GAGGTGSTAGGAGGAGGAGGI | AWSLGGAETLY | 0.463759 |
| GHGGTGAAGGNGGTGGAGGS( | AWSLGGAETLY | 0.6821548 |
| GNGGNGGAGGEGGPGGAGGA( | AWSLGGAETLY | 0.53186244 |
| GSTAGNGGNGGNATNGTIAG( | AWSLGGAETLY | 0.50926185 |
| GTGGDGGNGGNGGTGG | AWSLGGAETLY | 0.48924035 |
| GVRRPAGIPADAPV | AWSLGGAETLY | 0.6677704 |
| GYRQVTDAHLLTLARR | AWSLGGAETLY | 0.2916657 |
| IAKMGQDGHDRG | AWSLGGAETLY | 0.6355498 |
| KAEPSIEIQYT | AWSLGGAETLY | 0.43160653 |
| KPVRIRMAADEDTVAVT | AWSLGGAETLY | 0.73056096 |
| MPPRIAGMRLLVIKPEP | AWSLGGAETLY | 0.3651142 |
| MKFYAVALFRSRGVPPTR | AWSLGGAETLY | 0.6639485 |
| NAGGFGILGAVAHSP | AWSLGGAETLY | 0.5709059 |
| NEGREITLAMAEM | AWSLGGAETLY | 0.8074824 |
| NELVEWSHPLGHRWRWE | AWSLGGAETLY | 0.5021624 |
| NLGAGGTGGNGGNPG | AWSLGGAETLY | 0.63982654 |
| NTEWPKWETVMAADPVD | AWSLGGAETLY | 0.5738432 |
| PASAAPAETQP | AWSLGGAETLY | 0.7458211 |
| QQSSDGKNGVRAAEITGE | AWSLGGAETLY | 0.40877095 |
| RAPDAVTTSWWKE | AWSLGGAETLY | 0.49654394 |
| SGFGNVGAHTSGWFNQ | AWSLGGAETLY | 0.3027524 |
| SGVAGAVGQAASVGGLKV | AWSLGGAETLY | 0.6443817 |
| SNGGAGTGSAGGNGGT | AWSLGGAETLY | 0.6574705 |
| TGGAGGEGGAAG | AWSLGGAETLY | 0.3693965 |

| | | |
|---|---|---|
| TGGDGGAAGAGGAPGHG | AWSLGGAETLY | 0.60613585 |
| TMPPLLAAIANAGRHGV | AWSLGGAETLY | 0.6699121 |
| TSSTAGNSGGA | AWSLGGAETLY | 0.5102726 |
| VPATVPVGILC | AWSLGGAETLY | 0.7248836 |
| VQRPDSGGVVLTGRIS | AWSLGGAETLY | 0.44939768 |
| WTPNPVPATKGVPAGLVQAT | AWSLGGAETLY | 0.73707867 |
| YALKRALAKFSDRFDVVII | AWSLGGAETLY | 0.5878622 |
| YLIGSVIV | AWSLGGAETLY | 0.39070666 |
| YGQLVAVID | AWSLGGAETLY | 0.6655391 |
| VTVGGMTLPA | AWSLGGAETLY | 0.36577404 |
| VLALPAV | AWSLGGAETLY | 0.42294082 |
| VIPPTLNYET | AWSLGGAETLY | 0.7348631 |
| VGMGGMHPSSG | AWSLGGAETLY | 0.77890265 |
| VGLAGGAATA | AWSLGGAETLY | 0.7958977 |
| VGGAGGAGGSGP | AWSLGGAETLY | 0.6723653 |
| VFAAFMV | AWSLGGAETLY | 0.5674906 |
| TLYDKLR | AWSLGGAETLY | 0.69163525 |
| TLAVFTVL | AWSLGGAETLY | 0.7917924 |
| TGSVQPARV | AWSLGGAETLY | 0.8110999 |
| TGGGAGSGGT | AWSLGGAETLY | 0.7547299 |
| SHLETSL | AWSLGGAETLY | 0.49084663 |
| SFTTGMT | AWSLGGAETLY | 0.771542 |
| QVEVAKRMGVRQPTVS | AWSLGGAETLY | 0.6967282 |
| QGSGGPALAM | AWSLGGAETLY | 0.7286808 |
| MLAPSNP | AWSLGGAETLY | 0.75027597 |
| LVNNAAI | AWSLGGAETLY | 0.77196014 |
| LTGEINTLA | AWSLGGAETLY | 0.6911912 |
| LSIFGPV | AWSLGGAETLY | 0.6411891 |
| LLALLANL | AWSLGGAETLY | 0.74469554 |
| LISNASDALDKLR | AWSLGGAETLY | 0.7990627 |
| LINEAVP | AWSLGGAETLY | 0.7859197 |
| LGYAFSGL | AWSLGGAETLY | 0.81489635 |
| LDFVAVAL | AWSLGGAETLY | 0.81129265 |
| LAQPGAE | AWSLGGAETLY | 0.7959122 |
| ISNLGAVTGF | AWSLGGAETLY | 0.22928753 |
| HQEDPGRI | AWSLGGAETLY | 0.40551358 |
| GTPGEPTP | AWSLGGAETLY | 0.5929482 |
| GFFNSGS | AWSLGGAETLY | 0.70294374 |
| GAGGGGLTAGAN | AWSLGGAETLY | 0.27677238 |
| FVDALVNM | AWSLGGAETLY | 0.7424488 |
| FLTVAPEL | AWSLGGAETLY | 0.6407382 |
| FETLTAL | AWSLGGAETLY | 0.5826436 |
| DVINPIEL | AWSLGGAETLY | 0.80680335 |
| APVSGAPV | AWSLGGAETLY | 0.68828857 |
| ALVGGLAW | AWSLGGAETLY | 0.57104635 |
| ALAAAAATAI | AWSLGGAETLY | 0.5388803 |
| AFGLGLLL | AWSLGGAETLY | 0.80212766 |
| AFFAAASL | AWSLGGAETLY | 0.45385942 |
| AAAALPAFSPP | AWSLGGAETLY | 0.7985841 |
| VVLAPAL | AWSLGGAETLY | 0.5627283 |
| TIGHVDHGKTTL | AWSLGGAETLY | 0.8146547 |

| | | |
|---|---|---|
| LISNASDALDK | AWSLGGAETLY | 0.41303784 |
| LISNASDAL | AWSLGGAETLY | 0.7909936 |
| ISNASDALDK | AWSLGGAETLY | 0.72990036 |
| NQAETLVYQTE | AWSLGGAETLY | 0.84105384 |
| TGGTGGTAGAG | AWSLGGAETLY | 0.7019644 |
| TGSTAGGAGGA | AWSLGGAETLY | 0.742298 |
| DPLAVPVAAGPV | AWSLGGAETLY | 0.6859374 |
| GGTGGTPGNGGN | AWSLGGAETLY | 0.8076118 |
| PAGLAGVDPATL | AWSLGGAETLY | 0.41568792 |
| GNNLVVEAPHAGASV | AWSLGGAETLY | 0.7105004 |
| KVTRIMSASTLEQPA | AWSLGGAETLY | 0.38245454 |
| LGFALPIALPRRTVV | AWSLGGAETLY | 0.74536896 |
| TRLGNLTARKLVVLV | AWSLGGAETLY | 0.7742115 |
| WSQQRRLIYFPSAGP | AWSLGGAETLY | 0.28385875 |
| GPNGNPDPMAAAVDIR | AWSLGGAETLY | 0.79653347 |
| TQRCVSVADLRAAASR | AWSLGGAETLY | 0.65448743 |
| QFVLDVAEHPVEVLTETV | AWSLGGAETLY | 0.546937 |
| GGAGGTGGAGGNGGTQAGDG | AWSLGGAETLY | 0.79743165 |
| PLLATLRAIAVDVDAKPAQV | AWSLGGAETLY | 0.71164894 |
| APKFRTGGYMIDFWGVGYQV | AWSLGGAETLY | 0.81052566 |
| GAGGAGGAGGAAGGPGGTGG | AWSLGGAETLY | 0.63752353 |
| IGALGLLATDVAPAGSVAFG | AWSLGGAETLY | 0.6898767 |
| AGGAGGIGGNGGAGGAGGVL | AWSLGGAETLY | 0.76754403 |
| ADELIGLL | AWSLGGAETLY | 0.34708127 |
| ADRALKA | AWSLGGAETLY | 0.4625926 |
| AGAVGQVGGAGGA | AWSLGGAETLY | 0.8177812 |
| AGGGGGLGGAP | AWSLGGAETLY | 0.5714943 |
| AGGLLYGNGGAG | AWSLGGAETLY | 0.6315521 |
| AGLESIGL | AWSLGGAETLY | 0.5027931 |
| AGLKTGII | AWSLGGAETLY | 0.4630885 |
| ASLAALLL | AWSLGGAETLY | 0.54045475 |
| ASVINISSI | AWSLGGAETLY | 0.6190168 |
| ATPPGPA | AWSLGGAETLY | 0.6173167 |
| ATVASALV | AWSLGGAETLY | 0.5948849 |
| ATVASLIL | AWSLGGAETLY | 0.47229177 |
| AVNGPVLIRH | AWSLGGAETLY | 0.5594473 |
| DGAAGAVGI | AWSLGGAETLY | 0.50697696 |
| DPLLHGI | AWSLGGAETLY | 0.5405114 |
| ELGAALGAL | AWSLGGAETLY | 0.5821449 |
| ELGRTQAV | AWSLGGAETLY | 0.5767794 |
| FVAGGYLS | AWSLGGAETLY | 0.5442624 |
| GASGGAGGAGGW | AWSLGGAETLY | 0.5236009 |
| GGAGAGGDGGAGGI | AWSLGGAETLY | 0.58667994 |
| GTGHNGSQPGMGGNG | AWSLGGAETLY | 0.58767605 |
| GYAAASASAS | AWSLGGAETLY | 0.5532195 |
| HQVDVAAVQQG | AWSLGGAETLY | 0.64915246 |
| IGAAIAHT | AWSLGGAETLY | 0.60514194 |
| IGMIEVAQL | AWSLGGAETLY | 0.608959 |
| IRLAPEHTSAMD | AWSLGGAETLY | 0.33723825 |
| IYNVSVL | AWSLGGAETLY | 0.40430868 |
| KQGFLGM | AWSLGGAETLY | 0.5369042 |

| | | |
|---|---|---|
| LESVITQL | AWSLGGAETLY | 0.5209091 |
| LGLGALVVV | AWSLGGAETLY | 0.4019623 |
| LIGNGGNGGSGGI | AWSLGGAETLY | 0.59851384 |
| LNAINAPTL | AWSLGGAETLY | 0.5147222 |
| LQLLNTGV | AWSLGGAETLY | 0.4970958 |
| LQTSIEF | AWSLGGAETLY | 0.5984818 |
| MSFAVSEL | AWSLGGAETLY | 0.52927285 |
| NTGALGL | AWSLGGAETLY | 0.4975196 |
| PPGAGGAGGAGTT | AWSLGGAETLY | 0.5109689 |
| QLVGLGI | AWSLGGAETLY | 0.62380636 |
| RFLEQVHQL | AWSLGGAETLY | 0.41044706 |
| RGVGYVLR | AWSLGGAETLY | 0.5837768 |
| SATGVFAGVI | AWSLGGAETLY | 0.5189223 |
| SLPAITLP | AWSLGGAETLY | 0.61666673 |
| STALAFTV | AWSLGGAETLY | 0.4157549 |
| SVLTVGLGK | AWSLGGAETLY | 0.59248036 |
| TALAGAGSGI | AWSLGGAETLY | 0.5278168 |
| TGLPFTGI | AWSLGGAETLY | 0.65694517 |
| VAEHVTV | AWSLGGAETLY | 0.5121318 |
| VDIVLNSL | AWSLGGAETLY | 0.5577484 |
| VLTGGGAVL | AWSLGGAETLY | 0.5294342 |
| VNVGLAYGI | AWSLGGAETLY | 0.6165957 |
| VSSAAVGAV | AWSLGGAETLY | 0.41488835 |
| VTSVYEAGV | AWSLGGAETLY | 0.52201587 |
| WVALDRGA | AWSLGGAETLY | 0.4421568 |
| YLVGSVVI | AWSLGGAETLY | 0.5711353 |
| GPPGTGKTLL | AWSLGGAETLY | 0.58565116 |
| GVGMTKF | AWSLGGAETLY | 0.43365684 |
| IGIGGSDL | AWSLGGAETLY | 0.62931424 |
| IGTLLAEL | AWSLGGAETLY | 0.56152236 |
| IYTSGTT | AWSLGGAETLY | 0.47858012 |
| LSFPAGV | AWSLGGAETLY | 0.62309176 |
| LYTSGTT | AWSLGGAETLY | 0.5532625 |
| VGAFPFGF | AWSLGGAETLY | 0.62569904 |
| VGAGTVEF | AWSLGGAETLY | 0.5597651 |
| VGKSTLLL | AWSLGGAETLY | 0.537275 |
| AALAHAGGGIV | AWSLGGAETLY | 0.60298884 |
| AALVAVSLL | AWSLGGAETLY | 0.5140071 |
| AAVSRAVAAL | AWSLGGAETLY | 0.48793596 |
| AGGSSGIGGF | AWSLGGAETLY | 0.6195993 |
| AGLTVANANG | AWSLGGAETLY | 0.85891676 |
| AGLVGAVL | AWSLGGAETLY | 0.8854948 |
| ALSQAFTV | AWSLGGAETLY | 0.7943381 |
| ALTFSVSL | AWSLGGAETLY | 0.75302446 |
| ALTGAALVAL | AWSLGGAETLY | 0.81575286 |
| ASFSIGAGVL | AWSLGGAETLY | 0.88174754 |
| DGSVYKI | AWSLGGAETLY | 0.876758 |
| EQLAEIV | AWSLGGAETLY | 0.86375296 |
| FAVSELTVM | AWSLGGAETLY | 0.7299158 |
| GAAPLAAGL | AWSLGGAETLY | 0.8464284 |
| GADGGDGGSGGAGG | AWSLGGAETLY | 0.80831254 |

| | | |
|---|---|---|
| GFGLDTGGAGG | AWSLGGAETLY | 0.81474423 |
| GLASGGLL | AWSLGGAETLY | 0.8654416 |
| GRIAGAAVLL | AWSLGGAETLY | 0.8368879 |
| GSAGSLNTSF | AWSLGGAETLY | 0.80536765 |
| GTGQIGI | AWSLGGAETLY | 0.8203517 |
| GTLIALIL | AWSLGGAETLY | 0.83739865 |
| IAATMGFVVG | AWSLGGAETLY | 0.8564521 |
| IFAQALL | AWSLGGAETLY | 0.8435092 |
| IGAAVALGF | AWSLGGAETLY | 0.8960171 |
| IGLTPLAL | AWSLGGAETLY | 0.87743664 |
| ILGSNTSTL | AWSLGGAETLY | 0.8568765 |
| IVFSLTVL | AWSLGGAETLY | 0.37552357 |
| LASKATGF | AWSLGGAETLY | 0.7285737 |
| LASVALIL | AWSLGGAETLY | 0.8267275 |
| LFAVGLTL | AWSLGGAETLY | 0.828925 |
| LFNGGIGTYI | AWSLGGAETLY | 0.4731509 |
| LTLGAGII | AWSLGGAETLY | 0.86655486 |
| LVAAALGDI | AWSLGGAETLY | 0.81860757 |
| NTAFVQLGI | AWSLGGAETLY | 0.7519189 |
| QLFSGAEL | AWSLGGAETLY | 0.8682306 |
| SAATFITL | AWSLGGAETLY | 0.78371364 |
| SNYGLANLGV | AWSLGGAETLY | 0.7889589 |
| SYGGAYAVM | AWSLGGAETLY | 0.8229777 |
| TTGVGGLTLG | AWSLGGAETLY | 0.8774344 |
| TTQTIYI | AWSLGGAETLY | 0.74259716 |
| VAAGAGLGAVI | AWSLGGAETLY | 0.8701103 |
| VAAVVTLL | AWSLGGAETLY | 0.8289732 |
| VGFLGGLI | AWSLGGAETLY | 0.88461125 |
| VGVIPILL | AWSLGGAETLY | 0.6620824 |
| VIFASSLI | AWSLGGAETLY | 0.87509835 |
| VKGGDGGAAAG | AWSLGGAETLY | 0.83310854 |
| VTKTFTNI | AWSLGGAETLY | 0.8960333 |
| VVAAAILL | AWSLGGAETLY | 0.82083094 |
| YAALDVEL | AWSLGGAETLY | 0.84697497 |
| LASVALLL | AWSLGGAETLY | 0.8222196 |
| LDAVHRV | AWSLGGAETLY | 0.8740994 |
| QGLSSSIF | AWSLGGAETLY | 0.67920053 |
| RGTFANIRL | AWSLGGAETLY | 0.82733625 |
| RQLWWGHRI | AWSLGGAETLY | 0.73183465 |
| YGPPGTGKTLL | AWSLGGAETLY | 0.8528558 |
| AALVGAPANVA | AWSLGGAETLY | 0.8662149 |
| AATFRRTE | AWSLGGAETLY | 0.5815617 |
| AGALGGGI | AWSLGGAETLY | 0.8607426 |
| AGGAGGAGGIGAG | AWSLGGAETLY | 0.81702065 |
| AGSGALGL | AWSLGGAETLY | 0.7525817 |
| AQIADQI | AWSLGGAETLY | 0.8809989 |
| ASAVAVTYL | AWSLGGAETLY | 0.81402266 |
| ASTVLGGTGG | AWSLGGAETLY | 0.8784495 |
| AVGGEPAK | AWSLGGAETLY | 0.82425135 |
| CEQGTFGGTC | AWSLGGAETLY | 0.8211484 |
| DEGKAPAG | AWSLGGAETLY | 0.8687191 |

| | | |
|---|---|---|
| DTVVFPAL | AWSLGGAETLY | 0.70508814 |
| FEGITMAG | AWSLGGAETLY | 0.7795354 |
| FPLVVQD | AWSLGGAETLY | 0.88725996 |
| GAFTTPQ | AWSLGGAETLY | 0.6123185 |
| GFGGFGTAGGVGGA | AWSLGGAETLY | 0.81334186 |
| GFLAQFL | AWSLGGAETLY | 0.25722194 |
| GGAGGAGGNAGVA | AWSLGGAETLY | 0.25141016 |
| GGVGTAGAGGAGGAGG | AWSLGGAETLY | 0.4260937 |
| GNGGGGDAGNAGSGGN | AWSLGGAETLY | 0.7062457 |
| GTGGAGGAGNDAGS | AWSLGGAETLY | 0.8615347 |
| GTTNTGWFNS | AWSLGGAETLY | 0.6798138 |
| IFMGPLFTL | AWSLGGAETLY | 0.2713912 |
| ILAHAIEAI | AWSLGGAETLY | 0.5846808 |
| IRSAIREAL | AWSLGGAETLY | 0.3673743 |
| ITFAGGG | AWSLGGAETLY | 0.4394211 |
| ITGMSGHQEV | AWSLGGAETLY | 0.6258718 |
| IVAGLACTL | AWSLGGAETLY | 0.5433545 |
| IVGGGLAGA | AWSLGGAETLY | 0.37190986 |
| IVTGVLNI | AWSLGGAETLY | 0.29416725 |
| LAQQGCI | AWSLGGAETLY | 0.70037603 |
| LCLHVEN | AWSLGGAETLY | 0.7709515 |
| LDAISDGS | AWSLGGAETLY | 0.72464514 |
| LGGFAAML | AWSLGGAETLY | 0.817299 |
| LGGKNPMI | AWSLGGAETLY | 0.7179448 |
| LGPKGVAAIL | AWSLGGAETLY | 0.7165998 |
| LMVGVSI | AWSLGGAETLY | 0.11557934 |
| LVGAAFVL | AWSLGGAETLY | 0.18569055 |
| LVLAVAPL | AWSLGGAETLY | 0.36078763 |
| MAAIAVIDAA | AWSLGGAETLY | 0.45305854 |
| NNGINAELQ | AWSLGGAETLY | 0.18112113 |
| RGEPLVLVHG | AWSLGGAETLY | 0.68622994 |
| RGGAGGAAGLI | AWSLGGAETLY | 0.3414428 |
| RSYRMHPAV | AWSLGGAETLY | 0.28173727 |
| SAAHLLAGQLA | AWSLGGAETLY | 0.7309544 |
| SAAVGSY | AWSLGGAETLY | 0.32262194 |
| SPHAVLHRI | AWSLGGAETLY | 0.2746715 |
| SQIAKVL | AWSLGGAETLY | 0.37843293 |
| TALGHLVP | AWSLGGAETLY | 0.7148702 |
| TAPTTSGAA | AWSLGGAETLY | 0.22930185 |
| TDGDDGNGGSPGT | AWSLGGAETLY | 0.70597434 |
| TDGLGEN | AWSLGGAETLY | 0.52235544 |
| TGGPPLPGQA | AWSLGGAETLY | 0.70595646 |
| TIPAITI | AWSLGGAETLY | 0.20573449 |
| TIPGFTI | AWSLGGAETLY | 0.7046362 |
| TSIASGALA | AWSLGGAETLY | 0.46290255 |
| TVTLPGAAI | AWSLGGAETLY | 0.8407756 |
| VASGLSAALP | AWSLGGAETLY | 0.379744 |
| VGAGVIGI | AWSLGGAETLY | 0.560941 |
| VGPKRFGV | AWSLGGAETLY | 0.58333373 |
| VNVAFPD | AWSLGGAETLY | 0.81349957 |
| VSMTNSGVS | AWSLGGAETLY | 0.16346262 |

| | | |
|---|---|---|
| VTDVLGDLSV | AWSLGGAETLY | 0.3289979 |
| VVVVGAGFAGL | AWSLGGAETLY | 0.22762905 |
| VYTVGTVV | AWSLGGAETLY | 0.42755002 |
| WTTFSGP | AWSLGGAETLY | 0.64430165 |
| YDELVEAF | AWSLGGAETLY | 0.12858649 |
| YGTGGAGGNGG | AWSLGGAETLY | 0.57506675 |
| YTNVMAV | AWSLGGAETLY | 0.3980727 |
| FLEQVHQL | AWSLGGAETLY | 0.26553965 |
| GLSLLAVAV | AWSLGGAETLY | 0.704328 |
| LEQVHQL | AWSLGGAETLY | 0.56553614 |
| LNAAGAI | AWSLGGAETLY | 0.74481046 |
| PAVHASL | AWSLGGAETLY | 0.49759647 |
| TELEDDA | AWSLGGAETLY | 0.45139915 |
| TPPGPPN | AWSLGGAETLY | 0.46734333 |
| TPPPPPPP | AWSLGGAETLY | 0.23939638 |
| TQSPALI | ASSQVPGQGGEGQNTL | 0.35362315 |
| EFAAAFPQ | ASSQVPGQGGEGQNTL | 0.7695546 |
| LLSGNLDVL | ASSQVPGQGGEGQNTL | 0.53582025 |
| GPANLGNYNI | ASSQVPGQGGEGQNTL | 0.7175914 |
| TLGPTLP | ASSQVPGQGGEGQNTL | 0.22816649 |
| VYAYSSCCK | ASSQVPGQGGEGQNTL | 0.22971755 |
| APAPAPAPNI | ASSQVPGQGGEGQNTL | 0.37319887 |
| PGGPVPIP | ASSQVPGQGGEGQNTL | 0.6168556 |
| FLTVALEP | ASSQVPGQGGEGQNTL | 0.6912917 |
| SQYFGANL | ASSQVPGQGGEGQNTL | 0.6566492 |
| GGVKPAA | ASSQVPGQGGEGQNTL | 0.22884218 |
| GGTGGAGGAGGW | ASSQVPGQGGEGQNTL | 0.37414038 |
| GVGGVAGLGGAGG | ASSQVPGQGGEGQNTL | 0.248708 |
| VLLVLATTI | ASSQVPGQGGEGQNTL | 0.30187467 |
| IGIFAAML | ASSQVPGQGGEGQNTL | 0.40757325 |
| AVGDLID | ASSQVPGQGGEGQNTL | 0.2598353 |
| NISPLPL | ASSQVPGQGGEGQNTL | 0.2617308 |
| VTAVFVAI | ASSQVPGQGGEGQNTL | 0.2367298 |
| SGDDNDAAL | ASSQVPGQGGEGQNTL | 0.5746635 |
| AVAALAHS | ASSQVPGQGGEGQNTL | 0.6649606 |
| GGVTVMATP | ASSQVPGQGGEGQNTL | 0.5285393 |
| AGLAANVRD | ASSQVPGQGGEGQNTL | 0.7376867 |
| TNRPDLIDPALLRPG | ASSQVPGQGGEGQNTL | 0.5985345 |
| LANLGVTAL | ASSQVPGQGGEGQNTL | 0.53265905 |
| TGERIEL | ASSQVPGQGGEGQNTL | 0.1423712 |
| NGEELAA | ASSQVPGQGGEGQNTL | 0.1664084 |
| VLSLVWG | ASSQVPGQGGEGQNTL | 0.20515719 |
| IGYALWL | ASSQVPGQGGEGQNTL | 0.37328368 |
| LSLLLAPL | ASSQVPGQGGEGQNTL | 0.14869498 |
| TLDEALK | ASSQVPGQGGEGQNTL | 0.5065112 |
| TDGAGTAEGG | ASSQVPGQGGEGQNTL | 0.2900619 |
| NAPGGAGGNGG | ASSQVPGQGGEGQNTL | 0.20895113 |
| VSVFMGMGGNF | ASSQVPGQGGEGQNTL | 0.66257274 |
| LASYLAAL | ASSQVPGQGGEGQNTL | 0.2621417 |
| LVGGTPAL | ASSQVPGQGGEGQNTL | 0.21492153 |
| VRNADVL | ASSQVPGQGGEGQNTL | 0.24197073 |

| | | |
|---|---|---|
| TCSSPFLVSAM | ASSQVPGQGGEGQNTL | 0.68891543 |
| VHITDAE | ASSQVPGQGGEGQNTL | 0.17328438 |
| EYAAPSF | ASSQVPGQGGEGQNTL | 0.51816857 |
| VVDAFHGGL | ASSQVPGQGGEGQNTL | 0.38081372 |
| ASVVDPQ | ASSQVPGQGGEGQNTL | 0.48198164 |
| SYNEFGGLS | ASSQVPGQGGEGQNTL | 0.17690653 |
| DPDGSVY | ASSQVPGQGGEGQNTL | 0.598816 |
| MVSTIHATE | ASSQVPGQGGEGQNTL | 0.33358192 |
| LAAGVADPV | ASSQVPGQGGEGQNTL | 0.7827283 |
| TKEQIAA | ASSQVPGQGGEGQNTL | 0.23911954 |
| SLASPAAL | ASSQVPGQGGEGQNTL | 0.33432564 |
| IGLVFYGV | ASSQVPGQGGEGQNTL | 0.25018322 |
| FLLDASNL | ASSQVPGQGGEGQNTL | 0.5501617 |
| YQVGNFP | ASSQVPGQGGEGQNTL | 0.16288677 |
| ESLLSQST | ASSQVPGQGGEGQNTL | 0.25815362 |
| LFTVGFAAVG | ASSQVPGQGGEGQNTL | 0.18654083 |
| ATPPTPT | ASSQVPGQGGEGQNTL | 0.35280663 |
| TDLATLG | ASSQVPGQGGEGQNTL | 0.59473044 |
| SAGGTGR | ASSQVPGQGGEGQNTL | 0.15281416 |
| GFSNSGS | ASSQVPGQGGEGQNTL | 0.5837333 |
| GGNGGSGGTSV | ASSQVPGQGGEGQNTL | 0.26710543 |
| GTSVATGGAGN | ASSQVPGQGGEGQNTL | 0.21877958 |
| NGGNGGSGGTS | ASSQVPGQGGEGQNTL | 0.5269263 |
| AVVKFPH | ASSQVPGQGGEGQNTL | 0.20445934 |
| TPPPPPP | ASSQVPGQGGEGQNTL | 0.73269844 |
| AAPAAAAAPA | ASSQVPGQGGEGQNTL | 0.21052277 |
| IALNEDL | ASSQVPGQGGEGQNTL | 0.42590642 |
| IVNVSSM | ASSQVPGQGGEGQNTL | 0.45752162 |
| TGIPPAPR | ASSQVPGQGGEGQNTL | 0.1433429 |
| FDLGGGTF | ASSQVPGQGGEGQNTL | 0.20473793 |
| FDLGGGTFDVS | ASSQVPGQGGEGQNTL | 0.66609216 |
| NEPTAAA | ASSQVPGQGGEGQNTL | 0.74188817 |
| NEPTAAALA | ASSQVPGQGGEGQNTL | 0.7211041 |
| GSIINISSI | ASSQVPGQGGEGQNTL | 0.6866056 |
| VAPGNAGTA | ASSQVPGQGGEGQNTL | 0.6342169 |
| TGGGASLE | ASSQVPGQGGEGQNTL | 0.6939305 |
| VAVYNHYKRI | ASSQVPGQGGEGQNTL | 0.7407352 |
| TDPAPATT | ASSQVPGQGGEGQNTL | 0.75224876 |
| TNRPDILDPALLRPG | ASSQVPGQGGEGQNTL | 0.7406559 |
| LTTTTTS | ASSQVPGQGGEGQNTL | 0.6211107 |
| PTGSGKTL | ASSQVPGQGGEGQNTL | 0.74746203 |
| TGSGKTL | ASSQVPGQGGEGQNTL | 0.7109902 |
| TIGHVDHGK | ASSQVPGQGGEGQNTL | 0.71658623 |
| TIGHVDHGKTT | ASSQVPGQGGEGQNTL | 0.7329822 |
| GHVDHGKTT | ASSQVPGQGGEGQNTL | 0.7374888 |
| AQFEHTLL | ASSQVPGQGGEGQNTL | 0.71227396 |
| IVNQDEL | ASSQVPGQGGEGQNTL | 0.6720841 |
| TNGSQFF | ASSQVPGQGGEGQNTL | 0.74656963 |
| TGKGIVE | ASSQVPGQGGEGQNTL | 0.75507855 |
| LISNASDA | ASSQVPGQGGEGQNTL | 0.74960494 |
| AELRTLGELTKIAKAHGAQ | ASSQVPGQGGEGQNTL | 0.7650386 |

| | | |
|---|---|---|
| APNAVAAAEPSW | ASSQVPGQGGEGQNTL | 0.7504807 |
| APQVAGAATAVL | ASSQVPGQGGEGQNTL | 0.751577 |
| ARRDPAAAAAILPTDA | ASSQVPGQGGEGQNTL | 0.36414486 |
| DAAARDVLAEAGLAEYF | ASSQVPGQGGEGQNTL | 0.5905552 |
| DDQIVLSAKANLAKVEQ | ASSQVPGQGGEGQNTL | 0.66109645 |
| DELLPDTGEDSDEDGH | ASSQVPGQGGEGQNTL | 0.71525013 |
| ELPEAPANRATVPQD | ASSQVPGQGGEGQNTL | 0.39148492 |
| FGQSKIERVTGIG | ASSQVPGQGGEGQNTL | 0.7007332 |
| GAGGTGSTAGGAGGAGGAGGI | ASSQVPGQGGEGQNTL | 0.6992713 |
| GHGGTGAAGGNGGTGGAGGS( | ASSQVPGQGGEGQNTL | 0.65773773 |
| GNGGNGGAGGEGGPGGAGGA( | ASSQVPGQGGEGQNTL | 0.75992954 |
| GSTAGNGGNGGNATNGTIAG( | ASSQVPGQGGEGQNTL | 0.6988455 |
| GTGGDGGNGGNGGTGG | ASSQVPGQGGEGQNTL | 0.6444792 |
| GVRRPAGIPADAPV | ASSQVPGQGGEGQNTL | 0.6817932 |
| GYRQVTDAHLLTLARR | ASSQVPGQGGEGQNTL | 0.76172817 |
| IAKMGQDGHDRG | ASSQVPGQGGEGQNTL | 0.59462625 |
| KAEPSIEIQYT | ASSQVPGQGGEGQNTL | 0.75061667 |
| KPVRIRMAADEDTVAVT | ASSQVPGQGGEGQNTL | 0.6408479 |
| MPPRIAGMRLLVIKPEP | ASSQVPGQGGEGQNTL | 0.75555253 |
| MKFYAVALFRSRGVPPTR | ASSQVPGQGGEGQNTL | 0.5610323 |
| NAGGFGILGAVAHSP | ASSQVPGQGGEGQNTL | 0.74827087 |
| NEGREITLAMAEM | ASSQVPGQGGEGQNTL | 0.7090315 |
| NELVEWSHPLGHRWRWE | ASSQVPGQGGEGQNTL | 0.76765156 |
| NLGAGGTGGNGGNPG | ASSQVPGQGGEGQNTL | 0.6988675 |
| NTEWPKWETVMAADPVD | ASSQVPGQGGEGQNTL | 0.746978 |
| PASAAPAETQP | ASSQVPGQGGEGQNTL | 0.7168462 |
| QQSSDGKNGVRAAEITGE | ASSQVPGQGGEGQNTL | 0.7347107 |
| RAPDAVTTSWWKE | ASSQVPGQGGEGQNTL | 0.56694543 |
| SGFGNVGAHTSGWFNQ | ASSQVPGQGGEGQNTL | 0.7085861 |
| SGVAGAVGQAASVGGLKV | ASSQVPGQGGEGQNTL | 0.59691024 |
| SNGGAGTGSAGGNGGT | ASSQVPGQGGEGQNTL | 0.7278558 |
| TGGAGGEGGAAG | ASSQVPGQGGEGQNTL | 0.7395233 |
| TGGDGGAAGAGGAPGHG | ASSQVPGQGGEGQNTL | 0.3364006 |
| TMPPLLAAIANAGRHGV | ASSQVPGQGGEGQNTL | 0.75288016 |
| TSSTAGNSGGA | ASSQVPGQGGEGQNTL | 0.6437884 |
| VPATVPVGILC | ASSQVPGQGGEGQNTL | 0.6217249 |
| VQRPDSGGVVLTGRIS | ASSQVPGQGGEGQNTL | 0.7517011 |
| WTPNPVPATKGVPAGLVQAT | ASSQVPGQGGEGQNTL | 0.72730803 |
| YALKRALAKFSDRFDVVII | ASSQVPGQGGEGQNTL | 0.75048625 |
| YLIGSVIV | ASSQVPGQGGEGQNTL | 0.6981725 |
| YGQLVAVID | ASSQVPGQGGEGQNTL | 0.70892394 |
| VTVGGMTLPA | ASSQVPGQGGEGQNTL | 0.7312552 |
| VLALPAV | ASSQVPGQGGEGQNTL | 0.3826019 |
| VIPPTLNYET | ASSQVPGQGGEGQNTL | 0.6376947 |
| VGMGGMHPSSG | ASSQVPGQGGEGQNTL | 0.7574365 |
| VGLAGGAATA | ASSQVPGQGGEGQNTL | 0.78949845 |
| VGGAGGAGGSGP | ASSQVPGQGGEGQNTL | 0.80319697 |
| VFAAFMV | ASSQVPGQGGEGQNTL | 0.678115 |
| TLYDKLR | ASSQVPGQGGEGQNTL | 0.58304465 |
| TLAVFTVL | ASSQVPGQGGEGQNTL | 0.72265756 |
| TGSVQPARV | ASSQVPGQGGEGQNTL | 0.8334147 |

| | | |
|---|---|---|
| TGGGAGSGGT | ASSQVPGQGGEGQNTL | 0.8277485 |
| SHLETSL | ASSQVPGQGGEGQNTL | 0.7867265 |
| SFTTGMT | ASSQVPGQGGEGQNTL | 0.5871645 |
| QVEVAKRMGVRQPTVS | ASSQVPGQGGEGQNTL | 0.8056035 |
| QGSGGPALAM | ASSQVPGQGGEGQNTL | 0.7294314 |
| MLAPSNP | ASSQVPGQGGEGQNTL | 0.7355791 |
| LVNNAAI | ASSQVPGQGGEGQNTL | 0.8043339 |
| LTGEINTLA | ASSQVPGQGGEGQNTL | 0.8058868 |
| LSIFGPV | ASSQVPGQGGEGQNTL | 0.7516209 |
| LLALLANL | ASSQVPGQGGEGQNTL | 0.68651533 |
| LISNASDALDKLR | ASSQVPGQGGEGQNTL | 0.7930046 |
| LINEAVP | ASSQVPGQGGEGQNTL | 0.81374776 |
| LGYAFSGL | ASSQVPGQGGEGQNTL | 0.808723 |
| LDFVAVAL | ASSQVPGQGGEGQNTL | 0.85825026 |
| LAQPGAE | ASSQVPGQGGEGQNTL | 0.8319134 |
| ISNLGAVTGF | ASSQVPGQGGEGQNTL | 0.8034242 |
| HQEDPGRI | ASSQVPGQGGEGQNTL | 0.2373516 |
| GTPGEPTP | ASSQVPGQGGEGQNTL | 0.44487253 |
| GFFNSGS | ASSQVPGQGGEGQNTL | 0.68051696 |
| GAGGGGLTAGAN | ASSQVPGQGGEGQNTL | 0.72904944 |
| FVDALVNM | ASSQVPGQGGEGQNTL | 0.32764482 |
| FLTVAPEL | ASSQVPGQGGEGQNTL | 0.7907932 |
| FETLTAL | ASSQVPGQGGEGQNTL | 0.71746624 |
| DVINPIEL | ASSQVPGQGGEGQNTL | 0.6588409 |
| APVSGAPV | ASSQVPGQGGEGQNTL | 0.836715 |
| ALVGGLAW | ASSQVPGQGGEGQNTL | 0.7242187 |
| ALAAAAATAI | ASSQVPGQGGEGQNTL | 0.6936817 |
| AFGLGLLL | ASSQVPGQGGEGQNTL | 0.68240833 |
| AFFAAASL | ASSQVPGQGGEGQNTL | 0.86169755 |
| AAAALPAFSPP | ASSQVPGQGGEGQNTL | 0.5426024 |
| VVLAPAL | ASSQVPGQGGEGQNTL | 0.82321316 |
| TIGHVDHGKTTL | ASSQVPGQGGEGQNTL | 0.6969588 |
| LISNASDALDK | ASSQVPGQGGEGQNTL | 0.850247 |
| LISNASDAL | ASSQVPGQGGEGQNTL | 0.500505 |
| ISNASDALDK | ASSQVPGQGGEGQNTL | 0.817519 |
| NQAETLVYQTE | ASSQVPGQGGEGQNTL | 0.72862744 |
| TGGTGGTAGAG | ASSQVPGQGGEGQNTL | 0.87461734 |
| TGSTAGGAGGA | ASSQVPGQGGEGQNTL | 0.72852993 |
| DPLAVPVAAGPV | ASSQVPGQGGEGQNTL | 0.79056275 |
| GGTGGTPGNGGN | ASSQVPGQGGEGQNTL | 0.72756314 |
| PAGLAGVDPATL | ASSQVPGQGGEGQNTL | 0.80526954 |
| GNNLVVEAPHAGASV | ASSQVPGQGGEGQNTL | 0.47283024 |
| KVTRIMSASTLEQPA | ASSQVPGQGGEGQNTL | 0.7492224 |
| LGFALPIALPRRTVV | ASSQVPGQGGEGQNTL | 0.61189383 |
| TRLGNLTARKLVVLV | ASSQVPGQGGEGQNTL | 0.78454375 |
| WSQQRRLIYFPSAGP | ASSQVPGQGGEGQNTL | 0.7900461 |
| GPNGNPDPMAAAVDIR | ASSQVPGQGGEGQNTL | 0.3586459 |
| TQRCVSVADLRAAASR | ASSQVPGQGGEGQNTL | 0.8414644 |
| QFVLDVAEHPVEVLTETV | ASSQVPGQGGEGQNTL | 0.6995429 |
| GGAGGTGGAGGNGGTQAGDG | ASSQVPGQGGEGQNTL | 0.610828 |
| PLLATLRAIAVDVDAKPAQV | ASSQVPGQGGEGQNTL | 0.83671385 |

| | | |
|---|---|---|
| APKFRTGGYMIDFWGVGYQV/ | ASSQVPGQGGEGQNTL | 0.7548015 |
| GAGGAGGAGGAAGGPGGTGG | ASSQVPGQGGEGQNTL | 0.8381127 |
| IGALGLLATDVAPAGSVAFG | ASSQVPGQGGEGQNTL | 0.7427553 |
| AGGAGGIGGNGGAGGAGGVL | ASSQVPGQGGEGQNTL | 0.7574494 |
| ADELIGLL | ASSQVPGQGGEGQNTL | 0.8152034 |
| ADRALKA | ASSQVPGQGGEGQNTL | 0.49810007 |
| AGAVGQVGGAGGA | ASSQVPGQGGEGQNTL | 0.61037993 |
| AGGGGGLGGAP | ASSQVPGQGGEGQNTL | 0.8487328 |
| AGGLLYGNGGAG | ASSQVPGQGGEGQNTL | 0.764146 |
| AGLESIGL | ASSQVPGQGGEGQNTL | 0.79668045 |
| AGLKTGII | ASSQVPGQGGEGQNTL | 0.6942625 |
| ASLAALLL | ASSQVPGQGGEGQNTL | 0.619671 |
| ASVINISSI | ASSQVPGQGGEGQNTL | 0.7450213 |
| ATPPGPA | ASSQVPGQGGEGQNTL | 0.7703741 |
| ATVASALV | ASSQVPGQGGEGQNTL | 0.79635936 |
| ATVASLIL | ASSQVPGQGGEGQNTL | 0.77389574 |
| AVNGPVLIRH | ASSQVPGQGGEGQNTL | 0.5606788 |
| DGAAGAVGI | ASSQVPGQGGEGQNTL | 0.7695232 |
| DPLLHGI | ASSQVPGQGGEGQNTL | 0.70946395 |
| ELGAALGAL | ASSQVPGQGGEGQNTL | 0.7360357 |
| ELGRTQAV | ASSQVPGQGGEGQNTL | 0.7518517 |
| FVAGGYLS | ASSQVPGQGGEGQNTL | 0.69624376 |
| GASGGAGGAGGW | ASSQVPGQGGEGQNTL | 0.69941604 |
| GGAGAGGDGGAGGI | ASSQVPGQGGEGQNTL | 0.686314 |
| GTGHNGSQPGMGGNG | ASSQVPGQGGEGQNTL | 0.7459364 |
| GYAAASASAS | ASSQVPGQGGEGQNTL | 0.78088665 |
| HQVDVAAVQQG | ASSQVPGQGGEGQNTL | 0.77888596 |
| IGAAIAHT | ASSQVPGQGGEGQNTL | 0.78533316 |
| IGMIEVAQL | ASSQVPGQGGEGQNTL | 0.77863705 |
| IRLAPEHTSAMD | ASSQVPGQGGEGQNTL | 0.75842917 |
| IYNVSVL | ASSQVPGQGGEGQNTL | 0.34476098 |
| KQGFLGM | ASSQVPGQGGEGQNTL | 0.41173053 |
| LESVITQL | ASSQVPGQGGEGQNTL | 0.65252644 |
| LGLGALVVV | ASSQVPGQGGEGQNTL | 0.67475575 |
| LIGNGGNGGSGGI | ASSQVPGQGGEGQNTL | 0.3303984 |
| LNAINAPTL | ASSQVPGQGGEGQNTL | 0.7504773 |
| LQLLNTGV | ASSQVPGQGGEGQNTL | 0.69724125 |
| LQTSIEF | ASSQVPGQGGEGQNTL | 0.61493206 |
| MSFAVSEL | ASSQVPGQGGEGQNTL | 0.7871968 |
| NTGALGL | ASSQVPGQGGEGQNTL | 0.7118434 |
| PPGAGGAGGAGTT | ASSQVPGQGGEGQNTL | 0.5404367 |
| QLVGLGI | ASSQVPGQGGEGQNTL | 0.53285587 |
| RFLEQVHQL | ASSQVPGQGGEGQNTL | 0.72721076 |
| RGVGYVLR | ASSQVPGQGGEGQNTL | 0.45681095 |
| SATGVFAGVI | ASSQVPGQGGEGQNTL | 0.7733209 |
| SLPAITLP | ASSQVPGQGGEGQNTL | 0.53920877 |
| STALAFTV | ASSQVPGQGGEGQNTL | 0.7687136 |
| SVLTVGLGK | ASSQVPGQGGEGQNTL | 0.47979057 |
| TALAGAGSGI | ASSQVPGQGGEGQNTL | 0.7645606 |
| TGLPFTGI | ASSQVPGQGGEGQNTL | 0.7409129 |
| VAEHVTV | ASSQVPGQGGEGQNTL | 0.8057263 |

| | | |
|---|---|---|
| VDIVLNSL | ASSQVPGQGGEGQNTL | 0.7049931 |
| VLTGGGAVL | ASSQVPGQGGEGQNTL | 0.71781206 |
| VNVGLAYGI | ASSQVPGQGGEGQNTL | 0.5342998 |
| VSSAAVGAV | ASSQVPGQGGEGQNTL | 0.79436463 |
| VTSVYEAGV | ASSQVPGQGGEGQNTL | 0.5626885 |
| WVALDRGA | ASSQVPGQGGEGQNTL | 0.683608 |
| YLVGSVVI | ASSQVPGQGGEGQNTL | 0.46253318 |
| GPPGTGKTLL | ASSQVPGQGGEGQNTL | 0.7398577 |
| GVGMTKF | ASSQVPGQGGEGQNTL | 0.76628375 |
| IGIGGSDL | ASSQVPGQGGEGQNTL | 0.47508112 |
| IGTLLAEL | ASSQVPGQGGEGQNTL | 0.77300775 |
| IYTSGTT | ASSQVPGQGGEGQNTL | 0.71266115 |
| LSFPAGV | ASSQVPGQGGEGQNTL | 0.6548095 |
| LYTSGTT | ASSQVPGQGGEGQNTL | 0.7867379 |
| VGAFPFGF | ASSQVPGQGGEGQNTL | 0.71862406 |
| VGAGTVEF | ASSQVPGQGGEGQNTL | 0.79147583 |
| VGKSTLLL | ASSQVPGQGGEGQNTL | 0.59764934 |
| AALAHAGGGIV | ASSQVPGQGGEGQNTL | 0.64918727 |
| AALVAVSLL | ASSQVPGQGGEGQNTL | 0.72514194 |
| AAVSRAVAAL | ASSQVPGQGGEGQNTL | 0.5326789 |
| AGGSSGIGGF | ASSQVPGQGGEGQNTL | 0.48882687 |
| AGLTVANANG | ASSQVPGQGGEGQNTL | 0.7881141 |
| AGLVGAVL | ASSQVPGQGGEGQNTL | 0.7795154 |
| ALSQAFTV | ASSQVPGQGGEGQNTL | 0.82000923 |
| ALTFSVSL | ASSQVPGQGGEGQNTL | 0.72694844 |
| ALTGAALVAL | ASSQVPGQGGEGQNTL | 0.6604677 |
| ASFSIGAGVL | ASSQVPGQGGEGQNTL | 0.74579644 |
| DGSVYKI | ASSQVPGQGGEGQNTL | 0.8090003 |
| EQLAEIV | ASSQVPGQGGEGQNTL | 0.81459844 |
| FAVSELTVM | ASSQVPGQGGEGQNTL | 0.791405 |
| GAAPLAAGL | ASSQVPGQGGEGQNTL | 0.6105856 |
| GADGGDGGSGGAGG | ASSQVPGQGGEGQNTL | 0.74218297 |
| GFGLDTGGAGG | ASSQVPGQGGEGQNTL | 0.73793113 |
| GLASGGLL | ASSQVPGQGGEGQNTL | 0.72314 |
| GRIAGAAVLL | ASSQVPGQGGEGQNTL | 0.7769494 |
| GSAGSLNTSF | ASSQVPGQGGEGQNTL | 0.76668084 |
| GTGQIGI | ASSQVPGQGGEGQNTL | 0.6780504 |
| GTLIALIL | ASSQVPGQGGEGQNTL | 0.6974021 |
| IAATMGFVVG | ASSQVPGQGGEGQNTL | 0.7666203 |
| IFAQALL | ASSQVPGQGGEGQNTL | 0.79593945 |
| IGAAVALGF | ASSQVPGQGGEGQNTL | 0.7903404 |
| IGLTPLAL | ASSQVPGQGGEGQNTL | 0.8136127 |
| ILGSNTSTL | ASSQVPGQGGEGQNTL | 0.8043698 |
| IVFSLTVL | ASSQVPGQGGEGQNTL | 0.7865821 |
| LASKATGF | ASSQVPGQGGEGQNTL | 0.24426238 |
| LASVALIL | ASSQVPGQGGEGQNTL | 0.40077382 |
| LFAVGLTL | ASSQVPGQGGEGQNTL | 0.71537113 |
| LFNGGIGTYI | ASSQVPGQGGEGQNTL | 0.7244014 |
| LTLGAGII | ASSQVPGQGGEGQNTL | 0.47615978 |
| LVAAALGDI | ASSQVPGQGGEGQNTL | 0.7691942 |
| NTAFVQLGI | ASSQVPGQGGEGQNTL | 0.7047368 |

| | | |
|---|---|---|
| QLFSGAEL | ASSQVPGQGGEGQNTL | 0.66774887 |
| SAATFITL | ASSQVPGQGGEGQNTL | 0.80007267 |
| SNYGLANLGV | ASSQVPGQGGEGQNTL | 0.72136295 |
| SYGGAYAVM | ASSQVPGQGGEGQNTL | 0.6628072 |
| TTGVGGLTLG | ASSQVPGQGGEGQNTL | 0.6410297 |
| TTQTIYI | ASSQVPGQGGEGQNTL | 0.7950358 |
| VAAGAGLGAVI | ASSQVPGQGGEGQNTL | 0.47949937 |
| VAAVVTLL | ASSQVPGQGGEGQNTL | 0.78539443 |
| VGFLGGLI | ASSQVPGQGGEGQNTL | 0.58970016 |
| VGVIPILL | ASSQVPGQGGEGQNTL | 0.8074329 |
| VIFASSLI | ASSQVPGQGGEGQNTL | 0.5294595 |
| VKGGDGGAAAG | ASSQVPGQGGEGQNTL | 0.792934 |
| VTKTFTNI | ASSQVPGQGGEGQNTL | 0.7546661 |
| VVAAAILL | ASSQVPGQGGEGQNTL | 0.826125 |
| YAALDVEL | ASSQVPGQGGEGQNTL | 0.74040616 |
| LASVALLL | ASSQVPGQGGEGQNTL | 0.7683166 |
| LDAVHRV | ASSQVPGQGGEGQNTL | 0.6488496 |
| QGLSSSIF | ASSQVPGQGGEGQNTL | 0.80967534 |
| RGTFANIRL | ASSQVPGQGGEGQNTL | 0.6237945 |
| RQLWWGHRI | ASSQVPGQGGEGQNTL | 0.7458155 |
| YGPPGTGKTLL | ASSQVPGQGGEGQNTL | 0.4510346 |
| AALVGAPANVA | ASSQVPGQGGEGQNTL | 0.7799448 |
| AATFRRTE | ASSQVPGQGGEGQNTL | 0.7874923 |
| AGALGGGI | ASSQVPGQGGEGQNTL | 0.62242043 |
| AGGAGGAGGIGAG | ASSQVPGQGGEGQNTL | 0.7983676 |
| AGSGALGL | ASSQVPGQGGEGQNTL | 0.7782781 |
| AQIADQI | ASSQVPGQGGEGQNTL | 0.6964415 |
| ASAVAVTYL | ASSQVPGQGGEGQNTL | 0.8109937 |
| ASTVLGGTGG | ASSQVPGQGGEGQNTL | 0.7144002 |
| AVGGEPAK | ASSQVPGQGGEGQNTL | 0.8065703 |
| CEQGTFGGTC | ASSQVPGQGGEGQNTL | 0.7084347 |
| DEGKAPAG | ASSQVPGQGGEGQNTL | 0.6982112 |
| DTVVFPAL | ASSQVPGQGGEGQNTL | 0.7802149 |
| FEGITMAG | ASSQVPGQGGEGQNTL | 0.7414434 |
| FPLVVQD | ASSQVPGQGGEGQNTL | 0.64164424 |
| GAFTTPQ | ASSQVPGQGGEGQNTL | 0.80668044 |
| GFGGFGTAGGVGGA | ASSQVPGQGGEGQNTL | 0.8385132 |
| GFLAQFL | ASSQVPGQGGEGQNTL | 0.83471215 |
| GGAGGAGGNAGVA | ASSQVPGQGGEGQNTL | 0.7879031 |
| GGVGTAGAGGAGGAGG | ASSQVPGQGGEGQNTL | 0.7472036 |
| GNGGGGDAGNAGSGGN | ASSQVPGQGGEGQNTL | 0.78371054 |
| GTGGAGGAGNDAGS | ASSQVPGQGGEGQNTL | 0.82599145 |
| GTTNTGWFNS | ASSQVPGQGGEGQNTL | 0.849761 |
| IFMGPLFTL | ASSQVPGQGGEGQNTL | 0.8356924 |
| ILAHAIEAI | ASSQVPGQGGEGQNTL | 0.67141265 |
| IRSAIREAL | ASSQVPGQGGEGQNTL | 0.8453171 |
| ITFAGGG | ASSQVPGQGGEGQNTL | 0.803988 |
| ITGMSGHQEV | ASSQVPGQGGEGQNTL | 0.80165255 |
| IVAGLACTL | ASSQVPGQGGEGQNTL | 0.82730305 |
| IVGGGLAGA | ASSQVPGQGGEGQNTL | 0.8222266 |
| IVTGVLNI | ASSQVPGQGGEGQNTL | 0.7789384 |

| | | |
|---|---|---|
| LAQQGCI | ASSQVPGQGGEGQNTL | 0.6859153 |
| LCLHVEN | ASSQVPGQGGEGQNTL | 0.8171383 |
| LDAISDGS | ASSQVPGQGGEGQNTL | 0.84635246 |
| LGGFAAML | ASSQVPGQGGEGQNTL | 0.84538835 |
| LGGKNPMI | ASSQVPGQGGEGQNTL | 0.84514177 |
| LGPKGVAAIL | ASSQVPGQGGEGQNTL | 0.8489719 |
| LMVGVSI | ASSQVPGQGGEGQNTL | 0.83633727 |
| LVGAAFVL | ASSQVPGQGGEGQNTL | 0.34724498 |
| LVLAVAPL | ASSQVPGQGGEGQNTL | 0.5582156 |
| MAAIAVIDAA | ASSQVPGQGGEGQNTL | 0.6757338 |
| NNGINAELQ | ASSQVPGQGGEGQNTL | 0.7849109 |
| RGEPLVLVHG | ASSQVPGQGGEGQNTL | 0.30037564 |
| RGGAGGAAGLI | ASSQVPGQGGEGQNTL | 0.7351448 |
| RSYRMHPAV | ASSQVPGQGGEGQNTL | 0.7973113 |
| SAAHLLAGQLA | ASSQVPGQGGEGQNTL | 0.75090295 |
| SAAVGSY | ASSQVPGQGGEGQNTL | 0.84498006 |
| SPHAVLHRI | ASSQVPGQGGEGQNTL | 0.7974819 |
| SQIAKVL | ASSQVPGQGGEGQNTL | 0.6735511 |
| TALGHLVP | ASSQVPGQGGEGQNTL | 0.72441226 |
| TAPTTSGAA | ASSQVPGQGGEGQNTL | 0.8378267 |
| TDGDDGNGGSPGT | ASSQVPGQGGEGQNTL | 0.6394099 |
| TDGLGEN | ASSQVPGQGGEGQNTL | 0.8491002 |
| TGGPPLPGQA | ASSQVPGQGGEGQNTL | 0.47171527 |
| TIPAITI | ASSQVPGQGGEGQNTL | 0.84452975 |
| TIPGFTI | ASSQVPGQGGEGQNTL | 0.5983546 |
| TSIASGALA | ASSQVPGQGGEGQNTL | 0.8457448 |
| TVTLPGAAI | ASSQVPGQGGEGQNTL | 0.82601523 |
| VASGLSAALP | ASSQVPGQGGEGQNTL | 0.86055225 |
| VGAGVIGI | ASSQVPGQGGEGQNTL | 0.808453 |
| VGPKRFGV | ASSQVPGQGGEGQNTL | 0.84044266 |
| VNVAFPD | ASSQVPGQGGEGQNTL | 0.78015274 |
| VSMTNSGVS | ASSQVPGQGGEGQNTL | 0.8382292 |
| VTDVLGDLSV | ASSQVPGQGGEGQNTL | 0.6471081 |
| VVVVGAGFAGL | ASSQVPGQGGEGQNTL | 0.81185853 |
| VYTVGTVV | ASSQVPGQGGEGQNTL | 0.5682292 |
| WTTFSGP | ASSQVPGQGGEGQNTL | 0.8222121 |
| YDELVEAF | ASSQVPGQGGEGQNTL | 0.8385784 |
| YGTGGAGGNGG | ASSQVPGQGGEGQNTL | 0.25317022 |
| YTNVMAV | ASSQVPGQGGEGQNTL | 0.8366461 |
| FLEQVHQL | ASSQVPGQGGEGQNTL | 0.7164231 |
| GLSLLAVAV | ASSQVPGQGGEGQNTL | 0.6805556 |
| LEQVHQL | ASSQVPGQGGEGQNTL | 0.84918827 |
| LNAAGAI | ASSQVPGQGGEGQNTL | 0.81549925 |
| PAVHASL | ASSQVPGQGGEGQNTL | 0.8455405 |
| TELEDDA | ASSQVPGQGGEGQNTL | 0.758129 |
| TPPGPPN | ASSQVPGQGGEGQNTL | 0.76443124 |
| TPPPPPPP | ASSQVPGQGGEGQNTL | 0.813234 |
| TQSPALI | ASSQTGDSGNTLY | 0.36786497 |
| EFAAAFPQ | ASSQTGDSGNTLY | 0.6272416 |
| LLSGNLDVL | ASSQTGDSGNTLY | 0.8542174 |
| GPANLGNYNI | ASSQTGDSGNTLY | 0.51655024 |

| | | |
|---|---|---|
| TLGPTLP | ASSQTGDSGNTLY | 0.6447729 |
| VYAYSSCCK | ASSQTGDSGNTLY | 0.43980816 |
| APAPAPAPNI | ASSQTGDSGNTLY | 0.42251077 |
| PGGPVPIP | ASSQTGDSGNTLY | 0.4891985 |
| FLTVALEP | ASSQTGDSGNTLY | 0.57023287 |
| SQYFGANL | ASSQTGDSGNTLY | 0.6460241 |
| GGVKPAA | ASSQTGDSGNTLY | 0.5865379 |
| GGTGGAGGAGGW | ASSQTGDSGNTLY | 0.42901212 |
| GVGGVAGLGGAGG | ASSQTGDSGNTLY | 0.59265596 |
| VLLVLATTI | ASSQTGDSGNTLY | 0.50143206 |
| IGIFAAML | ASSQTGDSGNTLY | 0.4900331 |
| AVGDLID | ASSQTGDSGNTLY | 0.5169224 |
| NISPLPL | ASSQTGDSGNTLY | 0.4759203 |
| VTAVFVAI | ASSQTGDSGNTLY | 0.44914874 |
| SGDDNDAAL | ASSQTGDSGNTLY | 0.42331082 |
| AVAALAHS | ASSQTGDSGNTLY | 0.5528904 |
| GGVTVMATP | ASSQTGDSGNTLY | 0.5732404 |
| AGLAANVRD | ASSQTGDSGNTLY | 0.5601308 |
| TNRPDLIDPALLRPG | ASSQTGDSGNTLY | 0.6206504 |
| LANLGVTAL | ASSQTGDSGNTLY | 0.57430047 |
| TGERIEL | ASSQTGDSGNTLY | 0.46806163 |
| NGEELAA | ASSQTGDSGNTLY | 0.34740087 |
| VLSLVWG | ASSQTGDSGNTLY | 0.387724 |
| IGYALWL | ASSQTGDSGNTLY | 0.46283102 |
| LSLLLAPL | ASSQTGDSGNTLY | 0.5016596 |
| TLDEALK | ASSQTGDSGNTLY | 0.33857802 |
| TDGAGTAEGG | ASSQTGDSGNTLY | 0.58247197 |
| NAPGGAGGNGG | ASSQTGDSGNTLY | 0.49495733 |
| VSVFMGMGGNF | ASSQTGDSGNTLY | 0.47075903 |
| LASYLAAL | ASSQTGDSGNTLY | 0.6071409 |
| LVGGTPAL | ASSQTGDSGNTLY | 0.47373125 |
| VRNADVL | ASSQTGDSGNTLY | 0.41075322 |
| TCSSPFLVSAM | ASSQTGDSGNTLY | 0.43875447 |
| VHITDAE | ASSQTGDSGNTLY | 0.5871494 |
| EYAAPSF | ASSQTGDSGNTLY | 0.3815355 |
| VVDAFHGGL | ASSQTGDSGNTLY | 0.5914732 |
| ASVVDPQ | ASSQTGDSGNTLY | 0.42647296 |
| SYNEFGGLS | ASSQTGDSGNTLY | 0.5931573 |
| DPDGSVY | ASSQTGDSGNTLY | 0.39427072 |
| MVSTIHATE | ASSQTGDSGNTLY | 0.5642294 |
| LAAGVADPV | ASSQTGDSGNTLY | 0.5352534 |
| TKEQIAA | ASSQTGDSGNTLY | 0.6436329 |
| SLASPAAL | ASSQTGDSGNTLY | 0.4567489 |
| IGLVFYGV | ASSQTGDSGNTLY | 0.5404002 |
| FLLDASNL | ASSQTGDSGNTLY | 0.4684838 |
| YQVGNFP | ASSQTGDSGNTLY | 0.6410343 |
| ESLLSQST | ASSQTGDSGNTLY | 0.42115614 |
| LFTVGFAAVG | ASSQTGDSGNTLY | 0.43569288 |
| ATPPTPT | ASSQTGDSGNTLY | 0.4108104 |
| TDLATLG | ASSQTGDSGNTLY | 0.4832144 |
| SAGGTGR | ASSQTGDSGNTLY | 0.5693964 |

| | | |
|---|---|---|
| GFSNSGS | ASSQTGDSGNTLY | 0.4958111 |
| GGNGGSGGTSV | ASSQTGDSGNTLY | 0.52139705 |
| GTSVATGGAGN | ASSQTGDSGNTLY | 0.5864327 |
| NGGNGGSGGTS | ASSQTGDSGNTLY | 0.4265217 |
| AVVKFPH | ASSQTGDSGNTLY | 0.54608774 |
| TPPPPPP | ASSQTGDSGNTLY | 0.56456506 |
| AAPAAAAAPA | ASSQTGDSGNTLY | 0.5187462 |
| IALNEDL | ASSQTGDSGNTLY | 0.39702982 |
| IVNVSSM | ASSQTGDSGNTLY | 0.4735625 |
| TGIPPAPR | ASSQTGDSGNTLY | 0.5319963 |
| FDLGGGTF | ASSQTGDSGNTLY | 0.51862156 |
| FDLGGGTFDVS | ASSQTGDSGNTLY | 0.4381552 |
| NEPTAAA | ASSQTGDSGNTLY | 0.59914905 |
| NEPTAAALA | ASSQTGDSGNTLY | 0.8168879 |
| GSIINISSI | ASSQTGDSGNTLY | 0.83609384 |
| VAPGNAGTA | ASSQTGDSGNTLY | 0.71075463 |
| TGGGASLE | ASSQTGDSGNTLY | 0.6969827 |
| VAVYNHYKRI | ASSQTGDSGNTLY | 0.7458174 |
| TDPAPATT | ASSQTGDSGNTLY | 0.8206274 |
| TNRPDILDPALLRPG | ASSQTGDSGNTLY | 0.8483798 |
| LTTTTTS | ASSQTGDSGNTLY | 0.82579005 |
| PTGSGKTL | ASSQTGDSGNTLY | 0.73987126 |
| TGSGKTL | ASSQTGDSGNTLY | 0.8343625 |
| TIGHVDHGK | ASSQTGDSGNTLY | 0.80369353 |
| TIGHVDHGKTT | ASSQTGDSGNTLY | 0.7807332 |
| GHVDHGKTT | ASSQTGDSGNTLY | 0.8102079 |
| AQFEHTLL | ASSQTGDSGNTLY | 0.82009065 |
| IVNQDEL | ASSQTGDSGNTLY | 0.74851465 |
| TNGSQFF | ASSQTGDSGNTLY | 0.7705149 |
| TGKGIVE | ASSQTGDSGNTLY | 0.803931 |
| LISNASDA | ASSQTGDSGNTLY | 0.8240366 |
| AELRTLGELTKIAKAHGAQ | ASSQTGDSGNTLY | 0.8155518 |
| APNAVAAAEPSW | ASSQTGDSGNTLY | 0.8472978 |
| APQVAGAATAVL | ASSQTGDSGNTLY | 0.82792115 |
| ARRDPAAAAAILPTDA | ASSQTGDSGNTLY | 0.8230369 |
| DAAARDVLAEAGLAEYF | ASSQTGDSGNTLY | 0.5527532 |
| DDQIVLSAKANLAKVEQ | ASSQTGDSGNTLY | 0.6578875 |
| DELLPDTGEDSDEDGH | ASSQTGDSGNTLY | 0.81277686 |
| ELPEAPANRATVPQD | ASSQTGDSGNTLY | 0.7598816 |
| FGQSKIERVTGIG | ASSQTGDSGNTLY | 0.5132099 |
| GAGGTGSTAGGAGGAGGAGGI | ASSQTGDSGNTLY | 0.82817125 |
| GHGGTGAAGGNGGTGGAGGS( | ASSQTGDSGNTLY | 0.78963035 |
| GNGGNGGAGGEGGPGGAGGA( | ASSQTGDSGNTLY | 0.75782704 |
| GSTAGNGGNGGNATNGTIAG( | ASSQTGDSGNTLY | 0.8204978 |
| GTGGDGGNGGNGGTGG | ASSQTGDSGNTLY | 0.7774973 |
| GVRRPAGIPADAPV | ASSQTGDSGNTLY | 0.7568194 |
| GYRQVTDAHLLTLARR | ASSQTGDSGNTLY | 0.7995068 |
| IAKMGQDGHDRG | ASSQTGDSGNTLY | 0.83390576 |
| KAEPSIEIQYT | ASSQTGDSGNTLY | 0.74503744 |
| KPVRIRMAADEDTVAVT | ASSQTGDSGNTLY | 0.8343892 |
| MPPRIAGMRLLVIKPEP | ASSQTGDSGNTLY | 0.7559031 |

| | | |
|---|---|---|
| MKFYAVALFRSRGVPPTR | ASSQTGDSGNTLY | 0.8343424 |
| NAGGFGILGAVAHSP | ASSQTGDSGNTLY | 0.7017582 |
| NEGREITLAMAEM | ASSQTGDSGNTLY | 0.8352325 |
| NELVEWSHPLGHRWRWE | ASSQTGDSGNTLY | 0.78668225 |
| NLGAGGTGGNGGNPG | ASSQTGDSGNTLY | 0.84951675 |
| NTEWPKWETVMAADPVD | ASSQTGDSGNTLY | 0.77263314 |
| PASAAPAETQP | ASSQTGDSGNTLY | 0.8348598 |
| QQSSDGKNGVRAAEITGE | ASSQTGDSGNTLY | 0.82037187 |
| RAPDAVTTSWWKE | ASSQTGDSGNTLY | 0.8487447 |
| SGFGNVGAHTSGWFNQ | ASSQTGDSGNTLY | 0.7128058 |
| SGVAGAVGQAASVGGLKV | ASSQTGDSGNTLY | 0.77534294 |
| SNGGAGTGSAGGNGGT | ASSQTGDSGNTLY | 0.6951683 |
| TGGAGGEGGAAG | ASSQTGDSGNTLY | 0.8047054 |
| TGGDGGAAGAGGAPGHG | ASSQTGDSGNTLY | 0.8323984 |
| TMPPLLAAIANAGRHGV | ASSQTGDSGNTLY | 0.43423843 |
| TSSTAGNSGGA | ASSQTGDSGNTLY | 0.8363197 |
| VPATVPVGILC | ASSQTGDSGNTLY | 0.7012248 |
| VQRPDSGGVVLTGRIS | ASSQTGDSGNTLY | 0.7351258 |
| WTPNPVPATKGVPAGLVQAT | ASSQTGDSGNTLY | 0.83024967 |
| YALKRALAKFSDRFDVVII | ASSQTGDSGNTLY | 0.8175431 |
| YLIGSVIV | ASSQTGDSGNTLY | 0.81864595 |
| YGQLVAVID | ASSQTGDSGNTLY | 0.7841192 |
| VTVGGMTLPA | ASSQTGDSGNTLY | 0.7953353 |
| VLALPAV | ASSQTGDSGNTLY | 0.8017273 |
| VIPPTLNYET | ASSQTGDSGNTLY | 0.61751735 |
| VGMGGMHPSSG | ASSQTGDSGNTLY | 0.7609387 |
| VGLAGGAATA | ASSQTGDSGNTLY | 0.8456664 |
| VGGAGGAGGSGP | ASSQTGDSGNTLY | 0.764233 |
| VFAAFMV | ASSQTGDSGNTLY | 0.84218514 |
| TLYDKLR | ASSQTGDSGNTLY | 0.6499955 |
| TLAVFTVL | ASSQTGDSGNTLY | 0.55507904 |
| TGSVQPARV | ASSQTGDSGNTLY | 0.68864 |
| TGGGAGSGGT | ASSQTGDSGNTLY | 0.8323642 |
| SHLETSL | ASSQTGDSGNTLY | 0.82252264 |
| SFTTGMT | ASSQTGDSGNTLY | 0.78138494 |
| QVEVAKRMGVRQPTVS | ASSQTGDSGNTLY | 0.4913125 |
| QGSGGPALAM | ASSQTGDSGNTLY | 0.69978607 |
| MLAPSNP | ASSQTGDSGNTLY | 0.63297975 |
| LVNNAAI | ASSQTGDSGNTLY | 0.6598344 |
| LTGEINTLA | ASSQTGDSGNTLY | 0.760719 |
| LSIFGPV | ASSQTGDSGNTLY | 0.781 |
| LLALLANL | ASSQTGDSGNTLY | 0.6495424 |
| LISNASDALDKLR | ASSQTGDSGNTLY | 0.6321164 |
| LINEAVP | ASSQTGDSGNTLY | 0.7440797 |
| LGYAFSGL | ASSQTGDSGNTLY | 0.7844111 |
| LDFVAVAL | ASSQTGDSGNTLY | 0.7300619 |
| LAQPGAE | ASSQTGDSGNTLY | 0.85004544 |
| ISNLGAVTGF | ASSQTGDSGNTLY | 0.81839275 |
| HQEDPGRI | ASSQTGDSGNTLY | 0.811982 |
| GTPGEPTP | ASSQTGDSGNTLY | 0.18877512 |
| GFFNSGS | ASSQTGDSGNTLY | 0.35808787 |

| | | |
|---|---|---|
| GAGGGGLTAGAN | ASSQTGDSGNTLY | 0.6295289 |
| FVDALVNM | ASSQTGDSGNTLY | 0.67688155 |
| FLTVAPEL | ASSQTGDSGNTLY | 0.28587648 |
| FETLTAL | ASSQTGDSGNTLY | 0.7797698 |
| DVINPIEL | ASSQTGDSGNTLY | 0.5919846 |
| APVSGAPV | ASSQTGDSGNTLY | 0.55866385 |
| ALVGGLAW | ASSQTGDSGNTLY | 0.8057376 |
| ALAAAAATAI | ASSQTGDSGNTLY | 0.640617 |
| AFGLGLLL | ASSQTGDSGNTLY | 0.61559814 |
| AFFAAASL | ASSQTGDSGNTLY | 0.5667282 |
| AAAALPAFSPP | ASSQTGDSGNTLY | 0.81648004 |
| VVLAPAL | ASSQTGDSGNTLY | 0.3513991 |
| TIGHVDHGKTTL | ASSQTGDSGNTLY | 0.7590374 |
| LISNASDALDK | ASSQTGDSGNTLY | 0.60122484 |
| LISNASDAL | ASSQTGDSGNTLY | 0.8213239 |
| ISNASDALDK | ASSQTGDSGNTLY | 0.43564892 |
| NQAETLVYQTE | ASSQTGDSGNTLY | 0.80234706 |
| TGGTGGTAGAG | ASSQTGDSGNTLY | 0.7215219 |
| TGSTAGGAGGA | ASSQTGDSGNTLY | 0.86722803 |
| DPLAVPVAAGPV | ASSQTGDSGNTLY | 0.6756697 |
| GGTGGTPGNGGN | ASSQTGDSGNTLY | 0.7207606 |
| PAGLAGVDPATL | ASSQTGDSGNTLY | 0.6659054 |
| GNNLVVEAPHAGASV | ASSQTGDSGNTLY | 0.81945044 |
| KVTRIMSASTLEQPA | ASSQTGDSGNTLY | 0.4283749 |
| LGFALPIALPRRTVV | ASSQTGDSGNTLY | 0.6801449 |
| TRLGNLTARKLVVLV | ASSQTGDSGNTLY | 0.4410355 |
| WSQQRRLIYFPSAGP | ASSQTGDSGNTLY | 0.7362726 |
| GPNGNPDPMAAAVDIR | ASSQTGDSGNTLY | 0.7841354 |
| TQRCVSVADLRAAASR | ASSQTGDSGNTLY | 0.31654018 |
| QFVLDVAEHPVEVLTETV | ASSQTGDSGNTLY | 0.7508943 |
| GGAGGTGGAGGNGGTQAGDG | ASSQTGDSGNTLY | 0.7352337 |
| PLLATLRAIAVDVDAKPAQV | ASSQTGDSGNTLY | 0.5804024 |
| APKFRTGGYMIDFWGVGYQV | ASSQTGDSGNTLY | 0.82812655 |
| GAGGAGGAGGAAGGPGGTGG | ASSQTGDSGNTLY | 0.67333066 |
| IGALGLLATDVAPAGSVAFG | ASSQTGDSGNTLY | 0.833622 |
| AGGAGGIGGNGGAGGAGGVL | ASSQTGDSGNTLY | 0.67037606 |
| ADELIGLL | ASSQTGDSGNTLY | 0.6354747 |
| ADRALKA | ASSQTGDSGNTLY | 0.7769452 |
| AGAVGQVGGAGGA | ASSQTGDSGNTLY | 0.52191067 |
| AGGGGGLGGAP | ASSQTGDSGNTLY | 0.52036047 |
| AGGLLYGNGGAG | ASSQTGDSGNTLY | 0.837731 |
| AGLESIGL | ASSQTGDSGNTLY | 0.82012093 |
| AGLKTGII | ASSQTGDSGNTLY | 0.85257375 |
| ASLAALLL | ASSQTGDSGNTLY | 0.464282 |
| ASVINISSI | ASSQTGDSGNTLY | 0.5690968 |
| ATPPGPA | ASSQTGDSGNTLY | 0.63882506 |
| ATVASALV | ASSQTGDSGNTLY | 0.77271056 |
| ATVASLIL | ASSQTGDSGNTLY | 0.86324394 |
| AVNGPVLIRH | ASSQTGDSGNTLY | 0.8335147 |
| DGAAGAVGI | ASSQTGDSGNTLY | 0.5215517 |
| DPLLHGI | ASSQTGDSGNTLY | 0.745453 |

| | | |
|---|---|---|
| ELGAALGAL | ASSQTGDSGNTLY | 0.6368358 |
| ELGRTQAV | ASSQTGDSGNTLY | 0.7158308 |
| FVAGGYLS | ASSQTGDSGNTLY | 0.65298164 |
| GASGGAGGAGGW | ASSQTGDSGNTLY | 0.6665447 |
| GGAGAGGDGGAGGI | ASSQTGDSGNTLY | 0.5370493 |
| GTGHNGSQPGMGGNG | ASSQTGDSGNTLY | 0.45693472 |
| GYAAASASAS | ASSQTGDSGNTLY | 0.6616552 |
| HQVDVAAVQQG | ASSQTGDSGNTLY | 0.8063033 |
| IGAAIAHT | ASSQTGDSGNTLY | 0.79866207 |
| IGMIEVAQL | ASSQTGDSGNTLY | 0.81742704 |
| IRLAPEHTSAMD | ASSQTGDSGNTLY | 0.8072903 |
| IYNVSVL | ASSQTGDSGNTLY | 0.6966388 |
| KQGFLGM | ASSQTGDSGNTLY | 0.18176526 |
| LESVITQL | ASSQTGDSGNTLY | 0.3321168 |
| LGLGALVVV | ASSQTGDSGNTLY | 0.5304728 |
| LIGNGGNGGSGGI | ASSQTGDSGNTLY | 0.5784656 |
| LNAINAPTL | ASSQTGDSGNTLY | 0.19230719 |
| LQLLNTGV | ASSQTGDSGNTLY | 0.7134765 |
| LQTSIEF | ASSQTGDSGNTLY | 0.68701386 |
| MSFAVSEL | ASSQTGDSGNTLY | 0.59875154 |
| NTGALGL | ASSQTGDSGNTLY | 0.8082182 |
| PPGAGGAGGAGTT | ASSQTGDSGNTLY | 0.6292918 |
| QLVGLGI | ASSQTGDSGNTLY | 0.40366095 |
| RFLEQVHQL | ASSQTGDSGNTLY | 0.44768244 |
| RGVGYVLR | ASSQTGDSGNTLY | 0.80922425 |
| SATGVFAGVI | ASSQTGDSGNTLY | 0.39720148 |
| SLPAITLP | ASSQTGDSGNTLY | 0.81198895 |
| STALAFTV | ASSQTGDSGNTLY | 0.42398962 |
| SVLTVGLGK | ASSQTGDSGNTLY | 0.82705295 |
| TALAGAGSGI | ASSQTGDSGNTLY | 0.43385792 |
| TGLPFTGI | ASSQTGDSGNTLY | 0.7971679 |
| VAEHVTV | ASSQTGDSGNTLY | 0.7269908 |
| VDIVLNSL | ASSQTGDSGNTLY | 0.87228465 |
| VLTGGGAVL | ASSQTGDSGNTLY | 0.605652 |
| VNVGLAYGI | ASSQTGDSGNTLY | 0.73688394 |
| VSSAAVGAV | ASSQTGDSGNTLY | 0.48351204 |
| VTSVYEAGV | ASSQTGDSGNTLY | 0.83492076 |
| WVALDRGA | ASSQTGDSGNTLY | 0.4266941 |
| YLVGSVVI | ASSQTGDSGNTLY | 0.5369728 |
| GPPGTGKTLL | ASSQTGDSGNTLY | 0.34585604 |
| GVGMTKF | ASSQTGDSGNTLY | 0.5686121 |
| IGIGGSDL | ASSQTGDSGNTLY | 0.8431481 |
| IGTLLAEL | ASSQTGDSGNTLY | 0.2761051 |
| IYTSGTT | ASSQTGDSGNTLY | 0.6946434 |
| LSFPAGV | ASSQTGDSGNTLY | 0.60954756 |
| LYTSGTT | ASSQTGDSGNTLY | 0.49082404 |
| VGAFPFGF | ASSQTGDSGNTLY | 0.6975176 |
| VGAGTVEF | ASSQTGDSGNTLY | 0.6372023 |
| VGKSTLLL | ASSQTGDSGNTLY | 0.7836243 |
| AALAHAGGGIV | ASSQTGDSGNTLY | 0.45831794 |
| AALVAVSLL | ASSQTGDSGNTLY | 0.5789232 |

| | | |
|---|---|---|
| AAVSRAVAAL | ASSQTGDSGNTLY | 0.7129903 |
| AGGSSGIGGF | ASSQTGDSGNTLY | 0.42213678 |
| AGLTVANANG | ASSQTGDSGNTLY | 0.402218 |
| AGLVGAVL | ASSQTGDSGNTLY | 0.844651 |
| ALSQAFTV | ASSQTGDSGNTLY | 0.12102461 |
| ALTFSVSL | ASSQTGDSGNTLY | 0.120107874 |
| ALTGAALVAL | ASSQTGDSGNTLY | 0.11689159 |
| ASFSIGAGVL | ASSQTGDSGNTLY | 0.12022079 |
| DGSVYKI | ASSQTGDSGNTLY | 0.11785372 |
| EQLAEIV | ASSQTGDSGNTLY | 0.12722808 |
| FAVSELTVM | ASSQTGDSGNTLY | 0.12063443 |
| GAAPLAAGL | ASSQTGDSGNTLY | 0.120147124 |
| GADGGDGGSGGAGG | ASSQTGDSGNTLY | 0.12161384 |
| GFGLDTGGAGG | ASSQTGDSGNTLY | 0.1201645 |
| GLASGGLL | ASSQTGDSGNTLY | 0.11799664 |
| GRIAGAAVLL | ASSQTGDSGNTLY | 0.12074881 |
| GSAGSLNTSF | ASSQTGDSGNTLY | 0.119513474 |
| GTGQIGI | ASSQTGDSGNTLY | 0.12150877 |
| GTLIALIL | ASSQTGDSGNTLY | 0.11663465 |
| IAATMGFVVG | ASSQTGDSGNTLY | 0.12004847 |
| IFAQALL | ASSQTGDSGNTLY | 0.11958432 |
| IGAAVALGF | ASSQTGDSGNTLY | 0.12098101 |
| IGLTPLAL | ASSQTGDSGNTLY | 0.12006547 |
| ILGSNTSTL | ASSQTGDSGNTLY | 0.12836663 |
| IVFSLTVL | ASSQTGDSGNTLY | 0.123353295 |
| LASKATGF | ASSQTGDSGNTLY | 0.11797252 |
| LASVALIL | ASSQTGDSGNTLY | 0.11587732 |
| LFAVGLTL | ASSQTGDSGNTLY | 0.117791854 |
| LFNGGIGTYI | ASSQTGDSGNTLY | 0.11880723 |
| LTLGAGII | ASSQTGDSGNTLY | 0.12080458 |
| LVAAALGDI | ASSQTGDSGNTLY | 0.115052216 |
| NTAFVQLGI | ASSQTGDSGNTLY | 0.12406293 |
| QLFSGAEL | ASSQTGDSGNTLY | 0.12071767 |
| SAATFITL | ASSQTGDSGNTLY | 0.11907896 |
| SNYGLANLGV | ASSQTGDSGNTLY | 0.124352746 |
| SYGGAYAVM | ASSQTGDSGNTLY | 0.11734278 |
| TTGVGGLTLG | ASSQTGDSGNTLY | 0.11923845 |
| TTQTIYI | ASSQTGDSGNTLY | 0.11860249 |
| VAAGAGLGAVI | ASSQTGDSGNTLY | 0.124570265 |
| VAAVVTLL | ASSQTGDSGNTLY | 0.11803724 |
| VGFLGGLI | ASSQTGDSGNTLY | 0.12921199 |
| VGVIPILL | ASSQTGDSGNTLY | 0.11732788 |
| VIFASSLI | ASSQTGDSGNTLY | 0.13513978 |
| VKGGDGGAAAG | ASSQTGDSGNTLY | 0.1206302 |
| VTKTFTNI | ASSQTGDSGNTLY | 0.121175 |
| VVAAAILL | ASSQTGDSGNTLY | 0.12009845 |
| YAALDVEL | ASSQTGDSGNTLY | 0.13390407 |
| LASVALLL | ASSQTGDSGNTLY | 0.12338643 |
| LDAVHRV | ASSQTGDSGNTLY | 0.12665784 |
| QGLSSSIF | ASSQTGDSGNTLY | 0.11874008 |
| RGTFANIRL | ASSQTGDSGNTLY | 0.120641835 |

| | | |
|---|---|---|
| RQLWWGHRI | ASSQTGDSGNTLY | 0.11749982 |
| YGPPGTGKTLL | ASSQTGDSGNTLY | 0.12018728 |
| AALVGAPANVA | ASSQTGDSGNTLY | 0.11673095 |
| AATFRRTE | ASSQTGDSGNTLY | 0.12124827 |
| AGALGGGI | ASSQTGDSGNTLY | 0.1208088 |
| AGGAGGAGGIGAG | ASSQTGDSGNTLY | 0.11581913 |
| AGSGALGL | ASSQTGDSGNTLY | 0.11890824 |
| AQIADQI | ASSQTGDSGNTLY | 0.11609442 |
| ASAVAVTYL | ASSQTGDSGNTLY | 0.118432075 |
| ASTVLGGTGG | ASSQTGDSGNTLY | 0.119994886 |
| AVGGEPAK | ASSQTGDSGNTLY | 0.118850894 |
| CEQGTFGGTC | ASSQTGDSGNTLY | 0.11839665 |
| DEGKAPAG | ASSQTGDSGNTLY | 0.11610047 |
| DTVVFPAL | ASSQTGDSGNTLY | 0.11754788 |
| FEGITMAG | ASSQTGDSGNTLY | 0.12519868 |
| FPLVVQD | ASSQTGDSGNTLY | 0.11671409 |
| GAFTTPQ | ASSQTGDSGNTLY | 0.1176158 |
| GFGGFGTAGGVGGA | ASSQTGDSGNTLY | 0.12399458 |
| GFLAQFL | ASSQTGDSGNTLY | 0.8304093 |
| GGAGGAGGNAGVA | ASSQTGDSGNTLY | 0.79189515 |
| GGVGTAGAGGAGGAGG | ASSQTGDSGNTLY | 0.68736714 |
| GNGGGGDAGNAGSGGN | ASSQTGDSGNTLY | 0.6910156 |
| GTGGAGGAGNDAGS | ASSQTGDSGNTLY | 0.7391081 |
| GTTNTGWFNS | ASSQTGDSGNTLY | 0.84297454 |
| IFMGPLFTL | ASSQTGDSGNTLY | 0.83832973 |
| ILAHAIEAI | ASSQTGDSGNTLY | 0.8316825 |
| IRSAIREAL | ASSQTGDSGNTLY | 0.7048637 |
| ITFAGGG | ASSQTGDSGNTLY | 0.8585181 |
| ITGMSGHQEV | ASSQTGDSGNTLY | 0.79427767 |
| IVAGLACTL | ASSQTGDSGNTLY | 0.7770047 |
| IVGGGLAGA | ASSQTGDSGNTLY | 0.8320558 |
| IVTGVLNI | ASSQTGDSGNTLY | 0.812673 |
| LAQQGCI | ASSQTGDSGNTLY | 0.7581053 |
| LCLHVEN | ASSQTGDSGNTLY | 0.71607745 |
| LDAISDGS | ASSQTGDSGNTLY | 0.8258628 |
| LGGFAAML | ASSQTGDSGNTLY | 0.8425155 |
| LGGKNPMI | ASSQTGDSGNTLY | 0.8282237 |
| LGPKGVAAIL | ASSQTGDSGNTLY | 0.88794035 |
| LMVGVSI | ASSQTGDSGNTLY | 0.8503094 |
| LVGAAFVL | ASSQTGDSGNTLY | 0.8313726 |
| LVLAVAPL | ASSQTGDSGNTLY | 0.37372965 |
| MAAIAVIDAA | ASSQTGDSGNTLY | 0.63613987 |
| NNGINAELQ | ASSQTGDSGNTLY | 0.73893666 |
| RGEPLVLVHG | ASSQTGDSGNTLY | 0.77463233 |
| RGGAGGAAGLI | ASSQTGDSGNTLY | 0.30829772 |
| RSYRMHPAV | ASSQTGDSGNTLY | 0.8275384 |
| SAAHLLAGQLA | ASSQTGDSGNTLY | 0.81295145 |
| SAAVGSY | ASSQTGDSGNTLY | 0.7185477 |
| SPHAVLHRI | ASSQTGDSGNTLY | 0.84850836 |
| SQIAKVL | ASSQTGDSGNTLY | 0.7394276 |
| TALGHLVP | ASSQTGDSGNTLY | 0.7008573 |

| | | |
|---|---|---|
| TAPTTSGAA | ASSQTGDSGNTLY | 0.7748263 |
| TDGDDGNGGSPGT | ASSQTGDSGNTLY | 0.88122654 |
| TDGLGEN | ASSQTGDSGNTLY | 0.69520694 |
| TGGPPLPGQA | ASSQTGDSGNTLY | 0.856044 |
| TIPAITI | ASSQTGDSGNTLY | 0.76787835 |
| TIPGFTI | ASSQTGDSGNTLY | 0.87744987 |
| TSIASGALA | ASSQTGDSGNTLY | 0.6292223 |
| TVTLPGAAI | ASSQTGDSGNTLY | 0.84600985 |
| VASGLSAALP | ASSQTGDSGNTLY | 0.7574314 |
| VGAGVIGI | ASSQTGDSGNTLY | 0.88549936 |
| VGPKRFGV | ASSQTGDSGNTLY | 0.77346796 |
| VNVAFPD | ASSQTGDSGNTLY | 0.857882 |
| VSMTNSGVS | ASSQTGDSGNTLY | 0.78426933 |
| VTDVLGDLSV | ASSQTGDSGNTLY | 0.81961316 |
| VVVVGAGFAGL | ASSQTGDSGNTLY | 0.5823909 |
| VYTVGTVV | ASSQTGDSGNTLY | 0.7818407 |
| WTTFSGP | ASSQTGDSGNTLY | 0.72834706 |
| YDELVEAF | ASSQTGDSGNTLY | 0.8251536 |
| YGTGGAGGNGG | ASSQTGDSGNTLY | 0.8373503 |
| YTNVMAV | ASSQTGDSGNTLY | 0.23869726 |
| FLEQVHQL | ASSQTGDSGNTLY | 0.86340666 |
| GLSLLAVAV | ASSQTGDSGNTLY | 0.5646848 |
| LEQVHQL | ASSQTGDSGNTLY | 0.6824037 |
| LNAAGAI | ASSQTGDSGNTLY | 0.8456801 |
| PAVHASL | ASSQTGDSGNTLY | 0.8165354 |
| TELEDDA | ASSQTGDSGNTLY | 0.8471639 |
| TPPGPPN | ASSQTGDSGNTLY | 0.7396542 |
| TPPPPPPP | ASSQTGDSGNTLY | 0.81807387 |
| TQSPALI | ASSRADRGGNTEVF | 0.84844524 |
| EFAAAFPQ | ASSRADRGGNTEVF | 0.40826872 |
| LLSGNLDVL | ASSRADRGGNTEVF | 0.6968514 |
| GPANLGNYNI | ASSRADRGGNTEVF | 0.86566126 |
| TLGPTLP | ASSRADRGGNTEVF | 0.8753685 |
| VYAYSSCCK | ASSRADRGGNTEVF | 0.8823898 |
| APAPAPAPNI | ASSRADRGGNTEVF | 0.82560253 |
| PGGPVPIP | ASSRADRGGNTEVF | 0.7372794 |
| FLTVALEP | ASSRADRGGNTEVF | 0.8313168 |
| SQYFGANL | ASSRADRGGNTEVF | 0.88612115 |
| GGVKPAA | ASSRADRGGNTEVF | 0.8737154 |
| GGTGGAGGAGGW | ASSRADRGGNTEVF | 0.8660788 |
| GVGGVAGLGGAGG | ASSRADRGGNTEVF | 0.52862096 |
| VLLVLATTI | ASSRADRGGNTEVF | 0.8619423 |
| IGIFAAML | ASSRADRGGNTEVF | 0.8204664 |
| AVGDLID | ASSRADRGGNTEVF | 0.8248087 |
| NISPLPL | ASSRADRGGNTEVF | 0.871385 |
| VTAVFVAI | ASSRADRGGNTEVF | 0.82784647 |
| SGDDNDAAL | ASSRADRGGNTEVF | 0.8482288 |
| AVAALAHS | ASSRADRGGNTEVF | 0.8201817 |
| GGVTVMATP | ASSRADRGGNTEVF | 0.8636434 |
| AGLAANVRD | ASSRADRGGNTEVF | 0.8520571 |
| TNRPDLIDPALLRPG | ASSRADRGGNTEVF | 0.8112001 |

| | | |
|---|---|---|
| LANLGVTAL | ASSRADRGGNTEVF | 0.90159523 |
| TGERIEL | ASSRADRGGNTEVF | 0.8911965 |
| NGEELAA | ASSRADRGGNTEVF | 0.8832816 |
| VLSLVWG | ASSRADRGGNTEVF | 0.21093555 |
| IGYALWL | ASSRADRGGNTEVF | 0.6443517 |
| LSLLLAPL | ASSRADRGGNTEVF | 0.8129028 |
| TLDEALK | ASSRADRGGNTEVF | 0.84111404 |
| TDGAGTAEGG | ASSRADRGGNTEVF | 0.45761526 |
| NAPGGAGGNGG | ASSRADRGGNTEVF | 0.875567 |
| VSVFMGMGGNF | ASSRADRGGNTEVF | 0.8082098 |
| LASYLAAL | ASSRADRGGNTEVF | 0.7815179 |
| LVGGTPAL | ASSRADRGGNTEVF | 0.8910842 |
| VRNADVL | ASSRADRGGNTEVF | 0.8407624 |
| TCSSPFLVSAM | ASSRADRGGNTEVF | 0.7354474 |
| VHITDAE | ASSRADRGGNTEVF | 0.78669184 |
| EYAAPSF | ASSRADRGGNTEVF | 0.88575643 |
| VVDAFHGGL | ASSRADRGGNTEVF | 0.6738087 |
| ASVVDPQ | ASSRADRGGNTEVF | 0.8797157 |
| SYNEFGGLS | ASSRADRGGNTEVF | 0.81346655 |
| DPDGSVY | ASSRADRGGNTEVF | 0.8943334 |
| MVSTIHATE | ASSRADRGGNTEVF | 0.48645693 |
| LAAGVADPV | ASSRADRGGNTEVF | 0.8824016 |
| TKEQIAA | ASSRADRGGNTEVF | 0.8526734 |
| SLASPAAL | ASSRADRGGNTEVF | 0.9053217 |
| IGLVFYGV | ASSRADRGGNTEVF | 0.8246663 |
| FLLDASNL | ASSRADRGGNTEVF | 0.8151468 |
| YQVGNFP | ASSRADRGGNTEVF | 0.72437286 |
| ESLLSQST | ASSRADRGGNTEVF | 0.8857219 |
| LFTVGFAAVG | ASSRADRGGNTEVF | 0.66432184 |
| ATPPTPT | ASSRADRGGNTEVF | 0.79545605 |
| TDLATLG | ASSRADRGGNTEVF | 0.7292789 |
| SAGGTGR | ASSRADRGGNTEVF | 0.85654914 |
| GFSNSGS | ASSRADRGGNTEVF | 0.87009466 |
| GGNGGSGGTSV | ASSRADRGGNTEVF | 0.47152328 |
| GTSVATGGAGN | ASSRADRGGNTEVF | 0.8722595 |
| NGGNGGSGGTS | ASSRADRGGNTEVF | 0.808707 |
| AVVKFPH | ASSRADRGGNTEVF | 0.6393676 |
| TPPPPPP | ASSRADRGGNTEVF | 0.89351976 |
| AAPAAAAAPA | ASSRADRGGNTEVF | 0.78479916 |
| IALNEDL | ASSRADRGGNTEVF | 0.893686 |
| IVNVSSM | ASSRADRGGNTEVF | 0.82700527 |
| TGIPPAPR | ASSRADRGGNTEVF | 0.8175088 |
| FDLGGGTF | ASSRADRGGNTEVF | 0.8649866 |
| FDLGGGTFDVS | ASSRADRGGNTEVF | 0.7865262 |
| NEPTAAA | ASSRADRGGNTEVF | 0.75280225 |
| NEPTAAALA | ASSRADRGGNTEVF | 0.8946314 |
| GSIINISSI | ASSRADRGGNTEVF | 0.8277946 |
| VAPGNAGTA | ASSRADRGGNTEVF | 0.8494824 |
| TGGGASLE | ASSRADRGGNTEVF | 0.7729864 |
| VAVYNHYKRI | ASSRADRGGNTEVF | 0.6605923 |
| TDPAPATT | ASSRADRGGNTEVF | 0.77356017 |

| | | |
|---|---|---|
| TNRPDILDPALLRPG | ASSRADRGGNTEVF | 0.84907734 |
| LTTTTTS | ASSRADRGGNTEVF | 0.8447386 |
| PTGSGKTL | ASSRADRGGNTEVF | 0.84023166 |
| TGSGKTL | ASSRADRGGNTEVF | 0.46951827 |
| TIGHVDHGK | ASSRADRGGNTEVF | 0.8280173 |
| TIGHVDHGKTT | ASSRADRGGNTEVF | 0.77614707 |
| GHVDHGKTT | ASSRADRGGNTEVF | 0.8069501 |
| AQFEHTLL | ASSRADRGGNTEVF | 0.82048273 |
| IVNQDEL | ASSRADRGGNTEVF | 0.77274317 |
| TNGSQFF | ASSRADRGGNTEVF | 0.8044671 |
| TGKGIVE | ASSRADRGGNTEVF | 0.7125716 |
| LISNASDA | ASSRADRGGNTEVF | 0.8188478 |
| AELRTLGELTKIAKAHGAQ | ASSRADRGGNTEVF | 0.8077575 |
| APNAVAAAEPSW | ASSRADRGGNTEVF | 0.76148295 |
| APQVAGAATAVL | ASSRADRGGNTEVF | 0.866189 |
| ARRDPAAAAAILPTDA | ASSRADRGGNTEVF | 0.8510263 |
| DAAARDVLAEAGLAEYF | ASSRADRGGNTEVF | 0.83390784 |
| DDQIVLSAKANLAKVEQ | ASSRADRGGNTEVF | 0.18625787 |
| DELLPDTGEDSDEDGH | ASSRADRGGNTEVF | 0.62340426 |
| ELPEAPANRATVPQD | ASSRADRGGNTEVF | 0.7211869 |
| FGQSKIERVTGIG | ASSRADRGGNTEVF | 0.8090255 |
| GAGGTGSTAGGAGGAGGAGGI | ASSRADRGGNTEVF | 0.29507494 |
| GHGGTGAAGGNGGTGGAGGS( | ASSRADRGGNTEVF | 0.83988154 |
| GNGGNGGAGGEGGPGGAGGA( | ASSRADRGGNTEVF | 0.7506418 |
| GSTAGNGGNGGNATNGTIAG( | ASSRADRGGNTEVF | 0.76974416 |
| GTGGDGGNGGNGGTGG | ASSRADRGGNTEVF | 0.8547485 |
| GVRRPAGIPADAPV | ASSRADRGGNTEVF | 0.80080414 |
| GYRQVTDAHLLTLARR | ASSRADRGGNTEVF | 0.65371835 |
| IAKMGQDGHDRG | ASSRADRGGNTEVF | 0.6755356 |
| KAEPSIEIQYT | ASSRADRGGNTEVF | 0.8615314 |
| KPVRIRMAADEDTVAVT | ASSRADRGGNTEVF | 0.65247416 |
| MPPRIAGMRLLVIKPEP | ASSRADRGGNTEVF | 0.85714805 |
| MKFYAVALFRSRGVPPTR | ASSRADRGGNTEVF | 0.76593804 |
| NAGGFGILGAVAHSP | ASSRADRGGNTEVF | 0.8593504 |
| NEGREITLAMAEM | ASSRADRGGNTEVF | 0.36657485 |
| NELVEWSHPLGHRWRWE | ASSRADRGGNTEVF | 0.85167354 |
| NLGAGGTGGNGGNPG | ASSRADRGGNTEVF | 0.8139007 |
| NTEWPKWETVMAADPVD | ASSRADRGGNTEVF | 0.8662295 |
| PASAAPAETQP | ASSRADRGGNTEVF | 0.73984885 |
| QQSSDGKNGVRAAEITGE | ASSRADRGGNTEVF | 0.7931504 |
| RAPDAVTTSWWKE | ASSRADRGGNTEVF | 0.67543626 |
| SGFGNVGAHTSGWFNQ | ASSRADRGGNTEVF | 0.8574035 |
| SGVAGAVGQAASVGGLKV | ASSRADRGGNTEVF | 0.621907 |
| SNGGAGTGSAGGNGGT | ASSRADRGGNTEVF | 0.6330142 |
| TGGAGGEGGAAG | ASSRADRGGNTEVF | 0.6912583 |
| TGGDGGAAGAGGAPGHG | ASSRADRGGNTEVF | 0.7745153 |
| TMPPLLAAIANAGRHGV | ASSRADRGGNTEVF | 0.8427882 |
| TSSTAGNSGGA | ASSRADRGGNTEVF | 0.3539902 |
| VPATVPVGILC | ASSRADRGGNTEVF | 0.82809186 |
| VQRPDSGGVVLTGRIS | ASSRADRGGNTEVF | 0.75119734 |
| WTPNPVPATKGVPAGLVQAT | ASSRADRGGNTEVF | 0.48110276 |

| | | |
|---|---|---|
| YALKRALAKFSDRFDVVII | ASSRADRGGNTEVF | 0.8413135 |
| YLIGSVIV | ASSRADRGGNTEVF | 0.71170914 |
| YGQLVAVID | ASSRADRGGNTEVF | 0.84106326 |
| VTVGGMTLPA | ASSRADRGGNTEVF | 0.75704944 |
| VLALPAV | ASSRADRGGNTEVF | 0.7701987 |
| VIPPTLNYET | ASSRADRGGNTEVF | 0.808097 |
| VGMGGMHPSSG | ASSRADRGGNTEVF | 0.61972344 |
| VGLAGGAATA | ASSRADRGGNTEVF | 0.6695138 |
| VGGAGGAGGSGP | ASSRADRGGNTEVF | 0.8609259 |
| VFAAFMV | ASSRADRGGNTEVF | 0.72882104 |
| TLYDKLR | ASSRADRGGNTEVF | 0.7754083 |
| TLAVFTVL | ASSRADRGGNTEVF | 0.6385982 |
| TGSVQPARV | ASSRADRGGNTEVF | 0.57742137 |
| TGGGAGSGGT | ASSRADRGGNTEVF | 0.66154087 |
| SHLETSL | ASSRADRGGNTEVF | 0.7523367 |
| SFTTGMT | ASSRADRGGNTEVF | 0.7819594 |
| QVEVAKRMGVRQPTVS | ASSRADRGGNTEVF | 0.7390462 |
| QGSGGPALAM | ASSRADRGGNTEVF | 0.5739961 |
| MLAPSNP | ASSRADRGGNTEVF | 0.7079335 |
| LVNNAAI | ASSRADRGGNTEVF | 0.66973925 |
| LTGEINTLA | ASSRADRGGNTEVF | 0.65409815 |
| LSIFGPV | ASSRADRGGNTEVF | 0.7140287 |
| LLALLANL | ASSRADRGGNTEVF | 0.7203953 |
| LISNASDALDKLR | ASSRADRGGNTEVF | 0.6256006 |
| LINEAVP | ASSRADRGGNTEVF | 0.56068754 |
| LGYAFSGL | ASSRADRGGNTEVF | 0.7111033 |
| LDFVAVAL | ASSRADRGGNTEVF | 0.75526893 |
| LAQPGAE | ASSRADRGGNTEVF | 0.7326671 |
| ISNLGAVTGF | ASSRADRGGNTEVF | 0.7726604 |
| HQEDPGRI | ASSRADRGGNTEVF | 0.75039214 |
| GTPGEPTP | ASSRADRGGNTEVF | 0.73103166 |
| GFFNSGS | ASSRADRGGNTEVF | 0.25596523 |
| GAGGGGLTAGAN | ASSRADRGGNTEVF | 0.37063682 |
| FVDALVNM | ASSRADRGGNTEVF | 0.5694592 |
| FLTVAPEL | ASSRADRGGNTEVF | 0.6713735 |
| FETLTAL | ASSRADRGGNTEVF | 0.301056 |
| DVINPIEL | ASSRADRGGNTEVF | 0.6993832 |
| APVSGAPV | ASSRADRGGNTEVF | 0.62302655 |
| ALVGGLAW | ASSRADRGGNTEVF | 0.59022975 |
| ALAAAAATAI | ASSRADRGGNTEVF | 0.7534596 |
| AFGLGLLL | ASSRADRGGNTEVF | 0.6416824 |
| AFFAAASL | ASSRADRGGNTEVF | 0.5721837 |
| AAAALPAFSPP | ASSRADRGGNTEVF | 0.57583463 |
| VVLAPAL | ASSRADRGGNTEVF | 0.7726686 |
| TIGHVDHGKTTL | ASSRADRGGNTEVF | 0.37830418 |
| LISNASDALDK | ASSRADRGGNTEVF | 0.73442185 |
| LISNASDAL | ASSRADRGGNTEVF | 0.6352043 |
| ISNASDALDK | ASSRADRGGNTEVF | 0.7495463 |
| NQAETLVYQTE | ASSRADRGGNTEVF | 0.4695652 |
| TGGTGGTAGAG | ASSRADRGGNTEVF | 0.7453731 |
| TGSTAGGAGGA | ASSRADRGGNTEVF | 0.6744597 |

| | | |
|---|---|---|
| DPLAVPVAAGPV | ASSRADRGGNTEVF | 0.7843491 |
| GGTGGTPGNGGN | ASSRADRGGNTEVF | 0.6407797 |
| PAGLAGVDPATL | ASSRADRGGNTEVF | 0.71905744 |
| GNNLVVEAPHAGASV | ASSRADRGGNTEVF | 0.6040609 |
| KVTRIMSASTLEQPA | ASSRADRGGNTEVF | 0.76442397 |
| LGFALPIALPRRTVV | ASSRADRGGNTEVF | 0.44606504 |
| TRLGNLTARKLVVLV | ASSRADRGGNTEVF | 0.6535542 |
| WSQQRRLIYFPSAGP | ASSRADRGGNTEVF | 0.49624586 |
| GPNGNPDPMAAAVDIR | ASSRADRGGNTEVF | 0.7099868 |
| TQRCVSVADLRAAASR | ASSRADRGGNTEVF | 0.7366705 |
| QFVLDVAEHPVEVLTETV | ASSRADRGGNTEVF | 0.4041791 |
| GGAGGTGGAGGNGGTQAGDG | ASSRADRGGNTEVF | 0.76522684 |
| PLLATLRAIAVDVDAKPAQV | ASSRADRGGNTEVF | 0.6712643 |
| APKFRTGGYMIDFWGVGYQV | ASSRADRGGNTEVF | 0.5950847 |
| GAGGAGGAGGAAGGPGGTGG | ASSRADRGGNTEVF | 0.7487376 |
| IGALGLLATDVAPAGSVAFG | ASSRADRGGNTEVF | 0.6588055 |
| AGGAGGIGGNGGAGGAGGVL | ASSRADRGGNTEVF | 0.75616515 |
| ADELIGLL | ASSRADRGGNTEVF | 0.6169555 |
| ADRALKA | ASSRADRGGNTEVF | 0.7122124 |
| AGAVGQVGGAGGA | ASSRADRGGNTEVF | 0.73365545 |
| AGGGGGLGGAP | ASSRADRGGNTEVF | 0.5557227 |
| AGGLLYGNGGAG | ASSRADRGGNTEVF | 0.537812 |
| AGLESIGL | ASSRADRGGNTEVF | 0.7589937 |
| AGLKTGII | ASSRADRGGNTEVF | 0.88333285 |
| ASLAALLL | ASSRADRGGNTEVF | 0.90508187 |
| ASVINISSI | ASSRADRGGNTEVF | 0.7698089 |
| ATPPGPA | ASSRADRGGNTEVF | 0.70601606 |
| ATVASALV | ASSRADRGGNTEVF | 0.82531226 |
| ATVASLIL | ASSRADRGGNTEVF | 0.8930635 |
| AVNGPVLIRH | ASSRADRGGNTEVF | 0.91125715 |
| DGAAGAVGI | ASSRADRGGNTEVF | 0.87722194 |
| DPLLHGI | ASSRADRGGNTEVF | 0.71094245 |
| ELGAALGAL | ASSRADRGGNTEVF | 0.8617065 |
| ELGRTQAV | ASSRADRGGNTEVF | 0.83925545 |
| FVAGGYLS | ASSRADRGGNTEVF | 0.81292284 |
| GASGGAGGAGGW | ASSRADRGGNTEVF | 0.88158834 |
| GGAGAGGDGGAGGI | ASSRADRGGNTEVF | 0.8668472 |
| GTGHNGSQPGMGGNG | ASSRADRGGNTEVF | 0.8063998 |
| GYAAASASAS | ASSRADRGGNTEVF | 0.7994102 |
| HQVDVAAVQQG | ASSRADRGGNTEVF | 0.8737291 |
| IGAAIAHT | ASSRADRGGNTEVF | 0.8894237 |
| IGMIEVAQL | ASSRADRGGNTEVF | 0.8861476 |
| IRLAPEHTSAMD | ASSRADRGGNTEVF | 0.90429115 |
| IYNVSVL | ASSRADRGGNTEVF | 0.8929224 |
| KQGFLGM | ASSRADRGGNTEVF | 0.8832154 |
| LESVITQL | ASSRADRGGNTEVF | 0.25147182 |
| LGLGALVVV | ASSRADRGGNTEVF | 0.55233485 |
| LIGNGGNGGSGGI | ASSRADRGGNTEVF | 0.7641728 |
| LNAINAPTL | ASSRADRGGNTEVF | 0.82232577 |
| LQLLNTGV | ASSRADRGGNTEVF | 0.48276454 |
| LQTSIEF | ASSRADRGGNTEVF | 0.85818267 |

| | | |
|---|---|---|
| MSFAVSEL | ASSRADRGGNTEVF | 0.7800734 |
| NTGALGL | ASSRADRGGNTEVF | 0.72609353 |
| PPGAGGAGGAGTT | ASSRADRGGNTEVF | 0.8878304 |
| QLVGLGI | ASSRADRGGNTEVF | 0.8099816 |
| RFLEQVHQL | ASSRADRGGNTEVF | 0.7300416 |
| RGVGYVLR | ASSRADRGGNTEVF | 0.8134483 |
| SATGVFAGVI | ASSRADRGGNTEVF | 0.8990403 |
| SLPAITLP | ASSRADRGGNTEVF | 0.74336666 |
| STALAFTV | ASSRADRGGNTEVF | 0.883993 |
| SVLTVGLGK | ASSRADRGGNTEVF | 0.8244128 |
| TALAGAGSGI | ASSRADRGGNTEVF | 0.8993408 |
| TGLPFTGI | ASSRADRGGNTEVF | 0.591758 |
| VAEHVTV | ASSRADRGGNTEVF | 0.88810855 |
| VDIVLNSL | ASSRADRGGNTEVF | 0.8464309 |
| VLTGGGAVL | ASSRADRGGNTEVF | 0.9143264 |
| VNVGLAYGI | ASSRADRGGNTEVF | 0.8225534 |
| VSSAAVGAV | ASSRADRGGNTEVF | 0.875783 |
| VTSVYEAGV | ASSRADRGGNTEVF | 0.833221 |
| WVALDRGA | ASSRADRGGNTEVF | 0.9096416 |
| YLVGSVVI | ASSRADRGGNTEVF | 0.6400049 |
| GPPGTGKTLL | ASSRADRGGNTEVF | 0.83857477 |
| GVGMTKF | ASSRADRGGNTEVF | 0.5679311 |
| IGIGGSDL | ASSRADRGGNTEVF | 0.8554113 |
| IGTLLAEL | ASSRADRGGNTEVF | 0.8767334 |
| IYTSGTT | ASSRADRGGNTEVF | 0.36392957 |
| LSFPAGV | ASSRADRGGNTEVF | 0.8947476 |
| LYTSGTT | ASSRADRGGNTEVF | 0.79809815 |
| VGAFPFGF | ASSRADRGGNTEVF | 0.75746894 |
| VGAGTVEF | ASSRADRGGNTEVF | 0.89508903 |
| VGKSTLLL | ASSRADRGGNTEVF | 0.8221381 |
| AALAHAGGGIV | ASSRADRGGNTEVF | 0.8963816 |
| AALVAVSLL | ASSRADRGGNTEVF | 0.83235407 |
| AAVSRAVAAL | ASSRADRGGNTEVF | 0.8500222 |
| AGGSSGIGGF | ASSRADRGGNTEVF | 0.8628749 |
| AGLTVANANG | ASSRADRGGNTEVF | 0.6153341 |
| AGLVGAVL | ASSRADRGGNTEVF | 0.75200343 |
| ALSQAFTV | ASSRADRGGNTEVF | 0.9021814 |
| ALTFSVSL | ASSRADRGGNTEVF | 0.64069873 |
| ALTGAALVAL | ASSRADRGGNTEVF | 0.6600195 |
| ASFSIGAGVL | ASSRADRGGNTEVF | 0.5920738 |
| DGSVYKI | ASSRADRGGNTEVF | 0.5741616 |
| EQLAEIV | ASSRADRGGNTEVF | 0.59804726 |
| FAVSELTVM | ASSRADRGGNTEVF | 0.64969707 |
| GAAPLAAGL | ASSRADRGGNTEVF | 0.6653733 |
| GADGGDGGSGGAGG | ASSRADRGGNTEVF | 0.64820135 |
| GFGLDTGGAGG | ASSRADRGGNTEVF | 0.5943377 |
| GLASGGLL | ASSRADRGGNTEVF | 0.65059537 |
| GRIAGAAVLL | ASSRADRGGNTEVF | 0.63292044 |
| GSAGSLNTSF | ASSRADRGGNTEVF | 0.626769 |
| GTGQIGI | ASSRADRGGNTEVF | 0.6387842 |
| GTLIALIL | ASSRADRGGNTEVF | 0.6495865 |

| | | |
|---|---|---|
| IAATMGFVVG | ASSRADRGGNTEVF | 0.61154217 |
| IFAQALL | ASSRADRGGNTEVF | 0.6091511 |
| IGAAVALGF | ASSRADRGGNTEVF | 0.64005685 |
| IGLTPLAL | ASSRADRGGNTEVF | 0.64938444 |
| ILGSNTSTL | ASSRADRGGNTEVF | 0.6443868 |
| IVFSLTVL | ASSRADRGGNTEVF | 0.6661875 |
| LASKATGF | ASSRADRGGNTEVF | 0.6507175 |
| LASVALIL | ASSRADRGGNTEVF | 0.6489815 |
| LFAVGLTL | ASSRADRGGNTEVF | 0.4517712 |
| LFNGGIGTYI | ASSRADRGGNTEVF | 0.55281997 |
| LTLGAGII | ASSRADRGGNTEVF | 0.6318147 |
| LVAAALGDI | ASSRADRGGNTEVF | 0.6203679 |
| NTAFVQLGI | ASSRADRGGNTEVF | 0.43381026 |
| QLFSGAEL | ASSRADRGGNTEVF | 0.6512493 |
| SAATFITL | ASSRADRGGNTEVF | 0.6279936 |
| SNYGLANLGV | ASSRADRGGNTEVF | 0.6046524 |
| SYGGAYAVM | ASSRADRGGNTEVF | 0.6503657 |
| TTGVGGLTLG | ASSRADRGGNTEVF | 0.6199312 |
| TTQTIYI | ASSRADRGGNTEVF | 0.60180444 |
| VAAGAGLGAVI | ASSRADRGGNTEVF | 0.61737466 |
| VAAVVTLL | ASSRADRGGNTEVF | 0.6574185 |
| VGFLGGLI | ASSRADRGGNTEVF | 0.57014143 |
| VGVIPILL | ASSRADRGGNTEVF | 0.65690565 |
| VIFASSLI | ASSRADRGGNTEVF | 0.60800934 |
| VKGGDGGAAAG | ASSRADRGGNTEVF | 0.6595453 |
| VTKTFTNI | ASSRADRGGNTEVF | 0.56193554 |
| VVAAAILL | ASSRADRGGNTEVF | 0.6526848 |
| YAALDVEL | ASSRADRGGNTEVF | 0.6207646 |
| LASVALLL | ASSRADRGGNTEVF | 0.6682704 |
| LDAVHRV | ASSRADRGGNTEVF | 0.6098888 |
| QGLSSSIF | ASSRADRGGNTEVF | 0.6542145 |
| RGTFANIRL | ASSRADRGGNTEVF | 0.6483969 |
| RQLWWGHRI | ASSRADRGGNTEVF | 0.6625166 |
| YGPPGTGKTLL | ASSRADRGGNTEVF | 0.5675874 |
| AALVGAPANVA | ASSRADRGGNTEVF | 0.62227803 |
| AATFRRTE | ASSRADRGGNTEVF | 0.583016 |
| AGALGGGI | ASSRADRGGNTEVF | 0.6413397 |
| AGGAGGAGGIGAG | ASSRADRGGNTEVF | 0.6485724 |
| AGSGALGL | ASSRADRGGNTEVF | 0.41498518 |
| AQIADQI | ASSRADRGGNTEVF | 0.65937907 |
| ASAVAVTYL | ASSRADRGGNTEVF | 0.57400656 |
| ASTVLGGTGG | ASSRADRGGNTEVF | 0.59306186 |
| AVGGEPAK | ASSRADRGGNTEVF | 0.65373856 |
| CEQGTFGGTC | ASSRADRGGNTEVF | 0.63732123 |
| DEGKAPAG | ASSRADRGGNTEVF | 0.64888346 |
| DTVVFPAL | ASSRADRGGNTEVF | 0.62363786 |
| FEGITMAG | ASSRADRGGNTEVF | 0.6307346 |
| FPLVVQD | ASSRADRGGNTEVF | 0.64373523 |
| GAFTTPQ | ASSRADRGGNTEVF | 0.5429762 |
| GFGGFGTAGGVGGA | ASSRADRGGNTEVF | 0.5877981 |
| GFLAQFL | ASSRADRGGNTEVF | 0.6610985 |

| | | |
|---|---|---|
| GGAGGAGGNAGVA | ASSRADRGGNTEVF | 0.7668655 |
| GGVGTAGAGGAGGAGG | ASSRADRGGNTEVF | 0.78428006 |
| GNGGGGDAGNAGSGGN | ASSRADRGGNTEVF | 0.6793456 |
| GTGGAGGAGNDAGS | ASSRADRGGNTEVF | 0.61168295 |
| GTTNTGWFNS | ASSRADRGGNTEVF | 0.65944344 |
| IFMGPLFTL | ASSRADRGGNTEVF | 0.7718053 |
| ILAHAIEAI | ASSRADRGGNTEVF | 0.78449064 |
| IRSAIREAL | ASSRADRGGNTEVF | 0.78237224 |
| ITFAGGG | ASSRADRGGNTEVF | 0.5130834 |
| ITGMSGHQEV | ASSRADRGGNTEVF | 0.75778437 |
| IVAGLACTL | ASSRADRGGNTEVF | 0.7045882 |
| IVGGGLAGA | ASSRADRGGNTEVF | 0.71453154 |
| IVTGVLNI | ASSRADRGGNTEVF | 0.70806587 |
| LAQQGCI | ASSRADRGGNTEVF | 0.6157431 |
| LCLHVEN | ASSRADRGGNTEVF | 0.69724816 |
| LDAISDGS | ASSRADRGGNTEVF | 0.52954173 |
| LGGFAAML | ASSRADRGGNTEVF | 0.5209857 |
| LGGKNPMI | ASSRADRGGNTEVF | 0.73828804 |
| LGPKGVAAIL | ASSRADRGGNTEVF | 0.72041905 |
| LMVGVSI | ASSRADRGGNTEVF | 0.7765556 |
| LVGAAFVL | ASSRADRGGNTEVF | 0.77572227 |
| LVLAVAPL | ASSRADRGGNTEVF | 0.696236 |
| MAAIAVIDAA | ASSRADRGGNTEVF | 0.21698219 |
| NNGINAELQ | ASSRADRGGNTEVF | 0.52070284 |
| RGEPLVLVHG | ASSRADRGGNTEVF | 0.6272613 |
| RGGAGGAAGLI | ASSRADRGGNTEVF | 0.73180133 |
| RSYRMHPAV | ASSRADRGGNTEVF | 0.23103742 |
| SAAHLLAGQLA | ASSRADRGGNTEVF | 0.7378607 |
| SAAVGSY | ASSRADRGGNTEVF | 0.72857815 |
| SPHAVLHRI | ASSRADRGGNTEVF | 0.6933037 |
| SQIAKVL | ASSRADRGGNTEVF | 0.77047235 |
| TALGHLVP | ASSRADRGGNTEVF | 0.7125943 |
| TAPTTSGAA | ASSRADRGGNTEVF | 0.52717966 |
| TDGDDGNGGSPGT | ASSRADRGGNTEVF | 0.48485023 |
| TDGLGEN | ASSRADRGGNTEVF | 0.78045416 |
| TGGPPLPGQA | ASSRADRGGNTEVF | 0.42323864 |
| TIPAITI | ASSRADRGGNTEVF | 0.7714158 |
| TIPGFTI | ASSRADRGGNTEVF | 0.4947968 |
| TSIASGALA | ASSRADRGGNTEVF | 0.7716304 |
| TVTLPGAAI | ASSRADRGGNTEVF | 0.42465854 |
| VASGLSAALP | ASSRADRGGNTEVF | 0.76826733 |
| VGAGVIGI | ASSRADRGGNTEVF | 0.7400819 |
| VGPKRFGV | ASSRADRGGNTEVF | 0.78574955 |
| VNVAFPD | ASSRADRGGNTEVF | 0.6169749 |
| VSMTNSGVS | ASSRADRGGNTEVF | 0.72726214 |
| VTDVLGDLSV | ASSRADRGGNTEVF | 0.4119509 |
| VVVVGAGFAGL | ASSRADRGGNTEVF | 0.7836373 |
| VYTVGTVV | ASSRADRGGNTEVF | 0.5883913 |
| WTTFSGP | ASSRADRGGNTEVF | 0.6131142 |
| YDELVEAF | ASSRADRGGNTEVF | 0.4397849 |
| YGTGGAGGNGG | ASSRADRGGNTEVF | 0.7022298 |

| | | |
|---|---|---|
| YTNVMAV | ASSRADRGGNTEVF | 0.7824899 |
| FLEQVHQL | ASSRADRGGNTEVF | 0.5954226 |
| GLSLLAVAV | ASSRADRGGNTEVF | 0.7762227 |
| LEQVHQL | ASSRADRGGNTEVF | 0.7468877 |
| LNAAGAI | ASSRADRGGNTEVF | 0.4810232 |
| PAVHASL | ASSRADRGGNTEVF | 0.7372122 |
| TELEDDA | ASSRADRGGNTEVF | 0.51091987 |
| TPPGPPN | ASSRADRGGNTEVF | 0.76345026 |
| TPPPPPPP | ASSRADRGGNTEVF | 0.48755836 |
| TQSPALI | ASSSTGGALEQY | 0.7254584 |
| EFAAAFPQ | ASSSTGGALEQY | 0.741212 |
| LLSGNLDVL | ASSSTGGALEQY | 0.5822791 |
| GPANLGNYNI | ASSSTGGALEQY | 0.4413796 |
| TLGPTLP | ASSSTGGALEQY | 0.7822666 |
| VYAYSSCCK | ASSSTGGALEQY | 0.34343952 |
| APAPAPAPNI | ASSSTGGALEQY | 0.6921863 |
| PGGPVPIP | ASSSTGGALEQY | 0.14647067 |
| FLTVALEP | ASSSTGGALEQY | 0.15608715 |
| SQYFGANL | ASSSTGGALEQY | 0.21770915 |
| GGVKPAA | ASSSTGGALEQY | 0.33182198 |
| GGTGGAGGAGGW | ASSSTGGALEQY | 0.7769319 |
| GVGGVAGLGGAGG | ASSSTGGALEQY | 0.309201 |
| VLLVLATTI | ASSSTGGALEQY | 0.16014904 |
| IGIFAAML | ASSSTGGALEQY | 0.33466482 |
| AVGDLID | ASSSTGGALEQY | 0.17327303 |
| NISPLPL | ASSSTGGALEQY | 0.22706121 |
| VTAVFVAI | ASSSTGGALEQY | 0.21917179 |
| SGDDNDAAL | ASSSTGGALEQY | 0.20829815 |
| AVAALAHS | ASSSTGGALEQY | 0.21250364 |
| GGVTVMATP | ASSSTGGALEQY | 0.16264527 |
| AGLAANVRD | ASSSTGGALEQY | 0.32656455 |
| TNRPDLIDPALLRPG | ASSSTGGALEQY | 0.5162612 |
| LANLGVTAL | ASSSTGGALEQY | 0.43458277 |
| TGERIEL | ASSSTGGALEQY | 0.33212447 |
| NGEELAA | ASSSTGGALEQY | 0.37212053 |
| VLSLVWG | ASSSTGGALEQY | 0.2618695 |
| IGYALWL | ASSSTGGALEQY | 0.10510325 |
| LSLLLAPL | ASSSTGGALEQY | 0.121967055 |
| TLDEALK | ASSSTGGALEQY | 0.21402286 |
| TDGAGTAEGG | ASSSTGGALEQY | 0.17926699 |
| NAPGGAGGNGG | ASSSTGGALEQY | 0.11357724 |
| VSVFMGMGGNF | ASSSTGGALEQY | 0.26962572 |
| LASYLAAL | ASSSTGGALEQY | 0.19051248 |
| LVGGTPAL | ASSSTGGALEQY | 0.18698144 |
| VRNADVL | ASSSTGGALEQY | 0.47196627 |
| TCSSPFLVSAM | ASSSTGGALEQY | 0.18336001 |
| VHITDAE | ASSSTGGALEQY | 0.15692113 |
| EYAAPSF | ASSSTGGALEQY | 0.17217551 |
| VVDAFHGGL | ASSSTGGALEQY | 0.37002137 |
| ASVVDPQ | ASSSTGGALEQY | 0.14758211 |
| SYNEFGGLS | ASSSTGGALEQY | 0.32672477 |

| | | |
|---|---|---|
| DPDGSVY | ASSSTGGALEQY | 0.16876456 |
| MVSTIHATE | ASSSTGGALEQY | 0.42984456 |
| LAAGVADPV | ASSSTGGALEQY | 0.14353293 |
| TKEQIAA | ASSSTGGALEQY | 0.27926248 |
| SLASPAAL | ASSSTGGALEQY | 0.2081672 |
| IGLVFYGV | ASSSTGGALEQY | 0.727678 |
| FLLDASNL | ASSSTGGALEQY | 0.17393295 |
| YQVGNFP | ASSSTGGALEQY | 0.20240508 |
| ESLLSQST | ASSSTGGALEQY | 0.18467633 |
| LFTVGFAAVG | ASSSTGGALEQY | 0.80183107 |
| ATPPTPT | ASSSTGGALEQY | 0.124070436 |
| TDLATLG | ASSSTGGALEQY | 0.19327687 |
| SAGGTGR | ASSSTGGALEQY | 0.13565727 |
| GFSNSGS | ASSSTGGALEQY | 0.20727871 |
| GGNGGSGGTSV | ASSSTGGALEQY | 0.29518297 |
| GTSVATGGAGN | ASSSTGGALEQY | 0.10664797 |
| NGGNGGSGGTS | ASSSTGGALEQY | 0.25515142 |
| AVVKFPH | ASSSTGGALEQY | 0.13373776 |
| TPPPPPP | ASSSTGGALEQY | 0.14916033 |
| AAPAAAAAPA | ASSSTGGALEQY | 0.27470368 |
| IALNEDL | ASSSTGGALEQY | 0.21252847 |
| IVNVSSM | ASSSTGGALEQY | 0.37760943 |
| TGIPPAPR | ASSSTGGALEQY | 0.16804041 |
| FDLGGGTF | ASSSTGGALEQY | 0.18878391 |
| FDLGGGTFDVS | ASSSTGGALEQY | 0.24830844 |
| NEPTAAA | ASSSTGGALEQY | 0.14194727 |
| NEPTAAALA | ASSSTGGALEQY | 0.17225796 |
| GSIINISSI | ASSSTGGALEQY | 0.37020773 |
| VAPGNAGTA | ASSSTGGALEQY | 0.62795806 |
| TGGGASLE | ASSSTGGALEQY | 0.8350463 |
| VAVYNHYKRI | ASSSTGGALEQY | 0.42586476 |
| TDPAPATT | ASSSTGGALEQY | 0.3531052 |
| TNRPDILDPALLRPG | ASSSTGGALEQY | 0.63162553 |
| LTTTTTS | ASSSTGGALEQY | 0.76902515 |
| PTGSGKTL | ASSSTGGALEQY | 0.8595004 |
| TGSGKTL | ASSSTGGALEQY | 0.7590261 |
| TIGHVDHGK | ASSSTGGALEQY | 0.3541668 |
| TIGHVDHGKTT | ASSSTGGALEQY | 0.6300664 |
| GHVDHGKTT | ASSSTGGALEQY | 0.4477436 |
| AQFEHTLL | ASSSTGGALEQY | 0.5555355 |
| IVNQDEL | ASSSTGGALEQY | 0.7061622 |
| TNGSQFF | ASSSTGGALEQY | 0.6518786 |
| TGKGIVE | ASSSTGGALEQY | 0.40326375 |
| LISNASDA | ASSSTGGALEQY | 0.42461836 |
| AELRTLGELTKIAKAHGAQ | ASSSTGGALEQY | 0.76529235 |
| APNAVAAAEPSW | ASSSTGGALEQY | 0.81773263 |
| APQVAGAATAVL | ASSSTGGALEQY | 0.7962356 |
| ARRDPAAAAAILPTDA | ASSSTGGALEQY | 0.84671223 |
| DAAARDVLAEAGLAEYF | ASSSTGGALEQY | 0.7450706 |
| DDQIVLSAKANLAKVEQ | ASSSTGGALEQY | 0.784047 |
| DELLPDTGEDSDEDGH | ASSSTGGALEQY | 0.1241147 |

| | | |
|---|---|---|
| ELPEAPANRATVPQD | ASSSTGGALEQY | 0.21934704 |
| FGQSKIERVTGIG | ASSSTGGALEQY | 0.5307257 |
| GAGGTGSTAGGAGGAGGAGGI | ASSSTGGALEQY | 0.5732312 |
| GHGGTGAAGGNGGTGGAGGS( | ASSSTGGALEQY | 0.21193832 |
| GNGGNGGAGGEGGPGGAGGA( | ASSSTGGALEQY | 0.7447183 |
| GSTAGNGGNGGNATNGTIAG( | ASSSTGGALEQY | 0.43637428 |
| GTGGDGGNGGNGGTGG | ASSSTGGALEQY | 0.32720298 |
| GVRRPAGIPADAPV | ASSSTGGALEQY | 0.8035643 |
| GYRQVTDAHLLTLARR | ASSSTGGALEQY | 0.42588204 |
| IAKMGQDGHDRG | ASSSTGGALEQY | 0.3400371 |
| KAEPSIEIQYT | ASSSTGGALEQY | 0.47590187 |
| KPVRIRMAADEDTVAVT | ASSSTGGALEQY | 0.7242235 |
| MPPRIAGMRLLVIKPEP | ASSSTGGALEQY | 0.23291567 |
| MKFYAVALFRSRGVPPTR | ASSSTGGALEQY | 0.7513397 |
| NAGGFGILGAVAHSP | ASSSTGGALEQY | 0.5802332 |
| NEGREITLAMAEM | ASSSTGGALEQY | 0.7450379 |
| NELVEWSHPLGHRWRWE | ASSSTGGALEQY | 0.2751415 |
| NLGAGGTGGNGGNPG | ASSSTGGALEQY | 0.7453245 |
| NTEWPKWETVMAADPVD | ASSSTGGALEQY | 0.61936224 |
| PASAAPAETQP | ASSSTGGALEQY | 0.86873 |
| QQSSDGKNGVRAAEITGE | ASSSTGGALEQY | 0.4878406 |
| RAPDAVTTSWWKE | ASSSTGGALEQY | 0.6218999 |
| SGFGNVGAHTSGWFNQ | ASSSTGGALEQY | 0.6912099 |
| SGVAGAVGQAASVGGLKV | ASSSTGGALEQY | 0.83446264 |
| SNGGAGTGSAGGNGGT | ASSSTGGALEQY | 0.18698819 |
| TGGAGGEGGAAG | ASSSTGGALEQY | 0.5045378 |
| TGGDGGAAGAGGAPGHG | ASSSTGGALEQY | 0.21366665 |
| TMPPLLAAIANAGRHGV | ASSSTGGALEQY | 0.6056942 |
| TSSTAGNSGGA | ASSSTGGALEQY | 0.7163689 |
| VPATVPVGILC | ASSSTGGALEQY | 0.15865189 |
| VQRPDSGGVVLTGRIS | ASSSTGGALEQY | 0.6509111 |
| WTPNPVPATKGVPAGLVQAT | ASSSTGGALEQY | 0.55154467 |
| YALKRALAKFSDRFDVVII | ASSSTGGALEQY | 0.42601916 |
| YLIGSVIV | ASSSTGGALEQY | 0.79881614 |
| YGQLVAVID | ASSSTGGALEQY | 0.6479045 |
| VTVGGMTLPA | ASSSTGGALEQY | 0.793251 |
| VLALPAV | ASSSTGGALEQY | 0.6508714 |
| VIPPTLNYET | ASSSTGGALEQY | 0.5133883 |
| VGMGGMHPSSG | ASSSTGGALEQY | 0.6666239 |
| VGLAGGAATA | ASSSTGGALEQY | 0.32100523 |
| VGGAGGAGGSGP | ASSSTGGALEQY | 0.4691809 |
| VFAAFMV | ASSSTGGALEQY | 0.7961705 |
| TLYDKLR | ASSSTGGALEQY | 0.45117128 |
| TLAVFTVL | ASSSTGGALEQY | 0.71485233 |
| TGSVQPARV | ASSSTGGALEQY | 0.5164665 |
| TGGGAGSGGT | ASSSTGGALEQY | 0.4408146 |
| SHLETSL | ASSSTGGALEQY | 0.5788232 |
| SFTTGMT | ASSSTGGALEQY | 0.6462749 |
| QVEVAKRMGVRQPTVS | ASSSTGGALEQY | 0.7219794 |
| QGSGGPALAM | ASSSTGGALEQY | 0.55039597 |
| MLAPSNP | ASSSTGGALEQY | 0.42942357 |

| | | |
|---|---|---|
| LVNNAAI | ASSSTGGALEQY | 0.6057557 |
| LTGEINTLA | ASSSTGGALEQY | 0.41144148 |
| LSIFGPV | ASSSTGGALEQY | 0.5398391 |
| LLALLANL | ASSSTGGALEQY | 0.5755373 |
| LISNASDALDKLR | ASSSTGGALEQY | 0.6064321 |
| LINEAVP | ASSSTGGALEQY | 0.5516402 |
| LGYAFSGL | ASSSTGGALEQY | 0.39024135 |
| LDFVAVAL | ASSSTGGALEQY | 0.6479051 |
| LAQPGAE | ASSSTGGALEQY | 0.62363005 |
| ISNLGAVTGF | ASSSTGGALEQY | 0.48619992 |
| HQEDPGRI | ASSSTGGALEQY | 0.68057346 |
| GTPGEPTP | ASSSTGGALEQY | 0.6157713 |
| GFFNSGS | ASSSTGGALEQY | 0.51521975 |
| GAGGGGLTAGAN | ASSSTGGALEQY | 0.23379205 |
| FVDALVNM | ASSSTGGALEQY | 0.28657418 |
| FLTVAPEL | ASSSTGGALEQY | 0.44973594 |
| FETLTAL | ASSSTGGALEQY | 0.50840354 |
| DVINPIEL | ASSSTGGALEQY | 0.23445445 |
| APVSGAPV | ASSSTGGALEQY | 0.6703044 |
| ALVGGLAW | ASSSTGGALEQY | 0.450997 |
| ALAAAAATAI | ASSSTGGALEQY | 0.49345803 |
| AFGLGLLL | ASSSTGGALEQY | 0.6927952 |
| AFFAAASL | ASSSTGGALEQY | 0.4892508 |
| AAAALPAFSPP | ASSSTGGALEQY | 0.48664623 |
| VVLAPAL | ASSSTGGALEQY | 0.42522234 |
| TIGHVDHGKTTL | ASSSTGGALEQY | 0.68881667 |
| LISNASDALDK | ASSSTGGALEQY | 0.28579426 |
| LISNASDAL | ASSSTGGALEQY | 0.63578427 |
| ISNASDALDK | ASSSTGGALEQY | 0.42687774 |
| NQAETLVYQTE | ASSSTGGALEQY | 0.63024294 |
| TGGTGGTAGAG | ASSSTGGALEQY | 0.34628814 |
| TGSTAGGAGGA | ASSSTGGALEQY | 0.6028768 |
| DPLAVPVAAGPV | ASSSTGGALEQY | 0.5444692 |
| GGTGGTPGNGGN | ASSSTGGALEQY | 0.74255365 |
| PAGLAGVDPATL | ASSSTGGALEQY | 0.50262165 |
| GNNLVVEAPHAGASV | ASSSTGGALEQY | 0.56108963 |
| KVTRIMSASTLEQPA | ASSSTGGALEQY | 0.46890056 |
| LGFALPIALPRRTVV | ASSSTGGALEQY | 0.7165931 |
| TRLGNLTARKLVVLV | ASSSTGGALEQY | 0.32639614 |
| WSQQRRLIYFPSAGP | ASSSTGGALEQY | 0.4999152 |
| GPNGNPDPMAAAVDIR | ASSSTGGALEQY | 0.43132555 |
| TQRCVSVADLRAAASR | ASSSTGGALEQY | 0.6377592 |
| QFVLDVAEHPVEVLTETV | ASSSTGGALEQY | 0.49186972 |
| GGAGGTGGAGGNGGTQAGDG | ASSSTGGALEQY | 0.2825237 |
| PLLATLRAIAVDVDAKPAQV | ASSSTGGALEQY | 0.63774943 |
| APKFRTGGYMIDFWGVGYQV | ASSSTGGALEQY | 0.46841544 |
| GAGGAGGAGGAAGGPGGTGG | ASSSTGGALEQY | 0.38293964 |
| IGALGLLATDVAPAGSVAFG | ASSSTGGALEQY | 0.64320135 |
| AGGAGGIGGNGGAGGAGGVL | ASSSTGGALEQY | 0.5612562 |
| ADELIGLL | ASSSTGGALEQY | 0.6282276 |
| ADRALKA | ASSSTGGALEQY | 0.55847764 |

| | | |
|---|---|---|
| AGAVGQVGGAGGA | ASSSTGGALEQY | 0.5178752 |
| AGGGGGLGGAP | ASSSTGGALEQY | 0.66860324 |
| AGGLLYGNGGAG | ASSSTGGALEQY | 0.48831415 |
| AGLESIGL | ASSSTGGALEQY | 0.4008425 |
| AGLKTGII | ASSSTGGALEQY | 0.6391216 |
| ASLAALLL | ASSSTGGALEQY | 0.8201394 |
| ASVINISSI | ASSSTGGALEQY | 0.836749 |
| ATPPGPA | ASSSTGGALEQY | 0.7807907 |
| ATVASALV | ASSSTGGALEQY | 0.6996838 |
| ATVASLIL | ASSSTGGALEQY | 0.78084683 |
| AVNGPVLIRH | ASSSTGGALEQY | 0.8279007 |
| DGAAGAVGI | ASSSTGGALEQY | 0.84364086 |
| DPLLHGI | ASSSTGGALEQY | 0.8323867 |
| ELGAALGAL | ASSSTGGALEQY | 0.68348366 |
| ELGRTQAV | ASSSTGGALEQY | 0.83935255 |
| FVAGGYLS | ASSSTGGALEQY | 0.79473287 |
| GASGGAGGAGGW | ASSSTGGALEQY | 0.79237944 |
| GGAGAGGDGGAGGI | ASSSTGGALEQY | 0.79951894 |
| GTGHNGSQPGMGGNG | ASSSTGGALEQY | 0.815593 |
| GYAAASASAS | ASSSTGGALEQY | 0.7948537 |
| HQVDVAAVQQG | ASSSTGGALEQY | 0.7046471 |
| IGAAIAHT | ASSSTGGALEQY | 0.82609165 |
| IGMIEVAQL | ASSSTGGALEQY | 0.82878304 |
| IRLAPEHTSAMD | ASSSTGGALEQY | 0.826136 |
| IYNVSVL | ASSSTGGALEQY | 0.85226953 |
| KQGFLGM | ASSSTGGALEQY | 0.8398306 |
| LESVITQL | ASSSTGGALEQY | 0.8232555 |
| LGLGALVVV | ASSSTGGALEQY | 0.3366166 |
| LIGNGGNGGSGGI | ASSSTGGALEQY | 0.5720236 |
| LNAINAPTL | ASSSTGGALEQY | 0.734568 |
| LQLLNTGV | ASSSTGGALEQY | 0.7776204 |
| LQTSIEF | ASSSTGGALEQY | 0.3437971 |
| MSFAVSEL | ASSSTGGALEQY | 0.7807435 |
| NTGALGL | ASSSTGGALEQY | 0.8059686 |
| PPGAGGAGGAGTT | ASSSTGGALEQY | 0.7372861 |
| QLVGLGI | ASSSTGGALEQY | 0.839296 |
| RFLEQVHQL | ASSSTGGALEQY | 0.7881694 |
| RGVGYVLR | ASSSTGGALEQY | 0.6636455 |
| SATGVFAGVI | ASSSTGGALEQY | 0.6896748 |
| SLPAITLP | ASSSTGGALEQY | 0.84423935 |
| STALAFTV | ASSSTGGALEQY | 0.55066955 |
| SVLTVGLGK | ASSSTGGALEQY | 0.8372493 |
| TALAGAGSGI | ASSSTGGALEQY | 0.6921188 |
| TGLPFTGI | ASSSTGGALEQY | 0.837729 |
| VAEHVTV | ASSSTGGALEQY | 0.5633397 |
| VDIVLNSL | ASSSTGGALEQY | 0.83510584 |
| VLTGGGAVL | ASSSTGGALEQY | 0.797984 |
| VNVGLAYGI | ASSSTGGALEQY | 0.8564793 |
| VSSAAVGAV | ASSSTGGALEQY | 0.7587311 |
| VTSVYEAGV | ASSSTGGALEQY | 0.8273622 |
| WVALDRGA | ASSSTGGALEQY | 0.7502704 |

| | | |
|---|---|---|
| YLVGSVVI | ASSSTGGALEQY | 0.8339498 |
| GPPGTGKTLL | ASSSTGGALEQY | 0.6107923 |
| GVGMTKF | ASSSTGGALEQY | 0.7854743 |
| IGIGGSDL | ASSSTGGALEQY | 0.63470346 |
| IGTLLAEL | ASSSTGGALEQY | 0.8118118 |
| IYTSGTT | ASSSTGGALEQY | 0.83245873 |
| LSFPAGV | ASSSTGGALEQY | 0.4143532 |
| LYTSGTT | ASSSTGGALEQY | 0.8492254 |
| VGAFPFGF | ASSSTGGALEQY | 0.75078654 |
| VGAGTVEF | ASSSTGGALEQY | 0.70485824 |
| VGKSTLLL | ASSSTGGALEQY | 0.83976 |
| AALAHAGGGIV | ASSSTGGALEQY | 0.8093369 |
| AALVAVSLL | ASSSTGGALEQY | 0.8399562 |
| AAVSRAVAAL | ASSSTGGALEQY | 0.72843057 |
| AGGSSGIGGF | ASSSTGGALEQY | 0.792323 |
| AGLTVANANG | ASSSTGGALEQY | 0.8169285 |
| AGLVGAVL | ASSSTGGALEQY | 0.7248856 |
| ALSQAFTV | ASSSTGGALEQY | 0.64317423 |
| ALTFSVSL | ASSSTGGALEQY | 0.8454606 |
| ALTGAALVAL | ASSSTGGALEQY | 0.80398816 |
| ASFSIGAGVL | ASSSTGGALEQY | 0.865355 |
| DGSVYKI | ASSSTGGALEQY | 0.71448255 |
| EQLAEIV | ASSSTGGALEQY | 0.60203993 |
| FAVSELTVM | ASSSTGGALEQY | 0.7331431 |
| GAAPLAAGL | ASSSTGGALEQY | 0.8517974 |
| GADGGDGGSGGAGG | ASSSTGGALEQY | 0.8644489 |
| GFGLDTGGAGG | ASSSTGGALEQY | 0.8160369 |
| GLASGGLL | ASSSTGGALEQY | 0.5333375 |
| GRIAGAAVLL | ASSSTGGALEQY | 0.8052778 |
| GSAGSLNTSF | ASSSTGGALEQY | 0.7180468 |
| GTGQIGI | ASSSTGGALEQY | 0.7494934 |
| GTLIALIL | ASSSTGGALEQY | 0.8257201 |
| IAATMGFVVG | ASSSTGGALEQY | 0.76738966 |
| IFAQALL | ASSSTGGALEQY | 0.6899966 |
| IGAAVALGF | ASSSTGGALEQY | 0.65646803 |
| IGLTPLAL | ASSSTGGALEQY | 0.81268466 |
| ILGSNTSTL | ASSSTGGALEQY | 0.81508607 |
| IVFSLTVL | ASSSTGGALEQY | 0.81677437 |
| LASKATGF | ASSSTGGALEQY | 0.87613547 |
| LASVALIL | ASSSTGGALEQY | 0.8505055 |
| LFAVGLTL | ASSSTGGALEQY | 0.86260986 |
| LFNGGIGTYI | ASSSTGGALEQY | 0.21700442 |
| LTLGAGII | ASSSTGGALEQY | 0.41137582 |
| LVAAALGDI | ASSSTGGALEQY | 0.6871971 |
| NTAFVQLGI | ASSSTGGALEQY | 0.71326864 |
| QLFSGAEL | ASSSTGGALEQY | 0.44292897 |
| SAATFITL | ASSSTGGALEQY | 0.8053755 |
| SNYGLANLGV | ASSSTGGALEQY | 0.67799896 |
| SYGGAYAVM | ASSSTGGALEQY | 0.63691396 |
| TTGVGGLTLG | ASSSTGGALEQY | 0.8416542 |
| TTQTIYI | ASSSTGGALEQY | 0.7052062 |

| | | |
|---|---|---|
| VAAGAGLGAVI | ASSSTGGALEQY | 0.62673604 |
| VAAVVTLL | ASSSTGGALEQY | 0.61499393 |
| VGFLGGLI | ASSSTGGALEQY | 0.83218277 |
| VGVIPILL | ASSSTGGALEQY | 0.4514993 |
| VIFASSLI | ASSSTGGALEQY | 0.8256626 |
| VKGGDGGAAAG | ASSSTGGALEQY | 0.6233343 |
| VTKTFTNI | ASSSTGGALEQY | 0.8486093 |
| VVAAAILL | ASSSTGGALEQY | 0.43543833 |
| YAALDVEL | ASSSTGGALEQY | 0.8443857 |
| LASVALLL | ASSSTGGALEQY | 0.7712921 |
| LDAVHRV | ASSSTGGALEQY | 0.8860105 |
| QGLSSSIF | ASSSTGGALEQY | 0.7149317 |
| RGTFANIRL | ASSSTGGALEQY | 0.7548746 |
| RQLWWGHRI | ASSSTGGALEQY | 0.6154326 |
| YGPPGTGKTLL | ASSSTGGALEQY | 0.86130357 |
| AALVGAPANVA | ASSSTGGALEQY | 0.3889792 |
| AATFRRTE | ASSSTGGALEQY | 0.7543954 |
| AGALGGGI | ASSSTGGALEQY | 0.4259022 |
| AGGAGGAGGIGAG | ASSSTGGALEQY | 0.8139035 |
| AGSGALGL | ASSSTGGALEQY | 0.8194947 |
| AQIADQI | ASSSTGGALEQY | 0.3305482 |
| ASAVAVTYL | ASSSTGGALEQY | 0.82720995 |
| ASTVLGGTGG | ASSSTGGALEQY | 0.7819346 |
| AVGGEPAK | ASSSTGGALEQY | 0.6364188 |
| CEQGTFGGTC | ASSSTGGALEQY | 0.868148 |
| DEGKAPAG | ASSSTGGALEQY | 0.76874185 |
| DTVVFPAL | ASSSTGGALEQY | 0.8713173 |
| FEGITMAG | ASSSTGGALEQY | 0.7725078 |
| FPLVVQD | ASSSTGGALEQY | 0.67738163 |
| GAFTTPQ | ASSSTGGALEQY | 0.7945057 |
| GFGGFGTAGGVGGA | ASSSTGGALEQY | 0.5745384 |
| GFLAQFL | ASSSTGGALEQY | 0.5949546 |
| GGAGGAGGNAGVA | ASSSTGGALEQY | 0.8653481 |
| GGVGTAGAGGAGGAGG | ASSSTGGALEQY | 0.89551973 |
| GNGGGGDAGNAGSGGN | ASSSTGGALEQY | 0.8803779 |
| GTGGAGGAGNDAGS | ASSSTGGALEQY | 0.8027861 |
| GTTNTGWFNS | ASSSTGGALEQY | 0.8061657 |
| IFMGPLFTL | ASSSTGGALEQY | 0.83769166 |
| ILAHAIEAI | ASSSTGGALEQY | 0.89994824 |
| IRSAIREAL | ASSSTGGALEQY | 0.8882947 |
| ITFAGGG | ASSSTGGALEQY | 0.8908638 |
| ITGMSGHQEV | ASSSTGGALEQY | 0.8270882 |
| IVAGLACTL | ASSSTGGALEQY | 0.8643157 |
| IVGGGLAGA | ASSSTGGALEQY | 0.8584359 |
| IVTGVLNI | ASSSTGGALEQY | 0.82561886 |
| LAQQGCI | ASSSTGGALEQY | 0.88384444 |
| LCLHVEN | ASSSTGGALEQY | 0.8761042 |
| LDAISDGS | ASSSTGGALEQY | 0.82964844 |
| LGGFAAML | ASSSTGGALEQY | 0.8793577 |
| LGGKNPMI | ASSSTGGALEQY | 0.8575128 |
| LGPKGVAAIL | ASSSTGGALEQY | 0.88106537 |

| | | |
|---|---|---|
| LMVGVSI | ASSSTGGALEQY | 0.87438667 |
| LVGAAFVL | ASSSTGGALEQY | 0.91594714 |
| LVLAVAPL | ASSSTGGALEQY | 0.90051675 |
| MAAIAVIDAA | ASSSTGGALEQY | 0.89689267 |
| NNGINAELQ | ASSSTGGALEQY | 0.55482244 |
| RGEPLVLVHG | ASSSTGGALEQY | 0.751933 |
| RGGAGGAAGLI | ASSSTGGALEQY | 0.84806526 |
| RSYRMHPAV | ASSSTGGALEQY | 0.8508838 |
| SAAHLLAGQLA | ASSSTGGALEQY | 0.6719713 |
| SAAVGSY | ASSSTGGALEQY | 0.8724958 |
| SPHAVLHRI | ASSSTGGALEQY | 0.87014496 |
| SQIAKVL | ASSSTGGALEQY | 0.77651167 |
| TALGHLVP | ASSSTGGALEQY | 0.8918259 |
| TAPTTSGAA | ASSSTGGALEQY | 0.81963986 |
| TDGDDGNGGSPGT | ASSSTGGALEQY | 0.8355768 |
| TDGLGEN | ASSSTGGALEQY | 0.849262 |
| TGGPPLPGQA | ASSSTGGALEQY | 0.90300715 |
| TIPAITI | ASSSTGGALEQY | 0.80860883 |
| TIPGFTI | ASSSTGGALEQY | 0.8835772 |
| TSIASGALA | ASSSTGGALEQY | 0.8177042 |
| TVTLPGAAI | ASSSTGGALEQY | 0.90787446 |
| VASGLSAALP | ASSSTGGALEQY | 0.785499 |
| VGAGVIGI | ASSSTGGALEQY | 0.8975283 |
| VGPKRFGV | ASSSTGGALEQY | 0.8532553 |
| VNVAFPD | ASSSTGGALEQY | 0.91212046 |
| VSMTNSGVS | ASSSTGGALEQY | 0.8645927 |
| VTDVLGDLSV | ASSSTGGALEQY | 0.8898834 |
| VVVVGAGFAGL | ASSSTGGALEQY | 0.85180056 |
| VYTVGTVV | ASSSTGGALEQY | 0.8877002 |
| WTTFSGP | ASSSTGGALEQY | 0.7225396 |
| YDELVEAF | ASSSTGGALEQY | 0.8684397 |
| YGTGGAGGNGG | ASSSTGGALEQY | 0.6831974 |
| YTNVMAV | ASSSTGGALEQY | 0.87674034 |
| FLEQVHQL | ASSSTGGALEQY | 0.89094687 |
| GLSLLAVAV | ASSSTGGALEQY | 0.6595475 |
| LEQVHQL | ASSSTGGALEQY | 0.904239 |
| LNAAGAI | ASSSTGGALEQY | 0.8025901 |
| PAVHASL | ASSSTGGALEQY | 0.8518063 |
| TELEDDA | ASSSTGGALEQY | 0.8988043 |
| TPPGPPN | ASSSTGGALEQY | 0.8381145 |
| TPPPPPPP | ASSSTGGALEQY | 0.9078961 |
| TQSPALI | AWTGLANSDYT | 0.8325243 |
| EFAAAFPQ | AWTGLANSDYT | 0.82871103 |
| LLSGNLDVL | AWTGLANSDYT | 0.88230467 |
| GPANLGNYNI | AWTGLANSDYT | 0.6556657 |
| TLGPTLP | AWTGLANSDYT | 0.7932651 |
| VYAYSSCCK | AWTGLANSDYT | 0.9101958 |
| APAPAPAPNI | AWTGLANSDYT | 0.5603507 |
| PGGPVPIP | AWTGLANSDYT | 0.63633966 |
| FLTVALEP | AWTGLANSDYT | 0.5265975 |
| SQYFGANL | AWTGLANSDYT | 0.4361561 |

| | | |
|---|---|---|
| GGVKPAA | AWTGLANSDYT | 0.5811773 |
| GGTGGAGGAGGW | AWTGLANSDYT | 0.6038519 |
| GVGGVAGLGGAGG | AWTGLANSDYT | 0.64647734 |
| VLLVLATTI | AWTGLANSDYT | 0.5579753 |
| IGIFAAML | AWTGLANSDYT | 0.4178742 |
| AVGDLID | AWTGLANSDYT | 0.62577003 |
| NISPLPL | AWTGLANSDYT | 0.5917401 |
| VTAVFVAI | AWTGLANSDYT | 0.54871094 |
| SGDDNDAAL | AWTGLANSDYT | 0.598811 |
| AVAALAHS | AWTGLANSDYT | 0.59079456 |
| GGVTVMATP | AWTGLANSDYT | 0.5641097 |
| AGLAANVRD | AWTGLANSDYT | 0.43661824 |
| TNRPDLIDPALLRPG | AWTGLANSDYT | 0.5993708 |
| LANLGVTAL | AWTGLANSDYT | 0.61363685 |
| TGERIEL | AWTGLANSDYT | 0.61407065 |
| NGEELAA | AWTGLANSDYT | 0.64749146 |
| VLSLVWG | AWTGLANSDYT | 0.62223315 |
| IGYALWL | AWTGLANSDYT | 0.574176 |
| LSLLLAPL | AWTGLANSDYT | 0.320498 |
| TLDEALK | AWTGLANSDYT | 0.34976825 |
| TDGAGTAEGG | AWTGLANSDYT | 0.5615061 |
| NAPGGAGGNGG | AWTGLANSDYT | 0.51814735 |
| VSVFMGMGGNF | AWTGLANSDYT | 0.3521946 |
| LASYLAAL | AWTGLANSDYT | 0.6212133 |
| LVGGTPAL | AWTGLANSDYT | 0.5875485 |
| VRNADVL | AWTGLANSDYT | 0.5533294 |
| TCSSPFLVSAM | AWTGLANSDYT | 0.63484836 |
| VHITDAE | AWTGLANSDYT | 0.5441288 |
| EYAAPSF | AWTGLANSDYT | 0.47506088 |
| VVDAFHGGL | AWTGLANSDYT | 0.4916454 |
| ASVVDPQ | AWTGLANSDYT | 0.64153993 |
| SYNEFGGLS | AWTGLANSDYT | 0.3860854 |
| DPDGSVY | AWTGLANSDYT | 0.62679935 |
| MVSTIHATE | AWTGLANSDYT | 0.5355974 |
| LAAGVADPV | AWTGLANSDYT | 0.63706845 |
| TKEQIAA | AWTGLANSDYT | 0.38057676 |
| SLASPAAL | AWTGLANSDYT | 0.5871956 |
| IGLVFYGV | AWTGLANSDYT | 0.53858703 |
| FLLDASNL | AWTGLANSDYT | 0.6561881 |
| YQVGNFP | AWTGLANSDYT | 0.5111476 |
| ESLLSQST | AWTGLANSDYT | 0.5999977 |
| LFTVGFAAVG | AWTGLANSDYT | 0.5193824 |
| ATPPTPT | AWTGLANSDYT | 0.64238405 |
| TDLATLG | AWTGLANSDYT | 0.46143964 |
| SAGGTGR | AWTGLANSDYT | 0.55829906 |
| GFSNSGS | AWTGLANSDYT | 0.46475703 |
| GGNGGSGGTSV | AWTGLANSDYT | 0.59848213 |
| GTSVATGGAGN | AWTGLANSDYT | 0.5641214 |
| NGGNGGSGGTS | AWTGLANSDYT | 0.37668943 |
| AVVKFPH | AWTGLANSDYT | 0.64426935 |
| TPPPPPP | AWTGLANSDYT | 0.54253536 |

| | | |
|---|---|---|
| AAPAAAAAPA | AWTGLANSDYT | 0.44289848 |
| IALNEDL | AWTGLANSDYT | 0.6252803 |
| IVNVSSM | AWTGLANSDYT | 0.5969523 |
| TGIPPAPR | AWTGLANSDYT | 0.6086171 |
| FDLGGGTF | AWTGLANSDYT | 0.5467584 |
| FDLGGGTFDVS | AWTGLANSDYT | 0.60512245 |
| NEPTAAA | AWTGLANSDYT | 0.6147259 |
| NEPTAAALA | AWTGLANSDYT | 0.556671 |
| GSIINISSI | AWTGLANSDYT | 0.5067842 |
| VAPGNAGTA | AWTGLANSDYT | 0.611901 |
| TGGGASLE | AWTGLANSDYT | 0.74493974 |
| VAVYNHYKRI | AWTGLANSDYT | 0.7578368 |
| TDPAPATT | AWTGLANSDYT | 0.6641263 |
| TNRPDILDPALLRPG | AWTGLANSDYT | 0.63087666 |
| LTTTTTS | AWTGLANSDYT | 0.6396639 |
| PTGSGKTL | AWTGLANSDYT | 0.76646686 |
| TGSGKTL | AWTGLANSDYT | 0.7430135 |
| TIGHVDHGK | AWTGLANSDYT | 0.73235834 |
| TIGHVDHGKTT | AWTGLANSDYT | 0.65497464 |
| GHVDHGKTT | AWTGLANSDYT | 0.74212927 |
| AQFEHTLL | AWTGLANSDYT | 0.6980841 |
| IVNQDEL | AWTGLANSDYT | 0.66253674 |
| TNGSQFF | AWTGLANSDYT | 0.7396964 |
| TGKGIVE | AWTGLANSDYT | 0.74415517 |
| LISNASDA | AWTGLANSDYT | 0.6717638 |
| AELRTLGELTKIAKAHGAQ | AWTGLANSDYT | 0.72762954 |
| APNAVAAAEPSW | AWTGLANSDYT | 0.6647336 |
| APQVAGAATAVL | AWTGLANSDYT | 0.7246669 |
| ARRDPAAAAAILPTDA | AWTGLANSDYT | 0.7069872 |
| DAAARDVLAEAGLAEYF | AWTGLANSDYT | 0.7744003 |
| DDQIVLSAKANLAKVEQ | AWTGLANSDYT | 0.75636697 |
| DELLPDTGEDSDEDGH | AWTGLANSDYT | 0.75459963 |
| ELPEAPANRATVPQD | AWTGLANSDYT | 0.4473709 |
| FGQSKIERVTGIG | AWTGLANSDYT | 0.6127414 |
| GAGGTGSTAGGAGGAGGAGGI | AWTGLANSDYT | 0.71434176 |
| GHGGTGAAGGNGGTGGAGGS( | AWTGLANSDYT | 0.7001382 |
| GNGGNGGAGGEGGPGGAGGA( | AWTGLANSDYT | 0.5397418 |
| GSTAGNGGNGGNATNGTIAG( | AWTGLANSDYT | 0.742488 |
| GTGGDGGNGGNGGTGG | AWTGLANSDYT | 0.70049936 |
| GVRRPAGIPADAPV | AWTGLANSDYT | 0.6501996 |
| GYRQVTDAHLLTLARR | AWTGLANSDYT | 0.72872865 |
| IAKMGQDGHDRG | AWTGLANSDYT | 0.67484987 |
| KAEPSIEIQYT | AWTGLANSDYT | 0.71368814 |
| KPVRIRMAADEDTVAVT | AWTGLANSDYT | 0.68599164 |
| MPPRIAGMRLLVIKPEP | AWTGLANSDYT | 0.76195854 |
| MKFYAVALFRSRGVPPTR | AWTGLANSDYT | 0.63270235 |
| NAGGFGILGAVAHSP | AWTGLANSDYT | 0.7395959 |
| NEGREITLAMAEM | AWTGLANSDYT | 0.6513723 |
| NELVEWSHPLGHRWRWE | AWTGLANSDYT | 0.76394814 |
| NLGAGGTGGNGGNPG | AWTGLANSDYT | 0.61662966 |
| NTEWPKWETVMAADPVD | AWTGLANSDYT | 0.7436482 |

| | | |
|---|---|---|
| PASAAPAETQP | AWTGLANSDYT | 0.69661784 |
| QQSSDGKNGVRAAEITGE | AWTGLANSDYT | 0.77350146 |
| RAPDAVTTSWWKE | AWTGLANSDYT | 0.70767456 |
| SGFGNVGAHTSGWFNQ | AWTGLANSDYT | 0.7432094 |
| SGVAGAVGQAASVGGLKV | AWTGLANSDYT | 0.7159168 |
| SNGGAGTGSAGGNGGT | AWTGLANSDYT | 0.7523539 |
| TGGAGGEGGAAG | AWTGLANSDYT | 0.576213 |
| TGGDGGAAGAGGAPGHG | AWTGLANSDYT | 0.7151053 |
| TMPPLLAAIANAGRHGV | AWTGLANSDYT | 0.6055385 |
| TSSTAGNSGGA | AWTGLANSDYT | 0.74009657 |
| VPATVPVGILC | AWTGLANSDYT | 0.7431064 |
| VQRPDSGGVVLTGRIS | AWTGLANSDYT | 0.5830494 |
| WTPNPVPATKGVPAGLVQAT | AWTGLANSDYT | 0.7580287 |
| YALKRALAKFSDRFDVVII | AWTGLANSDYT | 0.69437647 |
| YLIGSVIV | AWTGLANSDYT | 0.69756085 |
| YGQLVAVID | AWTGLANSDYT | 0.7633471 |
| VTVGGMTLPA | AWTGLANSDYT | 0.7060522 |
| VLALPAV | AWTGLANSDYT | 0.7650477 |
| VIPPTLNYET | AWTGLANSDYT | 0.70289415 |
| VGMGGMHPSSG | AWTGLANSDYT | 0.69743425 |
| VGLAGGAATA | AWTGLANSDYT | 0.7428064 |
| VGGAGGAGGSGP | AWTGLANSDYT | 0.57909733 |
| VFAAFMV | AWTGLANSDYT | 0.6288827 |
| TLYDKLR | AWTGLANSDYT | 0.76290035 |
| TLAVFTVL | AWTGLANSDYT | 0.8270166 |
| TGSVQPARV | AWTGLANSDYT | 0.83214784 |
| TGGGAGSGGT | AWTGLANSDYT | 0.67621124 |
| SHLETSL | AWTGLANSDYT | 0.70689857 |
| SFTTGMT | AWTGLANSDYT | 0.7801948 |
| QVEVAKRMGVRQPTVS | AWTGLANSDYT | 0.84853506 |
| QGSGGPALAM | AWTGLANSDYT | 0.8607359 |
| MLAPSNP | AWTGLANSDYT | 0.83732545 |
| LVNNAAI | AWTGLANSDYT | 0.77371275 |
| LTGEINTLA | AWTGLANSDYT | 0.84085155 |
| LSIFGPV | AWTGLANSDYT | 0.8130913 |
| LLALLANL | AWTGLANSDYT | 0.79696476 |
| LISNASDALDKLR | AWTGLANSDYT | 0.85511154 |
| LINEAVP | AWTGLANSDYT | 0.8247261 |
| LGYAFSGL | AWTGLANSDYT | 0.7695812 |
| LDFVAVAL | AWTGLANSDYT | 0.7757759 |
| LAQPGAE | AWTGLANSDYT | 0.8315784 |
| ISNLGAVTGF | AWTGLANSDYT | 0.8499764 |
| HQEDPGRI | AWTGLANSDYT | 0.8437769 |
| GTPGEPTP | AWTGLANSDYT | 0.8775527 |
| GFFNSGS | AWTGLANSDYT | 0.84687525 |
| GAGGGGLTAGAN | AWTGLANSDYT | 0.83351487 |
| FVDALVNM | AWTGLANSDYT | 0.48295093 |
| FLTVAPEL | AWTGLANSDYT | 0.6114853 |
| FETLTAL | AWTGLANSDYT | 0.807143 |
| DVINPIEL | AWTGLANSDYT | 0.77108514 |
| APVSGAPV | AWTGLANSDYT | 0.46826893 |

| | | |
|---|---|---|
| ALVGGLAW | AWTGLANSDYT | 0.842957 |
| ALAAAAATAI | AWTGLANSDYT | 0.80601615 |
| AFGLGLLL | AWTGLANSDYT | 0.73894286 |
| AFFAAASL | AWTGLANSDYT | 0.85412055 |
| AAAALPAFSPP | AWTGLANSDYT | 0.7726957 |
| VVLAPAL | AWTGLANSDYT | 0.77351975 |
| TIGHVDHGKTTL | AWTGLANSDYT | 0.8209357 |
| LISNASDALDK | AWTGLANSDYT | 0.8737951 |
| LISNASDAL | AWTGLANSDYT | 0.7820576 |
| ISNASDALDK | AWTGLANSDYT | 0.86158895 |
| NQAETLVYQTE | AWTGLANSDYT | 0.81697285 |
| TGGTGGTAGAG | AWTGLANSDYT | 0.86148876 |
| TGSTAGGAGGA | AWTGLANSDYT | 0.7068014 |
| DPLAVPVAAGPV | AWTGLANSDYT | 0.8499966 |
| GGTGGTPGNGGN | AWTGLANSDYT | 0.79520476 |
| PAGLAGVDPATL | AWTGLANSDYT | 0.87971413 |
| GNNLVVEAPHAGASV | AWTGLANSDYT | 0.80031335 |
| KVTRIMSASTLEQPA | AWTGLANSDYT | 0.86425626 |
| LGFALPIALPRRTVV | AWTGLANSDYT | 0.83233404 |
| TRLGNLTARKLVVLV | AWTGLANSDYT | 0.8474118 |
| WSQQRRLIYFPSAGP | AWTGLANSDYT | 0.6795411 |
| GPNGNPDPMAAAVDIR | AWTGLANSDYT | 0.77831835 |
| TQRCVSVADLRAAASR | AWTGLANSDYT | 0.7329649 |
| QFVLDVAEHPVEVLTETV | AWTGLANSDYT | 0.8186287 |
| GGAGGTGGAGGNGGTQAGDG | AWTGLANSDYT | 0.84241664 |
| PLLATLRAIAVDVDAKPAQV | AWTGLANSDYT | 0.5101387 |
| APKFRTGGYMIDFWGVGYQV | AWTGLANSDYT | 0.8657744 |
| GAGGAGGAGGAAGGPGGTGG | AWTGLANSDYT | 0.7156817 |
| IGALGLLATDVAPAGSVAFG | AWTGLANSDYT | 0.77040553 |
| AGGAGGIGGNGGAGGAGGVL | AWTGLANSDYT | 0.8563846 |
| ADELIGLL | AWTGLANSDYT | 0.79591286 |
| ADRALKA | AWTGLANSDYT | 0.8309809 |
| AGAVGQVGGAGGA | AWTGLANSDYT | 0.7951727 |
| AGGGGGLGGAP | AWTGLANSDYT | 0.80372566 |
| AGGLLYGNGGAG | AWTGLANSDYT | 0.83354545 |
| AGLESIGL | AWTGLANSDYT | 0.54837865 |
| AGLKTGII | AWTGLANSDYT | 0.7630976 |
| ASLAALLL | AWTGLANSDYT | 0.8646828 |
| ASVINISSI | AWTGLANSDYT | 0.6207267 |
| ATPPGPA | AWTGLANSDYT | 0.74647677 |
| ATVASALV | AWTGLANSDYT | 0.39384872 |
| ATVASLIL | AWTGLANSDYT | 0.40707242 |
| AVNGPVLIRH | AWTGLANSDYT | 0.46265605 |
| DGAAGAVGI | AWTGLANSDYT | 0.5804882 |
| DPLLHGI | AWTGLANSDYT | 0.7533012 |
| ELGAALGAL | AWTGLANSDYT | 0.67550397 |
| ELGRTQAV | AWTGLANSDYT | 0.40512726 |
| FVAGGYLS | AWTGLANSDYT | 0.6070361 |
| GASGGAGGAGGW | AWTGLANSDYT | 0.50242984 |
| GGAGAGGDGGAGGI | AWTGLANSDYT | 0.51309687 |
| GTGHNGSQPGMGGNG | AWTGLANSDYT | 0.47577265 |

| | | |
|---|---|---|
| GYAAASASAS | AWTGLANSDYT | 0.46624184 |
| HQVDVAAVQQG | AWTGLANSDYT | 0.40541583 |
| IGAAIAHT | AWTGLANSDYT | 0.32594606 |
| IGMIEVAQL | AWTGLANSDYT | 0.5417818 |
| IRLAPEHTSAMD | AWTGLANSDYT | 0.608022 |
| IYNVSVL | AWTGLANSDYT | 0.5939566 |
| KQGFLGM | AWTGLANSDYT | 0.6284679 |
| LESVITQL | AWTGLANSDYT | 0.6746632 |
| LGLGALVVV | AWTGLANSDYT | 0.44661903 |
| LIGNGGNGGSGGI | AWTGLANSDYT | 0.2422729 |
| LNAINAPTL | AWTGLANSDYT | 0.28464004 |
| LQLLNTGV | AWTGLANSDYT | 0.41367736 |
| LQTSIEF | AWTGLANSDYT | 0.4794409 |
| MSFAVSEL | AWTGLANSDYT | 0.24278803 |
| NTGALGL | AWTGLANSDYT | 0.55997103 |
| PPGAGGAGGAGTT | AWTGLANSDYT | 0.4313384 |
| QLVGLGI | AWTGLANSDYT | 0.4771607 |
| RFLEQVHQL | AWTGLANSDYT | 0.6846508 |
| RGVGYVLR | AWTGLANSDYT | 0.4963867 |
| SATGVFAGVI | AWTGLANSDYT | 0.31986707 |
| SLPAITLP | AWTGLANSDYT | 0.36103576 |
| STALAFTV | AWTGLANSDYT | 0.64058137 |
| SVLTVGLGK | AWTGLANSDYT | 0.31188786 |
| TALAGAGSGI | AWTGLANSDYT | 0.6417503 |
| TGLPFTGI | AWTGLANSDYT | 0.34484744 |
| VAEHVTV | AWTGLANSDYT | 0.6389591 |
| VDIVLNSL | AWTGLANSDYT | 0.33428648 |
| VLTGGGAVL | AWTGLANSDYT | 0.61798716 |
| VNVGLAYGI | AWTGLANSDYT | 0.5978977 |
| VSSAAVGAV | AWTGLANSDYT | 0.73954064 |
| VTSVYEAGV | AWTGLANSDYT | 0.44336677 |
| WVALDRGA | AWTGLANSDYT | 0.61815923 |
| YLVGSVVI | AWTGLANSDYT | 0.35124922 |
| GPPGTGKTLL | AWTGLANSDYT | 0.74413717 |
| GVGMTKF | AWTGLANSDYT | 0.37987193 |
| IGIGGSDL | AWTGLANSDYT | 0.363203 |
| IGTLLAEL | AWTGLANSDYT | 0.30840623 |
| IYTSGTT | AWTGLANSDYT | 0.39853588 |
| LSFPAGV | AWTGLANSDYT | 0.6488044 |
| LYTSGTT | AWTGLANSDYT | 0.36203 |
| VGAFPFGF | AWTGLANSDYT | 0.5556075 |
| VGAGTVEF | AWTGLANSDYT | 0.5593362 |
| VGKSTLLL | AWTGLANSDYT | 0.36074317 |
| AALAHAGGGIV | AWTGLANSDYT | 0.51399386 |
| AALVAVSLL | AWTGLANSDYT | 0.4453048 |
| AAVSRAVAAL | AWTGLANSDYT | 0.5755459 |
| AGGSSGIGGF | AWTGLANSDYT | 0.35405746 |
| AGLTVANANG | AWTGLANSDYT | 0.48812497 |
| AGLVGAVL | AWTGLANSDYT | 0.47739866 |
| ALSQAFTV | AWTGLANSDYT | 0.38610664 |
| ALTFSVSL | AWTGLANSDYT | 0.3408718 |

| | | |
|---|---|---|
| ALTGAALVAL | AWTGLANSDYT | 0.6773366 |
| ASFSIGAGVL | AWTGLANSDYT | 0.62958324 |
| DGSVYKI | AWTGLANSDYT | 0.6651236 |
| EQLAEIV | AWTGLANSDYT | 0.59425247 |
| FAVSELTVM | AWTGLANSDYT | 0.56645864 |
| GAAPLAAGL | AWTGLANSDYT | 0.59487563 |
| GADGGDGGSGGAGG | AWTGLANSDYT | 0.65147585 |
| GFGLDTGGAGG | AWTGLANSDYT | 0.6572609 |
| GLASGGLL | AWTGLANSDYT | 0.63427603 |
| GRIAGAAVLL | AWTGLANSDYT | 0.5613963 |
| GSAGSLNTSF | AWTGLANSDYT | 0.6321483 |
| GTGQIGI | AWTGLANSDYT | 0.61337084 |
| GTLIALIL | AWTGLANSDYT | 0.60627776 |
| IAATMGFVVG | AWTGLANSDYT | 0.6273581 |
| IFAQALL | AWTGLANSDYT | 0.64079 |
| IGAAVALGF | AWTGLANSDYT | 0.59352046 |
| IGLTPLAL | AWTGLANSDYT | 0.60610783 |
| ILGSNTSTL | AWTGLANSDYT | 0.6077251 |
| IVFSLTVL | AWTGLANSDYT | 0.62886935 |
| LASKATGF | AWTGLANSDYT | 0.61602634 |
| LASVALIL | AWTGLANSDYT | 0.6626066 |
| LFAVGLTL | AWTGLANSDYT | 0.6481772 |
| LFNGGIGTYI | AWTGLANSDYT | 0.6373884 |
| LTLGAGII | AWTGLANSDYT | 0.3826941 |
| LVAAALGDI | AWTGLANSDYT | 0.5146867 |
| NTAFVQLGI | AWTGLANSDYT | 0.60229015 |
| QLFSGAEL | AWTGLANSDYT | 0.61918736 |
| SAATFITL | AWTGLANSDYT | 0.45411175 |
| SNYGLANLGV | AWTGLANSDYT | 0.6493201 |
| SYGGAYAVM | AWTGLANSDYT | 0.59211844 |
| TTGVGGLTLG | AWTGLANSDYT | 0.58456075 |
| TTQTIYI | AWTGLANSDYT | 0.64256966 |
| VAAGAGLGAVI | AWTGLANSDYT | 0.60002446 |
| VAAVVTLL | AWTGLANSDYT | 0.5995957 |
| VGFLGGLI | AWTGLANSDYT | 0.5757333 |
| VGVIPILL | AWTGLANSDYT | 0.64373326 |
| VIFASSLI | AWTGLANSDYT | 0.50841135 |
| VKGGDGGAAAG | AWTGLANSDYT | 0.6424089 |
| VTKTFTNI | AWTGLANSDYT | 0.5513493 |
| VVAAAILL | AWTGLANSDYT | 0.65331876 |
| YAALDVEL | AWTGLANSDYT | 0.5399327 |
| LASVALLL | AWTGLANSDYT | 0.6405663 |
| LDAVHRV | AWTGLANSDYT | 0.6081232 |
| QGLSSSIF | AWTGLANSDYT | 0.6679906 |
| RGTFANIRL | AWTGLANSDYT | 0.6060978 |
| RQLWWGHRI | AWTGLANSDYT | 0.6337796 |
| YGPPGTGKTLL | AWTGLANSDYT | 0.59410465 |
| AALVGAPANVA | AWTGLANSDYT | 0.65886533 |
| AATFRRTE | AWTGLANSDYT | 0.5256444 |
| AGALGGGI | AWTGLANSDYT | 0.6063807 |
| AGGAGGAGGIGAG | AWTGLANSDYT | 0.5020455 |

| | | |
|---|---|---|
| AGSGALGL | AWTGLANSDYT | 0.63339096 |
| AQIADQI | AWTGLANSDYT | 0.63460875 |
| ASAVAVTYL | AWTGLANSDYT | 0.50923747 |
| ASTVLGGTGG | AWTGLANSDYT | 0.6494329 |
| AVGGEPAK | AWTGLANSDYT | 0.60886896 |
| CEQGTFGGTC | AWTGLANSDYT | 0.58785665 |
| DEGKAPAG | AWTGLANSDYT | 0.6508997 |
| DTVVFPAL | AWTGLANSDYT | 0.61858517 |
| FEGITMAG | AWTGLANSDYT | 0.64883983 |
| FPLVVQD | AWTGLANSDYT | 0.6057092 |
| GAFTTPQ | AWTGLANSDYT | 0.60093045 |
| GFGGFGTAGGVGGA | AWTGLANSDYT | 0.6370271 |
| GFLAQFL | AWTGLANSDYT | 0.5845667 |
| GGAGGAGGNAGVA | AWTGLANSDYT | 0.5673962 |
| GGVGTAGAGGAGGAGG | AWTGLANSDYT | 0.6498449 |
| GNGGGGDAGNAGSGGN | AWTGLANSDYT | 0.7153511 |
| GTGGAGGAGNDAGS | AWTGLANSDYT | 0.8192878 |
| GTTNTGWFNS | AWTGLANSDYT | 0.5686207 |
| IFMGPLFTL | AWTGLANSDYT | 0.4880321 |
| ILAHAIEAI | AWTGLANSDYT | 0.62282836 |
| IRSAIREAL | AWTGLANSDYT | 0.8184556 |
| ITFAGGG | AWTGLANSDYT | 0.78541195 |
| ITGMSGHQEV | AWTGLANSDYT | 0.7599251 |
| IVAGLACTL | AWTGLANSDYT | 0.5057979 |
| IVGGGLAGA | AWTGLANSDYT | 0.622588 |
| IVTGVLNI | AWTGLANSDYT | 0.6047864 |
| LAQQGCI | AWTGLANSDYT | 0.642782 |
| LCLHVEN | AWTGLANSDYT | 0.7573621 |
| LDAISDGS | AWTGLANSDYT | 0.7334231 |
| LGGFAAML | AWTGLANSDYT | 0.6577574 |
| LGGKNPMI | AWTGLANSDYT | 0.5663845 |
| LGPKGVAAIL | AWTGLANSDYT | 0.66881186 |
| LMVGVSI | AWTGLANSDYT | 0.76002234 |
| LVGAAFVL | AWTGLANSDYT | 0.6873412 |
| LVLAVAPL | AWTGLANSDYT | 0.84619 |
| MAAIAVIDAA | AWTGLANSDYT | 0.79290414 |
| NNGINAELQ | AWTGLANSDYT | 0.7329831 |
| RGEPLVLVHG | AWTGLANSDYT | 0.13423647 |
| RGGAGGAAGLI | AWTGLANSDYT | 0.41135678 |
| RSYRMHPAV | AWTGLANSDYT | 0.70179725 |
| SAAHLLAGQLA | AWTGLANSDYT | 0.7158042 |
| SAAVGSY | AWTGLANSDYT | 0.34336734 |
| SPHAVLHRI | AWTGLANSDYT | 0.77626663 |
| SQIAKVL | AWTGLANSDYT | 0.6377086 |
| TALGHLVP | AWTGLANSDYT | 0.6165397 |
| TAPTTSGAA | AWTGLANSDYT | 0.8011224 |
| TDGDDGNGGSPGT | AWTGLANSDYT | 0.63906825 |
| TDGLGEN | AWTGLANSDYT | 0.60837626 |
| TGGPPLPGQA | AWTGLANSDYT | 0.624071 |
| TIPAITI | AWTGLANSDYT | 0.82496387 |
| TIPGFTI | AWTGLANSDYT | 0.4165223 |

| | | |
|---|---|---|
| TSIASGALA | AWTGLANSDYT | 0.76729417 |
| TVTLPGAAI | AWTGLANSDYT | 0.6914201 |
| VASGLSAALP | AWTGLANSDYT | 0.8146498 |
| VGAGVIGI | AWTGLANSDYT | 0.38909054 |
| VGPKRFGV | AWTGLANSDYT | 0.76093614 |
| VNVAFPD | AWTGLANSDYT | 0.6317352 |
| VSMTNSGVS | AWTGLANSDYT | 0.8696711 |
| VTDVLGDLSV | AWTGLANSDYT | 0.5834457 |
| VVVVGAGFAGL | AWTGLANSDYT | 0.7171172 |
| VYTVGTVV | AWTGLANSDYT | 0.5947381 |
| WTTFSGP | AWTGLANSDYT | 0.7427457 |
| YDELVEAF | AWTGLANSDYT | 0.43693805 |
| YGTGGAGGNGG | AWTGLANSDYT | 0.58320284 |
| YTNVMAV | AWTGLANSDYT | 0.43418738 |
| FLEQVHQL | AWTGLANSDYT | 0.69375145 |
| GLSLLAVAV | AWTGLANSDYT | 0.74655175 |
| LEQVHQL | AWTGLANSDYT | 0.43552312 |
| LNAAGAI | AWTGLANSDYT | 0.75735736 |
| PAVHASL | AWTGLANSDYT | 0.71434665 |
| TELEDDA | AWTGLANSDYT | 0.502953 |
| TPPGPPN | AWTGLANSDYT | 0.8075055 |
| TPPPPPPP | AWTGLANSDYT | 0.52768356 |
| TQSPALI | ASSLAWSSYEQY | 0.7726277 |
| EFAAAFPQ | ASSLAWSSYEQY | 0.6321442 |
| LLSGNLDVL | ASSLAWSSYEQY | 0.6648367 |
| GPANLGNYNI | ASSLAWSSYEQY | 0.7739785 |
| TLGPTLP | ASSLAWSSYEQY | 0.60160315 |
| VYAYSSCCK | ASSLAWSSYEQY | 0.5760133 |
| APAPAPAPNI | ASSLAWSSYEQY | 0.7964422 |
| PGGPVPIP | ASSLAWSSYEQY | 0.51056683 |
| FLTVALEP | ASSLAWSSYEQY | 0.65510935 |
| SQYFGANL | ASSLAWSSYEQY | 0.40532517 |
| GGVKPAA | ASSLAWSSYEQY | 0.43996978 |
| GGTGGAGGAGGW | ASSLAWSSYEQY | 0.5002721 |
| GVGGVAGLGGAGG | ASSLAWSSYEQY | 0.5820531 |
| VLLVLATTI | ASSLAWSSYEQY | 0.64991844 |
| IGIFAAML | ASSLAWSSYEQY | 0.5902528 |
| AVGDLID | ASSLAWSSYEQY | 0.45935476 |
| NISPLPL | ASSLAWSSYEQY | 0.53064144 |
| VTAVFVAI | ASSLAWSSYEQY | 0.50427514 |
| SGDDNDAAL | ASSLAWSSYEQY | 0.5107022 |
| AVAALAHS | ASSLAWSSYEQY | 0.52710724 |
| GGVTVMATP | ASSLAWSSYEQY | 0.50328326 |
| AGLAANVRD | ASSLAWSSYEQY | 0.4616976 |
| TNRPDLIDPALLRPG | ASSLAWSSYEQY | 0.41481227 |
| LANLGVTAL | ASSLAWSSYEQY | 0.51247513 |
| TGERIEL | ASSLAWSSYEQY | 0.58730197 |
| NGEELAA | ASSLAWSSYEQY | 0.571322 |
| VLSLVWG | ASSLAWSSYEQY | 0.6205109 |
| IGYALWL | ASSLAWSSYEQY | 0.5769794 |
| LSLLLAPL | ASSLAWSSYEQY | 0.46552357 |

| | | |
|---|---|---|
| TLDEALK | ASSLAWSSYEQY | 0.32783544 |
| TDGAGTAEGG | ASSLAWSSYEQY | 0.39353052 |
| NAPGGAGGNGG | ASSLAWSSYEQY | 0.4863725 |
| VSVFMGMGGNF | ASSLAWSSYEQY | 0.49774063 |
| LASYLAAL | ASSLAWSSYEQY | 0.33436182 |
| LVGGTPAL | ASSLAWSSYEQY | 0.5472826 |
| VRNADVL | ASSLAWSSYEQY | 0.5194148 |
| TCSSPFLVSAM | ASSLAWSSYEQY | 0.49147707 |
| VHITDAE | ASSLAWSSYEQY | 0.5854776 |
| EYAAPSF | ASSLAWSSYEQY | 0.48733813 |
| VVDAFHGGL | ASSLAWSSYEQY | 0.4178416 |
| ASVVDPQ | ASSLAWSSYEQY | 0.4059168 |
| SYNEFGGLS | ASSLAWSSYEQY | 0.59003484 |
| DPDGSVY | ASSLAWSSYEQY | 0.36572194 |
| MVSTIHATE | ASSLAWSSYEQY | 0.5755153 |
| LAAGVADPV | ASSLAWSSYEQY | 0.46719292 |
| TKEQIAA | ASSLAWSSYEQY | 0.5945352 |
| SLASPAAL | ASSLAWSSYEQY | 0.4063732 |
| IGLVFYGV | ASSLAWSSYEQY | 0.54032236 |
| FLLDASNL | ASSLAWSSYEQY | 0.50674987 |
| YQVGNFP | ASSLAWSSYEQY | 0.64346236 |
| ESLLSQST | ASSLAWSSYEQY | 0.48397467 |
| LFTVGFAAVG | ASSLAWSSYEQY | 0.5684173 |
| ATPPTPT | ASSLAWSSYEQY | 0.43825454 |
| TDLATLG | ASSLAWSSYEQY | 0.6429176 |
| SAGGTGR | ASSLAWSSYEQY | 0.45951265 |
| GFSNSGS | ASSLAWSSYEQY | 0.46684256 |
| GGNGGSGGTSV | ASSLAWSSYEQY | 0.39117897 |
| GTSVATGGAGN | ASSLAWSSYEQY | 0.49117514 |
| NGGNGGSGGTS | ASSLAWSSYEQY | 0.57735944 |
| AVVKFPH | ASSLAWSSYEQY | 0.5604414 |
| TPPPPPP | ASSLAWSSYEQY | 0.6106737 |
| AAPAAAAAPA | ASSLAWSSYEQY | 0.6125615 |
| IALNEDL | ASSLAWSSYEQY | 0.42612106 |
| IVNVSSM | ASSLAWSSYEQY | 0.57532716 |
| TGIPPAPR | ASSLAWSSYEQY | 0.48639008 |
| FDLGGGTF | ASSLAWSSYEQY | 0.52442104 |
| FDLGGGTFDVS | ASSLAWSSYEQY | 0.38629013 |
| NEPTAAA | ASSLAWSSYEQY | 0.55277133 |
| NEPTAAALA | ASSLAWSSYEQY | 0.58474517 |
| GSIINISSI | ASSLAWSSYEQY | 0.5524043 |
| VAPGNAGTA | ASSLAWSSYEQY | 0.40757927 |
| TGGGASLE | ASSLAWSSYEQY | 0.5845547 |
| VAVYNHYKRI | ASSLAWSSYEQY | 0.5684633 |
| TDPAPATT | ASSLAWSSYEQY | 0.82320064 |
| TNRPDILDPALLRPG | ASSLAWSSYEQY | 0.5597653 |
| LTTTTTS | ASSLAWSSYEQY | 0.29402953 |
| PTGSGKTL | ASSLAWSSYEQY | 0.4487123 |
| TGSGKTL | ASSLAWSSYEQY | 0.6187712 |
| TIGHVDHGK | ASSLAWSSYEQY | 0.83280647 |
| TIGHVDHGKTT | ASSLAWSSYEQY | 0.67655647 |

| | | |
|---|---|---|
| GHVDHGKTT | ASSLAWSSYEQY | 0.2270153 |
| AQFEHTLL | ASSLAWSSYEQY | 0.6166449 |
| IVNQDEL | ASSLAWSSYEQY | 0.452048 |
| TNGSQFF | ASSLAWSSYEQY | 0.47358382 |
| TGKGIVE | ASSLAWSSYEQY | 0.5919844 |
| LISNASDA | ASSLAWSSYEQY | 0.45148116 |
| AELRTLGELTKIAKAHGAQ | ASSLAWSSYEQY | 0.58420515 |
| APNAVAAAEPSW | ASSLAWSSYEQY | 0.36650532 |
| APQVAGAATAVL | ASSLAWSSYEQY | 0.64039534 |
| ARRDPAAAAAILPTDA | ASSLAWSSYEQY | 0.647884 |
| DAAARDVLAEAGLAEYF | ASSLAWSSYEQY | 0.5361674 |
| DDQIVLSAKANLAKVEQ | ASSLAWSSYEQY | 0.7186324 |
| DELLPDTGEDSDEDGH | ASSLAWSSYEQY | 0.71983325 |
| ELPEAPANRATVPQD | ASSLAWSSYEQY | 0.6866723 |
| FGQSKIERVTGIG | ASSLAWSSYEQY | 0.13093793 |
| GAGGTGSTAGGAGGAGGAGGI | ASSLAWSSYEQY | 0.20616098 |
| GHGGTGAAGGNGGTGGAGGS( | ASSLAWSSYEQY | 0.48410526 |
| GNGGNGGAGGEGGPGGAGGA( | ASSLAWSSYEQY | 0.49545094 |
| GSTAGNGGNGGNATNGTIAG( | ASSLAWSSYEQY | 0.17226277 |
| GTGGDGGNGGNGGTGG | ASSLAWSSYEQY | 0.4649912 |
| GVRRPAGIPADAPV | ASSLAWSSYEQY | 0.51842093 |
| GYRQVTDAHLLTLARR | ASSLAWSSYEQY | 0.41250357 |
| IAKMGQDGHDRG | ASSLAWSSYEQY | 0.75993156 |
| KAEPSIEIQYT | ASSLAWSSYEQY | 0.5917839 |
| KPVRIRMAADEDTVAVT | ASSLAWSSYEQY | 0.28984278 |
| MPPRIAGMRLLVIKPEP | ASSLAWSSYEQY | 0.29504186 |
| MKFYAVALFRSRGVPPTR | ASSLAWSSYEQY | 0.67265666 |
| NAGGFGILGAVAHSP | ASSLAWSSYEQY | 0.18551373 |
| NEGREITLAMAEM | ASSLAWSSYEQY | 0.74119085 |
| NELVEWSHPLGHRWRWE | ASSLAWSSYEQY | 0.31895176 |
| NLGAGGTGGNGGNPG | ASSLAWSSYEQY | 0.59720767 |
| NTEWPKWETVMAADPVD | ASSLAWSSYEQY | 0.18403062 |
| PASAAPAETQP | ASSLAWSSYEQY | 0.690004 |
| QQSSDGKNGVRAAEITGE | ASSLAWSSYEQY | 0.5511093 |
| RAPDAVTTSWWKE | ASSLAWSSYEQY | 0.8413597 |
| SGFGNVGAHTSGWFNQ | ASSLAWSSYEQY | 0.36071187 |
| SGVAGAVGQAASVGGLKV | ASSLAWSSYEQY | 0.4911904 |
| SNGGAGTGSAGGNGGT | ASSLAWSSYEQY | 0.324456 |
| TGGAGGEGGAAG | ASSLAWSSYEQY | 0.83640677 |
| TGGDGGAAGAGGAPGHG | ASSLAWSSYEQY | 0.25862196 |
| TMPPLLAAIANAGRHGV | ASSLAWSSYEQY | 0.41612792 |
| TSSTAGNSGGA | ASSLAWSSYEQY | 0.3198938 |
| VPATVPVGILC | ASSLAWSSYEQY | 0.60111237 |
| VQRPDSGGVVLTGRIS | ASSLAWSSYEQY | 0.68116355 |
| WTPNPVPATKGVPAGLVQAT | ASSLAWSSYEQY | 0.23851335 |
| YALKRALAKFSDRFDVVII | ASSLAWSSYEQY | 0.7677524 |
| YLIGSVIV | ASSLAWSSYEQY | 0.51821285 |
| YGQLVAVID | ASSLAWSSYEQY | 0.26244342 |
| VTVGGMTLPA | ASSLAWSSYEQY | 0.75375104 |
| VLALPAV | ASSLAWSSYEQY | 0.3658156 |
| VIPPTLNYET | ASSLAWSSYEQY | 0.7380785 |

| | | |
|---|---|---|
| VGMGGMHPSSG | ASSLAWSSYEQY | 0.3863157 |
| VGLAGGAATA | ASSLAWSSYEQY | 0.53252316 |
| VGGAGGAGGSGP | ASSLAWSSYEQY | 0.5688877 |
| VFAAFMV | ASSLAWSSYEQY | 0.5174403 |
| TLYDKLR | ASSLAWSSYEQY | 0.26669064 |
| TLAVFTVL | ASSLAWSSYEQY | 0.7562145 |
| TGSVQPARV | ASSLAWSSYEQY | 0.7288964 |
| TGGGAGSGGT | ASSLAWSSYEQY | 0.7298422 |
| SHLETSL | ASSLAWSSYEQY | 0.6895195 |
| SFTTGMT | ASSLAWSSYEQY | 0.68672776 |
| QVEVAKRMGVRQPTVS | ASSLAWSSYEQY | 0.67762136 |
| QGSGGPALAM | ASSLAWSSYEQY | 0.728658 |
| MLAPSNP | ASSLAWSSYEQY | 0.7341196 |
| LVNNAAI | ASSLAWSSYEQY | 0.73189676 |
| LTGEINTLA | ASSLAWSSYEQY | 0.6964954 |
| LSIFGPV | ASSLAWSSYEQY | 0.7270989 |
| LLALLANL | ASSLAWSSYEQY | 0.714504 |
| LISNASDALDKLR | ASSLAWSSYEQY | 0.70808136 |
| LINEAVP | ASSLAWSSYEQY | 0.7271996 |
| LGYAFSGL | ASSLAWSSYEQY | 0.7233583 |
| LDFVAVAL | ASSLAWSSYEQY | 0.68650687 |
| LAQPGAE | ASSLAWSSYEQY | 0.6976161 |
| ISNLGAVTGF | ASSLAWSSYEQY | 0.7154444 |
| HQEDPGRI | ASSLAWSSYEQY | 0.7317366 |
| GTPGEPTP | ASSLAWSSYEQY | 0.7275542 |
| GFFNSGS | ASSLAWSSYEQY | 0.7381611 |
| GAGGGGLTAGAN | ASSLAWSSYEQY | 0.7300699 |
| FVDALVNM | ASSLAWSSYEQY | 0.7333859 |
| FLTVAPEL | ASSLAWSSYEQY | 0.53471994 |
| FETLTAL | ASSLAWSSYEQY | 0.64704657 |
| DVINPIEL | ASSLAWSSYEQY | 0.6970035 |
| APVSGAPV | ASSLAWSSYEQY | 0.708332 |
| ALVGGLAW | ASSLAWSSYEQY | 0.48185027 |
| ALAAAAATAI | ASSLAWSSYEQY | 0.7269089 |
| AFGLGLLL | ASSLAWSSYEQY | 0.7159048 |
| AFFAAASL | ASSLAWSSYEQY | 0.6868182 |
| AAAALPAFSPP | ASSLAWSSYEQY | 0.72702986 |
| VVLAPAL | ASSLAWSSYEQY | 0.70239645 |
| TIGHVDHGKTTL | ASSLAWSSYEQY | 0.6807339 |
| LISNASDALDK | ASSLAWSSYEQY | 0.6999911 |
| LISNASDAL | ASSLAWSSYEQY | 0.73228014 |
| ISNASDALDK | ASSLAWSSYEQY | 0.6567292 |
| NQAETLVYQTE | ASSLAWSSYEQY | 0.7322124 |
| TGGTGGTAGAG | ASSLAWSSYEQY | 0.6558931 |
| TGSTAGGAGGA | ASSLAWSSYEQY | 0.7327005 |
| DPLAVPVAAGPV | ASSLAWSSYEQY | 0.67329264 |
| GGTGGTPGNGGN | ASSLAWSSYEQY | 0.7331534 |
| PAGLAGVDPATL | ASSLAWSSYEQY | 0.7160789 |
| GNNLVVEAPHAGASV | ASSLAWSSYEQY | 0.73849344 |
| KVTRIMSASTLEQPA | ASSLAWSSYEQY | 0.7117641 |
| LGFALPIALPRRTVV | ASSLAWSSYEQY | 0.7322742 |

| | | |
|---|---|---|
| TRLGNLTARKLVVLV | ASSLAWSSYEQY | 0.7289337 |
| WSQQRRLIYFPSAGP | ASSLAWSSYEQY | 0.7267894 |
| GPNGNPDPMAAAVDIR | ASSLAWSSYEQY | 0.6555357 |
| TQRCVSVADLRAAASR | ASSLAWSSYEQY | 0.71096504 |
| QFVLDVAEHPVEVLTETV | ASSLAWSSYEQY | 0.6619773 |
| GGAGGTGGAGGNGGTQAGDG | ASSLAWSSYEQY | 0.72089636 |
| PLLATLRAIAVDVDAKPAQV | ASSLAWSSYEQY | 0.7323744 |
| APKFRTGGYMIDFWGVGYQV/ | ASSLAWSSYEQY | 0.49295083 |
| GAGGAGGAGGAAGGPGGTGG' | ASSLAWSSYEQY | 0.73326135 |
| IGALGLLATDVAPAGSVAFG' | ASSLAWSSYEQY | 0.68527246 |
| AGGAGGIGGNGGAGGAGGVLI | ASSLAWSSYEQY | 0.7025001 |
| ADELIGLL | ASSLAWSSYEQY | 0.73310435 |
| ADRALKA | ASSLAWSSYEQY | 0.7180615 |
| AGAVGQVGGAGGA | ASSLAWSSYEQY | 0.728621 |
| AGGGGGLGGAP | ASSLAWSSYEQY | 0.70390314 |
| AGGLLYGNGGAG | ASSLAWSSYEQY | 0.70673203 |
| AGLESIGL | ASSLAWSSYEQY | 0.72135997 |
| AGLKTGII | ASSLAWSSYEQY | 0.5346817 |
| ASLAALLL | ASSLAWSSYEQY | 0.6546334 |
| ASVINISSI | ASSLAWSSYEQY | 0.73745394 |
| ATPPGPA | ASSLAWSSYEQY | 0.88369286 |
| ATVASALV | ASSLAWSSYEQY | 0.88523054 |
| ATVASLIL | ASSLAWSSYEQY | 0.83853674 |
| AVNGPVLIRH | ASSLAWSSYEQY | 0.82546663 |
| DGAAGAVGI | ASSLAWSSYEQY | 0.84867704 |
| DPLLHGI | ASSLAWSSYEQY | 0.89383364 |
| ELGAALGAL | ASSLAWSSYEQY | 0.8833417 |
| ELGRTQAV | ASSLAWSSYEQY | 0.8801952 |
| FVAGGYLS | ASSLAWSSYEQY | 0.7699003 |
| GASGGAGGAGGW | ASSLAWSSYEQY | 0.8876643 |
| GGAGAGGDGGAGGI | ASSLAWSSYEQY | 0.85331506 |
| GTGHNGSQPGMGGNG | ASSLAWSSYEQY | 0.85396826 |
| GYAAASASAS | ASSLAWSSYEQY | 0.87822956 |
| HQVDVAAVQQG | ASSLAWSSYEQY | 0.8562399 |
| IGAAIAHT | ASSLAWSSYEQY | 0.8694221 |
| IGMIEVAQL | ASSLAWSSYEQY | 0.86077964 |
| IRLAPEHTSAMD | ASSLAWSSYEQY | 0.87234616 |
| IYNVSVL | ASSLAWSSYEQY | 0.88324285 |
| KQGFLGM | ASSLAWSSYEQY | 0.8745226 |
| LESVITQL | ASSLAWSSYEQY | 0.9025887 |
| LGLGALVVV | ASSLAWSSYEQY | 0.8945358 |
| LIGNGGNGGSGGI | ASSLAWSSYEQY | 0.88807434 |
| LNAINAPTL | ASSLAWSSYEQY | 0.3295281 |
| LQLLNTGV | ASSLAWSSYEQY | 0.81432694 |
| LQTSIEF | ASSLAWSSYEQY | 0.8483112 |
| MSFAVSEL | ASSLAWSSYEQY | 0.8716795 |
| NTGALGL | ASSLAWSSYEQY | 0.60725105 |
| PPGAGGAGGAGTT | ASSLAWSSYEQY | 0.88099694 |
| QLVGLGI | ASSLAWSSYEQY | 0.87152267 |
| RFLEQVHQL | ASSLAWSSYEQY | 0.8368997 |
| RGVGYVLR | ASSLAWSSYEQY | 0.8957218 |

| | | |
|---|---|---|
| SATGVFAGVI | ASSLAWSSYEQY | 0.85647345 |
| SLPAITLP | ASSLAWSSYEQY | 0.8164696 |
| STALAFTV | ASSLAWSSYEQY | 0.8267305 |
| SVLTVGLGK | ASSLAWSSYEQY | 0.8963372 |
| TALAGAGSGI | ASSLAWSSYEQY | 0.7292186 |
| TGLPFTGI | ASSLAWSSYEQY | 0.8906002 |
| VAEHVTV | ASSLAWSSYEQY | 0.85388255 |
| VDIVLNSL | ASSLAWSSYEQY | 0.89943963 |
| VLTGGGAVL | ASSLAWSSYEQY | 0.71189076 |
| VNVGLAYGI | ASSLAWSSYEQY | 0.88671845 |
| VSSAAVGAV | ASSLAWSSYEQY | 0.8519617 |
| VTSVYEAGV | ASSLAWSSYEQY | 0.90326416 |
| WVALDRGA | ASSLAWSSYEQY | 0.85929835 |
| YLVGSVVI | ASSLAWSSYEQY | 0.87719446 |
| GPPGTGKTLL | ASSLAWSSYEQY | 0.8329215 |
| GVGMTKF | ASSLAWSSYEQY | 0.8878738 |
| IGIGGSDL | ASSLAWSSYEQY | 0.77141654 |
| IGTLLAEL | ASSLAWSSYEQY | 0.857128 |
| IYTSGTT | ASSLAWSSYEQY | 0.8350768 |
| LSFPAGV | ASSLAWSSYEQY | 0.87475085 |
| LYTSGTT | ASSLAWSSYEQY | 0.8837607 |
| VGAFPFGF | ASSLAWSSYEQY | 0.66499245 |
| VGAGTVEF | ASSLAWSSYEQY | 0.895771 |
| VGKSTLLL | ASSLAWSSYEQY | 0.8228829 |
| AALAHAGGGIV | ASSLAWSSYEQY | 0.8174937 |
| AALVAVSLL | ASSLAWSSYEQY | 0.8912505 |
| AAVSRAVAAL | ASSLAWSSYEQY | 0.8607359 |
| AGGSSGIGGF | ASSLAWSSYEQY | 0.8936919 |
| AGLTVANANG | ASSLAWSSYEQY | 0.82003593 |
| AGLVGAVL | ASSLAWSSYEQY | 0.87123466 |
| ALSQAFTV | ASSLAWSSYEQY | 0.882342 |
| ALTFSVSL | ASSLAWSSYEQY | 0.77623713 |
| ALTGAALVAL | ASSLAWSSYEQY | 0.7904416 |
| ASFSIGAGVL | ASSLAWSSYEQY | 0.8955539 |
| DGSVYKI | ASSLAWSSYEQY | 0.6460098 |
| EQLAEIV | ASSLAWSSYEQY | 0.65607417 |
| FAVSELTVM | ASSLAWSSYEQY | 0.6057929 |
| GAAPLAAGL | ASSLAWSSYEQY | 0.5629045 |
| GADGGDGGSGGAGG | ASSLAWSSYEQY | 0.61023664 |
| GFGLDTGGAGG | ASSLAWSSYEQY | 0.65099394 |
| GLASGGLL | ASSLAWSSYEQY | 0.65921557 |
| GRIAGAAVLL | ASSLAWSSYEQY | 0.6446609 |
| GSAGSLNTSF | ASSLAWSSYEQY | 0.53558517 |
| GTGQIGI | ASSLAWSSYEQY | 0.6541511 |
| GTLIALIL | ASSLAWSSYEQY | 0.638602 |
| IAATMGFVVG | ASSLAWSSYEQY | 0.6244178 |
| IFAQALL | ASSLAWSSYEQY | 0.64173037 |
| IGAAVALGF | ASSLAWSSYEQY | 0.64291954 |
| IGLTPLAL | ASSLAWSSYEQY | 0.5993856 |
| ILGSNTSTL | ASSLAWSSYEQY | 0.59861344 |
| IVFSLTVL | ASSLAWSSYEQY | 0.64288586 |

| | | |
|---|---|---|
| LASKATGF | ASSLAWSSYEQY | 0.656067 |
| LASVALIL | ASSLAWSSYEQY | 0.6574832 |
| LFAVGLTL | ASSLAWSSYEQY | 0.6665272 |
| LFNGGIGTYI | ASSLAWSSYEQY | 0.65932155 |
| LTLGAGII | ASSLAWSSYEQY | 0.64842695 |
| LVAAALGDI | ASSLAWSSYEQY | 0.39668506 |
| NTAFVQLGI | ASSLAWSSYEQY | 0.5128844 |
| QLFSGAEL | ASSLAWSSYEQY | 0.5885493 |
| SAATFITL | ASSLAWSSYEQY | 0.62427187 |
| SNYGLANLGV | ASSLAWSSYEQY | 0.4205253 |
| SYGGAYAVM | ASSLAWSSYEQY | 0.6426494 |
| TTGVGGLTLG | ASSLAWSSYEQY | 0.6079018 |
| TTQTIYI | ASSLAWSSYEQY | 0.59315515 |
| VAAGAGLGAVI | ASSLAWSSYEQY | 0.65595526 |
| VAAVVTLL | ASSLAWSSYEQY | 0.6081693 |
| VGFLGGLI | ASSLAWSSYEQY | 0.5655893 |
| VGVIPILL | ASSLAWSSYEQY | 0.5944881 |
| VIFASSLI | ASSLAWSSYEQY | 0.65208787 |
| VKGGDGGAAAG | ASSLAWSSYEQY | 0.5491497 |
| VTKTFTNI | ASSLAWSSYEQY | 0.65707195 |
| VVAAAILL | ASSLAWSSYEQY | 0.56687576 |
| YAALDVEL | ASSLAWSSYEQY | 0.6622816 |
| LASVALLL | ASSLAWSSYEQY | 0.5116151 |
| LDAVHRV | ASSLAWSSYEQY | 0.6571553 |
| QGLSSSIF | ASSLAWSSYEQY | 0.6307434 |
| RGTFANIRL | ASSLAWSSYEQY | 0.6705267 |
| RQLWWGHRI | ASSLAWSSYEQY | 0.6271507 |
| YGPPGTGKTLL | ASSLAWSSYEQY | 0.6512803 |
| AALVGAPANVA | ASSLAWSSYEQY | 0.6273678 |
| AATFRRTE | ASSLAWSSYEQY | 0.6592985 |
| AGALGGGI | ASSLAWSSYEQY | 0.52802753 |
| AGGAGGAGGIGAG | ASSLAWSSYEQY | 0.63067317 |
| AGSGALGL | ASSLAWSSYEQY | 0.5185791 |
| AQIADQI | ASSLAWSSYEQY | 0.63719285 |
| ASAVAVTYL | ASSLAWSSYEQY | 0.6499424 |
| ASTVLGGTGG | ASSLAWSSYEQY | 0.4278103 |
| AVGGEPAK | ASSLAWSSYEQY | 0.6575792 |
| CEQGTFGGTC | ASSLAWSSYEQY | 0.61131877 |
| DEGKAPAG | ASSLAWSSYEQY | 0.57763946 |
| DTVVFPAL | ASSLAWSSYEQY | 0.65784717 |
| FEGITMAG | ASSLAWSSYEQY | 0.6493577 |
| FPLVVQD | ASSLAWSSYEQY | 0.65033257 |
| GAFTTPQ | ASSLAWSSYEQY | 0.60133225 |
| GFGGFGTAGGVGGA | ASSLAWSSYEQY | 0.6174212 |
| GFLAQFL | ASSLAWSSYEQY | 0.63581157 |
| GGAGGAGGNAGVA | ASSLAWSSYEQY | 0.5929023 |
| GGVGTAGAGGAGGAGG | ASSLAWSSYEQY | 0.5696949 |
| GNGGGGDAGNAGSGGN | ASSLAWSSYEQY | 0.6620533 |
| GTGGAGGAGNDAGS | ASSLAWSSYEQY | 0.7966843 |
| GTTNTGWFNS | ASSLAWSSYEQY | 0.8378922 |
| IFMGPLFTL | ASSLAWSSYEQY | 0.64192474 |

| | | |
|---|---|---|
| ILAHAIEAI | ASSLAWSSYEQY | 0.62279403 |
| IRSAIREAL | ASSLAWSSYEQY | 0.7199834 |
| ITFAGGG | ASSLAWSSYEQY | 0.84558964 |
| ITGMSGHQEV | ASSLAWSSYEQY | 0.8588037 |
| IVAGLACTL | ASSLAWSSYEQY | 0.79940593 |
| IVGGGLAGA | ASSLAWSSYEQY | 0.67348254 |
| IVTGVLNI | ASSLAWSSYEQY | 0.7701889 |
| LAQQGCI | ASSLAWSSYEQY | 0.711612 |
| LCLHVEN | ASSLAWSSYEQY | 0.71142155 |
| LDAISDGS | ASSLAWSSYEQY | 0.8138058 |
| LGGFAAML | ASSLAWSSYEQY | 0.7850008 |
| LGGKNPMI | ASSLAWSSYEQY | 0.7094511 |
| LGPKGVAAIL | ASSLAWSSYEQY | 0.6939211 |
| LMVGVSI | ASSLAWSSYEQY | 0.76735723 |
| LVGAAFVL | ASSLAWSSYEQY | 0.8326062 |
| LVLAVAPL | ASSLAWSSYEQY | 0.81232333 |
| MAAIAVIDAA | ASSLAWSSYEQY | 0.8651092 |
| NNGINAELQ | ASSLAWSSYEQY | 0.82880104 |
| RGEPLVLVHG | ASSLAWSSYEQY | 0.81582713 |
| RGGAGGAAGLI | ASSLAWSSYEQY | 0.21950023 |
| RSYRMHPAV | ASSLAWSSYEQY | 0.492628 |
| SAAHLLAGQLA | ASSLAWSSYEQY | 0.69569373 |
| SAAVGSY | ASSLAWSSYEQY | 0.74375457 |
| SPHAVLHRI | ASSLAWSSYEQY | 0.34774813 |
| SQIAKVL | ASSLAWSSYEQY | 0.805464 |
| TALGHLVP | ASSLAWSSYEQY | 0.7144543 |
| TAPTTSGAA | ASSLAWSSYEQY | 0.6357999 |
| TDGDDGNGGSPGT | ASSLAWSSYEQY | 0.8316081 |
| TDGLGEN | ASSLAWSSYEQY | 0.7036903 |
| TGGPPLPGQA | ASSLAWSSYEQY | 0.68429387 |
| TIPAITI | ASSLAWSSYEQY | 0.7126007 |
| TIPGFTI | ASSLAWSSYEQY | 0.8461617 |
| TSIASGALA | ASSLAWSSYEQY | 0.5862947 |
| TVTLPGAAI | ASSLAWSSYEQY | 0.8102544 |
| VASGLSAALP | ASSLAWSSYEQY | 0.7686927 |
| VGAGVIGI | ASSLAWSSYEQY | 0.85159504 |
| VGPKRFGV | ASSLAWSSYEQY | 0.5626016 |
| VNVAFPD | ASSLAWSSYEQY | 0.8126645 |
| VSMTNSGVS | ASSLAWSSYEQY | 0.72359383 |
| VTDVLGDLSV | ASSLAWSSYEQY | 0.87467355 |
| VVVVGAGFAGL | ASSLAWSSYEQY | 0.71694815 |
| VYTVGTVV | ASSLAWSSYEQY | 0.80791426 |
| WTTFSGP | ASSLAWSSYEQY | 0.7722876 |
| YDELVEAF | ASSLAWSSYEQY | 0.8256272 |
| YGTGGAGGNGG | ASSLAWSSYEQY | 0.48911488 |
| YTNVMAV | ASSLAWSSYEQY | 0.7301135 |
| FLEQVHQL | ASSLAWSSYEQY | 0.62151754 |
| GLSLLAVAV | ASSLAWSSYEQY | 0.7686131 |
| LEQVHQL | ASSLAWSSYEQY | 0.7954054 |
| LNAAGAI | ASSLAWSSYEQY | 0.3484013 |
| PAVHASL | ASSLAWSSYEQY | 0.8207047 |

| | | |
|---|---|---|
| TELEDDA | ASSLAWSSYEQY | 0.6846601 |
| TPPGPPN | ASSLAWSSYEQY | 0.6528958 |
| TPPPPPPP | ASSLAWSSYEQY | 0.83090484 |
| TQSPALI | ASSWDLDSQNTLY | 0.7106352 |
| EFAAAFPQ | ASSWDLDSQNTLY | 0.8259825 |
| LLSGNLDVL | ASSWDLDSQNTLY | 0.7112768 |
| GPANLGNYNI | ASSWDLDSQNTLY | 0.763464 |
| TLGPTLP | ASSWDLDSQNTLY | 0.8128702 |
| VYAYSSCCK | ASSWDLDSQNTLY | 0.47139058 |
| APAPAPAPNI | ASSWDLDSQNTLY | 0.62022585 |
| PGGPVPIP | ASSWDLDSQNTLY | 0.8404244 |
| FLTVALEP | ASSWDLDSQNTLY | 0.6423429 |
| SQYFGANL | ASSWDLDSQNTLY | 0.65508974 |
| GGVKPAA | ASSWDLDSQNTLY | 0.58442646 |
| GGTGGAGGAGGW | ASSWDLDSQNTLY | 0.5609204 |
| GVGGVAGLGGAGG | ASSWDLDSQNTLY | 0.6233121 |
| VLLVLATTI | ASSWDLDSQNTLY | 0.65313005 |
| IGIFAAML | ASSWDLDSQNTLY | 0.6593433 |
| AVGDLID | ASSWDLDSQNTLY | 0.6384462 |
| NISPLPL | ASSWDLDSQNTLY | 0.5549129 |
| VTAVFVAI | ASSWDLDSQNTLY | 0.6335271 |
| SGDDNDAAL | ASSWDLDSQNTLY | 0.6242159 |
| AVAALAHS | ASSWDLDSQNTLY | 0.6111847 |
| GGVTVMATP | ASSWDLDSQNTLY | 0.64476174 |
| AGLAANVRD | ASSWDLDSQNTLY | 0.6229615 |
| TNRPDLIDPALLRPG | ASSWDLDSQNTLY | 0.57897294 |
| LANLGVTAL | ASSWDLDSQNTLY | 0.5929769 |
| TGERIEL | ASSWDLDSQNTLY | 0.6443359 |
| NGEELAA | ASSWDLDSQNTLY | 0.65775156 |
| VLSLVWG | ASSWDLDSQNTLY | 0.65730965 |
| IGYALWL | ASSWDLDSQNTLY | 0.66918266 |
| LSLLLAPL | ASSWDLDSQNTLY | 0.66017807 |
| TLDEALK | ASSWDLDSQNTLY | 0.6401484 |
| TDGAGTAEGG | ASSWDLDSQNTLY | 0.37934563 |
| NAPGGAGGNGG | ASSWDLDSQNTLY | 0.5334421 |
| VSVFMGMGGNF | ASSWDLDSQNTLY | 0.554208 |
| LASYLAAL | ASSWDLDSQNTLY | 0.63395584 |
| LVGGTPAL | ASSWDLDSQNTLY | 0.38824916 |
| VRNADVL | ASSWDLDSQNTLY | 0.6304708 |
| TCSSPFLVSAM | ASSWDLDSQNTLY | 0.6107993 |
| VHITDAE | ASSWDLDSQNTLY | 0.56756216 |
| EYAAPSF | ASSWDLDSQNTLY | 0.6669949 |
| VVDAFHGGL | ASSWDLDSQNTLY | 0.5890322 |
| ASVVDPQ | ASSWDLDSQNTLY | 0.54887044 |
| SYNEFGGLS | ASSWDLDSQNTLY | 0.56064504 |
| DPDGSVY | ASSWDLDSQNTLY | 0.66744804 |
| MVSTIHATE | ASSWDLDSQNTLY | 0.5250848 |
| LAAGVADPV | ASSWDLDSQNTLY | 0.657743 |
| TKEQIAA | ASSWDLDSQNTLY | 0.6026376 |
| SLASPAAL | ASSWDLDSQNTLY | 0.66521776 |
| IGLVFYGV | ASSWDLDSQNTLY | 0.52427435 |

| | | |
|---|---|---|
| FLLDASNL | ASSWDLDSQNTLY | 0.65530384 |
| YQVGNFP | ASSWDLDSQNTLY | 0.6170127 |
| ESLLSQST | ASSWDLDSQNTLY | 0.67521685 |
| LFTVGFAAVG | ASSWDLDSQNTLY | 0.62941414 |
| ATPPTPT | ASSWDLDSQNTLY | 0.65191406 |
| TDLATLG | ASSWDLDSQNTLY | 0.598773 |
| SAGGTGR | ASSWDLDSQNTLY | 0.6540607 |
| GFSNSGS | ASSWDLDSQNTLY | 0.52220017 |
| GGNGGSGGTSV | ASSWDLDSQNTLY | 0.62256986 |
| GTSVATGGAGN | ASSWDLDSQNTLY | 0.52362865 |
| NGGNGGSGGTS | ASSWDLDSQNTLY | 0.63331527 |
| AVVKFPH | ASSWDLDSQNTLY | 0.64631325 |
| TPPPPPP | ASSWDLDSQNTLY | 0.47251746 |
| AAPAAAAAPA | ASSWDLDSQNTLY | 0.6644861 |
| IALNEDL | ASSWDLDSQNTLY | 0.5971018 |
| IVNVSSM | ASSWDLDSQNTLY | 0.5873787 |
| TGIPPAPR | ASSWDLDSQNTLY | 0.6502843 |
| FDLGGGTF | ASSWDLDSQNTLY | 0.6123159 |
| FDLGGGTFDVS | ASSWDLDSQNTLY | 0.64755416 |
| NEPTAAA | ASSWDLDSQNTLY | 0.5346291 |
| NEPTAAALA | ASSWDLDSQNTLY | 0.63335764 |
| GSIINISSI | ASSWDLDSQNTLY | 0.64468133 |
| VAPGNAGTA | ASSWDLDSQNTLY | 0.5155418 |
| TGGGASLE | ASSWDLDSQNTLY | 0.5269084 |
| VAVYNHYKRI | ASSWDLDSQNTLY | 0.6618342 |
| TDPAPATT | ASSWDLDSQNTLY | 0.6428803 |
| TNRPDILDPALLRPG | ASSWDLDSQNTLY | 0.7014208 |
| LTTTTTS | ASSWDLDSQNTLY | 0.51033103 |
| PTGSGKTL | ASSWDLDSQNTLY | 0.47813445 |
| TGSGKTL | ASSWDLDSQNTLY | 0.5747032 |
| TIGHVDHGK | ASSWDLDSQNTLY | 0.6738625 |
| TIGHVDHGKTT | ASSWDLDSQNTLY | 0.69842815 |
| GHVDHGKTT | ASSWDLDSQNTLY | 0.65044206 |
| AQFEHTLL | ASSWDLDSQNTLY | 0.48570174 |
| IVNQDEL | ASSWDLDSQNTLY | 0.5720658 |
| TNGSQFF | ASSWDLDSQNTLY | 0.53714466 |
| TGKGIVE | ASSWDLDSQNTLY | 0.5667917 |
| LISNASDA | ASSWDLDSQNTLY | 0.6453116 |
| AELRTLGELTKIAKAHGAQ | ASSWDLDSQNTLY | 0.59749067 |
| APNAVAAAEPSW | ASSWDLDSQNTLY | 0.54177237 |
| APQVAGAATAVL | ASSWDLDSQNTLY | 0.5301045 |
| ARRDPAAAAAILPTDA | ASSWDLDSQNTLY | 0.6401231 |
| DAAARDVLAEAGLAEYF | ASSWDLDSQNTLY | 0.65377486 |
| DDQIVLSAKANLAKVEQ | ASSWDLDSQNTLY | 0.61630183 |
| DELLPDTGEDSDEDGH | ASSWDLDSQNTLY | 0.6954911 |
| ELPEAPANRATVPQD | ASSWDLDSQNTLY | 0.6614725 |
| FGQSKIERVTGIG | ASSWDLDSQNTLY | 0.6698871 |
| GAGGTGSTAGGAGGAGGAGGI | ASSWDLDSQNTLY | 0.29178062 |
| GHGGTGAAGGNGGTGGAGGS( | ASSWDLDSQNTLY | 0.37675297 |
| GNGGNGGAGGEGGPGGAGGA( | ASSWDLDSQNTLY | 0.5863321 |
| GSTAGNGGNGGNATNGTIAG( | ASSWDLDSQNTLY | 0.5459096 |

| | | |
|---|---|---|
| GTGGDGGNGGNGGTGG | ASSWDLDSQNTLY | 0.39601964 |
| GVRRPAGIPADAPV | ASSWDLDSQNTLY | 0.6243701 |
| GYRQVTDAHLLTLARR | ASSWDLDSQNTLY | 0.51482964 |
| IAKMGQDGHDRG | ASSWDLDSQNTLY | 0.5024864 |
| KAEPSIEIQYT | ASSWDLDSQNTLY | 0.65787977 |
| KPVRIRMAADEDTVAVT | ASSWDLDSQNTLY | 0.56255436 |
| MPPRIAGMRLLVIKPEP | ASSWDLDSQNTLY | 0.5085779 |
| MKFYAVALFRSRGVPPTR | ASSWDLDSQNTLY | 0.4913597 |
| NAGGFGILGAVAHSP | ASSWDLDSQNTLY | 0.6665372 |
| NEGREITLAMAEM | ASSWDLDSQNTLY | 0.3962221 |
| NELVEWSHPLGHRWRWE | ASSWDLDSQNTLY | 0.62869936 |
| NLGAGGTGGNGGNPG | ASSWDLDSQNTLY | 0.58755827 |
| NTEWPKWETVMAADPVD | ASSWDLDSQNTLY | 0.66405386 |
| PASAAPAETQP | ASSWDLDSQNTLY | 0.40806633 |
| QQSSDGKNGVRAAEITGE | ASSWDLDSQNTLY | 0.6434132 |
| RAPDAVTTSWWKE | ASSWDLDSQNTLY | 0.57816637 |
| SGFGNVGAHTSGWFNQ | ASSWDLDSQNTLY | 0.70796895 |
| SGVAGAVGQAASVGGLKV | ASSWDLDSQNTLY | 0.52343 |
| SNGGAGTGSAGGNGGT | ASSWDLDSQNTLY | 0.59412086 |
| TGGAGGEGGAAG | ASSWDLDSQNTLY | 0.58199376 |
| TGGDGGAAGAGGAPGHG | ASSWDLDSQNTLY | 0.6940706 |
| TMPPLLAAIANAGRHGV | ASSWDLDSQNTLY | 0.4250447 |
| TSSTAGNSGGA | ASSWDLDSQNTLY | 0.5551848 |
| VPATVPVGILC | ASSWDLDSQNTLY | 0.41010103 |
| VQRPDSGGVVLTGRIS | ASSWDLDSQNTLY | 0.60955846 |
| WTPNPVPATKGVPAGLVQAT | ASSWDLDSQNTLY | 0.64326644 |
| YALKRALAKFSDRFDVVII | ASSWDLDSQNTLY | 0.41274002 |
| YLIGSVIV | ASSWDLDSQNTLY | 0.6778799 |
| YGQLVAVID | ASSWDLDSQNTLY | 0.5946484 |
| VTVGGMTLPA | ASSWDLDSQNTLY | 0.5172259 |
| VLALPAV | ASSWDLDSQNTLY | 0.680495 |
| VIPPTLNYET | ASSWDLDSQNTLY | 0.5309961 |
| VGMGGMHPSSG | ASSWDLDSQNTLY | 0.6928959 |
| VGLAGGAATA | ASSWDLDSQNTLY | 0.6100333 |
| VGGAGGAGGSGP | ASSWDLDSQNTLY | 0.6085475 |
| VFAAFMV | ASSWDLDSQNTLY | 0.64075804 |
| TLYDKLR | ASSWDLDSQNTLY | 0.5757647 |
| TLAVFTVL | ASSWDLDSQNTLY | 0.5032423 |
| TGSVQPARV | ASSWDLDSQNTLY | 0.6773718 |
| TGGGAGSGGT | ASSWDLDSQNTLY | 0.60954845 |
| SHLETSL | ASSWDLDSQNTLY | 0.632885 |
| SFTTGMT | ASSWDLDSQNTLY | 0.56430763 |
| QVEVAKRMGVRQPTVS | ASSWDLDSQNTLY | 0.54169285 |
| QGSGGPALAM | ASSWDLDSQNTLY | 0.58348763 |
| MLAPSNP | ASSWDLDSQNTLY | 0.63258386 |
| LVNNAAI | ASSWDLDSQNTLY | 0.6425009 |
| LTGEINTLA | ASSWDLDSQNTLY | 0.6126163 |
| LSIFGPV | ASSWDLDSQNTLY | 0.5492991 |
| LLALLANL | ASSWDLDSQNTLY | 0.6354697 |
| LISNASDALDKLR | ASSWDLDSQNTLY | 0.5990522 |
| LINEAVP | ASSWDLDSQNTLY | 0.5801577 |

| | | |
|---|---|---|
| LGYAFSGL | ASSWDLDSQNTLY | 0.6195533 |
| LDFVAVAL | ASSWDLDSQNTLY | 0.6233144 |
| LAQPGAE | ASSWDLDSQNTLY | 0.5911113 |
| ISNLGAVTGF | ASSWDLDSQNTLY | 0.569676 |
| HQEDPGRI | ASSWDLDSQNTLY | 0.61976993 |
| GTPGEPTP | ASSWDLDSQNTLY | 0.63375396 |
| GFFNSGS | ASSWDLDSQNTLY | 0.62707096 |
| GAGGGGLTAGAN | ASSWDLDSQNTLY | 0.6485606 |
| FVDALVNM | ASSWDLDSQNTLY | 0.62415195 |
| FLTVAPEL | ASSWDLDSQNTLY | 0.6314736 |
| FETLTAL | ASSWDLDSQNTLY | 0.39215392 |
| DVINPIEL | ASSWDLDSQNTLY | 0.44410607 |
| APVSGAPV | ASSWDLDSQNTLY | 0.5930765 |
| ALVGGLAW | ASSWDLDSQNTLY | 0.59316087 |
| ALAAAAATAI | ASSWDLDSQNTLY | 0.44074216 |
| AFGLGLLL | ASSWDLDSQNTLY | 0.63531744 |
| AFFAAASL | ASSWDLDSQNTLY | 0.6061082 |
| AAAALPAFSPP | ASSWDLDSQNTLY | 0.5804687 |
| VVLAPAL | ASSWDLDSQNTLY | 0.6338433 |
| TIGHVDHGKTTL | ASSWDLDSQNTLY | 0.5793701 |
| LISNASDALDK | ASSWDLDSQNTLY | 0.5701699 |
| LISNASDAL | ASSWDLDSQNTLY | 0.58546495 |
| ISNASDALDK | ASSWDLDSQNTLY | 0.6434178 |
| NQAETLVYQTE | ASSWDLDSQNTLY | 0.4954214 |
| TGGTGGTAGAG | ASSWDLDSQNTLY | 0.6325889 |
| TGSTAGGAGGA | ASSWDLDSQNTLY | 0.5354996 |
| DPLAVPVAAGPV | ASSWDLDSQNTLY | 0.6400162 |
| GGTGGTPGNGGN | ASSWDLDSQNTLY | 0.50155944 |
| PAGLAGVDPATL | ASSWDLDSQNTLY | 0.6182649 |
| GNNLVVEAPHAGASV | ASSWDLDSQNTLY | 0.57506186 |
| KVTRIMSASTLEQPA | ASSWDLDSQNTLY | 0.65356284 |
| LGFALPIALPRRTVV | ASSWDLDSQNTLY | 0.57926255 |
| TRLGNLTARKLVVLV | ASSWDLDSQNTLY | 0.6289151 |
| WSQQRRLIYFPSAGP | ASSWDLDSQNTLY | 0.6122951 |
| GPNGNPDPMAAAVDIR | ASSWDLDSQNTLY | 0.63672847 |
| TQRCVSVADLRAAASR | ASSWDLDSQNTLY | 0.48682576 |
| QFVLDVAEHPVEVLTETV | ASSWDLDSQNTLY | 0.59892356 |
| GGAGGTGGAGGNGGTQAGDG | ASSWDLDSQNTLY | 0.49860582 |
| PLLATLRAIAVDVDAKPAQV | ASSWDLDSQNTLY | 0.61648303 |
| APKFRTGGYMIDFWGVGYQV | ASSWDLDSQNTLY | 0.6095544 |
| GAGGAGGAGGAAGGPGGTGG | ASSWDLDSQNTLY | 0.4478856 |
| IGALGLLATDVAPAGSVAFG | ASSWDLDSQNTLY | 0.6365004 |
| AGGAGGIGGNGGAGGAGGVL | ASSWDLDSQNTLY | 0.5824057 |
| ADELIGLL | ASSWDLDSQNTLY | 0.56361324 |
| ADRALKA | ASSWDLDSQNTLY | 0.6306725 |
| AGAVGQVGGAGGA | ASSWDLDSQNTLY | 0.6196143 |
| AGGGGGLGGAP | ASSWDLDSQNTLY | 0.6249155 |
| AGGLLYGNGGAG | ASSWDLDSQNTLY | 0.59727305 |
| AGLESIGL | ASSWDLDSQNTLY | 0.58433014 |
| AGLKTGII | ASSWDLDSQNTLY | 0.6263368 |
| ASLAALLL | ASSWDLDSQNTLY | 0.533877 |

| | | |
|---|---|---|
| ASVINISSI | ASSWDLDSQNTLY | 0.5625619 |
| ATPPGPA | ASSWDLDSQNTLY | 0.6217154 |
| ATVASALV | ASSWDLDSQNTLY | 0.6178915 |
| ATVASLIL | ASSWDLDSQNTLY | 0.7016816 |
| AVNGPVLIRH | ASSWDLDSQNTLY | 0.40877375 |
| DGAAGAVGI | ASSWDLDSQNTLY | 0.26882172 |
| DPLLHGI | ASSWDLDSQNTLY | 0.42921075 |
| ELGAALGAL | ASSWDLDSQNTLY | 0.6463525 |
| ELGRTQAV | ASSWDLDSQNTLY | 0.76060987 |
| FVAGGYLS | ASSWDLDSQNTLY | 0.6057182 |
| GASGGAGGAGGW | ASSWDLDSQNTLY | 0.2607872 |
| GGAGAGGDGGAGGI | ASSWDLDSQNTLY | 0.7042307 |
| GTGHNGSQPGMGGNG | ASSWDLDSQNTLY | 0.43067488 |
| GYAAASASAS | ASSWDLDSQNTLY | 0.4227211 |
| HQVDVAAVQQG | ASSWDLDSQNTLY | 0.5272596 |
| IGAAIAHT | ASSWDLDSQNTLY | 0.5300465 |
| IGMIEVAQL | ASSWDLDSQNTLY | 0.4400937 |
| IRLAPEHTSAMD | ASSWDLDSQNTLY | 0.32998583 |
| IYNVSVL | ASSWDLDSQNTLY | 0.5898049 |
| KQGFLGM | ASSWDLDSQNTLY | 0.71543765 |
| LESVITQL | ASSWDLDSQNTLY | 0.6848583 |
| LGLGALVVV | ASSWDLDSQNTLY | 0.72514796 |
| LIGNGGNGGSGGI | ASSWDLDSQNTLY | 0.7381511 |
| LNAINAPTL | ASSWDLDSQNTLY | 0.55533755 |
| LQLLNTGV | ASSWDLDSQNTLY | 0.11676362 |
| LQTSIEF | ASSWDLDSQNTLY | 0.21422212 |
| MSFAVSEL | ASSWDLDSQNTLY | 0.29655057 |
| NTGALGL | ASSWDLDSQNTLY | 0.414621 |
| PPGAGGAGGAGTT | ASSWDLDSQNTLY | 0.09819613 |
| QLVGLGI | ASSWDLDSQNTLY | 0.46326292 |
| RFLEQVHQL | ASSWDLDSQNTLY | 0.46230483 |
| RGVGYVLR | ASSWDLDSQNTLY | 0.29153067 |
| SATGVFAGVI | ASSWDLDSQNTLY | 0.746413 |
| SLPAITLP | ASSWDLDSQNTLY | 0.34652233 |
| STALAFTV | ASSWDLDSQNTLY | 0.32205415 |
| SVLTVGLGK | ASSWDLDSQNTLY | 0.32932937 |
| TALAGAGSGI | ASSWDLDSQNTLY | 0.7220594 |
| TGLPFTGI | ASSWDLDSQNTLY | 0.23345639 |
| VAEHVTV | ASSWDLDSQNTLY | 0.720613 |
| VDIVLNSL | ASSWDLDSQNTLY | 0.28026158 |
| VLTGGGAVL | ASSWDLDSQNTLY | 0.66853774 |
| VNVGLAYGI | ASSWDLDSQNTLY | 0.20814013 |
| VSSAAVGAV | ASSWDLDSQNTLY | 0.6854409 |
| VTSVYEAGV | ASSWDLDSQNTLY | 0.51614106 |
| WVALDRGA | ASSWDLDSQNTLY | 0.83355343 |
| YLVGSVVI | ASSWDLDSQNTLY | 0.4324515 |
| GPPGTGKTLL | ASSWDLDSQNTLY | 0.5539788 |
| GVGMTKF | ASSWDLDSQNTLY | 0.41199988 |
| IGIGGSDL | ASSWDLDSQNTLY | 0.7291313 |
| IGTLLAEL | ASSWDLDSQNTLY | 0.18868643 |
| IYTSGTT | ASSWDLDSQNTLY | 0.50694114 |

| | | |
|---|---|---|
| LSFPAGV | ASSWDLDSQNTLY | 0.2610261 |
| LYTSGTT | ASSWDLDSQNTLY | 0.59064937 |
| VGAFPFGF | ASSWDLDSQNTLY | 0.6265079 |
| VGAGTVEF | ASSWDLDSQNTLY | 0.10526252 |
| VGKSTLLL | ASSWDLDSQNTLY | 0.6991612 |
| AALAHAGGGIV | ASSWDLDSQNTLY | 0.24008983 |
| AALVAVSLL | ASSWDLDSQNTLY | 0.22685157 |
| AAVSRAVAAL | ASSWDLDSQNTLY | 0.6121305 |
| AGGSSGIGGF | ASSWDLDSQNTLY | 0.44775134 |
| AGLTVANANG | ASSWDLDSQNTLY | 0.75449383 |
| AGLVGAVL | ASSWDLDSQNTLY | 0.27963504 |
| ALSQAFTV | ASSWDLDSQNTLY | 0.5355685 |
| ALTFSVSL | ASSWDLDSQNTLY | 0.60603774 |
| ALTGAALVAL | ASSWDLDSQNTLY | 0.15677308 |
| ASFSIGAGVL | ASSWDLDSQNTLY | 0.25667858 |
| DGSVYKI | ASSWDLDSQNTLY | 0.75735295 |
| EQLAEIV | ASSWDLDSQNTLY | 0.8012141 |
| FAVSELTVM | ASSWDLDSQNTLY | 0.84949934 |
| GAAPLAAGL | ASSWDLDSQNTLY | 0.6980008 |
| GADGGDGGSGGAGG | ASSWDLDSQNTLY | 0.63508964 |
| GFGLDTGGAGG | ASSWDLDSQNTLY | 0.7258724 |
| GLASGGLL | ASSWDLDSQNTLY | 0.8471987 |
| GRIAGAAVLL | ASSWDLDSQNTLY | 0.8482716 |
| GSAGSLNTSF | ASSWDLDSQNTLY | 0.8041189 |
| GTGQIGI | ASSWDLDSQNTLY | 0.6166171 |
| GTLIALIL | ASSWDLDSQNTLY | 0.77855235 |
| IAATMGFVVG | ASSWDLDSQNTLY | 0.7111007 |
| IFAQALL | ASSWDLDSQNTLY | 0.7336458 |
| IGAAVALGF | ASSWDLDSQNTLY | 0.8222619 |
| IGLTPLAL | ASSWDLDSQNTLY | 0.7793181 |
| ILGSNTSTL | ASSWDLDSQNTLY | 0.69884837 |
| IVFSLTVL | ASSWDLDSQNTLY | 0.7356042 |
| LASKATGF | ASSWDLDSQNTLY | 0.7840488 |
| LASVALIL | ASSWDLDSQNTLY | 0.7973565 |
| LFAVGLTL | ASSWDLDSQNTLY | 0.79409754 |
| LFNGGIGTYI | ASSWDLDSQNTLY | 0.8688344 |
| LTLGAGII | ASSWDLDSQNTLY | 0.8318039 |
| LVAAALGDI | ASSWDLDSQNTLY | 0.8414707 |
| NTAFVQLGI | ASSWDLDSQNTLY | 0.2030027 |
| QLFSGAEL | ASSWDLDSQNTLY | 0.43435395 |
| SAATFITL | ASSWDLDSQNTLY | 0.7396255 |
| SNYGLANLGV | ASSWDLDSQNTLY | 0.71661425 |
| SYGGAYAVM | ASSWDLDSQNTLY | 0.5100758 |
| TTGVGGLTLG | ASSWDLDSQNTLY | 0.81454444 |
| TTQTIYI | ASSWDLDSQNTLY | 0.6907904 |
| VAAGAGLGAVI | ASSWDLDSQNTLY | 0.6474911 |
| VAAVVTLL | ASSWDLDSQNTLY | 0.8206476 |
| VGFLGGLI | ASSWDLDSQNTLY | 0.7123752 |
| VGVIPILL | ASSWDLDSQNTLY | 0.68619645 |
| VIFASSLI | ASSWDLDSQNTLY | 0.7047782 |
| VKGGDGGAAAG | ASSWDLDSQNTLY | 0.82790405 |

| | | |
|---|---|---|
| VTKTFTNI | ASSWDLDSQNTLY | 0.5102354 |
| VVAAAILL | ASSWDLDSQNTLY | 0.8106475 |
| YAALDVEL | ASSWDLDSQNTLY | 0.6701739 |
| LASVALLL | ASSWDLDSQNTLY | 0.84328747 |
| LDAVHRV | ASSWDLDSQNTLY | 0.538198 |
| QGLSSSIF | ASSWDLDSQNTLY | 0.81348956 |
| RGTFANIRL | ASSWDLDSQNTLY | 0.7443253 |
| RQLWWGHRI | ASSWDLDSQNTLY | 0.8728874 |
| YGPPGTGKTLL | ASSWDLDSQNTLY | 0.73236924 |
| AALVGAPANVA | ASSWDLDSQNTLY | 0.76902914 |
| AATFRRTE | ASSWDLDSQNTLY | 0.64517385 |
| AGALGGGI | ASSWDLDSQNTLY | 0.84390056 |
| AGGAGGAGGIGAG | ASSWDLDSQNTLY | 0.45227543 |
| AGSGALGL | ASSWDLDSQNTLY | 0.74125206 |
| AQIADQI | ASSWDLDSQNTLY | 0.44192356 |
| ASAVAVTYL | ASSWDLDSQNTLY | 0.803183 |
| ASTVLGGTGG | ASSWDLDSQNTLY | 0.79491854 |
| AVGGEPAK | ASSWDLDSQNTLY | 0.50512856 |
| CEQGTFGGTC | ASSWDLDSQNTLY | 0.8334174 |
| DEGKAPAG | ASSWDLDSQNTLY | 0.7614896 |
| DTVVFPAL | ASSWDLDSQNTLY | 0.6953628 |
| FEGITMAG | ASSWDLDSQNTLY | 0.8520639 |
| FPLVVQD | ASSWDLDSQNTLY | 0.7510553 |
| GAFTTPQ | ASSWDLDSQNTLY | 0.8455761 |
| GFGGFGTAGGVGGA | ASSWDLDSQNTLY | 0.7730007 |
| GFLAQFL | ASSWDLDSQNTLY | 0.6857425 |
| GGAGGAGGNAGVA | ASSWDLDSQNTLY | 0.81367284 |
| GGVGTAGAGGAGGAGG | ASSWDLDSQNTLY | 0.4993325 |
| GNGGGGDAGNAGSGGN | ASSWDLDSQNTLY | 0.6663895 |
| GTGGAGGAGNDAGS | ASSWDLDSQNTLY | 0.8412938 |
| GTTNTGWFNS | ASSWDLDSQNTLY | 0.68760324 |
| IFMGPLFTL | ASSWDLDSQNTLY | 0.6888435 |
| ILAHAIEAI | ASSWDLDSQNTLY | 0.53961027 |
| IRSAIREAL | ASSWDLDSQNTLY | 0.5775305 |
| ITFAGGG | ASSWDLDSQNTLY | 0.60521257 |
| ITGMSGHQEV | ASSWDLDSQNTLY | 0.67485714 |
| IVAGLACTL | ASSWDLDSQNTLY | 0.7150515 |
| IVGGGLAGA | ASSWDLDSQNTLY | 0.69055 |
| IVTGVLNI | ASSWDLDSQNTLY | 0.6215496 |
| LAQQGCI | ASSWDLDSQNTLY | 0.66760457 |
| LCLHVEN | ASSWDLDSQNTLY | 0.6134075 |
| LDAISDGS | ASSWDLDSQNTLY | 0.62443423 |
| LGGFAAML | ASSWDLDSQNTLY | 0.66676164 |
| LGGKNPMI | ASSWDLDSQNTLY | 0.65248525 |
| LGPKGVAAIL | ASSWDLDSQNTLY | 0.6063611 |
| LMVGVSI | ASSWDLDSQNTLY | 0.58494586 |
| LVGAAFVL | ASSWDLDSQNTLY | 0.6681461 |
| LVLAVAPL | ASSWDLDSQNTLY | 0.70522296 |
| MAAIAVIDAA | ASSWDLDSQNTLY | 0.6980792 |
| NNGINAELQ | ASSWDLDSQNTLY | 0.70955193 |
| RGEPLVLVHG | ASSWDLDSQNTLY | 0.6818226 |

| | | |
|---|---|---|
| RGGAGGAAGLI | ASSWDLDSQNTLY | 0.6910908 |
| RSYRMHPAV | ASSWDLDSQNTLY | 0.33493337 |
| SAAHLLAGQLA | ASSWDLDSQNTLY | 0.49721366 |
| SAAVGSY | ASSWDLDSQNTLY | 0.63819593 |
| SPHAVLHRI | ASSWDLDSQNTLY | 0.6265687 |
| SQIAKVL | ASSWDLDSQNTLY | 0.39354432 |
| TALGHLVP | ASSWDLDSQNTLY | 0.6722692 |
| TAPTTSGAA | ASSWDLDSQNTLY | 0.63019174 |
| TDGDDGNGGSPGT | ASSWDLDSQNTLY | 0.58264565 |
| TDGLGEN | ASSWDLDSQNTLY | 0.6874333 |
| TGGPPLPGQA | ASSWDLDSQNTLY | 0.6058403 |
| TIPAITI | ASSWDLDSQNTLY | 0.55960035 |
| TIPGFTI | ASSWDLDSQNTLY | 0.6242227 |
| TSIASGALA | ASSWDLDSQNTLY | 0.7085538 |
| TVTLPGAAI | ASSWDLDSQNTLY | 0.5531816 |
| VASGLSAALP | ASSWDLDSQNTLY | 0.6868482 |
| VGAGVIGI | ASSWDLDSQNTLY | 0.6158818 |
| VGPKRFGV | ASSWDLDSQNTLY | 0.6938858 |
| VNVAFPD | ASSWDLDSQNTLY | 0.54497683 |
| VSMTNSGVS | ASSWDLDSQNTLY | 0.6811133 |
| VTDVLGDLSV | ASSWDLDSQNTLY | 0.61591595 |
| VVVVGAGFAGL | ASSWDLDSQNTLY | 0.7218786 |
| VYTVGTVV | ASSWDLDSQNTLY | 0.6140786 |
| WTTFSGP | ASSWDLDSQNTLY | 0.6750751 |
| YDELVEAF | ASSWDLDSQNTLY | 0.6463376 |
| YGTGGAGGNGG | ASSWDLDSQNTLY | 0.704439 |
| YTNVMAV | ASSWDLDSQNTLY | 0.4870592 |
| FLEQVHQL | ASSWDLDSQNTLY | 0.6188436 |
| GLSLLAVAV | ASSWDLDSQNTLY | 0.53343874 |
| LEQVHQL | ASSWDLDSQNTLY | 0.6429105 |
| LNAAGAI | ASSWDLDSQNTLY | 0.6876873 |
| PAVHASL | ASSWDLDSQNTLY | 0.3755885 |
| TELEDDA | ASSWDLDSQNTLY | 0.6932363 |
| TPPGPPN | ASSWDLDSQNTLY | 0.5188095 |
| TPPPPPPP | ASSWDLDSQNTLY | 0.5944971 |
| TQSPALI | ASSDREVF | 0.68526363 |
| EFAAAFPQ | ASSDREVF | 0.6286382 |
| LLSGNLDVL | ASSDREVF | 0.69541466 |
| GPANLGNYNI | ASSDREVF | 0.6153713 |
| TLGPTLP | ASSDREVF | 0.64412796 |
| VYAYSSCCK | ASSDREVF | 0.68018997 |
| APAPAPAPNI | ASSDREVF | 0.50150645 |
| PGGPVPIP | ASSDREVF | 0.5780221 |
| FLTVALEP | ASSDREVF | 0.6982555 |
| SQYFGANL | ASSDREVF | 0.8151918 |
| GGVKPAA | ASSDREVF | 0.8266199 |
| GGTGGAGGAGGW | ASSDREVF | 0.7592749 |
| GVGGVAGLGGAGG | ASSDREVF | 0.6948126 |
| VLLVLATTI | ASSDREVF | 0.7635226 |
| IGIFAAML | ASSDREVF | 0.82618105 |
| AVGDLID | ASSDREVF | 0.81917524 |

| | | |
|---|---|---|
| NISPLPL | ASSDREVF | 0.8169698 |
| VTAVFVAI | ASSDREVF | 0.6880081 |
| SGDDNDAAL | ASSDREVF | 0.7947588 |
| AVAALAHS | ASSDREVF | 0.7818757 |
| GGVTVMATP | ASSDREVF | 0.7658719 |
| AGLAANVRD | ASSDREVF | 0.81795096 |
| TNRPDLIDPALLRPG | ASSDREVF | 0.7924695 |
| LANLGVTAL | ASSDREVF | 0.78291595 |
| TGERIEL | ASSDREVF | 0.7823208 |
| NGEELAA | ASSDREVF | 0.79303706 |
| VLSLVWG | ASSDREVF | 0.80759 |
| IGYALWL | ASSDREVF | 0.80205965 |
| LSLLLAPL | ASSDREVF | 0.8341925 |
| TLDEALK | ASSDREVF | 0.82249546 |
| TDGAGTAEGG | ASSDREVF | 0.8253857 |
| NAPGGAGGNGG | ASSDREVF | 0.27274248 |
| VSVFMGMGGNF | ASSDREVF | 0.57355756 |
| LASYLAAL | ASSDREVF | 0.8037982 |
| LVGGTPAL | ASSDREVF | 0.7655694 |
| VRNADVL | ASSDREVF | 0.7054247 |
| TCSSPFLVSAM | ASSDREVF | 0.81802803 |
| VHITDAE | ASSDREVF | 0.79237425 |
| EYAAPSF | ASSDREVF | 0.7320895 |
| VVDAFHGGL | ASSDREVF | 0.8061467 |
| ASVVDPQ | ASSDREVF | 0.76324195 |
| SYNEFGGLS | ASSDREVF | 0.75375676 |
| DPDGSVY | ASSDREVF | 0.778864 |
| MVSTIHATE | ASSDREVF | 0.82771635 |
| LAAGVADPV | ASSDREVF | 0.6970322 |
| TKEQIAA | ASSDREVF | 0.8049456 |
| SLASPAAL | ASSDREVF | 0.7970622 |
| IGLVFYGV | ASSDREVF | 0.8207824 |
| FLLDASNL | ASSDREVF | 0.60687435 |
| YQVGNFP | ASSDREVF | 0.8187009 |
| ESLLSQST | ASSDREVF | 0.78272486 |
| LFTVGFAAVG | ASSDREVF | 0.83181036 |
| ATPPTPT | ASSDREVF | 0.75528115 |
| TDLATLG | ASSDREVF | 0.77989054 |
| SAGGTGR | ASSDREVF | 0.74893224 |
| GFSNSGS | ASSDREVF | 0.81361794 |
| GGNGGSGGTSV | ASSDREVF | 0.51358014 |
| GTSVATGGAGN | ASSDREVF | 0.7848339 |
| NGGNGGSGGTS | ASSDREVF | 0.7169145 |
| AVVKFPH | ASSDREVF | 0.80042714 |
| TPPPPPP | ASSDREVF | 0.8149962 |
| AAPAAAAAPA | ASSDREVF | 0.6710596 |
| IALNEDL | ASSDREVF | 0.82073164 |
| IVNVSSM | ASSDREVF | 0.81124854 |
| TGIPPAPR | ASSDREVF | 0.7636448 |
| FDLGGGTF | ASSDREVF | 0.8264809 |
| FDLGGGTFDVS | ASSDREVF | 0.78868115 |

| | | |
|---|---|---|
| NEPTAAA | ASSDREVF | 0.82802534 |
| NEPTAAALA | ASSDREVF | 0.8009099 |
| GSIINISSI | ASSDREVF | 0.79634386 |
| VAPGNAGTA | ASSDREVF | 0.8120515 |
| TGGGASLE | ASSDREVF | 0.6430518 |
| VAVYNHYKRI | ASSDREVF | 0.76606613 |
| TDPAPATT | ASSDREVF | 0.82465184 |
| TNRPDILDPALLRPG | ASSDREVF | 0.75158584 |
| LTTTTTS | ASSDREVF | 0.81531507 |
| PTGSGKTL | ASSDREVF | 0.6563661 |
| TGSGKTL | ASSDREVF | 0.58337724 |
| TIGHVDHGK | ASSDREVF | 0.72387594 |
| TIGHVDHGKTT | ASSDREVF | 0.8069294 |
| GHVDHGKTT | ASSDREVF | 0.80649036 |
| AQFEHTLL | ASSDREVF | 0.78610885 |
| IVNQDEL | ASSDREVF | 0.58794326 |
| TNGSQFF | ASSDREVF | 0.7168572 |
| TGKGIVE | ASSDREVF | 0.72007805 |
| LISNASDA | ASSDREVF | 0.6685731 |
| AELRTLGELTKIAKAHGAQ | ASSDREVF | 0.7687465 |
| APNAVAAAEPSW | ASSDREVF | 0.72198087 |
| APQVAGAATAVL | ASSDREVF | 0.6093154 |
| ARRDPAAAAAILPTDA | ASSDREVF | 0.6303485 |
| DAAARDVLAEAGLAEYF | ASSDREVF | 0.75680786 |
| DDQIVLSAKANLAKVEQ | ASSDREVF | 0.785854 |
| DELLPDTGEDSDEDGH | ASSDREVF | 0.77115643 |
| ELPEAPANRATVPQD | ASSDREVF | 0.81807667 |
| FGQSKIERVTGIG | ASSDREVF | 0.7911675 |
| GAGGTGSTAGGAGGAGGAGGI | ASSDREVF | 0.75317943 |
| GHGGTGAAGGNGGTGGAGGS( | ASSDREVF | 0.2270671 |
| GNGGNGGAGGEGGPGGAGGA( | ASSDREVF | 0.4232739 |
| GSTAGNGGNGGNATNGTIAG( | ASSDREVF | 0.70756984 |
| GTGGDGGNGGNGGTGG | ASSDREVF | 0.72414994 |
| GVRRPAGIPADAPV | ASSDREVF | 0.53178984 |
| GYRQVTDAHLLTLARR | ASSDREVF | 0.78459823 |
| IAKMGQDGHDRG | ASSDREVF | 0.71045655 |
| KAEPSIEIQYT | ASSDREVF | 0.6474412 |
| KPVRIRMAADEDTVAVT | ASSDREVF | 0.78744674 |
| MPPRIAGMRLLVIKPEP | ASSDREVF | 0.61655223 |
| MKFYAVALFRSRGVPPTR | ASSDREVF | 0.61838526 |
| NAGGFGILGAVAHSP | ASSDREVF | 0.65964264 |
| NEGREITLAMAEM | ASSDREVF | 0.79944634 |
| NELVEWSHPLGHRWRWE | ASSDREVF | 0.46675915 |
| NLGAGGTGGNGGNPG | ASSDREVF | 0.7598578 |
| NTEWPKWETVMAADPVD | ASSDREVF | 0.6851473 |
| PASAAPAETQP | ASSDREVF | 0.796976 |
| QQSSDGKNGVRAAEITGE | ASSDREVF | 0.50211394 |
| RAPDAVTTSWWKE | ASSDREVF | 0.77794796 |
| SGFGNVGAHTSGWFNQ | ASSDREVF | 0.7160825 |
| SGVAGAVGQAASVGGLKV | ASSDREVF | 0.8205546 |
| SNGGAGTGSAGGNGGT | ASSDREVF | 0.68913656 |

| | | |
|---|---|---|
| TGGAGGEGGAAG | ASSDREVF | 0.7475828 |
| TGGDGGAAGAGGAPGHG | ASSDREVF | 0.62222314 |
| TMPPLLAAIANAGRHGV | ASSDREVF | 0.7963604 |
| TSSTAGNSGGA | ASSDREVF | 0.61064005 |
| VPATVPVGILC | ASSDREVF | 0.71108925 |
| VQRPDSGGVVLTGRIS | ASSDREVF | 0.50351244 |
| WTPNPVPATKGVPAGLVQAT | ASSDREVF | 0.75207996 |
| YALKRALAKFSDRFDVVII | ASSDREVF | 0.76609504 |
| YLIGSVIV | ASSDREVF | 0.70944214 |
| YGQLVAVID | ASSDREVF | 0.7768034 |
| VTVGGMTLPA | ASSDREVF | 0.7979323 |
| VLALPAV | ASSDREVF | 0.6427548 |
| VIPPTLNYET | ASSDREVF | 0.80439514 |
| VGMGGMHPSSG | ASSDREVF | 0.72144985 |
| VGLAGGAATA | ASSDREVF | 0.7949414 |
| VGGAGGAGGSGP | ASSDREVF | 0.6812235 |
| VFAAFMV | ASSDREVF | 0.7182349 |
| TLYDKLR | ASSDREVF | 0.78551424 |
| TLAVFTVL | ASSDREVF | 0.75422513 |
| TGSVQPARV | ASSDREVF | 0.68674004 |
| TGGGAGSGGT | ASSDREVF | 0.7851906 |
| SHLETSL | ASSDREVF | 0.39806667 |
| SFTTGMT | ASSDREVF | 0.7068958 |
| QVEVAKRMGVRQPTVS | ASSDREVF | 0.25013718 |
| QGSGGPALAM | ASSDREVF | 0.24810834 |
| MLAPSNP | ASSDREVF | 0.3274048 |
| LVNNAAI | ASSDREVF | 0.62757176 |
| LTGEINTLA | ASSDREVF | 0.6581973 |
| LSIFGPV | ASSDREVF | 0.5533401 |
| LLALLANL | ASSDREVF | 0.25983036 |
| LISNASDALDKLR | ASSDREVF | 0.28152585 |
| LINEAVP | ASSDREVF | 0.29029813 |
| LGYAFSGL | ASSDREVF | 0.30392334 |
| LDFVAVAL | ASSDREVF | 0.49012455 |
| LAQPGAE | ASSDREVF | 0.37982792 |
| ISNLGAVTGF | ASSDREVF | 0.25441 |
| HQEDPGRI | ASSDREVF | 0.33503035 |
| GTPGEPTP | ASSDREVF | 0.33255747 |
| GFFNSGS | ASSDREVF | 0.58285964 |
| GAGGGGLTAGAN | ASSDREVF | 0.5044477 |
| FVDALVNM | ASSDREVF | 0.6930821 |
| FLTVAPEL | ASSDREVF | 0.50714576 |
| FETLTAL | ASSDREVF | 0.5713617 |
| DVINPIEL | ASSDREVF | 0.12542386 |
| APVSGAPV | ASSDREVF | 0.1885332 |
| ALVGGLAW | ASSDREVF | 0.3563211 |
| ALAAAAATAI | ASSDREVF | 0.37172896 |
| AFGLGLLL | ASSDREVF | 0.20935836 |
| AFFAAASL | ASSDREVF | 0.51498896 |
| AAAALPAFSPP | ASSDREVF | 0.34834418 |
| VVLAPAL | ASSDREVF | 0.23141015 |

| | | |
|---|---|---|
| TIGHVDHGKTTL | ASSDREVF | 0.5475795 |
| LISNASDALDK | ASSDREVF | 0.23882611 |
| LISNASDAL | ASSDREVF | 0.3017301 |
| ISNASDALDK | ASSDREVF | 0.35996595 |
| NQAETLVYQTE | ASSDREVF | 0.49473283 |
| TGGTGGTAGAG | ASSDREVF | 0.2145767 |
| TGSTAGGAGGA | ASSDREVF | 0.44832137 |
| DPLAVPVAAGPV | ASSDREVF | 0.3728107 |
| GGTGGTPGNGGN | ASSDREVF | 0.5223038 |
| PAGLAGVDPATL | ASSDREVF | 0.23109344 |
| GNNLVVEAPHAGASV | ASSDREVF | 0.5110684 |
| KVTRIMSASTLEQPA | ASSDREVF | 0.29668015 |
| LGFALPIALPRRTVV | ASSDREVF | 0.73472273 |
| TRLGNLTARKLVVLV | ASSDREVF | 0.3201171 |
| WSQQRRLIYFPSAGP | ASSDREVF | 0.38107768 |
| GPNGNPDPMAAAVDIR | ASSDREVF | 0.34081632 |
| TQRCVSVADLRAAASR | ASSDREVF | 0.5970094 |
| QFVLDVAEHPVEVLTETV | ASSDREVF | 0.188696 |
| GGAGGTGGAGGNGGTQAGDG | ASSDREVF | 0.2777985 |
| PLLATLRAIAVDVDAKPAQV | ASSDREVF | 0.19050325 |
| APKFRTGGYMIDFWGVGYQV/ | ASSDREVF | 0.38396332 |
| GAGGAGGAGGAAGGPGGTGG' | ASSDREVF | 0.49603018 |
| IGALGLLATDVAPAGSVAFG' | ASSDREVF | 0.29590037 |
| AGGAGGIGGNGGAGGAGGVL! | ASSDREVF | 0.3982492 |
| ADELIGLL | ASSDREVF | 0.53205717 |
| ADRALKA | ASSDREVF | 0.31580964 |
| AGAVGQVGGAGGA | ASSDREVF | 0.60894394 |
| AGGGGGLGGAP | ASSDREVF | 0.25258803 |
| AGGLLYGNGGAG | ASSDREVF | 0.566542 |
| AGLESIGL | ASSDREVF | 0.31989527 |
| AGLKTGII | ASSDREVF | 0.27132782 |
| ASLAALLL | ASSDREVF | 0.46864498 |
| ASVINISSI | ASSDREVF | 0.3823201 |
| ATPPGPA | ASSDREVF | 0.28966352 |
| ATVASALV | ASSDREVF | 0.56920326 |
| ATVASLIL | ASSDREVF | 0.5505693 |
| AVNGPVLIRH | ASSDREVF | 0.81152594 |
| DGAAGAVGI | ASSDREVF | 0.28130347 |
| DPLLHGI | ASSDREVF | 0.2216151 |
| ELGAALGAL | ASSDREVF | 0.3670604 |
| ELGRTQAV | ASSDREVF | 0.68031925 |
| FVAGGYLS | ASSDREVF | 0.8268315 |
| GASGGAGGAGGW | ASSDREVF | 0.61601657 |
| GGAGAGGDGGAGGI | ASSDREVF | 0.21669991 |
| GTGHNGSQPGMGGNG | ASSDREVF | 0.5311009 |
| GYAAASASAS | ASSDREVF | 0.37205487 |
| HQVDVAAVQQG | ASSDREVF | 0.36656132 |
| IGAAIAHT | ASSDREVF | 0.43492728 |
| IGMIEVAQL | ASSDREVF | 0.5473344 |
| IRLAPEHTSAMD | ASSDREVF | 0.35151166 |
| IYNVSVL | ASSDREVF | 0.21827567 |

| | | |
|---|---|---|
| KQGFLGM | ASSDREVF | 0.5651838 |
| LESVITQL | ASSDREVF | 0.7228971 |
| LGLGALVVV | ASSDREVF | 0.676024 |
| LIGNGGNGGSGGI | ASSDREVF | 0.76563865 |
| LNAINAPTL | ASSDREVF | 0.7224028 |
| LQLLNTGV | ASSDREVF | 0.46647862 |
| LQTSIEF | ASSDREVF | 0.10505917 |
| MSFAVSEL | ASSDREVF | 0.16490468 |
| NTGALGL | ASSDREVF | 0.2682963 |
| PPGAGGAGGAGTT | ASSDREVF | 0.40908313 |
| QLVGLGI | ASSDREVF | 0.11401781 |
| RFLEQVHQL | ASSDREVF | 0.60225195 |
| RGVGYVLR | ASSDREVF | 0.29715037 |
| SATGVFAGVI | ASSDREVF | 0.29263505 |
| SLPAITLP | ASSDREVF | 0.7499331 |
| STALAFTV | ASSDREVF | 0.27733794 |
| SVLTVGLGK | ASSDREVF | 0.23690464 |
| TALAGAGSGI | ASSDREVF | 0.26990193 |
| TGLPFTGI | ASSDREVF | 0.720197 |
| VAEHVTV | ASSDREVF | 0.17801517 |
| VDIVLNSL | ASSDREVF | 0.60640013 |
| VLTGGGAVL | ASSDREVF | 0.36839274 |
| VNVGLAYGI | ASSDREVF | 0.7358516 |
| VSSAAVGAV | ASSDREVF | 0.1873872 |
| VTSVYEAGV | ASSDREVF | 0.65136063 |
| WVALDRGA | ASSDREVF | 0.41774178 |
| YLVGSVVI | ASSDREVF | 0.8615608 |
| GPPGTGKTLL | ASSDREVF | 0.31873062 |
| GVGMTKF | ASSDREVF | 0.49931538 |
| IGIGGSDL | ASSDREVF | 0.35163748 |
| IGTLLAEL | ASSDREVF | 0.79311323 |
| IYTSGTT | ASSDREVF | 0.18263158 |
| LSFPAGV | ASSDREVF | 0.3172989 |
| LYTSGTT | ASSDREVF | 0.18682301 |
| VGAFPFGF | ASSDREVF | 0.35773385 |
| VGAGTVEF | ASSDREVF | 0.62775326 |
| VGKSTLLL | ASSDREVF | 0.11291925 |
| AALAHAGGGIV | ASSDREVF | 0.510494 |
| AALVAVSLL | ASSDREVF | 0.3216581 |
| AAVSRAVAAL | ASSDREVF | 0.20783879 |
| AGGSSGIGGF | ASSDREVF | 0.5705362 |
| AGLTVANANG | ASSDREVF | 0.42917907 |
| AGLVGAVL | ASSDREVF | 0.74780095 |
| ALSQAFTV | ASSDREVF | 0.25734687 |
| ALTFSVSL | ASSDREVF | 0.4378428 |
| ALTGAALVAL | ASSDREVF | 0.4177174 |
| ASFSIGAGVL | ASSDREVF | 0.21531859 |
| DGSVYKI | ASSDREVF | 0.25244364 |
| EQLAEIV | ASSDREVF | 0.7189289 |
| FAVSELTVM | ASSDREVF | 0.40215433 |
| GAAPLAAGL | ASSDREVF | 0.8094425 |

| | | |
|---|---|---|
| GADGGDGGSGGAGG | ASSDREVF | 0.2680907 |
| GFGLDTGGAGG | ASSDREVF | 0.21214233 |
| GLASGGLL | ASSDREVF | 0.4041388 |
| GRIAGAAVLL | ASSDREVF | 0.71876323 |
| GSAGSLNTSF | ASSDREVF | 0.78572917 |
| GTGQIGI | ASSDREVF | 0.6127272 |
| GTLIALIL | ASSDREVF | 0.20961386 |
| IAATMGFVVG | ASSDREVF | 0.28559017 |
| IFAQALL | ASSDREVF | 0.26885176 |
| IGAAVALGF | ASSDREVF | 0.34416085 |
| IGLTPLAL | ASSDREVF | 0.51200104 |
| ILGSNTSTL | ASSDREVF | 0.2816908 |
| IVFSLTVL | ASSDREVF | 0.21628371 |
| LASKATGF | ASSDREVF | 0.27775276 |
| LASVALIL | ASSDREVF | 0.52025247 |
| LFAVGLTL | ASSDREVF | 0.6223257 |
| LFNGGIGTYI | ASSDREVF | 0.5655235 |
| LTLGAGII | ASSDREVF | 0.77550066 |
| LVAAALGDI | ASSDREVF | 0.5848147 |
| NTAFVQLGI | ASSDREVF | 0.6365129 |
| QLFSGAEL | ASSDREVF | 0.1257108 |
| SAATFITL | ASSDREVF | 0.15871692 |
| SNYGLANLGV | ASSDREVF | 0.29179704 |
| SYGGAYAVM | ASSDREVF | 0.34300235 |
| TTGVGGLTLG | ASSDREVF | 0.19265851 |
| TTQTIYI | ASSDREVF | 0.57355976 |
| VAAGAGLGAVI | ASSDREVF | 0.2919609 |
| VAAVVTLL | ASSDREVF | 0.20552419 |
| VGFLGGLI | ASSDREVF | 0.6574425 |
| VGVIPILL | ASSDREVF | 0.22686973 |
| VIFASSLI | ASSDREVF | 0.23540014 |
| VKGGDGGAAAG | ASSDREVF | 0.32747373 |
| VTKTFTNI | ASSDREVF | 0.55581737 |
| VVAAAILL | ASSDREVF | 0.1853045 |
| YAALDVEL | ASSDREVF | 0.4797773 |
| LASVALLL | ASSDREVF | 0.375929 |
| LDAVHRV | ASSDREVF | 0.57122934 |
| QGLSSSIF | ASSDREVF | 0.18378668 |
| RGTFANIRL | ASSDREVF | 0.61979085 |
| RQLWWGHRI | ASSDREVF | 0.38073158 |
| YGPPGTGKTLL | ASSDREVF | 0.81567746 |
| AALVGAPANVA | ASSDREVF | 0.26974353 |
| AATFRRTE | ASSDREVF | 0.30811515 |
| AGALGGGI | ASSDREVF | 0.27774164 |
| AGGAGGAGGIGAG | ASSDREVF | 0.76494795 |
| AGSGALGL | ASSDREVF | 0.17825705 |
| AQIADQI | ASSDREVF | 0.2906321 |
| ASAVAVTYL | ASSDREVF | 0.17189644 |
| ASTVLGGTGG | ASSDREVF | 0.44860968 |
| AVGGEPAK | ASSDREVF | 0.50996435 |
| CEQGTFGGTC | ASSDREVF | 0.24076301 |

| | | |
|---|---|---|
| DEGKAPAG | ASSDREVF | 0.46791565 |
| DTVVFPAL | ASSDREVF | 0.6782432 |
| FEGITMAG | ASSDREVF | 0.2702028 |
| FPLVVQD | ASSDREVF | 0.70765036 |
| GAFTTPQ | ASSDREVF | 0.2885976 |
| GFGGFGTAGGVGGA | ASSDREVF | 0.68147326 |
| GFLAQFL | ASSDREVF | 0.36996213 |
| GGAGGAGGNAGVA | ASSDREVF | 0.2717046 |
| GGVGTAGAGGAGGAGG | ASSDREVF | 0.56035465 |
| GNGGGGDAGNAGSGGN | ASSDREVF | 0.27880174 |
| GTGGAGGAGNDAGS | ASSDREVF | 0.30920708 |
| GTTNTGWFNS | ASSDREVF | 0.6387914 |
| IFMGPLFTL | ASSDREVF | 0.8243636 |
| ILAHAIEAI | ASSDREVF | 0.8431231 |
| IRSAIREAL | ASSDREVF | 0.7833861 |
| ITFAGGG | ASSDREVF | 0.7520683 |
| ITGMSGHQEV | ASSDREVF | 0.7962921 |
| IVAGLACTL | ASSDREVF | 0.8358687 |
| IVGGGLAGA | ASSDREVF | 0.83803535 |
| IVTGVLNI | ASSDREVF | 0.823612 |
| LAQQGCI | ASSDREVF | 0.73163456 |
| LCLHVEN | ASSDREVF | 0.8177979 |
| LDAISDGS | ASSDREVF | 0.796682 |
| LGGFAAML | ASSDREVF | 0.78809875 |
| LGGKNPMI | ASSDREVF | 0.82770944 |
| LGPKGVAAIL | ASSDREVF | 0.8069252 |
| LMVGVSI | ASSDREVF | 0.78638154 |
| LVGAAFVL | ASSDREVF | 0.8108361 |
| LVLAVAPL | ASSDREVF | 0.79991376 |
| MAAIAVIDAA | ASSDREVF | 0.81818545 |
| NNGINAELQ | ASSDREVF | 0.8107066 |
| RGEPLVLVHG | ASSDREVF | 0.84295547 |
| RGGAGGAAGLI | ASSDREVF | 0.83364433 |
| RSYRMHPAV | ASSDREVF | 0.8248955 |
| SAAHLLAGQLA | ASSDREVF | 0.44099653 |
| SAAVGSY | ASSDREVF | 0.7168714 |
| SPHAVLHRI | ASSDREVF | 0.80137444 |
| SQIAKVL | ASSDREVF | 0.8001789 |
| TALGHLVP | ASSDREVF | 0.5966538 |
| TAPTTSGAA | ASSDREVF | 0.82263184 |
| TDGDDGNGGSPGT | ASSDREVF | 0.79325235 |
| TDGLGEN | ASSDREVF | 0.74576545 |
| TGGPPLPGQA | ASSDREVF | 0.8249396 |
| TIPAITI | ASSDREVF | 0.78606844 |
| TIPGFTI | ASSDREVF | 0.77839315 |
| TSIASGALA | ASSDREVF | 0.7899913 |
| TVTLPGAAI | ASSDREVF | 0.82376766 |
| VASGLSAALP | ASSDREVF | 0.7414291 |
| VGAGVIGI | ASSDREVF | 0.823057 |
| VGPKRFGV | ASSDREVF | 0.77706265 |
| VNVAFPD | ASSDREVF | 0.8340243 |

| | | |
|---|---|---|
| VSMTNSGVS | ASSDREVF | 0.70269156 |
| VTDVLGDLSV | ASSDREVF | 0.828332 |
| VVVVGAGFAGL | ASSDREVF | 0.8037572 |
| VYTVGTVV | ASSDREVF | 0.84420645 |
| WTTFSGP | ASSDREVF | 0.7990974 |
| YDELVEAF | ASSDREVF | 0.8183197 |
| YGTGGAGGNGG | ASSDREVF | 0.787233 |
| YTNVMAV | ASSDREVF | 0.83924854 |
| FLEQVHQL | ASSDREVF | 0.69749016 |
| GLSLLAVAV | ASSDREVF | 0.81290305 |
| LEQVHQL | ASSDREVF | 0.72990793 |
| LNAAGAI | ASSDREVF | 0.8261212 |
| PAVHASL | ASSDREVF | 0.82377553 |
| TELEDDA | ASSDREVF | 0.7136997 |
| TPPGPPN | ASSDREVF | 0.8329265 |
| TPPPPPPP | ASSDREVF | 0.8138585 |
| TQSPALI | ASSPPTGGQEDTQY | 0.76791143 |
| EFAAAFPQ | ASSPPTGGQEDTQY | 0.8373014 |
| LLSGNLDVL | ASSPPTGGQEDTQY | 0.7892182 |
| GPANLGNYNI | ASSPPTGGQEDTQY | 0.83519757 |
| TLGPTLP | ASSPPTGGQEDTQY | 0.8051115 |
| VYAYSSCCK | ASSPPTGGQEDTQY | 0.81101114 |
| APAPAPAPNI | ASSPPTGGQEDTQY | 0.8220101 |
| PGGPVPIP | ASSPPTGGQEDTQY | 0.698753 |
| FLTVALEP | ASSPPTGGQEDTQY | 0.7370318 |
| SQYFGANL | ASSPPTGGQEDTQY | 0.8374647 |
| GGVKPAA | ASSPPTGGQEDTQY | 0.57681686 |
| GGTGGAGGAGGW | ASSPPTGGQEDTQY | 0.6855359 |
| GVGGVAGLGGAGG | ASSPPTGGQEDTQY | 0.4711763 |
| VLLVLATTI | ASSPPTGGQEDTQY | 0.4485498 |
| IGIFAAML | ASSPPTGGQEDTQY | 0.5664409 |
| AVGDLID | ASSPPTGGQEDTQY | 0.66843426 |
| NISPLPL | ASSPPTGGQEDTQY | 0.6748221 |
| VTAVFVAI | ASSPPTGGQEDTQY | 0.6161623 |
| SGDDNDAAL | ASSPPTGGQEDTQY | 0.46563822 |
| AVAALAHS | ASSPPTGGQEDTQY | 0.5417482 |
| GGVTVMATP | ASSPPTGGQEDTQY | 0.5024498 |
| AGLAANVRD | ASSPPTGGQEDTQY | 0.5519798 |
| TNRPDLIDPALLRPG | ASSPPTGGQEDTQY | 0.619918 |
| LANLGVTAL | ASSPPTGGQEDTQY | 0.54199433 |
| TGERIEL | ASSPPTGGQEDTQY | 0.4742432 |
| NGEELAA | ASSPPTGGQEDTQY | 0.5173409 |
| VLSLVWG | ASSPPTGGQEDTQY | 0.61790466 |
| IGYALWL | ASSPPTGGQEDTQY | 0.586208 |
| LSLLLAPL | ASSPPTGGQEDTQY | 0.5720712 |
| TLDEALK | ASSPPTGGQEDTQY | 0.68523633 |
| TDGAGTAEGG | ASSPPTGGQEDTQY | 0.62811697 |
| NAPGGAGGNGG | ASSPPTGGQEDTQY | 0.63089263 |
| VSVFMGMGGNF | ASSPPTGGQEDTQY | 0.30009004 |
| LASYLAAL | ASSPPTGGQEDTQY | 0.35151002 |
| LVGGTPAL | ASSPPTGGQEDTQY | 0.53735334 |

| | | |
|---|---|---|
| VRNADVL | ASSPPTGGQEDTQY | 0.5537927 |
| TCSSPFLVSAM | ASSPPTGGQEDTQY | 0.36666054 |
| VHITDAE | ASSPPTGGQEDTQY | 0.6357423 |
| EYAAPSF | ASSPPTGGQEDTQY | 0.49643183 |
| VVDAFHGGL | ASSPPTGGQEDTQY | 0.4710367 |
| ASVVDPQ | ASSPPTGGQEDTQY | 0.635674 |
| SYNEFGGLS | ASSPPTGGQEDTQY | 0.49771637 |
| DPDGSVY | ASSPPTGGQEDTQY | 0.48447943 |
| MVSTIHATE | ASSPPTGGQEDTQY | 0.5237168 |
| LAAGVADPV | ASSPPTGGQEDTQY | 0.60557616 |
| TKEQIAA | ASSPPTGGQEDTQY | 0.3763649 |
| SLASPAAL | ASSPPTGGQEDTQY | 0.6026575 |
| IGLVFYGV | ASSPPTGGQEDTQY | 0.5236872 |
| FLLDASNL | ASSPPTGGQEDTQY | 0.65583557 |
| YQVGNFP | ASSPPTGGQEDTQY | 0.40394375 |
| ESLLSQST | ASSPPTGGQEDTQY | 0.6128708 |
| LFTVGFAAVG | ASSPPTGGQEDTQY | 0.56556356 |
| ATPPTPT | ASSPPTGGQEDTQY | 0.69062346 |
| TDLATLG | ASSPPTGGQEDTQY | 0.50706416 |
| SAGGTGR | ASSPPTGGQEDTQY | 0.5735069 |
| GFSNSGS | ASSPPTGGQEDTQY | 0.55758005 |
| GGNGGSGGTSV | ASSPPTGGQEDTQY | 0.6765787 |
| GTSVATGGAGN | ASSPPTGGQEDTQY | 0.43854606 |
| NGGNGGSGGTS | ASSPPTGGQEDTQY | 0.50936264 |
| AVVKFPH | ASSPPTGGQEDTQY | 0.3863931 |
| TPPPPPP | ASSPPTGGQEDTQY | 0.6082996 |
| AAPAAAAAPA | ASSPPTGGQEDTQY | 0.5888025 |
| IALNEDL | ASSPPTGGQEDTQY | 0.4528386 |
| IVNVSSM | ASSPPTGGQEDTQY | 0.6322665 |
| TGIPPAPR | ASSPPTGGQEDTQY | 0.6177931 |
| FDLGGGTF | ASSPPTGGQEDTQY | 0.48452115 |
| FDLGGGTFDVS | ASSPPTGGQEDTQY | 0.65864694 |
| NEPTAAA | ASSPPTGGQEDTQY | 0.54306674 |
| NEPTAAALA | ASSPPTGGQEDTQY | 0.6442701 |
| GSIINISSI | ASSPPTGGQEDTQY | 0.58378434 |
| VAPGNAGTA | ASSPPTGGQEDTQY | 0.47903013 |
| TGGGASLE | ASSPPTGGQEDTQY | 0.62837744 |
| VAVYNHYKRI | ASSPPTGGQEDTQY | 0.55359876 |
| TDPAPATT | ASSPPTGGQEDTQY | 0.5101408 |
| TNRPDILDPALLRPG | ASSPPTGGQEDTQY | 0.6443224 |
| LTTTTTS | ASSPPTGGQEDTQY | 0.83768755 |
| PTGSGKTL | ASSPPTGGQEDTQY | 0.8144189 |
| TGSGKTL | ASSPPTGGQEDTQY | 0.6750434 |
| TIGHVDHGK | ASSPPTGGQEDTQY | 0.6729506 |
| TIGHVDHGKTT | ASSPPTGGQEDTQY | 0.71993107 |
| GHVDHGKTT | ASSPPTGGQEDTQY | 0.83328587 |
| AQFEHTLL | ASSPPTGGQEDTQY | 0.863611 |
| IVNQDEL | ASSPPTGGQEDTQY | 0.8361223 |
| TNGSQFF | ASSPPTGGQEDTQY | 0.588098 |
| TGKGIVE | ASSPPTGGQEDTQY | 0.860126 |
| LISNASDA | ASSPPTGGQEDTQY | 0.82299256 |

| | | |
|---|---|---|
| AELRTLGELTKIAKAHGAQ | ASSPPTGGQEDTQY | 0.8314125 |
| APNAVAAAEPSW | ASSPPTGGQEDTQY | 0.79579353 |
| APQVAGAATAVL | ASSPPTGGQEDTQY | 0.77321625 |
| ARRDPAAAAAILPTDA | ASSPPTGGQEDTQY | 0.698871 |
| DAAARDVLAEAGLAEYF | ASSPPTGGQEDTQY | 0.6137102 |
| DDQIVLSAKANLAKVEQ | ASSPPTGGQEDTQY | 0.7904241 |
| DELLPDTGEDSDEDGH | ASSPPTGGQEDTQY | 0.83296126 |
| ELPEAPANRATVPQD | ASSPPTGGQEDTQY | 0.82557404 |
| FGQSKIERVTGIG | ASSPPTGGQEDTQY | 0.8733572 |
| GAGGTGSTAGGAGGAGGAGGI | ASSPPTGGQEDTQY | 0.8579521 |
| GHGGTGAAGGNGGTGGAGGS( | ASSPPTGGQEDTQY | 0.8002444 |
| GNGGNGGAGGEGGPGGAGGA( | ASSPPTGGQEDTQY | 0.28055093 |
| GSTAGNGGNGGNATNGTIAG( | ASSPPTGGQEDTQY | 0.4770129 |
| GTGGDGGNGGNGGTGG | ASSPPTGGQEDTQY | 0.5808064 |
| GVRRPAGIPADAPV | ASSPPTGGQEDTQY | 0.7219707 |
| GYRQVTDAHLLTLARR | ASSPPTGGQEDTQY | 0.24431595 |
| IAKMGQDGHDRG | ASSPPTGGQEDTQY | 0.8028562 |
| KAEPSIEIQYT | ASSPPTGGQEDTQY | 0.75934595 |
| KPVRIRMAADEDTVAVT | ASSPPTGGQEDTQY | 0.72179115 |
| MPPRIAGMRLLVIKPEP | ASSPPTGGQEDTQY | 0.84343976 |
| MKFYAVALFRSRGVPPTR | ASSPPTGGQEDTQY | 0.7705598 |
| NAGGFGILGAVAHSP | ASSPPTGGQEDTQY | 0.55768365 |
| NEGREITLAMAEM | ASSPPTGGQEDTQY | 0.56562674 |
| NELVEWSHPLGHRWRWE | ASSPPTGGQEDTQY | 0.8642714 |
| NLGAGGTGGNGGNPG | ASSPPTGGQEDTQY | 0.6224908 |
| NTEWPKWETVMAADPVD | ASSPPTGGQEDTQY | 0.8752079 |
| PASAAPAETQP | ASSPPTGGQEDTQY | 0.5971439 |
| QQSSDGKNGVRAAEITGE | ASSPPTGGQEDTQY | 0.87393 |
| RAPDAVTTSWWKE | ASSPPTGGQEDTQY | 0.56857455 |
| SGFGNVGAHTSGWFNQ | ASSPPTGGQEDTQY | 0.86356723 |
| SGVAGAVGQAASVGGLKV | ASSPPTGGQEDTQY | 0.81656766 |
| SNGGAGTGSAGGNGGT | ASSPPTGGQEDTQY | 0.87718093 |
| TGGAGGEGGAAG | ASSPPTGGQEDTQY | 0.76485556 |
| TGGDGGAAGAGGAPGHG | ASSPPTGGQEDTQY | 0.840977 |
| TMPPLLAAIANAGRHGV | ASSPPTGGQEDTQY | 0.73657966 |
| TSSTAGNSGGA | ASSPPTGGQEDTQY | 0.8527988 |
| VPATVPVGILC | ASSPPTGGQEDTQY | 0.55394614 |
| VQRPDSGGVVLTGRIS | ASSPPTGGQEDTQY | 0.73156583 |
| WTPNPVPATKGVPAGLVQAT | ASSPPTGGQEDTQY | 0.5106665 |
| YALKRALAKFSDRFDVVII | ASSPPTGGQEDTQY | 0.7691641 |
| YLIGSVIV | ASSPPTGGQEDTQY | 0.8617468 |
| YGQLVAVID | ASSPPTGGQEDTQY | 0.1773041 |
| VTVGGMTLPA | ASSPPTGGQEDTQY | 0.8539883 |
| VLALPAV | ASSPPTGGQEDTQY | 0.56573784 |
| VIPPTLNYET | ASSPPTGGQEDTQY | 0.60224855 |
| VGMGGMHPSSG | ASSPPTGGQEDTQY | 0.83050245 |
| VGLAGGAATA | ASSPPTGGQEDTQY | 0.80293274 |
| VGGAGGAGGSGP | ASSPPTGGQEDTQY | 0.8306179 |
| VFAAFMV | ASSPPTGGQEDTQY | 0.61082 |
| TLYDKLR | ASSPPTGGQEDTQY | 0.7665731 |
| TLAVFTVL | ASSPPTGGQEDTQY | 0.7987565 |

| | | |
|---|---|---|
| TGSVQPARV | ASSPPTGGQEDTQY | 0.28473172 |
| TGGGAGSGGT | ASSPPTGGQEDTQY | 0.46718314 |
| SHLETSL | ASSPPTGGQEDTQY | 0.8789737 |
| SFTTGMT | ASSPPTGGQEDTQY | 0.79180765 |
| QVEVAKRMGVRQPTVS | ASSPPTGGQEDTQY | 0.77723897 |
| QGSGGPALAM | ASSPPTGGQEDTQY | 0.7103937 |
| MLAPSNP | ASSPPTGGQEDTQY | 0.69191974 |
| LVNNAAI | ASSPPTGGQEDTQY | 0.7459367 |
| LTGEINTLA | ASSPPTGGQEDTQY | 0.7981549 |
| LSIFGPV | ASSPPTGGQEDTQY | 0.7856244 |
| LLALLANL | ASSPPTGGQEDTQY | 0.781131 |
| LISNASDALDKLR | ASSPPTGGQEDTQY | 0.72627676 |
| LINEAVP | ASSPPTGGQEDTQY | 0.784004 |
| LGYAFSGL | ASSPPTGGQEDTQY | 0.7707494 |
| LDFVAVAL | ASSPPTGGQEDTQY | 0.73281497 |
| LAQPGAE | ASSPPTGGQEDTQY | 0.7938236 |
| ISNLGAVTGF | ASSPPTGGQEDTQY | 0.78254926 |
| HQEDPGRI | ASSPPTGGQEDTQY | 0.7355832 |
| GTPGEPTP | ASSPPTGGQEDTQY | 0.77102137 |
| GFFNSGS | ASSPPTGGQEDTQY | 0.7625747 |
| GAGGGGLTAGAN | ASSPPTGGQEDTQY | 0.7892247 |
| FVDALVNM | ASSPPTGGQEDTQY | 0.78610593 |
| FLTVAPEL | ASSPPTGGQEDTQY | 0.81708544 |
| FETLTAL | ASSPPTGGQEDTQY | 0.80093515 |
| DVINPIEL | ASSPPTGGQEDTQY | 0.7926072 |
| APVSGAPV | ASSPPTGGQEDTQY | 0.43759924 |
| ALVGGLAW | ASSPPTGGQEDTQY | 0.69415736 |
| ALAAAAATAI | ASSPPTGGQEDTQY | 0.76722324 |
| AFGLGLLL | ASSPPTGGQEDTQY | 0.76480883 |
| AFFAAASL | ASSPPTGGQEDTQY | 0.5752875 |
| AAAALPAFSPP | ASSPPTGGQEDTQY | 0.77537686 |
| VVLAPAL | ASSPPTGGQEDTQY | 0.75926054 |
| TIGHVDHGKTTL | ASSPPTGGQEDTQY | 0.6935812 |
| LISNASDALDK | ASSPPTGGQEDTQY | 0.8019883 |
| LISNASDAL | ASSPPTGGQEDTQY | 0.73685366 |
| ISNASDALDK | ASSPPTGGQEDTQY | 0.75273764 |
| NQAETLVYQTE | ASSPPTGGQEDTQY | 0.7721983 |
| TGGTGGTAGAG | ASSPPTGGQEDTQY | 0.8027824 |
| TGSTAGGAGGA | ASSPPTGGQEDTQY | 0.7183182 |
| DPLAVPVAAGPV | ASSPPTGGQEDTQY | 0.79423606 |
| GGTGGTPGNGGN | ASSPPTGGQEDTQY | 0.7420934 |
| PAGLAGVDPATL | ASSPPTGGQEDTQY | 0.8113487 |
| GNNLVVEAPHAGASV | ASSPPTGGQEDTQY | 0.6870307 |
| KVTRIMSASTLEQPA | ASSPPTGGQEDTQY | 0.79141223 |
| LGFALPIALPRRTVV | ASSPPTGGQEDTQY | 0.7538598 |
| TRLGNLTARKLVVLV | ASSPPTGGQEDTQY | 0.8164338 |
| WSQQRRLIYFPSAGP | ASSPPTGGQEDTQY | 0.7633405 |
| GPNGNPDPMAAAVDIR | ASSPPTGGQEDTQY | 0.80086136 |
| TQRCVSVADLRAAASR | ASSPPTGGQEDTQY | 0.7777573 |
| QFVLDVAEHPVEVLTETV | ASSPPTGGQEDTQY | 0.7855612 |
| GGAGGTGGAGGNGGTQAGDG | ASSPPTGGQEDTQY | 0.65192294 |

| | | |
|---|---|---|
| PLLATLRAIAVDVDAKPAQV | ASSPPTGGQEDTQY | 0.7746726 |
| APKFRTGGYMIDFWGVGYQV/ | ASSPPTGGQEDTQY | 0.68763274 |
| GAGGAGGAGGAAGGPGGTGG' | ASSPPTGGQEDTQY | 0.7874342 |
| IGALGLLATDVAPAGSVAFG' | ASSPPTGGQEDTQY | 0.78432053 |
| AGGAGGIGGNGGAGGAGGVLI | ASSPPTGGQEDTQY | 0.53965247 |
| ADELIGLL | ASSPPTGGQEDTQY | 0.79898536 |
| ADRALKA | ASSPPTGGQEDTQY | 0.69157517 |
| AGAVGQVGGAGGA | ASSPPTGGQEDTQY | 0.74754 |
| AGGGGGLGGAP | ASSPPTGGQEDTQY | 0.80579305 |
| AGGLLYGNGGAG | ASSPPTGGQEDTQY | 0.76130956 |
| AGLESIGL | ASSPPTGGQEDTQY | 0.8047513 |
| AGLKTGII | ASSPPTGGQEDTQY | 0.75917983 |
| ASLAALLL | ASSPPTGGQEDTQY | 0.7683448 |
| ASVINISSI | ASSPPTGGQEDTQY | 0.77599573 |
| ATPPGPA | ASSPPTGGQEDTQY | 0.64132315 |
| ATVASALV | ASSPPTGGQEDTQY | 0.7315736 |
| ATVASLIL | ASSPPTGGQEDTQY | 0.80528235 |
| AVNGPVLIRH | ASSPPTGGQEDTQY | 0.6459313 |
| DGAAGAVGI | ASSPPTGGQEDTQY | 0.7347442 |
| DPLLHGI | ASSPPTGGQEDTQY | 0.56669515 |
| ELGAALGAL | ASSPPTGGQEDTQY | 0.503768 |
| ELGRTQAV | ASSPPTGGQEDTQY | 0.6287564 |
| FVAGGYLS | ASSPPTGGQEDTQY | 0.7455999 |
| GASGGAGGAGGW | ASSPPTGGQEDTQY | 0.7224065 |
| GGAGAGGDGGAGGI | ASSPPTGGQEDTQY | 0.6855754 |
| GTGHNGSQPGMGGNG | ASSPPTGGQEDTQY | 0.54532915 |
| GYAAASASAS | ASSPPTGGQEDTQY | 0.7007196 |
| HQVDVAAVQQG | ASSPPTGGQEDTQY | 0.6162262 |
| IGAAIAHT | ASSPPTGGQEDTQY | 0.595928 |
| IGMIEVAQL | ASSPPTGGQEDTQY | 0.71759164 |
| IRLAPEHTSAMD | ASSPPTGGQEDTQY | 0.685348 |
| IYNVSVL | ASSPPTGGQEDTQY | 0.62281394 |
| KQGFLGM | ASSPPTGGQEDTQY | 0.62360257 |
| LESVITQL | ASSPPTGGQEDTQY | 0.64786947 |
| LGLGALVVV | ASSPPTGGQEDTQY | 0.66797066 |
| LIGNGGNGGSGGI | ASSPPTGGQEDTQY | 0.64817613 |
| LNAINAPTL | ASSPPTGGQEDTQY | 0.78558224 |
| LQLLNTGV | ASSPPTGGQEDTQY | 0.721173 |
| LQTSIEF | ASSPPTGGQEDTQY | 0.6977879 |
| MSFAVSEL | ASSPPTGGQEDTQY | 0.2650817 |
| NTGALGL | ASSPPTGGQEDTQY | 0.41089645 |
| PPGAGGAGGAGTT | ASSPPTGGQEDTQY | 0.61302966 |
| QLVGLGI | ASSPPTGGQEDTQY | 0.65240693 |
| RFLEQVHQL | ASSPPTGGQEDTQY | 0.35405365 |
| RGVGYVLR | ASSPPTGGQEDTQY | 0.7306988 |
| SATGVFAGVI | ASSPPTGGQEDTQY | 0.62916183 |
| SLPAITLP | ASSPPTGGQEDTQY | 0.5668365 |
| STALAFTV | ASSPPTGGQEDTQY | 0.74674916 |
| SVLTVGLGK | ASSPPTGGQEDTQY | 0.6018153 |
| TALAGAGSGI | ASSPPTGGQEDTQY | 0.6155956 |
| TGLPFTGI | ASSPPTGGQEDTQY | 0.6472118 |

| | | |
|---|---|---|
| VAEHVTV | ASSPPTGGQEDTQY | 0.7558956 |
| VDIVLNSL | ASSPPTGGQEDTQY | 0.49411762 |
| VLTGGGAVL | ASSPPTGGQEDTQY | 0.7140397 |
| VNVGLAYGI | ASSPPTGGQEDTQY | 0.58664834 |
| VSSAAVGAV | ASSPPTGGQEDTQY | 0.7477969 |
| VTSVYEAGV | ASSPPTGGQEDTQY | 0.47690743 |
| WVALDRGA | ASSPPTGGQEDTQY | 0.6994288 |
| YLVGSVVI | ASSPPTGGQEDTQY | 0.57534504 |
| GPPGTGKTLL | ASSPPTGGQEDTQY | 0.7915932 |
| GVGMTKF | ASSPPTGGQEDTQY | 0.61631835 |
| IGIGGSDL | ASSPPTGGQEDTQY | 0.6985635 |
| IGTLLAEL | ASSPPTGGQEDTQY | 0.6037808 |
| IYTSGTT | ASSPPTGGQEDTQY | 0.6999991 |
| LSFPAGV | ASSPPTGGQEDTQY | 0.43731785 |
| LYTSGTT | ASSPPTGGQEDTQY | 0.62647116 |
| VGAFPFGF | ASSPPTGGQEDTQY | 0.5215244 |
| VGAGTVEF | ASSPPTGGQEDTQY | 0.70596546 |
| VGKSTLLL | ASSPPTGGQEDTQY | 0.6672639 |
| AALAHAGGGIV | ASSPPTGGQEDTQY | 0.5021366 |
| AALVAVSLL | ASSPPTGGQEDTQY | 0.73998445 |
| AAVSRAVAAL | ASSPPTGGQEDTQY | 0.6492319 |
| AGGSSGIGGF | ASSPPTGGQEDTQY | 0.5668103 |
| AGLTVANANG | ASSPPTGGQEDTQY | 0.74945045 |
| AGLVGAVL | ASSPPTGGQEDTQY | 0.6075535 |
| ALSQAFTV | ASSPPTGGQEDTQY | 0.7049939 |
| ALTFSVSL | ASSPPTGGQEDTQY | 0.6279868 |
| ALTGAALVAL | ASSPPTGGQEDTQY | 0.6419657 |
| ASFSIGAGVL | ASSPPTGGQEDTQY | 0.72475797 |
| DGSVYKI | ASSPPTGGQEDTQY | 0.5269143 |
| EQLAEIV | ASSPPTGGQEDTQY | 0.5882136 |
| FAVSELTVM | ASSPPTGGQEDTQY | 0.7274939 |
| GAAPLAAGL | ASSPPTGGQEDTQY | 0.610113 |
| GADGGDGGSGGAGG | ASSPPTGGQEDTQY | 0.679588 |
| GFGLDTGGAGG | ASSPPTGGQEDTQY | 0.47081017 |
| GLASGGLL | ASSPPTGGQEDTQY | 0.42149597 |
| GRIAGAAVLL | ASSPPTGGQEDTQY | 0.53825796 |
| GSAGSLNTSF | ASSPPTGGQEDTQY | 0.679858 |
| GTGQIGI | ASSPPTGGQEDTQY | 0.6631521 |
| GTLIALIL | ASSPPTGGQEDTQY | 0.63464165 |
| IAATMGFVVG | ASSPPTGGQEDTQY | 0.4066108 |
| IFAQALL | ASSPPTGGQEDTQY | 0.5301194 |
| IGAAVALGF | ASSPPTGGQEDTQY | 0.51494455 |
| IGLTPLAL | ASSPPTGGQEDTQY | 0.5225499 |
| ILGSNTSTL | ASSPPTGGQEDTQY | 0.5711142 |
| IVFSLTVL | ASSPPTGGQEDTQY | 0.5233301 |
| LASKATGF | ASSPPTGGQEDTQY | 0.47327012 |
| LASVALIL | ASSPPTGGQEDTQY | 0.46595275 |
| LFAVGLTL | ASSPPTGGQEDTQY | 0.5796371 |
| LFNGGIGTYI | ASSPPTGGQEDTQY | 0.5615705 |
| LTLGAGII | ASSPPTGGQEDTQY | 0.52048314 |
| LVAAALGDI | ASSPPTGGQEDTQY | 0.69604254 |

| | | |
|---|---|---|
| NTAFVQLGI | ASSPPTGGQEDTQY | 0.65451854 |
| QLFSGAEL | ASSPPTGGQEDTQY | 0.6388509 |
| SAATFITL | ASSPPTGGQEDTQY | 0.28884155 |
| SNYGLANLGV | ASSPPTGGQEDTQY | 0.35660803 |
| SYGGAYAVM | ASSPPTGGQEDTQY | 0.43818414 |
| TTGVGGLTLG | ASSPPTGGQEDTQY | 0.49990982 |
| TTQTIYI | ASSPPTGGQEDTQY | 0.31426919 |
| VAAGAGLGAVI | ASSPPTGGQEDTQY | 0.6037523 |
| VAAVVTLL | ASSPPTGGQEDTQY | 0.54189306 |
| VGFLGGLI | ASSPPTGGQEDTQY | 0.43137017 |
| VGVIPILL | ASSPPTGGQEDTQY | 0.62544477 |
| VIFASSLI | ASSPPTGGQEDTQY | 0.43061537 |
| VKGGDGGAAAG | ASSPPTGGQEDTQY | 0.44051963 |
| VTKTFTNI | ASSPPTGGQEDTQY | 0.45057398 |
| VVAAAILL | ASSPPTGGQEDTQY | 0.641637 |
| YAALDVEL | ASSPPTGGQEDTQY | 0.37268606 |
| LASVALLL | ASSPPTGGQEDTQY | 0.60921407 |
| LDAVHRV | ASSPPTGGQEDTQY | 0.48721296 |
| QGLSSSIF | ASSPPTGGQEDTQY | 0.6765926 |
| RGTFANIRL | ASSPPTGGQEDTQY | 0.36780173 |
| RQLWWGHRI | ASSPPTGGQEDTQY | 0.6417462 |
| YGPPGTGKTLL | ASSPPTGGQEDTQY | 0.5543962 |
| AALVGAPANVA | ASSPPTGGQEDTQY | 0.6925106 |
| AATFRRTE | ASSPPTGGQEDTQY | 0.5027882 |
| AGALGGGI | ASSPPTGGQEDTQY | 0.5788854 |
| AGGAGGAGGIGAG | ASSPPTGGQEDTQY | 0.49463603 |
| AGSGALGL | ASSPPTGGQEDTQY | 0.6765693 |
| AQIADQI | ASSPPTGGQEDTQY | 0.40422946 |
| ASAVAVTYL | ASSPPTGGQEDTQY | 0.5656322 |
| ASTVLGGTGG | ASSPPTGGQEDTQY | 0.3992761 |
| AVGGEPAK | ASSPPTGGQEDTQY | 0.61865795 |
| CEQGTFGGTC | ASSPPTGGQEDTQY | 0.63084465 |
| DEGKAPAG | ASSPPTGGQEDTQY | 0.3252212 |
| DTVVFPAL | ASSPPTGGQEDTQY | 0.57419896 |
| FEGITMAG | ASSPPTGGQEDTQY | 0.5893949 |
| FPLVVQD | ASSPPTGGQEDTQY | 0.39792907 |
| GAFTTPQ | ASSPPTGGQEDTQY | 0.6410346 |
| GFGGFGTAGGVGGA | ASSPPTGGQEDTQY | 0.4682079 |
| GFLAQFL | ASSPPTGGQEDTQY | 0.67069364 |
| GGAGGAGGNAGVA | ASSPPTGGQEDTQY | 0.47696173 |
| GGVGTAGAGGAGGAGG | ASSPPTGGQEDTQY | 0.47681725 |
| GNGGGGDAGNAGSGGN | ASSPPTGGQEDTQY | 0.65040994 |
| GTGGAGGAGNDAGS | ASSPPTGGQEDTQY | 0.394634 |
| GTTNTGWFNS | ASSPPTGGQEDTQY | 0.40602478 |
| IFMGPLFTL | ASSPPTGGQEDTQY | 0.671087 |
| ILAHAIEAI | ASSPPTGGQEDTQY | 0.7938088 |
| IRSAIREAL | ASSPPTGGQEDTQY | 0.8176354 |
| ITFAGGG | ASSPPTGGQEDTQY | 0.6688396 |
| ITGMSGHQEV | ASSPPTGGQEDTQY | 0.60273296 |
| IVAGLACTL | ASSPPTGGQEDTQY | 0.7210402 |
| IVGGGLAGA | ASSPPTGGQEDTQY | 0.79981667 |

| | | |
|---|---|---|
| IVTGVLNI | ASSPPTGGQEDTQY | 0.82108176 |
| LAQQGCI | ASSPPTGGQEDTQY | 0.8038686 |
| LCLHVEN | ASSPPTGGQEDTQY | 0.577371 |
| LDAISDGS | ASSPPTGGQEDTQY | 0.76120776 |
| LGGFAAML | ASSPPTGGQEDTQY | 0.6839601 |
| LGGKNPMI | ASSPPTGGQEDTQY | 0.7104767 |
| LGPKGVAAIL | ASSPPTGGQEDTQY | 0.7982645 |
| LMVGVSI | ASSPPTGGQEDTQY | 0.75299156 |
| LVGAAFVL | ASSPPTGGQEDTQY | 0.7062577 |
| LVLAVAPL | ASSPPTGGQEDTQY | 0.6294873 |
| MAAIAVIDAA | ASSPPTGGQEDTQY | 0.80423045 |
| NNGINAELQ | ASSPPTGGQEDTQY | 0.8097619 |
| RGEPLVLVHG | ASSPPTGGQEDTQY | 0.8104306 |
| RGGAGGAAGLI | ASSPPTGGQEDTQY | 0.8140211 |
| RSYRMHPAV | ASSPPTGGQEDTQY | 0.7987559 |
| SAAHLLAGQLA | ASSPPTGGQEDTQY | 0.78448963 |
| SAAVGSY | ASSPPTGGQEDTQY | 0.24683179 |
| SPHAVLHRI | ASSPPTGGQEDTQY | 0.390513 |
| SQIAKVL | ASSPPTGGQEDTQY | 0.7679963 |
| TALGHLVP | ASSPPTGGQEDTQY | 0.6958958 |
| TAPTTSGAA | ASSPPTGGQEDTQY | 0.35547015 |
| TDGDDGNGGSPGT | ASSPPTGGQEDTQY | 0.7656942 |
| TDGLGEN | ASSPPTGGQEDTQY | 0.674646 |
| TGGPPLPGQA | ASSPPTGGQEDTQY | 0.66511923 |
| TIPAITI | ASSPPTGGQEDTQY | 0.8065875 |
| TIPGFTI | ASSPPTGGQEDTQY | 0.69626546 |
| TSIASGALA | ASSPPTGGQEDTQY | 0.5866667 |
| TVTLPGAAI | ASSPPTGGQEDTQY | 0.67940044 |
| VASGLSAALP | ASSPPTGGQEDTQY | 0.80512863 |
| VGAGVIGI | ASSPPTGGQEDTQY | 0.5566593 |
| VGPKRFGV | ASSPPTGGQEDTQY | 0.78353286 |
| VNVAFPD | ASSPPTGGQEDTQY | 0.7226227 |
| VSMTNSGVS | ASSPPTGGQEDTQY | 0.79856956 |
| VTDVLGDLSV | ASSPPTGGQEDTQY | 0.4810955 |
| VVVVGAGFAGL | ASSPPTGGQEDTQY | 0.7853683 |
| VYTVGTVV | ASSPPTGGQEDTQY | 0.73043346 |
| WTTFSGP | ASSPPTGGQEDTQY | 0.82860965 |
| YDELVEAF | ASSPPTGGQEDTQY | 0.6733457 |
| YGTGGAGGNGG | ASSPPTGGQEDTQY | 0.7393007 |
| YTNVMAV | ASSPPTGGQEDTQY | 0.72063243 |
| FLEQVHQL | ASSPPTGGQEDTQY | 0.81388545 |
| GLSLLAVAV | ASSPPTGGQEDTQY | 0.4730556 |
| LEQVHQL | ASSPPTGGQEDTQY | 0.7044843 |
| LNAAGAI | ASSPPTGGQEDTQY | 0.4660887 |
| PAVHASL | ASSPPTGGQEDTQY | 0.7442945 |
| TELEDDA | ASSPPTGGQEDTQY | 0.7832868 |
| TPPGPPN | ASSPPTGGQEDTQY | 0.43261343 |
| TPPPPPPP | ASSPPTGGQEDTQY | 0.8024184 |
| TQSPALI | ASSLSDRVNSDYT | 0.70939267 |
| EFAAAFPQ | ASSLSDRVNSDYT | 0.6498073 |
| LLSGNLDVL | ASSLSDRVNSDYT | 0.81602895 |

| | | |
|---|---|---|
| GPANLGNYNI | ASSLSDRVNSDYT | 0.75623596 |
| TLGPTLP | ASSLSDRVNSDYT | 0.8130195 |
| VYAYSSCCK | ASSLSDRVNSDYT | 0.75973374 |
| APAPAPAPNI | ASSLSDRVNSDYT | 0.7609923 |
| PGGPVPIP | ASSLSDRVNSDYT | 0.7737309 |
| FLTVALEP | ASSLSDRVNSDYT | 0.7296092 |
| SQYFGANL | ASSLSDRVNSDYT | 0.7116035 |
| GGVKPAA | ASSLSDRVNSDYT | 0.80110973 |
| GGTGGAGGAGGW | ASSLSDRVNSDYT | 0.34709483 |
| GVGGVAGLGGAGG | ASSLSDRVNSDYT | 0.73249614 |
| VLLVLATTI | ASSLSDRVNSDYT | 0.19597733 |
| IGIFAAML | ASSLSDRVNSDYT | 0.19753298 |
| AVGDLID | ASSLSDRVNSDYT | 0.2174945 |
| NISPLPL | ASSLSDRVNSDYT | 0.34425902 |
| VTAVFVAI | ASSLSDRVNSDYT | 0.7603595 |
| SGDDNDAAL | ASSLSDRVNSDYT | 0.40899783 |
| AVAALAHS | ASSLSDRVNSDYT | 0.21590191 |
| GGVTVMATP | ASSLSDRVNSDYT | 0.28832996 |
| AGLAANVRD | ASSLSDRVNSDYT | 0.22972107 |
| TNRPDLIDPALLRPG | ASSLSDRVNSDYT | 0.20968288 |
| LANLGVTAL | ASSLSDRVNSDYT | 0.3208393 |
| TGERIEL | ASSLSDRVNSDYT | 0.27940956 |
| NGEELAA | ASSLSDRVNSDYT | 0.22082907 |
| VLSLVWG | ASSLSDRVNSDYT | 0.22773348 |
| IGYALWL | ASSLSDRVNSDYT | 0.32369572 |
| LSLLLAPL | ASSLSDRVNSDYT | 0.37647313 |
| TLDEALK | ASSLSDRVNSDYT | 0.36584422 |
| TDGAGTAEGG | ASSLSDRVNSDYT | 0.41924036 |
| NAPGGAGGNGG | ASSLSDRVNSDYT | 0.36082977 |
| VSVFMGMGGNF | ASSLSDRVNSDYT | 0.3192637 |
| LASYLAAL | ASSLSDRVNSDYT | 0.118824914 |
| LVGGTPAL | ASSLSDRVNSDYT | 0.15494084 |
| VRNADVL | ASSLSDRVNSDYT | 0.3020109 |
| TCSSPFLVSAM | ASSLSDRVNSDYT | 0.22791451 |
| VHITDAE | ASSLSDRVNSDYT | 0.14259695 |
| EYAAPSF | ASSLSDRVNSDYT | 0.28948677 |
| VVDAFHGGL | ASSLSDRVNSDYT | 0.25968343 |
| ASVVDPQ | ASSLSDRVNSDYT | 0.21338116 |
| SYNEFGGLS | ASSLSDRVNSDYT | 0.42986596 |
| DPDGSVY | ASSLSDRVNSDYT | 0.2108353 |
| MVSTIHATE | ASSLSDRVNSDYT | 0.19518316 |
| LAAGVADPV | ASSLSDRVNSDYT | 0.22615461 |
| TKEQIAA | ASSLSDRVNSDYT | 0.3928237 |
| SLASPAAL | ASSLSDRVNSDYT | 0.18120113 |
| IGLVFYGV | ASSLSDRVNSDYT | 0.31626293 |
| FLLDASNL | ASSLSDRVNSDYT | 0.23900001 |
| YQVGNFP | ASSLSDRVNSDYT | 0.36558026 |
| ESLLSQST | ASSLSDRVNSDYT | 0.1747141 |
| LFTVGFAAVG | ASSLSDRVNSDYT | 0.30077448 |
| ATPPTPT | ASSLSDRVNSDYT | 0.22253008 |
| TDLATLG | ASSLSDRVNSDYT | 0.69510126 |

| | | |
|---|---|---|
| SAGGTGR | ASSLSDRVNSDYT | 0.22326237 |
| GFSNSGS | ASSLSDRVNSDYT | 0.29846993 |
| GGNGGSGGTSV | ASSLSDRVNSDYT | 0.25444362 |
| GTSVATGGAGN | ASSLSDRVNSDYT | 0.71812856 |
| NGGNGGSGGTS | ASSLSDRVNSDYT | 0.15174705 |
| AVVKFPH | ASSLSDRVNSDYT | 0.22220758 |
| TPPPPPP | ASSLSDRVNSDYT | 0.18744327 |
| AAPAAAAAPA | ASSLSDRVNSDYT | 0.25516295 |
| IALNEDL | ASSLSDRVNSDYT | 0.3534137 |
| IVNVSSM | ASSLSDRVNSDYT | 0.19315225 |
| TGIPPAPR | ASSLSDRVNSDYT | 0.40387183 |
| FDLGGGTF | ASSLSDRVNSDYT | 0.26573822 |
| FDLGGGTFDVS | ASSLSDRVNSDYT | 0.21453653 |
| NEPTAAA | ASSLSDRVNSDYT | 0.42264938 |
| NEPTAAALA | ASSLSDRVNSDYT | 0.26260582 |
| GSIINISSI | ASSLSDRVNSDYT | 0.37040204 |
| VAPGNAGTA | ASSLSDRVNSDYT | 0.22908708 |
| TGGGASLE | ASSLSDRVNSDYT | 0.29476145 |
| VAVYNHYKRI | ASSLSDRVNSDYT | 0.32815507 |
| TDPAPATT | ASSLSDRVNSDYT | 0.2869567 |
| TNRPDILDPALLRPG | ASSLSDRVNSDYT | 0.2142074 |
| LTTTTTS | ASSLSDRVNSDYT | 0.36144805 |
| PTGSGKTL | ASSLSDRVNSDYT | 0.8898705 |
| TGSGKTL | ASSLSDRVNSDYT | 0.8959831 |
| TIGHVDHGK | ASSLSDRVNSDYT | 0.84674734 |
| TIGHVDHGKTT | ASSLSDRVNSDYT | 0.82942027 |
| GHVDHGKTT | ASSLSDRVNSDYT | 0.85356647 |
| AQFEHTLL | ASSLSDRVNSDYT | 0.8972993 |
| IVNQDEL | ASSLSDRVNSDYT | 0.89743805 |
| TNGSQFF | ASSLSDRVNSDYT | 0.8943424 |
| TGKGIVE | ASSLSDRVNSDYT | 0.8105907 |
| LISNASDA | ASSLSDRVNSDYT | 0.8887106 |
| AELRTLGELTKIAKAHGAQ | ASSLSDRVNSDYT | 0.86140966 |
| APNAVAAAEPSW | ASSLSDRVNSDYT | 0.8739686 |
| APQVAGAATAVL | ASSLSDRVNSDYT | 0.88594353 |
| ARRDPAAAAAILPTDA | ASSLSDRVNSDYT | 0.8824413 |
| DAAARDVLAEAGLAEYF | ASSLSDRVNSDYT | 0.8662357 |
| DDQIVLSAKANLAKVEQ | ASSLSDRVNSDYT | 0.86796534 |
| DELLPDTGEDSDEDGH | ASSLSDRVNSDYT | 0.8790237 |
| ELPEAPANRATVPQD | ASSLSDRVNSDYT | 0.89335334 |
| FGQSKIERVTGIG | ASSLSDRVNSDYT | 0.8827517 |
| GAGGTGSTAGGAGGAGGAGGI | ASSLSDRVNSDYT | 0.908067 |
| GHGGTGAAGGNGGTGGAGGS( | ASSLSDRVNSDYT | 0.89970493 |
| GNGGNGGAGGEGGPGGAGGA( | ASSLSDRVNSDYT | 0.8944129 |
| GSTAGNGGNGGNATNGTIAG( | ASSLSDRVNSDYT | 0.48957524 |
| GTGGDGGNGGNGGTGG | ASSLSDRVNSDYT | 0.7793104 |
| GVRRPAGIPADAPV | ASSLSDRVNSDYT | 0.8293688 |
| GYRQVTDAHLLTLARR | ASSLSDRVNSDYT | 0.8718701 |
| IAKMGQDGHDRG | ASSLSDRVNSDYT | 0.53481287 |
| KAEPSIEIQYT | ASSLSDRVNSDYT | 0.8910432 |
| KPVRIRMAADEDTVAVT | ASSLSDRVNSDYT | 0.868245 |

| | | |
|---|---|---|
| MPPRIAGMRLLVIKPEP | ASSLSDRVNSDYT | 0.8333917 |
| MKFYAVALFRSRGVPPTR | ASSLSDRVNSDYT | 0.8962724 |
| NAGGFGILGAVAHSP | ASSLSDRVNSDYT | 0.86937064 |
| NEGREITLAMAEM | ASSLSDRVNSDYT | 0.83753264 |
| NELVEWSHPLGHRWRWE | ASSLSDRVNSDYT | 0.8226851 |
| NLGAGGTGGNGGNPG | ASSLSDRVNSDYT | 0.9047045 |
| NTEWPKWETVMAADPVD | ASSLSDRVNSDYT | 0.77931124 |
| PASAAPAETQP | ASSLSDRVNSDYT | 0.90073013 |
| QQSSDGKNGVRAAEITGE | ASSLSDRVNSDYT | 0.8331933 |
| RAPDAVTTSWWKE | ASSLSDRVNSDYT | 0.9026617 |
| SGFGNVGAHTSGWFNQ | ASSLSDRVNSDYT | 0.73061854 |
| SGVAGAVGQAASVGGLKV | ASSLSDRVNSDYT | 0.8969742 |
| SNGGAGTGSAGGNGGT | ASSLSDRVNSDYT | 0.87098616 |
| TGGAGGEGGAAG | ASSLSDRVNSDYT | 0.9099167 |
| TGGDGGAAGAGGAPGHG | ASSLSDRVNSDYT | 0.8669722 |
| TMPPLLAAIANAGRHGV | ASSLSDRVNSDYT | 0.8925111 |
| TSSTAGNSGGA | ASSLSDRVNSDYT | 0.8616634 |
| VPATVPVGILC | ASSLSDRVNSDYT | 0.90093255 |
| VQRPDSGGVVLTGRIS | ASSLSDRVNSDYT | 0.7643975 |
| WTPNPVPATKGVPAGLVQAT | ASSLSDRVNSDYT | 0.8591422 |
| YALKRALAKFSDRFDVVII | ASSLSDRVNSDYT | 0.73547965 |
| YLIGSVIV | ASSLSDRVNSDYT | 0.88060933 |
| YGQLVAVID | ASSLSDRVNSDYT | 0.8946553 |
| VTVGGMTLPA | ASSLSDRVNSDYT | 0.6548304 |
| VLALPAV | ASSLSDRVNSDYT | 0.9000635 |
| VIPPTLNYET | ASSLSDRVNSDYT | 0.84778345 |
| VGMGGMHPSSG | ASSLSDRVNSDYT | 0.8193381 |
| VGLAGGAATA | ASSLSDRVNSDYT | 0.8998878 |
| VGGAGGAGGSGP | ASSLSDRVNSDYT | 0.86781615 |
| VFAAFMV | ASSLSDRVNSDYT | 0.89987457 |
| TLYDKLR | ASSLSDRVNSDYT | 0.85542643 |
| TLAVFTVL | ASSLSDRVNSDYT | 0.8593245 |
| TGSVQPARV | ASSLSDRVNSDYT | 0.8893639 |
| TGGGAGSGGT | ASSLSDRVNSDYT | 0.6620028 |
| SHLETSL | ASSLSDRVNSDYT | 0.76311374 |
| SFTTGMT | ASSLSDRVNSDYT | 0.9051664 |
| QVEVAKRMGVRQPTVS | ASSLSDRVNSDYT | 0.88519955 |
| QGSGGPALAM | ASSLSDRVNSDYT | 0.88136333 |
| MLAPSNP | ASSLSDRVNSDYT | 0.8094177 |
| LVNNAAI | ASSLSDRVNSDYT | 0.7963191 |
| LTGEINTLA | ASSLSDRVNSDYT | 0.8219559 |
| LSIFGPV | ASSLSDRVNSDYT | 0.8886266 |
| LLALLANL | ASSLSDRVNSDYT | 0.89695525 |
| LISNASDALDKLR | ASSLSDRVNSDYT | 0.8911029 |
| LINEAVP | ASSLSDRVNSDYT | 0.7267517 |
| LGYAFSGL | ASSLSDRVNSDYT | 0.89198923 |
| LDFVAVAL | ASSLSDRVNSDYT | 0.8535853 |
| LAQPGAE | ASSLSDRVNSDYT | 0.8741766 |
| ISNLGAVTGF | ASSLSDRVNSDYT | 0.86086583 |
| HQEDPGRI | ASSLSDRVNSDYT | 0.84379524 |
| GTPGEPTP | ASSLSDRVNSDYT | 0.82992256 |

| | | |
|---|---|---|
| GFFNSGS | ASSLSDRVNSDYT | 0.79955363 |
| GAGGGGLTAGAN | ASSLSDRVNSDYT | 0.8696765 |
| FVDALVNM | ASSLSDRVNSDYT | 0.89083767 |
| FLTVAPEL | ASSLSDRVNSDYT | 0.8753753 |
| FETLTAL | ASSLSDRVNSDYT | 0.90933853 |
| DVINPIEL | ASSLSDRVNSDYT | 0.8997631 |
| APVSGAPV | ASSLSDRVNSDYT | 0.88304734 |
| ALVGGLAW | ASSLSDRVNSDYT | 0.45716733 |
| ALAAAAATAI | ASSLSDRVNSDYT | 0.7456612 |
| AFGLGLLL | ASSLSDRVNSDYT | 0.6935738 |
| AFFAAASL | ASSLSDRVNSDYT | 0.8526744 |
| AAAALPAFSPP | ASSLSDRVNSDYT | 0.29356956 |
| VVLAPAL | ASSLSDRVNSDYT | 0.87811613 |
| TIGHVDHGKTTL | ASSLSDRVNSDYT | 0.843682 |
| LISNASDALDK | ASSLSDRVNSDYT | 0.80669004 |
| LISNASDAL | ASSLSDRVNSDYT | 0.8928113 |
| ISNASDALDK | ASSLSDRVNSDYT | 0.8487021 |
| NQAETLVYQTE | ASSLSDRVNSDYT | 0.7319288 |
| TGGTGGTAGAG | ASSLSDRVNSDYT | 0.72631073 |
| TGSTAGGAGGA | ASSLSDRVNSDYT | 0.90579057 |
| DPLAVPVAAGPV | ASSLSDRVNSDYT | 0.7375692 |
| GGTGGTPGNGGN | ASSLSDRVNSDYT | 0.9061388 |
| PAGLAGVDPATL | ASSLSDRVNSDYT | 0.74697155 |
| GNNLVVEAPHAGASV | ASSLSDRVNSDYT | 0.9052192 |
| KVTRIMSASTLEQPA | ASSLSDRVNSDYT | 0.6133348 |
| LGFALPIALPRRTVV | ASSLSDRVNSDYT | 0.89747596 |
| TRLGNLTARKLVVLV | ASSLSDRVNSDYT | 0.86503726 |
| WSQQRRLIYFPSAGP | ASSLSDRVNSDYT | 0.9124634 |
| GPNGNPDPMAAAVDIR | ASSLSDRVNSDYT | 0.859645 |
| TQRCVSVADLRAAASR | ASSLSDRVNSDYT | 0.89198637 |
| QFVLDVAEHPVEVLTETV | ASSLSDRVNSDYT | 0.8221717 |
| GGAGGTGGAGGNGGTQAGDG | ASSLSDRVNSDYT | 0.89183867 |
| PLLATLRAIAVDVDAKPAQV | ASSLSDRVNSDYT | 0.7086366 |
| APKFRTGGYMIDFWGVGYQV | ASSLSDRVNSDYT | 0.82870764 |
| GAGGAGGAGGAAGGPGGTGG | ASSLSDRVNSDYT | 0.69063044 |
| IGALGLLATDVAPAGSVAFG | ASSLSDRVNSDYT | 0.8582829 |
| AGGAGGIGGNGGAGGAGGVL | ASSLSDRVNSDYT | 0.89423436 |
| ADELIGLL | ASSLSDRVNSDYT | 0.33191425 |
| ADRALKA | ASSLSDRVNSDYT | 0.89444304 |
| AGAVGQVGGAGGA | ASSLSDRVNSDYT | 0.75986457 |
| AGGGGGLGGAP | ASSLSDRVNSDYT | 0.70390755 |
| AGGLLYGNGGAG | ASSLSDRVNSDYT | 0.89051497 |
| AGLESIGL | ASSLSDRVNSDYT | 0.84868735 |
| AGLKTGII | ASSLSDRVNSDYT | 0.89161015 |
| ASLAALLL | ASSLSDRVNSDYT | 0.7676336 |
| ASVINISSI | ASSLSDRVNSDYT | 0.83287966 |
| ATPPGPA | ASSLSDRVNSDYT | 0.87501657 |
| ATVASALV | ASSLSDRVNSDYT | 0.44340372 |
| ATVASLIL | ASSLSDRVNSDYT | 0.61649036 |
| AVNGPVLIRH | ASSLSDRVNSDYT | 0.90781844 |
| DGAAGAVGI | ASSLSDRVNSDYT | 0.49378783 |

| | | |
|---|---|---|
| DPLLHGI | ASSLSDRVNSDYT | 0.6025562 |
| ELGAALGAL | ASSLSDRVNSDYT | 0.46938625 |
| ELGRTQAV | ASSLSDRVNSDYT | 0.4341663 |
| FVAGGYLS | ASSLSDRVNSDYT | 0.47463167 |
| GASGGAGGAGGW | ASSLSDRVNSDYT | 0.55275655 |
| GGAGAGGDGGAGGI | ASSLSDRVNSDYT | 0.6083844 |
| GTGHNGSQPGMGGNG | ASSLSDRVNSDYT | 0.50031865 |
| GYAAASASAS | ASSLSDRVNSDYT | 0.44767642 |
| HQVDVAAVQQG | ASSLSDRVNSDYT | 0.5305575 |
| IGAAIAHT | ASSLSDRVNSDYT | 0.45612264 |
| IGMIEVAQL | ASSLSDRVNSDYT | 0.4971642 |
| IRLAPEHTSAMD | ASSLSDRVNSDYT | 0.47823632 |
| IYNVSVL | ASSLSDRVNSDYT | 0.5268792 |
| KQGFLGM | ASSLSDRVNSDYT | 0.51329756 |
| LESVITQL | ASSLSDRVNSDYT | 0.4256399 |
| LGLGALVVV | ASSLSDRVNSDYT | 0.4541624 |
| LIGNGGNGGSGGI | ASSLSDRVNSDYT | 0.5245837 |
| LNAINAPTL | ASSLSDRVNSDYT | 0.45324612 |
| LQLLNTGV | ASSLSDRVNSDYT | 0.55353 |
| LQTSIEF | ASSLSDRVNSDYT | 0.5235066 |
| MSFAVSEL | ASSLSDRVNSDYT | 0.47195092 |
| NTGALGL | ASSLSDRVNSDYT | 0.32079646 |
| PPGAGGAGGAGTT | ASSLSDRVNSDYT | 0.34128907 |
| QLVGLGI | ASSLSDRVNSDYT | 0.45927694 |
| RFLEQVHQL | ASSLSDRVNSDYT | 0.43134305 |
| RGVGYVLR | ASSLSDRVNSDYT | 0.3322681 |
| SATGVFAGVI | ASSLSDRVNSDYT | 0.5779641 |
| SLPAITLP | ASSLSDRVNSDYT | 0.4897083 |
| STALAFTV | ASSLSDRVNSDYT | 0.49377185 |
| SVLTVGLGK | ASSLSDRVNSDYT | 0.5356087 |
| TALAGAGSGI | ASSLSDRVNSDYT | 0.5017379 |
| TGLPFTGI | ASSLSDRVNSDYT | 0.46285367 |
| VAEHVTV | ASSLSDRVNSDYT | 0.42447722 |
| VDIVLNSL | ASSLSDRVNSDYT | 0.55134284 |
| VLTGGGAVL | ASSLSDRVNSDYT | 0.35371047 |
| VNVGLAYGI | ASSLSDRVNSDYT | 0.5417174 |
| VSSAAVGAV | ASSLSDRVNSDYT | 0.37049207 |
| VTSVYEAGV | ASSLSDRVNSDYT | 0.53220755 |
| WVALDRGA | ASSLSDRVNSDYT | 0.41825402 |
| YLVGSVVI | ASSLSDRVNSDYT | 0.51528716 |
| GPPGTGKTLL | ASSLSDRVNSDYT | 0.4894286 |
| GVGMTKF | ASSLSDRVNSDYT | 0.62477446 |
| IGIGGSDL | ASSLSDRVNSDYT | 0.4577343 |
| IGTLLAEL | ASSLSDRVNSDYT | 0.5246477 |
| IYTSGTT | ASSLSDRVNSDYT | 0.42849112 |
| LSFPAGV | ASSLSDRVNSDYT | 0.6056256 |
| LYTSGTT | ASSLSDRVNSDYT | 0.39433807 |
| VGAFPFGF | ASSLSDRVNSDYT | 0.45307106 |
| VGAGTVEF | ASSLSDRVNSDYT | 0.38689685 |
| VGKSTLLL | ASSLSDRVNSDYT | 0.53017414 |
| AALAHAGGGIV | ASSLSDRVNSDYT | 0.50026095 |

| | | |
|---|---|---|
| AALVAVSLL | ASSLSDRVNSDYT | 0.4088307 |
| AAVSRAVAAL | ASSLSDRVNSDYT | 0.5010222 |
| AGGSSGIGGF | ASSLSDRVNSDYT | 0.51005113 |
| AGLTVANANG | ASSLSDRVNSDYT | 0.45278907 |
| AGLVGAVL | ASSLSDRVNSDYT | 0.55970913 |
| ALSQAFTV | ASSLSDRVNSDYT | 0.5048694 |
| ALTFSVSL | ASSLSDRVNSDYT | 0.54288876 |
| ALTGAALVAL | ASSLSDRVNSDYT | 0.4609315 |
| ASFSIGAGVL | ASSLSDRVNSDYT | 0.4192053 |
| DGSVYKI | ASSLSDRVNSDYT | 0.56045485 |
| EQLAEIV | ASSLSDRVNSDYT | 0.45808175 |
| FAVSELTVM | ASSLSDRVNSDYT | 0.40773976 |
| GAAPLAAGL | ASSLSDRVNSDYT | 0.54386 |
| GADGGDGGSGGAGG | ASSLSDRVNSDYT | 0.85320437 |
| GFGLDTGGAGG | ASSLSDRVNSDYT | 0.8586074 |
| GLASGGLL | ASSLSDRVNSDYT | 0.7331877 |
| GRIAGAAVLL | ASSLSDRVNSDYT | 0.69759274 |
| GSAGSLNTSF | ASSLSDRVNSDYT | 0.78975976 |
| GTGQIGI | ASSLSDRVNSDYT | 0.8657962 |
| GTLIALIL | ASSLSDRVNSDYT | 0.8792751 |
| IAATMGFVVG | ASSLSDRVNSDYT | 0.8531477 |
| IFAQALL | ASSLSDRVNSDYT | 0.6876631 |
| IGAAVALGF | ASSLSDRVNSDYT | 0.8567729 |
| IGLTPLAL | ASSLSDRVNSDYT | 0.7956254 |
| ILGSNTSTL | ASSLSDRVNSDYT | 0.8086922 |
| IVFSLTVL | ASSLSDRVNSDYT | 0.84603703 |
| LASKATGF | ASSLSDRVNSDYT | 0.82111824 |
| LASVALIL | ASSLSDRVNSDYT | 0.7782121 |
| LFAVGLTL | ASSLSDRVNSDYT | 0.6774466 |
| LFNGGIGTYI | ASSLSDRVNSDYT | 0.8622121 |
| LTLGAGII | ASSLSDRVNSDYT | 0.8760379 |
| LVAAALGDI | ASSLSDRVNSDYT | 0.86871135 |
| NTAFVQLGI | ASSLSDRVNSDYT | 0.8918922 |
| QLFSGAEL | ASSLSDRVNSDYT | 0.86961985 |
| SAATFITL | ASSLSDRVNSDYT | 0.86126924 |
| SNYGLANLGV | ASSLSDRVNSDYT | 0.27156037 |
| SYGGAYAVM | ASSLSDRVNSDYT | 0.5900458 |
| TTGVGGLTLG | ASSLSDRVNSDYT | 0.7164465 |
| TTQTIYI | ASSLSDRVNSDYT | 0.79775476 |
| VAAGAGLGAVI | ASSLSDRVNSDYT | 0.33725703 |
| VAAVVTLL | ASSLSDRVNSDYT | 0.8530356 |
| VGFLGGLI | ASSLSDRVNSDYT | 0.7368995 |
| VGVIPILL | ASSLSDRVNSDYT | 0.7315607 |
| VIFASSLI | ASSLSDRVNSDYT | 0.86974317 |
| VKGGDGGAAAG | ASSLSDRVNSDYT | 0.7664187 |
| VTKTFTNI | ASSLSDRVNSDYT | 0.6623813 |
| VVAAAILL | ASSLSDRVNSDYT | 0.7369833 |
| YAALDVEL | ASSLSDRVNSDYT | 0.8807597 |
| LASVALLL | ASSLSDRVNSDYT | 0.633222 |
| LDAVHRV | ASSLSDRVNSDYT | 0.86579645 |
| QGLSSSIF | ASSLSDRVNSDYT | 0.8068354 |

| | | |
|---|---|---|
| RGTFANIRL | ASSLSDRVNSDYT | 0.87891775 |
| RQLWWGHRI | ASSLSDRVNSDYT | 0.5534563 |
| YGPPGTGKTLL | ASSLSDRVNSDYT | 0.86615694 |
| AALVGAPANVA | ASSLSDRVNSDYT | 0.8101449 |
| AATFRRTE | ASSLSDRVNSDYT | 0.895844 |
| AGALGGGI | ASSLSDRVNSDYT | 0.798615 |
| AGGAGGAGGIGAG | ASSLSDRVNSDYT | 0.845332 |
| AGSGALGL | ASSLSDRVNSDYT | 0.82843494 |
| AQIADQI | ASSLSDRVNSDYT | 0.87767756 |
| ASAVAVTYL | ASSLSDRVNSDYT | 0.5445092 |
| ASTVLGGTGG | ASSLSDRVNSDYT | 0.7801037 |
| AVGGEPAK | ASSLSDRVNSDYT | 0.69505155 |
| CEQGTFGGTC | ASSLSDRVNSDYT | 0.8130335 |
| DEGKAPAG | ASSLSDRVNSDYT | 0.85505235 |
| DTVVFPAL | ASSLSDRVNSDYT | 0.23092684 |
| FEGITMAG | ASSLSDRVNSDYT | 0.85318595 |
| FPLVVQD | ASSLSDRVNSDYT | 0.70760983 |
| GAFTTPQ | ASSLSDRVNSDYT | 0.6804432 |
| GFGGFGTAGGVGGA | ASSLSDRVNSDYT | 0.86884093 |
| GFLAQFL | ASSLSDRVNSDYT | 0.8367765 |
| GGAGGAGGNAGVA | ASSLSDRVNSDYT | 0.87299746 |
| GGVGTAGAGGAGGAGG | ASSLSDRVNSDYT | 0.79117036 |
| GNGGGGDAGNAGSGGN | ASSLSDRVNSDYT | 0.8099386 |
| GTGGAGGAGNDAGS | ASSLSDRVNSDYT | 0.83047307 |
| GTTNTGWFNS | ASSLSDRVNSDYT | 0.5248393 |
| IFMGPLFTL | ASSLSDRVNSDYT | 0.72280896 |
| ILAHAIEAI | ASSLSDRVNSDYT | 0.8792938 |
| IRSAIREAL | ASSLSDRVNSDYT | 0.78917694 |
| ITFAGGG | ASSLSDRVNSDYT | 0.84060264 |
| ITGMSGHQEV | ASSLSDRVNSDYT | 0.62987345 |
| IVAGLACTL | ASSLSDRVNSDYT | 0.56770086 |
| IVGGGLAGA | ASSLSDRVNSDYT | 0.67543036 |
| IVTGVLNI | ASSLSDRVNSDYT | 0.85191584 |
| LAQQGCI | ASSLSDRVNSDYT | 0.8046305 |
| LCLHVEN | ASSLSDRVNSDYT | 0.7886249 |
| LDAISDGS | ASSLSDRVNSDYT | 0.54865634 |
| LGGFAAML | ASSLSDRVNSDYT | 0.7761178 |
| LGGKNPMI | ASSLSDRVNSDYT | 0.6968991 |
| LGPKGVAAIL | ASSLSDRVNSDYT | 0.7138284 |
| LMVGVSI | ASSLSDRVNSDYT | 0.8115309 |
| LVGAAFVL | ASSLSDRVNSDYT | 0.8015189 |
| LVLAVAPL | ASSLSDRVNSDYT | 0.6962371 |
| MAAIAVIDAA | ASSLSDRVNSDYT | 0.69827944 |
| NNGINAELQ | ASSLSDRVNSDYT | 0.73384225 |
| RGEPLVLVHG | ASSLSDRVNSDYT | 0.8005223 |
| RGGAGGAAGLI | ASSLSDRVNSDYT | 0.79122895 |
| RSYRMHPAV | ASSLSDRVNSDYT | 0.8797688 |
| SAAHLLAGQLA | ASSLSDRVNSDYT | 0.8321323 |
| SAAVGSY | ASSLSDRVNSDYT | 0.8511647 |
| SPHAVLHRI | ASSLSDRVNSDYT | 0.23577864 |
| SQIAKVL | ASSLSDRVNSDYT | 0.3945107 |

| | | |
|---|---|---|
| TALGHLVP | ASSLSDRVNSDYT | 0.7548842 |
| TAPTTSGAA | ASSLSDRVNSDYT | 0.71816033 |
| TDGDDGNGGSPGT | ASSLSDRVNSDYT | 0.5213794 |
| TDGLGEN | ASSLSDRVNSDYT | 0.8311156 |
| TGGPPLPGQA | ASSLSDRVNSDYT | 0.67856973 |
| TIPAITI | ASSLSDRVNSDYT | 0.6275735 |
| TIPGFTI | ASSLSDRVNSDYT | 0.82259405 |
| TSIASGALA | ASSLSDRVNSDYT | 0.6636284 |
| TVTLPGAAI | ASSLSDRVNSDYT | 0.69256306 |
| VASGLSAALP | ASSLSDRVNSDYT | 0.6808804 |
| VGAGVIGI | ASSLSDRVNSDYT | 0.8427936 |
| VGPKRFGV | ASSLSDRVNSDYT | 0.5410002 |
| VNVAFPD | ASSLSDRVNSDYT | 0.80958974 |
| VSMTNSGVS | ASSLSDRVNSDYT | 0.6991747 |
| VTDVLGDLSV | ASSLSDRVNSDYT | 0.8613452 |
| VVVVGAGFAGL | ASSLSDRVNSDYT | 0.46749154 |
| VYTVGTVV | ASSLSDRVNSDYT | 0.8254267 |
| WTTFSGP | ASSLSDRVNSDYT | 0.71947074 |
| YDELVEAF | ASSLSDRVNSDYT | 0.8860905 |
| YGTGGAGGNGG | ASSLSDRVNSDYT | 0.7097694 |
| YTNVMAV | ASSLSDRVNSDYT | 0.76029146 |
| FLEQVHQL | ASSLSDRVNSDYT | 0.6962978 |
| GLSLLAVAV | ASSLSDRVNSDYT | 0.80817443 |
| LEQVHQL | ASSLSDRVNSDYT | 0.4805519 |
| LNAAGAI | ASSLSDRVNSDYT | 0.6941844 |
| PAVHASL | ASSLSDRVNSDYT | 0.42184862 |
| TELEDDA | ASSLSDRVNSDYT | 0.77439415 |
| TPPGPPN | ASSLSDRVNSDYT | 0.796304 |
| TPPPPPPP | ASSLSDRVNSDYT | 0.5302065 |
| TQSPALI | ASSGTGENYAEQF | 0.7695389 |
| EFAAAFPQ | ASSGTGENYAEQF | 0.75859356 |
| LLSGNLDVL | ASSGTGENYAEQF | 0.61146784 |
| GPANLGNYNI | ASSGTGENYAEQF | 0.85574716 |
| TLGPTLP | ASSGTGENYAEQF | 0.7342994 |
| VYAYSSCCK | ASSGTGENYAEQF | 0.8414606 |
| APAPAPAPNI | ASSGTGENYAEQF | 0.74763 |
| PGGPVPIP | ASSGTGENYAEQF | 0.6560263 |
| FLTVALEP | ASSGTGENYAEQF | 0.81025076 |
| SQYFGANL | ASSGTGENYAEQF | 0.59347904 |
| GGVKPAA | ASSGTGENYAEQF | 0.6554926 |
| GGTGGAGGAGGW | ASSGTGENYAEQF | 0.85176206 |
| GVGGVAGLGGAGG | ASSGTGENYAEQF | 0.6597893 |
| VLLVLATTI | ASSGTGENYAEQF | 0.7085186 |
| IGIFAAML | ASSGTGENYAEQF | 0.5022931 |
| AVGDLID | ASSGTGENYAEQF | 0.51024663 |
| NISPLPL | ASSGTGENYAEQF | 0.5790013 |
| VTAVFVAI | ASSGTGENYAEQF | 0.6749903 |
| SGDDNDAAL | ASSGTGENYAEQF | 0.70855606 |
| AVAALAHS | ASSGTGENYAEQF | 0.6617442 |
| GGVTVMATP | ASSGTGENYAEQF | 0.45981854 |
| AGLAANVRD | ASSGTGENYAEQF | 0.6388833 |

| | | |
|---|---|---|
| TNRPDLIDPALLRPG | ASSGTGENYAEQF | 0.49793965 |
| LANLGVTAL | ASSGTGENYAEQF | 0.52889484 |
| TGERIEL | ASSGTGENYAEQF | 0.61437666 |
| NGEELAA | ASSGTGENYAEQF | 0.5297713 |
| VLSLVWG | ASSGTGENYAEQF | 0.48327214 |
| IGYALWL | ASSGTGENYAEQF | 0.5227229 |
| LSLLLAPL | ASSGTGENYAEQF | 0.5872165 |
| TLDEALK | ASSGTGENYAEQF | 0.67933416 |
| TDGAGTAEGG | ASSGTGENYAEQF | 0.6679724 |
| NAPGGAGGNGG | ASSGTGENYAEQF | 0.7128472 |
| VSVFMGMGGNF | ASSGTGENYAEQF | 0.69110286 |
| LASYLAAL | ASSGTGENYAEQF | 0.6571344 |
| LVGGTPAL | ASSGTGENYAEQF | 0.26252422 |
| VRNADVL | ASSGTGENYAEQF | 0.4084531 |
| TCSSPFLVSAM | ASSGTGENYAEQF | 0.3895061 |
| VHITDAE | ASSGTGENYAEQF | 0.5834087 |
| EYAAPSF | ASSGTGENYAEQF | 0.27054912 |
| VVDAFHGGL | ASSGTGENYAEQF | 0.6262245 |
| ASVVDPQ | ASSGTGENYAEQF | 0.505025 |
| SYNEFGGLS | ASSGTGENYAEQF | 0.4407314 |
| DPDGSVY | ASSGTGENYAEQF | 0.67452466 |
| MVSTIHATE | ASSGTGENYAEQF | 0.4799846 |
| LAAGVADPV | ASSGTGENYAEQF | 0.44303218 |
| TKEQIAA | ASSGTGENYAEQF | 0.42219457 |
| SLASPAAL | ASSGTGENYAEQF | 0.7091843 |
| IGLVFYGV | ASSGTGENYAEQF | 0.33636293 |
| FLLDASNL | ASSGTGENYAEQF | 0.66977787 |
| YQVGNFP | ASSGTGENYAEQF | 0.609115 |
| ESLLSQST | ASSGTGENYAEQF | 0.69741803 |
| LFTVGFAAVG | ASSGTGENYAEQF | 0.4027631 |
| ATPPTPT | ASSGTGENYAEQF | 0.6757504 |
| TDLATLG | ASSGTGENYAEQF | 0.5689939 |
| SAGGTGR | ASSGTGENYAEQF | 0.72474325 |
| GFSNSGS | ASSGTGENYAEQF | 0.5822951 |
| GGNGGSGGTSV | ASSGTGENYAEQF | 0.6030104 |
| GTSVATGGAGN | ASSGTGENYAEQF | 0.51518095 |
| NGGNGGSGGTS | ASSGTGENYAEQF | 0.7170322 |
| AVVKFPH | ASSGTGENYAEQF | 0.34763035 |
| TPPPPPP | ASSGTGENYAEQF | 0.60920954 |
| AAPAAAAAPA | ASSGTGENYAEQF | 0.43547624 |
| IALNEDL | ASSGTGENYAEQF | 0.63651246 |
| IVNVSSM | ASSGTGENYAEQF | 0.66843176 |
| TGIPPAPR | ASSGTGENYAEQF | 0.28257293 |
| FDLGGGTF | ASSGTGENYAEQF | 0.70704293 |
| FDLGGGTFDVS | ASSGTGENYAEQF | 0.53554285 |
| NEPTAAA | ASSGTGENYAEQF | 0.41967225 |
| NEPTAAALA | ASSGTGENYAEQF | 0.6660546 |
| GSIINISSI | ASSGTGENYAEQF | 0.5327988 |
| VAPGNAGTA | ASSGTGENYAEQF | 0.70789105 |
| TGGGASLE | ASSGTGENYAEQF | 0.40451202 |
| VAVYNHYKRI | ASSGTGENYAEQF | 0.6608926 |

| | | |
|---|---|---|
| TDPAPATT | ASSGTGENYAEQF | 0.65880144 |
| TNRPDILDPALLRPG | ASSGTGENYAEQF | 0.29670978 |
| LTTTTTS | ASSGTGENYAEQF | 0.37980756 |
| PTGSGKTL | ASSGTGENYAEQF | 0.7003566 |
| TGSGKTL | ASSGTGENYAEQF | 0.6580359 |
| TIGHVDHGK | ASSGTGENYAEQF | 0.6772131 |
| TIGHVDHGKTT | ASSGTGENYAEQF | 0.48013276 |
| GHVDHGKTT | ASSGTGENYAEQF | 0.46460158 |
| AQFEHTLL | ASSGTGENYAEQF | 0.58822453 |
| IVNQDEL | ASSGTGENYAEQF | 0.63966405 |
| TNGSQFF | ASSGTGENYAEQF | 0.68618023 |
| TGKGIVE | ASSGTGENYAEQF | 0.6478268 |
| LISNASDA | ASSGTGENYAEQF | 0.4495245 |
| AELRTLGELTKIAKAHGAQ | ASSGTGENYAEQF | 0.62814957 |
| APNAVAAAEPSW | ASSGTGENYAEQF | 0.5546803 |
| APQVAGAATAVL | ASSGTGENYAEQF | 0.5809212 |
| ARRDPAAAAAILPTDA | ASSGTGENYAEQF | 0.61514425 |
| DAAARDVLAEAGLAEYF | ASSGTGENYAEQF | 0.5957104 |
| DDQIVLSAKANLAKVEQ | ASSGTGENYAEQF | 0.5492641 |
| DELLPDTGEDSDEDGH | ASSGTGENYAEQF | 0.44821066 |
| ELPEAPANRATVPQD | ASSGTGENYAEQF | 0.6477155 |
| FGQSKIERVTGIG | ASSGTGENYAEQF | 0.6736994 |
| GAGGTGSTAGGAGGAGGAGGI | ASSGTGENYAEQF | 0.6721954 |
| GHGGTGAAGGNGGTGGAGGS( | ASSGTGENYAEQF | 0.65925837 |
| GNGGNGGAGGEGGPGGAGGA( | ASSGTGENYAEQF | 0.65806735 |
| GSTAGNGGNGGNATNGTIAG( | ASSGTGENYAEQF | 0.6039862 |
| GTGGDGGNGGNGGTGG | ASSGTGENYAEQF | 0.2906827 |
| GVRRPAGIPADAPV | ASSGTGENYAEQF | 0.33718726 |
| GYRQVTDAHLLTLARR | ASSGTGENYAEQF | 0.54539174 |
| IAKMGQDGHDRG | ASSGTGENYAEQF | 0.54414463 |
| KAEPSIEIQYT | ASSGTGENYAEQF | 0.3147377 |
| KPVRIRMAADEDTVAVT | ASSGTGENYAEQF | 0.5959731 |
| MPPRIAGMRLLVIKPEP | ASSGTGENYAEQF | 0.5257727 |
| MKFYAVALFRSRGVPPTR | ASSGTGENYAEQF | 0.53684825 |
| NAGGFGILGAVAHSP | ASSGTGENYAEQF | 0.66985565 |
| NEGREITLAMAEM | ASSGTGENYAEQF | 0.55415 |
| NELVEWSHPLGHRWRWE | ASSGTGENYAEQF | 0.44224888 |
| NLGAGGTGGNGGNPG | ASSGTGENYAEQF | 0.48473835 |
| NTEWPKWETVMAADPVD | ASSGTGENYAEQF | 0.662369 |
| PASAAPAETQP | ASSGTGENYAEQF | 0.39839447 |
| QQSSDGKNGVRAAEITGE | ASSGTGENYAEQF | 0.65024006 |
| RAPDAVTTSWWKE | ASSGTGENYAEQF | 0.42112976 |
| SGFGNVGAHTSGWFNQ | ASSGTGENYAEQF | 0.66662276 |
| SGVAGAVGQAASVGGLKV | ASSGTGENYAEQF | 0.4069676 |
| SNGGAGTGSAGGNGGT | ASSGTGENYAEQF | 0.6371063 |
| TGGAGGEGGAAG | ASSGTGENYAEQF | 0.5903884 |
| TGGDGGAAGAGGAPGHG | ASSGTGENYAEQF | 0.69865394 |
| TMPPLLAAIANAGRHGV | ASSGTGENYAEQF | 0.52141076 |
| TSSTAGNSGGA | ASSGTGENYAEQF | 0.60305876 |
| VPATVPVGILC | ASSGTGENYAEQF | 0.5374585 |
| VQRPDSGGVVLTGRIS | ASSGTGENYAEQF | 0.68710065 |

| | | |
|---|---|---|
| WTPNPVPATKGVPAGLVQAT | ASSGTGENYAEQF | 0.40676284 |
| YALKRALAKFSDRFDVVII | ASSGTGENYAEQF | 0.5454864 |
| YLIGSVIV | ASSGTGENYAEQF | 0.3461473 |
| YGQLVAVID | ASSGTGENYAEQF | 0.5382954 |
| VTVGGMTLPA | ASSGTGENYAEQF | 0.6410035 |
| VLALPAV | ASSGTGENYAEQF | 0.32885328 |
| VIPPTLNYET | ASSGTGENYAEQF | 0.6543234 |
| VGMGGMHPSSG | ASSGTGENYAEQF | 0.5030334 |
| VGLAGGAATA | ASSGTGENYAEQF | 0.489293 |
| VGGAGGAGGSGP | ASSGTGENYAEQF | 0.6529262 |
| VFAAFMV | ASSGTGENYAEQF | 0.5898833 |
| TLYDKLR | ASSGTGENYAEQF | 0.6659835 |
| TLAVFTVL | ASSGTGENYAEQF | 0.51135457 |
| TGSVQPARV | ASSGTGENYAEQF | 0.5144085 |
| TGGGAGSGGT | ASSGTGENYAEQF | 0.5804625 |
| SHLETSL | ASSGTGENYAEQF | 0.47765446 |
| SFTTGMT | ASSGTGENYAEQF | 0.5030135 |
| QVEVAKRMGVRQPTVS | ASSGTGENYAEQF | 0.6653285 |
| QGSGGPALAM | ASSGTGENYAEQF | 0.12626095 |
| MLAPSNP | ASSGTGENYAEQF | 0.12583214 |
| LVNNAAI | ASSGTGENYAEQF | 0.12314896 |
| LTGEINTLA | ASSGTGENYAEQF | 0.12597592 |
| LSIFGPV | ASSGTGENYAEQF | 0.12437014 |
| LLALLANL | ASSGTGENYAEQF | 0.13018417 |
| LISNASDALDKLR | ASSGTGENYAEQF | 0.12635522 |
| LINEAVP | ASSGTGENYAEQF | 0.12559313 |
| LGYAFSGL | ASSGTGENYAEQF | 0.12689763 |
| LDFVAVAL | ASSGTGENYAEQF | 0.12597069 |
| LAQPGAE | ASSGTGENYAEQF | 0.12387099 |
| ISNLGAVTGF | ASSGTGENYAEQF | 0.12588887 |
| HQEDPGRI | ASSGTGENYAEQF | 0.12523852 |
| GTPGEPTP | ASSGTGENYAEQF | 0.1270457 |
| GFFNSGS | ASSGTGENYAEQF | 0.122943185 |
| GAGGGGLTAGAN | ASSGTGENYAEQF | 0.12556607 |
| FVDALVNM | ASSGTGENYAEQF | 0.1257232 |
| FLTVAPEL | ASSGTGENYAEQF | 0.12669548 |
| FETLTAL | ASSGTGENYAEQF | 0.12559035 |
| DVINPIEL | ASSGTGENYAEQF | 0.13221663 |
| APVSGAPV | ASSGTGENYAEQF | 0.12797086 |
| ALVGGLAW | ASSGTGENYAEQF | 0.12398604 |
| ALAAAAATAI | ASSGTGENYAEQF | 0.122462526 |
| AFGLGLLL | ASSGTGENYAEQF | 0.12390473 |
| AFFAAASL | ASSGTGENYAEQF | 0.124349214 |
| AAAALPAFSPP | ASSGTGENYAEQF | 0.12657097 |
| VVLAPAL | ASSGTGENYAEQF | 0.12167575 |
| TIGHVDHGKTTL | ASSGTGENYAEQF | 0.12883146 |
| LISNASDALDK | ASSGTGENYAEQF | 0.12635711 |
| LISNASDAL | ASSGTGENYAEQF | 0.124906495 |
| ISNASDALDK | ASSGTGENYAEQF | 0.12925209 |
| NQAETLVYQTE | ASSGTGENYAEQF | 0.12361402 |
| TGGTGGTAGAG | ASSGTGENYAEQF | 0.12513496 |

| | | |
|---|---|---|
| TGSTAGGAGGA | ASSGTGENYAEQF | 0.124282256 |
| DPLAVPVAAGPV | ASSGTGENYAEQF | 0.12979138 |
| GGTGGTPGNGGN | ASSGTGENYAEQF | 0.123470336 |
| PAGLAGVDPATL | ASSGTGENYAEQF | 0.13270721 |
| GNNLVVEAPHAGASV | ASSGTGENYAEQF | 0.12357129 |
| KVTRIMSASTLEQPA | ASSGTGENYAEQF | 0.13628602 |
| LGFALPIALPRRTVV | ASSGTGENYAEQF | 0.12576187 |
| TRLGNLTARKLVVLV | ASSGTGENYAEQF | 0.12627499 |
| WSQQRRLIYFPSAGP | ASSGTGENYAEQF | 0.12560147 |
| GPNGNPDPMAAAVDIR | ASSGTGENYAEQF | 0.13505399 |
| TQRCVSVADLRAAASR | ASSGTGENYAEQF | 0.12847555 |
| QFVLDVAEHPVEVLTETV | ASSGTGENYAEQF | 0.13081309 |
| GGAGGTGGAGGNGGTQAGDG | ASSGTGENYAEQF | 0.12492477 |
| PLLATLRAIAVDVDAKPAQV | ASSGTGENYAEQF | 0.12647092 |
| APKFRTGGYMIDFWGVGYQVA | ASSGTGENYAEQF | 0.123595215 |
| GAGGAGGAGGAAGGPGGTGGT | ASSGTGENYAEQF | 0.1262417 |
| IGALGLLATDVAPAGSVAFGV | ASSGTGENYAEQF | 0.12318099 |
| AGGAGGIGGNGGAGGAGGVLM | ASSGTGENYAEQF | 0.12657706 |
| ADELIGLL | ASSGTGENYAEQF | 0.12612022 |
| ADRALKA | ASSGTGENYAEQF | 0.122361206 |
| AGAVGQVGGAGGA | ASSGTGENYAEQF | 0.124807134 |
| AGGGGGLGGAP | ASSGTGENYAEQF | 0.12255801 |
| AGGLLYGNGGAG | ASSGTGENYAEQF | 0.12438707 |
| AGLESIGL | ASSGTGENYAEQF | 0.12547049 |
| AGLKTGII | ASSGTGENYAEQF | 0.12499771 |
| ASLAALLL | ASSGTGENYAEQF | 0.12453287 |
| ASVINISSI | ASSGTGENYAEQF | 0.122547105 |
| ATPPGPA | ASSGTGENYAEQF | 0.12357445 |
| ATVASALV | ASSGTGENYAEQF | 0.12923746 |
| ATVASLIL | ASSGTGENYAEQF | 0.12311717 |
| AVNGPVLIRH | ASSGTGENYAEQF | 0.123594396 |
| DGAAGAVGI | ASSGTGENYAEQF | 0.12881228 |
| DPLLHGI | ASSGTGENYAEQF | 0.8244365 |
| ELGAALGAL | ASSGTGENYAEQF | 0.8542269 |
| ELGRTQAV | ASSGTGENYAEQF | 0.7559473 |
| FVAGGYLS | ASSGTGENYAEQF | 0.6600452 |
| GASGGAGGAGGW | ASSGTGENYAEQF | 0.748924 |
| GGAGAGGDGGAGGI | ASSGTGENYAEQF | 0.8700275 |
| GTGHNGSQPGMGGNG | ASSGTGENYAEQF | 0.8487354 |
| GYAAASASAS | ASSGTGENYAEQF | 0.8361734 |
| HQVDVAAVQQG | ASSGTGENYAEQF | 0.6321513 |
| IGAAIAHT | ASSGTGENYAEQF | 0.81921285 |
| IGMIEVAQL | ASSGTGENYAEQF | 0.75833035 |
| IRLAPEHTSAMD | ASSGTGENYAEQF | 0.77109975 |
| IYNVSVL | ASSGTGENYAEQF | 0.84735525 |
| KQGFLGM | ASSGTGENYAEQF | 0.7770721 |
| LESVITQL | ASSGTGENYAEQF | 0.77032995 |
| LGLGALVVV | ASSGTGENYAEQF | 0.7691039 |
| LIGNGGNGGSGGI | ASSGTGENYAEQF | 0.77944016 |
| LNAINAPTL | ASSGTGENYAEQF | 0.7840578 |
| LQLLNTGV | ASSGTGENYAEQF | 0.7844468 |

| | | |
|---|---|---|
| LQTSIEF | ASSGTGENYAEQF | 0.8819126 |
| MSFAVSEL | ASSGTGENYAEQF | 0.8612796 |
| NTGALGL | ASSGTGENYAEQF | 0.8598444 |
| PPGAGGAGGAGTT | ASSGTGENYAEQF | 0.20530835 |
| QLVGLGI | ASSGTGENYAEQF | 0.40972114 |
| RFLEQVHQL | ASSGTGENYAEQF | 0.8025297 |
| RGVGYVLR | ASSGTGENYAEQF | 0.731867 |
| SATGVFAGVI | ASSGTGENYAEQF | 0.5062564 |
| SLPAITLP | ASSGTGENYAEQF | 0.7875943 |
| STALAFTV | ASSGTGENYAEQF | 0.7469237 |
| SVLTVGLGK | ASSGTGENYAEQF | 0.6990836 |
| TALAGAGSGI | ASSGTGENYAEQF | 0.8433347 |
| TGLPFTGI | ASSGTGENYAEQF | 0.76655895 |
| VAEHVTV | ASSGTGENYAEQF | 0.69652736 |
| VDIVLNSL | ASSGTGENYAEQF | 0.72723615 |
| VLTGGGAVL | ASSGTGENYAEQF | 0.82452834 |
| VNVGLAYGI | ASSGTGENYAEQF | 0.5670424 |
| VSSAAVGAV | ASSGTGENYAEQF | 0.83680046 |
| VTSVYEAGV | ASSGTGENYAEQF | 0.6305089 |
| WVALDRGA | ASSGTGENYAEQF | 0.85978514 |
| YLVGSVVI | ASSGTGENYAEQF | 0.5513504 |
| GPPGTGKTLL | ASSGTGENYAEQF | 0.85053897 |
| GVGMTKF | ASSGTGENYAEQF | 0.790895 |
| IGIGGSDL | ASSGTGENYAEQF | 0.88844824 |
| IGTLLAEL | ASSGTGENYAEQF | 0.7554543 |
| IYTSGTT | ASSGTGENYAEQF | 0.7822225 |
| LSFPAGV | ASSGTGENYAEQF | 0.55464387 |
| LYTSGTT | ASSGTGENYAEQF | 0.84696656 |
| VGAFPFGF | ASSGTGENYAEQF | 0.5062959 |
| VGAGTVEF | ASSGTGENYAEQF | 0.76889277 |
| VGKSTLLL | ASSGTGENYAEQF | 0.47628927 |
| AALAHAGGGIV | ASSGTGENYAEQF | 0.83222586 |
| AALVAVSLL | ASSGTGENYAEQF | 0.8368424 |
| AAVSRAVAAL | ASSGTGENYAEQF | 0.60077894 |
| AGGSSGIGGF | ASSGTGENYAEQF | 0.85260797 |
| AGLTVANANG | ASSGTGENYAEQF | 0.8167865 |
| AGLVGAVL | ASSGTGENYAEQF | 0.7216438 |
| ALSQAFTV | ASSGTGENYAEQF | 0.877219 |
| ALTFSVSL | ASSGTGENYAEQF | 0.7673809 |
| ALTGAALVAL | ASSGTGENYAEQF | 0.8558809 |
| ASFSIGAGVL | ASSGTGENYAEQF | 0.8118212 |
| DGSVYKI | ASSGTGENYAEQF | 0.72217596 |
| EQLAEIV | ASSGTGENYAEQF | 0.8286688 |
| FAVSELTVM | ASSGTGENYAEQF | 0.42272806 |
| GAAPLAAGL | ASSGTGENYAEQF | 0.67765355 |
| GADGGDGGSGGAGG | ASSGTGENYAEQF | 0.8678147 |
| GFGLDTGGAGG | ASSGTGENYAEQF | 0.71063215 |
| GLASGGLL | ASSGTGENYAEQF | 0.79631805 |
| GRIAGAAVLL | ASSGTGENYAEQF | 0.56396365 |
| GSAGSLNTSF | ASSGTGENYAEQF | 0.5043216 |
| GTGQIGI | ASSGTGENYAEQF | 0.6340301 |

| | | |
|---|---|---|
| GTLIALIL | ASSGTGENYAEQF | 0.77612233 |
| IAATMGFVVG | ASSGTGENYAEQF | 0.77734566 |
| IFAQALL | ASSGTGENYAEQF | 0.724393 |
| IGAAVALGF | ASSGTGENYAEQF | 0.4949997 |
| IGLTPLAL | ASSGTGENYAEQF | 0.6507914 |
| ILGSNTSTL | ASSGTGENYAEQF | 0.58393455 |
| IVFSLTVL | ASSGTGENYAEQF | 0.6233173 |
| LASKATGF | ASSGTGENYAEQF | 0.75915766 |
| LASVALIL | ASSGTGENYAEQF | 0.6730431 |
| LFAVGLTL | ASSGTGENYAEQF | 0.57285595 |
| LFNGGIGTYI | ASSGTGENYAEQF | 0.62424296 |
| LTLGAGII | ASSGTGENYAEQF | 0.7007854 |
| LVAAALGDI | ASSGTGENYAEQF | 0.75585026 |
| NTAFVQLGI | ASSGTGENYAEQF | 0.76116794 |
| QLFSGAEL | ASSGTGENYAEQF | 0.81575024 |
| SAATFITL | ASSGTGENYAEQF | 0.75010145 |
| SNYGLANLGV | ASSGTGENYAEQF | 0.7830628 |
| SYGGAYAVM | ASSGTGENYAEQF | 0.24250321 |
| TTGVGGLTLG | ASSGTGENYAEQF | 0.38042825 |
| TTQTIYI | ASSGTGENYAEQF | 0.6757017 |
| VAAGAGLGAVI | ASSGTGENYAEQF | 0.6561314 |
| VAAVVTLL | ASSGTGENYAEQF | 0.4960161 |
| VGFLGGLI | ASSGTGENYAEQF | 0.7487867 |
| VGVIPILL | ASSGTGENYAEQF | 0.5834329 |
| VIFASSLI | ASSGTGENYAEQF | 0.5267846 |
| VKGGDGGAAAG | ASSGTGENYAEQF | 0.758777 |
| VTKTFTNI | ASSGTGENYAEQF | 0.5634259 |
| VVAAAILL | ASSGTGENYAEQF | 0.6146987 |
| YAALDVEL | ASSGTGENYAEQF | 0.6364372 |
| LASVALLL | ASSGTGENYAEQF | 0.7615855 |
| LDAVHRV | ASSGTGENYAEQF | 0.4584395 |
| QGLSSSIF | ASSGTGENYAEQF | 0.72211516 |
| RGTFANIRL | ASSGTGENYAEQF | 0.63420314 |
| RQLWWGHRI | ASSGTGENYAEQF | 0.7668991 |
| YGPPGTGKTLL | ASSGTGENYAEQF | 0.43470353 |
| AALVGAPANVA | ASSGTGENYAEQF | 0.7396348 |
| AATFRRTE | ASSGTGENYAEQF | 0.6123937 |
| AGALGGGI | ASSGTGENYAEQF | 0.81807625 |
| AGGAGGAGGIGAG | ASSGTGENYAEQF | 0.6351403 |
| AGSGALGL | ASSGTGENYAEQF | 0.660648 |
| AQIADQI | ASSGTGENYAEQF | 0.6284798 |
| ASAVAVTYL | ASSGTGENYAEQF | 0.7571598 |
| ASTVLGGTGG | ASSGTGENYAEQF | 0.3989888 |
| AVGGEPAK | ASSGTGENYAEQF | 0.6375909 |
| CEQGTFGGTC | ASSGTGENYAEQF | 0.40503064 |
| DEGKAPAG | ASSGTGENYAEQF | 0.7016749 |
| DTVVFPAL | ASSGTGENYAEQF | 0.6937542 |
| FEGITMAG | ASSGTGENYAEQF | 0.46393043 |
| FPLVVQD | ASSGTGENYAEQF | 0.74673647 |
| GAFTTPQ | ASSGTGENYAEQF | 0.6871749 |
| GFGGFGTAGGVGGA | ASSGTGENYAEQF | 0.58240604 |

| | | |
|---|---|---|
| GFLAQFL | ASSGTGENYAEQF | 0.80316365 |
| GGAGGAGGNAGVA | ASSGTGENYAEQF | 0.6760527 |
| GGVGTAGAGGAGGAGG | ASSGTGENYAEQF | 0.7704 |
| GNGGGGDAGNAGSGGN | ASSGTGENYAEQF | 0.69010866 |
| GTGGAGGAGNDAGS | ASSGTGENYAEQF | 0.58514047 |
| GTTNTGWFNS | ASSGTGENYAEQF | 0.7183403 |
| IFMGPLFTL | ASSGTGENYAEQF | 0.66514206 |
| ILAHAIEAI | ASSGTGENYAEQF | 0.62659556 |
| IRSAIREAL | ASSGTGENYAEQF | 0.7654147 |
| ITFAGGG | ASSGTGENYAEQF | 0.7851962 |
| ITGMSGHQEV | ASSGTGENYAEQF | 0.86352575 |
| IVAGLACTL | ASSGTGENYAEQF | 0.6902461 |
| IVGGGLAGA | ASSGTGENYAEQF | 0.6035217 |
| IVTGVLNI | ASSGTGENYAEQF | 0.7544544 |
| LAQQGCI | ASSGTGENYAEQF | 0.85287786 |
| LCLHVEN | ASSGTGENYAEQF | 0.8539977 |
| LDAISDGS | ASSGTGENYAEQF | 0.8197465 |
| LGGFAAML | ASSGTGENYAEQF | 0.54739785 |
| LGGKNPMI | ASSGTGENYAEQF | 0.76778316 |
| LGPKGVAAIL | ASSGTGENYAEQF | 0.7188345 |
| LMVGVSI | ASSGTGENYAEQF | 0.6825947 |
| LVGAAFVL | ASSGTGENYAEQF | 0.82013977 |
| LVLAVAPL | ASSGTGENYAEQF | 0.7801678 |
| MAAIAVIDAA | ASSGTGENYAEQF | 0.70218796 |
| NNGINAELQ | ASSGTGENYAEQF | 0.6163584 |
| RGEPLVLVHG | ASSGTGENYAEQF | 0.8049131 |
| RGGAGGAAGLI | ASSGTGENYAEQF | 0.82877606 |
| RSYRMHPAV | ASSGTGENYAEQF | 0.8183296 |
| SAAHLLAGQLA | ASSGTGENYAEQF | 0.86750484 |
| SAAVGSY | ASSGTGENYAEQF | 0.8345394 |
| SPHAVLHRI | ASSGTGENYAEQF | 0.821289 |
| SQIAKVL | ASSGTGENYAEQF | 0.17323586 |
| TALGHLVP | ASSGTGENYAEQF | 0.35553244 |
| TAPTTSGAA | ASSGTGENYAEQF | 0.7434983 |
| TDGDDGNGGSPGT | ASSGTGENYAEQF | 0.71202135 |
| TDGLGEN | ASSGTGENYAEQF | 0.49119663 |
| TGGPPLPGQA | ASSGTGENYAEQF | 0.8161018 |
| TIPAITI | ASSGTGENYAEQF | 0.71473575 |
| TIPGFTI | ASSGTGENYAEQF | 0.65593934 |
| TSIASGALA | ASSGTGENYAEQF | 0.8402871 |
| TVTLPGAAI | ASSGTGENYAEQF | 0.66295993 |
| VASGLSAALP | ASSGTGENYAEQF | 0.6027329 |
| VGAGVIGI | ASSGTGENYAEQF | 0.63981664 |
| VGPKRFGV | ASSGTGENYAEQF | 0.8250363 |
| VNVAFPD | ASSGTGENYAEQF | 0.44638854 |
| VSMTNSGVS | ASSGTGENYAEQF | 0.7934922 |
| VTDVLGDLSV | ASSGTGENYAEQF | 0.6259201 |
| VVVVGAGFAGL | ASSGTGENYAEQF | 0.85375273 |
| VYTVGTVV | ASSGTGENYAEQF | 0.4639702 |
| WTTFSGP | ASSGTGENYAEQF | 0.8190588 |
| YDELVEAF | ASSGTGENYAEQF | 0.7406721 |

| | | |
|---|---|---|
| YGTGGAGGNGG | ASSGTGENYAEQF | 0.87936676 |
| YTNVMAV | ASSGTGENYAEQF | 0.70703554 |
| FLEQVHQL | ASSGTGENYAEQF | 0.75426763 |
| GLSLLAVAV | ASSGTGENYAEQF | 0.5815222 |
| LEQVHQL | ASSGTGENYAEQF | 0.85064805 |
| LNAAGAI | ASSGTGENYAEQF | 0.54383856 |
| PAVHASL | ASSGTGENYAEQF | 0.7472036 |
| TELEDDA | ASSGTGENYAEQF | 0.42277905 |
| TPPGPPN | ASSGTGENYAEQF | 0.8045471 |
| TPPPPPPP | ASSGTGENYAEQF | 0.8012456 |
| TQSPALI | ASTDGGYEQY | 0.5146178 |
| EFAAAFPQ | ASTDGGYEQY | 0.79100764 |
| LLSGNLDVL | ASTDGGYEQY | 0.79438454 |
| GPANLGNYNI | ASTDGGYEQY | 0.62344223 |
| TLGPTLP | ASTDGGYEQY | 0.8647331 |
| VYAYSSCCK | ASTDGGYEQY | 0.7807915 |
| APAPAPAPNI | ASTDGGYEQY | 0.8503485 |
| PGGPVPIP | ASTDGGYEQY | 0.77468866 |
| FLTVALEP | ASTDGGYEQY | 0.63590103 |
| SQYFGANL | ASTDGGYEQY | 0.81170946 |
| GGVKPAA | ASTDGGYEQY | 0.7510661 |
| GGTGGAGGAGGW | ASTDGGYEQY | 0.6948333 |
| GVGGVAGLGGAGG | ASTDGGYEQY | 0.8346652 |
| VLLVLATTI | ASTDGGYEQY | 0.4345762 |
| IGIFAAML | ASTDGGYEQY | 0.8541141 |
| AVGDLID | ASTDGGYEQY | 0.24187744 |
| NISPLPL | ASTDGGYEQY | 0.2797653 |
| VTAVFVAI | ASTDGGYEQY | 0.46859467 |
| SGDDNDAAL | ASTDGGYEQY | 0.64755774 |
| AVAALAHS | ASTDGGYEQY | 0.8216857 |
| GGVTVMATP | ASTDGGYEQY | 0.57741344 |
| AGLAANVRD | ASTDGGYEQY | 0.25472796 |
| TNRPDLIDPALLRPG | ASTDGGYEQY | 0.37130517 |
| LANLGVTAL | ASTDGGYEQY | 0.23591945 |
| TGERIEL | ASTDGGYEQY | 0.36861098 |
| NGEELAA | ASTDGGYEQY | 0.5473231 |
| VLSLVWG | ASTDGGYEQY | 0.36924124 |
| IGYALWL | ASTDGGYEQY | 0.24085307 |
| LSLLLAPL | ASTDGGYEQY | 0.34890184 |
| TLDEALK | ASTDGGYEQY | 0.56582606 |
| TDGAGTAEGG | ASTDGGYEQY | 0.53686106 |
| NAPGGAGGNGG | ASTDGGYEQY | 0.40779546 |
| VSVFMGMGGNF | ASTDGGYEQY | 0.7464041 |
| LASYLAAL | ASTDGGYEQY | 0.6190555 |
| LVGGTPAL | ASTDGGYEQY | 0.59481835 |
| VRNADVL | ASTDGGYEQY | 0.11096532 |
| TCSSPFLVSAM | ASTDGGYEQY | 0.2183286 |
| VHITDAE | ASTDGGYEQY | 0.26914677 |
| EYAAPSF | ASTDGGYEQY | 0.44503438 |
| VVDAFHGGL | ASTDGGYEQY | 0.15135093 |
| ASVVDPQ | ASTDGGYEQY | 0.6311934 |

| | | |
|---|---|---|
| SYNEFGGLS | ASTDGGYEQY | 0.29530072 |
| DPDGSVY | ASTDGGYEQY | 0.30175444 |
| MVSTIHATE | ASTDGGYEQY | 0.675103 |
| LAAGVADPV | ASTDGGYEQY | 0.25545138 |
| TKEQIAA | ASTDGGYEQY | 0.30758262 |
| SLASPAAL | ASTDGGYEQY | 0.25807565 |
| IGLVFYGV | ASTDGGYEQY | 0.7059512 |
| FLLDASNL | ASTDGGYEQY | 0.15748262 |
| YQVGNFP | ASTDGGYEQY | 0.50994027 |
| ESLLSQST | ASTDGGYEQY | 0.26675683 |
| LFTVGFAAVG | ASTDGGYEQY | 0.58801615 |
| ATPPTPT | ASTDGGYEQY | 0.20612895 |
| TDLATLG | ASTDGGYEQY | 0.5997239 |
| SAGGTGR | ASTDGGYEQY | 0.35372984 |
| GFSNSGS | ASTDGGYEQY | 0.8632156 |
| GGNGGSGGTSV | ASTDGGYEQY | 0.38706774 |
| GTSVATGGAGN | ASTDGGYEQY | 0.51046586 |
| NGGNGGSGGTS | ASTDGGYEQY | 0.303494 |
| AVVKFPH | ASTDGGYEQY | 0.8269444 |
| TPPPPPP | ASTDGGYEQY | 0.18660758 |
| AAPAAAAAPA | ASTDGGYEQY | 0.4042661 |
| IALNEDL | ASTDGGYEQY | 0.19007084 |
| IVNVSSM | ASTDGGYEQY | 0.5148015 |
| TGIPPAPR | ASTDGGYEQY | 0.5612588 |
| FDLGGGTF | ASTDGGYEQY | 0.2426385 |
| FDLGGGTFDVS | ASTDGGYEQY | 0.5192975 |
| NEPTAAA | ASTDGGYEQY | 0.5128811 |
| NEPTAAALA | ASTDGGYEQY | 0.24980655 |
| GSIINISSI | ASTDGGYEQY | 0.7074621 |
| VAPGNAGTA | ASTDGGYEQY | 0.2672929 |
| TGGGASLE | ASTDGGYEQY | 0.77364725 |
| VAVYNHYKRI | ASTDGGYEQY | 0.36224118 |
| TDPAPATT | ASTDGGYEQY | 0.31743318 |
| TNRPDILDPALLRPG | ASTDGGYEQY | 0.54623044 |
| LTTTTTS | ASTDGGYEQY | 0.28126317 |
| PTGSGKTL | ASTDGGYEQY | 0.23835926 |
| TGSGKTL | ASTDGGYEQY | 0.6873375 |
| TIGHVDHGK | ASTDGGYEQY | 0.2784964 |
| TIGHVDHGKTT | ASTDGGYEQY | 0.8351393 |
| GHVDHGKTT | ASTDGGYEQY | 0.17038822 |
| AQFEHTLL | ASTDGGYEQY | 0.21010658 |
| IVNQDEL | ASTDGGYEQY | 0.31691742 |
| TNGSQFF | ASTDGGYEQY | 0.39974946 |
| TGKGIVE | ASTDGGYEQY | 0.81385386 |
| LISNASDA | ASTDGGYEQY | 0.3224656 |
| AELRTLGELTKIAKAHGAQ | ASTDGGYEQY | 0.19075857 |
| APNAVAAAEPSW | ASTDGGYEQY | 0.2400843 |
| APQVAGAATAVL | ASTDGGYEQY | 0.16881528 |
| ARRDPAAAAAILPTDA | ASTDGGYEQY | 0.26140442 |
| DAAARDVLAEAGLAEYF | ASTDGGYEQY | 0.3317143 |
| DDQIVLSAKANLAKVEQ | ASTDGGYEQY | 0.26819286 |

| | | |
|---|---|---|
| DELLPDTGEDSDEDGH | ASTDGGYEQY | 0.18007268 |
| ELPEAPANRATVPQD | ASTDGGYEQY | 0.21934305 |
| FGQSKIERVTGIG | ASTDGGYEQY | 0.33720067 |
| GAGGTGSTAGGAGGAGGAGGI | ASTDGGYEQY | 0.38082325 |
| GHGGTGAAGGNGGTGGAGGS( | ASTDGGYEQY | 0.30500773 |
| GNGGNGGAGGEGGPGGAGGA( | ASTDGGYEQY | 0.5312732 |
| GSTAGNGGNGGNATNGTIAG( | ASTDGGYEQY | 0.45116228 |
| GTGGDGGNGGNGGTGG | ASTDGGYEQY | 0.35926905 |
| GVRRPAGIPADAPV | ASTDGGYEQY | 0.09793778 |
| GYRQVTDAHLLTLARR | ASTDGGYEQY | 0.15907693 |
| IAKMGQDGHDRG | ASTDGGYEQY | 0.17134981 |
| KAEPSIEIQYT | ASTDGGYEQY | 0.2942505 |
| KPVRIRMAADEDTVAVT | ASTDGGYEQY | 0.10741605 |
| MPPRIAGMRLLVIKPEP | ASTDGGYEQY | 0.42558238 |
| MKFYAVALFRSRGVPPTR | ASTDGGYEQY | 0.20313281 |
| NAGGFGILGAVAHSP | ASTDGGYEQY | 0.20414592 |
| NEGREITLAMAEM | ASTDGGYEQY | 0.50747526 |
| NELVEWSHPLGHRWRWE | ASTDGGYEQY | 0.18555966 |
| NLGAGGTGGNGGNPG | ASTDGGYEQY | 0.22507189 |
| NTEWPKWETVMAADPVD | ASTDGGYEQY | 0.16819884 |
| PASAAPAETQP | ASTDGGYEQY | 0.5119649 |
| QQSSDGKNGVRAAEITGE | ASTDGGYEQY | 0.11982693 |
| RAPDAVTTSWWKE | ASTDGGYEQY | 0.38926184 |
| SGFGNVGAHTSGWFNQ | ASTDGGYEQY | 0.16410397 |
| SGVAGAVGQAASVGGLKV | ASTDGGYEQY | 0.37391168 |
| SNGGAGTGSAGGNGGT | ASTDGGYEQY | 0.15856989 |
| TGGAGGEGGAAG | ASTDGGYEQY | 0.4124849 |
| TGGDGGAAGAGGAPGHG | ASTDGGYEQY | 0.22772112 |
| TMPPLLAAIANAGRHGV | ASTDGGYEQY | 0.84152746 |
| TSSTAGNSGGA | ASTDGGYEQY | 0.31180888 |
| VPATVPVGILC | ASTDGGYEQY | 0.33988452 |
| VQRPDSGGVVLTGRIS | ASTDGGYEQY | 0.20934376 |
| WTPNPVPATKGVPAGLVQAT | ASTDGGYEQY | 0.82200193 |
| YALKRALAKFSDRFDVVII | ASTDGGYEQY | 0.12721878 |
| YLIGSVIV | ASTDGGYEQY | 0.27314034 |
| YGQLVAVID | ASTDGGYEQY | 0.13751012 |
| VTVGGMTLPA | ASTDGGYEQY | 0.3231556 |
| VLALPAV | ASTDGGYEQY | 0.36107266 |
| VIPPTLNYET | ASTDGGYEQY | 0.15714101 |
| VGMGGMHPSSG | ASTDGGYEQY | 0.3101855 |
| VGLAGGAATA | ASTDGGYEQY | 0.28316885 |
| VGGAGGAGGSGP | ASTDGGYEQY | 0.17240176 |
| VFAAFMV | ASTDGGYEQY | 0.4984989 |
| TLYDKLR | ASTDGGYEQY | 0.21293639 |
| TLAVFTVL | ASTDGGYEQY | 0.68536633 |
| TGSVQPARV | ASTDGGYEQY | 0.20674407 |
| TGGGAGSGGT | ASTDGGYEQY | 0.19559154 |
| SHLETSL | ASTDGGYEQY | 0.33621305 |
| SFTTGMT | ASTDGGYEQY | 0.17820156 |
| QVEVAKRMGVRQPTVS | ASTDGGYEQY | 0.15955392 |
| QGSGGPALAM | ASTDGGYEQY | 0.5384028 |

| | | |
|---|---|---|
| MLAPSNP | ASTDGGYEQY | 0.71279377 |
| LVNNAAI | ASTDGGYEQY | 0.73011655 |
| LTGEINTLA | ASTDGGYEQY | 0.6545268 |
| LSIFGPV | ASTDGGYEQY | 0.60826766 |
| LLALLANL | ASTDGGYEQY | 0.6907218 |
| LISNASDALDKLR | ASTDGGYEQY | 0.7266564 |
| LINEAVP | ASTDGGYEQY | 0.7234268 |
| LGYAFSGL | ASTDGGYEQY | 0.7162024 |
| LDFVAVAL | ASTDGGYEQY | 0.6106024 |
| LAQPGAE | ASTDGGYEQY | 0.68501675 |
| ISNLGAVTGF | ASTDGGYEQY | 0.66974664 |
| HQEDPGRI | ASTDGGYEQY | 0.64801496 |
| GTPGEPTP | ASTDGGYEQY | 0.7199166 |
| GFFNSGS | ASTDGGYEQY | 0.6677519 |
| GAGGGGLTAGAN | ASTDGGYEQY | 0.6599002 |
| FVDALVNM | ASTDGGYEQY | 0.6903466 |
| FLTVAPEL | ASTDGGYEQY | 0.69286764 |
| FETLTAL | ASTDGGYEQY | 0.716842 |
| DVINPIEL | ASTDGGYEQY | 0.7093734 |
| APVSGAPV | ASTDGGYEQY | 0.73383284 |
| ALVGGLAW | ASTDGGYEQY | 0.716661 |
| ALAAAAATAI | ASTDGGYEQY | 0.7185403 |
| AFGLGLLL | ASTDGGYEQY | 0.31618863 |
| AFFAAASL | ASTDGGYEQY | 0.5731528 |
| AAAALPAFSPP | ASTDGGYEQY | 0.6859693 |
| VVLAPAL | ASTDGGYEQY | 0.68784463 |
| TIGHVDHGKTTL | ASTDGGYEQY | 0.4795126 |
| LISNASDALDK | ASTDGGYEQY | 0.7084943 |
| LISNASDAL | ASTDGGYEQY | 0.69765556 |
| ISNASDALDK | ASTDGGYEQY | 0.59894717 |
| NQAETLVYQTE | ASTDGGYEQY | 0.7155766 |
| TGGTGGTAGAG | ASTDGGYEQY | 0.62182534 |
| TGSTAGGAGGA | ASTDGGYEQY | 0.6043289 |
| DPLAVPVAAGPV | ASTDGGYEQY | 0.6676345 |
| GGTGGTPGNGGN | ASTDGGYEQY | 0.72945136 |
| PAGLAGVDPATL | ASTDGGYEQY | 0.590425 |
| GNNLVVEAPHAGASV | ASTDGGYEQY | 0.70346564 |
| KVTRIMSASTLEQPA | ASTDGGYEQY | 0.70351124 |
| LGFALPIALPRRTVV | ASTDGGYEQY | 0.72062314 |
| TRLGNLTARKLVVLV | ASTDGGYEQY | 0.5439416 |
| WSQQRRLIYFPSAGP | ASTDGGYEQY | 0.714517 |
| GPNGNPDPMAAAVDIR | ASTDGGYEQY | 0.6838876 |
| TQRCVSVADLRAAASR | ASTDGGYEQY | 0.73235786 |
| QFVLDVAEHPVEVLTETV | ASTDGGYEQY | 0.6691754 |
| GGAGGTGGAGGNGGTQAGDG | ASTDGGYEQY | 0.69165987 |
| PLLATLRAIAVDVDAKPAQV | ASTDGGYEQY | 0.6413032 |
| APKFRTGGYMIDFWGVGYQV | ASTDGGYEQY | 0.7211778 |
| GAGGAGGAGGAAGGPGGTGG | ASTDGGYEQY | 0.56246233 |
| IGALGLLATDVAPAGSVAFG | ASTDGGYEQY | 0.68928915 |
| AGGAGGIGGNGGAGGAGGVL | ASTDGGYEQY | 0.62034214 |
| ADELIGLL | ASTDGGYEQY | 0.701686 |

| | | |
|---|---|---|
| ADRALKA | ASTDGGYEQY | 0.7149173 |
| AGAVGQVGGAGGA | ASTDGGYEQY | 0.6417601 |
| AGGGGGLGGAP | ASTDGGYEQY | 0.7240215 |
| AGGLLYGNGGAG | ASTDGGYEQY | 0.69954 |
| AGLESIGL | ASTDGGYEQY | 0.6553854 |
| AGLKTGII | ASTDGGYEQY | 0.7234243 |
| ASLAALLL | ASTDGGYEQY | 0.62854946 |
| ASVINISSI | ASTDGGYEQY | 0.7243834 |
| ATPPGPA | ASTDGGYEQY | 0.6633666 |
| ATVASALV | ASTDGGYEQY | 0.7056588 |
| ATVASLIL | ASTDGGYEQY | 0.71873564 |
| AVNGPVLIRH | ASTDGGYEQY | 0.57178 |
| DGAAGAVGI | ASTDGGYEQY | 0.6214185 |
| DPLLHGI | ASTDGGYEQY | 0.7223245 |
| ELGAALGAL | ASTDGGYEQY | 0.7731395 |
| ELGRTQAV | ASTDGGYEQY | 0.78875476 |
| FVAGGYLS | ASTDGGYEQY | 0.7350943 |
| GASGGAGGAGGW | ASTDGGYEQY | 0.588466 |
| GGAGAGGDGGAGGI | ASTDGGYEQY | 0.72166014 |
| GTGHNGSQPGMGGNG | ASTDGGYEQY | 0.78382325 |
| GYAAASASAS | ASTDGGYEQY | 0.78818053 |
| HQVDVAAVQQG | ASTDGGYEQY | 0.77179193 |
| IGAAIAHT | ASTDGGYEQY | 0.3860013 |
| IGMIEVAQL | ASTDGGYEQY | 0.7566035 |
| IRLAPEHTSAMD | ASTDGGYEQY | 0.70520175 |
| IYNVSVL | ASTDGGYEQY | 0.7361056 |
| KQGFLGM | ASTDGGYEQY | 0.7694627 |
| LESVITQL | ASTDGGYEQY | 0.72468406 |
| LGLGALVVV | ASTDGGYEQY | 0.7436047 |
| LIGNGGNGGSGGI | ASTDGGYEQY | 0.68293434 |
| LNAINAPTL | ASTDGGYEQY | 0.73366106 |
| LQLLNTGV | ASTDGGYEQY | 0.76262987 |
| LQTSIEF | ASTDGGYEQY | 0.7633101 |
| MSFAVSEL | ASTDGGYEQY | 0.78601575 |
| NTGALGL | ASTDGGYEQY | 0.78704906 |
| PPGAGGAGGAGTT | ASTDGGYEQY | 0.77700675 |
| QLVGLGI | ASTDGGYEQY | 0.20306914 |
| RFLEQVHQL | ASTDGGYEQY | 0.49871907 |
| RGVGYVLR | ASTDGGYEQY | 0.739208 |
| SATGVFAGVI | ASTDGGYEQY | 0.7363529 |
| SLPAITLP | ASTDGGYEQY | 0.4355882 |
| STALAFTV | ASTDGGYEQY | 0.7365632 |
| SVLTVGLGK | ASTDGGYEQY | 0.6781696 |
| TALAGAGSGI | ASTDGGYEQY | 0.7088754 |
| TGLPFTGI | ASTDGGYEQY | 0.78761584 |
| VAEHVTV | ASTDGGYEQY | 0.74376655 |
| VDIVLNSL | ASTDGGYEQY | 0.6487001 |
| VLTGGGAVL | ASTDGGYEQY | 0.6498462 |
| VNVGLAYGI | ASTDGGYEQY | 0.7645462 |
| VSSAAVGAV | ASTDGGYEQY | 0.59715796 |
| VTSVYEAGV | ASTDGGYEQY | 0.7759787 |

| | | |
|---|---|---|
| WVALDRGA | ASTDGGYEQY | 0.6173166 |
| YLVGSVVI | ASTDGGYEQY | 0.7821553 |
| GPPGTGKTLL | ASTDGGYEQY | 0.38638672 |
| GVGMTKF | ASTDGGYEQY | 0.77791715 |
| IGIGGSDL | ASTDGGYEQY | 0.7461493 |
| IGTLLAEL | ASTDGGYEQY | 0.8021999 |
| IYTSGTT | ASTDGGYEQY | 0.689619 |
| LSFPAGV | ASTDGGYEQY | 0.6847898 |
| LYTSGTT | ASTDGGYEQY | 0.5378964 |
| VGAFPFGF | ASTDGGYEQY | 0.78891504 |
| VGAGTVEF | ASTDGGYEQY | 0.5047318 |
| VGKSTLLL | ASTDGGYEQY | 0.72609353 |
| AALAHAGGGIV | ASTDGGYEQY | 0.41749012 |
| AALVAVSLL | ASTDGGYEQY | 0.76127815 |
| AAVSRAVAAL | ASTDGGYEQY | 0.7705921 |
| AGGSSGIGGF | ASTDGGYEQY | 0.38177302 |
| AGLTVANANG | ASTDGGYEQY | 0.78500384 |
| AGLVGAVL | ASTDGGYEQY | 0.73063815 |
| ALSQAFTV | ASTDGGYEQY | 0.5330603 |
| ALTFSVSL | ASTDGGYEQY | 0.79094166 |
| ALTGAALVAL | ASTDGGYEQY | 0.6394986 |
| ASFSIGAGVL | ASTDGGYEQY | 0.7910349 |
| DGSVYKI | ASTDGGYEQY | 0.7418699 |
| EQLAEIV | ASTDGGYEQY | 0.7405906 |
| FAVSELTVM | ASTDGGYEQY | 0.7460147 |
| GAAPLAAGL | ASTDGGYEQY | 0.7093896 |
| GADGGDGGSGGAGG | ASTDGGYEQY | 0.65952086 |
| GFGLDTGGAGG | ASTDGGYEQY | 0.78834295 |
| GLASGGLL | ASTDGGYEQY | 0.7937851 |
| GRIAGAAVLL | ASTDGGYEQY | 0.80254596 |
| GSAGSLNTSF | ASTDGGYEQY | 0.742207 |
| GTGQIGI | ASTDGGYEQY | 0.676242 |
| GTLIALIL | ASTDGGYEQY | 0.7647121 |
| IAATMGFVVG | ASTDGGYEQY | 0.8025919 |
| IFAQALL | ASTDGGYEQY | 0.8019359 |
| IGAAVALGF | ASTDGGYEQY | 0.78744996 |
| IGLTPLAL | ASTDGGYEQY | 0.5302096 |
| ILGSNTSTL | ASTDGGYEQY | 0.75691193 |
| IVFSLTVL | ASTDGGYEQY | 0.74423087 |
| LASKATGF | ASTDGGYEQY | 0.7094978 |
| LASVALIL | ASTDGGYEQY | 0.7961802 |
| LFAVGLTL | ASTDGGYEQY | 0.7278786 |
| LFNGGIGTYI | ASTDGGYEQY | 0.72846323 |
| LTLGAGII | ASTDGGYEQY | 0.7543584 |
| LVAAALGDI | ASTDGGYEQY | 0.76661646 |
| NTAFVQLGI | ASTDGGYEQY | 0.7993382 |
| QLFSGAEL | ASTDGGYEQY | 0.799942 |
| SAATFITL | ASTDGGYEQY | 0.8133594 |
| SNYGLANLGV | ASTDGGYEQY | 0.80526936 |
| SYGGAYAVM | ASTDGGYEQY | 0.8061463 |
| TTGVGGLTLG | ASTDGGYEQY | 0.24487448 |

| | | |
|---|---|---|
| TTQTIYI | ASTDGGYEQY | 0.5136744 |
| VAAGAGLGAVI | ASTDGGYEQY | 0.6929156 |
| VAAVVTLL | ASTDGGYEQY | 0.74017 |
| VGFLGGLI | ASTDGGYEQY | 0.5621846 |
| VGVIPILL | ASTDGGYEQY | 0.7282731 |
| VIFASSLI | ASTDGGYEQY | 0.7212615 |
| VKGGDGGAAAG | ASTDGGYEQY | 0.6385342 |
| VTKTFTNI | ASTDGGYEQY | 0.80221593 |
| VVAAAILL | ASTDGGYEQY | 0.70639515 |
| YAALDVEL | ASTDGGYEQY | 0.64553684 |
| LASVALLL | ASTDGGYEQY | 0.6286648 |
| LDAVHRV | ASTDGGYEQY | 0.79254216 |
| QGLSSSIF | ASTDGGYEQY | 0.5799719 |
| RGTFANIRL | ASTDGGYEQY | 0.78574383 |
| RQLWWGHRI | ASTDGGYEQY | 0.6225219 |
| YGPPGTGKTLL | ASTDGGYEQY | 0.7957047 |
| AALVGAPANVA | ASTDGGYEQY | 0.46995306 |
| AATFRRTE | ASTDGGYEQY | 0.79559064 |
| AGALGGGI | ASTDGGYEQY | 0.7633786 |
| AGGAGGAGGIGAG | ASTDGGYEQY | 0.8198547 |
| AGSGALGL | ASTDGGYEQY | 0.7449273 |
| AQIADQI | ASTDGGYEQY | 0.72265995 |
| ASAVAVTYL | ASTDGGYEQY | 0.5604128 |
| ASTVLGGTGG | ASTDGGYEQY | 0.79824454 |
| AVGGEPAK | ASTDGGYEQY | 0.52658373 |
| CEQGTFGGTC | ASTDGGYEQY | 0.7690546 |
| DEGKAPAG | ASTDGGYEQY | 0.50445604 |
| DTVVFPAL | ASTDGGYEQY | 0.78956467 |
| FEGITMAG | ASTDGGYEQY | 0.7864841 |
| FPLVVQD | ASTDGGYEQY | 0.59351456 |
| GAFTTPQ | ASTDGGYEQY | 0.8105767 |
| GFGGFGTAGGVGGA | ASTDGGYEQY | 0.75276613 |
| GFLAQFL | ASTDGGYEQY | 0.6562764 |
| GGAGGAGGNAGVA | ASTDGGYEQY | 0.81232864 |
| GGVGTAGAGGAGGAGG | ASTDGGYEQY | 0.6860145 |
| GNGGGGDAGNAGSGGN | ASTDGGYEQY | 0.8113161 |
| GTGGAGGAGNDAGS | ASTDGGYEQY | 0.7320516 |
| GTTNTGWFNS | ASTDGGYEQY | 0.7406052 |
| IFMGPLFTL | ASTDGGYEQY | 0.7710527 |
| ILAHAIEAI | ASTDGGYEQY | 0.5896025 |
| IRSAIREAL | ASTDGGYEQY | 0.5995984 |
| ITFAGGG | ASTDGGYEQY | 0.80491686 |
| ITGMSGHQEV | ASTDGGYEQY | 0.7796992 |
| IVAGLACTL | ASTDGGYEQY | 0.8192233 |
| IVGGGLAGA | ASTDGGYEQY | 0.62236613 |
| IVTGVLNI | ASTDGGYEQY | 0.60243505 |
| LAQQGCI | ASTDGGYEQY | 0.6848172 |
| LCLHVEN | ASTDGGYEQY | 0.8383903 |
| LDAISDGS | ASTDGGYEQY | 0.80332613 |
| LGGFAAML | ASTDGGYEQY | 0.7674022 |
| LGGKNPMI | ASTDGGYEQY | 0.5775912 |

| | | |
|---|---|---|
| LGPKGVAAIL | ASTDGGYEQY | 0.72065115 |
| LMVGVSI | ASTDGGYEQY | 0.7019299 |
| LVGAAFVL | ASTDGGYEQY | 0.648399 |
| LVLAVAPL | ASTDGGYEQY | 0.7949328 |
| MAAIAVIDAA | ASTDGGYEQY | 0.7757958 |
| NNGINAELQ | ASTDGGYEQY | 0.65416074 |
| RGEPLVLVHG | ASTDGGYEQY | 0.70787686 |
| RGGAGGAAGLI | ASTDGGYEQY | 0.714645 |
| RSYRMHPAV | ASTDGGYEQY | 0.79882485 |
| SAAHLLAGQLA | ASTDGGYEQY | 0.7879433 |
| SAAVGSY | ASTDGGYEQY | 0.85469955 |
| SPHAVLHRI | ASTDGGYEQY | 0.807135 |
| SQIAKVL | ASTDGGYEQY | 0.80315495 |
| TALGHLVP | ASTDGGYEQY | 0.23272263 |
| TAPTTSGAA | ASTDGGYEQY | 0.39830935 |
| TDGDDGNGGSPGT | ASTDGGYEQY | 0.72305715 |
| TDGLGEN | ASTDGGYEQY | 0.6921803 |
| TGGPPLPGQA | ASTDGGYEQY | 0.46819466 |
| TIPAITI | ASTDGGYEQY | 0.7899718 |
| TIPGFTI | ASTDGGYEQY | 0.70361865 |
| TSIASGALA | ASTDGGYEQY | 0.63107824 |
| TVTLPGAAI | ASTDGGYEQY | 0.7935761 |
| VASGLSAALP | ASTDGGYEQY | 0.63769007 |
| VGAGVIGI | ASTDGGYEQY | 0.69392025 |
| VGPKRFGV | ASTDGGYEQY | 0.642501 |
| VNVAFPD | ASTDGGYEQY | 0.79331875 |
| VSMTNSGVS | ASTDGGYEQY | 0.5697361 |
| VTDVLGDLSV | ASTDGGYEQY | 0.78269374 |
| VVVVGAGFAGL | ASTDGGYEQY | 0.6448096 |
| VYTVGTVV | ASTDGGYEQY | 0.8329564 |
| WTTFSGP | ASTDGGYEQY | 0.5527666 |
| YDELVEAF | ASTDGGYEQY | 0.77741265 |
| YGTGGAGGNGG | ASTDGGYEQY | 0.69230914 |
| YTNVMAV | ASTDGGYEQY | 0.85509694 |
| FLEQVHQL | ASTDGGYEQY | 0.7388253 |
| GLSLLAVAV | ASTDGGYEQY | 0.7712155 |
| LEQVHQL | ASTDGGYEQY | 0.5997584 |
| LNAAGAI | ASTDGGYEQY | 0.7660086 |
| PAVHASL | ASTDGGYEQY | 0.53628623 |
| TELEDDA | ASTDGGYEQY | 0.7202879 |
| TPPGPPN | ASTDGGYEQY | 0.4964617 |
| TPPPPPPP | ASTDGGYEQY | 0.76862 |
| TQSPALI | ASGEQRANSDYT | 0.7439829 |
| EFAAAFPQ | ASGEQRANSDYT | 0.5686975 |
| LLSGNLDVL | ASGEQRANSDYT | 0.8061243 |
| GPANLGNYNI | ASGEQRANSDYT | 0.75121653 |
| TLGPTLP | ASGEQRANSDYT | 0.66938055 |
| VYAYSSCCK | ASGEQRANSDYT | 0.8359865 |
| APAPAPAPNI | ASGEQRANSDYT | 0.72886765 |
| PGGPVPIP | ASGEQRANSDYT | 0.8148099 |
| FLTVALEP | ASGEQRANSDYT | 0.6944223 |

| | | |
|---|---|---|
| SQYFGANL | ASGEQRANSDYT | 0.66331226 |
| GGVKPAA | ASGEQRANSDYT | 0.7794553 |
| GGTGGAGGAGGW | ASGEQRANSDYT | 0.6889918 |
| GVGGVAGLGGAGG | ASGEQRANSDYT | 0.65400755 |
| VLLVLATTI | ASGEQRANSDYT | 0.8151432 |
| IGIFAAML | ASGEQRANSDYT | 0.80825824 |
| AVGDLID | ASGEQRANSDYT | 0.8540358 |
| NISPLPL | ASGEQRANSDYT | 0.6924657 |
| VTAVFVAI | ASGEQRANSDYT | 0.6830181 |
| SGDDNDAAL | ASGEQRANSDYT | 0.7489203 |
| AVAALAHS | ASGEQRANSDYT | 0.8482323 |
| GGVTVMATP | ASGEQRANSDYT | 0.81801116 |
| AGLAANVRD | ASGEQRANSDYT | 0.8071352 |
| TNRPDLIDPALLRPG | ASGEQRANSDYT | 0.58880055 |
| LANLGVTAL | ASGEQRANSDYT | 0.76065254 |
| TGERIEL | ASGEQRANSDYT | 0.67437696 |
| NGEELAA | ASGEQRANSDYT | 0.7283846 |
| VLSLVWG | ASGEQRANSDYT | 0.80742466 |
| IGYALWL | ASGEQRANSDYT | 0.752999 |
| LSLLLAPL | ASGEQRANSDYT | 0.7325177 |
| TLDEALK | ASGEQRANSDYT | 0.7728723 |
| TDGAGTAEGG | ASGEQRANSDYT | 0.77482575 |
| NAPGGAGGNGG | ASGEQRANSDYT | 0.7876494 |
| VSVFMGMGGNF | ASGEQRANSDYT | 0.7506847 |
| LASYLAAL | ASGEQRANSDYT | 0.86331177 |
| LVGGTPAL | ASGEQRANSDYT | 0.8316077 |
| VRNADVL | ASGEQRANSDYT | 0.842355 |
| TCSSPFLVSAM | ASGEQRANSDYT | 0.24371973 |
| VHITDAE | ASGEQRANSDYT | 0.50658673 |
| EYAAPSF | ASGEQRANSDYT | 0.7490147 |
| VVDAFHGGL | ASGEQRANSDYT | 0.7204051 |
| ASVVDPQ | ASGEQRANSDYT | 0.5263754 |
| SYNEFGGLS | ASGEQRANSDYT | 0.81770384 |
| DPDGSVY | ASGEQRANSDYT | 0.69344354 |
| MVSTIHATE | ASGEQRANSDYT | 0.64360166 |
| LAAGVADPV | ASGEQRANSDYT | 0.8198768 |
| TKEQIAA | ASGEQRANSDYT | 0.7468773 |
| SLASPAAL | ASGEQRANSDYT | 0.6706418 |
| IGLVFYGV | ASGEQRANSDYT | 0.59379053 |
| FLLDASNL | ASGEQRANSDYT | 0.81981397 |
| YQVGNFP | ASGEQRANSDYT | 0.4161266 |
| ESLLSQST | ASGEQRANSDYT | 0.77600396 |
| LFTVGFAAVG | ASGEQRANSDYT | 0.6511065 |
| ATPPTPT | ASGEQRANSDYT | 0.843852 |
| TDLATLG | ASGEQRANSDYT | 0.5168821 |
| SAGGTGR | ASGEQRANSDYT | 0.8155116 |
| GFSNSGS | ASGEQRANSDYT | 0.7635031 |
| GGNGGSGGTSV | ASGEQRANSDYT | 0.87202644 |
| GTSVATGGAGN | ASGEQRANSDYT | 0.7496728 |
| NGGNGGSGGTS | ASGEQRANSDYT | 0.75493085 |
| AVVKFPH | ASGEQRANSDYT | 0.6420542 |

| | | |
|---|---|---|
| TPPPPPP | ASGEQRANSDYT | 0.8280047 |
| AAPAAAAAPA | ASGEQRANSDYT | 0.53209406 |
| IALNEDL | ASGEQRANSDYT | 0.75309724 |
| IVNVSSM | ASGEQRANSDYT | 0.4561572 |
| TGIPPAPR | ASGEQRANSDYT | 0.79662234 |
| FDLGGGTF | ASGEQRANSDYT | 0.80505204 |
| FDLGGGTFDVS | ASGEQRANSDYT | 0.6561227 |
| NEPTAAA | ASGEQRANSDYT | 0.7931898 |
| NEPTAAALA | ASGEQRANSDYT | 0.7654671 |
| GSIINISSI | ASGEQRANSDYT | 0.7312958 |
| VAPGNAGTA | ASGEQRANSDYT | 0.8527845 |
| TGGGASLE | ASGEQRANSDYT | 0.7111987 |
| VAVYNHYKRI | ASGEQRANSDYT | 0.8608593 |
| TDPAPATT | ASGEQRANSDYT | 0.74647427 |
| TNRPDILDPALLRPG | ASGEQRANSDYT | 0.690658 |
| LTTTTTS | ASGEQRANSDYT | 0.80067444 |
| PTGSGKTL | ASGEQRANSDYT | 0.6123408 |
| TGSGKTL | ASGEQRANSDYT | 0.5705795 |
| TIGHVDHGK | ASGEQRANSDYT | 0.8550689 |
| TIGHVDHGKTT | ASGEQRANSDYT | 0.7455906 |
| GHVDHGKTT | ASGEQRANSDYT | 0.77456784 |
| AQFEHTLL | ASGEQRANSDYT | 0.6700896 |
| IVNQDEL | ASGEQRANSDYT | 0.59006697 |
| TNGSQFF | ASGEQRANSDYT | 0.6883118 |
| TGKGIVE | ASGEQRANSDYT | 0.7637352 |
| LISNASDA | ASGEQRANSDYT | 0.775805 |
| AELRTLGELTKIAKAHGAQ | ASGEQRANSDYT | 0.7459773 |
| APNAVAAAEPSW | ASGEQRANSDYT | 0.546883 |
| APQVAGAATAVL | ASGEQRANSDYT | 0.7475184 |
| ARRDPAAAAAILPTDA | ASGEQRANSDYT | 0.70902383 |
| DAAARDVLAEAGLAEYF | ASGEQRANSDYT | 0.6786695 |
| DDQIVLSAKANLAKVEQ | ASGEQRANSDYT | 0.71061504 |
| DELLPDTGEDSDEDGH | ASGEQRANSDYT | 0.7292514 |
| ELPEAPANRATVPQD | ASGEQRANSDYT | 0.6454543 |
| FGQSKIERVTGIG | ASGEQRANSDYT | 0.64674157 |
| GAGGTGSTAGGAGGAGGAGGI | ASGEQRANSDYT | 0.70552504 |
| GHGGTGAAGGNGGTGGAGGS( | ASGEQRANSDYT | 0.743822 |
| GNGGNGGAGGEGGPGGAGGA( | ASGEQRANSDYT | 0.74280775 |
| GSTAGNGGNGGNATNGTIAG( | ASGEQRANSDYT | 0.7779286 |
| GTGGDGGNGGNGGTGG | ASGEQRANSDYT | 0.7751962 |
| GVRRPAGIPADAPV | ASGEQRANSDYT | 0.73713344 |
| GYRQVTDAHLLTLARR | ASGEQRANSDYT | 0.25714538 |
| IAKMGQDGHDRG | ASGEQRANSDYT | 0.38781315 |
| KAEPSIEIQYT | ASGEQRANSDYT | 0.59772074 |
| KPVRIRMAADEDTVAVT | ASGEQRANSDYT | 0.6853361 |
| MPPRIAGMRLLVIKPEP | ASGEQRANSDYT | 0.32270548 |
| MKFYAVALFRSRGVPPTR | ASGEQRANSDYT | 0.7193601 |
| NAGGFGILGAVAHSP | ASGEQRANSDYT | 0.66722393 |
| NEGREITLAMAEM | ASGEQRANSDYT | 0.62457913 |
| NELVEWSHPLGHRWRWE | ASGEQRANSDYT | 0.76303434 |
| NLGAGGTGGNGGNPG | ASGEQRANSDYT | 0.666466 |

| | | |
|---|---|---|
| NTEWPKWETVMAADPVD | ASGEQRANSDYT | 0.5857275 |
| PASAAPAETQP | ASGEQRANSDYT | 0.57205755 |
| QQSSDGKNGVRAAEITGE | ASGEQRANSDYT | 0.75446296 |
| RAPDAVTTSWWKE | ASGEQRANSDYT | 0.4554386 |
| SGFGNVGAHTSGWFNQ | ASGEQRANSDYT | 0.76156616 |
| SGVAGAVGQAASVGGLKV | ASGEQRANSDYT | 0.44789296 |
| SNGGAGTGSAGGNGGT | ASGEQRANSDYT | 0.7731103 |
| TGGAGGEGGAAG | ASGEQRANSDYT | 0.4899338 |
| TGGDGGAAGAGGAPGHG | ASGEQRANSDYT | 0.7566782 |
| TMPPLLAAIANAGRHGV | ASGEQRANSDYT | 0.70097214 |
| TSSTAGNSGGA | ASGEQRANSDYT | 0.79660404 |
| VPATVPVGILC | ASGEQRANSDYT | 0.69240624 |
| VQRPDSGGVVLTGRIS | ASGEQRANSDYT | 0.74613845 |
| WTPNPVPATKGVPAGLVQAT | ASGEQRANSDYT | 0.57413584 |
| YALKRALAKFSDRFDVVII | ASGEQRANSDYT | 0.76934785 |
| YLIGSVIV | ASGEQRANSDYT | 0.5105353 |
| YGQLVAVID | ASGEQRANSDYT | 0.7006483 |
| VTVGGMTLPA | ASGEQRANSDYT | 0.36477578 |
| VLALPAV | ASGEQRANSDYT | 0.73620325 |
| VIPPTLNYET | ASGEQRANSDYT | 0.74983 |
| VGMGGMHPSSG | ASGEQRANSDYT | 0.49875605 |
| VGLAGGAATA | ASGEQRANSDYT | 0.7729603 |
| VGGAGGAGGSGP | ASGEQRANSDYT | 0.6985594 |
| VFAAFMV | ASGEQRANSDYT | 0.62131107 |
| TLYDKLR | ASGEQRANSDYT | 0.7691664 |
| TLAVFTVL | ASGEQRANSDYT | 0.69158506 |
| TGSVQPARV | ASGEQRANSDYT | 0.77167845 |
| TGGGAGSGGT | ASGEQRANSDYT | 0.6085603 |
| SHLETSL | ASGEQRANSDYT | 0.66259736 |
| SFTTGMT | ASGEQRANSDYT | 0.72269964 |
| QVEVAKRMGVRQPTVS | ASGEQRANSDYT | 0.6332469 |
| QGSGGPALAM | ASGEQRANSDYT | 0.5443468 |
| MLAPSNP | ASGEQRANSDYT | 0.7709713 |
| LVNNAAI | ASGEQRANSDYT | 0.0633583 |
| LTGEINTLA | ASGEQRANSDYT | 0.063407995 |
| LSIFGPV | ASGEQRANSDYT | 0.06300327 |
| LLALLANL | ASGEQRANSDYT | 0.06473357 |
| LISNASDALDKLR | ASGEQRANSDYT | 0.06334048 |
| LINEAVP | ASGEQRANSDYT | 0.06340712 |
| LGYAFSGL | ASGEQRANSDYT | 0.066350214 |
| LDFVAVAL | ASGEQRANSDYT | 0.063128024 |
| LAQPGAE | ASGEQRANSDYT | 0.06626932 |
| ISNLGAVTGF | ASGEQRANSDYT | 0.06971822 |
| HQEDPGRI | ASGEQRANSDYT | 0.0629121 |
| GTPGEPTP | ASGEQRANSDYT | 0.06536027 |
| GFFNSGS | ASGEQRANSDYT | 0.06336694 |
| GAGGGGLTAGAN | ASGEQRANSDYT | 0.06528536 |
| FVDALVNM | ASGEQRANSDYT | 0.062895045 |
| FLTVAPEL | ASGEQRANSDYT | 0.06353871 |
| FETLTAL | ASGEQRANSDYT | 0.06397965 |
| DVINPIEL | ASGEQRANSDYT | 0.0666474 |

| | | |
|---|---|---|
| APVSGAPV | ASGEQRANSDYT | 0.06533386 |
| ALVGGLAW | ASGEQRANSDYT | 0.0647971 |
| ALAAAAATAI | ASGEQRANSDYT | 0.06360256 |
| AFGLGLLL | ASGEQRANSDYT | 0.06331509 |
| AFFAAASL | ASGEQRANSDYT | 0.06292236 |
| AAAALPAFSPP | ASGEQRANSDYT | 0.06383026 |
| VVLAPAL | ASGEQRANSDYT | 0.06357202 |
| TIGHVDHGKTTL | ASGEQRANSDYT | 0.06512163 |
| LISNASDALDK | ASGEQRANSDYT | 0.06208238 |
| LISNASDAL | ASGEQRANSDYT | 0.0681125 |
| ISNASDALDK | ASGEQRANSDYT | 0.06315678 |
| NQAETLVYQTE | ASGEQRANSDYT | 0.06543457 |
| TGGTGGTAGAG | ASGEQRANSDYT | 0.06581018 |
| TGSTAGGAGGA | ASGEQRANSDYT | 0.06432378 |
| DPLAVPVAAGPV | ASGEQRANSDYT | 0.06467468 |
| GGTGGTPGNGGN | ASGEQRANSDYT | 0.063800484 |
| PAGLAGVDPATL | ASGEQRANSDYT | 0.06631427 |
| GNNLVVEAPHAGASV | ASGEQRANSDYT | 0.06316365 |
| KVTRIMSASTLEQPA | ASGEQRANSDYT | 0.07021062 |
| LGFALPIALPRRTVV | ASGEQRANSDYT | 0.06270528 |
| TRLGNLTARKLVVLV | ASGEQRANSDYT | 0.06640513 |
| WSQQRRLIYFPSAGP | ASGEQRANSDYT | 0.064651795 |
| GPNGNPDPMAAAVDIR | ASGEQRANSDYT | 0.06328105 |
| TQRCVSVADLRAAASR | ASGEQRANSDYT | 0.0635178 |
| QFVLDVAEHPVEVLTETV | ASGEQRANSDYT | 0.06907647 |
| GGAGGTGGAGGNGGTQAGDG | ASGEQRANSDYT | 0.06603172 |
| PLLATLRAIAVDVDAKPAQV | ASGEQRANSDYT | 0.067653105 |
| APKFRTGGYMIDFWGVGYQV | ASGEQRANSDYT | 0.06516248 |
| GAGGAGGAGGAAGGPGGTGG | ASGEQRANSDYT | 0.06497233 |
| IGALGLLATDVAPAGSVAFG | ASGEQRANSDYT | 0.06404616 |
| AGGAGGIGGNGGAGGAGGVL | ASGEQRANSDYT | 0.063353024 |
| ADELIGLL | ASGEQRANSDYT | 0.063099466 |
| ADRALKA | ASGEQRANSDYT | 0.06421669 |
| AGAVGQVGGAGGA | ASGEQRANSDYT | 0.0632912 |
| AGGGGGLGGAP | ASGEQRANSDYT | 0.07001294 |
| AGGLLYGNGGAG | ASGEQRANSDYT | 0.063194305 |
| AGLESIGL | ASGEQRANSDYT | 0.066965416 |
| AGLKTGII | ASGEQRANSDYT | 0.06354538 |
| ASLAALLL | ASGEQRANSDYT | 0.06360804 |
| ASVINISSI | ASGEQRANSDYT | 0.06789014 |
| ATPPGPA | ASGEQRANSDYT | 0.0627615 |
| ATVASALV | ASGEQRANSDYT | 0.06300521 |
| ATVASLIL | ASGEQRANSDYT | 0.062725134 |
| AVNGPVLIRH | ASGEQRANSDYT | 0.06516883 |
| DGAAGAVGI | ASGEQRANSDYT | 0.067849174 |
| DPLLHGI | ASGEQRANSDYT | 0.064050935 |
| ELGAALGAL | ASGEQRANSDYT | 0.06388358 |
| ELGRTQAV | ASGEQRANSDYT | 0.7670536 |
| FVAGGYLS | ASGEQRANSDYT | 0.78049815 |
| GASGGAGGAGGW | ASGEQRANSDYT | 0.566237 |
| GGAGAGGDGGAGGI | ASGEQRANSDYT | 0.47659755 |

| | | |
|---|---|---|
| GTGHNGSQPGMGGNG | ASGEQRANSDYT | 0.51854247 |
| GYAAASASAS | ASGEQRANSDYT | 0.7576303 |
| HQVDVAAVQQG | ASGEQRANSDYT | 0.802873 |
| IGAAIAHT | ASGEQRANSDYT | 0.7673055 |
| IGMIEVAQL | ASGEQRANSDYT | 0.38090318 |
| IRLAPEHTSAMD | ASGEQRANSDYT | 0.75067145 |
| IYNVSVL | ASGEQRANSDYT | 0.63901895 |
| KQGFLGM | ASGEQRANSDYT | 0.63487077 |
| LESVITQL | ASGEQRANSDYT | 0.6869676 |
| LGLGALVVV | ASGEQRANSDYT | 0.6018671 |
| LIGNGGNGGSGGI | ASGEQRANSDYT | 0.51098734 |
| LNAINAPTL | ASGEQRANSDYT | 0.5085046 |
| LQLLNTGV | ASGEQRANSDYT | 0.56033486 |
| LQTSIEF | ASGEQRANSDYT | 0.7366962 |
| MSFAVSEL | ASGEQRANSDYT | 0.6639123 |
| NTGALGL | ASGEQRANSDYT | 0.8059376 |
| PPGAGGAGGAGTT | ASGEQRANSDYT | 0.78628755 |
| QLVGLGI | ASGEQRANSDYT | 0.7639368 |
| RFLEQVHQL | ASGEQRANSDYT | 0.19609371 |
| RGVGYVLR | ASGEQRANSDYT | 0.3226022 |
| SATGVFAGVI | ASGEQRANSDYT | 0.40147984 |
| SLPAITLP | ASGEQRANSDYT | 0.6083466 |
| STALAFTV | ASGEQRANSDYT | 0.18844639 |
| SVLTVGLGK | ASGEQRANSDYT | 0.6473168 |
| TALAGAGSGI | ASGEQRANSDYT | 0.60214543 |
| TGLPFTGI | ASGEQRANSDYT | 0.4777793 |
| VAEHVTV | ASGEQRANSDYT | 0.7612331 |
| VDIVLNSL | ASGEQRANSDYT | 0.55432713 |
| VLTGGGAVL | ASGEQRANSDYT | 0.403561 |
| VNVGLAYGI | ASGEQRANSDYT | 0.49461752 |
| VSSAAVGAV | ASGEQRANSDYT | 0.7952728 |
| VTSVYEAGV | ASGEQRANSDYT | 0.35958332 |
| WVALDRGA | ASGEQRANSDYT | 0.79289573 |
| YLVGSVVI | ASGEQRANSDYT | 0.49308622 |
| GPPGTGKTLL | ASGEQRANSDYT | 0.77893525 |
| GVGMTKF | ASGEQRANSDYT | 0.32158604 |
| IGIGGSDL | ASGEQRANSDYT | 0.79234123 |
| IGTLLAEL | ASGEQRANSDYT | 0.7027613 |
| IYTSGTT | ASGEQRANSDYT | 0.8172472 |
| LSFPAGV | ASGEQRANSDYT | 0.6373967 |
| LYTSGTT | ASGEQRANSDYT | 0.715788 |
| VGAFPFGF | ASGEQRANSDYT | 0.5259644 |
| VGAGTVEF | ASGEQRANSDYT | 0.80281913 |
| VGKSTLLL | ASGEQRANSDYT | 0.30836207 |
| AALAHAGGGIV | ASGEQRANSDYT | 0.56748456 |
| AALVAVSLL | ASGEQRANSDYT | 0.35617155 |
| AAVSRAVAAL | ASGEQRANSDYT | 0.68102163 |
| AGGSSGIGGF | ASGEQRANSDYT | 0.78882396 |
| AGLTVANANG | ASGEQRANSDYT | 0.21107769 |
| AGLVGAVL | ASGEQRANSDYT | 0.8049469 |
| ALSQAFTV | ASGEQRANSDYT | 0.5424117 |

| | | |
|---|---|---|
| ALTFSVSL | ASGEQRANSDYT | 0.3972466 |
| ALTGAALVAL | ASGEQRANSDYT | 0.7481626 |
| ASFSIGAGVL | ASGEQRANSDYT | 0.56269175 |
| DGSVYKI | ASGEQRANSDYT | 0.7493329 |
| EQLAEIV | ASGEQRANSDYT | 0.4582436 |
| FAVSELTVM | ASGEQRANSDYT | 0.68178713 |
| GAAPLAAGL | ASGEQRANSDYT | 0.7242787 |
| GADGGDGGSGGAGG | ASGEQRANSDYT | 0.24081568 |
| GFGLDTGGAGG | ASGEQRANSDYT | 0.3589113 |
| GLASGGLL | ASGEQRANSDYT | 0.81943893 |
| GRIAGAAVLL | ASGEQRANSDYT | 0.6055401 |
| GSAGSLNTSF | ASGEQRANSDYT | 0.7333741 |
| GTGQIGI | ASGEQRANSDYT | 0.40995786 |
| GTLIALIL | ASGEQRANSDYT | 0.38519254 |
| IAATMGFVVG | ASGEQRANSDYT | 0.42532796 |
| IFAQALL | ASGEQRANSDYT | 0.54338115 |
| IGAAVALGF | ASGEQRANSDYT | 0.7382213 |
| IGLTPLAL | ASGEQRANSDYT | 0.5463805 |
| ILGSNTSTL | ASGEQRANSDYT | 0.29757342 |
| IVFSLTVL | ASGEQRANSDYT | 0.5610882 |
| LASKATGF | ASGEQRANSDYT | 0.49232805 |
| LASVALIL | ASGEQRANSDYT | 0.59753 |
| LFAVGLTL | ASGEQRANSDYT | 0.37330917 |
| LFNGGIGTYI | ASGEQRANSDYT | 0.37704274 |
| LTLGAGII | ASGEQRANSDYT | 0.34385216 |
| LVAAALGDI | ASGEQRANSDYT | 0.2924825 |
| NTAFVQLGI | ASGEQRANSDYT | 0.43853223 |
| QLFSGAEL | ASGEQRANSDYT | 0.6012568 |
| SAATFITL | ASGEQRANSDYT | 0.56212866 |
| SNYGLANLGV | ASGEQRANSDYT | 0.50621957 |
| SYGGAYAVM | ASGEQRANSDYT | 0.6849183 |
| TTGVGGLTLG | ASGEQRANSDYT | 0.3642048 |
| TTQTIYI | ASGEQRANSDYT | 0.22146384 |
| VAAGAGLGAVI | ASGEQRANSDYT | 0.27826005 |
| VAAVVTLL | ASGEQRANSDYT | 0.29926124 |
| VGFLGGLI | ASGEQRANSDYT | 0.43656695 |
| VGVIPILL | ASGEQRANSDYT | 0.21752004 |
| VIFASSLI | ASGEQRANSDYT | 0.3848671 |
| VKGGDGGAAAG | ASGEQRANSDYT | 0.3689537 |
| VTKTFTNI | ASGEQRANSDYT | 0.4244104 |
| VVAAAILL | ASGEQRANSDYT | 0.6364305 |
| YAALDVEL | ASGEQRANSDYT | 0.43547228 |
| LASVALLL | ASGEQRANSDYT | 0.28940684 |
| LDAVHRV | ASGEQRANSDYT | 0.28413817 |
| QGLSSSIF | ASGEQRANSDYT | 0.66535795 |
| RGTFANIRL | ASGEQRANSDYT | 0.26450193 |
| RQLWWGHRI | ASGEQRANSDYT | 0.63914764 |
| YGPPGTGKTLL | ASGEQRANSDYT | 0.3153182 |
| AALVGAPANVA | ASGEQRANSDYT | 0.6670594 |
| AATFRRTE | ASGEQRANSDYT | 0.29008853 |
| AGALGGGI | ASGEQRANSDYT | 0.5879922 |

| | | |
|---|---|---|
| AGGAGGAGGIGAG | ASGEQRANSDYT | 0.602919 |
| AGSGALGL | ASGEQRANSDYT | 0.7003834 |
| AQIADQI | ASGEQRANSDYT | 0.4222685 |
| ASAVAVTYL | ASGEQRANSDYT | 0.4299684 |
| ASTVLGGTGG | ASGEQRANSDYT | 0.29175815 |
| AVGGEPAK | ASGEQRANSDYT | 0.74383473 |
| CEQGTFGGTC | ASGEQRANSDYT | 0.28169772 |
| DEGKAPAG | ASGEQRANSDYT | 0.3962185 |
| DTVVFPAL | ASGEQRANSDYT | 0.2798415 |
| FEGITMAG | ASGEQRANSDYT | 0.37969935 |
| FPLVVQD | ASGEQRANSDYT | 0.6510755 |
| GAFTTPQ | ASGEQRANSDYT | 0.2244406 |
| GFGGFGTAGGVGGA | ASGEQRANSDYT | 0.5226487 |
| GFLAQFL | ASGEQRANSDYT | 0.33593118 |
| GGAGGAGGNAGVA | ASGEQRANSDYT | 0.30050293 |
| GGVGTAGAGGAGGAGG | ASGEQRANSDYT | 0.4441694 |
| GNGGGGDAGNAGSGGN | ASGEQRANSDYT | 0.37664545 |
| GTGGAGGAGNDAGS | ASGEQRANSDYT | 0.5028996 |
| GTTNTGWFNS | ASGEQRANSDYT | 0.2802631 |
| IFMGPLFTL | ASGEQRANSDYT | 0.52042335 |
| ILAHAIEAI | ASGEQRANSDYT | 0.4517377 |
| IRSAIREAL | ASGEQRANSDYT | 0.26914245 |
| ITFAGGG | ASGEQRANSDYT | 0.26512402 |
| ITGMSGHQEV | ASGEQRANSDYT | 0.66253185 |
| IVAGLACTL | ASGEQRANSDYT | 0.20897733 |
| IVGGGLAGA | ASGEQRANSDYT | 0.26244476 |
| IVTGVLNI | ASGEQRANSDYT | 0.11240978 |
| LAQQGCI | ASGEQRANSDYT | 0.12821306 |
| LCLHVEN | ASGEQRANSDYT | 0.118346736 |
| LDAISDGS | ASGEQRANSDYT | 0.15859483 |
| LGGFAAML | ASGEQRANSDYT | 0.17833194 |
| LGGKNPMI | ASGEQRANSDYT | 0.20360635 |
| LGPKGVAAIL | ASGEQRANSDYT | 0.12533869 |
| LMVGVSI | ASGEQRANSDYT | 0.13395071 |
| LVGAAFVL | ASGEQRANSDYT | 0.13337189 |
| LVLAVAPL | ASGEQRANSDYT | 0.14151351 |
| MAAIAVIDAA | ASGEQRANSDYT | 0.13697837 |
| NNGINAELQ | ASGEQRANSDYT | 0.115887545 |
| RGEPLVLVHG | ASGEQRANSDYT | 0.11150469 |
| RGGAGGAAGLI | ASGEQRANSDYT | 0.120954454 |
| RSYRMHPAV | ASGEQRANSDYT | 0.12025981 |
| SAAHLLAGQLA | ASGEQRANSDYT | 0.14943928 |
| SAAVGSY | ASGEQRANSDYT | 0.14153445 |
| SPHAVLHRI | ASGEQRANSDYT | 0.18499336 |
| SQIAKVL | ASGEQRANSDYT | 0.1808448 |
| TALGHLVP | ASGEQRANSDYT | 0.12745327 |
| TAPTTSGAA | ASGEQRANSDYT | 0.09896023 |
| TDGDDGNGGSPGT | ASGEQRANSDYT | 0.10821253 |
| TDGLGEN | ASGEQRANSDYT | 0.12247266 |
| TGGPPLPGQA | ASGEQRANSDYT | 0.117474854 |
| TIPAITI | ASGEQRANSDYT | 0.09659071 |

| | | |
|---|---|---|
| TIPGFTI | ASGEQRANSDYT | 0.13271718 |
| TSIASGALA | ASGEQRANSDYT | 0.16363564 |
| TVTLPGAAI | ASGEQRANSDYT | 0.11558879 |
| VASGLSAALP | ASGEQRANSDYT | 0.15706895 |
| VGAGVIGI | ASGEQRANSDYT | 0.11825252 |
| VGPKRFGV | ASGEQRANSDYT | 0.110496216 |
| VNVAFPD | ASGEQRANSDYT | 0.115005955 |
| VSMTNSGVS | ASGEQRANSDYT | 0.16927978 |
| VTDVLGDLSV | ASGEQRANSDYT | 0.110313915 |
| VVVVGAGFAGL | ASGEQRANSDYT | 0.17789242 |
| VYTVGTVV | ASGEQRANSDYT | 0.12576419 |
| WTTFSGP | ASGEQRANSDYT | 0.203366 |
| YDELVEAF | ASGEQRANSDYT | 0.11679621 |
| YGTGGAGGNGG | ASGEQRANSDYT | 0.16773386 |
| YTNVMAV | ASGEQRANSDYT | 0.13739996 |
| FLEQVHQL | ASGEQRANSDYT | 0.17142302 |
| GLSLLAVAV | ASGEQRANSDYT | 0.1318566 |
| LEQVHQL | ASGEQRANSDYT | 0.14811249 |
| LNAAGAI | ASGEQRANSDYT | 0.1129981 |
| PAVHASL | ASGEQRANSDYT | 0.15425253 |
| TELEDDA | ASGEQRANSDYT | 0.10809849 |
| TPPGPPN | ASGEQRANSDYT | 0.13077638 |
| TPPPPPPP | ASGEQRANSDYT | 0.1174875 |
| TQSPALI | ASSLEEGAGNTLY | 0.13112962 |
| EFAAAFPQ | ASSLEEGAGNTLY | 0.24583907 |
| LLSGNLDVL | ASSLEEGAGNTLY | 0.099447474 |
| GPANLGNYNI | ASSLEEGAGNTLY | 0.16320495 |
| TLGPTLP | ASSLEEGAGNTLY | 0.10483209 |
| VYAYSSCCK | ASSLEEGAGNTLY | 0.12392986 |
| APAPAPAPNI | ASSLEEGAGNTLY | 0.13753147 |
| PGGPVPIP | ASSLEEGAGNTLY | 0.11719578 |
| FLTVALEP | ASSLEEGAGNTLY | 0.1292534 |
| SQYFGANL | ASSLEEGAGNTLY | 0.10124514 |
| GGVKPAA | ASSLEEGAGNTLY | 0.13938174 |
| GGTGGAGGAGGW | ASSLEEGAGNTLY | 0.17032908 |
| GVGGVAGLGGAGG | ASSLEEGAGNTLY | 0.09930264 |
| VLLVLATTI | ASSLEEGAGNTLY | 0.105895236 |
| IGIFAAML | ASSLEEGAGNTLY | 0.20312251 |
| AVGDLID | ASSLEEGAGNTLY | 0.2958901 |
| NISPLPL | ASSLEEGAGNTLY | 0.288382 |
| VTAVFVAI | ASSLEEGAGNTLY | 0.2897724 |
| SGDDNDAAL | ASSLEEGAGNTLY | 0.2918704 |
| AVAALAHS | ASSLEEGAGNTLY | 0.29105508 |
| GGVTVMATP | ASSLEEGAGNTLY | 0.29419908 |
| AGLAANVRD | ASSLEEGAGNTLY | 0.29216594 |
| TNRPDLIDPALLRPG | ASSLEEGAGNTLY | 0.29917875 |
| LANLGVTAL | ASSLEEGAGNTLY | 0.2950649 |
| TGERIEL | ASSLEEGAGNTLY | 0.29915917 |
| NGEELAA | ASSLEEGAGNTLY | 0.29682738 |
| VLSLVWG | ASSLEEGAGNTLY | 0.2990711 |
| IGYALWL | ASSLEEGAGNTLY | 0.2926225 |

| | | |
|---|---|---|
| LSLLLAPL | ASSLEEGAGNTLY | 0.29133874 |
| TLDEALK | ASSLEEGAGNTLY | 0.2895813 |
| TDGAGTAEGG | ASSLEEGAGNTLY | 0.28997386 |
| NAPGGAGGNGG | ASSLEEGAGNTLY | 0.29068303 |
| VSVFMGMGGNF | ASSLEEGAGNTLY | 0.29271746 |
| LASYLAAL | ASSLEEGAGNTLY | 0.29206118 |
| LVGGTPAL | ASSLEEGAGNTLY | 0.3047265 |
| VRNADVL | ASSLEEGAGNTLY | 0.29859126 |
| TCSSPFLVSAM | ASSLEEGAGNTLY | 0.29287916 |
| VHITDAE | ASSLEEGAGNTLY | 0.28869715 |
| EYAAPSF | ASSLEEGAGNTLY | 0.2889785 |
| VVDAFHGGL | ASSLEEGAGNTLY | 0.2938054 |
| ASVVDPQ | ASSLEEGAGNTLY | 0.2907115 |
| SYNEFGGLS | ASSLEEGAGNTLY | 0.28646913 |
| DPDGSVY | ASSLEEGAGNTLY | 0.2936867 |
| MVSTIHATE | ASSLEEGAGNTLY | 0.29390076 |
| LAAGVADPV | ASSLEEGAGNTLY | 0.2931667 |
| TKEQIAA | ASSLEEGAGNTLY | 0.2988339 |
| SLASPAAL | ASSLEEGAGNTLY | 0.2923075 |
| IGLVFYGV | ASSLEEGAGNTLY | 0.2898129 |
| FLLDASNL | ASSLEEGAGNTLY | 0.29327065 |
| YQVGNFP | ASSLEEGAGNTLY | 0.30809098 |
| ESLLSQST | ASSLEEGAGNTLY | 0.29300764 |
| LFTVGFAAVG | ASSLEEGAGNTLY | 0.3111993 |
| ATPPTPT | ASSLEEGAGNTLY | 0.2903543 |
| TDLATLG | ASSLEEGAGNTLY | 0.30694824 |
| SAGGTGR | ASSLEEGAGNTLY | 0.29265332 |
| GFSNSGS | ASSLEEGAGNTLY | 0.29951432 |
| GGNGGSGGTSV | ASSLEEGAGNTLY | 0.29406434 |
| GTSVATGGAGN | ASSLEEGAGNTLY | 0.3009589 |
| NGGNGGSGGTS | ASSLEEGAGNTLY | 0.29421687 |
| AVVKFPH | ASSLEEGAGNTLY | 0.3050598 |
| TPPPPPP | ASSLEEGAGNTLY | 0.2907692 |
| AAPAAAAAPA | ASSLEEGAGNTLY | 0.29011756 |
| IALNEDL | ASSLEEGAGNTLY | 0.29044673 |
| IVNVSSM | ASSLEEGAGNTLY | 0.29081303 |
| TGIPPAPR | ASSLEEGAGNTLY | 0.2919121 |
| FDLGGGTF | ASSLEEGAGNTLY | 0.29308102 |
| FDLGGGTFDVS | ASSLEEGAGNTLY | 0.30062157 |
| NEPTAAA | ASSLEEGAGNTLY | 0.28757164 |
| NEPTAAALA | ASSLEEGAGNTLY | 0.29853255 |
| GSIINISSI | ASSLEEGAGNTLY | 0.2888288 |
| VAPGNAGTA | ASSLEEGAGNTLY | 0.29101798 |
| TGGGASLE | ASSLEEGAGNTLY | 0.29321375 |
| VAVYNHYKRI | ASSLEEGAGNTLY | 0.29462013 |
| TDPAPATT | ASSLEEGAGNTLY | 0.28981438 |
| TNRPDILDPALLRPG | ASSLEEGAGNTLY | 0.28861555 |
| LTTTTTS | ASSLEEGAGNTLY | 0.29471633 |
| PTGSGKTL | ASSLEEGAGNTLY | 0.2945977 |
| TGSGKTL | ASSLEEGAGNTLY | 0.28783122 |
| TIGHVDHGK | ASSLEEGAGNTLY | 0.29053387 |

| | | |
|---|---|---|
| TIGHVDHGKTT | ASSLEEGAGNTLY | 0.30335233 |
| GHVDHGKTT | ASSLEEGAGNTLY | 0.21955983 |
| AQFEHTLL | ASSLEEGAGNTLY | 0.21937476 |
| IVNQDEL | ASSLEEGAGNTLY | 0.21932712 |
| TNGSQFF | ASSLEEGAGNTLY | 0.21981928 |
| TGKGIVE | ASSLEEGAGNTLY | 0.21934389 |
| LISNASDA | ASSLEEGAGNTLY | 0.2194191 |
| AELRTLGELTKIAKAHGAQ | ASSLEEGAGNTLY | 0.21975894 |
| APNAVAAAEPSW | ASSLEEGAGNTLY | 0.21956944 |
| APQVAGAATAVL | ASSLEEGAGNTLY | 0.21981202 |
| ARRDPAAAAAILPTDA | ASSLEEGAGNTLY | 0.22003463 |
| DAAARDVLAEAGLAEYF | ASSLEEGAGNTLY | 0.2194775 |
| DDQIVLSAKANLAKVEQ | ASSLEEGAGNTLY | 0.21957237 |
| DELLPDTGEDSDEDGH | ASSLEEGAGNTLY | 0.21953534 |
| ELPEAPANRATVPQD | ASSLEEGAGNTLY | 0.22030044 |
| FGQSKIERVTGIG | ASSLEEGAGNTLY | 0.21941926 |
| GAGGTGSTAGGAGGAGGAGGI | ASSLEEGAGNTLY | 0.21932936 |
| GHGGTGAAGGNGGTGGAGGS( | ASSLEEGAGNTLY | 0.21985479 |
| GNGGNGGAGGEGGPGGAGGA( | ASSLEEGAGNTLY | 0.21995685 |
| GSTAGNGGNGGNATNGTIAG( | ASSLEEGAGNTLY | 0.21991089 |
| GTGGDGGNGGNGGTGG | ASSLEEGAGNTLY | 0.21946634 |
| GVRRPAGIPADAPV | ASSLEEGAGNTLY | 0.21957919 |
| GYRQVTDAHLLTLARR | ASSLEEGAGNTLY | 0.21966787 |
| IAKMGQDGHDRG | ASSLEEGAGNTLY | 0.21942177 |
| KAEPSIEIQYT | ASSLEEGAGNTLY | 0.21959785 |
| KPVRIRMAADEDTVAVT | ASSLEEGAGNTLY | 0.21957368 |
| MPPRIAGMRLLVIKPEP | ASSLEEGAGNTLY | 0.21967354 |
| MKFYAVALFRSRGVPPTR | ASSLEEGAGNTLY | 0.21892472 |
| NAGGFGILGAVAHSP | ASSLEEGAGNTLY | 0.22020803 |
| NEGREITLAMAEM | ASSLEEGAGNTLY | 0.22001374 |
| NELVEWSHPLGHRWRWE | ASSLEEGAGNTLY | 0.21973868 |
| NLGAGGTGGNGGNPG | ASSLEEGAGNTLY | 0.22002825 |
| NTEWPKWETVMAADPVD | ASSLEEGAGNTLY | 0.21951738 |
| PASAAPAETQP | ASSLEEGAGNTLY | 0.21978587 |
| QQSSDGKNGVRAAEITGE | ASSLEEGAGNTLY | 0.21947414 |
| RAPDAVTTSWWKE | ASSLEEGAGNTLY | 0.21968645 |
| SGFGNVGAHTSGWFNQ | ASSLEEGAGNTLY | 0.2196034 |
| SGVAGAVGQAASVGGLKV | ASSLEEGAGNTLY | 0.22027342 |
| SNGGAGTGSAGGNGGT | ASSLEEGAGNTLY | 0.21926092 |
| TGGAGGEGGAAG | ASSLEEGAGNTLY | 0.21998999 |
| TGGDGGAAGAGGAPGHG | ASSLEEGAGNTLY | 0.21970642 |
| TMPPLLAAIANAGRHGV | ASSLEEGAGNTLY | 0.21957025 |
| TSSTAGNSGGA | ASSLEEGAGNTLY | 0.21949011 |
| VPATVPVGILC | ASSLEEGAGNTLY | 0.2201705 |
| VQRPDSGGVVLTGRIS | ASSLEEGAGNTLY | 0.2198878 |
| WTPNPVPATKGVPAGLVQAT | ASSLEEGAGNTLY | 0.22028309 |
| YALKRALAKFSDRFDVVII | ASSLEEGAGNTLY | 0.21987471 |
| YLIGSVIV | ASSLEEGAGNTLY | 0.219788 |
| YGQLVAVID | ASSLEEGAGNTLY | 0.21954325 |
| VTVGGMTLPA | ASSLEEGAGNTLY | 0.21959767 |
| VLALPAV | ASSLEEGAGNTLY | 0.21957578 |

| | | |
|---|---|---|
| VIPPTLNYET | ASSLEEGAGNTLY | 0.21952012 |
| VGMGGMHPSSG | ASSLEEGAGNTLY | 0.21957424 |
| VGLAGGAATA | ASSLEEGAGNTLY | 0.21912025 |
| VGGAGGAGGSGP | ASSLEEGAGNTLY | 0.21933073 |
| VFAAFMV | ASSLEEGAGNTLY | 0.21909422 |
| TLYDKLR | ASSLEEGAGNTLY | 0.21965607 |
| TLAVFTVL | ASSLEEGAGNTLY | 0.21951443 |
| TGSVQPARV | ASSLEEGAGNTLY | 0.22029674 |
| TGGGAGSGGT | ASSLEEGAGNTLY | 0.2193037 |
| SHLETSL | ASSLEEGAGNTLY | 0.21941333 |
| SFTTGMT | ASSLEEGAGNTLY | 0.2193709 |
| QVEVAKRMGVRQPTVS | ASSLEEGAGNTLY | 0.21960871 |
| QGSGGPALAM | ASSLEEGAGNTLY | 0.21961051 |
| MLAPSNP | ASSLEEGAGNTLY | 0.21956381 |
| LVNNAAI | ASSLEEGAGNTLY | 0.21959147 |
| LTGEINTLA | ASSLEEGAGNTLY | 0.29641894 |
| LSIFGPV | ASSLEEGAGNTLY | 0.2939346 |
| LLALLANL | ASSLEEGAGNTLY | 0.26432735 |
| LISNASDALDKLR | ASSLEEGAGNTLY | 0.27705356 |
| LINEAVP | ASSLEEGAGNTLY | 0.2731553 |
| LGYAFSGL | ASSLEEGAGNTLY | 0.29347724 |
| LDFVAVAL | ASSLEEGAGNTLY | 0.29121342 |
| LAQPGAE | ASSLEEGAGNTLY | 0.29384893 |
| ISNLGAVTGF | ASSLEEGAGNTLY | 0.28695947 |
| HQEDPGRI | ASSLEEGAGNTLY | 0.27894443 |
| GTPGEPTP | ASSLEEGAGNTLY | 0.28354654 |
| GFFNSGS | ASSLEEGAGNTLY | 0.271675 |
| GAGGGGLTAGAN | ASSLEEGAGNTLY | 0.29097825 |
| FVDALVNM | ASSLEEGAGNTLY | 0.27446005 |
| FLTVAPEL | ASSLEEGAGNTLY | 0.2636755 |
| FETLTAL | ASSLEEGAGNTLY | 0.28060928 |
| DVINPIEL | ASSLEEGAGNTLY | 0.2683297 |
| APVSGAPV | ASSLEEGAGNTLY | 0.2954332 |
| ALVGGLAW | ASSLEEGAGNTLY | 0.29451814 |
| ALAAAAATAI | ASSLEEGAGNTLY | 0.33276817 |
| AFGLGLLL | ASSLEEGAGNTLY | 0.30191424 |
| AFFAAASL | ASSLEEGAGNTLY | 0.27662474 |
| AAAALPAFSPP | ASSLEEGAGNTLY | 0.2557976 |
| VVLAPAL | ASSLEEGAGNTLY | 0.2682376 |
| TIGHVDHGKTTL | ASSLEEGAGNTLY | 0.27613062 |
| LISNASDALDK | ASSLEEGAGNTLY | 0.29146618 |
| LISNASDAL | ASSLEEGAGNTLY | 0.25387058 |
| ISNASDALDK | ASSLEEGAGNTLY | 0.2838782 |
| NQAETLVYQTE | ASSLEEGAGNTLY | 0.2851179 |
| TGGTGGTAGAG | ASSLEEGAGNTLY | 0.2657761 |
| TGSTAGGAGGA | ASSLEEGAGNTLY | 0.31897452 |
| DPLAVPVAAGPV | ASSLEEGAGNTLY | 0.2659053 |
| GGTGGTPGNGGN | ASSLEEGAGNTLY | 0.2743426 |
| PAGLAGVDPATL | ASSLEEGAGNTLY | 0.27565414 |
| GNNLVVEAPHAGASV | ASSLEEGAGNTLY | 0.33714017 |
| KVTRIMSASTLEQPA | ASSLEEGAGNTLY | 0.27088594 |

| | | |
|---|---|---|
| LGFALPIALPRRTVV | ASSLEEGAGNTLY | 0.31488404 |
| TRLGNLTARKLVVLV | ASSLEEGAGNTLY | 0.26614776 |
| WSQQRRLIYFPSAGP | ASSLEEGAGNTLY | 0.3109058 |
| GPNGNPDPMAAAVDIR | ASSLEEGAGNTLY | 0.27491316 |
| TQRCVSVADLRAAASR | ASSLEEGAGNTLY | 0.29032224 |
| QFVLDVAEHPVEVLTETV | ASSLEEGAGNTLY | 0.28172067 |
| GGAGGTGGAGGNGGTQAGDG | ASSLEEGAGNTLY | 0.31177995 |
| PLLATLRAIAVDVDAKPAQV | ASSLEEGAGNTLY | 0.2914961 |
| APKFRTGGYMIDFWGVGYQV | ASSLEEGAGNTLY | 0.314437 |
| GAGGAGGAGGAAGGPGGTGG | ASSLEEGAGNTLY | 0.27173495 |
| IGALGLLATDVAPAGSVAFG | ASSLEEGAGNTLY | 0.27516788 |
| AGGAGGIGGNGGAGGAGGVL | ASSLEEGAGNTLY | 0.26439127 |
| ADELIGLL | ASSLEEGAGNTLY | 0.27965334 |
| ADRALKA | ASSLEEGAGNTLY | 0.2661926 |
| AGAVGQVGGAGGA | ASSLEEGAGNTLY | 0.27819797 |
| AGGGGGLGGAP | ASSLEEGAGNTLY | 0.2922882 |
| AGGLLYGNGGAG | ASSLEEGAGNTLY | 0.26197448 |
| AGLESIGL | ASSLEEGAGNTLY | 0.28779835 |
| AGLKTGII | ASSLEEGAGNTLY | 0.26212505 |
| ASLAALLL | ASSLEEGAGNTLY | 0.28037894 |
| ASVINISSI | ASSLEEGAGNTLY | 0.2906064 |
| ATPPGPA | ASSLEEGAGNTLY | 0.26770186 |
| ATVASALV | ASSLEEGAGNTLY | 0.28313738 |
| ATVASLIL | ASSLEEGAGNTLY | 0.260534 |
| AVNGPVLIRH | ASSLEEGAGNTLY | 0.27275544 |
| DGAAGAVGI | ASSLEEGAGNTLY | 0.29866922 |
| DPLLHGI | ASSLEEGAGNTLY | 0.25707352 |
| ELGAALGAL | ASSLEEGAGNTLY | 0.26757324 |
| ELGRTQAV | ASSLEEGAGNTLY | 0.3100786 |
| FVAGGYLS | ASSLEEGAGNTLY | 0.19119501 |
| GASGGAGGAGGW | ASSLEEGAGNTLY | 0.26928958 |
| GGAGAGGDGGAGGI | ASSLEEGAGNTLY | 0.13686633 |
| GTGHNGSQPGMGGNG | ASSLEEGAGNTLY | 0.1500416 |
| GYAAASASAS | ASSLEEGAGNTLY | 0.14442515 |
| HQVDVAAVQQG | ASSLEEGAGNTLY | 0.16034506 |
| IGAAIAHT | ASSLEEGAGNTLY | 0.2645093 |
| IGMIEVAQL | ASSLEEGAGNTLY | 0.2028234 |
| IRLAPEHTSAMD | ASSLEEGAGNTLY | 0.14969866 |
| IYNVSVL | ASSLEEGAGNTLY | 0.154504 |
| KQGFLGM | ASSLEEGAGNTLY | 0.13952185 |
| LESVITQL | ASSLEEGAGNTLY | 0.16207942 |
| LGLGALVVV | ASSLEEGAGNTLY | 0.15656549 |
| LIGNGGNGGSGGI | ASSLEEGAGNTLY | 0.14112759 |
| LNAINAPTL | ASSLEEGAGNTLY | 0.13884948 |
| LQLLNTGV | ASSLEEGAGNTLY | 0.14674015 |
| LQTSIEF | ASSLEEGAGNTLY | 0.14604275 |
| MSFAVSEL | ASSLEEGAGNTLY | 0.18750995 |
| NTGALGL | ASSLEEGAGNTLY | 0.17475115 |
| PPGAGGAGGAGTT | ASSLEEGAGNTLY | 0.178284 |
| QLVGLGI | ASSLEEGAGNTLY | 0.17863975 |
| RFLEQVHQL | ASSLEEGAGNTLY | 0.15393658 |

| | | |
|---|---|---|
| RGVGYVLR | ASSLEEGAGNTLY | 0.12869044 |
| SATGVFAGVI | ASSLEEGAGNTLY | 0.136214 |
| SLPAITLP | ASSLEEGAGNTLY | 0.13991876 |
| STALAFTV | ASSLEEGAGNTLY | 0.14599492 |
| SVLTVGLGK | ASSLEEGAGNTLY | 0.12787075 |
| TALAGAGSGI | ASSLEEGAGNTLY | 0.15526572 |
| TGLPFTGI | ASSLEEGAGNTLY | 0.1587728 |
| VAEHVTV | ASSLEEGAGNTLY | 0.13928826 |
| VDIVLNSL | ASSLEEGAGNTLY | 0.18082567 |
| VLTGGGAVL | ASSLEEGAGNTLY | 0.14479995 |
| VNVGLAYGI | ASSLEEGAGNTLY | 0.13934028 |
| VSSAAVGAV | ASSLEEGAGNTLY | 0.14357077 |
| VTSVYEAGV | ASSLEEGAGNTLY | 0.18232827 |
| WVALDRGA | ASSLEEGAGNTLY | 0.1363805 |
| YLVGSVVI | ASSLEEGAGNTLY | 0.19884728 |
| GPPGTGKTLL | ASSLEEGAGNTLY | 0.14464088 |
| GVGMTKF | ASSLEEGAGNTLY | 0.18218699 |
| IGIGGSDL | ASSLEEGAGNTLY | 0.13900824 |
| IGTLLAEL | ASSLEEGAGNTLY | 0.17503448 |
| IYTSGTT | ASSLEEGAGNTLY | 0.15198845 |
| LSFPAGV | ASSLEEGAGNTLY | 0.20041019 |
| LYTSGTT | ASSLEEGAGNTLY | 0.14838685 |
| VGAFPFGF | ASSLEEGAGNTLY | 0.15732366 |
| VGAGTVEF | ASSLEEGAGNTLY | 0.14375198 |
| VGKSTLLL | ASSLEEGAGNTLY | 0.25323942 |
| AALAHAGGGIV | ASSLEEGAGNTLY | 0.13139434 |
| AALVAVSLL | ASSLEEGAGNTLY | 0.14721358 |
| AAVSRAVAAL | ASSLEEGAGNTLY | 0.14187431 |
| AGGSSGIGGF | ASSLEEGAGNTLY | 0.15165645 |
| AGLTVANANG | ASSLEEGAGNTLY | 0.22886282 |
| AGLVGAVL | ASSLEEGAGNTLY | 0.13043642 |
| ALSQAFTV | ASSLEEGAGNTLY | 0.17880659 |
| ALTFSVSL | ASSLEEGAGNTLY | 0.13261622 |
| ALTGAALVAL | ASSLEEGAGNTLY | 0.14399074 |
| ASFSIGAGVL | ASSLEEGAGNTLY | 0.1623769 |
| DGSVYKI | ASSLEEGAGNTLY | 0.14098898 |
| EQLAEIV | ASSLEEGAGNTLY | 0.15749641 |
| FAVSELTVM | ASSLEEGAGNTLY | 0.13451894 |
| GAAPLAAGL | ASSLEEGAGNTLY | 0.14946488 |
| GADGGDGGSGGAGG | ASSLEEGAGNTLY | 0.17644894 |
| GFGLDTGGAGG | ASSLEEGAGNTLY | 0.13076131 |
| GLASGGLL | ASSLEEGAGNTLY | 0.13556804 |
| GRIAGAAVLL | ASSLEEGAGNTLY | 0.19540794 |
| GSAGSLNTSF | ASSLEEGAGNTLY | 0.18838994 |
| GTGQIGI | ASSLEEGAGNTLY | 0.3566116 |
| GTLIALIL | ASSLEEGAGNTLY | 0.16716956 |
| IAATMGFVVG | ASSLEEGAGNTLY | 0.17653541 |
| IFAQALL | ASSLEEGAGNTLY | 0.18077765 |
| IGAAVALGF | ASSLEEGAGNTLY | 0.19781294 |
| IGLTPLAL | ASSLEEGAGNTLY | 0.4471318 |
| ILGSNTSTL | ASSLEEGAGNTLY | 0.1898914 |

| | | |
|---|---|---|
| IVFSLTVL | ASSLEEGAGNTLY | 0.17783491 |
| LASKATGF | ASSLEEGAGNTLY | 0.17840324 |
| LASVALIL | ASSLEEGAGNTLY | 0.17010407 |
| LFAVGLTL | ASSLEEGAGNTLY | 0.1713194 |
| LFNGGIGTYI | ASSLEEGAGNTLY | 0.18192074 |
| LTLGAGII | ASSLEEGAGNTLY | 0.1783262 |
| LVAAALGDI | ASSLEEGAGNTLY | 0.17027576 |
| NTAFVQLGI | ASSLEEGAGNTLY | 0.17440942 |
| QLFSGAEL | ASSLEEGAGNTLY | 0.17497306 |
| SAATFITL | ASSLEEGAGNTLY | 0.20828357 |
| SNYGLANLGV | ASSLEEGAGNTLY | 0.19313297 |
| SYGGAYAVM | ASSLEEGAGNTLY | 0.21103387 |
| TTGVGGLTLG | ASSLEEGAGNTLY | 0.19100459 |
| TTQTIYI | ASSLEEGAGNTLY | 0.18055154 |
| VAAGAGLGAVI | ASSLEEGAGNTLY | 0.16169527 |
| VAAVVTLL | ASSLEEGAGNTLY | 0.1682747 |
| VGFLGGLI | ASSLEEGAGNTLY | 0.16934209 |
| VGVIPILL | ASSLEEGAGNTLY | 0.17999397 |
| VIFASSLI | ASSLEEGAGNTLY | 0.16192 |
| VKGGDGGAAAG | ASSLEEGAGNTLY | 0.18753411 |
| VTKTFTNI | ASSLEEGAGNTLY | 0.18336284 |
| VVAAAILL | ASSLEEGAGNTLY | 0.16926177 |
| YAALDVEL | ASSLEEGAGNTLY | 0.19872142 |
| LASVALLL | ASSLEEGAGNTLY | 0.16878274 |
| LDAVHRV | ASSLEEGAGNTLY | 0.17100853 |
| QGLSSSIF | ASSLEEGAGNTLY | 0.16698849 |
| RGTFANIRL | ASSLEEGAGNTLY | 0.19840066 |
| RQLWWGHRI | ASSLEEGAGNTLY | 0.16565175 |
| YGPPGTGKTLL | ASSLEEGAGNTLY | 0.20382494 |
| AALVGAPANVA | ASSLEEGAGNTLY | 0.17405364 |
| AATFRRTE | ASSLEEGAGNTLY | 0.19430801 |
| AGALGGGI | ASSLEEGAGNTLY | 0.16973546 |
| AGGAGGAGGIGAG | ASSLEEGAGNTLY | 0.18025826 |
| AGSGALGL | ASSLEEGAGNTLY | 0.17247003 |
| AQIADQI | ASSLEEGAGNTLY | 0.24082789 |
| ASAVAVTYL | ASSLEEGAGNTLY | 0.18520132 |
| ASTVLGGTGG | ASSLEEGAGNTLY | 0.19031724 |
| AVGGEPAK | ASSLEEGAGNTLY | 0.17180055 |
| CEQGTFGGTC | ASSLEEGAGNTLY | 0.35118166 |
| DEGKAPAG | ASSLEEGAGNTLY | 0.16602178 |
| DTVVFPAL | ASSLEEGAGNTLY | 0.17451301 |
| FEGITMAG | ASSLEEGAGNTLY | 0.17151088 |
| FPLVVQD | ASSLEEGAGNTLY | 0.17824003 |
| GAFTTPQ | ASSLEEGAGNTLY | 0.18630487 |
| GFGGFGTAGGVGGA | ASSLEEGAGNTLY | 0.16525802 |
| GFLAQFL | ASSLEEGAGNTLY | 0.18679647 |
| GGAGGAGGNAGVA | ASSLEEGAGNTLY | 0.17007257 |
| GGVGTAGAGGAGGAGG | ASSLEEGAGNTLY | 0.17113116 |
| GNGGGGDAGNAGSGGN | ASSLEEGAGNTLY | 0.18949117 |
| GTGGAGGAGNDAGS | ASSLEEGAGNTLY | 0.1710147 |
| GTTNTGWFNS | ASSLEEGAGNTLY | 0.18992133 |

| | | |
|---|---|---|
| IFMGPLFTL | ASSLEEGAGNTLY | 0.16497907 |
| ILAHAIEAI | ASSLEEGAGNTLY | 0.1760151 |
| IRSAIREAL | ASSLEEGAGNTLY | 0.1995275 |
| ITFAGGG | ASSLEEGAGNTLY | 0.16562584 |
| ITGMSGHQEV | ASSLEEGAGNTLY | 0.16566133 |
| IVAGLACTL | ASSLEEGAGNTLY | 0.19459097 |
| IVGGGLAGA | ASSLEEGAGNTLY | 0.10048373 |
| IVTGVLNI | ASSLEEGAGNTLY | 0.096643746 |
| LAQQGCI | ASSLEEGAGNTLY | 0.09178256 |
| LCLHVEN | ASSLEEGAGNTLY | 0.0992494 |
| LDAISDGS | ASSLEEGAGNTLY | 0.094942845 |
| LGGFAAML | ASSLEEGAGNTLY | 0.10055935 |
| LGGKNPMI | ASSLEEGAGNTLY | 0.101455204 |
| LGPKGVAAIL | ASSLEEGAGNTLY | 0.09878415 |
| LMVGVSI | ASSLEEGAGNTLY | 0.10484906 |
| LVGAAFVL | ASSLEEGAGNTLY | 0.10234936 |
| LVLAVAPL | ASSLEEGAGNTLY | 0.098950386 |
| MAAIAVIDAA | ASSLEEGAGNTLY | 0.10010351 |
| NNGINAELQ | ASSLEEGAGNTLY | 0.09707153 |
| RGEPLVLVHG | ASSLEEGAGNTLY | 0.09890477 |
| RGGAGGAAGLI | ASSLEEGAGNTLY | 0.09204482 |
| RSYRMHPAV | ASSLEEGAGNTLY | 0.092344455 |
| SAAHLLAGQLA | ASSLEEGAGNTLY | 0.100016415 |
| SAAVGSY | ASSLEEGAGNTLY | 0.10727678 |
| SPHAVLHRI | ASSLEEGAGNTLY | 0.11294752 |
| SQIAKVL | ASSLEEGAGNTLY | 0.105686314 |
| TALGHLVP | ASSLEEGAGNTLY | 0.106643304 |
| TAPTTSGAA | ASSLEEGAGNTLY | 0.09647399 |
| TDGDDGNGGSPGT | ASSLEEGAGNTLY | 0.09146428 |
| TDGLGEN | ASSLEEGAGNTLY | 0.09263008 |
| TGGPPLPGQA | ASSLEEGAGNTLY | 0.095455684 |
| TIPAITI | ASSLEEGAGNTLY | 0.09969045 |
| TIPGFTI | ASSLEEGAGNTLY | 0.08863141 |
| TSIASGALA | ASSLEEGAGNTLY | 0.100640945 |
| TVTLPGAAI | ASSLEEGAGNTLY | 0.099174626 |
| VASGLSAALP | ASSLEEGAGNTLY | 0.10057118 |
| VGAGVIGI | ASSLEEGAGNTLY | 0.10640677 |
| VGPKRFGV | ASSLEEGAGNTLY | 0.09405855 |
| VNVAFPD | ASSLEEGAGNTLY | 0.095724456 |
| VSMTNSGVS | ASSLEEGAGNTLY | 0.09668402 |
| VTDVLGDLSV | ASSLEEGAGNTLY | 0.099809654 |
| VVVVGAGFAGL | ASSLEEGAGNTLY | 0.094110675 |
| VYTVGTVV | ASSLEEGAGNTLY | 0.11735034 |
| WTTFSGP | ASSLEEGAGNTLY | 0.09362405 |
| YDELVEAF | ASSLEEGAGNTLY | 0.11792295 |
| YGTGGAGGNGG | ASSLEEGAGNTLY | 0.10192832 |
| YTNVMAV | ASSLEEGAGNTLY | 0.099872574 |
| FLEQVHQL | ASSLEEGAGNTLY | 0.09582558 |
| GLSLLAVAV | ASSLEEGAGNTLY | 0.1248452 |
| LEQVHQL | ASSLEEGAGNTLY | 0.10468687 |
| LNAAGAI | ASSLEEGAGNTLY | 0.11351326 |

| | | |
|---|---|---|
| PAVHASL | ASSLEEGAGNTLY | 0.096884124 |
| TELEDDA | ASSLEEGAGNTLY | 0.09833988 |
| TPPGPPN | ASSLEEGAGNTLY | 0.0966545 |
| TPPPPPPP | ASSLEEGAGNTLY | 0.0957119 |
| TQSPALI | ASRSRDKNTLY | 0.09275267 |
| EFAAAFPQ | ASRSRDKNTLY | 0.0955764 |
| LLSGNLDVL | ASRSRDKNTLY | 0.101743124 |
| GPANLGNYNI | ASRSRDKNTLY | 0.1157603 |
| TLGPTLP | ASRSRDKNTLY | 0.092705555 |
| VYAYSSCCK | ASRSRDKNTLY | 0.1435794 |
| APAPAPAPNI | ASRSRDKNTLY | 0.09348392 |
| PGGPVPIP | ASRSRDKNTLY | 0.09661172 |
| FLTVALEP | ASRSRDKNTLY | 0.10134276 |
| SQYFGANL | ASRSRDKNTLY | 0.09393698 |
| GGVKPAA | ASRSRDKNTLY | 0.0903493 |
| GGTGGAGGAGGW | ASRSRDKNTLY | 0.09289107 |
| GVGGVAGLGGAGG | ASRSRDKNTLY | 0.10548007 |
| VLLVLATTI | ASRSRDKNTLY | 0.10771416 |
| IGIFAAML | ASRSRDKNTLY | 0.09645071 |
| AVGDLID | ASRSRDKNTLY | 0.10280901 |
| NISPLPL | ASRSRDKNTLY | 0.64598626 |
| VTAVFVAI | ASRSRDKNTLY | 0.7155018 |
| SGDDNDAAL | ASRSRDKNTLY | 0.16675542 |
| AVAALAHS | ASRSRDKNTLY | 0.15921614 |
| GGVTVMATP | ASRSRDKNTLY | 0.4403436 |
| AGLAANVRD | ASRSRDKNTLY | 0.700684 |
| TNRPDLIDPALLRPG | ASRSRDKNTLY | 0.3432061 |
| LANLGVTAL | ASRSRDKNTLY | 0.65671676 |
| TGERIEL | ASRSRDKNTLY | 0.15239215 |
| NGEELAA | ASRSRDKNTLY | 0.1751139 |
| VLSLVWG | ASRSRDKNTLY | 0.17601162 |
| IGYALWL | ASRSRDKNTLY | 0.2120516 |
| LSLLLAPL | ASRSRDKNTLY | 0.51753247 |
| TLDEALK | ASRSRDKNTLY | 0.13086662 |
| TDGAGTAEGG | ASRSRDKNTLY | 0.12731308 |
| NAPGGAGGNGG | ASRSRDKNTLY | 0.28920412 |
| VSVFMGMGGNF | ASRSRDKNTLY | 0.47535563 |
| LASYLAAL | ASRSRDKNTLY | 0.727774 |
| LVGGTPAL | ASRSRDKNTLY | 0.68078387 |
| VRNADVL | ASRSRDKNTLY | 0.7575963 |
| TCSSPFLVSAM | ASRSRDKNTLY | 0.6678235 |
| VHITDAE | ASRSRDKNTLY | 0.56704473 |
| EYAAPSF | ASRSRDKNTLY | 0.09618607 |
| VVDAFHGGL | ASRSRDKNTLY | 0.15713915 |
| ASVVDPQ | ASRSRDKNTLY | 0.13772254 |
| SYNEFGGLS | ASRSRDKNTLY | 0.37315154 |
| DPDGSVY | ASRSRDKNTLY | 0.091506325 |
| MVSTIHATE | ASRSRDKNTLY | 0.25511062 |
| LAAGVADPV | ASRSRDKNTLY | 0.3134653 |
| TKEQIAA | ASRSRDKNTLY | 0.10648443 |
| SLASPAAL | ASRSRDKNTLY | 0.62469465 |

| | | |
|---|---|---|
| IGLVFYGV | ASRSRDKNTLY | 0.11999102 |
| FLLDASNL | ASRSRDKNTLY | 0.13340735 |
| YQVGNFP | ASRSRDKNTLY | 0.1680474 |
| ESLLSQST | ASRSRDKNTLY | 0.7241167 |
| LFTVGFAAVG | ASRSRDKNTLY | 0.14907277 |
| ATPPTPT | ASRSRDKNTLY | 0.51995146 |
| TDLATLG | ASRSRDKNTLY | 0.5039963 |
| SAGGTGR | ASRSRDKNTLY | 0.4352376 |
| GFSNSGS | ASRSRDKNTLY | 0.12221113 |
| GGNGGSGGTSV | ASRSRDKNTLY | 0.6110229 |
| GTSVATGGAGN | ASRSRDKNTLY | 0.3122452 |
| NGGNGGSGGTS | ASRSRDKNTLY | 0.76743305 |
| AVVKFPH | ASRSRDKNTLY | 0.30456945 |
| TPPPPPP | ASRSRDKNTLY | 0.23592257 |
| AAPAAAAAPA | ASRSRDKNTLY | 0.1738297 |
| IALNEDL | ASRSRDKNTLY | 0.15246175 |
| IVNVSSM | ASRSRDKNTLY | 0.10183822 |
| TGIPPAPR | ASRSRDKNTLY | 0.39501047 |
| FDLGGGTF | ASRSRDKNTLY | 0.17272861 |
| FDLGGGTFDVS | ASRSRDKNTLY | 0.31766832 |
| NEPTAAA | ASRSRDKNTLY | 0.61452365 |
| NEPTAAALA | ASRSRDKNTLY | 0.10966022 |
| GSIINISSI | ASRSRDKNTLY | 0.67433804 |
| VAPGNAGTA | ASRSRDKNTLY | 0.16347523 |
| TGGGASLE | ASRSRDKNTLY | 0.14746787 |
| VAVYNHYKRI | ASRSRDKNTLY | 0.65500605 |
| TDPAPATT | ASRSRDKNTLY | 0.117486246 |
| TNRPDILDPALLRPG | ASRSRDKNTLY | 0.72771704 |
| LTTTTTS | ASRSRDKNTLY | 0.10229373 |
| PTGSGKTL | ASRSRDKNTLY | 0.5251785 |
| TGSGKTL | ASRSRDKNTLY | 0.55438393 |
| TIGHVDHGK | ASRSRDKNTLY | 0.09057316 |
| TIGHVDHGKTT | ASRSRDKNTLY | 0.12150376 |
| GHVDHGKTT | ASRSRDKNTLY | 0.6751397 |
| AQFEHTLL | ASRSRDKNTLY | 0.21380739 |
| IVNQDEL | ASRSRDKNTLY | 0.5388138 |
| TNGSQFF | ASRSRDKNTLY | 0.106469065 |
| TGKGIVE | ASRSRDKNTLY | 0.116736084 |
| LISNASDA | ASRSRDKNTLY | 0.15552479 |
| AELRTLGELTKIAKAHGAQ | ASRSRDKNTLY | 0.2654817 |
| APNAVAAAEPSW | ASRSRDKNTLY | 0.14208715 |
| APQVAGAATAVL | ASRSRDKNTLY | 0.29750693 |
| ARRDPAAAAAILPTDA | ASRSRDKNTLY | 0.120270886 |
| DAAARDVLAEAGLAEYF | ASRSRDKNTLY | 0.11012417 |
| DDQIVLSAKANLAKVEQ | ASRSRDKNTLY | 0.11584417 |
| DELLPDTGEDSDEDGH | ASRSRDKNTLY | 0.13355829 |
| ELPEAPANRATVPQD | ASRSRDKNTLY | 0.183788 |
| FGQSKIERVTGIG | ASRSRDKNTLY | 0.11083106 |
| GAGGTGSTAGGAGGAGGAGG | ASRSRDKNTLY | 0.103048064 |
| GHGGTGAAGGNGGTGGAGGS( | ASRSRDKNTLY | 0.13310437 |
| GNGGNGGAGGEGGPGGAGGA( | ASRSRDKNTLY | 0.14731343 |

| | | |
|---|---|---|
| GSTAGNGGNGGNATNGTIAG( | ASRSRDKNTLY | 0.34501058 |
| GTGGDGGNGGNGGTGG | ASRSRDKNTLY | 0.33840215 |
| GVRRPAGIPADAPV | ASRSRDKNTLY | 0.3701688 |
| GYRQVTDAHLLTLARR | ASRSRDKNTLY | 0.28094065 |
| IAKMGQDGHDRG | ASRSRDKNTLY | 0.14533025 |
| KAEPSIEIQYT | ASRSRDKNTLY | 0.09302949 |
| KPVRIRMAADEDTVAVT | ASRSRDKNTLY | 0.11909805 |
| MPPRIAGMRLLVIKPEP | ASRSRDKNTLY | 0.109993674 |
| MKFYAVALFRSRGVPPTR | ASRSRDKNTLY | 0.19073918 |
| NAGGFGILGAVAHSP | ASRSRDKNTLY | 0.09060663 |
| NEGREITLAMAEM | ASRSRDKNTLY | 0.13612792 |
| NELVEWSHPLGHRWRWE | ASRSRDKNTLY | 0.15419696 |
| NLGAGGTGGNGGNPG | ASRSRDKNTLY | 0.098488525 |
| NTEWPKWETVMAADPVD | ASRSRDKNTLY | 0.3951534 |
| PASAAPAETQP | ASRSRDKNTLY | 0.10327485 |
| QQSSDGKNGVRAAEITGE | ASRSRDKNTLY | 0.11082547 |
| RAPDAVTTSWWKE | ASRSRDKNTLY | 0.11811496 |
| SGFGNVGAHTSGWFNQ | ASRSRDKNTLY | 0.2924067 |
| SGVAGAVGQAASVGGLKV | ASRSRDKNTLY | 0.11402987 |
| SNGGAGTGSAGGNGGT | ASRSRDKNTLY | 0.23969029 |
| TGGAGGEGGAAG | ASRSRDKNTLY | 0.16363695 |
| TGGDGGAAGAGGAPGHG | ASRSRDKNTLY | 0.17548099 |
| TMPPLLAAIANAGRHGV | ASRSRDKNTLY | 0.107774556 |
| TSSTAGNSGGA | ASRSRDKNTLY | 0.21837196 |
| VPATVPVGILC | ASRSRDKNTLY | 0.13750057 |
| VQRPDSGGVVLTGRIS | ASRSRDKNTLY | 0.69906545 |
| WTPNPVPATKGVPAGLVQAT | ASRSRDKNTLY | 0.15699959 |
| YALKRALAKFSDRFDVVII | ASRSRDKNTLY | 0.14779438 |
| YLIGSVIV | ASRSRDKNTLY | 0.11744326 |
| YGQLVAVID | ASRSRDKNTLY | 0.12695715 |
| VTVGGMTLPA | ASRSRDKNTLY | 0.098082565 |
| VLALPAV | ASRSRDKNTLY | 0.14691605 |
| VIPPTLNYET | ASRSRDKNTLY | 0.11017726 |
| VGMGGMHPSSG | ASRSRDKNTLY | 0.14064595 |
| VGLAGGAATA | ASRSRDKNTLY | 0.20946665 |
| VGGAGGAGGSGP | ASRSRDKNTLY | 0.10200342 |
| VFAAFMV | ASRSRDKNTLY | 0.2332821 |
| TLYDKLR | ASRSRDKNTLY | 0.10264733 |
| TLAVFTVL | ASRSRDKNTLY | 0.11568335 |
| TGSVQPARV | ASRSRDKNTLY | 0.23232368 |
| TGGGAGSGGT | ASRSRDKNTLY | 0.098869234 |
| SHLETSL | ASRSRDKNTLY | 0.53898925 |
| SFTTGMT | ASRSRDKNTLY | 0.09539784 |
| QVEVAKRMGVRQPTVS | ASRSRDKNTLY | 0.18028264 |
| QGSGGPALAM | ASRSRDKNTLY | 0.20251502 |
| MLAPSNP | ASRSRDKNTLY | 0.094396815 |
| LVNNAAI | ASRSRDKNTLY | 0.103233576 |
| LTGEINTLA | ASRSRDKNTLY | 0.29204857 |
| LSIFGPV | ASRSRDKNTLY | 0.17068176 |
| LLALLANL | ASRSRDKNTLY | 0.3139353 |
| LISNASDALDKLR | ASRSRDKNTLY | 0.118164 |

| | | |
|---|---|---|
| LINEAVP | ASRSRDKNTLY | 0.13705204 |
| LGYAFSGL | ASRSRDKNTLY | 0.16507417 |
| LDFVAVAL | ASRSRDKNTLY | 0.17085832 |
| LAQPGAE | ASRSRDKNTLY | 0.27429003 |
| ISNLGAVTGF | ASRSRDKNTLY | 0.16452268 |
| HQEDPGRI | ASRSRDKNTLY | 0.13609849 |
| GTPGEPTP | ASRSRDKNTLY | 0.1456303 |
| GFFNSGS | ASRSRDKNTLY | 0.1213408 |
| GAGGGGLTAGAN | ASRSRDKNTLY | 0.16574337 |
| FVDALVNM | ASRSRDKNTLY | 0.14793727 |
| FLTVAPEL | ASRSRDKNTLY | 0.15321769 |
| FETLTAL | ASRSRDKNTLY | 0.11216969 |
| DVINPIEL | ASRSRDKNTLY | 0.11568723 |
| APVSGAPV | ASRSRDKNTLY | 0.17035389 |
| ALVGGLAW | ASRSRDKNTLY | 0.38292944 |
| ALAAAAATAI | ASRSRDKNTLY | 0.3452711 |
| AFGLGLLL | ASRSRDKNTLY | 0.17370479 |
| AFFAAASL | ASRSRDKNTLY | 0.1943922 |
| AAAALPAFSPP | ASRSRDKNTLY | 0.117308654 |
| VVLAPAL | ASRSRDKNTLY | 0.10144714 |
| TIGHVDHGKTTL | ASRSRDKNTLY | 0.10828369 |
| LISNASDALDK | ASRSRDKNTLY | 0.13794184 |
| LISNASDAL | ASRSRDKNTLY | 0.15225047 |
| ISNASDALDK | ASRSRDKNTLY | 0.09744672 |
| NQAETLVYQTE | ASRSRDKNTLY | 0.16460034 |
| TGGTGGTAGAG | ASRSRDKNTLY | 0.13491859 |
| TGSTAGGAGGA | ASRSRDKNTLY | 0.124909714 |
| DPLAVPVAAGPV | ASRSRDKNTLY | 0.33509827 |
| GGTGGTPGNGGN | ASRSRDKNTLY | 0.11850537 |
| PAGLAGVDPATL | ASRSRDKNTLY | 0.12892953 |
| GNNLVVEAPHAGASV | ASRSRDKNTLY | 0.14463766 |
| KVTRIMSASTLEQPA | ASRSRDKNTLY | 0.13354026 |
| LGFALPIALPRRTVV | ASRSRDKNTLY | 0.12630196 |
| TRLGNLTARKLVVLV | ASRSRDKNTLY | 0.33683512 |
| WSQQRRLIYFPSAGP | ASRSRDKNTLY | 0.10933912 |
| GPNGNPDPMAAAVDIR | ASRSRDKNTLY | 0.19015065 |
| TQRCVSVADLRAAASR | ASRSRDKNTLY | 0.13026139 |
| QFVLDVAEHPVEVLTETV | ASRSRDKNTLY | 0.15940645 |
| GGAGGTGGAGGNGGTQAGDG | ASRSRDKNTLY | 0.195734 |
| PLLATLRAIAVDVDAKPAQV | ASRSRDKNTLY | 0.62619746 |
| APKFRTGGYMIDFWGVGYQV | ASRSRDKNTLY | 0.16788083 |
| GAGGAGGAGGAAGGPGGTGG | ASRSRDKNTLY | 0.16639256 |
| IGALGLLATDVAPAGSVAFG | ASRSRDKNTLY | 0.1230894 |
| AGGAGGIGGNGGAGGAGGVL | ASRSRDKNTLY | 0.20089357 |
| ADELIGLL | ASRSRDKNTLY | 0.112845495 |
| ADRALKA | ASRSRDKNTLY | 0.15113907 |
| AGAVGQVGGAGGA | ASRSRDKNTLY | 0.101701036 |
| AGGGGGLGGAP | ASRSRDKNTLY | 0.14816166 |
| AGGLLYGNGGAG | ASRSRDKNTLY | 0.15245706 |
| AGLESIGL | ASRSRDKNTLY | 0.11139767 |
| AGLKTGII | ASRSRDKNTLY | 0.11553242 |

| | | |
|---|---|---|
| ASLAALLL | ASRSRDKNTLY | 0.123638585 |
| ASVINISSI | ASRSRDKNTLY | 0.12951818 |
| ATPPGPA | ASRSRDKNTLY | 0.19716391 |
| ATVASALV | ASRSRDKNTLY | 0.1439199 |
| ATVASLIL | ASRSRDKNTLY | 0.10274558 |
| AVNGPVLIRH | ASRSRDKNTLY | 0.112567306 |
| DGAAGAVGI | ASRSRDKNTLY | 0.11202614 |
| DPLLHGI | ASRSRDKNTLY | 0.21870604 |
| ELGAALGAL | ASRSRDKNTLY | 0.10345838 |
| ELGRTQAV | ASRSRDKNTLY | 0.13150921 |
| FVAGGYLS | ASRSRDKNTLY | 0.17511262 |
| GASGGAGGAGGW | ASRSRDKNTLY | 0.46180895 |
| GGAGAGGDGGAGGI | ASRSRDKNTLY | 0.67177653 |
| GTGHNGSQPGMGGNG | ASRSRDKNTLY | 0.12467122 |
| GYAAASASAS | ASRSRDKNTLY | 0.14623368 |
| HQVDVAAVQQG | ASRSRDKNTLY | 0.3470293 |
| IGAAIAHT | ASRSRDKNTLY | 0.5916104 |
| IGMIEVAQL | ASRSRDKNTLY | 0.2664821 |
| IRLAPEHTSAMD | ASRSRDKNTLY | 0.56357884 |
| IYNVSVL | ASRSRDKNTLY | 0.14350303 |
| KQGFLGM | ASRSRDKNTLY | 0.13107781 |
| LESVITQL | ASRSRDKNTLY | 0.14124103 |
| LGLGALVVV | ASRSRDKNTLY | 0.17506601 |
| LIGNGGNGGSGGI | ASRSRDKNTLY | 0.42037654 |
| LNAINAPTL | ASRSRDKNTLY | 0.11940331 |
| LQLLNTGV | ASRSRDKNTLY | 0.109189466 |
| LQTSIEF | ASRSRDKNTLY | 0.22931574 |
| MSFAVSEL | ASRSRDKNTLY | 0.35949206 |
| NTGALGL | ASRSRDKNTLY | 0.6331213 |
| PPGAGGAGGAGTT | ASRSRDKNTLY | 0.61840934 |
| QLVGLGI | ASRSRDKNTLY | 0.6964078 |
| RFLEQVHQL | ASRSRDKNTLY | 0.4860309 |
| RGVGYVLR | ASRSRDKNTLY | 0.42770407 |
| SATGVFAGVI | ASRSRDKNTLY | 0.097119756 |
| SLPAITLP | ASRSRDKNTLY | 0.14524925 |
| STALAFTV | ASRSRDKNTLY | 0.123830855 |
| SVLTVGLGK | ASRSRDKNTLY | 0.29060206 |
| TALAGAGSGI | ASRSRDKNTLY | 0.094488755 |
| TGLPFTGI | ASRSRDKNTLY | 0.19539946 |
| VAEHVTV | ASRSRDKNTLY | 0.23135108 |
| VDIVLNSL | ASRSRDKNTLY | 0.10384313 |
| VLTGGGAVL | ASRSRDKNTLY | 0.552717 |
| VNVGLAYGI | ASRSRDKNTLY | 0.1074761 |
| VSSAAVGAV | ASRSRDKNTLY | 0.12575953 |
| VTSVYEAGV | ASRSRDKNTLY | 0.14213 |
| WVALDRGA | ASRSRDKNTLY | 0.54796207 |
| YLVGSVVI | ASRSRDKNTLY | 0.13834108 |
| GPPGTGKTLL | ASRSRDKNTLY | 0.37105668 |
| GVGMTKF | ASRSRDKNTLY | 0.37557715 |
| IGIGGSDL | ASRSRDKNTLY | 0.302166 |
| IGTLLAEL | ASRSRDKNTLY | 0.12085868 |

| | | |
|---|---|---|
| IYTSGTT | ASRSRDKNTLY | 0.40286058 |
| LSFPAGV | ASRSRDKNTLY | 0.20767623 |
| LYTSGTT | ASRSRDKNTLY | 0.71516794 |
| VGAFPFGF | ASRSRDKNTLY | 0.24840496 |
| VGAGTVEF | ASRSRDKNTLY | 0.1921508 |
| VGKSTLLL | ASRSRDKNTLY | 0.14468256 |
| AALAHAGGGIV | ASRSRDKNTLY | 0.14797224 |
| AALVAVSLL | ASRSRDKNTLY | 0.10186925 |
| AAVSRAVAAL | ASRSRDKNTLY | 0.22732173 |
| AGGSSGIGGF | ASRSRDKNTLY | 0.13533089 |
| AGLTVANANG | ASRSRDKNTLY | 0.22372097 |
| AGLVGAVL | ASRSRDKNTLY | 0.4210655 |
| ALSQAFTV | ASRSRDKNTLY | 0.11857872 |
| ALTFSVSL | ASRSRDKNTLY | 0.45596385 |
| ALTGAALVAL | ASRSRDKNTLY | 0.14361775 |
| ASFSIGAGVL | ASRSRDKNTLY | 0.14530542 |
| DGSVYKI | ASRSRDKNTLY | 0.54769015 |
| EQLAEIV | ASRSRDKNTLY | 0.112725824 |
| FAVSELTVM | ASRSRDKNTLY | 0.6693806 |
| GAAPLAAGL | ASRSRDKNTLY | 0.10112134 |
| GADGGDGGSGGAGG | ASRSRDKNTLY | 0.2764158 |
| GFGLDTGGAGG | ASRSRDKNTLY | 0.3961482 |
| GLASGGLL | ASRSRDKNTLY | 0.0916551 |
| GRIAGAAVLL | ASRSRDKNTLY | 0.11330758 |
| GSAGSLNTSF | ASRSRDKNTLY | 0.5230771 |
| GTGQIGI | ASRSRDKNTLY | 0.7362139 |
| GTLIALIL | ASRSRDKNTLY | 0.69605607 |
| IAATMGFVVG | ASRSRDKNTLY | 0.36224535 |
| IFAQALL | ASRSRDKNTLY | 0.3761687 |
| IGAAVALGF | ASRSRDKNTLY | 0.49668834 |
| IGLTPLAL | ASRSRDKNTLY | 0.7130529 |
| ILGSNTSTL | ASRSRDKNTLY | 0.7375077 |
| IVFSLTVL | ASRSRDKNTLY | 0.7218815 |
| LASKATGF | ASRSRDKNTLY | 0.36801282 |
| LASVALIL | ASRSRDKNTLY | 0.57638115 |
| LFAVGLTL | ASRSRDKNTLY | 0.49967575 |
| LFNGGIGTYI | ASRSRDKNTLY | 0.54743147 |
| LTLGAGII | ASRSRDKNTLY | 0.6064114 |
| LVAAALGDI | ASRSRDKNTLY | 0.3748774 |
| NTAFVQLGI | ASRSRDKNTLY | 0.37894103 |
| QLFSGAEL | ASRSRDKNTLY | 0.40613976 |
| SAATFITL | ASRSRDKNTLY | 0.5597061 |
| SNYGLANLGV | ASRSRDKNTLY | 0.7124596 |
| SYGGAYAVM | ASRSRDKNTLY | 0.68402874 |
| TTGVGGLTLG | ASRSRDKNTLY | 0.7898314 |
| TTQTIYI | ASRSRDKNTLY | 0.7614096 |
| VAAGAGLGAVI | ASRSRDKNTLY | 0.61378497 |
| VAAVVTLL | ASRSRDKNTLY | 0.15282373 |
| VGFLGGLI | ASRSRDKNTLY | 0.25741673 |
| VGVIPILL | ASRSRDKNTLY | 0.3529833 |
| VIFASSLI | ASRSRDKNTLY | 0.46830583 |

| | | |
|---|---|---|
| VKGGDGGAAAG | ASRSRDKNTLY | 0.18476957 |
| VTKTFTNI | ASRSRDKNTLY | 0.5482969 |
| VVAAAILL | ASRSRDKNTLY | 0.5590109 |
| YAALDVEL | ASRSRDKNTLY | 0.30810803 |
| LASVALLL | ASRSRDKNTLY | 0.72308254 |
| LDAVHRV | ASRSRDKNTLY | 0.33256415 |
| QGLSSSIF | ASRSRDKNTLY | 0.33946693 |
| RGTFANIRL | ASRSRDKNTLY | 0.3933589 |
| RQLWWGHRI | ASRSRDKNTLY | 0.74294317 |
| YGPPGTGKTLL | ASRSRDKNTLY | 0.38084972 |
| AALVGAPANVA | ASRSRDKNTLY | 0.70779914 |
| AATFRRTE | ASRSRDKNTLY | 0.4928192 |
| AGALGGGI | ASRSRDKNTLY | 0.77378786 |
| AGGAGGAGGIGAG | ASRSRDKNTLY | 0.30500397 |
| AGSGALGL | ASRSRDKNTLY | 0.7461934 |
| AQIADQI | ASRSRDKNTLY | 0.6164676 |
| ASAVAVTYL | ASRSRDKNTLY | 0.81726885 |
| ASTVLGGTGG | ASRSRDKNTLY | 0.48481315 |
| AVGGEPAK | ASRSRDKNTLY | 0.5268997 |
| CEQGTFGGTC | ASRSRDKNTLY | 0.33608723 |
| DEGKAPAG | ASRSRDKNTLY | 0.30369207 |
| DTVVFPAL | ASRSRDKNTLY | 0.23830177 |
| FEGITMAG | ASRSRDKNTLY | 0.55208254 |
| FPLVVQD | ASRSRDKNTLY | 0.29861322 |
| GAFTTPQ | ASRSRDKNTLY | 0.6011554 |
| GFGGFGTAGGVGGA | ASRSRDKNTLY | 0.74670744 |
| GFLAQFL | ASRSRDKNTLY | 0.19877973 |
| GGAGGAGGNAGVA | ASRSRDKNTLY | 0.69721127 |
| GGVGTAGAGGAGGAGG | ASRSRDKNTLY | 0.30468163 |
| GNGGGGDAGNAGSGGN | ASRSRDKNTLY | 0.36542943 |
| GTGGAGGAGNDAGS | ASRSRDKNTLY | 0.6522134 |
| GTTNTGWFNS | ASRSRDKNTLY | 0.36657774 |
| IFMGPLFTL | ASRSRDKNTLY | 0.7458481 |
| ILAHAIEAI | ASRSRDKNTLY | 0.3304994 |
| IRSAIREAL | ASRSRDKNTLY | 0.60441923 |
| ITFAGGG | ASRSRDKNTLY | 0.6985954 |
| ITGMSGHQEV | ASRSRDKNTLY | 0.15468949 |
| IVAGLACTL | ASRSRDKNTLY | 0.27728835 |
| IVGGGLAGA | ASRSRDKNTLY | 0.793347 |
| IVTGVLNI | ASRSRDKNTLY | 0.097359486 |
| LAQQGCI | ASRSRDKNTLY | 0.095927015 |
| LCLHVEN | ASRSRDKNTLY | 0.09600718 |
| LDAISDGS | ASRSRDKNTLY | 0.10242301 |
| LGGFAAML | ASRSRDKNTLY | 0.09681923 |
| LGGKNPMI | ASRSRDKNTLY | 0.096768 |
| LGPKGVAAIL | ASRSRDKNTLY | 0.097788185 |
| LMVGVSI | ASRSRDKNTLY | 0.09758431 |
| LVGAAFVL | ASRSRDKNTLY | 0.1049133 |
| LVLAVAPL | ASRSRDKNTLY | 0.10108026 |
| MAAIAVIDAA | ASRSRDKNTLY | 0.09652398 |
| NNGINAELQ | ASRSRDKNTLY | 0.10152556 |

| | | |
|---|---|---|
| RGEPLVLVHG | ASRSRDKNTLY | 0.09629015 |
| RGGAGGAAGLI | ASRSRDKNTLY | 0.10085558 |
| RSYRMHPAV | ASRSRDKNTLY | 0.095885985 |
| SAAHLLAGQLA | ASRSRDKNTLY | 0.09585853 |
| SAAVGSY | ASRSRDKNTLY | 0.09741974 |
| SPHAVLHRI | ASRSRDKNTLY | 0.10081574 |
| SQIAKVL | ASRSRDKNTLY | 0.09919756 |
| TALGHLVP | ASRSRDKNTLY | 0.096721955 |
| TAPTTSGAA | ASRSRDKNTLY | 0.09719895 |
| TDGDDGNGGSPGT | ASRSRDKNTLY | 0.095783696 |
| TDGLGEN | ASRSRDKNTLY | 0.09684462 |
| TGGPPLPGQA | ASRSRDKNTLY | 0.09727346 |
| TIPAITI | ASRSRDKNTLY | 0.0974345 |
| TIPGFTI | ASRSRDKNTLY | 0.09837051 |
| TSIASGALA | ASRSRDKNTLY | 0.09500214 |
| TVTLPGAAI | ASRSRDKNTLY | 0.09982203 |
| VASGLSAALP | ASRSRDKNTLY | 0.09797387 |
| VGAGVIGI | ASRSRDKNTLY | 0.105626956 |
| VGPKRFGV | ASRSRDKNTLY | 0.101229645 |
| VNVAFPD | ASRSRDKNTLY | 0.09760451 |
| VSMTNSGVS | ASRSRDKNTLY | 0.09857084 |
| VTDVLGDLSV | ASRSRDKNTLY | 0.096821986 |
| VVVVGAGFAGL | ASRSRDKNTLY | 0.09735276 |
| VYTVGTVV | ASRSRDKNTLY | 0.09621099 |
| WTTFSGP | ASRSRDKNTLY | 0.10349455 |
| YDELVEAF | ASRSRDKNTLY | 0.09548491 |
| YGTGGAGGNGG | ASRSRDKNTLY | 0.10389538 |
| YTNVMAV | ASRSRDKNTLY | 0.10402376 |
| FLEQVHQL | ASRSRDKNTLY | 0.096932 |
| GLSLLAVAV | ASRSRDKNTLY | 0.09737784 |
| LEQVHQL | ASRSRDKNTLY | 0.10300187 |
| LNAAGAI | ASRSRDKNTLY | 0.10114515 |
| PAVHASL | ASRSRDKNTLY | 0.10913055 |
| TELEDDA | ASRSRDKNTLY | 0.09657811 |
| TPPGPPN | ASRSRDKNTLY | 0.09676823 |
| TPPPPPPP | ASRSRDKNTLY | 0.10123264 |
| TQSPALI | ASGGGESQNTLY | 0.09682982 |
| EFAAAFPQ | ASGGGESQNTLY | 0.0965349 |
| LLSGNLDVL | ASGGGESQNTLY | 0.09694377 |
| GPANLGNYNI | ASGGGESQNTLY | 0.09833757 |
| TLGPTLP | ASGGGESQNTLY | 0.10190959 |
| VYAYSSCCK | ASGGGESQNTLY | 0.0958623 |
| APAPAPAPNI | ASGGGESQNTLY | 0.10289339 |
| PGGPVPIP | ASGGGESQNTLY | 0.09736547 |
| FLTVALEP | ASGGGESQNTLY | 0.09614646 |
| SQYFGANL | ASGGGESQNTLY | 0.10056302 |
| GGVKPAA | ASGGGESQNTLY | 0.095429264 |
| GGTGGAGGAGGW | ASGGGESQNTLY | 0.09540208 |
| GVGGVAGLGGAGG | ASGGGESQNTLY | 0.095876664 |
| VLLVLATTI | ASGGGESQNTLY | 0.10101723 |
| IGIFAAML | ASGGGESQNTLY | 0.10646643 |

| | | |
|---|---|---|
| AVGDLID | ASGGGESQNTLY | 0.097117886 |
| NISPLPL | ASGGGESQNTLY | 0.097369865 |
| VTAVFVAI | ASGGGESQNTLY | 0.76944983 |
| SGDDNDAAL | ASGGGESQNTLY | 0.78918165 |
| AVAALAHS | ASGGGESQNTLY | 0.4891146 |
| GGVTVMATP | ASGGGESQNTLY | 0.25240576 |
| AGLAANVRD | ASGGGESQNTLY | 0.6663124 |
| TNRPDLIDPALLRPG | ASGGGESQNTLY | 0.76277435 |
| LANLGVTAL | ASGGGESQNTLY | 0.8064282 |
| TGERIEL | ASGGGESQNTLY | 0.77546084 |
| NGEELAA | ASGGGESQNTLY | 0.2663224 |
| VLSLVWG | ASGGGESQNTLY | 0.52115846 |
| IGYALWL | ASGGGESQNTLY | 0.68099445 |
| LSLLLAPL | ASGGGESQNTLY | 0.42073926 |
| TLDEALK | ASGGGESQNTLY | 0.71201795 |
| TDGAGTAEGG | ASGGGESQNTLY | 0.24405685 |
| NAPGGAGGNGG | ASGGGESQNTLY | 0.6221057 |
| VSVFMGMGGNF | ASGGGESQNTLY | 0.4018857 |
| LASYLAAL | ASGGGESQNTLY | 0.5892919 |
| LVGGTPAL | ASGGGESQNTLY | 0.743486 |
| VRNADVL | ASGGGESQNTLY | 0.69969577 |
| TCSSPFLVSAM | ASGGGESQNTLY | 0.82859635 |
| VHITDAE | ASGGGESQNTLY | 0.79426193 |
| EYAAPSF | ASGGGESQNTLY | 0.64446914 |
| VVDAFHGGL | ASGGGESQNTLY | 0.15810418 |
| ASVVDPQ | ASGGGESQNTLY | 0.20720199 |
| SYNEFGGLS | ASGGGESQNTLY | 0.43534595 |
| DPDGSVY | ASGGGESQNTLY | 0.43017793 |
| MVSTIHATE | ASGGGESQNTLY | 0.18052892 |
| LAAGVADPV | ASGGGESQNTLY | 0.5693029 |
| TKEQIAA | ASGGGESQNTLY | 0.74055755 |
| SLASPAAL | ASGGGESQNTLY | 0.22829154 |
| IGLVFYGV | ASGGGESQNTLY | 0.70968604 |
| FLLDASNL | ASGGGESQNTLY | 0.38805905 |
| YQVGNFP | ASGGGESQNTLY | 0.24139817 |
| ESLLSQST | ASGGGESQNTLY | 0.3511552 |
| LFTVGFAAVG | ASGGGESQNTLY | 0.815985 |
| ATPPTPT | ASGGGESQNTLY | 0.27153894 |
| TDLATLG | ASGGGESQNTLY | 0.626843 |
| SAGGTGR | ASGGGESQNTLY | 0.698065 |
| GFSNSGS | ASGGGESQNTLY | 0.76196855 |
| GGNGGSGGTSV | ASGGGESQNTLY | 0.1913384 |
| GTSVATGGAGN | ASGGGESQNTLY | 0.7348963 |
| NGGNGGSGGTS | ASGGGESQNTLY | 0.5385004 |
| AVVKFPH | ASGGGESQNTLY | 0.80001444 |
| TPPPPPP | ASGGGESQNTLY | 0.31731886 |
| AAPAAAAAPA | ASGGGESQNTLY | 0.42407155 |
| IALNEDL | ASGGGESQNTLY | 0.3731457 |
| IVNVSSM | ASGGGESQNTLY | 0.7903051 |
| TGIPPAPR | ASGGGESQNTLY | 0.18909882 |
| FDLGGGTF | ASGGGESQNTLY | 0.68925905 |

| | | |
|---|---|---|
| FDLGGGTFDVS | ASGGGESQNTLY | 0.3652189 |
| NEPTAAA | ASGGGESQNTLY | 0.73595613 |
| NEPTAAALA | ASGGGESQNTLY | 0.76443547 |
| GSIINISSI | ASGGGESQNTLY | 0.22656342 |
| VAPGNAGTA | ASGGGESQNTLY | 0.80890036 |
| TGGGASLE | ASGGGESQNTLY | 0.4878326 |
| VAVYNHYKRI | ASGGGESQNTLY | 0.3149414 |
| TDPAPATT | ASGGGESQNTLY | 0.7013258 |
| TNRPDILDPALLRPG | ASGGGESQNTLY | 0.33775216 |
| LTTTTTS | ASGGGESQNTLY | 0.8007848 |
| PTGSGKTL | ASGGGESQNTLY | 0.2855752 |
| TGSGKTL | ASGGGESQNTLY | 0.7850669 |
| TIGHVDHGK | ASGGGESQNTLY | 0.78302205 |
| TIGHVDHGKTT | ASGGGESQNTLY | 0.20518492 |
| GHVDHGKTT | ASGGGESQNTLY | 0.3352607 |
| AQFEHTLL | ASGGGESQNTLY | 0.7794646 |
| IVNQDEL | ASGGGESQNTLY | 0.66984904 |
| TNGSQFF | ASGGGESQNTLY | 0.72998494 |
| TGKGIVE | ASGGGESQNTLY | 0.56525105 |
| LISNASDA | ASGGGESQNTLY | 0.47873425 |
| AELRTLGELTKIAKAHGAQ | ASGGGESQNTLY | 0.48700327 |
| APNAVAAAEPSW | ASGGGESQNTLY | 0.65037715 |
| APQVAGAATAVL | ASGGGESQNTLY | 0.7346356 |
| ARRDPAAAAAILPTDA | ASGGGESQNTLY | 0.66203034 |
| DAAARDVLAEAGLAEYF | ASGGGESQNTLY | 0.31235248 |
| DDQIVLSAKANLAKVEQ | ASGGGESQNTLY | 0.55454195 |
| DELLPDTGEDSDEDGH | ASGGGESQNTLY | 0.5502907 |
| ELPEAPANRATVPQD | ASGGGESQNTLY | 0.50092655 |
| FGQSKIERVTGIG | ASGGGESQNTLY | 0.6475247 |
| GAGGTGSTAGGAGGAGGAGGI | ASGGGESQNTLY | 0.5401084 |
| GHGGTGAAGGNGGTGGAGGS( | ASGGGESQNTLY | 0.558158 |
| GNGGNGGAGGEGGPGGAGGA( | ASGGGESQNTLY | 0.4721923 |
| GSTAGNGGNGGNATNGTIAG( | ASGGGESQNTLY | 0.45253572 |
| GTGGDGGNGGNGGTGG | ASGGGESQNTLY | 0.6315354 |
| GVRRPAGIPADAPV | ASGGGESQNTLY | 0.6625509 |
| GYRQVTDAHLLTLARR | ASGGGESQNTLY | 0.6807482 |
| IAKMGQDGHDRG | ASGGGESQNTLY | 0.7144505 |
| KAEPSIEIQYT | ASGGGESQNTLY | 0.5694152 |
| KPVRIRMAADEDTVAVT | ASGGGESQNTLY | 0.22149988 |
| MPPRIAGMRLLVIKPEP | ASGGGESQNTLY | 0.33615416 |
| MKFYAVALFRSRGVPPTR | ASGGGESQNTLY | 0.5197607 |
| NAGGFGILGAVAHSP | ASGGGESQNTLY | 0.5460726 |
| NEGREITLAMAEM | ASGGGESQNTLY | 0.25790924 |
| NELVEWSHPLGHRWRWE | ASGGGESQNTLY | 0.5394257 |
| NLGAGGTGGNGGNPG | ASGGGESQNTLY | 0.5784446 |
| NTEWPKWETVMAADPVD | ASGGGESQNTLY | 0.44237956 |
| PASAAPAETQP | ASGGGESQNTLY | 0.68904626 |
| QQSSDGKNGVRAAEITGE | ASGGGESQNTLY | 0.5636208 |
| RAPDAVTTSWWKE | ASGGGESQNTLY | 0.40467638 |
| SGFGNVGAHTSGWFNQ | ASGGGESQNTLY | 0.35048276 |
| SGVAGAVGQAASVGGLKV | ASGGGESQNTLY | 0.52636015 |

| | | |
|---|---|---|
| SNGGAGTGSAGGNGGT | ASGGGESQNTLY | 0.3478608 |
| TGGAGGEGGAAG | ASGGGESQNTLY | 0.71306205 |
| TGGDGGAAGAGGAPGHG | ASGGGESQNTLY | 0.3396602 |
| TMPPLLAAIANAGRHGV | ASGGGESQNTLY | 0.62753046 |
| TSSTAGNSGGA | ASGGGESQNTLY | 0.31275648 |
| VPATVPVGILC | ASGGGESQNTLY | 0.6769369 |
| VQRPDSGGVVLTGRIS | ASGGGESQNTLY | 0.6585382 |
| WTPNPVPATKGVPAGLVQAT | ASGGGESQNTLY | 0.7547952 |
| YALKRALAKFSDRFDVVII | ASGGGESQNTLY | 0.5934878 |
| YLIGSVIV | ASGGGESQNTLY | 0.5184586 |
| YGQLVAVID | ASGGGESQNTLY | 0.3138153 |
| VTVGGMTLPA | ASGGGESQNTLY | 0.7246554 |
| VLALPAV | ASGGGESQNTLY | 0.33689272 |
| VIPPTLNYET | ASGGGESQNTLY | 0.51122797 |
| VGMGGMHPSSG | ASGGGESQNTLY | 0.3000052 |
| VGLAGGAATA | ASGGGESQNTLY | 0.5615942 |
| VGGAGGAGGSGP | ASGGGESQNTLY | 0.67441726 |
| VFAAFMV | ASGGGESQNTLY | 0.41656673 |
| TLYDKLR | ASGGGESQNTLY | 0.6810818 |
| TLAVFTVL | ASGGGESQNTLY | 0.59942365 |
| TGSVQPARV | ASGGGESQNTLY | 0.4029992 |
| TGGGAGSGGT | ASGGGESQNTLY | 0.72038805 |
| SHLETSL | ASGGGESQNTLY | 0.4047262 |
| SFTTGMT | ASGGGESQNTLY | 0.73168623 |
| QVEVAKRMGVRQPTVS | ASGGGESQNTLY | 0.39616865 |
| QGSGGPALAM | ASGGGESQNTLY | 0.4562897 |
| MLAPSNP | ASGGGESQNTLY | 0.5011817 |
| LVNNAAI | ASGGGESQNTLY | 0.41655815 |
| LTGEINTLA | ASGGGESQNTLY | 0.3570184 |
| LSIFGPV | ASGGGESQNTLY | 0.7274527 |
| LLALLANL | ASGGGESQNTLY | 0.1652766 |
| LISNASDALDKLR | ASGGGESQNTLY | 0.13607465 |
| LINEAVP | ASGGGESQNTLY | 0.13778742 |
| LGYAFSGL | ASGGGESQNTLY | 0.16461287 |
| LDFVAVAL | ASGGGESQNTLY | 0.13940682 |
| LAQPGAE | ASGGGESQNTLY | 0.1509749 |
| ISNLGAVTGF | ASGGGESQNTLY | 0.14381734 |
| HQEDPGRI | ASGGGESQNTLY | 0.15749444 |
| GTPGEPTP | ASGGGESQNTLY | 0.17752817 |
| GFFNSGS | ASGGGESQNTLY | 0.15959239 |
| GAGGGGLTAGAN | ASGGGESQNTLY | 0.15605074 |
| FVDALVNM | ASGGGESQNTLY | 0.16096441 |
| FLTVAPEL | ASGGGESQNTLY | 0.14504321 |
| FETLTAL | ASGGGESQNTLY | 0.13863905 |
| DVINPIEL | ASGGGESQNTLY | 0.13620955 |
| APVSGAPV | ASGGGESQNTLY | 0.13659549 |
| ALVGGLAW | ASGGGESQNTLY | 0.13687919 |
| ALAAAAATAI | ASGGGESQNTLY | 0.1648161 |
| AFGLGLLL | ASGGGESQNTLY | 0.17027041 |
| AFFAAASL | ASGGGESQNTLY | 0.15994063 |
| AAAALPAFSPP | ASGGGESQNTLY | 0.16788696 |

| | | |
|---|---|---|
| VVLAPAL | ASGGGESQNTLY | 0.13613777 |
| TIGHVDHGKTTL | ASGGGESQNTLY | 0.13257852 |
| LISNASDALDK | ASGGGESQNTLY | 0.13844587 |
| LISNASDAL | ASGGGESQNTLY | 0.14700647 |
| ISNASDALDK | ASGGGESQNTLY | 0.15274917 |
| NQAETLVYQTE | ASGGGESQNTLY | 0.12640795 |
| TGGTGGTAGAG | ASGGGESQNTLY | 0.14780308 |
| TGSTAGGAGGA | ASGGGESQNTLY | 0.16168435 |
| DPLAVPVAAGPV | ASGGGESQNTLY | 0.16823801 |
| GGTGGTPGNGGN | ASGGGESQNTLY | 0.16888216 |
| PAGLAGVDPATL | ASGGGESQNTLY | 0.1420775 |
| GNNLVVEAPHAGASV | ASGGGESQNTLY | 0.14333303 |
| KVTRIMSASTLEQPA | ASGGGESQNTLY | 0.14215545 |
| LGFALPIALPRRTVV | ASGGGESQNTLY | 0.15206856 |
| TRLGNLTARKLVVLV | ASGGGESQNTLY | 0.13866131 |
| WSQQRRLIYFPSAGP | ASGGGESQNTLY | 0.1782976 |
| GPNGNPDPMAAAVDIR | ASGGGESQNTLY | 0.13261518 |
| TQRCVSVADLRAAASR | ASGGGESQNTLY | 0.20189212 |
| QFVLDVAEHPVEVLTETV | ASGGGESQNTLY | 0.16958708 |
| GGAGGTGGAGGNGGTQAGDG | ASGGGESQNTLY | 0.15109527 |
| PLLATLRAIAVDVDAKPAQV | ASGGGESQNTLY | 0.14878155 |
| APKFRTGGYMIDFWGVGYQV | ASGGGESQNTLY | 0.17814699 |
| GAGGAGGAGGAAGGPGGTGG | ASGGGESQNTLY | 0.16849469 |
| IGALGLLATDVAPAGSVAFG | ASGGGESQNTLY | 0.1849541 |
| AGGAGGIGGNGGAGGAGGVL | ASGGGESQNTLY | 0.1345299 |
| ADELIGLL | ASGGGESQNTLY | 0.13446417 |
| ADRALKA | ASGGGESQNTLY | 0.1518798 |
| AGAVGQVGGAGGA | ASGGGESQNTLY | 0.14816177 |
| AGGGGGLGGAP | ASGGGESQNTLY | 0.14154829 |
| AGGLLYGNGGAG | ASGGGESQNTLY | 0.14390914 |
| AGLESIGL | ASGGGESQNTLY | 0.16918543 |
| AGLKTGII | ASGGGESQNTLY | 0.22219953 |
| ASLAALLL | ASGGGESQNTLY | 0.14677882 |
| ASVINISSI | ASGGGESQNTLY | 0.2480611 |
| ATPPGPA | ASGGGESQNTLY | 0.14526607 |
| ATVASALV | ASGGGESQNTLY | 0.14487423 |
| ATVASLIL | ASGGGESQNTLY | 0.14534616 |
| AVNGPVLIRH | ASGGGESQNTLY | 0.1383513 |
| DGAAGAVGI | ASGGGESQNTLY | 0.12913862 |
| DPLLHGI | ASGGGESQNTLY | 0.14222412 |
| ELGAALGAL | ASGGGESQNTLY | 0.17714833 |
| ELGRTQAV | ASGGGESQNTLY | 0.16681004 |
| FVAGGYLS | ASGGGESQNTLY | 0.14061268 |
| GASGGAGGAGGW | ASGGGESQNTLY | 0.16544293 |
| GGAGAGGDGGAGGI | ASGGGESQNTLY | 0.11287062 |
| GTGHNGSQPGMGGNG | ASGGGESQNTLY | 0.11282811 |
| GYAAASASAS | ASGGGESQNTLY | 0.11280303 |
| HQVDVAAVQQG | ASGGGESQNTLY | 0.11322645 |
| IGAAIAHT | ASGGGESQNTLY | 0.11279924 |
| IGMIEVAQL | ASGGGESQNTLY | 0.11285537 |
| IRLAPEHTSAMD | ASGGGESQNTLY | 0.11346253 |

| | | |
|---|---|---|
| IYNVSVL | ASGGGESQNTLY | 0.112892665 |
| KQGFLGM | ASGGGESQNTLY | 0.11348482 |
| LESVITQL | ASGGGESQNTLY | 0.113483 |
| LGLGALVVV | ASGGGESQNTLY | 0.11290392 |
| LIGNGGNGGSGGI | ASGGGESQNTLY | 0.113102846 |
| LNAINAPTL | ASGGGESQNTLY | 0.11283529 |
| LQLLNTGV | ASGGGESQNTLY | 0.11376583 |
| LQTSIEF | ASGGGESQNTLY | 0.112799406 |
| MSFAVSEL | ASGGGESQNTLY | 0.11272233 |
| NTGALGL | ASGGGESQNTLY | 0.113122225 |
| PPGAGGAGGAGTT | ASGGGESQNTLY | 0.11316225 |
| QLVGLGI | ASGGGESQNTLY | 0.11313109 |
| RFLEQVHQL | ASGGGESQNTLY | 0.11289565 |
| RGVGYVLR | ASGGGESQNTLY | 0.11289997 |
| SATGVFAGVI | ASGGGESQNTLY | 0.11288799 |
| SLPAITLP | ASGGGESQNTLY | 0.1129338 |
| STALAFTV | ASGGGESQNTLY | 0.11292282 |
| SVLTVGLGK | ASGGGESQNTLY | 0.112928174 |
| TALAGAGSGI | ASGGGESQNTLY | 0.113091156 |
| TGLPFTGI | ASGGGESQNTLY | 0.11249877 |
| VAEHVTV | ASGGGESQNTLY | 0.11396624 |
| VDIVLNSL | ASGGGESQNTLY | 0.113103874 |
| VLTGGGAVL | ASGGGESQNTLY | 0.113780506 |
| VNVGLAYGI | ASGGGESQNTLY | 0.11334044 |
| VSSAAVGAV | ASGGGESQNTLY | 0.11305399 |
| VTSVYEAGV | ASGGGESQNTLY | 0.113236636 |
| WVALDRGA | ASGGGESQNTLY | 0.112897545 |
| YLVGSVVI | ASGGGESQNTLY | 0.11295234 |
| GPPGTGKTLL | ASGGGESQNTLY | 0.112914726 |
| GVGMTKF | ASGGGESQNTLY | 0.11352775 |
| IGIGGSDL | ASGGGESQNTLY | 0.11271978 |
| IGTLLAEL | ASGGGESQNTLY | 0.1133317 |
| IYTSGTT | ASGGGESQNTLY | 0.113267995 |
| LSFPAGV | ASGGGESQNTLY | 0.11288102 |
| LYTSGTT | ASGGGESQNTLY | 0.11284671 |
| VGAFPFGF | ASGGGESQNTLY | 0.11367 |
| VGAGTVEF | ASGGGESQNTLY | 0.113205925 |
| VGKSTLLL | ASGGGESQNTLY | 0.11377073 |
| AALAHAGGGIV | ASGGGESQNTLY | 0.11315102 |
| AALVAVSLL | ASGGGESQNTLY | 0.113259755 |
| AAVSRAVAAL | ASGGGESQNTLY | 0.1130974 |
| AGGSSGIGGF | ASGGGESQNTLY | 0.11285086 |
| AGLTVANANG | ASGGGESQNTLY | 0.112922944 |
| AGLVGAVL | ASGGGESQNTLY | 0.11288573 |
| ALSQAFTV | ASGGGESQNTLY | 0.11290426 |
| ALTFSVSL | ASGGGESQNTLY | 0.11262851 |
| ALTGAALVAL | ASGGGESQNTLY | 0.11273829 |
| ASFSIGAGVL | ASGGGESQNTLY | 0.11262131 |
| DGSVYKI | ASGGGESQNTLY | 0.113008425 |
| EQLAEIV | ASGGGESQNTLY | 0.11282774 |
| FAVSELTVM | ASGGGESQNTLY | 0.11389737 |

| | | |
|---|---|---|
| GAAPLAAGL | ASGGGESQNTLY | 0.11266615 |
| GADGGDGGSGGAGG | ASGGGESQNTLY | 0.11274795 |
| GFGLDTGGAGG | ASGGGESQNTLY | 0.11276868 |
| GLASGGLL | ASGGGESQNTLY | 0.11295935 |
| GRIAGAAVLL | ASGGGESQNTLY | 0.11293016 |
| GSAGSLNTSF | ASGGGESQNTLY | 0.11308809 |
| GTGQIGI | ASGGGESQNTLY | 0.11288394 |
| GTLIALIL | ASGGGESQNTLY | 0.70587015 |
| IAATMGFVVG | ASGGGESQNTLY | 0.5983963 |
| IFAQALL | ASGGGESQNTLY | 0.31683856 |
| IGAAVALGF | ASGGGESQNTLY | 0.25124124 |
| IGLTPLAL | ASGGGESQNTLY | 0.6162202 |
| ILGSNTSTL | ASGGGESQNTLY | 0.49144834 |
| IVFSLTVL | ASGGGESQNTLY | 0.23009439 |
| LASKATGF | ASGGGESQNTLY | 0.68741757 |
| LASVALIL | ASGGGESQNTLY | 0.18167645 |
| LFAVGLTL | ASGGGESQNTLY | 0.29969576 |
| LFNGGIGTYI | ASGGGESQNTLY | 0.26501435 |
| LTLGAGII | ASGGGESQNTLY | 0.2635424 |
| LVAAALGDI | ASGGGESQNTLY | 0.5535732 |
| NTAFVQLGI | ASGGGESQNTLY | 0.1596412 |
| QLFSGAEL | ASGGGESQNTLY | 0.25761995 |
| SAATFITL | ASGGGESQNTLY | 0.35337994 |
| SNYGLANLGV | ASGGGESQNTLY | 0.43700606 |
| SYGGAYAVM | ASGGGESQNTLY | 0.7345654 |
| TTGVGGLTLG | ASGGGESQNTLY | 0.699986 |
| TTQTIYI | ASGGGESQNTLY | 0.69570875 |
| VAAGAGLGAVI | ASGGGESQNTLY | 0.7142971 |
| VAAVVTLL | ASGGGESQNTLY | 0.71369946 |
| VGFLGGLI | ASGGGESQNTLY | 0.09783926 |
| VGVIPILL | ASGGGESQNTLY | 0.15401296 |
| VIFASSLI | ASGGGESQNTLY | 0.15988815 |
| VKGGDGGAAAG | ASGGGESQNTLY | 0.30951023 |
| VTKTFTNI | ASGGGESQNTLY | 0.12302018 |
| VVAAAILL | ASGGGESQNTLY | 0.2668375 |
| YAALDVEL | ASGGGESQNTLY | 0.494112 |
| LASVALLL | ASGGGESQNTLY | 0.1220109 |
| LDAVHRV | ASGGGESQNTLY | 0.68422043 |
| QGLSSSIF | ASGGGESQNTLY | 0.18103579 |
| RGTFANIRL | ASGGGESQNTLY | 0.17047411 |
| RQLWWGHRI | ASGGGESQNTLY | 0.23593442 |
| YGPPGTGKTLL | ASGGGESQNTLY | 0.7593061 |
| AALVGAPANVA | ASGGGESQNTLY | 0.19556293 |
| AATFRRTE | ASGGGESQNTLY | 0.5067548 |
| AGALGGGI | ASGGGESQNTLY | 0.34208244 |
| AGGAGGAGGIGAG | ASGGGESQNTLY | 0.5448696 |
| AGSGALGL | ASGGGESQNTLY | 0.13642095 |
| AQIADQI | ASGGGESQNTLY | 0.64345986 |
| ASAVAVTYL | ASGGGESQNTLY | 0.40691262 |
| ASTVLGGTGG | ASGGGESQNTLY | 0.77327424 |
| AVGGEPAK | ASGGGESQNTLY | 0.3863137 |

| | | |
|---|---|---|
| CEQGTFGGTC | ASGGGESQNTLY | 0.3052137 |
| DEGKAPAG | ASGGGESQNTLY | 0.19021155 |
| DTVVFPAL | ASGGGESQNTLY | 0.11874102 |
| FEGITMAG | ASGGGESQNTLY | 0.1114361 |
| FPLVVQD | ASGGGESQNTLY | 0.5947877 |
| GAFTTPQ | ASGGGESQNTLY | 0.21241874 |
| GFGGFGTAGGVGGA | ASGGGESQNTLY | 0.48438525 |
| GFLAQFL | ASGGGESQNTLY | 0.6895479 |
| GGAGGAGGNAGVA | ASGGGESQNTLY | 0.11263455 |
| GGVGTAGAGGAGGAGG | ASGGGESQNTLY | 0.76716125 |
| GNGGGGDAGNAGSGGN | ASGGGESQNTLY | 0.1455858 |
| GTGGAGGAGNDAGS | ASGGGESQNTLY | 0.19910508 |
| GTTNTGWFNS | ASGGGESQNTLY | 0.67812467 |
| IFMGPLFTL | ASGGGESQNTLY | 0.13043226 |
| ILAHAIEAI | ASGGGESQNTLY | 0.77733314 |
| IRSAIREAL | ASGGGESQNTLY | 0.16951765 |
| ITFAGGG | ASGGGESQNTLY | 0.63678414 |
| ITGMSGHQEV | ASGGGESQNTLY | 0.6773294 |
| IVAGLACTL | ASGGGESQNTLY | 0.09481536 |
| IVGGGLAGA | ASGGGESQNTLY | 0.13968939 |
| IVTGVLNI | ASGGGESQNTLY | 0.725965 |
| LAQQGCI | ASGGGESQNTLY | 0.43602538 |
| LCLHVEN | ASGGGESQNTLY | 0.5687683 |
| LDAISDGS | ASGGGESQNTLY | 0.15905055 |
| LGGFAAML | ASGGGESQNTLY | 0.18904714 |
| LGGKNPMI | ASGGGESQNTLY | 0.19839388 |
| LGPKGVAAIL | ASGGGESQNTLY | 0.40097553 |
| LMVGVSI | ASGGGESQNTLY | 0.59370375 |
| LVGAAFVL | ASGGGESQNTLY | 0.372638 |
| LVLAVAPL | ASGGGESQNTLY | 0.17835176 |
| MAAIAVIDAA | ASGGGESQNTLY | 0.32830405 |
| NNGINAELQ | ASGGGESQNTLY | 0.30835986 |
| RGEPLVLVHG | ASGGGESQNTLY | 0.29103187 |
| RGGAGGAAGLI | ASGGGESQNTLY | 0.26547217 |
| RSYRMHPAV | ASGGGESQNTLY | 0.32749325 |
| SAAHLLAGQLA | ASGGGESQNTLY | 0.2023226 |
| SAAVGSY | ASGGGESQNTLY | 0.1383137 |
| SPHAVLHRI | ASGGGESQNTLY | 0.26523113 |
| SQIAKVL | ASGGGESQNTLY | 0.6180482 |
| TALGHLVP | ASGGGESQNTLY | 0.4882905 |
| TAPTTSGAA | ASGGGESQNTLY | 0.4518023 |
| TDGDDGNGGSPGT | ASGGGESQNTLY | 0.5775901 |
| TDGLGEN | ASGGGESQNTLY | 0.24683136 |
| TGGPPLPGQA | ASGGGESQNTLY | 0.10460063 |
| TIPAITI | ASGGGESQNTLY | 0.11658567 |
| TIPGFTI | ASGGGESQNTLY | 0.17090636 |
| TSIASGALA | ASGGGESQNTLY | 0.22453932 |
| TVTLPGAAI | ASGGGESQNTLY | 0.09937545 |
| VASGLSAALP | ASGGGESQNTLY | 0.34707886 |
| VGAGVIGI | ASGGGESQNTLY | 0.23762041 |
| VGPKRFGV | ASGGGESQNTLY | 0.21503605 |

| | | |
|---|---|---|
| VNVAFPD | ASGGGESQNTLY | 0.5819396 |
| VSMTNSGVS | ASGGGESQNTLY | 0.20438787 |
| VTDVLGDLSV | ASGGGESQNTLY | 0.17543131 |
| VVVVGAGFAGL | ASGGGESQNTLY | 0.1823799 |
| VYTVGTVV | ASGGGESQNTLY | 0.39735457 |
| WTTFSGP | ASGGGESQNTLY | 0.14523354 |
| YDELVEAF | ASGGGESQNTLY | 0.5084545 |
| YGTGGAGGNGG | ASGGGESQNTLY | 0.15459053 |
| YTNVMAV | ASGGGESQNTLY | 0.50516546 |
| FLEQVHQL | ASGGGESQNTLY | 0.16381223 |
| GLSLLAVAV | ASGGGESQNTLY | 0.5255909 |
| LEQVHQL | ASGGGESQNTLY | 0.27913478 |
| LNAAGAI | ASGGGESQNTLY | 0.7082639 |
| PAVHASL | ASGGGESQNTLY | 0.27135795 |
| TELEDDA | ASGGGESQNTLY | 0.38360572 |
| TPPGPPN | ASGGGESQNTLY | 0.18937227 |
| TPPPPPPP | ASGGGESQNTLY | 0.41913858 |
| TQSPALI | ASGGGHSGNTLY | 0.13144676 |
| EFAAAFPQ | ASGGGHSGNTLY | 0.17757861 |
| LLSGNLDVL | ASGGGHSGNTLY | 0.119098976 |
| GPANLGNYNI | ASGGGHSGNTLY | 0.21496665 |
| TLGPTLP | ASGGGHSGNTLY | 0.3840697 |
| VYAYSSCCK | ASGGGHSGNTLY | 0.100956626 |
| APAPAPAPNI | ASGGGHSGNTLY | 0.27459064 |
| PGGPVPIP | ASGGGHSGNTLY | 0.13650015 |
| FLTVALEP | ASGGGHSGNTLY | 0.18006414 |
| SQYFGANL | ASGGGHSGNTLY | 0.34669703 |
| GGVKPAA | ASGGGHSGNTLY | 0.3174002 |
| GGTGGAGGAGGW | ASGGGHSGNTLY | 0.3190528 |
| GVGGVAGLGGAGG | ASGGGHSGNTLY | 0.1396148 |
| VLLVLATTI | ASGGGHSGNTLY | 0.20367527 |
| IGIFAAML | ASGGGHSGNTLY | 0.25626683 |
| AVGDLID | ASGGGHSGNTLY | 0.115903206 |
| NISPLPL | ASGGGHSGNTLY | 0.1600358 |
| VTAVFVAI | ASGGGHSGNTLY | 0.60534286 |
| SGDDNDAAL | ASGGGHSGNTLY | 0.74033785 |
| AVAALAHS | ASGGGHSGNTLY | 0.56797796 |
| GGVTVMATP | ASGGGHSGNTLY | 0.40650135 |
| AGLAANVRD | ASGGGHSGNTLY | 0.3783574 |
| TNRPDLIDPALLRPG | ASGGGHSGNTLY | 0.3162963 |
| LANLGVTAL | ASGGGHSGNTLY | 0.57874924 |
| TGERIEL | ASGGGHSGNTLY | 0.5617902 |
| NGEELAA | ASGGGHSGNTLY | 0.5506958 |
| VLSLVWG | ASGGGHSGNTLY | 0.27900693 |
| IGYALWL | ASGGGHSGNTLY | 0.35217083 |
| LSLLLAPL | ASGGGHSGNTLY | 0.5013586 |
| TLDEALK | ASGGGHSGNTLY | 0.48153913 |
| TDGAGTAEGG | ASGGGHSGNTLY | 0.3960191 |
| NAPGGAGGNGG | ASGGGHSGNTLY | 0.46267268 |
| VSVFMGMGGNF | ASGGGHSGNTLY | 0.3184271 |
| LASYLAAL | ASGGGHSGNTLY | 0.39837882 |

| | | |
|---|---|---|
| LVGGTPAL | ASGGGHSGNTLY | 0.30657023 |
| VRNADVL | ASGGGHSGNTLY | 0.61608934 |
| TCSSPFLVSAM | ASGGGHSGNTLY | 0.56449044 |
| VHITDAE | ASGGGHSGNTLY | 0.4933675 |
| EYAAPSF | ASGGGHSGNTLY | 0.7376599 |
| VVDAFHGGL | ASGGGHSGNTLY | 0.4091702 |
| ASVVDPQ | ASGGGHSGNTLY | 0.16763285 |
| SYNEFGGLS | ASGGGHSGNTLY | 0.25516933 |
| DPDGSVY | ASGGGHSGNTLY | 0.34785643 |
| MVSTIHATE | ASGGGHSGNTLY | 0.3957705 |
| LAAGVADPV | ASGGGHSGNTLY | 0.17509855 |
| TKEQIAA | ASGGGHSGNTLY | 0.294496 |
| SLASPAAL | ASGGGHSGNTLY | 0.27746457 |
| IGLVFYGV | ASGGGHSGNTLY | 0.27794713 |
| FLLDASNL | ASGGGHSGNTLY | 0.5693754 |
| YQVGNFP | ASGGGHSGNTLY | 0.40850258 |
| ESLLSQST | ASGGGHSGNTLY | 0.33909476 |
| LFTVGFAAVG | ASGGGHSGNTLY | 0.26724693 |
| ATPPTPT | ASGGGHSGNTLY | 0.3130122 |
| TDLATLG | ASGGGHSGNTLY | 0.3617588 |
| SAGGTGR | ASGGGHSGNTLY | 0.5948465 |
| GFSNSGS | ASGGGHSGNTLY | 0.21484219 |
| GGNGGSGGTSV | ASGGGHSGNTLY | 0.663028 |
| GTSVATGGAGN | ASGGGHSGNTLY | 0.30790925 |
| NGGNGGSGGTS | ASGGGHSGNTLY | 0.6467211 |
| AVVKFPH | ASGGGHSGNTLY | 0.627772 |
| TPPPPPP | ASGGGHSGNTLY | 0.7563015 |
| AAPAAAAAPA | ASGGGHSGNTLY | 0.62001735 |
| IALNEDL | ASGGGHSGNTLY | 0.52665925 |
| IVNVSSM | ASGGGHSGNTLY | 0.27806777 |
| TGIPPAPR | ASGGGHSGNTLY | 0.5595806 |
| FDLGGGTF | ASGGGHSGNTLY | 0.32826006 |
| FDLGGGTFDVS | ASGGGHSGNTLY | 0.40348727 |
| NEPTAAA | ASGGGHSGNTLY | 0.16472249 |
| NEPTAAALA | ASGGGHSGNTLY | 0.35541564 |
| GSIINISSI | ASGGGHSGNTLY | 0.6161094 |
| VAPGNAGTA | ASGGGHSGNTLY | 0.15958008 |
| TGGGASLE | ASGGGHSGNTLY | 0.33786476 |
| VAVYNHYKRI | ASGGGHSGNTLY | 0.26320392 |
| TDPAPATT | ASGGGHSGNTLY | 0.3300303 |
| TNRPDILDPALLRPG | ASGGGHSGNTLY | 0.5768569 |
| LTTTTTS | ASGGGHSGNTLY | 0.3032645 |
| PTGSGKTL | ASGGGHSGNTLY | 0.7591922 |
| TGSGKTL | ASGGGHSGNTLY | 0.3019417 |
| TIGHVDHGK | ASGGGHSGNTLY | 0.27676332 |
| TIGHVDHGKTT | ASGGGHSGNTLY | 0.28930432 |
| GHVDHGKTT | ASGGGHSGNTLY | 0.21039087 |
| AQFEHTLL | ASGGGHSGNTLY | 0.2511007 |
| IVNQDEL | ASGGGHSGNTLY | 0.7398397 |
| TNGSQFF | ASGGGHSGNTLY | 0.08321911 |
| TGKGIVE | ASGGGHSGNTLY | 0.082786836 |

| | | |
|---|---|---|
| LISNASDA | ASGGGHSGNTLY | 0.082916476 |
| AELRTLGELTKIAKAHGAQ | ASGGGHSGNTLY | 0.083999 |
| APNAVAAAEPSW | ASGGGHSGNTLY | 0.083043456 |
| APQVAGAATAVL | ASGGGHSGNTLY | 0.083152436 |
| ARRDPAAAAAILPTDA | ASGGGHSGNTLY | 0.083380364 |
| DAAARDVLAEAGLAEYF | ASGGGHSGNTLY | 0.083183356 |
| DDQIVLSAKANLAKVEQ | ASGGGHSGNTLY | 0.08521973 |
| DELLPDTGEDSDEDGH | ASGGGHSGNTLY | 0.08409613 |
| ELPEAPANRATVPQD | ASGGGHSGNTLY | 0.0832275 |
| FGQSKIERVTGIG | ASGGGHSGNTLY | 0.083777204 |
| GAGGTGSTAGGAGGAGGAGGI | ASGGGHSGNTLY | 0.08291894 |
| GHGGTGAAGGNGGTGGAGGS( | ASGGGHSGNTLY | 0.083809085 |
| GNGGNGGAGGEGGPGGAGGA( | ASGGGHSGNTLY | 0.082935795 |
| GSTAGNGGNGGNATNGTIAG( | ASGGGHSGNTLY | 0.08279351 |
| GTGGDGGNGGNGGTGG | ASGGGHSGNTLY | 0.08318172 |
| GVRRPAGIPADAPV | ASGGGHSGNTLY | 0.08361797 |
| GYRQVTDAHLLTLARR | ASGGGHSGNTLY | 0.083411865 |
| IAKMGQDGHDRG | ASGGGHSGNTLY | 0.08317973 |
| KAEPSIEIQYT | ASGGGHSGNTLY | 0.083282925 |
| KPVRIRMAADEDTVAVT | ASGGGHSGNTLY | 0.08288862 |
| MPPRIAGMRLLVIKPEP | ASGGGHSGNTLY | 0.0831127 |
| MKFYAVALFRSRGVPPTR | ASGGGHSGNTLY | 0.083040744 |
| NAGGFGILGAVAHSP | ASGGGHSGNTLY | 0.08326301 |
| NEGREITLAMAEM | ASGGGHSGNTLY | 0.08330351 |
| NELVEWSHPLGHRWRWE | ASGGGHSGNTLY | 0.08256797 |
| NLGAGGTGGNGGNPG | ASGGGHSGNTLY | 0.08371824 |
| NTEWPKWETVMAADPVD | ASGGGHSGNTLY | 0.08362071 |
| PASAAPAETQP | ASGGGHSGNTLY | 0.084512144 |
| QQSSDGKNGVRAAEITGE | ASGGGHSGNTLY | 0.08403128 |
| RAPDAVTTSWWKE | ASGGGHSGNTLY | 0.08323998 |
| SGFGNVGAHTSGWFNQ | ASGGGHSGNTLY | 0.08333699 |
| SGVAGAVGQAASVGGLKV | ASGGGHSGNTLY | 0.08322045 |
| SNGGAGTGSAGGNGGT | ASGGGHSGNTLY | 0.08338577 |
| TGGAGGEGGAAG | ASGGGHSGNTLY | 0.08301067 |
| TGGDGGAAGAGGAPGHG | ASGGGHSGNTLY | 0.08505653 |
| TMPPLLAAIANAGRHGV | ASGGGHSGNTLY | 0.08283122 |
| TSSTAGNSGGA | ASGGGHSGNTLY | 0.0847399 |
| VPATVPVGILC | ASGGGHSGNTLY | 0.08414497 |
| VQRPDSGGVVLTGRIS | ASGGGHSGNTLY | 0.083117515 |
| WTPNPVPATKGVPAGLVQAT | ASGGGHSGNTLY | 0.08312976 |
| YALKRALAKFSDRFDVVII | ASGGGHSGNTLY | 0.08400261 |
| YLIGSVIV | ASGGGHSGNTLY | 0.08368567 |
| YGQLVAVID | ASGGGHSGNTLY | 0.0860356 |
| VTVGGMTLPA | ASGGGHSGNTLY | 0.08325389 |
| VLALPAV | ASGGGHSGNTLY | 0.08316041 |
| VIPPTLNYET | ASGGGHSGNTLY | 0.08380972 |
| VGMGGMHPSSG | ASGGGHSGNTLY | 0.08307179 |
| VGLAGGAATA | ASGGGHSGNTLY | 0.0832408 |
| VGGAGGAGGSGP | ASGGGHSGNTLY | 0.08323519 |
| VFAAFMV | ASGGGHSGNTLY | 0.083494194 |
| TLYDKLR | ASGGGHSGNTLY | 0.083489664 |

| | | |
|---|---|---|
| TLAVFTVL | ASGGGHSGNTLY | 0.08296795 |
| TGSVQPARV | ASGGGHSGNTLY | 0.083672754 |
| TGGGAGSGGT | ASGGGHSGNTLY | 0.083143294 |
| SHLETSL | ASGGGHSGNTLY | 0.08286296 |
| SFTTGMT | ASGGGHSGNTLY | 0.08391226 |
| QVEVAKRMGVRQPTVS | ASGGGHSGNTLY | 0.08273284 |
| QGSGGPALAM | ASGGGHSGNTLY | 0.08274702 |
| MLAPSNP | ASGGGHSGNTLY | 0.08308341 |
| LVNNAAI | ASGGGHSGNTLY | 0.08438783 |
| LTGEINTLA | ASGGGHSGNTLY | 0.084339984 |
| LSIFGPV | ASGGGHSGNTLY | 0.08323383 |
| LLALLANL | ASGGGHSGNTLY | 0.08318845 |
| LISNASDALDKLR | ASGGGHSGNTLY | 0.5320349 |
| LINEAVP | ASGGGHSGNTLY | 0.7112256 |
| LGYAFSGL | ASGGGHSGNTLY | 0.12687996 |
| LDFVAVAL | ASGGGHSGNTLY | 0.15287986 |
| LAQPGAE | ASGGGHSGNTLY | 0.39336616 |
| ISNLGAVTGF | ASGGGHSGNTLY | 0.45712656 |
| HQEDPGRI | ASGGGHSGNTLY | 0.472106 |
| GTPGEPTP | ASGGGHSGNTLY | 0.50892836 |
| GFFNSGS | ASGGGHSGNTLY | 0.15525703 |
| GAGGGGLTAGAN | ASGGGHSGNTLY | 0.15322912 |
| FVDALVNM | ASGGGHSGNTLY | 0.15462402 |
| FLTVAPEL | ASGGGHSGNTLY | 0.2543281 |
| FETLTAL | ASGGGHSGNTLY | 0.4450921 |
| DVINPIEL | ASGGGHSGNTLY | 0.12670912 |
| APVSGAPV | ASGGGHSGNTLY | 0.1176547 |
| ALVGGLAW | ASGGGHSGNTLY | 0.17963794 |
| ALAAAAATAI | ASGGGHSGNTLY | 0.4780761 |
| AFGLGLLL | ASGGGHSGNTLY | 0.70360005 |
| AFFAAASL | ASGGGHSGNTLY | 0.69224066 |
| AAAALPAFSPP | ASGGGHSGNTLY | 0.6007905 |
| VVLAPAL | ASGGGHSGNTLY | 0.54382527 |
| TIGHVDHGKTTL | ASGGGHSGNTLY | 0.24454121 |
| LISNASDALDK | ASGGGHSGNTLY | 0.11161546 |
| LISNASDAL | ASGGGHSGNTLY | 0.13243343 |
| ISNASDALDK | ASGGGHSGNTLY | 0.12795694 |
| NQAETLVYQTE | ASGGGHSGNTLY | 0.33367354 |
| TGGTGGTAGAG | ASGGGHSGNTLY | 0.10708692 |
| TGSTAGGAGGA | ASGGGHSGNTLY | 0.19840337 |
| DPLAVPVAAGPV | ASGGGHSGNTLY | 0.18256131 |
| GGTGGTPGNGGN | ASGGGHSGNTLY | 0.11915395 |
| PAGLAGVDPATL | ASGGGHSGNTLY | 0.668437 |
| GNNLVVEAPHAGASV | ASGGGHSGNTLY | 0.12398388 |
| KVTRIMSASTLEQPA | ASGGGHSGNTLY | 0.12928966 |
| LGFALPIALPRRTVV | ASGGGHSGNTLY | 0.15189862 |
| TRLGNLTARKLVVLV | ASGGGHSGNTLY | 0.6550763 |
| WSQQRRLIYFPSAGP | ASGGGHSGNTLY | 0.14313677 |
| GPNGNPDPMAAAVDIR | ASGGGHSGNTLY | 0.62809575 |
| TQRCVSVADLRAAASR | ASGGGHSGNTLY | 0.2521192 |
| QFVLDVAEHPVEVLTETV | ASGGGHSGNTLY | 0.33752584 |

| | | |
|---|---|---|
| GGAGGTGGAGGNGGTQAGDG | ASGGGHSGNTLY | 0.13211824 |
| PLLATLRAIAVDVDAKPAQV | ASGGGHSGNTLY | 0.44810736 |
| APKFRTGGYMIDFWGVGYQV | ASGGGHSGNTLY | 0.30468482 |
| GAGGAGGAGGAAGGPGGTGG | ASGGGHSGNTLY | 0.7617966 |
| IGALGLLATDVAPAGSVAFG | ASGGGHSGNTLY | 0.30669555 |
| AGGAGGIGGNGGAGGAGGVL | ASGGGHSGNTLY | 0.2547641 |
| ADELIGLL | ASGGGHSGNTLY | 0.14876509 |
| ADRALKA | ASGGGHSGNTLY | 0.26788312 |
| AGAVGQVGGAGGA | ASGGGHSGNTLY | 0.117395006 |
| AGGGGGLGGAP | ASGGGHSGNTLY | 0.2712766 |
| AGGLLYGNGGAG | ASGGGHSGNTLY | 0.12852181 |
| AGLESIGL | ASGGGHSGNTLY | 0.18621084 |
| AGLKTGII | ASGGGHSGNTLY | 0.3960244 |
| ASLAALLL | ASGGGHSGNTLY | 0.1181978 |
| ASVINISSI | ASGGGHSGNTLY | 0.45578447 |
| ATPPGPA | ASGGGHSGNTLY | 0.14158583 |
| ATVASALV | ASGGGHSGNTLY | 0.14076826 |
| ATVASLIL | ASGGGHSGNTLY | 0.53025794 |
| AVNGPVLIRH | ASGGGHSGNTLY | 0.119279936 |
| DGAAGAVGI | ASGGGHSGNTLY | 0.44313908 |
| DPLLHGI | ASGGGHSGNTLY | 0.115182266 |
| ELGAALGAL | ASGGGHSGNTLY | 0.2971362 |
| ELGRTQAV | ASGGGHSGNTLY | 0.35094216 |
| FVAGGYLS | ASGGGHSGNTLY | 0.11111487 |
| GASGGAGGAGGW | ASGGGHSGNTLY | 0.12538476 |
| GGAGAGGDGGAGGI | ASGGGHSGNTLY | 0.5249674 |
| GTGHNGSQPGMGGNG | ASGGGHSGNTLY | 0.31662378 |
| GYAAASASAS | ASGGGHSGNTLY | 0.41747808 |
| HQVDVAAVQQG | ASGGGHSGNTLY | 0.18030821 |
| IGAAIAHT | ASGGGHSGNTLY | 0.18021923 |
| IGMIEVAQL | ASGGGHSGNTLY | 0.21230806 |
| IRLAPEHTSAMD | ASGGGHSGNTLY | 0.33755487 |
| IYNVSVL | ASGGGHSGNTLY | 0.50333714 |
| KQGFLGM | ASGGGHSGNTLY | 0.44482857 |
| LESVITQL | ASGGGHSGNTLY | 0.1751097 |
| LGLGALVVV | ASGGGHSGNTLY | 0.33072966 |
| LIGNGGNGGSGGI | ASGGGHSGNTLY | 0.20022878 |
| LNAINAPTL | ASGGGHSGNTLY | 0.2297241 |
| LQLLNTGV | ASGGGHSGNTLY | 0.25888577 |
| LQTSIEF | ASGGGHSGNTLY | 0.18629172 |
| MSFAVSEL | ASGGGHSGNTLY | 0.22620203 |
| NTGALGL | ASGGGHSGNTLY | 0.17081425 |
| PPGAGGAGGAGTT | ASGGGHSGNTLY | 0.20536076 |
| QLVGLGI | ASGGGHSGNTLY | 0.3947295 |
| RFLEQVHQL | ASGGGHSGNTLY | 0.26536506 |
| RGVGYVLR | ASGGGHSGNTLY | 0.55428684 |
| SATGVFAGVI | ASGGGHSGNTLY | 0.46015713 |
| SLPAITLP | ASGGGHSGNTLY | 0.1938126 |
| STALAFTV | ASGGGHSGNTLY | 0.14613354 |
| SVLTVGLGK | ASGGGHSGNTLY | 0.16841327 |
| TALAGAGSGI | ASGGGHSGNTLY | 0.17265135 |

| | | |
|---|---|---|
| TGLPFTGI | ASGGGHSGNTLY | 0.20858116 |
| VAEHVTV | ASGGGHSGNTLY | 0.1448317 |
| VDIVLNSL | ASGGGHSGNTLY | 0.23289213 |
| VLTGGGAVL | ASGGGHSGNTLY | 0.2403468 |
| VNVGLAYGI | ASGGGHSGNTLY | 0.20995343 |
| VSSAAVGAV | ASGGGHSGNTLY | 0.38928702 |
| VTSVYEAGV | ASGGGHSGNTLY | 0.21072929 |
| WVALDRGA | ASGGGHSGNTLY | 0.17806147 |
| YLVGSVVI | ASGGGHSGNTLY | 0.16483626 |
| GPPGTGKTLL | ASGGGHSGNTLY | 0.71527773 |
| GVGMTKF | ASGGGHSGNTLY | 0.16104029 |
| IGIGGSDL | ASGGGHSGNTLY | 0.5785427 |
| IGTLLAEL | ASGGGHSGNTLY | 0.31300423 |
| IYTSGTT | ASGGGHSGNTLY | 0.42576125 |
| LSFPAGV | ASGGGHSGNTLY | 0.16731818 |
| LYTSGTT | ASGGGHSGNTLY | 0.3381436 |
| VGAFPFGF | ASGGGHSGNTLY | 0.25193834 |
| VGAGTVEF | ASGGGHSGNTLY | 0.6123757 |
| VGKSTLLL | ASGGGHSGNTLY | 0.20035763 |
| AALAHAGGGIV | ASGGGHSGNTLY | 0.21442923 |
| AALVAVSLL | ASGGGHSGNTLY | 0.17504822 |
| AAVSRAVAAL | ASGGGHSGNTLY | 0.5031861 |
| AGGSSGIGGF | ASGGGHSGNTLY | 0.15435782 |
| AGLTVANANG | ASGGGHSGNTLY | 0.2184594 |
| AGLVGAVL | ASGGGHSGNTLY | 0.28603774 |
| ALSQAFTV | ASGGGHSGNTLY | 0.26359785 |
| ALTFSVSL | ASGGGHSGNTLY | 0.52611804 |
| ALTGAALVAL | ASGGGHSGNTLY | 0.14709553 |
| ASFSIGAGVL | ASGGGHSGNTLY | 0.59707606 |
| DGSVYKI | ASGGGHSGNTLY | 0.1547012 |
| EQLAEIV | ASGGGHSGNTLY | 0.164228 |
| FAVSELTVM | ASGGGHSGNTLY | 0.2880171 |
| GAAPLAAGL | ASGGGHSGNTLY | 0.1643562 |
| GADGGDGGSGGAGG | ASGGGHSGNTLY | 0.31248558 |
| GFGLDTGGAGG | ASGGGHSGNTLY | 0.16354372 |
| GLASGGLL | ASGGGHSGNTLY | 0.5267381 |
| GRIAGAAVLL | ASGGGHSGNTLY | 0.45558006 |
| GSAGSLNTSF | ASGGGHSGNTLY | 0.14921986 |
| GTGQIGI | ASGGGHSGNTLY | 0.15659405 |
| GTLIALIL | ASGGGHSGNTLY | 0.47409582 |
| IAATMGFVVG | ASGGGHSGNTLY | 0.67286 |
| IFAQALL | ASGGGHSGNTLY | 0.80068886 |
| IGAAVALGF | ASGGGHSGNTLY | 0.45967016 |
| IGLTPLAL | ASGGGHSGNTLY | 0.31003103 |
| ILGSNTSTL | ASGGGHSGNTLY | 0.50546867 |
| IVFSLTVL | ASGGGHSGNTLY | 0.6381349 |
| LASKATGF | ASGGGHSGNTLY | 0.8232639 |
| LASVALIL | ASGGGHSGNTLY | 0.70050746 |
| LFAVGLTL | ASGGGHSGNTLY | 0.2571268 |
| LFNGGIGTYI | ASGGGHSGNTLY | 0.6883671 |
| LTLGAGII | ASGGGHSGNTLY | 0.5454561 |

| | | |
|---|---|---|
| LVAAALGDI | ASGGGHSGNTLY | 0.6028364 |
| NTAFVQLGI | ASGGGHSGNTLY | 0.4029217 |
| QLFSGAEL | ASGGGHSGNTLY | 0.49045923 |
| SAATFITL | ASGGGHSGNTLY | 0.38438606 |
| SNYGLANLGV | ASGGGHSGNTLY | 0.25509962 |
| SYGGAYAVM | ASGGGHSGNTLY | 0.6143889 |
| TTGVGGLTLG | ASGGGHSGNTLY | 0.77180445 |
| TTQTIYI | ASGGGHSGNTLY | 0.76877487 |
| VAAGAGLGAVI | ASGGGHSGNTLY | 0.71342045 |
| VAAVVTLL | ASGGGHSGNTLY | 0.76299644 |
| VGFLGGLI | ASGGGHSGNTLY | 0.6459459 |
| VGVIPILL | ASGGGHSGNTLY | 0.157809 |
| VIFASSLI | ASGGGHSGNTLY | 0.20425598 |
| VKGGDGGAAAG | ASGGGHSGNTLY | 0.31540906 |
| VTKTFTNI | ASGGGHSGNTLY | 0.50891477 |
| VVAAAILL | ASGGGHSGNTLY | 0.14937785 |
| YAALDVEL | ASGGGHSGNTLY | 0.4784491 |
| LASVALLL | ASGGGHSGNTLY | 0.43485838 |
| LDAVHRV | ASGGGHSGNTLY | 0.37581134 |
| QGLSSSIF | ASGGGHSGNTLY | 0.7774179 |
| RGTFANIRL | ASGGGHSGNTLY | 0.4747457 |
| RQLWWGHRI | ASGGGHSGNTLY | 0.24722737 |
| YGPPGTGKTLL | ASGGGHSGNTLY | 0.2992149 |
| AALVGAPANVA | ASGGGHSGNTLY | 0.6638551 |
| AATFRRTE | ASGGGHSGNTLY | 0.21807514 |
| AGALGGGI | ASGGGHSGNTLY | 0.75726956 |
| AGGAGGAGGIGAG | ASGGGHSGNTLY | 0.28096953 |
| AGSGALGL | ASGGGHSGNTLY | 0.74490225 |
| AQIADQI | ASGGGHSGNTLY | 0.20550482 |
| ASAVAVTYL | ASGGGHSGNTLY | 0.7302325 |
| ASTVLGGTGG | ASGGGHSGNTLY | 0.5774416 |
| AVGGEPAK | ASGGGHSGNTLY | 0.81761056 |
| CEQGTFGGTC | ASGGGHSGNTLY | 0.43229258 |
| DEGKAPAG | ASGGGHSGNTLY | 0.56560636 |
| DTVVFPAL | ASGGGHSGNTLY | 0.38631397 |
| FEGITMAG | ASGGGHSGNTLY | 0.82262814 |
| FPLVVQD | ASGGGHSGNTLY | 0.28602767 |
| GAFTTPQ | ASGGGHSGNTLY | 0.3849955 |
| GFGGFGTAGGVGGA | ASGGGHSGNTLY | 0.19173977 |
| GFLAQFL | ASGGGHSGNTLY | 0.44724205 |
| GGAGGAGGNAGVA | ASGGGHSGNTLY | 0.71573615 |
| GGVGTAGAGGAGGAGG | ASGGGHSGNTLY | 0.16240147 |
| GNGGGGDAGNAGSGGN | ASGGGHSGNTLY | 0.6433733 |
| GTGGAGGAGNDAGS | ASGGGHSGNTLY | 0.3018825 |
| GTTNTGWFNS | ASGGGHSGNTLY | 0.24935189 |
| IFMGPLFTL | ASGGGHSGNTLY | 0.59214973 |
| ILAHAIEAI | ASGGGHSGNTLY | 0.5442395 |
| IRSAIREAL | ASGGGHSGNTLY | 0.65996575 |
| ITFAGGG | ASGGGHSGNTLY | 0.24768418 |
| ITGMSGHQEV | ASGGGHSGNTLY | 0.44500577 |
| IVAGLACTL | ASGGGHSGNTLY | 0.4749875 |

| | | |
|---|---|---|
| IVGGGLAGA | ASGGGHSGNTLY | 0.28984877 |
| IVTGVLNI | ASGGGHSGNTLY | 0.2643078 |
| LAQQGCI | ASGGGHSGNTLY | 0.77362657 |
| LCLHVEN | ASGGGHSGNTLY | 0.65561295 |
| LDAISDGS | ASGGGHSGNTLY | 0.62155515 |
| LGGFAAML | ASGGGHSGNTLY | 0.36836493 |
| LGGKNPMI | ASGGGHSGNTLY | 0.28751883 |
| LGPKGVAAIL | ASGGGHSGNTLY | 0.2928376 |
| LMVGVSI | ASGGGHSGNTLY | 0.60954297 |
| LVGAAFVL | ASGGGHSGNTLY | 0.6342263 |
| LVLAVAPL | ASGGGHSGNTLY | 0.5835346 |
| MAAIAVIDAA | ASGGGHSGNTLY | 0.2485406 |
| NNGINAELQ | ASGGGHSGNTLY | 0.27665856 |
| RGEPLVLVHG | ASGGGHSGNTLY | 0.4960382 |
| RGGAGGAAGLI | ASGGGHSGNTLY | 0.2624665 |
| RSYRMHPAV | ASGGGHSGNTLY | 0.32469374 |
| SAAHLLAGQLA | ASGGGHSGNTLY | 0.32059434 |
| SAAVGSY | ASGGGHSGNTLY | 0.34329504 |
| SPHAVLHRI | ASGGGHSGNTLY | 0.31761187 |
| SQIAKVL | ASGGGHSGNTLY | 0.23058678 |
| TALGHLVP | ASGGGHSGNTLY | 0.46990323 |
| TAPTTSGAA | ASGGGHSGNTLY | 0.42772526 |
| TDGDDGNGGSPGT | ASGGGHSGNTLY | 0.47257742 |
| TDGLGEN | ASGGGHSGNTLY | 0.6610224 |
| TGGPPLPGQA | ASGGGHSGNTLY | 0.3396782 |
| TIPAITI | ASGGGHSGNTLY | 0.21100216 |
| TIPGFTI | ASGGGHSGNTLY | 0.22640285 |
| TSIASGALA | ASGGGHSGNTLY | 0.28023553 |
| TVTLPGAAI | ASGGGHSGNTLY | 0.27183342 |
| VASGLSAALP | ASGGGHSGNTLY | 0.21305571 |
| VGAGVIGI | ASGGGHSGNTLY | 0.27285552 |
| VGPKRFGV | ASGGGHSGNTLY | 0.42871675 |
| VNVAFPD | ASGGGHSGNTLY | 0.2517056 |
| VSMTNSGVS | ASGGGHSGNTLY | 0.35536566 |
| VTDVLGDLSV | ASGGGHSGNTLY | 0.25426626 |
| VVVVGAGFAGL | ASGGGHSGNTLY | 0.27850422 |
| VYTVGTVV | ASGGGHSGNTLY | 0.2424458 |
| WTTFSGP | ASGGGHSGNTLY | 0.47160053 |
| YDELVEAF | ASGGGHSGNTLY | 0.2685159 |
| YGTGGAGGNGG | ASGGGHSGNTLY | 0.40918875 |
| YTNVMAV | ASGGGHSGNTLY | 0.2424337 |
| FLEQVHQL | ASGGGHSGNTLY | 0.5211998 |
| GLSLLAVAV | ASGGGHSGNTLY | 0.24180958 |
| LEQVHQL | ASGGGHSGNTLY | 0.55811673 |
| LNAAGAI | ASGGGHSGNTLY | 0.4783611 |
| PAVHASL | ASGGGHSGNTLY | 0.5795357 |
| TELEDDA | ASGGGHSGNTLY | 0.35176748 |
| TPPGPPN | ASGGGHSGNTLY | 0.3252135 |
| TPPPPPPP | ASGGGHSGNTLY | 0.22619733 |
| TQSPALI | ASSRDWVYAEQF | 0.6306178 |
| EFAAAFPQ | ASSRDWVYAEQF | 0.24540843 |

| | | |
|---|---|---|
| LLSGNLDVL | ASSRDWVYAEQF | 0.5095359 |
| GPANLGNYNI | ASSRDWVYAEQF | 0.21859382 |
| TLGPTLP | ASSRDWVYAEQF | 0.4836241 |
| VYAYSSCCK | ASSRDWVYAEQF | 0.6068663 |
| APAPAPAPNI | ASSRDWVYAEQF | 0.26351708 |
| PGGPVPIP | ASSRDWVYAEQF | 0.5512097 |
| FLTVALEP | ASSRDWVYAEQF | 0.504064 |
| SQYFGANL | ASSRDWVYAEQF | 0.24732211 |
| GGVKPAA | ASSRDWVYAEQF | 0.44915605 |
| GGTGGAGGAGGW | ASSRDWVYAEQF | 0.23773076 |
| GVGGVAGLGGAGG | ASSRDWVYAEQF | 0.666371 |
| VLLVLATTI | ASSRDWVYAEQF | 0.23488374 |
| IGIFAAML | ASSRDWVYAEQF | 0.479577 |
| AVGDLID | ASSRDWVYAEQF | 0.48985302 |
| NISPLPL | ASSRDWVYAEQF | 0.2572214 |
| VTAVFVAI | ASSRDWVYAEQF | 0.23165727 |
| SGDDNDAAL | ASSRDWVYAEQF | 0.6403047 |
| AVAALAHS | ASSRDWVYAEQF | 0.7117133 |
| GGVTVMATP | ASSRDWVYAEQF | 0.6974315 |
| AGLAANVRD | ASSRDWVYAEQF | 0.6379175 |
| TNRPDLIDPALLRPG | ASSRDWVYAEQF | 0.5675894 |
| LANLGVTAL | ASSRDWVYAEQF | 0.6494167 |
| TGERIEL | ASSRDWVYAEQF | 0.7000592 |
| NGEELAA | ASSRDWVYAEQF | 0.69514966 |
| VLSLVWG | ASSRDWVYAEQF | 0.6899392 |
| IGYALWL | ASSRDWVYAEQF | 0.4721626 |
| LSLLLAPL | ASSRDWVYAEQF | 0.6223984 |
| TLDEALK | ASSRDWVYAEQF | 0.6629272 |
| TDGAGTAEGG | ASSRDWVYAEQF | 0.5649258 |
| NAPGGAGGNGG | ASSRDWVYAEQF | 0.67610025 |
| VSVFMGMGGNF | ASSRDWVYAEQF | 0.65753555 |
| LASYLAAL | ASSRDWVYAEQF | 0.6184701 |
| LVGGTPAL | ASSRDWVYAEQF | 0.64427257 |
| VRNADVL | ASSRDWVYAEQF | 0.60706204 |
| TCSSPFLVSAM | ASSRDWVYAEQF | 0.69241446 |
| VHITDAE | ASSRDWVYAEQF | 0.685444 |
| EYAAPSF | ASSRDWVYAEQF | 0.70925033 |
| VVDAFHGGL | ASSRDWVYAEQF | 0.70095396 |
| ASVVDPQ | ASSRDWVYAEQF | 0.70017684 |
| SYNEFGGLS | ASSRDWVYAEQF | 0.29560545 |
| DPDGSVY | ASSRDWVYAEQF | 0.37397543 |
| MVSTIHATE | ASSRDWVYAEQF | 0.64932036 |
| LAAGVADPV | ASSRDWVYAEQF | 0.5959321 |
| TKEQIAA | ASSRDWVYAEQF | 0.39028704 |
| SLASPAAL | ASSRDWVYAEQF | 0.5808098 |
| IGLVFYGV | ASSRDWVYAEQF | 0.6449356 |
| FLLDASNL | ASSRDWVYAEQF | 0.5001385 |
| YQVGNFP | ASSRDWVYAEQF | 0.6789592 |
| ESLLSQST | ASSRDWVYAEQF | 0.6027254 |
| LFTVGFAAVG | ASSRDWVYAEQF | 0.5187839 |
| ATPPTPT | ASSRDWVYAEQF | 0.529647 |

| | | |
|---|---|---|
| TDLATLG | ASSRDWVYAEQF | 0.6822326 |
| SAGGTGR | ASSRDWVYAEQF | 0.50058645 |
| GFSNSGS | ASSRDWVYAEQF | 0.64910674 |
| GGNGGSGGTSV | ASSRDWVYAEQF | 0.50268376 |
| GTSVATGGAGN | ASSRDWVYAEQF | 0.69389373 |
| NGGNGGSGGTS | ASSRDWVYAEQF | 0.47251576 |
| AVVKFPH | ASSRDWVYAEQF | 0.6839427 |
| TPPPPPP | ASSRDWVYAEQF | 0.6662134 |
| AAPAAAAAPA | ASSRDWVYAEQF | 0.7135105 |
| IALNEDL | ASSRDWVYAEQF | 0.6326104 |
| IVNVSSM | ASSRDWVYAEQF | 0.66279936 |
| TGIPPAPR | ASSRDWVYAEQF | 0.54927385 |
| FDLGGGTF | ASSRDWVYAEQF | 0.68429387 |
| FDLGGGTFDVS | ASSRDWVYAEQF | 0.50299954 |
| NEPTAAA | ASSRDWVYAEQF | 0.67171645 |
| NEPTAAALA | ASSRDWVYAEQF | 0.3241223 |
| GSIINISSI | ASSRDWVYAEQF | 0.66274333 |
| VAPGNAGTA | ASSRDWVYAEQF | 0.69570553 |
| TGGGASLE | ASSRDWVYAEQF | 0.5039279 |
| VAVYNHYKRI | ASSRDWVYAEQF | 0.6948135 |
| TDPAPATT | ASSRDWVYAEQF | 0.6521756 |
| TNRPDILDPALLRPG | ASSRDWVYAEQF | 0.61013794 |
| LTTTTTS | ASSRDWVYAEQF | 0.70364094 |
| PTGSGKTL | ASSRDWVYAEQF | 0.5431751 |
| TGSGKTL | ASSRDWVYAEQF | 0.7182628 |
| TIGHVDHGK | ASSRDWVYAEQF | 0.5609138 |
| TIGHVDHGKTT | ASSRDWVYAEQF | 0.6425591 |
| GHVDHGKTT | ASSRDWVYAEQF | 0.6576616 |
| AQFEHTLL | ASSRDWVYAEQF | 0.5818536 |
| IVNQDEL | ASSRDWVYAEQF | 0.4718892 |
| TNGSQFF | ASSRDWVYAEQF | 0.7085104 |
| TGKGIVE | ASSRDWVYAEQF | 0.16217148 |
| LISNASDA | ASSRDWVYAEQF | 0.18253273 |
| AELRTLGELTKIAKAHGAQ | ASSRDWVYAEQF | 0.15515584 |
| APNAVAAAEPSW | ASSRDWVYAEQF | 0.16307095 |
| APQVAGAATAVL | ASSRDWVYAEQF | 0.1642482 |
| ARRDPAAAAAILPTDA | ASSRDWVYAEQF | 0.16579312 |
| DAAARDVLAEAGLAEYF | ASSRDWVYAEQF | 0.20361757 |
| DDQIVLSAKANLAKVEQ | ASSRDWVYAEQF | 0.16211386 |
| DELLPDTGEDSDEDGH | ASSRDWVYAEQF | 0.16989456 |
| ELPEAPANRATVPQD | ASSRDWVYAEQF | 0.16161685 |
| FGQSKIERVTGIG | ASSRDWVYAEQF | 0.15818694 |
| GAGGTGSTAGGAGGAGGAGGI | ASSRDWVYAEQF | 0.15949859 |
| GHGGTGAAGGNGGTGGAGGS( | ASSRDWVYAEQF | 0.17104907 |
| GNGGNGGAGGEGGPGGAGGA( | ASSRDWVYAEQF | 0.1795029 |
| GSTAGNGGNGGNATNGTIAG( | ASSRDWVYAEQF | 0.15228897 |
| GTGGDGGNGGNGGTGG | ASSRDWVYAEQF | 0.15984222 |
| GVRRPAGIPADAPV | ASSRDWVYAEQF | 0.1677389 |
| GYRQVTDAHLLTLARR | ASSRDWVYAEQF | 0.21279237 |
| IAKMGQDGHDRG | ASSRDWVYAEQF | 0.21066928 |
| KAEPSIEIQYT | ASSRDWVYAEQF | 0.17771623 |

| | | |
|---|---|---|
| KPVRIRMAADEDTVAVT | ASSRDWVYAEQF | 0.16993828 |
| MPPRIAGMRLLVIKPEP | ASSRDWVYAEQF | 0.1660222 |
| MKFYAVALFRSRGVPPTR | ASSRDWVYAEQF | 0.14996797 |
| NAGGFGILGAVAHSP | ASSRDWVYAEQF | 0.15636162 |
| NEGREITLAMAEM | ASSRDWVYAEQF | 0.16168523 |
| NELVEWSHPLGHRWRWE | ASSRDWVYAEQF | 0.17241924 |
| NLGAGGTGGNGGNPG | ASSRDWVYAEQF | 0.14652166 |
| NTEWPKWETVMAADPVD | ASSRDWVYAEQF | 0.16921724 |
| PASAAPAETQP | ASSRDWVYAEQF | 0.16499937 |
| QQSSDGKNGVRAAEITGE | ASSRDWVYAEQF | 0.15553351 |
| RAPDAVTTSWWKE | ASSRDWVYAEQF | 0.19615012 |
| SGFGNVGAHTSGWFNQ | ASSRDWVYAEQF | 0.15516955 |
| SGVAGAVGQAASVGGLKV | ASSRDWVYAEQF | 0.16534634 |
| SNGGAGTGSAGGNGGT | ASSRDWVYAEQF | 0.15981533 |
| TGGAGGEGGAAG | ASSRDWVYAEQF | 0.16660714 |
| TGGDGGAAGAGGAPGHG | ASSRDWVYAEQF | 0.1557406 |
| TMPPLLAAIANAGRHGV | ASSRDWVYAEQF | 0.18823676 |
| TSSTAGNSGGA | ASSRDWVYAEQF | 0.1547164 |
| VPATVPVGILC | ASSRDWVYAEQF | 0.1729494 |
| VQRPDSGGVVLTGRIS | ASSRDWVYAEQF | 0.15982947 |
| WTPNPVPATKGVPAGLVQAT | ASSRDWVYAEQF | 0.16373974 |
| YALKRALAKFSDRFDVVII | ASSRDWVYAEQF | 0.15649207 |
| YLIGSVIV | ASSRDWVYAEQF | 0.26311615 |
| YGQLVAVID | ASSRDWVYAEQF | 0.18793166 |
| VTVGGMTLPA | ASSRDWVYAEQF | 0.18207006 |
| VLALPAV | ASSRDWVYAEQF | 0.1676096 |
| VIPPTLNYET | ASSRDWVYAEQF | 0.16810395 |
| VGMGGMHPSSG | ASSRDWVYAEQF | 0.1528224 |
| VGLAGGAATA | ASSRDWVYAEQF | 0.16741644 |
| VGGAGGAGGSGP | ASSRDWVYAEQF | 0.15110087 |
| VFAAFMV | ASSRDWVYAEQF | 0.16474806 |
| TLYDKLR | ASSRDWVYAEQF | 0.15908743 |
| TLAVFTVL | ASSRDWVYAEQF | 0.1642093 |
| TGSVQPARV | ASSRDWVYAEQF | 0.15787672 |
| TGGGAGSGGT | ASSRDWVYAEQF | 0.1717907 |
| SHLETSL | ASSRDWVYAEQF | 0.16326837 |
| SFTTGMT | ASSRDWVYAEQF | 0.1746621 |
| QVEVAKRMGVRQPTVS | ASSRDWVYAEQF | 0.16946551 |
| QGSGGPALAM | ASSRDWVYAEQF | 0.15475462 |
| MLAPSNP | ASSRDWVYAEQF | 0.15154725 |
| LVNNAAI | ASSRDWVYAEQF | 0.15472077 |
| LTGEINTLA | ASSRDWVYAEQF | 0.18145052 |
| LSIFGPV | ASSRDWVYAEQF | 0.15262131 |
| LLALLANL | ASSRDWVYAEQF | 0.1581168 |
| LISNASDALDKLR | ASSRDWVYAEQF | 0.1673735 |
| LINEAVP | ASSRDWVYAEQF | 0.3503386 |
| LGYAFSGL | ASSRDWVYAEQF | 0.47036228 |
| LDFVAVAL | ASSRDWVYAEQF | 0.16657655 |
| LAQPGAE | ASSRDWVYAEQF | 0.19743522 |
| ISNLGAVTGF | ASSRDWVYAEQF | 0.2722066 |
| HQEDPGRI | ASSRDWVYAEQF | 0.3698298 |

| | | |
|---|---|---|
| GTPGEPTP | ASSRDWVYAEQF | 0.33021712 |
| GFFNSGS | ASSRDWVYAEQF | 0.4456815 |
| GAGGGGLTAGAN | ASSRDWVYAEQF | 0.2058211 |
| FVDALVNM | ASSRDWVYAEQF | 0.27110308 |
| FLTVAPEL | ASSRDWVYAEQF | 0.23808777 |
| FETLTAL | ASSRDWVYAEQF | 0.31838885 |
| DVINPIEL | ASSRDWVYAEQF | 0.2646581 |
| APVSGAPV | ASSRDWVYAEQF | 0.19217424 |
| ALVGGLAW | ASSRDWVYAEQF | 0.1570413 |
| ALAAAAATAI | ASSRDWVYAEQF | 0.19948374 |
| AFGLGLLL | ASSRDWVYAEQF | 0.2816117 |
| AFFAAASL | ASSRDWVYAEQF | 0.3474658 |
| AAAALPAFSPP | ASSRDWVYAEQF | 0.27312115 |
| VVLAPAL | ASSRDWVYAEQF | 0.399883 |
| TIGHVDHGKTTL | ASSRDWVYAEQF | 0.4586845 |
| LISNASDALDK | ASSRDWVYAEQF | 0.18184984 |
| LISNASDAL | ASSRDWVYAEQF | 0.12821084 |
| ISNASDALDK | ASSRDWVYAEQF | 0.15824564 |
| NQAETLVYQTE | ASSRDWVYAEQF | 0.19539091 |
| TGGTGGTAGAG | ASSRDWVYAEQF | 0.26122063 |
| TGSTAGGAGGA | ASSRDWVYAEQF | 0.11255159 |
| DPLAVPVAAGPV | ASSRDWVYAEQF | 0.28199774 |
| GGTGGTPGNGGN | ASSRDWVYAEQF | 0.2622906 |
| PAGLAGVDPATL | ASSRDWVYAEQF | 0.17028853 |
| GNNLVVEAPHAGASV | ASSRDWVYAEQF | 0.51282966 |
| KVTRIMSASTLEQPA | ASSRDWVYAEQF | 0.19383283 |
| LGFALPIALPRRTVV | ASSRDWVYAEQF | 0.18600342 |
| TRLGNLTARKLVVLV | ASSRDWVYAEQF | 0.22571944 |
| WSQQRRLIYFPSAGP | ASSRDWVYAEQF | 0.5339713 |
| GPNGNPDPMAAAVDIR | ASSRDWVYAEQF | 0.18859594 |
| TQRCVSVADLRAAASR | ASSRDWVYAEQF | 0.51972383 |
| QFVLDVAEHPVEVLTETV | ASSRDWVYAEQF | 0.19219005 |
| GGAGGTGGAGGNGGTQAGDG | ASSRDWVYAEQF | 0.40078604 |
| PLLATLRAIAVDVDAKPAQV | ASSRDWVYAEQF | 0.1749338 |
| APKFRTGGYMIDFWGVGYQV | ASSRDWVYAEQF | 0.44068414 |
| GAGGAGGAGGAAGGPGGTGG | ASSRDWVYAEQF | 0.31667256 |
| IGALGLLATDVAPAGSVAFG | ASSRDWVYAEQF | 0.5858443 |
| AGGAGGIGGNGGAGGAGGVL | ASSRDWVYAEQF | 0.2590354 |
| ADELIGLL | ASSRDWVYAEQF | 0.34199497 |
| ADRALKA | ASSRDWVYAEQF | 0.1896876 |
| AGAVGQVGGAGGA | ASSRDWVYAEQF | 0.20939076 |
| AGGGGGLGGAP | ASSRDWVYAEQF | 0.15288593 |
| AGGLLYGNGGAG | ASSRDWVYAEQF | 0.21941255 |
| AGLESIGL | ASSRDWVYAEQF | 0.16091558 |
| AGLKTGII | ASSRDWVYAEQF | 0.22655414 |
| ASLAALLL | ASSRDWVYAEQF | 0.4521738 |
| ASVINISSI | ASSRDWVYAEQF | 0.14478917 |
| ATPPGPA | ASSRDWVYAEQF | 0.3050847 |
| ATVASALV | ASSRDWVYAEQF | 0.16488267 |
| ATVASLIL | ASSRDWVYAEQF | 0.18560576 |
| AVNGPVLIRH | ASSRDWVYAEQF | 0.26844758 |

| | | |
|---|---|---|
| DGAAGAVGI | ASSRDWVYAEQF | 0.17301394 |
| DPLLHGI | ASSRDWVYAEQF | 0.23108438 |
| ELGAALGAL | ASSRDWVYAEQF | 0.13813628 |
| ELGRTQAV | ASSRDWVYAEQF | 0.24256046 |
| FVAGGYLS | ASSRDWVYAEQF | 0.302211 |
| GASGGAGGAGGW | ASSRDWVYAEQF | 0.120721795 |
| GGAGAGGDGGAGGI | ASSRDWVYAEQF | 0.169106 |
| GTGHNGSQPGMGGNG | ASSRDWVYAEQF | 0.5121622 |
| GYAAASASAS | ASSRDWVYAEQF | 0.47910732 |
| HQVDVAAVQQG | ASSRDWVYAEQF | 0.71227264 |
| IGAAIAHT | ASSRDWVYAEQF | 0.1283029 |
| IGMIEVAQL | ASSRDWVYAEQF | 0.1330807 |
| IRLAPEHTSAMD | ASSRDWVYAEQF | 0.2670841 |
| IYNVSVL | ASSRDWVYAEQF | 0.465946 |
| KQGFLGM | ASSRDWVYAEQF | 0.45029694 |
| LESVITQL | ASSRDWVYAEQF | 0.43407062 |
| LGLGALVVV | ASSRDWVYAEQF | 0.1361916 |
| LIGNGGNGGSGGI | ASSRDWVYAEQF | 0.17828068 |
| LNAINAPTL | ASSRDWVYAEQF | 0.17096357 |
| LQLLNTGV | ASSRDWVYAEQF | 0.2249687 |
| LQTSIEF | ASSRDWVYAEQF | 0.3359571 |
| MSFAVSEL | ASSRDWVYAEQF | 0.1793961 |
| NTGALGL | ASSRDWVYAEQF | 0.11915751 |
| PPGAGGAGGAGTT | ASSRDWVYAEQF | 0.17227796 |
| QLVGLGI | ASSRDWVYAEQF | 0.47715446 |
| RFLEQVHQL | ASSRDWVYAEQF | 0.66710186 |
| RGVGYVLR | ASSRDWVYAEQF | 0.6504518 |
| SATGVFAGVI | ASSRDWVYAEQF | 0.63443005 |
| SLPAITLP | ASSRDWVYAEQF | 0.5816658 |
| STALAFTV | ASSRDWVYAEQF | 0.39597672 |
| SVLTVGLGK | ASSRDWVYAEQF | 0.08970509 |
| TALAGAGSGI | ASSRDWVYAEQF | 0.119015835 |
| TGLPFTGI | ASSRDWVYAEQF | 0.14867364 |
| VAEHVTV | ASSRDWVYAEQF | 0.27644446 |
| VDIVLNSL | ASSRDWVYAEQF | 0.107456885 |
| VLTGGGAVL | ASSRDWVYAEQF | 0.25101995 |
| VNVGLAYGI | ASSRDWVYAEQF | 0.19809914 |
| VSSAAVGAV | ASSRDWVYAEQF | 0.11513618 |
| VTSVYEAGV | ASSRDWVYAEQF | 0.63971615 |
| WVALDRGA | ASSRDWVYAEQF | 0.116937324 |
| YLVGSVVI | ASSRDWVYAEQF | 0.13319296 |
| GPPGTGKTLL | ASSRDWVYAEQF | 0.16394004 |
| GVGMTKF | ASSRDWVYAEQF | 0.47670943 |
| IGIGGSDL | ASSRDWVYAEQF | 0.14072739 |
| IGTLLAEL | ASSRDWVYAEQF | 0.47248197 |
| IYTSGTT | ASSRDWVYAEQF | 0.2580996 |
| LSFPAGV | ASSRDWVYAEQF | 0.41299713 |
| LYTSGTT | ASSRDWVYAEQF | 0.12026488 |
| VGAFPFGF | ASSRDWVYAEQF | 0.5083115 |
| VGAGTVEF | ASSRDWVYAEQF | 0.27387553 |
| VGKSTLLL | ASSRDWVYAEQF | 0.7887707 |

| | | |
|---|---|---|
| AALAHAGGGIV | ASSRDWVYAEQF | 0.24977049 |
| AALVAVSLL | ASSRDWVYAEQF | 0.2596445 |
| AAVSRAVAAL | ASSRDWVYAEQF | 0.17106825 |
| AGGSSGIGGF | ASSRDWVYAEQF | 0.44685447 |
| AGLTVANANG | ASSRDWVYAEQF | 0.10746607 |
| AGLVGAVL | ASSRDWVYAEQF | 0.23035252 |
| ALSQAFTV | ASSRDWVYAEQF | 0.11326025 |
| ALTFSVSL | ASSRDWVYAEQF | 0.1838793 |
| ALTGAALVAL | ASSRDWVYAEQF | 0.42777747 |
| ASFSIGAGVL | ASSRDWVYAEQF | 0.12064058 |
| DGSVYKI | ASSRDWVYAEQF | 0.3595503 |
| EQLAEIV | ASSRDWVYAEQF | 0.18263066 |
| FAVSELTVM | ASSRDWVYAEQF | 0.13873112 |
| GAAPLAAGL | ASSRDWVYAEQF | 0.47077128 |
| GADGGDGGSGGAGG | ASSRDWVYAEQF | 0.14146332 |
| GFGLDTGGAGG | ASSRDWVYAEQF | 0.6464555 |
| GLASGGLL | ASSRDWVYAEQF | 0.13413146 |
| GRIAGAAVLL | ASSRDWVYAEQF | 0.24723385 |
| GSAGSLNTSF | ASSRDWVYAEQF | 0.2592913 |
| GTGQIGI | ASSRDWVYAEQF | 0.09981308 |
| GTLIALIL | ASSRDWVYAEQF | 0.13522904 |
| IAATMGFVVG | ASSRDWVYAEQF | 0.54235137 |
| IFAQALL | ASSRDWVYAEQF | 0.72827154 |
| IGAAVALGF | ASSRDWVYAEQF | 0.73714006 |
| IGLTPLAL | ASSRDWVYAEQF | 0.21451262 |
| ILGSNTSTL | ASSRDWVYAEQF | 0.27742118 |
| IVFSLTVL | ASSRDWVYAEQF | 0.4707011 |
| LASKATGF | ASSRDWVYAEQF | 0.6207867 |
| LASVALIL | ASSRDWVYAEQF | 0.8126042 |
| LFAVGLTL | ASSRDWVYAEQF | 0.7277734 |
| LFNGGIGTYI | ASSRDWVYAEQF | 0.24980699 |
| LTLGAGII | ASSRDWVYAEQF | 0.5881609 |
| LVAAALGDI | ASSRDWVYAEQF | 0.39976 |
| NTAFVQLGI | ASSRDWVYAEQF | 0.54676795 |
| QLFSGAEL | ASSRDWVYAEQF | 0.49105787 |
| SAATFITL | ASSRDWVYAEQF | 0.40757513 |
| SNYGLANLGV | ASSRDWVYAEQF | 0.33118445 |
| SYGGAYAVM | ASSRDWVYAEQF | 0.22331762 |
| TTGVGGLTLG | ASSRDWVYAEQF | 0.5903684 |
| TTQTIYI | ASSRDWVYAEQF | 0.70295537 |
| VAAGAGLGAVI | ASSRDWVYAEQF | 0.5747677 |
| VAAVVTLL | ASSRDWVYAEQF | 0.7429973 |
| VGFLGGLI | ASSRDWVYAEQF | 0.7317631 |
| VGVIPILL | ASSRDWVYAEQF | 0.40599638 |
| VIFASSLI | ASSRDWVYAEQF | 0.14037041 |
| VKGGDGGAAAG | ASSRDWVYAEQF | 0.16425478 |
| VTKTFTNI | ASSRDWVYAEQF | 0.24422807 |
| VVAAAILL | ASSRDWVYAEQF | 0.32876557 |
| YAALDVEL | ASSRDWVYAEQF | 0.13612482 |
| LASVALLL | ASSRDWVYAEQF | 0.561695 |
| LDAVHRV | ASSRDWVYAEQF | 0.3216415 |

| | | |
|---|---|---|
| QGLSSSIF | ASSRDWVYAEQF | 0.3178167 |
| RGTFANIRL | ASSRDWVYAEQF | 0.75062126 |
| RQLWWGHRI | ASSRDWVYAEQF | 0.31209826 |
| YGPPGTGKTLL | ASSRDWVYAEQF | 0.21769716 |
| AALVGAPANVA | ASSRDWVYAEQF | 0.27500764 |
| AATFRRTE | ASSRDWVYAEQF | 0.79767454 |
| AGALGGGI | ASSRDWVYAEQF | 0.24448827 |
| AGGAGGAGGIGAG | ASSRDWVYAEQF | 0.7544691 |
| AGSGALGL | ASSRDWVYAEQF | 0.37588298 |
| AQIADQI | ASSRDWVYAEQF | 0.74538934 |
| ASAVAVTYL | ASSRDWVYAEQF | 0.20016214 |
| ASTVLGGTGG | ASSRDWVYAEQF | 0.7001389 |
| AVGGEPAK | ASSRDWVYAEQF | 0.54429114 |
| CEQGTFGGTC | ASSRDWVYAEQF | 0.8084566 |
| DEGKAPAG | ASSRDWVYAEQF | 0.32875103 |
| DTVVFPAL | ASSRDWVYAEQF | 0.50058717 |
| FEGITMAG | ASSRDWVYAEQF | 0.36991918 |
| FPLVVQD | ASSRDWVYAEQF | 0.77405345 |
| GAFTTPQ | ASSRDWVYAEQF | 0.1990447 |
| GFGGFGTAGGVGGA | ASSRDWVYAEQF | 0.2879199 |
| GFLAQFL | ASSRDWVYAEQF | 0.2255683 |
| GGAGGAGGNAGVA | ASSRDWVYAEQF | 0.31126902 |
| GGVGTAGAGGAGGAGG | ASSRDWVYAEQF | 0.70884836 |
| GNGGGGDAGNAGSGGN | ASSRDWVYAEQF | 0.1339018 |
| GTGGAGGAGNDAGS | ASSRDWVYAEQF | 0.62985754 |
| GTTNTGWFNS | ASSRDWVYAEQF | 0.2333588 |
| IFMGPLFTL | ASSRDWVYAEQF | 0.21403658 |
| ILAHAIEAI | ASSRDWVYAEQF | 0.5190064 |
| IRSAIREAL | ASSRDWVYAEQF | 0.436305 |
| ITFAGGG | ASSRDWVYAEQF | 0.56787926 |
| ITGMSGHQEV | ASSRDWVYAEQF | 0.2456546 |
| IVAGLACTL | ASSRDWVYAEQF | 0.5245776 |
| IVGGGLAGA | ASSRDWVYAEQF | 0.47666457 |
| IVTGVLNI | ASSRDWVYAEQF | 0.13780074 |
| LAQQGCI | ASSRDWVYAEQF | 0.23178455 |
| LCLHVEN | ASSRDWVYAEQF | 0.74837434 |
| LDAISDGS | ASSRDWVYAEQF | 0.12427747 |
| LGGFAAML | ASSRDWVYAEQF | 0.15933439 |
| LGGKNPMI | ASSRDWVYAEQF | 0.10306741 |
| LGPKGVAAIL | ASSRDWVYAEQF | 0.10852855 |
| LMVGVSI | ASSRDWVYAEQF | 0.110435456 |
| LVGAAFVL | ASSRDWVYAEQF | 0.13067669 |
| LVLAVAPL | ASSRDWVYAEQF | 0.15517406 |
| MAAIAVIDAA | ASSRDWVYAEQF | 0.13130553 |
| NNGINAELQ | ASSRDWVYAEQF | 0.113274895 |
| RGEPLVLVHG | ASSRDWVYAEQF | 0.11123294 |
| RGGAGGAAGLI | ASSRDWVYAEQF | 0.11022494 |
| RSYRMHPAV | ASSRDWVYAEQF | 0.114702016 |
| SAAHLLAGQLA | ASSRDWVYAEQF | 0.118385494 |
| SAAVGSY | ASSRDWVYAEQF | 0.10704511 |
| SPHAVLHRI | ASSRDWVYAEQF | 0.10210954 |

| | | |
|---|---|---|
| SQIAKVL | ASSRDWVYAEQF | 0.10805421 |
| TALGHLVP | ASSRDWVYAEQF | 0.11164736 |
| TAPTTSGAA | ASSRDWVYAEQF | 0.12164227 |
| TDGDDGNGGSPGT | ASSRDWVYAEQF | 0.117900915 |
| TDGLGEN | ASSRDWVYAEQF | 0.1432266 |
| TGGPPLPGQA | ASSRDWVYAEQF | 0.1377111 |
| TIPAITI | ASSRDWVYAEQF | 0.10870596 |
| TIPGFTI | ASSRDWVYAEQF | 0.10009786 |
| TSIASGALA | ASSRDWVYAEQF | 0.103648886 |
| TVTLPGAAI | ASSRDWVYAEQF | 0.10969583 |
| VASGLSAALP | ASSRDWVYAEQF | 0.11318024 |
| VGAGVIGI | ASSRDWVYAEQF | 0.09818919 |
| VGPKRFGV | ASSRDWVYAEQF | 0.11470934 |
| VNVAFPD | ASSRDWVYAEQF | 0.11776623 |
| VSMTNSGVS | ASSRDWVYAEQF | 0.10470542 |
| VTDVLGDLSV | ASSRDWVYAEQF | 0.15256548 |
| VVVVGAGFAGL | ASSRDWVYAEQF | 0.103752784 |
| VYTVGTVV | ASSRDWVYAEQF | 0.104641065 |
| WTTFSGP | ASSRDWVYAEQF | 0.11101243 |
| YDELVEAF | ASSRDWVYAEQF | 0.13325967 |
| YGTGGAGGNGG | ASSRDWVYAEQF | 0.10928616 |
| YTNVMAV | ASSRDWVYAEQF | 0.13886614 |
| FLEQVHQL | ASSRDWVYAEQF | 0.111258164 |
| GLSLLAVAV | ASSRDWVYAEQF | 0.14946783 |
| LEQVHQL | ASSRDWVYAEQF | 0.10828415 |
| LNAAGAI | ASSRDWVYAEQF | 0.1256072 |
| PAVHASL | ASSRDWVYAEQF | 0.11354824 |
| TELEDDA | ASSRDWVYAEQF | 0.16490142 |
| TPPGPPN | ASSRDWVYAEQF | 0.11656981 |
| TPPPPPPP | ASSRDWVYAEQF | 0.12923971 |
| TQSPALI | ASSPGLGGYEQY | 0.10751216 |
| EFAAAFPQ | ASSPGLGGYEQY | 0.16123787 |
| LLSGNLDVL | ASSPGLGGYEQY | 0.10293241 |
| GPANLGNYNI | ASSPGLGGYEQY | 0.11213447 |
| TLGPTLP | ASSPGLGGYEQY | 0.10487768 |
| VYAYSSCCK | ASSPGLGGYEQY | 0.11293571 |
| APAPAPAPNI | ASSPGLGGYEQY | 0.13122182 |
| PGGPVPIP | ASSPGLGGYEQY | 0.10010602 |
| FLTVALEP | ASSPGLGGYEQY | 0.11840395 |
| SQYFGANL | ASSPGLGGYEQY | 0.10200823 |
| GGVKPAA | ASSPGLGGYEQY | 0.1079784 |
| GGTGGAGGAGGW | ASSPGLGGYEQY | 0.11586899 |
| GVGGVAGLGGAGG | ASSPGLGGYEQY | 0.10489576 |
| VLLVLATTI | ASSPGLGGYEQY | 0.10746064 |
| IGIFAAML | ASSPGLGGYEQY | 0.10079894 |
| AVGDLID | ASSPGLGGYEQY | 0.11483192 |
| NISPLPL | ASSPGLGGYEQY | 0.13047536 |
| VTAVFVAI | ASSPGLGGYEQY | 0.10038789 |
| SGDDNDAAL | ASSPGLGGYEQY | 0.10445466 |
| AVAALAHS | ASSPGLGGYEQY | 0.13629621 |
| GGVTVMATP | ASSPGLGGYEQY | 0.09084903 |

| | | |
|---|---|---|
| AGLAANVRD | ASSPGLGGYEQY | 0.090659484 |
| TNRPDLIDPALLRPG | ASSPGLGGYEQY | 0.09075812 |
| LANLGVTAL | ASSPGLGGYEQY | 0.09123977 |
| TGERIEL | ASSPGLGGYEQY | 0.09069403 |
| NGEELAA | ASSPGLGGYEQY | 0.09085042 |
| VLSLVWG | ASSPGLGGYEQY | 0.0909281 |
| IGYALWL | ASSPGLGGYEQY | 0.090842664 |
| LSLLLAPL | ASSPGLGGYEQY | 0.091460906 |
| TLDEALK | ASSPGLGGYEQY | 0.09127198 |
| TDGAGTAEGG | ASSPGLGGYEQY | 0.0909842 |
| NAPGGAGGNGG | ASSPGLGGYEQY | 0.09113577 |
| VSVFMGMGGNF | ASSPGLGGYEQY | 0.090712756 |
| LASYLAAL | ASSPGLGGYEQY | 0.09146167 |
| LVGGTPAL | ASSPGLGGYEQY | 0.09073785 |
| VRNADVL | ASSPGLGGYEQY | 0.09060975 |
| TCSSPFLVSAM | ASSPGLGGYEQY | 0.09097377 |
| VHITDAE | ASSPGLGGYEQY | 0.090963826 |
| EYAAPSF | ASSPGLGGYEQY | 0.090938784 |
| VVDAFHGGL | ASSPGLGGYEQY | 0.09084185 |
| ASVVDPQ | ASSPGLGGYEQY | 0.090888225 |
| SYNEFGGLS | ASSPGLGGYEQY | 0.090756595 |
| DPDGSVY | ASSPGLGGYEQY | 0.09097964 |
| MVSTIHATE | ASSPGLGGYEQY | 0.09075947 |
| LAAGVADPV | ASSPGLGGYEQY | 0.090992756 |
| TKEQIAA | ASSPGLGGYEQY | 0.0908243 |
| SLASPAAL | ASSPGLGGYEQY | 0.09041618 |
| IGLVFYGV | ASSPGLGGYEQY | 0.09132078 |
| FLLDASNL | ASSPGLGGYEQY | 0.09108768 |
| YQVGNFP | ASSPGLGGYEQY | 0.092517056 |
| ESLLSQST | ASSPGLGGYEQY | 0.09111562 |
| LFTVGFAAVG | ASSPGLGGYEQY | 0.09094477 |
| ATPPTPT | ASSPGLGGYEQY | 0.09097636 |
| TDLATLG | ASSPGLGGYEQY | 0.09089388 |
| SAGGTGR | ASSPGLGGYEQY | 0.09081728 |
| GFSNSGS | ASSPGLGGYEQY | 0.09086587 |
| GGNGGSGGTSV | ASSPGLGGYEQY | 0.09114261 |
| GTSVATGGAGN | ASSPGLGGYEQY | 0.09065384 |
| NGGNGGSGGTS | ASSPGLGGYEQY | 0.092459224 |
| AVVKFPH | ASSPGLGGYEQY | 0.09162126 |
| TPPPPPP | ASSPGLGGYEQY | 0.09083551 |
| AAPAAAAAPA | ASSPGLGGYEQY | 0.090848595 |
| IALNEDL | ASSPGLGGYEQY | 0.091411784 |
| IVNVSSM | ASSPGLGGYEQY | 0.091019355 |
| TGIPPAPR | ASSPGLGGYEQY | 0.091662236 |
| FDLGGGTF | ASSPGLGGYEQY | 0.09089807 |
| FDLGGGTFDVS | ASSPGLGGYEQY | 0.09091442 |
| NEPTAAA | ASSPGLGGYEQY | 0.0916359 |
| NEPTAAALA | ASSPGLGGYEQY | 0.09077433 |
| GSIINISSI | ASSPGLGGYEQY | 0.09087088 |
| VAPGNAGTA | ASSPGLGGYEQY | 0.09084406 |
| TGGGASLE | ASSPGLGGYEQY | 0.090995476 |

| | | |
|---|---|---|
| VAVYNHYKRI | ASSPGLGGYEQY | 0.091084436 |
| TDPAPATT | ASSPGLGGYEQY | 0.09063828 |
| TNRPDILDPALLRPG | ASSPGLGGYEQY | 0.091530986 |
| LTTTTTS | ASSPGLGGYEQY | 0.0908712 |
| PTGSGKTL | ASSPGLGGYEQY | 0.090688616 |
| TGSGKTL | ASSPGLGGYEQY | 0.09155907 |
| TIGHVDHGK | ASSPGLGGYEQY | 0.09055883 |
| TIGHVDHGKTT | ASSPGLGGYEQY | 0.09063359 |
| GHVDHGKTT | ASSPGLGGYEQY | 0.09068357 |
| AQFEHTLL | ASSPGLGGYEQY | 0.09118248 |
| IVNQDEL | ASSPGLGGYEQY | 0.09328786 |
| TNGSQFF | ASSPGLGGYEQY | 0.09100846 |
| TGKGIVE | ASSPGLGGYEQY | 0.09082863 |
| LISNASDA | ASSPGLGGYEQY | 0.5586102 |
| AELRTLGELTKIAKAHGAQ | ASSPGLGGYEQY | 0.68513083 |
| APNAVAAAEPSW | ASSPGLGGYEQY | 0.4341022 |
| APQVAGAATAVL | ASSPGLGGYEQY | 0.37569684 |
| ARRDPAAAAAILPTDA | ASSPGLGGYEQY | 0.5221379 |
| DAAARDVLAEAGLAEYF | ASSPGLGGYEQY | 0.6487373 |
| DDQIVLSAKANLAKVEQ | ASSPGLGGYEQY | 0.6988623 |
| DELLPDTGEDSDEDGH | ASSPGLGGYEQY | 0.6172309 |
| ELPEAPANRATVPQD | ASSPGLGGYEQY | 0.4273787 |
| FGQSKIERVTGIG | ASSPGLGGYEQY | 0.5323255 |
| GAGGTGSTAGGAGGAGGAGGI | ASSPGLGGYEQY | 0.4860593 |
| GHGGTGAAGGNGGTGGAGGS( | ASSPGLGGYEQY | 0.46734065 |
| GNGGNGGAGGEGGPGGAGGA( | ASSPGLGGYEQY | 0.50870687 |
| GSTAGNGGNGGNATNGTIAG( | ASSPGLGGYEQY | 0.57708883 |
| GTGGDGGNGGNGGTGG | ASSPGLGGYEQY | 0.45514643 |
| GVRRPAGIPADAPV | ASSPGLGGYEQY | 0.40973777 |
| GYRQVTDAHLLTLARR | ASSPGLGGYEQY | 0.51812565 |
| IAKMGQDGHDRG | ASSPGLGGYEQY | 0.62165403 |
| KAEPSIEIQYT | ASSPGLGGYEQY | 0.54434943 |
| KPVRIRMAADEDTVAVT | ASSPGLGGYEQY | 0.69028395 |
| MPPRIAGMRLLVIKPEP | ASSPGLGGYEQY | 0.63449913 |
| MKFYAVALFRSRGVPPTR | ASSPGLGGYEQY | 0.512627 |
| NAGGFGILGAVAHSP | ASSPGLGGYEQY | 0.20293927 |
| NEGREITLAMAEM | ASSPGLGGYEQY | 0.24238089 |
| NELVEWSHPLGHRWRWE | ASSPGLGGYEQY | 0.46665084 |
| NLGAGGTGGNGGNPG | ASSPGLGGYEQY | 0.49494505 |
| NTEWPKWETVMAADPVD | ASSPGLGGYEQY | 0.21706486 |
| PASAAPAETQP | ASSPGLGGYEQY | 0.5865054 |
| QQSSDGKNGVRAAEITGE | ASSPGLGGYEQY | 0.51383305 |
| RAPDAVTTSWWKE | ASSPGLGGYEQY | 0.4201688 |
| SGFGNVGAHTSGWFNQ | ASSPGLGGYEQY | 0.6404985 |
| SGVAGAVGQAASVGGLKV | ASSPGLGGYEQY | 0.4700215 |
| SNGGAGTGSAGGNGGT | ASSPGLGGYEQY | 0.36783636 |
| TGGAGGEGGAAG | ASSPGLGGYEQY | 0.42146277 |
| TGGDGGAAGAGGAPGHG | ASSPGLGGYEQY | 0.7171272 |
| TMPPLLAAIANAGRHGV | ASSPGLGGYEQY | 0.28556994 |
| TSSTAGNSGGA | ASSPGLGGYEQY | 0.63929045 |
| VPATVPVGILC | ASSPGLGGYEQY | 0.41456077 |

| | | |
|---|---|---|
| VQRPDSGGVVLTGRIS | ASSPGLGGYEQY | 0.6495333 |
| WTPNPVPATKGVPAGLVQAT | ASSPGLGGYEQY | 0.32856065 |
| YALKRALAKFSDRFDVVII | ASSPGLGGYEQY | 0.6177937 |
| YLIGSVIV | ASSPGLGGYEQY | 0.48814178 |
| YGQLVAVID | ASSPGLGGYEQY | 0.72187877 |
| VTVGGMTLPA | ASSPGLGGYEQY | 0.41603035 |
| VLALPAV | ASSPGLGGYEQY | 0.5931521 |
| VIPPTLNYET | ASSPGLGGYEQY | 0.4486286 |
| VGMGGMHPSSG | ASSPGLGGYEQY | 0.67081094 |
| VGLAGGAATA | ASSPGLGGYEQY | 0.30403528 |
| VGGAGGAGGSGP | ASSPGLGGYEQY | 0.4695018 |
| VFAAFMV | ASSPGLGGYEQY | 0.27960292 |
| TLYDKLR | ASSPGLGGYEQY | 0.53656745 |
| TLAVFTVL | ASSPGLGGYEQY | 0.6015042 |
| TGSVQPARV | ASSPGLGGYEQY | 0.272562 |
| TGGGAGSGGT | ASSPGLGGYEQY | 0.66095865 |
| SHLETSL | ASSPGLGGYEQY | 0.465846 |
| SFTTGMT | ASSPGLGGYEQY | 0.41183487 |
| QVEVAKRMGVRQPTVS | ASSPGLGGYEQY | 0.6200654 |
| QGSGGPALAM | ASSPGLGGYEQY | 0.46423656 |
| MLAPSNP | ASSPGLGGYEQY | 0.65288997 |
| LVNNAAI | ASSPGLGGYEQY | 0.37093425 |
| LTGEINTLA | ASSPGLGGYEQY | 0.54862165 |
| LSIFGPV | ASSPGLGGYEQY | 0.6025979 |
| LLALLANL | ASSPGLGGYEQY | 0.3219422 |
| LISNASDALDKLR | ASSPGLGGYEQY | 0.37054554 |
| LINEAVP | ASSPGLGGYEQY | 0.64533806 |
| LGYAFSGL | ASSPGLGGYEQY | 0.76534337 |
| LDFVAVAL | ASSPGLGGYEQY | 0.7648773 |
| LAQPGAE | ASSPGLGGYEQY | 0.69940233 |
| ISNLGAVTGF | ASSPGLGGYEQY | 0.6939366 |
| HQEDPGRI | ASSPGLGGYEQY | 0.69319284 |
| GTPGEPTP | ASSPGLGGYEQY | 0.7664094 |
| GFFNSGS | ASSPGLGGYEQY | 0.77302057 |
| GAGGGGLTAGAN | ASSPGLGGYEQY | 0.76303864 |
| FVDALVNM | ASSPGLGGYEQY | 0.6891217 |
| FLTVAPEL | ASSPGLGGYEQY | 0.75341505 |
| FETLTAL | ASSPGLGGYEQY | 0.7474902 |
| DVINPIEL | ASSPGLGGYEQY | 0.74096715 |
| APVSGAPV | ASSPGLGGYEQY | 0.7578969 |
| ALVGGLAW | ASSPGLGGYEQY | 0.7510879 |
| ALAAAAATAI | ASSPGLGGYEQY | 0.70286596 |
| AFGLGLLL | ASSPGLGGYEQY | 0.6781591 |
| AFFAAASL | ASSPGLGGYEQY | 0.7383522 |
| AAAALPAFSPP | ASSPGLGGYEQY | 0.76707053 |
| VVLAPAL | ASSPGLGGYEQY | 0.7606884 |
| TIGHVDHGKTTL | ASSPGLGGYEQY | 0.7733321 |
| LISNASDALDK | ASSPGLGGYEQY | 0.76669335 |
| LISNASDAL | ASSPGLGGYEQY | 0.76158744 |
| ISNASDALDK | ASSPGLGGYEQY | 0.37432662 |
| NQAETLVYQTE | ASSPGLGGYEQY | 0.58548087 |

| | | |
|---|---|---|
| TGGTGGTAGAG | ASSPGLGGYEQY | 0.70699584 |
| TGSTAGGAGGA | ASSPGLGGYEQY | 0.7282032 |
| DPLAVPVAAGPV | ASSPGLGGYEQY | 0.42913705 |
| GGTGGTPGNGGN | ASSPGLGGYEQY | 0.7500049 |
| PAGLAGVDPATL | ASSPGLGGYEQY | 0.7130656 |
| GNNLVVEAPHAGASV | ASSPGLGGYEQY | 0.7045611 |
| KVTRIMSASTLEQPA | ASSPGLGGYEQY | 0.7597846 |
| LGFALPIALPRRTVV | ASSPGLGGYEQY | 0.7300948 |
| TRLGNLTARKLVVLV | ASSPGLGGYEQY | 0.66906154 |
| WSQQRRLIYFPSAGP | ASSPGLGGYEQY | 0.66386294 |
| GPNGNPDPMAAAVDIR | ASSPGLGGYEQY | 0.767269 |
| TQRCVSVADLRAAASR | ASSPGLGGYEQY | 0.6764835 |
| QFVLDVAEHPVEVLTETV | ASSPGLGGYEQY | 0.76659584 |
| GGAGGTGGAGGNGGTQAGDG | ASSPGLGGYEQY | 0.6362076 |
| PLLATLRAIAVDVDAKPAQV | ASSPGLGGYEQY | 0.7702969 |
| APKFRTGGYMIDFWGVGYQV | ASSPGLGGYEQY | 0.6739768 |
| GAGGAGGAGGAAGGPGGTGG | ASSPGLGGYEQY | 0.7675574 |
| IGALGLLATDVAPAGSVAFG | ASSPGLGGYEQY | 0.7490436 |
| AGGAGGIGGNGGAGGAGGVL | ASSPGLGGYEQY | 0.7767129 |
| ADELIGLL | ASSPGLGGYEQY | 0.739572 |
| ADRALKA | ASSPGLGGYEQY | 0.76531804 |
| AGAVGQVGGAGGA | ASSPGLGGYEQY | 0.7372717 |
| AGGGGGLGGAP | ASSPGLGGYEQY | 0.75847054 |
| AGGLLYGNGGAG | ASSPGLGGYEQY | 0.6470283 |
| AGLESIGL | ASSPGLGGYEQY | 0.73185086 |
| AGLKTGII | ASSPGLGGYEQY | 0.6460613 |
| ASLAALLL | ASSPGLGGYEQY | 0.7427349 |
| ASVINISSI | ASSPGLGGYEQY | 0.766985 |
| ATPPGPA | ASSPGLGGYEQY | 0.3948384 |
| ATVASALV | ASSPGLGGYEQY | 0.7487697 |
| ATVASLIL | ASSPGLGGYEQY | 0.7124333 |
| AVNGPVLIRH | ASSPGLGGYEQY | 0.71518743 |
| DGAAGAVGI | ASSPGLGGYEQY | 0.76738274 |
| DPLLHGI | ASSPGLGGYEQY | 0.73358727 |
| ELGAALGAL | ASSPGLGGYEQY | 0.76148593 |
| ELGRTQAV | ASSPGLGGYEQY | 0.69202566 |
| FVAGGYLS | ASSPGLGGYEQY | 0.71650815 |
| GASGGAGGAGGW | ASSPGLGGYEQY | 0.74138206 |
| GGAGAGGDGGAGGI | ASSPGLGGYEQY | 0.39029488 |
| GTGHNGSQPGMGGNG | ASSPGLGGYEQY | 0.5988865 |
| GYAAASASAS | ASSPGLGGYEQY | 0.7744854 |
| HQVDVAAVQQG | ASSPGLGGYEQY | 0.20906162 |
| IGAAIAHT | ASSPGLGGYEQY | 0.20913279 |
| IGMIEVAQL | ASSPGLGGYEQY | 0.20896195 |
| IRLAPEHTSAMD | ASSPGLGGYEQY | 0.21017657 |
| IYNVSVL | ASSPGLGGYEQY | 0.20908621 |
| KQGFLGM | ASSPGLGGYEQY | 0.20920104 |
| LESVITQL | ASSPGLGGYEQY | 0.21394353 |
| LGLGALVVV | ASSPGLGGYEQY | 0.20909472 |
| LIGNGGNGGSGGI | ASSPGLGGYEQY | 0.21089593 |
| LNAINAPTL | ASSPGLGGYEQY | 0.21091773 |

| | | |
|---|---|---|
| LQLLNTGV | ASSPGLGGYEQY | 0.20907277 |
| LQTSIEF | ASSPGLGGYEQY | 0.20925468 |
| MSFAVSEL | ASSPGLGGYEQY | 0.20910068 |
| NTGALGL | ASSPGLGGYEQY | 0.21196932 |
| PPGAGGAGGAGTT | ASSPGLGGYEQY | 0.20898962 |
| QLVGLGI | ASSPGLGGYEQY | 0.20894475 |
| RFLEQVHQL | ASSPGLGGYEQY | 0.20939887 |
| RGVGYVLR | ASSPGLGGYEQY | 0.20952085 |
| SATGVFAGVI | ASSPGLGGYEQY | 0.20941602 |
| SLPAITLP | ASSPGLGGYEQY | 0.20933147 |
| STALAFTV | ASSPGLGGYEQY | 0.20922086 |
| SVLTVGLGK | ASSPGLGGYEQY | 0.20910521 |
| TALAGAGSGI | ASSPGLGGYEQY | 0.20925748 |
| TGLPFTGI | ASSPGLGGYEQY | 0.20949763 |
| VAEHVTV | ASSPGLGGYEQY | 0.20921102 |
| VDIVLNSL | ASSPGLGGYEQY | 0.21000299 |
| VLTGGGAVL | ASSPGLGGYEQY | 0.20841904 |
| VNVGLAYGI | ASSPGLGGYEQY | 0.21152778 |
| VSSAAVGAV | ASSPGLGGYEQY | 0.20940028 |
| VTSVYEAGV | ASSPGLGGYEQY | 0.21098287 |
| WVALDRGA | ASSPGLGGYEQY | 0.20972349 |
| YLVGSVVI | ASSPGLGGYEQY | 0.20915554 |
| GPPGTGKTLL | ASSPGLGGYEQY | 0.21022311 |
| GVGMTKF | ASSPGLGGYEQY | 0.20920295 |
| IGIGGSDL | ASSPGLGGYEQY | 0.20953844 |
| IGTLLAEL | ASSPGLGGYEQY | 0.20917663 |
| IYTSGTT | ASSPGLGGYEQY | 0.21086317 |
| LSFPAGV | ASSPGLGGYEQY | 0.20890963 |
| LYTSGTT | ASSPGLGGYEQY | 0.21023484 |
| VGAFPFGF | ASSPGLGGYEQY | 0.21069282 |
| VGAGTVEF | ASSPGLGGYEQY | 0.20911376 |
| VGKSTLLL | ASSPGLGGYEQY | 0.20908612 |
| AALAHAGGGIV | ASSPGLGGYEQY | 0.21004781 |
| AALVAVSLL | ASSPGLGGYEQY | 0.2100743 |
| AAVSRAVAAL | ASSPGLGGYEQY | 0.21151759 |
| AGGSSGIGGF | ASSPGLGGYEQY | 0.20975366 |
| AGLTVANANG | ASSPGLGGYEQY | 0.2149207 |
| AGLVGAVL | ASSPGLGGYEQY | 0.21003571 |
| ALSQAFTV | ASSPGLGGYEQY | 0.20917101 |
| ALTFSVSL | ASSPGLGGYEQY | 0.20920216 |
| ALTGAALVAL | ASSPGLGGYEQY | 0.20933709 |
| ASFSIGAGVL | ASSPGLGGYEQY | 0.20906541 |
| DGSVYKI | ASSPGLGGYEQY | 0.20884402 |
| EQLAEIV | ASSPGLGGYEQY | 0.20891184 |
| FAVSELTVM | ASSPGLGGYEQY | 0.20881316 |
| GAAPLAAGL | ASSPGLGGYEQY | 0.20915467 |
| GADGGDGGSGGAGG | ASSPGLGGYEQY | 0.20901835 |
| GFGLDTGGAGG | ASSPGLGGYEQY | 0.21097422 |
| GLASGGLL | ASSPGLGGYEQY | 0.2087616 |
| GRIAGAAVLL | ASSPGLGGYEQY | 0.20889574 |
| GSAGSLNTSF | ASSPGLGGYEQY | 0.20898041 |

| | | |
|---|---|---|
| GTGQIGI | ASSPGLGGYEQY | 0.20944801 |
| GTLIALIL | ASSPGLGGYEQY | 0.20939513 |
| IAATMGFVVG | ASSPGLGGYEQY | 0.20944536 |
| IFAQALL | ASSPGLGGYEQY | 0.20916235 |
| IGAAVALGF | ASSPGLGGYEQY | 0.08976493 |
| IGLTPLAL | ASSPGLGGYEQY | 0.08746554 |
| ILGSNTSTL | ASSPGLGGYEQY | 0.08344513 |
| IVFSLTVL | ASSPGLGGYEQY | 0.08734004 |
| LASKATGF | ASSPGLGGYEQY | 0.08565955 |
| LASVALIL | ASSPGLGGYEQY | 0.08869664 |
| LFAVGLTL | ASSPGLGGYEQY | 0.09080137 |
| LFNGGIGTYI | ASSPGLGGYEQY | 0.08830339 |
| LTLGAGII | ASSPGLGGYEQY | 0.090175115 |
| LVAAALGDI | ASSPGLGGYEQY | 0.08777653 |
| NTAFVQLGI | ASSPGLGGYEQY | 0.08572643 |
| QLFSGAEL | ASSPGLGGYEQY | 0.08835532 |
| SAATFITL | ASSPGLGGYEQY | 0.08615793 |
| SNYGLANLGV | ASSPGLGGYEQY | 0.08692683 |
| SYGGAYAVM | ASSPGLGGYEQY | 0.084468566 |
| TTGVGGLTLG | ASSPGLGGYEQY | 0.084189184 |
| TTQTIYI | ASSPGLGGYEQY | 0.08790796 |
| VAAGAGLGAVI | ASSPGLGGYEQY | 0.09548396 |
| VAAVVTLL | ASSPGLGGYEQY | 0.09326639 |
| VGFLGGLI | ASSPGLGGYEQY | 0.093252204 |
| VGVIPILL | ASSPGLGGYEQY | 0.093464546 |
| VIFASSLI | ASSPGLGGYEQY | 0.085503004 |
| VKGGDGGAAAG | ASSPGLGGYEQY | 0.08324056 |
| VTKTFTNI | ASSPGLGGYEQY | 0.08518783 |
| VVAAAILL | ASSPGLGGYEQY | 0.08489868 |
| YAALDVEL | ASSPGLGGYEQY | 0.087492615 |
| LASVALLL | ASSPGLGGYEQY | 0.08121122 |
| LDAVHRV | ASSPGLGGYEQY | 0.08749962 |
| QGLSSSIF | ASSPGLGGYEQY | 0.089794524 |
| RGTFANIRL | ASSPGLGGYEQY | 0.08438225 |
| RQLWWGHRI | ASSPGLGGYEQY | 0.092639536 |
| YGPPGTGKTLL | ASSPGLGGYEQY | 0.08406665 |
| AALVGAPANVA | ASSPGLGGYEQY | 0.08565849 |
| AATFRRTE | ASSPGLGGYEQY | 0.08528597 |
| AGALGGGI | ASSPGLGGYEQY | 0.08952398 |
| AGGAGGAGGIGAG | ASSPGLGGYEQY | 0.08380038 |
| AGSGALGL | ASSPGLGGYEQY | 0.09353793 |
| AQIADQI | ASSPGLGGYEQY | 0.08542385 |
| ASAVAVTYL | ASSPGLGGYEQY | 0.09913174 |
| ASTVLGGTGG | ASSPGLGGYEQY | 0.086000025 |
| AVGGEPAK | ASSPGLGGYEQY | 0.088149905 |
| CEQGTFGGTC | ASSPGLGGYEQY | 0.08487431 |
| DEGKAPAG | ASSPGLGGYEQY | 0.097799905 |
| DTVVFPAL | ASSPGLGGYEQY | 0.09110693 |
| FEGITMAG | ASSPGLGGYEQY | 0.095786065 |
| FPLVVQD | ASSPGLGGYEQY | 0.08738009 |
| GAFTTPQ | ASSPGLGGYEQY | 0.09340123 |

| | | |
|---|---|---|
| GFGGFGTAGGVGGA | ASSPGLGGYEQY | 0.08368642 |
| GFLAQFL | ASSPGLGGYEQY | 0.087361455 |
| GGAGGAGGNAGVA | ASSPGLGGYEQY | 0.08527246 |
| GGVGTAGAGGAGGAGG | ASSPGLGGYEQY | 0.087537944 |
| GNGGGGDAGNAGSGGN | ASSPGLGGYEQY | 0.08968675 |
| GTGGAGGAGNDAGS | ASSPGLGGYEQY | 0.08205488 |
| GTTNTGWFNS | ASSPGLGGYEQY | 0.087250344 |
| IFMGPLFTL | ASSPGLGGYEQY | 0.08212726 |
| ILAHAIEAI | ASSPGLGGYEQY | 0.084795356 |
| IRSAIREAL | ASSPGLGGYEQY | 0.08612691 |
| ITFAGGG | ASSPGLGGYEQY | 0.08600894 |
| ITGMSGHQEV | ASSPGLGGYEQY | 0.08683551 |
| IVAGLACTL | ASSPGLGGYEQY | 0.082560815 |
| IVGGGLAGA | ASSPGLGGYEQY | 0.08749307 |
| IVTGVLNI | ASSPGLGGYEQY | 0.09551452 |
| LAQQGCI | ASSPGLGGYEQY | 0.08307394 |
| LCLHVEN | ASSPGLGGYEQY | 0.08392016 |
| LDAISDGS | ASSPGLGGYEQY | 0.0914424 |
| LGGFAAML | ASSPGLGGYEQY | 0.21572353 |
| LGGKNPMI | ASSPGLGGYEQY | 0.22497278 |
| LGPKGVAAIL | ASSPGLGGYEQY | 0.21278486 |
| LMVGVSI | ASSPGLGGYEQY | 0.22101341 |
| LVGAAFVL | ASSPGLGGYEQY | 0.21445082 |
| LVLAVAPL | ASSPGLGGYEQY | 0.21629529 |
| MAAIAVIDAA | ASSPGLGGYEQY | 0.2582958 |
| NNGINAELQ | ASSPGLGGYEQY | 0.21822456 |
| RGEPLVLVHG | ASSPGLGGYEQY | 0.22641 |
| RGGAGGAAGLI | ASSPGLGGYEQY | 0.22102408 |
| RSYRMHPAV | ASSPGLGGYEQY | 0.21298982 |
| SAAHLLAGQLA | ASSPGLGGYEQY | 0.21718918 |
| SAAVGSY | ASSPGLGGYEQY | 0.21473974 |
| SPHAVLHRI | ASSPGLGGYEQY | 0.22122148 |
| SQIAKVL | ASSPGLGGYEQY | 0.21224444 |
| TALGHLVP | ASSPGLGGYEQY | 0.2154519 |
| TAPTTSGAA | ASSPGLGGYEQY | 0.21843088 |
| TDGDDGNGGSPGT | ASSPGLGGYEQY | 0.22941004 |
| TDGLGEN | ASSPGLGGYEQY | 0.23253554 |
| TGGPPLPGQA | ASSPGLGGYEQY | 0.22148824 |
| TIPAITI | ASSPGLGGYEQY | 0.21708398 |
| TIPGFTI | ASSPGLGGYEQY | 0.21783912 |
| TSIASGALA | ASSPGLGGYEQY | 0.2128978 |
| TVTLPGAAI | ASSPGLGGYEQY | 0.21651657 |
| VASGLSAALP | ASSPGLGGYEQY | 0.21479462 |
| VGAGVIGI | ASSPGLGGYEQY | 0.22074838 |
| VGPKRFGV | ASSPGLGGYEQY | 0.21073942 |
| VNVAFPD | ASSPGLGGYEQY | 0.22190498 |
| VSMTNSGVS | ASSPGLGGYEQY | 0.2149654 |
| VTDVLGDLSV | ASSPGLGGYEQY | 0.2173555 |
| VVVVGAGFAGL | ASSPGLGGYEQY | 0.22329701 |
| VYTVGTVV | ASSPGLGGYEQY | 0.21429417 |
| WTTFSGP | ASSPGLGGYEQY | 0.21789397 |

| | | |
|---|---|---|
| YDELVEAF | ASSPGLGGYEQY | 0.21495092 |
| YGTGGAGGNGG | ASSPGLGGYEQY | 0.22424449 |
| YTNVMAV | ASSPGLGGYEQY | 0.21347894 |
| FLEQVHQL | ASSPGLGGYEQY | 0.22952583 |
| GLSLLAVAV | ASSPGLGGYEQY | 0.21327469 |
| LEQVHQL | ASSPGLGGYEQY | 0.22496617 |
| LNAAGAI | ASSPGLGGYEQY | 0.2191466 |
| PAVHASL | ASSPGLGGYEQY | 0.2162478 |
| TELEDDA | ASSPGLGGYEQY | 0.21456248 |
| TPPGPPN | ASSPGLGGYEQY | 0.23929016 |
| TPPPPPPP | ASSPGLGGYEQY | 0.22273012 |
| TQSPALI | ASGEPGHQAPL | 0.23000042 |
| EFAAAFPQ | ASGEPGHQAPL | 0.21922387 |
| LLSGNLDVL | ASGEPGHQAPL | 0.23263176 |
| GPANLGNYNI | ASGEPGHQAPL | 0.21426108 |
| TLGPTLP | ASGEPGHQAPL | 0.2153624 |
| VYAYSSCCK | ASGEPGHQAPL | 0.21431348 |
| APAPAPAPNI | ASGEPGHQAPL | 0.21592417 |
| PGGPVPIP | ASGEPGHQAPL | 0.2175894 |
| FLTVALEP | ASGEPGHQAPL | 0.21178868 |
| SQYFGANL | ASGEPGHQAPL | 0.2184128 |
| GGVKPAA | ASGEPGHQAPL | 0.21160027 |
| GGTGGAGGAGGW | ASGEPGHQAPL | 0.21552123 |
| GVGGVAGLGGAGG | ASGEPGHQAPL | 0.21662869 |
| VLLVLATTI | ASGEPGHQAPL | 0.22081852 |
| IGIFAAML | ASGEPGHQAPL | 0.21424025 |
| AVGDLID | ASGEPGHQAPL | 0.21224657 |
| NISPLPL | ASGEPGHQAPL | 0.21371053 |
| VTAVFVAI | ASGEPGHQAPL | 0.2202901 |
| SGDDNDAAL | ASGEPGHQAPL | 0.21392459 |
| AVAALAHS | ASGEPGHQAPL | 0.21496381 |
| GGVTVMATP | ASGEPGHQAPL | 0.21874811 |
| AGLAANVRD | ASGEPGHQAPL | 0.72601664 |
| TNRPDLIDPALLRPG | ASGEPGHQAPL | 0.79135096 |
| LANLGVTAL | ASGEPGHQAPL | 0.64013064 |
| TGERIEL | ASGEPGHQAPL | 0.5276003 |
| NGEELAA | ASGEPGHQAPL | 0.6801814 |
| VLSLVWG | ASGEPGHQAPL | 0.77422184 |
| IGYALWL | ASGEPGHQAPL | 0.783016 |
| LSLLLAPL | ASGEPGHQAPL | 0.7472155 |
| TLDEALK | ASGEPGHQAPL | 0.52742106 |
| TDGAGTAEGG | ASGEPGHQAPL | 0.6948797 |
| NAPGGAGGNGG | ASGEPGHQAPL | 0.6723532 |
| VSVFMGMGGNF | ASGEPGHQAPL | 0.6691985 |
| LASYLAAL | ASGEPGHQAPL | 0.7595727 |
| LVGGTPAL | ASGEPGHQAPL | 0.7242404 |
| VRNADVL | ASGEPGHQAPL | 0.6567657 |
| TCSSPFLVSAM | ASGEPGHQAPL | 0.6403699 |
| VHITDAE | ASGEPGHQAPL | 0.7336767 |
| EYAAPSF | ASGEPGHQAPL | 0.7594627 |
| VVDAFHGGL | ASGEPGHQAPL | 0.74012065 |

| | | |
|---|---|---|
| ASVVDPQ | ASGEPGHQAPL | 0.8058381 |
| SYNEFGGLS | ASGEPGHQAPL | 0.7708598 |
| DPDGSVY | ASGEPGHQAPL | 0.7721589 |
| MVSTIHATE | ASGEPGHQAPL | 0.28309354 |
| LAAGVADPV | ASGEPGHQAPL | 0.40877128 |
| TKEQIAA | ASGEPGHQAPL | 0.6403048 |
| SLASPAAL | ASGEPGHQAPL | 0.64282066 |
| IGLVFYGV | ASGEPGHQAPL | 0.43071637 |
| FLLDASNL | ASGEPGHQAPL | 0.7132609 |
| YQVGNFP | ASGEPGHQAPL | 0.6462208 |
| ESLLSQST | ASGEPGHQAPL | 0.55715126 |
| LFTVGFAAVG | ASGEPGHQAPL | 0.77374315 |
| ATPPTPT | ASGEPGHQAPL | 0.6488277 |
| TDLATLG | ASGEPGHQAPL | 0.6152353 |
| SAGGTGR | ASGEPGHQAPL | 0.6281519 |
| GFSNSGS | ASGEPGHQAPL | 0.7897457 |
| GGNGGSGGTSV | ASGEPGHQAPL | 0.4959703 |
| GTSVATGGAGN | ASGEPGHQAPL | 0.72778285 |
| NGGNGGSGGTS | ASGEPGHQAPL | 0.71163464 |
| AVVKFPH | ASGEPGHQAPL | 0.77723557 |
| TPPPPPP | ASGEPGHQAPL | 0.47181156 |
| AAPAAAAAPA | ASGEPGHQAPL | 0.75455576 |
| IALNEDL | ASGEPGHQAPL | 0.6667428 |
| IVNVSSM | ASGEPGHQAPL | 0.8141651 |
| TGIPPAPR | ASGEPGHQAPL | 0.65425295 |
| FDLGGGTF | ASGEPGHQAPL | 0.72589266 |
| FDLGGGTFDVS | ASGEPGHQAPL | 0.6866211 |
| NEPTAAA | ASGEPGHQAPL | 0.7691666 |
| NEPTAAALA | ASGEPGHQAPL | 0.4797328 |
| GSIINISSI | ASGEPGHQAPL | 0.67577994 |
| VAPGNAGTA | ASGEPGHQAPL | 0.544893 |
| TGGGASLE | ASGEPGHQAPL | 0.72060627 |
| VAVYNHYKRI | ASGEPGHQAPL | 0.74450827 |
| TDPAPATT | ASGEPGHQAPL | 0.48741814 |
| TNRPDILDPALLRPG | ASGEPGHQAPL | 0.79102373 |
| LTTTTTS | ASGEPGHQAPL | 0.708872 |
| PTGSGKTL | ASGEPGHQAPL | 0.6071199 |
| TGSGKTL | ASGEPGHQAPL | 0.7874594 |
| TIGHVDHGK | ASGEPGHQAPL | 0.65844965 |
| TIGHVDHGKTT | ASGEPGHQAPL | 0.7809899 |
| GHVDHGKTT | ASGEPGHQAPL | 0.6986332 |
| AQFEHTLL | ASGEPGHQAPL | 0.7112937 |
| IVNQDEL | ASGEPGHQAPL | 0.7550063 |
| TNGSQFF | ASGEPGHQAPL | 0.60791165 |
| TGKGIVE | ASGEPGHQAPL | 0.56872195 |
| LISNASDA | ASGEPGHQAPL | 0.77774644 |
| AELRTLGELTKIAKAHGAQ | ASGEPGHQAPL | 0.63612556 |
| APNAVAAAEPSW | ASGEPGHQAPL | 0.84444475 |
| APQVAGAATAVL | ASGEPGHQAPL | 0.41759342 |
| ARRDPAAAAAILPTDA | ASGEPGHQAPL | 0.46584624 |
| DAAARDVLAEAGLAEYF | ASGEPGHQAPL | 0.51042706 |

| | | |
|---|---|---|
| DDQIVLSAKANLAKVEQ | ASGEPGHQAPL | 0.6917253 |
| DELLPDTGEDSDEDGH | ASGEPGHQAPL | 0.8364042 |
| ELPEAPANRATVPQD | ASGEPGHQAPL | 0.7681781 |
| FGQSKIERVTGIG | ASGEPGHQAPL | 0.4715892 |
| GAGGTGSTAGGAGGAGGAGGI | ASGEPGHQAPL | 0.6129892 |
| GHGGTGAAGGNGGTGGAGGS( | ASGEPGHQAPL | 0.5137621 |
| GNGGNGGAGGEGGPGGAGGA( | ASGEPGHQAPL | 0.5384046 |
| GSTAGNGGNGGNATNGTIAG( | ASGEPGHQAPL | 0.5885927 |
| GTGGDGGNGGNGGTGG | ASGEPGHQAPL | 0.5539274 |
| GVRRPAGIPADAPV | ASGEPGHQAPL | 0.52817947 |
| GYRQVTDAHLLTLARR | ASGEPGHQAPL | 0.37988573 |
| IAKMGQDGHDRG | ASGEPGHQAPL | 0.49992576 |
| KAEPSIEIQYT | ASGEPGHQAPL | 0.7127602 |
| KPVRIRMAADEDTVAVT | ASGEPGHQAPL | 0.6196394 |
| MPPRIAGMRLLVIKPEP | ASGEPGHQAPL | 0.76724195 |
| MKFYAVALFRSRGVPPTR | ASGEPGHQAPL | 0.694507 |
| NAGGFGILGAVAHSP | ASGEPGHQAPL | 0.50090736 |
| NEGREITLAMAEM | ASGEPGHQAPL | 0.15452503 |
| NELVEWSHPLGHRWRWE | ASGEPGHQAPL | 0.27909908 |
| NLGAGGTGGNGGNPG | ASGEPGHQAPL | 0.43625838 |
| NTEWPKWETVMAADPVD | ASGEPGHQAPL | 0.5605171 |
| PASAAPAETQP | ASGEPGHQAPL | 0.16332853 |
| QQSSDGKNGVRAAEITGE | ASGEPGHQAPL | 0.6271636 |
| RAPDAVTTSWWKE | ASGEPGHQAPL | 0.55336535 |
| SGFGNVGAHTSGWFNQ | ASGEPGHQAPL | 0.5257058 |
| SGVAGAVGQAASVGGLKV | ASGEPGHQAPL | 0.72271526 |
| SNGGAGTGSAGGNGGT | ASGEPGHQAPL | 0.58891624 |
| TGGAGGEGGAAG | ASGEPGHQAPL | 0.37084413 |
| TGGDGGAAGAGGAPGHG | ASGEPGHQAPL | 0.37127596 |
| TMPPLLAAIANAGRHGV | ASGEPGHQAPL | 0.7564964 |
| TSSTAGNSGGA | ASGEPGHQAPL | 0.23235063 |
| VPATVPVGILC | ASGEPGHQAPL | 0.67244947 |
| VQRPDSGGVVLTGRIS | ASGEPGHQAPL | 0.42088297 |
| WTPNPVPATKGVPAGLVQAT | ASGEPGHQAPL | 0.71410966 |
| YALKRALAKFSDRFDVVII | ASGEPGHQAPL | 0.35435435 |
| YLIGSVIV | ASGEPGHQAPL | 0.6994395 |
| YGQLVAVID | ASGEPGHQAPL | 0.5767812 |
| VTVGGMTLPA | ASGEPGHQAPL | 0.8079356 |
| VLALPAV | ASGEPGHQAPL | 0.46610776 |
| VIPPTLNYET | ASGEPGHQAPL | 0.66098547 |
| VGMGGMHPSSG | ASGEPGHQAPL | 0.4003726 |
| VGLAGGAATA | ASGEPGHQAPL | 0.82887244 |
| VGGAGGAGGSGP | ASGEPGHQAPL | 0.38499796 |
| VFAAFMV | ASGEPGHQAPL | 0.42404726 |
| TLYDKLR | ASGEPGHQAPL | 0.33685884 |
| TLAVFTVL | ASGEPGHQAPL | 0.52868783 |
| TGSVQPARV | ASGEPGHQAPL | 0.753232 |
| TGGGAGSGGT | ASGEPGHQAPL | 0.4765268 |
| SHLETSL | ASGEPGHQAPL | 0.7641401 |
| SFTTGMT | ASGEPGHQAPL | 0.62749517 |
| QVEVAKRMGVRQPTVS | ASGEPGHQAPL | 0.45483637 |

| | | |
|---|---|---|
| QGSGGPALAM | ASGEPGHQAPL | 0.6519576 |
| MLAPSNP | ASGEPGHQAPL | 0.46243387 |
| LVNNAAI | ASGEPGHQAPL | 0.6304015 |
| LTGEINTLA | ASGEPGHQAPL | 0.34079164 |
| LSIFGPV | ASGEPGHQAPL | 0.59617054 |
| LLALLANL | ASGEPGHQAPL | 0.678685 |
| LISNASDALDKLR | ASGEPGHQAPL | 0.44051793 |
| LINEAVP | ASGEPGHQAPL | 0.30522013 |
| LGYAFSGL | ASGEPGHQAPL | 0.76319164 |
| LDFVAVAL | ASGEPGHQAPL | 0.7683828 |
| LAQPGAE | ASGEPGHQAPL | 0.7220106 |
| ISNLGAVTGF | ASGEPGHQAPL | 0.54787606 |
| HQEDPGRI | ASGEPGHQAPL | 0.46637365 |
| GTPGEPTP | ASGEPGHQAPL | 0.6211793 |
| GFFNSGS | ASGEPGHQAPL | 0.7892545 |
| GAGGGGLTAGAN | ASGEPGHQAPL | 0.7891704 |
| FVDALVNM | ASGEPGHQAPL | 0.7207992 |
| FLTVAPEL | ASGEPGHQAPL | 0.4313956 |
| FETLTAL | ASGEPGHQAPL | 0.7551557 |
| DVINPIEL | ASGEPGHQAPL | 0.68191004 |
| APVSGAPV | ASGEPGHQAPL | 0.6577992 |
| ALVGGLAW | ASGEPGHQAPL | 0.75961435 |
| ALAAAAATAI | ASGEPGHQAPL | 0.7082001 |
| AFGLGLLL | ASGEPGHQAPL | 0.6099115 |
| AFFAAASL | ASGEPGHQAPL | 0.5025393 |
| AAAALPAFSPP | ASGEPGHQAPL | 0.7474998 |
| VVLAPAL | ASGEPGHQAPL | 0.77819675 |
| TIGHVDHGKTTL | ASGEPGHQAPL | 0.7704936 |
| LISNASDALDK | ASGEPGHQAPL | 0.8274871 |
| LISNASDAL | ASGEPGHQAPL | 0.8039962 |
| ISNASDALDK | ASGEPGHQAPL | 0.7092058 |
| NQAETLVYQTE | ASGEPGHQAPL | 0.1941903 |
| TGGTGGTAGAG | ASGEPGHQAPL | 0.33607164 |
| TGSTAGGAGGA | ASGEPGHQAPL | 0.5981327 |
| DPLAVPVAAGPV | ASGEPGHQAPL | 0.6233594 |
| GGTGGTPGNGGN | ASGEPGHQAPL | 0.24913727 |
| PAGLAGVDPATL | ASGEPGHQAPL | 0.7219402 |
| GNNLVVEAPHAGASV | ASGEPGHQAPL | 0.59610385 |
| KVTRIMSASTLEQPA | ASGEPGHQAPL | 0.5365108 |
| LGFALPIALPRRTVV | ASGEPGHQAPL | 0.80301964 |
| TRLGNLTARKLVVLV | ASGEPGHQAPL | 0.5863643 |
| WSQQRRLIYFPSAGP | ASGEPGHQAPL | 0.54227775 |
| GPNGNPDPMAAAVDIR | ASGEPGHQAPL | 0.59513557 |
| TQRCVSVADLRAAASR | ASGEPGHQAPL | 0.7868546 |
| QFVLDVAEHPVEVLTETV | ASGEPGHQAPL | 0.4901568 |
| GGAGGTGGAGGNGGTQAGDG | ASGEPGHQAPL | 0.78440875 |
| PLLATLRAIAVDVDAKPAQV | ASGEPGHQAPL | 0.5482248 |
| APKFRTGGYMIDFWGVGYQV | ASGEPGHQAPL | 0.83363223 |
| GAGGAGGAGGAAGGPGGTGG | ASGEPGHQAPL | 0.39160466 |
| IGALGLLATDVAPAGSVAFG | ASGEPGHQAPL | 0.7814807 |
| AGGAGGIGGNGGAGGAGGVL | ASGEPGHQAPL | 0.684871 |

| | | |
|---|---|---|
| ADELIGLL | ASGEPGHQAPL | 0.86677605 |
| ADRALKA | ASGEPGHQAPL | 0.6389865 |
| AGAVGQVGGAGGA | ASGEPGHQAPL | 0.7242651 |
| AGGGGGLGGAP | ASGEPGHQAPL | 0.60948265 |
| AGGLLYGNGGAG | ASGEPGHQAPL | 0.7672098 |
| AGLESIGL | ASGEPGHQAPL | 0.36058593 |
| AGLKTGII | ASGEPGHQAPL | 0.65485185 |
| ASLAALLL | ASGEPGHQAPL | 0.32555893 |
| ASVINISSI | ASGEPGHQAPL | 0.6942206 |
| ATPPGPA | ASGEPGHQAPL | 0.7420863 |
| ATVASALV | ASGEPGHQAPL | 0.19961345 |
| ATVASLIL | ASGEPGHQAPL | 0.75813353 |
| AVNGPVLIRH | ASGEPGHQAPL | 0.47409835 |
| DGAAGAVGI | ASGEPGHQAPL | 0.4868011 |
| DPLLHGI | ASGEPGHQAPL | 0.79669726 |
| ELGAALGAL | ASGEPGHQAPL | 0.70214486 |
| ELGRTQAV | ASGEPGHQAPL | 0.76809895 |
| FVAGGYLS | ASGEPGHQAPL | 0.61072946 |
| GASGGAGGAGGW | ASGEPGHQAPL | 0.6508914 |
| GGAGAGGDGGAGGI | ASGEPGHQAPL | 0.6916313 |
| GTGHNGSQPGMGGNG | ASGEPGHQAPL | 0.35482168 |
| GYAAASASAS | ASGEPGHQAPL | 0.55096793 |
| HQVDVAAVQQG | ASGEPGHQAPL | 0.8177536 |
| IGAAIAHT | ASGEPGHQAPL | 0.66448855 |
| IGMIEVAQL | ASGEPGHQAPL | 0.6687147 |
| IRLAPEHTSAMD | ASGEPGHQAPL | 0.6354489 |
| IYNVSVL | ASGEPGHQAPL | 0.6343331 |
| KQGFLGM | ASGEPGHQAPL | 0.652681 |
| LESVITQL | ASGEPGHQAPL | 0.66745853 |
| LGLGALVVV | ASGEPGHQAPL | 0.6753318 |
| LIGNGGNGGSGGI | ASGEPGHQAPL | 0.6699095 |
| LNAINAPTL | ASGEPGHQAPL | 0.64053774 |
| LQLLNTGV | ASGEPGHQAPL | 0.6739737 |
| LQTSIEF | ASGEPGHQAPL | 0.661789 |
| MSFAVSEL | ASGEPGHQAPL | 0.6607357 |
| NTGALGL | ASGEPGHQAPL | 0.66874605 |
| PPGAGGAGGAGTT | ASGEPGHQAPL | 0.66218984 |
| QLVGLGI | ASGEPGHQAPL | 0.6489133 |
| RFLEQVHQL | ASGEPGHQAPL | 0.6470775 |
| RGVGYVLR | ASGEPGHQAPL | 0.6701729 |
| SATGVFAGVI | ASGEPGHQAPL | 0.66964746 |
| SLPAITLP | ASGEPGHQAPL | 0.66930073 |
| STALAFTV | ASGEPGHQAPL | 0.6774758 |
| SVLTVGLGK | ASGEPGHQAPL | 0.6704359 |
| TALAGAGSGI | ASGEPGHQAPL | 0.66828376 |
| TGLPFTGI | ASGEPGHQAPL | 0.5269045 |
| VAEHVTV | ASGEPGHQAPL | 0.6203187 |
| VDIVLNSL | ASGEPGHQAPL | 0.6412426 |
| VLTGGGAVL | ASGEPGHQAPL | 0.6565564 |
| VNVGLAYGI | ASGEPGHQAPL | 0.49554676 |
| VSSAAVGAV | ASGEPGHQAPL | 0.66657835 |

| | | |
|---|---|---|
| VTSVYEAGV | ASGEPGHQAPL | 0.6606804 |
| WVALDRGA | ASGEPGHQAPL | 0.64374995 |
| YLVGSVVI | ASGEPGHQAPL | 0.6732558 |
| GPPGTGKTLL | ASGEPGHQAPL | 0.6536375 |
| GVGMTKF | ASGEPGHQAPL | 0.6370537 |
| IGIGGSDL | ASGEPGHQAPL | 0.6534637 |
| IGTLLAEL | ASGEPGHQAPL | 0.6753884 |
| IYTSGTT | ASGEPGHQAPL | 0.642816 |
| LSFPAGV | ASGEPGHQAPL | 0.6741693 |
| LYTSGTT | ASGEPGHQAPL | 0.64954364 |
| VGAFPFGF | ASGEPGHQAPL | 0.6744313 |
| VGAGTVEF | ASGEPGHQAPL | 0.61576545 |
| VGKSTLLL | ASGEPGHQAPL | 0.67196935 |
| AALAHAGGGIV | ASGEPGHQAPL | 0.6575401 |
| AALVAVSLL | ASGEPGHQAPL | 0.67832 |
| AAVSRAVAAL | ASGEPGHQAPL | 0.6560021 |
| AGGSSGIGGF | ASGEPGHQAPL | 0.6737319 |
| AGLTVANANG | ASGEPGHQAPL | 0.66422623 |
| AGLVGAVL | ASGEPGHQAPL | 0.6720996 |
| ALSQAFTV | ASGEPGHQAPL | 0.60579526 |
| ALTFSVSL | ASGEPGHQAPL | 0.6553543 |
| ALTGAALVAL | ASGEPGHQAPL | 0.64164114 |
| ASFSIGAGVL | ASGEPGHQAPL | 0.66618645 |
| DGSVYKI | ASGEPGHQAPL | 0.6694038 |
| EQLAEIV | ASGEPGHQAPL | 0.4677014 |
| FAVSELTVM | ASGEPGHQAPL | 0.6754456 |
| GAAPLAAGL | ASGEPGHQAPL | 0.6256297 |
| GADGGDGGSGGAGG | ASGEPGHQAPL | 0.63761103 |
| GFGLDTGGAGG | ASGEPGHQAPL | 0.67256 |
| GLASGGLL | ASGEPGHQAPL | 0.6635829 |
| GRIAGAAVLL | ASGEPGHQAPL | 0.6678705 |
| GSAGSLNTSF | ASGEPGHQAPL | 0.65326846 |
| GTGQIGI | ASGEPGHQAPL | 0.66206586 |
| GTLIALIL | ASGEPGHQAPL | 0.6667777 |
| IAATMGFVVG | ASGEPGHQAPL | 0.5692543 |
| IFAQALL | ASGEPGHQAPL | 0.62901187 |
| IGAAVALGF | ASGEPGHQAPL | 0.67588276 |
| IGLTPLAL | ASGEPGHQAPL | 0.45834306 |
| ILGSNTSTL | ASGEPGHQAPL | 0.8072641 |
| IVFSLTVL | ASGEPGHQAPL | 0.31071663 |
| LASKATGF | ASGEPGHQAPL | 0.2047937 |
| LASVALIL | ASGEPGHQAPL | 0.256368 |
| LFAVGLTL | ASGEPGHQAPL | 0.61239034 |
| LFNGGIGTYI | ASGEPGHQAPL | 0.8228689 |
| LTLGAGII | ASGEPGHQAPL | 0.60951966 |
| LVAAALGDI | ASGEPGHQAPL | 0.20080829 |
| NTAFVQLGI | ASGEPGHQAPL | 0.3528263 |
| QLFSGAEL | ASGEPGHQAPL | 0.25947398 |
| SAATFITL | ASGEPGHQAPL | 0.31808463 |
| SNYGLANLGV | ASGEPGHQAPL | 0.45962298 |
| SYGGAYAVM | ASGEPGHQAPL | 0.345443 |

| | | |
|---|---|---|
| TTGVGGLTLG | ASGEPGHQAPL | 0.29826653 |
| TTQTIYI | ASGEPGHQAPL | 0.25996792 |
| VAAGAGLGAVI | ASGEPGHQAPL | 0.37117338 |
| VAAVVTLL | ASGEPGHQAPL | 0.54140073 |
| VGFLGGLI | ASGEPGHQAPL | 0.35982373 |
| VGVIPILL | ASGEPGHQAPL | 0.6555648 |
| VIFASSLI | ASGEPGHQAPL | 0.65519094 |
| VKGGDGGAAAG | ASGEPGHQAPL | 0.64427054 |
| VTKTFTNI | ASGEPGHQAPL | 0.11196446 |
| VVAAAILL | ASGEPGHQAPL | 0.19413587 |
| YAALDVEL | ASGEPGHQAPL | 0.2423331 |
| LASVALLL | ASGEPGHQAPL | 0.384581 |
| LDAVHRV | ASGEPGHQAPL | 0.13914509 |
| QGLSSSIF | ASGEPGHQAPL | 0.43115553 |
| RGTFANIRL | ASGEPGHQAPL | 0.27548602 |
| RQLWWGHRI | ASGEPGHQAPL | 0.242762 |
| YGPPGTGKTLL | ASGEPGHQAPL | 0.62232584 |
| AALVGAPANVA | ASGEPGHQAPL | 0.31736934 |
| AATFRRTE | ASGEPGHQAPL | 0.21425834 |
| AGALGGGI | ASGEPGHQAPL | 0.20324242 |
| AGGAGGAGGIGAG | ASGEPGHQAPL | 0.7101972 |
| AGSGALGL | ASGEPGHQAPL | 0.16840382 |
| AQIADQI | ASGEPGHQAPL | 0.66564554 |
| ASAVAVTYL | ASGEPGHQAPL | 0.2506013 |
| ASTVLGGTGG | ASGEPGHQAPL | 0.56589043 |
| AVGGEPAK | ASGEPGHQAPL | 0.16678308 |
| CEQGTFGGTC | ASGEPGHQAPL | 0.65201 |
| DEGKAPAG | ASGEPGHQAPL | 0.4642517 |
| DTVVFPAL | ASGEPGHQAPL | 0.8484546 |
| FEGITMAG | ASGEPGHQAPL | 0.27826446 |
| FPLVVQD | ASGEPGHQAPL | 0.35061327 |
| GAFTTPQ | ASGEPGHQAPL | 0.24559535 |
| GFGGFGTAGGVGGA | ASGEPGHQAPL | 0.8107679 |
| GFLAQFL | ASGEPGHQAPL | 0.15354456 |
| GGAGGAGGNAGVA | ASGEPGHQAPL | 0.22301763 |
| GGVGTAGAGGAGGAGG | ASGEPGHQAPL | 0.17526394 |
| GNGGGGDAGNAGSGGN | ASGEPGHQAPL | 0.32482493 |
| GTGGAGGAGNDAGS | ASGEPGHQAPL | 0.6055753 |
| GTTNTGWFNS | ASGEPGHQAPL | 0.14279653 |
| IFMGPLFTL | ASGEPGHQAPL | 0.6388012 |
| ILAHAIEAI | ASGEPGHQAPL | 0.32897532 |
| IRSAIREAL | ASGEPGHQAPL | 0.19876762 |
| ITFAGGG | ASGEPGHQAPL | 0.69431925 |
| ITGMSGHQEV | ASGEPGHQAPL | 0.2320078 |
| IVAGLACTL | ASGEPGHQAPL | 0.72306186 |
| IVGGGLAGA | ASGEPGHQAPL | 0.2469278 |
| IVTGVLNI | ASGEPGHQAPL | 0.29755884 |
| LAQQGCI | ASGEPGHQAPL | 0.4328117 |
| LCLHVEN | ASGEPGHQAPL | 0.1808921 |
| LDAISDGS | ASGEPGHQAPL | 0.18359144 |
| LGGFAAML | ASGEPGHQAPL | 0.7451863 |

| | | |
|---|---|---|
| LGGKNPMI | ASGEPGHQAPL | 0.32866332 |
| LGPKGVAAIL | ASGEPGHQAPL | 0.72254324 |
| LMVGVSI | ASGEPGHQAPL | 0.192102 |
| LVGAAFVL | ASGEPGHQAPL | 0.16769919 |
| LVLAVAPL | ASGEPGHQAPL | 0.24069847 |
| MAAIAVIDAA | ASGEPGHQAPL | 0.5310337 |
| NNGINAELQ | ASGEPGHQAPL | 0.63721186 |
| RGEPLVLVHG | ASGEPGHQAPL | 0.3572514 |
| RGGAGGAAGLI | ASGEPGHQAPL | 0.19146861 |
| RSYRMHPAV | ASGEPGHQAPL | 0.27814353 |
| SAAHLLAGQLA | ASGEPGHQAPL | 0.23676822 |
| SAAVGSY | ASGEPGHQAPL | 0.20274921 |
| SPHAVLHRI | ASGEPGHQAPL | 0.36319578 |
| SQIAKVL | ASGEPGHQAPL | 0.33674166 |
| TALGHLVP | ASGEPGHQAPL | 0.20778675 |
| TAPTTSGAA | ASGEPGHQAPL | 0.24415123 |
| TDGDDGNGGSPGT | ASGEPGHQAPL | 0.3002922 |
| TDGLGEN | ASGEPGHQAPL | 0.42907575 |
| TGGPPLPGQA | ASGEPGHQAPL | 0.36417246 |
| TIPAITI | ASGEPGHQAPL | 0.5855933 |
| TIPGFTI | ASGEPGHQAPL | 0.42323002 |
| TSIASGALA | ASGEPGHQAPL | 0.38500455 |
| TVTLPGAAI | ASGEPGHQAPL | 0.10677173 |
| VASGLSAALP | ASGEPGHQAPL | 0.13369742 |
| VGAGVIGI | ASGEPGHQAPL | 0.29882252 |
| VGPKRFGV | ASGEPGHQAPL | 0.23103787 |
| VNVAFPD | ASGEPGHQAPL | 0.1505366 |
| VSMTNSGVS | ASGEPGHQAPL | 0.43096796 |
| VTDVLGDLSV | ASGEPGHQAPL | 0.2538471 |
| VVVVGAGFAGL | ASGEPGHQAPL | 0.1948863 |
| VYTVGTVV | ASGEPGHQAPL | 0.41845232 |
| WTTFSGP | ASGEPGHQAPL | 0.18768919 |
| YDELVEAF | ASGEPGHQAPL | 0.21520565 |
| YGTGGAGGNGG | ASGEPGHQAPL | 0.23557709 |
| YTNVMAV | ASGEPGHQAPL | 0.38279432 |
| FLEQVHQL | ASGEPGHQAPL | 0.17528637 |
| GLSLLAVAV | ASGEPGHQAPL | 0.32090908 |
| LEQVHQL | ASGEPGHQAPL | 0.25704032 |
| LNAAGAI | ASGEPGHQAPL | 0.4491511 |
| PAVHASL | ASGEPGHQAPL | 0.16363741 |
| TELEDDA | ASGEPGHQAPL | 0.34772113 |
| TPPGPPN | ASGEPGHQAPL | 0.21271145 |
| TPPPPPPP | ASGEPGHQAPL | 0.71679455 |
| TQSPALI | ASSQVPRDKNSDYT | 0.23462918 |
| EFAAAFPQ | ASSQVPRDKNSDYT | 0.32876948 |
| LLSGNLDVL | ASSQVPRDKNSDYT | 0.29729882 |
| GPANLGNYNI | ASSQVPRDKNSDYT | 0.642367 |
| TLGPTLP | ASSQVPRDKNSDYT | 0.14168142 |
| VYAYSSCCK | ASSQVPRDKNSDYT | 0.24479958 |
| APAPAPAPNI | ASSQVPRDKNSDYT | 0.1546956 |
| PGGPVPIP | ASSQVPRDKNSDYT | 0.32752842 |

| | | |
|---|---|---|
| FLTVALEP | ASSQVPRDKNSDYT | 0.3215693 |
| SQYFGANL | ASSQVPRDKNSDYT | 0.17963782 |
| GGVKPAA | ASSQVPRDKNSDYT | 0.36044735 |
| GGTGGAGGAGGW | ASSQVPRDKNSDYT | 0.38889694 |
| GVGGVAGLGGAGG | ASSQVPRDKNSDYT | 0.19249798 |
| VLLVLATTI | ASSQVPRDKNSDYT | 0.53916985 |
| IGIFAAML | ASSQVPRDKNSDYT | 0.3015038 |
| AVGDLID | ASSQVPRDKNSDYT | 0.5346406 |
| NISPLPL | ASSQVPRDKNSDYT | 0.28177965 |
| VTAVFVAI | ASSQVPRDKNSDYT | 0.22623557 |
| SGDDNDAAL | ASSQVPRDKNSDYT | 0.35714877 |
| AVAALAHS | ASSQVPRDKNSDYT | 0.30820903 |
| GGVTVMATP | ASSQVPRDKNSDYT | 0.21390805 |
| AGLAANVRD | ASSQVPRDKNSDYT | 0.4205402 |
| TNRPDLIDPALLRPG | ASSQVPRDKNSDYT | 0.8328234 |
| LANLGVTAL | ASSQVPRDKNSDYT | 0.8205913 |
| TGERIEL | ASSQVPRDKNSDYT | 0.6322391 |
| NGEELAA | ASSQVPRDKNSDYT | 0.6682781 |
| VLSLVWG | ASSQVPRDKNSDYT | 0.74733937 |
| IGYALWL | ASSQVPRDKNSDYT | 0.82700074 |
| LSLLLAPL | ASSQVPRDKNSDYT | 0.8513248 |
| TLDEALK | ASSQVPRDKNSDYT | 0.8303665 |
| TDGAGTAEGG | ASSQVPRDKNSDYT | 0.6187122 |
| NAPGGAGGNGG | ASSQVPRDKNSDYT | 0.8344149 |
| VSVFMGMGGNF | ASSQVPRDKNSDYT | 0.79016423 |
| LASYLAAL | ASSQVPRDKNSDYT | 0.81630504 |
| LVGGTPAL | ASSQVPRDKNSDYT | 0.7906556 |
| VRNADVL | ASSQVPRDKNSDYT | 0.73726296 |
| TCSSPFLVSAM | ASSQVPRDKNSDYT | 0.6806142 |
| VHITDAE | ASSQVPRDKNSDYT | 0.48549446 |
| EYAAPSF | ASSQVPRDKNSDYT | 0.8296759 |
| VVDAFHGGL | ASSQVPRDKNSDYT | 0.83119625 |
| ASVVDPQ | ASSQVPRDKNSDYT | 0.81483227 |
| SYNEFGGLS | ASSQVPRDKNSDYT | 0.8532541 |
| DPDGSVY | ASSQVPRDKNSDYT | 0.8448127 |
| MVSTIHATE | ASSQVPRDKNSDYT | 0.7844186 |
| LAAGVADPV | ASSQVPRDKNSDYT | 0.23707528 |
| TKEQIAA | ASSQVPRDKNSDYT | 0.38458622 |
| SLASPAAL | ASSQVPRDKNSDYT | 0.56889975 |
| IGLVFYGV | ASSQVPRDKNSDYT | 0.70670885 |
| FLLDASNL | ASSQVPRDKNSDYT | 0.2121729 |
| YQVGNFP | ASSQVPRDKNSDYT | 0.78627473 |
| ESLLSQST | ASSQVPRDKNSDYT | 0.6839144 |
| LFTVGFAAVG | ASSQVPRDKNSDYT | 0.7336097 |
| ATPPTPT | ASSQVPRDKNSDYT | 0.84588593 |
| TDLATLG | ASSQVPRDKNSDYT | 0.7145176 |
| SAGGTGR | ASSQVPRDKNSDYT | 0.4635339 |
| GFSNSGS | ASSQVPRDKNSDYT | 0.5881252 |
| GGNGGSGGTSV | ASSQVPRDKNSDYT | 0.8556005 |
| GTSVATGGAGN | ASSQVPRDKNSDYT | 0.6280991 |
| NGGNGGSGGTS | ASSQVPRDKNSDYT | 0.850331 |

| | | |
|---|---|---|
| AVVKFPH | ASSQVPRDKNSDYT | 0.7048979 |
| TPPPPPP | ASSQVPRDKNSDYT | 0.85656375 |
| AAPAAAAAPA | ASSQVPRDKNSDYT | 0.53990316 |
| IALNEDL | ASSQVPRDKNSDYT | 0.8441758 |
| IVNVSSM | ASSQVPRDKNSDYT | 0.81434476 |
| TGIPPAPR | ASSQVPRDKNSDYT | 0.8624784 |
| FDLGGGTF | ASSQVPRDKNSDYT | 0.7343161 |
| FDLGGGTFDVS | ASSQVPRDKNSDYT | 0.80597997 |
| NEPTAAA | ASSQVPRDKNSDYT | 0.6570418 |
| NEPTAAALA | ASSQVPRDKNSDYT | 0.8326225 |
| GSIINISSI | ASSQVPRDKNSDYT | 0.5873032 |
| VAPGNAGTA | ASSQVPRDKNSDYT | 0.6886854 |
| TGGGASLE | ASSQVPRDKNSDYT | 0.5639427 |
| VAVYNHYKRI | ASSQVPRDKNSDYT | 0.71940684 |
| TDPAPATT | ASSQVPRDKNSDYT | 0.8389956 |
| TNRPDILDPALLRPG | ASSQVPRDKNSDYT | 0.16741852 |
| LTTTTTS | ASSQVPRDKNSDYT | 0.8245373 |
| PTGSGKTL | ASSQVPRDKNSDYT | 0.54303634 |
| TGSGKTL | ASSQVPRDKNSDYT | 0.508229 |
| TIGHVDHGK | ASSQVPRDKNSDYT | 0.8144536 |
| TIGHVDHGKTT | ASSQVPRDKNSDYT | 0.7857305 |
| GHVDHGKTT | ASSQVPRDKNSDYT | 0.80575573 |
| AQFEHTLL | ASSQVPRDKNSDYT | 0.63805836 |
| IVNQDEL | ASSQVPRDKNSDYT | 0.78926456 |
| TNGSQFF | ASSQVPRDKNSDYT | 0.79481196 |
| TGKGIVE | ASSQVPRDKNSDYT | 0.25968647 |
| LISNASDA | ASSQVPRDKNSDYT | 0.55787146 |
| AELRTLGELTKIAKAHGAQ | ASSQVPRDKNSDYT | 0.85423565 |
| APNAVAAAEPSW | ASSQVPRDKNSDYT | 0.508184 |
| APQVAGAATAVL | ASSQVPRDKNSDYT | 0.7890878 |
| ARRDPAAAAAILPTDA | ASSQVPRDKNSDYT | 0.17116539 |
| DAAARDVLAEAGLAEYF | ASSQVPRDKNSDYT | 0.16721441 |
| DDQIVLSAKANLAKVEQ | ASSQVPRDKNSDYT | 0.3144819 |
| DELLPDTGEDSDEDGH | ASSQVPRDKNSDYT | 0.557057 |
| ELPEAPANRATVPQD | ASSQVPRDKNSDYT | 0.81142366 |
| FGQSKIERVTGIG | ASSQVPRDKNSDYT | 0.5579696 |
| GAGGTGSTAGGAGGAGGAGGI | ASSQVPRDKNSDYT | 0.16989037 |
| GHGGTGAAGGNGGTGGAGGS( | ASSQVPRDKNSDYT | 0.3844866 |
| GNGGNGGAGGEGGPGGAGGA( | ASSQVPRDKNSDYT | 0.25421345 |
| GSTAGNGGNGGNATNGTIAG( | ASSQVPRDKNSDYT | 0.27810973 |
| GTGGDGGNGGNGGTGG | ASSQVPRDKNSDYT | 0.40242866 |
| GVRRPAGIPADAPV | ASSQVPRDKNSDYT | 0.3569422 |
| GYRQVTDAHLLTLARR | ASSQVPRDKNSDYT | 0.1917029 |
| IAKMGQDGHDRG | ASSQVPRDKNSDYT | 0.17661333 |
| KAEPSIEIQYT | ASSQVPRDKNSDYT | 0.60945696 |
| KPVRIRMAADEDTVAVT | ASSQVPRDKNSDYT | 0.7312291 |
| MPPRIAGMRLLVIKPEP | ASSQVPRDKNSDYT | 0.6725136 |
| MKFYAVALFRSRGVPPTR | ASSQVPRDKNSDYT | 0.7517729 |
| NAGGFGILGAVAHSP | ASSQVPRDKNSDYT | 0.55950224 |
| NEGREITLAMAEM | ASSQVPRDKNSDYT | 0.56091195 |
| NELVEWSHPLGHRWRWE | ASSQVPRDKNSDYT | 0.09609616 |

| | | |
|---|---|---|
| NLGAGGTGGNGGNPG | ASSQVPRDKNSDYT | 0.11330595 |
| NTEWPKWETVMAADPVD | ASSQVPRDKNSDYT | 0.21065046 |
| PASAAPAETQP | ASSQVPRDKNSDYT | 0.2983306 |
| QQSSDGKNGVRAAEITGE | ASSQVPRDKNSDYT | 0.12953447 |
| RAPDAVTTSWWKE | ASSQVPRDKNSDYT | 0.5253136 |
| SGFGNVGAHTSGWFNQ | ASSQVPRDKNSDYT | 0.22376868 |
| SGVAGAVGQAASVGGLKV | ASSQVPRDKNSDYT | 0.19073895 |
| SNGGAGTGSAGGNGGT | ASSQVPRDKNSDYT | 0.6833879 |
| TGGAGGEGGAAG | ASSQVPRDKNSDYT | 0.17579071 |
| TGGDGGAAGAGGAPGHG | ASSQVPRDKNSDYT | 0.17102863 |
| TMPPLLAAIANAGRHGV | ASSQVPRDKNSDYT | 0.23467281 |
| TSSTAGNSGGA | ASSQVPRDKNSDYT | 0.6570407 |
| VPATVPVGILC | ASSQVPRDKNSDYT | 0.15808551 |
| VQRPDSGGVVLTGRIS | ASSQVPRDKNSDYT | 0.5061941 |
| WTPNPVPATKGVPAGLVQAT | ASSQVPRDKNSDYT | 0.26833254 |
| YALKRALAKFSDRFDVVII | ASSQVPRDKNSDYT | 0.5964358 |
| YLIGSVIV | ASSQVPRDKNSDYT | 0.13930121 |
| YGQLVAVID | ASSQVPRDKNSDYT | 0.5250353 |
| VTVGGMTLPA | ASSQVPRDKNSDYT | 0.29493326 |
| VLALPAV | ASSQVPRDKNSDYT | 0.8450303 |
| VIPPTLNYET | ASSQVPRDKNSDYT | 0.2082892 |
| VGMGGMHPSSG | ASSQVPRDKNSDYT | 0.34417927 |
| VGLAGGAATA | ASSQVPRDKNSDYT | 0.24012226 |
| VGGAGGAGGSGP | ASSQVPRDKNSDYT | 0.8000002 |
| VFAAFMV | ASSQVPRDKNSDYT | 0.12803045 |
| TLYDKLR | ASSQVPRDKNSDYT | 0.22753188 |
| TLAVFTVL | ASSQVPRDKNSDYT | 0.12369781 |
| TGSVQPARV | ASSQVPRDKNSDYT | 0.2806363 |
| TGGGAGSGGT | ASSQVPRDKNSDYT | 0.49822348 |
| SHLETSL | ASSQVPRDKNSDYT | 0.10882071 |
| SFTTGMT | ASSQVPRDKNSDYT | 0.44761252 |
| QVEVAKRMGVRQPTVS | ASSQVPRDKNSDYT | 0.26608163 |
| QGSGGPALAM | ASSQVPRDKNSDYT | 0.16361131 |
| MLAPSNP | ASSQVPRDKNSDYT | 0.5425071 |
| LVNNAAI | ASSQVPRDKNSDYT | 0.305058 |
| LTGEINTLA | ASSQVPRDKNSDYT | 0.7245499 |
| LSIFGPV | ASSQVPRDKNSDYT | 0.31472874 |
| LLALLANL | ASSQVPRDKNSDYT | 0.27418303 |
| LISNASDALDKLR | ASSQVPRDKNSDYT | 0.36092603 |
| LINEAVP | ASSQVPRDKNSDYT | 0.24834919 |
| LGYAFSGL | ASSQVPRDKNSDYT | 0.21540244 |
| LDFVAVAL | ASSQVPRDKNSDYT | 0.5834728 |
| LAQPGAE | ASSQVPRDKNSDYT | 0.90386796 |
| ISNLGAVTGF | ASSQVPRDKNSDYT | 0.88475835 |
| HQEDPGRI | ASSQVPRDKNSDYT | 0.8441825 |
| GTPGEPTP | ASSQVPRDKNSDYT | 0.8333093 |
| GFFNSGS | ASSQVPRDKNSDYT | 0.8710356 |
| GAGGGGLTAGAN | ASSQVPRDKNSDYT | 0.9047679 |
| FVDALVNM | ASSQVPRDKNSDYT | 0.9132674 |
| FLTVAPEL | ASSQVPRDKNSDYT | 0.9028518 |
| FETLTAL | ASSQVPRDKNSDYT | 0.8391089 |

| | | |
|---|---|---|
| DVINPIEL | ASSQVPRDKNSDYT | 0.9070462 |
| APVSGAPV | ASSQVPRDKNSDYT | 0.8952085 |
| ALVGGLAW | ASSQVPRDKNSDYT | 0.87513375 |
| ALAAAAATAI | ASSQVPRDKNSDYT | 0.9052949 |
| AFGLGLLL | ASSQVPRDKNSDYT | 0.8997728 |
| AFFAAASL | ASSQVPRDKNSDYT | 0.85586494 |
| AAAALPAFSPP | ASSQVPRDKNSDYT | 0.86100245 |
| VVLAPAL | ASSQVPRDKNSDYT | 0.8926157 |
| TIGHVDHGKTTL | ASSQVPRDKNSDYT | 0.902913 |
| LISNASDALDK | ASSQVPRDKNSDYT | 0.9032513 |
| LISNASDAL | ASSQVPRDKNSDYT | 0.9137963 |
| ISNASDALDK | ASSQVPRDKNSDYT | 0.91084254 |
| NQAETLVYQTE | ASSQVPRDKNSDYT | 0.9022935 |
| TGGTGGTAGAG | ASSQVPRDKNSDYT | 0.47202185 |
| TGSTAGGAGGA | ASSQVPRDKNSDYT | 0.74113697 |
| DPLAVPVAAGPV | ASSQVPRDKNSDYT | 0.86863875 |
| GGTGGTPGNGGN | ASSQVPRDKNSDYT | 0.85957146 |
| PAGLAGVDPATL | ASSQVPRDKNSDYT | 0.48079818 |
| GNNLVVEAPHAGASV | ASSQVPRDKNSDYT | 0.8618407 |
| KVTRIMSASTLEQPA | ASSQVPRDKNSDYT | 0.8794167 |
| LGFALPIALPRRTVV | ASSQVPRDKNSDYT | 0.840619 |
| TRLGNLTARKLVVLV | ASSQVPRDKNSDYT | 0.90647256 |
| WSQQRRLIYFPSAGP | ASSQVPRDKNSDYT | 0.87113744 |
| GPNGNPDPMAAAVDIR | ASSQVPRDKNSDYT | 0.8283565 |
| TQRCVSVADLRAAASR | ASSQVPRDKNSDYT | 0.8739947 |
| QFVLDVAEHPVEVLTETV | ASSQVPRDKNSDYT | 0.9099835 |
| GGAGGTGGAGGNGGTQAGDG | ASSQVPRDKNSDYT | 0.85534704 |
| PLLATLRAIAVDVDAKPAQV | ASSQVPRDKNSDYT | 0.91130143 |
| APKFRTGGYMIDFWGVGYQV | ASSQVPRDKNSDYT | 0.77604043 |
| GAGGAGGAGGAAGGPGGTGG | ASSQVPRDKNSDYT | 0.914163 |
| IGALGLLATDVAPAGSVAFG | ASSQVPRDKNSDYT | 0.81275874 |
| AGGAGGIGGNGGAGGAGGVL | ASSQVPRDKNSDYT | 0.9092585 |
| ADELIGLL | ASSQVPRDKNSDYT | 0.8904078 |
| ADRALKA | ASSQVPRDKNSDYT | 0.91946006 |
| AGAVGQVGGAGGA | ASSQVPRDKNSDYT | 0.88786995 |
| AGGGGGLGGAP | ASSQVPRDKNSDYT | 0.91269183 |
| AGGLLYGNGGAG | ASSQVPRDKNSDYT | 0.88780004 |
| AGLESIGL | ASSQVPRDKNSDYT | 0.9001361 |
| AGLKTGII | ASSQVPRDKNSDYT | 0.8148105 |
| ASLAALLL | ASSQVPRDKNSDYT | 0.88361865 |
| ASVINISSI | ASSQVPRDKNSDYT | 0.6995574 |
| ATPPGPA | ASSQVPRDKNSDYT | 0.8933343 |
| ATVASALV | ASSQVPRDKNSDYT | 0.9043556 |
| ATVASLIL | ASSQVPRDKNSDYT | 0.35178676 |
| AVNGPVLIRH | ASSQVPRDKNSDYT | 0.9086434 |
| DGAAGAVGI | ASSQVPRDKNSDYT | 0.8071586 |
| DPLLHGI | ASSQVPRDKNSDYT | 0.84521776 |
| ELGAALGAL | ASSQVPRDKNSDYT | 0.9128983 |
| ELGRTQAV | ASSQVPRDKNSDYT | 0.8897544 |
| FVAGGYLS | ASSQVPRDKNSDYT | 0.910043 |
| GASGGAGGAGGW | ASSQVPRDKNSDYT | 0.866852 |

| | | |
|---|---|---|
| GGAGAGGDGGAGGI | ASSQVPRDKNSDYT | 0.8857378 |
| GTGHNGSQPGMGGNG | ASSQVPRDKNSDYT | 0.88526165 |
| GYAAASASAS | ASSQVPRDKNSDYT | 0.6408527 |
| HQVDVAAVQQG | ASSQVPRDKNSDYT | 0.8335744 |
| IGAAIAHT | ASSQVPRDKNSDYT | 0.91719574 |
| IGMIEVAQL | ASSQVPRDKNSDYT | 0.6365719 |
| IRLAPEHTSAMD | ASSQVPRDKNSDYT | 0.6252891 |
| IYNVSVL | ASSQVPRDKNSDYT | 0.53952384 |
| KQGFLGM | ASSQVPRDKNSDYT | 0.52779746 |
| LESVITQL | ASSQVPRDKNSDYT | 0.57502407 |
| LGLGALVVV | ASSQVPRDKNSDYT | 0.6475499 |
| LIGNGGNGGSGGI | ASSQVPRDKNSDYT | 0.62159395 |
| LNAINAPTL | ASSQVPRDKNSDYT | 0.6313691 |
| LQLLNTGV | ASSQVPRDKNSDYT | 0.54089725 |
| LQTSIEF | ASSQVPRDKNSDYT | 0.60221726 |
| MSFAVSEL | ASSQVPRDKNSDYT | 0.59240675 |
| NTGALGL | ASSQVPRDKNSDYT | 0.5685209 |
| PPGAGGAGGAGTT | ASSQVPRDKNSDYT | 0.6276256 |
| QLVGLGI | ASSQVPRDKNSDYT | 0.58745885 |
| RFLEQVHQL | ASSQVPRDKNSDYT | 0.5745838 |
| RGVGYVLR | ASSQVPRDKNSDYT | 0.5725895 |
| SATGVFAGVI | ASSQVPRDKNSDYT | 0.59941787 |
| SLPAITLP | ASSQVPRDKNSDYT | 0.63061035 |
| STALAFTV | ASSQVPRDKNSDYT | 0.6225828 |
| SVLTVGLGK | ASSQVPRDKNSDYT | 0.65818256 |
| TALAGAGSGI | ASSQVPRDKNSDYT | 0.64442945 |
| TGLPFTGI | ASSQVPRDKNSDYT | 0.64312166 |
| VAEHVTV | ASSQVPRDKNSDYT | 0.35992384 |
| VDIVLNSL | ASSQVPRDKNSDYT | 0.48022354 |
| VLTGGGAVL | ASSQVPRDKNSDYT | 0.56718826 |
| VNVGLAYGI | ASSQVPRDKNSDYT | 0.5878806 |
| VSSAAVGAV | ASSQVPRDKNSDYT | 0.4338918 |
| VTSVYEAGV | ASSQVPRDKNSDYT | 0.6107856 |
| WVALDRGA | ASSQVPRDKNSDYT | 0.61784667 |
| YLVGSVVI | ASSQVPRDKNSDYT | 0.53078663 |
| GPPGTGKTLL | ASSQVPRDKNSDYT | 0.62754357 |
| GVGMTKF | ASSQVPRDKNSDYT | 0.54649824 |
| IGIGGSDL | ASSQVPRDKNSDYT | 0.5262515 |
| IGTLLAEL | ASSQVPRDKNSDYT | 0.538394 |
| IYTSGTT | ASSQVPRDKNSDYT | 0.64919084 |
| LSFPAGV | ASSQVPRDKNSDYT | 0.476947 |
| LYTSGTT | ASSQVPRDKNSDYT | 0.62248576 |
| VGAFPFGF | ASSQVPRDKNSDYT | 0.5778891 |
| VGAGTVEF | ASSQVPRDKNSDYT | 0.6442222 |
| VGKSTLLL | ASSQVPRDKNSDYT | 0.4939665 |
| AALAHAGGGIV | ASSQVPRDKNSDYT | 0.63421595 |
| AALVAVSLL | ASSQVPRDKNSDYT | 0.57251734 |
| AAVSRAVAAL | ASSQVPRDKNSDYT | 0.6516863 |
| AGGSSGIGGF | ASSQVPRDKNSDYT | 0.58735096 |
| AGLTVANANG | ASSQVPRDKNSDYT | 0.610786 |
| AGLVGAVL | ASSQVPRDKNSDYT | 0.5634247 |

| | | |
|---|---|---|
| ALSQAFTV | ASSQVPRDKNSDYT | 0.61122966 |
| ALTFSVSL | ASSQVPRDKNSDYT | 0.46092257 |
| ALTGAALVAL | ASSQVPRDKNSDYT | 0.5888804 |
| ASFSIGAGVL | ASSQVPRDKNSDYT | 0.491786 |
| DGSVYKI | ASSQVPRDKNSDYT | 0.6058928 |
| EQLAEIV | ASSQVPRDKNSDYT | 0.63119453 |
| FAVSELTVM | ASSQVPRDKNSDYT | 0.4848364 |
| GAAPLAAGL | ASSQVPRDKNSDYT | 0.6312188 |
| GADGGDGGSGGAGG | ASSQVPRDKNSDYT | 0.5880988 |
| GFGLDTGGAGG | ASSQVPRDKNSDYT | 0.5695321 |
| GLASGGLL | ASSQVPRDKNSDYT | 0.6363077 |
| GRIAGAAVLL | ASSQVPRDKNSDYT | 0.57793975 |
| GSAGSLNTSF | ASSQVPRDKNSDYT | 0.6482345 |
| GTGQIGI | ASSQVPRDKNSDYT | 0.5277737 |
| GTLIALIL | ASSQVPRDKNSDYT | 0.5961972 |
| IAATMGFVVG | ASSQVPRDKNSDYT | 0.6338161 |
| IFAQALL | ASSQVPRDKNSDYT | 0.45360267 |
| IGAAVALGF | ASSQVPRDKNSDYT | 0.5280438 |
| IGLTPLAL | ASSQVPRDKNSDYT | 0.64136624 |
| ILGSNTSTL | ASSQVPRDKNSDYT | 0.8249103 |
| IVFSLTVL | ASSQVPRDKNSDYT | 0.81599903 |
| LASKATGF | ASSQVPRDKNSDYT | 0.7681354 |
| LASVALIL | ASSQVPRDKNSDYT | 0.74771416 |
| LFAVGLTL | ASSQVPRDKNSDYT | 0.7787399 |
| LFNGGIGTYI | ASSQVPRDKNSDYT | 0.8272662 |
| LTLGAGII | ASSQVPRDKNSDYT | 0.8323919 |
| LVAAALGDI | ASSQVPRDKNSDYT | 0.82163405 |
| NTAFVQLGI | ASSQVPRDKNSDYT | 0.7417598 |
| QLFSGAEL | ASSQVPRDKNSDYT | 0.8273691 |
| SAATFITL | ASSQVPRDKNSDYT | 0.80855465 |
| SNYGLANLGV | ASSQVPRDKNSDYT | 0.79297227 |
| SYGGAYAVM | ASSQVPRDKNSDYT | 0.8288356 |
| TTGVGGLTLG | ASSQVPRDKNSDYT | 0.8196088 |
| TTQTIYI | ASSQVPRDKNSDYT | 0.77764887 |
| VAAGAGLGAVI | ASSQVPRDKNSDYT | 0.7736896 |
| VAAVVTLL | ASSQVPRDKNSDYT | 0.81808317 |
| VGFLGGLI | ASSQVPRDKNSDYT | 0.8310906 |
| VGVIPILL | ASSQVPRDKNSDYT | 0.8294818 |
| VIFASSLI | ASSQVPRDKNSDYT | 0.8409846 |
| VKGGDGGAAAG | ASSQVPRDKNSDYT | 0.8357301 |
| VTKTFTNI | ASSQVPRDKNSDYT | 0.8245708 |
| VVAAAILL | ASSQVPRDKNSDYT | 0.4484933 |
| YAALDVEL | ASSQVPRDKNSDYT | 0.6938134 |
| LASVALLL | ASSQVPRDKNSDYT | 0.7726544 |
| LDAVHRV | ASSQVPRDKNSDYT | 0.79637104 |
| QGLSSSIF | ASSQVPRDKNSDYT | 0.46123695 |
| RGTFANIRL | ASSQVPRDKNSDYT | 0.7948595 |
| RQLWWGHRI | ASSQVPRDKNSDYT | 0.79738855 |
| YGPPGTGKTLL | ASSQVPRDKNSDYT | 0.75429946 |
| AALVGAPANVA | ASSQVPRDKNSDYT | 0.8349565 |
| AATFRRTE | ASSQVPRDKNSDYT | 0.7843766 |

| | | |
|---|---|---|
| AGALGGGI | ASSQVPRDKNSDYT | 0.754914 |
| AGGAGGAGGIGAG | ASSQVPRDKNSDYT | 0.791193 |
| AGSGALGL | ASSQVPRDKNSDYT | 0.83790195 |
| AQIADQI | ASSQVPRDKNSDYT | 0.7422253 |
| ASAVAVTYL | ASSQVPRDKNSDYT | 0.8334527 |
| ASTVLGGTGG | ASSQVPRDKNSDYT | 0.7058042 |
| AVGGEPAK | ASSQVPRDKNSDYT | 0.84030217 |
| CEQGTFGGTC | ASSQVPRDKNSDYT | 0.6971104 |
| DEGKAPAG | ASSQVPRDKNSDYT | 0.8307353 |
| DTVVFPAL | ASSQVPRDKNSDYT | 0.8019784 |
| FEGITMAG | ASSQVPRDKNSDYT | 0.8477894 |
| FPLVVQD | ASSQVPRDKNSDYT | 0.8088053 |
| GAFTTPQ | ASSQVPRDKNSDYT | 0.8355826 |
| GFGGFGTAGGVGGA | ASSQVPRDKNSDYT | 0.811525 |
| GFLAQFL | ASSQVPRDKNSDYT | 0.81397176 |
| GGAGGAGGNAGVA | ASSQVPRDKNSDYT | 0.6746887 |
| GGVGTAGAGGAGGAGG | ASSQVPRDKNSDYT | 0.81030554 |
| GNGGGGDAGNAGSGGN | ASSQVPRDKNSDYT | 0.67905843 |
| GTGGAGGAGNDAGS | ASSQVPRDKNSDYT | 0.8177356 |
| GTTNTGWFNS | ASSQVPRDKNSDYT | 0.8208655 |
| IFMGPLFTL | ASSQVPRDKNSDYT | 0.40107888 |
| ILAHAIEAI | ASSQVPRDKNSDYT | 0.8313112 |
| IRSAIREAL | ASSQVPRDKNSDYT | 0.73718536 |
| ITFAGGG | ASSQVPRDKNSDYT | 0.7561924 |
| ITGMSGHQEV | ASSQVPRDKNSDYT | 0.8374129 |
| IVAGLACTL | ASSQVPRDKNSDYT | 0.81068915 |
| IVGGGLAGA | ASSQVPRDKNSDYT | 0.8291844 |
| IVTGVLNI | ASSQVPRDKNSDYT | 0.78941274 |
| LAQQGCI | ASSQVPRDKNSDYT | 0.7982839 |
| LCLHVEN | ASSQVPRDKNSDYT | 0.81143856 |
| LDAISDGS | ASSQVPRDKNSDYT | 0.48866326 |
| LGGFAAML | ASSQVPRDKNSDYT | 0.74747276 |
| LGGKNPMI | ASSQVPRDKNSDYT | 0.8397677 |
| LGPKGVAAIL | ASSQVPRDKNSDYT | 0.58867884 |
| LMVGVSI | ASSQVPRDKNSDYT | 0.64038086 |
| LVGAAFVL | ASSQVPRDKNSDYT | 0.5017922 |
| LVLAVAPL | ASSQVPRDKNSDYT | 0.47105822 |
| MAAIAVIDAA | ASSQVPRDKNSDYT | 0.52430624 |
| NNGINAELQ | ASSQVPRDKNSDYT | 0.6337943 |
| RGEPLVLVHG | ASSQVPRDKNSDYT | 0.63462555 |
| RGGAGGAAGLI | ASSQVPRDKNSDYT | 0.6060767 |
| RSYRMHPAV | ASSQVPRDKNSDYT | 0.4516042 |
| SAAHLLAGQLA | ASSQVPRDKNSDYT | 0.54245615 |
| SAAVGSY | ASSQVPRDKNSDYT | 0.5243488 |
| SPHAVLHRI | ASSQVPRDKNSDYT | 0.49647865 |
| SQIAKVL | ASSQVPRDKNSDYT | 0.53729343 |
| TALGHLVP | ASSQVPRDKNSDYT | 0.56834006 |
| TAPTTSGAA | ASSQVPRDKNSDYT | 0.48778915 |
| TDGDDGNGGSPGT | ASSQVPRDKNSDYT | 0.4654743 |
| TDGLGEN | ASSQVPRDKNSDYT | 0.5259519 |
| TGGPPLPGQA | ASSQVPRDKNSDYT | 0.6065302 |

| | | |
|---|---|---|
| TIPAITI | ASSQVPRDKNSDYT | 0.5897811 |
| TIPGFTI | ASSQVPRDKNSDYT | 0.6492791 |
| TSIASGALA | ASSQVPRDKNSDYT | 0.6228831 |
| TVTLPGAAI | ASSQVPRDKNSDYT | 0.5971241 |
| VASGLSAALP | ASSQVPRDKNSDYT | 0.32406592 |
| VGAGVIGI | ASSQVPRDKNSDYT | 0.38876235 |
| VGPKRFGV | ASSQVPRDKNSDYT | 0.46053436 |
| VNVAFPD | ASSQVPRDKNSDYT | 0.53392196 |
| VSMTNSGVS | ASSQVPRDKNSDYT | 0.3467606 |
| VTDVLGDLSV | ASSQVPRDKNSDYT | 0.59472424 |
| VVVVGAGFAGL | ASSQVPRDKNSDYT | 0.546826 |
| VYTVGTVV | ASSQVPRDKNSDYT | 0.48546478 |
| WTTFSGP | ASSQVPRDKNSDYT | 0.6161158 |
| YDELVEAF | ASSQVPRDKNSDYT | 0.4667135 |
| YGTGGAGGNGG | ASSQVPRDKNSDYT | 0.47816914 |
| YTNVMAV | ASSQVPRDKNSDYT | 0.47126603 |
| FLEQVHQL | ASSQVPRDKNSDYT | 0.61736643 |
| GLSLLAVAV | ASSQVPRDKNSDYT | 0.40295208 |
| LEQVHQL | ASSQVPRDKNSDYT | 0.6014581 |
| LNAAGAI | ASSQVPRDKNSDYT | 0.44820714 |
| PAVHASL | ASSQVPRDKNSDYT | 0.6355751 |
| TELEDDA | ASSQVPRDKNSDYT | 0.42351308 |
| TPPGPPN | ASSQVPRDKNSDYT | 0.6123442 |
| TPPPPPPP | ASSQVPRDKNSDYT | 0.5297767 |
| TQSPALI | ASSFYRENTGQLY | 0.6558275 |
| EFAAAFPQ | ASSFYRENTGQLY | 0.52695334 |
| LLSGNLDVL | ASSFYRENTGQLY | 0.55833745 |
| GPANLGNYNI | ASSFYRENTGQLY | 0.46036276 |
| TLGPTLP | ASSFYRENTGQLY | 0.62184054 |
| VYAYSSCCK | ASSFYRENTGQLY | 0.43436113 |
| APAPAPAPNI | ASSFYRENTGQLY | 0.5401916 |
| PGGPVPIP | ASSFYRENTGQLY | 0.39093783 |
| FLTVALEP | ASSFYRENTGQLY | 0.5910973 |
| SQYFGANL | ASSFYRENTGQLY | 0.60648465 |
| GGVKPAA | ASSFYRENTGQLY | 0.3662529 |
| GGTGGAGGAGGW | ASSFYRENTGQLY | 0.56239265 |
| GVGGVAGLGGAGG | ASSFYRENTGQLY | 0.55939686 |
| VLLVLATTI | ASSFYRENTGQLY | 0.44381747 |
| IGIFAAML | ASSFYRENTGQLY | 0.599332 |
| AVGDLID | ASSFYRENTGQLY | 0.496972 |
| NISPLPL | ASSFYRENTGQLY | 0.63012373 |
| VTAVFVAI | ASSFYRENTGQLY | 0.44665867 |
| SGDDNDAAL | ASSFYRENTGQLY | 0.47423527 |
| AVAALAHS | ASSFYRENTGQLY | 0.61018467 |
| GGVTVMATP | ASSFYRENTGQLY | 0.46394897 |
| AGLAANVRD | ASSFYRENTGQLY | 0.46300247 |
| TNRPDLIDPALLRPG | ASSFYRENTGQLY | 0.6255457 |
| LANLGVTAL | ASSFYRENTGQLY | 0.67566544 |
| TGERIEL | ASSFYRENTGQLY | 0.8465973 |
| NGEELAA | ASSFYRENTGQLY | 0.5187333 |
| VLSLVWG | ASSFYRENTGQLY | 0.44029933 |

| | | |
|---|---|---|
| IGYALWL | ASSFYRENTGQLY | 0.6612565 |
| LSLLLAPL | ASSFYRENTGQLY | 0.8073539 |
| TLDEALK | ASSFYRENTGQLY | 0.82590324 |
| TDGAGTAEGG | ASSFYRENTGQLY | 0.74035954 |
| NAPGGAGGNGG | ASSFYRENTGQLY | 0.38596204 |
| VSVFMGMGGNF | ASSFYRENTGQLY | 0.6182493 |
| LASYLAAL | ASSFYRENTGQLY | 0.508867 |
| LVGGTPAL | ASSFYRENTGQLY | 0.6184959 |
| VRNADVL | ASSFYRENTGQLY | 0.7293216 |
| TCSSPFLVSAM | ASSFYRENTGQLY | 0.65428233 |
| VHITDAE | ASSFYRENTGQLY | 0.42523974 |
| EYAAPSF | ASSFYRENTGQLY | 0.45854312 |
| VVDAFHGGL | ASSFYRENTGQLY | 0.73099816 |
| ASVVDPQ | ASSFYRENTGQLY | 0.77308214 |
| SYNEFGGLS | ASSFYRENTGQLY | 0.76831543 |
| DPDGSVY | ASSFYRENTGQLY | 0.83896434 |
| MVSTIHATE | ASSFYRENTGQLY | 0.79359686 |
| LAAGVADPV | ASSFYRENTGQLY | 0.76324874 |
| TKEQIAA | ASSFYRENTGQLY | 0.1405038 |
| SLASPAAL | ASSFYRENTGQLY | 0.2276568 |
| IGLVFYGV | ASSFYRENTGQLY | 0.44673103 |
| FLLDASNL | ASSFYRENTGQLY | 0.63964903 |
| YQVGNFP | ASSFYRENTGQLY | 0.21249884 |
| ESLLSQST | ASSFYRENTGQLY | 0.75474834 |
| LFTVGFAAVG | ASSFYRENTGQLY | 0.49941558 |
| ATPPTPT | ASSFYRENTGQLY | 0.41795665 |
| TDLATLG | ASSFYRENTGQLY | 0.80377704 |
| SAGGTGR | ASSFYRENTGQLY | 0.40197057 |
| GFSNSGS | ASSFYRENTGQLY | 0.44483584 |
| GGNGGSGGTSV | ASSFYRENTGQLY | 0.48380184 |
| GTSVATGGAGN | ASSFYRENTGQLY | 0.777449 |
| NGGNGGSGGTS | ASSFYRENTGQLY | 0.28282148 |
| AVVKFPH | ASSFYRENTGQLY | 0.76590544 |
| TPPPPPP | ASSFYRENTGQLY | 0.5790833 |
| AAPAAAAAPA | ASSFYRENTGQLY | 0.79933655 |
| IALNEDL | ASSFYRENTGQLY | 0.30986613 |
| IVNVSSM | ASSFYRENTGQLY | 0.79069346 |
| TGIPPAPR | ASSFYRENTGQLY | 0.67409605 |
| FDLGGGTF | ASSFYRENTGQLY | 0.8583217 |
| FDLGGGTFDVS | ASSFYRENTGQLY | 0.6053251 |
| NEPTAAA | ASSFYRENTGQLY | 0.651219 |
| NEPTAAALA | ASSFYRENTGQLY | 0.46749696 |
| GSIINISSI | ASSFYRENTGQLY | 0.82368374 |
| VAPGNAGTA | ASSFYRENTGQLY | 0.30610037 |
| TGGGASLE | ASSFYRENTGQLY | 0.58865607 |
| VAVYNHYKRI | ASSFYRENTGQLY | 0.2303389 |
| TDPAPATT | ASSFYRENTGQLY | 0.6546293 |
| TNRPDILDPALLRPG | ASSFYRENTGQLY | 0.73419535 |
| LTTTTTS | ASSFYRENTGQLY | 0.26158205 |
| PTGSGKTL | ASSFYRENTGQLY | 0.7112897 |
| TGSGKTL | ASSFYRENTGQLY | 0.7676581 |

| | | |
|---|---|---|
| TIGHVDHGK | ASSFYRENTGQLY | 0.42361033 |
| TIGHVDHGKTT | ASSFYRENTGQLY | 0.80836517 |
| GHVDHGKTT | ASSFYRENTGQLY | 0.56944364 |
| AQFEHTLL | ASSFYRENTGQLY | 0.8125135 |
| IVNQDEL | ASSFYRENTGQLY | 0.5925264 |
| TNGSQFF | ASSFYRENTGQLY | 0.57688797 |
| TGKGIVE | ASSFYRENTGQLY | 0.7365687 |
| LISNASDA | ASSFYRENTGQLY | 0.36387622 |
| AELRTLGELTKIAKAHGAQ | ASSFYRENTGQLY | 0.4734825 |
| APNAVAAAEPSW | ASSFYRENTGQLY | 0.8109169 |
| APQVAGAATAVL | ASSFYRENTGQLY | 0.68847203 |
| ARRDPAAAAAILPTDA | ASSFYRENTGQLY | 0.71409684 |
| DAAARDVLAEAGLAEYF | ASSFYRENTGQLY | 0.6383213 |
| DDQIVLSAKANLAKVEQ | ASSFYRENTGQLY | 0.5931009 |
| DELLPDTGEDSDEDGH | ASSFYRENTGQLY | 0.6455022 |
| ELPEAPANRATVPQD | ASSFYRENTGQLY | 0.709102 |
| FGQSKIERVTGIG | ASSFYRENTGQLY | 0.70718575 |
| GAGGTGSTAGGAGGAGGAGGI | ASSFYRENTGQLY | 0.6883161 |
| GHGGTGAAGGNGGTGGAGGS( | ASSFYRENTGQLY | 0.5689345 |
| GNGGNGGAGGEGGPGGAGGA( | ASSFYRENTGQLY | 0.70160896 |
| GSTAGNGGNGGNATNGTIAG( | ASSFYRENTGQLY | 0.66679144 |
| GTGGDGGNGGNGGTGG | ASSFYRENTGQLY | 0.65085566 |
| GVRRPAGIPADAPV | ASSFYRENTGQLY | 0.6967385 |
| GYRQVTDAHLLTLARR | ASSFYRENTGQLY | 0.6847089 |
| IAKMGQDGHDRG | ASSFYRENTGQLY | 0.6669854 |
| KAEPSIEIQYT | ASSFYRENTGQLY | 0.6566862 |
| KPVRIRMAADEDTVAVT | ASSFYRENTGQLY | 0.64724284 |
| MPPRIAGMRLLVIKPEP | ASSFYRENTGQLY | 0.6868458 |
| MKFYAVALFRSRGVPPTR | ASSFYRENTGQLY | 0.6688565 |
| NAGGFGILGAVAHSP | ASSFYRENTGQLY | 0.718662 |
| NEGREITLAMAEM | ASSFYRENTGQLY | 0.7055701 |
| NELVEWSHPLGHRWRWE | ASSFYRENTGQLY | 0.69105697 |
| NLGAGGTGGNGGNPG | ASSFYRENTGQLY | 0.30050528 |
| NTEWPKWETVMAADPVD | ASSFYRENTGQLY | 0.57919353 |
| PASAAPAETQP | ASSFYRENTGQLY | 0.6816454 |
| QQSSDGKNGVRAAEITGE | ASSFYRENTGQLY | 0.67396736 |
| RAPDAVTTSWWKE | ASSFYRENTGQLY | 0.47128502 |
| SGFGNVGAHTSGWFNQ | ASSFYRENTGQLY | 0.7109808 |
| SGVAGAVGQAASVGGLKV | ASSFYRENTGQLY | 0.66106164 |
| SNGGAGTGSAGGNGGT | ASSFYRENTGQLY | 0.6665572 |
| TGGAGGEGGAAG | ASSFYRENTGQLY | 0.7088126 |
| TGGDGGAAGAGGAPGHG | ASSFYRENTGQLY | 0.66450524 |
| TMPPLLAAIANAGRHGV | ASSFYRENTGQLY | 0.65950036 |
| TSSTAGNSGGA | ASSFYRENTGQLY | 0.6710334 |
| VPATVPVGILC | ASSFYRENTGQLY | 0.70986235 |
| VQRPDSGGVVLTGRIS | ASSFYRENTGQLY | 0.6100904 |
| WTPNPVPATKGVPAGLVQAT | ASSFYRENTGQLY | 0.6996299 |
| YALKRALAKFSDRFDVVII | ASSFYRENTGQLY | 0.63897985 |
| YLIGSVIV | ASSFYRENTGQLY | 0.71508765 |
| YGQLVAVID | ASSFYRENTGQLY | 0.5354142 |
| VTVGGMTLPA | ASSFYRENTGQLY | 0.69873095 |

| | | |
|---|---|---|
| VLALPAV | ASSFYRENTGQLY | 0.65714157 |
| VIPPTLNYET | ASSFYRENTGQLY | 0.7238651 |
| VGMGGMHPSSG | ASSFYRENTGQLY | 0.64639974 |
| VGLAGGAATA | ASSFYRENTGQLY | 0.6920691 |
| VGGAGGAGGSGP | ASSFYRENTGQLY | 0.6511118 |
| VFAAFMV | ASSFYRENTGQLY | 0.71610034 |
| TLYDKLR | ASSFYRENTGQLY | 0.56329185 |
| TLAVFTVL | ASSFYRENTGQLY | 0.65739584 |
| TGSVQPARV | ASSFYRENTGQLY | 0.6091161 |
| TGGGAGSGGT | ASSFYRENTGQLY | 0.68420726 |
| SHLETSL | ASSFYRENTGQLY | 0.69353354 |
| SFTTGMT | ASSFYRENTGQLY | 0.5051547 |
| QVEVAKRMGVRQPTVS | ASSFYRENTGQLY | 0.7085969 |
| QGSGGPALAM | ASSFYRENTGQLY | 0.6703083 |
| MLAPSNP | ASSFYRENTGQLY | 0.61794186 |
| LVNNAAI | ASSFYRENTGQLY | 0.7075496 |
| LTGEINTLA | ASSFYRENTGQLY | 0.6671064 |
| LSIFGPV | ASSFYRENTGQLY | 0.68819773 |
| LLALLANL | ASSFYRENTGQLY | 0.6582526 |
| LISNASDALDKLR | ASSFYRENTGQLY | 0.6734047 |
| LINEAVP | ASSFYRENTGQLY | 0.695532 |
| LGYAFSGL | ASSFYRENTGQLY | 0.5963532 |
| LDFVAVAL | ASSFYRENTGQLY | 0.6371942 |
| LAQPGAE | ASSFYRENTGQLY | 0.70658743 |
| ISNLGAVTGF | ASSFYRENTGQLY | 0.7784643 |
| HQEDPGRI | ASSFYRENTGQLY | 0.82858825 |
| GTPGEPTP | ASSFYRENTGQLY | 0.6742182 |
| GFFNSGS | ASSFYRENTGQLY | 0.54870623 |
| GAGGGGLTAGAN | ASSFYRENTGQLY | 0.7123692 |
| FVDALVNM | ASSFYRENTGQLY | 0.8444578 |
| FLTVAPEL | ASSFYRENTGQLY | 0.82514226 |
| FETLTAL | ASSFYRENTGQLY | 0.7911922 |
| DVINPIEL | ASSFYRENTGQLY | 0.5752426 |
| APVSGAPV | ASSFYRENTGQLY | 0.7453858 |
| ALVGGLAW | ASSFYRENTGQLY | 0.6883304 |
| ALAAAAATAI | ASSFYRENTGQLY | 0.64379764 |
| AFGLGLLL | ASSFYRENTGQLY | 0.8172373 |
| AFFAAASL | ASSFYRENTGQLY | 0.7952299 |
| AAAALPAFSPP | ASSFYRENTGQLY | 0.6963498 |
| VVLAPAL | ASSFYRENTGQLY | 0.7037376 |
| TIGHVDHGKTTL | ASSFYRENTGQLY | 0.7723182 |
| LISNASDALDK | ASSFYRENTGQLY | 0.72734433 |
| LISNASDAL | ASSFYRENTGQLY | 0.7069032 |
| ISNASDALDK | ASSFYRENTGQLY | 0.8647214 |
| NQAETLVYQTE | ASSFYRENTGQLY | 0.82424086 |
| TGGTGGTAGAG | ASSFYRENTGQLY | 0.83245116 |
| TGSTAGGAGGA | ASSFYRENTGQLY | 0.19706915 |
| DPLAVPVAAGPV | ASSFYRENTGQLY | 0.3524658 |
| GGTGGTPGNGGN | ASSFYRENTGQLY | 0.75960606 |
| PAGLAGVDPATL | ASSFYRENTGQLY | 0.69342583 |
| GNNLVVEAPHAGASV | ASSFYRENTGQLY | 0.46561423 |

| | | |
|---|---|---|
| KVTRIMSASTLEQPA | ASSFYRENTGQLY | 0.7490734 |
| LGFALPIALPRRTVV | ASSFYRENTGQLY | 0.67283356 |
| TRLGNLTARKLVVLV | ASSFYRENTGQLY | 0.6193712 |
| WSQQRRLIYFPSAGP | ASSFYRENTGQLY | 0.80789244 |
| GPNGNPDPMAAAVDIR | ASSFYRENTGQLY | 0.6578739 |
| TQRCVSVADLRAAASR | ASSFYRENTGQLY | 0.6555998 |
| QFVLDVAEHPVEVLTETV | ASSFYRENTGQLY | 0.6785562 |
| GGAGGTGGAGGNGGTQAGDG | ASSFYRENTGQLY | 0.82070124 |
| PLLATLRAIAVDVDAKPAQV | ASSFYRENTGQLY | 0.48382038 |
| APKFRTGGYMIDFWGVGYQV[illegible] | ASSFYRENTGQLY | 0.7665905 |
| GAGGAGGAGGAAGGPGGTGG[illegible] | ASSFYRENTGQLY | 0.6480133 |
| IGALGLLATDVAPAGSVAFG[illegible] | ASSFYRENTGQLY | 0.82018626 |
| AGGAGGIGGNGGAGGAGGVL[illegible] | ASSFYRENTGQLY | 0.5050044 |
| ADELIGLL | ASSFYRENTGQLY | 0.8102572 |
| ADRALKA | ASSFYRENTGQLY | 0.7125919 |
| AGAVGQVGGAGGA | ASSFYRENTGQLY | 0.8674048 |
| AGGGGGLGGAP | ASSFYRENTGQLY | 0.694507 |
| AGGLLYGNGGAG | ASSFYRENTGQLY | 0.76182675 |
| AGLESIGL | ASSFYRENTGQLY | 0.719833 |
| AGLKTGII | ASSFYRENTGQLY | 0.8140408 |
| ASLAALLL | ASSFYRENTGQLY | 0.49395007 |
| ASVINISSI | ASSFYRENTGQLY | 0.6971809 |
| ATPPGPA | ASSFYRENTGQLY | 0.44640034 |
| ATVASALV | ASSFYRENTGQLY | 0.7821188 |
| ATVASLIL | ASSFYRENTGQLY | 0.7758883 |
| AVNGPVLIRH | ASSFYRENTGQLY | 0.48260304 |
| DGAAGAVGI | ASSFYRENTGQLY | 0.82261086 |
| DPLLHGI | ASSFYRENTGQLY | 0.79813015 |
| ELGAALGAL | ASSFYRENTGQLY | 0.65257764 |
| ELGRTQAV | ASSFYRENTGQLY | 0.8529817 |
| FVAGGYLS | ASSFYRENTGQLY | 0.7456503 |
| GASGGAGGAGGW | ASSFYRENTGQLY | 0.83809984 |
| GGAGAGGDGGAGGI | ASSFYRENTGQLY | 0.78361696 |
| GTGHNGSQPGMGGNG | ASSFYRENTGQLY | 0.7213114 |
| GYAAASASAS | ASSFYRENTGQLY | 0.78549373 |
| HQVDVAAVQQG | ASSFYRENTGQLY | 0.6281831 |
| IGAAIAHT | ASSFYRENTGQLY | 0.65214145 |
| IGMIEVAQL | ASSFYRENTGQLY | 0.8327202 |
| IRLAPEHTSAMD | ASSFYRENTGQLY | 0.7882079 |
| IYNVSVL | ASSFYRENTGQLY | 0.6903665 |
| KQGFLGM | ASSFYRENTGQLY | 0.5064126 |
| LESVITQL | ASSFYRENTGQLY | 0.35280263 |
| LGLGALVVV | ASSFYRENTGQLY | 0.53193694 |
| LIGNGGNGGSGGI | ASSFYRENTGQLY | 0.73508453 |
| LNAINAPTL | ASSFYRENTGQLY | 0.64800555 |
| LQLLNTGV | ASSFYRENTGQLY | 0.7374679 |
| LQTSIEF | ASSFYRENTGQLY | 0.239481 |
| MSFAVSEL | ASSFYRENTGQLY | 0.5419426 |
| NTGALGL | ASSFYRENTGQLY | 0.5173855 |
| PPGAGGAGGAGTT | ASSFYRENTGQLY | 0.5036912 |
| QLVGLGI | ASSFYRENTGQLY | 0.64370775 |

| | | |
|---|---|---|
| RFLEQVHQL | ASSFYRENTGQLY | 0.23746787 |
| RGVGYVLR | ASSFYRENTGQLY | 0.4909425 |
| SATGVFAGVI | ASSFYRENTGQLY | 0.4712872 |
| SLPAITLP | ASSFYRENTGQLY | 0.5213142 |
| STALAFTV | ASSFYRENTGQLY | 0.7343876 |
| SVLTVGLGK | ASSFYRENTGQLY | 0.7142843 |
| TALAGAGSGI | ASSFYRENTGQLY | 0.77703744 |
| TGLPFTGI | ASSFYRENTGQLY | 0.77444804 |
| VAEHVTV | ASSFYRENTGQLY | 0.7419205 |
| VDIVLNSL | ASSFYRENTGQLY | 0.117276266 |
| VLTGGGAVL | ASSFYRENTGQLY | 0.2806483 |
| VNVGLAYGI | ASSFYRENTGQLY | 0.21467537 |
| VSSAAVGAV | ASSFYRENTGQLY | 0.5896192 |
| VTSVYEAGV | ASSFYRENTGQLY | 0.13975799 |
| WVALDRGA | ASSFYRENTGQLY | 0.5719429 |
| YLVGSVVI | ASSFYRENTGQLY | 0.68972725 |
| GPPGTGKTLL | ASSFYRENTGQLY | 0.23573153 |
| GVGMTKF | ASSFYRENTGQLY | 0.7383987 |
| IGIGGSDL | ASSFYRENTGQLY | 0.3951106 |
| IGTLLAEL | ASSFYRENTGQLY | 0.24268812 |
| IYTSGTT | ASSFYRENTGQLY | 0.2566663 |
| LSFPAGV | ASSFYRENTGQLY | 0.78158927 |
| LYTSGTT | ASSFYRENTGQLY | 0.3228954 |
| VGAFPFGF | ASSFYRENTGQLY | 0.7666528 |
| VGAGTVEF | ASSFYRENTGQLY | 0.40468496 |
| VGKSTLLL | ASSFYRENTGQLY | 0.7081085 |
| AALAHAGGGIV | ASSFYRENTGQLY | 0.17277397 |
| AALVAVSLL | ASSFYRENTGQLY | 0.74259937 |
| AAVSRAVAAL | ASSFYRENTGQLY | 0.64157695 |
| AGGSSGIGGF | ASSFYRENTGQLY | 0.77794635 |
| AGLTVANANG | ASSFYRENTGQLY | 0.5627495 |
| AGLVGAVL | ASSFYRENTGQLY | 0.49740085 |
| ALSQAFTV | ASSFYRENTGQLY | 0.30345565 |
| ALTFSVSL | ASSFYRENTGQLY | 0.4780392 |
| ALTGAALVAL | ASSFYRENTGQLY | 0.17806576 |
| ASFSIGAGVL | ASSFYRENTGQLY | 0.59452957 |
| DGSVYKI | ASSFYRENTGQLY | 0.24300359 |
| EQLAEIV | ASSFYRENTGQLY | 0.5926051 |
| FAVSELTVM | ASSFYRENTGQLY | 0.76161766 |
| GAAPLAAGL | ASSFYRENTGQLY | 0.1402745 |
| GADGGDGGSGGAGG | ASSFYRENTGQLY | 0.70639217 |
| GFGLDTGGAGG | ASSFYRENTGQLY | 0.42552805 |
| GLASGGLL | ASSFYRENTGQLY | 0.21309818 |
| GRIAGAAVLL | ASSFYRENTGQLY | 0.7080564 |
| GSAGSLNTSF | ASSFYRENTGQLY | 0.20300694 |
| GTGQIGI | ASSFYRENTGQLY | 0.770465 |
| GTLIALIL | ASSFYRENTGQLY | 0.2130693 |
| IAATMGFVVG | ASSFYRENTGQLY | 0.64162123 |
| IFAQALL | ASSFYRENTGQLY | 0.71346617 |
| IGAAVALGF | ASSFYRENTGQLY | 0.11110756 |
| IGLTPLAL | ASSFYRENTGQLY | 0.19331278 |

| | | |
|---|---|---|
| ILGSNTSTL | ASSFYRENTGQLY | 0.7763268 |
| IVFSLTVL | ASSFYRENTGQLY | 0.60342604 |
| LASKATGF | ASSFYRENTGQLY | 0.70951307 |
| LASVALIL | ASSFYRENTGQLY | 0.49986488 |
| LFAVGLTL | ASSFYRENTGQLY | 0.4411685 |
| LFNGGIGTYI | ASSFYRENTGQLY | 0.51990044 |
| LTLGAGII | ASSFYRENTGQLY | 0.6651159 |
| LVAAALGDI | ASSFYRENTGQLY | 0.704748 |
| NTAFVQLGI | ASSFYRENTGQLY | 0.648046 |
| QLFSGAEL | ASSFYRENTGQLY | 0.42340094 |
| SAATFITL | ASSFYRENTGQLY | 0.6359798 |
| SNYGLANLGV | ASSFYRENTGQLY | 0.53782326 |
| SYGGAYAVM | ASSFYRENTGQLY | 0.5924134 |
| TTGVGGLTLG | ASSFYRENTGQLY | 0.59656155 |
| TTQTIYI | ASSFYRENTGQLY | 0.6106931 |
| VAAGAGLGAVI | ASSFYRENTGQLY | 0.47394246 |
| VAAVVTLL | ASSFYRENTGQLY | 0.41948834 |
| VGFLGGLI | ASSFYRENTGQLY | 0.6793401 |
| VGVIPILL | ASSFYRENTGQLY | 0.68029374 |
| VIFASSLI | ASSFYRENTGQLY | 0.6805159 |
| VKGGDGGAAAG | ASSFYRENTGQLY | 0.68719417 |
| VTKTFTNI | ASSFYRENTGQLY | 0.6776942 |
| VVAAAILL | ASSFYRENTGQLY | 0.61509997 |
| YAALDVEL | ASSFYRENTGQLY | 0.2998041 |
| LASVALLL | ASSFYRENTGQLY | 0.3601993 |
| LDAVHRV | ASSFYRENTGQLY | 0.487575 |
| QGLSSSIF | ASSFYRENTGQLY | 0.5699919 |
| RGTFANIRL | ASSFYRENTGQLY | 0.3156778 |
| RQLWWGHRI | ASSFYRENTGQLY | 0.63445735 |
| YGPPGTGKTLL | ASSFYRENTGQLY | 0.44439057 |
| AALVGAPANVA | ASSFYRENTGQLY | 0.4892969 |
| AATFRRTE | ASSFYRENTGQLY | 0.6835743 |
| AGALGGGI | ASSFYRENTGQLY | 0.48384613 |
| AGGAGGAGGIGAG | ASSFYRENTGQLY | 0.43135503 |
| AGSGALGL | ASSFYRENTGQLY | 0.41821003 |
| AQIADQI | ASSFYRENTGQLY | 0.5880758 |
| ASAVAVTYL | ASSFYRENTGQLY | 0.37142828 |
| ASTVLGGTGG | ASSFYRENTGQLY | 0.6411906 |
| AVGGEPAK | ASSFYRENTGQLY | 0.42692614 |
| CEQGTFGGTC | ASSFYRENTGQLY | 0.6856309 |
| DEGKAPAG | ASSFYRENTGQLY | 0.3855947 |
| DTVVFPAL | ASSFYRENTGQLY | 0.6637711 |
| FEGITMAG | ASSFYRENTGQLY | 0.5907662 |
| FPLVVQD | ASSFYRENTGQLY | 0.72008336 |
| GAFTTPQ | ASSFYRENTGQLY | 0.536595 |
| GFGGFGTAGGVGGA | ASSFYRENTGQLY | 0.60040367 |
| GFLAQFL | ASSFYRENTGQLY | 0.5324752 |
| GGAGGAGGNAGVA | ASSFYRENTGQLY | 0.7025075 |
| GGVGTAGAGGAGGAGG | ASSFYRENTGQLY | 0.38360202 |
| GNGGGGDAGNAGSGGN | ASSFYRENTGQLY | 0.51692665 |
| GTGGAGGAGNDAGS | ASSFYRENTGQLY | 0.3476948 |

| | | |
|---|---|---|
| GTTNTGWFNS | ASSFYRENTGQLY | 0.55993485 |
| IFMGPLFTL | ASSFYRENTGQLY | 0.6364841 |
| ILAHAIEAI | ASSFYRENTGQLY | 0.31780913 |
| IRSAIREAL | ASSFYRENTGQLY | 0.5791849 |
| ITFAGGG | ASSFYRENTGQLY | 0.52316284 |
| ITGMSGHQEV | ASSFYRENTGQLY | 0.4250316 |
| IVAGLACTL | ASSFYRENTGQLY | 0.67653984 |
| IVGGGLAGA | ASSFYRENTGQLY | 0.624933 |
| IVTGVLNI | ASSFYRENTGQLY | 0.67575634 |
| LAQQGCI | ASSFYRENTGQLY | 0.5246297 |
| LCLHVEN | ASSFYRENTGQLY | 0.48610526 |
| LDAISDGS | ASSFYRENTGQLY | 0.6040883 |
| LGGFAAML | ASSFYRENTGQLY | 0.60099465 |
| LGGKNPMI | ASSFYRENTGQLY | 0.4622013 |
| LGPKGVAAIL | ASSFYRENTGQLY | 0.6793232 |
| LMVGVSI | ASSFYRENTGQLY | 0.7096214 |
| LVGAAFVL | ASSFYRENTGQLY | 0.7236608 |
| LVLAVAPL | ASSFYRENTGQLY | 0.6458995 |
| MAAIAVIDAA | ASSFYRENTGQLY | 0.6240716 |
| NNGINAELQ | ASSFYRENTGQLY | 0.68638605 |
| RGEPLVLVHG | ASSFYRENTGQLY | 0.71864 |
| RGGAGGAAGLI | ASSFYRENTGQLY | 0.7294352 |
| RSYRMHPAV | ASSFYRENTGQLY | 0.71026397 |
| SAAHLLAGQLA | ASSFYRENTGQLY | 0.5877889 |
| SAAVGSY | ASSFYRENTGQLY | 0.71381515 |
| SPHAVLHRI | ASSFYRENTGQLY | 0.6882496 |
| SQIAKVL | ASSFYRENTGQLY | 0.6843306 |
| TALGHLVP | ASSFYRENTGQLY | 0.7149606 |
| TAPTTSGAA | ASSFYRENTGQLY | 0.69078565 |
| TDGDDGNGGSPGT | ASSFYRENTGQLY | 0.6578765 |
| TDGLGEN | ASSFYRENTGQLY | 0.62058794 |
| TGGPPLPGQA | ASSFYRENTGQLY | 0.7175218 |
| TIPAITI | ASSFYRENTGQLY | 0.7214596 |
| TIPGFTI | ASSFYRENTGQLY | 0.719413 |
| TSIASGALA | ASSFYRENTGQLY | 0.7308049 |
| TVTLPGAAI | ASSFYRENTGQLY | 0.71988106 |
| VASGLSAALP | ASSFYRENTGQLY | 0.7040734 |
| VGAGVIGI | ASSFYRENTGQLY | 0.3392108 |
| VGPKRFGV | ASSFYRENTGQLY | 0.5030504 |
| VNVAFPD | ASSFYRENTGQLY | 0.5974144 |
| VSMTNSGVS | ASSFYRENTGQLY | 0.6825675 |
| VTDVLGDLSV | ASSFYRENTGQLY | 0.3509218 |
| VVVVGAGFAGL | ASSFYRENTGQLY | 0.7113538 |
| VYTVGTVV | ASSFYRENTGQLY | 0.6670018 |
| WTTFSGP | ASSFYRENTGQLY | 0.6350208 |
| YDELVEAF | ASSFYRENTGQLY | 0.72290754 |
| YGTGGAGGNGG | ASSFYRENTGQLY | 0.651314 |
| YTNVMAV | ASSFYRENTGQLY | 0.58034617 |
| FLEQVHQL | ASSFYRENTGQLY | 0.6525385 |
| GLSLLAVAV | ASSFYRENTGQLY | 0.7289463 |
| LEQVHQL | ASSFYRENTGQLY | 0.54258394 |

| | | |
|---|---|---|
| LNAAGAI | ASSFYRENTGQLY | 0.7214774 |
| PAVHASL | ASSFYRENTGQLY | 0.6875005 |
| TELEDDA | ASSFYRENTGQLY | 0.7263755 |
| TPPGPPN | ASSFYRENTGQLY | 0.5062287 |
| TPPPPPPP | ASSFYRENTGQLY | 0.7168945 |
| TQSPALI | ASSQDWGGDAEQF | 0.6914587 |
| EFAAAFPQ | ASSQDWGGDAEQF | 0.7343125 |
| LLSGNLDVL | ASSQDWGGDAEQF | 0.6763225 |
| GPANLGNYNI | ASSQDWGGDAEQF | 0.710562 |
| TLGPTLP | ASSQDWGGDAEQF | 0.7024591 |
| VYAYSSCCK | ASSQDWGGDAEQF | 0.7252232 |
| APAPAPAPNI | ASSQDWGGDAEQF | 0.5491657 |
| PGGPVPIP | ASSQDWGGDAEQF | 0.684358 |
| FLTVALEP | ASSQDWGGDAEQF | 0.6426929 |
| SQYFGANL | ASSQDWGGDAEQF | 0.7008153 |
| GGVKPAA | ASSQDWGGDAEQF | 0.7082449 |
| GGTGGAGGAGGW | ASSQDWGGDAEQF | 0.3871123 |
| GVGGVAGLGGAGG | ASSQDWGGDAEQF | 0.72533834 |
| VLLVLATTI | ASSQDWGGDAEQF | 0.66144264 |
| IGIFAAML | ASSQDWGGDAEQF | 0.6029136 |
| AVGDLID | ASSQDWGGDAEQF | 0.71880984 |
| NISPLPL | ASSQDWGGDAEQF | 0.6965061 |
| VTAVFVAI | ASSQDWGGDAEQF | 0.7108933 |
| SGDDNDAAL | ASSQDWGGDAEQF | 0.6401477 |
| AVAALAHS | ASSQDWGGDAEQF | 0.6951689 |
| GGVTVMATP | ASSQDWGGDAEQF | 0.7135755 |
| AGLAANVRD | ASSQDWGGDAEQF | 0.5576404 |
| TNRPDLIDPALLRPG | ASSQDWGGDAEQF | 0.6163434 |
| LANLGVTAL | ASSQDWGGDAEQF | 0.7227732 |
| TGERIEL | ASSQDWGGDAEQF | 0.8850968 |
| NGEELAA | ASSQDWGGDAEQF | 0.8869593 |
| VLSLVWG | ASSQDWGGDAEQF | 0.82932687 |
| IGYALWL | ASSQDWGGDAEQF | 0.7738749 |
| LSLLLAPL | ASSQDWGGDAEQF | 0.8408111 |
| TLDEALK | ASSQDWGGDAEQF | 0.8928704 |
| TDGAGTAEGG | ASSQDWGGDAEQF | 0.88935554 |
| NAPGGAGGNGG | ASSQDWGGDAEQF | 0.88133657 |
| VSVFMGMGGNF | ASSQDWGGDAEQF | 0.75575256 |
| LASYLAAL | ASSQDWGGDAEQF | 0.8772957 |
| LVGGTPAL | ASSQDWGGDAEQF | 0.85245126 |
| VRNADVL | ASSQDWGGDAEQF | 0.84757376 |
| TCSSPFLVSAM | ASSQDWGGDAEQF | 0.8834747 |
| VHITDAE | ASSQDWGGDAEQF | 0.87008023 |
| EYAAPSF | ASSQDWGGDAEQF | 0.8520795 |
| VVDAFHGGL | ASSQDWGGDAEQF | 0.8566781 |
| ASVVDPQ | ASSQDWGGDAEQF | 0.8627155 |
| SYNEFGGLS | ASSQDWGGDAEQF | 0.88080657 |
| DPDGSVY | ASSQDWGGDAEQF | 0.8815242 |
| MVSTIHATE | ASSQDWGGDAEQF | 0.90179294 |
| LAAGVADPV | ASSQDWGGDAEQF | 0.8966008 |
| TKEQIAA | ASSQDWGGDAEQF | 0.8879125 |

| | | |
|---|---|---|
| SLASPAAL | ASSQDWGGDAEQF | 0.2822454 |
| IGLVFYGV | ASSQDWGGDAEQF | 0.6141459 |
| FLLDASNL | ASSQDWGGDAEQF | 0.8120856 |
| YQVGNFP | ASSQDWGGDAEQF | 0.8382994 |
| ESLLSQST | ASSQDWGGDAEQF | 0.6486273 |
| LFTVGFAAVG | ASSQDWGGDAEQF | 0.8619763 |
| ATPPTPT | ASSQDWGGDAEQF | 0.8441584 |
| TDLATLG | ASSQDWGGDAEQF | 0.7811817 |
| SAGGTGR | ASSQDWGGDAEQF | 0.8895037 |
| GFSNSGS | ASSQDWGGDAEQF | 0.84884423 |
| GGNGGSGGTSV | ASSQDWGGDAEQF | 0.8067285 |
| GTSVATGGAGN | ASSQDWGGDAEQF | 0.8262471 |
| NGGNGGSGGTS | ASSQDWGGDAEQF | 0.8919685 |
| AVVKFPH | ASSQDWGGDAEQF | 0.7716963 |
| TPPPPPP | ASSQDWGGDAEQF | 0.88662934 |
| AAPAAAAAPA | ASSQDWGGDAEQF | 0.80237377 |
| IALNEDL | ASSQDWGGDAEQF | 0.89738077 |
| IVNVSSM | ASSQDWGGDAEQF | 0.6816463 |
| TGIPPAPR | ASSQDWGGDAEQF | 0.88997793 |
| FDLGGGTF | ASSQDWGGDAEQF | 0.8602352 |
| FDLGGGTFDVS | ASSQDWGGDAEQF | 0.907742 |
| NEPTAAA | ASSQDWGGDAEQF | 0.848441 |
| NEPTAAALA | ASSQDWGGDAEQF | 0.86880845 |
| GSIINISSI | ASSQDWGGDAEQF | 0.7942451 |
| VAPGNAGTA | ASSQDWGGDAEQF | 0.88433874 |
| TGGGASLE | ASSQDWGGDAEQF | 0.6660143 |
| VAVYNHYKRI | ASSQDWGGDAEQF | 0.8649322 |
| TDPAPATT | ASSQDWGGDAEQF | 0.6704071 |
| TNRPDILDPALLRPG | ASSQDWGGDAEQF | 0.8757483 |
| LTTTTTS | ASSQDWGGDAEQF | 0.87977934 |
| PTGSGKTL | ASSQDWGGDAEQF | 0.62886184 |
| TGSGKTL | ASSQDWGGDAEQF | 0.8984678 |
| TIGHVDHGK | ASSQDWGGDAEQF | 0.8542702 |
| TIGHVDHGKTT | ASSQDWGGDAEQF | 0.8030288 |
| GHVDHGKTT | ASSQDWGGDAEQF | 0.8987242 |
| AQFEHTLL | ASSQDWGGDAEQF | 0.8423952 |
| IVNQDEL | ASSQDWGGDAEQF | 0.90144813 |
| TNGSQFF | ASSQDWGGDAEQF | 0.8546988 |
| TGKGIVE | ASSQDWGGDAEQF | 0.85187215 |
| LISNASDA | ASSQDWGGDAEQF | 0.87283665 |
| AELRTLGELTKIAKAHGAQ | ASSQDWGGDAEQF | 0.60247374 |
| APNAVAAAEPSW | ASSQDWGGDAEQF | 0.7702341 |
| APQVAGAATAVL | ASSQDWGGDAEQF | 0.9000548 |
| ARRDPAAAAAILPTDA | ASSQDWGGDAEQF | 0.83144397 |
| DAAARDVLAEAGLAEYF | ASSQDWGGDAEQF | 0.8188144 |
| DDQIVLSAKANLAKVEQ | ASSQDWGGDAEQF | 0.68899953 |
| DELLPDTGEDSDEDGH | ASSQDWGGDAEQF | 0.5719845 |
| ELPEAPANRATVPQD | ASSQDWGGDAEQF | 0.694401 |
| FGQSKIERVTGIG | ASSQDWGGDAEQF | 0.8546617 |
| GAGGTGSTAGGAGGAGGAGGI | ASSQDWGGDAEQF | 0.8433417 |
| GHGGTGAAGGNGGTGGAGGS( | ASSQDWGGDAEQF | 0.81131697 |

| | | |
|---|---|---|
| GNGGNGGAGGEGGPGGAGGA( | ASSQDWGGDAEQF | 0.47394195 |
| GSTAGNGGNGGNATNGTIAG( | ASSQDWGGDAEQF | 0.8303608 |
| GTGGDGGNGGNGGTGG | ASSQDWGGDAEQF | 0.75273913 |
| GVRRPAGIPADAPV | ASSQDWGGDAEQF | 0.7717047 |
| GYRQVTDAHLLTLARR | ASSQDWGGDAEQF | 0.81051594 |
| IAKMGQDGHDRG | ASSQDWGGDAEQF | 0.74511236 |
| KAEPSIEIQYT | ASSQDWGGDAEQF | 0.7567662 |
| KPVRIRMAADEDTVAVT | ASSQDWGGDAEQF | 0.5813118 |
| MPPRIAGMRLLVIKPEP | ASSQDWGGDAEQF | 0.7819754 |
| MKFYAVALFRSRGVPPTR | ASSQDWGGDAEQF | 0.82608044 |
| NAGGFGILGAVAHSP | ASSQDWGGDAEQF | 0.8038925 |
| NEGREITLAMAEM | ASSQDWGGDAEQF | 0.8755344 |
| NELVEWSHPLGHRWRWE | ASSQDWGGDAEQF | 0.8677516 |
| NLGAGGTGGNGGNPG | ASSQDWGGDAEQF | 0.82089424 |
| NTEWPKWETVMAADPVD | ASSQDWGGDAEQF | 0.17499492 |
| PASAAPAETQP | ASSQDWGGDAEQF | 0.47385317 |
| QQSSDGKNGVRAAEITGE | ASSQDWGGDAEQF | 0.63031256 |
| RAPDAVTTSWWKE | ASSQDWGGDAEQF | 0.7627354 |
| SGFGNVGAHTSGWFNQ | ASSQDWGGDAEQF | 0.25167328 |
| SGVAGAVGQAASVGGLKV | ASSQDWGGDAEQF | 0.7906655 |
| SNGGAGTGSAGGNGGT | ASSQDWGGDAEQF | 0.68190265 |
| TGGAGGEGGAAG | ASSQDWGGDAEQF | 0.69897026 |
| TGGDGGAAGAGGAPGHG | ASSQDWGGDAEQF | 0.87356174 |
| TMPPLLAAIANAGRHGV | ASSQDWGGDAEQF | 0.7447785 |
| TSSTAGNSGGA | ASSQDWGGDAEQF | 0.6296363 |
| VPATVPVGILC | ASSQDWGGDAEQF | 0.6523504 |
| VQRPDSGGVVLTGRIS | ASSQDWGGDAEQF | 0.8669435 |
| WTPNPVPATKGVPAGLVQAT | ASSQDWGGDAEQF | 0.4627136 |
| YALKRALAKFSDRFDVVII | ASSQDWGGDAEQF | 0.8607271 |
| YLIGSVIV | ASSQDWGGDAEQF | 0.6175271 |
| YGQLVAVID | ASSQDWGGDAEQF | 0.8838934 |
| VTVGGMTLPA | ASSQDWGGDAEQF | 0.41710535 |
| VLALPAV | ASSQDWGGDAEQF | 0.847942 |
| VIPPTLNYET | ASSQDWGGDAEQF | 0.77923644 |
| VGMGGMHPSSG | ASSQDWGGDAEQF | 0.89645016 |
| VGLAGGAATA | ASSQDWGGDAEQF | 0.7321819 |
| VGGAGGAGGSGP | ASSQDWGGDAEQF | 0.79920363 |
| VFAAFMV | ASSQDWGGDAEQF | 0.6154618 |
| TLYDKLR | ASSQDWGGDAEQF | 0.83662045 |
| TLAVFTVL | ASSQDWGGDAEQF | 0.48261845 |
| TGSVQPARV | ASSQDWGGDAEQF | 0.702151 |
| TGGGAGSGGT | ASSQDWGGDAEQF | 0.5165793 |
| SHLETSL | ASSQDWGGDAEQF | 0.78242284 |
| SFTTGMT | ASSQDWGGDAEQF | 0.8271514 |
| QVEVAKRMGVRQPTVS | ASSQDWGGDAEQF | 0.20486774 |
| QGSGGPALAM | ASSQDWGGDAEQF | 0.8290626 |
| MLAPSNP | ASSQDWGGDAEQF | 0.58858585 |
| LVNNAAI | ASSQDWGGDAEQF | 0.5123261 |
| LTGEINTLA | ASSQDWGGDAEQF | 0.8477446 |
| LSIFGPV | ASSQDWGGDAEQF | 0.70874965 |
| LLALLANL | ASSQDWGGDAEQF | 0.8394878 |

| | | |
|---|---|---|
| LISNASDALDKLR | ASSQDWGGDAEQF | 0.6651174 |
| LINEAVP | ASSQDWGGDAEQF | 0.74297726 |
| LGYAFSGL | ASSQDWGGDAEQF | 0.7988929 |
| LDFVAVAL | ASSQDWGGDAEQF | 0.3607319 |
| LAQPGAE | ASSQDWGGDAEQF | 0.58204085 |
| ISNLGAVTGF | ASSQDWGGDAEQF | 0.87276995 |
| HQEDPGRI | ASSQDWGGDAEQF | 0.6159184 |
| GTPGEPTP | ASSQDWGGDAEQF | 0.69778645 |
| GFFNSGS | ASSQDWGGDAEQF | 0.5982221 |
| GAGGGGLTAGAN | ASSQDWGGDAEQF | 0.53270483 |
| FVDALVNM | ASSQDWGGDAEQF | 0.6069864 |
| FLTVAPEL | ASSQDWGGDAEQF | 0.6868813 |
| FETLTAL | ASSQDWGGDAEQF | 0.6919879 |
| DVINPIEL | ASSQDWGGDAEQF | 0.66229475 |
| APVSGAPV | ASSQDWGGDAEQF | 0.51044554 |
| ALVGGLAW | ASSQDWGGDAEQF | 0.65538156 |
| ALAAAAATAI | ASSQDWGGDAEQF | 0.5801457 |
| AFGLGLLL | ASSQDWGGDAEQF | 0.62864697 |
| AFFAAASL | ASSQDWGGDAEQF | 0.65358126 |
| AAAALPAFSPP | ASSQDWGGDAEQF | 0.63019276 |
| VVLAPAL | ASSQDWGGDAEQF | 0.6107397 |
| TIGHVDHGKTTL | ASSQDWGGDAEQF | 0.5622582 |
| LISNASDALDK | ASSQDWGGDAEQF | 0.651285 |
| LISNASDAL | ASSQDWGGDAEQF | 0.6240181 |
| ISNASDALDK | ASSQDWGGDAEQF | 0.58566797 |
| NQAETLVYQTE | ASSQDWGGDAEQF | 0.69448006 |
| TGGTGGTAGAG | ASSQDWGGDAEQF | 0.6711705 |
| TGSTAGGAGGA | ASSQDWGGDAEQF | 0.6569737 |
| DPLAVPVAAGPV | ASSQDWGGDAEQF | 0.31387812 |
| GGTGGTPGNGGN | ASSQDWGGDAEQF | 0.39750692 |
| PAGLAGVDPATL | ASSQDWGGDAEQF | 0.606987 |
| GNNLVVEAPHAGASV | ASSQDWGGDAEQF | 0.58605146 |
| KVTRIMSASTLEQPA | ASSQDWGGDAEQF | 0.39640006 |
| LGFALPIALPRRTVV | ASSQDWGGDAEQF | 0.66674936 |
| TRLGNLTARKLVVLV | ASSQDWGGDAEQF | 0.5991474 |
| WSQQRRLIYFPSAGP | ASSQDWGGDAEQF | 0.59585994 |
| GPNGNPDPMAAAVDIR | ASSQDWGGDAEQF | 0.6720736 |
| TQRCVSVADLRAAASR | ASSQDWGGDAEQF | 0.6119386 |
| QFVLDVAEHPVEVLTETV | ASSQDWGGDAEQF | 0.575801 |
| GGAGGTGGAGGNGGTQAGDG | ASSQDWGGDAEQF | 0.5533545 |
| PLLATLRAIAVDVDAKPAQV | ASSQDWGGDAEQF | 0.67673945 |
| APKFRTGGYMIDFWGVGYQV | ASSQDWGGDAEQF | 0.45322287 |
| GAGGAGGAGGAAGGPGGTGG | ASSQDWGGDAEQF | 0.6634927 |
| IGALGLLATDVAPAGSVAFG | ASSQDWGGDAEQF | 0.53205985 |
| AGGAGGIGGNGGAGGAGGVL | ASSQDWGGDAEQF | 0.6761796 |
| ADELIGLL | ASSQDWGGDAEQF | 0.46777117 |
| ADRALKA | ASSQDWGGDAEQF | 0.66660315 |
| AGAVGQVGGAGGA | ASSQDWGGDAEQF | 0.622186 |
| AGGGGGLGGAP | ASSQDWGGDAEQF | 0.7092668 |
| AGGLLYGNGGAG | ASSQDWGGDAEQF | 0.5691693 |
| AGLESIGL | ASSQDWGGDAEQF | 0.6142029 |

| | | |
|---|---|---|
| AGLKTGII | ASSQDWGGDAEQF | 0.5753541 |
| ASLAALLL | ASSQDWGGDAEQF | 0.6888971 |
| ASVINISSI | ASSQDWGGDAEQF | 0.462427 |
| ATPPGPA | ASSQDWGGDAEQF | 0.5893919 |
| ATVASALV | ASSQDWGGDAEQF | 0.51344544 |
| ATVASLIL | ASSQDWGGDAEQF | 0.664577 |
| AVNGPVLIRH | ASSQDWGGDAEQF | 0.6512657 |
| DGAAGAVGI | ASSQDWGGDAEQF | 0.4485189 |
| DPLLHGI | ASSQDWGGDAEQF | 0.6748232 |
| ELGAALGAL | ASSQDWGGDAEQF | 0.6380235 |
| ELGRTQAV | ASSQDWGGDAEQF | 0.55388176 |
| FVAGGYLS | ASSQDWGGDAEQF | 0.6873622 |
| GASGGAGGAGGW | ASSQDWGGDAEQF | 0.62480414 |
| GGAGAGGDGGAGGI | ASSQDWGGDAEQF | 0.6686701 |
| GTGHNGSQPGMGGNG | ASSQDWGGDAEQF | 0.62999064 |
| GYAAASASAS | ASSQDWGGDAEQF | 0.5991941 |
| HQVDVAAVQQG | ASSQDWGGDAEQF | 0.6672595 |
| IGAAIAHT | ASSQDWGGDAEQF | 0.61540484 |
| IGMIEVAQL | ASSQDWGGDAEQF | 0.54082 |
| IRLAPEHTSAMD | ASSQDWGGDAEQF | 0.6824316 |
| IYNVSVL | ASSQDWGGDAEQF | 0.7406752 |
| KQGFLGM | ASSQDWGGDAEQF | 0.7431843 |
| LESVITQL | ASSQDWGGDAEQF | 0.6625484 |
| LGLGALVVV | ASSQDWGGDAEQF | 0.6531613 |
| LIGNGGNGGSGGI | ASSQDWGGDAEQF | 0.69479966 |
| LNAINAPTL | ASSQDWGGDAEQF | 0.7448077 |
| LQLLNTGV | ASSQDWGGDAEQF | 0.7673827 |
| LQTSIEF | ASSQDWGGDAEQF | 0.74955535 |
| MSFAVSEL | ASSQDWGGDAEQF | 0.68903655 |
| NTGALGL | ASSQDWGGDAEQF | 0.75604993 |
| PPGAGGAGGAGTT | ASSQDWGGDAEQF | 0.73123294 |
| QLVGLGI | ASSQDWGGDAEQF | 0.72857714 |
| RFLEQVHQL | ASSQDWGGDAEQF | 0.74609536 |
| RGVGYVLR | ASSQDWGGDAEQF | 0.73423773 |
| SATGVFAGVI | ASSQDWGGDAEQF | 0.6893362 |
| SLPAITLP | ASSQDWGGDAEQF | 0.69475424 |
| STALAFTV | ASSQDWGGDAEQF | 0.743945 |
| SVLTVGLGK | ASSQDWGGDAEQF | 0.7524338 |
| TALAGAGSGI | ASSQDWGGDAEQF | 0.74636054 |
| TGLPFTGI | ASSQDWGGDAEQF | 0.77409583 |
| VAEHVTV | ASSQDWGGDAEQF | 0.75569975 |
| VDIVLNSL | ASSQDWGGDAEQF | 0.7448987 |
| VLTGGGAVL | ASSQDWGGDAEQF | 0.4949079 |
| VNVGLAYGI | ASSQDWGGDAEQF | 0.64222395 |
| VSSAAVGAV | ASSQDWGGDAEQF | 0.697323 |
| VTSVYEAGV | ASSQDWGGDAEQF | 0.711596 |
| WVALDRGA | ASSQDWGGDAEQF | 0.44323885 |
| YLVGSVVI | ASSQDWGGDAEQF | 0.7476826 |
| GPPGTGKTLL | ASSQDWGGDAEQF | 0.7327773 |
| GVGMTKF | ASSQDWGGDAEQF | 0.66898465 |
| IGIGGSDL | ASSQDWGGDAEQF | 0.75731003 |

| | | |
|---|---|---|
| IGTLLAEL | ASSQDWGGDAEQF | 0.70301944 |
| IYTSGTT | ASSQDWGGDAEQF | 0.6821174 |
| LSFPAGV | ASSQDWGGDAEQF | 0.7175421 |
| LYTSGTT | ASSQDWGGDAEQF | 0.77117586 |
| VGAFPFGF | ASSQDWGGDAEQF | 0.66871727 |
| VGAGTVEF | ASSQDWGGDAEQF | 0.76198655 |
| VGKSTLLL | ASSQDWGGDAEQF | 0.7250344 |
| AALAHAGGGIV | ASSQDWGGDAEQF | 0.7651269 |
| AALVAVSLL | ASSQDWGGDAEQF | 0.63974047 |
| AAVSRAVAAL | ASSQDWGGDAEQF | 0.7597969 |
| AGGSSGIGGF | ASSQDWGGDAEQF | 0.7193315 |
| AGLTVANANG | ASSQDWGGDAEQF | 0.7737818 |
| AGLVGAVL | ASSQDWGGDAEQF | 0.71886563 |
| ALSQAFTV | ASSQDWGGDAEQF | 0.76317215 |
| ALTFSVSL | ASSQDWGGDAEQF | 0.73998547 |
| ALTGAALVAL | ASSQDWGGDAEQF | 0.75517714 |
| ASFSIGAGVL | ASSQDWGGDAEQF | 0.61202025 |
| DGSVYKI | ASSQDWGGDAEQF | 0.7096624 |
| EQLAEIV | ASSQDWGGDAEQF | 0.6851551 |
| FAVSELTVM | ASSQDWGGDAEQF | 0.7359307 |
| GAAPLAAGL | ASSQDWGGDAEQF | 0.75410867 |
| GADGGDGGSGGAGG | ASSQDWGGDAEQF | 0.3767089 |
| GFGLDTGGAGG | ASSQDWGGDAEQF | 0.76610774 |
| GLASGGLL | ASSQDWGGDAEQF | 0.6125766 |
| GRIAGAAVLL | ASSQDWGGDAEQF | 0.67227405 |
| GSAGSLNTSF | ASSQDWGGDAEQF | 0.7562154 |
| GTGQIGI | ASSQDWGGDAEQF | 0.7275351 |
| GTLIALIL | ASSQDWGGDAEQF | 0.7451066 |
| IAATMGFVVG | ASSQDWGGDAEQF | 0.706902 |
| IFAQALL | ASSQDWGGDAEQF | 0.72846127 |
| IGAAVALGF | ASSQDWGGDAEQF | 0.74354553 |
| IGLTPLAL | ASSQDWGGDAEQF | 0.4677795 |
| ILGSNTSTL | ASSQDWGGDAEQF | 0.64319146 |
| IVFSLTVL | ASSQDWGGDAEQF | 0.7696336 |
| LASKATGF | ASSQDWGGDAEQF | 0.25932705 |
| LASVALIL | ASSQDWGGDAEQF | 0.40798962 |
| LFAVGLTL | ASSQDWGGDAEQF | 0.25260288 |
| LFNGGIGTYI | ASSQDWGGDAEQF | 0.24816576 |
| LTLGAGII | ASSQDWGGDAEQF | 0.26213646 |
| LVAAALGDI | ASSQDWGGDAEQF | 0.26550692 |
| NTAFVQLGI | ASSQDWGGDAEQF | 0.53042793 |
| QLFSGAEL | ASSQDWGGDAEQF | 0.25299543 |
| SAATFITL | ASSQDWGGDAEQF | 0.24894655 |
| SNYGLANLGV | ASSQDWGGDAEQF | 0.34671307 |
| SYGGAYAVM | ASSQDWGGDAEQF | 0.23946068 |
| TTGVGGLTLG | ASSQDWGGDAEQF | 0.24331231 |
| TTQTIYI | ASSQDWGGDAEQF | 0.24415958 |
| VAAGAGLGAVI | ASSQDWGGDAEQF | 0.2581008 |
| VAAVVTLL | ASSQDWGGDAEQF | 0.22860658 |
| VGFLGGLI | ASSQDWGGDAEQF | 0.2479089 |
| VGVIPILL | ASSQDWGGDAEQF | 0.23828813 |

| | | |
|---|---|---|
| VIFASSLI | ASSQDWGGDAEQF | 0.31310433 |
| VKGGDGGAAAG | ASSQDWGGDAEQF | 0.27400354 |
| VTKTFTNI | ASSQDWGGDAEQF | 0.2829488 |
| VVAAAILL | ASSQDWGGDAEQF | 0.27589008 |
| YAALDVEL | ASSQDWGGDAEQF | 0.23897888 |
| LASVALLL | ASSQDWGGDAEQF | 0.21956383 |
| LDAVHRV | ASSQDWGGDAEQF | 0.23252739 |
| QGLSSSIF | ASSQDWGGDAEQF | 0.23453963 |
| RGTFANIRL | ASSQDWGGDAEQF | 0.26221848 |
| RQLWWGHRI | ASSQDWGGDAEQF | 0.21257424 |
| YGPPGTGKTLL | ASSQDWGGDAEQF | 0.28920224 |
| AALVGAPANVA | ASSQDWGGDAEQF | 0.24357569 |
| AATFRRTE | ASSQDWGGDAEQF | 0.24269791 |
| AGALGGGI | ASSQDWGGDAEQF | 0.2784728 |
| AGGAGGAGGIGAG | ASSQDWGGDAEQF | 0.23291993 |
| AGSGALGL | ASSQDWGGDAEQF | 0.23718157 |
| AQIADQI | ASSQDWGGDAEQF | 0.24566272 |
| ASAVAVTYL | ASSQDWGGDAEQF | 0.31780788 |
| ASTVLGGTGG | ASSQDWGGDAEQF | 0.23270436 |
| AVGGEPAK | ASSQDWGGDAEQF | 0.38801846 |
| CEQGTFGGTC | ASSQDWGGDAEQF | 0.22320601 |
| DEGKAPAG | ASSQDWGGDAEQF | 0.305086 |
| DTVVFPAL | ASSQDWGGDAEQF | 0.23641373 |
| FEGITMAG | ASSQDWGGDAEQF | 0.25557396 |
| FPLVVQD | ASSQDWGGDAEQF | 0.29265445 |
| GAFTTPQ | ASSQDWGGDAEQF | 0.3171633 |
| GFGGFGTAGGVGGA | ASSQDWGGDAEQF | 0.2562645 |
| GFLAQFL | ASSQDWGGDAEQF | 0.29597867 |
| GGAGGAGGNAGVA | ASSQDWGGDAEQF | 0.24819562 |
| GGVGTAGAGGAGGAGG | ASSQDWGGDAEQF | 0.5443058 |
| GNGGGGDAGNAGSGGN | ASSQDWGGDAEQF | 0.24604535 |
| GTGGAGGAGNDAGS | ASSQDWGGDAEQF | 0.27753648 |
| GTTNTGWFNS | ASSQDWGGDAEQF | 0.22584133 |
| IFMGPLFTL | ASSQDWGGDAEQF | 0.31706256 |
| ILAHAIEAI | ASSQDWGGDAEQF | 0.2756744 |
| IRSAIREAL | ASSQDWGGDAEQF | 0.21537514 |
| ITFAGGG | ASSQDWGGDAEQF | 0.2602666 |
| ITGMSGHQEV | ASSQDWGGDAEQF | 0.22046724 |
| IVAGLACTL | ASSQDWGGDAEQF | 0.22702263 |
| IVGGGLAGA | ASSQDWGGDAEQF | 0.24964964 |
| IVTGVLNI | ASSQDWGGDAEQF | 0.25520244 |
| LAQQGCI | ASSQDWGGDAEQF | 0.23966138 |
| LCLHVEN | ASSQDWGGDAEQF | 0.22474863 |
| LDAISDGS | ASSQDWGGDAEQF | 0.23194501 |
| LGGFAAML | ASSQDWGGDAEQF | 0.34618783 |
| LGGKNPMI | ASSQDWGGDAEQF | 0.2261756 |
| LGPKGVAAIL | ASSQDWGGDAEQF | 0.2287011 |
| LMVGVSI | ASSQDWGGDAEQF | 0.2805368 |
| LVGAAFVL | ASSQDWGGDAEQF | 0.8431548 |
| LVLAVAPL | ASSQDWGGDAEQF | 0.87345904 |
| MAAIAVIDAA | ASSQDWGGDAEQF | 0.754871 |

| | | |
|---|---|---|
| NNGINAELQ | ASSQDWGGDAEQF | 0.6151357 |
| RGEPLVLVHG | ASSQDWGGDAEQF | 0.76107633 |
| RGGAGGAAGLI | ASSQDWGGDAEQF | 0.86446726 |
| RSYRMHPAV | ASSQDWGGDAEQF | 0.8779261 |
| SAAHLLAGQLA | ASSQDWGGDAEQF | 0.84814894 |
| SAAVGSY | ASSQDWGGDAEQF | 0.4908029 |
| SPHAVLHRI | ASSQDWGGDAEQF | 0.83830285 |
| SQIAKVL | ASSQDWGGDAEQF | 0.7619724 |
| TALGHLVP | ASSQDWGGDAEQF | 0.8190006 |
| TAPTTSGAA | ASSQDWGGDAEQF | 0.8408486 |
| TDGDDGNGGSPGT | ASSQDWGGDAEQF | 0.79068315 |
| TDGLGEN | ASSQDWGGDAEQF | 0.7909896 |
| TGGPPLPGQA | ASSQDWGGDAEQF | 0.66845363 |
| TIPAITI | ASSQDWGGDAEQF | 0.7951543 |
| TIPGFTI | ASSQDWGGDAEQF | 0.836195 |
| TSIASGALA | ASSQDWGGDAEQF | 0.78157794 |
| TVTLPGAAI | ASSQDWGGDAEQF | 0.8813574 |
| VASGLSAALP | ASSQDWGGDAEQF | 0.875327 |
| VGAGVIGI | ASSQDWGGDAEQF | 0.8468901 |
| VGPKRFGV | ASSQDWGGDAEQF | 0.1476251 |
| VNVAFPD | ASSQDWGGDAEQF | 0.47401303 |
| VSMTNSGVS | ASSQDWGGDAEQF | 0.7106314 |
| VTDVLGDLSV | ASSQDWGGDAEQF | 0.7916452 |
| VVVVGAGFAGL | ASSQDWGGDAEQF | 0.23375778 |
| VYTVGTVV | ASSQDWGGDAEQF | 0.83988476 |
| WTTFSGP | ASSQDWGGDAEQF | 0.74054694 |
| YDELVEAF | ASSQDWGGDAEQF | 0.73977184 |
| YGTGGAGGNGG | ASSQDWGGDAEQF | 0.8759749 |
| YTNVMAV | ASSQDWGGDAEQF | 0.80322504 |
| FLEQVHQL | ASSQDWGGDAEQF | 0.6412506 |
| GLSLLAVAV | ASSQDWGGDAEQF | 0.6139994 |
| LEQVHQL | ASSQDWGGDAEQF | 0.87558484 |
| LNAAGAI | ASSQDWGGDAEQF | 0.51800615 |
| PAVHASL | ASSQDWGGDAEQF | 0.8696373 |
| TELEDDA | ASSQDWGGDAEQF | 0.6768339 |
| TPPGPPN | ASSQDWGGDAEQF | 0.88035405 |
| TPPPPPPP | ASSQDWGGDAEQF | 0.397713 |
| TQSPALI | ASSRGAINTGQLY | 0.8627932 |
| EFAAAFPQ | ASSRGAINTGQLY | 0.80798846 |
| LLSGNLDVL | ASSRGAINTGQLY | 0.8996291 |
| GPANLGNYNI | ASSRGAINTGQLY | 0.69242084 |
| TLGPTLP | ASSRGAINTGQLY | 0.785167 |
| VYAYSSCCK | ASSRGAINTGQLY | 0.6564686 |
| APAPAPAPNI | ASSRGAINTGQLY | 0.87278676 |
| PGGPVPIP | ASSRGAINTGQLY | 0.48465583 |
| FLTVALEP | ASSRGAINTGQLY | 0.7165189 |
| SQYFGANL | ASSRGAINTGQLY | 0.5445781 |
| GGVKPAA | ASSRGAINTGQLY | 0.81119204 |
| GGTGGAGGAGGW | ASSRGAINTGQLY | 0.8562032 |
| GVGGVAGLGGAGG | ASSRGAINTGQLY | 0.27565905 |
| VLLVLATTI | ASSRGAINTGQLY | 0.8682285 |

| | | |
|---|---|---|
| IGIFAAML | ASSRGAINTGQLY | 0.7392535 |
| AVGDLID | ASSRGAINTGQLY | 0.5588523 |
| NISPLPL | ASSRGAINTGQLY | 0.87259334 |
| VTAVFVAI | ASSRGAINTGQLY | 0.700853 |
| SGDDNDAAL | ASSRGAINTGQLY | 0.86320317 |
| AVAALAHS | ASSRGAINTGQLY | 0.68429947 |
| GGVTVMATP | ASSRGAINTGQLY | 0.7491404 |
| AGLAANVRD | ASSRGAINTGQLY | 0.8451032 |
| TNRPDLIDPALLRPG | ASSRGAINTGQLY | 0.39584303 |
| LANLGVTAL | ASSRGAINTGQLY | 0.480688 |
| TGERIEL | ASSRGAINTGQLY | 0.8864334 |
| NGEELAA | ASSRGAINTGQLY | 0.8226683 |
| VLSLVWG | ASSRGAINTGQLY | 0.8495092 |
| IGYALWL | ASSRGAINTGQLY | 0.6710056 |
| LSLLLAPL | ASSRGAINTGQLY | 0.6584012 |
| TLDEALK | ASSRGAINTGQLY | 0.7691241 |
| TDGAGTAEGG | ASSRGAINTGQLY | 0.85579705 |
| NAPGGAGGNGG | ASSRGAINTGQLY | 0.8566751 |
| VSVFMGMGGNF | ASSRGAINTGQLY | 0.8300174 |
| LASYLAAL | ASSRGAINTGQLY | 0.73278314 |
| LVGGTPAL | ASSRGAINTGQLY | 0.8075385 |
| VRNADVL | ASSRGAINTGQLY | 0.78511786 |
| TCSSPFLVSAM | ASSRGAINTGQLY | 0.75503886 |
| VHITDAE | ASSRGAINTGQLY | 0.8481855 |
| EYAAPSF | ASSRGAINTGQLY | 0.8201352 |
| VVDAFHGGL | ASSRGAINTGQLY | 0.7403467 |
| ASVVDPQ | ASSRGAINTGQLY | 0.79184365 |
| SYNEFGGLS | ASSRGAINTGQLY | 0.8067976 |
| DPDGSVY | ASSRGAINTGQLY | 0.84151185 |
| MVSTIHATE | ASSRGAINTGQLY | 0.82979614 |
| LAAGVADPV | ASSRGAINTGQLY | 0.88217866 |
| TKEQIAA | ASSRGAINTGQLY | 0.85014004 |
| SLASPAAL | ASSRGAINTGQLY | 0.82937324 |
| IGLVFYGV | ASSRGAINTGQLY | 0.33046794 |
| FLLDASNL | ASSRGAINTGQLY | 0.59862703 |
| YQVGNFP | ASSRGAINTGQLY | 0.7851713 |
| ESLLSQST | ASSRGAINTGQLY | 0.7926674 |
| LFTVGFAAVG | ASSRGAINTGQLY | 0.5005194 |
| ATPPTPT | ASSRGAINTGQLY | 0.8440013 |
| TDLATLG | ASSRGAINTGQLY | 0.77390695 |
| SAGGTGR | ASSRGAINTGQLY | 0.6856588 |
| GFSNSGS | ASSRGAINTGQLY | 0.84750056 |
| GGNGGSGGTSV | ASSRGAINTGQLY | 0.7364359 |
| GTSVATGGAGN | ASSRGAINTGQLY | 0.75383675 |
| NGGNGGSGGTS | ASSRGAINTGQLY | 0.8009334 |
| AVVKFPH | ASSRGAINTGQLY | 0.8774014 |
| TPPPPPP | ASSRGAINTGQLY | 0.68977815 |
| AAPAAAAAPA | ASSRGAINTGQLY | 0.8473596 |
| IALNEDL | ASSRGAINTGQLY | 0.8018497 |
| IVNVSSM | ASSRGAINTGQLY | 0.86252046 |
| TGIPPAPR | ASSRGAINTGQLY | 0.62595874 |

| | | |
|---|---|---|
| FDLGGGTF | ASSRGAINTGQLY | 0.8428493 |
| FDLGGGTFDVS | ASSRGAINTGQLY | 0.77910435 |
| NEPTAAA | ASSRGAINTGQLY | 0.88225746 |
| NEPTAAALA | ASSRGAINTGQLY | 0.7871913 |
| GSIINISSI | ASSRGAINTGQLY | 0.84579176 |
| VAPGNAGTA | ASSRGAINTGQLY | 0.8037914 |
| TGGGASLE | ASSRGAINTGQLY | 0.8245969 |
| VAVYNHYKRI | ASSRGAINTGQLY | 0.58573264 |
| TDPAPATT | ASSRGAINTGQLY | 0.78947014 |
| TNRPDILDPALLRPG | ASSRGAINTGQLY | 0.61806846 |
| LTTTTTS | ASSRGAINTGQLY | 0.81763303 |
| PTGSGKTL | ASSRGAINTGQLY | 0.8283209 |
| TGSGKTL | ASSRGAINTGQLY | 0.58954793 |
| TIGHVDHGK | ASSRGAINTGQLY | 0.8429936 |
| TIGHVDHGKTT | ASSRGAINTGQLY | 0.75404537 |
| GHVDHGKTT | ASSRGAINTGQLY | 0.73904 |
| AQFEHTLL | ASSRGAINTGQLY | 0.8564247 |
| IVNQDEL | ASSRGAINTGQLY | 0.74900794 |
| TNGSQFF | ASSRGAINTGQLY | 0.8443177 |
| TGKGIVE | ASSRGAINTGQLY | 0.7585426 |
| LISNASDA | ASSRGAINTGQLY | 0.7692782 |
| AELRTLGELTKIAKAHGAQ | ASSRGAINTGQLY | 0.83285594 |
| APNAVAAAEPSW | ASSRGAINTGQLY | 0.5375984 |
| APQVAGAATAVL | ASSRGAINTGQLY | 0.72166383 |
| ARRDPAAAAAILPTDA | ASSRGAINTGQLY | 0.8617036 |
| DAAARDVLAEAGLAEYF | ASSRGAINTGQLY | 0.6203607 |
| DDQIVLSAKANLAKVEQ | ASSRGAINTGQLY | 0.69547987 |
| DELLPDTGEDSDEDGH | ASSRGAINTGQLY | 0.53664684 |
| ELPEAPANRATVPQD | ASSRGAINTGQLY | 0.4570393 |
| FGQSKIERVTGIG | ASSRGAINTGQLY | 0.54543734 |
| GAGGTGSTAGGAGGAGGAGGI | ASSRGAINTGQLY | 0.57248664 |
| GHGGTGAAGGNGGTGGAGGS( | ASSRGAINTGQLY | 0.70151114 |
| GNGGNGGAGGEGGPGGAGGA( | ASSRGAINTGQLY | 0.62014 |
| GSTAGNGGNGGNATNGTIAG( | ASSRGAINTGQLY | 0.38416177 |
| GTGGDGGNGGNGGTGG | ASSRGAINTGQLY | 0.64286315 |
| GVRRPAGIPADAPV | ASSRGAINTGQLY | 0.5648594 |
| GYRQVTDAHLLTLARR | ASSRGAINTGQLY | 0.5672697 |
| IAKMGQDGHDRG | ASSRGAINTGQLY | 0.49952158 |
| KAEPSIEIQYT | ASSRGAINTGQLY | 0.59109116 |
| KPVRIRMAADEDTVAVT | ASSRGAINTGQLY | 0.56988156 |
| MPPRIAGMRLLVIKPEP | ASSRGAINTGQLY | 0.4096276 |
| MKFYAVALFRSRGVPPTR | ASSRGAINTGQLY | 0.5422611 |
| NAGGFGILGAVAHSP | ASSRGAINTGQLY | 0.6048736 |
| NEGREITLAMAEM | ASSRGAINTGQLY | 0.5779394 |
| NELVEWSHPLGHRWRWE | ASSRGAINTGQLY | 0.5939951 |
| NLGAGGTGGNGGNPG | ASSRGAINTGQLY | 0.6647936 |
| NTEWPKWETVMAADPVD | ASSRGAINTGQLY | 0.59127665 |
| PASAAPAETQP | ASSRGAINTGQLY | 0.27556202 |
| QQSSDGKNGVRAAEITGE | ASSRGAINTGQLY | 0.323915 |
| RAPDAVTTSWWKE | ASSRGAINTGQLY | 0.46511376 |
| SGFGNVGAHTSGWFNQ | ASSRGAINTGQLY | 0.48654264 |

| | | |
|---|---|---|
| SGVAGAVGQAASVGGLKV | ASSRGAINTGQLY | 0.2834879 |
| SNGGAGTGSAGGNGGT | ASSRGAINTGQLY | 0.51793385 |
| TGGAGGEGGAAG | ASSRGAINTGQLY | 0.5244149 |
| TGGDGGAAGAGGAPGHG | ASSRGAINTGQLY | 0.5217835 |
| TMPPLLAAIANAGRHGV | ASSRGAINTGQLY | 0.6488581 |
| TSSTAGNSGGA | ASSRGAINTGQLY | 0.56223214 |
| VPATVPVGILC | ASSRGAINTGQLY | 0.39219743 |
| VQRPDSGGVVLTGRIS | ASSRGAINTGQLY | 0.3602021 |
| WTPNPVPATKGVPAGLVQAT | ASSRGAINTGQLY | 0.5606248 |
| YALKRALAKFSDRFDVVII | ASSRGAINTGQLY | 0.35398245 |
| YLIGSVIV | ASSRGAINTGQLY | 0.64076763 |
| YGQLVAVID | ASSRGAINTGQLY | 0.32380283 |
| VTVGGMTLPA | ASSRGAINTGQLY | 0.6119253 |
| VLALPAV | ASSRGAINTGQLY | 0.39026973 |
| VIPPTLNYET | ASSRGAINTGQLY | 0.61389846 |
| VGMGGMHPSSG | ASSRGAINTGQLY | 0.5933385 |
| VGLAGGAATA | ASSRGAINTGQLY | 0.688991 |
| VGGAGGAGGSGP | ASSRGAINTGQLY | 0.51794434 |
| VFAAFMV | ASSRGAINTGQLY | 0.55356336 |
| TLYDKLR | ASSRGAINTGQLY | 0.38766682 |
| TLAVFTVL | ASSRGAINTGQLY | 0.6960993 |
| TGSVQPARV | ASSRGAINTGQLY | 0.39885792 |
| TGGGAGSGGT | ASSRGAINTGQLY | 0.52233964 |
| SHLETSL | ASSRGAINTGQLY | 0.329869 |
| SFTTGMT | ASSRGAINTGQLY | 0.530061 |
| QVEVAKRMGVRQPTVS | ASSRGAINTGQLY | 0.6427331 |
| QGSGGPALAM | ASSRGAINTGQLY | 0.31394553 |
| MLAPSNP | ASSRGAINTGQLY | 0.61021006 |
| LVNNAAI | ASSRGAINTGQLY | 0.46274874 |
| LTGEINTLA | ASSRGAINTGQLY | 0.42809194 |
| LSIFGPV | ASSRGAINTGQLY | 0.5912926 |
| LLALLANL | ASSRGAINTGQLY | 0.57622284 |
| LISNASDALDKLR | ASSRGAINTGQLY | 0.6745705 |
| LINEAVP | ASSRGAINTGQLY | 0.40111458 |
| LGYAFSGL | ASSRGAINTGQLY | 0.46720064 |
| LDFVAVAL | ASSRGAINTGQLY | 0.56282794 |
| LAQPGAE | ASSRGAINTGQLY | 0.46492654 |
| ISNLGAVTGF | ASSRGAINTGQLY | 0.3855139 |
| HQEDPGRI | ASSRGAINTGQLY | 0.66985816 |
| GTPGEPTP | ASSRGAINTGQLY | 0.7521541 |
| GFFNSGS | ASSRGAINTGQLY | 0.7649729 |
| GAGGGGLTAGAN | ASSRGAINTGQLY | 0.7128737 |
| FVDALVNM | ASSRGAINTGQLY | 0.71223044 |
| FLTVAPEL | ASSRGAINTGQLY | 0.7178465 |
| FETLTAL | ASSRGAINTGQLY | 0.76058036 |
| DVINPIEL | ASSRGAINTGQLY | 0.7617612 |
| APVSGAPV | ASSRGAINTGQLY | 0.75223625 |
| ALVGGLAW | ASSRGAINTGQLY | 0.63999 |
| ALAAAAATAI | ASSRGAINTGQLY | 0.75072706 |
| AFGLGLLL | ASSRGAINTGQLY | 0.7225337 |
| AFFAAASL | ASSRGAINTGQLY | 0.7223333 |

| | | |
|---|---|---|
| AAAALPAFSPP | ASSRGAINTGQLY | 0.74831134 |
| VVLAPAL | ASSRGAINTGQLY | 0.739371 |
| TIGHVDHGKTTL | ASSRGAINTGQLY | 0.709082 |
| LISNASDALDK | ASSRGAINTGQLY | 0.7148914 |
| LISNASDAL | ASSRGAINTGQLY | 0.7428588 |
| ISNASDALDK | ASSRGAINTGQLY | 0.7564918 |
| NQAETLVYQTE | ASSRGAINTGQLY | 0.7538806 |
| TGGTGGTAGAG | ASSRGAINTGQLY | 0.76646984 |
| TGSTAGGAGGA | ASSRGAINTGQLY | 0.7602192 |
| DPLAVPVAAGPV | ASSRGAINTGQLY | 0.75248194 |
| GGTGGTPGNGGN | ASSRGAINTGQLY | 0.3882687 |
| PAGLAGVDPATL | ASSRGAINTGQLY | 0.64900696 |
| GNNLVVEAPHAGASV | ASSRGAINTGQLY | 0.7198076 |
| KVTRIMSASTLEQPA | ASSRGAINTGQLY | 0.7295177 |
| LGFALPIALPRRTVV | ASSRGAINTGQLY | 0.38555574 |
| TRLGNLTARKLVVLV | ASSRGAINTGQLY | 0.74592125 |
| WSQQRRLIYFPSAGP | ASSRGAINTGQLY | 0.7286517 |
| GPNGNPDPMAAAVDIR | ASSRGAINTGQLY | 0.70273834 |
| TQRCVSVADLRAAASR | ASSRGAINTGQLY | 0.75608444 |
| QFVLDVAEHPVEVLTETV | ASSRGAINTGQLY | 0.71630096 |
| GGAGGTGGAGGNGGTQAGDG | ASSRGAINTGQLY | 0.6818322 |
| PLLATLRAIAVDVDAKPAQV | ASSRGAINTGQLY | 0.6321955 |
| APKFRTGGYMIDFWGVGYQV | ASSRGAINTGQLY | 0.76005995 |
| GAGGAGGAGGAAGGPGGTGG | ASSRGAINTGQLY | 0.6237596 |
| IGALGLLATDVAPAGSVAFG | ASSRGAINTGQLY | 0.75627947 |
| AGGAGGIGGNGGAGGAGGVL | ASSRGAINTGQLY | 0.6786973 |
| ADELIGLL | ASSRGAINTGQLY | 0.7624098 |
| ADRALKA | ASSRGAINTGQLY | 0.6046219 |
| AGAVGQVGGAGGA | ASSRGAINTGQLY | 0.75524724 |
| AGGGGGLGGAP | ASSRGAINTGQLY | 0.73776853 |
| AGGLLYGNGGAG | ASSRGAINTGQLY | 0.77187175 |
| AGLESIGL | ASSRGAINTGQLY | 0.7348131 |
| AGLKTGII | ASSRGAINTGQLY | 0.74548805 |
| ASLAALLL | ASSRGAINTGQLY | 0.67176825 |
| ASVINISSI | ASSRGAINTGQLY | 0.7620268 |
| ATPPGPA | ASSRGAINTGQLY | 0.62410045 |
| ATVASALV | ASSRGAINTGQLY | 0.7294402 |
| ATVASLIL | ASSRGAINTGQLY | 0.5878793 |
| AVNGPVLIRH | ASSRGAINTGQLY | 0.7436956 |
| DGAAGAVGI | ASSRGAINTGQLY | 0.7543597 |
| DPLLHGI | ASSRGAINTGQLY | 0.65933454 |
| ELGAALGAL | ASSRGAINTGQLY | 0.75816476 |
| ELGRTQAV | ASSRGAINTGQLY | 0.72554815 |
| FVAGGYLS | ASSRGAINTGQLY | 0.68248576 |
| GASGGAGGAGGW | ASSRGAINTGQLY | 0.7616614 |
| GGAGAGGDGGAGGI | ASSRGAINTGQLY | 0.73446804 |
| GTGHNGSQPGMGGNG | ASSRGAINTGQLY | 0.7610389 |
| GYAAASASAS | ASSRGAINTGQLY | 0.6451423 |
| HQVDVAAVQQG | ASSRGAINTGQLY | 0.7416415 |
| IGAAIAHT | ASSRGAINTGQLY | 0.747347 |
| IGMIEVAQL | ASSRGAINTGQLY | 0.6970328 |

| | | |
|---|---|---|
| IRLAPEHTSAMD | ASSRGAINTGQLY | 0.6556901 |
| IYNVSVL | ASSRGAINTGQLY | 0.76438546 |
| KQGFLGM | ASSRGAINTGQLY | 0.72635436 |
| LESVITQL | ASSRGAINTGQLY | 0.75403637 |
| LGLGALVVV | ASSRGAINTGQLY | 0.6585809 |
| LIGNGGNGGSGGI | ASSRGAINTGQLY | 0.53398144 |
| LNAINAPTL | ASSRGAINTGQLY | 0.6567863 |
| LQLLNTGV | ASSRGAINTGQLY | 0.75316834 |
| LQTSIEF | ASSRGAINTGQLY | 0.7557317 |
| MSFAVSEL | ASSRGAINTGQLY | 0.7371081 |
| NTGALGL | ASSRGAINTGQLY | 0.5185759 |
| PPGAGGAGGAGTT | ASSRGAINTGQLY | 0.6896812 |
| QLVGLGI | ASSRGAINTGQLY | 0.6635215 |
| RFLEQVHQL | ASSRGAINTGQLY | 0.64574856 |
| RGVGYVLR | ASSRGAINTGQLY | 0.72555053 |
| SATGVFAGVI | ASSRGAINTGQLY | 0.7242619 |
| SLPAITLP | ASSRGAINTGQLY | 0.6773429 |
| STALAFTV | ASSRGAINTGQLY | 0.62203366 |
| SVLTVGLGK | ASSRGAINTGQLY | 0.70520926 |
| TALAGAGSGI | ASSRGAINTGQLY | 0.74205726 |
| TGLPFTGI | ASSRGAINTGQLY | 0.7311096 |
| VAEHVTV | ASSRGAINTGQLY | 0.76880753 |
| VDIVLNSL | ASSRGAINTGQLY | 0.74577093 |
| VLTGGGAVL | ASSRGAINTGQLY | 0.7373991 |
| VNVGLAYGI | ASSRGAINTGQLY | 0.22361931 |
| VSSAAVGAV | ASSRGAINTGQLY | 0.28713083 |
| VTSVYEAGV | ASSRGAINTGQLY | 0.66586405 |
| WVALDRGA | ASSRGAINTGQLY | 0.59551895 |
| YLVGSVVI | ASSRGAINTGQLY | 0.3063954 |
| GPPGTGKTLL | ASSRGAINTGQLY | 0.70370543 |
| GVGMTKF | ASSRGAINTGQLY | 0.66615486 |
| IGIGGSDL | ASSRGAINTGQLY | 0.5641361 |
| IGTLLAEL | ASSRGAINTGQLY | 0.74321467 |
| IYTSGTT | ASSRGAINTGQLY | 0.64460623 |
| LSFPAGV | ASSRGAINTGQLY | 0.55538726 |
| LYTSGTT | ASSRGAINTGQLY | 0.5699103 |
| VGAFPFGF | ASSRGAINTGQLY | 0.7434951 |
| VGAGTVEF | ASSRGAINTGQLY | 0.514452 |
| VGKSTLLL | ASSRGAINTGQLY | 0.7326612 |
| AALAHAGGGIV | ASSRGAINTGQLY | 0.50934786 |
| AALVAVSLL | ASSRGAINTGQLY | 0.7503226 |
| AAVSRAVAAL | ASSRGAINTGQLY | 0.45193478 |
| AGGSSGIGGF | ASSRGAINTGQLY | 0.73790014 |
| AGLTVANANG | ASSRGAINTGQLY | 0.6961626 |
| AGLVGAVL | ASSRGAINTGQLY | 0.78147423 |
| ALSQAFTV | ASSRGAINTGQLY | 0.62051314 |
| ALTFSVSL | ASSRGAINTGQLY | 0.68183184 |
| ALTGAALVAL | ASSRGAINTGQLY | 0.60884446 |
| ASFSIGAGVL | ASSRGAINTGQLY | 0.73942053 |
| DGSVYKI | ASSRGAINTGQLY | 0.4057293 |
| EQLAEIV | ASSRGAINTGQLY | 0.67877924 |

| | | |
|---|---|---|
| FAVSELTVM | ASSRGAINTGQLY | 0.32817784 |
| GAAPLAAGL | ASSRGAINTGQLY | 0.71665704 |
| GADGGDGGSGGAGG | ASSRGAINTGQLY | 0.7247523 |
| GFGLDTGGAGG | ASSRGAINTGQLY | 0.30633044 |
| GLASGGLL | ASSRGAINTGQLY | 0.74656415 |
| GRIAGAAVLL | ASSRGAINTGQLY | 0.6198524 |
| GSAGSLNTSF | ASSRGAINTGQLY | 0.5780339 |
| GTGQIGI | ASSRGAINTGQLY | 0.7603397 |
| GTLIALIL | ASSRGAINTGQLY | 0.6874611 |
| IAATMGFVVG | ASSRGAINTGQLY | 0.75011045 |
| IFAQALL | ASSRGAINTGQLY | 0.69262373 |
| IGAAVALGF | ASSRGAINTGQLY | 0.6562574 |
| IGLTPLAL | ASSRGAINTGQLY | 0.7132778 |
| ILGSNTSTL | ASSRGAINTGQLY | 0.49646562 |
| IVFSLTVL | ASSRGAINTGQLY | 0.54283607 |
| LASKATGF | ASSRGAINTGQLY | 0.75456774 |
| LASVALIL | ASSRGAINTGQLY | 0.71260434 |
| LFAVGLTL | ASSRGAINTGQLY | 0.7449527 |
| LFNGGIGTYI | ASSRGAINTGQLY | 0.61548346 |
| LTLGAGII | ASSRGAINTGQLY | 0.50959593 |
| LVAAALGDI | ASSRGAINTGQLY | 0.66091764 |
| NTAFVQLGI | ASSRGAINTGQLY | 0.7284862 |
| QLFSGAEL | ASSRGAINTGQLY | 0.7373576 |
| SAATFITL | ASSRGAINTGQLY | 0.72241855 |
| SNYGLANLGV | ASSRGAINTGQLY | 0.40086502 |
| SYGGAYAVM | ASSRGAINTGQLY | 0.6868712 |
| TTGVGGLTLG | ASSRGAINTGQLY | 0.61340296 |
| TTQTIYI | ASSRGAINTGQLY | 0.6471479 |
| VAAGAGLGAVI | ASSRGAINTGQLY | 0.7238858 |
| VAAVVTLL | ASSRGAINTGQLY | 0.5863191 |
| VGFLGGLI | ASSRGAINTGQLY | 0.6404812 |
| VGVIPILL | ASSRGAINTGQLY | 0.62133926 |
| VIFASSLI | ASSRGAINTGQLY | 0.6899814 |
| VKGGDGGAAAG | ASSRGAINTGQLY | 0.71613777 |
| VTKTFTNI | ASSRGAINTGQLY | 0.7029294 |
| VVAAAILL | ASSRGAINTGQLY | 0.739499 |
| YAALDVEL | ASSRGAINTGQLY | 0.7220073 |
| LASVALLL | ASSRGAINTGQLY | 0.70902884 |
| LDAVHRV | ASSRGAINTGQLY | 0.24107769 |
| QGLSSSIF | ASSRGAINTGQLY | 0.5062002 |
| RGTFANIRL | ASSRGAINTGQLY | 0.66031194 |
| RQLWWGHRI | ASSRGAINTGQLY | 0.69583666 |
| YGPPGTGKTLL | ASSRGAINTGQLY | 0.39639384 |
| AALVGAPANVA | ASSRGAINTGQLY | 0.7216961 |
| AATFRRTE | ASSRGAINTGQLY | 0.63168967 |
| AGALGGGI | ASSRGAINTGQLY | 0.6005386 |
| AGGAGGAGGIGAG | ASSRGAINTGQLY | 0.7325816 |
| AGSGALGL | ASSRGAINTGQLY | 0.6270896 |
| AQIADQI | ASSRGAINTGQLY | 0.5023535 |
| ASAVAVTYL | ASSRGAINTGQLY | 0.58576685 |
| ASTVLGGTGG | ASSRGAINTGQLY | 0.7313783 |

| | | |
|---|---|---|
| AVGGEPAK | ASSRGAINTGQLY | 0.4762236 |
| CEQGTFGGTC | ASSRGAINTGQLY | 0.71161956 |
| DEGKAPAG | ASSRGAINTGQLY | 0.6823392 |
| DTVVFPAL | ASSRGAINTGQLY | 0.7176068 |
| FEGITMAG | ASSRGAINTGQLY | 0.35096022 |
| FPLVVQD | ASSRGAINTGQLY | 0.7174807 |
| GAFTTPQ | ASSRGAINTGQLY | 0.6760824 |
| GFGGFGTAGGVGGA | ASSRGAINTGQLY | 0.74518985 |
| GFLAQFL | ASSRGAINTGQLY | 0.5750004 |
| GGAGGAGGNAGVA | ASSRGAINTGQLY | 0.6366271 |
| GGVGTAGAGGAGGAGG | ASSRGAINTGQLY | 0.46323872 |
| GNGGGGDAGNAGSGGN | ASSRGAINTGQLY | 0.7357478 |
| GTGGAGGAGNDAGS | ASSRGAINTGQLY | 0.49580717 |
| GTTNTGWFNS | ASSRGAINTGQLY | 0.59647316 |
| IFMGPLFTL | ASSRGAINTGQLY | 0.53572786 |
| ILAHAIEAI | ASSRGAINTGQLY | 0.67511255 |
| IRSAIREAL | ASSRGAINTGQLY | 0.7151582 |
| ITFAGGG | ASSRGAINTGQLY | 0.57971066 |
| ITGMSGHQEV | ASSRGAINTGQLY | 0.7237205 |
| IVAGLACTL | ASSRGAINTGQLY | 0.7083554 |
| IVGGGLAGA | ASSRGAINTGQLY | 0.4640824 |
| IVTGVLNI | ASSRGAINTGQLY | 0.7323691 |
| LAQQGCI | ASSRGAINTGQLY | 0.54655886 |
| LCLHVEN | ASSRGAINTGQLY | 0.72813666 |
| LDAISDGS | ASSRGAINTGQLY | 0.6541373 |
| LGGFAAML | ASSRGAINTGQLY | 0.6925993 |
| LGGKNPMI | ASSRGAINTGQLY | 0.71121 |
| LGPKGVAAIL | ASSRGAINTGQLY | 0.66653407 |
| LMVGVSI | ASSRGAINTGQLY | 0.58828896 |
| LVGAAFVL | ASSRGAINTGQLY | 0.72721195 |
| LVLAVAPL | ASSRGAINTGQLY | 0.7314851 |
| MAAIAVIDAA | ASSRGAINTGQLY | 0.84630144 |
| NNGINAELQ | ASSRGAINTGQLY | 0.5455905 |
| RGEPLVLVHG | ASSRGAINTGQLY | 0.4930207 |
| RGGAGGAAGLI | ASSRGAINTGQLY | 0.6139846 |
| RSYRMHPAV | ASSRGAINTGQLY | 0.824034 |
| SAAHLLAGQLA | ASSRGAINTGQLY | 0.82064366 |
| SAAVGSY | ASSRGAINTGQLY | 0.75707304 |
| SPHAVLHRI | ASSRGAINTGQLY | 0.53619695 |
| SQIAKVL | ASSRGAINTGQLY | 0.6096264 |
| TALGHLVP | ASSRGAINTGQLY | 0.5709045 |
| TAPTTSGAA | ASSRGAINTGQLY | 0.61318856 |
| TDGDDGNGGSPGT | ASSRGAINTGQLY | 0.78722334 |
| TDGLGEN | ASSRGAINTGQLY | 0.71406054 |
| TGGPPLPGQA | ASSRGAINTGQLY | 0.59647644 |
| TIPAITI | ASSRGAINTGQLY | 0.6361521 |
| TIPGFTI | ASSRGAINTGQLY | 0.65969235 |
| TSIASGALA | ASSRGAINTGQLY | 0.7442499 |
| TVTLPGAAI | ASSRGAINTGQLY | 0.7023603 |
| VASGLSAALP | ASSRGAINTGQLY | 0.8659171 |
| VGAGVIGI | ASSRGAINTGQLY | 0.7781353 |

| | | |
|---|---|---|
| VGPKRFGV | ASSRGAINTGQLY | 0.80477786 |
| VNVAFPD | ASSRGAINTGQLY | 0.14420514 |
| VSMTNSGVS | ASSRGAINTGQLY | 0.35955626 |
| VTDVLGDLSV | ASSRGAINTGQLY | 0.70905775 |
| VVVVGAGFAGL | ASSRGAINTGQLY | 0.6815995 |
| VYTVGTVV | ASSRGAINTGQLY | 0.43884975 |
| WTTFSGP | ASSRGAINTGQLY | 0.7873925 |
| YDELVEAF | ASSRGAINTGQLY | 0.6119042 |
| YGTGGAGGNGG | ASSRGAINTGQLY | 0.5195394 |
| YTNVMAV | ASSRGAINTGQLY | 0.7819849 |
| FLEQVHQL | ASSRGAINTGQLY | 0.58159614 |
| GLSLLAVAV | ASSRGAINTGQLY | 0.5928855 |
| LEQVHQL | ASSRGAINTGQLY | 0.63595825 |
| LNAAGAI | ASSRGAINTGQLY | 0.8064344 |
| PAVHASL | ASSRGAINTGQLY | 0.41233248 |
| TELEDDA | ASSRGAINTGQLY | 0.7353856 |
| TPPGPPN | ASSRGAINTGQLY | 0.68071866 |
| TPPPPPPP | ASSRGAINTGQLY | 0.8061884 |
| TQSPALI | ASGERTGGAANTGQLY | 0.43004057 |
| EFAAAFPQ | ASGERTGGAANTGQLY | 0.7597553 |
| LLSGNLDVL | ASGERTGGAANTGQLY | 0.64135706 |
| GPANLGNYNI | ASGERTGGAANTGQLY | 0.8606769 |
| TLGPTLP | ASGERTGGAANTGQLY | 0.5904613 |
| VYAYSSCCK | ASGERTGGAANTGQLY | 0.70017356 |
| APAPAPAPNI | ASGERTGGAANTGQLY | 0.66260993 |
| PGGPVPIP | ASGERTGGAANTGQLY | 0.7872783 |
| FLTVALEP | ASGERTGGAANTGQLY | 0.42116582 |
| SQYFGANL | ASGERTGGAANTGQLY | 0.5948241 |
| GGVKPAA | ASGERTGGAANTGQLY | 0.38970163 |
| GGTGGAGGAGGW | ASGERTGGAANTGQLY | 0.7134354 |
| GVGGVAGLGGAGG | ASGERTGGAANTGQLY | 0.7313732 |
| VLLVLATTI | ASGERTGGAANTGQLY | 0.4352048 |
| IGIFAAML | ASGERTGGAANTGQLY | 0.7472054 |
| AVGDLID | ASGERTGGAANTGQLY | 0.70068777 |
| NISPLPL | ASGERTGGAANTGQLY | 0.5874085 |
| VTAVFVAI | ASGERTGGAANTGQLY | 0.82774985 |
| SGDDNDAAL | ASGERTGGAANTGQLY | 0.5067011 |
| AVAALAHS | ASGERTGGAANTGQLY | 0.8109057 |
| GGVTVMATP | ASGERTGGAANTGQLY | 0.70186156 |
| AGLAANVRD | ASGERTGGAANTGQLY | 0.59715056 |
| TNRPDLIDPALLRPG | ASGERTGGAANTGQLY | 0.7690387 |
| LANLGVTAL | ASGERTGGAANTGQLY | 0.5290592 |
| TGERIEL | ASGERTGGAANTGQLY | 0.5701187 |
| NGEELAA | ASGERTGGAANTGQLY | 0.8146448 |
| VLSLVWG | ASGERTGGAANTGQLY | 0.82137096 |
| IGYALWL | ASGERTGGAANTGQLY | 0.85386705 |
| LSLLLAPL | ASGERTGGAANTGQLY | 0.7717583 |
| TLDEALK | ASGERTGGAANTGQLY | 0.71541977 |
| TDGAGTAEGG | ASGERTGGAANTGQLY | 0.8064978 |
| NAPGGAGGNGG | ASGERTGGAANTGQLY | 0.85262567 |
| VSVFMGMGGNF | ASGERTGGAANTGQLY | 0.8443624 |

| | | |
|---|---|---|
| LASYLAAL | ASGERTGGAANTGQLY | 0.83677936 |
| LVGGTPAL | ASGERTGGAANTGQLY | 0.7034793 |
| VRNADVL | ASGERTGGAANTGQLY | 0.79524916 |
| TCSSPFLVSAM | ASGERTGGAANTGQLY | 0.79644406 |
| VHITDAE | ASGERTGGAANTGQLY | 0.7737024 |
| EYAAPSF | ASGERTGGAANTGQLY | 0.8423901 |
| VVDAFHGGL | ASGERTGGAANTGQLY | 0.8008162 |
| ASVVDPQ | ASGERTGGAANTGQLY | 0.74503213 |
| SYNEFGGLS | ASGERTGGAANTGQLY | 0.7976966 |
| DPDGSVY | ASGERTGGAANTGQLY | 0.81715804 |
| MVSTIHATE | ASGERTGGAANTGQLY | 0.8204486 |
| LAAGVADPV | ASGERTGGAANTGQLY | 0.81565094 |
| TKEQIAA | ASGERTGGAANTGQLY | 0.86073065 |
| SLASPAAL | ASGERTGGAANTGQLY | 0.8438524 |
| IGLVFYGV | ASGERTGGAANTGQLY | 0.83381677 |
| FLLDASNL | ASGERTGGAANTGQLY | 0.24459454 |
| YQVGNFP | ASGERTGGAANTGQLY | 0.61318475 |
| ESLLSQST | ASGERTGGAANTGQLY | 0.7805058 |
| LFTVGFAAVG | ASGERTGGAANTGQLY | 0.80852294 |
| ATPPTPT | ASGERTGGAANTGQLY | 0.67694104 |
| TDLATLG | ASGERTGGAANTGQLY | 0.84090245 |
| SAGGTGR | ASGERTGGAANTGQLY | 0.78180796 |
| GFSNSGS | ASGERTGGAANTGQLY | 0.72767115 |
| GGNGGSGGTSV | ASGERTGGAANTGQLY | 0.8380284 |
| GTSVATGGAGN | ASGERTGGAANTGQLY | 0.73165166 |
| NGGNGGSGGTS | ASGERTGGAANTGQLY | 0.75987226 |
| AVVKFPH | ASGERTGGAANTGQLY | 0.78099346 |
| TPPPPPP | ASGERTGGAANTGQLY | 0.8499695 |
| AAPAAAAAPA | ASGERTGGAANTGQLY | 0.66801775 |
| IALNEDL | ASGERTGGAANTGQLY | 0.827391 |
| IVNVSSM | ASGERTGGAANTGQLY | 0.7996919 |
| TGIPPAPR | ASGERTGGAANTGQLY | 0.84790325 |
| FDLGGGTF | ASGERTGGAANTGQLY | 0.6255245 |
| FDLGGGTFDVS | ASGERTGGAANTGQLY | 0.8389855 |
| NEPTAAA | ASGERTGGAANTGQLY | 0.81009287 |
| NEPTAAALA | ASGERTGGAANTGQLY | 0.8593672 |
| GSIINISSI | ASGERTGGAANTGQLY | 0.7955215 |
| VAPGNAGTA | ASGERTGGAANTGQLY | 0.81412804 |
| TGGGASLE | ASGERTGGAANTGQLY | 0.7435664 |
| VAVYNHYKRI | ASGERTGGAANTGQLY | 0.83945626 |
| TDPAPATT | ASGERTGGAANTGQLY | 0.70407206 |
| TNRPDILDPALLRPG | ASGERTGGAANTGQLY | 0.80086464 |
| LTTTTTS | ASGERTGGAANTGQLY | 0.68421364 |
| PTGSGKTL | ASGERTGGAANTGQLY | 0.82369685 |
| TGSGKTL | ASGERTGGAANTGQLY | 0.8283404 |
| TIGHVDHGK | ASGERTGGAANTGQLY | 0.77402914 |
| TIGHVDHGKTT | ASGERTGGAANTGQLY | 0.8415256 |
| GHVDHGKTT | ASGERTGGAANTGQLY | 0.84746754 |
| AQFEHTLL | ASGERTGGAANTGQLY | 0.7493289 |
| IVNQDEL | ASGERTGGAANTGQLY | 0.8539975 |
| TNGSQFF | ASGERTGGAANTGQLY | 0.77916473 |

| | | |
|---|---|---|
| TGKGIVE | ASGERTGGAANTGQLY | 0.8426222 |
| LISNASDA | ASGERTGGAANTGQLY | 0.8089814 |
| AELRTLGELTKIAKAHGAQ | ASGERTGGAANTGQLY | 0.79378736 |
| APNAVAAAEPSW | ASGERTGGAANTGQLY | 0.8401381 |
| APQVAGAATAVL | ASGERTGGAANTGQLY | 0.76715547 |
| ARRDPAAAAAILPTDA | ASGERTGGAANTGQLY | 0.7768857 |
| DAAARDVLAEAGLAEYF | ASGERTGGAANTGQLY | 0.8458519 |
| DDQIVLSAKANLAKVEQ | ASGERTGGAANTGQLY | 0.7322502 |
| DELLPDTGEDSDEDGH | ASGERTGGAANTGQLY | 0.7817036 |
| ELPEAPANRATVPQD | ASGERTGGAANTGQLY | 0.3337406 |
| FGQSKIERVTGIG | ASGERTGGAANTGQLY | 0.31687576 |
| GAGGTGSTAGGAGGAGGAGGI | ASGERTGGAANTGQLY | 0.53756356 |
| GHGGTGAAGGNGGTGGAGGS( | ASGERTGGAANTGQLY | 0.67181116 |
| GNGGNGGAGGEGGPGGAGGA( | ASGERTGGAANTGQLY | 0.83651805 |
| GSTAGNGGNGGNATNGTIAG( | ASGERTGGAANTGQLY | 0.76663095 |
| GTGGDGGNGGNGGTGG | ASGERTGGAANTGQLY | 0.32376742 |
| GVRRPAGIPADAPV | ASGERTGGAANTGQLY | 0.6474843 |
| GYRQVTDAHLLTLARR | ASGERTGGAANTGQLY | 0.4583309 |
| IAKMGQDGHDRG | ASGERTGGAANTGQLY | 0.5351371 |
| KAEPSIEIQYT | ASGERTGGAANTGQLY | 0.57323444 |
| KPVRIRMAADEDTVAVT | ASGERTGGAANTGQLY | 0.43644637 |
| MPPRIAGMRLLVIKPEP | ASGERTGGAANTGQLY | 0.39222303 |
| MKFYAVALFRSRGVPPTR | ASGERTGGAANTGQLY | 0.23943368 |
| NAGGFGILGAVAHSP | ASGERTGGAANTGQLY | 0.6868752 |
| NEGREITLAMAEM | ASGERTGGAANTGQLY | 0.7985358 |
| NELVEWSHPLGHRWRWE | ASGERTGGAANTGQLY | 0.74344766 |
| NLGAGGTGGNGGNPG | ASGERTGGAANTGQLY | 0.81138146 |
| NTEWPKWETVMAADPVD | ASGERTGGAANTGQLY | 0.7773945 |
| PASAAPAETQP | ASGERTGGAANTGQLY | 0.64138234 |
| QQSSDGKNGVRAAEITGE | ASGERTGGAANTGQLY | 0.13074492 |
| RAPDAVTTSWWKE | ASGERTGGAANTGQLY | 0.19912477 |
| SGFGNVGAHTSGWFNQ | ASGERTGGAANTGQLY | 0.26428303 |
| SGVAGAVGQAASVGGLKV | ASGERTGGAANTGQLY | 0.5029506 |
| SNGGAGTGSAGGNGGT | ASGERTGGAANTGQLY | 0.13657922 |
| TGGAGGEGGAAG | ASGERTGGAANTGQLY | 0.6157776 |
| TGGDGGAAGAGGAPGHG | ASGERTGGAANTGQLY | 0.4036641 |
| TMPPLLAAIANAGRHGV | ASGERTGGAANTGQLY | 0.3391145 |
| TSSTAGNSGGA | ASGERTGGAANTGQLY | 0.7863888 |
| VPATVPVGILC | ASGERTGGAANTGQLY | 0.36081666 |
| VQRPDSGGVVLTGRIS | ASGERTGGAANTGQLY | 0.22666423 |
| WTPNPVPATKGVPAGLVQAT | ASGERTGGAANTGQLY | 0.34306598 |
| YALKRALAKFSDRFDVVII | ASGERTGGAANTGQLY | 0.8250667 |
| YLIGSVIV | ASGERTGGAANTGQLY | 0.24578938 |
| YGQLVAVID | ASGERTGGAANTGQLY | 0.77903533 |
| VTVGGMTLPA | ASGERTGGAANTGQLY | 0.48809502 |
| VLALPAV | ASGERTGGAANTGQLY | 0.7794628 |
| VIPPTLNYET | ASGERTGGAANTGQLY | 0.207032 |
| VGMGGMHPSSG | ASGERTGGAANTGQLY | 0.7502301 |
| VGLAGGAATA | ASGERTGGAANTGQLY | 0.59314495 |
| VGGAGGAGGSGP | ASGERTGGAANTGQLY | 0.84132326 |
| VFAAFMV | ASGERTGGAANTGQLY | 0.4051421 |

| | | |
|---|---|---|
| TLYDKLR | ASGERTGGAANTGQLY | 0.55554104 |
| TLAVFTVL | ASGERTGGAANTGQLY | 0.45222333 |
| TGSVQPARV | ASGERTGGAANTGQLY | 0.80524886 |
| TGGGAGSGGT | ASGERTGGAANTGQLY | 0.20753819 |
| SHLETSL | ASGERTGGAANTGQLY | 0.39444876 |
| SFTTGMT | ASGERTGGAANTGQLY | 0.28178447 |
| QVEVAKRMGVRQPTVS | ASGERTGGAANTGQLY | 0.4476561 |
| QGSGGPALAM | ASGERTGGAANTGQLY | 0.7294877 |
| MLAPSNP | ASGERTGGAANTGQLY | 0.12435269 |
| LVNNAAI | ASGERTGGAANTGQLY | 0.71091735 |
| LTGEINTLA | ASGERTGGAANTGQLY | 0.24890846 |
| LSIFGPV | ASGERTGGAANTGQLY | 0.25000468 |
| LLALLANL | ASGERTGGAANTGQLY | 0.6425928 |
| LISNASDALDKLR | ASGERTGGAANTGQLY | 0.49480587 |
| LINEAVP | ASGERTGGAANTGQLY | 0.73639643 |
| LGYAFSGL | ASGERTGGAANTGQLY | 0.27450854 |
| LDFVAVAL | ASGERTGGAANTGQLY | 0.64882755 |
| LAQPGAE | ASGERTGGAANTGQLY | 0.57207274 |
| ISNLGAVTGF | ASGERTGGAANTGQLY | 0.14585589 |
| HQEDPGRI | ASGERTGGAANTGQLY | 0.32863343 |
| GTPGEPTP | ASGERTGGAANTGQLY | 0.7833193 |
| GFFNSGS | ASGERTGGAANTGQLY | 0.7985352 |
| GAGGGGLTAGAN | ASGERTGGAANTGQLY | 0.84196365 |
| FVDALVNM | ASGERTGGAANTGQLY | 0.66811275 |
| FLTVAPEL | ASGERTGGAANTGQLY | 0.61692524 |
| FETLTAL | ASGERTGGAANTGQLY | 0.71218526 |
| DVINPIEL | ASGERTGGAANTGQLY | 0.7784606 |
| APVSGAPV | ASGERTGGAANTGQLY | 0.84772277 |
| ALVGGLAW | ASGERTGGAANTGQLY | 0.8238622 |
| ALAAAAATAI | ASGERTGGAANTGQLY | 0.5494672 |
| AFGLGLLL | ASGERTGGAANTGQLY | 0.79219717 |
| AFFAAASL | ASGERTGGAANTGQLY | 0.74593806 |
| AAAALPAFSPP | ASGERTGGAANTGQLY | 0.7144115 |
| VVLAPAL | ASGERTGGAANTGQLY | 0.6831256 |
| TIGHVDHGKTTL | ASGERTGGAANTGQLY | 0.7148647 |
| LISNASDALDK | ASGERTGGAANTGQLY | 0.6368791 |
| LISNASDAL | ASGERTGGAANTGQLY | 0.4305424 |
| ISNASDALDK | ASGERTGGAANTGQLY | 0.6520928 |
| NQAETLVYQTE | ASGERTGGAANTGQLY | 0.78708655 |
| TGGTGGTAGAG | ASGERTGGAANTGQLY | 0.77337503 |
| TGSTAGGAGGA | ASGERTGGAANTGQLY | 0.7880646 |
| DPLAVPVAAGPV | ASGERTGGAANTGQLY | 0.8171464 |
| GGTGGTPGNGGN | ASGERTGGAANTGQLY | 0.593989 |
| PAGLAGVDPATL | ASGERTGGAANTGQLY | 0.18547048 |
| GNNLVVEAPHAGASV | ASGERTGGAANTGQLY | 0.307646 |
| KVTRIMSASTLEQPA | ASGERTGGAANTGQLY | 0.6226057 |
| LGFALPIALPRRTVV | ASGERTGGAANTGQLY | 0.665962 |
| TRLGNLTARKLVVLV | ASGERTGGAANTGQLY | 0.18001956 |
| WSQQRRLIYFPSAGP | ASGERTGGAANTGQLY | 0.66929024 |
| GPNGNPDPMAAAVDIR | ASGERTGGAANTGQLY | 0.7005599 |
| TQRCVSVADLRAAASR | ASGERTGGAANTGQLY | 0.6997118 |

| | | |
|---|---|---|
| QFVLDVAEHPVEVLTETV | ASGERTGGAANTGQLY | 0.80830437 |
| GGAGGTGGAGGNGGTQAGDG | ASGERTGGAANTGQLY | 0.7523979 |
| PLLATLRAIAVDVDAKPAQV | ASGERTGGAANTGQLY | 0.38763118 |
| APKFRTGGYMIDFWGVGYQV/ | ASGERTGGAANTGQLY | 0.44347033 |
| GAGGAGGAGGAAGGPGGTGG' | ASGERTGGAANTGQLY | 0.81393623 |
| IGALGLLATDVAPAGSVAFG' | ASGERTGGAANTGQLY | 0.37342185 |
| AGGAGGIGGNGGAGGAGGVLI | ASGERTGGAANTGQLY | 0.812354 |
| ADELIGLL | ASGERTGGAANTGQLY | 0.3845557 |
| ADRALKA | ASGERTGGAANTGQLY | 0.8122263 |
| AGAVGQVGGAGGA | ASGERTGGAANTGQLY | 0.50600004 |
| AGGGGGLGGAP | ASGERTGGAANTGQLY | 0.79253757 |
| AGGLLYGNGGAG | ASGERTGGAANTGQLY | 0.7795631 |
| AGLESIGL | ASGERTGGAANTGQLY | 0.84148926 |
| AGLKTGII | ASGERTGGAANTGQLY | 0.6496308 |
| ASLAALLL | ASGERTGGAANTGQLY | 0.7921929 |
| ASVINISSI | ASGERTGGAANTGQLY | 0.46502328 |
| ATPPGPA | ASGERTGGAANTGQLY | 0.8415718 |
| ATVASALV | ASGERTGGAANTGQLY | 0.6277091 |
| ATVASLIL | ASGERTGGAANTGQLY | 0.6116816 |
| AVNGPVLIRH | ASGERTGGAANTGQLY | 0.34449092 |
| DGAAGAVGI | ASGERTGGAANTGQLY | 0.66297054 |
| DPLLHGI | ASGERTGGAANTGQLY | 0.82497567 |
| ELGAALGAL | ASGERTGGAANTGQLY | 0.52030045 |
| ELGRTQAV | ASGERTGGAANTGQLY | 0.8113585 |
| FVAGGYLS | ASGERTGGAANTGQLY | 0.6932359 |
| GASGGAGGAGGW | ASGERTGGAANTGQLY | 0.58702666 |
| GGAGAGGDGGAGGI | ASGERTGGAANTGQLY | 0.7621919 |
| GTGHNGSQPGMGGNG | ASGERTGGAANTGQLY | 0.6527092 |
| GYAAASASAS | ASGERTGGAANTGQLY | 0.7924843 |
| HQVDVAAVQQG | ASGERTGGAANTGQLY | 0.47740844 |
| IGAAIAHT | ASGERTGGAANTGQLY | 0.68779457 |
| IGMIEVAQL | ASGERTGGAANTGQLY | 0.7458254 |
| IRLAPEHTSAMD | ASGERTGGAANTGQLY | 0.5510196 |
| IYNVSVL | ASGERTGGAANTGQLY | 0.4057005 |
| KQGFLGM | ASGERTGGAANTGQLY | 0.82776606 |
| LESVITQL | ASGERTGGAANTGQLY | 0.77267516 |
| LGLGALVVV | ASGERTGGAANTGQLY | 0.79282063 |
| LIGNGGNGGSGGI | ASGERTGGAANTGQLY | 0.6893804 |
| LNAINAPTL | ASGERTGGAANTGQLY | 0.5759932 |
| LQLLNTGV | ASGERTGGAANTGQLY | 0.7311725 |
| LQTSIEF | ASGERTGGAANTGQLY | 0.7826228 |
| MSFAVSEL | ASGERTGGAANTGQLY | 0.8064275 |
| NTGALGL | ASGERTGGAANTGQLY | 0.75558853 |
| PPGAGGAGGAGTT | ASGERTGGAANTGQLY | 0.48368293 |
| QLVGLGI | ASGERTGGAANTGQLY | 0.7901501 |
| RFLEQVHQL | ASGERTGGAANTGQLY | 0.7122191 |
| RGVGYVLR | ASGERTGGAANTGQLY | 0.7376646 |
| SATGVFAGVI | ASGERTGGAANTGQLY | 0.7617611 |
| SLPAITLP | ASGERTGGAANTGQLY | 0.7441962 |
| STALAFTV | ASGERTGGAANTGQLY | 0.7101075 |
| SVLTVGLGK | ASGERTGGAANTGQLY | 0.620743 |

| | | |
|---|---|---|
| TALAGAGSGI | ASGERTGGAANTGQLY | 0.77762973 |
| TGLPFTGI | ASGERTGGAANTGQLY | 0.8009842 |
| VAEHVTV | ASGERTGGAANTGQLY | 0.7957999 |
| VDIVLNSL | ASGERTGGAANTGQLY | 0.8187336 |
| VLTGGGAVL | ASGERTGGAANTGQLY | 0.8107281 |
| VNVGLAYGI | ASGERTGGAANTGQLY | 0.7918032 |
| VSSAAVGAV | ASGERTGGAANTGQLY | 0.2298812 |
| VTSVYEAGV | ASGERTGGAANTGQLY | 0.43024832 |
| WVALDRGA | ASGERTGGAANTGQLY | 0.514804 |
| YLVGSVVI | ASGERTGGAANTGQLY | 0.7202009 |
| GPPGTGKTLL | ASGERTGGAANTGQLY | 0.26981753 |
| GVGMTKF | ASGERTGGAANTGQLY | 0.7025173 |
| IGIGGSDL | ASGERTGGAANTGQLY | 0.6670283 |
| IGTLLAEL | ASGERTGGAANTGQLY | 0.5985666 |
| IYTSGTT | ASGERTGGAANTGQLY | 0.8199284 |
| LSFPAGV | ASGERTGGAANTGQLY | 0.68099666 |
| LYTSGTT | ASGERTGGAANTGQLY | 0.55496013 |
| VGAFPFGF | ASGERTGGAANTGQLY | 0.57687014 |
| VGAGTVEF | ASGERTGGAANTGQLY | 0.8131021 |
| VGKSTLLL | ASGERTGGAANTGQLY | 0.4566792 |
| AALAHAGGGIV | ASGERTGGAANTGQLY | 0.80496305 |
| AALVAVSLL | ASGERTGGAANTGQLY | 0.6442253 |
| AAVSRAVAAL | ASGERTGGAANTGQLY | 0.8133358 |
| AGGSSGIGGF | ASGERTGGAANTGQLY | 0.3909107 |
| AGLTVANANG | ASGERTGGAANTGQLY | 0.7949593 |
| AGLVGAVL | ASGERTGGAANTGQLY | 0.7221551 |
| ALSQAFTV | ASGERTGGAANTGQLY | 0.83957946 |
| ALTFSVSL | ASGERTGGAANTGQLY | 0.7067441 |
| ALTGAALVAL | ASGERTGGAANTGQLY | 0.74613905 |
| ASFSIGAGVL | ASGERTGGAANTGQLY | 0.7039757 |
| DGSVYKI | ASGERTGGAANTGQLY | 0.8049786 |
| EQLAEIV | ASGERTGGAANTGQLY | 0.39960837 |
| FAVSELTVM | ASGERTGGAANTGQLY | 0.73353785 |
| GAAPLAAGL | ASGERTGGAANTGQLY | 0.44654018 |
| GADGGDGGSGGAGG | ASGERTGGAANTGQLY | 0.7640406 |
| GFGLDTGGAGG | ASGERTGGAANTGQLY | 0.77256286 |
| GLASGGLL | ASGERTGGAANTGQLY | 0.26999465 |
| GRIAGAAVLL | ASGERTGGAANTGQLY | 0.8059125 |
| GSAGSLNTSF | ASGERTGGAANTGQLY | 0.6331943 |
| GTGQIGI | ASGERTGGAANTGQLY | 0.5032264 |
| GTLIALIL | ASGERTGGAANTGQLY | 0.7956905 |
| IAATMGFVVG | ASGERTGGAANTGQLY | 0.71347123 |
| IFAQALL | ASGERTGGAANTGQLY | 0.8073815 |
| IGAAVALGF | ASGERTGGAANTGQLY | 0.632339 |
| IGLTPLAL | ASGERTGGAANTGQLY | 0.73647714 |
| ILGSNTSTL | ASGERTGGAANTGQLY | 0.78426635 |
| IVFSLTVL | ASGERTGGAANTGQLY | 0.3796726 |
| LASKATGF | ASGERTGGAANTGQLY | 0.49954414 |
| LASVALIL | ASGERTGGAANTGQLY | 0.8114679 |
| LFAVGLTL | ASGERTGGAANTGQLY | 0.87837 |
| LFNGGIGTYI | ASGERTGGAANTGQLY | 0.8753225 |

| | | |
|---|---|---|
| LTLGAGII | ASGERTGGAANTGQLY | 0.8113562 |
| LVAAALGDI | ASGERTGGAANTGQLY | 0.7534952 |
| NTAFVQLGI | ASGERTGGAANTGQLY | 0.8277576 |
| QLFSGAEL | ASGERTGGAANTGQLY | 0.8863958 |
| SAATFITL | ASGERTGGAANTGQLY | 0.88723695 |
| SNYGLANLGV | ASGERTGGAANTGQLY | 0.88183033 |
| SYGGAYAVM | ASGERTGGAANTGQLY | 0.65050846 |
| TTGVGGLTLG | ASGERTGGAANTGQLY | 0.88885295 |
| TTQTIYI | ASGERTGGAANTGQLY | 0.84829646 |
| VAAGAGLGAVI | ASGERTGGAANTGQLY | 0.839235 |
| VAAVVTLL | ASGERTGGAANTGQLY | 0.87835383 |
| VGFLGGLI | ASGERTGGAANTGQLY | 0.8506974 |
| VGVIPILL | ASGERTGGAANTGQLY | 0.8486885 |
| VIFASSLI | ASGERTGGAANTGQLY | 0.8279867 |
| VKGGDGGAAAG | ASGERTGGAANTGQLY | 0.86470336 |
| VTKTFTNI | ASGERTGGAANTGQLY | 0.8849289 |
| VVAAAILL | ASGERTGGAANTGQLY | 0.87855196 |
| YAALDVEL | ASGERTGGAANTGQLY | 0.90743744 |
| LASVALLL | ASGERTGGAANTGQLY | 0.8960991 |
| LDAVHRV | ASGERTGGAANTGQLY | 0.8900602 |
| QGLSSSIF | ASGERTGGAANTGQLY | 0.32043236 |
| RGTFANIRL | ASGERTGGAANTGQLY | 0.5923625 |
| RQLWWGHRI | ASGERTGGAANTGQLY | 0.791617 |
| YGPPGTGKTLL | ASGERTGGAANTGQLY | 0.83301055 |
| AALVGAPANVA | ASGERTGGAANTGQLY | 0.40754732 |
| AATFRRTE | ASGERTGGAANTGQLY | 0.835339 |
| AGALGGGI | ASGERTGGAANTGQLY | 0.842453 |
| AGGAGGAGGIGAG | ASGERTGGAANTGQLY | 0.77491295 |
| AGSGALGL | ASGERTGGAANTGQLY | 0.89571416 |
| AQIADQI | ASGERTGGAANTGQLY | 0.83461106 |
| ASAVAVTYL | ASGERTGGAANTGQLY | 0.7592181 |
| ASTVLGGTGG | ASGERTGGAANTGQLY | 0.79481685 |
| AVGGEPAK | ASGERTGGAANTGQLY | 0.9021274 |
| CEQGTFGGTC | ASGERTGGAANTGQLY | 0.73398936 |
| DEGKAPAG | ASGERTGGAANTGQLY | 0.89412093 |
| DTVVFPAL | ASGERTGGAANTGQLY | 0.7995922 |
| FEGITMAG | ASGERTGGAANTGQLY | 0.9016146 |
| FPLVVQD | ASGERTGGAANTGQLY | 0.55427724 |
| GAFTTPQ | ASGERTGGAANTGQLY | 0.8895304 |
| GFGGFGTAGGVGGA | ASGERTGGAANTGQLY | 0.84355724 |
| GFLAQFL | ASGERTGGAANTGQLY | 0.91214454 |
| GGAGGAGGNAGVA | ASGERTGGAANTGQLY | 0.8335166 |
| GGVGTAGAGGAGGAGG | ASGERTGGAANTGQLY | 0.8698429 |
| GNGGGGDAGNAGSGGN | ASGERTGGAANTGQLY | 0.8160208 |
| GTGGAGGAGNDAGS | ASGERTGGAANTGQLY | 0.89119595 |
| GTTNTGWFNS | ASGERTGGAANTGQLY | 0.61067927 |
| IFMGPLFTL | ASGERTGGAANTGQLY | 0.8315792 |
| ILAHAIEAI | ASGERTGGAANTGQLY | 0.69568324 |
| IRSAIREAL | ASGERTGGAANTGQLY | 0.8697566 |
| ITFAGGG | ASGERTGGAANTGQLY | 0.88273025 |
| ITGMSGHQEV | ASGERTGGAANTGQLY | 0.43121338 |

| | | |
|---|---|---|
| IVAGLACTL | ASGERTGGAANTGQLY | 0.89884317 |
| IVGGGLAGA | ASGERTGGAANTGQLY | 0.77414715 |
| IVTGVLNI | ASGERTGGAANTGQLY | 0.699445 |
| LAQQGCI | ASGERTGGAANTGQLY | 0.89602065 |
| LCLHVEN | ASGERTGGAANTGQLY | 0.8465635 |
| LDAISDGS | ASGERTGGAANTGQLY | 0.89632547 |
| LGGFAAML | ASGERTGGAANTGQLY | 0.8285246 |
| LGGKNPMI | ASGERTGGAANTGQLY | 0.8547653 |
| LGPKGVAAIL | ASGERTGGAANTGQLY | 0.87390137 |
| LMVGVSI | ASGERTGGAANTGQLY | 0.5187585 |
| LVGAAFVL | ASGERTGGAANTGQLY | 0.7438246 |
| LVLAVAPL | ASGERTGGAANTGQLY | 0.9017532 |
| MAAIAVIDAA | ASGERTGGAANTGQLY | 0.6967344 |
| NNGINAELQ | ASGERTGGAANTGQLY | 0.7109275 |
| RGEPLVLVHG | ASGERTGGAANTGQLY | 0.6519848 |
| RGGAGGAAGLI | ASGERTGGAANTGQLY | 0.61982137 |
| RSYRMHPAV | ASGERTGGAANTGQLY | 0.6549074 |
| SAAHLLAGQLA | ASGERTGGAANTGQLY | 0.7107619 |
| SAAVGSY | ASGERTGGAANTGQLY | 0.7074574 |
| SPHAVLHRI | ASGERTGGAANTGQLY | 0.70007765 |
| SQIAKVL | ASGERTGGAANTGQLY | 0.533369 |
| TALGHLVP | ASGERTGGAANTGQLY | 0.6894938 |
| TAPTTSGAA | ASGERTGGAANTGQLY | 0.66111076 |
| TDGDDGNGGSPGT | ASGERTGGAANTGQLY | 0.6486411 |
| TDGLGEN | ASGERTGGAANTGQLY | 0.6979734 |
| TGGPPLPGQA | ASGERTGGAANTGQLY | 0.6844143 |
| TIPAITI | ASGERTGGAANTGQLY | 0.6530905 |
| TIPGFTI | ASGERTGGAANTGQLY | 0.67570436 |
| TSIASGALA | ASGERTGGAANTGQLY | 0.67803067 |
| TVTLPGAAI | ASGERTGGAANTGQLY | 0.67944866 |
| VASGLSAALP | ASGERTGGAANTGQLY | 0.68057394 |
| VGAGVIGI | ASGERTGGAANTGQLY | 0.7148018 |
| VGPKRFGV | ASGERTGGAANTGQLY | 0.70792365 |
| VNVAFPD | ASGERTGGAANTGQLY | 0.705446 |
| VSMTNSGVS | ASGERTGGAANTGQLY | 0.3333112 |
| VTDVLGDLSV | ASGERTGGAANTGQLY | 0.45687598 |
| VVVVGAGFAGL | ASGERTGGAANTGQLY | 0.6463312 |
| VYTVGTVV | ASGERTGGAANTGQLY | 0.63300735 |
| WTTFSGP | ASGERTGGAANTGQLY | 0.455923 |
| YDELVEAF | ASGERTGGAANTGQLY | 0.6761099 |
| YGTGGAGGNGG | ASGERTGGAANTGQLY | 0.6621082 |
| YTNVMAV | ASGERTGGAANTGQLY | 0.6227503 |
| FLEQVHQL | ASGERTGGAANTGQLY | 0.6990725 |
| GLSLLAVAV | ASGERTGGAANTGQLY | 0.65722954 |
| LEQVHQL | ASGERTGGAANTGQLY | 0.6436284 |
| LNAAGAI | ASGERTGGAANTGQLY | 0.59670126 |
| PAVHASL | ASGERTGGAANTGQLY | 0.67059565 |
| TELEDDA | ASGERTGGAANTGQLY | 0.5148684 |
| TPPGPPN | ASGERTGGAANTGQLY | 0.70284057 |
| TPPPPPPP | ASGERTGGAANTGQLY | 0.52137 |
| TQSPALI | ASSDNAETLY | 0.704461 |

| | | |
|---|---|---|
| EFAAAFPQ | ASSDNAETLY | 0.5443265 |
| LLSGNLDVL | ASSDNAETLY | 0.7005426 |
| GPANLGNYNI | ASSDNAETLY | 0.68089867 |
| TLGPTLP | ASSDNAETLY | 0.7260579 |
| VYAYSSCCK | ASSDNAETLY | 0.6730181 |
| APAPAPAPNI | ASSDNAETLY | 0.6819825 |
| PGGPVPIP | ASSDNAETLY | 0.59739804 |
| FLTVALEP | ASSDNAETLY | 0.7101628 |
| SQYFGANL | ASSDNAETLY | 0.5314698 |
| GGVKPAA | ASSDNAETLY | 0.67337203 |
| GGTGGAGGAGGW | ASSDNAETLY | 0.46357474 |
| GVGGVAGLGGAGG | ASSDNAETLY | 0.7010739 |
| VLLVLATTI | ASSDNAETLY | 0.7004268 |
| IGIFAAML | ASSDNAETLY | 0.54778856 |
| AVGDLID | ASSDNAETLY | 0.710266 |
| NISPLPL | ASSDNAETLY | 0.67935073 |
| VTAVFVAI | ASSDNAETLY | 0.63208055 |
| SGDDNDAAL | ASSDNAETLY | 0.7174506 |
| AVAALAHS | ASSDNAETLY | 0.6581236 |
| GGVTVMATP | ASSDNAETLY | 0.71600366 |
| AGLAANVRD | ASSDNAETLY | 0.6618277 |
| TNRPDLIDPALLRPG | ASSDNAETLY | 0.6236214 |
| LANLGVTAL | ASSDNAETLY | 0.68104875 |
| TGERIEL | ASSDNAETLY | 0.5192974 |
| NGEELAA | ASSDNAETLY | 0.5569233 |
| VLSLVWG | ASSDNAETLY | 0.71542853 |
| IGYALWL | ASSDNAETLY | 0.6553277 |
| LSLLLAPL | ASSDNAETLY | 0.68756366 |
| TLDEALK | ASSDNAETLY | 0.6179413 |
| TDGAGTAEGG | ASSDNAETLY | 0.56834626 |
| NAPGGAGGNGG | ASSDNAETLY | 0.6250619 |
| VSVFMGMGGNF | ASSDNAETLY | 0.6736698 |
| LASYLAAL | ASSDNAETLY | 0.68257666 |
| LVGGTPAL | ASSDNAETLY | 0.66415644 |
| VRNADVL | ASSDNAETLY | 0.5438884 |
| TCSSPFLVSAM | ASSDNAETLY | 0.64130867 |
| VHITDAE | ASSDNAETLY | 0.6193471 |
| EYAAPSF | ASSDNAETLY | 0.63025707 |
| VVDAFHGGL | ASSDNAETLY | 0.6604821 |
| ASVVDPQ | ASSDNAETLY | 0.64576095 |
| SYNEFGGLS | ASSDNAETLY | 0.59422725 |
| DPDGSVY | ASSDNAETLY | 0.617519 |
| MVSTIHATE | ASSDNAETLY | 0.65262127 |
| LAAGVADPV | ASSDNAETLY | 0.6681719 |
| TKEQIAA | ASSDNAETLY | 0.6657611 |
| SLASPAAL | ASSDNAETLY | 0.68576926 |
| IGLVFYGV | ASSDNAETLY | 0.67393374 |
| FLLDASNL | ASSDNAETLY | 0.6756952 |
| YQVGNFP | ASSDNAETLY | 0.3683279 |
| ESLLSQST | ASSDNAETLY | 0.47924116 |
| LFTVGFAAVG | ASSDNAETLY | 0.6070286 |

| | | |
|---|---|---|
| ATPPTPT | ASSDNAETLY | 0.6335532 |
| TDLATLG | ASSDNAETLY | 0.49597356 |
| SAGGTGR | ASSDNAETLY | 0.6569706 |
| GFSNSGS | ASSDNAETLY | 0.5901502 |
| GGNGGSGGTSV | ASSDNAETLY | 0.56422263 |
| GTSVATGGAGN | ASSDNAETLY | 0.6714796 |
| NGGNGGSGGTS | ASSDNAETLY | 0.6173048 |
| AVVKFPH | ASSDNAETLY | 0.58998734 |
| TPPPPPP | ASSDNAETLY | 0.59770167 |
| AAPAAAAAPA | ASSDNAETLY | 0.67155325 |
| IALNEDL | ASSDNAETLY | 0.49504802 |
| IVNVSSM | ASSDNAETLY | 0.66504663 |
| TGIPPAPR | ASSDNAETLY | 0.5912142 |
| FDLGGGTF | ASSDNAETLY | 0.6769586 |
| FDLGGGTFDVS | ASSDNAETLY | 0.4934997 |
| NEPTAAA | ASSDNAETLY | 0.6709173 |
| NEPTAAALA | ASSDNAETLY | 0.6395546 |
| GSIINISSI | ASSDNAETLY | 0.6897933 |
| VAPGNAGTA | ASSDNAETLY | 0.6247324 |
| TGGGASLE | ASSDNAETLY | 0.6438289 |
| VAVYNHYKRI | ASSDNAETLY | 0.60486925 |
| TDPAPATT | ASSDNAETLY | 0.68124837 |
| TNRPDILDPALLRPG | ASSDNAETLY | 0.4971617 |
| LTTTTTS | ASSDNAETLY | 0.6339171 |
| PTGSGKTL | ASSDNAETLY | 0.5123471 |
| TGSGKTL | ASSDNAETLY | 0.6540006 |
| TIGHVDHGK | ASSDNAETLY | 0.6621593 |
| TIGHVDHGKTT | ASSDNAETLY | 0.511656 |
| GHVDHGKTT | ASSDNAETLY | 0.6755209 |
| AQFEHTLL | ASSDNAETLY | 0.6468827 |
| IVNQDEL | ASSDNAETLY | 0.5957302 |
| TNGSQFF | ASSDNAETLY | 0.68214095 |
| TGKGIVE | ASSDNAETLY | 0.6216796 |
| LISNASDA | ASSDNAETLY | 0.6727862 |
| AELRTLGELTKIAKAHGAQ | ASSDNAETLY | 0.6383802 |
| APNAVAAAEPSW | ASSDNAETLY | 0.6300009 |
| APQVAGAATAVL | ASSDNAETLY | 0.6578462 |
| ARRDPAAAAAILPTDA | ASSDNAETLY | 0.6346841 |
| DAAARDVLAEAGLAEYF | ASSDNAETLY | 0.56578034 |
| DDQIVLSAKANLAKVEQ | ASSDNAETLY | 0.68007326 |
| DELLPDTGEDSDEDGH | ASSDNAETLY | 0.7705086 |
| ELPEAPANRATVPQD | ASSDNAETLY | 0.79017913 |
| FGQSKIERVTGIG | ASSDNAETLY | 0.72745264 |
| GAGGTGSTAGGAGGAGGAGGI | ASSDNAETLY | 0.6803163 |
| GHGGTGAAGGNGGTGGAGGS( | ASSDNAETLY | 0.7281528 |
| GNGGNGGAGGEGGPGGAGGA( | ASSDNAETLY | 0.7809356 |
| GSTAGNGGNGGNATNGTIAG( | ASSDNAETLY | 0.78926814 |
| GTGGDGGNGGNGGTGG | ASSDNAETLY | 0.772733 |
| GVRRPAGIPADAPV | ASSDNAETLY | 0.66180456 |
| GYRQVTDAHLLTLARR | ASSDNAETLY | 0.76289475 |
| IAKMGQDGHDRG | ASSDNAETLY | 0.746405 |

| | | |
|---|---|---|
| KAEPSIEIQYT | ASSDNAETLY | 0.7462768 |
| KPVRIRMAADEDTVAVT | ASSDNAETLY | 0.77496105 |
| MPPRIAGMRLLVIKPEP | ASSDNAETLY | 0.76207674 |
| MKFYAVALFRSRGVPPTR | ASSDNAETLY | 0.72352517 |
| NAGGFGILGAVAHSP | ASSDNAETLY | 0.7306819 |
| NEGREITLAMAEM | ASSDNAETLY | 0.7574605 |
| NELVEWSHPLGHRWRWE | ASSDNAETLY | 0.77119565 |
| NLGAGGTGGNGGNPG | ASSDNAETLY | 0.76592404 |
| NTEWPKWETVMAADPVD | ASSDNAETLY | 0.79111934 |
| PASAAPAETQP | ASSDNAETLY | 0.7822021 |
| QQSSDGKNGVRAAEITGE | ASSDNAETLY | 0.7790361 |
| RAPDAVTTSWWKE | ASSDNAETLY | 0.38503513 |
| SGFGNVGAHTSGWFNQ | ASSDNAETLY | 0.65218073 |
| SGVAGAVGQAASVGGLKV | ASSDNAETLY | 0.74026537 |
| SNGGAGTGSAGGNGGT | ASSDNAETLY | 0.74239707 |
| TGGAGGEGGAAG | ASSDNAETLY | 0.5610684 |
| TGGDGGAAGAGGAPGHG | ASSDNAETLY | 0.7595529 |
| TMPPLLAAIANAGRHGV | ASSDNAETLY | 0.7316693 |
| TSSTAGNSGGA | ASSDNAETLY | 0.68052435 |
| VPATVPVGILC | ASSDNAETLY | 0.77487445 |
| VQRPDSGGVVLTGRIS | ASSDNAETLY | 0.727141 |
| WTPNPVPATKGVPAGLVQAT | ASSDNAETLY | 0.7164687 |
| YALKRALAKFSDRFDVVII | ASSDNAETLY | 0.7243418 |
| YLIGSVIV | ASSDNAETLY | 0.78403366 |
| YGQLVAVID | ASSDNAETLY | 0.6591885 |
| VTVGGMTLPA | ASSDNAETLY | 0.77288866 |
| VLALPAV | ASSDNAETLY | 0.74945074 |
| VIPPTLNYET | ASSDNAETLY | 0.7839439 |
| VGMGGMHPSSG | ASSDNAETLY | 0.61520123 |
| VGLAGGAATA | ASSDNAETLY | 0.7784791 |
| VGGAGGAGGSGP | ASSDNAETLY | 0.7493335 |
| VFAAFMV | ASSDNAETLY | 0.7930189 |
| TLYDKLR | ASSDNAETLY | 0.73912394 |
| TLAVFTVL | ASSDNAETLY | 0.7649159 |
| TGSVQPARV | ASSDNAETLY | 0.7578399 |
| TGGGAGSGGT | ASSDNAETLY | 0.78823596 |
| SHLETSL | ASSDNAETLY | 0.5982895 |
| SFTTGMT | ASSDNAETLY | 0.75641733 |
| QVEVAKRMGVRQPTVS | ASSDNAETLY | 0.6795467 |
| QGSGGPALAM | ASSDNAETLY | 0.7693895 |
| MLAPSNP | ASSDNAETLY | 0.7743148 |
| LVNNAAI | ASSDNAETLY | 0.5101038 |
| LTGEINTLA | ASSDNAETLY | 0.78340733 |
| LSIFGPV | ASSDNAETLY | 0.7379336 |
| LLALLANL | ASSDNAETLY | 0.7068851 |
| LISNASDALDKLR | ASSDNAETLY | 0.7842276 |
| LINEAVP | ASSDNAETLY | 0.7405056 |
| LGYAFSGL | ASSDNAETLY | 0.7827681 |
| LDFVAVAL | ASSDNAETLY | 0.75430655 |
| LAQPGAE | ASSDNAETLY | 0.7659366 |
| ISNLGAVTGF | ASSDNAETLY | 0.77279496 |

| | | |
|---|---|---|
| HQEDPGRI | ASSDNAETLY | 0.6201763 |
| GTPGEPTP | ASSDNAETLY | 0.6999433 |
| GFFNSGS | ASSDNAETLY | 0.7870252 |
| GAGGGGLTAGAN | ASSDNAETLY | 0.7806827 |
| FVDALVNM | ASSDNAETLY | 0.7635346 |
| FLTVAPEL | ASSDNAETLY | 0.7131655 |
| FETLTAL | ASSDNAETLY | 0.6958398 |
| DVINPIEL | ASSDNAETLY | 0.72296786 |
| APVSGAPV | ASSDNAETLY | 0.7741674 |
| ALVGGLAW | ASSDNAETLY | 0.7838799 |
| ALAAAAATAI | ASSDNAETLY | 0.7827981 |
| AFGLGLLL | ASSDNAETLY | 0.59210014 |
| AFFAAASL | ASSDNAETLY | 0.78669155 |
| AAAALPAFSPP | ASSDNAETLY | 0.76887685 |
| VVLAPAL | ASSDNAETLY | 0.77584976 |
| TIGHVDHGKTTL | ASSDNAETLY | 0.7508725 |
| LISNASDALDK | ASSDNAETLY | 0.7537154 |
| LISNASDAL | ASSDNAETLY | 0.7246587 |
| ISNASDALDK | ASSDNAETLY | 0.6563297 |
| NQAETLVYQTE | ASSDNAETLY | 0.7687334 |
| TGGTGGTAGAG | ASSDNAETLY | 0.77681744 |
| TGSTAGGAGGA | ASSDNAETLY | 0.77520716 |
| DPLAVPVAAGPV | ASSDNAETLY | 0.7890511 |
| GGTGGTPGNGGN | ASSDNAETLY | 0.7878952 |
| PAGLAGVDPATL | ASSDNAETLY | 0.77702343 |
| GNNLVVEAPHAGASV | ASSDNAETLY | 0.3745508 |
| KVTRIMSASTLEQPA | ASSDNAETLY | 0.5670281 |
| LGFALPIALPRRTVV | ASSDNAETLY | 0.69718975 |
| TRLGNLTARKLVVLV | ASSDNAETLY | 0.71987474 |
| WSQQRRLIYFPSAGP | ASSDNAETLY | 0.31676218 |
| GPNGNPDPMAAAVDIR | ASSDNAETLY | 0.70105606 |
| TQRCVSVADLRAAASR | ASSDNAETLY | 0.70692134 |
| QFVLDVAEHPVEVLTETV | ASSDNAETLY | 0.73021436 |
| GGAGGTGGAGGNGGTQAGDG | ASSDNAETLY | 0.7815542 |
| PLLATLRAIAVDVDAKPAQV | ASSDNAETLY | 0.7535839 |
| APKFRTGGYMIDFWGVGYQV/ | ASSDNAETLY | 0.575783 |
| GAGGAGGAGGAAGGPGGTGG' | ASSDNAETLY | 0.6533764 |
| IGALGLLATDVAPAGSVAFG' | ASSDNAETLY | 0.7869724 |
| AGGAGGIGGNGGAGGAGGVLI | ASSDNAETLY | 0.67484456 |
| ADELIGLL | ASSDNAETLY | 0.7900008 |
| ADRALKA | ASSDNAETLY | 0.59816444 |
| AGAVGQVGGAGGA | ASSDNAETLY | 0.78847605 |
| AGGGGGLGGAP | ASSDNAETLY | 0.6100174 |
| AGGLLYGNGGAG | ASSDNAETLY | 0.78914595 |
| AGLESIGL | ASSDNAETLY | 0.77542365 |
| AGLKTGII | ASSDNAETLY | 0.79172444 |
| ASLAALLL | ASSDNAETLY | 0.7423856 |
| ASVINISSI | ASSDNAETLY | 0.7766516 |
| ATPPGPA | ASSDNAETLY | 0.72493154 |
| ATVASALV | ASSDNAETLY | 0.76948726 |
| ATVASLIL | ASSDNAETLY | 0.68241835 |

| | | |
|---|---|---|
| AVNGPVLIRH | ASSDNAETLY | 0.72840273 |
| DGAAGAVGI | ASSDNAETLY | 0.5896797 |
| DPLLHGI | ASSDNAETLY | 0.7489133 |
| ELGAALGAL | ASSDNAETLY | 0.78643614 |
| ELGRTQAV | ASSDNAETLY | 0.24161342 |
| FVAGGYLS | ASSDNAETLY | 0.7811875 |
| GASGGAGGAGGW | ASSDNAETLY | 0.67275214 |
| GGAGAGGDGGAGGI | ASSDNAETLY | 0.60893536 |
| GTGHNGSQPGMGGNG | ASSDNAETLY | 0.781392 |
| GYAAASASAS | ASSDNAETLY | 0.7639065 |
| HQVDVAAVQQG | ASSDNAETLY | 0.77471435 |
| IGAAIAHT | ASSDNAETLY | 0.7047124 |
| IGMIEVAQL | ASSDNAETLY | 0.73632526 |
| IRLAPEHTSAMD | ASSDNAETLY | 0.7464268 |
| IYNVSVL | ASSDNAETLY | 0.38310054 |
| KQGFLGM | ASSDNAETLY | 0.56948197 |
| LESVITQL | ASSDNAETLY | 0.79275846 |
| LGLGALVVV | ASSDNAETLY | 0.82281613 |
| LIGNGGNGGSGGI | ASSDNAETLY | 0.82955086 |
| LNAINAPTL | ASSDNAETLY | 0.69636804 |
| LQLLNTGV | ASSDNAETLY | 0.6357162 |
| LQTSIEF | ASSDNAETLY | 0.7302134 |
| MSFAVSEL | ASSDNAETLY | 0.8464637 |
| NTGALGL | ASSDNAETLY | 0.8592987 |
| PPGAGGAGGAGTT | ASSDNAETLY | 0.8331909 |
| QLVGLGI | ASSDNAETLY | 0.6620181 |
| RFLEQVHQL | ASSDNAETLY | 0.8369359 |
| RGVGYVLR | ASSDNAETLY | 0.7772602 |
| SATGVFAGVI | ASSDNAETLY | 0.75135016 |
| SLPAITLP | ASSDNAETLY | 0.8314943 |
| STALAFTV | ASSDNAETLY | 0.81533486 |
| SVLTVGLGK | ASSDNAETLY | 0.7396306 |
| TALAGAGSGI | ASSDNAETLY | 0.726312 |
| TGLPFTGI | ASSDNAETLY | 0.8249864 |
| VAEHVTV | ASSDNAETLY | 0.8400667 |
| VDIVLNSL | ASSDNAETLY | 0.83231056 |
| VLTGGGAVL | ASSDNAETLY | 0.8892005 |
| VNVGLAYGI | ASSDNAETLY | 0.8506604 |
| VSSAAVGAV | ASSDNAETLY | 0.8524314 |
| VTSVYEAGV | ASSDNAETLY | 0.25649345 |
| WVALDRGA | ASSDNAETLY | 0.5024384 |
| YLVGSVVI | ASSDNAETLY | 0.72312856 |
| GPPGTGKTLL | ASSDNAETLY | 0.76426923 |
| GVGMTKF | ASSDNAETLY | 0.40072757 |
| IGIGGSDL | ASSDNAETLY | 0.8265337 |
| IGTLLAEL | ASSDNAETLY | 0.7684821 |
| IYTSGTT | ASSDNAETLY | 0.67349553 |
| LSFPAGV | ASSDNAETLY | 0.8480034 |
| LYTSGTT | ASSDNAETLY | 0.72149086 |
| VGAFPFGF | ASSDNAETLY | 0.69799614 |
| VGAGTVEF | ASSDNAETLY | 0.74175763 |

| | | |
|---|---|---|
| VGKSTLLL | ASSDNAETLY | 0.8712295 |
| AALAHAGGGIV | ASSDNAETLY | 0.6221439 |
| AALVAVSLL | ASSDNAETLY | 0.8446294 |
| AAVSRAVAAL | ASSDNAETLY | 0.7200496 |
| AGGSSGIGGF | ASSDNAETLY | 0.86616635 |
| AGLTVANANG | ASSDNAETLY | 0.56620246 |
| AGLVGAVL | ASSDNAETLY | 0.8453102 |
| ALSQAFTV | ASSDNAETLY | 0.7652929 |
| ALTFSVSL | ASSDNAETLY | 0.8920814 |
| ALTGAALVAL | ASSDNAETLY | 0.7493057 |
| ASFSIGAGVL | ASSDNAETLY | 0.8332523 |
| DGSVYKI | ASSDNAETLY | 0.7903498 |
| EQLAEIV | ASSDNAETLY | 0.8366104 |
| FAVSELTVM | ASSDNAETLY | 0.48985526 |
| GAAPLAAGL | ASSDNAETLY | 0.77938086 |
| GADGGDGGSGGAGG | ASSDNAETLY | 0.5656887 |
| GFGLDTGGAGG | ASSDNAETLY | 0.8133775 |
| GLASGGLL | ASSDNAETLY | 0.826749 |
| GRIAGAAVLL | ASSDNAETLY | 0.31905252 |
| GSAGSLNTSF | ASSDNAETLY | 0.8609717 |
| GTGQIGI | ASSDNAETLY | 0.6922296 |
| GTLIALIL | ASSDNAETLY | 0.680385 |
| IAATMGFVVG | ASSDNAETLY | 0.86055815 |
| IFAQALL | ASSDNAETLY | 0.7942519 |
| IGAAVALGF | ASSDNAETLY | 0.8558184 |
| IGLTPLAL | ASSDNAETLY | 0.7740373 |
| ILGSNTSTL | ASSDNAETLY | 0.7814173 |
| IVFSLTVL | ASSDNAETLY | 0.8275834 |
| LASKATGF | ASSDNAETLY | 0.5197904 |
| LASVALIL | ASSDNAETLY | 0.70034945 |
| LFAVGLTL | ASSDNAETLY | 0.8686148 |
| LFNGGIGTYI | ASSDNAETLY | 0.8819182 |
| LTLGAGII | ASSDNAETLY | 0.88018346 |
| LVAAALGDI | ASSDNAETLY | 0.7923728 |
| NTAFVQLGI | ASSDNAETLY | 0.7588358 |
| QLFSGAEL | ASSDNAETLY | 0.82018965 |
| SAATFITL | ASSDNAETLY | 0.88706434 |
| SNYGLANLGV | ASSDNAETLY | 0.903713 |
| SYGGAYAVM | ASSDNAETLY | 0.8921672 |
| TTGVGGLTLG | ASSDNAETLY | 0.8015455 |
| TTQTIYI | ASSDNAETLY | 0.8969798 |
| VAAGAGLGAVI | ASSDNAETLY | 0.87491226 |
| VAAVVTLL | ASSDNAETLY | 0.8507749 |
| VGFLGGLI | ASSDNAETLY | 0.8841146 |
| VGVIPILL | ASSDNAETLY | 0.8894803 |
| VIFASSLI | ASSDNAETLY | 0.8219351 |
| VKGGDGGAAAG | ASSDNAETLY | 0.8426266 |
| VTKTFTNI | ASSDNAETLY | 0.87042785 |
| VVAAAILL | ASSDNAETLY | 0.88465166 |
| YAALDVEL | ASSDNAETLY | 0.8844489 |
| LASVALLL | ASSDNAETLY | 0.9149115 |

| | | |
|---|---|---|
| LDAVHRV | ASSDNAETLY | 0.89648503 |
| QGLSSSIF | ASSDNAETLY | 0.88317436 |
| RGTFANIRL | ASSDNAETLY | 0.40555748 |
| RQLWWGHRI | ASSDNAETLY | 0.73407006 |
| YGPPGTGKTLL | ASSDNAETLY | 0.842292 |
| AALVGAPANVA | ASSDNAETLY | 0.8531152 |
| AATFRRTE | ASSDNAETLY | 0.47438633 |
| AGALGGGI | ASSDNAETLY | 0.87659 |
| AGGAGGAGGIGAG | ASSDNAETLY | 0.8489592 |
| AGSGALGL | ASSDNAETLY | 0.81418514 |
| AQIADQI | ASSDNAETLY | 0.89808416 |
| ASAVAVTYL | ASSDNAETLY | 0.84802043 |
| ASTVLGGTGG | ASSDNAETLY | 0.8243295 |
| AVGGEPAK | ASSDNAETLY | 0.8494524 |
| CEQGTFGGTC | ASSDNAETLY | 0.9060188 |
| DEGKAPAG | ASSDNAETLY | 0.792572 |
| DTVVFPAL | ASSDNAETLY | 0.9047278 |
| FEGITMAG | ASSDNAETLY | 0.7734524 |
| FPLVVQD | ASSDNAETLY | 0.9024608 |
| GAFTTPQ | ASSDNAETLY | 0.76353586 |
| GFGGFGTAGGVGGA | ASSDNAETLY | 0.90053594 |
| GFLAQFL | ASSDNAETLY | 0.86212444 |
| GGAGGAGGNAGVA | ASSDNAETLY | 0.9179178 |
| GGVGTAGAGGAGGAGG | ASSDNAETLY | 0.8467869 |
| GNGGGGDAGNAGSGGN | ASSDNAETLY | 0.9053477 |
| GTGGAGGAGNDAGS | ASSDNAETLY | 0.86034834 |
| GTTNTGWFNS | ASSDNAETLY | 0.90091574 |
| IFMGPLFTL | ASSDNAETLY | 0.7583711 |
| ILAHAIEAI | ASSDNAETLY | 0.85503775 |
| IRSAIREAL | ASSDNAETLY | 0.67858857 |
| ITFAGGG | ASSDNAETLY | 0.8726038 |
| ITGMSGHQEV | ASSDNAETLY | 0.89206886 |
| IVAGLACTL | ASSDNAETLY | 0.5880377 |
| IVGGGLAGA | ASSDNAETLY | 0.9069263 |
| IVTGVLNI | ASSDNAETLY | 0.79763275 |
| LAQQGCI | ASSDNAETLY | 0.8117422 |
| LCLHVEN | ASSDNAETLY | 0.9032272 |
| LDAISDGS | ASSDNAETLY | 0.87323964 |
| LGGFAAML | ASSDNAETLY | 0.89044315 |
| LGGKNPMI | ASSDNAETLY | 0.844967 |
| LGPKGVAAIL | ASSDNAETLY | 0.85532427 |
| LMVGVSI | ASSDNAETLY | 0.8687498 |
| LVGAAFVL | ASSDNAETLY | 0.61973196 |
| LVLAVAPL | ASSDNAETLY | 0.8027109 |
| MAAIAVIDAA | ASSDNAETLY | 0.91337466 |
| NNGINAELQ | ASSDNAETLY | 0.78687435 |
| RGEPLVLVHG | ASSDNAETLY | 0.77068615 |
| RGGAGGAAGLI | ASSDNAETLY | 0.59266424 |
| RSYRMHPAV | ASSDNAETLY | 0.51745427 |
| SAAHLLAGQLA | ASSDNAETLY | 0.62874013 |
| SAAVGSY | ASSDNAETLY | 0.7844429 |

| | | |
|---|---|---|
| SPHAVLHRI | ASSDNAETLY | 0.8237687 |
| SQIAKVL | ASSDNAETLY | 0.7761165 |
| TALGHLVP | ASSDNAETLY | 0.5105365 |
| TAPTTSGAA | ASSDNAETLY | 0.79424345 |
| TDGDDGNGGSPGT | ASSDNAETLY | 0.74624777 |
| TDGLGEN | ASSDNAETLY | 0.69187766 |
| TGGPPLPGQA | ASSDNAETLY | 0.7399232 |
| TIPAITI | ASSDNAETLY | 0.7342997 |
| TIPGFTI | ASSDNAETLY | 0.5654894 |
| TSIASGALA | ASSDNAETLY | 0.57097864 |
| TVTLPGAAI | ASSDNAETLY | 0.7385186 |
| VASGLSAALP | ASSDNAETLY | 0.7951589 |
| VGAGVIGI | ASSDNAETLY | 0.79514503 |
| VGPKRFGV | ASSDNAETLY | 0.8443732 |
| VNVAFPD | ASSDNAETLY | 0.8054512 |
| VSMTNSGVS | ASSDNAETLY | 0.784714 |
| VTDVLGDLSV | ASSDNAETLY | 0.25527886 |
| VVVVGAGFAGL | ASSDNAETLY | 0.35028362 |
| VYTVGTVV | ASSDNAETLY | 0.6892165 |
| WTTFSGP | ASSDNAETLY | 0.5983083 |
| YDELVEAF | ASSDNAETLY | 0.2614277 |
| YGTGGAGGNGG | ASSDNAETLY | 0.7145189 |
| YTNVMAV | ASSDNAETLY | 0.6695485 |
| FLEQVHQL | ASSDNAETLY | 0.57455826 |
| GLSLLAVAV | ASSDNAETLY | 0.79074144 |
| LEQVHQL | ASSDNAETLY | 0.63923895 |
| LNAAGAI | ASSDNAETLY | 0.48633057 |
| PAVHASL | ASSDNAETLY | 0.6750959 |
| TELEDDA | ASSDNAETLY | 0.8016054 |
| TPPGPPN | ASSDNAETLY | 0.5868779 |
| TPPPPPPP | ASSDNAETLY | 0.8117688 |
| TQSPALI | ASSETNSDYT | 0.5880806 |
| EFAAAFPQ | ASSETNSDYT | 0.8333153 |
| LLSGNLDVL | ASSETNSDYT | 0.45330745 |
| GPANLGNYNI | ASSETNSDYT | 0.8091304 |
| TLGPTLP | ASSETNSDYT | 0.71837544 |
| VYAYSSCCK | ASSETNSDYT | 0.86327773 |
| APAPAPAPNI | ASSETNSDYT | 0.6598215 |
| PGGPVPIP | ASSETNSDYT | 0.7992902 |
| FLTVALEP | ASSETNSDYT | 0.76072305 |
| SQYFGANL | ASSETNSDYT | 0.8212291 |
| GGVKPAA | ASSETNSDYT | 0.5127115 |
| GGTGGAGGAGGW | ASSETNSDYT | 0.6894626 |
| GVGGVAGLGGAGG | ASSETNSDYT | 0.41641438 |
| VLLVLATTI | ASSETNSDYT | 0.73383296 |
| IGIFAAML | ASSETNSDYT | 0.8049656 |
| AVGDLID | ASSETNSDYT | 0.19782963 |
| NISPLPL | ASSETNSDYT | 0.82154197 |
| VTAVFVAI | ASSETNSDYT | 0.4464711 |
| SGDDNDAAL | ASSETNSDYT | 0.56011784 |
| AVAALAHS | ASSETNSDYT | 0.80050236 |

| | | |
|---|---|---|
| GGVTVMATP | ASSETNSDYT | 0.73499966 |
| AGLAANVRD | ASSETNSDYT | 0.8065933 |
| TNRPDLIDPALLRPG | ASSETNSDYT | 0.6403287 |
| LANLGVTAL | ASSETNSDYT | 0.7230592 |
| TGERIEL | ASSETNSDYT | 0.73601687 |
| NGEELAA | ASSETNSDYT | 0.4093616 |
| VLSLVWG | ASSETNSDYT | 0.56195146 |
| IGYALWL | ASSETNSDYT | 0.8375016 |
| LSLLLAPL | ASSETNSDYT | 0.7162702 |
| TLDEALK | ASSETNSDYT | 0.7186885 |
| TDGAGTAEGG | ASSETNSDYT | 0.68987805 |
| NAPGGAGGNGG | ASSETNSDYT | 0.6785298 |
| VSVFMGMGGNF | ASSETNSDYT | 0.70159054 |
| LASYLAAL | ASSETNSDYT | 0.7177342 |
| LVGGTPAL | ASSETNSDYT | 0.7218553 |
| VRNADVL | ASSETNSDYT | 0.71796024 |
| TCSSPFLVSAM | ASSETNSDYT | 0.67117816 |
| VHITDAE | ASSETNSDYT | 0.72062874 |
| EYAAPSF | ASSETNSDYT | 0.7111363 |
| VVDAFHGGL | ASSETNSDYT | 0.7088547 |
| ASVVDPQ | ASSETNSDYT | 0.7150098 |
| SYNEFGGLS | ASSETNSDYT | 0.70864725 |
| DPDGSVY | ASSETNSDYT | 0.6928467 |
| MVSTIHATE | ASSETNSDYT | 0.6871267 |
| LAAGVADPV | ASSETNSDYT | 0.7146232 |
| TKEQIAA | ASSETNSDYT | 0.7170025 |
| SLASPAAL | ASSETNSDYT | 0.7182723 |
| IGLVFYGV | ASSETNSDYT | 0.7242626 |
| FLLDASNL | ASSETNSDYT | 0.7202146 |
| YQVGNFP | ASSETNSDYT | 0.7162408 |
| ESLLSQST | ASSETNSDYT | 0.4789902 |
| LFTVGFAAVG | ASSETNSDYT | 0.6467613 |
| ATPPTPT | ASSETNSDYT | 0.6811478 |
| TDLATLG | ASSETNSDYT | 0.70385176 |
| SAGGTGR | ASSETNSDYT | 0.468994 |
| GFSNSGS | ASSETNSDYT | 0.7069551 |
| GGNGGSGGTSV | ASSETNSDYT | 0.70225406 |
| GTSVATGGAGN | ASSETNSDYT | 0.69113463 |
| NGGNGGSGGTS | ASSETNSDYT | 0.72056365 |
| AVVKFPH | ASSETNSDYT | 0.70286655 |
| TPPPPPP | ASSETNSDYT | 0.66804034 |
| AAPAAAAAPA | ASSETNSDYT | 0.6961681 |
| IALNEDL | ASSETNSDYT | 0.72209656 |
| IVNVSSM | ASSETNSDYT | 0.69010293 |
| TGIPPAPR | ASSETNSDYT | 0.72161376 |
| FDLGGGTF | ASSETNSDYT | 0.6710106 |
| FDLGGGTFDVS | ASSETNSDYT | 0.722731 |
| NEPTAAA | ASSETNSDYT | 0.65113926 |
| NEPTAAALA | ASSETNSDYT | 0.72104025 |
| GSIINISSI | ASSETNSDYT | 0.71087295 |
| VAPGNAGTA | ASSETNSDYT | 0.7261211 |

| | | |
|---|---|---|
| TGGGASLE | ASSETNSDYT | 0.70512104 |
| VAVYNHYKRI | ASSETNSDYT | 0.72000855 |
| TDPAPATT | ASSETNSDYT | 0.7016523 |
| TNRPDILDPALLRPG | ASSETNSDYT | 0.7210783 |
| LTTTTS | ASSETNSDYT | 0.6610085 |
| PTGSGKTL | ASSETNSDYT | 0.70568085 |
| TGSGKTL | ASSETNSDYT | 0.6421454 |
| TIGHVDHGK | ASSETNSDYT | 0.7138531 |
| TIGHVDHGKTT | ASSETNSDYT | 0.7193161 |
| GHVDHGKTT | ASSETNSDYT | 0.45949122 |
| AQFEHTLL | ASSETNSDYT | 0.7221847 |
| IVNQDEL | ASSETNSDYT | 0.68414503 |
| TNGSQFF | ASSETNSDYT | 0.68277776 |
| TGKGIVE | ASSETNSDYT | 0.72107005 |
| LISNASDA | ASSETNSDYT | 0.7096151 |
| AELRTLGELTKIAKAHGAQ | ASSETNSDYT | 0.71760714 |
| APNAVAAAEPSW | ASSETNSDYT | 0.691105 |
| APQVAGAATAVL | ASSETNSDYT | 0.7083818 |
| ARRDPAAAAAILPTDA | ASSETNSDYT | 0.7137836 |
| DAAARDVLAEAGLAEYF | ASSETNSDYT | 0.62130815 |
| DDQIVLSAKANLAKVEQ | ASSETNSDYT | 0.6767906 |
| DELLPDTGEDSDEDGH | ASSETNSDYT | 0.72427213 |
| ELPEAPANRATVPQD | ASSETNSDYT | 0.28193805 |
| FGQSKIERVTGIG | ASSETNSDYT | 0.82775944 |
| GAGGTGSTAGGAGGAGGAGGI | ASSETNSDYT | 0.185258 |
| GHGGTGAAGGNGGTGGAGGS( | ASSETNSDYT | 0.22171402 |
| GNGGNGGAGGEGGPGGAGGA( | ASSETNSDYT | 0.30210277 |
| GSTAGNGGNGGNATNGTIAG( | ASSETNSDYT | 0.38289618 |
| GTGGDGGNGGNGGTGG | ASSETNSDYT | 0.8043747 |
| GVRRPAGIPADAPV | ASSETNSDYT | 0.3279071 |
| GYRQVTDAHLLTLARR | ASSETNSDYT | 0.18572472 |
| IAKMGQDGHDRG | ASSETNSDYT | 0.2380941 |
| KAEPSIEIQYT | ASSETNSDYT | 0.17068824 |
| KPVRIRMAADEDTVAVT | ASSETNSDYT | 0.26729074 |
| MPPRIAGMRLLVIKPEP | ASSETNSDYT | 0.3259269 |
| MKFYAVALFRSRGVPPTR | ASSETNSDYT | 0.26535505 |
| NAGGFGILGAVAHSP | ASSETNSDYT | 0.17680553 |
| NEGREITLAMAEM | ASSETNSDYT | 0.21865778 |
| NELVEWSHPLGHRWRWE | ASSETNSDYT | 0.3365026 |
| NLGAGGTGGNGGNPG | ASSETNSDYT | 0.36884022 |
| NTEWPKWETVMAADPVD | ASSETNSDYT | 0.29878134 |
| PASAAPAETQP | ASSETNSDYT | 0.53015065 |
| QQSSDGKNGVRAAEITGE | ASSETNSDYT | 0.446697 |
| RAPDAVTTSWWKE | ASSETNSDYT | 0.36747938 |
| SGFGNVGAHTSGWFNQ | ASSETNSDYT | 0.11231359 |
| SGVAGAVGQAASVGGLKV | ASSETNSDYT | 0.15591212 |
| SNGGAGTGSAGGNGGT | ASSETNSDYT | 0.1692062 |
| TGGAGGEGGAAG | ASSETNSDYT | 0.27375424 |
| TGGDGGAAGAGGAPGHG | ASSETNSDYT | 0.11735942 |
| TMPPLLAAIANAGRHGV | ASSETNSDYT | 0.3753379 |
| TSSTAGNSGGA | ASSETNSDYT | 0.19767037 |

| | | |
|---|---|---|
| VPATVPVGILC | ASSETNSDYT | 0.20754966 |
| VQRPDSGGVVLTGRIS | ASSETNSDYT | 0.50725263 |
| WTPNPVPATKGVPAGLVQAT | ASSETNSDYT | 0.19805947 |
| YALKRALAKFSDRFDVVII | ASSETNSDYT | 0.22169492 |
| YLIGSVIV | ASSETNSDYT | 0.16896641 |
| YGQLVAVID | ASSETNSDYT | 0.48949897 |
| VTVGGMTLPA | ASSETNSDYT | 0.12820907 |
| VLALPAV | ASSETNSDYT | 0.37589824 |
| VIPPTLNYET | ASSETNSDYT | 0.16135766 |
| VGMGGMHPSSG | ASSETNSDYT | 0.36546692 |
| VGLAGGAATA | ASSETNSDYT | 0.16445954 |
| VGGAGGAGGSGP | ASSETNSDYT | 0.4279968 |
| VFAAFMV | ASSETNSDYT | 0.24504869 |
| TLYDKLR | ASSETNSDYT | 0.8307725 |
| TLAVFTVL | ASSETNSDYT | 0.30746862 |
| TGSVQPARV | ASSETNSDYT | 0.3329857 |
| TGGGAGSGGT | ASSETNSDYT | 0.20788026 |
| SHLETSL | ASSETNSDYT | 0.8148204 |
| SFTTGMT | ASSETNSDYT | 0.13684069 |
| QVEVAKRMGVRQPTVS | ASSETNSDYT | 0.28390896 |
| QGSGGPALAM | ASSETNSDYT | 0.14223237 |
| MLAPSNP | ASSETNSDYT | 0.3280301 |
| LVNNAAI | ASSETNSDYT | 0.37857494 |
| LTGEINTLA | ASSETNSDYT | 0.15796256 |
| LSIFGPV | ASSETNSDYT | 0.31637788 |
| LLALLANL | ASSETNSDYT | 0.28710634 |
| LISNASDALDKLR | ASSETNSDYT | 0.17705403 |
| LINEAVP | ASSETNSDYT | 0.51330477 |
| LGYAFSGL | ASSETNSDYT | 0.21735385 |
| LDFVAVAL | ASSETNSDYT | 0.67515934 |
| LAQPGAE | ASSETNSDYT | 0.2099445 |
| ISNLGAVTGF | ASSETNSDYT | 0.19919536 |
| HQEDPGRI | ASSETNSDYT | 0.30792844 |
| GTPGEPTP | ASSETNSDYT | 0.16818455 |
| GFFNSGS | ASSETNSDYT | 0.16148868 |
| GAGGGGLTAGAN | ASSETNSDYT | 0.5537092 |
| FVDALVNM | ASSETNSDYT | 0.8071692 |
| FLTVAPEL | ASSETNSDYT | 0.7545202 |
| FETLTAL | ASSETNSDYT | 0.71693593 |
| DVINPIEL | ASSETNSDYT | 0.6083436 |
| APVSGAPV | ASSETNSDYT | 0.71530735 |
| ALVGGLAW | ASSETNSDYT | 0.7939217 |
| ALAAAAATAI | ASSETNSDYT | 0.7768028 |
| AFGLGLLL | ASSETNSDYT | 0.7771853 |
| AFFAAASL | ASSETNSDYT | 0.52452284 |
| AAAALPAFSPP | ASSETNSDYT | 0.7804935 |
| VVLAPAL | ASSETNSDYT | 0.75461787 |
| TIGHVDHGKTTL | ASSETNSDYT | 0.61211276 |
| LISNASDALDK | ASSETNSDYT | 0.7768855 |
| LISNASDAL | ASSETNSDYT | 0.67784977 |
| ISNASDALDK | ASSETNSDYT | 0.67381203 |

| | | |
|---|---|---|
| NQAETLVYQTE | ASSETNSDYT | 0.7270324 |
| TGGTGGTAGAG | ASSETNSDYT | 0.6242664 |
| TGSTAGGAGGA | ASSETNSDYT | 0.77327275 |
| DPLAVPVAAGPV | ASSETNSDYT | 0.74683 |
| GGTGGTPGNGGN | ASSETNSDYT | 0.8199513 |
| PAGLAGVDPATL | ASSETNSDYT | 0.8086106 |
| GNNLVVEAPHAGASV | ASSETNSDYT | 0.7644292 |
| KVTRIMSASTLEQPA | ASSETNSDYT | 0.27715564 |
| LGFALPIALPRRTVV | ASSETNSDYT | 0.45567894 |
| TRLGNLTARKLVVLV | ASSETNSDYT | 0.7250886 |
| WSQQRRLIYFPSAGP | ASSETNSDYT | 0.6665901 |
| GPNGNPDPMAAAVDIR | ASSETNSDYT | 0.28390345 |
| TQRCVSVADLRAAASR | ASSETNSDYT | 0.6942078 |
| QFVLDVAEHPVEVLTETV | ASSETNSDYT | 0.7636634 |
| GGAGGTGGAGGNGGTQAGDG | ASSETNSDYT | 0.5714227 |
| PLLATLRAIAVDVDAKPAQV | ASSETNSDYT | 0.7485189 |
| APKFRTGGYMIDFWGVGYQV | ASSETNSDYT | 0.5973662 |
| GAGGAGGAGGAAGGPGGTGG | ASSETNSDYT | 0.49741334 |
| IGALGLLATDVAPAGSVAFG | ASSETNSDYT | 0.6596327 |
| AGGAGGIGGNGGAGGAGGVL | ASSETNSDYT | 0.8032606 |
| ADELIGLL | ASSETNSDYT | 0.6184815 |
| ADRALKA | ASSETNSDYT | 0.79808706 |
| AGAVGQVGGAGGA | ASSETNSDYT | 0.59882766 |
| AGGGGGLGGAP | ASSETNSDYT | 0.7967051 |
| AGGLLYGNGGAG | ASSETNSDYT | 0.4831375 |
| AGLESIGL | ASSETNSDYT | 0.7862759 |
| AGLKTGII | ASSETNSDYT | 0.750366 |
| ASLAALLL | ASSETNSDYT | 0.80519485 |
| ASVINISSI | ASSETNSDYT | 0.67803836 |
| ATPPGPA | ASSETNSDYT | 0.75723875 |
| ATVASALV | ASSETNSDYT | 0.5887269 |
| ATVASLIL | ASSETNSDYT | 0.77908033 |
| AVNGPVLIRH | ASSETNSDYT | 0.60682875 |
| DGAAGAVGI | ASSETNSDYT | 0.7824712 |
| DPLLHGI | ASSETNSDYT | 0.44284105 |
| ELGAALGAL | ASSETNSDYT | 0.79125655 |
| ELGRTQAV | ASSETNSDYT | 0.8050587 |
| FVAGGYLS | ASSETNSDYT | 0.417598 |
| GASGGAGGAGGW | ASSETNSDYT | 0.8238621 |
| GGAGAGGDGGAGGI | ASSETNSDYT | 0.681346 |
| GTGHNGSQPGMGGNG | ASSETNSDYT | 0.5811111 |
| GYAAASASAS | ASSETNSDYT | 0.80229616 |
| HQVDVAAVQQG | ASSETNSDYT | 0.6441554 |
| IGAAIAHT | ASSETNSDYT | 0.8092116 |
| IGMIEVAQL | ASSETNSDYT | 0.6110709 |
| IRLAPEHTSAMD | ASSETNSDYT | 0.7591196 |
| IYNVSVL | ASSETNSDYT | 0.80532527 |
| KQGFLGM | ASSETNSDYT | 0.5605575 |
| LESVITQL | ASSETNSDYT | 0.47036242 |
| LGLGALVVV | ASSETNSDYT | 0.8036371 |
| LIGNGGNGGSGGI | ASSETNSDYT | 0.73426104 |

| | | |
|---|---|---|
| LNAINAPTL | ASSETNSDYT | 0.7590821 |
| LQLLNTGV | ASSETNSDYT | 0.6521671 |
| LQTSIEF | ASSETNSDYT | 0.57596076 |
| MSFAVSEL | ASSETNSDYT | 0.6608937 |
| NTGALGL | ASSETNSDYT | 0.74826837 |
| PPGAGGAGGAGTT | ASSETNSDYT | 0.7639844 |
| QLVGLGI | ASSETNSDYT | 0.73004746 |
| RFLEQVHQL | ASSETNSDYT | 0.52036035 |
| RGVGYVLR | ASSETNSDYT | 0.7452328 |
| SATGVFAGVI | ASSETNSDYT | 0.70789236 |
| SLPAITLP | ASSETNSDYT | 0.6737167 |
| STALAFTV | ASSETNSDYT | 0.6952267 |
| SVLTVGLGK | ASSETNSDYT | 0.72010225 |
| TALAGAGSGI | ASSETNSDYT | 0.6273794 |
| TGLPFTGI | ASSETNSDYT | 0.6133511 |
| VAEHVTV | ASSETNSDYT | 0.7002338 |
| VDIVLNSL | ASSETNSDYT | 0.7377145 |
| VLTGGGAVL | ASSETNSDYT | 0.7436739 |
| VNVGLAYGI | ASSETNSDYT | 0.7670692 |
| VSSAAVGAV | ASSETNSDYT | 0.767455 |
| VTSVYEAGV | ASSETNSDYT | 0.72306156 |
| WVALDRGA | ASSETNSDYT | 0.2909665 |
| YLVGSVVI | ASSETNSDYT | 0.42804232 |
| GPPGTGKTLL | ASSETNSDYT | 0.5869127 |
| GVGMTKF | ASSETNSDYT | 0.67296684 |
| IGIGGSDL | ASSETNSDYT | 0.3133512 |
| IGTLLAEL | ASSETNSDYT | 0.70466995 |
| IYTSGTT | ASSETNSDYT | 0.6545714 |
| LSFPAGV | ASSETNSDYT | 0.6197815 |
| LYTSGTT | ASSETNSDYT | 0.75273293 |
| VGAFPFGF | ASSETNSDYT | 0.6533067 |
| VGAGTVEF | ASSETNSDYT | 0.57697254 |
| VGKSTLLL | ASSETNSDYT | 0.57446104 |
| AALAHAGGGIV | ASSETNSDYT | 0.7410512 |
| AALVAVSLL | ASSETNSDYT | 0.4828782 |
| AAVSRAVAAL | ASSETNSDYT | 0.7568493 |
| AGGSSGIGGF | ASSETNSDYT | 0.4543152 |
| AGLTVANANG | ASSETNSDYT | 0.76961076 |
| AGLVGAVL | ASSETNSDYT | 0.48588073 |
| ALSQAFTV | ASSETNSDYT | 0.74912554 |
| ALTFSVSL | ASSETNSDYT | 0.6947282 |
| ALTGAALVAL | ASSETNSDYT | 0.7884835 |
| ASFSIGAGVL | ASSETNSDYT | 0.68154144 |
| DGSVYKI | ASSETNSDYT | 0.7408378 |
| EQLAEIV | ASSETNSDYT | 0.5858224 |
| FAVSELTVM | ASSETNSDYT | 0.757145 |
| GAAPLAAGL | ASSETNSDYT | 0.51978767 |
| GADGGDGGSGGAGG | ASSETNSDYT | 0.69291687 |
| GFGLDTGGAGG | ASSETNSDYT | 0.39929295 |
| GLASGGLL | ASSETNSDYT | 0.7167174 |
| GRIAGAAVLL | ASSETNSDYT | 0.7377207 |

| | | |
|---|---|---|
| GSAGSLNTSF | ASSETNSDYT | 0.41933084 |
| GTGQIGI | ASSETNSDYT | 0.7588787 |
| GTLIALIL | ASSETNSDYT | 0.65621185 |
| IAATMGFVVG | ASSETNSDYT | 0.59014255 |
| IFAQALL | ASSETNSDYT | 0.75719506 |
| IGAAVALGF | ASSETNSDYT | 0.7018416 |
| IGLTPLAL | ASSETNSDYT | 0.7597647 |
| ILGSNTSTL | ASSETNSDYT | 0.5851555 |
| IVFSLTVL | ASSETNSDYT | 0.6514809 |
| LASKATGF | ASSETNSDYT | 0.7093418 |
| LASVALIL | ASSETNSDYT | 0.61531377 |
| LFAVGLTL | ASSETNSDYT | 0.5449941 |
| LFNGGIGTYI | ASSETNSDYT | 0.76389235 |
| LTLGAGII | ASSETNSDYT | 0.5105047 |
| LVAAALGDI | ASSETNSDYT | 0.60779804 |
| NTAFVQLGI | ASSETNSDYT | 0.32946712 |
| QLFSGAEL | ASSETNSDYT | 0.33755383 |
| SAATFITL | ASSETNSDYT | 0.39187512 |
| SNYGLANLGV | ASSETNSDYT | 0.5302627 |
| SYGGAYAVM | ASSETNSDYT | 0.57535183 |
| TTGVGGLTLG | ASSETNSDYT | 0.5078817 |
| TTQTIYI | ASSETNSDYT | 0.34830537 |
| VAAGAGLGAVI | ASSETNSDYT | 0.49537316 |
| VAAVVTLL | ASSETNSDYT | 0.39988288 |
| VGFLGGLI | ASSETNSDYT | 0.3412215 |
| VGVIPILL | ASSETNSDYT | 0.45757315 |
| VIFASSLI | ASSETNSDYT | 0.4433841 |
| VKGGDGGAAAG | ASSETNSDYT | 0.35632032 |
| VTKTFTNI | ASSETNSDYT | 0.4370508 |
| VVAAAILL | ASSETNSDYT | 0.41069266 |
| YAALDVEL | ASSETNSDYT | 0.49702525 |
| LASVALLL | ASSETNSDYT | 0.45299533 |
| LDAVHRV | ASSETNSDYT | 0.5966674 |
| QGLSSSIF | ASSETNSDYT | 0.54951614 |
| RGTFANIRL | ASSETNSDYT | 0.4699468 |
| RQLWWGHRI | ASSETNSDYT | 0.22380014 |
| YGPPGTGKTLL | ASSETNSDYT | 0.29658636 |
| AALVGAPANVA | ASSETNSDYT | 0.41866323 |
| AATFRRTE | ASSETNSDYT | 0.37809426 |
| AGALGGGI | ASSETNSDYT | 0.25942862 |
| AGGAGGAGGIGAG | ASSETNSDYT | 0.47533837 |
| AGSGALGL | ASSETNSDYT | 0.45036 |
| AQIADQI | ASSETNSDYT | 0.32703912 |
| ASAVAVTYL | ASSETNSDYT | 0.47369325 |
| ASTVLGGTGG | ASSETNSDYT | 0.30811745 |
| AVGGEPAK | ASSETNSDYT | 0.33579454 |
| CEQGTFGGTC | ASSETNSDYT | 0.37617293 |
| DEGKAPAG | ASSETNSDYT | 0.5299654 |
| DTVVFPAL | ASSETNSDYT | 0.3389202 |
| FEGITMAG | ASSETNSDYT | 0.4863589 |
| FPLVVQD | ASSETNSDYT | 0.39906198 |

| | | |
|---|---|---|
| GAFTTPQ | ASSETNSDYT | 0.57149935 |
| GFGGFGTAGGVGGA | ASSETNSDYT | 0.3116407 |
| GFLAQFL | ASSETNSDYT | 0.49050066 |
| GGAGGAGGNAGVA | ASSETNSDYT | 0.39513615 |
| GGVGTAGAGGAGGAGG | ASSETNSDYT | 0.5998132 |
| GNGGGGDAGNAGSGGN | ASSETNSDYT | 0.38363913 |
| GTGGAGGAGNDAGS | ASSETNSDYT | 0.47366565 |
| GTTNTGWFNS | ASSETNSDYT | 0.40760833 |
| IFMGPLFTL | ASSETNSDYT | 0.6080845 |
| ILAHAIEAI | ASSETNSDYT | 0.27618477 |
| IRSAIREAL | ASSETNSDYT | 0.418577 |
| ITFAGGG | ASSETNSDYT | 0.31920594 |
| ITGMSGHQEV | ASSETNSDYT | 0.45482635 |
| IVAGLACTL | ASSETNSDYT | 0.5303326 |
| IVGGGLAGA | ASSETNSDYT | 0.2750701 |
| IVTGVLNI | ASSETNSDYT | 0.57697797 |
| LAQQGCI | ASSETNSDYT | 0.35701454 |
| LCLHVEN | ASSETNSDYT | 0.34834644 |
| LDAISDGS | ASSETNSDYT | 0.50153095 |
| LGGFAAML | ASSETNSDYT | 0.38051343 |
| LGGKNPMI | ASSETNSDYT | 0.57932985 |
| LGPKGVAAIL | ASSETNSDYT | 0.3727733 |
| LMVGVSI | ASSETNSDYT | 0.43425813 |
| LVGAAFVL | ASSETNSDYT | 0.49103028 |
| LVLAVAPL | ASSETNSDYT | 0.31065434 |
| MAAIAVIDAA | ASSETNSDYT | 0.315539 |
| NNGINAELQ | ASSETNSDYT | 0.5604008 |
| RGEPLVLVHG | ASSETNSDYT | 0.45261112 |
| RGGAGGAAGLI | ASSETNSDYT | 0.65393466 |
| RSYRMHPAV | ASSETNSDYT | 0.31240627 |
| SAAHLLAGQLA | ASSETNSDYT | 0.30764455 |
| SAAVGSY | ASSETNSDYT | 0.3159543 |
| SPHAVLHRI | ASSETNSDYT | 0.40234607 |
| SQIAKVL | ASSETNSDYT | 0.65808254 |
| TALGHLVP | ASSETNSDYT | 0.51494324 |
| TAPTTSGAA | ASSETNSDYT | 0.3163631 |
| TDGDDGNGGSPGT | ASSETNSDYT | 0.3751402 |
| TDGLGEN | ASSETNSDYT | 0.32111174 |
| TGGPPLPGQA | ASSETNSDYT | 0.4566291 |
| TIPAITI | ASSETNSDYT | 0.36008316 |
| TIPGFTI | ASSETNSDYT | 0.33705127 |
| TSIASGALA | ASSETNSDYT | 0.36865497 |
| TVTLPGAAI | ASSETNSDYT | 0.30154672 |
| VASGLSAALP | ASSETNSDYT | 0.3286053 |
| VGAGVIGI | ASSETNSDYT | 0.5506991 |
| VGPKRFGV | ASSETNSDYT | 0.49063304 |
| VNVAFPD | ASSETNSDYT | 0.4141951 |
| VSMTNSGVS | ASSETNSDYT | 0.5707581 |
| VTDVLGDLSV | ASSETNSDYT | 0.3438839 |
| VVVVGAGFAGL | ASSETNSDYT | 0.2544088 |
| VYTVGTVV | ASSETNSDYT | 0.30376038 |

| | | |
|---|---|---|
| WTTFSGP | ASSETNSDYT | 0.31025514 |
| YDELVEAF | ASSETNSDYT | 0.3647511 |
| YGTGGAGGNGG | ASSETNSDYT | 0.25913057 |
| YTNVMAV | ASSETNSDYT | 0.35754722 |
| FLEQVHQL | ASSETNSDYT | 0.32927638 |
| GLSLLAVAV | ASSETNSDYT | 0.35741666 |
| LEQVHQL | ASSETNSDYT | 0.50954616 |
| LNAAGAI | ASSETNSDYT | 0.366286 |
| PAVHASL | ASSETNSDYT | 0.30746606 |
| TELEDDA | ASSETNSDYT | 0.3014319 |
| TPPGPPN | ASSETNSDYT | 0.5489613 |
| TPPPPPPP | ASSETNSDYT | 0.28587788 |
| TQSPALI | ASSDPGANSDYT | 0.5139159 |
| EFAAAFPQ | ASSDPGANSDYT | 0.31348073 |
| LLSGNLDVL | ASSDPGANSDYT | 0.4669928 |
| GPANLGNYNI | ASSDPGANSDYT | 0.2920704 |
| TLGPTLP | ASSDPGANSDYT | 0.52192265 |
| VYAYSSCCK | ASSDPGANSDYT | 0.4078501 |
| APAPAPAPNI | ASSDPGANSDYT | 0.6135888 |
| PGGPVPIP | ASSDPGANSDYT | 0.3300827 |
| FLTVALEP | ASSDPGANSDYT | 0.371778 |
| SQYFGANL | ASSDPGANSDYT | 0.3077086 |
| GGVKPAA | ASSDPGANSDYT | 0.66309655 |
| GGTGGAGGAGGW | ASSDPGANSDYT | 0.2843745 |
| GVGGVAGLGGAGG | ASSDPGANSDYT | 0.31369147 |
| VLLVLATTI | ASSDPGANSDYT | 0.29861537 |
| IGIFAAML | ASSDPGANSDYT | 0.31998274 |
| AVGDLID | ASSDPGANSDYT | 0.53366095 |
| NISPLPL | ASSDPGANSDYT | 0.2604514 |
| VTAVFVAI | ASSDPGANSDYT | 0.47578037 |
| SGDDNDAAL | ASSDPGANSDYT | 0.30031642 |
| AVAALAHS | ASSDPGANSDYT | 0.30048594 |
| GGVTVMATP | ASSDPGANSDYT | 0.39292735 |
| AGLAANVRD | ASSDPGANSDYT | 0.31210417 |
| TNRPDLIDPALLRPG | ASSDPGANSDYT | 0.47230715 |
| LANLGVTAL | ASSDPGANSDYT | 0.3000131 |
| TGERIEL | ASSDPGANSDYT | 0.37293175 |
| NGEELAA | ASSDPGANSDYT | 0.36460105 |
| VLSLVWG | ASSDPGANSDYT | 0.28517303 |
| IGYALWL | ASSDPGANSDYT | 0.29334095 |
| LSLLLAPL | ASSDPGANSDYT | 0.6095306 |
| TLDEALK | ASSDPGANSDYT | 0.63571227 |
| TDGAGTAEGG | ASSDPGANSDYT | 0.7569201 |
| NAPGGAGGNGG | ASSDPGANSDYT | 0.23619357 |
| VSVFMGMGGNF | ASSDPGANSDYT | 0.22484793 |
| LASYLAAL | ASSDPGANSDYT | 0.36100549 |
| LVGGTPAL | ASSDPGANSDYT | 0.6844816 |
| VRNADVL | ASSDPGANSDYT | 0.8048206 |
| TCSSPFLVSAM | ASSDPGANSDYT | 0.6449256 |
| VHITDAE | ASSDPGANSDYT | 0.27038342 |
| EYAAPSF | ASSDPGANSDYT | 0.5680988 |

| | | |
|---|---|---|
| VVDAFHGGL | ASSDPGANSDYT | 0.3750863 |
| ASVVDPQ | ASSDPGANSDYT | 0.44281152 |
| SYNEFGGLS | ASSDPGANSDYT | 0.5911813 |
| DPDGSVY | ASSDPGANSDYT | 0.54789996 |
| MVSTIHATE | ASSDPGANSDYT | 0.3691621 |
| LAAGVADPV | ASSDPGANSDYT | 0.26874045 |
| TKEQIAA | ASSDPGANSDYT | 0.67632747 |
| SLASPAAL | ASSDPGANSDYT | 0.75154406 |
| IGLVFYGV | ASSDPGANSDYT | 0.7286701 |
| FLLDASNL | ASSDPGANSDYT | 0.8060165 |
| YQVGNFP | ASSDPGANSDYT | 0.7131854 |
| ESLLSQST | ASSDPGANSDYT | 0.7077347 |
| LFTVGFAAVG | ASSDPGANSDYT | 0.118864276 |
| ATPPTPT | ASSDPGANSDYT | 0.16569884 |
| TDLATLG | ASSDPGANSDYT | 0.38463104 |
| SAGGTGR | ASSDPGANSDYT | 0.3776747 |
| GFSNSGS | ASSDPGANSDYT | 0.17087081 |
| GGNGGSGGTSV | ASSDPGANSDYT | 0.6279226 |
| GTSVATGGAGN | ASSDPGANSDYT | 0.32353875 |
| NGGNGGSGGTS | ASSDPGANSDYT | 0.3072342 |
| AVVKFPH | ASSDPGANSDYT | 0.73824215 |
| TPPPPPP | ASSDPGANSDYT | 0.29087242 |
| AAPAAAAAPA | ASSDPGANSDYT | 0.28586242 |
| IALNEDL | ASSDPGANSDYT | 0.3783764 |
| IVNVSSM | ASSDPGANSDYT | 0.7045127 |
| TGIPPAPR | ASSDPGANSDYT | 0.273813 |
| FDLGGGTF | ASSDPGANSDYT | 0.64902544 |
| FDLGGGTFDVS | ASSDPGANSDYT | 0.41942286 |
| NEPTAAA | ASSDPGANSDYT | 0.75335175 |
| NEPTAAALA | ASSDPGANSDYT | 0.22353256 |
| GSIINISSI | ASSDPGANSDYT | 0.6971365 |
| VAPGNAGTA | ASSDPGANSDYT | 0.45309988 |
| TGGGASLE | ASSDPGANSDYT | 0.84596777 |
| VAVYNHYKRI | ASSDPGANSDYT | 0.32292178 |
| TDPAPATT | ASSDPGANSDYT | 0.49981034 |
| TNRPDILDPALLRPG | ASSDPGANSDYT | 0.4423477 |
| LTTTTTS | ASSDPGANSDYT | 0.79699576 |
| PTGSGKTL | ASSDPGANSDYT | 0.19466601 |
| TGSGKTL | ASSDPGANSDYT | 0.31121323 |
| TIGHVDHGK | ASSDPGANSDYT | 0.19528793 |
| TIGHVDHGKTT | ASSDPGANSDYT | 0.40023634 |
| GHVDHGKTT | ASSDPGANSDYT | 0.60549664 |
| AQFEHTLL | ASSDPGANSDYT | 0.12190325 |
| IVNQDEL | ASSDPGANSDYT | 0.57699096 |
| TNGSQFF | ASSDPGANSDYT | 0.36280274 |
| TGKGIVE | ASSDPGANSDYT | 0.2502443 |
| LISNASDA | ASSDPGANSDYT | 0.7257298 |
| AELRTLGELTKIAKAHGAQ | ASSDPGANSDYT | 0.5617279 |
| APNAVAAAEPSW | ASSDPGANSDYT | 0.74612373 |
| APQVAGAATAVL | ASSDPGANSDYT | 0.528021 |
| ARRDPAAAAAILPTDA | ASSDPGANSDYT | 0.3855236 |

| | | |
|---|---|---|
| DAAARDVLAEAGLAEYF | ASSDPGANSDYT | 0.452458 |
| DDQIVLSAKANLAKVEQ | ASSDPGANSDYT | 0.22264889 |
| DELLPDTGEDSDEDGH | ASSDPGANSDYT | 0.36192554 |
| ELPEAPANRATVPQD | ASSDPGANSDYT | 0.752712 |
| FGQSKIERVTGIG | ASSDPGANSDYT | 0.56734395 |
| GAGGTGSTAGGAGGAGGAGGI | ASSDPGANSDYT | 0.7688517 |
| GHGGTGAAGGNGGTGGAGGS( | ASSDPGANSDYT | 0.35022178 |
| GNGGNGGAGGEGGPGGAGGA( | ASSDPGANSDYT | 0.29685712 |
| GSTAGNGGNGGNATNGTIAG( | ASSDPGANSDYT | 0.48447347 |
| GTGGDGGNGGNGGTGG | ASSDPGANSDYT | 0.7011262 |
| GVRRPAGIPADAPV | ASSDPGANSDYT | 0.7934268 |
| GYRQVTDAHLLTLARR | ASSDPGANSDYT | 0.69651055 |
| IAKMGQDGHDRG | ASSDPGANSDYT | 0.32676807 |
| KAEPSIEIQYT | ASSDPGANSDYT | 0.5569627 |
| KPVRIRMAADEDTVAVT | ASSDPGANSDYT | 0.4360842 |
| MPPRIAGMRLLVIKPEP | ASSDPGANSDYT | 0.44268844 |
| MKFYAVALFRSRGVPPTR | ASSDPGANSDYT | 0.5141505 |
| NAGGFGILGAVAHSP | ASSDPGANSDYT | 0.5700786 |
| NEGREITLAMAEM | ASSDPGANSDYT | 0.40486157 |
| NELVEWSHPLGHRWRWE | ASSDPGANSDYT | 0.3213334 |
| NLGAGGTGGNGGNPG | ASSDPGANSDYT | 0.5679467 |
| NTEWPKWETVMAADPVD | ASSDPGANSDYT | 0.6937648 |
| PASAAPAETQP | ASSDPGANSDYT | 0.633638 |
| QQSSDGKNGVRAAEITGE | ASSDPGANSDYT | 0.7972022 |
| RAPDAVTTSWWKE | ASSDPGANSDYT | 0.691764 |
| SGFGNVGAHTSGWFNQ | ASSDPGANSDYT | 0.53560764 |
| SGVAGAVGQAASVGGLKV | ASSDPGANSDYT | 0.12802319 |
| SNGGAGTGSAGGNGGT | ASSDPGANSDYT | 0.19357637 |
| TGGAGGEGGAAG | ASSDPGANSDYT | 0.38167095 |
| TGGDGGAAGAGGAPGHG | ASSDPGANSDYT | 0.4767515 |
| TMPPLLAAIANAGRHGV | ASSDPGANSDYT | 0.14408118 |
| TSSTAGNSGGA | ASSDPGANSDYT | 0.6447021 |
| VPATVPVGILC | ASSDPGANSDYT | 0.42372805 |
| VQRPDSGGVVLTGRIS | ASSDPGANSDYT | 0.36244744 |
| WTPNPVPATKGVPAGLVQAT | ASSDPGANSDYT | 0.7245115 |
| YALKRALAKFSDRFDVVII | ASSDPGANSDYT | 0.39083475 |
| YLIGSVIV | ASSDPGANSDYT | 0.33372498 |
| YGQLVAVID | ASSDPGANSDYT | 0.40834674 |
| VTVGGMTLPA | ASSDPGANSDYT | 0.78871596 |
| VLALPAV | ASSDPGANSDYT | 0.2516461 |
| VIPPTLNYET | ASSDPGANSDYT | 0.6831629 |
| VGMGGMHPSSG | ASSDPGANSDYT | 0.43833694 |
| VGLAGGAATA | ASSDPGANSDYT | 0.723348 |
| VGGAGGAGGSGP | ASSDPGANSDYT | 0.24944918 |
| VFAAFMV | ASSDPGANSDYT | 0.66801274 |
| TLYDKLR | ASSDPGANSDYT | 0.45632517 |
| TLAVFTVL | ASSDPGANSDYT | 0.8386569 |
| TGSVQPARV | ASSDPGANSDYT | 0.35932767 |
| TGGGAGSGGT | ASSDPGANSDYT | 0.5694697 |
| SHLETSL | ASSDPGANSDYT | 0.38759 |
| SFTTGMT | ASSDPGANSDYT | 0.768687 |

| | | |
|---|---|---|
| QVEVAKRMGVRQPTVS | ASSDPGANSDYT | 0.22781 |
| QGSGGPALAM | ASSDPGANSDYT | 0.39883038 |
| MLAPSNP | ASSDPGANSDYT | 0.22825962 |
| LVNNAAI | ASSDPGANSDYT | 0.4840831 |
| LTGEINTLA | ASSDPGANSDYT | 0.66375387 |
| LSIFGPV | ASSDPGANSDYT | 0.15484838 |
| LLALLANL | ASSDPGANSDYT | 0.69979703 |
| LISNASDALDKLR | ASSDPGANSDYT | 0.38694155 |
| LINEAVP | ASSDPGANSDYT | 0.31202948 |
| LGYAFSGL | ASSDPGANSDYT | 0.66151494 |
| LDFVAVAL | ASSDPGANSDYT | 0.4502911 |
| LAQPGAE | ASSDPGANSDYT | 0.7477187 |
| ISNLGAVTGF | ASSDPGANSDYT | 0.35132974 |
| HQEDPGRI | ASSDPGANSDYT | 0.5445037 |
| GTPGEPTP | ASSDPGANSDYT | 0.5884874 |
| GFFNSGS | ASSDPGANSDYT | 0.2671815 |
| GAGGGGLTAGAN | ASSDPGANSDYT | 0.3624279 |
| FVDALVNM | ASSDPGANSDYT | 0.7261378 |
| FLTVAPEL | ASSDPGANSDYT | 0.6393907 |
| FETLTAL | ASSDPGANSDYT | 0.64479256 |
| DVINPIEL | ASSDPGANSDYT | 0.47110778 |
| APVSGAPV | ASSDPGANSDYT | 0.51818675 |
| ALVGGLAW | ASSDPGANSDYT | 0.44426215 |
| ALAAAAATAI | ASSDPGANSDYT | 0.6135242 |
| AFGLGLLL | ASSDPGANSDYT | 0.66362095 |
| AFFAAASL | ASSDPGANSDYT | 0.63797957 |
| AAAALPAFSPP | ASSDPGANSDYT | 0.4990192 |
| VVLAPAL | ASSDPGANSDYT | 0.6336505 |
| TIGHVDHGKTTL | ASSDPGANSDYT | 0.57857037 |
| LISNASDALDK | ASSDPGANSDYT | 0.5864245 |
| LISNASDAL | ASSDPGANSDYT | 0.5681373 |
| ISNASDALDK | ASSDPGANSDYT | 0.5822459 |
| NQAETLVYQTE | ASSDPGANSDYT | 0.4447552 |
| TGGTGGTAGAG | ASSDPGANSDYT | 0.43143728 |
| TGSTAGGAGGA | ASSDPGANSDYT | 0.5355525 |
| DPLAVPVAAGPV | ASSDPGANSDYT | 0.6387723 |
| GGTGGTPGNGGN | ASSDPGANSDYT | 0.6318641 |
| PAGLAGVDPATL | ASSDPGANSDYT | 0.64777523 |
| GNNLVVEAPHAGASV | ASSDPGANSDYT | 0.64367485 |
| KVTRIMSASTLEQPA | ASSDPGANSDYT | 0.6170456 |
| LGFALPIALPRRTVV | ASSDPGANSDYT | 0.326252 |
| TRLGNLTARKLVVLV | ASSDPGANSDYT | 0.3734552 |
| WSQQRRLIYFPSAGP | ASSDPGANSDYT | 0.43745536 |
| GPNGNPDPMAAAVDIR | ASSDPGANSDYT | 0.5211402 |
| TQRCVSVADLRAAASR | ASSDPGANSDYT | 0.30396134 |
| QFVLDVAEHPVEVLTETV | ASSDPGANSDYT | 0.59134984 |
| GGAGGTGGAGGNGGTQAGDG | ASSDPGANSDYT | 0.5113482 |
| PLLATLRAIAVDVDAKPAQV | ASSDPGANSDYT | 0.53629243 |
| APKFRTGGYMIDFWGVGYQV[illegible] | ASSDPGANSDYT | 0.61805254 |
| GAGGAGGAGGAAGGPGGTGG[illegible] | ASSDPGANSDYT | 0.53879535 |
| IGALGLLATDVAPAGSVAFG[illegible] | ASSDPGANSDYT | 0.42070147 |

| | | |
|---|---|---|
| AGGAGGIGGNGGAGGAGGVLM | ASSDPGANSDYT | 0.46408966 |
| ADELIGLL | ASSDPGANSDYT | 0.64317065 |
| ADRALKA | ASSDPGANSDYT | 0.43715292 |
| AGAVGQVGGAGGA | ASSDPGANSDYT | 0.6566618 |
| AGGGGGLGGAP | ASSDPGANSDYT | 0.37724456 |
| AGGLLYGNGGAG | ASSDPGANSDYT | 0.645045 |
| AGLESIGL | ASSDPGANSDYT | 0.47160292 |
| AGLKTGII | ASSDPGANSDYT | 0.6479907 |
| ASLAALLL | ASSDPGANSDYT | 0.613831 |
| ASVINISSI | ASSDPGANSDYT | 0.66885936 |
| ATPPGPA | ASSDPGANSDYT | 0.547035 |
| ATVASALV | ASSDPGANSDYT | 0.6457032 |
| ATVASLIL | ASSDPGANSDYT | 0.55671084 |
| AVNGPVLIRH | ASSDPGANSDYT | 0.63773245 |
| DGAAGAVGI | ASSDPGANSDYT | 0.45146137 |
| DPLLHGI | ASSDPGANSDYT | 0.5093715 |
| ELGAALGAL | ASSDPGANSDYT | 0.37948292 |
| ELGRTQAV | ASSDPGANSDYT | 0.5592989 |
| FVAGGYLS | ASSDPGANSDYT | 0.6477424 |
| GASGGAGGAGGW | ASSDPGANSDYT | 0.30585998 |
| GGAGAGGDGGAGGI | ASSDPGANSDYT | 0.61690456 |
| GTGHNGSQPGMGGNG | ASSDPGANSDYT | 0.40287948 |
| GYAAASASAS | ASSDPGANSDYT | 0.4675422 |
| HQVDVAAVQQG | ASSDPGANSDYT | 0.6121452 |
| IGAAIAHT | ASSDPGANSDYT | 0.5889696 |
| IGMIEVAQL | ASSDPGANSDYT | 0.6026721 |
| IRLAPEHTSAMD | ASSDPGANSDYT | 0.4320809 |
| IYNVSVL | ASSDPGANSDYT | 0.5041275 |
| KQGFLGM | ASSDPGANSDYT | 0.5915509 |
| LESVITQL | ASSDPGANSDYT | 0.31581303 |
| LGLGALVVV | ASSDPGANSDYT | 0.41108853 |
| LIGNGGNGGSGGI | ASSDPGANSDYT | 0.66454655 |
| LNAINAPTL | ASSDPGANSDYT | 0.25711894 |
| LQLLNTGV | ASSDPGANSDYT | 0.80580145 |
| LQTSIEF | ASSDPGANSDYT | 0.17761964 |
| MSFAVSEL | ASSDPGANSDYT | 0.15926471 |
| NTGALGL | ASSDPGANSDYT | 0.30041122 |
| PPGAGGAGGAGTT | ASSDPGANSDYT | 0.54589194 |
| QLVGLGI | ASSDPGANSDYT | 0.81028825 |
| RFLEQVHQL | ASSDPGANSDYT | 0.48113188 |
| RGVGYVLR | ASSDPGANSDYT | 0.15933013 |
| SATGVFAGVI | ASSDPGANSDYT | 0.20709525 |
| SLPAITLP | ASSDPGANSDYT | 0.16438028 |
| STALAFTV | ASSDPGANSDYT | 0.23958355 |
| SVLTVGLGK | ASSDPGANSDYT | 0.45744467 |
| TALAGAGSGI | ASSDPGANSDYT | 0.21043412 |
| TGLPFTGI | ASSDPGANSDYT | 0.17674954 |
| VAEHVTV | ASSDPGANSDYT | 0.21138744 |
| VDIVLNSL | ASSDPGANSDYT | 0.48309782 |
| VLTGGGAVL | ASSDPGANSDYT | 0.44823673 |
| VNVGLAYGI | ASSDPGANSDYT | 0.33621126 |

| | | |
|---|---|---|
| VSSAAVGAV | ASSDPGANSDYT | 0.6878265 |
| VTSVYEAGV | ASSDPGANSDYT | 0.3339789 |
| WVALDRGA | ASSDPGANSDYT | 0.5897088 |
| YLVGSVVI | ASSDPGANSDYT | 0.1072717 |
| GPPGTGKTLL | ASSDPGANSDYT | 0.1277392 |
| GVGMTKF | ASSDPGANSDYT | 0.2562593 |
| IGIGGSDL | ASSDPGANSDYT | 0.23154944 |
| IGTLLAEL | ASSDPGANSDYT | 0.15578382 |
| IYTSGTT | ASSDPGANSDYT | 0.43663546 |
| LSFPAGV | ASSDPGANSDYT | 0.18947971 |
| LYTSGTT | ASSDPGANSDYT | 0.1473891 |
| VGAFPFGF | ASSDPGANSDYT | 0.59976447 |
| VGAGTVEF | ASSDPGANSDYT | 0.17296718 |
| VGKSTLLL | ASSDPGANSDYT | 0.17463747 |
| AALAHAGGGIV | ASSDPGANSDYT | 0.21491474 |
| AALVAVSLL | ASSDPGANSDYT | 0.29921874 |
| AAVSRAVAAL | ASSDPGANSDYT | 0.1464798 |
| AGGSSGIGGF | ASSDPGANSDYT | 0.33210346 |
| AGLTVANANG | ASSDPGANSDYT | 0.26451853 |
| AGLVGAVL | ASSDPGANSDYT | 0.36675388 |
| ALSQAFTV | ASSDPGANSDYT | 0.14075276 |
| ALTFSVSL | ASSDPGANSDYT | 0.39846155 |
| ALTGAALVAL | ASSDPGANSDYT | 0.2687237 |
| ASFSIGAGVL | ASSDPGANSDYT | 0.8265874 |
| DGSVYKI | ASSDPGANSDYT | 0.18359311 |
| EQLAEIV | ASSDPGANSDYT | 0.21414013 |
| FAVSELTVM | ASSDPGANSDYT | 0.23957601 |
| GAAPLAAGL | ASSDPGANSDYT | 0.79728913 |
| GADGGDGGSGGAGG | ASSDPGANSDYT | 0.13033023 |
| GFGLDTGGAGG | ASSDPGANSDYT | 0.183849 |
| GLASGGLL | ASSDPGANSDYT | 0.12835714 |
| GRIAGAAVLL | ASSDPGANSDYT | 0.30030802 |
| GSAGSLNTSF | ASSDPGANSDYT | 0.3398151 |
| GTGQIGI | ASSDPGANSDYT | 0.14402738 |
| GTLIALIL | ASSDPGANSDYT | 0.26916745 |
| IAATMGFVVG | ASSDPGANSDYT | 0.31369984 |
| IFAQALL | ASSDPGANSDYT | 0.19468081 |
| IGAAVALGF | ASSDPGANSDYT | 0.67546594 |
| IGLTPLAL | ASSDPGANSDYT | 0.21020412 |
| ILGSNTSTL | ASSDPGANSDYT | 0.5123846 |
| IVFSLTVL | ASSDPGANSDYT | 0.36318618 |
| LASKATGF | ASSDPGANSDYT | 0.1498452 |
| LASVALIL | ASSDPGANSDYT | 0.38245195 |
| LFAVGLTL | ASSDPGANSDYT | 0.17158256 |
| LFNGGIGTYI | ASSDPGANSDYT | 0.18610747 |
| LTLGAGII | ASSDPGANSDYT | 0.5006949 |
| LVAAALGDI | ASSDPGANSDYT | 0.81908166 |
| NTAFVQLGI | ASSDPGANSDYT | 0.8146767 |
| QLFSGAEL | ASSDPGANSDYT | 0.61026 |
| SAATFITL | ASSDPGANSDYT | 0.63168335 |
| SNYGLANLGV | ASSDPGANSDYT | 0.6914201 |

| | | |
|---|---|---|
| SYGGAYAVM | ASSDPGANSDYT | 0.80704236 |
| TTGVGGLTLG | ASSDPGANSDYT | 0.84674585 |
| TTQTIYI | ASSDPGANSDYT | 0.8040555 |
| VAAGAGLGAVI | ASSDPGANSDYT | 0.6216736 |
| VAAVVTLL | ASSDPGANSDYT | 0.78784513 |
| VGFLGGLI | ASSDPGANSDYT | 0.7545498 |
| VGVIPILL | ASSDPGANSDYT | 0.75689685 |
| VIFASSLI | ASSDPGANSDYT | 0.7732617 |
| VKGGDGGAAAG | ASSDPGANSDYT | 0.7819164 |
| VTKTFTNI | ASSDPGANSDYT | 0.6580212 |
| VVAAAILL | ASSDPGANSDYT | 0.6028991 |
| YAALDVEL | ASSDPGANSDYT | 0.78133744 |
| LASVALLL | ASSDPGANSDYT | 0.81953007 |
| LDAVHRV | ASSDPGANSDYT | 0.8095101 |
| QGLSSSIF | ASSDPGANSDYT | 0.849829 |
| RGTFANIRL | ASSDPGANSDYT | 0.83369875 |
| RQLWWGHRI | ASSDPGANSDYT | 0.77396065 |
| YGPPGTGKTLL | ASSDPGANSDYT | 0.23263505 |
| AALVGAPANVA | ASSDPGANSDYT | 0.48222226 |
| AATFRRTE | ASSDPGANSDYT | 0.6437 |
| AGALGGGI | ASSDPGANSDYT | 0.7245882 |
| AGGAGGAGGIGAG | ASSDPGANSDYT | 0.22123766 |
| AGSGALGL | ASSDPGANSDYT | 0.7924035 |
| AQIADQI | ASSDPGANSDYT | 0.71865344 |
| ASAVAVTYL | ASSDPGANSDYT | 0.67637515 |
| ASTVLGGTGG | ASSDPGANSDYT | 0.82044613 |
| AVGGEPAK | ASSDPGANSDYT | 0.707746 |
| CEQGTFGGTC | ASSDPGANSDYT | 0.57214886 |
| DEGKAPAG | ASSDPGANSDYT | 0.60801893 |
| DTVVFPAL | ASSDPGANSDYT | 0.8527322 |
| FEGITMAG | ASSDPGANSDYT | 0.5791068 |
| FPLVVQD | ASSDPGANSDYT | 0.8404932 |
| GAFTTPQ | ASSDPGANSDYT | 0.66592425 |
| GFGGFGTAGGVGGA | ASSDPGANSDYT | 0.84690535 |
| GFLAQFL | ASSDPGANSDYT | 0.5375723 |
| GGAGGAGGNAGVA | ASSDPGANSDYT | 0.8270706 |
| GGVGTAGAGGAGGAGG | ASSDPGANSDYT | 0.7584703 |
| GNGGGGDAGNAGSGGN | ASSDPGANSDYT | 0.86263525 |
| GTGGAGGAGNDAGS | ASSDPGANSDYT | 0.74042964 |
| GTTNTGWFNS | ASSDPGANSDYT | 0.81694824 |
| IFMGPLFTL | ASSDPGANSDYT | 0.7425551 |
| ILAHAIEAI | ASSDPGANSDYT | 0.81542903 |
| IRSAIREAL | ASSDPGANSDYT | 0.5361464 |
| ITFAGGG | ASSDPGANSDYT | 0.73487574 |
| ITGMSGHQEV | ASSDPGANSDYT | 0.47817028 |
| IVAGLACTL | ASSDPGANSDYT | 0.73215544 |
| IVGGGLAGA | ASSDPGANSDYT | 0.8214531 |
| IVTGVLNI | ASSDPGANSDYT | 0.19396399 |
| LAQQGCI | ASSDPGANSDYT | 0.82627606 |
| LCLHVEN | ASSDPGANSDYT | 0.5420685 |
| LDAISDGS | ASSDPGANSDYT | 0.6202237 |

| | | |
|---|---|---|
| LGGFAAML | ASSDPGANSDYT | 0.807682 |
| LGGKNPMI | ASSDPGANSDYT | 0.74920744 |
| LGPKGVAAIL | ASSDPGANSDYT | 0.82648766 |
| LMVGVSI | ASSDPGANSDYT | 0.62583905 |
| LVGAAFVL | ASSDPGANSDYT | 0.7667405 |
| LVLAVAPL | ASSDPGANSDYT | 0.7594565 |
| MAAIAVIDAA | ASSDPGANSDYT | 0.32294714 |
| NNGINAELQ | ASSDPGANSDYT | 0.513669 |
| RGEPLVLVHG | ASSDPGANSDYT | 0.8532007 |
| RGGAGGAAGLI | ASSDPGANSDYT | 0.24134031 |
| RSYRMHPAV | ASSDPGANSDYT | 0.372374 |
| SAAHLLAGQLA | ASSDPGANSDYT | 0.13277659 |
| SAAVGSY | ASSDPGANSDYT | 0.15274628 |
| SPHAVLHRI | ASSDPGANSDYT | 0.17318937 |
| SQIAKVL | ASSDPGANSDYT | 0.22337839 |
| TALGHLVP | ASSDPGANSDYT | 0.373631 |
| TAPTTSGAA | ASSDPGANSDYT | 0.3131187 |
| TDGDDGNGGSPGT | ASSDPGANSDYT | 0.14869148 |
| TDGLGEN | ASSDPGANSDYT | 0.18637629 |
| TGGPPLPGQA | ASSDPGANSDYT | 0.15919891 |
| TIPAITI | ASSDPGANSDYT | 0.17797421 |
| TIPGFTI | ASSDPGANSDYT | 0.15785542 |
| TSIASGALA | ASSDPGANSDYT | 0.17258993 |
| TVTLPGAAI | ASSDPGANSDYT | 0.15377815 |
| VASGLSAALP | ASSDPGANSDYT | 0.13863467 |
| VGAGVIGI | ASSDPGANSDYT | 0.15753739 |
| VGPKRFGV | ASSDPGANSDYT | 0.39694244 |
| VNVAFPD | ASSDPGANSDYT | 0.23858446 |
| VSMTNSGVS | ASSDPGANSDYT | 0.29929802 |
| VTDVLGDLSV | ASSDPGANSDYT | 0.28044876 |
| VVVVGAGFAGL | ASSDPGANSDYT | 0.15115555 |
| VYTVGTVV | ASSDPGANSDYT | 0.11923188 |
| WTTFSGP | ASSDPGANSDYT | 0.12612638 |
| YDELVEAF | ASSDPGANSDYT | 0.13816653 |
| YGTGGAGGNGG | ASSDPGANSDYT | 0.15589367 |
| YTNVMAV | ASSDPGANSDYT | 0.11786224 |
| FLEQVHQL | ASSDPGANSDYT | 0.17324491 |
| GLSLLAVAV | ASSDPGANSDYT | 0.16109174 |
| LEQVHQL | ASSDPGANSDYT | 0.13908629 |
| LNAAGAI | ASSDPGANSDYT | 0.2558692 |
| PAVHASL | ASSDPGANSDYT | 0.14503628 |
| TELEDDA | ASSDPGANSDYT | 0.14372516 |
| TPPGPPN | ASSDPGANSDYT | 0.13416032 |
| TPPPPPPP | ASSDPGANSDYT | 0.46829778 |
| TQSPALI | ASSRGQKNTLY | 0.13008352 |
| EFAAAFPQ | ASSRGQKNTLY | 0.2779851 |
| LLSGNLDVL | ASSRGQKNTLY | 0.13657945 |
| GPANLGNYNI | ASSRGQKNTLY | 0.22209175 |
| TLGPTLP | ASSRGQKNTLY | 0.13784155 |
| VYAYSSCCK | ASSRGQKNTLY | 0.24307156 |
| APAPAPAPNI | ASSRGQKNTLY | 0.19888881 |

| | | |
|---|---|---|
| PGGPVPIP | ASSRGQKNTLY | 0.5327859 |
| FLTVALEP | ASSRGQKNTLY | 0.15835641 |
| SQYFGANL | ASSRGQKNTLY | 0.17548089 |
| GGVKPAA | ASSRGQKNTLY | 0.1317876 |
| GGTGGAGGAGGW | ASSRGQKNTLY | 0.53230447 |
| GVGGVAGLGGAGG | ASSRGQKNTLY | 0.12645988 |
| VLLVLATTI | ASSRGQKNTLY | 0.15861775 |
| IGIFAAML | ASSRGQKNTLY | 0.12968622 |
| AVGDLID | ASSRGQKNTLY | 0.16229671 |
| NISPLPL | ASSRGQKNTLY | 0.27919102 |
| VTAVFVAI | ASSRGQKNTLY | 0.12240176 |
| SGDDNDAAL | ASSRGQKNTLY | 0.25177303 |
| AVAALAHS | ASSRGQKNTLY | 0.12798235 |
| GGVTVMATP | ASSRGQKNTLY | 0.13830015 |
| AGLAANVRD | ASSRGQKNTLY | 0.18014537 |
| TNRPDLIDPALLRPG | ASSRGQKNTLY | 0.14584288 |
| LANLGVTAL | ASSRGQKNTLY | 0.3228861 |
| TGERIEL | ASSRGQKNTLY | 0.12814276 |
| NGEELAA | ASSRGQKNTLY | 0.19564281 |
| VLSLVWG | ASSRGQKNTLY | 0.21650112 |
| IGYALWL | ASSRGQKNTLY | 0.120551206 |
| LSLLLAPL | ASSRGQKNTLY | 0.12863341 |
| TLDEALK | ASSRGQKNTLY | 0.4028054 |
| TDGAGTAEGG | ASSRGQKNTLY | 0.15114798 |
| NAPGGAGGNGG | ASSRGQKNTLY | 0.0772749 |
| VSVFMGMGGNF | ASSRGQKNTLY | 0.09344259 |
| LASYLAAL | ASSRGQKNTLY | 0.11345403 |
| LVGGTPAL | ASSRGQKNTLY | 0.08616181 |
| VRNADVL | ASSRGQKNTLY | 0.10548389 |
| TCSSPFLVSAM | ASSRGQKNTLY | 0.11593512 |
| VHITDAE | ASSRGQKNTLY | 0.12567626 |
| EYAAPSF | ASSRGQKNTLY | 0.13712588 |
| VVDAFHGGL | ASSRGQKNTLY | 0.1506448 |
| ASVVDPQ | ASSRGQKNTLY | 0.12554875 |
| SYNEFGGLS | ASSRGQKNTLY | 0.12075166 |
| DPDGSVY | ASSRGQKNTLY | 0.08963964 |
| MVSTIHATE | ASSRGQKNTLY | 0.10191171 |
| LAAGVADPV | ASSRGQKNTLY | 0.09632639 |
| TKEQIAA | ASSRGQKNTLY | 0.078549854 |
| SLASPAAL | ASSRGQKNTLY | 0.08040389 |
| IGLVFYGV | ASSRGQKNTLY | 0.12932989 |
| FLLDASNL | ASSRGQKNTLY | 0.12631445 |
| YQVGNFP | ASSRGQKNTLY | 0.11023108 |
| ESLLSQST | ASSRGQKNTLY | 0.16025409 |
| LFTVGFAAVG | ASSRGQKNTLY | 0.07737995 |
| ATPPTPT | ASSRGQKNTLY | 0.07266308 |
| TDLATLG | ASSRGQKNTLY | 0.08205918 |
| SAGGTGR | ASSRGQKNTLY | 0.10014918 |
| GFSNSGS | ASSRGQKNTLY | 0.09618706 |
| GGNGGSGGTSV | ASSRGQKNTLY | 0.063903466 |
| GTSVATGGAGN | ASSRGQKNTLY | 0.10244371 |

| | | |
|---|---|---|
| NGGNGGSGGTS | ASSRGQKNTLY | 0.15218565 |
| AVVKFPH | ASSRGQKNTLY | 0.106714234 |
| TPPPPPP | ASSRGQKNTLY | 0.11287677 |
| AAPAAAAAPA | ASSRGQKNTLY | 0.10035377 |
| IALNEDL | ASSRGQKNTLY | 0.10122358 |
| IVNVSSM | ASSRGQKNTLY | 0.095935516 |
| TGIPPAPR | ASSRGQKNTLY | 0.10475988 |
| FDLGGGTF | ASSRGQKNTLY | 0.09059294 |
| FDLGGGTFDVS | ASSRGQKNTLY | 0.20273125 |
| NEPTAAA | ASSRGQKNTLY | 0.07282372 |
| NEPTAAALA | ASSRGQKNTLY | 0.18199416 |
| GSIINISSI | ASSRGQKNTLY | 0.109263845 |
| VAPGNAGTA | ASSRGQKNTLY | 0.11323129 |
| TGGGASLE | ASSRGQKNTLY | 0.10952329 |
| VAVYNHYKRI | ASSRGQKNTLY | 0.13128883 |
| TDPAPATT | ASSRGQKNTLY | 0.13732186 |
| TNRPDILDPALLRPG | ASSRGQKNTLY | 0.16610485 |
| LTTTTTS | ASSRGQKNTLY | 0.08039068 |
| PTGSGKTL | ASSRGQKNTLY | 0.09893153 |
| TGSGKTL | ASSRGQKNTLY | 0.08926733 |
| TIGHVDHGK | ASSRGQKNTLY | 0.12987685 |
| TIGHVDHGKTT | ASSRGQKNTLY | 0.09800995 |
| GHVDHGKTT | ASSRGQKNTLY | 0.11179644 |
| AQFEHTLL | ASSRGQKNTLY | 0.13680638 |
| IVNQDEL | ASSRGQKNTLY | 0.14141057 |
| TNGSQFF | ASSRGQKNTLY | 0.10960481 |
| TGKGIVE | ASSRGQKNTLY | 0.20205773 |
| LISNASDA | ASSRGQKNTLY | 0.10280636 |
| AELRTLGELTKIAKAHGAQ | ASSRGQKNTLY | 0.0934499 |
| APNAVAAAEPSW | ASSRGQKNTLY | 0.10449009 |
| APQVAGAATAVL | ASSRGQKNTLY | 0.083790034 |
| ARRDPAAAAAILPTDA | ASSRGQKNTLY | 0.06979422 |
| DAAARDVLAEAGLAEYF | ASSRGQKNTLY | 0.10183517 |
| DDQIVLSAKANLAKVEQ | ASSRGQKNTLY | 0.17381139 |
| DELLPDTGEDSDEDGH | ASSRGQKNTLY | 0.17562087 |
| ELPEAPANRATVPQD | ASSRGQKNTLY | 0.09102441 |
| FGQSKIERVTGIG | ASSRGQKNTLY | 0.13541569 |
| GAGGTGSTAGGAGGAGGAGGI | ASSRGQKNTLY | 0.8381317 |
| GHGGTGAAGGNGGTGGAGGS( | ASSRGQKNTLY | 0.70140904 |
| GNGGNGGAGGEGGPGGAGGA( | ASSRGQKNTLY | 0.51274073 |
| GSTAGNGGNGGNATNGTIAG( | ASSRGQKNTLY | 0.406613 |
| GTGGDGGNGGNGGTGG | ASSRGQKNTLY | 0.5432927 |
| GVRRPAGIPADAPV | ASSRGQKNTLY | 0.8010143 |
| GYRQVTDAHLLTLARR | ASSRGQKNTLY | 0.70754135 |
| IAKMGQDGHDRG | ASSRGQKNTLY | 0.8131527 |
| KAEPSIEIQYT | ASSRGQKNTLY | 0.2954916 |
| KPVRIRMAADEDTVAVT | ASSRGQKNTLY | 0.6804323 |
| MPPRIAGMRLLVIKPEP | ASSRGQKNTLY | 0.63112015 |
| MKFYAVALFRSRGVPPTR | ASSRGQKNTLY | 0.6086476 |
| NAGGFGILGAVAHSP | ASSRGQKNTLY | 0.7027892 |
| NEGREITLAMAEM | ASSRGQKNTLY | 0.33210492 |

| | | |
|---|---|---|
| NELVEWSHPLGHRWRWE | ASSRGQKNTLY | 0.44277728 |
| NLGAGGTGGNGGNPG | ASSRGQKNTLY | 0.39043272 |
| NTEWPKWETVMAADPVD | ASSRGQKNTLY | 0.54687524 |
| PASAAPAETQP | ASSRGQKNTLY | 0.76004803 |
| QQSSDGKNGVRAAEITGE | ASSRGQKNTLY | 0.7697079 |
| RAPDAVTTSWWKE | ASSRGQKNTLY | 0.80857664 |
| SGFGNVGAHTSGWFNQ | ASSRGQKNTLY | 0.81475455 |
| SGVAGAVGQAASVGGLKV | ASSRGQKNTLY | 0.71562684 |
| SNGGAGTGSAGGNGGT | ASSRGQKNTLY | 0.090432696 |
| TGGAGGEGGAAG | ASSRGQKNTLY | 0.17746209 |
| TGGDGGAAGAGGAPGHG | ASSRGQKNTLY | 0.3529977 |
| TMPPLLAAIANAGRHGV | ASSRGQKNTLY | 0.4567324 |
| TSSTAGNSGGA | ASSRGQKNTLY | 0.10339891 |
| VPATVPVGILC | ASSRGQKNTLY | 0.5473201 |
| VQRPDSGGVVLTGRIS | ASSRGQKNTLY | 0.69129467 |
| WTPNPVPATKGVPAGLVQAT | ASSRGQKNTLY | 0.35333937 |
| YALKRALAKFSDRFDVVII | ASSRGQKNTLY | 0.7290957 |
| YLIGSVIV | ASSRGQKNTLY | 0.43053892 |
| YGQLVAVID | ASSRGQKNTLY | 0.29248872 |
| VTVGGMTLPA | ASSRGQKNTLY | 0.41959512 |
| VLALPAV | ASSRGQKNTLY | 0.7932712 |
| VIPPTLNYET | ASSRGQKNTLY | 0.5893189 |
| VGMGGMHPSSG | ASSRGQKNTLY | 0.78337944 |
| VGLAGGAATA | ASSRGQKNTLY | 0.34482932 |
| VGGAGGAGGSGP | ASSRGQKNTLY | 0.8094975 |
| VFAAFMV | ASSRGQKNTLY | 0.26136184 |
| TLYDKLR | ASSRGQKNTLY | 0.7919642 |
| TLAVFTVL | ASSRGQKNTLY | 0.7400803 |
| TGSVQPARV | ASSRGQKNTLY | 0.8455491 |
| TGGGAGSGGT | ASSRGQKNTLY | 0.5931361 |
| SHLETSL | ASSRGQKNTLY | 0.58077234 |
| SFTTGMT | ASSRGQKNTLY | 0.21886498 |
| QVEVAKRMGVRQPTVS | ASSRGQKNTLY | 0.5221638 |
| QGSGGPALAM | ASSRGQKNTLY | 0.23591922 |
| MLAPSNP | ASSRGQKNTLY | 0.68429023 |
| LVNNAAI | ASSRGQKNTLY | 0.23221686 |
| LTGEINTLA | ASSRGQKNTLY | 0.6733655 |
| LSIFGPV | ASSRGQKNTLY | 0.8308334 |
| LLALLANL | ASSRGQKNTLY | 0.098832436 |
| LISNASDALDKLR | ASSRGQKNTLY | 0.7980939 |
| LINEAVP | ASSRGQKNTLY | 0.32921174 |
| LGYAFSGL | ASSRGQKNTLY | 0.30970627 |
| LDFVAVAL | ASSRGQKNTLY | 0.7835457 |
| LAQPGAE | ASSRGQKNTLY | 0.29190505 |
| ISNLGAVTGF | ASSRGQKNTLY | 0.78772 |
| HQEDPGRI | ASSRGQKNTLY | 0.33593217 |
| GTPGEPTP | ASSRGQKNTLY | 0.645678 |
| GFFNSGS | ASSRGQKNTLY | 0.79603434 |
| GAGGGGLTAGAN | ASSRGQKNTLY | 0.08432314 |
| FVDALVNM | ASSRGQKNTLY | 0.29522517 |
| FLTVAPEL | ASSRGQKNTLY | 0.83214295 |

| | | |
|---|---|---|
| FETLTAL | ASSRGQKNTLY | 0.24714822 |
| DVINPIEL | ASSRGQKNTLY | 0.16423175 |
| APVSGAPV | ASSRGQKNTLY | 0.15358692 |
| ALVGGLAW | ASSRGQKNTLY | 0.16648561 |
| ALAAAAATAI | ASSRGQKNTLY | 0.16465782 |
| AFGLGLLL | ASSRGQKNTLY | 0.18419327 |
| AFFAAASL | ASSRGQKNTLY | 0.19404513 |
| AAAALPAFSPP | ASSRGQKNTLY | 0.27830926 |
| VVLAPAL | ASSRGQKNTLY | 0.16719808 |
| TIGHVDHGKTTL | ASSRGQKNTLY | 0.17557222 |
| LISNASDALDK | ASSRGQKNTLY | 0.15933391 |
| LISNASDAL | ASSRGQKNTLY | 0.15721719 |
| ISNASDALDK | ASSRGQKNTLY | 0.17846629 |
| NQAETLVYQTE | ASSRGQKNTLY | 0.16472659 |
| TGGTGGTAGAG | ASSRGQKNTLY | 0.1581322 |
| TGSTAGGAGGA | ASSRGQKNTLY | 0.16235521 |
| DPLAVPVAAGPV | ASSRGQKNTLY | 0.16411741 |
| GGTGGTPGNGGN | ASSRGQKNTLY | 0.2802997 |
| PAGLAGVDPATL | ASSRGQKNTLY | 0.21731757 |
| GNNLVVEAPHAGASV | ASSRGQKNTLY | 0.23560058 |
| KVTRIMSASTLEQPA | ASSRGQKNTLY | 0.21444826 |
| LGFALPIALPRRTVV | ASSRGQKNTLY | 0.20152944 |
| TRLGNLTARKLVVLV | ASSRGQKNTLY | 0.13975552 |
| WSQQRRLIYFPSAGP | ASSRGQKNTLY | 0.15207456 |
| GPNGNPDPMAAAVDIR | ASSRGQKNTLY | 0.15859993 |
| TQRCVSVADLRAAASR | ASSRGQKNTLY | 0.1589813 |
| QFVLDVAEHPVEVLTETV | ASSRGQKNTLY | 0.13923566 |
| GGAGGTGGAGGNGGTQAGDG | ASSRGQKNTLY | 0.17010324 |
| PLLATLRAIAVDVDAKPAQV | ASSRGQKNTLY | 0.2065317 |
| APKFRTGGYMIDFWGVGYQV | ASSRGQKNTLY | 0.15351976 |
| GAGGAGGAGGAAGGPGGTGG | ASSRGQKNTLY | 0.20293032 |
| IGALGLLATDVAPAGSVAFG | ASSRGQKNTLY | 0.15160464 |
| AGGAGGIGGNGGAGGAGGVL | ASSRGQKNTLY | 0.15396744 |
| ADELIGLL | ASSRGQKNTLY | 0.1596646 |
| ADRALKA | ASSRGQKNTLY | 0.27252042 |
| AGAVGQVGGAGGA | ASSRGQKNTLY | 0.15411516 |
| AGGGGGLGGAP | ASSRGQKNTLY | 0.18907 |
| AGGLLYGNGGAG | ASSRGQKNTLY | 0.15939124 |
| AGLESIGL | ASSRGQKNTLY | 0.2322523 |
| AGLKTGII | ASSRGQKNTLY | 0.1572981 |
| ASLAALLL | ASSRGQKNTLY | 0.19573775 |
| ASVINISSI | ASSRGQKNTLY | 0.16480269 |
| ATPPGPA | ASSRGQKNTLY | 0.2728145 |
| ATVASALV | ASSRGQKNTLY | 0.16917382 |
| ATVASLIL | ASSRGQKNTLY | 0.18596525 |
| AVNGPVLIRH | ASSRGQKNTLY | 0.1611382 |
| DGAAGAVGI | ASSRGQKNTLY | 0.16622116 |
| DPLLHGI | ASSRGQKNTLY | 0.14522037 |
| ELGAALGAL | ASSRGQKNTLY | 0.17175193 |
| ELGRTQAV | ASSRGQKNTLY | 0.15996951 |
| FVAGGYLS | ASSRGQKNTLY | 0.17162311 |

| | | |
|---|---|---|
| GASGGAGGAGGW | ASSRGQKNTLY | 0.2742671 |
| GGAGAGGDGGAGGI | ASSRGQKNTLY | 0.1383678 |
| GTGHNGSQPGMGGNG | ASSRGQKNTLY | 0.23038486 |
| GYAAASASAS | ASSRGQKNTLY | 0.13950363 |
| HQVDVAAVQQG | ASSRGQKNTLY | 0.15983076 |
| IGAAIAHT | ASSRGQKNTLY | 0.18892387 |
| IGMIEVAQL | ASSRGQKNTLY | 0.16192086 |
| IRLAPEHTSAMD | ASSRGQKNTLY | 0.236704 |
| IYNVSVL | ASSRGQKNTLY | 0.15212338 |
| KQGFLGM | ASSRGQKNTLY | 0.17425266 |
| LESVITQL | ASSRGQKNTLY | 0.20596454 |
| LGLGALVVV | ASSRGQKNTLY | 0.14063996 |
| LIGNGGNGGSGGI | ASSRGQKNTLY | 0.14773571 |
| LNAINAPTL | ASSRGQKNTLY | 0.22084072 |
| LQLLNTGV | ASSRGQKNTLY | 0.24298117 |
| LQTSIEF | ASSRGQKNTLY | 0.4499309 |
| MSFAVSEL | ASSRGQKNTLY | 0.21305144 |
| NTGALGL | ASSRGQKNTLY | 0.20971577 |
| PPGAGGAGGAGTT | ASSRGQKNTLY | 0.23693532 |
| QLVGLGI | ASSRGQKNTLY | 0.26265034 |
| RFLEQVHQL | ASSRGQKNTLY | 0.43532372 |
| RGVGYVLR | ASSRGQKNTLY | 0.2634863 |
| SATGVFAGVI | ASSRGQKNTLY | 0.2137309 |
| SLPAITLP | ASSRGQKNTLY | 0.21807542 |
| STALAFTV | ASSRGQKNTLY | 0.21947247 |
| SVLTVGLGK | ASSRGQKNTLY | 0.22006148 |
| TALAGAGSGI | ASSRGQKNTLY | 0.24350381 |
| TGLPFTGI | ASSRGQKNTLY | 0.21271352 |
| VAEHVTV | ASSRGQKNTLY | 0.20822266 |
| VDIVLNSL | ASSRGQKNTLY | 0.21381488 |
| VLTGGGAVL | ASSRGQKNTLY | 0.23617738 |
| VNVGLAYGI | ASSRGQKNTLY | 0.30378476 |
| VSSAAVGAV | ASSRGQKNTLY | 0.28404066 |
| VTSVYEAGV | ASSRGQKNTLY | 0.30215836 |
| WVALDRGA | ASSRGQKNTLY | 0.26456764 |
| YLVGSVVI | ASSRGQKNTLY | 0.23583438 |
| GPPGTGKTLL | ASSRGQKNTLY | 0.19510959 |
| GVGMTKF | ASSRGQKNTLY | 0.20472851 |
| IGIGGSDL | ASSRGQKNTLY | 0.21610332 |
| IGTLLAEL | ASSRGQKNTLY | 0.22462013 |
| IYTSGTT | ASSRGQKNTLY | 0.19423321 |
| LSFPAGV | ASSRGQKNTLY | 0.22844526 |
| LYTSGTT | ASSRGQKNTLY | 0.24262354 |
| VGAFPFGF | ASSRGQKNTLY | 0.20043948 |
| VGAGTVEF | ASSRGQKNTLY | 0.3059256 |
| VGKSTLLL | ASSRGQKNTLY | 0.20722161 |
| AALAHAGGGIV | ASSRGQKNTLY | 0.20867883 |
| AALVAVSLL | ASSRGQKNTLY | 0.21209608 |
| AAVSRAVAAL | ASSRGQKNTLY | 0.27663344 |
| AGGSSGIGGF | ASSRGQKNTLY | 0.20415668 |
| AGLTVANANG | ASSRGQKNTLY | 0.2583346 |

| | | |
|---|---|---|
| AGLVGAVL | ASSRGQKNTLY | 0.22122921 |
| ALSQAFTV | ASSRGQKNTLY | 0.25515172 |
| ALTFSVSL | ASSRGQKNTLY | 0.2026539 |
| ALTGAALVAL | ASSRGQKNTLY | 0.251498 |
| ASFSIGAGVL | ASSRGQKNTLY | 0.22700918 |
| DGSVYKI | ASSRGQKNTLY | 0.43597135 |
| EQLAEIV | ASSRGQKNTLY | 0.22340512 |
| FAVSELTVM | ASSRGQKNTLY | 0.24184965 |
| GAAPLAAGL | ASSRGQKNTLY | 0.22562735 |
| GADGGDGGSGGAGG | ASSRGQKNTLY | 0.30185214 |
| GFGLDTGGAGG | ASSRGQKNTLY | 0.19959751 |
| GLASGGLL | ASSRGQKNTLY | 0.24082363 |
| GRIAGAAVLL | ASSRGQKNTLY | 0.20896813 |
| GSAGSLNTSF | ASSRGQKNTLY | 0.25184163 |
| GTGQIGI | ASSRGQKNTLY | 0.2695303 |
| GTLIALIL | ASSRGQKNTLY | 0.20513497 |
| IAATMGFVVG | ASSRGQKNTLY | 0.27306768 |
| IFAQALL | ASSRGQKNTLY | 0.21341938 |
| IGAAVALGF | ASSRGQKNTLY | 0.2098933 |
| IGLTPLAL | ASSRGQKNTLY | 0.25847688 |
| ILGSNTSTL | ASSRGQKNTLY | 0.20427163 |
| IVFSLTVL | ASSRGQKNTLY | 0.24607684 |
| LASKATGF | ASSRGQKNTLY | 0.20349501 |
| LASVALIL | ASSRGQKNTLY | 0.23697375 |
| LFAVGLTL | ASSRGQKNTLY | 0.31474698 |
| LFNGGIGTYI | ASSRGQKNTLY | 0.19908848 |
| LTLGAGII | ASSRGQKNTLY | 0.20448487 |
| LVAAALGDI | ASSRGQKNTLY | 0.26958507 |
| NTAFVQLGI | ASSRGQKNTLY | 0.5922461 |
| QLFSGAEL | ASSRGQKNTLY | 0.5670562 |
| SAATFITL | ASSRGQKNTLY | 0.23100127 |
| SNYGLANLGV | ASSRGQKNTLY | 0.23994046 |
| SYGGAYAVM | ASSRGQKNTLY | 0.27292898 |
| TTGVGGLTLG | ASSRGQKNTLY | 0.40005055 |
| TTQTIYI | ASSRGQKNTLY | 0.6563253 |
| VAAGAGLGAVI | ASSRGQKNTLY | 0.52635056 |
| VAAVVTLL | ASSRGQKNTLY | 0.23111537 |
| VGFLGGLI | ASSRGQKNTLY | 0.32153583 |
| VGVIPILL | ASSRGQKNTLY | 0.32661533 |
| VIFASSLI | ASSRGQKNTLY | 0.33423674 |
| VKGGDGGAAAG | ASSRGQKNTLY | 0.29023048 |
| VTKTFTNI | ASSRGQKNTLY | 0.28991035 |
| VVAAAILL | ASSRGQKNTLY | 0.25304368 |
| YAALDVEL | ASSRGQKNTLY | 0.22832729 |
| LASVALLL | ASSRGQKNTLY | 0.2821899 |
| LDAVHRV | ASSRGQKNTLY | 0.60266685 |
| QGLSSSIF | ASSRGQKNTLY | 0.6028131 |
| RGTFANIRL | ASSRGQKNTLY | 0.3664043 |
| RQLWWGHRI | ASSRGQKNTLY | 0.5811616 |
| YGPPGTGKTLL | ASSRGQKNTLY | 0.38235372 |
| AALVGAPANVA | ASSRGQKNTLY | 0.19439048 |

| | | |
|---|---|---|
| AATFRRTE | ASSRGQKNTLY | 0.20217495 |
| AGALGGGI | ASSRGQKNTLY | 0.22033443 |
| AGGAGGAGGIGAG | ASSRGQKNTLY | 0.24060364 |
| AGSGALGL | ASSRGQKNTLY | 0.19785334 |
| AQIADQI | ASSRGQKNTLY | 0.29533565 |
| ASAVAVTYL | ASSRGQKNTLY | 0.24783775 |
| ASTVLGGTGG | ASSRGQKNTLY | 0.23717493 |
| AVGGEPAK | ASSRGQKNTLY | 0.41727 |
| CEQGTFGGTC | ASSRGQKNTLY | 0.27056208 |
| DEGKAPAG | ASSRGQKNTLY | 0.22505306 |
| DTVVFPAL | ASSRGQKNTLY | 0.22509769 |
| FEGITMAG | ASSRGQKNTLY | 0.50202584 |
| FPLVVQD | ASSRGQKNTLY | 0.21932733 |
| GAFTTPQ | ASSRGQKNTLY | 0.50791335 |
| GFGGFGTAGGVGGA | ASSRGQKNTLY | 0.21607295 |
| GFLAQFL | ASSRGQKNTLY | 0.4700709 |
| GGAGGAGGNAGVA | ASSRGQKNTLY | 0.21764472 |
| GGVGTAGAGGAGGAGG | ASSRGQKNTLY | 0.50087 |
| GNGGGGDAGNAGSGGN | ASSRGQKNTLY | 0.41142997 |
| GTGGAGGAGNDAGS | ASSRGQKNTLY | 0.6928675 |
| GTTNTGWFNS | ASSRGQKNTLY | 0.27775818 |
| IFMGPLFTL | ASSRGQKNTLY | 0.3460297 |
| ILAHAIEAI | ASSRGQKNTLY | 0.22893302 |
| IRSAIREAL | ASSRGQKNTLY | 0.7005782 |
| ITFAGGG | ASSRGQKNTLY | 0.20423517 |
| ITGMSGHQEV | ASSRGQKNTLY | 0.2889744 |
| IVAGLACTL | ASSRGQKNTLY | 0.198811 |
| IVGGGLAGA | ASSRGQKNTLY | 0.26392394 |
| IVTGVLNI | ASSRGQKNTLY | 0.5372302 |
| LAQQGCI | ASSRGQKNTLY | 0.20535854 |
| LCLHVEN | ASSRGQKNTLY | 0.35728619 |
| LDAISDGS | ASSRGQKNTLY | 0.2542708 |
| LGGFAAML | ASSRGQKNTLY | 0.23490007 |
| LGGKNPMI | ASSRGQKNTLY | 0.36611515 |
| LGPKGVAAIL | ASSRGQKNTLY | 0.25871375 |
| LMVGVSI | ASSRGQKNTLY | 0.61272496 |
| LVGAAFVL | ASSRGQKNTLY | 0.22036809 |
| LVLAVAPL | ASSRGQKNTLY | 0.28271547 |
| MAAIAVIDAA | ASSRGQKNTLY | 0.28735903 |
| NNGINAELQ | ASSRGQKNTLY | 0.2086653 |
| RGEPLVLVHG | ASSRGQKNTLY | 0.21550724 |
| RGGAGGAAGLI | ASSRGQKNTLY | 0.62426996 |
| RSYRMHPAV | ASSRGQKNTLY | 0.24666111 |
| SAAHLLAGQLA | ASSRGQKNTLY | 0.19670318 |
| SAAVGSY | ASSRGQKNTLY | 0.15678251 |
| SPHAVLHRI | ASSRGQKNTLY | 0.16115154 |
| SQIAKVL | ASSRGQKNTLY | 0.14390118 |
| TALGHLVP | ASSRGQKNTLY | 0.1911897 |
| TAPTTSGAA | ASSRGQKNTLY | 0.22461744 |
| TDGDDGNGGSPGT | ASSRGQKNTLY | 0.20875731 |
| TDGLGEN | ASSRGQKNTLY | 0.15790632 |

| | | |
|---|---|---|
| TGGPPLPGQA | ASSRGQKNTLY | 0.17800795 |
| TIPAITI | ASSRGQKNTLY | 0.17459334 |
| TIPGFTI | ASSRGQKNTLY | 0.15605088 |
| TSIASGALA | ASSRGQKNTLY | 0.165882 |
| TVTLPGAAI | ASSRGQKNTLY | 0.16521382 |
| VASGLSAALP | ASSRGQKNTLY | 0.15543625 |
| VGAGVIGI | ASSRGQKNTLY | 0.16074935 |
| VGPKRFGV | ASSRGQKNTLY | 0.14475563 |
| VNVAFPD | ASSRGQKNTLY | 0.1885019 |
| VSMTNSGVS | ASSRGQKNTLY | 0.17498916 |
| VTDVLGDLSV | ASSRGQKNTLY | 0.20601729 |
| VVVVGAGFAGL | ASSRGQKNTLY | 0.24058163 |
| VYTVGTVV | ASSRGQKNTLY | 0.17320432 |
| WTTFSGP | ASSRGQKNTLY | 0.13407314 |
| YDELVEAF | ASSRGQKNTLY | 0.14420648 |
| YGTGGAGGNGG | ASSRGQKNTLY | 0.17170754 |
| YTNVMAV | ASSRGQKNTLY | 0.15473306 |
| FLEQVHQL | ASSRGQKNTLY | 0.1291632 |
| GLSLLAVAV | ASSRGQKNTLY | 0.16565214 |
| LEQVHQL | ASSRGQKNTLY | 0.1882875 |
| LNAAGAI | ASSRGQKNTLY | 0.15724646 |
| PAVHASL | ASSRGQKNTLY | 0.16271658 |
| TELEDDA | ASSRGQKNTLY | 0.14657007 |
| TPPGPPN | ASSRGQKNTLY | 0.15077698 |
| TPPPPPPP | ASSRGQKNTLY | 0.15999454 |
| TQSPALI | ASSLARGNSDYT | 0.176677 |
| EFAAAFPQ | ASSLARGNSDYT | 0.15659352 |
| LLSGNLDVL | ASSLARGNSDYT | 0.18450393 |
| GPANLGNYNI | ASSLARGNSDYT | 0.14289963 |
| TLGPTLP | ASSLARGNSDYT | 0.22783531 |
| VYAYSSCCK | ASSLARGNSDYT | 0.1552018 |
| APAPAPAPNI | ASSLARGNSDYT | 0.19653687 |
| PGGPVPIP | ASSLARGNSDYT | 0.1714059 |
| FLTVALEP | ASSLARGNSDYT | 0.19493318 |
| SQYFGANL | ASSLARGNSDYT | 0.16703518 |
| GGVKPAA | ASSLARGNSDYT | 0.18637426 |
| GGTGGAGGAGGW | ASSLARGNSDYT | 0.15651178 |
| GVGGVAGLGGAGG | ASSLARGNSDYT | 0.24411407 |
| VLLVLATTI | ASSLARGNSDYT | 0.14373626 |
| IGIFAAML | ASSLARGNSDYT | 0.18969879 |
| AVGDLID | ASSLARGNSDYT | 0.14535874 |
| NISPLPL | ASSLARGNSDYT | 0.18824038 |
| VTAVFVAI | ASSLARGNSDYT | 0.2531343 |
| SGDDNDAAL | ASSLARGNSDYT | 0.13344571 |
| AVAALAHS | ASSLARGNSDYT | 0.20364764 |
| GGVTVMATP | ASSLARGNSDYT | 0.14163885 |
| AGLAANVRD | ASSLARGNSDYT | 0.14944147 |
| TNRPDLIDPALLRPG | ASSLARGNSDYT | 0.18755208 |
| LANLGVTAL | ASSLARGNSDYT | 0.16269438 |
| TGERIEL | ASSLARGNSDYT | 0.21123421 |
| NGEELAA | ASSLARGNSDYT | 0.14606582 |

| | | |
|---|---|---|
| VLSLVWG | ASSLARGNSDYT | 0.17766964 |
| IGYALWL | ASSLARGNSDYT | 0.23515035 |
| LSLLLAPL | ASSLARGNSDYT | 0.13847783 |
| TLDEALK | ASSLARGNSDYT | 0.14794123 |
| TDGAGTAEGG | ASSLARGNSDYT | 0.23095618 |
| NAPGGAGGNGG | ASSLARGNSDYT | 0.15413983 |
| VSVFMGMGGNF | ASSLARGNSDYT | 0.14963296 |
| LASYLAAL | ASSLARGNSDYT | 0.09459994 |
| LVGGTPAL | ASSLARGNSDYT | 0.09876027 |
| VRNADVL | ASSLARGNSDYT | 0.09282075 |
| TCSSPFLVSAM | ASSLARGNSDYT | 0.108965434 |
| VHITDAE | ASSLARGNSDYT | 0.16983631 |
| EYAAPSF | ASSLARGNSDYT | 0.14025863 |
| VVDAFHGGL | ASSLARGNSDYT | 0.09542643 |
| ASVVDPQ | ASSLARGNSDYT | 0.14736451 |
| SYNEFGGLS | ASSLARGNSDYT | 0.09692602 |
| DPDGSVY | ASSLARGNSDYT | 0.10748243 |
| MVSTIHATE | ASSLARGNSDYT | 0.10021877 |
| LAAGVADPV | ASSLARGNSDYT | 0.09846401 |
| TKEQIAA | ASSLARGNSDYT | 0.08845548 |
| SLASPAAL | ASSLARGNSDYT | 0.09230242 |
| IGLVFYGV | ASSLARGNSDYT | 0.08645801 |
| FLLDASNL | ASSLARGNSDYT | 0.29341394 |
| YQVGNFP | ASSLARGNSDYT | 0.20815982 |
| ESLLSQST | ASSLARGNSDYT | 0.10923402 |
| LFTVGFAAVG | ASSLARGNSDYT | 0.14779472 |
| ATPPTPT | ASSLARGNSDYT | 0.114158876 |
| TDLATLG | ASSLARGNSDYT | 0.07925427 |
| SAGGTGR | ASSLARGNSDYT | 0.085385494 |
| GFSNSGS | ASSLARGNSDYT | 0.0877208 |
| GGNGGSGGTSV | ASSLARGNSDYT | 0.096586846 |
| GTSVATGGAGN | ASSLARGNSDYT | 0.07761235 |
| NGGNGGSGGTS | ASSLARGNSDYT | 0.099754445 |
| AVVKFPH | ASSLARGNSDYT | 0.10637251 |
| TPPPPPP | ASSLARGNSDYT | 0.09013712 |
| AAPAAAAAPA | ASSLARGNSDYT | 0.14133492 |
| IALNEDL | ASSLARGNSDYT | 0.09443466 |
| IVNVSSM | ASSLARGNSDYT | 0.09456706 |
| TGIPPAPR | ASSLARGNSDYT | 0.08982818 |
| FDLGGGTF | ASSLARGNSDYT | 0.11495706 |
| FDLGGGTFDVS | ASSLARGNSDYT | 0.08526177 |
| NEPTAAA | ASSLARGNSDYT | 0.16231507 |
| NEPTAAALA | ASSLARGNSDYT | 0.08190635 |
| GSIINISSI | ASSLARGNSDYT | 0.12010866 |
| VAPGNAGTA | ASSLARGNSDYT | 0.089975685 |
| TGGGASLE | ASSLARGNSDYT | 0.13468067 |
| VAVYNHYKRI | ASSLARGNSDYT | 0.12331584 |
| TDPAPATT | ASSLARGNSDYT | 0.28547025 |
| TNRPDILDPALLRPG | ASSLARGNSDYT | 0.11501089 |
| LTTTTTS | ASSLARGNSDYT | 0.120342486 |
| PTGSGKTL | ASSLARGNSDYT | 0.08684588 |

| | | |
|---|---|---|
| TGSGKTL | ASSLARGNSDYT | 0.11979697 |
| TIGHVDHGK | ASSLARGNSDYT | 0.08348636 |
| TIGHVDHGKTT | ASSLARGNSDYT | 0.11134779 |
| GHVDHGKTT | ASSLARGNSDYT | 0.08375798 |
| AQFEHTLL | ASSLARGNSDYT | 0.1173724 |
| IVNQDEL | ASSLARGNSDYT | 0.15918694 |
| TNGSQFF | ASSLARGNSDYT | 0.08643293 |
| TGKGIVE | ASSLARGNSDYT | 0.12622981 |
| LISNASDA | ASSLARGNSDYT | 0.09091025 |
| AELRTLGELTKIAKAHGAQ | ASSLARGNSDYT | 0.0917131 |
| APNAVAAAEPSW | ASSLARGNSDYT | 0.1231894 |
| APQVAGAATAVL | ASSLARGNSDYT | 0.09545769 |
| ARRDPAAAAAILPTDA | ASSLARGNSDYT | 0.1282628 |
| DAAARDVLAEAGLAEYF | ASSLARGNSDYT | 0.08289527 |
| DDQIVLSAKANLAKVEQ | ASSLARGNSDYT | 0.100030124 |
| DELLPDTGEDSDEDGH | ASSLARGNSDYT | 0.15774949 |
| ELPEAPANRATVPQD | ASSLARGNSDYT | 0.08262493 |
| FGQSKIERVTGIG | ASSLARGNSDYT | 0.08634465 |
| GAGGTGSTAGGAGGAGGAGGI | ASSLARGNSDYT | 0.1762743 |
| GHGGTGAAGGNGGTGGAGGS( | ASSLARGNSDYT | 0.28224438 |
| GNGGNGGAGGEGGPGGAGGA( | ASSLARGNSDYT | 0.18429509 |
| GSTAGNGGNGGNATNGTIAG( | ASSLARGNSDYT | 0.090914614 |
| GTGGDGGNGGNGGTGG | ASSLARGNSDYT | 0.112717874 |
| GVRRPAGIPADAPV | ASSLARGNSDYT | 0.13340504 |
| GYRQVTDAHLLTLARR | ASSLARGNSDYT | 0.17838974 |
| IAKMGQDGHDRG | ASSLARGNSDYT | 0.40283552 |
| KAEPSIEIQYT | ASSLARGNSDYT | 0.29960966 |
| KPVRIRMAADEDTVAVT | ASSLARGNSDYT | 0.11780619 |
| MPPRIAGMRLLVIKPEP | ASSLARGNSDYT | 0.15837926 |
| MKFYAVALFRSRGVPPTR | ASSLARGNSDYT | 0.10679744 |
| NAGGFGILGAVAHSP | ASSLARGNSDYT | 0.16324233 |
| NEGREITLAMAEM | ASSLARGNSDYT | 0.15869604 |
| NELVEWSHPLGHRWRWE | ASSLARGNSDYT | 0.106647946 |
| NLGAGGTGGNGGNPG | ASSLARGNSDYT | 0.09173756 |
| NTEWPKWETVMAADPVD | ASSLARGNSDYT | 0.092789516 |
| PASAAPAETQP | ASSLARGNSDYT | 0.18371505 |
| QQSSDGKNGVRAAEITGE | ASSLARGNSDYT | 0.3473579 |
| RAPDAVTTSWWKE | ASSLARGNSDYT | 0.18094017 |
| SGFGNVGAHTSGWFNQ | ASSLARGNSDYT | 0.21611422 |
| SGVAGAVGQAASVGGLKV | ASSLARGNSDYT | 0.28667176 |
| SNGGAGTGSAGGNGGT | ASSLARGNSDYT | 0.10493038 |
| TGGAGGEGGAAG | ASSLARGNSDYT | 0.08249729 |
| TGGDGGAAGAGGAPGHG | ASSLARGNSDYT | 0.08755323 |
| TMPPLLAAIANAGRHGV | ASSLARGNSDYT | 0.10041963 |
| TSSTAGNSGGA | ASSLARGNSDYT | 0.11828075 |
| VPATVPVGILC | ASSLARGNSDYT | 0.077116005 |
| VQRPDSGGVVLTGRIS | ASSLARGNSDYT | 0.16495891 |
| WTPNPVPATKGVPAGLVQAT | ASSLARGNSDYT | 0.11562412 |
| YALKRALAKFSDRFDVVII | ASSLARGNSDYT | 0.09962285 |
| YLIGSVIV | ASSLARGNSDYT | 0.31879854 |
| YGQLVAVID | ASSLARGNSDYT | 0.09715763 |

| | | |
|---|---|---|
| VTVGGMTLPA | ASSLARGNSDYT | 0.09709155 |
| VLALPAV | ASSLARGNSDYT | 0.11432247 |
| VIPPTLNYET | ASSLARGNSDYT | 0.18629491 |
| VGMGGMHPSSG | ASSLARGNSDYT | 0.10485962 |
| VGLAGGAATA | ASSLARGNSDYT | 0.38036907 |
| VGGAGGAGGSGP | ASSLARGNSDYT | 0.10737911 |
| VFAAFMV | ASSLARGNSDYT | 0.2357132 |
| TLYDKLR | ASSLARGNSDYT | 0.105124265 |
| TLAVFTVL | ASSLARGNSDYT | 0.22852299 |
| TGSVQPARV | ASSLARGNSDYT | 0.1640946 |
| TGGGAGSGGT | ASSLARGNSDYT | 0.61090946 |
| SHLETSL | ASSLARGNSDYT | 0.14208269 |
| SFTTGMT | ASSLARGNSDYT | 0.18508838 |
| QVEVAKRMGVRQPTVS | ASSLARGNSDYT | 0.11062644 |
| QGSGGPALAM | ASSLARGNSDYT | 0.10586913 |
| MLAPSNP | ASSLARGNSDYT | 0.09011519 |
| LVNNAAI | ASSLARGNSDYT | 0.1001269 |
| LTGEINTLA | ASSLARGNSDYT | 0.09147369 |
| LSIFGPV | ASSLARGNSDYT | 0.10427941 |
| LLALLANL | ASSLARGNSDYT | 0.23289631 |
| LISNASDALDKLR | ASSLARGNSDYT | 0.0881836 |
| LINEAVP | ASSLARGNSDYT | 0.10414998 |
| LGYAFSGL | ASSLARGNSDYT | 0.103117876 |
| LDFVAVAL | ASSLARGNSDYT | 0.10116395 |
| LAQPGAE | ASSLARGNSDYT | 0.17815845 |
| ISNLGAVTGF | ASSLARGNSDYT | 0.118368156 |
| HQEDPGRI | ASSLARGNSDYT | 0.086115085 |
| GTPGEPTP | ASSLARGNSDYT | 0.08625935 |
| GFFNSGS | ASSLARGNSDYT | 0.10559935 |
| GAGGGGLTAGAN | ASSLARGNSDYT | 0.1449177 |
| FVDALVNM | ASSLARGNSDYT | 0.081664324 |
| FLTVAPEL | ASSLARGNSDYT | 0.09894023 |
| FETLTAL | ASSLARGNSDYT | 0.27437207 |
| DVINPIEL | ASSLARGNSDYT | 0.28710476 |
| APVSGAPV | ASSLARGNSDYT | 0.28484973 |
| ALVGGLAW | ASSLARGNSDYT | 0.2853654 |
| ALAAAAATAI | ASSLARGNSDYT | 0.28694823 |
| AFGLGLLL | ASSLARGNSDYT | 0.2851384 |
| AFFAAASL | ASSLARGNSDYT | 0.28666845 |
| AAAALPAFSPP | ASSLARGNSDYT | 0.2875327 |
| VVLAPAL | ASSLARGNSDYT | 0.2864607 |
| TIGHVDHGKTTL | ASSLARGNSDYT | 0.28742468 |
| LISNASDALDK | ASSLARGNSDYT | 0.28936103 |
| LISNASDAL | ASSLARGNSDYT | 0.2884549 |
| ISNASDALDK | ASSLARGNSDYT | 0.29025427 |
| NQAETLVYQTE | ASSLARGNSDYT | 0.28589413 |
| TGGTGGTAGAG | ASSLARGNSDYT | 0.2873801 |
| TGSTAGGAGGA | ASSLARGNSDYT | 0.28495398 |
| DPLAVPVAAGPV | ASSLARGNSDYT | 0.28476045 |
| GGTGGTPGNGGN | ASSLARGNSDYT | 0.28795275 |
| PAGLAGVDPATL | ASSLARGNSDYT | 0.28903168 |

| | | |
|---|---|---|
| GNNLVVEAPHAGASV | ASSLARGNSDYT | 0.2902372 |
| KVTRIMSASTLEQPA | ASSLARGNSDYT | 0.289615 |
| LGFALPIALPRRTVV | ASSLARGNSDYT | 0.28981248 |
| TRLGNLTARKLVVLV | ASSLARGNSDYT | 0.28666422 |
| WSQQRRLIYFPSAGP | ASSLARGNSDYT | 0.28511274 |
| GPNGNPDPMAAAVDIR | ASSLARGNSDYT | 0.28511605 |
| TQRCVSVADLRAAASR | ASSLARGNSDYT | 0.28584808 |
| QFVLDVAEHPVEVLTETV | ASSLARGNSDYT | 0.28661984 |
| GGAGGTGGAGGNGGTQAGDG | ASSLARGNSDYT | 0.28328508 |
| PLLATLRAIAVDVDAKPAQV | ASSLARGNSDYT | 0.28882897 |
| APKFRTGGYMIDFWGVGYQV | ASSLARGNSDYT | 0.28768787 |
| GAGGAGGAGGAAGGPGGTGG | ASSLARGNSDYT | 0.2874288 |
| IGALGLLATDVAPAGSVAFG | ASSLARGNSDYT | 0.29217988 |
| AGGAGGIGGNGGAGGAGGVL | ASSLARGNSDYT | 0.28594586 |
| ADELIGLL | ASSLARGNSDYT | 0.28572112 |
| ADRALKA | ASSLARGNSDYT | 0.28631145 |
| AGAVGQVGGAGGA | ASSLARGNSDYT | 0.28738308 |
| AGGGGGLGGAP | ASSLARGNSDYT | 0.28592545 |
| AGGLLYGNGGAG | ASSLARGNSDYT | 0.29228356 |
| AGLESIGL | ASSLARGNSDYT | 0.28474343 |
| AGLKTGII | ASSLARGNSDYT | 0.2956404 |
| ASLAALLL | ASSLARGNSDYT | 0.28733194 |
| ASVINISSI | ASSLARGNSDYT | 0.28827012 |
| ATPPGPA | ASSLARGNSDYT | 0.28651834 |
| ATVASALV | ASSLARGNSDYT | 0.29104885 |
| ATVASLIL | ASSLARGNSDYT | 0.290388 |
| AVNGPVLIRH | ASSLARGNSDYT | 0.2934488 |
| DGAAGAVGI | ASSLARGNSDYT | 0.28766203 |
| DPLLHGI | ASSLARGNSDYT | 0.2865127 |
| ELGAALGAL | ASSLARGNSDYT | 0.28671587 |
| ELGRTQAV | ASSLARGNSDYT | 0.28607166 |
| FVAGGYLS | ASSLARGNSDYT | 0.2855183 |
| GASGGAGGAGGW | ASSLARGNSDYT | 0.2865431 |
| GGAGAGGDGGAGGI | ASSLARGNSDYT | 0.2879535 |
| GTGHNGSQPGMGGNG | ASSLARGNSDYT | 0.29029682 |
| GYAAASASAS | ASSLARGNSDYT | 0.28544158 |
| HQVDVAAVQQG | ASSLARGNSDYT | 0.30191058 |
| IGAAIAHT | ASSLARGNSDYT | 0.2858142 |
| IGMIEVAQL | ASSLARGNSDYT | 0.28583127 |
| IRLAPEHTSAMD | ASSLARGNSDYT | 0.289468 |
| IYNVSVL | ASSLARGNSDYT | 0.28538123 |
| KQGFLGM | ASSLARGNSDYT | 0.28455696 |
| LESVITQL | ASSLARGNSDYT | 0.28536642 |
| LGLGALVVV | ASSLARGNSDYT | 0.28771058 |
| LIGNGGNGGSGGI | ASSLARGNSDYT | 0.2934272 |
| LNAINAPTL | ASSLARGNSDYT | 0.28581128 |
| LQLLNTGV | ASSLARGNSDYT | 0.28952217 |
| LQTSIEF | ASSLARGNSDYT | 0.25966582 |
| MSFAVSEL | ASSLARGNSDYT | 0.360862 |
| NTGALGL | ASSLARGNSDYT | 0.2081496 |
| PPGAGGAGGAGTT | ASSLARGNSDYT | 0.21678941 |

| | | |
|---|---|---|
| QLVGLGI | ASSLARGNSDYT | 0.23338862 |
| RFLEQVHQL | ASSLARGNSDYT | 0.26566413 |
| RGVGYVLR | ASSLARGNSDYT | 0.24491194 |
| SATGVFAGVI | ASSLARGNSDYT | 0.2500484 |
| SLPAITLP | ASSLARGNSDYT | 0.22556467 |
| STALAFTV | ASSLARGNSDYT | 0.21421437 |
| SVLTVGLGK | ASSLARGNSDYT | 0.21289186 |
| TALAGAGSGI | ASSLARGNSDYT | 0.22797091 |
| TGLPFTGI | ASSLARGNSDYT | 0.24670753 |
| VAEHVTV | ASSLARGNSDYT | 0.21241719 |
| VDIVLNSL | ASSLARGNSDYT | 0.20870756 |
| VLTGGGAVL | ASSLARGNSDYT | 0.22763783 |
| VNVGLAYGI | ASSLARGNSDYT | 0.22746864 |
| VSSAAVGAV | ASSLARGNSDYT | 0.29146582 |
| VTSVYEAGV | ASSLARGNSDYT | 0.26605275 |
| WVALDRGA | ASSLARGNSDYT | 0.2744922 |
| YLVGSVVI | ASSLARGNSDYT | 0.26694614 |
| GPPGTGKTLL | ASSLARGNSDYT | 0.23048548 |
| GVGMTKF | ASSLARGNSDYT | 0.199825 |
| IGIGGSDL | ASSLARGNSDYT | 0.21898934 |
| IGTLLAEL | ASSLARGNSDYT | 0.21120107 |
| IYTSGTT | ASSLARGNSDYT | 0.23747678 |
| LSFPAGV | ASSLARGNSDYT | 0.19634804 |
| LYTSGTT | ASSLARGNSDYT | 0.22678585 |
| VGAFPFGF | ASSLARGNSDYT | 0.23773496 |
| VGAGTVEF | ASSLARGNSDYT | 0.20451686 |
| VGKSTLLL | ASSLARGNSDYT | 0.2973421 |
| AALAHAGGGIV | ASSLARGNSDYT | 0.21135512 |
| AALVAVSLL | ASSLARGNSDYT | 0.21645069 |
| AAVSRAVAAL | ASSLARGNSDYT | 0.21349166 |
| AGGSSGIGGF | ASSLARGNSDYT | 0.27293095 |
| AGLTVANANG | ASSLARGNSDYT | 0.20807919 |
| AGLVGAVL | ASSLARGNSDYT | 0.25865877 |
| ALSQAFTV | ASSLARGNSDYT | 0.21732716 |
| ALTFSVSL | ASSLARGNSDYT | 0.25181705 |
| ALTGAALVAL | ASSLARGNSDYT | 0.21341372 |
| ASFSIGAGVL | ASSLARGNSDYT | 0.24369147 |
| DGSVYKI | ASSLARGNSDYT | 0.22146928 |
| EQLAEIV | ASSLARGNSDYT | 0.46820474 |
| FAVSELTVM | ASSLARGNSDYT | 0.24055289 |
| GAAPLAAGL | ASSLARGNSDYT | 0.25234753 |
| GADGGDGGSGGAGG | ASSLARGNSDYT | 0.22708094 |
| GFGLDTGGAGG | ASSLARGNSDYT | 0.20780793 |
| GLASGGLL | ASSLARGNSDYT | 0.2032982 |
| GRIAGAAVLL | ASSLARGNSDYT | 0.2365795 |
| GSAGSLNTSF | ASSLARGNSDYT | 0.21169093 |
| GTGQIGI | ASSLARGNSDYT | 0.23263298 |
| GTLIALIL | ASSLARGNSDYT | 0.24824616 |
| IAATMGFVVG | ASSLARGNSDYT | 0.2053833 |
| IFAQALL | ASSLARGNSDYT | 0.24947216 |
| IGAAVALGF | ASSLARGNSDYT | 0.20493627 |

| | | |
|---|---|---|
| IGLTPLAL | ASSLARGNSDYT | 0.22268786 |
| ILGSNTSTL | ASSLARGNSDYT | 0.2570778 |
| IVFSLTVL | ASSLARGNSDYT | 0.2040321 |
| LASKATGF | ASSLARGNSDYT | 0.27925497 |
| LASVALIL | ASSLARGNSDYT | 0.20306614 |
| LFAVGLTL | ASSLARGNSDYT | 0.22909105 |
| LFNGGIGTYI | ASSLARGNSDYT | 0.2771275 |
| LTLGAGII | ASSLARGNSDYT | 0.19802056 |
| LVAAALGDI | ASSLARGNSDYT | 0.20539212 |
| NTAFVQLGI | ASSLARGNSDYT | 0.2850615 |
| QLFSGAEL | ASSLARGNSDYT | 0.6984951 |
| SAATFITL | ASSLARGNSDYT | 0.7073464 |
| SNYGLANLGV | ASSLARGNSDYT | 0.19449154 |
| SYGGAYAVM | ASSLARGNSDYT | 0.19607157 |
| TTGVGGLTLG | ASSLARGNSDYT | 0.5178281 |
| TTQTIYI | ASSLARGNSDYT | 0.6621803 |
| VAAGAGLGAVI | ASSLARGNSDYT | 0.41027737 |
| VAAVVTLL | ASSLARGNSDYT | 0.68550086 |
| VGFLGGLI | ASSLARGNSDYT | 0.1870303 |
| VGVIPILL | ASSLARGNSDYT | 0.31709555 |
| VIFASSLI | ASSLARGNSDYT | 0.27526623 |
| VKGGDGGAAAG | ASSLARGNSDYT | 0.373844 |
| VTKTFTNI | ASSLARGNSDYT | 0.47614086 |
| VVAAAILL | ASSLARGNSDYT | 0.1606183 |
| YAALDVEL | ASSLARGNSDYT | 0.15846132 |
| LASVALLL | ASSLARGNSDYT | 0.28356463 |
| LDAVHRV | ASSLARGNSDYT | 0.53297347 |
| QGLSSSIF | ASSLARGNSDYT | 0.7153716 |
| RGTFANIRL | ASSLARGNSDYT | 0.70374656 |
| RQLWWGHRI | ASSLARGNSDYT | 0.7479937 |
| YGPPGTGKTLL | ASSLARGNSDYT | 0.72904265 |
| AALVGAPANVA | ASSLARGNSDYT | 0.57780445 |
| AATFRRTE | ASSLARGNSDYT | 0.10331896 |
| AGALGGGI | ASSLARGNSDYT | 0.19704148 |
| AGGAGGAGGIGAG | ASSLARGNSDYT | 0.16371217 |
| AGSGALGL | ASSLARGNSDYT | 0.50440323 |
| AQIADQI | ASSLARGNSDYT | 0.11241645 |
| ASAVAVTYL | ASSLARGNSDYT | 0.36742854 |
| ASTVLGGTGG | ASSLARGNSDYT | 0.3519835 |
| AVGGEPAK | ASSLARGNSDYT | 0.13914472 |
| CEQGTFGGTC | ASSLARGNSDYT | 0.71308506 |
| DEGKAPAG | ASSLARGNSDYT | 0.18002628 |
| DTVVFPAL | ASSLARGNSDYT | 0.17106721 |
| FEGITMAG | ASSLARGNSDYT | 0.22389399 |
| FPLVVQD | ASSLARGNSDYT | 0.75389624 |
| GAFTTPQ | ASSLARGNSDYT | 0.18112536 |
| GFGGFGTAGGVGGA | ASSLARGNSDYT | 0.71048164 |
| GFLAQFL | ASSLARGNSDYT | 0.48835897 |
| GGAGGAGGNAGVA | ASSLARGNSDYT | 0.6266811 |
| GGVGTAGAGGAGGAGG | ASSLARGNSDYT | 0.14450352 |
| GNGGGGDAGNAGSGGN | ASSLARGNSDYT | 0.68028104 |

| | | |
|---|---|---|
| GTGGAGGAGNDAGS | ASSLARGNSDYT | 0.4604581 |
| GTTNTGWFNS | ASSLARGNSDYT | 0.78750306 |
| IFMGPLFTL | ASSLARGNSDYT | 0.40265876 |
| ILAHAIEAI | ASSLARGNSDYT | 0.40118507 |
| IRSAIREAL | ASSLARGNSDYT | 0.20875838 |
| ITFAGGG | ASSLARGNSDYT | 0.19149548 |
| ITGMSGHQEV | ASSLARGNSDYT | 0.124949984 |
| IVAGLACTL | ASSLARGNSDYT | 0.41891706 |
| IVGGGLAGA | ASSLARGNSDYT | 0.18722671 |
| IVTGVLNI | ASSLARGNSDYT | 0.3287047 |
| LAQQGCI | ASSLARGNSDYT | 0.67124844 |
| LCLHVEN | ASSLARGNSDYT | 0.10434757 |
| LDAISDGS | ASSLARGNSDYT | 0.64308214 |
| LGGFAAML | ASSLARGNSDYT | 0.16947909 |
| LGGKNPMI | ASSLARGNSDYT | 0.17709036 |
| LGPKGVAAIL | ASSLARGNSDYT | 0.51370466 |
| LMVGVSI | ASSLARGNSDYT | 0.13737835 |
| LVGAAFVL | ASSLARGNSDYT | 0.7359224 |
| LVLAVAPL | ASSLARGNSDYT | 0.114088066 |
| MAAIAVIDAA | ASSLARGNSDYT | 0.6002546 |
| NNGINAELQ | ASSLARGNSDYT | 0.48793584 |
| RGEPLVLVHG | ASSLARGNSDYT | 0.089660786 |
| RGGAGGAAGLI | ASSLARGNSDYT | 0.15230067 |
| RSYRMHPAV | ASSLARGNSDYT | 0.74492705 |
| SAAHLLAGQLA | ASSLARGNSDYT | 0.098434664 |
| SAAVGSY | ASSLARGNSDYT | 0.09319131 |
| SPHAVLHRI | ASSLARGNSDYT | 0.08740299 |
| SQIAKVL | ASSLARGNSDYT | 0.09457206 |
| TALGHLVP | ASSLARGNSDYT | 0.088889904 |
| TAPTTSGAA | ASSLARGNSDYT | 0.097569056 |
| TDGDDGNGGSPGT | ASSLARGNSDYT | 0.099893205 |
| TDGLGEN | ASSLARGNSDYT | 0.10525514 |
| TGGPPLPGQA | ASSLARGNSDYT | 0.10172528 |
| TIPAITI | ASSLARGNSDYT | 0.096500374 |
| TIPGFTI | ASSLARGNSDYT | 0.09264553 |
| TSIASGALA | ASSLARGNSDYT | 0.09881852 |
| TVTLPGAAI | ASSLARGNSDYT | 0.093646675 |
| VASGLSAALP | ASSLARGNSDYT | 0.08999593 |
| VGAGVIGI | ASSLARGNSDYT | 0.086665414 |
| VGPKRFGV | ASSLARGNSDYT | 0.08819676 |
| VNVAFPD | ASSLARGNSDYT | 0.09385758 |
| VSMTNSGVS | ASSLARGNSDYT | 0.10161339 |
| VTDVLGDLSV | ASSLARGNSDYT | 0.09818582 |
| VVVVGAGFAGL | ASSLARGNSDYT | 0.10868882 |
| VYTVGTVV | ASSLARGNSDYT | 0.100361645 |
| WTTFSGP | ASSLARGNSDYT | 0.09034267 |
| YDELVEAF | ASSLARGNSDYT | 0.086035214 |
| YGTGGAGGNGG | ASSLARGNSDYT | 0.08790776 |
| YTNVMAV | ASSLARGNSDYT | 0.09058775 |
| FLEQVHQL | ASSLARGNSDYT | 0.09442581 |
| GLSLLAVAV | ASSLARGNSDYT | 0.08342212 |

| | | |
|---|---|---|
| LEQVHQL | ASSLARGNSDYT | 0.093920276 |
| LNAAGAI | ASSLARGNSDYT | 0.09640878 |
| PAVHASL | ASSLARGNSDYT | 0.09224815 |
| TELEDDA | ASSLARGNSDYT | 0.105437 |
| TPPGPPN | ASSLARGNSDYT | 0.08892627 |
| TPPPPPPP | ASSLARGNSDYT | 0.08890128 |
| TQSPALI | ASSDGTGWNTLY | 0.09325921 |
| EFAAAFPQ | ASSDGTGWNTLY | 0.103606746 |
| LLSGNLDVL | ASSDGTGWNTLY | 0.08897972 |
| GPANLGNYNI | ASSDGTGWNTLY | 0.10945842 |
| TLGPTLP | ASSDGTGWNTLY | 0.08870882 |
| VYAYSSCCK | ASSDGTGWNTLY | 0.11138498 |
| APAPAPAPNI | ASSDGTGWNTLY | 0.09541096 |
| PGGPVPIP | ASSDGTGWNTLY | 0.101080514 |
| FLTVALEP | ASSDGTGWNTLY | 0.09280402 |
| SQYFGANL | ASSDGTGWNTLY | 0.109305985 |
| GGVKPAA | ASSDGTGWNTLY | 0.09692706 |
| GGTGGAGGAGGW | ASSDGTGWNTLY | 0.10775339 |
| GVGGVAGLGGAGG | ASSDGTGWNTLY | 0.08964827 |
| VLLVLATTI | ASSDGTGWNTLY | 0.09113029 |
| IGIFAAML | ASSDGTGWNTLY | 0.088075586 |
| AVGDLID | ASSDGTGWNTLY | 0.09184424 |
| NISPLPL | ASSDGTGWNTLY | 0.09197643 |
| VTAVFVAI | ASSDGTGWNTLY | 0.09409986 |
| SGDDNDAAL | ASSDGTGWNTLY | 0.1076579 |
| AVAALAHS | ASSDGTGWNTLY | 0.0846262 |
| GGVTVMATP | ASSDGTGWNTLY | 0.09419365 |
| AGLAANVRD | ASSDGTGWNTLY | 0.085370734 |
| TNRPDLIDPALLRPG | ASSDGTGWNTLY | 0.09127324 |
| LANLGVTAL | ASSDGTGWNTLY | 0.09348778 |
| TGERIEL | ASSDGTGWNTLY | 0.090213746 |
| NGEELAA | ASSDGTGWNTLY | 0.08764315 |
| VLSLVWG | ASSDGTGWNTLY | 0.08529923 |
| IGYALWL | ASSDGTGWNTLY | 0.09149445 |
| LSLLLAPL | ASSDGTGWNTLY | 0.11541373 |
| TLDEALK | ASSDGTGWNTLY | 0.08600291 |
| TDGAGTAEGG | ASSDGTGWNTLY | 0.088185124 |
| NAPGGAGGNGG | ASSDGTGWNTLY | 0.10604029 |
| VSVFMGMGGNF | ASSDGTGWNTLY | 0.21745215 |
| LASYLAAL | ASSDGTGWNTLY | 0.32281548 |
| LVGGTPAL | ASSDGTGWNTLY | 0.08895821 |
| VRNADVL | ASSDGTGWNTLY | 0.10775094 |
| TCSSPFLVSAM | ASSDGTGWNTLY | 0.14632092 |
| VHITDAE | ASSDGTGWNTLY | 0.2152614 |
| EYAAPSF | ASSDGTGWNTLY | 0.17131618 |
| VVDAFHGGL | ASSDGTGWNTLY | 0.23980379 |
| ASVVDPQ | ASSDGTGWNTLY | 0.11091534 |
| SYNEFGGLS | ASSDGTGWNTLY | 0.11240429 |
| DPDGSVY | ASSDGTGWNTLY | 0.10482277 |
| MVSTIHATE | ASSDGTGWNTLY | 0.13826683 |
| LAAGVADPV | ASSDGTGWNTLY | 0.15109923 |

| | | |
|---|---|---|
| TKEQIAA | ASSDGTGWNTLY | 0.096404135 |
| SLASPAAL | ASSDGTGWNTLY | 0.084428415 |
| IGLVFYGV | ASSDGTGWNTLY | 0.09882845 |
| FLLDASNL | ASSDGTGWNTLY | 0.19481292 |
| YQVGNFP | ASSDGTGWNTLY | 0.42762262 |
| ESLLSQST | ASSDGTGWNTLY | 0.40963218 |
| LFTVGFAAVG | ASSDGTGWNTLY | 0.2519307 |
| ATPPTPT | ASSDGTGWNTLY | 0.24682248 |
| TDLATLG | ASSDGTGWNTLY | 0.09820209 |
| SAGGTGR | ASSDGTGWNTLY | 0.07868137 |
| GFSNSGS | ASSDGTGWNTLY | 0.08675622 |
| GGNGGSGGTSV | ASSDGTGWNTLY | 0.09598992 |
| GTSVATGGAGN | ASSDGTGWNTLY | 0.13501194 |
| NGGNGGSGGTS | ASSDGTGWNTLY | 0.07401819 |
| AVVKFPH | ASSDGTGWNTLY | 0.13955043 |
| TPPPPPP | ASSDGTGWNTLY | 0.12702693 |
| AAPAAAAAPA | ASSDGTGWNTLY | 0.08663405 |
| IALNEDL | ASSDGTGWNTLY | 0.35762924 |
| IVNVSSM | ASSDGTGWNTLY | 0.088851124 |
| TGIPPAPR | ASSDGTGWNTLY | 0.092675135 |
| FDLGGGTF | ASSDGTGWNTLY | 0.10676422 |
| FDLGGGTFDVS | ASSDGTGWNTLY | 0.25688076 |
| NEPTAAA | ASSDGTGWNTLY | 0.101277635 |
| NEPTAAALA | ASSDGTGWNTLY | 0.2724836 |
| GSIINISSI | ASSDGTGWNTLY | 0.10605761 |
| VAPGNAGTA | ASSDGTGWNTLY | 0.20684284 |
| TGGGASLE | ASSDGTGWNTLY | 0.099186465 |
| VAVYNHYKRI | ASSDGTGWNTLY | 0.19008146 |
| TDPAPATT | ASSDGTGWNTLY | 0.13810632 |
| TNRPDILDPALLRPG | ASSDGTGWNTLY | 0.67134273 |
| LTTTTTS | ASSDGTGWNTLY | 0.13681063 |
| PTGSGKTL | ASSDGTGWNTLY | 0.15867884 |
| TGSGKTL | ASSDGTGWNTLY | 0.10641371 |
| TIGHVDHGK | ASSDGTGWNTLY | 0.09561555 |
| TIGHVDHGKTT | ASSDGTGWNTLY | 0.08381426 |
| GHVDHGKTT | ASSDGTGWNTLY | 0.12173483 |
| AQFEHTLL | ASSDGTGWNTLY | 0.08790661 |
| IVNQDEL | ASSDGTGWNTLY | 0.119095534 |
| TNGSQFF | ASSDGTGWNTLY | 0.21375895 |
| TGKGIVE | ASSDGTGWNTLY | 0.08230339 |
| LISNASDA | ASSDGTGWNTLY | 0.13789949 |
| AELRTLGELTKIAKAHGAQ | ASSDGTGWNTLY | 0.08298928 |
| APNAVAAAEPSW | ASSDGTGWNTLY | 0.100966565 |
| APQVAGAATAVL | ASSDGTGWNTLY | 0.17085102 |
| ARRDPAAAAAILPTDA | ASSDGTGWNTLY | 0.08981809 |
| DAAARDVLAEAGLAEYF | ASSDGTGWNTLY | 0.14689696 |
| DDQIVLSAKANLAKVEQ | ASSDGTGWNTLY | 0.082209006 |
| DELLPDTGEDSDEDGH | ASSDGTGWNTLY | 0.11853793 |
| ELPEAPANRATVPQD | ASSDGTGWNTLY | 0.18992108 |
| FGQSKIERVTGIG | ASSDGTGWNTLY | 0.07600553 |
| GAGGTGSTAGGAGGAGGAGGI | ASSDGTGWNTLY | 0.089916624 |

| | | |
|---|---|---|
| GHGGTGAAGGNGGTGGAGGS( | ASSDGTGWNTLY | 0.22925028 |
| GNGGNGGAGGEGGPGGAGGA( | ASSDGTGWNTLY | 0.8086355 |
| GSTAGNGGNGGNATNGTIAG( | ASSDGTGWNTLY | 0.82198346 |
| GTGGDGGNGGNGGTGG | ASSDGTGWNTLY | 0.7419383 |
| GVRRPAGIPADAPV | ASSDGTGWNTLY | 0.6685958 |
| GYRQVTDAHLLTLARR | ASSDGTGWNTLY | 0.74287164 |
| IAKMGQDGHDRG | ASSDGTGWNTLY | 0.7983905 |
| KAEPSIEIQYT | ASSDGTGWNTLY | 0.8243041 |
| KPVRIRMAADEDTVAVT | ASSDGTGWNTLY | 0.81087077 |
| MPPRIAGMRLLVIKPEP | ASSDGTGWNTLY | 0.46981388 |
| MKFYAVALFRSRGVPPTR | ASSDGTGWNTLY | 0.80692077 |
| NAGGFGILGAVAHSP | ASSDGTGWNTLY | 0.7410727 |
| NEGREITLAMAEM | ASSDGTGWNTLY | 0.76314676 |
| NELVEWSHPLGHRWRWE | ASSDGTGWNTLY | 0.7622167 |
| NLGAGGTGGNGGNPG | ASSDGTGWNTLY | 0.6964826 |
| NTEWPKWETVMAADPVD | ASSDGTGWNTLY | 0.7682648 |
| PASAAPAETQP | ASSDGTGWNTLY | 0.61994195 |
| QQSSDGKNGVRAAEITGE | ASSDGTGWNTLY | 0.76104486 |
| RAPDAVTTSWWKE | ASSDGTGWNTLY | 0.80602753 |
| SGFGNVGAHTSGWFNQ | ASSDGTGWNTLY | 0.7931895 |
| SGVAGAVGQAASVGGLKV | ASSDGTGWNTLY | 0.80654144 |
| SNGGAGTGSAGGNGGT | ASSDGTGWNTLY | 0.81597245 |
| TGGAGGEGGAAG | ASSDGTGWNTLY | 0.78767943 |
| TGGDGGAAGAGGAPGHG | ASSDGTGWNTLY | 0.21099882 |
| TMPPLLAAIANAGRHGV | ASSDGTGWNTLY | 0.45906174 |
| TSSTAGNSGGA | ASSDGTGWNTLY | 0.7173236 |
| VPATVPVGILC | ASSDGTGWNTLY | 0.7243123 |
| VQRPDSGGVVLTGRIS | ASSDGTGWNTLY | 0.22706547 |
| WTPNPVPATKGVPAGLVQAT | ASSDGTGWNTLY | 0.7677917 |
| YALKRALAKFSDRFDVVII | ASSDGTGWNTLY | 0.72331643 |
| YLIGSVIV | ASSDGTGWNTLY | 0.75123405 |
| YGQLVAVID | ASSDGTGWNTLY | 0.81379235 |
| VTVGGMTLPA | ASSDGTGWNTLY | 0.77952254 |
| VLALPAV | ASSDGTGWNTLY | 0.5831201 |
| VIPPTLNYET | ASSDGTGWNTLY | 0.5654725 |
| VGMGGMHPSSG | ASSDGTGWNTLY | 0.8224231 |
| VGLAGGAATA | ASSDGTGWNTLY | 0.46339697 |
| VGGAGGAGGSGP | ASSDGTGWNTLY | 0.8150313 |
| VFAAFMV | ASSDGTGWNTLY | 0.6696775 |
| TLYDKLR | ASSDGTGWNTLY | 0.80660236 |
| TLAVFTVL | ASSDGTGWNTLY | 0.40574878 |
| TGSVQPARV | ASSDGTGWNTLY | 0.8066 |
| TGGGAGSGGT | ASSDGTGWNTLY | 0.780629 |
| SHLETSL | ASSDGTGWNTLY | 0.82984 |
| SFTTGMT | ASSDGTGWNTLY | 0.7006986 |
| QVEVAKRMGVRQPTVS | ASSDGTGWNTLY | 0.739423 |
| QGSGGPALAM | ASSDGTGWNTLY | 0.5616439 |
| MLAPSNP | ASSDGTGWNTLY | 0.8239558 |
| LVNNAAI | ASSDGTGWNTLY | 0.5824496 |
| LTGEINTLA | ASSDGTGWNTLY | 0.7055697 |
| LSIFGPV | ASSDGTGWNTLY | 0.58472973 |

| | | |
|---|---|---|
| LLALLANL | ASSDGTGWNTLY | 0.7606272 |
| LISNASDALDKLR | ASSDGTGWNTLY | 0.812281 |
| LINEAVP | ASSDGTGWNTLY | 0.37230018 |
| LGYAFSGL | ASSDGTGWNTLY | 0.8145087 |
| LDFVAVAL | ASSDGTGWNTLY | 0.71253574 |
| LAQPGAE | ASSDGTGWNTLY | 0.5351728 |
| ISNLGAVTGF | ASSDGTGWNTLY | 0.8048783 |
| HQEDPGRI | ASSDGTGWNTLY | 0.73207086 |
| GTPGEPTP | ASSDGTGWNTLY | 0.8083751 |
| GFFNSGS | ASSDGTGWNTLY | 0.64000165 |
| GAGGGGLTAGAN | ASSDGTGWNTLY | 0.79639125 |
| FVDALVNM | ASSDGTGWNTLY | 0.7741983 |
| FLTVAPEL | ASSDGTGWNTLY | 0.66009605 |
| FETLTAL | ASSDGTGWNTLY | 0.589785 |
| DVINPIEL | ASSDGTGWNTLY | 0.8223142 |
| APVSGAPV | ASSDGTGWNTLY | 0.8053011 |
| ALVGGLAW | ASSDGTGWNTLY | 0.801969 |
| ALAAAAATAI | ASSDGTGWNTLY | 0.7331611 |
| AFGLGLLL | ASSDGTGWNTLY | 0.73486435 |
| AFFAAASL | ASSDGTGWNTLY | 0.76859796 |
| AAAALPAFSPP | ASSDGTGWNTLY | 0.7949637 |
| VVLAPAL | ASSDGTGWNTLY | 0.80721486 |
| TIGHVDHGKTTL | ASSDGTGWNTLY | 0.80462563 |
| LISNASDALDK | ASSDGTGWNTLY | 0.6959962 |
| LISNASDAL | ASSDGTGWNTLY | 0.7910963 |
| ISNASDALDK | ASSDGTGWNTLY | 0.75084925 |
| NQAETLVYQTE | ASSDGTGWNTLY | 0.7652377 |
| TGGTGGTAGAG | ASSDGTGWNTLY | 0.78829104 |
| TGSTAGGAGGA | ASSDGTGWNTLY | 0.7520722 |
| DPLAVPVAAGPV | ASSDGTGWNTLY | 0.75299555 |
| GGTGGTPGNGGN | ASSDGTGWNTLY | 0.7470161 |
| PAGLAGVDPATL | ASSDGTGWNTLY | 0.788496 |
| GNNLVVEAPHAGASV | ASSDGTGWNTLY | 0.80566 |
| KVTRIMSASTLEQPA | ASSDGTGWNTLY | 0.8007414 |
| LGFALPIALPRRTVV | ASSDGTGWNTLY | 0.8096071 |
| TRLGNLTARKLVVLV | ASSDGTGWNTLY | 0.8032217 |
| WSQQRRLIYFPSAGP | ASSDGTGWNTLY | 0.80406046 |
| GPNGNPDPMAAAVDIR | ASSDGTGWNTLY | 0.3397009 |
| TQRCVSVADLRAAASR | ASSDGTGWNTLY | 0.6309471 |
| QFVLDVAEHPVEVLTETV | ASSDGTGWNTLY | 0.7615359 |
| GGAGGTGGAGGNGGTQAGDG | ASSDGTGWNTLY | 0.7611981 |
| PLLATLRAIAVDVDAKPAQV | ASSDGTGWNTLY | 0.39957473 |
| APKFRTGGYMIDFWGVGYQV | ASSDGTGWNTLY | 0.7839274 |
| GAGGAGGAGGAAGGPGGTGG | ASSDGTGWNTLY | 0.78398985 |
| IGALGLLATDVAPAGSVAFG | ASSDGTGWNTLY | 0.72747993 |
| AGGAGGIGGNGGAGGAGGVL | ASSDGTGWNTLY | 0.8023583 |
| ADELIGLL | ASSDGTGWNTLY | 0.7612017 |
| ADRALKA | ASSDGTGWNTLY | 0.69515014 |
| AGAVGQVGGAGGA | ASSDGTGWNTLY | 0.71608174 |
| AGGGGGLGGAP | ASSDGTGWNTLY | 0.8017403 |
| AGGLLYGNGGAG | ASSDGTGWNTLY | 0.68132144 |

| | | |
|---|---|---|
| AGLESIGL | ASSDGTGWNTLY | 0.8045639 |
| AGLKTGII | ASSDGTGWNTLY | 0.6350993 |
| ASLAALLL | ASSDGTGWNTLY | 0.8057349 |
| ASVINISSI | ASSDGTGWNTLY | 0.68049526 |
| ATPPGPA | ASSDGTGWNTLY | 0.803643 |
| ATVASALV | ASSDGTGWNTLY | 0.77591944 |
| ATVASLIL | ASSDGTGWNTLY | 0.8151387 |
| AVNGPVLIRH | ASSDGTGWNTLY | 0.75873077 |
| DGAAGAVGI | ASSDGTGWNTLY | 0.78582966 |
| DPLLHGI | ASSDGTGWNTLY | 0.7410626 |
| ELGAALGAL | ASSDGTGWNTLY | 0.80805045 |
| ELGRTQAV | ASSDGTGWNTLY | 0.64417464 |
| FVAGGYLS | ASSDGTGWNTLY | 0.77703285 |
| GASGGAGGAGGW | ASSDGTGWNTLY | 0.5235721 |
| GGAGAGGDGGAGGI | ASSDGTGWNTLY | 0.77812034 |
| GTGHNGSQPGMGGNG | ASSDGTGWNTLY | 0.80618775 |
| GYAAASASAS | ASSDGTGWNTLY | 0.4739198 |
| HQVDVAAVQQG | ASSDGTGWNTLY | 0.8041577 |
| IGAAIAHT | ASSDGTGWNTLY | 0.7043688 |
| IGMIEVAQL | ASSDGTGWNTLY | 0.7514723 |
| IRLAPEHTSAMD | ASSDGTGWNTLY | 0.8037988 |
| IYNVSVL | ASSDGTGWNTLY | 0.76518965 |
| KQGFLGM | ASSDGTGWNTLY | 0.80794775 |
| LESVITQL | ASSDGTGWNTLY | 0.73697174 |
| LGLGALVVV | ASSDGTGWNTLY | 0.76497674 |
| LIGNGGNGGSGGI | ASSDGTGWNTLY | 0.7771849 |
| LNAINAPTL | ASSDGTGWNTLY | 0.65501857 |
| LQLLNTGV | ASSDGTGWNTLY | 0.7073167 |
| LQTSIEF | ASSDGTGWNTLY | 0.81174135 |
| MSFAVSEL | ASSDGTGWNTLY | 0.5073705 |
| NTGALGL | ASSDGTGWNTLY | 0.5091988 |
| PPGAGGAGGAGTT | ASSDGTGWNTLY | 0.168221 |
| QLVGLGI | ASSDGTGWNTLY | 0.18200684 |
| RFLEQVHQL | ASSDGTGWNTLY | 0.20163268 |
| RGVGYVLR | ASSDGTGWNTLY | 0.29586905 |
| SATGVFAGVI | ASSDGTGWNTLY | 0.7223499 |
| SLPAITLP | ASSDGTGWNTLY | 0.48858804 |
| STALAFTV | ASSDGTGWNTLY | 0.18813346 |
| SVLTVGLGK | ASSDGTGWNTLY | 0.32585785 |
| TALAGAGSGI | ASSDGTGWNTLY | 0.23951837 |
| TGLPFTGI | ASSDGTGWNTLY | 0.2984853 |
| VAEHVTV | ASSDGTGWNTLY | 0.2230251 |
| VDIVLNSL | ASSDGTGWNTLY | 0.20433617 |
| VLTGGGAVL | ASSDGTGWNTLY | 0.17419292 |
| VNVGLAYGI | ASSDGTGWNTLY | 0.16067332 |
| VSSAAVGAV | ASSDGTGWNTLY | 0.32349956 |
| VTSVYEAGV | ASSDGTGWNTLY | 0.4448997 |
| WVALDRGA | ASSDGTGWNTLY | 0.31877816 |
| YLVGSVVI | ASSDGTGWNTLY | 0.39168245 |
| GPPGTGKTLL | ASSDGTGWNTLY | 0.5296291 |
| GVGMTKF | ASSDGTGWNTLY | 0.2424323 |

| | | |
|---|---|---|
| IGIGGSDL | ASSDGTGWNTLY | 0.14670485 |
| IGTLLAEL | ASSDGTGWNTLY | 0.15084231 |
| IYTSGTT | ASSDGTGWNTLY | 0.183447 |
| LSFPAGV | ASSDGTGWNTLY | 0.18058285 |
| LYTSGTT | ASSDGTGWNTLY | 0.14426693 |
| VGAFPFGF | ASSDGTGWNTLY | 0.25130075 |
| VGAGTVEF | ASSDGTGWNTLY | 0.19679311 |
| VGKSTLLL | ASSDGTGWNTLY | 0.18526152 |
| AALAHAGGGIV | ASSDGTGWNTLY | 0.5001341 |
| AALVAVSLL | ASSDGTGWNTLY | 0.1841715 |
| AAVSRAVAAL | ASSDGTGWNTLY | 0.1629563 |
| AGGSSGIGGF | ASSDGTGWNTLY | 0.18816647 |
| AGLTVANANG | ASSDGTGWNTLY | 0.43062103 |
| AGLVGAVL | ASSDGTGWNTLY | 0.17915067 |
| ALSQAFTV | ASSDGTGWNTLY | 0.43048415 |
| ALTFSVSL | ASSDGTGWNTLY | 0.18500052 |
| ALTGAALVAL | ASSDGTGWNTLY | 0.48400992 |
| ASFSIGAGVL | ASSDGTGWNTLY | 0.16988775 |
| DGSVYKI | ASSDGTGWNTLY | 0.44603392 |
| EQLAEIV | ASSDGTGWNTLY | 0.30580282 |
| FAVSELTVM | ASSDGTGWNTLY | 0.7112589 |
| GAAPLAAGL | ASSDGTGWNTLY | 0.20198318 |
| GADGGDGGSGGAGG | ASSDGTGWNTLY | 0.30465326 |
| GFGLDTGGAGG | ASSDGTGWNTLY | 0.200452 |
| GLASGGLL | ASSDGTGWNTLY | 0.64427054 |
| GRIAGAAVLL | ASSDGTGWNTLY | 0.16252393 |
| GSAGSLNTSF | ASSDGTGWNTLY | 0.18259108 |
| GTGQIGI | ASSDGTGWNTLY | 0.16034743 |
| GTLIALIL | ASSDGTGWNTLY | 0.19072036 |
| IAATMGFVVG | ASSDGTGWNTLY | 0.5312884 |
| IFAQALL | ASSDGTGWNTLY | 0.1440071 |
| IGAAVALGF | ASSDGTGWNTLY | 0.23018585 |
| IGLTPLAL | ASSDGTGWNTLY | 0.16225488 |
| ILGSNTSTL | ASSDGTGWNTLY | 0.166374 |
| IVFSLTVL | ASSDGTGWNTLY | 0.2617087 |
| LASKATGF | ASSDGTGWNTLY | 0.24479362 |
| LASVALIL | ASSDGTGWNTLY | 0.23161763 |
| LFAVGLTL | ASSDGTGWNTLY | 0.16931662 |
| LFNGGIGTYI | ASSDGTGWNTLY | 0.22657435 |
| LTLGAGII | ASSDGTGWNTLY | 0.2417578 |
| LVAAALGDI | ASSDGTGWNTLY | 0.1507854 |
| NTAFVQLGI | ASSDGTGWNTLY | 0.17019558 |
| QLFSGAEL | ASSDGTGWNTLY | 0.50066566 |
| SAATFITL | ASSDGTGWNTLY | 0.75842273 |
| SNYGLANLGV | ASSDGTGWNTLY | 0.82425874 |
| SYGGAYAVM | ASSDGTGWNTLY | 0.36372763 |
| TTGVGGLTLG | ASSDGTGWNTLY | 0.3097506 |
| TTQTIYI | ASSDGTGWNTLY | 0.5526321 |
| VAAGAGLGAVI | ASSDGTGWNTLY | 0.78939444 |
| VAAVVTLL | ASSDGTGWNTLY | 0.8612047 |
| VGFLGGLI | ASSDGTGWNTLY | 0.79534996 |

| | | |
|---|---|---|
| VGVIPILL | ASSDGTGWNTLY | 0.3025236 |
| VIFASSLI | ASSDGTGWNTLY | 0.6706388 |
| VKGGDGGAAAG | ASSDGTGWNTLY | 0.46995524 |
| VTKTFTNI | ASSDGTGWNTLY | 0.62881637 |
| VVAAAILL | ASSDGTGWNTLY | 0.71914834 |
| YAALDVEL | ASSDGTGWNTLY | 0.665587 |
| LASVALLL | ASSDGTGWNTLY | 0.3995666 |
| LDAVHRV | ASSDGTGWNTLY | 0.33555868 |
| QGLSSSIF | ASSDGTGWNTLY | 0.7635522 |
| RGTFANIRL | ASSDGTGWNTLY | 0.8165127 |
| RQLWWGHRI | ASSDGTGWNTLY | 0.7859191 |
| YGPPGTGKTLL | ASSDGTGWNTLY | 0.85097307 |
| AALVGAPANVA | ASSDGTGWNTLY | 0.7917741 |
| AATFRRTE | ASSDGTGWNTLY | 0.76921856 |
| AGALGGGI | ASSDGTGWNTLY | 0.12165257 |
| AGGAGGAGGIGAG | ASSDGTGWNTLY | 0.18255568 |
| AGSGALGL | ASSDGTGWNTLY | 0.4186835 |
| AQIADQI | ASSDGTGWNTLY | 0.56101865 |
| ASAVAVTYL | ASSDGTGWNTLY | 0.16225925 |
| ASTVLGGTGG | ASSDGTGWNTLY | 0.7822888 |
| AVGGEPAK | ASSDGTGWNTLY | 0.43434802 |
| CEQGTFGGTC | ASSDGTGWNTLY | 0.3339457 |
| DEGKAPAG | ASSDGTGWNTLY | 0.81255335 |
| DTVVFPAL | ASSDGTGWNTLY | 0.36800754 |
| FEGITMAG | ASSDGTGWNTLY | 0.29681495 |
| FPLVVQD | ASSDGTGWNTLY | 0.45244288 |
| GAFTTPQ | ASSDGTGWNTLY | 0.78172046 |
| GFGGFGTAGGVGGA | ASSDGTGWNTLY | 0.27535647 |
| GFLAQFL | ASSDGTGWNTLY | 0.76737297 |
| GGAGGAGGNAGVA | ASSDGTGWNTLY | 0.5614425 |
| GGVGTAGAGGAGGAGG | ASSDGTGWNTLY | 0.8026116 |
| GNGGGGDAGNAGSGGN | ASSDGTGWNTLY | 0.23601727 |
| GTGGAGGAGNDAGS | ASSDGTGWNTLY | 0.80140877 |
| GTTNTGWFNS | ASSDGTGWNTLY | 0.6464129 |
| IFMGPLFTL | ASSDGTGWNTLY | 0.8715776 |
| ILAHAIEAI | ASSDGTGWNTLY | 0.44712028 |
| IRSAIREAL | ASSDGTGWNTLY | 0.6183802 |
| ITFAGGG | ASSDGTGWNTLY | 0.5761305 |
| ITGMSGHQEV | ASSDGTGWNTLY | 0.83952236 |
| IVAGLACTL | ASSDGTGWNTLY | 0.22092211 |
| IVGGGLAGA | ASSDGTGWNTLY | 0.43176752 |
| IVTGVLNI | ASSDGTGWNTLY | 0.19312444 |
| LAQQGCI | ASSDGTGWNTLY | 0.50545174 |
| LCLHVEN | ASSDGTGWNTLY | 0.77320194 |
| LDAISDGS | ASSDGTGWNTLY | 0.14103538 |
| LGGFAAML | ASSDGTGWNTLY | 0.68723726 |
| LGGKNPMI | ASSDGTGWNTLY | 0.542789 |
| LGPKGVAAIL | ASSDGTGWNTLY | 0.33767393 |
| LMVGVSI | ASSDGTGWNTLY | 0.79137135 |
| LVGAAFVL | ASSDGTGWNTLY | 0.6805428 |
| LVLAVAPL | ASSDGTGWNTLY | 0.76990867 |

| | | |
|---|---|---|
| MAAIAVIDAA | ASSDGTGWNTLY | 0.577086 |
| NNGINAELQ | ASSDGTGWNTLY | 0.5267717 |
| RGEPLVLVHG | ASSDGTGWNTLY | 0.5983143 |
| RGGAGGAAGLI | ASSDGTGWNTLY | 0.22223383 |
| RSYRMHPAV | ASSDGTGWNTLY | 0.47529685 |
| SAAHLLAGQLA | ASSDGTGWNTLY | 0.82825756 |
| SAAVGSY | ASSDGTGWNTLY | 0.67944074 |
| SPHAVLHRI | ASSDGTGWNTLY | 0.73914427 |
| SQIAKVL | ASSDGTGWNTLY | 0.20931278 |
| TALGHLVP | ASSDGTGWNTLY | 0.23416436 |
| TAPTTSGAA | ASSDGTGWNTLY | 0.49493092 |
| TDGDDGNGGSPGT | ASSDGTGWNTLY | 0.6801435 |
| TDGLGEN | ASSDGTGWNTLY | 0.817569 |
| TGGPPLPGQA | ASSDGTGWNTLY | 0.7083862 |
| TIPAITI | ASSDGTGWNTLY | 0.26837212 |
| TIPGFTI | ASSDGTGWNTLY | 0.45688623 |
| TSIASGALA | ASSDGTGWNTLY | 0.37250108 |
| TVTLPGAAI | ASSDGTGWNTLY | 0.37662596 |
| VASGLSAALP | ASSDGTGWNTLY | 0.49639133 |
| VGAGVIGI | ASSDGTGWNTLY | 0.5006186 |
| VGPKRFGV | ASSDGTGWNTLY | 0.3629994 |
| VNVAFPD | ASSDGTGWNTLY | 0.25157872 |
| VSMTNSGVS | ASSDGTGWNTLY | 0.5001256 |
| VTDVLGDLSV | ASSDGTGWNTLY | 0.7469111 |
| VVVVGAGFAGL | ASSDGTGWNTLY | 0.6213665 |
| VYTVGTVV | ASSDGTGWNTLY | 0.78551894 |
| WTTFSGP | ASSDGTGWNTLY | 0.68244064 |
| YDELVEAF | ASSDGTGWNTLY | 0.46000925 |
| YGTGGAGGNGG | ASSDGTGWNTLY | 0.10880606 |
| YTNVMAV | ASSDGTGWNTLY | 0.15593769 |
| FLEQVHQL | ASSDGTGWNTLY | 0.22354867 |
| GLSLLAVAV | ASSDGTGWNTLY | 0.48164693 |
| LEQVHQL | ASSDGTGWNTLY | 0.11350567 |
| LNAAGAI | ASSDGTGWNTLY | 0.5617151 |
| PAVHASL | ASSDGTGWNTLY | 0.36293375 |
| TELEDDA | ASSDGTGWNTLY | 0.24191512 |
| TPPGPPN | ASSDGTGWNTLY | 0.7173116 |
| TPPPPPPP | ASSDGTGWNTLY | 0.30103123 |
| TQSPALI | ASSQEGQNNQAPL | 0.21601236 |
| EFAAAFPQ | ASSQEGQNNQAPL | 0.28881162 |
| LLSGNLDVL | ASSQEGQNNQAPL | 0.7880068 |
| GPANLGNYNI | ASSQEGQNNQAPL | 0.19752884 |
| TLGPTLP | ASSQEGQNNQAPL | 0.70201087 |
| VYAYSSCCK | ASSQEGQNNQAPL | 0.4471822 |
| APAPAPAPNI | ASSQEGQNNQAPL | 0.6757195 |
| PGGPVPIP | ASSQEGQNNQAPL | 0.17880307 |
| FLTVALEP | ASSQEGQNNQAPL | 0.6491276 |
| SQYFGANL | ASSQEGQNNQAPL | 0.41629064 |
| GGVKPAA | ASSQEGQNNQAPL | 0.80145335 |
| GGTGGAGGAGGW | ASSQEGQNNQAPL | 0.28658217 |
| GVGGVAGLGGAGG | ASSQEGQNNQAPL | 0.51918656 |

| | | |
|---|---|---|
| VLLVLATTI | ASSQEGQNNQAPL | 0.40694183 |
| IGIFAAML | ASSQEGQNNQAPL | 0.7529936 |
| AVGDLID | ASSQEGQNNQAPL | 0.14699174 |
| NISPLPL | ASSQEGQNNQAPL | 0.3611713 |
| VTAVFVAI | ASSQEGQNNQAPL | 0.19032063 |
| SGDDNDAAL | ASSQEGQNNQAPL | 0.41662902 |
| AVAALAHS | ASSQEGQNNQAPL | 0.6272579 |
| GGVTVMATP | ASSQEGQNNQAPL | 0.112996206 |
| AGLAANVRD | ASSQEGQNNQAPL | 0.71619606 |
| TNRPDLIDPALLRPG | ASSQEGQNNQAPL | 0.3126017 |
| LANLGVTAL | ASSQEGQNNQAPL | 0.25325182 |
| TGERIEL | ASSQEGQNNQAPL | 0.558485 |
| NGEELAA | ASSQEGQNNQAPL | 0.38258728 |
| VLSLVWG | ASSQEGQNNQAPL | 0.7035702 |
| IGYALWL | ASSQEGQNNQAPL | 0.24338366 |
| LSLLLAPL | ASSQEGQNNQAPL | 0.5743238 |
| TLDEALK | ASSQEGQNNQAPL | 0.56526077 |
| TDGAGTAEGG | ASSQEGQNNQAPL | 0.11513954 |
| NAPGGAGGNGG | ASSQEGQNNQAPL | 0.24258381 |
| VSVFMGMGGNF | ASSQEGQNNQAPL | 0.7033738 |
| LASYLAAL | ASSQEGQNNQAPL | 0.76439583 |
| LVGGTPAL | ASSQEGQNNQAPL | 0.81169677 |
| VRNADVL | ASSQEGQNNQAPL | 0.6154644 |
| TCSSPFLVSAM | ASSQEGQNNQAPL | 0.520699 |
| VHITDAE | ASSQEGQNNQAPL | 0.6891286 |
| EYAAPSF | ASSQEGQNNQAPL | 0.8019416 |
| VVDAFHGGL | ASSQEGQNNQAPL | 0.81821704 |
| ASVVDPQ | ASSQEGQNNQAPL | 0.77042913 |
| SYNEFGGLS | ASSQEGQNNQAPL | 0.5075708 |
| DPDGSVY | ASSQEGQNNQAPL | 0.75176 |
| MVSTIHATE | ASSQEGQNNQAPL | 0.6865943 |
| LAAGVADPV | ASSQEGQNNQAPL | 0.6969403 |
| TKEQIAA | ASSQEGQNNQAPL | 0.7959819 |
| SLASPAAL | ASSQEGQNNQAPL | 0.72510344 |
| IGLVFYGV | ASSQEGQNNQAPL | 0.685761 |
| FLLDASNL | ASSQEGQNNQAPL | 0.6189638 |
| YQVGNFP | ASSQEGQNNQAPL | 0.7272008 |
| ESLLSQST | ASSQEGQNNQAPL | 0.7872044 |
| LFTVGFAAVG | ASSQEGQNNQAPL | 0.74444485 |
| ATPPTPT | ASSQEGQNNQAPL | 0.839559 |
| TDLATLG | ASSQEGQNNQAPL | 0.80558515 |
| SAGGTGR | ASSQEGQNNQAPL | 0.776422 |
| GFSNSGS | ASSQEGQNNQAPL | 0.1740664 |
| GGNGGSGGTSV | ASSQEGQNNQAPL | 0.49804 |
| GTSVATGGAGN | ASSQEGQNNQAPL | 0.6607716 |
| NGGNGGSGGTS | ASSQEGQNNQAPL | 0.73935056 |
| AVVKFPH | ASSQEGQNNQAPL | 0.3028759 |
| TPPPPPP | ASSQEGQNNQAPL | 0.7931385 |
| AAPAAAAAPA | ASSQEGQNNQAPL | 0.6907555 |
| IALNEDL | ASSQEGQNNQAPL | 0.6209384 |
| IVNVSSM | ASSQEGQNNQAPL | 0.82982874 |

| | | |
|---|---|---|
| TGIPPAPR | ASSQEGQNNQAPL | 0.6683373 |
| FDLGGGTF | ASSQEGQNNQAPL | 0.63682455 |
| FDLGGGTFDVS | ASSQEGQNNQAPL | 0.6598667 |
| NEPTAAA | ASSQEGQNNQAPL | 0.8399559 |
| NEPTAAALA | ASSQEGQNNQAPL | 0.50573176 |
| GSIINISSI | ASSQEGQNNQAPL | 0.8013976 |
| VAPGNAGTA | ASSQEGQNNQAPL | 0.7144031 |
| TGGGASLE | ASSQEGQNNQAPL | 0.8245537 |
| VAVYNHYKRI | ASSQEGQNNQAPL | 0.397245 |
| TDPAPATT | ASSQEGQNNQAPL | 0.7921263 |
| TNRPDILDPALLRPG | ASSQEGQNNQAPL | 0.68367875 |
| LTTTTTS | ASSQEGQNNQAPL | 0.8544105 |
| PTGSGKTL | ASSQEGQNNQAPL | 0.673364 |
| TGSGKTL | ASSQEGQNNQAPL | 0.7529582 |
| TIGHVDHGK | ASSQEGQNNQAPL | 0.65738434 |
| TIGHVDHGKTT | ASSQEGQNNQAPL | 0.79857856 |
| GHVDHGKTT | ASSQEGQNNQAPL | 0.42792314 |
| AQFEHTLL | ASSQEGQNNQAPL | 0.6632006 |
| IVNQDEL | ASSQEGQNNQAPL | 0.59282005 |
| TNGSQFF | ASSQEGQNNQAPL | 0.724316 |
| TGKGIVE | ASSQEGQNNQAPL | 0.7728714 |
| LISNASDA | ASSQEGQNNQAPL | 0.31720942 |
| AELRTLGELTKIAKAHGAQ | ASSQEGQNNQAPL | 0.7821378 |
| APNAVAAAEPSW | ASSQEGQNNQAPL | 0.6329056 |
| APQVAGAATAVL | ASSQEGQNNQAPL | 0.51323235 |
| ARRDPAAAAAILPTDA | ASSQEGQNNQAPL | 0.8089293 |
| DAAARDVLAEAGLAEYF | ASSQEGQNNQAPL | 0.66297704 |
| DDQIVLSAKANLAKVEQ | ASSQEGQNNQAPL | 0.78542644 |
| DELLPDTGEDSDEDGH | ASSQEGQNNQAPL | 0.6799278 |
| ELPEAPANRATVPQD | ASSQEGQNNQAPL | 0.71169776 |
| FGQSKIERVTGIG | ASSQEGQNNQAPL | 0.7808934 |
| GAGGTGSTAGGAGGAGGAGGI | ASSQEGQNNQAPL | 0.5094272 |
| GHGGTGAAGGNGGTGGAGGS( | ASSQEGQNNQAPL | 0.56345594 |
| GNGGNGGAGGEGGPGGAGGA( | ASSQEGQNNQAPL | 0.81612754 |
| GSTAGNGGNGGNATNGTIAG( | ASSQEGQNNQAPL | 0.8961103 |
| GTGGDGGNGGNGGTGG | ASSQEGQNNQAPL | 0.8922885 |
| GVRRPAGIPADAPV | ASSQEGQNNQAPL | 0.83259577 |
| GYRQVTDAHLLTLARR | ASSQEGQNNQAPL | 0.8114618 |
| IAKMGQDGHDRG | ASSQEGQNNQAPL | 0.85400224 |
| KAEPSIEIQYT | ASSQEGQNNQAPL | 0.9006929 |
| KPVRIRMAADEDTVAVT | ASSQEGQNNQAPL | 0.899453 |
| MPPRIAGMRLLVIKPEP | ASSQEGQNNQAPL | 0.8884702 |
| MKFYAVALFRSRGVPPTR | ASSQEGQNNQAPL | 0.81849 |
| NAGGFGILGAVAHSP | ASSQEGQNNQAPL | 0.8849178 |
| NEGREITLAMAEM | ASSQEGQNNQAPL | 0.85408515 |
| NELVEWSHPLGHRWRWE | ASSQEGQNNQAPL | 0.85321134 |
| NLGAGGTGGNGGNPG | ASSQEGQNNQAPL | 0.8990192 |
| NTEWPKWETVMAADPVD | ASSQEGQNNQAPL | 0.87091935 |
| PASAAPAETQP | ASSQEGQNNQAPL | 0.861524 |
| QQSSDGKNGVRAAEITGE | ASSQEGQNNQAPL | 0.8656218 |
| RAPDAVTTSWWKE | ASSQEGQNNQAPL | 0.88347423 |

| | | |
|---|---|---|
| SGFGNVGAHTSGWFNQ | ASSQEGQNNQAPL | 0.89907897 |
| SGVAGAVGQAASVGGLKV | ASSQEGQNNQAPL | 0.8928354 |
| SNGGAGTGSAGGNGGT | ASSQEGQNNQAPL | 0.9108742 |
| TGGAGGEGGAAG | ASSQEGQNNQAPL | 0.89873075 |
| TGGDGGAAGAGGAPGHG | ASSQEGQNNQAPL | 0.8947561 |
| TMPPLLAAIANAGRHGV | ASSQEGQNNQAPL | 0.41413885 |
| TSSTAGNSGGA | ASSQEGQNNQAPL | 0.7415917 |
| VPATVPVGILC | ASSQEGQNNQAPL | 0.8618859 |
| VQRPDSGGVVLTGRIS | ASSQEGQNNQAPL | 0.8550743 |
| WTPNPVPATKGVPAGLVQAT | ASSQEGQNNQAPL | 0.61288154 |
| YALKRALAKFSDRFDVVII | ASSQEGQNNQAPL | 0.89005625 |
| YLIGSVIV | ASSQEGQNNQAPL | 0.8746965 |
| YGQLVAVID | ASSQEGQNNQAPL | 0.80249286 |
| VTVGGMTLPA | ASSQEGQNNQAPL | 0.8960264 |
| VLALPAV | ASSQEGQNNQAPL | 0.8360694 |
| VIPPTLNYET | ASSQEGQNNQAPL | 0.8233483 |
| VGMGGMHPSSG | ASSQEGQNNQAPL | 0.86314476 |
| VGLAGGAATA | ASSQEGQNNQAPL | 0.90599924 |
| VGGAGGAGGSGP | ASSQEGQNNQAPL | 0.8323252 |
| VFAAFMV | ASSQEGQNNQAPL | 0.894122 |
| TLYDKLR | ASSQEGQNNQAPL | 0.87150455 |
| TLAVFTVL | ASSQEGQNNQAPL | 0.9034567 |
| TGSVQPARV | ASSQEGQNNQAPL | 0.7494848 |
| TGGGAGSGGT | ASSQEGQNNQAPL | 0.8929764 |
| SHLETSL | ASSQEGQNNQAPL | 0.864071 |
| SFTTGMT | ASSQEGQNNQAPL | 0.91266143 |
| QVEVAKRMGVRQPTVS | ASSQEGQNNQAPL | 0.8621848 |
| QGSGGPALAM | ASSQEGQNNQAPL | 0.8833546 |
| MLAPSNP | ASSQEGQNNQAPL | 0.8325701 |
| LVNNAAI | ASSQEGQNNQAPL | 0.8879424 |
| LTGEINTLA | ASSQEGQNNQAPL | 0.6797211 |
| LSIFGPV | ASSQEGQNNQAPL | 0.8790709 |
| LLALLANL | ASSQEGQNNQAPL | 0.8111496 |
| LISNASDALDKLR | ASSQEGQNNQAPL | 0.88572836 |
| LINEAVP | ASSQEGQNNQAPL | 0.8867937 |
| LGYAFSGL | ASSQEGQNNQAPL | 0.5473764 |
| LDFVAVAL | ASSQEGQNNQAPL | 0.9012088 |
| LAQPGAE | ASSQEGQNNQAPL | 0.82221514 |
| ISNLGAVTGF | ASSQEGQNNQAPL | 0.81505656 |
| HQEDPGRI | ASSQEGQNNQAPL | 0.9053288 |
| GTPGEPTP | ASSQEGQNNQAPL | 0.8692751 |
| GFFNSGS | ASSQEGQNNQAPL | 0.9013729 |
| GAGGGGLTAGAN | ASSQEGQNNQAPL | 0.87946844 |
| FVDALVNM | ASSQEGQNNQAPL | 0.8750316 |
| FLTVAPEL | ASSQEGQNNQAPL | 0.8974601 |
| FETLTAL | ASSQEGQNNQAPL | 0.64191025 |
| DVINPIEL | ASSQEGQNNQAPL | 0.8086719 |
| APVSGAPV | ASSQEGQNNQAPL | 0.9025544 |
| ALVGGLAW | ASSQEGQNNQAPL | 0.6549635 |
| ALAAAAATAI | ASSQEGQNNQAPL | 0.6561897 |
| AFGLGLLL | ASSQEGQNNQAPL | 0.6177372 |

| | | |
|---|---|---|
| AFFAAASL | ASSQEGQNNQAPL | 0.5939647 |
| AAAALPAFSPP | ASSQEGQNNQAPL | 0.6183606 |
| VVLAPAL | ASSQEGQNNQAPL | 0.65247715 |
| TIGHVDHGKTTL | ASSQEGQNNQAPL | 0.66472393 |
| LISNASDALDK | ASSQEGQNNQAPL | 0.6555257 |
| LISNASDAL | ASSQEGQNNQAPL | 0.60330915 |
| ISNASDALDK | ASSQEGQNNQAPL | 0.6619066 |
| NQAETLVYQTE | ASSQEGQNNQAPL | 0.647895 |
| TGGTGGTAGAG | ASSQEGQNNQAPL | 0.6402973 |
| TGSTAGGAGGA | ASSQEGQNNQAPL | 0.6457334 |
| DPLAVPVAAGPV | ASSQEGQNNQAPL | 0.6574079 |
| GGTGGTPGNGGN | ASSQEGQNNQAPL | 0.62913436 |
| PAGLAGVDPATL | ASSQEGQNNQAPL | 0.61302453 |
| GNNLVVEAPHAGASV | ASSQEGQNNQAPL | 0.6483181 |
| KVTRIMSASTLEQPA | ASSQEGQNNQAPL | 0.6556591 |
| LGFALPIALPRRTVV | ASSQEGQNNQAPL | 0.65478975 |
| TRLGNLTARKLVVLV | ASSQEGQNNQAPL | 0.66645575 |
| WSQQRRLIYFPSAGP | ASSQEGQNNQAPL | 0.66008365 |
| GPNGNPDPMAAAVDIR | ASSQEGQNNQAPL | 0.6567222 |
| TQRCVSVADLRAAASR | ASSQEGQNNQAPL | 0.44113848 |
| QFVLDVAEHPVEVLTETV | ASSQEGQNNQAPL | 0.5712104 |
| GGAGGTGGAGGNGGTQAGDG | ASSQEGQNNQAPL | 0.62915653 |
| PLLATLRAIAVDVDAKPAQV | ASSQEGQNNQAPL | 0.63427097 |
| APKFRTGGYMIDFWGVGYQV | ASSQEGQNNQAPL | 0.43138215 |
| GAGGAGGAGGAAGGPGGTGG | ASSQEGQNNQAPL | 0.65473455 |
| IGALGLLATDVAPAGSVAFG | ASSQEGQNNQAPL | 0.64028835 |
| AGGAGGIGGNGGAGGAGGVL | ASSQEGQNNQAPL | 0.6262135 |
| ADELIGLL | ASSQEGQNNQAPL | 0.65841955 |
| ADRALKA | ASSQEGQNNQAPL | 0.6352563 |
| AGAVGQVGGAGGA | ASSQEGQNNQAPL | 0.60196614 |
| AGGGGGLGGAP | ASSQEGQNNQAPL | 0.6281465 |
| AGGLLYGNGGAG | ASSQEGQNNQAPL | 0.6626615 |
| AGLESIGL | ASSQEGQNNQAPL | 0.60039055 |
| AGLKTGII | ASSQEGQNNQAPL | 0.6629239 |
| ASLAALLL | ASSQEGQNNQAPL | 0.5936548 |
| ASVINISSI | ASSQEGQNNQAPL | 0.6639925 |
| ATPPGPA | ASSQEGQNNQAPL | 0.57687294 |
| ATVASALV | ASSQEGQNNQAPL | 0.6607851 |
| ATVASLIL | ASSQEGQNNQAPL | 0.64112496 |
| AVNGPVLIRH | ASSQEGQNNQAPL | 0.66903275 |
| DGAAGAVGI | ASSQEGQNNQAPL | 0.6303136 |
| DPLLHGI | ASSQEGQNNQAPL | 0.6605148 |
| ELGAALGAL | ASSQEGQNNQAPL | 0.6480975 |
| ELGRTQAV | ASSQEGQNNQAPL | 0.6637445 |
| FVAGGYLS | ASSQEGQNNQAPL | 0.591748 |
| GASGGAGGAGGW | ASSQEGQNNQAPL | 0.6377914 |
| GGAGAGGDGGAGGI | ASSQEGQNNQAPL | 0.56273335 |
| GTGHNGSQPGMGGNG | ASSQEGQNNQAPL | 0.64729244 |
| GYAAASASAS | ASSQEGQNNQAPL | 0.6577225 |
| HQVDVAAVQQG | ASSQEGQNNQAPL | 0.38703135 |
| IGAAIAHT | ASSQEGQNNQAPL | 0.66257066 |

| | | |
|---|---|---|
| IGMIEVAQL | ASSQEGQNNQAPL | 0.57690185 |
| IRLAPEHTSAMD | ASSQEGQNNQAPL | 0.6045441 |
| IYNVSVL | ASSQEGQNNQAPL | 0.658371 |
| KQGFLGM | ASSQEGQNNQAPL | 0.65652764 |
| LESVITQL | ASSQEGQNNQAPL | 0.65720487 |
| LGLGALVVV | ASSQEGQNNQAPL | 0.63523877 |
| LIGNGGNGGSGGI | ASSQEGQNNQAPL | 0.64061356 |
| LNAINAPTL | ASSQEGQNNQAPL | 0.64785427 |
| LQLLNTGV | ASSQEGQNNQAPL | 0.5567763 |
| LQTSIEF | ASSQEGQNNQAPL | 0.6097768 |
| MSFAVSEL | ASSQEGQNNQAPL | 0.66529036 |
| NTGALGL | ASSQEGQNNQAPL | 0.50199133 |
| PPGAGGAGGAGTT | ASSQEGQNNQAPL | 0.62211823 |
| QLVGLGI | ASSQEGQNNQAPL | 0.49982074 |
| RFLEQVHQL | ASSQEGQNNQAPL | 0.4277776 |
| RGVGYVLR | ASSQEGQNNQAPL | 0.4848321 |
| SATGVFAGVI | ASSQEGQNNQAPL | 0.5483941 |
| SLPAITLP | ASSQEGQNNQAPL | 0.623692 |
| STALAFTV | ASSQEGQNNQAPL | 0.51251316 |
| SVLTVGLGK | ASSQEGQNNQAPL | 0.41867635 |
| TALAGAGSGI | ASSQEGQNNQAPL | 0.5636178 |
| TGLPFTGI | ASSQEGQNNQAPL | 0.4762559 |
| VAEHVTV | ASSQEGQNNQAPL | 0.52830833 |
| VDIVLNSL | ASSQEGQNNQAPL | 0.49379367 |
| VLTGGGAVL | ASSQEGQNNQAPL | 0.5558358 |
| VNVGLAYGI | ASSQEGQNNQAPL | 0.54484105 |
| VSSAAVGAV | ASSQEGQNNQAPL | 0.44058996 |
| VTSVYEAGV | ASSQEGQNNQAPL | 0.48579204 |
| WVALDRGA | ASSQEGQNNQAPL | 0.45101917 |
| YLVGSVVI | ASSQEGQNNQAPL | 0.44542047 |
| GPPGTGKTLL | ASSQEGQNNQAPL | 0.5374174 |
| GVGMTKF | ASSQEGQNNQAPL | 0.5553671 |
| IGIGGSDL | ASSQEGQNNQAPL | 0.4772211 |
| IGTLLAEL | ASSQEGQNNQAPL | 0.31854293 |
| IYTSGTT | ASSQEGQNNQAPL | 0.33607095 |
| LSFPAGV | ASSQEGQNNQAPL | 0.4687208 |
| LYTSGTT | ASSQEGQNNQAPL | 0.4247163 |
| VGAFPFGF | ASSQEGQNNQAPL | 0.331051 |
| VGAGTVEF | ASSQEGQNNQAPL | 0.57079244 |
| VGKSTLLL | ASSQEGQNNQAPL | 0.5082775 |
| AALAHAGGGIV | ASSQEGQNNQAPL | 0.50472504 |
| AALVAVSLL | ASSQEGQNNQAPL | 0.55251795 |
| AAVSRAVAAL | ASSQEGQNNQAPL | 0.52612495 |
| AGGSSGIGGF | ASSQEGQNNQAPL | 0.44334194 |
| AGLTVANANG | ASSQEGQNNQAPL | 0.4257726 |
| AGLVGAVL | ASSQEGQNNQAPL | 0.43139955 |
| ALSQAFTV | ASSQEGQNNQAPL | 0.35129368 |
| ALTFSVSL | ASSQEGQNNQAPL | 0.58374834 |
| ALTGAALVAL | ASSQEGQNNQAPL | 0.3534319 |
| ASFSIGAGVL | ASSQEGQNNQAPL | 0.5327685 |
| DGSVYKI | ASSQEGQNNQAPL | 0.41077 |

| | | |
|---|---|---|
| EQLAEIV | ASSQEGQNNQAPL | 0.54965615 |
| FAVSELTVM | ASSQEGQNNQAPL | 0.5213333 |
| GAAPLAAGL | ASSQEGQNNQAPL | 0.63931715 |
| GADGGDGGSGGAGG | ASSQEGQNNQAPL | 0.47153914 |
| GFGLDTGGAGG | ASSQEGQNNQAPL | 0.5337795 |
| GLASGGLL | ASSQEGQNNQAPL | 0.42012778 |
| GRIAGAAVLL | ASSQEGQNNQAPL | 0.62872326 |
| GSAGSLNTSF | ASSQEGQNNQAPL | 0.37727284 |
| GTGQIGI | ASSQEGQNNQAPL | 0.46986324 |
| GTLIALIL | ASSQEGQNNQAPL | 0.37205485 |
| IAATMGFVVG | ASSQEGQNNQAPL | 0.5479181 |
| IFAQALL | ASSQEGQNNQAPL | 0.52149296 |
| IGAAVALGF | ASSQEGQNNQAPL | 0.37605035 |
| IGLTPLAL | ASSQEGQNNQAPL | 0.5518663 |
| ILGSNTSTL | ASSQEGQNNQAPL | 0.5022108 |
| IVFSLTVL | ASSQEGQNNQAPL | 0.4379828 |
| LASKATGF | ASSQEGQNNQAPL | 0.58929545 |
| LASVALIL | ASSQEGQNNQAPL | 0.5423622 |
| LFAVGLTL | ASSQEGQNNQAPL | 0.58212936 |
| LFNGGIGTYI | ASSQEGQNNQAPL | 0.4933375 |
| LTLGAGII | ASSQEGQNNQAPL | 0.4242112 |
| LVAAALGDI | ASSQEGQNNQAPL | 0.47666356 |
| NTAFVQLGI | ASSQEGQNNQAPL | 0.46086136 |
| QLFSGAEL | ASSQEGQNNQAPL | 0.41262722 |
| SAATFITL | ASSQEGQNNQAPL | 0.58140504 |
| SNYGLANLGV | ASSQEGQNNQAPL | 0.7455771 |
| SYGGAYAVM | ASSQEGQNNQAPL | 0.82449853 |
| TTGVGGLTLG | ASSQEGQNNQAPL | 0.68895733 |
| TTQTIYI | ASSQEGQNNQAPL | 0.6040516 |
| VAAGAGLGAVI | ASSQEGQNNQAPL | 0.72580326 |
| VAAVVTLL | ASSQEGQNNQAPL | 0.8141297 |
| VGFLGGLI | ASSQEGQNNQAPL | 0.8187985 |
| VGVIPILL | ASSQEGQNNQAPL | 0.7856928 |
| VIFASSLI | ASSQEGQNNQAPL | 0.52686834 |
| VKGGDGGAAAG | ASSQEGQNNQAPL | 0.7633811 |
| VTKTFTNI | ASSQEGQNNQAPL | 0.6685916 |
| VVAAAILL | ASSQEGQNNQAPL | 0.733431 |
| YAALDVEL | ASSQEGQNNQAPL | 0.7870371 |
| LASVALLL | ASSQEGQNNQAPL | 0.71721935 |
| LDAVHRV | ASSQEGQNNQAPL | 0.6514547 |
| QGLSSSIF | ASSQEGQNNQAPL | 0.6705409 |
| RGTFANIRL | ASSQEGQNNQAPL | 0.798133 |
| RQLWWGHRI | ASSQEGQNNQAPL | 0.75559694 |
| YGPPGTGKTLL | ASSQEGQNNQAPL | 0.762805 |
| AALVGAPANVA | ASSQEGQNNQAPL | 0.83077437 |
| AATFRRTE | ASSQEGQNNQAPL | 0.79921883 |
| AGALGGGI | ASSQEGQNNQAPL | 0.80726045 |
| AGGAGGAGGIGAG | ASSQEGQNNQAPL | 0.2471476 |
| AGSGALGL | ASSQEGQNNQAPL | 0.43491116 |
| AQIADQI | ASSQEGQNNQAPL | 0.67390466 |
| ASAVAVTYL | ASSQEGQNNQAPL | 0.70845556 |

| | | |
|---|---|---|
| ASTVLGGTGG | ASSQEGQNNQAPL | 0.50514954 |
| AVGGEPAK | ASSQEGQNNQAPL | 0.7789583 |
| CEQGTFGGTC | ASSQEGQNNQAPL | 0.62885153 |
| DEGKAPAG | ASSQEGQNNQAPL | 0.6061137 |
| DTVVFPAL | ASSQEGQNNQAPL | 0.8050592 |
| FEGITMAG | ASSQEGQNNQAPL | 0.66975474 |
| FPLVVQD | ASSQEGQNNQAPL | 0.61783427 |
| GAFTTPQ | ASSQEGQNNQAPL | 0.6585967 |
| GFGGFGTAGGVGGA | ASSQEGQNNQAPL | 0.78851867 |
| GFLAQFL | ASSQEGQNNQAPL | 0.45968145 |
| GGAGGAGGNAGVA | ASSQEGQNNQAPL | 0.78151983 |
| GGVGTAGAGGAGGAGG | ASSQEGQNNQAPL | 0.6905291 |
| GNGGGGDAGNAGSGGN | ASSQEGQNNQAPL | 0.81478846 |
| GTGGAGGAGNDAGS | ASSQEGQNNQAPL | 0.47987175 |
| GTTNTGWFNS | ASSQEGQNNQAPL | 0.7943087 |
| IFMGPLFTL | ASSQEGQNNQAPL | 0.7448075 |
| ILAHAIEAI | ASSQEGQNNQAPL | 0.8374138 |
| IRSAIREAL | ASSQEGQNNQAPL | 0.67479694 |
| ITFAGGG | ASSQEGQNNQAPL | 0.7014823 |
| ITGMSGHQEV | ASSQEGQNNQAPL | 0.6581354 |
| IVAGLACTL | ASSQEGQNNQAPL | 0.8214572 |
| IVGGGLAGA | ASSQEGQNNQAPL | 0.46748698 |
| IVTGVLNI | ASSQEGQNNQAPL | 0.69760686 |
| LAQQGCI | ASSQEGQNNQAPL | 0.4877437 |
| LCLHVEN | ASSQEGQNNQAPL | 0.7722814 |
| LDAISDGS | ASSQEGQNNQAPL | 0.77496296 |
| LGGFAAML | ASSQEGQNNQAPL | 0.4305076 |
| LGGKNPMI | ASSQEGQNNQAPL | 0.7925204 |
| LGPKGVAAIL | ASSQEGQNNQAPL | 0.75159365 |
| LMVGVSI | ASSQEGQNNQAPL | 0.61851716 |
| LVGAAFVL | ASSQEGQNNQAPL | 0.818433 |
| LVLAVAPL | ASSQEGQNNQAPL | 0.7366768 |
| MAAIAVIDAA | ASSQEGQNNQAPL | 0.81222206 |
| NNGINAELQ | ASSQEGQNNQAPL | 0.760638 |
| RGEPLVLVHG | ASSQEGQNNQAPL | 0.7019702 |
| RGGAGGAAGLI | ASSQEGQNNQAPL | 0.7832135 |
| RSYRMHPAV | ASSQEGQNNQAPL | 0.65129775 |
| SAAHLLAGQLA | ASSQEGQNNQAPL | 0.6658436 |
| SAAVGSY | ASSQEGQNNQAPL | 0.8120474 |
| SPHAVLHRI | ASSQEGQNNQAPL | 0.87334776 |
| SQIAKVL | ASSQEGQNNQAPL | 0.8747381 |
| TALGHLVP | ASSQEGQNNQAPL | 0.8044629 |
| TAPTTSGAA | ASSQEGQNNQAPL | 0.7308177 |
| TDGDDGNGGSPGT | ASSQEGQNNQAPL | 0.8253859 |
| TDGLGEN | ASSQEGQNNQAPL | 0.87802976 |
| TGGPPLPGQA | ASSQEGQNNQAPL | 0.89338624 |
| TIPAITI | ASSQEGQNNQAPL | 0.87075317 |
| TIPGFTI | ASSQEGQNNQAPL | 0.6523923 |
| TSIASGALA | ASSQEGQNNQAPL | 0.87963474 |
| TVTLPGAAI | ASSQEGQNNQAPL | 0.86141264 |
| VASGLSAALP | ASSQEGQNNQAPL | 0.8416452 |

| | | |
|---|---|---|
| VGAGVIGI | ASSQEGQNNQAPL | 0.87863934 |
| VGPKRFGV | ASSQEGQNNQAPL | 0.85927445 |
| VNVAFPD | ASSQEGQNNQAPL | 0.8048593 |
| VSMTNSGVS | ASSQEGQNNQAPL | 0.8161115 |
| VTDVLGDLSV | ASSQEGQNNQAPL | 0.8701831 |
| VVVVGAGFAGL | ASSQEGQNNQAPL | 0.8806952 |
| VYTVGTVV | ASSQEGQNNQAPL | 0.8860028 |
| WTTFSGP | ASSQEGQNNQAPL | 0.90958476 |
| YDELVEAF | ASSQEGQNNQAPL | 0.8958434 |
| YGTGGAGGNGG | ASSQEGQNNQAPL | 0.88516676 |
| YTNVMAV | ASSQEGQNNQAPL | 0.34484375 |
| FLEQVHQL | ASSQEGQNNQAPL | 0.65394676 |
| GLSLLAVAV | ASSQEGQNNQAPL | 0.79881346 |
| LEQVHQL | ASSQEGQNNQAPL | 0.8446926 |
| LNAAGAI | ASSQEGQNNQAPL | 0.487949 |
| PAVHASL | ASSQEGQNNQAPL | 0.8376697 |
| TELEDDA | ASSQEGQNNQAPL | 0.8010013 |
| TPPGPPN | ASSQEGQNNQAPL | 0.7723984 |
| TPPPPPPP | ASSQEGQNNQAPL | 0.8965782 |
| TQSPALI | ASSTIANTEVF | 0.8248821 |
| EFAAAFPQ | ASSTIANTEVF | 0.7722717 |
| LLSGNLDVL | ASSTIANTEVF | 0.8279915 |
| GPANLGNYNI | ASSTIANTEVF | 0.8999644 |
| TLGPTLP | ASSTIANTEVF | 0.76636577 |
| VYAYSSCCK | ASSTIANTEVF | 0.88962203 |
| APAPAPAPNI | ASSTIANTEVF | 0.7383288 |
| PGGPVPIP | ASSTIANTEVF | 0.90110403 |
| FLTVALEP | ASSTIANTEVF | 0.63698953 |
| SQYFGANL | ASSTIANTEVF | 0.89618397 |
| GGVKPAA | ASSTIANTEVF | 0.8562787 |
| GGTGGAGGAGGW | ASSTIANTEVF | 0.9166807 |
| GVGGVAGLGGAGG | ASSTIANTEVF | 0.84240806 |
| VLLVLATTI | ASSTIANTEVF | 0.87596184 |
| IGIFAAML | ASSTIANTEVF | 0.7910674 |
| AVGDLID | ASSTIANTEVF | 0.8910432 |
| NISPLPL | ASSTIANTEVF | 0.6690939 |
| VTAVFVAI | ASSTIANTEVF | 0.8553791 |
| SGDDNDAAL | ASSTIANTEVF | 0.5824327 |
| AVAALAHS | ASSTIANTEVF | 0.86913013 |
| GGVTVMATP | ASSTIANTEVF | 0.8775531 |
| AGLAANVRD | ASSTIANTEVF | 0.45180434 |
| TNRPDLIDPALLRPG | ASSTIANTEVF | 0.9011178 |
| LANLGVTAL | ASSTIANTEVF | 0.780774 |
| TGERIEL | ASSTIANTEVF | 0.7431823 |
| NGEELAA | ASSTIANTEVF | 0.9026673 |
| VLSLVWG | ASSTIANTEVF | 0.8540987 |
| IGYALWL | ASSTIANTEVF | 0.88912 |
| LSLLLAPL | ASSTIANTEVF | 0.83677113 |
| TLDEALK | ASSTIANTEVF | 0.83026445 |
| TDGAGTAEGG | ASSTIANTEVF | 0.8572606 |
| NAPGGAGGNGG | ASSTIANTEVF | 0.7035261 |

| | | |
|---|---|---|
| VSVFMGMGGNF | ASSTIANTEVF | 0.7847651 |
| LASYLAAL | ASSTIANTEVF | 0.90681386 |
| LVGGTPAL | ASSTIANTEVF | 0.82849836 |
| VRNADVL | ASSTIANTEVF | 0.82262343 |
| TCSSPFLVSAM | ASSTIANTEVF | 0.7240554 |
| VHITDAE | ASSTIANTEVF | 0.66618186 |
| EYAAPSF | ASSTIANTEVF | 0.760116 |
| VVDAFHGGL | ASSTIANTEVF | 0.836787 |
| ASVVDPQ | ASSTIANTEVF | 0.8536272 |
| SYNEFGGLS | ASSTIANTEVF | 0.8256807 |
| DPDGSVY | ASSTIANTEVF | 0.66785765 |
| MVSTIHATE | ASSTIANTEVF | 0.83703685 |
| LAAGVADPV | ASSTIANTEVF | 0.8156891 |
| TKEQIAA | ASSTIANTEVF | 0.7687515 |
| SLASPAAL | ASSTIANTEVF | 0.8491781 |
| IGLVFYGV | ASSTIANTEVF | 0.8268108 |
| FLLDASNL | ASSTIANTEVF | 0.7487327 |
| YQVGNFP | ASSTIANTEVF | 0.790752 |
| ESLLSQST | ASSTIANTEVF | 0.81948674 |
| LFTVGFAAVG | ASSTIANTEVF | 0.8387991 |
| ATPPTPT | ASSTIANTEVF | 0.84531415 |
| TDLATLG | ASSTIANTEVF | 0.8769562 |
| SAGGTGR | ASSTIANTEVF | 0.8552889 |
| GFSNSGS | ASSTIANTEVF | 0.8339629 |
| GGNGGSGGTSV | ASSTIANTEVF | 0.36659318 |
| GTSVATGGAGN | ASSTIANTEVF | 0.63843143 |
| NGGNGGSGGTS | ASSTIANTEVF | 0.78642064 |
| AVVKFPH | ASSTIANTEVF | 0.79957235 |
| TPPPPPP | ASSTIANTEVF | 0.47505808 |
| AAPAAAAAPA | ASSTIANTEVF | 0.81146526 |
| IALNEDL | ASSTIANTEVF | 0.7777326 |
| IVNVSSM | ASSTIANTEVF | 0.71003854 |
| TGIPPAPR | ASSTIANTEVF | 0.85929614 |
| FDLGGGTF | ASSTIANTEVF | 0.76197195 |
| FDLGGGTFDVS | ASSTIANTEVF | 0.7735772 |
| NEPTAAA | ASSTIANTEVF | 0.81291693 |
| NEPTAAALA | ASSTIANTEVF | 0.8677169 |
| GSIINISSI | ASSTIANTEVF | 0.733706 |
| VAPGNAGTA | ASSTIANTEVF | 0.85174924 |
| TGGGASLE | ASSTIANTEVF | 0.7315839 |
| VAVYNHYKRI | ASSTIANTEVF | 0.8655969 |
| TDPAPATT | ASSTIANTEVF | 0.639608 |
| TNRPDILDPALLRPG | ASSTIANTEVF | 0.8485076 |
| LTTTTTS | ASSTIANTEVF | 0.7865153 |
| PTGSGKTL | ASSTIANTEVF | 0.88442147 |
| TGSGKTL | ASSTIANTEVF | 0.80403554 |
| TIGHVDHGK | ASSTIANTEVF | 0.85266906 |
| TIGHVDHGKTT | ASSTIANTEVF | 0.77639914 |
| GHVDHGKTT | ASSTIANTEVF | 0.82254416 |
| AQFEHTLL | ASSTIANTEVF | 0.60954094 |
| IVNQDEL | ASSTIANTEVF | 0.8177467 |

| | | |
|---|---|---|
| TNGSQFF | ASSTIANTEVF | 0.6084024 |
| TGKGIVE | ASSTIANTEVF | 0.8300764 |
| LISNASDA | ASSTIANTEVF | 0.8226617 |
| AELRTLGELTKIAKAHGAQ | ASSTIANTEVF | 0.46350515 |
| APNAVAAAEPSW | ASSTIANTEVF | 0.863532 |
| APQVAGAATAVL | ASSTIANTEVF | 0.6910434 |
| ARRDPAAAAAILPTDA | ASSTIANTEVF | 0.73625654 |
| DAAARDVLAEAGLAEYF | ASSTIANTEVF | 0.8666211 |
| DDQIVLSAKANLAKVEQ | ASSTIANTEVF | 0.8049509 |
| DELLPDTGEDSDEDGH | ASSTIANTEVF | 0.8507746 |
| ELPEAPANRATVPQD | ASSTIANTEVF | 0.8013958 |
| FGQSKIERVTGIG | ASSTIANTEVF | 0.7932848 |
| GAGGTGSTAGGAGGAGGAGGI | ASSTIANTEVF | 0.81367195 |
| GHGGTGAAGGNGGTGGAGGS( | ASSTIANTEVF | 0.6045797 |
| GNGGNGGAGGEGGPGGAGGA( | ASSTIANTEVF | 0.766499 |
| GSTAGNGGNGGNATNGTIAG( | ASSTIANTEVF | 0.86558 |
| GTGGDGGNGGNGGTGG | ASSTIANTEVF | 0.3822161 |
| GVRRPAGIPADAPV | ASSTIANTEVF | 0.77196634 |
| GYRQVTDAHLLTLARR | ASSTIANTEVF | 0.275167 |
| IAKMGQDGHDRG | ASSTIANTEVF | 0.3523206 |
| KAEPSIEIQYT | ASSTIANTEVF | 0.33771157 |
| KPVRIRMAADEDTVAVT | ASSTIANTEVF | 0.5021174 |
| MPPRIAGMRLLVIKPEP | ASSTIANTEVF | 0.74446476 |
| MKFYAVALFRSRGVPPTR | ASSTIANTEVF | 0.5480075 |
| NAGGFGILGAVAHSP | ASSTIANTEVF | 0.2713869 |
| NEGREITLAMAEM | ASSTIANTEVF | 0.48455447 |
| NELVEWSHPLGHRWRWE | ASSTIANTEVF | 0.26816982 |
| NLGAGGTGGNGGNPG | ASSTIANTEVF | 0.32126504 |
| NTEWPKWETVMAADPVD | ASSTIANTEVF | 0.47599065 |
| PASAAPAETQP | ASSTIANTEVF | 0.36245885 |
| QQSSDGKNGVRAAEITGE | ASSTIANTEVF | 0.27495408 |
| RAPDAVTTSWWKE | ASSTIANTEVF | 0.30735597 |
| SGFGNVGAHTSGWFNQ | ASSTIANTEVF | 0.37449124 |
| SGVAGAVGQAASVGGLKV | ASSTIANTEVF | 0.43576306 |
| SNGGAGTGSAGGNGGT | ASSTIANTEVF | 0.43044823 |
| TGGAGGEGGAAG | ASSTIANTEVF | 0.6755514 |
| TGGDGGAAGAGGAPGHG | ASSTIANTEVF | 0.5626007 |
| TMPPLLAAIANAGRHGV | ASSTIANTEVF | 0.59734666 |
| TSSTAGNSGGA | ASSTIANTEVF | 0.1772949 |
| VPATVPVGILC | ASSTIANTEVF | 0.2353794 |
| VQRPDSGGVVLTGRIS | ASSTIANTEVF | 0.25877634 |
| WTPNPVPATKGVPAGLVQAT | ASSTIANTEVF | 0.3458785 |
| YALKRALAKFSDRFDVVII | ASSTIANTEVF | 0.19925688 |
| YLIGSVIV | ASSTIANTEVF | 0.4678107 |
| YGQLVAVID | ASSTIANTEVF | 0.29337612 |
| VTVGGMTLPA | ASSTIANTEVF | 0.30980843 |
| VLALPAV | ASSTIANTEVF | 0.5764293 |
| VIPPTLNYET | ASSTIANTEVF | 0.2689417 |
| VGMGGMHPSSG | ASSTIANTEVF | 0.26118132 |
| VGLAGGAATA | ASSTIANTEVF | 0.23098381 |
| VGGAGGAGGSGP | ASSTIANTEVF | 0.41709387 |

| | | |
|---|---|---|
| VFAAFMV | ASSTIANTEVF | 0.21324039 |
| TLYDKLR | ASSTIANTEVF | 0.51880705 |
| TLAVFTVL | ASSTIANTEVF | 0.25743636 |
| TGSVQPARV | ASSTIANTEVF | 0.5464333 |
| TGGGAGSGGT | ASSTIANTEVF | 0.259194 |
| SHLETSL | ASSTIANTEVF | 0.562533 |
| SFTTGMT | ASSTIANTEVF | 0.36435986 |
| QVEVAKRMGVRQPTVS | ASSTIANTEVF | 0.7970088 |
| QGSGGPALAM | ASSTIANTEVF | 0.41922957 |
| MLAPSNP | ASSTIANTEVF | 0.33329156 |
| LVNNAAI | ASSTIANTEVF | 0.23240806 |
| LTGEINTLA | ASSTIANTEVF | 0.76751643 |
| LSIFGPV | ASSTIANTEVF | 0.21381323 |
| LLALLANL | ASSTIANTEVF | 0.3105998 |
| LISNASDALDKLR | ASSTIANTEVF | 0.23095287 |
| LINEAVP | ASSTIANTEVF | 0.43773302 |
| LGYAFSGL | ASSTIANTEVF | 0.5144259 |
| LDFVAVAL | ASSTIANTEVF | 0.18939124 |
| LAQPGAE | ASSTIANTEVF | 0.49739432 |
| ISNLGAVTGF | ASSTIANTEVF | 0.432267 |
| HQEDPGRI | ASSTIANTEVF | 0.2378872 |
| GTPGEPTP | ASSTIANTEVF | 0.681013 |
| GFFNSGS | ASSTIANTEVF | 0.38246867 |
| GAGGGGLTAGAN | ASSTIANTEVF | 0.69675744 |
| FVDALVNM | ASSTIANTEVF | 0.29940233 |
| FLTVAPEL | ASSTIANTEVF | 0.27219003 |
| FETLTAL | ASSTIANTEVF | 0.37419242 |
| DVINPIEL | ASSTIANTEVF | 0.21740565 |
| APVSGAPV | ASSTIANTEVF | 0.2166947 |
| ALVGGLAW | ASSTIANTEVF | 0.6503484 |
| ALAAAAATAI | ASSTIANTEVF | 0.74660957 |
| AFGLGLLL | ASSTIANTEVF | 0.7463409 |
| AFFAAASL | ASSTIANTEVF | 0.6887418 |
| AAAALPAFSPP | ASSTIANTEVF | 0.6965883 |
| VVLAPAL | ASSTIANTEVF | 0.7265148 |
| TIGHVDHGKTTL | ASSTIANTEVF | 0.74969256 |
| LISNASDALDK | ASSTIANTEVF | 0.75719756 |
| LISNASDAL | ASSTIANTEVF | 0.75224257 |
| ISNASDALDK | ASSTIANTEVF | 0.71595675 |
| NQAETLVYQTE | ASSTIANTEVF | 0.7500782 |
| TGGTGGTAGAG | ASSTIANTEVF | 0.7332319 |
| TGSTAGGAGGA | ASSTIANTEVF | 0.7358624 |
| DPLAVPVAAGPV | ASSTIANTEVF | 0.745201 |
| GGTGGTPGNGGN | ASSTIANTEVF | 0.73592526 |
| PAGLAGVDPATL | ASSTIANTEVF | 0.7143845 |
| GNNLVVEAPHAGASV | ASSTIANTEVF | 0.6873481 |
| KVTRIMSASTLEQPA | ASSTIANTEVF | 0.7455596 |
| LGFALPIALPRRTVV | ASSTIANTEVF | 0.7543703 |
| TRLGNLTARKLVVLV | ASSTIANTEVF | 0.7507322 |
| WSQQRRLIYFPSAGP | ASSTIANTEVF | 0.7595822 |
| GPNGNPDPMAAAVDIR | ASSTIANTEVF | 0.75032 |

| | | |
|---|---|---|
| TQRCVSVADLRAAASR | ASSTIANTEVF | 0.74956465 |
| QFVLDVAEHPVEVLTETV | ASSTIANTEVF | 0.407682 |
| GGAGGTGGAGGNGGTQAGDG | ASSTIANTEVF | 0.60031927 |
| PLLATLRAIAVDVDAKPAQV | ASSTIANTEVF | 0.7108824 |
| APKFRTGGYMIDFWGVGYQV | ASSTIANTEVF | 0.7210764 |
| GAGGAGGAGGAAGGPGGTGG | ASSTIANTEVF | 0.39556134 |
| IGALGLLATDVAPAGSVAFG | ASSTIANTEVF | 0.7503885 |
| AGGAGGIGGNGGAGGAGGVL | ASSTIANTEVF | 0.72928977 |
| ADELIGLL | ASSTIANTEVF | 0.7101649 |
| ADRALKA | ASSTIANTEVF | 0.75251806 |
| AGAVGQVGGAGGA | ASSTIANTEVF | 0.7254181 |
| AGGGGGLGGAP | ASSTIANTEVF | 0.6385442 |
| AGGLLYGNGGAG | ASSTIANTEVF | 0.7120238 |
| AGLESIGL | ASSTIANTEVF | 0.7576982 |
| AGLKTGII | ASSTIANTEVF | 0.7049326 |
| ASLAALLL | ASSTIANTEVF | 0.7552258 |
| ASVINISSI | ASSTIANTEVF | 0.70112765 |
| ATPPGPA | ASSTIANTEVF | 0.75508034 |
| ATVASALV | ASSTIANTEVF | 0.66963506 |
| ATVASLIL | ASSTIANTEVF | 0.75310075 |
| AVNGPVLIRH | ASSTIANTEVF | 0.7363437 |
| DGAAGAVGI | ASSTIANTEVF | 0.7609941 |
| DPLLHGI | ASSTIANTEVF | 0.7272823 |
| ELGAALGAL | ASSTIANTEVF | 0.7540735 |
| ELGRTQAV | ASSTIANTEVF | 0.742416 |
| FVAGGYLS | ASSTIANTEVF | 0.75066996 |
| GASGGAGGAGGW | ASSTIANTEVF | 0.6735739 |
| GGAGAGGDGGAGGI | ASSTIANTEVF | 0.7201698 |
| GTGHNGSQPGMGGNG | ASSTIANTEVF | 0.6554604 |
| GYAAASASAS | ASSTIANTEVF | 0.7306175 |
| HQVDVAAVQQG | ASSTIANTEVF | 0.7528108 |
| IGAAIAHT | ASSTIANTEVF | 0.31763506 |
| IGMIEVAQL | ASSTIANTEVF | 0.74626315 |
| IRLAPEHTSAMD | ASSTIANTEVF | 0.657228 |
| IYNVSVL | ASSTIANTEVF | 0.7051512 |
| KQGFLGM | ASSTIANTEVF | 0.75048745 |
| LESVITQL | ASSTIANTEVF | 0.7343826 |
| LGLGALVVV | ASSTIANTEVF | 0.74670374 |
| LIGNGGNGGSGGI | ASSTIANTEVF | 0.7028083 |
| LNAINAPTL | ASSTIANTEVF | 0.7256992 |
| LQLLNTGV | ASSTIANTEVF | 0.7411357 |
| LQTSIEF | ASSTIANTEVF | 0.40694577 |
| MSFAVSEL | ASSTIANTEVF | 0.67862463 |
| NTGALGL | ASSTIANTEVF | 0.7579284 |
| PPGAGGAGGAGTT | ASSTIANTEVF | 0.6672772 |
| QLVGLGI | ASSTIANTEVF | 0.67985743 |
| RFLEQVHQL | ASSTIANTEVF | 0.6222793 |
| RGVGYVLR | ASSTIANTEVF | 0.620692 |
| SATGVFAGVI | ASSTIANTEVF | 0.6492667 |
| SLPAITLP | ASSTIANTEVF | 0.6782114 |
| STALAFTV | ASSTIANTEVF | 0.68460953 |

| | | |
|---|---|---|
| SVLTVGLGK | ASSTIANTEVF | 0.67181337 |
| TALAGAGSGI | ASSTIANTEVF | 0.6371008 |
| TGLPFTGI | ASSTIANTEVF | 0.6725698 |
| VAEHVTV | ASSTIANTEVF | 0.6544353 |
| VDIVLNSL | ASSTIANTEVF | 0.6506624 |
| VLTGGGAVL | ASSTIANTEVF | 0.67323166 |
| VNVGLAYGI | ASSTIANTEVF | 0.6672802 |
| VSSAAVGAV | ASSTIANTEVF | 0.64563906 |
| VTSVYEAGV | ASSTIANTEVF | 0.649548 |
| WVALDRGA | ASSTIANTEVF | 0.6717973 |
| YLVGSVVI | ASSTIANTEVF | 0.67607415 |
| GPPGTGKTLL | ASSTIANTEVF | 0.67322046 |
| GVGMTKF | ASSTIANTEVF | 0.6864633 |
| IGIGGSDL | ASSTIANTEVF | 0.67404723 |
| IGTLLAEL | ASSTIANTEVF | 0.67209625 |
| IYTSGTT | ASSTIANTEVF | 0.45734295 |
| LSFPAGV | ASSTIANTEVF | 0.57467353 |
| LYTSGTT | ASSTIANTEVF | 0.65591943 |
| VGAFPFGF | ASSTIANTEVF | 0.655619 |
| VGAGTVEF | ASSTIANTEVF | 0.5140811 |
| VGKSTLLL | ASSTIANTEVF | 0.67806077 |
| AALAHAGGGIV | ASSTIANTEVF | 0.65304905 |
| AALVAVSLL | ASSTIANTEVF | 0.63615847 |
| AAVSRAVAAL | ASSTIANTEVF | 0.67517316 |
| AGGSSGIGGF | ASSTIANTEVF | 0.6461792 |
| AGLTVANANG | ASSTIANTEVF | 0.6407046 |
| AGLVGAVL | ASSTIANTEVF | 0.6558019 |
| ALSQAFTV | ASSTIANTEVF | 0.6839869 |
| ALTFSVSL | ASSTIANTEVF | 0.62556785 |
| ALTGAALVAL | ASSTIANTEVF | 0.67646646 |
| ASFSIGAGVL | ASSTIANTEVF | 0.65667003 |
| DGSVYKI | ASSTIANTEVF | 0.6812333 |
| EQLAEIV | ASSTIANTEVF | 0.60088503 |
| FAVSELTVM | ASSTIANTEVF | 0.67105985 |
| GAAPLAAGL | ASSTIANTEVF | 0.64810866 |
| GADGGDGGSGGAGG | ASSTIANTEVF | 0.68876183 |
| GFGLDTGGAGG | ASSTIANTEVF | 0.65072393 |
| GLASGGLL | ASSTIANTEVF | 0.67635685 |
| GRIAGAAVLL | ASSTIANTEVF | 0.6686554 |
| GSAGSLNTSF | ASSTIANTEVF | 0.68165517 |
| GTGQIGI | ASSTIANTEVF | 0.587806 |
| GTLIALIL | ASSTIANTEVF | 0.6557 |
| IAATMGFVVG | ASSTIANTEVF | 0.6297242 |
| IFAQALL | ASSTIANTEVF | 0.66797644 |
| IGAAVALGF | ASSTIANTEVF | 0.66690195 |
| IGLTPLAL | ASSTIANTEVF | 0.48865214 |
| ILGSNTSTL | ASSTIANTEVF | 0.68010604 |
| IVFSLTVL | ASSTIANTEVF | 0.6229404 |
| LASKATGF | ASSTIANTEVF | 0.6366501 |
| LASVALIL | ASSTIANTEVF | 0.67805564 |
| LFAVGLTL | ASSTIANTEVF | 0.6575233 |

| | | |
|---|---|---|
| LFNGGIGTYI | ASSTIANTEVF | 0.6744503 |
| LTLGAGII | ASSTIANTEVF | 0.65768325 |
| LVAAALGDI | ASSTIANTEVF | 0.66426873 |
| NTAFVQLGI | ASSTIANTEVF | 0.6733747 |
| QLFSGAEL | ASSTIANTEVF | 0.5791364 |
| SAATFITL | ASSTIANTEVF | 0.637828 |
| SNYGLANLGV | ASSTIANTEVF | 0.67907107 |
| SYGGAYAVM | ASSTIANTEVF | 0.79131913 |
| TTGVGGLTLG | ASSTIANTEVF | 0.7720578 |
| TTQTIYI | ASSTIANTEVF | 0.6204008 |
| VAAGAGLGAVI | ASSTIANTEVF | 0.5574992 |
| VAAVVTLL | ASSTIANTEVF | 0.7254506 |
| VGFLGGLI | ASSTIANTEVF | 0.78055096 |
| VGVIPILL | ASSTIANTEVF | 0.825023 |
| VIFASSLI | ASSTIANTEVF | 0.772128 |
| VKGGDGGAAAG | ASSTIANTEVF | 0.5716733 |
| VTKTFTNI | ASSTIANTEVF | 0.80122334 |
| VVAAAILL | ASSTIANTEVF | 0.7346301 |
| YAALDVEL | ASSTIANTEVF | 0.75747335 |
| LASVALLL | ASSTIANTEVF | 0.74863625 |
| LDAVHRV | ASSTIANTEVF | 0.7173334 |
| QGLSSSIF | ASSTIANTEVF | 0.66643786 |
| RGTFANIRL | ASSTIANTEVF | 0.5761295 |
| RQLWWGHRI | ASSTIANTEVF | 0.7933316 |
| YGPPGTGKTLL | ASSTIANTEVF | 0.8093959 |
| AALVGAPANVA | ASSTIANTEVF | 0.8008929 |
| AATFRRTE | ASSTIANTEVF | 0.8439351 |
| AGALGGGI | ASSTIANTEVF | 0.82207465 |
| AGGAGGAGGIGAG | ASSTIANTEVF | 0.7902186 |
| AGSGALGL | ASSTIANTEVF | 0.32243693 |
| AQIADQI | ASSTIANTEVF | 0.4772978 |
| ASAVAVTYL | ASSTIANTEVF | 0.6124774 |
| ASTVLGGTGG | ASSTIANTEVF | 0.70240366 |
| AVGGEPAK | ASSTIANTEVF | 0.3263825 |
| CEQGTFGGTC | ASSTIANTEVF | 0.7710079 |
| DEGKAPAG | ASSTIANTEVF | 0.67513835 |
| DTVVFPAL | ASSTIANTEVF | 0.6310589 |
| FEGITMAG | ASSTIANTEVF | 0.82420355 |
| FPLVVQD | ASSTIANTEVF | 0.6744733 |
| GAFTTPQ | ASSTIANTEVF | 0.5625277 |
| GFGGFGTAGGVGGA | ASSTIANTEVF | 0.6989149 |
| GFLAQFL | ASSTIANTEVF | 0.842948 |
| GGAGGAGGNAGVA | ASSTIANTEVF | 0.57524765 |
| GGVGTAGAGGAGGAGG | ASSTIANTEVF | 0.8194361 |
| GNGGGDAGNAGSGGN | ASSTIANTEVF | 0.72286665 |
| GTGGAGGAGNDAGS | ASSTIANTEVF | 0.83979744 |
| GTTNTGWFNS | ASSTIANTEVF | 0.46639508 |
| IFMGPLFTL | ASSTIANTEVF | 0.8222829 |
| ILAHAIEAI | ASSTIANTEVF | 0.7377281 |
| IRSAIREAL | ASSTIANTEVF | 0.85573536 |
| ITFAGGG | ASSTIANTEVF | 0.70422214 |

| | | |
|---|---|---|
| ITGMSGHQEV | ASSTIANTEVF | 0.778423 |
| IVAGLACTL | ASSTIANTEVF | 0.7281841 |
| IVGGGLAGA | ASSTIANTEVF | 0.8088708 |
| IVTGVLNI | ASSTIANTEVF | 0.47316542 |
| LAQQGCI | ASSTIANTEVF | 0.6934896 |
| LCLHVEN | ASSTIANTEVF | 0.5753092 |
| LDAISDGS | ASSTIANTEVF | 0.74197996 |
| LGGFAAML | ASSTIANTEVF | 0.79816014 |
| LGGKNPMI | ASSTIANTEVF | 0.23578708 |
| LGPKGVAAIL | ASSTIANTEVF | 0.80676347 |
| LMVGVSI | ASSTIANTEVF | 0.4653826 |
| LVGAAFVL | ASSTIANTEVF | 0.5077422 |
| LVLAVAPL | ASSTIANTEVF | 0.7910328 |
| MAAIAVIDAA | ASSTIANTEVF | 0.7484534 |
| NNGINAELQ | ASSTIANTEVF | 0.81036466 |
| RGEPLVLVHG | ASSTIANTEVF | 0.65339106 |
| RGGAGGAAGLI | ASSTIANTEVF | 0.74894667 |
| RSYRMHPAV | ASSTIANTEVF | 0.77438986 |
| SAAHLLAGQLA | ASSTIANTEVF | 0.4077735 |
| SAAVGSY | ASSTIANTEVF | 0.61478025 |
| SPHAVLHRI | ASSTIANTEVF | 0.8353093 |
| SQIAKVL | ASSTIANTEVF | 0.7961829 |
| TALGHLVP | ASSTIANTEVF | 0.79502964 |
| TAPTTSGAA | ASSTIANTEVF | 0.6823118 |
| TDGDDGNGGSPGT | ASSTIANTEVF | 0.5476958 |
| TDGLGEN | ASSTIANTEVF | 0.7187421 |
| TGGPPLPGQA | ASSTIANTEVF | 0.8250203 |
| TIPAITI | ASSTIANTEVF | 0.82301867 |
| TIPGFTI | ASSTIANTEVF | 0.7758182 |
| TSIASGALA | ASSTIANTEVF | 0.49985576 |
| TVTLPGAAI | ASSTIANTEVF | 0.7959473 |
| VASGLSAALP | ASSTIANTEVF | 0.71569777 |
| VGAGVIGI | ASSTIANTEVF | 0.6987404 |
| VGPKRFGV | ASSTIANTEVF | 0.80845964 |
| VNVAFPD | ASSTIANTEVF | 0.7236665 |
| VSMTNSGVS | ASSTIANTEVF | 0.72758555 |
| VTDVLGDLSV | ASSTIANTEVF | 0.6888564 |
| VVVVGAGFAGL | ASSTIANTEVF | 0.7648817 |
| VYTVGTVV | ASSTIANTEVF | 0.809034 |
| WTTFSGP | ASSTIANTEVF | 0.80913436 |
| YDELVEAF | ASSTIANTEVF | 0.87148786 |
| YGTGGAGGNGG | ASSTIANTEVF | 0.8421235 |
| YTNVMAV | ASSTIANTEVF | 0.86255467 |
| FLEQVHQL | ASSTIANTEVF | 0.26591858 |
| GLSLLAVAV | ASSTIANTEVF | 0.4690276 |
| LEQVHQL | ASSTIANTEVF | 0.6890181 |
| LNAAGAI | ASSTIANTEVF | 0.69480324 |
| PAVHASL | ASSTIANTEVF | 0.42428568 |
| TELEDDA | ASSTIANTEVF | 0.746971 |
| TPPGPPN | ASSTIANTEVF | 0.73378307 |
| TPPPPPPP | ASSTIANTEVF | 0.5431036 |

| | | |
|---|---|---|
| TQSPALI | ASSQASTGGHEQY | 0.8199889 |
| EFAAAFPQ | ASSQASTGGHEQY | 0.6602068 |
| LLSGNLDVL | ASSQASTGGHEQY | 0.59618974 |
| GPANLGNYNI | ASSQASTGGHEQY | 0.6891992 |
| TLGPTLP | ASSQASTGGHEQY | 0.82423806 |
| VYAYSSCCK | ASSQASTGGHEQY | 0.52927816 |
| APAPAPAPNI | ASSQASTGGHEQY | 0.79112554 |
| PGGPVPIP | ASSQASTGGHEQY | 0.67108583 |
| FLTVALEP | ASSQASTGGHEQY | 0.84701955 |
| SQYFGANL | ASSQASTGGHEQY | 0.41305476 |
| GGVKPAA | ASSQASTGGHEQY | 0.8225516 |
| GGTGGAGGAGGW | ASSQASTGGHEQY | 0.6966144 |
| GVGGVAGLGGAGG | ASSQASTGGHEQY | 0.87956715 |
| VLLVLATTI | ASSQASTGGHEQY | 0.70528734 |
| IGIFAAML | ASSQASTGGHEQY | 0.73463845 |
| AVGDLID | ASSQASTGGHEQY | 0.6769006 |
| NISPLPL | ASSQASTGGHEQY | 0.82904565 |
| VTAVFVAI | ASSQASTGGHEQY | 0.35187823 |
| SGDDNDAAL | ASSQASTGGHEQY | 0.7886294 |
| AVAALAHS | ASSQASTGGHEQY | 0.5175043 |
| GGVTVMATP | ASSQASTGGHEQY | 0.8086839 |
| AGLAANVRD | ASSQASTGGHEQY | 0.7916343 |
| TNRPDLIDPALLRPG | ASSQASTGGHEQY | 0.34256607 |
| LANLGVTAL | ASSQASTGGHEQY | 0.8329185 |
| TGERIEL | ASSQASTGGHEQY | 0.6567317 |
| NGEELAA | ASSQASTGGHEQY | 0.5683829 |
| VLSLVWG | ASSQASTGGHEQY | 0.85484254 |
| IGYALWL | ASSQASTGGHEQY | 0.7275549 |
| LSLLLAPL | ASSQASTGGHEQY | 0.8654715 |
| TLDEALK | ASSQASTGGHEQY | 0.7569254 |
| TDGAGTAEGG | ASSQASTGGHEQY | 0.71214557 |
| NAPGGAGGNGG | ASSQASTGGHEQY | 0.81001437 |
| VSVFMGMGGNF | ASSQASTGGHEQY | 0.54174435 |
| LASYLAAL | ASSQASTGGHEQY | 0.6335303 |
| LVGGTPAL | ASSQASTGGHEQY | 0.8402709 |
| VRNADVL | ASSQASTGGHEQY | 0.86025536 |
| TCSSPFLVSAM | ASSQASTGGHEQY | 0.8655181 |
| VHITDAE | ASSQASTGGHEQY | 0.7756722 |
| EYAAPSF | ASSQASTGGHEQY | 0.7150774 |
| VVDAFHGGL | ASSQASTGGHEQY | 0.8079818 |
| ASVVDPQ | ASSQASTGGHEQY | 0.86822397 |
| SYNEFGGLS | ASSQASTGGHEQY | 0.8778027 |
| DPDGSVY | ASSQASTGGHEQY | 0.85976505 |
| MVSTIHATE | ASSQASTGGHEQY | 0.71767336 |
| LAAGVADPV | ASSQASTGGHEQY | 0.8443054 |
| TKEQIAA | ASSQASTGGHEQY | 0.82183266 |
| SLASPAAL | ASSQASTGGHEQY | 0.811091 |
| IGLVFYGV | ASSQASTGGHEQY | 0.865373 |
| FLLDASNL | ASSQASTGGHEQY | 0.86158895 |
| YQVGNFP | ASSQASTGGHEQY | 0.8122317 |
| ESLLSQST | ASSQASTGGHEQY | 0.7633674 |

| | | |
|---|---|---|
| LFTVGFAAVG | ASSQASTGGHEQY | 0.8643469 |
| ATPPTPT | ASSQASTGGHEQY | 0.8704823 |
| TDLATLG | ASSQASTGGHEQY | 0.86748713 |
| SAGGTGR | ASSQASTGGHEQY | 0.8804605 |
| GFSNSGS | ASSQASTGGHEQY | 0.8672186 |
| GGNGGSGGTSV | ASSQASTGGHEQY | 0.86121583 |
| GTSVATGGAGN | ASSQASTGGHEQY | 0.30748367 |
| NGGNGGSGGTS | ASSQASTGGHEQY | 0.56629723 |
| AVVKFPH | ASSQASTGGHEQY | 0.8270975 |
| TPPPPPP | ASSQASTGGHEQY | 0.8078559 |
| AAPAAAAAPA | ASSQASTGGHEQY | 0.561967 |
| IALNEDL | ASSQASTGGHEQY | 0.84749377 |
| IVNVSSM | ASSQASTGGHEQY | 0.80967325 |
| TGIPPAPR | ASSQASTGGHEQY | 0.75184405 |
| FDLGGGTF | ASSQASTGGHEQY | 0.8671929 |
| FDLGGGTFDVS | ASSQASTGGHEQY | 0.7927839 |
| NEPTAAA | ASSQASTGGHEQY | 0.736253 |
| NEPTAAALA | ASSQASTGGHEQY | 0.80298173 |
| GSIINISSI | ASSQASTGGHEQY | 0.87555826 |
| VAPGNAGTA | ASSQASTGGHEQY | 0.7674278 |
| TGGGASLE | ASSQASTGGHEQY | 0.8631962 |
| VAVYNHYKRI | ASSQASTGGHEQY | 0.84678876 |
| TDPAPATT | ASSQASTGGHEQY | 0.8716924 |
| TNRPDILDPALLRPG | ASSQASTGGHEQY | 0.62729186 |
| LTTTTTS | ASSQASTGGHEQY | 0.86378217 |
| PTGSGKTL | ASSQASTGGHEQY | 0.82021844 |
| TGSGKTL | ASSQASTGGHEQY | 0.8855477 |
| TIGHVDHGK | ASSQASTGGHEQY | 0.80036485 |
| TIGHVDHGKTT | ASSQASTGGHEQY | 0.8546009 |
| GHVDHGKTT | ASSQASTGGHEQY | 0.85807276 |
| AQFEHTLL | ASSQASTGGHEQY | 0.8687732 |
| IVNQDEL | ASSQASTGGHEQY | 0.6165578 |
| TNGSQFF | ASSQASTGGHEQY | 0.8135 |
| TGKGIVE | ASSQASTGGHEQY | 0.6397934 |
| LISNASDA | ASSQASTGGHEQY | 0.830586 |
| AELRTLGELTKIAKAHGAQ | ASSQASTGGHEQY | 0.8546495 |
| APNAVAAAEPSW | ASSQASTGGHEQY | 0.38216347 |
| APQVAGAATAVL | ASSQASTGGHEQY | 0.87507886 |
| ARRDPAAAAAILPTDA | ASSQASTGGHEQY | 0.7740599 |
| DAAARDVLAEAGLAEYF | ASSQASTGGHEQY | 0.77041376 |
| DDQIVLSAKANLAKVEQ | ASSQASTGGHEQY | 0.8724914 |
| DELLPDTGEDSDEDGH | ASSQASTGGHEQY | 0.85133016 |
| ELPEAPANRATVPQD | ASSQASTGGHEQY | 0.8642989 |
| FGQSKIERVTGIG | ASSQASTGGHEQY | 0.8445953 |
| GAGGTGSTAGGAGGAGGAGGI | ASSQASTGGHEQY | 0.84991145 |
| GHGGTGAAGGNGGTGGAGGS( | ASSQASTGGHEQY | 0.84785247 |
| GNGGNGGAGGEGGPGGAGGA( | ASSQASTGGHEQY | 0.7565439 |
| GSTAGNGGNGGNATNGTIAG( | ASSQASTGGHEQY | 0.7891512 |
| GTGGDGGNGGNGGTGG | ASSQASTGGHEQY | 0.86852014 |
| GVRRPAGIPADAPV | ASSQASTGGHEQY | 0.6439897 |
| GYRQVTDAHLLTLARR | ASSQASTGGHEQY | 0.83388543 |

| | | |
|---|---|---|
| IAKMGQDGHDRG | ASSQASTGGHEQY | 0.38124406 |
| KAEPSIEIQYT | ASSQASTGGHEQY | 0.28280818 |
| KPVRIRMAADEDTVAVT | ASSQASTGGHEQY | 0.49394906 |
| MPPRIAGMRLLVIKPEP | ASSQASTGGHEQY | 0.70678926 |
| MKFYAVALFRSRGVPPTR | ASSQASTGGHEQY | 0.85520124 |
| NAGGFGILGAVAHSP | ASSQASTGGHEQY | 0.7041836 |
| NEGREITLAMAEM | ASSQASTGGHEQY | 0.2883935 |
| NELVEWSHPLGHRWRWE | ASSQASTGGHEQY | 0.6672289 |
| NLGAGGTGGNGGNPG | ASSQASTGGHEQY | 0.51461804 |
| NTEWPKWETVMAADPVD | ASSQASTGGHEQY | 0.49179864 |
| PASAAPAETQP | ASSQASTGGHEQY | 0.6322382 |
| QQSSDGKNGVRAAEITGE | ASSQASTGGHEQY | 0.6229628 |
| RAPDAVTTSWWKE | ASSQASTGGHEQY | 0.41115755 |
| SGFGNVGAHTSGWFNQ | ASSQASTGGHEQY | 0.26748493 |
| SGVAGAVGQAASVGGLKV | ASSQASTGGHEQY | 0.7478827 |
| SNGGAGTGSAGGNGGT | ASSQASTGGHEQY | 0.7851982 |
| TGGAGGEGGAAG | ASSQASTGGHEQY | 0.7782149 |
| TGGDGGAAGAGGAPGHG | ASSQASTGGHEQY | 0.8091588 |
| TMPPLLAAIANAGRHGV | ASSQASTGGHEQY | 0.7609391 |
| TSSTAGNSGGA | ASSQASTGGHEQY | 0.6122097 |
| VPATVPVGILC | ASSQASTGGHEQY | 0.14335588 |
| VQRPDSGGVVLTGRIS | ASSQASTGGHEQY | 0.21550936 |
| WTPNPVPATKGVPAGLVQAT | ASSQASTGGHEQY | 0.44457132 |
| YALKRALAKFSDRFDVVII | ASSQASTGGHEQY | 0.52566457 |
| YLIGSVIV | ASSQASTGGHEQY | 0.17782156 |
| YGQLVAVID | ASSQASTGGHEQY | 0.696612 |
| VTVGGMTLPA | ASSQASTGGHEQY | 0.36021554 |
| VLALPAV | ASSQASTGGHEQY | 0.36397552 |
| VIPPTLNYET | ASSQASTGGHEQY | 0.7848809 |
| VGMGGMHPSSG | ASSQASTGGHEQY | 0.38781664 |
| VGLAGGAATA | ASSQASTGGHEQY | 0.2843398 |
| VGGAGGAGGSGP | ASSQASTGGHEQY | 0.4053073 |
| VFAAFMV | ASSQASTGGHEQY | 0.7541501 |
| TLYDKLR | ASSQASTGGHEQY | 0.23775701 |
| TLAVFTVL | ASSQASTGGHEQY | 0.7336119 |
| TGSVQPARV | ASSQASTGGHEQY | 0.5711044 |
| TGGGAGSGGT | ASSQASTGGHEQY | 0.76477504 |
| SHLETSL | ASSQASTGGHEQY | 0.22431727 |
| SFTTGMT | ASSQASTGGHEQY | 0.72308385 |
| QVEVAKRMGVRQPTVS | ASSQASTGGHEQY | 0.54779017 |
| QGSGGPALAM | ASSQASTGGHEQY | 0.8521786 |
| MLAPSNP | ASSQASTGGHEQY | 0.42163807 |
| LVNNAAI | ASSQASTGGHEQY | 0.63073736 |
| LTGEINTLA | ASSQASTGGHEQY | 0.6086462 |
| LSIFGPV | ASSQASTGGHEQY | 0.825752 |
| LLALLANL | ASSQASTGGHEQY | 0.2325927 |
| LISNASDALDKLR | ASSQASTGGHEQY | 0.42259508 |
| LINEAVP | ASSQASTGGHEQY | 0.27166742 |
| LGYAFSGL | ASSQASTGGHEQY | 0.46832573 |
| LDFVAVAL | ASSQASTGGHEQY | 0.6861027 |
| LAQPGAE | ASSQASTGGHEQY | 0.14835861 |

| | | |
|---|---|---|
| ISNLGAVTGF | ASSQASTGGHEQY | 0.6705941 |
| HQEDPGRI | ASSQASTGGHEQY | 0.41595477 |
| GTPGEPTP | ASSQASTGGHEQY | 0.2826699 |
| GFFNSGS | ASSQASTGGHEQY | 0.7230646 |
| GAGGGGLTAGAN | ASSQASTGGHEQY | 0.6369895 |
| FVDALVNM | ASSQASTGGHEQY | 0.74940157 |
| FLTVAPEL | ASSQASTGGHEQY | 0.46703953 |
| FETLTAL | ASSQASTGGHEQY | 0.6011752 |
| DVINPIEL | ASSQASTGGHEQY | 0.51313007 |
| APVSGAPV | ASSQASTGGHEQY | 0.4313265 |
| ALVGGLAW | ASSQASTGGHEQY | 0.45846397 |
| ALAAAAATAI | ASSQASTGGHEQY | 0.758286 |
| AFGLGLLL | ASSQASTGGHEQY | 0.66578066 |
| AFFAAASL | ASSQASTGGHEQY | 0.65710616 |
| AAAALPAFSPP | ASSQASTGGHEQY | 0.16396576 |
| VVLAPAL | ASSQASTGGHEQY | 0.19744183 |
| TIGHVDHGKTTL | ASSQASTGGHEQY | 0.2776993 |
| LISNASDALDK | ASSQASTGGHEQY | 0.48125952 |
| LISNASDAL | ASSQASTGGHEQY | 0.6748444 |
| ISNASDALDK | ASSQASTGGHEQY | 0.672265 |
| NQAETLVYQTE | ASSQASTGGHEQY | 0.21141098 |
| TGGTGGTAGAG | ASSQASTGGHEQY | 0.3171699 |
| TGSTAGGAGGA | ASSQASTGGHEQY | 0.27483445 |
| DPLAVPVAAGPV | ASSQASTGGHEQY | 0.30912167 |
| GGTGGTPGNGGN | ASSQASTGGHEQY | 0.38414556 |
| PAGLAGVDPATL | ASSQASTGGHEQY | 0.19594803 |
| GNNLVVEAPHAGASV | ASSQASTGGHEQY | 0.1849798 |
| KVTRIMSASTLEQPA | ASSQASTGGHEQY | 0.20920768 |
| LGFALPIALPRRTVV | ASSQASTGGHEQY | 0.24150158 |
| TRLGNLTARKLVVLV | ASSQASTGGHEQY | 0.64746296 |
| WSQQRRLIYFPSAGP | ASSQASTGGHEQY | 0.65497124 |
| GPNGNPDPMAAAVDIR | ASSQASTGGHEQY | 0.51493955 |
| TQRCVSVADLRAAASR | ASSQASTGGHEQY | 0.6407759 |
| QFVLDVAEHPVEVLTETV | ASSQASTGGHEQY | 0.40982282 |
| GGAGGTGGAGGNGGTQAGDG | ASSQASTGGHEQY | 0.11772815 |
| PLLATLRAIAVDVDAKPAQV | ASSQASTGGHEQY | 0.15444078 |
| APKFRTGGYMIDFWGVGYQV | ASSQASTGGHEQY | 0.20274214 |
| GAGGAGGAGGAAGGPGGTGG | ASSQASTGGHEQY | 0.2517476 |
| IGALGLLATDVAPAGSVAFG | ASSQASTGGHEQY | 0.12372327 |
| AGGAGGIGGNGGAGGAGGVL | ASSQASTGGHEQY | 0.36076042 |
| ADELIGLL | ASSQASTGGHEQY | 0.30795577 |
| ADRALKA | ASSQASTGGHEQY | 0.17883228 |
| AGAVGQVGGAGGA | ASSQASTGGHEQY | 0.5763649 |
| AGGGGGLGGAP | ASSQASTGGHEQY | 0.21582244 |
| AGGLLYGNGGAG | ASSQASTGGHEQY | 0.17405844 |
| AGLESIGL | ASSQASTGGHEQY | 0.2140066 |
| AGLKTGII | ASSQASTGGHEQY | 0.65591645 |
| ASLAALLL | ASSQASTGGHEQY | 0.19678013 |
| ASVINISSI | ASSQASTGGHEQY | 0.6134513 |
| ATPPGPA | ASSQASTGGHEQY | 0.26170066 |
| ATVASALV | ASSQASTGGHEQY | 0.5288796 |

| | | |
|---|---|---|
| ATVASLIL | ASSQASTGGHEQY | 0.16541703 |
| AVNGPVLIRH | ASSQASTGGHEQY | 0.5762807 |
| DGAAGAVGI | ASSQASTGGHEQY | 0.3735397 |
| DPLLHGI | ASSQASTGGHEQY | 0.74778545 |
| ELGAALGAL | ASSQASTGGHEQY | 0.28441155 |
| ELGRTQAV | ASSQASTGGHEQY | 0.41432285 |
| FVAGGYLS | ASSQASTGGHEQY | 0.17575073 |
| GASGGAGGAGGW | ASSQASTGGHEQY | 0.4058059 |
| GGAGAGGDGGAGGI | ASSQASTGGHEQY | 0.1470831 |
| GTGHNGSQPGMGGNG | ASSQASTGGHEQY | 0.28213367 |
| GYAAASASAS | ASSQASTGGHEQY | 0.15748182 |
| HQVDVAAVQQG | ASSQASTGGHEQY | 0.22886272 |
| IGAAIAHT | ASSQASTGGHEQY | 0.6340769 |
| IGMIEVAQL | ASSQASTGGHEQY | 0.12898263 |
| IRLAPEHTSAMD | ASSQASTGGHEQY | 0.5194086 |
| IYNVSVL | ASSQASTGGHEQY | 0.17451751 |
| KQGFLGM | ASSQASTGGHEQY | 0.20113397 |
| LESVITQL | ASSQASTGGHEQY | 0.4548526 |
| LGLGALVVV | ASSQASTGGHEQY | 0.18074651 |
| LIGNGGNGGSGGI | ASSQASTGGHEQY | 0.55920786 |
| LNAINAPTL | ASSQASTGGHEQY | 0.15201479 |
| LQLLNTGV | ASSQASTGGHEQY | 0.43103653 |
| LQTSIEF | ASSQASTGGHEQY | 0.33838862 |
| MSFAVSEL | ASSQASTGGHEQY | 0.1187674 |
| NTGALGL | ASSQASTGGHEQY | 0.18220648 |
| PPGAGGAGGAGTT | ASSQASTGGHEQY | 0.6595128 |
| QLVGLGI | ASSQASTGGHEQY | 0.2006039 |
| RFLEQVHQL | ASSQASTGGHEQY | 0.51453066 |
| RGVGYVLR | ASSQASTGGHEQY | 0.10030307 |
| SATGVFAGVI | ASSQASTGGHEQY | 0.11235358 |
| SLPAITLP | ASSQASTGGHEQY | 0.16913639 |
| STALAFTV | ASSQASTGGHEQY | 0.2589967 |
| SVLTVGLGK | ASSQASTGGHEQY | 0.14524943 |
| TALAGAGSGI | ASSQASTGGHEQY | 0.25721923 |
| TGLPFTGI | ASSQASTGGHEQY | 0.113047436 |
| VAEHVTV | ASSQASTGGHEQY | 0.10898444 |
| VDIVLNSL | ASSQASTGGHEQY | 0.10498274 |
| VLTGGGAVL | ASSQASTGGHEQY | 0.117401585 |
| VNVGLAYGI | ASSQASTGGHEQY | 0.17359164 |
| VSSAAVGAV | ASSQASTGGHEQY | 0.10261629 |
| VTSVYEAGV | ASSQASTGGHEQY | 0.10022671 |
| WVALDRGA | ASSQASTGGHEQY | 0.14226718 |
| YLVGSVVI | ASSQASTGGHEQY | 0.2105616 |
| GPPGTGKTLL | ASSQASTGGHEQY | 0.27259797 |
| GVGMTKF | ASSQASTGGHEQY | 0.23569444 |
| IGIGGSDL | ASSQASTGGHEQY | 0.42512003 |
| IGTLLAEL | ASSQASTGGHEQY | 0.20810404 |
| IYTSGTT | ASSQASTGGHEQY | 0.1458513 |
| LSFPAGV | ASSQASTGGHEQY | 0.08846388 |
| LYTSGTT | ASSQASTGGHEQY | 0.113387555 |
| VGAFPFGF | ASSQASTGGHEQY | 0.10206158 |

| | | |
|---|---|---|
| VGAGTVEF | ASSQASTGGHEQY | 0.14828254 |
| VGKSTLLL | ASSQASTGGHEQY | 0.085004486 |
| AALAHAGGGIV | ASSQASTGGHEQY | 0.13138106 |
| AALVAVSLL | ASSQASTGGHEQY | 0.14931352 |
| AAVSRAVAAL | ASSQASTGGHEQY | 0.093646586 |
| AGGSSGIGGF | ASSQASTGGHEQY | 0.243315 |
| AGLTVANANG | ASSQASTGGHEQY | 0.09548701 |
| AGLVGAVL | ASSQASTGGHEQY | 0.10706208 |
| ALSQAFTV | ASSQASTGGHEQY | 0.112718195 |
| ALTFSVSL | ASSQASTGGHEQY | 0.48062697 |
| ALTGAALVAL | ASSQASTGGHEQY | 0.10694203 |
| ASFSIGAGVL | ASSQASTGGHEQY | 0.21592495 |
| DGSVYKI | ASSQASTGGHEQY | 0.18882158 |
| EQLAEIV | ASSQASTGGHEQY | 0.15345523 |
| FAVSELTVM | ASSQASTGGHEQY | 0.0989684 |
| GAAPLAAGL | ASSQASTGGHEQY | 0.18127058 |
| GADGGDGGSGGAGG | ASSQASTGGHEQY | 0.12519008 |
| GFGLDTGGAGG | ASSQASTGGHEQY | 0.661234 |
| GLASGGLL | ASSQASTGGHEQY | 0.15343708 |
| GRIAGAAVLL | ASSQASTGGHEQY | 0.12759574 |
| GSAGSLNTSF | ASSQASTGGHEQY | 0.108194456 |
| GTGQIGI | ASSQASTGGHEQY | 0.105282374 |
| GTLIALIL | ASSQASTGGHEQY | 0.09019643 |
| IAATMGFVVG | ASSQASTGGHEQY | 0.15734129 |
| IFAQALL | ASSQASTGGHEQY | 0.13188353 |
| IGAAVALGF | ASSQASTGGHEQY | 0.1418171 |
| IGLTPLAL | ASSQASTGGHEQY | 0.19175819 |
| ILGSNTSTL | ASSQASTGGHEQY | 0.10587545 |
| IVFSLTVL | ASSQASTGGHEQY | 0.31900257 |
| LASKATGF | ASSQASTGGHEQY | 0.10997834 |
| LASVALIL | ASSQASTGGHEQY | 0.10629217 |
| LFAVGLTL | ASSQASTGGHEQY | 0.23884925 |
| LFNGGIGTYI | ASSQASTGGHEQY | 0.098727286 |
| LTLGAGII | ASSQASTGGHEQY | 0.4097014 |
| LVAAALGDI | ASSQASTGGHEQY | 0.09075022 |
| NTAFVQLGI | ASSQASTGGHEQY | 0.20186435 |
| QLFSGAEL | ASSQASTGGHEQY | 0.24644321 |
| SAATFITL | ASSQASTGGHEQY | 0.08705834 |
| SNYGLANLGV | ASSQASTGGHEQY | 0.098607734 |
| SYGGAYAVM | ASSQASTGGHEQY | 0.20374526 |
| TTGVGGLTLG | ASSQASTGGHEQY | 0.7056521 |
| TTQTIYI | ASSQASTGGHEQY | 0.7462895 |
| VAAGAGLGAVI | ASSQASTGGHEQY | 0.43909982 |
| VAAVVTLL | ASSQASTGGHEQY | 0.3883875 |
| VGFLGGLI | ASSQASTGGHEQY | 0.5444604 |
| VGVIPILL | ASSQASTGGHEQY | 0.71846217 |
| VIFASSLI | ASSQASTGGHEQY | 0.79344356 |
| VKGGDGGAAAG | ASSQASTGGHEQY | 0.71239823 |
| VTKTFTNI | ASSQASTGGHEQY | 0.45171466 |
| VVAAAILL | ASSQASTGGHEQY | 0.70281714 |
| YAALDVEL | ASSQASTGGHEQY | 0.6599009 |

| | | |
|---|---|---|
| LASVALLL | ASSQASTGGHEQY | 0.6399629 |
| LDAVHRV | ASSQASTGGHEQY | 0.642132 |
| QGLSSSIF | ASSQASTGGHEQY | 0.6794232 |
| RGTFANIRL | ASSQASTGGHEQY | 0.49714932 |
| RQLWWGHRI | ASSQASTGGHEQY | 0.43544734 |
| YGPPGTGKTLL | ASSQASTGGHEQY | 0.7098121 |
| AALVGAPANVA | ASSQASTGGHEQY | 0.7609026 |
| AATFRRTE | ASSQASTGGHEQY | 0.749568 |
| AGALGGGI | ASSQASTGGHEQY | 0.8065412 |
| AGGAGGAGGIGAG | ASSQASTGGHEQY | 0.76243174 |
| AGSGALGL | ASSQASTGGHEQY | 0.6514126 |
| AQIADQI | ASSQASTGGHEQY | 0.2277494 |
| ASAVAVTYL | ASSQASTGGHEQY | 0.31032544 |
| ASTVLGGTGG | ASSQASTGGHEQY | 0.54716384 |
| AVGGEPAK | ASSQASTGGHEQY | 0.5623941 |
| CEQGTFGGTC | ASSQASTGGHEQY | 0.20847987 |
| DEGKAPAG | ASSQASTGGHEQY | 0.6662041 |
| DTVVFPAL | ASSQASTGGHEQY | 0.5616425 |
| FEGITMAG | ASSQASTGGHEQY | 0.52477396 |
| FPLVVQD | ASSQASTGGHEQY | 0.77188045 |
| GAFTTPQ | ASSQASTGGHEQY | 0.535745 |
| GFGGFGTAGGVGGA | ASSQASTGGHEQY | 0.45632273 |
| GFLAQFL | ASSQASTGGHEQY | 0.56681633 |
| GGAGGAGGNAGVA | ASSQASTGGHEQY | 0.79244035 |
| GGVGTAGAGGAGGAGG | ASSQASTGGHEQY | 0.4328633 |
| GNGGGGDAGNAGSGGN | ASSQASTGGHEQY | 0.7869278 |
| GTGGAGGAGNDAGS | ASSQASTGGHEQY | 0.5547415 |
| GTTNTGWFNS | ASSQASTGGHEQY | 0.79886746 |
| IFMGPLFTL | ASSQASTGGHEQY | 0.3634207 |
| ILAHAIEAI | ASSQASTGGHEQY | 0.77227056 |
| IRSAIREAL | ASSQASTGGHEQY | 0.65472865 |
| ITFAGGG | ASSQASTGGHEQY | 0.84235054 |
| ITGMSGHQEV | ASSQASTGGHEQY | 0.5942682 |
| IVAGLACTL | ASSQASTGGHEQY | 0.7590094 |
| IVGGGLAGA | ASSQASTGGHEQY | 0.6213447 |
| IVTGVLNI | ASSQASTGGHEQY | 0.7639066 |
| LAQQGCI | ASSQASTGGHEQY | 0.45307815 |
| LCLHVEN | ASSQASTGGHEQY | 0.52759075 |
| LDAISDGS | ASSQASTGGHEQY | 0.3801447 |
| LGGFAAML | ASSQASTGGHEQY | 0.5491907 |
| LGGKNPMI | ASSQASTGGHEQY | 0.747188 |
| LGPKGVAAIL | ASSQASTGGHEQY | 0.23040915 |
| LMVGVSI | ASSQASTGGHEQY | 0.6913629 |
| LVGAAFVL | ASSQASTGGHEQY | 0.4416951 |
| LVLAVAPL | ASSQASTGGHEQY | 0.41521996 |
| MAAIAVIDAA | ASSQASTGGHEQY | 0.7015583 |
| NNGINAELQ | ASSQASTGGHEQY | 0.66599494 |
| RGEPLVLVHG | ASSQASTGGHEQY | 0.6751404 |
| RGGAGGAAGLI | ASSQASTGGHEQY | 0.46830073 |
| RSYRMHPAV | ASSQASTGGHEQY | 0.5737929 |
| SAAHLLAGQLA | ASSQASTGGHEQY | 0.62245566 |

| | | |
|---|---|---|
| SAAVGSY | ASSQASTGGHEQY | 0.33769774 |
| SPHAVLHRI | ASSQASTGGHEQY | 0.4739622 |
| SQIAKVL | ASSQASTGGHEQY | 0.7982183 |
| TALGHLVP | ASSQASTGGHEQY | 0.32161546 |
| TAPTTSGAA | ASSQASTGGHEQY | 0.793194 |
| TDGDDGNGGSPGT | ASSQASTGGHEQY | 0.20342363 |
| TDGLGEN | ASSQASTGGHEQY | 0.19581994 |
| TGGPPLPGQA | ASSQASTGGHEQY | 0.18326041 |
| TIPAITI | ASSQASTGGHEQY | 0.34419805 |
| TIPGFTI | ASSQASTGGHEQY | 0.7986528 |
| TSIASGALA | ASSQASTGGHEQY | 0.45351064 |
| TVTLPGAAI | ASSQASTGGHEQY | 0.1815479 |
| VASGLSAALP | ASSQASTGGHEQY | 0.30894 |
| VGAGVIGI | ASSQASTGGHEQY | 0.20549738 |
| VGPKRFGV | ASSQASTGGHEQY | 0.22051635 |
| VNVAFPD | ASSQASTGGHEQY | 0.26828197 |
| VSMTNSGVS | ASSQASTGGHEQY | 0.22941606 |
| VTDVLGDLSV | ASSQASTGGHEQY | 0.20533988 |
| VVVVGAGFAGL | ASSQASTGGHEQY | 0.19104369 |
| VYTVGTVV | ASSQASTGGHEQY | 0.19288331 |
| WTTFSGP | ASSQASTGGHEQY | 0.40322164 |
| YDELVEAF | ASSQASTGGHEQY | 0.3484763 |
| YGTGGAGGNGG | ASSQASTGGHEQY | 0.40278417 |
| YTNVMAV | ASSQASTGGHEQY | 0.3958213 |
| FLEQVHQL | ASSQASTGGHEQY | 0.25865388 |
| GLSLLAVAV | ASSQASTGGHEQY | 0.1406133 |
| LEQVHQL | ASSQASTGGHEQY | 0.16475259 |
| LNAAGAI | ASSQASTGGHEQY | 0.17923619 |
| PAVHASL | ASSQASTGGHEQY | 0.25378636 |
| TELEDDA | ASSQASTGGHEQY | 0.14139697 |
| TPPGPPN | ASSQASTGGHEQY | 0.33446613 |
| TPPPPPPP | ASSQASTGGHEQY | 0.2242401 |
| TQSPALI | ASSIQNTLY | 0.21304946 |
| EFAAAFPQ | ASSIQNTLY | 0.36438265 |
| LLSGNLDVL | ASSIQNTLY | 0.23159693 |
| GPANLGNYNI | ASSIQNTLY | 0.17653768 |
| TLGPTLP | ASSIQNTLY | 0.17619452 |
| VYAYSSCCK | ASSIQNTLY | 0.29967406 |
| APAPAPAPNI | ASSIQNTLY | 0.15191375 |
| PGGPVPIP | ASSIQNTLY | 0.36936432 |
| FLTVALEP | ASSIQNTLY | 0.1587043 |
| SQYFGANL | ASSIQNTLY | 0.25541347 |
| GGVKPAA | ASSIQNTLY | 0.16606028 |
| GGTGGAGGAGGW | ASSIQNTLY | 0.41842324 |
| GVGGVAGLGGAGG | ASSIQNTLY | 0.2448657 |
| VLLVLATTI | ASSIQNTLY | 0.5460245 |
| IGIFAAML | ASSIQNTLY | 0.21630608 |
| AVGDLID | ASSIQNTLY | 0.26480645 |
| NISPLPL | ASSIQNTLY | 0.194506 |
| VTAVFVAI | ASSIQNTLY | 0.7821009 |
| SGDDNDAAL | ASSIQNTLY | 0.1560242 |

| | | |
|---|---|---|
| AVAALAHS | ASSIQNTLY | 0.19696511 |
| GGVTVMATP | ASSIQNTLY | 0.1617488 |
| AGLAANVRD | ASSIQNTLY | 0.23675671 |
| TNRPDLIDPALLRPG | ASSIQNTLY | 0.4315827 |
| LANLGVTAL | ASSIQNTLY | 0.18320438 |
| TGERIEL | ASSIQNTLY | 0.3136345 |
| NGEELAA | ASSIQNTLY | 0.28105462 |
| VLSLVWG | ASSIQNTLY | 0.1871022 |
| IGYALWL | ASSIQNTLY | 0.3406274 |
| LSLLLAPL | ASSIQNTLY | 0.2458147 |
| TLDEALK | ASSIQNTLY | 0.3445724 |
| TDGAGTAEGG | ASSIQNTLY | 0.17349896 |
| NAPGGAGGNGG | ASSIQNTLY | 0.21912381 |
| VSVFMGMGGNF | ASSIQNTLY | 0.24391231 |
| LASYLAAL | ASSIQNTLY | 0.2052235 |
| LVGGTPAL | ASSIQNTLY | 0.16825789 |
| VRNADVL | ASSIQNTLY | 0.5229678 |
| TCSSPFLVSAM | ASSIQNTLY | 0.8741358 |
| VHITDAE | ASSIQNTLY | 0.8785157 |
| EYAAPSF | ASSIQNTLY | 0.7613068 |
| VVDAFHGGL | ASSIQNTLY | 0.77042705 |
| ASVVDPQ | ASSIQNTLY | 0.8260385 |
| SYNEFGGLS | ASSIQNTLY | 0.88193595 |
| DPDGSVY | ASSIQNTLY | 0.89953065 |
| MVSTIHATE | ASSIQNTLY | 0.8759469 |
| LAAGVADPV | ASSIQNTLY | 0.81353974 |
| TKEQIAA | ASSIQNTLY | 0.88387465 |
| SLASPAAL | ASSIQNTLY | 0.85143733 |
| IGLVFYGV | ASSIQNTLY | 0.83888036 |
| FLLDASNL | ASSIQNTLY | 0.8746865 |
| YQVGNFP | ASSIQNTLY | 0.8673719 |
| ESLLSQST | ASSIQNTLY | 0.82177496 |
| LFTVGFAAVG | ASSIQNTLY | 0.8052095 |
| ATPPTPT | ASSIQNTLY | 0.872971 |
| TDLATLG | ASSIQNTLY | 0.8857272 |
| SAGGTGR | ASSIQNTLY | 0.87474597 |
| GFSNSGS | ASSIQNTLY | 0.902039 |
| GGNGGSGGTSV | ASSIQNTLY | 0.8833715 |
| GTSVATGGAGN | ASSIQNTLY | 0.87055385 |
| NGGNGGSGGTS | ASSIQNTLY | 0.47801515 |
| AVVKFPH | ASSIQNTLY | 0.6546848 |
| TPPPPPP | ASSIQNTLY | 0.8300097 |
| AAPAAAAAPA | ASSIQNTLY | 0.8238032 |
| IALNEDL | ASSIQNTLY | 0.40384024 |
| IVNVSSM | ASSIQNTLY | 0.8798643 |
| TGIPPAPR | ASSIQNTLY | 0.84542644 |
| FDLGGGTF | ASSIQNTLY | 0.7997391 |
| FDLGGGTFDVS | ASSIQNTLY | 0.8833835 |
| NEPTAAA | ASSIQNTLY | 0.8310185 |
| NEPTAAALA | ASSIQNTLY | 0.7751691 |
| GSIINISSI | ASSIQNTLY | 0.82837105 |

| | | |
|---|---|---|
| VAPGNAGTA | ASSIQNTLY | 0.89464784 |
| TGGGASLE | ASSIQNTLY | 0.7635571 |
| VAVYNHYKRI | ASSIQNTLY | 0.8870477 |
| TDPAPATT | ASSIQNTLY | 0.80505216 |
| TNRPDILDPALLRPG | ASSIQNTLY | 0.8920795 |
| LTTTTTS | ASSIQNTLY | 0.7602298 |
| PTGSGKTL | ASSIQNTLY | 0.8839022 |
| TGSGKTL | ASSIQNTLY | 0.84330845 |
| TIGHVDHGK | ASSIQNTLY | 0.9015944 |
| TIGHVDHGKTT | ASSIQNTLY | 0.8402114 |
| GHVDHGKTT | ASSIQNTLY | 0.8903055 |
| AQFEHTLL | ASSIQNTLY | 0.86666703 |
| IVNQDEL | ASSIQNTLY | 0.89023864 |
| TNGSQFF | ASSIQNTLY | 0.7448729 |
| TGKGIVE | ASSIQNTLY | 0.8297859 |
| LISNASDA | ASSIQNTLY | 0.7609676 |
| AELRTLGELTKIAKAHGAQ | ASSIQNTLY | 0.8549905 |
| APNAVAAAEPSW | ASSIQNTLY | 0.87947476 |
| APQVAGAATAVL | ASSIQNTLY | 0.3676181 |
| ARRDPAAAAAILPTDA | ASSIQNTLY | 0.8824762 |
| DAAARDVLAEAGLAEYF | ASSIQNTLY | 0.759396 |
| DDQIVLSAKANLAKVEQ | ASSIQNTLY | 0.80252653 |
| DELLPDTGEDSDEDGH | ASSIQNTLY | 0.8823235 |
| ELPEAPANRATVPQD | ASSIQNTLY | 0.85547984 |
| FGQSKIERVTGIG | ASSIQNTLY | 0.87701917 |
| GAGGTGSTAGGAGGAGGAGGI | ASSIQNTLY | 0.8306266 |
| GHGGTGAAGGNGGTGGAGGS( | ASSIQNTLY | 0.8384495 |
| GNGGNGGAGGEGGPGGAGGA( | ASSIQNTLY | 0.86471266 |
| GSTAGNGGNGGNATNGTIAG( | ASSIQNTLY | 0.45194376 |
| GTGGDGGNGGNGGTGG | ASSIQNTLY | 0.75055796 |
| GVRRPAGIPADAPV | ASSIQNTLY | 0.8967161 |
| GYRQVTDAHLLTLARR | ASSIQNTLY | 0.7268688 |
| IAKMGQDGHDRG | ASSIQNTLY | 0.7793846 |
| KAEPSIEIQYT | ASSIQNTLY | 0.5829827 |
| KPVRIRMAADEDTVAVT | ASSIQNTLY | 0.52745247 |
| MPPRIAGMRLLVIKPEP | ASSIQNTLY | 0.58993626 |
| MKFYAVALFRSRGVPPTR | ASSIQNTLY | 0.76058125 |
| NAGGFGILGAVAHSP | ASSIQNTLY | 0.7718819 |
| NEGREITLAMAEM | ASSIQNTLY | 0.7234616 |
| NELVEWSHPLGHRWRWE | ASSIQNTLY | 0.51461184 |
| NLGAGGTGGNGGNPG | ASSIQNTLY | 0.70431966 |
| NTEWPKWETVMAADPVD | ASSIQNTLY | 0.60913926 |
| PASAAPAETQP | ASSIQNTLY | 0.6278391 |
| QQSSDGKNGVRAAEITGE | ASSIQNTLY | 0.693637 |
| RAPDAVTTSWWKE | ASSIQNTLY | 0.7153457 |
| SGFGNVGAHTSGWFNQ | ASSIQNTLY | 0.6476097 |
| SGVAGAVGQAASVGGLKV | ASSIQNTLY | 0.566279 |
| SNGGAGTGSAGGNGGT | ASSIQNTLY | 0.67507374 |
| TGGAGGEGGAAG | ASSIQNTLY | 0.7568587 |
| TGGDGGAAGAGGAPGHG | ASSIQNTLY | 0.7366141 |
| TMPPLLAAIANAGRHGV | ASSIQNTLY | 0.819468 |

| | | |
|---|---|---|
| TSSTAGNSGGA | ASSIQNTLY | 0.7682513 |
| VPATVPVGILC | ASSIQNTLY | 0.74058676 |
| VQRPDSGGVVLTGRIS | ASSIQNTLY | 0.17435671 |
| WTPNPVPATKGVPAGLVQAT | ASSIQNTLY | 0.3992331 |
| YALKRALAKFSDRFDVVII | ASSIQNTLY | 0.56947356 |
| YLIGSVIV | ASSIQNTLY | 0.64159036 |
| YGQLVAVID | ASSIQNTLY | 0.23958756 |
| VTVGGMTLPA | ASSIQNTLY | 0.7149543 |
| VLALPAV | ASSIQNTLY | 0.62132514 |
| VIPPTLNYET | ASSIQNTLY | 0.54380673 |
| VGMGGMHPSSG | ASSIQNTLY | 0.7716714 |
| VGLAGGAATA | ASSIQNTLY | 0.63255274 |
| VGGAGGAGGSGP | ASSIQNTLY | 0.5561019 |
| VFAAFMV | ASSIQNTLY | 0.5406816 |
| TLYDKLR | ASSIQNTLY | 0.7799361 |
| TLAVFTVL | ASSIQNTLY | 0.41158006 |
| TGSVQPARV | ASSIQNTLY | 0.7544342 |
| TGGGAGSGGT | ASSIQNTLY | 0.45251286 |
| SHLETSL | ASSIQNTLY | 0.7797026 |
| SFTTGMT | ASSIQNTLY | 0.4097754 |
| QVEVAKRMGVRQPTVS | ASSIQNTLY | 0.74550736 |
| QGSGGPALAM | ASSIQNTLY | 0.65277857 |
| MLAPSNP | ASSIQNTLY | 0.8531529 |
| LVNNAAI | ASSIQNTLY | 0.61415994 |
| LTGEINTLA | ASSIQNTLY | 0.70680726 |
| LSIFGPV | ASSIQNTLY | 0.5158629 |
| LLALLANL | ASSIQNTLY | 0.76019 |
| LISNASDALDKLR | ASSIQNTLY | 0.39665228 |
| LINEAVP | ASSIQNTLY | 0.6461696 |
| LGYAFSGL | ASSIQNTLY | 0.43591332 |
| LDFVAVAL | ASSIQNTLY | 0.7077106 |
| LAQPGAE | ASSIQNTLY | 0.72841 |
| ISNLGAVTGF | ASSIQNTLY | 0.2634146 |
| HQEDPGRI | ASSIQNTLY | 0.7483362 |
| GTPGEPTP | ASSIQNTLY | 0.5179297 |
| GFFNSGS | ASSIQNTLY | 0.4682864 |
| GAGGGGLTAGAN | ASSIQNTLY | 0.7859641 |
| FVDALVNM | ASSIQNTLY | 0.6068796 |
| FLTVAPEL | ASSIQNTLY | 0.80981934 |
| FETLTAL | ASSIQNTLY | 0.59261197 |
| DVINPIEL | ASSIQNTLY | 0.5957442 |
| APVSGAPV | ASSIQNTLY | 0.73300207 |
| ALVGGLAW | ASSIQNTLY | 0.4607775 |
| ALAAAAATAI | ASSIQNTLY | 0.48229393 |
| AFGLGLLL | ASSIQNTLY | 0.802834 |
| AFFAAASL | ASSIQNTLY | 0.7681709 |
| AAAALPAFSPP | ASSIQNTLY | 0.7750802 |
| VVLAPAL | ASSIQNTLY | 0.57466775 |
| TIGHVDHGKTTL | ASSIQNTLY | 0.57325464 |
| LISNASDALDK | ASSIQNTLY | 0.6994059 |
| LISNASDAL | ASSIQNTLY | 0.78260416 |

| | | |
|---|---|---|
| ISNASDALDK | ASSIQNTLY | 0.8172345 |
| NQAETLVYQTE | ASSIQNTLY | 0.740675 |
| TGGTGGTAGAG | ASSIQNTLY | 0.59235144 |
| TGSTAGGAGGA | ASSIQNTLY | 0.790807 |
| DPLAVPVAAGPV | ASSIQNTLY | 0.7340725 |
| GGTGGTPGNGGN | ASSIQNTLY | 0.66121924 |
| PAGLAGVDPATL | ASSIQNTLY | 0.77930796 |
| GNNLVVEAPHAGASV | ASSIQNTLY | 0.7525934 |
| KVTRIMSASTLEQPA | ASSIQNTLY | 0.6870193 |
| LGFALPIALPRRTVV | ASSIQNTLY | 0.66459113 |
| TRLGNLTARKLVVLV | ASSIQNTLY | 0.75555694 |
| WSQQRRLIYFPSAGP | ASSIQNTLY | 0.7738278 |
| GPNGNPDPMAAAVDIR | ASSIQNTLY | 0.76946867 |
| TQRCVSVADLRAAASR | ASSIQNTLY | 0.84683746 |
| QFVLDVAEHPVEVLTETV | ASSIQNTLY | 0.7925342 |
| GGAGGTGGAGGNGGTQAGDG | ASSIQNTLY | 0.7341102 |
| PLLATLRAIAVDVDAKPAQV | ASSIQNTLY | 0.28882882 |
| APKFRTGGYMIDFWGVGYQV[illegible] | ASSIQNTLY | 0.48912624 |
| GAGGAGGAGGAAGGPGGTGG[illegible] | ASSIQNTLY | 0.72685623 |
| IGALGLLATDVAPAGSVAFG[illegible] | ASSIQNTLY | 0.6898506 |
| AGGAGGIGGNGGAGGAGGVL[illegible] | ASSIQNTLY | 0.3239429 |
| ADELIGLL | ASSIQNTLY | 0.7858083 |
| ADRALKA | ASSIQNTLY | 0.7453991 |
| AGAVGQVGGAGGA | ASSIQNTLY | 0.64420164 |
| AGGGGGLGGAP | ASSIQNTLY | 0.79745334 |
| AGGLLYGNGGAG | ASSIQNTLY | 0.6684559 |
| AGLESIGL | ASSIQNTLY | 0.6951793 |
| AGLKTGII | ASSIQNTLY | 0.6789948 |
| ASLAALLL | ASSIQNTLY | 0.8390211 |
| ASVINISSI | ASSIQNTLY | 0.5932371 |
| ATPPGPA | ASSIQNTLY | 0.8008574 |
| ATVASALV | ASSIQNTLY | 0.7365403 |
| ATVASLIL | ASSIQNTLY | 0.8114154 |
| AVNGPVLIRH | ASSIQNTLY | 0.5512303 |
| DGAAGAVGI | ASSIQNTLY | 0.7779983 |
| DPLLHGI | ASSIQNTLY | 0.7056645 |
| ELGAALGAL | ASSIQNTLY | 0.8334791 |
| ELGRTQAV | ASSIQNTLY | 0.7178018 |
| FVAGGYLS | ASSIQNTLY | 0.79661006 |
| GASGGAGGAGGW | ASSIQNTLY | 0.69994247 |
| GGAGAGGDGGAGGI | ASSIQNTLY | 0.8220867 |
| GTGHNGSQPGMGGNG | ASSIQNTLY | 0.56236964 |
| GYAAASASAS | ASSIQNTLY | 0.73009974 |
| HQVDVAAVQQG | ASSIQNTLY | 0.672233 |
| IGAAIAHT | ASSIQNTLY | 0.7566663 |
| IGMIEVAQL | ASSIQNTLY | 0.7475418 |
| IRLAPEHTSAMD | ASSIQNTLY | 0.34144366 |
| IYNVSVL | ASSIQNTLY | 0.8349478 |
| KQGFLGM | ASSIQNTLY | 0.5702364 |
| LESVITQL | ASSIQNTLY | 0.63749367 |
| LGLGALVVV | ASSIQNTLY | 0.7881867 |

| | | |
|---|---|---|
| LIGNGGNGGSGGI | ASSIQNTLY | 0.71723044 |
| LNAINAPTL | ASSIQNTLY | 0.79969376 |
| LQLLNTGV | ASSIQNTLY | 0.69425166 |
| LQTSIEF | ASSIQNTLY | 0.768633 |
| MSFAVSEL | ASSIQNTLY | 0.77021885 |
| NTGALGL | ASSIQNTLY | 0.5041417 |
| PPGAGGAGGAGTT | ASSIQNTLY | 0.61011 |
| QLVGLGI | ASSIQNTLY | 0.8110774 |
| RFLEQVHQL | ASSIQNTLY | 0.7877562 |
| RGVGYVLR | ASSIQNTLY | 0.8685679 |
| SATGVFAGVI | ASSIQNTLY | 0.69230413 |
| SLPAITLP | ASSIQNTLY | 0.566386 |
| STALAFTV | ASSIQNTLY | 0.7433208 |
| SVLTVGLGK | ASSIQNTLY | 0.8502805 |
| TALAGAGSGI | ASSIQNTLY | 0.85549575 |
| TGLPFTGI | ASSIQNTLY | 0.8279853 |
| VAEHVTV | ASSIQNTLY | 0.5109592 |
| VDIVLNSL | ASSIQNTLY | 0.7383479 |
| VLTGGGAVL | ASSIQNTLY | 0.6746142 |
| VNVGLAYGI | ASSIQNTLY | 0.7528195 |
| VSSAAVGAV | ASSIQNTLY | 0.80594057 |
| VTSVYEAGV | ASSIQNTLY | 0.76051754 |
| WVALDRGA | ASSIQNTLY | 0.72650754 |
| YLVGSVVI | ASSIQNTLY | 0.63694656 |
| GPPGTGKTLL | ASSIQNTLY | 0.77136225 |
| GVGMTKF | ASSIQNTLY | 0.84054077 |
| IGIGGSDL | ASSIQNTLY | 0.81415546 |
| IGTLLAEL | ASSIQNTLY | 0.88193244 |
| IYTSGTT | ASSIQNTLY | 0.8441221 |
| LSFPAGV | ASSIQNTLY | 0.84748757 |
| LYTSGTT | ASSIQNTLY | 0.13978735 |
| VGAFPFGF | ASSIQNTLY | 0.30350035 |
| VGAGTVEF | ASSIQNTLY | 0.5427772 |
| VGKSTLLL | ASSIQNTLY | 0.70382357 |
| AALAHAGGGIV | ASSIQNTLY | 0.30607072 |
| AALVAVSLL | ASSIQNTLY | 0.8040174 |
| AAVSRAVAAL | ASSIQNTLY | 0.7029296 |
| AGGSSGIGGF | ASSIQNTLY | 0.59761006 |
| AGLTVANANG | ASSIQNTLY | 0.86348367 |
| AGLVGAVL | ASSIQNTLY | 0.6979214 |
| ALSQAFTV | ASSIQNTLY | 0.5906601 |
| ALTFSVSL | ASSIQNTLY | 0.59534013 |
| ALTGAALVAL | ASSIQNTLY | 0.86517656 |
| ASFSIGAGVL | ASSIQNTLY | 0.3955372 |
| DGSVYKI | ASSIQNTLY | 0.8262126 |
| EQLAEIV | ASSIQNTLY | 0.5812793 |
| FAVSELTVM | ASSIQNTLY | 0.8424485 |
| GAAPLAAGL | ASSIQNTLY | 0.41302383 |
| GADGGDGGSGGAGG | ASSIQNTLY | 0.8362671 |
| GFGLDTGGAGG | ASSIQNTLY | 0.7397441 |
| GLASGGLL | ASSIQNTLY | 0.90199566 |

| | | |
|---|---|---|
| GRIAGAAVLL | ASSIQNTLY | 0.6933809 |
| GSAGSLNTSF | ASSIQNTLY | 0.75250506 |
| GTGQIGI | ASSIQNTLY | 0.54425085 |
| GTLIALIL | ASSIQNTLY | 0.84379226 |
| IAATMGFVVG | ASSIQNTLY | 0.42977366 |
| IFAQALL | ASSIQNTLY | 0.7075467 |
| IGAAVALGF | ASSIQNTLY | 0.44235402 |
| IGLTPLAL | ASSIQNTLY | 0.80068237 |
| ILGSNTSTL | ASSIQNTLY | 0.8225772 |
| IVFSLTVL | ASSIQNTLY | 0.47816044 |
| LASKATGF | ASSIQNTLY | 0.86596346 |
| LASVALIL | ASSIQNTLY | 0.7839047 |
| LFAVGLTL | ASSIQNTLY | 0.61799735 |
| LFNGGIGTYI | ASSIQNTLY | 0.85863286 |
| LTLGAGII | ASSIQNTLY | 0.58965194 |
| LVAAALGDI | ASSIQNTLY | 0.85389066 |
| NTAFVQLGI | ASSIQNTLY | 0.6940665 |
| QLFSGAEL | ASSIQNTLY | 0.7190522 |
| SAATFITL | ASSIQNTLY | 0.85283095 |
| SNYGLANLGV | ASSIQNTLY | 0.479962 |
| SYGGAYAVM | ASSIQNTLY | 0.48990336 |
| TTGVGGLTLG | ASSIQNTLY | 0.85927665 |
| TTQTIYI | ASSIQNTLY | 0.86338294 |
| VAAGAGLGAVI | ASSIQNTLY | 0.87830055 |
| VAAVVTLL | ASSIQNTLY | 0.76395375 |
| VGFLGGLI | ASSIQNTLY | 0.6803691 |
| VGVIPILL | ASSIQNTLY | 0.7833709 |
| VIFASSLI | ASSIQNTLY | 0.87834495 |
| VKGGDGGAAAG | ASSIQNTLY | 0.86150014 |
| VTKTFTNI | ASSIQNTLY | 0.8580954 |
| VVAAAILL | ASSIQNTLY | 0.6952508 |
| YAALDVEL | ASSIQNTLY | 0.79559225 |
| LASVALLL | ASSIQNTLY | 0.773931 |
| LDAVHRV | ASSIQNTLY | 0.76721203 |
| QGLSSSIF | ASSIQNTLY | 0.8600633 |
| RGTFANIRL | ASSIQNTLY | 0.7860979 |
| RQLWWGHRI | ASSIQNTLY | 0.820786 |
| YGPPGTGKTLL | ASSIQNTLY | 0.8249449 |
| AALVGAPANVA | ASSIQNTLY | 0.8384863 |
| AATFRRTE | ASSIQNTLY | 0.8531424 |
| AGALGGGI | ASSIQNTLY | 0.8316461 |
| AGGAGGAGGIGAG | ASSIQNTLY | 0.8921292 |
| AGSGALGL | ASSIQNTLY | 0.8673326 |
| AQIADQI | ASSIQNTLY | 0.8729142 |
| ASAVAVTYL | ASSIQNTLY | 0.20763057 |
| ASTVLGGTGG | ASSIQNTLY | 0.5070532 |
| AVGGEPAK | ASSIQNTLY | 0.81970906 |
| CEQGTFGGTC | ASSIQNTLY | 0.76478213 |
| DEGKAPAG | ASSIQNTLY | 0.68730134 |
| DTVVFPAL | ASSIQNTLY | 0.8460231 |
| FEGITMAG | ASSIQNTLY | 0.8275902 |

| | | |
|---|---|---|
| FPLVVQD | ASSIQNTLY | 0.70410514 |
| GAFTTPQ | ASSIQNTLY | 0.8574977 |
| GFGGFGTAGGVGGA | ASSIQNTLY | 0.7739365 |
| GFLAQFL | ASSIQNTLY | 0.7335804 |
| GGAGGAGGNAGVA | ASSIQNTLY | 0.7956476 |
| GGVGTAGAGGAGGAGG | ASSIQNTLY | 0.87968683 |
| GNGGGGDAGNAGSGGN | ASSIQNTLY | 0.69680834 |
| GTGGAGGAGNDAGS | ASSIQNTLY | 0.81690466 |
| GTTNTGWFNS | ASSIQNTLY | 0.84118986 |
| IFMGPLFTL | ASSIQNTLY | 0.8672664 |
| ILAHAIEAI | ASSIQNTLY | 0.5511651 |
| IRSAIREAL | ASSIQNTLY | 0.85590005 |
| ITFAGGG | ASSIQNTLY | 0.7752731 |
| ITGMSGHQEV | ASSIQNTLY | 0.89376366 |
| IVAGLACTL | ASSIQNTLY | 0.7397084 |
| IVGGGLAGA | ASSIQNTLY | 0.7921653 |
| IVTGVLNI | ASSIQNTLY | 0.78011554 |
| LAQQGCI | ASSIQNTLY | 0.8493974 |
| LCLHVEN | ASSIQNTLY | 0.5128554 |
| LDAISDGS | ASSIQNTLY | 0.81068987 |
| LGGFAAML | ASSIQNTLY | 0.73394966 |
| LGGKNPMI | ASSIQNTLY | 0.83903927 |
| LGPKGVAAIL | ASSIQNTLY | 0.85204005 |
| LMVGVSI | ASSIQNTLY | 0.6884835 |
| LVGAAFVL | ASSIQNTLY | 0.88427484 |
| LVLAVAPL | ASSIQNTLY | 0.80240774 |
| MAAIAVIDAA | ASSIQNTLY | 0.7843723 |
| NNGINAELQ | ASSIQNTLY | 0.87718195 |
| RGEPLVLVHG | ASSIQNTLY | 0.7386524 |
| RGGAGGAAGLI | ASSIQNTLY | 0.88676566 |
| RSYRMHPAV | ASSIQNTLY | 0.8282317 |
| SAAHLLAGQLA | ASSIQNTLY | 0.84332097 |
| SAAVGSY | ASSIQNTLY | 0.86240214 |
| SPHAVLHRI | ASSIQNTLY | 0.7491573 |
| SQIAKVL | ASSIQNTLY | 0.7287278 |
| TALGHLVP | ASSIQNTLY | 0.8705636 |
| TAPTTSGAA | ASSIQNTLY | 0.8071138 |
| TDGDDGNGGSPGT | ASSIQNTLY | 0.84234476 |
| TDGLGEN | ASSIQNTLY | 0.7693294 |
| TGGPPLPGQA | ASSIQNTLY | 0.71532637 |
| TIPAITI | ASSIQNTLY | 0.7385875 |
| TIPGFTI | ASSIQNTLY | 0.82396746 |
| TSIASGALA | ASSIQNTLY | 0.83970606 |
| TVTLPGAAI | ASSIQNTLY | 0.81727934 |
| VASGLSAALP | ASSIQNTLY | 0.5165471 |
| VGAGVIGI | ASSIQNTLY | 0.8192401 |
| VGPKRFGV | ASSIQNTLY | 0.76884466 |
| VNVAFPD | ASSIQNTLY | 0.75838184 |
| VSMTNSGVS | ASSIQNTLY | 0.80094177 |
| VTDVLGDLSV | ASSIQNTLY | 0.79031706 |
| VVVVGAGFAGL | ASSIQNTLY | 0.7611197 |

| | | |
|---|---|---|
| VYTVGTVV | ASSIQNTLY | 0.73946947 |
| WTTFSGP | ASSIQNTLY | 0.74772656 |
| YDELVEAF | ASSIQNTLY | 0.8166218 |
| YGTGGAGGNGG | ASSIQNTLY | 0.8062469 |
| YTNVMAV | ASSIQNTLY | 0.8381344 |
| FLEQVHQL | ASSIQNTLY | 0.8373547 |
| GLSLLAVAV | ASSIQNTLY | 0.7875771 |
| LEQVHQL | ASSIQNTLY | 0.25153744 |
| LNAAGAI | ASSIQNTLY | 0.4590704 |
| PAVHASL | ASSIQNTLY | 0.70845485 |
| TELEDDA | ASSIQNTLY | 0.76409674 |
| TPPGPPN | ASSIQNTLY | 0.28728428 |
| TPPPPPPP | ASSIQNTLY | 0.78521186 |
| TQSPALI | ASSPRERTAEQF | 0.77423155 |
| EFAAAFPQ | ASSPRERTAEQF | 0.7371932 |
| LLSGNLDVL | ASSPRERTAEQF | 0.8313815 |
| GPANLGNYNI | ASSPRERTAEQF | 0.78270245 |
| TLGPTLP | ASSPRERTAEQF | 0.69358075 |
| VYAYSSCCK | ASSPRERTAEQF | 0.61444575 |
| APAPAPAPNI | ASSPRERTAEQF | 0.78945374 |
| PGGPVPIP | ASSPRERTAEQF | 0.42593545 |
| FLTVALEP | ASSPRERTAEQF | 0.8358432 |
| SQYFGANL | ASSPRERTAEQF | 0.42806077 |
| GGVKPAA | ASSPRERTAEQF | 0.8179455 |
| GGTGGAGGAGGW | ASSPRERTAEQF | 0.50324833 |
| GVGGVAGLGGAGG | ASSPRERTAEQF | 0.82530713 |
| VLLVLATTI | ASSPRERTAEQF | 0.78904927 |
| IGIFAAML | ASSPRERTAEQF | 0.86227834 |
| AVGDLID | ASSPRERTAEQF | 0.77287227 |
| NISPLPL | ASSPRERTAEQF | 0.79167825 |
| VTAVFVAI | ASSPRERTAEQF | 0.5923431 |
| SGDDNDAAL | ASSPRERTAEQF | 0.8336908 |
| AVAALAHS | ASSPRERTAEQF | 0.6318768 |
| GGVTVMATP | ASSPRERTAEQF | 0.7671016 |
| AGLAANVRD | ASSPRERTAEQF | 0.4389553 |
| TNRPDLIDPALLRPG | ASSPRERTAEQF | 0.81004524 |
| LANLGVTAL | ASSPRERTAEQF | 0.82165825 |
| TGERIEL | ASSPRERTAEQF | 0.5997814 |
| NGEELAA | ASSPRERTAEQF | 0.82312685 |
| VLSLVWG | ASSPRERTAEQF | 0.77988267 |
| IGYALWL | ASSPRERTAEQF | 0.6582083 |
| LSLLLAPL | ASSPRERTAEQF | 0.8422953 |
| TLDEALK | ASSPRERTAEQF | 0.7087392 |
| TDGAGTAEGG | ASSPRERTAEQF | 0.83826 |
| NAPGGAGGNGG | ASSPRERTAEQF | 0.7033758 |
| VSVFMGMGGNF | ASSPRERTAEQF | 0.72820675 |
| LASYLAAL | ASSPRERTAEQF | 0.77873546 |
| LVGGTPAL | ASSPRERTAEQF | 0.63913465 |
| VRNADVL | ASSPRERTAEQF | 0.5654825 |
| TCSSPFLVSAM | ASSPRERTAEQF | 0.84092784 |
| VHITDAE | ASSPRERTAEQF | 0.80721676 |

| | | |
|---|---|---|
| EYAAPSF | ASSPRERTAEQF | 0.8236487 |
| VVDAFHGGL | ASSPRERTAEQF | 0.6648462 |
| ASVVDPQ | ASSPRERTAEQF | 0.58298737 |
| SYNEFGGLS | ASSPRERTAEQF | 0.6722972 |
| DPDGSVY | ASSPRERTAEQF | 0.8130511 |
| MVSTIHATE | ASSPRERTAEQF | 0.8385378 |
| LAAGVADPV | ASSPRERTAEQF | 0.800501 |
| TKEQIAA | ASSPRERTAEQF | 0.5910754 |
| SLASPAAL | ASSPRERTAEQF | 0.7815531 |
| IGLVFYGV | ASSPRERTAEQF | 0.7321483 |
| FLLDASNL | ASSPRERTAEQF | 0.73735857 |
| YQVGNFP | ASSPRERTAEQF | 0.7848255 |
| ESLLSQST | ASSPRERTAEQF | 0.7784054 |
| LFTVGFAAVG | ASSPRERTAEQF | 0.69214916 |
| ATPPTPT | ASSPRERTAEQF | 0.6544169 |
| TDLATLG | ASSPRERTAEQF | 0.7696872 |
| SAGGTGR | ASSPRERTAEQF | 0.82474434 |
| GFSNSGS | ASSPRERTAEQF | 0.8147707 |
| GGNGGSGGTSV | ASSPRERTAEQF | 0.8501551 |
| GTSVATGGAGN | ASSPRERTAEQF | 0.8255787 |
| NGGNGGSGGTS | ASSPRERTAEQF | 0.82943225 |
| AVVKFPH | ASSPRERTAEQF | 0.29100198 |
| TPPPPPP | ASSPRERTAEQF | 0.47961846 |
| AAPAAAAAPA | ASSPRERTAEQF | 0.6551676 |
| IALNEDL | ASSPRERTAEQF | 0.7275099 |
| IVNVSSM | ASSPRERTAEQF | 0.35658723 |
| TGIPPAPR | ASSPRERTAEQF | 0.7921363 |
| FDLGGGTF | ASSPRERTAEQF | 0.71327657 |
| FDLGGGTFDVS | ASSPRERTAEQF | 0.61043674 |
| NEPTAAA | ASSPRERTAEQF | 0.8155407 |
| NEPTAAALA | ASSPRERTAEQF | 0.69019455 |
| GSIINISSI | ASSPRERTAEQF | 0.60897213 |
| VAPGNAGTA | ASSPRERTAEQF | 0.65737426 |
| TGGGASLE | ASSPRERTAEQF | 0.836313 |
| VAVYNHYKRI | ASSPRERTAEQF | 0.5907043 |
| TDPAPATT | ASSPRERTAEQF | 0.80926573 |
| TNRPDILDPALLRPG | ASSPRERTAEQF | 0.62827295 |
| LTTTTTS | ASSPRERTAEQF | 0.8352434 |
| PTGSGKTL | ASSPRERTAEQF | 0.50150275 |
| TGSGKTL | ASSPRERTAEQF | 0.8262581 |
| TIGHVDHGK | ASSPRERTAEQF | 0.75093514 |
| TIGHVDHGKTT | ASSPRERTAEQF | 0.8636539 |
| GHVDHGKTT | ASSPRERTAEQF | 0.7024999 |
| AQFEHTLL | ASSPRERTAEQF | 0.78296185 |
| IVNQDEL | ASSPRERTAEQF | 0.73302555 |
| TNGSQFF | ASSPRERTAEQF | 0.81968564 |
| TGKGIVE | ASSPRERTAEQF | 0.47996804 |
| LISNASDA | ASSPRERTAEQF | 0.70450914 |
| AELRTLGELTKIAKAHGAQ | ASSPRERTAEQF | 0.5058109 |
| APNAVAAAEPSW | ASSPRERTAEQF | 0.76534176 |
| APQVAGAATAVL | ASSPRERTAEQF | 0.8094629 |

| | | |
|---|---|---|
| ARRDPAAAAAILPTDA | ASSPRERTAEQF | 0.33589014 |
| DAAARDVLAEAGLAEYF | ASSPRERTAEQF | 0.8039416 |
| DDQIVLSAKANLAKVEQ | ASSPRERTAEQF | 0.676878 |
| DELLPDTGEDSDEDGH | ASSPRERTAEQF | 0.6195265 |
| ELPEAPANRATVPQD | ASSPRERTAEQF | 0.8299041 |
| FGQSKIERVTGIG | ASSPRERTAEQF | 0.76458013 |
| GAGGTGSTAGGAGGAGGAGGI | ASSPRERTAEQF | 0.82377326 |
| GHGGTGAAGGNGGTGGAGGS( | ASSPRERTAEQF | 0.692686 |
| GNGGNGGAGGEGGPGGAGGA( | ASSPRERTAEQF | 0.7102201 |
| GSTAGNGGNGGNATNGTIAG( | ASSPRERTAEQF | 0.77637035 |
| GTGGDGGNGGNGGTGG | ASSPRERTAEQF | 0.49884665 |
| GVRRPAGIPADAPV | ASSPRERTAEQF | 0.5934404 |
| GYRQVTDAHLLTLARR | ASSPRERTAEQF | 0.8469155 |
| IAKMGQDGHDRG | ASSPRERTAEQF | 0.35279697 |
| KAEPSIEIQYT | ASSPRERTAEQF | 0.75167966 |
| KPVRIRMAADEDTVAVT | ASSPRERTAEQF | 0.15168166 |
| MPPRIAGMRLLVIKPEP | ASSPRERTAEQF | 0.1459253 |
| MKFYAVALFRSRGVPPTR | ASSPRERTAEQF | 0.2530283 |
| NAGGFGILGAVAHSP | ASSPRERTAEQF | 0.5738739 |
| NEGREITLAMAEM | ASSPRERTAEQF | 0.7597558 |
| NELVEWSHPLGHRWRWE | ASSPRERTAEQF | 0.49217275 |
| NLGAGGTGGNGGNPG | ASSPRERTAEQF | 0.15872198 |
| NTEWPKWETVMAADPVD | ASSPRERTAEQF | 0.1958057 |
| PASAAPAETQP | ASSPRERTAEQF | 0.19583982 |
| QQSSDGKNGVRAAEITGE | ASSPRERTAEQF | 0.18047956 |
| RAPDAVTTSWWKE | ASSPRERTAEQF | 0.32284325 |
| SGFGNVGAHTSGWFNQ | ASSPRERTAEQF | 0.22118604 |
| SGVAGAVGQAASVGGLKV | ASSPRERTAEQF | 0.17106831 |
| SNGGAGTGSAGGNGGT | ASSPRERTAEQF | 0.16454193 |
| TGGAGGEGGAAG | ASSPRERTAEQF | 0.30609006 |
| TGGDGGAAGAGGAPGHG | ASSPRERTAEQF | 0.55872667 |
| TMPPLLAAIANAGRHGV | ASSPRERTAEQF | 0.45106742 |
| TSSTAGNSGGA | ASSPRERTAEQF | 0.59960824 |
| VPATVPVGILC | ASSPRERTAEQF | 0.41083318 |
| VQRPDSGGVVLTGRIS | ASSPRERTAEQF | 0.3692571 |
| WTPNPVPATKGVPAGLVQAT | ASSPRERTAEQF | 0.10522713 |
| YALKRALAKFSDRFDVVII | ASSPRERTAEQF | 0.117935576 |
| YLIGSVIV | ASSPRERTAEQF | 0.21985278 |
| YGQLVAVID | ASSPRERTAEQF | 0.17188297 |
| VTVGGMTLPA | ASSPRERTAEQF | 0.13238588 |
| VLALPAV | ASSPRERTAEQF | 0.39201447 |
| VIPPTLNYET | ASSPRERTAEQF | 0.20839033 |
| VGMGGMHPSSG | ASSPRERTAEQF | 0.15625067 |
| VGLAGGAATA | ASSPRERTAEQF | 0.4578601 |
| VGGAGGAGGSGP | ASSPRERTAEQF | 0.15944588 |
| VFAAFMV | ASSPRERTAEQF | 0.1565742 |
| TLYDKLR | ASSPRERTAEQF | 0.16402902 |
| TLAVFTVL | ASSPRERTAEQF | 0.4443804 |
| TGSVQPARV | ASSPRERTAEQF | 0.1401367 |
| TGGGAGSGGT | ASSPRERTAEQF | 0.278018 |
| SHLETSL | ASSPRERTAEQF | 0.24057212 |

| | | |
|---|---|---|
| SFTTGMT | ASSPRERTAEQF | 0.37939674 |
| QVEVAKRMGVRQPTVS | ASSPRERTAEQF | 0.13622063 |
| QGSGGPALAM | ASSPRERTAEQF | 0.32580602 |
| MLAPSNP | ASSPRERTAEQF | 0.1838226 |
| LVNNAAI | ASSPRERTAEQF | 0.79660344 |
| LTGEINTLA | ASSPRERTAEQF | 0.17081507 |
| LSIFGPV | ASSPRERTAEQF | 0.2309428 |
| LLALLANL | ASSPRERTAEQF | 0.16958642 |
| LISNASDALDKLR | ASSPRERTAEQF | 0.71332026 |
| LINEAVP | ASSPRERTAEQF | 0.12858711 |
| LGYAFSGL | ASSPRERTAEQF | 0.20406587 |
| LDFVAVAL | ASSPRERTAEQF | 0.13274458 |
| LAQPGAE | ASSPRERTAEQF | 0.29235962 |
| ISNLGAVTGF | ASSPRERTAEQF | 0.36815354 |
| HQEDPGRI | ASSPRERTAEQF | 0.1220419 |
| GTPGEPTP | ASSPRERTAEQF | 0.39401335 |
| GFFNSGS | ASSPRERTAEQF | 0.27367634 |
| GAGGGGLTAGAN | ASSPRERTAEQF | 0.17335947 |
| FVDALVNM | ASSPRERTAEQF | 0.5010315 |
| FLTVAPEL | ASSPRERTAEQF | 0.19090745 |
| FETLTAL | ASSPRERTAEQF | 0.5998368 |
| DVINPIEL | ASSPRERTAEQF | 0.20489794 |
| APVSGAPV | ASSPRERTAEQF | 0.22407681 |
| ALVGGLAW | ASSPRERTAEQF | 0.4272016 |
| ALAAAAATAI | ASSPRERTAEQF | 0.23832679 |
| AFGLGLLL | ASSPRERTAEQF | 0.15657729 |
| AFFAAASL | ASSPRERTAEQF | 0.39690995 |
| AAAALPAFSPP | ASSPRERTAEQF | 0.7443172 |
| VVLAPAL | ASSPRERTAEQF | 0.7424415 |
| TIGHVDHGKTTL | ASSPRERTAEQF | 0.56733185 |
| LISNASDALDK | ASSPRERTAEQF | 0.5712809 |
| LISNASDAL | ASSPRERTAEQF | 0.62747717 |
| ISNASDALDK | ASSPRERTAEQF | 0.76071024 |
| NQAETLVYQTE | ASSPRERTAEQF | 0.78212297 |
| TGGTGGTAGAG | ASSPRERTAEQF | 0.74857426 |
| TGSTAGGAGGA | ASSPRERTAEQF | 0.6543036 |
| DPLAVPVAAGPV | ASSPRERTAEQF | 0.74980026 |
| GGTGGTPGNGGN | ASSPRERTAEQF | 0.6881769 |
| PAGLAGVDPATL | ASSPRERTAEQF | 0.6943326 |
| GNNLVVEAPHAGASV | ASSPRERTAEQF | 0.7340497 |
| KVTRIMSASTLEQPA | ASSPRERTAEQF | 0.7228313 |
| LGFALPIALPRRTVV | ASSPRERTAEQF | 0.63902426 |
| TRLGNLTARKLVVLV | ASSPRERTAEQF | 0.60596514 |
| WSQQRRLIYFPSAGP | ASSPRERTAEQF | 0.7521112 |
| GPNGNPDPMAAAVDIR | ASSPRERTAEQF | 0.77586913 |
| TQRCVSVADLRAAASR | ASSPRERTAEQF | 0.76293635 |
| QFVLDVAEHPVEVLTETV | ASSPRERTAEQF | 0.80703354 |
| GGAGGTGGAGGNGGTQAGDG | ASSPRERTAEQF | 0.7629433 |
| PLLATLRAIAVDVDAKPAQV | ASSPRERTAEQF | 0.7526389 |
| APKFRTGGYMIDFWGVGYQV[illegible] | ASSPRERTAEQF | 0.34899303 |
| GAGGAGGAGGAAGGPGGTGG[illegible] | ASSPRERTAEQF | 0.49418992 |

| | | |
|---|---|---|
| IGALGLLATDVAPAGSVAFG' | ASSPRERTAEQF | 0.6404592 |
| AGGAGGIGGNGGAGGAGGVLI | ASSPRERTAEQF | 0.66151005 |
| ADELIGLL | ASSPRERTAEQF | 0.33138403 |
| ADRALKA | ASSPRERTAEQF | 0.7480441 |
| AGAVGQVGGAGGA | ASSPRERTAEQF | 0.7043556 |
| AGGGGGLGGAP | ASSPRERTAEQF | 0.6071527 |
| AGGLLYGNGGAG | ASSPRERTAEQF | 0.7659582 |
| AGLESIGL | ASSPRERTAEQF | 0.64870363 |
| AGLKTGII | ASSPRERTAEQF | 0.5993577 |
| ASLAALLL | ASSPRERTAEQF | 0.64066905 |
| ASVINISSI | ASSPRERTAEQF | 0.7899536 |
| ATPPGPA | ASSPRERTAEQF | 0.5116351 |
| ATVASALV | ASSPRERTAEQF | 0.7790803 |
| ATVASLIL | ASSPRERTAEQF | 0.6497807 |
| AVNGPVLIRH | ASSPRERTAEQF | 0.78614414 |
| DGAAGAVGI | ASSPRERTAEQF | 0.54676974 |
| DPLLHGI | ASSPRERTAEQF | 0.7619213 |
| ELGAALGAL | ASSPRERTAEQF | 0.6757367 |
| ELGRTQAV | ASSPRERTAEQF | 0.8086823 |
| FVAGGYLS | ASSPRERTAEQF | 0.68075776 |
| GASGGAGGAGGW | ASSPRERTAEQF | 0.77368855 |
| GGAGAGGDGGAGGI | ASSPRERTAEQF | 0.73035216 |
| GTGHNGSQPGMGGNG | ASSPRERTAEQF | 0.77183473 |
| GYAAASASAS | ASSPRERTAEQF | 0.486637 |
| HQVDVAAVQQG | ASSPRERTAEQF | 0.6583618 |
| IGAAIAHT | ASSPRERTAEQF | 0.62246084 |
| IGMIEVAQL | ASSPRERTAEQF | 0.70678455 |
| IRLAPEHTSAMD | ASSPRERTAEQF | 0.752913 |
| IYNVSVL | ASSPRERTAEQF | 0.2876968 |
| KQGFLGM | ASSPRERTAEQF | 0.7686057 |
| LESVITQL | ASSPRERTAEQF | 0.518245 |
| LGLGALVVV | ASSPRERTAEQF | 0.6078896 |
| LIGNGGNGGSGGI | ASSPRERTAEQF | 0.7598121 |
| LNAINAPTL | ASSPRERTAEQF | 0.701804 |
| LQLLNTGV | ASSPRERTAEQF | 0.75456154 |
| LQTSIEF | ASSPRERTAEQF | 0.6234094 |
| MSFAVSEL | ASSPRERTAEQF | 0.6558703 |
| NTGALGL | ASSPRERTAEQF | 0.7479171 |
| PPGAGGAGGAGTT | ASSPRERTAEQF | 0.43198916 |
| QLVGLGI | ASSPRERTAEQF | 0.5290822 |
| RFLEQVHQL | ASSPRERTAEQF | 0.79112005 |
| RGVGYVLR | ASSPRERTAEQF | 0.6348158 |
| SATGVFAGVI | ASSPRERTAEQF | 0.64469147 |
| SLPAITLP | ASSPRERTAEQF | 0.56604326 |
| STALAFTV | ASSPRERTAEQF | 0.57190955 |
| SVLTVGLGK | ASSPRERTAEQF | 0.59923226 |
| TALAGAGSGI | ASSPRERTAEQF | 0.6445512 |
| TGLPFTGI | ASSPRERTAEQF | 0.65651846 |
| VAEHVTV | ASSPRERTAEQF | 0.63563305 |
| VDIVLNSL | ASSPRERTAEQF | 0.5965724 |
| VLTGGGAVL | ASSPRERTAEQF | 0.632705 |

| | | |
|---|---|---|
| VNVGLAYGI | ASSPRERTAEQF | 0.6012613 |
| VSSAAVGAV | ASSPRERTAEQF | 0.6074505 |
| VTSVYEAGV | ASSPRERTAEQF | 0.63769054 |
| WVALDRGA | ASSPRERTAEQF | 0.61959386 |
| YLVGSVVI | ASSPRERTAEQF | 0.5963706 |
| GPPGTGKTLL | ASSPRERTAEQF | 0.59246176 |
| GVGMTKF | ASSPRERTAEQF | 0.63478065 |
| IGIGGSDL | ASSPRERTAEQF | 0.6471343 |
| IGTLLAEL | ASSPRERTAEQF | 0.6425255 |
| IYTSGTT | ASSPRERTAEQF | 0.66234386 |
| LSFPAGV | ASSPRERTAEQF | 0.64316916 |
| LYTSGTT | ASSPRERTAEQF | 0.64218897 |
| VGAFPFGF | ASSPRERTAEQF | 0.38597327 |
| VGAGTVEF | ASSPRERTAEQF | 0.52863705 |
| VGKSTLLL | ASSPRERTAEQF | 0.60473686 |
| AALAHAGGGIV | ASSPRERTAEQF | 0.6128519 |
| AALVAVSLL | ASSPRERTAEQF | 0.4318249 |
| AAVSRAVAAL | ASSPRERTAEQF | 0.6381425 |
| AGGSSGIGGF | ASSPRERTAEQF | 0.6090456 |
| AGLTVANANG | ASSPRERTAEQF | 0.5750129 |
| AGLVGAVL | ASSPRERTAEQF | 0.6447984 |
| ALSQAFTV | ASSPRERTAEQF | 0.5896946 |
| ALTFSVSL | ASSPRERTAEQF | 0.5779877 |
| ALTGAALVAL | ASSPRERTAEQF | 0.6031072 |
| ASFSIGAGVL | ASSPRERTAEQF | 0.6582854 |
| DGSVYKI | ASSPRERTAEQF | 0.5599818 |
| EQLAEIV | ASSPRERTAEQF | 0.6428937 |
| FAVSELTVM | ASSPRERTAEQF | 0.61281 |
| GAAPLAAGL | ASSPRERTAEQF | 0.6530819 |
| GADGGDGGSGGAGG | ASSPRERTAEQF | 0.53775764 |
| GFGLDTGGAGG | ASSPRERTAEQF | 0.6389391 |
| GLASGGLL | ASSPRERTAEQF | 0.6009395 |
| GRIAGAAVLL | ASSPRERTAEQF | 0.66774136 |
| GSAGSLNTSF | ASSPRERTAEQF | 0.6018501 |
| GTGQIGI | ASSPRERTAEQF | 0.63496625 |
| GTLIALIL | ASSPRERTAEQF | 0.61342657 |
| IAATMGFVVG | ASSPRERTAEQF | 0.64485747 |
| IFAQALL | ASSPRERTAEQF | 0.48960906 |
| IGAAVALGF | ASSPRERTAEQF | 0.6103722 |
| IGLTPLAL | ASSPRERTAEQF | 0.5679758 |
| ILGSNTSTL | ASSPRERTAEQF | 0.6284555 |
| IVFSLTVL | ASSPRERTAEQF | 0.6323446 |
| LASKATGF | ASSPRERTAEQF | 0.4213323 |
| LASVALIL | ASSPRERTAEQF | 0.6498376 |
| LFAVGLTL | ASSPRERTAEQF | 0.5484733 |
| LFNGGIGTYI | ASSPRERTAEQF | 0.5812812 |
| LTLGAGII | ASSPRERTAEQF | 0.64869213 |
| LVAAALGDI | ASSPRERTAEQF | 0.6117517 |
| NTAFVQLGI | ASSPRERTAEQF | 0.6487751 |
| QLFSGAEL | ASSPRERTAEQF | 0.60494924 |
| SAATFITL | ASSPRERTAEQF | 0.61496156 |

| | | |
|---|---|---|
| SNYGLANLGV | ASSPRERTAEQF | 0.6432136 |
| SYGGAYAVM | ASSPRERTAEQF | 0.5285848 |
| TTGVGGLTLG | ASSPRERTAEQF | 0.5768625 |
| TTQTIYI | ASSPRERTAEQF | 0.6506953 |
| VAAGAGLGAVI | ASSPRERTAEQF | 0.79659206 |
| VAAVVTLL | ASSPRERTAEQF | 0.8237735 |
| VGFLGGLI | ASSPRERTAEQF | 0.7303037 |
| VGVIPILL | ASSPRERTAEQF | 0.6481035 |
| VIFASSLI | ASSPRERTAEQF | 0.73972386 |
| VKGGDGGAAAG | ASSPRERTAEQF | 0.79708076 |
| VTKTFTNI | ASSPRERTAEQF | 0.82001 |
| VVAAAILL | ASSPRERTAEQF | 0.7454113 |
| YAALDVEL | ASSPRERTAEQF | 0.45073166 |
| LASVALLL | ASSPRERTAEQF | 0.72921485 |
| LDAVHRV | ASSPRERTAEQF | 0.7332376 |
| QGLSSSIF | ASSPRERTAEQF | 0.71528924 |
| RGTFANIRL | ASSPRERTAEQF | 0.7567748 |
| RQLWWGHRI | ASSPRERTAEQF | 0.74227947 |
| YGPPGTGKTLL | ASSPRERTAEQF | 0.71164167 |
| AALVGAPANVA | ASSPRERTAEQF | 0.73271245 |
| AATFRRTE | ASSPRERTAEQF | 0.6672172 |
| AGALGGGI | ASSPRERTAEQF | 0.7923025 |
| AGGAGGAGGIGAG | ASSPRERTAEQF | 0.8055235 |
| AGSGALGL | ASSPRERTAEQF | 0.7952574 |
| AQIADQI | ASSPRERTAEQF | 0.8126342 |
| ASAVAVTYL | ASSPRERTAEQF | 0.78417206 |
| ASTVLGGTGG | ASSPRERTAEQF | 0.20004463 |
| AVGGEPAK | ASSPRERTAEQF | 0.37156093 |
| CEQGTFGGTC | ASSPRERTAEQF | 0.6979921 |
| DEGKAPAG | ASSPRERTAEQF | 0.661899 |
| DTVVFPAL | ASSPRERTAEQF | 0.35154122 |
| FEGITMAG | ASSPRERTAEQF | 0.63756895 |
| FPLVVQD | ASSPRERTAEQF | 0.6704619 |
| GAFTTPQ | ASSPRERTAEQF | 0.63623416 |
| GFGGFGTAGGVGGA | ASSPRERTAEQF | 0.787611 |
| GFLAQFL | ASSPRERTAEQF | 0.725037 |
| GGAGGAGGNAGVA | ASSPRERTAEQF | 0.67904276 |
| GGVGTAGAGGAGGAGG | ASSPRERTAEQF | 0.55179787 |
| GNGGGGDAGNAGSGGN | ASSPRERTAEQF | 0.6701519 |
| GTGGAGGAGNDAGS | ASSPRERTAEQF | 0.37633955 |
| GTTNTGWFNS | ASSPRERTAEQF | 0.790437 |
| IFMGPLFTL | ASSPRERTAEQF | 0.38172552 |
| ILAHAIEAI | ASSPRERTAEQF | 0.78709173 |
| IRSAIREAL | ASSPRERTAEQF | 0.50145584 |
| ITFAGGG | ASSPRERTAEQF | 0.78961754 |
| ITGMSGHQEV | ASSPRERTAEQF | 0.77134967 |
| IVAGLACTL | ASSPRERTAEQF | 0.8404765 |
| IVGGGLAGA | ASSPRERTAEQF | 0.7640733 |
| IVTGVLNI | ASSPRERTAEQF | 0.691877 |
| LAQQGCI | ASSPRERTAEQF | 0.42476904 |
| LCLHVEN | ASSPRERTAEQF | 0.81314826 |

| | | |
|---|---|---|
| LDAISDGS | ASSPRERTAEQF | 0.63743824 |
| LGGFAAML | ASSPRERTAEQF | 0.76249737 |
| LGGKNPMI | ASSPRERTAEQF | 0.32702765 |
| LGPKGVAAIL | ASSPRERTAEQF | 0.79424554 |
| LMVGVSI | ASSPRERTAEQF | 0.7569945 |
| LVGAAFVL | ASSPRERTAEQF | 0.52935994 |
| LVLAVAPL | ASSPRERTAEQF | 0.7973007 |
| MAAIAVIDAA | ASSPRERTAEQF | 0.7752192 |
| NNGINAELQ | ASSPRERTAEQF | 0.6266187 |
| RGEPLVLVHG | ASSPRERTAEQF | 0.8220271 |
| RGGAGGAAGLI | ASSPRERTAEQF | 0.64564204 |
| RSYRMHPAV | ASSPRERTAEQF | 0.8299544 |
| SAAHLLAGQLA | ASSPRERTAEQF | 0.68668914 |
| SAAVGSY | ASSPRERTAEQF | 0.66935617 |
| SPHAVLHRI | ASSPRERTAEQF | 0.70809555 |
| SQIAKVL | ASSPRERTAEQF | 0.73104036 |
| TALGHLVP | ASSPRERTAEQF | 0.5740593 |
| TAPTTSGAA | ASSPRERTAEQF | 0.82250273 |
| TDGDDGNGGSPGT | ASSPRERTAEQF | 0.6917815 |
| TDGLGEN | ASSPRERTAEQF | 0.69786155 |
| TGGPPLPGQA | ASSPRERTAEQF | 0.62238896 |
| TIPAITI | ASSPRERTAEQF | 0.5948111 |
| TIPGFTI | ASSPRERTAEQF | 0.6266783 |
| TSIASGALA | ASSPRERTAEQF | 0.710282 |
| TVTLPGAAI | ASSPRERTAEQF | 0.71385366 |
| VASGLSAALP | ASSPRERTAEQF | 0.69225407 |
| VGAGVIGI | ASSPRERTAEQF | 0.61492455 |
| VGPKRFGV | ASSPRERTAEQF | 0.7171665 |
| VNVAFPD | ASSPRERTAEQF | 0.6717925 |
| VSMTNSGVS | ASSPRERTAEQF | 0.63908046 |
| VTDVLGDLSV | ASSPRERTAEQF | 0.69360614 |
| VVVVGAGFAGL | ASSPRERTAEQF | 0.7081341 |
| VYTVGTVV | ASSPRERTAEQF | 0.64800817 |
| WTTFSGP | ASSPRERTAEQF | 0.6497853 |
| YDELVEAF | ASSPRERTAEQF | 0.6698588 |
| YGTGGAGGNGG | ASSPRERTAEQF | 0.70373523 |
| YTNVMAV | ASSPRERTAEQF | 0.69139993 |
| FLEQVHQL | ASSPRERTAEQF | 0.7280427 |
| GLSLLAVAV | ASSPRERTAEQF | 0.70831794 |
| LEQVHQL | ASSPRERTAEQF | 0.70281243 |
| LNAAGAI | ASSPRERTAEQF | 0.36374497 |
| PAVHASL | ASSPRERTAEQF | 0.4816036 |
| TELEDDA | ASSPRERTAEQF | 0.68593425 |
| TPPGPPN | ASSPRERTAEQF | 0.65501195 |
| TPPPPPPP | ASSPRERTAEQF | 0.41347688 |
| TQSPALI | ASSSGTGGEQY | 0.7034779 |
| EFAAAFPQ | ASSSGTGGEQY | 0.6607249 |
| LLSGNLDVL | ASSSGTGGEQY | 0.66104364 |
| GPANLGNYNI | ASSSGTGGEQY | 0.71009403 |
| TLGPTLP | ASSSGTGGEQY | 0.6623079 |
| VYAYSSCCK | ASSSGTGGEQY | 0.6461688 |

| | | |
|---|---|---|
| APAPAPAPNI | ASSSGTGGEQY | 0.66071635 |
| PGGPVPIP | ASSSGTGGEQY | 0.72967124 |
| FLTVALEP | ASSSGTGGEQY | 0.5506974 |
| SQYFGANL | ASSSGTGGEQY | 0.70778966 |
| GGVKPAA | ASSSGTGGEQY | 0.57884854 |
| GGTGGAGGAGGW | ASSSGTGGEQY | 0.722291 |
| GVGGVAGLGGAGG | ASSSGTGGEQY | 0.56102306 |
| VLLVLATTI | ASSSGTGGEQY | 0.7014216 |
| IGIFAAML | ASSSGTGGEQY | 0.65084046 |
| AVGDLID | ASSSGTGGEQY | 0.74005735 |
| NISPLPL | ASSSGTGGEQY | 0.6525351 |
| VTAVFVAI | ASSSGTGGEQY | 0.71266377 |
| SGDDNDAAL | ASSSGTGGEQY | 0.68468887 |
| AVAALAHS | ASSSGTGGEQY | 0.7050591 |
| GGVTVMATP | ASSSGTGGEQY | 0.56865454 |
| AGLAANVRD | ASSSGTGGEQY | 0.6747066 |
| TNRPDLIDPALLRPG | ASSSGTGGEQY | 0.5451222 |
| LANLGVTAL | ASSSGTGGEQY | 0.6962842 |
| TGERIEL | ASSSGTGGEQY | 0.6935351 |
| NGEELAA | ASSSGTGGEQY | 0.47091335 |
| VLSLVWG | ASSSGTGGEQY | 0.7270281 |
| IGYALWL | ASSSGTGGEQY | 0.62991184 |
| LSLLLAPL | ASSSGTGGEQY | 0.63078636 |
| TLDEALK | ASSSGTGGEQY | 0.7127496 |
| TDGAGTAEGG | ASSSGTGGEQY | 0.69352454 |
| NAPGGAGGNGG | ASSSGTGGEQY | 0.70532846 |
| VSVFMGMGGNF | ASSSGTGGEQY | 0.66250503 |
| LASYLAAL | ASSSGTGGEQY | 0.66854376 |
| LVGGTPAL | ASSSGTGGEQY | 0.70443577 |
| VRNADVL | ASSSGTGGEQY | 0.5873144 |
| TCSSPFLVSAM | ASSSGTGGEQY | 0.6263318 |
| VHITDAE | ASSSGTGGEQY | 0.7163672 |
| EYAAPSF | ASSSGTGGEQY | 0.88858235 |
| VVDAFHGGL | ASSSGTGGEQY | 0.8938867 |
| ASVVDPQ | ASSSGTGGEQY | 0.7971779 |
| SYNEFGGLS | ASSSGTGGEQY | 0.7775067 |
| DPDGSVY | ASSSGTGGEQY | 0.8419514 |
| MVSTIHATE | ASSSGTGGEQY | 0.8992063 |
| LAAGVADPV | ASSSGTGGEQY | 0.89901245 |
| TKEQIAA | ASSSGTGGEQY | 0.88888943 |
| SLASPAAL | ASSSGTGGEQY | 0.80080116 |
| IGLVFYGV | ASSSGTGGEQY | 0.874948 |
| FLLDASNL | ASSSGTGGEQY | 0.8554095 |
| YQVGNFP | ASSSGTGGEQY | 0.85467947 |
| ESLLSQST | ASSSGTGGEQY | 0.89040947 |
| LFTVGFAAVG | ASSSGTGGEQY | 0.8769429 |
| ATPPTPT | ASSSGTGGEQY | 0.8306756 |
| TDLATLG | ASSSGTGGEQY | 0.8361063 |
| SAGGTGR | ASSSGTGGEQY | 0.87871444 |
| GFSNSGS | ASSSGTGGEQY | 0.89453447 |
| GGNGGSGGTSV | ASSSGTGGEQY | 0.8869965 |

| | | |
|---|---|---|
| GTSVATGGAGN | ASSSGTGGEQY | 0.9089113 |
| NGGNGGSGGTS | ASSSGTGGEQY | 0.894917 |
| AVVKFPH | ASSSGTGGEQY | 0.8893714 |
| TPPPPPP | ASSSGTGGEQY | 0.29436868 |
| AAPAAAAAPA | ASSSGTGGEQY | 0.6637019 |
| IALNEDL | ASSSGTGGEQY | 0.8367661 |
| IVNVSSM | ASSSGTGGEQY | 0.8488767 |
| TGIPPAPR | ASSSGTGGEQY | 0.603721 |
| FDLGGGTF | ASSSGTGGEQY | 0.8883605 |
| FDLGGGTFDVS | ASSSGTGGEQY | 0.83803946 |
| NEPTAAA | ASSSGTGGEQY | 0.80542815 |
| NEPTAAALA | ASSSGTGGEQY | 0.8937001 |
| GSIINISSI | ASSSGTGGEQY | 0.8326098 |
| VAPGNAGTA | ASSSGTGGEQY | 0.8086856 |
| TGGGASLE | ASSSGTGGEQY | 0.8583605 |
| VAVYNHYKRI | ASSSGTGGEQY | 0.90503824 |
| TDPAPATT | ASSSGTGGEQY | 0.7759763 |
| TNRPDILDPALLRPG | ASSSGTGGEQY | 0.89397424 |
| LTTTTTS | ASSSGTGGEQY | 0.863576 |
| PTGSGKTL | ASSSGTGGEQY | 0.9020073 |
| TGSGKTL | ASSSGTGGEQY | 0.7258091 |
| TIGHVDHGK | ASSSGTGGEQY | 0.8954127 |
| TIGHVDHGKTT | ASSSGTGGEQY | 0.8596572 |
| GHVDHGKTT | ASSSGTGGEQY | 0.91184765 |
| AQFEHTLL | ASSSGTGGEQY | 0.8493639 |
| IVNQDEL | ASSSGTGGEQY | 0.8852561 |
| TNGSQFF | ASSSGTGGEQY | 0.8667681 |
| TGKGIVE | ASSSGTGGEQY | 0.89832866 |
| LISNASDA | ASSSGTGGEQY | 0.70580435 |
| AELRTLGELTKIAKAHGAQ | ASSSGTGGEQY | 0.85152364 |
| APNAVAAAEPSW | ASSSGTGGEQY | 0.69431657 |
| APQVAGAATAVL | ASSSGTGGEQY | 0.87327284 |
| ARRDPAAAAAILPTDA | ASSSGTGGEQY | 0.8898997 |
| DAAARDVLAEAGLAEYF | ASSSGTGGEQY | 0.45170468 |
| DDQIVLSAKANLAKVEQ | ASSSGTGGEQY | 0.8958918 |
| DELLPDTGEDSDEDGH | ASSSGTGGEQY | 0.8116082 |
| ELPEAPANRATVPQD | ASSSGTGGEQY | 0.8127245 |
| FGQSKIERVTGIG | ASSSGTGGEQY | 0.8993346 |
| GAGGTGSTAGGAGGAGGAGGI | ASSSGTGGEQY | 0.8532776 |
| GHGGTGAAGGNGGTGGAGGS( | ASSSGTGGEQY | 0.8945087 |
| GNGGNGGAGGEGGPGGAGGA( | ASSSGTGGEQY | 0.85459447 |
| GSTAGNGGNGGNATNGTIAG( | ASSSGTGGEQY | 0.86260664 |
| GTGGDGGNGGNGGTGG | ASSSGTGGEQY | 0.8811146 |
| GVRRPAGIPADAPV | ASSSGTGGEQY | 0.66074944 |
| GYRQVTDAHLLTLARR | ASSSGTGGEQY | 0.82509553 |
| IAKMGQDGHDRG | ASSSGTGGEQY | 0.90405196 |
| KAEPSIEIQYT | ASSSGTGGEQY | 0.72940505 |
| KPVRIRMAADEDTVAVT | ASSSGTGGEQY | 0.7470779 |
| MPPRIAGMRLLVIKPEP | ASSSGTGGEQY | 0.28729138 |
| MKFYAVALFRSRGVPPTR | ASSSGTGGEQY | 0.305837 |
| NAGGFGILGAVAHSP | ASSSGTGGEQY | 0.5408467 |

| | | |
|---|---|---|
| NEGREITLAMAEM | ASSSGTGGEQY | 0.69202 |
| NELVEWSHPLGHRWRWE | ASSSGTGGEQY | 0.76582456 |
| NLGAGGTGGNGGNPG | ASSSGTGGEQY | 0.7266269 |
| NTEWPKWETVMAADPVD | ASSSGTGGEQY | 0.27652854 |
| PASAAPAETQP | ASSSGTGGEQY | 0.65783155 |
| QQSSDGKNGVRAAEITGE | ASSSGTGGEQY | 0.4550232 |
| RAPDAVTTSWWKE | ASSSGTGGEQY | 0.58039165 |
| SGFGNVGAHTSGWFNQ | ASSSGTGGEQY | 0.65929747 |
| SGVAGAVGQAASVGGLKV | ASSSGTGGEQY | 0.51092285 |
| SNGGAGTGSAGGNGGT | ASSSGTGGEQY | 0.43971384 |
| TGGAGGEGGAAG | ASSSGTGGEQY | 0.2748511 |
| TGGDGGAAGAGGAPGHG | ASSSGTGGEQY | 0.68945855 |
| TMPPLLAAIANAGRHGV | ASSSGTGGEQY | 0.73129916 |
| TSSTAGNSGGA | ASSSGTGGEQY | 0.7094954 |
| VPATVPVGILC | ASSSGTGGEQY | 0.7590928 |
| VQRPDSGGVVLTGRIS | ASSSGTGGEQY | 0.7357152 |
| WTPNPVPATKGVPAGLVQAT | ASSSGTGGEQY | 0.59927994 |
| YALKRALAKFSDRFDVVII | ASSSGTGGEQY | 0.16910735 |
| YLIGSVIV | ASSSGTGGEQY | 0.2029571 |
| YGQLVAVID | ASSSGTGGEQY | 0.3050011 |
| VTVGGMTLPA | ASSSGTGGEQY | 0.4446857 |
| VLALPAV | ASSSGTGGEQY | 0.18262158 |
| VIPPTLNYET | ASSSGTGGEQY | 0.67143196 |
| VGMGGMHPSSG | ASSSGTGGEQY | 0.3856305 |
| VGLAGGAATA | ASSSGTGGEQY | 0.3903076 |
| VGGAGGAGGSGP | ASSSGTGGEQY | 0.7386888 |
| VFAAFMV | ASSSGTGGEQY | 0.39357156 |
| TLYDKLR | ASSSGTGGEQY | 0.2604142 |
| TLAVFTVL | ASSSGTGGEQY | 0.3800085 |
| TGSVQPARV | ASSSGTGGEQY | 0.74051416 |
| TGGGAGSGGT | ASSSGTGGEQY | 0.33401465 |
| SHLETSL | ASSSGTGGEQY | 0.7332194 |
| SFTTGMT | ASSSGTGGEQY | 0.48525518 |
| QVEVAKRMGVRQPTVS | ASSSGTGGEQY | 0.7322074 |
| QGSGGPALAM | ASSSGTGGEQY | 0.22310048 |
| MLAPSNP | ASSSGTGGEQY | 0.7180271 |
| LVNNAAI | ASSSGTGGEQY | 0.6378289 |
| LTGEINTLA | ASSSGTGGEQY | 0.7873006 |
| LSIFGPV | ASSSGTGGEQY | 0.4473703 |
| LLALLANL | ASSSGTGGEQY | 0.6201908 |
| LISNASDALDKLR | ASSSGTGGEQY | 0.424825 |
| LINEAVP | ASSSGTGGEQY | 0.7248242 |
| LGYAFSGL | ASSSGTGGEQY | 0.24605192 |
| LDFVAVAL | ASSSGTGGEQY | 0.4399238 |
| LAQPGAE | ASSSGTGGEQY | 0.24308565 |
| ISNLGAVTGF | ASSSGTGGEQY | 0.40522128 |
| HQEDPGRI | ASSSGTGGEQY | 0.70774686 |
| GTPGEPTP | ASSSGTGGEQY | 0.17410506 |
| GFFNSGS | ASSSGTGGEQY | 0.6633033 |
| GAGGGGLTAGAN | ASSSGTGGEQY | 0.4067924 |
| FVDALVNM | ASSSGTGGEQY | 0.26966316 |

| | | |
|---|---|---|
| FLTVAPEL | ASSSGTGGEQY | 0.720436 |
| FETLTAL | ASSSGTGGEQY | 0.5371296 |
| DVINPIEL | ASSSGTGGEQY | 0.6958046 |
| APVSGAPV | ASSSGTGGEQY | 0.55286634 |
| ALVGGLAW | ASSSGTGGEQY | 0.54570687 |
| ALAAAAATAI | ASSSGTGGEQY | 0.52453333 |
| AFGLGLLL | ASSSGTGGEQY | 0.17431386 |
| AFFAAASL | ASSSGTGGEQY | 0.32224685 |
| AAAALPAFSPP | ASSSGTGGEQY | 0.7500216 |
| VVLAPAL | ASSSGTGGEQY | 0.82832247 |
| TIGHVDHGKTTL | ASSSGTGGEQY | 0.83997667 |
| LISNASDALDK | ASSSGTGGEQY | 0.78429246 |
| LISNASDAL | ASSSGTGGEQY | 0.7297349 |
| ISNASDALDK | ASSSGTGGEQY | 0.7813482 |
| NQAETLVYQTE | ASSSGTGGEQY | 0.8368359 |
| TGGTGGTAGAG | ASSSGTGGEQY | 0.83863544 |
| TGSTAGGAGGA | ASSSGTGGEQY | 0.82769287 |
| DPLAVPVAAGPV | ASSSGTGGEQY | 0.72159755 |
| GGTGGTPGNGGN | ASSSGTGGEQY | 0.81750286 |
| PAGLAGVDPATL | ASSSGTGGEQY | 0.80014133 |
| GNNLVVEAPHAGASV | ASSSGTGGEQY | 0.77904254 |
| KVTRIMSASTLEQPA | ASSSGTGGEQY | 0.82722116 |
| LGFALPIALPRRTVV | ASSSGTGGEQY | 0.8213159 |
| TRLGNLTARKLVVLV | ASSSGTGGEQY | 0.7823912 |
| WSQQRRLIYFPSAGP | ASSSGTGGEQY | 0.7844323 |
| GPNGNPDPMAAAVDIR | ASSSGTGGEQY | 0.8112342 |
| TQRCVSVADLRAAASR | ASSSGTGGEQY | 0.8275659 |
| QFVLDVAEHPVEVLTETV | ASSSGTGGEQY | 0.8235832 |
| GGAGGTGGAGGNGGTQAGDG | ASSSGTGGEQY | 0.8413274 |
| PLLATLRAIAVDVDAKPAQV | ASSSGTGGEQY | 0.83551514 |
| APKFRTGGYMIDFWGVGYQV | ASSSGTGGEQY | 0.8335756 |
| GAGGAGGAGGAAGGPGGTGG | ASSSGTGGEQY | 0.30685213 |
| IGALGLLATDVAPAGSVAFG | ASSSGTGGEQY | 0.5592093 |
| AGGAGGIGGNGGAGGAGGVL | ASSSGTGGEQY | 0.8083002 |
| ADELIGLL | ASSSGTGGEQY | 0.7805036 |
| ADRALKA | ASSSGTGGEQY | 0.60348064 |
| AGAVGQVGGAGGA | ASSSGTGGEQY | 0.8186104 |
| AGGGGGLGGAP | ASSSGTGGEQY | 0.7793782 |
| AGGLLYGNGGAG | ASSSGTGGEQY | 0.7602884 |
| AGLESIGL | ASSSGTGGEQY | 0.83017254 |
| AGLKTGII | ASSSGTGGEQY | 0.7884915 |
| ASLAALLL | ASSSGTGGEQY | 0.776901 |
| ASVINISSI | ASSSGTGGEQY | 0.7858926 |
| ATPPGPA | ASSSGTGGEQY | 0.8367556 |
| ATVASALV | ASSSGTGGEQY | 0.72396654 |
| ATVASLIL | ASSSGTGGEQY | 0.82271075 |
| AVNGPVLIRH | ASSSGTGGEQY | 0.7653983 |
| DGAAGAVGI | ASSSGTGGEQY | 0.83704656 |
| DPLLHGI | ASSSGTGGEQY | 0.6757543 |
| ELGAALGAL | ASSSGTGGEQY | 0.8299098 |
| ELGRTQAV | ASSSGTGGEQY | 0.80339694 |

| | | |
|---|---|---|
| FVAGGYLS | ASSSGTGGEQY | 0.84742856 |
| GASGGAGGAGGW | ASSSGTGGEQY | 0.79808724 |
| GGAGAGGDGGAGGI | ASSSGTGGEQY | 0.8113116 |
| GTGHNGSQPGMGGNG | ASSSGTGGEQY | 0.77997255 |
| GYAAASASAS | ASSSGTGGEQY | 0.8332104 |
| HQVDVAAVQQG | ASSSGTGGEQY | 0.6718842 |
| IGAAIAHT | ASSSGTGGEQY | 0.80582386 |
| IGMIEVAQL | ASSSGTGGEQY | 0.63961166 |
| IRLAPEHTSAMD | ASSSGTGGEQY | 0.81899405 |
| IYNVSVL | ASSSGTGGEQY | 0.82381225 |
| KQGFLGM | ASSSGTGGEQY | 0.56625515 |
| LESVITQL | ASSSGTGGEQY | 0.8346268 |
| LGLGALVVV | ASSSGTGGEQY | 0.7981511 |
| LIGNGGNGGSGGI | ASSSGTGGEQY | 0.77563995 |
| LNAINAPTL | ASSSGTGGEQY | 0.8400668 |
| LQLLNTGV | ASSSGTGGEQY | 0.804883 |
| LQTSIEF | ASSSGTGGEQY | 0.83906025 |
| MSFAVSEL | ASSSGTGGEQY | 0.8096035 |
| NTGALGL | ASSSGTGGEQY | 0.81209856 |
| PPGAGGAGGAGTT | ASSSGTGGEQY | 0.8227915 |
| QLVGLGI | ASSSGTGGEQY | 0.65147007 |
| RFLEQVHQL | ASSSGTGGEQY | 0.7741449 |
| RGVGYVLR | ASSSGTGGEQY | 0.837363 |
| SATGVFAGVI | ASSSGTGGEQY | 0.807462 |
| SLPAITLP | ASSSGTGGEQY | 0.8334614 |
| STALAFTV | ASSSGTGGEQY | 0.6906574 |
| SVLTVGLGK | ASSSGTGGEQY | 0.6654241 |
| TALAGAGSGI | ASSSGTGGEQY | 0.69316053 |
| TGLPFTGI | ASSSGTGGEQY | 0.8195762 |
| VAEHVTV | ASSSGTGGEQY | 0.8255014 |
| VDIVLNSL | ASSSGTGGEQY | 0.8023997 |
| VLTGGGAVL | ASSSGTGGEQY | 0.66128576 |
| VNVGLAYGI | ASSSGTGGEQY | 0.77547187 |
| VSSAAVGAV | ASSSGTGGEQY | 0.7460666 |
| VTSVYEAGV | ASSSGTGGEQY | 0.73754644 |
| WVALDRGA | ASSSGTGGEQY | 0.80593693 |
| YLVGSVVI | ASSSGTGGEQY | 0.78025806 |
| GPPGTGKTLL | ASSSGTGGEQY | 0.7384695 |
| GVGMTKF | ASSSGTGGEQY | 0.7535642 |
| IGIGGSDL | ASSSGTGGEQY | 0.7434331 |
| IGTLLAEL | ASSSGTGGEQY | 0.8081974 |
| IYTSGTT | ASSSGTGGEQY | 0.7920519 |
| LSFPAGV | ASSSGTGGEQY | 0.837558 |
| LYTSGTT | ASSSGTGGEQY | 0.8181323 |
| VGAFPFGF | ASSSGTGGEQY | 0.80550164 |
| VGAGTVEF | ASSSGTGGEQY | 0.28985146 |
| VGKSTLLL | ASSSGTGGEQY | 0.66659594 |
| AALAHAGGGIV | ASSSGTGGEQY | 0.7449617 |
| AALVAVSLL | ASSSGTGGEQY | 0.7695246 |
| AAVSRAVAAL | ASSSGTGGEQY | 0.5042335 |
| AGGSSGIGGF | ASSSGTGGEQY | 0.80562294 |

| | | |
|---|---|---|
| AGLTVANANG | ASSSGTGGEQY | 0.76871645 |
| AGLVGAVL | ASSSGTGGEQY | 0.6842714 |
| ALSQAFTV | ASSSGTGGEQY | 0.8147702 |
| ALTFSVSL | ASSSGTGGEQY | 0.7215925 |
| ALTGAALVAL | ASSSGTGGEQY | 0.74599874 |
| ASFSIGAGVL | ASSSGTGGEQY | 0.72950524 |
| DGSVYKI | ASSSGTGGEQY | 0.81331503 |
| EQLAEIV | ASSSGTGGEQY | 0.67511874 |
| FAVSELTVM | ASSSGTGGEQY | 0.80427057 |
| GAAPLAAGL | ASSSGTGGEQY | 0.70324117 |
| GADGGDGGSGGAGG | ASSSGTGGEQY | 0.8243873 |
| GFGLDTGGAGG | ASSSGTGGEQY | 0.61822295 |
| GLASGGLL | ASSSGTGGEQY | 0.8185635 |
| GRIAGAAVLL | ASSSGTGGEQY | 0.7579005 |
| GSAGSLNTSF | ASSSGTGGEQY | 0.84672964 |
| GTGQIGI | ASSSGTGGEQY | 0.76064765 |
| GTLIALIL | ASSSGTGGEQY | 0.8032795 |
| IAATMGFVVG | ASSSGTGGEQY | 0.725748 |
| IFAQALL | ASSSGTGGEQY | 0.82084167 |
| IGAAVALGF | ASSSGTGGEQY | 0.5830674 |
| IGLTPLAL | ASSSGTGGEQY | 0.74393165 |
| ILGSNTSTL | ASSSGTGGEQY | 0.6427955 |
| IVFSLTVL | ASSSGTGGEQY | 0.7727848 |
| LASKATGF | ASSSGTGGEQY | 0.8079446 |
| LASVALIL | ASSSGTGGEQY | 0.54634595 |
| LFAVGLTL | ASSSGTGGEQY | 0.8149766 |
| LFNGGIGTYI | ASSSGTGGEQY | 0.7210352 |
| LTLGAGII | ASSSGTGGEQY | 0.7204198 |
| LVAAALGDI | ASSSGTGGEQY | 0.82116866 |
| NTAFVQLGI | ASSSGTGGEQY | 0.74158424 |
| QLFSGAEL | ASSSGTGGEQY | 0.82298946 |
| SAATFITL | ASSSGTGGEQY | 0.7509526 |
| SNYGLANLGV | ASSSGTGGEQY | 0.74531674 |
| SYGGAYAVM | ASSSGTGGEQY | 0.7929573 |
| TTGVGGLTLG | ASSSGTGGEQY | 0.6386124 |
| TTQTIYI | ASSSGTGGEQY | 0.67456913 |
| VAAGAGLGAVI | ASSSGTGGEQY | 0.833445 |
| VAAVVTLL | ASSSGTGGEQY | 0.8079841 |
| VGFLGGLI | ASSSGTGGEQY | 0.8027962 |
| VGVIPILL | ASSSGTGGEQY | 0.7192091 |
| VIFASSLI | ASSSGTGGEQY | 0.57480276 |
| VKGGDGGAAAG | ASSSGTGGEQY | 0.7459679 |
| VTKTFTNI | ASSSGTGGEQY | 0.8098614 |
| VVAAAILL | ASSSGTGGEQY | 0.7939953 |
| YAALDVEL | ASSSGTGGEQY | 0.7985301 |
| LASVALLL | ASSSGTGGEQY | 0.44422308 |
| LDAVHRV | ASSSGTGGEQY | 0.6557488 |
| QGLSSSIF | ASSSGTGGEQY | 0.7518665 |
| RGTFANIRL | ASSSGTGGEQY | 0.67054343 |
| RQLWWGHRI | ASSSGTGGEQY | 0.7899031 |
| YGPPGTGKTLL | ASSSGTGGEQY | 0.65042305 |

| | | |
|---|---|---|
| AALVGAPANVA | ASSSGTGGEQY | 0.7164203 |
| AATFRRTE | ASSSGTGGEQY | 0.7307918 |
| AGALGGGI | ASSSGTGGEQY | 0.74669373 |
| AGGAGGAGGIGAG | ASSSGTGGEQY | 0.77994764 |
| AGSGALGL | ASSSGTGGEQY | 0.76513386 |
| AQIADQI | ASSSGTGGEQY | 0.8182651 |
| ASAVAVTYL | ASSSGTGGEQY | 0.8095342 |
| ASTVLGGTGG | ASSSGTGGEQY | 0.80650365 |
| AVGGEPAK | ASSSGTGGEQY | 0.21634041 |
| CEQGTFGGTC | ASSSGTGGEQY | 0.41242242 |
| DEGKAPAG | ASSSGTGGEQY | 0.6767848 |
| DTVVFPAL | ASSSGTGGEQY | 0.62429947 |
| FEGITMAG | ASSSGTGGEQY | 0.6055068 |
| FPLVVQD | ASSSGTGGEQY | 0.7361488 |
| GAFTTPQ | ASSSGTGGEQY | 0.7925495 |
| GFGGFGTAGGVGGA | ASSSGTGGEQY | 0.4828919 |
| GFLAQFL | ASSSGTGGEQY | 0.7874215 |
| GGAGGAGGNAGVA | ASSSGTGGEQY | 0.6331993 |
| GGVGTAGAGGAGGAGG | ASSSGTGGEQY | 0.5781998 |
| GNGGGGDAGNAGSGGN | ASSSGTGGEQY | 0.6498646 |
| GTGGAGGAGNDAGS | ASSSGTGGEQY | 0.8031769 |
| GTTNTGWFNS | ASSSGTGGEQY | 0.5609817 |
| IFMGPLFTL | ASSSGTGGEQY | 0.7452465 |
| ILAHAIEAI | ASSSGTGGEQY | 0.7392317 |
| IRSAIREAL | ASSSGTGGEQY | 0.8029921 |
| ITFAGGG | ASSSGTGGEQY | 0.41435874 |
| ITGMSGHQEV | ASSSGTGGEQY | 0.80074215 |
| IVAGLACTL | ASSSGTGGEQY | 0.7502845 |
| IVGGGLAGA | ASSSGTGGEQY | 0.81341183 |
| IVTGVLNI | ASSSGTGGEQY | 0.6738285 |
| LAQQGCI | ASSSGTGGEQY | 0.67001164 |
| LCLHVEN | ASSSGTGGEQY | 0.6000898 |
| LDAISDGS | ASSSGTGGEQY | 0.78527004 |
| LGGFAAML | ASSSGTGGEQY | 0.41944218 |
| LGGKNPMI | ASSSGTGGEQY | 0.76723945 |
| LGPKGVAAIL | ASSSGTGGEQY | 0.61364996 |
| LMVGVSI | ASSSGTGGEQY | 0.77350086 |
| LVGAAFVL | ASSSGTGGEQY | 0.79795873 |
| LVLAVAPL | ASSSGTGGEQY | 0.5134306 |
| MAAIAVIDAA | ASSSGTGGEQY | 0.793684 |
| NNGINAELQ | ASSSGTGGEQY | 0.7741605 |
| RGEPLVLVHG | ASSSGTGGEQY | 0.68184555 |
| RGGAGGAAGLI | ASSSGTGGEQY | 0.8040184 |
| RSYRMHPAV | ASSSGTGGEQY | 0.55533314 |
| SAAHLLAGQLA | ASSSGTGGEQY | 0.81648463 |
| SAAVGSY | ASSSGTGGEQY | 0.7167396 |
| SPHAVLHRI | ASSSGTGGEQY | 0.7373233 |
| SQIAKVL | ASSSGTGGEQY | 0.7790657 |
| TALGHLVP | ASSSGTGGEQY | 0.3796503 |
| TAPTTSGAA | ASSSGTGGEQY | 0.53019893 |
| TDGDDGNGGSPGT | ASSSGTGGEQY | 0.8101023 |

| | | |
|---|---|---|
| TDGLGEN | ASSSGTGGEQY | 0.7642282 |
| TGGPPLPGQA | ASSSGTGGEQY | 0.7697899 |
| TIPAITI | ASSSGTGGEQY | 0.71424866 |
| TIPGFTI | ASSSGTGGEQY | 0.67357206 |
| TSIASGALA | ASSSGTGGEQY | 0.7004223 |
| TVTLPGAAI | ASSSGTGGEQY | 0.7659281 |
| VASGLSAALP | ASSSGTGGEQY | 0.7699125 |
| VGAGVIGI | ASSSGTGGEQY | 0.7533245 |
| VGPKRFGV | ASSSGTGGEQY | 0.6551904 |
| VNVAFPD | ASSSGTGGEQY | 0.7407783 |
| VSMTNSGVS | ASSSGTGGEQY | 0.73567593 |
| VTDVLGDLSV | ASSSGTGGEQY | 0.7174735 |
| VVVVGAGFAGL | ASSSGTGGEQY | 0.7525909 |
| VYTVGTVV | ASSSGTGGEQY | 0.7663727 |
| WTTFSGP | ASSSGTGGEQY | 0.7220433 |
| YDELVEAF | ASSSGTGGEQY | 0.7065878 |
| YGTGGAGGNGG | ASSSGTGGEQY | 0.7180434 |
| YTNVMAV | ASSSGTGGEQY | 0.7662427 |
| FLEQVHQL | ASSSGTGGEQY | 0.75437087 |
| GLSLLAVAV | ASSSGTGGEQY | 0.7785791 |
| LEQVHQL | ASSSGTGGEQY | 0.77002 |
| LNAAGAI | ASSSGTGGEQY | 0.7675115 |
| PAVHASL | ASSSGTGGEQY | 0.3562955 |
| TELEDDA | ASSSGTGGEQY | 0.49263018 |
| TPPGPPN | ASSSGTGGEQY | 0.73600346 |
| TPPPPPPP | ASSSGTGGEQY | 0.7079483 |
| TQSPALI | ASSQSGYSQNTLY | 0.49811286 |
| EFAAAFPQ | ASSQSGYSQNTLY | 0.74591553 |
| LLSGNLDVL | ASSQSGYSQNTLY | 0.7135526 |
| GPANLGNYNI | ASSQSGYSQNTLY | 0.6880866 |
| TLGPTLP | ASSQSGYSQNTLY | 0.7583878 |
| VYAYSSCCK | ASSQSGYSQNTLY | 0.72499025 |
| APAPAPAPNI | ASSQSGYSQNTLY | 0.68886304 |
| PGGPVPIP | ASSQSGYSQNTLY | 0.653878 |
| FLTVALEP | ASSQSGYSQNTLY | 0.7643089 |
| SQYFGANL | ASSQSGYSQNTLY | 0.6084603 |
| GGVKPAA | ASSQSGYSQNTLY | 0.75632495 |
| GGTGGAGGAGGW | ASSQSGYSQNTLY | 0.5991243 |
| GVGGVAGLGGAGG | ASSQSGYSQNTLY | 0.7627151 |
| VLLVLATTI | ASSQSGYSQNTLY | 0.6144717 |
| IGIFAAML | ASSQSGYSQNTLY | 0.76753473 |
| AVGDLID | ASSQSGYSQNTLY | 0.73697567 |
| NISPLPL | ASSQSGYSQNTLY | 0.78368497 |
| VTAVFVAI | ASSQSGYSQNTLY | 0.7229974 |
| SGDDNDAAL | ASSQSGYSQNTLY | 0.75548035 |
| AVAALAHS | ASSQSGYSQNTLY | 0.7219363 |
| GGVTVMATP | ASSQSGYSQNTLY | 0.7575705 |
| AGLAANVRD | ASSQSGYSQNTLY | 0.61498773 |
| TNRPDLIDPALLRPG | ASSQSGYSQNTLY | 0.726645 |
| LANLGVTAL | ASSQSGYSQNTLY | 0.506038 |
| TGERIEL | ASSQSGYSQNTLY | 0.7469208 |

| | | |
|---|---|---|
| NGEELAA | ASSQSGYSQNTLY | 0.76084363 |
| VLSLVWG | ASSQSGYSQNTLY | 0.48054683 |
| IGYALWL | ASSQSGYSQNTLY | 0.7513652 |
| LSLLLAPL | ASSQSGYSQNTLY | 0.73827463 |
| TLDEALK | ASSQSGYSQNTLY | 0.70747435 |
| TDGAGTAEGG | ASSQSGYSQNTLY | 0.7751447 |
| NAPGGAGGNGG | ASSQSGYSQNTLY | 0.73733056 |
| VSVFMGMGGNF | ASSQSGYSQNTLY | 0.77692676 |
| LASYLAAL | ASSQSGYSQNTLY | 0.7247454 |
| LVGGTPAL | ASSQSGYSQNTLY | 0.7083351 |
| VRNADVL | ASSQSGYSQNTLY | 0.73405397 |
| TCSSPFLVSAM | ASSQSGYSQNTLY | 0.45729858 |
| VHITDAE | ASSQSGYSQNTLY | 0.5961325 |
| EYAAPSF | ASSQSGYSQNTLY | 0.7755549 |
| VVDAFHGGL | ASSQSGYSQNTLY | 0.82973444 |
| ASVVDPQ | ASSQSGYSQNTLY | 0.83603 |
| SYNEFGGLS | ASSQSGYSQNTLY | 0.78309494 |
| DPDGSVY | ASSQSGYSQNTLY | 0.75892186 |
| MVSTIHATE | ASSQSGYSQNTLY | 0.7963279 |
| LAAGVADPV | ASSQSGYSQNTLY | 0.8362615 |
| TKEQIAA | ASSQSGYSQNTLY | 0.8403534 |
| SLASPAAL | ASSQSGYSQNTLY | 0.82412314 |
| IGLVFYGV | ASSQSGYSQNTLY | 0.730902 |
| FLLDASNL | ASSQSGYSQNTLY | 0.82564986 |
| YQVGNFP | ASSQSGYSQNTLY | 0.8089216 |
| ESLLSQST | ASSQSGYSQNTLY | 0.8029028 |
| LFTVGFAAVG | ASSQSGYSQNTLY | 0.8238244 |
| ATPPTPT | ASSQSGYSQNTLY | 0.82955414 |
| TDLATLG | ASSQSGYSQNTLY | 0.79204494 |
| SAGGTGR | ASSQSGYSQNTLY | 0.7932852 |
| GFSNSGS | ASSQSGYSQNTLY | 0.8140791 |
| GGNGGSGGTSV | ASSQSGYSQNTLY | 0.83243406 |
| GTSVATGGAGN | ASSQSGYSQNTLY | 0.8289908 |
| NGGNGGSGGTS | ASSQSGYSQNTLY | 0.8411256 |
| AVVKFPH | ASSQSGYSQNTLY | 0.8355609 |
| TPPPPPP | ASSQSGYSQNTLY | 0.82739866 |
| AAPAAAAAPA | ASSQSGYSQNTLY | 0.3303368 |
| IALNEDL | ASSQSGYSQNTLY | 0.52515304 |
| IVNVSSM | ASSQSGYSQNTLY | 0.7931641 |
| TGIPPAPR | ASSQSGYSQNTLY | 0.7977141 |
| FDLGGGTF | ASSQSGYSQNTLY | 0.45448875 |
| FDLGGGTFDVS | ASSQSGYSQNTLY | 0.8281276 |
| NEPTAAA | ASSQSGYSQNTLY | 0.78849554 |
| NEPTAAALA | ASSQSGYSQNTLY | 0.7731371 |
| GSIINISSI | ASSQSGYSQNTLY | 0.831419 |
| VAPGNAGTA | ASSQSGYSQNTLY | 0.80535233 |
| TGGGASLE | ASSQSGYSQNTLY | 0.7475585 |
| VAVYNHYKRI | ASSQSGYSQNTLY | 0.7810308 |
| TDPAPATT | ASSQSGYSQNTLY | 0.8245958 |
| TNRPDILDPALLRPG | ASSQSGYSQNTLY | 0.6507071 |
| LTTTTTS | ASSQSGYSQNTLY | 0.8339952 |

| | | |
|---|---|---|
| PTGSGKTL | ASSQSGYSQNTLY | 0.69890237 |
| TGSGKTL | ASSQSGYSQNTLY | 0.8376148 |
| TIGHVDHGK | ASSQSGYSQNTLY | 0.7006374 |
| TIGHVDHGKTT | ASSQSGYSQNTLY | 0.8305865 |
| GHVDHGKTT | ASSQSGYSQNTLY | 0.8123088 |
| AQFEHTLL | ASSQSGYSQNTLY | 0.848988 |
| IVNQDEL | ASSQSGYSQNTLY | 0.8021155 |
| TNGSQFF | ASSQSGYSQNTLY | 0.8281315 |
| TGKGIVE | ASSQSGYSQNTLY | 0.8135873 |
| LISNASDA | ASSQSGYSQNTLY | 0.83937085 |
| AELRTLGELTKIAKAHGAQ | ASSQSGYSQNTLY | 0.7039353 |
| APNAVAAAEPSW | ASSQSGYSQNTLY | 0.81094694 |
| APQVAGAATAVL | ASSQSGYSQNTLY | 0.52542615 |
| ARRDPAAAAAILPTDA | ASSQSGYSQNTLY | 0.82045627 |
| DAAARDVLAEAGLAEYF | ASSQSGYSQNTLY | 0.8282886 |
| DDQIVLSAKANLAKVEQ | ASSQSGYSQNTLY | 0.47990012 |
| DELLPDTGEDSDEDGH | ASSQSGYSQNTLY | 0.8322411 |
| ELPEAPANRATVPQD | ASSQSGYSQNTLY | 0.79540825 |
| FGQSKIERVTGIG | ASSQSGYSQNTLY | 0.78371465 |
| GAGGTGSTAGGAGGAGGAGGI | ASSQSGYSQNTLY | 0.8356573 |
| GHGGTGAAGGNGGTGGAGGS( | ASSQSGYSQNTLY | 0.81870526 |
| GNGGNGGAGGEGGPGGAGGA( | ASSQSGYSQNTLY | 0.83326334 |
| GSTAGNGGNGGNATNGTIAG( | ASSQSGYSQNTLY | 0.8002053 |
| GTGGDGGNGGNGGTGG | ASSQSGYSQNTLY | 0.7917495 |
| GVRRPAGIPADAPV | ASSQSGYSQNTLY | 0.80791247 |
| GYRQVTDAHLLTLARR | ASSQSGYSQNTLY | 0.42437202 |
| IAKMGQDGHDRG | ASSQSGYSQNTLY | 0.719163 |
| KAEPSIEIQYT | ASSQSGYSQNTLY | 0.84066516 |
| KPVRIRMAADEDTVAVT | ASSQSGYSQNTLY | 0.769474 |
| MPPRIAGMRLLVIKPEP | ASSQSGYSQNTLY | 0.75607693 |
| MKFYAVALFRSRGVPPTR | ASSQSGYSQNTLY | 0.6695523 |
| NAGGFGILGAVAHSP | ASSQSGYSQNTLY | 0.6151923 |
| NEGREITLAMAEM | ASSQSGYSQNTLY | 0.722034 |
| NELVEWSHPLGHRWRWE | ASSQSGYSQNTLY | 0.7653239 |
| NLGAGGTGGNGGNPG | ASSQSGYSQNTLY | 0.79036343 |
| NTEWPKWETVMAADPVD | ASSQSGYSQNTLY | 0.76178974 |
| PASAAPAETQP | ASSQSGYSQNTLY | 0.6281718 |
| QQSSDGKNGVRAAEITGE | ASSQSGYSQNTLY | 0.783941 |
| RAPDAVTTSWWKE | ASSQSGYSQNTLY | 0.75044507 |
| SGFGNVGAHTSGWFNQ | ASSQSGYSQNTLY | 0.7241222 |
| SGVAGAVGQAASVGGLKV | ASSQSGYSQNTLY | 0.76758987 |
| SNGGAGTGSAGGNGGT | ASSQSGYSQNTLY | 0.7685679 |
| TGGAGGEGGAAG | ASSQSGYSQNTLY | 0.72263587 |
| TGGDGGAAGAGGAPGHG | ASSQSGYSQNTLY | 0.6199826 |
| TMPPLLAAIANAGRHGV | ASSQSGYSQNTLY | 0.7716956 |
| TSSTAGNSGGA | ASSQSGYSQNTLY | 0.7832451 |
| VPATVPVGILC | ASSQSGYSQNTLY | 0.7810971 |
| VQRPDSGGVVLTGRIS | ASSQSGYSQNTLY | 0.7927317 |
| WTPNPVPATKGVPAGLVQAT | ASSQSGYSQNTLY | 0.7813346 |
| YALKRALAKFSDRFDVVII | ASSQSGYSQNTLY | 0.77210486 |
| YLIGSVIV | ASSQSGYSQNTLY | 0.31055123 |

| | | |
|---|---|---|
| YGQLVAVID | ASSQSGYSQNTLY | 0.5133668 |
| VTVGGMTLPA | ASSQSGYSQNTLY | 0.67787534 |
| VLALPAV | ASSQSGYSQNTLY | 0.71837795 |
| VIPPTLNYET | ASSQSGYSQNTLY | 0.33724537 |
| VGMGGMHPSSG | ASSQSGYSQNTLY | 0.74572533 |
| VGLAGGAATA | ASSQSGYSQNTLY | 0.72470176 |
| VGGAGGAGGSGP | ASSQSGYSQNTLY | 0.6943903 |
| VFAAFMV | ASSQSGYSQNTLY | 0.78865457 |
| TLYDKLR | ASSQSGYSQNTLY | 0.71667755 |
| TLAVFTVL | ASSQSGYSQNTLY | 0.63565946 |
| TGSVQPARV | ASSQSGYSQNTLY | 0.70992184 |
| TGGGAGSGGT | ASSQSGYSQNTLY | 0.79058635 |
| SHLETSL | ASSQSGYSQNTLY | 0.63279086 |
| SFTTGMT | ASSQSGYSQNTLY | 0.78645587 |
| QVEVAKRMGVRQPTVS | ASSQSGYSQNTLY | 0.6807536 |
| QGSGGPALAM | ASSQSGYSQNTLY | 0.7891623 |
| MLAPSNP | ASSQSGYSQNTLY | 0.5507134 |
| LVNNAAI | ASSQSGYSQNTLY | 0.7780675 |
| LTGEINTLA | ASSQSGYSQNTLY | 0.7326857 |
| LSIFGPV | ASSQSGYSQNTLY | 0.80429274 |
| LLALLANL | ASSQSGYSQNTLY | 0.7225373 |
| LISNASDALDKLR | ASSQSGYSQNTLY | 0.7796917 |
| LINEAVP | ASSQSGYSQNTLY | 0.74929416 |
| LGYAFSGL | ASSQSGYSQNTLY | 0.78220445 |
| LDFVAVAL | ASSQSGYSQNTLY | 0.58272433 |
| LAQPGAE | ASSQSGYSQNTLY | 0.723365 |
| ISNLGAVTGF | ASSQSGYSQNTLY | 0.6134008 |
| HQEDPGRI | ASSQSGYSQNTLY | 0.7386348 |
| GTPGEPTP | ASSQSGYSQNTLY | 0.7609197 |
| GFFNSGS | ASSQSGYSQNTLY | 0.2917299 |
| GAGGGGLTAGAN | ASSQSGYSQNTLY | 0.7858021 |
| FVDALVNM | ASSQSGYSQNTLY | 0.6286169 |
| FLTVAPEL | ASSQSGYSQNTLY | 0.65118706 |
| FETLTAL | ASSQSGYSQNTLY | 0.7823186 |
| DVINPIEL | ASSQSGYSQNTLY | 0.7687696 |
| APVSGAPV | ASSQSGYSQNTLY | 0.7684742 |
| ALVGGLAW | ASSQSGYSQNTLY | 0.71978307 |
| ALAAAAATAI | ASSQSGYSQNTLY | 0.74089414 |
| AFGLGLLL | ASSQSGYSQNTLY | 0.7527664 |
| AFFAAASL | ASSQSGYSQNTLY | 0.440528 |
| AAAALPAFSPP | ASSQSGYSQNTLY | 0.64670247 |
| VVLAPAL | ASSQSGYSQNTLY | 0.7874639 |
| TIGHVDHGKTTL | ASSQSGYSQNTLY | 0.71777457 |
| LISNASDALDK | ASSQSGYSQNTLY | 0.78525513 |
| LISNASDAL | ASSQSGYSQNTLY | 0.5781001 |
| ISNASDALDK | ASSQSGYSQNTLY | 0.6294471 |
| NQAETLVYQTE | ASSQSGYSQNTLY | 0.68846476 |
| TGGTGGTAGAG | ASSQSGYSQNTLY | 0.74497235 |
| TGSTAGGAGGA | ASSQSGYSQNTLY | 0.78017056 |
| DPLAVPVAAGPV | ASSQSGYSQNTLY | 0.75429595 |
| GGTGGTPGNGGN | ASSQSGYSQNTLY | 0.5884869 |

| | | |
|---|---|---|
| PAGLAGVDPATL | ASSQSGYSQNTLY | 0.73576367 |
| GNNLVVEAPHAGASV | ASSQSGYSQNTLY | 0.55046535 |
| KVTRIMSASTLEQPA | ASSQSGYSQNTLY | 0.6983955 |
| LGFALPIALPRRTVV | ASSQSGYSQNTLY | 0.7301216 |
| TRLGNLTARKLVVLV | ASSQSGYSQNTLY | 0.6909367 |
| WSQQRRLIYFPSAGP | ASSQSGYSQNTLY | 0.57968897 |
| GPNGNPDPMAAAVDIR | ASSQSGYSQNTLY | 0.6060821 |
| TQRCVSVADLRAAASR | ASSQSGYSQNTLY | 0.7490927 |
| QFVLDVAEHPVEVLTETV | ASSQSGYSQNTLY | 0.77063036 |
| GGAGGTGGAGGNGGTQAGDG | ASSQSGYSQNTLY | 0.7513854 |
| PLLATLRAIAVDVDAKPAQV | ASSQSGYSQNTLY | 0.7819774 |
| APKFRTGGYMIDFWGVGYQV | ASSQSGYSQNTLY | 0.75946057 |
| GAGGAGGAGGAAGGPGGTGG | ASSQSGYSQNTLY | 0.7408282 |
| IGALGLLATDVAPAGSVAFG | ASSQSGYSQNTLY | 0.24259277 |
| AGGAGGIGGNGGAGGAGGVL | ASSQSGYSQNTLY | 0.5030941 |
| ADELIGLL | ASSQSGYSQNTLY | 0.55178237 |
| ADRALKA | ASSQSGYSQNTLY | 0.7191343 |
| AGAVGQVGGAGGA | ASSQSGYSQNTLY | 0.23748615 |
| AGGGGGLGGAP | ASSQSGYSQNTLY | 0.7561051 |
| AGGLLYGNGGAG | ASSQSGYSQNTLY | 0.61558217 |
| AGLESIGL | ASSQSGYSQNTLY | 0.61504245 |
| AGLKTGII | ASSQSGYSQNTLY | 0.7745763 |
| ASLAALLL | ASSQSGYSQNTLY | 0.6452037 |
| ASVINISSI | ASSQSGYSQNTLY | 0.56681705 |
| ATPPGPA | ASSQSGYSQNTLY | 0.54758537 |
| ATVASALV | ASSQSGYSQNTLY | 0.7820169 |
| ATVASLIL | ASSQSGYSQNTLY | 0.33246115 |
| AVNGPVLIRH | ASSQSGYSQNTLY | 0.7563372 |
| DGAAGAVGI | ASSQSGYSQNTLY | 0.5710857 |
| DPLLHGI | ASSQSGYSQNTLY | 0.7570942 |
| ELGAALGAL | ASSQSGYSQNTLY | 0.47659826 |
| ELGRTQAV | ASSQSGYSQNTLY | 0.76057196 |
| FVAGGYLS | ASSQSGYSQNTLY | 0.6923216 |
| GASGGAGGAGGW | ASSQSGYSQNTLY | 0.79585004 |
| GGAGAGGDGGAGGI | ASSQSGYSQNTLY | 0.67188823 |
| GTGHNGSQPGMGGNG | ASSQSGYSQNTLY | 0.72300535 |
| GYAAASASAS | ASSQSGYSQNTLY | 0.5962502 |
| HQVDVAAVQQG | ASSQSGYSQNTLY | 0.772999 |
| IGAAIAHT | ASSQSGYSQNTLY | 0.39739037 |
| IGMIEVAQL | ASSQSGYSQNTLY | 0.6506717 |
| IRLAPEHTSAMD | ASSQSGYSQNTLY | 0.51603377 |
| IYNVSVL | ASSQSGYSQNTLY | 0.6901804 |
| KQGFLGM | ASSQSGYSQNTLY | 0.74680185 |
| LESVITQL | ASSQSGYSQNTLY | 0.36393777 |
| LGLGALVVV | ASSQSGYSQNTLY | 0.7595924 |
| LIGNGGNGGSGGI | ASSQSGYSQNTLY | 0.6543725 |
| LNAINAPTL | ASSQSGYSQNTLY | 0.5446657 |
| LQLLNTGV | ASSQSGYSQNTLY | 0.76669747 |
| LQTSIEF | ASSQSGYSQNTLY | 0.70373297 |
| MSFAVSEL | ASSQSGYSQNTLY | 0.7673074 |
| NTGALGL | ASSQSGYSQNTLY | 0.5914355 |

| | | |
|---|---|---|
| PPGAGGAGGAGTT | ASSQSGYSQNTLY | 0.6949954 |
| QLVGLGI | ASSQSGYSQNTLY | 0.7069634 |
| RFLEQVHQL | ASSQSGYSQNTLY | 0.389901 |
| RGVGYVLR | ASSQSGYSQNTLY | 0.5266017 |
| SATGVFAGVI | ASSQSGYSQNTLY | 0.7730408 |
| SLPAITLP | ASSQSGYSQNTLY | 0.76993483 |
| STALAFTV | ASSQSGYSQNTLY | 0.77265674 |
| SVLTVGLGK | ASSQSGYSQNTLY | 0.6096229 |
| TALAGAGSGI | ASSQSGYSQNTLY | 0.6138727 |
| TGLPFTGI | ASSQSGYSQNTLY | 0.6754899 |
| VAEHVTV | ASSQSGYSQNTLY | 0.7930038 |
| VDIVLNSL | ASSQSGYSQNTLY | 0.8139577 |
| VLTGGGAVL | ASSQSGYSQNTLY | 0.78460354 |
| VNVGLAYGI | ASSQSGYSQNTLY | 0.71019876 |
| VSSAAVGAV | ASSQSGYSQNTLY | 0.7753551 |
| VTSVYEAGV | ASSQSGYSQNTLY | 0.73582643 |
| WVALDRGA | ASSQSGYSQNTLY | 0.7157923 |
| YLVGSVVI | ASSQSGYSQNTLY | 0.7729347 |
| GPPGTGKTLL | ASSQSGYSQNTLY | 0.78489125 |
| GVGMTKF | ASSQSGYSQNTLY | 0.6860312 |
| IGIGGSDL | ASSQSGYSQNTLY | 0.69997805 |
| IGTLLAEL | ASSQSGYSQNTLY | 0.7512305 |
| IYTSGTT | ASSQSGYSQNTLY | 0.78696084 |
| LSFPAGV | ASSQSGYSQNTLY | 0.7684211 |
| LYTSGTT | ASSQSGYSQNTLY | 0.83121514 |
| VGAFPFGF | ASSQSGYSQNTLY | 0.7833331 |
| VGAGTVEF | ASSQSGYSQNTLY | 0.7664054 |
| VGKSTLLL | ASSQSGYSQNTLY | 0.39213264 |
| AALAHAGGGIV | ASSQSGYSQNTLY | 0.5622989 |
| AALVAVSLL | ASSQSGYSQNTLY | 0.77606666 |
| AAVSRAVAAL | ASSQSGYSQNTLY | 0.707018 |
| AGGSSGIGGF | ASSQSGYSQNTLY | 0.30836195 |
| AGLTVANANG | ASSQSGYSQNTLY | 0.79488146 |
| AGLVGAVL | ASSQSGYSQNTLY | 0.72679347 |
| ALSQAFTV | ASSQSGYSQNTLY | 0.69071424 |
| ALTFSVSL | ASSQSGYSQNTLY | 0.78152895 |
| ALTGAALVAL | ASSQSGYSQNTLY | 0.7052027 |
| ASFSIGAGVL | ASSQSGYSQNTLY | 0.6951894 |
| DGSVYKI | ASSQSGYSQNTLY | 0.7514384 |
| EQLAEIV | ASSQSGYSQNTLY | 0.81835985 |
| FAVSELTVM | ASSQSGYSQNTLY | 0.6522026 |
| GAAPLAAGL | ASSQSGYSQNTLY | 0.7981367 |
| GADGGDGGSGGAGG | ASSQSGYSQNTLY | 0.7237425 |
| GFGLDTGGAGG | ASSQSGYSQNTLY | 0.8103025 |
| GLASGGLL | ASSQSGYSQNTLY | 0.64482063 |
| GRIAGAAVLL | ASSQSGYSQNTLY | 0.7911358 |
| GSAGSLNTSF | ASSQSGYSQNTLY | 0.71260476 |
| GTGQIGI | ASSQSGYSQNTLY | 0.8361261 |
| GTLIALIL | ASSQSGYSQNTLY | 0.70321816 |
| IAATMGFVVG | ASSQSGYSQNTLY | 0.8047778 |
| IFAQALL | ASSQSGYSQNTLY | 0.7737772 |

| | | |
|---|---|---|
| IGAAVALGF | ASSQSGYSQNTLY | 0.7867569 |
| IGLTPLAL | ASSQSGYSQNTLY | 0.61822486 |
| ILGSNTSTL | ASSQSGYSQNTLY | 0.7024996 |
| IVFSLTVL | ASSQSGYSQNTLY | 0.6195839 |
| LASKATGF | ASSQSGYSQNTLY | 0.7467538 |
| LASVALIL | ASSQSGYSQNTLY | 0.7844997 |
| LFAVGLTL | ASSQSGYSQNTLY | 0.34829855 |
| LFNGGIGTYI | ASSQSGYSQNTLY | 0.78770113 |
| LTLGAGII | ASSQSGYSQNTLY | 0.5898271 |
| LVAAALGDI | ASSQSGYSQNTLY | 0.68070936 |
| NTAFVQLGI | ASSQSGYSQNTLY | 0.7952225 |
| QLFSGAEL | ASSQSGYSQNTLY | 0.7544336 |
| SAATFITL | ASSQSGYSQNTLY | 0.76991236 |
| SNYGLANLGV | ASSQSGYSQNTLY | 0.70504135 |
| SYGGAYAVM | ASSQSGYSQNTLY | 0.7164858 |
| TTGVGGLTLG | ASSQSGYSQNTLY | 0.7697579 |
| TTQTIYI | ASSQSGYSQNTLY | 0.48172146 |
| VAAGAGLGAVI | ASSQSGYSQNTLY | 0.69795233 |
| VAAVVTLL | ASSQSGYSQNTLY | 0.8121021 |
| VGFLGGLI | ASSQSGYSQNTLY | 0.8192137 |
| VGVIPILL | ASSQSGYSQNTLY | 0.83321536 |
| VIFASSLI | ASSQSGYSQNTLY | 0.7293093 |
| VKGGDGGAAAG | ASSQSGYSQNTLY | 0.7186222 |
| VTKTFTNI | ASSQSGYSQNTLY | 0.77654624 |
| VVAAAILL | ASSQSGYSQNTLY | 0.8448665 |
| YAALDVEL | ASSQSGYSQNTLY | 0.83748144 |
| LASVALLL | ASSQSGYSQNTLY | 0.8172542 |
| LDAVHRV | ASSQSGYSQNTLY | 0.69910586 |
| QGLSSSIF | ASSQSGYSQNTLY | 0.8098668 |
| RGTFANIRL | ASSQSGYSQNTLY | 0.7628461 |
| RQLWWGHRI | ASSQSGYSQNTLY | 0.7762266 |
| YGPPGTGKTLL | ASSQSGYSQNTLY | 0.83226776 |
| AALVGAPANVA | ASSQSGYSQNTLY | 0.81438076 |
| AATFRRTE | ASSQSGYSQNTLY | 0.77257174 |
| AGALGGGI | ASSQSGYSQNTLY | 0.75167835 |
| AGGAGGAGGIGAG | ASSQSGYSQNTLY | 0.8091355 |
| AGSGALGL | ASSQSGYSQNTLY | 0.83812696 |
| AQIADQI | ASSQSGYSQNTLY | 0.82909596 |
| ASAVAVTYL | ASSQSGYSQNTLY | 0.8568926 |
| ASTVLGGTGG | ASSQSGYSQNTLY | 0.83504057 |
| AVGGEPAK | ASSQSGYSQNTLY | 0.8306722 |
| CEQGTFGGTC | ASSQSGYSQNTLY | 0.32809997 |
| DEGKAPAG | ASSQSGYSQNTLY | 0.56370896 |
| DTVVFPAL | ASSQSGYSQNTLY | 0.762486 |
| FEGITMAG | ASSQSGYSQNTLY | 0.7803023 |
| FPLVVQD | ASSQSGYSQNTLY | 0.51504433 |
| GAFTTPQ | ASSQSGYSQNTLY | 0.83390474 |
| GFGGFGTAGGVGGA | ASSQSGYSQNTLY | 0.7587143 |
| GFLAQFL | ASSQSGYSQNTLY | 0.7197473 |
| GGAGGAGGNAGVA | ASSQSGYSQNTLY | 0.8354681 |
| GGVGTAGAGGAGGAGG | ASSQSGYSQNTLY | 0.75368893 |

| | | |
|---|---|---|
| GNGGGGDAGNAGSGGN | ASSQSGYSQNTLY | 0.75206375 |
| GTGGAGGAGNDAGS | ASSQSGYSQNTLY | 0.7503115 |
| GTTNTGWFNS | ASSQSGYSQNTLY | 0.84466314 |
| IFMGPLFTL | ASSQSGYSQNTLY | 0.6442447 |
| ILAHAIEAI | ASSQSGYSQNTLY | 0.8244165 |
| IRSAIREAL | ASSQSGYSQNTLY | 0.78703046 |
| ITFAGGG | ASSQSGYSQNTLY | 0.8459556 |
| ITGMSGHQEV | ASSQSGYSQNTLY | 0.59830743 |
| IVAGLACTL | ASSQSGYSQNTLY | 0.8279413 |
| IVGGGLAGA | ASSQSGYSQNTLY | 0.7843425 |
| IVTGVLNI | ASSQSGYSQNTLY | 0.85999775 |
| LAQQGCI | ASSQSGYSQNTLY | 0.7814558 |
| LCLHVEN | ASSQSGYSQNTLY | 0.8143296 |
| LDAISDGS | ASSQSGYSQNTLY | 0.78071463 |
| LGGFAAML | ASSQSGYSQNTLY | 0.8271249 |
| LGGKNPMI | ASSQSGYSQNTLY | 0.5749177 |
| LGPKGVAAIL | ASSQSGYSQNTLY | 0.78618395 |
| LMVGVSI | ASSQSGYSQNTLY | 0.7016804 |
| LVGAAFVL | ASSQSGYSQNTLY | 0.80368435 |
| LVLAVAPL | ASSQSGYSQNTLY | 0.8131288 |
| MAAIAVIDAA | ASSQSGYSQNTLY | 0.508506 |
| NNGINAELQ | ASSQSGYSQNTLY | 0.8176392 |
| RGEPLVLVHG | ASSQSGYSQNTLY | 0.75594723 |
| RGGAGGAAGLI | ASSQSGYSQNTLY | 0.7240078 |
| RSYRMHPAV | ASSQSGYSQNTLY | 0.8427793 |
| SAAHLLAGQLA | ASSQSGYSQNTLY | 0.79583347 |
| SAAVGSY | ASSQSGYSQNTLY | 0.83223903 |
| SPHAVLHRI | ASSQSGYSQNTLY | 0.7874822 |
| SQIAKVL | ASSQSGYSQNTLY | 0.76630986 |
| TALGHLVP | ASSQSGYSQNTLY | 0.83019805 |
| TAPTTSGAA | ASSQSGYSQNTLY | 0.59681404 |
| TDGDDGNGGSPGT | ASSQSGYSQNTLY | 0.7177378 |
| TDGLGEN | ASSQSGYSQNTLY | 0.843611 |
| TGGPPLPGQA | ASSQSGYSQNTLY | 0.86783683 |
| TIPAITI | ASSQSGYSQNTLY | 0.8682888 |
| TIPGFTI | ASSQSGYSQNTLY | 0.84244484 |
| TSIASGALA | ASSQSGYSQNTLY | 0.82516456 |
| TVTLPGAAI | ASSQSGYSQNTLY | 0.82600296 |
| VASGLSAALP | ASSQSGYSQNTLY | 0.8733197 |
| VGAGVIGI | ASSQSGYSQNTLY | 0.86834997 |
| VGPKRFGV | ASSQSGYSQNTLY | 0.8687725 |
| VNVAFPD | ASSQSGYSQNTLY | 0.77610695 |
| VSMTNSGVS | ASSQSGYSQNTLY | 0.8696611 |
| VTDVLGDLSV | ASSQSGYSQNTLY | 0.8530184 |
| VVVVGAGFAGL | ASSQSGYSQNTLY | 0.8466617 |
| VYTVGTVV | ASSQSGYSQNTLY | 0.8687783 |
| WTTFSGP | ASSQSGYSQNTLY | 0.8572507 |
| YDELVEAF | ASSQSGYSQNTLY | 0.8495322 |
| YGTGGAGGNGG | ASSQSGYSQNTLY | 0.8542159 |
| YTNVMAV | ASSQSGYSQNTLY | 0.85203433 |
| FLEQVHQL | ASSQSGYSQNTLY | 0.85770065 |

| | | |
|---|---|---|
| GLSLLAVAV | ASSQSGYSQNTLY | 0.8465495 |
| LEQVHQL | ASSQSGYSQNTLY | 0.87785685 |
| LNAAGAI | ASSQSGYSQNTLY | 0.872025 |
| PAVHASL | ASSQSGYSQNTLY | 0.86981326 |
| TELEDDA | ASSQSGYSQNTLY | 0.38900572 |
| TPPGPPN | ASSQSGYSQNTLY | 0.80699265 |
| TPPPPPPP | ASSQSGYSQNTLY | 0.85625464 |
| TQSPALI | ASSLPGQNYEQY | 0.8539888 |
| EFAAAFPQ | ASSLPGQNYEQY | 0.6694205 |
| LLSGNLDVL | ASSLPGQNYEQY | 0.8580933 |
| GPANLGNYNI | ASSLPGQNYEQY | 0.8475196 |
| TLGPTLP | ASSLPGQNYEQY | 0.83735955 |
| VYAYSSCCK | ASSLPGQNYEQY | 0.8711449 |
| APAPAPAPNI | ASSLPGQNYEQY | 0.8528031 |
| PGGPVPIP | ASSLPGQNYEQY | 0.8380804 |
| FLTVALEP | ASSLPGQNYEQY | 0.84895 |
| SQYFGANL | ASSLPGQNYEQY | 0.8747542 |
| GGVKPAA | ASSLPGQNYEQY | 0.8261142 |
| GGTGGAGGAGGW | ASSLPGQNYEQY | 0.8703747 |
| GVGGVAGLGGAGG | ASSLPGQNYEQY | 0.8106763 |
| VLLVLATTI | ASSLPGQNYEQY | 0.8753408 |
| IGIFAAML | ASSLPGQNYEQY | 0.7466782 |
| AVGDLID | ASSLPGQNYEQY | 0.8706232 |
| NISPLPL | ASSLPGQNYEQY | 0.85463566 |
| VTAVFVAI | ASSLPGQNYEQY | 0.8799714 |
| SGDDNDAAL | ASSLPGQNYEQY | 0.8503076 |
| AVAALAHS | ASSLPGQNYEQY | 0.8676711 |
| GGVTVMATP | ASSLPGQNYEQY | 0.8374431 |
| AGLAANVRD | ASSLPGQNYEQY | 0.86374533 |
| TNRPDLIDPALLRPG | ASSLPGQNYEQY | 0.79362327 |
| LANLGVTAL | ASSLPGQNYEQY | 0.8434427 |
| TGERIEL | ASSLPGQNYEQY | 0.81953776 |
| NGEELAA | ASSLPGQNYEQY | 0.86219823 |
| VLSLVWG | ASSLPGQNYEQY | 0.86923754 |
| IGYALWL | ASSLPGQNYEQY | 0.74667835 |
| LSLLLAPL | ASSLPGQNYEQY | 0.87339824 |
| TLDEALK | ASSLPGQNYEQY | 0.83350325 |
| TDGAGTAEGG | ASSLPGQNYEQY | 0.8337598 |
| NAPGGAGGNGG | ASSLPGQNYEQY | 0.8755543 |
| VSVFMGMGGNF | ASSLPGQNYEQY | 0.8465221 |
| LASYLAAL | ASSLPGQNYEQY | 0.8701042 |
| LVGGTPAL | ASSLPGQNYEQY | 0.8498442 |
| VRNADVL | ASSLPGQNYEQY | 0.84818405 |
| TCSSPFLVSAM | ASSLPGQNYEQY | 0.8639038 |
| VHITDAE | ASSLPGQNYEQY | 0.78634804 |
| EYAAPSF | ASSLPGQNYEQY | 0.83412755 |
| VVDAFHGGL | ASSLPGQNYEQY | 0.87649405 |
| ASVVDPQ | ASSLPGQNYEQY | 0.64863485 |
| SYNEFGGLS | ASSLPGQNYEQY | 0.70279956 |
| DPDGSVY | ASSLPGQNYEQY | 0.5763879 |
| MVSTIHATE | ASSLPGQNYEQY | 0.51651156 |

| | | |
|---|---|---|
| LAAGVADPV | ASSLPGQNYEQY | 0.5814961 |
| TKEQIAA | ASSLPGQNYEQY | 0.65755534 |
| SLASPAAL | ASSLPGQNYEQY | 0.6962905 |
| IGLVFYGV | ASSLPGQNYEQY | 0.6487963 |
| FLLDASNL | ASSLPGQNYEQY | 0.45513755 |
| YQVGNFP | ASSLPGQNYEQY | 0.6193014 |
| ESLLSQST | ASSLPGQNYEQY | 0.57299757 |
| LFTVGFAAVG | ASSLPGQNYEQY | 0.6007874 |
| ATPPTPT | ASSLPGQNYEQY | 0.58967394 |
| TDLATLG | ASSLPGQNYEQY | 0.61048776 |
| SAGGTGR | ASSLPGQNYEQY | 0.5798594 |
| GFSNSGS | ASSLPGQNYEQY | 0.49281022 |
| GGNGGSGGTSV | ASSLPGQNYEQY | 0.6068709 |
| GTSVATGGAGN | ASSLPGQNYEQY | 0.61032283 |
| NGGNGGSGGTS | ASSLPGQNYEQY | 0.5863682 |
| AVVKFPH | ASSLPGQNYEQY | 0.66067344 |
| TPPPPPP | ASSLPGQNYEQY | 0.6820579 |
| AAPAAAAAPA | ASSLPGQNYEQY | 0.6478331 |
| IALNEDL | ASSLPGQNYEQY | 0.28517535 |
| IVNVSSM | ASSLPGQNYEQY | 0.3530892 |
| TGIPPAPR | ASSLPGQNYEQY | 0.5462495 |
| FDLGGGTF | ASSLPGQNYEQY | 0.575364 |
| FDLGGGTFDVS | ASSLPGQNYEQY | 0.30088916 |
| NEPTAAA | ASSLPGQNYEQY | 0.60418844 |
| NEPTAAALA | ASSLPGQNYEQY | 0.5158369 |
| GSIINISSI | ASSLPGQNYEQY | 0.55695534 |
| VAPGNAGTA | ASSLPGQNYEQY | 0.6722222 |
| TGGGASLE | ASSLPGQNYEQY | 0.60351455 |
| VAVYNHYKRI | ASSLPGQNYEQY | 0.4822948 |
| TDPAPATT | ASSLPGQNYEQY | 0.4516772 |
| TNRPDILDPALLRPG | ASSLPGQNYEQY | 0.6280236 |
| LTTTTTS | ASSLPGQNYEQY | 0.3742176 |
| PTGSGKTL | ASSLPGQNYEQY | 0.647628 |
| TGSGKTL | ASSLPGQNYEQY | 0.38787302 |
| TIGHVDHGK | ASSLPGQNYEQY | 0.65694964 |
| TIGHVDHGKTT | ASSLPGQNYEQY | 0.4232168 |
| GHVDHGKTT | ASSLPGQNYEQY | 0.6576207 |
| AQFEHTLL | ASSLPGQNYEQY | 0.6239679 |
| IVNQDEL | ASSLPGQNYEQY | 0.70691156 |
| TNGSQFF | ASSLPGQNYEQY | 0.56999874 |
| TGKGIVE | ASSLPGQNYEQY | 0.5839622 |
| LISNASDA | ASSLPGQNYEQY | 0.46988684 |
| AELRTLGELTKIAKAHGAQ | ASSLPGQNYEQY | 0.70075196 |
| APNAVAAAEPSW | ASSLPGQNYEQY | 0.46308053 |
| APQVAGAATAVL | ASSLPGQNYEQY | 0.5749091 |
| ARRDPAAAAAILPTDA | ASSLPGQNYEQY | 0.35560805 |
| DAAARDVLAEAGLAEYF | ASSLPGQNYEQY | 0.6022544 |
| DDQIVLSAKANLAKVEQ | ASSLPGQNYEQY | 0.65742815 |
| DELLPDTGEDSDEDGH | ASSLPGQNYEQY | 0.47076094 |
| ELPEAPANRATVPQD | ASSLPGQNYEQY | 0.65506256 |
| FGQSKIERVTGIG | ASSLPGQNYEQY | 0.6209067 |

| | | |
|---|---|---|
| GAGGTGSTAGGAGGAGGAGGI | ASSLPGQNYEQY | 0.51761615 |
| GHGGTGAAGGNGGTGGAGGS( | ASSLPGQNYEQY | 0.6721453 |
| GNGGNGGAGGEGGPGGAGGA( | ASSLPGQNYEQY | 0.58744216 |
| GSTAGNGGNGGNATNGTIAG( | ASSLPGQNYEQY | 0.68452513 |
| GTGGDGGNGGNGGTGG | ASSLPGQNYEQY | 0.5170044 |
| GVRRPAGIPADAPV | ASSLPGQNYEQY | 0.53048915 |
| GYRQVTDAHLLTLARR | ASSLPGQNYEQY | 0.60174567 |
| IAKMGQDGHDRG | ASSLPGQNYEQY | 0.55008894 |
| KAEPSIEIQYT | ASSLPGQNYEQY | 0.47644383 |
| KPVRIRMAADEDTVAVT | ASSLPGQNYEQY | 0.68508327 |
| MPPRIAGMRLLVIKPEP | ASSLPGQNYEQY | 0.8548862 |
| MKFYAVALFRSRGVPPTR | ASSLPGQNYEQY | 0.87760943 |
| NAGGFGILGAVAHSP | ASSLPGQNYEQY | 0.79273814 |
| NEGREITLAMAEM | ASSLPGQNYEQY | 0.6787941 |
| NELVEWSHPLGHRWRWE | ASSLPGQNYEQY | 0.8055661 |
| NLGAGGTGGNGGNPG | ASSLPGQNYEQY | 0.874665 |
| NTEWPKWETVMAADPVD | ASSLPGQNYEQY | 0.8808887 |
| PASAAPAETQP | ASSLPGQNYEQY | 0.8655908 |
| QQSSDGKNGVRAAEITGE | ASSLPGQNYEQY | 0.5770446 |
| RAPDAVTTSWWKE | ASSLPGQNYEQY | 0.8535869 |
| SGFGNVGAHTSGWFNQ | ASSLPGQNYEQY | 0.79831064 |
| SGVAGAVGQAASVGGLKV | ASSLPGQNYEQY | 0.81539613 |
| SNGGAGTGSAGGNGGT | ASSLPGQNYEQY | 0.8659394 |
| TGGAGGEGGAAG | ASSLPGQNYEQY | 0.8026801 |
| TGGDGGAAGAGGAPGHG | ASSLPGQNYEQY | 0.7676302 |
| TMPPLLAAIANAGRHGV | ASSLPGQNYEQY | 0.72544575 |
| TSSTAGNSGGA | ASSLPGQNYEQY | 0.86783236 |
| VPATVPVGILC | ASSLPGQNYEQY | 0.87316 |
| VQRPDSGGVVLTGRIS | ASSLPGQNYEQY | 0.872455 |
| WTPNPVPATKGVPAGLVQAT | ASSLPGQNYEQY | 0.888319 |
| YALKRALAKFSDRFDVVII | ASSLPGQNYEQY | 0.87901294 |
| YLIGSVIV | ASSLPGQNYEQY | 0.8770791 |
| YGQLVAVID | ASSLPGQNYEQY | 0.28363836 |
| VTVGGMTLPA | ASSLPGQNYEQY | 0.528621 |
| VLALPAV | ASSLPGQNYEQY | 0.7511785 |
| VIPPTLNYET | ASSLPGQNYEQY | 0.79184586 |
| VGMGGMHPSSG | ASSLPGQNYEQY | 0.48171064 |
| VGLAGGAATA | ASSLPGQNYEQY | 0.8195354 |
| VGGAGGAGGSGP | ASSLPGQNYEQY | 0.7296062 |
| VFAAFMV | ASSLPGQNYEQY | 0.69760114 |
| TLYDKLR | ASSLPGQNYEQY | 0.8745292 |
| TLAVFTVL | ASSLPGQNYEQY | 0.7411786 |
| TGSVQPARV | ASSLPGQNYEQY | 0.64596266 |
| TGGGAGSGGT | ASSLPGQNYEQY | 0.720065 |
| SHLETSL | ASSLPGQNYEQY | 0.8807161 |
| SFTTGMT | ASSLPGQNYEQY | 0.658356 |
| QVEVAKRMGVRQPTVS | ASSLPGQNYEQY | 0.8629396 |
| QGSGGPALAM | ASSLPGQNYEQY | 0.81401885 |
| MLAPSNP | ASSLPGQNYEQY | 0.8779997 |
| LVNNAAI | ASSLPGQNYEQY | 0.49120462 |
| LTGEINTLA | ASSLPGQNYEQY | 0.87528825 |

| | | |
|---|---|---|
| LSIFGPV | ASSLPGQNYEQY | 0.84014285 |
| LLALLANL | ASSLPGQNYEQY | 0.893311 |
| LISNASDALDKLR | ASSLPGQNYEQY | 0.7832681 |
| LINEAVP | ASSLPGQNYEQY | 0.7970224 |
| LGYAFSGL | ASSLPGQNYEQY | 0.72755015 |
| LDFVAVAL | ASSLPGQNYEQY | 0.8755454 |
| LAQPGAE | ASSLPGQNYEQY | 0.47005135 |
| ISNLGAVTGF | ASSLPGQNYEQY | 0.81762815 |
| HQEDPGRI | ASSLPGQNYEQY | 0.68955046 |
| GTPGEPTP | ASSLPGQNYEQY | 0.83887416 |
| GFFNSGS | ASSLPGQNYEQY | 0.8614943 |
| GAGGGGLTAGAN | ASSLPGQNYEQY | 0.32187396 |
| FVDALVNM | ASSLPGQNYEQY | 0.87559974 |
| FLTVAPEL | ASSLPGQNYEQY | 0.8137484 |
| FETLTAL | ASSLPGQNYEQY | 0.61718595 |
| DVINPIEL | ASSLPGQNYEQY | 0.8837461 |
| APVSGAPV | ASSLPGQNYEQY | 0.82435733 |
| ALVGGLAW | ASSLPGQNYEQY | 0.8821453 |
| ALAAAAATAI | ASSLPGQNYEQY | 0.8290746 |
| AFGLGLLL | ASSLPGQNYEQY | 0.85495394 |
| AFFAAASL | ASSLPGQNYEQY | 0.8570185 |
| AAAALPAFSPP | ASSLPGQNYEQY | 0.66596925 |
| VVLAPAL | ASSLPGQNYEQY | 0.7390077 |
| TIGHVDHGKTTL | ASSLPGQNYEQY | 0.8834102 |
| LISNASDALDK | ASSLPGQNYEQY | 0.77791154 |
| LISNASDAL | ASSLPGQNYEQY | 0.83402824 |
| ISNASDALDK | ASSLPGQNYEQY | 0.5920626 |
| NQAETLVYQTE | ASSLPGQNYEQY | 0.5047146 |
| TGGTGGTAGAG | ASSLPGQNYEQY | 0.6652746 |
| TGSTAGGAGGA | ASSLPGQNYEQY | 0.82032746 |
| DPLAVPVAAGPV | ASSLPGQNYEQY | 0.83171546 |
| GGTGGTPGNGGN | ASSLPGQNYEQY | 0.7960962 |
| PAGLAGVDPATL | ASSLPGQNYEQY | 0.5555742 |
| GNNLVVEAPHAGASV | ASSLPGQNYEQY | 0.7000238 |
| KVTRIMSASTLEQPA | ASSLPGQNYEQY | 0.6613935 |
| LGFALPIALPRRTVV | ASSLPGQNYEQY | 0.6804852 |
| TRLGNLTARKLVVLV | ASSLPGQNYEQY | 0.76733667 |
| WSQQRRLIYFPSAGP | ASSLPGQNYEQY | 0.758629 |
| GPNGNPDPMAAAVDIR | ASSLPGQNYEQY | 0.6584399 |
| TQRCVSVADLRAAASR | ASSLPGQNYEQY | 0.60371023 |
| QFVLDVAEHPVEVLTETV | ASSLPGQNYEQY | 0.73380446 |
| GGAGGTGGAGGNGGTQAGDG | ASSLPGQNYEQY | 0.8065729 |
| PLLATLRAIAVDVDAKPAQV | ASSLPGQNYEQY | 0.7871957 |
| APKFRTGGYMIDFWGVGYQV | ASSLPGQNYEQY | 0.8477782 |
| GAGGAGGAGGAAGGPGGTGG | ASSLPGQNYEQY | 0.8126292 |
| IGALGLLATDVAPAGSVAFG | ASSLPGQNYEQY | 0.81880975 |
| AGGAGGIGGNGGAGGAGGVL | ASSLPGQNYEQY | 0.1929799 |
| ADELIGLL | ASSLPGQNYEQY | 0.36561704 |
| ADRALKA | ASSLPGQNYEQY | 0.6219213 |
| AGAVGQVGGAGGA | ASSLPGQNYEQY | 0.68322676 |
| AGGGGGLGGAP | ASSLPGQNYEQY | 0.33671996 |

| | | |
|---|---|---|
| AGGLLYGNGGAG | ASSLPGQNYEQY | 0.76484823 |
| AGLESIGL | ASSLPGQNYEQY | 0.60098195 |
| AGLKTGII | ASSLPGQNYEQY | 0.55985373 |
| ASLAALLL | ASSLPGQNYEQY | 0.80754256 |
| ASVINISSI | ASSLPGQNYEQY | 0.670367 |
| ATPPGPA | ASSLPGQNYEQY | 0.58771145 |
| ATVASALV | ASSLPGQNYEQY | 0.5834059 |
| ATVASLIL | ASSLPGQNYEQY | 0.8142408 |
| AVNGPVLIRH | ASSLPGQNYEQY | 0.41851372 |
| DGAAGAVGI | ASSLPGQNYEQY | 0.78101593 |
| DPLLHGI | ASSLPGQNYEQY | 0.7227495 |
| ELGAALGAL | ASSLPGQNYEQY | 0.8147427 |
| ELGRTQAV | ASSLPGQNYEQY | 0.434406 |
| FVAGGYLS | ASSLPGQNYEQY | 0.8064782 |
| GASGGAGGAGGW | ASSLPGQNYEQY | 0.7013848 |
| GGAGAGGDGGAGGI | ASSLPGQNYEQY | 0.8565371 |
| GTGHNGSQPGMGGNG | ASSLPGQNYEQY | 0.61766005 |
| GYAAASASAS | ASSLPGQNYEQY | 0.74344003 |
| HQVDVAAVQQG | ASSLPGQNYEQY | 0.70776737 |
| IGAAIAHT | ASSLPGQNYEQY | 0.8265196 |
| IGMIEVAQL | ASSLPGQNYEQY | 0.39194053 |
| IRLAPEHTSAMD | ASSLPGQNYEQY | 0.6310991 |
| IYNVSVL | ASSLPGQNYEQY | 0.4685663 |
| KQGFLGM | ASSLPGQNYEQY | 0.7171906 |
| LESVITQL | ASSLPGQNYEQY | 0.7966962 |
| LGLGALVVV | ASSLPGQNYEQY | 0.2953386 |
| LIGNGGNGGSGGI | ASSLPGQNYEQY | 0.80394197 |
| LNAINAPTL | ASSLPGQNYEQY | 0.72878706 |
| LQLLNTGV | ASSLPGQNYEQY | 0.57080996 |
| LQTSIEF | ASSLPGQNYEQY | 0.8210057 |
| MSFAVSEL | ASSLPGQNYEQY | 0.6398239 |
| NTGALGL | ASSLPGQNYEQY | 0.8273668 |
| PPGAGGAGGAGTT | ASSLPGQNYEQY | 0.6517692 |
| QLVGLGI | ASSLPGQNYEQY | 0.7012651 |
| RFLEQVHQL | ASSLPGQNYEQY | 0.767605 |
| RGVGYVLR | ASSLPGQNYEQY | 0.33750924 |
| SATGVFAGVI | ASSLPGQNYEQY | 0.50860703 |
| SLPAITLP | ASSLPGQNYEQY | 0.83407485 |
| STALAFTV | ASSLPGQNYEQY | 0.62255603 |
| SVLTVGLGK | ASSLPGQNYEQY | 0.6478248 |
| TALAGAGSGI | ASSLPGQNYEQY | 0.5390429 |
| TGLPFTGI | ASSLPGQNYEQY | 0.4945155 |
| VAEHVTV | ASSLPGQNYEQY | 0.53685904 |
| VDIVLNSL | ASSLPGQNYEQY | 0.62360257 |
| VLTGGGAVL | ASSLPGQNYEQY | 0.66075367 |
| VNVGLAYGI | ASSLPGQNYEQY | 0.6224741 |
| VSSAAVGAV | ASSLPGQNYEQY | 0.50166506 |
| VTSVYEAGV | ASSLPGQNYEQY | 0.62506604 |
| WVALDRGA | ASSLPGQNYEQY | 0.59036577 |
| YLVGSVVI | ASSLPGQNYEQY | 0.5679831 |
| GPPGTGKTLL | ASSLPGQNYEQY | 0.5788387 |

| | | |
|---|---|---|
| GVGMTKF | ASSLPGQNYEQY | 0.6341329 |
| IGIGGSDL | ASSLPGQNYEQY | 0.55818284 |
| IGTLLAEL | ASSLPGQNYEQY | 0.4945029 |
| IYTSGTT | ASSLPGQNYEQY | 0.580731 |
| LSFPAGV | ASSLPGQNYEQY | 0.64068425 |
| LYTSGTT | ASSLPGQNYEQY | 0.6248931 |
| VGAFPFGF | ASSLPGQNYEQY | 0.6587771 |
| VGAGTVEF | ASSLPGQNYEQY | 0.6444229 |
| VGKSTLLL | ASSLPGQNYEQY | 0.60924053 |
| AALAHAGGGIV | ASSLPGQNYEQY | 0.33671522 |
| AALVAVSLL | ASSLPGQNYEQY | 0.40036795 |
| AAVSRAVAAL | ASSLPGQNYEQY | 0.56517744 |
| AGGSSGIGGF | ASSLPGQNYEQY | 0.5615662 |
| AGLTVANANG | ASSLPGQNYEQY | 0.33751488 |
| AGLVGAVL | ASSLPGQNYEQY | 0.61786664 |
| ALSQAFTV | ASSLPGQNYEQY | 0.5520415 |
| ALTFSVSL | ASSLPGQNYEQY | 0.5514038 |
| ALTGAALVAL | ASSLPGQNYEQY | 0.6345632 |
| ASFSIGAGVL | ASSLPGQNYEQY | 0.5634073 |
| DGSVYKI | ASSLPGQNYEQY | 0.4990095 |
| EQLAEIV | ASSLPGQNYEQY | 0.53155404 |
| FAVSELTVM | ASSLPGQNYEQY | 0.65831494 |
| GAAPLAAGL | ASSLPGQNYEQY | 0.45856377 |
| GADGGDGGSGGAGG | ASSLPGQNYEQY | 0.63211644 |
| GFGLDTGGAGG | ASSLPGQNYEQY | 0.5041348 |
| GLASGGLL | ASSLPGQNYEQY | 0.651328 |
| GRIAGAAVLL | ASSLPGQNYEQY | 0.45075333 |
| GSAGSLNTSF | ASSLPGQNYEQY | 0.6342992 |
| GTGQIGI | ASSLPGQNYEQY | 0.5817656 |
| GTLIALIL | ASSLPGQNYEQY | 0.6738961 |
| IAATMGFVVG | ASSLPGQNYEQY | 0.544062 |
| IFAQALL | ASSLPGQNYEQY | 0.6230024 |
| IGAAVALGF | ASSLPGQNYEQY | 0.52946454 |
| IGLTPLAL | ASSLPGQNYEQY | 0.64125794 |
| ILGSNTSTL | ASSLPGQNYEQY | 0.4732751 |
| IVFSLTVL | ASSLPGQNYEQY | 0.5555464 |
| LASKATGF | ASSLPGQNYEQY | 0.43003017 |
| LASVALIL | ASSLPGQNYEQY | 0.58181363 |
| LFAVGLTL | ASSLPGQNYEQY | 0.6313237 |
| LFNGGIGTYI | ASSLPGQNYEQY | 0.34461534 |
| LTLGAGII | ASSLPGQNYEQY | 0.639504 |
| LVAAALGDI | ASSLPGQNYEQY | 0.4864811 |
| NTAFVQLGI | ASSLPGQNYEQY | 0.48981762 |
| QLFSGAEL | ASSLPGQNYEQY | 0.63097304 |
| SAATFITL | ASSLPGQNYEQY | 0.61118996 |
| SNYGLANLGV | ASSLPGQNYEQY | 0.6412397 |
| SYGGAYAVM | ASSLPGQNYEQY | 0.53959316 |
| TTGVGGLTLG | ASSLPGQNYEQY | 0.5827333 |
| TTQTIYI | ASSLPGQNYEQY | 0.6083605 |
| VAAGAGLGAVI | ASSLPGQNYEQY | 0.439083 |
| VAAVVTLL | ASSLPGQNYEQY | 0.5108472 |

| | | |
|---|---|---|
| VGFLGGLI | ASSLPGQNYEQY | 0.6495633 |
| VGVIPILL | ASSLPGQNYEQY | 0.5903413 |
| VIFASSLI | ASSLPGQNYEQY | 0.82256913 |
| VKGGDGGAAAG | ASSLPGQNYEQY | 0.46988085 |
| VTKTFTNI | ASSLPGQNYEQY | 0.3463663 |
| VVAAAILL | ASSLPGQNYEQY | 0.49772412 |
| YAALDVEL | ASSLPGQNYEQY | 0.6692985 |
| LASVALLL | ASSLPGQNYEQY | 0.8182197 |
| LDAVHRV | ASSLPGQNYEQY | 0.6440402 |
| QGLSSSIF | ASSLPGQNYEQY | 0.27098194 |
| RGTFANIRL | ASSLPGQNYEQY | 0.534414 |
| RQLWWGHRI | ASSLPGQNYEQY | 0.40287983 |
| YGPPGTGKTLL | ASSLPGQNYEQY | 0.45403206 |
| AALVGAPANVA | ASSLPGQNYEQY | 0.54007506 |
| AATFRRTE | ASSLPGQNYEQY | 0.57621706 |
| AGALGGGI | ASSLPGQNYEQY | 0.5023989 |
| AGGAGGAGGIGAG | ASSLPGQNYEQY | 0.32602125 |
| AGSGALGL | ASSLPGQNYEQY | 0.6118237 |
| AQIADQI | ASSLPGQNYEQY | 0.7090875 |
| ASAVAVTYL | ASSLPGQNYEQY | 0.638395 |
| ASTVLGGTGG | ASSLPGQNYEQY | 0.72887087 |
| AVGGEPAK | ASSLPGQNYEQY | 0.70647657 |
| CEQGTFGGTC | ASSLPGQNYEQY | 0.558462 |
| DEGKAPAG | ASSLPGQNYEQY | 0.14804703 |
| DTVVFPAL | ASSLPGQNYEQY | 0.20536782 |
| FEGITMAG | ASSLPGQNYEQY | 0.41290274 |
| FPLVVQD | ASSLPGQNYEQY | 0.44552726 |
| GAFTTPQ | ASSLPGQNYEQY | 0.17984442 |
| GFGGFGTAGGVGGA | ASSLPGQNYEQY | 0.53653234 |
| GFLAQFL | ASSLPGQNYEQY | 0.418499 |
| GGAGGAGGNAGVA | ASSLPGQNYEQY | 0.40775698 |
| GGVGTAGAGGAGGAGG | ASSLPGQNYEQY | 0.7661288 |
| GNGGGGDAGNAGSGGN | ASSLPGQNYEQY | 0.46661496 |
| GTGGAGGAGNDAGS | ASSLPGQNYEQY | 0.37395287 |
| GTTNTGWFNS | ASSLPGQNYEQY | 0.30645698 |
| IFMGPLFTL | ASSLPGQNYEQY | 0.6928618 |
| ILAHAIEAI | ASSLPGQNYEQY | 0.23008063 |
| IRSAIREAL | ASSLPGQNYEQY | 0.6143204 |
| ITFAGGG | ASSLPGQNYEQY | 0.29541004 |
| ITGMSGHQEV | ASSLPGQNYEQY | 0.6561856 |
| IVAGLACTL | ASSLPGQNYEQY | 0.23027432 |
| IVGGGLAGA | ASSLPGQNYEQY | 0.65825665 |
| IVTGVLNI | ASSLPGQNYEQY | 0.5453342 |
| LAQQGCI | ASSLPGQNYEQY | 0.84023863 |
| LCLHVEN | ASSLPGQNYEQY | 0.42126906 |
| LDAISDGS | ASSLPGQNYEQY | 0.49192038 |
| LGGFAAML | ASSLPGQNYEQY | 0.2966033 |
| LGGKNPMI | ASSLPGQNYEQY | 0.8039876 |
| LGPKGVAAIL | ASSLPGQNYEQY | 0.27805862 |
| LMVGVSI | ASSLPGQNYEQY | 0.45127618 |
| LVGAAFVL | ASSLPGQNYEQY | 0.23071085 |

| | | |
|---|---|---|
| LVLAVAPL | ASSLPGQNYEQY | 0.5869109 |
| MAAIAVIDAA | ASSLPGQNYEQY | 0.623925 |
| NNGINAELQ | ASSLPGQNYEQY | 0.21939151 |
| RGEPLVLVHG | ASSLPGQNYEQY | 0.6298608 |
| RGGAGGAAGLI | ASSLPGQNYEQY | 0.5568047 |
| RSYRMHPAV | ASSLPGQNYEQY | 0.27913165 |
| SAAHLLAGQLA | ASSLPGQNYEQY | 0.7176839 |
| SAAVGSY | ASSLPGQNYEQY | 0.3393683 |
| SPHAVLHRI | ASSLPGQNYEQY | 0.75437653 |
| SQIAKVL | ASSLPGQNYEQY | 0.39673787 |
| TALGHLVP | ASSLPGQNYEQY | 0.3988364 |
| TAPTTSGAA | ASSLPGQNYEQY | 0.6176634 |
| TDGDDGNGGSPGT | ASSLPGQNYEQY | 0.4341058 |
| TDGLGEN | ASSLPGQNYEQY | 0.29648328 |
| TGGPPLPGQA | ASSLPGQNYEQY | 0.72480583 |
| TIPAITI | ASSLPGQNYEQY | 0.76422215 |
| TIPGFTI | ASSLPGQNYEQY | 0.7900232 |
| TSIASGALA | ASSLPGQNYEQY | 0.6838526 |
| TVTLPGAAI | ASSLPGQNYEQY | 0.69546485 |
| VASGLSAALP | ASSLPGQNYEQY | 0.75254405 |
| VGAGVIGI | ASSLPGQNYEQY | 0.7776358 |
| VGPKRFGV | ASSLPGQNYEQY | 0.8035506 |
| VNVAFPD | ASSLPGQNYEQY | 0.7720388 |
| VSMTNSGVS | ASSLPGQNYEQY | 0.72953284 |
| VTDVLGDLSV | ASSLPGQNYEQY | 0.76980805 |
| VVVVGAGFAGL | ASSLPGQNYEQY | 0.7452383 |
| VYTVGTVV | ASSLPGQNYEQY | 0.74672735 |
| WTTFSGP | ASSLPGQNYEQY | 0.7866634 |
| YDELVEAF | ASSLPGQNYEQY | 0.7462762 |
| YGTGGAGGNGG | ASSLPGQNYEQY | 0.6985204 |
| YTNVMAV | ASSLPGQNYEQY | 0.7134302 |
| FLEQVHQL | ASSLPGQNYEQY | 0.783154 |
| GLSLLAVAV | ASSLPGQNYEQY | 0.7839906 |
| LEQVHQL | ASSLPGQNYEQY | 0.78116924 |
| LNAAGAI | ASSLPGQNYEQY | 0.8046493 |
| PAVHASL | ASSLPGQNYEQY | 0.78065056 |
| TELEDDA | ASSLPGQNYEQY | 0.7659244 |
| TPPGPPN | ASSLPGQNYEQY | 0.46799773 |
| TPPPPPPP | ASSLPGQNYEQY | 0.66070104 |
| TQSPALI | ASSLEGWGGSYEQY | 0.7065722 |
| EFAAAFPQ | ASSLEGWGGSYEQY | 0.7562039 |
| LLSGNLDVL | ASSLEGWGGSYEQY | 0.4367991 |
| GPANLGNYNI | ASSLEGWGGSYEQY | 0.76750016 |
| TLGPTLP | ASSLEGWGGSYEQY | 0.74945766 |
| VYAYSSCCK | ASSLEGWGGSYEQY | 0.68604434 |
| APAPAPAPNI | ASSLEGWGGSYEQY | 0.7904002 |
| PGGPVPIP | ASSLEGWGGSYEQY | 0.70025754 |
| FLTVALEP | ASSLEGWGGSYEQY | 0.69963485 |
| SQYFGANL | ASSLEGWGGSYEQY | 0.7409246 |
| GGVKPAA | ASSLEGWGGSYEQY | 0.8033032 |
| GGTGGAGGAGGW | ASSLEGWGGSYEQY | 0.6848504 |

| | | |
|---|---|---|
| GVGGVAGLGGAGG | ASSLEGWGGSYEQY | 0.7913029 |
| VLLVLATTI | ASSLEGWGGSYEQY | 0.7714249 |
| IGIFAAML | ASSLEGWGGSYEQY | 0.7930279 |
| AVGDLID | ASSLEGWGGSYEQY | 0.6586453 |
| NISPLPL | ASSLEGWGGSYEQY | 0.78401595 |
| VTAVFVAI | ASSLEGWGGSYEQY | 0.73927176 |
| SGDDNDAAL | ASSLEGWGGSYEQY | 0.804137 |
| AVAALAHS | ASSLEGWGGSYEQY | 0.7556399 |
| GGVTVMATP | ASSLEGWGGSYEQY | 0.7869462 |
| AGLAANVRD | ASSLEGWGGSYEQY | 0.7703667 |
| TNRPDLIDPALLRPG | ASSLEGWGGSYEQY | 0.7962251 |
| LANLGVTAL | ASSLEGWGGSYEQY | 0.62191296 |
| TGERIEL | ASSLEGWGGSYEQY | 0.7440963 |
| NGEELAA | ASSLEGWGGSYEQY | 0.7314024 |
| VLSLVWG | ASSLEGWGGSYEQY | 0.7639285 |
| IGYALWL | ASSLEGWGGSYEQY | 0.7748683 |
| LSLLLAPL | ASSLEGWGGSYEQY | 0.40455696 |
| TLDEALK | ASSLEGWGGSYEQY | 0.79016197 |
| TDGAGTAEGG | ASSLEGWGGSYEQY | 0.64279413 |
| NAPGGAGGNGG | ASSLEGWGGSYEQY | 0.7061882 |
| VSVFMGMGGNF | ASSLEGWGGSYEQY | 0.7844388 |
| LASYLAAL | ASSLEGWGGSYEQY | 0.73097646 |
| LVGGTPAL | ASSLEGWGGSYEQY | 0.7763717 |
| VRNADVL | ASSLEGWGGSYEQY | 0.71804 |
| TCSSPFLVSAM | ASSLEGWGGSYEQY | 0.7637228 |
| VHITDAE | ASSLEGWGGSYEQY | 0.77807164 |
| EYAAPSF | ASSLEGWGGSYEQY | 0.46154767 |
| VVDAFHGGL | ASSLEGWGGSYEQY | 0.67322075 |
| ASVVDPQ | ASSLEGWGGSYEQY | 0.7939248 |
| SYNEFGGLS | ASSLEGWGGSYEQY | 0.7240579 |
| DPDGSVY | ASSLEGWGGSYEQY | 0.75124323 |
| MVSTIHATE | ASSLEGWGGSYEQY | 0.6833058 |
| LAAGVADPV | ASSLEGWGGSYEQY | 0.59798586 |
| TKEQIAA | ASSLEGWGGSYEQY | 0.69856364 |
| SLASPAAL | ASSLEGWGGSYEQY | 0.74072754 |
| IGLVFYGV | ASSLEGWGGSYEQY | 0.74958026 |
| FLLDASNL | ASSLEGWGGSYEQY | 0.7320871 |
| YQVGNFP | ASSLEGWGGSYEQY | 0.57584167 |
| ESLLSQST | ASSLEGWGGSYEQY | 0.70636827 |
| LFTVGFAAVG | ASSLEGWGGSYEQY | 0.6845957 |
| ATPPTPT | ASSLEGWGGSYEQY | 0.695336 |
| TDLATLG | ASSLEGWGGSYEQY | 0.7335075 |
| SAGGTGR | ASSLEGWGGSYEQY | 0.71832776 |
| GFSNSGS | ASSLEGWGGSYEQY | 0.67485976 |
| GGNGGSGGTSV | ASSLEGWGGSYEQY | 0.6494095 |
| GTSVATGGAGN | ASSLEGWGGSYEQY | 0.7164078 |
| NGGNGGSGGTS | ASSLEGWGGSYEQY | 0.7414663 |
| AVVKFPH | ASSLEGWGGSYEQY | 0.7336248 |
| TPPPPPP | ASSLEGWGGSYEQY | 0.75302243 |
| AAPAAAAAPA | ASSLEGWGGSYEQY | 0.743669 |
| IALNEDL | ASSLEGWGGSYEQY | 0.7429516 |

| | | |
|---|---|---|
| IVNVSSM | ASSLEGWGGSYEQY | 0.3091372 |
| TGIPPAPR | ASSLEGWGGSYEQY | 0.47876203 |
| FDLGGGTF | ASSLEGWGGSYEQY | 0.6424962 |
| FDLGGGTFDVS | ASSLEGWGGSYEQY | 0.6820208 |
| NEPTAAA | ASSLEGWGGSYEQY | 0.4721133 |
| NEPTAAALA | ASSLEGWGGSYEQY | 0.708112 |
| GSIINISSI | ASSLEGWGGSYEQY | 0.6567234 |
| VAPGNAGTA | ASSLEGWGGSYEQY | 0.6202407 |
| TGGGASLE | ASSLEGWGGSYEQY | 0.7438549 |
| VAVYNHYKRI | ASSLEGWGGSYEQY | 0.6780631 |
| TDPAPATT | ASSLEGWGGSYEQY | 0.6388664 |
| TNRPDILDPALLRPG | ASSLEGWGGSYEQY | 0.643033 |
| LTTTTTS | ASSLEGWGGSYEQY | 0.75067735 |
| PTGSGKTL | ASSLEGWGGSYEQY | 0.5144386 |
| TGSGKTL | ASSLEGWGGSYEQY | 0.7258377 |
| TIGHVDHGK | ASSLEGWGGSYEQY | 0.69899035 |
| TIGHVDHGKTT | ASSLEGWGGSYEQY | 0.7413599 |
| GHVDHGKTT | ASSLEGWGGSYEQY | 0.51267624 |
| AQFEHTLL | ASSLEGWGGSYEQY | 0.7390623 |
| IVNQDEL | ASSLEGWGGSYEQY | 0.70387936 |
| TNGSQFF | ASSLEGWGGSYEQY | 0.7582219 |
| TGKGIVE | ASSLEGWGGSYEQY | 0.68323094 |
| LISNASDA | ASSLEGWGGSYEQY | 0.7149599 |
| AELRTLGELTKIAKAHGAQ | ASSLEGWGGSYEQY | 0.6819924 |
| APNAVAAAEPSW | ASSLEGWGGSYEQY | 0.74524033 |
| APQVAGAATAVL | ASSLEGWGGSYEQY | 0.53045607 |
| ARRDPAAAAAILPTDA | ASSLEGWGGSYEQY | 0.6928277 |
| DAAARDVLAEAGLAEYF | ASSLEGWGGSYEQY | 0.57015276 |
| DDQIVLSAKANLAKVEQ | ASSLEGWGGSYEQY | 0.71837455 |
| DELLPDTGEDSDEDGH | ASSLEGWGGSYEQY | 0.7318072 |
| ELPEAPANRATVPQD | ASSLEGWGGSYEQY | 0.5281389 |
| FGQSKIERVTGIG | ASSLEGWGGSYEQY | 0.748922 |
| GAGGTGSTAGGAGGAGGAGGI | ASSLEGWGGSYEQY | 0.7199062 |
| GHGGTGAAGGNGGTGGAGGS( | ASSLEGWGGSYEQY | 0.63610375 |
| GNGGNGGAGGEGGPGGAGGA( | ASSLEGWGGSYEQY | 0.7484413 |
| GSTAGNGGNGGNATNGTIAG( | ASSLEGWGGSYEQY | 0.67673916 |
| GTGGDGGNGGNGGTGG | ASSLEGWGGSYEQY | 0.74737227 |
| GVRRPAGIPADAPV | ASSLEGWGGSYEQY | 0.69350725 |
| GYRQVTDAHLLTLARR | ASSLEGWGGSYEQY | 0.71484375 |
| IAKMGQDGHDRG | ASSLEGWGGSYEQY | 0.7333519 |
| KAEPSIEIQYT | ASSLEGWGGSYEQY | 0.57914186 |
| KPVRIRMAADEDTVAVT | ASSLEGWGGSYEQY | 0.6071218 |
| MPPRIAGMRLLVIKPEP | ASSLEGWGGSYEQY | 0.7467748 |
| MKFYAVALFRSRGVPPTR | ASSLEGWGGSYEQY | 0.8108503 |
| NAGGFGILGAVAHSP | ASSLEGWGGSYEQY | 0.82570976 |
| NEGREITLAMAEM | ASSLEGWGGSYEQY | 0.6796074 |
| NELVEWSHPLGHRWRWE | ASSLEGWGGSYEQY | 0.59299016 |
| NLGAGGTGGNGGNPG | ASSLEGWGGSYEQY | 0.7393143 |
| NTEWPKWETVMAADPVD | ASSLEGWGGSYEQY | 0.8226238 |
| PASAAPAETQP | ASSLEGWGGSYEQY | 0.8221078 |
| QQSSDGKNGVRAAEITGE | ASSLEGWGGSYEQY | 0.8130109 |

| | | |
|---|---|---|
| RAPDAVTTSWWKE | ASSLEGWGGSYEQY | 0.51406676 |
| SGFGNVGAHTSGWFNQ | ASSLEGWGGSYEQY | 0.7722828 |
| SGVAGAVGQAASVGGLKV | ASSLEGWGGSYEQY | 0.72673535 |
| SNGGAGTGSAGGNGGT | ASSLEGWGGSYEQY | 0.7571169 |
| TGGAGGEGGAAG | ASSLEGWGGSYEQY | 0.80509126 |
| TGGDGGAAGAGGAPGHG | ASSLEGWGGSYEQY | 0.75669175 |
| TMPPLLAAIANAGRHGV | ASSLEGWGGSYEQY | 0.70634955 |
| TSSTAGNSGGA | ASSLEGWGGSYEQY | 0.68245023 |
| VPATVPVGILC | ASSLEGWGGSYEQY | 0.80922127 |
| VQRPDSGGVVLTGRIS | ASSLEGWGGSYEQY | 0.8180202 |
| WTPNPVPATKGVPAGLVQAT | ASSLEGWGGSYEQY | 0.8137852 |
| YALKRALAKFSDRFDVVII | ASSLEGWGGSYEQY | 0.83185637 |
| YLIGSVIV | ASSLEGWGGSYEQY | 0.81521094 |
| YGQLVAVID | ASSLEGWGGSYEQY | 0.80874765 |
| VTVGGMTLPA | ASSLEGWGGSYEQY | 0.21617323 |
| VLALPAV | ASSLEGWGGSYEQY | 0.4456926 |
| VIPPTLNYET | ASSLEGWGGSYEQY | 0.6826452 |
| VGMGGMHPSSG | ASSLEGWGGSYEQY | 0.733794 |
| VGLAGGAATA | ASSLEGWGGSYEQY | 0.35850054 |
| VGGAGGAGGSGP | ASSLEGWGGSYEQY | 0.8021184 |
| VFAAFMV | ASSLEGWGGSYEQY | 0.7158743 |
| TLYDKLR | ASSLEGWGGSYEQY | 0.59042865 |
| TLAVFTVL | ASSLEGWGGSYEQY | 0.8131739 |
| TGSVQPARV | ASSLEGWGGSYEQY | 0.66542304 |
| TGGGAGSGGT | ASSLEGWGGSYEQY | 0.60939324 |
| SHLETSL | ASSLEGWGGSYEQY | 0.69850314 |
| SFTTGMT | ASSLEGWGGSYEQY | 0.81415176 |
| QVEVAKRMGVRQPTVS | ASSLEGWGGSYEQY | 0.57092965 |
| QGSGGPALAM | ASSLEGWGGSYEQY | 0.8073551 |
| MLAPSNP | ASSLEGWGGSYEQY | 0.748253 |
| LVNNAAI | ASSLEGWGGSYEQY | 0.8204309 |
| LTGEINTLA | ASSLEGWGGSYEQY | 0.45930895 |
| LSIFGPV | ASSLEGWGGSYEQY | 0.81557673 |
| LLALLANL | ASSLEGWGGSYEQY | 0.778213 |
| LISNASDALDKLR | ASSLEGWGGSYEQY | 0.8376475 |
| LINEAVP | ASSLEGWGGSYEQY | 0.72856534 |
| LGYAFSGL | ASSLEGWGGSYEQY | 0.7553425 |
| LDFVAVAL | ASSLEGWGGSYEQY | 0.64343643 |
| LAQPGAE | ASSLEGWGGSYEQY | 0.82285845 |
| ISNLGAVTGF | ASSLEGWGGSYEQY | 0.39152712 |
| HQEDPGRI | ASSLEGWGGSYEQY | 0.7496722 |
| GTPGEPTP | ASSLEGWGGSYEQY | 0.4874946 |
| GFFNSGS | ASSLEGWGGSYEQY | 0.7585776 |
| GAGGGGLTAGAN | ASSLEGWGGSYEQY | 0.8108032 |
| FVDALVNM | ASSLEGWGGSYEQY | 0.32201385 |
| FLTVAPEL | ASSLEGWGGSYEQY | 0.8073838 |
| FETLTAL | ASSLEGWGGSYEQY | 0.7433772 |
| DVINPIEL | ASSLEGWGGSYEQY | 0.62946635 |
| APVSGAPV | ASSLEGWGGSYEQY | 0.8218368 |
| ALVGGLAW | ASSLEGWGGSYEQY | 0.7487309 |
| ALAAAAATAI | ASSLEGWGGSYEQY | 0.8266442 |

| | | |
|---|---|---|
| AFGLGLLL | ASSLEGWGGSYEQY | 0.76131225 |
| AFFAAASL | ASSLEGWGGSYEQY | 0.76587653 |
| AAAALPAFSPP | ASSLEGWGGSYEQY | 0.7892812 |
| VVLAPAL | ASSLEGWGGSYEQY | 0.55059856 |
| TIGHVDHGKTTL | ASSLEGWGGSYEQY | 0.64959586 |
| LISNASDALDK | ASSLEGWGGSYEQY | 0.8268808 |
| LISNASDAL | ASSLEGWGGSYEQY | 0.7660732 |
| ISNASDALDK | ASSLEGWGGSYEQY | 0.8427002 |
| NQAETLVYQTE | ASSLEGWGGSYEQY | 0.6340623 |
| TGGTGGTAGAG | ASSLEGWGGSYEQY | 0.60001576 |
| TGSTAGGAGGA | ASSLEGWGGSYEQY | 0.6835052 |
| DPLAVPVAAGPV | ASSLEGWGGSYEQY | 0.8563969 |
| GGTGGTPGNGGN | ASSLEGWGGSYEQY | 0.83146477 |
| PAGLAGVDPATL | ASSLEGWGGSYEQY | 0.7838264 |
| GNNLVVEAPHAGASV | ASSLEGWGGSYEQY | 0.61823314 |
| KVTRIMSASTLEQPA | ASSLEGWGGSYEQY | 0.7849112 |
| LGFALPIALPRRTVV | ASSLEGWGGSYEQY | 0.67383075 |
| TRLGNLTARKLVVLV | ASSLEGWGGSYEQY | 0.6842206 |
| WSQQRRLIYFPSAGP | ASSLEGWGGSYEQY | 0.80350286 |
| GPNGNPDPMAAAVDIR | ASSLEGWGGSYEQY | 0.77912045 |
| TQRCVSVADLRAAASR | ASSLEGWGGSYEQY | 0.7127427 |
| QFVLDVAEHPVEVLTETV | ASSLEGWGGSYEQY | 0.68855053 |
| GGAGGTGGAGGNGGTQAGDG | ASSLEGWGGSYEQY | 0.76340485 |
| PLLATLRAIAVDVDAKPAQV | ASSLEGWGGSYEQY | 0.7912745 |
| APKFRTGGYMIDFWGVGYQV[illegible] | ASSLEGWGGSYEQY | 0.73499703 |
| GAGGAGGAGGAAGGPGGTGG[illegible] | ASSLEGWGGSYEQY | 0.88406885 |
| IGALGLLATDVAPAGSVAFG[illegible] | ASSLEGWGGSYEQY | 0.8200668 |
| AGGAGGIGGNGGAGGAGGVL[illegible] | ASSLEGWGGSYEQY | 0.813853 |
| ADELIGLL | ASSLEGWGGSYEQY | 0.21704996 |
| ADRALKA | ASSLEGWGGSYEQY | 0.44659352 |
| AGAVGQVGGAGGA | ASSLEGWGGSYEQY | 0.7202971 |
| AGGGGGLGGAP | ASSLEGWGGSYEQY | 0.6957304 |
| AGGLLYGNGGAG | ASSLEGWGGSYEQY | 0.41825622 |
| AGLESIGL | ASSLEGWGGSYEQY | 0.8306222 |
| AGLKTGII | ASSLEGWGGSYEQY | 0.7115924 |
| ASLAALLL | ASSLEGWGGSYEQY | 0.64084756 |
| ASVINISSI | ASSLEGWGGSYEQY | 0.80107135 |
| ATPPGPA | ASSLEGWGGSYEQY | 0.678449 |
| ATVASALV | ASSLEGWGGSYEQY | 0.70000136 |
| ATVASLIL | ASSLEGWGGSYEQY | 0.6830119 |
| AVNGPVLIRH | ASSLEGWGGSYEQY | 0.86124265 |
| DGAAGAVGI | ASSLEGWGGSYEQY | 0.5271617 |
| DPLLHGI | ASSLEGWGGSYEQY | 0.7948236 |
| ELGAALGAL | ASSLEGWGGSYEQY | 0.7226882 |
| ELGRTQAV | ASSLEGWGGSYEQY | 0.85110956 |
| FVAGGYLS | ASSLEGWGGSYEQY | 0.51137066 |
| GASGGAGGAGGW | ASSLEGWGGSYEQY | 0.7886516 |
| GGAGAGGDGGAGGI | ASSLEGWGGSYEQY | 0.68989146 |
| GTGHNGSQPGMGGNG | ASSLEGWGGSYEQY | 0.8812079 |
| GYAAASASAS | ASSLEGWGGSYEQY | 0.6887784 |
| HQVDVAAVQQG | ASSLEGWGGSYEQY | 0.76932895 |

| | | |
|---|---|---|
| IGAAIAHT | ASSLEGWGGSYEQY | 0.66988796 |
| IGMIEVAQL | ASSLEGWGGSYEQY | 0.8337654 |
| IRLAPEHTSAMD | ASSLEGWGGSYEQY | 0.49414814 |
| IYNVSVL | ASSLEGWGGSYEQY | 0.71481454 |
| KQGFLGM | ASSLEGWGGSYEQY | 0.61810434 |
| LESVITQL | ASSLEGWGGSYEQY | 0.79980254 |
| LGLGALVVV | ASSLEGWGGSYEQY | 0.78034836 |
| LIGNGGNGGSGGI | ASSLEGWGGSYEQY | 0.46241698 |
| LNAINAPTL | ASSLEGWGGSYEQY | 0.822893 |
| LQLLNTGV | ASSLEGWGGSYEQY | 0.72656953 |
| LQTSIEF | ASSLEGWGGSYEQY | 0.64081573 |
| MSFAVSEL | ASSLEGWGGSYEQY | 0.83910435 |
| NTGALGL | ASSLEGWGGSYEQY | 0.71173906 |
| PPGAGGAGGAGTT | ASSLEGWGGSYEQY | 0.8456746 |
| QLVGLGI | ASSLEGWGGSYEQY | 0.76136005 |
| RFLEQVHQL | ASSLEGWGGSYEQY | 0.70818055 |
| RGVGYVLR | ASSLEGWGGSYEQY | 0.83962023 |
| SATGVFAGVI | ASSLEGWGGSYEQY | 0.6022332 |
| SLPAITLP | ASSLEGWGGSYEQY | 0.63323057 |
| STALAFTV | ASSLEGWGGSYEQY | 0.8326764 |
| SVLTVGLGK | ASSLEGWGGSYEQY | 0.7110336 |
| TALAGAGSGI | ASSLEGWGGSYEQY | 0.77219653 |
| TGLPFTGI | ASSLEGWGGSYEQY | 0.5882097 |
| VAEHVTV | ASSLEGWGGSYEQY | 0.5124856 |
| VDIVLNSL | ASSLEGWGGSYEQY | 0.65538645 |
| VLTGGGAVL | ASSLEGWGGSYEQY | 0.78504264 |
| VNVGLAYGI | ASSLEGWGGSYEQY | 0.7580941 |
| VSSAAVGAV | ASSLEGWGGSYEQY | 0.73599315 |
| VTSVYEAGV | ASSLEGWGGSYEQY | 0.52858156 |
| WVALDRGA | ASSLEGWGGSYEQY | 0.6875638 |
| YLVGSVVI | ASSLEGWGGSYEQY | 0.63349026 |
| GPPGTGKTLL | ASSLEGWGGSYEQY | 0.6281729 |
| GVGMTKF | ASSLEGWGGSYEQY | 0.75794053 |
| IGIGGSDL | ASSLEGWGGSYEQY | 0.69349265 |
| IGTLLAEL | ASSLEGWGGSYEQY | 0.60610497 |
| IYTSGTT | ASSLEGWGGSYEQY | 0.6534637 |
| LSFPAGV | ASSLEGWGGSYEQY | 0.73115367 |
| LYTSGTT | ASSLEGWGGSYEQY | 0.72075236 |
| VGAFPFGF | ASSLEGWGGSYEQY | 0.70384914 |
| VGAGTVEF | ASSLEGWGGSYEQY | 0.8170425 |
| VGKSTLLL | ASSLEGWGGSYEQY | 0.75933194 |
| AALAHAGGGIV | ASSLEGWGGSYEQY | 0.7598978 |
| AALVAVSLL | ASSLEGWGGSYEQY | 0.28910854 |
| AAVSRAVAAL | ASSLEGWGGSYEQY | 0.40162376 |
| AGGSSGIGGF | ASSLEGWGGSYEQY | 0.67121637 |
| AGLTVANANG | ASSLEGWGGSYEQY | 0.6471828 |
| AGLVGAVL | ASSLEGWGGSYEQY | 0.43464255 |
| ALSQAFTV | ASSLEGWGGSYEQY | 0.76577485 |
| ALTFSVSL | ASSLEGWGGSYEQY | 0.62210906 |
| ALTGAALVAL | ASSLEGWGGSYEQY | 0.5643612 |
| ASFSIGAGVL | ASSLEGWGGSYEQY | 0.7593861 |

| | | |
|---|---|---|
| DGSVYKI | ASSLEGWGGSYEQY | 0.6048638 |
| EQLAEIV | ASSLEGWGGSYEQY | 0.6114863 |
| FAVSELTVM | ASSLEGWGGSYEQY | 0.63396466 |
| GAAPLAAGL | ASSLEGWGGSYEQY | 0.7751081 |
| GADGGDGGSGGAGG | ASSLEGWGGSYEQY | 0.51576096 |
| GFGLDTGGAGG | ASSLEGWGGSYEQY | 0.7179975 |
| GLASGGLL | ASSLEGWGGSYEQY | 0.62500614 |
| GRIAGAAVLL | ASSLEGWGGSYEQY | 0.77803206 |
| GSAGSLNTSF | ASSLEGWGGSYEQY | 0.46229702 |
| GTGQIGI | ASSLEGWGGSYEQY | 0.7460079 |
| GTLIALIL | ASSLEGWGGSYEQY | 0.64664286 |
| IAATMGFVVG | ASSLEGWGGSYEQY | 0.8193252 |
| IFAQALL | ASSLEGWGGSYEQY | 0.6451191 |
| IGAAVALGF | ASSLEGWGGSYEQY | 0.68799883 |
| IGLTPLAL | ASSLEGWGGSYEQY | 0.6337468 |
| ILGSNTSTL | ASSLEGWGGSYEQY | 0.7493211 |
| IVFSLTVL | ASSLEGWGGSYEQY | 0.43838894 |
| LASKATGF | ASSLEGWGGSYEQY | 0.6400566 |
| LASVALIL | ASSLEGWGGSYEQY | 0.48683783 |
| LFAVGLTL | ASSLEGWGGSYEQY | 0.7117278 |
| LFNGGIGTYI | ASSLEGWGGSYEQY | 0.7207475 |
| LTLGAGII | ASSLEGWGGSYEQY | 0.5137577 |
| LVAAALGDI | ASSLEGWGGSYEQY | 0.7555234 |
| NTAFVQLGI | ASSLEGWGGSYEQY | 0.7035999 |
| QLFSGAEL | ASSLEGWGGSYEQY | 0.5940075 |
| SAATFITL | ASSLEGWGGSYEQY | 0.7930502 |
| SNYGLANLGV | ASSLEGWGGSYEQY | 0.66332704 |
| SYGGAYAVM | ASSLEGWGGSYEQY | 0.779971 |
| TTGVGGLTLG | ASSLEGWGGSYEQY | 0.69173086 |
| TTQTIYI | ASSLEGWGGSYEQY | 0.6100942 |
| VAAGAGLGAVI | ASSLEGWGGSYEQY | 0.7352129 |
| VAAVVTLL | ASSLEGWGGSYEQY | 0.6403317 |
| VGFLGGLI | ASSLEGWGGSYEQY | 0.60498583 |
| VGVIPILL | ASSLEGWGGSYEQY | 0.7735719 |
| VIFASSLI | ASSLEGWGGSYEQY | 0.8078283 |
| VKGGDGGAAAG | ASSLEGWGGSYEQY | 0.78815126 |
| VTKTFTNI | ASSLEGWGGSYEQY | 0.68525606 |
| VVAAAILL | ASSLEGWGGSYEQY | 0.633817 |
| YAALDVEL | ASSLEGWGGSYEQY | 0.7297261 |
| LASVALLL | ASSLEGWGGSYEQY | 0.82308096 |
| LDAVHRV | ASSLEGWGGSYEQY | 0.82132655 |
| QGLSSSIF | ASSLEGWGGSYEQY | 0.7896832 |
| RGTFANIRL | ASSLEGWGGSYEQY | 0.59688324 |
| RQLWWGHRI | ASSLEGWGGSYEQY | 0.8150863 |
| YGPPGTGKTLL | ASSLEGWGGSYEQY | 0.76799816 |
| AALVGAPANVA | ASSLEGWGGSYEQY | 0.7565291 |
| AATFRRTE | ASSLEGWGGSYEQY | 0.8017106 |
| AGALGGGI | ASSLEGWGGSYEQY | 0.7969477 |
| AGGAGGAGGIGAG | ASSLEGWGGSYEQY | 0.75207204 |
| AGSGALGL | ASSLEGWGGSYEQY | 0.71761143 |
| AQIADQI | ASSLEGWGGSYEQY | 0.8062544 |

| | | |
|---|---|---|
| ASAVAVTYL | ASSLEGWGGSYEQY | 0.830817 |
| ASTVLGGTGG | ASSLEGWGGSYEQY | 0.82041264 |
| AVGGEPAK | ASSLEGWGGSYEQY | 0.8531635 |
| CEQGTFGGTC | ASSLEGWGGSYEQY | 0.82949555 |
| DEGKAPAG | ASSLEGWGGSYEQY | 0.8211926 |
| DTVVFPAL | ASSLEGWGGSYEQY | 0.2930462 |
| FEGITMAG | ASSLEGWGGSYEQY | 0.5108007 |
| FPLVVQD | ASSLEGWGGSYEQY | 0.7105148 |
| GAFTTPQ | ASSLEGWGGSYEQY | 0.7455335 |
| GFGGFGTAGGVGGA | ASSLEGWGGSYEQY | 0.36630324 |
| GFLAQFL | ASSLEGWGGSYEQY | 0.7987313 |
| GGAGGAGGNAGVA | ASSLEGWGGSYEQY | 0.7474277 |
| GGVGTAGAGGAGGAGG | ASSLEGWGGSYEQY | 0.6681361 |
| GNGGGGDAGNAGSGGN | ASSLEGWGGSYEQY | 0.82684577 |
| GTGGAGGAGNDAGS | ASSLEGWGGSYEQY | 0.7288569 |
| GTTNTGWFNS | ASSLEGWGGSYEQY | 0.70081115 |
| IFMGPLFTL | ASSLEGWGGSYEQY | 0.68360883 |
| ILAHAIEAI | ASSLEGWGGSYEQY | 0.84460926 |
| IRSAIREAL | ASSLEGWGGSYEQY | 0.6167837 |
| ITFAGGG | ASSLEGWGGSYEQY | 0.8249757 |
| ITGMSGHQEV | ASSLEGWGGSYEQY | 0.7031009 |
| IVAGLACTL | ASSLEGWGGSYEQY | 0.8495946 |
| IVGGGLAGA | ASSLEGWGGSYEQY | 0.50532055 |
| IVTGVLNI | ASSLEGWGGSYEQY | 0.8200763 |
| LAQQGCI | ASSLEGWGGSYEQY | 0.75719124 |
| LCLHVEN | ASSLEGWGGSYEQY | 0.86138844 |
| LDAISDGS | ASSLEGWGGSYEQY | 0.7547761 |
| LGGFAAML | ASSLEGWGGSYEQY | 0.801314 |
| LGGKNPMI | ASSLEGWGGSYEQY | 0.72845745 |
| LGPKGVAAIL | ASSLEGWGGSYEQY | 0.8100916 |
| LMVGVSI | ASSLEGWGGSYEQY | 0.50616884 |
| LVGAAFVL | ASSLEGWGGSYEQY | 0.7575513 |
| LVLAVAPL | ASSLEGWGGSYEQY | 0.5950465 |
| MAAIAVIDAA | ASSLEGWGGSYEQY | 0.7944058 |
| NNGINAELQ | ASSLEGWGGSYEQY | 0.8031553 |
| RGEPLVLVHG | ASSLEGWGGSYEQY | 0.33228385 |
| RGGAGGAAGLI | ASSLEGWGGSYEQY | 0.8167648 |
| RSYRMHPAV | ASSLEGWGGSYEQY | 0.6625477 |
| SAAHLLAGQLA | ASSLEGWGGSYEQY | 0.63164276 |
| SAAVGSY | ASSLEGWGGSYEQY | 0.8304186 |
| SPHAVLHRI | ASSLEGWGGSYEQY | 0.77959985 |
| SQIAKVL | ASSLEGWGGSYEQY | 0.8338125 |
| TALGHLVP | ASSLEGWGGSYEQY | 0.73016655 |
| TAPTTSGAA | ASSLEGWGGSYEQY | 0.7406033 |
| TDGDDGNGGSPGT | ASSLEGWGGSYEQY | 0.80950195 |
| TDGLGEN | ASSLEGWGGSYEQY | 0.48678234 |
| TGGPPLPGQA | ASSLEGWGGSYEQY | 0.6147719 |
| TIPAITI | ASSLEGWGGSYEQY | 0.84248304 |
| TIPGFTI | ASSLEGWGGSYEQY | 0.81605613 |
| TSIASGALA | ASSLEGWGGSYEQY | 0.86072934 |
| TVTLPGAAI | ASSLEGWGGSYEQY | 0.6942942 |

| | | |
|---|---|---|
| VASGLSAALP | ASSLEGWGGSYEQY | 0.60471094 |
| VGAGVIGI | ASSLEGWGGSYEQY | 0.76441324 |
| VGPKRFGV | ASSLEGWGGSYEQY | 0.87376666 |
| VNVAFPD | ASSLEGWGGSYEQY | 0.8538001 |
| VSMTNSGVS | ASSLEGWGGSYEQY | 0.8287346 |
| VTDVLGDLSV | ASSLEGWGGSYEQY | 0.5602973 |
| VVVVGAGFAGL | ASSLEGWGGSYEQY | 0.7631265 |
| VYTVGTVV | ASSLEGWGGSYEQY | 0.71879673 |
| WTTFSGP | ASSLEGWGGSYEQY | 0.7301804 |
| YDELVEAF | ASSLEGWGGSYEQY | 0.84839237 |
| YGTGGAGGNGG | ASSLEGWGGSYEQY | 0.7918881 |
| YTNVMAV | ASSLEGWGGSYEQY | 0.7158745 |
| FLEQVHQL | ASSLEGWGGSYEQY | 0.692916 |
| GLSLLAVAV | ASSLEGWGGSYEQY | 0.8051808 |
| LEQVHQL | ASSLEGWGGSYEQY | 0.863377 |
| LNAAGAI | ASSLEGWGGSYEQY | 0.84974706 |
| PAVHASL | ASSLEGWGGSYEQY | 0.8922652 |
| TELEDDA | ASSLEGWGGSYEQY | 0.85352933 |
| TPPGPPN | ASSLEGWGGSYEQY | 0.8638672 |
| TPPPPPPP | ASSLEGWGGSYEQY | 0.18238789 |
| TQSPALI | ASLSYSGNTLY | 0.3707407 |
| EFAAAFPQ | ASLSYSGNTLY | 0.7573265 |
| LLSGNLDVL | ASLSYSGNTLY | 0.7129737 |
| GPANLGNYNI | ASLSYSGNTLY | 0.5262456 |
| TLGPTLP | ASLSYSGNTLY | 0.80195475 |
| VYAYSSCCK | ASLSYSGNTLY | 0.6988411 |
| APAPAPAPNI | ASLSYSGNTLY | 0.64038885 |
| PGGPVPIP | ASLSYSGNTLY | 0.85567594 |
| FLTVALEP | ASLSYSGNTLY | 0.6973125 |
| SQYFGANL | ASLSYSGNTLY | 0.6761123 |
| GGVKPAA | ASLSYSGNTLY | 0.6701687 |
| GGTGGAGGAGGW | ASLSYSGNTLY | 0.8630835 |
| GVGGVAGLGGAGG | ASLSYSGNTLY | 0.5388725 |
| VLLVLATTI | ASLSYSGNTLY | 0.81839085 |
| IGIFAAML | ASLSYSGNTLY | 0.73536193 |
| AVGDLID | ASLSYSGNTLY | 0.874272 |
| NISPLPL | ASLSYSGNTLY | 0.477127 |
| VTAVFVAI | ASLSYSGNTLY | 0.83661085 |
| SGDDNDAAL | ASLSYSGNTLY | 0.7378016 |
| AVAALAHS | ASLSYSGNTLY | 0.89775467 |
| GGVTVMATP | ASLSYSGNTLY | 0.7353758 |
| AGLAANVRD | ASLSYSGNTLY | 0.79010487 |
| TNRPDLIDPALLRPG | ASLSYSGNTLY | 0.6691943 |
| LANLGVTAL | ASLSYSGNTLY | 0.83797944 |
| TGERIEL | ASLSYSGNTLY | 0.5357293 |
| NGEELAA | ASLSYSGNTLY | 0.74581206 |
| VLSLVWG | ASLSYSGNTLY | 0.514968 |
| IGYALWL | ASLSYSGNTLY | 0.8118762 |
| LSLLLAPL | ASLSYSGNTLY | 0.814013 |
| TLDEALK | ASLSYSGNTLY | 0.47053713 |
| TDGAGTAEGG | ASLSYSGNTLY | 0.86116016 |

| | | |
|---|---|---|
| NAPGGAGGNGG | ASLSYSGNTLY | 0.77658975 |
| VSVFMGMGGNF | ASLSYSGNTLY | 0.6902235 |
| LASYLAAL | ASLSYSGNTLY | 0.88191265 |
| LVGGTPAL | ASLSYSGNTLY | 0.7618797 |
| VRNADVL | ASLSYSGNTLY | 0.85921323 |
| TCSSPFLVSAM | ASLSYSGNTLY | 0.79343945 |
| VHITDAE | ASLSYSGNTLY | 0.7262641 |
| EYAAPSF | ASLSYSGNTLY | 0.85103667 |
| VVDAFHGGL | ASLSYSGNTLY | 0.7482039 |
| ASVVDPQ | ASLSYSGNTLY | 0.67769337 |
| SYNEFGGLS | ASLSYSGNTLY | 0.85939205 |
| DPDGSVY | ASLSYSGNTLY | 0.73000526 |
| MVSTIHATE | ASLSYSGNTLY | 0.7606522 |
| LAAGVADPV | ASLSYSGNTLY | 0.7015122 |
| TKEQIAA | ASLSYSGNTLY | 0.63376564 |
| SLASPAAL | ASLSYSGNTLY | 0.70263827 |
| IGLVFYGV | ASLSYSGNTLY | 0.7540598 |
| FLLDASNL | ASLSYSGNTLY | 0.7577981 |
| YQVGNFP | ASLSYSGNTLY | 0.7354691 |
| ESLLSQST | ASLSYSGNTLY | 0.61259735 |
| LFTVGFAAVG | ASLSYSGNTLY | 0.72490776 |
| ATPPTPT | ASLSYSGNTLY | 0.70381534 |
| TDLATLG | ASLSYSGNTLY | 0.70342696 |
| SAGGTGR | ASLSYSGNTLY | 0.7216072 |
| GFSNSGS | ASLSYSGNTLY | 0.7409178 |
| GGNGGSGGTSV | ASLSYSGNTLY | 0.6958247 |
| GTSVATGGAGN | ASLSYSGNTLY | 0.6694861 |
| NGGNGGSGGTS | ASLSYSGNTLY | 0.72978586 |
| AVVKFPH | ASLSYSGNTLY | 0.7192942 |
| TPPPPPP | ASLSYSGNTLY | 0.7027317 |
| AAPAAAAAPA | ASLSYSGNTLY | 0.7601072 |
| IALNEDL | ASLSYSGNTLY | 0.7523836 |
| IVNVSSM | ASLSYSGNTLY | 0.7429684 |
| TGIPPAPR | ASLSYSGNTLY | 0.28326085 |
| FDLGGGTF | ASLSYSGNTLY | 0.40280914 |
| FDLGGGTFDVS | ASLSYSGNTLY | 0.70207644 |
| NEPTAAA | ASLSYSGNTLY | 0.692144 |
| NEPTAAALA | ASLSYSGNTLY | 0.4375451 |
| GSIINISSI | ASLSYSGNTLY | 0.7310958 |
| VAPGNAGTA | ASLSYSGNTLY | 0.6589631 |
| TGGGASLE | ASLSYSGNTLY | 0.6798246 |
| VAVYNHYKRI | ASLSYSGNTLY | 0.7480401 |
| TDPAPATT | ASLSYSGNTLY | 0.7026115 |
| TNRPDILDPALLRPG | ASLSYSGNTLY | 0.6574186 |
| LTTTTTS | ASLSYSGNTLY | 0.6670843 |
| PTGSGKTL | ASLSYSGNTLY | 0.75124955 |
| TGSGKTL | ASLSYSGNTLY | 0.5835899 |
| TIGHVDHGK | ASLSYSGNTLY | 0.7353991 |
| TIGHVDHGKTT | ASLSYSGNTLY | 0.6349775 |
| GHVDHGKTT | ASLSYSGNTLY | 0.7549797 |
| AQFEHTLL | ASLSYSGNTLY | 0.5773848 |

| | | |
|---|---|---|
| IVNQDEL | ASLSYSGNTLY | 0.7462766 |
| TNGSQFF | ASLSYSGNTLY | 0.71866727 |
| TGKGIVE | ASLSYSGNTLY | 0.7705089 |
| LISNASDA | ASLSYSGNTLY | 0.6945158 |
| AELRTLGELTKIAKAHGAQ | ASLSYSGNTLY | 0.7203884 |
| APNAVAAAEPSW | ASLSYSGNTLY | 0.65370625 |
| APQVAGAATAVL | ASLSYSGNTLY | 0.75665855 |
| ARRDPAAAAAILPTDA | ASLSYSGNTLY | 0.587419 |
| DAAARDVLAEAGLAEYF | ASLSYSGNTLY | 0.69372976 |
| DDQIVLSAKANLAKVEQ | ASLSYSGNTLY | 0.4604475 |
| DELLPDTGEDSDEDGH | ASLSYSGNTLY | 0.7301938 |
| ELPEAPANRATVPQD | ASLSYSGNTLY | 0.73206776 |
| FGQSKIERVTGIG | ASLSYSGNTLY | 0.463876 |
| GAGGTGSTAGGAGGAGGAGGI | ASLSYSGNTLY | 0.74912256 |
| GHGGTGAAGGNGGTGGAGGS( | ASLSYSGNTLY | 0.70532745 |
| GNGGNGGAGGEGGPGGAGGA( | ASLSYSGNTLY | 0.67128706 |
| GSTAGNGGNGGNATNGTIAG( | ASLSYSGNTLY | 0.7541213 |
| GTGGDGGNGGNGGTGG | ASLSYSGNTLY | 0.71984434 |
| GVRRPAGIPADAPV | ASLSYSGNTLY | 0.7527781 |
| GYRQVTDAHLLTLARR | ASLSYSGNTLY | 0.7203342 |
| IAKMGQDGHDRG | ASLSYSGNTLY | 0.71327245 |
| KAEPSIEIQYT | ASLSYSGNTLY | 0.73345184 |
| KPVRIRMAADEDTVAVT | ASLSYSGNTLY | 0.69083375 |
| MPPRIAGMRLLVIKPEP | ASLSYSGNTLY | 0.6833258 |
| MKFYAVALFRSRGVPPTR | ASLSYSGNTLY | 0.75382316 |
| NAGGFGILGAVAHSP | ASLSYSGNTLY | 0.83232224 |
| NEGREITLAMAEM | ASLSYSGNTLY | 0.8725646 |
| NELVEWSHPLGHRWRWE | ASLSYSGNTLY | 0.73268116 |
| NLGAGGTGGNGGNPG | ASLSYSGNTLY | 0.62804544 |
| NTEWPKWETVMAADPVD | ASLSYSGNTLY | 0.78675306 |
| PASAAPAETQP | ASLSYSGNTLY | 0.8810846 |
| QQSSDGKNGVRAAEITGE | ASLSYSGNTLY | 0.86729205 |
| RAPDAVTTSWWKE | ASLSYSGNTLY | 0.8423817 |
| SGFGNVGAHTSGWFNQ | ASLSYSGNTLY | 0.5842799 |
| SGVAGAVGQAASVGGLKV | ASLSYSGNTLY | 0.7853371 |
| SNGGAGTGSAGGNGGT | ASLSYSGNTLY | 0.75056064 |
| TGGAGGEGGAAG | ASLSYSGNTLY | 0.7592816 |
| TGGDGGAAGAGGAPGHG | ASLSYSGNTLY | 0.85615486 |
| TMPPLLAAIANAGRHGV | ASLSYSGNTLY | 0.8106003 |
| TSSTAGNSGGA | ASLSYSGNTLY | 0.73266757 |
| VPATVPVGILC | ASLSYSGNTLY | 0.7201004 |
| VQRPDSGGVVLTGRIS | ASLSYSGNTLY | 0.8289284 |
| WTPNPVPATKGVPAGLVQAT | ASLSYSGNTLY | 0.87376153 |
| YALKRALAKFSDRFDVVII | ASLSYSGNTLY | 0.8650766 |
| YLIGSVIV | ASLSYSGNTLY | 0.89640534 |
| YGQLVAVID | ASLSYSGNTLY | 0.8663657 |
| VTVGGMTLPA | ASLSYSGNTLY | 0.8754271 |
| VLALPAV | ASLSYSGNTLY | 0.19330056 |
| VIPPTLNYET | ASLSYSGNTLY | 0.4280934 |
| VGMGGMHPSSG | ASLSYSGNTLY | 0.76499635 |
| VGLAGGAATA | ASLSYSGNTLY | 0.758029 |

| | | |
|---|---|---|
| VGGAGGAGGSGP | ASLSYSGNTLY | 0.5653057 |
| VFAAFMV | ASLSYSGNTLY | 0.81624293 |
| TLYDKLR | ASLSYSGNTLY | 0.7153386 |
| TLAVFTVL | ASLSYSGNTLY | 0.6668186 |
| TGSVQPARV | ASLSYSGNTLY | 0.8692694 |
| TGGGAGSGGT | ASLSYSGNTLY | 0.717582 |
| SHLETSL | ASLSYSGNTLY | 0.69877344 |
| SFTTGMT | ASLSYSGNTLY | 0.70100856 |
| QVEVAKRMGVRQPTVS | ASLSYSGNTLY | 0.8762952 |
| QGSGGPALAM | ASLSYSGNTLY | 0.537079 |
| MLAPSNP | ASLSYSGNTLY | 0.8376981 |
| LVNNAAI | ASLSYSGNTLY | 0.754846 |
| LTGEINTLA | ASLSYSGNTLY | 0.8832059 |
| LSIFGPV | ASLSYSGNTLY | 0.4967381 |
| LLALLANL | ASLSYSGNTLY | 0.85228616 |
| LISNASDALDKLR | ASLSYSGNTLY | 0.7709388 |
| LINEAVP | ASLSYSGNTLY | 0.90202284 |
| LGYAFSGL | ASLSYSGNTLY | 0.76256764 |
| LDFVAVAL | ASLSYSGNTLY | 0.8006739 |
| LAQPGAE | ASLSYSGNTLY | 0.6852807 |
| ISNLGAVTGF | ASLSYSGNTLY | 0.8557011 |
| HQEDPGRI | ASLSYSGNTLY | 0.5714998 |
| GTPGEPTP | ASLSYSGNTLY | 0.77717924 |
| GFFNSGS | ASLSYSGNTLY | 0.55309236 |
| GAGGGGLTAGAN | ASLSYSGNTLY | 0.8252671 |
| FVDALVNM | ASLSYSGNTLY | 0.8338514 |
| FLTVAPEL | ASLSYSGNTLY | 0.50865024 |
| FETLTAL | ASLSYSGNTLY | 0.8700095 |
| DVINPIEL | ASLSYSGNTLY | 0.8049493 |
| APVSGAPV | ASLSYSGNTLY | 0.71269107 |
| ALVGGLAW | ASLSYSGNTLY | 0.8889208 |
| ALAAAAATAI | ASLSYSGNTLY | 0.7959161 |
| AFGLGLLL | ASLSYSGNTLY | 0.86926115 |
| AFFAAASL | ASLSYSGNTLY | 0.8092867 |
| AAAALPAFSPP | ASLSYSGNTLY | 0.7491385 |
| VVLAPAL | ASLSYSGNTLY | 0.8602463 |
| TIGHVDHGKTTL | ASLSYSGNTLY | 0.7749514 |
| LISNASDALDK | ASLSYSGNTLY | 0.7090825 |
| LISNASDAL | ASLSYSGNTLY | 0.87046695 |
| ISNASDALDK | ASLSYSGNTLY | 0.7466692 |
| NQAETLVYQTE | ASLSYSGNTLY | 0.78002304 |
| TGGTGGTAGAG | ASLSYSGNTLY | 0.54379636 |
| TGSTAGGAGGA | ASLSYSGNTLY | 0.5753027 |
| DPLAVPVAAGPV | ASLSYSGNTLY | 0.6700016 |
| GGTGGTPGNGGN | ASLSYSGNTLY | 0.8120929 |
| PAGLAGVDPATL | ASLSYSGNTLY | 0.8230274 |
| GNNLVVEAPHAGASV | ASLSYSGNTLY | 0.77206993 |
| KVTRIMSASTLEQPA | ASLSYSGNTLY | 0.6544363 |
| LGFALPIALPRRTVV | ASLSYSGNTLY | 0.77031016 |
| TRLGNLTARKLVVLV | ASLSYSGNTLY | 0.6668431 |
| WSQQRRLIYFPSAGP | ASLSYSGNTLY | 0.693275 |

| | | |
|---|---|---|
| GPNGNPDPMAAAVDIR | ASLSYSGNTLY | 0.7822745 |
| TQRCVSVADLRAAASR | ASLSYSGNTLY | 0.74130917 |
| QFVLDVAEHPVEVLTETV | ASLSYSGNTLY | 0.6891848 |
| GGAGGTGGAGGNGGTQAGDG | ASLSYSGNTLY | 0.62497675 |
| PLLATLRAIAVDVDAKPAQV | ASLSYSGNTLY | 0.77586555 |
| APKFRTGGYMIDFWGVGYQV | ASLSYSGNTLY | 0.8069384 |
| GAGGAGGAGGAAGGPGGTGG | ASLSYSGNTLY | 0.7772835 |
| IGALGLLATDVAPAGSVAFG | ASLSYSGNTLY | 0.8502234 |
| AGGAGGIGGNGGAGGAGGVL | ASLSYSGNTLY | 0.77600324 |
| ADELIGLL | ASLSYSGNTLY | 0.7830546 |
| ADRALKA | ASLSYSGNTLY | 0.26522204 |
| AGAVGQVGGAGGA | ASLSYSGNTLY | 0.44954917 |
| AGGGGGLGGAP | ASLSYSGNTLY | 0.69750655 |
| AGGLLYGNGGAG | ASLSYSGNTLY | 0.69931185 |
| AGLESIGL | ASLSYSGNTLY | 0.33014894 |
| AGLKTGII | ASLSYSGNTLY | 0.7974175 |
| ASLAALLL | ASLSYSGNTLY | 0.6900737 |
| ASVINISSI | ASLSYSGNTLY | 0.62876713 |
| ATPPGPA | ASLSYSGNTLY | 0.7955122 |
| ATVASALV | ASLSYSGNTLY | 0.65853995 |
| ATVASLIL | ASLSYSGNTLY | 0.6370257 |
| AVNGPVLIRH | ASLSYSGNTLY | 0.6819626 |
| DGAAGAVGI | ASLSYSGNTLY | 0.8302088 |
| DPLLHGI | ASLSYSGNTLY | 0.58699346 |
| ELGAALGAL | ASLSYSGNTLY | 0.78103125 |
| ELGRTQAV | ASLSYSGNTLY | 0.7383407 |
| FVAGGYLS | ASLSYSGNTLY | 0.81802404 |
| GASGGAGGAGGW | ASLSYSGNTLY | 0.5194495 |
| GGAGAGGDGGAGGI | ASLSYSGNTLY | 0.7728783 |
| GTGHNGSQPGMGGNG | ASLSYSGNTLY | 0.6592144 |
| GYAAASASAS | ASLSYSGNTLY | 0.8498448 |
| HQVDVAAVQQG | ASLSYSGNTLY | 0.65977573 |
| IGAAIAHT | ASLSYSGNTLY | 0.77965117 |
| IGMIEVAQL | ASLSYSGNTLY | 0.76627004 |
| IRLAPEHTSAMD | ASLSYSGNTLY | 0.7756188 |
| IYNVSVL | ASLSYSGNTLY | 0.47216648 |
| KQGFLGM | ASLSYSGNTLY | 0.6487036 |
| LESVITQL | ASLSYSGNTLY | 0.679129 |
| LGLGALVVV | ASLSYSGNTLY | 0.74005705 |
| LIGNGGNGGSGGI | ASLSYSGNTLY | 0.7608912 |
| LNAINAPTL | ASLSYSGNTLY | 0.31899703 |
| LQLLNTGV | ASLSYSGNTLY | 0.7818762 |
| LQTSIEF | ASLSYSGNTLY | 0.57914764 |
| MSFAVSEL | ASLSYSGNTLY | 0.62821585 |
| NTGALGL | ASLSYSGNTLY | 0.79587084 |
| PPGAGGAGGAGTT | ASLSYSGNTLY | 0.71773565 |
| QLVGLGI | ASLSYSGNTLY | 0.7668326 |
| RFLEQVHQL | ASLSYSGNTLY | 0.6969608 |
| RGVGYVLR | ASLSYSGNTLY | 0.69357896 |
| SATGVFAGVI | ASLSYSGNTLY | 0.7869023 |
| SLPAITLP | ASLSYSGNTLY | 0.4588181 |

| | | |
|---|---|---|
| STALAFTV | ASLSYSGNTLY | 0.5950651 |
| SVLTVGLGK | ASLSYSGNTLY | 0.80996007 |
| TALAGAGSGI | ASLSYSGNTLY | 0.84394485 |
| TGLPFTGI | ASLSYSGNTLY | 0.846887 |
| VAEHVTV | ASLSYSGNTLY | 0.7769255 |
| VDIVLNSL | ASLSYSGNTLY | 0.7341244 |
| VLTGGGAVL | ASLSYSGNTLY | 0.79474384 |
| VNVGLAYGI | ASLSYSGNTLY | 0.8495743 |
| VSSAAVGAV | ASLSYSGNTLY | 0.8511207 |
| VTSVYEAGV | ASLSYSGNTLY | 0.8384596 |
| WVALDRGA | ASLSYSGNTLY | 0.63019943 |
| YLVGSVVI | ASLSYSGNTLY | 0.837951 |
| GPPGTGKTLL | ASLSYSGNTLY | 0.80407 |
| GVGMTKF | ASLSYSGNTLY | 0.80664515 |
| IGIGGSDL | ASLSYSGNTLY | 0.8408482 |
| IGTLLAEL | ASLSYSGNTLY | 0.82539845 |
| IYTSGTT | ASLSYSGNTLY | 0.80154604 |
| LSFPAGV | ASLSYSGNTLY | 0.7464863 |
| LYTSGTT | ASLSYSGNTLY | 0.8368881 |
| VGAFPFGF | ASLSYSGNTLY | 0.8517713 |
| VGAGTVEF | ASLSYSGNTLY | 0.8490716 |
| VGKSTLLL | ASLSYSGNTLY | 0.854639 |
| AALAHAGGGIV | ASLSYSGNTLY | 0.8515056 |
| AALVAVSLL | ASLSYSGNTLY | 0.849586 |
| AAVSRAVAAL | ASLSYSGNTLY | 0.27762017 |
| AGGSSGIGGF | ASLSYSGNTLY | 0.5136151 |
| AGLTVANANG | ASLSYSGNTLY | 0.7805384 |
| AGLVGAVL | ASLSYSGNTLY | 0.7749748 |
| ALSQAFTV | ASLSYSGNTLY | 0.44060338 |
| ALTFSVSL | ASLSYSGNTLY | 0.8301096 |
| ALTGAALVAL | ASLSYSGNTLY | 0.7757712 |
| ASFSIGAGVL | ASLSYSGNTLY | 0.75433254 |
| DGSVYKI | ASLSYSGNTLY | 0.8449449 |
| EQLAEIV | ASLSYSGNTLY | 0.79668385 |
| FAVSELTVM | ASLSYSGNTLY | 0.7460287 |
| GAAPLAAGL | ASLSYSGNTLY | 0.7412132 |
| GADGGDGGSGGAGG | ASLSYSGNTLY | 0.85239565 |
| GFGLDTGGAGG | ASLSYSGNTLY | 0.7043835 |
| GLASGGLL | ASLSYSGNTLY | 0.8461292 |
| GRIAGAAVLL | ASLSYSGNTLY | 0.7015732 |
| GSAGSLNTSF | ASLSYSGNTLY | 0.8533099 |
| GTGQIGI | ASLSYSGNTLY | 0.5564853 |
| GTLIALIL | ASLSYSGNTLY | 0.84965503 |
| IAATMGFVVG | ASLSYSGNTLY | 0.83254725 |
| IFAQALL | ASLSYSGNTLY | 0.8622308 |
| IGAAVALGF | ASLSYSGNTLY | 0.8140015 |
| IGLTPLAL | ASLSYSGNTLY | 0.8282155 |
| ILGSNTSTL | ASLSYSGNTLY | 0.7832625 |
| IVFSLTVL | ASLSYSGNTLY | 0.8443432 |
| LASKATGF | ASLSYSGNTLY | 0.6059692 |
| LASVALIL | ASLSYSGNTLY | 0.81186676 |

| | | |
|---|---|---|
| LFAVGLTL | ASLSYSGNTLY | 0.5799125 |
| LFNGGIGTYI | ASLSYSGNTLY | 0.829813 |
| LTLGAGII | ASLSYSGNTLY | 0.8389893 |
| LVAAALGDI | ASLSYSGNTLY | 0.40564242 |
| NTAFVQLGI | ASLSYSGNTLY | 0.83897924 |
| QLFSGAEL | ASLSYSGNTLY | 0.808337 |
| SAATFITL | ASLSYSGNTLY | 0.7115915 |
| SNYGLANLGV | ASLSYSGNTLY | 0.85510373 |
| SYGGAYAVM | ASLSYSGNTLY | 0.81750876 |
| TTGVGGLTLG | ASLSYSGNTLY | 0.8517722 |
| TTQTIYI | ASLSYSGNTLY | 0.7984836 |
| VAAGAGLGAVI | ASLSYSGNTLY | 0.78734756 |
| VAAVVTLL | ASLSYSGNTLY | 0.82484114 |
| VGFLGGLI | ASLSYSGNTLY | 0.53664494 |
| VGVIPILL | ASLSYSGNTLY | 0.7046311 |
| VIFASSLI | ASLSYSGNTLY | 0.8586493 |
| VKGGDGGAAAG | ASLSYSGNTLY | 0.7053169 |
| VTKTFTNI | ASLSYSGNTLY | 0.7867321 |
| VVAAAILL | ASLSYSGNTLY | 0.57678354 |
| YAALDVEL | ASLSYSGNTLY | 0.5413074 |
| LASVALLL | ASLSYSGNTLY | 0.6583668 |
| LDAVHRV | ASLSYSGNTLY | 0.73304945 |
| QGLSSSIF | ASLSYSGNTLY | 0.78289837 |
| RGTFANIRL | ASLSYSGNTLY | 0.7417509 |
| RQLWWGHRI | ASLSYSGNTLY | 0.4814608 |
| YGPPGTGKTLL | ASLSYSGNTLY | 0.6887078 |
| AALVGAPANVA | ASLSYSGNTLY | 0.53316313 |
| AATFRRTE | ASLSYSGNTLY | 0.62286747 |
| AGALGGGI | ASLSYSGNTLY | 0.6979842 |
| AGGAGGAGGIGAG | ASLSYSGNTLY | 0.5703161 |
| AGSGALGL | ASLSYSGNTLY | 0.5559625 |
| AQIADQI | ASLSYSGNTLY | 0.5447256 |
| ASAVAVTYL | ASLSYSGNTLY | 0.67667747 |
| ASTVLGGTGG | ASLSYSGNTLY | 0.7771367 |
| AVGGEPAK | ASLSYSGNTLY | 0.7567909 |
| CEQGTFGGTC | ASLSYSGNTLY | 0.7852025 |
| DEGKAPAG | ASLSYSGNTLY | 0.76291555 |
| DTVVFPAL | ASLSYSGNTLY | 0.76679134 |
| FEGITMAG | ASLSYSGNTLY | 0.22251126 |
| FPLVVQD | ASLSYSGNTLY | 0.38771528 |
| GAFTTPQ | ASLSYSGNTLY | 0.4764651 |
| GFGGFGTAGGVGGA | ASLSYSGNTLY | 0.6257987 |
| GFLAQFL | ASLSYSGNTLY | 0.27183297 |
| GGAGGAGGNAGVA | ASLSYSGNTLY | 0.7026311 |
| GGVGTAGAGGAGGAGG | ASLSYSGNTLY | 0.6000829 |
| GNGGGGDAGNAGSGGN | ASLSYSGNTLY | 0.48556072 |
| GTGGAGGAGNDAGS | ASLSYSGNTLY | 0.7603085 |
| GTTNTGWFNS | ASLSYSGNTLY | 0.52686703 |
| IFMGPLFTL | ASLSYSGNTLY | 0.4302796 |
| ILAHAIEAI | ASLSYSGNTLY | 0.48486644 |
| IRSAIREAL | ASLSYSGNTLY | 0.7541121 |

| | | |
|---|---|---|
| ITFAGGG | ASLSYSGNTLY | 0.33327413 |
| ITGMSGHQEV | ASLSYSGNTLY | 0.7201604 |
| IVAGLACTL | ASLSYSGNTLY | 0.40012938 |
| IVGGGLAGA | ASLSYSGNTLY | 0.7602246 |
| IVTGVLNI | ASLSYSGNTLY | 0.378579 |
| LAQQGCI | ASLSYSGNTLY | 0.74843335 |
| LCLHVEN | ASLSYSGNTLY | 0.61396337 |
| LDAISDGS | ASLSYSGNTLY | 0.81041014 |
| LGGFAAML | ASLSYSGNTLY | 0.61561537 |
| LGGKNPMI | ASLSYSGNTLY | 0.6538148 |
| LGPKGVAAIL | ASLSYSGNTLY | 0.41132963 |
| LMVGVSI | ASLSYSGNTLY | 0.77638817 |
| LVGAAFVL | ASLSYSGNTLY | 0.32953057 |
| LVLAVAPL | ASLSYSGNTLY | 0.6612861 |
| MAAIAVIDAA | ASLSYSGNTLY | 0.34945786 |
| NNGINAELQ | ASLSYSGNTLY | 0.7042471 |
| RGEPLVLVHG | ASLSYSGNTLY | 0.73720634 |
| RGGAGGAAGLI | ASLSYSGNTLY | 0.32478112 |
| RSYRMHPAV | ASLSYSGNTLY | 0.6875071 |
| SAAHLLAGQLA | ASLSYSGNTLY | 0.5531538 |
| SAAVGSY | ASLSYSGNTLY | 0.4950401 |
| SPHAVLHRI | ASLSYSGNTLY | 0.7719363 |
| SQIAKVL | ASLSYSGNTLY | 0.64007306 |
| TALGHLVP | ASLSYSGNTLY | 0.7725946 |
| TAPTTSGAA | ASLSYSGNTLY | 0.5198142 |
| TDGDDGNGGSPGT | ASLSYSGNTLY | 0.5024605 |
| TDGLGEN | ASLSYSGNTLY | 0.72097605 |
| TGGPPLPGQA | ASLSYSGNTLY | 0.5063852 |
| TIPAITI | ASLSYSGNTLY | 0.42063892 |
| TIPGFTI | ASLSYSGNTLY | 0.7687309 |
| TSIASGALA | ASLSYSGNTLY | 0.68965614 |
| TVTLPGAAI | ASLSYSGNTLY | 0.7038357 |
| VASGLSAALP | ASLSYSGNTLY | 0.63055074 |
| VGAGVIGI | ASLSYSGNTLY | 0.63475084 |
| VGPKRFGV | ASLSYSGNTLY | 0.64586747 |
| VNVAFPD | ASLSYSGNTLY | 0.6996262 |
| VSMTNSGVS | ASLSYSGNTLY | 0.71048295 |
| VTDVLGDLSV | ASLSYSGNTLY | 0.6999519 |
| VVVVGAGFAGL | ASLSYSGNTLY | 0.65555274 |
| VYTVGTVV | ASLSYSGNTLY | 0.6959441 |
| WTTFSGP | ASLSYSGNTLY | 0.6696617 |
| YDELVEAF | ASLSYSGNTLY | 0.67234504 |
| YGTGGAGGNGG | ASLSYSGNTLY | 0.69149524 |
| YTNVMAV | ASLSYSGNTLY | 0.6914693 |
| FLEQVHQL | ASLSYSGNTLY | 0.64219666 |
| GLSLLAVAV | ASLSYSGNTLY | 0.64047873 |
| LEQVHQL | ASLSYSGNTLY | 0.67940223 |
| LNAAGAI | ASLSYSGNTLY | 0.69828296 |
| PAVHASL | ASLSYSGNTLY | 0.6854899 |
| TELEDDA | ASLSYSGNTLY | 0.7135769 |
| TPPGPPN | ASLSYSGNTLY | 0.6960253 |

| | | |
|---|---|---|
| TPPPPPPP | ASLSYSGNTLY | 0.6907909 |
| TQSPALI | ASSPPGGTGNTLY | 0.40889794 |
| EFAAAFPQ | ASSPPGGTGNTLY | 0.5855614 |
| LLSGNLDVL | ASSPPGGTGNTLY | 0.6458594 |
| GPANLGNYNI | ASSPPGGTGNTLY | 0.67397624 |
| TLGPTLP | ASSPPGGTGNTLY | 0.3313627 |
| VYAYSSCCK | ASSPPGGTGNTLY | 0.6985253 |
| APAPAPAPNI | ASSPPGGTGNTLY | 0.672687 |
| PGGPVPIP | ASSPPGGTGNTLY | 0.64677095 |
| FLTVALEP | ASSPPGGTGNTLY | 0.69768983 |
| SQYFGANL | ASSPPGGTGNTLY | 0.6600529 |
| GGVKPAA | ASSPPGGTGNTLY | 0.6409649 |
| GGTGGAGGAGGW | ASSPPGGTGNTLY | 0.6650092 |
| GVGGVAGLGGAGG | ASSPPGGTGNTLY | 0.70882726 |
| VLLVLATTI | ASSPPGGTGNTLY | 0.59545434 |
| IGIFAAML | ASSPPGGTGNTLY | 0.7032026 |
| AVGDLID | ASSPPGGTGNTLY | 0.6538987 |
| NISPLPL | ASSPPGGTGNTLY | 0.70373106 |
| VTAVFVAI | ASSPPGGTGNTLY | 0.6238822 |
| SGDDNDAAL | ASSPPGGTGNTLY | 0.70291317 |
| AVAALAHS | ASSPPGGTGNTLY | 0.67135215 |
| GGVTVMATP | ASSPPGGTGNTLY | 0.71399456 |
| AGLAANVRD | ASSPPGGTGNTLY | 0.6680263 |
| TNRPDLIDPALLRPG | ASSPPGGTGNTLY | 0.701779 |
| LANLGVTAL | ASSPPGGTGNTLY | 0.6804953 |
| TGERIEL | ASSPPGGTGNTLY | 0.6988889 |
| NGEELAA | ASSPPGGTGNTLY | 0.59578764 |
| VLSLVWG | ASSPPGGTGNTLY | 0.66053784 |
| IGYALWL | ASSPPGGTGNTLY | 0.6198679 |
| LSLLLAPL | ASSPPGGTGNTLY | 0.6824154 |
| TLDEALK | ASSPPGGTGNTLY | 0.70028377 |
| TDGAGTAEGG | ASSPPGGTGNTLY | 0.43053862 |
| NAPGGAGGNGG | ASSPPGGTGNTLY | 0.69287133 |
| VSVFMGMGGNF | ASSPPGGTGNTLY | 0.63311404 |
| LASYLAAL | ASSPPGGTGNTLY | 0.6442698 |
| LVGGTPAL | ASSPPGGTGNTLY | 0.69910896 |
| VRNADVL | ASSPPGGTGNTLY | 0.67524624 |
| TCSSPFLVSAM | ASSPPGGTGNTLY | 0.6902895 |
| VHITDAE | ASSPPGGTGNTLY | 0.6489423 |
| EYAAPSF | ASSPPGGTGNTLY | 0.6663712 |
| VVDAFHGGL | ASSPPGGTGNTLY | 0.69089675 |
| ASVVDPQ | ASSPPGGTGNTLY | 0.42390785 |
| SYNEFGGLS | ASSPPGGTGNTLY | 0.6014148 |
| DPDGSVY | ASSPPGGTGNTLY | 0.70976937 |
| MVSTIHATE | ASSPPGGTGNTLY | 0.6864321 |
| LAAGVADPV | ASSPPGGTGNTLY | 0.6963798 |
| TKEQIAA | ASSPPGGTGNTLY | 0.6320643 |
| SLASPAAL | ASSPPGGTGNTLY | 0.6030957 |
| IGLVFYGV | ASSPPGGTGNTLY | 0.6237032 |
| FLLDASNL | ASSPPGGTGNTLY | 0.6813648 |
| YQVGNFP | ASSPPGGTGNTLY | 0.7060694 |

| | | |
|---|---|---|
| ESLLSQST | ASSPPGGTGNTLY | 0.6891247 |
| LFTVGFAAVG | ASSPPGGTGNTLY | 0.5574498 |
| ATPPTPT | ASSPPGGTGNTLY | 0.7083995 |
| TDLATLG | ASSPPGGTGNTLY | 0.67677575 |
| SAGGTGR | ASSPPGGTGNTLY | 0.6685795 |
| GFSNSGS | ASSPPGGTGNTLY | 0.67076445 |
| GGNGGSGGTSV | ASSPPGGTGNTLY | 0.6735225 |
| GTSVATGGAGN | ASSPPGGTGNTLY | 0.6356356 |
| NGGNGGSGGTS | ASSPPGGTGNTLY | 0.57132345 |
| AVVKFPH | ASSPPGGTGNTLY | 0.664885 |
| TPPPPPP | ASSPPGGTGNTLY | 0.69421685 |
| AAPAAAAAPA | ASSPPGGTGNTLY | 0.6891557 |
| IALNEDL | ASSPPGGTGNTLY | 0.70890707 |
| IVNVSSM | ASSPPGGTGNTLY | 0.70221406 |
| TGIPPAPR | ASSPPGGTGNTLY | 0.67957103 |
| FDLGGGTF | ASSPPGGTGNTLY | 0.37871873 |
| FDLGGGTFDVS | ASSPPGGTGNTLY | 0.567718 |
| NEPTAAA | ASSPPGGTGNTLY | 0.54320794 |
| NEPTAAALA | ASSPPGGTGNTLY | 0.6659939 |
| GSIINISSI | ASSPPGGTGNTLY | 0.33603144 |
| VAPGNAGTA | ASSPPGGTGNTLY | 0.66898376 |
| TGGGASLE | ASSPPGGTGNTLY | 0.6608859 |
| VAVYNHYKRI | ASSPPGGTGNTLY | 0.63142085 |
| TDPAPATT | ASSPPGGTGNTLY | 0.7017748 |
| TNRPDILDPALLRPG | ASSPPGGTGNTLY | 0.64720756 |
| LTTTTTS | ASSPPGGTGNTLY | 0.57068074 |
| PTGSGKTL | ASSPPGGTGNTLY | 0.57775813 |
| TGSGKTL | ASSPPGGTGNTLY | 0.7028699 |
| TIGHVDHGK | ASSPPGGTGNTLY | 0.56987715 |
| TIGHVDHGKTT | ASSPPGGTGNTLY | 0.709118 |
| GHVDHGKTT | ASSPPGGTGNTLY | 0.51024336 |
| AQFEHTLL | ASSPPGGTGNTLY | 0.7064242 |
| IVNQDEL | ASSPPGGTGNTLY | 0.5384741 |
| TNGSQFF | ASSPPGGTGNTLY | 0.7015927 |
| TGKGIVE | ASSPPGGTGNTLY | 0.6705228 |
| LISNASDA | ASSPPGGTGNTLY | 0.715512 |
| AELRTLGELTKIAKAHGAQ | ASSPPGGTGNTLY | 0.65831393 |
| APNAVAAAEPSW | ASSPPGGTGNTLY | 0.69695145 |
| APQVAGAATAVL | ASSPPGGTGNTLY | 0.611897 |
| ARRDPAAAAAILPTDA | ASSPPGGTGNTLY | 0.7034731 |
| DAAARDVLAEAGLAEYF | ASSPPGGTGNTLY | 0.55163366 |
| DDQIVLSAKANLAKVEQ | ASSPPGGTGNTLY | 0.6626179 |
| DELLPDTGEDSDEDGH | ASSPPGGTGNTLY | 0.5549928 |
| ELPEAPANRATVPQD | ASSPPGGTGNTLY | 0.6735828 |
| FGQSKIERVTGIG | ASSPPGGTGNTLY | 0.6988446 |
| GAGGTGSTAGGAGGAGGAGGI | ASSPPGGTGNTLY | 0.36172935 |
| GHGGTGAAGGNGGTGGAGGS( | ASSPPGGTGNTLY | 0.69800234 |
| GNGGNGGAGGEGGPGGAGGA( | ASSPPGGTGNTLY | 0.5864806 |
| GSTAGNGGNGGNATNGTIAG( | ASSPPGGTGNTLY | 0.5780807 |
| GTGGDGGNGGNGGTGG | ASSPPGGTGNTLY | 0.6942197 |
| GVRRPAGIPADAPV | ASSPPGGTGNTLY | 0.6847378 |

| | | |
|---|---|---|
| GYRQVTDAHLLTLARR | ASSPPGGTGNTLY | 0.687986 |
| IAKMGQDGHDRG | ASSPPGGTGNTLY | 0.5693698 |
| KAEPSIEIQYT | ASSPPGGTGNTLY | 0.6636125 |
| KPVRIRMAADEDTVAVT | ASSPPGGTGNTLY | 0.6836895 |
| MPPRIAGMRLLVIKPEP | ASSPPGGTGNTLY | 0.40883982 |
| MKFYAVALFRSRGVPPTR | ASSPPGGTGNTLY | 0.51881826 |
| NAGGFGILGAVAHSP | ASSPPGGTGNTLY | 0.71075815 |
| NEGREITLAMAEM | ASSPPGGTGNTLY | 0.6808887 |
| NELVEWSHPLGHRWRWE | ASSPPGGTGNTLY | 0.6807864 |
| NLGAGGTGGNGGNPG | ASSPPGGTGNTLY | 0.6399187 |
| NTEWPKWETVMAADPVD | ASSPPGGTGNTLY | 0.6168888 |
| PASAAPAETQP | ASSPPGGTGNTLY | 0.61441416 |
| QQSSDGKNGVRAAEITGE | ASSPPGGTGNTLY | 0.6783133 |
| RAPDAVTTSWWKE | ASSPPGGTGNTLY | 0.68703663 |
| SGFGNVGAHTSGWFNQ | ASSPPGGTGNTLY | 0.6823983 |
| SGVAGAVGQAASVGGLKV | ASSPPGGTGNTLY | 0.5295108 |
| SNGGAGTGSAGGNGGT | ASSPPGGTGNTLY | 0.6826575 |
| TGGAGGEGGAAG | ASSPPGGTGNTLY | 0.66756594 |
| TGGDGGAAGAGGAPGHG | ASSPPGGTGNTLY | 0.6678679 |
| TMPPLLAAIANAGRHGV | ASSPPGGTGNTLY | 0.6665077 |
| TSSTAGNSGGA | ASSPPGGTGNTLY | 0.6564237 |
| VPATVPVGILC | ASSPPGGTGNTLY | 0.6004938 |
| VQRPDSGGVVLTGRIS | ASSPPGGTGNTLY | 0.59710103 |
| WTPNPVPATKGVPAGLVQAT | ASSPPGGTGNTLY | 0.6550491 |
| YALKRALAKFSDRFDVVII | ASSPPGGTGNTLY | 0.68692046 |
| YLIGSVIV | ASSPPGGTGNTLY | 0.6867837 |
| YGQLVAVID | ASSPPGGTGNTLY | 0.6904751 |
| VTVGGMTLPA | ASSPPGGTGNTLY | 0.6897433 |
| VLALPAV | ASSPPGGTGNTLY | 0.68158746 |
| VIPPTLNYET | ASSPPGGTGNTLY | 0.38680312 |
| VGMGGMHPSSG | ASSPPGGTGNTLY | 0.55888206 |
| VGLAGGAATA | ASSPPGGTGNTLY | 0.58697486 |
| VGGAGGAGGSGP | ASSPPGGTGNTLY | 0.6530935 |
| VFAAFMV | ASSPPGGTGNTLY | 0.40901145 |
| TLYDKLR | ASSPPGGTGNTLY | 0.6104648 |
| TLAVFTVL | ASSPPGGTGNTLY | 0.6380918 |
| TGSVQPARV | ASSPPGGTGNTLY | 0.624529 |
| TGGGAGSGGT | ASSPPGGTGNTLY | 0.6843306 |
| SHLETSL | ASSPPGGTGNTLY | 0.64746964 |
| SFTTGMT | ASSPPGGTGNTLY | 0.5782487 |
| QVEVAKRMGVRQPTVS | ASSPPGGTGNTLY | 0.558609 |
| QGSGGPALAM | ASSPPGGTGNTLY | 0.68072695 |
| MLAPSNP | ASSPPGGTGNTLY | 0.5972194 |
| LVNNAAI | ASSPPGGTGNTLY | 0.6902494 |
| LTGEINTLA | ASSPPGGTGNTLY | 0.49532312 |
| LSIFGPV | ASSPPGGTGNTLY | 0.69314647 |
| LLALLANL | ASSPPGGTGNTLY | 0.55231166 |
| LISNASDALDKLR | ASSPPGGTGNTLY | 0.69024765 |
| LINEAVP | ASSPPGGTGNTLY | 0.6705762 |
| LGYAFSGL | ASSPPGGTGNTLY | 0.69899726 |
| LDFVAVAL | ASSPPGGTGNTLY | 0.6637258 |

| | | |
|---|---|---|
| LAQPGAE | ASSPPGGTGNTLY | 0.67613006 |
| ISNLGAVTGF | ASSPPGGTGNTLY | 0.5663971 |
| HQEDPGRI | ASSPPGGTGNTLY | 0.68229073 |
| GTPGEPTP | ASSPPGGTGNTLY | 0.55282307 |
| GFFNSGS | ASSPPGGTGNTLY | 0.65290165 |
| GAGGGGLTAGAN | ASSPPGGTGNTLY | 0.53718233 |
| FVDALVNM | ASSPPGGTGNTLY | 0.66020566 |
| FLTVAPEL | ASSPPGGTGNTLY | 0.6855241 |
| FETLTAL | ASSPPGGTGNTLY | 0.3905962 |
| DVINPIEL | ASSPPGGTGNTLY | 0.6825502 |
| APVSGAPV | ASSPPGGTGNTLY | 0.63429177 |
| ALVGGLAW | ASSPPGGTGNTLY | 0.6037326 |
| ALAAAAATAI | ASSPPGGTGNTLY | 0.6893513 |
| AFGLGLLL | ASSPPGGTGNTLY | 0.6646494 |
| AFFAAASL | ASSPPGGTGNTLY | 0.68161714 |
| AAAALPAFSPP | ASSPPGGTGNTLY | 0.57409614 |
| VVLAPAL | ASSPPGGTGNTLY | 0.6335316 |
| TIGHVDHGKTTL | ASSPPGGTGNTLY | 0.6621448 |
| LISNASDALDK | ASSPPGGTGNTLY | 0.6087922 |
| LISNASDAL | ASSPPGGTGNTLY | 0.5404495 |
| ISNASDALDK | ASSPPGGTGNTLY | 0.69469273 |
| NQAETLVYQTE | ASSPPGGTGNTLY | 0.6982275 |
| TGGTGGTAGAG | ASSPPGGTGNTLY | 0.7771275 |
| TGSTAGGAGGA | ASSPPGGTGNTLY | 0.49612415 |
| DPLAVPVAAGPV | ASSPPGGTGNTLY | 0.43727678 |
| GGTGGTPGNGGN | ASSPPGGTGNTLY | 0.6290843 |
| PAGLAGVDPATL | ASSPPGGTGNTLY | 0.7773807 |
| GNNLVVEAPHAGASV | ASSPPGGTGNTLY | 0.7754373 |
| KVTRIMSASTLEQPA | ASSPPGGTGNTLY | 0.7274918 |
| LGFALPIALPRRTVV | ASSPPGGTGNTLY | 0.4090879 |
| TRLGNLTARKLVVLV | ASSPPGGTGNTLY | 0.6085856 |
| WSQQRRLIYFPSAGP | ASSPPGGTGNTLY | 0.5738784 |
| GPNGNPDPMAAAVDIR | ASSPPGGTGNTLY | 0.5648127 |
| TQRCVSVADLRAAASR | ASSPPGGTGNTLY | 0.70601475 |
| QFVLDVAEHPVEVLTETV | ASSPPGGTGNTLY | 0.6122515 |
| GGAGGTGGAGGNGGTQAGDG | ASSPPGGTGNTLY | 0.47819257 |
| PLLATLRAIAVDVDAKPAQV | ASSPPGGTGNTLY | 0.5094434 |
| APKFRTGGYMIDFWGVGYQV | ASSPPGGTGNTLY | 0.6261629 |
| GAGGAGGAGGAAGGPGGTGG | ASSPPGGTGNTLY | 0.75820136 |
| IGALGLLATDVAPAGSVAFG | ASSPPGGTGNTLY | 0.73227227 |
| AGGAGGIGGNGGAGGAGGVL | ASSPPGGTGNTLY | 0.7948258 |
| ADELIGLL | ASSPPGGTGNTLY | 0.74362123 |
| ADRALKA | ASSPPGGTGNTLY | 0.7387997 |
| AGAVGQVGGAGGA | ASSPPGGTGNTLY | 0.18468039 |
| AGGGGGLGGAP | ASSPPGGTGNTLY | 0.2654702 |
| AGGLLYGNGGAG | ASSPPGGTGNTLY | 0.5554941 |
| AGLESIGL | ASSPPGGTGNTLY | 0.60121363 |
| AGLKTGII | ASSPPGGTGNTLY | 0.35431093 |
| ASLAALLL | ASSPPGGTGNTLY | 0.7382946 |
| ASVINISSI | ASSPPGGTGNTLY | 0.5593434 |
| ATPPGPA | ASSPPGGTGNTLY | 0.46785718 |

| | | |
|---|---|---|
| ATVASALV | ASSPPGGTGNTLY | 0.75149345 |
| ATVASLIL | ASSPPGGTGNTLY | 0.48605168 |
| AVNGPVLIRH | ASSPPGGTGNTLY | 0.46903458 |
| DGAAGAVGI | ASSPPGGTGNTLY | 0.49084878 |
| DPLLHGI | ASSPPGGTGNTLY | 0.7571917 |
| ELGAALGAL | ASSPPGGTGNTLY | 0.34972197 |
| ELGRTQAV | ASSPPGGTGNTLY | 0.691479 |
| FVAGGYLS | ASSPPGGTGNTLY | 0.53416246 |
| GASGGAGGAGGW | ASSPPGGTGNTLY | 0.744313 |
| GGAGAGGDGGAGGI | ASSPPGGTGNTLY | 0.3570828 |
| GTGHNGSQPGMGGNG | ASSPPGGTGNTLY | 0.7184001 |
| GYAAASASAS | ASSPPGGTGNTLY | 0.59047705 |
| HQVDVAAVQQG | ASSPPGGTGNTLY | 0.8000536 |
| IGAAIAHT | ASSPPGGTGNTLY | 0.5706029 |
| IGMIEVAQL | ASSPPGGTGNTLY | 0.6567751 |
| IRLAPEHTSAMD | ASSPPGGTGNTLY | 0.46347594 |
| IYNVSVL | ASSPPGGTGNTLY | 0.7368882 |
| KQGFLGM | ASSPPGGTGNTLY | 0.42560565 |
| LESVITQL | ASSPPGGTGNTLY | 0.597702 |
| LGLGALVVV | ASSPPGGTGNTLY | 0.33935246 |
| LIGNGGNGGSGGI | ASSPPGGTGNTLY | 0.67619574 |
| LNAINAPTL | ASSPPGGTGNTLY | 0.6938801 |
| LQLLNTGV | ASSPPGGTGNTLY | 0.39027685 |
| LQTSIEF | ASSPPGGTGNTLY | 0.6675509 |
| MSFAVSEL | ASSPPGGTGNTLY | 0.69727457 |
| NTGALGL | ASSPPGGTGNTLY | 0.5217482 |
| PPGAGGAGGAGTT | ASSPPGGTGNTLY | 0.76555586 |
| QLVGLGI | ASSPPGGTGNTLY | 0.5823768 |
| RFLEQVHQL | ASSPPGGTGNTLY | 0.7469362 |
| RGVGYVLR | ASSPPGGTGNTLY | 0.53577435 |
| SATGVFAGVI | ASSPPGGTGNTLY | 0.52891016 |
| SLPAITLP | ASSPPGGTGNTLY | 0.7335538 |
| STALAFTV | ASSPPGGTGNTLY | 0.55652785 |
| SVLTVGLGK | ASSPPGGTGNTLY | 0.48503712 |
| TALAGAGSGI | ASSPPGGTGNTLY | 0.7336965 |
| TGLPFTGI | ASSPPGGTGNTLY | 0.6566634 |
| VAEHVTV | ASSPPGGTGNTLY | 0.7059827 |
| VDIVLNSL | ASSPPGGTGNTLY | 0.39428818 |
| VLTGGGAVL | ASSPPGGTGNTLY | 0.35263902 |
| VNVGLAYGI | ASSPPGGTGNTLY | 0.39419731 |
| VSSAAVGAV | ASSPPGGTGNTLY | 0.7120633 |
| VTSVYEAGV | ASSPPGGTGNTLY | 0.73969626 |
| WVALDRGA | ASSPPGGTGNTLY | 0.6683389 |
| YLVGSVVI | ASSPPGGTGNTLY | 0.36858177 |
| GPPGTGKTLL | ASSPPGGTGNTLY | 0.5927409 |
| GVGMTKF | ASSPPGGTGNTLY | 0.43931046 |
| IGIGGSDL | ASSPPGGTGNTLY | 0.51130265 |
| IGTLLAEL | ASSPPGGTGNTLY | 0.52259403 |
| IYTSGTT | ASSPPGGTGNTLY | 0.6205448 |
| LSFPAGV | ASSPPGGTGNTLY | 0.43492717 |
| LYTSGTT | ASSPPGGTGNTLY | 0.38821915 |

| | | |
|---|---|---|
| VGAFPFGF | ASSPPGGTGNTLY | 0.56431293 |
| VGAGTVEF | ASSPPGGTGNTLY | 0.7204174 |
| VGKSTLLL | ASSPPGGTGNTLY | 0.65218794 |
| AALAHAGGGIV | ASSPPGGTGNTLY | 0.75847214 |
| AALVAVSLL | ASSPPGGTGNTLY | 0.70210147 |
| AAVSRAVAAL | ASSPPGGTGNTLY | 0.63010347 |
| AGGSSGIGGF | ASSPPGGTGNTLY | 0.17497063 |
| AGLTVANANG | ASSPPGGTGNTLY | 0.27692595 |
| AGLVGAVL | ASSPPGGTGNTLY | 0.3958197 |
| ALSQAFTV | ASSPPGGTGNTLY | 0.4896242 |
| ALTFSVSL | ASSPPGGTGNTLY | 0.16823155 |
| ALTGAALVAL | ASSPPGGTGNTLY | 0.61920846 |
| ASFSIGAGVL | ASSPPGGTGNTLY | 0.46681568 |
| DGSVYKI | ASSPPGGTGNTLY | 0.3802337 |
| EQLAEIV | ASSPPGGTGNTLY | 0.6605854 |
| FAVSELTVM | ASSPPGGTGNTLY | 0.44409198 |
| GAAPLAAGL | ASSPPGGTGNTLY | 0.4208318 |
| GADGGDGGSGGAGG | ASSPPGGTGNTLY | 0.41682407 |
| GFGLDTGGAGG | ASSPPGGTGNTLY | 0.7127309 |
| GLASGGLL | ASSPPGGTGNTLY | 0.27320728 |
| GRIAGAAVLL | ASSPPGGTGNTLY | 0.6834508 |
| GSAGSLNTSF | ASSPPGGTGNTLY | 0.33003044 |
| GTGQIGI | ASSPPGGTGNTLY | 0.7121761 |
| GTLIALIL | ASSPPGGTGNTLY | 0.31937382 |
| IAATMGFVVG | ASSPPGGTGNTLY | 0.7068533 |
| IFAQALL | ASSPPGGTGNTLY | 0.56184435 |
| IGAAVALGF | ASSPPGGTGNTLY | 0.80571735 |
| IGLTPLAL | ASSPPGGTGNTLY | 0.46124822 |
| ILGSNTSTL | ASSPPGGTGNTLY | 0.587439 |
| IVFSLTVL | ASSPPGGTGNTLY | 0.504511 |
| LASKATGF | ASSPPGGTGNTLY | 0.7224369 |
| LASVALIL | ASSPPGGTGNTLY | 0.25467902 |
| LFAVGLTL | ASSPPGGTGNTLY | 0.4807561 |
| LFNGGIGTYI | ASSPPGGTGNTLY | 0.26864737 |
| LTLGAGII | ASSPPGGTGNTLY | 0.5714999 |
| LVAAALGDI | ASSPPGGTGNTLY | 0.69047356 |
| NTAFVQLGI | ASSPPGGTGNTLY | 0.1728208 |
| QLFSGAEL | ASSPPGGTGNTLY | 0.5343037 |
| SAATFITL | ASSPPGGTGNTLY | 0.3765684 |
| SNYGLANLGV | ASSPPGGTGNTLY | 0.3519846 |
| SYGGAYAVM | ASSPPGGTGNTLY | 0.6360818 |
| TTGVGGLTLG | ASSPPGGTGNTLY | 0.54338735 |
| TTQTIYI | ASSPPGGTGNTLY | 0.7003857 |
| VAAGAGLGAVI | ASSPPGGTGNTLY | 0.40336582 |
| VAAVVTLL | ASSPPGGTGNTLY | 0.38813084 |
| VGFLGGLI | ASSPPGGTGNTLY | 0.62397826 |
| VGVIPILL | ASSPPGGTGNTLY | 0.2527951 |
| VIFASSLI | ASSPPGGTGNTLY | 0.32996288 |
| VKGGDGGAAAG | ASSPPGGTGNTLY | 0.7623122 |
| VTKTFTNI | ASSPPGGTGNTLY | 0.80085444 |
| VVAAAILL | ASSPPGGTGNTLY | 0.83826387 |

| | | |
|---|---|---|
| YAALDVEL | ASSPPGGTGNTLY | 0.6606102 |
| LASVALLL | ASSPPGGTGNTLY | 0.60722816 |
| LDAVHRV | ASSPPGGTGNTLY | 0.7069081 |
| QGLSSSIF | ASSPPGGTGNTLY | 0.8255177 |
| RGTFANIRL | ASSPPGGTGNTLY | 0.85003674 |
| RQLWWGHRI | ASSPPGGTGNTLY | 0.8089099 |
| YGPPGTGKTLL | ASSPPGGTGNTLY | 0.6353762 |
| AALVGAPANVA | ASSPPGGTGNTLY | 0.80459166 |
| AATFRRTE | ASSPPGGTGNTLY | 0.74944466 |
| AGALGGGI | ASSPPGGTGNTLY | 0.73997027 |
| AGGAGGAGGIGAG | ASSPPGGTGNTLY | 0.8048745 |
| AGSGALGL | ASSPPGGTGNTLY | 0.78664887 |
| AQIADQI | ASSPPGGTGNTLY | 0.70752376 |
| ASAVAVTYL | ASSPPGGTGNTLY | 0.6972053 |
| ASTVLGGTGG | ASSPPGGTGNTLY | 0.7875532 |
| AVGGEPAK | ASSPPGGTGNTLY | 0.8358016 |
| CEQGTFGGTC | ASSPPGGTGNTLY | 0.82360333 |
| DEGKAPAG | ASSPPGGTGNTLY | 0.8709428 |
| DTVVFPAL | ASSPPGGTGNTLY | 0.833648 |
| FEGITMAG | ASSPPGGTGNTLY | 0.83819824 |
| FPLVVQD | ASSPPGGTGNTLY | 0.28734314 |
| GAFTTPQ | ASSPPGGTGNTLY | 0.5597978 |
| GFGGFGTAGGVGGA | ASSPPGGTGNTLY | 0.68622017 |
| GFLAQFL | ASSPPGGTGNTLY | 0.74823886 |
| GGAGGAGGNAGVA | ASSPPGGTGNTLY | 0.37664348 |
| GGVGTAGAGGAGGAGG | ASSPPGGTGNTLY | 0.82258475 |
| GNGGGGDAGNAGSGGN | ASSPPGGTGNTLY | 0.7408105 |
| GTGGAGGAGNDAGS | ASSPPGGTGNTLY | 0.6292292 |
| GTTNTGWFNS | ASSPPGGTGNTLY | 0.8369411 |
| IFMGPLFTL | ASSPPGGTGNTLY | 0.7051998 |
| ILAHAIEAI | ASSPPGGTGNTLY | 0.67196923 |
| IRSAIREAL | ASSPPGGTGNTLY | 0.716079 |
| ITFAGGG | ASSPPGGTGNTLY | 0.85414326 |
| ITGMSGHQEV | ASSPPGGTGNTLY | 0.60258996 |
| IVAGLACTL | ASSPPGGTGNTLY | 0.8263064 |
| IVGGGLAGA | ASSPPGGTGNTLY | 0.70169234 |
| IVTGVLNI | ASSPPGGTGNTLY | 0.85515416 |
| LAQQGCI | ASSPPGGTGNTLY | 0.5310998 |
| LCLHVEN | ASSPPGGTGNTLY | 0.83354706 |
| LDAISDGS | ASSPPGGTGNTLY | 0.74442697 |
| LGGFAAML | ASSPPGGTGNTLY | 0.8884933 |
| LGGKNPMI | ASSPPGGTGNTLY | 0.7354602 |
| LGPKGVAAIL | ASSPPGGTGNTLY | 0.8064052 |
| LMVGVSI | ASSPPGGTGNTLY | 0.7467818 |
| LVGAAFVL | ASSPPGGTGNTLY | 0.84110296 |
| LVLAVAPL | ASSPPGGTGNTLY | 0.4719211 |
| MAAIAVIDAA | ASSPPGGTGNTLY | 0.7522724 |
| NNGINAELQ | ASSPPGGTGNTLY | 0.61590934 |
| RGEPLVLVHG | ASSPPGGTGNTLY | 0.7866812 |
| RGGAGGAAGLI | ASSPPGGTGNTLY | 0.81276655 |
| RSYRMHPAV | ASSPPGGTGNTLY | 0.33145532 |

| | | |
|---|---|---|
| SAAHLLAGQLA | ASSPPGGTGNTLY | 0.8313956 |
| SAAVGSY | ASSPPGGTGNTLY | 0.670461 |
| SPHAVLHRI | ASSPPGGTGNTLY | 0.6328299 |
| SQIAKVL | ASSPPGGTGNTLY | 0.84895205 |
| TALGHLVP | ASSPPGGTGNTLY | 0.7709651 |
| TAPTTSGAA | ASSPPGGTGNTLY | 0.8367017 |
| TDGDDGNGGSPGT | ASSPPGGTGNTLY | 0.7518332 |
| TDGLGEN | ASSPPGGTGNTLY | 0.7589407 |
| TGGPPLPGQA | ASSPPGGTGNTLY | 0.81115806 |
| TIPAITI | ASSPPGGTGNTLY | 0.58329415 |
| TIPGFTI | ASSPPGGTGNTLY | 0.649343 |
| TSIASGALA | ASSPPGGTGNTLY | 0.8613116 |
| TVTLPGAAI | ASSPPGGTGNTLY | 0.72449124 |
| VASGLSAALP | ASSPPGGTGNTLY | 0.8086207 |
| VGAGVIGI | ASSPPGGTGNTLY | 0.62381434 |
| VGPKRFGV | ASSPPGGTGNTLY | 0.49324393 |
| VNVAFPD | ASSPPGGTGNTLY | 0.72204447 |
| VSMTNSGVS | ASSPPGGTGNTLY | 0.71559834 |
| VTDVLGDLSV | ASSPPGGTGNTLY | 0.81970143 |
| VVVVGAGFAGL | ASSPPGGTGNTLY | 0.7463875 |
| VYTVGTVV | ASSPPGGTGNTLY | 0.39038154 |
| WTTFSGP | ASSPPGGTGNTLY | 0.766663 |
| YDELVEAF | ASSPPGGTGNTLY | 0.67163396 |
| YGTGGAGGNGG | ASSPPGGTGNTLY | 0.61554545 |
| YTNVMAV | ASSPPGGTGNTLY | 0.66534996 |
| FLEQVHQL | ASSPPGGTGNTLY | 0.6939564 |
| GLSLLAVAV | ASSPPGGTGNTLY | 0.63291055 |
| LEQVHQL | ASSPPGGTGNTLY | 0.4566459 |
| LNAAGAI | ASSPPGGTGNTLY | 0.73723304 |
| PAVHASL | ASSPPGGTGNTLY | 0.78202957 |
| TELEDDA | ASSPPGGTGNTLY | 0.7862562 |
| TPPGPPN | ASSPPGGTGNTLY | 0.7595595 |
| TPPPPPPP | ASSPPGGTGNTLY | 0.7698528 |
| TQSPALI | ASSPPVYEQY | 0.72698283 |
| EFAAAFPQ | ASSPPVYEQY | 0.19032271 |
| LLSGNLDVL | ASSPPVYEQY | 0.2559687 |
| GPANLGNYNI | ASSPPVYEQY | 0.48171538 |
| TLGPTLP | ASSPPVYEQY | 0.56116915 |
| VYAYSSCCK | ASSPPVYEQY | 0.23870316 |
| APAPAPAPNI | ASSPPVYEQY | 0.59826523 |
| PGGPVPIP | ASSPPVYEQY | 0.6155391 |
| FLTVALEP | ASSPPVYEQY | 0.5028228 |
| SQYFGANL | ASSPPVYEQY | 0.8029889 |
| GGVKPAA | ASSPPVYEQY | 0.6034353 |
| GGTGGAGGAGGW | ASSPPVYEQY | 0.4121909 |
| GVGGVAGLGGAGG | ASSPPVYEQY | 0.45390898 |
| VLLVLATTI | ASSPPVYEQY | 0.72299266 |
| IGIFAAML | ASSPPVYEQY | 0.331874 |
| AVGDLID | ASSPPVYEQY | 0.76443756 |
| NISPLPL | ASSPPVYEQY | 0.42014468 |
| VTAVFVAI | ASSPPVYEQY | 0.755088 |

| | | |
|---|---|---|
| SGDDNDAAL | ASSPPVYEQY | 0.35083005 |
| AVAALAHS | ASSPPVYEQY | 0.7343366 |
| GGVTVMATP | ASSPPVYEQY | 0.6625501 |
| AGLAANVRD | ASSPPVYEQY | 0.83261085 |
| TNRPDLIDPALLRPG | ASSPPVYEQY | 0.59720206 |
| LANLGVTAL | ASSPPVYEQY | 0.68196636 |
| TGERIEL | ASSPPVYEQY | 0.44730186 |
| NGEELAA | ASSPPVYEQY | 0.8076408 |
| VLSLVWG | ASSPPVYEQY | 0.36680442 |
| IGYALWL | ASSPPVYEQY | 0.6397881 |
| LSLLLAPL | ASSPPVYEQY | 0.29683864 |
| TLDEALK | ASSPPVYEQY | 0.67668223 |
| TDGAGTAEGG | ASSPPVYEQY | 0.71479475 |
| NAPGGAGGNGG | ASSPPVYEQY | 0.2857421 |
| VSVFMGMGGNF | ASSPPVYEQY | 0.79364777 |
| LASYLAAL | ASSPPVYEQY | 0.5793926 |
| LVGGTPAL | ASSPPVYEQY | 0.49794504 |
| VRNADVL | ASSPPVYEQY | 0.77253896 |
| TCSSPFLVSAM | ASSPPVYEQY | 0.65633404 |
| VHITDAE | ASSPPVYEQY | 0.79042244 |
| EYAAPSF | ASSPPVYEQY | 0.5093461 |
| VVDAFHGGL | ASSPPVYEQY | 0.6312035 |
| ASVVDPQ | ASSPPVYEQY | 0.68641675 |
| SYNEFGGLS | ASSPPVYEQY | 0.5359895 |
| DPDGSVY | ASSPPVYEQY | 0.45309776 |
| MVSTIHATE | ASSPPVYEQY | 0.7644124 |
| LAAGVADPV | ASSPPVYEQY | 0.6399102 |
| TKEQIAA | ASSPPVYEQY | 0.6859475 |
| SLASPAAL | ASSPPVYEQY | 0.6052374 |
| IGLVFYGV | ASSPPVYEQY | 0.54873663 |
| FLLDASNL | ASSPPVYEQY | 0.62548816 |
| YQVGNFP | ASSPPVYEQY | 0.67315257 |
| ESLLSQST | ASSPPVYEQY | 0.68805915 |
| LFTVGFAAVG | ASSPPVYEQY | 0.6659177 |
| ATPPTPT | ASSPPVYEQY | 0.51268363 |
| TDLATLG | ASSPPVYEQY | 0.68140924 |
| SAGGTGR | ASSPPVYEQY | 0.61105406 |
| GFSNSGS | ASSPPVYEQY | 0.62684107 |
| GGNGGSGGTSV | ASSPPVYEQY | 0.66371554 |
| GTSVATGGAGN | ASSPPVYEQY | 0.6414934 |
| NGGNGGSGGTS | ASSPPVYEQY | 0.62151337 |
| AVVKFPH | ASSPPVYEQY | 0.5821637 |
| TPPPPPP | ASSPPVYEQY | 0.65864325 |
| AAPAAAAAPA | ASSPPVYEQY | 0.6711874 |
| IALNEDL | ASSPPVYEQY | 0.6650293 |
| IVNVSSM | ASSPPVYEQY | 0.7012781 |
| TGIPPAPR | ASSPPVYEQY | 0.6770625 |
| FDLGGGTF | ASSPPVYEQY | 0.6779543 |
| FDLGGGTFDVS | ASSPPVYEQY | 0.34646857 |
| NEPTAAA | ASSPPVYEQY | 0.4405673 |
| NEPTAAALA | ASSPPVYEQY | 0.56021076 |

| | | |
|---|---|---|
| GSIINISSI | ASSPPVYEQY | 0.61838657 |
| VAPGNAGTA | ASSPPVYEQY | 0.37350106 |
| TGGGASLE | ASSPPVYEQY | 0.6548409 |
| VAVYNHYKRI | ASSPPVYEQY | 0.6271919 |
| TDPAPATT | ASSPPVYEQY | 0.5641892 |
| TNRPDILDPALLRPG | ASSPPVYEQY | 0.6843027 |
| LTTTTTS | ASSPPVYEQY | 0.6070509 |
| PTGSGKTL | ASSPPVYEQY | 0.5639137 |
| TGSGKTL | ASSPPVYEQY | 0.5627834 |
| TIGHVDHGK | ASSPPVYEQY | 0.6809126 |
| TIGHVDHGKTT | ASSPPVYEQY | 0.47010148 |
| GHVDHGKTT | ASSPPVYEQY | 0.6789773 |
| AQFEHTLL | ASSPPVYEQY | 0.53038895 |
| IVNQDEL | ASSPPVYEQY | 0.67296326 |
| TNGSQFF | ASSPPVYEQY | 0.46939886 |
| TGKGIVE | ASSPPVYEQY | 0.67584 |
| LISNASDA | ASSPPVYEQY | 0.61681 |
| AELRTLGELTKIAKAHGAQ | ASSPPVYEQY | 0.7076917 |
| APNAVAAAEPSW | ASSPPVYEQY | 0.613196 |
| APQVAGAATAVL | ASSPPVYEQY | 0.6487403 |
| ARRDPAAAAAILPTDA | ASSPPVYEQY | 0.53806925 |
| DAAARDVLAEAGLAEYF | ASSPPVYEQY | 0.68753934 |
| DDQIVLSAKANLAKVEQ | ASSPPVYEQY | 0.42984393 |
| DELLPDTGEDSDEDGH | ASSPPVYEQY | 0.6263168 |
| ELPEAPANRATVPQD | ASSPPVYEQY | 0.5279294 |
| FGQSKIERVTGIG | ASSPPVYEQY | 0.66527784 |
| GAGGTGSTAGGAGGAGGAGGI | ASSPPVYEQY | 0.66236407 |
| GHGGTGAAGGNGGTGGAGGS( | ASSPPVYEQY | 0.4706412 |
| GNGGNGGAGGEGGPGGAGGA( | ASSPPVYEQY | 0.69172096 |
| GSTAGNGGNGGNATNGTIAG( | ASSPPVYEQY | 0.6370813 |
| GTGGDGGNGGNGGTGG | ASSPPVYEQY | 0.557545 |
| GVRRPAGIPADAPV | ASSPPVYEQY | 0.68783057 |
| GYRQVTDAHLLTLARR | ASSPPVYEQY | 0.6558173 |
| IAKMGQDGHDRG | ASSPPVYEQY | 0.66968024 |
| KAEPSIEIQYT | ASSPPVYEQY | 0.62181085 |
| KPVRIRMAADEDTVAVT | ASSPPVYEQY | 0.62510026 |
| MPPRIAGMRLLVIKPEP | ASSPPVYEQY | 0.6706731 |
| MKFYAVALFRSRGVPPTR | ASSPPVYEQY | 0.51891375 |
| NAGGFGILGAVAHSP | ASSPPVYEQY | 0.5087594 |
| NEGREITLAMAEM | ASSPPVYEQY | 0.68549156 |
| NELVEWSHPLGHRWRWE | ASSPPVYEQY | 0.8240542 |
| NLGAGGTGGNGGNPG | ASSPPVYEQY | 0.8123746 |
| NTEWPKWETVMAADPVD | ASSPPVYEQY | 0.6836258 |
| PASAAPAETQP | ASSPPVYEQY | 0.60374767 |
| QQSSDGKNGVRAAEITGE | ASSPPVYEQY | 0.6700609 |
| RAPDAVTTSWWKE | ASSPPVYEQY | 0.80766046 |
| SGFGNVGAHTSGWFNQ | ASSPPVYEQY | 0.8310634 |
| SGVAGAVGQAASVGGLKV | ASSPPVYEQY | 0.8131359 |
| SNGGAGTGSAGGNGGT | ASSPPVYEQY | 0.4853835 |
| TGGAGGEGGAAG | ASSPPVYEQY | 0.82213664 |
| TGGDGGAAGAGGAPGHG | ASSPPVYEQY | 0.7833061 |

| | | |
|---|---|---|
| TMPPLLAAIANAGRHGV | ASSPPVYEQY | 0.7940182 |
| TSSTAGNSGGA | ASSPPVYEQY | 0.74134207 |
| VPATVPVGILC | ASSPPVYEQY | 0.7253995 |
| VQRPDSGGVVLTGRIS | ASSPPVYEQY | 0.65433466 |
| WTPNPVPATKGVPAGLVQAT | ASSPPVYEQY | 0.5368632 |
| YALKRALAKFSDRFDVVII | ASSPPVYEQY | 0.775398 |
| YLIGSVIV | ASSPPVYEQY | 0.8182506 |
| YGQLVAVID | ASSPPVYEQY | 0.82050633 |
| VTVGGMTLPA | ASSPPVYEQY | 0.84861887 |
| VLALPAV | ASSPPVYEQY | 0.8463613 |
| VIPPTLNYET | ASSPPVYEQY | 0.80867773 |
| VGMGGMHPSSG | ASSPPVYEQY | 0.26758277 |
| VGLAGGAATA | ASSPPVYEQY | 0.48169023 |
| VGGAGGAGGSGP | ASSPPVYEQY | 0.56237113 |
| VFAAFMV | ASSPPVYEQY | 0.71586436 |
| TLYDKLR | ASSPPVYEQY | 0.2868298 |
| TLAVFTVL | ASSPPVYEQY | 0.62844896 |
| TGSVQPARV | ASSPPVYEQY | 0.659631 |
| TGGGAGSGGT | ASSPPVYEQY | 0.6513599 |
| SHLETSL | ASSPPVYEQY | 0.826306 |
| SFTTGMT | ASSPPVYEQY | 0.7131239 |
| QVEVAKRMGVRQPTVS | ASSPPVYEQY | 0.5002342 |
| QGSGGPALAM | ASSPPVYEQY | 0.5141657 |
| MLAPSNP | ASSPPVYEQY | 0.82593596 |
| LVNNAAI | ASSPPVYEQY | 0.58023006 |
| LTGEINTLA | ASSPPVYEQY | 0.84444547 |
| LSIFGPV | ASSPPVYEQY | 0.4980754 |
| LLALLANL | ASSPPVYEQY | 0.8503773 |
| LISNASDALDKLR | ASSPPVYEQY | 0.46711725 |
| LINEAVP | ASSPPVYEQY | 0.8433403 |
| LGYAFSGL | ASSPPVYEQY | 0.79305553 |
| LDFVAVAL | ASSPPVYEQY | 0.8659695 |
| LAQPGAE | ASSPPVYEQY | 0.739226 |
| ISNLGAVTGF | ASSPPVYEQY | 0.78282696 |
| HQEDPGRI | ASSPPVYEQY | 0.6289658 |
| GTPGEPTP | ASSPPVYEQY | 0.8082176 |
| GFFNSGS | ASSPPVYEQY | 0.47979563 |
| GAGGGGLTAGAN | ASSPPVYEQY | 0.7100328 |
| FVDALVNM | ASSPPVYEQY | 0.48330265 |
| FLTVAPEL | ASSPPVYEQY | 0.74210715 |
| FETLTAL | ASSPPVYEQY | 0.8302803 |
| DVINPIEL | ASSPPVYEQY | 0.20872785 |
| APVSGAPV | ASSPPVYEQY | 0.8015147 |
| ALVGGLAW | ASSPPVYEQY | 0.6463256 |
| ALAAAAATAI | ASSPPVYEQY | 0.5146315 |
| AFGLGLLL | ASSPPVYEQY | 0.8284125 |
| AFFAAASL | ASSPPVYEQY | 0.7675277 |
| AAAALPAFSPP | ASSPPVYEQY | 0.8285369 |
| VVLAPAL | ASSPPVYEQY | 0.6050191 |
| TIGHVDHGKTTL | ASSPPVYEQY | 0.7046876 |
| LISNASDALDK | ASSPPVYEQY | 0.75281197 |

| | | |
|---|---|---|
| LISNASDAL | ASSPPVYEQY | 0.35357457 |
| ISNASDALDK | ASSPPVYEQY | 0.45831177 |
| NQAETLVYQTE | ASSPPVYEQY | 0.85849 |
| TGGTGGTAGAG | ASSPPVYEQY | 0.72384983 |
| TGSTAGGAGGA | ASSPPVYEQY | 0.75135034 |
| DPLAVPVAAGPV | ASSPPVYEQY | 0.6316393 |
| GGTGGTPGNGGN | ASSPPVYEQY | 0.5669028 |
| PAGLAGVDPATL | ASSPPVYEQY | 0.60725546 |
| GNNLVVEAPHAGASV | ASSPPVYEQY | 0.7392103 |
| KVTRIMSASTLEQPA | ASSPPVYEQY | 0.7479011 |
| LGFALPIALPRRTVV | ASSPPVYEQY | 0.7044464 |
| TRLGNLTARKLVVLV | ASSPPVYEQY | 0.52678806 |
| WSQQRRLIYFPSAGP | ASSPPVYEQY | 0.7020075 |
| GPNGNPDPMAAAVDIR | ASSPPVYEQY | 0.66238475 |
| TQRCVSVADLRAAASR | ASSPPVYEQY | 0.66222596 |
| QFVLDVAEHPVEVLTETV | ASSPPVYEQY | 0.6569675 |
| GGAGGTGGAGGNGGTQAGDG | ASSPPVYEQY | 0.7276271 |
| PLLATLRAIAVDVDAKPAQV | ASSPPVYEQY | 0.6407167 |
| APKFRTGGYMIDFWGVGYQV | ASSPPVYEQY | 0.5503197 |
| GAGGAGGAGGAAGGPGGTGG | ASSPPVYEQY | 0.6802695 |
| IGALGLLATDVAPAGSVAFG | ASSPPVYEQY | 0.7402125 |
| AGGAGGIGGNGGAGGAGGVL | ASSPPVYEQY | 0.7182497 |
| ADELIGLL | ASSPPVYEQY | 0.7472931 |
| ADRALKA | ASSPPVYEQY | 0.74189466 |
| AGAVGQVGGAGGA | ASSPPVYEQY | 0.71061677 |
| AGGGGGLGGAP | ASSPPVYEQY | 0.2699734 |
| AGGLLYGNGGAG | ASSPPVYEQY | 0.33120692 |
| AGLESIGL | ASSPPVYEQY | 0.5873642 |
| AGLKTGII | ASSPPVYEQY | 0.6070058 |
| ASLAALLL | ASSPPVYEQY | 0.3082769 |
| ASVINISSI | ASSPPVYEQY | 0.6981876 |
| ATPPGPA | ASSPPVYEQY | 0.5941292 |
| ATVASALV | ASSPPVYEQY | 0.6146878 |
| ATVASLIL | ASSPPVYEQY | 0.7318317 |
| AVNGPVLIRH | ASSPPVYEQY | 0.6280519 |
| DGAAGAVGI | ASSPPVYEQY | 0.5730315 |
| DPLLHGI | ASSPPVYEQY | 0.49570185 |
| ELGAALGAL | ASSPPVYEQY | 0.73384774 |
| ELGRTQAV | ASSPPVYEQY | 0.37289977 |
| FVAGGYLS | ASSPPVYEQY | 0.7233329 |
| GASGGAGGAGGW | ASSPPVYEQY | 0.46692318 |
| GGAGAGGDGGAGGI | ASSPPVYEQY | 0.7388717 |
| GTGHNGSQPGMGGNG | ASSPPVYEQY | 0.51025045 |
| GYAAASASAS | ASSPPVYEQY | 0.73181397 |
| HQVDVAAVQQG | ASSPPVYEQY | 0.6800772 |
| IGAAIAHT | ASSPPVYEQY | 0.76735306 |
| IGMIEVAQL | ASSPPVYEQY | 0.64487165 |
| IRLAPEHTSAMD | ASSPPVYEQY | 0.68598807 |
| IYNVSVL | ASSPPVYEQY | 0.5594491 |
| KQGFLGM | ASSPPVYEQY | 0.7361541 |
| LESVITQL | ASSPPVYEQY | 0.48204035 |

| | | |
|---|---|---|
| LGLGALVVV | ASSPPVYEQY | 0.6505843 |
| LIGNGGNGGSGGI | ASSPPVYEQY | 0.42849767 |
| LNAINAPTL | ASSPPVYEQY | 0.68224347 |
| LQLLNTGV | ASSPPVYEQY | 0.7188262 |
| LQTSIEF | ASSPPVYEQY | 0.31330305 |
| MSFAVSEL | ASSPPVYEQY | 0.7036028 |
| NTGALGL | ASSPPVYEQY | 0.6266874 |
| PPGAGGAGGAGTT | ASSPPVYEQY | 0.5632696 |
| QLVGLGI | ASSPPVYEQY | 0.7323668 |
| RFLEQVHQL | ASSPPVYEQY | 0.70413715 |
| RGVGYVLR | ASSPPVYEQY | 0.7419578 |
| SATGVFAGVI | ASSPPVYEQY | 0.6263846 |
| SLPAITLP | ASSPPVYEQY | 0.64345014 |
| STALAFTV | ASSPPVYEQY | 0.7062103 |
| SVLTVGLGK | ASSPPVYEQY | 0.5777303 |
| TALAGAGSGI | ASSPPVYEQY | 0.4670992 |
| TGLPFTGI | ASSPPVYEQY | 0.7435944 |
| VAEHVTV | ASSPPVYEQY | 0.63857645 |
| VDIVLNSL | ASSPPVYEQY | 0.72939545 |
| VLTGGGAVL | ASSPPVYEQY | 0.5130689 |
| VNVGLAYGI | ASSPPVYEQY | 0.4868098 |
| VSSAAVGAV | ASSPPVYEQY | 0.5691519 |
| VTSVYEAGV | ASSPPVYEQY | 0.72391284 |
| WVALDRGA | ASSPPVYEQY | 0.7218535 |
| YLVGSVVI | ASSPPVYEQY | 0.65418863 |
| GPPGTGKTLL | ASSPPVYEQY | 0.50510395 |
| GVGMTKF | ASSPPVYEQY | 0.59871376 |
| IGIGGSDL | ASSPPVYEQY | 0.53730595 |
| IGTLLAEL | ASSPPVYEQY | 0.57115376 |
| IYTSGTT | ASSPPVYEQY | 0.6758828 |
| LSFPAGV | ASSPPVYEQY | 0.6628011 |
| LYTSGTT | ASSPPVYEQY | 0.57463026 |
| VGAFPFGF | ASSPPVYEQY | 0.57312787 |
| VGAGTVEF | ASSPPVYEQY | 0.5925486 |
| VGKSTLLL | ASSPPVYEQY | 0.6964342 |
| AALAHAGGGIV | ASSPPVYEQY | 0.66492456 |
| AALVAVSLL | ASSPPVYEQY | 0.73572874 |
| AAVSRAVAAL | ASSPPVYEQY | 0.68356866 |
| AGGSSGIGGF | ASSPPVYEQY | 0.64411676 |
| AGLTVANANG | ASSPPVYEQY | 0.23854436 |
| AGLVGAVL | ASSPPVYEQY | 0.3420984 |
| ALSQAFTV | ASSPPVYEQY | 0.6041597 |
| ALTFSVSL | ASSPPVYEQY | 0.583831 |
| ALTGAALVAL | ASSPPVYEQY | 0.33590677 |
| ASFSIGAGVL | ASSPPVYEQY | 0.70981044 |
| DGSVYKI | ASSPPVYEQY | 0.56399506 |
| EQLAEIV | ASSPPVYEQY | 0.55641514 |
| FAVSELTVM | ASSPPVYEQY | 0.70716476 |
| GAAPLAAGL | ASSPPVYEQY | 0.59391785 |
| GADGGDGGSGGAGG | ASSPPVYEQY | 0.576016 |
| GFGLDTGGAGG | ASSPPVYEQY | 0.5415188 |

| | | |
|---|---|---|
| GLASGGLL | ASSPPVYEQY | 0.69030243 |
| GRIAGAAVLL | ASSPPVYEQY | 0.42834195 |
| GSAGSLNTSF | ASSPPVYEQY | 0.6632147 |
| GTGQIGI | ASSPPVYEQY | 0.46899223 |
| GTLIALIL | ASSPPVYEQY | 0.70590746 |
| IAATMGFVVG | ASSPPVYEQY | 0.43968835 |
| IFAQALL | ASSPPVYEQY | 0.65006006 |
| IGAAVALGF | ASSPPVYEQY | 0.5592993 |
| IGLTPLAL | ASSPPVYEQY | 0.77967215 |
| ILGSNTSTL | ASSPPVYEQY | 0.58934915 |
| IVFSLTVL | ASSPPVYEQY | 0.64100266 |
| LASKATGF | ASSPPVYEQY | 0.52088815 |
| LASVALIL | ASSPPVYEQY | 0.69639015 |
| LFAVGLTL | ASSPPVYEQY | 0.4091601 |
| LFNGGIGTYI | ASSPPVYEQY | 0.5656066 |
| LTLGAGII | ASSPPVYEQY | 0.39244452 |
| LVAAALGDI | ASSPPVYEQY | 0.64597344 |
| NTAFVQLGI | ASSPPVYEQY | 0.6211038 |
| QLFSGAEL | ASSPPVYEQY | 0.44053987 |
| SAATFITL | ASSPPVYEQY | 0.69341934 |
| SNYGLANLGV | ASSPPVYEQY | 0.61647546 |
| SYGGAYAVM | ASSPPVYEQY | 0.55856586 |
| TTGVGGLTLG | ASSPPVYEQY | 0.7274307 |
| TTQTIYI | ASSPPVYEQY | 0.540954 |
| VAAGAGLGAVI | ASSPPVYEQY | 0.7223413 |
| VAAVVTLL | ASSPPVYEQY | 0.60260254 |
| VGFLGGLI | ASSPPVYEQY | 0.49955764 |
| VGVIPILL | ASSPPVYEQY | 0.65606534 |
| VIFASSLI | ASSPPVYEQY | 0.5565613 |
| VKGGDGGAAAG | ASSPPVYEQY | 0.5522249 |
| VTKTFTNI | ASSPPVYEQY | 0.6948892 |
| VVAAAILL | ASSPPVYEQY | 0.7663597 |
| YAALDVEL | ASSPPVYEQY | 0.75793767 |
| LASVALLL | ASSPPVYEQY | 0.7138401 |
| LDAVHRV | ASSPPVYEQY | 0.69446754 |
| QGLSSSIF | ASSPPVYEQY | 0.74409264 |
| RGTFANIRL | ASSPPVYEQY | 0.7555703 |
| RQLWWGHRI | ASSPPVYEQY | 0.77077746 |
| YGPPGTGKTLL | ASSPPVYEQY | 0.76129436 |
| AALVGAPANVA | ASSPPVYEQY | 0.63370645 |
| AATFRRTE | ASSPPVYEQY | 0.7673156 |
| AGALGGGI | ASSPPVYEQY | 0.74497664 |
| AGGAGGAGGIGAG | ASSPPVYEQY | 0.7457898 |
| AGSGALGL | ASSPPVYEQY | 0.752519 |
| AQIADQI | ASSPPVYEQY | 0.74632055 |
| ASAVAVTYL | ASSPPVYEQY | 0.7366606 |
| ASTVLGGTGG | ASSPPVYEQY | 0.71147203 |
| AVGGEPAK | ASSPPVYEQY | 0.75534534 |
| CEQGTFGGTC | ASSPPVYEQY | 0.7684597 |
| DEGKAPAG | ASSPPVYEQY | 0.7671788 |
| DTVVFPAL | ASSPPVYEQY | 0.77023697 |

| | | |
|---|---|---|
| FEGITMAG | ASSPPVYEQY | 0.7701453 |
| FPLVVQD | ASSPPVYEQY | 0.7655766 |
| GAFTTPQ | ASSPPVYEQY | 0.4406567 |
| GFGGFGTAGGVGGA | ASSPPVYEQY | 0.61224353 |
| GFLAQFL | ASSPPVYEQY | 0.7244942 |
| GGAGGAGGNAGVA | ASSPPVYEQY | 0.7285783 |
| GGVGTAGAGGAGGAGG | ASSPPVYEQY | 0.41083154 |
| GNGGGGDAGNAGSGGN | ASSPPVYEQY | 0.7346519 |
| GTGGAGGAGNDAGS | ASSPPVYEQY | 0.7410753 |
| GTTNTGWFNS | ASSPPVYEQY | 0.70099527 |
| IFMGPLFTL | ASSPPVYEQY | 0.7698648 |
| ILAHAIEAI | ASSPPVYEQY | 0.7392447 |
| IRSAIREAL | ASSPPVYEQY | 0.64555734 |
| ITFAGGG | ASSPPVYEQY | 0.71349144 |
| ITGMSGHQEV | ASSPPVYEQY | 0.7590722 |
| IVAGLACTL | ASSPPVYEQY | 0.6833006 |
| IVGGGLAGA | ASSPPVYEQY | 0.77110934 |
| IVTGVLNI | ASSPPVYEQY | 0.6619925 |
| LAQQGCI | ASSPPVYEQY | 0.7714788 |
| LCLHVEN | ASSPPVYEQY | 0.57367945 |
| LDAISDGS | ASSPPVYEQY | 0.7640078 |
| LGGFAAML | ASSPPVYEQY | 0.7462521 |
| LGGKNPMI | ASSPPVYEQY | 0.781924 |
| LGPKGVAAIL | ASSPPVYEQY | 0.7336806 |
| LMVGVSI | ASSPPVYEQY | 0.76372707 |
| LVGAAFVL | ASSPPVYEQY | 0.7023001 |
| LVLAVAPL | ASSPPVYEQY | 0.7701566 |
| MAAIAVIDAA | ASSPPVYEQY | 0.6677331 |
| NNGINAELQ | ASSPPVYEQY | 0.7425953 |
| RGEPLVLVHG | ASSPPVYEQY | 0.5772141 |
| RGGAGGAAGLI | ASSPPVYEQY | 0.74768865 |
| RSYRMHPAV | ASSPPVYEQY | 0.7625295 |
| SAAHLLAGQLA | ASSPPVYEQY | 0.3410398 |
| SAAVGSY | ASSPPVYEQY | 0.77169776 |
| SPHAVLHRI | ASSPPVYEQY | 0.63434 |
| SQIAKVL | ASSPPVYEQY | 0.6619583 |
| TALGHLVP | ASSPPVYEQY | 0.7707369 |
| TAPTTSGAA | ASSPPVYEQY | 0.73285466 |
| TDGDDGNGGSPGT | ASSPPVYEQY | 0.77567035 |
| TDGLGEN | ASSPPVYEQY | 0.71718997 |
| TGGPPLPGQA | ASSPPVYEQY | 0.74760497 |
| TIPAITI | ASSPPVYEQY | 0.7414406 |
| TIPGFTI | ASSPPVYEQY | 0.6767672 |
| TSIASGALA | ASSPPVYEQY | 0.68155676 |
| TVTLPGAAI | ASSPPVYEQY | 0.7752363 |
| VASGLSAALP | ASSPPVYEQY | 0.74047947 |
| VGAGVIGI | ASSPPVYEQY | 0.7272793 |
| VGPKRFGV | ASSPPVYEQY | 0.66792893 |
| VNVAFPD | ASSPPVYEQY | 0.5600354 |
| VSMTNSGVS | ASSPPVYEQY | 0.649297 |
| VTDVLGDLSV | ASSPPVYEQY | 0.7372305 |

| | | |
|---|---|---|
| VVVVGAGFAGL | ASSPPVYEQY | 0.7451096 |
| VYTVGTVV | ASSPPVYEQY | 0.73202646 |
| WTTFSGP | ASSPPVYEQY | 0.5098226 |
| YDELVEAF | ASSPPVYEQY | 0.71509385 |
| YGTGGAGGNGG | ASSPPVYEQY | 0.67236745 |
| YTNVMAV | ASSPPVYEQY | 0.6578369 |
| FLEQVHQL | ASSPPVYEQY | 0.7202619 |
| GLSLLAVAV | ASSPPVYEQY | 0.71685815 |
| LEQVHQL | ASSPPVYEQY | 0.652742 |
| LNAAGAI | ASSPPVYEQY | 0.60925066 |
| PAVHASL | ASSPPVYEQY | 0.7207625 |
| TELEDDA | ASSPPVYEQY | 0.7567526 |
| TPPGPPN | ASSPPVYEQY | 0.7524147 |
| TPPPPPPP | ASSPPVYEQY | 0.7662001 |
| TQSPALI | ASRVQGAEQY | 0.7613473 |
| EFAAAFPQ | ASRVQGAEQY | 0.7568029 |
| LLSGNLDVL | ASRVQGAEQY | 0.23933397 |
| GPANLGNYNI | ASRVQGAEQY | 0.46808603 |
| TLGPTLP | ASRVQGAEQY | 0.6050813 |
| VYAYSSCCK | ASRVQGAEQY | 0.69974804 |
| APAPAPAPNI | ASRVQGAEQY | 0.32774767 |
| PGGPVPIP | ASRVQGAEQY | 0.62030077 |
| FLTVALEP | ASRVQGAEQY | 0.6605984 |
| SQYFGANL | ASRVQGAEQY | 0.53722525 |
| GGVKPAA | ASRVQGAEQY | 0.7737623 |
| GGTGGAGGAGGW | ASRVQGAEQY | 0.6406605 |
| GVGGVAGLGGAGG | ASRVQGAEQY | 0.5354241 |
| VLLVLATTI | ASRVQGAEQY | 0.6193241 |
| IGIFAAML | ASRVQGAEQY | 0.74490964 |
| AVGDLID | ASRVQGAEQY | 0.52394664 |
| NISPLPL | ASRVQGAEQY | 0.7354197 |
| VTAVFVAI | ASRVQGAEQY | 0.5434852 |
| SGDDNDAAL | ASRVQGAEQY | 0.7536831 |
| AVAALAHS | ASRVQGAEQY | 0.43671975 |
| GGVTVMATP | ASRVQGAEQY | 0.7479671 |
| AGLAANVRD | ASRVQGAEQY | 0.6764411 |
| TNRPDLIDPALLRPG | ASRVQGAEQY | 0.7844727 |
| LANLGVTAL | ASRVQGAEQY | 0.6603422 |
| TGERIEL | ASRVQGAEQY | 0.7333073 |
| NGEELAA | ASRVQGAEQY | 0.62816757 |
| VLSLVWG | ASRVQGAEQY | 0.7379359 |
| IGYALWL | ASRVQGAEQY | 0.47361663 |
| LSLLLAPL | ASRVQGAEQY | 0.6631394 |
| TLDEALK | ASRVQGAEQY | 0.38198888 |
| TDGAGTAEGG | ASRVQGAEQY | 0.7046392 |
| NAPGGAGGNGG | ASRVQGAEQY | 0.73415804 |
| VSVFMGMGGNF | ASRVQGAEQY | 0.31585118 |
| LASYLAAL | ASRVQGAEQY | 0.75014985 |
| LVGGTPAL | ASRVQGAEQY | 0.6069936 |
| VRNADVL | ASRVQGAEQY | 0.52341586 |
| TCSSPFLVSAM | ASRVQGAEQY | 0.7650168 |

| | | |
|---|---|---|
| VHITDAE | ASRVQGAEQY | 0.6377209 |
| EYAAPSF | ASRVQGAEQY | 0.7621766 |
| VVDAFHGGL | ASRVQGAEQY | 0.6219053 |
| ASVVDPQ | ASRVQGAEQY | 0.6567447 |
| SYNEFGGLS | ASRVQGAEQY | 0.7134609 |
| DPDGSVY | ASRVQGAEQY | 0.6059226 |
| MVSTIHATE | ASRVQGAEQY | 0.5809558 |
| LAAGVADPV | ASRVQGAEQY | 0.7583158 |
| TKEQIAA | ASRVQGAEQY | 0.7309264 |
| SLASPAAL | ASRVQGAEQY | 0.7340982 |
| IGLVFYGV | ASRVQGAEQY | 0.6654199 |
| FLLDASNL | ASRVQGAEQY | 0.6650933 |
| YQVGNFP | ASRVQGAEQY | 0.69597846 |
| ESLLSQST | ASRVQGAEQY | 0.73031116 |
| LFTVGFAAVG | ASRVQGAEQY | 0.74238294 |
| ATPPTPT | ASRVQGAEQY | 0.73378277 |
| TDLATLG | ASRVQGAEQY | 0.6700072 |
| SAGGTGR | ASRVQGAEQY | 0.73231494 |
| GFSNSGS | ASRVQGAEQY | 0.7127214 |
| GGNGGSGGTSV | ASRVQGAEQY | 0.7163782 |
| GTSVATGGAGN | ASRVQGAEQY | 0.7138356 |
| NGGNGGSGGTS | ASRVQGAEQY | 0.7024669 |
| AVVKFPH | ASRVQGAEQY | 0.67382175 |
| TPPPPPP | ASRVQGAEQY | 0.595886 |
| AAPAAAAAPA | ASRVQGAEQY | 0.7273 |
| IALNEDL | ASRVQGAEQY | 0.73478127 |
| IVNVSSM | ASRVQGAEQY | 0.72856057 |
| TGIPPAPR | ASRVQGAEQY | 0.7448033 |
| FDLGGGTF | ASRVQGAEQY | 0.73522466 |
| FDLGGGTFDVS | ASRVQGAEQY | 0.7238021 |
| NEPTAAA | ASRVQGAEQY | 0.42703182 |
| NEPTAAALA | ASRVQGAEQY | 0.5713005 |
| GSIINISSI | ASRVQGAEQY | 0.6209115 |
| VAPGNAGTA | ASRVQGAEQY | 0.6943139 |
| TGGGASLE | ASRVQGAEQY | 0.332024 |
| VAVYNHYKRI | ASRVQGAEQY | 0.72518766 |
| TDPAPATT | ASRVQGAEQY | 0.6964376 |
| TNRPDILDPALLRPG | ASRVQGAEQY | 0.6770508 |
| LTTTTTS | ASRVQGAEQY | 0.7350215 |
| PTGSGKTL | ASRVQGAEQY | 0.6899955 |
| TGSGKTL | ASRVQGAEQY | 0.5706551 |
| TIGHVDHGK | ASRVQGAEQY | 0.66308594 |
| TIGHVDHGKTT | ASRVQGAEQY | 0.73872876 |
| GHVDHGKTT | ASRVQGAEQY | 0.62470794 |
| AQFEHTLL | ASRVQGAEQY | 0.7388521 |
| IVNQDEL | ASRVQGAEQY | 0.69654244 |
| TNGSQFF | ASRVQGAEQY | 0.74040794 |
| TGKGIVE | ASRVQGAEQY | 0.61003447 |
| LISNASDA | ASRVQGAEQY | 0.73769975 |
| AELRTLGELTKIAKAHGAQ | ASRVQGAEQY | 0.715346 |
| APNAVAAAEPSW | ASRVQGAEQY | 0.7460982 |

| | | |
|---|---|---|
| APQVAGAATAVL | ASRVQGAEQY | 0.7004709 |
| ARRDPAAAAAILPTDA | ASRVQGAEQY | 0.7331207 |
| DAAARDVLAEAGLAEYF | ASRVQGAEQY | 0.716463 |
| DDQIVLSAKANLAKVEQ | ASRVQGAEQY | 0.7362893 |
| DELLPDTGEDSDEDGH | ASRVQGAEQY | 0.6204609 |
| ELPEAPANRATVPQD | ASRVQGAEQY | 0.6789518 |
| FGQSKIERVTGIG | ASRVQGAEQY | 0.6694559 |
| GAGGTGSTAGGAGGAGGAGGI | ASRVQGAEQY | 0.69870144 |
| GHGGTGAAGGNGGTGGAGGS( | ASRVQGAEQY | 0.7365804 |
| GNGGNGGAGGEGGPGGAGGA( | ASRVQGAEQY | 0.30176967 |
| GSTAGNGGNGGNATNGTIAG( | ASRVQGAEQY | 0.717859 |
| GTGGDGGNGGNGGTGG | ASRVQGAEQY | 0.64751697 |
| GVRRPAGIPADAPV | ASRVQGAEQY | 0.60835034 |
| GYRQVTDAHLLTLARR | ASRVQGAEQY | 0.7278013 |
| IAKMGQDGHDRG | ASRVQGAEQY | 0.7136471 |
| KAEPSIEIQYT | ASRVQGAEQY | 0.72829205 |
| KPVRIRMAADEDTVAVT | ASRVQGAEQY | 0.633463 |
| MPPRIAGMRLLVIKPEP | ASRVQGAEQY | 0.70291877 |
| MKFYAVALFRSRGVPPTR | ASRVQGAEQY | 0.718626 |
| NAGGFGILGAVAHSP | ASRVQGAEQY | 0.45256957 |
| NEGREITLAMAEM | ASRVQGAEQY | 0.6283988 |
| NELVEWSHPLGHRWRWE | ASRVQGAEQY | 0.7429888 |
| NLGAGGTGGNGGNPG | ASRVQGAEQY | 0.17724496 |
| NTEWPKWETVMAADPVD | ASRVQGAEQY | 0.403885 |
| PASAAPAETQP | ASRVQGAEQY | 0.13197376 |
| QQSSDGKNGVRAAEITGE | ASRVQGAEQY | 0.14308424 |
| RAPDAVTTSWWKE | ASRVQGAEQY | 0.14740156 |
| SGFGNVGAHTSGWFNQ | ASRVQGAEQY | 0.19837004 |
| SGVAGAVGQAASVGGLKV | ASRVQGAEQY | 0.3009539 |
| SNGGAGTGSAGGNGGT | ASRVQGAEQY | 0.22159664 |
| TGGAGGEGGAAG | ASRVQGAEQY | 0.1458876 |
| TGGDGGAAGAGGAPGHG | ASRVQGAEQY | 0.14261672 |
| TMPPLLAAIANAGRHGV | ASRVQGAEQY | 0.13787907 |
| TSSTAGNSGGA | ASRVQGAEQY | 0.14559962 |
| VPATVPVGILC | ASRVQGAEQY | 0.16338438 |
| VQRPDSGGVVLTGRIS | ASRVQGAEQY | 0.1391631 |
| WTPNPVPATKGVPAGLVQAT | ASRVQGAEQY | 0.13390908 |
| YALKRALAKFSDRFDVVII | ASRVQGAEQY | 0.14988047 |
| YLIGSVIV | ASRVQGAEQY | 0.14767358 |
| YGQLVAVID | ASRVQGAEQY | 0.18109551 |
| VTVGGMTLPA | ASRVQGAEQY | 0.1755303 |
| VLALPAV | ASRVQGAEQY | 0.22441444 |
| VIPPTLNYET | ASRVQGAEQY | 0.17463148 |
| VGMGGMHPSSG | ASRVQGAEQY | 0.15895844 |
| VGLAGGAATA | ASRVQGAEQY | 0.12484053 |
| VGGAGGAGGSGP | ASRVQGAEQY | 0.13229564 |
| VFAAFMV | ASRVQGAEQY | 0.14833236 |
| TLYDKLR | ASRVQGAEQY | 0.14980835 |
| TLAVFTVL | ASRVQGAEQY | 0.1262833 |
| TGSVQPARV | ASRVQGAEQY | 0.15950197 |
| TGGGAGSGGT | ASRVQGAEQY | 0.16358736 |

| | | |
|---|---|---|
| SHLETSL | ASRVQGAEQY | 0.13187784 |
| SFTTGMT | ASRVQGAEQY | 0.17497371 |
| QVEVAKRMGVRQPTVS | ASRVQGAEQY | 0.13472012 |
| QGSGGPALAM | ASRVQGAEQY | 0.1348049 |
| MLAPSNP | ASRVQGAEQY | 0.14257677 |
| LVNNAAI | ASRVQGAEQY | 0.18540004 |
| LTGEINTLA | ASRVQGAEQY | 0.13719061 |
| LSIFGPV | ASRVQGAEQY | 0.1739271 |
| LLALLANL | ASRVQGAEQY | 0.16281681 |
| LISNASDALDKLR | ASRVQGAEQY | 0.1808471 |
| LINEAVP | ASRVQGAEQY | 0.13613501 |
| LGYAFSGL | ASRVQGAEQY | 0.16292754 |
| LDFVAVAL | ASRVQGAEQY | 0.13947731 |
| LAQPGAE | ASRVQGAEQY | 0.23161857 |
| ISNLGAVTGF | ASRVQGAEQY | 0.15002814 |
| HQEDPGRI | ASRVQGAEQY | 0.16647881 |
| GTPGEPTP | ASRVQGAEQY | 0.14507504 |
| GFFNSGS | ASRVQGAEQY | 0.2053203 |
| GAGGGGLTAGAN | ASRVQGAEQY | 0.12821974 |
| FVDALVNM | ASRVQGAEQY | 0.14021158 |
| FLTVAPEL | ASRVQGAEQY | 0.14107452 |
| FETLTAL | ASRVQGAEQY | 0.14512852 |
| DVINPIEL | ASRVQGAEQY | 0.20223169 |
| APVSGAPV | ASRVQGAEQY | 0.13951068 |
| ALVGGLAW | ASRVQGAEQY | 0.16454679 |
| ALAAAAATAI | ASRVQGAEQY | 0.14908186 |
| AFGLGLLL | ASRVQGAEQY | 0.1476243 |
| AFFAAASL | ASRVQGAEQY | 0.16629265 |
| AAAALPAFSPP | ASRVQGAEQY | 0.13223225 |
| VVLAPAL | ASRVQGAEQY | 0.16091281 |
| TIGHVDHGKTTL | ASRVQGAEQY | 0.12822658 |
| LISNASDALDK | ASRVQGAEQY | 0.15065357 |
| LISNASDAL | ASRVQGAEQY | 0.18407156 |
| ISNASDALDK | ASRVQGAEQY | 0.13076854 |
| NQAETLVYQTE | ASRVQGAEQY | 0.13170932 |
| TGGTGGTAGAG | ASRVQGAEQY | 0.18388967 |
| TGSTAGGAGGA | ASRVQGAEQY | 0.69704336 |
| DPLAVPVAAGPV | ASRVQGAEQY | 0.7537371 |
| GGTGGTPGNGGN | ASRVQGAEQY | 0.3158864 |
| PAGLAGVDPATL | ASRVQGAEQY | 0.38192046 |
| GNNLVVEAPHAGASV | ASRVQGAEQY | 0.5329218 |
| KVTRIMSASTLEQPA | ASRVQGAEQY | 0.7534667 |
| LGFALPIALPRRTVV | ASRVQGAEQY | 0.8041611 |
| TRLGNLTARKLVVLV | ASRVQGAEQY | 0.73067796 |
| WSQQRRLIYFPSAGP | ASRVQGAEQY | 0.39346805 |
| GPNGNPDPMAAAVDIR | ASRVQGAEQY | 0.66069126 |
| TQRCVSVADLRAAASR | ASRVQGAEQY | 0.5407789 |
| QFVLDVAEHPVEVLTETV | ASRVQGAEQY | 0.54342324 |
| GGAGGTGGAGGNGGTQAGDG | ASRVQGAEQY | 0.6550198 |
| PLLATLRAIAVDVDAKPAQV | ASRVQGAEQY | 0.61912674 |
| APKFRTGGYMIDFWGVGYQV/ | ASRVQGAEQY | 0.41525888 |

| | | |
|---|---|---|
| GAGGAGGAGGAAGGPGGTGG[illegible] | ASRVQGAEQY | 0.34652367 |
| IGALGLLATDVAPAGSVAFG[illegible] | ASRVQGAEQY | 0.68009657 |
| AGGAGGIGGNGGAGGAGGVL[illegible] | ASRVQGAEQY | 0.75923645 |
| ADELIGLL | ASRVQGAEQY | 0.73387474 |
| ADRALKA | ASRVQGAEQY | 0.8365363 |
| AGAVGQVGGAGGA | ASRVQGAEQY | 0.74540627 |
| AGGGGGLGGAP | ASRVQGAEQY | 0.6594515 |
| AGGLLYGNGGAG | ASRVQGAEQY | 0.1363601 |
| AGLESIGL | ASRVQGAEQY | 0.26470083 |
| AGLKTGII | ASRVQGAEQY | 0.3993513 |
| ASLAALLL | ASRVQGAEQY | 0.6103987 |
| ASVINISSI | ASRVQGAEQY | 0.16860941 |
| ATPPGPA | ASRVQGAEQY | 0.71044934 |
| ATVASALV | ASRVQGAEQY | 0.4372394 |
| ATVASLIL | ASRVQGAEQY | 0.41791397 |
| AVNGPVLIRH | ASRVQGAEQY | 0.758111 |
| DGAAGAVGI | ASRVQGAEQY | 0.36593965 |
| DPLLHGI | ASRVQGAEQY | 0.39491528 |
| ELGAALGAL | ASRVQGAEQY | 0.4177376 |
| ELGRTQAV | ASRVQGAEQY | 0.7742648 |
| FVAGGYLS | ASRVQGAEQY | 0.32160872 |
| GASGGAGGAGGW | ASRVQGAEQY | 0.7557324 |
| GGAGAGGDGGAGGI | ASRVQGAEQY | 0.4781521 |
| GTGHNGSQPGMGGNG | ASRVQGAEQY | 0.7729663 |
| GYAAASASAS | ASRVQGAEQY | 0.3190722 |
| HQVDVAAVQQG | ASRVQGAEQY | 0.7233387 |
| IGAAIAHT | ASRVQGAEQY | 0.58568037 |
| IGMIEVAQL | ASRVQGAEQY | 0.83359766 |
| IRLAPEHTSAMD | ASRVQGAEQY | 0.56530035 |
| IYNVSVL | ASRVQGAEQY | 0.733873 |
| KQGFLGM | ASRVQGAEQY | 0.5395723 |
| LESVITQL | ASRVQGAEQY | 0.76052123 |
| LGLGALVVV | ASRVQGAEQY | 0.27954942 |
| LIGNGGNGGSGGI | ASRVQGAEQY | 0.44267708 |
| LNAINAPTL | ASRVQGAEQY | 0.2764622 |
| LQLLNTGV | ASRVQGAEQY | 0.5166168 |
| LQTSIEF | ASRVQGAEQY | 0.68382585 |
| MSFAVSEL | ASRVQGAEQY | 0.14099514 |
| NTGALGL | ASRVQGAEQY | 0.54499507 |
| PPGAGGAGGAGTT | ASRVQGAEQY | 0.41520828 |
| QLVGLGI | ASRVQGAEQY | 0.3578717 |
| RFLEQVHQL | ASRVQGAEQY | 0.71729565 |
| RGVGYVLR | ASRVQGAEQY | 0.5883963 |
| SATGVFAGVI | ASRVQGAEQY | 0.7470813 |
| SLPAITLP | ASRVQGAEQY | 0.44096762 |
| STALAFTV | ASRVQGAEQY | 0.4651043 |
| SVLTVGLGK | ASRVQGAEQY | 0.5794187 |
| TALAGAGSGI | ASRVQGAEQY | 0.20498759 |
| TGLPFTGI | ASRVQGAEQY | 0.38553274 |
| VAEHVTV | ASRVQGAEQY | 0.7885637 |
| VDIVLNSL | ASRVQGAEQY | 0.7506167 |

| | | |
|---|---|---|
| VLTGGGAVL | ASRVQGAEQY | 0.7607467 |
| VNVGLAYGI | ASRVQGAEQY | 0.6996937 |
| VSSAAVGAV | ASRVQGAEQY | 0.6669005 |
| VTSVYEAGV | ASRVQGAEQY | 0.73027205 |
| WVALDRGA | ASRVQGAEQY | 0.75654817 |
| YLVGSVVI | ASRVQGAEQY | 0.764769 |
| GPPGTGKTLL | ASRVQGAEQY | 0.7554903 |
| GVGMTKF | ASRVQGAEQY | 0.6668549 |
| IGIGGSDL | ASRVQGAEQY | 0.74788904 |
| IGTLLAEL | ASRVQGAEQY | 0.73311543 |
| IYTSGTT | ASRVQGAEQY | 0.7283835 |
| LSFPAGV | ASRVQGAEQY | 0.7554516 |
| LYTSGTT | ASRVQGAEQY | 0.73207414 |
| VGAFPFGF | ASRVQGAEQY | 0.6913922 |
| VGAGTVEF | ASRVQGAEQY | 0.6918837 |
| VGKSTLLL | ASRVQGAEQY | 0.75339794 |
| AALAHAGGGIV | ASRVQGAEQY | 0.76078546 |
| AALVAVSLL | ASRVQGAEQY | 0.75948817 |
| AAVSRAVAAL | ASRVQGAEQY | 0.76765597 |
| AGGSSGIGGF | ASRVQGAEQY | 0.75913465 |
| AGLTVANANG | ASRVQGAEQY | 0.7531878 |
| AGLVGAVL | ASRVQGAEQY | 0.3490082 |
| ALSQAFTV | ASRVQGAEQY | 0.5961502 |
| ALTFSVSL | ASRVQGAEQY | 0.6982256 |
| ALTGAALVAL | ASRVQGAEQY | 0.7346056 |
| ASFSIGAGVL | ASRVQGAEQY | 0.49898595 |
| DGSVYKI | ASRVQGAEQY | 0.75098103 |
| EQLAEIV | ASRVQGAEQY | 0.7002466 |
| FAVSELTVM | ASRVQGAEQY | 0.67373013 |
| GAAPLAAGL | ASRVQGAEQY | 0.7604668 |
| GADGGDGGSGGAGG | ASRVQGAEQY | 0.6880137 |
| GFGLDTGGAGG | ASRVQGAEQY | 0.6311445 |
| GLASGGLL | ASRVQGAEQY | 0.71896124 |
| GRIAGAAVLL | ASRVQGAEQY | 0.7653239 |
| GSAGSLNTSF | ASRVQGAEQY | 0.64228106 |
| GTGQIGI | ASRVQGAEQY | 0.7580211 |
| GTLIALIL | ASRVQGAEQY | 0.73896575 |
| IAATMGFVVG | ASRVQGAEQY | 0.76300395 |
| IFAQALL | ASRVQGAEQY | 0.5793042 |
| IGAAVALGF | ASRVQGAEQY | 0.75770193 |
| IGLTPLAL | ASRVQGAEQY | 0.7321171 |
| ILGSNTSTL | ASRVQGAEQY | 0.76887363 |
| IVFSLTVL | ASRVQGAEQY | 0.72923446 |
| LASKATGF | ASRVQGAEQY | 0.74821997 |
| LASVALIL | ASRVQGAEQY | 0.73004943 |
| LFAVGLTL | ASRVQGAEQY | 0.7599387 |
| LFNGGIGTYI | ASRVQGAEQY | 0.59910333 |
| LTLGAGII | ASRVQGAEQY | 0.7249166 |
| LVAAALGDI | ASRVQGAEQY | 0.6687514 |
| NTAFVQLGI | ASRVQGAEQY | 0.73455894 |
| QLFSGAEL | ASRVQGAEQY | 0.75222933 |

| | | |
|---|---|---|
| SAATFITL | ASRVQGAEQY | 0.4175507 |
| SNYGLANLGV | ASRVQGAEQY | 0.7584412 |
| SYGGAYAVM | ASRVQGAEQY | 0.7095041 |
| TTGVGGLTLG | ASRVQGAEQY | 0.67202985 |
| TTQTIYI | ASRVQGAEQY | 0.7605312 |
| VAAGAGLGAVI | ASRVQGAEQY | 0.739638 |
| VAAVVTLL | ASRVQGAEQY | 0.75445235 |
| VGFLGGLI | ASRVQGAEQY | 0.71601605 |
| VGVIPILL | ASRVQGAEQY | 0.7426014 |
| VIFASSLI | ASRVQGAEQY | 0.7455508 |
| VKGGDGGAAAG | ASRVQGAEQY | 0.69846547 |
| VTKTFTNI | ASRVQGAEQY | 0.7038306 |
| VVAAAILL | ASRVQGAEQY | 0.7611935 |
| YAALDVEL | ASRVQGAEQY | 0.8631104 |
| LASVALLL | ASRVQGAEQY | 0.8545021 |
| LDAVHRV | ASRVQGAEQY | 0.7424189 |
| QGLSSSIF | ASRVQGAEQY | 0.7573533 |
| RGTFANIRL | ASRVQGAEQY | 0.7956196 |
| RQLWWGHRI | ASRVQGAEQY | 0.8666177 |
| YGPPGTGKTLL | ASRVQGAEQY | 0.8700917 |
| AALVGAPANVA | ASRVQGAEQY | 0.8663807 |
| AATFRRTE | ASRVQGAEQY | 0.7856051 |
| AGALGGGI | ASRVQGAEQY | 0.8486651 |
| AGGAGGAGGIGAG | ASRVQGAEQY | 0.8157471 |
| AGSGALGL | ASRVQGAEQY | 0.8350063 |
| AQIADQI | ASRVQGAEQY | 0.85436666 |
| ASAVAVTYL | ASRVQGAEQY | 0.8318474 |
| ASTVLGGTGG | ASRVQGAEQY | 0.7587979 |
| AVGGEPAK | ASRVQGAEQY | 0.7851418 |
| CEQGTFGGTC | ASRVQGAEQY | 0.8464323 |
| DEGKAPAG | ASRVQGAEQY | 0.86154604 |
| DTVVFPAL | ASRVQGAEQY | 0.8603006 |
| FEGITMAG | ASRVQGAEQY | 0.8919674 |
| FPLVVQD | ASRVQGAEQY | 0.8708335 |
| GAFTTPQ | ASRVQGAEQY | 0.8520761 |
| GFGGFGTAGGVGGA | ASRVQGAEQY | 0.449856 |
| GFLAQFL | ASRVQGAEQY | 0.6846186 |
| GGAGGAGGNAGVA | ASRVQGAEQY | 0.79157186 |
| GGVGTAGAGGAGGAGG | ASRVQGAEQY | 0.8016174 |
| GNGGGDAGNAGSGGN | ASRVQGAEQY | 0.40506506 |
| GTGGAGGAGNDAGS | ASRVQGAEQY | 0.854323 |
| GTTNTGWFNS | ASRVQGAEQY | 0.8254664 |
| IFMGPLFTL | ASRVQGAEQY | 0.7502639 |
| ILAHAIEAI | ASRVQGAEQY | 0.8709449 |
| IRSAIREAL | ASRVQGAEQY | 0.7902397 |
| ITFAGGG | ASRVQGAEQY | 0.75461906 |
| ITGMSGHQEV | ASRVQGAEQY | 0.80548173 |
| IVAGLACTL | ASRVQGAEQY | 0.8768512 |
| IVGGGLAGA | ASRVQGAEQY | 0.7950933 |
| IVTGVLNI | ASRVQGAEQY | 0.8754411 |
| LAQQGCI | ASRVQGAEQY | 0.7878588 |

| | | |
|---|---|---|
| LCLHVEN | ASRVQGAEQY | 0.8839302 |
| LDAISDGS | ASRVQGAEQY | 0.7421801 |
| LGGFAAML | ASRVQGAEQY | 0.8789816 |
| LGGKNPMI | ASRVQGAEQY | 0.83655083 |
| LGPKGVAAIL | ASRVQGAEQY | 0.8995493 |
| LMVGVSI | ASRVQGAEQY | 0.83560777 |
| LVGAAFVL | ASRVQGAEQY | 0.8714634 |
| LVLAVAPL | ASRVQGAEQY | 0.78578025 |
| MAAIAVIDAA | ASRVQGAEQY | 0.84776735 |
| NNGINAELQ | ASRVQGAEQY | 0.69525075 |
| RGEPLVLVHG | ASRVQGAEQY | 0.8011405 |
| RGGAGGAAGLI | ASRVQGAEQY | 0.6912154 |
| RSYRMHPAV | ASRVQGAEQY | 0.82892513 |
| SAAHLLAGQLA | ASRVQGAEQY | 0.8727213 |
| SAAVGSY | ASRVQGAEQY | 0.35615245 |
| SPHAVLHRI | ASRVQGAEQY | 0.8583391 |
| SQIAKVL | ASRVQGAEQY | 0.6762444 |
| TALGHLVP | ASRVQGAEQY | 0.7708323 |
| TAPTTSGAA | ASRVQGAEQY | 0.8721012 |
| TDGDDGNGGSPGT | ASRVQGAEQY | 0.8229933 |
| TDGLGEN | ASRVQGAEQY | 0.8653111 |
| TGGPPLPGQA | ASRVQGAEQY | 0.7679058 |
| TIPAITI | ASRVQGAEQY | 0.80752134 |
| TIPGFTI | ASRVQGAEQY | 0.8459475 |
| TSIASGALA | ASRVQGAEQY | 0.5494733 |
| TVTLPGAAI | ASRVQGAEQY | 0.735957 |
| VASGLSAALP | ASRVQGAEQY | 0.8920015 |
| VGAGVIGI | ASRVQGAEQY | 0.6035302 |
| VGPKRFGV | ASRVQGAEQY | 0.64350027 |
| VNVAFPD | ASRVQGAEQY | 0.5133322 |
| VSMTNSGVS | ASRVQGAEQY | 0.49615827 |
| VTDVLGDLSV | ASRVQGAEQY | 0.5305543 |
| VVVVGAGFAGL | ASRVQGAEQY | 0.63722605 |
| VYTVGTVV | ASRVQGAEQY | 0.6373571 |
| WTTFSGP | ASRVQGAEQY | 0.6177756 |
| YDELVEAF | ASRVQGAEQY | 0.52817047 |
| YGTGGAGGNGG | ASRVQGAEQY | 0.59896445 |
| YTNVMAV | ASRVQGAEQY | 0.5595378 |
| FLEQVHQL | ASRVQGAEQY | 0.55689406 |
| GLSLLAVAV | ASRVQGAEQY | 0.61671865 |
| LEQVHQL | ASRVQGAEQY | 0.62797946 |
| LNAAGAI | ASRVQGAEQY | 0.5632502 |
| PAVHASL | ASRVQGAEQY | 0.5305393 |
| TELEDDA | ASRVQGAEQY | 0.6031493 |
| TPPGPPN | ASRVQGAEQY | 0.6336779 |
| TPPPPPPP | ASRVQGAEQY | 0.6190819 |
| TQSPALI | AWGDNERLF | 0.6607329 |
| EFAAAFPQ | AWGDNERLF | 0.62398386 |
| LLSGNLDVL | AWGDNERLF | 0.62497044 |
| GPANLGNYNI | AWGDNERLF | 0.34892914 |
| TLGPTLP | AWGDNERLF | 0.41217563 |

| | | |
|---|---|---|
| VYAYSSCCK | AWGDNERLF | 0.60494524 |
| APAPAPAPNI | AWGDNERLF | 0.5679616 |
| PGGPVPIP | AWGDNERLF | 0.44227505 |
| FLTVALEP | AWGDNERLF | 0.6186236 |
| SQYFGANL | AWGDNERLF | 0.55638194 |
| GGVKPAA | AWGDNERLF | 0.5559349 |
| GGTGGAGGAGGW | AWGDNERLF | 0.6199318 |
| GVGGVAGLGGAGG | AWGDNERLF | 0.54037654 |
| VLLVLATTI | AWGDNERLF | 0.5316157 |
| IGIFAAML | AWGDNERLF | 0.54921407 |
| AVGDLID | AWGDNERLF | 0.641281 |
| NISPLPL | AWGDNERLF | 0.45324692 |
| VTAVFVAI | AWGDNERLF | 0.61099863 |
| SGDDNDAAL | AWGDNERLF | 0.5611012 |
| AVAALAHS | AWGDNERLF | 0.6410101 |
| GGVTVMATP | AWGDNERLF | 0.4850783 |
| AGLAANVRD | AWGDNERLF | 0.6236454 |
| TNRPDLIDPALLRPG | AWGDNERLF | 0.54699725 |
| LANLGVTAL | AWGDNERLF | 0.6641288 |
| TGERIEL | AWGDNERLF | 0.5527817 |
| NGEELAA | AWGDNERLF | 0.61612236 |
| VLSLVWG | AWGDNERLF | 0.60826695 |
| IGYALWL | AWGDNERLF | 0.61275315 |
| LSLLLAPL | AWGDNERLF | 0.44051957 |
| TLDEALK | AWGDNERLF | 0.5554902 |
| TDGAGTAEGG | AWGDNERLF | 0.46356764 |
| NAPGGAGGNGG | AWGDNERLF | 0.5863971 |
| VSVFMGMGGNF | AWGDNERLF | 0.616469 |
| LASYLAAL | AWGDNERLF | 0.4247301 |
| LVGGTPAL | AWGDNERLF | 0.6135075 |
| VRNADVL | AWGDNERLF | 0.56658643 |
| TCSSPFLVSAM | AWGDNERLF | 0.54246795 |
| VHITDAE | AWGDNERLF | 0.6416095 |
| EYAAPSF | AWGDNERLF | 0.5973924 |
| VVDAFHGGL | AWGDNERLF | 0.630427 |
| ASVVDPQ | AWGDNERLF | 0.5870303 |
| SYNEFGGLS | AWGDNERLF | 0.5456931 |
| DPDGSVY | AWGDNERLF | 0.6133441 |
| MVSTIHATE | AWGDNERLF | 0.5416766 |
| LAAGVADPV | AWGDNERLF | 0.5318643 |
| TKEQIAA | AWGDNERLF | 0.63149315 |
| SLASPAAL | AWGDNERLF | 0.83787334 |
| IGLVFYGV | AWGDNERLF | 0.8853466 |
| FLLDASNL | AWGDNERLF | 0.7266039 |
| YQVGNFP | AWGDNERLF | 0.70863813 |
| ESLLSQST | AWGDNERLF | 0.78696096 |
| LFTVGFAAVG | AWGDNERLF | 0.8824435 |
| ATPPTPT | AWGDNERLF | 0.873338 |
| TDLATLG | AWGDNERLF | 0.84775615 |
| SAGGTGR | AWGDNERLF | 0.7122784 |
| GFSNSGS | AWGDNERLF | 0.8124205 |

| | | |
|---|---|---|
| GGNGGSGGTSV | AWGDNERLF | 0.7448185 |
| GTSVATGGAGN | AWGDNERLF | 0.7810229 |
| NGGNGGSGGTS | AWGDNERLF | 0.85912955 |
| AVVKFPH | AWGDNERLF | 0.8002881 |
| TPPPPPP | AWGDNERLF | 0.76157755 |
| AAPAAAAAPA | AWGDNERLF | 0.787822 |
| IALNEDL | AWGDNERLF | 0.82431674 |
| IVNVSSM | AWGDNERLF | 0.86548316 |
| TGIPPAPR | AWGDNERLF | 0.83923674 |
| FDLGGGTF | AWGDNERLF | 0.90008366 |
| FDLGGGTFDVS | AWGDNERLF | 0.85757315 |
| NEPTAAA | AWGDNERLF | 0.87052906 |
| NEPTAAALA | AWGDNERLF | 0.22161528 |
| GSIINISSI | AWGDNERLF | 0.54331267 |
| VAPGNAGTA | AWGDNERLF | 0.7758118 |
| TGGGASLE | AWGDNERLF | 0.79800874 |
| VAVYNHYKRI | AWGDNERLF | 0.5591163 |
| TDPAPATT | AWGDNERLF | 0.8615457 |
| TNRPDILDPALLRPG | AWGDNERLF | 0.77423733 |
| LTTTTTS | AWGDNERLF | 0.7186084 |
| PTGSGKTL | AWGDNERLF | 0.85968506 |
| TGSGKTL | AWGDNERLF | 0.7627029 |
| TIGHVDHGK | AWGDNERLF | 0.7428822 |
| TIGHVDHGKTT | AWGDNERLF | 0.73643154 |
| GHVDHGKTT | AWGDNERLF | 0.8809961 |
| AQFEHTLL | AWGDNERLF | 0.5173832 |
| IVNQDEL | AWGDNERLF | 0.83995354 |
| TNGSQFF | AWGDNERLF | 0.81434685 |
| TGKGIVE | AWGDNERLF | 0.87223464 |
| LISNASDA | AWGDNERLF | 0.58907664 |
| AELRTLGELTKIAKAHGAQ | AWGDNERLF | 0.8440306 |
| APNAVAAAEPSW | AWGDNERLF | 0.7737844 |
| APQVAGAATAVL | AWGDNERLF | 0.9003626 |
| ARRDPAAAAAILPTDA | AWGDNERLF | 0.7727257 |
| DAAARDVLAEAGLAEYF | AWGDNERLF | 0.81828 |
| DDQIVLSAKANLAKVEQ | AWGDNERLF | 0.73324394 |
| DELLPDTGEDSDEDGH | AWGDNERLF | 0.85444975 |
| ELPEAPANRATVPQD | AWGDNERLF | 0.60824066 |
| FGQSKIERVTGIG | AWGDNERLF | 0.787194 |
| GAGGTGSTAGGAGGAGGAGGI | AWGDNERLF | 0.7213267 |
| GHGGTGAAGGNGGTGGAGGS( | AWGDNERLF | 0.8378858 |
| GNGGNGGAGGEGGPGGAGGA( | AWGDNERLF | 0.8389201 |
| GSTAGNGGNGGNATNGTIAG( | AWGDNERLF | 0.6725862 |
| GTGGDGGNGGNGGTGG | AWGDNERLF | 0.86432624 |
| GVRRPAGIPADAPV | AWGDNERLF | 0.80398834 |
| GYRQVTDAHLLTLARR | AWGDNERLF | 0.7523798 |
| IAKMGQDGHDRG | AWGDNERLF | 0.879946 |
| KAEPSIEIQYT | AWGDNERLF | 0.76407886 |
| KPVRIRMAADEDTVAVT | AWGDNERLF | 0.8680863 |
| MPPRIAGMRLLVIKPEP | AWGDNERLF | 0.789069 |
| MKFYAVALFRSRGVPPTR | AWGDNERLF | 0.8073316 |

| | | |
|---|---|---|
| NAGGFGILGAVAHSP | AWGDNERLF | 0.8702364 |
| NEGREITLAMAEM | AWGDNERLF | 0.6719084 |
| NELVEWSHPLGHRWRWE | AWGDNERLF | 0.6698421 |
| NLGAGGTGGNGGNPG | AWGDNERLF | 0.8711555 |
| NTEWPKWETVMAADPVD | AWGDNERLF | 0.811105 |
| PASAAPAETQP | AWGDNERLF | 0.8280312 |
| QQSSDGKNGVRAAEITGE | AWGDNERLF | 0.67250586 |
| RAPDAVTTSWWKE | AWGDNERLF | 0.61981344 |
| SGFGNVGAHTSGWFNQ | AWGDNERLF | 0.70846343 |
| SGVAGAVGQAASVGGLKV | AWGDNERLF | 0.8253773 |
| SNGGAGTGSAGGNGGT | AWGDNERLF | 0.8523207 |
| TGGAGGEGGAAG | AWGDNERLF | 0.7955735 |
| TGGDGGAAGAGGAPGHG | AWGDNERLF | 0.66797674 |
| TMPPLLAAIANAGRHGV | AWGDNERLF | 0.80161285 |
| TSSTAGNSGGA | AWGDNERLF | 0.76039106 |
| VPATVPVGILC | AWGDNERLF | 0.74688995 |
| VQRPDSGGVVLTGRIS | AWGDNERLF | 0.80567706 |
| WTPNPVPATKGVPAGLVQAT | AWGDNERLF | 0.798611 |
| YALKRALAKFSDRFDVVII | AWGDNERLF | 0.72982526 |
| YLIGSVIV | AWGDNERLF | 0.69249284 |
| YGQLVAVID | AWGDNERLF | 0.79326 |
| VTVGGMTLPA | AWGDNERLF | 0.83566 |
| VLALPAV | AWGDNERLF | 0.8294561 |
| VIPPTLNYET | AWGDNERLF | 0.86337644 |
| VGMGGMHPSSG | AWGDNERLF | 0.8396311 |
| VGLAGGAATA | AWGDNERLF | 0.81502545 |
| VGGAGGAGGSGP | AWGDNERLF | 0.26812288 |
| VFAAFMV | AWGDNERLF | 0.55392194 |
| TLYDKLR | AWGDNERLF | 0.65220356 |
| TLAVFTVL | AWGDNERLF | 0.7593178 |
| TGSVQPARV | AWGDNERLF | 0.32734516 |
| TGGGAGSGGT | AWGDNERLF | 0.80149496 |
| SHLETSL | AWGDNERLF | 0.73819816 |
| SFTTGMT | AWGDNERLF | 0.6564407 |
| QVEVAKRMGVRQPTVS | AWGDNERLF | 0.8383957 |
| QGSGGPALAM | AWGDNERLF | 0.721617 |
| MLAPSNP | AWGDNERLF | 0.69512594 |
| LVNNAAI | AWGDNERLF | 0.70587486 |
| LTGEINTLA | AWGDNERLF | 0.8622483 |
| LSIFGPV | AWGDNERLF | 0.56644475 |
| LLALLANL | AWGDNERLF | 0.83108294 |
| LISNASDALDKLR | AWGDNERLF | 0.71238536 |
| LINEAVP | AWGDNERLF | 0.85480535 |
| LGYAFSGL | AWGDNERLF | 0.5466235 |
| LDFVAVAL | AWGDNERLF | 0.82598114 |
| LAQPGAE | AWGDNERLF | 0.7524444 |
| ISNLGAVTGF | AWGDNERLF | 0.8773296 |
| HQEDPGRI | AWGDNERLF | 0.76012254 |
| GTPGEPTP | AWGDNERLF | 0.81604743 |
| GFFNSGS | AWGDNERLF | 0.7248019 |
| GAGGGGLTAGAN | AWGDNERLF | 0.8292012 |

| | | |
|---|---|---|
| FVDALVNM | AWGDNERLF | 0.5166856 |
| FLTVAPEL | AWGDNERLF | 0.768972 |
| FETLTAL | AWGDNERLF | 0.64600456 |
| DVINPIEL | AWGDNERLF | 0.7877177 |
| APVSGAPV | AWGDNERLF | 0.80278575 |
| ALVGGLAW | AWGDNERLF | 0.33414525 |
| ALAAAAATAI | AWGDNERLF | 0.83024514 |
| AFGLGLLL | AWGDNERLF | 0.66315174 |
| AFFAAASL | AWGDNERLF | 0.6504177 |
| AAAALPAFSPP | AWGDNERLF | 0.8383935 |
| VVLAPAL | AWGDNERLF | 0.7627124 |
| TIGHVDHGKTTL | AWGDNERLF | 0.8365042 |
| LISNASDALDK | AWGDNERLF | 0.7104378 |
| LISNASDAL | AWGDNERLF | 0.74981505 |
| ISNASDALDK | AWGDNERLF | 0.81367123 |
| NQAETLVYQTE | AWGDNERLF | 0.4694814 |
| TGGTGGTAGAG | AWGDNERLF | 0.6223936 |
| TGSTAGGAGGA | AWGDNERLF | 0.85285556 |
| DPLAVPVAAGPV | AWGDNERLF | 0.8023646 |
| GGTGGTPGNGGN | AWGDNERLF | 0.7374368 |
| PAGLAGVDPATL | AWGDNERLF | 0.6175033 |
| GNNLVVEAPHAGASV | AWGDNERLF | 0.57017654 |
| KVTRIMSASTLEQPA | AWGDNERLF | 0.6968051 |
| LGFALPIALPRRTVV | AWGDNERLF | 0.80364645 |
| TRLGNLTARKLVVLV | AWGDNERLF | 0.80941033 |
| WSQQRRLIYFPSAGP | AWGDNERLF | 0.7938384 |
| GPNGNPDPMAAAVDIR | AWGDNERLF | 0.601475 |
| TQRCVSVADLRAAASR | AWGDNERLF | 0.798736 |
| QFVLDVAEHPVEVLTETV | AWGDNERLF | 0.71756244 |
| GGAGGTGGAGGNGGTQAGDG | AWGDNERLF | 0.7095967 |
| PLLATLRAIAVDVDAKPAQV | AWGDNERLF | 0.787048 |
| APKFRTGGYMIDFWGVGYQV[illegible] | AWGDNERLF | 0.7258125 |
| GAGGAGGAGGAAGGPGGTGG[illegible] | AWGDNERLF | 0.65565765 |
| IGALGLLATDVAPAGSVAFG[illegible] | AWGDNERLF | 0.5715036 |
| AGGAGGIGGNGGAGGAGGVL[illegible] | AWGDNERLF | 0.80372417 |
| ADELIGLL | AWGDNERLF | 0.8120315 |
| ADRALKA | AWGDNERLF | 0.783425 |
| AGAVGQVGGAGGA | AWGDNERLF | 0.8509238 |
| AGGGGGLGGAP | AWGDNERLF | 0.8156738 |
| AGGLLYGNGGAG | AWGDNERLF | 0.7781347 |
| AGLESIGL | AWGDNERLF | 0.29160744 |
| AGLKTGII | AWGDNERLF | 0.41202202 |
| ASLAALLL | AWGDNERLF | 0.6485174 |
| ASVINISSI | AWGDNERLF | 0.6681099 |
| ATPPGPA | AWGDNERLF | 0.19398206 |
| ATVASALV | AWGDNERLF | 0.691155 |
| ATVASLIL | AWGDNERLF | 0.72054017 |
| AVNGPVLIRH | AWGDNERLF | 0.5746068 |
| DGAAGAVGI | AWGDNERLF | 0.82407403 |
| DPLLHGI | AWGDNERLF | 0.65274113 |
| ELGAALGAL | AWGDNERLF | 0.5316381 |

| | | |
|---|---|---|
| ELGRTQAV | AWGDNERLF | 0.7062504 |
| FVAGGYLS | AWGDNERLF | 0.84592736 |
| GASGGAGGAGGW | AWGDNERLF | 0.6336872 |
| GGAGAGGDGGAGGI | AWGDNERLF | 0.806113 |
| GTGHNGSQPGMGGNG | AWGDNERLF | 0.68971 |
| GYAAASASAS | AWGDNERLF | 0.837188 |
| HQVDVAAVQQG | AWGDNERLF | 0.47059274 |
| IGAAIAHT | AWGDNERLF | 0.8058748 |
| IGMIEVAQL | AWGDNERLF | 0.70280874 |
| IRLAPEHTSAMD | AWGDNERLF | 0.85466903 |
| IYNVSVL | AWGDNERLF | 0.6871073 |
| KQGFLGM | AWGDNERLF | 0.812901 |
| LESVITQL | AWGDNERLF | 0.78078234 |
| LGLGALVVV | AWGDNERLF | 0.78827137 |
| LIGNGGNGGSGGI | AWGDNERLF | 0.43350762 |
| LNAINAPTL | AWGDNERLF | 0.7003914 |
| LQLLNTGV | AWGDNERLF | 0.6111219 |
| LQTSIEF | AWGDNERLF | 0.75349385 |
| MSFAVSEL | AWGDNERLF | 0.79662067 |
| NTGALGL | AWGDNERLF | 0.14707105 |
| PPGAGGAGGAGTT | AWGDNERLF | 0.8209061 |
| QLVGLGI | AWGDNERLF | 0.40480968 |
| RFLEQVHQL | AWGDNERLF | 0.5616684 |
| RGVGYVLR | AWGDNERLF | 0.81151843 |
| SATGVFAGVI | AWGDNERLF | 0.7358825 |
| SLPAITLP | AWGDNERLF | 0.79110646 |
| STALAFTV | AWGDNERLF | 0.63428295 |
| SVLTVGLGK | AWGDNERLF | 0.7584423 |
| TALAGAGSGI | AWGDNERLF | 0.7839277 |
| TGLPFTGI | AWGDNERLF | 0.28357342 |
| VAEHVTV | AWGDNERLF | 0.5784867 |
| VDIVLNSL | AWGDNERLF | 0.8288878 |
| VLTGGGAVL | AWGDNERLF | 0.88763016 |
| VNVGLAYGI | AWGDNERLF | 0.8884444 |
| VSSAAVGAV | AWGDNERLF | 0.83541906 |
| VTSVYEAGV | AWGDNERLF | 0.8065988 |
| WVALDRGA | AWGDNERLF | 0.84996957 |
| YLVGSVVI | AWGDNERLF | 0.8887982 |
| GPPGTGKTLL | AWGDNERLF | 0.8968053 |
| GVGMTKF | AWGDNERLF | 0.8853563 |
| IGIGGSDL | AWGDNERLF | 0.7948253 |
| IGTLLAEL | AWGDNERLF | 0.8924609 |
| IYTSGTT | AWGDNERLF | 0.8712228 |
| LSFPAGV | AWGDNERLF | 0.86172295 |
| LYTSGTT | AWGDNERLF | 0.8842392 |
| VGAFPFGF | AWGDNERLF | 0.87502205 |
| VGAGTVEF | AWGDNERLF | 0.8420211 |
| VGKSTLLL | AWGDNERLF | 0.81937784 |
| AALAHAGGGIV | AWGDNERLF | 0.8803349 |
| AALVAVSLL | AWGDNERLF | 0.89273745 |
| AAVSRAVAAL | AWGDNERLF | 0.89236015 |

| | | |
|---|---|---|
| AGGSSGIGGF | AWGDNERLF | 0.9009731 |
| AGLTVANANG | AWGDNERLF | 0.89521277 |
| AGLVGAVL | AWGDNERLF | 0.8812518 |
| ALSQAFTV | AWGDNERLF | 0.3454075 |
| ALTFSVSL | AWGDNERLF | 0.76067746 |
| ALTGAALVAL | AWGDNERLF | 0.7914964 |
| ASFSIGAGVL | AWGDNERLF | 0.86438924 |
| DGSVYKI | AWGDNERLF | 0.35868412 |
| EQLAEIV | AWGDNERLF | 0.853804 |
| FAVSELTVM | AWGDNERLF | 0.86434907 |
| GAAPLAAGL | AWGDNERLF | 0.8268039 |
| GADGGDGGSGGAGG | AWGDNERLF | 0.8938595 |
| GFGLDTGGAGG | AWGDNERLF | 0.84189624 |
| GLASGGLL | AWGDNERLF | 0.76988864 |
| GRIAGAAVLL | AWGDNERLF | 0.8029394 |
| GSAGSLNTSF | AWGDNERLF | 0.8993434 |
| GTGQIGI | AWGDNERLF | 0.7957418 |
| GTLIALIL | AWGDNERLF | 0.8954608 |
| IAATMGFVVG | AWGDNERLF | 0.7928321 |
| IFAQALL | AWGDNERLF | 0.8990758 |
| IGAAVALGF | AWGDNERLF | 0.7435688 |
| IGLTPLAL | AWGDNERLF | 0.8913127 |
| ILGSNTSTL | AWGDNERLF | 0.870912 |
| IVFSLTVL | AWGDNERLF | 0.9048422 |
| LASKATGF | AWGDNERLF | 0.8616455 |
| LASVALIL | AWGDNERLF | 0.8923459 |
| LFAVGLTL | AWGDNERLF | 0.8503678 |
| LFNGGIGTYI | AWGDNERLF | 0.89022315 |
| LTLGAGII | AWGDNERLF | 0.76012814 |
| LVAAALGDI | AWGDNERLF | 0.8720082 |
| NTAFVQLGI | AWGDNERLF | 0.7653395 |
| QLFSGAEL | AWGDNERLF | 0.88063264 |
| SAATFITL | AWGDNERLF | 0.88803905 |
| SNYGLANLGV | AWGDNERLF | 0.405586 |
| SYGGAYAVM | AWGDNERLF | 0.89706165 |
| TTGVGGLTLG | AWGDNERLF | 0.81458044 |
| TTQTIYI | AWGDNERLF | 0.81087005 |
| VAAGAGLGAVI | AWGDNERLF | 0.8930244 |
| VAAVVTLL | AWGDNERLF | 0.8657515 |
| VGFLGGLI | AWGDNERLF | 0.8919221 |
| VGVIPILL | AWGDNERLF | 0.79784834 |
| VIFASSLI | AWGDNERLF | 0.8737204 |
| VKGGDGGAAAG | AWGDNERLF | 0.8818852 |
| VTKTFTNI | AWGDNERLF | 0.6086717 |
| VVAAAILL | AWGDNERLF | 0.7062478 |
| YAALDVEL | AWGDNERLF | 0.8989881 |
| LASVALLL | AWGDNERLF | 0.8390775 |
| LDAVHRV | AWGDNERLF | 0.8268855 |
| QGLSSSIF | AWGDNERLF | 0.7274985 |
| RGTFANIRL | AWGDNERLF | 0.7480564 |
| RQLWWGHRI | AWGDNERLF | 0.7796418 |

| | | |
|---|---|---|
| YGPPGTGKTLL | AWGDNERLF | 0.8415675 |
| AALVGAPANVA | AWGDNERLF | 0.8516067 |
| AATFRRTE | AWGDNERLF | 0.84298086 |
| AGALGGGI | AWGDNERLF | 0.7730721 |
| AGGAGGAGGIGAG | AWGDNERLF | 0.8370563 |
| AGSGALGL | AWGDNERLF | 0.80347717 |
| AQIADQI | AWGDNERLF | 0.80271983 |
| ASAVAVTYL | AWGDNERLF | 0.82835364 |
| ASTVLGGTGG | AWGDNERLF | 0.8249465 |
| AVGGEPAK | AWGDNERLF | 0.76296544 |
| CEQGTFGGTC | AWGDNERLF | 0.696752 |
| DEGKAPAG | AWGDNERLF | 0.8202548 |
| DTVVFPAL | AWGDNERLF | 0.8463888 |
| FEGITMAG | AWGDNERLF | 0.8306309 |
| FPLVVQD | AWGDNERLF | 0.8567627 |
| GAFTTPQ | AWGDNERLF | 0.8417938 |
| GFGGFGTAGGVGGA | AWGDNERLF | 0.8360613 |
| GFLAQFL | AWGDNERLF | 0.30338287 |
| GGAGGAGGNAGVA | AWGDNERLF | 0.54728484 |
| GGVGTAGAGGAGGAGG | AWGDNERLF | 0.73902977 |
| GNGGGGDAGNAGSGGN | AWGDNERLF | 0.77738106 |
| GTGGAGGAGNDAGS | AWGDNERLF | 0.27934116 |
| GTTNTGWFNS | AWGDNERLF | 0.8253733 |
| IFMGPLFTL | AWGDNERLF | 0.7840142 |
| ILAHAIEAI | AWGDNERLF | 0.77301955 |
| IRSAIREAL | AWGDNERLF | 0.8413749 |
| ITFAGGG | AWGDNERLF | 0.78741443 |
| ITGMSGHQEV | AWGDNERLF | 0.661945 |
| IVAGLACTL | AWGDNERLF | 0.7597825 |
| IVGGGLAGA | AWGDNERLF | 0.8512481 |
| IVTGVLNI | AWGDNERLF | 0.69953376 |
| LAQQGCI | AWGDNERLF | 0.8455628 |
| LCLHVEN | AWGDNERLF | 0.6886308 |
| LDAISDGS | AWGDNERLF | 0.8507979 |
| LGGFAAML | AWGDNERLF | 0.7291305 |
| LGGKNPMI | AWGDNERLF | 0.84663695 |
| LGPKGVAAIL | AWGDNERLF | 0.81404114 |
| LMVGVSI | AWGDNERLF | 0.8593309 |
| LVGAAFVL | AWGDNERLF | 0.7970549 |
| LVLAVAPL | AWGDNERLF | 0.84727037 |
| MAAIAVIDAA | AWGDNERLF | 0.81689525 |
| NNGINAELQ | AWGDNERLF | 0.8351998 |
| RGEPLVLVHG | AWGDNERLF | 0.6947468 |
| RGGAGGAAGLI | AWGDNERLF | 0.78004235 |
| RSYRMHPAV | AWGDNERLF | 0.66083646 |
| SAAHLLAGQLA | AWGDNERLF | 0.80615723 |
| SAAVGSY | AWGDNERLF | 0.84635633 |
| SPHAVLHRI | AWGDNERLF | 0.24316922 |
| SQIAKVL | AWGDNERLF | 0.82203925 |
| TALGHLVP | AWGDNERLF | 0.70810056 |
| TAPTTSGAA | AWGDNERLF | 0.7456893 |

| | | |
|---|---|---|
| TDGDDGNGGSPGT | AWGDNERLF | 0.8402976 |
| TDGLGEN | AWGDNERLF | 0.8089472 |
| TGGPPLPGQA | AWGDNERLF | 0.8349547 |
| TIPAITI | AWGDNERLF | 0.72859526 |
| TIPGFTI | AWGDNERLF | 0.77500427 |
| TSIASGALA | AWGDNERLF | 0.8215667 |
| TVTLPGAAI | AWGDNERLF | 0.33513427 |
| VASGLSAALP | AWGDNERLF | 0.6391802 |
| VGAGVIGI | AWGDNERLF | 0.8567246 |
| VGPKRFGV | AWGDNERLF | 0.8022922 |
| VNVAFPD | AWGDNERLF | 0.81382537 |
| VSMTNSGVS | AWGDNERLF | 0.67606694 |
| VTDVLGDLSV | AWGDNERLF | 0.64398843 |
| VVVVGAGFAGL | AWGDNERLF | 0.70165217 |
| VYTVGTVV | AWGDNERLF | 0.8290642 |
| WTTFSGP | AWGDNERLF | 0.82578087 |
| YDELVEAF | AWGDNERLF | 0.8041004 |
| YGTGGAGGNGG | AWGDNERLF | 0.6577074 |
| YTNVMAV | AWGDNERLF | 0.8098055 |
| FLEQVHQL | AWGDNERLF | 0.74915445 |
| GLSLLAVAV | AWGDNERLF | 0.7387856 |
| LEQVHQL | AWGDNERLF | 0.8152314 |
| LNAAGAI | AWGDNERLF | 0.79189575 |
| PAVHASL | AWGDNERLF | 0.72502005 |
| TELEDDA | AWGDNERLF | 0.73231137 |
| TPPGPPN | AWGDNERLF | 0.80003595 |
| TPPPPPPP | AWGDNERLF | 0.8189042 |
| TQSPALI | ASSRDRGWDSYEQY | 0.8120136 |
| EFAAAFPQ | ASSRDRGWDSYEQY | 0.8576797 |
| LLSGNLDVL | ASSRDRGWDSYEQY | 0.8235564 |
| GPANLGNYNI | ASSRDRGWDSYEQY | 0.81186163 |
| TLGPTLP | ASSRDRGWDSYEQY | 0.3527778 |
| VYAYSSCCK | ASSRDRGWDSYEQY | 0.5608741 |
| APAPAPAPNI | ASSRDRGWDSYEQY | 0.72589153 |
| PGGPVPIP | ASSRDRGWDSYEQY | 0.7470012 |
| FLTVALEP | ASSRDRGWDSYEQY | 0.3863725 |
| SQYFGANL | ASSRDRGWDSYEQY | 0.8156187 |
| GGVKPAA | ASSRDRGWDSYEQY | 0.75440085 |
| GGTGGAGGAGGW | ASSRDRGWDSYEQY | 0.6644412 |
| GVGGVAGLGGAGG | ASSRDRGWDSYEQY | 0.81933004 |
| VLLVLATTI | ASSRDRGWDSYEQY | 0.71342444 |
| IGIFAAML | ASSRDRGWDSYEQY | 0.71346724 |
| AVGDLID | ASSRDRGWDSYEQY | 0.7438303 |
| NISPLPL | ASSRDRGWDSYEQY | 0.84210825 |
| VTAVFVAI | ASSRDRGWDSYEQY | 0.648313 |
| SGDDNDAAL | ASSRDRGWDSYEQY | 0.81906915 |
| AVAALAHS | ASSRDRGWDSYEQY | 0.7154132 |
| GGVTVMATP | ASSRDRGWDSYEQY | 0.8419478 |
| AGLAANVRD | ASSRDRGWDSYEQY | 0.585252 |
| TNRPDLIDPALLRPG | ASSRDRGWDSYEQY | 0.8190256 |
| LANLGVTAL | ASSRDRGWDSYEQY | 0.7496959 |

| | | |
|---|---|---|
| TGERIEL | ASSRDRGWDSYEQY | 0.86298877 |
| NGEELAA | ASSRDRGWDSYEQY | 0.7397311 |
| VLSLVWG | ASSRDRGWDSYEQY | 0.8066055 |
| IGYALWL | ASSRDRGWDSYEQY | 0.7745303 |
| LSLLLAPL | ASSRDRGWDSYEQY | 0.80142945 |
| TLDEALK | ASSRDRGWDSYEQY | 0.5022861 |
| TDGAGTAEGG | ASSRDRGWDSYEQY | 0.7650907 |
| NAPGGAGGNGG | ASSRDRGWDSYEQY | 0.6609948 |
| VSVFMGMGGNF | ASSRDRGWDSYEQY | 0.79629743 |
| LASYLAAL | ASSRDRGWDSYEQY | 0.8097545 |
| LVGGTPAL | ASSRDRGWDSYEQY | 0.3994943 |
| VRNADVL | ASSRDRGWDSYEQY | 0.8255805 |
| TCSSPFLVSAM | ASSRDRGWDSYEQY | 0.69150543 |
| VHITDAE | ASSRDRGWDSYEQY | 0.6929041 |
| EYAAPSF | ASSRDRGWDSYEQY | 0.8373194 |
| VVDAFHGGL | ASSRDRGWDSYEQY | 0.77492213 |
| ASVVDPQ | ASSRDRGWDSYEQY | 0.8240151 |
| SYNEFGGLS | ASSRDRGWDSYEQY | 0.75116813 |
| DPDGSVY | ASSRDRGWDSYEQY | 0.73313236 |
| MVSTIHATE | ASSRDRGWDSYEQY | 0.8149264 |
| LAAGVADPV | ASSRDRGWDSYEQY | 0.63451916 |
| TKEQIAA | ASSRDRGWDSYEQY | 0.6723323 |
| SLASPAAL | ASSRDRGWDSYEQY | 0.8402616 |
| IGLVFYGV | ASSRDRGWDSYEQY | 0.7609097 |
| FLLDASNL | ASSRDRGWDSYEQY | 0.7722386 |
| YQVGNFP | ASSRDRGWDSYEQY | 0.64934015 |
| ESLLSQST | ASSRDRGWDSYEQY | 0.5812464 |
| LFTVGFAAVG | ASSRDRGWDSYEQY | 0.7127515 |
| ATPPTPT | ASSRDRGWDSYEQY | 0.7771622 |
| TDLATLG | ASSRDRGWDSYEQY | 0.785242 |
| SAGGTGR | ASSRDRGWDSYEQY | 0.74710923 |
| GFSNSGS | ASSRDRGWDSYEQY | 0.5658523 |
| GGNGGSGGTSV | ASSRDRGWDSYEQY | 0.75423586 |
| GTSVATGGAGN | ASSRDRGWDSYEQY | 0.7099925 |
| NGGNGGSGGTS | ASSRDRGWDSYEQY | 0.6899864 |
| AVVKFPH | ASSRDRGWDSYEQY | 0.74980336 |
| TPPPPPP | ASSRDRGWDSYEQY | 0.73089933 |
| AAPAAAAAPA | ASSRDRGWDSYEQY | 0.67655057 |
| IALNEDL | ASSRDRGWDSYEQY | 0.6086482 |
| IVNVSSM | ASSRDRGWDSYEQY | 0.74944615 |
| TGIPPAPR | ASSRDRGWDSYEQY | 0.77806026 |
| FDLGGGTF | ASSRDRGWDSYEQY | 0.7757176 |
| FDLGGGTFDVS | ASSRDRGWDSYEQY | 0.79207146 |
| NEPTAAA | ASSRDRGWDSYEQY | 0.77947927 |
| NEPTAAALA | ASSRDRGWDSYEQY | 0.76852477 |
| GSIINISSI | ASSRDRGWDSYEQY | 0.2665566 |
| VAPGNAGTA | ASSRDRGWDSYEQY | 0.35329613 |
| TGGGASLE | ASSRDRGWDSYEQY | 0.68682975 |
| VAVYNHYKRI | ASSRDRGWDSYEQY | 0.6783583 |
| TDPAPATT | ASSRDRGWDSYEQY | 0.35943875 |
| TNRPDILDPALLRPG | ASSRDRGWDSYEQY | 0.740391 |

| | | |
|---|---|---|
| LTTTTTS | ASSRDRGWDSYEQY | 0.65658456 |
| PTGSGKTL | ASSRDRGWDSYEQY | 0.6238008 |
| TGSGKTL | ASSRDRGWDSYEQY | 0.7793846 |
| TIGHVDHGK | ASSRDRGWDSYEQY | 0.66389394 |
| TIGHVDHGKTT | ASSRDRGWDSYEQY | 0.61457825 |
| GHVDHGKTT | ASSRDRGWDSYEQY | 0.6389048 |
| AQFEHTLL | ASSRDRGWDSYEQY | 0.7607441 |
| IVNQDEL | ASSRDRGWDSYEQY | 0.5387223 |
| TNGSQFF | ASSRDRGWDSYEQY | 0.7660752 |
| TGKGIVE | ASSRDRGWDSYEQY | 0.5353904 |
| LISNASDA | ASSRDRGWDSYEQY | 0.7794079 |
| AELRTLGELTKIAKAHGAQ | ASSRDRGWDSYEQY | 0.4948343 |
| APNAVAAAEPSW | ASSRDRGWDSYEQY | 0.7601899 |
| APQVAGAATAVL | ASSRDRGWDSYEQY | 0.70292056 |
| ARRDPAAAAAILPTDA | ASSRDRGWDSYEQY | 0.80117714 |
| DAAARDVLAEAGLAEYF | ASSRDRGWDSYEQY | 0.7014191 |
| DDQIVLSAKANLAKVEQ | ASSRDRGWDSYEQY | 0.739563 |
| DELLPDTGEDSDEDGH | ASSRDRGWDSYEQY | 0.6153693 |
| ELPEAPANRATVPQD | ASSRDRGWDSYEQY | 0.7706076 |
| FGQSKIERVTGIG | ASSRDRGWDSYEQY | 0.44525892 |
| GAGGTGSTAGGAGGAGGAGGI | ASSRDRGWDSYEQY | 0.723786 |
| GHGGTGAAGGNGGTGGAGGS( | ASSRDRGWDSYEQY | 0.4194451 |
| GNGGNGGAGGEGGPGGAGGA( | ASSRDRGWDSYEQY | 0.73290366 |
| GSTAGNGGNGGNATNGTIAG( | ASSRDRGWDSYEQY | 0.73757607 |
| GTGGDGGNGGNGGTGG | ASSRDRGWDSYEQY | 0.42548627 |
| GVRRPAGIPADAPV | ASSRDRGWDSYEQY | 0.7564603 |
| GYRQVTDAHLLTLARR | ASSRDRGWDSYEQY | 0.6869836 |
| IAKMGQDGHDRG | ASSRDRGWDSYEQY | 0.62972546 |
| KAEPSIEIQYT | ASSRDRGWDSYEQY | 0.7851195 |
| KPVRIRMAADEDTVAVT | ASSRDRGWDSYEQY | 0.7517273 |
| MPPRIAGMRLLVIKPEP | ASSRDRGWDSYEQY | 0.7759659 |
| MKFYAVALFRSRGVPPTR | ASSRDRGWDSYEQY | 0.68465126 |
| NAGGFGILGAVAHSP | ASSRDRGWDSYEQY | 0.6262115 |
| NEGREITLAMAEM | ASSRDRGWDSYEQY | 0.74077916 |
| NELVEWSHPLGHRWRWE | ASSRDRGWDSYEQY | 0.61246693 |
| NLGAGGTGGNGGNPG | ASSRDRGWDSYEQY | 0.64869964 |
| NTEWPKWETVMAADPVD | ASSRDRGWDSYEQY | 0.772261 |
| PASAAPAETQP | ASSRDRGWDSYEQY | 0.8030705 |
| QQSSDGKNGVRAAEITGE | ASSRDRGWDSYEQY | 0.819315 |
| RAPDAVTTSWWKE | ASSRDRGWDSYEQY | 0.7489351 |
| SGFGNVGAHTSGWFNQ | ASSRDRGWDSYEQY | 0.65531313 |
| SGVAGAVGQAASVGGLKV | ASSRDRGWDSYEQY | 0.7519624 |
| SNGGAGTGSAGGNGGT | ASSRDRGWDSYEQY | 0.81544024 |
| TGGAGGEGGAAG | ASSRDRGWDSYEQY | 0.81990457 |
| TGGDGGAAGAGGAPGHG | ASSRDRGWDSYEQY | 0.7887044 |
| TMPPLLAAIANAGRHGV | ASSRDRGWDSYEQY | 0.521513 |
| TSSTAGNSGGA | ASSRDRGWDSYEQY | 0.79016346 |
| VPATVPVGILC | ASSRDRGWDSYEQY | 0.7589958 |
| VQRPDSGGVVLTGRIS | ASSRDRGWDSYEQY | 0.71236956 |
| WTPNPVPATKGVPAGLVQAT | ASSRDRGWDSYEQY | 0.79549724 |
| YALKRALAKFSDRFDVVII | ASSRDRGWDSYEQY | 0.7739067 |

| | | |
|---|---|---|
| YLIGSVIV | ASSRDRGWDSYEQY | 0.7012541 |
| YGQLVAVID | ASSRDRGWDSYEQY | 0.7000041 |
| VTVGGMTLPA | ASSRDRGWDSYEQY | 0.7707535 |
| VLALPAV | ASSRDRGWDSYEQY | 0.8120324 |
| VIPPTLNYET | ASSRDRGWDSYEQY | 0.8165318 |
| VGMGGMHPSSG | ASSRDRGWDSYEQY | 0.82694274 |
| VGLAGGAATA | ASSRDRGWDSYEQY | 0.8304497 |
| VGGAGGAGGSGP | ASSRDRGWDSYEQY | 0.814834 |
| VFAAFMV | ASSRDRGWDSYEQY | 0.248002 |
| TLYDKLR | ASSRDRGWDSYEQY | 0.35057884 |
| TLAVFTVL | ASSRDRGWDSYEQY | 0.7126531 |
| TGSVQPARV | ASSRDRGWDSYEQY | 0.69642043 |
| TGGGAGSGGT | ASSRDRGWDSYEQY | 0.4134899 |
| SHLETSL | ASSRDRGWDSYEQY | 0.63107955 |
| SFTTGMT | ASSRDRGWDSYEQY | 0.7017901 |
| QVEVAKRMGVRQPTVS | ASSRDRGWDSYEQY | 0.62104666 |
| QGSGGPALAM | ASSRDRGWDSYEQY | 0.8221223 |
| MLAPSNP | ASSRDRGWDSYEQY | 0.7175813 |
| LVNNAAI | ASSRDRGWDSYEQY | 0.6478412 |
| LTGEINTLA | ASSRDRGWDSYEQY | 0.67302644 |
| LSIFGPV | ASSRDRGWDSYEQY | 0.7755446 |
| LLALLANL | ASSRDRGWDSYEQY | 0.58175474 |
| LISNASDALDKLR | ASSRDRGWDSYEQY | 0.80390584 |
| LINEAVP | ASSRDRGWDSYEQY | 0.49528745 |
| LGYAFSGL | ASSRDRGWDSYEQY | 0.8246388 |
| LDFVAVAL | ASSRDRGWDSYEQY | 0.52621776 |
| LAQPGAE | ASSRDRGWDSYEQY | 0.8116622 |
| ISNLGAVTGF | ASSRDRGWDSYEQY | 0.7691449 |
| HQEDPGRI | ASSRDRGWDSYEQY | 0.85053796 |
| GTPGEPTP | ASSRDRGWDSYEQY | 0.77200127 |
| GFFNSGS | ASSRDRGWDSYEQY | 0.7779788 |
| GAGGGGLTAGAN | ASSRDRGWDSYEQY | 0.59758615 |
| FVDALVNM | ASSRDRGWDSYEQY | 0.8044125 |
| FLTVAPEL | ASSRDRGWDSYEQY | 0.55714417 |
| FETLTAL | ASSRDRGWDSYEQY | 0.7776356 |
| DVINPIEL | ASSRDRGWDSYEQY | 0.3922315 |
| APVSGAPV | ASSRDRGWDSYEQY | 0.7996991 |
| ALVGGLAW | ASSRDRGWDSYEQY | 0.79233503 |
| ALAAAAATAI | ASSRDRGWDSYEQY | 0.4454275 |
| AFGLGLLL | ASSRDRGWDSYEQY | 0.8114325 |
| AFFAAASL | ASSRDRGWDSYEQY | 0.7571164 |
| AAAALPAFSPP | ASSRDRGWDSYEQY | 0.66602373 |
| VVLAPAL | ASSRDRGWDSYEQY | 0.83686066 |
| TIGHVDHGKTTL | ASSRDRGWDSYEQY | 0.7449399 |
| LISNASDALDK | ASSRDRGWDSYEQY | 0.8330229 |
| LISNASDAL | ASSRDRGWDSYEQY | 0.7323001 |
| ISNASDALDK | ASSRDRGWDSYEQY | 0.70216954 |
| NQAETLVYQTE | ASSRDRGWDSYEQY | 0.76840895 |
| TGGTGGTAGAG | ASSRDRGWDSYEQY | 0.7019884 |
| TGSTAGGAGGA | ASSRDRGWDSYEQY | 0.6676258 |
| DPLAVPVAAGPV | ASSRDRGWDSYEQY | 0.82636 |

| | | |
|---|---|---|
| GGTGGTPGNGGN | ASSRDRGWDSYEQY | 0.8876395 |
| PAGLAGVDPATL | ASSRDRGWDSYEQY | 0.8890732 |
| GNNLVVEAPHAGASV | ASSRDRGWDSYEQY | 0.8491287 |
| KVTRIMSASTLEQPA | ASSRDRGWDSYEQY | 0.8231973 |
| LGFALPIALPRRTVV | ASSRDRGWDSYEQY | 0.86152315 |
| TRLGNLTARKLVVLV | ASSRDRGWDSYEQY | 0.89161146 |
| WSQQRRLIYFPSAGP | ASSRDRGWDSYEQY | 0.8920661 |
| GPNGNPDPMAAAVDIR | ASSRDRGWDSYEQY | 0.8819215 |
| TQRCVSVADLRAAASR | ASSRDRGWDSYEQY | 0.82712036 |
| QFVLDVAEHPVEVLTETV | ASSRDRGWDSYEQY | 0.878559 |
| GGAGGTGGAGGNGGTQAGDG | ASSRDRGWDSYEQY | 0.86401254 |
| PLLATLRAIAVDVDAKPAQV | ASSRDRGWDSYEQY | 0.8582964 |
| APKFRTGGYMIDFWGVGYQV | ASSRDRGWDSYEQY | 0.8853505 |
| GAGGAGGAGGAAGGPGGTGG | ASSRDRGWDSYEQY | 0.878599 |
| IGALGLLATDVAPAGSVAFG | ASSRDRGWDSYEQY | 0.860078 |
| AGGAGGIGGNGGAGGAGGVL | ASSRDRGWDSYEQY | 0.84742653 |
| ADELIGLL | ASSRDRGWDSYEQY | 0.873448 |
| ADRALKA | ASSRDRGWDSYEQY | 0.8904858 |
| AGAVGQVGGAGGA | ASSRDRGWDSYEQY | 0.8849456 |
| AGGGGGLGGAP | ASSRDRGWDSYEQY | 0.90169823 |
| AGGLLYGNGGAG | ASSRDRGWDSYEQY | 0.89311147 |
| AGLESIGL | ASSRDRGWDSYEQY | 0.88807714 |
| AGLKTGII | ASSRDRGWDSYEQY | 0.38609773 |
| ASLAALLL | ASSRDRGWDSYEQY | 0.7410623 |
| ASVINISSI | ASSRDRGWDSYEQY | 0.8717608 |
| ATPPGPA | ASSRDRGWDSYEQY | 0.8661494 |
| ATVASALV | ASSRDRGWDSYEQY | 0.5955576 |
| ATVASLIL | ASSRDRGWDSYEQY | 0.87934625 |
| AVNGPVLIRH | ASSRDRGWDSYEQY | 0.86786526 |
| DGAAGAVGI | ASSRDRGWDSYEQY | 0.8318322 |
| DPLLHGI | ASSRDRGWDSYEQY | 0.89266443 |
| ELGAALGAL | ASSRDRGWDSYEQY | 0.852669 |
| ELGRTQAV | ASSRDRGWDSYEQY | 0.8236588 |
| FVAGGYLS | ASSRDRGWDSYEQY | 0.85804564 |
| GASGGAGGAGGW | ASSRDRGWDSYEQY | 0.89773 |
| GGAGAGGDGGAGGI | ASSRDRGWDSYEQY | 0.8162148 |
| GTGHNGSQPGMGGNG | ASSRDRGWDSYEQY | 0.8878511 |
| GYAAASASAS | ASSRDRGWDSYEQY | 0.8716843 |
| HQVDVAAVQQG | ASSRDRGWDSYEQY | 0.8972868 |
| IGAAIAHT | ASSRDRGWDSYEQY | 0.772506 |
| IGMIEVAQL | ASSRDRGWDSYEQY | 0.8866161 |
| IRLAPEHTSAMD | ASSRDRGWDSYEQY | 0.8595937 |
| IYNVSVL | ASSRDRGWDSYEQY | 0.90287244 |
| KQGFLGM | ASSRDRGWDSYEQY | 0.8610946 |
| LESVITQL | ASSRDRGWDSYEQY | 0.8848027 |
| LGLGALVVV | ASSRDRGWDSYEQY | 0.8505573 |
| LIGNGGNGGSGGI | ASSRDRGWDSYEQY | 0.88058406 |
| LNAINAPTL | ASSRDRGWDSYEQY | 0.7914572 |
| LQLLNTGV | ASSRDRGWDSYEQY | 0.87290883 |
| LQTSIEF | ASSRDRGWDSYEQY | 0.7915009 |
| MSFAVSEL | ASSRDRGWDSYEQY | 0.88004243 |

| | | |
|---|---|---|
| NTGALGL | ASSRDRGWDSYEQY | 0.88428533 |
| PPGAGGAGGAGTT | ASSRDRGWDSYEQY | 0.6627829 |
| QLVGLGI | ASSRDRGWDSYEQY | 0.89345413 |
| RFLEQVHQL | ASSRDRGWDSYEQY | 0.837305 |
| RGVGYVLR | ASSRDRGWDSYEQY | 0.8445279 |
| SATGVFAGVI | ASSRDRGWDSYEQY | 0.89496917 |
| SLPAITLP | ASSRDRGWDSYEQY | 0.8690249 |
| STALAFTV | ASSRDRGWDSYEQY | 0.89253885 |
| SVLTVGLGK | ASSRDRGWDSYEQY | 0.85584694 |
| TALAGAGSGI | ASSRDRGWDSYEQY | 0.87431437 |
| TGLPFTGI | ASSRDRGWDSYEQY | 0.88885516 |
| VAEHVTV | ASSRDRGWDSYEQY | 0.7897564 |
| VDIVLNSL | ASSRDRGWDSYEQY | 0.84229565 |
| VLTGGGAVL | ASSRDRGWDSYEQY | 0.89276123 |
| VNVGLAYGI | ASSRDRGWDSYEQY | 0.82176566 |
| VSSAAVGAV | ASSRDRGWDSYEQY | 0.86208665 |
| VTSVYEAGV | ASSRDRGWDSYEQY | 0.7362114 |
| WVALDRGA | ASSRDRGWDSYEQY | 0.64275455 |
| YLVGSVVI | ASSRDRGWDSYEQY | 0.73253953 |
| GPPGTGKTLL | ASSRDRGWDSYEQY | 0.82926905 |
| GVGMTKF | ASSRDRGWDSYEQY | 0.8708527 |
| IGIGGSDL | ASSRDRGWDSYEQY | 0.8359331 |
| IGTLLAEL | ASSRDRGWDSYEQY | 0.6298584 |
| IYTSGTT | ASSRDRGWDSYEQY | 0.85396874 |
| LSFPAGV | ASSRDRGWDSYEQY | 0.7911105 |
| LYTSGTT | ASSRDRGWDSYEQY | 0.7665721 |
| VGAFPFGF | ASSRDRGWDSYEQY | 0.80807745 |
| VGAGTVEF | ASSRDRGWDSYEQY | 0.83556795 |
| VGKSTLLL | ASSRDRGWDSYEQY | 0.7650143 |
| AALAHAGGGIV | ASSRDRGWDSYEQY | 0.6698077 |
| AALVAVSLL | ASSRDRGWDSYEQY | 0.7819307 |
| AAVSRAVAAL | ASSRDRGWDSYEQY | 0.8416033 |
| AGGSSGIGGF | ASSRDRGWDSYEQY | 0.8324003 |
| AGLTVANANG | ASSRDRGWDSYEQY | 0.8829396 |
| AGLVGAVL | ASSRDRGWDSYEQY | 0.86372936 |
| ALSQAFTV | ASSRDRGWDSYEQY | 0.8062264 |
| ALTFSVSL | ASSRDRGWDSYEQY | 0.2283912 |
| ALTGAALVAL | ASSRDRGWDSYEQY | 0.49648076 |
| ASFSIGAGVL | ASSRDRGWDSYEQY | 0.68452215 |
| DGSVYKI | ASSRDRGWDSYEQY | 0.76593804 |
| EQLAEIV | ASSRDRGWDSYEQY | 0.27745977 |
| FAVSELTVM | ASSRDRGWDSYEQY | 0.7892848 |
| GAAPLAAGL | ASSRDRGWDSYEQY | 0.79017985 |
| GADGGDGGSGGAGG | ASSRDRGWDSYEQY | 0.69688267 |
| GFGLDTGGAGG | ASSRDRGWDSYEQY | 0.86026484 |
| GLASGGLL | ASSRDRGWDSYEQY | 0.7445456 |
| GRIAGAAVLL | ASSRDRGWDSYEQY | 0.65027773 |
| GSAGSLNTSF | ASSRDRGWDSYEQY | 0.6810279 |
| GTGQIGI | ASSRDRGWDSYEQY | 0.8640008 |
| GTLIALIL | ASSRDRGWDSYEQY | 0.5455451 |
| IAATMGFVVG | ASSRDRGWDSYEQY | 0.85737085 |

| | | |
|---|---|---|
| IFAQALL | ASSRDRGWDSYEQY | 0.60525095 |
| IGAAVALGF | ASSRDRGWDSYEQY | 0.86770236 |
| IGLTPLAL | ASSRDRGWDSYEQY | 0.5322778 |
| ILGSNTSTL | ASSRDRGWDSYEQY | 0.8478397 |
| IVFSLTVL | ASSRDRGWDSYEQY | 0.76430726 |
| LASKATGF | ASSRDRGWDSYEQY | 0.90107083 |
| LASVALIL | ASSRDRGWDSYEQY | 0.7489635 |
| LFAVGLTL | ASSRDRGWDSYEQY | 0.8469678 |
| LFNGGIGTYI | ASSRDRGWDSYEQY | 0.6887572 |
| LTLGAGII | ASSRDRGWDSYEQY | 0.84831774 |
| LVAAALGDI | ASSRDRGWDSYEQY | 0.5348826 |
| NTAFVQLGI | ASSRDRGWDSYEQY | 0.77693343 |
| QLFSGAEL | ASSRDRGWDSYEQY | 0.51509166 |
| SAATFITL | ASSRDRGWDSYEQY | 0.8107736 |
| SNYGLANLGV | ASSRDRGWDSYEQY | 0.8368144 |
| SYGGAYAVM | ASSRDRGWDSYEQY | 0.35410586 |
| TTGVGGLTLG | ASSRDRGWDSYEQY | 0.87097406 |
| TTQTIYI | ASSRDRGWDSYEQY | 0.6945216 |
| VAAGAGLGAVI | ASSRDRGWDSYEQY | 0.64018893 |
| VAAVVTLL | ASSRDRGWDSYEQY | 0.86191684 |
| VGFLGGLI | ASSRDRGWDSYEQY | 0.7975726 |
| VGVIPILL | ASSRDRGWDSYEQY | 0.86710095 |
| VIFASSLI | ASSRDRGWDSYEQY | 0.70654714 |
| VKGGDGGAAAG | ASSRDRGWDSYEQY | 0.7890098 |
| VTKTFTNI | ASSRDRGWDSYEQY | 0.8185315 |
| VVAAAILL | ASSRDRGWDSYEQY | 0.58098984 |
| YAALDVEL | ASSRDRGWDSYEQY | 0.61081785 |
| LASVALLL | ASSRDRGWDSYEQY | 0.8712092 |
| LDAVHRV | ASSRDRGWDSYEQY | 0.7959257 |
| QGLSSSIF | ASSRDRGWDSYEQY | 0.7885774 |
| RGTFANIRL | ASSRDRGWDSYEQY | 0.630787 |
| RQLWWGHRI | ASSRDRGWDSYEQY | 0.618227 |
| YGPPGTGKTLL | ASSRDRGWDSYEQY | 0.6920613 |
| AALVGAPANVA | ASSRDRGWDSYEQY | 0.8181279 |
| AATFRRTE | ASSRDRGWDSYEQY | 0.83932966 |
| AGALGGGI | ASSRDRGWDSYEQY | 0.80561435 |
| AGGAGGAGGIGAG | ASSRDRGWDSYEQY | 0.6905183 |
| AGSGALGL | ASSRDRGWDSYEQY | 0.82408625 |
| AQIADQI | ASSRDRGWDSYEQY | 0.76089233 |
| ASAVAVTYL | ASSRDRGWDSYEQY | 0.74866253 |
| ASTVLGGTGG | ASSRDRGWDSYEQY | 0.81909406 |
| AVGGEPAK | ASSRDRGWDSYEQY | 0.7967746 |
| CEQGTFGGTC | ASSRDRGWDSYEQY | 0.7116247 |
| DEGKAPAG | ASSRDRGWDSYEQY | 0.69271183 |
| DTVVFPAL | ASSRDRGWDSYEQY | 0.7975714 |
| FEGITMAG | ASSRDRGWDSYEQY | 0.8244489 |
| FPLVVQD | ASSRDRGWDSYEQY | 0.81135464 |
| GAFTTPQ | ASSRDRGWDSYEQY | 0.86792445 |
| GFGGFGTAGGVGGA | ASSRDRGWDSYEQY | 0.8254779 |
| GFLAQFL | ASSRDRGWDSYEQY | 0.79496145 |
| GGAGGAGGNAGVA | ASSRDRGWDSYEQY | 0.32222062 |

| | | |
|---|---|---|
| GGVGTAGAGGAGGAGG | ASSRDRGWDSYEQY | 0.5999819 |
| GNGGGGDAGNAGSGGN | ASSRDRGWDSYEQY | 0.70876956 |
| GTGGAGGAGNDAGS | ASSRDRGWDSYEQY | 0.75615036 |
| GTTNTGWFNS | ASSRDRGWDSYEQY | 0.32222468 |
| IFMGPLFTL | ASSRDRGWDSYEQY | 0.7999417 |
| ILAHAIEAI | ASSRDRGWDSYEQY | 0.76086235 |
| IRSAIREAL | ASSRDRGWDSYEQY | 0.6649225 |
| ITFAGGG | ASSRDRGWDSYEQY | 0.8301542 |
| ITGMSGHQEV | ASSRDRGWDSYEQY | 0.69058037 |
| IVAGLACTL | ASSRDRGWDSYEQY | 0.7047493 |
| IVGGGLAGA | ASSRDRGWDSYEQY | 0.7476633 |
| IVTGVLNI | ASSRDRGWDSYEQY | 0.85705787 |
| LAQQGCI | ASSRDRGWDSYEQY | 0.644497 |
| LCLHVEN | ASSRDRGWDSYEQY | 0.8353037 |
| LDAISDGS | ASSRDRGWDSYEQY | 0.71171486 |
| LGGFAAML | ASSRDRGWDSYEQY | 0.84930515 |
| LGGKNPMI | ASSRDRGWDSYEQY | 0.597996 |
| LGPKGVAAIL | ASSRDRGWDSYEQY | 0.82287735 |
| LMVGVSI | ASSRDRGWDSYEQY | 0.73478544 |
| LVGAAFVL | ASSRDRGWDSYEQY | 0.87163115 |
| LVLAVAPL | ASSRDRGWDSYEQY | 0.7454888 |
| MAAIAVIDAA | ASSRDRGWDSYEQY | 0.83985126 |
| NNGINAELQ | ASSRDRGWDSYEQY | 0.78858614 |
| RGEPLVLVHG | ASSRDRGWDSYEQY | 0.7910052 |
| RGGAGGAAGLI | ASSRDRGWDSYEQY | 0.5229714 |
| RSYRMHPAV | ASSRDRGWDSYEQY | 0.7439842 |
| SAAHLLAGQLA | ASSRDRGWDSYEQY | 0.64878 |
| SAAVGSY | ASSRDRGWDSYEQY | 0.79168165 |
| SPHAVLHRI | ASSRDRGWDSYEQY | 0.8025833 |
| SQIAKVL | ASSRDRGWDSYEQY | 0.29958 |
| TALGHLVP | ASSRDRGWDSYEQY | 0.8309397 |
| TAPTTSGAA | ASSRDRGWDSYEQY | 0.588029 |
| TDGDDGNGGSPGT | ASSRDRGWDSYEQY | 0.6590823 |
| TDGLGEN | ASSRDRGWDSYEQY | 0.83331805 |
| TGGPPLPGQA | ASSRDRGWDSYEQY | 0.7694929 |
| TIPAITI | ASSRDRGWDSYEQY | 0.8058634 |
| TIPGFTI | ASSRDRGWDSYEQY | 0.73622453 |
| TSIASGALA | ASSRDRGWDSYEQY | 0.7652168 |
| TVTLPGAAI | ASSRDRGWDSYEQY | 0.8057839 |
| VASGLSAALP | ASSRDRGWDSYEQY | 0.47022563 |
| VGAGVIGI | ASSRDRGWDSYEQY | 0.6576367 |
| VGPKRFGV | ASSRDRGWDSYEQY | 0.8468223 |
| VNVAFPD | ASSRDRGWDSYEQY | 0.79594177 |
| VSMTNSGVS | ASSRDRGWDSYEQY | 0.84466696 |
| VTDVLGDLSV | ASSRDRGWDSYEQY | 0.64530957 |
| VVVVGAGFAGL | ASSRDRGWDSYEQY | 0.5409898 |
| VYTVGTVV | ASSRDRGWDSYEQY | 0.73561996 |
| WTTFSGP | ASSRDRGWDSYEQY | 0.84058094 |
| YDELVEAF | ASSRDRGWDSYEQY | 0.845034 |
| YGTGGAGGNGG | ASSRDRGWDSYEQY | 0.81297743 |
| YTNVMAV | ASSRDRGWDSYEQY | 0.56062317 |

| | | |
|---|---|---|
| FLEQVHQL | ASSRDRGWDSYEQY | 0.6574276 |
| GLSLLAVAV | ASSRDRGWDSYEQY | 0.6576971 |
| LEQVHQL | ASSRDRGWDSYEQY | 0.6581849 |
| LNAAGAI | ASSRDRGWDSYEQY | 0.809676 |
| PAVHASL | ASSRDRGWDSYEQY | 0.7724196 |
| TELEDDA | ASSRDRGWDSYEQY | 0.6690042 |
| TPPGPPN | ASSRDRGWDSYEQY | 0.6607671 |
| TPPPPPPP | ASSRDRGWDSYEQY | 0.79217416 |
| TQSPALI | ASSFGGGGNTEVF | 0.8188647 |
| EFAAAFPQ | ASSFGGGGNTEVF | 0.7582655 |
| LLSGNLDVL | ASSFGGGGNTEVF | 0.8679771 |
| GPANLGNYNI | ASSFGGGGNTEVF | 0.82004845 |
| TLGPTLP | ASSFGGGGNTEVF | 0.843874 |
| VYAYSSCCK | ASSFGGGGNTEVF | 0.14580274 |
| APAPAPAPNI | ASSFGGGGNTEVF | 0.2748559 |
| PGGPVPIP | ASSFGGGGNTEVF | 0.65391237 |
| FLTVALEP | ASSFGGGGNTEVF | 0.63870066 |
| SQYFGANL | ASSFGGGGNTEVF | 0.37851068 |
| GGVKPAA | ASSFGGGGNTEVF | 0.7618283 |
| GGTGGAGGAGGW | ASSFGGGGNTEVF | 0.6068623 |
| GVGGVAGLGGAGG | ASSFGGGGNTEVF | 0.54481214 |
| VLLVLATTI | ASSFGGGGNTEVF | 0.8320391 |
| IGIFAAML | ASSFGGGGNTEVF | 0.66517407 |
| AVGDLID | ASSFGGGGNTEVF | 0.5884641 |
| NISPLPL | ASSFGGGGNTEVF | 0.5160493 |
| VTAVFVAI | ASSFGGGGNTEVF | 0.836508 |
| SGDDNDAAL | ASSFGGGGNTEVF | 0.29962912 |
| AVAALAHS | ASSFGGGGNTEVF | 0.77566826 |
| GGVTVMATP | ASSFGGGGNTEVF | 0.63081014 |
| AGLAANVRD | ASSFGGGGNTEVF | 0.8189858 |
| TNRPDLIDPALLRPG | ASSFGGGGNTEVF | 0.45696968 |
| LANLGVTAL | ASSFGGGGNTEVF | 0.81239253 |
| TGERIEL | ASSFGGGGNTEVF | 0.7389034 |
| NGEELAA | ASSFGGGGNTEVF | 0.87895596 |
| VLSLVWG | ASSFGGGGNTEVF | 0.67323434 |
| IGYALWL | ASSFGGGGNTEVF | 0.7593671 |
| LSLLLAPL | ASSFGGGGNTEVF | 0.6411346 |
| TLDEALK | ASSFGGGGNTEVF | 0.8270628 |
| TDGAGTAEGG | ASSFGGGGNTEVF | 0.48622978 |
| NAPGGAGGNGG | ASSFGGGGNTEVF | 0.67738396 |
| VSVFMGMGGNF | ASSFGGGGNTEVF | 0.3634798 |
| LASYLAAL | ASSFGGGGNTEVF | 0.7658727 |
| LVGGTPAL | ASSFGGGGNTEVF | 0.79384184 |
| VRNADVL | ASSFGGGGNTEVF | 0.36564207 |
| TCSSPFLVSAM | ASSFGGGGNTEVF | 0.8290771 |
| VHITDAE | ASSFGGGGNTEVF | 0.76818883 |
| EYAAPSF | ASSFGGGGNTEVF | 0.65660477 |
| VVDAFHGGL | ASSFGGGGNTEVF | 0.85633767 |
| ASVVDPQ | ASSFGGGGNTEVF | 0.6265865 |
| SYNEFGGLS | ASSFGGGGNTEVF | 0.8632554 |
| DPDGSVY | ASSFGGGGNTEVF | 0.71580595 |

| | | |
|---|---|---|
| MVSTIHATE | ASSFGGGGNTEVF | 0.66954684 |
| LAAGVADPV | ASSFGGGGNTEVF | 0.80878633 |
| TKEQIAA | ASSFGGGGNTEVF | 0.38334072 |
| SLASPAAL | ASSFGGGGNTEVF | 0.4737602 |
| IGLVFYGV | ASSFGGGGNTEVF | 0.84555733 |
| FLLDASNL | ASSFGGGGNTEVF | 0.8801351 |
| YQVGNFP | ASSFGGGGNTEVF | 0.87886316 |
| ESLLSQST | ASSFGGGGNTEVF | 0.7990379 |
| LFTVGFAAVG | ASSFGGGGNTEVF | 0.7032191 |
| ATPPTPT | ASSFGGGGNTEVF | 0.8324062 |
| TDLATLG | ASSFGGGGNTEVF | 0.8830007 |
| SAGGTGR | ASSFGGGGNTEVF | 0.88845956 |
| GFSNSGS | ASSFGGGGNTEVF | 0.87065566 |
| GGNGGSGGTSV | ASSFGGGGNTEVF | 0.6837654 |
| GTSVATGGAGN | ASSFGGGGNTEVF | 0.8650815 |
| NGGNGGSGGTS | ASSFGGGGNTEVF | 0.8329467 |
| AVVKFPH | ASSFGGGGNTEVF | 0.8251706 |
| TPPPPPP | ASSFGGGGNTEVF | 0.88096255 |
| AAPAAAAAPA | ASSFGGGGNTEVF | 0.8636128 |
| IALNEDL | ASSFGGGGNTEVF | 0.82243514 |
| IVNVSSM | ASSFGGGGNTEVF | 0.7793476 |
| TGIPPAPR | ASSFGGGGNTEVF | 0.8714489 |
| FDLGGGTF | ASSFGGGGNTEVF | 0.88801277 |
| FDLGGGTFDVS | ASSFGGGGNTEVF | 0.8869709 |
| NEPTAAA | ASSFGGGGNTEVF | 0.8968379 |
| NEPTAAALA | ASSFGGGGNTEVF | 0.8890219 |
| GSIINISSI | ASSFGGGGNTEVF | 0.8804276 |
| VAPGNAGTA | ASSFGGGGNTEVF | 0.24586023 |
| TGGGASLE | ASSFGGGGNTEVF | 0.5481368 |
| VAVYNHYKRI | ASSFGGGGNTEVF | 0.7942602 |
| TDPAPATT | ASSFGGGGNTEVF | 0.8263534 |
| TNRPDILDPALLRPG | ASSFGGGGNTEVF | 0.49464947 |
| LTTTTTS | ASSFGGGGNTEVF | 0.8354475 |
| PTGSGKTL | ASSFGGGGNTEVF | 0.8073109 |
| TGSGKTL | ASSFGGGGNTEVF | 0.7422097 |
| TIGHVDHGK | ASSFGGGGNTEVF | 0.8895809 |
| TIGHVDHGKTT | ASSFGGGGNTEVF | 0.8056767 |
| GHVDHGKTT | ASSFGGGGNTEVF | 0.747406 |
| AQFEHTLL | ASSFGGGGNTEVF | 0.78976357 |
| IVNQDEL | ASSFGGGGNTEVF | 0.89529324 |
| TNGSQFF | ASSFGGGGNTEVF | 0.7223049 |
| TGKGIVE | ASSFGGGGNTEVF | 0.87868404 |
| LISNASDA | ASSFGGGGNTEVF | 0.80665004 |
| AELRTLGELTKIAKAHGAQ | ASSFGGGGNTEVF | 0.89154136 |
| APNAVAAAEPSW | ASSFGGGGNTEVF | 0.57950634 |
| APQVAGAATAVL | ASSFGGGGNTEVF | 0.8820301 |
| ARRDPAAAAAILPTDA | ASSFGGGGNTEVF | 0.84195006 |
| DAAARDVLAEAGLAEYF | ASSFGGGGNTEVF | 0.90580714 |
| DDQIVLSAKANLAKVEQ | ASSFGGGGNTEVF | 0.825567 |
| DELLPDTGEDSDEDGH | ASSFGGGGNTEVF | 0.8622948 |
| ELPEAPANRATVPQD | ASSFGGGGNTEVF | 0.8324347 |

| | | |
|---|---|---|
| FGQSKIERVTGIG | ASSFGGGGNTEVF | 0.87955666 |
| GAGGTGSTAGGAGGAGGAGGI | ASSFGGGGNTEVF | 0.6220447 |
| GHGGTGAAGGNGGTGGAGGS( | ASSFGGGGNTEVF | 0.844832 |
| GNGGNGGAGGEGGPGGAGGA( | ASSFGGGGNTEVF | 0.6058756 |
| GSTAGNGGNGGNATNGTIAG( | ASSFGGGGNTEVF | 0.85336876 |
| GTGGDGGNGGNGGTGG | ASSFGGGGNTEVF | 0.8679158 |
| GVRRPAGIPADAPV | ASSFGGGGNTEVF | 0.35596693 |
| GYRQVTDAHLLTLARR | ASSFGGGGNTEVF | 0.88594615 |
| IAKMGQDGHDRG | ASSFGGGGNTEVF | 0.79514754 |
| KAEPSIEIQYT | ASSFGGGGNTEVF | 0.75769377 |
| KPVRIRMAADEDTVAVT | ASSFGGGGNTEVF | 0.89368105 |
| MPPRIAGMRLLVIKPEP | ASSFGGGGNTEVF | 0.84833944 |
| MKFYAVALFRSRGVPPTR | ASSFGGGGNTEVF | 0.89066476 |
| NAGGFGILGAVAHSP | ASSFGGGGNTEVF | 0.84239566 |
| NEGREITLAMAEM | ASSFGGGGNTEVF | 0.85173124 |
| NELVEWSHPLGHRWRWE | ASSFGGGGNTEVF | 0.86131436 |
| NLGAGGTGGNGGNPG | ASSFGGGGNTEVF | 0.6037434 |
| NTEWPKWETVMAADPVD | ASSFGGGGNTEVF | 0.7634481 |
| PASAAPAETQP | ASSFGGGGNTEVF | 0.8920232 |
| QQSSDGKNGVRAAEITGE | ASSFGGGGNTEVF | 0.82265824 |
| RAPDAVTTSWWKE | ASSFGGGGNTEVF | 0.81023824 |
| SGFGNVGAHTSGWFNQ | ASSFGGGGNTEVF | 0.681461 |
| SGVAGAVGQAASVGGLKV | ASSFGGGGNTEVF | 0.6923649 |
| SNGGAGTGSAGGNGGT | ASSFGGGGNTEVF | 0.753959 |
| TGGAGGEGGAAG | ASSFGGGGNTEVF | 0.80660653 |
| TGGDGGAAGAGGAPGHG | ASSFGGGGNTEVF | 0.845391 |
| TMPPLLAAIANAGRHGV | ASSFGGGGNTEVF | 0.81925523 |
| TSSTAGNSGGA | ASSFGGGGNTEVF | 0.735947 |
| VPATVPVGILC | ASSFGGGGNTEVF | 0.81221604 |
| VQRPDSGGVVLTGRIS | ASSFGGGGNTEVF | 0.78625274 |
| WTPNPVPATKGVPAGLVQAT | ASSFGGGGNTEVF | 0.7782563 |
| YALKRALAKFSDRFDVVII | ASSFGGGGNTEVF | 0.80149716 |
| YLIGSVIV | ASSFGGGGNTEVF | 0.78917205 |
| YGQLVAVID | ASSFGGGGNTEVF | 0.7028337 |
| VTVGGMTLPA | ASSFGGGGNTEVF | 0.69249296 |
| VLALPAV | ASSFGGGGNTEVF | 0.81307834 |
| VIPPTLNYET | ASSFGGGGNTEVF | 0.8301492 |
| VGMGGMHPSSG | ASSFGGGGNTEVF | 0.82840335 |
| VGLAGGAATA | ASSFGGGGNTEVF | 0.84585536 |
| VGGAGGAGGSGP | ASSFGGGGNTEVF | 0.83486676 |
| VFAAFMV | ASSFGGGGNTEVF | 0.8156667 |
| TLYDKLR | ASSFGGGGNTEVF | 0.40023333 |
| TLAVFTVL | ASSFGGGGNTEVF | 0.60115784 |
| TGSVQPARV | ASSFGGGGNTEVF | 0.74091494 |
| TGGGAGSGGT | ASSFGGGGNTEVF | 0.75299203 |
| SHLETSL | ASSFGGGGNTEVF | 0.39354667 |
| SFTTGMT | ASSFGGGGNTEVF | 0.8014803 |
| QVEVAKRMGVRQPTVS | ASSFGGGGNTEVF | 0.7651403 |
| QGSGGPALAM | ASSFGGGGNTEVF | 0.7022776 |
| MLAPSNP | ASSFGGGGNTEVF | 0.82993925 |
| LVNNAAI | ASSFGGGGNTEVF | 0.7438519 |

| | | |
|---|---|---|
| LTGEINTLA | ASSFGGGGNTEVF | 0.6454469 |
| LSIFGPV | ASSFGGGGNTEVF | 0.73606753 |
| LLALLANL | ASSFGGGGNTEVF | 0.8470882 |
| LISNASDALDKLR | ASSFGGGGNTEVF | 0.71176314 |
| LINEAVP | ASSFGGGGNTEVF | 0.83718723 |
| LGYAFSGL | ASSFGGGGNTEVF | 0.74967897 |
| LDFVAVAL | ASSFGGGGNTEVF | 0.83800054 |
| LAQPGAE | ASSFGGGGNTEVF | 0.66428703 |
| ISNLGAVTGF | ASSFGGGGNTEVF | 0.83684057 |
| HQEDPGRI | ASSFGGGGNTEVF | 0.79243934 |
| GTPGEPTP | ASSFGGGGNTEVF | 0.85323155 |
| GFFNSGS | ASSFGGGGNTEVF | 0.7753986 |
| GAGGGGLTAGAN | ASSFGGGGNTEVF | 0.8246399 |
| FVDALVNM | ASSFGGGGNTEVF | 0.7811485 |
| FLTVAPEL | ASSFGGGGNTEVF | 0.83046657 |
| FETLTAL | ASSFGGGGNTEVF | 0.6473143 |
| DVINPIEL | ASSFGGGGNTEVF | 0.7584748 |
| APVSGAPV | ASSFGGGGNTEVF | 0.6616224 |
| ALVGGLAW | ASSFGGGGNTEVF | 0.76548016 |
| ALAAAAATAI | ASSFGGGGNTEVF | 0.8279277 |
| AFGLGLLL | ASSFGGGGNTEVF | 0.3074366 |
| AFFAAASL | ASSFGGGGNTEVF | 0.8307981 |
| AAAALPAFSPP | ASSFGGGGNTEVF | 0.5906359 |
| VVLAPAL | ASSFGGGGNTEVF | 0.6930384 |
| TIGHVDHGKTTL | ASSFGGGGNTEVF | 0.8270521 |
| LISNASDALDK | ASSFGGGGNTEVF | 0.7873206 |
| LISNASDAL | ASSFGGGGNTEVF | 0.8304179 |
| ISNASDALDK | ASSFGGGGNTEVF | 0.7286551 |
| NQAETLVYQTE | ASSFGGGGNTEVF | 0.7869556 |
| TGGTGGTAGAG | ASSFGGGGNTEVF | 0.79050195 |
| TGSTAGGAGGA | ASSFGGGGNTEVF | 0.54328644 |
| DPLAVPVAAGPV | ASSFGGGGNTEVF | 0.67636853 |
| GGTGGTPGNGGN | ASSFGGGGNTEVF | 0.84880614 |
| PAGLAGVDPATL | ASSFGGGGNTEVF | 0.7614557 |
| GNNLVVEAPHAGASV | ASSFGGGGNTEVF | 0.76860344 |
| KVTRIMSASTLEQPA | ASSFGGGGNTEVF | 0.64882743 |
| LGFALPIALPRRTVV | ASSFGGGGNTEVF | 0.64601487 |
| TRLGNLTARKLVVLV | ASSFGGGGNTEVF | 0.6824434 |
| WSQQRRLIYFPSAGP | ASSFGGGGNTEVF | 0.7716928 |
| GPNGNPDPMAAAVDIR | ASSFGGGGNTEVF | 0.7817229 |
| TQRCVSVADLRAAASR | ASSFGGGGNTEVF | 0.7635981 |
| QFVLDVAEHPVEVLTETV | ASSFGGGGNTEVF | 0.62437797 |
| GGAGGTGGAGGNGGTQAGDG | ASSFGGGGNTEVF | 0.7390926 |
| PLLATLRAIAVDVDAKPAQV | ASSFGGGGNTEVF | 0.69567347 |
| APKFRTGGYMIDFWGVGYQV | ASSFGGGGNTEVF | 0.70613796 |
| GAGGAGGAGGAAGGPGGTGG | ASSFGGGGNTEVF | 0.74918437 |
| IGALGLLATDVAPAGSVAFG | ASSFGGGGNTEVF | 0.73149955 |
| AGGAGGIGGNGGAGGAGGVL | ASSFGGGGNTEVF | 0.7007828 |
| ADELIGLL | ASSFGGGGNTEVF | 0.5830108 |
| ADRALKA | ASSFGGGGNTEVF | 0.74822307 |
| AGAVGQVGGAGGA | ASSFGGGGNTEVF | 0.78611267 |

| | | |
|---|---|---|
| AGGGGGLGGAP | ASSFGGGGNTEVF | 0.772164 |
| AGGLLYGNGGAG | ASSFGGGGNTEVF | 0.7877131 |
| AGLESIGL | ASSFGGGGNTEVF | 0.7713301 |
| AGLKTGII | ASSFGGGGNTEVF | 0.7679243 |
| ASLAALLL | ASSFGGGGNTEVF | 0.26347774 |
| ASVINISSI | ASSFGGGGNTEVF | 0.43641132 |
| ATPPGPA | ASSFGGGGNTEVF | 0.55785286 |
| ATVASALV | ASSFGGGGNTEVF | 0.69660723 |
| ATVASLIL | ASSFGGGGNTEVF | 0.30449274 |
| AVNGPVLIRH | ASSFGGGGNTEVF | 0.7472564 |
| DGAAGAVGI | ASSFGGGGNTEVF | 0.6907202 |
| DPLLHGI | ASSFGGGGNTEVF | 0.6607335 |
| ELGAALGAL | ASSFGGGGNTEVF | 0.77378035 |
| ELGRTQAV | ASSFGGGGNTEVF | 0.6905016 |
| FVAGGYLS | ASSFGGGGNTEVF | 0.5951497 |
| GASGGAGGAGGW | ASSFGGGGNTEVF | 0.59996414 |
| GGAGAGGDGGAGGI | ASSFGGGGNTEVF | 0.7877831 |
| GTGHNGSQPGMGGNG | ASSFGGGGNTEVF | 0.44310308 |
| GYAAASASAS | ASSFGGGGNTEVF | 0.7671089 |
| HQVDVAAVQQG | ASSFGGGGNTEVF | 0.59472203 |
| IGAAIAHT | ASSFGGGGNTEVF | 0.7780735 |
| IGMIEVAQL | ASSFGGGGNTEVF | 0.55655193 |
| IRLAPEHTSAMD | ASSFGGGGNTEVF | 0.772691 |
| IYNVSVL | ASSFGGGGNTEVF | 0.7176153 |
| KQGFLGM | ASSFGGGGNTEVF | 0.7986499 |
| LESVITQL | ASSFGGGGNTEVF | 0.70054865 |
| LGLGALVVV | ASSFGGGGNTEVF | 0.7533779 |
| LIGNGGNGGSGGI | ASSFGGGGNTEVF | 0.7167245 |
| LNAINAPTL | ASSFGGGGNTEVF | 0.7669697 |
| LQLLNTGV | ASSFGGGGNTEVF | 0.46672332 |
| LQTSIEF | ASSFGGGGNTEVF | 0.6933658 |
| MSFAVSEL | ASSFGGGGNTEVF | 0.59367895 |
| NTGALGL | ASSFGGGGNTEVF | 0.72254884 |
| PPGAGGAGGAGTT | ASSFGGGGNTEVF | 0.7675371 |
| QLVGLGI | ASSFGGGGNTEVF | 0.30004507 |
| RFLEQVHQL | ASSFGGGGNTEVF | 0.74810183 |
| RGVGYVLR | ASSFGGGGNTEVF | 0.65192246 |
| SATGVFAGVI | ASSFGGGGNTEVF | 0.65072227 |
| SLPAITLP | ASSFGGGGNTEVF | 0.76606023 |
| STALAFTV | ASSFGGGGNTEVF | 0.7182095 |
| SVLTVGLGK | ASSFGGGGNTEVF | 0.7619152 |
| TALAGAGSGI | ASSFGGGGNTEVF | 0.61384153 |
| TGLPFTGI | ASSFGGGGNTEVF | 0.6638422 |
| VAEHVTV | ASSFGGGGNTEVF | 0.7517582 |
| VDIVLNSL | ASSFGGGGNTEVF | 0.37444696 |
| VLTGGGAVL | ASSFGGGGNTEVF | 0.5043089 |
| VNVGLAYGI | ASSFGGGGNTEVF | 0.7845073 |
| VSSAAVGAV | ASSFGGGGNTEVF | 0.6707293 |
| VTSVYEAGV | ASSFGGGGNTEVF | 0.6800804 |
| WVALDRGA | ASSFGGGGNTEVF | 0.63691556 |
| YLVGSVVI | ASSFGGGGNTEVF | 0.6030546 |

| | | |
|---|---|---|
| GPPGTGKTLL | ASSFGGGGNTEVF | 0.6343422 |
| GVGMTKF | ASSFGGGGNTEVF | 0.6824764 |
| IGIGGSDL | ASSFGGGGNTEVF | 0.67636573 |
| IGTLLAEL | ASSFGGGGNTEVF | 0.66458666 |
| IYTSGTT | ASSFGGGGNTEVF | 0.5166646 |
| LSFPAGV | ASSFGGGGNTEVF | 0.66196847 |
| LYTSGTT | ASSFGGGGNTEVF | 0.6494815 |
| VGAFPFGF | ASSFGGGGNTEVF | 0.6326117 |
| VGAGTVEF | ASSFGGGGNTEVF | 0.6614986 |
| VGKSTLLL | ASSFGGGGNTEVF | 0.6670786 |
| AALAHAGGGIV | ASSFGGGGNTEVF | 0.6169703 |
| AALVAVSLL | ASSFGGGGNTEVF | 0.640151 |
| AAVSRAVAAL | ASSFGGGGNTEVF | 0.65706915 |
| AGGSSGIGGF | ASSFGGGGNTEVF | 0.67602843 |
| AGLTVANANG | ASSFGGGGNTEVF | 0.6761329 |
| AGLVGAVL | ASSFGGGGNTEVF | 0.68776256 |
| ALSQAFTV | ASSFGGGGNTEVF | 0.68359375 |
| ALTFSVSL | ASSFGGGGNTEVF | 0.6815003 |
| ALTGAALVAL | ASSFGGGGNTEVF | 0.37495598 |
| ASFSIGAGVL | ASSFGGGGNTEVF | 0.49249464 |
| DGSVYKI | ASSFGGGGNTEVF | 0.6327317 |
| EQLAEIV | ASSFGGGGNTEVF | 0.6355009 |
| FAVSELTVM | ASSFGGGGNTEVF | 0.507735 |
| GAAPLAAGL | ASSFGGGGNTEVF | 0.64794326 |
| GADGGDGGSGGAGG | ASSFGGGGNTEVF | 0.6192772 |
| GFGLDTGGAGG | ASSFGGGGNTEVF | 0.5962961 |
| GLASGGLL | ASSFGGGGNTEVF | 0.677199 |
| GRIAGAAVLL | ASSFGGGGNTEVF | 0.6282279 |
| GSAGSLNTSF | ASSFGGGGNTEVF | 0.6203972 |
| GTGQIGI | ASSFGGGGNTEVF | 0.58518195 |
| GTLIALIL | ASSFGGGGNTEVF | 0.67556465 |
| IAATMGFVVG | ASSFGGGGNTEVF | 0.54276943 |
| IFAQALL | ASSFGGGGNTEVF | 0.67074394 |
| IGAAVALGF | ASSFGGGGNTEVF | 0.53706545 |
| IGLTPLAL | ASSFGGGGNTEVF | 0.68531835 |
| ILGSNTSTL | ASSFGGGGNTEVF | 0.5147404 |
| IVFSLTVL | ASSFGGGGNTEVF | 0.67574656 |
| LASKATGF | ASSFGGGGNTEVF | 0.64888805 |
| LASVALIL | ASSFGGGGNTEVF | 0.6955316 |
| LFAVGLTL | ASSFGGGGNTEVF | 0.6568489 |
| LFNGGIGTYI | ASSFGGGGNTEVF | 0.6568847 |
| LTLGAGII | ASSFGGGGNTEVF | 0.58683074 |
| LVAAALGDI | ASSFGGGGNTEVF | 0.6763808 |
| NTAFVQLGI | ASSFGGGGNTEVF | 0.5228194 |
| QLFSGAEL | ASSFGGGGNTEVF | 0.65637964 |
| SAATFITL | ASSFGGGGNTEVF | 0.47861353 |
| SNYGLANLGV | ASSFGGGGNTEVF | 0.6660656 |
| SYGGAYAVM | ASSFGGGGNTEVF | 0.6677685 |
| TTGVGGLTLG | ASSFGGGGNTEVF | 0.5201019 |
| TTQTIYI | ASSFGGGGNTEVF | 0.6746088 |
| VAAGAGLGAVI | ASSFGGGGNTEVF | 0.6465232 |

| | | |
|---|---|---|
| VAAVVTLL | ASSFGGGGNTEVF | 0.606468 |
| VGFLGGLI | ASSFGGGGNTEVF | 0.687456 |
| VGVIPILL | ASSFGGGGNTEVF | 0.65453637 |
| VIFASSLI | ASSFGGGGNTEVF | 0.6828472 |
| VKGGDGGAAAG | ASSFGGGGNTEVF | 0.6407182 |
| VTKTFTNI | ASSFGGGGNTEVF | 0.6380374 |
| VVAAAILL | ASSFGGGGNTEVF | 0.6632572 |
| YAALDVEL | ASSFGGGGNTEVF | 0.6433865 |
| LASVALLL | ASSFGGGGNTEVF | 0.5970509 |
| LDAVHRV | ASSFGGGGNTEVF | 0.68385017 |
| QGLSSSIF | ASSFGGGGNTEVF | 0.73855054 |
| RGTFANIRL | ASSFGGGGNTEVF | 0.727961 |
| RQLWWGHRI | ASSFGGGGNTEVF | 0.69838446 |
| YGPPGTGKTLL | ASSFGGGGNTEVF | 0.6775757 |
| AALVGAPANVA | ASSFGGGGNTEVF | 0.7061318 |
| AATFRRTE | ASSFGGGGNTEVF | 0.7370863 |
| AGALGGGI | ASSFGGGGNTEVF | 0.73978937 |
| AGGAGGAGGIGAG | ASSFGGGGNTEVF | 0.73173076 |
| AGSGALGL | ASSFGGGGNTEVF | 0.68784773 |
| AQIADQI | ASSFGGGGNTEVF | 0.7332895 |
| ASAVAVTYL | ASSFGGGGNTEVF | 0.7289 |
| ASTVLGGTGG | ASSFGGGGNTEVF | 0.700601 |
| AVGGEPAK | ASSFGGGGNTEVF | 0.73735905 |
| CEQGTFGGTC | ASSFGGGGNTEVF | 0.7375266 |
| DEGKAPAG | ASSFGGGGNTEVF | 0.7081604 |
| DTVVFPAL | ASSFGGGGNTEVF | 0.7170004 |
| FEGITMAG | ASSFGGGGNTEVF | 0.7292826 |
| FPLVVQD | ASSFGGGGNTEVF | 0.73833966 |
| GAFTTPQ | ASSFGGGGNTEVF | 0.7370552 |
| GFGGFGTAGGVGGA | ASSFGGGGNTEVF | 0.7442024 |
| GFLAQFL | ASSFGGGGNTEVF | 0.7413875 |
| GGAGGAGGNAGVA | ASSFGGGGNTEVF | 0.7361435 |
| GGVGTAGAGGAGGAGG | ASSFGGGGNTEVF | 0.5830363 |
| GNGGGGDAGNAGSGGN | ASSFGGGGNTEVF | 0.6313139 |
| GTGGAGGAGNDAGS | ASSFGGGGNTEVF | 0.73138326 |
| GTTNTGWFNS | ASSFGGGGNTEVF | 0.7110082 |
| IFMGPLFTL | ASSFGGGGNTEVF | 0.5877509 |
| ILAHAIEAI | ASSFGGGGNTEVF | 0.72439003 |
| IRSAIREAL | ASSFGGGGNTEVF | 0.7174382 |
| ITFAGGG | ASSFGGGGNTEVF | 0.6906378 |
| ITGMSGHQEV | ASSFGGGGNTEVF | 0.7365813 |
| IVAGLACTL | ASSFGGGGNTEVF | 0.71321297 |
| IVGGGLAGA | ASSFGGGGNTEVF | 0.7032317 |
| IVTGVLNI | ASSFGGGGNTEVF | 0.7196647 |
| LAQQGCI | ASSFGGGGNTEVF | 0.73803115 |
| LCLHVEN | ASSFGGGGNTEVF | 0.6929378 |
| LDAISDGS | ASSFGGGGNTEVF | 0.7365327 |
| LGGFAAML | ASSFGGGGNTEVF | 0.6903069 |
| LGGKNPMI | ASSFGGGGNTEVF | 0.74311274 |
| LGPKGVAAIL | ASSFGGGGNTEVF | 0.6602991 |
| LMVGVSI | ASSFGGGGNTEVF | 0.73612505 |

| | | |
|---|---|---|
| LVGAAFVL | ASSFGGGGNTEVF | 0.7176029 |
| LVLAVAPL | ASSFGGGGNTEVF | 0.74867904 |
| MAAIAVIDAA | ASSFGGGGNTEVF | 0.7219423 |
| NNGINAELQ | ASSFGGGGNTEVF | 0.7400578 |
| RGEPLVLVHG | ASSFGGGGNTEVF | 0.7302996 |
| RGGAGGAAGLI | ASSFGGGGNTEVF | 0.7312084 |
| RSYRMHPAV | ASSFGGGGNTEVF | 0.66894454 |
| SAAHLLAGQLA | ASSFGGGGNTEVF | 0.72946405 |
| SAAVGSY | ASSFGGGGNTEVF | 0.6664772 |
| SPHAVLHRI | ASSFGGGGNTEVF | 0.7301698 |
| SQIAKVL | ASSFGGGGNTEVF | 0.7334143 |
| TALGHLVP | ASSFGGGGNTEVF | 0.5106019 |
| TAPTTSGAA | ASSFGGGGNTEVF | 0.7375958 |
| TDGDDGNGGSPGT | ASSFGGGGNTEVF | 0.6917077 |
| TDGLGEN | ASSFGGGGNTEVF | 0.7079031 |
| TGGPPLPGQA | ASSFGGGGNTEVF | 0.744582 |
| TIPAITI | ASSFGGGGNTEVF | 0.7346847 |
| TIPGFTI | ASSFGGGGNTEVF | 0.7423638 |
| TSIASGALA | ASSFGGGGNTEVF | 0.72410834 |
| TVTLPGAAI | ASSFGGGGNTEVF | 0.71913636 |
| VASGLSAALP | ASSFGGGGNTEVF | 0.7230583 |
| VGAGVIGI | ASSFGGGGNTEVF | 0.6042977 |
| VGPKRFGV | ASSFGGGGNTEVF | 0.7035001 |
| VNVAFPD | ASSFGGGGNTEVF | 0.7430242 |
| VSMTNSGVS | ASSFGGGGNTEVF | 0.6875708 |
| VTDVLGDLSV | ASSFGGGGNTEVF | 0.75526965 |
| VVVVGAGFAGL | ASSFGGGGNTEVF | 0.5028872 |
| VYTVGTVV | ASSFGGGGNTEVF | 0.4215807 |
| WTTFSGP | ASSFGGGGNTEVF | 0.47513002 |
| YDELVEAF | ASSFGGGGNTEVF | 0.7714149 |
| YGTGGAGGNGG | ASSFGGGGNTEVF | 0.7138519 |
| YTNVMAV | ASSFGGGGNTEVF | 0.7261342 |
| FLEQVHQL | ASSFGGGGNTEVF | 0.4388564 |
| GLSLLAVAV | ASSFGGGGNTEVF | 0.5791461 |
| LEQVHQL | ASSFGGGGNTEVF | 0.5075742 |
| LNAAGAI | ASSFGGGGNTEVF | 0.5888474 |
| PAVHASL | ASSFGGGGNTEVF | 0.66446537 |
| TELEDDA | ASSFGGGGNTEVF | 0.7239369 |
| TPPGPPN | ASSFGGGGNTEVF | 0.5432215 |
| TPPPPPPP | ASSFGGGGNTEVF | 0.48522562 |
| TQSPALI | ASSSRISNERLF | 0.57939583 |
| EFAAAFPQ | ASSSRISNERLF | 0.6935901 |
| LLSGNLDVL | ASSSRISNERLF | 0.64342666 |
| GPANLGNYNI | ASSSRISNERLF | 0.7980677 |
| TLGPTLP | ASSSRISNERLF | 0.7324655 |
| VYAYSSCCK | ASSSRISNERLF | 0.724317 |
| APAPAPAPNI | ASSSRISNERLF | 0.18464863 |
| PGGPVPIP | ASSSRISNERLF | 0.2853673 |
| FLTVALEP | ASSSRISNERLF | 0.5063839 |
| SQYFGANL | ASSSRISNERLF | 0.5804137 |
| GGVKPAA | ASSSRISNERLF | 0.2476841 |

| | | |
|---|---|---|
| GGTGGAGGAGGW | ASSSRISNERLF | 0.6893182 |
| GVGGVAGLGGAGG | ASSSRISNERLF | 0.53856635 |
| VLLVLATTI | ASSSRISNERLF | 0.50984204 |
| IGIFAAML | ASSSRISNERLF | 0.7126704 |
| AVGDLID | ASSSRISNERLF | 0.54699826 |
| NISPLPL | ASSSRISNERLF | 0.52778363 |
| VTAVFVAI | ASSSRISNERLF | 0.509606 |
| SGDDNDAAL | ASSSRISNERLF | 0.71933436 |
| AVAALAHS | ASSSRISNERLF | 0.3039616 |
| GGVTVMATP | ASSSRISNERLF | 0.6977759 |
| AGLAANVRD | ASSSRISNERLF | 0.46572632 |
| TNRPDLIDPALLRPG | ASSSRISNERLF | 0.7488506 |
| LANLGVTAL | ASSSRISNERLF | 0.39139116 |
| TGERIEL | ASSSRISNERLF | 0.74399364 |
| NGEELAA | ASSSRISNERLF | 0.61132264 |
| VLSLVWG | ASSSRISNERLF | 0.80812025 |
| IGYALWL | ASSSRISNERLF | 0.5203875 |
| LSLLLAPL | ASSSRISNERLF | 0.6302289 |
| TLDEALK | ASSSRISNERLF | 0.56457967 |
| TDGAGTAEGG | ASSSRISNERLF | 0.71702635 |
| NAPGGAGGNGG | ASSSRISNERLF | 0.321457 |
| VSVFMGMGGNF | ASSSRISNERLF | 0.5403531 |
| LASYLAAL | ASSSRISNERLF | 0.28998932 |
| LVGGTPAL | ASSSRISNERLF | 0.65057915 |
| VRNADVL | ASSSRISNERLF | 0.7385122 |
| TCSSPFLVSAM | ASSSRISNERLF | 0.3004903 |
| VHITDAE | ASSSRISNERLF | 0.67122436 |
| EYAAPSF | ASSSRISNERLF | 0.65707856 |
| VVDAFHGGL | ASSSRISNERLF | 0.4935724 |
| ASVVDPQ | ASSSRISNERLF | 0.74467444 |
| SYNEFGGLS | ASSSRISNERLF | 0.5661391 |
| DPDGSVY | ASSSRISNERLF | 0.7559959 |
| MVSTIHATE | ASSSRISNERLF | 0.6030171 |
| LAAGVADPV | ASSSRISNERLF | 0.47783473 |
| TKEQIAA | ASSSRISNERLF | 0.7028737 |
| SLASPAAL | ASSSRISNERLF | 0.28446257 |
| IGLVFYGV | ASSSRISNERLF | 0.428036 |
| FLLDASNL | ASSSRISNERLF | 0.77123004 |
| YQVGNFP | ASSSRISNERLF | 0.7800793 |
| ESLLSQST | ASSSRISNERLF | 0.8601353 |
| LFTVGFAAVG | ASSSRISNERLF | 0.6078423 |
| ATPPTPT | ASSSRISNERLF | 0.4777396 |
| TDLATLG | ASSSRISNERLF | 0.6666405 |
| SAGGTGR | ASSSRISNERLF | 0.84513414 |
| GFSNSGS | ASSSRISNERLF | 0.8522328 |
| GGNGGSGGTSV | ASSSRISNERLF | 0.8041494 |
| GTSVATGGAGN | ASSSRISNERLF | 0.44037914 |
| NGGNGGSGGTS | ASSSRISNERLF | 0.7334945 |
| AVVKFPH | ASSSRISNERLF | 0.6485585 |
| TPPPPPP | ASSSRISNERLF | 0.6969279 |
| AAPAAAAAPA | ASSSRISNERLF | 0.76535076 |

| | | |
|---|---|---|
| IALNEDL | ASSSRISNERLF | 0.78512424 |
| IVNVSSM | ASSSRISNERLF | 0.65194786 |
| TGIPPAPR | ASSSRISNERLF | 0.5245749 |
| FDLGGGTF | ASSSRISNERLF | 0.78897536 |
| FDLGGGTFDVS | ASSSRISNERLF | 0.7956822 |
| NEPTAAA | ASSSRISNERLF | 0.8008537 |
| NEPTAAALA | ASSSRISNERLF | 0.8574185 |
| GSIINISSI | ASSSRISNERLF | 0.8406116 |
| VAPGNAGTA | ASSSRISNERLF | 0.8265466 |
| TGGGASLE | ASSSRISNERLF | 0.14436096 |
| VAVYNHYKRI | ASSSRISNERLF | 0.25010014 |
| TDPAPATT | ASSSRISNERLF | 0.55952626 |
| TNRPDILDPALLRPG | ASSSRISNERLF | 0.6826279 |
| LTTTTTS | ASSSRISNERLF | 0.33946553 |
| PTGSGKTL | ASSSRISNERLF | 0.7544155 |
| TGSGKTL | ASSSRISNERLF | 0.5458276 |
| TIGHVDHGK | ASSSRISNERLF | 0.564796 |
| TIGHVDHGKTT | ASSSRISNERLF | 0.83149797 |
| GHVDHGKTT | ASSSRISNERLF | 0.6254846 |
| AQFEHTLL | ASSSRISNERLF | 0.54733944 |
| IVNQDEL | ASSSRISNERLF | 0.575474 |
| TNGSQFF | ASSSRISNERLF | 0.7887654 |
| TGKGIVE | ASSSRISNERLF | 0.47946516 |
| LISNASDA | ASSSRISNERLF | 0.7863668 |
| AELRTLGELTKIAKAHGAQ | ASSSRISNERLF | 0.65770316 |
| APNAVAAAEPSW | ASSSRISNERLF | 0.8418365 |
| APQVAGAATAVL | ASSSRISNERLF | 0.37768513 |
| ARRDPAAAAAILPTDA | ASSSRISNERLF | 0.8295011 |
| DAAARDVLAEAGLAEYF | ASSSRISNERLF | 0.73896074 |
| DDQIVLSAKANLAKVEQ | ASSSRISNERLF | 0.87788653 |
| DELLPDTGEDSDEDGH | ASSSRISNERLF | 0.647812 |
| ELPEAPANRATVPQD | ASSSRISNERLF | 0.73615557 |
| FGQSKIERVTGIG | ASSSRISNERLF | 0.65342104 |
| GAGGTGSTAGGAGGAGGAGGI | ASSSRISNERLF | 0.85107803 |
| GHGGTGAAGGNGGTGGAGGS( | ASSSRISNERLF | 0.4080158 |
| GNGGNGGAGGEGGPGGAGGA( | ASSSRISNERLF | 0.5939894 |
| GSTAGNGGNGGNATNGTIAG( | ASSSRISNERLF | 0.22402641 |
| GTGGDGGNGGNGGTGG | ASSSRISNERLF | 0.71970534 |
| GVRRPAGIPADAPV | ASSSRISNERLF | 0.7942321 |
| GYRQVTDAHLLTLARR | ASSSRISNERLF | 0.27050024 |
| IAKMGQDGHDRG | ASSSRISNERLF | 0.8337923 |
| KAEPSIEIQYT | ASSSRISNERLF | 0.8052981 |
| KPVRIRMAADEDTVAVT | ASSSRISNERLF | 0.5324062 |
| MPPRIAGMRLLVIKPEP | ASSSRISNERLF | 0.8409415 |
| MKFYAVALFRSRGVPPTR | ASSSRISNERLF | 0.68792117 |
| NAGGFGILGAVAHSP | ASSSRISNERLF | 0.85376346 |
| NEGREITLAMAEM | ASSSRISNERLF | 0.7542951 |
| NELVEWSHPLGHRWRWE | ASSSRISNERLF | 0.7105894 |
| NLGAGGTGGNGGNPG | ASSSRISNERLF | 0.77362067 |
| NTEWPKWETVMAADPVD | ASSSRISNERLF | 0.4779705 |
| PASAAPAETQP | ASSSRISNERLF | 0.5665345 |

| | | |
|---|---|---|
| QQSSDGKNGVRAAEITGE | ASSSRISNERLF | 0.84668064 |
| RAPDAVTTSWWKE | ASSSRISNERLF | 0.48520896 |
| SGFGNVGAHTSGWFNQ | ASSSRISNERLF | 0.7271036 |
| SGVAGAVGQAASVGGLKV | ASSSRISNERLF | 0.24215722 |
| SNGGAGTGSAGGNGGT | ASSSRISNERLF | 0.18704256 |
| TGGAGGEGGAAG | ASSSRISNERLF | 0.29000807 |
| TGGDGGAAGAGGAPGHG | ASSSRISNERLF | 0.61533195 |
| TMPPLLAAIANAGRHGV | ASSSRISNERLF | 0.78753626 |
| TSSTAGNSGGA | ASSSRISNERLF | 0.5368476 |
| VPATVPVGILC | ASSSRISNERLF | 0.20286983 |
| VQRPDSGGVVLTGRIS | ASSSRISNERLF | 0.4106164 |
| WTPNPVPATKGVPAGLVQAT | ASSSRISNERLF | 0.26132348 |
| YALKRALAKFSDRFDVVII | ASSSRISNERLF | 0.3186865 |
| YLIGSVIV | ASSSRISNERLF | 0.43311167 |
| YGQLVAVID | ASSSRISNERLF | 0.38026989 |
| VTVGGMTLPA | ASSSRISNERLF | 0.3067427 |
| VLALPAV | ASSSRISNERLF | 0.2091034 |
| VIPPTLNYET | ASSSRISNERLF | 0.63644314 |
| VGMGGMHPSSG | ASSSRISNERLF | 0.7025545 |
| VGLAGGAATA | ASSSRISNERLF | 0.661314 |
| VGGAGGAGGSGP | ASSSRISNERLF | 0.72656745 |
| VFAAFMV | ASSSRISNERLF | 0.6281769 |
| TLYDKLR | ASSSRISNERLF | 0.52613056 |
| TLAVFTVL | ASSSRISNERLF | 0.09452147 |
| TGSVQPARV | ASSSRISNERLF | 0.14347261 |
| TGGGAGSGGT | ASSSRISNERLF | 0.26439437 |
| SHLETSL | ASSSRISNERLF | 0.32774782 |
| SFTTGMT | ASSSRISNERLF | 0.103265315 |
| QVEVAKRMGVRQPTVS | ASSSRISNERLF | 0.44739014 |
| QGSGGPALAM | ASSSRISNERLF | 0.2644514 |
| MLAPSNP | ASSSRISNERLF | 0.21353742 |
| LVNNAAI | ASSSRISNERLF | 0.71078396 |
| LTGEINTLA | ASSSRISNERLF | 0.23004654 |
| LSIFGPV | ASSSRISNERLF | 0.22492144 |
| LLALLANL | ASSSRISNERLF | 0.2583034 |
| LISNASDALDKLR | ASSSRISNERLF | 0.66113114 |
| LINEAVP | ASSSRISNERLF | 0.17009057 |
| LGYAFSGL | ASSSRISNERLF | 0.5224962 |
| LDFVAVAL | ASSSRISNERLF | 0.30218366 |
| LAQPGAE | ASSSRISNERLF | 0.6466288 |
| ISNLGAVTGF | ASSSRISNERLF | 0.15583588 |
| HQEDPGRI | ASSSRISNERLF | 0.58108974 |
| GTPGEPTP | ASSSRISNERLF | 0.35296062 |
| GFFNSGS | ASSSRISNERLF | 0.8400731 |
| GAGGGGLTAGAN | ASSSRISNERLF | 0.2445027 |
| FVDALVNM | ASSSRISNERLF | 0.3115735 |
| FLTVAPEL | ASSSRISNERLF | 0.2754047 |
| FETLTAL | ASSSRISNERLF | 0.754325 |
| DVINPIEL | ASSSRISNERLF | 0.13461053 |
| APVSGAPV | ASSSRISNERLF | 0.29787198 |
| ALVGGLAW | ASSSRISNERLF | 0.17128311 |

| | | |
|---|---|---|
| ALAAAAATAI | ASSSRISNERLF | 0.37312186 |
| AFGLGLLL | ASSSRISNERLF | 0.47931513 |
| AFFAAASL | ASSSRISNERLF | 0.096700475 |
| AAAALPAFSPP | ASSSRISNERLF | 0.5570848 |
| VVLAPAL | ASSSRISNERLF | 0.1836583 |
| TIGHVDHGKTTL | ASSSRISNERLF | 0.17870046 |
| LISNASDALDK | ASSSRISNERLF | 0.62541807 |
| LISNASDAL | ASSSRISNERLF | 0.31532001 |
| ISNASDALDK | ASSSRISNERLF | 0.71840554 |
| NQAETLVYQTE | ASSSRISNERLF | 0.2715667 |
| TGGTGGTAGAG | ASSSRISNERLF | 0.4082821 |
| TGSTAGGAGGA | ASSSRISNERLF | 0.4712289 |
| DPLAVPVAAGPV | ASSSRISNERLF | 0.13477412 |
| GGTGGTPGNGGN | ASSSRISNERLF | 0.2572508 |
| PAGLAGVDPATL | ASSSRISNERLF | 0.6399452 |
| GNNLVVEAPHAGASV | ASSSRISNERLF | 0.8364189 |
| KVTRIMSASTLEQPA | ASSSRISNERLF | 0.8861209 |
| LGFALPIALPRRTVV | ASSSRISNERLF | 0.746783 |
| TRLGNLTARKLVVLV | ASSSRISNERLF | 0.7039113 |
| WSQQRRLIYFPSAGP | ASSSRISNERLF | 0.7807615 |
| GPNGNPDPMAAAVDIR | ASSSRISNERLF | 0.8748285 |
| TQRCVSVADLRAAASR | ASSSRISNERLF | 0.8810869 |
| QFVLDVAEHPVEVLTETV | ASSSRISNERLF | 0.84806764 |
| GGAGGTGGAGGNGGTQAGDG | ASSSRISNERLF | 0.6006308 |
| PLLATLRAIAVDVDAKPAQV | ASSSRISNERLF | 0.7818201 |
| APKFRTGGYMIDFWGVGYQV | ASSSRISNERLF | 0.7603608 |
| GAGGAGGAGGAAGGPGGTGG | ASSSRISNERLF | 0.7435876 |
| IGALGLLATDVAPAGSVAFG | ASSSRISNERLF | 0.8654419 |
| AGGAGGIGGNGGAGGAGGVL | ASSSRISNERLF | 0.8069917 |
| ADELIGLL | ASSSRISNERLF | 0.7398952 |
| ADRALKA | ASSSRISNERLF | 0.78870916 |
| AGAVGQVGGAGGA | ASSSRISNERLF | 0.7942022 |
| AGGGGGLGGAP | ASSSRISNERLF | 0.8262322 |
| AGGLLYGNGGAG | ASSSRISNERLF | 0.8389945 |
| AGLESIGL | ASSSRISNERLF | 0.8803865 |
| AGLKTGII | ASSSRISNERLF | 0.86604893 |
| ASLAALLL | ASSSRISNERLF | 0.8232904 |
| ASVINISSI | ASSSRISNERLF | 0.1868131 |
| ATPPGPA | ASSSRISNERLF | 0.4962911 |
| ATVASALV | ASSSRISNERLF | 0.81259847 |
| ATVASLIL | ASSSRISNERLF | 0.7427511 |
| AVNGPVLIRH | ASSSRISNERLF | 0.3371398 |
| DGAAGAVGI | ASSSRISNERLF | 0.7527462 |
| DPLLHGI | ASSSRISNERLF | 0.75205743 |
| ELGAALGAL | ASSSRISNERLF | 0.6681946 |
| ELGRTQAV | ASSSRISNERLF | 0.8600286 |
| FVAGGYLS | ASSSRISNERLF | 0.7856136 |
| GASGGAGGAGGW | ASSSRISNERLF | 0.65951645 |
| GGAGAGGDGGAGGI | ASSSRISNERLF | 0.6825719 |
| GTGHNGSQPGMGGNG | ASSSRISNERLF | 0.86814356 |
| GYAAASASAS | ASSSRISNERLF | 0.47823673 |

| | | |
|---|---|---|
| HQVDVAAVQQG | ASSSRISNERLF | 0.8370261 |
| IGAAIAHT | ASSSRISNERLF | 0.7742605 |
| IGMIEVAQL | ASSSRISNERLF | 0.8697318 |
| IRLAPEHTSAMD | ASSSRISNERLF | 0.5645278 |
| IYNVSVL | ASSSRISNERLF | 0.843343 |
| KQGFLGM | ASSSRISNERLF | 0.7988236 |
| LESVITQL | ASSSRISNERLF | 0.9025738 |
| LGLGALVVV | ASSSRISNERLF | 0.789044 |
| LIGNGGNGGSGGI | ASSSRISNERLF | 0.79391754 |
| LNAINAPTL | ASSSRISNERLF | 0.7123216 |
| LQLLNTGV | ASSSRISNERLF | 0.8756899 |
| LQTSIEF | ASSSRISNERLF | 0.61469233 |
| MSFAVSEL | ASSSRISNERLF | 0.8032304 |
| NTGALGL | ASSSRISNERLF | 0.46103242 |
| PPGAGGAGGAGTT | ASSSRISNERLF | 0.83179367 |
| QLVGLGI | ASSSRISNERLF | 0.8468995 |
| RFLEQVHQL | ASSSRISNERLF | 0.7317204 |
| RGVGYVLR | ASSSRISNERLF | 0.8856612 |
| SATGVFAGVI | ASSSRISNERLF | 0.81493133 |
| SLPAITLP | ASSSRISNERLF | 0.75771683 |
| STALAFTV | ASSSRISNERLF | 0.89194655 |
| SVLTVGLGK | ASSSRISNERLF | 0.7064717 |
| TALAGAGSGI | ASSSRISNERLF | 0.88966495 |
| TGLPFTGI | ASSSRISNERLF | 0.75823015 |
| VAEHVTV | ASSSRISNERLF | 0.8361681 |
| VDIVLNSL | ASSSRISNERLF | 0.85054976 |
| VLTGGGAVL | ASSSRISNERLF | 0.70877326 |
| VNVGLAYGI | ASSSRISNERLF | 0.66550314 |
| VSSAAVGAV | ASSSRISNERLF | 0.876521 |
| VTSVYEAGV | ASSSRISNERLF | 0.82240176 |
| WVALDRGA | ASSSRISNERLF | 0.8185445 |
| YLVGSVVI | ASSSRISNERLF | 0.74366623 |
| GPPGTGKTLL | ASSSRISNERLF | 0.7513376 |
| GVGMTKF | ASSSRISNERLF | 0.74992466 |
| IGIGGSDL | ASSSRISNERLF | 0.81495714 |
| IGTLLAEL | ASSSRISNERLF | 0.82912695 |
| IYTSGTT | ASSSRISNERLF | 0.82295954 |
| LSFPAGV | ASSSRISNERLF | 0.69775975 |
| LYTSGTT | ASSSRISNERLF | 0.8225522 |
| VGAFPFGF | ASSSRISNERLF | 0.78369343 |
| VGAGTVEF | ASSSRISNERLF | 0.8045338 |
| VGKSTLLL | ASSSRISNERLF | 0.79610014 |
| AALAHAGGGIV | ASSSRISNERLF | 0.79497534 |
| AALVAVSLL | ASSSRISNERLF | 0.7484195 |
| AAVSRAVAAL | ASSSRISNERLF | 0.6117284 |
| AGGSSGIGGF | ASSSRISNERLF | 0.81126153 |
| AGLTVANANG | ASSSRISNERLF | 0.8231367 |
| AGLVGAVL | ASSSRISNERLF | 0.8124094 |
| ALSQAFTV | ASSSRISNERLF | 0.8295922 |
| ALTFSVSL | ASSSRISNERLF | 0.8265372 |
| ALTGAALVAL | ASSSRISNERLF | 0.81334436 |

| | | |
|---|---|---|
| ASFSIGAGVL | ASSSRISNERLF | 0.32694858 |
| DGSVYKI | ASSSRISNERLF | 0.5583631 |
| EQLAEIV | ASSSRISNERLF | 0.6064211 |
| FAVSELTVM | ASSSRISNERLF | 0.7681693 |
| GAAPLAAGL | ASSSRISNERLF | 0.25044727 |
| GADGGDGGSGGAGG | ASSSRISNERLF | 0.80121595 |
| GFGLDTGGAGG | ASSSRISNERLF | 0.7360031 |
| GLASGGLL | ASSSRISNERLF | 0.76521707 |
| GRIAGAAVLL | ASSSRISNERLF | 0.82421106 |
| GSAGSLNTSF | ASSSRISNERLF | 0.7779795 |
| GTGQIGI | ASSSRISNERLF | 0.6059648 |
| GTLIALIL | ASSSRISNERLF | 0.6771656 |
| IAATMGFVVG | ASSSRISNERLF | 0.830743 |
| IFAQALL | ASSSRISNERLF | 0.5794879 |
| IGAAVALGF | ASSSRISNERLF | 0.8271397 |
| IGLTPLAL | ASSSRISNERLF | 0.7202391 |
| ILGSNTSTL | ASSSRISNERLF | 0.8267734 |
| IVFSLTVL | ASSSRISNERLF | 0.65487814 |
| LASKATGF | ASSSRISNERLF | 0.82745385 |
| LASVALIL | ASSSRISNERLF | 0.8064943 |
| LFAVGLTL | ASSSRISNERLF | 0.83423555 |
| LFNGGIGTYI | ASSSRISNERLF | 0.78225446 |
| LTLGAGII | ASSSRISNERLF | 0.8129083 |
| LVAAALGDI | ASSSRISNERLF | 0.76414084 |
| NTAFVQLGI | ASSSRISNERLF | 0.81672674 |
| QLFSGAEL | ASSSRISNERLF | 0.62164515 |
| SAATFITL | ASSSRISNERLF | 0.7633679 |
| SNYGLANLGV | ASSSRISNERLF | 0.72437614 |
| SYGGAYAVM | ASSSRISNERLF | 0.7801487 |
| TTGVGGLTLG | ASSSRISNERLF | 0.82608116 |
| TTQTIYI | ASSSRISNERLF | 0.2244038 |
| VAAGAGLGAVI | ASSSRISNERLF | 0.81526566 |
| VAAVVTLL | ASSSRISNERLF | 0.7046515 |
| VGFLGGLI | ASSSRISNERLF | 0.63076794 |
| VGVIPILL | ASSSRISNERLF | 0.81907654 |
| VIFASSLI | ASSSRISNERLF | 0.7978141 |
| VKGGDGGAAAG | ASSSRISNERLF | 0.81632286 |
| VTKTFTNI | ASSSRISNERLF | 0.70768005 |
| VVAAAILL | ASSSRISNERLF | 0.79451275 |
| YAALDVEL | ASSSRISNERLF | 0.7904042 |
| LASVALLL | ASSSRISNERLF | 0.3027192 |
| LDAVHRV | ASSSRISNERLF | 0.5851068 |
| QGLSSSIF | ASSSRISNERLF | 0.83220387 |
| RGTFANIRL | ASSSRISNERLF | 0.846033 |
| RQLWWGHRI | ASSSRISNERLF | 0.8377698 |
| YGPPGTGKTLL | ASSSRISNERLF | 0.80878556 |
| AALVGAPANVA | ASSSRISNERLF | 0.803731 |
| AATFRRTE | ASSSRISNERLF | 0.8084642 |
| AGALGGGI | ASSSRISNERLF | 0.84445065 |
| AGGAGGAGGIGAG | ASSSRISNERLF | 0.8519807 |
| AGSGALGL | ASSSRISNERLF | 0.8495599 |

| | | |
|---|---|---|
| AQIADQI | ASSSRISNERLF | 0.80469126 |
| ASAVAVTYL | ASSSRISNERLF | 0.85367775 |
| ASTVLGGTGG | ASSSRISNERLF | 0.8343334 |
| AVGGEPAK | ASSSRISNERLF | 0.8307294 |
| CEQGTFGGTC | ASSSRISNERLF | 0.8463975 |
| DEGKAPAG | ASSSRISNERLF | 0.84043634 |
| DTVVFPAL | ASSSRISNERLF | 0.81847084 |
| FEGITMAG | ASSSRISNERLF | 0.8057213 |
| FPLVVQD | ASSSRISNERLF | 0.8377733 |
| GAFTTPQ | ASSSRISNERLF | 0.84981155 |
| GFGGFGTAGGVGGA | ASSSRISNERLF | 0.8449788 |
| GFLAQFL | ASSSRISNERLF | 0.85710645 |
| GGAGGAGGNAGVA | ASSSRISNERLF | 0.85027087 |
| GGVGTAGAGGAGGAGG | ASSSRISNERLF | 0.84467065 |
| GNGGGGDAGNAGSGGN | ASSSRISNERLF | 0.6300626 |
| GTGGAGGAGNDAGS | ASSSRISNERLF | 0.7803677 |
| GTTNTGWFNS | ASSSRISNERLF | 0.804046 |
| IFMGPLFTL | ASSSRISNERLF | 0.829558 |
| ILAHAIEAI | ASSSRISNERLF | 0.44308564 |
| IRSAIREAL | ASSSRISNERLF | 0.83511996 |
| ITFAGGG | ASSSRISNERLF | 0.83743954 |
| ITGMSGHQEV | ASSSRISNERLF | 0.8121513 |
| IVAGLACTL | ASSSRISNERLF | 0.85119957 |
| IVGGGLAGA | ASSSRISNERLF | 0.8232708 |
| IVTGVLNI | ASSSRISNERLF | 0.7935711 |
| LAQQGCI | ASSSRISNERLF | 0.8238127 |
| LCLHVEN | ASSSRISNERLF | 0.85553837 |
| LDAISDGS | ASSSRISNERLF | 0.7908066 |
| LGGFAAML | ASSSRISNERLF | 0.85387254 |
| LGGKNPMI | ASSSRISNERLF | 0.7453203 |
| LGPKGVAAIL | ASSSRISNERLF | 0.8536251 |
| LMVGVSI | ASSSRISNERLF | 0.7749433 |
| LVGAAFVL | ASSSRISNERLF | 0.8515769 |
| LVLAVAPL | ASSSRISNERLF | 0.831725 |
| MAAIAVIDAA | ASSSRISNERLF | 0.8589149 |
| NNGINAELQ | ASSSRISNERLF | 0.8304238 |
| RGEPLVLVHG | ASSSRISNERLF | 0.85330236 |
| RGGAGGAAGLI | ASSSRISNERLF | 0.83236 |
| RSYRMHPAV | ASSSRISNERLF | 0.831502 |
| SAAHLLAGQLA | ASSSRISNERLF | 0.76207066 |
| SAAVGSY | ASSSRISNERLF | 0.8319342 |
| SPHAVLHRI | ASSSRISNERLF | 0.78258216 |
| SQIAKVL | ASSSRISNERLF | 0.8407732 |
| TALGHLVP | ASSSRISNERLF | 0.84978104 |
| TAPTTSGAA | ASSSRISNERLF | 0.44635108 |
| TDGDDGNGGSPGT | ASSSRISNERLF | 0.8526044 |
| TDGLGEN | ASSSRISNERLF | 0.7796011 |
| TGGPPLPGQA | ASSSRISNERLF | 0.7955137 |
| TIPAITI | ASSSRISNERLF | 0.852121 |
| TIPGFTI | ASSSRISNERLF | 0.8366449 |
| TSIASGALA | ASSSRISNERLF | 0.8463588 |

| | | |
|---|---|---|
| TVTLPGAAI | ASSSRISNERLF | 0.80788875 |
| VASGLSAALP | ASSSRISNERLF | 0.8282543 |
| VGAGVIGI | ASSSRISNERLF | 0.8432463 |
| VGPKRFGV | ASSSRISNERLF | 0.4406905 |
| VNVAFPD | ASSSRISNERLF | 0.77903414 |
| VSMTNSGVS | ASSSRISNERLF | 0.85644865 |
| VTDVLGDLSV | ASSSRISNERLF | 0.85209703 |
| VVVVGAGFAGL | ASSSRISNERLF | 0.8730028 |
| VYTVGTVV | ASSSRISNERLF | 0.78655064 |
| WTTFSGP | ASSSRISNERLF | 0.74235505 |
| YDELVEAF | ASSSRISNERLF | 0.8240534 |
| YGTGGAGGNGG | ASSSRISNERLF | 0.8742226 |
| YTNVMAV | ASSSRISNERLF | 0.87643015 |
| FLEQVHQL | ASSSRISNERLF | 0.86164117 |
| GLSLLAVAV | ASSSRISNERLF | 0.72839403 |
| LEQVHQL | ASSSRISNERLF | 0.84985083 |
| LNAAGAI | ASSSRISNERLF | 0.78807926 |
| PAVHASL | ASSSRISNERLF | 0.82688034 |
| TELEDDA | ASSSRISNERLF | 0.86475813 |
| TPPGPPN | ASSSRISNERLF | 0.8421918 |
| TPPPPPPP | ASSSRISNERLF | 0.8112409 |
| TQSPALI | ASSQGQGANTEVF | 0.7740438 |
| EFAAAFPQ | ASSQGQGANTEVF | 0.8581001 |
| LLSGNLDVL | ASSQGQGANTEVF | 0.8698724 |
| GPANLGNYNI | ASSQGQGANTEVF | 0.8674022 |
| TLGPTLP | ASSQGQGANTEVF | 0.884231 |
| VYAYSSCCK | ASSQGQGANTEVF | 0.8692306 |
| APAPAPAPNI | ASSQGQGANTEVF | 0.8641177 |
| PGGPVPIP | ASSQGQGANTEVF | 0.2412515 |
| FLTVALEP | ASSQGQGANTEVF | 0.49747765 |
| SQYFGANL | ASSQGQGANTEVF | 0.7896916 |
| GGVKPAA | ASSQGQGANTEVF | 0.8237528 |
| GGTGGAGGAGGW | ASSQGQGANTEVF | 0.52759904 |
| GVGGVAGLGGAGG | ASSQGQGANTEVF | 0.8622909 |
| VLLVLATTI | ASSQGQGANTEVF | 0.7778443 |
| IGIFAAML | ASSQGQGANTEVF | 0.7670852 |
| AVGDLID | ASSQGQGANTEVF | 0.8715182 |
| NISPLPL | ASSQGQGANTEVF | 0.7986602 |
| VTAVFVAI | ASSQGQGANTEVF | 0.76736736 |
| SGDDNDAAL | ASSQGQGANTEVF | 0.78851914 |
| AVAALAHS | ASSQGQGANTEVF | 0.88200444 |
| GGVTVMATP | ASSQGQGANTEVF | 0.695531 |
| AGLAANVRD | ASSQGQGANTEVF | 0.86332476 |
| TNRPDLIDPALLRPG | ASSQGQGANTEVF | 0.8154316 |
| LANLGVTAL | ASSQGQGANTEVF | 0.8749472 |
| TGERIEL | ASSQGQGANTEVF | 0.6058923 |
| NGEELAA | ASSQGQGANTEVF | 0.86422443 |
| VLSLVWG | ASSQGQGANTEVF | 0.8335254 |
| IGYALWL | ASSQGQGANTEVF | 0.88819104 |
| LSLLLAPL | ASSQGQGANTEVF | 0.7986135 |
| TLDEALK | ASSQGQGANTEVF | 0.8287306 |

| | | |
|---|---|---|
| TDGAGTAEGG | ASSQGQGANTEVF | 0.76555026 |
| NAPGGAGGNGG | ASSQGQGANTEVF | 0.8668362 |
| VSVFMGMGGNF | ASSQGQGANTEVF | 0.5739981 |
| LASYLAAL | ASSQGQGANTEVF | 0.8193289 |
| LVGGTPAL | ASSQGQGANTEVF | 0.6969171 |
| VRNADVL | ASSQGQGANTEVF | 0.84444857 |
| TCSSPFLVSAM | ASSQGQGANTEVF | 0.8516941 |
| VHITDAE | ASSQGQGANTEVF | 0.43608987 |
| EYAAPSF | ASSQGQGANTEVF | 0.8697586 |
| VVDAFHGGL | ASSQGQGANTEVF | 0.8079701 |
| ASVVDPQ | ASSQGQGANTEVF | 0.72917485 |
| SYNEFGGLS | ASSQGQGANTEVF | 0.8779964 |
| DPDGSVY | ASSQGQGANTEVF | 0.8161286 |
| MVSTIHATE | ASSQGQGANTEVF | 0.87227666 |
| LAAGVADPV | ASSQGQGANTEVF | 0.8384725 |
| TKEQIAA | ASSQGQGANTEVF | 0.8302026 |
| SLASPAAL | ASSQGQGANTEVF | 0.86585534 |
| IGLVFYGV | ASSQGQGANTEVF | 0.63373613 |
| FLLDASNL | ASSQGQGANTEVF | 0.7728726 |
| YQVGNFP | ASSQGQGANTEVF | 0.87457615 |
| ESLLSQST | ASSQGQGANTEVF | 0.777974 |
| LFTVGFAAVG | ASSQGQGANTEVF | 0.8687608 |
| ATPPTPT | ASSQGQGANTEVF | 0.7236886 |
| TDLATLG | ASSQGQGANTEVF | 0.603614 |
| SAGGTGR | ASSQGQGANTEVF | 0.7371705 |
| GFSNSGS | ASSQGQGANTEVF | 0.8192587 |
| GGNGGSGGTSV | ASSQGQGANTEVF | 0.86474144 |
| GTSVATGGAGN | ASSQGQGANTEVF | 0.79416984 |
| NGGNGGSGGTS | ASSQGQGANTEVF | 0.48930496 |
| AVVKFPH | ASSQGQGANTEVF | 0.77983344 |
| TPPPPPP | ASSQGQGANTEVF | 0.6834872 |
| AAPAAAAAPA | ASSQGQGANTEVF | 0.7154751 |
| IALNEDL | ASSQGQGANTEVF | 0.79416275 |
| IVNVSSM | ASSQGQGANTEVF | 0.7593492 |
| TGIPPAPR | ASSQGQGANTEVF | 0.7114803 |
| FDLGGGTF | ASSQGQGANTEVF | 0.6271465 |
| FDLGGGTFDVS | ASSQGQGANTEVF | 0.78907853 |
| NEPTAAA | ASSQGQGANTEVF | 0.8155348 |
| NEPTAAALA | ASSQGQGANTEVF | 0.80002296 |
| GSIINISSI | ASSQGQGANTEVF | 0.85334975 |
| VAPGNAGTA | ASSQGQGANTEVF | 0.8353587 |
| TGGGASLE | ASSQGQGANTEVF | 0.82901424 |
| VAVYNHYKRI | ASSQGQGANTEVF | 0.1757617 |
| TDPAPATT | ASSQGQGANTEVF | 0.29698986 |
| TNRPDILDPALLRPG | ASSQGQGANTEVF | 0.58001506 |
| LTTTTTS | ASSQGQGANTEVF | 0.65344286 |
| PTGSGKTL | ASSQGQGANTEVF | 0.25502768 |
| TGSGKTL | ASSQGQGANTEVF | 0.78603536 |
| TIGHVDHGK | ASSQGQGANTEVF | 0.665122 |
| TIGHVDHGKTT | ASSQGQGANTEVF | 0.6269494 |
| GHVDHGKTT | ASSQGQGANTEVF | 0.83955634 |

| | | |
|---|---|---|
| AQFEHTLL | ASSQGQGANTEVF | 0.7263112 |
| IVNQDEL | ASSQGQGANTEVF | 0.6126517 |
| TNGSQFF | ASSQGQGANTEVF | 0.5795368 |
| TGKGIVE | ASSQGQGANTEVF | 0.8265334 |
| LISNASDA | ASSQGQGANTEVF | 0.3418274 |
| AELRTLGELTKIAKAHGAQ | ASSQGQGANTEVF | 0.8120917 |
| APNAVAAAEPSW | ASSQGQGANTEVF | 0.45024496 |
| APQVAGAATAVL | ASSQGQGANTEVF | 0.8212471 |
| ARRDPAAAAAILPTDA | ASSQGQGANTEVF | 0.41145492 |
| DAAARDVLAEAGLAEYF | ASSQGQGANTEVF | 0.8210776 |
| DDQIVLSAKANLAKVEQ | ASSQGQGANTEVF | 0.7725072 |
| DELLPDTGEDSDEDGH | ASSQGQGANTEVF | 0.8898517 |
| ELPEAPANRATVPQD | ASSQGQGANTEVF | 0.70496917 |
| FGQSKIERVTGIG | ASSQGQGANTEVF | 0.7132479 |
| GAGGTGSTAGGAGGAGGAGGI | ASSQGQGANTEVF | 0.54381216 |
| GHGGTGAAGGNGGTGGAGGS( | ASSQGQGANTEVF | 0.85962886 |
| GNGGNGGAGGEGGPGGAGGA( | ASSQGQGANTEVF | 0.3685267 |
| GSTAGNGGNGGNATNGTIAG( | ASSQGQGANTEVF | 0.73658514 |
| GTGGDGGNGGNGGTGG | ASSQGQGANTEVF | 0.4739328 |
| GVRRPAGIPADAPV | ASSQGQGANTEVF | 0.80750334 |
| GYRQVTDAHLLTLARR | ASSQGQGANTEVF | 0.79354334 |
| IAKMGQDGHDRG | ASSQGQGANTEVF | 0.3133566 |
| KAEPSIEIQYT | ASSQGQGANTEVF | 0.8264088 |
| KPVRIRMAADEDTVAVT | ASSQGQGANTEVF | 0.75384784 |
| MPPRIAGMRLLVIKPEP | ASSQGQGANTEVF | 0.57138574 |
| MKFYAVALFRSRGVPPTR | ASSQGQGANTEVF | 0.85666025 |
| NAGGFGILGAVAHSP | ASSQGQGANTEVF | 0.70662564 |
| NEGREITLAMAEM | ASSQGQGANTEVF | 0.86361647 |
| NELVEWSHPLGHRWRWE | ASSQGQGANTEVF | 0.677752 |
| NLGAGGTGGNGGNPG | ASSQGQGANTEVF | 0.5981033 |
| NTEWPKWETVMAADPVD | ASSQGQGANTEVF | 0.7916217 |
| PASAAPAETQP | ASSQGQGANTEVF | 0.41516024 |
| QQSSDGKNGVRAAEITGE | ASSQGQGANTEVF | 0.50094426 |
| RAPDAVTTSWWKE | ASSQGQGANTEVF | 0.85685873 |
| SGFGNVGAHTSGWFNQ | ASSQGQGANTEVF | 0.7815529 |
| SGVAGAVGQAASVGGLKV | ASSQGQGANTEVF | 0.7896231 |
| SNGGAGTGSAGGNGGT | ASSQGQGANTEVF | 0.7342276 |
| TGGAGGEGGAAG | ASSQGQGANTEVF | 0.69857204 |
| TGGDGGAAGAGGAPGHG | ASSQGQGANTEVF | 0.7619347 |
| TMPPLLAAIANAGRHGV | ASSQGQGANTEVF | 0.79617083 |
| TSSTAGNSGGA | ASSQGQGANTEVF | 0.79939365 |
| VPATVPVGILC | ASSQGQGANTEVF | 0.7886948 |
| VQRPDSGGVVLTGRIS | ASSQGQGANTEVF | 0.7018588 |
| WTPNPVPATKGVPAGLVQAT | ASSQGQGANTEVF | 0.7936914 |
| YALKRALAKFSDRFDVVII | ASSQGQGANTEVF | 0.7722195 |
| YLIGSVIV | ASSQGQGANTEVF | 0.7578467 |
| YGQLVAVID | ASSQGQGANTEVF | 0.79738843 |
| VTVGGMTLPA | ASSQGQGANTEVF | 0.7830376 |
| VLALPAV | ASSQGQGANTEVF | 0.7457919 |
| VIPPTLNYET | ASSQGQGANTEVF | 0.74837184 |
| VGMGGMHPSSG | ASSQGQGANTEVF | 0.7805214 |

| | | |
|---|---|---|
| VGLAGGAATA | ASSQGQGANTEVF | 0.7910551 |
| VGGAGGAGGSGP | ASSQGQGANTEVF | 0.7942983 |
| VFAAFMV | ASSQGQGANTEVF | 0.8110354 |
| TLYDKLR | ASSQGQGANTEVF | 0.79817545 |
| TLAVFTVL | ASSQGQGANTEVF | 0.7870759 |
| TGSVQPARV | ASSQGQGANTEVF | 0.3334057 |
| TGGGAGSGGT | ASSQGQGANTEVF | 0.5473958 |
| SHLETSL | ASSQGQGANTEVF | 0.75473595 |
| SFTTGMT | ASSQGQGANTEVF | 0.76782995 |
| QVEVAKRMGVRQPTVS | ASSQGQGANTEVF | 0.47447997 |
| QGSGGPALAM | ASSQGQGANTEVF | 0.7746735 |
| MLAPSNP | ASSQGQGANTEVF | 0.7542713 |
| LVNNAAI | ASSQGQGANTEVF | 0.72925234 |
| LTGEINTLA | ASSQGQGANTEVF | 0.8032302 |
| LSIFGPV | ASSQGQGANTEVF | 0.75506544 |
| LLALLANL | ASSQGQGANTEVF | 0.7222134 |
| LISNASDALDKLR | ASSQGQGANTEVF | 0.75984144 |
| LINEAVP | ASSQGQGANTEVF | 0.80506563 |
| LGYAFSGL | ASSQGQGANTEVF | 0.71425307 |
| LDFVAVAL | ASSQGQGANTEVF | 0.79849845 |
| LAQPGAE | ASSQGQGANTEVF | 0.67120314 |
| ISNLGAVTGF | ASSQGQGANTEVF | 0.80167764 |
| HQEDPGRI | ASSQGQGANTEVF | 0.65596026 |
| GTPGEPTP | ASSQGQGANTEVF | 0.7953743 |
| GFFNSGS | ASSQGQGANTEVF | 0.7653731 |
| GAGGGGLTAGAN | ASSQGQGANTEVF | 0.8146949 |
| FVDALVNM | ASSQGQGANTEVF | 0.7620336 |
| FLTVAPEL | ASSQGQGANTEVF | 0.7968762 |
| FETLTAL | ASSQGQGANTEVF | 0.7460341 |
| DVINPIEL | ASSQGQGANTEVF | 0.7926984 |
| APVSGAPV | ASSQGQGANTEVF | 0.644328 |
| ALVGGLAW | ASSQGQGANTEVF | 0.7748352 |
| ALAAAAATAI | ASSQGQGANTEVF | 0.5340202 |
| AFGLGLLL | ASSQGQGANTEVF | 0.78279424 |
| AFFAAASL | ASSQGQGANTEVF | 0.7849359 |
| AAAALPAFSPP | ASSQGQGANTEVF | 0.62763315 |
| VVLAPAL | ASSQGQGANTEVF | 0.8013825 |
| TIGHVDHGKTTL | ASSQGQGANTEVF | 0.76220596 |
| LISNASDALDK | ASSQGQGANTEVF | 0.72689444 |
| LISNASDAL | ASSQGQGANTEVF | 0.8056929 |
| ISNASDALDK | ASSQGQGANTEVF | 0.7665893 |
| NQAETLVYQTE | ASSQGQGANTEVF | 0.7929241 |
| TGGTGGTAGAG | ASSQGQGANTEVF | 0.7587341 |
| TGSTAGGAGGA | ASSQGQGANTEVF | 0.7461031 |
| DPLAVPVAAGPV | ASSQGQGANTEVF | 0.7835093 |
| GGTGGTPGNGGN | ASSQGQGANTEVF | 0.6477242 |
| PAGLAGVDPATL | ASSQGQGANTEVF | 0.7334838 |
| GNNLVVEAPHAGASV | ASSQGQGANTEVF | 0.8039199 |
| KVTRIMSASTLEQPA | ASSQGQGANTEVF | 0.83501387 |
| LGFALPIALPRRTVV | ASSQGQGANTEVF | 0.8424077 |
| TRLGNLTARKLVVLV | ASSQGQGANTEVF | 0.63942885 |

| | | |
|---|---|---|
| WSQQRRLIYFPSAGP | ASSQGQGANTEVF | 0.6554326 |
| GPNGNPDPMAAAVDIR | ASSQGQGANTEVF | 0.7345764 |
| TQRCVSVADLRAAASR | ASSQGQGANTEVF | 0.8423105 |
| QFVLDVAEHPVEVLTETV | ASSQGQGANTEVF | 0.8613353 |
| GGAGGTGGAGGNGGTQAGDG | ASSQGQGANTEVF | 0.8410311 |
| PLLATLRAIAVDVDAKPAQV | ASSQGQGANTEVF | 0.683336 |
| APKFRTGGYMIDFWGVGYQV/ | ASSQGQGANTEVF | 0.818035 |
| GAGGAGGAGGAAGGPGGTGG | ASSQGQGANTEVF | 0.75427294 |
| IGALGLLATDVAPAGSVAFG | ASSQGQGANTEVF | 0.75253224 |
| AGGAGGIGGNGGAGGAGGVL | ASSQGQGANTEVF | 0.8468558 |
| ADELIGLL | ASSQGQGANTEVF | 0.78748 |
| ADRALKA | ASSQGQGANTEVF | 0.6838498 |
| AGAVGQVGGAGGA | ASSQGQGANTEVF | 0.68891037 |
| AGGGGGLGGAP | ASSQGQGANTEVF | 0.80198205 |
| AGGLLYGNGGAG | ASSQGQGANTEVF | 0.8322988 |
| AGLESIGL | ASSQGQGANTEVF | 0.82497984 |
| AGLKTGII | ASSQGQGANTEVF | 0.89272153 |
| ASLAALLL | ASSQGQGANTEVF | 0.85197634 |
| ASVINISSI | ASSQGQGANTEVF | 0.8413956 |
| ATPPGPA | ASSQGQGANTEVF | 0.25369602 |
| ATVASALV | ASSQGQGANTEVF | 0.5027964 |
| ATVASLIL | ASSQGQGANTEVF | 0.74445295 |
| AVNGPVLIRH | ASSQGQGANTEVF | 0.74919486 |
| DGAAGAVGI | ASSQGQGANTEVF | 0.38284636 |
| DPLLHGI | ASSQGQGANTEVF | 0.8240682 |
| ELGAALGAL | ASSQGQGANTEVF | 0.75363505 |
| ELGRTQAV | ASSQGQGANTEVF | 0.6463332 |
| FVAGGYLS | ASSQGQGANTEVF | 0.8397206 |
| GASGGAGGAGGW | ASSQGQGANTEVF | 0.6276411 |
| GGAGAGGDGGAGGI | ASSQGQGANTEVF | 0.5825874 |
| GTGHNGSQPGMGGNG | ASSQGQGANTEVF | 0.7538248 |
| GYAAASASAS | ASSQGQGANTEVF | 0.8708639 |
| HQVDVAAVQQG | ASSQGQGANTEVF | 0.68234503 |
| IGAAIAHT | ASSQGQGANTEVF | 0.8476131 |
| IGMIEVAQL | ASSQGQGANTEVF | 0.79519254 |
| IRLAPEHTSAMD | ASSQGQGANTEVF | 0.86180484 |
| IYNVSVL | ASSQGQGANTEVF | 0.5564997 |
| KQGFLGM | ASSQGQGANTEVF | 0.8521369 |
| LESVITQL | ASSQGQGANTEVF | 0.75671804 |
| LGLGALVVV | ASSQGQGANTEVF | 0.88142425 |
| LIGNGGNGGSGGI | ASSQGQGANTEVF | 0.7687032 |
| LNAINAPTL | ASSQGQGANTEVF | 0.8421147 |
| LQLLNTGV | ASSQGQGANTEVF | 0.7821053 |
| LQTSIEF | ASSQGQGANTEVF | 0.858544 |
| MSFAVSEL | ASSQGQGANTEVF | 0.5297564 |
| NTGALGL | ASSQGQGANTEVF | 0.74699533 |
| PPGAGGAGGAGTT | ASSQGQGANTEVF | 0.5430554 |
| QLVGLGI | ASSQGQGANTEVF | 0.77833223 |
| RFLEQVHQL | ASSQGQGANTEVF | 0.8339212 |
| RGVGYVLR | ASSQGQGANTEVF | 0.2158774 |
| SATGVFAGVI | ASSQGQGANTEVF | 0.8493932 |

| | | |
|---|---|---|
| SLPAITLP | ASSQGQGANTEVF | 0.59684336 |
| STALAFTV | ASSQGQGANTEVF | 0.631421 |
| SVLTVGLGK | ASSQGQGANTEVF | 0.8565305 |
| TALAGAGSGI | ASSQGQGANTEVF | 0.7546563 |
| TGLPFTGI | ASSQGQGANTEVF | 0.86619973 |
| VAEHVTV | ASSQGQGANTEVF | 0.74842167 |
| VDIVLNSL | ASSQGQGANTEVF | 0.78697526 |
| VLTGGGAVL | ASSQGQGANTEVF | 0.7965555 |
| VNVGLAYGI | ASSQGQGANTEVF | 0.37624094 |
| VSSAAVGAV | ASSQGQGANTEVF | 0.6550975 |
| VTSVYEAGV | ASSQGQGANTEVF | 0.8712437 |
| WVALDRGA | ASSQGQGANTEVF | 0.85033 |
| YLVGSVVI | ASSQGQGANTEVF | 0.8708602 |
| GPPGTGKTLL | ASSQGQGANTEVF | 0.77049637 |
| GVGMTKF | ASSQGQGANTEVF | 0.74370444 |
| IGIGGSDL | ASSQGQGANTEVF | 0.7926408 |
| IGTLLAEL | ASSQGQGANTEVF | 0.8740273 |
| IYTSGTT | ASSQGQGANTEVF | 0.8550163 |
| LSFPAGV | ASSQGQGANTEVF | 0.8445 |
| LYTSGTT | ASSQGQGANTEVF | 0.6927849 |
| VGAFPFGF | ASSQGQGANTEVF | 0.78303623 |
| VGAGTVEF | ASSQGQGANTEVF | 0.7568228 |
| VGKSTLLL | ASSQGQGANTEVF | 0.77288544 |
| AALAHAGGGIV | ASSQGQGANTEVF | 0.8628739 |
| AALVAVSLL | ASSQGQGANTEVF | 0.77617764 |
| AAVSRAVAAL | ASSQGQGANTEVF | 0.7678627 |
| AGGSSGIGGF | ASSQGQGANTEVF | 0.8208624 |
| AGLTVANANG | ASSQGQGANTEVF | 0.7728629 |
| AGLVGAVL | ASSQGQGANTEVF | 0.8123256 |
| ALSQAFTV | ASSQGQGANTEVF | 0.79594505 |
| ALTFSVSL | ASSQGQGANTEVF | 0.8877015 |
| ALTGAALVAL | ASSQGQGANTEVF | 0.8663299 |
| ASFSIGAGVL | ASSQGQGANTEVF | 0.8628695 |
| DGSVYKI | ASSQGQGANTEVF | 0.20122328 |
| EQLAEIV | ASSQGQGANTEVF | 0.57091445 |
| FAVSELTVM | ASSQGQGANTEVF | 0.82335734 |
| GAAPLAAGL | ASSQGQGANTEVF | 0.77561116 |
| GADGGDGGSGGAGG | ASSQGQGANTEVF | 0.51296926 |
| GFGLDTGGAGG | ASSQGQGANTEVF | 0.8236039 |
| GLASGGLL | ASSQGQGANTEVF | 0.790177 |
| GRIAGAAVLL | ASSQGQGANTEVF | 0.6765158 |
| GSAGSLNTSF | ASSQGQGANTEVF | 0.854252 |
| GTGQIGI | ASSQGQGANTEVF | 0.7815216 |
| GTLIALIL | ASSQGQGANTEVF | 0.67588556 |
| IAATMGFVVG | ASSQGQGANTEVF | 0.76221377 |
| IFAQALL | ASSQGQGANTEVF | 0.8534378 |
| IGAAVALGF | ASSQGQGANTEVF | 0.6187563 |
| IGLTPLAL | ASSQGQGANTEVF | 0.8350195 |
| ILGSNTSTL | ASSQGQGANTEVF | 0.7243892 |
| IVFSLTVL | ASSQGQGANTEVF | 0.8764777 |
| LASKATGF | ASSQGQGANTEVF | 0.634432 |

| | | |
|---|---|---|
| LASVALIL | ASSQGQGANTEVF | 0.85326046 |
| LFAVGLTL | ASSQGQGANTEVF | 0.80540866 |
| LFNGGIGTYI | ASSQGQGANTEVF | 0.89595973 |
| LTLGAGII | ASSQGQGANTEVF | 0.7984614 |
| LVAAALGDI | ASSQGQGANTEVF | 0.8251352 |
| NTAFVQLGI | ASSQGQGANTEVF | 0.6988735 |
| QLFSGAEL | ASSQGQGANTEVF | 0.82382905 |
| SAATFITL | ASSQGQGANTEVF | 0.5783073 |
| SNYGLANLGV | ASSQGQGANTEVF | 0.81715953 |
| SYGGAYAVM | ASSQGQGANTEVF | 0.56661797 |
| TTGVGGLTLG | ASSQGQGANTEVF | 0.8473693 |
| TTQTIYI | ASSQGQGANTEVF | 0.84033245 |
| VAAGAGLGAVI | ASSQGQGANTEVF | 0.60192955 |
| VAAVVTLL | ASSQGQGANTEVF | 0.8656379 |
| VGFLGGLI | ASSQGQGANTEVF | 0.77269316 |
| VGVIPILL | ASSQGQGANTEVF | 0.7730485 |
| VIFASSLI | ASSQGQGANTEVF | 0.8869681 |
| VKGGDGGAAAG | ASSQGQGANTEVF | 0.74531335 |
| VTKTFTNI | ASSQGQGANTEVF | 0.8819843 |
| VVAAAILL | ASSQGQGANTEVF | 0.7831853 |
| YAALDVEL | ASSQGQGANTEVF | 0.7814408 |
| LASVALLL | ASSQGQGANTEVF | 0.840158 |
| LDAVHRV | ASSQGQGANTEVF | 0.3287851 |
| QGLSSSIF | ASSQGQGANTEVF | 0.68286765 |
| RGTFANIRL | ASSQGQGANTEVF | 0.8773202 |
| RQLWWGHRI | ASSQGQGANTEVF | 0.8363265 |
| YGPPGTGKTLL | ASSQGQGANTEVF | 0.8242358 |
| AALVGAPANVA | ASSQGQGANTEVF | 0.7904419 |
| AATFRRTE | ASSQGQGANTEVF | 0.7716768 |
| AGALGGGI | ASSQGQGANTEVF | 0.8124334 |
| AGGAGGAGGIGAG | ASSQGQGANTEVF | 0.819973 |
| AGSGALGL | ASSQGQGANTEVF | 0.8460573 |
| AQIADQI | ASSQGQGANTEVF | 0.8363167 |
| ASAVAVTYL | ASSQGQGANTEVF | 0.73380494 |
| ASTVLGGTGG | ASSQGQGANTEVF | 0.8444343 |
| AVGGEPAK | ASSQGQGANTEVF | 0.8273438 |
| CEQGTFGGTC | ASSQGQGANTEVF | 0.8229453 |
| DEGKAPAG | ASSQGQGANTEVF | 0.82109725 |
| DTVVFPAL | ASSQGQGANTEVF | 0.80725217 |
| FEGITMAG | ASSQGQGANTEVF | 0.79951715 |
| FPLVVQD | ASSQGQGANTEVF | 0.6947533 |
| GAFTTPQ | ASSQGQGANTEVF | 0.8290493 |
| GFGGFGTAGGVGGA | ASSQGQGANTEVF | 0.8383851 |
| GFLAQFL | ASSQGQGANTEVF | 0.83749974 |
| GGAGGAGGNAGVA | ASSQGQGANTEVF | 0.8406671 |
| GGVGTAGAGGAGGAGG | ASSQGQGANTEVF | 0.8426676 |
| GNGGGGDAGNAGSGGN | ASSQGQGANTEVF | 0.8293768 |
| GTGGAGGAGNDAGS | ASSQGQGANTEVF | 0.44142112 |
| GTTNTGWFNS | ASSQGQGANTEVF | 0.611576 |
| IFMGPLFTL | ASSQGQGANTEVF | 0.71327704 |
| ILAHAIEAI | ASSQGQGANTEVF | 0.79432654 |

| | | |
|---|---|---|
| IRSAIREAL | ASSQGQGANTEVF | 0.2994244 |
| ITFAGGG | ASSQGQGANTEVF | 0.7723057 |
| ITGMSGHQEV | ASSQGQGANTEVF | 0.810522 |
| IVAGLACTL | ASSQGQGANTEVF | 0.7853631 |
| IVGGGLAGA | ASSQGQGANTEVF | 0.84178567 |
| IVTGVLNI | ASSQGQGANTEVF | 0.8095275 |
| LAQQGCI | ASSQGQGANTEVF | 0.5995157 |
| LCLHVEN | ASSQGQGANTEVF | 0.77000344 |
| LDAISDGS | ASSQGQGANTEVF | 0.83649665 |
| LGGFAAML | ASSQGQGANTEVF | 0.75573224 |
| LGGKNPMI | ASSQGQGANTEVF | 0.84551793 |
| LGPKGVAAIL | ASSQGQGANTEVF | 0.6839272 |
| LMVGVSI | ASSQGQGANTEVF | 0.8431561 |
| LVGAAFVL | ASSQGQGANTEVF | 0.686543 |
| LVLAVAPL | ASSQGQGANTEVF | 0.84210706 |
| MAAIAVIDAA | ASSQGQGANTEVF | 0.8267784 |
| NNGINAELQ | ASSQGQGANTEVF | 0.8480808 |
| RGEPLVLVHG | ASSQGQGANTEVF | 0.8054214 |
| RGGAGGAAGLI | ASSQGQGANTEVF | 0.83781826 |
| RSYRMHPAV | ASSQGQGANTEVF | 0.8025862 |
| SAAHLLAGQLA | ASSQGQGANTEVF | 0.8308674 |
| SAAVGSY | ASSQGQGANTEVF | 0.72546947 |
| SPHAVLHRI | ASSQGQGANTEVF | 0.80340695 |
| SQIAKVL | ASSQGQGANTEVF | 0.6921153 |
| TALGHLVP | ASSQGQGANTEVF | 0.8093563 |
| TAPTTSGAA | ASSQGQGANTEVF | 0.83972746 |
| TDGDDGNGGSPGT | ASSQGQGANTEVF | 0.22246356 |
| TDGLGEN | ASSQGQGANTEVF | 0.84050626 |
| TGGPPLPGQA | ASSQGQGANTEVF | 0.68173957 |
| TIPAITI | ASSQGQGANTEVF | 0.7002871 |
| TIPGFTI | ASSQGQGANTEVF | 0.8390084 |
| TSIASGALA | ASSQGQGANTEVF | 0.82589 |
| TVTLPGAAI | ASSQGQGANTEVF | 0.8328053 |
| VASGLSAALP | ASSQGQGANTEVF | 0.7379397 |
| VGAGVIGI | ASSQGQGANTEVF | 0.8068572 |
| VGPKRFGV | ASSQGQGANTEVF | 0.81776327 |
| VNVAFPD | ASSQGQGANTEVF | 0.3376701 |
| VSMTNSGVS | ASSQGQGANTEVF | 0.6692543 |
| VTDVLGDLSV | ASSQGQGANTEVF | 0.8469643 |
| VVVVGAGFAGL | ASSQGQGANTEVF | 0.7147647 |
| VYTVGTVV | ASSQGQGANTEVF | 0.7183598 |
| WTTFSGP | ASSQGQGANTEVF | 0.66543174 |
| YDELVEAF | ASSQGQGANTEVF | 0.66618055 |
| YGTGGAGGNGG | ASSQGQGANTEVF | 0.69791734 |
| YTNVMAV | ASSQGQGANTEVF | 0.717305 |
| FLEQVHQL | ASSQGQGANTEVF | 0.72960544 |
| GLSLLAVAV | ASSQGQGANTEVF | 0.7160327 |
| LEQVHQL | ASSQGQGANTEVF | 0.69003975 |
| LNAAGAI | ASSQGQGANTEVF | 0.71650064 |
| PAVHASL | ASSQGQGANTEVF | 0.7081834 |
| TELEDDA | ASSQGQGANTEVF | 0.6963301 |

| | | |
|---|---|---|
| TPPGPPN | ASSQGQGANTEVF | 0.72228837 |
| TPPPPPPP | ASSQGQGANTEVF | 0.7095514 |
| TQSPALI | ASSLEGSSYEQY | 0.67779905 |
| EFAAAFPQ | ASSLEGSSYEQY | 0.6970644 |
| LLSGNLDVL | ASSLEGSSYEQY | 0.71722347 |
| GPANLGNYNI | ASSLEGSSYEQY | 0.7219424 |
| TLGPTLP | ASSLEGSSYEQY | 0.7183008 |
| VYAYSSCCK | ASSLEGSSYEQY | 0.7329986 |
| APAPAPAPNI | ASSLEGSSYEQY | 0.72289586 |
| PGGPVPIP | ASSLEGSSYEQY | 0.7174016 |
| FLTVALEP | ASSLEGSSYEQY | 0.5738892 |
| SQYFGANL | ASSLEGSSYEQY | 0.66190606 |
| GGVKPAA | ASSLEGSSYEQY | 0.6911775 |
| GGTGGAGGAGGW | ASSLEGSSYEQY | 0.70077443 |
| GVGGVAGLGGAGG | ASSLEGSSYEQY | 0.50932884 |
| VLLVLATTI | ASSLEGSSYEQY | 0.71116555 |
| IGIFAAML | ASSLEGSSYEQY | 0.7091317 |
| AVGDLID | ASSLEGSSYEQY | 0.6691677 |
| NISPLPL | ASSLEGSSYEQY | 0.7235092 |
| VTAVFVAI | ASSLEGSSYEQY | 0.6855692 |
| SGDDNDAAL | ASSLEGSSYEQY | 0.6908212 |
| AVAALAHS | ASSLEGSSYEQY | 0.69415754 |
| GGVTVMATP | ASSLEGSSYEQY | 0.7328424 |
| AGLAANVRD | ASSLEGSSYEQY | 0.6745868 |
| TNRPDLIDPALLRPG | ASSLEGSSYEQY | 0.72614235 |
| LANLGVTAL | ASSLEGSSYEQY | 0.7149196 |
| TGERIEL | ASSLEGSSYEQY | 0.7276059 |
| NGEELAA | ASSLEGSSYEQY | 0.65847737 |
| VLSLVWG | ASSLEGSSYEQY | 0.72295445 |
| IGYALWL | ASSLEGSSYEQY | 0.7033106 |
| LSLLLAPL | ASSLEGSSYEQY | 0.73207426 |
| TLDEALK | ASSLEGSSYEQY | 0.7091561 |
| TDGAGTAEGG | ASSLEGSSYEQY | 0.72773015 |
| NAPGGAGGNGG | ASSLEGSSYEQY | 0.71113783 |
| VSVFMGMGGNF | ASSLEGSSYEQY | 0.7200997 |
| LASYLAAL | ASSLEGSSYEQY | 0.64549965 |
| LVGGTPAL | ASSLEGSSYEQY | 0.7039357 |
| VRNADVL | ASSLEGSSYEQY | 0.68804324 |
| TCSSPFLVSAM | ASSLEGSSYEQY | 0.71196187 |
| VHITDAE | ASSLEGSSYEQY | 0.71688825 |
| EYAAPSF | ASSLEGSSYEQY | 0.496024 |
| VVDAFHGGL | ASSLEGSSYEQY | 0.7299098 |
| ASVVDPQ | ASSLEGSSYEQY | 0.6424004 |
| SYNEFGGLS | ASSLEGSSYEQY | 0.6897303 |
| DPDGSVY | ASSLEGSSYEQY | 0.7242596 |
| MVSTIHATE | ASSLEGSSYEQY | 0.69814616 |
| LAAGVADPV | ASSLEGSSYEQY | 0.7190942 |
| TKEQIAA | ASSLEGSSYEQY | 0.68644893 |
| SLASPAAL | ASSLEGSSYEQY | 0.71320367 |
| IGLVFYGV | ASSLEGSSYEQY | 0.71526355 |
| FLLDASNL | ASSLEGSSYEQY | 0.5259188 |

| | | |
|---|---|---|
| YQVGNFP | ASSLEGSSYEQY | 0.6512214 |
| ESLLSQST | ASSLEGSSYEQY | 0.7296735 |
| LFTVGFAAVG | ASSLEGSSYEQY | 0.7511704 |
| ATPPTPT | ASSLEGSSYEQY | 0.78017664 |
| TDLATLG | ASSLEGSSYEQY | 0.7173872 |
| SAGGTGR | ASSLEGSSYEQY | 0.63716936 |
| GFSNSGS | ASSLEGSSYEQY | 0.70350593 |
| GGNGGSGGTSV | ASSLEGSSYEQY | 0.7528637 |
| GTSVATGGAGN | ASSLEGSSYEQY | 0.78094953 |
| NGGNGGSGGTS | ASSLEGSSYEQY | 0.76728046 |
| AVVKFPH | ASSLEGSSYEQY | 0.57305574 |
| TPPPPPP | ASSLEGSSYEQY | 0.77429324 |
| AAPAAAAAPA | ASSLEGSSYEQY | 0.72576326 |
| IALNEDL | ASSLEGSSYEQY | 0.731839 |
| IVNVSSM | ASSLEGSSYEQY | 0.7158908 |
| TGIPPAPR | ASSLEGSSYEQY | 0.748476 |
| FDLGGGTF | ASSLEGSSYEQY | 0.7204011 |
| FDLGGGTFDVS | ASSLEGSSYEQY | 0.594874 |
| NEPTAAA | ASSLEGSSYEQY | 0.7264607 |
| NEPTAAALA | ASSLEGSSYEQY | 0.7637473 |
| GSIINISSI | ASSLEGSSYEQY | 0.7577132 |
| VAPGNAGTA | ASSLEGSSYEQY | 0.78362054 |
| TGGGASLE | ASSLEGSSYEQY | 0.77559936 |
| VAVYNHYKRI | ASSLEGSSYEQY | 0.7543124 |
| TDPAPATT | ASSLEGSSYEQY | 0.27388868 |
| TNRPDILDPALLRPG | ASSLEGSSYEQY | 0.4885625 |
| LTTTTTS | ASSLEGSSYEQY | 0.6052111 |
| PTGSGKTL | ASSLEGSSYEQY | 0.72243035 |
| TGSGKTL | ASSLEGSSYEQY | 0.2958534 |
| TIGHVDHGK | ASSLEGSSYEQY | 0.71815574 |
| TIGHVDHGKTT | ASSLEGSSYEQY | 0.721298 |
| GHVDHGKTT | ASSLEGSSYEQY | 0.6725994 |
| AQFEHTLL | ASSLEGSSYEQY | 0.7733947 |
| IVNQDEL | ASSLEGSSYEQY | 0.7126993 |
| TNGSQFF | ASSLEGSSYEQY | 0.5757497 |
| TGKGIVE | ASSLEGSSYEQY | 0.5519657 |
| LISNASDA | ASSLEGSSYEQY | 0.7713419 |
| AELRTLGELTKIAKAHGAQ | ASSLEGSSYEQY | 0.45542008 |
| APNAVAAAEPSW | ASSLEGSSYEQY | 0.7728789 |
| APQVAGAATAVL | ASSLEGSSYEQY | 0.53220105 |
| ARRDPAAAAAILPTDA | ASSLEGSSYEQY | 0.76969856 |
| DAAARDVLAEAGLAEYF | ASSLEGSSYEQY | 0.50500995 |
| DDQIVLSAKANLAKVEQ | ASSLEGSSYEQY | 0.7717092 |
| DELLPDTGEDSDEDGH | ASSLEGSSYEQY | 0.73293424 |
| ELPEAPANRATVPQD | ASSLEGSSYEQY | 0.78994864 |
| FGQSKIERVTGIG | ASSLEGSSYEQY | 0.67988527 |
| GAGGTGSTAGGAGGAGGAGGI | ASSLEGSSYEQY | 0.7473499 |
| GHGGTGAAGGNGGTGGAGGS( | ASSLEGSSYEQY | 0.60034513 |
| GNGGNGGAGGEGGPGGAGGA( | ASSLEGSSYEQY | 0.7761222 |
| GSTAGNGGNGGNATNGTIAG( | ASSLEGSSYEQY | 0.5319872 |
| GTGGDGGNGGNGGTGG | ASSLEGSSYEQY | 0.70346653 |

| | | |
|---|---|---|
| GVRRPAGIPADAPV | ASSLEGSSYEQY | 0.46919844 |
| GYRQVTDAHLLTLARR | ASSLEGSSYEQY | 0.7385167 |
| IAKMGQDGHDRG | ASSLEGSSYEQY | 0.7689327 |
| KAEPSIEIQYT | ASSLEGSSYEQY | 0.37614238 |
| KPVRIRMAADEDTVAVT | ASSLEGSSYEQY | 0.77600646 |
| MPPRIAGMRLLVIKPEP | ASSLEGSSYEQY | 0.6776658 |
| MKFYAVALFRSRGVPPTR | ASSLEGSSYEQY | 0.5963482 |
| NAGGFGILGAVAHSP | ASSLEGSSYEQY | 0.76479626 |
| NEGREITLAMAEM | ASSLEGSSYEQY | 0.73277795 |
| NELVEWSHPLGHRWRWE | ASSLEGSSYEQY | 0.7741613 |
| NLGAGGTGGNGGNPG | ASSLEGSSYEQY | 0.59988296 |
| NTEWPKWETVMAADPVD | ASSLEGSSYEQY | 0.71725965 |
| PASAAPAETQP | ASSLEGSSYEQY | 0.74707365 |
| QQSSDGKNGVRAAEITGE | ASSLEGSSYEQY | 0.596856 |
| RAPDAVTTSWWKE | ASSLEGSSYEQY | 0.5310316 |
| SGFGNVGAHTSGWFNQ | ASSLEGSSYEQY | 0.78072464 |
| SGVAGAVGQAASVGGLKV | ASSLEGSSYEQY | 0.66801906 |
| SNGGAGTGSAGGNGGT | ASSLEGSSYEQY | 0.67681825 |
| TGGAGGEGGAAG | ASSLEGSSYEQY | 0.6002606 |
| TGGDGGAAGAGGAPGHG | ASSLEGSSYEQY | 0.61594635 |
| TMPPLLAAIANAGRHGV | ASSLEGSSYEQY | 0.6146263 |
| TSSTAGNSGGA | ASSLEGSSYEQY | 0.66494787 |
| VPATVPVGILC | ASSLEGSSYEQY | 0.6799613 |
| VQRPDSGGVVLTGRIS | ASSLEGSSYEQY | 0.6698197 |
| WTPNPVPATKGVPAGLVQAT | ASSLEGSSYEQY | 0.592726 |
| YALKRALAKFSDRFDVVII | ASSLEGSSYEQY | 0.6637076 |
| YLIGSVIV | ASSLEGSSYEQY | 0.630503 |
| YGQLVAVID | ASSLEGSSYEQY | 0.63884115 |
| VTVGGMTLPA | ASSLEGSSYEQY | 0.65065837 |
| VLALPAV | ASSLEGSSYEQY | 0.64648265 |
| VIPPTLNYET | ASSLEGSSYEQY | 0.6234951 |
| VGMGGMHPSSG | ASSLEGSSYEQY | 0.57037973 |
| VGLAGGAATA | ASSLEGSSYEQY | 0.6561919 |
| VGGAGGAGGSGP | ASSLEGSSYEQY | 0.67439926 |
| VFAAFMV | ASSLEGSSYEQY | 0.66867 |
| TLYDKLR | ASSLEGSSYEQY | 0.6730373 |
| TLAVFTVL | ASSLEGSSYEQY | 0.66914827 |
| TGSVQPARV | ASSLEGSSYEQY | 0.65916723 |
| TGGGAGSGGT | ASSLEGSSYEQY | 0.38838136 |
| SHLETSL | ASSLEGSSYEQY | 0.52947676 |
| SFTTGMT | ASSLEGSSYEQY | 0.5976205 |
| QVEVAKRMGVRQPTVS | ASSLEGSSYEQY | 0.64051926 |
| QGSGGPALAM | ASSLEGSSYEQY | 0.3607441 |
| MLAPSNP | ASSLEGSSYEQY | 0.6629832 |
| LVNNAAI | ASSLEGSSYEQY | 0.6352879 |
| LTGEINTLA | ASSLEGSSYEQY | 0.6244232 |
| LSIFGPV | ASSLEGSSYEQY | 0.66897 |
| LLALLANL | ASSLEGSSYEQY | 0.62921053 |
| LISNASDALDKLR | ASSLEGSSYEQY | 0.5801146 |
| LINEAVP | ASSLEGSSYEQY | 0.54920334 |
| LGYAFSGL | ASSLEGSSYEQY | 0.6729721 |

| | | |
|---|---|---|
| LDFVAVAL | ASSLEGSSYEQY | 0.52486 |
| LAQPGAE | ASSLEGSSYEQY | 0.6712211 |
| ISNLGAVTGF | ASSLEGSSYEQY | 0.4918077 |
| HQEDPGRI | ASSLEGSSYEQY | 0.67057806 |
| GTPGEPTP | ASSLEGSSYEQY | 0.5682971 |
| GFFNSGS | ASSLEGSSYEQY | 0.6684933 |
| GAGGGGLTAGAN | ASSLEGSSYEQY | 0.63964033 |
| FVDALVNM | ASSLEGSSYEQY | 0.68404347 |
| FLTVAPEL | ASSLEGSSYEQY | 0.6318979 |
| FETLTAL | ASSLEGSSYEQY | 0.6603422 |
| DVINPIEL | ASSLEGSSYEQY | 0.6225199 |
| APVSGAPV | ASSLEGSSYEQY | 0.6785193 |
| ALVGGLAW | ASSLEGSSYEQY | 0.5327505 |
| ALAAAAATAI | ASSLEGSSYEQY | 0.634591 |
| AFGLGLLL | ASSLEGSSYEQY | 0.51213974 |
| AFFAAASL | ASSLEGSSYEQY | 0.64256185 |
| AAAALPAFSPP | ASSLEGSSYEQY | 0.66904914 |
| VVLAPAL | ASSLEGSSYEQY | 0.4416244 |
| TIGHVDHGKTTL | ASSLEGSSYEQY | 0.65791595 |
| LISNASDALDK | ASSLEGSSYEQY | 0.6029277 |
| LISNASDAL | ASSLEGSSYEQY | 0.6062282 |
| ISNASDALDK | ASSLEGSSYEQY | 0.66427237 |
| NQAETLVYQTE | ASSLEGSSYEQY | 0.64933074 |
| TGGTGGTAGAG | ASSLEGSSYEQY | 0.6663817 |
| TGSTAGGAGGA | ASSLEGSSYEQY | 0.57490146 |
| DPLAVPVAAGPV | ASSLEGSSYEQY | 0.6094 |
| GGTGGTPGNGGN | ASSLEGSSYEQY | 0.65010357 |
| PAGLAGVDPATL | ASSLEGSSYEQY | 0.57787675 |
| GNNLVVEAPHAGASV | ASSLEGSSYEQY | 0.57358325 |
| KVTRIMSASTLEQPA | ASSLEGSSYEQY | 0.67601293 |
| LGFALPIALPRRTVV | ASSLEGSSYEQY | 0.6240363 |
| TRLGNLTARKLVVLV | ASSLEGSSYEQY | 0.64047796 |
| WSQQRRLIYFPSAGP | ASSLEGSSYEQY | 0.56705713 |
| GPNGNPDPMAAAVDIR | ASSLEGSSYEQY | 0.554867 |
| TQRCVSVADLRAAASR | ASSLEGSSYEQY | 0.5884847 |
| QFVLDVAEHPVEVLTETV | ASSLEGSSYEQY | 0.6451005 |
| GGAGGTGGAGGNGGTQAGDG | ASSLEGSSYEQY | 0.64110297 |
| PLLATLRAIAVDVDAKPAQV | ASSLEGSSYEQY | 0.61787826 |
| APKFRTGGYMIDFWGVGYQV | ASSLEGSSYEQY | 0.5454469 |
| GAGGAGGAGGAAGGPGGTGG | ASSLEGSSYEQY | 0.61962694 |
| IGALGLLATDVAPAGSVAFG | ASSLEGSSYEQY | 0.6106169 |
| AGGAGGIGGNGGAGGAGGVL | ASSLEGSSYEQY | 0.5857735 |
| ADELIGLL | ASSLEGSSYEQY | 0.63353753 |
| ADRALKA | ASSLEGSSYEQY | 0.62269235 |
| AGAVGQVGGAGGA | ASSLEGSSYEQY | 0.5801691 |
| AGGGGGLGGAP | ASSLEGSSYEQY | 0.59041476 |
| AGGLLYGNGGAG | ASSLEGSSYEQY | 0.61095655 |
| AGLESIGL | ASSLEGSSYEQY | 0.63715464 |
| AGLKTGII | ASSLEGSSYEQY | 0.6356119 |
| ASLAALLL | ASSLEGSSYEQY | 0.65622646 |
| ASVINISSI | ASSLEGSSYEQY | 0.6402879 |

| | | |
|---|---|---|
| ATPPGPA | ASSLEGSSYEQY | 0.62623703 |
| ATVASALV | ASSLEGSSYEQY | 0.38962904 |
| ATVASLIL | ASSLEGSSYEQY | 0.44593096 |
| AVNGPVLIRH | ASSLEGSSYEQY | 0.5909265 |
| DGAAGAVGI | ASSLEGSSYEQY | 0.60457945 |
| DPLLHGI | ASSLEGSSYEQY | 0.47563475 |
| ELGAALGAL | ASSLEGSSYEQY | 0.6366458 |
| ELGRTQAV | ASSLEGSSYEQY | 0.59338087 |
| FVAGGYLS | ASSLEGSSYEQY | 0.57316005 |
| GASGGAGGAGGW | ASSLEGSSYEQY | 0.6375945 |
| GGAGAGGDGGAGGI | ASSLEGSSYEQY | 0.5762016 |
| GTGHNGSQPGMGGNG | ASSLEGSSYEQY | 0.58887815 |
| GYAAASASAS | ASSLEGSSYEQY | 0.5846813 |
| HQVDVAAVQQG | ASSLEGSSYEQY | 0.6430709 |
| IGAAIAHT | ASSLEGSSYEQY | 0.54364485 |
| IGMIEVAQL | ASSLEGSSYEQY | 0.6324011 |
| IRLAPEHTSAMD | ASSLEGSSYEQY | 0.54501545 |
| IYNVSVL | ASSLEGSSYEQY | 0.6484166 |
| KQGFLGM | ASSLEGSSYEQY | 0.52015483 |
| LESVITQL | ASSLEGSSYEQY | 0.62848324 |
| LGLGALVVV | ASSLEGSSYEQY | 0.589695 |
| LIGNGGNGGSGGI | ASSLEGSSYEQY | 0.6571611 |
| LNAINAPTL | ASSLEGSSYEQY | 0.6119645 |
| LQLLNTGV | ASSLEGSSYEQY | 0.63225406 |
| LQTSIEF | ASSLEGSSYEQY | 0.5894487 |
| MSFAVSEL | ASSLEGSSYEQY | 0.6274494 |
| NTGALGL | ASSLEGSSYEQY | 0.5188415 |
| PPGAGGAGGAGTT | ASSLEGSSYEQY | 0.6115277 |
| QLVGLGI | ASSLEGSSYEQY | 0.51794785 |
| RFLEQVHQL | ASSLEGSSYEQY | 0.6208925 |
| RGVGYVLR | ASSLEGSSYEQY | 0.61611444 |
| SATGVFAGVI | ASSLEGSSYEQY | 0.54005307 |
| SLPAITLP | ASSLEGSSYEQY | 0.6307268 |
| STALAFTV | ASSLEGSSYEQY | 0.6127994 |
| SVLTVGLGK | ASSLEGSSYEQY | 0.5762979 |
| TALAGAGSGI | ASSLEGSSYEQY | 0.64464384 |
| TGLPFTGI | ASSLEGSSYEQY | 0.6129761 |
| VAEHVTV | ASSLEGSSYEQY | 0.63354486 |
| VDIVLNSL | ASSLEGSSYEQY | 0.5887003 |
| VLTGGGAVL | ASSLEGSSYEQY | 0.5603953 |
| VNVGLAYGI | ASSLEGSSYEQY | 0.62746036 |
| VSSAAVGAV | ASSLEGSSYEQY | 0.58801484 |
| VTSVYEAGV | ASSLEGSSYEQY | 0.58275753 |
| WVALDRGA | ASSLEGSSYEQY | 0.6360308 |
| YLVGSVVI | ASSLEGSSYEQY | 0.8259618 |
| GPPGTGKTLL | ASSLEGSSYEQY | 0.8161638 |
| GVGMTKF | ASSLEGSSYEQY | 0.6490574 |
| IGIGGSDL | ASSLEGSSYEQY | 0.605734 |
| IGTLLAEL | ASSLEGSSYEQY | 0.6998174 |
| IYTSGTT | ASSLEGSSYEQY | 0.8369626 |
| LSFPAGV | ASSLEGSSYEQY | 0.7974782 |

| | | |
|---|---|---|
| LYTSGTT | ASSLEGSSYEQY | 0.80749446 |
| VGAFPFGF | ASSLEGSSYEQY | 0.54690534 |
| VGAGTVEF | ASSLEGSSYEQY | 0.73929703 |
| VGKSTLLL | ASSLEGSSYEQY | 0.724452 |
| AALAHAGGGIV | ASSLEGSSYEQY | 0.725253 |
| AALVAVSLL | ASSLEGSSYEQY | 0.80782604 |
| AAVSRAVAAL | ASSLEGSSYEQY | 0.7115915 |
| AGGSSGIGGF | ASSLEGSSYEQY | 0.7260536 |
| AGLTVANANG | ASSLEGSSYEQY | 0.6875653 |
| AGLVGAVL | ASSLEGSSYEQY | 0.7488916 |
| ALSQAFTV | ASSLEGSSYEQY | 0.8077034 |
| ALTFSVSL | ASSLEGSSYEQY | 0.79180884 |
| ALTGAALVAL | ASSLEGSSYEQY | 0.86336696 |
| ASFSIGAGVL | ASSLEGSSYEQY | 0.82598567 |
| DGSVYKI | ASSLEGSSYEQY | 0.81571585 |
| EQLAEIV | ASSLEGSSYEQY | 0.20188843 |
| FAVSELTVM | ASSLEGSSYEQY | 0.43791318 |
| GAAPLAAGL | ASSLEGSSYEQY | 0.6698458 |
| GADGGDGGSGGAGG | ASSLEGSSYEQY | 0.6678585 |
| GFGLDTGGAGG | ASSLEGSSYEQY | 0.3365206 |
| GLASGGLL | ASSLEGSSYEQY | 0.7895744 |
| GRIAGAAVLL | ASSLEGSSYEQY | 0.7920893 |
| GSAGSLNTSF | ASSLEGSSYEQY | 0.56763995 |
| GTGQIGI | ASSLEGSSYEQY | 0.8178123 |
| GTLIALIL | ASSLEGSSYEQY | 0.6630906 |
| IAATMGFVVG | ASSLEGSSYEQY | 0.6333694 |
| IFAQALL | ASSLEGSSYEQY | 0.65242386 |
| IGAAVALGF | ASSLEGSSYEQY | 0.8303557 |
| IGLTPLAL | ASSLEGSSYEQY | 0.5960307 |
| ILGSNTSTL | ASSLEGSSYEQY | 0.8106732 |
| IVFSLTVL | ASSLEGSSYEQY | 0.6721468 |
| LASKATGF | ASSLEGSSYEQY | 0.8449408 |
| LASVALIL | ASSLEGSSYEQY | 0.5133722 |
| LFAVGLTL | ASSLEGSSYEQY | 0.81320745 |
| LFNGGIGTYI | ASSLEGSSYEQY | 0.7329895 |
| LTLGAGII | ASSLEGSSYEQY | 0.8717099 |
| LVAAALGDI | ASSLEGSSYEQY | 0.7117355 |
| NTAFVQLGI | ASSLEGSSYEQY | 0.71861404 |
| QLFSGAEL | ASSLEGSSYEQY | 0.5810812 |
| SAATFITL | ASSLEGSSYEQY | 0.76385266 |
| SNYGLANLGV | ASSLEGSSYEQY | 0.39646286 |
| SYGGAYAVM | ASSLEGSSYEQY | 0.7304627 |
| TTGVGGLTLG | ASSLEGSSYEQY | 0.54600084 |
| TTQTIYI | ASSLEGSSYEQY | 0.7559173 |
| VAAGAGLGAVI | ASSLEGSSYEQY | 0.8065322 |
| VAAVVTLL | ASSLEGSSYEQY | 0.3638129 |
| VGFLGGLI | ASSLEGSSYEQY | 0.7873777 |
| VGVIPILL | ASSLEGSSYEQY | 0.668756 |
| VIFASSLI | ASSLEGSSYEQY | 0.7031888 |
| VKGGDGGAAAG | ASSLEGSSYEQY | 0.8275673 |
| VTKTFTNI | ASSLEGSSYEQY | 0.66015255 |

| | | |
|---|---|---|
| VVAAAILL | ASSLEGSSYEQY | 0.83475363 |
| YAALDVEL | ASSLEGSSYEQY | 0.6623126 |
| LASVALLL | ASSLEGSSYEQY | 0.6954243 |
| LDAVHRV | ASSLEGSSYEQY | 0.8063022 |
| QGLSSSIF | ASSLEGSSYEQY | 0.3311783 |
| RGTFANIRL | ASSLEGSSYEQY | 0.5540809 |
| RQLWWGHRI | ASSLEGSSYEQY | 0.8419863 |
| YGPPGTGKTLL | ASSLEGSSYEQY | 0.70931506 |
| AALVGAPANVA | ASSLEGSSYEQY | 0.7222361 |
| AATFRRTE | ASSLEGSSYEQY | 0.60108775 |
| AGALGGGI | ASSLEGSSYEQY | 0.5566684 |
| AGGAGGAGGIGAG | ASSLEGSSYEQY | 0.65806735 |
| AGSGALGL | ASSLEGSSYEQY | 0.71948266 |
| AQIADQI | ASSLEGSSYEQY | 0.71695274 |
| ASAVAVTYL | ASSLEGSSYEQY | 0.7104775 |
| ASTVLGGTGG | ASSLEGSSYEQY | 0.5427189 |
| AVGGEPAK | ASSLEGSSYEQY | 0.6681348 |
| CEQGTFGGTC | ASSLEGSSYEQY | 0.65393806 |
| DEGKAPAG | ASSLEGSSYEQY | 0.6462669 |
| DTVVFPAL | ASSLEGSSYEQY | 0.7006246 |
| FEGITMAG | ASSLEGSSYEQY | 0.59395623 |
| FPLVVQD | ASSLEGSSYEQY | 0.65610677 |
| GAFTTPQ | ASSLEGSSYEQY | 0.6301261 |
| GFGGFGTAGGVGGA | ASSLEGSSYEQY | 0.6935338 |
| GFLAQFL | ASSLEGSSYEQY | 0.7036106 |
| GGAGGAGGNAGVA | ASSLEGSSYEQY | 0.695153 |
| GGVGTAGAGGAGGAGG | ASSLEGSSYEQY | 0.72526807 |
| GNGGGGDAGNAGSGGN | ASSLEGSSYEQY | 0.7128497 |
| GTGGAGGAGNDAGS | ASSLEGSSYEQY | 0.70904267 |
| GTTNTGWFNS | ASSLEGSSYEQY | 0.29939473 |
| IFMGPLFTL | ASSLEGSSYEQY | 0.44461426 |
| ILAHAIEAI | ASSLEGSSYEQY | 0.6400287 |
| IRSAIREAL | ASSLEGSSYEQY | 0.6212431 |
| ITFAGGG | ASSLEGSSYEQY | 0.49150127 |
| ITGMSGHQEV | ASSLEGSSYEQY | 0.6847235 |
| IVAGLACTL | ASSLEGSSYEQY | 0.69195735 |
| IVGGGLAGA | ASSLEGSSYEQY | 0.5703325 |
| IVTGVLNI | ASSLEGSSYEQY | 0.69564867 |
| LAQQGCI | ASSLEGSSYEQY | 0.5990815 |
| LCLHVEN | ASSLEGSSYEQY | 0.5604688 |
| LDAISDGS | ASSLEGSSYEQY | 0.5816257 |
| LGGFAAML | ASSLEGSSYEQY | 0.71484685 |
| LGGKNPMI | ASSLEGSSYEQY | 0.5015965 |
| LGPKGVAAIL | ASSLEGSSYEQY | 0.67934835 |
| LMVGVSI | ASSLEGSSYEQY | 0.68901885 |
| LVGAAFVL | ASSLEGSSYEQY | 0.71183425 |
| LVLAVAPL | ASSLEGSSYEQY | 0.4354691 |
| MAAIAVIDAA | ASSLEGSSYEQY | 0.7053871 |
| NNGINAELQ | ASSLEGSSYEQY | 0.6440845 |
| RGEPLVLVHG | ASSLEGSSYEQY | 0.7215553 |
| RGGAGGAAGLI | ASSLEGSSYEQY | 0.5978613 |

| | | |
|---|---|---|
| RSYRMHPAV | ASSLEGSSYEQY | 0.6342364 |
| SAAHLLAGQLA | ASSLEGSSYEQY | 0.60799944 |
| SAAVGSY | ASSLEGSSYEQY | 0.70996594 |
| SPHAVLHRI | ASSLEGSSYEQY | 0.40164274 |
| SQIAKVL | ASSLEGSSYEQY | 0.6710683 |
| TALGHLVP | ASSLEGSSYEQY | 0.6389636 |
| TAPTTSGAA | ASSLEGSSYEQY | 0.68925524 |
| TDGDDGNGGSPGT | ASSLEGSSYEQY | 0.70730776 |
| TDGLGEN | ASSLEGSSYEQY | 0.50580597 |
| TGGPPLPGQA | ASSLEGSSYEQY | 0.7158979 |
| TIPAITI | ASSLEGSSYEQY | 0.66597366 |
| TIPGFTI | ASSLEGSSYEQY | 0.6040354 |
| TSIASGALA | ASSLEGSSYEQY | 0.70910543 |
| TVTLPGAAI | ASSLEGSSYEQY | 0.62810135 |
| VASGLSAALP | ASSLEGSSYEQY | 0.7194401 |
| VGAGVIGI | ASSLEGSSYEQY | 0.6613872 |
| VGPKRFGV | ASSLEGSSYEQY | 0.6954044 |
| VNVAFPD | ASSLEGSSYEQY | 0.7040782 |
| VSMTNSGVS | ASSLEGSSYEQY | 0.5954162 |
| VTDVLGDLSV | ASSLEGSSYEQY | 0.5517366 |
| VVVVGAGFAGL | ASSLEGSSYEQY | 0.7131957 |
| VYTVGTVV | ASSLEGSSYEQY | 0.75791377 |
| WTTFSGP | ASSLEGSSYEQY | 0.8301445 |
| YDELVEAF | ASSLEGSSYEQY | 0.4605295 |
| YGTGGAGGNGG | ASSLEGSSYEQY | 0.4390933 |
| YTNVMAV | ASSLEGSSYEQY | 0.6604688 |
| FLEQVHQL | ASSLEGSSYEQY | 0.8110375 |
| GLSLLAVAV | ASSLEGSSYEQY | 0.8283931 |
| LEQVHQL | ASSLEGSSYEQY | 0.7859501 |
| LNAAGAI | ASSLEGSSYEQY | 0.38144535 |
| PAVHASL | ASSLEGSSYEQY | 0.6713631 |
| TELEDDA | ASSLEGSSYEQY | 0.52789223 |
| TPPGPPN | ASSLEGSSYEQY | 0.6740843 |
| TPPPPPPP | ASSLEGSSYEQY | 0.7525892 |
| TQSPALI | TCSALGISGNTLY | 0.67448825 |
| EFAAAFPQ | TCSALGISGNTLY | 0.5692072 |
| LLSGNLDVL | TCSALGISGNTLY | 0.5004093 |
| GPANLGNYNI | TCSALGISGNTLY | 0.78376025 |
| TLGPTLP | TCSALGISGNTLY | 0.8073205 |
| VYAYSSCCK | TCSALGISGNTLY | 0.7828656 |
| APAPAPAPNI | TCSALGISGNTLY | 0.8413899 |
| PGGPVPIP | TCSALGISGNTLY | 0.7889291 |
| FLTVALEP | TCSALGISGNTLY | 0.7632724 |
| SQYFGANL | TCSALGISGNTLY | 0.1655879 |
| GGVKPAA | TCSALGISGNTLY | 0.2831906 |
| GGTGGAGGAGGW | TCSALGISGNTLY | 0.4867212 |
| GVGGVAGLGGAGG | TCSALGISGNTLY | 0.6202153 |
| VLLVLATTI | TCSALGISGNTLY | 0.22469765 |
| IGIFAAML | TCSALGISGNTLY | 0.74649024 |
| AVGDLID | TCSALGISGNTLY | 0.5080895 |
| NISPLPL | TCSALGISGNTLY | 0.45805776 |

| | | |
|---|---|---|
| VTAVFVAI | TCSALGISGNTLY | 0.7972959 |
| SGDDNDAAL | TCSALGISGNTLY | 0.49169248 |
| AVAALAHS | TCSALGISGNTLY | 0.46663415 |
| GGVTVMATP | TCSALGISGNTLY | 0.54094565 |
| AGLAANVRD | TCSALGISGNTLY | 0.7852779 |
| TNRPDLIDPALLRPG | TCSALGISGNTLY | 0.36128134 |
| LANLGVTAL | TCSALGISGNTLY | 0.75962734 |
| TGERIEL | TCSALGISGNTLY | 0.6743243 |
| NGEELAA | TCSALGISGNTLY | 0.818434 |
| VLSLVWG | TCSALGISGNTLY | 0.3343578 |
| IGYALWL | TCSALGISGNTLY | 0.7811265 |
| LSLLLAPL | TCSALGISGNTLY | 0.6808964 |
| TLDEALK | TCSALGISGNTLY | 0.8523028 |
| TDGAGTAEGG | TCSALGISGNTLY | 0.5403815 |
| NAPGGAGGNGG | TCSALGISGNTLY | 0.64374137 |
| VSVFMGMGGNF | TCSALGISGNTLY | 0.5240553 |
| LASYLAAL | TCSALGISGNTLY | 0.81879115 |
| LVGGTPAL | TCSALGISGNTLY | 0.28114837 |
| VRNADVL | TCSALGISGNTLY | 0.59921956 |
| TCSSPFLVSAM | TCSALGISGNTLY | 0.35768017 |
| VHITDAE | TCSALGISGNTLY | 0.6463824 |
| EYAAPSF | TCSALGISGNTLY | 0.772606 |
| VVDAFHGGL | TCSALGISGNTLY | 0.21074504 |
| ASVVDPQ | TCSALGISGNTLY | 0.77013505 |
| SYNEFGGLS | TCSALGISGNTLY | 0.57625437 |
| DPDGSVY | TCSALGISGNTLY | 0.43369496 |
| MVSTIHATE | TCSALGISGNTLY | 0.8070208 |
| LAAGVADPV | TCSALGISGNTLY | 0.59391433 |
| TKEQIAA | TCSALGISGNTLY | 0.81667113 |
| SLASPAAL | TCSALGISGNTLY | 0.6720547 |
| IGLVFYGV | TCSALGISGNTLY | 0.64582825 |
| FLLDASNL | TCSALGISGNTLY | 0.738799 |
| YQVGNFP | TCSALGISGNTLY | 0.29542583 |
| ESLLSQST | TCSALGISGNTLY | 0.49130565 |
| LFTVGFAAVG | TCSALGISGNTLY | 0.8178297 |
| ATPPTPT | TCSALGISGNTLY | 0.6890776 |
| TDLATLG | TCSALGISGNTLY | 0.6990124 |
| SAGGTGR | TCSALGISGNTLY | 0.62850416 |
| GFSNSGS | TCSALGISGNTLY | 0.6184245 |
| GGNGGSGGTSV | TCSALGISGNTLY | 0.648644 |
| GTSVATGGAGN | TCSALGISGNTLY | 0.68651915 |
| NGGNGGSGGTS | TCSALGISGNTLY | 0.70993984 |
| AVVKFPH | TCSALGISGNTLY | 0.6960666 |
| TPPPPPP | TCSALGISGNTLY | 0.5908926 |
| AAPAAAAAPA | TCSALGISGNTLY | 0.6942397 |
| IALNEDL | TCSALGISGNTLY | 0.66762495 |
| IVNVSSM | TCSALGISGNTLY | 0.6847892 |
| TGIPPAPR | TCSALGISGNTLY | 0.6731077 |
| FDLGGGTF | TCSALGISGNTLY | 0.6378038 |
| FDLGGGTFDVS | TCSALGISGNTLY | 0.6217258 |
| NEPTAAA | TCSALGISGNTLY | 0.57216483 |

| | | |
|---|---|---|
| NEPTAAALA | TCSALGISGNTLY | 0.66300046 |
| GSIINISSI | TCSALGISGNTLY | 0.6954836 |
| VAPGNAGTA | TCSALGISGNTLY | 0.6857538 |
| TGGGASLE | TCSALGISGNTLY | 0.7082125 |
| VAVYNHYKRI | TCSALGISGNTLY | 0.69857883 |
| TDPAPATT | TCSALGISGNTLY | 0.6733433 |
| TNRPDILDPALLRPG | TCSALGISGNTLY | 0.33793217 |
| LTTTTTS | TCSALGISGNTLY | 0.5573358 |
| PTGSGKTL | TCSALGISGNTLY | 0.51907754 |
| TGSGKTL | TCSALGISGNTLY | 0.6646896 |
| TIGHVDHGK | TCSALGISGNTLY | 0.2980296 |
| TIGHVDHGKTT | TCSALGISGNTLY | 0.6693046 |
| GHVDHGKTT | TCSALGISGNTLY | 0.66272247 |
| AQFEHTLL | TCSALGISGNTLY | 0.6212652 |
| IVNQDEL | TCSALGISGNTLY | 0.6977482 |
| TNGSQFF | TCSALGISGNTLY | 0.656523 |
| TGKGIVE | TCSALGISGNTLY | 0.5318545 |
| LISNASDA | TCSALGISGNTLY | 0.5362233 |
| AELRTLGELTKIAKAHGAQ | TCSALGISGNTLY | 0.71319926 |
| APNAVAAAEPSW | TCSALGISGNTLY | 0.53091407 |
| APQVAGAATAVL | TCSALGISGNTLY | 0.7097037 |
| ARRDPAAAAAILPTDA | TCSALGISGNTLY | 0.57870483 |
| DAAARDVLAEAGLAEYF | TCSALGISGNTLY | 0.70448756 |
| DDQIVLSAKANLAKVEQ | TCSALGISGNTLY | 0.5193489 |
| DELLPDTGEDSDEDGH | TCSALGISGNTLY | 0.698614 |
| ELPEAPANRATVPQD | TCSALGISGNTLY | 0.6771953 |
| FGQSKIERVTGIG | TCSALGISGNTLY | 0.7124501 |
| GAGGTGSTAGGAGGAGGAGGI | TCSALGISGNTLY | 0.6463524 |
| GHGGTGAAGGNGGTGGAGGS( | TCSALGISGNTLY | 0.69202447 |
| GNGGNGGAGGEGGPGGAGGA( | TCSALGISGNTLY | 0.5748162 |
| GSTAGNGGNGGNATNGTIAG( | TCSALGISGNTLY | 0.71020895 |
| GTGGDGGNGGNGGTGG | TCSALGISGNTLY | 0.5300108 |
| GVRRPAGIPADAPV | TCSALGISGNTLY | 0.65668535 |
| GYRQVTDAHLLTLARR | TCSALGISGNTLY | 0.5955026 |
| IAKMGQDGHDRG | TCSALGISGNTLY | 0.6707736 |
| KAEPSIEIQYT | TCSALGISGNTLY | 0.6981081 |
| KPVRIRMAADEDTVAVT | TCSALGISGNTLY | 0.3380031 |
| MPPRIAGMRLLVIKPEP | TCSALGISGNTLY | 0.7048032 |
| MKFYAVALFRSRGVPPTR | TCSALGISGNTLY | 0.61233264 |
| NAGGFGILGAVAHSP | TCSALGISGNTLY | 0.57467467 |
| NEGREITLAMAEM | TCSALGISGNTLY | 0.69321144 |
| NELVEWSHPLGHRWRWE | TCSALGISGNTLY | 0.62245506 |
| NLGAGGTGGNGGNPG | TCSALGISGNTLY | 0.68871886 |
| NTEWPKWETVMAADPVD | TCSALGISGNTLY | 0.53481483 |
| PASAAPAETQP | TCSALGISGNTLY | 0.68000805 |
| QQSSDGKNGVRAAEITGE | TCSALGISGNTLY | 0.6932076 |
| RAPDAVTTSWWKE | TCSALGISGNTLY | 0.38641226 |
| SGFGNVGAHTSGWFNQ | TCSALGISGNTLY | 0.42237556 |
| SGVAGAVGQAASVGGLKV | TCSALGISGNTLY | 0.7086135 |
| SNGGAGTGSAGGNGGT | TCSALGISGNTLY | 0.8490422 |
| TGGAGGEGGAAG | TCSALGISGNTLY | 0.8441926 |

| | | |
|---|---|---|
| TGGDGGAAGAGGAPGHG | TCSALGISGNTLY | 0.7507941 |
| TMPPLLAAIANAGRHGV | TCSALGISGNTLY | 0.71996117 |
| TSSTAGNSGGA | TCSALGISGNTLY | 0.79173064 |
| VPATVPVGILC | TCSALGISGNTLY | 0.8727806 |
| VQRPDSGGVVLTGRIS | TCSALGISGNTLY | 0.8504261 |
| WTPNPVPATKGVPAGLVQAT | TCSALGISGNTLY | 0.8447359 |
| YALKRALAKFSDRFDVVII | TCSALGISGNTLY | 0.71859425 |
| YLIGSVIV | TCSALGISGNTLY | 0.86763954 |
| YGQLVAVID | TCSALGISGNTLY | 0.78579855 |
| VTVGGMTLPA | TCSALGISGNTLY | 0.8034773 |
| VLALPAV | TCSALGISGNTLY | 0.8688279 |
| VIPPTLNYET | TCSALGISGNTLY | 0.8339709 |
| VGMGGMHPSSG | TCSALGISGNTLY | 0.81885874 |
| VGLAGGAATA | TCSALGISGNTLY | 0.8149046 |
| VGGAGGAGGSGP | TCSALGISGNTLY | 0.8488732 |
| VFAAFMV | TCSALGISGNTLY | 0.8637816 |
| TLYDKLR | TCSALGISGNTLY | 0.8410573 |
| TLAVFTVL | TCSALGISGNTLY | 0.9027633 |
| TGSVQPARV | TCSALGISGNTLY | 0.86029905 |
| TGGGAGSGGT | TCSALGISGNTLY | 0.88549924 |
| SHLETSL | TCSALGISGNTLY | 0.38190365 |
| SFTTGMT | TCSALGISGNTLY | 0.6744721 |
| QVEVAKRMGVRQPTVS | TCSALGISGNTLY | 0.84771717 |
| QGSGGPALAM | TCSALGISGNTLY | 0.8052671 |
| MLAPSNP | TCSALGISGNTLY | 0.55096745 |
| LVNNAAI | TCSALGISGNTLY | 0.8492981 |
| LTGEINTLA | TCSALGISGNTLY | 0.8119275 |
| LSIFGPV | TCSALGISGNTLY | 0.7312244 |
| LLALLANL | TCSALGISGNTLY | 0.87241495 |
| LISNASDALDKLR | TCSALGISGNTLY | 0.78893137 |
| LINEAVP | TCSALGISGNTLY | 0.8162115 |
| LGYAFSGL | TCSALGISGNTLY | 0.8061305 |
| LDFVAVAL | TCSALGISGNTLY | 0.88822424 |
| LAQPGAE | TCSALGISGNTLY | 0.7326386 |
| ISNLGAVTGF | TCSALGISGNTLY | 0.85583246 |
| HQEDPGRI | TCSALGISGNTLY | 0.8236504 |
| GTPGEPTP | TCSALGISGNTLY | 0.8816247 |
| GFFNSGS | TCSALGISGNTLY | 0.6380888 |
| GAGGGGLTAGAN | TCSALGISGNTLY | 0.86009604 |
| FVDALVNM | TCSALGISGNTLY | 0.78960073 |
| FLTVAPEL | TCSALGISGNTLY | 0.90257275 |
| FETLTAL | TCSALGISGNTLY | 0.80023646 |
| DVINPIEL | TCSALGISGNTLY | 0.8372762 |
| APVSGAPV | TCSALGISGNTLY | 0.84334266 |
| ALVGGLAW | TCSALGISGNTLY | 0.8458404 |
| ALAAAAATAI | TCSALGISGNTLY | 0.5928575 |
| AFGLGLLL | TCSALGISGNTLY | 0.8024607 |
| AFFAAASL | TCSALGISGNTLY | 0.7654747 |
| AAAALPAFSPP | TCSALGISGNTLY | 0.8470913 |
| VVLAPAL | TCSALGISGNTLY | 0.8431599 |
| TIGHVDHGKTTL | TCSALGISGNTLY | 0.4788466 |

| | | |
|---|---|---|
| LISNASDALDK | TCSALGISGNTLY | 0.87406313 |
| LISNASDAL | TCSALGISGNTLY | 0.7494582 |
| ISNASDALDK | TCSALGISGNTLY | 0.74660003 |
| NQAETLVYQTE | TCSALGISGNTLY | 0.8889121 |
| TGGTGGTAGAG | TCSALGISGNTLY | 0.84819347 |
| TGSTAGGAGGA | TCSALGISGNTLY | 0.8780869 |
| DPLAVPVAAGPV | TCSALGISGNTLY | 0.83238333 |
| GGTGGTPGNGGN | TCSALGISGNTLY | 0.8002378 |
| PAGLAGVDPATL | TCSALGISGNTLY | 0.86725855 |
| GNNLVVEAPHAGASV | TCSALGISGNTLY | 0.625387 |
| KVTRIMSASTLEQPA | TCSALGISGNTLY | 0.76099706 |
| LGFALPIALPRRTVV | TCSALGISGNTLY | 0.88320094 |
| TRLGNLTARKLVVLV | TCSALGISGNTLY | 0.88794136 |
| WSQQRRLIYFPSAGP | TCSALGISGNTLY | 0.9040675 |
| GPNGNPDPMAAAVDIR | TCSALGISGNTLY | 0.7951285 |
| TQRCVSVADLRAAASR | TCSALGISGNTLY | 0.7957024 |
| QFVLDVAEHPVEVLTETV | TCSALGISGNTLY | 0.8603012 |
| GGAGGTGGAGGNGGTQAGDG | TCSALGISGNTLY | 0.90310293 |
| PLLATLRAIAVDVDAKPAQV | TCSALGISGNTLY | 0.91451794 |
| APKFRTGGYMIDFWGVGYQV | TCSALGISGNTLY | 0.8978603 |
| GAGGAGGAGGAAGGPGGTGG | TCSALGISGNTLY | 0.8231739 |
| IGALGLLATDVAPAGSVAFG | TCSALGISGNTLY | 0.89107627 |
| AGGAGGIGGNGGAGGAGGVL | TCSALGISGNTLY | 0.86457956 |
| ADELIGLL | TCSALGISGNTLY | 0.86506927 |
| ADRALKA | TCSALGISGNTLY | 0.9008415 |
| AGAVGQVGGAGGA | TCSALGISGNTLY | 0.8731791 |
| AGGGGGLGGAP | TCSALGISGNTLY | 0.8353637 |
| AGGLLYGNGGAG | TCSALGISGNTLY | 0.8111174 |
| AGLESIGL | TCSALGISGNTLY | 0.891436 |
| AGLKTGII | TCSALGISGNTLY | 0.90100056 |
| ASLAALLL | TCSALGISGNTLY | 0.8967466 |
| ASVINISSI | TCSALGISGNTLY | 0.9165994 |
| ATPPGPA | TCSALGISGNTLY | 0.89857256 |
| ATVASALV | TCSALGISGNTLY | 0.893388 |
| ATVASLIL | TCSALGISGNTLY | 0.36865452 |
| AVNGPVLIRH | TCSALGISGNTLY | 0.74473023 |
| DGAAGAVGI | TCSALGISGNTLY | 0.8319268 |
| DPLLHGI | TCSALGISGNTLY | 0.8663162 |
| ELGAALGAL | TCSALGISGNTLY | 0.58571965 |
| ELGRTQAV | TCSALGISGNTLY | 0.8963752 |
| FVAGGYLS | TCSALGISGNTLY | 0.8562724 |
| GASGGAGGAGGW | TCSALGISGNTLY | 0.8107064 |
| GGAGAGGDGGAGGI | TCSALGISGNTLY | 0.90146214 |
| GTGHNGSQPGMGGNG | TCSALGISGNTLY | 0.8339 |
| GYAAASASAS | TCSALGISGNTLY | 0.7856175 |
| HQVDVAAVQQG | TCSALGISGNTLY | 0.85918033 |
| IGAAIAHT | TCSALGISGNTLY | 0.9129161 |
| IGMIEVAQL | TCSALGISGNTLY | 0.8171679 |
| IRLAPEHTSAMD | TCSALGISGNTLY | 0.90388346 |
| IYNVSVL | TCSALGISGNTLY | 0.8837478 |
| KQGFLGM | TCSALGISGNTLY | 0.90808266 |

| | | |
|---|---|---|
| LESVITQL | TCSALGISGNTLY | 0.70248413 |
| LGLGALVVV | TCSALGISGNTLY | 0.9002384 |
| LIGNGGNGGSGGI | TCSALGISGNTLY | 0.8640212 |
| LNAINAPTL | TCSALGISGNTLY | 0.9183649 |
| LQLLNTGV | TCSALGISGNTLY | 0.8619567 |
| LQTSIEF | TCSALGISGNTLY | 0.8974955 |
| MSFAVSEL | TCSALGISGNTLY | 0.88233876 |
| NTGALGL | TCSALGISGNTLY | 0.90523154 |
| PPGAGGAGGAGTT | TCSALGISGNTLY | 0.7137959 |
| QLVGLGI | TCSALGISGNTLY | 0.8513645 |
| RFLEQVHQL | TCSALGISGNTLY | 0.84110564 |
| RGVGYVLR | TCSALGISGNTLY | 0.87218654 |
| SATGVFAGVI | TCSALGISGNTLY | 0.8922887 |
| SLPAITLP | TCSALGISGNTLY | 0.3819977 |
| STALAFTV | TCSALGISGNTLY | 0.8992049 |
| SVLTVGLGK | TCSALGISGNTLY | 0.80259633 |
| TALAGAGSGI | TCSALGISGNTLY | 0.80172896 |
| TGLPFTGI | TCSALGISGNTLY | 0.90479434 |
| VAEHVTV | TCSALGISGNTLY | 0.86754024 |
| VDIVLNSL | TCSALGISGNTLY | 0.8933565 |
| VLTGGGAVL | TCSALGISGNTLY | 0.8703286 |
| VNVGLAYGI | TCSALGISGNTLY | 0.8723695 |
| VSSAAVGAV | TCSALGISGNTLY | 0.890916 |
| VTSVYEAGV | TCSALGISGNTLY | 0.61861706 |
| WVALDRGA | TCSALGISGNTLY | 0.821612 |
| YLVGSVVI | TCSALGISGNTLY | 0.90850544 |
| GPPGTGKTLL | TCSALGISGNTLY | 0.71306634 |
| GVGMTKF | TCSALGISGNTLY | 0.80200016 |
| IGIGGSDL | TCSALGISGNTLY | 0.26760522 |
| IGTLLAEL | TCSALGISGNTLY | 0.24173059 |
| IYTSGTT | TCSALGISGNTLY | 0.5272599 |
| LSFPAGV | TCSALGISGNTLY | 0.7245579 |
| LYTSGTT | TCSALGISGNTLY | 0.78445715 |
| VGAFPFGF | TCSALGISGNTLY | 0.7318877 |
| VGAGTVEF | TCSALGISGNTLY | 0.22019969 |
| VGKSTLLL | TCSALGISGNTLY | 0.5291601 |
| AALAHAGGGIV | TCSALGISGNTLY | 0.308136 |
| AALVAVSLL | TCSALGISGNTLY | 0.50466335 |
| AAVSRAVAAL | TCSALGISGNTLY | 0.5923339 |
| AGGSSGIGGF | TCSALGISGNTLY | 0.33754975 |
| AGLTVANANG | TCSALGISGNTLY | 0.306256 |
| AGLVGAVL | TCSALGISGNTLY | 0.24700849 |
| ALSQAFTV | TCSALGISGNTLY | 0.75431865 |
| ALTFSVSL | TCSALGISGNTLY | 0.78144175 |
| ALTGAALVAL | TCSALGISGNTLY | 0.753131 |
| ASFSIGAGVL | TCSALGISGNTLY | 0.80063343 |
| DGSVYKI | TCSALGISGNTLY | 0.7383926 |
| EQLAEIV | TCSALGISGNTLY | 0.7155815 |
| FAVSELTVM | TCSALGISGNTLY | 0.12308827 |
| GAAPLAAGL | TCSALGISGNTLY | 0.16184108 |
| GADGGDGGSGGAGG | TCSALGISGNTLY | 0.21603149 |

| | | |
|---|---|---|
| GFGLDTGGAGG | TCSALGISGNTLY | 0.48639333 |
| GLASGGLL | TCSALGISGNTLY | 0.1526405 |
| GRIAGAAVLL | TCSALGISGNTLY | 0.60046875 |
| GSAGSLNTSF | TCSALGISGNTLY | 0.3066104 |
| GTGQIGI | TCSALGISGNTLY | 0.22319527 |
| GTLIALIL | TCSALGISGNTLY | 0.7761216 |
| IAATMGFVVG | TCSALGISGNTLY | 0.27684003 |
| IFAQALL | TCSALGISGNTLY | 0.19708039 |
| IGAAVALGF | TCSALGISGNTLY | 0.25710085 |
| IGLTPLAL | TCSALGISGNTLY | 0.768471 |
| ILGSNTSTL | TCSALGISGNTLY | 0.17545608 |
| IVFSLTVL | TCSALGISGNTLY | 0.7084673 |
| LASKATGF | TCSALGISGNTLY | 0.48093098 |
| LASVALIL | TCSALGISGNTLY | 0.6739395 |
| LFAVGLTL | TCSALGISGNTLY | 0.16432202 |
| LFNGGIGTYI | TCSALGISGNTLY | 0.73649514 |
| LTLGAGII | TCSALGISGNTLY | 0.5651394 |
| LVAAALGDI | TCSALGISGNTLY | 0.83807135 |
| NTAFVQLGI | TCSALGISGNTLY | 0.3220495 |
| QLFSGAEL | TCSALGISGNTLY | 0.44006965 |
| SAATFITL | TCSALGISGNTLY | 0.3302714 |
| SNYGLANLGV | TCSALGISGNTLY | 0.74344885 |
| SYGGAYAVM | TCSALGISGNTLY | 0.15965937 |
| TTGVGGLTLG | TCSALGISGNTLY | 0.36754665 |
| TTQTIYI | TCSALGISGNTLY | 0.19423433 |
| VAAGAGLGAVI | TCSALGISGNTLY | 0.39625508 |
| VAAVVTLL | TCSALGISGNTLY | 0.720459 |
| VGFLGGLI | TCSALGISGNTLY | 0.13339864 |
| VGVIPILL | TCSALGISGNTLY | 0.66570556 |
| VIFASSLI | TCSALGISGNTLY | 0.5472287 |
| VKGGDGGAAAG | TCSALGISGNTLY | 0.21262442 |
| VTKTFTNI | TCSALGISGNTLY | 0.7195045 |
| VVAAAILL | TCSALGISGNTLY | 0.34403366 |
| YAALDVEL | TCSALGISGNTLY | 0.78225374 |
| LASVALLL | TCSALGISGNTLY | 0.41742045 |
| LDAVHRV | TCSALGISGNTLY | 0.5488627 |
| QGLSSSIF | TCSALGISGNTLY | 0.54753125 |
| RGTFANIRL | TCSALGISGNTLY | 0.12686251 |
| RQLWWGHRI | TCSALGISGNTLY | 0.21416438 |
| YGPPGTGKTLL | TCSALGISGNTLY | 0.78103876 |
| AALVGAPANVA | TCSALGISGNTLY | 0.70953065 |
| AATFRRTE | TCSALGISGNTLY | 0.8163595 |
| AGALGGGI | TCSALGISGNTLY | 0.5398284 |
| AGGAGGAGGIGAG | TCSALGISGNTLY | 0.45262164 |
| AGSGALGL | TCSALGISGNTLY | 0.60017174 |
| AQIADQI | TCSALGISGNTLY | 0.76828325 |
| ASAVAVTYL | TCSALGISGNTLY | 0.8146244 |
| ASTVLGGTGG | TCSALGISGNTLY | 0.73755795 |
| AVGGEPAK | TCSALGISGNTLY | 0.48902723 |
| CEQGTFGGTC | TCSALGISGNTLY | 0.65565896 |
| DEGKAPAG | TCSALGISGNTLY | 0.5968723 |

| | | |
|---|---|---|
| DTVVFPAL | TCSALGISGNTLY | 0.58763146 |
| FEGITMAG | TCSALGISGNTLY | 0.6864337 |
| FPLVVQD | TCSALGISGNTLY | 0.71298856 |
| GAFTTPQ | TCSALGISGNTLY | 0.5419451 |
| GFGGFGTAGGVGGA | TCSALGISGNTLY | 0.5362865 |
| GFLAQFL | TCSALGISGNTLY | 0.6789552 |
| GGAGGAGGNAGVA | TCSALGISGNTLY | 0.76622653 |
| GGVGTAGAGGAGGAGG | TCSALGISGNTLY | 0.7315036 |
| GNGGGGDAGNAGSGGN | TCSALGISGNTLY | 0.82899153 |
| GTGGAGGAGNDAGS | TCSALGISGNTLY | 0.77364135 |
| GTTNTGWFNS | TCSALGISGNTLY | 0.7558356 |
| IFMGPLFTL | TCSALGISGNTLY | 0.20173942 |
| ILAHAIEAI | TCSALGISGNTLY | 0.33948785 |
| IRSAIREAL | TCSALGISGNTLY | 0.5959834 |
| ITFAGGG | TCSALGISGNTLY | 0.62396944 |
| ITGMSGHQEV | TCSALGISGNTLY | 0.32230723 |
| IVAGLACTL | TCSALGISGNTLY | 0.72933435 |
| IVGGGLAGA | TCSALGISGNTLY | 0.6065748 |
| IVTGVLNI | TCSALGISGNTLY | 0.48378283 |
| LAQQGCI | TCSALGISGNTLY | 0.7685585 |
| LCLHVEN | TCSALGISGNTLY | 0.5281735 |
| LDAISDGS | TCSALGISGNTLY | 0.5100039 |
| LGGFAAML | TCSALGISGNTLY | 0.5651975 |
| LGGKNPMI | TCSALGISGNTLY | 0.7862369 |
| LGPKGVAAIL | TCSALGISGNTLY | 0.39555672 |
| LMVGVSI | TCSALGISGNTLY | 0.7352345 |
| LVGAAFVL | TCSALGISGNTLY | 0.6020781 |
| LVLAVAPL | TCSALGISGNTLY | 0.7951856 |
| MAAIAVIDAA | TCSALGISGNTLY | 0.3950452 |
| NNGINAELQ | TCSALGISGNTLY | 0.7605905 |
| RGEPLVLVHG | TCSALGISGNTLY | 0.60071826 |
| RGGAGGAAGLI | TCSALGISGNTLY | 0.857135 |
| RSYRMHPAV | TCSALGISGNTLY | 0.59603107 |
| SAAHLLAGQLA | TCSALGISGNTLY | 0.7010923 |
| SAAVGSY | TCSALGISGNTLY | 0.6177857 |
| SPHAVLHRI | TCSALGISGNTLY | 0.7731418 |
| SQIAKVL | TCSALGISGNTLY | 0.3582633 |
| TALGHLVP | TCSALGISGNTLY | 0.5844609 |
| TAPTTSGAA | TCSALGISGNTLY | 0.35963216 |
| TDGDDGNGGSPGT | TCSALGISGNTLY | 0.64392126 |
| TDGLGEN | TCSALGISGNTLY | 0.7384778 |
| TGGPPLPGQA | TCSALGISGNTLY | 0.31762195 |
| TIPAITI | TCSALGISGNTLY | 0.7300074 |
| TIPGFTI | TCSALGISGNTLY | 0.613108 |
| TSIASGALA | TCSALGISGNTLY | 0.5008322 |
| TVTLPGAAI | TCSALGISGNTLY | 0.789433 |
| VASGLSAALP | TCSALGISGNTLY | 0.63799894 |
| VGAGVIGI | TCSALGISGNTLY | 0.7864896 |
| VGPKRFGV | TCSALGISGNTLY | 0.6034565 |
| VNVAFPD | TCSALGISGNTLY | 0.60658956 |
| VSMTNSGVS | TCSALGISGNTLY | 0.7175772 |

| | | |
|---|---|---|
| VTDVLGDLSV | TCSALGISGNTLY | 0.53252286 |
| VVVVGAGFAGL | TCSALGISGNTLY | 0.5189885 |
| VYTVGTVV | TCSALGISGNTLY | 0.7942712 |
| WTTFSGP | TCSALGISGNTLY | 0.7544784 |
| YDELVEAF | TCSALGISGNTLY | 0.7627583 |
| YGTGGAGGNGG | TCSALGISGNTLY | 0.6538918 |
| YTNVMAV | TCSALGISGNTLY | 0.6521198 |
| FLEQVHQL | TCSALGISGNTLY | 0.68374074 |
| GLSLLAVAV | TCSALGISGNTLY | 0.764602 |
| LEQVHQL | TCSALGISGNTLY | 0.7741111 |
| LNAAGAI | TCSALGISGNTLY | 0.7559281 |
| PAVHASL | TCSALGISGNTLY | 0.68692416 |
| TELEDDA | TCSALGISGNTLY | 0.7476851 |
| TPPGPPN | TCSALGISGNTLY | 0.7067569 |
| TPPPPPPP | TCSALGISGNTLY | 0.69987124 |
| TQSPALI | ASSLEHPNTEVF | 0.7546431 |
| EFAAAFPQ | ASSLEHPNTEVF | 0.73822343 |
| LLSGNLDVL | ASSLEHPNTEVF | 0.6695824 |
| GPANLGNYNI | ASSLEHPNTEVF | 0.72953403 |
| TLGPTLP | ASSLEHPNTEVF | 0.7273512 |
| VYAYSSCCK | ASSLEHPNTEVF | 0.75659823 |
| APAPAPAPNI | ASSLEHPNTEVF | 0.7573931 |
| PGGPVPIP | ASSLEHPNTEVF | 0.78112864 |
| FLTVALEP | ASSLEHPNTEVF | 0.7652966 |
| SQYFGANL | ASSLEHPNTEVF | 0.75345075 |
| GGVKPAA | ASSLEHPNTEVF | 0.4067863 |
| GGTGGAGGAGGW | ASSLEHPNTEVF | 0.59851205 |
| GVGGVAGLGGAGG | ASSLEHPNTEVF | 0.72023225 |
| VLLVLATTI | ASSLEHPNTEVF | 0.7214023 |
| IGIFAAML | ASSLEHPNTEVF | 0.51361966 |
| AVGDLID | ASSLEHPNTEVF | 0.7475867 |
| NISPLPL | ASSLEHPNTEVF | 0.72457933 |
| VTAVFVAI | ASSLEHPNTEVF | 0.66317064 |
| SGDDNDAAL | ASSLEHPNTEVF | 0.7622093 |
| AVAALAHS | ASSLEHPNTEVF | 0.678647 |
| GGVTVMATP | ASSLEHPNTEVF | 0.6974895 |
| AGLAANVRD | ASSLEHPNTEVF | 0.7101941 |
| TNRPDLIDPALLRPG | ASSLEHPNTEVF | 0.7702579 |
| LANLGVTAL | ASSLEHPNTEVF | 0.6687498 |
| TGERIEL | ASSLEHPNTEVF | 0.76072097 |
| NGEELAA | ASSLEHPNTEVF | 0.6614878 |
| VLSLVWG | ASSLEHPNTEVF | 0.7709408 |
| IGYALWL | ASSLEHPNTEVF | 0.64882267 |
| LSLLLAPL | ASSLEHPNTEVF | 0.75522816 |
| TLDEALK | ASSLEHPNTEVF | 0.71169734 |
| TDGAGTAEGG | ASSLEHPNTEVF | 0.79219973 |
| NAPGGAGGNGG | ASSLEHPNTEVF | 0.7248858 |
| VSVFMGMGGNF | ASSLEHPNTEVF | 0.75802565 |
| LASYLAAL | ASSLEHPNTEVF | 0.7224198 |
| LVGGTPAL | ASSLEHPNTEVF | 0.76795137 |
| VRNADVL | ASSLEHPNTEVF | 0.5726261 |

| | | |
|---|---|---|
| TCSSPFLVSAM | ASSLEHPNTEVF | 0.728371 |
| VHITDAE | ASSLEHPNTEVF | 0.57747734 |
| EYAAPSF | ASSLEHPNTEVF | 0.74439585 |
| VVDAFHGGL | ASSLEHPNTEVF | 0.75367093 |
| ASVVDPQ | ASSLEHPNTEVF | 0.5319505 |
| SYNEFGGLS | ASSLEHPNTEVF | 0.7767266 |
| DPDGSVY | ASSLEHPNTEVF | 0.654986 |
| MVSTIHATE | ASSLEHPNTEVF | 0.7067758 |
| LAAGVADPV | ASSLEHPNTEVF | 0.77252495 |
| TKEQIAA | ASSLEHPNTEVF | 0.71001256 |
| SLASPAAL | ASSLEHPNTEVF | 0.77743435 |
| IGLVFYGV | ASSLEHPNTEVF | 0.7077459 |
| FLLDASNL | ASSLEHPNTEVF | 0.7254018 |
| YQVGNFP | ASSLEHPNTEVF | 0.75273496 |
| ESLLSQST | ASSLEHPNTEVF | 0.60308313 |
| LFTVGFAAVG | ASSLEHPNTEVF | 0.6714279 |
| ATPPTPT | ASSLEHPNTEVF | 0.77548075 |
| TDLATLG | ASSLEHPNTEVF | 0.6501505 |
| SAGGTGR | ASSLEHPNTEVF | 0.6487778 |
| GFSNSGS | ASSLEHPNTEVF | 0.5908096 |
| GGNGGSGGTSV | ASSLEHPNTEVF | 0.5436729 |
| GTSVATGGAGN | ASSLEHPNTEVF | 0.5804347 |
| NGGNGGSGGTS | ASSLEHPNTEVF | 0.6450441 |
| AVVKFPH | ASSLEHPNTEVF | 0.6588094 |
| TPPPPPP | ASSLEHPNTEVF | 0.643333 |
| AAPAAAAAPA | ASSLEHPNTEVF | 0.5305302 |
| IALNEDL | ASSLEHPNTEVF | 0.6460884 |
| IVNVSSM | ASSLEHPNTEVF | 0.62947917 |
| TGIPPAPR | ASSLEHPNTEVF | 0.6225965 |
| FDLGGGTF | ASSLEHPNTEVF | 0.6250595 |
| FDLGGGTFDVS | ASSLEHPNTEVF | 0.6529967 |
| NEPTAAA | ASSLEHPNTEVF | 0.6165066 |
| NEPTAAALA | ASSLEHPNTEVF | 0.50646454 |
| GSIINISSI | ASSLEHPNTEVF | 0.63687867 |
| VAPGNAGTA | ASSLEHPNTEVF | 0.65774554 |
| TGGGASLE | ASSLEHPNTEVF | 0.651071 |
| VAVYNHYKRI | ASSLEHPNTEVF | 0.6596842 |
| TDPAPATT | ASSLEHPNTEVF | 0.6569009 |
| TNRPDILDPALLRPG | ASSLEHPNTEVF | 0.6344882 |
| LTTTTTS | ASSLEHPNTEVF | 0.3659472 |
| PTGSGKTL | ASSLEHPNTEVF | 0.49204192 |
| TGSGKTL | ASSLEHPNTEVF | 0.55693734 |
| TIGHVDHGK | ASSLEHPNTEVF | 0.6076618 |
| TIGHVDHGKTT | ASSLEHPNTEVF | 0.36219287 |
| GHVDHGKTT | ASSLEHPNTEVF | 0.6332324 |
| AQFEHTLL | ASSLEHPNTEVF | 0.5994806 |
| IVNQDEL | ASSLEHPNTEVF | 0.6045153 |
| TNGSQFF | ASSLEHPNTEVF | 0.65435135 |
| TGKGIVE | ASSLEHPNTEVF | 0.6132136 |
| LISNASDA | ASSLEHPNTEVF | 0.55068934 |
| AELRTLGELTKIAKAHGAQ | ASSLEHPNTEVF | 0.5672715 |

| | | |
|---|---|---|
| APNAVAAAEPSW | ASSLEHPNTEVF | 0.6615084 |
| APQVAGAATAVL | ASSLEHPNTEVF | 0.5122048 |
| ARRDPAAAAAILPTDA | ASSLEHPNTEVF | 0.6564586 |
| DAAARDVLAEAGLAEYF | ASSLEHPNTEVF | 0.5702296 |
| DDQIVLSAKANLAKVEQ | ASSLEHPNTEVF | 0.6609505 |
| DELLPDTGEDSDEDGH | ASSLEHPNTEVF | 0.5090166 |
| ELPEAPANRATVPQD | ASSLEHPNTEVF | 0.65500236 |
| FGQSKIERVTGIG | ASSLEHPNTEVF | 0.6279779 |
| GAGGTGSTAGGAGGAGGAGGI | ASSLEHPNTEVF | 0.66882986 |
| GHGGTGAAGGNGGTGGAGGS( | ASSLEHPNTEVF | 0.6054972 |
| GNGGNGGAGGEGGPGGAGGA( | ASSLEHPNTEVF | 0.65098774 |
| GSTAGNGGNGGNATNGTIAG( | ASSLEHPNTEVF | 0.62080884 |
| GTGGDGGNGGNGGTGG | ASSLEHPNTEVF | 0.64876753 |
| GVRRPAGIPADAPV | ASSLEHPNTEVF | 0.5380174 |
| GYRQVTDAHLLTLARR | ASSLEHPNTEVF | 0.6052338 |
| IAKMGQDGHDRG | ASSLEHPNTEVF | 0.51324046 |
| KAEPSIEIQYT | ASSLEHPNTEVF | 0.6162042 |
| KPVRIRMAADEDTVAVT | ASSLEHPNTEVF | 0.6507776 |
| MPPRIAGMRLLVIKPEP | ASSLEHPNTEVF | 0.341928 |
| MKFYAVALFRSRGVPPTR | ASSLEHPNTEVF | 0.64875317 |
| NAGGFGILGAVAHSP | ASSLEHPNTEVF | 0.51124656 |
| NEGREITLAMAEM | ASSLEHPNTEVF | 0.5392126 |
| NELVEWSHPLGHRWRWE | ASSLEHPNTEVF | 0.6466259 |
| NLGAGGTGGNGGNPG | ASSLEHPNTEVF | 0.64451504 |
| NTEWPKWETVMAADPVD | ASSLEHPNTEVF | 0.6480605 |
| PASAAPAETQP | ASSLEHPNTEVF | 0.59329295 |
| QQSSDGKNGVRAAEITGE | ASSLEHPNTEVF | 0.6150607 |
| RAPDAVTTSWWKE | ASSLEHPNTEVF | 0.6327965 |
| SGFGNVGAHTSGWFNQ | ASSLEHPNTEVF | 0.4930222 |
| SGVAGAVGQAASVGGLKV | ASSLEHPNTEVF | 0.53103286 |
| SNGGAGTGSAGGNGGT | ASSLEHPNTEVF | 0.66007674 |
| TGGAGGEGGAAG | ASSLEHPNTEVF | 0.61778176 |
| TGGDGGAAGAGGAPGHG | ASSLEHPNTEVF | 0.7477107 |
| TMPPLLAAIANAGRHGV | ASSLEHPNTEVF | 0.4758304 |
| TSSTAGNSGGA | ASSLEHPNTEVF | 0.34969223 |
| VPATVPVGILC | ASSLEHPNTEVF | 0.4917567 |
| VQRPDSGGVVLTGRIS | ASSLEHPNTEVF | 0.7197052 |
| WTPNPVPATKGVPAGLVQAT | ASSLEHPNTEVF | 0.7517665 |
| YALKRALAKFSDRFDVVII | ASSLEHPNTEVF | 0.6697637 |
| YLIGSVIV | ASSLEHPNTEVF | 0.3779037 |
| YGQLVAVID | ASSLEHPNTEVF | 0.54157597 |
| VTVGGMTLPA | ASSLEHPNTEVF | 0.4656504 |
| VLALPAV | ASSLEHPNTEVF | 0.50786686 |
| VIPPTLNYET | ASSLEHPNTEVF | 0.591089 |
| VGMGGMHPSSG | ASSLEHPNTEVF | 0.592631 |
| VGLAGGAATA | ASSLEHPNTEVF | 0.51012266 |
| VGGAGGAGGSGP | ASSLEHPNTEVF | 0.407588 |
| VFAAFMV | ASSLEHPNTEVF | 0.54514647 |
| TLYDKLR | ASSLEHPNTEVF | 0.6592275 |
| TLAVFTVL | ASSLEHPNTEVF | 0.55909324 |
| TGSVQPARV | ASSLEHPNTEVF | 0.7547602 |

| | | |
|---|---|---|
| TGGGAGSGGT | ASSLEHPNTEVF | 0.6968855 |
| SHLETSL | ASSLEHPNTEVF | 0.63443017 |
| SFTTGMT | ASSLEHPNTEVF | 0.1951896 |
| QVEVAKRMGVRQPTVS | ASSLEHPNTEVF | 0.25425565 |
| QGSGGPALAM | ASSLEHPNTEVF | 0.47767514 |
| MLAPSNP | ASSLEHPNTEVF | 0.5006506 |
| LVNNAAI | ASSLEHPNTEVF | 0.2109731 |
| LTGEINTLA | ASSLEHPNTEVF | 0.64153564 |
| LSIFGPV | ASSLEHPNTEVF | 0.4676407 |
| LLALLANL | ASSLEHPNTEVF | 0.40423527 |
| LISNASDALDKLR | ASSLEHPNTEVF | 0.70004535 |
| LINEAVP | ASSLEHPNTEVF | 0.438967 |
| LGYAFSGL | ASSLEHPNTEVF | 0.41651422 |
| LDFVAVAL | ASSLEHPNTEVF | 0.44230926 |
| LAQPGAE | ASSLEHPNTEVF | 0.7025311 |
| ISNLGAVTGF | ASSLEHPNTEVF | 0.27863052 |
| HQEDPGRI | ASSLEHPNTEVF | 0.6439231 |
| GTPGEPTP | ASSLEHPNTEVF | 0.4978604 |
| GFFNSGS | ASSLEHPNTEVF | 0.6898191 |
| GAGGGGLTAGAN | ASSLEHPNTEVF | 0.299971 |
| FVDALVNM | ASSLEHPNTEVF | 0.696027 |
| FLTVAPEL | ASSLEHPNTEVF | 0.55485183 |
| FETLTAL | ASSLEHPNTEVF | 0.7701951 |
| DVINPIEL | ASSLEHPNTEVF | 0.4356236 |
| APVSGAPV | ASSLEHPNTEVF | 0.50491107 |
| ALVGGLAW | ASSLEHPNTEVF | 0.45843506 |
| ALAAAAATAI | ASSLEHPNTEVF | 0.73249614 |
| AFGLGLLL | ASSLEHPNTEVF | 0.268851 |
| AFFAAASL | ASSLEHPNTEVF | 0.4964801 |
| AAAALPAFSPP | ASSLEHPNTEVF | 0.27964053 |
| VVLAPAL | ASSLEHPNTEVF | 0.5964848 |
| TIGHVDHGKTTL | ASSLEHPNTEVF | 0.65461487 |
| LISNASDALDK | ASSLEHPNTEVF | 0.23114724 |
| LISNASDAL | ASSLEHPNTEVF | 0.6707559 |
| ISNASDALDK | ASSLEHPNTEVF | 0.469397 |
| NQAETLVYQTE | ASSLEHPNTEVF | 0.3841862 |
| TGGTGGTAGAG | ASSLEHPNTEVF | 0.70351464 |
| TGSTAGGAGGA | ASSLEHPNTEVF | 0.48693436 |
| DPLAVPVAAGPV | ASSLEHPNTEVF | 0.7276345 |
| GGTGGTPGNGGN | ASSLEHPNTEVF | 0.5029408 |
| PAGLAGVDPATL | ASSLEHPNTEVF | 0.56980914 |
| GNNLVVEAPHAGASV | ASSLEHPNTEVF | 0.61370814 |
| KVTRIMSASTLEQPA | ASSLEHPNTEVF | 0.310778 |
| LGFALPIALPRRTVV | ASSLEHPNTEVF | 0.43845314 |
| TRLGNLTARKLVVLV | ASSLEHPNTEVF | 0.71028787 |
| WSQQRRLIYFPSAGP | ASSLEHPNTEVF | 0.8633489 |
| GPNGNPDPMAAAVDIR | ASSLEHPNTEVF | 0.88851756 |
| TQRCVSVADLRAAASR | ASSLEHPNTEVF | 0.7944646 |
| QFVLDVAEHPVEVLTETV | ASSLEHPNTEVF | 0.74332577 |
| GGAGGTGGAGGNGGTQAGDG | ASSLEHPNTEVF | 0.80522066 |
| PLLATLRAIAVDVDAKPAQV | ASSLEHPNTEVF | 0.8835522 |

| | | |
|---|---|---|
| APKFRTGGYMIDFWGVGYQV/ | ASSLEHPNTEVF | 0.87434304 |
| GAGGAGGAGGAAGGPGGTGG | ASSLEHPNTEVF | 0.8611075 |
| IGALGLLATDVAPAGSVAFG | ASSLEHPNTEVF | 0.60450804 |
| AGGAGGIGGNGGAGGAGGVL | ASSLEHPNTEVF | 0.8103168 |
| ADELIGLL | ASSLEHPNTEVF | 0.77967864 |
| ADRALKA | ASSLEHPNTEVF | 0.7873806 |
| AGAVGQVGGAGGA | ASSLEHPNTEVF | 0.8615856 |
| AGGGGGLGGAP | ASSLEHPNTEVF | 0.83058214 |
| AGGLLYGNGGAG | ASSLEHPNTEVF | 0.769764 |
| AGLESIGL | ASSLEHPNTEVF | 0.7766438 |
| AGLKTGII | ASSLEHPNTEVF | 0.82475597 |
| ASLAALLL | ASSLEHPNTEVF | 0.8506037 |
| ASVINISSI | ASSLEHPNTEVF | 0.84454143 |
| ATPPGPA | ASSLEHPNTEVF | 0.9001321 |
| ATVASALV | ASSLEHPNTEVF | 0.8798444 |
| ATVASLIL | ASSLEHPNTEVF | 0.8824086 |
| AVNGPVLIRH | ASSLEHPNTEVF | 0.15482679 |
| DGAAGAVGI | ASSLEHPNTEVF | 0.60333514 |
| DPLLHGI | ASSLEHPNTEVF | 0.7641026 |
| ELGAALGAL | ASSLEHPNTEVF | 0.8317281 |
| ELGRTQAV | ASSLEHPNTEVF | 0.56125116 |
| FVAGGYLS | ASSLEHPNTEVF | 0.85129964 |
| GASGGAGGAGGW | ASSLEHPNTEVF | 0.79902905 |
| GGAGAGGDGGAGGI | ASSLEHPNTEVF | 0.70887876 |
| GTGHNGSQPGMGGNG | ASSLEHPNTEVF | 0.8769468 |
| GYAAASASAS | ASSLEHPNTEVF | 0.76012826 |
| HQVDVAAVQQG | ASSLEHPNTEVF | 0.6996776 |
| IGAAIAHT | ASSLEHPNTEVF | 0.68004847 |
| IGMIEVAQL | ASSLEHPNTEVF | 0.87394875 |
| IRLAPEHTSAMD | ASSLEHPNTEVF | 0.597816 |
| IYNVSVL | ASSLEHPNTEVF | 0.8664721 |
| KQGFLGM | ASSLEHPNTEVF | 0.7226876 |
| LESVITQL | ASSLEHPNTEVF | 0.88745373 |
| LGLGALVVV | ASSLEHPNTEVF | 0.55545616 |
| LIGNGGNGGSGGI | ASSLEHPNTEVF | 0.8756435 |
| LNAINAPTL | ASSLEHPNTEVF | 0.83252645 |
| LQLLNTGV | ASSLEHPNTEVF | 0.9002955 |
| LQTSIEF | ASSLEHPNTEVF | 0.7927234 |
| MSFAVSEL | ASSLEHPNTEVF | 0.80991244 |
| NTGALGL | ASSLEHPNTEVF | 0.6347537 |
| PPGAGGAGGAGTT | ASSLEHPNTEVF | 0.87968475 |
| QLVGLGI | ASSLEHPNTEVF | 0.61433387 |
| RFLEQVHQL | ASSLEHPNTEVF | 0.82997835 |
| RGVGYVLR | ASSLEHPNTEVF | 0.52253115 |
| SATGVFAGVI | ASSLEHPNTEVF | 0.8608328 |
| SLPAITLP | ASSLEHPNTEVF | 0.86519074 |
| STALAFTV | ASSLEHPNTEVF | 0.32713598 |
| SVLTVGLGK | ASSLEHPNTEVF | 0.8647747 |
| TALAGAGSGI | ASSLEHPNTEVF | 0.8145534 |
| TGLPFTGI | ASSLEHPNTEVF | 0.5957204 |
| VAEHVTV | ASSLEHPNTEVF | 0.8867749 |

| | | |
|---|---|---|
| VDIVLNSL | ASSLEHPNTEVF | 0.7028059 |
| VLTGGGAVL | ASSLEHPNTEVF | 0.89337516 |
| VNVGLAYGI | ASSLEHPNTEVF | 0.7967899 |
| VSSAAVGAV | ASSLEHPNTEVF | 0.7957515 |
| VTSVYEAGV | ASSLEHPNTEVF | 0.85136354 |
| WVALDRGA | ASSLEHPNTEVF | 0.6053947 |
| YLVGSVVI | ASSLEHPNTEVF | 0.64059913 |
| GPPGTGKTLL | ASSLEHPNTEVF | 0.8889667 |
| GVGMTKF | ASSLEHPNTEVF | 0.7190069 |
| IGIGGSDL | ASSLEHPNTEVF | 0.85912824 |
| IGTLLAEL | ASSLEHPNTEVF | 0.5534112 |
| IYTSGTT | ASSLEHPNTEVF | 0.5313479 |
| LSFPAGV | ASSLEHPNTEVF | 0.64601445 |
| LYTSGTT | ASSLEHPNTEVF | 0.7984232 |
| VGAFPFGF | ASSLEHPNTEVF | 0.86149216 |
| VGAGTVEF | ASSLEHPNTEVF | 0.79815495 |
| VGKSTLLL | ASSLEHPNTEVF | 0.42030287 |
| AALAHAGGGIV | ASSLEHPNTEVF | 0.6803256 |
| AALVAVSLL | ASSLEHPNTEVF | 0.5917839 |
| AAVSRAVAAL | ASSLEHPNTEVF | 0.6623401 |
| AGGSSGIGGF | ASSLEHPNTEVF | 0.77643144 |
| AGLTVANANG | ASSLEHPNTEVF | 0.6176204 |
| AGLVGAVL | ASSLEHPNTEVF | 0.58103687 |
| ALSQAFTV | ASSLEHPNTEVF | 0.46526685 |
| ALTFSVSL | ASSLEHPNTEVF | 0.7605833 |
| ALTGAALVAL | ASSLEHPNTEVF | 0.79622316 |
| ASFSIGAGVL | ASSLEHPNTEVF | 0.7686393 |
| DGSVYKI | ASSLEHPNTEVF | 0.83505416 |
| EQLAEIV | ASSLEHPNTEVF | 0.80527437 |
| FAVSELTVM | ASSLEHPNTEVF | 0.665298 |
| GAAPLAAGL | ASSLEHPNTEVF | 0.15617591 |
| GADGGDGGSGGAGG | ASSLEHPNTEVF | 0.361418 |
| GFGLDTGGAGG | ASSLEHPNTEVF | 0.6650317 |
| GLASGGLL | ASSLEHPNTEVF | 0.68208265 |
| GRIAGAAVLL | ASSLEHPNTEVF | 0.19105504 |
| GSAGSLNTSF | ASSLEHPNTEVF | 0.78183746 |
| GTGQIGI | ASSLEHPNTEVF | 0.57768035 |
| GTLIALIL | ASSLEHPNTEVF | 0.59328157 |
| IAATMGFVVG | ASSLEHPNTEVF | 0.82216996 |
| IFAQALL | ASSLEHPNTEVF | 0.64306945 |
| IGAAVALGF | ASSLEHPNTEVF | 0.4488621 |
| IGLTPLAL | ASSLEHPNTEVF | 0.4013995 |
| ILGSNTSTL | ASSLEHPNTEVF | 0.78506285 |
| IVFSLTVL | ASSLEHPNTEVF | 0.33530694 |
| LASKATGF | ASSLEHPNTEVF | 0.7727372 |
| LASVALIL | ASSLEHPNTEVF | 0.5598272 |
| LFAVGLTL | ASSLEHPNTEVF | 0.78085816 |
| LFNGGIGTYI | ASSLEHPNTEVF | 0.30509835 |
| LTLGAGII | ASSLEHPNTEVF | 0.7763925 |
| LVAAALGDI | ASSLEHPNTEVF | 0.7036991 |
| NTAFVQLGI | ASSLEHPNTEVF | 0.859452 |

| | | |
|---|---|---|
| QLFSGAEL | ASSLEHPNTEVF | 0.5992515 |
| SAATFITL | ASSLEHPNTEVF | 0.69372284 |
| SNYGLANLGV | ASSLEHPNTEVF | 0.5387001 |
| SYGGAYAVM | ASSLEHPNTEVF | 0.8539568 |
| TTGVGGLTLG | ASSLEHPNTEVF | 0.48356804 |
| TTQTIYI | ASSLEHPNTEVF | 0.49601737 |
| VAAGAGLGAVI | ASSLEHPNTEVF | 0.33013967 |
| VAAVVTLL | ASSLEHPNTEVF | 0.661738 |
| VGFLGGLI | ASSLEHPNTEVF | 0.77565813 |
| VGVIPILL | ASSLEHPNTEVF | 0.6219449 |
| VIFASSLI | ASSLEHPNTEVF | 0.7378197 |
| VKGGDGGAAAG | ASSLEHPNTEVF | 0.72490704 |
| VTKTFTNI | ASSLEHPNTEVF | 0.44550294 |
| VVAAAILL | ASSLEHPNTEVF | 0.8220508 |
| YAALDVEL | ASSLEHPNTEVF | 0.53945017 |
| LASVALLL | ASSLEHPNTEVF | 0.7949089 |
| LDAVHRV | ASSLEHPNTEVF | 0.5184027 |
| QGLSSSIF | ASSLEHPNTEVF | 0.6519936 |
| RGTFANIRL | ASSLEHPNTEVF | 0.7102572 |
| RQLWWGHRI | ASSLEHPNTEVF | 0.6752311 |
| YGPPGTGKTLL | ASSLEHPNTEVF | 0.45156917 |
| AALVGAPANVA | ASSLEHPNTEVF | 0.8189601 |
| AATFRRTE | ASSLEHPNTEVF | 0.8441154 |
| AGALGGGI | ASSLEHPNTEVF | 0.85242367 |
| AGGAGGAGGIGAG | ASSLEHPNTEVF | 0.7551836 |
| AGSGALGL | ASSLEHPNTEVF | 0.70404506 |
| AQIADQI | ASSLEHPNTEVF | 0.7908044 |
| ASAVAVTYL | ASSLEHPNTEVF | 0.84496087 |
| ASTVLGGTGG | ASSLEHPNTEVF | 0.85388625 |
| AVGGEPAK | ASSLEHPNTEVF | 0.8369232 |
| CEQGTFGGTC | ASSLEHPNTEVF | 0.64698744 |
| DEGKAPAG | ASSLEHPNTEVF | 0.81245947 |
| DTVVFPAL | ASSLEHPNTEVF | 0.79515326 |
| FEGITMAG | ASSLEHPNTEVF | 0.7869054 |
| FPLVVQD | ASSLEHPNTEVF | 0.81899774 |
| GAFTTPQ | ASSLEHPNTEVF | 0.7981158 |
| GFGGFGTAGGVGGA | ASSLEHPNTEVF | 0.7654202 |
| GFLAQFL | ASSLEHPNTEVF | 0.7532056 |
| GGAGGAGGNAGVA | ASSLEHPNTEVF | 0.81468296 |
| GGVGTAGAGGAGGAGG | ASSLEHPNTEVF | 0.8435006 |
| GNGGGGDAGNAGSGGN | ASSLEHPNTEVF | 0.8338404 |
| GTGGAGGAGNDAGS | ASSLEHPNTEVF | 0.86181426 |
| GTTNTGWFNS | ASSLEHPNTEVF | 0.8519009 |
| IFMGPLFTL | ASSLEHPNTEVF | 0.8369653 |
| ILAHAIEAI | ASSLEHPNTEVF | 0.259707 |
| IRSAIREAL | ASSLEHPNTEVF | 0.56986856 |
| ITFAGGG | ASSLEHPNTEVF | 0.72367 |
| ITGMSGHQEV | ASSLEHPNTEVF | 0.80879915 |
| IVAGLACTL | ASSLEHPNTEVF | 0.3609478 |
| IVGGGLAGA | ASSLEHPNTEVF | 0.8226466 |
| IVTGVLNI | ASSLEHPNTEVF | 0.7851611 |

| | | |
|---|---|---|
| LAQQGCI | ASSLEHPNTEVF | 0.73584735 |
| LCLHVEN | ASSLEHPNTEVF | 0.84808546 |
| LDAISDGS | ASSLEHPNTEVF | 0.77397406 |
| LGGFAAML | ASSLEHPNTEVF | 0.6765748 |
| LGGKNPMI | ASSLEHPNTEVF | 0.6308551 |
| LGPKGVAAIL | ASSLEHPNTEVF | 0.8476801 |
| LMVGVSI | ASSLEHPNTEVF | 0.5854148 |
| LVGAAFVL | ASSLEHPNTEVF | 0.8480641 |
| LVLAVAPL | ASSLEHPNTEVF | 0.69232345 |
| MAAIAVIDAA | ASSLEHPNTEVF | 0.85109365 |
| NNGINAELQ | ASSLEHPNTEVF | 0.54616296 |
| RGEPLVLVHG | ASSLEHPNTEVF | 0.84789705 |
| RGGAGGAAGLI | ASSLEHPNTEVF | 0.8185625 |
| RSYRMHPAV | ASSLEHPNTEVF | 0.8644775 |
| SAAHLLAGQLA | ASSLEHPNTEVF | 0.790367 |
| SAAVGSY | ASSLEHPNTEVF | 0.81985885 |
| SPHAVLHRI | ASSLEHPNTEVF | 0.7884527 |
| SQIAKVL | ASSLEHPNTEVF | 0.8503449 |
| TALGHLVP | ASSLEHPNTEVF | 0.70867085 |
| TAPTTSGAA | ASSLEHPNTEVF | 0.7822065 |
| TDGDDGNGGSPGT | ASSLEHPNTEVF | 0.41395172 |
| TDGLGEN | ASSLEHPNTEVF | 0.7998364 |
| TGGPPLPGQA | ASSLEHPNTEVF | 0.8419467 |
| TIPAITI | ASSLEHPNTEVF | 0.3921064 |
| TIPGFTI | ASSLEHPNTEVF | 0.8077023 |
| TSIASGALA | ASSLEHPNTEVF | 0.76858294 |
| TVTLPGAAI | ASSLEHPNTEVF | 0.6706296 |
| VASGLSAALP | ASSLEHPNTEVF | 0.84709775 |
| VGAGVIGI | ASSLEHPNTEVF | 0.7763233 |
| VGPKRFGV | ASSLEHPNTEVF | 0.8577679 |
| VNVAFPD | ASSLEHPNTEVF | 0.72385246 |
| VSMTNSGVS | ASSLEHPNTEVF | 0.76940495 |
| VTDVLGDLSV | ASSLEHPNTEVF | 0.8090148 |
| VVVVGAGFAGL | ASSLEHPNTEVF | 0.55838907 |
| VYTVGTVV | ASSLEHPNTEVF | 0.66854745 |
| WTTFSGP | ASSLEHPNTEVF | 0.8561341 |
| YDELVEAF | ASSLEHPNTEVF | 0.6847094 |
| YGTGGAGGNGG | ASSLEHPNTEVF | 0.710145 |
| YTNVMAV | ASSLEHPNTEVF | 0.65027297 |
| FLEQVHQL | ASSLEHPNTEVF | 0.6021583 |
| GLSLLAVAV | ASSLEHPNTEVF | 0.65831316 |
| LEQVHQL | ASSLEHPNTEVF | 0.70034885 |
| LNAAGAI | ASSLEHPNTEVF | 0.7072078 |
| PAVHASL | ASSLEHPNTEVF | 0.69022185 |
| TELEDDA | ASSLEHPNTEVF | 0.5488726 |
| TPPGPPN | ASSLEHPNTEVF | 0.677323 |
| TPPPPPPP | ASSLEHPNTEVF | 0.6572546 |
| TQSPALI | TCSAGTGLYEQY | 0.6586392 |
| EFAAAFPQ | TCSAGTGLYEQY | 0.68465984 |
| LLSGNLDVL | TCSAGTGLYEQY | 0.6783078 |
| GPANLGNYNI | TCSAGTGLYEQY | 0.6378776 |

| | | |
|---|---|---|
| TLGPTLP | TCSAGTGLYEQY | 0.61619973 |
| VYAYSSCCK | TCSAGTGLYEQY | 0.6760801 |
| APAPAPAPNI | TCSAGTGLYEQY | 0.68364096 |
| PGGPVPIP | TCSAGTGLYEQY | 0.6820973 |
| FLTVALEP | TCSAGTGLYEQY | 0.7110306 |
| SQYFGANL | TCSAGTGLYEQY | 0.70696783 |
| GGVKPAA | TCSAGTGLYEQY | 0.69526565 |
| GGTGGAGGAGGW | TCSAGTGLYEQY | 0.3230405 |
| GVGGVAGLGGAGG | TCSAGTGLYEQY | 0.4876435 |
| VLLVLATTI | TCSAGTGLYEQY | 0.6206869 |
| IGIFAAML | TCSAGTGLYEQY | 0.6575248 |
| AVGDLID | TCSAGTGLYEQY | 0.3737976 |
| NISPLPL | TCSAGTGLYEQY | 0.6618103 |
| VTAVFVAI | TCSAGTGLYEQY | 0.61327577 |
| SGDDNDAAL | TCSAGTGLYEQY | 0.62027085 |
| AVAALAHS | TCSAGTGLYEQY | 0.7028153 |
| GGVTVMATP | TCSAGTGLYEQY | 0.65565056 |
| AGLAANVRD | TCSAGTGLYEQY | 0.6134778 |
| TNRPDLIDPALLRPG | TCSAGTGLYEQY | 0.5851147 |
| LANLGVTAL | TCSAGTGLYEQY | 0.6859602 |
| TGERIEL | TCSAGTGLYEQY | 0.48641312 |
| NGEELAA | TCSAGTGLYEQY | 0.69877386 |
| VLSLVWG | TCSAGTGLYEQY | 0.51977175 |
| IGYALWL | TCSAGTGLYEQY | 0.70423406 |
| LSLLLAPL | TCSAGTGLYEQY | 0.521975 |
| TLDEALK | TCSAGTGLYEQY | 0.69992733 |
| TDGAGTAEGG | TCSAGTGLYEQY | 0.67657536 |
| NAPGGAGGNGG | TCSAGTGLYEQY | 0.72139347 |
| VSVFMGMGGNF | TCSAGTGLYEQY | 0.65412176 |
| LASYLAAL | TCSAGTGLYEQY | 0.6753119 |
| LVGGTPAL | TCSAGTGLYEQY | 0.5868416 |
| VRNADVL | TCSAGTGLYEQY | 0.7106994 |
| TCSSPFLVSAM | TCSAGTGLYEQY | 0.52274704 |
| VHITDAE | TCSAGTGLYEQY | 0.66576886 |
| EYAAPSF | TCSAGTGLYEQY | 0.45518357 |
| VVDAFHGGL | TCSAGTGLYEQY | 0.68786365 |
| ASVVDPQ | TCSAGTGLYEQY | 0.6919962 |
| SYNEFGGLS | TCSAGTGLYEQY | 0.39688513 |
| DPDGSVY | TCSAGTGLYEQY | 0.69905865 |
| MVSTIHATE | TCSAGTGLYEQY | 0.64895475 |
| LAAGVADPV | TCSAGTGLYEQY | 0.59623677 |
| TKEQIAA | TCSAGTGLYEQY | 0.7085035 |
| SLASPAAL | TCSAGTGLYEQY | 0.6412219 |
| IGLVFYGV | TCSAGTGLYEQY | 0.70887434 |
| FLLDASNL | TCSAGTGLYEQY | 0.6333097 |
| YQVGNFP | TCSAGTGLYEQY | 0.6526258 |
| ESLLSQST | TCSAGTGLYEQY | 0.67392206 |
| LFTVGFAAVG | TCSAGTGLYEQY | 0.5237349 |
| ATPPTPT | TCSAGTGLYEQY | 0.55863094 |
| TDLATLG | TCSAGTGLYEQY | 0.71060723 |
| SAGGTGR | TCSAGTGLYEQY | 0.71852934 |

| | | |
|---|---|---|
| GFSNSGS | TCSAGTGLYEQY | 0.82399225 |
| GGNGGSGGTSV | TCSAGTGLYEQY | 0.5269391 |
| GTSVATGGAGN | TCSAGTGLYEQY | 0.38978127 |
| NGGNGGSGGTS | TCSAGTGLYEQY | 0.6026206 |
| AVVKFPH | TCSAGTGLYEQY | 0.7589744 |
| TPPPPPP | TCSAGTGLYEQY | 0.85851586 |
| AAPAAAAAPA | TCSAGTGLYEQY | 0.721267 |
| IALNEDL | TCSAGTGLYEQY | 0.31289423 |
| IVNVSSM | TCSAGTGLYEQY | 0.69761944 |
| TGIPPAPR | TCSAGTGLYEQY | 0.49394706 |
| FDLGGGTF | TCSAGTGLYEQY | 0.54297864 |
| FDLGGGTFDVS | TCSAGTGLYEQY | 0.65256596 |
| NEPTAAA | TCSAGTGLYEQY | 0.60788035 |
| NEPTAAALA | TCSAGTGLYEQY | 0.62462187 |
| GSIINISSI | TCSAGTGLYEQY | 0.44919485 |
| VAPGNAGTA | TCSAGTGLYEQY | 0.6690722 |
| TGGGASLE | TCSAGTGLYEQY | 0.76252294 |
| VAVYNHYKRI | TCSAGTGLYEQY | 0.57427853 |
| TDPAPATT | TCSAGTGLYEQY | 0.8282787 |
| TNRPDILDPALLRPG | TCSAGTGLYEQY | 0.7648009 |
| LTTTTTS | TCSAGTGLYEQY | 0.76669943 |
| PTGSGKTL | TCSAGTGLYEQY | 0.11856085 |
| TGSGKTL | TCSAGTGLYEQY | 0.21361238 |
| TIGHVDHGK | TCSAGTGLYEQY | 0.42498112 |
| TIGHVDHGKTT | TCSAGTGLYEQY | 0.52801436 |
| GHVDHGKTT | TCSAGTGLYEQY | 0.16291992 |
| AQFEHTLL | TCSAGTGLYEQY | 0.7027022 |
| IVNQDEL | TCSAGTGLYEQY | 0.5084363 |
| TNGSQFF | TCSAGTGLYEQY | 0.45532188 |
| TGKGIVE | TCSAGTGLYEQY | 0.80072296 |
| LISNASDA | TCSAGTGLYEQY | 0.5788586 |
| AELRTLGELTKIAKAHGAQ | TCSAGTGLYEQY | 0.37027705 |
| APNAVAAAEPSW | TCSAGTGLYEQY | 0.32109624 |
| APQVAGAATAVL | TCSAGTGLYEQY | 0.7768881 |
| ARRDPAAAAAILPTDA | TCSAGTGLYEQY | 0.22210926 |
| DAAARDVLAEAGLAEYF | TCSAGTGLYEQY | 0.75643957 |
| DDQIVLSAKANLAKVEQ | TCSAGTGLYEQY | 0.40008032 |
| DELLPDTGEDSDEDGH | TCSAGTGLYEQY | 0.7187774 |
| ELPEAPANRATVPQD | TCSAGTGLYEQY | 0.22732124 |
| FGQSKIERVTGIG | TCSAGTGLYEQY | 0.7362454 |
| GAGGTGSTAGGAGGAGGAGGI | TCSAGTGLYEQY | 0.61668146 |
| GHGGTGAAGGNGGTGGAGGS( | TCSAGTGLYEQY | 0.8721311 |
| GNGGNGGAGGEGGPGGAGGA( | TCSAGTGLYEQY | 0.4652368 |
| GSTAGNGGNGGNATNGTIAG( | TCSAGTGLYEQY | 0.5788396 |
| GTGGDGGNGGNGGTGG | TCSAGTGLYEQY | 0.58219266 |
| GVRRPAGIPADAPV | TCSAGTGLYEQY | 0.8444998 |
| GYRQVTDAHLLTLARR | TCSAGTGLYEQY | 0.3011777 |
| IAKMGQDGHDRG | TCSAGTGLYEQY | 0.40784806 |
| KAEPSIEIQYT | TCSAGTGLYEQY | 0.29281956 |
| KPVRIRMAADEDTVAVT | TCSAGTGLYEQY | 0.62326366 |
| MPPRIAGMRLLVIKPEP | TCSAGTGLYEQY | 0.7215551 |

| | | |
|---|---|---|
| MKFYAVALFRSRGVPPTR | TCSAGTGLYEQY | 0.16714925 |
| NAGGFGILGAVAHSP | TCSAGTGLYEQY | 0.68131673 |
| NEGREITLAMAEM | TCSAGTGLYEQY | 0.53053916 |
| NELVEWSHPLGHRWRWE | TCSAGTGLYEQY | 0.328121 |
| NLGAGGTGGNGGNPG | TCSAGTGLYEQY | 0.78872097 |
| NTEWPKWETVMAADPVD | TCSAGTGLYEQY | 0.5074887 |
| PASAAPAETQP | TCSAGTGLYEQY | 0.84874964 |
| QQSSDGKNGVRAAEITGE | TCSAGTGLYEQY | 0.4459006 |
| RAPDAVTTSWWKE | TCSAGTGLYEQY | 0.41196698 |
| SGFGNVGAHTSGWFNQ | TCSAGTGLYEQY | 0.6599388 |
| SGVAGAVGQAASVGGLKV | TCSAGTGLYEQY | 0.3072953 |
| SNGGAGTGSAGGNGGT | TCSAGTGLYEQY | 0.29409635 |
| TGGAGGEGGAAG | TCSAGTGLYEQY | 0.80247533 |
| TGGDGGAAGAGGAPGHG | TCSAGTGLYEQY | 0.63342965 |
| TMPPLLAAIANAGRHGV | TCSAGTGLYEQY | 0.65546656 |
| TSSTAGNSGGA | TCSAGTGLYEQY | 0.58796346 |
| VPATVPVGILC | TCSAGTGLYEQY | 0.57549965 |
| VQRPDSGGVVLTGRIS | TCSAGTGLYEQY | 0.58848274 |
| WTPNPVPATKGVPAGLVQAT | TCSAGTGLYEQY | 0.6486712 |
| YALKRALAKFSDRFDVVII | TCSAGTGLYEQY | 0.6542046 |
| YLIGSVIV | TCSAGTGLYEQY | 0.6398804 |
| YGQLVAVID | TCSAGTGLYEQY | 0.58818376 |
| VTVGGMTLPA | TCSAGTGLYEQY | 0.6322051 |
| VLALPAV | TCSAGTGLYEQY | 0.60800254 |
| VIPPTLNYET | TCSAGTGLYEQY | 0.6018986 |
| VGMGGMHPSSG | TCSAGTGLYEQY | 0.6345078 |
| VGLAGGAATA | TCSAGTGLYEQY | 0.6351009 |
| VGGAGGAGGSGP | TCSAGTGLYEQY | 0.5904804 |
| VFAAFMV | TCSAGTGLYEQY | 0.59814537 |
| TLYDKLR | TCSAGTGLYEQY | 0.621657 |
| TLAVFTVL | TCSAGTGLYEQY | 0.64352185 |
| TGSVQPARV | TCSAGTGLYEQY | 0.63419366 |
| TGGGAGSGGT | TCSAGTGLYEQY | 0.6583384 |
| SHLETSL | TCSAGTGLYEQY | 0.6437617 |
| SFTTGMT | TCSAGTGLYEQY | 0.63897634 |
| QVEVAKRMGVRQPTVS | TCSAGTGLYEQY | 0.3753829 |
| QGSGGPALAM | TCSAGTGLYEQY | 0.49647307 |
| MLAPSNP | TCSAGTGLYEQY | 0.56362194 |
| LVNNAAI | TCSAGTGLYEQY | 0.6171691 |
| LTGEINTLA | TCSAGTGLYEQY | 0.4137642 |
| LSIFGPV | TCSAGTGLYEQY | 0.644338 |
| LLALLANL | TCSAGTGLYEQY | 0.6093781 |
| LISNASDALDKLR | TCSAGTGLYEQY | 0.5811843 |
| LINEAVP | TCSAGTGLYEQY | 0.64538497 |
| LGYAFSGL | TCSAGTGLYEQY | 0.6030697 |
| LDFVAVAL | TCSAGTGLYEQY | 0.5885581 |
| LAQPGAE | TCSAGTGLYEQY | 0.5811091 |
| ISNLGAVTGF | TCSAGTGLYEQY | 0.6476972 |
| HQEDPGRI | TCSAGTGLYEQY | 0.49013105 |
| GTPGEPTP | TCSAGTGLYEQY | 0.6449483 |
| GFFNSGS | TCSAGTGLYEQY | 0.5286585 |

| | | |
|---|---|---|
| GAGGGGLTAGAN | TCSAGTGLYEQY | 0.649752 |
| FVDALVNM | TCSAGTGLYEQY | 0.5507798 |
| FLTVAPEL | TCSAGTGLYEQY | 0.6420227 |
| FETLTAL | TCSAGTGLYEQY | 0.6120818 |
| DVINPIEL | TCSAGTGLYEQY | 0.6629648 |
| APVSGAPV | TCSAGTGLYEQY | 0.6079853 |
| ALVGGLAW | TCSAGTGLYEQY | 0.6427035 |
| ALAAAAATAI | TCSAGTGLYEQY | 0.6234361 |
| AFGLGLLL | TCSAGTGLYEQY | 0.653399 |
| AFFAAASL | TCSAGTGLYEQY | 0.5076215 |
| AAAALPAFSPP | TCSAGTGLYEQY | 0.61039925 |
| VVLAPAL | TCSAGTGLYEQY | 0.49574423 |
| TIGHVDHGKTTL | TCSAGTGLYEQY | 0.6320534 |
| LISNASDALDK | TCSAGTGLYEQY | 0.63968885 |
| LISNASDAL | TCSAGTGLYEQY | 0.4673751 |
| ISNASDALDK | TCSAGTGLYEQY | 0.65218014 |
| NQAETLVYQTE | TCSAGTGLYEQY | 0.60132945 |
| TGGTGGTAGAG | TCSAGTGLYEQY | 0.5966657 |
| TGSTAGGAGGA | TCSAGTGLYEQY | 0.6447755 |
| DPLAVPVAAGPV | TCSAGTGLYEQY | 0.6116998 |
| GGTGGTPGNGGN | TCSAGTGLYEQY | 0.64361596 |
| PAGLAGVDPATL | TCSAGTGLYEQY | 0.58780867 |
| GNNLVVEAPHAGASV | TCSAGTGLYEQY | 0.6021018 |
| KVTRIMSASTLEQPA | TCSAGTGLYEQY | 0.6400184 |
| LGFALPIALPRRTVV | TCSAGTGLYEQY | 0.5172744 |
| TRLGNLTARKLVVLV | TCSAGTGLYEQY | 0.53490806 |
| WSQQRRLIYFPSAGP | TCSAGTGLYEQY | 0.65120465 |
| GPNGNPDPMAAAVDIR | TCSAGTGLYEQY | 0.78297746 |
| TQRCVSVADLRAAASR | TCSAGTGLYEQY | 0.80281687 |
| QFVLDVAEHPVEVLTETV | TCSAGTGLYEQY | 0.727704 |
| GGAGGTGGAGGNGGTQAGDG | TCSAGTGLYEQY | 0.71108645 |
| PLLATLRAIAVDVDAKPAQV | TCSAGTGLYEQY | 0.7399055 |
| APKFRTGGYMIDFWGVGYQV | TCSAGTGLYEQY | 0.80037665 |
| GAGGAGGAGGAAGGPGGTGG | TCSAGTGLYEQY | 0.799337 |
| IGALGLLATDVAPAGSVAFG | TCSAGTGLYEQY | 0.7833119 |
| AGGAGGIGGNGGAGGAGGVL | TCSAGTGLYEQY | 0.73451614 |
| ADELIGLL | TCSAGTGLYEQY | 0.7776682 |
| ADRALKA | TCSAGTGLYEQY | 0.7578634 |
| AGAVGQVGGAGGA | TCSAGTGLYEQY | 0.7483005 |
| AGGGGGLGGAP | TCSAGTGLYEQY | 0.78918326 |
| AGGLLYGNGGAG | TCSAGTGLYEQY | 0.78031486 |
| AGLESIGL | TCSAGTGLYEQY | 0.7450376 |
| AGLKTGII | TCSAGTGLYEQY | 0.75783825 |
| ASLAALLL | TCSAGTGLYEQY | 0.7698647 |
| ASVINISSI | TCSAGTGLYEQY | 0.7856402 |
| ATPPGPA | TCSAGTGLYEQY | 0.7792214 |
| ATVASALV | TCSAGTGLYEQY | 0.81256795 |
| ATVASLIL | TCSAGTGLYEQY | 0.794155 |
| AVNGPVLIRH | TCSAGTGLYEQY | 0.79352826 |
| DGAAGAVGI | TCSAGTGLYEQY | 0.4295174 |
| DPLLHGI | TCSAGTGLYEQY | 0.6668085 |

| | | |
|---|---|---|
| ELGAALGAL | TCSAGTGLYEQY | 0.770038 |
| ELGRTQAV | TCSAGTGLYEQY | 0.759569 |
| FVAGGYLS | TCSAGTGLYEQY | 0.5897239 |
| GASGGAGGAGGW | TCSAGTGLYEQY | 0.7900049 |
| GGAGAGGDGGAGGI | TCSAGTGLYEQY | 0.746004 |
| GTGHNGSQPGMGGNG | TCSAGTGLYEQY | 0.7189245 |
| GYAAASASAS | TCSAGTGLYEQY | 0.78675747 |
| HQVDVAAVQQG | TCSAGTGLYEQY | 0.74679255 |
| IGAAIAHT | TCSAGTGLYEQY | 0.74841714 |
| IGMIEVAQL | TCSAGTGLYEQY | 0.7556692 |
| IRLAPEHTSAMD | TCSAGTGLYEQY | 0.80188316 |
| IYNVSVL | TCSAGTGLYEQY | 0.6894028 |
| KQGFLGM | TCSAGTGLYEQY | 0.7858592 |
| LESVITQL | TCSAGTGLYEQY | 0.7612418 |
| LGLGALVVV | TCSAGTGLYEQY | 0.8019837 |
| LIGNGGNGGSGGI | TCSAGTGLYEQY | 0.68938565 |
| LNAINAPTL | TCSAGTGLYEQY | 0.788412 |
| LQLLNTGV | TCSAGTGLYEQY | 0.75469744 |
| LQTSIEF | TCSAGTGLYEQY | 0.81177074 |
| MSFAVSEL | TCSAGTGLYEQY | 0.758553 |
| NTGALGL | TCSAGTGLYEQY | 0.78627133 |
| PPGAGGAGGAGTT | TCSAGTGLYEQY | 0.7795246 |
| QLVGLGI | TCSAGTGLYEQY | 0.79756093 |
| RFLEQVHQL | TCSAGTGLYEQY | 0.6285013 |
| RGVGYVLR | TCSAGTGLYEQY | 0.76103985 |
| SATGVFAGVI | TCSAGTGLYEQY | 0.70949435 |
| SLPAITLP | TCSAGTGLYEQY | 0.77732503 |
| STALAFTV | TCSAGTGLYEQY | 0.7843773 |
| SVLTVGLGK | TCSAGTGLYEQY | 0.6479969 |
| TALAGAGSGI | TCSAGTGLYEQY | 0.7879152 |
| TGLPFTGI | TCSAGTGLYEQY | 0.7506459 |
| VAEHVTV | TCSAGTGLYEQY | 0.7462305 |
| VDIVLNSL | TCSAGTGLYEQY | 0.80196565 |
| VLTGGGAVL | TCSAGTGLYEQY | 0.7614904 |
| VNVGLAYGI | TCSAGTGLYEQY | 0.7953944 |
| VSSAAVGAV | TCSAGTGLYEQY | 0.75971824 |
| VTSVYEAGV | TCSAGTGLYEQY | 0.7531828 |
| WVALDRGA | TCSAGTGLYEQY | 0.78992087 |
| YLVGSVVI | TCSAGTGLYEQY | 0.68762535 |
| GPPGTGKTLL | TCSAGTGLYEQY | 0.7265228 |
| GVGMTKF | TCSAGTGLYEQY | 0.8002924 |
| IGIGGSDL | TCSAGTGLYEQY | 0.12994418 |
| IGTLLAEL | TCSAGTGLYEQY | 0.12967339 |
| IYTSGTT | TCSAGTGLYEQY | 0.10440951 |
| LSFPAGV | TCSAGTGLYEQY | 0.113226086 |
| LYTSGTT | TCSAGTGLYEQY | 0.10850064 |
| VGAFPFGF | TCSAGTGLYEQY | 0.13352638 |
| VGAGTVEF | TCSAGTGLYEQY | 0.2385433 |
| VGKSTLLL | TCSAGTGLYEQY | 0.1303725 |
| AALAHAGGGIV | TCSAGTGLYEQY | 0.1185917 |
| AALVAVSLL | TCSAGTGLYEQY | 0.11740026 |

| | | |
|---|---|---|
| AAVSRAVAAL | TCSAGTGLYEQY | 0.11692645 |
| AGGSSGIGGF | TCSAGTGLYEQY | 0.11399704 |
| AGLTVANANG | TCSAGTGLYEQY | 0.114180796 |
| AGLVGAVL | TCSAGTGLYEQY | 0.11815121 |
| ALSQAFTV | TCSAGTGLYEQY | 0.10950926 |
| ALTFSVSL | TCSAGTGLYEQY | 0.104347 |
| ALTGAALVAL | TCSAGTGLYEQY | 0.11144094 |
| ASFSIGAGVL | TCSAGTGLYEQY | 0.14051504 |
| DGSVYKI | TCSAGTGLYEQY | 0.13374528 |
| EQLAEIV | TCSAGTGLYEQY | 0.1399588 |
| FAVSELTVM | TCSAGTGLYEQY | 0.1551428 |
| GAAPLAAGL | TCSAGTGLYEQY | 0.107107416 |
| GADGGDGGSGGAGG | TCSAGTGLYEQY | 0.098407164 |
| GFGLDTGGAGG | TCSAGTGLYEQY | 0.10446049 |
| GLASGGLL | TCSAGTGLYEQY | 0.110075094 |
| GRIAGAAVLL | TCSAGTGLYEQY | 0.114507645 |
| GSAGSLNTSF | TCSAGTGLYEQY | 0.09568254 |
| GTGQIGI | TCSAGTGLYEQY | 0.1249313 |
| GTLIALIL | TCSAGTGLYEQY | 0.11978366 |
| IAATMGFVVG | TCSAGTGLYEQY | 0.10859871 |
| IFAQALL | TCSAGTGLYEQY | 0.14007986 |
| IGAAVALGF | TCSAGTGLYEQY | 0.1057263 |
| IGLTPLAL | TCSAGTGLYEQY | 0.11389127 |
| ILGSNTSTL | TCSAGTGLYEQY | 0.11251818 |
| IVFSLTVL | TCSAGTGLYEQY | 0.13297425 |
| LASKATGF | TCSAGTGLYEQY | 0.10684852 |
| LASVALIL | TCSAGTGLYEQY | 0.13527742 |
| LFAVGLTL | TCSAGTGLYEQY | 0.108865775 |
| LFNGGIGTYI | TCSAGTGLYEQY | 0.15788382 |
| LTLGAGII | TCSAGTGLYEQY | 0.11025465 |
| LVAAALGDI | TCSAGTGLYEQY | 0.12762824 |
| NTAFVQLGI | TCSAGTGLYEQY | 0.11125557 |
| QLFSGAEL | TCSAGTGLYEQY | 0.26010785 |
| SAATFITL | TCSAGTGLYEQY | 0.12313135 |
| SNYGLANLGV | TCSAGTGLYEQY | 0.13674082 |
| SYGGAYAVM | TCSAGTGLYEQY | 0.11152033 |
| TTGVGGLTLG | TCSAGTGLYEQY | 0.14801645 |
| TTQTIYI | TCSAGTGLYEQY | 0.10296678 |
| VAAGAGLGAVI | TCSAGTGLYEQY | 0.11598107 |
| VAAVVTLL | TCSAGTGLYEQY | 0.10471068 |
| VGFLGGLI | TCSAGTGLYEQY | 0.116492316 |
| VGVIPILL | TCSAGTGLYEQY | 0.1279048 |
| VIFASSLI | TCSAGTGLYEQY | 0.098304905 |
| VKGGDGGAAAG | TCSAGTGLYEQY | 0.114501394 |
| VTKTFTNI | TCSAGTGLYEQY | 0.10032659 |
| VVAAAILL | TCSAGTGLYEQY | 0.106467515 |
| YAALDVEL | TCSAGTGLYEQY | 0.11489135 |
| LASVALLL | TCSAGTGLYEQY | 0.10881627 |
| LDAVHRV | TCSAGTGLYEQY | 0.12431959 |
| QGLSSSIF | TCSAGTGLYEQY | 0.10200246 |
| RGTFANIRL | TCSAGTGLYEQY | 0.11593861 |

| | | |
|---|---|---|
| RQLWWGHRI | TCSAGTGLYEQY | 0.13804501 |
| YGPPGTGKTLL | TCSAGTGLYEQY | 0.10013273 |
| AALVGAPANVA | TCSAGTGLYEQY | 0.10694495 |
| AATFRRTE | TCSAGTGLYEQY | 0.13963471 |
| AGALGGGI | TCSAGTGLYEQY | 0.7987964 |
| AGGAGGAGGIGAG | TCSAGTGLYEQY | 0.8001354 |
| AGSGALGL | TCSAGTGLYEQY | 0.76674974 |
| AQIADQI | TCSAGTGLYEQY | 0.76027477 |
| ASAVAVTYL | TCSAGTGLYEQY | 0.78354895 |
| ASTVLGGTGG | TCSAGTGLYEQY | 0.79997534 |
| AVGGEPAK | TCSAGTGLYEQY | 0.80483174 |
| CEQGTFGGTC | TCSAGTGLYEQY | 0.7977426 |
| DEGKAPAG | TCSAGTGLYEQY | 0.7663908 |
| DTVVFPAL | TCSAGTGLYEQY | 0.79417366 |
| FEGITMAG | TCSAGTGLYEQY | 0.7931117 |
| FPLVVQD | TCSAGTGLYEQY | 0.7814084 |
| GAFTTPQ | TCSAGTGLYEQY | 0.79895294 |
| GFGGFGTAGGVGGA | TCSAGTGLYEQY | 0.79438925 |
| GFLAQFL | TCSAGTGLYEQY | 0.77294135 |
| GGAGGAGGNAGVA | TCSAGTGLYEQY | 0.77546775 |
| GGVGTAGAGGAGGAGG | TCSAGTGLYEQY | 0.7956592 |
| GNGGGGDAGNAGSGGN | TCSAGTGLYEQY | 0.8000118 |
| GTGGAGGAGNDAGS | TCSAGTGLYEQY | 0.7999367 |
| GTTNTGWFNS | TCSAGTGLYEQY | 0.8053037 |
| IFMGPLFTL | TCSAGTGLYEQY | 0.80172527 |
| ILAHAIEAI | TCSAGTGLYEQY | 0.79746807 |
| IRSAIREAL | TCSAGTGLYEQY | 0.49445567 |
| ITFAGGG | TCSAGTGLYEQY | 0.72898614 |
| ITGMSGHQEV | TCSAGTGLYEQY | 0.7735021 |
| IVAGLACTL | TCSAGTGLYEQY | 0.78793985 |
| IVGGGLAGA | TCSAGTGLYEQY | 0.5862602 |
| IVTGVLNI | TCSAGTGLYEQY | 0.7945709 |
| LAQQGCI | TCSAGTGLYEQY | 0.78292084 |
| LCLHVEN | TCSAGTGLYEQY | 0.76147956 |
| LDAISDGS | TCSAGTGLYEQY | 0.80086315 |
| LGGFAAML | TCSAGTGLYEQY | 0.7776679 |
| LGGKNPMI | TCSAGTGLYEQY | 0.76742303 |
| LGPKGVAAIL | TCSAGTGLYEQY | 0.7788121 |
| LMVGVSI | TCSAGTGLYEQY | 0.80432343 |
| LVGAAFVL | TCSAGTGLYEQY | 0.77285314 |
| LVLAVAPL | TCSAGTGLYEQY | 0.80085194 |
| MAAIAVIDAA | TCSAGTGLYEQY | 0.7752714 |
| NNGINAELQ | TCSAGTGLYEQY | 0.80362296 |
| RGEPLVLVHG | TCSAGTGLYEQY | 0.7441441 |
| RGGAGGAAGLI | TCSAGTGLYEQY | 0.80060524 |
| RSYRMHPAV | TCSAGTGLYEQY | 0.7910204 |
| SAAHLLAGQLA | TCSAGTGLYEQY | 0.8074851 |
| SAAVGSY | TCSAGTGLYEQY | 0.7914963 |
| SPHAVLHRI | TCSAGTGLYEQY | 0.8012785 |
| SQIAKVL | TCSAGTGLYEQY | 0.77716637 |
| TALGHLVP | TCSAGTGLYEQY | 0.8017299 |

| | | |
|---|---|---|
| TAPTTSGAA | TCSAGTGLYEQY | 0.7440871 |
| TDGDDGNGGSPGT | TCSAGTGLYEQY | 0.78865504 |
| TDGLGEN | TCSAGTGLYEQY | 0.7147159 |
| TGGPPLPGQA | TCSAGTGLYEQY | 0.790621 |
| TIPAITI | TCSAGTGLYEQY | 0.7967975 |
| TIPGFTI | TCSAGTGLYEQY | 0.57281554 |
| TSIASGALA | TCSAGTGLYEQY | 0.8012675 |
| TVTLPGAAI | TCSAGTGLYEQY | 0.76555425 |
| VASGLSAALP | TCSAGTGLYEQY | 0.7746362 |
| VGAGVIGI | TCSAGTGLYEQY | 0.80269325 |
| VGPKRFGV | TCSAGTGLYEQY | 0.7841176 |
| VNVAFPD | TCSAGTGLYEQY | 0.8018173 |
| VSMTNSGVS | TCSAGTGLYEQY | 0.7784698 |
| VTDVLGDLSV | TCSAGTGLYEQY | 0.7860725 |
| VVVVGAGFAGL | TCSAGTGLYEQY | 0.7930168 |
| VYTVGTVV | TCSAGTGLYEQY | 0.6896783 |
| WTTFSGP | TCSAGTGLYEQY | 0.7680054 |
| YDELVEAF | TCSAGTGLYEQY | 0.8038172 |
| YGTGGAGGNGG | TCSAGTGLYEQY | 0.59041363 |
| YTNVMAV | TCSAGTGLYEQY | 0.52295953 |
| FLEQVHQL | TCSAGTGLYEQY | 0.16588236 |
| GLSLLAVAV | TCSAGTGLYEQY | 0.20533898 |
| LEQVHQL | TCSAGTGLYEQY | 0.24381138 |
| LNAAGAI | TCSAGTGLYEQY | 0.31525058 |
| PAVHASL | TCSAGTGLYEQY | 0.7428265 |
| TELEDDA | TCSAGTGLYEQY | 0.5881265 |
| TPPGPPN | TCSAGTGLYEQY | 0.20436586 |
| TPPPPPPP | TCSAGTGLYEQY | 0.39099473 |
| TQSPALI | ASSATGGSQNTLY | 0.24525094 |
| EFAAAFPQ | ASSATGGSQNTLY | 0.33317578 |
| LLSGNLDVL | ASSATGGSQNTLY | 0.25756246 |
| GPANLGNYNI | ASSATGGSQNTLY | 0.2828998 |
| TLGPTLP | ASSATGGSQNTLY | 0.1923244 |
| VYAYSSCCK | ASSATGGSQNTLY | 0.1627945 |
| APAPAPAPNI | ASSATGGSQNTLY | 0.44207007 |
| PGGPVPIP | ASSATGGSQNTLY | 0.6485646 |
| FLTVALEP | ASSATGGSQNTLY | 0.48939887 |
| SQYFGANL | ASSATGGSQNTLY | 0.4208616 |
| GGVKPAA | ASSATGGSQNTLY | 0.56821746 |
| GGTGGAGGAGGW | ASSATGGSQNTLY | 0.3271485 |
| GVGGVAGLGGAGG | ASSATGGSQNTLY | 0.13736993 |
| VLLVLATTI | ASSATGGSQNTLY | 0.14491232 |
| IGIFAAML | ASSATGGSQNTLY | 0.19401602 |
| AVGDLID | ASSATGGSQNTLY | 0.19793877 |
| NISPLPL | ASSATGGSQNTLY | 0.13524099 |
| VTAVFVAI | ASSATGGSQNTLY | 0.33071956 |
| SGDDNDAAL | ASSATGGSQNTLY | 0.23021641 |
| AVAALAHS | ASSATGGSQNTLY | 0.20857856 |
| GGVTVMATP | ASSATGGSQNTLY | 0.5884636 |
| AGLAANVRD | ASSATGGSQNTLY | 0.20372915 |
| TNRPDLIDPALLRPG | ASSATGGSQNTLY | 0.16887677 |

| | | |
|---|---|---|
| LANLGVTAL | ASSATGGSQNTLY | 0.20188612 |
| TGERIEL | ASSATGGSQNTLY | 0.40331966 |
| NGEELAA | ASSATGGSQNTLY | 0.17574027 |
| VLSLVWG | ASSATGGSQNTLY | 0.54417896 |
| IGYALWL | ASSATGGSQNTLY | 0.19943677 |
| LSLLLAPL | ASSATGGSQNTLY | 0.49513623 |
| TLDEALK | ASSATGGSQNTLY | 0.17811081 |
| TDGAGTAEGG | ASSATGGSQNTLY | 0.52289176 |
| NAPGGAGGNGG | ASSATGGSQNTLY | 0.32988358 |
| VSVFMGMGGNF | ASSATGGSQNTLY | 0.75368947 |
| LASYLAAL | ASSATGGSQNTLY | 0.24596903 |
| LVGGTPAL | ASSATGGSQNTLY | 0.3989371 |
| VRNADVL | ASSATGGSQNTLY | 0.25553802 |
| TCSSPFLVSAM | ASSATGGSQNTLY | 0.6779671 |
| VHITDAE | ASSATGGSQNTLY | 0.16171359 |
| EYAAPSF | ASSATGGSQNTLY | 0.18358462 |
| VVDAFHGGL | ASSATGGSQNTLY | 0.15044932 |
| ASVVDPQ | ASSATGGSQNTLY | 0.19282678 |
| SYNEFGGLS | ASSATGGSQNTLY | 0.5964363 |
| DPDGSVY | ASSATGGSQNTLY | 0.1328952 |
| MVSTIHATE | ASSATGGSQNTLY | 0.2225259 |
| LAAGVADPV | ASSATGGSQNTLY | 0.15696216 |
| TKEQIAA | ASSATGGSQNTLY | 0.18504196 |
| SLASPAAL | ASSATGGSQNTLY | 0.31882104 |
| IGLVFYGV | ASSATGGSQNTLY | 0.35344684 |
| FLLDASNL | ASSATGGSQNTLY | 0.35310137 |
| YQVGNFP | ASSATGGSQNTLY | 0.19176883 |
| ESLLSQST | ASSATGGSQNTLY | 0.21282238 |
| LFTVGFAAVG | ASSATGGSQNTLY | 0.23901743 |
| ATPPTPT | ASSATGGSQNTLY | 0.14635755 |
| TDLATLG | ASSATGGSQNTLY | 0.17888014 |
| SAGGTGR | ASSATGGSQNTLY | 0.59662664 |
| GFSNSGS | ASSATGGSQNTLY | 0.122973904 |
| GGNGGSGGTSV | ASSATGGSQNTLY | 0.11205649 |
| GTSVATGGAGN | ASSATGGSQNTLY | 0.110982865 |
| NGGNGGSGGTS | ASSATGGSQNTLY | 0.11749422 |
| AVVKFPH | ASSATGGSQNTLY | 0.11314094 |
| TPPPPPP | ASSATGGSQNTLY | 0.1335657 |
| AAPAAAAAPA | ASSATGGSQNTLY | 0.12553093 |
| IALNEDL | ASSATGGSQNTLY | 0.12695819 |
| IVNVSSM | ASSATGGSQNTLY | 0.11994866 |
| TGIPPAPR | ASSATGGSQNTLY | 0.12294388 |
| FDLGGGTF | ASSATGGSQNTLY | 0.119730815 |
| FDLGGGTFDVS | ASSATGGSQNTLY | 0.12134001 |
| NEPTAAA | ASSATGGSQNTLY | 0.114657186 |
| NEPTAAALA | ASSATGGSQNTLY | 0.122287154 |
| GSIINISSI | ASSATGGSQNTLY | 0.1131874 |
| VAPGNAGTA | ASSATGGSQNTLY | 0.11005595 |
| TGGGASLE | ASSATGGSQNTLY | 0.114533365 |
| VAVYNHYKRI | ASSATGGSQNTLY | 0.12427865 |
| TDPAPATT | ASSATGGSQNTLY | 0.12159832 |

| | | |
|---|---|---|
| TNRPDILDPALLRPG | ASSATGGSQNTLY | 0.13367014 |
| LTTTTTS | ASSATGGSQNTLY | 0.13425748 |
| PTGSGKTL | ASSATGGSQNTLY | 0.11071362 |
| TGSGKTL | ASSATGGSQNTLY | 0.107068375 |
| TIGHVDHGK | ASSATGGSQNTLY | 0.1099197 |
| TIGHVDHGKTT | ASSATGGSQNTLY | 0.11599819 |
| GHVDHGKTT | ASSATGGSQNTLY | 0.118131414 |
| AQFEHTLL | ASSATGGSQNTLY | 0.10367084 |
| IVNQDEL | ASSATGGSQNTLY | 0.12656173 |
| TNGSQFF | ASSATGGSQNTLY | 0.12099999 |
| TGKGIVE | ASSATGGSQNTLY | 0.115433514 |
| LISNASDA | ASSATGGSQNTLY | 0.12705183 |
| AELRTLGELTKIAKAHGAQ | ASSATGGSQNTLY | 0.11196284 |
| APNAVAAAEPSW | ASSATGGSQNTLY | 0.11746017 |
| APQVAGAATAVL | ASSATGGSQNTLY | 0.12323956 |
| ARRDPAAAAAILPTDA | ASSATGGSQNTLY | 0.12612174 |
| DAAARDVLAEAGLAEYF | ASSATGGSQNTLY | 0.115440786 |
| DDQIVLSAKANLAKVEQ | ASSATGGSQNTLY | 0.13738327 |
| DELLPDTGEDSDEDGH | ASSATGGSQNTLY | 0.112909585 |
| ELPEAPANRATVPQD | ASSATGGSQNTLY | 0.1561137 |
| FGQSKIERVTGIG | ASSATGGSQNTLY | 0.11722618 |
| GAGGTGSTAGGAGGAGGAGGI | ASSATGGSQNTLY | 0.12428282 |
| GHGGTGAAGGNGGTGGAGGS | ASSATGGSQNTLY | 0.11620755 |
| GNGGNGGAGGEGGPGGAGGA | ASSATGGSQNTLY | 0.17926641 |
| GSTAGNGGNGGNATNGTIAG | ASSATGGSQNTLY | 0.12183114 |
| GTGGDGGNGGNGGTGG | ASSATGGSQNTLY | 0.13356288 |
| GVRRPAGIPADAPV | ASSATGGSQNTLY | 0.11453234 |
| GYRQVTDAHLLTLARR | ASSATGGSQNTLY | 0.120740056 |
| IAKMGQDGHDRG | ASSATGGSQNTLY | 0.110053204 |
| KAEPSIEIQYT | ASSATGGSQNTLY | 0.11494644 |
| KPVRIRMAADEDTVAVT | ASSATGGSQNTLY | 0.11017052 |
| MPPRIAGMRLLVIKPEP | ASSATGGSQNTLY | 0.120449394 |
| MKFYAVALFRSRGVPPTR | ASSATGGSQNTLY | 0.12719235 |
| NAGGFGILGAVAHSP | ASSATGGSQNTLY | 0.10531631 |
| NEGREITLAMAEM | ASSATGGSQNTLY | 0.113801114 |
| NELVEWSHPLGHRWRWE | ASSATGGSQNTLY | 0.105247445 |
| NLGAGGTGGNGGNPG | ASSATGGSQNTLY | 0.11173576 |
| NTEWPKWETVMAADPVD | ASSATGGSQNTLY | 0.11356909 |
| PASAAPAETQP | ASSATGGSQNTLY | 0.118399076 |
| QQSSDGKNGVRAAEITGE | ASSATGGSQNTLY | 0.11520922 |
| RAPDAVTTSWWKE | ASSATGGSQNTLY | 0.1091454 |
| SGFGNVGAHTSGWFNQ | ASSATGGSQNTLY | 0.11387752 |
| SGVAGAVGQAASVGGLKV | ASSATGGSQNTLY | 0.13306536 |
| SNGGAGTGSAGGNGGT | ASSATGGSQNTLY | 0.10690231 |
| TGGAGGEGGAAG | ASSATGGSQNTLY | 0.116035484 |
| TGGDGGAAGAGGAPGHG | ASSATGGSQNTLY | 0.1341117 |
| TMPPLLAAIANAGRHGV | ASSATGGSQNTLY | 0.2519747 |
| TSSTAGNSGGA | ASSATGGSQNTLY | 0.6727566 |
| VPATVPVGILC | ASSATGGSQNTLY | 0.107106395 |
| VQRPDSGGVVLTGRIS | ASSATGGSQNTLY | 0.13598981 |
| WTPNPVPATKGVPAGLVQAT | ASSATGGSQNTLY | 0.3045619 |

| | | |
|---|---|---|
| YALKRALAKFSDRFDVVII | ASSATGGSQNTLY | 0.33633327 |
| YLIGSVIV | ASSATGGSQNTLY | 0.12896307 |
| YGQLVAVID | ASSATGGSQNTLY | 0.35938746 |
| VTVGGMTLPA | ASSATGGSQNTLY | 0.12767985 |
| VLALPAV | ASSATGGSQNTLY | 0.1246252 |
| VIPPTLNYET | ASSATGGSQNTLY | 0.12219638 |
| VGMGGMHPSSG | ASSATGGSQNTLY | 0.18938848 |
| VGLAGGAATA | ASSATGGSQNTLY | 0.27388847 |
| VGGAGGAGGSGP | ASSATGGSQNTLY | 0.1121241 |
| VFAAFMV | ASSATGGSQNTLY | 0.09407067 |
| TLYDKLR | ASSATGGSQNTLY | 0.17220421 |
| TLAVFTVL | ASSATGGSQNTLY | 0.41001832 |
| TGSVQPARV | ASSATGGSQNTLY | 0.64858496 |
| TGGGAGSGGT | ASSATGGSQNTLY | 0.65009904 |
| SHLETSL | ASSATGGSQNTLY | 0.5058579 |
| SFTTGMT | ASSATGGSQNTLY | 0.3588603 |
| QVEVAKRMGVRQPTVS | ASSATGGSQNTLY | 0.25776938 |
| QGSGGPALAM | ASSATGGSQNTLY | 0.084451854 |
| MLAPSNP | ASSATGGSQNTLY | 0.12350085 |
| LVNNAAI | ASSATGGSQNTLY | 0.11012101 |
| LTGEINTLA | ASSATGGSQNTLY | 0.27136916 |
| LSIFGPV | ASSATGGSQNTLY | 0.07623825 |
| LLALLANL | ASSATGGSQNTLY | 0.16912092 |
| LISNASDALDKLR | ASSATGGSQNTLY | 0.18053086 |
| LINEAVP | ASSATGGSQNTLY | 0.09177969 |
| LGYAFSGL | ASSATGGSQNTLY | 0.55217147 |
| LDFVAVAL | ASSATGGSQNTLY | 0.098530374 |
| LAQPGAE | ASSATGGSQNTLY | 0.11598812 |
| ISNLGAVTGF | ASSATGGSQNTLY | 0.13416547 |
| HQEDPGRI | ASSATGGSQNTLY | 0.53972954 |
| GTPGEPTP | ASSATGGSQNTLY | 0.1257482 |
| GFFNSGS | ASSATGGSQNTLY | 0.41167596 |
| GAGGGGLTAGAN | ASSATGGSQNTLY | 0.18482153 |
| FVDALVNM | ASSATGGSQNTLY | 0.2580961 |
| FLTVAPEL | ASSATGGSQNTLY | 0.106025316 |
| FETLTAL | ASSATGGSQNTLY | 0.3330261 |
| DVINPIEL | ASSATGGSQNTLY | 0.1969957 |
| APVSGAPV | ASSATGGSQNTLY | 0.76985884 |
| ALVGGLAW | ASSATGGSQNTLY | 0.24119045 |
| ALAAAAATAI | ASSATGGSQNTLY | 0.18304041 |
| AFGLGLLL | ASSATGGSQNTLY | 0.11451744 |
| AFFAAASL | ASSATGGSQNTLY | 0.098504305 |
| AAAALPAFSPP | ASSATGGSQNTLY | 0.09235587 |
| VVLAPAL | ASSATGGSQNTLY | 0.20708062 |
| TIGHVDHGKTTL | ASSATGGSQNTLY | 0.116166234 |
| LISNASDALDK | ASSATGGSQNTLY | 0.16885552 |
| LISNASDAL | ASSATGGSQNTLY | 0.27781814 |
| ISNASDALDK | ASSATGGSQNTLY | 0.109619096 |
| NQAETLVYQTE | ASSATGGSQNTLY | 0.3172078 |
| TGGTGGTAGAG | ASSATGGSQNTLY | 0.12407233 |
| TGSTAGGAGGA | ASSATGGSQNTLY | 0.12162883 |

| | | |
|---|---|---|
| DPLAVPVAAGPV | ASSATGGSQNTLY | 0.3572644 |
| GGTGGTPGNGGN | ASSATGGSQNTLY | 0.097069286 |
| PAGLAGVDPATL | ASSATGGSQNTLY | 0.62030476 |
| GNNLVVEAPHAGASV | ASSATGGSQNTLY | 0.085258074 |
| KVTRIMSASTLEQPA | ASSATGGSQNTLY | 0.20591417 |
| LGFALPIALPRRTVV | ASSATGGSQNTLY | 0.31264794 |
| TRLGNLTARKLVVLV | ASSATGGSQNTLY | 0.07899773 |
| WSQQRRLIYFPSAGP | ASSATGGSQNTLY | 0.10416396 |
| GPNGNPDPMAAAVDIR | ASSATGGSQNTLY | 0.38996166 |
| TQRCVSVADLRAAASR | ASSATGGSQNTLY | 0.38860548 |
| QFVLDVAEHPVEVLTETV | ASSATGGSQNTLY | 0.7535172 |
| GGAGGTGGAGGNGGTQAGDG | ASSATGGSQNTLY | 0.17346638 |
| PLLATLRAIAVDVDAKPAQV | ASSATGGSQNTLY | 0.16987267 |
| APKFRTGGYMIDFWGVGYQV | ASSATGGSQNTLY | 0.3258764 |
| GAGGAGGAGGAAGGPGGTGG | ASSATGGSQNTLY | 0.4799708 |
| IGALGLLATDVAPAGSVAFG | ASSATGGSQNTLY | 0.46372163 |
| AGGAGGIGGNGGAGGAGGVL | ASSATGGSQNTLY | 0.44614312 |
| ADELIGLL | ASSATGGSQNTLY | 0.15929933 |
| ADRALKA | ASSATGGSQNTLY | 0.26415038 |
| AGAVGQVGGAGGA | ASSATGGSQNTLY | 0.18660755 |
| AGGGGGLGGAP | ASSATGGSQNTLY | 0.24450547 |
| AGGLLYGNGGAG | ASSATGGSQNTLY | 0.35151786 |
| AGLESIGL | ASSATGGSQNTLY | 0.22351262 |
| AGLKTGII | ASSATGGSQNTLY | 0.16183923 |
| ASLAALLL | ASSATGGSQNTLY | 0.21012454 |
| ASVINISSI | ASSATGGSQNTLY | 0.5685452 |
| ATPPGPA | ASSATGGSQNTLY | 0.6197078 |
| ATVASALV | ASSATGGSQNTLY | 0.55991733 |
| ATVASLIL | ASSATGGSQNTLY | 0.6620221 |
| AVNGPVLIRH | ASSATGGSQNTLY | 0.5503885 |
| DGAAGAVGI | ASSATGGSQNTLY | 0.48018092 |
| DPLLHGI | ASSATGGSQNTLY | 0.111095496 |
| ELGAALGAL | ASSATGGSQNTLY | 0.14667591 |
| ELGRTQAV | ASSATGGSQNTLY | 0.17580783 |
| FVAGGYLS | ASSATGGSQNTLY | 0.30701917 |
| GASGGAGGAGGW | ASSATGGSQNTLY | 0.12493417 |
| GGAGAGGDGGAGGI | ASSATGGSQNTLY | 0.3627755 |
| GTGHNGSQPGMGGNG | ASSATGGSQNTLY | 0.20913967 |
| GYAAASASAS | ASSATGGSQNTLY | 0.14723802 |
| HQVDVAAVQQG | ASSATGGSQNTLY | 0.66498166 |
| IGAAIAHT | ASSATGGSQNTLY | 0.14936274 |
| IGMIEVAQL | ASSATGGSQNTLY | 0.17360708 |
| IRLAPEHTSAMD | ASSATGGSQNTLY | 0.19824527 |
| IYNVSVL | ASSATGGSQNTLY | 0.5033996 |
| KQGFLGM | ASSATGGSQNTLY | 0.15313442 |
| LESVITQL | ASSATGGSQNTLY | 0.48403555 |
| LGLGALVVV | ASSATGGSQNTLY | 0.32917476 |
| LIGNGGNGGSGGI | ASSATGGSQNTLY | 0.5281047 |
| LNAINAPTL | ASSATGGSQNTLY | 0.14118282 |
| LQLLNTGV | ASSATGGSQNTLY | 0.5214486 |
| LQTSIEF | ASSATGGSQNTLY | 0.31080082 |

| | | |
|---|---|---|
| MSFAVSEL | ASSATGGSQNTLY | 0.8172618 |
| NTGALGL | ASSATGGSQNTLY | 0.26664048 |
| PPGAGGAGGAGTT | ASSATGGSQNTLY | 0.2747066 |
| QLVGLGI | ASSATGGSQNTLY | 0.20407808 |
| RFLEQVHQL | ASSATGGSQNTLY | 0.51966345 |
| RGVGYVLR | ASSATGGSQNTLY | 0.13099308 |
| SATGVFAGVI | ASSATGGSQNTLY | 0.2364704 |
| SLPAITLP | ASSATGGSQNTLY | 0.14175102 |
| STALAFTV | ASSATGGSQNTLY | 0.24513716 |
| SVLTVGLGK | ASSATGGSQNTLY | 0.42735967 |
| TALAGAGSGI | ASSATGGSQNTLY | 0.1332037 |
| TGLPFTGI | ASSATGGSQNTLY | 0.41187495 |
| VAEHVTV | ASSATGGSQNTLY | 0.20917521 |
| VDIVLNSL | ASSATGGSQNTLY | 0.16399945 |
| VLTGGGAVL | ASSATGGSQNTLY | 0.4694205 |
| VNVGLAYGI | ASSATGGSQNTLY | 0.18978323 |
| VSSAAVGAV | ASSATGGSQNTLY | 0.68873143 |
| VTSVYEAGV | ASSATGGSQNTLY | 0.19044296 |
| WVALDRGA | ASSATGGSQNTLY | 0.32929185 |
| YLVGSVVI | ASSATGGSQNTLY | 0.36001366 |
| GPPGTGKTLL | ASSATGGSQNTLY | 0.12113218 |
| GVGMTKF | ASSATGGSQNTLY | 0.17314483 |
| IGIGGSDL | ASSATGGSQNTLY | 0.5858325 |
| IGTLLAEL | ASSATGGSQNTLY | 0.45791382 |
| IYTSGTT | ASSATGGSQNTLY | 0.80719006 |
| LSFPAGV | ASSATGGSQNTLY | 0.20632583 |
| LYTSGTT | ASSATGGSQNTLY | 0.18144795 |
| VGAFPFGF | ASSATGGSQNTLY | 0.26251122 |
| VGAGTVEF | ASSATGGSQNTLY | 0.61027265 |
| VGKSTLLL | ASSATGGSQNTLY | 0.79699826 |
| AALAHAGGGIV | ASSATGGSQNTLY | 0.55415785 |
| AALVAVSLL | ASSATGGSQNTLY | 0.20448588 |
| AAVSRAVAAL | ASSATGGSQNTLY | 0.24351679 |
| AGGSSGIGGF | ASSATGGSQNTLY | 0.24639452 |
| AGLTVANANG | ASSATGGSQNTLY | 0.21726802 |
| AGLVGAVL | ASSATGGSQNTLY | 0.5152718 |
| ALSQAFTV | ASSATGGSQNTLY | 0.28736874 |
| ALTFSVSL | ASSATGGSQNTLY | 0.19626534 |
| ALTGAALVAL | ASSATGGSQNTLY | 0.26064748 |
| ASFSIGAGVL | ASSATGGSQNTLY | 0.36983716 |
| DGSVYKI | ASSATGGSQNTLY | 0.6751071 |
| EQLAEIV | ASSATGGSQNTLY | 0.6248282 |
| FAVSELTVM | ASSATGGSQNTLY | 0.7045187 |
| GAAPLAAGL | ASSATGGSQNTLY | 0.54331464 |
| GADGGDGGSGGAGG | ASSATGGSQNTLY | 0.62482744 |
| GFGLDTGGAGG | ASSATGGSQNTLY | 0.09857003 |
| GLASGGLL | ASSATGGSQNTLY | 0.14449081 |
| GRIAGAAVLL | ASSATGGSQNTLY | 0.2810398 |
| GSAGSLNTSF | ASSATGGSQNTLY | 0.24799676 |
| GTGQIGI | ASSATGGSQNTLY | 0.13943726 |
| GTLIALIL | ASSATGGSQNTLY | 0.38101885 |

| | | |
|---|---|---|
| IAATMGFVVG | ASSATGGSQNTLY | 0.264104 |
| IFAQALL | ASSATGGSQNTLY | 0.15593758 |
| IGAAVALGF | ASSATGGSQNTLY | 0.52750075 |
| IGLTPLAL | ASSATGGSQNTLY | 0.17980559 |
| ILGSNTSTL | ASSATGGSQNTLY | 0.219012 |
| IVFSLTVL | ASSATGGSQNTLY | 0.22670484 |
| LASKATGF | ASSATGGSQNTLY | 0.59491247 |
| LASVALIL | ASSATGGSQNTLY | 0.16977498 |
| LFAVGLTL | ASSATGGSQNTLY | 0.38664663 |
| LFNGGIGTYI | ASSATGGSQNTLY | 0.31553283 |
| LTLGAGII | ASSATGGSQNTLY | 0.52930826 |
| LVAAALGDI | ASSATGGSQNTLY | 0.14981638 |
| NTAFVQLGI | ASSATGGSQNTLY | 0.4993226 |
| QLFSGAEL | ASSATGGSQNTLY | 0.256149 |
| SAATFITL | ASSATGGSQNTLY | 0.807376 |
| SNYGLANLGV | ASSATGGSQNTLY | 0.24870068 |
| SYGGAYAVM | ASSATGGSQNTLY | 0.2888589 |
| TTGVGGLTLG | ASSATGGSQNTLY | 0.24002561 |
| TTQTIYI | ASSATGGSQNTLY | 0.75998867 |
| VAAGAGLGAVI | ASSATGGSQNTLY | 0.1211395 |
| VAAVVTLL | ASSATGGSQNTLY | 0.2777785 |
| VGFLGGLI | ASSATGGSQNTLY | 0.17940155 |
| VGVIPILL | ASSATGGSQNTLY | 0.40282845 |
| VIFASSLI | ASSATGGSQNTLY | 0.48266336 |
| VKGGDGGAAAG | ASSATGGSQNTLY | 0.15874083 |
| VTKTFTNI | ASSATGGSQNTLY | 0.5765507 |
| VVAAAILL | ASSATGGSQNTLY | 0.39655408 |
| YAALDVEL | ASSATGGSQNTLY | 0.217009 |
| LASVALLL | ASSATGGSQNTLY | 0.6893213 |
| LDAVHRV | ASSATGGSQNTLY | 0.23109199 |
| QGLSSSIF | ASSATGGSQNTLY | 0.65819865 |
| RGTFANIRL | ASSATGGSQNTLY | 0.28808594 |
| RQLWWGHRI | ASSATGGSQNTLY | 0.32469216 |
| YGPPGTGKTLL | ASSATGGSQNTLY | 0.53326935 |
| AALVGAPANVA | ASSATGGSQNTLY | 0.22338073 |
| AATFRRTE | ASSATGGSQNTLY | 0.18984914 |
| AGALGGGI | ASSATGGSQNTLY | 0.60979724 |
| AGGAGGAGGIGAG | ASSATGGSQNTLY | 0.8250977 |
| AGSGALGL | ASSATGGSQNTLY | 0.8203572 |
| AQIADQI | ASSATGGSQNTLY | 0.67665464 |
| ASAVAVTYL | ASSATGGSQNTLY | 0.58630943 |
| ASTVLGGTGG | ASSATGGSQNTLY | 0.68041164 |
| AVGGEPAK | ASSATGGSQNTLY | 0.8592303 |
| CEQGTFGGTC | ASSATGGSQNTLY | 0.835126 |
| DEGKAPAG | ASSATGGSQNTLY | 0.8153113 |
| DTVVFPAL | ASSATGGSQNTLY | 0.5520238 |
| FEGITMAG | ASSATGGSQNTLY | 0.78456646 |
| FPLVVQD | ASSATGGSQNTLY | 0.7626519 |
| GAFTTPQ | ASSATGGSQNTLY | 0.68783444 |
| GFGGFGTAGGVGGA | ASSATGGSQNTLY | 0.84609437 |
| GFLAQFL | ASSATGGSQNTLY | 0.82278347 |

| | | |
|---|---|---|
| GGAGGAGGNAGVA | ASSATGGSQNTLY | 0.70890385 |
| GGVGTAGAGGAGGAGG | ASSATGGSQNTLY | 0.7399124 |
| GNGGGGDAGNAGSGGN | ASSATGGSQNTLY | 0.80751383 |
| GTGGAGGAGNDAGS | ASSATGGSQNTLY | 0.8285167 |
| GTTNTGWFNS | ASSATGGSQNTLY | 0.82784 |
| IFMGPLFTL | ASSATGGSQNTLY | 0.8822603 |
| ILAHAIEAI | ASSATGGSQNTLY | 0.8513589 |
| IRSAIREAL | ASSATGGSQNTLY | 0.8354436 |
| ITFAGGG | ASSATGGSQNTLY | 0.23759931 |
| ITGMSGHQEV | ASSATGGSQNTLY | 0.44554186 |
| IVAGLACTL | ASSATGGSQNTLY | 0.76241153 |
| IVGGGLAGA | ASSATGGSQNTLY | 0.70258224 |
| IVTGVLNI | ASSATGGSQNTLY | 0.40592283 |
| LAQQGCI | ASSATGGSQNTLY | 0.74448586 |
| LCLHVEN | ASSATGGSQNTLY | 0.7080457 |
| LDAISDGS | ASSATGGSQNTLY | 0.6073267 |
| LGGFAAML | ASSATGGSQNTLY | 0.8412074 |
| LGGKNPMI | ASSATGGSQNTLY | 0.70250016 |
| LGPKGVAAIL | ASSATGGSQNTLY | 0.7266027 |
| LMVGVSI | ASSATGGSQNTLY | 0.686599 |
| LVGAAFVL | ASSATGGSQNTLY | 0.83715296 |
| LVLAVAPL | ASSATGGSQNTLY | 0.62084097 |
| MAAIAVIDAA | ASSATGGSQNTLY | 0.8127471 |
| NNGINAELQ | ASSATGGSQNTLY | 0.66172355 |
| RGEPLVLVHG | ASSATGGSQNTLY | 0.8577502 |
| RGGAGGAAGLI | ASSATGGSQNTLY | 0.5412791 |
| RSYRMHPAV | ASSATGGSQNTLY | 0.82925606 |
| SAAHLLAGQLA | ASSATGGSQNTLY | 0.7516556 |
| SAAVGSY | ASSATGGSQNTLY | 0.8892622 |
| SPHAVLHRI | ASSATGGSQNTLY | 0.75007796 |
| SQIAKVL | ASSATGGSQNTLY | 0.8130994 |
| TALGHLVP | ASSATGGSQNTLY | 0.7586576 |
| TAPTTSGAA | ASSATGGSQNTLY | 0.793771 |
| TDGDDGNGGSPGT | ASSATGGSQNTLY | 0.4695073 |
| TDGLGEN | ASSATGGSQNTLY | 0.76855534 |
| TGGPPLPGQA | ASSATGGSQNTLY | 0.4642906 |
| TIPAITI | ASSATGGSQNTLY | 0.7921728 |
| TIPGFTI | ASSATGGSQNTLY | 0.79821616 |
| TSIASGALA | ASSATGGSQNTLY | 0.43044272 |
| TVTLPGAAI | ASSATGGSQNTLY | 0.845899 |
| VASGLSAALP | ASSATGGSQNTLY | 0.7238596 |
| VGAGVIGI | ASSATGGSQNTLY | 0.69563377 |
| VGPKRFGV | ASSATGGSQNTLY | 0.87025756 |
| VNVAFPD | ASSATGGSQNTLY | 0.76070917 |
| VSMTNSGVS | ASSATGGSQNTLY | 0.8629291 |
| VTDVLGDLSV | ASSATGGSQNTLY | 0.78689665 |
| VVVVGAGFAGL | ASSATGGSQNTLY | 0.7215258 |
| VYTVGTVV | ASSATGGSQNTLY | 0.7786221 |
| WTTFSGP | ASSATGGSQNTLY | 0.55999994 |
| YDELVEAF | ASSATGGSQNTLY | 0.6880086 |
| YGTGGAGGNGG | ASSATGGSQNTLY | 0.8591637 |

| | | |
|---|---|---|
| YTNVMAV | ASSATGGSQNTLY | 0.8105607 |
| FLEQVHQL | ASSATGGSQNTLY | 0.7961374 |
| GLSLLAVAV | ASSATGGSQNTLY | 0.7665565 |
| LEQVHQL | ASSATGGSQNTLY | 0.74596953 |
| LNAAGAI | ASSATGGSQNTLY | 0.7746484 |
| PAVHASL | ASSATGGSQNTLY | 0.8067184 |
| TELEDDA | ASSATGGSQNTLY | 0.81245005 |
| TPPGPPN | ASSATGGSQNTLY | 0.8008877 |
| TPPPPPPP | ASSATGGSQNTLY | 0.6288229 |
| TQSPALI | ASSLGWGNYAEQF | 0.8132657 |
| EFAAAFPQ | ASSLGWGNYAEQF | 0.7978046 |
| LLSGNLDVL | ASSLGWGNYAEQF | 0.7933433 |
| GPANLGNYNI | ASSLGWGNYAEQF | 0.80456024 |
| TLGPTLP | ASSLGWGNYAEQF | 0.80044186 |
| VYAYSSCCK | ASSLGWGNYAEQF | 0.7742446 |
| APAPAPAPNI | ASSLGWGNYAEQF | 0.76494074 |
| PGGPVPIP | ASSLGWGNYAEQF | 0.7950822 |
| FLTVALEP | ASSLGWGNYAEQF | 0.81208694 |
| SQYFGANL | ASSLGWGNYAEQF | 0.80988204 |
| GGVKPAA | ASSLGWGNYAEQF | 0.8132752 |
| GGTGGAGGAGGW | ASSLGWGNYAEQF | 0.81542504 |
| GVGGVAGLGGAGG | ASSLGWGNYAEQF | 0.8037834 |
| VLLVLATTI | ASSLGWGNYAEQF | 0.50762326 |
| IGIFAAML | ASSLGWGNYAEQF | 0.6513268 |
| AVGDLID | ASSLGWGNYAEQF | 0.74439776 |
| NISPLPL | ASSLGWGNYAEQF | 0.77690285 |
| VTAVFVAI | ASSLGWGNYAEQF | 0.43413228 |
| SGDDNDAAL | ASSLGWGNYAEQF | 0.73947597 |
| AVAALAHS | ASSLGWGNYAEQF | 0.7864746 |
| GGVTVMATP | ASSLGWGNYAEQF | 0.7525974 |
| AGLAANVRD | ASSLGWGNYAEQF | 0.8115325 |
| TNRPDLIDPALLRPG | ASSLGWGNYAEQF | 0.7771976 |
| LANLGVTAL | ASSLGWGNYAEQF | 0.75158536 |
| TGERIEL | ASSLGWGNYAEQF | 0.73339427 |
| NGEELAA | ASSLGWGNYAEQF | 0.8071641 |
| VLSLVWG | ASSLGWGNYAEQF | 0.731714 |
| IGYALWL | ASSLGWGNYAEQF | 0.8155237 |
| LSLLLAPL | ASSLGWGNYAEQF | 0.7040742 |
| TLDEALK | ASSLGWGNYAEQF | 0.81754124 |
| TDGAGTAEGG | ASSLGWGNYAEQF | 0.6381029 |
| NAPGGAGGNGG | ASSLGWGNYAEQF | 0.8121043 |
| VSVFMGMGGNF | ASSLGWGNYAEQF | 0.7935792 |
| LASYLAAL | ASSLGWGNYAEQF | 0.82245034 |
| LVGGTPAL | ASSLGWGNYAEQF | 0.7974397 |
| VRNADVL | ASSLGWGNYAEQF | 0.8102423 |
| TCSSPFLVSAM | ASSLGWGNYAEQF | 0.79047024 |
| VHITDAE | ASSLGWGNYAEQF | 0.80429626 |
| EYAAPSF | ASSLGWGNYAEQF | 0.68080306 |
| VVDAFHGGL | ASSLGWGNYAEQF | 0.7958826 |
| ASVVDPQ | ASSLGWGNYAEQF | 0.6326044 |
| SYNEFGGLS | ASSLGWGNYAEQF | 0.80171615 |

| | | |
|---|---|---|
| DPDGSVY | ASSLGWGNYAEQF | 0.80661815 |
| MVSTIHATE | ASSLGWGNYAEQF | 0.39408407 |
| LAAGVADPV | ASSLGWGNYAEQF | 0.8091849 |
| TKEQIAA | ASSLGWGNYAEQF | 0.73123294 |
| SLASPAAL | ASSLGWGNYAEQF | 0.7178244 |
| IGLVFYGV | ASSLGWGNYAEQF | 0.81557286 |
| FLLDASNL | ASSLGWGNYAEQF | 0.7939894 |
| YQVGNFP | ASSLGWGNYAEQF | 0.8135522 |
| ESLLSQST | ASSLGWGNYAEQF | 0.7634571 |
| LFTVGFAAVG | ASSLGWGNYAEQF | 0.7881739 |
| ATPPTPT | ASSLGWGNYAEQF | 0.7940712 |
| TDLATLG | ASSLGWGNYAEQF | 0.4739694 |
| SAGGTGR | ASSLGWGNYAEQF | 0.67657495 |
| GFSNSGS | ASSLGWGNYAEQF | 0.81920457 |
| GGNGGSGGTSV | ASSLGWGNYAEQF | 0.8538384 |
| GTSVATGGAGN | ASSLGWGNYAEQF | 0.8306439 |
| NGGNGGSGGTS | ASSLGWGNYAEQF | 0.7191482 |
| AVVKFPH | ASSLGWGNYAEQF | 0.7207781 |
| TPPPPPP | ASSLGWGNYAEQF | 0.76516545 |
| AAPAAAAAPA | ASSLGWGNYAEQF | 0.85899985 |
| IALNEDL | ASSLGWGNYAEQF | 0.84389067 |
| IVNVSSM | ASSLGWGNYAEQF | 0.84162927 |
| TGIPPAPR | ASSLGWGNYAEQF | 0.68841565 |
| FDLGGGTF | ASSLGWGNYAEQF | 0.8451743 |
| FDLGGGTFDVS | ASSLGWGNYAEQF | 0.8023238 |
| NEPTAAA | ASSLGWGNYAEQF | 0.7699896 |
| NEPTAAALA | ASSLGWGNYAEQF | 0.82092464 |
| GSIINISSI | ASSLGWGNYAEQF | 0.83262837 |
| VAPGNAGTA | ASSLGWGNYAEQF | 0.79754674 |
| TGGGASLE | ASSLGWGNYAEQF | 0.75127906 |
| VAVYNHYKRI | ASSLGWGNYAEQF | 0.80985963 |
| TDPAPATT | ASSLGWGNYAEQF | 0.8601848 |
| TNRPDILDPALLRPG | ASSLGWGNYAEQF | 0.8411794 |
| LTTTTTS | ASSLGWGNYAEQF | 0.8822058 |
| PTGSGKTL | ASSLGWGNYAEQF | 0.86236537 |
| TGSGKTL | ASSLGWGNYAEQF | 0.83204454 |
| TIGHVDHGK | ASSLGWGNYAEQF | 0.35959095 |
| TIGHVDHGKTT | ASSLGWGNYAEQF | 0.5115325 |
| GHVDHGKTT | ASSLGWGNYAEQF | 0.7421512 |
| AQFEHTLL | ASSLGWGNYAEQF | 0.7666228 |
| IVNQDEL | ASSLGWGNYAEQF | 0.32765895 |
| TNGSQFF | ASSLGWGNYAEQF | 0.83675504 |
| TGKGIVE | ASSLGWGNYAEQF | 0.83733994 |
| LISNASDA | ASSLGWGNYAEQF | 0.7321196 |
| AELRTLGELTKIAKAHGAQ | ASSLGWGNYAEQF | 0.8563607 |
| APNAVAAAEPSW | ASSLGWGNYAEQF | 0.7560564 |
| APQVAGAATAVL | ASSLGWGNYAEQF | 0.7332086 |
| ARRDPAAAAAILPTDA | ASSLGWGNYAEQF | 0.6846639 |
| DAAARDVLAEAGLAEYF | ASSLGWGNYAEQF | 0.8542544 |
| DDQIVLSAKANLAKVEQ | ASSLGWGNYAEQF | 0.5851179 |
| DELLPDTGEDSDEDGH | ASSLGWGNYAEQF | 0.8583467 |

| | | |
|---|---|---|
| ELPEAPANRATVPQD | ASSLGWGNYAEQF | 0.635137 |
| FGQSKIERVTGIG | ASSLGWGNYAEQF | 0.8796226 |
| GAGGTGSTAGGAGGAGGAGGI | ASSLGWGNYAEQF | 0.6385883 |
| GHGGTGAAGGNGGTGGAGGS( | ASSLGWGNYAEQF | 0.84769535 |
| GNGGNGGAGGEGGPGGAGGA( | ASSLGWGNYAEQF | 0.7743906 |
| GSTAGNGGNGGNATNGTIAG( | ASSLGWGNYAEQF | 0.88770425 |
| GTGGDGGNGGNGGTGG | ASSLGWGNYAEQF | 0.80214745 |
| GVRRPAGIPADAPV | ASSLGWGNYAEQF | 0.85424745 |
| GYRQVTDAHLLTLARR | ASSLGWGNYAEQF | 0.7315787 |
| IAKMGQDGHDRG | ASSLGWGNYAEQF | 0.832281 |
| KAEPSIEIQYT | ASSLGWGNYAEQF | 0.63344234 |
| KPVRIRMAADEDTVAVT | ASSLGWGNYAEQF | 0.81535006 |
| MPPRIAGMRLLVIKPEP | ASSLGWGNYAEQF | 0.51248527 |
| MKFYAVALFRSRGVPPTR | ASSLGWGNYAEQF | 0.8203063 |
| NAGGFGILGAVAHSP | ASSLGWGNYAEQF | 0.84769326 |
| NEGREITLAMAEM | ASSLGWGNYAEQF | 0.3429807 |
| NELVEWSHPLGHRWRWE | ASSLGWGNYAEQF | 0.8545382 |
| NLGAGGTGGNGGNPG | ASSLGWGNYAEQF | 0.69219923 |
| NTEWPKWETVMAADPVD | ASSLGWGNYAEQF | 0.72465384 |
| PASAAPAETQP | ASSLGWGNYAEQF | 0.8471501 |
| QQSSDGKNGVRAAEITGE | ASSLGWGNYAEQF | 0.820959 |
| RAPDAVTTSWWKE | ASSLGWGNYAEQF | 0.87473506 |
| SGFGNVGAHTSGWFNQ | ASSLGWGNYAEQF | 0.70112467 |
| SGVAGAVGQAASVGGLKV | ASSLGWGNYAEQF | 0.7613215 |
| SNGGAGTGSAGGNGGT | ASSLGWGNYAEQF | 0.8194107 |
| TGGAGGEGGAAG | ASSLGWGNYAEQF | 0.51289946 |
| TGGDGGAAGAGGAPGHG | ASSLGWGNYAEQF | 0.61959565 |
| TMPPLLAAIANAGRHGV | ASSLGWGNYAEQF | 0.87502944 |
| TSSTAGNSGGA | ASSLGWGNYAEQF | 0.87154865 |
| VPATVPVGILC | ASSLGWGNYAEQF | 0.8924351 |
| VQRPDSGGVVLTGRIS | ASSLGWGNYAEQF | 0.7526959 |
| WTPNPVPATKGVPAGLVQAT | ASSLGWGNYAEQF | 0.756711 |
| YALKRALAKFSDRFDVVII | ASSLGWGNYAEQF | 0.8281102 |
| YLIGSVIV | ASSLGWGNYAEQF | 0.8994713 |
| YGQLVAVID | ASSLGWGNYAEQF | 0.9042145 |
| VTVGGMTLPA | ASSLGWGNYAEQF | 0.8845097 |
| VLALPAV | ASSLGWGNYAEQF | 0.80953455 |
| VIPPTLNYET | ASSLGWGNYAEQF | 0.8648995 |
| VGMGGMHPSSG | ASSLGWGNYAEQF | 0.82737863 |
| VGLAGGAATA | ASSLGWGNYAEQF | 0.83310276 |
| VGGAGGAGGSGP | ASSLGWGNYAEQF | 0.8897145 |
| VFAAFMV | ASSLGWGNYAEQF | 0.86549914 |
| TLYDKLR | ASSLGWGNYAEQF | 0.8077528 |
| TLAVFTVL | ASSLGWGNYAEQF | 0.81955326 |
| TGSVQPARV | ASSLGWGNYAEQF | 0.86554146 |
| TGGGAGSGGT | ASSLGWGNYAEQF | 0.88583994 |
| SHLETSL | ASSLGWGNYAEQF | 0.8800423 |
| SFTTGMT | ASSLGWGNYAEQF | 0.9142599 |
| QVEVAKRMGVRQPTVS | ASSLGWGNYAEQF | 0.88521767 |
| QGSGGPALAM | ASSLGWGNYAEQF | 0.88291514 |
| MLAPSNP | ASSLGWGNYAEQF | 0.29113045 |

| | | |
|---|---|---|
| LVNNAAI | ASSLGWGNYAEQF | 0.6270751 |
| LTGEINTLA | ASSLGWGNYAEQF | 0.85814595 |
| LSIFGPV | ASSLGWGNYAEQF | 0.83781624 |
| LLALLANL | ASSLGWGNYAEQF | 0.5578368 |
| LISNASDALDKLR | ASSLGWGNYAEQF | 0.88365006 |
| LINEAVP | ASSLGWGNYAEQF | 0.8205267 |
| LGYAFSGL | ASSLGWGNYAEQF | 0.78415966 |
| LDFVAVAL | ASSLGWGNYAEQF | 0.8860509 |
| LAQPGAE | ASSLGWGNYAEQF | 0.8129351 |
| ISNLGAVTGF | ASSLGWGNYAEQF | 0.7985765 |
| HQEDPGRI | ASSLGWGNYAEQF | 0.8561974 |
| GTPGEPTP | ASSLGWGNYAEQF | 0.9093428 |
| GFFNSGS | ASSLGWGNYAEQF | 0.8019879 |
| GAGGGGLTAGAN | ASSLGWGNYAEQF | 0.88862616 |
| FVDALVNM | ASSLGWGNYAEQF | 0.856632 |
| FLTVAPEL | ASSLGWGNYAEQF | 0.90023446 |
| FETLTAL | ASSLGWGNYAEQF | 0.7272397 |
| DVINPIEL | ASSLGWGNYAEQF | 0.88099277 |
| APVSGAPV | ASSLGWGNYAEQF | 0.8286856 |
| ALVGGLAW | ASSLGWGNYAEQF | 0.9154111 |
| ALAAAAATAI | ASSLGWGNYAEQF | 0.82110614 |
| AFGLGLLL | ASSLGWGNYAEQF | 0.880949 |
| AFFAAASL | ASSLGWGNYAEQF | 0.84989566 |
| AAAALPAFSPP | ASSLGWGNYAEQF | 0.8940848 |
| VVLAPAL | ASSLGWGNYAEQF | 0.68633306 |
| TIGHVDHGKTTL | ASSLGWGNYAEQF | 0.8221278 |
| LISNASDALDK | ASSLGWGNYAEQF | 0.7251538 |
| LISNASDAL | ASSLGWGNYAEQF | 0.86458755 |
| ISNASDALDK | ASSLGWGNYAEQF | 0.87233984 |
| NQAETLVYQTE | ASSLGWGNYAEQF | 0.46413484 |
| TGGTGGTAGAG | ASSLGWGNYAEQF | 0.8906612 |
| TGSTAGGAGGA | ASSLGWGNYAEQF | 0.7756007 |
| DPLAVPVAAGPV | ASSLGWGNYAEQF | 0.7918302 |
| GGTGGTPGNGGN | ASSLGWGNYAEQF | 0.8994719 |
| PAGLAGVDPATL | ASSLGWGNYAEQF | 0.8265035 |
| GNNLVVEAPHAGASV | ASSLGWGNYAEQF | 0.88475245 |
| KVTRIMSASTLEQPA | ASSLGWGNYAEQF | 0.8594958 |
| LGFALPIALPRRTVV | ASSLGWGNYAEQF | 0.84166706 |
| TRLGNLTARKLVVLV | ASSLGWGNYAEQF | 0.88586587 |
| WSQQRRLIYFPSAGP | ASSLGWGNYAEQF | 0.68484604 |
| GPNGNPDPMAAAVDIR | ASSLGWGNYAEQF | 0.81474304 |
| TQRCVSVADLRAAASR | ASSLGWGNYAEQF | 0.89840984 |
| QFVLDVAEHPVEVLTETV | ASSLGWGNYAEQF | 0.6187817 |
| GGAGGTGGAGGNGGTQAGDG | ASSLGWGNYAEQF | 0.8409827 |
| PLLATLRAIAVDVDAKPAQV | ASSLGWGNYAEQF | 0.38357294 |
| APKFRTGGYMIDFWGVGYQV | ASSLGWGNYAEQF | 0.31512538 |
| GAGGAGGAGGAAGGPGGTGG | ASSLGWGNYAEQF | 0.34028193 |
| IGALGLLATDVAPAGSVAFG | ASSLGWGNYAEQF | 0.7798412 |
| AGGAGGIGGNGGAGGAGGVL | ASSLGWGNYAEQF | 0.8044403 |
| ADELIGLL | ASSLGWGNYAEQF | 0.7406864 |
| ADRALKA | ASSLGWGNYAEQF | 0.32340008 |

| | | |
|---|---|---|
| AGAVGQVGGAGGA | ASSLGWGNYAEQF | 0.52247804 |
| AGGGGGLGGAP | ASSLGWGNYAEQF | 0.31599218 |
| AGGLLYGNGGAG | ASSLGWGNYAEQF | 0.44751227 |
| AGLESIGL | ASSLGWGNYAEQF | 0.7110387 |
| AGLKTGII | ASSLGWGNYAEQF | 0.46098155 |
| ASLAALLL | ASSLGWGNYAEQF | 0.4149118 |
| ASVINISSI | ASSLGWGNYAEQF | 0.42324835 |
| ATPPGPA | ASSLGWGNYAEQF | 0.62351286 |
| ATVASALV | ASSLGWGNYAEQF | 0.64838636 |
| ATVASLIL | ASSLGWGNYAEQF | 0.48207378 |
| AVNGPVLIRH | ASSLGWGNYAEQF | 0.83509004 |
| DGAAGAVGI | ASSLGWGNYAEQF | 0.69568086 |
| DPLLHGI | ASSLGWGNYAEQF | 0.76356065 |
| ELGAALGAL | ASSLGWGNYAEQF | 0.11199025 |
| ELGRTQAV | ASSLGWGNYAEQF | 0.25435287 |
| FVAGGYLS | ASSLGWGNYAEQF | 0.42423174 |
| GASGGAGGAGGW | ASSLGWGNYAEQF | 0.5674814 |
| GGAGAGGDGGAGGI | ASSLGWGNYAEQF | 0.21643981 |
| GTGHNGSQPGMGGNG | ASSLGWGNYAEQF | 0.6398731 |
| GYAAASASAS | ASSLGWGNYAEQF | 0.45002395 |
| HQVDVAAVQQG | ASSLGWGNYAEQF | 0.3804611 |
| IGAAIAHT | ASSLGWGNYAEQF | 0.744074 |
| IGMIEVAQL | ASSLGWGNYAEQF | 0.38805014 |
| IRLAPEHTSAMD | ASSLGWGNYAEQF | 0.3983999 |
| IYNVSVL | ASSLGWGNYAEQF | 0.42188585 |
| KQGFLGM | ASSLGWGNYAEQF | 0.8114121 |
| LESVITQL | ASSLGWGNYAEQF | 0.25898433 |
| LGLGALVVV | ASSLGWGNYAEQF | 0.69279695 |
| LIGNGGNGGSGGI | ASSLGWGNYAEQF | 0.43615907 |
| LNAINAPTL | ASSLGWGNYAEQF | 0.64305955 |
| LQLLNTGV | ASSLGWGNYAEQF | 0.25669548 |
| LQTSIEF | ASSLGWGNYAEQF | 0.72888124 |
| MSFAVSEL | ASSLGWGNYAEQF | 0.46604818 |
| NTGALGL | ASSLGWGNYAEQF | 0.8650935 |
| PPGAGGAGGAGTT | ASSLGWGNYAEQF | 0.36179748 |
| QLVGLGI | ASSLGWGNYAEQF | 0.47755018 |
| RFLEQVHQL | ASSLGWGNYAEQF | 0.37103134 |
| RGVGYVLR | ASSLGWGNYAEQF | 0.8042048 |
| SATGVFAGVI | ASSLGWGNYAEQF | 0.19109191 |
| SLPAITLP | ASSLGWGNYAEQF | 0.37243053 |
| STALAFTV | ASSLGWGNYAEQF | 0.37173194 |
| SVLTVGLGK | ASSLGWGNYAEQF | 0.5807352 |
| TALAGAGSGI | ASSLGWGNYAEQF | 0.7017719 |
| TGLPFTGI | ASSLGWGNYAEQF | 0.32463536 |
| VAEHVTV | ASSLGWGNYAEQF | 0.77853703 |
| VDIVLNSL | ASSLGWGNYAEQF | 0.6433346 |
| VLTGGGAVL | ASSLGWGNYAEQF | 0.31608328 |
| VNVGLAYGI | ASSLGWGNYAEQF | 0.79893035 |
| VSSAAVGAV | ASSLGWGNYAEQF | 0.31472883 |
| VTSVYEAGV | ASSLGWGNYAEQF | 0.77577925 |
| WVALDRGA | ASSLGWGNYAEQF | 0.58688384 |

| | | |
|---|---|---|
| YLVGSVVI | ASSLGWGNYAEQF | 0.52969193 |
| GPPGTGKTLL | ASSLGWGNYAEQF | 0.632698 |
| GVGMTKF | ASSLGWGNYAEQF | 0.43435085 |
| IGIGGSDL | ASSLGWGNYAEQF | 0.3334152 |
| IGTLLAEL | ASSLGWGNYAEQF | 0.7553269 |
| IYTSGTT | ASSLGWGNYAEQF | 0.86980045 |
| LSFPAGV | ASSLGWGNYAEQF | 0.87844336 |
| LYTSGTT | ASSLGWGNYAEQF | 0.7845111 |
| VGAFPFGF | ASSLGWGNYAEQF | 0.7221703 |
| VGAGTVEF | ASSLGWGNYAEQF | 0.8089578 |
| VGKSTLLL | ASSLGWGNYAEQF | 0.86234915 |
| AALAHAGGGIV | ASSLGWGNYAEQF | 0.8886625 |
| AALVAVSLL | ASSLGWGNYAEQF | 0.8654456 |
| AAVSRAVAAL | ASSLGWGNYAEQF | 0.6941442 |
| AGGSSGIGGF | ASSLGWGNYAEQF | 0.8632282 |
| AGLTVANANG | ASSLGWGNYAEQF | 0.8399246 |
| AGLVGAVL | ASSLGWGNYAEQF | 0.8309152 |
| ALSQAFTV | ASSLGWGNYAEQF | 0.86123276 |
| ALTFSVSL | ASSLGWGNYAEQF | 0.8370966 |
| ALTGAALVAL | ASSLGWGNYAEQF | 0.7824009 |
| ASFSIGAGVL | ASSLGWGNYAEQF | 0.7168283 |
| DGSVYKI | ASSLGWGNYAEQF | 0.8643828 |
| EQLAEIV | ASSLGWGNYAEQF | 0.8745333 |
| FAVSELTVM | ASSLGWGNYAEQF | 0.87241983 |
| GAAPLAAGL | ASSLGWGNYAEQF | 0.8841858 |
| GADGGDGGSGGAGG | ASSLGWGNYAEQF | 0.8768113 |
| GFGLDTGGAGG | ASSLGWGNYAEQF | 0.86978793 |
| GLASGGLL | ASSLGWGNYAEQF | 0.38830906 |
| GRIAGAAVLL | ASSLGWGNYAEQF | 0.6072742 |
| GSAGSLNTSF | ASSLGWGNYAEQF | 0.78537536 |
| GTGQIGI | ASSLGWGNYAEQF | 0.8018073 |
| GTLIALIL | ASSLGWGNYAEQF | 0.43746847 |
| IAATMGFVVG | ASSLGWGNYAEQF | 0.8126353 |
| IFAQALL | ASSLGWGNYAEQF | 0.80217886 |
| IGAAVALGF | ASSLGWGNYAEQF | 0.75326914 |
| IGLTPLAL | ASSLGWGNYAEQF | 0.873793 |
| ILGSNTSTL | ASSLGWGNYAEQF | 0.80346954 |
| IVFSLTVL | ASSLGWGNYAEQF | 0.67697495 |
| LASKATGF | ASSLGWGNYAEQF | 0.79432124 |
| LASVALIL | ASSLGWGNYAEQF | 0.8741459 |
| LFAVGLTL | ASSLGWGNYAEQF | 0.77046907 |
| LFNGGIGTYI | ASSLGWGNYAEQF | 0.87090766 |
| LTLGAGII | ASSLGWGNYAEQF | 0.802399 |
| LVAAALGDI | ASSLGWGNYAEQF | 0.87856394 |
| NTAFVQLGI | ASSLGWGNYAEQF | 0.63393724 |
| QLFSGAEL | ASSLGWGNYAEQF | 0.875072 |
| SAATFITL | ASSLGWGNYAEQF | 0.83988494 |
| SNYGLANLGV | ASSLGWGNYAEQF | 0.88834906 |
| SYGGAYAVM | ASSLGWGNYAEQF | 0.8297926 |
| TTGVGGLTLG | ASSLGWGNYAEQF | 0.8646843 |
| TTQTIYI | ASSLGWGNYAEQF | 0.8358389 |

| | | |
|---|---|---|
| VAAGAGLGAVI | ASSLGWGNYAEQF | 0.88310087 |
| VAAVVTLL | ASSLGWGNYAEQF | 0.6793351 |
| VGFLGGLI | ASSLGWGNYAEQF | 0.8198403 |
| VGVIPILL | ASSLGWGNYAEQF | 0.7536031 |
| VIFASSLI | ASSLGWGNYAEQF | 0.83740646 |
| VKGGDGGAAAG | ASSLGWGNYAEQF | 0.8658399 |
| VTKTFTNI | ASSLGWGNYAEQF | 0.29308578 |
| VVAAAILL | ASSLGWGNYAEQF | 0.87077415 |
| YAALDVEL | ASSLGWGNYAEQF | 0.7400522 |
| LASVALLL | ASSLGWGNYAEQF | 0.7314253 |
| LDAVHRV | ASSLGWGNYAEQF | 0.87838936 |
| QGLSSSIF | ASSLGWGNYAEQF | 0.84417886 |
| RGTFANIRL | ASSLGWGNYAEQF | 0.87418014 |
| RQLWWGHRI | ASSLGWGNYAEQF | 0.82298076 |
| YGPPGTGKTLL | ASSLGWGNYAEQF | 0.8502071 |
| AALVGAPANVA | ASSLGWGNYAEQF | 0.8401519 |
| AATFRRTE | ASSLGWGNYAEQF | 0.50683594 |
| AGALGGGI | ASSLGWGNYAEQF | 0.73675966 |
| AGGAGGAGGIGAG | ASSLGWGNYAEQF | 0.8833583 |
| AGSGALGL | ASSLGWGNYAEQF | 0.7852913 |
| AQIADQI | ASSLGWGNYAEQF | 0.8457414 |
| ASAVAVTYL | ASSLGWGNYAEQF | 0.61327606 |
| ASTVLGGTGG | ASSLGWGNYAEQF | 0.53880703 |
| AVGGEPAK | ASSLGWGNYAEQF | 0.67399275 |
| CEQGTFGGTC | ASSLGWGNYAEQF | 0.8257041 |
| DEGKAPAG | ASSLGWGNYAEQF | 0.853263 |
| DTVVFPAL | ASSLGWGNYAEQF | 0.80999553 |
| FEGITMAG | ASSLGWGNYAEQF | 0.4662999 |
| FPLVVQD | ASSLGWGNYAEQF | 0.81576526 |
| GAFTTPQ | ASSLGWGNYAEQF | 0.628963 |
| GFGGFGTAGGVGGA | ASSLGWGNYAEQF | 0.73068726 |
| GFLAQFL | ASSLGWGNYAEQF | 0.7848562 |
| GGAGGAGGNAGVA | ASSLGWGNYAEQF | 0.74048066 |
| GGVGTAGAGGAGGAGG | ASSLGWGNYAEQF | 0.6725377 |
| GNGGGGDAGNAGSGGN | ASSLGWGNYAEQF | 0.46401352 |
| GTGGAGGAGNDAGS | ASSLGWGNYAEQF | 0.84146625 |
| GTTNTGWFNS | ASSLGWGNYAEQF | 0.847314 |
| IFMGPLFTL | ASSLGWGNYAEQF | 0.83741343 |
| ILAHAIEAI | ASSLGWGNYAEQF | 0.8506318 |
| IRSAIREAL | ASSLGWGNYAEQF | 0.8347877 |
| ITFAGGG | ASSLGWGNYAEQF | 0.81838155 |
| ITGMSGHQEV | ASSLGWGNYAEQF | 0.1952425 |
| IVAGLACTL | ASSLGWGNYAEQF | 0.3564548 |
| IVGGGLAGA | ASSLGWGNYAEQF | 0.49017122 |
| IVTGVLNI | ASSLGWGNYAEQF | 0.7027442 |
| LAQQGCI | ASSLGWGNYAEQF | 0.2565359 |
| LCLHVEN | ASSLGWGNYAEQF | 0.81346154 |
| LDAISDGS | ASSLGWGNYAEQF | 0.57674176 |
| LGGFAAML | ASSLGWGNYAEQF | 0.624823 |
| LGGKNPMI | ASSLGWGNYAEQF | 0.840047 |
| LGPKGVAAIL | ASSLGWGNYAEQF | 0.61569124 |

| | | |
|---|---|---|
| LMVGVSI | ASSLGWGNYAEQF | 0.49689156 |
| LVGAAFVL | ASSLGWGNYAEQF | 0.55993474 |
| LVLAVAPL | ASSLGWGNYAEQF | 0.8391862 |
| MAAIAVIDAA | ASSLGWGNYAEQF | 0.43621755 |
| NNGINAELQ | ASSLGWGNYAEQF | 0.8242334 |
| RGEPLVLVHG | ASSLGWGNYAEQF | 0.5695261 |
| RGGAGGAAGLI | ASSLGWGNYAEQF | 0.83483255 |
| RSYRMHPAV | ASSLGWGNYAEQF | 0.3840583 |
| SAAHLLAGQLA | ASSLGWGNYAEQF | 0.8297463 |
| SAAVGSY | ASSLGWGNYAEQF | 0.7499393 |
| SPHAVLHRI | ASSLGWGNYAEQF | 0.86727035 |
| SQIAKVL | ASSLGWGNYAEQF | 0.6687133 |
| TALGHLVP | ASSLGWGNYAEQF | 0.7327135 |
| TAPTTSGAA | ASSLGWGNYAEQF | 0.6614797 |
| TDGDDGNGGSPGT | ASSLGWGNYAEQF | 0.8469339 |
| TDGLGEN | ASSLGWGNYAEQF | 0.33736894 |
| TGGPPLPGQA | ASSLGWGNYAEQF | 0.664479 |
| TIPAITI | ASSLGWGNYAEQF | 0.47453764 |
| TIPGFTI | ASSLGWGNYAEQF | 0.72698855 |
| TSIASGALA | ASSLGWGNYAEQF | 0.79952496 |
| TVTLPGAAI | ASSLGWGNYAEQF | 0.18387339 |
| VASGLSAALP | ASSLGWGNYAEQF | 0.7736442 |
| VGAGVIGI | ASSLGWGNYAEQF | 0.6436531 |
| VGPKRFGV | ASSLGWGNYAEQF | 0.4476014 |
| VNVAFPD | ASSLGWGNYAEQF | 0.83109117 |
| VSMTNSGVS | ASSLGWGNYAEQF | 0.7835763 |
| VTDVLGDLSV | ASSLGWGNYAEQF | 0.8407215 |
| VVVVGAGFAGL | ASSLGWGNYAEQF | 0.7135738 |
| VYTVGTVV | ASSLGWGNYAEQF | 0.6673323 |
| WTTFSGP | ASSLGWGNYAEQF | 0.76005375 |
| YDELVEAF | ASSLGWGNYAEQF | 0.42053658 |
| YGTGGAGGNGG | ASSLGWGNYAEQF | 0.5720812 |
| YTNVMAV | ASSLGWGNYAEQF | 0.8437594 |
| FLEQVHQL | ASSLGWGNYAEQF | 0.8252498 |
| GLSLLAVAV | ASSLGWGNYAEQF | 0.8617344 |
| LEQVHQL | ASSLGWGNYAEQF | 0.7470842 |
| LNAAGAI | ASSLGWGNYAEQF | 0.60950214 |
| PAVHASL | ASSLGWGNYAEQF | 0.73854244 |
| TELEDDA | ASSLGWGNYAEQF | 0.85718226 |
| TPPGPPN | ASSLGWGNYAEQF | 0.8650634 |
| TPPPPPPP | ASSLGWGNYAEQF | 0.84333825 |
| TQSPALI | ASGDRGDTQY | 0.5780281 |
| EFAAAFPQ | ASGDRGDTQY | 0.74977636 |
| LLSGNLDVL | ASGDRGDTQY | 0.70510244 |
| GPANLGNYNI | ASGDRGDTQY | 0.7253902 |
| TLGPTLP | ASGDRGDTQY | 0.85155916 |
| VYAYSSCCK | ASGDRGDTQY | 0.79903656 |
| APAPAPAPNI | ASGDRGDTQY | 0.7378111 |
| PGGPVPIP | ASGDRGDTQY | 0.7511823 |
| FLTVALEP | ASGDRGDTQY | 0.75496995 |
| SQYFGANL | ASGDRGDTQY | 0.820485 |

| | | |
|---|---|---|
| GGVKPAA | ASGDRGDTQY | 0.80025125 |
| GGTGGAGGAGGW | ASGDRGDTQY | 0.88270056 |
| GVGGVAGLGGAGG | ASGDRGDTQY | 0.85190296 |
| VLLVLATTI | ASGDRGDTQY | 0.8662224 |
| IGIFAAML | ASGDRGDTQY | 0.18580958 |
| AVGDLID | ASGDRGDTQY | 0.37197524 |
| NISPLPL | ASGDRGDTQY | 0.7530412 |
| VTAVFVAI | ASGDRGDTQY | 0.7021842 |
| SGDDNDAAL | ASGDRGDTQY | 0.6035569 |
| AVAALAHS | ASGDRGDTQY | 0.78660023 |
| GGVTVMATP | ASGDRGDTQY | 0.6719513 |
| AGLAANVRD | ASGDRGDTQY | 0.6327081 |
| TNRPDLIDPALLRPG | ASGDRGDTQY | 0.84890616 |
| LANLGVTAL | ASGDRGDTQY | 0.7516142 |
| TGERIEL | ASGDRGDTQY | 0.6786825 |
| NGEELAA | ASGDRGDTQY | 0.6992842 |
| VLSLVWG | ASGDRGDTQY | 0.8358947 |
| IGYALWL | ASGDRGDTQY | 0.48214895 |
| LSLLLAPL | ASGDRGDTQY | 0.80011594 |
| TLDEALK | ASGDRGDTQY | 0.64059037 |
| TDGAGTAEGG | ASGDRGDTQY | 0.8345079 |
| NAPGGAGGNGG | ASGDRGDTQY | 0.4871431 |
| VSVFMGMGGNF | ASGDRGDTQY | 0.8513674 |
| LASYLAAL | ASGDRGDTQY | 0.7823827 |
| LVGGTPAL | ASGDRGDTQY | 0.89230794 |
| VRNADVL | ASGDRGDTQY | 0.71073794 |
| TCSSPFLVSAM | ASGDRGDTQY | 0.742398 |
| VHITDAE | ASGDRGDTQY | 0.64023465 |
| EYAAPSF | ASGDRGDTQY | 0.84358615 |
| VVDAFHGGL | ASGDRGDTQY | 0.49580175 |
| ASVVDPQ | ASGDRGDTQY | 0.73759544 |
| SYNEFGGLS | ASGDRGDTQY | 0.42264968 |
| DPDGSVY | ASGDRGDTQY | 0.8154207 |
| MVSTIHATE | ASGDRGDTQY | 0.83294773 |
| LAAGVADPV | ASGDRGDTQY | 0.50515467 |
| TKEQIAA | ASGDRGDTQY | 0.86274314 |
| SLASPAAL | ASGDRGDTQY | 0.80332613 |
| IGLVFYGV | ASGDRGDTQY | 0.6823282 |
| FLLDASNL | ASGDRGDTQY | 0.8808644 |
| YQVGNFP | ASGDRGDTQY | 0.6790432 |
| ESLLSQST | ASGDRGDTQY | 0.8785099 |
| LFTVGFAAVG | ASGDRGDTQY | 0.8176054 |
| ATPPTPT | ASGDRGDTQY | 0.68288267 |
| TDLATLG | ASGDRGDTQY | 0.8104633 |
| SAGGTGR | ASGDRGDTQY | 0.46448773 |
| GFSNSGS | ASGDRGDTQY | 0.61812705 |
| GGNGGSGGTSV | ASGDRGDTQY | 0.8734304 |
| GTSVATGGAGN | ASGDRGDTQY | 0.78955704 |
| NGGNGGSGGTS | ASGDRGDTQY | 0.8139707 |
| AVVKFPH | ASGDRGDTQY | 0.7390105 |
| TPPPPPP | ASGDRGDTQY | 0.7148831 |

| | | |
|---|---|---|
| AAPAAAAAPA | ASGDRGDTQY | 0.7743425 |
| IALNEDL | ASGDRGDTQY | 0.79817295 |
| IVNVSSM | ASGDRGDTQY | 0.81463367 |
| TGIPPAPR | ASGDRGDTQY | 0.79274976 |
| FDLGGGTF | ASGDRGDTQY | 0.69563115 |
| FDLGGGTFDVS | ASGDRGDTQY | 0.7862893 |
| NEPTAAA | ASGDRGDTQY | 0.752854 |
| NEPTAAALA | ASGDRGDTQY | 0.74772406 |
| GSIINISSI | ASGDRGDTQY | 0.7981447 |
| VAPGNAGTA | ASGDRGDTQY | 0.74170196 |
| TGGGASLE | ASGDRGDTQY | 0.74957204 |
| VAVYNHYKRI | ASGDRGDTQY | 0.75816524 |
| TDPAPATT | ASGDRGDTQY | 0.7854121 |
| TNRPDILDPALLRPG | ASGDRGDTQY | 0.80309296 |
| LTTTTTS | ASGDRGDTQY | 0.80109864 |
| PTGSGKTL | ASGDRGDTQY | 0.8114569 |
| TGSGKTL | ASGDRGDTQY | 0.79551196 |
| TIGHVDHGK | ASGDRGDTQY | 0.7811295 |
| TIGHVDHGKTT | ASGDRGDTQY | 0.38444942 |
| GHVDHGKTT | ASGDRGDTQY | 0.6657226 |
| AQFEHTLL | ASGDRGDTQY | 0.76634926 |
| IVNQDEL | ASGDRGDTQY | 0.75758916 |
| TNGSQFF | ASGDRGDTQY | 0.3958097 |
| TGKGIVE | ASGDRGDTQY | 0.7800034 |
| LISNASDA | ASGDRGDTQY | 0.77256715 |
| AELRTLGELTKIAKAHGAQ | ASGDRGDTQY | 0.70935214 |
| APNAVAAAEPSW | ASGDRGDTQY | 0.7961962 |
| APQVAGAATAVL | ASGDRGDTQY | 0.74225104 |
| ARRDPAAAAAILPTDA | ASGDRGDTQY | 0.6640738 |
| DAAARDVLAEAGLAEYF | ASGDRGDTQY | 0.74473006 |
| DDQIVLSAKANLAKVEQ | ASGDRGDTQY | 0.80782676 |
| DELLPDTGEDSDEDGH | ASGDRGDTQY | 0.694381 |
| ELPEAPANRATVPQD | ASGDRGDTQY | 0.7922307 |
| FGQSKIERVTGIG | ASGDRGDTQY | 0.7723954 |
| GAGGTGSTAGGAGGAGGAGGI | ASGDRGDTQY | 0.800298 |
| GHGGTGAAGGNGGTGGAGGS( | ASGDRGDTQY | 0.6147362 |
| GNGGNGGAGGEGGPGGAGGA( | ASGDRGDTQY | 0.78356063 |
| GSTAGNGGNGGNATNGTIAG( | ASGDRGDTQY | 0.75803155 |
| GTGGDGGNGGNGGTGG | ASGDRGDTQY | 0.81597614 |
| GVRRPAGIPADAPV | ASGDRGDTQY | 0.746794 |
| GYRQVTDAHLLTLARR | ASGDRGDTQY | 0.78349805 |
| IAKMGQDGHDRG | ASGDRGDTQY | 0.7560066 |
| KAEPSIEIQYT | ASGDRGDTQY | 0.8127527 |
| KPVRIRMAADEDTVAVT | ASGDRGDTQY | 0.6333168 |
| MPPRIAGMRLLVIKPEP | ASGDRGDTQY | 0.7809384 |
| MKFYAVALFRSRGVPPTR | ASGDRGDTQY | 0.73326766 |
| NAGGFGILGAVAHSP | ASGDRGDTQY | 0.78977966 |
| NEGREITLAMAEM | ASGDRGDTQY | 0.79040265 |
| NELVEWSHPLGHRWRWE | ASGDRGDTQY | 0.69587725 |
| NLGAGGTGGNGGNPG | ASGDRGDTQY | 0.80877614 |
| NTEWPKWETVMAADPVD | ASGDRGDTQY | 0.7371642 |

| | | |
|---|---|---|
| PASAAPAETQP | ASGDRGDTQY | 0.7277211 |
| QQSSDGKNGVRAAEITGE | ASGDRGDTQY | 0.80287635 |
| RAPDAVTTSWWKE | ASGDRGDTQY | 0.75659287 |
| SGFGNVGAHTSGWFNQ | ASGDRGDTQY | 0.79780185 |
| SGVAGAVGQAASVGGLKV | ASGDRGDTQY | 0.73243594 |
| SNGGAGTGSAGGNGGT | ASGDRGDTQY | 0.7989535 |
| TGGAGGEGGAAG | ASGDRGDTQY | 0.8002257 |
| TGGDGGAAGAGGAPGHG | ASGDRGDTQY | 0.7265545 |
| TMPPLLAAIANAGRHGV | ASGDRGDTQY | 0.7022066 |
| TSSTAGNSGGA | ASGDRGDTQY | 0.796308 |
| VPATVPVGILC | ASGDRGDTQY | 0.77645195 |
| VQRPDSGGVVLTGRIS | ASGDRGDTQY | 0.8002258 |
| WTPNPVPATKGVPAGLVQAT | ASGDRGDTQY | 0.69554555 |
| YALKRALAKFSDRFDVVII | ASGDRGDTQY | 0.6709542 |
| YLIGSVIV | ASGDRGDTQY | 0.7073859 |
| YGQLVAVID | ASGDRGDTQY | 0.79148966 |
| VTVGGMTLPA | ASGDRGDTQY | 0.80433637 |
| VLALPAV | ASGDRGDTQY | 0.7680533 |
| VIPPTLNYET | ASGDRGDTQY | 0.6996249 |
| VGMGGMHPSSG | ASGDRGDTQY | 0.7575666 |
| VGLAGGAATA | ASGDRGDTQY | 0.74050474 |
| VGGAGGAGGSGP | ASGDRGDTQY | 0.7220503 |
| VFAAFMV | ASGDRGDTQY | 0.7738745 |
| TLYDKLR | ASGDRGDTQY | 0.77063197 |
| TLAVFTVL | ASGDRGDTQY | 0.7160337 |
| TGSVQPARV | ASGDRGDTQY | 0.7330657 |
| TGGGAGSGGT | ASGDRGDTQY | 0.7439339 |
| SHLETSL | ASGDRGDTQY | 0.7871131 |
| SFTTGMT | ASGDRGDTQY | 0.77141565 |
| QVEVAKRMGVRQPTVS | ASGDRGDTQY | 0.80789435 |
| QGSGGPALAM | ASGDRGDTQY | 0.7871049 |
| MLAPSNP | ASGDRGDTQY | 0.78685427 |
| LVNNAAI | ASGDRGDTQY | 0.39579776 |
| LTGEINTLA | ASGDRGDTQY | 0.5964816 |
| LSIFGPV | ASGDRGDTQY | 0.72410834 |
| LLALLANL | ASGDRGDTQY | 0.73894155 |
| LISNASDALDKLR | ASGDRGDTQY | 0.48514563 |
| LINEAVP | ASGDRGDTQY | 0.77363765 |
| LGYAFSGL | ASGDRGDTQY | 0.73752517 |
| LDFVAVAL | ASGDRGDTQY | 0.67155993 |
| LAQPGAE | ASGDRGDTQY | 0.78345 |
| ISNLGAVTGF | ASGDRGDTQY | 0.7185155 |
| HQEDPGRI | ASGDRGDTQY | 0.7182706 |
| GTPGEPTP | ASGDRGDTQY | 0.7259124 |
| GFFNSGS | ASGDRGDTQY | 0.800825 |
| GAGGGGLTAGAN | ASGDRGDTQY | 0.6004987 |
| FVDALVNM | ASGDRGDTQY | 0.7806907 |
| FLTVAPEL | ASGDRGDTQY | 0.7137533 |
| FETLTAL | ASGDRGDTQY | 0.7950603 |
| DVINPIEL | ASGDRGDTQY | 0.6247492 |
| APVSGAPV | ASGDRGDTQY | 0.7808442 |

| | | |
|---|---|---|
| ALVGGLAW | ASGDRGDTQY | 0.73537135 |
| ALAAAAATAI | ASGDRGDTQY | 0.8123573 |
| AFGLGLLL | ASGDRGDTQY | 0.7326223 |
| AFFAAASL | ASGDRGDTQY | 0.7835455 |
| AAAALPAFSPP | ASGDRGDTQY | 0.7636366 |
| VVLAPAL | ASGDRGDTQY | 0.79289556 |
| TIGHVDHGKTTL | ASGDRGDTQY | 0.61582744 |
| LISNASDALDK | ASGDRGDTQY | 0.74897456 |
| LISNASDAL | ASGDRGDTQY | 0.6756879 |
| ISNASDALDK | ASGDRGDTQY | 0.77167714 |
| NQAETLVYQTE | ASGDRGDTQY | 0.77432406 |
| TGGTGGTAGAG | ASGDRGDTQY | 0.53083223 |
| TGSTAGGAGGA | ASGDRGDTQY | 0.796818 |
| DPLAVPVAAGPV | ASGDRGDTQY | 0.69624823 |
| GGTGGTPGNGGN | ASGDRGDTQY | 0.70250916 |
| PAGLAGVDPATL | ASGDRGDTQY | 0.7903372 |
| GNNLVVEAPHAGASV | ASGDRGDTQY | 0.7292117 |
| KVTRIMSASTLEQPA | ASGDRGDTQY | 0.78850603 |
| LGFALPIALPRRTVV | ASGDRGDTQY | 0.7224145 |
| TRLGNLTARKLVVLV | ASGDRGDTQY | 0.74898565 |
| WSQQRRLIYFPSAGP | ASGDRGDTQY | 0.77944255 |
| GPNGNPDPMAAAVDIR | ASGDRGDTQY | 0.6280527 |
| TQRCVSVADLRAAASR | ASGDRGDTQY | 0.6585612 |
| QFVLDVAEHPVEVLTETV | ASGDRGDTQY | 0.79644144 |
| GGAGGTGGAGGNGGTQAGDG | ASGDRGDTQY | 0.6270125 |
| PLLATLRAIAVDVDAKPAQV | ASGDRGDTQY | 0.6849451 |
| APKFRTGGYMIDFWGVGYQV | ASGDRGDTQY | 0.48458982 |
| GAGGAGGAGGAAGGPGGTGG | ASGDRGDTQY | 0.49384376 |
| IGALGLLATDVAPAGSVAFG | ASGDRGDTQY | 0.55222356 |
| AGGAGGIGGNGGAGGAGGVL | ASGDRGDTQY | 0.65566325 |
| ADELIGLL | ASGDRGDTQY | 0.69561434 |
| ADRALKA | ASGDRGDTQY | 0.64495295 |
| AGAVGQVGGAGGA | ASGDRGDTQY | 0.5363006 |
| AGGGGGLGGAP | ASGDRGDTQY | 0.63627774 |
| AGGLLYGNGGAG | ASGDRGDTQY | 0.55826336 |
| AGLESIGL | ASGDRGDTQY | 0.57532775 |
| AGLKTGII | ASGDRGDTQY | 0.6168478 |
| ASLAALLL | ASGDRGDTQY | 0.6127356 |
| ASVINISSI | ASGDRGDTQY | 0.5423067 |
| ATPPGPA | ASGDRGDTQY | 0.5388055 |
| ATVASALV | ASGDRGDTQY | 0.64180636 |
| ATVASLIL | ASGDRGDTQY | 0.67187494 |
| AVNGPVLIRH | ASGDRGDTQY | 0.6527229 |
| DGAAGAVGI | ASGDRGDTQY | 0.702964 |
| DPLLHGI | ASGDRGDTQY | 0.6489314 |
| ELGAALGAL | ASGDRGDTQY | 0.649546 |
| ELGRTQAV | ASGDRGDTQY | 0.319731 |
| FVAGGYLS | ASGDRGDTQY | 0.4159998 |
| GASGGAGGAGGW | ASGDRGDTQY | 0.56183106 |
| GGAGAGGDGGAGGI | ASGDRGDTQY | 0.59011877 |
| GTGHNGSQPGMGGNG | ASGDRGDTQY | 0.35518208 |

| | | |
|---|---|---|
| GYAAASASAS | ASGDRGDTQY | 0.660603 |
| HQVDVAAVQQG | ASGDRGDTQY | 0.576964 |
| IGAAIAHT | ASGDRGDTQY | 0.52899134 |
| IGMIEVAQL | ASGDRGDTQY | 0.65713155 |
| IRLAPEHTSAMD | ASGDRGDTQY | 0.54156476 |
| IYNVSVL | ASGDRGDTQY | 0.51141506 |
| KQGFLGM | ASGDRGDTQY | 0.54051614 |
| LESVITQL | ASGDRGDTQY | 0.69999444 |
| LGLGALVVV | ASGDRGDTQY | 0.42338622 |
| LIGNGGNGGSGGI | ASGDRGDTQY | 0.66043323 |
| LNAINAPTL | ASGDRGDTQY | 0.55364835 |
| LQLLNTGV | ASGDRGDTQY | 0.6677295 |
| LQTSIEF | ASGDRGDTQY | 0.45861676 |
| MSFAVSEL | ASGDRGDTQY | 0.644182 |
| NTGALGL | ASGDRGDTQY | 0.5446702 |
| PPGAGGAGGAGTT | ASGDRGDTQY | 0.71081406 |
| QLVGLGI | ASGDRGDTQY | 0.54997593 |
| RFLEQVHQL | ASGDRGDTQY | 0.63290715 |
| RGVGYVLR | ASGDRGDTQY | 0.63133395 |
| SATGVFAGVI | ASGDRGDTQY | 0.6663486 |
| SLPAITLP | ASGDRGDTQY | 0.38803285 |
| STALAFTV | ASGDRGDTQY | 0.54669887 |
| SVLTVGLGK | ASGDRGDTQY | 0.5160012 |
| TALAGAGSGI | ASGDRGDTQY | 0.5958889 |
| TGLPFTGI | ASGDRGDTQY | 0.64245677 |
| VAEHVTV | ASGDRGDTQY | 0.436558 |
| VDIVLNSL | ASGDRGDTQY | 0.65961593 |
| VLTGGGAVL | ASGDRGDTQY | 0.550362 |
| VNVGLAYGI | ASGDRGDTQY | 0.5344929 |
| VSSAAVGAV | ASGDRGDTQY | 0.6505194 |
| VTSVYEAGV | ASGDRGDTQY | 0.600963 |
| WVALDRGA | ASGDRGDTQY | 0.65990543 |
| YLVGSVVI | ASGDRGDTQY | 0.5394145 |
| GPPGTGKTLL | ASGDRGDTQY | 0.57833683 |
| GVGMTKF | ASGDRGDTQY | 0.64200747 |
| IGIGGSDL | ASGDRGDTQY | 0.53918886 |
| IGTLLAEL | ASGDRGDTQY | 0.5003303 |
| IYTSGTT | ASGDRGDTQY | 0.66781175 |
| LSFPAGV | ASGDRGDTQY | 0.7873436 |
| LYTSGTT | ASGDRGDTQY | 0.8004557 |
| VGAFPFGF | ASGDRGDTQY | 0.64724636 |
| VGAGTVEF | ASGDRGDTQY | 0.5710213 |
| VGKSTLLL | ASGDRGDTQY | 0.6927567 |
| AALAHAGGGIV | ASGDRGDTQY | 0.8310144 |
| AALVAVSLL | ASGDRGDTQY | 0.8375889 |
| AAVSRAVAAL | ASGDRGDTQY | 0.7761969 |
| AGGSSGIGGF | ASGDRGDTQY | 0.5871892 |
| AGLTVANANG | ASGDRGDTQY | 0.78079045 |
| AGLVGAVL | ASGDRGDTQY | 0.7198213 |
| ALSQAFTV | ASGDRGDTQY | 0.71426785 |
| ALTFSVSL | ASGDRGDTQY | 0.78742635 |

| | | |
|---|---|---|
| ALTGAALVAL | ASGDRGDTQY | 0.78528714 |
| ASFSIGAGVL | ASGDRGDTQY | 0.713184 |
| DGSVYKI | ASGDRGDTQY | 0.66132367 |
| EQLAEIV | ASGDRGDTQY | 0.75964355 |
| FAVSELTVM | ASGDRGDTQY | 0.8209617 |
| GAAPLAAGL | ASGDRGDTQY | 0.7943028 |
| GADGGDGGSGGAGG | ASGDRGDTQY | 0.86249006 |
| GFGLDTGGAGG | ASGDRGDTQY | 0.8256166 |
| GLASGGLL | ASGDRGDTQY | 0.8063048 |
| GRIAGAAVLL | ASGDRGDTQY | 0.2206478 |
| GSAGSLNTSF | ASGDRGDTQY | 0.43266577 |
| GTGQIGI | ASGDRGDTQY | 0.6311871 |
| GTLIALIL | ASGDRGDTQY | 0.7260565 |
| IAATMGFVVG | ASGDRGDTQY | 0.3181336 |
| IFAQALL | ASGDRGDTQY | 0.8057218 |
| IGAAVALGF | ASGDRGDTQY | 0.7108621 |
| IGLTPLAL | ASGDRGDTQY | 0.6289643 |
| ILGSNTSTL | ASGDRGDTQY | 0.81340724 |
| IVFSLTVL | ASGDRGDTQY | 0.69362986 |
| LASKATGF | ASGDRGDTQY | 0.6746984 |
| LASVALIL | ASGDRGDTQY | 0.6341305 |
| LFAVGLTL | ASGDRGDTQY | 0.8523103 |
| LFNGGIGTYI | ASGDRGDTQY | 0.48431626 |
| LTLGAGII | ASGDRGDTQY | 0.81605613 |
| LVAAALGDI | ASGDRGDTQY | 0.656355 |
| NTAFVQLGI | ASGDRGDTQY | 0.84848344 |
| QLFSGAEL | ASGDRGDTQY | 0.4932671 |
| SAATFITL | ASGDRGDTQY | 0.8124567 |
| SNYGLANLGV | ASGDRGDTQY | 0.7151463 |
| SYGGAYAVM | ASGDRGDTQY | 0.8721342 |
| TTGVGGLTLG | ASGDRGDTQY | 0.71433604 |
| TTQTIYI | ASGDRGDTQY | 0.7800243 |
| VAAGAGLGAVI | ASGDRGDTQY | 0.7244544 |
| VAAVVTLL | ASGDRGDTQY | 0.79953444 |
| VGFLGGLI | ASGDRGDTQY | 0.42193738 |
| VGVIPILL | ASGDRGDTQY | 0.7439182 |
| VIFASSLI | ASGDRGDTQY | 0.5276252 |
| VKGGDGGAAAG | ASGDRGDTQY | 0.7784806 |
| VTKTFTNI | ASGDRGDTQY | 0.78880185 |
| VVAAAILL | ASGDRGDTQY | 0.3464521 |
| YAALDVEL | ASGDRGDTQY | 0.82172847 |
| LASVALLL | ASGDRGDTQY | 0.6822235 |
| LDAVHRV | ASGDRGDTQY | 0.627591 |
| QGLSSSIF | ASGDRGDTQY | 0.8219962 |
| RGTFANIRL | ASGDRGDTQY | 0.72858256 |
| RQLWWGHRI | ASGDRGDTQY | 0.8314686 |
| YGPPGTGKTLL | ASGDRGDTQY | 0.69282246 |
| AALVGAPANVA | ASGDRGDTQY | 0.71905434 |
| AATFRRTE | ASGDRGDTQY | 0.8048762 |
| AGALGGGI | ASGDRGDTQY | 0.44532675 |
| AGGAGGAGGIGAG | ASGDRGDTQY | 0.553171 |

| | | |
|---|---|---|
| AGSGALGL | ASGDRGDTQY | 0.84351134 |
| AQIADQI | ASGDRGDTQY | 0.7719759 |
| ASAVAVTYL | ASGDRGDTQY | 0.8357891 |
| ASTVLGGTGG | ASGDRGDTQY | 0.5866165 |
| AVGGEPAK | ASGDRGDTQY | 0.47987026 |
| CEQGTFGGTC | ASGDRGDTQY | 0.6750095 |
| DEGKAPAG | ASGDRGDTQY | 0.8411747 |
| DTVVFPAL | ASGDRGDTQY | 0.84630954 |
| FEGITMAG | ASGDRGDTQY | 0.78061116 |
| FPLVVQD | ASGDRGDTQY | 0.49258685 |
| GAFTTPQ | ASGDRGDTQY | 0.72262144 |
| GFGGFGTAGGVGGA | ASGDRGDTQY | 0.61869615 |
| GFLAQFL | ASGDRGDTQY | 0.6446907 |
| GGAGGAGGNAGVA | ASGDRGDTQY | 0.817258 |
| GGVGTAGAGGAGGAGG | ASGDRGDTQY | 0.6355474 |
| GNGGGGDAGNAGSGGN | ASGDRGDTQY | 0.6484158 |
| GTGGAGGAGNDAGS | ASGDRGDTQY | 0.639772 |
| GTTNTGWFNS | ASGDRGDTQY | 0.73517025 |
| IFMGPLFTL | ASGDRGDTQY | 0.813383 |
| ILAHAIEAI | ASGDRGDTQY | 0.7739377 |
| IRSAIREAL | ASGDRGDTQY | 0.889498 |
| ITFAGGG | ASGDRGDTQY | 0.8214948 |
| ITGMSGHQEV | ASGDRGDTQY | 0.83890474 |
| IVAGLACTL | ASGDRGDTQY | 0.17361233 |
| IVGGGLAGA | ASGDRGDTQY | 0.3760076 |
| IVTGVLNI | ASGDRGDTQY | 0.6705269 |
| LAQQGCI | ASGDRGDTQY | 0.64570427 |
| LCLHVEN | ASGDRGDTQY | 0.39764434 |
| LDAISDGS | ASGDRGDTQY | 0.80003035 |
| LGGFAAML | ASGDRGDTQY | 0.67150784 |
| LGGKNPMI | ASGDRGDTQY | 0.45625514 |
| LGPKGVAAIL | ASGDRGDTQY | 0.79250646 |
| LMVGVSI | ASGDRGDTQY | 0.5742228 |
| LVGAAFVL | ASGDRGDTQY | 0.53617436 |
| LVLAVAPL | ASGDRGDTQY | 0.63303244 |
| MAAIAVIDAA | ASGDRGDTQY | 0.82641554 |
| NNGINAELQ | ASGDRGDTQY | 0.46165788 |
| RGEPLVLVHG | ASGDRGDTQY | 0.7835187 |
| RGGAGGAAGLI | ASGDRGDTQY | 0.6820024 |
| RSYRMHPAV | ASGDRGDTQY | 0.8365935 |
| SAAHLLAGQLA | ASGDRGDTQY | 0.35079548 |
| SAAVGSY | ASGDRGDTQY | 0.80378413 |
| SPHAVLHRI | ASGDRGDTQY | 0.67833066 |
| SQIAKVL | ASGDRGDTQY | 0.8844639 |
| TALGHLVP | ASGDRGDTQY | 0.6182222 |
| TAPTTSGAA | ASGDRGDTQY | 0.66951436 |
| TDGDDGNGGSPGT | ASGDRGDTQY | 0.59257805 |
| TDGLGEN | ASGDRGDTQY | 0.8250805 |
| TGGPPLPGQA | ASGDRGDTQY | 0.3553948 |
| TIPAITI | ASGDRGDTQY | 0.69741154 |
| TIPGFTI | ASGDRGDTQY | 0.45550022 |

| | | |
|---|---|---|
| TSIASGALA | ASGDRGDTQY | 0.77232563 |
| TVTLPGAAI | ASGDRGDTQY | 0.75200367 |
| VASGLSAALP | ASGDRGDTQY | 0.42328307 |
| VGAGVIGI | ASGDRGDTQY | 0.80523324 |
| VGPKRFGV | ASGDRGDTQY | 0.7139919 |
| VNVAFPD | ASGDRGDTQY | 0.55113876 |
| VSMTNSGVS | ASGDRGDTQY | 0.85684574 |
| VTDVLGDLSV | ASGDRGDTQY | 0.6298269 |
| VVVVGAGFAGL | ASGDRGDTQY | 0.8564377 |
| VYTVGTVV | ASGDRGDTQY | 0.75160605 |
| WTTFSGP | ASGDRGDTQY | 0.62083673 |
| YDELVEAF | ASGDRGDTQY | 0.8193251 |
| YGTGGAGGNGG | ASGDRGDTQY | 0.38491762 |
| YTNVMAV | ASGDRGDTQY | 0.5878608 |
| FLEQVHQL | ASGDRGDTQY | 0.83722126 |
| GLSLLAVAV | ASGDRGDTQY | 0.86041427 |
| LEQVHQL | ASGDRGDTQY | 0.8496752 |
| LNAAGAI | ASGDRGDTQY | 0.7326335 |
| PAVHASL | ASGDRGDTQY | 0.64443815 |
| TELEDDA | ASGDRGDTQY | 0.71591216 |
| TPPGPPN | ASGDRGDTQY | 0.85701376 |
| TPPPPPPP | ASGDRGDTQY | 0.8603426 |
| TQSPALI | ASGEDWGGTETLY | 0.84197134 |
| EFAAAFPQ | ASGEDWGGTETLY | 0.4657923 |
| LLSGNLDVL | ASGEDWGGTETLY | 0.8141912 |
| GPANLGNYNI | ASGEDWGGTETLY | 0.749465 |
| TLGPTLP | ASGEDWGGTETLY | 0.7929244 |
| VYAYSSCCK | ASGEDWGGTETLY | 0.7943555 |
| APAPAPAPNI | ASGEDWGGTETLY | 0.7189107 |
| PGGPVPIP | ASGEDWGGTETLY | 0.69629294 |
| FLTVALEP | ASGEDWGGTETLY | 0.5901838 |
| SQYFGANL | ASGEDWGGTETLY | 0.8405249 |
| GGVKPAA | ASGEDWGGTETLY | 0.87089646 |
| GGTGGAGGAGGW | ASGEDWGGTETLY | 0.8615444 |
| GVGGVAGLGGAGG | ASGEDWGGTETLY | 0.8650869 |
| VLLVLATTI | ASGEDWGGTETLY | 0.86988294 |
| IGIFAAML | ASGEDWGGTETLY | 0.84776527 |
| AVGDLID | ASGEDWGGTETLY | 0.20777354 |
| NISPLPL | ASGEDWGGTETLY | 0.37032002 |
| VTAVFVAI | ASGEDWGGTETLY | 0.579438 |
| SGDDNDAAL | ASGEDWGGTETLY | 0.7206539 |
| AVAALAHS | ASGEDWGGTETLY | 0.22195321 |
| GGVTVMATP | ASGEDWGGTETLY | 0.769935 |
| AGLAANVRD | ASGEDWGGTETLY | 0.69445443 |
| TNRPDLIDPALLRPG | ASGEDWGGTETLY | 0.59892434 |
| LANLGVTAL | ASGEDWGGTETLY | 0.8574964 |
| TGERIEL | ASGEDWGGTETLY | 0.70442355 |
| NGEELAA | ASGEDWGGTETLY | 0.559134 |
| VLSLVWG | ASGEDWGGTETLY | 0.50328493 |
| IGYALWL | ASGEDWGGTETLY | 0.83930916 |
| LSLLLAPL | ASGEDWGGTETLY | 0.5479396 |

| | | |
|---|---|---|
| TLDEALK | ASGEDWGGTETLY | 0.84851575 |
| TDGAGTAEGG | ASGEDWGGTETLY | 0.4634732 |
| NAPGGAGGNGG | ASGEDWGGTETLY | 0.86798424 |
| VSVFMGMGGNF | ASGEDWGGTETLY | 0.48671123 |
| LASYLAAL | ASGEDWGGTETLY | 0.8648196 |
| LVGGTPAL | ASGEDWGGTETLY | 0.8144553 |
| VRNADVL | ASGEDWGGTETLY | 0.8937436 |
| TCSSPFLVSAM | ASGEDWGGTETLY | 0.7548558 |
| VHITDAE | ASGEDWGGTETLY | 0.74485576 |
| EYAAPSF | ASGEDWGGTETLY | 0.54203737 |
| VVDAFHGGL | ASGEDWGGTETLY | 0.84478474 |
| ASVVDPQ | ASGEDWGGTETLY | 0.42115635 |
| SYNEFGGLS | ASGEDWGGTETLY | 0.7726065 |
| DPDGSVY | ASGEDWGGTETLY | 0.4120681 |
| MVSTIHATE | ASGEDWGGTETLY | 0.787811 |
| LAAGVADPV | ASGEDWGGTETLY | 0.8568934 |
| TKEQIAA | ASGEDWGGTETLY | 0.20236538 |
| SLASPAAL | ASGEDWGGTETLY | 0.7942474 |
| IGLVFYGV | ASGEDWGGTETLY | 0.63958776 |
| FLLDASNL | ASGEDWGGTETLY | 0.58320427 |
| YQVGNFP | ASGEDWGGTETLY | 0.8539773 |
| ESLLSQST | ASGEDWGGTETLY | 0.7848512 |
| LFTVGFAAVG | ASGEDWGGTETLY | 0.8713908 |
| ATPPTPT | ASGEDWGGTETLY | 0.6352715 |
| TDLATLG | ASGEDWGGTETLY | 0.71578944 |
| SAGGTGR | ASGEDWGGTETLY | 0.81244075 |
| GFSNSGS | ASGEDWGGTETLY | 0.33376187 |
| GGNGGSGGTSV | ASGEDWGGTETLY | 0.4973167 |
| GTSVATGGAGN | ASGEDWGGTETLY | 0.8794471 |
| NGGNGGSGGTS | ASGEDWGGTETLY | 0.8360085 |
| AVVKFPH | ASGEDWGGTETLY | 0.85494167 |
| TPPPPPP | ASGEDWGGTETLY | 0.74530214 |
| AAPAAAAAPA | ASGEDWGGTETLY | 0.68775785 |
| IALNEDL | ASGEDWGGTETLY | 0.74565494 |
| IVNVSSM | ASGEDWGGTETLY | 0.8704632 |
| TGIPPAPR | ASGEDWGGTETLY | 0.8433532 |
| FDLGGGTF | ASGEDWGGTETLY | 0.83807755 |
| FDLGGGTFDVS | ASGEDWGGTETLY | 0.7237435 |
| NEPTAAA | ASGEDWGGTETLY | 0.82515633 |
| NEPTAAALA | ASGEDWGGTETLY | 0.81175596 |
| GSIINISSI | ASGEDWGGTETLY | 0.75870466 |
| VAPGNAGTA | ASGEDWGGTETLY | 0.8579532 |
| TGGGASLE | ASGEDWGGTETLY | 0.8436005 |
| VAVYNHYKRI | ASGEDWGGTETLY | 0.7817321 |
| TDPAPATT | ASGEDWGGTETLY | 0.81198823 |
| TNRPDILDPALLRPG | ASGEDWGGTETLY | 0.7750211 |
| LTTTTTS | ASGEDWGGTETLY | 0.8107695 |
| PTGSGKTL | ASGEDWGGTETLY | 0.78597933 |
| TGSGKTL | ASGEDWGGTETLY | 0.89050305 |
| TIGHVDHGK | ASGEDWGGTETLY | 0.8656077 |
| TIGHVDHGKTT | ASGEDWGGTETLY | 0.83841765 |

| | | |
|---|---|---|
| GHVDHGKTT | ASGEDWGGTETLY | 0.3872267 |
| AQFEHTLL | ASGEDWGGTETLY | 0.6285857 |
| IVNQDEL | ASGEDWGGTETLY | 0.8189353 |
| TNGSQFF | ASGEDWGGTETLY | 0.7970921 |
| TGKGIVE | ASGEDWGGTETLY | 0.55982095 |
| LISNASDA | ASGEDWGGTETLY | 0.8278402 |
| AELRTLGELTKIAKAHGAQ | ASGEDWGGTETLY | 0.7893214 |
| APNAVAAAEPSW | ASGEDWGGTETLY | 0.7455634 |
| APQVAGAATAVL | ASGEDWGGTETLY | 0.85193706 |
| ARRDPAAAAAILPTDA | ASGEDWGGTETLY | 0.79208755 |
| DAAARDVLAEAGLAEYF | ASGEDWGGTETLY | 0.80086684 |
| DDQIVLSAKANLAKVEQ | ASGEDWGGTETLY | 0.8043164 |
| DELLPDTGEDSDEDGH | ASGEDWGGTETLY | 0.87264776 |
| ELPEAPANRATVPQD | ASGEDWGGTETLY | 0.7000543 |
| FGQSKIERVTGIG | ASGEDWGGTETLY | 0.8330873 |
| GAGGTGSTAGGAGGAGGAGGI | ASGEDWGGTETLY | 0.7378347 |
| GHGGTGAAGGNGGTGGAGGS( | ASGEDWGGTETLY | 0.8759754 |
| GNGGNGGAGGEGGPGGAGGA( | ASGEDWGGTETLY | 0.682943 |
| GSTAGNGGNGGNATNGTIAG( | ASGEDWGGTETLY | 0.8520843 |
| GTGGDGGNGGNGGTGG | ASGEDWGGTETLY | 0.77361166 |
| GVRRPAGIPADAPV | ASGEDWGGTETLY | 0.89643025 |
| GYRQVTDAHLLTLARR | ASGEDWGGTETLY | 0.7905222 |
| IAKMGQDGHDRG | ASGEDWGGTETLY | 0.8512598 |
| KAEPSIEIQYT | ASGEDWGGTETLY | 0.81775117 |
| KPVRIRMAADEDTVAVT | ASGEDWGGTETLY | 0.83442503 |
| MPPRIAGMRLLVIKPEP | ASGEDWGGTETLY | 0.6637906 |
| MKFYAVALFRSRGVPPTR | ASGEDWGGTETLY | 0.804765 |
| NAGGFGILGAVAHSP | ASGEDWGGTETLY | 0.7082094 |
| NEGREITLAMAEM | ASGEDWGGTETLY | 0.84142876 |
| NELVEWSHPLGHRWRWE | ASGEDWGGTETLY | 0.8347382 |
| NLGAGGTGGNGGNPG | ASGEDWGGTETLY | 0.6767186 |
| NTEWPKWETVMAADPVD | ASGEDWGGTETLY | 0.8664181 |
| PASAAPAETQP | ASGEDWGGTETLY | 0.79801667 |
| QQSSDGKNGVRAAEITGE | ASGEDWGGTETLY | 0.78026843 |
| RAPDAVTTSWWKE | ASGEDWGGTETLY | 0.87635016 |
| SGFGNVGAHTSGWFNQ | ASGEDWGGTETLY | 0.7898191 |
| SGVAGAVGQAASVGGLKV | ASGEDWGGTETLY | 0.8576536 |
| SNGGAGTGSAGGNGGT | ASGEDWGGTETLY | 0.8081959 |
| TGGAGGEGGAAG | ASGEDWGGTETLY | 0.81043005 |
| TGGDGGAAGAGGAPGHG | ASGEDWGGTETLY | 0.83447033 |
| TMPPLLAAIANAGRHGV | ASGEDWGGTETLY | 0.6902997 |
| TSSTAGNSGGA | ASGEDWGGTETLY | 0.75289893 |
| VPATVPVGILC | ASGEDWGGTETLY | 0.87049353 |
| VQRPDSGGVVLTGRIS | ASGEDWGGTETLY | 0.8687553 |
| WTPNPVPATKGVPAGLVQAT | ASGEDWGGTETLY | 0.859249 |
| YALKRALAKFSDRFDVVII | ASGEDWGGTETLY | 0.756312 |
| YLIGSVIV | ASGEDWGGTETLY | 0.63161755 |
| YGQLVAVID | ASGEDWGGTETLY | 0.8154468 |
| VTVGGMTLPA | ASGEDWGGTETLY | 0.8502668 |
| VLALPAV | ASGEDWGGTETLY | 0.88975877 |
| VIPPTLNYET | ASGEDWGGTETLY | 0.8583374 |

| | | |
|---|---|---|
| VGMGGMHPSSG | ASGEDWGGTETLY | 0.5840185 |
| VGLAGGAATA | ASGEDWGGTETLY | 0.87943614 |
| VGGAGGAGGSGP | ASGEDWGGTETLY | 0.81898403 |
| VFAAFMV | ASGEDWGGTETLY | 0.8118776 |
| TLYDKLR | ASGEDWGGTETLY | 0.8299327 |
| TLAVFTVL | ASGEDWGGTETLY | 0.80649036 |
| TGSVQPARV | ASGEDWGGTETLY | 0.7889045 |
| TGGGAGSGGT | ASGEDWGGTETLY | 0.6242516 |
| SHLETSL | ASGEDWGGTETLY | 0.8631247 |
| SFTTGMT | ASGEDWGGTETLY | 0.881294 |
| QVEVAKRMGVRQPTVS | ASGEDWGGTETLY | 0.87685907 |
| QGSGGPALAM | ASGEDWGGTETLY | 0.87700343 |
| MLAPSNP | ASGEDWGGTETLY | 0.881654 |
| LVNNAAI | ASGEDWGGTETLY | 0.8682864 |
| LTGEINTLA | ASGEDWGGTETLY | 0.20261855 |
| LSIFGPV | ASGEDWGGTETLY | 0.4167775 |
| LLALLANL | ASGEDWGGTETLY | 0.5662115 |
| LISNASDALDKLR | ASGEDWGGTETLY | 0.7800591 |
| LINEAVP | ASGEDWGGTETLY | 0.2697125 |
| LGYAFSGL | ASGEDWGGTETLY | 0.7961843 |
| LDFVAVAL | ASGEDWGGTETLY | 0.7576499 |
| LAQPGAE | ASGEDWGGTETLY | 0.70025766 |
| ISNLGAVTGF | ASGEDWGGTETLY | 0.8857914 |
| HQEDPGRI | ASGEDWGGTETLY | 0.7864177 |
| GTPGEPTP | ASGEDWGGTETLY | 0.554029 |
| GFFNSGS | ASGEDWGGTETLY | 0.7128449 |
| GAGGGGLTAGAN | ASGEDWGGTETLY | 0.8797524 |
| FVDALVNM | ASGEDWGGTETLY | 0.6154507 |
| FLTVAPEL | ASGEDWGGTETLY | 0.8785995 |
| FETLTAL | ASGEDWGGTETLY | 0.5992545 |
| DVINPIEL | ASGEDWGGTETLY | 0.8817849 |
| APVSGAPV | ASGEDWGGTETLY | 0.4854777 |
| ALVGGLAW | ASGEDWGGTETLY | 0.8769231 |
| ALAAAAATAI | ASGEDWGGTETLY | 0.8309696 |
| AFGLGLLL | ASGEDWGGTETLY | 0.90369564 |
| AFFAAASL | ASGEDWGGTETLY | 0.7669256 |
| AAAALPAFSPP | ASGEDWGGTETLY | 0.84807336 |
| VVLAPAL | ASGEDWGGTETLY | 0.741964 |
| TIGHVDHGKTTL | ASGEDWGGTETLY | 0.8807639 |
| LISNASDALDK | ASGEDWGGTETLY | 0.51858693 |
| LISNASDAL | ASGEDWGGTETLY | 0.7908825 |
| ISNASDALDK | ASGEDWGGTETLY | 0.43835598 |
| NQAETLVYQTE | ASGEDWGGTETLY | 0.8160826 |
| TGGTGGTAGAG | ASGEDWGGTETLY | 0.86010873 |
| TGSTAGGAGGA | ASGEDWGGTETLY | 0.20699398 |
| DPLAVPVAAGPV | ASGEDWGGTETLY | 0.883664 |
| GGTGGTPGNGGN | ASGEDWGGTETLY | 0.6659149 |
| PAGLAGVDPATL | ASGEDWGGTETLY | 0.6036308 |
| GNNLVVEAPHAGASV | ASGEDWGGTETLY | 0.8720191 |
| KVTRIMSASTLEQPA | ASGEDWGGTETLY | 0.82347316 |
| LGFALPIALPRRTVV | ASGEDWGGTETLY | 0.8818247 |

| | | |
|---|---|---|
| TRLGNLTARKLVVLV | ASGEDWGGTETLY | 0.71060467 |
| WSQQRRLIYFPSAGP | ASGEDWGGTETLY | 0.7943403 |
| GPNGNPDPMAAAVDIR | ASGEDWGGTETLY | 0.8256396 |
| TQRCVSVADLRAAASR | ASGEDWGGTETLY | 0.39414844 |
| QFVLDVAEHPVEVLTETV | ASGEDWGGTETLY | 0.61714345 |
| GGAGGTGGAGGNGGTQAGDG | ASGEDWGGTETLY | 0.8903084 |
| PLLATLRAIAVDVDAKPAQV | ASGEDWGGTETLY | 0.76646924 |
| APKFRTGGYMIDFWGVGYQV | ASGEDWGGTETLY | 0.7774519 |
| GAGGAGGAGGAAGGPGGTGG | ASGEDWGGTETLY | 0.70223844 |
| IGALGLLATDVAPAGSVAFG | ASGEDWGGTETLY | 0.6970403 |
| AGGAGGIGGNGGAGGAGGVL | ASGEDWGGTETLY | 0.70138067 |
| ADELIGLL | ASGEDWGGTETLY | 0.7706433 |
| ADRALKA | ASGEDWGGTETLY | 0.78792214 |
| AGAVGQVGGAGGA | ASGEDWGGTETLY | 0.7750896 |
| AGGGGGLGGAP | ASGEDWGGTETLY | 0.7310225 |
| AGGLLYGNGGAG | ASGEDWGGTETLY | 0.77362025 |
| AGLESIGL | ASGEDWGGTETLY | 0.75033367 |
| AGLKTGII | ASGEDWGGTETLY | 0.745905 |
| ASLAALLL | ASGEDWGGTETLY | 0.7689573 |
| ASVINISSI | ASGEDWGGTETLY | 0.76011515 |
| ATPPGPA | ASGEDWGGTETLY | 0.6924111 |
| ATVASALV | ASGEDWGGTETLY | 0.7239227 |
| ATVASLIL | ASGEDWGGTETLY | 0.7501893 |
| AVNGPVLIRH | ASGEDWGGTETLY | 0.7727023 |
| DGAAGAVGI | ASGEDWGGTETLY | 0.7667633 |
| DPLLHGI | ASGEDWGGTETLY | 0.7895912 |
| ELGAALGAL | ASGEDWGGTETLY | 0.7737036 |
| ELGRTQAV | ASGEDWGGTETLY | 0.7753244 |
| FVAGGYLS | ASGEDWGGTETLY | 0.4689388 |
| GASGGAGGAGGW | ASGEDWGGTETLY | 0.6707204 |
| GGAGAGGDGGAGGI | ASGEDWGGTETLY | 0.7219599 |
| GTGHNGSQPGMGGNG | ASGEDWGGTETLY | 0.7444084 |
| GYAAASASAS | ASGEDWGGTETLY | 0.41423285 |
| HQVDVAAVQQG | ASGEDWGGTETLY | 0.7701401 |
| IGAAIAHT | ASGEDWGGTETLY | 0.7505243 |
| IGMIEVAQL | ASGEDWGGTETLY | 0.7067046 |
| IRLAPEHTSAMD | ASGEDWGGTETLY | 0.7753067 |
| IYNVSVL | ASGEDWGGTETLY | 0.7264618 |
| KQGFLGM | ASGEDWGGTETLY | 0.71093655 |
| LESVITQL | ASGEDWGGTETLY | 0.75124615 |
| LGLGALVVV | ASGEDWGGTETLY | 0.78297234 |
| LIGNGGNGGSGGI | ASGEDWGGTETLY | 0.6798325 |
| LNAINAPTL | ASGEDWGGTETLY | 0.7822703 |
| LQLLNTGV | ASGEDWGGTETLY | 0.672271 |
| LQTSIEF | ASGEDWGGTETLY | 0.7795245 |
| MSFAVSEL | ASGEDWGGTETLY | 0.6906135 |
| NTGALGL | ASGEDWGGTETLY | 0.7806124 |
| PPGAGGAGGAGTT | ASGEDWGGTETLY | 0.7507367 |
| QLVGLGI | ASGEDWGGTETLY | 0.7888243 |
| RFLEQVHQL | ASGEDWGGTETLY | 0.7457669 |
| RGVGYVLR | ASGEDWGGTETLY | 0.7827232 |

| | | |
|---|---|---|
| SATGVFAGVI | ASGEDWGGTETLY | 0.7648177 |
| SLPAITLP | ASGEDWGGTETLY | 0.7808155 |
| STALAFTV | ASGEDWGGTETLY | 0.65941954 |
| SVLTVGLGK | ASGEDWGGTETLY | 0.7419475 |
| TALAGAGSGI | ASGEDWGGTETLY | 0.6565198 |
| TGLPFTGI | ASGEDWGGTETLY | 0.76372975 |
| VAEHVTV | ASGEDWGGTETLY | 0.77694774 |
| VDIVLNSL | ASGEDWGGTETLY | 0.4578293 |
| VLTGGGAVL | ASGEDWGGTETLY | 0.78388417 |
| VNVGLAYGI | ASGEDWGGTETLY | 0.67773616 |
| VSSAAVGAV | ASGEDWGGTETLY | 0.71727717 |
| VTSVYEAGV | ASGEDWGGTETLY | 0.7793986 |
| WVALDRGA | ASGEDWGGTETLY | 0.7480525 |
| YLVGSVVI | ASGEDWGGTETLY | 0.7681495 |
| GPPGTGKTLL | ASGEDWGGTETLY | 0.73217356 |
| GVGMTKF | ASGEDWGGTETLY | 0.74451005 |
| IGIGGSDL | ASGEDWGGTETLY | 0.76517284 |
| IGTLLAEL | ASGEDWGGTETLY | 0.4904893 |
| IYTSGTT | ASGEDWGGTETLY | 0.68050325 |
| LSFPAGV | ASGEDWGGTETLY | 0.7879208 |
| LYTSGTT | ASGEDWGGTETLY | 0.72819054 |
| VGAFPFGF | ASGEDWGGTETLY | 0.6732563 |
| VGAGTVEF | ASGEDWGGTETLY | 0.37985247 |
| VGKSTLLL | ASGEDWGGTETLY | 0.4249626 |
| AALAHAGGGIV | ASGEDWGGTETLY | 0.48291785 |
| AALVAVSLL | ASGEDWGGTETLY | 0.57354146 |
| AAVSRAVAAL | ASGEDWGGTETLY | 0.81551147 |
| AGGSSGIGGF | ASGEDWGGTETLY | 0.68787634 |
| AGLTVANANG | ASGEDWGGTETLY | 0.41442788 |
| AGLVGAVL | ASGEDWGGTETLY | 0.81997514 |
| ALSQAFTV | ASGEDWGGTETLY | 0.64313245 |
| ALTFSVSL | ASGEDWGGTETLY | 0.6822609 |
| ALTGAALVAL | ASGEDWGGTETLY | 0.58524215 |
| ASFSIGAGVL | ASGEDWGGTETLY | 0.5627235 |
| DGSVYKI | ASGEDWGGTETLY | 0.5241303 |
| EQLAEIV | ASGEDWGGTETLY | 0.4213599 |
| FAVSELTVM | ASGEDWGGTETLY | 0.65460825 |
| GAAPLAAGL | ASGEDWGGTETLY | 0.75568056 |
| GADGGDGGSGGAGG | ASGEDWGGTETLY | 0.7429395 |
| GFGLDTGGAGG | ASGEDWGGTETLY | 0.6856131 |
| GLASGGLL | ASGEDWGGTETLY | 0.7686871 |
| GRIAGAAVLL | ASGEDWGGTETLY | 0.5401311 |
| GSAGSLNTSF | ASGEDWGGTETLY | 0.21316099 |
| GTGQIGI | ASGEDWGGTETLY | 0.3400591 |
| GTLIALIL | ASGEDWGGTETLY | 0.44070137 |
| IAATMGFVVG | ASGEDWGGTETLY | 0.46046874 |
| IFAQALL | ASGEDWGGTETLY | 0.18900214 |
| IGAAVALGF | ASGEDWGGTETLY | 0.53966933 |
| IGLTPLAL | ASGEDWGGTETLY | 0.5991864 |
| ILGSNTSTL | ASGEDWGGTETLY | 0.5353185 |
| IVFSLTVL | ASGEDWGGTETLY | 0.73384404 |

| | | |
|---|---|---|
| LASKATGF | ASGEDWGGTETLY | 0.53923595 |
| LASVALIL | ASGEDWGGTETLY | 0.41257346 |
| LFAVGLTL | ASGEDWGGTETLY | 0.4457096 |
| LFNGGIGTYI | ASGEDWGGTETLY | 0.7911866 |
| LTLGAGII | ASGEDWGGTETLY | 0.40857744 |
| LVAAALGDI | ASGEDWGGTETLY | 0.83664626 |
| NTAFVQLGI | ASGEDWGGTETLY | 0.44618887 |
| QLFSGAEL | ASGEDWGGTETLY | 0.757759 |
| SAATFITL | ASGEDWGGTETLY | 0.3530868 |
| SNYGLANLGV | ASGEDWGGTETLY | 0.71541095 |
| SYGGAYAVM | ASGEDWGGTETLY | 0.6406297 |
| TTGVGGLTLG | ASGEDWGGTETLY | 0.7616019 |
| TTQTIYI | ASGEDWGGTETLY | 0.5101678 |
| VAAGAGLGAVI | ASGEDWGGTETLY | 0.6959438 |
| VAAVVTLL | ASGEDWGGTETLY | 0.53767985 |
| VGFLGGLI | ASGEDWGGTETLY | 0.74316776 |
| VGVIPILL | ASGEDWGGTETLY | 0.3403512 |
| VIFASSLI | ASGEDWGGTETLY | 0.5340775 |
| VKGGDGGAAAG | ASGEDWGGTETLY | 0.45089373 |
| VTKTFTNI | ASGEDWGGTETLY | 0.5352726 |
| VVAAAILL | ASGEDWGGTETLY | 0.7692157 |
| YAALDVEL | ASGEDWGGTETLY | 0.1346237 |
| LASVALLL | ASGEDWGGTETLY | 0.7290245 |
| LDAVHRV | ASGEDWGGTETLY | 0.25129598 |
| QGLSSSIF | ASGEDWGGTETLY | 0.39078987 |
| RGTFANIRL | ASGEDWGGTETLY | 0.6223817 |
| RQLWWGHRI | ASGEDWGGTETLY | 0.6380504 |
| YGPPGTGKTLL | ASGEDWGGTETLY | 0.6302625 |
| AALVGAPANVA | ASGEDWGGTETLY | 0.42183945 |
| AATFRRTE | ASGEDWGGTETLY | 0.5786 |
| AGALGGGI | ASGEDWGGTETLY | 0.6152878 |
| AGGAGGAGGIGAG | ASGEDWGGTETLY | 0.21718344 |
| AGSGALGL | ASGEDWGGTETLY | 0.35962898 |
| AQIADQI | ASGEDWGGTETLY | 0.7882467 |
| ASAVAVTYL | ASGEDWGGTETLY | 0.6221355 |
| ASTVLGGTGG | ASGEDWGGTETLY | 0.77515584 |
| AVGGEPAK | ASGEDWGGTETLY | 0.4417462 |
| CEQGTFGGTC | ASGEDWGGTETLY | 0.47304946 |
| DEGKAPAG | ASGEDWGGTETLY | 0.51893336 |
| DTVVFPAL | ASGEDWGGTETLY | 0.7154637 |
| FEGITMAG | ASGEDWGGTETLY | 0.72306174 |
| FPLVVQD | ASGEDWGGTETLY | 0.63782954 |
| GAFTTPQ | ASGEDWGGTETLY | 0.4489585 |
| GFGGFGTAGGVGGA | ASGEDWGGTETLY | 0.5365796 |
| GFLAQFL | ASGEDWGGTETLY | 0.45643672 |
| GGAGGAGGNAGVA | ASGEDWGGTETLY | 0.5175576 |
| GGVGTAGAGGAGGAGG | ASGEDWGGTETLY | 0.59972245 |
| GNGGGGDAGNAGSGGN | ASGEDWGGTETLY | 0.62664247 |
| GTGGAGGAGNDAGS | ASGEDWGGTETLY | 0.47213995 |
| GTTNTGWFNS | ASGEDWGGTETLY | 0.52270204 |
| IFMGPLFTL | ASGEDWGGTETLY | 0.60617864 |

| | | |
|---|---|---|
| ILAHAIEAI | ASGEDWGGTETLY | 0.7011797 |
| IRSAIREAL | ASGEDWGGTETLY | 0.6752967 |
| ITFAGGG | ASGEDWGGTETLY | 0.75737226 |
| ITGMSGHQEV | ASGEDWGGTETLY | 0.7192904 |
| IVAGLACTL | ASGEDWGGTETLY | 0.6732652 |
| IVGGGLAGA | ASGEDWGGTETLY | 0.19340031 |
| IVTGVLNI | ASGEDWGGTETLY | 0.3825662 |
| LAQQGCI | ASGEDWGGTETLY | 0.45409122 |
| LCLHVEN | ASGEDWGGTETLY | 0.5517191 |
| LDAISDGS | ASGEDWGGTETLY | 0.2792623 |
| LGGFAAML | ASGEDWGGTETLY | 0.64207065 |
| LGGKNPMI | ASGEDWGGTETLY | 0.5392824 |
| LGPKGVAAIL | ASGEDWGGTETLY | 0.43590888 |
| LMVGVSI | ASGEDWGGTETLY | 0.7035987 |
| LVGAAFVL | ASGEDWGGTETLY | 0.48460665 |
| LVLAVAPL | ASGEDWGGTETLY | 0.4950199 |
| MAAIAVIDAA | ASGEDWGGTETLY | 0.38512635 |
| NNGINAELQ | ASGEDWGGTETLY | 0.674194 |
| RGEPLVLVHG | ASGEDWGGTETLY | 0.3165618 |
| RGGAGGAAGLI | ASGEDWGGTETLY | 0.68052226 |
| RSYRMHPAV | ASGEDWGGTETLY | 0.44581822 |
| SAAHLLAGQLA | ASGEDWGGTETLY | 0.7171625 |
| SAAVGSY | ASGEDWGGTETLY | 0.38148388 |
| SPHAVLHRI | ASGEDWGGTETLY | 0.6781986 |
| SQIAKVL | ASGEDWGGTETLY | 0.5548179 |
| TALGHLVP | ASGEDWGGTETLY | 0.82067776 |
| TAPTTSGAA | ASGEDWGGTETLY | 0.58551264 |
| TDGDDGNGGSPGT | ASGEDWGGTETLY | 0.5858667 |
| TDGLGEN | ASGEDWGGTETLY | 0.49587044 |
| TGGPPLPGQA | ASGEDWGGTETLY | 0.73565555 |
| TIPAITI | ASGEDWGGTETLY | 0.30783182 |
| TIPGFTI | ASGEDWGGTETLY | 0.5169508 |
| TSIASGALA | ASGEDWGGTETLY | 0.334083 |
| TVTLPGAAI | ASGEDWGGTETLY | 0.5592233 |
| VASGLSAALP | ASGEDWGGTETLY | 0.66273 |
| VGAGVIGI | ASGEDWGGTETLY | 0.34990925 |
| VGPKRFGV | ASGEDWGGTETLY | 0.6119807 |
| VNVAFPD | ASGEDWGGTETLY | 0.5588639 |
| VSMTNSGVS | ASGEDWGGTETLY | 0.47730038 |
| VTDVLGDLSV | ASGEDWGGTETLY | 0.71334887 |
| VVVVGAGFAGL | ASGEDWGGTETLY | 0.51395845 |
| VYTVGTVV | ASGEDWGGTETLY | 0.7646094 |
| WTTFSGP | ASGEDWGGTETLY | 0.44440538 |
| YDELVEAF | ASGEDWGGTETLY | 0.45383054 |
| YGTGGAGGNGG | ASGEDWGGTETLY | 0.6403949 |
| YTNVMAV | ASGEDWGGTETLY | 0.4681041 |
| FLEQVHQL | ASGEDWGGTETLY | 0.3426197 |
| GLSLLAVAV | ASGEDWGGTETLY | 0.7681476 |
| LEQVHQL | ASGEDWGGTETLY | 0.6043469 |
| LNAAGAI | ASGEDWGGTETLY | 0.78195137 |
| PAVHASL | ASGEDWGGTETLY | 0.3342084 |

| | | |
|---|---|---|
| TELEDDA | ASGEDWGGTETLY | 0.25545377 |
| TPPGPPN | ASGEDWGGTETLY | 0.3874013 |
| TPPPPPPP | ASGEDWGGTETLY | 0.67671263 |
| TQSPALI | ASGDRGENYAEQF | 0.77385736 |
| EFAAAFPQ | ASGDRGENYAEQF | 0.625722 |
| LLSGNLDVL | ASGDRGENYAEQF | 0.20951346 |
| GPANLGNYNI | ASGDRGENYAEQF | 0.39787152 |
| TLGPTLP | ASGDRGENYAEQF | 0.3681296 |
| VYAYSSCCK | ASGDRGENYAEQF | 0.38088673 |
| APAPAPAPNI | ASGDRGENYAEQF | 0.48360187 |
| PGGPVPIP | ASGDRGENYAEQF | 0.39925563 |
| FLTVALEP | ASGDRGENYAEQF | 0.3552224 |
| SQYFGANL | ASGDRGENYAEQF | 0.26959413 |
| GGVKPAA | ASGDRGENYAEQF | 0.52254695 |
| GGTGGAGGAGGW | ASGDRGENYAEQF | 0.5760521 |
| GVGGVAGLGGAGG | ASGDRGENYAEQF | 0.41099188 |
| VLLVLATTI | ASGDRGENYAEQF | 0.7166269 |
| IGIFAAML | ASGDRGENYAEQF | 0.73739773 |
| AVGDLID | ASGDRGENYAEQF | 0.6142617 |
| NISPLPL | ASGDRGENYAEQF | 0.13764158 |
| VTAVFVAI | ASGDRGENYAEQF | 0.21701764 |
| SGDDNDAAL | ASGDRGENYAEQF | 0.27013934 |
| AVAALAHS | ASGDRGENYAEQF | 0.46692917 |
| GGVTVMATP | ASGDRGENYAEQF | 0.15780494 |
| AGLAANVRD | ASGDRGENYAEQF | 0.54206705 |
| TNRPDLIDPALLRPG | ASGDRGENYAEQF | 0.3959725 |
| LANLGVTAL | ASGDRGENYAEQF | 0.25387406 |
| TGERIEL | ASGDRGENYAEQF | 0.72017527 |
| NGEELAA | ASGDRGENYAEQF | 0.34960198 |
| VLSLVWG | ASGDRGENYAEQF | 0.23744464 |
| IGYALWL | ASGDRGENYAEQF | 0.23727441 |
| LSLLLAPL | ASGDRGENYAEQF | 0.6130477 |
| TLDEALK | ASGDRGENYAEQF | 0.21278007 |
| TDGAGTAEGG | ASGDRGENYAEQF | 0.6391295 |
| NAPGGAGGNGG | ASGDRGENYAEQF | 0.23498824 |
| VSVFMGMGGNF | ASGDRGENYAEQF | 0.64205545 |
| LASYLAAL | ASGDRGENYAEQF | 0.19116847 |
| LVGGTPAL | ASGDRGENYAEQF | 0.70016646 |
| VRNADVL | ASGDRGENYAEQF | 0.48664948 |
| TCSSPFLVSAM | ASGDRGENYAEQF | 0.82045186 |
| VHITDAE | ASGDRGENYAEQF | 0.37135017 |
| EYAAPSF | ASGDRGENYAEQF | 0.3903317 |
| VVDAFHGGL | ASGDRGENYAEQF | 0.30698797 |
| ASVVDPQ | ASGDRGENYAEQF | 0.77714825 |
| SYNEFGGLS | ASGDRGENYAEQF | 0.18063822 |
| DPDGSVY | ASGDRGENYAEQF | 0.25795007 |
| MVSTIHATE | ASGDRGENYAEQF | 0.18859029 |
| LAAGVADPV | ASGDRGENYAEQF | 0.35291815 |
| TKEQIAA | ASGDRGENYAEQF | 0.62531 |
| SLASPAAL | ASGDRGENYAEQF | 0.16633862 |
| IGLVFYGV | ASGDRGENYAEQF | 0.30249807 |

| | | |
|---|---|---|
| FLLDASNL | ASGDRGENYAEQF | 0.4692588 |
| YQVGNFP | ASGDRGENYAEQF | 0.22469589 |
| ESLLSQST | ASGDRGENYAEQF | 0.64168453 |
| LFTVGFAAVG | ASGDRGENYAEQF | 0.31077588 |
| ATPPTPT | ASGDRGENYAEQF | 0.7629839 |
| TDLATLG | ASGDRGENYAEQF | 0.3044616 |
| SAGGTGR | ASGDRGENYAEQF | 0.30005318 |
| GFSNSGS | ASGDRGENYAEQF | 0.34315068 |
| GGNGGSGGTSV | ASGDRGENYAEQF | 0.2636733 |
| GTSVATGGAGN | ASGDRGENYAEQF | 0.22833748 |
| NGGNGGSGGTS | ASGDRGENYAEQF | 0.7597989 |
| AVVKFPH | ASGDRGENYAEQF | 0.87240463 |
| TPPPPPP | ASGDRGENYAEQF | 0.86438555 |
| AAPAAAAAPA | ASGDRGENYAEQF | 0.7718772 |
| IALNEDL | ASGDRGENYAEQF | 0.7867143 |
| IVNVSSM | ASGDRGENYAEQF | 0.8302649 |
| TGIPPAPR | ASGDRGENYAEQF | 0.8809817 |
| FDLGGGTF | ASGDRGENYAEQF | 0.88369185 |
| FDLGGGTFDVS | ASGDRGENYAEQF | 0.87696993 |
| NEPTAAA | ASGDRGENYAEQF | 0.7746179 |
| NEPTAAALA | ASGDRGENYAEQF | 0.8614074 |
| GSIINISSI | ASGDRGENYAEQF | 0.8350614 |
| VAPGNAGTA | ASGDRGENYAEQF | 0.8503576 |
| TGGGASLE | ASGDRGENYAEQF | 0.85943747 |
| VAVYNHYKRI | ASGDRGENYAEQF | 0.85156894 |
| TDPAPATT | ASGDRGENYAEQF | 0.80879176 |
| TNRPDILDPALLRPG | ASGDRGENYAEQF | 0.76579 |
| LTTTTTS | ASGDRGENYAEQF | 0.8616305 |
| PTGSGKTL | ASGDRGENYAEQF | 0.8759196 |
| TGSGKTL | ASGDRGENYAEQF | 0.8654338 |
| TIGHVDHGK | ASGDRGENYAEQF | 0.8902953 |
| TIGHVDHGKTT | ASGDRGENYAEQF | 0.8747358 |
| GHVDHGKTT | ASGDRGENYAEQF | 0.86547446 |
| AQFEHTLL | ASGDRGENYAEQF | 0.34864557 |
| IVNQDEL | ASGDRGENYAEQF | 0.65334857 |
| TNGSQFF | ASGDRGENYAEQF | 0.7772148 |
| TGKGIVE | ASGDRGENYAEQF | 0.83302855 |
| LISNASDA | ASGDRGENYAEQF | 0.3863088 |
| AELRTLGELTKIAKAHGAQ | ASGDRGENYAEQF | 0.87120146 |
| APNAVAAAEPSW | ASGDRGENYAEQF | 0.84253025 |
| APQVAGAATAVL | ASGDRGENYAEQF | 0.7842018 |
| ARRDPAAAAAILPTDA | ASGDRGENYAEQF | 0.87699175 |
| DAAARDVLAEAGLAEYF | ASGDRGENYAEQF | 0.8134732 |
| DDQIVLSAKANLAKVEQ | ASGDRGENYAEQF | 0.7553576 |
| DELLPDTGEDSDEDGH | ASGDRGENYAEQF | 0.8046337 |
| ELPEAPANRATVPQD | ASGDRGENYAEQF | 0.8837173 |
| FGQSKIERVTGIG | ASGDRGENYAEQF | 0.7100202 |
| GAGGTGSTAGGAGGAGGAGGI | ASGDRGENYAEQF | 0.8808093 |
| GHGGTGAAGGNGGTGGAGGS( | ASGDRGENYAEQF | 0.79627895 |
| GNGGNGGAGGEGGPGGAGGA( | ASGDRGENYAEQF | 0.8834672 |
| GSTAGNGGNGGNATNGTIAG( | ASGDRGENYAEQF | 0.74149877 |

| | | |
|---|---|---|
| GTGGDGGNGGNGGTGG | ASGDRGENYAEQF | 0.877718 |
| GVRRPAGIPADAPV | ASGDRGENYAEQF | 0.8459863 |
| GYRQVTDAHLLTLARR | ASGDRGENYAEQF | 0.8933672 |
| IAKMGQDGHDRG | ASGDRGENYAEQF | 0.84013855 |
| KAEPSIEIQYT | ASGDRGENYAEQF | 0.86965656 |
| KPVRIRMAADEDTVAVT | ASGDRGENYAEQF | 0.8340034 |
| MPPRIAGMRLLVIKPEP | ASGDRGENYAEQF | 0.86839163 |
| MKFYAVALFRSRGVPPTR | ASGDRGENYAEQF | 0.7080076 |
| NAGGFGILGAVAHSP | ASGDRGENYAEQF | 0.82866144 |
| NEGREITLAMAEM | ASGDRGENYAEQF | 0.68489516 |
| NELVEWSHPLGHRWRWE | ASGDRGENYAEQF | 0.8424885 |
| NLGAGGTGGNGGNPG | ASGDRGENYAEQF | 0.8782885 |
| NTEWPKWETVMAADPVD | ASGDRGENYAEQF | 0.3206706 |
| PASAAPAETQP | ASGDRGENYAEQF | 0.86230177 |
| QQSSDGKNGVRAAEITGE | ASGDRGENYAEQF | 0.7707865 |
| RAPDAVTTSWWKE | ASGDRGENYAEQF | 0.79541916 |
| SGFGNVGAHTSGWFNQ | ASGDRGENYAEQF | 0.8728123 |
| SGVAGAVGQAASVGGLKV | ASGDRGENYAEQF | 0.84545964 |
| SNGGAGTGSAGGNGGT | ASGDRGENYAEQF | 0.86945534 |
| TGGAGGEGGAAG | ASGDRGENYAEQF | 0.7980022 |
| TGGDGGAAGAGGAPGHG | ASGDRGENYAEQF | 0.8395363 |
| TMPPLLAAIANAGRHGV | ASGDRGENYAEQF | 0.8583025 |
| TSSTAGNSGGA | ASGDRGENYAEQF | 0.5382629 |
| VPATVPVGILC | ASGDRGENYAEQF | 0.7387661 |
| VQRPDSGGVVLTGRIS | ASGDRGENYAEQF | 0.88757837 |
| WTPNPVPATKGVPAGLVQAT | ASGDRGENYAEQF | 0.744383 |
| YALKRALAKFSDRFDVVII | ASGDRGENYAEQF | 0.7602763 |
| YLIGSVIV | ASGDRGENYAEQF | 0.57584924 |
| YGQLVAVID | ASGDRGENYAEQF | 0.48446816 |
| VTVGGMTLPA | ASGDRGENYAEQF | 0.5854204 |
| VLALPAV | ASGDRGENYAEQF | 0.7555516 |
| VIPPTLNYET | ASGDRGENYAEQF | 0.73814493 |
| VGMGGMHPSSG | ASGDRGENYAEQF | 0.7273103 |
| VGLAGGAATA | ASGDRGENYAEQF | 0.5145704 |
| VGGAGGAGGSGP | ASGDRGENYAEQF | 0.6483034 |
| VFAAFMV | ASGDRGENYAEQF | 0.66356426 |
| TLYDKLR | ASGDRGENYAEQF | 0.57711476 |
| TLAVFTVL | ASGDRGENYAEQF | 0.7137725 |
| TGSVQPARV | ASGDRGENYAEQF | 0.66826844 |
| TGGGAGSGGT | ASGDRGENYAEQF | 0.6569127 |
| SHLETSL | ASGDRGENYAEQF | 0.6354946 |
| SFTTGMT | ASGDRGENYAEQF | 0.6670479 |
| QVEVAKRMGVRQPTVS | ASGDRGENYAEQF | 0.70731914 |
| QGSGGPALAM | ASGDRGENYAEQF | 0.673465 |
| MLAPSNP | ASGDRGENYAEQF | 0.7793486 |
| LVNNAAI | ASGDRGENYAEQF | 0.7515174 |
| LTGEINTLA | ASGDRGENYAEQF | 0.73893535 |
| LSIFGPV | ASGDRGENYAEQF | 0.31554383 |
| LLALLANL | ASGDRGENYAEQF | 0.399116 |
| LISNASDALDKLR | ASGDRGENYAEQF | 0.70107126 |
| LINEAVP | ASGDRGENYAEQF | 0.5628422 |

| | | |
|---|---|---|
| LGYAFSGL | ASGDRGENYAEQF | 0.41501158 |
| LDFVAVAL | ASGDRGENYAEQF | 0.691113 |
| LAQPGAE | ASGDRGENYAEQF | 0.71430445 |
| ISNLGAVTGF | ASGDRGENYAEQF | 0.5356767 |
| HQEDPGRI | ASGDRGENYAEQF | 0.70306367 |
| GTPGEPTP | ASGDRGENYAEQF | 0.5541584 |
| GFFNSGS | ASGDRGENYAEQF | 0.5650388 |
| GAGGGGLTAGAN | ASGDRGENYAEQF | 0.6145202 |
| FVDALVNM | ASGDRGENYAEQF | 0.7394811 |
| FLTVAPEL | ASGDRGENYAEQF | 0.51440537 |
| FETLTAL | ASGDRGENYAEQF | 0.6987613 |
| DVINPIEL | ASGDRGENYAEQF | 0.6474038 |
| APVSGAPV | ASGDRGENYAEQF | 0.7487362 |
| ALVGGLAW | ASGDRGENYAEQF | 0.4605391 |
| ALAAAAATAI | ASGDRGENYAEQF | 0.73198366 |
| AFGLGLLL | ASGDRGENYAEQF | 0.6337882 |
| AFFAAASL | ASGDRGENYAEQF | 0.76694393 |
| AAAALPAFSPP | ASGDRGENYAEQF | 0.5819577 |
| VVLAPAL | ASGDRGENYAEQF | 0.6534227 |
| TIGHVDHGKTTL | ASGDRGENYAEQF | 0.6617465 |
| LISNASDALDK | ASGDRGENYAEQF | 0.7183431 |
| LISNASDAL | ASGDRGENYAEQF | 0.44985297 |
| ISNASDALDK | ASGDRGENYAEQF | 0.64857197 |
| NQAETLVYQTE | ASGDRGENYAEQF | 0.55783165 |
| TGGTGGTAGAG | ASGDRGENYAEQF | 0.68593514 |
| TGSTAGGAGGA | ASGDRGENYAEQF | 0.7265899 |
| DPLAVPVAAGPV | ASGDRGENYAEQF | 0.43331626 |
| GGTGGTPGNGGN | ASGDRGENYAEQF | 0.73920643 |
| PAGLAGVDPATL | ASGDRGENYAEQF | 0.6724774 |
| GNNLVVEAPHAGASV | ASGDRGENYAEQF | 0.5925613 |
| KVTRIMSASTLEQPA | ASGDRGENYAEQF | 0.74865824 |
| LGFALPIALPRRTVV | ASGDRGENYAEQF | 0.64422035 |
| TRLGNLTARKLVVLV | ASGDRGENYAEQF | 0.76247674 |
| WSQQRRLIYFPSAGP | ASGDRGENYAEQF | 0.6472742 |
| GPNGNPDPMAAAVDIR | ASGDRGENYAEQF | 0.6685403 |
| TQRCVSVADLRAAASR | ASGDRGENYAEQF | 0.72619903 |
| QFVLDVAEHPVEVLTETV | ASGDRGENYAEQF | 0.54000956 |
| GGAGGTGGAGGNGGTQAGDG | ASGDRGENYAEQF | 0.53970647 |
| PLLATLRAIAVDVDAKPAQV | ASGDRGENYAEQF | 0.75098675 |
| APKFRTGGYMIDFWGVGYQV | ASGDRGENYAEQF | 0.71112275 |
| GAGGAGGAGGAAGGPGGTGG | ASGDRGENYAEQF | 0.7098652 |
| IGALGLLATDVAPAGSVAFG | ASGDRGENYAEQF | 0.67246014 |
| AGGAGGIGGNGGAGGAGGVL | ASGDRGENYAEQF | 0.67217803 |
| ADELIGLL | ASGDRGENYAEQF | 0.6771481 |
| ADRALKA | ASGDRGENYAEQF | 0.7063551 |
| AGAVGQVGGAGGA | ASGDRGENYAEQF | 0.717612 |
| AGGGGGLGGAP | ASGDRGENYAEQF | 0.7105721 |
| AGGLLYGNGGAG | ASGDRGENYAEQF | 0.67246825 |
| AGLESIGL | ASGDRGENYAEQF | 0.71232665 |
| AGLKTGII | ASGDRGENYAEQF | 0.7050434 |
| ASLAALLL | ASGDRGENYAEQF | 0.69368994 |

| | | |
|---|---|---|
| ASVINISSI | ASGDRGENYAEQF | 0.7000456 |
| ATPPGPA | ASGDRGENYAEQF | 0.69420063 |
| ATVASALV | ASGDRGENYAEQF | 0.6791363 |
| ATVASLIL | ASGDRGENYAEQF | 0.63382494 |
| AVNGPVLIRH | ASGDRGENYAEQF | 0.6951997 |
| DGAAGAVGI | ASGDRGENYAEQF | 0.71276784 |
| DPLLHGI | ASGDRGENYAEQF | 0.7085391 |
| ELGAALGAL | ASGDRGENYAEQF | 0.72011757 |
| ELGRTQAV | ASGDRGENYAEQF | 0.7128782 |
| FVAGGYLS | ASGDRGENYAEQF | 0.70258343 |
| GASGGAGGAGGW | ASGDRGENYAEQF | 0.49157834 |
| GGAGAGGDGGAGGI | ASGDRGENYAEQF | 0.61923593 |
| GTGHNGSQPGMGGNG | ASGDRGENYAEQF | 0.6569618 |
| GYAAASASAS | ASGDRGENYAEQF | 0.6908114 |
| HQVDVAAVQQG | ASGDRGENYAEQF | 0.38847816 |
| IGAAIAHT | ASGDRGENYAEQF | 0.70416653 |
| IGMIEVAQL | ASGDRGENYAEQF | 0.6992378 |
| IRLAPEHTSAMD | ASGDRGENYAEQF | 0.67961323 |
| IYNVSVL | ASGDRGENYAEQF | 0.71004957 |
| KQGFLGM | ASGDRGENYAEQF | 0.69153154 |
| LESVITQL | ASGDRGENYAEQF | 0.6410854 |
| LGLGALVVV | ASGDRGENYAEQF | 0.6730521 |
| LIGNGGNGGSGGI | ASGDRGENYAEQF | 0.7208203 |
| LNAINAPTL | ASGDRGENYAEQF | 0.6304686 |
| LQLLNTGV | ASGDRGENYAEQF | 0.718239 |
| LQTSIEF | ASGDRGENYAEQF | 0.6270757 |
| MSFAVSEL | ASGDRGENYAEQF | 0.7175357 |
| NTGALGL | ASGDRGENYAEQF | 0.65009093 |
| PPGAGGAGGAGTT | ASGDRGENYAEQF | 0.7149204 |
| QLVGLGI | ASGDRGENYAEQF | 0.6979307 |
| RFLEQVHQL | ASGDRGENYAEQF | 0.722353 |
| RGVGYVLR | ASGDRGENYAEQF | 0.68826675 |
| SATGVFAGVI | ASGDRGENYAEQF | 0.71397877 |
| SLPAITLP | ASGDRGENYAEQF | 0.68412024 |
| STALAFTV | ASGDRGENYAEQF | 0.71086884 |
| SVLTVGLGK | ASGDRGENYAEQF | 0.64002514 |
| TALAGAGSGI | ASGDRGENYAEQF | 0.68754506 |
| TGLPFTGI | ASGDRGENYAEQF | 0.6530646 |
| VAEHVTV | ASGDRGENYAEQF | 0.6951789 |
| VDIVLNSL | ASGDRGENYAEQF | 0.7152791 |
| VLTGGGAVL | ASGDRGENYAEQF | 0.5273316 |
| VNVGLAYGI | ASGDRGENYAEQF | 0.71321714 |
| VSSAAVGAV | ASGDRGENYAEQF | 0.6444056 |
| VTSVYEAGV | ASGDRGENYAEQF | 0.6672225 |
| WVALDRGA | ASGDRGENYAEQF | 0.70808595 |
| YLVGSVVI | ASGDRGENYAEQF | 0.7001618 |
| GPPGTGKTLL | ASGDRGENYAEQF | 0.71076 |
| GVGMTKF | ASGDRGENYAEQF | 0.6468985 |
| IGIGGSDL | ASGDRGENYAEQF | 0.69651496 |
| IGTLLAEL | ASGDRGENYAEQF | 0.7032124 |
| IYTSGTT | ASGDRGENYAEQF | 0.5115578 |

| | | |
|---|---|---|
| LSFPAGV | ASGDRGENYAEQF | 0.6443173 |
| LYTSGTT | ASGDRGENYAEQF | 0.7186033 |
| VGAFPFGF | ASGDRGENYAEQF | 0.67977405 |
| VGAGTVEF | ASGDRGENYAEQF | 0.6988323 |
| VGKSTLLL | ASGDRGENYAEQF | 0.6047004 |
| AALAHAGGGIV | ASGDRGENYAEQF | 0.5891649 |
| AALVAVSLL | ASGDRGENYAEQF | 0.6401571 |
| AAVSRAVAAL | ASGDRGENYAEQF | 0.6925545 |
| AGGSSGIGGF | ASGDRGENYAEQF | 0.70948774 |
| AGLTVANANG | ASGDRGENYAEQF | 0.68488455 |
| AGLVGAVL | ASGDRGENYAEQF | 0.6307212 |
| ALSQAFTV | ASGDRGENYAEQF | 0.6743438 |
| ALTFSVSL | ASGDRGENYAEQF | 0.66419315 |
| ALTGAALVAL | ASGDRGENYAEQF | 0.64717436 |
| ASFSIGAGVL | ASGDRGENYAEQF | 0.69319165 |
| DGSVYKI | ASGDRGENYAEQF | 0.6817407 |
| EQLAEIV | ASGDRGENYAEQF | 0.6323949 |
| FAVSELTVM | ASGDRGENYAEQF | 0.64620745 |
| GAAPLAAGL | ASGDRGENYAEQF | 0.67936695 |
| GADGGDGGSGGAGG | ASGDRGENYAEQF | 0.69300765 |
| GFGLDTGGAGG | ASGDRGENYAEQF | 0.6867801 |
| GLASGGLL | ASGDRGENYAEQF | 0.7139555 |
| GRIAGAAVLL | ASGDRGENYAEQF | 0.6930665 |
| GSAGSLNTSF | ASGDRGENYAEQF | 0.6800732 |
| GTGQIGI | ASGDRGENYAEQF | 0.440384 |
| GTLIALIL | ASGDRGENYAEQF | 0.5766453 |
| IAATMGFVVG | ASGDRGENYAEQF | 0.6755901 |
| IFAQALL | ASGDRGENYAEQF | 0.658005 |
| IGAAVALGF | ASGDRGENYAEQF | 0.44897583 |
| IGLTPLAL | ASGDRGENYAEQF | 0.6834724 |
| ILGSNTSTL | ASGDRGENYAEQF | 0.6526722 |
| IVFSLTVL | ASGDRGENYAEQF | 0.62432295 |
| LASKATGF | ASGDRGENYAEQF | 0.69638395 |
| LASVALIL | ASGDRGENYAEQF | 0.63866544 |
| LFAVGLTL | ASGDRGENYAEQF | 0.6414491 |
| LFNGGIGTYI | ASGDRGENYAEQF | 0.6714055 |
| LTLGAGII | ASGDRGENYAEQF | 0.7093487 |
| LVAAALGDI | ASGDRGENYAEQF | 0.62159795 |
| NTAFVQLGI | ASGDRGENYAEQF | 0.69749284 |
| QLFSGAEL | ASGDRGENYAEQF | 0.6835888 |
| SAATFITL | ASGDRGENYAEQF | 0.7023939 |
| SNYGLANLGV | ASGDRGENYAEQF | 0.587747 |
| SYGGAYAVM | ASGDRGENYAEQF | 0.691893 |
| TTGVGGLTLG | ASGDRGENYAEQF | 0.64792573 |
| TTQTIYI | ASGDRGENYAEQF | 0.71592253 |
| VAAGAGLGAVI | ASGDRGENYAEQF | 0.65098685 |
| VAAVVTLL | ASGDRGENYAEQF | 0.69930756 |
| VGFLGGLI | ASGDRGENYAEQF | 0.6820286 |
| VGVIPILL | ASGDRGENYAEQF | 0.6957588 |
| VIFASSLI | ASGDRGENYAEQF | 0.58883023 |
| VKGGDGGAAAG | ASGDRGENYAEQF | 0.6530942 |

| | | |
|---|---|---|
| VTKTFTNI | ASGDRGENYAEQF | 0.62049735 |
| VVAAAILL | ASGDRGENYAEQF | 0.67020667 |
| YAALDVEL | ASGDRGENYAEQF | 0.6874991 |
| LASVALLL | ASGDRGENYAEQF | 0.48248845 |
| LDAVHRV | ASGDRGENYAEQF | 0.70262665 |
| QGLSSSIF | ASGDRGENYAEQF | 0.6491705 |
| RGTFANIRL | ASGDRGENYAEQF | 0.63025117 |
| RQLWWGHRI | ASGDRGENYAEQF | 0.6972454 |
| YGPPGTGKTLL | ASGDRGENYAEQF | 0.6560751 |
| AALVGAPANVA | ASGDRGENYAEQF | 0.67667896 |
| AATFRRTE | ASGDRGENYAEQF | 0.64843994 |
| AGALGGGI | ASGDRGENYAEQF | 0.67261684 |
| AGGAGGAGGIGAG | ASGDRGENYAEQF | 0.6810634 |
| AGSGALGL | ASGDRGENYAEQF | 0.5414211 |
| AQIADQI | ASGDRGENYAEQF | 0.6362073 |
| ASAVAVTYL | ASGDRGENYAEQF | 0.7003849 |
| ASTVLGGTGG | ASGDRGENYAEQF | 0.6866073 |
| AVGGEPAK | ASGDRGENYAEQF | 0.7046331 |
| CEQGTFGGTC | ASGDRGENYAEQF | 0.5831138 |
| DEGKAPAG | ASGDRGENYAEQF | 0.56687677 |
| DTVVFPAL | ASGDRGENYAEQF | 0.6333773 |
| FEGITMAG | ASGDRGENYAEQF | 0.69715166 |
| FPLVVQD | ASGDRGENYAEQF | 0.70013505 |
| GAFTTPQ | ASGDRGENYAEQF | 0.6876793 |
| GFGGFGTAGGVGGA | ASGDRGENYAEQF | 0.5587013 |
| GFLAQFL | ASGDRGENYAEQF | 0.6610376 |
| GGAGGAGGNAGVA | ASGDRGENYAEQF | 0.6378875 |
| GGVGTAGAGGAGGAGG | ASGDRGENYAEQF | 0.62985647 |
| GNGGGGDAGNAGSGGN | ASGDRGENYAEQF | 0.66601104 |
| GTGGAGGAGNDAGS | ASGDRGENYAEQF | 0.66823685 |
| GTTNTGWFNS | ASGDRGENYAEQF | 0.612273 |
| IFMGPLFTL | ASGDRGENYAEQF | 0.61136854 |
| ILAHAIEAI | ASGDRGENYAEQF | 0.63089395 |
| IRSAIREAL | ASGDRGENYAEQF | 0.68314844 |
| ITFAGGG | ASGDRGENYAEQF | 0.6664124 |
| ITGMSGHQEV | ASGDRGENYAEQF | 0.7058152 |
| IVAGLACTL | ASGDRGENYAEQF | 0.6917934 |
| IVGGGLAGA | ASGDRGENYAEQF | 0.682366 |
| IVTGVLNI | ASGDRGENYAEQF | 0.30738622 |
| LAQQGCI | ASGDRGENYAEQF | 0.37413728 |
| LCLHVEN | ASGDRGENYAEQF | 0.5953397 |
| LDAISDGS | ASGDRGENYAEQF | 0.62933075 |
| LGGFAAML | ASGDRGENYAEQF | 0.3399885 |
| LGGKNPMI | ASGDRGENYAEQF | 0.66827846 |
| LGPKGVAAIL | ASGDRGENYAEQF | 0.6435405 |
| LMVGVSI | ASGDRGENYAEQF | 0.58683705 |
| LVGAAFVL | ASGDRGENYAEQF | 0.6725851 |
| LVLAVAPL | ASGDRGENYAEQF | 0.6162895 |
| MAAIAVIDAA | ASGDRGENYAEQF | 0.5720689 |
| NNGINAELQ | ASGDRGENYAEQF | 0.5554486 |
| RGEPLVLVHG | ASGDRGENYAEQF | 0.680781 |

| | | |
|---|---|---|
| RGGAGGAAGLI | ASGDRGENYAEQF | 0.46178287 |
| RSYRMHPAV | ASGDRGENYAEQF | 0.679917 |
| SAAHLLAGQLA | ASGDRGENYAEQF | 0.49973893 |
| SAAVGSY | ASGDRGENYAEQF | 0.6870768 |
| SPHAVLHRI | ASGDRGENYAEQF | 0.5453683 |
| SQIAKVL | ASGDRGENYAEQF | 0.6917978 |
| TALGHLVP | ASGDRGENYAEQF | 0.651227 |
| TAPTTSGAA | ASGDRGENYAEQF | 0.7064502 |
| TDGDDGNGGSPGT | ASGDRGENYAEQF | 0.6102319 |
| TDGLGEN | ASGDRGENYAEQF | 0.668821 |
| TGGPPLPGQA | ASGDRGENYAEQF | 0.6247522 |
| TIPAITI | ASGDRGENYAEQF | 0.70426977 |
| TIPGFTI | ASGDRGENYAEQF | 0.47042045 |
| TSIASGALA | ASGDRGENYAEQF | 0.64646137 |
| TVTLPGAAI | ASGDRGENYAEQF | 0.39729828 |
| VASGLSAALP | ASGDRGENYAEQF | 0.6681588 |
| VGAGVIGI | ASGDRGENYAEQF | 0.69195783 |
| VGPKRFGV | ASGDRGENYAEQF | 0.39031908 |
| VNVAFPD | ASGDRGENYAEQF | 0.6809194 |
| VSMTNSGVS | ASGDRGENYAEQF | 0.66264427 |
| VTDVLGDLSV | ASGDRGENYAEQF | 0.60744375 |
| VVVVGAGFAGL | ASGDRGENYAEQF | 0.687806 |
| VYTVGTVV | ASGDRGENYAEQF | 0.63078415 |
| WTTFSGP | ASGDRGENYAEQF | 0.69532585 |
| YDELVEAF | ASGDRGENYAEQF | 0.5979988 |
| YGTGGAGGNGG | ASGDRGENYAEQF | 0.6164659 |
| YTNVMAV | ASGDRGENYAEQF | 0.66564715 |
| FLEQVHQL | ASGDRGENYAEQF | 0.39852 |
| GLSLLAVAV | ASGDRGENYAEQF | 0.47981137 |
| LEQVHQL | ASGDRGENYAEQF | 0.7023237 |
| LNAAGAI | ASGDRGENYAEQF | 0.672751 |
| PAVHASL | ASGDRGENYAEQF | 0.68003416 |
| TELEDDA | ASGDRGENYAEQF | 0.6287337 |
| TPPGPPN | ASGDRGENYAEQF | 0.6158809 |
| TPPPPPPP | ASGDRGENYAEQF | 0.6350242 |
| TQSPALI | ASSHHREGTEVF | 0.6736811 |
| EFAAAFPQ | ASSHHREGTEVF | 0.6802085 |
| LLSGNLDVL | ASSHHREGTEVF | 0.67364806 |
| GPANLGNYNI | ASSHHREGTEVF | 0.6149526 |
| TLGPTLP | ASSHHREGTEVF | 0.6730459 |
| VYAYSSCCK | ASSHHREGTEVF | 0.6479692 |
| APAPAPAPNI | ASSHHREGTEVF | 0.64784646 |
| PGGPVPIP | ASSHHREGTEVF | 0.66554165 |
| FLTVALEP | ASSHHREGTEVF | 0.6713237 |
| SQYFGANL | ASSHHREGTEVF | 0.64256424 |
| GGVKPAA | ASSHHREGTEVF | 0.6424652 |
| GGTGGAGGAGGW | ASSHHREGTEVF | 0.66514283 |
| GVGGVAGLGGAGG | ASSHHREGTEVF | 0.6726284 |
| VLLVLATTI | ASSHHREGTEVF | 0.6698396 |
| IGIFAAML | ASSHHREGTEVF | 0.68463063 |
| AVGDLID | ASSHHREGTEVF | 0.6758796 |

| | | |
|---|---|---|
| NISPLPL | ASSHHREGTEVF | 0.6759187 |
| VTAVFVAI | ASSHHREGTEVF | 0.40748197 |
| SGDDNDAAL | ASSHHREGTEVF | 0.524082 |
| AVAALAHS | ASSHHREGTEVF | 0.6536822 |
| GGVTVMATP | ASSHHREGTEVF | 0.6494779 |
| AGLAANVRD | ASSHHREGTEVF | 0.46363306 |
| TNRPDLIDPALLRPG | ASSHHREGTEVF | 0.67594594 |
| LANLGVTAL | ASSHHREGTEVF | 0.6453759 |
| TGERIEL | ASSHHREGTEVF | 0.6379559 |
| NGEELAA | ASSHHREGTEVF | 0.6759218 |
| VLSLVWG | ASSHHREGTEVF | 0.65174115 |
| IGYALWL | ASSHHREGTEVF | 0.6272358 |
| LSLLLAPL | ASSHHREGTEVF | 0.640148 |
| TLDEALK | ASSHHREGTEVF | 0.6685965 |
| TDGAGTAEGG | ASSHHREGTEVF | 0.60327375 |
| NAPGGAGGNGG | ASSHHREGTEVF | 0.6745554 |
| VSVFMGMGGNF | ASSHHREGTEVF | 0.55608124 |
| LASYLAAL | ASSHHREGTEVF | 0.67817605 |
| LVGGTPAL | ASSHHREGTEVF | 0.5945107 |
| VRNADVL | ASSHHREGTEVF | 0.6744217 |
| TCSSPFLVSAM | ASSHHREGTEVF | 0.6523257 |
| VHITDAE | ASSHHREGTEVF | 0.691373 |
| EYAAPSF | ASSHHREGTEVF | 0.6458194 |
| VVDAFHGGL | ASSHHREGTEVF | 0.6706402 |
| ASVVDPQ | ASSHHREGTEVF | 0.6535892 |
| SYNEFGGLS | ASSHHREGTEVF | 0.68148136 |
| DPDGSVY | ASSHHREGTEVF | 0.57112217 |
| MVSTIHATE | ASSHHREGTEVF | 0.65144193 |
| LAAGVADPV | ASSHHREGTEVF | 0.47240072 |
| TKEQIAA | ASSHHREGTEVF | 0.66399276 |
| SLASPAAL | ASSHHREGTEVF | 0.6739383 |
| IGLVFYGV | ASSHHREGTEVF | 0.53997767 |
| FLLDASNL | ASSHHREGTEVF | 0.6792928 |
| YQVGNFP | ASSHHREGTEVF | 0.63534534 |
| ESLLSQST | ASSHHREGTEVF | 0.63214314 |
| LFTVGFAAVG | ASSHHREGTEVF | 0.6800208 |
| ATPPTPT | ASSHHREGTEVF | 0.6680866 |
| TDLATLG | ASSHHREGTEVF | 0.67969966 |
| SAGGTGR | ASSHHREGTEVF | 0.65412927 |
| GFSNSGS | ASSHHREGTEVF | 0.6406921 |
| GGNGGSGGTSV | ASSHHREGTEVF | 0.66533804 |
| GTSVATGGAGN | ASSHHREGTEVF | 0.6193322 |
| NGGNGGSGGTS | ASSHHREGTEVF | 0.63141483 |
| AVVKFPH | ASSHHREGTEVF | 0.68286926 |
| TPPPPPP | ASSHHREGTEVF | 0.6514127 |
| AAPAAAAAPA | ASSHHREGTEVF | 0.6650094 |
| IALNEDL | ASSHHREGTEVF | 0.5704985 |
| IVNVSSM | ASSHHREGTEVF | 0.5445176 |
| TGIPPAPR | ASSHHREGTEVF | 0.5945717 |
| FDLGGGTF | ASSHHREGTEVF | 0.66134423 |
| FDLGGGTFDVS | ASSHHREGTEVF | 0.6776346 |

| | | |
|---|---|---|
| NEPTAAA | ASSHHREGTEVF | 0.64784086 |
| NEPTAAALA | ASSHHREGTEVF | 0.5707849 |
| GSIINISSI | ASSHHREGTEVF | 0.643504 |
| VAPGNAGTA | ASSHHREGTEVF | 0.6182276 |
| TGGGASLE | ASSHHREGTEVF | 0.6078464 |
| VAVYNHYKRI | ASSHHREGTEVF | 0.65750796 |
| TDPAPATT | ASSHHREGTEVF | 0.6485435 |
| TNRPDILDPALLRPG | ASSHHREGTEVF | 0.6010527 |
| LTTTTTS | ASSHHREGTEVF | 0.5985832 |
| PTGSGKTL | ASSHHREGTEVF | 0.6523748 |
| TGSGKTL | ASSHHREGTEVF | 0.66572034 |
| TIGHVDHGK | ASSHHREGTEVF | 0.6571696 |
| TIGHVDHGKTT | ASSHHREGTEVF | 0.68671465 |
| GHVDHGKTT | ASSHHREGTEVF | 0.660479 |
| AQFEHTLL | ASSHHREGTEVF | 0.67338085 |
| IVNQDEL | ASSHHREGTEVF | 0.40314204 |
| TNGSQFF | ASSHHREGTEVF | 0.5029149 |
| TGKGIVE | ASSHHREGTEVF | 0.62136376 |
| LISNASDA | ASSHHREGTEVF | 0.6115974 |
| AELRTLGELTKIAKAHGAQ | ASSHHREGTEVF | 0.47050506 |
| APNAVAAAEPSW | ASSHHREGTEVF | 0.6540618 |
| APQVAGAATAVL | ASSHHREGTEVF | 0.61822534 |
| ARRDPAAAAAILPTDA | ASSHHREGTEVF | 0.55356944 |
| DAAARDVLAEAGLAEYF | ASSHHREGTEVF | 0.656614 |
| DDQIVLSAKANLAKVEQ | ASSHHREGTEVF | 0.5882364 |
| DELLPDTGEDSDEDGH | ASSHHREGTEVF | 0.56960857 |
| ELPEAPANRATVPQD | ASSHHREGTEVF | 0.6054876 |
| FGQSKIERVTGIG | ASSHHREGTEVF | 0.6705893 |
| GAGGTGSTAGGAGGAGGAGGI | ASSHHREGTEVF | 0.5473568 |
| GHGGTGAAGGNGGTGGAGGS( | ASSHHREGTEVF | 0.6537672 |
| GNGGNGGAGGEGGPGGAGGA( | ASSHHREGTEVF | 0.6166469 |
| GSTAGNGGNGGNATNGTIAG( | ASSHHREGTEVF | 0.6702582 |
| GTGGDGGNGGNGGTGG | ASSHHREGTEVF | 0.51689327 |
| GVRRPAGIPADAPV | ASSHHREGTEVF | 0.6586145 |
| GYRQVTDAHLLTLARR | ASSHHREGTEVF | 0.60763466 |
| IAKMGQDGHDRG | ASSHHREGTEVF | 0.689327 |
| KAEPSIEIQYT | ASSHHREGTEVF | 0.6046802 |
| KPVRIRMAADEDTVAVT | ASSHHREGTEVF | 0.6563736 |
| MPPRIAGMRLLVIKPEP | ASSHHREGTEVF | 0.65906787 |
| MKFYAVALFRSRGVPPTR | ASSHHREGTEVF | 0.66830736 |
| NAGGFGILGAVAHSP | ASSHHREGTEVF | 0.48360416 |
| NEGREITLAMAEM | ASSHHREGTEVF | 0.6178435 |
| NELVEWSHPLGHRWRWE | ASSHHREGTEVF | 0.5552081 |
| NLGAGGTGGNGGNPG | ASSHHREGTEVF | 0.63489425 |
| NTEWPKWETVMAADPVD | ASSHHREGTEVF | 0.64722204 |
| PASAAPAETQP | ASSHHREGTEVF | 0.41736814 |
| QQSSDGKNGVRAAEITGE | ASSHHREGTEVF | 0.66737217 |
| RAPDAVTTSWWKE | ASSHHREGTEVF | 0.5758759 |
| SGFGNVGAHTSGWFNQ | ASSHHREGTEVF | 0.5872484 |
| SGVAGAVGQAASVGGLKV | ASSHHREGTEVF | 0.6716988 |
| SNGGAGTGSAGGNGGT | ASSHHREGTEVF | 0.64009887 |

| | | |
|---|---|---|
| TGGAGGEGGAAG | ASSHHREGTEVF | 0.6672723 |
| TGGDGGAAGAGGAPGHG | ASSHHREGTEVF | 0.63424736 |
| TMPPLLAAIANAGRHGV | ASSHHREGTEVF | 0.6175639 |
| TSSTAGNSGGA | ASSHHREGTEVF | 0.64224887 |
| VPATVPVGILC | ASSHHREGTEVF | 0.51189834 |
| VQRPDSGGVVLTGRIS | ASSHHREGTEVF | 0.56903243 |
| WTPNPVPATKGVPAGLVQAT | ASSHHREGTEVF | 0.67324394 |
| YALKRALAKFSDRFDVVII | ASSHHREGTEVF | 0.80679435 |
| YLIGSVIV | ASSHHREGTEVF | 0.8135743 |
| YGQLVAVID | ASSHHREGTEVF | 0.6857181 |
| VTVGGMTLPA | ASSHHREGTEVF | 0.67835283 |
| VLALPAV | ASSHHREGTEVF | 0.7330575 |
| VIPPTLNYET | ASSHHREGTEVF | 0.8332647 |
| VGMGGMHPSSG | ASSHHREGTEVF | 0.83705705 |
| VGLAGGAATA | ASSHHREGTEVF | 0.7917341 |
| VGGAGGAGGSGP | ASSHHREGTEVF | 0.730999 |
| VFAAFMV | ASSHHREGTEVF | 0.8313016 |
| TLYDKLR | ASSHHREGTEVF | 0.7932043 |
| TLAVFTVL | ASSHHREGTEVF | 0.7637096 |
| TGSVQPARV | ASSHHREGTEVF | 0.82589424 |
| TGGGAGSGGT | ASSHHREGTEVF | 0.8168389 |
| SHLETSL | ASSHHREGTEVF | 0.7513244 |
| SFTTGMT | ASSHHREGTEVF | 0.7310306 |
| QVEVAKRMGVRQPTVS | ASSHHREGTEVF | 0.807632 |
| QGSGGPALAM | ASSHHREGTEVF | 0.8262649 |
| MLAPSNP | ASSHHREGTEVF | 0.821087 |
| LVNNAAI | ASSHHREGTEVF | 0.8713072 |
| LTGEINTLA | ASSHHREGTEVF | 0.8371308 |
| LSIFGPV | ASSHHREGTEVF | 0.8084323 |
| LLALLANL | ASSHHREGTEVF | 0.43758503 |
| LISNASDALDKLR | ASSHHREGTEVF | 0.64944845 |
| LINEAVP | ASSHHREGTEVF | 0.77204573 |
| LGYAFSGL | ASSHHREGTEVF | 0.7824459 |
| LDFVAVAL | ASSHHREGTEVF | 0.3895123 |
| LAQPGAE | ASSHHREGTEVF | 0.8329415 |
| ISNLGAVTGF | ASSHHREGTEVF | 0.75656533 |
| HQEDPGRI | ASSHHREGTEVF | 0.7064004 |
| GTPGEPTP | ASSHHREGTEVF | 0.83388424 |
| GFFNSGS | ASSHHREGTEVF | 0.72593844 |
| GAGGGGLTAGAN | ASSHHREGTEVF | 0.74976134 |
| FVDALVNM | ASSHHREGTEVF | 0.78246266 |
| FLTVAPEL | ASSHHREGTEVF | 0.8556305 |
| FETLTAL | ASSHHREGTEVF | 0.69495463 |
| DVINPIEL | ASSHHREGTEVF | 0.83985853 |
| APVSGAPV | ASSHHREGTEVF | 0.78622866 |
| ALVGGLAW | ASSHHREGTEVF | 0.8596865 |
| ALAAAAATAI | ASSHHREGTEVF | 0.6616149 |
| AFGLGLLL | ASSHHREGTEVF | 0.82654023 |
| AFFAAASL | ASSHHREGTEVF | 0.76150453 |
| AAAALPAFSPP | ASSHHREGTEVF | 0.8685305 |
| VVLAPAL | ASSHHREGTEVF | 0.7832981 |

| | | |
|---|---|---|
| TIGHVDHGKTTL | ASSHHREGTEVF | 0.8421099 |
| LISNASDALDK | ASSHHREGTEVF | 0.8115844 |
| LISNASDAL | ASSHHREGTEVF | 0.8389766 |
| ISNASDALDK | ASSHHREGTEVF | 0.6023352 |
| NQAETLVYQTE | ASSHHREGTEVF | 0.7723847 |
| TGGTGGTAGAG | ASSHHREGTEVF | 0.72251856 |
| TGSTAGGAGGA | ASSHHREGTEVF | 0.80603784 |
| DPLAVPVAAGPV | ASSHHREGTEVF | 0.8107908 |
| GGTGGTPGNGGN | ASSHHREGTEVF | 0.4208482 |
| PAGLAGVDPATL | ASSHHREGTEVF | 0.82265806 |
| GNNLVVEAPHAGASV | ASSHHREGTEVF | 0.6723529 |
| KVTRIMSASTLEQPA | ASSHHREGTEVF | 0.70140326 |
| LGFALPIALPRRTVV | ASSHHREGTEVF | 0.8337306 |
| TRLGNLTARKLVVLV | ASSHHREGTEVF | 0.80950934 |
| WSQQRRLIYFPSAGP | ASSHHREGTEVF | 0.8295408 |
| GPNGNPDPMAAAVDIR | ASSHHREGTEVF | 0.76184994 |
| TQRCVSVADLRAAASR | ASSHHREGTEVF | 0.7674371 |
| QFVLDVAEHPVEVLTETV | ASSHHREGTEVF | 0.81784594 |
| GGAGGTGGAGGNGGTQAGDG | ASSHHREGTEVF | 0.6228603 |
| PLLATLRAIAVDVDAKPAQV | ASSHHREGTEVF | 0.72508264 |
| APKFRTGGYMIDFWGVGYQV | ASSHHREGTEVF | 0.8466505 |
| GAGGAGGAGGAAGGPGGTGG | ASSHHREGTEVF | 0.43418995 |
| IGALGLLATDVAPAGSVAFG | ASSHHREGTEVF | 0.69170064 |
| AGGAGGIGGNGGAGGAGGVL | ASSHHREGTEVF | 0.23379469 |
| ADELIGLL | ASSHHREGTEVF | 0.24561217 |
| ADRALKA | ASSHHREGTEVF | 0.34160116 |
| AGAVGQVGGAGGA | ASSHHREGTEVF | 0.5159604 |
| AGGGGGLGGAP | ASSHHREGTEVF | 0.6905513 |
| AGGLLYGNGGAG | ASSHHREGTEVF | 0.508533 |
| AGLESIGL | ASSHHREGTEVF | 0.25233358 |
| AGLKTGII | ASSHHREGTEVF | 0.33886912 |
| ASLAALLL | ASSHHREGTEVF | 0.29223314 |
| ASVINISSI | ASSHHREGTEVF | 0.2876674 |
| ATPPGPA | ASSHHREGTEVF | 0.43616745 |
| ATVASALV | ASSHHREGTEVF | 0.2780222 |
| ATVASLIL | ASSHHREGTEVF | 0.2567578 |
| AVNGPVLIRH | ASSHHREGTEVF | 0.26440367 |
| DGAAGAVGI | ASSHHREGTEVF | 0.3395388 |
| DPLLHGI | ASSHHREGTEVF | 0.53736395 |
| ELGAALGAL | ASSHHREGTEVF | 0.55301946 |
| ELGRTQAV | ASSHHREGTEVF | 0.59860754 |
| FVAGGYLS | ASSHHREGTEVF | 0.44596624 |
| GASGGAGGAGGW | ASSHHREGTEVF | 0.32914668 |
| GGAGAGGDGGAGGI | ASSHHREGTEVF | 0.2008322 |
| GTGHNGSQPGMGGNG | ASSHHREGTEVF | 0.22370481 |
| GYAAASASAS | ASSHHREGTEVF | 0.2957955 |
| HQVDVAAVQQG | ASSHHREGTEVF | 0.30760795 |
| IGAAIAHT | ASSHHREGTEVF | 0.20632686 |
| IGMIEVAQL | ASSHHREGTEVF | 0.4613921 |
| IRLAPEHTSAMD | ASSHHREGTEVF | 0.3378417 |
| IYNVSVL | ASSHHREGTEVF | 0.28337952 |

| | | |
|---|---|---|
| KQGFLGM | ASSHHREGTEVF | 0.5255813 |
| LESVITQL | ASSHHREGTEVF | 0.26106 |
| LGLGALVVV | ASSHHREGTEVF | 0.24397284 |
| LIGNGGNGGSGGI | ASSHHREGTEVF | 0.24900475 |
| LNAINAPTL | ASSHHREGTEVF | 0.50313526 |
| LQLLNTGV | ASSHHREGTEVF | 0.22494216 |
| LQTSIEF | ASSHHREGTEVF | 0.45885372 |
| MSFAVSEL | ASSHHREGTEVF | 0.31748465 |
| NTGALGL | ASSHHREGTEVF | 0.48732018 |
| PPGAGGAGGAGTT | ASSHHREGTEVF | 0.23213054 |
| QLVGLGI | ASSHHREGTEVF | 0.37344247 |
| RFLEQVHQL | ASSHHREGTEVF | 0.29314482 |
| RGVGYVLR | ASSHHREGTEVF | 0.6754411 |
| SATGVFAGVI | ASSHHREGTEVF | 0.30031186 |
| SLPAITLP | ASSHHREGTEVF | 0.3670531 |
| STALAFTV | ASSHHREGTEVF | 0.23658876 |
| SVLTVGLGK | ASSHHREGTEVF | 0.64845717 |
| TALAGAGSGI | ASSHHREGTEVF | 0.2410501 |
| TGLPFTGI | ASSHHREGTEVF | 0.32305527 |
| VAEHVTV | ASSHHREGTEVF | 0.22379658 |
| VDIVLNSL | ASSHHREGTEVF | 0.3396014 |
| VLTGGGAVL | ASSHHREGTEVF | 0.41159898 |
| VNVGLAYGI | ASSHHREGTEVF | 0.26428583 |
| VSSAAVGAV | ASSHHREGTEVF | 0.4202162 |
| VTSVYEAGV | ASSHHREGTEVF | 0.3836758 |
| WVALDRGA | ASSHHREGTEVF | 0.26709378 |
| YLVGSVVI | ASSHHREGTEVF | 0.5324369 |
| GPPGTGKTLL | ASSHHREGTEVF | 0.31399983 |
| GVGMTKF | ASSHHREGTEVF | 0.43330285 |
| IGIGGSDL | ASSHHREGTEVF | 0.2648803 |
| IGTLLAEL | ASSHHREGTEVF | 0.3431154 |
| IYTSGTT | ASSHHREGTEVF | 0.4643743 |
| LSFPAGV | ASSHHREGTEVF | 0.25134444 |
| LYTSGTT | ASSHHREGTEVF | 0.26417518 |
| VGAFPFGF | ASSHHREGTEVF | 0.42092663 |
| VGAGTVEF | ASSHHREGTEVF | 0.62333727 |
| VGKSTLLL | ASSHHREGTEVF | 0.6643058 |
| AALAHAGGGIV | ASSHHREGTEVF | 0.43032813 |
| AALVAVSLL | ASSHHREGTEVF | 0.453192 |
| AAVSRAVAAL | ASSHHREGTEVF | 0.49983016 |
| AGGSSGIGGF | ASSHHREGTEVF | 0.6456525 |
| AGLTVANANG | ASSHHREGTEVF | 0.6715802 |
| AGLVGAVL | ASSHHREGTEVF | 0.62601495 |
| ALSQAFTV | ASSHHREGTEVF | 0.47549534 |
| ALTFSVSL | ASSHHREGTEVF | 0.55712306 |
| ALTGAALVAL | ASSHHREGTEVF | 0.47354364 |
| ASFSIGAGVL | ASSHHREGTEVF | 0.5174502 |
| DGSVYKI | ASSHHREGTEVF | 0.5991117 |
| EQLAEIV | ASSHHREGTEVF | 0.5853789 |
| FAVSELTVM | ASSHHREGTEVF | 0.5213883 |
| GAAPLAAGL | ASSHHREGTEVF | 0.49688715 |

| | | |
|---|---|---|
| GADGGDGGSGGAGG | ASSHHREGTEVF | 0.6015849 |
| GFGLDTGGAGG | ASSHHREGTEVF | 0.65382487 |
| GLASGGLL | ASSHHREGTEVF | 0.630633 |
| GRIAGAAVLL | ASSHHREGTEVF | 0.6766019 |
| GSAGSLNTSF | ASSHHREGTEVF | 0.6289061 |
| GTGQIGI | ASSHHREGTEVF | 0.6257608 |
| GTLIALIL | ASSHHREGTEVF | 0.27827793 |
| IAATMGFVVG | ASSHHREGTEVF | 0.3653158 |
| IFAQALL | ASSHHREGTEVF | 0.55971044 |
| IGAAVALGF | ASSHHREGTEVF | 0.51608986 |
| IGLTPLAL | ASSHHREGTEVF | 0.34969938 |
| ILGSNTSTL | ASSHHREGTEVF | 0.61717665 |
| IVFSLTVL | ASSHHREGTEVF | 0.5234199 |
| LASKATGF | ASSHHREGTEVF | 0.47796947 |
| LASVALIL | ASSHHREGTEVF | 0.63546735 |
| LFAVGLTL | ASSHHREGTEVF | 0.488761 |
| LFNGGIGTYI | ASSHHREGTEVF | 0.47321087 |
| LTLGAGII | ASSHHREGTEVF | 0.47260904 |
| LVAAALGDI | ASSHHREGTEVF | 0.66091275 |
| NTAFVQLGI | ASSHHREGTEVF | 0.4008259 |
| QLFSGAEL | ASSHHREGTEVF | 0.61540806 |
| SAATFITL | ASSHHREGTEVF | 0.49618486 |
| SNYGLANLGV | ASSHHREGTEVF | 0.65155756 |
| SYGGAYAVM | ASSHHREGTEVF | 0.41232097 |
| TTGVGGLTLG | ASSHHREGTEVF | 0.61004007 |
| TTQTIYI | ASSHHREGTEVF | 0.50676763 |
| VAAGAGLGAVI | ASSHHREGTEVF | 0.71001583 |
| VAAVVTLL | ASSHHREGTEVF | 0.50005776 |
| VGFLGGLI | ASSHHREGTEVF | 0.5589318 |
| VGVIPILL | ASSHHREGTEVF | 0.53170854 |
| VIFASSLI | ASSHHREGTEVF | 0.66782445 |
| VKGGDGGAAAG | ASSHHREGTEVF | 0.36343938 |
| VTKTFTNI | ASSHHREGTEVF | 0.5134194 |
| VVAAAILL | ASSHHREGTEVF | 0.40800568 |
| YAALDVEL | ASSHHREGTEVF | 0.54925805 |
| LASVALLL | ASSHHREGTEVF | 0.61359847 |
| LDAVHRV | ASSHHREGTEVF | 0.34055513 |
| QGLSSSIF | ASSHHREGTEVF | 0.6304697 |
| RGTFANIRL | ASSHHREGTEVF | 0.49049 |
| RQLWWGHRI | ASSHHREGTEVF | 0.47354034 |
| YGPPGTGKTLL | ASSHHREGTEVF | 0.64374477 |
| AALVGAPANVA | ASSHHREGTEVF | 0.51398224 |
| AATFRRTE | ASSHHREGTEVF | 0.67139584 |
| AGALGGGI | ASSHHREGTEVF | 0.5181526 |
| AGGAGGAGGIGAG | ASSHHREGTEVF | 0.5145892 |
| AGSGALGL | ASSHHREGTEVF | 0.61606675 |
| AQIADQI | ASSHHREGTEVF | 0.47240418 |
| ASAVAVTYL | ASSHHREGTEVF | 0.45243225 |
| ASTVLGGTGG | ASSHHREGTEVF | 0.6442578 |
| AVGGEPAK | ASSHHREGTEVF | 0.47823057 |
| CEQGTFGGTC | ASSHHREGTEVF | 0.6953125 |

| | | |
|---|---|---|
| DEGKAPAG | ASSHHREGTEVF | 0.2597333 |
| DTVVFPAL | ASSHHREGTEVF | 0.31259876 |
| FEGITMAG | ASSHHREGTEVF | 0.36920616 |
| FPLVVQD | ASSHHREGTEVF | 0.57872564 |
| GAFTTPQ | ASSHHREGTEVF | 0.71622646 |
| GFGGFGTAGGVGGA | ASSHHREGTEVF | 0.5000724 |
| GFLAQFL | ASSHHREGTEVF | 0.32651195 |
| GGAGGAGGNAGVA | ASSHHREGTEVF | 0.4179151 |
| GGVGTAGAGGAGGAGG | ASSHHREGTEVF | 0.37896088 |
| GNGGGGDAGNAGSGGN | ASSHHREGTEVF | 0.43694887 |
| GTGGAGGAGNDAGS | ASSHHREGTEVF | 0.45647234 |
| GTTNTGWFNS | ASSHHREGTEVF | 0.4465106 |
| IFMGPLFTL | ASSHHREGTEVF | 0.32053185 |
| ILAHAIEAI | ASSHHREGTEVF | 0.33636186 |
| IRSAIREAL | ASSHHREGTEVF | 0.5353488 |
| ITFAGGG | ASSHHREGTEVF | 0.6223177 |
| ITGMSGHQEV | ASSHHREGTEVF | 0.60419226 |
| IVAGLACTL | ASSHHREGTEVF | 0.63368285 |
| IVGGGLAGA | ASSHHREGTEVF | 0.57653564 |
| IVTGVLNI | ASSHHREGTEVF | 0.41743883 |
| LAQQGCI | ASSHHREGTEVF | 0.18457812 |
| LCLHVEN | ASSHHREGTEVF | 0.24643975 |
| LDAISDGS | ASSHHREGTEVF | 0.37559018 |
| LGGFAAML | ASSHHREGTEVF | 0.4059991 |
| LGGKNPMI | ASSHHREGTEVF | 0.22969915 |
| LGPKGVAAIL | ASSHHREGTEVF | 0.5079232 |
| LMVGVSI | ASSHHREGTEVF | 0.3978151 |
| LVGAAFVL | ASSHHREGTEVF | 0.33762744 |
| LVLAVAPL | ASSHHREGTEVF | 0.6277224 |
| MAAIAVIDAA | ASSHHREGTEVF | 0.29877424 |
| NNGINAELQ | ASSHHREGTEVF | 0.31280494 |
| RGEPLVLVHG | ASSHHREGTEVF | 0.33144602 |
| RGGAGGAAGLI | ASSHHREGTEVF | 0.50353944 |
| RSYRMHPAV | ASSHHREGTEVF | 0.28774744 |
| SAAHLLAGQLA | ASSHHREGTEVF | 0.58855075 |
| SAAVGSY | ASSHHREGTEVF | 0.36009222 |
| SPHAVLHRI | ASSHHREGTEVF | 0.6389139 |
| SQIAKVL | ASSHHREGTEVF | 0.29355487 |
| TALGHLVP | ASSHHREGTEVF | 0.53989756 |
| TAPTTSGAA | ASSHHREGTEVF | 0.38945535 |
| TDGDDGNGGSPGT | ASSHHREGTEVF | 0.7457244 |
| TDGLGEN | ASSHHREGTEVF | 0.39460486 |
| TGGPPLPGQA | ASSHHREGTEVF | 0.48630962 |
| TIPAITI | ASSHHREGTEVF | 0.39405861 |
| TIPGFTI | ASSHHREGTEVF | 0.68421173 |
| TSIASGALA | ASSHHREGTEVF | 0.24549638 |
| TVTLPGAAI | ASSHHREGTEVF | 0.34234294 |
| VASGLSAALP | ASSHHREGTEVF | 0.23625898 |
| VGAGVIGI | ASSHHREGTEVF | 0.37645268 |
| VGPKRFGV | ASSHHREGTEVF | 0.50939965 |
| VNVAFPD | ASSHHREGTEVF | 0.2445142 |

| | | |
|---|---|---|
| VSMTNSGVS | ASSHHREGTEVF | 0.383221 |
| VTDVLGDLSV | ASSHHREGTEVF | 0.34997487 |
| VVVVGAGFAGL | ASSHHREGTEVF | 0.3216653 |
| VYTVGTVV | ASSHHREGTEVF | 0.48588544 |
| WTTFSGP | ASSHHREGTEVF | 0.41580302 |
| YDELVEAF | ASSHHREGTEVF | 0.56353587 |
| YGTGGAGGNGG | ASSHHREGTEVF | 0.3029511 |
| YTNVMAV | ASSHHREGTEVF | 0.30979136 |
| FLEQVHQL | ASSHHREGTEVF | 0.44263956 |
| GLSLLAVAV | ASSHHREGTEVF | 0.34181064 |
| LEQVHQL | ASSHHREGTEVF | 0.28103906 |
| LNAAGAI | ASSHHREGTEVF | 0.6161623 |
| PAVHASL | ASSHHREGTEVF | 0.8847209 |
| TELEDDA | ASSHHREGTEVF | 0.8715813 |
| TPPGPPN | ASSHHREGTEVF | 0.7960794 |
| TPPPPPPP | ASSHHREGTEVF | 0.77561754 |
| TQSPALI | ASSSGQSANTEVF | 0.8235667 |
| EFAAAFPQ | ASSSGQSANTEVF | 0.90043056 |
| LLSGNLDVL | ASSSGQSANTEVF | 0.8940417 |
| GPANLGNYNI | ASSSGQSANTEVF | 0.8758688 |
| TLGPTLP | ASSSGQSANTEVF | 0.7753447 |
| VYAYSSCCK | ASSSGQSANTEVF | 0.87710714 |
| APAPAPAPNI | ASSSGQSANTEVF | 0.8553628 |
| PGGPVPIP | ASSSGQSANTEVF | 0.83843327 |
| FLTVALEP | ASSSGQSANTEVF | 0.88953817 |
| SQYFGANL | ASSSGQSANTEVF | 0.8819343 |
| GGVKPAA | ASSSGQSANTEVF | 0.83725536 |
| GGTGGAGGAGGW | ASSSGQSANTEVF | 0.84787947 |
| GVGGVAGLGGAGG | ASSSGQSANTEVF | 0.86479115 |
| VLLVLATTI | ASSSGQSANTEVF | 0.8853267 |
| IGIFAAML | ASSSGQSANTEVF | 0.88574016 |
| AVGDLID | ASSSGQSANTEVF | 0.90273786 |
| NISPLPL | ASSSGQSANTEVF | 0.8960791 |
| VTAVFVAI | ASSSGQSANTEVF | 0.88038874 |
| SGDDNDAAL | ASSSGQSANTEVF | 0.31335545 |
| AVAALAHS | ASSSGQSANTEVF | 0.5715643 |
| GGVTVMATP | ASSSGQSANTEVF | 0.84726906 |
| AGLAANVRD | ASSSGQSANTEVF | 0.83013046 |
| TNRPDLIDPALLRPG | ASSSGQSANTEVF | 0.5296365 |
| LANLGVTAL | ASSSGQSANTEVF | 0.8650506 |
| TGERIEL | ASSSGQSANTEVF | 0.8377262 |
| NGEELAA | ASSSGQSANTEVF | 0.79653907 |
| VLSLVWG | ASSSGQSANTEVF | 0.88922244 |
| IGYALWL | ASSSGQSANTEVF | 0.84032094 |
| LSLLLAPL | ASSSGQSANTEVF | 0.8174465 |
| TLDEALK | ASSSGQSANTEVF | 0.830428 |
| TDGAGTAEGG | ASSSGQSANTEVF | 0.8976821 |
| NAPGGAGGNGG | ASSSGQSANTEVF | 0.7563901 |
| VSVFMGMGGNF | ASSSGQSANTEVF | 0.88973886 |
| LASYLAAL | ASSSGQSANTEVF | 0.7667242 |
| LVGGTPAL | ASSSGQSANTEVF | 0.9052114 |

| | | |
|---|---|---|
| VRNADVL | ASSSGQSANTEVF | 0.739028 |
| TCSSPFLVSAM | ASSSGQSANTEVF | 0.8860073 |
| VHITDAE | ASSSGQSANTEVF | 0.84452784 |
| EYAAPSF | ASSSGQSANTEVF | 0.9164956 |
| VVDAFHGGL | ASSSGQSANTEVF | 0.8527192 |
| ASVVDPQ | ASSSGQSANTEVF | 0.8913827 |
| SYNEFGGLS | ASSSGQSANTEVF | 0.8517605 |
| DPDGSVY | ASSSGQSANTEVF | 0.8821175 |
| MVSTIHATE | ASSSGQSANTEVF | 0.7097447 |
| LAAGVADPV | ASSSGQSANTEVF | 0.8662604 |
| TKEQIAA | ASSSGQSANTEVF | 0.5760914 |
| SLASPAAL | ASSSGQSANTEVF | 0.8757647 |
| IGLVFYGV | ASSSGQSANTEVF | 0.8760824 |
| FLLDASNL | ASSSGQSANTEVF | 0.5039102 |
| YQVGNFP | ASSSGQSANTEVF | 0.8922622 |
| ESLLSQST | ASSSGQSANTEVF | 0.7965944 |
| LFTVGFAAVG | ASSSGQSANTEVF | 0.82685244 |
| ATPPTPT | ASSSGQSANTEVF | 0.9014653 |
| TDLATLG | ASSSGQSANTEVF | 0.852788 |
| SAGGTGR | ASSSGQSANTEVF | 0.8971822 |
| GFSNSGS | ASSSGQSANTEVF | 0.85624796 |
| GGNGGSGGTSV | ASSSGQSANTEVF | 0.8488833 |
| GTSVATGGAGN | ASSSGQSANTEVF | 0.8698678 |
| NGGNGGSGGTS | ASSSGQSANTEVF | 0.5345924 |
| AVVKFPH | ASSSGQSANTEVF | 0.8004726 |
| TPPPPPP | ASSSGQSANTEVF | 0.904638 |
| AAPAAAAAPA | ASSSGQSANTEVF | 0.8270389 |
| IALNEDL | ASSSGQSANTEVF | 0.86478305 |
| IVNVSSM | ASSSGQSANTEVF | 0.7324175 |
| TGIPPAPR | ASSSGQSANTEVF | 0.66532356 |
| FDLGGGTF | ASSSGQSANTEVF | 0.75699174 |
| FDLGGGTFDVS | ASSSGQSANTEVF | 0.88129807 |
| NEPTAAA | ASSSGQSANTEVF | 0.8590038 |
| NEPTAAALA | ASSSGQSANTEVF | 0.82981217 |
| GSIINISSI | ASSSGQSANTEVF | 0.6089027 |
| VAPGNAGTA | ASSSGQSANTEVF | 0.78864795 |
| TGGGASLE | ASSSGQSANTEVF | 0.7487521 |
| VAVYNHYKRI | ASSSGQSANTEVF | 0.7297262 |
| TDPAPATT | ASSSGQSANTEVF | 0.86234295 |
| TNRPDILDPALLRPG | ASSSGQSANTEVF | 0.8222798 |
| LTTTTTS | ASSSGQSANTEVF | 0.7404567 |
| PTGSGKTL | ASSSGQSANTEVF | 0.7873821 |
| TGSGKTL | ASSSGQSANTEVF | 0.80663526 |
| TIGHVDHGK | ASSSGQSANTEVF | 0.7850243 |
| TIGHVDHGKTT | ASSSGQSANTEVF | 0.7754822 |
| GHVDHGKTT | ASSSGQSANTEVF | 0.8839151 |
| AQFEHTLL | ASSSGQSANTEVF | 0.8538171 |
| IVNQDEL | ASSSGQSANTEVF | 0.86188555 |
| TNGSQFF | ASSSGQSANTEVF | 0.19623071 |
| TGKGIVE | ASSSGQSANTEVF | 0.38300318 |
| LISNASDA | ASSSGQSANTEVF | 0.8145001 |

| | | |
|---|---|---|
| AELRTLGELTKIAKAHGAQ | ASSSGQSANTEVF | 0.6966783 |
| APNAVAAAEPSW | ASSSGQSANTEVF | 0.5778968 |
| APQVAGAATAVL | ASSSGQSANTEVF | 0.8084458 |
| ARRDPAAAAAILPTDA | ASSSGQSANTEVF | 0.73105407 |
| DAAARDVLAEAGLAEYF | ASSSGQSANTEVF | 0.65953815 |
| DDQIVLSAKANLAKVEQ | ASSSGQSANTEVF | 0.8418531 |
| DELLPDTGEDSDEDGH | ASSSGQSANTEVF | 0.7573241 |
| ELPEAPANRATVPQD | ASSSGQSANTEVF | 0.712311 |
| FGQSKIERVTGIG | ASSSGQSANTEVF | 0.7128581 |
| GAGGTGSTAGGAGGAGGAGGI | ASSSGQSANTEVF | 0.82610106 |
| GHGGTGAAGGNGGTGGAGGS( | ASSSGQSANTEVF | 0.58700943 |
| GNGGNGGAGGEGGPGGAGGA( | ASSSGQSANTEVF | 0.8160259 |
| GSTAGNGGNGGNATNGTIAG( | ASSSGQSANTEVF | 0.6646991 |
| GTGGDGGNGGNGGTGG | ASSSGQSANTEVF | 0.864059 |
| GVRRPAGIPADAPV | ASSSGQSANTEVF | 0.5706485 |
| GYRQVTDAHLLTLARR | ASSSGQSANTEVF | 0.8328479 |
| IAKMGQDGHDRG | ASSSGQSANTEVF | 0.76714903 |
| KAEPSIEIQYT | ASSSGQSANTEVF | 0.90543675 |
| KPVRIRMAADEDTVAVT | ASSSGQSANTEVF | 0.76543677 |
| MPPRIAGMRLLVIKPEP | ASSSGQSANTEVF | 0.79898834 |
| MKFYAVALFRSRGVPPTR | ASSSGQSANTEVF | 0.70840824 |
| NAGGFGILGAVAHSP | ASSSGQSANTEVF | 0.8440887 |
| NEGREITLAMAEM | ASSSGQSANTEVF | 0.5893122 |
| NELVEWSHPLGHRWRWE | ASSSGQSANTEVF | 0.7718576 |
| NLGAGGTGGNGGNPG | ASSSGQSANTEVF | 0.44928578 |
| NTEWPKWETVMAADPVD | ASSSGQSANTEVF | 0.82952106 |
| PASAAPAETQP | ASSSGQSANTEVF | 0.8050004 |
| QQSSDGKNGVRAAEITGE | ASSSGQSANTEVF | 0.5983362 |
| RAPDAVTTSWWKE | ASSSGQSANTEVF | 0.8505837 |
| SGFGNVGAHTSGWFNQ | ASSSGQSANTEVF | 0.8071341 |
| SGVAGAVGQAASVGGLKV | ASSSGQSANTEVF | 0.7597625 |
| SNGGAGTGSAGGNGGT | ASSSGQSANTEVF | 0.8915774 |
| TGGAGGEGGAAG | ASSSGQSANTEVF | 0.7725147 |
| TGGDGGAAGAGGAPGHG | ASSSGQSANTEVF | 0.87977016 |
| TMPPLLAAIANAGRHGV | ASSSGQSANTEVF | 0.83286786 |
| TSSTAGNSGGA | ASSSGQSANTEVF | 0.71862555 |
| VPATVPVGILC | ASSSGQSANTEVF | 0.81836426 |
| VQRPDSGGVVLTGRIS | ASSSGQSANTEVF | 0.64692396 |
| WTPNPVPATKGVPAGLVQAT | ASSSGQSANTEVF | 0.72380006 |
| YALKRALAKFSDRFDVVII | ASSSGQSANTEVF | 0.86710525 |
| YLIGSVIV | ASSSGQSANTEVF | 0.68145454 |
| YGQLVAVID | ASSSGQSANTEVF | 0.69667614 |
| VTVGGMTLPA | ASSSGQSANTEVF | 0.6194936 |
| VLALPAV | ASSSGQSANTEVF | 0.5636716 |
| VIPPTLNYET | ASSSGQSANTEVF | 0.63315254 |
| VGMGGMHPSSG | ASSSGQSANTEVF | 0.6946279 |
| VGLAGGAATA | ASSSGQSANTEVF | 0.7046442 |
| VGGAGGAGGSGP | ASSSGQSANTEVF | 0.68385863 |
| VFAAFMV | ASSSGQSANTEVF | 0.5544163 |
| TLYDKLR | ASSSGQSANTEVF | 0.6942713 |
| TLAVFTVL | ASSSGQSANTEVF | 0.6631888 |

| | | |
|---|---|---|
| TGSVQPARV | ASSSGQSANTEVF | 0.66448236 |
| TGGGAGSGGT | ASSSGQSANTEVF | 0.6777804 |
| SHLETSL | ASSSGQSANTEVF | 0.6839311 |
| SFTTGMT | ASSSGQSANTEVF | 0.63998246 |
| QVEVAKRMGVRQPTVS | ASSSGQSANTEVF | 0.5766572 |
| QGSGGPALAM | ASSSGQSANTEVF | 0.6849796 |
| MLAPSNP | ASSSGQSANTEVF | 0.6941627 |
| LVNNAAI | ASSSGQSANTEVF | 0.69225276 |
| LTGEINTLA | ASSSGQSANTEVF | 0.70871204 |
| LSIFGPV | ASSSGQSANTEVF | 0.6989732 |
| LLALLANL | ASSSGQSANTEVF | 0.6735513 |
| LISNASDALDKLR | ASSSGQSANTEVF | 0.34599227 |
| LINEAVP | ASSSGQSANTEVF | 0.4811231 |
| LGYAFSGL | ASSSGQSANTEVF | 0.61581206 |
| LDFVAVAL | ASSSGQSANTEVF | 0.6512788 |
| LAQPGAE | ASSSGQSANTEVF | 0.34994665 |
| ISNLGAVTGF | ASSSGQSANTEVF | 0.6812781 |
| HQEDPGRI | ASSSGQSANTEVF | 0.61526585 |
| GTPGEPTP | ASSSGQSANTEVF | 0.6368742 |
| GFFNSGS | ASSSGQSANTEVF | 0.69684386 |
| GAGGGGLTAGAN | ASSSGQSANTEVF | 0.63115346 |
| FVDALVNM | ASSSGQSANTEVF | 0.573477 |
| FLTVAPEL | ASSSGQSANTEVF | 0.6331726 |
| FETLTAL | ASSSGQSANTEVF | 0.70746195 |
| DVINPIEL | ASSSGQSANTEVF | 0.5654259 |
| APVSGAPV | ASSSGQSANTEVF | 0.70035064 |
| ALVGGLAW | ASSSGQSANTEVF | 0.62224287 |
| ALAAAAATAI | ASSSGQSANTEVF | 0.7063011 |
| AFGLGLLL | ASSSGQSANTEVF | 0.5187254 |
| AFFAAASL | ASSSGQSANTEVF | 0.696323 |
| AAAALPAFSPP | ASSSGQSANTEVF | 0.66793 |
| VVLAPAL | ASSSGQSANTEVF | 0.7161263 |
| TIGHVDHGKTTL | ASSSGQSANTEVF | 0.6362622 |
| LISNASDALDK | ASSSGQSANTEVF | 0.68437976 |
| LISNASDAL | ASSSGQSANTEVF | 0.64849347 |
| ISNASDALDK | ASSSGQSANTEVF | 0.700451 |
| NQAETLVYQTE | ASSSGQSANTEVF | 0.5424567 |
| TGGTGGTAGAG | ASSSGQSANTEVF | 0.6398706 |
| TGSTAGGAGGA | ASSSGQSANTEVF | 0.52844 |
| DPLAVPVAAGPV | ASSSGQSANTEVF | 0.6646433 |
| GGTGGTPGNGGN | ASSSGQSANTEVF | 0.6825969 |
| PAGLAGVDPATL | ASSSGQSANTEVF | 0.34921488 |
| GNNLVVEAPHAGASV | ASSSGQSANTEVF | 0.69843984 |
| KVTRIMSASTLEQPA | ASSSGQSANTEVF | 0.55443853 |
| LGFALPIALPRRTVV | ASSSGQSANTEVF | 0.55181944 |
| TRLGNLTARKLVVLV | ASSSGQSANTEVF | 0.69583267 |
| WSQQRRLIYFPSAGP | ASSSGQSANTEVF | 0.68097186 |
| GPNGNPDPMAAAVDIR | ASSSGQSANTEVF | 0.6891516 |
| TQRCVSVADLRAAASR | ASSSGQSANTEVF | 0.6452322 |
| QFVLDVAEHPVEVLTETV | ASSSGQSANTEVF | 0.66889125 |
| GGAGGTGGAGGNGGTQAGDG | ASSSGQSANTEVF | 0.6783948 |

| | | |
|---|---|---|
| PLLATLRAIAVDVDAKPAQV | ASSSGQSANTEVF | 0.5578664 |
| APKFRTGGYMIDFWGVGYQV | ASSSGQSANTEVF | 0.6163697 |
| GAGGAGGAGGAAGGPGGTGG | ASSSGQSANTEVF | 0.70265126 |
| IGALGLLATDVAPAGSVAFG | ASSSGQSANTEVF | 0.78956383 |
| AGGAGGIGGNGGAGGAGGVL | ASSSGQSANTEVF | 0.8069701 |
| ADELIGLL | ASSSGQSANTEVF | 0.73615456 |
| ADRALKA | ASSSGQSANTEVF | 0.6436909 |
| AGAVGQVGGAGGA | ASSSGQSANTEVF | 0.7595349 |
| AGGGGGLGGAP | ASSSGQSANTEVF | 0.8105406 |
| AGGLLYGNGGAG | ASSSGQSANTEVF | 0.8184098 |
| AGLESIGL | ASSSGQSANTEVF | 0.8015004 |
| AGLKTGII | ASSSGQSANTEVF | 0.5635207 |
| ASLAALLL | ASSSGQSANTEVF | 0.80314755 |
| ASVINISSI | ASSSGQSANTEVF | 0.747193 |
| ATPPGPA | ASSSGQSANTEVF | 0.7612785 |
| ATVASALV | ASSSGQSANTEVF | 0.7963044 |
| ATVASLIL | ASSSGQSANTEVF | 0.75163954 |
| AVNGPVLIRH | ASSSGQSANTEVF | 0.74835044 |
| DGAAGAVGI | ASSSGQSANTEVF | 0.66702855 |
| DPLLHGI | ASSSGQSANTEVF | 0.80957735 |
| ELGAALGAL | ASSSGQSANTEVF | 0.81334096 |
| ELGRTQAV | ASSSGQSANTEVF | 0.810264 |
| FVAGGYLS | ASSSGQSANTEVF | 0.8270384 |
| GASGGAGGAGGW | ASSSGQSANTEVF | 0.8128664 |
| GGAGAGGDGGAGGI | ASSSGQSANTEVF | 0.79217595 |
| GTGHNGSQPGMGGNG | ASSSGQSANTEVF | 0.26069376 |
| GYAAASASAS | ASSSGQSANTEVF | 0.4892745 |
| HQVDVAAVQQG | ASSSGQSANTEVF | 0.662607 |
| IGAAIAHT | ASSSGQSANTEVF | 0.74563324 |
| IGMIEVAQL | ASSSGQSANTEVF | 0.26724294 |
| IRLAPEHTSAMD | ASSSGQSANTEVF | 0.75812185 |
| IYNVSVL | ASSSGQSANTEVF | 0.7242665 |
| KQGFLGM | ASSSGQSANTEVF | 0.66262555 |
| LESVITQL | ASSSGQSANTEVF | 0.8169848 |
| LGLGALVVV | ASSSGQSANTEVF | 0.70673555 |
| LIGNGGNGGSGGI | ASSSGQSANTEVF | 0.64931107 |
| LNAINAPTL | ASSSGQSANTEVF | 0.7108785 |
| LQLLNTGV | ASSSGQSANTEVF | 0.8250281 |
| LQTSIEF | ASSSGQSANTEVF | 0.6045431 |
| MSFAVSEL | ASSSGQSANTEVF | 0.8103266 |
| NTGALGL | ASSSGQSANTEVF | 0.74870574 |
| PPGAGGAGGAGTT | ASSSGQSANTEVF | 0.81942284 |
| QLVGLGI | ASSSGQSANTEVF | 0.46064615 |
| RFLEQVHQL | ASSSGQSANTEVF | 0.8081353 |
| RGVGYVLR | ASSSGQSANTEVF | 0.7758826 |
| SATGVFAGVI | ASSSGQSANTEVF | 0.8325274 |
| SLPAITLP | ASSSGQSANTEVF | 0.733134 |
| STALAFTV | ASSSGQSANTEVF | 0.7620063 |
| SVLTVGLGK | ASSSGQSANTEVF | 0.64644754 |
| TALAGAGSGI | ASSSGQSANTEVF | 0.8131939 |
| TGLPFTGI | ASSSGQSANTEVF | 0.48025835 |

| | | |
|---|---|---|
| VAEHVTV | ASSSGQSANTEVF | 0.76190054 |
| VDIVLNSL | ASSSGQSANTEVF | 0.61797786 |
| VLTGGGAVL | ASSSGQSANTEVF | 0.7892322 |
| VNVGLAYGI | ASSSGQSANTEVF | 0.7976952 |
| VSSAAVGAV | ASSSGQSANTEVF | 0.28082263 |
| VTSVYEAGV | ASSSGQSANTEVF | 0.8188132 |
| WVALDRGA | ASSSGQSANTEVF | 0.6629578 |
| YLVGSVVI | ASSSGQSANTEVF | 0.5953784 |
| GPPGTGKTLL | ASSSGQSANTEVF | 0.8126638 |
| GVGMTKF | ASSSGQSANTEVF | 0.7504606 |
| IGIGGSDL | ASSSGQSANTEVF | 0.8100307 |
| IGTLLAEL | ASSSGQSANTEVF | 0.71411467 |
| IYTSGTT | ASSSGQSANTEVF | 0.7896172 |
| LSFPAGV | ASSSGQSANTEVF | 0.8068723 |
| LYTSGTT | ASSSGQSANTEVF | 0.4422556 |
| VGAFPFGF | ASSSGQSANTEVF | 0.675468 |
| VGAGTVEF | ASSSGQSANTEVF | 0.81912315 |
| VGKSTLLL | ASSSGQSANTEVF | 0.6523516 |
| AALAHAGGGIV | ASSSGQSANTEVF | 0.6815292 |
| AALVAVSLL | ASSSGQSANTEVF | 0.5521404 |
| AAVSRAVAAL | ASSSGQSANTEVF | 0.42828318 |
| AGGSSGIGGF | ASSSGQSANTEVF | 0.5215076 |
| AGLTVANANG | ASSSGQSANTEVF | 0.6724617 |
| AGLVGAVL | ASSSGQSANTEVF | 0.69912744 |
| ALSQAFTV | ASSSGQSANTEVF | 0.6338416 |
| ALTFSVSL | ASSSGQSANTEVF | 0.40361288 |
| ALTGAALVAL | ASSSGQSANTEVF | 0.66928554 |
| ASFSIGAGVL | ASSSGQSANTEVF | 0.62395686 |
| DGSVYKI | ASSSGQSANTEVF | 0.5392238 |
| EQLAEIV | ASSSGQSANTEVF | 0.6020764 |
| FAVSELTVM | ASSSGQSANTEVF | 0.64085656 |
| GAAPLAAGL | ASSSGQSANTEVF | 0.5063051 |
| GADGGDGGSGGAGG | ASSSGQSANTEVF | 0.4874832 |
| GFGLDTGGAGG | ASSSGQSANTEVF | 0.633838 |
| GLASGGLL | ASSSGQSANTEVF | 0.6756834 |
| GRIAGAAVLL | ASSSGQSANTEVF | 0.6771222 |
| GSAGSLNTSF | ASSSGQSANTEVF | 0.73279405 |
| GTGQIGI | ASSSGQSANTEVF | 0.70837027 |
| GTLIALIL | ASSSGQSANTEVF | 0.68055975 |
| IAATMGFVVG | ASSSGQSANTEVF | 0.2864595 |
| IFAQALL | ASSSGQSANTEVF | 0.35863757 |
| IGAAVALGF | ASSSGQSANTEVF | 0.5652228 |
| IGLTPLAL | ASSSGQSANTEVF | 0.5381856 |
| ILGSNTSTL | ASSSGQSANTEVF | 0.30064562 |
| IVFSLTVL | ASSSGQSANTEVF | 0.6047143 |
| LASKATGF | ASSSGQSANTEVF | 0.5387881 |
| LASVALIL | ASSSGQSANTEVF | 0.46611142 |
| LFAVGLTL | ASSSGQSANTEVF | 0.68816626 |
| LFNGGIGTYI | ASSSGQSANTEVF | 0.53048146 |
| LTLGAGII | ASSSGQSANTEVF | 0.46259922 |
| LVAAALGDI | ASSSGQSANTEVF | 0.51290596 |

| | | |
|---|---|---|
| NTAFVQLGI | ASSSGQSANTEVF | 0.6549436 |
| QLFSGAEL | ASSSGQSANTEVF | 0.41869578 |
| SAATFITL | ASSSGQSANTEVF | 0.6724365 |
| SNYGLANLGV | ASSSGQSANTEVF | 0.4455424 |
| SYGGAYAVM | ASSSGQSANTEVF | 0.72196305 |
| TTGVGGLTLG | ASSSGQSANTEVF | 0.37636703 |
| TTQTIYI | ASSSGQSANTEVF | 0.68877435 |
| VAAGAGLGAVI | ASSSGQSANTEVF | 0.5862928 |
| VAAVVTLL | ASSSGQSANTEVF | 0.7615862 |
| VGFLGGLI | ASSSGQSANTEVF | 0.56960773 |
| VGVIPILL | ASSSGQSANTEVF | 0.6508961 |
| VIFASSLI | ASSSGQSANTEVF | 0.57193744 |
| VKGGDGGAAAG | ASSSGQSANTEVF | 0.6804067 |
| VTKTFTNI | ASSSGQSANTEVF | 0.3770104 |
| VVAAAILL | ASSSGQSANTEVF | 0.5943209 |
| YAALDVEL | ASSSGQSANTEVF | 0.37454838 |
| LASVALLL | ASSSGQSANTEVF | 0.6153087 |
| LDAVHRV | ASSSGQSANTEVF | 0.66233075 |
| QGLSSSIF | ASSSGQSANTEVF | 0.2757411 |
| RGTFANIRL | ASSSGQSANTEVF | 0.667461 |
| RQLWWGHRI | ASSSGQSANTEVF | 0.46399426 |
| YGPPGTGKTLL | ASSSGQSANTEVF | 0.47453505 |
| AALVGAPANVA | ASSSGQSANTEVF | 0.70595145 |
| AATFRRTE | ASSSGQSANTEVF | 0.65594137 |
| AGALGGGI | ASSSGQSANTEVF | 0.6955248 |
| AGGAGGAGGIGAG | ASSSGQSANTEVF | 0.52371895 |
| AGSGALGL | ASSSGQSANTEVF | 0.551407 |
| AQIADQI | ASSSGQSANTEVF | 0.5846334 |
| ASAVAVTYL | ASSSGQSANTEVF | 0.49314165 |
| ASTVLGGTGG | ASSSGQSANTEVF | 0.49726453 |
| AVGGEPAK | ASSSGQSANTEVF | 0.7164818 |
| CEQGTFGGTC | ASSSGQSANTEVF | 0.8622898 |
| DEGKAPAG | ASSSGQSANTEVF | 0.8672913 |
| DTVVFPAL | ASSSGQSANTEVF | 0.7576283 |
| FEGITMAG | ASSSGQSANTEVF | 0.7240299 |
| FPLVVQD | ASSSGQSANTEVF | 0.8177974 |
| GAFTTPQ | ASSSGQSANTEVF | 0.86564827 |
| GFGGFGTAGGVGGA | ASSSGQSANTEVF | 0.8792687 |
| GFLAQFL | ASSSGQSANTEVF | 0.85692585 |
| GGAGGAGGNAGVA | ASSSGQSANTEVF | 0.7371692 |
| GGVGTAGAGGAGGAGG | ASSSGQSANTEVF | 0.8464432 |
| GNGGGGDAGNAGSGGN | ASSSGQSANTEVF | 0.8182548 |
| GTGGAGGAGNDAGS | ASSSGQSANTEVF | 0.8083863 |
| GTTNTGWFNS | ASSSGQSANTEVF | 0.8620286 |
| IFMGPLFTL | ASSSGQSANTEVF | 0.84411204 |
| ILAHAIEAI | ASSSGQSANTEVF | 0.8136685 |
| IRSAIREAL | ASSSGQSANTEVF | 0.79000926 |
| ITFAGGG | ASSSGQSANTEVF | 0.84751356 |
| ITGMSGHQEV | ASSSGQSANTEVF | 0.875147 |
| IVAGLACTL | ASSSGQSANTEVF | 0.86584646 |
| IVGGGLAGA | ASSSGQSANTEVF | 0.8827202 |

| | | |
|---|---|---|
| IVTGVLNI | ASSSGQSANTEVF | 0.8685711 |
| LAQQGCI | ASSSGQSANTEVF | 0.8646339 |
| LCLHVEN | ASSSGQSANTEVF | 0.35889408 |
| LDAISDGS | ASSSGQSANTEVF | 0.6073711 |
| LGGFAAML | ASSSGQSANTEVF | 0.78910005 |
| LGGKNPMI | ASSSGQSANTEVF | 0.81081116 |
| LGPKGVAAIL | ASSSGQSANTEVF | 0.5813632 |
| LMVGVSI | ASSSGQSANTEVF | 0.8593956 |
| LVGAAFVL | ASSSGQSANTEVF | 0.8083576 |
| LVLAVAPL | ASSSGQSANTEVF | 0.73201233 |
| MAAIAVIDAA | ASSSGQSANTEVF | 0.8679831 |
| NNGINAELQ | ASSSGQSANTEVF | 0.80791557 |
| RGEPLVLVHG | ASSSGQSANTEVF | 0.7414142 |
| RGGAGGAAGLI | ASSSGQSANTEVF | 0.77164364 |
| RSYRMHPAV | ASSSGQSANTEVF | 0.8775022 |
| SAAHLLAGQLA | ASSSGQSANTEVF | 0.6900008 |
| SAAVGSY | ASSSGQSANTEVF | 0.8591502 |
| SPHAVLHRI | ASSSGQSANTEVF | 0.8141775 |
| SQIAKVL | ASSSGQSANTEVF | 0.8700688 |
| TALGHLVP | ASSSGQSANTEVF | 0.59356666 |
| TAPTTSGAA | ASSSGQSANTEVF | 0.8646716 |
| TDGDDGNGGSPGT | ASSSGQSANTEVF | 0.8240964 |
| TDGLGEN | ASSSGQSANTEVF | 0.8868344 |
| TGGPPLPGQA | ASSSGQSANTEVF | 0.8138244 |
| TIPAITI | ASSSGQSANTEVF | 0.85605776 |
| TIPGFTI | ASSSGQSANTEVF | 0.83865446 |
| TSIASGALA | ASSSGQSANTEVF | 0.86007017 |
| TVTLPGAAI | ASSSGQSANTEVF | 0.63307846 |
| VASGLSAALP | ASSSGQSANTEVF | 0.8168434 |
| VGAGVIGI | ASSSGQSANTEVF | 0.72870266 |
| VGPKRFGV | ASSSGQSANTEVF | 0.8344865 |
| VNVAFPD | ASSSGQSANTEVF | 0.85565734 |
| VSMTNSGVS | ASSSGQSANTEVF | 0.4290874 |
| VTDVLGDLSV | ASSSGQSANTEVF | 0.86050063 |
| VVVVGAGFAGL | ASSSGQSANTEVF | 0.77896965 |
| VYTVGTVV | ASSSGQSANTEVF | 0.7647516 |
| WTTFSGP | ASSSGQSANTEVF | 0.8729643 |
| YDELVEAF | ASSSGQSANTEVF | 0.82774687 |
| YGTGGAGGNGG | ASSSGQSANTEVF | 0.8719442 |
| YTNVMAV | ASSSGQSANTEVF | 0.823015 |
| FLEQVHQL | ASSSGQSANTEVF | 0.8196659 |
| GLSLLAVAV | ASSSGQSANTEVF | 0.8347033 |
| LEQVHQL | ASSSGQSANTEVF | 0.6081101 |
| LNAAGAI | ASSSGQSANTEVF | 0.7195597 |
| PAVHASL | ASSSGQSANTEVF | 0.8783474 |
| TELEDDA | ASSSGQSANTEVF | 0.7345206 |
| TPPGPPN | ASSSGQSANTEVF | 0.8510379 |
| TPPPPPPP | ASSSGQSANTEVF | 0.54099166 |
| TQSPALI | GGRDSSGNTLY | 0.4662999 |
| EFAAAFPQ | GGRDSSGNTLY | 0.69473225 |
| LLSGNLDVL | GGRDSSGNTLY | 0.7997631 |

| | | |
|---|---|---|
| GPANLGNYNI | GGRDSSGNTLY | 0.8502567 |
| TLGPTLP | GGRDSSGNTLY | 0.773456 |
| VYAYSSCCK | GGRDSSGNTLY | 0.4269557 |
| APAPAPAPNI | GGRDSSGNTLY | 0.6896876 |
| PGGPVPIP | GGRDSSGNTLY | 0.5312536 |
| FLTVALEP | GGRDSSGNTLY | 0.66007864 |
| SQYFGANL | GGRDSSGNTLY | 0.78102016 |
| GGVKPAA | GGRDSSGNTLY | 0.5832628 |
| GGTGGAGGAGGW | GGRDSSGNTLY | 0.61298275 |
| GVGGVAGLGGAGG | GGRDSSGNTLY | 0.56591815 |
| VLLVLATTI | GGRDSSGNTLY | 0.8133169 |
| IGIFAAML | GGRDSSGNTLY | 0.81061274 |
| AVGDLID | GGRDSSGNTLY | 0.7704153 |
| NISPLPL | GGRDSSGNTLY | 0.8736528 |
| VTAVFVAI | GGRDSSGNTLY | 0.79344845 |
| SGDDNDAAL | GGRDSSGNTLY | 0.8325628 |
| AVAALAHS | GGRDSSGNTLY | 0.17572784 |
| GGVTVMATP | GGRDSSGNTLY | 0.31751162 |
| AGLAANVRD | GGRDSSGNTLY | 0.60381305 |
| TNRPDLIDPALLRPG | GGRDSSGNTLY | 0.62452286 |
| LANLGVTAL | GGRDSSGNTLY | 0.33447623 |
| TGERIEL | GGRDSSGNTLY | 0.79158247 |
| NGEELAA | GGRDSSGNTLY | 0.6212628 |
| VLSLVWG | GGRDSSGNTLY | 0.42000538 |
| IGYALWL | GGRDSSGNTLY | 0.8208426 |
| LSLLLAPL | GGRDSSGNTLY | 0.6219301 |
| TLDEALK | GGRDSSGNTLY | 0.5500271 |
| TDGAGTAEGG | GGRDSSGNTLY | 0.6010778 |
| NAPGGAGGNGG | GGRDSSGNTLY | 0.8087754 |
| VSVFMGMGGNF | GGRDSSGNTLY | 0.38386166 |
| LASYLAAL | GGRDSSGNTLY | 0.7292118 |
| LVGGTPAL | GGRDSSGNTLY | 0.6408323 |
| VRNADVL | GGRDSSGNTLY | 0.7953379 |
| TCSSPFLVSAM | GGRDSSGNTLY | 0.30938116 |
| VHITDAE | GGRDSSGNTLY | 0.7679592 |
| EYAAPSF | GGRDSSGNTLY | 0.6199069 |
| VVDAFHGGL | GGRDSSGNTLY | 0.8893311 |
| ASVVDPQ | GGRDSSGNTLY | 0.5603938 |
| SYNEFGGLS | GGRDSSGNTLY | 0.6114676 |
| DPDGSVY | GGRDSSGNTLY | 0.632564 |
| MVSTIHATE | GGRDSSGNTLY | 0.8295618 |
| LAAGVADPV | GGRDSSGNTLY | 0.28852487 |
| TKEQIAA | GGRDSSGNTLY | 0.63345885 |
| SLASPAAL | GGRDSSGNTLY | 0.4152615 |
| IGLVFYGV | GGRDSSGNTLY | 0.73295987 |
| FLLDASNL | GGRDSSGNTLY | 0.7315481 |
| YQVGNFP | GGRDSSGNTLY | 0.37857017 |
| ESLLSQST | GGRDSSGNTLY | 0.7755904 |
| LFTVGFAAVG | GGRDSSGNTLY | 0.61362267 |
| ATPPTPT | GGRDSSGNTLY | 0.6049717 |
| TDLATLG | GGRDSSGNTLY | 0.8396074 |

| | | |
|---|---|---|
| SAGGTGR | GGRDSSGNTLY | 0.6440314 |
| GFSNSGS | GGRDSSGNTLY | 0.861371 |
| GGNGGSGGTSV | GGRDSSGNTLY | 0.7300496 |
| GTSVATGGAGN | GGRDSSGNTLY | 0.640712 |
| NGGNGGSGGTS | GGRDSSGNTLY | 0.7921032 |
| AVVKFPH | GGRDSSGNTLY | 0.5325293 |
| TPPPPPP | GGRDSSGNTLY | 0.61424613 |
| AAPAAAAAPA | GGRDSSGNTLY | 0.8052368 |
| IALNEDL | GGRDSSGNTLY | 0.80836105 |
| IVNVSSM | GGRDSSGNTLY | 0.81785 |
| TGIPPAPR | GGRDSSGNTLY | 0.7537738 |
| FDLGGGTF | GGRDSSGNTLY | 0.74773914 |
| FDLGGGTFDVS | GGRDSSGNTLY | 0.7762395 |
| NEPTAAA | GGRDSSGNTLY | 0.81944776 |
| NEPTAAALA | GGRDSSGNTLY | 0.818506 |
| GSIINISSI | GGRDSSGNTLY | 0.81127733 |
| VAPGNAGTA | GGRDSSGNTLY | 0.7556443 |
| TGGGASLE | GGRDSSGNTLY | 0.80359364 |
| VAVYNHYKRI | GGRDSSGNTLY | 0.7943574 |
| TDPAPATT | GGRDSSGNTLY | 0.7816359 |
| TNRPDILDPALLRPG | GGRDSSGNTLY | 0.81140393 |
| LTTTTTS | GGRDSSGNTLY | 0.8056673 |
| PTGSGKTL | GGRDSSGNTLY | 0.75950426 |
| TGSGKTL | GGRDSSGNTLY | 0.7917903 |
| TIGHVDHGK | GGRDSSGNTLY | 0.8013803 |
| TIGHVDHGKTT | GGRDSSGNTLY | 0.80832756 |
| GHVDHGKTT | GGRDSSGNTLY | 0.8077114 |
| AQFEHTLL | GGRDSSGNTLY | 0.82694924 |
| IVNQDEL | GGRDSSGNTLY | 0.81693625 |
| TNGSQFF | GGRDSSGNTLY | 0.80980945 |
| TGKGIVE | GGRDSSGNTLY | 0.46248794 |
| LISNASDA | GGRDSSGNTLY | 0.6758868 |
| AELRTLGELTKIAKAHGAQ | GGRDSSGNTLY | 0.7843691 |
| APNAVAAAEPSW | GGRDSSGNTLY | 0.7875279 |
| APQVAGAATAVL | GGRDSSGNTLY | 0.5744494 |
| ARRDPAAAAAILPTDA | GGRDSSGNTLY | 0.810233 |
| DAAARDVLAEAGLAEYF | GGRDSSGNTLY | 0.7690987 |
| DDQIVLSAKANLAKVEQ | GGRDSSGNTLY | 0.7593578 |
| DELLPDTGEDSDEDGH | GGRDSSGNTLY | 0.8150569 |
| ELPEAPANRATVPQD | GGRDSSGNTLY | 0.78574 |
| FGQSKIERVTGIG | GGRDSSGNTLY | 0.77843255 |
| GAGGTGSTAGGAGGAGGAGGI | GGRDSSGNTLY | 0.78988487 |
| GHGGTGAAGGNGGTGGAGGS( | GGRDSSGNTLY | 0.8166915 |
| GNGGNGGAGGEGGPGGAGGA( | GGRDSSGNTLY | 0.74158424 |
| GSTAGNGGNGGNATNGTIAG( | GGRDSSGNTLY | 0.8163246 |
| GTGGDGGNGGNGGTGG | GGRDSSGNTLY | 0.7376387 |
| GVRRPAGIPADAPV | GGRDSSGNTLY | 0.82310075 |
| GYRQVTDAHLLTLARR | GGRDSSGNTLY | 0.72918236 |
| IAKMGQDGHDRG | GGRDSSGNTLY | 0.81487054 |
| KAEPSIEIQYT | GGRDSSGNTLY | 0.79489744 |
| KPVRIRMAADEDTVAVT | GGRDSSGNTLY | 0.8304879 |

| | | |
|---|---|---|
| MPPRIAGMRLLVIKPEP | GGRDSSGNTLY | 0.789098 |
| MKFYAVALFRSRGVPPTR | GGRDSSGNTLY | 0.81497586 |
| NAGGFGILGAVAHSP | GGRDSSGNTLY | 0.78567827 |
| NEGREITLAMAEM | GGRDSSGNTLY | 0.8196875 |
| NELVEWSHPLGHRWRWE | GGRDSSGNTLY | 0.71608305 |
| NLGAGGTGGNGGNPG | GGRDSSGNTLY | 0.7943652 |
| NTEWPKWETVMAADPVD | GGRDSSGNTLY | 0.6435678 |
| PASAAPAETQP | GGRDSSGNTLY | 0.8081175 |
| QQSSDGKNGVRAAEITGE | GGRDSSGNTLY | 0.8107865 |
| RAPDAVTTSWWKE | GGRDSSGNTLY | 0.57254696 |
| SGFGNVGAHTSGWFNQ | GGRDSSGNTLY | 0.8199941 |
| SGVAGAVGQAASVGGLKV | GGRDSSGNTLY | 0.7569953 |
| SNGGAGTGSAGGNGGT | GGRDSSGNTLY | 0.77291274 |
| TGGAGGEGGAAG | GGRDSSGNTLY | 0.8208202 |
| TGGDGGAAGAGGAPGHG | GGRDSSGNTLY | 0.7844745 |
| TMPPLLAAIANAGRHGV | GGRDSSGNTLY | 0.81416166 |
| TSSTAGNSGGA | GGRDSSGNTLY | 0.77324754 |
| VPATVPVGILC | GGRDSSGNTLY | 0.7809706 |
| VQRPDSGGVVLTGRIS | GGRDSSGNTLY | 0.80249554 |
| WTPNPVPATKGVPAGLVQAT | GGRDSSGNTLY | 0.66557956 |
| YALKRALAKFSDRFDVVII | GGRDSSGNTLY | 0.76323366 |
| YLIGSVIV | GGRDSSGNTLY | 0.82472134 |
| YGQLVAVID | GGRDSSGNTLY | 0.63859373 |
| VTVGGMTLPA | GGRDSSGNTLY | 0.7066731 |
| VLALPAV | GGRDSSGNTLY | 0.53934824 |
| VIPPTLNYET | GGRDSSGNTLY | 0.51135314 |
| VGMGGMHPSSG | GGRDSSGNTLY | 0.5369369 |
| VGLAGGAATA | GGRDSSGNTLY | 0.6994611 |
| VGGAGGAGGSGP | GGRDSSGNTLY | 0.6882129 |
| VFAAFMV | GGRDSSGNTLY | 0.6405535 |
| TLYDKLR | GGRDSSGNTLY | 0.54893225 |
| TLAVFTVL | GGRDSSGNTLY | 0.62103623 |
| TGSVQPARV | GGRDSSGNTLY | 0.5919423 |
| TGGGAGSGGT | GGRDSSGNTLY | 0.540781 |
| SHLETSL | GGRDSSGNTLY | 0.6419468 |
| SFTTGMT | GGRDSSGNTLY | 0.66011745 |
| QVEVAKRMGVRQPTVS | GGRDSSGNTLY | 0.5675555 |
| QGSGGPALAM | GGRDSSGNTLY | 0.5846289 |
| MLAPSNP | GGRDSSGNTLY | 0.58386725 |
| LVNNAAI | GGRDSSGNTLY | 0.66563904 |
| LTGEINTLA | GGRDSSGNTLY | 0.63416475 |
| LSIFGPV | GGRDSSGNTLY | 0.73027396 |
| LLALLANL | GGRDSSGNTLY | 0.67424786 |
| LISNASDALDKLR | GGRDSSGNTLY | 0.65567034 |
| LINEAVP | GGRDSSGNTLY | 0.30877885 |
| LGYAFSGL | GGRDSSGNTLY | 0.4141206 |
| LDFVAVAL | GGRDSSGNTLY | 0.6587751 |
| LAQPGAE | GGRDSSGNTLY | 0.57634544 |
| ISNLGAVTGF | GGRDSSGNTLY | 0.37913546 |
| HQEDPGRI | GGRDSSGNTLY | 0.6815744 |
| GTPGEPTP | GGRDSSGNTLY | 0.5951924 |

| | | |
|---|---|---|
| GFFNSGS | GGRDSSGNTLY | 0.56959337 |
| GAGGGGLTAGAN | GGRDSSGNTLY | 0.6535737 |
| FVDALVNM | GGRDSSGNTLY | 0.5660152 |
| FLTVAPEL | GGRDSSGNTLY | 0.5709308 |
| FETLTAL | GGRDSSGNTLY | 0.57901245 |
| DVINPIEL | GGRDSSGNTLY | 0.7008119 |
| APVSGAPV | GGRDSSGNTLY | 0.4817023 |
| ALVGGLAW | GGRDSSGNTLY | 0.6454488 |
| ALAAAAATAI | GGRDSSGNTLY | 0.5545499 |
| AFGLGLLL | GGRDSSGNTLY | 0.7055867 |
| AFFAAASL | GGRDSSGNTLY | 0.48547798 |
| AAAALPAFSPP | GGRDSSGNTLY | 0.64134467 |
| VVLAPAL | GGRDSSGNTLY | 0.5592492 |
| TIGHVDHGKTTL | GGRDSSGNTLY | 0.74263686 |
| LISNASDALDK | GGRDSSGNTLY | 0.58738995 |
| LISNASDAL | GGRDSSGNTLY | 0.6537807 |
| ISNASDALDK | GGRDSSGNTLY | 0.59330255 |
| NQAETLVYQTE | GGRDSSGNTLY | 0.6563734 |
| TGGTGGTAGAG | GGRDSSGNTLY | 0.4501627 |
| TGSTAGGAGGA | GGRDSSGNTLY | 0.57533824 |
| DPLAVPVAAGPV | GGRDSSGNTLY | 0.4700703 |
| GGTGGTPGNGGN | GGRDSSGNTLY | 0.6368904 |
| PAGLAGVDPATL | GGRDSSGNTLY | 0.63820606 |
| GNNLVVEAPHAGASV | GGRDSSGNTLY | 0.52560747 |
| KVTRIMSASTLEQPA | GGRDSSGNTLY | 0.663756 |
| LGFALPIALPRRTVV | GGRDSSGNTLY | 0.63317525 |
| TRLGNLTARKLVVLV | GGRDSSGNTLY | 0.5643101 |
| WSQQRRLIYFPSAGP | GGRDSSGNTLY | 0.7045861 |
| GPNGNPDPMAAAVDIR | GGRDSSGNTLY | 0.6112889 |
| TQRCVSVADLRAAASR | GGRDSSGNTLY | 0.6759122 |
| QFVLDVAEHPVEVLTETV | GGRDSSGNTLY | 0.5692768 |
| GGAGGTGGAGGNGGTQAGDG | GGRDSSGNTLY | 0.5564949 |
| PLLATLRAIAVDVDAKPAQV | GGRDSSGNTLY | 0.6841655 |
| APKFRTGGYMIDFWGVGYQV | GGRDSSGNTLY | 0.59401524 |
| GAGGAGGAGGAAGGPGGTGG | GGRDSSGNTLY | 0.5691834 |
| IGALGLLATDVAPAGSVAFG | GGRDSSGNTLY | 0.6658397 |
| AGGAGGIGGNGGAGGAGGVL | GGRDSSGNTLY | 0.5968363 |
| ADELIGLL | GGRDSSGNTLY | 0.8670577 |
| ADRALKA | GGRDSSGNTLY | 0.48196697 |
| AGAVGQVGGAGGA | GGRDSSGNTLY | 0.49566874 |
| AGGGGGLGGAP | GGRDSSGNTLY | 0.39343694 |
| AGGLLYGNGGAG | GGRDSSGNTLY | 0.7034321 |
| AGLESIGL | GGRDSSGNTLY | 0.8609435 |
| AGLKTGII | GGRDSSGNTLY | 0.7753258 |
| ASLAALLL | GGRDSSGNTLY | 0.4060226 |
| ASVINISSI | GGRDSSGNTLY | 0.6320378 |
| ATPPGPA | GGRDSSGNTLY | 0.4943774 |
| ATVASALV | GGRDSSGNTLY | 0.49883163 |
| ATVASLIL | GGRDSSGNTLY | 0.59702015 |
| AVNGPVLIRH | GGRDSSGNTLY | 0.5834893 |
| DGAAGAVGI | GGRDSSGNTLY | 0.4065319 |

| | | |
|---|---|---|
| DPLLHGI | GGRDSSGNTLY | 0.31464317 |
| ELGAALGAL | GGRDSSGNTLY | 0.3875007 |
| ELGRTQAV | GGRDSSGNTLY | 0.6191589 |
| FVAGGYLS | GGRDSSGNTLY | 0.5302321 |
| GASGGAGGAGGW | GGRDSSGNTLY | 0.76767254 |
| GGAGAGGDGGAGGI | GGRDSSGNTLY | 0.71651465 |
| GTGHNGSQPGMGGNG | GGRDSSGNTLY | 0.41026348 |
| GYAAASASAS | GGRDSSGNTLY | 0.11351741 |
| HQVDVAAVQQG | GGRDSSGNTLY | 0.23013648 |
| IGAAIAHT | GGRDSSGNTLY | 0.42349184 |
| IGMIEVAQL | GGRDSSGNTLY | 0.59853053 |
| IRLAPEHTSAMD | GGRDSSGNTLY | 0.10873074 |
| IYNVSVL | GGRDSSGNTLY | 0.57885665 |
| KQGFLGM | GGRDSSGNTLY | 0.5250913 |
| LESVITQL | GGRDSSGNTLY | 0.58310026 |
| LGLGALVVV | GGRDSSGNTLY | 0.7609468 |
| LIGNGGNGGSGGI | GGRDSSGNTLY | 0.55110186 |
| LNAINAPTL | GGRDSSGNTLY | 0.3418971 |
| LQLLNTGV | GGRDSSGNTLY | 0.3001687 |
| LQTSIEF | GGRDSSGNTLY | 0.77901757 |
| MSFAVSEL | GGRDSSGNTLY | 0.18834415 |
| NTGALGL | GGRDSSGNTLY | 0.7170092 |
| PPGAGGAGGAGTT | GGRDSSGNTLY | 0.27006966 |
| QLVGLGI | GGRDSSGNTLY | 0.67446065 |
| RFLEQVHQL | GGRDSSGNTLY | 0.34155852 |
| RGVGYVLR | GGRDSSGNTLY | 0.715095 |
| SATGVFAGVI | GGRDSSGNTLY | 0.6224187 |
| SLPAITLP | GGRDSSGNTLY | 0.8481328 |
| STALAFTV | GGRDSSGNTLY | 0.5055952 |
| SVLTVGLGK | GGRDSSGNTLY | 0.65869415 |
| TALAGAGSGI | GGRDSSGNTLY | 0.2642865 |
| TGLPFTGI | GGRDSSGNTLY | 0.8482654 |
| VAEHVTV | GGRDSSGNTLY | 0.40684468 |
| VDIVLNSL | GGRDSSGNTLY | 0.4145398 |
| VLTGGGAVL | GGRDSSGNTLY | 0.2130522 |
| VNVGLAYGI | GGRDSSGNTLY | 0.5357413 |
| VSSAAVGAV | GGRDSSGNTLY | 0.74895835 |
| VTSVYEAGV | GGRDSSGNTLY | 0.49142253 |
| WVALDRGA | GGRDSSGNTLY | 0.7548742 |
| YLVGSVVI | GGRDSSGNTLY | 0.6987387 |
| GPPGTGKTLL | GGRDSSGNTLY | 0.3934164 |
| GVGMTKF | GGRDSSGNTLY | 0.7174196 |
| IGIGGSDL | GGRDSSGNTLY | 0.43710038 |
| IGTLLAEL | GGRDSSGNTLY | 0.58049905 |
| IYTSGTT | GGRDSSGNTLY | 0.24108396 |
| LSFPAGV | GGRDSSGNTLY | 0.5974168 |
| LYTSGTT | GGRDSSGNTLY | 0.6165242 |
| VGAFPFGF | GGRDSSGNTLY | 0.4259809 |
| VGAGTVEF | GGRDSSGNTLY | 0.3124134 |
| VGKSTLLL | GGRDSSGNTLY | 0.77310944 |
| AALAHAGGGIV | GGRDSSGNTLY | 0.5002352 |

| | | |
|---|---|---|
| AALVAVSLL | GGRDSSGNTLY | 0.5939226 |
| AAVSRAVAAL | GGRDSSGNTLY | 0.3550333 |
| AGGSSGIGGF | GGRDSSGNTLY | 0.31877512 |
| AGLTVANANG | GGRDSSGNTLY | 0.39286754 |
| AGLVGAVL | GGRDSSGNTLY | 0.52021885 |
| ALSQAFTV | GGRDSSGNTLY | 0.6485386 |
| ALTFSVSL | GGRDSSGNTLY | 0.5150421 |
| ALTGAALVAL | GGRDSSGNTLY | 0.3415407 |
| ASFSIGAGVL | GGRDSSGNTLY | 0.5443119 |
| DGSVYKI | GGRDSSGNTLY | 0.4184314 |
| EQLAEIV | GGRDSSGNTLY | 0.44310513 |
| FAVSELTVM | GGRDSSGNTLY | 0.40766487 |
| GAAPLAAGL | GGRDSSGNTLY | 0.4277507 |
| GADGGDGGSGGAGG | GGRDSSGNTLY | 0.3852361 |
| GFGLDTGGAGG | GGRDSSGNTLY | 0.325133 |
| GLASGGLL | GGRDSSGNTLY | 0.48473784 |
| GRIAGAAVLL | GGRDSSGNTLY | 0.6017959 |
| GSAGSLNTSF | GGRDSSGNTLY | 0.54342544 |
| GTGQIGI | GGRDSSGNTLY | 0.59792936 |
| GTLIALIL | GGRDSSGNTLY | 0.59371805 |
| IAATMGFVVG | GGRDSSGNTLY | 0.49719983 |
| IFAQALL | GGRDSSGNTLY | 0.25930157 |
| IGAAVALGF | GGRDSSGNTLY | 0.30266896 |
| IGLTPLAL | GGRDSSGNTLY | 0.34557033 |
| ILGSNTSTL | GGRDSSGNTLY | 0.42041823 |
| IVFSLTVL | GGRDSSGNTLY | 0.25800055 |
| LASKATGF | GGRDSSGNTLY | 0.43752187 |
| LASVALIL | GGRDSSGNTLY | 0.36805683 |
| LFAVGLTL | GGRDSSGNTLY | 0.36020577 |
| LFNGGIGTYI | GGRDSSGNTLY | 0.5989622 |
| LTLGAGII | GGRDSSGNTLY | 0.38142097 |
| LVAAALGDI | GGRDSSGNTLY | 0.33103767 |
| NTAFVQLGI | GGRDSSGNTLY | 0.36442646 |
| QLFSGAEL | GGRDSSGNTLY | 0.6389533 |
| SAATFITL | GGRDSSGNTLY | 0.3167229 |
| SNYGLANLGV | GGRDSSGNTLY | 0.57134247 |
| SYGGAYAVM | GGRDSSGNTLY | 0.3890827 |
| TTGVGGLTLG | GGRDSSGNTLY | 0.6025926 |
| TTQTIYI | GGRDSSGNTLY | 0.3008021 |
| VAAGAGLGAVI | GGRDSSGNTLY | 0.5546494 |
| VAAVVTLL | GGRDSSGNTLY | 0.4456821 |
| VGFLGGLI | GGRDSSGNTLY | 0.6681363 |
| VGVIPILL | GGRDSSGNTLY | 0.3699721 |
| VIFASSLI | GGRDSSGNTLY | 0.48337498 |
| VKGGDGGAAAG | GGRDSSGNTLY | 0.391968 |
| VTKTFTNI | GGRDSSGNTLY | 0.61732405 |
| VVAAAILL | GGRDSSGNTLY | 0.29558325 |
| YAALDVEL | GGRDSSGNTLY | 0.36988795 |
| LASVALLL | GGRDSSGNTLY | 0.353422 |
| LDAVHRV | GGRDSSGNTLY | 0.43595755 |
| QGLSSSIF | GGRDSSGNTLY | 0.52820396 |

| | | |
|---|---|---|
| RGTFANIRL | GGRDSSGNTLY | 0.25175565 |
| RQLWWGHRI | GGRDSSGNTLY | 0.52420324 |
| YGPPGTGKTLL | GGRDSSGNTLY | 0.30342746 |
| AALVGAPANVA | GGRDSSGNTLY | 0.30668807 |
| AATFRRTE | GGRDSSGNTLY | 0.46598205 |
| AGALGGGI | GGRDSSGNTLY | 0.4129975 |
| AGGAGGAGGIGAG | GGRDSSGNTLY | 0.52104646 |
| AGSGALGL | GGRDSSGNTLY | 0.3433593 |
| AQIADQI | GGRDSSGNTLY | 0.49637604 |
| ASAVAVTYL | GGRDSSGNTLY | 0.47992745 |
| ASTVLGGTGG | GGRDSSGNTLY | 0.2772623 |
| AVGGEPAK | GGRDSSGNTLY | 0.32447398 |
| CEQGTFGGTC | GGRDSSGNTLY | 0.6005623 |
| DEGKAPAG | GGRDSSGNTLY | 0.7687031 |
| DTVVFPAL | GGRDSSGNTLY | 0.86254275 |
| FEGITMAG | GGRDSSGNTLY | 0.7138238 |
| FPLVVQD | GGRDSSGNTLY | 0.5148189 |
| GAFTTPQ | GGRDSSGNTLY | 0.7290443 |
| GFGGFGTAGGVGGA | GGRDSSGNTLY | 0.8569086 |
| GFLAQFL | GGRDSSGNTLY | 0.8586076 |
| GGAGGAGGNAGVA | GGRDSSGNTLY | 0.826719 |
| GGVGTAGAGGAGGAGG | GGRDSSGNTLY | 0.4686929 |
| GNGGGGDAGNAGSGGN | GGRDSSGNTLY | 0.7269993 |
| GTGGAGGAGNDAGS | GGRDSSGNTLY | 0.6215966 |
| GTTNTGWFNS | GGRDSSGNTLY | 0.75122327 |
| IFMGPLFTL | GGRDSSGNTLY | 0.8254081 |
| ILAHAIEAI | GGRDSSGNTLY | 0.7270398 |
| IRSAIREAL | GGRDSSGNTLY | 0.72269344 |
| ITFAGGG | GGRDSSGNTLY | 0.6290529 |
| ITGMSGHQEV | GGRDSSGNTLY | 0.7947868 |
| IVAGLACTL | GGRDSSGNTLY | 0.8125603 |
| IVGGGLAGA | GGRDSSGNTLY | 0.7774875 |
| IVTGVLNI | GGRDSSGNTLY | 0.88592005 |
| LAQQGCI | GGRDSSGNTLY | 0.8502103 |
| LCLHVEN | GGRDSSGNTLY | 0.8440911 |
| LDAISDGS | GGRDSSGNTLY | 0.14480293 |
| LGGFAAML | GGRDSSGNTLY | 0.31923667 |
| LGGKNPMI | GGRDSSGNTLY | 0.66639054 |
| LGPKGVAAIL | GGRDSSGNTLY | 0.6815178 |
| LMVGVSI | GGRDSSGNTLY | 0.30582726 |
| LVGAAFVL | GGRDSSGNTLY | 0.7750217 |
| LVLAVAPL | GGRDSSGNTLY | 0.61787295 |
| MAAIAVIDAA | GGRDSSGNTLY | 0.55533105 |
| NNGINAELQ | GGRDSSGNTLY | 0.86364293 |
| RGEPLVLVHG | GGRDSSGNTLY | 0.7023604 |
| RGGAGGAAGLI | GGRDSSGNTLY | 0.5864179 |
| RSYRMHPAV | GGRDSSGNTLY | 0.65116596 |
| SAAHLLAGQLA | GGRDSSGNTLY | 0.86713284 |
| SAAVGSY | GGRDSSGNTLY | 0.4324367 |
| SPHAVLHRI | GGRDSSGNTLY | 0.810211 |
| SQIAKVL | GGRDSSGNTLY | 0.69167215 |

| | | |
|---|---|---|
| TALGHLVP | GGRDSSGNTLY | 0.85148954 |
| TAPTTSGAA | GGRDSSGNTLY | 0.34955847 |
| TDGDDGNGGSPGT | GGRDSSGNTLY | 0.8422146 |
| TDGLGEN | GGRDSSGNTLY | 0.765625 |
| TGGPPLPGQA | GGRDSSGNTLY | 0.9041168 |
| TIPAITI | GGRDSSGNTLY | 0.6248155 |
| TIPGFTI | GGRDSSGNTLY | 0.686668 |
| TSIASGALA | GGRDSSGNTLY | 0.51515526 |
| TVTLPGAAI | GGRDSSGNTLY | 0.85652274 |
| VASGLSAALP | GGRDSSGNTLY | 0.37532556 |
| VGAGVIGI | GGRDSSGNTLY | 0.66839075 |
| VGPKRFGV | GGRDSSGNTLY | 0.434385 |
| VNVAFPD | GGRDSSGNTLY | 0.79079676 |
| VSMTNSGVS | GGRDSSGNTLY | 0.81702745 |
| VTDVLGDLSV | GGRDSSGNTLY | 0.32311347 |
| VVVVGAGFAGL | GGRDSSGNTLY | 0.863858 |
| VYTVGTVV | GGRDSSGNTLY | 0.7388717 |
| WTTFSGP | GGRDSSGNTLY | 0.5252397 |
| YDELVEAF | GGRDSSGNTLY | 0.87497926 |
| YGTGGAGGNGG | GGRDSSGNTLY | 0.5413351 |
| YTNVMAV | GGRDSSGNTLY | 0.8634163 |
| FLEQVHQL | GGRDSSGNTLY | 0.75603366 |
| GLSLLAVAV | GGRDSSGNTLY | 0.72623837 |
| LEQVHQL | GGRDSSGNTLY | 0.8339721 |
| LNAAGAI | GGRDSSGNTLY | 0.391879 |
| PAVHASL | GGRDSSGNTLY | 0.5673782 |
| TELEDDA | GGRDSSGNTLY | 0.8723125 |
| TPPGPPN | GGRDSSGNTLY | 0.7509851 |
| TPPPPPPP | GGRDSSGNTLY | 0.8818009 |
| TQSPALI | ASSDGQGNNQAPL | 0.6147772 |
| EFAAAFPQ | ASSDGQGNNQAPL | 0.47455564 |
| LLSGNLDVL | ASSDGQGNNQAPL | 0.7348915 |
| GPANLGNYNI | ASSDGQGNNQAPL | 0.85108507 |
| TLGPTLP | ASSDGQGNNQAPL | 0.87524986 |
| VYAYSSCCK | ASSDGQGNNQAPL | 0.81001335 |
| APAPAPAPNI | ASSDGQGNNQAPL | 0.48841318 |
| PGGPVPIP | ASSDGQGNNQAPL | 0.6263893 |
| FLTVALEP | ASSDGQGNNQAPL | 0.61619395 |
| SQYFGANL | ASSDGQGNNQAPL | 0.67804325 |
| GGVKPAA | ASSDGQGNNQAPL | 0.83371353 |
| GGTGGAGGAGGW | ASSDGQGNNQAPL | 0.59313315 |
| GVGGVAGLGGAGG | ASSDGQGNNQAPL | 0.55060923 |
| VLLVLATTI | ASSDGQGNNQAPL | 0.59996516 |
| IGIFAAML | ASSDGQGNNQAPL | 0.79815125 |
| AVGDLID | ASSDGQGNNQAPL | 0.81634057 |
| NISPLPL | ASSDGQGNNQAPL | 0.7935512 |
| VTAVFVAI | ASSDGQGNNQAPL | 0.8835287 |
| SGDDNDAAL | ASSDGQGNNQAPL | 0.8253979 |
| AVAALAHS | ASSDGQGNNQAPL | 0.837796 |
| GGVTVMATP | ASSDGQGNNQAPL | 0.15955406 |
| AGLAANVRD | ASSDGQGNNQAPL | 0.31695879 |

| | | |
|---|---|---|
| TNRPDLIDPALLRPG | ASSDGQGNNQAPL | 0.6372213 |
| LANLGVTAL | ASSDGQGNNQAPL | 0.6815464 |
| TGERIEL | ASSDGQGNNQAPL | 0.32134277 |
| NGEELAA | ASSDGQGNNQAPL | 0.7495215 |
| VLSLVWG | ASSDGQGNNQAPL | 0.56894445 |
| IGYALWL | ASSDGQGNNQAPL | 0.4295327 |
| LSLLLAPL | ASSDGQGNNQAPL | 0.84126574 |
| TLDEALK | ASSDGQGNNQAPL | 0.5485612 |
| TDGAGTAEGG | ASSDGQGNNQAPL | 0.51098853 |
| NAPGGAGGNGG | ASSDGQGNNQAPL | 0.554209 |
| VSVFMGMGGNF | ASSDGQGNNQAPL | 0.8575967 |
| LASYLAAL | ASSDGQGNNQAPL | 0.31071126 |
| LVGGTPAL | ASSDGQGNNQAPL | 0.7784552 |
| VRNADVL | ASSDGQGNNQAPL | 0.76817065 |
| TCSSPFLVSAM | ASSDGQGNNQAPL | 0.8255147 |
| VHITDAE | ASSDGQGNNQAPL | 0.32148346 |
| EYAAPSF | ASSDGQGNNQAPL | 0.81637776 |
| VVDAFHGGL | ASSDGQGNNQAPL | 0.6917279 |
| ASVVDPQ | ASSDGQGNNQAPL | 0.88649595 |
| SYNEFGGLS | ASSDGQGNNQAPL | 0.6199291 |
| DPDGSVY | ASSDGQGNNQAPL | 0.67798615 |
| MVSTIHATE | ASSDGQGNNQAPL | 0.59477 |
| LAAGVADPV | ASSDGQGNNQAPL | 0.8591861 |
| TKEQIAA | ASSDGQGNNQAPL | 0.3797533 |
| SLASPAAL | ASSDGQGNNQAPL | 0.6464299 |
| IGLVFYGV | ASSDGQGNNQAPL | 0.49350435 |
| FLLDASNL | ASSDGQGNNQAPL | 0.76993215 |
| YQVGNFP | ASSDGQGNNQAPL | 0.7792338 |
| ESLLSQST | ASSDGQGNNQAPL | 0.512184 |
| LFTVGFAAVG | ASSDGQGNNQAPL | 0.8485577 |
| ATPPTPT | ASSDGQGNNQAPL | 0.7958476 |
| TDLATLG | ASSDGQGNNQAPL | 0.56548214 |
| SAGGTGR | ASSDGQGNNQAPL | 0.86258197 |
| GFSNSGS | ASSDGQGNNQAPL | 0.52273256 |
| GGNGGSGGTSV | ASSDGQGNNQAPL | 0.83331245 |
| GTSVATGGAGN | ASSDGQGNNQAPL | 0.7316958 |
| NGGNGGSGGTS | ASSDGQGNNQAPL | 0.75770015 |
| AVVKFPH | ASSDGQGNNQAPL | 0.8333044 |
| TPPPPPP | ASSDGQGNNQAPL | 0.44231254 |
| AAPAAAAAPA | ASSDGQGNNQAPL | 0.49547687 |
| IALNEDL | ASSDGQGNNQAPL | 0.84952366 |
| IVNVSSM | ASSDGQGNNQAPL | 0.7651234 |
| TGIPPAPR | ASSDGQGNNQAPL | 0.7810062 |
| FDLGGGTF | ASSDGQGNNQAPL | 0.67768717 |
| FDLGGGTFDVS | ASSDGQGNNQAPL | 0.6021056 |
| NEPTAAA | ASSDGQGNNQAPL | 0.6653386 |
| NEPTAAALA | ASSDGQGNNQAPL | 0.7586569 |
| GSIINISSI | ASSDGQGNNQAPL | 0.7861588 |
| VAPGNAGTA | ASSDGQGNNQAPL | 0.76881754 |
| TGGGASLE | ASSDGQGNNQAPL | 0.4508772 |
| VAVYNHYKRI | ASSDGQGNNQAPL | 0.75678 |

| | | |
|---|---|---|
| TDPAPATT | ASSDGQGNNQAPL | 0.67875373 |
| TNRPDILDPALLRPG | ASSDGQGNNQAPL | 0.7020153 |
| LTTTTTS | ASSDGQGNNQAPL | 0.6970652 |
| PTGSGKTL | ASSDGQGNNQAPL | 0.67322755 |
| TGSGKTL | ASSDGQGNNQAPL | 0.72683585 |
| TIGHVDHGK | ASSDGQGNNQAPL | 0.53896725 |
| TIGHVDHGKTT | ASSDGQGNNQAPL | 0.67346394 |
| GHVDHGKTT | ASSDGQGNNQAPL | 0.77090144 |
| AQFEHTLL | ASSDGQGNNQAPL | 0.7455431 |
| IVNQDEL | ASSDGQGNNQAPL | 0.7762754 |
| TNGSQFF | ASSDGQGNNQAPL | 0.77283585 |
| TGKGIVE | ASSDGQGNNQAPL | 0.7469046 |
| LISNASDA | ASSDGQGNNQAPL | 0.22440062 |
| AELRTLGELTKIAKAHGAQ | ASSDGQGNNQAPL | 0.4556299 |
| APNAVAAAEPSW | ASSDGQGNNQAPL | 0.6403841 |
| APQVAGAATAVL | ASSDGQGNNQAPL | 0.6969607 |
| ARRDPAAAAAILPTDA | ASSDGQGNNQAPL | 0.24479088 |
| DAAARDVLAEAGLAEYF | ASSDGQGNNQAPL | 0.7049563 |
| DDQIVLSAKANLAKVEQ | ASSDGQGNNQAPL | 0.65113753 |
| DELLPDTGEDSDEDGH | ASSDGQGNNQAPL | 0.7015467 |
| ELPEAPANRATVPQD | ASSDGQGNNQAPL | 0.77214086 |
| FGQSKIERVTGIG | ASSDGQGNNQAPL | 0.7286267 |
| GAGGTGSTAGGAGGAGGAGGI | ASSDGQGNNQAPL | 0.5340988 |
| GHGGTGAAGGNGGTGGAGGS( | ASSDGQGNNQAPL | 0.5141465 |
| GNGGNGGAGGEGGPGGAGGA( | ASSDGQGNNQAPL | 0.78876287 |
| GSTAGNGGNGGNATNGTIAG( | ASSDGQGNNQAPL | 0.4106396 |
| GTGGDGGNGGNGGTGG | ASSDGQGNNQAPL | 0.7718686 |
| GVRRPAGIPADAPV | ASSDGQGNNQAPL | 0.6634919 |
| GYRQVTDAHLLTLARR | ASSDGQGNNQAPL | 0.7699286 |
| IAKMGQDGHDRG | ASSDGQGNNQAPL | 0.37230182 |
| KAEPSIEIQYT | ASSDGQGNNQAPL | 0.76564413 |
| KPVRIRMAADEDTVAVT | ASSDGQGNNQAPL | 0.7238636 |
| MPPRIAGMRLLVIKPEP | ASSDGQGNNQAPL | 0.7895576 |
| MKFYAVALFRSRGVPPTR | ASSDGQGNNQAPL | 0.63759595 |
| NAGGFGILGAVAHSP | ASSDGQGNNQAPL | 0.7160995 |
| NEGREITLAMAEM | ASSDGQGNNQAPL | 0.5794906 |
| NELVEWSHPLGHRWRWE | ASSDGQGNNQAPL | 0.78511155 |
| NLGAGGTGGNGGNPG | ASSDGQGNNQAPL | 0.538211 |
| NTEWPKWETVMAADPVD | ASSDGQGNNQAPL | 0.61643934 |
| PASAAPAETQP | ASSDGQGNNQAPL | 0.5713454 |
| QQSSDGKNGVRAAEITGE | ASSDGQGNNQAPL | 0.7049202 |
| RAPDAVTTSWWKE | ASSDGQGNNQAPL | 0.7703009 |
| SGFGNVGAHTSGWFNQ | ASSDGQGNNQAPL | 0.34938812 |
| SGVAGAVGQAASVGGLKV | ASSDGQGNNQAPL | 0.785364 |
| SNGGAGTGSAGGNGGT | ASSDGQGNNQAPL | 0.67403424 |
| TGGAGGEGGAAG | ASSDGQGNNQAPL | 0.506886 |
| TGGDGGAAGAGGAPGHG | ASSDGQGNNQAPL | 0.7459332 |
| TMPPLLAAIANAGRHGV | ASSDGQGNNQAPL | 0.64184284 |
| TSSTAGNSGGA | ASSDGQGNNQAPL | 0.7674097 |
| VPATVPVGILC | ASSDGQGNNQAPL | 0.52270067 |
| VQRPDSGGVVLTGRIS | ASSDGQGNNQAPL | 0.74981046 |

| | | |
|---|---|---|
| WTPNPVPATKGVPAGLVQAT | ASSDGQGNNQAPL | 0.71153724 |
| YALKRALAKFSDRFDVVII | ASSDGQGNNQAPL | 0.584773 |
| YLIGSVIV | ASSDGQGNNQAPL | 0.52781093 |
| YGQLVAVID | ASSDGQGNNQAPL | 0.78074133 |
| VTVGGMTLPA | ASSDGQGNNQAPL | 0.36308455 |
| VLALPAV | ASSDGQGNNQAPL | 0.7834575 |
| VIPPTLNYET | ASSDGQGNNQAPL | 0.2114245 |
| VGMGGMHPSSG | ASSDGQGNNQAPL | 0.19586726 |
| VGLAGGAATA | ASSDGQGNNQAPL | 0.36418593 |
| VGGAGGAGGSGP | ASSDGQGNNQAPL | 0.64572334 |
| VFAAFMV | ASSDGQGNNQAPL | 0.8056152 |
| TLYDKLR | ASSDGQGNNQAPL | 0.5958442 |
| TLAVFTVL | ASSDGQGNNQAPL | 0.21029794 |
| TGSVQPARV | ASSDGQGNNQAPL | 0.3123396 |
| TGGGAGSGGT | ASSDGQGNNQAPL | 0.2257632 |
| SHLETSL | ASSDGQGNNQAPL | 0.32335627 |
| SFTTGMT | ASSDGQGNNQAPL | 0.53315485 |
| QVEVAKRMGVRQPTVS | ASSDGQGNNQAPL | 0.4011984 |
| QGSGGPALAM | ASSDGQGNNQAPL | 0.27363133 |
| MLAPSNP | ASSDGQGNNQAPL | 0.28909114 |
| LVNNAAI | ASSDGQGNNQAPL | 0.5781073 |
| LTGEINTLA | ASSDGQGNNQAPL | 0.57574165 |
| LSIFGPV | ASSDGQGNNQAPL | 0.47814885 |
| LLALLANL | ASSDGQGNNQAPL | 0.76933086 |
| LISNASDALDKLR | ASSDGQGNNQAPL | 0.489067 |
| LINEAVP | ASSDGQGNNQAPL | 0.64721596 |
| LGYAFSGL | ASSDGQGNNQAPL | 0.11039659 |
| LDFVAVAL | ASSDGQGNNQAPL | 0.1485112 |
| LAQPGAE | ASSDGQGNNQAPL | 0.34687567 |
| ISNLGAVTGF | ASSDGQGNNQAPL | 0.3252458 |
| HQEDPGRI | ASSDGQGNNQAPL | 0.17509143 |
| GTPGEPTP | ASSDGQGNNQAPL | 0.57483715 |
| GFFNSGS | ASSDGQGNNQAPL | 0.24410126 |
| GAGGGGLTAGAN | ASSDGQGNNQAPL | 0.20184195 |
| FVDALVNM | ASSDGQGNNQAPL | 0.65436184 |
| FLTVAPEL | ASSDGQGNNQAPL | 0.25789735 |
| FETLTAL | ASSDGQGNNQAPL | 0.25302762 |
| DVINPIEL | ASSDGQGNNQAPL | 0.30901837 |
| APVSGAPV | ASSDGQGNNQAPL | 0.4105459 |
| ALVGGLAW | ASSDGQGNNQAPL | 0.18381815 |
| ALAAAAATAI | ASSDGQGNNQAPL | 0.47061613 |
| AFGLGLLL | ASSDGQGNNQAPL | 0.40516296 |
| AFFAAASL | ASSDGQGNNQAPL | 0.51498806 |
| AAAALPAFSPP | ASSDGQGNNQAPL | 0.18110593 |
| VVLAPAL | ASSDGQGNNQAPL | 0.54377323 |
| TIGHVDHGKTTL | ASSDGQGNNQAPL | 0.34033725 |
| LISNASDALDK | ASSDGQGNNQAPL | 0.8219107 |
| LISNASDAL | ASSDGQGNNQAPL | 0.24532814 |
| ISNASDALDK | ASSDGQGNNQAPL | 0.32686958 |
| NQAETLVYQTE | ASSDGQGNNQAPL | 0.3908757 |
| TGGTGGTAGAG | ASSDGQGNNQAPL | 0.76705235 |

| | | |
|---|---|---|
| TGSTAGGAGGA | ASSDGQGNNQAPL | 0.14768983 |
| DPLAVPVAAGPV | ASSDGQGNNQAPL | 0.21913671 |
| GGTGGTPGNGGN | ASSDGQGNNQAPL | 0.15477154 |
| PAGLAGVDPATL | ASSDGQGNNQAPL | 0.36111048 |
| GNNLVVEAPHAGASV | ASSDGQGNNQAPL | 0.4929309 |
| KVTRIMSASTLEQPA | ASSDGQGNNQAPL | 0.15152258 |
| LGFALPIALPRRTVV | ASSDGQGNNQAPL | 0.42935604 |
| TRLGNLTARKLVVLV | ASSDGQGNNQAPL | 0.41177407 |
| WSQQRRLIYFPSAGP | ASSDGQGNNQAPL | 0.27359602 |
| GPNGNPDPMAAAVDIR | ASSDGQGNNQAPL | 0.69278735 |
| TQRCVSVADLRAAASR | ASSDGQGNNQAPL | 0.30960307 |
| QFVLDVAEHPVEVLTETV | ASSDGQGNNQAPL | 0.66008174 |
| GGAGGTGGAGGNGGTQAGDG | ASSDGQGNNQAPL | 0.50056213 |
| PLLATLRAIAVDVDAKPAQV | ASSDGQGNNQAPL | 0.22706398 |
| APKFRTGGYMIDFWGVGYQV | ASSDGQGNNQAPL | 0.43996587 |
| GAGGAGGAGGAAGGPGGTGG | ASSDGQGNNQAPL | 0.23106316 |
| IGALGLLATDVAPAGSVAFG | ASSDGQGNNQAPL | 0.2906385 |
| AGGAGGIGGNGGAGGAGGVL | ASSDGQGNNQAPL | 0.64259905 |
| ADELIGLL | ASSDGQGNNQAPL | 0.7959014 |
| ADRALKA | ASSDGQGNNQAPL | 0.86511314 |
| AGAVGQVGGAGGA | ASSDGQGNNQAPL | 0.69282097 |
| AGGGGGLGGAP | ASSDGQGNNQAPL | 0.58977926 |
| AGGLLYGNGGAG | ASSDGQGNNQAPL | 0.73762524 |
| AGLESIGL | ASSDGQGNNQAPL | 0.85208803 |
| AGLKTGII | ASSDGQGNNQAPL | 0.8599804 |
| ASLAALLL | ASSDGQGNNQAPL | 0.8165174 |
| ASVINISSI | ASSDGQGNNQAPL | 0.61443007 |
| ATPPGPA | ASSDGQGNNQAPL | 0.7698362 |
| ATVASALV | ASSDGQGNNQAPL | 0.6990297 |
| ATVASLIL | ASSDGQGNNQAPL | 0.7621386 |
| AVNGPVLIRH | ASSDGQGNNQAPL | 0.8133398 |
| DGAAGAVGI | ASSDGQGNNQAPL | 0.7762289 |
| DPLLHGI | ASSDGQGNNQAPL | 0.7475008 |
| ELGAALGAL | ASSDGQGNNQAPL | 0.699247 |
| ELGRTQAV | ASSDGQGNNQAPL | 0.78966665 |
| FVAGGYLS | ASSDGQGNNQAPL | 0.82636 |
| GASGGAGGAGGW | ASSDGQGNNQAPL | 0.7908653 |
| GGAGAGGDGGAGGI | ASSDGQGNNQAPL | 0.8807472 |
| GTGHNGSQPGMGGNG | ASSDGQGNNQAPL | 0.84437245 |
| GYAAASASAS | ASSDGQGNNQAPL | 0.8389014 |
| HQVDVAAVQQG | ASSDGQGNNQAPL | 0.18779337 |
| IGAAIAHT | ASSDGQGNNQAPL | 0.45717573 |
| IGMIEVAQL | ASSDGQGNNQAPL | 0.68234026 |
| IRLAPEHTSAMD | ASSDGQGNNQAPL | 0.75031555 |
| IYNVSVL | ASSDGQGNNQAPL | 0.31759837 |
| KQGFLGM | ASSDGQGNNQAPL | 0.80890584 |
| LESVITQL | ASSDGQGNNQAPL | 0.698274 |
| LGLGALVVV | ASSDGQGNNQAPL | 0.64451635 |
| LIGNGGNGGSGGI | ASSDGQGNNQAPL | 0.8494513 |
| LNAINAPTL | ASSDGQGNNQAPL | 0.7485274 |
| LQLLNTGV | ASSDGQGNNQAPL | 0.6712485 |

| | | |
|---|---|---|
| LQTSIEF | ASSDGQGNNQAPL | 0.66189814 |
| MSFAVSEL | ASSDGQGNNQAPL | 0.8698656 |
| NTGALGL | ASSDGQGNNQAPL | 0.4446714 |
| PPGAGGAGGAGTT | ASSDGQGNNQAPL | 0.8299943 |
| QLVGLGI | ASSDGQGNNQAPL | 0.77099055 |
| RFLEQVHQL | ASSDGQGNNQAPL | 0.85418713 |
| RGVGYVLR | ASSDGQGNNQAPL | 0.48664123 |
| SATGVFAGVI | ASSDGQGNNQAPL | 0.835102 |
| SLPAITLP | ASSDGQGNNQAPL | 0.7509128 |
| STALAFTV | ASSDGQGNNQAPL | 0.8896071 |
| SVLTVGLGK | ASSDGQGNNQAPL | 0.7002239 |
| TALAGAGSGI | ASSDGQGNNQAPL | 0.77221775 |
| TGLPFTGI | ASSDGQGNNQAPL | 0.7171755 |
| VAEHVTV | ASSDGQGNNQAPL | 0.85320604 |
| VDIVLNSL | ASSDGQGNNQAPL | 0.50080585 |
| VLTGGGAVL | ASSDGQGNNQAPL | 0.7117144 |
| VNVGLAYGI | ASSDGQGNNQAPL | 0.5575244 |
| VSSAAVGAV | ASSDGQGNNQAPL | 0.7921951 |
| VTSVYEAGV | ASSDGQGNNQAPL | 0.8184364 |
| WVALDRGA | ASSDGQGNNQAPL | 0.41211742 |
| YLVGSVVI | ASSDGQGNNQAPL | 0.8589593 |
| GPPGTGKTLL | ASSDGQGNNQAPL | 0.7582171 |
| GVGMTKF | ASSDGQGNNQAPL | 0.6528851 |
| IGIGGSDL | ASSDGQGNNQAPL | 0.8543893 |
| IGTLLAEL | ASSDGQGNNQAPL | 0.6674595 |
| IYTSGTT | ASSDGQGNNQAPL | 0.84911776 |
| LSFPAGV | ASSDGQGNNQAPL | 0.72187793 |
| LYTSGTT | ASSDGQGNNQAPL | 0.757761 |
| VGAFPFGF | ASSDGQGNNQAPL | 0.8381802 |
| VGAGTVEF | ASSDGQGNNQAPL | 0.5208273 |
| VGKSTLLL | ASSDGQGNNQAPL | 0.5635699 |
| AALAHAGGGIV | ASSDGQGNNQAPL | 0.8624867 |
| AALVAVSLL | ASSDGQGNNQAPL | 0.75085247 |
| AAVSRAVAAL | ASSDGQGNNQAPL | 0.772689 |
| AGGSSGIGGF | ASSDGQGNNQAPL | 0.7068511 |
| AGLTVANANG | ASSDGQGNNQAPL | 0.639616 |
| AGLVGAVL | ASSDGQGNNQAPL | 0.6846918 |
| ALSQAFTV | ASSDGQGNNQAPL | 0.7639737 |
| ALTFSVSL | ASSDGQGNNQAPL | 0.77323985 |
| ALTGAALVAL | ASSDGQGNNQAPL | 0.75447875 |
| ASFSIGAGVL | ASSDGQGNNQAPL | 0.508932 |
| DGSVYKI | ASSDGQGNNQAPL | 0.7550856 |
| EQLAEIV | ASSDGQGNNQAPL | 0.6806916 |
| FAVSELTVM | ASSDGQGNNQAPL | 0.6917294 |
| GAAPLAAGL | ASSDGQGNNQAPL | 0.7447926 |
| GADGGDGGSGGAGG | ASSDGQGNNQAPL | 0.666558 |
| GFGLDTGGAGG | ASSDGQGNNQAPL | 0.6953546 |
| GLASGGLL | ASSDGQGNNQAPL | 0.6827419 |
| GRIAGAAVLL | ASSDGQGNNQAPL | 0.7288052 |
| GSAGSLNTSF | ASSDGQGNNQAPL | 0.7593364 |
| GTGQIGI | ASSDGQGNNQAPL | 0.74521226 |

| | | |
|---|---|---|
| GTLIALIL | ASSDGQGNNQAPL | 0.77118266 |
| IAATMGFVVG | ASSDGQGNNQAPL | 0.76281786 |
| IFAQALL | ASSDGQGNNQAPL | 0.7609161 |
| IGAAVALGF | ASSDGQGNNQAPL | 0.23339729 |
| IGLTPLAL | ASSDGQGNNQAPL | 0.5314555 |
| ILGSNTSTL | ASSDGQGNNQAPL | 0.6073122 |
| IVFSLTVL | ASSDGQGNNQAPL | 0.7320056 |
| LASKATGF | ASSDGQGNNQAPL | 0.32672995 |
| LASVALIL | ASSDGQGNNQAPL | 0.74126345 |
| LFAVGLTL | ASSDGQGNNQAPL | 0.7116256 |
| LFNGGIGTYI | ASSDGQGNNQAPL | 0.66436446 |
| LTLGAGII | ASSDGQGNNQAPL | 0.769773 |
| LVAAALGDI | ASSDGQGNNQAPL | 0.6977955 |
| NTAFVQLGI | ASSDGQGNNQAPL | 0.5628216 |
| QLFSGAEL | ASSDGQGNNQAPL | 0.66155255 |
| SAATFITL | ASSDGQGNNQAPL | 0.7680719 |
| SNYGLANLGV | ASSDGQGNNQAPL | 0.49089193 |
| SYGGAYAVM | ASSDGQGNNQAPL | 0.7620525 |
| TTGVGGLTLG | ASSDGQGNNQAPL | 0.6401764 |
| TTQTIYI | ASSDGQGNNQAPL | 0.7526778 |
| VAAGAGLGAVI | ASSDGQGNNQAPL | 0.38679472 |
| VAAVVTLL | ASSDGQGNNQAPL | 0.7594081 |
| VGFLGGLI | ASSDGQGNNQAPL | 0.71924245 |
| VGVIPILL | ASSDGQGNNQAPL | 0.7798502 |
| VIFASSLI | ASSDGQGNNQAPL | 0.68453765 |
| VKGGDGGAAAG | ASSDGQGNNQAPL | 0.72809696 |
| VTKTFTNI | ASSDGQGNNQAPL | 0.6616129 |
| VVAAAILL | ASSDGQGNNQAPL | 0.7712376 |
| YAALDVEL | ASSDGQGNNQAPL | 0.47718197 |
| LASVALLL | ASSDGQGNNQAPL | 0.66080904 |
| LDAVHRV | ASSDGQGNNQAPL | 0.6051638 |
| QGLSSSIF | ASSDGQGNNQAPL | 0.7359276 |
| RGTFANIRL | ASSDGQGNNQAPL | 0.7560259 |
| RQLWWGHRI | ASSDGQGNNQAPL | 0.5213349 |
| YGPPGTGKTLL | ASSDGQGNNQAPL | 0.7710884 |
| AALVGAPANVA | ASSDGQGNNQAPL | 0.70996106 |
| AATFRRTE | ASSDGQGNNQAPL | 0.5624713 |
| AGALGGGI | ASSDGQGNNQAPL | 0.7596133 |
| AGGAGGAGGIGAG | ASSDGQGNNQAPL | 0.67494357 |
| AGSGALGL | ASSDGQGNNQAPL | 0.7619244 |
| AQIADQI | ASSDGQGNNQAPL | 0.6417955 |
| ASAVAVTYL | ASSDGQGNNQAPL | 0.72950625 |
| ASTVLGGTGG | ASSDGQGNNQAPL | 0.7523885 |
| AVGGEPAK | ASSDGQGNNQAPL | 0.5545542 |
| CEQGTFGGTC | ASSDGQGNNQAPL | 0.58543754 |
| DEGKAPAG | ASSDGQGNNQAPL | 0.766796 |
| DTVVFPAL | ASSDGQGNNQAPL | 0.72220004 |
| FEGITMAG | ASSDGQGNNQAPL | 0.7529223 |
| FPLVVQD | ASSDGQGNNQAPL | 0.6420103 |
| GAFTTPQ | ASSDGQGNNQAPL | 0.57314 |
| GFGGFGTAGGVGGA | ASSDGQGNNQAPL | 0.6375551 |

| | | |
|---|---|---|
| GFLAQFL | ASSDGQGNNQAPL | 0.7384808 |
| GGAGGAGGNAGVA | ASSDGQGNNQAPL | 0.7638352 |
| GGVGTAGAGGAGGAGG | ASSDGQGNNQAPL | 0.73810863 |
| GNGGGGDAGNAGSGGN | ASSDGQGNNQAPL | 0.5858998 |
| GTGGAGGAGNDAGS | ASSDGQGNNQAPL | 0.7281915 |
| GTTNTGWFNS | ASSDGQGNNQAPL | 0.6937406 |
| IFMGPLFTL | ASSDGQGNNQAPL | 0.69267803 |
| ILAHAIEAI | ASSDGQGNNQAPL | 0.696538 |
| IRSAIREAL | ASSDGQGNNQAPL | 0.70522 |
| ITFAGGG | ASSDGQGNNQAPL | 0.6455718 |
| ITGMSGHQEV | ASSDGQGNNQAPL | 0.58713126 |
| IVAGLACTL | ASSDGQGNNQAPL | 0.69541943 |
| IVGGGLAGA | ASSDGQGNNQAPL | 0.75137484 |
| IVTGVLNI | ASSDGQGNNQAPL | 0.7433883 |
| LAQQGCI | ASSDGQGNNQAPL | 0.772017 |
| LCLHVEN | ASSDGQGNNQAPL | 0.754112 |
| LDAISDGS | ASSDGQGNNQAPL | 0.7269002 |
| LGGFAAML | ASSDGQGNNQAPL | 0.31276008 |
| LGGKNPMI | ASSDGQGNNQAPL | 0.45981556 |
| LGPKGVAAIL | ASSDGQGNNQAPL | 0.60682327 |
| LMVGVSI | ASSDGQGNNQAPL | 0.6769933 |
| LVGAAFVL | ASSDGQGNNQAPL | 0.31066677 |
| LVLAVAPL | ASSDGQGNNQAPL | 0.71079624 |
| MAAIAVIDAA | ASSDGQGNNQAPL | 0.6687508 |
| NNGINAELQ | ASSDGQGNNQAPL | 0.594658 |
| RGEPLVLVHG | ASSDGQGNNQAPL | 0.748471 |
| RGGAGGAAGLI | ASSDGQGNNQAPL | 0.65775746 |
| RSYRMHPAV | ASSDGQGNNQAPL | 0.56201756 |
| SAAHLLAGQLA | ASSDGQGNNQAPL | 0.59523976 |
| SAAVGSY | ASSDGQGNNQAPL | 0.76680744 |
| SPHAVLHRI | ASSDGQGNNQAPL | 0.53608954 |
| SQIAKVL | ASSDGQGNNQAPL | 0.74361414 |
| TALGHLVP | ASSDGQGNNQAPL | 0.6228597 |
| TAPTTSGAA | ASSDGQGNNQAPL | 0.7600846 |
| TDGDDGNGGSPGT | ASSDGQGNNQAPL | 0.47916168 |
| TDGLGEN | ASSDGQGNNQAPL | 0.74930894 |
| TGGPPLPGQA | ASSDGQGNNQAPL | 0.6894096 |
| TIPAITI | ASSDGQGNNQAPL | 0.78237057 |
| TIPGFTI | ASSDGQGNNQAPL | 0.644651 |
| TSIASGALA | ASSDGQGNNQAPL | 0.7235102 |
| TVTLPGAAI | ASSDGQGNNQAPL | 0.6011426 |
| VASGLSAALP | ASSDGQGNNQAPL | 0.7419224 |
| VGAGVIGI | ASSDGQGNNQAPL | 0.4835814 |
| VGPKRFGV | ASSDGQGNNQAPL | 0.6587108 |
| VNVAFPD | ASSDGQGNNQAPL | 0.57804775 |
| VSMTNSGVS | ASSDGQGNNQAPL | 0.70353425 |
| VTDVLGDLSV | ASSDGQGNNQAPL | 0.7422267 |
| VVVVGAGFAGL | ASSDGQGNNQAPL | 0.35169432 |
| VYTVGTVV | ASSDGQGNNQAPL | 0.74976295 |
| WTTFSGP | ASSDGQGNNQAPL | 0.6203856 |
| YDELVEAF | ASSDGQGNNQAPL | 0.5538183 |

| | | |
|---|---|---|
| YGTGGAGGNGG | ASSDGQGNNQAPL | 0.7480775 |
| YTNVMAV | ASSDGQGNNQAPL | 0.67647576 |
| FLEQVHQL | ASSDGQGNNQAPL | 0.7477108 |
| GLSLLAVAV | ASSDGQGNNQAPL | 0.57776535 |
| LEQVHQL | ASSDGQGNNQAPL | 0.70013106 |
| LNAAGAI | ASSDGQGNNQAPL | 0.7343972 |
| PAVHASL | ASSDGQGNNQAPL | 0.53204423 |
| TELEDDA | ASSDGQGNNQAPL | 0.5470457 |
| TPPGPPN | ASSDGQGNNQAPL | 0.7655481 |
| TPPPPPPP | ASSDGQGNNQAPL | 0.6945363 |
| TQSPALI | ASSLWDWGYEQY | 0.7310566 |
| EFAAAFPQ | ASSLWDWGYEQY | 0.6280527 |
| LLSGNLDVL | ASSLWDWGYEQY | 0.6386279 |
| GPANLGNYNI | ASSLWDWGYEQY | 0.6308051 |
| TLGPTLP | ASSLWDWGYEQY | 0.718351 |
| VYAYSSCCK | ASSLWDWGYEQY | 0.7267865 |
| APAPAPAPNI | ASSLWDWGYEQY | 0.71153414 |
| PGGPVPIP | ASSLWDWGYEQY | 0.53644615 |
| FLTVALEP | ASSLWDWGYEQY | 0.6974236 |
| SQYFGANL | ASSLWDWGYEQY | 0.6392648 |
| GGVKPAA | ASSLWDWGYEQY | 0.66564566 |
| GGTGGAGGAGGW | ASSLWDWGYEQY | 0.69296384 |
| GVGGVAGLGGAGG | ASSLWDWGYEQY | 0.64490515 |
| VLLVLATTI | ASSLWDWGYEQY | 0.6423137 |
| IGIFAAML | ASSLWDWGYEQY | 0.61670834 |
| AVGDLID | ASSLWDWGYEQY | 0.66401374 |
| NISPLPL | ASSLWDWGYEQY | 0.70770645 |
| VTAVFVAI | ASSLWDWGYEQY | 0.69847125 |
| SGDDNDAAL | ASSLWDWGYEQY | 0.7316128 |
| AVAALAHS | ASSLWDWGYEQY | 0.7194495 |
| GGVTVMATP | ASSLWDWGYEQY | 0.68448406 |
| AGLAANVRD | ASSLWDWGYEQY | 0.30949056 |
| TNRPDLIDPALLRPG | ASSLWDWGYEQY | 0.5538776 |
| LANLGVTAL | ASSLWDWGYEQY | 0.591636 |
| TGERIEL | ASSLWDWGYEQY | 0.6930741 |
| NGEELAA | ASSLWDWGYEQY | 0.33072057 |
| VLSLVWG | ASSLWDWGYEQY | 0.69445074 |
| IGYALWL | ASSLWDWGYEQY | 0.66769457 |
| LSLLLAPL | ASSLWDWGYEQY | 0.6398132 |
| TLDEALK | ASSLWDWGYEQY | 0.720209 |
| TDGAGTAEGG | ASSLWDWGYEQY | 0.6603973 |
| NAPGGAGGNGG | ASSLWDWGYEQY | 0.5732454 |
| VSVFMGMGGNF | ASSLWDWGYEQY | 0.4901518 |
| LASYLAAL | ASSLWDWGYEQY | 0.707001 |
| LVGGTPAL | ASSLWDWGYEQY | 0.46871096 |
| VRNADVL | ASSLWDWGYEQY | 0.715017 |
| TCSSPFLVSAM | ASSLWDWGYEQY | 0.49469894 |
| VHITDAE | ASSLWDWGYEQY | 0.7167212 |
| EYAAPSF | ASSLWDWGYEQY | 0.47851276 |
| VVDAFHGGL | ASSLWDWGYEQY | 0.7122867 |
| ASVVDPQ | ASSLWDWGYEQY | 0.66419846 |

| | | |
|---|---|---|
| SYNEFGGLS | ASSLWDWGYEQY | 0.7427433 |
| DPDGSVY | ASSLWDWGYEQY | 0.6588596 |
| MVSTIHATE | ASSLWDWGYEQY | 0.67073834 |
| LAAGVADPV | ASSLWDWGYEQY | 0.5390314 |
| TKEQIAA | ASSLWDWGYEQY | 0.72442055 |
| SLASPAAL | ASSLWDWGYEQY | 0.5260877 |
| IGLVFYGV | ASSLWDWGYEQY | 0.6215647 |
| FLLDASNL | ASSLWDWGYEQY | 0.49651557 |
| YQVGNFP | ASSLWDWGYEQY | 0.6848678 |
| ESLLSQST | ASSLWDWGYEQY | 0.7114836 |
| LFTVGFAAVG | ASSLWDWGYEQY | 0.62555444 |
| ATPPTPT | ASSLWDWGYEQY | 0.71301246 |
| TDLATLG | ASSLWDWGYEQY | 0.66253847 |
| SAGGTGR | ASSLWDWGYEQY | 0.5740439 |
| GFSNSGS | ASSLWDWGYEQY | 0.71832114 |
| GGNGGSGGTSV | ASSLWDWGYEQY | 0.6060953 |
| GTSVATGGAGN | ASSLWDWGYEQY | 0.71393186 |
| NGGNGGSGGTS | ASSLWDWGYEQY | 0.53432536 |
| AVVKFPH | ASSLWDWGYEQY | 0.6555902 |
| TPPPPPP | ASSLWDWGYEQY | 0.6959622 |
| AAPAAAAAPA | ASSLWDWGYEQY | 0.63286734 |
| IALNEDL | ASSLWDWGYEQY | 0.5327657 |
| IVNVSSM | ASSLWDWGYEQY | 0.722902 |
| TGIPPAPR | ASSLWDWGYEQY | 0.8904679 |
| FDLGGGTF | ASSLWDWGYEQY | 0.88278544 |
| FDLGGGTFDVS | ASSLWDWGYEQY | 0.79729724 |
| NEPTAAA | ASSLWDWGYEQY | 0.8044902 |
| NEPTAAALA | ASSLWDWGYEQY | 0.84560204 |
| GSIINISSI | ASSLWDWGYEQY | 0.8977016 |
| VAPGNAGTA | ASSLWDWGYEQY | 0.90679336 |
| TGGGASLE | ASSLWDWGYEQY | 0.8957406 |
| VAVYNHYKRI | ASSLWDWGYEQY | 0.8411892 |
| TDPAPATT | ASSLWDWGYEQY | 0.8999605 |
| TNRPDILDPALLRPG | ASSLWDWGYEQY | 0.87569433 |
| LTTTTTS | ASSLWDWGYEQY | 0.86683136 |
| PTGSGKTL | ASSLWDWGYEQY | 0.8988507 |
| TGSGKTL | ASSLWDWGYEQY | 0.8859799 |
| TIGHVDHGK | ASSLWDWGYEQY | 0.83954203 |
| TIGHVDHGKTT | ASSLWDWGYEQY | 0.85493875 |
| GHVDHGKTT | ASSLWDWGYEQY | 0.8869153 |
| AQFEHTLL | ASSLWDWGYEQY | 0.89639175 |
| IVNQDEL | ASSLWDWGYEQY | 0.89129555 |
| TNGSQFF | ASSLWDWGYEQY | 0.9138197 |
| TGKGIVE | ASSLWDWGYEQY | 0.90026534 |
| LISNASDA | ASSLWDWGYEQY | 0.8855715 |
| AELRTLGELTKIAKAHGAQ | ASSLWDWGYEQY | 0.5614169 |
| APNAVAAAEPSW | ASSLWDWGYEQY | 0.7437563 |
| APQVAGAATAVL | ASSLWDWGYEQY | 0.8735 |
| ARRDPAAAAAILPTDA | ASSLWDWGYEQY | 0.84623873 |
| DAAARDVLAEAGLAEYF | ASSLWDWGYEQY | 0.54093105 |
| DDQIVLSAKANLAKVEQ | ASSLWDWGYEQY | 0.896844 |

| | | |
|---|---|---|
| DELLPDTGEDSDEDGH | ASSLWDWGYEQY | 0.87226033 |
| ELPEAPANRATVPQD | ASSLWDWGYEQY | 0.8323202 |
| FGQSKIERVTGIG | ASSLWDWGYEQY | 0.8997642 |
| GAGGTGSTAGGAGGAGGAGGI | ASSLWDWGYEQY | 0.8575971 |
| GHGGTGAAGGNGGTGGAGGS( | ASSLWDWGYEQY | 0.8426982 |
| GNGGNGGAGGEGGPGGAGGA( | ASSLWDWGYEQY | 0.8798366 |
| GSTAGNGGNGGNATNGTIAG( | ASSLWDWGYEQY | 0.9109698 |
| GTGGDGGNGGNGGTGG | ASSLWDWGYEQY | 0.8478985 |
| GVRRPAGIPADAPV | ASSLWDWGYEQY | 0.9050166 |
| GYRQVTDAHLLTLARR | ASSLWDWGYEQY | 0.8626131 |
| IAKMGQDGHDRG | ASSLWDWGYEQY | 0.90925264 |
| KAEPSIEIQYT | ASSLWDWGYEQY | 0.80037904 |
| KPVRIRMAADEDTVAVT | ASSLWDWGYEQY | 0.90138805 |
| MPPRIAGMRLLVIKPEP | ASSLWDWGYEQY | 0.8709369 |
| MKFYAVALFRSRGVPPTR | ASSLWDWGYEQY | 0.9164759 |
| NAGGFGILGAVAHSP | ASSLWDWGYEQY | 0.8692533 |
| NEGREITLAMAEM | ASSLWDWGYEQY | 0.9062683 |
| NELVEWSHPLGHRWRWE | ASSLWDWGYEQY | 0.88073224 |
| NLGAGGTGGNGGNPG | ASSLWDWGYEQY | 0.88904065 |
| NTEWPKWETVMAADPVD | ASSLWDWGYEQY | 0.77484167 |
| PASAAPAETQP | ASSLWDWGYEQY | 0.85977846 |
| QQSSDGKNGVRAAEITGE | ASSLWDWGYEQY | 0.8040397 |
| RAPDAVTTSWWKE | ASSLWDWGYEQY | 0.88035816 |
| SGFGNVGAHTSGWFNQ | ASSLWDWGYEQY | 0.8952243 |
| SGVAGAVGQAASVGGLKV | ASSLWDWGYEQY | 0.42044163 |
| SNGGAGTGSAGGNGGT | ASSLWDWGYEQY | 0.9068824 |
| TGGAGGEGGAAG | ASSLWDWGYEQY | 0.7583503 |
| TGGDGGAAGAGGAPGHG | ASSLWDWGYEQY | 0.83397675 |
| TMPPLLAAIANAGRHGV | ASSLWDWGYEQY | 0.9049634 |
| TSSTAGNSGGA | ASSLWDWGYEQY | 0.87406594 |
| VPATVPVGILC | ASSLWDWGYEQY | 0.8970165 |
| VQRPDSGGVVLTGRIS | ASSLWDWGYEQY | 0.86615056 |
| WTPNPVPATKGVPAGLVQAT | ASSLWDWGYEQY | 0.8722886 |
| YALKRALAKFSDRFDVVII | ASSLWDWGYEQY | 0.88647515 |
| YLIGSVIV | ASSLWDWGYEQY | 0.58935547 |
| YGQLVAVID | ASSLWDWGYEQY | 0.83630776 |
| VTVGGMTLPA | ASSLWDWGYEQY | 0.91145825 |
| VLALPAV | ASSLWDWGYEQY | 0.6760603 |
| VIPPTLNYET | ASSLWDWGYEQY | 0.6860228 |
| VGMGGMHPSSG | ASSLWDWGYEQY | 0.65104425 |
| VGLAGGAATA | ASSLWDWGYEQY | 0.6308166 |
| VGGAGGAGGSGP | ASSLWDWGYEQY | 0.65179163 |
| VFAAFMV | ASSLWDWGYEQY | 0.6793834 |
| TLYDKLR | ASSLWDWGYEQY | 0.68568116 |
| TLAVFTVL | ASSLWDWGYEQY | 0.674351 |
| TGSVQPARV | ASSLWDWGYEQY | 0.6407708 |
| TGGGAGSGGT | ASSLWDWGYEQY | 0.67229795 |
| SHLETSL | ASSLWDWGYEQY | 0.66658044 |
| SFTTGMT | ASSLWDWGYEQY | 0.65684354 |
| QVEVAKRMGVRQPTVS | ASSLWDWGYEQY | 0.675866 |
| QGSGGPALAM | ASSLWDWGYEQY | 0.67157346 |

| | | |
|---|---|---|
| MLAPSNP | ASSLWDWGYEQY | 0.64439535 |
| LVNNAAI | ASSLWDWGYEQY | 0.6615309 |
| LTGEINTLA | ASSLWDWGYEQY | 0.66296023 |
| LSIFGPV | ASSLWDWGYEQY | 0.6753479 |
| LLALLANL | ASSLWDWGYEQY | 0.6729543 |
| LISNASDALDKLR | ASSLWDWGYEQY | 0.68780404 |
| LINEAVP | ASSLWDWGYEQY | 0.6811111 |
| LGYAFSGL | ASSLWDWGYEQY | 0.67880964 |
| LDFVAVAL | ASSLWDWGYEQY | 0.47735396 |
| LAQPGAE | ASSLWDWGYEQY | 0.608721 |
| ISNLGAVTGF | ASSLWDWGYEQY | 0.6596132 |
| HQEDPGRI | ASSLWDWGYEQY | 0.6636882 |
| GTPGEPTP | ASSLWDWGYEQY | 0.54118216 |
| GFFNSGS | ASSLWDWGYEQY | 0.6771153 |
| GAGGGGLTAGAN | ASSLWDWGYEQY | 0.65883374 |
| FVDALVNM | ASSLWDWGYEQY | 0.6359434 |
| FLTVAPEL | ASSLWDWGYEQY | 0.6782901 |
| FETLTAL | ASSLWDWGYEQY | 0.65453815 |
| DVINPIEL | ASSLWDWGYEQY | 0.65130514 |
| APVSGAPV | ASSLWDWGYEQY | 0.66330457 |
| ALVGGLAW | ASSLWDWGYEQY | 0.6831871 |
| ALAAAAATAI | ASSLWDWGYEQY | 0.6277636 |
| AFGLGLLL | ASSLWDWGYEQY | 0.6804807 |
| AFFAAASL | ASSLWDWGYEQY | 0.646799 |
| AAAALPAFSPP | ASSLWDWGYEQY | 0.68446904 |
| VVLAPAL | ASSLWDWGYEQY | 0.61669093 |
| TIGHVDHGKTTL | ASSLWDWGYEQY | 0.6785097 |
| LISNASDALDK | ASSLWDWGYEQY | 0.665742 |
| LISNASDAL | ASSLWDWGYEQY | 0.6896681 |
| ISNASDALDK | ASSLWDWGYEQY | 0.66209376 |
| NQAETLVYQTE | ASSLWDWGYEQY | 0.68011975 |
| TGGTGGTAGAG | ASSLWDWGYEQY | 0.672983 |
| TGSTAGGAGGA | ASSLWDWGYEQY | 0.68473107 |
| DPLAVPVAAGPV | ASSLWDWGYEQY | 0.61807895 |
| GGTGGTPGNGGN | ASSLWDWGYEQY | 0.6686282 |
| PAGLAGVDPATL | ASSLWDWGYEQY | 0.60366476 |
| GNNLVVEAPHAGASV | ASSLWDWGYEQY | 0.67381346 |
| KVTRIMSASTLEQPA | ASSLWDWGYEQY | 0.67661625 |
| LGFALPIALPRRTVV | ASSLWDWGYEQY | 0.5987184 |
| TRLGNLTARKLVVLV | ASSLWDWGYEQY | 0.6848823 |
| WSQQRRLIYFPSAGP | ASSLWDWGYEQY | 0.6574013 |
| GPNGNPDPMAAAVDIR | ASSLWDWGYEQY | 0.64974564 |
| TQRCVSVADLRAAASR | ASSLWDWGYEQY | 0.6832071 |
| QFVLDVAEHPVEVLTETV | ASSLWDWGYEQY | 0.66075736 |
| GGAGGTGGAGGNGGTQAGDG | ASSLWDWGYEQY | 0.67878485 |
| PLLATLRAIAVDVDAKPAQV | ASSLWDWGYEQY | 0.65694964 |
| APKFRTGGYMIDFWGVGYQV | ASSLWDWGYEQY | 0.666664 |
| GAGGAGGAGGAAGGPGGTGG | ASSLWDWGYEQY | 0.67387676 |
| IGALGLLATDVAPAGSVAFG | ASSLWDWGYEQY | 0.611378 |
| AGGAGGIGGNGGAGGAGGVL | ASSLWDWGYEQY | 0.63245976 |
| ADELIGLL | ASSLWDWGYEQY | 0.68431526 |

| | | |
|---|---|---|
| ADRALKA | ASSLWDWGYEQY | 0.7291608 |
| AGAVGQVGGAGGA | ASSLWDWGYEQY | 0.7251965 |
| AGGGGGLGGAP | ASSLWDWGYEQY | 0.6829567 |
| AGGLLYGNGGAG | ASSLWDWGYEQY | 0.668116 |
| AGLESIGL | ASSLWDWGYEQY | 0.69904035 |
| AGLKTGII | ASSLWDWGYEQY | 0.74269366 |
| ASLAALLL | ASSLWDWGYEQY | 0.7320564 |
| ASVINISSI | ASSLWDWGYEQY | 0.7327266 |
| ATPPGPA | ASSLWDWGYEQY | 0.6549307 |
| ATVASALV | ASSLWDWGYEQY | 0.7425878 |
| ATVASLIL | ASSLWDWGYEQY | 0.71875536 |
| AVNGPVLIRH | ASSLWDWGYEQY | 0.71864265 |
| DGAAGAVGI | ASSLWDWGYEQY | 0.7424201 |
| DPLLHGI | ASSLWDWGYEQY | 0.7133093 |
| ELGAALGAL | ASSLWDWGYEQY | 0.70459574 |
| ELGRTQAV | ASSLWDWGYEQY | 0.713603 |
| FVAGGYLS | ASSLWDWGYEQY | 0.71057034 |
| GASGGAGGAGGW | ASSLWDWGYEQY | 0.72629076 |
| GGAGAGGDGGAGGI | ASSLWDWGYEQY | 0.7188105 |
| GTGHNGSQPGMGGNG | ASSLWDWGYEQY | 0.7549001 |
| GYAAASASAS | ASSLWDWGYEQY | 0.74354863 |
| HQVDVAAVQQG | ASSLWDWGYEQY | 0.7309449 |
| IGAAIAHT | ASSLWDWGYEQY | 0.36251527 |
| IGMIEVAQL | ASSLWDWGYEQY | 0.67261547 |
| IRLAPEHTSAMD | ASSLWDWGYEQY | 0.7115122 |
| IYNVSVL | ASSLWDWGYEQY | 0.7256297 |
| KQGFLGM | ASSLWDWGYEQY | 0.48487407 |
| LESVITQL | ASSLWDWGYEQY | 0.7348596 |
| LGLGALVVV | ASSLWDWGYEQY | 0.71107507 |
| LIGNGGNGGSGGI | ASSLWDWGYEQY | 0.6960273 |
| LNAINAPTL | ASSLWDWGYEQY | 0.74917233 |
| LQLLNTGV | ASSLWDWGYEQY | 0.69949305 |
| LQTSIEF | ASSLWDWGYEQY | 0.7080468 |
| MSFAVSEL | ASSLWDWGYEQY | 0.70044976 |
| NTGALGL | ASSLWDWGYEQY | 0.75589323 |
| PPGAGGAGGAGTT | ASSLWDWGYEQY | 0.68547714 |
| QLVGLGI | ASSLWDWGYEQY | 0.74818283 |
| RFLEQVHQL | ASSLWDWGYEQY | 0.71178967 |
| RGVGYVLR | ASSLWDWGYEQY | 0.7514921 |
| SATGVFAGVI | ASSLWDWGYEQY | 0.6209769 |
| SLPAITLP | ASSLWDWGYEQY | 0.7414462 |
| STALAFTV | ASSLWDWGYEQY | 0.7112774 |
| SVLTVGLGK | ASSLWDWGYEQY | 0.7553241 |
| TALAGAGSGI | ASSLWDWGYEQY | 0.7183751 |
| TGLPFTGI | ASSLWDWGYEQY | 0.73825234 |
| VAEHVTV | ASSLWDWGYEQY | 0.70349395 |
| VDIVLNSL | ASSLWDWGYEQY | 0.73537564 |
| VLTGGGAVL | ASSLWDWGYEQY | 0.6330972 |
| VNVGLAYGI | ASSLWDWGYEQY | 0.70377606 |
| VSSAAVGAV | ASSLWDWGYEQY | 0.6623784 |
| VTSVYEAGV | ASSLWDWGYEQY | 0.72873783 |

| | | |
|---|---|---|
| WVALDRGA | ASSLWDWGYEQY | 0.73502004 |
| YLVGSVVI | ASSLWDWGYEQY | 0.5489893 |
| GPPGTGKTLL | ASSLWDWGYEQY | 0.74673915 |
| GVGMTKF | ASSLWDWGYEQY | 0.65681684 |
| IGIGGSDL | ASSLWDWGYEQY | 0.6842575 |
| IGTLLAEL | ASSLWDWGYEQY | 0.74362385 |
| IYTSGTT | ASSLWDWGYEQY | 0.71040493 |
| LSFPAGV | ASSLWDWGYEQY | 0.7384902 |
| LYTSGTT | ASSLWDWGYEQY | 0.69652414 |
| VGAFPFGF | ASSLWDWGYEQY | 0.7146784 |
| VGAGTVEF | ASSLWDWGYEQY | 0.7348946 |
| VGKSTLLL | ASSLWDWGYEQY | 0.56551546 |
| AALAHAGGGIV | ASSLWDWGYEQY | 0.6748334 |
| AALVAVSLL | ASSLWDWGYEQY | 0.7486707 |
| AAVSRAVAAL | ASSLWDWGYEQY | 0.58272266 |
| AGGSSGIGGF | ASSLWDWGYEQY | 0.67921287 |
| AGLTVANANG | ASSLWDWGYEQY | 0.4681733 |
| AGLVGAVL | ASSLWDWGYEQY | 0.44979873 |
| ALSQAFTV | ASSLWDWGYEQY | 0.5465036 |
| ALTFSVSL | ASSLWDWGYEQY | 0.6647162 |
| ALTGAALVAL | ASSLWDWGYEQY | 0.6560343 |
| ASFSIGAGVL | ASSLWDWGYEQY | 0.6175797 |
| DGSVYKI | ASSLWDWGYEQY | 0.4775219 |
| EQLAEIV | ASSLWDWGYEQY | 0.53317386 |
| FAVSELTVM | ASSLWDWGYEQY | 0.49987906 |
| GAAPLAAGL | ASSLWDWGYEQY | 0.5424735 |
| GADGGDGGSGGAGG | ASSLWDWGYEQY | 0.6271751 |
| GFGLDTGGAGG | ASSLWDWGYEQY | 0.5447812 |
| GLASGGLL | ASSLWDWGYEQY | 0.49100322 |
| GRIAGAAVLL | ASSLWDWGYEQY | 0.52802414 |
| GSAGSLNTSF | ASSLWDWGYEQY | 0.60243213 |
| GTGQIGI | ASSLWDWGYEQY | 0.5891412 |
| GTLIALIL | ASSLWDWGYEQY | 0.57910395 |
| IAATMGFVVG | ASSLWDWGYEQY | 0.6840191 |
| IFAQALL | ASSLWDWGYEQY | 0.62216103 |
| IGAAVALGF | ASSLWDWGYEQY | 0.63763857 |
| IGLTPLAL | ASSLWDWGYEQY | 0.31805456 |
| ILGSNTSTL | ASSLWDWGYEQY | 0.37198442 |
| IVFSLTVL | ASSLWDWGYEQY | 0.5684751 |
| LASKATGF | ASSLWDWGYEQY | 0.54276955 |
| LASVALIL | ASSLWDWGYEQY | 0.42379686 |
| LFAVGLTL | ASSLWDWGYEQY | 0.6295377 |
| LFNGGIGTYI | ASSLWDWGYEQY | 0.5229564 |
| LTLGAGII | ASSLWDWGYEQY | 0.47206196 |
| LVAAALGDI | ASSLWDWGYEQY | 0.6209562 |
| NTAFVQLGI | ASSLWDWGYEQY | 0.49253973 |
| QLFSGAEL | ASSLWDWGYEQY | 0.4952452 |
| SAATFITL | ASSLWDWGYEQY | 0.5393616 |
| SNYGLANLGV | ASSLWDWGYEQY | 0.62381184 |
| SYGGAYAVM | ASSLWDWGYEQY | 0.41857514 |
| TTGVGGLTLG | ASSLWDWGYEQY | 0.593746 |

| | | |
|---|---|---|
| TTQTIYI | ASSLWDWGYEQY | 0.5561899 |
| VAAGAGLGAVI | ASSLWDWGYEQY | 0.6476304 |
| VAAVVTLL | ASSLWDWGYEQY | 0.426403 |
| VGFLGGLI | ASSLWDWGYEQY | 0.6142088 |
| VGVIPILL | ASSLWDWGYEQY | 0.54622436 |
| VIFASSLI | ASSLWDWGYEQY | 0.68623304 |
| VKGGDGGAAAG | ASSLWDWGYEQY | 0.50614005 |
| VTKTFTNI | ASSLWDWGYEQY | 0.56857294 |
| VVAAAILL | ASSLWDWGYEQY | 0.56708306 |
| YAALDVEL | ASSLWDWGYEQY | 0.656091 |
| LASVALLL | ASSLWDWGYEQY | 0.42950636 |
| LDAVHRV | ASSLWDWGYEQY | 0.50859934 |
| QGLSSSIF | ASSLWDWGYEQY | 0.41117287 |
| RGTFANIRL | ASSLWDWGYEQY | 0.59643596 |
| RQLWWGHRI | ASSLWDWGYEQY | 0.59528995 |
| YGPPGTGKTLL | ASSLWDWGYEQY | 0.45441422 |
| AALVGAPANVA | ASSLWDWGYEQY | 0.6260751 |
| AATFRRTE | ASSLWDWGYEQY | 0.6168192 |
| AGALGGGI | ASSLWDWGYEQY | 0.5000126 |
| AGGAGGAGGIGAG | ASSLWDWGYEQY | 0.65920097 |
| AGSGALGL | ASSLWDWGYEQY | 0.5411442 |
| AQIADQI | ASSLWDWGYEQY | 0.64308715 |
| ASAVAVTYL | ASSLWDWGYEQY | 0.57786494 |
| ASTVLGGTGG | ASSLWDWGYEQY | 0.49497557 |
| AVGGEPAK | ASSLWDWGYEQY | 0.63173723 |
| CEQGTFGGTC | ASSLWDWGYEQY | 0.5620845 |
| DEGKAPAG | ASSLWDWGYEQY | 0.5103554 |
| DTVVFPAL | ASSLWDWGYEQY | 0.64537245 |
| FEGITMAG | ASSLWDWGYEQY | 0.53448975 |
| FPLVVQD | ASSLWDWGYEQY | 0.6650639 |
| GAFTTPQ | ASSLWDWGYEQY | 0.4131938 |
| GFGGFGTAGGVGGA | ASSLWDWGYEQY | 0.40112102 |
| GFLAQFL | ASSLWDWGYEQY | 0.52647775 |
| GGAGGAGGNAGVA | ASSLWDWGYEQY | 0.64220107 |
| GGVGTAGAGGAGGAGG | ASSLWDWGYEQY | 0.656551 |
| GNGGGGDAGNAGSGGN | ASSLWDWGYEQY | 0.58351296 |
| GTGGAGGAGNDAGS | ASSLWDWGYEQY | 0.4221254 |
| GTTNTGWFNS | ASSLWDWGYEQY | 0.4780531 |
| IFMGPLFTL | ASSLWDWGYEQY | 0.44903195 |
| ILAHAIEAI | ASSLWDWGYEQY | 0.49677497 |
| IRSAIREAL | ASSLWDWGYEQY | 0.5792501 |
| ITFAGGG | ASSLWDWGYEQY | 0.49485344 |
| ITGMSGHQEV | ASSLWDWGYEQY | 0.41697067 |
| IVAGLACTL | ASSLWDWGYEQY | 0.4662541 |
| IVGGGLAGA | ASSLWDWGYEQY | 0.5881052 |
| IVTGVLNI | ASSLWDWGYEQY | 0.55588764 |
| LAQQGCI | ASSLWDWGYEQY | 0.53446484 |
| LCLHVEN | ASSLWDWGYEQY | 0.6651536 |
| LDAISDGS | ASSLWDWGYEQY | 0.5907676 |
| LGGFAAML | ASSLWDWGYEQY | 0.60561717 |
| LGGKNPMI | ASSLWDWGYEQY | 0.29875827 |

| | | |
|---|---|---|
| LGPKGVAAIL | ASSLWDWGYEQY | 0.32920322 |
| LMVGVSI | ASSLWDWGYEQY | 0.47605464 |
| LVGAAFVL | ASSLWDWGYEQY | 0.5029959 |
| LVLAVAPL | ASSLWDWGYEQY | 0.3527145 |
| MAAIAVIDAA | ASSLWDWGYEQY | 0.59957653 |
| NNGINAELQ | ASSLWDWGYEQY | 0.45555276 |
| RGEPLVLVHG | ASSLWDWGYEQY | 0.4154493 |
| RGGAGGAAGLI | ASSLWDWGYEQY | 0.6071199 |
| RSYRMHPAV | ASSLWDWGYEQY | 0.43948567 |
| SAAHLLAGQLA | ASSLWDWGYEQY | 0.42860353 |
| SAAVGSY | ASSLWDWGYEQY | 0.45972323 |
| SPHAVLHRI | ASSLWDWGYEQY | 0.5518865 |
| SQIAKVL | ASSLWDWGYEQY | 0.34767795 |
| TALGHLVP | ASSLWDWGYEQY | 0.55883145 |
| TAPTTSGAA | ASSLWDWGYEQY | 0.46701264 |
| TDGDDGNGGSPGT | ASSLWDWGYEQY | 0.6228748 |
| TDGLGEN | ASSLWDWGYEQY | 0.37116805 |
| TGGPPLPGQA | ASSLWDWGYEQY | 0.5735376 |
| TIPAITI | ASSLWDWGYEQY | 0.5225595 |
| TIPGFTI | ASSLWDWGYEQY | 0.6728622 |
| TSIASGALA | ASSLWDWGYEQY | 0.45333278 |
| TVTLPGAAI | ASSLWDWGYEQY | 0.5316546 |
| VASGLSAALP | ASSLWDWGYEQY | 0.5236849 |
| VGAGVIGI | ASSLWDWGYEQY | 0.6550946 |
| VGPKRFGV | ASSLWDWGYEQY | 0.40088618 |
| VNVAFPD | ASSLWDWGYEQY | 0.4566579 |
| VSMTNSGVS | ASSLWDWGYEQY | 0.3501779 |
| VTDVLGDLSV | ASSLWDWGYEQY | 0.5615404 |
| VVVVGAGFAGL | ASSLWDWGYEQY | 0.5482038 |
| VYTVGTVV | ASSLWDWGYEQY | 0.4104722 |
| WTTFSGP | ASSLWDWGYEQY | 0.5911664 |
| YDELVEAF | ASSLWDWGYEQY | 0.58371353 |
| YGTGGAGGNGG | ASSLWDWGYEQY | 0.4402955 |
| YTNVMAV | ASSLWDWGYEQY | 0.6309664 |
| FLEQVHQL | ASSLWDWGYEQY | 0.49257398 |
| GLSLLAVAV | ASSLWDWGYEQY | 0.6177246 |
| LEQVHQL | ASSLWDWGYEQY | 0.54205316 |
| LNAAGAI | ASSLWDWGYEQY | 0.41719776 |
| PAVHASL | ASSLWDWGYEQY | 0.5853611 |
| TELEDDA | ASSLWDWGYEQY | 0.48673436 |
| TPPGPPN | ASSLWDWGYEQY | 0.45007092 |
| TPPPPPPP | ASSLWDWGYEQY | 0.60737586 |
